# Analysis of the Dynamics and Distribution of SARS-CoV-2 Mutations and its Possible Structural and Functional Implications

**DOI:** 10.1101/2020.11.13.381228

**Authors:** Santiago Justo Arévalo, Daniela Zapata Sifuentes, César Huallpa Robles, Gianfranco Landa Bianchi, Adriana Castillo Chávez, Romina Garavito-Salini Casas, Roberto Pineda Chavarría, Guillermo Uceda-Campos

**Author notes:** These authors contributed equally to this work (Daniela Zapata Sifuentes, César Huallpa Robles, Gianfranco Landa Bianchi, Adriana Castillo Chávez, Romina Garavito-Salini Casas, Roberto Pineda Chavarría).

## Abstract

After eight months of the pandemic declaration, COVID-19 has not been globally controlled. Several efforts to control SARS-CoV-2 dissemination are still running including vaccines and drug treatments. The effectiveness of these procedures depends, in part, that the regions to which these treatments are directed do not vary considerably. Although, it is known that the mutation rate of SARS-CoV-2 is relatively low it is necessary to monitor the adaptation and evolution of the virus in the different stages of the pandemic. Thus, identification, analysis of the dynamics, and possible functional and structural implication of mutations are relevant. Here, we first estimate the number of COVID-19 cases with a virus with a specific mutation and then calculate its global relative frequency (NRFp). Using this approach in a dataset of 100 924 genomes from GISAID, we identified 41 mutations to be present in viruses in an estimated number of 750 000 global COVID-19 cases (0.03 NRFp). We classified these mutations into three groups: high-frequent, low-frequent non-synonymous, and low-frequent synonymous. Analysis of the dynamics of these mutations by month and continent showed that high-frequent mutations appeared early in the pandemic, all are present in all continents and some of them are almost fixed in the global population. On the other hand, low-frequent mutations (non-synonymous and synonymous) appear late in the pandemic and seems to be at least partially continent-specific. This could be due to that high-frequent mutation appeared early when lockdown policies had not yet been applied and low-frequent mutations appeared after lockdown policies. Thus, preventing global dissemination of them. Finally, we present a brief structural and functional review of the analyzed ORFs and the possible implications of the 25 identified non-synonymous mutations.

## INTRODUCTION

COVID-19 cases reached approximately 52 million and 1.3 million deaths (as of November 12th) (WHO. 2020), several countries have reactivated their lockdown policies to control a second-wave of infections (Looi. 2020) showing that the COVID-19 pandemic is not yet controlled. Efforts to develop and license treatments are still running with few of them in the final stages (Krammer. 2020).

In this context, is still important to track the adaptation and evolution of SARS-CoV-2 all around the world. Efforts to sequence genomes are different in each region causing regions with few sequenced genomes to be left out of global analyses. For this reason, we use an approach that takes into account the difference in the number of sequenced genomes in each continent and each month and normalized them by the number of COVID-19 cases resulting in a less biased procedure to determine which are the most frequent and then the most relevant mutation in all around the world.

The study of the dynamics of these relevant mutations and their possible implications in the structure and function of the virus is important to understand the adaptation and evolution process of SARS-CoV-2. Thus, we analyze 100 924 genomes from the GISAID database to identify the most relevant mutations in the world and then explore their changes in frequencies during the pandemic. After that, we performed a revision of the possible structural and functional implications of the non-synonymous mutations identified.

## RESULTS AND DISCUSSION

### Identification of Relevant Mutations (RM)

Our normalized by COVID-19 cases relative frequency analysis (see material and methods for details) performed in each of 26 protein sequences from 100 924 SARS-CoV-2 genomes showed 41 mutations with an NRFp greater than 0.03 (representing an estimation to be in SARS-CoV-2 viruses from ~750 000 cases) (Fig. S1 and Table 1). Those mutations are named Relevant Mutations (RM) from here.

**Table 1.** Summary of the 41 relevant mutations identified by our NRFp analysis.

| Mutation |  | Region | ID | NRFp |
| --- | --- | --- | --- | --- |
| nt | AA |  |  |  |
| High-Frequency (HF) |  |  |  |  |
| C241T | --- | 5'UTR | C241T_5'UTR | 0.922 |
| C1059T | T85I | nsp2 | T85I_nsp2 | 0.217 |
| C3037T | Syn | nsp3 | C3037_nsp3(S) | 0.938 |
| G11083T | L37F | nsp6 | L37F_nsp6 | 0.072 |
| C14408T | P323L | nsp12 | P323L_nsp12 | 0.933 |
| A23403G | D614G | S | D614G_S | 0.935 |
| G25563T | Q57H | orf3a | Q57H_orf3a | 0.291 |
| G28881A | R203K | N | G28881A_N | 0.430 |
| G28882A | R203K | N | G28882A_N | 0.428 |
| G28883C | G204R | N | G204R_N | 0.429 |
| Low-Frequency non-Synonymous (LF_nS) |  |  |  |  |
| C4002T | T428I | nsp3 | T428I_nsp3 | 0.056 |
| C5700A | A994D | nsp3 | A994D_nsp3 | 0.044 |
| C6573T | S1285F | nsp3 | S1285F_nsp3 | 0.042 |
| G10097A | G15S | nsp5 | G15S_nsp5 | 0.064 |
| C10319T | L89F | nsp5 | L89F_nsp5 | 0.047 |
| C11916T | S25L | nsp7 | S25L_nsp7 | 0.046 |
| C13730T | A97V | nsp12 | A97V_nsp12 | 0.031 |
| A18424G | N129D | nsp14 | N129D_nsp14 | 0.032 |
| C21304T | R216C | nsp16 | R216C_nsp16 | 0.032 |
| C22227T | A222V | S | A222V_S | 0.032 |
| C25528T | L46F | orf3a | L46F_orf3a | 0.041 |
| G25907T | G172V | orf3a | G172V_orf3a | 0.031 |
| C27964T | S24L | orf8 | S24L_orf8 | 0.064 |
| C28311T | P13L | N | P13L_N | 0.030 |
| C28854T | S194L | N | S194L_N | 0.075 |
| C28869T | P199L | N | P199L_N | 0.034 |
| C28932T | A220V | N | A220V_N | 0.030 |
| G29645T | V30L | orf10 | V30L_orf10 | 0.030 |
| Low-Frequency Synonymous (LF_S) |  |  |  |  |
| C313T | Syn | nsp1 | C313T_nsp1(S) | 0.057 |
| G4354A | Syn | nsp3 | G4354A_nsp3(S) | 0.040 |
| C6286T | Syn | nsp3 | C6286T_nsp3(S) | 0.031 |
| C13536T | Syn | nsp12 | C13536T_nsp12(S) | 0.058 |
| C15324T | Syn | nsp12 | C15324T_nsp12(S) | 0.034 |
| C18877T | Syn | nsp14 | C18877T_nsp14(S) | 0.057 |
| C19524T | Syn | nsp14 | C19524T_nsp14(S) | 0.031 |
| T19839C | Syn | nsp15 | T19839C_nsp15(S) | 0.039 |
| A20268G | Syn | nsp15 | A20268G_nsp15(S) | 0.069 |
| G21255C | Syn | nsp16 | G21255C_nsp16(S) | 0.031 |
| C23731T | Syn | S | C23731T_S(S) | 0.062 |
| C26735T | Syn | M | C26735T_M(S) | 0.046 |
| G29179T | Syn | N | G29179T_N(S) | 0.040 |

From those 41, 10 exceed the 0.3 globally NRFp in at least one month. Those were named high-frequency (HF) mutations. The other 31 were classified as low-frequency (LF) mutations and divided into synonymous (LF_S) and non-synonymous (LF_nS) to perform the analysis (Fig. 1).

**Figure 1.**
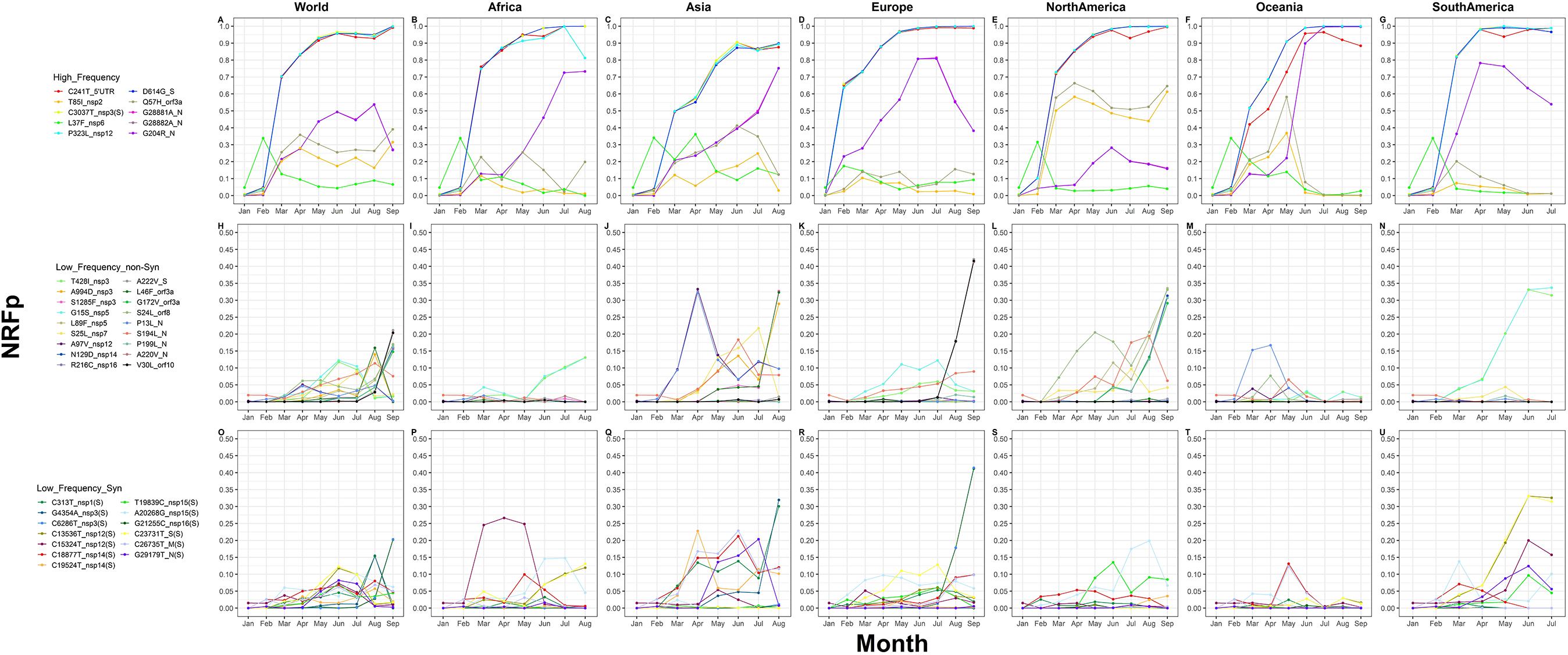
Analysis of temporal and geographical dynamics analysis of the 41 relevant mutations shows different patterns. Panels are divided by continents in x-axis and by mutation groups in y-axis (High-frequent (HF), Low-Frequent non-Synonymous (LF_nS) and Low-Frequent Synonymous (LF_S)). Each panel shows the variation by month of relative frequency of each mutation. A-G) HF mutations in general present high frequencies since February or March and in some cases are almost fixed in the continent. H-N) Most of LF_nS mutations begin to increase their frequencies after February and seems to be continente-specific. O-U) A pattern very similar to the LF_nS mutations are observed for the LF_S mutations.

### High-Frequency (HF) mutations

Of the 10 HF mutations, 4 (C241T_5’UTR, C3037T_nsp3(S), P323L_nsp12, D614G_S) appear almost fixed in the world (more than ~0.9 NRFp since June). In all continents except Asia and Africa, those mutations have reached 1 NRFp in the last months analyzed. Some months have lower frequencies of C241_5’UTR compared to the other three (Fig. 1.B, E, F, G). This is mainly due to the absence of information of the 5’UTR in some sequenced genomes. In Africa, in May, June, and August an NRFp of ~0.9, 0.9, and 0.8, respectively, is reported for the mutation P323L_nsp12. This is mainly due to the presence of genomes from Egypt with P323_nsp12 but with the other 3 HF mutations (C241T_5’UTR, C3037T_nsp3(S), D614G_S). Those were isolated from samples collected between June and August probably indicating a mutant reversion (Fig. S2).

Mutations G28881A_N, G28882A_N, and G204R_N (or G28883C_N) showed a rapid global increase from February to June. In July and August, they stabilized its frequencies but in September they showed a global decreasing (Fig. 1.A). This drop in frequency seems to be predominantly due to the decrease in frequency in Europe during September (Fig. 1.D) and the absence of information for Africa, Asia, and South America on this month. On the other hand, Africa, Asia, and Oceania showed a constant increase of these mutations since March (Fig. 1.B, C, F). In the case of Oceania those mutations are apparently fixed since July (Fig. 1.F). Intriguingly, North America showed lower frequencies of theses mutations despite having also been detected since February (Fig. 1.E).

In North America, mutations T85I_nsp2 and Q57H_orf3a had frequencies between 0.4 and 0.7 since March (Fig. 1.E). In contrast, other continents show lower frequencies (up to 0.3). Except for June and July in Asia (Fig. 1.C) and May in Oceania (Fig. 1.F). Why those mutations are more frequent than Nucleocapsid mutations (G28881A_N, G28882A_N, and G204R_N) in North America, and why T85I_nsp2 and Q57H_orf3a mutations are in lower frequencies in the other continents are still open questions.

Mutation L37F_nsp6 showed a global peak in February (Fig 1.A) that corresponds with a peak in all the continents (Fig. 1.B-G). However, since February the frequency of this mutation was below 0.2 in all months and continents except in Asia-April (Fig. 1.B-G). Furthermore; in Africa, Europe, North America, Oceania, and South America this mutation seems to be near to extinction on the last analyzed months (Fig. 1.B, D-G).

### Low-Frequency non-Synonymous (LF_nS) mutations

Respect to the LF mutations, we identified 15 LF_nS mutations (T428I_nsp3, A994D_nsp3, S1285F_nsp3, G15S_nsp5, L89F_nsp5, N129D_nsp14, R216C_nsp16, A222V_S, L46F_orf3a, G172V_orf3a, S24L_orf8, S194L_N, P199L_N, A220V_N, V30L_orf10) that reach a global NRFp greater than 0.1 in at least one month (Fig 1.H).

Analyses by continents showed that most of those 15 LF_nS have a continent-specific distribution; for example, in Asia, A994D_nsp3, S1285F_nsp3, L46F_orf3a are maintained with ~0.05 relative frequency from May to July and showed a high peak of ~0.3 in August (Fig 1.J); in other continents, those mutations have frequencies near to 0 (Fig 1.I, K-N). In Europe, three LF_nS seems to be specific for this continent, A222V_S, A220V_N, and V30L_orf10 that showed a rapid increasing relative frequency in August (~0.18) and September (~0.42) (Fig 1.K). Those mutations appear in genomes that do not present HF nucleocapsid mutations (G28881A_N, G28882A_N, G204R_N) explaining the decrease in the relative frequency of these mutations in August and September (Fig 1.D). Similarly, just in North America, L89F_nsp5, N129D_nsp14, R216C_nsp16, G172V_orf3a, S24L_orf8, P199L_N showed increasing frequencies since August (Fig 1.L). One of them (S24L_orf8) with relatively high frequencies already reported in previous months (April, May, and June) (Fig 1.L).

Interestingly, T428I_nsp3 and G15S_nsp5 appear with frequencies up to ~0.12 in Europe (Fig 1.K). In Africa and South America, a constant increase of relative frequency (up to ~0.14 in Africa and ~0.33 in South America) is reported from June to August in Africa and from March to June in South America (July is the last month analyzed in South America and with 89 analyzed genomes the frequencies of those mutations are similar to those in June) (Fig 1.I, N). Mutation S194L_N showed a variable relative frequency in Asia, Europe, North America, and Oceania (Fig 1.J-M). In Africa and South America its frequency is near to 0 in all the months analyzed (Fig 1.I, N).

### Low-Frequency Synonymous (LF_S) mutations

Thirteen LF but synonymous mutations (LF_S) were identified in our analysis and 6 of them reaching 0.1 global NRFp in at least one month. Those mutations follow a similar pattern that those described by the LF_nS mutations. Thus, C313T_nsp1(S) and G4354A_nsp3(S) accompany the three LF_nS mutations specific to Asia (Fig 1.J, Q). In the same way, C6286T_nsp3(S) and G21255C_nsp16(S) shows the same pattern as LF_nS mutations A222V_S, A220V_N, and V30L_orf10 in Europe (Fig. 1.K, R). LF_nS mutations T428I_nsp3 and G15S_nsp5 (from Africa, Europe, and South America) are accompanied by LF_S mutations C13536T_nsp12(S) and C23731T_S(S) (Fig. 1.I, K, N, P, R, U). Is interesting to note that in Africa and South America these four mutations have similar frequencies in all the months. However, in Europe T428I_nsp3 and C13536T_nsp12(S) have similar frequencies but different from G15S_nsp5 and C23731T_S(S) (Fig 1.K, R). LF_nS mutations in NorthAmerica (L89F_nsp5, N129D_nsp14, R216C_nsp16, G172V_orf3a, S24L_orf8, P199L_N) are not accompanied by LF_S mutations (Fig. 1.S).

Oceania is the only continent that did not show LF mutations with a tendency to increase in the last months analyzed (Fig. 1.M, T). Instead (and already previously mentioned), HF nucleocapsid mutations (G28881A_N, G28882A_N, G204R_N) appear near to fixation (Fig. 1.F).

### Functional and Structural implications of non-Synonymous Relevant Mutations (RM)

#### Nsp1

Non-structural protein 1 (nsp1) is conserved in the Coronaviridae family and has a sequence homology of up to 85% with SARS-CoV (Schubert et al. 2020). Nsp1 has two regions: the globular N-terminal domain (1-128 aa) (Almeida et al. 2007) and the C-terminal domain (residues 148-180) (Thoms et al. 2020, Schubert et al. 2020) linked by a flexible region of 20 residues (Fig. S3). Nsp1 forms complexes with the host translational machinery, sequestering the 40S minor ribosomal subunit and positioning its C-terminal domain in the mRNA binding site, thus preventing the host’s mRNA translation (Thoms et al. 2020, Schubert et al. 2020). It also interferes with the innate immune response and the expression levels of IFN type I and III by preventing the translation of antiviral response factors mediated by IFN-stimulated genes (ISGs) (Thoms et al. 2020). Our analysis did not show relevant non-synonymous mutations in nsp1.

#### Nsp2

Nsp2 function in SARS-CoV-2 is unknown. Pfam (El-Gebali et al. 2019) analysis did not show any predicted domain, SWISS-MODEL (Waterhouse et al. 2018) template search found as the best template Ubiquitin-40S ribosomal protein S31 (PDB ID: 6I7O) with a very bad homology (0.26 similarity in an alignment coverage of 0.09) and BLAST search (Altschul et al. 1990) (excluding sequences from Coronaviridae) found 0.21 identity with 0.81 coverage to a hypothetical protein from E. coli (accession code: WP_162751855.1) and to an RNA-dependent RNA polymerase domain protein from E. coli (accession code: EYE07746.1). The deletion of Nsp2 from SARS-CoV has little effect on viral titers (Graham et al. 2005). Also, In SARS-CoV, was showed that nsp2 can interact with prohibitin 1 and 2 (PBH1 and PBH2) (Cornillez-Ty et al. 2009). These proteins are involved in several cellular functions including cell cycle progression (Wang et al. 1999), cell migration (Rajalingam et al. 2005), cellular differentiation (Sun et al. 2004), apoptosis (Fusaro et al. 2003), and mitochondrial biogenesis (Merkwirth et al. 2008). The absence of structural and functional information of nsp2 impede us to hypothesize the effects of the HF mutation T85I that our analysis showed with an NRFp of 0.22.

#### Nsp3

The nsp3 protein is the longest encoded in coronaviruses, to which different functions are attributed due to its multiple domains. Nsp3 is best known for its protease activity and for being essential in the replication/transcription complex (RTC). Structurally, Nsp3 is a 1945 amino acid protein; there is no consensus on the number of domains of nsp3 in SARS-CoV and SARS-Cov-2. Lei 2018 mentions 14 domains in SARS-CoV (Ubl-1, Ac, X, SUD, Ubl-2, PL2pro, NAB, βSM, TM1, 3Ecto, TM2, AH1, Y1, and CoV-Y). In nsp3 we identified three non-synonymous relevant mutations.

The SARS Unique Domain (SUD) is composed of 3 subdomains: Mac2, Mac3, and DPUP. The function of these is binding to RNA to form G-quadruplexes (Lei 2018). Besides, Mac3 has been described as an essential region for the RdRp complex (Kusov 2015). According to Tan 2009, the positive interface of Mac2-Mac3 is the possible region for nucleic acid binding (Fig. 2B). It is unlikely that the LF_nS mutation T428I (NRFp 0.056) varies the electrostatic potential in this region because it is located far from this interface (Fig. 2A). In the T428 variant, there are no hydrogen bonds between this residue and the nearby residues. Instead, we observed several hydrophobic atoms from residues L431, T423, Y536, and E427 (Fig. 2C). An in silico model of T428I showed that isoleucine could accommodate at this site (Fig. 2D).

**Figure 2.**
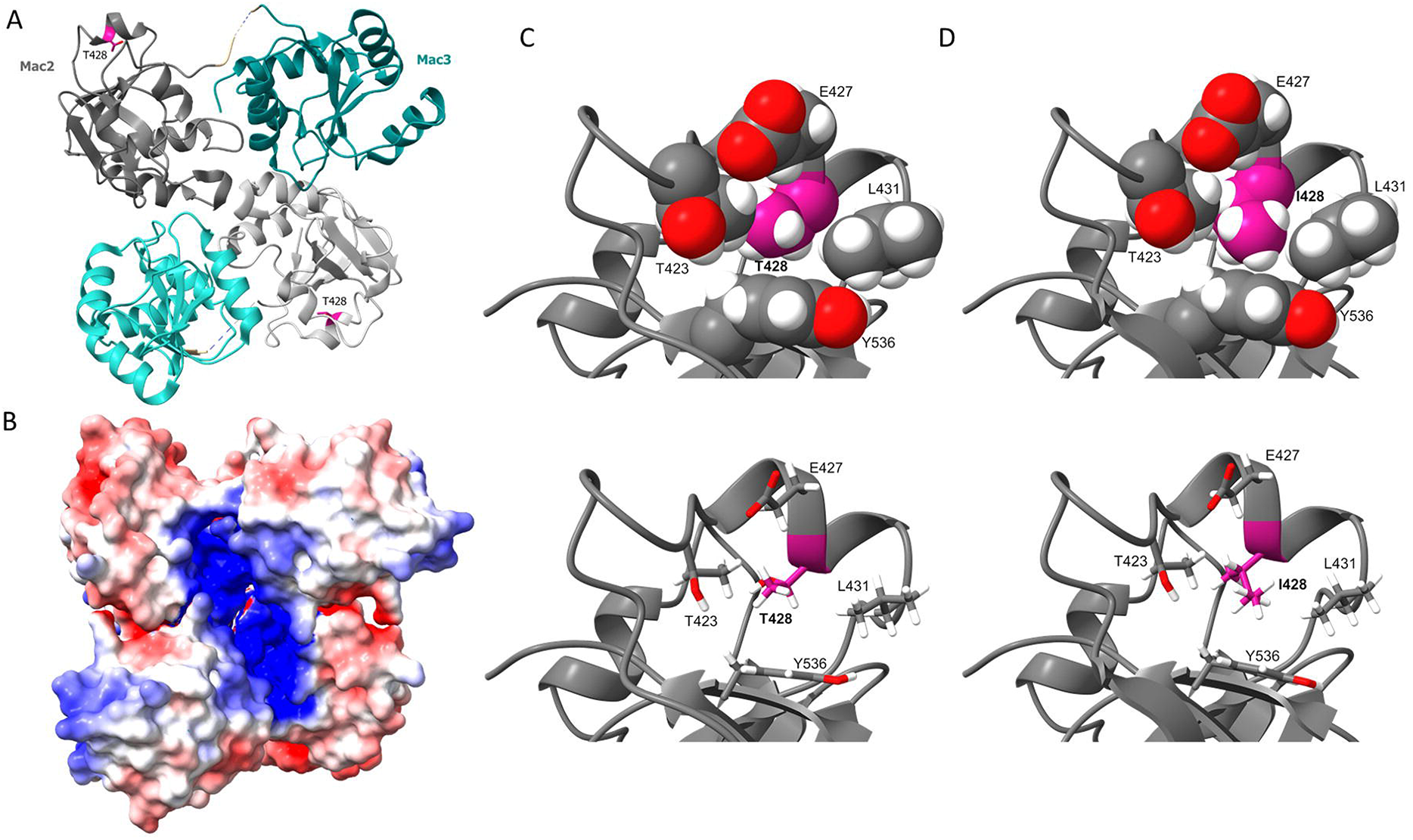
T428I mutation in domain of nsp3 could modify hydrophobic packaging in mac2 domain. A) Structure of the mac2-mac3 dimer (PDB: 2W2G) showing the residue T428 in magenta. B) Electrostatic Surface potential representation of A showing the positive patch in the middle that could binds nucleic acids. C) Zoom in the region near to the mutation, showing residues 2 Å close to T428 side chain in spheres (above) or sticks (below) representation, no hydrogen bonds were observed involving this threonine. D) Model of the T428I variant in the same view of C, after a potential energy minimization we noted that internal spaces are better filled.

The PL2pro domain processes the amino-terminal of polyprotein 1a/ab to generate 2 or 3 products: nsp1, nsp2, and nsp3, through to its protease activity mediated by the catalytic triad Cys-His-Asp (Barretto 2005) (Fig. S4). Also, Pl2pro cleave Interferon Stimulated Gene 15 (ISG-15) causing the loss of ISGylation from interferon responsive factor 3 (IRF3), an important component in the Interferon I pathway (Shin 2020). Structurally, it is made up of 3 subdomains: thumb, palm, and fingers. It also has a Ubl-2 domain linked by an alpha helix (Clasman 2017) (Fig. S4) that determines the substrate specificity for ISG (Shin 2020). The subdomain corresponding to fingers contains a zinc-binding region (Fig. S4) and has been described as essential for the maintenance of the structure of Pl2pro (Baez-Santos 2014). The A994D LF_nS mutation (NRFp 0.044) is located on a beta-sheet of the palm subdomain, far away from the catalytic site (Fig. S4). This residue is exposed to the surface, where D could favor solvent interactions.

The last mutation observed in the nsp3 protein corresponds to S1285F, located in the βSM domain, for which, to our knowledge, there is no structural or functional information.

#### Nsp4

Nsp4 has four transmembrane domains and both termini on the cytoplasmic side (Oostra et al. 2007). In SARS-CoV, together with Nsp3, it causes membrane rearrangement in HEK293T cells (Sakai et al. 2017). Amino acids from positions 112–164 in the large luminal loop of nsp4 are responsible for binding to the c-terminal region of nsp3 (Sakai et al. 2017). H120 and F121 in nsp4 are crucial for binding with nsp3 and the induction of the membrane rearrangements in HEK293T cells (Sakai et al. 2017). In MERS, nsp3-nsp4 has a significant effect on the membrane conformation. When expressed in HuH-7 cells they show induction of perinuclear double-membrane bodies (Oudshoorn et al. 2017). No relevant mutations were identified on Nsp4.

#### Nsp5

The nsp5 encodes the main protease or 3CLPro that cleaves the polyprotein 1a/ab in 11 sites generating nsp4 to nsp16. SARS-CoV-2 nsp5 is a homodimer with three domains: domain I (DI) (8-101), domain II (DII) or N-terminal domain (102-184), and domain III (DIII) or C-terminal domain (201-303) (Fig 3A). Residues 1-7 called N-finger are located between DII and DIII (Fig 3A) (Ulrich and Nitsche 2020). Nsp5 cleaves through a Cys-His catalytic dyad located between DI and DII, where a pocket (S1, F140, and E166) is formed to accommodate the substrate (Fig 3A) (Zhang et al. 2020).

**Figure 3.**
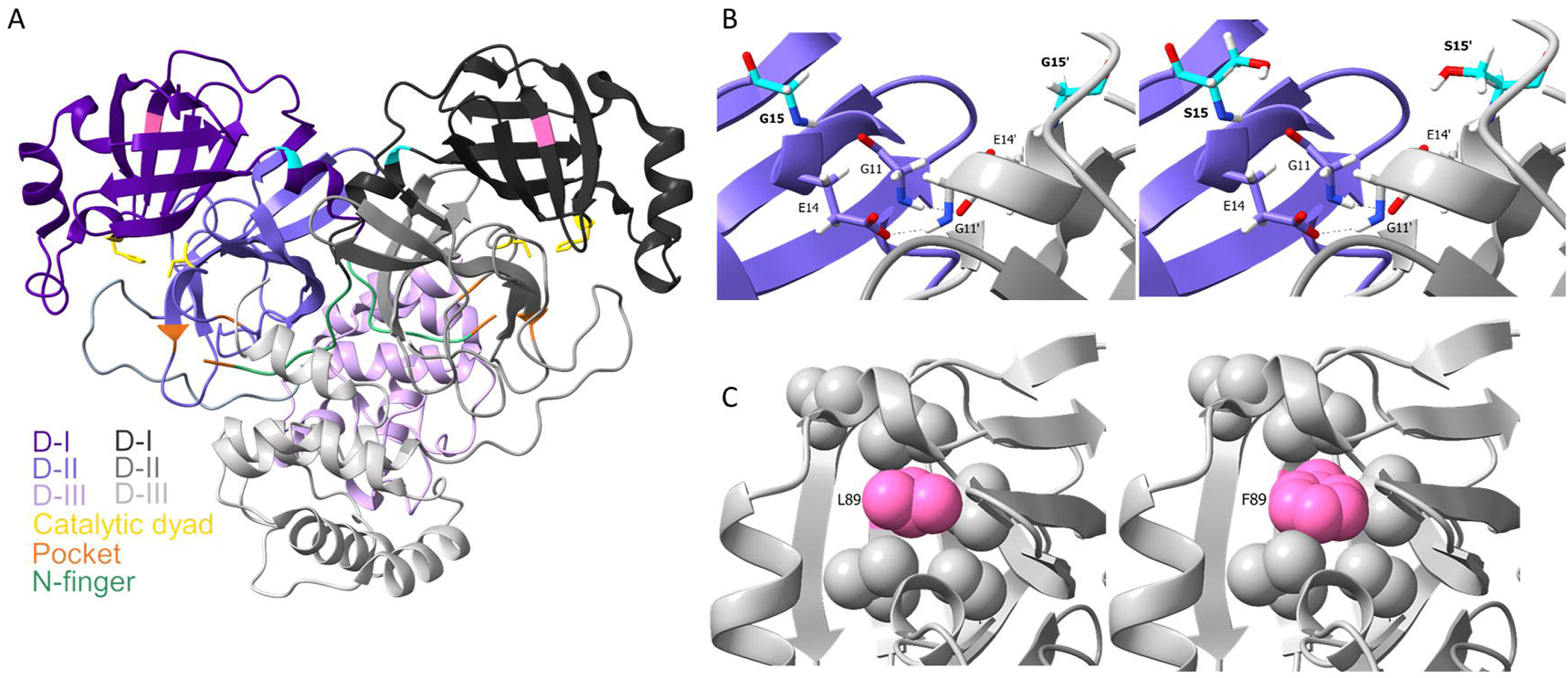
Mutations in nsp5 G15S and L89F probably have minor effects in structure and function. A) nsp5 homodimer (PDB: 6Y2G) colored by domains and highlighting regions such as substrate pocket, N-finger and catalytic dyad. B) G15S mutation in domain II is present just after E14 that forms a hydrogen bond with G11 in the dimer interface. S15 could confer solubility and less flexibility due to its location. C) Packing of the side chains near to L89 in domain I showing that mutations L89F slightly changes the packaging in this region (a potential energy minimization was done after the mutation was modelled).

DII, DIII, and the N-finger stabilize the homodimer through interactions at the monomers interface (Xia 2011). We have found two LF_nS mutations (G15S and L89F) located in DI (Fig. 3A). G15S (NRFp: 0.064) is close to residue E14, described by Xia et al (2011) as being important for dimerization since it forms a hydrogen bond with G11 of the opposite monomer (Fig 3B). Due to the S15 orientation, it would not favor the formation of inter- or intra-monomer hydrogen bonds. Instead, it could favor solubility through solvent interactions. L89F mutation is located in a beta-sheet of DII with the side chain points towards the center of the protein. The bigger hydrophobic chain of phenylalanine could favor the packaging between the beta-sheets of this domain (Fig 3C).

#### Nsp6

The NSP6 of coronaviruses are proteins with 6 transmembrane domains (Baliji et al. 2009). It is located in the endoplasmic reticulum participating in viral autophagic regulation (Cottam et al. 2014). The HF L37F mutation (NRFp 0.07), there are no experimental studies that allow us to hypothesize the effects of this mutation on protein structure or function. Benvenuto et al. (2020) speculate through bioinformatic analysis the possible effect of this mutation.

#### Nsp7

Nsp7 is an 83 amino acids formed by four helices: α1 (K2-S25), α2 (K27-A42), α3 (D44-S61), and α4 (D67-N78) (Fig. 4A). Together with nsp8 is involved in RNA polymerase processivity (Subissi L et al. 2014). Nsp7 confers processivity through direct RNA-binding (K7, H36, and N37) (Subissi L et al. 2014). The LF_nS S25L had 0.05 of NRFp is at the end of α1 along with S24 and S26 (Fig. 4B). This mutation is near to the interface with nsp8 (Fig. 4C). Our structural in silico analysis shows that this mutation could increase the hydrophobic packaging in the region (Fig. 4D).

**Figure 4.**
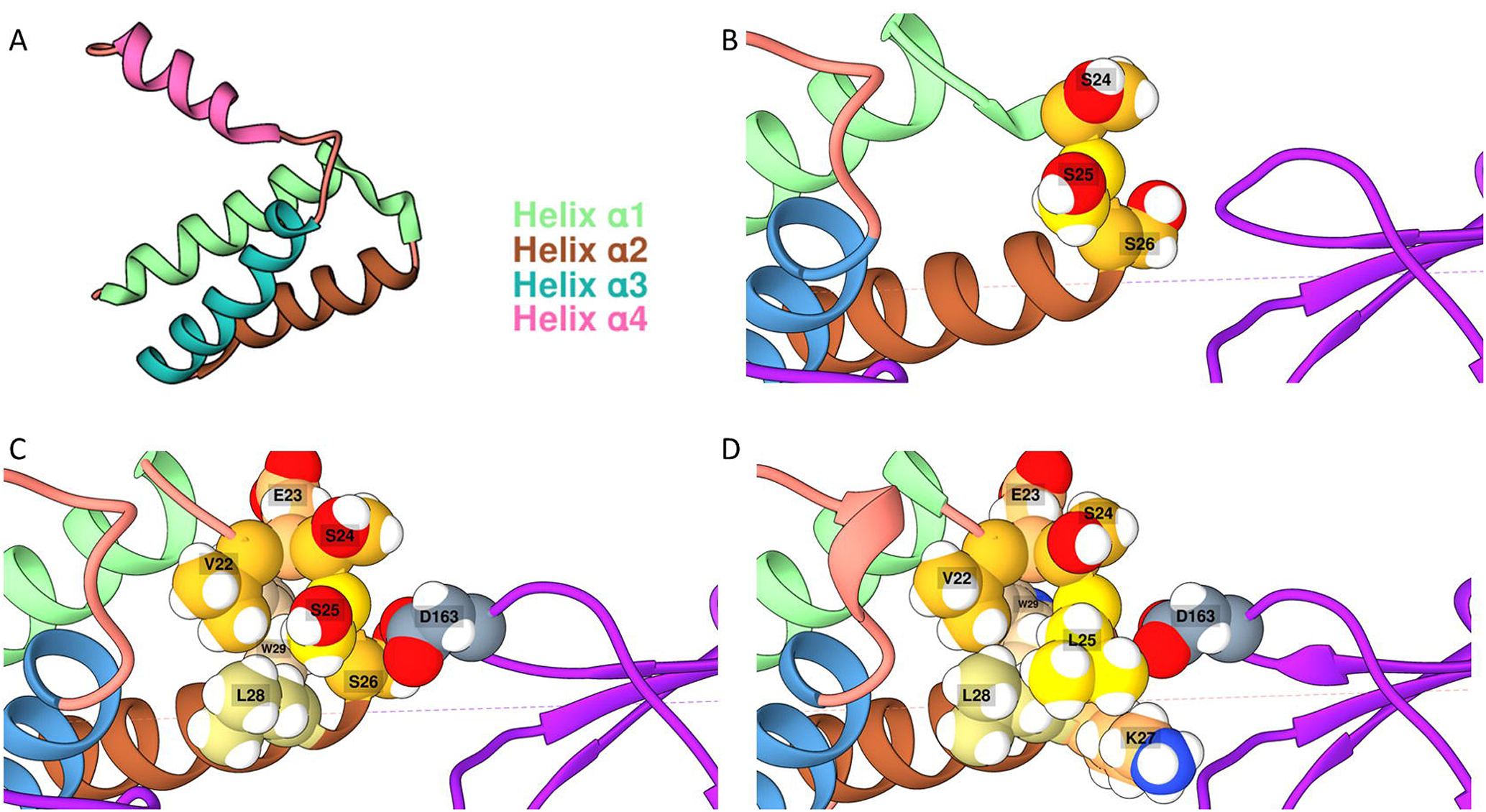
Mutation S25L confers hydrophocity in the connection between α1 and α2. A) nsp7 is formed by 4 alpha helix (PDB: 6M5I). B) Connection between α1 and α2 is formed by three serines (S24, S25 and S26). C) S26 is forming a hydrogen bond with D163 from nsp8 and hydroxyl of S25 is pointing to the solvent. D) Computational modelling of S25L shows a gaining in hydrophobicity of the region with the three serines.

#### Nsp8

Nsp8 is a 198 amino acid protein showed to have a key role in forming cytoplasmic complexes (nsp7/nsp8/nsp12) for viral RNA synthesis (Kumar et al. 2007). Nsp8 has an N-terminal motif important for RNA polymerase activity since these residues are part of the Mg2 binding site of nsp12 (Te Velthuis et al. 2011). Evidence has been presented that nsp8 can generate radiolabeled RNA products in a template-dependent manner, with sizes that exceed those of the template, indicative of both de novo and primer-dependent RNA polymerase activities. (Te Velthuis et al. 2011). On our frequency analysis, we did not find relevant mutations in nsp8.

#### Nsp9

SARS-CoV-2 Nsp9 is a 113 amino acid RNA-binding homodimeric protein (Littler, 2020). The monomer is composed of seven β-strands (β1-β7), an N-terminal β7 extension, and a C-terminal α-helix with a conserved “GxxxG” motif (Fig. S5) (Littler, 2020; Zhang, 2020). Its dimerization occurs via the “GxxxG” motif present also in SARS-CoV Nsp9 (Sutton, 2004; Littler, 2020) and by the N-terminal β7 extension (Fig. S5) (Littler, 2020; Zhang, 2020). Zhang et al. (2020) reported a SARS-CoV-2 nsp9 homotetramer structure with an interface composed mainly of β5 and three connection loops. It is proposed that the nsp9 function is critical to the replication and transcription machinery since mutations in the SARS-CoV Nsp9 gene prevent viral replication (Miknis, 2009). Nearly 97% sequence identity is found between SARS-CoV and SARS-CoV-2 Nsp9 protomers, suggesting a related function (Littler, 2020). We did not find relevant mutations in nsp9.

#### Nsp10

The SARS-CoV-2 Nsp10 is divided into two subdomains: the alpha helix subdomain (α1-α4 and α6) and a beta subdomain (two antiparallel sheets, a short alpha-helix, and coiled-coil regions) (Fig. S6A) (Viswanathan, 2020). Nsp10 has two zinc-binding sites: i) C74, C77, C90, and H83 (between α2 and α3) (Fig. S6B), ii) C117, C120, C128, and C130; both with stabilizing effects (Krafcikova, 2020) (Fig. S6C). Nsp10 interacts with the Nsp14 and Nsp16 methyltransferases, activating them (Krafcikova 2020). The SARS-CoV-2 Nsp10 helices α2, α3, α4, and a coiled-coil region between α1 and β1 (N40 to T49) interact with Nsp16 through hydrophobic interactions or water molecules. Two important Nsp10 residues are immersed into hydrophobic pockets from Nsp16: V42 (pocket 1: M41, V44, V78, A79 and P8O from Nsp16) and L45 (pocket 2: P37, I40, V44, T48, L244 and M247 from Nsp16) (Fig. S7A) (Krafcikova, 2020). In SARS-CoV, the Nsp10-Nsp14 is mediated by the N-terminal loop, the α1 helix (1-20), the loop following α2, and residues around zinc finger 1 (Fig. S7B) (Ma, 2015). We did not find relevant mutations in nsp10.

#### Nsp12

Nsp12, in complex with nsp7-nsp8, forms the RdRp complex (Pachetti M et al. 2020). Nsp12 is a 932 amino acid protein formed by an N-terminal extension domain that adopts a nidovirus RdRp-associated nucleotidyltransferase (NiRAN) (60-49), a polymerase domain (367-920), and an interface domain (250-365) that connects the polymerase and NiRAN domains (Fig. 5A) (Gao et al. 2020). Furthermore, the polymerase domain has three subdomains: finger subdomain (366-A581 and 621-679), palm subdomain (582-620 and 680-815), and a thumb subdomain (816-920) (Fig. 5B) (Gao et al. 2020). Our analysis identified two relevant mutations in nsp12: P323L and A97V, with 0.88 and 0.06 NRFp, respectively. P323, together with P322, in the interface domain, ends the helix 10 generating a turn that precedes a beta-sheet (Fig. 5C). The P323L mutation would be disadvantaged due to the loosening of the turn at the end of the helix 10. Thus, the presence of P322 could allow that P323 mutate to another residue avoiding a reasonable impact on the overall structure. L323 could change hydrophobic packaging in the interface between the polymerase domain and interface domain (Fig. 5D). A97 in the NiRAN domain falls in a loop exposed to the solvent (Fig. 5E). In the A97V variant, additional methyl groups of valine gain interactions with V96 increasing hydrophobic contacts (Fig. 5F).

**Figure 5.**
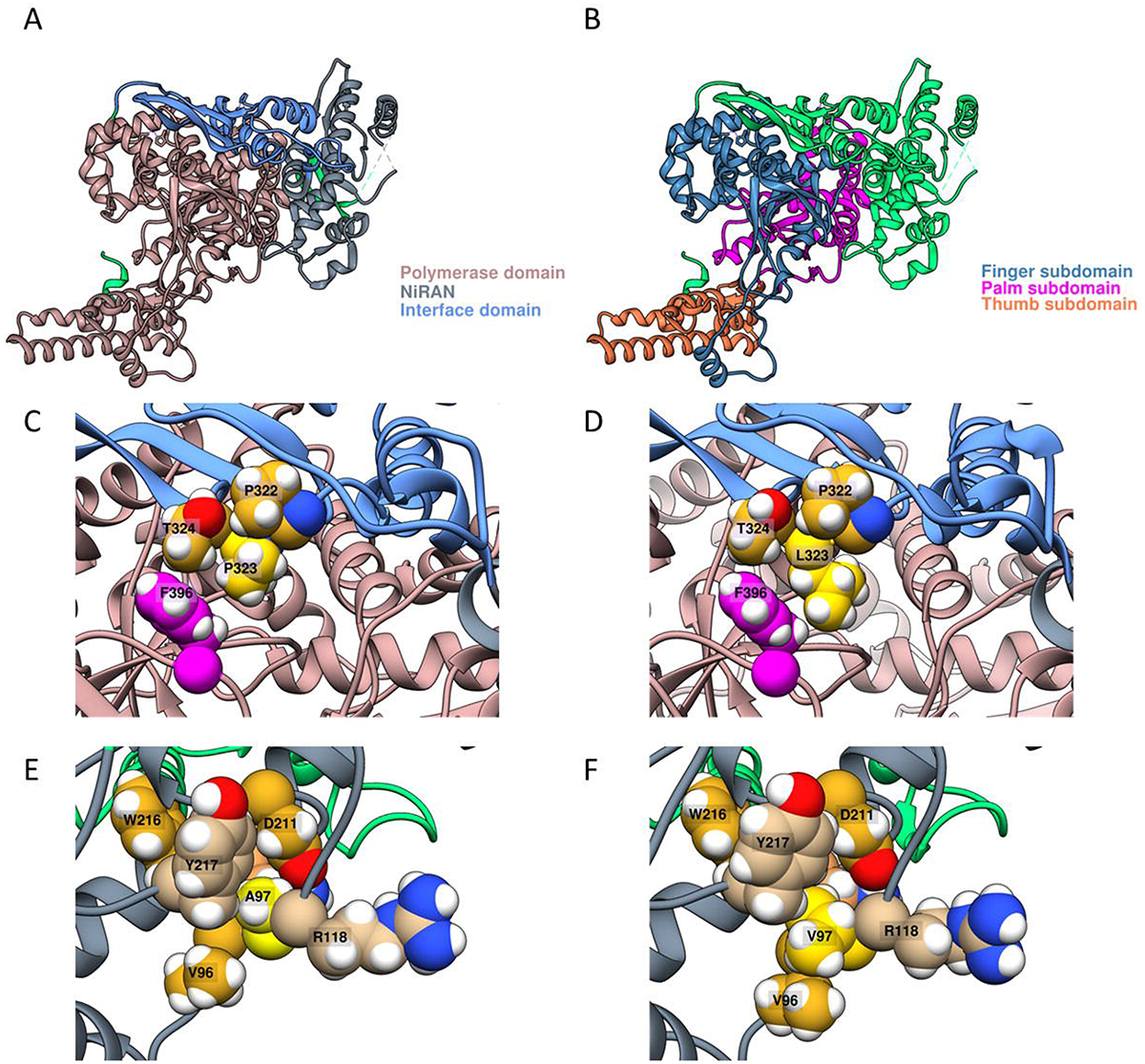
Two mutations in nsp12 (A97L and P323L) modify the hydrophobic packaging. A) nsp12 (PDB: 6YYT) is formed by three domains showed in different colors. B) Polymerase domain of nsp12 is divided in three subdomains colored in blue, magenta and orange. C-D) Mutation P323L could affect the rigidity of the loop, but another proline is present in position P322. Thus, P323L increases the internal hydrophobicity probably modifying the stability of nsp12. E-F) Presence of A97V mutation allows that a methyl group from valine accomodates near to V96 increasing the contacts between methyl groups in that region.

#### Nsp13

Nsp13 is a dsRNA and dsDNA helicase (Adedeji 2012, Ivanov 2004) of 603 amino acids. Nsp13 (PDB: 6ZSL) and its homolog in SARS-CoV (PDB: 6JYT) has 5 domains: ZBD (Zinc binding domain), Stem (connects ZBD and 1B), 1B, 1A, and 2A (RecA-like domains) (Fig. S8A) (Jia 2019). Additionally, it has a phosphate-binding site at the interface of 1A and 2A (P284, K288, Q404, R443, R567) (Fig. S8B) and three Zinc Fingers (ZnF): ZnF1 (C5, C8, C26, C29), ZnF2 (C16, C19, H33, H39) and ZnF3 (C50, C55, C72, H75) (Fig. S8C). No relevant mutations were found in nsp3.

#### Nsp14

The SARS-CoV-2 nsp14 is conserved in Coronaviridae (Ma et al. 2015, Robson et al. 2020) and comprises two domains: the N-terminal exonuclease (1-287, ExoN) and the C-terminal N7-methyltransferase (288-527, N7-MTase) (Selvaraj et al. 2020). ExoN domain functions as a proofreading exoribonuclease (Ma et al. 2015). It removes mismatched nucleotides from the 3’ end of growing RNAs from RdRp RNA synthesis (Ferron, 2018). Additionally, nsp14 shows (guanine-N7) methyltransferase activity of the cap using S-adenosyl-L-methionine (SAM) as the methyl donor (Robson, 2020). Our analysis showed that the region comprised residues 420 to 503 within the N7-MTase domain seems to be difficult to sequence as shown by the relatively high frequency of Ns in this region (Fig. S9). Also, we identified the LF_nS N129D mutation in the ExoN domain (NRFp: 0.03). In the SARS-CoV (PDB ID: 5C8U) N129 side chain is exposed to the protein surface and far from interaction surfaces with nsp10 indicating little probability to impact the structure or function of nsp14.

#### Nsp15

Nsp15 is a nidoviral RNA uridine-specific endoribonuclease (NendoU) with 347 amino acids in SARS-CoV-2. It is conserved among all coronaviruses (Deng et al. 2017, Kim et al. 2020). It processes viral RNA to evade detection by the innate immune system by cleaving RNA substrates at the 3’ of uridines (Deng, 2017; Pillon, 2020). Nsp15 monomer has three domains: the N-terminal (NTD), the middle (MD), and the C-terminal (CTD) (Fig. S10A) (Kim et al. 2020, Pillon et al. 2020). In its active form, SARS-CoV-2 Nsp15 is a hexamer made from a dimer of trimers (Fig. S10B) (Kim et al. 2020, Pillon et al. 2020). The NTD is responsible for the hexamer stability, the CTD has the catalytic NendoU activity and the MD connects NTD and CTD (Kim et al. 2020, Pillon et al. 2020). H235, H250, and K290 conserved in SARS-CoV, MERS-CoV, and SARS-CoV-2 form the catalytic triad (Kim et al. 2020). We did not find relevant mutations in this protein.

#### Nsp16

Nsp16 methylates the 5’-end of viral mRNAs to mimic cellular mRNAs (Viswanathan et al. 2020). As Nsp14, Nsp16 requires Nsp10 as a cofactor (Krafcikova, 2020). Specifically, it catalyzes a ribose 2’-O methyltransferase activity, being the methyl donor the S-adenosyl-L-methionine (SAM) (Viswanathan et al. 2020, Krafcikova et al. 2020). The SAM binding site has two important regions: a nucleoside pocket and a methionine pocket (Fig. 6A, B). The nucleoside forms hydrogen bonds with D99 and D114 side chains, L100, C115, and Y132 main chains, and through water molecules with N101 (Fig. 6A) (PDB: 6W4H, Viswanathan et al. 2020). Methionine moiety forms hydrogen bonds with N43, Y47, and D130 side chains and G71 and G81 main chains (Fig. 6B) (PDB: 6W4H, Viswanathan, 2020). We found the LF_nS R216C mutation (0.03 NRFp). This residue seems not to be involved in the catalytic or dimerization functionalities. R216 forms a salt bridge with E217 (PDB: 7JHE, 2.872 A) (Fig. 6C), however in other structure (PDB: 6W4H) these two residues are further away from each other (4.930 A) (Fig. 6C), indicating that probably this salt bridge is not enough strong to contribute significantly to the stability. Anyway, R216C mutation removes the possibility to form this salt bridge. None cysteines were observed near C216 to speculate the formation of a disulfuric bridge.

**Figure 6.**
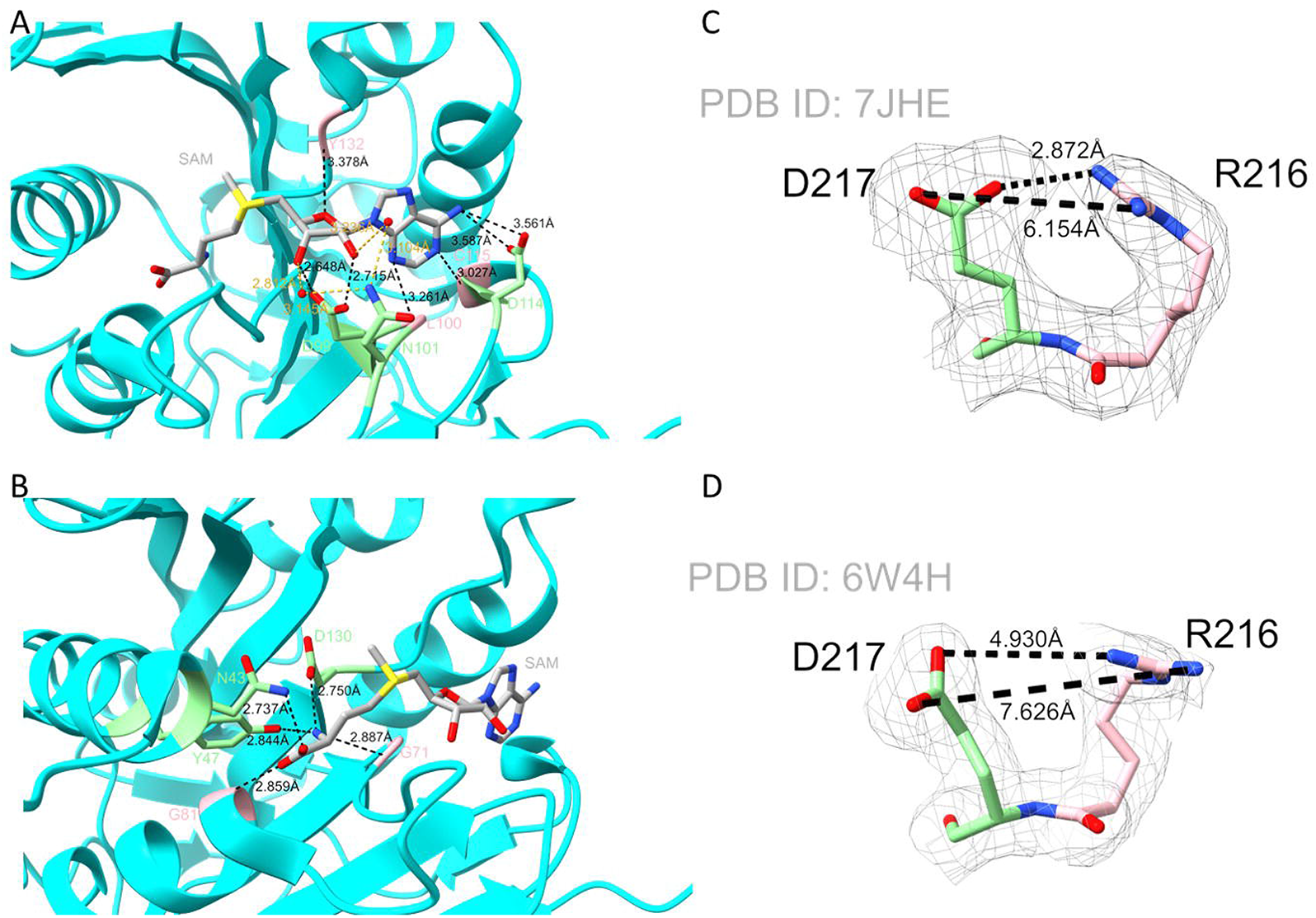
A weak salt bridge is interrupted by R216C mutation in nsp16. A-B) nsp16 binds SAM through several residues, one group of residues stabilized the binding of the nucleoside (A) and another group stabilized the mehtionine moiety (B). C) PDB: 7JHE shows a well established salt-bridge between D217 and R216. D) however in PDB: 6W4H salt-bridge seems not well established due to the greater distance between residues. These dofferences are probably due to the competence for interactions with the solvent.

#### Spike

Spike (S) protein is a trimer that expresses in the virion surface. It is recognized by the human receptor ACE2 (Tortorici and Vessler. 2019) and before that recognition, a process of viral fusion with the host cell begins (Bosch et al. 2003). Due to its superficial expression and its immunogenicity is the main protein used as a target of vaccines (Krammer 2020). Structurally, the spike is divided into two domains: S1 (1-681) and S2 (686-1213). Between these two domains, exist the cleavage region S1/S2 (682-685) (Fig. 7A). S1 domain has two subdomains: the N-terminal domain (NTD: 13-303) and the receptor-binding domain (RBD: 319-541). RBD presents the receptor-binding motif (RBM: 437-508) that has direct contacts with the ACE2 receptor (Fig. 7B). S2 also presents the fusion peptide (FP: 816-833), heptad repeat 1 (HR: 908-985), central helix (CH: 986-1035), and connector domain (CD: 1076-1141) (Shang et al. 2020) (Fig. 7C). Two relevant mutations and a region seemingly difficult to sequence between amino acids 260 and 321 were identified. The highly discussed mutation D614G with 0.93 NRFp presents in the S1 domain is now apparently fixed in the world. Discussions about its possible structural implications are described elsewhere (Justo et al. 2020). Mutation A222V, in the NTD of S1, showed 0.032 NRFp. The side chain of alanine is immersed between side chains of V36, Y38, F220, and I285 (Fig. 7D). The presence of a bigger side chain could fill better the spaces probably affecting the stability of this region.

**Figure 7.**
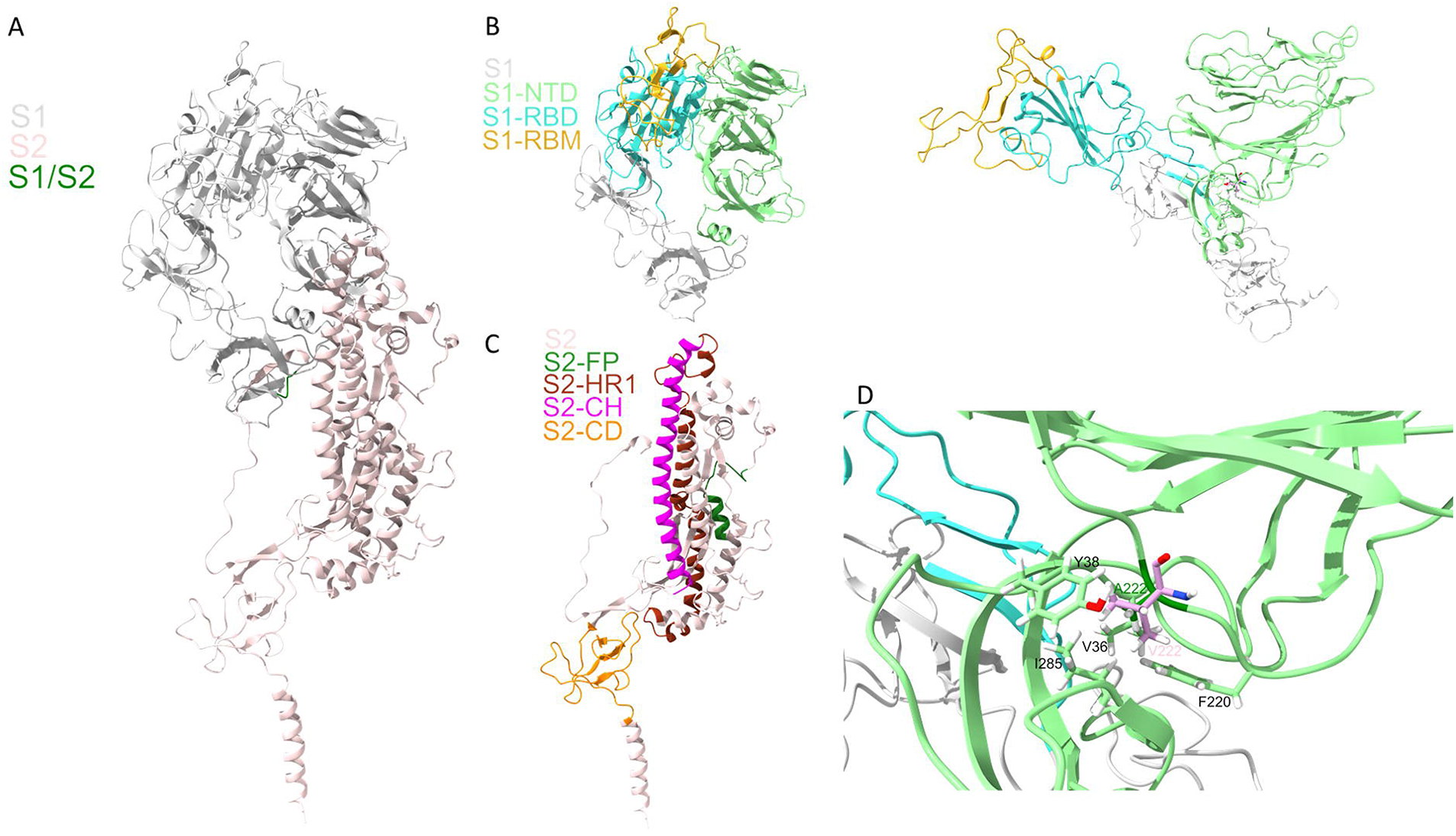
A bigger hydrophobic residue in A222V variant could confer slightly higher stability on spike protein. A) The monomer of spike protein (PDB 6VYB, chain A) is formed by two domains named S1 and S2. B) S1 domains is subdivided in and N-terminal domain (NTD) and in a Receptor Binding Domain (RBD), inside RBD can be found the region responsible for ACE2 receptor recognition called Receptor Binding Motif (RBM). C) S2 is formed for a small region called Fusion Peptide (FP), two helix subdomains involved in intermonomer interactions (HR1 and CH) and a C-terminal domain. D) Mutation A222V in the NTD domain of spike interacts better with Y38, F220 and I285.

#### Orf3a

Orf3a functions as a potassium ion channel (Lu et al. 2006, Kern et al. 2020), induces apoptotic (Chan et al. 2009) and necrotic cell death (Yue et al. 2018), and is involved in the activation of the inflammasome (Siu et al. 2019). It has 275 residues and two regions. The transmembrane domain (TMD) towards the extracellular face contains three transmembrane α-helix that form the ion channel. The cytosolic domain (CD) with several β-folded sheets found on the cytosolic face (Fig. 8A) (Zeng et al. 2004, Lu et al. 2006, Kern et al. 2020). Orf3a is expressed mainly in the endoplasmic reticulum (ER)/Golgi apparatus and to a lesser extent in the cell membrane (Padhan et al 2007).

**Figure 8.**
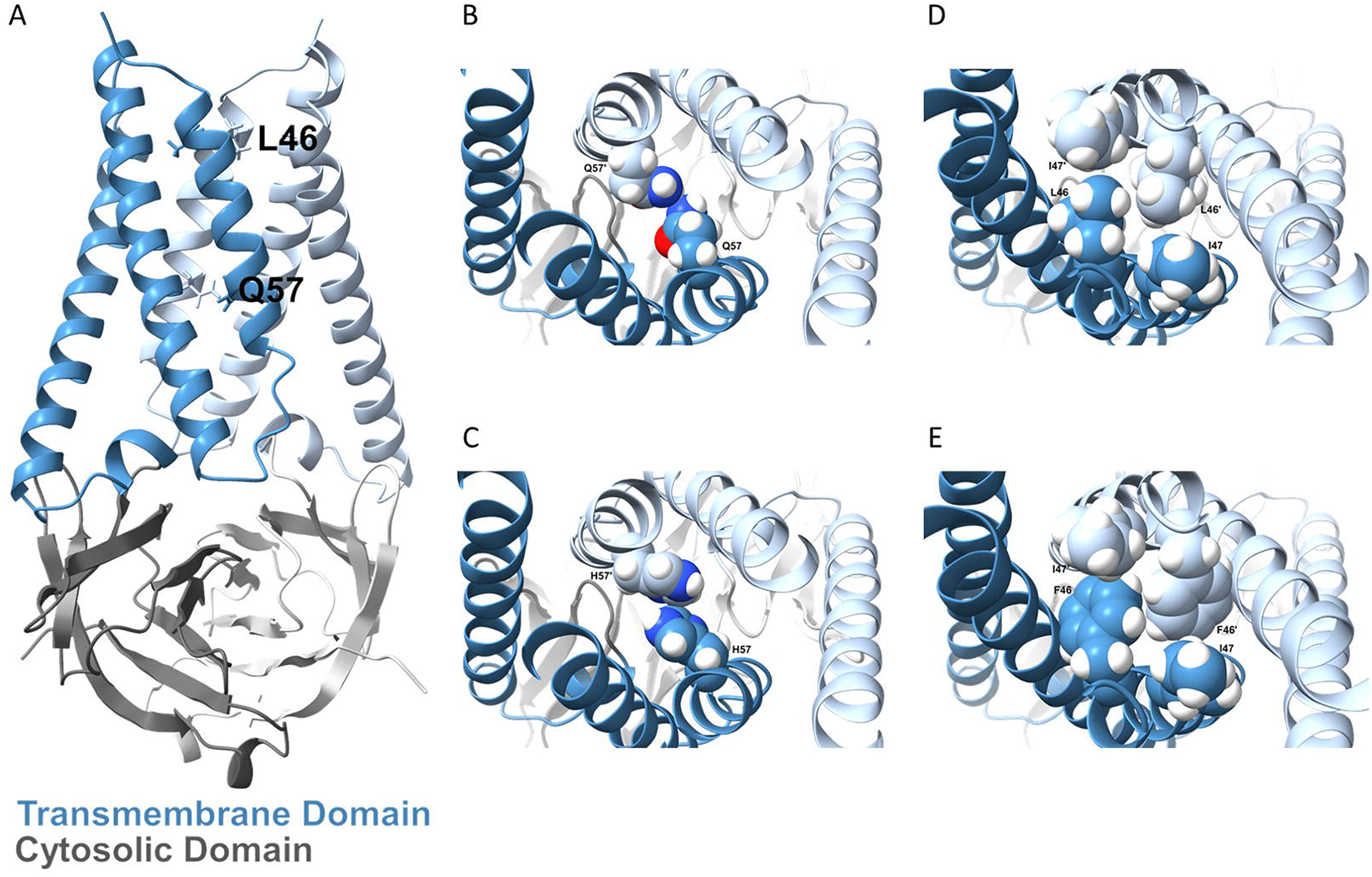
Two mutations (Q57H and L46F) in the pore of orf3a could change transportation properties. A) Orf3a dimer forms a ion-channel composed by two domains (PDB: 6XDC). B) Q57 forms a constriction in the central region of the ion-channel. C) The mutation Q57H apparently does not change the diameter of the pore but could confer different electrostatic properties. D) L46 together with I47 forms another constriction in the ion-channel. E) L46F could affect the diameter of the pore due to the bigger side chain of Phenylalanine.

In our analysis, we found three relevant mutations (L46F (0.04), Q57H (0.29), and G172V(0.03)). Two of them in the ion channel pore. The ion channel shows six constrictions along the pore, the fifth corresponding to the side chain of Q57 (Fig. 8B). Mutational studies do not show differences in the expression, stability, conductance, selectivity, or gating behavior in the Q57H variant (Kern et al. 2020). Our in-silico analysis of Q578H did not show a reduction of pore size on this constriction (Fig. 8C). Due to the difference in the pKa of the glutamine and histidine is interesting to speculate that in different pHs functional differences could be observed. The second constriction of the pore corresponds to L46 and I47 side chains (Fig. 8D) (Kern et al. 2020). L46F mutation could produce a steric effect in the pore impacting its functionality (Fig. 8E). The G172 on the CD is part of a putative di-acidic EXD motif (171-173) described as a canonical cellular traffic signal responsible for proteins export from the endoplasmic reticulum to the cell membrane in viruses, yeasts, and plants. (Nishimura et al. 1997, Votsmeier et al. 2001, Hanton et al. 2005). However, mutational studies in this region do not show significant differences in the role of the intracellular trafficking of orf3a in SARS-CoV (Minakshi et al. 2014).

#### Envelope

Envelope (E) protein is a small 75 amino acid viroporin with functions described in several stages of viral infection including viral entry, replication, virion assembly, and virion release (Schoeman and Fielding. 2019). Recently, the structure of the transmembrane domain in ERGIC-mimetic lipid bilayers has been solved by solid-state NMR (Mandala et al. 2020). The structure shows the homopentameric helices of ~35 Å vertical length (AA 14-34) with 7 aminoacids (N15, L18, L21, V25, L28, A32, and T35) pointing toward the center of the pore (Fig. S11A) and F23, F26, V29, L31, and I33 stabilizing the helix-helix interfaces (Fig. S11B). This homopentameric conformation differs from that described for SARS-CoV (8-64 AA) (Fig. S11C) (Surya et al. 2018). We did not identify any relevant mutation in E protein.

#### Membrane

The membrane (M) protein of SARS-CoV-2 consists of 222 residues and is predicted to has three transmembrane domains for SARS-CoV (Ma et al. 2008) and SARS-CoV-2 (Wang et al. 2020). It is localized in the Endoplasmic Reticulum (ER)/Golgi apparatus complex (Wang et al. 2020). M protein from SARS-CoV-2 is necessary for the maturation, assembly, and release of virus-like particles when co-expressed with E, N, or S proteins (Boson et al. 2020). M protein inhibits the production of type I (α/β) and III (λ) IFNs induced by the RIG-I/MDA-5-MAVS signaling pathway by preventing the formation of multiprotein complexes that allow translocation of transcription factors to the nucleus (Wang et al. 2020).

#### Orf6

Orf6 is a 61 amino acid residue protein that is localized in membranous vesicles in the endoplasmic reticulum (Frieman et al. 2007). Orf6 exhibits the highest suppression of IFN synthesis and signaling among all viral proteins (Yuen et al. 2020). Orf6 inhibits the phosphorylation and translocation of IRF-3 to the nucleus, a transcription factor necessary for IFN synthesis (Kopecky-Bromberg et al. 2007, Li et al. 2020, Yuen et al. 2020). It interferes with the expression from the Interferon-sensitive response element (ISRE) promoter inhibiting IFN cellular response (Kopecky-Bromberg et al. 2007). It also prevents the translocation of STAT1 as a result of the sequestration of the nuclear translocation factors carioferin-α-2 and carioferin-β-1, which are retained in the ER membrane (Frieman et al. 2007). Our analysis does not show a relevant mutation in orf6.

#### Orf7a

Orf7a consists of 121 amino acids and shows a compact seven-stranded beta-sandwich (PDB: 6W37) (Fig. S12A) similar to members of the Ig superfamily (Fig. S12B). SARS-CoV orf7a presents % identity and 95.9% similarity with SARS-CoV-2 (Fig. S12C) and a similar 3D structure(Fig. S12D). Orf7a of SARS-Cov can be located in the Golgi apparatus exported by the COPII protein (118-120 are critical for this function) (Pekosz et al. 2006, Kopecky-Bromberg et al. 2006, Schaecher et al. 2007a). Also, it induces apoptosis through a caspase-dependent pathway (Tan et al. 2004, Schaecher et al. 2007b), activates the mitogen-activated protein kinase signaling pathway, inhibits translation (Kopecky-Bromberg et al 2006), suppresses progression of cell growth-inducing the arrest of the G0 / G1 cell phase through the cyclin D3/pRb pathway (Yuang et al. 2006) and is identified as a structural protein of the virion (Huang et al 2006). No relevant mutations were identified in orf7a.

#### Orf7b

Orf7b consists of 43 amino acids and does not have a described function or structure. It presents an 85.4 % identity and a 97.2 % sequence similarity with SARS-CoV (Fig. S13). Orf7b in SARS-CoV can be located in the Golgi apparatus (Schaecher et al. 2008), contributes to cell apoptosis (Schaecher et al. 2007b), and is a structural component of the virion (Schaecher et al. 2007a). The SARS-CoV homolog is an integral membrane protein (Schaecher et al. 2007a) extremely hydrophobic between amino acids 9 to 28 (Pekosz et al. 2006, Schaecher et al. 2007a). It has an eight residue N-terminus, a 22-residue transmembrane domain (TMD), and a 14-residue C-terminal tail exposed to the cytosol. TMD is essential for the localization of the protein, with residues 13-15 and 19-22 being the most critical for its localization on the Golgi complex (Schaecher et al. 2008). No relevant mutations were found in this protein.

#### Orf8

Orf8 from SARS-CoV-2 can directly interact with MHC-I molecules and reduce their surface expression (Zhang et al. 2020). In protein-protein interactions mapping between SARS-CoV-2 and human proteins using affinity-purification mass spectrometry, ORF8 shows physical interaction with 11 endoplasmic reticulum proteins (Gordon et al. 2020). Recently, its structure was solved (PDB: 7JTL) (Flower et al. 2020). SARS-CoV-2 orf8 is a 121 amino acid protein with an N-terminal signal sequence. The structure has two different dimerization interfaces: i) a covalent interface with a C20 disulfide bridge and other interactions through a specific N-terminal sequence (115-120 of SARS-CoV-2) and ii) a non-covalent interface formed by another specific motif of SARS-CoV-2 (73-YIDI-76) (Fig. 9A). These interfaces could allow SARS-CoV-2 orf8 to form oligomers that are not seen in other coronaviruses. S24L in the interface-1 showed 0.06 NRFp. In the crystallographic model, S24 forms a hydrogen bond with K53 of the other monomer, which stabilizes the dimerization interface. However, this only occurs at one end of the oligomer and not at the other showing that is not a stable h-bond (Fig. 9B). The presence of leucine in this position cannot stabilize the oligomer through hydrogen bonds.

**Figure 9.**
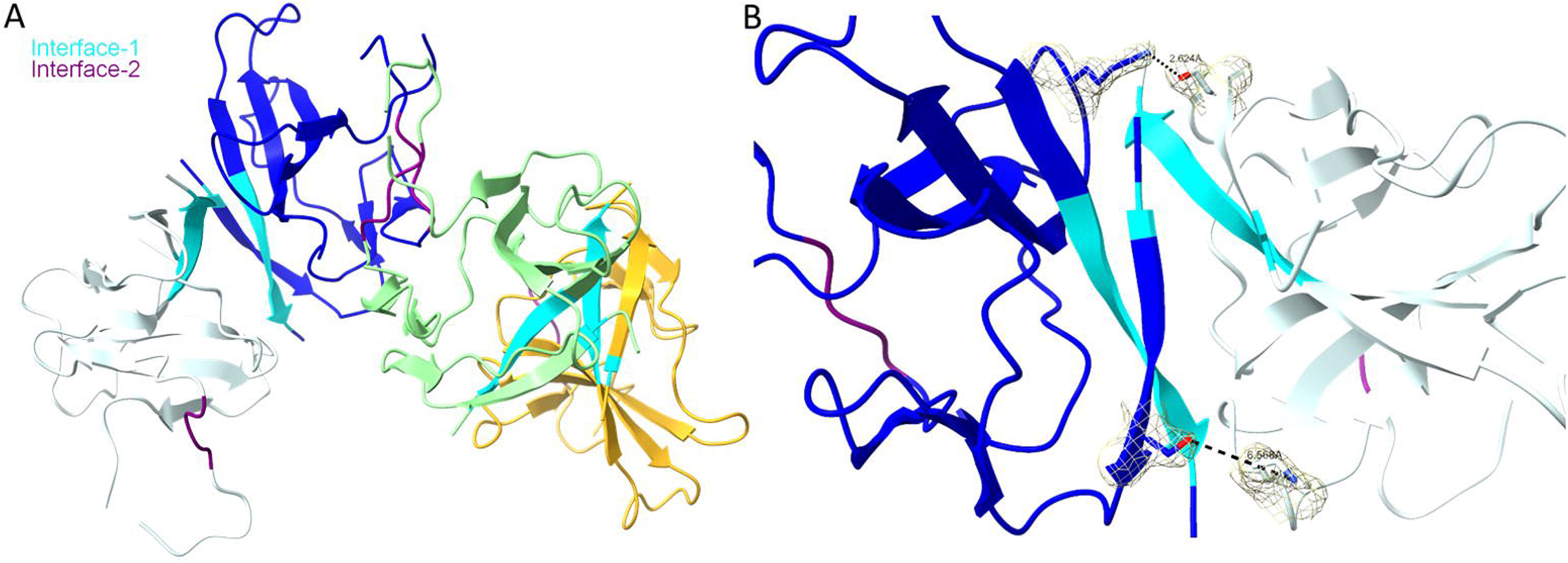
Mutation S24L would prevent the formation of a hydrogen bond a intermonomer interface of orf8. A) orf8 presents two interfaces of dimerization showed in cyan and magenta. B) Residue S24 is capable to form a hydrogen bond with K53, however, just one of the possible interactions is visualized in the structure indicating that is not sufficient stable. Thus, the mutation S25L would not significantly affects dimer stability.

#### Nucleocapsid

The nucleocapsid (N) is a highly conserved structural protein in the Coronavirus family (Lu et al. 2020). It has two well-differentiated domains: N-terminal (NTD), which is the main domain of interaction with RNA (Kang et al. 2020), and C-terminal (CTD), which allows protein dimerization (Zeng et al. 2020). The N protein is positively charged and contains disordered regions flanking NTD and CTD to facilitates its interaction with nucleic acids (Chang et al. 2009).

We found two HF (R203K and G204R) and four LF_nS (P13L, S194L, P199L, and A220V) mutations in N. Five of the six mutations are found in the central disordered region which contains serine/arginine (S/R) rich motifs prone to being phosphorylated (Surjit et al. 2005, Peng et al. 2008, Wu et al. 2009). N phosphorylation modulates its interaction with RNA and prevents its multimerization (Peng et al. 2008). Furthermore, S/R motifs have transcriptional regulation functions by recruiting DDX1 RNA helicase to the replication/transcription complex facilitating the processing of subgenomic mRNAs (Wu et al. 2014, Cong et al. 2020).

Using NetPhos 3.1 (Blom et al. 2004) we predicted the phosphorylation potential (PP) of eight phosphorylation sites (PS) (193S, 194S, 197S, 198T, 201S, 202S, 205T, and 206S) by different kinases (including protein kinase A (PKA), protein kinase B (PKB), protein kinase C (PKC), ribosomal S6 Kinase (RSK), cyclin-dependent kinase 1 (cdk1/CDC-2), casein kinase 1 (CKI), cyclin-dependent kinase 5 (CDK-5), p38 mitogen-activated protein kinase (p38MAPK) and glycogen synthase kinase 3 (GSK-3)) (Fig. 10) and evaluate the effects of ten identified genomic variants (Table. 2).

**Figure 10.**
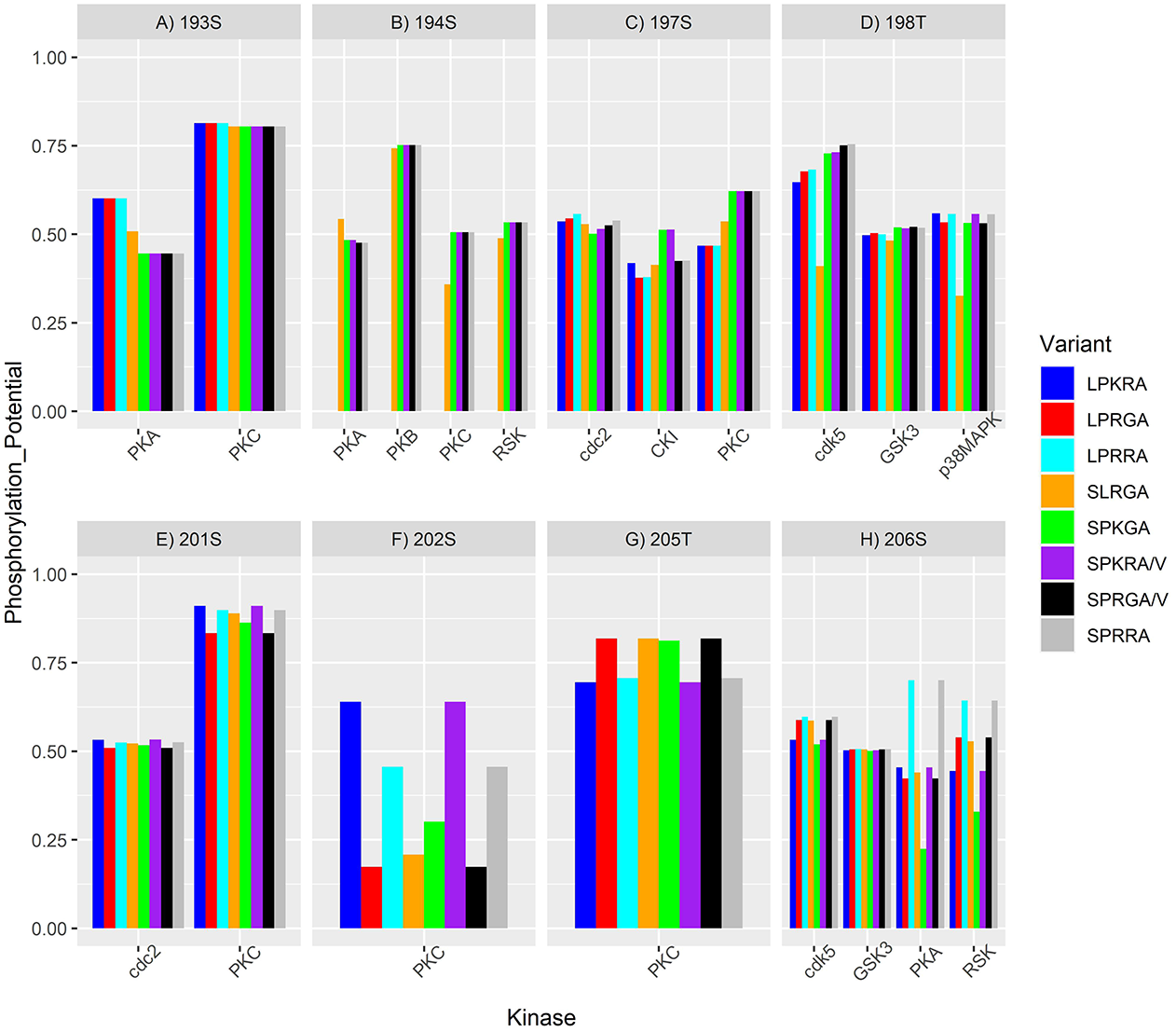
Nucleocapsid mutation affects the phosphorylation potential (PP) predicted by NetPhos in the central disordered region. A-H) Each panel shows the results of PP in each predicted phosphorylation site. X-axis in each panel shows the kinases predicted to phosphorylate the site in at least one variant with 0.5 or more PP. Variants are named based in the combination of the 5 positions where we identify mutations. i.e: Variant LPKRA means; L194,P199,K203,R204,A220.

**Table 2.**
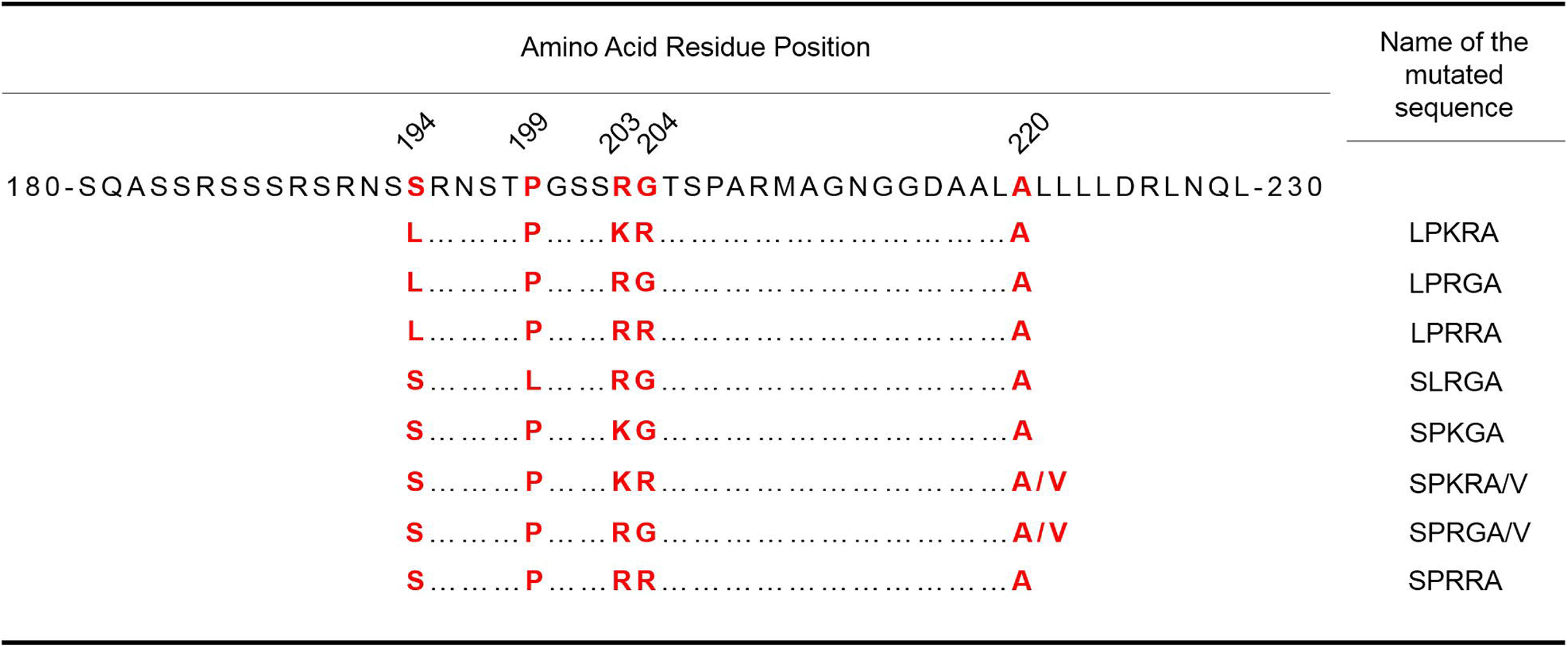
Description of the 10 nucleocapsid variants identified in the 100 924 genomes.

In the PS S193, PKA kinase showed differences between variants (Fig. 10A). PKA recognizes sequences where arginine is preferred at positions −3 and −2, whereas hydrophobic residues are preferred at the C-terminal to the phosphorylatable serine/threonine (Fig. S14) (Songyang et al 1994). Thus, the S194L and P199L mutations confer a hydrophobic amino acid at position +1 and +6, respectively, (Table. 2) increasing the PP on those variants (Fig. 10A). The distance of the hydrophobic residue from the PS seems to affects the PP (Fig. 10A).

At position S194 we observed PKA, PKC, and RSK with a different PP for the variant SLRGA (Fig. 10B). Similar to site S193, PKA has a slightly higher PP in the SLRGA variant (due to a Leucine in position +5). PKC shows the opposite pattern indicating that proline is preferred in position +5 for this kinase (Fig. 10B). RSK is a highly selective kinase that preferentially phosphorylates serine with basic amino acid residues at positions −5 and −3 (Fig. S14) (Leighton et al. 1995), with high selectivity toward arginine over any other basic residues (Galan et al. 2014). All variants for this kinase have the RSRNSS sequence (Table. 2), which matches the consensus phosphorylation sequence (Fig. S14). Despite this, the SLRGA variant had a slightly lower phosphorylation potential in comparison with variants with proline in position +5. S194L excluded the LPKRA, LPRGA, and LPRRA variants as possible targets due to the loss of the PS.

At residue S197 NetPhos identified cdc2, CKI, and PKC as possible kinases (Fig. 10C). For CKI, the consensus sequence S(P)-Xn-S has been proposed, where prior phosphorylation (S(P)) is a critical determinant of kinase action and two residues spacing (Xn = 2) is the best recognition site (Fig. S14) (Flotow et al. 1990). LPKRA, LPRGA, and LPRRA do not have the PS S194 reducing the PP for these variants (Fig. 10C). Also, the absence of K or the presence of R in position 203 seems to impact the PP of S197. For PKC, variants with S194 and P199 show the greatest PP. For cdc-2, subtle differences are predicted between the variants.

At PS T198, three kinases were identified, cdk-5, GSK-3, and p38MAPK (Fig. 10D). CDK family and MAPKs require a proline just after the phospho-acceptor residue (S/T-P motif) (Fig. S14) (Sheridan et al. 2008, Songyang et al. 1996). Thus, in the SLRGA variant, the loss of proline significantly decreases its PP (Fig. 10D). The greatest difference between the other variants may be due to differences in position 194 (−4) for cdk-5 and position 204 (+6) for p38MAPK. GSK-3 recognizes the sequence S-X-X-X-S(P), where prior phosphorylation is required for the recognition site (Fig. S14) (Fiol et al. 1987). All the variants present this sequence and show little differences among their PPs (Fig. 10D).

PKC and cdc2 were predicted for PS S201 (Fig. 10E). Cdc2 has shown slight differences between the variants. Woodgett et al., (1986) reported that the sequence S/T-X-K/R, where X is usually an uncharged residue, is a target sequence for PKC (Fig. S14). Additionally, basic residues at both the N and C termini of the PS enhance the PP. We observed that all the variants have a high PP due to the presence of the uncharged serine at position +1 and the basic lysine/arginine at position +2. The differences between the variants are due to the difference in the basic residue (lysine or arginine) and proline or leucine in position −2 (Fig. 10E).

Similar results for PKC are observed in the PS S202, except for the variant SLRGA that opposite to the S201 it showed lesser PP than SPKGA (Fig. 10F). The PS T205 shows a higher PP for PKC in variants where residue G204 is present (Fig. 10G). G204 creates the PKC consensus sequence described by Nishikawa et al., (1997) (Fig. S14).

NetPhos predicted cdk5, PKA, and RSK as potential kinases for S206 with variable PP for the variants (Fig. 10H). All variants show a proline at the position +1 necessary for cdk5 recognition (Fig. S14). Thus, subtle differences appear due to differences in the c-terminal of S206 with variants with RR (−2, −3) as those with the highest PP (Fig. 10H). For PKA and RSK, LPRRA and SPRRA variants showed the highest PP. Again, this can be explained by the presence of RR. When just one R is present the PP goes down and when there is none it goes down even more. Also, we noted that NetPhos predicts a higher PP where R is presented in position 204 than when it is present at position 203 for PKA and the opposite for RSK.

The P13L and A220V mutation do not generate differences in kinase PPs predicted by NetPhos due to its distance from the predicted phosphorylation sites.

#### Orf10

ORF10 is a 38 amino acid with no known function or with homologous proteins in other coronaviruses, not relevant results in Pfam (El-Gebali et al. 2019), BLAST search (Altschul et al. 1990) or SWISS-MODEL (Waterhouse et al. 2018) was found. V30L mutations showed 0.03 NRFp, but we cannot speculate its function due to the lack of information.

## MATERIAL AND METHODS

### Global normalized relative frequency analysis of each base in each ORF from SARS-CoV-2

To perform the mutation frequency analysis in each ORF, we first downloaded a total of 100 924 complete and high coverage genomes from the GISAID database (as of October 05^th^, 2020). After that, we randomly divided the genomes in 4 groups of 25 231 genomes each. Each set of genomes was divided in genomes with at least one ambiguity and without ambiguities (any nucleotide different to A, C, T or G was identified as ambiguous). The 8 sets of genomes were aligned using MAFFT with FFT-NS-2 strategy and default parameter settings (Katoh et al. 2002). In each alignment, we removed columns that do not correspond to the region from nt 203 to nt 29674, and insertions respect to the genome EPI_ISL_402125 (Thus, our analysis does not cover the frequency of insertions respect to the reference genome EPI_ISL_402125). After that, we repeat the alignment with MAFFT (Katoh et al. 2002) (same setting that the previous one) and the columns removal iteratively until we do not obtain difference in the number of columns after the alignment (indicating that any other insertion was detected in the region comprised nt 203 to nt 29674 in the reference genome). Finally, we bound the 8 alignments using cat function in Linux and use this to extract regions corresponding to each of the ORFs and nsp regions of SARS-CoV-2 (regions as annotated in the NCBI database of the Wuhan-Hu-1 reference genome). These regions were translated to obtain the AA sequences. After that, sequences were divided by continent-month combinations, aligned using MAFFT with FFT-NS-2 strategy and default parameter settings (Katoh et al. 2002), columns with more than 98 % gaps were removed and relative frequencies of each base or gap in each position were calculated (***RF***_*p,m*−*c*_). To obtain Normalized Relative Frequencies by COVID-19 cases (***NRF***_*p*_) we follow similar steps as described in (Justo et al. 2020). Thus, relative frequencies were multiplied by the number of cases reported in the respective continent-month combination (***CN***_*m*−*c*_) obtaining an estimation of the number of cases that present a virus with a specific base or gap in a specific genomic position (***RF***_*p*_***CN***_*m*−*c*_). Finally, we added the ***RF***_*p*_***CN***_*m*−*c*_ of each subalignment and divided it by the total number of cases in the world (Σ_*m*−*c*_ ***RF***_*o*_***CN***_*m*−*c*1_)/***TCN***_*w*_. This procedure allows us to obtain a relative frequency normalized by cases of each base or gap in each genomic position (***NRF***_*p*_). The number of cases of each country was obtained from the European Centre for Disease Prevention and Control: https://www.ecdc.europa.eu/en/publications-data/download-todays-data-geographic-distribution-covid-19-cases-worldwide. We used the number of cases of countries with at least one genome sequenced and deposited in GISAID database. Also, we just consider in the analysis month-continent combinations with at least 90 genomes sequenced. Plot figures were done using R (v.4.0.3) (R core team. 2014) with the ggplot2 package (Wickham 2016).

### Frequency analysis by continent and month

***RF***_*p,m*−*c*_ from the previous calculations were used to analyze the variation of relative frequencies of the most frequents mutations in each continent by month. Plot figures were done using R (v.4.0.3) (R core team. 2014) with the ggplot2 package (Wickham 2016).

### Energy minimization of structural models of mutations

Protein structures models of proteins … were downloaded from Protein Data Bank (PDB) (Berman et al. 2000), open in UCSF ChimeraX (v.1.1) (Pettersen et al. 2020) and mutations were modelled using swapaa command. Potential energy of mutational and wild-type models was minimized using Gromacs (v.2018.8) (Berendsen et al. 1995). Briefly, models were solvated and ionized using gmx solvate and gmx genion commands. Finally, 10 000 steps of energy minimization were performed using steepest descent integrator. All structural images were produced using ChimeraX (v.1.1) (Pettersen et al. 2020) or Chimera (v.1.15) (Pettersen et al. 2004).

### Predictions of Phosphorylation Potential (PP)

SARS-CoV-2 nucleocapsid sequences from AA 180 to 229 of the ten different identified variants were extracted and submitted to NetPhosK (Blom et al 2004) to perform the phosphorylation potential predictions. Plot figures were done using R (v.4.0.3) (R core team. 2014) with the ggplot2 package (Wickham 2016).

## Supporting information

Figure_S1_interactive

GISAID_acknowledgements_part_1

GISAID_acknowledgements_part_2

GISAID_acknowledgements_part_3

GISAID_acknowledgements_part_4

GISAID_acknowledgements_part_5

GISAID_acknowledgements_part_6

GISAID_acknowledgements_part_7

GISAID_acknowledgements_part_8

GISAID_acknowledgements_part_9

GISAID_acknowledgements_part_10

GISAID_acknowledgements_part_11

GISAID_acknowledgements_part_12

## DATA AVAILABILITY

The data that support the findings of this study comes from the GISAID initiative (Shu and McCaluey. 2017) (gisaid.org).

## Acknowledgements

We are very grateful to the GISAID Initiative and all its data contributors, i.e. the Authors from the Originating laboratories responsible for obtaining the specimens and the Submitting laboratories where genetic sequence data were generated and shared via the GISAID Initiative, on which this research is based. Complete acknowledgements of the 100 924 genomes used are available in supplementary file (S-). To the Ricardo Palma University High-Performance Computational Cluster (URPHPC) managers Gustavo Adolfo Abarca Valdiviezo and Roxana Paola Mier Hermoza at the Ricardo Palma Informatic Department (OFICIC) for their contribution in programs and remote use configuration of URPHPC.

## REFERENCES

1. Adedeji A, Marchand B, Te Velthuis A, Snijder E, Weiss S, Eoff R, Singh K, Sarafianos S. 2012. Mechanism of nucleic acid unwinding by SARS-CoV helicase. Vol. 7(5): 1–11. PloS One.

2. Altschul S, Gish W, Miller W, Myers E, Lipman D. 1990. Basic Local Alignment Search Tool. Vol. 215, 403–410. Journal of Molecular Biology.

3. Baez-Santos Y, John S, Mesecar A. 2014. The SARS-coronavirus papain-like protease: Structure, function and inhibition by designed antiviral compounds. Vol. 115: 21–38. Antiviral Research.

4. Baliji S, Cammer S A, Sobral B, Baker S. 2009. Detection of Nonstructural Protein 6 in Murine Coronavirus-Infected Cells and Analysis of the Transmembrane Topology by Using Bioinformatics and Molecular Approaches. Vol. 83(13), 6957–6962. Journal of Virology.

5. Barretto N, Jukneliene D, Ratia K, Chen Z, Mesecar A, Baker S. 2005. The Papain-Like Protease of Severe Acute Respiratory Syndrome Coronavirus Has Deubiquitinating Activity. Vol. 79(24): 15189–15198. Journal of Virology.

6. Benvenuto D, Angeletti S, Giovanetti M, Bianchi M, Pascarella S, Cauda R, Ciccozzi M, Cassone, A. 2020. Evolutionary analysis of SARS-CoV-2: how mutation of Non-Structural Protein 6 (NSP6) could affect viral autophagy. Vol. 81(1): e24–e27. Journal of Infection.

7. Berendsen H, van der Spoel D, van Drunen R. 1995. GROMACS: A message-passing parallel molecular dynamics implementation. Vol. 91(1-3): 43–56. Computer Physics Communications.

8. Berman H, Westbrook J, Feng Z, Gilliland G, Bhat T, Weissig H, Shindyalov I, Bourne P. 2000. The Protein Data Bank. Vol. 28(1): 235–242. Nucleic Acids Research.

9. Blom N, Sicheritz-Pontén T, Gupta R, Gammeltoft S, Brunak S. 2004. Prediction of post-translational glycosylation and phosphorylation of proteins from the amino acid sequence. Vol. 4(6):1633–1649. Proteomics.

10. Bosch B, van der Zee R, de Haan C, Rottier P. 2003. The Coronavirus Spike Protein is a Class I Virus Fusion Protein: Structural and Functional Characterization of the fusion core complex. Vol. 77(16), 8801–8811. Journal of Virology.

11. Chang CK, Hsu YL, Chang YH, Chao FA, Wu MC, Huang YS, Hu CK, Huang TH. 2009. Multiple nucleic acid binding sites and intrinsic disorder of severe acute respiratory syndrome coronavirus nucleocapsid protein: implications for ribonucleocapsid protein packaging. Vol. 83(5): 2255–2264. Journal of virology.

12. Clasman J, Báez-Santos Y, Mettelman R, O’Brien A, Baker S, Mesecar A. 2017. X-ray Structure and Enzymatic Activity Profile of a Core Papain-like Protease of MERS Coronavirus with utility for structure-based drug design. Vol. 7: 40292. Scientific Reports.

13. Cong Y, Ulasli M, Schepers H, Mauthe M, V’kovski P, Kriegenburg F, Thiel V, de Haan CAM, Reggiori F. 2020. Nucleocapsid Protein Recruitment to Replication-Transcription Complexes Plays a Crucial Role in Coronaviral Life Cycle. Vol. 94(4): e01925–19. Journal of virology.

14. Cornillez-Ty C, Liao L, Yates J, Kuhn P, Buchmeier M. 2009. Severe acute respiratory syndrome coronavirus nonstructural protein 2 interacts with a host protein complex involved in mitochondrial biogenesis and intracellular signaling. Vol. 83(19), 10314–10318. Journal of Virology.

15. Cottam E, Whelband M, Wileman T. 2014. Coronavirus NSP6 restricts autophagosome expansion. Vol. 10(8): 1426–1441. Autophagy.

16. Deng X, Hackbart M, Mettelman R, O’Brien A, Mielech A, Yi G, Kao C, Baker S. 2017. Coronavirus nonstructural protein 15 mediates evasion of dsRNA sensors and limits apoptosis in macrophages. Vol. 114(21): 4251–4260. Proceeding of the National Academy of Sciences of the United States of America.

17. El-Gebali S, Mistry J, Bateman A, Eddy S, Luciani A, Potter S, Qureshi M, Richardson L, Salazar G, Smart A, Sonnhammer E, Hirsh L, Paladin L, Piovesan D, Tosatto S, Finn R. 2019. The Pfam protein families database in 2019. Vol. 47(D1), D427–D432. Nucleic Acids Research.

18. Ferron F, Subissi L, Silveira A, Le N, Sevajol M, Gluais L, Decroly E, Vonrhein C, Bricogne G, Canard B, Imbert I. 2018. Structural and molecular basis of mismatch correction and ribavirin excision from coronavirus RNA. Vol. 115(2): 162–171. Proceeding of the National Academy of Sciences of the United States of America.

19. Fiol CJ, Mahrenholz AM, Wang Y, Roeske RW, Roach PJ. 1987. Formation of protein kinase recognition sites by covalent modification of the substrate. Molecular mechanism for the synergistic action of casein kinase II and glycogen synthase kinase 3. Vol. 262(29): 14042–14048. The Journal of biological chemistry.

20. Flotow H, Graves PR, Wang AQ, Fiol CJ, Roeske RW, Roach PJ. 1990. Phosphate groups as substrate determinants for casein kinase I action. Vol. 265(24):14264–14269. The Journal of biological chemistry.

21. Flower T, Buffalo C, Hooy R, Allaire M, Ren X, Hurley J. 2020. Structure of SARS-CoV-2 ORF8, a rapidly evolving coronavirus protein implicated in immune evasion. bioRxiv preprint DOI: https://doi.org/10.1101/2020.08.27.270637

22. Fusaro G, Dasgupta P, Rastogi S, Joshi B, Chellappan S. 2003. Prohibitin induces the transcriptional activity of p53 and is exported from the nucleus upon apoptotic signaling. Vol. 278(48), 47853–47861. The Journal of Biological Chemistry.

23. Galan JA, Geraghty KM, Lavoie G, Kanshin E, Tcherkezian J, Calabrese V, Jeschke GR, Turk BE, Ballif BA, Blenis J, Thibault P, Roux PP. 2014. Phosphoproteomic analysis identifies the tumor suppressor PDCD4 as a RSK substrate negatively regulated by 14-3-3. Vol. 111(29):E2918–E2927. Proceedings of the National Academy of Sciences of the United States of America.

24. Gao Y, Yan L, Huang Y, Liu F, Zhao Y, Cao L, Wang T, Sun Q, Ming Z, Zhang L, Ge J, Zheng L, Zhang Y, Wang H, Zhu Y, Zhu C, Hu T, Hua T, Zhang B, Yang X, Li J, Yang H, Liu Zhijie, Xu W, Guddat L, Wang Q, Zhiyong, Rao Z. 2020. Structure of the RNA-dependent RNA polymerase from COVID-19 virus. Vol. 368: 779–782. Science.

25. Gordon D, Jang G, Bouhaddou M, Xu J, Obernier K, White K et al. 2020. A SARS-CoV-2 protein interaction map reveals targets for drug repurposing. Vol. 583(7816): 459–468. Nature.

26. Graham R, Sims A, Brockway S, Baric S, Denison M. 2005. The nsp2 replicase protein of murine hepatitis virus and severe acute respiratory syndrome coronavirus is dispensable for viral replication. Vol. 79(21), 13399–13411. Journal of Virology.

27. Huang C, Ito N, Tseng C, Makino S. 2006. Severe Acute Respiratory Syndrome Coronavirus 7a Accessory Protein Is a Viral Structural Protein. Vol. 80(15): 7287–7294. Journal of Virology.

28. Ivanov K, Thiel V, Dobbe J, Van der Meer Y, Snijder E, Ziebuhr J. 2004. Multiple Enzymatic Activities Associated with Severe Acute Respiratory Syndrome Coronavirus Helicase. Vol. 78(11): 5619–5632. Journal of Virology.

29. Jia Z, Yan L, Ren Z, Wu L, Wang J, Guo J, Zheng L, Ming Z, Zhang L, Lou Z, Rao Z. 2019. Delicate structural coordination of the Severe Acute Respiratory Syndrome coronavirus Nsp13 upon ATP hydrolysis. Vol. 47(12): 6538–6550. Nucleic Acids Research.

30. Justo S, Zapata D, Huallpa C, Landa G, Castillo A, Garavito-Salini R, Uceda-Campos G, Pineda R. 2020. Global geographic and temporal analysis of SARS-CoV-2 haplotypes normalized by COVID-19 cases during the first seven months of the pandemic. bioRxiv preprint doi: https://doi.org/10.1101/2020.07.12.199414.

31. Kang S, Yang M, Hong Z, Zhang L, Huang Z, Chen X, He S, Zhou Z, Zhou Z, Chen Q, Yan Y, Zhang C, Shan H, Chen S. 2020. Crystal structure of SARS-CoV-2 nucleocapsid protein RNA binding domain reveals potential unique drug targeting sites. Vol. 10(7): 1228–1238. Acta pharmaceutica Sinica. B.

32. Katoh K, Misawa K, Kuma K, Miyata T. 2002. MAFFT: a novel method for rapid multiple sequence alignment based on fast Fourier transform. Vol. 30(14): 3059–3066. Nucleic Acids Research.

33. Kim Y, Jedrzejczak R, Maltseva N, Wilamowski M, Endres M, Godzik A, Michalska K, Joachimiak A. 2020. Crystal structure of Nsp15 endoribonuclease NendoU from SARS-CoV-2. Vol. 29: 1596–1605. Protein Science.

34. Kopecky-Bromberg S, Martinez-Sobrido L, Palese, P. 2006. 7a Protein of Severe Acute Respiratory Syndrome Coronavirus Inhibits Cellular Protein Synthesis and Activates p38 Mitogen-Activated Protein Kinase. Vol. 80(2): 785–793. Journal of Virology.

35. Krafcikova P, Silhan J, Nencka R, Boura E. 2020. Structural analysis of the SARS-CoV-2 methyltransferase complex involved in RNA cap creation bound to sinefungin. Vol. 11:3717 Nature Communications.

36. Krammer F. 2020. SARS-CoV-2 vaccines in development. Vol. 586, 516–527. Nature.

37. Kumar P, Gunalan V, Liu B, Chow V, Druce J, Birch C, Catton M, Fielding B, Tan Y, Lal S. 2007 The nonstructural protein 8 (nsp8) of the SARS coronavirus interacts with its ORF6 accessory protein. Vol. 366: 293–303. Virology.

38. Kusov Y, Tan J, Alvarez E, Enjuanes L, Hilgenfeld R. 2015. A G-quadruplex-binding macrodomain within the “SARS-unique domain” is essential for the activity of the SARS-coronavirus replication–transcription complex. Vol. 484: 313–322. Virology.

39. Leighton IA, Dalby KN, Caudwell FB, Cohen PT, Cohen P. 1995. Comparison of the specificities of p70 S6 kinase and MAPKAP kinase-1 identifies a relatively specific substrate for p70 S6 kinase: the N-terminal kinase domain of MAPKAP kinase-1 is essential for peptide phosphorylation. Vol. 375(3): 289–293. FEBS letters.

40. Lei J, Kusov Y, Hilgenfeld R. 2018. Nsp3 of coronaviruses: Structures and functions of a large multi-domain protein. Vol. 149: 58–74. Antiviral Research.

41. Littler D, Gully B, Colson R, Rossjohn J. 2020. Crystal Structure of the SARS-CoV-2 Non-structural Protein 9, Nsp9. Vol. 23(7): 101258. iScience.

42. Looi M. 2020. Covid-19: Is a second wave hitting Europe?. Vol. 371: m4113. BMJ. DOI: https://doi.org/10.1136/bmj.m4113

43. Lu R, Zhao X, Li J, Niu P, Yang B, Wu H, Wang W, Song H, Huang B, Zhu N, Bi Y, Ma X, Zhan F, Wang L, Hu T, Zhou H, Hu Z, Zhou W, Zhao L, Chen J, Meng Y, Wang J, Lin Y, Yuan J, Xie Z, Ma J, Liu WJ, Wang D, Xu W, Holmes EC, Gao GF, Wu G, Chen W, Shi W, Tan W. 2020. Genomic characterization and epidemiology of 2019 novel coronavirus: implications for virus origins and receptor binding. Vol. 395(10224): 565–574. Lancet.

44. Ma Y, Wu L, Shaw N, Gao Y, Wang J, Sun Y, Lou Z, Yan L, Zhang R, Rao Z. 2015. Structural basis and functional analysis of the SARS coronavirus nsp14–nsp10 complex. Vol. 112(30): 9436–9441. Proceeding of the National Academy of Sciences of the United States of America.

45. Merkwirth C and Langer T. 2008. Prohibitin function within mitochondria: essential roles for cell proliferation and cristae morphogenesis. Vol. 1793: 27–32. Biochimica et Biophysica Acta.

46. Miknis Z, Donaldson E, Umland T, Rimmer R, Baric R, Schultz, Wayne L. 2009. Severe Acute Respiratory Syndrome Coronavirus nsp9 Dimerization Is Essential for Efficient Viral Growth. Vol. 83(7): 3007–3018. Journal of Virology.

47. Nelson C, Pekosz A, Lee C, Diamond M, Fremont D. 2005. Structure and intracellular targeting of the SARS-coronavirus orf7a accessory protein. Vol. 13(1): 75–85. Structure.

48. Nishikawa K, Toker A, Johannes FJ, Songyang Z, Cantley LC. 1997. Determination of the specific substrate sequence motifs of protein kinase C isozymes. Vol. 272(2):952–960. The Journal of biological chemistry. https://doi.org/10.1074/jbc.272.2.952

49. Oostra M, te Lintelo E, Deijs M, Verheije M H, Rottier P, de Haan M. 2007. Localization and Membrane Topology of Coronavirus Nonstructural Protein 4: Involvement of the Early Secretory Pathway in Replication. Vol. 81(22): 12323–12336. Journal of Virology.

50. Oudshoorn D, Rijs K, Limpens R, Groen K, Koster A, Snijder E, Kikkert M, Barcena M. 2017. Expression and cleavage of middle east respiratory syndrome coronavirus nsp3-4 polyprotein induce the formation of double-membrane vesicles that mimic those associated with coronaviral RNA replication. Vol. 8(6): e01658–17. mBio.

51. Pachetti M, Marini B, Benedetti F, Giudici F, Mauro E, Storici P, Masciovecchio C, Angeletti S, Ciccozzi M, Gallo R, Zella D, Ippodrino R. 2020. Emerging SARS-CoV-2 mutation hot spots include a novel RNA-dependent-RNA polymerase variant. Vol. 18(179): 1–9. Journal of Translational Medicine.

52. Pekosz A, Schaecher S, Diamond M, Fremont D, Sims A, Baric R. 2006. Structure, expression, and intracellular localization of the SARS-CoV accessory proteins 7a and 7b. Vol 581: 115–120. Advances in Experimental Medicine and Biology.

53. Pettersen E, Goddard T, Huang C, Meng E, Couch G, Croll T, Morris J, Ferrin T. 2020. UCSF ChimeraX: structure visualization for researchers, educators, and developers. Protein Science. DOI: 10.1002/pro.3943

54. Pettersen E, Goddard T, Huang C, Couch G, Greenblatt D, Meng E, Ferrin R. 2004. UCSF Chimera—A visualization sustem for exploratory research and analysis. Vol. 25(13): 1605–1612. Journal of Computational Chemistry.

55. Pillon M, Frazier M, Dillard L, Williams J, Kocaman S, Krahn J, Perera L, Hayne C, Gordon J, Stewart Z, Sobhany M, Deterding L, Hsu A, Dandey V, Borgnia M, Stanley R. 2020. Cryo-EM Structures of the SARS-CoV-2 Endoribonuclease Nsp15. BioRxiv preprint DOI: https://doi.org/10.1101/2020.08.11.244863

56. Peng TY, Lee KR, Tarn WY. 2008. Phosphorylation of the arginine/serine dipeptide-rich motif of the severe acute respiratory syndrome coronavirus nucleocapsid protein modulates its multimerization, translation inhibitory activity and cellular localization. Vol. 275(16): 4152–4163. The FEBS journal.

57. Rajalingam K, Wunder C, Brinkmann V, Churin Y, Hekman M, Sievers C, Rapp U, Rudel T. 2005. Prohibitin is required for RAS-induced RAF-MEK-ERK activation and epithelial cell migration. Vol. 7(8), 837–843. Nature Cell Biology.

58. R core team. 2014. R: A language and environment for statistical computing. R Foundation for statistical computing, Vienna, Austria. URL: http://www.R-project.org/.

59. Robson F, Khan K, Le T, Paris C, Demirbag S, Barfuss P, Rocchi P, Ng Wl. 2020. Coronavirus RNA Proofreading: Molecular Basis and Therapeutic Targeting. Vol. 79(5): 710–727. Molecular Cell.

60. Sakai Y, Kawachi K, Terada Y, Omori H, Matsuura Y, Kamitani W. 2017. Two-amino acids change in the nsp4 of SARS coronavirus abolishes viral replication. Vol. 510: 165–174. Virology.

61. Schaecher S, Diamond M, Pekosz A. 2008. The Transmembrane Domain of the Severe Acute Respiratory Syndrome Coronavirus ORF7b Protein Is Necessary and Sufficient for Its Retention in the Golgi Complex. Vol. 82(19): 9477–9491. Journal of Virology.

62. Schaecher S, Mackenzie J, Pekosz A. 2007a. The ORF7b Protein of Severe Acute Respiratory Syndrome Coronavirus (SARS-CoV) Is Expressed in Virus-Infected Cells and Incorporated into SARS-CoV Particles. Vol. 81(2): 718–731. Journal of Virology.

63. Schaecher S, Touchette E, Schriewer J, Buller R, Pekosz A. 2007b. Severe Acute Respiratory Syndrome Coronavirus Gene 7 Products Contribute to Virus-Induced Apoptosis. Vol. 81(20): 11054–11068. Journal of Virology.

64. Selvaraj C, Dinesh D, Panwar U, Abhirami R, Boura E, Singh S. 2020. Structure-based virtual screening and molecular dynamics simulation of SARS-CoV-2 Guanine-N7 methyltransferase (nsp14) for identifying antiviral inhibitors against COVID-19. Journal of Biomolecular Structure & Dynamics. DOI: doi.org/10.1080/07391102.2020.1778535

65. Sheridan DL, Kong Y, Parker SA, Dalby KN, Turk BE. 2008. Substrate discrimination among mitogen-activated protein kinases through distinct docking sequence motifs. Vol. 283(28):19511–19520. The Journal of biological chemistry.

66. Shin D, Mukherjee R, Grewe D, Bojkova D, Baek K, Bhattacharya A, Schulz L, Widera M, Mehdipour A, Tascher G, Geurink P, Wilhelm A, van Noort G, Ovaa H, Müller S, Knobeloch K-P, Rajalingam K, Schulman B, Cinatl J, Hummer G, Clesek S, Dikic I. 2020. Papain-like protease regulates SARS-CoV-2 viral spread and innate immunity. https://doi.org/10.1038/s41586-020-2601-5. Nature.

67. Shu Y and McCauley J. 2017. GISAID: Global initiative on sharing all influenza data – from vision to reality. Vol. 22(13). Eurosurveillance.

68. Songyang Z, Blechner S, Hoagland N, Hoekstra MF, Piwnica-Worms H, Cantley LC. 1994. Use of an oriented peptide library to determine the optimal substrates of protein kinases. Vol. 4(11):973–982. Current biology.

69. Songyang Z, Lu KP, Kwon YT, Tsai LH, Filhol O, Cochet C, Brickey DA, Soderling TR, Bartleson C, Graves DJ, DeMaggio AJ, Hoekstra MF, Blenis J, Hunter T, Cantley LC. 1996. A structural basis for substrate specificities of protein Ser/Thr kinases: primary sequence preference of casein kinases I and II, NIMA, phosphorylase kinase, calmodulin-dependent kinase II, CDK5, and Erk1. Vol. 16(11):6486–6493. Molecular and cellular biology.

70. Subissi L, Posthuma C, Collet A, Zevenhoven-Dobbe J, Gorbalenya A, Decroly E, Snijder E, Canard B, Imbert I. 2014. One severe acute respiratory syndrome coronavirus protein complex integrates processive RNA polymerase and exonuclease activities. Vol. 111(37): E3900–E3909. PNAS.

71. Sun L, Liu L, Yang X, Wu Z. 2004. Akt binds prohibitin 2 and relieves its repression of MyoD and muscle differentiation. Vol. 117(14), 3021–3029. Journal of Cell Science.

72. Surjit M, Kumar R, Mishra RN, Reddy MK, Chow VT, Lal SK. 2005. The severe acute respiratory syndrome coronavirus nucleocapsid protein is phosphorylated and localizes in the cytoplasm by 14-3-3-mediated translocation. Vol. 79(17): 11476–11486. Journal of virology.

73. Sutton G, Fry E, Carter L, Sainsbury S, Walter T, Nettleship J, Berrow N, Owens R, Gilbert R, Davidson A, Siddell S, Poon L, Diprose J, Alderton D, Walsh, Grimes J, Stuart D. 2004. The nsp9 Replicase Protein of SARS-Coronavirus, Structure and Functional Insights. Vol. 12(2): 341–353. Structure.

74. Tan J, Vonrhein C, Smart O, Bricogne G, Bollati M, Kusov Y, Hansen G, Mesters J, Schmidt C, Hilgenfeld R. 2009. The SARS-Unique Domain (SUD) of SARS Coronavirus Contains Two Macrodomains That Bind G-Quadruplexes. Vol. 5(5): e1000428. Plos Pathogens.

75. Tan Y, Fielding B, Goh P, Shen S, Tan T, Lim S, Hong W. 2004. Overexpression of 7a, a Protein Specifically Encoded by the Severe Acute Respiratory Syndrome Coronavirus, Induces Apoptosis via a Caspase-Dependent Pathway. Vol. 78(24): 14043–14047. Journal of Virology.

76. Te Velthuis A, van den Worm S, Snijder E. 2011. The SARS-coronavirus nsp7+nsp8 complex is a unique multimeric RNA polymerase capable of both de novo initiation and primer extension. Vol. 40(4): 1737–1747. Nucleic Acids Research.

77. Tortorici A, Vessler D. 2019. Structural insights into coronavirus entry. Vol. 105, 93–116. Advances in Virus Research.

78. Ulrich S and Nitsche C. 2020. The SARS-CoV-2 main protease as drug target. Vol. 30: 127377. Bioorganic & Medicinal Chemistry Letters.

79. Viswanathan T, Arya S, Chan S, Qi S, Dai N, Misra A, Park J, Oladunni F, Kovalskyy D, Hromas R, Martinez-Sobrido L, Gupta Y. 2020. Structural basis of RNA cap modification by SARS-CoV-2. Vol. 11: 3718. Nature Communications.

80. Wang S, Nath N, Adlam M, Chellappan S. 1999. Prohibitin, a potential tumor suppressor, interacts with RB and regulates E2F function. Vol. 18, 3501–3510. Oncogene.

81. Waterhouse A, Bertoni M, Bienert S, Studer G, Tauriello G, Gumienny R, Heer F, de Beer T, Rempfer C, Bardoli L, Lepore R, Schwede T. 2018. SWISS-MODEL: homology modelling of a protein structures and complexes. Vol. 46(W1), W296–W303. Nucleic Acids Research.

82. WHO. 2020. https://www.who.int/emergencies/diseases/novel-coronavirus-2019. Retrieved on 12 November 2020.

83. Wickham. 2016. Ggplot2: Elegant graphics for data analysis. Springer-Verlag New York. ISBN 978-3-319-24277-4. https://ggplot2.tidyverse.org.

84. Woodgett JR, Gould KL, Hunter T. 1986. Substrate specificity of protein kinase C. Use of synthetic peptides corresponding to physiological sites as probes for substrate recognition requirements. Vol. 161(1):177–184. European journal of biochemistry.

85. Wu CH, Chen PJ, Yeh SH. 2014. Nucleocapsid phosphorylation and RNA helicase DDX1 recruitment enables coronavirus transition from discontinuous to continuous transcription. Vol. 16(4): 462–472. Cell host & microbe.

86. Wu CH, Yeh SH, Tsay YG, Shieh YH, Kao CL, Chen YS, Wang SH, Kuo TJ, Chen DS, Chen PJ. 2009. Glycogen synthase kinase-3 regulates the phosphorylation of severe acute respiratory syndrome coronavirus nucleocapsid protein and viral replication. Vol. 284(8): 5229–5239. The Journal of biological chemistry.

87. Xia B and Kang X. 2011. Activation and maturation of SARS-CoV main protease. Vol. 2(4): 282–290. Protein Cell.

88. Yuan X, Wu J, Shan Y, Yao Z, Dong B, Chen B et al. 2006. SARS coronavirus 7a protein blocks cell cycle progression at G0/G1 phase via the cyclin D3/pRb pathway. Vol. 346(1): 74–85. Virology.

89. Zeng W, Liu G, Ma H, Zhao D, Yang Y, Liu M, Mohammed A, Zhao C, Yang Y, Xie J, Ding C, Ma X, Weng J, Gao Y, He H, Jin T. 2020. Biochemical characterization of SARS-CoV-2 nucleocapsid protein. Vol. 527(3): 618–623. Biochemical and biophysical research communications.

90. Zhang C, Chen Y, Li L, Yang Y, He J, Chen C, Su D. 2020. Structural basis for the multimerization of nonstructural protein nsp9 from SARS-CoV-2. Vol. 1(1):5. Molecular Biomedicine.

91. Zhang L, Lin D, Sun X, Curth U, Drosten C, Sauerhering L, Becker S, Rox K, Hilgenfeld R. 2020. Crystal structure of SARS-CoV-2 main protease provides a basis for design of improved a-ketoamide inhibitors. Vol. 368: 409–412. Science.

92. Zhang Y, Zhang J, Chen Y, Luo B, Yuan Y, Huang F, Yang T, Yu F, Liu J, Liu B, Song Z, Chen J, Pan T, Zhang X, Li Y, Li R, Huang W, Xiao F, Zhang H. 2020. The ORF8 Protein of SARS-CoV-2 Mediates Immune Evasion through Potently Downregulating MHC-I. bioRxiv preprint DOI: https://doi.org/10.1101/2020.05.24.111823.

