## Supplementary material for "Analysis of the Dynamics and Distribution of SARS-CoV-2 Mutations and its Possible Structural and Functional Implications": GISAID_acknowledgements_part_2

We gratefully acknowledge the following Authors from the Originating laboratories responsible for obtaining the specimens, as well as the Submitting laboratories where the genome data were generated and shared via GISAID, on which this research is based.

All Submitters of data may be contacted directly via www.gisaid.org

| Accession ID | Originating Laboratory | Submitting Laboratory | Authors |
|---|---|---|---|
| EPI_ISL_431013 | Alaska State Virology Laboratory | Alaska State Virology Laboratory | Jack Chen |
| EPI_ISL_431014, EPI_ISL_431015, EPI_ISL_431016, EPI_ISL_431017, EPI_ISL_431018, EPI_ISL_431019 | Alaska State Virology Laboratory | Alaska State Virology Laboratory | Jack Chen, Ph.D. |
| EPI_ISL_431080, EPI_ISL_431085 | Yale COVID-19 Biorepository | Grubaugh Lab - Yale School of Public Health | Joseph Fauver, Tara Alpert, Anderson Brito, Anne Wyllie, Chantal Vogels, Mary Petrone, Cole Jensen, Chaney Kalinich, Isabel Ott, Arnau Casanovas, Catherine Muenker, Adam Moore, Alice Lu, Maria Tokuyama, Patrick Wong, Peiwen Lu, Saad Omer, Richard Martinello, Allison Nelson, Shelli Farhadian, Akiko Iwasaki, Charlese Dela Cruz, Albert Ko, Nathan Grubaugh |
| EPI_ISL_431101 | Department of Microbiology,Gandhi Medical College and Hospital | Virus Research Laboratory, Department of Zoology, Osmania University,Hyderabad,India | Muttineni Radhakrishna, Nagamani K, Thrilok Chander B, Raja Rao M, Kalyani Putty, Ravikumar P, Sunitha P, Pankaj Singh D, Anand Kumar K, Amit A. Upadhyay Steven E. Bosinger, Rama Amara |
| EPI_ISL_431102 | Department of MicroBiology,Gandhi Medical College and Hospital,Secendrabad,Hyderabad,India | Department of Microbiology, Gandhi Medical College and Hospital, Secendrabad, Hyderabad | Nagamani K, Muttineni Radhakrishna, Thrilok Chander B, Raja Rao M, Kalyani Putty, Ravikumar P, Sunitha P, Pankaj Singh D, Anand Kumar K, Amit A. Upadhyay, Steven E. Bosinger, Rama Amara |
| EPI_ISL_431103 | Department of Microbiology, Gandhi Medical College and Hospital, Secendrabad, Hyderabad, India | Department of Microbiology, Gandhi Medical College and Hospital, Secendrabad, Hyderabad, India | Nagamani K, Muttineni Radhakrishna, Thrilok Chander B, Raja Rao M, Kalyani Putty, Ravikumar P, Sunitha P, Pankaj Singh D, Anand Kumar K, Amit A. Upadhyay, Steven E. Bosinger, Rama Amara |
| EPI_ISL_431117 | Department of Microbiology, Gandhi Medical College and Hospital, Secendrabad, Hyderabad, India | Department of Microbiology, Gandhi Medical College and Hospital, Secendrabad, Hyderabad, India | Thrilok Chander B, Muttineni Radhakrishna, Nagamani K, Raja Rao M, Kalyani Putty, Ravikumar P, Sunitha P, Pankaj Singh D, Anand Kumar K, Amit A. Upadhyay, Steven E.Bosinger, Rama Amara |
| EPI_ISL_431118, EPI_ISL_431180, EPI_ISL_431240, EPI_ISL_431782, EPI_ISL_431783, EPI_ISL_431784, EPI_ISL_431785 | Fujian Center for Disease Control and Prevention | Fujian Center for Disease Control and Prevention | Lin Qi, Huang Zhimiao, Zhang Yanhua, Weng Yuwei |
| EPI_ISL_431944, EPI_ISL_432176, EPI_ISL_432177, EPI_ISL_432178, EPI_ISL_432179, EPI_ISL_432180, EPI_ISL_432181, EPI_ISL_432182, EPI_ISL_432183, EPI_ISL_432184, EPI_ISL_432185, EPI_ISL_432186, EPI_ISL_432187, EPI_ISL_432188, EPI_ISL_432189, EPI_ISL_432190, EPI_ISL_432191, EPI_ISL_432192, EPI_ISL_432193, EPI_ISL_432194, EPI_ISL_432195, EPI_ISL_432196, EPI_ISL_432197, EPI_ISL_432198, EPI_ISL_432199, EPI_ISL_432200, EPI_ISL_432201, EPI_ISL_432202, EPI_ISL_432203, EPI_ISL_432204, EPI_ISL_432205, EPI_ISL_432206, EPI_ISL_432207, EPI_ISL_432208, EPI_ISL_432209, EPI_ISL_432210, EPI_ISL_432211, EPI_ISL_432212, EPI_ISL_432213, EPI_ISL_432214, EPI_ISL_432215, EPI_ISL_432216, EPI_ISL_432217, EPI_ISL_432218, EPI_ISL_432219, EPI_ISL_432220, EPI_ISL_432221, EPI_ISL_432222, EPI_ISL_432223, EPI_ISL_432224, EPI_ISL_432225, EPI_ISL_432226, EPI_ISL_432227, EPI_ISL_432228, EPI_ISL_432229, EPI_ISL_432230, EPI_ISL_432231, EPI_ISL_432232, EPI_ISL_432233, EPI_ISL_432234, EPI_ISL_432235, EPI_ISL_432236, EPI_ISL_432237, EPI_ISL_432238, EPI_ISL_432239, EPI_ISL_432240, EPI_ISL_432241, EPI_ISL_432242, EPI_ISL_432243, EPI_ISL_432244, EPI_ISL_432245, EPI_ISL_432246, EPI_ISL_432247, EPI_ISL_432248, EPI_ISL_432249, EPI_ISL_432250, EPI_ISL_432251, EPI_ISL_432252, EPI_ISL_432253, EPI_ISL_432254, EPI_ISL_432255, EPI_ISL_432256, EPI_ISL_432257, EPI_ISL_432258, EPI_ISL_432259, EPI_ISL_432260, EPI_ISL_432261, EPI_ISL_432262, EPI_ISL_432263, EPI_ISL_432264, EPI_ISL_432265, EPI_ISL_432266, EPI_ISL_432267, EPI_ISL_432268, EPI_ISL_432269, EPI_ISL_432270, EPI_ISL_432271, EPI_ISL_432272, EPI_ISL_432273, EPI_ISL_432274, EPI_ISL_432275, EPI_ISL_432276, EPI_ISL_432277, EPI_ISL_432278, EPI_ISL_432279, EPI_ISL_432280, EPI_ISL_432281, EPI_ISL_432282, EPI_ISL_432283, EPI_ISL_432284, EPI_ISL_432285, EPI_ISL_432286, EPI_ISL_432287, EPI_ISL_432288, EPI_ISL_432289, EPI_ISL_432290, EPI_ISL_432291, EPI_ISL_432292, EPI_ISL_432293, EPI_ISL_432294, EPI_ISL_432295, EPI_ISL_432296, EPI_ISL_432297, EPI_ISL_432298, EPI_ISL_432299, EPI_ISL_432300, EPI_ISL_432301, EPI_ISL_432302, EPI_ISL_432303, EPI_ISL_432304, EPI_ISL_432305, EPI_ISL_432306, EPI_ISL_432307, EPI_ISL_432308, EPI_ISL_432309, EPI_ISL_432310, EPI_ISL_432311, EPI_ISL_432312, EPI_ISL_432313, EPI_ISL_432314, EPI_ISL_432315, EPI_ISL_432316, EPI_ISL_432317, EPI_ISL_432318, EPI_ISL_432319, EPI_ISL_432320, EPI_ISL_432321, EPI_ISL_432322, EPI_ISL_432323, EPI_ISL_432325, EPI_ISL_432326, EPI_ISL_432327, EPI_ISL_432328, EPI_ISL_432329, EPI_ISL_432330, EPI_ISL_432331, EPI_ISL_432332, EPI_ISL_432334, EPI_ISL_432335, EPI_ISL_432336, EPI_ISL_432337, EPI_ISL_432338, EPI_ISL_432339, EPI_ISL_432340, EPI_ISL_432342, EPI_ISL_432343, EPI_ISL_432344, EPI_ISL_432345, EPI_ISL_432346, EPI_ISL_432347, EPI_ISL_432349, EPI_ISL_432350, EPI_ISL_432351, EPI_ISL_432352, EPI_ISL_432353, EPI_ISL_432354, EPI_ISL_432355, EPI_ISL_432356, EPI_ISL_432357, EPI_ISL_432358, EPI_ISL_432359, EPI_ISL_432360, EPI_ISL_432361, EPI_ISL_432362, EPI_ISL_432365, EPI_ISL_432366, EPI_ISL_432367, EPI_ISL_432369, EPI_ISL_432370, EPI_ISL_432371, EPI_ISL_432374, EPI_ISL_432375, EPI_ISL_432376, EPI_ISL_432377, EPI_ISL_432378, EPI_ISL_432379, EPI_ISL_432382, EPI_ISL_432383, EPI_ISL_432385, EPI_ISL_432387, EPI_ISL_432389, EPI_ISL_432390, EPI_ISL_432393, EPI_ISL_432395, EPI_ISL_432396, EPI_ISL_432397, EPI_ISL_432399, EPI_ISL_432400, EPI_ISL_432401, EPI_ISL_432403, EPI_ISL_432405, EPI_ISL_432406, EPI_ISL_432407, EPI_ISL_432416, EPI_ISL_432417, EPI_ISL_432418, EPI_ISL_432419, EPI_ISL_432421, EPI_ISL_432423, EPI_ISL_432425, EPI_ISL_432428, EPI_ISL_432430 | | | |
| see above | Wales Specialist Virology Centre | Public Health Wales Microbiology Cardiff | Catherine Moore, Johnathan Evans, Malorie Perry, Simon Cottrell, Alec Birchley, Alexander Adams, Amy Gaskin, Bree Gatica-Wilcox, Jason Coombes, Lauren Gilbert, Lee Graham, Nicole Pacchiarini, Sara Kumziene-Summerhayes, Sarah Taylor, Sophie Jones, Sara Rey, Matthew Bull, Joanne Watkins, Sally Corden, Tom Connor |
| EPI_ISL_432458, EPI_ISL_432466, EPI_ISL_432478, EPI_ISL_432480, EPI_ISL_432481, EPI_ISL_432487, EPI_ISL_432502, EPI_ISL_432505, EPI_ISL_432510, EPI_ISL_432517, EPI_ISL_432519, EPI_ISL_432529, EPI_ISL_432533, EPI_ISL_432539, EPI_ISL_432541, EPI_ISL_432542, EPI_ISL_432547, EPI_ISL_432548, EPI_ISL_432549, EPI_ISL_432551, EPI_ISL_432570, EPI_ISL_432578, EPI_ISL_432579, EPI_ISL_432588, EPI_ISL_432601, EPI_ISL_432602, EPI_ISL_432614, EPI_ISL_432620, EPI_ISL_432622, EPI_ISL_432630, EPI_ISL_432642, EPI_ISL_432644 | | | |
| see above | Virology Department, Sheffield Teaching Hospitals NHS Foundation Trust / Virology Department, Sheffield Teaching Hospitals NHS Foundation Trust | COVID-19 Genomics UK (COG-UK) Consortium | Thushan de Silva, Matthew Parker,Adri Angyal, Rebecca Brown, Luke Green, Rachel Tucker, Paul Parsons, Danielle Groves, Alex Keeley, Dave Partridge, Matthew Wyles, Benjamin Lindsey, Mehmet Yavuz, Mohammad Raza, Cariad Evans |
| EPI_ISL_432675, EPI_ISL_432691, EPI_ISL_432706 | Queens Medical Centre, Clinical Microbiology Department / DeepSeq Nottingham | COVID-19 Genomics UK (COG-UK) Consortium | Gemma Clark, Wendy Smith, Manjinder Khakh, Hannah Howson-Wells, Jonathan Ball, Patrick McClure, Joseph Chappell, Theocharis Tsoleridis, Nadine Holmes, Matthew Carlisle, Christopher Moore, Fei Sang, Johnny Debebe, Victoria Wright, Matthew Loose |
| EPI_ISL_432712, EPI_ISL_432713, EPI_ISL_432716, EPI_ISL_432720, EPI_ISL_432722, EPI_ISL_432724, EPI_ISL_432727, EPI_ISL_432732, EPI_ISL_432734, EPI_ISL_432742, EPI_ISL_432744, EPI_ISL_432746, EPI_ISL_432753, EPI_ISL_432762, EPI_ISL_432766, EPI_ISL_432768, EPI_ISL_432777, EPI_ISL_432780, EPI_ISL_432786, EPI_ISL_432788, EPI_ISL_432790, EPI_ISL_432799, EPI_ISL_432802, EPI_ISL_432804, EPI_ISL_432812, EPI_ISL_432813, EPI_ISL_432825, EPI_ISL_432828, EPI_ISL_432831, EPI_ISL_432836, EPI_ISL_432837, EPI_ISL_432850, EPI_ISL_432859, EPI_ISL_432861, EPI_ISL_432864, EPI_ISL_432867 | | | |
| see above | Virology Department, Sheffield Teaching Hospitals NHS Foundation Trust / Virology Department, Sheffield Teaching Hospitals NHS Foundation Trust | COVID-19 Genomics UK (COG-UK) Consortium | Thushan de Silva, Matthew Parker,Adri Angyal, Rebecca Brown, Luke Green, Rachel Tucker, Paul Parsons, Danielle Groves, Alex Keeley, Dave Partridge, Matthew Wyles, Benjamin Lindsey, Mehmet Yavuz, Mohammad Raza, Cariad Evans |
| EPI_ISL_432868, EPI_ISL_432869, EPI_ISL_432870, EPI_ISL_432871, EPI_ISL_432872, EPI_ISL_432873, EPI_ISL_432874, EPI_ISL_432875, EPI_ISL_432876, EPI_ISL_432877, EPI_ISL_432878, EPI_ISL_432880, EPI_ISL_432881, EPI_ISL_432883, EPI_ISL_432885, EPI_ISL_432886, EPI_ISL_432888, EPI_ISL_432889, EPI_ISL_432890, EPI_ISL_432891, EPI_ISL_432892, EPI_ISL_432893, EPI_ISL_432894, EPI_ISL_432895, EPI_ISL_432897, EPI_ISL_432898, EPI_ISL_432899 | | | |
| see above | Virology Department, Royal Infirmary of Edinburgh, NHS Lothian / School of Biological Sciences, University of Edinburgh / Institute of Genetics and Molecular Medicine, University of Edinburgh | COVID-19 Genomics UK (COG-UK) Consortium | McHugh M, Dewar R, Rooke S, Gallagher M, Balcaza C, O'Toole A, Hill V, McCrone JT, Colquhoun R, Yu X, Jackson B, Rambaut A, Williams TC, Templeton K |
| EPI_ISL_432900, EPI_ISL_432916, EPI_ISL_432921, EPI_ISL_432925, EPI_ISL_432930, EPI_ISL_432935, EPI_ISL_432938, EPI_ISL_432942, EPI_ISL_432944, EPI_ISL_432951, EPI_ISL_432954, EPI_ISL_432955, EPI_ISL_432956, EPI_ISL_432961, EPI_ISL_432963, EPI_ISL_432965, EPI_ISL_432967, EPI_ISL_432970, EPI_ISL_432979, EPI_ISL_432984, EPI_ISL_432987, EPI_ISL_432988, EPI_ISL_432990, EPI_ISL_432992, EPI_ISL_432993, EPI_ISL_432998, EPI_ISL_433000, EPI_ISL_433001, EPI_ISL_433002, EPI_ISL_433012, EPI_ISL_433016, EPI_ISL_433034, EPI_ISL_433049, EPI_ISL_433066 | | | |
| see above | Queens Medical Centre, Clinical Microbiology Department / DeepSeq Nottingham | COVID-19 Genomics UK (COG-UK) Consortium | Gemma Clark, Wendy Smith, Manjinder Khakh, Hannah Howson-Wells, Jonathan Ball, Patrick McClure, Joseph Chappell, Theocharis Tsoleridis, Nadine Holmes, Matthew Carlisle, Christopher Moore, Fei Sang, Johnny Debebe, Victoria Wright, Matthew Loose |
| EPI_ISL_433068, EPI_ISL_433069, EPI_ISL_433070, EPI_ISL_433071, EPI_ISL_433072, EPI_ISL_433073, EPI_ISL_433074, EPI_ISL_433075, EPI_ISL_433076, EPI_ISL_433077, EPI_ISL_433078, EPI_ISL_433079, EPI_ISL_433080, EPI_ISL_433081, EPI_ISL_433082, EPI_ISL_433083, EPI_ISL_433084, EPI_ISL_433085, EPI_ISL_433086, EPI_ISL_433087, EPI_ISL_433088, EPI_ISL_433089, EPI_ISL_433090, EPI_ISL_433091, EPI_ISL_433092, EPI_ISL_433093, EPI_ISL_433094, EPI_ISL_433095, EPI_ISL_433096, EPI_ISL_433097, EPI_ISL_433098, EPI_ISL_433099, EPI_ISL_433100, EPI_ISL_433101, EPI_ISL_433102, EPI_ISL_433104, EPI_ISL_433105, EPI_ISL_433106, EPI_ISL_433107, EPI_ISL_433108, EPI_ISL_433109, EPI_ISL_433110, EPI_ISL_433111, EPI_ISL_433112, EPI_ISL_433113, EPI_ISL_433114, EPI_ISL_433115, EPI_ISL_433116, EPI_ISL_433117, EPI_ISL_433118, EPI_ISL_433119, EPI_ISL_433120, EPI_ISL_433121, EPI_ISL_433122, EPI_ISL_433123, EPI_ISL_433124, EPI_ISL_433125, EPI_ISL_433126, EPI_ISL_433127, EPI_ISL_433129, EPI_ISL_433130, EPI_ISL_433131, EPI_ISL_433132, EPI_ISL_433133, EPI_ISL_433135, EPI_ISL_433136, EPI_ISL_433137, EPI_ISL_433139, EPI_ISL_433140, EPI_ISL_433141, EPI_ISL_433142, EPI_ISL_433143, EPI_ISL_433145, EPI_ISL_433147, EPI_ISL_433148, EPI_ISL_433149, EPI_ISL_433150, EPI_ISL_433153, EPI_ISL_433154, EPI_ISL_433155, EPI_ISL_433156, EPI_ISL_433162, EPI_ISL_433164, EPI_ISL_433165, EPI_ISL_433166, EPI_ISL_433167, EPI_ISL_433168, EPI_ISL_433169, EPI_ISL_433170, EPI_ISL_433171, EPI_ISL_433172, EPI_ISL_433173, EPI_ISL_433174, EPI_ISL_433175, EPI_ISL_433176, EPI_ISL_433177, EPI_ISL_433178, EPI_ISL_433180, EPI_ISL_433181, EPI_ISL_433182, EPI_ISL_433183, EPI_ISL_433184, EPI_ISL_433185, EPI_ISL_433186, EPI_ISL_433187, EPI_ISL_433188, EPI_ISL_433190, EPI_ISL_433192, EPI_ISL_433193, EPI_ISL_433194, EPI_ISL_433195, EPI_ISL_433196, EPI_ISL_433197, EPI_ISL_433198, EPI_ISL_433200, EPI_ISL_433201, EPI_ISL_433202, EPI_ISL_433203, EPI_ISL_433204, EPI_ISL_433205, EPI_ISL_433206, EPI_ISL_433207, EPI_ISL_433208, EPI_ISL_433210, EPI_ISL_433211, EPI_ISL_433213, EPI_ISL_433214, EPI_ISL_433215, EPI_ISL_433216, EPI_ISL_433217, EPI_ISL_433218, EPI_ISL_433220, EPI_ISL_433221, EPI_ISL_433222, EPI_ISL_433223, EPI_ISL_433224, EPI_ISL_433225, EPI_ISL_433226, EPI_ISL_433227, EPI_ISL_433228, EPI_ISL_433229, EPI_ISL_433233, EPI_ISL_433234, EPI_ISL_433236, EPI_ISL_433237, EPI_ISL_433238, EPI_ISL_433240, EPI_ISL_433241, EPI_ISL_433242, EPI_ISL_433243, EPI_ISL_433244, EPI_ISL_433245, EPI_ISL_433246, EPI_ISL_433247, EPI_ISL_433248, EPI_ISL_433249, EPI_ISL_433250, EPI_ISL_433253, EPI_ISL_433254, EPI_ISL_433255, EPI_ISL_433256, EPI_ISL_433257, EPI_ISL_433260, EPI_ISL_433261, EPI_ISL_433262, EPI_ISL_433263, EPI_ISL_433265, EPI_ISL_433266, EPI_ISL_433267 | | | |
| see above | Virology Department, Royal Infirmary of Edinburgh, NHS Lothian / School of Biological Sciences, University of Edinburgh / Institute of Genetics and Molecular Medicine, University of Edinburgh | COVID-19 Genomics UK (COG-UK) Consortium | McHugh M, Dewar R, Rooke S, Gallagher M, Balcaza C, O'Toole A, Hill V, McCrone JT, Colquhoun R, Yu X, Jackson B, Rambaut A, Williams TC, Templeton K |
| EPI_ISL_433270, EPI_ISL_433274, EPI_ISL_433276, EPI_ISL_433277, EPI_ISL_433278, EPI_ISL_433279, EPI_ISL_433280, EPI_ISL_433281, EPI_ISL_433282, EPI_ISL_433283, EPI_ISL_433284, EPI_ISL_433285, EPI_ISL_433286, EPI_ISL_433287, EPI_ISL_433288, EPI_ISL_433289, EPI_ISL_433291, EPI_ISL_433293, EPI_ISL_433294, EPI_ISL_433295, EPI_ISL_433298, EPI_ISL_433299, EPI_ISL_433302, EPI_ISL_433303, EPI_ISL_433304, EPI_ISL_433305, EPI_ISL_433307, EPI_ISL_433308, EPI_ISL_433313, EPI_ISL_433314, EPI_ISL_433316, EPI_ISL_433321, EPI_ISL_433323, EPI_ISL_433324, EPI_ISL_433326, EPI_ISL_433327, EPI_ISL_433329, EPI_ISL_433330, EPI_ISL_433333, EPI_ISL_433334, EPI_ISL_433335, EPI_ISL_433338, EPI_ISL_433339, EPI_ISL_433340, EPI_ISL_433341, EPI_ISL_433342, EPI_ISL_433344, EPI_ISL_433346, EPI_ISL_433347, EPI_ISL_433348, EPI_ISL_433350, EPI_ISL_433351, EPI_ISL_433353, EPI_ISL_433355, EPI_ISL_433356, EPI_ISL_433357, EPI_ISL_433358, EPI_ISL_433359, EPI_ISL_433360, EPI_ISL_433361, EPI_ISL_433362, EPI_ISL_433364, EPI_ISL_433366, EPI_ISL_433367, EPI_ISL_433368, EPI_ISL_433370, EPI_ISL_433371, EPI_ISL_433372 | | | |
| see above | West of Scotland Specialist Virology Centre, NHSGGC / MRC-University of Glasgow Centre for Virus Research | COVID-19 Genomics UK (COG-UK) Consortium | Ana da Silva Filipe, Natasha Johnson, Kathy Smollett, Daniel Mair, Stephen Carmichael, Lily Tong, Jenna Nichols, Elihu Aranday-Cortes, Kirstyn Brunker, Yasmin Parr, Kyriaki Nomikou; Sarah McDonald, Marc Niebel, Patawee Asamaphan; Richard Orton, Joseph Hughes, Sreenu Vattipally, David L Robertson; Alasdair MacLean, Rory Gunson; Kathy Li, Natasha Jesudason, Rajiv Shah, James Shepherd, Antonia Ho, Emma Thomson |
| EPI_ISL_433374, EPI_ISL_433375, EPI_ISL_433376, EPI_ISL_433377, EPI_ISL_433378, EPI_ISL_433379, EPI_ISL_433380, EPI_ISL_433381, EPI_ISL_433382, EPI_ISL_433383, EPI_ISL_433384, EPI_ISL_433385, EPI_ISL_433386, EPI_ISL_433387, EPI_ISL_433388, EPI_ISL_433389, EPI_ISL_433390, EPI_ISL_433391, EPI_ISL_433392, EPI_ISL_433393, EPI_ISL_433394, EPI_ISL_433395, EPI_ISL_433396, EPI_ISL_433397, EPI_ISL_433398, EPI_ISL_433399, EPI_ISL_433400, EPI_ISL_433401, EPI_ISL_433402, EPI_ISL_433403, EPI_ISL_433405, EPI_ISL_433406, EPI_ISL_433407, EPI_ISL_433408, EPI_ISL_433409, EPI_ISL_433410, EPI_ISL_433411, EPI_ISL_433412, EPI_ISL_433413, EPI_ISL_433414, EPI_ISL_433415, EPI_ISL_433416, EPI_ISL_433417, EPI_ISL_433418, EPI_ISL_433419, EPI_ISL_433420, EPI_ISL_433421, EPI_ISL_433423, EPI_ISL_433424, EPI_ISL_433425, EPI_ISL_433426, EPI_ISL_433428, EPI_ISL_433429, EPI_ISL_433430, EPI_ISL_433431, EPI_ISL_433432, EPI_ISL_433433, EPI_ISL_433434, EPI_ISL_433435, EPI_ISL_433436, EPI_ISL_433437, EPI_ISL_433439, EPI_ISL_433440, EPI_ISL_433441, EPI_ISL_433442, EPI_ISL_433443, EPI_ISL_433444, EPI_ISL_433446, EPI_ISL_433447, EPI_ISL_433448, EPI_ISL_433449, EPI_ISL_433450, EPI_ISL_433451, EPI_ISL_433452, EPI_ISL_433453, EPI_ISL_433454, EPI_ISL_433455, EPI_ISL_433456, EPI_ISL_433457, EPI_ISL_433458, EPI_ISL_433459, EPI_ISL_433460, EPI_ISL_433461, EPI_ISL_433462, EPI_ISL_433463, EPI_ISL_433464 | | | |
| see above | Virology Department, Royal Infirmary of Edinburgh, NHS Lothian / School of Biological Sciences, University of Edinburgh / Institute of Genetics and Molecular Medicine, University of Edinburgh | COVID-19 Genomics UK (COG-UK) Consortium | McHugh M, Dewar R, Rooke S, Gallagher M, Balcaza C, O'Toole A, Hill V, McCrone JT, Colquhoun R, Yu X, Jackson B, Rambaut A, Williams TC, Templeton K |
| EPI_ISL_433467, EPI_ISL_433469, EPI_ISL_433470, EPI_ISL_433471, EPI_ISL_433472, EPI_ISL_433478, EPI_ISL_433479, EPI_ISL_433480, EPI_ISL_433481, EPI_ISL_433482, EPI_ISL_433483, EPI_ISL_433484, EPI_ISL_433485, EPI_ISL_433486, EPI_ISL_433487, EPI_ISL_433488, EPI_ISL_433489, EPI_ISL_433490, EPI_ISL_433491, EPI_ISL_433492, EPI_ISL_433493, EPI_ISL_433494, EPI_ISL_433495, EPI_ISL_433496, EPI_ISL_433497 | | | |
| see above | Department of Pathology, University of Cambridge | COVID-19 Genomics UK (COG-UK) Consortium | Luke W Meredith, M. Estee Torok , Myra Hosmillo, William L. Hamilton, Martin D. Curran, Theresa Feltwell, Grant Hall, Anna Yakovleva, Fahad A Khokhar, Charlotte J. Houldcroft, Laura G Caller, Aminu S. Jahun, Sarah L. Caddy, Ian Goodfellow |
| EPI_ISL_433498, EPI_ISL_433499, EPI_ISL_433500, EPI_ISL_433501, EPI_ISL_433502, EPI_ISL_433505, EPI_ISL_433507, EPI_ISL_433508, EPI_ISL_433509, EPI_ISL_433510, EPI_ISL_433511, EPI_ISL_433514, EPI_ISL_433515, EPI_ISL_433517, EPI_ISL_433518, EPI_ISL_433519, EPI_ISL_433521, EPI_ISL_433522, EPI_ISL_433525, EPI_ISL_433527, EPI_ISL_433529, EPI_ISL_433534, EPI_ISL_433535, EPI_ISL_433536, EPI_ISL_433537, EPI_ISL_433538, EPI_ISL_433539, EPI_ISL_433541, EPI_ISL_433542, EPI_ISL_433543, EPI_ISL_433544, EPI_ISL_433545, EPI_ISL_433546, EPI_ISL_433547, EPI_ISL_433548, EPI_ISL_433549, EPI_ISL_433550, EPI_ISL_433551, EPI_ISL_433552, EPI_ISL_433554, EPI_ISL_433557, EPI_ISL_433559, EPI_ISL_433560, EPI_ISL_433561, EPI_ISL_433562, EPI_ISL_433563, EPI_ISL_433565, EPI_ISL_433566, EPI_ISL_433567, EPI_ISL_433568, EPI_ISL_433569, EPI_ISL_433570, EPI_ISL_433571, EPI_ISL_433572, EPI_ISL_433574, EPI_ISL_433575, EPI_ISL_433576, EPI_ISL_433580, EPI_ISL_433581, EPI_ISL_433582, EPI_ISL_433583, EPI_ISL_433584, EPI_ISL_433585, EPI_ISL_433586, EPI_ISL_433587, EPI_ISL_433588, EPI_ISL_433589, EPI_ISL_433590, EPI_ISL_433591, EPI_ISL_433593, EPI_ISL_433594, EPI_ISL_433595, EPI_ISL_433597, EPI_ISL_433598, EPI_ISL_433600, EPI_ISL_433602, EPI_ISL_433603, EPI_ISL_433607, EPI_ISL_433608, EPI_ISL_433609, EPI_ISL_433614, EPI_ISL_433615, EPI_ISL_433617, EPI_ISL_433621, EPI_ISL_433622, EPI_ISL_433626, EPI_ISL_433630, EPI_ISL_433632, EPI_ISL_433633, EPI_ISL_433636, EPI_ISL_433638, EPI_ISL_433640, EPI_ISL_433641, EPI_ISL_433642, EPI_ISL_433643, EPI_ISL_433644, | | | |

| | | | |
|---|---|---|---|
| EPI_ISL_433646, EPI_ISL_433647, EPI_ISL_433648, EPI_ISL_433649, EPI_ISL_433650, EPI_ISL_433651, EPI_ISL_433653, EPI_ISL_433655, EPI_ISL_433656, EPI_ISL_433657, EPI_ISL_433658, EPI_ISL_433660, EPI_ISL_433663, EPI_ISL_433664, EPI_ISL_433665 | | | |
| see above | West of Scotland Specialist Virology Centre, NHSGGC / MRC-University of Glasgow Centre for Virus Research | COVID-19 Genomics UK (COG-UK) Consortium | Ana da Silva Filipe, Natasha Johnson, Kathy Smollett, Daniel Mair, Stephen Carmichael, Lily Tong, Jenna Nichols, Elihu Aranday-Cortes, Kirstyn Brunker, Yasmin Parr, Kyriaki Nomikou; Sarah McDonald, Marc Niebel, Patawee Asamaphan; Richard Orton, Joseph Hughes, Sreenu Vattipally, David L Robertson; Alasdair MacLean, Rory Gunson; Kathy Li, Natasha Jesudason, Rajiv Shah, James Shepherd, Antonia Ho, Emma Thomson |
| EPI_ISL_433666, EPI_ISL_433667, EPI_ISL_433668, EPI_ISL_433669, EPI_ISL_433671, EPI_ISL_433672, EPI_ISL_433673, EPI_ISL_433674, EPI_ISL_433675, EPI_ISL_433676, EPI_ISL_433677, EPI_ISL_433678, EPI_ISL_433679, EPI_ISL_433680, EPI_ISL_433681, EPI_ISL_433682, EPI_ISL_433683, EPI_ISL_433684, EPI_ISL_433685, EPI_ISL_433686, EPI_ISL_433687, EPI_ISL_433688, EPI_ISL_433689, EPI_ISL_433690, EPI_ISL_433691, EPI_ISL_433693, EPI_ISL_433694, EPI_ISL_433695, EPI_ISL_433697, EPI_ISL_433698, EPI_ISL_433699, EPI_ISL_433701, EPI_ISL_433702, EPI_ISL_433704, EPI_ISL_433705, EPI_ISL_433706, EPI_ISL_433707, EPI_ISL_433709, EPI_ISL_433710, EPI_ISL_433711, EPI_ISL_433712, EPI_ISL_433713, EPI_ISL_433714, EPI_ISL_433716, EPI_ISL_433717, EPI_ISL_433718, EPI_ISL_433719, EPI_ISL_433720, EPI_ISL_433721, EPI_ISL_433722, EPI_ISL_433724, EPI_ISL_433728, EPI_ISL_433729, EPI_ISL_433730, EPI_ISL_433731, EPI_ISL_433732, EPI_ISL_433733, EPI_ISL_433734, EPI_ISL_433735, EPI_ISL_433736, EPI_ISL_433737, EPI_ISL_433738, EPI_ISL_433739, EPI_ISL_433740, EPI_ISL_433741, EPI_ISL_433742, EPI_ISL_433743, EPI_ISL_433744, EPI_ISL_433745, EPI_ISL_433746, EPI_ISL_433747, EPI_ISL_433748, EPI_ISL_433749, EPI_ISL_433750, EPI_ISL_433751, EPI_ISL_433752, EPI_ISL_433753, EPI_ISL_433755, EPI_ISL_433757, EPI_ISL_433758, EPI_ISL_433759, EPI_ISL_433760, EPI_ISL_433761, EPI_ISL_433762, EPI_ISL_433763, EPI_ISL_433764, EPI_ISL_433765, EPI_ISL_433766, EPI_ISL_433767, EPI_ISL_433768, EPI_ISL_433770, EPI_ISL_433771, EPI_ISL_433772, EPI_ISL_433775, EPI_ISL_433776, EPI_ISL_433777, EPI_ISL_433778, EPI_ISL_433779, EPI_ISL_433780, EPI_ISL_433781, EPI_ISL_433782, EPI_ISL_433783, EPI_ISL_433784, EPI_ISL_433785, EPI_ISL_433786, EPI_ISL_433787, EPI_ISL_433789, EPI_ISL_433790, EPI_ISL_433791, EPI_ISL_433792, EPI_ISL_433793, EPI_ISL_433794, EPI_ISL_433795, EPI_ISL_433796, EPI_ISL_433797, EPI_ISL_433798, EPI_ISL_433799, EPI_ISL_433800, EPI_ISL_433801, EPI_ISL_433803, EPI_ISL_433805, EPI_ISL_433806, EPI_ISL_433807, EPI_ISL_433808, EPI_ISL_433809, EPI_ISL_433811, EPI_ISL_433812, EPI_ISL_433813, EPI_ISL_433814, EPI_ISL_433815, EPI_ISL_433816, EPI_ISL_433817, EPI_ISL_433818, EPI_ISL_433819, EPI_ISL_433820, EPI_ISL_433821, EPI_ISL_433823, EPI_ISL_433824, EPI_ISL_433825, EPI_ISL_433826, EPI_ISL_433827, EPI_ISL_433828, EPI_ISL_433829, EPI_ISL_433833, EPI_ISL_433834, EPI_ISL_433835, EPI_ISL_433837, EPI_ISL_433838, EPI_ISL_433839, EPI_ISL_433840, EPI_ISL_433842, EPI_ISL_433843, EPI_ISL_433844, EPI_ISL_433846, EPI_ISL_433847, EPI_ISL_433848, EPI_ISL_433849, EPI_ISL_433851, EPI_ISL_433852, EPI_ISL_433854, EPI_ISL_433855, EPI_ISL_433856, EPI_ISL_433857, EPI_ISL_433858, EPI_ISL_433859, EPI_ISL_433860, EPI_ISL_433861, EPI_ISL_433862, EPI_ISL_433863, EPI_ISL_433864, EPI_ISL_433865, EPI_ISL_433866, EPI_ISL_433868, EPI_ISL_433870, EPI_ISL_433871, EPI_ISL_433872, EPI_ISL_433873, EPI_ISL_433875, EPI_ISL_433876, EPI_ISL_433877, EPI_ISL_433878, EPI_ISL_433880, EPI_ISL_433882, EPI_ISL_433883, EPI_ISL_433884, EPI_ISL_433885, EPI_ISL_433886, EPI_ISL_433888, EPI_ISL_433890, EPI_ISL_433891, EPI_ISL_433892, EPI_ISL_433893, EPI_ISL_433894, EPI_ISL_433895, EPI_ISL_433896, EPI_ISL_433897, EPI_ISL_433898, EPI_ISL_433899, EPI_ISL_433900, EPI_ISL_433902, EPI_ISL_433904, EPI_ISL_433905, EPI_ISL_433906, EPI_ISL_433907, EPI_ISL_433909, EPI_ISL_433910, EPI_ISL_433911, EPI_ISL_433913, EPI_ISL_433914, EPI_ISL_433915, EPI_ISL_433918, EPI_ISL_433919, EPI_ISL_433920, EPI_ISL_433922, EPI_ISL_433923, EPI_ISL_433926, EPI_ISL_433927, EPI_ISL_433928, EPI_ISL_433929, EPI_ISL_433930, EPI_ISL_433931, EPI_ISL_433933, EPI_ISL_433934, EPI_ISL_433935, EPI_ISL_433936, EPI_ISL_433937, EPI_ISL_433938, EPI_ISL_433939, EPI_ISL_433940, EPI_ISL_433942, EPI_ISL_433944, EPI_ISL_433945, EPI_ISL_433948, EPI_ISL_433949, EPI_ISL_433950, EPI_ISL_433951, EPI_ISL_433952, EPI_ISL_433953, EPI_ISL_433954, EPI_ISL_433955, EPI_ISL_433956, EPI_ISL_433957, EPI_ISL_433958, EPI_ISL_433959, EPI_ISL_433961, EPI_ISL_433962, EPI_ISL_433963, EPI_ISL_433964, EPI_ISL_433965, EPI_ISL_433966, EPI_ISL_433967, EPI_ISL_433968, EPI_ISL_433969, EPI_ISL_433970, EPI_ISL_433971, EPI_ISL_433972, EPI_ISL_433973, EPI_ISL_433974, EPI_ISL_433975, EPI_ISL_433976, EPI_ISL_433982, EPI_ISL_433992, EPI_ISL_433999, EPI_ISL_434001, EPI_ISL_434002, EPI_ISL_434007, EPI_ISL_434009, EPI_ISL_434010, EPI_ISL_434012, EPI_ISL_434013, EPI_ISL_434014, EPI_ISL_434015, EPI_ISL_434020, EPI_ISL_434021, EPI_ISL_434022, EPI_ISL_434024, EPI_ISL_434027, EPI_ISL_434029, EPI_ISL_434032, EPI_ISL_434034, EPI_ISL_434035, EPI_ISL_434036, EPI_ISL_434038, EPI_ISL_434039, EPI_ISL_434041, EPI_ISL_434045, EPI_ISL_434046, EPI_ISL_434047, EPI_ISL_434049, EPI_ISL_434051, EPI_ISL_434054 | | | |
| see above | Department of Pathology, University of Cambridge | COVID-19 Genomics UK (COG-UK) Consortium | Luke W Meredith, M. Estee Torok , Myra Hosmillo, William L. Hamilton, Martin D. Curran, Theresa Feltwell, Grant Hall, Anna Yakovleva, Fahad A Khokhar, Charlotte J. Houldcroft, Laura G Caller, Aminu S. Jahun, Sarah L. Caddy, Ian Goodfellow |
| EPI_ISL_434063, EPI_ISL_434064, EPI_ISL_434065, EPI_ISL_434066, EPI_ISL_434068, EPI_ISL_434069, EPI_ISL_434070, EPI_ISL_434071, EPI_ISL_434072, EPI_ISL_434073, EPI_ISL_434074, EPI_ISL_434075, EPI_ISL_434076, EPI_ISL_434077, EPI_ISL_434079, EPI_ISL_434080, EPI_ISL_434081, EPI_ISL_434082, EPI_ISL_434083, EPI_ISL_434084, EPI_ISL_434085, EPI_ISL_434086, EPI_ISL_434087, EPI_ISL_434088, EPI_ISL_434089, EPI_ISL_434090, EPI_ISL_434091, EPI_ISL_434092, EPI_ISL_434093, EPI_ISL_434094, EPI_ISL_434095, EPI_ISL_434096, EPI_ISL_434097, EPI_ISL_434098, EPI_ISL_434099, EPI_ISL_434100, EPI_ISL_434101, EPI_ISL_434102, EPI_ISL_434103, EPI_ISL_434104, EPI_ISL_434105, EPI_ISL_434106, EPI_ISL_434107, EPI_ISL_434108, EPI_ISL_434109, EPI_ISL_434110, EPI_ISL_434111, EPI_ISL_434112, EPI_ISL_434113, EPI_ISL_434114, EPI_ISL_434115, EPI_ISL_434116, EPI_ISL_434117, EPI_ISL_434118, EPI_ISL_434119, EPI_ISL_434120, EPI_ISL_434121, EPI_ISL_434122, EPI_ISL_434123, EPI_ISL_434124, EPI_ISL_434128, EPI_ISL_434129, EPI_ISL_434130, EPI_ISL_434131, EPI_ISL_434132, EPI_ISL_434133, EPI_ISL_434134, EPI_ISL_434135, EPI_ISL_434136, EPI_ISL_434137, EPI_ISL_434138, EPI_ISL_434139, EPI_ISL_434140, EPI_ISL_434141, EPI_ISL_434142, EPI_ISL_434143, EPI_ISL_434144, EPI_ISL_434145, EPI_ISL_434146, EPI_ISL_434147, EPI_ISL_434148, EPI_ISL_434149, EPI_ISL_434150, EPI_ISL_434151, EPI_ISL_434152, EPI_ISL_434153, EPI_ISL_434154, EPI_ISL_434155, EPI_ISL_434156, EPI_ISL_434157, EPI_ISL_434158, EPI_ISL_434159, EPI_ISL_434160, EPI_ISL_434161, EPI_ISL_434162, EPI_ISL_434163, EPI_ISL_434164, EPI_ISL_434165, EPI_ISL_434166, EPI_ISL_434167, EPI_ISL_434169, EPI_ISL_434170, EPI_ISL_434171, EPI_ISL_434172, EPI_ISL_434173, EPI_ISL_434175, EPI_ISL_434176, EPI_ISL_434177, EPI_ISL_434178, EPI_ISL_434179, EPI_ISL_434180, EPI_ISL_434182, EPI_ISL_434183, EPI_ISL_434184, EPI_ISL_434185, EPI_ISL_434186, EPI_ISL_434187, EPI_ISL_434188, EPI_ISL_434189, EPI_ISL_434190, EPI_ISL_434191, EPI_ISL_434192, EPI_ISL_434193, EPI_ISL_434194, EPI_ISL_434196, EPI_ISL_434197, EPI_ISL_434199, EPI_ISL_434200, EPI_ISL_434201, EPI_ISL_434202, EPI_ISL_434204, EPI_ISL_434205, EPI_ISL_434206, EPI_ISL_434207, EPI_ISL_434208, EPI_ISL_434209, EPI_ISL_434211, EPI_ISL_434212, EPI_ISL_434213, EPI_ISL_434214, EPI_ISL_434215, EPI_ISL_434216, EPI_ISL_434217, EPI_ISL_434218, EPI_ISL_434219, EPI_ISL_434220, EPI_ISL_434221, EPI_ISL_434222, EPI_ISL_434223, EPI_ISL_434224, EPI_ISL_434225, EPI_ISL_434226, EPI_ISL_434227, EPI_ISL_434228, EPI_ISL_434229, EPI_ISL_434230, EPI_ISL_434231, EPI_ISL_434232, EPI_ISL_434233, EPI_ISL_434234, EPI_ISL_434235, EPI_ISL_434236, EPI_ISL_434237, EPI_ISL_434238, EPI_ISL_434239, EPI_ISL_434240, EPI_ISL_434241, EPI_ISL_434242, EPI_ISL_434243, EPI_ISL_434244, EPI_ISL_434245, EPI_ISL_434246, EPI_ISL_434247, EPI_ISL_434248, EPI_ISL_434249, EPI_ISL_434250, EPI_ISL_434251, EPI_ISL_434252, EPI_ISL_434253, EPI_ISL_434254, EPI_ISL_434255, EPI_ISL_434256, EPI_ISL_434257, EPI_ISL_434258, EPI_ISL_434259, EPI_ISL_434260, EPI_ISL_434262, EPI_ISL_434263, EPI_ISL_434264, EPI_ISL_434265, EPI_ISL_434266, EPI_ISL_434268, EPI_ISL_434269, EPI_ISL_434270, EPI_ISL_434271, EPI_ISL_434272, EPI_ISL_434273, EPI_ISL_434274, EPI_ISL_434276, EPI_ISL_434277, EPI_ISL_434279, EPI_ISL_434280, EPI_ISL_434281, EPI_ISL_434282, EPI_ISL_434283, EPI_ISL_434284, EPI_ISL_434285, EPI_ISL_434286, EPI_ISL_434287, EPI_ISL_434288, EPI_ISL_434289, EPI_ISL_434290, EPI_ISL_434291, EPI_ISL_434293, EPI_ISL_434294, EPI_ISL_434296, EPI_ISL_434298, EPI_ISL_434299, EPI_ISL_434300, EPI_ISL_434301, EPI_ISL_434302, EPI_ISL_434303, EPI_ISL_434305, EPI_ISL_434306, EPI_ISL_434307, EPI_ISL_434308, EPI_ISL_434309, EPI_ISL_434310, EPI_ISL_434311, EPI_ISL_434313, EPI_ISL_434314, EPI_ISL_434315, EPI_ISL_434316, EPI_ISL_434317, EPI_ISL_434319, EPI_ISL_434320, EPI_ISL_434321, EPI_ISL_434322, EPI_ISL_434323, EPI_ISL_434324, EPI_ISL_434327, EPI_ISL_434328, EPI_ISL_434329, EPI_ISL_434330, EPI_ISL_434331, EPI_ISL_434332, EPI_ISL_434334, EPI_ISL_434335, EPI_ISL_434336, EPI_ISL_434337, EPI_ISL_434338, EPI_ISL_434339, EPI_ISL_434341, EPI_ISL_434342, EPI_ISL_434343, EPI_ISL_434345, EPI_ISL_434346 | | | |
| see above | Washington State Department of Health | Seattle Flu Study | Chu et al |
| EPI_ISL_434347, EPI_ISL_434348, EPI_ISL_434349, EPI_ISL_434351, EPI_ISL_434352, EPI_ISL_434354, EPI_ISL_434355 | Lab voor klinische biologie | Onderzoeksgroep Virologie | Laurens Lambrechts, Nick Vereecke, Marthe Pauwels, Jozefien De Clercq, Bruno Verhasselt, Linos Vandekerckhove, Hans Nauwynck, Sebastiaan Theuns |
| EPI_ISL_434357, EPI_ISL_434360, EPI_ISL_434361, EPI_ISL_434362 | Lab voor klinische biologie | Onderzoeksgroep Virologie | Nick Vereecke, Laurens Lambrechts, Marthe Pauwels, Jozefien De Clercq, Bruno Verhasselt, Linos Vandekerckhove, Hans Nauwynck, Sebastiaan Theuns |
| EPI_ISL_434366, EPI_ISL_434368, EPI_ISL_434369, EPI_ISL_434370, EPI_ISL_434371 | Hospital AZ Rivierenland | Institute of Tropical Medicine | Philippe Selhorst, Colin Anthony |
| EPI_ISL_434372, EPI_ISL_434373, EPI_ISL_434374, EPI_ISL_434375, EPI_ISL_434376, EPI_ISL_434377, EPI_ISL_434378, EPI_ISL_434379, EPI_ISL_434380, EPI_ISL_434381, EPI_ISL_434382, EPI_ISL_434383 | | | |
| see above | Hospital AZ Rivierenland | Institute of Tropical Medicine | Philippe Selhorst, Colin Anthony, |
| EPI_ISL_434384 | Hospital AZ Rivierenland | Institute of Tropical Medicine | Philippe Selhorst, Colin Anthony |
| EPI_ISL_434455, EPI_ISL_434456, EPI_ISL_434457, EPI_ISL_434458, EPI_ISL_434459, EPI_ISL_434460, EPI_ISL_434461, EPI_ISL_434462, EPI_ISL_434463, EPI_ISL_434464, EPI_ISL_434465, EPI_ISL_434466, EPI_ISL_434467, EPI_ISL_434468, EPI_ISL_434469, EPI_ISL_434470, EPI_ISL_434471, EPI_ISL_434472, EPI_ISL_434473, EPI_ISL_434474, EPI_ISL_434475, EPI_ISL_434476, EPI_ISL_434477, EPI_ISL_434478, EPI_ISL_434479, EPI_ISL_434480, EPI_ISL_434481, EPI_ISL_434482, EPI_ISL_434483, EPI_ISL_434484, EPI_ISL_434485, EPI_ISL_434486 | | | |
| see above | Laboratory of Microbiology, Medical School, National and Kapodistrian University of Athens | Laboratory of Biology, Department of Medicine, Democritus University of Thrace | Kassela K., Bampali,M., Dovrolis,N., Gatzidou,E., Froukala,E., Stavropoulou,A., Veletza,S., Tsakris,A., Spanakis,N. and Karakasiliotis,I. |
| EPI_ISL_434487, EPI_ISL_434488, EPI_ISL_434489, EPI_ISL_434491, EPI_ISL_434492, EPI_ISL_434493, EPI_ISL_434495, EPI_ISL_434496, EPI_ISL_434497, EPI_ISL_434498, EPI_ISL_434500, EPI_ISL_434502, EPI_ISL_434503, EPI_ISL_434504, EPI_ISL_434505, EPI_ISL_434506, EPI_ISL_434507, EPI_ISL_434511, EPI_ISL_434512, EPI_ISL_434514, EPI_ISL_434515 | | | |
| see above | Laboratoire National de Sante, Microbiology, Virology | Laboratoire National de Sante, Microbiology, Epidemiology and Microbial Genomics | Anke Wienecke-Baldacchino, Ardashel Latsuzbaia, Jessica Tapp, Catherine Ragimbeau, Guillaume Fournier, Tamir Abdelrahman, Trung Nguyen Nguyen, Joel Mossong |
| EPI_ISL_434516 | Biolab Diagnostic Laboratories | Andersen lab at Scripps Research | Issa Abu-Dayyeh, Ahmad Tibi, Lama Hussein, Lina Mohammad, Zein Naber, Amid Abdelnour with SEARCH Alliance San Diego |
| EPI_ISL_434517, EPI_ISL_434518, EPI_ISL_434519, EPI_ISL_434526, EPI_ISL_434527, EPI_ISL_434530 | Robert Garry lab | Andersen lab at Scripps Research | Allison Smither, Gilberto Sabino-Santos, Patricia Snarski, Lilia Melnik, Antoinette Bell, Kaylynn Genemaras, Arnaud Drouin, Dahlene Fusco, Robert Garry with SEARCH Alliance San Diego |
| EPI_ISL_434533, EPI_ISL_434535 | Area de Salud Alajuela Sur | Inciensa, Instituto Costarricense de Investigación y Enseñanza en Nutrición y Salud | Francisco Duarte, Hebleen Porras, Claudio Soto-Garita, Estela Cordero, Adriana Godinez & Melany Calderon |
| EPI_ISL_434538 | COOPESAIN | Inciensa, Instituto Costarricense de Investigación y Enseñanza en Nutrición y Salud | Francisco Duarte, Hebleen Porras, Claudio Soto-Garita, Estela Cordero, Adriana Godinez & Melany Calderon |
| EPI_ISL_434539 | Area de Salud Orotina | Inciensa, Instituto Costarricense de Investigación y Enseñanza en Nutrición y Salud | Francisco Duarte, Hebleen Porras, Claudio Soto-Garita, Estela Cordero, Adriana Godinez & Melany Calderon |
| EPI_ISL_434541, EPI_ISL_434543, EPI_ISL_434545, EPI_ISL_434546, EPI_ISL_434547, EPI_ISL_434548, EPI_ISL_434549, EPI_ISL_434551 | Puerto Rico Department of Health | Centers for Disease Control and Prevention, Dengue Branch | Gilberto A. Santiago, Glenda Gonzalez, Betzabel Flores, Keyla Charriez, Fabiola Cruz, Chaney Kalinich, Joseph Fauver, Jessica I. Falcon, Nathan Grubaugh, Jorge L. Munoz-Jordan |
| EPI_ISL_434555, EPI_ISL_434558 | National Institutes of Health, University of the Philippines Manila | Philippine Genome Center | Carlo M. Lapid, Francis A. Tablizo, Benedict A. Maralit, Jan Michael C. Yap, Raul V. Destura, Marissa M. Alejandria, El King D. Morado, Joshua Gregor A. Dizon, Jo-Hannah S. Llames, Shiela Mae M. Araiza, Kris P. Punayan, Kristianne Arielle D. Gabriel, Shebna Rose D. Fabilloren, Shana F. Genavia, Jarvin E. Nipales, Alessandra C. Sanchez, Haifa L.Gaza, Joy Ann Petronio-Santos, Julius Aaron Mejia, Maribell Dollete, Sonia Salamat, Christina Tan, Bernard Demot, John Mark Velasco, Eva Maria Cutiongco-de la Paz, and Cynthia P. Saloma |
| EPI_ISL_434572 | The National Institute of Public Health Center for Epidemiology and Microbiology | The National Institute of Public Health Center for Epidemiology and Microbiology | Alexander Nagy, Helena Jirincova, Ludmila Novakova, Dusan Trnka, Jaromira Vecerova |
| EPI_ISL_434586, EPI_ISL_434587, EPI_ISL_434588 | Johns Hopkins Hospital Department of Pathology | Johns Hopkins Hospital Department of Pathology | Peter M. Thielen, Thomas Mehoke, Shirlee Wohl, Srividya Ramakrishnan, Oluwaseun Nwulia-Falade, Amanda Ernlund, Melanie Kirsche, Paul Morris, Norah Sadowski, Nídiá Trovao, Victoria Gniazdowski, Michael Schatz, Stuart C. Ray, Winston Timp, Heba Mostafa |
| EPI_ISL_434590, EPI_ISL_434592, EPI_ISL_434593, EPI_ISL_434594, EPI_ISL_434595, EPI_ISL_434596, EPI_ISL_434597, EPI_ISL_434598, EPI_ISL_434599, EPI_ISL_434600, EPI_ISL_434601, EPI_ISL_434602, EPI_ISL_434603, EPI_ISL_434604, EPI_ISL_434605 | | | |
| see above | Virginia DCLS | Virginia DCLS | Virginia DCLS |
| EPI_ISL_434607, EPI_ISL_434608, EPI_ISL_434609, EPI_ISL_434610, EPI_ISL_434611, EPI_ISL_434612, EPI_ISL_434613, EPI_ISL_434614, EPI_ISL_434615 | University of Wisconsin-Madison AIDS Vaccine Research Laboratories | University of Wisconsin-Madison AIDS Vaccine Research Laboratories | Gage Moreno, Katarina Braun, et al. AIDS Vaccine Research Laboratories |
| EPI_ISL_434616, EPI_ISL_434617, EPI_ISL_434618, EPI_ISL_434619, EPI_ISL_434620, EPI_ISL_434621, EPI_ISL_434622, EPI_ISL_434623, EPI_ISL_434624, EPI_ISL_434625, EPI_ISL_434626, EPI_ISL_434627, EPI_ISL_434628, EPI_ISL_434629, EPI_ISL_434630, EPI_ISL_434631, EPI_ISL_434632, EPI_ISL_434633, EPI_ISL_434634, EPI_ISL_434635 | | | |
| see above | CHU Purpan - Laboratoire de Virologie - Institut Fédératif de Biologie | Laboratoire de virologie - École Nationale Vétérinaire de Toulouse | Guillaume Croville, Jean-Luc Guérin, Jacques Izopet |
| EPI_ISL_434636 | Lednicky Laboratory, Emerging Pathogens Institute, University of Florida | Lednicky Laboratory at Emerging Pathogens Institute, University of Florida | Elbadry,M.A., Subramaniam,K., Waltzek,T.B., Stephenson,C.J.,Gibson,J.C., Alam,M., Morris,J.G. Jr. and Lednicky,J.A. |
| EPI_ISL_434637 | Lednicky Laboratory, Emerging Pathogens Institute, University of Florida. | Lednicky Laboratory, Emerging Pathogens Institute, University of Florida. | Elbadry,M.A.; Subramaniam,K.; Waltzek,T.B.; Gibson,J.C.; Stephenson,C.J.; Morris,J.G. Jr. and Lednicky,J.A. |
| EPI_ISL_434638, EPI_ISL_434639, EPI_ISL_434640 | Johns Hopkins Hospital Department of Pathology | Johns Hopkins Hospital Department of Pathology | Peter M. Thielen, Thomas Mehoke, Shirlee Wohl, Srividya Ramakrishnan, Oluwaseun Nwulia-Falade, Amanda Ernlund, Melanie Kirsche, Paul Morris, Norah Sadowski, Nídiá Trovao, Victoria Gniazdowski, Michael Schatz, Stuart C. Ray, Winston Timp, Heba Mostafa |
| EPI_ISL_434641, EPI_ISL_434642 | Laboratoriemedicin | The Public Health Agency of Sweden | Oskar Karlsson Lindsjo, Maria Lind Karlberg, Anna-Malin Linde, Olov Svartstrom, Anna Risberg, Shaman Muradrasoli, Karin Tegmark-Wisell |
| EPI_ISL_434643 | Uppsala Narakut Aleris | The Public Health Agency of Sweden | Annika Nilsson, Oskar Karlsson Lindsjo, Maria Lind Karlberg, Anna-Malin Linde, Olov Svartstrom, Anna Risberg, Theresa Enkirch, Mia Brytting, Karin Tegmark-Wisell |
| EPI_ISL_434644 | Kungsholmsdoktorn | The Public Health Agency of Sweden | Linus Hammar, Oskar Karlsson Lindsjo, Maria Lind Karlberg, Anna-Malin Linde, Olov Svartstrom, Anna Risberg, Theresa Enkirch, Mia Brytting, Karin Tegmark-Wisell |
| EPI_ISL_434645 | Svardsjo VC | The Public Health Agency of Sweden | Tommy Janers, Oskar Karlsson Lindsjo, Maria Lind Karlberg, Anna-Malin Linde, Olov Svartstrom, Anna Risberg, Theresa Enkirch, Mia Brytting, Karin Tegmark-Wisell |
| EPI_ISL_434646 | Ulltuna Vardcentral | The Public Health Agency of Sweden | Heidi Lindback, Oskar Karlsson Lindsjo, Maria Lind Karlberg, Anna-Malin Linde, Olov Svartstrom, Anna Risberg, Theresa Enkirch, Mia Brytting, Karin Tegmark-Wisell |
| EPI_ISL_434647, EPI_ISL_434648 | Victoria Vard och Halsa | The Public Health Agency of Sweden | Sarah Henriksson, Oskar Karlsson Lindsjo, Maria Lind Karlberg, Anna-Malin Linde, Olov Svartstrom, Anna Risberg, Theresa Enkirch, Mia Brytting, Karin Tegmark-Wisell |
| EPI_ISL_434649 | Svardsjo VC | The Public Health Agency of Sweden | Tommy Janers, Oskar Karlsson Lindsjo, Maria Lind Karlberg, Anna-Malin Linde, Olov Svartstrom, Anna Risberg, Theresa Enkirch, Mia Brytting, Karin Tegmark-Wisell |
| EPI_ISL_434650 | Follinge Halsocentral | The Public Health Agency of Sweden | Kerstin Persson Moberg, Oskar Karlsson Lindsjo, Maria Lind Karlberg, Anna-Malin Linde, Olov Svartstrom, Anna Risberg, Theresa Enkirch, Mia Brytting, Karin Tegmark-Wisell |
| EPI_ISL_434651 | Krokoms Halsocentral | The Public Health Agency of Sweden | Martin Ersson, Oskar Karlsson Lindsjo, Maria Lind Karlberg, Anna-Malin Linde, Olov Svartstrom, Anna Risberg, Theresa Enkirch, Mia Brytting, Karin Tegmark-Wisell |
| EPI_ISL_434652 | Ektorps Vardcentral | The Public Health Agency of Sweden | Eva Espmark, Oskar Karlsson Lindsjo, Maria Lind Karlberg, Anna-Malin Linde, Olov Svartstrom, Anna Risberg, Theresa Enkirch, Mia Brytting, Karin Tegmark-Wisell |

| | | | |
|---|---|---|---|
| EPI_ISL_434653 | Trollbackens VC | The Public Health Agency of Sweden | Amelie Holmqvist, Oskar Karlsson Lindsjo, Maria Lind Karlberg, Anna-Malin Linde, Olov Svartstrom, Anna Risberg, Theresa Enkirch, Mia Brytting, Karin Tegmark-Wisell |
| EPI_ISL_434654 | Saroledens Familjelakare | The Public Health Agency of Sweden | Katarina Jarbur, Oskar Karlsson Lindsjo, Maria Lind Karlberg, Anna-Malin Linde, Olov Svartstrom, Anna Risberg, Theresa Enkirch, Mia Brytting, Karin Tegmark-Wisell |
| EPI_ISL_434655 | Uppsala Narakut Aleris | The Public Health Agency of Sweden | Annika Nilsson, Oskar Karlsson Lindsjo, Maria Lind Karlberg, Anna-Malin Linde, Olov Svartstrom, Anna Risberg, Theresa Enkirch, Mia Brytting, Karin Tegmark-Wisell |
| EPI_ISL_434656 | Trollbackens VC | The Public Health Agency of Sweden | Amelie Holmqvist, Oskar Karlsson Lindsjo, Maria Lind Karlberg, Anna-Malin Linde, Olov Svartstrom, Anna Risberg, Theresa Enkirch, Mia Brytting, Karin Tegmark-Wisell |
| EPI_ISL_434657 | Kristianstadkliniken | The Public Health Agency of Sweden | Mia Settergren Hammer, Oskar Karlsson Lindsjo, Maria Lind Karlberg, Anna-Malin Linde, Olov Svartstrom, Anna Risberg, Theresa Enkirch, Mia Brytting, Karin Tegmark-Wisell |
| EPI_ISL_434658 | Lundens VC | The Public Health Agency of Sweden | Marita Dagner, Oskar Karlsson Lindsjo, Maria Lind Karlberg, Anna-Malin Linde, Olov Svartstrom, Anna Risberg, Theresa Enkirch, Mia Brytting, Karin Tegmark-Wisell |
| EPI_ISL_434659 | Knivsta VC | The Public Health Agency of Sweden | Johanna Carlson, Oskar Karlsson Lindsjo, Maria Lind Karlberg, Anna-Malin Linde, Olov Svartstrom, Anna Risberg, Theresa Enkirch, Mia Brytting, Karin Tegmark-Wisell |
| EPI_ISL_434660 | Narhalsan Backa vardcentral | The Public Health Agency of Sweden | Mats Olsson, Oskar Karlsson Lindsjo, Maria Lind Karlberg, Anna-Malin Linde, Olov Svartstrom, Anna Risberg, Theresa Enkirch, Mia Brytting, Karin Tegmark-Wisell |
| EPI_ISL_434661, EPI_ISL_434662 | Omtanken Grimmered | The Public Health Agency of Sweden | Bernd Sengpiel, Oskar Karlsson Lindsjo, Maria Lind Karlberg, Anna-Malin Linde, Olov Svartstrom, Anna Risberg, Theresa Enkirch, Mia Brytting, Karin Tegmark-Wisell |
| EPI_ISL_434663 | Surbrunns VC | The Public Health Agency of Sweden | Erik Embring, Oskar Karlsson Lindsjo, Maria Lind Karlberg, Anna-Malin Linde, Olov Svartstrom, Anna Risberg, Theresa Enkirch, Mia Brytting, Karin Tegmark-Wisell |
| EPI_ISL_434664 | Omtanken Grimmered | The Public Health Agency of Sweden | Bernd Sengpiel, Oskar Karlsson Lindsjo, Maria Lind Karlberg, Anna-Malin Linde, Olov Svartstrom, Anna Risberg, Theresa Enkirch, Mia Brytting, Karin Tegmark-Wisell |
| EPI_ISL_434665 | Uppsala Narakut Aleris | The Public Health Agency of Sweden | Annika Nilsson, Oskar Karlsson Lindsjo, Maria Lind Karlberg, Anna-Malin Linde, Olov Svartstrom, Anna Risberg, Theresa Enkirch, Mia Brytting, Karin Tegmark-Wisell |
| EPI_ISL_434666 | Hornefors Halsocentral | The Public Health Agency of Sweden | Camilla Eiback, Oskar Karlsson Lindsjo, Maria Lind Karlberg, Anna-Malin Linde, Olov Svartstrom, Anna Risberg, Theresa Enkirch, Mia Brytting, Karin Tegmark-Wisell |
| EPI_ISL_434667 | Ulltuna Vardcentral | The Public Health Agency of Sweden | Heidi Lindback, Oskar Karlsson Lindsjo, Maria Lind Karlberg, Anna-Malin Linde, Olov Svartstrom, Anna Risberg, Theresa Enkirch, Mia Brytting, Karin Tegmark-Wisell |
| EPI_ISL_434668 | Kungsors VC | The Public Health Agency of Sweden | Jessica Karlsson, Oskar Karlsson Lindsjo, Maria Lind Karlberg, Anna-Malin Linde, Olov Svartstrom, Anna Risberg, Theresa Enkirch, Mia Brytting, Karin Tegmark-Wisell |
| EPI_ISL_434669 | Lakargruppen | The Public Health Agency of Sweden | Boris Klanger, Oskar Karlsson Lindsjo, Maria Lind Karlberg, Anna-Malin Linde, Olov Svartstrom, Anna Risberg, Theresa Enkirch, Mia Brytting, Karin Tegmark-Wisell |
| EPI_ISL_434670 | Narhalsan Molnlycke, Barn och ungdomsmedicin | The Public Health Agency of Sweden | Mats Reimer, Oskar Karlsson Lindsjo, Maria Lind Karlberg, Anna-Malin Linde, Olov Svartstrom, Anna Risberg, Theresa Enkirch, Mia Brytting, Karin Tegmark-Wisell |
| EPI_ISL_434671 | Narhalsan Sjobo vardcentral | The Public Health Agency of Sweden | Lovisa Hjerten, Oskar Karlsson Lindsjo, Maria Lind Karlberg, Anna-Malin Linde, Olov Svartstrom, Anna Risberg, Theresa Enkirch, Mia Brytting, Karin Tegmark-Wisell |
| EPI_ISL_434672 | Surbrunns VC | The Public Health Agency of Sweden | Erik Embring, Oskar Karlsson Lindsjo, Maria Lind Karlberg, Anna-Malin Linde, Olov Svartstrom, Anna Risberg, Theresa Enkirch, Mia Brytting, Karin Tegmark-Wisell |
| EPI_ISL_434673 | Omtanken Grimmered | The Public Health Agency of Sweden | Bernd Sengpiel, Oskar Karlsson Lindsjo, Maria Lind Karlberg, Anna-Malin Linde, Olov Svartstrom, Anna Risberg, Theresa Enkirch, Mia Brytting, Karin Tegmark-Wisell |
| EPI_ISL_434674 | Narhalsan Backa vardcentral | The Public Health Agency of Sweden | Mats Olsson, Oskar Karlsson Lindsjo, Maria Lind Karlberg, Anna-Malin Linde, Olov Svartstrom, Anna Risberg, Theresa Enkirch, Mia Brytting, Karin Tegmark-Wisell |
| EPI_ISL_434675, EPI_ISL_434676 | Surbrunns VC | The Public Health Agency of Sweden | Erik Embring, Oskar Karlsson Lindsjo, Maria Lind Karlberg, Anna-Malin Linde, Olov Svartstrom, Anna Risberg, Theresa Enkirch, Mia Brytting, Karin Tegmark-Wisell |
| EPI_ISL_434678 | Viral Respiratory Lab, National Institute for Biomedical Research (INRB) | Pathogen Sequencing Lab, National Institute for Biomedical Research (INRB) | Placide Mbala-Kingebeni; Edith Nkwembe; Eddy Kinganda-Lusamaki; Amuri Aziza; Francisca Muyembe Mawete; Catherine Pratt; Matthias Pauthner; Josh Quick; Allison Black; James Hadfield; Trevor Bedford; Ian Goodfellow; Andrew Rambaut; Nick Loman; Kristian Andersen; Michael Wiley; Steve Ahuka-Mundeke; Jean-Jacques Muyembe Tamfum |
| EPI_ISL_434682, EPI_ISL_434683, EPI_ISL_434684, EPI_ISL_434685, EPI_ISL_434686, EPI_ISL_434687, EPI_ISL_434688, EPI_ISL_434689, EPI_ISL_434690, EPI_ISL_434691 | Johns Hopkins Hospital Department of Pathology | Johns Hopkins Hospital Department of Pathology | Peter M. Thielen, Thomas Mehoke, Shirlee Wohl, Srividya Ramakrishnan, Melanie Kirsche, Amanda Ernlund, Oluwaseun Falade-Nwulia, Timothy Gilpatrick, Paul Morris, Norah Sadowski, Nídiá Trovao, Victoria Gniazdowski, Michael Schatz, Stuart C. Ray, Winston Timp, Heba Mostafa |
| EPI_ISL_434693, EPI_ISL_434694 | Bamrasnaradura hospital | National Institute of Health. Department of medical Sciences, Ministry of Public Health, Thailand | Pilailuk,Okada; Siripaporn,Phuygun; Thanutsapa,Thanadachakul; Sittiporn,Parnmen;Warawan,Wongboot; Sunthareeya,Waicharoen; Malinee,Chittaganpitch |
| EPI_ISL_434697 | unknown | National Institute of Health. Department of medical Sciences, Ministry of Public Health, Thailand | Pilailuk,Okada; Siripaporn,Phuygun; Thanutsapa,Thanadachakul; Sittiporn,Parnmen;Warawan,Wongboot; Sunthareeya,Waicharoen; Malinee,Chittaganpitch |
| EPI_ISL_434699 | Praram 9 Hospital | National Institute of Health. Department of medical Sciences, Ministry of Public Health, Thailand | Pilailuk,Okada; Siripaporn,Phuygun; Thanutsapa,Thanadachakul; Sittiporn,Parnmen;Warawan,Wongboot; Sunthareeya,Waicharoen; Malinee,Chittaganpitch |
| EPI_ISL_434701 | Panyanunthaphikkhu Chonprathan Medical Center (PCMC) | National Institute of Health. Department of medical Sciences, Ministry of Public Health, Thailand | Pilailuk,Okada; Siripaporn,Phuygun; Thanutsapa,Thanadachakul; Sittiporn,Parnmen;Warawan,Wongboot; Sunthareeya,Waicharoen; Malinee,Chittaganpitch |
| EPI_ISL_434703, EPI_ISL_434705, EPI_ISL_434706 | Praram 9 Hospital | National Institute of Health. Department of medical Sciences, Ministry of Public Health, Thailand | Pilailuk,Okada; Siripaporn,Phuygun; Thanutsapa,Thanadachakul; Sittiporn,Parnmen;Warawan,Wongboot; Sunthareeya,Waicharoen; Malinee,Chittaganpitch |
| EPI_ISL_434708, EPI_ISL_434709 | unknown | National Institute of Health. Department of medical Sciences, Ministry of Public Health, Thailand | Pilailuk,Okada; Siripaporn,Phuygun; Thanutsapa,Thanadachakul; Sittiporn,Parnmen;Warawan,Wongboot; Sunthareeya,Waicharoen; Malinee,Chittaganpitch |
| EPI_ISL_434710 | Viral Respiratory Lab, National Institute for Biomedical Research (INRB) | Pathogen Sequencing Lab, National Institute for Biomedical Research (INRB) | Placide Mbala-Kingebeni, Edith Nkwembe, Eddy Kinganda-Lusamaki, Adrienne Amuri Aziza, Francisca Muyembe Mawete, Catherine Pratt, Matthias Pauthner, Josh Quick, Allison Black, James Hadfield, Trevor Bedford, Ian Goodfellow, Andrew Rambaut, Nick Loman, Kristian Andersen, Michael Wiley, Steve Ahuka-Mundeke, Jean-Jacques Muyembe Tamfum |
| EPI_ISL_434712, EPI_ISL_434713, EPI_ISL_434714, EPI_ISL_434715, EPI_ISL_434716, EPI_ISL_434717, EPI_ISL_434718, EPI_ISL_434720, EPI_ISL_434721, EPI_ISL_434722, EPI_ISL_434724, EPI_ISL_434725, EPI_ISL_434726, EPI_ISL_434727, EPI_ISL_434728, EPI_ISL_434729, EPI_ISL_434730, EPI_ISL_434731, EPI_ISL_434732, EPI_ISL_434735, EPI_ISL_434737, EPI_ISL_434744, EPI_ISL_434745, EPI_ISL_434746, EPI_ISL_434753, EPI_ISL_434755, EPI_ISL_434759, EPI_ISL_434760, EPI_ISL_434764, EPI_ISL_434766, EPI_ISL_434768, EPI_ISL_434772, EPI_ISL_434776, EPI_ISL_434777, EPI_ISL_434780, EPI_ISL_434781, EPI_ISL_434783, EPI_ISL_434784, EPI_ISL_434785, EPI_ISL_434787, EPI_ISL_434788, EPI_ISL_434790, EPI_ISL_434791, EPI_ISL_434792, EPI_ISL_434795, EPI_ISL_434797, EPI_ISL_434798, EPI_ISL_434799, EPI_ISL_434800, EPI_ISL_434801, EPI_ISL_434802, EPI_ISL_434803, EPI_ISL_434805, EPI_ISL_434807, EPI_ISL_434808, EPI_ISL_434810, EPI_ISL_434811, EPI_ISL_434812, EPI_ISL_434813, EPI_ISL_434815, EPI_ISL_434821, EPI_ISL_434822, EPI_ISL_434824, EPI_ISL_434825, EPI_ISL_434827, EPI_ISL_434828, EPI_ISL_434829, EPI_ISL_434831, EPI_ISL_434832, EPI_ISL_434833, EPI_ISL_434834, EPI_ISL_434837, EPI_ISL_434839, EPI_ISL_434840, EPI_ISL_434841, EPI_ISL_434843, EPI_ISL_434844, EPI_ISL_434845, EPI_ISL_434846, EPI_ISL_434847, EPI_ISL_434849, EPI_ISL_434850, EPI_ISL_434851, EPI_ISL_434852, EPI_ISL_434853, EPI_ISL_434854, EPI_ISL_434855, EPI_ISL_434856, EPI_ISL_434857, EPI_ISL_434858, EPI_ISL_434859, EPI_ISL_434860, EPI_ISL_434861, EPI_ISL_434863, EPI_ISL_434864, EPI_ISL_434865, EPI_ISL_434866, EPI_ISL_434867, EPI_ISL_434868, EPI_ISL_434870, EPI_ISL_434871, EPI_ISL_434873, EPI_ISL_434875, EPI_ISL_434876, EPI_ISL_434877, EPI_ISL_434878, EPI_ISL_434879, EPI_ISL_434880, EPI_ISL_434881, EPI_ISL_434882, EPI_ISL_434883, EPI_ISL_434884, EPI_ISL_434885, EPI_ISL_434887, EPI_ISL_434888, EPI_ISL_434889, EPI_ISL_434890, EPI_ISL_434891, EPI_ISL_434892, EPI_ISL_434893, EPI_ISL_434894, EPI_ISL_434895, EPI_ISL_434896, EPI_ISL_434897, EPI_ISL_434898, EPI_ISL_434899, EPI_ISL_434900, EPI_ISL_434901, EPI_ISL_434902, EPI_ISL_434903, EPI_ISL_434904, EPI_ISL_434905, EPI_ISL_434906, EPI_ISL_434907, EPI_ISL_434908, EPI_ISL_434909, EPI_ISL_434910, EPI_ISL_434911, EPI_ISL_434912, EPI_ISL_434913, EPI_ISL_434914, EPI_ISL_434915, EPI_ISL_434916, EPI_ISL_434917, EPI_ISL_434918, EPI_ISL_434919, EPI_ISL_434920, EPI_ISL_434921, EPI_ISL_434922, EPI_ISL_434923, EPI_ISL_434924, EPI_ISL_434925, EPI_ISL_434926, EPI_ISL_434927, EPI_ISL_434928, EPI_ISL_434930, EPI_ISL_434931, EPI_ISL_434933, EPI_ISL_434934, EPI_ISL_434935, EPI_ISL_434936, EPI_ISL_434937, EPI_ISL_434938, EPI_ISL_434939, EPI_ISL_434940, EPI_ISL_434941, EPI_ISL_434942, EPI_ISL_434943, EPI_ISL_434945, EPI_ISL_434946, EPI_ISL_434947, EPI_ISL_434948, EPI_ISL_434949, EPI_ISL_434950, EPI_ISL_434951, EPI_ISL_434952, EPI_ISL_434953, EPI_ISL_434954, EPI_ISL_434955, EPI_ISL_434956, EPI_ISL_434957, EPI_ISL_434958, EPI_ISL_434959, EPI_ISL_434960, EPI_ISL_434961, EPI_ISL_434962, EPI_ISL_434963, EPI_ISL_434964, EPI_ISL_434965, EPI_ISL_434966, EPI_ISL_434967, EPI_ISL_434969, EPI_ISL_434970, EPI_ISL_434971, EPI_ISL_434972, EPI_ISL_434973, EPI_ISL_434974, EPI_ISL_434975, EPI_ISL_434976, EPI_ISL_434977, EPI_ISL_434978, EPI_ISL_434979, EPI_ISL_434980, EPI_ISL_434981, EPI_ISL_434982, EPI_ISL_434983, EPI_ISL_434984, EPI_ISL_434985, EPI_ISL_434986, EPI_ISL_434987, EPI_ISL_434988, EPI_ISL_434989, EPI_ISL_434990, EPI_ISL_434991, EPI_ISL_434992, EPI_ISL_434993, EPI_ISL_434994, EPI_ISL_434995, EPI_ISL_434996, EPI_ISL_434997, EPI_ISL_434998, EPI_ISL_434999, EPI_ISL_435000, EPI_ISL_435001, EPI_ISL_435002, EPI_ISL_435003, EPI_ISL_435005, EPI_ISL_435006, EPI_ISL_435007, EPI_ISL_435008, EPI_ISL_435009, EPI_ISL_435010, EPI_ISL_435011, EPI_ISL_435012, EPI_ISL_435013, EPI_ISL_435014, EPI_ISL_435015, EPI_ISL_435016, EPI_ISL_435017, EPI_ISL_435018, EPI_ISL_435019, EPI_ISL_435020, EPI_ISL_435021, EPI_ISL_435022, EPI_ISL_435023, EPI_ISL_435024, EPI_ISL_435028, EPI_ISL_435029, EPI_ISL_435030, EPI_ISL_435031 | | | |
| see above | Houston Methodist Hospital | Houston Methodist Hospital | S. Wesley Long, Randall J. Olsen, Paul A. Christensen, David W. Bernard, James J. Davis, Maulik Shukla, Marcus Nguyen, Matthew Ojeda Saavedra, Concepcion C. Cantu, Prasanti Yerramilli, Layne Pruitt, Sishir Subedi, Heather Hendrickson, Ghazaleh Eskandari, Muthiah Kumaraswami, Jason S. McLellan, Hakon Jonsson, Kari Stefansson, and James M. Musser |
| EPI_ISL_435032, EPI_ISL_435033 | Viral Respiratory Lab, National Institute for Biomedical Research (INRB) | Pathogen Sequencing Lab, National Institute for Biomedical Research (INRB) | Placide Mbala-Kingebeni, Edith Nkwembe, Eddy Kinganda-Lusamaki, Adrienne Amuri Aziza, Francisca Muyembe Mawete, Catherine Pratt, Matthias Pauthner, Josh Quick, Allison Black, James Hadfield, Trevor Bedford, Ian Goodfellow, Andrew Rambaut, Nick Loman, Kristian Andersen, Michael Wiley, Steve Ahuka-Mundeke, Jean-Jacques Muyembe Tamfum |
| EPI_ISL_435034 | LSUHS Emerging Viral Threat Laboratory | Microbial Genome Sequencing Center | Jeremy P. Kamil, John A. Vanchiere, Rona S. Scott, Camille F. Abshire, Abida Siddiqa, Byeong-Jae Lee, Chan-ki Min, Md Maksudul Alam, Monica Gestal-Cartele, Edna Ondari, Adam Greer, Malgorzata Bienkowska-Haba, Katarzyna Zwolinska, Michelle M. Arnold, Jason M. Bodily, Andrew D. Yurochko, Paul M. Weinberger, Christopher G. Kevil, Martin J. Sapp, Daniel J. Snyder, Vaughn S. Cooper |
| EPI_ISL_435035, EPI_ISL_435036, EPI_ISL_435037, EPI_ISL_435038, EPI_ISL_435039, EPI_ISL_435040, EPI_ISL_435041, EPI_ISL_435042, EPI_ISL_435043, EPI_ISL_435044 | LSUHS Emerging Viral Threat Laboratory | Microbial Genome Sequencing Center | Jeremy P. Kamil, John A. Vanchiere, Rona S. Scott, Camille F. Abshire, Abida Siddiqa, Byeong-Jae Lee, Chan-ki Min, Md Maksudul Alam, Monica Gestal-Cartele, Edna Ondari, Adam Greer, Malgorzata Bienkowska-Haba, Katarzyna Zwolinska, Jason M. Bodily, Andrew D. Yurochko, Paul M. Weinberger, Christopher G. Kevil, Martin J. Sapp, Daniel J. Snyder, Vaughn S. Cooper |
| EPI_ISL_435045, EPI_ISL_435046, EPI_ISL_435047, EPI_ISL_435048 | Laboratory of Applied Genetics | RSE "National Center for Biotechnology" | Alexandr Shevtsov, Ilyas Akhmetollayev, Viktoriya Lutsay, Asylulan Amirgazin, Ruslan Kalendar, Yerlan Ramanculov |
| EPI_ISL_435049 | B.J. Medical College and Civil hospital | Gujarat Biotechnology Research Centre | Pinal Trivedi, Maharshi Pandya, Amit Kanani, Akanksha Verma, Nitin Savaliya, Raghawendra Kumar, Dinesh Kumar, Zuber Saiyed, Dipa Kinariwala, Disha Patel, Binita Aring, Geeta Vaghela, Sonia Barve, Bhavesh Modi, Kairavi Joshi, Gaurishankar Shrimali, Nidhi Sood, Pranay Shah, R D Dixit, Snehal Bagatharia, Kamlesh J Upadhyay, Ramesh Pandit, Tejas Shah, Ankit Hinsu, Pritesh Sabara, Apurvasinh Puvar, Janvi Raval, Monika Gandhi, Neha Rajpara, Chaitanya Joshi, Madhvi Joshi |
| EPI_ISL_435050 | B.J. Medical College and Civil hospital | Gujarat Biotechnology Research Centre | Ankit Hinsu, Pritesh Sabara, Apurvasinh Puvar, Janvi Raval, Monika Gandhi, Pinal Trivedi, Maharshi Pandya, Amit Kanani, Akanksha Verma, Nitin Savaliya, Raghawendra Kumar, Dinesh Kumar, Zuber Saiyed, Dipa Kinariwala, Disha Patel, Binita Aring, Geeta Vaghela, Sonia Barve, Bhavesh Modi, Kairavi Joshi, Gaurishankar Shrimali, Nidhi Sood, Pranay Shah, R D Dixit, Snehal Bagatharia, Kamlesh J Upadhyay, Ramesh Pandit, Tejas Shah, Dipeshwari Shewale, Chaitanya Joshi, Madhvi Joshi |
| EPI_ISL_435051 | B.J. Medical College and Civil hospital | Gujarat Biotechnology Research Centre | Pritesh Sabara, Apurvasinh Puvar, Janvi Raval, Monika Gandhi, Pinal Trivedi, Maharshi Pandya, Amit Kanani, Akanksha Verma, Nitin Savaliya, Raghawendra Kumar, Dinesh Kumar, Zuber Saiyed, Dipa Kinariwala, Disha Patel, Binita Aring, Geeta Vaghela, Sonia Barve, Bhavesh Modi, Kairavi Joshi, Gaurishankar Shrimali, Nidhi Sood, Pranay Shah, R D Dixit, Snehal Bagatharia, Kamlesh J Upadhyay, Ramesh Pandit, Tejas Shah, Ankit Hinsu, Vasudha Sharma, Chaitanya Joshi, Madhvi Joshi |
| EPI_ISL_435052 | B.J. Medical College and Civil hospital | Gujarat Biotechnology Research Centre | Apurvasinh Puvar, Janvi Raval, Monika Gandhi, Pinal Trivedi, Maharshi Pandya, Amit Kanani, Akanksha Verma, Nitin Savaliya, Raghawendra Kumar, Dinesh Kumar, Zuber Saiyed, Dipa Kinariwala, Disha Patel, Binita Aring, Geeta Vaghela, Sonia Barve, Bhavesh Modi, Kairavi Joshi, Gaurishankar Shrimali, Nidhi Sood, Pranay Shah, R D Dixit, Snehal Bagatharia, Kamlesh J Upadhyay, Ramesh Pandit, Tejas Shah, Ankit Hinsu, Pritesh Sabara, Pooja P Doshi, Chaitanya Joshi, Madhvi Joshi |
| EPI_ISL_435053 | B.J. Medical College and Civil hospital | Gujarat Biotechnology Research Centre | Janvi Raval, Monika Gandhi, Pinal Trivedi, Maharshi Pandya, Amit Kanani, Akanksha Verma, Nitin Savaliya, Raghawendra Kumar, Dinesh Kumar, Zuber Saiyed, Dipa Kinariwala, Disha Patel, Binita Aring, Geeta Vaghela, Sonia Barve, Bhavesh Modi, Kairavi Joshi, Gaurishankar Shrimali, Nidhi Sood, Pranay Shah, R D Dixit, Snehal Bagatharia, Kamlesh J Upadhyay, Ramesh Pandit, Tejas Shah, Ankit Hinsu, Pritesh Sabara, Apurvasinh Puvar, Nidhi Patel, Chaitanya Joshi, Madhvi Joshi |
| EPI_ISL_435054 | B.J. Medical College and Civil hospital | Gujarat Biotechnology Research Centre | Monika Gandhi, Pinal Trivedi, Maharshi Pandya, Amit Kanani, Akanksha Verma, Nitin Savaliya, Raghawendra Kumar, Dinesh Kumar, Zuber Saiyed, Dipa Kinariwala, Disha Patel, Binita Aring, Geeta Vaghela, Sonia Barve, Bhavesh Modi, Kairavi Joshi, Gaurishankar Shrimali, Nidhi Sood, Pranay Shah, R D Dixit, Snehal Bagatharia, Kamlesh J Upadhyay, Ramesh Pandit, Tejas Shah, Ankit Hinsu, Pritesh Sabara, Apurvasinh Puvar, Janvi Raval, Priti Pandita, Chaitanya Joshi, Madhvi Joshi |
| EPI_ISL_435055 | Gujarat Biotechnology Research Centre | Gujarat Biotechnology Research Centre | Tejas Shah, Ankit Hinsu, Pritesh Sabara, Apurvasinh Puvar, Janvi Raval, Monika Gandhi, Pinal Trivedi, Maharshi Pandya, Amit Kanani, Akanksha Verma, Nitin Savaliya, Raghawendra Kumar, Dinesh Kumar, Zuber Saiyed, Dipa Kinariwala, Disha Patel, Binita Aring, Geeta Vaghela, Sonia Barve, Bhavesh Modi, Kairavi Joshi, Gaurishankar Shrimali, Nidhi Sood, Pranay Shah, R D Dixit, Snehal Bagatharia, Kamlesh J Upadhyay, Ramesh Pandit, Anjali Rajwal, Chaitanya Joshi, Madhvi Joshi |
| EPI_ISL_435056 | Gujarat Biotechnology Research Centre | Gujarat Biotechnology Research Centre | Maharshi Pandya, Amit Kanani, Akanksha Verma, Nitin Savaliya, Raghawendra Kumar, Dinesh Kumar, Zuber Saiyed, Dipa Kinariwala, Disha Patel, Binita Aring, Geeta Vaghela, Sonia Barve, Bhavesh Modi, Kairavi Joshi, Gaurishankar Shrimali, Nidhi Sood, Pranay Shah, R D Dixit, Snehal Bagatharia, Kamlesh J Upadhyay, Ramesh Pandit, Tejas Shah, Ankit Hinsu, Pritesh Sabara, Apurvasinh Puvar, Janvi Raval, Monika Gandhi, Pinal Trivedi, Afzal Ansari, Chaitanya Joshi, Madhvi Joshi |
| EPI_ISL_435060, EPI_ISL_435061, EPI_ISL_435062, EPI_ISL_435063, EPI_ISL_435064, EPI_ISL_435065, EPI_ISL_435066, EPI_ISL_435067, EPI_ISL_435068, EPI_ISL_435069, EPI_ISL_435070, EPI_ISL_435071, EPI_ISL_435072, EPI_ISL_435074, EPI_ISL_435078, EPI_ISL_435080, EPI_ISL_435081, EPI_ISL_435082, EPI_ISL_435083, EPI_ISL_435084, EPI_ISL_435085, EPI_ISL_435086, EPI_ISL_435087, EPI_ISL_435088, | | | |

| | | | |
|---|---|---|---|
| EPI_ISL_435090, EPI_ISL_435091, EPI_ISL_435092, EPI_ISL_435093, EPI_ISL_435094, EPI_ISL_435095, EPI_ISL_435096, EPI_ISL_435097, EPI_ISL_435098, EPI_ISL_435099, EPI_ISL_435100, EPI_ISL_435101, EPI_ISL_435102, EPI_ISL_435103, EPI_ISL_435104, EPI_ISL_435105, EPI_ISL_435106, EPI_ISL_435108, EPI_ISL_435109, EPI_ISL_435110, EPI_ISL_435111, EPI_ISL_435112 | | | |
| see above | National Centre for Disease control (NCDC), CSIR-Institute of Genomics and Integrative Biology (CSIR-IGIB) | NCDC/CSIR-IGIB | Pramod Kumar, Rajesh Pandey, Pooja Sharma, Mahesh Dhar, Vivekanand A, Bharathram Uppili, Himanshu Vashisht, Saruchi Wadhwa, Nishu Tyagi, Uma Sharma, Priyanka Singh, Hemlata Lall, Meena Datta, Poonam Gupta, Nidhi Saini, Aarti Tewari, Bibhash Nandi, Dhirendra Kumar, Satyabrata Bag, Varun Jaiswal, Hema Gogia, Preeti Madan, Simrita Singh, Prateek Singh, Debasis Dash, Mitali Mukerji, Manju Bala, Sandhya Kabra, Sujeet Singh, Mohammed Faruq, Anurag Agrawal, Partha Rakshit |
| EPI_ISL_435114 | Viral Respiratory Lab, National Institute for Biomedical Research (INRB) | Pathogen Sequencing Lab, National Institute for Biomedical Research (INRB) | Placide Mbala-Kingebeni, Edith Nkwembe, Eddy Kinganda-Lusamaki, Adrienne Amuri Aziza, Francisca Muyembe Mawete, Catherine Pratt, Matthias Pauthner, Josh Quick, Allison Black, James Hadfield, Trevor Bedford, Ian Goodfellow, Andrew Rambaut, Nick Loman, Kristian Andersen, Michael Wiley, Steve Ahuka-Mundeke, Jean-Jacques Muyembe Tamfum |
| EPI_ISL_435119 | Mohammed Bin Rashid University of Medicine and Health Sciences | Al Jalila Children's Hospital | Ahmad Abou Tayoun, Tom Loney, Hamda Khansaheb, Sathishkumar Ramaswamy, Divinlal Harilal, Zulfa Omar Deesi, Rupa Murthy Varghese, Hanan Al Suwaidi, Abdulmajeed Alkhaja, Mohammed Uddin, Rifat Hamoudi, Rabih Halwani, Abiola Catherine Senok, Qutayba Hamid, Norbert Nowotny, Alawi Alsheikh-Ali |
| EPI_ISL_435120, EPI_ISL_435121, EPI_ISL_435122, EPI_ISL_435123, EPI_ISL_435124, EPI_ISL_435125, EPI_ISL_435126, EPI_ISL_435127, EPI_ISL_435128, EPI_ISL_435129, EPI_ISL_435130, EPI_ISL_435131, EPI_ISL_435132, EPI_ISL_435133, EPI_ISL_435134, EPI_ISL_435135, EPI_ISL_435136, EPI_ISL_435137, EPI_ISL_435138, EPI_ISL_435139, EPI_ISL_435140, EPI_ISL_435141, EPI_ISL_435142, EPI_ISL_435143 | | | |
| see above | Mohammed Bin Rashid University of Medicine and Health Sciences | Al Jalila Genomics Center | Ahmad Abou Tayoun, Tom Loney, Hamda Khansaheb, Sathishkumar Ramaswamy, Divinlal Harilal, Zulfa Omar Deesi, Rupa Murthy Varghese, Hanan Al Suwaidi, Abdulmajeed Alkhaja, Mohammed Uddin, Rifat Hamoudi, Rabih Halwani, Abiola Catherine Senok, Qutayba Hamid, Norbert Nowotny, Alawi Alsheikh-Ali |
| EPI_ISL_435144 | Hospital Universitario La Paz | Hospital Universitario 12 de Octubre | Elias Dahdouh, Sara González, Raúl Recio, Fernando Lázaro, Esther Viedma, Natalia Stella, Julio García, Juan Carlos Galán, Rafael Cantón, Mª Dolores Folgueira, Rafael Delgado, Jesús Mingorance |
| EPI_ISL_435145 | Ospedale Civile Giuseppe Mazzini | Istituto Zooprofilattico Sperimentale dell'Abruzzo e Molise "G.Caporale" | Lorusso A, Marcacci M, Di Domenico M, Ancora M, Curini V, Mangone I, Rinaldi A, Di Pasquale A, Cammà C, Puglia I, Savini G |
| EPI_ISL_435146, EPI_ISL_435147 | Villa Serena del Dr. Leonardo Petruzzi | Istituto Zooprofilattico Sperimentale dell'Abruzzo e Molise "G.Caporale" | Lorusso A, Marcacci M, Di Domenico M, Ancora M, Curini V, Mangone I, Rinaldi A, Di Pasquale A, Cammà C, Puglia I, Savini G |
| EPI_ISL_435148 | Ospedale SS Annunziata | Istituto Zooprofilattico Sperimentale dell'Abruzzo e Molise "G.Caporale" | Lorusso A, Marcacci M, Di Domenico M, Ancora M, Curini V, Mangone I, Rinaldi A, Di Pasquale A, Cammà C, Puglia I, Savini G |
| EPI_ISL_435149 | SERVIZIO DI IGIENE E SANITÀ PUBBLICA ASL Teramo | Istituto Zooprofilattico Sperimentale dell'Abruzzo e Molise "G.Caporale" | Lorusso A, Marcacci M, Di Domenico M, Ancora M, Curini V, Mangone I, Rinaldi A, Di Pasquale A, Cammà C, Puglia I, Savini G |
| EPI_ISL_435150, EPI_ISL_435151 | Ospedale SS Annunziata | Istituto Zooprofilattico Sperimentale dell'Abruzzo e Molise "G.Caporale" | Lorusso A, Marcacci M, Di Domenico M, Ancora M, Curini V, Mangone I, Rinaldi A, Di Pasquale A, Cammà C, Puglia I, Savini G |
| EPI_ISL_435152 | Servizio di Igiene, Epidemiologia e Sanità Pubblica (SIESP) Avezzano | Istituto Zooprofilattico Sperimentale dell'Abruzzo e Molise "G.Caporale" | Lorusso A, Marcacci M, Di Domenico M, Ancora M, Curini V, Mangone I, Rinaldi A, Di Pasquale A, Cammà C, Puglia I, Savini G |
| EPI_ISL_435153, EPI_ISL_435154, EPI_ISL_435155 | SERVIZIO DI IGIENE E SANITÀ PUBBLICA ASL Teramo | Istituto Zooprofilattico Sperimentale dell'Abruzzo e Molise "G.Caporale" | Lorusso A, Marcacci M, Di Domenico M, Ancora M, Curini V, Mangone I, Rinaldi A, Di Pasquale A, Cammà C, Puglia I, Savini G |
| EPI_ISL_435156, EPI_ISL_435163 | Viral Respiratory Lab, National Institute for Biomedical Research (INRB) | Pathogen Sequencing Lab, National Institute for Biomedical Research (INRB) | Placide Mbala-Kingebeni, Edith Nkwembe, Eddy Kinganda-Lusamaki, Amuri Aziza, Francisca Muyembe Mawete, Catherine Pratt, Matthias Pauthner, Josh Quick, Allison Black, James Hadfield, Trevor Bedford, Ian Goodfellow, Andrew Rambaut, Nick Loman, Kristian Andersen, Michael Wiley, Steve Ahuka-Mundeke, Jean-Jacques Muyembe Tamfum |
| EPI_ISL_435281 | Medistra Hospital Jakarta | Eijkman Institute for Molecular Biology, Ministry of Research and Technology/National Agency for Research and Innovation | Edison Johar, Frilasita A Yudhaputri, Hidayat Trimarsanto, David H Muljono, Safarina G Malik, Khin Saw Myint, Amin Soebandrio |
| EPI_ISL_435282, EPI_ISL_435283 | RS Pondok Indah Hospital – Pondok Indah | Eijkman Institute for Molecular Biology, Ministry of Research and Technology/National Agency for Research and Innovation | Edison Johar, Frilasita A Yudhaputri, Hidayat Trimarsanto, David H Muljono, Safarina G Malik, Khin Saw Myint, Amin Soebandrio |
| EPI_ISL_435284 | Central Virology Laboratory, Israel Ministry of Health | Central Virology Laboratory, Israel Ministry of Health | Neta Zuckerman, Efrat Bucris, Oran Erster, Danit Sofer, Orna Mor, Ella Mendelson, Michal Mandelboim |
| EPI_ISL_435286 | Central Virology Laboratory, Israel Ministry of Health | Central Virology Laboratory, Israel Ministry of Health | eta Zuckerman, Efrat Bucris, Oran Erster, Orna Mor, Ella Mendelson, Michal Mandelboim, Danit Sofer |
| EPI_ISL_435289 | Central Virology Laboratory, Israel Ministry of Health | Central Virology Laboratory, Israel Ministry of Health | Neta Zuckerman, Efrat Bucris, Oran Erster, Danit Sofer, Orna Mor, Ella Mendelson, Michal Mandelboim |
| EPI_ISL_435291 | Central Virology Laboratory, Israel Ministry of Health | Central Virology Laboratory, Israel Ministry of Health | Neta Zuckerman, Efrat Bucris, Oran Erster, Danit Sofer, Orna Mor, Ella Mendelson, Michal Mandelboim |
| EPI_ISL_435292 | Central Virology Laboratory, Israel Ministry of Health | Central Virology Laboratory, Israel Ministry of Health | Neta Zuckerman, Efrat Bucris, Oran Erster, Danit Sofer, Ella Mendelson, Michal Mandelboim, Orna Mor |
| EPI_ISL_435303 | National Hospital of Tropical Diseases | Oxford University Clinical Research Unit, Hanoi, Vietnam | Nguyen Thi Tam, Van Dinh Trang, Nguyen Thu Trang, Nguyen Thi Ngoc Diep, Le Nguyen Minh Hoa, Pham Ngoc Thach, H.Rogier van Doorn, on behalf of the OUCRU COVID-19 research group |
| EPI_ISL_435305 | National Hospital of Tropical Diseases | Oxford University Clinical Research Unit, Hanoi, Vietnam | Nguyen Thi Tam, Van Dinh Trang, Nguyen Thu Trang, Nguyen Thi Ngoc Diep, Le Nguyen Minh Hoa, Pham Ngoc Thach, H. Rogier van Doorn, on behalf of the OUCRU COVID-19 research group |
| EPI_ISL_435308 | National Hospital of Tropical Diseases | Oxford University Clinical Research Unit, Hanoi, Vietnam | Nguyen Thi Tam, Van Dinh Trang, Nguyen Thu Trang, Nguyen Thi Ngoc Diep, Le Nguyen Minh Hoa, Pham Ngoc Thach, H. Rogier van Doorn, on behalf of the OUCRU COVID-19 research group |
| EPI_ISL_435310 | National Hospital of Tropical Diseases | Oxford University Clinical Research Unit, Hanoi, Vietnam | Nguyen Thi Tam, Van Dinh Trang, Nguyen Thu Trang, Nguyen Thi Ngoc Diep, Le Nguyen Minh Hoa, Pham Ngoc Thach, H. Rogier van Doorn, on behalf of the OUCRU COVID-19 research group |
| EPI_ISL_435311 | National Hospital of Tropical Diseases | Oxford University Clinical Research Unit, Hanoi, Vietnam | Nguyen Thi Tam, Van Dinh Trang, Nguyen Thu Trang, Nguyen Thi Ngoc Diep, Le Nguyen Minh Hoa, Pham Ngoc Thach, H. Rogier van Doorn, on behalf of the OUCRU COVID-19 research group |
| EPI_ISL_435312, EPI_ISL_435313 | National Hospital of Tropical Diseases | Oxford University Clinical Research Unit, Hanoi, Vietnam | Nguyen Thi Tam, Van Dinh Trang, Nguyen Thu Trang, Nguyen Thi Ngoc Diep, Le Nguyen Minh Hoa, Pham Ngoc Thach, H. Rogier van Doorn, on behalf of the OUCRU COVID-19 research group |
| EPI_ISL_435314 | National Hospital of Tropical Diseases | Oxford University Clinical Research Unit, Hanoi, Vietnam | Nguyen Thi Tam, Van Dinh Trang, Nguyen Thu Trang, Nguyen Thi Ngoc Diep, Le Nguyen Minh Hoa, Pham Ngoc Thach, H. Rogier van Doorn, on behalf of the OUCRU COVID-19 research group |
| EPI_ISL_435315, EPI_ISL_435316, EPI_ISL_435317 | National Hospital of Tropical Diseases | Oxford University Clinical Research Unit, Hanoi, Vietnam | Nguyen Thi Tam, Van Dinh Trang, Nguyen Thu Trang, Nguyen Thi Ngoc Diep, Le Nguyen Minh Hoa, Pham Ngoc Thach, H. Rogier van Doorn, on behalf of the OUCRU COVID-19 research group |
| EPI_ISL_435350, EPI_ISL_435353, EPI_ISL_435356, EPI_ISL_435358, EPI_ISL_435359, EPI_ISL_435360, EPI_ISL_435362, EPI_ISL_435363, EPI_ISL_435368, EPI_ISL_435369, EPI_ISL_435372, EPI_ISL_435375, EPI_ISL_435376, EPI_ISL_435377, EPI_ISL_435379, EPI_ISL_435384, EPI_ISL_435389, EPI_ISL_435390, EPI_ISL_435391, EPI_ISL_435393 | | | |
| see above | Utah Public Health Laboratory | Utah Public Health Laboratory | Erin Young, Kelly Oakeson |
| EPI_ISL_435394, EPI_ISL_435395, EPI_ISL_435397, EPI_ISL_435398, EPI_ISL_435399, EPI_ISL_435400, EPI_ISL_435401, EPI_ISL_435402 | Gundersen Molecular Diagnostics Laboratory | Kabara Cancer Research Institute | Craig S. Richmond, Paraic A. Kenny |
| EPI_ISL_435403, EPI_ISL_435404, EPI_ISL_435405, EPI_ISL_435406, EPI_ISL_435407, EPI_ISL_435408, EPI_ISL_435409, EPI_ISL_435410, EPI_ISL_435411, EPI_ISL_435412, EPI_ISL_435413, EPI_ISL_435414, EPI_ISL_435415, EPI_ISL_435416, EPI_ISL_435417, EPI_ISL_435418, EPI_ISL_435419, EPI_ISL_435420, EPI_ISL_435421, EPI_ISL_435422, EPI_ISL_435423, EPI_ISL_435424, EPI_ISL_435425, EPI_ISL_435426, EPI_ISL_435427, EPI_ISL_435428, EPI_ISL_435429, EPI_ISL_435430, EPI_ISL_435431 | | | |
| see above | Virological Research Group, Szentágothai Research Centre | Bioinformatics Research Group, Szentágothai Research Centre | Péter Urbán, Endre Gábor Tóth, Gábor Kemenesi, Róbert Herczeg, Attila Gyenesei, Ferenc Jakab |
| EPI_ISL_435441, EPI_ISL_435442, EPI_ISL_435443, EPI_ISL_435444 | Alaska State Virology Laboratory | Alaska State Virology Laboratory | Jack Chen, Ph.D. |
| EPI_ISL_435445, EPI_ISL_435446, EPI_ISL_435447, EPI_ISL_435448, EPI_ISL_435449, EPI_ISL_435450, EPI_ISL_435451, EPI_ISL_435452, EPI_ISL_435453, EPI_ISL_435454, EPI_ISL_435456, EPI_ISL_435457, EPI_ISL_435458, EPI_ISL_435459, EPI_ISL_435460, EPI_ISL_435461, EPI_ISL_435462, EPI_ISL_435464, EPI_ISL_435465, EPI_ISL_435466, EPI_ISL_435467, EPI_ISL_435468, EPI_ISL_435469, EPI_ISL_435470, EPI_ISL_435471 | | | |
| see above | Robert Garry lab | Andersen lab at Scripps Research | Allison Smither, Gilberto Sabino-Santos, Patricia Snarski, Lilia Melnik, Antoinette Bell, Kaylynn Genemaras, Arnaud Drouin, Dahlene Fusco, Robert Garry with SEARCH Alliance San Diego |
| EPI_ISL_435473, EPI_ISL_435474 | Rady's Childrens Hospital | Andersen lab at Scripps Research | SEARCH Alliance San Diego |
| EPI_ISL_435475, EPI_ISL_435476, EPI_ISL_435477, EPI_ISL_435478, EPI_ISL_435479, EPI_ISL_435480, EPI_ISL_435481, EPI_ISL_435482, EPI_ISL_435483, EPI_ISL_435484, EPI_ISL_435485, EPI_ISL_435486, EPI_ISL_435487, EPI_ISL_435488, EPI_ISL_435489, EPI_ISL_435490, EPI_ISL_435491, EPI_ISL_435492, EPI_ISL_435493, EPI_ISL_435494, EPI_ISL_435495, EPI_ISL_435496, EPI_ISL_435497, EPI_ISL_435498, EPI_ISL_435499, EPI_ISL_435500, EPI_ISL_435501, EPI_ISL_435502, EPI_ISL_435503, EPI_ISL_435504, EPI_ISL_435505, EPI_ISL_435506, EPI_ISL_435507, EPI_ISL_435508, EPI_ISL_435509, EPI_ISL_435510, EPI_ISL_435511, EPI_ISL_435512, EPI_ISL_435513, EPI_ISL_435514, EPI_ISL_435515, EPI_ISL_435516, EPI_ISL_435517, EPI_ISL_435518, EPI_ISL_435519, EPI_ISL_435520, EPI_ISL_435521, EPI_ISL_435522, EPI_ISL_435523, EPI_ISL_435524, EPI_ISL_435525, EPI_ISL_435526, EPI_ISL_435527, EPI_ISL_435528, EPI_ISL_435529, EPI_ISL_435530, EPI_ISL_435531, EPI_ISL_435532, EPI_ISL_435533, EPI_ISL_435534, EPI_ISL_435535, EPI_ISL_435536, EPI_ISL_435537, EPI_ISL_435538, EPI_ISL_435539, EPI_ISL_435540, EPI_ISL_435541, EPI_ISL_435542, EPI_ISL_435543, EPI_ISL_435544, EPI_ISL_435545, EPI_ISL_435546, EPI_ISL_435547, EPI_ISL_435548, EPI_ISL_435549 | | | |
| see above | NYU Langone Health | Departments of Pathology and Medicine, New York University School of Medicine | Maria Aguero-Rosenfeld, Brendan Belovarac, Margaret Black, Ludovic Boytard, John Cadley, Paolo Cotzia, John Chen, Dacia Dimartino, Xiaojun Feng, Tatyana Gindin, Emily Guzman, Adriana Heguy, Megan Hogan, Emily Huang, George Jour, Lawrence H. Lin, Raven Luther, Andrew Lytle, Christian Marier, Matthew T. Maurano, Mark J. Mulligan, Peter Meyn, Raquel Ordonez Ciriza, Iman Osman, Jared Pinnell, Vanessa Raabe, Sitharam Ramaswami, Amy Rapkiewicz, Andre M. Ribeiro-dos-Santos, Marie Samanovic-Golden, Antonio Serrano, Guomiao Shen, Matija Snuderl, Theodore Vougiouklakis, Nick Vulpescu, Gael Westby, Paul Zappile, Yutong Zhang |
| EPI_ISL_435550, EPI_ISL_435551, EPI_ISL_435552, EPI_ISL_435553, EPI_ISL_435554 | LSUHS Emerging Viral Threat Laboratory | Microbial Genome Sequencing Center | Rona S. Scott, Jeremy P. Kamil, John A. Vanchiere, Camille F. Abshire, Abida Siddiqa, Byeong-Jae Lee, Chan-ki Min, Md Maksudul Alam, Monica Gestal-Cartele, Edna Ondari, Adam Greer, Malgorzata Bienkowska-Haba, Katarzyna Zwolinska, Jason M. Bodily, Andrew D. Yurochko, Paul M. Weinberger, Christopher G. Kevil, Martin J. Sapp, Daniel J. Snyder, Vaughn S. Cooper |
| EPI_ISL_435555, EPI_ISL_435556, EPI_ISL_435557, EPI_ISL_435558, EPI_ISL_435559, EPI_ISL_435560, EPI_ISL_435561, EPI_ISL_435562, EPI_ISL_435563, EPI_ISL_435564, EPI_ISL_435565, EPI_ISL_435566, EPI_ISL_435567, EPI_ISL_435568 | | | |
| see above | LSUHS Emerging Viral Threat Laboratory | Microbial Genome Sequencing Center | John A. Vanchiere, Jeremy P. Kamil, Rona S. Scott, Camille F. Abshire, Abida Siddiqa, Byeong-Jae Lee, Chan-ki Min, Md Maksudul Alam, Monica Gestal-Cartele, Edna Ondari, Adam Greer, Malgorzata Bienkowska-Haba, Katarzyna Zwolinska, Jason M. Bodily, Andrew D. Yurochko, Paul M. Weinberger, Christopher G. Kevil, Martin J. Sapp, Daniel J. Snyder, Vaughn S. Cooper |
| EPI_ISL_435569, EPI_ISL_435570, EPI_ISL_435571, EPI_ISL_435572, EPI_ISL_435573, EPI_ISL_435574, EPI_ISL_435575, EPI_ISL_435576, EPI_ISL_435577, EPI_ISL_435578, EPI_ISL_435579 | | | |
| see above | LSUHS Emerging Viral Threat Laboratory | Microbial Genome Sequencing Center | Jeremy P. Kamil, John A. Vanchiere, Rona S. Scott, Camille F. Abshire, Abida Siddiqa, Byeong-Jae Lee, Chan-ki Min, Md Maksudul Alam, Monica Gestal-Cartele, Edna Ondari, Adam Greer, Malgorzata Bienkowska-Haba, Katarzyna Zwolinska, Jason M. Bodily, Andrew D. Yurochko, Paul M. Weinberger, Christopher G. Kevil, Martin J. Sapp, Daniel J. Snyder, Vaughn S. Cooper |
| EPI_ISL_435580, EPI_ISL_435583, EPI_ISL_435586, EPI_ISL_435587, EPI_ISL_435594, EPI_ISL_435596, EPI_ISL_435601, EPI_ISL_435608, EPI_ISL_435609, EPI_ISL_435612, EPI_ISL_435615, EPI_ISL_435621, EPI_ISL_435627, EPI_ISL_435631, EPI_ISL_435634, EPI_ISL_435636, EPI_ISL_435637, EPI_ISL_435639, EPI_ISL_435641, EPI_ISL_435643, EPI_ISL_435645, EPI_ISL_435649, EPI_ISL_435650, EPI_ISL_435654, EPI_ISL_435657, EPI_ISL_435662, EPI_ISL_435663, EPI_ISL_435670, EPI_ISL_435671 | | | |
| see above | Santa Clara County Public Health Department | Chiu Laboratory, University of California, San Francisco | Xianding Deng, Scot Federman, Wei Gu, Elsa Villarino, Brandon Bonin, Debra A. Wadford, and Charles Y. Chiu |
| EPI_ISL_435674, EPI_ISL_435675, EPI_ISL_435676, EPI_ISL_435677 | National Virology Reference Laboratory | National Public Health Laboratory, National Centre for Infectious Diseases | Mak Tze Minn, Octavia Sophie, Chavatte Jean-Marc, Zaini Zainun, Taib Surita, Cui Lin, Lin Raymond Tzer Pin |
| EPI_ISL_435678, EPI_ISL_435679, EPI_ISL_435680, EPI_ISL_435681, EPI_ISL_435682, EPI_ISL_435683, EPI_ISL_435684, EPI_ISL_435685, EPI_ISL_435686, EPI_ISL_435687, EPI_ISL_435688, EPI_ISL_435689, EPI_ISL_435690, EPI_ISL_435691, EPI_ISL_435692, EPI_ISL_435693, EPI_ISL_435694, EPI_ISL_435695, EPI_ISL_435696, EPI_ISL_435697, EPI_ISL_435698, EPI_ISL_435699, EPI_ISL_435700 | | | |
| see above | National Public Health Laboratory, National Centre for Infectious Diseases | National Public Health Laboratory, National Centre for Infectious Diseases | Mak Tze Minn, Octavia Sophie, Chavatte Jean-Marc, Cui Lin, Lin Raymond Tzer Pin |
| EPI_ISL_435702, EPI_ISL_435703, EPI_ISL_435704, EPI_ISL_435705, EPI_ISL_435706, EPI_ISL_435708, EPI_ISL_435709 | Yale COVID-19 Biorepository | Grubaugh Lab - Yale School of Public Health | Joseph Fauver, Tara Alpert, Anderson Brito, Anne Wyllie, Chantal Vogels, Mary Petrone, Cole Jensen, Chaney Kalinich, Isabel Ott, Arnau Casanovas, Catherine Muenker, Adam Moore, Alice Lu, Maria Tokuyama, Patrick Wong, Peiwen Lu, Saad Omer, Richard Martinello, Allison Nelson, Shelli Farhadian, Akiko Iwasaki, Charlese Dela Cruz, Albert Ko, Nathan Grubaugh |
| EPI_ISL_435716 | Connecticut State Department of Public Health | Grubaugh Lab - Yale School of Public Health | Joseph Fauver, Tara Alpert, Anderson Brito, Anne Wyllie, Chantal Vogels, Mary Petrone, Cole Jensen, Chaney Kalinich, Isabel Ott, Arnau Casanovas, Catherine Muenker, Adam Moore, Alice Lu, Maria Tokuyama, Patrick Wong, Peiwen Lu, Saad Omer, Richard Martinello, Allison Nelson, Shelli Farhadian, Akiko Iwasaki, Charlese Dela Cruz, Albert Ko, Nathan Grubaugh |
| EPI_ISL_435720 | Yale Clinical virology | Grubaugh Lab - Yale School of Public Health | Joseph Fauver, Tara Alpert, Anderson Brito, Anne Wyllie, Chantal Vogels, Mary Petrone, Cole Jensen, Chaney Kalinich, Isabel Ott, Arnau Casanovas, Catherine Muenker, Adam Moore, Alice Lu, Maria Tokuyama, Patrick Wong, Peiwen Lu, Saad Omer, Richard Martinello, Allison Nelson, Shelli Farhadian, Akiko Iwasaki, Charlese Dela Cruz, Albert Ko, Nathan Grubaugh |
| EPI_ISL_436040, EPI_ISL_436041, EPI_ISL_436042, | DC Public Health Lab Dept of Forensic Science | Pathogen Discovery, Respiratory Viruses Branch, Division of Viral Diseases, | Ying Tao, Jing Zhang, Krista Queen, Yan Li, Anna Uehara, Clinton R. Paden, Haibin Wang, Zachary Weiner, Bettina Bankamp, Suxiang Tong |

| | | | |
|---|---|---|---|
| EPI_ISL_436043 | | Centers for Disease Control and Prevention | |
| EPI_ISL_436044 | Louisiana Office of Public Health Laboratories | Pathogen Discovery, Respiratory Viruses Branch, Division of Viral Diseases, Centers for Disease Control and Prevention | Ying Tao, Jing Zhang, Krista Queen, Yan Li, Anna Uehara, Clinton R. Paden, Haibin Wang, Zachary Weiner, Bettina Bankamp, Suxiang Tong |
| EPI_ISL_436045, EPI_ISL_436046 | US VI Department of Health | Pathogen Discovery, Respiratory Viruses Branch, Division of Viral Diseases, Centers for Disease Control and Prevention | Ying Tao, Jing Zhang, Krista Queen, Yan Li, Anna Uehara, Clinton R. Paden, Haibin Wang, Zachary Weiner, Bettina Bankamp, Suxiang Tong |
| EPI_ISL_436047, EPI_ISL_436048, EPI_ISL_436049, EPI_ISL_436050, EPI_ISL_436051, EPI_ISL_436052, EPI_ISL_436053, EPI_ISL_436054, EPI_ISL_436055, EPI_ISL_436056, EPI_ISL_436057, EPI_ISL_436058, EPI_ISL_436059, EPI_ISL_436060, EPI_ISL_436061, EPI_ISL_436062, EPI_ISL_436063, EPI_ISL_436064, EPI_ISL_436065, EPI_ISL_436066, EPI_ISL_436067, EPI_ISL_436068, EPI_ISL_436069, EPI_ISL_436070, EPI_ISL_436071, EPI_ISL_436072, EPI_ISL_436073, EPI_ISL_436074, EPI_ISL_436075, EPI_ISL_436076, EPI_ISL_436077, EPI_ISL_436078, EPI_ISL_436079, EPI_ISL_436080, EPI_ISL_436081, EPI_ISL_436082 | | | |
| see above | NYC Department of Health and Mental Hygiene | Pathogen Discovery, Respiratory Viruses Branch, Division of Viral Diseases, Centers for Disease Control and Prevention | Ying Tao, Krista Queen, Christy Harrison, Jennifer Rakeman, Clinton R. Paden, Jing Zhang, Anna Uehara, Yan Li, Haibin Wang, Jasmine Padilla, Justin Lee, Bettina Bankamp, Zachary Weiner, Suxiang Tong |
| EPI_ISL_436097 | Prince Charles Hospital | Public Health Virology Laboratory, Forensics and Scientific Services, Queensland Health | Alyssa Pyke, Neelima Nair, Natalie Simpson, Lisa Leckie, Jamie McMahon, Jean Barcelon, Amanda De Jong, Sean Moody, Doris Genge, Glen Hewitson, Peter Burtonclay, Judy Northill, Ian Maxwell Mackay, Carmel Taylor, Bixing Huang, David Warrilow, Mitchell Finger, Peter Moore, Sarah Wheatley, Sonja Hall-Mendelin, Andrew Van Den Hurk, Elisabeth Gamez, Inga Sultana and Frederick Moore |
| EPI_ISL_436098 | Royal Brisbane and Women's Hospital | Public Health Virology Laboratory, Forensic and Scientific Services, Queensland Health | Alyssa Pyke, Neelima Nair, Natalie Simpson, Lisa Leckie, Jamie McMahon, Jean Barcelon, Amanda De Jong, Sean Moody, Doris Genge, Glen Hewitson, Peter Burtonclay, Judy Northill, Ian Maxwell Mackay, Carmel Taylor, Bixing Huang, David Warrilow, Mitchell Finger, Peter Moore, Sarah Wheatley, Sonja Hall-Mendelin, Andrew Van Den Hurk, Elisabeth Gamez, Inga Sultana and Frederick Moore |
| EPI_ISL_436099, EPI_ISL_436101, EPI_ISL_436102, EPI_ISL_436104, EPI_ISL_436106, EPI_ISL_436107, EPI_ISL_436108 | TSGH-CP molecular lab | TSGH-CP molecular lab | Cherng-Lih Perng, Ming-Jr JIAN, Chih-Kai Chang, Jung-Chung Lin, Kuo-Ming Yeh, Chien-Wen Chen, Sheng-Kang Chiu, Hsing-Yi Chung, Shih-Hung Tsai, Kuo-Sheng Hung, Tien-Yao Chang, Feng-Yee Chang, Hung-Sheng Shang |
| EPI_ISL_436113, EPI_ISL_436114, EPI_ISL_436119, EPI_ISL_436120, EPI_ISL_436126, EPI_ISL_436131 | Victorian Infectious Diseases Reference Laboratory (VIDRL) | Microbiological Diagnostic Unit Public Health Laboratory and Victorian Infectious Diseases Reference Laboratory, Doherty Institute | Caly L., Seemann T., Sait, M., Schultz M., Druce J., Sherry, N. |
| EPI_ISL_436157 | District Surveillance Unit | Department of Neurovirology, National Institute of Mental Health and Neuroscience (NIMHANS) | Chitra Pattabiraman, Vijayalakshmi Reddy, Harsha PK, Risha Rasheed, Shafeeq S Hameed, Manjunatha Venkataswamy, Anita Desai, Ravi Vasanthapuram |
| EPI_ISL_436197 | Servicio de Microbiología. Consorcio Hospital General Universitario de Valencia | Sequencing and Bioinformatics Service and Molecular Epidemiology Research Group. FISABIO-Public Health | Beatriz Beamud, Lidia Ruiz Roldan, Marta Pla Diaz,Neris Garcia-Gonzalez, Loreto Ferrús Abad, Maria Dolores Ocete, Inma Galán Vendrell, Paula Ruiz-Hueso, Mariana Reyes-Prieto, Vicente Soriano Chirona, Maria Alma Bracho, Griselda De Marco, Llúcia Martínez-Priego, Concepcion Gimeno, Giuseppe D'Auria, Fernando Gonzalez-Candelas |
| EPI_ISL_436200 | Servicio de Microbiología. Consorcio Hospital General Universitario de Valencia | Sequencing and Bioinformatics Service and Molecular Epidemiology Research Group. FISABIO-Public Health | Neris Garcia-Gonzalez, Loreto Ferrús Abad, Maria Dolores Ocete, Inma Galán Vendrell, Paula Ruiz-Hueso, Mariana Reyes-Prieto, Vicente Soriano Chirona, Maria Alma Bracho, Griselda De Marco, Beatriz Beamud, Lidia Ruiz Roldan, Marta Pla Diaz, Llúcia Martínez-Priego, Concepcion Gimeno, Giuseppe D'Auria, Fernando Gonzalez-Candelas |
| EPI_ISL_436201 | Servicio de Microbiología. Consorcio Hospital General Universitario de Valencia | Sequencing and Bioinformatics Service and Molecular Epidemiology Research Group. FISABIO-Public Health | Loreto Ferrús Abad, Maria Dolores Ocete, Inma Galán Vendrell, Paula Ruiz-Hueso, Mariana Reyes-Prieto, Vicente Soriano Chirona, Maria Alma Bracho, Griselda De Marco, Beatriz Beamud, Lidia Ruiz Roldan, Marta Pla Diaz,Neris Garcia-Gonzalez, Llúcia Martínez-Priego, Concepcion Gimeno, Giuseppe D'Auria, Fernando Gonzalez-Candelas |
| EPI_ISL_436202 | Servicio de Microbiología. Consorcio Hospital General Universitario de Valencia | Sequencing and Bioinformatics Service and Molecular Epidemiology Research Group. FISABIO-Public Health | Loreto Ferrús Abad, Maria Dolores Ocete,Inma Galán Vendrell, Paula Ruiz-Hueso, Mariana Reyes-Prieto, Vicente Soriano Chirona, Maria Alma Bracho, Griselda De Marco, Beatriz Beamud, Lidia Ruiz Roldan, Marta Pla Diaz,Neris Garcia-Gonzalez, Llúcia Martínez-Priego, Concepcion Gimeno, Giuseppe D'Auria, Fernando Gonzalez-Candelas |
| EPI_ISL_436203 | Servicio de Microbiología. Consorcio Hospital General Universitario de Valencia | Sequencing and Bioinformatics Service and Molecular Epidemiology Research Group. FISABIO-Public Health | Maria Dolores Ocete, Inma Galán Vendrell, Paula Ruiz-Hueso, Mariana Reyes-Prieto, Vicente Soriano Chirona, Maria Alma Bracho, Griselda De Marco, Beatriz Beamud, Lidia Ruiz Roldan, Marta Pla Diaz,Neris Garcia-Gonzalez, Loreto Ferrús Abad, Llúcia Martínez-Priego, Concepcion Gimeno, Giuseppe D'Auria, Fernando Gonzalez-Candelas |
| EPI_ISL_436205 | Servicio de Microbiología. Consorcio Hospital General Universitario de Valencia | Sequencing and Bioinformatics Service and Molecular Epidemiology Research Group. FISABIO-Public Health | Beatriz Beamud, Lidia Ruiz Roldan, Marta Pla Diaz,Neris Garcia-Gonzalez, Loreto Ferrús Abad, Maria Dolores Ocete, Inma Galán Vendrell, Paula Ruiz-Hueso, Mariana Reyes-Prieto, Vicente Soriano Chirona, Maria Alma Bracho, Griselda De Marco, Llúcia Martínez-Priego, Concepcion Gimeno, Giuseppe D'Auria, Fernando Gonzalez-Candelas |
| EPI_ISL_436206 | Servicio de Microbiología. Consorcio Hospital General Universitario de Valencia | Sequencing and Bioinformatics Service and Molecular Epidemiology Research Group. FISABIO-Public Health | Lidia Ruiz Roldan, Marta Pla Diaz,Neris Garcia-Gonzalez, Loreto Ferrús Abad, Maria Dolores Ocete, Inma Galán Vendrell, Paula Ruiz-Hueso, Mariana Reyes-Prieto, Vicente Soriano Chirona, Maria Alma Bracho, Griselda De Marco, Beatriz Beamud, Llúcia Martínez-Priego, Concepcion Gimeno, Giuseppe D'Auria, Fernando Gonzalez-Candelas |
| EPI_ISL_436207 | Servicio de Microbiología. Consorcio Hospital General Universitario de Valencia | Sequencing and Bioinformatics Service and Molecular Epidemiology Research Group. FISABIO-Public Health | Marta Pla Diaz,Neris Garcia-Gonzalez, Loreto Ferrús Abad, Maria Dolores Ocete, Inma Galán Vendrell, Paula Ruiz-Hueso, Mariana Reyes-Prieto, Vicente Soriano Chirona, Maria Alma Bracho, Griselda De Marco, Beatriz Beamud, Lidia Ruiz Roldan, Llúcia Martínez-Priego, Concepcion Gimeno, Giuseppe D'Auria, Fernando Gonzalez-Candelas |
| EPI_ISL_436209 | Servicio de Microbiología. Consorcio Hospital General Universitario de Valencia | Sequencing and Bioinformatics Service and Molecular Epidemiology Research Group. FISABIO-Public Health | Loreto Ferrús Abad, Maria Dolores Ocete, Inma Galán Vendrell, Paula Ruiz-Hueso, Mariana Reyes-Prieto, Vicente Soriano Chirona, Maria Alma Bracho, Griselda De Marco, Beatriz Beamud, Lidia Ruiz Roldan, Marta Pla Diaz,Neris Garcia-Gonzalez, Llúcia Martínez-Priego, Concepcion Gimeno, Giuseppe D'Auria, Fernando Gonzalez-Candelas |
| EPI_ISL_436210 | Servicio de Microbiología. Consorcio Hospital General Universitario de Valencia | Sequencing and Bioinformatics Service and Molecular Epidemiology Research Group. FISABIO-Public Health | Loreto Ferrús Abad, Maria Dolores Ocete,Inma Galán Vendrell, Paula Ruiz-Hueso, Mariana Reyes-Prieto, Vicente Soriano Chirona, Maria Alma Bracho, Griselda De Marco, Beatriz Beamud, Lidia Ruiz Roldan, Marta Pla Diaz,Neris Garcia-Gonzalez, Llúcia Martínez-Priego, Concepcion Gimeno, Giuseppe D'Auria, Fernando Gonzalez-Candelas |
| EPI_ISL_436211 | Servicio de Microbiología. Consorcio Hospital General Universitario de Valencia | Sequencing and Bioinformatics Service and Molecular Epidemiology Research Group. FISABIO-Public Health | Maria Dolores Ocete, Inma Galán Vendrell, Paula Ruiz-Hueso, Mariana Reyes-Prieto, Vicente Soriano Chirona, Maria Alma Bracho, Griselda De Marco, Beatriz Beamud, Lidia Ruiz Roldan, Marta Pla Diaz,Neris Garcia-Gonzalez, Loreto Ferrús Abad, Llúcia Martínez-Priego, Concepcion Gimeno, Giuseppe D'Auria, Fernando Gonzalez-Candelas |
| EPI_ISL_436212 | Servicio de Microbiología. Consorcio Hospital General Universitario de Valencia | Sequencing and Bioinformatics Service and Molecular Epidemiology Research Group. FISABIO-Public Health | Griselda De Marco, Beatriz Beamud, Lidia Ruiz Roldan, Marta Pla Diaz,Neris Garcia-Gonzalez, Loreto Ferrús Abad, Maria Dolores Ocete, Inma Galán Vendrell, Paula Ruiz-Hueso, Mariana Reyes-Prieto, Vicente Soriano Chirona, Maria Alma Bracho, Llúcia Martínez-Priego, Concepcion Gimeno, Giuseppe D'Auria, Fernando Gonzalez-Candelas |
| EPI_ISL_436213 | Servicio de Microbiología. Consorcio Hospital General Universitario de Valencia | Sequencing and Bioinformatics Service and Molecular Epidemiology Research Group. FISABIO-Public Health | Beatriz Beamud, Lidia Ruiz Roldan, Marta Pla Diaz,Neris Garcia-Gonzalez, Loreto Ferrús Abad, Maria Dolores Ocete, Inma Galán Vendrell, Paula Ruiz-Hueso, Mariana Reyes-Prieto, Vicente Soriano Chirona, Maria Alma Bracho, Griselda De Marco, Llúcia Martínez-Priego, Concepcion Gimeno, Giuseppe D'Auria, Fernando Gonzalez-Candelas |
| EPI_ISL_436214 | Servicio de Microbiología. Consorcio Hospital General Universitario de Valencia | Sequencing and Bioinformatics Service and Molecular Epidemiology Research Group. FISABIO-Public Health | Lidia Ruiz Roldan, Marta Pla Diaz,Neris Garcia-Gonzalez, Loreto Ferrús Abad, Maria Dolores Ocete, Inma Galán Vendrell, Paula Ruiz-Hueso, Mariana Reyes-Prieto, Vicente Soriano Chirona, Maria Alma Bracho, Griselda De Marco, Beatriz Beamud, Llúcia Martínez-Priego, Concepcion Gimeno, Giuseppe D'Auria, Fernando Gonzalez-Candelas |
| EPI_ISL_436217 | Servicio de Microbiología. Consorcio Hospital General Universitario de Valencia | Sequencing and Bioinformatics Service and Molecular Epidemiology Research Group. FISABIO-Public Health | Loreto Ferrús Abad, Maria Dolores Ocete, Inma Galán Vendrell, Paula Ruiz-Hueso, Mariana Reyes-Prieto, Vicente Soriano Chirona, Maria Alma Bracho, Griselda De Marco, Beatriz Beamud, Lidia Ruiz Roldan, Marta Pla Diaz,Neris Garcia-Gonzalez, Llúcia Martínez-Priego, Concepcion Gimeno, Giuseppe D'Auria, Fernando Gonzalez-Candelas |
| EPI_ISL_436219 | Servicio de Microbiología. Consorcio Hospital General Universitario de Valencia | Sequencing and Bioinformatics Service and Molecular Epidemiology Research Group. FISABIO-Public Health | Maria Dolores Ocete, Inma Galán Vendrell, Paula Ruiz-Hueso, Mariana Reyes-Prieto, Vicente Soriano Chirona, Maria Alma Bracho, Griselda De Marco, Beatriz Beamud, Lidia Ruiz Roldan, Marta Pla Diaz,Neris Garcia-Gonzalez, Loreto Ferrús Abad, Llúcia Martínez-Priego, Concepcion Gimeno, Giuseppe D'Auria, Fernando Gonzalez-Candelas |
| EPI_ISL_436220 | Servicio de Microbiología. Consorcio Hospital General Universitario de Valencia | Sequencing and Bioinformatics Service and Molecular Epidemiology Research Group. FISABIO-Public Health | Griselda De Marco, Beatriz Beamud, Lidia Ruiz Roldan, Marta Pla Diaz,Neris Garcia-Gonzalez, Loreto Ferrús Abad, Maria Dolores Ocete, Inma Galán Vendrell, Paula Ruiz-Hueso, Mariana Reyes-Prieto, Vicente Soriano Chirona, Maria Alma Bracho, Llúcia Martínez-Priego, Concepcion Gimeno, Giuseppe D'Auria, Fernando Gonzalez-Candelas |
| EPI_ISL_436223 | Servicio de Microbiología. Consorcio Hospital General Universitario de Valencia | Sequencing and Bioinformatics Service and Molecular Epidemiology Research Group. FISABIO-Public Health | Marta Pla Diaz,Neris Garcia-Gonzalez, Loreto Ferrús Abad, Maria Dolores Ocete, Inma Galán Vendrell, Paula Ruiz-Hueso, Mariana Reyes-Prieto, Vicente Soriano Chirona, Maria Alma Bracho, Griselda De Marco, Beatriz Beamud, Lidia Ruiz Roldan, Llúcia Martínez-Priego, Concepcion Gimeno, Giuseppe D'Auria, Fernando Gonzalez-Candelas |
| EPI_ISL_436224 | Servicio de Microbiología. Consorcio Hospital General Universitario de Valencia | Sequencing and Bioinformatics Service and Molecular Epidemiology Research Group. FISABIO-Public Health | Neris Garcia-Gonzalez, Loreto Ferrús Abad, Maria Dolores Ocete, Inma Galán Vendrell, Paula Ruiz-Hueso, Mariana Reyes-Prieto, Vicente Soriano Chirona, Maria Alma Bracho, Griselda De Marco, Beatriz Beamud, Lidia Ruiz Roldan, Marta Pla Diaz, Llúcia Martínez-Priego, Concepcion Gimeno, Giuseppe D'Auria, Fernando Gonzalez-Candelas |
| EPI_ISL_436225 | Servicio de Microbiología. Consorcio Hospital General Universitario de Valencia | Sequencing and Bioinformatics Service and Molecular Epidemiology Research Group. FISABIO-Public Health | Loreto Ferrús Abad, Maria Dolores Ocete, Inma Galán Vendrell, Paula Ruiz-Hueso, Mariana Reyes-Prieto, Vicente Soriano Chirona, Maria Alma Bracho, Griselda De Marco, Beatriz Beamud, Lidia Ruiz Roldan, Marta Pla Diaz,Neris Garcia-Gonzalez, Llúcia Martínez-Priego, Concepcion Gimeno, Giuseppe D'Auria, Fernando Gonzalez-Candelas |
| EPI_ISL_436227 | Servicio de Microbiología. Consorcio Hospital General Universitario de Valencia | Sequencing and Bioinformatics Service and Molecular Epidemiology Research Group. FISABIO-Public Health | Griselda De Marco, Beatriz Beamud, Lidia Ruiz Roldan, Marta Pla Diaz,Neris Garcia-Gonzalez, Loreto Ferrús Abad, Maria Dolores Ocete, Inma Galán Vendrell, Paula Ruiz-Hueso, Mariana Reyes-Prieto, Vicente Soriano Chirona, Maria Alma Bracho, Llúcia Martínez-Priego, Concepcion Gimeno, Giuseppe D'Auria, Fernando Gonzalez-Candelas |
| EPI_ISL_436229 | Servicio de Microbiología. Consorcio Hospital General Universitario de Valencia | Sequencing and Bioinformatics Service and Molecular Epidemiology Research Group. FISABIO-Public Health | Lidia Ruiz Roldan, Marta Pla Diaz,Neris Garcia-Gonzalez, Loreto Ferrús Abad, Maria Dolores Ocete, Inma Galán Vendrell, Paula Ruiz-Hueso, Mariana Reyes-Prieto, Vicente Soriano Chirona, Maria Alma Bracho, Griselda De Marco, Beatriz Beamud, Llúcia Martínez-Priego, Concepcion Gimeno, Giuseppe D'Auria, Fernando Gonzalez-Candelas |
| EPI_ISL_436230 | Servicio de Microbiología. Consorcio Hospital General Universitario de Valencia | Sequencing and Bioinformatics Service and Molecular Epidemiology Research Group. FISABIO-Public Health | Marta Pla Diaz,Neris Garcia-Gonzalez, Loreto Ferrús Abad, Maria Dolores Ocete, Inma Galán Vendrell, Paula Ruiz-Hueso, Mariana Reyes-Prieto, Vicente Soriano Chirona, Maria Alma Bracho, Griselda De Marco, Beatriz Beamud, Lidia Ruiz Roldan, Llúcia Martínez-Priego, Concepcion Gimeno, Giuseppe D'Auria, Fernando Gonzalez-Candelas |
| EPI_ISL_436231 | Servicio de Microbiología. Consorcio Hospital General Universitario de Valencia | Sequencing and Bioinformatics Service and Molecular Epidemiology Research Group. FISABIO-Public Health | Neris Garcia-Gonzalez, Loreto Ferrús Abad, Maria Dolores Ocete, Inma Galán Vendrell, Paula Ruiz-Hueso, Mariana Reyes-Prieto, Vicente Soriano Chirona, Maria Alma Bracho, Griselda De Marco, Beatriz Beamud, Lidia Ruiz Roldan, Marta Pla Diaz, Llúcia Martínez-Priego, Concepcion Gimeno, Giuseppe D'Auria, Fernando Gonzalez-Candelas |
| EPI_ISL_436233 | Servicio de Microbiología. Consorcio Hospital General Universitario de Valencia | Sequencing and Bioinformatics Service and Molecular Epidemiology Research Group. FISABIO-Public Health | Loreto Ferrús Abad, Maria Dolores Ocete,Inma Galán Vendrell, Paula Ruiz-Hueso, Mariana Reyes-Prieto, Vicente Soriano Chirona, Maria Alma Bracho, Griselda De Marco, Beatriz Beamud, Lidia Ruiz Roldan, Marta Pla Diaz,Neris Garcia-Gonzalez, Llúcia Martínez-Priego, Concepcion Gimeno, Giuseppe D'Auria, Fernando Gonzalez-Candelas |
| EPI_ISL_436234 | Servicio de Microbiología. Consorcio Hospital General Universitario de Valencia | Sequencing and Bioinformatics Service and Molecular Epidemiology Research Group. FISABIO-Public Health | Maria Dolores Ocete, Inma Galán Vendrell, Paula Ruiz-Hueso, Mariana Reyes-Prieto, Vicente Soriano Chirona, Maria Alma Bracho, Griselda De Marco, Beatriz Beamud, Lidia Ruiz Roldan, Marta Pla Diaz,Neris Garcia-Gonzalez, Loreto Ferrús Abad, Llúcia Martínez-Priego, Concepcion Gimeno, Giuseppe D'Auria, Fernando Gonzalez-Candelas |
| EPI_ISL_436235, EPI_ISL_436236, EPI_ISL_436237, EPI_ISL_436238, EPI_ISL_436239, EPI_ISL_436240, EPI_ISL_436241, EPI_ISL_436242, EPI_ISL_436244, EPI_ISL_436245, EPI_ISL_436246, EPI_ISL_436247, EPI_ISL_436248, EPI_ISL_436249, EPI_ISL_436250, EPI_ISL_436251, EPI_ISL_436252, EPI_ISL_436254, EPI_ISL_436255, EPI_ISL_436258, EPI_ISL_436261, EPI_ISL_436262, EPI_ISL_436263, EPI_ISL_436264, EPI_ISL_436265, EPI_ISL_436266, EPI_ISL_436267, EPI_ISL_436268, EPI_ISL_436269 | | | |
| see above | Servicio de Microbiologia. Hospital Universitario Doctor Peset | Sequencing and Bioinformatics Service and Molecular Epidemiology Research Group. FISABIO-Public Health | Juan Alberola Enguídanos, Juan Jose Camarena Miñana, Rosa González Pellicer, Neris Garcia-Gonzalez, Inma Galán Vendrell, Sandra Carbo, Loreto Ferrús Abad, Paula Ruiz-Hueso, Mariana Reyes-Prieto, Vicente Soriano Chirona, Ivan Ansari, Maria Alma Bracho, Griselda De Marco, Beatriz Beamud, Lidia Ruiz Roldan, Marta Pla Diaz, Llúcia Martínez-Priego, Giuseppe D'Auria, Jose Miguel Nogueira Coito, Fernando Gonzalez-Candelas |
| EPI_ISL_436271 | Servicio de Microbiología. Consorcio Hospital General Universitario de Valencia | Sequencing and Bioinformatics Service and Molecular Epidemiology Research Group. FISABIO-Public Health | Griselda De Marco, Beatriz Beamud, Lidia Ruiz Roldan, Marta Pla Diaz,Neris Garcia-Gonzalez, Loreto Ferrús Abad, Maria Dolores Ocete, Inma Galán Vendrell, Paula Ruiz-Hueso, Mariana Reyes-Prieto, Vicente Soriano Chirona, Maria Alma Bracho, Llúcia Martínez-Priego, Concepcion Gimeno, Giuseppe D'Auria, Fernando Gonzalez-Candelas |
| EPI_ISL_436272 | Servicio de Microbiología. Consorcio Hospital General Universitario de Valencia | Sequencing and Bioinformatics Service and Molecular Epidemiology Research Group. FISABIO-Public Health | Beatriz Beamud, Lidia Ruiz Roldan, Marta Pla Diaz,Neris Garcia-Gonzalez, Loreto Ferrús Abad, Maria Dolores Ocete, Inma Galán Vendrell, Paula Ruiz-Hueso, Mariana Reyes-Prieto, Vicente Soriano Chirona, Maria Alma Bracho, Griselda De Marco, Llúcia Martínez-Priego, Concepcion Gimeno, Giuseppe D'Auria, Fernando Gonzalez-Candelas |
| EPI_ISL_436282 | Servicio de Microbiologia. Hospital Clinico Universitario de Valencia | Sequencing and Bioinformatics Service and Molecular Epidemiology Research Group. FISABIO-Public Health | Marta Pla Diaz, Neris Garcia-Gonzalez, Inma Galán Vendrell, Sandra Carbo, Loreto Ferrús Abad, Paula Ruiz-Hueso, Mariana Reyes-Prieto, Vicente Soriano Chirona, Ivan Ansari, David Navarro, Maria Alma Bracho, Griselda De Marco, Beatriz Beamud, Lidia Ruiz Roldan, Llúcia Martínez-Priego, Giuseppe D'Auria, Fernando Gonzalez-Candelas |
| EPI_ISL_436283 | Servicio de Microbiologia. Hospital Clinico Universitario de Valencia | Sequencing and Bioinformatics Service and Molecular Epidemiology Research Group. FISABIO-Public Health | Neris Garcia-Gonzalez, Inma Galán Vendrell, Sandra Carbo, Loreto Ferrús Abad, Paula Ruiz-Hueso, Mariana Reyes-Prieto, Vicente Soriano Chirona, Ivan Ansari, David Navarro, Maria Alma Bracho, Griselda De Marco, Beatriz Beamud, Lidia Ruiz Roldan, Marta Pla Diaz, Llúcia Martínez-Priego, Giuseppe D'Auria, Fernando Gonzalez-Candelas |
| EPI_ISL_436284 | Servicio de Microbiologia. Hospital Clinico Universitario de Valencia | Sequencing and Bioinformatics Service and Molecular Epidemiology Research Group. FISABIO-Public Health | Inma Galán Vendrell, Sandra Carbo, Loreto Ferrús Abad, Paula Ruiz-Hueso, Mariana Reyes-Prieto, Vicente Soriano Chirona, Ivan Ansari, David Navarro, Maria Alma Bracho, Griselda De Marco, Beatriz Beamud, Lidia Ruiz Roldan, Marta Pla Diaz, Neris Garcia-Gonzalez, Llúcia Martínez-Priego, Giuseppe D'Auria, Fernando Gonzalez-Candelas |
| EPI_ISL_436285 | Servicio de Microbiologia. Hospital Clinico Universitario de Valencia | Sequencing and Bioinformatics Service and Molecular Epidemiology Research Group. FISABIO-Public Health | Sandra Carbo, Loreto Ferrús Abad, Paula Ruiz-Hueso, Mariana Reyes-Prieto, Vicente Soriano Chirona, Ivan Ansari, David Navarro, Maria Alma Bracho, Griselda De Marco, Beatriz Beamud, Lidia Ruiz Roldan, Marta Pla Diaz, Neris Garcia-Gonzalez, Inma Galán Vendrell, Llúcia Martínez-Priego, Giuseppe D'Auria, Fernando Gonzalez-Candelas |
| EPI_ISL_436287 | Servicio de Microbiologia. Hospital Clinico Universitario de | Sequencing and Bioinformatics Service and Molecular Epidemiology Research | Paula Ruiz-Hueso, Mariana Reyes-Prieto, Vicente Soriano Chirona, Ivan Ansari, David Navarro, Maria Alma Bracho, Griselda De Marco, Beatriz Beamud, Lidia Ruiz Roldan, Marta Pla Diaz, Neris Garcia-Gonzalez, |

| | | | |
|---|---|---|---|
| | Valencia | Group. FISABIO-Public Health | Inma Galán Vendrell, Sandra Carbo, Loreto Ferrús Abad, Llúcia Martínez-Priego, Giuseppe D'Auria, Fernando Gonzalez-Candelas |
| EPI_ISL_436288 | Servicio de Microbiologia. Hospital Clinico Universitario de Valencia | Sequencing and Bioinformatics Service and Molecular Epidemiology Research Group. FISABIO-Public Health | Mariana Reyes-Prieto, Vicente Soriano Chirona, Ivan Ansari, David Navarro, Maria Alma Bracho, Griselda De Marco, Beatriz Beamud, Lidia Ruiz Roldan, Marta Pla Diaz, Neris Garcia-Gonzalez, Inma Galán Vendrell, Sandra Carbo, Loreto Ferrús Abad, Paula Ruiz-Hueso, Llúcia Martínez-Priego, Giuseppe D'Auria, Fernando Gonzalez-Candelas |
| EPI_ISL_436290 | Servicio de Microbiologia. Hospital Clinico Universitario de Valencia | Sequencing and Bioinformatics Service and Molecular Epidemiology Research Group. FISABIO-Public Health | Ivan Ansari, David Navarro, Maria Alma Bracho, Griselda De Marco, Beatriz Beamud, Lidia Ruiz Roldan, Marta Pla Diaz, Neris Garcia-Gonzalez, Inma Galán Vendrell, Sandra Carbo, Loreto Ferrús Abad, Paula Ruiz-Hueso, Mariana Reyes-Prieto, Vicente Soriano Chirona, Llúcia Martínez-Priego, Giuseppe D'Auria, Fernando Gonzalez-Candelas |
| EPI_ISL_436291 | Servicio de Microbiologia. Hospital Clinico Universitario de Valencia | Sequencing and Bioinformatics Service and Molecular Epidemiology Research Group. FISABIO-Public Health | David Navarro, Maria Alma Bracho, Griselda De Marco, Beatriz Beamud, Lidia Ruiz Roldan, Marta Pla Diaz, Neris Garcia-Gonzalez, Inma Galán Vendrell, Sandra Carbo, Loreto Ferrús Abad, Paula Ruiz-Hueso, Mariana Reyes-Prieto, Vicente Soriano Chirona, Ivan Ansari, Llúcia Martínez-Priego, Giuseppe D'Auria, Fernando Gonzalez-Candelas |
| EPI_ISL_436292 | Servicio de Microbiologia. Hospital Clinico Universitario de Valencia | Sequencing and Bioinformatics Service and Molecular Epidemiology Research Group. FISABIO-Public Health | Maria Alma Bracho, Griselda De Marco, Beatriz Beamud, Lidia Ruiz Roldan, Marta Pla Diaz, Neris Garcia-Gonzalez, Inma Galán Vendrell, Sandra Carbo, Loreto Ferrús Abad, Paula Ruiz-Hueso, Mariana Reyes-Prieto, Vicente Soriano Chirona, Ivan Ansari, David Navarro, Llúcia Martínez-Priego, Giuseppe D'Auria, Fernando Gonzalez-Candelas |
| EPI_ISL_436295 | Servicio de Microbiologia. Hospital Clinico Universitario de Valencia | Sequencing and Bioinformatics Service and Molecular Epidemiology Research Group. FISABIO-Public Health | Lidia Ruiz Roldan, Marta Pla Diaz, Neris Garcia-Gonzalez, Inma Galán Vendrell, Sandra Carbo, Loreto Ferrús Abad, Paula Ruiz-Hueso, Mariana Reyes-Prieto, Vicente Soriano Chirona, Ivan Ansari, David Navarro, Maria Alma Bracho, Griselda De Marco, Beatriz Beamud, Llúcia Martínez-Priego, Giuseppe D'Auria, Fernando Gonzalez-Candelas |
| EPI_ISL_436297 | Servicio de Microbiologia. Hospital Clinico Universitario de Valencia | Sequencing and Bioinformatics Service and Molecular Epidemiology Research Group. FISABIO-Public Health | Neris Garcia-Gonzalez, Inma Galán Vendrell, Sandra Carbo, Loreto Ferrús Abad, Paula Ruiz-Hueso, Mariana Reyes-Prieto, Vicente Soriano Chirona, Ivan Ansari, David Navarro, Maria Alma Bracho, Griselda De Marco, Beatriz Beamud, Lidia Ruiz Roldan, Marta Pla Diaz, Llúcia Martínez-Priego, Giuseppe D'Auria, Fernando Gonzalez-Candelas |
| EPI_ISL_436301 | Servicio de Microbiologia. Hospital Clinico Universitario de Valencia | Sequencing and Bioinformatics Service and Molecular Epidemiology Research Group. FISABIO-Public Health | Lidia Ruiz Roldan, Marta Pla Diaz, Neris Garcia-Gonzalez, Inma Galán Vendrell, Sandra Carbo, Loreto Ferrús Abad, Paula Ruiz-Hueso, Mariana Reyes-Prieto, Vicente Soriano Chirona, Ivan Ansari, David Navarro, Maria Alma Bracho, Griselda De Marco, Beatriz Beamud, Llúcia Martínez-Priego, Giuseppe D'Auria, Fernando Gonzalez-Candelas |
| EPI_ISL_436303 | Servicio de Microbiologia. Hospital Clinico Universitario de Valencia | Sequencing and Bioinformatics Service and Molecular Epidemiology Research Group. FISABIO-Public Health | Neris Garcia-Gonzalez, Inma Galán Vendrell, Sandra Carbo, Loreto Ferrús Abad, Paula Ruiz-Hueso, Mariana Reyes-Prieto, Vicente Soriano Chirona, Ivan Ansari, David Navarro, Maria Alma Bracho, Griselda De Marco, Beatriz Beamud, Lidia Ruiz Roldan, Marta Pla Diaz, Llúcia Martínez-Priego, Giuseppe D'Auria, Fernando Gonzalez-Candelas |
| EPI_ISL_436304 | Servicio de Microbiologia. Hospital Clinico Universitario de Valencia | Sequencing and Bioinformatics Service and Molecular Epidemiology Research Group. FISABIO-Public Health | Inma Galán Vendrell, Sandra Carbo, Loreto Ferrús Abad, Paula Ruiz-Hueso, Mariana Reyes-Prieto, Vicente Soriano Chirona, Ivan Ansari, David Navarro, Maria Alma Bracho, Griselda De Marco, Beatriz Beamud, Lidia Ruiz Roldan, Marta Pla Diaz, Neris Garcia-Gonzalez, Llúcia Martínez-Priego, Giuseppe D'Auria, Fernando Gonzalez-Candelas |
| EPI_ISL_436307 | Servicio de Microbiologia. Hospital Clinico Universitario de Valencia | Sequencing and Bioinformatics Service and Molecular Epidemiology Research Group. FISABIO-Public Health | Paula Ruiz-Hueso, Mariana Reyes-Prieto, Vicente Soriano Chirona, Ivan Ansari, David Navarro, Maria Alma Bracho, Griselda De Marco, Beatriz Beamud, Lidia Ruiz Roldan, Marta Pla Diaz, Neris Garcia-Gonzalez, Inma Galán Vendrell, Sandra Carbo, Loreto Ferrús Abad, Llúcia Martínez-Priego, Giuseppe D'Auria, Fernando Gonzalez-Candelas |
| EPI_ISL_436310 | Servicio de Microbiologia. Hospital Clinico Universitario de Valencia | Sequencing and Bioinformatics Service and Molecular Epidemiology Research Group. FISABIO-Public Health | Ivan Ansari, David Navarro, Maria Alma Bracho, Griselda De Marco, Beatriz Beamud, Lidia Ruiz Roldan, Marta Pla Diaz, Neris Garcia-Gonzalez, Inma Galán Vendrell, Sandra Carbo, Loreto Ferrús Abad, Paula Ruiz-Hueso, Mariana Reyes-Prieto, Vicente Soriano Chirona, Llúcia Martínez-Priego, Giuseppe D'Auria, Fernando Gonzalez-Candelas |
| EPI_ISL_436311 | Servicio de Microbiologia. Hospital Clinico Universitario de Valencia | Sequencing and Bioinformatics Service and Molecular Epidemiology Research Group. FISABIO-Public Health | David Navarro, Maria Alma Bracho, Griselda De Marco, Beatriz Beamud, Lidia Ruiz Roldan, Marta Pla Diaz, Neris Garcia-Gonzalez, Inma Galán Vendrell, Sandra Carbo, Loreto Ferrús Abad, Paula Ruiz-Hueso, Mariana Reyes-Prieto, Vicente Soriano Chirona, Ivan Ansari, Llúcia Martínez-Priego, Giuseppe D'Auria, Fernando Gonzalez-Candelas |
| EPI_ISL_436314 | Servicio de Microbiologia. Hospital Clinico Universitario de Valencia | Sequencing and Bioinformatics Service and Molecular Epidemiology Research Group. FISABIO-Public Health | Beatriz Beamud, Lidia Ruiz Roldan, Marta Pla Diaz, Neris Garcia-Gonzalez, Inma Galán Vendrell, Sandra Carbo, Loreto Ferrús Abad, Paula Ruiz-Hueso, Mariana Reyes-Prieto, Vicente Soriano Chirona, Ivan Ansari, David Navarro, Maria Alma Bracho, Griselda De Marco, Llúcia Martínez-Priego, Giuseppe D'Auria, Fernando Gonzalez-Candelas |
| EPI_ISL_436316 | Servicio de Microbiologia. Hospital Clinico Universitario de Valencia | Sequencing and Bioinformatics Service and Molecular Epidemiology Research Group. FISABIO-Public Health | Marta Pla Diaz, Neris Garcia-Gonzalez, Inma Galán Vendrell, Sandra Carbo, Loreto Ferrús Abad, Paula Ruiz-Hueso, Mariana Reyes-Prieto, Vicente Soriano Chirona, Ivan Ansari, David Navarro, Maria Alma Bracho, Griselda De Marco, Beatriz Beamud, Lidia Ruiz Roldan, Llúcia Martínez-Priego, Giuseppe D'Auria, Fernando Gonzalez-Candelas |
| EPI_ISL_436317 | Servicio de Microbiologia. Hospital Clinico Universitario de Valencia | Sequencing and Bioinformatics Service and Molecular Epidemiology Research Group. FISABIO-Public Health | Neris Garcia-Gonzalez, Inma Galán Vendrell, Sandra Carbo, Loreto Ferrús Abad, Paula Ruiz-Hueso, Mariana Reyes-Prieto, Vicente Soriano Chirona, Ivan Ansari, David Navarro, Maria Alma Bracho, Griselda De Marco, Beatriz Beamud, Lidia Ruiz Roldan, Marta Pla Diaz, Llúcia Martínez-Priego, Giuseppe D'Auria, Fernando Gonzalez-Candelas |
| EPI_ISL_436318 | Servicio de Microbiologia. Hospital Clinico Universitario de Valencia | Sequencing and Bioinformatics Service and Molecular Epidemiology Research Group. FISABIO-Public Health | David Navarro, Maria Alma Bracho, Griselda De Marco, Beatriz Beamud, Lidia Ruiz Roldan, Marta Pla Diaz, Neris Garcia-Gonzalez, Inma Galán Vendrell, Sandra Carbo, Loreto Ferrús Abad, Paula Ruiz-Hueso, Mariana Reyes-Prieto, Vicente Soriano Chirona, Ivan Ansari, David Navarro, Llúcia Martínez-Priego, Giuseppe D'Auria, Fernando Gonzalez-Candelas |
| EPI_ISL_436325 | Servicio de Microbiologia. Hospital Clinico Universitario de Valencia | Sequencing and Bioinformatics Service and Molecular Epidemiology Research Group. FISABIO-Public Health | Sandra Carbo, Loreto Ferrús Abad, Paula Ruiz-Hueso, Mariana Reyes-Prieto, Vicente Soriano Chirona, Ivan Ansari, David Navarro, Maria Alma Bracho, Griselda De Marco, Beatriz Beamud, Lidia Ruiz Roldan, Marta Pla Diaz, Neris Garcia-Gonzalez, Inma Galán Vendrell, Llúcia Martínez-Priego, Giuseppe D'Auria, Fernando Gonzalez-Candelas |
| EPI_ISL_436326 | Servicio de Microbiologia. Hospital Clinico Universitario de Valencia | Sequencing and Bioinformatics Service and Molecular Epidemiology Research Group. FISABIO-Public Health | Loreto Ferrús Abad, Paula Ruiz-Hueso, Mariana Reyes-Prieto, Vicente Soriano Chirona, Ivan Ansari, David Navarro, Maria Alma Bracho, Griselda De Marco, Beatriz Beamud, Lidia Ruiz Roldan, Marta Pla Diaz, Neris Garcia-Gonzalez, Inma Galán Vendrell, Sandra Carbo, Llúcia Martínez-Priego, Giuseppe D'Auria, Fernando Gonzalez-Candelas |
| EPI_ISL_436327 | Servicio de Microbiologia. Hospital Clinico Universitario de Valencia | Sequencing and Bioinformatics Service and Molecular Epidemiology Research Group. FISABIO-Public Health | Paula Ruiz-Hueso, Mariana Reyes-Prieto, Vicente Soriano Chirona, Ivan Ansari, David Navarro, Maria Alma Bracho, Griselda De Marco, Beatriz Beamud, Lidia Ruiz Roldan, Marta Pla Diaz, Neris Garcia-Gonzalez, Inma Galán Vendrell, Sandra Carbo, Loreto Ferrús Abad, Llúcia Martínez-Priego, Giuseppe D'Auria, Fernando Gonzalez-Candelas |
| EPI_ISL_436328 | Servicio de Microbiologia. Hospital Clinico Universitario de Valencia | Sequencing and Bioinformatics Service and Molecular Epidemiology Research Group. FISABIO-Public Health | Mariana Reyes-Prieto, Vicente Soriano Chirona, Ivan Ansari, David Navarro, Maria Alma Bracho, Griselda De Marco, Beatriz Beamud, Lidia Ruiz Roldan, Marta Pla Diaz, Neris Garcia-Gonzalez, Inma Galán Vendrell, Sandra Carbo, Loreto Ferrús Abad, Paula Ruiz-Hueso, Llúcia Martínez-Priego, Giuseppe D'Auria, Fernando Gonzalez-Candelas |
| EPI_ISL_436330 | Servicio de Microbiologia. Hospital Clinico Universitario de Valencia | Sequencing and Bioinformatics Service and Molecular Epidemiology Research Group. FISABIO-Public Health | Ivan Ansari, David Navarro, Maria Alma Bracho, Griselda De Marco, Beatriz Beamud, Lidia Ruiz Roldan, Marta Pla Diaz, Neris Garcia-Gonzalez, Inma Galán Vendrell, Sandra Carbo, Loreto Ferrús Abad, Paula Ruiz-Hueso, Mariana Reyes-Prieto, Vicente Soriano Chirona, Llúcia Martínez-Priego, Giuseppe D'Auria, Fernando Gonzalez-Candelas |
| EPI_ISL_436335 | Servicio de Microbiologia. Hospital Clinico Universitario de Valencia | Sequencing and Bioinformatics Service and Molecular Epidemiology Research Group. FISABIO-Public Health | Lidia Ruiz Roldan, Marta Pla Diaz, Neris Garcia-Gonzalez, Inma Galán Vendrell, Sandra Carbo, Loreto Ferrús Abad, Paula Ruiz-Hueso, Mariana Reyes-Prieto, Vicente Soriano Chirona, Ivan Ansari, David Navarro, Maria Alma Bracho, Griselda De Marco, Beatriz Beamud, Llúcia Martínez-Priego, Giuseppe D'Auria, Fernando Gonzalez-Candelas |
| EPI_ISL_436336 | Servicio de Microbiologia. Hospital Clinico Universitario de Valencia | Sequencing and Bioinformatics Service and Molecular Epidemiology Research Group. FISABIO-Public Health | Marta Pla Diaz, Neris Garcia-Gonzalez, Inma Galán Vendrell, Sandra Carbo, Loreto Ferrús Abad, Paula Ruiz-Hueso, Mariana Reyes-Prieto, Vicente Soriano Chirona, Ivan Ansari, David Navarro, Maria Alma Bracho, Griselda De Marco, Beatriz Beamud, Lidia Ruiz Roldan, Llúcia Martínez-Priego, Giuseppe D'Auria, Fernando Gonzalez-Candelas |
| EPI_ISL_436337 | Servicio de Microbiologia. Hospital Clinico Universitario de Valencia | Sequencing and Bioinformatics Service and Molecular Epidemiology Research Group. FISABIO-Public Health | Neris Garcia-Gonzalez, Inma Galán Vendrell, Sandra Carbo, Loreto Ferrús Abad, Paula Ruiz-Hueso, Mariana Reyes-Prieto, Vicente Soriano Chirona, Ivan Ansari, David Navarro, Maria Alma Bracho, Griselda De Marco, Beatriz Beamud, Lidia Ruiz Roldan, Marta Pla Diaz, Llúcia Martínez-Priego, Giuseppe D'Auria, Fernando Gonzalez-Candelas |
| EPI_ISL_436341 | Servicio de Microbiologia. Hospital Clinico Universitario de Valencia | Sequencing and Bioinformatics Service and Molecular Epidemiology Research Group. FISABIO-Public Health | Lidia Ruiz Roldan, Marta Pla Diaz, Neris Garcia-Gonzalez, Inma Galán Vendrell, Sandra Carbo, Loreto Ferrús Abad, Paula Ruiz-Hueso, Mariana Reyes-Prieto, Vicente Soriano Chirona, Ivan Ansari, David Navarro, Maria Alma Bracho, Griselda De Marco, Beatriz Beamud, Llúcia Martínez-Priego, Giuseppe D'Auria, Fernando Gonzalez-Candelas |
| EPI_ISL_436342 | Servicio de Microbiologia. Hospital Clinico Universitario de Valencia | Sequencing and Bioinformatics Service and Molecular Epidemiology Research Group. FISABIO-Public Health | Marta Pla Diaz, Neris Garcia-Gonzalez, Inma Galán Vendrell, Sandra Carbo, Loreto Ferrús Abad, Paula Ruiz-Hueso, Mariana Reyes-Prieto, Vicente Soriano Chirona, Ivan Ansari, David Navarro, Maria Alma Bracho, Griselda De Marco, Beatriz Beamud, Lidia Ruiz Roldan, Llúcia Martínez-Priego, Giuseppe D'Auria, Fernando Gonzalez-Candelas |
| EPI_ISL_436345 | Servicio de Microbiologia. Hospital Clinico Universitario de Valencia | Sequencing and Bioinformatics Service and Molecular Epidemiology Research Group. FISABIO-Public Health | Sandra Carbo, Loreto Ferrús Abad, Paula Ruiz-Hueso, Mariana Reyes-Prieto, Vicente Soriano Chirona, Ivan Ansari, David Navarro, Maria Alma Bracho, Griselda De Marco, Beatriz Beamud, Lidia Ruiz Roldan, Marta Pla Diaz, Neris Garcia-Gonzalez, Inma Galán Vendrell, Llúcia Martínez-Priego, Giuseppe D'Auria, Fernando Gonzalez-Candelas |
| EPI_ISL_436346 | Servicio de Microbiologia. Hospital Clinico Universitario de Valencia | Sequencing and Bioinformatics Service and Molecular Epidemiology Research Group. FISABIO-Public Health | Loreto Ferrús Abad, Paula Ruiz-Hueso, Mariana Reyes-Prieto, Vicente Soriano Chirona, Ivan Ansari, David Navarro, Maria Alma Bracho, Griselda De Marco, Beatriz Beamud, Lidia Ruiz Roldan, Marta Pla Diaz, Neris Garcia-Gonzalez, Inma Galán Vendrell, Sandra Carbo, Llúcia Martínez-Priego, Giuseppe D'Auria, Fernando Gonzalez-Candelas |
| EPI_ISL_436353 | Servicio de Microbiologia. Hospital Clinico Universitario de Valencia | Sequencing and Bioinformatics Service and Molecular Epidemiology Research Group. FISABIO-Public Health | Griselda De Marco, Beatriz Beamud, Lidia Ruiz Roldan, Marta Pla Diaz, Neris Garcia-Gonzalez, Inma Galán Vendrell, Sandra Carbo, Loreto Ferrús Abad, Paula Ruiz-Hueso, Mariana Reyes-Prieto, Vicente Soriano Chirona, Ivan Ansari, David Navarro, Maria Alma Bracho, Llúcia Martínez-Priego, Giuseppe D'Auria, Fernando Gonzalez-Candelas |
| EPI_ISL_436355 | Servicio de Microbiologia. Hospital Clinico Universitario de Valencia | Sequencing and Bioinformatics Service and Molecular Epidemiology Research Group. FISABIO-Public Health | Lidia Ruiz Roldan, Marta Pla Diaz, Neris Garcia-Gonzalez, Inma Galán Vendrell, Sandra Carbo, Loreto Ferrús Abad, Paula Ruiz-Hueso, Mariana Reyes-Prieto, Vicente Soriano Chirona, Ivan Ansari, David Navarro, Maria Alma Bracho, Griselda De Marco, Beatriz Beamud, Llúcia Martínez-Priego, Giuseppe D'Auria, Fernando Gonzalez-Candelas |
| EPI_ISL_436356 | Servicio de Microbiologia. Hospital Clinico Universitario de Valencia | Sequencing and Bioinformatics Service and Molecular Epidemiology Research Group. FISABIO-Public Health | Marta Pla Diaz, Neris Garcia-Gonzalez, Inma Galán Vendrell, Sandra Carbo, Loreto Ferrús Abad, Paula Ruiz-Hueso, Mariana Reyes-Prieto, Vicente Soriano Chirona, Ivan Ansari, David Navarro, Maria Alma Bracho, Griselda De Marco, Beatriz Beamud, Lidia Ruiz Roldan, Llúcia Martínez-Priego, Giuseppe D'Auria, Fernando Gonzalez-Candelas |
| EPI_ISL_436361 | Servicio de Microbiología. Consorcio Hospital General Universitario de Valencia | Sequencing and Bioinformatics Service and Molecular Epidemiology Research Group. FISABIO-Public Health | Neris Garcia-Gonzalez, Loreto Ferrús Abad, Maria Dolores Ocete, Inma Galán Vendrell, Paula Ruiz-Hueso, Mariana Reyes-Prieto, Vicente Soriano Chirona, Maria Alma Bracho, Griselda De Marco, Beatriz Beamud, Lidia Ruiz Roldan, Marta Pla Diaz, Llúcia Martínez-Priego, Concepcion Gimeno, Giuseppe D'Auria, Fernando Gonzalez-Candelas |
| EPI_ISL_436364 | Servicio de Microbiologia. Hospital Clinico Universitario de Valencia | Sequencing and Bioinformatics Service and Molecular Epidemiology Research Group. FISABIO-Public Health | Beatriz Beamud, Lidia Ruiz Roldan, Marta Pla Diaz, Neris Garcia-Gonzalez, Inma Galán Vendrell, Sandra Carbo, Loreto Ferrús Abad, Paula Ruiz-Hueso, Mariana Reyes-Prieto, Vicente Soriano Chirona, Ivan Ansari, David Navarro, Maria Alma Bracho, Griselda De Marco, Llúcia Martínez-Priego, Giuseppe D'Auria, Fernando Gonzalez-Candelas |
| EPI_ISL_436366 | Servicio de Microbiologia. Hospital Clinico Universitario de Valencia | Sequencing and Bioinformatics Service and Molecular Epidemiology Research Group. FISABIO-Public Health | Marta Pla Diaz, Neris Garcia-Gonzalez, Inma Galán Vendrell, Sandra Carbo, Loreto Ferrús Abad, Paula Ruiz-Hueso, Mariana Reyes-Prieto, Vicente Soriano Chirona, Ivan Ansari, David Navarro, Maria Alma Bracho, Griselda De Marco, Beatriz Beamud, Lidia Ruiz Roldan, Llúcia Martínez-Priego, Giuseppe D'Auria, Fernando Gonzalez-Candelas |
| EPI_ISL_436367 | Servicio de Microbiologia. Hospital Clinico Universitario de Valencia | Sequencing and Bioinformatics Service and Molecular Epidemiology Research Group. FISABIO-Public Health | Neris Garcia-Gonzalez, Inma Galán Vendrell, Sandra Carbo, Loreto Ferrús Abad, Paula Ruiz-Hueso, Mariana Reyes-Prieto, Vicente Soriano Chirona, Ivan Ansari, David Navarro, Maria Alma Bracho, Griselda De Marco, Beatriz Beamud, Lidia Ruiz Roldan, Marta Pla Diaz, Llúcia Martínez-Priego, Giuseppe D'Auria, Fernando Gonzalez-Candelas |
| EPI_ISL_436368 | Servicio de Microbiologia. Hospital Clinico Universitario de Valencia | Sequencing and Bioinformatics Service and Molecular Epidemiology Research Group. FISABIO-Public Health | Inma Galán Vendrell, Sandra Carbo, Loreto Ferrús Abad, Paula Ruiz-Hueso, Mariana Reyes-Prieto, Vicente Soriano Chirona, Ivan Ansari, David Navarro, Maria Alma Bracho, Griselda De Marco, Beatriz Beamud, Lidia Ruiz Roldan, Marta Pla Diaz, Neris Garcia-Gonzalez, Llúcia Martínez-Priego, Giuseppe D'Auria, Fernando Gonzalez-Candelas |
| EPI_ISL_436369 | Servicio de Microbiologia. Hospital Clinico Universitario de Valencia | Sequencing and Bioinformatics Service and Molecular Epidemiology Research Group. FISABIO-Public Health | Sandra Carbo, Loreto Ferrús Abad, Paula Ruiz-Hueso, Mariana Reyes-Prieto, Vicente Soriano Chirona, Ivan Ansari, David Navarro, Maria Alma Bracho, Griselda De Marco, Beatriz Beamud, Lidia Ruiz Roldan, Marta Pla Diaz, Neris Garcia-Gonzalez, Inma Galán Vendrell, Llúcia Martínez-Priego, Giuseppe D'Auria, Fernando Gonzalez-Candelas |
| EPI_ISL_436371 | Servicio de Microbiologia. Hospital Clinico Universitario de Valencia | Sequencing and Bioinformatics Service and Molecular Epidemiology Research Group. FISABIO-Public Health | Paula Ruiz-Hueso, Mariana Reyes-Prieto, Vicente Soriano Chirona, Ivan Ansari, David Navarro, Maria Alma Bracho, Griselda De Marco, Beatriz Beamud, Lidia Ruiz Roldan, Marta Pla Diaz, Neris Garcia-Gonzalez, Inma Galán Vendrell, Sandra Carbo, Loreto Ferrús Abad, Llúcia Martínez-Priego, Giuseppe D'Auria, Fernando Gonzalez-Candelas |
| EPI_ISL_436373, EPI_ISL_436374, EPI_ISL_436376, EPI_ISL_436377, EPI_ISL_436378, EPI_ISL_436379, EPI_ISL_436380 | Servicio de Microbiologia. Hospital Universitario Doctor Peset | Sequencing and Bioinformatics Service and Molecular Epidemiology Research Group. FISABIO-Public Health | Juan Alberola Enguídanos, Juan Jose Camarena Miñana, Rosa González Pellicer, Neris Garcia-Gonzalez, Inma Galán Vendrell, Sandra Carbo, Loreto Ferrús Abad, Paula Ruiz-Hueso, Mariana Reyes-Prieto, Vicente Soriano Chirona, Ivan Ansari, Maria Alma Bracho, Griselda De Marco, Beatriz Beamud, Lidia Ruiz Roldan, Marta Pla Diaz, Llúcia Martínez-Priego, Giuseppe D'Auria, Jose Miguel Nogueira Coito, Fernando Gonzalez-Candelas |
| EPI_ISL_436382 | Servicio de Microbiología. Consorcio Hospital General Universitario de Valencia | Sequencing and Bioinformatics Service and Molecular Epidemiology Research Group. FISABIO-Public Health | Loreto Ferrús Abad, Maria Dolores Ocete,Inma Galán Vendrell, Paula Ruiz-Hueso, Mariana Reyes-Prieto, Vicente Soriano Chirona, Maria Alma Bracho, Griselda De Marco, Beatriz Beamud, Lidia Ruiz Roldan, Marta Pla Diaz,Neris Garcia-Gonzalez, Llúcia Martínez-Priego, Concepcion Gimeno, Giuseppe D'Auria, Fernando Gonzalez-Candelas |
| EPI_ISL_436383 | Servicio de Microbiología. Consorcio Hospital General Universitario de Valencia | Sequencing and Bioinformatics Service and Molecular Epidemiology Research Group. FISABIO-Public Health | Maria Dolores Ocete, Inma Galán Vendrell, Paula Ruiz-Hueso, Mariana Reyes-Prieto, Vicente Soriano Chirona, Maria Alma Bracho, Griselda De Marco, Beatriz Beamud, Lidia Ruiz Roldan, Marta Pla Diaz,Neris Garcia-Gonzalez, Loreto Ferrús Abad, Llúcia Martínez-Priego, Concepcion Gimeno, Giuseppe D'Auria, Fernando Gonzalez-Candelas |
| EPI_ISL_436384 | Servicio de Microbiología. Consorcio Hospital General Universitario de Valencia | Sequencing and Bioinformatics Service and Molecular Epidemiology Research Group. FISABIO-Public Health | Griselda De Marco, Beatriz Beamud, Lidia Ruiz Roldan, Marta Pla Diaz,Neris Garcia-Gonzalez, Loreto Ferrús Abad, Maria Dolores Ocete, Inma Galán Vendrell, Paula Ruiz-Hueso, Mariana Reyes-Prieto, Vicente Soriano Chirona, Maria Alma Bracho, Llúcia Martínez-Priego, Concepcion Gimeno, Giuseppe D'Auria, Fernando Gonzalez-Candelas |
| EPI_ISL_436386 | Servicio de Microbiología. Consorcio Hospital General Universitario de Valencia | Sequencing and Bioinformatics Service and Molecular Epidemiology Research Group. FISABIO-Public Health | Lidia Ruiz Roldan, Marta Pla Diaz,Neris Garcia-Gonzalez, Loreto Ferrús Abad, Maria Dolores Ocete, Inma Galán Vendrell, Paula Ruiz-Hueso, Mariana Reyes-Prieto, Vicente Soriano Chirona, Maria Alma Bracho, Griselda De Marco, Beatriz Beamud, Llúcia Martínez-Priego, Concepcion Gimeno, Giuseppe D'Auria, Fernando Gonzalez-Candelas |
| EPI_ISL_436387 | Servicio de Microbiología. Consorcio Hospital General Universitario de Valencia | Sequencing and Bioinformatics Service and Molecular Epidemiology Research Group. FISABIO-Public Health | Marta Pla Diaz,Neris Garcia-Gonzalez, Loreto Ferrús Abad, Maria Dolores Ocete, Inma Galán Vendrell, Paula Ruiz-Hueso, Mariana Reyes-Prieto, Vicente Soriano Chirona, Maria Alma Bracho, Griselda De Marco, Beatriz Beamud, Lidia Ruiz Roldan, Llúcia Martínez-Priego, Concepcion Gimeno, Giuseppe D'Auria, Fernando Gonzalez-Candelas |

| | | | |
|---|---|---|---|
| EPI_ISL_436388 | Servicio de Microbiología. Consorcio Hospital General Universitario de Valencia | Sequencing and Bioinformatics Service and Molecular Epidemiology Research Group. FISABIO-Public Health | Neris Garcia-Gonzalez, Loreto Ferrús Abad, Maria Dolores Ocete, Inma Galán Vendrell, Paula Ruiz-Hueso, Mariana Reyes-Prieto, Vicente Soriano Chirona, Maria Alma Bracho, Griselda De Marco, Beatriz Beamud, Lidia Ruiz Roldan, Marta Pla Diaz, Llúcia Martínez-Priego, Concepcion Gimeno, Giuseppe D'Auria, Fernando Gonzalez-Candelas |
| EPI_ISL_436389 | Servicio de Microbiología. Consorcio Hospital General Universitario de Valencia | Sequencing and Bioinformatics Service and Molecular Epidemiology Research Group. FISABIO-Public Health | Loreto Ferrús Abad, Maria Dolores Ocete, Inma Galán Vendrell, Paula Ruiz-Hueso, Mariana Reyes-Prieto, Vicente Soriano Chirona, Maria Alma Bracho, Griselda De Marco, Beatriz Beamud, Lidia Ruiz Roldan, Marta Pla Diaz,Neris Garcia-Gonzalez, Llúcia Martínez-Priego, Concepcion Gimeno, Giuseppe D'Auria, Fernando Gonzalez-Candelas |
| EPI_ISL_436390 | Servicio de Microbiología. Consorcio Hospital General Universitario de Valencia | Sequencing and Bioinformatics Service and Molecular Epidemiology Research Group. FISABIO-Public Health | Maria Dolores Ocete, Inma Galán Vendrell, Paula Ruiz-Hueso, Mariana Reyes-Prieto, Vicente Soriano Chirona, Maria Alma Bracho, Griselda De Marco, Beatriz Beamud, Lidia Ruiz Roldan, Marta Pla Diaz,Neris Garcia-Gonzalez, Loreto Ferrús Abad, Llúcia Martínez-Priego, Concepcion Gimeno, Giuseppe D'Auria, Fernando Gonzalez-Candelas |
| EPI_ISL_436391 | Servicio de Microbiología. Consorcio Hospital General Universitario de Valencia | Sequencing and Bioinformatics Service and Molecular Epidemiology Research Group. FISABIO-Public Health | Griselda De Marco, Beatriz Beamud, Lidia Ruiz Roldan, Marta Pla Diaz,Neris Garcia-Gonzalez, Loreto Ferrús Abad, Maria Dolores Ocete, Inma Galán Vendrell, Paula Ruiz-Hueso, Mariana Reyes-Prieto, Vicente Soriano Chirona, Maria Alma Bracho, Llúcia Martínez-Priego, Concepcion Gimeno, Giuseppe D'Auria, Fernando Gonzalez-Candelas |
| EPI_ISL_436397 | Servicio de Microbiologia. Hospital Clinico Universitario de Valencia | Sequencing and Bioinformatics Service and Molecular Epidemiology Research Group. FISABIO-Public Health | Loreto Ferrús Abad, Paula Ruiz-Hueso, Mariana Reyes-Prieto, Vicente Soriano Chirona, Ivan Ansari, David Navarro, Maria Alma Bracho, Griselda De Marco, Beatriz Beamud, Lidia Ruiz Roldan, Marta Pla Diaz, Neris Garcia-Gonzalez, Inma Galán Vendrell, Sandra Carbo, Llúcia Martínez-Priego, Giuseppe D'Auria, Fernando Gonzalez-Candelas |
| EPI_ISL_436411 | Servicio de Microbiologia. Hospital Clinico Universitario de Valencia | Sequencing and Bioinformatics Service and Molecular Epidemiology Research Group. FISABIO-Public Health | Neris Garcia-Gonzalez, Inma Galán Vendrell, Sandra Carbo, Loreto Ferrús Abad, Paula Ruiz-Hueso, Mariana Reyes-Prieto, Vicente Soriano Chirona, Ivan Ansari, David Navarro, Maria Alma Bracho, Griselda De Marco, Beatriz Beamud, Lidia Ruiz Roldan, Marta Pla Diaz, Llúcia Martínez-Priego, Giuseppe D'Auria, Fernando Gonzalez-Candelas |
| EPI_ISL_436412 | Viral Respiratory Lab, National Institute for Biomedical Research (INRB) | Pathogen Sequencing Lab, National Institute for Biomedical Research (INRB) | Placide Mbala-Kingebeni, Edith Nkwembe, Eddy Kinganda-Lusamaki, Amuri Aziza, Francisca Muyembe Mawete, Catherine Pratt, Matthias Pauthner, Josh Quick, Allison Black, James Hadfield, Trevor Bedford, Ian Goodfellow, Andrew Rambaut, Nick Loman, Kristian Andersen, Michael Wiley, Steve Ahuka-Mundeke, Jean-Jacques Muyembe Tamfum |
| EPI_ISL_436413, EPI_ISL_436414, EPI_ISL_436415, EPI_ISL_436417, EPI_ISL_436418, EPI_ISL_436419, EPI_ISL_436420, EPI_ISL_436421, EPI_ISL_436422, EPI_ISL_436424, EPI_ISL_436425, EPI_ISL_436426, EPI_ISL_436428, EPI_ISL_436429, EPI_ISL_436430, EPI_ISL_436431, EPI_ISL_436432, EPI_ISL_436433, EPI_ISL_436434, EPI_ISL_436435, EPI_ISL_436436, EPI_ISL_436437, EPI_ISL_436440, EPI_ISL_436444, EPI_ISL_436445, EPI_ISL_436447, EPI_ISL_436448, EPI_ISL_436449, EPI_ISL_436450, EPI_ISL_436451, EPI_ISL_436452, EPI_ISL_436453, EPI_ISL_436454, EPI_ISL_436455, EPI_ISL_436456, EPI_ISL_436457, EPI_ISL_436458, EPI_ISL_436459, EPI_ISL_436460, EPI_ISL_436461, EPI_ISL_436462, EPI_ISL_436463 | | | |
| see above | National Centre for Disease control (NCDC) | NCDC/CSIR-IGIB | Pramod Kumar#, Rajesh Pandey#, Pooja Sharma, Mahesh S Dhar, Vivekanand A, Bharathram Uppili, Himanshu Vashisht, Saruchi Wadhwa, Nishu Tyagi, Uma Sharma, Priyanka Singh, Hemlata Lall, Meena Datta, Poonam Gupta, Nidhi Saini, Aarti Tewari, Bibhash Nandi, Dhirendra Kumar, Satyabrata Bag, Varun Jaiswal, Hema Gogia, Preeti Madan, Simrita Singh, Prateek Singh, Debasis Dash, Mitali Mukerji, Manju Bala, Sandhya Kabra, Sujeet Singh, Mohammed Faruq, Anurag Agrawal*, Partha Rakshit* |
| EPI_ISL_436464 | Alaska State Virology Laboratory | Alaska State Virology Laboratory | Jack Chen |
| EPI_ISL_436466, EPI_ISL_436467, EPI_ISL_436468, EPI_ISL_436469, EPI_ISL_436470, EPI_ISL_436471, EPI_ISL_436472, EPI_ISL_436473, EPI_ISL_436474, EPI_ISL_436475, EPI_ISL_436476, EPI_ISL_436477, EPI_ISL_436478, EPI_ISL_436479, EPI_ISL_436480, EPI_ISL_436481, EPI_ISL_436482, EPI_ISL_436483, EPI_ISL_436484, EPI_ISL_436485, EPI_ISL_436486, EPI_ISL_436487, EPI_ISL_436488, EPI_ISL_436489, EPI_ISL_436490, EPI_ISL_436491, EPI_ISL_436492, EPI_ISL_436493, EPI_ISL_436494, EPI_ISL_436495, EPI_ISL_436496, EPI_ISL_436497, EPI_ISL_436498, EPI_ISL_436499, EPI_ISL_436500, EPI_ISL_436501, EPI_ISL_436502, EPI_ISL_436503, EPI_ISL_436504 | | | |
| see above | UPMC Clinical Laboratory | Microbial Genome Sequencing Center, Microbial Genomic Epidemiological Laboratory | Dan Snyder, Stephanie L Mitchell, Mustapha M Mustapha, Marissa P Griffith, Vatsala R Srinivasa, Kady D Waggle, Chinelo Ezeonwuku, Jane W. Marsh, Lee H. Harrison, Vaughn S. Cooper |
| EPI_ISL_436512, EPI_ISL_436549 | Florida Bureau of Public Health Laboratories | Florida Bureau of Public Health Laboratories | Sarah Schmedes, Jason Blanton |
| EPI_ISL_436564, EPI_ISL_436565, EPI_ISL_436566, EPI_ISL_436567, EPI_ISL_436568, EPI_ISL_436569, EPI_ISL_436570, EPI_ISL_436571, EPI_ISL_436572, EPI_ISL_436573, EPI_ISL_436574, EPI_ISL_436575, EPI_ISL_436576, EPI_ISL_436577, EPI_ISL_436578, EPI_ISL_436579, EPI_ISL_436580, EPI_ISL_436581, EPI_ISL_436582, EPI_ISL_436583, EPI_ISL_436584, EPI_ISL_436585, EPI_ISL_436586, EPI_ISL_436587, EPI_ISL_436588, EPI_ISL_436589, EPI_ISL_436590, EPI_ISL_436591, EPI_ISL_436592, EPI_ISL_436593, EPI_ISL_436594, EPI_ISL_436595, EPI_ISL_436596, EPI_ISL_436597, EPI_ISL_436598, EPI_ISL_436599, EPI_ISL_436600, EPI_ISL_436601, EPI_ISL_436602, EPI_ISL_436603, EPI_ISL_436604, EPI_ISL_436605, EPI_ISL_436606, EPI_ISL_436607, EPI_ISL_436608, EPI_ISL_436609, EPI_ISL_436610, EPI_ISL_436612, EPI_ISL_436613, EPI_ISL_436614, EPI_ISL_436615, EPI_ISL_436616, EPI_ISL_436617, EPI_ISL_436618, EPI_ISL_436619, EPI_ISL_436620, EPI_ISL_436621, EPI_ISL_436622, EPI_ISL_436623, EPI_ISL_436625, EPI_ISL_436626, EPI_ISL_436627, EPI_ISL_436628, EPI_ISL_436629, EPI_ISL_436630, EPI_ISL_436631, EPI_ISL_436632, EPI_ISL_436634, EPI_ISL_436635, EPI_ISL_436636, EPI_ISL_436637, EPI_ISL_436638, EPI_ISL_436640 | | | |
| see above | University of Wisconsin-Madison AIDS Vaccine Research Laboratories | University of Wisconsin-Madison AIDS Vaccine Research Laboratories | Gage Moreno, Katarina Braun, et al. AIDS Vaccine Research Laboratories |
| EPI_ISL_436641, EPI_ISL_436642, EPI_ISL_436643, EPI_ISL_436644, EPI_ISL_436645, EPI_ISL_436646, EPI_ISL_436647, EPI_ISL_436648, EPI_ISL_436650, EPI_ISL_436651, EPI_ISL_436652, EPI_ISL_436653, EPI_ISL_436654, EPI_ISL_436655, EPI_ISL_436656, EPI_ISL_436657, EPI_ISL_436658, EPI_ISL_436659, EPI_ISL_436660, EPI_ISL_436661, EPI_ISL_436662, EPI_ISL_436663, EPI_ISL_436664, EPI_ISL_436665, EPI_ISL_436666, EPI_ISL_436667, EPI_ISL_436669, EPI_ISL_436670, EPI_ISL_436671, EPI_ISL_436672, EPI_ISL_436673, EPI_ISL_436674, EPI_ISL_436675, EPI_ISL_436676, EPI_ISL_436677, EPI_ISL_436678, EPI_ISL_436679, EPI_ISL_436680, EPI_ISL_436681, EPI_ISL_436682, EPI_ISL_436683 | | | |
| see above | County of Santa Clara Public Health Department | Chan-Zuckerberg Biohub | CZB Cliahub Consortium |
| EPI_ISL_436684, EPI_ISL_436686 | KRISP, KZN Research Innovation and Sequencing Platform | KRISP, KZN Research Innovation and Sequencing Platform | Giandhari J, Pillay S, Lessells R, Chimukangara B, Deforche K, Tegally H, Wilkinson E, de Oliveira T |
| EPI_ISL_436688, EPI_ISL_436689 | Victorian Infectious Diseases Reference Laboratory (VIDRL) | Microbiological Diagnostic Unit Public Health Laboratory and Victorian Infectious Diseases Reference Laboratory, The Peter Doherty Institute for Infection & Immunity | Caly L., Seemann T., Sait, M., Schultz M., Druce J., Sherry, N. |
| EPI_ISL_436715 | Genomics and Computational Biology Lab, Scientific Research Institute of Physical-Chemical Medicine, FMBA of Russia | Genomics and Computational Biology Lab, Scientific Research Institute of Physical-Chemical Medicine, FMBA of Russia | A. Pavlenko, O. Guskova, K. Klimina, V. Veselovsky, A. Manolov, D. Fedorov, V. Govorun and E. Ilina |
| EPI_ISL_436716 | Genomics and Computational Biology Lab, Scientific Research Institute of Physical-Chemical Medicine, FMBA of Russia | Genomics and Computational Biology Lab, Scientific Research Institute of Physical-Chemical Medicine, FMBA of Russia | A. Pavlenko, O. Guskova, K. Klimina, V. Veselovsky, A. Manolov, D. Fedorov, V. Govorun and E. Ilina |
| EPI_ISL_436717 | Genomics and Computational Biology Lab, Scientific Research Institute of Physical-Chemical Medicine, FMBA of Russia | Genomics and Computational Biology Lab, Scientific Research Institute of Physical-Chemical Medicine, FMBA of Russia | A. Pavlenko, O. Guskova, K. Klimina, V. Veselovsky, A. Manolov, D. Fedorov, V. Govorun and E. Ilina |
| EPI_ISL_436718 | Ospedale Regionale San Salvatore | Istituto Zooprofilattico Sperimentale dell'Abruzzo e Molise "G.Caporale" | Lorusso A, Marcacci M, Di Domenico M, Ancora M, Curini V, Mangone I, Rinaldi A, Di Pasquale A, Cammà C, Puglia I, Savini G |
| EPI_ISL_436719, EPI_ISL_436720, EPI_ISL_436721, EPI_ISL_436722 | Ospedale Civile S. Liberatore di Atri | Istituto Zooprofilattico Sperimentale dell'Abruzzo e Molise "G.Caporale" | Lorusso A, Marcacci M, Di Domenico M, Ancora M, Curini V, Mangone I, Rinaldi A, Di Pasquale A, Cammà C, Puglia I, Savini G |
| EPI_ISL_436723 | Ospedale Civile Giuseppe Mazzini | Istituto Zooprofilattico Sperimentale dell'Abruzzo e Molise "G.Caporale" | Lorusso A, Marcacci M, Di Domenico M, Ancora M, Curini V, Mangone I, Rinaldi A, Di Pasquale A, Cammà C, Puglia I, Savini G |
| EPI_ISL_436724 | Ospedale Civile S. Liberatore di Atri | Istituto Zooprofilattico Sperimentale dell'Abruzzo e Molise "G.Caporale" | Lorusso A, Marcacci M, Di Domenico M, Ancora M, Curini V, Mangone I, Rinaldi A, Di Pasquale A, Cammà C, Puglia I, Savini G |
| EPI_ISL_436725 | RSA/RP Villa San Giovanni - Gruppo Edos | Istituto Zooprofilattico Sperimentale dell'Abruzzo e Molise "G.Caporale" | Lorusso A, Marcacci M, Di Domenico M, Ancora M, Curini V, Mangone I, Rinaldi A, Di Pasquale A, Cammà C, Puglia I, Savini G |
| EPI_ISL_436726, EPI_ISL_436727, EPI_ISL_436729 | SERVIZIO DI IGIENE E SANITÀ PUBBLICA ASL Teramo | Istituto Zooprofilattico Sperimentale dell'Abruzzo e Molise "G.Caporale" | Lorusso A, Marcacci M, Di Domenico M, Ancora M, Curini V, Mangone I, Rinaldi A, Di Pasquale A, Cammà C, Puglia I, Savini G |
| EPI_ISL_436730 | Servizio di igiene epidemiologia e sanità pubblica (Siesp) Chieti | Istituto Zooprofilattico Sperimentale dell'Abruzzo e Molise "G.Caporale" | Lorusso A, Marcacci M, Di Domenico M, Ancora M, Curini V, Mangone I, Rinaldi A, Di Pasquale A, Cammà C, Puglia I, Savini G |
| EPI_ISL_436731, EPI_ISL_436732 | Ospedale Civile S. Liberatore di Atri | Istituto Zooprofilattico Sperimentale dell'Abruzzo e Molise "G.Caporale" | Lorusso A, Marcacci M, Di Domenico M, Ancora M, Curini V, Mangone I, Rinaldi A, Di Pasquale A, Cammà C, Puglia I, Savini G |
| EPI_ISL_436800, EPI_ISL_436801, EPI_ISL_436802, EPI_ISL_436803, EPI_ISL_436804, EPI_ISL_436807, EPI_ISL_436808, EPI_ISL_436812, EPI_ISL_436814, EPI_ISL_436815, EPI_ISL_436818, EPI_ISL_436819, EPI_ISL_436820, EPI_ISL_436821, EPI_ISL_436822, EPI_ISL_436823, EPI_ISL_436824, EPI_ISL_436825, EPI_ISL_436827, EPI_ISL_436829, EPI_ISL_436830, EPI_ISL_436831, EPI_ISL_436833, EPI_ISL_436834, EPI_ISL_436836, EPI_ISL_436838, EPI_ISL_436841, EPI_ISL_436842, EPI_ISL_436843, EPI_ISL_436846, EPI_ISL_436847, EPI_ISL_436848, EPI_ISL_436850, EPI_ISL_436852, EPI_ISL_436853, EPI_ISL_436854, EPI_ISL_436855, EPI_ISL_436856, EPI_ISL_436857, EPI_ISL_436859, EPI_ISL_436861, EPI_ISL_436863, EPI_ISL_436864, EPI_ISL_436865, EPI_ISL_436867, EPI_ISL_436868, EPI_ISL_436869, EPI_ISL_436870, EPI_ISL_436874, EPI_ISL_436875, EPI_ISL_436876, EPI_ISL_436877, EPI_ISL_436878, EPI_ISL_436879, EPI_ISL_436880, EPI_ISL_436884, EPI_ISL_436885, EPI_ISL_436886, EPI_ISL_436887, EPI_ISL_436889 | | | |
| see above | Michigan Department of Health and Human Services, Bureau of Laboratories | Michigan Department of Health and Human Services, Bureau of Laboratories | Blankenship HM, Riner D, Soehnlen MK |
| EPI_ISL_436892, EPI_ISL_436893, EPI_ISL_436894, EPI_ISL_436895, EPI_ISL_436896, EPI_ISL_436897, EPI_ISL_436898, EPI_ISL_436899, EPI_ISL_436900 | Gundersen Molecular Diagnostics Laboratory | Kabara Cancer Research Institute | Craig S. Richmond, Paraic A. Kenny |
| EPI_ISL_436903, EPI_ISL_436905, EPI_ISL_436907, EPI_ISL_436914, EPI_ISL_436915, EPI_ISL_436917, EPI_ISL_436921, EPI_ISL_436922, EPI_ISL_436923, EPI_ISL_436924, EPI_ISL_436925 | | | |
| see above | Utah Public Health Laboratory | Utah Public Health Laboratory | Erin Young, Kelly Oakeson |
| EPI_ISL_436926 | $x^2$ | Utah Public Health Laboratory | Erin Young, Kelly Oakeson |
| EPI_ISL_436927, EPI_ISL_436928, EPI_ISL_436930, EPI_ISL_436931, EPI_ISL_436934, EPI_ISL_436935, EPI_ISL_436937, EPI_ISL_436938 | Utah Public Health Laboratory | Utah Public Health Laboratory | Erin Young, Kelly Oakeson |
| EPI_ISL_436939, EPI_ISL_436942, EPI_ISL_436943, EPI_ISL_436944, EPI_ISL_436945, EPI_ISL_436946, EPI_ISL_436947, EPI_ISL_436948, EPI_ISL_436949, EPI_ISL_436950, EPI_ISL_436952, EPI_ISL_436953, EPI_ISL_436954, EPI_ISL_436956, EPI_ISL_436957, EPI_ISL_436958, EPI_ISL_436959, EPI_ISL_436960, EPI_ISL_436961 | | | |
| see above | Ochsner Health | Bioinfoexperts, LLC | Amy Feehan, David J. Nolan, Rebecca Rose, Sissy Cross, David Moraga Amador, Tong Yang, Luke Caruso, Wayra Navia, Lydia Von Borstel, Xiao Hui Zhou, Julia-Garcia-Diaz, Susanna L. Lamers |
| EPI_ISL_436962, EPI_ISL_436963, EPI_ISL_436965, EPI_ISL_436967, EPI_ISL_436969, EPI_ISL_436973, EPI_ISL_436975, EPI_ISL_436978, EPI_ISL_436979, EPI_ISL_436980, EPI_ISL_436981, EPI_ISL_436982, EPI_ISL_436984, EPI_ISL_436985, EPI_ISL_436986, EPI_ISL_436987, EPI_ISL_436988, EPI_ISL_436989, EPI_ISL_436990, EPI_ISL_436991, EPI_ISL_436993, EPI_ISL_436995, EPI_ISL_436996, EPI_ISL_436997, EPI_ISL_436998, EPI_ISL_436999, EPI_ISL_437000, EPI_ISL_437001, EPI_ISL_437003, EPI_ISL_437004, EPI_ISL_437006, EPI_ISL_437007, EPI_ISL_437011, EPI_ISL_437013, EPI_ISL_437014, EPI_ISL_437015, EPI_ISL_437016, EPI_ISL_437017, EPI_ISL_437018, EPI_ISL_437019, EPI_ISL_437020, EPI_ISL_437021, EPI_ISL_437023, EPI_ISL_437024, EPI_ISL_437025, EPI_ISL_437026, EPI_ISL_437027, EPI_ISL_437028, EPI_ISL_437029, EPI_ISL_437030, EPI_ISL_437031, EPI_ISL_437032, EPI_ISL_437033, EPI_ISL_437035, EPI_ISL_437036, EPI_ISL_437037, EPI_ISL_437038, EPI_ISL_437039, EPI_ISL_437040, EPI_ISL_437041, EPI_ISL_437042 | | | |
| see above | Department of Virus and Microbiological Special Diagnostics, Statens Serum Institut, Copenhagen, Denmark, Artillerivej 5, 2300 Copenahgen S | Albertsen lab, Department of Chemistry and Bioscience, Aalborg University, Denmark | Rasmus Kirkegaard |
| EPI_ISL_437044, EPI_ISL_437045, EPI_ISL_437046, EPI_ISL_437047, EPI_ISL_437048, EPI_ISL_437049, EPI_ISL_437050, EPI_ISL_437051, EPI_ISL_437053, EPI_ISL_437054, EPI_ISL_437055, EPI_ISL_437056, EPI_ISL_437057, EPI_ISL_437059, EPI_ISL_437060, EPI_ISL_437061, EPI_ISL_437063, EPI_ISL_437064, EPI_ISL_437065, EPI_ISL_437066, EPI_ISL_437067, EPI_ISL_437068, EPI_ISL_437071, EPI_ISL_437072, EPI_ISL_437073, EPI_ISL_437074, EPI_ISL_437075, EPI_ISL_437076, EPI_ISL_437077, EPI_ISL_437079, EPI_ISL_437081, EPI_ISL_437082, EPI_ISL_437083, EPI_ISL_437084, EPI_ISL_437085, EPI_ISL_437087, EPI_ISL_437088 | | | |
| see above | County of Santa Clara Public Health | Chan-Zuckerberg Biohub | CZB Cliahub Consortium |
| EPI_ISL_437089, EPI_ISL_437090, EPI_ISL_437091, EPI_ISL_437092, EPI_ISL_437093, EPI_ISL_437094, EPI_ISL_437095, EPI_ISL_437096 | Latvijas Infektoloģijas centrs | Latvian Biomedical Research and Study Centre | Ivars Silamiķelis, Kaspars Megnis, Monta Ustinova, Ņikita Zrelovs, Vita Rovīte, Jeļena Storoženko, Tatjana Kolupajeva, Oksana Savicka, Uga Dumpis, Jānis Kloviņš |
| EPI_ISL_437099, EPI_ISL_437100, EPI_ISL_437101, EPI_ISL_437107, EPI_ISL_437109, EPI_ISL_437111, EPI_ISL_437113, EPI_ISL_437115, EPI_ISL_437117, EPI_ISL_437120, EPI_ISL_437123, EPI_ISL_437128, EPI_ISL_437131, EPI_ISL_437132, EPI_ISL_437135, EPI_ISL_437137, EPI_ISL_437138, EPI_ISL_437139, EPI_ISL_437140, EPI_ISL_437142, EPI_ISL_437145, EPI_ISL_437146, EPI_ISL_437147, EPI_ISL_437150, EPI_ISL_437152, EPI_ISL_437153, EPI_ISL_437154, EPI_ISL_437155, EPI_ISL_437157, EPI_ISL_437162, EPI_ISL_437163, EPI_ISL_437164, EPI_ISL_437166, EPI_ISL_437170, EPI_ISL_437174, EPI_ISL_437176, EPI_ISL_437180 | | | |
| see above | Michigan Department of Health and Human Services, Bureau of Laboratories | Michigan Department of Health and Human Services, Bureau of Laboratories | Blankenship HM, Riner D, Soehnlen MK |
| EPI_ISL_437187 | Siloam Hospitals | Institute of Tropical Disease, Universitas Airlangga | Kazufumi Shimizu, Krisnoadi Rahardjo, Aldise M Nastri, Jezzy R Dewantari, Rima R Prasetya, Maria M Padmidewi, Gatot Soegiarto, Laksmi Wulandari, Retno A Setyoningrum, Resti Y Meliana, Yohko K Shimizu, Mitsuhiro Nishimura, Yasuko Mori, Soetjipto, Maria I Lusida |

| | | | |
|---|---|---|---|
| EPI_ISL_437188 | RSUD Dr. Soetomo | Institute of Tropical Disease, Universitas Airlangga | Krisnoadi Rahardjo, Aldise M Nastri, Jezzy R Dewantari, Rima R Prasetya, Joni Wahyuhadi, Gatot Soegiarto, Laksmi Wulandari, Retno A Setyoningrum, Resti Y Meliana, Yohko K Shimizu, Mitsuhiro Nishimura, Yasuko Mori, Soetjipto, Kazufumi Shimizu, Maria I Lusida |
| EPI_ISL_437189 | Pusat Pertamina Hospital | Eijkman Institute for Molecular Biology, Ministry of Research and Technology/National Agency for Research and Innovation | Edison Johar, Frilasita A Yudhaputri, Hidayat Trimarsanto, David H Muljono, Safarina G Malik, Khin Saw Myint, Amin Soebandrio |
| EPI_ISL_437190, EPI_ISL_437191 | RS Pondok Indah Hospital – Pondok Indah | Eijkman Institute for Molecular Biology, Ministry of Research and Technology/National Agency for Research and Innovation | Edison Johar, Frilasita A Yudhaputri, Hidayat Trimarsanto, David H Muljono, Safarina G Malik, Khin Saw Myint, Amin Soebandrio |
| EPI_ISL_437192 | Mitra Keluarga Kelapa Gading Hospital | Eijkman Institute for Molecular Biology, Ministry of Research and Technology/National Agency for Research and Innovation | Edison Johar, Frilasita A Yudhaputri, Hidayat Trimarsanto, David H Muljono, Safarina G Malik, Khin Saw Myint, Amin Soebandrio |
| EPI_ISL_437194 | Viral Respiratory Lab, National Institute for Biomedical Research (INRB) | Pathogen Sequencing Lab, National Institute for Biomedical Research (INRB) | Placide Mbala-Kingebeni, Edith Nkwembe, Eddy Kinganda-Lusamaki, Amuri Aziza, Francisca Muyembe Mawete, Catherine Pratt, Matthias Pauthner, Josh Quick, Allison Black, James Hadfield, Trevor Bedford, Ian Goodfellow, Andrew Rambaut, Nick Loman, Kristian Andersen, Michael Wiley, Steve Ahuka-Mundeke, Jean-Jacques Muyembe Tamfum |
| EPI_ISL_437197 | Diagnostic- and Research Institute of Pathology, Medical University of Graz | Diagnostic- and Research Institute of Pathology, Medical University of Graz | Karl Kashofer, Peter Regitnig, Martin Zacharias, Gregor Gorkiewicz |
| EPI_ISL_437198, EPI_ISL_437199, EPI_ISL_437200, EPI_ISL_437201, EPI_ISL_437202 | Diagnostic- and Research Institute of Pathology, Medical University of Graz | Diagnostic- and Research Institute of Pathology, Medical University of Graz | Karl Kashofer, Peter Regitnig, Martin Zacharias, Gregor Gorkiewicz |
| EPI_ISL_437203 | Diagnostic- and Research Institute of Pathology, Medical University of Graz | Diagnostic- and Research Institute of Pathology, Medical University of Graz | Karl Kashofer, Peter Regitnig, Martin Zacharias, Gregor Gorkiewicz |
| EPI_ISL_437204, EPI_ISL_437205, EPI_ISL_437206, EPI_ISL_437207, EPI_ISL_437208, EPI_ISL_437209, EPI_ISL_437210, EPI_ISL_437211, EPI_ISL_437212, EPI_ISL_437213, EPI_ISL_437214, EPI_ISL_437215, EPI_ISL_437216, EPI_ISL_437217, EPI_ISL_437218, EPI_ISL_437219, EPI_ISL_437220, EPI_ISL_437221, EPI_ISL_437222, EPI_ISL_437223, EPI_ISL_437224, EPI_ISL_437225, EPI_ISL_437227, EPI_ISL_437228, EPI_ISL_437229, EPI_ISL_437230, EPI_ISL_437231, EPI_ISL_437232, EPI_ISL_437233, EPI_ISL_437234, EPI_ISL_437235, EPI_ISL_437236, EPI_ISL_437237, EPI_ISL_437238, EPI_ISL_437239, EPI_ISL_437241, EPI_ISL_437242, EPI_ISL_437243, EPI_ISL_437244, EPI_ISL_437245, EPI_ISL_437246, EPI_ISL_437247, EPI_ISL_437248, EPI_ISL_437249, EPI_ISL_437250, EPI_ISL_437251, EPI_ISL_437252, EPI_ISL_437253, EPI_ISL_437254, EPI_ISL_437255, EPI_ISL_437256, EPI_ISL_437257, EPI_ISL_437258, EPI_ISL_437259, EPI_ISL_437260, EPI_ISL_437261, EPI_ISL_437262, EPI_ISL_437263, EPI_ISL_437264, EPI_ISL_437265, EPI_ISL_437266, EPI_ISL_437267, EPI_ISL_437269, EPI_ISL_437270, EPI_ISL_437271, EPI_ISL_437272, EPI_ISL_437273, EPI_ISL_437274, EPI_ISL_437275, EPI_ISL_437276, EPI_ISL_437277, EPI_ISL_437278, EPI_ISL_437279, EPI_ISL_437280, EPI_ISL_437281, EPI_ISL_437282, EPI_ISL_437283, EPI_ISL_437284, EPI_ISL_437285, EPI_ISL_437286, EPI_ISL_437287, EPI_ISL_437288, EPI_ISL_437289, EPI_ISL_437290, EPI_ISL_437291, EPI_ISL_437292, EPI_ISL_437294, EPI_ISL_437295, EPI_ISL_437296, EPI_ISL_437297 | | | |
| see above | Max von Pettenkofer Institute, Virology, National Reference Center for Retroviruses, LMU München | Laboratory for Functional Genome Analysis, Dept. Genomics, Gene Center of the LMU Munich | Max Muenchhoff, Stefan Krebs, Alexander Graf, Oliver Keppler, Helmut Blum |
| EPI_ISL_437298, EPI_ISL_437299, EPI_ISL_437300, EPI_ISL_437301, EPI_ISL_437302, EPI_ISL_437303 | Diagnostic- and Research Institute of Pathology, Medical University of Graz | Diagnostic- and Research Institute of Pathology, Medical University of Graz | Karl Kashofer, Peter Regitnig, Martin Zacharias, Gregor Gorkiewicz |
| EPI_ISL_437304, EPI_ISL_437305, EPI_ISL_437306, EPI_ISL_437307, EPI_ISL_437308, EPI_ISL_437309, EPI_ISL_437310, EPI_ISL_437311, EPI_ISL_437312, EPI_ISL_437313, EPI_ISL_437314, EPI_ISL_437315, EPI_ISL_437316, EPI_ISL_437317, EPI_ISL_437318 | | | |
| see above | Ministry of Health Turkey | Ministry of Health Turkey | Fatma Bayrakdar,Tülin Demir,Süleyman Yalçın, Selçuk Kılıç |
| EPI_ISL_437319, EPI_ISL_437320, EPI_ISL_437321 | Ministry of Health Turkey | Ministry of Health Turkey | Fatma Bayrakdar,Ayşe Başak Altaş,Yasemin Coşgun,Süleyman Yalçın, Gülay Korukluoğlu,Selçuk Kılıç |
| EPI_ISL_437322 | Ministry of Health Turkey | Ministry of Health Turkey | Fatma Bayrakdar,Tülin Demir,Süleyman Yalçın, Selçuk Kılıç |
| EPI_ISL_437323, EPI_ISL_437324, EPI_ISL_437325, EPI_ISL_437326, EPI_ISL_437327, EPI_ISL_437328, EPI_ISL_437329, EPI_ISL_437330 | Ministry of Health Turkey | Ministry of Health Turkey | Fatma Bayrakdar,Ayşe Başak Altaş,Yasemin Coşgun,Süleyman Yalçın, Gülay Korukluoğlu,Selçuk Kılıç |
| EPI_ISL_437331 | Ministry of Health Turkey | Ministry of Health Turkey | Fatma Bayrakdar,Tülin Demir,Süleyman Yalçın, Selçuk Kılıç |
| EPI_ISL_437332, EPI_ISL_437333, EPI_ISL_437334, EPI_ISL_437335 | Ministry of Health Turkey | Ministry of Health Turkey | Fatma Bayrakdar,Ayşe Başak Altaş,Yasemin Coşgun,Süleyman Yalçın, Gülay Korukluoğlu,Selçuk Kılıç |
| EPI_ISL_437337, EPI_ISL_437338, EPI_ISL_437339, EPI_ISL_437340, EPI_ISL_437341, EPI_ISL_437343, EPI_ISL_437346, EPI_ISL_437348, EPI_ISL_437350, EPI_ISL_437351, EPI_ISL_437352, EPI_ISL_437354, EPI_ISL_437356, EPI_ISL_437357, EPI_ISL_437358 | | | |
| see above | Viral Respiratory Lab, National Institute for Biomedical Research (INRB) | Pathogen Sequencing Lab, National Institute for Biomedical Research (INRB) | Placide Mbala-Kingebeni, Edith Nkwembe, Eddy Kinganda-Lusamaki, Amuri Aziza, Francisca Muyembe Mawete, Catherine Pratt, Matthias Pauthner, Josh Quick, Allison Black, James Hadfield, Trevor Bedford, Ian Goodfellow, Andrew Rambaut, Nick Loman, Kristian Andersen, Michael Wiley, Steve Ahuka-Mundeke, Jean-Jacques Muyembe Tamfum |
| EPI_ISL_437359 | Max von Pettenkofer Institute, Virology, National Reference Center for Retroviruses, LMU München | Laboratory for Functional Genome Analysis, Dept. Genomics, Gene Center of the LMU Munich | Max Muenchhoff, Stefan Krebs, Alexander Graf, Oliver Keppler, Helmut Blum |
| EPI_ISL_437361, EPI_ISL_437362, EPI_ISL_437363, EPI_ISL_437364, EPI_ISL_437365, EPI_ISL_437366, EPI_ISL_437367, EPI_ISL_437368, EPI_ISL_437369, EPI_ISL_437370, EPI_ISL_437371, EPI_ISL_437372, EPI_ISL_437373, EPI_ISL_437374, EPI_ISL_437375, EPI_ISL_437376, EPI_ISL_437377, EPI_ISL_437378, EPI_ISL_437379, EPI_ISL_437380, EPI_ISL_437381, EPI_ISL_437382, EPI_ISL_437383, EPI_ISL_437384, EPI_ISL_437385, EPI_ISL_437386 | | | |
| see above | Minnesota Department of Health, Public Health Laboratory | Minnesota Department of Health, Public Health Laboratory | Matt Plumb, Jacob Garfin, and Xiong Wang |
| EPI_ISL_437387, EPI_ISL_437388, EPI_ISL_437389, EPI_ISL_437390, EPI_ISL_437391, EPI_ISL_437392, EPI_ISL_437393, EPI_ISL_437394, EPI_ISL_437395, EPI_ISL_437396, EPI_ISL_437397, EPI_ISL_437398, EPI_ISL_437399, EPI_ISL_437400, EPI_ISL_437401, EPI_ISL_437403, EPI_ISL_437404, EPI_ISL_437405, EPI_ISL_437406, EPI_ISL_437407, EPI_ISL_437408, EPI_ISL_437409, EPI_ISL_437410, EPI_ISL_437411, EPI_ISL_437412, EPI_ISL_437413, EPI_ISL_437414, EPI_ISL_437415, EPI_ISL_437416, EPI_ISL_437417, EPI_ISL_437418, EPI_ISL_437419, EPI_ISL_437420, EPI_ISL_437421, EPI_ISL_437422, EPI_ISL_437423, EPI_ISL_437424, EPI_ISL_437425, EPI_ISL_437426, EPI_ISL_437427, EPI_ISL_437428, EPI_ISL_437429, EPI_ISL_437431, EPI_ISL_437432 | | | |
| see above | Virginia DCLS | Virginia DCLS | Virginia DCLS |
| EPI_ISL_437435, EPI_ISL_437436 | Veterinary Specialized Institue Kraljevo | Veterinary Specialized Institue Kraljevo | Dejan Vidanovic. Bojana Tesovic, Milanko Sekler, Marko Dmitric, Kazimir Matovic, Zoran Debeljak, Nikola Vaskovic, Tamas Petrovic, Jeremy Volkening, Claudio L Afonso |
| EPI_ISL_437437 | Alaska State Virology Laboratory | Alaska State Virology Laboratory | Jack Chen, Ph.D. |
| EPI_ISL_437438 | Department of MicroBiology, Government Medical College, Surat | Gujarat Biotechnology Research Centre | Amit Kanani, Akanksha Verma, Nitin Savaliya, Raghawendra Kumar, Dinesh Kumar, Zuber Saiyed, Dipa Kinariwala, Disha Patel, Binita Aring, Neeta Khandelwal, Geeta Vaghela, Sonia Barve, Bhavesh Modi, Kairavi Joshi, Gaurishankar Shrimali, Nidhi Sood, Pranay Shah, R D Dixit, Snehal Bagatharia, Kamlesh J Upadhyay, Ramesh Pandit, Tejas Shah, Ankit Hinsu, Pritesh Sabara, Apurvasinh Puvar, Janvi Raval, Monika Gandhi, Pinal Trivedi, Maharshi Pandya, Neelam Nathani, Chaitanya Joshi, Madhvi Joshi |
| EPI_ISL_437440 | Department of MicroBiology, Government Medical College, Surat | Gujarat Biotechnology Research Centre | Nitin Savaliya, Raghawendra Kumar, Dinesh Kumar, Zuber Saiyed, Dipa Kinariwala, Disha Patel, Binita Aring, Neeta Khandelwal, Geeta Vaghela, Sonia Barve, Bhavesh Modi, Kairavi Joshi, Gaurishankar Shrimali, Nidhi Sood, Pranay Shah, R D Dixit, Snehal Bagatharia, Kamlesh J Upadhyay, Ramesh Pandit, Tejas Shah, Ankit Hinsu, Pritesh Sabara, Apurvasinh Puvar, Janvi Raval, Monika Gandhi, Pinal Trivedi, Maharshi Pandya, Amit Kanani, Akanksha Verma, Bhavya Jindal, Chaitanya Joshi, Madhvi Joshi |
| EPI_ISL_437441 | Department of MicroBiology, Government Medical College, Surat | Gujarat Biotechnology Research Centre | Raghawendra Kumar, Dinesh Kumar, Zuber Saiyed, Dipa Kinariwala, Disha Patel, Binita Aring, Neeta Khandelwal, Geeta Vaghela, Sonia Barve, Bhavesh Modi, Kairavi Joshi, Gaurishankar Shrimali, Nidhi Sood, Pranay Shah, R D Dixit, Snehal Bagatharia, Kamlesh J Upadhyay, Ramesh Pandit, Tejas Shah, Ankit Hinsu, Pritesh Sabara, Apurvasinh Puvar, Janvi Raval, Monika Gandhi, Pinal Trivedi, Maharshi Pandya, Amit Kanani, Akanksha Verma, Nitin Savaliya, Anjali Rajwar, Chaitanya Joshi, Madhvi Joshi |
| EPI_ISL_437442 | Department of MicroBiology, Government Medical College, Surat | Gujarat Biotechnology Research Centre | Dinesh Kumar, Zuber Saiyed, Dipa Kinariwala, Disha Patel, Binita Aring, Neeta Khandelwal, Geeta Vaghela, Sonia Barve, Bhavesh Modi, Kairavi Joshi, Gaurishankar Shrimali, Nidhi Sood, Pranay Shah, R D Dixit, Snehal Bagatharia, Kamlesh J Upadhyay, Ramesh Pandit, Tejas Shah, Ankit Hinsu, Pritesh Sabara, Apurvasinh Puvar, Janvi Raval, Monika Gandhi, Pinal Trivedi, Maharshi Pandya, Amit Kanani, Akanksha Verma, Nitin Savaliya, Raghawendra Kumar, Dipeshwari Shewale, Chaitanya Joshi, Madhvi Joshi |
| EPI_ISL_437444 | Department of MicroBiology, Government Medical College, Surat | Gujarat Biotechnology Research Centre | Dipa Kinariwala, Disha Patel, Binita Aring, Neeta Khandelwal, Geeta Vaghela, Sonia Barve, Bhavesh Modi, Kairavi Joshi, Gaurishankar Shrimali, Nidhi Sood, Pranay Shah, R D Dixit, Snehal Bagatharia, Kamlesh J Upadhyay, Ramesh Pandit, Tejas Shah, Ankit Hinsu, Pritesh Sabara, Apurvasinh Puvar, Janvi Raval, Monika Gandhi, Pinal Trivedi, Maharshi Pandya, Amit Kanani, Akanksha Verma, Nitin Savaliya, Raghawendra Kumar, Dinesh Kumar, Zuber Saiyed, Pooja P Doshi, Chaitanya Joshi, Madhvi Joshi |
| EPI_ISL_437445 | B.J. Medical College and Civil hospital | Gujarat Biotechnology Research Centre | Disha Patel, Binita Aring, Neeta Khandelwal, Geeta Vaghela, Sonia Barve, Bhavesh Modi, Kairavi Joshi, Gaurishankar Shrimali, Nidhi Sood, Pranay Shah, R D Dixit, Snehal Bagatharia, Kamlesh J Upadhyay, Ramesh Pandit, Tejas Shah, Ankit Hinsu, Pritesh Sabara, Apurvasinh Puvar, Janvi Raval, Monika Gandhi, Pinal Trivedi, Maharshi Pandya, Amit Kanani, Akanksha Verma, Nitin Savaliya, Raghawendra Kumar, Dinesh Kumar, Zuber Saiyed, Dipa Kinariwala, Nidhi Patel, Chaitanya Joshi, Madhvi Joshi |
| EPI_ISL_437446 | B.J. Medical College and Civil hospital | Gujarat Biotechnology Research Centre | Binita Aring, Neeta Khandelwal, Geeta Vaghela, Sonia Barve, Bhavesh Modi, Kairavi Joshi, Gaurishankar Shrimali, Nidhi Sood, Pranay Shah, R D Dixit, Snehal Bagatharia, Kamlesh J Upadhyay, Ramesh Pandit, Tejas Shah, Ankit Hinsu, Pritesh Sabara, Apurvasinh Puvar, Janvi Raval, Monika Gandhi, Pinal Trivedi, Maharshi Pandya, Amit Kanani, Akanksha Verma, Nitin Savaliya, Raghawendra Kumar, Dinesh Kumar, Zuber Saiyed, Dipa Kinariwala, Disha Patel, Priti Pandita, Chaitanya Joshi, Madhvi Joshi |
| EPI_ISL_437447 | B.J. Medical College and Civil hospital | Gujarat Biotechnology Research Centre | Neeta Khandelwal, Geeta Vaghela, Sonia Barve, Bhavesh Modi, Kairavi Joshi, Gaurishankar Shrimali, Nidhi Sood, Pranay Shah, R D Dixit, Snehal Bagatharia, Kamlesh J Upadhyay, Ramesh Pandit, Tejas Shah, Ankit Hinsu, Pritesh Sabara, Apurvasinh Puvar, Janvi Raval, Monika Gandhi, Pinal Trivedi, Maharshi Pandya, Amit Kanani, Akanksha Verma, Nitin Savaliya, Raghawendra Kumar, Dinesh Kumar, Zuber Saiyed, Dipa Kinariwala, Disha Patel, Binita Aring, Neha Rajpara, Chaitanya Joshi, Madhvi Joshi |
| EPI_ISL_437448 | B.J. Medical College and Civil hospital | Gujarat Biotechnology Research Centre | Geeta Vaghela, Sonia Barve, Bhavesh Modi, Kairavi Joshi, Gaurishankar Shrimali, Nidhi Sood, Pranay Shah, R D Dixit, Snehal Bagatharia, Kamlesh J Upadhyay, Ramesh Pandit, Tejas Shah, Ankit Hinsu, Pritesh Sabara, Apurvasinh Puvar, Janvi Raval, Monika Gandhi, Pinal Trivedi, Maharshi Pandya, Amit Kanani, Akanksha Verma, Nitin Savaliya, Raghawendra Kumar, Dinesh Kumar, Zuber Saiyed, Dipa Kinariwala, Disha Patel, Binita Aring, Neeta Khandelwal, Afzal Ansari, Chaitanya Joshi, Madhvi Joshi |
| EPI_ISL_437449 | B.J. Medical College and Civil hospital | Gujarat Biotechnology Research Centre | Sonia Barve, Bhavesh Modi, Kairavi Joshi, Gaurishankar Shrimali, Nidhi Sood, Pranay Shah, R D Dixit, Snehal Bagatharia, Kamlesh J Upadhyay, Ramesh Pandit, Tejas Shah, Ankit Hinsu, Pritesh Sabara, Apurvasinh Puvar, Janvi Raval, Monika Gandhi, Pinal Trivedi, Maharshi Pandya, Amit Kanani, Akanksha Verma, Nitin Savaliya, Raghawendra Kumar, Dinesh Kumar, Zuber Saiyed, Dipa Kinariwala, Disha Patel, Binita Aring, Neeta Khandelwal, Geeta Vaghela, Neelam Nathani, Chaitanya Joshi, Madhvi Joshi |
| EPI_ISL_437450 | B.J. Medical College and Civil hospital | Gujarat Biotechnology Research Centre | Bhavesh Modi, Kairavi Joshi, Gaurishankar Shrimali, Nidhi Sood, Pranay Shah, R D Dixit, Snehal Bagatharia, Kamlesh J Upadhyay, Ramesh Pandit, Tejas Shah, Ankit Hinsu, Pritesh Sabara, Apurvasinh Puvar, Janvi Raval, Monika Gandhi, Pinal Trivedi, Maharshi Pandya, Amit Kanani, Akanksha Verma, Nitin Savaliya, Raghawendra Kumar, Dinesh Kumar, Zuber Saiyed, Dipa Kinariwala, Disha Patel, Binita Aring, Neeta Khandelwal, Geeta Vaghela, Sonia Barve, Armi Chaudhari, Chaitanya Joshi, Madhvi Joshi |
| EPI_ISL_437451 | B.J. Medical College and Civil hospital | Gujarat Biotechnology Research Centre | Kairavi Joshi, Gaurishankar Shrimali, Nidhi Sood, Pranay Shah, R D Dixit, Snehal Bagatharia, Kamlesh J Upadhyay, Ramesh Pandit, Tejas Shah, Ankit Hinsu, Pritesh Sabara, Apurvasinh Puvar, Janvi Raval, Monika Gandhi, Pinal Trivedi, Maharshi Pandya, Amit Kanani, Akanksha Verma, Nitin Savaliya, Raghawendra Kumar, Dinesh Kumar, Zuber Saiyed, Dipa Kinariwala, Disha Patel, Binita Aring, Neeta Khandelwal, Geeta Vaghela, Sonia Barve, Bhavesh Modi, Bhavya Jindal, Chaitanya Joshi, Madhvi Joshi |
| EPI_ISL_437452 | B.J. Medical College and Civil hospital | Gujarat Biotechnology Research Centre | Gaurishankar Shrimali, Nidhi Sood, Pranay Shah, R D Dixit, Snehal Bagatharia, Kamlesh J Upadhyay, Ramesh Pandit, Tejas Shah, Ankit Hinsu, Pritesh Sabara, Apurvasinh Puvar, Janvi Raval, Monika Gandhi, Pinal Trivedi, Maharshi Pandya, Amit Kanani, Akanksha Verma, Nitin Savaliya, Raghawendra Kumar, Dinesh Kumar, Zuber Saiyed, Dipa Kinariwala, Disha Patel, Binita Aring, Neeta Khandelwal, Geeta Vaghela, Sonia Barve, Bhavesh Modi, Kairavi Joshi, Chaitanya Joshi, Anjali Rajwar, Madhvi Joshi |
| EPI_ISL_437453 | B.J. Medical College and Civil hospital | Gujarat Biotechnology Research Centre | Nidhi Sood, Pranay Shah, R D Dixit, Snehal Bagatharia, Kamlesh J Upadhyay, Ramesh Pandit, Tejas Shah, Ankit Hinsu, Pritesh Sabara, Apurvasinh Puvar, Janvi Raval, Monika Gandhi, Pinal Trivedi, Maharshi Pandya, Amit Kanani, Akanksha Verma, Nitin Savaliya, Raghawendra Kumar, Dinesh Kumar, Zuber Saiyed, Dipa Kinariwala, Disha Patel, Binita Aring, Neeta Khandelwal, Geeta Vaghela, Sonia Barve, Bhavesh Modi, Kairavi Joshi, Gaurishankar Shrimali, Chaitanya Joshi, Dipeshwari Shewale, Madhvi Joshi |
| EPI_ISL_437454 | B.J. Medical College and Civil hospital | Gujarat Biotechnology Research Centre | Pranay Shah, R D Dixit, Snehal Bagatharia, Kamlesh J Upadhyay, Ramesh Pandit, Tejas Shah, Ankit Hinsu, Pritesh Sabara, Apurvasinh Puvar, Janvi Raval, Monika Gandhi, Pinal Trivedi, Maharshi Pandya, Amit Kanani, Akanksha Verma, Nitin Savaliya, Raghawendra Kumar, Dinesh Kumar, Zuber Saiyed, Dipa Kinariwala, Disha Patel, Binita Aring, Neeta Khandelwal, Geeta Vaghela, Sonia Barve, Bhavesh Modi, Kairavi |

| | | | |
|---|---|---|---|
| | | | Joshi, Gaurishankar Shrimali, Nidhi Sood, Chaitanya Joshi, Sharmistha Majumdar, Madhvi Joshi |
| EPI_ISL_437455, EPI_ISL_437456, EPI_ISL_437457, EPI_ISL_437458 | Clinical Diagnostics Laboratory, Diagnostic & Experimental Pathology, Lilly Research Laboratories | Clinical Diagnostics Laboratory, Diagnostic & Experimental Pathology, Lilly Research Laboratories | Tim Holzer, Mayuri Vaidya, Angie Fulford, Sam McNeely, Rachael Redmond, Phil Ebert, John Calley, Leslie ONeill Reising, Pat Finnegan, Erin Wray, John McElwee, Jeff Fill, Joe Oakley, Andrew Schade |
| EPI_ISL_437459, EPI_ISL_437460, EPI_ISL_437461, EPI_ISL_437462, EPI_ISL_437463, EPI_ISL_437464, EPI_ISL_437465, EPI_ISL_437466, EPI_ISL_437468, EPI_ISL_437469, EPI_ISL_437470, EPI_ISL_437471, EPI_ISL_437472, EPI_ISL_437473, EPI_ISL_437474, EPI_ISL_437475, EPI_ISL_437477, EPI_ISL_437479 | | | |
| see above | Pathogen Genomics Lab King Abdullah University of Science and Technology(KAUST) | Pathogen Genomics Lab King Abdullah University of Science and Technology(KAUST) | Sharif Hala,Raeece Naeem,Sara Mfarrej,Arnab Pain |
| EPI_ISL_437481, EPI_ISL_437482, EPI_ISL_437483, EPI_ISL_437484, EPI_ISL_437485, EPI_ISL_437486, EPI_ISL_437487, EPI_ISL_437488, EPI_ISL_437489, EPI_ISL_437490, EPI_ISL_437491, EPI_ISL_437492, EPI_ISL_437493, EPI_ISL_437494, EPI_ISL_437495, EPI_ISL_437496, EPI_ISL_437497 | | | |
| see above | Pathogen Genomics Lab King Abdullah University of Science and Technology(KAUST) | Pathogen Genomics Lab King Abdullah University of Science and Technology(KAUST) | Sara Mfarrej,Raeece Naeem,Sharif Hala,Amit Subudhi,Fathia Rached,Arnab Pain |
| EPI_ISL_437512 | Human Genetic Research Center, Kawsar Biotech Company | Human Genetic Research Center, Kawsar Biotech Company | Khosravi,M.A., Abbasalipour,M., Zeinali,S., Sabeghi,S., Kehsvar,Y., Hosseini,F. and Haghdoost,Y. |
| EPI_ISL_437513, EPI_ISL_437514, EPI_ISL_437515, EPI_ISL_437516, EPI_ISL_437517, EPI_ISL_437518 | Alaska State Virology Laboratory | Alaska State Virology Laboratory | Jack Chen, Ph.D. |
| EPI_ISL_437519 | The National Institute of Public Health Center for Epidemiology and Microbiology | The National Institute of Public Health Center for Epidemiology and Microbiology | Alexander Nagy, Helena Jirincova, Ludmila Novakova, Dusan Trnka, Jaromira Vecerova |
| EPI_ISL_437521, EPI_ISL_437523, EPI_ISL_437524, EPI_ISL_437525, EPI_ISL_437528, EPI_ISL_437529, EPI_ISL_437531, EPI_ISL_437532, EPI_ISL_437535 | OHSU Lab Services Molecular Microbiology Lab | Oregon SARS-CoV-2 Genome Sequencing Center | Brendan L. O'Connell, Ruth V. Nichols, Alec J. Hirsch, Guang Fan, Daniel N. Streblow, William B. Messer, Andrew C. Adey, Benjamin N. Bimber, Brian J. O'Roak |
| EPI_ISL_437539 | ICMR-National Institute of Cholera and Enteric Diseases | National Institute of Biomedical Genomics | Arindam Maitra, Mamta Chawla Sarkar, Sreedhar Chinnaswamy, Hasina Banu, Ananya Chatterjee, Shanta Dutta, Saumitra Das |
| EPI_ISL_437541, EPI_ISL_437542, EPI_ISL_437546, EPI_ISL_437547 | Robert Garry lab | Andersen lab at Scripps Research | Allison Smither, Gilberto Sabino-Santos, Patricia Snarski, Lilia Melnik, Antoinette Bell, Kaylynn Genemaras, Arnaud Drouin, Dahlene Fusco, Robert Garry with SEARCH Alliance San Diego |
| EPI_ISL_437549, EPI_ISL_437550, EPI_ISL_437551, EPI_ISL_437552, EPI_ISL_437553, EPI_ISL_437554, EPI_ISL_437555, EPI_ISL_437556, EPI_ISL_437559, EPI_ISL_437560, EPI_ISL_437561, EPI_ISL_437562, EPI_ISL_437563, EPI_ISL_437564, EPI_ISL_437566, EPI_ISL_437567, EPI_ISL_437568, EPI_ISL_437569, EPI_ISL_437570, EPI_ISL_437571, EPI_ISL_437572, EPI_ISL_437574, EPI_ISL_437575, EPI_ISL_437576, EPI_ISL_437580, EPI_ISL_437582, EPI_ISL_437583, EPI_ISL_437584, EPI_ISL_437585, EPI_ISL_437586, EPI_ISL_437587, EPI_ISL_437588, EPI_ISL_437589, EPI_ISL_437590, EPI_ISL_437591, EPI_ISL_437592, EPI_ISL_437594, EPI_ISL_437595, EPI_ISL_437596, EPI_ISL_437598, EPI_ISL_437599, EPI_ISL_437600 | | | |
| see above | Scripps Medical Laboratory | Andersen lab at Scripps Research | SEARCH Alliance San Diego with Michael Quigley, Ellen Stefanski, Ian Mchardy |
| EPI_ISL_437601 | Keio University School of Medicine | Keio University School of Medicine | Kenjiro Kosaki, Yuka Iwasaki, Toshiki Takenouchi, Haruhiko Siomi, |
| EPI_ISL_437602, EPI_ISL_437603, EPI_ISL_437604, EPI_ISL_437605, EPI_ISL_437606, EPI_ISL_437607, EPI_ISL_437608, EPI_ISL_437609, EPI_ISL_437610, EPI_ISL_437611, EPI_ISL_437612, EPI_ISL_437613, EPI_ISL_437614, EPI_ISL_437615, EPI_ISL_437616, EPI_ISL_437617, EPI_ISL_437618, EPI_ISL_437619, EPI_ISL_437620, EPI_ISL_437621, EPI_ISL_437622, EPI_ISL_437623, EPI_ISL_437624 | | | |
| see above | unknown | Faculty of Medicine | Rodpan,A., Joyjinda,Y., Wacharapluesadee,S., Buathong,R., Ghai,S., Petcharat,S., Bunprakob,S., Sirichan,N., Prasithsirikul,W., Mungaomklang,A., Plipat,T. and Hemachudha,T. |
| EPI_ISL_437625 | Laboratory of Genomics & Bioinformatics, Institute of Immunology and Experimental Therapy, Polish Academy of Sciences Oddzial Mikrobiologii Wojewodzkiej Stacji Sanitarno-Epidemiologicznej. | Laboratory of Genomics & Bioinformatics, Institute of Immunology and Experimental Therapy, Polish Academy of Sciences | Dorota Kujawa, Aleksandra Herud, Dariusz Martynowski, Krzysztof Jakub Pawlik, Joanna Sikorska, Paulina Zebrowska, Grazyna Zalewska, Oskar Karpinski and Lukasz Laczmanski |
| EPI_ISL_437626 | Department of Microbiology,Gandhi Medical College and Hospital | Department of Veterinary Biotechnalogy, College of Veterinary Science, Rajendranagar, PV Narsimha Rao Telengana Veterinary University | Kalyani Putty, Muttineni Radhakrishna, Nagamani K, Thrilok Chander B, Raja Rao M, Ravikumar P, Sunitha P, Pankaj Singh D, Anand Kumar K, Amit A. Upadhyay, Steven Bosinger, Rama Amara |
| EPI_ISL_437628, EPI_ISL_437629, EPI_ISL_437630, EPI_ISL_437631, EPI_ISL_437632, EPI_ISL_437633, EPI_ISL_437634, EPI_ISL_437635, EPI_ISL_437636, EPI_ISL_437637, EPI_ISL_437639, EPI_ISL_437640, EPI_ISL_437642, EPI_ISL_437643, EPI_ISL_437644, EPI_ISL_437645, EPI_ISL_437646, EPI_ISL_437647, EPI_ISL_437648, EPI_ISL_437649, EPI_ISL_437650, EPI_ISL_437651, EPI_ISL_437653, EPI_ISL_437654, EPI_ISL_437655, EPI_ISL_437656, EPI_ISL_437657, EPI_ISL_437658, EPI_ISL_437659, EPI_ISL_437660, EPI_ISL_437661, EPI_ISL_437663, EPI_ISL_437664, EPI_ISL_437667, EPI_ISL_437668, EPI_ISL_437669, EPI_ISL_437670, EPI_ISL_437672, EPI_ISL_437674, EPI_ISL_437676, EPI_ISL_437677, EPI_ISL_437678, EPI_ISL_437679, EPI_ISL_437683 | | | |
| see above | Department of Virus and Microbiological Special Diagnostics, Statens Serum Institut, Copenhagen, Denmark, Artillerivej 5, 2300 Copenahgen S | Albertsen lab, Department of Chemistry and Bioscience, Aalborg University, Denmark | Rasmus Kirkegaard |
| EPI_ISL_437684, EPI_ISL_437685, EPI_ISL_437686, EPI_ISL_437687, EPI_ISL_437688 | UCD National Virus Reference Laboratory | UCD National Virus Reference Laboratory | Michael J. Carr, Gabriel Gonzalez, Brendan Crowley, Cillian F De Gascun |
| EPI_ISL_437689 | Laboratory for Urgent Response to Biological Threats | Institut Pasteur CIBU / ERI | V. Caro, A. Kwasiborski, V. Hourdel, C. Balière, J. Vanhomwegen, C. Batéjat, JC. Manuguerra |
| EPI_ISL_437690 | Laboratory for Urgent Response to Biological Threats | Institut Pasteur CIBU /ERI | V. Caro, A. Kwasiborski, H. Hourdel, C. Balière, J. Vanhomwegen, C. Batéjat, JC. Manuguerra |
| EPI_ISL_437691, EPI_ISL_437692, EPI_ISL_437693, EPI_ISL_437694, EPI_ISL_437695, EPI_ISL_437696, EPI_ISL_437697, EPI_ISL_437698, EPI_ISL_437699, EPI_ISL_437700, EPI_ISL_437701, EPI_ISL_437702, EPI_ISL_437703, EPI_ISL_437704, EPI_ISL_437705, EPI_ISL_437706, EPI_ISL_437707, EPI_ISL_437708, EPI_ISL_437709, EPI_ISL_437710, EPI_ISL_437711, EPI_ISL_437712, EPI_ISL_437713, EPI_ISL_437714, EPI_ISL_437715, EPI_ISL_437716, EPI_ISL_437717, EPI_ISL_437718, EPI_ISL_437719, EPI_ISL_437720, EPI_ISL_437721, EPI_ISL_437722, EPI_ISL_437723, EPI_ISL_437724, EPI_ISL_437725, EPI_ISL_437726, EPI_ISL_437727, EPI_ISL_437728, EPI_ISL_437729, EPI_ISL_437730, EPI_ISL_437731, EPI_ISL_437732, EPI_ISL_437733, EPI_ISL_437734, EPI_ISL_437735, EPI_ISL_437736, EPI_ISL_437737, EPI_ISL_437738, EPI_ISL_437739, EPI_ISL_437740, EPI_ISL_437741, EPI_ISL_437742, EPI_ISL_437743, EPI_ISL_437744, EPI_ISL_437745, EPI_ISL_437746, EPI_ISL_437747, EPI_ISL_437748, EPI_ISL_437749, EPI_ISL_437750, EPI_ISL_437751, EPI_ISL_437752, EPI_ISL_437753, EPI_ISL_437754, EPI_ISL_437755, EPI_ISL_437756, EPI_ISL_437757, EPI_ISL_437758, EPI_ISL_437759, EPI_ISL_437760, EPI_ISL_437761, EPI_ISL_437762 | | | |
| see above | Pathogen Genomics Lab King Abdullah University of Science and Technology(KAUST) | Pathogen Genomics Lab King Abdullah University of Science and Technology(KAUST) | Sharif Hala,Fadwa Alofi,Afrah Alsomali, Asim Khogeer, Sara Mfarrej, Khaled Algithami,Raeece Naeem, Amit Kumar Subudhi,Fathia Ben-Rached, Rahul Salunke, Anwar Hashem, Naif Almontashiri, Arnab Pain |
| EPI_ISL_437763, EPI_ISL_437764, EPI_ISL_437765, EPI_ISL_437766, EPI_ISL_437767, EPI_ISL_437768, EPI_ISL_437769, EPI_ISL_437770, EPI_ISL_437771, EPI_ISL_437772, EPI_ISL_437773, EPI_ISL_437774, EPI_ISL_437775, EPI_ISL_437776, EPI_ISL_437777, EPI_ISL_437778, EPI_ISL_437779, EPI_ISL_437780, EPI_ISL_437781, EPI_ISL_437782, EPI_ISL_437783, EPI_ISL_437784, EPI_ISL_437785, EPI_ISL_437786, EPI_ISL_437788, EPI_ISL_437789, EPI_ISL_437790, EPI_ISL_437791, EPI_ISL_437792, EPI_ISL_437793, EPI_ISL_437794, EPI_ISL_437795, EPI_ISL_437796, EPI_ISL_437797, EPI_ISL_437798, EPI_ISL_437799, EPI_ISL_437800, EPI_ISL_437801, EPI_ISL_437802 | | | |
| see above | Virginia DCLS | Virginia DCLS | Virginia DCLS |
| EPI_ISL_437803, EPI_ISL_437804, EPI_ISL_437805, EPI_ISL_437806, EPI_ISL_437807, EPI_ISL_437808, EPI_ISL_437809, EPI_ISL_437810, EPI_ISL_437811, EPI_ISL_437812, EPI_ISL_437813, EPI_ISL_437814, EPI_ISL_437815, EPI_ISL_437816, EPI_ISL_437817, EPI_ISL_437818, EPI_ISL_437819, EPI_ISL_437820, EPI_ISL_437821, EPI_ISL_437822, EPI_ISL_437824, EPI_ISL_437825, EPI_ISL_437826, EPI_ISL_437827, EPI_ISL_437828, EPI_ISL_437829, EPI_ISL_437830, EPI_ISL_437831, EPI_ISL_437832, EPI_ISL_437833, EPI_ISL_437835, EPI_ISL_437836, EPI_ISL_437837, EPI_ISL_437838, EPI_ISL_437839, EPI_ISL_437840, EPI_ISL_437841, EPI_ISL_437842, EPI_ISL_437843, EPI_ISL_437844, EPI_ISL_437845, EPI_ISL_437848, EPI_ISL_437850, EPI_ISL_437851, EPI_ISL_437852, EPI_ISL_437853, EPI_ISL_437854, EPI_ISL_437855, EPI_ISL_437856, EPI_ISL_437857, EPI_ISL_437858, EPI_ISL_437859, EPI_ISL_437860, EPI_ISL_437861, EPI_ISL_437862, EPI_ISL_437863, EPI_ISL_437864, EPI_ISL_437865, EPI_ISL_437866, EPI_ISL_437868, EPI_ISL_437869, EPI_ISL_437870, EPI_ISL_437871, EPI_ISL_437872 | | | |
| see above | UW Virology Lab | UW Virology Lab | Pavitra Roychoudhury, Hong Xie, Keith Jerome, Alexander Greninger |
| EPI_ISL_437873 | Alaska State Virology Laboratory | Alaska State Virology Laboratory | Jack Chen, Ph.D. |
| EPI_ISL_437874, EPI_ISL_437875, EPI_ISL_437876, EPI_ISL_437877, EPI_ISL_437878, EPI_ISL_437879, EPI_ISL_437880, EPI_ISL_437881, EPI_ISL_437882, EPI_ISL_437883, EPI_ISL_437884, EPI_ISL_437885, EPI_ISL_437886, EPI_ISL_437887, EPI_ISL_437888, EPI_ISL_437889, EPI_ISL_437890, EPI_ISL_437891, EPI_ISL_437892, EPI_ISL_437893, EPI_ISL_437894, EPI_ISL_437895, EPI_ISL_437896, EPI_ISL_437897, EPI_ISL_437898, EPI_ISL_437899, EPI_ISL_437900, EPI_ISL_437901, EPI_ISL_437902, EPI_ISL_437903, EPI_ISL_437904, EPI_ISL_437905, EPI_ISL_437906, EPI_ISL_437907, EPI_ISL_437908, EPI_ISL_437909, EPI_ISL_437910, EPI_ISL_437911 | | | |
| see above | Laboratory of Microbiology, Medical School, National and Kapodistrian University of Athens | Laboratory of Biology, Department of Medicine, Democritus University of Thrace | Kassela K., Dovrolis,N., Bampali,M., Gatzidou,E., Froukala,E., Stavropoulou,A., Veletza,S., Tsakiris,A., Spanakis,N. and Karakasiliotis,I. |
| EPI_ISL_437912 | Child Health Research Foundation | Child Health Research Lab | Senjuti Saha, Roly Malaker, Md Saiful Islam Sajib, Md Hasanuzzaman, Md Hafizur Rahman, Md Shahidul Islam, Zabed B Ahmed, Maksuda Islam, Samir K Saha |
| EPI_ISL_437913, EPI_ISL_437914, EPI_ISL_437915, EPI_ISL_437916, EPI_ISL_437917, EPI_ISL_437918, EPI_ISL_437919, EPI_ISL_437920, EPI_ISL_437921, EPI_ISL_437922, EPI_ISL_437923, EPI_ISL_437924, EPI_ISL_437925, EPI_ISL_437926, EPI_ISL_437927, EPI_ISL_437928, EPI_ISL_437929, EPI_ISL_437930, EPI_ISL_437931, EPI_ISL_437932 | | | |
| see above | Institut für Virologie am Department für Hygiene, Mikrobiologie und Public Health | Bergthaler laboratory, CeMM Research Center for Molecular Medicine of the Austrian Academy of Sciences | Alexandra Popa, Benedikt Agerer, Henrique Colaco, Lukas Endler, Jakob-Wendelin Genger, Alexander Lercher, Mark Smyth, Thomas Penz, Michael Schuster, Jan Laine, Martin Senekowitsch, Judith Aberle, Stephan Aberle, Elisabeth Puchhammer-Stoeckl, Manfred Nairz, Guenter Weiss, Wegene Borena, Dorothee von Laer, Christoph Bock, Andreas Bergthaler |
| EPI_ISL_437933, EPI_ISL_437934, EPI_ISL_437935, EPI_ISL_437936, EPI_ISL_437937, EPI_ISL_437938, EPI_ISL_437939, EPI_ISL_437940, EPI_ISL_437941, EPI_ISL_437942, EPI_ISL_437943, EPI_ISL_437944, EPI_ISL_437945, EPI_ISL_437946, EPI_ISL_437947, EPI_ISL_437948, EPI_ISL_437949, EPI_ISL_437950, EPI_ISL_437951, EPI_ISL_437952, EPI_ISL_437953, EPI_ISL_437954, EPI_ISL_437955, EPI_ISL_437956, EPI_ISL_437957, EPI_ISL_437958, EPI_ISL_437959, EPI_ISL_437960, EPI_ISL_437961, EPI_ISL_437962, EPI_ISL_437963, EPI_ISL_437964, EPI_ISL_437965, EPI_ISL_437966, EPI_ISL_437967, EPI_ISL_437968, EPI_ISL_437969, EPI_ISL_437970, EPI_ISL_437971, EPI_ISL_437972, EPI_ISL_437973 | | | |
| see above | Universitaetsklinik für Innere Medizin II Innsbruck | Bergthaler laboratory, CeMM Research Center for Molecular Medicine of the Austrian Academy of Sciences | Alexandra Popa, Benedikt Agerer, Henrique Colaco, Lukas Endler, Jakob-Wendelin Genger, Alexander Lercher, Mark Smyth, Thomas Penz, Michael Schuster, Jan Laine, Martin Senekowitsch, Judith Aberle, Stephan Aberle, Elisabeth Puchhammer-Stoeckl, Manfred Nairz, Guenter Weiss, Wegene Borena, Dorothee von Laer, Christoph Bock, Andreas Bergthaler |
| EPI_ISL_437974, EPI_ISL_437975, EPI_ISL_437976, EPI_ISL_437977, EPI_ISL_437978, EPI_ISL_437979, EPI_ISL_437980, EPI_ISL_437981, EPI_ISL_437982, EPI_ISL_437983, EPI_ISL_437984, EPI_ISL_437985, EPI_ISL_437986, EPI_ISL_437987, EPI_ISL_437988, EPI_ISL_437989, EPI_ISL_437990, EPI_ISL_437991, EPI_ISL_437992 | | | |
| see above | Institut für Virologie am Department für Hygiene, Mikrobiologie und Public Health | Bergthaler laboratory, CeMM Research Center for Molecular Medicine of the Austrian Academy of Sciences | Alexandra Popa, Benedikt Agerer, Henrique Colaco, Lukas Endler, Jakob-Wendelin Genger, Alexander Lercher, Mark Smyth, Thomas Penz, Michael Schuster, Jan Laine, Martin Senekowitsch, Judith Aberle, Stephan Aberle, Elisabeth Puchhammer-Stoeckl, Manfred Nairz, Guenter Weiss, Wegene Borena, Dorothee von Laer, Christoph Bock, Andreas Bergthaler |
| EPI_ISL_437993, EPI_ISL_437994, EPI_ISL_437995, EPI_ISL_437996, EPI_ISL_437997, EPI_ISL_437998, EPI_ISL_437999, EPI_ISL_438000, EPI_ISL_438001, EPI_ISL_438002, EPI_ISL_438003, EPI_ISL_438004, EPI_ISL_438005, EPI_ISL_438006, EPI_ISL_438007, EPI_ISL_438008, EPI_ISL_438009, EPI_ISL_438010, EPI_ISL_438011, EPI_ISL_438012, EPI_ISL_438013, EPI_ISL_438014, EPI_ISL_438015, EPI_ISL_438016, EPI_ISL_438017, EPI_ISL_438018, EPI_ISL_438019, EPI_ISL_438020, EPI_ISL_438021, EPI_ISL_438022, EPI_ISL_438023, EPI_ISL_438024, EPI_ISL_438025, EPI_ISL_438026, EPI_ISL_438027, EPI_ISL_438028, EPI_ISL_438029, EPI_ISL_438030, EPI_ISL_438031, EPI_ISL_438032, EPI_ISL_438033, EPI_ISL_438034, EPI_ISL_438035, EPI_ISL_438036, EPI_ISL_438037, EPI_ISL_438038, EPI_ISL_438039, EPI_ISL_438040, EPI_ISL_438041, EPI_ISL_438042, EPI_ISL_438043, EPI_ISL_438044, EPI_ISL_438045, EPI_ISL_438046, EPI_ISL_438047, EPI_ISL_438048, EPI_ISL_438049, EPI_ISL_438050, EPI_ISL_438051, EPI_ISL_438052, EPI_ISL_438053, EPI_ISL_438054, EPI_ISL_438055, EPI_ISL_438056, EPI_ISL_438057, EPI_ISL_438058, EPI_ISL_438059, EPI_ISL_438060, EPI_ISL_438061, EPI_ISL_438062, EPI_ISL_438063, EPI_ISL_438064, EPI_ISL_438065, EPI_ISL_438066, EPI_ISL_438067, EPI_ISL_438068, EPI_ISL_438069, EPI_ISL_438070, EPI_ISL_438071, EPI_ISL_438072, EPI_ISL_438073, EPI_ISL_438074, EPI_ISL_438075, EPI_ISL_438076, EPI_ISL_438077, EPI_ISL_438078, EPI_ISL_438079, EPI_ISL_438080, EPI_ISL_438081, EPI_ISL_438082, EPI_ISL_438083, EPI_ISL_438084, EPI_ISL_438085, EPI_ISL_438086, EPI_ISL_438087, EPI_ISL_438088, EPI_ISL_438089, EPI_ISL_438090, EPI_ISL_438091, EPI_ISL_438092, EPI_ISL_438093, EPI_ISL_438094, EPI_ISL_438095, EPI_ISL_438096, EPI_ISL_438097, EPI_ISL_438098, EPI_ISL_438099, EPI_ISL_438100, EPI_ISL_438101, EPI_ISL_438102, EPI_ISL_438103, EPI_ISL_438104, EPI_ISL_438105, EPI_ISL_438106, EPI_ISL_438107, EPI_ISL_438108, EPI_ISL_438109, EPI_ISL_438110, EPI_ISL_438111, EPI_ISL_438112, EPI_ISL_438113, EPI_ISL_438114, EPI_ISL_438115, EPI_ISL_438116, EPI_ISL_438117, EPI_ISL_438118, EPI_ISL_438119, EPI_ISL_438120, EPI_ISL_438121, EPI_ISL_438122, EPI_ISL_438123, EPI_ISL_438124, EPI_ISL_438125, EPI_ISL_438126, EPI_ISL_438127, EPI_ISL_438128 | | | |
| see above | Center for Virology, Medical University of Vienna | Bergthaler laboratory, CeMM Research Center for Molecular Medicine of the Austrian Academy of Sciences | Alexandra Popa, Benedikt Agerer, Henrique Colaco, Lukas Endler, Jakob-Wendelin Genger, Alexander Lercher, Mark Smyth, Thomas Penz, Michael Schuster, Jan Laine, Martin Senekowitsch, Judith Aberle, Stephan Aberle, Elisabeth Puchhammer-Stoeckl, Manfred Nairz, Guenter Weiss, Wegene Borena, Dorothee von Laer, Christoph Bock, Andreas Bergthaler |
| EPI_ISL_438138 | Department of Microbiology,Gandhi Medical College and Hospital | Department of Microbiology, Gandhi Medical College and Hospital Secendrabad, Hyderbad, India | Raja Rao Mesipogu, Muttineni Radhakrishna, Nagamani K, Thrilok Chander B, Kalyani Putty, Ravikumar P, Sunitha P, Pankaj Singh D, Anand Kumar K, Amit A. Upadhyay, Steven Bosinger, Rama Amara |
| EPI_ISL_438139 | Department of Microbiology,Gandhi Medical College and Hospital,Hyderabad | Virus Research Laboratory, Department of Zoology, Osmania University, Hyderabad, India | Muttineni Radhakrishna, Nagamani K, Thrilok Chander B, Raja Rao M, Kalyani Putty, Ravikumar P, Sunitha P, Pankaj Singh D, Anand Kumar K, Amit A. Upadhyay, Steven Bosinger, Rama Amara |
| EPI_ISL_438140, EPI_ISL_438141, EPI_ISL_438142, EPI_ISL_438143, EPI_ISL_438144, EPI_ISL_438145, EPI_ISL_438147, EPI_ISL_438148, EPI_ISL_438149, EPI_ISL_438150, EPI_ISL_438151, EPI_ISL_438152, EPI_ISL_438153, EPI_ISL_438154, EPI_ISL_438155, EPI_ISL_438156, EPI_ISL_438157, EPI_ISL_438158, EPI_ISL_438159, EPI_ISL_438160, EPI_ISL_438161, EPI_ISL_438162, EPI_ISL_438163, EPI_ISL_438164, EPI_ISL_438165, EPI_ISL_438166, EPI_ISL_438167, EPI_ISL_438168, EPI_ISL_438169, EPI_ISL_438170, EPI_ISL_438171, EPI_ISL_438172, EPI_ISL_438173, EPI_ISL_438174, EPI_ISL_438175 | | | |
| see above | Seattle Flu Study | Seattle Flu Study | Chu et al |
| EPI_ISL_438176, EPI_ISL_438177, EPI_ISL_438179, EPI_ISL_438180, EPI_ISL_438181, EPI_ISL_438182, EPI_ISL_438183, EPI_ISL_438184, EPI_ISL_438185, EPI_ISL_438186, EPI_ISL_438187, EPI_ISL_438188, EPI_ISL_438189, EPI_ISL_438190, EPI_ISL_438191, EPI_ISL_438192, EPI_ISL_438193, EPI_ISL_438194, EPI_ISL_438195, EPI_ISL_438196, EPI_ISL_438197, EPI_ISL_438198, EPI_ISL_438199, EPI_ISL_438200, EPI_ISL_438201, EPI_ISL_438202, EPI_ISL_438203, EPI_ISL_438204, EPI_ISL_438205, EPI_ISL_438206, EPI_ISL_438207, EPI_ISL_438208, EPI_ISL_438209, EPI_ISL_438210, EPI_ISL_438211, EPI_ISL_438212, EPI_ISL_438213, EPI_ISL_438214, EPI_ISL_438215, EPI_ISL_438216, EPI_ISL_438217, EPI_ISL_438218, EPI_ISL_438219, EPI_ISL_438220, EPI_ISL_438221 | | | |
| see above | Washington State Department of Health | Seattle Flu Study | Chu et al |
| EPI_ISL_438222, EPI_ISL_438223, EPI_ISL_438224, EPI_ISL_438225, EPI_ISL_438226, EPI_ISL_438228, EPI_ISL_438229, EPI_ISL_438230, EPI_ISL_438231, EPI_ISL_438233, EPI_ISL_438234 | | | |
| see above | Johns Hopkins Hospital Department of Pathology | Johns Hopkins Hospital Department of Pathology | Peter M. Thielen, Thomas Mehoke, Shirlee Wohl, Srividya Ramakrishnan, Melanie Kirsche,_Amanda Ernlund,_Oluwaseun Falade-Nwulia, Timothy Gilpatrick, Paul Morris, Norah Sadowski, N_di_ Trovao, Victoria |

| | | | |
|---|---|---|---|
| | | | Gniazdowski, Michael Schatz, Stuart C. Ray, Winston Timp, Heba Mostafa |
| EPI_ISL_438235, EPI_ISL_438237, EPI_ISL_438240, EPI_ISL_438241, EPI_ISL_438242, EPI_ISL_438243, EPI_ISL_438244, EPI_ISL_438245, EPI_ISL_438246, EPI_ISL_438247 | Johns Hopkins Hospital Department of Pathology | Johns Hopkins Hospital Department of Pathology | Peter M. Thielen, Thomas Mehoke, Shirlee Wohl, Srividya Ramakrishnan, Melanie Kirsche, Amanda Ernlund, Oluwaseun Falade-Nwulia, Timothy Gilpatrick, Paul Morris, Norah Sadowski, Nidia Trovao, Victoria Gniazdowski, Michael Schatz, Stuart C. Ray, Winston Timp, Heba Mostafa |
| EPI_ISL_438257, EPI_ISL_438259, EPI_ISL_438260, EPI_ISL_438262, EPI_ISL_438265, EPI_ISL_438266, EPI_ISL_438267, EPI_ISL_438268, EPI_ISL_438269, EPI_ISL_438270, EPI_ISL_438272, EPI_ISL_438273, EPI_ISL_438274, EPI_ISL_438276, EPI_ISL_438277, EPI_ISL_438278, EPI_ISL_438279, EPI_ISL_438280, EPI_ISL_438282, EPI_ISL_438284, EPI_ISL_438286, EPI_ISL_438287, EPI_ISL_438288, EPI_ISL_438289, EPI_ISL_438290, EPI_ISL_438291, EPI_ISL_438292, EPI_ISL_438293, EPI_ISL_438294, EPI_ISL_438295, EPI_ISL_438296, EPI_ISL_438297, EPI_ISL_438299, EPI_ISL_438300, EPI_ISL_438301, EPI_ISL_438302, EPI_ISL_438303, EPI_ISL_438305, EPI_ISL_438306, EPI_ISL_438307, EPI_ISL_438308, EPI_ISL_438309, EPI_ISL_438310, EPI_ISL_438311, EPI_ISL_438312, EPI_ISL_438315, EPI_ISL_438316, EPI_ISL_438318, EPI_ISL_438319, EPI_ISL_438321, EPI_ISL_438322, EPI_ISL_438323, EPI_ISL_438324, EPI_ISL_438325, EPI_ISL_438326, EPI_ISL_438327, EPI_ISL_438329, EPI_ISL_438330, EPI_ISL_438332, EPI_ISL_438333, EPI_ISL_438334, EPI_ISL_438337, EPI_ISL_438338, EPI_ISL_438340, EPI_ISL_438342, EPI_ISL_438343, EPI_ISL_438344, EPI_ISL_438345, EPI_ISL_438346, EPI_ISL_438347, EPI_ISL_438349, EPI_ISL_438350, EPI_ISL_438351, EPI_ISL_438352, EPI_ISL_438354, EPI_ISL_438355, EPI_ISL_438356, EPI_ISL_438358, EPI_ISL_438359, EPI_ISL_438360, EPI_ISL_438361, EPI_ISL_438362, EPI_ISL_438363, EPI_ISL_438365, EPI_ISL_438366, EPI_ISL_438367, EPI_ISL_438368, EPI_ISL_438369, EPI_ISL_438370, EPI_ISL_438371, EPI_ISL_438372, EPI_ISL_438375, EPI_ISL_438376, EPI_ISL_438377, EPI_ISL_438378, EPI_ISL_438380, EPI_ISL_438381, EPI_ISL_438382, EPI_ISL_438383, EPI_ISL_438384, EPI_ISL_438385, EPI_ISL_438387, EPI_ISL_438388, EPI_ISL_438389, EPI_ISL_438390, EPI_ISL_438391, EPI_ISL_438392, EPI_ISL_438394, EPI_ISL_438396, EPI_ISL_438398, EPI_ISL_438399, EPI_ISL_438400, EPI_ISL_438401, EPI_ISL_438403, EPI_ISL_438404, EPI_ISL_438406, EPI_ISL_438407, EPI_ISL_438408, EPI_ISL_438409, EPI_ISL_438411, EPI_ISL_438412, EPI_ISL_438413, EPI_ISL_438414, EPI_ISL_438415, EPI_ISL_438416, EPI_ISL_438419, EPI_ISL_438420, EPI_ISL_438421, EPI_ISL_438422, EPI_ISL_438423, EPI_ISL_438424, EPI_ISL_438425, EPI_ISL_438426, EPI_ISL_438427, EPI_ISL_438428, EPI_ISL_438429, EPI_ISL_438430, EPI_ISL_438431, EPI_ISL_438434, EPI_ISL_438436, EPI_ISL_438437, EPI_ISL_438440, EPI_ISL_438441, EPI_ISL_438442, EPI_ISL_438443, EPI_ISL_438446, EPI_ISL_438447, EPI_ISL_438448, EPI_ISL_438451, EPI_ISL_438452, EPI_ISL_438453, EPI_ISL_438454, EPI_ISL_438455, EPI_ISL_438456, EPI_ISL_438457, EPI_ISL_438458, EPI_ISL_438459, EPI_ISL_438460, EPI_ISL_438462, EPI_ISL_438463, EPI_ISL_438464, EPI_ISL_438465, EPI_ISL_438466, EPI_ISL_438467, EPI_ISL_438468, EPI_ISL_438469, EPI_ISL_438470, EPI_ISL_438471, EPI_ISL_438472, EPI_ISL_438474, EPI_ISL_438475, EPI_ISL_438476, EPI_ISL_438477, EPI_ISL_438479, EPI_ISL_438480, EPI_ISL_438481, EPI_ISL_438482, EPI_ISL_438484, EPI_ISL_438485, EPI_ISL_438486, EPI_ISL_438488, EPI_ISL_438489, EPI_ISL_438491, EPI_ISL_438492, EPI_ISL_438493, EPI_ISL_438496, EPI_ISL_438497, EPI_ISL_438498, EPI_ISL_438499, EPI_ISL_438500, EPI_ISL_438501, EPI_ISL_438502, EPI_ISL_438504, EPI_ISL_438505, EPI_ISL_438506, EPI_ISL_438507, EPI_ISL_438509, EPI_ISL_438510, EPI_ISL_438511, EPI_ISL_438512, EPI_ISL_438513, EPI_ISL_438514, EPI_ISL_438515, EPI_ISL_438516, EPI_ISL_438517, EPI_ISL_438518, EPI_ISL_438519, EPI_ISL_438520, EPI_ISL_438521, EPI_ISL_438523, EPI_ISL_438524, EPI_ISL_438525, EPI_ISL_438527, EPI_ISL_438528, EPI_ISL_438529, EPI_ISL_438531, EPI_ISL_438532, EPI_ISL_438534, EPI_ISL_438535, EPI_ISL_438536, EPI_ISL_438537, EPI_ISL_438538, EPI_ISL_438539, EPI_ISL_438540, EPI_ISL_438541, EPI_ISL_438543, EPI_ISL_438544, EPI_ISL_438545 | | | |
| see above | Department of Pathology, University of Cambridge | Wellcome Sanger Institute for the COVID-19 Genomics UK Consortium | Luke W Meredith, M. Estée Török , Myra Hosmillo, William L. Hamilton, Martin D. Curran, Theresa Feltwell, Grant Hall, Anna Yakovleva, Fahad A Khokhar, Charlotte J. Houldcroft, Laura G Caller, Aminu S. Jahun, Sarah L. Caddy, Ian Goodfellow, Alex Alderton, Roberto Amato, Sonia Goncalves, Ewan Harrison, David K. Jackson, Ian Johnston, Dominic Kwiatkowski, Cordelia Langford, John Sillitoe on behalf of the Wellcome Sanger Institute COVID-19 Surveillance Team (http://www.sanger.ac.uk/covid-team) |
| EPI_ISL_438550, EPI_ISL_438551, EPI_ISL_438554, EPI_ISL_438557, EPI_ISL_438558, EPI_ISL_438559, EPI_ISL_438560, EPI_ISL_438561, EPI_ISL_438562, EPI_ISL_438563, EPI_ISL_438564, EPI_ISL_438565, EPI_ISL_438566, EPI_ISL_438567, EPI_ISL_438568, EPI_ISL_438571, EPI_ISL_438572, EPI_ISL_438573, EPI_ISL_438574, EPI_ISL_438575, EPI_ISL_438576, EPI_ISL_438577, EPI_ISL_438578, EPI_ISL_438579, EPI_ISL_438580, EPI_ISL_438581, EPI_ISL_438583, EPI_ISL_438587, EPI_ISL_438588, EPI_ISL_438590, EPI_ISL_438592, EPI_ISL_438593, EPI_ISL_438594, EPI_ISL_438597, EPI_ISL_438599, EPI_ISL_438600, EPI_ISL_438601, EPI_ISL_438602, EPI_ISL_438604, EPI_ISL_438605, EPI_ISL_438608, EPI_ISL_438610, EPI_ISL_438618, EPI_ISL_438619, EPI_ISL_438620, EPI_ISL_438621, EPI_ISL_438622, EPI_ISL_438623, EPI_ISL_438624, EPI_ISL_438625, EPI_ISL_438626, EPI_ISL_438628, EPI_ISL_438630, EPI_ISL_438631, EPI_ISL_438632, EPI_ISL_438633, EPI_ISL_438644, EPI_ISL_438645, EPI_ISL_438647, EPI_ISL_438649, EPI_ISL_438650, EPI_ISL_438652, EPI_ISL_438656, EPI_ISL_438658, EPI_ISL_438660, EPI_ISL_438662, EPI_ISL_438663, EPI_ISL_438664, EPI_ISL_438665, EPI_ISL_438666, EPI_ISL_438667, EPI_ISL_438669, EPI_ISL_438670, EPI_ISL_438671, EPI_ISL_438672, EPI_ISL_438676, EPI_ISL_438677, EPI_ISL_438678, EPI_ISL_438679, EPI_ISL_438681, EPI_ISL_438682, EPI_ISL_438683, EPI_ISL_438684, EPI_ISL_438686, EPI_ISL_438687, EPI_ISL_438688, EPI_ISL_438689, EPI_ISL_438690, EPI_ISL_438691, EPI_ISL_438692, EPI_ISL_438693, EPI_ISL_438696, EPI_ISL_438697, EPI_ISL_438698, EPI_ISL_438699, EPI_ISL_438700, EPI_ISL_438701, EPI_ISL_438702, EPI_ISL_438703, EPI_ISL_438704, EPI_ISL_438705, EPI_ISL_438707, EPI_ISL_438708, EPI_ISL_438709, EPI_ISL_438710, EPI_ISL_438711, EPI_ISL_438712, EPI_ISL_438713, EPI_ISL_438714, EPI_ISL_438715, EPI_ISL_438716, EPI_ISL_438717, EPI_ISL_438719, EPI_ISL_438720, EPI_ISL_438721, EPI_ISL_438722, EPI_ISL_438725, EPI_ISL_438726, EPI_ISL_438729, EPI_ISL_438733, EPI_ISL_438735, EPI_ISL_438736, EPI_ISL_438737, EPI_ISL_438740, EPI_ISL_438744, EPI_ISL_438745, EPI_ISL_438746, EPI_ISL_438747, EPI_ISL_438748 | | | |
| see above | Department of Pathology, University of Cambridge | COVID-19 Genomics UK (COG-UK) Consortium | Luke W Meredith, M. Est√©e T√∂r√∂k , Myra Hosmillo, William L. Hamilton, Martin D. Curran, Theresa Feltwell, Grant Hall, Anna Yakovleva, Fahad A Khokhar, Charlotte J. Houldcroft, Laura G Caller, Aminu S. Jahun, Sarah L. Caddy, Ian Goodfellow |
| EPI_ISL_438749, EPI_ISL_438754, EPI_ISL_438757, EPI_ISL_438760, EPI_ISL_438762, EPI_ISL_438763, EPI_ISL_438764, EPI_ISL_438766, EPI_ISL_438775, EPI_ISL_438776, EPI_ISL_438779, EPI_ISL_438783, EPI_ISL_438784, EPI_ISL_438798, EPI_ISL_438806, EPI_ISL_438815, EPI_ISL_438824, EPI_ISL_438827, EPI_ISL_438829, EPI_ISL_438830, EPI_ISL_438831, EPI_ISL_438836, EPI_ISL_438837, EPI_ISL_438839, EPI_ISL_438840, EPI_ISL_438841, EPI_ISL_438842, EPI_ISL_438844, EPI_ISL_438845, EPI_ISL_438849, EPI_ISL_438850, EPI_ISL_438860, EPI_ISL_438868, EPI_ISL_438870, EPI_ISL_438875, EPI_ISL_438877, EPI_ISL_438878, EPI_ISL_438879, EPI_ISL_438883, EPI_ISL_438884, EPI_ISL_438889, EPI_ISL_438890, EPI_ISL_438891, EPI_ISL_438892, EPI_ISL_438893, EPI_ISL_438894, EPI_ISL_438895, EPI_ISL_438896, EPI_ISL_438900, EPI_ISL_438901, EPI_ISL_438902, EPI_ISL_438903, EPI_ISL_438905, EPI_ISL_438907, EPI_ISL_438908, EPI_ISL_438910, EPI_ISL_438912, EPI_ISL_438913, EPI_ISL_438914, EPI_ISL_438915, EPI_ISL_438916, EPI_ISL_438917, EPI_ISL_438920, EPI_ISL_438921, EPI_ISL_438923, EPI_ISL_438925, EPI_ISL_438926, EPI_ISL_438928, EPI_ISL_438929, EPI_ISL_438930, EPI_ISL_438931, EPI_ISL_438933, EPI_ISL_438937, EPI_ISL_438941, EPI_ISL_438944 | | | |
| see above | West of Scotland Specialist Virology Centre, NHSGGC / MRC-University of Glasgow Centre for Virus Research | COVID-19 Genomics UK (COG-UK) Consortium | Ana da Silva Filipe, Natasha Johnson, Kathy Smollett, Daniel Mair, Stephen Carmichael, Lily Tong, Jenna Nichols, Elihu Aranday-Cortes, Kirstyn Brunker, Yasmin Parr, Kyriaki Nomikou; Sarah McDonald, Marc Niebel, Patawee Asamaphan; Richard Orton, Joseph Hughes, Sreenu Vattipally, David L Robertson; Alasdair MacLean, Rory Gunson; Kathy Li, Natasha Jesudason, Rajiv Shah, James Shepherd, Antonia Ho, Emma Thomson |
| EPI_ISL_438947, EPI_ISL_438948, EPI_ISL_438949, EPI_ISL_438951, EPI_ISL_438952, EPI_ISL_438953, EPI_ISL_438954, EPI_ISL_438955, EPI_ISL_438956, EPI_ISL_438958, EPI_ISL_438959, EPI_ISL_438961, EPI_ISL_438965, EPI_ISL_438966, EPI_ISL_438967, EPI_ISL_438969, EPI_ISL_438970, EPI_ISL_438971, EPI_ISL_438972 | | | |
| see above | Keio University School of Medicine | Keio University School of Medicine | Kenjiro Kosaki |
| EPI_ISL_438973, EPI_ISL_438974, EPI_ISL_438975, EPI_ISL_438976, EPI_ISL_438977, EPI_ISL_438978, EPI_ISL_438979, EPI_ISL_438980, EPI_ISL_438981, EPI_ISL_438982, EPI_ISL_438983, EPI_ISL_438984, EPI_ISL_438985, EPI_ISL_438986, EPI_ISL_438995, EPI_ISL_438998, EPI_ISL_438999, EPI_ISL_439000, EPI_ISL_439003, EPI_ISL_439004, EPI_ISL_439005, EPI_ISL_439006, EPI_ISL_439007, EPI_ISL_439009, EPI_ISL_439012, EPI_ISL_439013, EPI_ISL_439014, EPI_ISL_439015, EPI_ISL_439016, EPI_ISL_439018, EPI_ISL_439020, EPI_ISL_439022, EPI_ISL_439024, EPI_ISL_439026, EPI_ISL_439027, EPI_ISL_439028, EPI_ISL_439029, EPI_ISL_439030, EPI_ISL_439031, EPI_ISL_439032, EPI_ISL_439033, EPI_ISL_439034, EPI_ISL_439035, EPI_ISL_439036, EPI_ISL_439037, EPI_ISL_439038, EPI_ISL_439041, EPI_ISL_439043, EPI_ISL_439045, EPI_ISL_439046, EPI_ISL_439048, EPI_ISL_439049, EPI_ISL_439052, EPI_ISL_439056, EPI_ISL_439057, EPI_ISL_439062, EPI_ISL_439063, EPI_ISL_439065, EPI_ISL_439066, EPI_ISL_439067, EPI_ISL_439070, EPI_ISL_439072, EPI_ISL_439076, EPI_ISL_439080, EPI_ISL_439081, EPI_ISL_439082, EPI_ISL_439084, EPI_ISL_439085, EPI_ISL_439087, EPI_ISL_439090, EPI_ISL_439091, EPI_ISL_439092, EPI_ISL_439093, EPI_ISL_439095, EPI_ISL_439098, EPI_ISL_439101, EPI_ISL_439103, EPI_ISL_439105, EPI_ISL_439106, EPI_ISL_439116, EPI_ISL_439117, EPI_ISL_439118, EPI_ISL_439120, EPI_ISL_439123, EPI_ISL_439125, EPI_ISL_439126, EPI_ISL_439127, EPI_ISL_439130, EPI_ISL_439132, EPI_ISL_439134, EPI_ISL_439135, EPI_ISL_439140, EPI_ISL_439142 | | | |
| see above | West of Scotland Specialist Virology Centre, NHSGGC / MRC-University of Glasgow Centre for Virus Research | COVID-19 Genomics UK (COG-UK) Consortium | Ana da Silva Filipe, Natasha Johnson, Kathy Smollett, Daniel Mair, Stephen Carmichael, Lily Tong, Jenna Nichols, Elihu Aranday-Cortes, Kirstyn Brunker, Yasmin Parr, Kyriaki Nomikou; Sarah McDonald, Marc Niebel, Patawee Asamaphan; Richard Orton, Joseph Hughes, Sreenu Vattipally, David L Robertson; Alasdair MacLean, Rory Gunson; Kathy Li, Natasha Jesudason, Rajiv Shah, James Shepherd, Antonia Ho, Emma Thomson |
| EPI_ISL_439144, EPI_ISL_439145, EPI_ISL_439146, EPI_ISL_439147, EPI_ISL_439148, EPI_ISL_439150, EPI_ISL_439151, EPI_ISL_439152, EPI_ISL_439153, EPI_ISL_439154, EPI_ISL_439155, EPI_ISL_439157, EPI_ISL_439159, EPI_ISL_439160, EPI_ISL_439161, EPI_ISL_439162, EPI_ISL_439164, EPI_ISL_439166, EPI_ISL_439167, EPI_ISL_439168, EPI_ISL_439170, EPI_ISL_439171, EPI_ISL_439172, EPI_ISL_439173, EPI_ISL_439174, EPI_ISL_439175, EPI_ISL_439176, EPI_ISL_439177, EPI_ISL_439178, EPI_ISL_439179, EPI_ISL_439180, EPI_ISL_439181, EPI_ISL_439182, EPI_ISL_439183, EPI_ISL_439184, EPI_ISL_439185, EPI_ISL_439187, EPI_ISL_439188, EPI_ISL_439189, EPI_ISL_439190, EPI_ISL_439191, EPI_ISL_439193, EPI_ISL_439194, EPI_ISL_439195, EPI_ISL_439196, EPI_ISL_439197, EPI_ISL_439198, EPI_ISL_439199, EPI_ISL_439200, EPI_ISL_439201, EPI_ISL_439202, EPI_ISL_439203, EPI_ISL_439204, EPI_ISL_439205, EPI_ISL_439206, EPI_ISL_439207, EPI_ISL_439208, EPI_ISL_439209, EPI_ISL_439210, EPI_ISL_439211, EPI_ISL_439213, EPI_ISL_439214, EPI_ISL_439215, EPI_ISL_439216, EPI_ISL_439217, EPI_ISL_439218, EPI_ISL_439219, EPI_ISL_439220, EPI_ISL_439221, EPI_ISL_439222, EPI_ISL_439223, EPI_ISL_439224, EPI_ISL_439225, EPI_ISL_439226, EPI_ISL_439227, EPI_ISL_439228, EPI_ISL_439229, EPI_ISL_439230, EPI_ISL_439231, EPI_ISL_439232, EPI_ISL_439233, EPI_ISL_439234, EPI_ISL_439235, EPI_ISL_439236, EPI_ISL_439237, EPI_ISL_439238, EPI_ISL_439239, EPI_ISL_439240, EPI_ISL_439241, EPI_ISL_439242, EPI_ISL_439243, EPI_ISL_439244, EPI_ISL_439245, EPI_ISL_439246, EPI_ISL_439247, EPI_ISL_439248, EPI_ISL_439249, EPI_ISL_439250, EPI_ISL_439251, EPI_ISL_439253, EPI_ISL_439254, EPI_ISL_439255, EPI_ISL_439256, EPI_ISL_439257, EPI_ISL_439258, EPI_ISL_439259, EPI_ISL_439261, EPI_ISL_439262, EPI_ISL_439263, EPI_ISL_439264, EPI_ISL_439265, EPI_ISL_439266, EPI_ISL_439267, EPI_ISL_439268, EPI_ISL_439269, EPI_ISL_439270, EPI_ISL_439271, EPI_ISL_439272, EPI_ISL_439273, EPI_ISL_439274, EPI_ISL_439275, EPI_ISL_439276, EPI_ISL_439277, EPI_ISL_439278, EPI_ISL_439279, EPI_ISL_439280, EPI_ISL_439281, EPI_ISL_439282, EPI_ISL_439283, EPI_ISL_439284, EPI_ISL_439285, EPI_ISL_439286, EPI_ISL_439287, EPI_ISL_439288, EPI_ISL_439289, EPI_ISL_439290, EPI_ISL_439291, EPI_ISL_439292, EPI_ISL_439293, EPI_ISL_439294, EPI_ISL_439296, EPI_ISL_439297, EPI_ISL_439298, EPI_ISL_439299, EPI_ISL_439300, EPI_ISL_439301, EPI_ISL_439302, EPI_ISL_439303, EPI_ISL_439304, EPI_ISL_439305, EPI_ISL_439306, EPI_ISL_439307, EPI_ISL_439308, EPI_ISL_439309, EPI_ISL_439310, EPI_ISL_439311, EPI_ISL_439312, EPI_ISL_439313, EPI_ISL_439314, EPI_ISL_439315, EPI_ISL_439316, EPI_ISL_439317, EPI_ISL_439318, EPI_ISL_439319, EPI_ISL_439320, EPI_ISL_439322, EPI_ISL_439323, EPI_ISL_439324, EPI_ISL_439326, EPI_ISL_439327, EPI_ISL_439328, EPI_ISL_439329, EPI_ISL_439330, EPI_ISL_439331, EPI_ISL_439332, EPI_ISL_439333, EPI_ISL_439335, EPI_ISL_439336, EPI_ISL_439337, EPI_ISL_439338, EPI_ISL_439340, EPI_ISL_439341, EPI_ISL_439342, EPI_ISL_439343, EPI_ISL_439344, EPI_ISL_439345, EPI_ISL_439346, EPI_ISL_439347, EPI_ISL_439348, EPI_ISL_439349, EPI_ISL_439350, EPI_ISL_439351, EPI_ISL_439352, EPI_ISL_439353, EPI_ISL_439354, EPI_ISL_439355, EPI_ISL_439357, EPI_ISL_439358, EPI_ISL_439359, EPI_ISL_439360, EPI_ISL_439361, EPI_ISL_439363, EPI_ISL_439364, EPI_ISL_439365, EPI_ISL_439366, EPI_ISL_439367 | | | |
| see above | Virology Department, Royal Infirmary of Edinburgh, NHS Lothian / School of Biological Sciences, University of Edinburgh / Institute of Genetics and Molecular Medicine, University of Edinburgh | COVID-19 Genomics UK (COG-UK) Consortium | McHugh M, Dewar R, Rooke S, Gallagher M, Balcaza C, O,ÄôToole √Å, Scher E, Hill V, McCrone JT, Colquhoun R, Yu X, Jackson B, Rambaut A, Williams TC, Templeton K |
| EPI_ISL_439368, EPI_ISL_439369, EPI_ISL_439371, EPI_ISL_439372, EPI_ISL_439373, EPI_ISL_439374, EPI_ISL_439375, EPI_ISL_439376, EPI_ISL_439377, EPI_ISL_439378, EPI_ISL_439379, EPI_ISL_439381, EPI_ISL_439382, EPI_ISL_439383, EPI_ISL_439384, EPI_ISL_439385, EPI_ISL_439386, EPI_ISL_439387, EPI_ISL_439388, EPI_ISL_439389, EPI_ISL_439391, EPI_ISL_439393, EPI_ISL_439396, EPI_ISL_439397, EPI_ISL_439399, EPI_ISL_439400, EPI_ISL_439401, EPI_ISL_439402, EPI_ISL_439403, EPI_ISL_439404, EPI_ISL_439405, EPI_ISL_439406, EPI_ISL_439407, EPI_ISL_439408, EPI_ISL_439409, EPI_ISL_439410, EPI_ISL_439411, EPI_ISL_439414, EPI_ISL_439415, EPI_ISL_439416, EPI_ISL_439417, EPI_ISL_439418, EPI_ISL_439419, EPI_ISL_439420, EPI_ISL_439421, EPI_ISL_439422, EPI_ISL_439423, EPI_ISL_439424, EPI_ISL_439426, EPI_ISL_439428, EPI_ISL_439429, EPI_ISL_439430, EPI_ISL_439431, EPI_ISL_439432, EPI_ISL_439433, EPI_ISL_439434, EPI_ISL_439436, EPI_ISL_439437, EPI_ISL_439438, EPI_ISL_439439, EPI_ISL_439440, EPI_ISL_439441, EPI_ISL_439442, EPI_ISL_439443, EPI_ISL_439444, EPI_ISL_439446, EPI_ISL_439447, EPI_ISL_439448, EPI_ISL_439449, EPI_ISL_439450, EPI_ISL_439451, EPI_ISL_439452, EPI_ISL_439453, EPI_ISL_439454, EPI_ISL_439455, EPI_ISL_439456, EPI_ISL_439457, EPI_ISL_439459, EPI_ISL_439460, EPI_ISL_439461, EPI_ISL_439463, EPI_ISL_439464, EPI_ISL_439465, EPI_ISL_439466, EPI_ISL_439468, EPI_ISL_439469, EPI_ISL_439470, EPI_ISL_439471, EPI_ISL_439473, EPI_ISL_439475, EPI_ISL_439476, EPI_ISL_439478, EPI_ISL_439479, EPI_ISL_439481, EPI_ISL_439482, EPI_ISL_439484, EPI_ISL_439485, EPI_ISL_439486, EPI_ISL_439487, EPI_ISL_439489, EPI_ISL_439490, EPI_ISL_439491, EPI_ISL_439492, EPI_ISL_439494, EPI_ISL_439495, EPI_ISL_439496, EPI_ISL_439498, EPI_ISL_439499, EPI_ISL_439501, EPI_ISL_439502, EPI_ISL_439503, EPI_ISL_439504, EPI_ISL_439505, EPI_ISL_439507, EPI_ISL_439508, EPI_ISL_439509, EPI_ISL_439510, EPI_ISL_439511, EPI_ISL_439514, EPI_ISL_439516, EPI_ISL_439517, EPI_ISL_439518, EPI_ISL_439519, EPI_ISL_439520, EPI_ISL_439521, EPI_ISL_439522, EPI_ISL_439523, EPI_ISL_439524, EPI_ISL_439525, EPI_ISL_439526, EPI_ISL_439527, EPI_ISL_439528, EPI_ISL_439532, EPI_ISL_439533, EPI_ISL_439534, EPI_ISL_439535, EPI_ISL_439536, EPI_ISL_439538, EPI_ISL_439540, EPI_ISL_439541, EPI_ISL_439542, EPI_ISL_439545, EPI_ISL_439547, EPI_ISL_439548, EPI_ISL_439550, EPI_ISL_439555, EPI_ISL_439557, EPI_ISL_439559, EPI_ISL_439562, EPI_ISL_439564, EPI_ISL_439565, EPI_ISL_439567, EPI_ISL_439568, EPI_ISL_439570, EPI_ISL_439571, EPI_ISL_439572, EPI_ISL_439574, EPI_ISL_439575, EPI_ISL_439576, EPI_ISL_439582, EPI_ISL_439585, EPI_ISL_439587, EPI_ISL_439588, EPI_ISL_439589, EPI_ISL_439590, EPI_ISL_439591, EPI_ISL_439592, EPI_ISL_439594, EPI_ISL_439596, EPI_ISL_439598, EPI_ISL_439599, EPI_ISL_439601, EPI_ISL_439602, EPI_ISL_439604, EPI_ISL_439606, EPI_ISL_439607, EPI_ISL_439608, EPI_ISL_439609, EPI_ISL_439610, EPI_ISL_439611, EPI_ISL_439612, EPI_ISL_439613, EPI_ISL_439614, EPI_ISL_439615, EPI_ISL_439618, EPI_ISL_439619, EPI_ISL_439620, EPI_ISL_439621, EPI_ISL_439622, EPI_ISL_439623, EPI_ISL_439625, EPI_ISL_439626, EPI_ISL_439627, EPI_ISL_439628, EPI_ISL_439630, EPI_ISL_439631, EPI_ISL_439633, EPI_ISL_439634, EPI_ISL_439635, EPI_ISL_439636, EPI_ISL_439637, EPI_ISL_439638, EPI_ISL_439639, EPI_ISL_439640, EPI_ISL_439641, EPI_ISL_439642, EPI_ISL_439643, EPI_ISL_439644, EPI_ISL_439645, EPI_ISL_439646, EPI_ISL_439647, EPI_ISL_439648, EPI_ISL_439649, EPI_ISL_439652, EPI_ISL_439653, EPI_ISL_439655, EPI_ISL_439656, EPI_ISL_439657, EPI_ISL_439659, EPI_ISL_439660, EPI_ISL_439662, EPI_ISL_439664 | | | |
| see above | Department of Pathology, University of Cambridge | Wellcome Sanger Institute for the COVID-19 Genomics UK Consortium | Luke W Meredith, M. Estée Török , Myra Hosmillo, William L. Hamilton, Martin D. Curran, Theresa Feltwell, Grant Hall, Anna Yakovleva, Fahad A Khokhar, Charlotte J. Houldcroft, Laura G Caller, Aminu S. Jahun, Sarah L. Caddy, Ian Goodfellow, Alex Alderton, Roberto Amato, Sonia Goncalves, Ewan Harrison, David K. Jackson, Ian Johnston, Dominic Kwiatkowski, Cordelia Langford, John Sillitoe on behalf of the Wellcome Sanger Institute COVID-19 Surveillance Team (http://www.sanger.ac.uk/covid-team) |
| EPI_ISL_439665, EPI_ISL_439666, EPI_ISL_439668, EPI_ISL_439669, EPI_ISL_439671 | Virology Department, Royal Infirmary of Edinburgh, NHS Lothian / School of Biological Sciences, University of Edinburgh / Institute of Genetics and Molecular Medicine, University of Edinburgh | COVID-19 Genomics UK (COG-UK) Consortium | McHugh M, Dewar R, Rooke S, Gallagher M, Balcaza C, O,ÄôToole √Å, Scher E, Hill V, McCrone JT, Colquhoun R, Yu X, Jackson B, Rambaut A, Williams TC, Templeton K |
| EPI_ISL_439672, EPI_ISL_439673, EPI_ISL_439677, EPI_ISL_439681, EPI_ISL_439682, EPI_ISL_439683, EPI_ISL_439684, EPI_ISL_439687, EPI_ISL_439697, EPI_ISL_439704, EPI_ISL_439706, EPI_ISL_439707, EPI_ISL_439708, EPI_ISL_439709, EPI_ISL_439710, EPI_ISL_439712, EPI_ISL_439714, EPI_ISL_439715, EPI_ISL_439718, EPI_ISL_439719, EPI_ISL_439720, EPI_ISL_439721, EPI_ISL_439722, EPI_ISL_439724, EPI_ISL_439725, EPI_ISL_439728, EPI_ISL_439730, EPI_ISL_439731, EPI_ISL_439733, EPI_ISL_439737, EPI_ISL_439744, EPI_ISL_439747, EPI_ISL_439748, EPI_ISL_439751, EPI_ISL_439752, EPI_ISL_439753, EPI_ISL_439754, EPI_ISL_439756, EPI_ISL_439757, EPI_ISL_439758, EPI_ISL_439762, EPI_ISL_439763, EPI_ISL_439765, EPI_ISL_439767, EPI_ISL_439770, EPI_ISL_439775, EPI_ISL_439777, EPI_ISL_439780, EPI_ISL_439782, EPI_ISL_439784, EPI_ISL_439785, EPI_ISL_439787, EPI_ISL_439788, EPI_ISL_439790, EPI_ISL_439791, EPI_ISL_439793, EPI_ISL_439795, EPI_ISL_439796, EPI_ISL_439798, EPI_ISL_439799, EPI_ISL_439804, EPI_ISL_439806, EPI_ISL_439807, EPI_ISL_439808, EPI_ISL_439810, EPI_ISL_439812, EPI_ISL_439815, EPI_ISL_439817, EPI_ISL_439820, EPI_ISL_439821, EPI_ISL_439824, EPI_ISL_439825, EPI_ISL_439827, EPI_ISL_439832, EPI_ISL_439834, EPI_ISL_439836, EPI_ISL_439837, EPI_ISL_439848, EPI_ISL_439849, EPI_ISL_439855, EPI_ISL_439857, EPI_ISL_439861 | | | |
| see above | Liverpool Clinical Laboratories | COVID-19 Genomics UK (COG-UK) Consortium | Sam Haldenby, Anita Lucaci, Steve Paterson, Julian Hiscox, Alistair Darby, M Almsaud, A Alrezaihi, Muhannad Alruwaili, Stuart D Armstrong, Jones Benjamin , Eleanor G Bentley, Anu Chawla, Jordan J Clark, Angela Cowell, Richard Eccles, Isabel Garc√≠a-Dorival, Matthew Gemmell, Alessandro Gerada, PKF Gilmore, Richard Gregory, Ximeng Han, Catherine Hartley, Margaret Hughes, Miren Iturriza-Gomara, James Johnson, L Luu, Jenifer Manson , Charlotte Nelson, Elaine O,ÄôToole, Cassie Olateju, Rebekah Penrice-Randal¬†, Lucille Rainbow, N.P Randle, Trevor Ian Robinson, Parul Sharma, Ghada T Shawli, James P Stewart , Neil Swainston, Ecaterina Vamos, Joanne Watts, Mark Whitehead |
| EPI_ISL_439863, EPI_ISL_439865, EPI_ISL_439867, EPI_ISL_439868, EPI_ISL_439872, EPI_ISL_439874, EPI_ISL_439875, EPI_ISL_439876, EPI_ISL_439877, EPI_ISL_439880, EPI_ISL_439882, EPI_ISL_439883, EPI_ISL_439884, EPI_ISL_439886, EPI_ISL_439888, EPI_ISL_439889, EPI_ISL_439891, EPI_ISL_439892, EPI_ISL_439895, EPI_ISL_439898, EPI_ISL_439900, EPI_ISL_439902, EPI_ISL_439903, EPI_ISL_439904, EPI_ISL_439905, EPI_ISL_439907, EPI_ISL_439909, EPI_ISL_439911, EPI_ISL_439914, EPI_ISL_439917, EPI_ISL_439918, EPI_ISL_439919, EPI_ISL_439920, EPI_ISL_439921, EPI_ISL_439923, EPI_ISL_439926, EPI_ISL_439927, EPI_ISL_439930, EPI_ISL_439931, EPI_ISL_439932, EPI_ISL_439933, EPI_ISL_439934, EPI_ISL_439935, EPI_ISL_439936, EPI_ISL_439937, EPI_ISL_439938, EPI_ISL_439941, EPI_ISL_439943, EPI_ISL_439945, EPI_ISL_439947, EPI_ISL_439949, EPI_ISL_439954, EPI_ISL_439956 | | | |
| see above | Department of Pathology, University of Cambridge | Wellcome Sanger Institute for the COVID-19 Genomics UK Consortium | Luke W Meredith, M. Estée Török , Myra Hosmillo, William L. Hamilton, Martin D. Curran, Theresa Feltwell, Grant Hall, Anna Yakovleva, Fahad A Khokhar, Charlotte J. Houldcroft, Laura G Caller, Aminu S. Jahun, Sarah L. Caddy, Ian Goodfellow, Alex Alderton, Roberto Amato, Sonia Goncalves, Ewan Harrison, David K. Jackson, Ian Johnston, Dominic Kwiatkowski, Cordelia Langford, John Sillitoe on behalf of the Wellcome Sanger Institute COVID-19 Surveillance Team (http://www.sanger.ac.uk/covid-team) |
| EPI_ISL_439957 | PHE South West Regional Laboratory, National Infection Service | Wellcome Sanger Institute for the COVID-19 Genomics UK Consortium | Stephanie Hutchings, Hannah Pymont, Dr Peter Muir, Barry Vipond, Rich Hopes, Alex Alderton, Roberto Amato, Sonia Goncalves, Ewan Harrison, David K. Jackson, Ian Johnston, Dominic Kwiatkowski, Cordelia Langford, John Sillitoe on behalf of the Wellcome Sanger Institute COVID-19 Surveillance Team (http://www.sanger.ac.uk/covid-team) |
| EPI_ISL_439958 | Department of Pathology, University of Cambridge | Wellcome Sanger Institute for the COVID-19 Genomics UK Consortium | Luke W Meredith, M. Estée Török , Myra Hosmillo, William L. Hamilton, Martin D. Curran, Theresa Feltwell, Grant Hall, Anna Yakovleva, Fahad A Khokhar, Charlotte J. Houldcroft, Laura G Caller, Aminu S. Jahun, Sarah L. Caddy, Ian Goodfellow, Alex Alderton, Roberto Amato, Sonia Goncalves, Ewan Harrison, David K. Jackson, Ian Johnston, Dominic Kwiatkowski, Cordelia Langford, John Sillitoe on behalf of the Wellcome Sanger Institute COVID-19 Surveillance Team (http://www.sanger.ac.uk/covid-team) |
| EPI_ISL_439959 | PHE South West Regional Laboratory, National Infection Service | Wellcome Sanger Institute for the COVID-19 Genomics UK Consortium | Stephanie Hutchings, Hannah Pymont, Dr Peter Muir, Barry Vipond, Rich Hopes, Alex Alderton, Roberto Amato, Sonia Goncalves, Ewan Harrison, David K. Jackson, Ian Johnston, Dominic Kwiatkowski, Cordelia Langford, John Sillitoe on behalf of the Wellcome Sanger Institute COVID-19 Surveillance Team (http://www.sanger.ac.uk/covid-team) |
| EPI_ISL_439961, EPI_ISL_439963 | Department of Pathology, University of Cambridge | Wellcome Sanger Institute for the COVID-19 Genomics UK Consortium | Luke W Meredith, M. Estée Török , Myra Hosmillo, William L. Hamilton, Martin D. Curran, Theresa Feltwell, Grant Hall, Anna Yakovleva, Fahad A Khokhar, Charlotte J. Houldcroft, Laura G Caller, Aminu S. Jahun, Sarah L. Caddy, Ian Goodfellow, Alex Alderton, Roberto Amato, Sonia Goncalves, Ewan Harrison, David K. Jackson, Ian Johnston, Dominic Kwiatkowski, Cordelia Langford, John Sillitoe on behalf of the Wellcome Sanger Institute COVID-19 Surveillance Team (http://www.sanger.ac.uk/covid-team) |
| EPI_ISL_439964, EPI_ISL_439965, EPI_ISL_439966, EPI_ISL_439967 | PHE South West Regional Laboratory, National Infection Service | Wellcome Sanger Institute for the COVID-19 Genomics UK Consortium | Stephanie Hutchings, Hannah Pymont, Dr Peter Muir, Barry Vipond, Rich Hopes, Alex Alderton, Roberto Amato, Sonia Goncalves, Ewan Harrison, David K. Jackson, Ian Johnston, Dominic Kwiatkowski, Cordelia Langford, John Sillitoe on behalf of the Wellcome Sanger Institute COVID-19 Surveillance Team (http://www.sanger.ac.uk/covid-team) |
| EPI_ISL_439968 | Department of Pathology, University of Cambridge | Wellcome Sanger Institute for the COVID-19 Genomics UK Consortium | Luke W Meredith, M. Estée Török , Myra Hosmillo, William L. Hamilton, Martin D. Curran, Theresa Feltwell, Grant Hall, Anna Yakovleva, Fahad A Khokhar, Charlotte J. Houldcroft, Laura G Caller, Aminu S. Jahun, |

| | | | |
|---|---|---|---|
| | | | Sarah L. Caddy, Ian Goodfellow, Alex Alderton, Roberto Amato, Sonia Goncalves, Ewan Harrison, David K. Jackson, Ian Johnston, Dominic Kwiatkowski, Cordelia Langford, John Sillitoe on behalf of the Wellcome Sanger Institute COVID-19 Surveillance Team (http://www.sanger.ac.uk/covid-team) |
| EPI_ISL_439969 | PHE South West Regional Laboratory, National Infection Service | Wellcome Sanger Institute for the COVID-19 Genomics UK Consortium | Stephanie Hutchings, Hannah Pymont, Dr Peter Muir, Barry Vipond, Rich Hopes, Alex Alderton, Roberto Amato, Sonia Goncalves, Ewan Harrison, David K. Jackson, Ian Johnston, Dominic Kwiatkowski, Cordelia Langford, John Sillitoe on behalf of the Wellcome Sanger Institute COVID-19 Surveillance Team (http://www.sanger.ac.uk/covid-team) |
| EPI_ISL_439970 | Department of Pathology, University of Cambridge | Wellcome Sanger Institute for the COVID-19 Genomics UK Consortium | Luke W Meredith, M. Estée Török , Myra Hosmillo, William L. Hamilton, Martin D. Curran, Theresa Feltwell, Grant Hall, Anna Yakovleva, Fahad A Khokhar, Charlotte J. Houldcroft, Laura G Caller, Aminu S. Jahun, Sarah L. Caddy, Ian Goodfellow, Alex Alderton, Roberto Amato, Sonia Goncalves, Ewan Harrison, David K. Jackson, Ian Johnston, Dominic Kwiatkowski, Cordelia Langford, John Sillitoe on behalf of the Wellcome Sanger Institute COVID-19 Surveillance Team (http://www.sanger.ac.uk/covid-team) |
| EPI_ISL_439971, EPI_ISL_439972, EPI_ISL_439973, EPI_ISL_439974 | PHE South West Regional Laboratory, National Infection Service | Wellcome Sanger Institute for the COVID-19 Genomics UK Consortium | Stephanie Hutchings, Hannah Pymont, Dr Peter Muir, Barry Vipond, Rich Hopes, Alex Alderton, Roberto Amato, Sonia Goncalves, Ewan Harrison, David K. Jackson, Ian Johnston, Dominic Kwiatkowski, Cordelia Langford, John Sillitoe on behalf of the Wellcome Sanger Institute COVID-19 Surveillance Team (http://www.sanger.ac.uk/covid-team) |
| EPI_ISL_439975 | Department of Pathology, University of Cambridge | Wellcome Sanger Institute for the COVID-19 Genomics UK Consortium | Luke W Meredith, M. Estée Török , Myra Hosmillo, William L. Hamilton, Martin D. Curran, Theresa Feltwell, Grant Hall, Anna Yakovleva, Fahad A Khokhar, Charlotte J. Houldcroft, Laura G Caller, Aminu S. Jahun, Sarah L. Caddy, Ian Goodfellow, Alex Alderton, Roberto Amato, Sonia Goncalves, Ewan Harrison, David K. Jackson, Ian Johnston, Dominic Kwiatkowski, Cordelia Langford, John Sillitoe on behalf of the Wellcome Sanger Institute COVID-19 Surveillance Team (http://www.sanger.ac.uk/covid-team) |
| EPI_ISL_439978, EPI_ISL_439979 | PHE South West Regional Laboratory, National Infection Service | Wellcome Sanger Institute for the COVID-19 Genomics UK Consortium | Stephanie Hutchings, Hannah Pymont, Dr Peter Muir, Barry Vipond, Rich Hopes, Alex Alderton, Roberto Amato, Sonia Goncalves, Ewan Harrison, David K. Jackson, Ian Johnston, Dominic Kwiatkowski, Cordelia Langford, John Sillitoe on behalf of the Wellcome Sanger Institute COVID-19 Surveillance Team (http://www.sanger.ac.uk/covid-team) |
| EPI_ISL_439980 | Department of Pathology, University of Cambridge | Wellcome Sanger Institute for the COVID-19 Genomics UK Consortium | Luke W Meredith, M. Estée Török , Myra Hosmillo, William L. Hamilton, Martin D. Curran, Theresa Feltwell, Grant Hall, Anna Yakovleva, Fahad A Khokhar, Charlotte J. Houldcroft, Laura G Caller, Aminu S. Jahun, Sarah L. Caddy, Ian Goodfellow, Alex Alderton, Roberto Amato, Sonia Goncalves, Ewan Harrison, David K. Jackson, Ian Johnston, Dominic Kwiatkowski, Cordelia Langford, John Sillitoe on behalf of the Wellcome Sanger Institute COVID-19 Surveillance Team (http://www.sanger.ac.uk/covid-team) |
| EPI_ISL_439981, EPI_ISL_439982, EPI_ISL_439983, EPI_ISL_439984, EPI_ISL_439985, EPI_ISL_439986 | PHE South West Regional Laboratory, National Infection Service | Wellcome Sanger Institute for the COVID-19 Genomics UK Consortium | Stephanie Hutchings, Hannah Pymont, Dr Peter Muir, Barry Vipond, Rich Hopes, Alex Alderton, Roberto Amato, Sonia Goncalves, Ewan Harrison, David K. Jackson, Ian Johnston, Dominic Kwiatkowski, Cordelia Langford, John Sillitoe on behalf of the Wellcome Sanger Institute COVID-19 Surveillance Team (http://www.sanger.ac.uk/covid-team) |
| EPI_ISL_439987 | Department of Pathology, University of Cambridge | Wellcome Sanger Institute for the COVID-19 Genomics UK Consortium | Luke W Meredith, M. Estée Török , Myra Hosmillo, William L. Hamilton, Martin D. Curran, Theresa Feltwell, Grant Hall, Anna Yakovleva, Fahad A Khokhar, Charlotte J. Houldcroft, Laura G Caller, Aminu S. Jahun, Sarah L. Caddy, Ian Goodfellow, Alex Alderton, Roberto Amato, Sonia Goncalves, Ewan Harrison, David K. Jackson, Ian Johnston, Dominic Kwiatkowski, Cordelia Langford, John Sillitoe on behalf of the Wellcome Sanger Institute COVID-19 Surveillance Team (http://www.sanger.ac.uk/covid-team) |
| EPI_ISL_439988 | PHE South West Regional Laboratory, National Infection Service | Wellcome Sanger Institute for the COVID-19 Genomics UK Consortium | Stephanie Hutchings, Hannah Pymont, Dr Peter Muir, Barry Vipond, Rich Hopes, Alex Alderton, Roberto Amato, Sonia Goncalves, Ewan Harrison, David K. Jackson, Ian Johnston, Dominic Kwiatkowski, Cordelia Langford, John Sillitoe on behalf of the Wellcome Sanger Institute COVID-19 Surveillance Team (http://www.sanger.ac.uk/covid-team) |
| EPI_ISL_439989, EPI_ISL_439990, EPI_ISL_439991 | Department of Pathology, University of Cambridge | Wellcome Sanger Institute for the COVID-19 Genomics UK Consortium | Luke W Meredith, M. Estée Török , Myra Hosmillo, William L. Hamilton, Martin D. Curran, Theresa Feltwell, Grant Hall, Anna Yakovleva, Fahad A Khokhar, Charlotte J. Houldcroft, Laura G Caller, Aminu S. Jahun, Sarah L. Caddy, Ian Goodfellow, Alex Alderton, Roberto Amato, Sonia Goncalves, Ewan Harrison, David K. Jackson, Ian Johnston, Dominic Kwiatkowski, Cordelia Langford, John Sillitoe on behalf of the Wellcome Sanger Institute COVID-19 Surveillance Team (http://www.sanger.ac.uk/covid-team) |
| EPI_ISL_439992 | PHE South West Regional Laboratory, National Infection Service | Wellcome Sanger Institute for the COVID-19 Genomics UK Consortium | Stephanie Hutchings, Hannah Pymont, Dr Peter Muir, Barry Vipond, Rich Hopes, Alex Alderton, Roberto Amato, Sonia Goncalves, Ewan Harrison, David K. Jackson, Ian Johnston, Dominic Kwiatkowski, Cordelia Langford, John Sillitoe on behalf of the Wellcome Sanger Institute COVID-19 Surveillance Team (http://www.sanger.ac.uk/covid-team) |
| EPI_ISL_439993, EPI_ISL_439997 | Department of Pathology, University of Cambridge | Wellcome Sanger Institute for the COVID-19 Genomics UK Consortium | Luke W Meredith, M. Estée Török , Myra Hosmillo, William L. Hamilton, Martin D. Curran, Theresa Feltwell, Grant Hall, Anna Yakovleva, Fahad A Khokhar, Charlotte J. Houldcroft, Laura G Caller, Aminu S. Jahun, Sarah L. Caddy, Ian Goodfellow, Alex Alderton, Roberto Amato, Sonia Goncalves, Ewan Harrison, David K. Jackson, Ian Johnston, Dominic Kwiatkowski, Cordelia Langford, John Sillitoe on behalf of the Wellcome Sanger Institute COVID-19 Surveillance Team (http://www.sanger.ac.uk/covid-team) |
| EPI_ISL_439998 | PHE South West Regional Laboratory, National Infection Service | Wellcome Sanger Institute for the COVID-19 Genomics UK Consortium | Stephanie Hutchings, Hannah Pymont, Dr Peter Muir, Barry Vipond, Rich Hopes, Alex Alderton, Roberto Amato, Sonia Goncalves, Ewan Harrison, David K. Jackson, Ian Johnston, Dominic Kwiatkowski, Cordelia Langford, John Sillitoe on behalf of the Wellcome Sanger Institute COVID-19 Surveillance Team (http://www.sanger.ac.uk/covid-team) |
| EPI_ISL_439999 | Department of Pathology, University of Cambridge | Wellcome Sanger Institute for the COVID-19 Genomics UK Consortium | Luke W Meredith, M. Estée Török , Myra Hosmillo, William L. Hamilton, Martin D. Curran, Theresa Feltwell, Grant Hall, Anna Yakovleva, Fahad A Khokhar, Charlotte J. Houldcroft, Laura G Caller, Aminu S. Jahun, Sarah L. Caddy, Ian Goodfellow, Alex Alderton, Roberto Amato, Sonia Goncalves, Ewan Harrison, David K. Jackson, Ian Johnston, Dominic Kwiatkowski, Cordelia Langford, John Sillitoe on behalf of the Wellcome Sanger Institute COVID-19 Surveillance Team (http://www.sanger.ac.uk/covid-team) |
| EPI_ISL_440001, EPI_ISL_440002 | PHE South West Regional Laboratory, National Infection Service | Wellcome Sanger Institute for the COVID-19 Genomics UK Consortium | Stephanie Hutchings, Hannah Pymont, Dr Peter Muir, Barry Vipond, Rich Hopes, Alex Alderton, Roberto Amato, Sonia Goncalves, Ewan Harrison, David K. Jackson, Ian Johnston, Dominic Kwiatkowski, Cordelia Langford, John Sillitoe on behalf of the Wellcome Sanger Institute COVID-19 Surveillance Team (http://www.sanger.ac.uk/covid-team) |
| EPI_ISL_440004 | Department of Pathology, University of Cambridge | Wellcome Sanger Institute for the COVID-19 Genomics UK Consortium | Luke W Meredith, M. Estée Török , Myra Hosmillo, William L. Hamilton, Martin D. Curran, Theresa Feltwell, Grant Hall, Anna Yakovleva, Fahad A Khokhar, Charlotte J. Houldcroft, Laura G Caller, Aminu S. Jahun, Sarah L. Caddy, Ian Goodfellow, Alex Alderton, Roberto Amato, Sonia Goncalves, Ewan Harrison, David K. Jackson, Ian Johnston, Dominic Kwiatkowski, Cordelia Langford, John Sillitoe on behalf of the Wellcome Sanger Institute COVID-19 Surveillance Team (http://www.sanger.ac.uk/covid-team) |
| EPI_ISL_440005, EPI_ISL_440006 | PHE South West Regional Laboratory, National Infection Service | Wellcome Sanger Institute for the COVID-19 Genomics UK Consortium | Stephanie Hutchings, Hannah Pymont, Dr Peter Muir, Barry Vipond, Rich Hopes, Alex Alderton, Roberto Amato, Sonia Goncalves, Ewan Harrison, David K. Jackson, Ian Johnston, Dominic Kwiatkowski, Cordelia Langford, John Sillitoe on behalf of the Wellcome Sanger Institute COVID-19 Surveillance Team (http://www.sanger.ac.uk/covid-team) |
| EPI_ISL_440007 | Department of Pathology, University of Cambridge | Wellcome Sanger Institute for the COVID-19 Genomics UK Consortium | Luke W Meredith, M. Estée Török , Myra Hosmillo, William L. Hamilton, Martin D. Curran, Theresa Feltwell, Grant Hall, Anna Yakovleva, Fahad A Khokhar, Charlotte J. Houldcroft, Laura G Caller, Aminu S. Jahun, Sarah L. Caddy, Ian Goodfellow, Alex Alderton, Roberto Amato, Sonia Goncalves, Ewan Harrison, David K. Jackson, Ian Johnston, Dominic Kwiatkowski, Cordelia Langford, John Sillitoe on behalf of the Wellcome Sanger Institute COVID-19 Surveillance Team (http://www.sanger.ac.uk/covid-team) |
| EPI_ISL_440008 | PHE South West Regional Laboratory, National Infection Service | Wellcome Sanger Institute for the COVID-19 Genomics UK Consortium | Stephanie Hutchings, Hannah Pymont, Dr Peter Muir, Barry Vipond, Rich Hopes, Alex Alderton, Roberto Amato, Sonia Goncalves, Ewan Harrison, David K. Jackson, Ian Johnston, Dominic Kwiatkowski, Cordelia Langford, John Sillitoe on behalf of the Wellcome Sanger Institute COVID-19 Surveillance Team (http://www.sanger.ac.uk/covid-team) |
| EPI_ISL_440009, EPI_ISL_440010, EPI_ISL_440011 | Department of Pathology, University of Cambridge | Wellcome Sanger Institute for the COVID-19 Genomics UK Consortium | Luke W Meredith, M. Estée Török , Myra Hosmillo, William L. Hamilton, Martin D. Curran, Theresa Feltwell, Grant Hall, Anna Yakovleva, Fahad A Khokhar, Charlotte J. Houldcroft, Laura G Caller, Aminu S. Jahun, Sarah L. Caddy, Ian Goodfellow, Alex Alderton, Roberto Amato, Sonia Goncalves, Ewan Harrison, David K. Jackson, Ian Johnston, Dominic Kwiatkowski, Cordelia Langford, John Sillitoe on behalf of the Wellcome Sanger Institute COVID-19 Surveillance Team (http://www.sanger.ac.uk/covid-team) |
| EPI_ISL_440013 | PHE South West Regional Laboratory, National Infection Service | Wellcome Sanger Institute for the COVID-19 Genomics UK Consortium | Stephanie Hutchings, Hannah Pymont, Dr Peter Muir, Barry Vipond, Rich Hopes, Alex Alderton, Roberto Amato, Sonia Goncalves, Ewan Harrison, David K. Jackson, Ian Johnston, Dominic Kwiatkowski, Cordelia Langford, John Sillitoe on behalf of the Wellcome Sanger Institute COVID-19 Surveillance Team (http://www.sanger.ac.uk/covid-team) |
| EPI_ISL_440014, EPI_ISL_440017 | Department of Pathology, University of Cambridge | Wellcome Sanger Institute for the COVID-19 Genomics UK Consortium | Luke W Meredith, M. Estée Török , Myra Hosmillo, William L. Hamilton, Martin D. Curran, Theresa Feltwell, Grant Hall, Anna Yakovleva, Fahad A Khokhar, Charlotte J. Houldcroft, Laura G Caller, Aminu S. Jahun, Sarah L. Caddy, Ian Goodfellow, Alex Alderton, Roberto Amato, Sonia Goncalves, Ewan Harrison, David K. Jackson, Ian Johnston, Dominic Kwiatkowski, Cordelia Langford, John Sillitoe on behalf of the Wellcome Sanger Institute COVID-19 Surveillance Team (http://www.sanger.ac.uk/covid-team) |
| EPI_ISL_440018, EPI_ISL_440020, EPI_ISL_440021, EPI_ISL_440022, EPI_ISL_440023 | PHE South West Regional Laboratory, National Infection Service | Wellcome Sanger Institute for the COVID-19 Genomics UK Consortium | Stephanie Hutchings, Hannah Pymont, Dr Peter Muir, Barry Vipond, Rich Hopes, Alex Alderton, Roberto Amato, Sonia Goncalves, Ewan Harrison, David K. Jackson, Ian Johnston, Dominic Kwiatkowski, Cordelia Langford, John Sillitoe on behalf of the Wellcome Sanger Institute COVID-19 Surveillance Team (http://www.sanger.ac.uk/covid-team) |
| EPI_ISL_440025 | Department of Pathology, University of Cambridge | Wellcome Sanger Institute for the COVID-19 Genomics UK Consortium | Luke W Meredith, M. Estée Török , Myra Hosmillo, William L. Hamilton, Martin D. Curran, Theresa Feltwell, Grant Hall, Anna Yakovleva, Fahad A Khokhar, Charlotte J. Houldcroft, Laura G Caller, Aminu S. Jahun, Sarah L. Caddy, Ian Goodfellow, Alex Alderton, Roberto Amato, Sonia Goncalves, Ewan Harrison, David K. Jackson, Ian Johnston, Dominic Kwiatkowski, Cordelia Langford, John Sillitoe on behalf of the Wellcome Sanger Institute COVID-19 Surveillance Team (http://www.sanger.ac.uk/covid-team) |
| EPI_ISL_440027, EPI_ISL_440029 | PHE South West Regional Laboratory, National Infection Service | Wellcome Sanger Institute for the COVID-19 Genomics UK Consortium | Stephanie Hutchings, Hannah Pymont, Dr Peter Muir, Barry Vipond, Rich Hopes, Alex Alderton, Roberto Amato, Sonia Goncalves, Ewan Harrison, David K. Jackson, Ian Johnston, Dominic Kwiatkowski, Cordelia Langford, John Sillitoe on behalf of the Wellcome Sanger Institute COVID-19 Surveillance Team (http://www.sanger.ac.uk/covid-team) |
| EPI_ISL_440030 | Department of Pathology, University of Cambridge | Wellcome Sanger Institute for the COVID-19 Genomics UK Consortium | Luke W Meredith, M. Estée Török , Myra Hosmillo, William L. Hamilton, Martin D. Curran, Theresa Feltwell, Grant Hall, Anna Yakovleva, Fahad A Khokhar, Charlotte J. Houldcroft, Laura G Caller, Aminu S. Jahun, Sarah L. Caddy, Ian Goodfellow, Alex Alderton, Roberto Amato, Sonia Goncalves, Ewan Harrison, David K. Jackson, Ian Johnston, Dominic Kwiatkowski, Cordelia Langford, John Sillitoe on behalf of the Wellcome Sanger Institute COVID-19 Surveillance Team (http://www.sanger.ac.uk/covid-team) |
| EPI_ISL_440032, EPI_ISL_440033 | PHE South West Regional Laboratory, National Infection Service | Wellcome Sanger Institute for the COVID-19 Genomics UK Consortium | Stephanie Hutchings, Hannah Pymont, Dr Peter Muir, Barry Vipond, Rich Hopes, Alex Alderton, Roberto Amato, Sonia Goncalves, Ewan Harrison, David K. Jackson, Ian Johnston, Dominic Kwiatkowski, Cordelia Langford, John Sillitoe on behalf of the Wellcome Sanger Institute COVID-19 Surveillance Team (http://www.sanger.ac.uk/covid-team) |
| EPI_ISL_440034 | Department of Pathology, University of Cambridge | Wellcome Sanger Institute for the COVID-19 Genomics UK Consortium | Luke W Meredith, M. Estée Török , Myra Hosmillo, William L. Hamilton, Martin D. Curran, Theresa Feltwell, Grant Hall, Anna Yakovleva, Fahad A Khokhar, Charlotte J. Houldcroft, Laura G Caller, Aminu S. Jahun, Sarah L. Caddy, Ian Goodfellow, Alex Alderton, Roberto Amato, Sonia Goncalves, Ewan Harrison, David K. Jackson, Ian Johnston, Dominic Kwiatkowski, Cordelia Langford, John Sillitoe on behalf of the Wellcome Sanger Institute COVID-19 Surveillance Team (http://www.sanger.ac.uk/covid-team) |
| EPI_ISL_440037, EPI_ISL_440038, EPI_ISL_440039 | PHE South West Regional Laboratory, National Infection Service | Wellcome Sanger Institute for the COVID-19 Genomics UK Consortium | Stephanie Hutchings, Hannah Pymont, Dr Peter Muir, Barry Vipond, Rich Hopes, Alex Alderton, Roberto Amato, Sonia Goncalves, Ewan Harrison, David K. Jackson, Ian Johnston, Dominic Kwiatkowski, Cordelia Langford, John Sillitoe on behalf of the Wellcome Sanger Institute COVID-19 Surveillance Team (http://www.sanger.ac.uk/covid-team) |
| EPI_ISL_440040, EPI_ISL_440041 | Department of Pathology, University of Cambridge | Wellcome Sanger Institute for the COVID-19 Genomics UK Consortium | Luke W Meredith, M. Estée Török , Myra Hosmillo, William L. Hamilton, Martin D. Curran, Theresa Feltwell, Grant Hall, Anna Yakovleva, Fahad A Khokhar, Charlotte J. Houldcroft, Laura G Caller, Aminu S. Jahun, Sarah L. Caddy, Ian Goodfellow, Alex Alderton, Roberto Amato, Sonia Goncalves, Ewan Harrison, David K. Jackson, Ian Johnston, Dominic Kwiatkowski, Cordelia Langford, John Sillitoe on behalf of the Wellcome Sanger Institute COVID-19 Surveillance Team (http://www.sanger.ac.uk/covid-team) |
| EPI_ISL_440042 | PHE South West Regional Laboratory, National Infection Service | Wellcome Sanger Institute for the COVID-19 Genomics UK Consortium | Stephanie Hutchings, Hannah Pymont, Dr Peter Muir, Barry Vipond, Rich Hopes, Alex Alderton, Roberto Amato, Sonia Goncalves, Ewan Harrison, David K. Jackson, Ian Johnston, Dominic Kwiatkowski, Cordelia Langford, John Sillitoe on behalf of the Wellcome Sanger Institute COVID-19 Surveillance Team (http://www.sanger.ac.uk/covid-team) |
| EPI_ISL_440044, EPI_ISL_440045, EPI_ISL_440046, EPI_ISL_440047 | Department of Pathology, University of Cambridge | Wellcome Sanger Institute for the COVID-19 Genomics UK Consortium | Luke W Meredith, M. Estée Török , Myra Hosmillo, William L. Hamilton, Martin D. Curran, Theresa Feltwell, Grant Hall, Anna Yakovleva, Fahad A Khokhar, Charlotte J. Houldcroft, Laura G Caller, Aminu S. Jahun, Sarah L. Caddy, Ian Goodfellow, Alex Alderton, Roberto Amato, Sonia Goncalves, Ewan Harrison, David K. Jackson, Ian Johnston, Dominic Kwiatkowski, Cordelia Langford, John Sillitoe on behalf of the Wellcome Sanger Institute COVID-19 Surveillance Team (http://www.sanger.ac.uk/covid-team) |
| EPI_ISL_440050, EPI_ISL_440052 | PHE South West Regional Laboratory, National Infection Service | Wellcome Sanger Institute for the COVID-19 Genomics UK Consortium | Stephanie Hutchings, Hannah Pymont, Dr Peter Muir, Barry Vipond, Rich Hopes, Alex Alderton, Roberto Amato, Sonia Goncalves, Ewan Harrison, David K. Jackson, Ian Johnston, Dominic Kwiatkowski, Cordelia Langford, John Sillitoe on behalf of the Wellcome Sanger Institute COVID-19 Surveillance Team (http://www.sanger.ac.uk/covid-team) |
| EPI_ISL_440053 | Department of Pathology, University of Cambridge | Wellcome Sanger Institute for the COVID-19 Genomics UK Consortium | Luke W Meredith, M. Estée Török , Myra Hosmillo, William L. Hamilton, Martin D. Curran, Theresa Feltwell, Grant Hall, Anna Yakovleva, Fahad A Khokhar, Charlotte J. Houldcroft, Laura G Caller, Aminu S. Jahun, Sarah L. Caddy, Ian Goodfellow, Alex Alderton, Roberto Amato, Sonia Goncalves, Ewan Harrison, David K. Jackson, Ian Johnston, Dominic Kwiatkowski, Cordelia Langford, John Sillitoe on behalf of the Wellcome Sanger Institute COVID-19 Surveillance Team (http://www.sanger.ac.uk/covid-team) |
| EPI_ISL_440054 | PHE South West Regional Laboratory, National Infection Service | Wellcome Sanger Institute for the COVID-19 Genomics UK Consortium | Stephanie Hutchings, Hannah Pymont, Dr Peter Muir, Barry Vipond, Rich Hopes, Alex Alderton, Roberto Amato, Sonia Goncalves, Ewan Harrison, David K. Jackson, Ian Johnston, Dominic Kwiatkowski, Cordelia Langford, John Sillitoe on behalf of the Wellcome Sanger Institute COVID-19 Surveillance Team (http://www.sanger.ac.uk/covid-team) |
| EPI_ISL_440055 | Department of Pathology, University of Cambridge | Wellcome Sanger Institute for the COVID-19 Genomics UK Consortium | Luke W Meredith, M. Estée Török , Myra Hosmillo, William L. Hamilton, Martin D. Curran, Theresa Feltwell, Grant Hall, Anna Yakovleva, Fahad A Khokhar, Charlotte J. Houldcroft, Laura G Caller, Aminu S. Jahun, Sarah L. Caddy, Ian Goodfellow, Alex Alderton, Roberto Amato, Sonia Goncalves, Ewan Harrison, David K. Jackson, Ian Johnston, Dominic Kwiatkowski, Cordelia Langford, John Sillitoe on behalf of the Wellcome Sanger Institute COVID-19 Surveillance Team (http://www.sanger.ac.uk/covid-team) |
| EPI_ISL_440056 | PHE South West Regional Laboratory, National Infection | Wellcome Sanger Institute for the COVID-19 Genomics UK Consortium | Stephanie Hutchings, Hannah Pymont, Dr Peter Muir, Barry Vipond, Rich Hopes, Alex Alderton, Roberto Amato, Sonia Goncalves, Ewan Harrison, David K. Jackson, Ian Johnston, Dominic Kwiatkowski, Cordelia |

| | | | |
|---|---|---|---|
| | Service | | Langford, John Sillitoe on behalf of the Wellcome Sanger Institute COVID-19 Surveillance Team (http://www.sanger.ac.uk/covid-team) |
| EPI_ISL_440057, EPI_ISL_440059, EPI_ISL_440060, EPI_ISL_440061, EPI_ISL_440062 | Department of Pathology, University of Cambridge | Wellcome Sanger Institute for the COVID-19 Genomics UK Consortium | Luke W Meredith, M. Estée Török , Myra Hosmillo, William L. Hamilton, Martin D. Curran, Theresa Feltwell, Grant Hall, Anna Yakovleva, Fahad A Khokhar, Charlotte J. Houldcroft, Laura G Caller, Aminu S. Jahun, Sarah L. Caddy, Ian Goodfellow, Alex Alderton, Roberto Amato, Sonia Goncalves, Ewan Harrison, David K. Jackson, Ian Johnston, Dominic Kwiatkowski, Cordelia Langford, John Sillitoe on behalf of the Wellcome Sanger Institute COVID-19 Surveillance Team (http://www.sanger.ac.uk/covid-team) |
| EPI_ISL_440063 | PHE South West Regional Laboratory, National Infection Service | Wellcome Sanger Institute for the COVID-19 Genomics UK Consortium | Stephanie Hutchings, Hannah Pymont, Dr Peter Muir, Barry Vipond, Rich Hopes, Alex Alderton, Roberto Amato, Sonia Goncalves, Ewan Harrison, David K. Jackson, Ian Johnston, Dominic Kwiatkowski, Cordelia Langford, John Sillitoe on behalf of the Wellcome Sanger Institute COVID-19 Surveillance Team (http://www.sanger.ac.uk/covid-team) |
| EPI_ISL_440064 | Department of Pathology, University of Cambridge | Wellcome Sanger Institute for the COVID-19 Genomics UK Consortium | Luke W Meredith, M. Estée Török , Myra Hosmillo, William L. Hamilton, Martin D. Curran, Theresa Feltwell, Grant Hall, Anna Yakovleva, Fahad A Khokhar, Charlotte J. Houldcroft, Laura G Caller, Aminu S. Jahun, Sarah L. Caddy, Ian Goodfellow, Alex Alderton, Roberto Amato, Sonia Goncalves, Ewan Harrison, David K. Jackson, Ian Johnston, Dominic Kwiatkowski, Cordelia Langford, John Sillitoe on behalf of the Wellcome Sanger Institute COVID-19 Surveillance Team (http://www.sanger.ac.uk/covid-team) |
| EPI_ISL_440065 | PHE South West Regional Laboratory, National Infection Service | Wellcome Sanger Institute for the COVID-19 Genomics UK Consortium | Stephanie Hutchings, Hannah Pymont, Dr Peter Muir, Barry Vipond, Rich Hopes, Alex Alderton, Roberto Amato, Sonia Goncalves, Ewan Harrison, David K. Jackson, Ian Johnston, Dominic Kwiatkowski, Cordelia Langford, John Sillitoe on behalf of the Wellcome Sanger Institute COVID-19 Surveillance Team (http://www.sanger.ac.uk/covid-team) |
| EPI_ISL_440066 | Department of Pathology, University of Cambridge | Wellcome Sanger Institute for the COVID-19 Genomics UK Consortium | Luke W Meredith, M. Estée Török , Myra Hosmillo, William L. Hamilton, Martin D. Curran, Theresa Feltwell, Grant Hall, Anna Yakovleva, Fahad A Khokhar, Charlotte J. Houldcroft, Laura G Caller, Aminu S. Jahun, Sarah L. Caddy, Ian Goodfellow, Alex Alderton, Roberto Amato, Sonia Goncalves, Ewan Harrison, David K. Jackson, Ian Johnston, Dominic Kwiatkowski, Cordelia Langford, John Sillitoe on behalf of the Wellcome Sanger Institute COVID-19 Surveillance Team (http://www.sanger.ac.uk/covid-team) |
| EPI_ISL_440068 | PHE South West Regional Laboratory, National Infection Service | Wellcome Sanger Institute for the COVID-19 Genomics UK Consortium | Stephanie Hutchings, Hannah Pymont, Dr Peter Muir, Barry Vipond, Rich Hopes, Alex Alderton, Roberto Amato, Sonia Goncalves, Ewan Harrison, David K. Jackson, Ian Johnston, Dominic Kwiatkowski, Cordelia Langford, John Sillitoe on behalf of the Wellcome Sanger Institute COVID-19 Surveillance Team (http://www.sanger.ac.uk/covid-team) |
| EPI_ISL_440069 | Department of Pathology, University of Cambridge | Wellcome Sanger Institute for the COVID-19 Genomics UK Consortium | Luke W Meredith, M. Estée Török , Myra Hosmillo, William L. Hamilton, Martin D. Curran, Theresa Feltwell, Grant Hall, Anna Yakovleva, Fahad A Khokhar, Charlotte J. Houldcroft, Laura G Caller, Aminu S. Jahun, Sarah L. Caddy, Ian Goodfellow, Alex Alderton, Roberto Amato, Sonia Goncalves, Ewan Harrison, David K. Jackson, Ian Johnston, Dominic Kwiatkowski, Cordelia Langford, John Sillitoe on behalf of the Wellcome Sanger Institute COVID-19 Surveillance Team (http://www.sanger.ac.uk/covid-team) |
| EPI_ISL_440070, EPI_ISL_440071 | PHE South West Regional Laboratory, National Infection Service | Wellcome Sanger Institute for the COVID-19 Genomics UK Consortium | Stephanie Hutchings, Hannah Pymont, Dr Peter Muir, Barry Vipond, Rich Hopes, Alex Alderton, Roberto Amato, Sonia Goncalves, Ewan Harrison, David K. Jackson, Ian Johnston, Dominic Kwiatkowski, Cordelia Langford, John Sillitoe on behalf of the Wellcome Sanger Institute COVID-19 Surveillance Team (http://www.sanger.ac.uk/covid-team) |
| EPI_ISL_440072, EPI_ISL_440073, EPI_ISL_440074, EPI_ISL_440075, EPI_ISL_440076, EPI_ISL_440077, EPI_ISL_440079, EPI_ISL_440080 | Department of Pathology, University of Cambridge | Wellcome Sanger Institute for the COVID-19 Genomics UK Consortium | Luke W Meredith, M. Estée Török , Myra Hosmillo, William L. Hamilton, Martin D. Curran, Theresa Feltwell, Grant Hall, Anna Yakovleva, Fahad A Khokhar, Charlotte J. Houldcroft, Laura G Caller, Aminu S. Jahun, Sarah L. Caddy, Ian Goodfellow, Alex Alderton, Roberto Amato, Sonia Goncalves, Ewan Harrison, David K. Jackson, Ian Johnston, Dominic Kwiatkowski, Cordelia Langford, John Sillitoe on behalf of the Wellcome Sanger Institute COVID-19 Surveillance Team (http://www.sanger.ac.uk/covid-team) |
| EPI_ISL_440081 | PHE South West Regional Laboratory, National Infection Service | Wellcome Sanger Institute for the COVID-19 Genomics UK Consortium | Stephanie Hutchings, Hannah Pymont, Dr Peter Muir, Barry Vipond, Rich Hopes, Alex Alderton, Roberto Amato, Sonia Goncalves, Ewan Harrison, David K. Jackson, Ian Johnston, Dominic Kwiatkowski, Cordelia Langford, John Sillitoe on behalf of the Wellcome Sanger Institute COVID-19 Surveillance Team (http://www.sanger.ac.uk/covid-team) |
| EPI_ISL_440085, EPI_ISL_440086 | Department of Pathology, University of Cambridge | Wellcome Sanger Institute for the COVID-19 Genomics UK Consortium | Luke W Meredith, M. Estée Török , Myra Hosmillo, William L. Hamilton, Martin D. Curran, Theresa Feltwell, Grant Hall, Anna Yakovleva, Fahad A Khokhar, Charlotte J. Houldcroft, Laura G Caller, Aminu S. Jahun, Sarah L. Caddy, Ian Goodfellow, Alex Alderton, Roberto Amato, Sonia Goncalves, Ewan Harrison, David K. Jackson, Ian Johnston, Dominic Kwiatkowski, Cordelia Langford, John Sillitoe on behalf of the Wellcome Sanger Institute COVID-19 Surveillance Team (http://www.sanger.ac.uk/covid-team) |
| EPI_ISL_440087 | PHE South West Regional Laboratory, National Infection Service | Wellcome Sanger Institute for the COVID-19 Genomics UK Consortium | Stephanie Hutchings, Hannah Pymont, Dr Peter Muir, Barry Vipond, Rich Hopes, Alex Alderton, Roberto Amato, Sonia Goncalves, Ewan Harrison, David K. Jackson, Ian Johnston, Dominic Kwiatkowski, Cordelia Langford, John Sillitoe on behalf of the Wellcome Sanger Institute COVID-19 Surveillance Team (http://www.sanger.ac.uk/covid-team) |
| EPI_ISL_440088, EPI_ISL_440090 | Department of Pathology, University of Cambridge | Wellcome Sanger Institute for the COVID-19 Genomics UK Consortium | Luke W Meredith, M. Estée Török , Myra Hosmillo, William L. Hamilton, Martin D. Curran, Theresa Feltwell, Grant Hall, Anna Yakovleva, Fahad A Khokhar, Charlotte J. Houldcroft, Laura G Caller, Aminu S. Jahun, Sarah L. Caddy, Ian Goodfellow, Alex Alderton, Roberto Amato, Sonia Goncalves, Ewan Harrison, David K. Jackson, Ian Johnston, Dominic Kwiatkowski, Cordelia Langford, John Sillitoe on behalf of the Wellcome Sanger Institute COVID-19 Surveillance Team (http://www.sanger.ac.uk/covid-team) |
| EPI_ISL_440091, EPI_ISL_440092, EPI_ISL_440093, EPI_ISL_440094, EPI_ISL_440095 | PHE South West Regional Laboratory, National Infection Service | Wellcome Sanger Institute for the COVID-19 Genomics UK Consortium | Stephanie Hutchings, Hannah Pymont, Dr Peter Muir, Barry Vipond, Rich Hopes, Alex Alderton, Roberto Amato, Sonia Goncalves, Ewan Harrison, David K. Jackson, Ian Johnston, Dominic Kwiatkowski, Cordelia Langford, John Sillitoe on behalf of the Wellcome Sanger Institute COVID-19 Surveillance Team (http://www.sanger.ac.uk/covid-team) |
| EPI_ISL_440096, EPI_ISL_440097 | Department of Pathology, University of Cambridge | Wellcome Sanger Institute for the COVID-19 Genomics UK Consortium | Luke W Meredith, M. Estée Török , Myra Hosmillo, William L. Hamilton, Martin D. Curran, Theresa Feltwell, Grant Hall, Anna Yakovleva, Fahad A Khokhar, Charlotte J. Houldcroft, Laura G Caller, Aminu S. Jahun, Sarah L. Caddy, Ian Goodfellow, Alex Alderton, Roberto Amato, Sonia Goncalves, Ewan Harrison, David K. Jackson, Ian Johnston, Dominic Kwiatkowski, Cordelia Langford, John Sillitoe on behalf of the Wellcome Sanger Institute COVID-19 Surveillance Team (http://www.sanger.ac.uk/covid-team) |
| EPI_ISL_440098 | PHE South West Regional Laboratory, National Infection Service | Wellcome Sanger Institute for the COVID-19 Genomics UK Consortium | Stephanie Hutchings, Hannah Pymont, Dr Peter Muir, Barry Vipond, Rich Hopes, Alex Alderton, Roberto Amato, Sonia Goncalves, Ewan Harrison, David K. Jackson, Ian Johnston, Dominic Kwiatkowski, Cordelia Langford, John Sillitoe on behalf of the Wellcome Sanger Institute COVID-19 Surveillance Team (http://www.sanger.ac.uk/covid-team) |
| EPI_ISL_440099 | Department of Pathology, University of Cambridge | Wellcome Sanger Institute for the COVID-19 Genomics UK Consortium | Luke W Meredith, M. Estée Török , Myra Hosmillo, William L. Hamilton, Martin D. Curran, Theresa Feltwell, Grant Hall, Anna Yakovleva, Fahad A Khokhar, Charlotte J. Houldcroft, Laura G Caller, Aminu S. Jahun, Sarah L. Caddy, Ian Goodfellow, Alex Alderton, Roberto Amato, Sonia Goncalves, Ewan Harrison, David K. Jackson, Ian Johnston, Dominic Kwiatkowski, Cordelia Langford, John Sillitoe on behalf of the Wellcome Sanger Institute COVID-19 Surveillance Team (http://www.sanger.ac.uk/covid-team) |
| EPI_ISL_440100, EPI_ISL_440101, EPI_ISL_440102, EPI_ISL_440103 | PHE South West Regional Laboratory, National Infection Service | Wellcome Sanger Institute for the COVID-19 Genomics UK Consortium | Stephanie Hutchings, Hannah Pymont, Dr Peter Muir, Barry Vipond, Rich Hopes, Alex Alderton, Roberto Amato, Sonia Goncalves, Ewan Harrison, David K. Jackson, Ian Johnston, Dominic Kwiatkowski, Cordelia Langford, John Sillitoe on behalf of the Wellcome Sanger Institute COVID-19 Surveillance Team (http://www.sanger.ac.uk/covid-team) |
| EPI_ISL_440104 | Department of Pathology, University of Cambridge | Wellcome Sanger Institute for the COVID-19 Genomics UK Consortium | Luke W Meredith, M. Estée Török , Myra Hosmillo, William L. Hamilton, Martin D. Curran, Theresa Feltwell, Grant Hall, Anna Yakovleva, Fahad A Khokhar, Charlotte J. Houldcroft, Laura G Caller, Aminu S. Jahun, Sarah L. Caddy, Ian Goodfellow, Alex Alderton, Roberto Amato, Sonia Goncalves, Ewan Harrison, David K. Jackson, Ian Johnston, Dominic Kwiatkowski, Cordelia Langford, John Sillitoe on behalf of the Wellcome Sanger Institute COVID-19 Surveillance Team (http://www.sanger.ac.uk/covid-team) |
| EPI_ISL_440105, EPI_ISL_440106 | PHE South West Regional Laboratory, National Infection Service | Wellcome Sanger Institute for the COVID-19 Genomics UK Consortium | Stephanie Hutchings, Hannah Pymont, Dr Peter Muir, Barry Vipond, Rich Hopes, Alex Alderton, Roberto Amato, Sonia Goncalves, Ewan Harrison, David K. Jackson, Ian Johnston, Dominic Kwiatkowski, Cordelia Langford, John Sillitoe on behalf of the Wellcome Sanger Institute COVID-19 Surveillance Team (http://www.sanger.ac.uk/covid-team) |
| EPI_ISL_440107, EPI_ISL_440108, EPI_ISL_440109, EPI_ISL_440110, EPI_ISL_440111, EPI_ISL_440112 | Department of Pathology, University of Cambridge | Wellcome Sanger Institute for the COVID-19 Genomics UK Consortium | Luke W Meredith, M. Estée Török , Myra Hosmillo, William L. Hamilton, Martin D. Curran, Theresa Feltwell, Grant Hall, Anna Yakovleva, Fahad A Khokhar, Charlotte J. Houldcroft, Laura G Caller, Aminu S. Jahun, Sarah L. Caddy, Ian Goodfellow, Alex Alderton, Roberto Amato, Sonia Goncalves, Ewan Harrison, David K. Jackson, Ian Johnston, Dominic Kwiatkowski, Cordelia Langford, John Sillitoe on behalf of the Wellcome Sanger Institute COVID-19 Surveillance Team (http://www.sanger.ac.uk/covid-team) |
| EPI_ISL_440114, EPI_ISL_440115, EPI_ISL_440116 | PHE South West Regional Laboratory, National Infection Service | Wellcome Sanger Institute for the COVID-19 Genomics UK Consortium | Stephanie Hutchings, Hannah Pymont, Dr Peter Muir, Barry Vipond, Rich Hopes, Alex Alderton, Roberto Amato, Sonia Goncalves, Ewan Harrison, David K. Jackson, Ian Johnston, Dominic Kwiatkowski, Cordelia Langford, John Sillitoe on behalf of the Wellcome Sanger Institute COVID-19 Surveillance Team (http://www.sanger.ac.uk/covid-team) |
| EPI_ISL_440117, EPI_ISL_440120, EPI_ISL_440121, EPI_ISL_440122 | Department of Pathology, University of Cambridge | Wellcome Sanger Institute for the COVID-19 Genomics UK Consortium | Luke W Meredith, M. Estée Török , Myra Hosmillo, William L. Hamilton, Martin D. Curran, Theresa Feltwell, Grant Hall, Anna Yakovleva, Fahad A Khokhar, Charlotte J. Houldcroft, Laura G Caller, Aminu S. Jahun, Sarah L. Caddy, Ian Goodfellow, Alex Alderton, Roberto Amato, Sonia Goncalves, Ewan Harrison, David K. Jackson, Ian Johnston, Dominic Kwiatkowski, Cordelia Langford, John Sillitoe on behalf of the Wellcome Sanger Institute COVID-19 Surveillance Team (http://www.sanger.ac.uk/covid-team) |
| EPI_ISL_440123, EPI_ISL_440124 | PHE South West Regional Laboratory, National Infection Service | Wellcome Sanger Institute for the COVID-19 Genomics UK Consortium | Stephanie Hutchings, Hannah Pymont, Dr Peter Muir, Barry Vipond, Rich Hopes, Alex Alderton, Roberto Amato, Sonia Goncalves, Ewan Harrison, David K. Jackson, Ian Johnston, Dominic Kwiatkowski, Cordelia Langford, John Sillitoe on behalf of the Wellcome Sanger Institute COVID-19 Surveillance Team (http://www.sanger.ac.uk/covid-team) |
| EPI_ISL_440125 | Department of Pathology, University of Cambridge | Wellcome Sanger Institute for the COVID-19 Genomics UK Consortium | Luke W Meredith, M. Estée Török , Myra Hosmillo, William L. Hamilton, Martin D. Curran, Theresa Feltwell, Grant Hall, Anna Yakovleva, Fahad A Khokhar, Charlotte J. Houldcroft, Laura G Caller, Aminu S. Jahun, Sarah L. Caddy, Ian Goodfellow, Alex Alderton, Roberto Amato, Sonia Goncalves, Ewan Harrison, David K. Jackson, Ian Johnston, Dominic Kwiatkowski, Cordelia Langford, John Sillitoe on behalf of the Wellcome Sanger Institute COVID-19 Surveillance Team (http://www.sanger.ac.uk/covid-team) |
| EPI_ISL_440126, EPI_ISL_440127, EPI_ISL_440128 | PHE South West Regional Laboratory, National Infection Service | Wellcome Sanger Institute for the COVID-19 Genomics UK Consortium | Stephanie Hutchings, Hannah Pymont, Dr Peter Muir, Barry Vipond, Rich Hopes, Alex Alderton, Roberto Amato, Sonia Goncalves, Ewan Harrison, David K. Jackson, Ian Johnston, Dominic Kwiatkowski, Cordelia Langford, John Sillitoe on behalf of the Wellcome Sanger Institute COVID-19 Surveillance Team (http://www.sanger.ac.uk/covid-team) |
| EPI_ISL_440129, EPI_ISL_440132 | Department of Pathology, University of Cambridge | Wellcome Sanger Institute for the COVID-19 Genomics UK Consortium | Luke W Meredith, M. Estée Török , Myra Hosmillo, William L. Hamilton, Martin D. Curran, Theresa Feltwell, Grant Hall, Anna Yakovleva, Fahad A Khokhar, Charlotte J. Houldcroft, Laura G Caller, Aminu S. Jahun, Sarah L. Caddy, Ian Goodfellow, Alex Alderton, Roberto Amato, Sonia Goncalves, Ewan Harrison, David K. Jackson, Ian Johnston, Dominic Kwiatkowski, Cordelia Langford, John Sillitoe on behalf of the Wellcome Sanger Institute COVID-19 Surveillance Team (http://www.sanger.ac.uk/covid-team) |
| EPI_ISL_440135 | PHE South West Regional Laboratory, National Infection Service | Wellcome Sanger Institute for the COVID-19 Genomics UK Consortium | Stephanie Hutchings, Hannah Pymont, Dr Peter Muir, Barry Vipond, Rich Hopes, Alex Alderton, Roberto Amato, Sonia Goncalves, Ewan Harrison, David K. Jackson, Ian Johnston, Dominic Kwiatkowski, Cordelia Langford, John Sillitoe on behalf of the Wellcome Sanger Institute COVID-19 Surveillance Team (http://www.sanger.ac.uk/covid-team) |
| EPI_ISL_440138 | Department of Pathology, University of Cambridge | Wellcome Sanger Institute for the COVID-19 Genomics UK Consortium | Luke W Meredith, M. Estée Török , Myra Hosmillo, William L. Hamilton, Martin D. Curran, Theresa Feltwell, Grant Hall, Anna Yakovleva, Fahad A Khokhar, Charlotte J. Houldcroft, Laura G Caller, Aminu S. Jahun, Sarah L. Caddy, Ian Goodfellow, Alex Alderton, Roberto Amato, Sonia Goncalves, Ewan Harrison, David K. Jackson, Ian Johnston, Dominic Kwiatkowski, Cordelia Langford, John Sillitoe on behalf of the Wellcome Sanger Institute COVID-19 Surveillance Team (http://www.sanger.ac.uk/covid-team) |
| EPI_ISL_440139, EPI_ISL_440141 | PHE South West Regional Laboratory, National Infection Service | Wellcome Sanger Institute for the COVID-19 Genomics UK Consortium | Stephanie Hutchings, Hannah Pymont, Dr Peter Muir, Barry Vipond, Rich Hopes, Alex Alderton, Roberto Amato, Sonia Goncalves, Ewan Harrison, David K. Jackson, Ian Johnston, Dominic Kwiatkowski, Cordelia Langford, John Sillitoe on behalf of the Wellcome Sanger Institute COVID-19 Surveillance Team (http://www.sanger.ac.uk/covid-team) |
| EPI_ISL_440143, EPI_ISL_440144, EPI_ISL_440146, EPI_ISL_440147 | Department of Pathology, University of Cambridge | Wellcome Sanger Institute for the COVID-19 Genomics UK Consortium | Luke W Meredith, M. Estée Török , Myra Hosmillo, William L. Hamilton, Martin D. Curran, Theresa Feltwell, Grant Hall, Anna Yakovleva, Fahad A Khokhar, Charlotte J. Houldcroft, Laura G Caller, Aminu S. Jahun, Sarah L. Caddy, Ian Goodfellow, Alex Alderton, Roberto Amato, Sonia Goncalves, Ewan Harrison, David K. Jackson, Ian Johnston, Dominic Kwiatkowski, Cordelia Langford, John Sillitoe on behalf of the Wellcome Sanger Institute COVID-19 Surveillance Team (http://www.sanger.ac.uk/covid-team) |
| EPI_ISL_440149 | PHE South West Regional Laboratory, National Infection Service | Wellcome Sanger Institute for the COVID-19 Genomics UK Consortium | Stephanie Hutchings, Hannah Pymont, Dr Peter Muir, Barry Vipond, Rich Hopes, Alex Alderton, Roberto Amato, Sonia Goncalves, Ewan Harrison, David K. Jackson, Ian Johnston, Dominic Kwiatkowski, Cordelia Langford, John Sillitoe on behalf of the Wellcome Sanger Institute COVID-19 Surveillance Team (http://www.sanger.ac.uk/covid-team) |
| EPI_ISL_440150, EPI_ISL_440152, EPI_ISL_440155 | Department of Pathology, University of Cambridge | Wellcome Sanger Institute for the COVID-19 Genomics UK Consortium | Luke W Meredith, M. Estée Török , Myra Hosmillo, William L. Hamilton, Martin D. Curran, Theresa Feltwell, Grant Hall, Anna Yakovleva, Fahad A Khokhar, Charlotte J. Houldcroft, Laura G Caller, Aminu S. Jahun, Sarah L. Caddy, Ian Goodfellow, Alex Alderton, Roberto Amato, Sonia Goncalves, Ewan Harrison, David K. Jackson, Ian Johnston, Dominic Kwiatkowski, Cordelia Langford, John Sillitoe on behalf of the Wellcome Sanger Institute COVID-19 Surveillance Team (http://www.sanger.ac.uk/covid-team) |
| EPI_ISL_440156, EPI_ISL_440158 | PHE South West Regional Laboratory, National Infection Service | Wellcome Sanger Institute for the COVID-19 Genomics UK Consortium | Stephanie Hutchings, Hannah Pymont, Dr Peter Muir, Barry Vipond, Rich Hopes, Alex Alderton, Roberto Amato, Sonia Goncalves, Ewan Harrison, David K. Jackson, Ian Johnston, Dominic Kwiatkowski, Cordelia Langford, John Sillitoe on behalf of the Wellcome Sanger Institute COVID-19 Surveillance Team (http://www.sanger.ac.uk/covid-team) |
| EPI_ISL_440159, EPI_ISL_440161 | Department of Pathology, University of Cambridge | Wellcome Sanger Institute for the COVID-19 Genomics UK Consortium | Luke W Meredith, M. Estée Török , Myra Hosmillo, William L. Hamilton, Martin D. Curran, Theresa Feltwell, Grant Hall, Anna Yakovleva, Fahad A Khokhar, Charlotte J. Houldcroft, Laura G Caller, Aminu S. Jahun, Sarah L. Caddy, Ian Goodfellow, Alex Alderton, Roberto Amato, Sonia Goncalves, Ewan Harrison, David K. Jackson, Ian Johnston, Dominic Kwiatkowski, Cordelia Langford, John Sillitoe on behalf of the Wellcome Sanger Institute COVID-19 Surveillance Team (http://www.sanger.ac.uk/covid-team) |
| EPI_ISL_440162 | PHE South West Regional Laboratory, National Infection Service | Wellcome Sanger Institute for the COVID-19 Genomics UK Consortium | Stephanie Hutchings, Hannah Pymont, Dr Peter Muir, Barry Vipond, Rich Hopes, Alex Alderton, Roberto Amato, Sonia Goncalves, Ewan Harrison, David K. Jackson, Ian Johnston, Dominic Kwiatkowski, Cordelia Langford, John Sillitoe on behalf of the Wellcome Sanger Institute COVID-19 Surveillance Team (http://www.sanger.ac.uk/covid-team) |
| EPI_ISL_440163, EPI_ISL_440164, EPI_ISL_440166 | Department of Pathology, University of Cambridge | Wellcome Sanger Institute for the COVID-19 Genomics UK Consortium | Luke W Meredith, M. Estée Török , Myra Hosmillo, William L. Hamilton, Martin D. Curran, Theresa Feltwell, Grant Hall, Anna Yakovleva, Fahad A Khokhar, Charlotte J. Houldcroft, Laura G Caller, Aminu S. Jahun, Sarah L. Caddy, Ian Goodfellow, Alex Alderton, Roberto Amato, Sonia Goncalves, Ewan Harrison, David K. Jackson, Ian Johnston, Dominic Kwiatkowski, Cordelia Langford, John Sillitoe on behalf of the Wellcome Sanger Institute COVID-19 Surveillance Team (http://www.sanger.ac.uk/covid-team) |

| | | | |
|---|---|---|---|
| EPI_ISL_440167, EPI_ISL_440168 | PHE South West Regional Laboratory, National Infection Service | Wellcome Sanger Institute for the COVID-19 Genomics UK Consortium | Stephanie Hutchings, Hannah Pymont, Dr Peter Muir, Barry Vipond, Rich Hopes, Alex Alderton, Roberto Amato, Sonia Goncalves, Ewan Harrison, David K. Jackson, Ian Johnston, Dominic Kwiatkowski, Cordelia Langford, John Sillitoe on behalf of the Wellcome Sanger Institute COVID-19 Surveillance Team (http://www.sanger.ac.uk/covid-team) |
| EPI_ISL_440169, EPI_ISL_440170, EPI_ISL_440176, EPI_ISL_440177 | Department of Pathology, University of Cambridge | Wellcome Sanger Institute for the COVID-19 Genomics UK Consortium | Luke W Meredith, M. Estée Török, Myra Hosmillo, William L. Hamilton, Martin D. Curran, Theresa Feltwell, Grant Hall, Anna Yakovleva, Fahad A Khokhar, Charlotte J. Houldcroft, Laura G Caller, Aminu S. Jahun, Sarah L. Caddy, Ian Goodfellow, Alex Alderton, Roberto Amato, Sonia Goncalves, Ewan Harrison, David K. Jackson, Ian Johnston, Dominic Kwiatkowski, Cordelia Langford, John Sillitoe on behalf of the Wellcome Sanger Institute COVID-19 Surveillance Team (http://www.sanger.ac.uk/covid-team) |
| EPI_ISL_440178 | PHE South West Regional Laboratory, National Infection Service | Wellcome Sanger Institute for the COVID-19 Genomics UK Consortium | Stephanie Hutchings, Hannah Pymont, Dr Peter Muir, Barry Vipond, Rich Hopes, Alex Alderton, Roberto Amato, Sonia Goncalves, Ewan Harrison, David K. Jackson, Ian Johnston, Dominic Kwiatkowski, Cordelia Langford, John Sillitoe on behalf of the Wellcome Sanger Institute COVID-19 Surveillance Team (http://www.sanger.ac.uk/covid-team) |
| EPI_ISL_440179, EPI_ISL_440181, EPI_ISL_440182, EPI_ISL_440183, EPI_ISL_440184, EPI_ISL_440185, EPI_ISL_440186 | Department of Pathology, University of Cambridge | Wellcome Sanger Institute for the COVID-19 Genomics UK Consortium | Luke W Meredith, M. Estée Török , Myra Hosmillo, William L. Hamilton, Martin D. Curran, Theresa Feltwell, Grant Hall, Anna Yakovleva, Fahad A Khokhar, Charlotte J. Houldcroft, Laura G Caller, Aminu S. Jahun, Sarah L. Caddy, Ian Goodfellow, Alex Alderton, Roberto Amato, Sonia Goncalves, Ewan Harrison, David K. Jackson, Ian Johnston, Dominic Kwiatkowski, Cordelia Langford, John Sillitoe on behalf of the Wellcome Sanger Institute COVID-19 Surveillance Team (http://www.sanger.ac.uk/covid-team) |
| EPI_ISL_440187 | PHE South West Regional Laboratory, National Infection Service | Wellcome Sanger Institute for the COVID-19 Genomics UK Consortium | Stephanie Hutchings, Hannah Pymont, Dr Peter Muir, Barry Vipond, Rich Hopes, Alex Alderton, Roberto Amato, Sonia Goncalves, Ewan Harrison, David K. Jackson, Ian Johnston, Dominic Kwiatkowski, Cordelia Langford, John Sillitoe on behalf of the Wellcome Sanger Institute COVID-19 Surveillance Team (http://www.sanger.ac.uk/covid-team) |
| EPI_ISL_440188 | Department of Pathology, University of Cambridge | Wellcome Sanger Institute for the COVID-19 Genomics UK Consortium | Luke W Meredith, M. Estée Török , Myra Hosmillo, William L. Hamilton, Martin D. Curran, Theresa Feltwell, Grant Hall, Anna Yakovleva, Fahad A Khokhar, Charlotte J. Houldcroft, Laura G Caller, Aminu S. Jahun, Sarah L. Caddy, Ian Goodfellow, Alex Alderton, Roberto Amato, Sonia Goncalves, Ewan Harrison, David K. Jackson, Ian Johnston, Dominic Kwiatkowski, Cordelia Langford, John Sillitoe on behalf of the Wellcome Sanger Institute COVID-19 Surveillance Team (http://www.sanger.ac.uk/covid-team) |
| EPI_ISL_440189, EPI_ISL_440194 | PHE South West Regional Laboratory, National Infection Service | Wellcome Sanger Institute for the COVID-19 Genomics UK Consortium | Stephanie Hutchings, Hannah Pymont, Dr Peter Muir, Barry Vipond, Rich Hopes, Alex Alderton, Roberto Amato, Sonia Goncalves, Ewan Harrison, David K. Jackson, Ian Johnston, Dominic Kwiatkowski, Cordelia Langford, John Sillitoe on behalf of the Wellcome Sanger Institute COVID-19 Surveillance Team (http://www.sanger.ac.uk/covid-team) |
| EPI_ISL_440195 | Department of Pathology, University of Cambridge | Wellcome Sanger Institute for the COVID-19 Genomics UK Consortium | Luke W Meredith, M. Estée Török , Myra Hosmillo, William L. Hamilton, Martin D. Curran, Theresa Feltwell, Grant Hall, Anna Yakovleva, Fahad A Khokhar, Charlotte J. Houldcroft, Laura G Caller, Aminu S. Jahun, Sarah L. Caddy, Ian Goodfellow, Alex Alderton, Roberto Amato, Sonia Goncalves, Ewan Harrison, David K. Jackson, Ian Johnston, Dominic Kwiatkowski, Cordelia Langford, John Sillitoe on behalf of the Wellcome Sanger Institute COVID-19 Surveillance Team (http://www.sanger.ac.uk/covid-team) |
| EPI_ISL_440196, EPI_ISL_440198, EPI_ISL_440201 | PHE South West Regional Laboratory, National Infection Service | Wellcome Sanger Institute for the COVID-19 Genomics UK Consortium | Stephanie Hutchings, Hannah Pymont, Dr Peter Muir, Barry Vipond, Rich Hopes, Alex Alderton, Roberto Amato, Sonia Goncalves, Ewan Harrison, David K. Jackson, Ian Johnston, Dominic Kwiatkowski, Cordelia Langford, John Sillitoe on behalf of the Wellcome Sanger Institute COVID-19 Surveillance Team (http://www.sanger.ac.uk/covid-team) |
| EPI_ISL_440202, EPI_ISL_440203 | Department of Pathology, University of Cambridge | Wellcome Sanger Institute for the COVID-19 Genomics UK Consortium | Luke W Meredith, M. Estée Török , Myra Hosmillo, William L. Hamilton, Martin D. Curran, Theresa Feltwell, Grant Hall, Anna Yakovleva, Fahad A Khokhar, Charlotte J. Houldcroft, Laura G Caller, Aminu S. Jahun, Sarah L. Caddy, Ian Goodfellow, Alex Alderton, Roberto Amato, Sonia Goncalves, Ewan Harrison, David K. Jackson, Ian Johnston, Dominic Kwiatkowski, Cordelia Langford, John Sillitoe on behalf of the Wellcome Sanger Institute COVID-19 Surveillance Team (http://www.sanger.ac.uk/covid-team) |
| EPI_ISL_440206 | PHE South West Regional Laboratory, National Infection Service | Wellcome Sanger Institute for the COVID-19 Genomics UK Consortium | Stephanie Hutchings, Hannah Pymont, Dr Peter Muir, Barry Vipond, Rich Hopes, Alex Alderton, Roberto Amato, Sonia Goncalves, Ewan Harrison, David K. Jackson, Ian Johnston, Dominic Kwiatkowski, Cordelia Langford, John Sillitoe on behalf of the Wellcome Sanger Institute COVID-19 Surveillance Team (http://www.sanger.ac.uk/covid-team) |
| EPI_ISL_440207, EPI_ISL_440208, EPI_ISL_440213 | Department of Pathology, University of Cambridge | Wellcome Sanger Institute for the COVID-19 Genomics UK Consortium | Luke W Meredith, M. Estée Török , Myra Hosmillo, William L. Hamilton, Martin D. Curran, Theresa Feltwell, Grant Hall, Anna Yakovleva, Fahad A Khokhar, Charlotte J. Houldcroft, Laura G Caller, Aminu S. Jahun, Sarah L. Caddy, Ian Goodfellow, Alex Alderton, Roberto Amato, Sonia Goncalves, Ewan Harrison, David K. Jackson, Ian Johnston, Dominic Kwiatkowski, Cordelia Langford, John Sillitoe on behalf of the Wellcome Sanger Institute COVID-19 Surveillance Team (http://www.sanger.ac.uk/covid-team) |
| EPI_ISL_440214 | PHE South West Regional Laboratory, National Infection Service | Wellcome Sanger Institute for the COVID-19 Genomics UK Consortium | Stephanie Hutchings, Hannah Pymont, Dr Peter Muir, Barry Vipond, Rich Hopes, Alex Alderton, Roberto Amato, Sonia Goncalves, Ewan Harrison, David K. Jackson, Ian Johnston, Dominic Kwiatkowski, Cordelia Langford, John Sillitoe on behalf of the Wellcome Sanger Institute COVID-19 Surveillance Team (http://www.sanger.ac.uk/covid-team) |
| EPI_ISL_440217 | Department of Pathology, University of Cambridge | Wellcome Sanger Institute for the COVID-19 Genomics UK Consortium | Luke W Meredith, M. Estée Török , Myra Hosmillo, William L. Hamilton, Martin D. Curran, Theresa Feltwell, Grant Hall, Anna Yakovleva, Fahad A Khokhar, Charlotte J. Houldcroft, Laura G Caller, Aminu S. Jahun, Sarah L. Caddy, Ian Goodfellow, Alex Alderton, Roberto Amato, Sonia Goncalves, Ewan Harrison, David K. Jackson, Ian Johnston, Dominic Kwiatkowski, Cordelia Langford, John Sillitoe on behalf of the Wellcome Sanger Institute COVID-19 Surveillance Team (http://www.sanger.ac.uk/covid-team) |
| EPI_ISL_440218, EPI_ISL_440219, EPI_ISL_440222 | PHE South West Regional Laboratory, National Infection Service | Wellcome Sanger Institute for the COVID-19 Genomics UK Consortium | Stephanie Hutchings, Hannah Pymont, Dr Peter Muir, Barry Vipond, Rich Hopes, Alex Alderton, Roberto Amato, Sonia Goncalves, Ewan Harrison, David K. Jackson, Ian Johnston, Dominic Kwiatkowski, Cordelia Langford, John Sillitoe on behalf of the Wellcome Sanger Institute COVID-19 Surveillance Team (http://www.sanger.ac.uk/covid-team) |
| EPI_ISL_440223 | Department of Pathology, University of Cambridge | Wellcome Sanger Institute for the COVID-19 Genomics UK Consortium | Luke W Meredith, M. Estée Török , Myra Hosmillo, William L. Hamilton, Martin D. Curran, Theresa Feltwell, Grant Hall, Anna Yakovleva, Fahad A Khokhar, Charlotte J. Houldcroft, Laura G Caller, Aminu S. Jahun, Sarah L. Caddy, Ian Goodfellow, Alex Alderton, Roberto Amato, Sonia Goncalves, Ewan Harrison, David K. Jackson, Ian Johnston, Dominic Kwiatkowski, Cordelia Langford, John Sillitoe on behalf of the Wellcome Sanger Institute COVID-19 Surveillance Team (http://www.sanger.ac.uk/covid-team) |
| EPI_ISL_440224, EPI_ISL_440225 | PHE South West Regional Laboratory, National Infection Service | Wellcome Sanger Institute for the COVID-19 Genomics UK Consortium | Stephanie Hutchings, Hannah Pymont, Dr Peter Muir, Barry Vipond, Rich Hopes, Alex Alderton, Roberto Amato, Sonia Goncalves, Ewan Harrison, David K. Jackson, Ian Johnston, Dominic Kwiatkowski, Cordelia Langford, John Sillitoe on behalf of the Wellcome Sanger Institute COVID-19 Surveillance Team (http://www.sanger.ac.uk/covid-team) |
| EPI_ISL_440226, EPI_ISL_440231 | Department of Pathology, University of Cambridge | Wellcome Sanger Institute for the COVID-19 Genomics UK Consortium | Luke W Meredith, M. Estée Török , Myra Hosmillo, William L. Hamilton, Martin D. Curran, Theresa Feltwell, Grant Hall, Anna Yakovleva, Fahad A Khokhar, Charlotte J. Houldcroft, Laura G Caller, Aminu S. Jahun, Sarah L. Caddy, Ian Goodfellow, Alex Alderton, Roberto Amato, Sonia Goncalves, Ewan Harrison, David K. Jackson, Ian Johnston, Dominic Kwiatkowski, Cordelia Langford, John Sillitoe on behalf of the Wellcome Sanger Institute COVID-19 Surveillance Team (http://www.sanger.ac.uk/covid-team) |
| EPI_ISL_440232, EPI_ISL_440233 | PHE South West Regional Laboratory, National Infection Service | Wellcome Sanger Institute for the COVID-19 Genomics UK Consortium | Stephanie Hutchings, Hannah Pymont, Dr Peter Muir, Barry Vipond, Rich Hopes, Alex Alderton, Roberto Amato, Sonia Goncalves, Ewan Harrison, David K. Jackson, Ian Johnston, Dominic Kwiatkowski, Cordelia Langford, John Sillitoe on behalf of the Wellcome Sanger Institute COVID-19 Surveillance Team (http://www.sanger.ac.uk/covid-team) |
| EPI_ISL_440234 | Department of Pathology, University of Cambridge | Wellcome Sanger Institute for the COVID-19 Genomics UK Consortium | Luke W Meredith, M. Estée Török , Myra Hosmillo, William L. Hamilton, Martin D. Curran, Theresa Feltwell, Grant Hall, Anna Yakovleva, Fahad A Khokhar, Charlotte J. Houldcroft, Laura G Caller, Aminu S. Jahun, Sarah L. Caddy, Ian Goodfellow, Alex Alderton, Roberto Amato, Sonia Goncalves, Ewan Harrison, David K. Jackson, Ian Johnston, Dominic Kwiatkowski, Cordelia Langford, John Sillitoe on behalf of the Wellcome Sanger Institute COVID-19 Surveillance Team (http://www.sanger.ac.uk/covid-team) |
| EPI_ISL_440236 | PHE South West Regional Laboratory, National Infection Service | Wellcome Sanger Institute for the COVID-19 Genomics UK Consortium | Stephanie Hutchings, Hannah Pymont, Dr Peter Muir, Barry Vipond, Rich Hopes, Alex Alderton, Roberto Amato, Sonia Goncalves, Ewan Harrison, David K. Jackson, Ian Johnston, Dominic Kwiatkowski, Cordelia Langford, John Sillitoe on behalf of the Wellcome Sanger Institute COVID-19 Surveillance Team (http://www.sanger.ac.uk/covid-team) |
| EPI_ISL_440237, EPI_ISL_440239 | Department of Pathology, University of Cambridge | Wellcome Sanger Institute for the COVID-19 Genomics UK Consortium | Luke W Meredith, M. Estée Török , Myra Hosmillo, William L. Hamilton, Martin D. Curran, Theresa Feltwell, Grant Hall, Anna Yakovleva, Fahad A Khokhar, Charlotte J. Houldcroft, Laura G Caller, Aminu S. Jahun, Sarah L. Caddy, Ian Goodfellow, Alex Alderton, Roberto Amato, Sonia Goncalves, Ewan Harrison, David K. Jackson, Ian Johnston, Dominic Kwiatkowski, Cordelia Langford, John Sillitoe on behalf of the Wellcome Sanger Institute COVID-19 Surveillance Team (http://www.sanger.ac.uk/covid-team) |
| EPI_ISL_440241, EPI_ISL_440243 | PHE South West Regional Laboratory, National Infection Service | Wellcome Sanger Institute for the COVID-19 Genomics UK Consortium | Stephanie Hutchings, Hannah Pymont, Dr Peter Muir, Barry Vipond, Rich Hopes, Alex Alderton, Roberto Amato, Sonia Goncalves, Ewan Harrison, David K. Jackson, Ian Johnston, Dominic Kwiatkowski, Cordelia Langford, John Sillitoe on behalf of the Wellcome Sanger Institute COVID-19 Surveillance Team (http://www.sanger.ac.uk/covid-team) |
| EPI_ISL_440244, EPI_ISL_440246 | Department of Pathology, University of Cambridge | Wellcome Sanger Institute for the COVID-19 Genomics UK Consortium | Luke W Meredith, M. Estée Török , Myra Hosmillo, William L. Hamilton, Martin D. Curran, Theresa Feltwell, Grant Hall, Anna Yakovleva, Fahad A Khokhar, Charlotte J. Houldcroft, Laura G Caller, Aminu S. Jahun, Sarah L. Caddy, Ian Goodfellow, Alex Alderton, Roberto Amato, Sonia Goncalves, Ewan Harrison, David K. Jackson, Ian Johnston, Dominic Kwiatkowski, Cordelia Langford, John Sillitoe on behalf of the Wellcome Sanger Institute COVID-19 Surveillance Team (http://www.sanger.ac.uk/covid-team) |
| EPI_ISL_440248, EPI_ISL_440250 | PHE South West Regional Laboratory, National Infection Service | Wellcome Sanger Institute for the COVID-19 Genomics UK Consortium | Stephanie Hutchings, Hannah Pymont, Dr Peter Muir, Barry Vipond, Rich Hopes, Alex Alderton, Roberto Amato, Sonia Goncalves, Ewan Harrison, David K. Jackson, Ian Johnston, Dominic Kwiatkowski, Cordelia Langford, John Sillitoe on behalf of the Wellcome Sanger Institute COVID-19 Surveillance Team (http://www.sanger.ac.uk/covid-team) |
| EPI_ISL_440254 | Department of Pathology, University of Cambridge | Wellcome Sanger Institute for the COVID-19 Genomics UK Consortium | Luke W Meredith, M. Estée Török , Myra Hosmillo, William L. Hamilton, Martin D. Curran, Theresa Feltwell, Grant Hall, Anna Yakovleva, Fahad A Khokhar, Charlotte J. Houldcroft, Laura G Caller, Aminu S. Jahun, Sarah L. Caddy, Ian Goodfellow, Alex Alderton, Roberto Amato, Sonia Goncalves, Ewan Harrison, David K. Jackson, Ian Johnston, Dominic Kwiatkowski, Cordelia Langford, John Sillitoe on behalf of the Wellcome Sanger Institute COVID-19 Surveillance Team (http://www.sanger.ac.uk/covid-team) |
| EPI_ISL_440255, EPI_ISL_440256 | PHE South West Regional Laboratory, National Infection Service | Wellcome Sanger Institute for the COVID-19 Genomics UK Consortium | Stephanie Hutchings, Hannah Pymont, Dr Peter Muir, Barry Vipond, Rich Hopes, Alex Alderton, Roberto Amato, Sonia Goncalves, Ewan Harrison, David K. Jackson, Ian Johnston, Dominic Kwiatkowski, Cordelia Langford, John Sillitoe on behalf of the Wellcome Sanger Institute COVID-19 Surveillance Team (http://www.sanger.ac.uk/covid-team) |
| EPI_ISL_440257, EPI_ISL_440258, EPI_ISL_440259, EPI_ISL_440260, EPI_ISL_440261, EPI_ISL_440264, EPI_ISL_440265, EPI_ISL_440267, EPI_ISL_440268 | Department of Pathology, University of Cambridge | Wellcome Sanger Institute for the COVID-19 Genomics UK Consortium | Luke W Meredith, M. Estée Török , Myra Hosmillo, William L. Hamilton, Martin D. Curran, Theresa Feltwell, Grant Hall, Anna Yakovleva, Fahad A Khokhar, Charlotte J. Houldcroft, Laura G Caller, Aminu S. Jahun, Sarah L. Caddy, Ian Goodfellow, Alex Alderton, Roberto Amato, Sonia Goncalves, Ewan Harrison, David K. Jackson, Ian Johnston, Dominic Kwiatkowski, Cordelia Langford, John Sillitoe on behalf of the Wellcome Sanger Institute COVID-19 Surveillance Team (http://www.sanger.ac.uk/covid-team) |
| EPI_ISL_440269 | PHE South West Regional Laboratory, National Infection Service | Wellcome Sanger Institute for the COVID-19 Genomics UK Consortium | Stephanie Hutchings, Hannah Pymont, Dr Peter Muir, Barry Vipond, Rich Hopes, Alex Alderton, Roberto Amato, Sonia Goncalves, Ewan Harrison, David K. Jackson, Ian Johnston, Dominic Kwiatkowski, Cordelia Langford, John Sillitoe on behalf of the Wellcome Sanger Institute COVID-19 Surveillance Team (http://www.sanger.ac.uk/covid-team) |
| EPI_ISL_440270, EPI_ISL_440271 | Department of Pathology, University of Cambridge | Wellcome Sanger Institute for the COVID-19 Genomics UK Consortium | Luke W Meredith, M. Estée Török , Myra Hosmillo, William L. Hamilton, Martin D. Curran, Theresa Feltwell, Grant Hall, Anna Yakovleva, Fahad A Khokhar, Charlotte J. Houldcroft, Laura G Caller, Aminu S. Jahun, Sarah L. Caddy, Ian Goodfellow, Alex Alderton, Roberto Amato, Sonia Goncalves, Ewan Harrison, David K. Jackson, Ian Johnston, Dominic Kwiatkowski, Cordelia Langford, John Sillitoe on behalf of the Wellcome Sanger Institute COVID-19 Surveillance Team (http://www.sanger.ac.uk/covid-team) |
| EPI_ISL_440272, EPI_ISL_440275 | PHE South West Regional Laboratory, National Infection Service | Wellcome Sanger Institute for the COVID-19 Genomics UK Consortium | Stephanie Hutchings, Hannah Pymont, Dr Peter Muir, Barry Vipond, Rich Hopes, Alex Alderton, Roberto Amato, Sonia Goncalves, Ewan Harrison, David K. Jackson, Ian Johnston, Dominic Kwiatkowski, Cordelia Langford, John Sillitoe on behalf of the Wellcome Sanger Institute COVID-19 Surveillance Team (http://www.sanger.ac.uk/covid-team) |
| EPI_ISL_440278, EPI_ISL_440279, EPI_ISL_440280, EPI_ISL_440282, EPI_ISL_440283, EPI_ISL_440284, EPI_ISL_440285 | Department of Pathology, University of Cambridge | Wellcome Sanger Institute for the COVID-19 Genomics UK Consortium | Luke W Meredith, M. Estée Török , Myra Hosmillo, William L. Hamilton, Martin D. Curran, Theresa Feltwell, Grant Hall, Anna Yakovleva, Fahad A Khokhar, Charlotte J. Houldcroft, Laura G Caller, Aminu S. Jahun, Sarah L. Caddy, Ian Goodfellow, Alex Alderton, Roberto Amato, Sonia Goncalves, Ewan Harrison, David K. Jackson, Ian Johnston, Dominic Kwiatkowski, Cordelia Langford, John Sillitoe on behalf of the Wellcome Sanger Institute COVID-19 Surveillance Team (http://www.sanger.ac.uk/covid-team) |
| EPI_ISL_440286 | PHE South West Regional Laboratory, National Infection Service | Wellcome Sanger Institute for the COVID-19 Genomics UK Consortium | Stephanie Hutchings, Hannah Pymont, Dr Peter Muir, Barry Vipond, Rich Hopes, Alex Alderton, Roberto Amato, Sonia Goncalves, Ewan Harrison, David K. Jackson, Ian Johnston, Dominic Kwiatkowski, Cordelia Langford, John Sillitoe on behalf of the Wellcome Sanger Institute COVID-19 Surveillance Team (http://www.sanger.ac.uk/covid-team) |
| EPI_ISL_440287, EPI_ISL_440288, EPI_ISL_440289, EPI_ISL_440290 | Department of Pathology, University of Cambridge | Wellcome Sanger Institute for the COVID-19 Genomics UK Consortium | Luke W Meredith, M. Estée Török , Myra Hosmillo, William L. Hamilton, Martin D. Curran, Theresa Feltwell, Grant Hall, Anna Yakovleva, Fahad A Khokhar, Charlotte J. Houldcroft, Laura G Caller, Aminu S. Jahun, Sarah L. Caddy, Ian Goodfellow, Alex Alderton, Roberto Amato, Sonia Goncalves, Ewan Harrison, David K. Jackson, Ian Johnston, Dominic Kwiatkowski, Cordelia Langford, John Sillitoe on behalf of the Wellcome Sanger Institute COVID-19 Surveillance Team (http://www.sanger.ac.uk/covid-team) |
| EPI_ISL_440291 | PHE South West Regional Laboratory, National Infection Service | Wellcome Sanger Institute for the COVID-19 Genomics UK Consortium | Stephanie Hutchings, Hannah Pymont, Dr Peter Muir, Barry Vipond, Rich Hopes, Alex Alderton, Roberto Amato, Sonia Goncalves, Ewan Harrison, David K. Jackson, Ian Johnston, Dominic Kwiatkowski, Cordelia Langford, John Sillitoe on behalf of the Wellcome Sanger Institute COVID-19 Surveillance Team (http://www.sanger.ac.uk/covid-team) |
| EPI_ISL_440293, EPI_ISL_440295, EPI_ISL_440296, EPI_ISL_440297 | Department of Pathology, University of Cambridge | Wellcome Sanger Institute for the COVID-19 Genomics UK Consortium | Luke W Meredith, M. Estée Török , Myra Hosmillo, William L. Hamilton, Martin D. Curran, Theresa Feltwell, Grant Hall, Anna Yakovleva, Fahad A Khokhar, Charlotte J. Houldcroft, Laura G Caller, Aminu S. Jahun, Sarah L. Caddy, Ian Goodfellow, Alex Alderton, Roberto Amato, Sonia Goncalves, Ewan Harrison, David K. Jackson, Ian Johnston, Dominic Kwiatkowski, Cordelia Langford, John Sillitoe on behalf of the Wellcome Sanger Institute COVID-19 Surveillance Team (http://www.sanger.ac.uk/covid-team) |
| EPI_ISL_440298, EPI_ISL_440299 | PHE South West Regional Laboratory, National Infection Service | Wellcome Sanger Institute for the COVID-19 Genomics UK Consortium | Stephanie Hutchings, Hannah Pymont, Dr Peter Muir, Barry Vipond, Rich Hopes, Alex Alderton, Roberto Amato, Sonia Goncalves, Ewan Harrison, David K. Jackson, Ian Johnston, Dominic Kwiatkowski, Cordelia Langford, John Sillitoe on behalf of the Wellcome Sanger Institute COVID-19 Surveillance Team (http://www.sanger.ac.uk/covid-team) |
| EPI_ISL_440300, EPI_ISL_440301, EPI_ISL_440302, EPI_ISL_440303, EPI_ISL_440304, EPI_ISL_440305, EPI_ISL_440306, EPI_ISL_440307, EPI_ISL_440309, EPI_ISL_440310, EPI_ISL_440311, EPI_ISL_440313, EPI_ISL_440314, EPI_ISL_440315, EPI_ISL_440318, EPI_ISL_440319, EPI_ISL_440321, EPI_ISL_440323, EPI_ISL_440328, EPI_ISL_440329, EPI_ISL_440330, EPI_ISL_440333, EPI_ISL_440335, EPI_ISL_440337, EPI_ISL_440338, EPI_ISL_440340, EPI_ISL_440341, EPI_ISL_440342, EPI_ISL_440343, EPI_ISL_440344, EPI_ISL_440345, EPI_ISL_440347, EPI_ISL_440348, EPI_ISL_440351, EPI_ISL_440352, EPI_ISL_440354, EPI_ISL_440356, EPI_ISL_440357, EPI_ISL_440358, EPI_ISL_440359, EPI_ISL_440361, EPI_ISL_440364, EPI_ISL_440365, EPI_ISL_440367, EPI_ISL_440369, EPI_ISL_440373, EPI_ISL_440374, EPI_ISL_440375, | | | |

| | | | |
|---|---|---|---|
| EPI_ISL_440376, EPI_ISL_440378, EPI_ISL_440379, EPI_ISL_440380, EPI_ISL_440381, EPI_ISL_440382, EPI_ISL_440384, EPI_ISL_440386, EPI_ISL_440387, EPI_ISL_440389, EPI_ISL_440391, EPI_ISL_440393, EPI_ISL_440395, EPI_ISL_440396, EPI_ISL_440397, EPI_ISL_440399, EPI_ISL_440400, EPI_ISL_440401, EPI_ISL_440402, EPI_ISL_440403, EPI_ISL_440404, EPI_ISL_440405, EPI_ISL_440407, EPI_ISL_440409, EPI_ISL_440411, EPI_ISL_440413, EPI_ISL_440414, EPI_ISL_440417, EPI_ISL_440418, EPI_ISL_440419, EPI_ISL_440420, EPI_ISL_440421, EPI_ISL_440422, EPI_ISL_440426, EPI_ISL_440428, EPI_ISL_440429, EPI_ISL_440430, EPI_ISL_440431, EPI_ISL_440432, EPI_ISL_440433, EPI_ISL_440434, EPI_ISL_440436, EPI_ISL_440437, EPI_ISL_440438, EPI_ISL_440439, EPI_ISL_440445, EPI_ISL_440451, EPI_ISL_440459, EPI_ISL_440460, EPI_ISL_440464, EPI_ISL_440480, EPI_ISL_440483, EPI_ISL_440486, EPI_ISL_440487, EPI_ISL_440489, EPI_ISL_440491, EPI_ISL_440492, EPI_ISL_440494, EPI_ISL_440501, EPI_ISL_440503, EPI_ISL_440506, EPI_ISL_440507, EPI_ISL_440510, EPI_ISL_440511, EPI_ISL_440515, EPI_ISL_440516, EPI_ISL_440518, EPI_ISL_440519, EPI_ISL_440521, EPI_ISL_440525, EPI_ISL_440526, EPI_ISL_440532, EPI_ISL_440533, EPI_ISL_440534, EPI_ISL_440538, EPI_ISL_440539, EPI_ISL_440541, EPI_ISL_440542, EPI_ISL_440547, EPI_ISL_440551, EPI_ISL_440552, EPI_ISL_440553, EPI_ISL_440555, EPI_ISL_440557, EPI_ISL_440558, EPI_ISL_440560, EPI_ISL_440561, EPI_ISL_440562, EPI_ISL_440563, EPI_ISL_440564, EPI_ISL_440565, EPI_ISL_440566, EPI_ISL_440567, EPI_ISL_440568, EPI_ISL_440569, EPI_ISL_440570, EPI_ISL_440571, EPI_ISL_440573, EPI_ISL_440574, EPI_ISL_440576, EPI_ISL_440577, EPI_ISL_440579, EPI_ISL_440581, EPI_ISL_440582, EPI_ISL_440583, EPI_ISL_440584, EPI_ISL_440586, EPI_ISL_440589, EPI_ISL_440590, EPI_ISL_440591, EPI_ISL_440592, EPI_ISL_440593, EPI_ISL_440595, EPI_ISL_440597, EPI_ISL_440598, EPI_ISL_440599, EPI_ISL_440600, EPI_ISL_440601, EPI_ISL_440603, EPI_ISL_440604, EPI_ISL_440605, EPI_ISL_440606, EPI_ISL_440607, EPI_ISL_440608, EPI_ISL_440610, EPI_ISL_440611, EPI_ISL_440612, EPI_ISL_440613, EPI_ISL_440615, EPI_ISL_440616, EPI_ISL_440617, EPI_ISL_440618, EPI_ISL_440619, EPI_ISL_440620, EPI_ISL_440621, EPI_ISL_440622 | | | |
| see above | Department of Pathology, University of Cambridge | Wellcome Sanger Institute for the COVID-19 Genomics UK Consortium | Luke W Meredith, M. Estée Török , Myra Hosmillo, William L. Hamilton, Martin D. Curran, Theresa Feltwell, Grant Hall, Anna Yakovleva, Fahad A Khokhar, Charlotte J. Houldcroft, Laura G Caller, Aminu S. Jahun, Sarah L. Caddy, Ian Goodfellow, Alex Alderton, Roberto Amato, Sonia Goncalves, Ewan Harrison, David K. Jackson, Ian Johnston, Dominic Kwiatkowski, Cordelia Langford, John Sillitoe on behalf of the Wellcome Sanger Institute COVID-19 Surveillance Team (http://www.sanger.ac.uk/covid-team) |
| EPI_ISL_440624, EPI_ISL_440626, EPI_ISL_440628, EPI_ISL_440632, EPI_ISL_440634, EPI_ISL_440635, EPI_ISL_440636, EPI_ISL_440639, EPI_ISL_440640, EPI_ISL_440641, EPI_ISL_440642, EPI_ISL_440643, EPI_ISL_440645, EPI_ISL_440646, EPI_ISL_440647, EPI_ISL_440649, EPI_ISL_440650, EPI_ISL_440653, EPI_ISL_440655, EPI_ISL_440656, EPI_ISL_440657, EPI_ISL_440658, EPI_ISL_440659, EPI_ISL_440660, EPI_ISL_440662, EPI_ISL_440663, EPI_ISL_440664, EPI_ISL_440665, EPI_ISL_440666, EPI_ISL_440667, EPI_ISL_440670, EPI_ISL_440671, EPI_ISL_440673, EPI_ISL_440674, EPI_ISL_440676, EPI_ISL_440679, EPI_ISL_440681, EPI_ISL_440682, EPI_ISL_440683, EPI_ISL_440684, EPI_ISL_440685, EPI_ISL_440688, EPI_ISL_440689, EPI_ISL_440690, EPI_ISL_440691, EPI_ISL_440692, EPI_ISL_440694, EPI_ISL_440695, EPI_ISL_440697, EPI_ISL_440698, EPI_ISL_440699, EPI_ISL_440700, EPI_ISL_440701, EPI_ISL_440702, EPI_ISL_440703, EPI_ISL_440704, EPI_ISL_440705, EPI_ISL_440707, EPI_ISL_440710, EPI_ISL_440711, EPI_ISL_440712, EPI_ISL_440715, EPI_ISL_440716, EPI_ISL_440717, EPI_ISL_440718, EPI_ISL_440719, EPI_ISL_440721, EPI_ISL_440723, EPI_ISL_440724, EPI_ISL_440726, EPI_ISL_440728, EPI_ISL_440729, EPI_ISL_440730, EPI_ISL_440732, EPI_ISL_440733, EPI_ISL_440734, EPI_ISL_440735, EPI_ISL_440736, EPI_ISL_440737, EPI_ISL_440738, EPI_ISL_440739, EPI_ISL_440740, EPI_ISL_440741, EPI_ISL_440742, EPI_ISL_440744, EPI_ISL_440745, EPI_ISL_440746, EPI_ISL_440747, EPI_ISL_440751, EPI_ISL_440752, EPI_ISL_440753, EPI_ISL_440754, EPI_ISL_440755, EPI_ISL_440757, EPI_ISL_440758, EPI_ISL_440759, EPI_ISL_440761, EPI_ISL_440762, EPI_ISL_440763, EPI_ISL_440764, EPI_ISL_440765, EPI_ISL_440766, EPI_ISL_440767, EPI_ISL_440768, EPI_ISL_440770, EPI_ISL_440773, EPI_ISL_440774, EPI_ISL_440775, EPI_ISL_440776, EPI_ISL_440777, EPI_ISL_440779, EPI_ISL_440781, EPI_ISL_440783, EPI_ISL_440784, EPI_ISL_440785, EPI_ISL_440787, EPI_ISL_440788, EPI_ISL_440789, EPI_ISL_440790, EPI_ISL_440791, EPI_ISL_440793, EPI_ISL_440794, EPI_ISL_440795, EPI_ISL_440796, EPI_ISL_440799, EPI_ISL_440800, EPI_ISL_440801, EPI_ISL_440802, EPI_ISL_440803, EPI_ISL_440804, EPI_ISL_440806, EPI_ISL_440807, EPI_ISL_440808, EPI_ISL_440809 | | | |
| see above | PHE South West Regional Laboratory, National Infection Service | Wellcome Sanger Institute for the COVID-19 Genomics UK Consortium | Stephanie Hutchings, Hannah Pymont, Dr Peter Muir, Barry Vipond, Rich Hopes, Alex Alderton, Roberto Amato, Sonia Goncalves, Ewan Harrison, David K. Jackson, Ian Johnston, Dominic Kwiatkowski, Cordelia Langford, John Sillitoe on behalf of the Wellcome Sanger Institute COVID-19 Surveillance Team (http://www.sanger.ac.uk/covid-team) |
| EPI_ISL_440814, EPI_ISL_440832, EPI_ISL_440839 | Department of Pathology, University of Cambridge | Wellcome Sanger Institute for the COVID-19 Genomics UK Consortium | Luke W Meredith, M. Estée Török , Myra Hosmillo, William L. Hamilton, Martin D. Curran, Theresa Feltwell, Grant Hall, Anna Yakovleva, Fahad A Khokhar, Charlotte J. Houldcroft, Laura G Caller, Aminu S. Jahun, Sarah L. Caddy, Ian Goodfellow, Alex Alderton, Roberto Amato, Sonia Goncalves, Ewan Harrison, David K. Jackson, Ian Johnston, Dominic Kwiatkowski, Cordelia Langford, John Sillitoe on behalf of the Wellcome Sanger Institute COVID-19 Surveillance Team (http://www.sanger.ac.uk/covid-team) |
| EPI_ISL_440853, EPI_ISL_440863, EPI_ISL_440864, EPI_ISL_440866, EPI_ISL_440867, EPI_ISL_440873, EPI_ISL_440875, EPI_ISL_440876, EPI_ISL_440878, EPI_ISL_440881, EPI_ISL_440883, EPI_ISL_440884, EPI_ISL_440886, EPI_ISL_440887, EPI_ISL_440892, EPI_ISL_440894, EPI_ISL_440895, EPI_ISL_440897, EPI_ISL_440898, EPI_ISL_440900, EPI_ISL_440903, EPI_ISL_440904, EPI_ISL_440905, EPI_ISL_440910, EPI_ISL_440912, EPI_ISL_440917, EPI_ISL_440922, EPI_ISL_440923, EPI_ISL_440924, EPI_ISL_440925, EPI_ISL_440927, EPI_ISL_440930, EPI_ISL_440935, EPI_ISL_440936, EPI_ISL_440939, EPI_ISL_440946, EPI_ISL_440948, EPI_ISL_440949 | | | |
| see above | Liverpool Clinical Laboratories | COVID-19 Genomics UK (COG-UK) Consortium | Sam Haldenby, Anita Lucaci, Steve Paterson, Julian Hiscox, Alistair Darby, M Almsaud, A Alrezaihi, Muhannad Alruwaili, Stuart D Armstrong, Jones Benjamin , Eleanor G Bentley, Anu Chawla, Jordan J Clark, Angela Cowell, Richard Eccles, Isabel Garc√≠a-Dorival, Matthew Gemmell, Alessandro Gerada, PKF Gilmore, Richard Gregory, Ximeng Han, Catherine Hartley, Margaret Hughes, Miren Iturriza-Gomara, James Johnson, L Luu, Jenifer Manson , Charlotte Nelson, Elaine O‚ÄôToole, Cassie Olateju, Rebekah Penrice-Randal¬†, Lucille Rainbow, N.P Randle, Trevor Ian Robinson, Parul Sharma, Ghada T Shawli, James P Stewart , Neil Swainston, Ecaterina Vamos, Joanne Watts, Mark Whitehead |
| EPI_ISL_440951, EPI_ISL_440952, EPI_ISL_440953, EPI_ISL_440954, EPI_ISL_440956, EPI_ISL_440957, EPI_ISL_440958, EPI_ISL_440960, EPI_ISL_440962, EPI_ISL_440964, EPI_ISL_440965, EPI_ISL_440966, EPI_ISL_440967, EPI_ISL_440968, EPI_ISL_440969, EPI_ISL_440970, EPI_ISL_440971, EPI_ISL_440972, EPI_ISL_440973, EPI_ISL_440975, EPI_ISL_440976, EPI_ISL_440980, EPI_ISL_440981, EPI_ISL_440983, EPI_ISL_440984, EPI_ISL_440985, EPI_ISL_440986, EPI_ISL_440987, EPI_ISL_440988, EPI_ISL_440989, EPI_ISL_440990, EPI_ISL_440991, EPI_ISL_440993, EPI_ISL_440994, EPI_ISL_440995, EPI_ISL_440996, EPI_ISL_440997, EPI_ISL_440998, EPI_ISL_440999, EPI_ISL_441000, EPI_ISL_441001, EPI_ISL_441002, EPI_ISL_441003, EPI_ISL_441004, EPI_ISL_441005, EPI_ISL_441006, EPI_ISL_441007, EPI_ISL_441008, EPI_ISL_441009, EPI_ISL_441010, EPI_ISL_441011, EPI_ISL_441012, EPI_ISL_441013, EPI_ISL_441014, EPI_ISL_441015, EPI_ISL_441016, EPI_ISL_441017, EPI_ISL_441018, EPI_ISL_441019, EPI_ISL_441020, EPI_ISL_441021, EPI_ISL_441022, EPI_ISL_441023, EPI_ISL_441024, EPI_ISL_441025, EPI_ISL_441026, EPI_ISL_441027, EPI_ISL_441028, EPI_ISL_441029, EPI_ISL_441030, EPI_ISL_441031, EPI_ISL_441032, EPI_ISL_441033, EPI_ISL_441034, EPI_ISL_441035, EPI_ISL_441036, EPI_ISL_441037, EPI_ISL_441038, EPI_ISL_441039, EPI_ISL_441040, EPI_ISL_441041, EPI_ISL_441042, EPI_ISL_441043, EPI_ISL_441045, EPI_ISL_441046, EPI_ISL_441047, EPI_ISL_441049, EPI_ISL_441050, EPI_ISL_441051 | | | |
| see above | University College London, Great Ormond Street Hospital for Children NHS Foundation Trust, Imperial College Healthcare NHS Trust | COVID-19 Genomics UK (COG-UK) Consortium | Sergi Castellano, Rachel Williams, Mark Kristiansen, Paola Resende Silva, Sunando Roy, Tony Brooks, Helena Tutill, Paola Niola, Patricia Dyal, Charlotte Williams, Leysa Forrest, Yasmin Panchbhaya, Jacqueline Findlay, Sam Weeks, Julianne Brown, Kathryn Harris, Paul Randell, James Price, Alison Holmes, Judith Breuer |
| EPI_ISL_441053, EPI_ISL_441054, EPI_ISL_441055, EPI_ISL_441057, EPI_ISL_441058, EPI_ISL_441059, EPI_ISL_441062, EPI_ISL_441063, EPI_ISL_441064, EPI_ISL_441066, EPI_ISL_441068, EPI_ISL_441070, EPI_ISL_441072, EPI_ISL_441073, EPI_ISL_441074, EPI_ISL_441075, EPI_ISL_441076, EPI_ISL_441077, EPI_ISL_441079, EPI_ISL_441081, EPI_ISL_441082, EPI_ISL_441083, EPI_ISL_441084, EPI_ISL_441085, EPI_ISL_441086, EPI_ISL_441088, EPI_ISL_441089, EPI_ISL_441090, EPI_ISL_441091, EPI_ISL_441092, EPI_ISL_441093, EPI_ISL_441095, EPI_ISL_441096, EPI_ISL_441098, EPI_ISL_441099, EPI_ISL_441100, EPI_ISL_441101, EPI_ISL_441102, EPI_ISL_441103, EPI_ISL_441104, EPI_ISL_441107, EPI_ISL_441108, EPI_ISL_441110, EPI_ISL_441111, EPI_ISL_441112, EPI_ISL_441114, EPI_ISL_441115, EPI_ISL_441118, EPI_ISL_441119, EPI_ISL_441120, EPI_ISL_441122, EPI_ISL_441123, EPI_ISL_441124, EPI_ISL_441125, EPI_ISL_441126, EPI_ISL_441127, EPI_ISL_441128, EPI_ISL_441130, EPI_ISL_441131, EPI_ISL_441132, EPI_ISL_441135, EPI_ISL_441136, EPI_ISL_441137, EPI_ISL_441138, EPI_ISL_441139, EPI_ISL_441140, EPI_ISL_441141, EPI_ISL_441142, EPI_ISL_441143, EPI_ISL_441144, EPI_ISL_441145, EPI_ISL_441146, EPI_ISL_441147, EPI_ISL_441148, EPI_ISL_441150, EPI_ISL_441152, EPI_ISL_441153, EPI_ISL_441154, EPI_ISL_441155, EPI_ISL_441156, EPI_ISL_441157, EPI_ISL_441158, EPI_ISL_441160, EPI_ISL_441161, EPI_ISL_441162, EPI_ISL_441164, EPI_ISL_441165, EPI_ISL_441166, EPI_ISL_441167, EPI_ISL_441169, EPI_ISL_441170, EPI_ISL_441171, EPI_ISL_441172, EPI_ISL_441173, EPI_ISL_441174, EPI_ISL_441175, EPI_ISL_441176, EPI_ISL_441177, EPI_ISL_441178, EPI_ISL_441179, EPI_ISL_441180, EPI_ISL_441183, EPI_ISL_441187, EPI_ISL_441188, EPI_ISL_441191, EPI_ISL_441192, EPI_ISL_441194, EPI_ISL_441195, EPI_ISL_441196, EPI_ISL_441198, EPI_ISL_441200, EPI_ISL_441201, EPI_ISL_441202, EPI_ISL_441203, EPI_ISL_441204, EPI_ISL_441205, EPI_ISL_441206, EPI_ISL_441207, EPI_ISL_441208, EPI_ISL_441209, EPI_ISL_441210, EPI_ISL_441211, EPI_ISL_441213, EPI_ISL_441214, EPI_ISL_441215, EPI_ISL_441216, EPI_ISL_441217, EPI_ISL_441218, EPI_ISL_441220, EPI_ISL_441221, EPI_ISL_441222, EPI_ISL_441223, EPI_ISL_441224, EPI_ISL_441225, EPI_ISL_441226, EPI_ISL_441227, EPI_ISL_441228, EPI_ISL_441229, EPI_ISL_441231, EPI_ISL_441232, EPI_ISL_441234, EPI_ISL_441236, EPI_ISL_441237, EPI_ISL_441239, EPI_ISL_441240, EPI_ISL_441242, EPI_ISL_441243, EPI_ISL_441244, EPI_ISL_441245, EPI_ISL_441247, EPI_ISL_441249, EPI_ISL_441250, EPI_ISL_441251, EPI_ISL_441253, EPI_ISL_441254, EPI_ISL_441255, EPI_ISL_441256, EPI_ISL_441259, EPI_ISL_441260, EPI_ISL_441262, EPI_ISL_441266, EPI_ISL_441268, EPI_ISL_441269, EPI_ISL_441270, EPI_ISL_441271, EPI_ISL_441272, EPI_ISL_441274, EPI_ISL_441278, EPI_ISL_441280, EPI_ISL_441281, EPI_ISL_441282, EPI_ISL_441283, EPI_ISL_441284, EPI_ISL_441286, EPI_ISL_441288, EPI_ISL_441289, EPI_ISL_441290, EPI_ISL_441292, EPI_ISL_441293, EPI_ISL_441294, EPI_ISL_441295, EPI_ISL_441296, EPI_ISL_441297, EPI_ISL_441298, EPI_ISL_441299, EPI_ISL_441300, EPI_ISL_441301, EPI_ISL_441302, EPI_ISL_441303, EPI_ISL_441304, EPI_ISL_441305, EPI_ISL_441307, EPI_ISL_441308, EPI_ISL_441309, EPI_ISL_441310, EPI_ISL_441311, EPI_ISL_441312, EPI_ISL_441313, EPI_ISL_441314, EPI_ISL_441315, EPI_ISL_441317, EPI_ISL_441319, EPI_ISL_441320, EPI_ISL_441321, EPI_ISL_441322, EPI_ISL_441325, EPI_ISL_441326, EPI_ISL_441328, EPI_ISL_441331, EPI_ISL_441332, EPI_ISL_441333, EPI_ISL_441334, EPI_ISL_441335, EPI_ISL_441337, EPI_ISL_441338, EPI_ISL_441339, EPI_ISL_441340, EPI_ISL_441341, EPI_ISL_441342, EPI_ISL_441344, EPI_ISL_441345, EPI_ISL_441346, EPI_ISL_441347, EPI_ISL_441349 | | | |
| see above | Department of Pathology, University of Cambridge | Wellcome Sanger Institute for the COVID-19 Genomics UK Consortium | Luke W Meredith, M. Estée Török , Myra Hosmillo, William L. Hamilton, Martin D. Curran, Theresa Feltwell, Grant Hall, Anna Yakovleva, Fahad A Khokhar, Charlotte J. Houldcroft, Laura G Caller, Aminu S. Jahun, Sarah L. Caddy, Ian Goodfellow, Alex Alderton, Roberto Amato, Sonia Goncalves, Ewan Harrison, David K. Jackson, Ian Johnston, Dominic Kwiatkowski, Cordelia Langford, John Sillitoe on behalf of the Wellcome Sanger Institute COVID-19 Surveillance Team (http://www.sanger.ac.uk/covid-team) |
| EPI_ISL_441355, EPI_ISL_441356, EPI_ISL_441357, EPI_ISL_441358, EPI_ISL_441359, EPI_ISL_441360, EPI_ISL_441361, EPI_ISL_441363, EPI_ISL_441364, EPI_ISL_441365, EPI_ISL_441366, EPI_ISL_441367, EPI_ISL_441368, EPI_ISL_441369, EPI_ISL_441370, EPI_ISL_441371, EPI_ISL_441372, EPI_ISL_441373, EPI_ISL_441375, EPI_ISL_441376, EPI_ISL_441378, EPI_ISL_441379, EPI_ISL_441380, EPI_ISL_441381, EPI_ISL_441382, EPI_ISL_441397, EPI_ISL_441398, EPI_ISL_441399, EPI_ISL_441401, EPI_ISL_441402, EPI_ISL_441403, EPI_ISL_441404, EPI_ISL_441405, EPI_ISL_441406, EPI_ISL_441407, EPI_ISL_441408, EPI_ISL_441409, EPI_ISL_441410, EPI_ISL_441411, EPI_ISL_441412, EPI_ISL_441414, EPI_ISL_441415, EPI_ISL_441416, EPI_ISL_441417, EPI_ISL_441418, EPI_ISL_441419, EPI_ISL_441420, EPI_ISL_441421, EPI_ISL_441422, EPI_ISL_441423, EPI_ISL_441424, EPI_ISL_441425, EPI_ISL_441426, EPI_ISL_441427, EPI_ISL_441428, EPI_ISL_441429, EPI_ISL_441430, EPI_ISL_441431, EPI_ISL_441432, EPI_ISL_441433, EPI_ISL_441434, EPI_ISL_441435, EPI_ISL_441436 | | | |
| see above | Regional Virus Laboratory, Belfast Health and Social Care Trust | COVID-19 Genomics UK (COG-UK) Consortium | Conall McCaughey, James McKenna, Tanya Curran, Susan Feeney, Alison Watt, Ciara Cox, Mairead Connor, Zoltan Molnar, David Simpson, Derek Fairley |
| EPI_ISL_441465, EPI_ISL_441478, EPI_ISL_441479 | Queens Medical Centre, Clinical Microbiology Department / DeepSeq Nottingham | COVID-19 Genomics UK (COG-UK) Consortium | Gemma Clark, Wendy Smith, Manjinder Khakh, Hannah Howson-Wells, Jonathan Ball, Patrick McClure, Joseph Chappell, Theocharis Tsoleridis, Nadine Holmes, Matthew Carlisle, Christopher Moore, Fei Sang, Johnny Debebe, Victoria Wright, Matthew Loose |
| EPI_ISL_441547, EPI_ISL_441549, EPI_ISL_441550, EPI_ISL_441551, EPI_ISL_441552, EPI_ISL_441554, EPI_ISL_441555, EPI_ISL_441556, EPI_ISL_441557, EPI_ISL_441558, EPI_ISL_441559, EPI_ISL_441560, EPI_ISL_441561, EPI_ISL_441562, EPI_ISL_441563, EPI_ISL_441564, EPI_ISL_441565, EPI_ISL_441566, EPI_ISL_441569, EPI_ISL_441571, EPI_ISL_441572, EPI_ISL_441573, EPI_ISL_441574, EPI_ISL_441575, EPI_ISL_441576, EPI_ISL_441577, EPI_ISL_441578, EPI_ISL_441579, EPI_ISL_441582, EPI_ISL_441583, EPI_ISL_441584, EPI_ISL_441585, EPI_ISL_441586, EPI_ISL_441587, EPI_ISL_441588, EPI_ISL_441590, EPI_ISL_441591, EPI_ISL_441593, EPI_ISL_441594, EPI_ISL_441595, EPI_ISL_441596, EPI_ISL_441597, EPI_ISL_441599, EPI_ISL_441600, EPI_ISL_441602, EPI_ISL_441603, EPI_ISL_441604, EPI_ISL_441605, EPI_ISL_441606, EPI_ISL_441607, EPI_ISL_441609, EPI_ISL_441610, EPI_ISL_441611, EPI_ISL_441612, EPI_ISL_441613, EPI_ISL_441614, EPI_ISL_441615, EPI_ISL_441616, EPI_ISL_441617, EPI_ISL_441618, EPI_ISL_441619, EPI_ISL_441621, EPI_ISL_441623, EPI_ISL_441624, EPI_ISL_441626, EPI_ISL_441627, EPI_ISL_441628, EPI_ISL_441629, EPI_ISL_441630, EPI_ISL_441631, EPI_ISL_441632, EPI_ISL_441633, EPI_ISL_441634, EPI_ISL_441635, EPI_ISL_441636, EPI_ISL_441637, EPI_ISL_441638, EPI_ISL_441639, EPI_ISL_441640, EPI_ISL_441641, EPI_ISL_441642, EPI_ISL_441643, EPI_ISL_441644, EPI_ISL_441645, EPI_ISL_441647, EPI_ISL_441648, EPI_ISL_441649, EPI_ISL_441650, EPI_ISL_441651, EPI_ISL_441653, EPI_ISL_441654, EPI_ISL_441656, EPI_ISL_441657, EPI_ISL_441658 | | | |
| see above | Department of Pathology, University of Cambridge | Wellcome Sanger Institute for the COVID-19 Genomics UK Consortium | Luke W Meredith, M. Estée Török , Myra Hosmillo, William L. Hamilton, Martin D. Curran, Theresa Feltwell, Grant Hall, Anna Yakovleva, Fahad A Khokhar, Charlotte J. Houldcroft, Laura G Caller, Aminu S. Jahun, Sarah L. Caddy, Ian Goodfellow, Alex Alderton, Roberto Amato, Sonia Goncalves, Ewan Harrison, David K. Jackson, Ian Johnston, Dominic Kwiatkowski, Cordelia Langford, John Sillitoe on behalf of the Wellcome Sanger Institute COVID-19 Surveillance Team (http://www.sanger.ac.uk/covid-team) |
| EPI_ISL_441659 | Regional Virus Laboratory, Belfast Health and Social Care Trust | Wellcome Sanger Institute for the COVID-19 Genomics UK Consortium | Conall McCaughey, James McKenna, Tanya Curran, Susan Feeney, Alison Watt, Ciara Cox, Mairead Connor, Zoltan Molnar, David Simpson, Derek Fairley, Alex Alderton, Roberto Amato, Sonia Goncalves, Ewan Harrison, David K. Jackson, Ian Johnston, Dominic Kwiatkowski, Cordelia Langford, John Sillitoe on behalf of the Wellcome Sanger Institute COVID-19 Surveillance Team (http://www.sanger.ac.uk/covid-team) |
| EPI_ISL_441660 | Department of Pathology, University of Cambridge | Wellcome Sanger Institute for the COVID-19 Genomics UK Consortium | Luke W Meredith, M. Estée Török , Myra Hosmillo, William L. Hamilton, Martin D. Curran, Theresa Feltwell, Grant Hall, Anna Yakovleva, Fahad A Khokhar, Charlotte J. Houldcroft, Laura G Caller, Aminu S. Jahun, Sarah L. Caddy, Ian Goodfellow, Alex Alderton, Roberto Amato, Sonia Goncalves, Ewan Harrison, David K. Jackson, Ian Johnston, Dominic Kwiatkowski, Cordelia Langford, John Sillitoe on behalf of the Wellcome Sanger Institute COVID-19 Surveillance Team (http://www.sanger.ac.uk/covid-team) |
| EPI_ISL_441662, EPI_ISL_441663 | Regional Virus Laboratory, Belfast Health and Social Care Trust | Wellcome Sanger Institute for the COVID-19 Genomics UK Consortium | Conall McCaughey, James McKenna, Tanya Curran, Susan Feeney, Alison Watt, Ciara Cox, Mairead Connor, Zoltan Molnar, David Simpson, Derek Fairley, Alex Alderton, Roberto Amato, Sonia Goncalves, Ewan Harrison, David K. Jackson, Ian Johnston, Dominic Kwiatkowski, Cordelia Langford, John Sillitoe on behalf of the Wellcome Sanger Institute COVID-19 Surveillance Team (http://www.sanger.ac.uk/covid-team) |
| EPI_ISL_441664, EPI_ISL_441665, EPI_ISL_441666, EPI_ISL_441667, EPI_ISL_441668, EPI_ISL_441669 | Department of Pathology, University of Cambridge | Wellcome Sanger Institute for the COVID-19 Genomics UK Consortium | Luke W Meredith, M. Estée Török , Myra Hosmillo, William L. Hamilton, Martin D. Curran, Theresa Feltwell, Grant Hall, Anna Yakovleva, Fahad A Khokhar, Charlotte J. Houldcroft, Laura G Caller, Aminu S. Jahun, Sarah L. Caddy, Ian Goodfellow, Alex Alderton, Roberto Amato, Sonia Goncalves, Ewan Harrison, David K. Jackson, Ian Johnston, Dominic Kwiatkowski, Cordelia Langford, John Sillitoe on behalf of the Wellcome Sanger Institute COVID-19 Surveillance Team (http://www.sanger.ac.uk/covid-team) |
| EPI_ISL_441670 | Regional Virus Laboratory, Belfast Health and Social Care Trust | Wellcome Sanger Institute for the COVID-19 Genomics UK Consortium | Conall McCaughey, James McKenna, Tanya Curran, Susan Feeney, Alison Watt, Ciara Cox, Mairead Connor, Zoltan Molnar, David Simpson, Derek Fairley, Alex Alderton, Roberto Amato, Sonia Goncalves, Ewan Harrison, David K. Jackson, Ian Johnston, Dominic Kwiatkowski, Cordelia Langford, John Sillitoe on behalf of the Wellcome Sanger Institute COVID-19 Surveillance Team (http://www.sanger.ac.uk/covid-team) |
| EPI_ISL_441671, EPI_ISL_441672, EPI_ISL_441673, EPI_ISL_441674 | Department of Pathology, University of Cambridge | Wellcome Sanger Institute for the COVID-19 Genomics UK Consortium | Luke W Meredith, M. Estée Török , Myra Hosmillo, William L. Hamilton, Martin D. Curran, Theresa Feltwell, Grant Hall, Anna Yakovleva, Fahad A Khokhar, Charlotte J. Houldcroft, Laura G Caller, Aminu S. Jahun, Sarah L. Caddy, Ian Goodfellow, Alex Alderton, Roberto Amato, Sonia Goncalves, Ewan Harrison, David K. Jackson, Ian Johnston, Dominic Kwiatkowski, Cordelia Langford, John Sillitoe on behalf of the Wellcome Sanger Institute COVID-19 Surveillance Team (http://www.sanger.ac.uk/covid-team) |
| EPI_ISL_441675, EPI_ISL_441676 | Regional Virus Laboratory, Belfast Health and Social Care Trust | Wellcome Sanger Institute for the COVID-19 Genomics UK Consortium | Conall McCaughey, James McKenna, Tanya Curran, Susan Feeney, Alison Watt, Ciara Cox, Mairead Connor, Zoltan Molnar, David Simpson, Derek Fairley, Alex Alderton, Roberto Amato, Sonia Goncalves, Ewan Harrison, David K. Jackson, Ian Johnston, Dominic Kwiatkowski, Cordelia Langford, John Sillitoe on behalf of the Wellcome Sanger Institute COVID-19 Surveillance Team (http://www.sanger.ac.uk/covid-team) |
| EPI_ISL_441677, EPI_ISL_441678, EPI_ISL_441679, EPI_ISL_441680, EPI_ISL_441681, EPI_ISL_441682, EPI_ISL_441683, EPI_ISL_441684, EPI_ISL_441686, EPI_ISL_441687, EPI_ISL_441688, EPI_ISL_441689 | | | |
| see above | Department of Pathology, University of Cambridge | Wellcome Sanger Institute for the COVID-19 Genomics UK Consortium | Luke W Meredith, M. Estée Török , Myra Hosmillo, William L. Hamilton, Martin D. Curran, Theresa Feltwell, Grant Hall, Anna Yakovleva, Fahad A Khokhar, Charlotte J. Houldcroft, Laura G Caller, Aminu S. Jahun, Sarah L. Caddy, Ian Goodfellow, Alex Alderton, Roberto Amato, Sonia Goncalves, Ewan Harrison, David K. Jackson, Ian Johnston, Dominic Kwiatkowski, Cordelia Langford, John Sillitoe on behalf of the Wellcome Sanger Institute COVID-19 Surveillance Team (http://www.sanger.ac.uk/covid-team) |
| EPI_ISL_441690, EPI_ISL_441691 | Regional Virus Laboratory, Belfast Health and Social Care Trust | Wellcome Sanger Institute for the COVID-19 Genomics UK Consortium | Conall McCaughey, James McKenna, Tanya Curran, Susan Feeney, Alison Watt, Ciara Cox, Mairead Connor, Zoltan Molnar, David Simpson, Derek Fairley, Alex Alderton, Roberto Amato, Sonia Goncalves, Ewan Harrison, David K. Jackson, Ian Johnston, Dominic Kwiatkowski, Cordelia Langford, John Sillitoe on behalf of the Wellcome Sanger Institute COVID-19 Surveillance Team (http://www.sanger.ac.uk/covid-team) |
| EPI_ISL_441692 | Department of Pathology, University of Cambridge | Wellcome Sanger Institute for the COVID-19 Genomics UK Consortium | Luke W Meredith, M. Estée Török , Myra Hosmillo, William L. Hamilton, Martin D. Curran, Theresa Feltwell, Grant Hall, Anna Yakovleva, Fahad A Khokhar, Charlotte J. Houldcroft, Laura G Caller, Aminu S. Jahun, Sarah L. Caddy, Ian Goodfellow, Alex Alderton, Roberto Amato, Sonia Goncalves, Ewan Harrison, David K. Jackson, Ian Johnston, Dominic Kwiatkowski, Cordelia Langford, John Sillitoe on behalf of the Wellcome Sanger Institute COVID-19 Surveillance Team (http://www.sanger.ac.uk/covid-team) |
| EPI_ISL_441693, EPI_ISL_441694 | Regional Virus Laboratory, Belfast Health and Social Care Trust | Wellcome Sanger Institute for the COVID-19 Genomics UK Consortium | Conall McCaughey, James McKenna, Tanya Curran, Susan Feeney, Alison Watt, Ciara Cox, Mairead Connor, Zoltan Molnar, David Simpson, Derek Fairley, Alex Alderton, Roberto Amato, Sonia Goncalves, Ewan Harrison, David K. Jackson, Ian Johnston, Dominic Kwiatkowski, Cordelia Langford, John Sillitoe on behalf of the Wellcome Sanger Institute COVID-19 Surveillance Team (http://www.sanger.ac.uk/covid-team) |
| EPI_ISL_441695 | Department of Pathology, University of Cambridge | Wellcome Sanger Institute for the COVID-19 Genomics UK Consortium | Luke W Meredith, M. Estée Török , Myra Hosmillo, William L. Hamilton, Martin D. Curran, Theresa Feltwell, Grant Hall, Anna Yakovleva, Fahad A Khokhar, Charlotte J. Houldcroft, Laura G Caller, Aminu S. Jahun, Sarah L. Caddy, Ian Goodfellow, Alex Alderton, Roberto Amato, Sonia Goncalves, Ewan Harrison, David K. Jackson, Ian Johnston, Dominic Kwiatkowski, Cordelia Langford, John Sillitoe on behalf of the Wellcome Sanger Institute COVID-19 Surveillance Team (http://www.sanger.ac.uk/covid-team) |
| EPI_ISL_441696, EPI_ISL_441697, EPI_ISL_441698, EPI_ISL_441699 | Regional Virus Laboratory, Belfast Health and Social Care Trust | Wellcome Sanger Institute for the COVID-19 Genomics UK Consortium | Conall McCaughey, James McKenna, Tanya Curran, Susan Feeney, Alison Watt, Ciara Cox, Mairead Connor, Zoltan Molnar, David Simpson, Derek Fairley, Alex Alderton, Roberto Amato, Sonia Goncalves, Ewan Harrison, David K. Jackson, Ian Johnston, Dominic Kwiatkowski, Cordelia Langford, John Sillitoe on behalf of the Wellcome Sanger Institute COVID-19 Surveillance Team (http://www.sanger.ac.uk/covid-team) |
| EPI_ISL_441700, EPI_ISL_441701, EPI_ISL_441702, EPI_ISL_441703, EPI_ISL_441707 | Department of Pathology, University of Cambridge | Wellcome Sanger Institute for the COVID-19 Genomics UK Consortium | Luke W Meredith, M. Estée Török , Myra Hosmillo, William L. Hamilton, Martin D. Curran, Theresa Feltwell, Grant Hall, Anna Yakovleva, Fahad A Khokhar, Charlotte J. Houldcroft, Laura G Caller, Aminu S. Jahun, Sarah L. Caddy, Ian Goodfellow, Alex Alderton, Roberto Amato, Sonia Goncalves, Ewan Harrison, David K. Jackson, Ian Johnston, Dominic Kwiatkowski, Cordelia Langford, John Sillitoe on behalf of the Wellcome Sanger Institute COVID-19 Surveillance Team (http://www.sanger.ac.uk/covid-team) |

| | | | |
|---|---|---|---|
| EPI_ISL_441708 | Regional Virus Laboratory, Belfast Health and Social Care Trust | Wellcome Sanger Institute for the COVID-19 Genomics UK Consortium | Conall McCaughey, James McKenna, Tanya Curran, Susan Feeney, Alison Watt, Ciara Cox, Mairead Connor, Zoltan Molnar, David Simpson, Derek Fairley, Alex Alderton, Roberto Amato, Sonia Goncalves, Ewan Harrison, David K. Jackson, Ian Johnston, Dominic Kwiatkowski, Cordelia Langford, John Sillitoe on behalf of the Wellcome Sanger Institute COVID-19 Surveillance Team (http://www.sanger.ac.uk/covid-team) |
| EPI_ISL_441709, EPI_ISL_441711, EPI_ISL_441713 | Department of Pathology, University of Cambridge | Wellcome Sanger Institute for the COVID-19 Genomics UK Consortium | Luke W Meredith, M. Estée Török , Myra Hosmillo, William L. Hamilton, Martin D. Curran, Theresa Feltwell, Grant Hall, Anna Yakovleva, Fahad A Khokhar, Charlotte J. Houldcroft, Laura G Caller, Aminu S. Jahun, Sarah L. Caddy, Ian Goodfellow, Alex Alderton, Roberto Amato, Sonia Goncalves, Ewan Harrison, David K. Jackson, Ian Johnston, Dominic Kwiatkowski, Cordelia Langford, John Sillitoe on behalf of the Wellcome Sanger Institute COVID-19 Surveillance Team (http://www.sanger.ac.uk/covid-team) |
| EPI_ISL_441714 | Regional Virus Laboratory, Belfast Health and Social Care Trust | Wellcome Sanger Institute for the COVID-19 Genomics UK Consortium | Conall McCaughey, James McKenna, Tanya Curran, Susan Feeney, Alison Watt, Ciara Cox, Mairead Connor, Zoltan Molnar, David Simpson, Derek Fairley, Alex Alderton, Roberto Amato, Sonia Goncalves, Ewan Harrison, David K. Jackson, Ian Johnston, Dominic Kwiatkowski, Cordelia Langford, John Sillitoe on behalf of the Wellcome Sanger Institute COVID-19 Surveillance Team (http://www.sanger.ac.uk/covid-team) |
| EPI_ISL_441715 | Department of Pathology, University of Cambridge | Wellcome Sanger Institute for the COVID-19 Genomics UK Consortium | Luke W Meredith, M. Estée Török , Myra Hosmillo, William L. Hamilton, Martin D. Curran, Theresa Feltwell, Grant Hall, Anna Yakovleva, Fahad A Khokhar, Charlotte J. Houldcroft, Laura G Caller, Aminu S. Jahun, Sarah L. Caddy, Ian Goodfellow, Alex Alderton, Roberto Amato, Sonia Goncalves, Ewan Harrison, David K. Jackson, Ian Johnston, Dominic Kwiatkowski, Cordelia Langford, John Sillitoe on behalf of the Wellcome Sanger Institute COVID-19 Surveillance Team (http://www.sanger.ac.uk/covid-team) |
| EPI_ISL_441717 | Regional Virus Laboratory, Belfast Health and Social Care Trust | Wellcome Sanger Institute for the COVID-19 Genomics UK Consortium | Conall McCaughey, James McKenna, Tanya Curran, Susan Feeney, Alison Watt, Ciara Cox, Mairead Connor, Zoltan Molnar, David Simpson, Derek Fairley, Alex Alderton, Roberto Amato, Sonia Goncalves, Ewan Harrison, David K. Jackson, Ian Johnston, Dominic Kwiatkowski, Cordelia Langford, John Sillitoe on behalf of the Wellcome Sanger Institute COVID-19 Surveillance Team (http://www.sanger.ac.uk/covid-team) |
| EPI_ISL_441718, EPI_ISL_441720, EPI_ISL_441721, EPI_ISL_441723, EPI_ISL_441724, EPI_ISL_441725, EPI_ISL_441726, EPI_ISL_441727, EPI_ISL_441728, EPI_ISL_441729 | Department of Pathology, University of Cambridge | Wellcome Sanger Institute for the COVID-19 Genomics UK Consortium | Luke W Meredith, M. Estée Török , Myra Hosmillo, William L. Hamilton, Martin D. Curran, Theresa Feltwell, Grant Hall, Anna Yakovleva, Fahad A Khokhar, Charlotte J. Houldcroft, Laura G Caller, Aminu S. Jahun, Sarah L. Caddy, Ian Goodfellow, Alex Alderton, Roberto Amato, Sonia Goncalves, Ewan Harrison, David K. Jackson, Ian Johnston, Dominic Kwiatkowski, Cordelia Langford, John Sillitoe on behalf of the Wellcome Sanger Institute COVID-19 Surveillance Team (http://www.sanger.ac.uk/covid-team) |
| EPI_ISL_441730 | Regional Virus Laboratory, Belfast Health and Social Care Trust | Wellcome Sanger Institute for the COVID-19 Genomics UK Consortium | Conall McCaughey, James McKenna, Tanya Curran, Susan Feeney, Alison Watt, Ciara Cox, Mairead Connor, Zoltan Molnar, David Simpson, Derek Fairley, Alex Alderton, Roberto Amato, Sonia Goncalves, Ewan Harrison, David K. Jackson, Ian Johnston, Dominic Kwiatkowski, Cordelia Langford, John Sillitoe on behalf of the Wellcome Sanger Institute COVID-19 Surveillance Team (http://www.sanger.ac.uk/covid-team) |
| EPI_ISL_441731, EPI_ISL_441732 | Department of Pathology, University of Cambridge | Wellcome Sanger Institute for the COVID-19 Genomics UK Consortium | Luke W Meredith, M. Estée Török , Myra Hosmillo, William L. Hamilton, Martin D. Curran, Theresa Feltwell, Grant Hall, Anna Yakovleva, Fahad A Khokhar, Charlotte J. Houldcroft, Laura G Caller, Aminu S. Jahun, Sarah L. Caddy, Ian Goodfellow, Alex Alderton, Roberto Amato, Sonia Goncalves, Ewan Harrison, David K. Jackson, Ian Johnston, Dominic Kwiatkowski, Cordelia Langford, John Sillitoe on behalf of the Wellcome Sanger Institute COVID-19 Surveillance Team (http://www.sanger.ac.uk/covid-team) |
| EPI_ISL_441733 | Regional Virus Laboratory, Belfast Health and Social Care Trust | Wellcome Sanger Institute for the COVID-19 Genomics UK Consortium | Conall McCaughey, James McKenna, Tanya Curran, Susan Feeney, Alison Watt, Ciara Cox, Mairead Connor, Zoltan Molnar, David Simpson, Derek Fairley, Alex Alderton, Roberto Amato, Sonia Goncalves, Ewan Harrison, David K. Jackson, Ian Johnston, Dominic Kwiatkowski, Cordelia Langford, John Sillitoe on behalf of the Wellcome Sanger Institute COVID-19 Surveillance Team (http://www.sanger.ac.uk/covid-team) |
| EPI_ISL_441734, EPI_ISL_441735, EPI_ISL_441736 | Department of Pathology, University of Cambridge | Wellcome Sanger Institute for the COVID-19 Genomics UK Consortium | Luke W Meredith, M. Estée Török , Myra Hosmillo, William L. Hamilton, Martin D. Curran, Theresa Feltwell, Grant Hall, Anna Yakovleva, Fahad A Khokhar, Charlotte J. Houldcroft, Laura G Caller, Aminu S. Jahun, Sarah L. Caddy, Ian Goodfellow, Alex Alderton, Roberto Amato, Sonia Goncalves, Ewan Harrison, David K. Jackson, Ian Johnston, Dominic Kwiatkowski, Cordelia Langford, John Sillitoe on behalf of the Wellcome Sanger Institute COVID-19 Surveillance Team (http://www.sanger.ac.uk/covid-team) |
| EPI_ISL_441737 | Regional Virus Laboratory, Belfast Health and Social Care Trust | Wellcome Sanger Institute for the COVID-19 Genomics UK Consortium | Conall McCaughey, James McKenna, Tanya Curran, Susan Feeney, Alison Watt, Ciara Cox, Mairead Connor, Zoltan Molnar, David Simpson, Derek Fairley, Alex Alderton, Roberto Amato, Sonia Goncalves, Ewan Harrison, David K. Jackson, Ian Johnston, Dominic Kwiatkowski, Cordelia Langford, John Sillitoe on behalf of the Wellcome Sanger Institute COVID-19 Surveillance Team (http://www.sanger.ac.uk/covid-team) |
| EPI_ISL_441738, EPI_ISL_441739, EPI_ISL_441740, EPI_ISL_441741, EPI_ISL_441742, EPI_ISL_441743, EPI_ISL_441744, EPI_ISL_441745, EPI_ISL_441746 | Department of Pathology, University of Cambridge | Wellcome Sanger Institute for the COVID-19 Genomics UK Consortium | Luke W Meredith, M. Estée Török , Myra Hosmillo, William L. Hamilton, Martin D. Curran, Theresa Feltwell, Grant Hall, Anna Yakovleva, Fahad A Khokhar, Charlotte J. Houldcroft, Laura G Caller, Aminu S. Jahun, Sarah L. Caddy, Ian Goodfellow, Alex Alderton, Roberto Amato, Sonia Goncalves, Ewan Harrison, David K. Jackson, Ian Johnston, Dominic Kwiatkowski, Cordelia Langford, John Sillitoe on behalf of the Wellcome Sanger Institute COVID-19 Surveillance Team (http://www.sanger.ac.uk/covid-team) |
| EPI_ISL_441747 | Regional Virus Laboratory, Belfast Health and Social Care Trust | Wellcome Sanger Institute for the COVID-19 Genomics UK Consortium | Conall McCaughey, James McKenna, Tanya Curran, Susan Feeney, Alison Watt, Ciara Cox, Mairead Connor, Zoltan Molnar, David Simpson, Derek Fairley, Alex Alderton, Roberto Amato, Sonia Goncalves, Ewan Harrison, David K. Jackson, Ian Johnston, Dominic Kwiatkowski, Cordelia Langford, John Sillitoe on behalf of the Wellcome Sanger Institute COVID-19 Surveillance Team (http://www.sanger.ac.uk/covid-team) |
| EPI_ISL_441748, EPI_ISL_441749, EPI_ISL_441750 | Department of Pathology, University of Cambridge | Wellcome Sanger Institute for the COVID-19 Genomics UK Consortium | Luke W Meredith, M. Estée Török , Myra Hosmillo, William L. Hamilton, Martin D. Curran, Theresa Feltwell, Grant Hall, Anna Yakovleva, Fahad A Khokhar, Charlotte J. Houldcroft, Laura G Caller, Aminu S. Jahun, Sarah L. Caddy, Ian Goodfellow, Alex Alderton, Roberto Amato, Sonia Goncalves, Ewan Harrison, David K. Jackson, Ian Johnston, Dominic Kwiatkowski, Cordelia Langford, John Sillitoe on behalf of the Wellcome Sanger Institute COVID-19 Surveillance Team (http://www.sanger.ac.uk/covid-team) |
| EPI_ISL_441751 | Regional Virus Laboratory, Belfast Health and Social Care Trust | Wellcome Sanger Institute for the COVID-19 Genomics UK Consortium | Conall McCaughey, James McKenna, Tanya Curran, Susan Feeney, Alison Watt, Ciara Cox, Mairead Connor, Zoltan Molnar, David Simpson, Derek Fairley, Alex Alderton, Roberto Amato, Sonia Goncalves, Ewan Harrison, David K. Jackson, Ian Johnston, Dominic Kwiatkowski, Cordelia Langford, John Sillitoe on behalf of the Wellcome Sanger Institute COVID-19 Surveillance Team (http://www.sanger.ac.uk/covid-team) |
| EPI_ISL_441752, EPI_ISL_441754, EPI_ISL_441755, EPI_ISL_441758, EPI_ISL_441759, EPI_ISL_441760, EPI_ISL_441762, EPI_ISL_441764 | Department of Pathology, University of Cambridge | Wellcome Sanger Institute for the COVID-19 Genomics UK Consortium | Luke W Meredith, M. Estée Török , Myra Hosmillo, William L. Hamilton, Martin D. Curran, Theresa Feltwell, Grant Hall, Anna Yakovleva, Fahad A Khokhar, Charlotte J. Houldcroft, Laura G Caller, Aminu S. Jahun, Sarah L. Caddy, Ian Goodfellow, Alex Alderton, Roberto Amato, Sonia Goncalves, Ewan Harrison, David K. Jackson, Ian Johnston, Dominic Kwiatkowski, Cordelia Langford, John Sillitoe on behalf of the Wellcome Sanger Institute COVID-19 Surveillance Team (http://www.sanger.ac.uk/covid-team) |
| EPI_ISL_441765, EPI_ISL_441766 | Regional Virus Laboratory, Belfast Health and Social Care Trust | Wellcome Sanger Institute for the COVID-19 Genomics UK Consortium | Conall McCaughey, James McKenna, Tanya Curran, Susan Feeney, Alison Watt, Ciara Cox, Mairead Connor, Zoltan Molnar, David Simpson, Derek Fairley, Alex Alderton, Roberto Amato, Sonia Goncalves, Ewan Harrison, David K. Jackson, Ian Johnston, Dominic Kwiatkowski, Cordelia Langford, John Sillitoe on behalf of the Wellcome Sanger Institute COVID-19 Surveillance Team (http://www.sanger.ac.uk/covid-team) |
| EPI_ISL_441767, EPI_ISL_441768, EPI_ISL_441769 | Department of Pathology, University of Cambridge | Wellcome Sanger Institute for the COVID-19 Genomics UK Consortium | Luke W Meredith, M. Estée Török , Myra Hosmillo, William L. Hamilton, Martin D. Curran, Theresa Feltwell, Grant Hall, Anna Yakovleva, Fahad A Khokhar, Charlotte J. Houldcroft, Laura G Caller, Aminu S. Jahun, Sarah L. Caddy, Ian Goodfellow, Alex Alderton, Roberto Amato, Sonia Goncalves, Ewan Harrison, David K. Jackson, Ian Johnston, Dominic Kwiatkowski, Cordelia Langford, John Sillitoe on behalf of the Wellcome Sanger Institute COVID-19 Surveillance Team (http://www.sanger.ac.uk/covid-team) |
| EPI_ISL_441770, EPI_ISL_441771 | Regional Virus Laboratory, Belfast Health and Social Care Trust | Wellcome Sanger Institute for the COVID-19 Genomics UK Consortium | Conall McCaughey, James McKenna, Tanya Curran, Susan Feeney, Alison Watt, Ciara Cox, Mairead Connor, Zoltan Molnar, David Simpson, Derek Fairley, Alex Alderton, Roberto Amato, Sonia Goncalves, Ewan Harrison, David K. Jackson, Ian Johnston, Dominic Kwiatkowski, Cordelia Langford, John Sillitoe on behalf of the Wellcome Sanger Institute COVID-19 Surveillance Team (http://www.sanger.ac.uk/covid-team) |
| EPI_ISL_441772, EPI_ISL_441773, EPI_ISL_441774 | Department of Pathology, University of Cambridge | Wellcome Sanger Institute for the COVID-19 Genomics UK Consortium | Luke W Meredith, M. Estée Török , Myra Hosmillo, William L. Hamilton, Martin D. Curran, Theresa Feltwell, Grant Hall, Anna Yakovleva, Fahad A Khokhar, Charlotte J. Houldcroft, Laura G Caller, Aminu S. Jahun, Sarah L. Caddy, Ian Goodfellow, Alex Alderton, Roberto Amato, Sonia Goncalves, Ewan Harrison, David K. Jackson, Ian Johnston, Dominic Kwiatkowski, Cordelia Langford, John Sillitoe on behalf of the Wellcome Sanger Institute COVID-19 Surveillance Team (http://www.sanger.ac.uk/covid-team) |
| EPI_ISL_441775 | Regional Virus Laboratory, Belfast Health and Social Care Trust | Wellcome Sanger Institute for the COVID-19 Genomics UK Consortium | Conall McCaughey, James McKenna, Tanya Curran, Susan Feeney, Alison Watt, Ciara Cox, Mairead Connor, Zoltan Molnar, David Simpson, Derek Fairley, Alex Alderton, Roberto Amato, Sonia Goncalves, Ewan Harrison, David K. Jackson, Ian Johnston, Dominic Kwiatkowski, Cordelia Langford, John Sillitoe on behalf of the Wellcome Sanger Institute COVID-19 Surveillance Team (http://www.sanger.ac.uk/covid-team) |
| EPI_ISL_441777, EPI_ISL_441778, EPI_ISL_441779, EPI_ISL_441780 | Department of Pathology, University of Cambridge | Wellcome Sanger Institute for the COVID-19 Genomics UK Consortium | Luke W Meredith, M. Estée Török , Myra Hosmillo, William L. Hamilton, Martin D. Curran, Theresa Feltwell, Grant Hall, Anna Yakovleva, Fahad A Khokhar, Charlotte J. Houldcroft, Laura G Caller, Aminu S. Jahun, Sarah L. Caddy, Ian Goodfellow, Alex Alderton, Roberto Amato, Sonia Goncalves, Ewan Harrison, David K. Jackson, Ian Johnston, Dominic Kwiatkowski, Cordelia Langford, John Sillitoe on behalf of the Wellcome Sanger Institute COVID-19 Surveillance Team (http://www.sanger.ac.uk/covid-team) |
| EPI_ISL_441781 | Regional Virus Laboratory, Belfast Health and Social Care Trust | Wellcome Sanger Institute for the COVID-19 Genomics UK Consortium | Conall McCaughey, James McKenna, Tanya Curran, Susan Feeney, Alison Watt, Ciara Cox, Mairead Connor, Zoltan Molnar, David Simpson, Derek Fairley, Alex Alderton, Roberto Amato, Sonia Goncalves, Ewan Harrison, David K. Jackson, Ian Johnston, Dominic Kwiatkowski, Cordelia Langford, John Sillitoe on behalf of the Wellcome Sanger Institute COVID-19 Surveillance Team (http://www.sanger.ac.uk/covid-team) |
| EPI_ISL_441782, EPI_ISL_441784, EPI_ISL_441785, EPI_ISL_441786, EPI_ISL_441787, EPI_ISL_441788, EPI_ISL_441789, EPI_ISL_441790, EPI_ISL_441791, EPI_ISL_441792, EPI_ISL_441793, EPI_ISL_441795, EPI_ISL_441797, EPI_ISL_441798, EPI_ISL_441800, EPI_ISL_441801, EPI_ISL_441802, EPI_ISL_441803, EPI_ISL_441804, EPI_ISL_441806, EPI_ISL_441809, EPI_ISL_441811, EPI_ISL_441813, EPI_ISL_441816, EPI_ISL_441817, EPI_ISL_441818, EPI_ISL_441820, EPI_ISL_441824, EPI_ISL_441826, EPI_ISL_441827, EPI_ISL_441830, EPI_ISL_441831, EPI_ISL_441832, EPI_ISL_441833, EPI_ISL_441835, EPI_ISL_441836, EPI_ISL_441837, EPI_ISL_441841, EPI_ISL_441842, EPI_ISL_441843, EPI_ISL_441844, EPI_ISL_441845 | | | |
| see above | Department of Pathology, University of Cambridge | Wellcome Sanger Institute for the COVID-19 Genomics UK Consortium | Luke W Meredith, M. Estée Török , Myra Hosmillo, William L. Hamilton, Martin D. Curran, Theresa Feltwell, Grant Hall, Anna Yakovleva, Fahad A Khokhar, Charlotte J. Houldcroft, Laura G Caller, Aminu S. Jahun, Sarah L. Caddy, Ian Goodfellow, Alex Alderton, Roberto Amato, Sonia Goncalves, Ewan Harrison, David K. Jackson, Ian Johnston, Dominic Kwiatkowski, Cordelia Langford, John Sillitoe on behalf of the Wellcome Sanger Institute COVID-19 Surveillance Team (http://www.sanger.ac.uk/covid-team) |
| EPI_ISL_441863, EPI_ISL_441864, EPI_ISL_441865, EPI_ISL_441866, EPI_ISL_441867, EPI_ISL_441868, EPI_ISL_441869, EPI_ISL_441870, EPI_ISL_441871, EPI_ISL_441872, EPI_ISL_441873, EPI_ISL_441878, EPI_ISL_441879, EPI_ISL_441880, EPI_ISL_441881, EPI_ISL_441882, EPI_ISL_441883, EPI_ISL_441885, EPI_ISL_441886, EPI_ISL_441888, EPI_ISL_441890, EPI_ISL_441891, EPI_ISL_441892, EPI_ISL_441894, EPI_ISL_441895, EPI_ISL_441896, EPI_ISL_441897 | | | |
| see above | Queens Medical Centre, Clinical Microbiology Department / DeepSeq Nottingham | COVID-19 Genomics UK (COG-UK) Consortium | Gemma Clark, Wendy Smith, Manjinder Khakh, Hannah Howson-Wells, Jonathan Ball, Patrick McClure, Joseph Chappell, Theocharis Tsoleridis, Nadine Holmes, Matthew Carlisle, Christopher Moore, Fei Sang, Johnny Debebe, Victoria Wright, Matthew Loose |
| EPI_ISL_441898, EPI_ISL_441913, EPI_ISL_441916, EPI_ISL_441932, EPI_ISL_441935, EPI_ISL_441942, EPI_ISL_441952, EPI_ISL_441955, EPI_ISL_441975, EPI_ISL_441981, EPI_ISL_441984, EPI_ISL_441991, EPI_ISL_441999, EPI_ISL_442001, EPI_ISL_442013, EPI_ISL_442032, EPI_ISL_442043 | | | |
| see above | Virology Department, Sheffield Teaching Hospitals NHS Foundation Trust/Department of Infection, Immunity and Cardiovascular Disease, The Medical School, University of Sheffield | COVID-19 Genomics UK (COG-UK) Consortium | Thushan de Silva, Matthew Parker, Nikki Smith, Adri Angyal, Rebecca Brown, Luke Green, Rachel Tucker, Paul Parsons, Danielle Groves, Katie Johnson, Laura Carrilero, Alex Keeley, Dave Partridge, Matthew Wyles, Benjamin Lindsey, Mehmet Yavuz, Mohammad Raza, Cariad Evans |
| EPI_ISL_442044 | Kawsar Human Genetic Research Center | Kawsar Human Genetic Research Center | Mohammad Ali Khosravi, Maryam Abbasalipour Bashash, Sirous Zeinali, Solmaz Sabeghi, Yeganeh Keshvar, Fatemeh Hosseini, Yeganeh Haghdoost |
| EPI_ISL_442045, EPI_ISL_442046, EPI_ISL_442047, EPI_ISL_442048, EPI_ISL_442049, EPI_ISL_442050, EPI_ISL_442051, EPI_ISL_442053, EPI_ISL_442054, EPI_ISL_442055, EPI_ISL_442056, EPI_ISL_442057, EPI_ISL_442058, EPI_ISL_442059, EPI_ISL_442060, EPI_ISL_442061, EPI_ISL_442062, EPI_ISL_442064, EPI_ISL_442066, EPI_ISL_442067, EPI_ISL_442068, EPI_ISL_442070, EPI_ISL_442071, EPI_ISL_442073, EPI_ISL_442074, EPI_ISL_442075, EPI_ISL_442076, EPI_ISL_442077, EPI_ISL_442078, EPI_ISL_442079, EPI_ISL_442080, EPI_ISL_442081, EPI_ISL_442082, EPI_ISL_442083, EPI_ISL_442084, EPI_ISL_442085, EPI_ISL_442086, EPI_ISL_442088, EPI_ISL_442089, EPI_ISL_442090, EPI_ISL_442091, EPI_ISL_442092, EPI_ISL_442093, EPI_ISL_442094, EPI_ISL_442095, EPI_ISL_442097, EPI_ISL_442098, EPI_ISL_442101, EPI_ISL_442102, EPI_ISL_442103, EPI_ISL_442104, EPI_ISL_442105, EPI_ISL_442106, EPI_ISL_442107, EPI_ISL_442108, EPI_ISL_442109, EPI_ISL_442111, EPI_ISL_442112, EPI_ISL_442113, EPI_ISL_442114, EPI_ISL_442116, EPI_ISL_442117, EPI_ISL_442120, EPI_ISL_442121, EPI_ISL_442122, EPI_ISL_442123, EPI_ISL_442124, EPI_ISL_442125, EPI_ISL_442126, EPI_ISL_442129, EPI_ISL_442130, EPI_ISL_442133, EPI_ISL_442134, EPI_ISL_442136, EPI_ISL_442139, EPI_ISL_442140, EPI_ISL_442142, EPI_ISL_442143, EPI_ISL_442144, EPI_ISL_442146, EPI_ISL_442147, EPI_ISL_442149, EPI_ISL_442150, EPI_ISL_442154, EPI_ISL_442157, EPI_ISL_442158, EPI_ISL_442160, EPI_ISL_442161, EPI_ISL_442163, EPI_ISL_442164, EPI_ISL_442165, EPI_ISL_442167, EPI_ISL_442168, EPI_ISL_442169, EPI_ISL_442170, EPI_ISL_442171, EPI_ISL_442172, EPI_ISL_442174, EPI_ISL_442175, EPI_ISL_442176, EPI_ISL_442179, EPI_ISL_442180, EPI_ISL_442182, EPI_ISL_442183, EPI_ISL_442184, EPI_ISL_442186, EPI_ISL_442187, EPI_ISL_442188, EPI_ISL_442189, EPI_ISL_442190, EPI_ISL_442192, EPI_ISL_442193, EPI_ISL_442194, EPI_ISL_442196, EPI_ISL_442197, EPI_ISL_442198, EPI_ISL_442199, EPI_ISL_442200, EPI_ISL_442201, EPI_ISL_442202, EPI_ISL_442203, EPI_ISL_442204, EPI_ISL_442206, EPI_ISL_442207, EPI_ISL_442208, EPI_ISL_442210, EPI_ISL_442211, EPI_ISL_442213, EPI_ISL_442214, EPI_ISL_442215, EPI_ISL_442216, EPI_ISL_442217, EPI_ISL_442220, EPI_ISL_442221, EPI_ISL_442222, EPI_ISL_442223, EPI_ISL_442224, EPI_ISL_442225, EPI_ISL_442226, EPI_ISL_442229, EPI_ISL_442230, EPI_ISL_442231, EPI_ISL_442232, EPI_ISL_442234, EPI_ISL_442235, EPI_ISL_442236, EPI_ISL_442237, EPI_ISL_442240, EPI_ISL_442241, EPI_ISL_442242, EPI_ISL_442244, EPI_ISL_442246, EPI_ISL_442247, EPI_ISL_442248, EPI_ISL_442250, EPI_ISL_442251, EPI_ISL_442252, EPI_ISL_442253, EPI_ISL_442254, EPI_ISL_442255, EPI_ISL_442256, EPI_ISL_442257, EPI_ISL_442258, EPI_ISL_442259, EPI_ISL_442260, EPI_ISL_442261, EPI_ISL_442262, EPI_ISL_442263, EPI_ISL_442265, EPI_ISL_442266, EPI_ISL_442268, EPI_ISL_442269, EPI_ISL_442270, EPI_ISL_442272, EPI_ISL_442273, EPI_ISL_442274, EPI_ISL_442275, EPI_ISL_442276, EPI_ISL_442277, EPI_ISL_442278, EPI_ISL_442281, EPI_ISL_442282, EPI_ISL_442283, EPI_ISL_442284, EPI_ISL_442285, EPI_ISL_442286, EPI_ISL_442287, EPI_ISL_442288, EPI_ISL_442289, EPI_ISL_442291, EPI_ISL_442292, EPI_ISL_442294, EPI_ISL_442295, EPI_ISL_442296, EPI_ISL_442297, EPI_ISL_442299, EPI_ISL_442301, EPI_ISL_442302, EPI_ISL_442305, EPI_ISL_442306, EPI_ISL_442307, EPI_ISL_442308, EPI_ISL_442309, EPI_ISL_442310, EPI_ISL_442312, EPI_ISL_442313, EPI_ISL_442315, EPI_ISL_442316, EPI_ISL_442317, EPI_ISL_442318, EPI_ISL_442319, EPI_ISL_442320, EPI_ISL_442321, EPI_ISL_442323, EPI_ISL_442324, EPI_ISL_442325, EPI_ISL_442326, EPI_ISL_442327, EPI_ISL_442329, EPI_ISL_442330, EPI_ISL_442331, EPI_ISL_442332, EPI_ISL_442333, EPI_ISL_442334, EPI_ISL_442335, EPI_ISL_442336, EPI_ISL_442337, EPI_ISL_442338, EPI_ISL_442339, EPI_ISL_442340, EPI_ISL_442341, EPI_ISL_442342, EPI_ISL_442343 | | | |
| see above | Department of Pathology, University of Cambridge | Wellcome Sanger Institute for the COVID-19 Genomics UK Consortium | Luke W Meredith, M. Estée Török , Myra Hosmillo, William L. Hamilton, Martin D. Curran, Theresa Feltwell, Grant Hall, Anna Yakovleva, Fahad A Khokhar, Charlotte J. Houldcroft, Laura G Caller, Aminu S. Jahun, Sarah L. Caddy, Ian Goodfellow, Alex Alderton, Roberto Amato, Sonia Goncalves, Ewan Harrison, David K. Jackson, Ian Johnston, Dominic Kwiatkowski, Cordelia Langford, John Sillitoe on behalf of the Wellcome Sanger Institute COVID-19 Surveillance Team (http://www.sanger.ac.uk/covid-team) |
| EPI_ISL_442350, EPI_ISL_442353, EPI_ISL_442354, EPI_ISL_442357, EPI_ISL_442358, EPI_ISL_442370, EPI_ISL_442372, EPI_ISL_442380, EPI_ISL_442387, EPI_ISL_442390, EPI_ISL_442397, EPI_ISL_442398, EPI_ISL_442404, EPI_ISL_442407, EPI_ISL_442413, EPI_ISL_442418, EPI_ISL_442432, EPI_ISL_442438, EPI_ISL_442448, EPI_ISL_442468, EPI_ISL_442472, EPI_ISL_442478, EPI_ISL_442479, EPI_ISL_442481, EPI_ISL_442487, EPI_ISL_442489, EPI_ISL_442502, EPI_ISL_442506, EPI_ISL_442510 | | | |
| see above | Virology Department, Sheffield Teaching Hospitals NHS Foundation Trust/Department of Infection, Immunity and Cardiovascular Disease, The Medical School, University of Sheffield | COVID-19 Genomics UK (COG-UK) Consortium | Thushan de Silva, Matthew Parker, Nikki Smith, Adri Angyal, Rebecca Brown, Luke Green, Rachel Tucker, Paul Parsons, Danielle Groves, Katie Johnson, Laura Carrilero, Alex Keeley, Dave Partridge, Matthew Wyles, Benjamin Lindsey, Mehmet Yavuz, Mohammad Raza, Cariad Evans |
| EPI_ISL_442523 | Pasteur Institute of Iran | Kawsar Human Genetic Research Company | Sirous Zeinali, Mohammad Ali Khosravi,Maryam Abbasalipour Bashash, Sanaz Mostafavi Jabbari, Maraym Firoozi, Sormeh Pourtavakoli, Elmira Khateri, Razieh Zeinali and Fahimeh Hoseini |
| EPI_ISL_442524, EPI_ISL_442526, EPI_ISL_442527, EPI_ISL_442528, EPI_ISL_442529, EPI_ISL_442531, EPI_ISL_442532, EPI_ISL_442533, EPI_ISL_442534, EPI_ISL_442535, EPI_ISL_442536, EPI_ISL_442537, EPI_ISL_442538, EPI_ISL_442539, EPI_ISL_442541, EPI_ISL_442542, EPI_ISL_442543, EPI_ISL_442545, EPI_ISL_442546, EPI_ISL_442548, EPI_ISL_442553, EPI_ISL_442556, EPI_ISL_442558, EPI_ISL_442559, | | | |

| | | | |
|---|---|---|---|
| EPI_ISL_442560, EPI_ISL_442561, EPI_ISL_442562, EPI_ISL_442563, EPI_ISL_442564, EPI_ISL_442565, EPI_ISL_442567, EPI_ISL_442569, EPI_ISL_442570, EPI_ISL_442572, EPI_ISL_442573, EPI_ISL_442574, EPI_ISL_442575, EPI_ISL_442577, EPI_ISL_442578, EPI_ISL_442581, EPI_ISL_442582, EPI_ISL_442583, EPI_ISL_442584, EPI_ISL_442586, EPI_ISL_442587, EPI_ISL_442588, EPI_ISL_442589, EPI_ISL_442592, EPI_ISL_442593, EPI_ISL_442594, EPI_ISL_442595, EPI_ISL_442596, EPI_ISL_442597, EPI_ISL_442599, EPI_ISL_442601, EPI_ISL_442602, EPI_ISL_442603, EPI_ISL_442604, EPI_ISL_442605, EPI_ISL_442606, EPI_ISL_442607, EPI_ISL_442610, EPI_ISL_442611, EPI_ISL_442613, EPI_ISL_442614, EPI_ISL_442615, EPI_ISL_442616, EPI_ISL_442618, EPI_ISL_442620, EPI_ISL_442622, EPI_ISL_442623 | | | |
| see above | Department of Pathology, University of Cambridge | Wellcome Sanger Institute for the COVID-19 Genomics UK Consortium | Luke W Meredith, M. Estée Török , Myra Hosmillo, William L. Hamilton, Martin D. Curran, Theresa Feltwell, Grant Hall, Anna Yakovleva, Fahad A Khokhar, Charlotte J. Houldcroft, Laura G Caller, Aminu S. Jahun, Sarah L. Caddy, Ian Goodfellow, Alex Alderton, Roberto Amato, Sonia Goncalves, Ewan Harrison, David K. Jackson, Ian Johnston, Dominic Kwiatkowski, Cordelia Langford, John Sillitoe on behalf of the Wellcome Sanger Institute COVID-19 Surveillance Team (http://www.sanger.ac.uk/covid-team) |
| EPI_ISL_442624, EPI_ISL_442625, EPI_ISL_442626, EPI_ISL_442627, EPI_ISL_442628, EPI_ISL_442629, EPI_ISL_442630, EPI_ISL_442632, EPI_ISL_442633, EPI_ISL_442634, EPI_ISL_442636, EPI_ISL_442637, EPI_ISL_442638, EPI_ISL_442639, EPI_ISL_442640, EPI_ISL_442641, EPI_ISL_442643, EPI_ISL_442644, EPI_ISL_442645, EPI_ISL_442646, EPI_ISL_442647, EPI_ISL_442648, EPI_ISL_442649, EPI_ISL_442650, EPI_ISL_442652, EPI_ISL_442653, EPI_ISL_442654, EPI_ISL_442655, EPI_ISL_442656, EPI_ISL_442657, EPI_ISL_442658, EPI_ISL_442660, EPI_ISL_442662, EPI_ISL_442663, EPI_ISL_442664, EPI_ISL_442665, EPI_ISL_442667, EPI_ISL_442668, EPI_ISL_442669, EPI_ISL_442670, EPI_ISL_442671, EPI_ISL_442672, EPI_ISL_442673, EPI_ISL_442674, EPI_ISL_442675, EPI_ISL_442676, EPI_ISL_442679, EPI_ISL_442680, EPI_ISL_442681, EPI_ISL_442682, EPI_ISL_442683, EPI_ISL_442684, EPI_ISL_442685, EPI_ISL_442686, EPI_ISL_442688, EPI_ISL_442689, EPI_ISL_442690, EPI_ISL_442691, EPI_ISL_442692, EPI_ISL_442694, EPI_ISL_442695, EPI_ISL_442696, EPI_ISL_442697, EPI_ISL_442699, EPI_ISL_442700, EPI_ISL_442701, EPI_ISL_442702, EPI_ISL_442704, EPI_ISL_442705, EPI_ISL_442706, EPI_ISL_442707, EPI_ISL_442708, EPI_ISL_442709, EPI_ISL_442710, EPI_ISL_442711, EPI_ISL_442712, EPI_ISL_442713, EPI_ISL_442714, EPI_ISL_442718, EPI_ISL_442719, EPI_ISL_442720, EPI_ISL_442722, EPI_ISL_442724, EPI_ISL_442725, EPI_ISL_442726, EPI_ISL_442727, EPI_ISL_442728, EPI_ISL_442729, EPI_ISL_442730, EPI_ISL_442731, EPI_ISL_442732, EPI_ISL_442734, EPI_ISL_442735, EPI_ISL_442736, EPI_ISL_442737, EPI_ISL_442739, EPI_ISL_442740, EPI_ISL_442741, EPI_ISL_442742, EPI_ISL_442743, EPI_ISL_442744, EPI_ISL_442745, EPI_ISL_442747, EPI_ISL_442748, EPI_ISL_442749, EPI_ISL_442751, EPI_ISL_442752, EPI_ISL_442753, EPI_ISL_442754, EPI_ISL_442755, EPI_ISL_442756, EPI_ISL_442757, EPI_ISL_442758, EPI_ISL_442759, EPI_ISL_442760, EPI_ISL_442761, EPI_ISL_442762, EPI_ISL_442763, EPI_ISL_442764, EPI_ISL_442765, EPI_ISL_442767, EPI_ISL_442768, EPI_ISL_442769, EPI_ISL_442770, EPI_ISL_442771, EPI_ISL_442772, EPI_ISL_442773, EPI_ISL_442774, EPI_ISL_442775, EPI_ISL_442777, EPI_ISL_442778, EPI_ISL_442779, EPI_ISL_442780, EPI_ISL_442781, EPI_ISL_442782, EPI_ISL_442783, EPI_ISL_442784, EPI_ISL_442785, EPI_ISL_442786, EPI_ISL_442787, EPI_ISL_442788, EPI_ISL_442789, EPI_ISL_442790, EPI_ISL_442791, EPI_ISL_442792, EPI_ISL_442793, EPI_ISL_442794, EPI_ISL_442795, EPI_ISL_442796, EPI_ISL_442797, EPI_ISL_442799, EPI_ISL_442800, EPI_ISL_442801, EPI_ISL_442802, EPI_ISL_442804, EPI_ISL_442805, EPI_ISL_442807, EPI_ISL_442808, EPI_ISL_442809, EPI_ISL_442810 | | | |
| see above | PHE South West Regional Laboratory, National Infection Service | Wellcome Sanger Institute for the COVID-19 Genomics UK Consortium | Stephanie Hutchings, Hannah Pymont, Dr Peter Muir, Barry Vipond, Rich Hopes, Alex Alderton, Roberto Amato, Sonia Goncalves, Ewan Harrison, David K. Jackson, Ian Johnston, Dominic Kwiatkowski, Cordelia Langford, John Sillitoe on behalf of the Wellcome Sanger Institute COVID-19 Surveillance Team (http://www.sanger.ac.uk/covid-team) |
| EPI_ISL_442811, EPI_ISL_442812, EPI_ISL_442813, EPI_ISL_442814, EPI_ISL_442815, EPI_ISL_442816, EPI_ISL_442817, EPI_ISL_442818, EPI_ISL_442819, EPI_ISL_442820, EPI_ISL_442822, EPI_ISL_442823, EPI_ISL_442824, EPI_ISL_442825, EPI_ISL_442826, EPI_ISL_442828, EPI_ISL_442829, EPI_ISL_442830, EPI_ISL_442831, EPI_ISL_442832, EPI_ISL_442833, EPI_ISL_442834, EPI_ISL_442835, EPI_ISL_442839, EPI_ISL_442841, EPI_ISL_442842, EPI_ISL_442844, EPI_ISL_442845, EPI_ISL_442846, EPI_ISL_442847, EPI_ISL_442848, EPI_ISL_442849, EPI_ISL_442850, EPI_ISL_442851, EPI_ISL_442852, EPI_ISL_442853, EPI_ISL_442854, EPI_ISL_442856, EPI_ISL_442857, EPI_ISL_442858, EPI_ISL_442859, EPI_ISL_442860, EPI_ISL_442862, EPI_ISL_442863, EPI_ISL_442864, EPI_ISL_442865, EPI_ISL_442866, EPI_ISL_442867, EPI_ISL_442868, EPI_ISL_442869, EPI_ISL_442870, EPI_ISL_442871, EPI_ISL_442872, EPI_ISL_442873, EPI_ISL_442874, EPI_ISL_442875, EPI_ISL_442876, EPI_ISL_442878, EPI_ISL_442879, EPI_ISL_442880, EPI_ISL_442881, EPI_ISL_442882, EPI_ISL_442883, EPI_ISL_442884, EPI_ISL_442885, EPI_ISL_442886, EPI_ISL_442888, EPI_ISL_442889, EPI_ISL_442890, EPI_ISL_442891, EPI_ISL_442892, EPI_ISL_442894, EPI_ISL_442895, EPI_ISL_442897, EPI_ISL_442898, EPI_ISL_442899, EPI_ISL_442900, EPI_ISL_442901, EPI_ISL_442903, EPI_ISL_442904, EPI_ISL_442905, EPI_ISL_442906, EPI_ISL_442907, EPI_ISL_442908, EPI_ISL_442909, EPI_ISL_442910, EPI_ISL_442912, EPI_ISL_442913, EPI_ISL_442914, EPI_ISL_442915, EPI_ISL_442916, EPI_ISL_442917, EPI_ISL_442918, EPI_ISL_442919, EPI_ISL_442920, EPI_ISL_442922, EPI_ISL_442923, EPI_ISL_442926, EPI_ISL_442927, EPI_ISL_442930, EPI_ISL_442932, EPI_ISL_442934, EPI_ISL_442935, EPI_ISL_442936, EPI_ISL_442937, EPI_ISL_442938, EPI_ISL_442939, EPI_ISL_442940, EPI_ISL_442943, EPI_ISL_442944, EPI_ISL_442945, EPI_ISL_442947, EPI_ISL_442952, EPI_ISL_442953, EPI_ISL_442954, EPI_ISL_442956, EPI_ISL_442957, EPI_ISL_442959, EPI_ISL_442961, EPI_ISL_442962, EPI_ISL_442964, EPI_ISL_442965, EPI_ISL_442966, EPI_ISL_442967, EPI_ISL_442968, EPI_ISL_442969, EPI_ISL_442970, EPI_ISL_442973, EPI_ISL_442974, EPI_ISL_442975, EPI_ISL_442976, EPI_ISL_442977, EPI_ISL_442978, EPI_ISL_442979, EPI_ISL_442980, EPI_ISL_442981, EPI_ISL_442982, EPI_ISL_442983, EPI_ISL_442984, EPI_ISL_442985, EPI_ISL_442987, EPI_ISL_442988, EPI_ISL_442989, EPI_ISL_442991, EPI_ISL_442993, EPI_ISL_442994, EPI_ISL_442995, EPI_ISL_442996, EPI_ISL_442997, EPI_ISL_442998, EPI_ISL_442999, EPI_ISL_443000, EPI_ISL_443001, EPI_ISL_443002, EPI_ISL_443003, EPI_ISL_443004, EPI_ISL_443007, EPI_ISL_443009, EPI_ISL_443010, EPI_ISL_443011, EPI_ISL_443015, EPI_ISL_443017, EPI_ISL_443020, EPI_ISL_443021, EPI_ISL_443022, EPI_ISL_443023, EPI_ISL_443024, EPI_ISL_443025, EPI_ISL_443026, EPI_ISL_443027, EPI_ISL_443029, EPI_ISL_443030, EPI_ISL_443031, EPI_ISL_443032, EPI_ISL_443033, EPI_ISL_443035, EPI_ISL_443036, EPI_ISL_443037, EPI_ISL_443038, EPI_ISL_443039, EPI_ISL_443041, EPI_ISL_443042, EPI_ISL_443043, EPI_ISL_443044, EPI_ISL_443045, EPI_ISL_443046, EPI_ISL_443047, EPI_ISL_443049, EPI_ISL_443050, EPI_ISL_443051, EPI_ISL_443053, EPI_ISL_443055, EPI_ISL_443058, EPI_ISL_443059, EPI_ISL_443060, EPI_ISL_443061, EPI_ISL_443062, EPI_ISL_443063, EPI_ISL_443065, EPI_ISL_443066, EPI_ISL_443068, EPI_ISL_443070, EPI_ISL_443072, EPI_ISL_443074, EPI_ISL_443075, EPI_ISL_443076, EPI_ISL_443077, EPI_ISL_443078, EPI_ISL_443080, EPI_ISL_443081, EPI_ISL_443083, EPI_ISL_443084, EPI_ISL_443086, EPI_ISL_443087, EPI_ISL_443089, EPI_ISL_443090, EPI_ISL_443091, EPI_ISL_443092, EPI_ISL_443093, EPI_ISL_443094, EPI_ISL_443095, EPI_ISL_443096, EPI_ISL_443097, EPI_ISL_443098, EPI_ISL_443099, EPI_ISL_443100, EPI_ISL_443101, EPI_ISL_443102, EPI_ISL_443103, EPI_ISL_443104, EPI_ISL_443105, EPI_ISL_443111, EPI_ISL_443112, EPI_ISL_443113, EPI_ISL_443114, EPI_ISL_443116 | | | |
| see above | Department of Pathology, University of Cambridge | Wellcome Sanger Institute for the COVID-19 Genomics UK Consortium | Luke W Meredith, M. Estée Török , Myra Hosmillo, William L. Hamilton, Martin D. Curran, Theresa Feltwell, Grant Hall, Anna Yakovleva, Fahad A Khokhar, Charlotte J. Houldcroft, Laura G Caller, Aminu S. Jahun, Sarah L. Caddy, Ian Goodfellow, Alex Alderton, Roberto Amato, Sonia Goncalves, Ewan Harrison, David K. Jackson, Ian Johnston, Dominic Kwiatkowski, Cordelia Langford, John Sillitoe on behalf of the Wellcome Sanger Institute COVID-19 Surveillance Team (http://www.sanger.ac.uk/covid-team) |
| EPI_ISL_443183 | M Health Fairview | University of Minnesota Genomics Center | Daryl M. Gohl, John Garbe, Patrick Grady, Jerry Daniel, Ray Watson, Benjamin Auch, Andrew Nelson, Sophia Yohe, and Kenneth B. Beckman |
| EPI_ISL_443184, EPI_ISL_443185, EPI_ISL_443186 | UW Virology Lab | UW Virology Lab | Pavitra Roychoudhury, Hong Xie, Keith Jerome, Alexander Greninger |
| EPI_ISL_443187 | National Virology Reference Laboratory | National Public Health Laboratory, National Centre for Infectious Diseases | Mak Tze Minn, Octavia Sophie, Chavatte Jean-Marc, Zaini Zainun, Taib Surita, Cui Lin, Lin Raymond Tzer Pin |
| EPI_ISL_443188, EPI_ISL_443189, EPI_ISL_443190, EPI_ISL_443191, EPI_ISL_443192, EPI_ISL_443193, EPI_ISL_443195, EPI_ISL_443197, EPI_ISL_443198, EPI_ISL_443199, EPI_ISL_443200, EPI_ISL_443201, EPI_ISL_443202, EPI_ISL_443203, EPI_ISL_443204, EPI_ISL_443205, EPI_ISL_443206, EPI_ISL_443207, EPI_ISL_443208, EPI_ISL_443209, EPI_ISL_443210, EPI_ISL_443211, EPI_ISL_443212, EPI_ISL_443213, EPI_ISL_443214, EPI_ISL_443215, EPI_ISL_443216, EPI_ISL_443217, EPI_ISL_443218, EPI_ISL_443219, EPI_ISL_443220, EPI_ISL_443221, EPI_ISL_443222, EPI_ISL_443223, EPI_ISL_443224, EPI_ISL_443225, EPI_ISL_443226, EPI_ISL_443227, EPI_ISL_443228, EPI_ISL_443229, EPI_ISL_443230, EPI_ISL_443231, EPI_ISL_443232, EPI_ISL_443233, EPI_ISL_443234, EPI_ISL_443235, EPI_ISL_443236, EPI_ISL_443237, EPI_ISL_443238, EPI_ISL_443239, EPI_ISL_443240, EPI_ISL_443241, EPI_ISL_443242, EPI_ISL_443243, EPI_ISL_443244, EPI_ISL_443245, EPI_ISL_443246, EPI_ISL_443247, EPI_ISL_443248, EPI_ISL_443249 | | | |
| see above | National Public Health Laboratory, National Centre for Infectious Diseases | National Public Health Laboratory, National Centre for Infectious Diseases | Mak Tze Minn, Octavia Sophie, Chavatte Jean-Marc, Cui Lin, Lin Raymond Tzer Pin |
| EPI_ISL_443253, EPI_ISL_443254, EPI_ISL_443255, EPI_ISL_443256 | M Health Fairview | University of Minnesota Genomics Center | Daryl M. Gohl, John Garbe, Patrick Grady, Jerry Daniel, Ray Watson, Benjamin Auch, Andrew Nelson, Sophia Yohe, and Kenneth B. Beckman |
| EPI_ISL_443258, EPI_ISL_443259 | Résidence Ornano | National Reference Center for Viruses of Respiratory Infections, Institut Pasteur, Paris | Mélanie Albert, Marion Barbet, Sylvie Behillil, Méline Bizard, Angela Brisebarre, Flora Donati, Etienne Simon-Lorière, Vincent Enouf, Maud Vanpeene, Sylvie van der Werf |
| EPI_ISL_443260 | LABM GH nord Essonne de Longjumeau - BP 125 | National Reference Center for Viruses of Respiratory Infections, Institut Pasteur, Paris | Mélanie Albert, Marion Barbet, Sylvie Behillil, Méline Bizard, Angela Brisebarre, Flora Donati, Etienne Simon-Lorière, Vincent Enouf, Maud Vanpeene, Sylvie van der Werf |
| EPI_ISL_443261, EPI_ISL_443262, EPI_ISL_443263, EPI_ISL_443264 | CHU de Dijon - Laboratoire de Virologie | National Reference Center for Viruses of Respiratory Infections, Institut Pasteur, Paris | Mélanie Albert, Marion Barbet, Sylvie Behillil, Méline Bizard, Angela Brisebarre, Flora Donati, Etienne Simon-Lorière, Vincent Enouf, Maud Vanpeene, Sylvie van der Werf, Jean-Baptiste Bour |
| EPI_ISL_443265, EPI_ISL_443266, EPI_ISL_443267, EPI_ISL_443268, EPI_ISL_443269, EPI_ISL_443270, EPI_ISL_443271, EPI_ISL_443272, EPI_ISL_443273, EPI_ISL_443274, EPI_ISL_443275, EPI_ISL_443276, EPI_ISL_443277, EPI_ISL_443278, EPI_ISL_443279, EPI_ISL_443280, EPI_ISL_443281, EPI_ISL_443282, EPI_ISL_443283 | | | |
| see above | CHU - Hôpital Cavale Blanche - Labo. de Virologie | National Reference Center for Viruses of Respiratory Infections, Institut Pasteur, Paris | Mélanie Albert, Marion Barbet, Sylvie Behillil, Méline Bizard, Angela Brisebarre, Flora Donati, Etienne Simon-Lorière, Vincent Enouf, Maud Vanpeene, Sylvie van der Werf, Léa Pilorge |
| EPI_ISL_443284, EPI_ISL_443285, EPI_ISL_443286, EPI_ISL_443287, EPI_ISL_443288 | Laboratoire de Microbiologie - Bât A - CH René Dubois | National Reference Center for Viruses of Respiratory Infections, Institut Pasteur, Paris | Mélanie Albert, Marion Barbet, Sylvie Behillil, Méline Bizard, Angela Brisebarre, Flora Donati, Etienne Simon-Lorière, Vincent Enouf, Maud Vanpeene, Sylvie van der Werf, Pascale Martres |
| EPI_ISL_443289, EPI_ISL_443290, EPI_ISL_443291, EPI_ISL_443292, EPI_ISL_443293, EPI_ISL_443294 | CHRU Pontchaillou - Laboratoire de Virologie | National Reference Center for Viruses of Respiratory Infections, Institut Pasteur, Paris | Mélanie Albert, Marion Barbet, Sylvie Behillil, Méline Bizard, Angela Brisebarre, Flora Donati, Etienne Simon-Lorière, Vincent Enouf, Maud Vanpeene, Sylvie van der Werf, Gisèle Lagathu |
| EPI_ISL_443295, EPI_ISL_443296, EPI_ISL_443298, EPI_ISL_443299 | Hôpital Necker - Enfants - Malades Laboratoire de Virologie | National Reference Center for Viruses of Respiratory Infections, Institut Pasteur, Paris | Mélanie Albert, Marion Barbet, Sylvie Behillil, Méline Bizard, Angela Brisebarre, Flora Donati, Etienne Simon-Lorière, Vincent Enouf, Maud Vanpeene, Sylvie van der Werf, Marianne Leruez-Ville |
| EPI_ISL_443300, EPI_ISL_443301, EPI_ISL_443302 | Cabinet Médical | National Reference Center for Viruses of Respiratory Infections, Institut Pasteur, Paris | Mélanie Albert, Marion Barbet, Sylvie Behillil, Méline Bizard, Angela Brisebarre, Flora Donati, Etienne Simon-Lorière, Vincent Enouf, Maud Vanpeene, Sylvie van der Werf |
| EPI_ISL_443303 | Résidence Les Marines | National Reference Center for Viruses of Respiratory Infections, Institut Pasteur, Paris | Mélanie Albert, Marion Barbet, Sylvie Behillil, Méline Bizard, Angela Brisebarre, Flora Donati, Etienne Simon-Lorière, Vincent Enouf, Maud Vanpeene, Sylvie van der Werf |
| EPI_ISL_443305 | LABM GH nord Essonne de Longjumeau - BP 125 | National Reference Center for Viruses of Respiratory Infections, Institut Pasteur, Paris | Mélanie Albert, Marion Barbet, Sylvie Behillil, Méline Bizard, Angela Brisebarre, Flora Donati, Etienne Simon-Lorière, Vincent Enouf, Maud Vanpeene, Sylvie van der Werf |
| EPI_ISL_443306 | Cabinet Médical | National Reference Center for Viruses of Respiratory Infections, Institut Pasteur, Paris | Mélanie Albert, Marion Barbet, Sylvie Behillil, Méline Bizard, Angela Brisebarre, Flora Donati, Etienne Simon-Lorière, Vincent Enouf, Maud Vanpeene, Sylvie van der Werf |
| EPI_ISL_443307 | La Villa Papyri | National Reference Center for Viruses of Respiratory Infections, Institut Pasteur, Paris | Mélanie Albert, Marion Barbet, Sylvie Behillil, Méline Bizard, Angela Brisebarre, Flora Donati, Etienne Simon-Lorière, Vincent Enouf, Maud Vanpeene, Sylvie van der Werf |
| EPI_ISL_443308 | Plaisance | National Reference Center for Viruses of Respiratory Infections, Institut Pasteur, Paris | Mélanie Albert, Marion Barbet, Sylvie Behillil, Méline Bizard, Angela Brisebarre, Flora Donati, Etienne Simon-Lorière, Vincent Enouf, Maud Vanpeene, Sylvie van der Werf |
| EPI_ISL_443309 | CH Compiègne Laboratoire de Biologie | National Reference Center for Viruses of Respiratory Infections, Institut Pasteur, Paris | Mélanie Albert, Marion Barbet, Sylvie Behillil, Méline Bizard, Angela Brisebarre, Flora Donati, Etienne Simon-Lorière, Vincent Enouf, Maud Vanpeene, Sylvie van der Werf |
| EPI_ISL_443310 | Centre de santé Filieris | National Reference Center for Viruses of Respiratory Infections, Institut Pasteur, Paris | Mélanie Albert, Marion Barbet, Sylvie Behillil, Méline Bizard, Angela Brisebarre, Flora Donati, Etienne Simon-Lorière, Vincent Enouf, Maud Vanpeene, Sylvie van der Werf |
| EPI_ISL_443311, EPI_ISL_443312, EPI_ISL_443313 | Cabinet Médical | National Reference Center for Viruses of Respiratory Infections, Institut Pasteur, Paris | Mélanie Albert, Marion Barbet, Sylvie Behillil, Méline Bizard, Angela Brisebarre, Flora Donati, Etienne Simon-Lorière, Vincent Enouf, Maud Vanpeene, Sylvie van der Werf |
| EPI_ISL_443314 | LABM GH nord Essonne de Longjumeau - BP 125 | National Reference Center for Viruses of Respiratory Infections, Institut Pasteur, Paris | Mélanie Albert, Marion Barbet, Sylvie Behillil, Méline Bizard, Angela Brisebarre, Flora Donati, Etienne Simon-Lorière, Vincent Enouf, Maud Vanpeene, Sylvie van der Werf |
| EPI_ISL_443315 | Château de la Source | National Reference Center for Viruses of Respiratory Infections, Institut Pasteur, Paris | Mélanie Albert, Marion Barbet, Sylvie Behillil, Méline Bizard, Angela Brisebarre, Flora Donati, Etienne Simon-Lorière, Vincent Enouf, Maud Vanpeene, Sylvie van der Werf |
| EPI_ISL_443316 | CH Compiègne Laboratoire de Biologie | National Reference Center for Viruses of Respiratory Infections, Institut Pasteur, Paris | Mélanie Albert, Marion Barbet, Sylvie Behillil, Méline Bizard, Angela Brisebarre, Flora Donati, Etienne Simon-Lorière, Vincent Enouf, Maud Vanpeene, Sylvie van der Werf, Olivia Raulin |
| EPI_ISL_443317 | Cabinet Médical | National Reference Center for Viruses of Respiratory Infections, Institut Pasteur, Paris | Mélanie Albert, Marion Barbet, Sylvie Behillil, Méline Bizard, Angela Brisebarre, Flora Donati, Etienne Simon-Lorière, Vincent Enouf, Maud Vanpeene, Sylvie van der Werf |
| EPI_ISL_443319, EPI_ISL_443332, EPI_ISL_443335, EPI_ISL_443336, EPI_ISL_443338, EPI_ISL_443340, EPI_ISL_443342, EPI_ISL_443343, EPI_ISL_443345, EPI_ISL_443349, EPI_ISL_443353, EPI_ISL_443359, EPI_ISL_443360, EPI_ISL_443371, EPI_ISL_443372, EPI_ISL_443375, EPI_ISL_443378, EPI_ISL_443383, EPI_ISL_443384, EPI_ISL_443385, EPI_ISL_443386, EPI_ISL_443387, EPI_ISL_443388, EPI_ISL_443389, EPI_ISL_443390, EPI_ISL_443391, EPI_ISL_443392, EPI_ISL_443393, EPI_ISL_443394, EPI_ISL_443395, EPI_ISL_443396, EPI_ISL_443398, EPI_ISL_443399, EPI_ISL_443402, EPI_ISL_443403, EPI_ISL_443404, EPI_ISL_443406, EPI_ISL_443408, EPI_ISL_443409, EPI_ISL_443410, EPI_ISL_443411, EPI_ISL_443414, EPI_ISL_443415, EPI_ISL_443416, EPI_ISL_443417, EPI_ISL_443419, EPI_ISL_443421, EPI_ISL_443425, EPI_ISL_443426, EPI_ISL_443427, EPI_ISL_443428, EPI_ISL_443429, EPI_ISL_443434, EPI_ISL_443436, EPI_ISL_443437, EPI_ISL_443440, EPI_ISL_443441, EPI_ISL_443444, EPI_ISL_443445, EPI_ISL_443446, EPI_ISL_443447, EPI_ISL_443448, EPI_ISL_443450, EPI_ISL_443452, EPI_ISL_443453, EPI_ISL_443454, EPI_ISL_443455, EPI_ISL_443456, EPI_ISL_443457, EPI_ISL_443458, EPI_ISL_443459, EPI_ISL_443460, EPI_ISL_443461, EPI_ISL_443463, EPI_ISL_443464, EPI_ISL_443465, EPI_ISL_443466, EPI_ISL_443467, EPI_ISL_443469, EPI_ISL_443470, EPI_ISL_443472, EPI_ISL_443473, EPI_ISL_443474, EPI_ISL_443476, EPI_ISL_443477, EPI_ISL_443478, EPI_ISL_443481, EPI_ISL_443482, EPI_ISL_443483, EPI_ISL_443487, EPI_ISL_443488, EPI_ISL_443489, EPI_ISL_443490, EPI_ISL_443493, EPI_ISL_443494, EPI_ISL_443495, EPI_ISL_443496, EPI_ISL_443498, EPI_ISL_443500, EPI_ISL_443501, EPI_ISL_443502, EPI_ISL_443503, EPI_ISL_443504, EPI_ISL_443505, EPI_ISL_443507, EPI_ISL_443509, EPI_ISL_443510, EPI_ISL_443511, EPI_ISL_443512, EPI_ISL_443514, EPI_ISL_443515, EPI_ISL_443517, EPI_ISL_443519, EPI_ISL_443522, EPI_ISL_443523, EPI_ISL_443524, EPI_ISL_443525, EPI_ISL_443526, EPI_ISL_443527, EPI_ISL_443528, EPI_ISL_443529, EPI_ISL_443530, EPI_ISL_443531, EPI_ISL_443532, EPI_ISL_443533, EPI_ISL_443534, EPI_ISL_443535, EPI_ISL_443536, EPI_ISL_443537, EPI_ISL_443538, EPI_ISL_443540, EPI_ISL_443545, EPI_ISL_443546, EPI_ISL_443548, EPI_ISL_443549, EPI_ISL_443552, EPI_ISL_443554, EPI_ISL_443557, EPI_ISL_443558, EPI_ISL_443559, EPI_ISL_443560, EPI_ISL_443561, EPI_ISL_443562, EPI_ISL_443564, EPI_ISL_443566, EPI_ISL_443567, EPI_ISL_443568, EPI_ISL_443569, EPI_ISL_443571, EPI_ISL_443572, EPI_ISL_443574, EPI_ISL_443575, EPI_ISL_443576, EPI_ISL_443577, EPI_ISL_443578, EPI_ISL_443579, EPI_ISL_443581, EPI_ISL_443582, EPI_ISL_443583, EPI_ISL_443585, EPI_ISL_443586, EPI_ISL_443587, EPI_ISL_443588, EPI_ISL_443589, EPI_ISL_443590, EPI_ISL_443591, EPI_ISL_443592, EPI_ISL_443593, EPI_ISL_443594, EPI_ISL_443595, EPI_ISL_443598, EPI_ISL_443599, EPI_ISL_443600, EPI_ISL_443602, EPI_ISL_443604, EPI_ISL_443605, EPI_ISL_443607, EPI_ISL_443608, EPI_ISL_443609, EPI_ISL_443612, EPI_ISL_443613, EPI_ISL_443614, EPI_ISL_443616, EPI_ISL_443617, EPI_ISL_443618, EPI_ISL_443621, EPI_ISL_443622, EPI_ISL_443623, EPI_ISL_443625, EPI_ISL_443626, EPI_ISL_443628, EPI_ISL_443629, EPI_ISL_443630, EPI_ISL_443631, EPI_ISL_443632, EPI_ISL_443633, EPI_ISL_443634, EPI_ISL_443636, EPI_ISL_443637, EPI_ISL_443639, EPI_ISL_443640, EPI_ISL_443644, EPI_ISL_443646, EPI_ISL_443648, EPI_ISL_443649, EPI_ISL_443651, EPI_ISL_443653, EPI_ISL_443654, EPI_ISL_443656, EPI_ISL_443657, EPI_ISL_443658, EPI_ISL_443661, EPI_ISL_443662, EPI_ISL_443663, EPI_ISL_443667, EPI_ISL_443668, EPI_ISL_443670, EPI_ISL_443672, EPI_ISL_443673, EPI_ISL_443674, EPI_ISL_443676, EPI_ISL_443677, EPI_ISL_443680, EPI_ISL_443681, EPI_ISL_443682, EPI_ISL_443685 | | | |
| see above | Department of Pathology, University of Cambridge | Wellcome Sanger Institute for the COVID-19 Genomics UK Consortium | Luke W Meredith, M. Estée Török , Myra Hosmillo, William L. Hamilton, Martin D. Curran, Theresa Feltwell, Grant Hall, Anna Yakovleva, Fahad A Khokhar, Charlotte J. Houldcroft, Laura G Caller, Aminu S. Jahun, Sarah L. Caddy, Ian Goodfellow; and Alex Alderton, Roberto Amato, Sonia Goncalves, Ewan Harrison, David K. Jackson, Ian Johnston, Dominic Kwiatkowski, Cordelia Langford, John Sillitoe on behalf of the Wellcome Sanger Institute COVID-19 Surveillance Team (http://www.sanger.ac.uk/covid-team) |
| EPI_ISL_443686, EPI_ISL_443687, EPI_ISL_443690, EPI_ISL_443691, EPI_ISL_443692, EPI_ISL_443697, EPI_ISL_443698, EPI_ISL_443699, EPI_ISL_443701, EPI_ISL_443702, EPI_ISL_443704, EPI_ISL_443706, EPI_ISL_443707, EPI_ISL_443709, EPI_ISL_443710, EPI_ISL_443711, EPI_ISL_443714, EPI_ISL_443715, EPI_ISL_443716, EPI_ISL_443717, EPI_ISL_443718, EPI_ISL_443719, EPI_ISL_443720, EPI_ISL_443726, EPI_ISL_443729, EPI_ISL_443730, EPI_ISL_443733, EPI_ISL_443738, EPI_ISL_443739, EPI_ISL_443740, EPI_ISL_443741, EPI_ISL_443742, EPI_ISL_443743, EPI_ISL_443744, EPI_ISL_443745, EPI_ISL_443746, EPI_ISL_443747, EPI_ISL_443748, EPI_ISL_443749, EPI_ISL_443751, EPI_ISL_443752, EPI_ISL_443757, EPI_ISL_443758, EPI_ISL_443759, EPI_ISL_443760, EPI_ISL_443763, EPI_ISL_443764, EPI_ISL_443765, EPI_ISL_443768, EPI_ISL_443769, EPI_ISL_443770, EPI_ISL_443772, EPI_ISL_443773, EPI_ISL_443774, EPI_ISL_443775, EPI_ISL_443776, EPI_ISL_443777, EPI_ISL_443778, EPI_ISL_443780, EPI_ISL_443781, EPI_ISL_443782, EPI_ISL_443783, EPI_ISL_443784, EPI_ISL_443786, EPI_ISL_443787, EPI_ISL_443788, EPI_ISL_443789, EPI_ISL_443790, EPI_ISL_443791, EPI_ISL_443793, EPI_ISL_443795, EPI_ISL_443796, | | | |

| | | | |
|---|---|---|---|
| EPI_ISL_443797, EPI_ISL_443798, EPI_ISL_443800, EPI_ISL_443801, EPI_ISL_443802, EPI_ISL_443803, EPI_ISL_443804, EPI_ISL_443805, EPI_ISL_443807, EPI_ISL_443808, EPI_ISL_443809, EPI_ISL_443811, EPI_ISL_443812, EPI_ISL_443813, EPI_ISL_443815, EPI_ISL_443816, EPI_ISL_443817, EPI_ISL_443819, EPI_ISL_443820, EPI_ISL_443821, EPI_ISL_443825, EPI_ISL_443826, EPI_ISL_443827, EPI_ISL_443828, EPI_ISL_443830, EPI_ISL_443831, EPI_ISL_443832, EPI_ISL_443833, EPI_ISL_443834, EPI_ISL_443835, EPI_ISL_443836, EPI_ISL_443838, EPI_ISL_443839, EPI_ISL_443840, EPI_ISL_443841, EPI_ISL_443842, EPI_ISL_443843, EPI_ISL_443844, EPI_ISL_443846, EPI_ISL_443847, EPI_ISL_443851, EPI_ISL_443852, EPI_ISL_443853, EPI_ISL_443854, EPI_ISL_443855, EPI_ISL_443856, EPI_ISL_443857, EPI_ISL_443858, EPI_ISL_443859, EPI_ISL_443860, EPI_ISL_443862, EPI_ISL_443864, EPI_ISL_443865, EPI_ISL_443866, EPI_ISL_443867, EPI_ISL_443869, EPI_ISL_443870, EPI_ISL_443871, EPI_ISL_443872, EPI_ISL_443873, EPI_ISL_443876, EPI_ISL_443877, EPI_ISL_443879, EPI_ISL_443880, EPI_ISL_443881, EPI_ISL_443882, EPI_ISL_443883, EPI_ISL_443884, EPI_ISL_443885, EPI_ISL_443888, EPI_ISL_443891, EPI_ISL_443892, EPI_ISL_443893, EPI_ISL_443894, EPI_ISL_443895, EPI_ISL_443896, EPI_ISL_443897, EPI_ISL_443898, EPI_ISL_443899, EPI_ISL_443901, EPI_ISL_443902, EPI_ISL_443904, EPI_ISL_443906, EPI_ISL_443907, EPI_ISL_443909, EPI_ISL_443910, EPI_ISL_443911, EPI_ISL_443912, EPI_ISL_443914, EPI_ISL_443915, EPI_ISL_443916, EPI_ISL_443918, EPI_ISL_443919, EPI_ISL_443922, EPI_ISL_443924, EPI_ISL_443925, EPI_ISL_443927, EPI_ISL_443928, EPI_ISL_443929, EPI_ISL_443933, EPI_ISL_443935, EPI_ISL_443936, EPI_ISL_443938, EPI_ISL_443939, EPI_ISL_443940, EPI_ISL_443941, EPI_ISL_443942, EPI_ISL_443943, EPI_ISL_443944, EPI_ISL_443945, EPI_ISL_443946, EPI_ISL_443947, EPI_ISL_443948, EPI_ISL_443949, EPI_ISL_443950, EPI_ISL_443951, EPI_ISL_443952, EPI_ISL_443953, EPI_ISL_443954, EPI_ISL_443955, EPI_ISL_443956, EPI_ISL_443957, EPI_ISL_443958, EPI_ISL_443959, EPI_ISL_443961, EPI_ISL_443962, EPI_ISL_443963, EPI_ISL_443964, EPI_ISL_443967, EPI_ISL_443968, EPI_ISL_443969, EPI_ISL_443970, EPI_ISL_443971, EPI_ISL_443972, EPI_ISL_443973, EPI_ISL_443975, EPI_ISL_443976, EPI_ISL_443978, EPI_ISL_443980, EPI_ISL_443981, EPI_ISL_443982, EPI_ISL_443983, EPI_ISL_443985, EPI_ISL_443986, EPI_ISL_443987, EPI_ISL_443988, EPI_ISL_443989, EPI_ISL_443990, EPI_ISL_443991, EPI_ISL_443992, EPI_ISL_443996, EPI_ISL_443997, EPI_ISL_443998, EPI_ISL_443999, EPI_ISL_444000, EPI_ISL_444001, EPI_ISL_444002, EPI_ISL_444003, EPI_ISL_444006, EPI_ISL_444007, EPI_ISL_444009, EPI_ISL_444010, EPI_ISL_444011, EPI_ISL_444012, EPI_ISL_444013, EPI_ISL_444015, EPI_ISL_444016, EPI_ISL_444018, EPI_ISL_444019, EPI_ISL_444020, EPI_ISL_444021 | | | |
| see above | PHE South West Regional Laboratory, National Infection Service | Wellcome Sanger Institute for the COVID-19 Genomics UK Consortium | Stephanie Hutchings, Hannah Pymont, Dr Peter Muir, Barry Vipond, Rich Hopes; and Alex Alderton, Roberto Amato, Sonia Goncalves, Ewan Harrison, David K. Jackson, Ian Johnston, Dominic Kwiatkowski, Cordelia Langford, John Sillitoe on behalf of the Wellcome Sanger Institute COVID-19 Surveillance Team (http://www.sanger.ac.uk/covid-team) |
| EPI_ISL_444022 | Baylor College of Medicine | Baylor College of Medicine: HGSC | Vasanthi Avadhanula, Erin Nicholson, David Henke, Pedro Piedra, Harsha Doddapaneni, Donna Muzny, Qingchang Meng, Hsu Chao, Zeineen Momin, Hua Shen, George Weissenberger, Kavya Kottapalli, Yimiti Meiheerguli, Sejal Salvi, Ginger Metcalf, Vipin Menon, Sara J.J. Cregeen, Matthew C. Ross, Tulin Ayvaz, Richard Sucgang, Kristi L. Hoffman, Matthew Wong, Joseph F. Petrosino |
| EPI_ISL_444023, EPI_ISL_444024, EPI_ISL_444025, EPI_ISL_444026 | County of Santa Clara Public Health | Chan-Zuckerberg Biohub | CZB Cliahub Consortium |
| EPI_ISL_444051, EPI_ISL_444052, EPI_ISL_444053, EPI_ISL_444054, EPI_ISL_444055, EPI_ISL_444057, EPI_ISL_444058, EPI_ISL_444059, EPI_ISL_444063, EPI_ISL_444065, EPI_ISL_444066, EPI_ISL_444067, EPI_ISL_444068, EPI_ISL_444069, EPI_ISL_444070, EPI_ISL_444071, EPI_ISL_444073, EPI_ISL_444074, EPI_ISL_444075, EPI_ISL_444076, EPI_ISL_444077, EPI_ISL_444078 | | | |
| see above | UCSF Clinical Microbiology Laboratory | Chan-Zuckerberg Biohub | CZB Cliahub Consortium |
| EPI_ISL_444079, EPI_ISL_444080, EPI_ISL_444081, EPI_ISL_444082, EPI_ISL_444083, EPI_ISL_444084, EPI_ISL_444085, EPI_ISL_444086, EPI_ISL_444087, EPI_ISL_444088, EPI_ISL_444089, EPI_ISL_444090, EPI_ISL_444091, EPI_ISL_444092, EPI_ISL_444093, EPI_ISL_444094, EPI_ISL_444095, EPI_ISL_444096, EPI_ISL_444097, EPI_ISL_444098, EPI_ISL_444099, EPI_ISL_444100, EPI_ISL_444101, EPI_ISL_444102, EPI_ISL_444103, EPI_ISL_444104, EPI_ISL_444105, EPI_ISL_444106, EPI_ISL_444108, EPI_ISL_444109, EPI_ISL_444110, EPI_ISL_444111, EPI_ISL_444112, EPI_ISL_444114, EPI_ISL_444116, EPI_ISL_444117, EPI_ISL_444120, EPI_ISL_444122, EPI_ISL_444123, EPI_ISL_444124, EPI_ISL_444125, EPI_ISL_444126, EPI_ISL_444127, EPI_ISL_444128, EPI_ISL_444129, EPI_ISL_444130, EPI_ISL_444133, EPI_ISL_444134, EPI_ISL_444137, EPI_ISL_444139, EPI_ISL_444141, EPI_ISL_444143, EPI_ISL_444144, EPI_ISL_444146, EPI_ISL_444147, EPI_ISL_444149, EPI_ISL_444150, EPI_ISL_444151, EPI_ISL_444152, EPI_ISL_444153, EPI_ISL_444155, EPI_ISL_444156, EPI_ISL_444159, EPI_ISL_444160, EPI_ISL_444161, EPI_ISL_444162, EPI_ISL_444163, EPI_ISL_444164, EPI_ISL_444165, EPI_ISL_444166, EPI_ISL_444167, EPI_ISL_444168, EPI_ISL_444169, EPI_ISL_444170, EPI_ISL_444171, EPI_ISL_444172, EPI_ISL_444173, EPI_ISL_444174, EPI_ISL_444175, EPI_ISL_444178, EPI_ISL_444179, EPI_ISL_444180, EPI_ISL_444181, EPI_ISL_444182, EPI_ISL_444185, EPI_ISL_444188, EPI_ISL_444189, EPI_ISL_444190, EPI_ISL_444191, EPI_ISL_444192, EPI_ISL_444193, EPI_ISL_444194, EPI_ISL_444195, EPI_ISL_444196, EPI_ISL_444197, EPI_ISL_444198, EPI_ISL_444199, EPI_ISL_444201, EPI_ISL_444202, EPI_ISL_444203, EPI_ISL_444204, EPI_ISL_444205, EPI_ISL_444206, EPI_ISL_444207, EPI_ISL_444208, EPI_ISL_444209, EPI_ISL_444210, EPI_ISL_444211, EPI_ISL_444212, EPI_ISL_444213, EPI_ISL_444214, EPI_ISL_444215, EPI_ISL_444216, EPI_ISL_444217, EPI_ISL_444218, EPI_ISL_444219, EPI_ISL_444220, EPI_ISL_444221, EPI_ISL_444222, EPI_ISL_444223, EPI_ISL_444224, EPI_ISL_444225, EPI_ISL_444226, EPI_ISL_444227, EPI_ISL_444228, EPI_ISL_444229, EPI_ISL_444230, EPI_ISL_444231, EPI_ISL_444232, EPI_ISL_444233, EPI_ISL_444235, EPI_ISL_444236, EPI_ISL_444238, EPI_ISL_444239, EPI_ISL_444241, EPI_ISL_444242, EPI_ISL_444243, EPI_ISL_444244, EPI_ISL_444245, EPI_ISL_444246, EPI_ISL_444247, EPI_ISL_444248, EPI_ISL_444249, EPI_ISL_444250, EPI_ISL_444251, EPI_ISL_444252, EPI_ISL_444253, EPI_ISL_444254, EPI_ISL_444256, EPI_ISL_444257, EPI_ISL_444258, EPI_ISL_444259, EPI_ISL_444260, EPI_ISL_444261, EPI_ISL_444262, EPI_ISL_444263, EPI_ISL_444265, EPI_ISL_444268, EPI_ISL_444269, EPI_ISL_444270, EPI_ISL_444272 | | | |
| see above | University College London, Great Ormond Street Hospital for Children NHS Foundation Trust, Imperial College Healthcare NHS Trust | COVID-19 Genomics UK (COG-UK) Consortium | Sergi Castellano, Rachel Williams, Mark Kristiansen, Paola Resende Silva, Sunando Roy, Tony Brooks, Helena Tutill, Paola Niola, Patricia Dyal, Charlotte Williams, Leysa Forrest, Yasmin Panchbhaya, Jacqueline Findlay, Sam Weeks, Julianne Brown, Kathryn Harris, Paul Randell, James Price, Alison Holmes, Judith Breuer |
| EPI_ISL_444273 | State Key Laboratory of Respiratory Disease, National Clinical Research Center for Respiratory Disease, Guangzhou Institute of Respiratory Health, the First Affiliated Hospital of Guangzhou Medical University | State Key Laboratory of Respiratory Disease, National Clinical Research Center for Respiratory Disease, Guangzhou Institute of Respiratory Health, the First Affiliated Hospital of Guangzhou Medical University | Sun,J., Shi,Y., Zheng,K., Huang,J. and Zhao,J. |
| EPI_ISL_444274, EPI_ISL_444276, EPI_ISL_444277, EPI_ISL_444278 | Laboratory Medicine | Department of Laboratory Medicine, Lin-Kou Chang Gung Memorial Hospital, Taoyuan, Taiwan | Kuo-Chien Tsao, Yu-Nong Gong, Shu-Li Yang, Yi-Chun Liu, Chung-Guei Huang, Mei-Jen Hsiao, Po-Wei Huang, Cheng-Ta Yang, Cheng-Hsun Chiu, Peng-Nien Huang, Kuo-Ming Lee, Guang-Wu Chen, Shin-Ru Shih |
| EPI_ISL_444279, EPI_ISL_444280, EPI_ISL_444281, EPI_ISL_444283, EPI_ISL_444284, EPI_ISL_444285, EPI_ISL_444286, EPI_ISL_444288, EPI_ISL_444289, EPI_ISL_444290, EPI_ISL_444291, EPI_ISL_444292, EPI_ISL_444293, EPI_ISL_444299, EPI_ISL_444300 | | | |
| see above | University of Birmingham | COVID-19 Genomics UK (COG-UK) Consortium | Loman Lab: Claire McMurray, Joanne Stockton, Samuel Nicholls, Radoslaw Poplawski, Will Rowe, Josh Quick, Nicholas Loman // UHB Lab: Celina M Whalley, Andrew Bosworth, Charlotte Poxon, Kasun Wanigasooriya, Oliver Pickles, Mike Kidd, Alex Richter, Andrew D Beggs // PHE Heartlands Lab: Husam Osman, Andrew Bosworth |
| EPI_ISL_444320, EPI_ISL_444323, EPI_ISL_444328, EPI_ISL_444329, EPI_ISL_444334, EPI_ISL_444335, EPI_ISL_444336, EPI_ISL_444337, EPI_ISL_444338, EPI_ISL_444339, EPI_ISL_444340, EPI_ISL_444342, EPI_ISL_444343, EPI_ISL_444345, EPI_ISL_444346, EPI_ISL_444347, EPI_ISL_444348, EPI_ISL_444349, EPI_ISL_444351, EPI_ISL_444352, EPI_ISL_444353, EPI_ISL_444354, EPI_ISL_444355, EPI_ISL_444356, EPI_ISL_444357, EPI_ISL_444358, EPI_ISL_444359, EPI_ISL_444360, EPI_ISL_444361, EPI_ISL_444362, EPI_ISL_444363, EPI_ISL_444364, EPI_ISL_444365, EPI_ISL_444366, EPI_ISL_444368, EPI_ISL_444370, EPI_ISL_444371, EPI_ISL_444372, EPI_ISL_444374, EPI_ISL_444375, EPI_ISL_444376, EPI_ISL_444377, EPI_ISL_444384, EPI_ISL_444393, EPI_ISL_444394, EPI_ISL_444395, EPI_ISL_444397, EPI_ISL_444398, EPI_ISL_444400, EPI_ISL_444402, EPI_ISL_444407, EPI_ISL_444413, EPI_ISL_444415, EPI_ISL_444416, EPI_ISL_444417, EPI_ISL_444418, EPI_ISL_444419, EPI_ISL_444420, EPI_ISL_444421, EPI_ISL_444422, EPI_ISL_444423, EPI_ISL_444430, EPI_ISL_444432, EPI_ISL_444434, EPI_ISL_444435, EPI_ISL_444436, EPI_ISL_444437, EPI_ISL_444438, EPI_ISL_444439, EPI_ISL_444440, EPI_ISL_444441, EPI_ISL_444442, EPI_ISL_444443, EPI_ISL_444445, EPI_ISL_444446, EPI_ISL_444447, EPI_ISL_444448, EPI_ISL_444449, EPI_ISL_444450, EPI_ISL_444451, EPI_ISL_444452, EPI_ISL_444453 | | | |
| see above | Department of Pathology, University of Cambridge | COVID-19 Genomics UK (COG-UK) Consortium | Luke W Meredith, M. Estée Török , Myra Hosmillo, William L. Hamilton, Martin D. Curran, Theresa Feltwell, Grant Hall, Anna Yakovleva, Fahad A Khokhar, Charlotte J. Houldcroft, Laura G Caller, Aminu S. Jahun, Sarah L. Caddy, Ian Goodfellow |
| EPI_ISL_444456 | B.J. Medical College and Civil hospital | Gujarat Biotechnology Research Centre | R D Dixit, Snehal Bagatharia, Kamlesh J Upadhyay, Ramesh Pandit, Tejas Shah, Ankit Hinsu, Pritesh Sabara, Apurvasinh Puvar, Janvi Raval, Monika Gandhi, Pinal Trivedi, Maharshi Pandya, Amit Kanani, Akanksha Verma, Nitin Savaliya, Raghawendra Kumar, Dinesh Kumar, Zuber Saiyed, Dipa Kinariwala, Disha Patel, Binita Aring, Neeta Khandelwal, Geeta Vaghela, Sonia Barve, Bhavesh Modi, Kairavi Joshi, Gaurishankar Shrimali, Nidhi Sood, Pranay Shah, Pooja P Doshi, Chaitanya Joshi, Madhvi Joshi |
| EPI_ISL_444457 | B.J. Medical College and Civil hospital | Gujarat Biotechnology Research Centre | Snehal Bagatharia, Kamlesh J Upadhyay, Ramesh Pandit, Tejas Shah, Ankit Hinsu, Pritesh Sabara, Apurvasinh Puvar, Janvi Raval, Monika Gandhi, Pinal Trivedi, Maharshi Pandya, Amit Kanani, Akanksha Verma, Nitin Savaliya, Raghawendra Kumar, Dinesh Kumar, Zuber Saiyed, Dipa Kinariwala, Disha Patel, Binita Aring, Neeta Khandelwal, Geeta Vaghela, Sonia Barve, Bhavesh Modi, Kairavi Joshi, Gaurishankar Shrimali, Nidhi Sood, Pranay Shah, R D Dixit, Nidhi Patel, Chaitanya Joshi, Madhvi Joshi |
| EPI_ISL_444458 | B.J. Medical College and Civil hospital | Gujarat Biotechnology Research Centre | Kamlesh J Upadhyay, Ramesh Pandit, Tejas Shah, Ankit Hinsu, Pritesh Sabara, Apurvasinh Puvar, Janvi Raval, Monika Gandhi, Pinal Trivedi, Maharshi Pandya, Amit Kanani, Akanksha Verma, Nitin Savaliya, Raghawendra Kumar, Dinesh Kumar, Zuber Saiyed, Dipa Kinariwala, Disha Patel, Binita Aring, Neeta Khandelwal, Geeta Vaghela, Sonia Barve, Bhavesh Modi, Kairavi Joshi, Gaurishankar Shrimali, Nidhi Sood, Pranay Shah, R D Dixit, Snehal Bagatharia, Priti Pandita, Chaitanya Joshi, Madhvi Joshi |
| EPI_ISL_444459 | B.J. Medical College and Civil hospital | Gujarat Biotechnology Research Centre | Ramesh Pandit, Tejas Shah, Ankit Hinsu, Pritesh Sabara, Apurvasinh Puvar, Janvi Raval, Monika Gandhi, Pinal Trivedi, Maharshi Pandya, Amit Kanani, Akanksha Verma, Nitin Savaliya, Raghawendra Kumar, Dinesh Kumar, Zuber Saiyed, Dipa Kinariwala, Disha Patel, Binita Aring, Neeta Khandelwal, Geeta Vaghela, Sonia Barve, Bhavesh Modi, Kairavi Joshi, Gaurishankar Shrimali, Nidhi Sood, Pranay Shah, R D Dixit, Snehal Bagatharia, Kamlesh J Upadhyay, Neha Rajpara, Chaitanya Joshi, Madhvi Joshi |
| EPI_ISL_444460 | B.J. Medical College and Civil hospital | Gujarat Biotechnology Research Centre | Tejas Shah, Ankit Hinsu, Pritesh Sabara, Apurvasinh Puvar, Janvi Raval, Monika Gandhi, Pinal Trivedi, Maharshi Pandya, Amit Kanani, Akanksha Verma, Nitin Savaliya, Raghawendra Kumar, Dinesh Kumar, Zuber Saiyed, Dipa Kinariwala, Disha Patel, Binita Aring, Neeta Khandelwal, Geeta Vaghela, Sonia Barve, Bhavesh Modi, Kairavi Joshi, Gaurishankar Shrimali, Nidhi Sood, Pranay Shah, R D Dixit, Snehal Bagatharia, Kamlesh J Upadhyay, Ramesh Pandit, Afzal Ansari, Chaitanya Joshi, Madhvi Joshi |
| EPI_ISL_444461 | B.J. Medical College and Civil hospital | Gujarat Biotechnology Research Centre | Ankit Hinsu, Pritesh Sabara, Apurvasinh Puvar, Janvi Raval, Monika Gandhi, Pinal Trivedi, Maharshi Pandya, Amit Kanani, Akanksha Verma, Nitin Savaliya, Raghawendra Kumar, Dinesh Kumar, Zuber Saiyed, Dipa Kinariwala, Disha Patel, Binita Aring, Neeta Khandelwal, Geeta Vaghela, Sonia Barve, Bhavesh Modi, Kairavi Joshi, Gaurishankar Shrimali, Nidhi Sood, Pranay Shah, R D Dixit, Snehal Bagatharia, Kamlesh J Upadhyay, Ramesh Pandit, Neelam Nathani, Chaitanya Joshi, Madhvi Joshi, Tejas Shah |
| EPI_ISL_444462 | B.J. Medical College and Civil hospital | Gujarat Biotechnology Research Centre | Pritesh Sabara, Apurvasinh Puvar, Janvi Raval, Monika Gandhi, Pinal Trivedi, Maharshi Pandya, Amit Kanani, Akanksha Verma, Nitin Savaliya, Raghawendra Kumar, Dinesh Kumar, Zuber Saiyed, Dipa Kinariwala, Disha Patel, Binita Aring, Neeta Khandelwal, Geeta Vaghela, Sonia Barve, Bhavesh Modi, Kairavi Joshi, Gaurishankar Shrimali, Nidhi Sood, Pranay Shah, R D Dixit, Snehal Bagatharia, Kamlesh J Upadhyay, Ramesh Pandit, Tejas Shah, Ankit Hinsu, Armi Chaudhari, Chaitanya Joshi, Madhvi Joshi |
| EPI_ISL_444463 | B.J. Medical College and Civil hospital | Gujarat Biotechnology Research Centre | Apurvasinh Puvar, Janvi Raval, Monika Gandhi, Pinal Trivedi, Maharshi Pandya, Amit Kanani, Akanksha Verma, Nitin Savaliya, Raghawendra Kumar, Dinesh Kumar, Zuber Saiyed, Dipa Kinariwala, Disha Patel, Binita Aring, Neeta Khandelwal, Geeta Vaghela, Sonia Barve, Bhavesh Modi, Kairavi Joshi, Gaurishankar Shrimali, Nidhi Sood, Pranay Shah, R D Dixit, Snehal Bagatharia, Kamlesh J Upadhyay, Ramesh Pandit, Tejas Shah, Ankit Hinsu, Pritesh Sabara, Bhavya Jindal, Chaitanya Joshi, Madhvi Joshi |
| EPI_ISL_444464 | B.J. Medical College and Civil hospital | Gujarat Biotechnology Research Centre | Janvi Raval, Monika Gandhi, Pinal Trivedi, Maharshi Pandya, Amit Kanani, Akanksha Verma, Nitin Savaliya, Raghawendra Kumar, Dinesh Kumar, Zuber Saiyed, Dipa Kinariwala, Disha Patel, Binita Aring, Neeta Khandelwal, Geeta Vaghela, Sonia Barve, Bhavesh Modi, Kairavi Joshi, Gaurishankar Shrimali, Nidhi Sood, Pranay Shah, R D Dixit, Snehal Bagatharia, Kamlesh J Upadhyay, Ramesh Pandit, Tejas Shah, Ankit Hinsu, Pritesh Sabara, Apurvasinh Puvar, Dipeshwari Shewale, Chaitanya Joshi, Madhvi Joshi |
| EPI_ISL_444465 | B.J. Medical College and Civil hospital | Gujarat Biotechnology Research Centre | Monika Gandhi, Pinal Trivedi, Maharshi Pandya, Amit Kanani, Akanksha Verma, Nitin Savaliya, Raghawendra Kumar, Dinesh Kumar, Zuber Saiyed, Dipa Kinariwala, Disha Patel, Binita Aring, Neeta Khandelwal, Geeta Vaghela, Sonia Barve, Bhavesh Modi, Kairavi Joshi, Gaurishankar Shrimali, Nidhi Sood, Pranay Shah, R D Dixit, Snehal Bagatharia, Kamlesh J Upadhyay, Ramesh Pandit, Tejas Shah, Ankit Hinsu, Pritesh Sabara, Apurvasinh Puvar, Janvi Raval, Anjali Rajwar, Chaitanya Joshi, Madhvi Joshi |
| EPI_ISL_444466 | B.J. Medical College and Civil hospital | Gujarat Biotechnology Research Centre | Pinal Trivedi, Maharshi Pandya, Amit Kanani, Akanksha Verma, Nitin Savaliya, Raghawendra Kumar, Dinesh Kumar, Zuber Saiyed, Dipa Kinariwala, Disha Patel, Binita Aring, Neeta Khandelwal, Geeta Vaghela, Sonia Barve, Bhavesh Modi, Kairavi Joshi, Gaurishankar Shrimali, Nidhi Sood, Pranay Shah, R D Dixit, Snehal Bagatharia, Kamlesh J Upadhyay, Ramesh Pandit, Tejas Shah, Ankit Hinsu, Pritesh Sabara, Apurvasinh Puvar, Janvi Raval, Monika Gandhi, Sharmistha Majumdar, Chaitanya Joshi, Madhvi Joshi |
| EPI_ISL_444467 | B.J. Medical College and Civil hospital | Gujarat Biotechnology Research Centre | Maharshi Pandya, Amit Kanani, Akanksha Verma, Nitin Savaliya, Raghawendra Kumar, Dinesh Kumar, Zuber Saiyed, Dipa Kinariwala, Disha Patel, Binita Aring, Neeta Khandelwal, Geeta Vaghela, Sonia Barve, Bhavesh Modi, Kairavi Joshi, Gaurishankar Shrimali, Nidhi Sood, Pranay Shah, R D Dixit, Snehal Bagatharia, Kamlesh J Upadhyay, Ramesh Pandit, Tejas Shah, Ankit Hinsu, Pritesh Sabara, Apurvasinh Puvar, Janvi Raval, Monika Gandhi, Pinal Trivedi, Pooja P Doshi, Chaitanya Joshi, Madhvi Joshi |
| EPI_ISL_444468 | B.J. Medical College and Civil hospital | Gujarat Biotechnology Research Centre | Amit Kanani, Akanksha Verma, Nitin Savaliya, Raghawendra Kumar, Dinesh Kumar, Zuber Saiyed, Dipa Kinariwala, Disha Patel, Binita Aring, Neeta Khandelwal, Geeta Vaghela, Sonia Barve, Bhavesh Modi, Kairavi Joshi, Gaurishankar Shrimali, Nidhi Sood, Pranay Shah, R D Dixit, Snehal Bagatharia, Kamlesh J Upadhyay, Ramesh Pandit, Tejas Shah, Ankit Hinsu, Pritesh Sabara, Apurvasinh Puvar, Janvi Raval, Monika Gandhi, Pinal Trivedi, Maharshi Pandya, Nidhi Patel, Chaitanya Joshi, Madhvi Joshi |
| EPI_ISL_444469 | B.J. Medical College and Civil hospital | Gujarat Biotechnology Research Centre | Akanksha Verma, Nitin Savaliya, Raghawendra Kumar, Dinesh Kumar, Zuber Saiyed, Dipa Kinariwala, Disha Patel, Binita Aring, Neeta Khandelwal, Geeta Vaghela, Sonia Barve, Bhavesh Modi, Kairavi Joshi, Gaurishankar Shrimali, Nidhi Sood, Pranay Shah, R D Dixit, Snehal Bagatharia, Kamlesh J Upadhyay, Ramesh Pandit, Tejas Shah, Ankit Hinsu, Pritesh Sabara, Apurvasinh Puvar, Janvi Raval, Monika Gandhi, Pinal Trivedi, Maharshi Pandya, Amit Kanani, Priti Pandita, Chaitanya Joshi, Madhvi Joshi |
| EPI_ISL_444470 | B.J. Medical College and Civil hospital | Gujarat Biotechnology Research Centre | Nitin Savaliya, Raghawendra Kumar, Dinesh Kumar, Zuber Saiyed, Dipa Kinariwala, Disha Patel, Binita Aring, Neeta Khandelwal, Geeta Vaghela, Sonia Barve, Bhavesh Modi, Kairavi Joshi, Gaurishankar Shrimali, Nidhi Sood, Pranay Shah, R D Dixit, Snehal Bagatharia, Kamlesh J Upadhyay, Ramesh Pandit, Tejas Shah, Ankit Hinsu, Pritesh Sabara, Apurvasinh Puvar, Janvi Raval, Monika Gandhi, Pinal Trivedi, Maharshi Pandya, Amit Kanani, Akanksha Verma, Neha Rajpara, Chaitanya Joshi, Madhvi Joshi |
| EPI_ISL_444471 | B.J. Medical College and Civil hospital | Gujarat Biotechnology Research Centre | Raghawendra Kumar, Dinesh Kumar, Zuber Saiyed, Dipa Kinariwala, Disha Patel, Binita Aring, Neeta Khandelwal, Geeta Vaghela, Sonia Barve, Bhavesh Modi, Kairavi Joshi, Gaurishankar Shrimali, Nidhi Sood, Pranay Shah, R D Dixit, Snehal Bagatharia, Kamlesh J Upadhyay, Ramesh Pandit, Tejas Shah, Ankit Hinsu, Pritesh Sabara, Apurvasinh Puvar, Janvi Raval, Monika Gandhi, Pinal Trivedi, Maharshi Pandya, Amit Kanani, Akanksha Verma, Nitin Savaliya, Afzal Ansari, Chaitanya Joshi, Madhvi Joshi |
| EPI_ISL_444472 | B.J. Medical College and Civil hospital | Gujarat Biotechnology Research Centre | Dinesh Kumar, Zuber Saiyed, Dipa Kinariwala, Disha Patel, Binita Aring, Neeta Khandelwal, Geeta Vaghela, Sonia Barve, Bhavesh Modi, Kairavi Joshi, Gaurishankar Shrimali, Nidhi Sood, Pranay Shah, R D Dixit, Snehal Bagatharia, Kamlesh J Upadhyay, Ramesh Pandit, Tejas Shah, Ankit Hinsu, Pritesh Sabara, Apurvasinh Puvar, Janvi Raval, Monika Gandhi, Pinal Trivedi, Maharshi Pandya, Amit Kanani, Akanksha Verma, Nitin Savaliya, Raghawendra Kumar, Neelam Nathani, Chaitanya Joshi, Madhvi Joshi |
| EPI_ISL_444473 | B.J. Medical College and Civil hospital | Gujarat Biotechnology Research Centre | Zuber Saiyed, Dipa Kinariwala, Disha Patel, Binita Aring, Neeta Khandelwal, Geeta Vaghela, Sonia Barve, Bhavesh Modi, Kairavi Joshi, Gaurishankar Shrimali, Nidhi Sood, Pranay Shah, R D Dixit, Snehal Bagatharia, Kamlesh J Upadhyay, Ramesh Pandit, Tejas Shah, Ankit Hinsu, Pritesh Sabara, Apurvasinh Puvar, Janvi Raval, Monika Gandhi, Pinal Trivedi, Maharshi Pandya, Amit Kanani, Akanksha Verma, Nitin Savaliya, Raghawendra Kumar, Dinesh Kumar, Armi Chaudhari, Chaitanya Joshi, Madhvi Joshi |
| EPI_ISL_444474 | B.J. Medical College and Civil hospital | Gujarat Biotechnology Research Centre | Dipa Kinariwala, Disha Patel, Binita Aring, Neeta Khandelwal, Geeta Vaghela, Sonia Barve, Bhavesh Modi, Kairavi Joshi, Gaurishankar Shrimali, Nidhi Sood, Pranay Shah, R D Dixit, Snehal Bagatharia, Kamlesh J |

| | | | |
|---|---|---|---|
| | | | Upadhyay, Ramesh Pandit, Tejas Shah, Ankit Hinsu, Pritesh Sabara, Apurvasinh Puvar, Janvi Raval, Monika Gandhi, Pinal Trivedi, Maharshi Pandya, Amit Kanani, Akanksha Verma, Nitin Savaliya, Raghawendra Kumar, Dinesh Kumar, Zuber Saiyed, Bhavya Jindal, Chaitanya Joshi, Madhvi Joshi |
| EPI_ISL_444475 | B.J. Medical College and Civil hospital | Gujarat Biotechnology Research Centre | Disha Patel, Binita Aring, Neeta Khandelwal, Geeta Vaghela, Sonia Barve, Bhavesh Modi, Kairavi Joshi, Gaurishankar Shrimali, Nidhi Sood, Pranay Shah, R D Dixit, Snehal Bagatharia, Kamlesh J Upadhyay, Ramesh Pandit, Tejas Shah, Ankit Hinsu, Pritesh Sabara, Apurvasinh Puvar, Janvi Raval, Monika Gandhi, Pinal Trivedi, Maharshi Pandya, Amit Kanani, Akanksha Verma, Nitin Savaliya, Raghawendra Kumar, Dinesh Kumar, Zuber Saiyed, Dipa Kinariwala, Dipeshwari Shewale, Chaitanya Joshi, Madhvi Joshi |
| EPI_ISL_444476 | B.J. Medical College and Civil hospital | Gujarat Biotechnology Research Centre | Binita Aring, Neeta Khandelwal, Geeta Vaghela, Sonia Barve, Bhavesh Modi, Kairavi Joshi, Gaurishankar Shrimali, Nidhi Sood, Pranay Shah, R D Dixit, Snehal Bagatharia, Kamlesh J Upadhyay, Ramesh Pandit, Tejas Shah, Ankit Hinsu, Pritesh Sabara, Apurvasinh Puvar, Janvi Raval, Monika Gandhi, Pinal Trivedi, Maharshi Pandya, Amit Kanani, Akanksha Verma, Nitin Savaliya, Raghawendra Kumar, Dinesh Kumar, Zuber Saiyed, Dipa Kinariwala, Disha Patel, Chaitanya Joshi, Madhvi Joshi, Dipeshwari Shewale |
| EPI_ISL_444477 | B.J. Medical College and Civil hospital | Gujarat Biotechnology Research Centre | Neeta Khandelwal, Geeta Vaghela, Sonia Barve, Bhavesh Modi, Kairavi Joshi, Gaurishankar Shrimali, Nidhi Sood, Pranay Shah, R D Dixit, Snehal Bagatharia, Kamlesh J Upadhyay, Ramesh Pandit, Tejas Shah, Ankit Hinsu, Pritesh Sabara, Apurvasinh Puvar, Janvi Raval, Monika Gandhi, Pinal Trivedi, Maharshi Pandya, Amit Kanani, Akanksha Verma, Nitin Savaliya, Raghawendra Kumar, Dinesh Kumar, Zuber Saiyed, Dipa Kinariwala, Disha Patel, Binita Aring, Sharmistha Majumdar, Chaitanya Joshi, Madhvi Joshi |
| EPI_ISL_444478 | B.J. Medical College and Civil hospital | Gujarat Biotechnology Research Centre | Geeta Vaghela, Sonia Barve, Bhavesh Modi, Kairavi Joshi, Gaurishankar Shrimali, Nidhi Sood, Pranay Shah, R D Dixit, Snehal Bagatharia, Kamlesh J Upadhyay, Ramesh Pandit, Tejas Shah, Ankit Hinsu, Pritesh Sabara, Apurvasinh Puvar, Janvi Raval, Monika Gandhi, Pinal Trivedi, Maharshi Pandya, Amit Kanani, Akanksha Verma, Nitin Savaliya, Raghawendra Kumar, Dinesh Kumar, Zuber Saiyed, Dipa Kinariwala, Disha Patel, Binita Aring, Neeta Khandelwal, Pooja P Doshi, Chaitanya Joshi, Madhvi Joshi |
| EPI_ISL_444479 | B.J. Medical College and Civil hospital | Gujarat Biotechnology Research Centre | Sonia Barve, Bhavesh Modi, Kairavi Joshi, Gaurishankar Shrimali, Nidhi Sood, Pranay Shah, R D Dixit, Snehal Bagatharia, Kamlesh J Upadhyay, Ramesh Pandit, Tejas Shah, Ankit Hinsu, Pritesh Sabara, Apurvasinh Puvar, Janvi Raval, Monika Gandhi, Pinal Trivedi, Maharshi Pandya, Amit Kanani, Akanksha Verma, Nitin Savaliya, Raghawendra Kumar, Dinesh Kumar, Zuber Saiyed, Dipa Kinariwala, Disha Patel, Binita Aring, Neeta Khandelwal, Geeta Vaghela, Nidhi Patel, Chaitanya Joshi, Madhvi Joshi |
| EPI_ISL_444480 | B.J. Medical College and Civil hospital | Gujarat Biotechnology Research Centre | Bhavesh Modi, Kairavi Joshi, Gaurishankar Shrimali, Nidhi Sood, Pranay Shah, R D Dixit, Snehal Bagatharia, Kamlesh J Upadhyay, Ramesh Pandit, Tejas Shah, Ankit Hinsu, Pritesh Sabara, Apurvasinh Puvar, Janvi Raval, Monika Gandhi, Pinal Trivedi, Maharshi Pandya, Amit Kanani, Akanksha Verma, Nitin Savaliya, Raghawendra Kumar, Dinesh Kumar, Zuber Saiyed, Dipa Kinariwala, Disha Patel, Binita Aring, Neeta Khandelwal, Geeta Vaghela, Sonia Barve, Priti Pandita, Chaitanya Joshi, Madhvi Joshi |
| EPI_ISL_444481 | B.J. Medical College and Civil hospital | Gujarat Biotechnology Research Centre | Kairavi Joshi, Gaurishankar Shrimali, Nidhi Sood, Pranay Shah, R D Dixit, Snehal Bagatharia, Kamlesh J Upadhyay, Ramesh Pandit, Tejas Shah, Ankit Hinsu, Pritesh Sabara, Apurvasinh Puvar, Janvi Raval, Monika Gandhi, Pinal Trivedi, Maharshi Pandya, Amit Kanani, Akanksha Verma, Nitin Savaliya, Raghawendra Kumar, Dinesh Kumar, Zuber Saiyed, Dipa Kinariwala, Disha Patel, Binita Aring, Neeta Khandelwal, Geeta Vaghela, Sonia Barve, Bhavesh Modi, Neha Rajpara, Chaitanya Joshi, Madhvi Joshi |
| EPI_ISL_444482 | Gujarat Biotechnology Research Centre | Gujarat Biotechnology Research Centre | Gaurishankar Shrimali, Nidhi Sood, Pranay Shah, R D Dixit, Snehal Bagatharia, Kamlesh J Upadhyay, Ramesh Pandit, Tejas Shah, Ankit Hinsu, Pritesh Sabara, Apurvasinh Puvar, Janvi Raval, Monika Gandhi, Pinal Trivedi, Maharshi Pandya, Amit Kanani, Akanksha Verma, Nitin Savaliya, Raghawendra Kumar, Dinesh Kumar, Zuber Saiyed, Dipa Kinariwala, Disha Patel, Binita Aring, Neeta Khandelwal, Geeta Vaghela, Sonia Barve, Bhavesh Modi, Kairavi Joshi, Afzal Ansari, Chaitanya Joshi, Madhvi Joshi |
| EPI_ISL_444483 | Gujarat Biotechnology Research Centre | Gujarat Biotechnology Research Centre | Nidhi Sood, Pranay Shah, R D Dixit, Snehal Bagatharia, Kamlesh J Upadhyay, Ramesh Pandit, Tejas Shah, Ankit Hinsu, Pritesh Sabara, Apurvasinh Puvar, Janvi Raval, Monika Gandhi, Pinal Trivedi, Maharshi Pandya, Amit Kanani, Akanksha Verma, Nitin Savaliya, Raghawendra Kumar, Dinesh Kumar, Zuber Saiyed, Dipa Kinariwala, Disha Patel, Binita Aring, Neeta Khandelwal, Geeta Vaghela, Sonia Barve, Bhavesh Modi, Kairavi Joshi, Gaurishankar Shrimali, Neelam Nathani, Chaitanya Joshi, Madhvi Joshi |
| EPI_ISL_444484 | Gujarat Biotechnology Research Centre | Gujarat Biotechnology Research Centre | Pranay Shah, R D Dixit, Snehal Bagatharia, Kamlesh J Upadhyay, Ramesh Pandit, Tejas Shah, Ankit Hinsu, Pritesh Sabara, Apurvasinh Puvar, Janvi Raval, Monika Gandhi, Pinal Trivedi, Maharshi Pandya, Amit Kanani, Akanksha Verma, Nitin Savaliya, Raghawendra Kumar, Dinesh Kumar, Zuber Saiyed, Dipa Kinariwala, Disha Patel, Binita Aring, Neeta Khandelwal, Geeta Vaghela, Sonia Barve, Bhavesh Modi, Kairavi Joshi, Gaurishankar Shrimali, Nidhi Sood, Armi Chaudhari, Chaitanya Joshi, Madhvi Joshi |
| EPI_ISL_444485 | Gujarat Biotechnology Research Centre | Gujarat Biotechnology Research Centre | R D Dixit, Snehal Bagatharia, Kamlesh J Upadhyay, Ramesh Pandit, Tejas Shah, Ankit Hinsu, Pritesh Sabara, Apurvasinh Puvar, Janvi Raval, Monika Gandhi, Pinal Trivedi, Maharshi Pandya, Amit Kanani, Akanksha Verma, Nitin Savaliya, Raghawendra Kumar, Dinesh Kumar, Zuber Saiyed, Dipa Kinariwala, Disha Patel, Binita Aring, Neeta Khandelwal, Geeta Vaghela, Sonia Barve, Bhavesh Modi, Kairavi Joshi, Gaurishankar Shrimali, Nidhi Sood, Pranay Shah, Bhavya Jindal, Chaitanya Joshi, Madhvi Joshi |
| EPI_ISL_444486 | Gujarat Biotechnology Research Centre | Gujarat Biotechnology Research Centre | Snehal Bagatharia, Kamlesh J Upadhyay, Ramesh Pandit, Tejas Shah, Ankit Hinsu, Pritesh Sabara, Apurvasinh Puvar, Janvi Raval, Monika Gandhi, Pinal Trivedi, Maharshi Pandya, Amit Kanani, Akanksha Verma, Nitin Savaliya, Raghawendra Kumar, Dinesh Kumar, Zuber Saiyed, Dipa Kinariwala, Disha Patel, Binita Aring, Neeta Khandelwal, Geeta Vaghela, Sonia Barve, Bhavesh Modi, Kairavi Joshi, Gaurishankar Shrimali, Nidhi Sood, Pranay Shah, R D Dixit, Dipeshwari Shewale, Chaitanya Joshi, Madhvi Joshi |
| EPI_ISL_444488, EPI_ISL_444489, EPI_ISL_444492 | Karolinska Universitetslaboratoriet | CTMR, Karolinska Institutet, Stockholm, Sweden | Yue Hu, Stefanie Prast-Nielsen, Jingkai Ji, Fredrik Boulund, Jing Wang, Shuiqin Li, Yinghua Zha, Caroline Bjurnemark, Linnéa Påvénius, Marica Hamsten, Vivien Lan Yang Swartz, Lars Engstrand |
| EPI_ISL_444493 | Departamento de Laboratorios de Salud Publica (DLSP, Division Epidemiologia, Ministerio de Salud Publica) | Facultad de Ciencias (Sección Genética Evolutiva, Sección Virología). | Panzera,Y., Delfraro,A., Ramos,N., Frabasile,S., Calleros,L., Techera,C., Grecco,S., Fuques,E., Goni,N., Coppola,L., Ramos,V., Chiparelli,H., Arbiza,J. and Perez,R. |
| EPI_ISL_444495, EPI_ISL_444500 | Laboratoire de microbiologie, Hopital de Verdun | Smith Laboratory, Centre de Recherche CHU Sainte-Justine | Martin Smith, Marieke Rozendaal, Ivan Pavlov |
| EPI_ISL_444518 | unknown | Molecular Infectious Disease | Anderson,B.P., Rosenthal,S.H., Gerasimova,A., Kagan,R.M. and Owen,R. |
| EPI_ISL_444521, EPI_ISL_444522, EPI_ISL_444523, EPI_ISL_444524, EPI_ISL_444525, EPI_ISL_444527, EPI_ISL_444528, EPI_ISL_444530, EPI_ISL_444531, EPI_ISL_444532, EPI_ISL_444533, EPI_ISL_444534, EPI_ISL_444535, EPI_ISL_444537, EPI_ISL_444538, EPI_ISL_444540, EPI_ISL_444544, EPI_ISL_444545, EPI_ISL_444546, EPI_ISL_444547, EPI_ISL_444549, EPI_ISL_444550, EPI_ISL_444552, EPI_ISL_444553, EPI_ISL_444555, EPI_ISL_444556, EPI_ISL_444558, EPI_ISL_444560, EPI_ISL_444561, EPI_ISL_444562, EPI_ISL_444564, EPI_ISL_444566, EPI_ISL_444567, EPI_ISL_444568, EPI_ISL_444569, EPI_ISL_444570, EPI_ISL_444571, EPI_ISL_444572, EPI_ISL_444573, EPI_ISL_444574, EPI_ISL_444575, EPI_ISL_444576, EPI_ISL_444577, EPI_ISL_444578, EPI_ISL_444579, EPI_ISL_444580, EPI_ISL_444581, EPI_ISL_444583, EPI_ISL_444585, EPI_ISL_444586, EPI_ISL_444587, EPI_ISL_444588, EPI_ISL_444589, EPI_ISL_444590, EPI_ISL_444591, EPI_ISL_444592, EPI_ISL_444593, EPI_ISL_444594, EPI_ISL_444595, EPI_ISL_444596, EPI_ISL_444597, EPI_ISL_444598, EPI_ISL_444599, EPI_ISL_444600, EPI_ISL_444601, EPI_ISL_444603, EPI_ISL_444604, EPI_ISL_444605, EPI_ISL_444606, EPI_ISL_444607, EPI_ISL_444608, EPI_ISL_444609 | | | |
| see above | Northwestern Memorial Hospital | Ozer Lab | Ramon Lorenzo-Redondo, Hannah H. Nam, Scott C. Roberts, Lacy M. Simons, Chad J. Achenbach, Lawrence J. Jennings, Chao Qi, Alan R. Hauser, Michael G. Ison, Judd F. Hultquist, Egon A. Ozer |
| EPI_ISL_444611 | Pathology Queensland | Public Health Virology Laboratory | Bixing Huang, Alyssa Pyke, Amanda De Jong, Andrew Van Den Hurk, Carmel Taylor, David Warrilow, Doris Genge, Elisabeth Gamez, Glen Hewitson, Ian Maxwell Mackay, Inga Sultana, Jamie McMahon, Jean Barcelon, Judy Northill, Mitchell Finger, Natalie Simpson, Neelima Nair, Peter Burtonclay, Peter Moore, Sarah Wheatley, Sean Moody, Sonja Hall-Mendelin, Timothy Gardam, and Frederick Moore |
| EPI_ISL_444612 | QML Pathology | Public Health Virology Laboratory | Bixing Huang, Alyssa Pyke, Amanda De Jong, Andrew Van Den Hurk, Carmel Taylor, David Warrilow, Doris Genge, Elisabeth Gamez, Glen Hewitson, Ian Maxwell Mackay, Inga Sultana, Jamie McMahon, Jean Barcelon, Judy Northill, Mitchell Finger, Natalie Simpson, Neelima Nair, Peter Burtonclay, Peter Moore, Sarah Wheatley, Sean Moody, Sonja Hall-Mendelin, Timothy Gardam, and Frederick Moore |
| EPI_ISL_444613, EPI_ISL_444615, EPI_ISL_444616, EPI_ISL_444617, EPI_ISL_444618, EPI_ISL_444619, EPI_ISL_444620, EPI_ISL_444621, EPI_ISL_444622, EPI_ISL_444623, EPI_ISL_444624, EPI_ISL_444625, EPI_ISL_444626, EPI_ISL_444627, EPI_ISL_444628, EPI_ISL_444629, EPI_ISL_444630, EPI_ISL_444631, EPI_ISL_444632, EPI_ISL_444633, EPI_ISL_444634, EPI_ISL_444635, EPI_ISL_444636, EPI_ISL_444637, EPI_ISL_444638, EPI_ISL_444639, EPI_ISL_444640, EPI_ISL_444641, EPI_ISL_444642, EPI_ISL_444643, EPI_ISL_444644, EPI_ISL_444645, EPI_ISL_444646, EPI_ISL_444647, EPI_ISL_444648, EPI_ISL_444650, EPI_ISL_444651, EPI_ISL_444652, EPI_ISL_444653, EPI_ISL_444654, EPI_ISL_444655, EPI_ISL_444656, EPI_ISL_444657, EPI_ISL_444658, EPI_ISL_444659, EPI_ISL_444660, EPI_ISL_444661, EPI_ISL_444662, EPI_ISL_444663, EPI_ISL_444664, EPI_ISL_444665, EPI_ISL_444666, EPI_ISL_444667, EPI_ISL_444668, EPI_ISL_444669, EPI_ISL_444670, EPI_ISL_444671, EPI_ISL_444673, EPI_ISL_444675, EPI_ISL_444677, EPI_ISL_444678, EPI_ISL_444679, EPI_ISL_444680, EPI_ISL_444681, EPI_ISL_444682, EPI_ISL_444683, EPI_ISL_444684, EPI_ISL_444685, EPI_ISL_444686, EPI_ISL_444687, EPI_ISL_444688, EPI_ISL_444690, EPI_ISL_444691, EPI_ISL_444692, EPI_ISL_444693, EPI_ISL_444694, EPI_ISL_444695, EPI_ISL_444696, EPI_ISL_444697, EPI_ISL_444698, EPI_ISL_444699, EPI_ISL_444700, EPI_ISL_444701, EPI_ISL_444702, EPI_ISL_444703, EPI_ISL_444704, EPI_ISL_444705, EPI_ISL_444706, EPI_ISL_444707, EPI_ISL_444708, EPI_ISL_444710, EPI_ISL_444711, EPI_ISL_444712, EPI_ISL_444713, EPI_ISL_444714, EPI_ISL_444715, EPI_ISL_444716, EPI_ISL_444717, EPI_ISL_444718, EPI_ISL_444719, EPI_ISL_444720, EPI_ISL_444721, EPI_ISL_444722, EPI_ISL_444723, EPI_ISL_444724, EPI_ISL_444725, EPI_ISL_444726, EPI_ISL_444727, EPI_ISL_444728, EPI_ISL_444729, EPI_ISL_444730, EPI_ISL_444731, EPI_ISL_444732, EPI_ISL_444733, EPI_ISL_444734, EPI_ISL_444735, EPI_ISL_444736, EPI_ISL_444737, EPI_ISL_444738, EPI_ISL_444739, EPI_ISL_444740, EPI_ISL_444741, EPI_ISL_444742, EPI_ISL_444743, EPI_ISL_444744, EPI_ISL_444745, EPI_ISL_444746, EPI_ISL_444747, EPI_ISL_444748, EPI_ISL_444749, EPI_ISL_444750, EPI_ISL_444751, EPI_ISL_444752, EPI_ISL_444753, EPI_ISL_444754, EPI_ISL_444755, EPI_ISL_444756, EPI_ISL_444757, EPI_ISL_444758, EPI_ISL_444759, EPI_ISL_444760, EPI_ISL_444761, EPI_ISL_444762, EPI_ISL_444763, EPI_ISL_444764, EPI_ISL_444765, EPI_ISL_444766, EPI_ISL_444767, EPI_ISL_444768, EPI_ISL_444769, EPI_ISL_444770, EPI_ISL_444771, EPI_ISL_444772, EPI_ISL_444773, EPI_ISL_444774, EPI_ISL_444775, EPI_ISL_444776, EPI_ISL_444777, EPI_ISL_444780, EPI_ISL_444781, EPI_ISL_444782, EPI_ISL_444784, EPI_ISL_444786, EPI_ISL_444787, EPI_ISL_444788, EPI_ISL_444789, EPI_ISL_444790, EPI_ISL_444791 | | | |
| see above | NYU Langone Health | Departments of Pathology and Medicine, New York University School of Medicine | Maria Aguero-Rosenfeld, Brendan Belovarac, Margaret Black, Ludovic Boytard, John Cadley, Paolo Cotzia, John Chen, Dacia Dimartino, Xiaojun Feng, Tatyana Gindin, Emily Guzman, Adriana Heguy, Megan Hogan, Emily Huang, George Jour, Alireza Khodadadi-Jamayran, Lawrence H. Lin, Raven Luther, Andrew Lytle, Christian Marier, Matthew T. Maurano, Mark J. Mulligan, Peter Meyn, Raquel Ordonez Ciriza, Iman Osman, Jared Pinnell, Vanessa Raabe, Sitharam Ramaswami, Amy Rapkiewicz, Andre M. Ribeiro-dos-Santos, Marie Samanovic-Golden, Antonio Serrano, Guomiao Shen, Matija Snuderl, Theodore Vougiouklakis, Nick Vulpescu, Gael Westby, Paul Zappile, Yutong Zhang |
| EPI_ISL_444793 | Pathology Queensland | Public Health Virology Laboratory | Bixing Huang, Alyssa Pyke, Amanda De Jong, Andrew Van Den Hurk, Carmel Taylor, David Warrilow, Doris Genge, Elisabeth Gamez, Glen Hewitson, Ian Maxwell Mackay, Inga Sultana, Jamie McMahon, Jean Barcelon, Judy Northill, Mitchell Finger, Natalie Simpson, Neelima Nair, Peter Burtonclay, Peter Moore, Sarah Wheatley, Sean Moody, Sonja Hall-Mendelin, Timothy Gardam, and Frederick Moore |
| EPI_ISL_444794 | Cairns Hospital | Public Health Virology Laboratory | Bixing Huang, Alyssa Pyke, Amanda De Jong, Andrew Van Den Hurk, Carmel Taylor, David Warrilow, Doris Genge, Elisabeth Gamez, Glen Hewitson, Ian Maxwell Mackay, Inga Sultana, Jamie McMahon, Jean Barcelon, Judy Northill, Mitchell Finger, Natalie Simpson, Neelima Nair, Peter Burtonclay, Peter Moore, Sarah Wheatley, Sean Moody, Sonja Hall-Mendelin, Timothy Gardam, and Frederick Moore |
| EPI_ISL_444817, EPI_ISL_444818, EPI_ISL_444819, EPI_ISL_444820, EPI_ISL_444822, EPI_ISL_444823, EPI_ISL_444824, EPI_ISL_444825, EPI_ISL_444826, EPI_ISL_444827, EPI_ISL_444828, EPI_ISL_444829, EPI_ISL_444831, EPI_ISL_444832, EPI_ISL_444833, EPI_ISL_444834, EPI_ISL_444835, EPI_ISL_444836, EPI_ISL_444837, EPI_ISL_444838, EPI_ISL_444839, EPI_ISL_444840, EPI_ISL_444841, EPI_ISL_444844, EPI_ISL_444846, EPI_ISL_444848, EPI_ISL_444849, EPI_ISL_444850, EPI_ISL_444852, EPI_ISL_444853, EPI_ISL_444854, EPI_ISL_444855, EPI_ISL_444856, EPI_ISL_444857, EPI_ISL_444859, EPI_ISL_444860, EPI_ISL_444861, EPI_ISL_444862, EPI_ISL_444863, EPI_ISL_444864, EPI_ISL_444866, EPI_ISL_444867, EPI_ISL_444869, EPI_ISL_444870, EPI_ISL_444872, EPI_ISL_444873, EPI_ISL_444874, EPI_ISL_444875, EPI_ISL_444877, EPI_ISL_444879, EPI_ISL_444880, EPI_ISL_444881, EPI_ISL_444883, EPI_ISL_444884, EPI_ISL_444885, EPI_ISL_444887, EPI_ISL_444888, EPI_ISL_444890, EPI_ISL_444891, EPI_ISL_444892, EPI_ISL_444893, EPI_ISL_444894, EPI_ISL_444895, EPI_ISL_444896, EPI_ISL_444897, EPI_ISL_444898, EPI_ISL_444899, EPI_ISL_444900, EPI_ISL_444901, EPI_ISL_444902, EPI_ISL_444903, EPI_ISL_444904, EPI_ISL_444905, EPI_ISL_444906, EPI_ISL_444908, EPI_ISL_444909, EPI_ISL_444910, EPI_ISL_444911, EPI_ISL_444912, EPI_ISL_444915, EPI_ISL_444916, EPI_ISL_444917, EPI_ISL_444918, EPI_ISL_444920, EPI_ISL_444921, EPI_ISL_444922, EPI_ISL_444923, EPI_ISL_444924, EPI_ISL_444925, EPI_ISL_444926, EPI_ISL_444927, EPI_ISL_444928, EPI_ISL_444929, EPI_ISL_444930, EPI_ISL_444931, EPI_ISL_444933, EPI_ISL_444934, EPI_ISL_444935, EPI_ISL_444937, EPI_ISL_444938, EPI_ISL_444939, EPI_ISL_444940, EPI_ISL_444941, EPI_ISL_444942, EPI_ISL_444944, EPI_ISL_444945, EPI_ISL_444946, EPI_ISL_444947, EPI_ISL_444948, EPI_ISL_444949, EPI_ISL_444950, EPI_ISL_444952, EPI_ISL_444954, EPI_ISL_444955, EPI_ISL_444956, EPI_ISL_444957, EPI_ISL_444958, EPI_ISL_444959, EPI_ISL_444960, EPI_ISL_444961, EPI_ISL_444963, EPI_ISL_444964, EPI_ISL_444965, EPI_ISL_444966, EPI_ISL_444967, EPI_ISL_444968 | | | |
| see above | Department of Virus and Microbiological Special Diagnostics, Statens Serum Institut, Copenhagen, Denmark, Artillerivej 5, 2300 Copenahgen S | Albertsen lab, Department of Chemistry and Bioscience, Aalborg University, Denmark | Rasmus Kirkegaard |
| EPI_ISL_444969 | Guangzhou Eighth People's Hospital (Jiahe Sector) | Institute of Human Virology, Zhongshan School of Medicine, Sun Yat-sen University | Junsong Zhang, Fei Yu, Jun Liu, Huimin Fan, Ruosu Ying, Feng Huang, Ting Pan, Bingfeng Liu,Yiwen Zhang, Xu Zhang, Mang Shi, Fengyu Hu, Fang Li, Kai Deng, Hui Zhang |
| EPI_ISL_444971 | Hospital Universitari Vall d'Hebron - Vall d'Hebron Institut de Recerca | Hospital Universitari Vall d'Hebron | Cristina Andrés, Maria Piñana, Damir Garcia-Cehic, Mercedes Guerrero-Murillo, Ariadna Rando, Juliana Esperalba, Maria Gema Codina, Tomàs Pumarola, Josep Quer, Andrés Antón |
| EPI_ISL_444972 | Hospital Universitari Vall d'Hebron - Vall d'Hebron Institut de Recerca | Hospital Universitari Vall d'Hebron | Cristina Andrés, Maria Piñana, DAmir Garcia-Cehic, Mercedes Guerrero-Murillo, Ariadna Rando, Juliana Esperalba, Maria Gema Codina, Tomàs Pumarola, Josep Quer, Andrés Antón |
| EPI_ISL_444973 | Hospital Universitari Vall d`Hebron - Vall d'Hebron Institut de Recerca | Hospital Universitari Vall d'Hebron | Cristina Andrés, Maria Piñana, DAmir Garcia-Cehic, Mercedes Guerrero-Murillo, Ariadna Rando, Juliana Esperalba, Maria Gema Codina, Tomàs Pumarola, Josep Quer, Andrés Antón |
| EPI_ISL_444974, EPI_ISL_444975, EPI_ISL_444976, EPI_ISL_444977, EPI_ISL_444978, EPI_ISL_444979 | Hospital Universitari Vall d'Hebron - Vall d'Hebron Institut de Recerca | Hospital Universitari Vall d'Hebron | Cristina Andrés, Maria Piñana, Damir Garcia-Cehic, Mercedes Guerrero-Murillo, Ariadna Rando, Juliana Esperalba, Maria Gema Codina, Tomàs Pumarola, Josep Quer, Andrés Antón |
| EPI_ISL_444980, EPI_ISL_444981, EPI_ISL_444982, EPI_ISL_444983 | Hospital Universitari Vall d'Hebron - Vall d"hebron Institut de Recerca | Hospital Universitari Vall d'Hebron | Cristina Andrés, Maria Piñana, Damir Garcia-Cehic, Mercedes Guerrero-Murillo, Ariadna Rando, Juliana Esperalba, Maria Gema Codina, Tomàs Pumarola, Josep Quer, Andrés Antón |
| EPI_ISL_444984, EPI_ISL_444985, EPI_ISL_444986, EPI_ISL_444987, EPI_ISL_444988, EPI_ISL_444989, EPI_ISL_444990 | Hospital Universitari Vall d'Hebron - Vall d'Hebron Institut de Recerca | Hospital Universitari Vall d'Hebron | Cristina Andrés, Maria Piñana, Damir Garcia-Cehic, Mercedes Guerrero-Murillo, Ariadna Rando, Juliana Esperalba, Maria Gema Codina, Tomàs Pumarola, Josep Quer, Andrés Antón |
| EPI_ISL_444994, EPI_ISL_444995, EPI_ISL_444996, EPI_ISL_444997, EPI_ISL_444998, EPI_ISL_444999, EPI_ISL_445000 | Naval Health Research Center | Naval Medical Research Center Biological Defense Research Directorate | Logan Voegtly, Regina Cer, Dessiree Pena-Gomez, Adrian Paskey,Kyle Long, Roger Pan, Melinda Balansay-Ames, Chris Myers, Ewell Hollis, Nathaniel Christy, Kimberly Bishop-Lilly |
| EPI_ISL_445043 | Florida Bureau of Public Health Laboratories | Florida Bureau of Public Health Laboratories | Sarah Schmedes, Jason Blanton |

| | | | |
|---|---|---|---|
| EPI_ISL_445054, EPI_ISL_445055, EPI_ISL_445056, EPI_ISL_445057, EPI_ISL_445058, EPI_ISL_445060, EPI_ISL_445061, EPI_ISL_445064, EPI_ISL_445065, EPI_ISL_445066, EPI_ISL_445067, EPI_ISL_445069, EPI_ISL_445070, EPI_ISL_445071, EPI_ISL_445072, EPI_ISL_445074, EPI_ISL_445075, EPI_ISL_445076 | | | |
| see above | Laboratoire National de Sante, Microbiology, Virology | Laboratoire National de Sante, Microbiology, Epidemiology and Microbial Genomics | Anke Wienecke-Baldacchino, Ardashel Latsuzbaia, Jessica Tapp, Catherine Ragimbeau, Guillaume Fournier, Tamir Abdelrahman, Trung Nguyen Nguyen, Joel Mossong |
| EPI_ISL_445077 | M Health Fairview | University of Minnesota Genomics Center | Daryl M. Gohl, John Garbe, Patrick Grady, Jerry Daniel, Ray Watson, Benjamin Auch, Andrew Nelson, Sophia Yohe, and Kenneth B. Beckman |
| EPI_ISL_445078, EPI_ISL_445079, EPI_ISL_445080, EPI_ISL_445081, EPI_ISL_445082, EPI_ISL_445083, EPI_ISL_445084 | Baylor College of Medicine | Baylor College of Medicine: HGSC | Vasanthi Avadhanula, Erin Nicholson, David Henke, Pedro Piedra, Harsha Doddapaneni, Donna Muzny, Qingchang Meng, Hsu Chao, Zeineen Momin, Hua Shen, George Weissenberger, Kavya Kottapalli, Yimiti Meiheerguli, Sejal Salvi, Ginger Metcalf, Vipin Menon, Sara J.J. Cregeen, Matthew C. Ross, Tulin Ayvaz, Richard Sucgang, Kristi L. Hoffman, Matthew Wong, Joseph F. Petrosino |
| EPI_ISL_445085 | Virology Unit, Agrobiodiversity and Biotechnology Project, CIAT - International Center for Tropical Agriculture | Virology Unit, Agrobiodiversity and Biotechnology Project, CIAT - International Center for Tropical Agriculture | Lopez,D., Parra,B. and Cuellar,W.J. |
| EPI_ISL_445086 | Laboratory Diagnostic, Veterinary Specialized Institute Kraljevo | Laboratory Diagnostic, Veterinary Specialized Institute Kraljevo | Vidanovic,D., Tesovic,B., Sekler,M., Dmitric,M., Debeljak,Z., Matovic,K., Vaskovic,N., Petrovic,T., Volkening,J. and Alfonso,C.L. |
| EPI_ISL_445087 | Laboratory Diagnostic, Veterinary Specialized Institute Kraljevo | Laboratory Diagnostic, Veterinary Specialized Institute Kraljevo | Vidanovic,D., Tesovic,B., Sekler,M., Dmitric,M., Debeljak,Z., Matovic,K., Vaskovic,N., Petrovic,T., Volkening,J. and Alfonso,C. |
| EPI_ISL_445094, EPI_ISL_445096, EPI_ISL_445097, EPI_ISL_445098, EPI_ISL_445101, EPI_ISL_445102, EPI_ISL_445105, EPI_ISL_445107, EPI_ISL_445108, EPI_ISL_445109, EPI_ISL_445110, EPI_ISL_445111, EPI_ISL_445113, EPI_ISL_445114, EPI_ISL_445115, EPI_ISL_445116, EPI_ISL_445117 | | | |
| see above | UC San Diego Center for Advanced Laboratory Medicine | Andersen lab at Scripps Research | SEARCH Alliance San Diego with David Pride, Ji H Shin |
| EPI_ISL_445118 | Rady's Childrens Hospital | Andersen lab at Scripps Research | SEARCH Alliance San Diego |
| EPI_ISL_445119, EPI_ISL_445120, EPI_ISL_445121, EPI_ISL_445122, EPI_ISL_445123, EPI_ISL_445124, EPI_ISL_445125, EPI_ISL_445126, EPI_ISL_445127, EPI_ISL_445128, EPI_ISL_445129, EPI_ISL_445130, EPI_ISL_445132, EPI_ISL_445133, EPI_ISL_445134, EPI_ISL_445135, EPI_ISL_445136, EPI_ISL_445137, EPI_ISL_445138, EPI_ISL_445139, EPI_ISL_445140, EPI_ISL_445141, EPI_ISL_445143, EPI_ISL_445144, EPI_ISL_445145, EPI_ISL_445146, EPI_ISL_445147, EPI_ISL_445148, EPI_ISL_445149, EPI_ISL_445150, EPI_ISL_445152, EPI_ISL_445153, EPI_ISL_445154, EPI_ISL_445155, EPI_ISL_445156, EPI_ISL_445157, EPI_ISL_445159, EPI_ISL_445161, EPI_ISL_445162 | | | |
| see above | Robert Garry lab | Andersen lab at Scripps Research | Allison Smither, Gilberto Sabino-Santos, Patricia Snarski, Lilia Melnik, Antoinette Bell, Kaylynn Genemaras, Arnaud Drouin, Dahlene Fusco, Robert Garry with SEARCH Alliance San Diego |
| EPI_ISL_445164, EPI_ISL_445166, EPI_ISL_445167 | Scripps Medical Laboratory | Andersen lab at Scripps Research | SEARCH Alliance San Diego with Michael Quigley, Ellen Stefanski, Ian Mchardy |
| EPI_ISL_445169, EPI_ISL_445171, EPI_ISL_445172, EPI_ISL_445173, EPI_ISL_445175, EPI_ISL_445176, EPI_ISL_445177, EPI_ISL_445179, EPI_ISL_445180, EPI_ISL_445181, EPI_ISL_445182 | | | |
| see above | UCSF Clinical Microbiology Laboratory | Chan-Zuckerberg Biohub | CZB Cliahub Consortium |
| EPI_ISL_445183 | Takayuki Hishiki Kanagawa Prefectural Institute of Public Health | Takayuki Hishiki Kanagawa Prefectural Institute of Public Health | Hishiki,T., Suzuki,R., Sakuragi,J., Usui,K., Tanaka,Y., Kawai,J., Kogo,Y., Matsuki,Y., An,T., Hayashizaki,Y. and Takasaki,T. |
| EPI_ISL_445214, EPI_ISL_445215, EPI_ISL_445216 | DNA Solution Ltd. | DNA Solution Ltd. | Md. Imran Khan, Kazi Nadim Hasan, Abu Sufian, Mohammed Nafiz Imtiaz Polol, Abdul Khaleque, Mizanur Rahman, MSM Chowdhury, Hasan Ul Haider, Mamudul Hasan Razu, Mala Khan, Mohammad Fazle Alam Rabbi |
| EPI_ISL_445219 | Universidad del Valle, Laboratorio de Microbiologia, VIREM | Universidad del Valle, Universidad Nacional de Colombia-Sede Palmira, International Center for Tropical Agriculture | Beatriz Parra, Diana López-Alvarez, Wilmer J. Cuellar |
| EPI_ISL_445220 | Laboratory for Respiratory Viruses, "Cantacuzino" National Military-Medical Institute for Resararch and Development | Cantacuzino Institute | M.Lazar, L.Ustea, A.Cretu |
| EPI_ISL_445221 | Wasterlakarna | The Public Health Agency of Sweden | Frida Ahlfors, Oskar Karlsson Lindsjo, Maria Lind Karlberg, Anna-Malin Linde, Olov Svartstrom, Anna Risberg, Theresa Enkirch, Mia Brytting, Karin Tegmark-Wisell |
| EPI_ISL_445222 | Saroledens Familjelakare | The Public Health Agency of Sweden | Katarina Jarbur, Oskar Karlsson Lindsjo, Maria Lind Karlberg, Anna-Malin Linde, Olov Svartstrom, Anna Risberg, Theresa Enkirch, Mia Brytting, Karin Tegmark-Wisell |
| EPI_ISL_445223 | Victoria Vard och Halsa | The Public Health Agency of Sweden | Sarah Henriksson, Oskar Karlsson Lindsjo, Maria Lind Karlberg, Anna-Malin Linde, Olov Svartstrom, Anna Risberg, Theresa Enkirch, Mia Brytting, Karin Tegmark-Wisell |
| EPI_ISL_445224 | Narhalsan Olskroken VC | The Public Health Agency of Sweden | Mahin Ghoroghi, Oskar Karlsson Lindsjo, Maria Lind Karlberg, Anna-Malin Linde, Olov Svartstrom, Anna Risberg, Theresa Enkirch, Mia Brytting, Karin Tegmark-Wisell |
| EPI_ISL_445225 | Surbrunns VC | The Public Health Agency of Sweden | Erik Embring, Oskar Karlsson Lindsjo, Maria Lind Karlberg, Anna-Malin Linde, Olov Svartstrom, Anna Risberg, Theresa Enkirch, Mia Brytting, Karin Tegmark-Wisell |
| EPI_ISL_445226 | Saroledens Familjelakare | The Public Health Agency of Sweden | Katarina Jarbur, Oskar Karlsson Lindsjo, Maria Lind Karlberg, Anna-Malin Linde, Olov Svartstrom, Anna Risberg, Theresa Enkirch, Mia Brytting, Karin Tegmark-Wisell |
| EPI_ISL_445227 | Uppsala Narakut Aleris | The Public Health Agency of Sweden | Annika Nilsson, Oskar Karlsson Lindsjo, Maria Lind Karlberg, Anna-Malin Linde, Olov Svartstrom, Anna Risberg, Theresa Enkirch, Mia Brytting, Karin Tegmark-Wisell |
| EPI_ISL_445228 | Ulltuna Vardcentral | The Public Health Agency of Sweden | Heidi Lindback, Oskar Karlsson Lindsjo, Maria Lind Karlberg, Anna-Malin Linde, Olov Svartstrom, Anna Risberg, Theresa Enkirch, Mia Brytting, Karin Tegmark-Wisell |
| EPI_ISL_445229 | Narhalsan Backa vardcentral | The Public Health Agency of Sweden | Mats Olsson, Oskar Karlsson Lindsjo, Maria Lind Karlberg, Anna-Malin Linde, Olov Svartstrom, Anna Risberg, Theresa Enkirch, Mia Brytting, Karin Tegmark-Wisell |
| EPI_ISL_445230, EPI_ISL_445231 | Uppsala Narakut Aleris | The Public Health Agency of Sweden | Annika Nilsson, Oskar Karlsson Lindsjo, Maria Lind Karlberg, Anna-Malin Linde, Olov Svartstrom, Anna Risberg, Theresa Enkirch, Mia Brytting, Karin Tegmark-Wisell |
| EPI_ISL_445232 | Kungsors VC | The Public Health Agency of Sweden | Jessica Karlsson, Oskar Karlsson Lindsjo, Maria Lind Karlberg, Anna-Malin Linde, Olov Svartstrom, Anna Risberg, Theresa Enkirch, Mia Brytting, Karin Tegmark-Wisell |
| EPI_ISL_445233 | Vardcentralen Brinken | The Public Health Agency of Sweden | Agnes Wigh, Oskar Karlsson Lindsjo, Maria Lind Karlberg, Anna-Malin Linde, Olov Svartstrom, Anna Risberg, Theresa Enkirch, Mia Brytting, Karin Tegmark-Wisell |
| EPI_ISL_445234, EPI_ISL_445235 | Wasterlakarna | The Public Health Agency of Sweden | Frida Ahlfors, Oskar Karlsson Lindsjo, Maria Lind Karlberg, Anna-Malin Linde, Olov Svartstrom, Anna Risberg, Theresa Enkirch, Mia Brytting, Karin Tegmark-Wisell |
| EPI_ISL_445236 | Narhalsan Backa vardcentral | The Public Health Agency of Sweden | Mats Olsson, Oskar Karlsson Lindsjo, Maria Lind Karlberg, Anna-Malin Linde, Olov Svartstrom, Anna Risberg, Theresa Enkirch, Mia Brytting, Karin Tegmark-Wisell |
| EPI_ISL_445237 | Narhalsan Molnlycke, Barn och ungdomsmedicin | The Public Health Agency of Sweden | Mats Reimer, Oskar Karlsson Lindsjo, Maria Lind Karlberg, Anna-Malin Linde, Olov Svartstrom, Anna Risberg, Theresa Enkirch, Mia Brytting, Karin Tegmark-Wisell |
| EPI_ISL_445238 | Ã–resundslakarna | The Public Health Agency of Sweden | Del Akrawi, Oskar Karlsson Lindsjo, Maria Lind Karlberg, Anna-Malin Linde, Olov Svartstrom, Anna Risberg, Theresa Enkirch, Mia Brytting, Karin Tegmark-Wisell |
| EPI_ISL_445239 | Uppsala Narakut Aleris | The Public Health Agency of Sweden | Annika Nilsson, Oskar Karlsson Lindsjo, Maria Lind Karlberg, Anna-Malin Linde, Olov Svartstrom, Anna Risberg, Theresa Enkirch, Mia Brytting, Karin Tegmark-Wisell |
| EPI_ISL_445240 | Ulltuna Vardcentral | The Public Health Agency of Sweden | Heidi Lindback, Oskar Karlsson Lindsjo, Maria Lind Karlberg, Anna-Malin Linde, Olov Svartstrom, Anna Risberg, Theresa Enkirch, Mia Brytting, Karin Tegmark-Wisell |
| EPI_ISL_445241 | Ã–restadsklinikens VC | The Public Health Agency of Sweden | Lisa Kjellberg / Laura Plavitu, Oskar Karlsson Lindsjo, Maria Lind Karlberg, Anna-Malin Linde, Olov Svartstrom, Anna Risberg, Theresa Enkirch, Mia Brytting, Karin Tegmark-Wisell |
| EPI_ISL_445242 | Jokkmokks Halsocentral | The Public Health Agency of Sweden | Markus Beland, Oskar Karlsson Lindsjo, Maria Lind Karlberg, Anna-Malin Linde, Olov Svartstrom, Anna Risberg, Theresa Enkirch, Mia Brytting, Karin Tegmark-Wisell |
| EPI_ISL_445243 | Laboratory for Respiratory Viruses, Cantacuzino National Military-Medical Institute for Research and Development | Cantacuzino Institute | M.Lazar, L.Ustea, A.Cretu |
| EPI_ISL_445244 | Akbiomed lab | Tejgaon College bmb lab | Md.Abdul kaium,Md.Easin Arafat |
| EPI_ISL_445245 | CLINICA ALEMANA DE SANTIAGO S.A. | Instituto de Salud Publica de Chile | Andrés E Castillo, Bárbara Parra,Paz Tapia, Jaime Lagos, Loredana Arata, Alejandra Acevedo, Winston Andrade, Gabriel Leal, Carolina Tambley, Patricia Bustos, Rodrigo Fasce, Jorge Fernandez |
| EPI_ISL_445247 | UNIVERSIDAD DE LOS ANDES | Instituto de Salud Publica de Chile | Andrés E Castillo, Bárbara Parra,Paz Tapia, Jaime Lagos, Loredana Arata, Alejandra Acevedo, Winston Andrade, Gabriel Leal, Carolina Tambley, Patricia Bustos, Rodrigo Fasce, Jorge Fernandez |
| EPI_ISL_445248 | CLINICA ALEMANA DE SANTIAGO S.A. | Instituto de Salud Publica de Chile | Andrés E Castillo, Bárbara Parra,Paz Tapia, Jaime Lagos, Loredana Arata, Alejandra Acevedo, Winston Andrade, Gabriel Leal, Carolina Tambley, Patricia Bustos, Rodrigo Fasce, Jorge Fernandez |
| EPI_ISL_445249 | CLINICA SANTA MARIA S.A. | Instituto de Salud Publica de Chile | Andrés E Castillo, Bárbara Parra,Paz Tapia, Jaime Lagos, Loredana Arata, Alejandra Acevedo, Winston Andrade, Gabriel Leal, Carolina Tambley, Patricia Bustos, Rodrigo Fasce, Jorge Fernandez |
| EPI_ISL_445250 | CLINICA ALEMANA DE SANTIAGO S.A. | Instituto de Salud Publica de Chile | Andrés E Castillo, Bárbara Parra,Paz Tapia, Jaime Lagos, Loredana Arata, Alejandra Acevedo, Winston Andrade, Gabriel Leal, Carolina Tambley, Patricia Bustos, Rodrigo Fasce, Jorge Fernandez |
| EPI_ISL_445251 | HOSPITAL DE CARABINEROS | Instituto de Salud Publica de Chile | Andrés E Castillo, Bárbara Parra,Paz Tapia, Jaime Lagos, Loredana Arata, Alejandra Acevedo, Winston Andrade, Gabriel Leal, Carolina Tambley, Patricia Bustos, Rodrigo Fasce, Jorge Fernandez |
| EPI_ISL_445254, EPI_ISL_445255, EPI_ISL_445258 | CLINICA ALEMANA DE SANTIAGO S.A. | Instituto de Salud Publica de Chile | Andrés E Castillo, Bárbara Parra,Paz Tapia, Jaime Lagos, Loredana Arata, Alejandra Acevedo, Winston Andrade, Gabriel Leal, Carolina Tambley, Patricia Bustos, Rodrigo Fasce, Jorge Fernandez |
| EPI_ISL_445259 | CLINICA LAS CONDES S.A. | Instituto de Salud Publica de Chile | Andrés E Castillo, Bárbara Parra,Paz Tapia, Jaime Lagos, Loredana Arata, Alejandra Acevedo, Winston Andrade, Gabriel Leal, Carolina Tambley, Patricia Bustos, Rodrigo Fasce, Jorge Fernandez |
| EPI_ISL_445260 | CLINICA ALEMANA DE SANTIAGO S.A. | Instituto de Salud Publica de Chile | Andrés E Castillo, Bárbara Parra,Paz Tapia, Jaime Lagos, Loredana Arata, Alejandra Acevedo, Winston Andrade, Gabriel Leal, Carolina Tambley, Patricia Bustos, Rodrigo Fasce, Jorge Fernandez |
| EPI_ISL_445261 | INTEGRAMEDICA LAB. CLINICO LTDA. | Instituto de Salud Publica de Chile | Andrés E Castillo, Bárbara Parra,Paz Tapia, Jaime Lagos, Loredana Arata, Alejandra Acevedo, Winston Andrade, Gabriel Leal, Carolina Tambley, Patricia Bustos, Rodrigo Fasce, Jorge Fernandez |
| EPI_ISL_445262 | CLINICA TABANCURA | Instituto de Salud Publica de Chile | Andrés E Castillo, Bárbara Parra,Paz Tapia, Jaime Lagos, Loredana Arata, Alejandra Acevedo, Winston Andrade, Gabriel Leal, Carolina Tambley, Patricia Bustos, Rodrigo Fasce, Jorge Fernandez |
| EPI_ISL_445263 | MEGASALUD SPA. | Instituto de Salud Publica de Chile | Andrés E Castillo, Bárbara Parra,Paz Tapia, Jaime Lagos, Loredana Arata, Alejandra Acevedo, Winston Andrade, Gabriel Leal, Carolina Tambley, Patricia Bustos, Rodrigo Fasce, Jorge Fernandez |
| EPI_ISL_445264 | CLINICA REDSALUD VITACURA. | Instituto de Salud Publica de Chile | Andrés E Castillo, Bárbara Parra,Paz Tapia, Jaime Lagos, Loredana Arata, Alejandra Acevedo, Winston Andrade, Gabriel Leal, Carolina Tambley, Patricia Bustos, Rodrigo Fasce, Jorge Fernandez |
| EPI_ISL_445266, EPI_ISL_445267 | CENTRO ONCOLOGICO DEL NORTE | Instituto de Salud Publica de Chile | Andrés E Castillo, Bárbara Parra,Paz Tapia, Jaime Lagos, Loredana Arata, Alejandra Acevedo, Winston Andrade, Gabriel Leal, Carolina Tambley, Patricia Bustos, Rodrigo Fasce, Jorge Fernandez |
| EPI_ISL_445268, EPI_ISL_445269 | HOSPITAL REG.LAUTARO NAVARRO AVARIA | Instituto de Salud Publica de Chile | Andrés E Castillo, Bárbara Parra,Paz Tapia, Jaime Lagos, Loredana Arata, Alejandra Acevedo, Winston Andrade, Gabriel Leal, Carolina Tambley, Patricia Bustos, Rodrigo Fasce, Jorge Fernandez |
| EPI_ISL_445270 | HOSPITAL DR.HERNAN HENRIQUEZ ARAVENA | Instituto de Salud Publica de Chile | Andrés E Castillo, Bárbara Parra,Paz Tapia, Jaime Lagos, Loredana Arata, Alejandra Acevedo, Winston Andrade, Gabriel Leal, Carolina Tambley, Patricia Bustos, Rodrigo Fasce, Jorge Fernandez |
| EPI_ISL_445271 | LABORATORIO CLINICA CHILLAN | Instituto de Salud Publica de Chile | Andrés E Castillo, Bárbara Parra,Paz Tapia, Jaime Lagos, Loredana Arata, Alejandra Acevedo, Winston Andrade, Gabriel Leal, Carolina Tambley, Patricia Bustos, Rodrigo Fasce, Jorge Fernandez |
| EPI_ISL_445272 | CLINICA CIUDAD DEL MAR | Instituto de Salud Publica de Chile | Andrés E Castillo, Bárbara Parra,Paz Tapia, Jaime Lagos, Loredana Arata, Alejandra Acevedo, Winston Andrade, Gabriel Leal, Carolina Tambley, Patricia Bustos, Rodrigo Fasce, Jorge Fernandez |
| EPI_ISL_445273, EPI_ISL_445274 | LABORATORIO TORRE MEDICA LTDA. | Instituto de Salud Publica de Chile | Andrés E Castillo, Bárbara Parra,Paz Tapia, Jaime Lagos, Loredana Arata, Alejandra Acevedo, Winston Andrade, Gabriel Leal, Carolina Tambley, Patricia Bustos, Rodrigo Fasce, Jorge Fernandez |
| EPI_ISL_445275 | HOSPITAL CLINICO FUSAT | Instituto de Salud Publica de Chile | Andrés E Castillo, Bárbara Parra,Paz Tapia, Jaime Lagos, Loredana Arata, Alejandra Acevedo, Winston Andrade, Gabriel Leal, Carolina Tambley, Patricia Bustos, Rodrigo Fasce, Jorge Fernandez |
| EPI_ISL_445277 | FUNDACION DE SALUD EL TENIENTE | Instituto de Salud Publica de Chile | Andrés E Castillo, Bárbara Parra,Paz Tapia, Jaime Lagos, Loredana Arata, Alejandra Acevedo, Winston Andrade, Gabriel Leal, Carolina Tambley, Patricia Bustos, Rodrigo Fasce, Jorge Fernandez |
| EPI_ISL_445278 | LABORATORIO TORRE MEDICA LTDA. | Instituto de Salud Publica de Chile | Andrés E Castillo, Bárbara Parra,Paz Tapia, Jaime Lagos, Loredana Arata, Alejandra Acevedo, Winston Andrade, Gabriel Leal, Carolina Tambley, Patricia Bustos, Rodrigo Fasce, Jorge Fernandez |
| EPI_ISL_445279 | LABORATORIO INMUNOLAB SPA | Instituto de Salud Publica de Chile | Andrés E Castillo, Bárbara Parra,Paz Tapia, Jaime Lagos, Loredana Arata, Alejandra Acevedo, Winston Andrade, Gabriel Leal, Carolina Tambley, Patricia Bustos, Rodrigo Fasce, Jorge Fernandez |
| EPI_ISL_445280 | HOSPITAL REG.LAUTARO NAVARRO AVARIA | Instituto de Salud Publica de Chile | Andrés E Castillo, Bárbara Parra,Paz Tapia, Jaime Lagos, Loredana Arata, Alejandra Acevedo, Winston Andrade, Gabriel Leal, Carolina Tambley, Patricia Bustos, Rodrigo Fasce, Jorge Fernandez |
| EPI_ISL_445281 | HOSPITAL CLINICO DEL SUR | Instituto de Salud Publica de Chile | Andrés E Castillo, Bárbara Parra,Paz Tapia, Jaime Lagos, Loredana Arata, Alejandra Acevedo, Winston Andrade, Gabriel Leal, Carolina Tambley, Patricia Bustos, Rodrigo Fasce, Jorge Fernandez |
| EPI_ISL_445282, EPI_ISL_445283 | CLINICA MAGALLANES S.A. | Instituto de Salud Publica de Chile | Andrés E Castillo, Bárbara Parra,Paz Tapia, Jaime Lagos, Loredana Arata, Alejandra Acevedo, Winston Andrade, Gabriel Leal, Carolina Tambley, Patricia Bustos, Rodrigo Fasce, Jorge Fernandez |
| EPI_ISL_445284 | HOSPITAL REG.LAUTARO NAVARRO AVARIA | Instituto de Salud Publica de Chile | Andrés E Castillo, Bárbara Parra,Paz Tapia, Jaime Lagos, Loredana Arata, Alejandra Acevedo, Winston Andrade, Gabriel Leal, Carolina Tambley, Patricia Bustos, Rodrigo Fasce, Jorge Fernandez |

| | | | |
|---|---|---|---|
| EPI_ISL_445285 | CLINICA CIUDAD DEL MAR | Instituto de Salud Publica de Chile | Andrés E Castillo, Bárbara Parra,Paz Tapia, Jaime Lagos, Loredana Arata, Alejandra Acevedo, Winston Andrade, Gabriel Leal, Carolina Tambley, Patricia Bustos, Rodrigo Fasce, Jorge Fernandez |
| EPI_ISL_445286 | HOSPITAL HANGA ROA | Instituto de Salud Publica de Chile | Andrés E Castillo, Bárbara Parra,Paz Tapia, Jaime Lagos, Loredana Arata, Alejandra Acevedo, Winston Andrade, Gabriel Leal, Carolina Tambley, Patricia Bustos, Rodrigo Fasce, Jorge Fernandez |
| EPI_ISL_445287 | CLINICA CIUDAD DEL MAR | Instituto de Salud Publica de Chile | Andrés E Castillo, Bárbara Parra,Paz Tapia, Jaime Lagos, Loredana Arata, Alejandra Acevedo, Winston Andrade, Gabriel Leal, Carolina Tambley, Patricia Bustos, Rodrigo Fasce, Jorge Fernandez |
| EPI_ISL_445288 | HOSPITAL REG.LAUTARO NAVARRO AVARIA | Instituto de Salud Publica de Chile | Andrés E Castillo, Bárbara Parra,Paz Tapia, Jaime Lagos, Loredana Arata, Alejandra Acevedo, Winston Andrade, Gabriel Leal, Carolina Tambley, Patricia Bustos, Rodrigo Fasce, Jorge Fernandez |
| EPI_ISL_445289 | HOSPITAL NAVAL PUERTO WILLIAMS | Instituto de Salud Publica de Chile | Andrés E Castillo, Bárbara Parra,Paz Tapia, Jaime Lagos, Loredana Arata, Alejandra Acevedo, Winston Andrade, Gabriel Leal, Carolina Tambley, Patricia Bustos, Rodrigo Fasce, Jorge Fernandez |
| EPI_ISL_445290, EPI_ISL_445292 | CLINICA MAGALLANES S.A. | Instituto de Salud Publica de Chile | Andrés E Castillo, Bárbara Parra,Paz Tapia, Jaime Lagos, Loredana Arata, Alejandra Acevedo, Winston Andrade, Gabriel Leal, Carolina Tambley, Patricia Bustos, Rodrigo Fasce, Jorge Fernandez |
| EPI_ISL_445293, EPI_ISL_445294 | HOSPITAL REG.LAUTARO NAVARRO AVARIA | Instituto de Salud Publica de Chile | Andrés E Castillo, Bárbara Parra,Paz Tapia, Jaime Lagos, Loredana Arata, Alejandra Acevedo, Winston Andrade, Gabriel Leal, Carolina Tambley, Patricia Bustos, Rodrigo Fasce, Jorge Fernandez |
| EPI_ISL_445297 | CLINICA INTEGRAL S.A. | Instituto de Salud Publica de Chile | Andrés E Castillo, Bárbara Parra,Paz Tapia, Jaime Lagos, Loredana Arata, Alejandra Acevedo, Winston Andrade, Gabriel Leal, Carolina Tambley, Patricia Bustos, Rodrigo Fasce, Jorge Fernandez |
| EPI_ISL_445300 | HOSPITAL DE RANCAGUA | Instituto de Salud Publica de Chile | Andrés E Castillo, Bárbara Parra,Paz Tapia, Jaime Lagos, Loredana Arata, Alejandra Acevedo, Winston Andrade, Gabriel Leal, Carolina Tambley, Patricia Bustos, Rodrigo Fasce, Jorge Fernandez |
| EPI_ISL_445302 | INSTITUTO MEDICO LEGAL | Instituto de Salud Publica de Chile | Andrés E Castillo, Bárbara Parra,Paz Tapia, Jaime Lagos, Loredana Arata, Alejandra Acevedo, Winston Andrade, Gabriel Leal, Carolina Tambley, Patricia Bustos, Rodrigo Fasce, Jorge Fernandez |
| EPI_ISL_445303 | CTRO.DE SALUD FAMILIAR DR. RAUL YAZIGI | Instituto de Salud Publica de Chile | Andrés E Castillo, Bárbara Parra,Paz Tapia, Jaime Lagos, Loredana Arata, Alejandra Acevedo, Winston Andrade, Gabriel Leal, Carolina Tambley, Patricia Bustos, Rodrigo Fasce, Jorge Fernandez |
| EPI_ISL_445304 | HOSPITAL SAN JUAN DE DIOS | Instituto de Salud Publica de Chile | Andrés E Castillo, Bárbara Parra,Paz Tapia, Jaime Lagos, Loredana Arata, Alejandra Acevedo, Winston Andrade, Gabriel Leal, Carolina Tambley, Patricia Bustos, Rodrigo Fasce, Jorge Fernandez |
| EPI_ISL_445305 | HOSPITAL DE CARABINEROS | Instituto de Salud Publica de Chile | Andrés E Castillo, Bárbara Parra,Paz Tapia, Jaime Lagos, Loredana Arata, Alejandra Acevedo, Winston Andrade, Gabriel Leal, Carolina Tambley, Patricia Bustos, Rodrigo Fasce, Jorge Fernandez |
| EPI_ISL_445306 | CLINICA UC SAN CARLOS DE APOQUINDO | Instituto de Salud Publica de Chile | Andrés E Castillo, Bárbara Parra,Paz Tapia, Jaime Lagos, Loredana Arata, Alejandra Acevedo, Winston Andrade, Gabriel Leal, Carolina Tambley, Patricia Bustos, Rodrigo Fasce, Jorge Fernandez |
| EPI_ISL_445307 | HOSPITAL SAN JOSE DE MAIPO | Instituto de Salud Publica de Chile | Andrés E Castillo, Bárbara Parra,Paz Tapia, Jaime Lagos, Loredana Arata, Alejandra Acevedo, Winston Andrade, Gabriel Leal, Carolina Tambley, Patricia Bustos, Rodrigo Fasce, Jorge Fernandez |
| EPI_ISL_445309 | MUTUAL DE SEGURIDAD C.CH.C. | Instituto de Salud Publica de Chile | Andrés E Castillo, Bárbara Parra,Paz Tapia, Jaime Lagos, Loredana Arata, Alejandra Acevedo, Winston Andrade, Gabriel Leal, Carolina Tambley, Patricia Bustos, Rodrigo Fasce, Jorge Fernandez |
| EPI_ISL_445310 | HOSPITAL DR.SOTERO DEL RIO | Instituto de Salud Publica de Chile | Andrés E Castillo, Bárbara Parra,Paz Tapia, Jaime Lagos, Loredana Arata, Alejandra Acevedo, Winston Andrade, Gabriel Leal, Carolina Tambley, Patricia Bustos, Rodrigo Fasce, Jorge Fernandez |
| EPI_ISL_445311 | MEGASALUD S.A. | Instituto de Salud Publica de Chile | Andrés E Castillo, Bárbara Parra,Paz Tapia, Jaime Lagos, Loredana Arata, Alejandra Acevedo, Winston Andrade, Gabriel Leal, Carolina Tambley, Patricia Bustos, Rodrigo Fasce, Jorge Fernandez |
| EPI_ISL_445312 | CLINICA UC SAN CARLOS DE APOQUINDO | Instituto de Salud Publica de Chile | Andrés E Castillo, Bárbara Parra,Paz Tapia, Jaime Lagos, Loredana Arata, Alejandra Acevedo, Winston Andrade, Gabriel Leal, Carolina Tambley, Patricia Bustos, Rodrigo Fasce, Jorge Fernandez |
| EPI_ISL_445313, EPI_ISL_445314 | HOSPITAL EL CARMEN DR.LUIS VALENTIN F. | Instituto de Salud Publica de Chile | Andrés E Castillo, Bárbara Parra,Paz Tapia, Jaime Lagos, Loredana Arata, Alejandra Acevedo, Winston Andrade, Gabriel Leal, Carolina Tambley, Patricia Bustos, Rodrigo Fasce, Jorge Fernandez |
| EPI_ISL_445315 | CLINICA UC SAN CARLOS DE APOQUINDO | Instituto de Salud Publica de Chile | Andrés E Castillo, Bárbara Parra,Paz Tapia, Jaime Lagos, Loredana Arata, Alejandra Acevedo, Winston Andrade, Gabriel Leal, Carolina Tambley, Patricia Bustos, Rodrigo Fasce, Jorge Fernandez |
| EPI_ISL_445316 | CESFAM BALMACEDA DE RENCA | Instituto de Salud Publica de Chile | Andrés E Castillo, Bárbara Parra,Paz Tapia, Jaime Lagos, Loredana Arata, Alejandra Acevedo, Winston Andrade, Gabriel Leal, Carolina Tambley, Patricia Bustos, Rodrigo Fasce, Jorge Fernandez |
| EPI_ISL_445317 | UNIV.DE CHILE HOSP.CLINICO | Instituto de Salud Publica de Chile | Andrés E Castillo, Bárbara Parra,Paz Tapia, Jaime Lagos, Loredana Arata, Alejandra Acevedo, Winston Andrade, Gabriel Leal, Carolina Tambley, Patricia Bustos, Rodrigo Fasce, Jorge Fernandez |
| EPI_ISL_445318 | C.DE SALUD FAMILIAR PABLO NERUDA | Instituto de Salud Publica de Chile | Andrés E Castillo, Bárbara Parra,Paz Tapia, Jaime Lagos, Loredana Arata, Alejandra Acevedo, Winston Andrade, Gabriel Leal, Carolina Tambley, Patricia Bustos, Rodrigo Fasce, Jorge Fernandez |
| EPI_ISL_445320 | C.C.SALUD FAMILIAR PADRE FELIX DONOSO G. | Instituto de Salud Publica de Chile | Andrés E Castillo, Bárbara Parra,Paz Tapia, Jaime Lagos, Loredana Arata, Alejandra Acevedo, Winston Andrade, Gabriel Leal, Carolina Tambley, Patricia Bustos, Rodrigo Fasce, Jorge Fernandez |
| EPI_ISL_445322 | UNIV.DE CHILE HOSP.CLINICO | Instituto de Salud Publica de Chile | Andrés E Castillo, Bárbara Parra,Paz Tapia, Jaime Lagos, Loredana Arata, Alejandra Acevedo, Winston Andrade, Gabriel Leal, Carolina Tambley, Patricia Bustos, Rodrigo Fasce, Jorge Fernandez |
| EPI_ISL_445323 | HOSP.ENFERMEDADES INFECCIOSAS | Instituto de Salud Publica de Chile | Andrés E Castillo, Bárbara Parra,Paz Tapia, Jaime Lagos, Loredana Arata, Alejandra Acevedo, Winston Andrade, Gabriel Leal, Carolina Tambley, Patricia Bustos, Rodrigo Fasce, Jorge Fernandez |
| EPI_ISL_445326 | ASISTENCIA PUBLICA DR.ALEJANDRO DEL RIO | Instituto de Salud Publica de Chile | Andrés E Castillo, Bárbara Parra,Paz Tapia, Jaime Lagos, Loredana Arata, Alejandra Acevedo, Winston Andrade, Gabriel Leal, Carolina Tambley, Patricia Bustos, Rodrigo Fasce, Jorge Fernandez |
| EPI_ISL_445327 | HOSPITAL PADRE HURTADO | Instituto de Salud Publica de Chile | Andrés E Castillo, Bárbara Parra,Paz Tapia, Jaime Lagos, Loredana Arata, Alejandra Acevedo, Winston Andrade, Gabriel Leal, Carolina Tambley, Patricia Bustos, Rodrigo Fasce, Jorge Fernandez |
| EPI_ISL_445328 | MEGASALUD S.A. | Instituto de Salud Publica de Chile | Andrés E Castillo, Bárbara Parra,Paz Tapia, Jaime Lagos, Loredana Arata, Alejandra Acevedo, Winston Andrade, Gabriel Leal, Carolina Tambley, Patricia Bustos, Rodrigo Fasce, Jorge Fernandez |
| EPI_ISL_445329 | HOSPITAL DR.SOTERO DEL RIO | Instituto de Salud Publica de Chile | Andrés E Castillo, Bárbara Parra,Paz Tapia, Jaime Lagos, Loredana Arata, Alejandra Acevedo, Winston Andrade, Gabriel Leal, Carolina Tambley, Patricia Bustos, Rodrigo Fasce, Jorge Fernandez |
| EPI_ISL_445330 | CLINICA VESPUCIO S. A. | Instituto de Salud Publica de Chile | Andrés E Castillo, Bárbara Parra,Paz Tapia, Jaime Lagos, Loredana Arata, Alejandra Acevedo, Winston Andrade, Gabriel Leal, Carolina Tambley, Patricia Bustos, Rodrigo Fasce, Jorge Fernandez |
| EPI_ISL_445331, EPI_ISL_445332 | HOSPITAL HERMINDA MARTIN CHILLAN | Instituto de Salud Publica de Chile | Andrés E Castillo, Bárbara Parra,Paz Tapia, Jaime Lagos, Loredana Arata, Alejandra Acevedo, Winston Andrade, Gabriel Leal, Carolina Tambley, Patricia Bustos, Rodrigo Fasce, Jorge Fernandez |
| EPI_ISL_445333 | LABORATORIO CLINICA UNIVERSITARIA DE CONCEPCION | Instituto de Salud Publica de Chile | Andrés E Castillo, Bárbara Parra,Paz Tapia, Jaime Lagos, Loredana Arata, Alejandra Acevedo, Winston Andrade, Gabriel Leal, Carolina Tambley, Patricia Bustos, Rodrigo Fasce, Jorge Fernandez |
| EPI_ISL_445334 | CLINICA UNIVERSITARIA DE PUERTO MONTT S.A. | Instituto de Salud Publica de Chile | Andrés E Castillo, Bárbara Parra,Paz Tapia, Jaime Lagos, Loredana Arata, Alejandra Acevedo, Winston Andrade, Gabriel Leal, Carolina Tambley, Patricia Bustos, Rodrigo Fasce, Jorge Fernandez |
| EPI_ISL_445335 | HOSPITAL DE CALBUCO | Instituto de Salud Publica de Chile | Andrés E Castillo, Bárbara Parra,Paz Tapia, Jaime Lagos, Loredana Arata, Alejandra Acevedo, Winston Andrade, Gabriel Leal, Carolina Tambley, Patricia Bustos, Rodrigo Fasce, Jorge Fernandez |
| EPI_ISL_445340, EPI_ISL_445342, EPI_ISL_445343, EPI_ISL_445344, EPI_ISL_445345, EPI_ISL_445346, EPI_ISL_445347, EPI_ISL_445348 | HOSPITAL DR.HERNAN HENRIQUEZ ARAVENA | Instituto de Salud Publica de Chile | Andrés E Castillo, Bárbara Parra,Paz Tapia, Jaime Lagos, Loredana Arata, Alejandra Acevedo, Winston Andrade, Gabriel Leal, Carolina Tambley, Patricia Bustos, Rodrigo Fasce, Jorge Fernandez |
| EPI_ISL_445349, EPI_ISL_445350, EPI_ISL_445351 | HOSPITAL SAN JUAN DE DIOS | Instituto de Salud Publica de Chile | Andrés E Castillo, Bárbara Parra,Paz Tapia, Jaime Lagos, Loredana Arata, Alejandra Acevedo, Winston Andrade, Gabriel Leal, Carolina Tambley, Patricia Bustos, Rodrigo Fasce, Jorge Fernandez |
| EPI_ISL_445352 | HOSPITAL DEL PROFESOR | Instituto de Salud Publica de Chile | Andrés E Castillo, Bárbara Parra,Paz Tapia, Jaime Lagos, Loredana Arata, Alejandra Acevedo, Winston Andrade, Gabriel Leal, Carolina Tambley, Patricia Bustos, Rodrigo Fasce, Jorge Fernandez |
| EPI_ISL_445353 | HOSPITAL PADRE HURTADO | Instituto de Salud Publica de Chile | Andrés E Castillo, Bárbara Parra,Paz Tapia, Jaime Lagos, Loredana Arata, Alejandra Acevedo, Winston Andrade, Gabriel Leal, Carolina Tambley, Patricia Bustos, Rodrigo Fasce, Jorge Fernandez |
| EPI_ISL_445355 | MUTUAL DE SEGURIDAD C.CH.C. | Instituto de Salud Publica de Chile | Andrés E Castillo, Bárbara Parra,Paz Tapia, Jaime Lagos, Loredana Arata, Alejandra Acevedo, Winston Andrade, Gabriel Leal, Carolina Tambley, Patricia Bustos, Rodrigo Fasce, Jorge Fernandez |
| EPI_ISL_445356 | PONTIFICIA U. CATOLICA SERV. LABORATORIO | Instituto de Salud Publica de Chile | Andrés E Castillo, Bárbara Parra,Paz Tapia, Jaime Lagos, Loredana Arata, Alejandra Acevedo, Winston Andrade, Gabriel Leal, Carolina Tambley, Patricia Bustos, Rodrigo Fasce, Jorge Fernandez |
| EPI_ISL_445358 | MEGASALUD SPA. | Instituto de Salud Publica de Chile | Andrés E Castillo, Bárbara Parra,Paz Tapia, Jaime Lagos, Loredana Arata, Alejandra Acevedo, Winston Andrade, Gabriel Leal, Carolina Tambley, Patricia Bustos, Rodrigo Fasce, Jorge Fernandez |
| EPI_ISL_445360 | HOSPITAL DEL PROFESOR | Instituto de Salud Publica de Chile | Andrés E Castillo, Bárbara Parra,Paz Tapia, Jaime Lagos, Loredana Arata, Alejandra Acevedo, Winston Andrade, Gabriel Leal, Carolina Tambley, Patricia Bustos, Rodrigo Fasce, Jorge Fernandez |
| EPI_ISL_445361 | CLINICA UC SAN CARLOS DE APOQUINDO | Instituto de Salud Publica de Chile | Andrés E Castillo, Bárbara Parra,Paz Tapia, Jaime Lagos, Loredana Arata, Alejandra Acevedo, Winston Andrade, Gabriel Leal, Carolina Tambley, Patricia Bustos, Rodrigo Fasce, Jorge Fernandez |
| EPI_ISL_445364 | HOSPITAL EL CARMEN DR.LUIS VALENTIN F. | Instituto de Salud Publica de Chile | Andrés E Castillo, Bárbara Parra,Paz Tapia, Jaime Lagos, Loredana Arata, Alejandra Acevedo, Winston Andrade, Gabriel Leal, Carolina Tambley, Patricia Bustos, Rodrigo Fasce, Jorge Fernandez |
| EPI_ISL_445365, EPI_ISL_445366 | HOSPITAL DR.SOTERO DEL RIO | Instituto de Salud Publica de Chile | Andrés E Castillo, Bárbara Parra,Paz Tapia, Jaime Lagos, Loredana Arata, Alejandra Acevedo, Winston Andrade, Gabriel Leal, Carolina Tambley, Patricia Bustos, Rodrigo Fasce, Jorge Fernandez |
| EPI_ISL_445367 | ASISTENCIA PUBLICA DR.ALEJANDRO DEL RIO | Instituto de Salud Publica de Chile | Andrés E Castillo, Bárbara Parra,Paz Tapia, Jaime Lagos, Loredana Arata, Alejandra Acevedo, Winston Andrade, Gabriel Leal, Carolina Tambley, Patricia Bustos, Rodrigo Fasce, Jorge Fernandez |
| EPI_ISL_445368 | HOSPITAL DEL PROFESOR | Instituto de Salud Publica de Chile | Andrés E Castillo, Bárbara Parra,Paz Tapia, Jaime Lagos, Loredana Arata, Alejandra Acevedo, Winston Andrade, Gabriel Leal, Carolina Tambley, Patricia Bustos, Rodrigo Fasce, Jorge Fernandez |
| EPI_ISL_445371 | HOSPITAL DR.SOTERO DEL RIO | Instituto de Salud Publica de Chile | Andrés E Castillo, Bárbara Parra,Paz Tapia, Jaime Lagos, Loredana Arata, Alejandra Acevedo, Winston Andrade, Gabriel Leal, Carolina Tambley, Patricia Bustos, Rodrigo Fasce, Jorge Fernandez |
| EPI_ISL_445372 | HOSPITAL FF.AA. "CIRUJANO C. GUZMAN | Instituto de Salud Publica de Chile | Andrés E Castillo, Bárbara Parra,Paz Tapia, Jaime Lagos, Loredana Arata, Alejandra Acevedo, Winston Andrade, Gabriel Leal, Carolina Tambley, Patricia Bustos, Rodrigo Fasce, Jorge Fernandez |
| EPI_ISL_445373, EPI_ISL_445375, EPI_ISL_445376, EPI_ISL_445377 | HOSPITAL SAN JUAN DE DIOS | Instituto de Salud Publica de Chile | Andrés E Castillo, Bárbara Parra,Paz Tapia, Jaime Lagos, Loredana Arata, Alejandra Acevedo, Winston Andrade, Gabriel Leal, Carolina Tambley, Patricia Bustos, Rodrigo Fasce, Jorge Fernandez |
| EPI_ISL_445378 | HOSPITAL DE BULNES | Instituto de Salud Publica de Chile | Andrés E Castillo, Bárbara Parra,Paz Tapia, Jaime Lagos, Loredana Arata, Alejandra Acevedo, Winston Andrade, Gabriel Leal, Carolina Tambley, Patricia Bustos, Rodrigo Fasce, Jorge Fernandez |

EPI_ISL_445385, EPI_ISL_445387, EPI_ISL_445388, EPI_ISL_445392, EPI_ISL_445398, EPI_ISL_445405, EPI_ISL_445406, EPI_ISL_445408, EPI_ISL_445409, EPI_ISL_445412, EPI_ISL_445413, EPI_ISL_445414, EPI_ISL_445416, EPI_ISL_445417, EPI_ISL_445419, EPI_ISL_445420, EPI_ISL_445422, EPI_ISL_445423, EPI_ISL_445427, EPI_ISL_445428, EPI_ISL_445433, EPI_ISL_445434, EPI_ISL_445438, EPI_ISL_445439, EPI_ISL_445448, EPI_ISL_445452, EPI_ISL_445453, EPI_ISL_445454, EPI_ISL_445457, EPI_ISL_445458, EPI_ISL_445459, EPI_ISL_445462, EPI_ISL_445470, EPI_ISL_445475, EPI_ISL_445478, EPI_ISL_445481, EPI_ISL_445485, EPI_ISL_445489, EPI_ISL_445494, EPI_ISL_445496, EPI_ISL_445500, EPI_ISL_445507, EPI_ISL_445510, EPI_ISL_445531, EPI_ISL_445541, EPI_ISL_445545, EPI_ISL_445552, EPI_ISL_445553, EPI_ISL_445564, EPI_ISL_445567, EPI_ISL_445571, EPI_ISL_445572, EPI_ISL_445573, EPI_ISL_445576, EPI_ISL_445583, EPI_ISL_445585, EPI_ISL_445589, EPI_ISL_445591, EPI_ISL_445594, EPI_ISL_445595, EPI_ISL_445597, EPI_ISL_445601, EPI_ISL_445602, EPI_ISL_445606, EPI_ISL_445614, EPI_ISL_445617, EPI_ISL_445619, EPI_ISL_445625, EPI_ISL_445628, EPI_ISL_445631, EPI_ISL_445632, EPI_ISL_445637, EPI_ISL_445638, EPI_ISL_445639, EPI_ISL_445640, EPI_ISL_445641, EPI_ISL_445642, EPI_ISL_445644, EPI_ISL_445649, EPI_ISL_445650, EPI_ISL_445651, EPI_ISL_445654, EPI_ISL_445662, EPI_ISL_445671, EPI_ISL_445672, EPI_ISL_445673, EPI_ISL_445674, EPI_ISL_445675, EPI_ISL_445677, EPI_ISL_445678, EPI_ISL_445680, EPI_ISL_445681, EPI_ISL_445682, EPI_ISL_445683, EPI_ISL_445686, EPI_ISL_445688, EPI_ISL_445689, EPI_ISL_445691, EPI_ISL_445692, EPI_ISL_445696, EPI_ISL_445697, EPI_ISL_445698, EPI_ISL_445701, EPI_ISL_445702, EPI_ISL_445703, EPI_ISL_445705, EPI_ISL_445708, EPI_ISL_445709, EPI_ISL_445716, EPI_ISL_445719, EPI_ISL_445720, EPI_ISL_445721, EPI_ISL_445723, EPI_ISL_445724, EPI_ISL_445725, EPI_ISL_445726, EPI_ISL_445727, EPI_ISL_445734, EPI_ISL_445735, EPI_ISL_445737, EPI_ISL_445741, EPI_ISL_445745, EPI_ISL_445746, EPI_ISL_445747, EPI_ISL_445748, EPI_ISL_445753, EPI_ISL_445754, EPI_ISL_445755, EPI_ISL_445756, EPI_ISL_445760, EPI_ISL_445761, EPI_ISL_445762, EPI_ISL_445763, EPI_ISL_445766, EPI_ISL_445767, EPI_ISL_445769, EPI_ISL_445775, EPI_ISL_445782, EPI_ISL_445786, EPI_ISL_445794, EPI_ISL_445823, EPI_ISL_445824, EPI_ISL_445831, EPI_ISL_445833, EPI_ISL_445842, EPI_ISL_445846, EPI_ISL_445849, EPI_ISL_445850, EPI_ISL_445856, EPI_ISL_445862, EPI_ISL_445863, EPI_ISL_445866, EPI_ISL_445870, EPI_ISL_445878, EPI_ISL_445885, EPI_ISL_445887, EPI_ISL_445895, EPI_ISL_445910, EPI_ISL_445915, EPI_ISL_445916, EPI_ISL_445931, EPI_ISL_445938, EPI_ISL_445944, EPI_ISL_445945, EPI_ISL_445946, EPI_ISL_445950, EPI_ISL_445951, EPI_ISL_445980, EPI_ISL_446014, EPI_ISL_446025, EPI_ISL_446027, EPI_ISL_446031, EPI_ISL_446038, EPI_ISL_446039, EPI_ISL_446040, EPI_ISL_446042, EPI_ISL_446043, EPI_ISL_446044, EPI_ISL_446047, EPI_ISL_446050, EPI_ISL_446060, EPI_ISL_446061, EPI_ISL_446064, EPI_ISL_446067, EPI_ISL_446068, EPI_ISL_446069, EPI_ISL_446072, EPI_ISL_446079, EPI_ISL_446080, EPI_ISL_446083, EPI_ISL_446086, EPI_ISL_446098, EPI_ISL_446101, EPI_ISL_446102, EPI_ISL_446103, EPI_ISL_446115, EPI_ISL_446121, EPI_ISL_446124, EPI_ISL_446125, EPI_ISL_446126, EPI_ISL_446130, EPI_ISL_446131, EPI_ISL_446139, EPI_ISL_446143, EPI_ISL_446144, EPI_ISL_446146, EPI_ISL_446147, EPI_ISL_446151, EPI_ISL_446153, EPI_ISL_446156, EPI_ISL_446160, EPI_ISL_446162, EPI_ISL_446164, EPI_ISL_446167, EPI_ISL_446171, EPI_ISL_446172, EPI_ISL_446177, EPI_ISL_446182, EPI_ISL_446183, EPI_ISL_446184, EPI_ISL_446186, EPI_ISL_446187, EPI_ISL_446190, EPI_ISL_446192, EPI_ISL_446193, EPI_ISL_446199, EPI_ISL_446200, EPI_ISL_446203, EPI_ISL_446206, EPI_ISL_446207, EPI_ISL_446210, EPI_ISL_446211, EPI_ISL_446212, EPI_ISL_446213, EPI_ISL_446214, EPI_ISL_446216, EPI_ISL_446217, EPI_ISL_446219, EPI_ISL_446220, EPI_ISL_446221, EPI_ISL_446222, EPI_ISL_446223, EPI_ISL_446224, EPI_ISL_446227, EPI_ISL_446230, EPI_ISL_446232, EPI_ISL_446234, EPI_ISL_446236, EPI_ISL_446237, EPI_ISL_446239, EPI_ISL_446242, EPI_ISL_446243, EPI_ISL_446257, EPI_ISL_446269, EPI_ISL_446270, EPI_ISL_446271, EPI_ISL_446280, EPI_ISL_446282, EPI_ISL_446283, EPI_ISL_446285, EPI_ISL_446287, EPI_ISL_446288, EPI_ISL_446294, EPI_ISL_446301, EPI_ISL_446305, EPI_ISL_446307, EPI_ISL_446309, EPI_ISL_446310, EPI_ISL_446317, EPI_ISL_446320, EPI_ISL_446321, EPI_ISL_446328, EPI_ISL_446332, EPI_ISL_446335, EPI_ISL_446336, EPI_ISL_446339, EPI_ISL_446340, EPI_ISL_446343, EPI_ISL_446347, EPI_ISL_446352, EPI_ISL_446363, EPI_ISL_446388, EPI_ISL_446423, EPI_ISL_446425, EPI_ISL_446435, EPI_ISL_446441, EPI_ISL_446442, EPI_ISL_446455, EPI_ISL_446461, EPI_ISL_446464, EPI_ISL_446466, EPI_ISL_446473, EPI_ISL_446474, EPI_ISL_446475, EPI_ISL_446476, EPI_ISL_446477, EPI_ISL_446483, EPI_ISL_446484, EPI_ISL_446486, EPI_ISL_446493, EPI_ISL_446494, EPI_ISL_446496, EPI_ISL_446497, EPI_ISL_446498, EPI_ISL_446501, EPI_ISL_446505, EPI_ISL_446514, EPI_ISL_446516, EPI_ISL_446522, EPI_ISL_446528, EPI_ISL_446535, EPI_ISL_446580, EPI_ISL_446589, EPI_ISL_446590, EPI_ISL_446594, EPI_ISL_446600, EPI_ISL_446601, EPI_ISL_446609, EPI_ISL_446612, EPI_ISL_446614, EPI_ISL_446621, EPI_ISL_446623, EPI_ISL_446626, EPI_ISL_446628, EPI_ISL_446629, EPI_ISL_446634, EPI_ISL_446636, EPI_ISL_446638, EPI_ISL_446639, EPI_ISL_446643, EPI_ISL_446644, EPI_ISL_446646, EPI_ISL_446647, EPI_ISL_446649, EPI_ISL_446679, EPI_ISL_446691, EPI_ISL_446698, EPI_ISL_446699, EPI_ISL_446706, EPI_ISL_446714, EPI_ISL_446715, EPI_ISL_446718, EPI_ISL_446723, EPI_ISL_446727, EPI_ISL_446733, EPI_ISL_446741, EPI_ISL_446744, EPI_ISL_446749, EPI_ISL_446758, EPI_ISL_446762, EPI_ISL_446767, EPI_ISL_446771, EPI_ISL_446774, EPI_ISL_446780, EPI_ISL_446788, EPI_ISL_446794, EPI_ISL_446795, EPI_ISL_446796, EPI_ISL_446802, EPI_ISL_446811, EPI_ISL_446823, EPI_ISL_446835, EPI_ISL_446851, EPI_ISL_446862, EPI_ISL_446866, EPI_ISL_446873, EPI_ISL_446877, EPI_ISL_446887, EPI_ISL_446894, EPI_ISL_446920, EPI_ISL_446923, EPI_ISL_446926, EPI_ISL_446934, EPI_ISL_446943, EPI_ISL_446949, EPI_ISL_446957, EPI_ISL_446959, EPI_ISL_446968, EPI_ISL_446979, EPI_ISL_446988, EPI_ISL_446992, EPI_ISL_446994, EPI_ISL_446995

| | | | |
|---|---|---|---|
| see above | Wales Specialist Virology Centre | Public Health Wales Microbiology Cardiff | Catherine Moore, Johnathan Evans, Laura Gifford, Malorie Perry, Simon Cottrell, Alec Birchley, Alexander Adams, Amy Gaskin, Bree Gatica-Wilcox, Jason Coombes, Lauren Gilbert, Lee Graham, Nicole Pacchiarini, Sara Kumziene-Summerhayes, Sarah Taylor, Sophie Jones, Sara Rey, Matthew Bull, Joanne Watkins, Sally Corden, Tom Connor |
| EPI_ISL_446996, EPI_ISL_447011, EPI_ISL_447015, EPI_ISL_447017, EPI_ISL_447019, EPI_ISL_447020, EPI_ISL_447025, EPI_ISL_447026 | Ramathibodi Hospital | COVID-19 Network Investigations (CONI) Alliance | Elizabeth Batty, Wasun Chantratita, Thanat Chookajorn, Stefan Fernandez, Angkana Huang, Anthony R. Jones, Khajohn Joonsalak, Chonticha Klungtong, Theerarat Kochakarn, Namfon Kotanan, Krittikorn Kumpornsin, Wudtichai Manasatienkij, Bhakbhoom Panthan, Ekawat Pasomsub, Kingkan Rakmanee, Insee Sensorn, Janjira Thaipadungpanit, Arporn Wangwiwatsin,Treewat Watthanachockchai |
| EPI_ISL_447030 | B.J. Medical College and Civil hospital | Gujarat Biotechnology Research Centre | Kamlesh J Upadhyay, Ramesh Pandit, Tejas Shah, Ankit Hinsu, Pritesh Sabara, Apurvasinh Puvar, Janvi Raval, Monika Gandhi, Pinal Trivedi, Maharshi Pandya, Amit Kanani, Akanksha Verma, Nitin Savaliya, Raghawendra Kumar, Dinesh Kumar, Zuber Saiyed, Dipa Kinariwala, Disha Patel, Binita Aring, Neeta Khandelwal, Geeta Vaghela, Sonia Barve, Bhavesh Modi, Kairavi Joshi, Gaurishankar Shrimali, Nidhi Sood, Pranay Shah, R D Dixit, Snehal Bagatharia, Anjali Rajwar, Chaitanya Joshi, Madhvi Joshi |
| EPI_ISL_447031 | B.J. Medical College and Civil hospital | Gujarat Biotechnology Research Centre | Ramesh Pandit, Tejas Shah, Ankit Hinsu, Pritesh Sabara, Apurvasinh Puvar, Janvi Raval, Monika Gandhi, Pinal Trivedi, Maharshi Pandya, Amit Kanani, Akanksha Verma, Nitin Savaliya, Raghawendra Kumar, Dinesh Kumar, Zuber Saiyed, Dipa Kinariwala, Disha Patel, Binita Aring, Neeta Khandelwal, Geeta Vaghela, Sonia Barve, Bhavesh Modi, Kairavi Joshi, Gaurishankar Shrimali, Nidhi Sood, Pranay Shah, R D Dixit, Snehal Bagatharia, Kamlesh J Upadhyay, Sharmistha Majumdar, Chaitanya Joshi, Madhvi Joshi |
| EPI_ISL_447032 | B.J. Medical College and Civil hospital | Gujarat Biotechnology Research Centre | Tejas Shah, Ankit Hinsu, Pritesh Sabara, Apurvasinh Puvar, Janvi Raval, Monika Gandhi, Pinal Trivedi, Maharshi Pandya, Amit Kanani, Akanksha Verma, Nitin Savaliya, Raghawendra Kumar, Dinesh Kumar, |

| | | | |
|---|---|---|---|
| | | | Zuber Saiyed, Dipa Kinariwala, Disha Patel, Binita Aring, Neeta Khandelwal, Geeta Vaghela, Sonia Barve, Bhavesh Modi, Kairavi Joshi, Gaurishankar Shrimali, Nidhi Sood, Pranay Shah, R D Dixit, Snehal Bagatharia, Ramesh Pandit, Kamlesh J Upadhyay, Pooja P Doshi, Chaitanya Joshi, Madhvi Joshi |
| EPI_ISL_447033 | B.J. Medical College and Civil hospital | Gujarat Biotechnology Research Centre | Ankit Hinsu, Pritesh Sabara, Apurvasinh Puvar, Janvi Raval, Monika Gandhi, Pinal Trivedi, Maharshi Pandya, Amit Kanani, Akanksha Verma, Nitin Savaliya, Raghawendra Kumar, Dinesh Kumar, Zuber Saiyed, Dipa Kinariwala, Disha Patel, Binita Aring, Neeta Khandelwal, Geeta Vaghela, Sonia Barve, Bhavesh Modi, Kairavi Joshi, Gaurishankar Shrimali, Nidhi Sood, Pranay Shah, R D Dixit, Snehal Bagatharia, Kamlesh J Upadhyay, Ramesh Pandit, Tejas Shah, Nidhi Patel, Chaitanya Joshi, Madhvi Joshi |
| EPI_ISL_447034 | B.J. Medical College and Civil hospital | Gujarat Biotechnology Research Centre | Pritesh Sabara, Apurvasinh Puvar, Janvi Raval, Monika Gandhi, Pinal Trivedi, Maharshi Pandya, Amit Kanani, Akanksha Verma, Nitin Savaliya, Raghawendra Kumar, Dinesh Kumar, Zuber Saiyed, Dipa Kinariwala, Disha Patel, Binita Aring, Neeta Khandelwal, Geeta Vaghela, Sonia Barve, Bhavesh Modi, Kairavi Joshi, Gaurishankar Shrimali, Nidhi Sood, Pranay Shah, R D Dixit, Snehal Bagatharia, Kamlesh J Upadhyay, Ramesh Pandit, Tejas Shah, Ankit Hinsu, Priti Pandita, Chaitanya Joshi, Madhvi Joshi |
| EPI_ISL_447035 | B.J. Medical College and Civil hospital | Gujarat Biotechnology Research Centre | Apurvasinh Puvar, Janvi Raval, Monika Gandhi, Pinal Trivedi, Maharshi Pandya, Amit Kanani, Akanksha Verma, Nitin Savaliya, Raghawendra Kumar, Dinesh Kumar, Zuber Saiyed, Dipa Kinariwala, Disha Patel, Binita Aring, Neeta Khandelwal, Geeta Vaghela, Sonia Barve, Bhavesh Modi, Kairavi Joshi, Gaurishankar Shrimali, Nidhi Sood, Pranay Shah, R D Dixit, Snehal Bagatharia, Kamlesh J Upadhyay, Ramesh Pandit, Tejas Shah, Ankit Hinsu, Pritesh Sabara, Neha Rajpara, Chaitanya Joshi, Madhvi Joshi |
| EPI_ISL_447036 | B.J. Medical College and Civil hospital | Gujarat Biotechnology Research Centre | Janvi Raval, Monika Gandhi, Pinal Trivedi, Maharshi Pandya, Amit Kanani, Akanksha Verma, Nitin Savaliya, Raghawendra Kumar, Dinesh Kumar, Zuber Saiyed, Dipa Kinariwala, Disha Patel, Binita Aring, Neeta Khandelwal, Geeta Vaghela, Sonia Barve, Bhavesh Modi, Kairavi Joshi, Gaurishankar Shrimali, Nidhi Sood, Pranay Shah, R D Dixit, Snehal Bagatharia, Kamlesh J Upadhyay, Ramesh Pandit, Tejas Shah, Ankit Hinsu, Pritesh Sabara, Apurvasinh Puvar, Afzal Ansari, Chaitanya Joshi, Madhvi Joshi |
| EPI_ISL_447037 | B.J. Medical College and Civil hospital | Gujarat Biotechnology Research Centre | Monika Gandhi, Pinal Trivedi, Maharshi Pandya, Amit Kanani, Akanksha Verma, Nitin Savaliya, Raghawendra Kumar, Dinesh Kumar, Zuber Saiyed, Dipa Kinariwala, Disha Patel, Binita Aring, Neeta Khandelwal, Geeta Vaghela, Sonia Barve, Bhavesh Modi, Kairavi Joshi, Gaurishankar Shrimali, Nidhi Sood, Pranay Shah, R D Dixit, Snehal Bagatharia, Kamlesh J Upadhyay, Ramesh Pandit, Tejas Shah, Ankit Hinsu, Pritesh Sabara, Apurvasinh Puvar, Janvi Raval, Neelam Nathani, Chaitanya Joshi, Madhvi Joshi |
| EPI_ISL_447038 | B.J. Medical College and Civil hospital | Gujarat Biotechnology Research Centre | Pinal Trivedi, Maharshi Pandya, Amit Kanani, Akanksha Verma, Nitin Savaliya, Raghawendra Kumar, Dinesh Kumar, Zuber Saiyed, Dipa Kinariwala, Disha Patel, Binita Aring, Neeta Khandelwal, Geeta Vaghela, Sonia Barve, Bhavesh Modi, Kairavi Joshi, Gaurishankar Shrimali, Nidhi Sood, Pranay Shah, R D Dixit, Snehal Bagatharia, Kamlesh J Upadhyay, Ramesh Pandit, Tejas Shah, Ankit Hinsu, Pritesh Sabara, Apurvasinh Puvar, Janvi Raval, Monika Gandhi, Armi Chaudhari, Chaitanya Joshi, Madhvi Joshi |
| EPI_ISL_447039 | B.J. Medical College and Civil hospital | Gujarat Biotechnology Research Centre | Maharshi Pandya, Amit Kanani, Akanksha Verma, Nitin Savaliya, Raghawendra Kumar, Dinesh Kumar, Zuber Saiyed, Dipa Kinariwala, Disha Patel, Binita Aring, Neeta Khandelwal, Geeta Vaghela, Sonia Barve, Bhavesh Modi, Kairavi Joshi, Gaurishankar Shrimali, Nidhi Sood, Pranay Shah, R D Dixit, Snehal Bagatharia, Kamlesh J Upadhyay, Ramesh Pandit, Tejas Shah, Ankit Hinsu, Pritesh Sabara, Apurvasinh Puvar, Janvi Raval, Monika Gandhi, Pinal Trivedi, Bhavya Jindal, Chaitanya Joshi, Madhvi Joshi |
| EPI_ISL_447040 | B.J. Medical College and Civil hospital | Gujarat Biotechnology Research Centre | Amit Kanani, Akanksha Verma, Nitin Savaliya, Raghawendra Kumar, Dinesh Kumar, Zuber Saiyed, Dipa Kinariwala, Disha Patel, Binita Aring, Neeta Khandelwal, Geeta Vaghela, Sonia Barve, Bhavesh Modi, Kairavi Joshi, Gaurishankar Shrimali, Nidhi Sood, Pranay Shah, R D Dixit, Snehal Bagatharia, Kamlesh J Upadhyay, Ramesh Pandit, Tejas Shah, Ankit Hinsu, Pritesh Sabara, Apurvasinh Puvar, Janvi Raval, Monika Gandhi, Pinal Trivedi, Maharshi Pandya, Anjali Rajwar, Chaitanya Joshi, Madhvi Joshi |
| EPI_ISL_447041 | B.J. Medical College and Civil hospital | Gujarat Biotechnology Research Centre | Akanksha Verma, Nitin Savaliya, Raghawendra Kumar, Dinesh Kumar, Zuber Saiyed, Dipa Kinariwala, Disha Patel, Binita Aring, Neeta Khandelwal, Geeta Vaghela, Sonia Barve, Bhavesh Modi, Kairavi Joshi, Gaurishankar Shrimali, Nidhi Sood, Pranay Shah, R D Dixit, Snehal Bagatharia, Kamlesh J Upadhyay, Ramesh Pandit, Tejas Shah, Ankit Hinsu, Pritesh Sabara, Apurvasinh Puvar, Janvi Raval, Monika Gandhi, Pinal Trivedi, Maharshi Pandya, Amit Kanani, Sharmistha Majumdar, Chaitanya Joshi, Madhvi Joshi |
| EPI_ISL_447042 | B.J. Medical College and Civil hospital | Gujarat Biotechnology Research Centre | Nitin Savaliya, Raghawendra Kumar, Dinesh Kumar, Zuber Saiyed, Dipa Kinariwala, Disha Patel, Binita Aring, Neeta Khandelwal, Geeta Vaghela, Sonia Barve, Bhavesh Modi, Kairavi Joshi, Gaurishankar Shrimali, Nidhi Sood, Pranay Shah, R D Dixit, Snehal Bagatharia, Kamlesh J Upadhyay, Ramesh Pandit, Tejas Shah, Ankit Hinsu, Pritesh Sabara, Apurvasinh Puvar, Janvi Raval, Monika Gandhi, Pinal Trivedi, Maharshi Pandya, Amit Kanani, Akanksha Verma, Pooja P Doshi, Chaitanya Joshi, Madhvi Joshi |
| EPI_ISL_447043 | B.J. Medical College and Civil hospital | Gujarat Biotechnology Research Centre | Raghawendra Kumar, Dinesh Kumar, Zuber Saiyed, Dipa Kinariwala, Disha Patel, Binita Aring, Neeta Khandelwal, Geeta Vaghela, Sonia Barve, Bhavesh Modi, Kairavi Joshi, Gaurishankar Shrimali, Nidhi Sood, Pranay Shah, R D Dixit, Snehal Bagatharia, Kamlesh J Upadhyay, Ramesh Pandit, Tejas Shah, Ankit Hinsu, Pritesh Sabara, Apurvasinh Puvar, Janvi Raval, Monika Gandhi, Pinal Trivedi, Maharshi Pandya, Amit Kanani, Akanksha Verma, Nitin Savaliya, Nidhi Patel, Chaitanya Joshi, Madhvi Joshi |
| EPI_ISL_447044 | B.J. Medical College and Civil hospital | Gujarat Biotechnology Research Centre | Dinesh Kumar, Zuber Saiyed, Dipa Kinariwala, Disha Patel, Binita Aring, Neeta Khandelwal, Geeta Vaghela, Sonia Barve, Bhavesh Modi, Kairavi Joshi, Gaurishankar Shrimali, Nidhi Sood, Pranay Shah, R D Dixit, Snehal Bagatharia, Kamlesh J Upadhyay, Ramesh Pandit, Tejas Shah, Ankit Hinsu, Pritesh Sabara, Apurvasinh Puvar, Janvi Raval, Monika Gandhi, Pinal Trivedi, Maharshi Pandya, Amit Kanani, Akanksha Verma, Nitin Savaliya, Raghawendra Kumar, Priti Pandita, Chaitanya Joshi, Madhvi Joshi |
| EPI_ISL_447045 | B.J. Medical College and Civil hospital | Gujarat Biotechnology Research Centre | Zuber Saiyed, Dipa Kinariwala, Disha Patel, Binita Aring, Neeta Khandelwal, Geeta Vaghela, Sonia Barve, Bhavesh Modi, Kairavi Joshi, Gaurishankar Shrimali, Nidhi Sood, Pranay Shah, R D Dixit, Snehal Bagatharia, Kamlesh J Upadhyay, Ramesh Pandit, Tejas Shah, Ankit Hinsu, Pritesh Sabara, Apurvasinh Puvar, Janvi Raval, Monika Gandhi, Pinal Trivedi, Maharshi Pandya, Amit Kanani, Akanksha Verma, Nitin Savaliya, Raghawendra Kumar, Dinesh Kumar, Neha Rajpara, Chaitanya Joshi, Madhvi Joshi |
| EPI_ISL_447046 | B.J. Medical College and Civil hospital | Gujarat Biotechnology Research Centre | Dipa Kinariwala, Disha Patel, Binita Aring, Neeta Khandelwal, Geeta Vaghela, Sonia Barve, Bhavesh Modi, Kairavi Joshi, Gaurishankar Shrimali, Nidhi Sood, Pranay Shah, R D Dixit, Snehal Bagatharia, Kamlesh J Upadhyay, Ramesh Pandit, Tejas Shah, Ankit Hinsu, Pritesh Sabara, Apurvasinh Puvar, Janvi Raval, Monika Gandhi, Pinal Trivedi, Maharshi Pandya, Amit Kanani, Akanksha Verma, Nitin Savaliya, Raghawendra Kumar, Dinesh Kumar, Zuber Saiyed, Afzal Ansari, Chaitanya Joshi, Madhvi Joshi |
| EPI_ISL_447047 | GMERS Medical College and Hospital, Gandhinagar | Gujarat Biotechnology Research Centre | Disha Patel, Binita Aring, Neeta Khandelwal, Geeta Vaghela, Sonia Barve, Bhavesh Modi, Kairavi Joshi, Gaurishankar Shrimali, Nidhi Sood, Pranay Shah, R D Dixit, Snehal Bagatharia, Kamlesh J Upadhyay, Ramesh Pandit, Tejas Shah, Ankit Hinsu, Pritesh Sabara, Apurvasinh Puvar, Janvi Raval, Monika Gandhi, Pinal Trivedi, Maharshi Pandya, Amit Kanani, Akanksha Verma, Nitin Savaliya, Raghawendra Kumar, Dinesh Kumar, Zuber Saiyed, Dipa Kinariwala, Neelam Nathani, Chaitanya Joshi, Madhvi Joshi |
| EPI_ISL_447048 | GMERS Medical College and Hospital, Gandhinagar | Gujarat Biotechnology Research Centre | Binita Aring, Neeta Khandelwal, Geeta Vaghela, Sonia Barve, Bhavesh Modi, Kairavi Joshi, Gaurishankar Shrimali, Nidhi Sood, Pranay Shah, R D Dixit, Snehal Bagatharia, Kamlesh J Upadhyay, Ramesh Pandit, Tejas Shah, Ankit Hinsu, Pritesh Sabara, Apurvasinh Puvar, Janvi Raval, Monika Gandhi, Pinal Trivedi, Maharshi Pandya, Amit Kanani, Akanksha Verma, Nitin Savaliya, Raghawendra Kumar, Dinesh Kumar, Zuber Saiyed, Dipa Kinariwala, Disha Patel, Armi Chaudhari, Chaitanya Joshi, Madhvi Joshi |
| EPI_ISL_447049 | GMERS Medical College and Hospital, Gandhinagar | Gujarat Biotechnology Research Centre | Neeta Khandelwal, Geeta Vaghela, Sonia Barve, Bhavesh Modi, Kairavi Joshi, Gaurishankar Shrimali, Nidhi Sood, Pranay Shah, R D Dixit, Snehal Bagatharia, Kamlesh J Upadhyay, Ramesh Pandit, Tejas Shah, Ankit Hinsu, Pritesh Sabara, Apurvasinh Puvar, Janvi Raval, Monika Gandhi, Pinal Trivedi, Maharshi Pandya, Amit Kanani, Akanksha Verma, Nitin Savaliya, Raghawendra Kumar, Dinesh Kumar, Zuber Saiyed, Dipa Kinariwala, Disha Patel, Binita Aring, Bhavya Jindal, Chaitanya Joshi, Madhvi Joshi |
| EPI_ISL_447050 | GMERS Medical College and Hospital, Gandhinagar | Gujarat Biotechnology Research Centre | Geeta Vaghela, Sonia Barve, Bhavesh Modi, Kairavi Joshi, Gaurishankar Shrimali, Nidhi Sood, Pranay Shah, R D Dixit, Snehal Bagatharia, Kamlesh J Upadhyay, Ramesh Pandit, Tejas Shah, Ankit Hinsu, Pritesh Sabara, Apurvasinh Puvar, Janvi Raval, Monika Gandhi, Pinal Trivedi, Maharshi Pandya, Amit Kanani, Akanksha Verma, Nitin Savaliya, Raghawendra Kumar, Dinesh Kumar, Zuber Saiyed, Dipa Kinariwala, Disha Patel, Binita Aring, Neeta Khandelwal, Dipeshwari Shewale, Chaitanya Joshi, Madhvi Joshi |
| EPI_ISL_447051 | GMERS Medical College and Hospital, Gandhinagar | Gujarat Biotechnology Research Centre | Sonia Barve, Bhavesh Modi, Kairavi Joshi, Gaurishankar Shrimali, Nidhi Sood, Pranay Shah, R D Dixit, Snehal Bagatharia, Kamlesh J Upadhyay, Ramesh Pandit, Tejas Shah, Ankit Hinsu, Pritesh Sabara, Apurvasinh Puvar, Janvi Raval, Monika Gandhi, Pinal Trivedi, Maharshi Pandya, Amit Kanani, Akanksha Verma, Nitin Savaliya, Raghawendra Kumar, Dinesh Kumar, Zuber Saiyed, Dipa Kinariwala, Disha Patel, Binita Aring, Neeta Khandelwal, Geeta Vaghela, Anjali Rajwar, Chaitanya Joshi, Madhvi Joshi |
| EPI_ISL_447052 | GMERS Medical College and Hospital, Gandhinagar | Gujarat Biotechnology Research Centre | Bhavesh Modi, Kairavi Joshi, Gaurishankar Shrimali, Nidhi Sood, Pranay Shah, R D Dixit, Snehal Bagatharia, Kamlesh J Upadhyay, Ramesh Pandit, Tejas Shah, Ankit Hinsu, Pritesh Sabara, Apurvasinh Puvar, Janvi Raval, Monika Gandhi, Pinal Trivedi, Maharshi Pandya, Amit Kanani, Akanksha Verma, Nitin Savaliya, Raghawendra Kumar, Dinesh Kumar, Zuber Saiyed, Dipa Kinariwala, Disha Patel, Binita Aring, Neeta Khandelwal, Geeta Vaghela, Sonia Barve, Sharmistha Majumdar, Chaitanya Joshi, Madhvi Joshi |
| EPI_ISL_447053 | GMERS Medical College and Hospital, Gandhinagar | Gujarat Biotechnology Research Centre | Kairavi Joshi, Gaurishankar Shrimali, Nidhi Sood, Pranay Shah, R D Dixit, Snehal Bagatharia, Kamlesh J Upadhyay, Ramesh Pandit, Tejas Shah, Ankit Hinsu, Pritesh Sabara, Apurvasinh Puvar, Janvi Raval, Monika Gandhi, Pinal Trivedi, Maharshi Pandya, Amit Kanani, Akanksha Verma, Nitin Savaliya, Raghawendra Kumar, Dinesh Kumar, Zuber Saiyed, Dipa Kinariwala, Disha Patel, Binita Aring, Neeta Khandelwal, Geeta Vaghela, Sonia Barve, Bhavesh Modi, Pooja P Doshi, Chaitanya Joshi, Madhvi Joshi |
| EPI_ISL_447054 | Cantacuzino National Military-Medical Institute for Research and Development | Cantacuzino Institute | M.Lazar, L.Ustea, A.Cretu |
| EPI_ISL_447055 | Department for Virology, Molecular Biology and Genome Research, R. G. Lugar Center for Public Health Research, National Center for Disease Control and Public Health (NCDC) of Georgia. | Department for Virology, Molecular Biology and Genome Research, R. G. Lugar Center for Public Health Research, National Center for Disease Control and Public Health (NCDC) of Georgia. | Meri Pantsulaia, Gvantsa Brachveli, Giorgi Tomashvili, Gvantsa Chanturia, Ann Machablishvili, Nato Kotaria, Marine Murtskhvaladze, Lela Sabadze, Mari Gavashelidze, Ana Papkiauri, Tata Imnadze, Tamar Jashiashvili, Tea Tevdoradze, Ketevan Sidamonidze, Ekaterine Khmaladze, Ekaterine Zhghenti, Roena Sukhiashvili, Mariam Zakalashvili, Lela Urushadze, Magda Dgebuadze, Davit Tsaguria, Ekaterine Zangaladze, Nino Berishvili, Adam Kotorashvili, Maia Alkhazashvili, Irma Burjanadze, Anna Kasradze, Khatuna Zakhashvili, Paata Imnadze, Amiran Gamkrelidze. |
| EPI_ISL_447056 | Department for Virology, Molecular Biology and Genome Research, R. G. Lugar Center for Public Health Research, National Center for Disease Control and Public Health (NCDC) of Georgia. | Department for Virology, Molecular Biology and Genome Research, R. G. Lugar Center for Public Health Research, National Center for Disease Control and Public Health (NCDC) of Georgia. | Gvantsa Brachveli, Meri Pantsulaia, Giorgi Tomashvili, Gvantsa Chanturia, Ann Machablishvili, Nato Kotaria, Marine Murtskhvaladze, Lela Sabadze, Mari Gavashelidze, Ana Papkiauri, Gvantsa Brachveli, Tata Imnadze, Tamar Jashiashvili, Tea Tevdoradze, Ketevan Sidamonidze, Ekaterine Khmaladze, Ekaterine Zhghenti, Roena Sukhiashvili, Mariam Zakalashvili, Lela Urushadze, Magda Dgebuadze, Davit Tsaguria, Ekaterine Zangaladze, Nino Berishvili, Adam Kotorashvili, Maia Alkhazashvili, Irma Burjanadze, Anna Kasradze, Khatuna Zakhashvili, Paata Imnadze, Amiran Gamkrelidze. |
| EPI_ISL_447057, EPI_ISL_447059, EPI_ISL_447063, EPI_ISL_447064, EPI_ISL_447068, EPI_ISL_447069, EPI_ISL_447071, EPI_ISL_447072, EPI_ISL_447077, EPI_ISL_447081, EPI_ISL_447082, EPI_ISL_447083, EPI_ISL_447088, EPI_ISL_447089, EPI_ISL_447090, EPI_ISL_447091, EPI_ISL_447092, EPI_ISL_447093, EPI_ISL_447094, EPI_ISL_447095, EPI_ISL_447096, EPI_ISL_447098, EPI_ISL_447099, EPI_ISL_447101, EPI_ISL_447102, EPI_ISL_447104, EPI_ISL_447105, EPI_ISL_447106, EPI_ISL_447107, EPI_ISL_447111, EPI_ISL_447113, EPI_ISL_447115, EPI_ISL_447116, EPI_ISL_447117, EPI_ISL_447118 | | | |
| see above | Michigan Department of Health and Human Services, Bureau of Laboratories | Michigan Department of Health and Human Services, Bureau of Laboratories | Blankenship HM, Riner D, Soehnlen MK |
| EPI_ISL_447119 | HOSPITAL DR.HERNAN HENRIQUEZ ARAVENA | Instituto de Salud Publica de Chile | Andrés E Castillo, Bárbara Parra,Paz Tapia, Jaime Lagos, Loredana Arata, Alejandra Acevedo, Winston Andrade, Gabriel Leal, Carolina Tambley, Patricia Bustos, Rodrigo Fasce, Jorge Fernandez |
| EPI_ISL_447120, EPI_ISL_447121, EPI_ISL_447122, EPI_ISL_447123, EPI_ISL_447124, EPI_ISL_447127, EPI_ISL_447128, EPI_ISL_447129, EPI_ISL_447130, EPI_ISL_447133, EPI_ISL_447134, EPI_ISL_447135, EPI_ISL_447136, EPI_ISL_447137, EPI_ISL_447138, EPI_ISL_447139, EPI_ISL_447140, EPI_ISL_447141, EPI_ISL_447148, EPI_ISL_447152, EPI_ISL_447153, EPI_ISL_447155 | | | |
| see above | Department of Clinical Microbiology | GIGA Medical Genomics | Keith Durkin, Maria Artesi, Sébastien Bontems, Raphaël Boreux, Cécile Meex, Pierrette Melin, Marie-Pierre Hayette, Vincent Bours. |
| EPI_ISL_447163, EPI_ISL_447165, EPI_ISL_447167, EPI_ISL_447168, EPI_ISL_447172, EPI_ISL_447173, EPI_ISL_447174, EPI_ISL_447175, EPI_ISL_447177, EPI_ISL_447178, EPI_ISL_447179, EPI_ISL_447183, EPI_ISL_447184, EPI_ISL_447189, EPI_ISL_447190, EPI_ISL_447191, EPI_ISL_447193, EPI_ISL_447194, EPI_ISL_447196, EPI_ISL_447197, EPI_ISL_447200, EPI_ISL_447201, EPI_ISL_447204, EPI_ISL_447205, EPI_ISL_447206, EPI_ISL_447207, EPI_ISL_447209, EPI_ISL_447210, EPI_ISL_447211, EPI_ISL_447212, EPI_ISL_447215, EPI_ISL_447216, EPI_ISL_447218, EPI_ISL_447219, EPI_ISL_447220, EPI_ISL_447221, EPI_ISL_447224, EPI_ISL_447228, EPI_ISL_447229 | | | |
| see above | Michigan Department of Health and Human Services, Bureau of Laboratories | Michigan Department of Health and Human Services, Bureau of Laboratories | Blankenship HM, Riner D, Soehnlen MK |
| EPI_ISL_447231, EPI_ISL_447232, EPI_ISL_447233, EPI_ISL_447234, EPI_ISL_447235, EPI_ISL_447236, EPI_ISL_447237, EPI_ISL_447239, EPI_ISL_447240, EPI_ISL_447245, EPI_ISL_447246, EPI_ISL_447248, EPI_ISL_447249 | | | |
| see above | Viral Respiratory Lab, National Institute for Biomedical Research (INRB) | Pathogen Sequencing Lab, National Institute for Biomedical Research (INRB) | Placide Mbala-Kingebeni, Edith Nkwembe, Eddy Kinganda-Lusamaki, Amuri Aziza, Francisca Muyembe Mawete, Catherine Pratt, Matthias Pauthner, Josh Quick, Allison Black, James Hadfield, Trevor Bedford, Ian Goodfellow, Andrew Rambaut, Nick Loman, Kristian Andersen, Michael Wiley, Steve Ahuka-Mundeke, Jean-Jacques Muyembe Tamfum |
| EPI_ISL_447251 | Central Virology Laboratory | Central Virology Laboratory | Neta Zuckerman, Efrat Bucris, Oran Erster, Danit Sofer, Orna Mor, Ella Mendelson, Michal Mandelboim |
| EPI_ISL_447252, EPI_ISL_447253, EPI_ISL_447255, EPI_ISL_447257 | TSGH-CP molecular lab | TSGH-CP molecular lab | Cherng-Lih Perng, Ming-Jr JIAN, Chih-Kai Chang, Jung-Chung Lin, Kuo-Ming Yeh, Chien-Wen Chen, Sheng-Kang Chiu, Hsing-Yi Chung, Shih-Hung Tsai, Kuo-Sheng Hung, Tien-Yao Chang, Feng-Yee Chang, Hung-Sheng Shang |
| EPI_ISL_447260, EPI_ISL_447263, EPI_ISL_447264, EPI_ISL_447265, EPI_ISL_447267, EPI_ISL_447268, EPI_ISL_447269, EPI_ISL_447271, EPI_ISL_447272, EPI_ISL_447273, EPI_ISL_447274, EPI_ISL_447275, EPI_ISL_447277, EPI_ISL_447279, EPI_ISL_447280 | | | |
| see above | Microbiology laboratory, Assuta Ashdod University-Affiliated | Stern Lab | Stern Lab |

| | | | |
|---|---|---|---|
| | Hospital | | |
| EPI_ISL_447281, EPI_ISL_447282, EPI_ISL_447283, EPI_ISL_447284, EPI_ISL_447285, EPI_ISL_447286, EPI_ISL_447287, EPI_ISL_447288, EPI_ISL_447289, EPI_ISL_447290, EPI_ISL_447291, EPI_ISL_447292, EPI_ISL_447293, EPI_ISL_447294, EPI_ISL_447295, EPI_ISL_447296, EPI_ISL_447297, EPI_ISL_447299, EPI_ISL_447300, EPI_ISL_447301, EPI_ISL_447302, EPI_ISL_447303, EPI_ISL_447305, EPI_ISL_447306, EPI_ISL_447307, EPI_ISL_447308, EPI_ISL_447309, EPI_ISL_447310 | | | |
| see above | Microbiology Division, Barzilai University Medical Center | Stern Lab | Stern Lab |
| EPI_ISL_447312, EPI_ISL_447313, EPI_ISL_447314, EPI_ISL_447315, EPI_ISL_447316, EPI_ISL_447317, EPI_ISL_447319, EPI_ISL_447320, EPI_ISL_447321, EPI_ISL_447323, EPI_ISL_447324, EPI_ISL_447327, EPI_ISL_447328, EPI_ISL_447330 | | | |
| see above | Clinical Virology Laboratory, Soroka Medical Center and the Faculty of Health Sciences, Ben-Gurion University of the Negev | Stern Lab | Stern Lab |
| EPI_ISL_447331, EPI_ISL_447332, EPI_ISL_447334, EPI_ISL_447337, EPI_ISL_447338, EPI_ISL_447339, EPI_ISL_447340, EPI_ISL_447341, EPI_ISL_447342, EPI_ISL_447343, EPI_ISL_447344, EPI_ISL_447345, EPI_ISL_447346, EPI_ISL_447347, EPI_ISL_447348, EPI_ISL_447349, EPI_ISL_447350, EPI_ISL_447351, EPI_ISL_447352, EPI_ISL_447353, EPI_ISL_447355, EPI_ISL_447356, EPI_ISL_447357, EPI_ISL_447359, EPI_ISL_447360, EPI_ISL_447361, EPI_ISL_447364, EPI_ISL_447365, EPI_ISL_447366, EPI_ISL_447367, EPI_ISL_447369, EPI_ISL_447370, EPI_ISL_447372, EPI_ISL_447374, EPI_ISL_447375, EPI_ISL_447379, EPI_ISL_447380, EPI_ISL_447381, EPI_ISL_447382 | | | |
| see above | Clinical Virology Unit, Hadassah Hebrew University Medical Center | Stern Lab | Stern Lab |
| EPI_ISL_447384, EPI_ISL_447385, EPI_ISL_447386, EPI_ISL_447387, EPI_ISL_447388, EPI_ISL_447389, EPI_ISL_447391, EPI_ISL_447393, EPI_ISL_447394, EPI_ISL_447395, EPI_ISL_447396, EPI_ISL_447397, EPI_ISL_447399, EPI_ISL_447400, EPI_ISL_447401, EPI_ISL_447402, EPI_ISL_447403, EPI_ISL_447404, EPI_ISL_447406 | | | |
| see above | Clinical Microbiology Laboratory, The Baruch Padeh Medical Center, Poriya | Stern Lab | Stern Lab |
| EPI_ISL_447408, EPI_ISL_447409, EPI_ISL_447410, EPI_ISL_447411, EPI_ISL_447412, EPI_ISL_447416 | Clinical Virology Unit, Hadassah Hebrew University Medical Center | Stern Lab | Stern Lab |
| EPI_ISL_447417, EPI_ISL_447418 | Clinical Microbiology Laboratory, The Baruch Padeh Medical Center, Poriya | Stern Lab | Stern Lab |
| EPI_ISL_447419, EPI_ISL_447420, EPI_ISL_447422, EPI_ISL_447423, EPI_ISL_447424, EPI_ISL_447426, EPI_ISL_447427, EPI_ISL_447428, EPI_ISL_447429, EPI_ISL_447430, EPI_ISL_447432, EPI_ISL_447434, EPI_ISL_447435, EPI_ISL_447436, EPI_ISL_447438, EPI_ISL_447440, EPI_ISL_447441, EPI_ISL_447442, EPI_ISL_447443, EPI_ISL_447444, EPI_ISL_447445, EPI_ISL_447446, EPI_ISL_447447, EPI_ISL_447448, EPI_ISL_447449, EPI_ISL_447450, EPI_ISL_447452, EPI_ISL_447453, EPI_ISL_447454, EPI_ISL_447455, EPI_ISL_447456, EPI_ISL_447457, EPI_ISL_447460, EPI_ISL_447461, EPI_ISL_447462, EPI_ISL_447463, EPI_ISL_447464, EPI_ISL_447465, EPI_ISL_447467, EPI_ISL_447468, EPI_ISL_447469 | | | |
| see above | Clinical Microbiology Laboratory, Sheba Medical Center | Stern Lab | Stern Lab |
| EPI_ISL_447471 | Servicio de Microbiologia. Hospital Clinico Universitario de Valencia | Sequencing and Bioinformatics Service and Molecular Epidemiology Research Group. FISABIO-Public Health | Eliseo Albert, Maria Alma Bracho, Griselda De Marco, Lidia Ruiz Roldan, Neris Garcia-Gonzalez, Inma Galán Vendrell, Sandra Carbo, Loreto Ferrús Abad, Paula Ruiz-Hueso, Mariana Reyes-Prieto, Vicente Soriano Chirona, Ivan Ansari, Llúcia Martínez-Priego, Giuseppe 'Auria, David Navarro, Fernando Gonzalez-Candelas |
| EPI_ISL_447472 | Servicio de Microbiologia. Hospital Clinico Universitario de Valencia | Sequencing and Bioinformatics Service and Molecular Epidemiology Research Group. FISABIO-Public Health | Maria Alma Bracho, Griselda De Marco, Lidia Ruiz Roldan, Neris Garcia-Gonzalez, Inma Galán Vendrell, Sandra Carbo, Loreto Ferrús Abad, Paula Ruiz-Hueso, Mariana Reyes-Prieto, Vicente Soriano Chirona, Ivan Ansari, Llúcia Martínez-Priego, Giuseppe 'Auria, David Navarro, Eliseo Albert, Fernando Gonzalez-Candelas |
| EPI_ISL_447473 | Servicio de Microbiologia. Hospital Clinico Universitario de Valencia | Sequencing and Bioinformatics Service and Molecular Epidemiology Research Group. FISABIO-Public Health | Griselda De Marco, Lidia Ruiz Roldan, Neris Garcia-Gonzalez, Inma Galán Vendrell, Sandra Carbo, Loreto Ferrús Abad, Paula Ruiz-Hueso, Mariana Reyes-Prieto, Vicente Soriano Chirona, Ivan Ansari, Llúcia Martínez-Priego, Giuseppe 'Auria, David Navarro, Eliseo Albert, Maria Alma Bracho, Fernando Gonzalez-Candelas |
| EPI_ISL_447474 | Servicio de Microbiologia. Hospital Clinico Universitario de Valencia | Sequencing and Bioinformatics Service and Molecular Epidemiology Research Group. FISABIO-Public Health | Lidia Ruiz Roldan, Neris Garcia-Gonzalez, Inma Galán Vendrell, Sandra Carbo, Loreto Ferrús Abad, Paula Ruiz-Hueso, Mariana Reyes-Prieto, Vicente Soriano Chirona, Ivan Ansari, Llúcia Martínez-Priego, Giuseppe 'Auria, David Navarro, Eliseo Albert, Maria Alma Bracho, Fernando Gonzalez-Candelas |
| EPI_ISL_447477 | Servicio de Microbiologia. Hospital Clinico Universitario de Valencia | Sequencing and Bioinformatics Service and Molecular Epidemiology Research Group. FISABIO-Public Health | Sandra Carbo, Loreto Ferrús Abad, Paula Ruiz-Hueso, Mariana Reyes-Prieto, Vicente Soriano Chirona, Ivan Ansari, Llúcia Martínez-Priego, Giuseppe 'Auria, David Navarro, Eliseo Albert, Maria Alma Bracho, Lidia Ruiz Roldan, Neris Garcia-Gonzalez, Inma Galán Vendrell, Fernando Gonzalez-Candelas |
| EPI_ISL_447478 | Servicio de Microbiologia. Hospital Clinico Universitario de Valencia | Sequencing and Bioinformatics Service and Molecular Epidemiology Research Group. FISABIO-Public Health | Loreto Ferrús Abad, Paula Ruiz-Hueso, Mariana Reyes-Prieto, Vicente Soriano Chirona, Ivan Ansari, Llúcia Martínez-Priego, Giuseppe 'Auria, David Navarro, Eliseo Albert, Maria Alma Bracho, Lidia Ruiz Roldan, Neris Garcia-Gonzalez, Inma Galán Vendrell, Sandra Carbo, Fernando Gonzalez-Candelas |
| EPI_ISL_447481 | Servicio de Microbiologia. Hospital Clinico Universitario de Valencia | Sequencing and Bioinformatics Service and Molecular Epidemiology Research Group. FISABIO-Public Health | Vicente Soriano Chirona, Ivan Ansari, Llúcia Martínez-Priego, Giuseppe 'Auria, David Navarro, Eliseo Albert, Maria Alma Bracho, Lidia Ruiz Roldan, Neris Garcia-Gonzalez, Inma Galán Vendrell, Sandra Carbo, Loreto Ferrús Abad, Paula Ruiz-Hueso, Mariana Reyes-Prieto, Fernando Gonzalez-Candelas |
| EPI_ISL_447482 | Servicio de Microbiologia. Hospital Clinico Universitario de Valencia | Sequencing and Bioinformatics Service and Molecular Epidemiology Research Group. FISABIO-Public Health | Giuseppe 'Auria, David Navarro, Eliseo Albert, Maria Alma Bracho, Lidia Ruiz Roldan, Neris Garcia-Gonzalez, Inma Galán Vendrell, Sandra Carbo, Loreto Ferrús Abad, Paula Ruiz-Hueso, Mariana Reyes-Prieto, Vicente Soriano Chirona, Ivan Ansari, Llúcia Martínez-Priego, Fernando Gonzalez-Candelas |
| EPI_ISL_447483 | Servicio de Microbiologia. Hospital Clinico Universitario de Valencia | Sequencing and Bioinformatics Service and Molecular Epidemiology Research Group. FISABIO-Public Health | Llúcia Martínez-Priego, Giuseppe 'Auria, David Navarro, Eliseo Albert, Maria Alma Bracho, Lidia Ruiz Roldan, Neris Garcia-Gonzalez, Inma Galán Vendrell, Sandra Carbo, Loreto Ferrús Abad, Paula Ruiz-Hueso, Mariana Reyes-Prieto, Vicente Soriano Chirona, Ivan Ansari, Fernando Gonzalez-Candelas |
| EPI_ISL_447485 | Servicio de Microbiologia. Hospital Clinico Universitario de Valencia | Sequencing and Bioinformatics Service and Molecular Epidemiology Research Group. FISABIO-Public Health | Eliseo Albert, Maria Alma Bracho, Griselda De Marco, Lidia Ruiz Roldan, Neris Garcia-Gonzalez, Inma Galán Vendrell, Sandra Carbo, Loreto Ferrús Abad, Paula Ruiz-Hueso, Mariana Reyes-Prieto, Vicente Soriano Chirona, Ivan Ansari, Llúcia Martínez-Priego, Giuseppe 'Auria, David Navarro, Fernando Gonzalez-Candelas |
| EPI_ISL_447493 | Servicio de Microbiologia. Hospital Clinico Universitario de Valencia | Sequencing and Bioinformatics Service and Molecular Epidemiology Research Group. FISABIO-Public Health | Paula Ruiz-Hueso, Mariana Reyes-Prieto, Vicente Soriano Chirona, Ivan Ansari, Llúcia Martínez-Priego, Giuseppe 'Auria, David Navarro, Eliseo Albert, Maria Alma Bracho, Lidia Ruiz Roldan, Neris Garcia-Gonzalez, Inma Galán Vendrell, Sandra Carbo, Loreto Ferrús Abad, Fernando Gonzalez-Candelas |
| EPI_ISL_447497 | Servicio de Microbiologia. Hospital Clinico Universitario de Valencia | Sequencing and Bioinformatics Service and Molecular Epidemiology Research Group. FISABIO-Public Health | Llúcia Martínez-Priego, Giuseppe 'Auria, David Navarro, Eliseo Albert, Maria Alma Bracho, Lidia Ruiz Roldan, Neris Garcia-Gonzalez, Inma Galán Vendrell, Sandra Carbo, Loreto Ferrús Abad, Paula Ruiz-Hueso, Mariana Reyes-Prieto, Vicente Soriano Chirona, Ivan Ansari, Fernando Gonzalez-Candelas |
| EPI_ISL_447498 | Servicio de Microbiologia. Hospital Clinico Universitario de Valencia | Sequencing and Bioinformatics Service and Molecular Epidemiology Research Group. FISABIO-Public Health | David Navarro, Eliseo Albert, Maria Alma Bracho, Griselda De Marco, Lidia Ruiz Roldan, Neris Garcia-Gonzalez, Inma Galán Vendrell, Sandra Carbo, Loreto Ferrús Abad, Paula Ruiz-Hueso, Mariana Reyes-Prieto, Vicente Soriano Chirona, Ivan Ansari, Llúcia Martínez-Priego, Giuseppe 'Auria, Fernando Gonzalez-Candelas |
| EPI_ISL_447500 | Servicio de Microbiologia. Hospital Clinico Universitario de Valencia | Sequencing and Bioinformatics Service and Molecular Epidemiology Research Group. FISABIO-Public Health | Maria Alma Bracho, Griselda De Marco, Lidia Ruiz Roldan, Neris Garcia-Gonzalez, Inma Galán Vendrell, Sandra Carbo, Loreto Ferrús Abad, Paula Ruiz-Hueso, Mariana Reyes-Prieto, Vicente Soriano Chirona, Ivan Ansari, Llúcia Martínez-Priego, Giuseppe 'Auria, David Navarro, Eliseo Albert, Fernando Gonzalez-Candelas |
| EPI_ISL_447501 | Servicio de Microbiologia. Hospital Clinico Universitario de Valencia | Sequencing and Bioinformatics Service and Molecular Epidemiology Research Group. FISABIO-Public Health | Griselda De Marco, Lidia Ruiz Roldan, Neris Garcia-Gonzalez, Inma Galán Vendrell, Sandra Carbo, Loreto Ferrús Abad, Paula Ruiz-Hueso, Mariana Reyes-Prieto, Vicente Soriano Chirona, Ivan Ansari, Llúcia Martínez-Priego, Giuseppe 'Auria, David Navarro, Eliseo Albert, Maria Alma Bracho, Fernando Gonzalez-Candelas |
| EPI_ISL_447502 | Servicio de Microbiologia. Hospital Clinico Universitario de Valencia | Sequencing and Bioinformatics Service and Molecular Epidemiology Research Group. FISABIO-Public Health | Lidia Ruiz Roldan, Neris Garcia-Gonzalez, Inma Galán Vendrell, Sandra Carbo, Loreto Ferrús Abad, Paula Ruiz-Hueso, Mariana Reyes-Prieto, Vicente Soriano Chirona, Ivan Ansari, Llúcia Martínez-Priego, Giuseppe 'Auria, David Navarro, Eliseo Albert, Maria Alma Bracho, Fernando Gonzalez-Candelas |
| EPI_ISL_447503 | Servicio de Microbiologia. Hospital Clinico Universitario de Valencia | Sequencing and Bioinformatics Service and Molecular Epidemiology Research Group. FISABIO-Public Health | Neris Garcia-Gonzalez, Inma Galán Vendrell, Sandra Carbo, Loreto Ferrús Abad, Paula Ruiz-Hueso, Mariana Reyes-Prieto, Vicente Soriano Chirona, Ivan Ansari, Llúcia Martínez-Priego, Giuseppe 'Auria, David Navarro, Eliseo Albert, Maria Alma Bracho, Lidia Ruiz Roldan, Fernando Gonzalez-Candelas |
| EPI_ISL_447505 | Servicio de Microbiologia. Hospital Clinico Universitario de Valencia | Sequencing and Bioinformatics Service and Molecular Epidemiology Research Group. FISABIO-Public Health | Sandra Carbo, Loreto Ferrús Abad, Paula Ruiz-Hueso, Mariana Reyes-Prieto, Vicente Soriano Chirona, Ivan Ansari, Llúcia Martínez-Priego, Giuseppe 'Auria, David Navarro, Eliseo Albert, Maria Alma Bracho, Lidia Ruiz Roldan, Neris Garcia-Gonzalez, Inma Galán Vendrell, Fernando Gonzalez-Candelas |
| EPI_ISL_447506 | Servicio de Microbiologia. Hospital Clinico Universitario de Valencia | Sequencing and Bioinformatics Service and Molecular Epidemiology Research Group. FISABIO-Public Health | Loreto Ferrús Abad, Paula Ruiz-Hueso, Mariana Reyes-Prieto, Vicente Soriano Chirona, Ivan Ansari, Llúcia Martínez-Priego, Giuseppe 'Auria, David Navarro, Eliseo Albert, Maria Alma Bracho, Lidia Ruiz Roldan, Neris Garcia-Gonzalez, Inma Galán Vendrell, Sandra Carbo, Fernando Gonzalez-Candelas |
| EPI_ISL_447510 | Servicio de Microbiologia. Hospital Clinico Universitario de Valencia | Sequencing and Bioinformatics Service and Molecular Epidemiology Research Group. FISABIO-Public Health | Giuseppe 'Auria, David Navarro, Eliseo Albert, Maria Alma Bracho, Lidia Ruiz Roldan, Neris Garcia-Gonzalez, Inma Galán Vendrell, Sandra Carbo, Loreto Ferrús Abad, Paula Ruiz-Hueso, Mariana Reyes-Prieto, Vicente Soriano Chirona, Ivan Ansari, Llúcia Martínez-Priego, Fernando Gonzalez-Candelas |
| EPI_ISL_447511 | Servicio de Microbiologia. Hospital Clinico Universitario de Valencia | Sequencing and Bioinformatics Service and Molecular Epidemiology Research Group. FISABIO-Public Health | Llúcia Martínez-Priego, Giuseppe 'Auria, David Navarro, Eliseo Albert, Maria Alma Bracho, Lidia Ruiz Roldan, Neris Garcia-Gonzalez, Inma Galán Vendrell, Sandra Carbo, Loreto Ferrús Abad, Paula Ruiz-Hueso, Mariana Reyes-Prieto, Vicente Soriano Chirona, Ivan Ansari, Fernando Gonzalez-Candelas |
| EPI_ISL_447512 | Servicio de Microbiologia. Hospital Clinico Universitario de Valencia | Sequencing and Bioinformatics Service and Molecular Epidemiology Research Group. FISABIO-Public Health | David Navarro, Eliseo Albert, Maria Alma Bracho, Griselda De Marco, Lidia Ruiz Roldan, Neris Garcia-Gonzalez, Inma Galán Vendrell, Sandra Carbo, Loreto Ferrús Abad, Paula Ruiz-Hueso, Mariana Reyes-Prieto, Vicente Soriano Chirona, Ivan Ansari, Llúcia Martínez-Priego, Giuseppe 'Auria, Fernando Gonzalez-Candelas |
| EPI_ISL_447513 | Servicio de Microbiologia. Hospital Clinico Universitario de Valencia | Sequencing and Bioinformatics Service and Molecular Epidemiology Research Group. FISABIO-Public Health | Eliseo Albert, Maria Alma Bracho, Griselda De Marco, Lidia Ruiz Roldan, Neris Garcia-Gonzalez, Inma Galán Vendrell, Sandra Carbo, Loreto Ferrús Abad, Paula Ruiz-Hueso, Mariana Reyes-Prieto, Vicente Soriano Chirona, Ivan Ansari, Llúcia Martínez-Priego, Giuseppe 'Auria, David Navarro, Fernando Gonzalez-Candelas |
| EPI_ISL_447514 | Servicio de Microbiologia. Hospital Clinico Universitario de Valencia | Sequencing and Bioinformatics Service and Molecular Epidemiology Research Group. FISABIO-Public Health | Maria Alma Bracho, Griselda De Marco, Lidia Ruiz Roldan, Neris Garcia-Gonzalez, Inma Galán Vendrell, Sandra Carbo, Loreto Ferrús Abad, Paula Ruiz-Hueso, Mariana Reyes-Prieto, Vicente Soriano Chirona, Ivan Ansari, Llúcia Martínez-Priego, Giuseppe 'Auria, David Navarro, Eliseo Albert, Fernando Gonzalez-Candelas |
| EPI_ISL_447515 | Servicio de Microbiologia. Hospital Clinico Universitario de Valencia | Sequencing and Bioinformatics Service and Molecular Epidemiology Research Group. FISABIO-Public Health | Griselda De Marco, Lidia Ruiz Roldan, Neris Garcia-Gonzalez, Inma Galán Vendrell, Sandra Carbo, Loreto Ferrús Abad, Paula Ruiz-Hueso, Mariana Reyes-Prieto, Vicente Soriano Chirona, Ivan Ansari, Llúcia Martínez-Priego, Giuseppe 'Auria, David Navarro, Eliseo Albert, Maria Alma Bracho, Fernando Gonzalez-Candelas |
| EPI_ISL_447517 | Servicio de Microbiologia. Hospital Clinico Universitario de Valencia | Sequencing and Bioinformatics Service and Molecular Epidemiology Research Group. FISABIO-Public Health | Neris Garcia-Gonzalez, Inma Galán Vendrell, Sandra Carbo, Loreto Ferrús Abad, Paula Ruiz-Hueso, Mariana Reyes-Prieto, Vicente Soriano Chirona, Ivan Ansari, Llúcia Martínez-Priego, Giuseppe 'Auria, David Navarro, Eliseo Albert, Maria Alma Bracho, Lidia Ruiz Roldan, Fernando Gonzalez-Candelas |
| EPI_ISL_447519 | Servicio de Microbiologia. Hospital Clinico Universitario de Valencia | Sequencing and Bioinformatics Service and Molecular Epidemiology Research Group. FISABIO-Public Health | Sandra Carbo, Loreto Ferrús Abad, Paula Ruiz-Hueso, Mariana Reyes-Prieto, Vicente Soriano Chirona, Ivan Ansari, Llúcia Martínez-Priego, Giuseppe 'Auria, David Navarro, Eliseo Albert, Maria Alma Bracho, Lidia Ruiz Roldan, Neris Garcia-Gonzalez, Inma Galán Vendrell, Fernando Gonzalez-Candelas |
| EPI_ISL_447520 | Servicio de Microbiologia. Hospital Clinico Universitario de Valencia | Sequencing and Bioinformatics Service and Molecular Epidemiology Research Group. FISABIO-Public Health | Loreto Ferrús Abad, Paula Ruiz-Hueso, Mariana Reyes-Prieto, Vicente Soriano Chirona, Ivan Ansari, Llúcia Martínez-Priego, Giuseppe 'Auria, David Navarro, Eliseo Albert, Maria Alma Bracho, Lidia Ruiz Roldan, Neris Garcia-Gonzalez, Inma Galán Vendrell, Sandra Carbo, Fernando Gonzalez-Candelas |
| EPI_ISL_447522 | Servicio de Microbiologia. Hospital Clinico Universitario de Valencia | Sequencing and Bioinformatics Service and Molecular Epidemiology Research Group. FISABIO-Public Health | Vicente Soriano Chirona, Ivan Ansari, Llúcia Martínez-Priego, Giuseppe 'Auria, David Navarro, Eliseo Albert, Maria Alma Bracho, Lidia Ruiz Roldan, Neris Garcia-Gonzalez, Inma Galán Vendrell, Sandra Carbo, Loreto Ferrús Abad, Paula Ruiz-Hueso, Mariana Reyes-Prieto, Fernando Gonzalez-Candelas |
| EPI_ISL_447523 | Servicio de Microbiologia. Hospital Clinico Universitario de Valencia | Sequencing and Bioinformatics Service and Molecular Epidemiology Research Group. FISABIO-Public Health | Giuseppe 'Auria, David Navarro, Eliseo Albert, Maria Alma Bracho, Lidia Ruiz Roldan, Neris Garcia-Gonzalez, Inma Galán Vendrell, Sandra Carbo, Loreto Ferrús Abad, Paula Ruiz-Hueso, Mariana Reyes-Prieto, Vicente Soriano Chirona, Ivan Ansari, Llúcia Martínez-Priego, Fernando Gonzalez-Candelas |
| EPI_ISL_447527 | Servicio de Microbiologia. Hospital Clinico Universitario de Valencia | Sequencing and Bioinformatics Service and Molecular Epidemiology Research Group. FISABIO-Public Health | Maria Alma Bracho, Griselda De Marco, Lidia Ruiz Roldan, Neris Garcia-Gonzalez, Inma Galán Vendrell, Sandra Carbo, Loreto Ferrús Abad, Paula Ruiz-Hueso, Mariana Reyes-Prieto, Vicente Soriano Chirona, Ivan Ansari, Llúcia Martínez-Priego, Giuseppe 'Auria, David Navarro, Eliseo Albert, Fernando Gonzalez-Candelas |
| EPI_ISL_447528 | Servicio de Microbiologia. Hospital Clinico Universitario de Valencia | Sequencing and Bioinformatics Service and Molecular Epidemiology Research Group. FISABIO-Public Health | Griselda De Marco, Lidia Ruiz Roldan, Neris Garcia-Gonzalez, Inma Galán Vendrell, Sandra Carbo, Loreto Ferrús Abad, Paula Ruiz-Hueso, Mariana Reyes-Prieto, Vicente Soriano Chirona, Ivan Ansari, Llúcia Martínez-Priego, Giuseppe 'Auria, David Navarro, Eliseo Albert, Maria Alma Bracho, Fernando Gonzalez-Candelas |
| EPI_ISL_447529 | Servicio de Microbiologia. Hospital Clinico Universitario de Valencia | Sequencing and Bioinformatics Service and Molecular Epidemiology Research Group. FISABIO-Public Health | Lidia Ruiz Roldan, Neris Garcia-Gonzalez, Inma Galán Vendrell, Sandra Carbo, Loreto Ferrús Abad, Paula Ruiz-Hueso, Mariana Reyes-Prieto, Vicente Soriano Chirona, Ivan Ansari, Llúcia Martínez-Priego, Giuseppe 'Auria, David Navarro, Eliseo Albert, Maria Alma Bracho, Fernando Gonzalez-Candelas |
| EPI_ISL_447530 | Servicio de Microbiologia. Hospital Clinico Universitario de | Sequencing and Bioinformatics Service and Molecular Epidemiology Research | Neris Garcia-Gonzalez, Inma Galán Vendrell, Sandra Carbo, Loreto Ferrús Abad, Paula Ruiz-Hueso, Mariana Reyes-Prieto, Vicente Soriano Chirona, Ivan Ansari, Llúcia Martínez-Priego, Giuseppe 'Auria, David |

| | | | |
|---|---|---|---|
| | Valencia | Group. FISABIO-Public Health | Navarro, Eliseo Albert, Maria Alma Bracho, Lidia Ruiz Roldan, Fernando Gonzalez-Candelas |
| EPI_ISL_447531 | Servicio de Microbiologia. Hospital Clinico Universitario de Valencia | Sequencing and Bioinformatics Service and Molecular Epidemiology Research Group. FISABIO-Public Health | Inma Galán Vendrell, Sandra Carbo, Loreto Ferrús Abad, Paula Ruiz-Hueso, Mariana Reyes-Prieto, Vicente Soriano Chirona, Ivan Ansari, Llúcia Martínez-Priego, Giuseppe 'Auria, David Navarro, Eliseo Albert, Maria Alma Bracho, Lidia Ruiz Roldan, Neris Garcia-Gonzalez, Fernando Gonzalez-Candelas |
| EPI_ISL_447532, EPI_ISL_447533 | Hospital Universitari Vall d'Hebron - Vall d'Hebron Institut de Recerca | Hospital Universitari Vall d'Hebron | Cristina Andrés, Maria Piñana, Damir Garcia-Cehic, Mercedes Guerrero-Murillo, Ariadna Rando, Juliana Esperalba, Maria Gema Codina, Tomàs Pumarola, Josep Quer, Andrés Antón |
| EPI_ISL_447534 | Gujarat Biotechnology Research Centre | Gujarat Biotechnology Research Centre | Gaurishankar Shrimali, Nidhi Sood, Pranay Shah, R D Dixit, Snehal Bagatharia, Kamlesh J Upadhyay, Ramesh Pandit, Tejas Shah, Ankit Hinsu, Pritesh Sabara, Apurvasinh Puvar, Janvi Raval, Monika Gandhi, Pinal Trivedi, Maharshi Pandya, Amit Kanani, Akanksha Verma, Nitin Savaliya, Raghawendra Kumar, Dinesh Kumar, Zuber Saiyed, Dipa Kinariwala, Disha Patel, Binita Aring, Neeta Khandelwal, Geeta Vaghela, Sonia Barve, Bhavesh Modi, Kairavi Joshi, Nidhi Patel, Chaitanya Joshi, Madhvi Joshi |
| EPI_ISL_447535 | Gujarat Biotechnology Research Centre | Gujarat Biotechnology Research Centre | Nidhi Sood, Pranay Shah, R D Dixit, Snehal Bagatharia, Kamlesh J Upadhyay, Ramesh Pandit, Tejas Shah, Ankit Hinsu, Pritesh Sabara, Apurvasinh Puvar, Janvi Raval, Monika Gandhi, Pinal Trivedi, Maharshi Pandya, Amit Kanani, Akanksha Verma, Nitin Savaliya, Raghawendra Kumar, Dinesh Kumar, Zuber Saiyed, Dipa Kinariwala, Disha Patel, Binita Aring, Neeta Khandelwal, Geeta Vaghela, Sonia Barve, Bhavesh Modi, Kairavi Joshi, Gaurishankar Shrimali, Priti Pandita, Chaitanya Joshi, Madhvi Joshi |
| EPI_ISL_447536 | Gujarat Biotechnology Research Centre | Gujarat Biotechnology Research Centre | Pranay Shah, R D Dixit, Snehal Bagatharia, Kamlesh J Upadhyay, Ramesh Pandit, Tejas Shah, Ankit Hinsu, Pritesh Sabara, Apurvasinh Puvar, Janvi Raval, Monika Gandhi, Pinal Trivedi, Maharshi Pandya, Amit Kanani, Akanksha Verma, Nitin Savaliya, Raghawendra Kumar, Dinesh Kumar, Zuber Saiyed, Dipa Kinariwala, Disha Patel, Binita Aring, Neeta Khandelwal, Geeta Vaghela, Sonia Barve, Bhavesh Modi, Kairavi Joshi, Gaurishankar Shrimali, Nidhi Sood, Neha Rajpara, Chaitanya Joshi, Madhvi Joshi |
| EPI_ISL_447537 | Gujarat Biotechnology Research Centre | Gujarat Biotechnology Research Centre | R D Dixit, Snehal Bagatharia, Kamlesh J Upadhyay, Ramesh Pandit, Tejas Shah, Ankit Hinsu, Pritesh Sabara, Apurvasinh Puvar, Janvi Raval, Monika Gandhi, Pinal Trivedi, Maharshi Pandya, Amit Kanani, Akanksha Verma, Nitin Savaliya, Raghawendra Kumar, Dinesh Kumar, Zuber Saiyed, Dipa Kinariwala, Disha Patel, Binita Aring, Neeta Khandelwal, Geeta Vaghela, Sonia Barve, Bhavesh Modi, Kairavi Joshi, Gaurishankar Shrimali, Nidhi Sood, Pranay Shah, Afzal Ansari, Chaitanya Joshi, Madhvi Joshi |
| EPI_ISL_447538 | Gujarat Biotechnology Research Centre | Gujarat Biotechnology Research Centre | Snehal Bagatharia, Kamlesh J Upadhyay, Ramesh Pandit, Tejas Shah, Ankit Hinsu, Pritesh Sabara, Apurvasinh Puvar, Janvi Raval, Monika Gandhi, Pinal Trivedi, Maharshi Pandya, Amit Kanani, Akanksha Verma, Nitin Savaliya, Raghawendra Kumar, Dinesh Kumar, Zuber Saiyed, Dipa Kinariwala, Disha Patel, Binita Aring, Neeta Khandelwal, Geeta Vaghela, Sonia Barve, Bhavesh Modi, Kairavi Joshi, Gaurishankar Shrimali, Nidhi Sood, Pranay Shah, R D Dixit, Neelam Nathani, Chaitanya Joshi, Madhvi Joshi |
| EPI_ISL_447539 | Gujarat Biotechnology Research Centre | Gujarat Biotechnology Research Centre | Kamlesh J Upadhyay, Ramesh Pandit, Tejas Shah, Ankit Hinsu, Pritesh Sabara, Apurvasinh Puvar, Janvi Raval, Monika Gandhi, Pinal Trivedi, Maharshi Pandya, Amit Kanani, Akanksha Verma, Nitin Savaliya, Raghawendra Kumar, Dinesh Kumar, Zuber Saiyed, Dipa Kinariwala, Disha Patel, Binita Aring, Neeta Khandelwal, Geeta Vaghela, Sonia Barve, Bhavesh Modi, Kairavi Joshi, Gaurishankar Shrimali, Nidhi Sood, Pranay Shah, R D Dixit, Snehal Bagatharia, Armi Chaudhari, Chaitanya Joshi, Madhvi Joshi |
| EPI_ISL_447540 | Gujarat Biotechnology Research Centre | Gujarat Biotechnology Research Centre | Ramesh Pandit, Tejas Shah, Ankit Hinsu, Pritesh Sabara, Apurvasinh Puvar, Janvi Raval, Monika Gandhi, Pinal Trivedi, Maharshi Pandya, Amit Kanani, Akanksha Verma, Nitin Savaliya, Raghawendra Kumar, Dinesh Kumar, Zuber Saiyed, Dipa Kinariwala, Disha Patel, Binita Aring, Neeta Khandelwal, Geeta Vaghela, Sonia Barve, Bhavesh Modi, Kairavi Joshi, Gaurishankar Shrimali, Nidhi Sood, Pranay Shah, R D Dixit, Snehal Bagatharia, Kamlesh J Upadhyay, Bhavya Jindal, Chaitanya Joshi, Madhvi Joshi |
| EPI_ISL_447541 | Gujarat Biotechnology Research Centre | Gujarat Biotechnology Research Centre | Tejas Shah, Ankit Hinsu, Pritesh Sabara, Apurvasinh Puvar, Janvi Raval, Monika Gandhi, Pinal Trivedi, Maharshi Pandya, Amit Kanani, Akanksha Verma, Nitin Savaliya, Raghawendra Kumar, Dinesh Kumar, Zuber Saiyed, Dipa Kinariwala, Disha Patel, Binita Aring, Neeta Khandelwal, Geeta Vaghela, Sonia Barve, Bhavesh Modi, Kairavi Joshi, Gaurishankar Shrimali, Nidhi Sood, Pranay Shah, R D Dixit, Snehal Bagatharia, Kamlesh J Upadhyay, Ramesh Pandit, Anjali Rajwar, Chaitanya Joshi, Madhvi Joshi |
| EPI_ISL_447542 | Gujarat Biotechnology Research Centre | Gujarat Biotechnology Research Centre | Ankit Hinsu, Pritesh Sabara, Apurvasinh Puvar, Janvi Raval, Monika Gandhi, Pinal Trivedi, Maharshi Pandya, Amit Kanani, Akanksha Verma, Nitin Savaliya, Raghawendra Kumar, Dinesh Kumar, Zuber Saiyed, Dipa Kinariwala, Disha Patel, Binita Aring, Neeta Khandelwal, Geeta Vaghela, Sonia Barve, Bhavesh Modi, Kairavi Joshi, Gaurishankar Shrimali, Nidhi Sood, Pranay Shah, R D Dixit, Snehal Bagatharia, Kamlesh J Upadhyay, Ramesh Pandit, Tejas Shah, Dipeshwari Shewale, Chaitanya Joshi, Madhvi Joshi |
| EPI_ISL_447543 | Gujarat Biotechnology Research Centre | Gujarat Biotechnology Research Centre | Pritesh Sabara, Apurvasinh Puvar, Janvi Raval, Monika Gandhi, Pinal Trivedi, Maharshi Pandya, Amit Kanani, Akanksha Verma, Nitin Savaliya, Raghawendra Kumar, Dinesh Kumar, Zuber Saiyed, Dipa Kinariwala, Disha Patel, Binita Aring, Neeta Khandelwal, Geeta Vaghela, Sonia Barve, Bhavesh Modi, Kairavi Joshi, Gaurishankar Shrimali, Nidhi Sood, Pranay Shah, R D Dixit, Snehal Bagatharia, Kamlesh J Upadhyay, Ramesh Pandit, Tejas Shah, Ankit Hinsu, Sharmistha Majumdar, Chaitanya Joshi, Madhvi Joshi |
| EPI_ISL_447544 | Gujarat Biotechnology Research Centre | Gujarat Biotechnology Research Centre | Apurvasinh Puvar, Janvi Raval, Monika Gandhi, Pinal Trivedi, Maharshi Pandya, Amit Kanani, Akanksha Verma, Nitin Savaliya, Raghawendra Kumar, Dinesh Kumar, Zuber Saiyed, Dipa Kinariwala, Disha Patel, Binita Aring, Neeta Khandelwal, Geeta Vaghela, Sonia Barve, Bhavesh Modi, Kairavi Joshi, Gaurishankar Shrimali, Nidhi Sood, Pranay Shah, R D Dixit, Snehal Bagatharia, Kamlesh J Upadhyay, Ramesh Pandit, Tejas Shah, Ankit Hinsu, Pritesh Sabara, Pooja P Doshi, Chaitanya Joshi, Madhvi Joshi |
| EPI_ISL_447545 | Gujarat Biotechnology Research Centre | Gujarat Biotechnology Research Centre | Janvi Raval, Monika Gandhi, Pinal Trivedi, Maharshi Pandya, Amit Kanani, Akanksha Verma, Nitin Savaliya, Raghawendra Kumar, Dinesh Kumar, Zuber Saiyed, Dipa Kinariwala, Disha Patel, Binita Aring, Neeta Khandelwal, Geeta Vaghela, Sonia Barve, Bhavesh Modi, Kairavi Joshi, Gaurishankar Shrimali, Nidhi Sood, Pranay Shah, R D Dixit, Snehal Bagatharia, Kamlesh J Upadhyay, Ramesh Pandit, Tejas Shah, Ankit Hinsu, Pritesh Sabara, Apurvasinh Puvar, Nidhi Patel, Chaitanya Joshi, Madhvi Joshi |
| EPI_ISL_447546 | Gujarat Biotechnology Research Centre | Gujarat Biotechnology Research Centre | Monika Gandhi, Pinal Trivedi, Maharshi Pandya, Amit Kanani, Akanksha Verma, Nitin Savaliya, Raghawendra Kumar, Dinesh Kumar, Zuber Saiyed, Dipa Kinariwala, Disha Patel, Binita Aring, Neeta Khandelwal, Geeta Vaghela, Sonia Barve, Bhavesh Modi, Kairavi Joshi, Gaurishankar Shrimali, Nidhi Sood, Pranay Shah, R D Dixit, Snehal Bagatharia, Kamlesh J Upadhyay, Ramesh Pandit, Tejas Shah, Ankit Hinsu, Pritesh Sabara, Apurvasinh Puvar, Janvi Raval, Priti Pandita, Chaitanya Joshi, Madhvi Joshi |
| EPI_ISL_447547 | GMERS Medical College and Hospital, Gandhinagar | Gujarat Biotechnology Research Centre | Pinal Trivedi, Maharshi Pandya, Amit Kanani, Akanksha Verma, Nitin Savaliya, Raghawendra Kumar, Dinesh Kumar, Zuber Saiyed, Dipa Kinariwala, Disha Patel, Binita Aring, Neeta Khandelwal, Geeta Vaghela, Sonia Barve, Bhavesh Modi, Kairavi Joshi, Gaurishankar Shrimali, Nidhi Sood, Pranay Shah, R D Dixit, Snehal Bagatharia, Kamlesh J Upadhyay, Ramesh Pandit, Tejas Shah, Ankit Hinsu, Pritesh Sabara, Apurvasinh Puvar, Janvi Raval, Monika Gandhi, Neha Rajpara, Chaitanya Joshi, Madhvi Joshi |
| EPI_ISL_447548 | GMERS Medical College and Hospital, Gandhinagar | Gujarat Biotechnology Research Centre | Maharshi Pandya, Amit Kanani, Akanksha Verma, Nitin Savaliya, Raghawendra Kumar, Dinesh Kumar, Zuber Saiyed, Dipa Kinariwala, Disha Patel, Binita Aring, Neeta Khandelwal, Geeta Vaghela, Sonia Barve, Bhavesh Modi, Kairavi Joshi, Gaurishankar Shrimali, Nidhi Sood, Pranay Shah, R D Dixit, Snehal Bagatharia, Kamlesh J Upadhyay, Ramesh Pandit, Tejas Shah, Ankit Hinsu, Pritesh Sabara, Apurvasinh Puvar, Janvi Raval, Monika Gandhi, Pinal Trivedi, Afzal Ansari, Chaitanya Joshi, Madhvi Joshi |
| EPI_ISL_447549 | GMERS Medical College and Hospital, Gandhinagar | Gujarat Biotechnology Research Centre | Amit Kanani, Akanksha Verma, Nitin Savaliya, Raghawendra Kumar, Dinesh Kumar, Zuber Saiyed, Dipa Kinariwala, Disha Patel, Binita Aring, Neeta Khandelwal, Geeta Vaghela, Sonia Barve, Bhavesh Modi, Kairavi Joshi, Gaurishankar Shrimali, Nidhi Sood, Pranay Shah, R D Dixit, Snehal Bagatharia, Kamlesh J Upadhyay, Ramesh Pandit, Tejas Shah, Ankit Hinsu, Pritesh Sabara, Apurvasinh Puvar, Janvi Raval, Monika Gandhi, Pinal Trivedi, Maharshi Pandya, Neelam Nathani, Chaitanya Joshi, Madhvi Joshi |
| EPI_ISL_447550 | GMERS Medical College and Hospital, Gandhinagar | Gujarat Biotechnology Research Centre | Akanksha Verma, Nitin Savaliya, Raghawendra Kumar, Dinesh Kumar, Zuber Saiyed, Dipa Kinariwala, Disha Patel, Binita Aring, Neeta Khandelwal, Geeta Vaghela, Sonia Barve, Bhavesh Modi, Kairavi Joshi, Gaurishankar Shrimali, Nidhi Sood, Pranay Shah, R D Dixit, Snehal Bagatharia, Kamlesh J Upadhyay, Ramesh Pandit, Tejas Shah, Ankit Hinsu, Pritesh Sabara, Apurvasinh Puvar, Janvi Raval, Monika Gandhi, Pinal Trivedi, Maharshi Pandya, Amit Kanani, Armi Chaudhari, Chaitanya Joshi, Madhvi Joshi |
| EPI_ISL_447551 | GMERS Medical College and Hospital, Gandhinagar | Gujarat Biotechnology Research Centre | Nitin Savaliya, Raghawendra Kumar, Dinesh Kumar, Zuber Saiyed, Dipa Kinariwala, Disha Patel, Binita Aring, Neeta Khandelwal, Geeta Vaghela, Sonia Barve, Bhavesh Modi, Kairavi Joshi, Gaurishankar Shrimali, Nidhi Sood, Pranay Shah, R D Dixit, Snehal Bagatharia, Kamlesh J Upadhyay, Ramesh Pandit, Tejas Shah, Ankit Hinsu, Pritesh Sabara, Apurvasinh Puvar, Janvi Raval, Monika Gandhi, Pinal Trivedi, Maharshi Pandya, Amit Kanani, Akanksha Verma, Bhavya Jindal, Chaitanya Joshi, Madhvi Joshi |
| EPI_ISL_447552 | GMERS Medical College and Hospital, Gandhinagar | Gujarat Biotechnology Research Centre | Raghawendra Kumar, Dinesh Kumar, Zuber Saiyed, Dipa Kinariwala, Disha Patel, Binita Aring, Neeta Khandelwal, Geeta Vaghela, Sonia Barve, Bhavesh Modi, Kairavi Joshi, Gaurishankar Shrimali, Nidhi Sood, Pranay Shah, R D Dixit, Snehal Bagatharia, Kamlesh J Upadhyay, Ramesh Pandit, Tejas Shah, Ankit Hinsu, Pritesh Sabara, Apurvasinh Puvar, Janvi Raval, Monika Gandhi, Pinal Trivedi, Maharshi Pandya, Amit Kanani, Akanksha Verma, Nitin Savaliya, Anjali Rajwar, Chaitanya Joshi, Madhvi Joshi |
| EPI_ISL_447553 | GMERS Medical College and Hospital, Gandhinagar | Gujarat Biotechnology Research Centre | Dinesh Kumar, Zuber Saiyed, Dipa Kinariwala, Disha Patel, Binita Aring, Neeta Khandelwal, Geeta Vaghela, Sonia Barve, Bhavesh Modi, Kairavi Joshi, Gaurishankar Shrimali, Nidhi Sood, Pranay Shah, R D Dixit, Snehal Bagatharia, Kamlesh J Upadhyay, Ramesh Pandit, Tejas Shah, Ankit Hinsu, Pritesh Sabara, Apurvasinh Puvar, Janvi Raval, Monika Gandhi, Pinal Trivedi, Maharshi Pandya, Amit Kanani, Akanksha Verma, Nitin Savaliya, Raghawendra Kumar, Dipeshwari Shewale, Chaitanya Joshi, Madhvi Joshi |
| EPI_ISL_447554 | GMERS Medical College and Hospital, Gandhinagar | Gujarat Biotechnology Research Centre | Zuber Saiyed, Dipa Kinariwala, Disha Patel, Binita Aring, Neeta Khandelwal, Geeta Vaghela, Sonia Barve, Bhavesh Modi, Kairavi Joshi, Gaurishankar Shrimali, Nidhi Sood, Pranay Shah, R D Dixit, Snehal Bagatharia, Kamlesh J Upadhyay, Ramesh Pandit, Tejas Shah, Ankit Hinsu, Pritesh Sabara, Apurvasinh Puvar, Janvi Raval, Monika Gandhi, Pinal Trivedi, Maharshi Pandya, Amit Kanani, Akanksha Verma, Nitin Savaliya, Raghawendra Kumar, Dinesh Kumar, Sharmistha Majumdar, Chaitanya Joshi, Madhvi Joshi |
| EPI_ISL_447555 | GMERS Medical College and Hospital, Gandhinagar | Gujarat Biotechnology Research Centre | Dipa Kinariwala, Disha Patel, Binita Aring, Neeta Khandelwal, Geeta Vaghela, Sonia Barve, Bhavesh Modi, Kairavi Joshi, Gaurishankar Shrimali, Nidhi Sood, Pranay Shah, R D Dixit, Snehal Bagatharia, Kamlesh J Upadhyay, Ramesh Pandit, Tejas Shah, Ankit Hinsu, Pritesh Sabara, Apurvasinh Puvar, Janvi Raval, Monika Gandhi, Pinal Trivedi, Maharshi Pandya, Amit Kanani, Akanksha Verma, Nitin Savaliya, Raghawendra Kumar, Dinesh Kumar, Zuber Saiyed, Pooja P Doshi, Chaitanya Joshi, Madhvi Joshi |
| EPI_ISL_447556 | CSIR-Centre for Cellular and Molecular Biology | CSIR-Centre for Cellular and Molecular Biology | Sofia Banu, Payel Mukherjee, Priya Singh, Dhiviya Vedagiri, Divya Gupta, Vishal Sah, Santosh Kumar Kuncha, Krishnan Harinivas Harshan, Archana Bharadwaj Siva, Karthik Bharadwaj Tallapaka, Shagufta Khan, Lamuk Zaveri, Namami Gaur, Sakshi Shambhavi, Tulasi Nagabandi, Purushotham Vodnala, Rakesh K Mishra, Divya Tej Sowpati |
| EPI_ISL_447557, EPI_ISL_447558 | CSIR-Centre for Cellular and Molecular Biology | CSIR-Centre for Cellular and Molecular Biology | Payel Mukherjee, Sofia Banu, Priya Singh, Dhiviya Vedagiri, Divya Gupta, Vishal Sah, Santosh Kumar Kuncha, Krishnan Harinivas Harshan, Archana Bharadwaj Siva, Karthik Bharadwaj Tallapaka, Shagufta Khan, Lamuk Zaveri, Namami Gaur, Sakshi Shambhavi, Tulasi Nagabandi, Purushotham Vodnala, Rakesh K Mishra, Divya Tej Sowpati |
| EPI_ISL_447559 | CSIR-Centre for Cellular and Molecular Biology | CSIR-Centre for Cellular and Molecular Biology | Sofia Banu, Payel Mukherjee, Priya Singh, Dhiviya Vedagiri, Divya Gupta, Vishal Sah, Santosh Kumar Kuncha, Krishnan Harinivas Harshan, Archana Bharadwaj Siva, Karthik Bharadwaj Tallapaka, Shagufta Khan, Lamuk Zaveri, Namami Gaur, Sakshi Shambhavi, Tulasi Nagabandi, Purushotham Vodnala, Rakesh K Mishra, Divya Tej Sowpati |
| EPI_ISL_447560 | CSIR-Centre for Cellular and Molecular Biology | CSIR-Centre for Cellular and Molecular Biology | Payel Mukherjee, Sofia Banu, Priya Singh, Dhiviya Vedagiri, Divya Gupta, Vishal Sah, Santosh Kumar Kuncha, Krishnan Harinivas Harshan, Archana Bharadwaj Siva, Karthik Bharadwaj Tallapaka, Shagufta Khan, Lamuk Zaveri, Namami Gaur, Sakshi Shambhavi, Tulasi Nagabandi, Purushotham Vodnala, Rakesh K Mishra, Divya Tej Sowpati |
| EPI_ISL_447561, EPI_ISL_447562 | CSIR-Centre for Cellular and Molecular Biology | CSIR-Centre for Cellular and Molecular Biology | Shagufta Khan, Lamuk Zaveri, Namami Gaur, Sakshi Shambhavi, Tulasi Nagabandi, Purushotham Vodnala, Payel Mukherjee, Sofia Banu, Priya Singh, Dhiviya Vedagiri, Divya Gupta, Vishal Sah, Santosh Kumar Kuncha, Krishnan Harinivas Harshan, Archana Bharadwaj Siva, Karthik Bharadwaj Tallapaka, Rakesh K Mishra, Divya Tej Sowpati |
| EPI_ISL_447563 | CSIR-Centre for Cellular and Molecular Biology | CSIR-Centre for Cellular and Molecular Biology | Sakshi Shambhavi, Lamuk Zaveri, Shagufta Khan, Namami Gaur, Tulasi Nagabandi, Purushotham Vodnala, Payel Mukherjee, Sofia Banu, Priya Singh, Dhiviya Vedagiri, Divya Gupta, Vishal Sah, Santosh Kumar Kuncha, Krishnan Harinivas Harshan, Archana Bharadwaj Siva, Karthik Bharadwaj Tallapaka, Rakesh K Mishra, Divya Tej Sowpati |
| EPI_ISL_447564, EPI_ISL_447565, EPI_ISL_447566 | CSIR-Centre for Cellular and Molecular Biology | CSIR-Centre for Cellular and Molecular Biology | Sofia Banu, Payel Mukherjee, Priya Singh, Dhiviya Vedagiri, Divya Gupta, Vishal Sah, Santosh Kumar Kuncha, Krishnan Harinivas Harshan, Archana Bharadwaj Siva, Karthik Bharadwaj Tallapaka, Shagufta Khan, Lamuk Zaveri, Namami Gaur, Sakshi Shambhavi, Tulasi Nagabandi, Purushotham Vodnala, Rakesh K Mishra, Divya Tej Sowpati |
| EPI_ISL_447567 | CSIR-Centre for Cellular and Molecular Biology | CSIR-Centre for Cellular and Molecular Biology | Payel Mukherjee, Sofia Banu, Priya Singh, Dhiviya Vedagiri, Divya Gupta, Vishal Sah, Santosh Kumar Kuncha, Krishnan Harinivas Harshan, Archana Bharadwaj Siva, Karthik Bharadwaj Tallapaka, Shagufta Khan, Lamuk Zaveri, Namami Gaur, Sakshi Shambhavi, Tulasi Nagabandi, Purushotham Vodnala, Rakesh K Mishra, Divya Tej Sowpati |
| EPI_ISL_447568, EPI_ISL_447569 | CSIR-Centre for Cellular and Molecular Biology | CSIR-Centre for Cellular and Molecular Biology | Shagufta Khan, Lamuk Zaveri, Namami Gaur, Sakshi Shambhavi, Tulasi Nagabandi, Purushotham Vodnala, Payel Mukherjee, Sofia Banu, Priya Singh, Dhiviya Vedagiri, Divya Gupta, Vishal Sah, Santosh Kumar Kuncha, Krishnan Harinivas Harshan, Archana Bharadwaj Siva, Karthik Bharadwaj Tallapaka, Rakesh K Mishra, Divya Tej Sowpati |
| EPI_ISL_447570, EPI_ISL_447571, EPI_ISL_447572 | CSIR-Centre for Cellular and Molecular Biology | CSIR-Centre for Cellular and Molecular Biology | Lamuk Zaveri, Shagufta Khan, Namami Gaur, Sakshi Shambhavi, Tulasi Nagabandi, Purushotham Vodnala, Payel Mukherjee, Sofia Banu, Priya Singh, Dhiviya Vedagiri, Divya Gupta, Vishal Sah, Santosh Kumar Kuncha, Krishnan Harinivas Harshan, Archana Bharadwaj Siva, Karthik Bharadwaj Tallapaka, Rakesh K Mishra, Divya Tej Sowpati |
| EPI_ISL_447573 | CSIR-Centre for Cellular and Molecular Biology | CSIR-Centre for Cellular and Molecular Biology | Sakshi Shambhavi, Lamuk Zaveri, Shagufta Khan, Namami Gaur, Tulasi Nagabandi, Purushotham Vodnala, Payel Mukherjee, Sofia Banu, Priya Singh, Dhiviya Vedagiri, Divya Gupta, Vishal Sah, Santosh Kumar Kuncha, Krishnan Harinivas Harshan, Archana Bharadwaj Siva, Karthik Bharadwaj Tallapaka, Rakesh K Mishra, Divya Tej Sowpati |
| EPI_ISL_447574 | CSIR-Centre for Cellular and Molecular Biology | CSIR-Centre for Cellular and Molecular Biology | Namami Gaur, Sakshi Shambhavi, Lamuk Zaveri, Shagufta Khan, Tulasi Nagabandi, Purushotham Vodnala, Payel Mukherjee, Sofia Banu, Priya Singh, Dhiviya Vedagiri, Divya Gupta, Vishal Sah, Santosh Kumar |

| | | | |
|---|---|---|---|
| | | | Kuncha, Krishnan Harinivas Harshan, Archana Bharadwaj Siva, Karthik Bharadwaj Tallapaka, Rakesh K Mishra, Divya Tej Sowpati |
| EPI_ISL_447575 | CSIR-Centre for Cellular and Molecular Biology | CSIR-Centre for Cellular and Molecular Biology | Sofia Banu, Payel Mukherjee, Priya Singh, Dhiviya Vedagiri, Divya Gupta, Vishal Sah, Santosh Kumar Kuncha, Krishnan Harinivas Harshan, Archana Bharadwaj Siva, Karthik Bharadwaj Tallapaka, Shagufta Khan, Lamuk Zaveri, Namami Gaur, Sakshi Shambhavi, Tulasi Nagabandi, Purushotham Vodnala, Rakesh K Mishra, Divya Tej Sowpati |
| EPI_ISL_447576, EPI_ISL_447577, EPI_ISL_447578 | CSIR-Centre for Cellular and Molecular Biology | CSIR-Centre for Cellular and Molecular Biology | Namami Gaur, Sakshi Shambhavi, Lamuk Zaveri, Shagufta Khan, Tulasi Nagabandi, Purushotham Vodnala, Payel Mukherjee, Sofia Banu, Priya Singh, Dhiviya Vedagiri, Divya Gupta, Vishal Sah, Santosh Kumar Kuncha, Krishnan Harinivas Harshan, Archana Bharadwaj Siva, Karthik Bharadwaj Tallapaka, Rakesh K Mishra, Divya Tej Sowpati |
| EPI_ISL_447579, EPI_ISL_447580 | CSIR-Centre for Cellular and Molecular Biology | CSIR-Centre for Cellular and Molecular Biology | Tulasi Nagabandi, Namami Gaur, Sakshi Shambhavi, Lamuk Zaveri, Shagufta Khan, Purushotham Vodnala, Payel Mukherjee, Sofia Banu, Priya Singh, Dhiviya Vedagiri, Divya Gupta, Vishal Sah, Santosh Kumar Kuncha, Krishnan Harinivas Harshan, Archana Bharadwaj Siva, Karthik Bharadwaj Tallapaka, Rakesh K Mishra, Divya Tej Sowpati |
| EPI_ISL_447581 | CSIR-Centre for Cellular and Molecular Biology | CSIR-Centre for Cellular and Molecular Biology | Sakshi Shambhavi, Lamuk Zaveri, Shagufta Khan, Namami Gaur, Tulasi Nagabandi, Purushotham Vodnala, Payel Mukherjee, Sofia Banu, Priya Singh, Dhiviya Vedagiri, Divya Gupta, Vishal Sah, Santosh Kumar Kuncha, Krishnan Harinivas Harshan, Archana Bharadwaj Siva, Karthik Bharadwaj Tallapaka, Rakesh K Mishra, Divya Tej Sowpati |
| EPI_ISL_447582, EPI_ISL_447583 | CSIR-Centre for Cellular and Molecular Biology | CSIR-Centre for Cellular and Molecular Biology | Tulasi Nagabandi, Namami Gaur, Sakshi Shambhavi, Lamuk Zaveri, Shagufta Khan, Purushotham Vodnala, Payel Mukherjee, Sofia Banu, Priya Singh, Dhiviya Vedagiri, Divya Gupta, Vishal Sah, Santosh Kumar Kuncha, Krishnan Harinivas Harshan, Archana Bharadwaj Siva, Karthik Bharadwaj Tallapaka, Rakesh K Mishra, Divya Tej Sowpati |
| EPI_ISL_447584, EPI_ISL_447585, EPI_ISL_447586, EPI_ISL_447587 | Tamil Nadu Veterinary and Animal Sciences University | CSIR-Centre for Cellular and Molecular Biology | K Kaveri, S Sivasubramanian, S Vennila, P Padmapriya, R Kiruba, S Magesh, G Dhinakar Raj, G Ravi Kumar, Payel Mukherjee, Tulasi Nagabandi, Namami Gaur, Sakshi Shambhavi, Lamuk Zaveri, Shagufta Khan, Purushotham Vodnala, Sofia Banu, Priya Singh, Dhiviya Vedagiri, Divya Gupta, Vishal Sah, Santosh Kumar Kuncha, Krishnan Harinivas Harshan, Archana Bharadwaj Siva, Karthik Bharadwaj Tallapaka, Kumarasamy Thangaraj, Rakesh K Mishra, Divya Tej Sowpati |
| EPI_ISL_447588 | Lednicky Lab | Lednicky lab | Elbadry,M.A., Subramaniam,K., Waltzek,T.B., Gibson,J.C., Stephenson,C.J., Alam,M.M., Morris,J.G. Jr. and Lednicky,J.A. |
| EPI_ISL_447589 | University of Florida, Lednicky Lab | University of Florida, Lednicky Lab | Elbadry,M.A., Subramaniam,K., Waltzek,T.B., Gibson,J.C., Stephenson,C.J., Alam,M.M., Morris,J.G. Jr. and Lednicky,J.A. |
| EPI_ISL_447592, EPI_ISL_447593 | TSGH-CP molecular lab | TSGH-CP molecular lab | Cherng-Lih Perng, Ming-Jr JIAN, Chih-Kai Chang, Jung-Chung Lin, Kuo-Ming Yeh, Chien-Wen Chen, Sheng-Kang Chiu, Hsing-Yi Chung, Shih-Hung Tsai, Kuo-Sheng Hung, Tien-Yao Chang, Feng-Yee Chang, Hung-Sheng Shang |
| EPI_ISL_447594 | Caloundra Hospital | Public Health Virology Laboratory | Bixing Huang, Alyssa Pyke, Amanda De Jong, Andrew Van Den Hurk, Carmel Taylor, David Warrilow, Doris Genge, Elisabeth Gamez, Glen Hewitson, Ian Maxwell Mackay, Inga Sultana, Jamie McMahon, Jean Barcelon, Judy Northill, Mitchell Finger, Natalie Simpson, Neelima Nair, Peter Burtonclay, Peter Moore, Sarah Wheatley, Sean Moody, Sonja Hall-Mendelin, Timothy Gardam, and Frederick Moore |
| EPI_ISL_447595 | Pathology Queensland, Sunshine Coast University Hospital | Public Health Virology Laboratory | Bixing Huang, Alyssa Pyke, Amanda De Jong, Andrew Van Den Hurk, Carmel Taylor, David Warrilow, Doris Genge, Elisabeth Gamez, Glen Hewitson, Ian Maxwell Mackay, Inga Sultana, Jamie McMahon, Jean Barcelon, Judy Northill, Mitchell Finger, Natalie Simpson, Neelima Nair, Peter Burtonclay, Peter Moore, Sarah Wheatley, Sean Moody, Sonja Hall-Mendelin, Timothy Gardam, and Frederick Moore |
| EPI_ISL_447596, EPI_ISL_447597, EPI_ISL_447598, EPI_ISL_447599, EPI_ISL_447606, EPI_ISL_447607 | Viral Respiratory Lab, National Institute for Biomedical Research (INRB) | Pathogen Sequencing Lab, National Institute for Biomedical Research (INRB) | Placide Mbala-Kingebeni, Edith Nkwembe, Eddy Kinganda-Lusamaki, Amuri Aziza, Francisca Muyembe Mawete, Catherine Pratt, Matthias Pauthner, Josh Quick, Allison Black, James Hadfield, Trevor Bedford, Ian Goodfellow, Andrew Rambaut, Nick Loman, Kristian Andersen, Michael Wiley, Steve Ahuka-Mundeke, Jean-Jacques Muyembe Tamfum |
| EPI_ISL_447608, EPI_ISL_447609, EPI_ISL_447610, EPI_ISL_447611, EPI_ISL_447612, EPI_ISL_447613 | Goethe University Hospital Frankfurt | Institute for Medical Virology, Goethe University Hospital Frankfurt | Tuna Toptan, Sebastian Hoehl, Sandra Westhaus, Denisa Bojkova, Annemarie Berger, Björn Rotter, Klaus Hoffmeier, Jindrich Cinatl, Sandra Ciesek, and Marek Widera |
| EPI_ISL_447614, EPI_ISL_447615, EPI_ISL_447616, EPI_ISL_447617, EPI_ISL_447618, EPI_ISL_447619, EPI_ISL_447620, EPI_ISL_447621, EPI_ISL_447622 | Department of Laboratory Medicine, National Taiwan University Hospital | Microbial Genomics Core Lab, National Taiwan University Centers of Genomic and Precision Medicine | Shiou-Hwei Yeh, You-Yu Lin, Ya-Yun Lai, Chiao-Ling Li, Shan-Chwen Chang, Pei-Jer Chen, Sui-Yuan Chang |
| EPI_ISL_447635, EPI_ISL_447636, EPI_ISL_447637, EPI_ISL_447638, EPI_ISL_447639, EPI_ISL_447640, EPI_ISL_447641, EPI_ISL_447642, EPI_ISL_447643, EPI_ISL_447644, EPI_ISL_447645, EPI_ISL_447646, EPI_ISL_447647, EPI_ISL_447648, EPI_ISL_447649, EPI_ISL_447650, EPI_ISL_447651, EPI_ISL_447652, EPI_ISL_447653 | | | |
| see above | unknown | Department of Medicine | Kassela,K., Dovrolis,N., Bampali,M., Gatzidou,E., Froukala,E., Stavropoulou,A., Veletza,S., Tsakris,A., Spanakis,N. and Karakasiliotis,I. |
| EPI_ISL_447654, EPI_ISL_447655 | Hôpital Henri-Mondor Ap-Hp | Hôpital Henri-Mondor Ap-Hp | Rodriguez,C., De Prost,N., Fourati,S., Lamoureux,C., Schmitz,D., Deveaux,I., Picard,O., Lepeule,R., Surgers,L., Mekontso-Dessap,A., Woerther,P.-L., Canoui-Poitrine,F., Pawlotsky,J.-M., Clinical Study Group,C., Gricourt,G., N'debi,M., Demontant,V., Trawinski,E. |
| EPI_ISL_447656 | unknown | Genomic platform | De Prost,N., Fourati,S., Lamoureux,C., Schmitz,D., Deveaux,I., Picard,O., Lepeule,R., Surgers,L., Mekontso-Dessap,A., Woerther,P.-L., Canoui-Poitrine,F., Pawlotsky,J.-M., Clinical Study Group,C., Rodrigue,C., Gricourt,G., N'debi,M., Demontant,V., Trawinski,E. |
| EPI_ISL_447657, EPI_ISL_447658, EPI_ISL_447659, EPI_ISL_447660, EPI_ISL_447661, EPI_ISL_447662, EPI_ISL_447663, EPI_ISL_447664, EPI_ISL_447665, EPI_ISL_447666, EPI_ISL_447667, EPI_ISL_447668, EPI_ISL_447669, EPI_ISL_447670, EPI_ISL_447671, EPI_ISL_447672, EPI_ISL_447673, EPI_ISL_447674, EPI_ISL_447675, EPI_ISL_447676, EPI_ISL_447677, EPI_ISL_447678, EPI_ISL_447679, EPI_ISL_447680, EPI_ISL_447681, EPI_ISL_447682, EPI_ISL_447683, EPI_ISL_447684, EPI_ISL_447685, EPI_ISL_447686, EPI_ISL_447687, EPI_ISL_447688, EPI_ISL_447689, EPI_ISL_447690, EPI_ISL_447691, EPI_ISL_447692, EPI_ISL_447693, EPI_ISL_447694, EPI_ISL_447695, EPI_ISL_447696, EPI_ISL_447697, EPI_ISL_447698, EPI_ISL_447699, EPI_ISL_447700, EPI_ISL_447701, EPI_ISL_447702, EPI_ISL_447703, EPI_ISL_447704, EPI_ISL_447705, EPI_ISL_447706, EPI_ISL_447707, EPI_ISL_447708, EPI_ISL_447709, EPI_ISL_447710, EPI_ISL_447711, EPI_ISL_447712, EPI_ISL_447713, EPI_ISL_447714, EPI_ISL_447715, EPI_ISL_447716, EPI_ISL_447717, EPI_ISL_447718, EPI_ISL_447719, EPI_ISL_447720, EPI_ISL_447721, EPI_ISL_447722, EPI_ISL_447723, EPI_ISL_447724, EPI_ISL_447725, EPI_ISL_447726, EPI_ISL_447727, EPI_ISL_447728, EPI_ISL_447729, EPI_ISL_447730, EPI_ISL_447731, EPI_ISL_447732, EPI_ISL_447733 | | | |
| see above | Hôpital Henri-Mondor Ap-Hp | Hôpital Henri-Mondor Ap-Hp | Rodriguez,C., De Prost,N., Fourati,S., Lamoureux,C., Schmitz,D., Deveaux,I., Picard,O., Lepeule,R., Surgers,L., Mekontso-Dessap,A., Woerther,P.-L., Canoui-Poitrine,F., Pawlotsky,J.-M., Clinical Study Group,C., Gricourt,G., N'debi,M., Demontant,V., Trawinski,E. |
| EPI_ISL_447734, EPI_ISL_447735, EPI_ISL_447736, EPI_ISL_447738, EPI_ISL_447739, EPI_ISL_447740, EPI_ISL_447741, EPI_ISL_447742, EPI_ISL_447743, EPI_ISL_447744, EPI_ISL_447745, EPI_ISL_447746, EPI_ISL_447747, EPI_ISL_447748, EPI_ISL_447750, EPI_ISL_447754 | | | |
| see above | Grupo de Investigaciones Microbiológicas-UR (GIMUR), Departamento de Biología, Facultad de Ciencias Naturales, Universidad del Rosario, Bogotá, Colombia | Grupo de Investigaciones Microbiológicas-UR (GIMUR), Departamento de Biología, Facultad de Ciencias Naturales, Universidad del Rosario, Bogotá, Colombia Instituto Nacional de Salud, Bogotá, Colombia Icahn School of Medicine at Mount Sinai, New York, USA | Juan David Ramírez, Carolina Florez, Marina Muñoz, Carolina Hernandez, Adriana Castillo, Sergio Castañeda, Nathalia Ballesteros, David Martínez, Laura Vega, Jesús E. Jaimes, Sergio Gomez, Angelica Rico, Lisseth Pardo, Esther C. Barros, Martha L. Ospina, Aníbal A. Teherán, Ana S. Gonzalez-Reiche, Matthew M. Hernandez, Emilia Mia Sordillo, Viviana Simon, Harm van Bakel, Alberto Paniz-Mondolfi |
| EPI_ISL_447755, EPI_ISL_447756, EPI_ISL_447757, EPI_ISL_447759, EPI_ISL_447760, EPI_ISL_447761, EPI_ISL_447762, EPI_ISL_447763, EPI_ISL_447765, EPI_ISL_447766, EPI_ISL_447767, EPI_ISL_447768, EPI_ISL_447769, EPI_ISL_447771, EPI_ISL_447774, EPI_ISL_447775, EPI_ISL_447777, EPI_ISL_447778, EPI_ISL_447779, EPI_ISL_447780, EPI_ISL_447781, EPI_ISL_447785, EPI_ISL_447786, EPI_ISL_447789, EPI_ISL_447790, EPI_ISL_447791, EPI_ISL_447792, EPI_ISL_447793, EPI_ISL_447794, EPI_ISL_447795, EPI_ISL_447796, EPI_ISL_447797, EPI_ISL_447798, EPI_ISL_447800, EPI_ISL_447802, EPI_ISL_447803, EPI_ISL_447804, EPI_ISL_447805, EPI_ISL_447806, EPI_ISL_447807, EPI_ISL_447808, EPI_ISL_447809, EPI_ISL_447810, EPI_ISL_447812, EPI_ISL_447813, EPI_ISL_447816, EPI_ISL_447817 | | | |
| see above | Instituto Nacional de Salud, Bogotá, Colombia | Grupo de Investigaciones Microbiológicas-UR (GIMUR), Departamento de Biología, Facultad de Ciencias Naturales, Universidad del Rosario, Bogotá, Colombia Instituto Nacional de Salud, Bogotá, Colombia Icahn School of Medicine at Mount Sinai, New York, USA | Juan David Ramírez, Carolina Florez, Marina Muñoz, Carolina Hernandez, Adriana Castillo, Sergio Castañeda, Nathalia Ballesteros, David Martínez, Laura Vega, Jesús E. Jaimes, Sergio Gomez, Angelica Rico, Lisseth Pardo, Esther C. Barros, Martha L. Ospina, Aníbal A. Teherán, Ana S. Gonzalez-Reiche, Matthew M. Hernandez, Emilia Mia Sordillo, Viviana Simon, Harm van Bakel, Alberto Paniz-Mondolfi |
| EPI_ISL_447832, EPI_ISL_447833, EPI_ISL_447834, EPI_ISL_447835, EPI_ISL_447836 | unknown | Department of Medicine | Kassela,K., Dovrolis,N., Bampali,M., Gatzidou,E., Froukala,E., Stavropoulou,A., Veletza,S., Tsakris,A., Spanakis,N. and Karakasiliotis,I. |
| EPI_ISL_447837 | Dept. of Medical Microbiology, Stavanger University Hospital, Helse Stavanger HF, | Norwegian Institute of Public Health, Department of Virology | Kathrine Stene-Johansen, Kamilla Heddeland Instefjord, Hilde Elshaug, Rasmus Riis Kopperud, Karoline Bragstad, Olav Hungnes |
| EPI_ISL_447840 | DC Public Health Lab/ Dept. of Forensic Sciences | Pathogen Discovery, Respiratory Viruses Branch, Division of Viral Diseases, Centers for Disease Control and Prevention | Krista Queen, Yan Li, Anna Uehara, Jing Zhang, Ying Tao, Clinton R. Paden, Haibin Wang, Jasmine Padilla, Mary S. Keckler, Alison S. Laufer Halpin, Justin Lee, Christopher A. Elkins, Suxiang Tong |
| EPI_ISL_447841 | FL Bureau of Public Health Laboratories-Tampa | Pathogen Discovery, Respiratory Viruses Branch, Division of Viral Diseases, Centers for Disease Control and Prevention | Krista Queen, Yan Li, Anna Uehara, Jing Zhang, Ying Tao, Clinton R. Paden, Haibin Wang, Jasmine Padilla, Mary S. Keckler, Alison S. Laufer Halpin, Justin Lee, Christopher A. Elkins, Suxiang Tong |
| EPI_ISL_447842 | IA State Hygienic Laboratory | Pathogen Discovery, Respiratory Viruses Branch, Division of Viral Diseases, Centers for Disease Control and Prevention | Krista Queen, Yan Li, Anna Uehara, Jing Zhang, Ying Tao, Clinton R. Paden, Haibin Wang, Jasmine Padilla, Mary S. Keckler, Alison S. Laufer Halpin, Justin Lee, Christopher A. Elkins, Suxiang Tong |
| EPI_ISL_447843 | MD DOH Laboratories Administration | Pathogen Discovery, Respiratory Viruses Branch, Division of Viral Diseases, Centers for Disease Control and Prevention | Krista Queen, Yan Li, Anna Uehara, Jing Zhang, Ying Tao, Clinton R. Paden, Haibin Wang, Jasmine Padilla, Mary S. Keckler, Alison S. Laufer Halpin, Justin Lee, Christopher A. Elkins, Suxiang Tong |
| EPI_ISL_447844 | PA Department of Health, Bureau of Laboratories | Pathogen Discovery, Respiratory Viruses Branch, Division of Viral Diseases, Centers for Disease Control and Prevention | Krista Queen, Yan Li, Anna Uehara, Jing Zhang, Ying Tao, Clinton R. Paden, Haibin Wang, Jasmine Padilla, Mary S. Keckler, Alison S. Laufer Halpin, Justin Lee, Christopher A. Elkins, Suxiang Tong |
| EPI_ISL_447845 | PR - Biological and Chemical Emergencies Lab Office of Public Health Preparedness and Response | Pathogen Discovery, Respiratory Viruses Branch, Division of Viral Diseases, Centers for Disease Control and Prevention | Krista Queen, Yan Li, Anna Uehara, Jing Zhang, Ying Tao, Clinton R. Paden, Haibin Wang, Jasmine Padilla, Mary S. Keckler, Alison S. Laufer Halpin, Justin Lee, Christopher A. Elkins, Suxiang Tong |
| EPI_ISL_447846 | VT Dept. of Health Laboratory | Pathogen Discovery, Respiratory Viruses Branch, Division of Viral Diseases, Centers for Disease Control and Prevention | Krista Queen, Yan Li, Anna Uehara, Jing Zhang, Ying Tao, Clinton R. Paden, Haibin Wang, Jasmine Padilla, Mary S. Keckler, Alison S. Laufer Halpin, Justin Lee, Christopher A. Elkins, Suxiang Tong |
| EPI_ISL_447847 | CSIR-Centre for Cellular and Molecular Biology | CSIR-Centre for Cellular and Molecular Biology | Payel Mukherjee, Sofia Banu, Priya Singh, Dhiviya Vedagiri, Divya Gupta, Vishal Sah, Santosh Kumar Kuncha, Krishnan Harinivas Harshan, Archana Bharadwaj Siva, Karthik Bharadwaj Tallapaka, Shagufta Khan, Lamuk Zaveri, Namami Gaur, Sakshi Shambhavi, Tulasi Nagabandi, Purushotham Vodnala, Rakesh K Mishra, Divya Tej Sowpati |
| EPI_ISL_447848 | CSIR-Centre for Cellular and Molecular Biology | CSIR-Centre for Cellular and Molecular Biology | Sofia Banu, Payel Mukherjee, Priya Singh, Dhiviya Vedagiri, Divya Gupta, Vishal Sah, Santosh Kumar Kuncha, Krishnan Harinivas Harshan, Archana Bharadwaj Siva, Karthik Bharadwaj Tallapaka, Shagufta Khan, Lamuk Zaveri, Namami Gaur, Sakshi Shambhavi, Tulasi Nagabandi, Purushotham Vodnala, Rakesh K Mishra, Divya Tej Sowpati |
| EPI_ISL_447849, EPI_ISL_447850 | CSIR-Centre for Cellular and Molecular Biology | CSIR-Centre for Cellular and Molecular Biology | Shagufta Khan, Lamuk Zaveri, Namami Gaur, Sakshi Shambhavi, Tulasi Nagabandi, Purushotham Vodnala, Payel Mukherjee, Sofia Banu, Priya Singh, Dhiviya Vedagiri, Divya Gupta, Vishal Sah, Santosh Kumar Kuncha, Krishnan Harinivas Harshan, Archana Bharadwaj Siva, Karthik Bharadwaj Tallapaka, Rakesh K Mishra, Divya Tej Sowpati |
| EPI_ISL_447851, EPI_ISL_447852 | CSIR-Centre for Cellular and Molecular Biology | CSIR-Centre for Cellular and Molecular Biology | Lamuk Zaveri, Shagufta Khan, Namami Gaur, Sakshi Shambhavi, Tulasi Nagabandi, Purushotham Vodnala, Payel Mukherjee, Sofia Banu, Priya Singh, Dhiviya Vedagiri, Divya Gupta, Vishal Sah, Santosh Kumar Kuncha, Krishnan Harinivas Harshan, Archana Bharadwaj Siva, Karthik Bharadwaj Tallapaka, Rakesh K Mishra, Divya Tej Sowpati |
| EPI_ISL_447853 | CSIR-Centre for Cellular and Molecular Biology | CSIR-Centre for Cellular and Molecular Biology | Namami Gaur, Sakshi Shambhavi, Lamuk Zaveri, Shagufta Khan, Tulasi Nagabandi, Purushotham Vodnala, Payel Mukherjee, Sofia Banu, Priya Singh, Dhiviya Vedagiri, Divya Gupta, Vishal Sah, Santosh Kumar Kuncha, Krishnan Harinivas Harshan, Archana Bharadwaj Siva, Karthik Bharadwaj Tallapaka, Rakesh K Mishra, Divya Tej Sowpati |
| EPI_ISL_447854 | CSIR-Centre for Cellular and Molecular Biology | CSIR-Centre for Cellular and Molecular Biology | Payel Mukherjee, Sofia Banu, Priya Singh, Dhiviya Vedagiri, Divya Gupta, Vishal Sah, Santosh Kumar Kuncha, Krishnan Harinivas Harshan, Archana Bharadwaj Siva, Karthik Bharadwaj Tallapaka, Shagufta Khan, Lamuk Zaveri, Namami Gaur, Sakshi Shambhavi, Tulasi Nagabandi, Purushotham Vodnala, Rakesh K Mishra, Divya Tej Sowpati |
| EPI_ISL_447855 | CSIR-Centre for Cellular and Molecular Biology | CSIR-Centre for Cellular and Molecular Biology | Lamuk Zaveri, Shagufta Khan, Namami Gaur, Sakshi Shambhavi, Tulasi Nagabandi, Purushotham Vodnala, Payel Mukherjee, Sofia Banu, Priya Singh, Dhiviya Vedagiri, Divya Gupta, Vishal Sah, Santosh Kumar Kuncha, Krishnan Harinivas Harshan, Archana Bharadwaj Siva, Karthik Bharadwaj Tallapaka, Rakesh K Mishra, Divya Tej Sowpati |
| EPI_ISL_447856, EPI_ISL_447857, EPI_ISL_447858 | CSIR-Centre for Cellular and Molecular Biology | CSIR-Centre for Cellular and Molecular Biology | Sakshi Shambhavi, Lamuk Zaveri, Shagufta Khan, Namami Gaur, Tulasi Nagabandi, Purushotham Vodnala, Payel Mukherjee, Sofia Banu, Priya Singh, Dhiviya Vedagiri, Divya Gupta, Vishal Sah, Santosh Kumar Kuncha, Krishnan Harinivas Harshan, Archana Bharadwaj Siva, Karthik Bharadwaj Tallapaka, Rakesh K Mishra, Divya Tej Sowpati |
| EPI_ISL_447859 | CSIR-Centre for Cellular and Molecular Biology | CSIR-Centre for Cellular and Molecular Biology | Payel Mukherjee, Sofia Banu, Priya Singh, Dhiviya Vedagiri, Divya Gupta, Vishal Sah, Santosh Kumar Kuncha, Krishnan Harinivas Harshan, Archana Bharadwaj Siva, Karthik Bharadwaj Tallapaka, Shagufta Khan, Lamuk Zaveri, Namami Gaur, Sakshi Shambhavi, Tulasi Nagabandi, Purushotham Vodnala, Rakesh K Mishra, Divya Tej Sowpati |
| EPI_ISL_447860, EPI_ISL_447861 | CSIR-Centre for Cellular and Molecular Biology | CSIR-Centre for Cellular and Molecular Biology | Tulasi Nagabandi, Namami Gaur, Sakshi Shambhavi, Lamuk Zaveri, Shagufta Khan, Purushotham Vodnala, Payel Mukherjee, Sofia Banu, Priya Singh, Dhiviya Vedagiri, Divya Gupta, Vishal Sah, Santosh Kumar Kuncha, Krishnan Harinivas Harshan, Archana Bharadwaj Siva, Karthik Bharadwaj Tallapaka, Rakesh K Mishra, Divya Tej Sowpati |
| EPI_ISL_447862, EPI_ISL_447863, EPI_ISL_447864 | CSIR-Centre for Cellular and Molecular Biology | CSIR-Centre for Cellular and Molecular Biology | Payel Mukherjee, Sofia Banu, Priya Singh, Dhiviya Vedagiri, Divya Gupta, Vishal Sah, Santosh Kumar Kuncha, Krishnan Harinivas Harshan, Archana Bharadwaj Siva, Karthik Bharadwaj Tallapaka, Shagufta Khan, |

| | | | |
|---|---|---|---|
| | | | Lamuk Zaveri, Namami Gaur, Sakshi Shambhavi, Tulasi Nagabandi, Purushotham Vodnala, Rakesh K Mishra, Divya Tej Sowpati |
| EPI_ISL_447865, EPI_ISL_447866 | CSIR-Centre for Cellular and Molecular Biology | CSIR-Centre for Cellular and Molecular Biology | Sofia Banu, Payel Mukherjee, Priya Singh, Dhiviya Vedagiri, Divya Gupta, Vishal Sah, Santosh Kumar Kuncha, Krishnan Harinivas Harshan, Archana Bharadwaj Siva, Karthik Bharadwaj Tallapaka, Shagufta Khan, Lamuk Zaveri, Namami Gaur, Sakshi Shambhavi, Tulasi Nagabandi, Purushotham Vodnala, Rakesh K Mishra, Divya Tej Sowpati |
| EPI_ISL_447886 | unknown | Pathogen Discovery | Ying Tao, Yan Li, Jing Zhang, Clinton R. Paden, Krista Queen, Anna Uehara, Haibin Wang, Julu Bhatnagar, Suxiang Tong |
| EPI_ISL_447887, EPI_ISL_447888, EPI_ISL_447889, EPI_ISL_447890, EPI_ISL_447891, EPI_ISL_447892, EPI_ISL_447893, EPI_ISL_447894, EPI_ISL_447895, EPI_ISL_447896 | University of California, Davis | Chan-Zuckerberg Biohub | CZB Cliahub Consortium |
| EPI_ISL_447903 | University of Florida | University of Florida | Elbadry,M.A., Subramaniam,K., Waltzek,T.B., Lauzardo,M., Gibson,J.C., Stephenson,C.J., Alam,M.M., Morris,J.G. Jr. and Lednicky,J.A. |
| EPI_ISL_447905 | University of Florida | University of Florida | Elbadry,M.A., Subramaniam,K., Waltzek,T.B., Gibson,J.C., Stephenson,C.J., Alam,M.M., Morris,J.G. Jr. and Lednicky,J.A. |
| EPI_ISL_449480, EPI_ISL_449481, EPI_ISL_449484, EPI_ISL_449486 | unknown | Department of Respiratory and Critical Care | Wang,X., Zhou,Q., He,Y., Liu,L., Ma,X., Wei,X., Jiang,N., Liang,L., Zheng,Y., Ma,L., Xu,Y., Yang,D., Zhang,J., Yang,B., Jiang,N., Zheng,Y., Ma,L., Xu,Y., Yang,D., Zhang,J., Yang,B., Jiang,N., Deng,T., Zhai,B., Gao,Y., Liu,W., Bai,X., Pan,T., Wang,G., Chang,Y., Zhang,Z., Shi,H., Ma,W.L. and Gao,Z. |
| EPI_ISL_450212, EPI_ISL_450213, EPI_ISL_450214 | unknown | Microbiological Diagnostic Unit Public Health Laboratory (MDU-PHL) and Victorian Infectious Disease Reference Laboratory (VIDRL) | Seemann,T., Lane,C.R., Sherry,N.L., Duchene,S., Goncalves da Silva,A., Caly,L., Sait,M., Ballard,S.A., Horan,K., Schultz,M.B., Hoang,T., Easton,M., Dougal,S., Stinear,T.P., Druce,J., Catton,M., Sutton,B., van Diemen,A., Alpren,C., Williamson,D.A., Howden,B.P. |
| EPI_ISL_450408, EPI_ISL_450409, EPI_ISL_450410, EPI_ISL_450411, EPI_ISL_450412 | unknown | Microbiology | To,K.K.W., Yuen,K.-Y. |
| EPI_ISL_450442 | The Department of Infectious Disease Prevention and Control, Henan Provincial Center for Disease Control and Prevention | The Department of Infectious Disease Prevention and Control, Henan Provincial Center for Disease Control and Prevention | Li,X., Lu,S., Wu,B., Hu,X., Li,D., Huang,X. and Guo,W. |
| EPI_ISL_450484, EPI_ISL_450485, EPI_ISL_450486, EPI_ISL_450487 | unknown | Data Science | Carroll,T.D., Tran,N.K., Cohen,S.H., Miller,C.J. |
| EPI_ISL_454692, EPI_ISL_454693 | Quest Diagnostics | Quest Diagnostics | Anderson,B.P., Rosenthal,S.H., Gerasimova,A., Kagan,R.M. and Owen, R. |
| EPI_ISL_524433, EPI_ISL_524434 | Environmental and Global Health, University of Florida - Gainesville | University of Florida | Elbadry,M.A., Subramaniam,K., Waltzek,T.B., Gibson,J.C., Stephenson,C.J., Alam,M.M., Morris,J.G. Jr., Lednicky,J.A. |
| EPI_ISL_529147, EPI_ISL_529148 | Democritus University of Thrace, Department of Medicine | Democritus University of Thrace, Department of Medicine | Kassela,K., Dovrolis,N., Bampali,M., Gatzidou,E., Froukala,E., Stavropoulou,A., Veletza,S., Tsakris,A., Spanakis,N., Karakasiliotis,I. |
| EPI_ISL_605929, EPI_ISL_605930 | Department of Infectious Disease Prevention and Control, Henan Provincial Center for Disease Control and Prevention | Department of Infectious Disease Prevention and Control, Henan Provincial Center for Disease Control and Prevention | Li,X., Lu,S., Wu,B., Hu,X., Li,D., Ye,Y., Huang,X., Guo,W. |
