## Supplementary material for "Analysis of the Dynamics and Distribution of SARS-CoV-2 Mutations and its Possible Structural and Functional Implications": GISAID_acknowledgements_part_3

We gratefully acknowledge the following Authors from the Originating laboratories responsible for obtaining the specimens, as well as the Submitting laboratories where the genome data were generated and shared via GISAID, on which this research is based.

All Submitters of data may be contacted directly via www.gisaid.org

| Accession ID | Originating Laboratory | Submitting Laboratory | Authors |
|---|---|---|---|
| EPI_ISL_447904 | National Institute of Biotechnology | National Institute of Biotechnology | Md. Moniruzzaman, Mohammad Uzzal Hossain, Md. Nazrul Islam, Md. Hadisur Rahman, Irfan Ahmed, Tahia Anan Rahman, Arittra Bhattacharjee, Md. Ruhul Amin, Asif Rashid, Chaman Ara Keya, Keshob Chandra Das, Md. Salimullah |
| EPI_ISL_447906, EPI_ISL_447907, EPI_ISL_447908 | Siriraj hospital | National Institute of Health. Department of medical Sciences, Ministry of Public Health, Thailand | Pilailuk,Okada; Navin Horthongkham, Siripaporn,Phuygun; Thanutsapa,Thanadachakul; Sittiporn,Parnmen;Warawan,Wongboot; Sunthareeya,Waicharoen; Malinee,Chittaganpitch |
| EPI_ISL_447909, EPI_ISL_447910, EPI_ISL_447911, EPI_ISL_447912, EPI_ISL_447913, EPI_ISL_447914, EPI_ISL_447915, EPI_ISL_447916, EPI_ISL_447917, EPI_ISL_447918, EPI_ISL_447919, EPI_ISL_447920, EPI_ISL_447921 | | | |
| see above | n/a | National Institute of Health. Department of medical Sciences, Ministry of Public Health, Thailand | Pilailuk,Okada; Siripaporn,Phuygun; Thanutsapa,Thanadachakul; Sittiporn,Parnmen;Warawan,Wongboot; Sunthareeya,Waicharoen; Malinee,Chittaganpitch |
| EPI_ISL_447927, EPI_ISL_447928, EPI_ISL_447931, EPI_ISL_447933, EPI_ISL_447936, EPI_ISL_447939, EPI_ISL_447940 | University of Birmingham | COVID-19 Genomics UK (COG-UK) Consortium | Claire McMurray, Joanne Stockton, Samuel Nicholls, Radoslaw Poplawski, Will Rowe, Josh Quick, Nicholas Loman, Celina M Whalley, Andrew Bosworth, Charlotte Poxon, Kasun Wanigasooriya, Oliver Pickles, Mike Kidd, Alex Richter, Andrew D Beggs, Husam Osman, Andrew Bosworth |
| EPI_ISL_447953, EPI_ISL_447954, EPI_ISL_447955, EPI_ISL_447956, EPI_ISL_447959, EPI_ISL_447960, EPI_ISL_447963, EPI_ISL_447964, EPI_ISL_447966, EPI_ISL_447967, EPI_ISL_447969, EPI_ISL_447970, EPI_ISL_447971, EPI_ISL_447972, EPI_ISL_447973, EPI_ISL_447976, EPI_ISL_447977, EPI_ISL_447978, EPI_ISL_447979, EPI_ISL_447980, EPI_ISL_447981, EPI_ISL_447983, EPI_ISL_447984, EPI_ISL_447985, EPI_ISL_447986, EPI_ISL_447987, EPI_ISL_447988, EPI_ISL_447989, EPI_ISL_447990, EPI_ISL_447994, EPI_ISL_447995, EPI_ISL_447997, EPI_ISL_447998, EPI_ISL_447999, EPI_ISL_448000, EPI_ISL_448001, EPI_ISL_448003, EPI_ISL_448007, EPI_ISL_448008, EPI_ISL_448010, EPI_ISL_448011, EPI_ISL_448012, EPI_ISL_448013, EPI_ISL_448014, EPI_ISL_448015, EPI_ISL_448016, EPI_ISL_448017, EPI_ISL_448018, EPI_ISL_448019, EPI_ISL_448021, EPI_ISL_448022, EPI_ISL_448023, EPI_ISL_448024, EPI_ISL_448028, EPI_ISL_448029, EPI_ISL_448030, EPI_ISL_448033, EPI_ISL_448034, EPI_ISL_448035, EPI_ISL_448036, EPI_ISL_448037, EPI_ISL_448038, EPI_ISL_448039, EPI_ISL_448040, EPI_ISL_448041, EPI_ISL_448042, EPI_ISL_448043, EPI_ISL_448044, EPI_ISL_448046, EPI_ISL_448047, EPI_ISL_448049, EPI_ISL_448051, EPI_ISL_448053, EPI_ISL_448056, EPI_ISL_448057, EPI_ISL_448059, EPI_ISL_448060, EPI_ISL_448063, EPI_ISL_448064, EPI_ISL_448065, EPI_ISL_448066, EPI_ISL_448068, EPI_ISL_448072, EPI_ISL_448073, EPI_ISL_448077, EPI_ISL_448078, EPI_ISL_448079, EPI_ISL_448080, EPI_ISL_448081, EPI_ISL_448082, EPI_ISL_448083, EPI_ISL_448084, EPI_ISL_448085, EPI_ISL_448086, EPI_ISL_448087, EPI_ISL_448088, EPI_ISL_448089, EPI_ISL_448090, EPI_ISL_448091, EPI_ISL_448092, EPI_ISL_448095, EPI_ISL_448096, EPI_ISL_448097, EPI_ISL_448098, EPI_ISL_448100, EPI_ISL_448101, EPI_ISL_448102, EPI_ISL_448103, EPI_ISL_448105, EPI_ISL_448106, EPI_ISL_448107, EPI_ISL_448108, EPI_ISL_448109, EPI_ISL_448111, EPI_ISL_448114 | | | |
| see above | Department of Pathology, University of Cambridge | COVID-19 Genomics UK (COG-UK) Consortium | Luke W Meredith, M. Estée Török , Myra Hosmillo, William L. Hamilton, Martin D. Curran, Theresa Feltwell, Grant Hall, Anna Yakovleva, Fahad A Khokhar, Charlotte J. Houldcroft, Laura G Caller, Aminu S. Jahun, Sarah L. Caddy, Ian Goodfellow |
| EPI_ISL_448117, EPI_ISL_448118, EPI_ISL_448119, EPI_ISL_448120, EPI_ISL_448121, EPI_ISL_448122, EPI_ISL_448123, EPI_ISL_448124, EPI_ISL_448126, EPI_ISL_448127, EPI_ISL_448128, EPI_ISL_448129, EPI_ISL_448130, EPI_ISL_448131, EPI_ISL_448132, EPI_ISL_448133, EPI_ISL_448134, EPI_ISL_448136, EPI_ISL_448138, EPI_ISL_448139, EPI_ISL_448140, EPI_ISL_448141, EPI_ISL_448142, EPI_ISL_448143, EPI_ISL_448144, EPI_ISL_448145, EPI_ISL_448147, EPI_ISL_448148, EPI_ISL_448149, EPI_ISL_448151, EPI_ISL_448152, EPI_ISL_448153, EPI_ISL_448154, EPI_ISL_448155, EPI_ISL_448156, EPI_ISL_448157, EPI_ISL_448158, EPI_ISL_448161, EPI_ISL_448162, EPI_ISL_448163, EPI_ISL_448164, EPI_ISL_448165, EPI_ISL_448166, EPI_ISL_448167, EPI_ISL_448168, EPI_ISL_448169, EPI_ISL_448170, EPI_ISL_448171, EPI_ISL_448172, EPI_ISL_448175, EPI_ISL_448176, EPI_ISL_448177, EPI_ISL_448178, EPI_ISL_448179, EPI_ISL_448180, EPI_ISL_448182, EPI_ISL_448183, EPI_ISL_448184, EPI_ISL_448185, EPI_ISL_448187, EPI_ISL_448188, EPI_ISL_448189, EPI_ISL_448190, EPI_ISL_448191, EPI_ISL_448192, EPI_ISL_448193, EPI_ISL_448195, EPI_ISL_448196, EPI_ISL_448197, EPI_ISL_448198, EPI_ISL_448199, EPI_ISL_448200, EPI_ISL_448201, EPI_ISL_448204, EPI_ISL_448205, EPI_ISL_448206, EPI_ISL_448207, EPI_ISL_448209, EPI_ISL_448210, EPI_ISL_448211, EPI_ISL_448212, EPI_ISL_448213, EPI_ISL_448214, EPI_ISL_448215, EPI_ISL_448216, EPI_ISL_448217, EPI_ISL_448218, EPI_ISL_448219, EPI_ISL_448220, EPI_ISL_448221 | | | |
| see above | West of Scotland Specialist Virology Centre, NHSGGC / MRC-University of Glasgow Centre for Virus Research | COVID-19 Genomics UK (COG-UK) Consortium | Ana da Silva Filipe, Natasha Johnson, Kathy Smollett, Daniel Mair, Stephen Carmichael, Lily Tong, Jenna Nichols, Elihu Aranday-Cortes, Kirstyn Brunker, Yasmin Parr, Kyriaki Nomikou, Sarah McDonald, Marc Niebel, Patawee Asamaphan, Richard Orton, Joseph Hughes, Sreenu Vattipally, David L Robertson, Alasdair MacLean, Rory Gunson, Kathy Li, Natasha Jesudason, Rajiv Shah, James Shepherd, Antonia Ho, Emma Thomson |
| EPI_ISL_448222 | Pasteur Insitute Ho Chi Minh City | National Key Laboratory of Gene Technology, Institute of Biotechnology, Vietnam Academy of Science and Technology | Le Tung Lam, Nguyen Trung Hieu, Nguyen Hong Trang, Ho Thi Thuong, Nguyen Thi Ngoc Thao, Huynh Thi Kim Loan, Luu Thuy Tien, Tran Huyen Linh, Pham Duy Quang, Luong Chan Quang, Cao Minh Thang, Nguyen Vu Thuong, Hoang Ha, Chu Hoang Ha, Phan Trong Lan, Truong Nam Hai |
| EPI_ISL_448223, EPI_ISL_448224, EPI_ISL_448225, EPI_ISL_448226, EPI_ISL_448227, EPI_ISL_448228, EPI_ISL_448229, EPI_ISL_448230, EPI_ISL_448232, EPI_ISL_448233, EPI_ISL_448234, EPI_ISL_448235, EPI_ISL_448236, EPI_ISL_448237, EPI_ISL_448238, EPI_ISL_448239, EPI_ISL_448240, EPI_ISL_448241, EPI_ISL_448242, EPI_ISL_448243, EPI_ISL_448244, EPI_ISL_448245, EPI_ISL_448246, EPI_ISL_448247, EPI_ISL_448248, EPI_ISL_448249, EPI_ISL_448250, EPI_ISL_448251, EPI_ISL_448253, EPI_ISL_448254, EPI_ISL_448255, EPI_ISL_448256, EPI_ISL_448258, EPI_ISL_448259, EPI_ISL_448260, EPI_ISL_448261, EPI_ISL_448262, EPI_ISL_448263, EPI_ISL_448264, EPI_ISL_448265, EPI_ISL_448266, EPI_ISL_448267, EPI_ISL_448268, EPI_ISL_448269, EPI_ISL_448270, EPI_ISL_448271, EPI_ISL_448272, EPI_ISL_448273, EPI_ISL_448274, EPI_ISL_448276, EPI_ISL_448277, EPI_ISL_448278, EPI_ISL_448279, EPI_ISL_448280, EPI_ISL_448281, EPI_ISL_448282, EPI_ISL_448283, EPI_ISL_448285, EPI_ISL_448286, EPI_ISL_448287, EPI_ISL_448288, EPI_ISL_448289, EPI_ISL_448290, EPI_ISL_448291, EPI_ISL_448292, EPI_ISL_448293, EPI_ISL_448294, EPI_ISL_448295, EPI_ISL_448296, EPI_ISL_448297, EPI_ISL_448299, EPI_ISL_448300, EPI_ISL_448301, EPI_ISL_448304, EPI_ISL_448305, EPI_ISL_448306, EPI_ISL_448307, EPI_ISL_448308, EPI_ISL_448309, EPI_ISL_448311, EPI_ISL_448312, EPI_ISL_448313, EPI_ISL_448314, EPI_ISL_448315, EPI_ISL_448316, EPI_ISL_448317, EPI_ISL_448320, EPI_ISL_448321, EPI_ISL_448322, EPI_ISL_448323, EPI_ISL_448324, EPI_ISL_448325, EPI_ISL_448326, EPI_ISL_448327, EPI_ISL_448328, EPI_ISL_448329, EPI_ISL_448330, EPI_ISL_448331, EPI_ISL_448332, EPI_ISL_448335, EPI_ISL_448336, EPI_ISL_448338, EPI_ISL_448339, EPI_ISL_448340, EPI_ISL_448341, EPI_ISL_448342, EPI_ISL_448344, EPI_ISL_448345, EPI_ISL_448346, EPI_ISL_448348, EPI_ISL_448350, EPI_ISL_448351, EPI_ISL_448352, EPI_ISL_448355, EPI_ISL_448357, EPI_ISL_448358, EPI_ISL_448359, EPI_ISL_448360, EPI_ISL_448361, EPI_ISL_448362, EPI_ISL_448363, EPI_ISL_448364, EPI_ISL_448365, EPI_ISL_448366, EPI_ISL_448369, EPI_ISL_448370, EPI_ISL_448371, EPI_ISL_448372, EPI_ISL_448374, EPI_ISL_448375, EPI_ISL_448376, EPI_ISL_448377, EPI_ISL_448379, EPI_ISL_448380, EPI_ISL_448383, EPI_ISL_448384, EPI_ISL_448385, EPI_ISL_448387, EPI_ISL_448388 | | | |
| see above | Quadram Institute Bioscience | COVID-19 Genomics UK (COG-UK) Consortium | Dave J. Baker, Gemma L. Kay, Alp Aydin, Thanh Le-Viet, Steven Rudder, Ana P. Tedim, Anastasia Kolyva, Maria Diaz, Leonardo de Oliveira Martins, Nabil-Fareed Alikhan, Lizzie Meadows, Rachael Stanley, Ngozi Elumogo, Muhammed Yasir, Nicholas M. Thomson, Alexander J Trotter, Rachel Gilroy, Samuel Bloomfield, Claire Stuart, Andrew Bell, Reenesh Prakash, Samir Dervisevic, Alison E. Mather, John Wain, Mark Webber, Andrew J. Page, Justin O'Grady |
| EPI_ISL_448389, EPI_ISL_448391, EPI_ISL_448393, EPI_ISL_448395, EPI_ISL_448397, EPI_ISL_448398, EPI_ISL_448399, EPI_ISL_448400, EPI_ISL_448401, EPI_ISL_448402, EPI_ISL_448404, EPI_ISL_448405, EPI_ISL_448406, EPI_ISL_448407, EPI_ISL_448408, EPI_ISL_448409, EPI_ISL_448410, EPI_ISL_448412, EPI_ISL_448413, EPI_ISL_448414, EPI_ISL_448415, EPI_ISL_448416, EPI_ISL_448417, EPI_ISL_448418, EPI_ISL_448420, EPI_ISL_448421, EPI_ISL_448430, EPI_ISL_448432, EPI_ISL_448436, EPI_ISL_448438, EPI_ISL_448440, EPI_ISL_448441, EPI_ISL_448443, EPI_ISL_448444, EPI_ISL_448445, EPI_ISL_448446, EPI_ISL_448448 | | | |
| see above | Queens Medical Centre, Clinical Microbiology Department / DeepSeq Nottingham | COVID-19 Genomics UK (COG-UK) Consortium | Gemma Clark, Wendy Smith, Manjinder Khakh, Hannah Howson-Wells, Jonathan Ball, Patrick McClure, Joseph Chappell, Theocharis Tsoleridis, Nadine Holmes, Matthew Carlisle, Christopher Moore, Fei Sang, Johnny Debebe, Victoria Wright, Matthew Loose |
| EPI_ISL_448449, EPI_ISL_448450, EPI_ISL_448451, EPI_ISL_448453, EPI_ISL_448454, EPI_ISL_448455, EPI_ISL_448456, EPI_ISL_448457, EPI_ISL_448458, EPI_ISL_448459, EPI_ISL_448460, EPI_ISL_448461, EPI_ISL_448462, EPI_ISL_448463, EPI_ISL_448465, EPI_ISL_448467, EPI_ISL_448468, EPI_ISL_448469, EPI_ISL_448470, EPI_ISL_448471, EPI_ISL_448472, EPI_ISL_448473, EPI_ISL_448476, EPI_ISL_448477, EPI_ISL_448478, EPI_ISL_448479, EPI_ISL_448480, EPI_ISL_448481, EPI_ISL_448482, EPI_ISL_448483, EPI_ISL_448485, EPI_ISL_448486, EPI_ISL_448487, EPI_ISL_448488, EPI_ISL_448490, EPI_ISL_448491, EPI_ISL_448492, EPI_ISL_448493, EPI_ISL_448494, EPI_ISL_448495, EPI_ISL_448496, EPI_ISL_448497, EPI_ISL_448498, EPI_ISL_448499, EPI_ISL_448500, EPI_ISL_448501, EPI_ISL_448502, EPI_ISL_448503, EPI_ISL_448504, EPI_ISL_448505, EPI_ISL_448506, EPI_ISL_448507, EPI_ISL_448508, EPI_ISL_448510, EPI_ISL_448511, EPI_ISL_448512, EPI_ISL_448513, EPI_ISL_448514, EPI_ISL_448515, EPI_ISL_448516, EPI_ISL_448517, EPI_ISL_448518, EPI_ISL_448519, EPI_ISL_448520, EPI_ISL_448521, EPI_ISL_448522, EPI_ISL_448525, EPI_ISL_448526, EPI_ISL_448527, EPI_ISL_448529, EPI_ISL_448530, EPI_ISL_448531, EPI_ISL_448532, EPI_ISL_448533, EPI_ISL_448534, EPI_ISL_448535, EPI_ISL_448536, EPI_ISL_448537, EPI_ISL_448538, EPI_ISL_448540, EPI_ISL_448541, EPI_ISL_448542, EPI_ISL_448545, EPI_ISL_448546, EPI_ISL_448547, EPI_ISL_448548, EPI_ISL_448549, EPI_ISL_448550, EPI_ISL_448551, EPI_ISL_448552, EPI_ISL_448553, EPI_ISL_448555, EPI_ISL_448556, EPI_ISL_448557, EPI_ISL_448559, EPI_ISL_448561, EPI_ISL_448563, EPI_ISL_448564, EPI_ISL_448565, EPI_ISL_448566, EPI_ISL_448567, EPI_ISL_448568, EPI_ISL_448570, EPI_ISL_448571, EPI_ISL_448572, EPI_ISL_448573, EPI_ISL_448575, EPI_ISL_448577, EPI_ISL_448579, EPI_ISL_448580, EPI_ISL_448582, EPI_ISL_448584, EPI_ISL_448585, EPI_ISL_448587, EPI_ISL_448588, EPI_ISL_448590, EPI_ISL_448591, EPI_ISL_448592, EPI_ISL_448593, EPI_ISL_448594, EPI_ISL_448595, EPI_ISL_448596, EPI_ISL_448597, EPI_ISL_448598, EPI_ISL_448599, EPI_ISL_448600, EPI_ISL_448601, EPI_ISL_448602, EPI_ISL_448603, EPI_ISL_448604, EPI_ISL_448605, EPI_ISL_448606, EPI_ISL_448607, EPI_ISL_448608, EPI_ISL_448609, EPI_ISL_448610, EPI_ISL_448611, EPI_ISL_448612, EPI_ISL_448613, EPI_ISL_448614, EPI_ISL_448616, EPI_ISL_448617, EPI_ISL_448618, EPI_ISL_448619, EPI_ISL_448620, EPI_ISL_448621, EPI_ISL_448622, EPI_ISL_448623, EPI_ISL_448624, EPI_ISL_448625, EPI_ISL_448626, EPI_ISL_448627, EPI_ISL_448628, EPI_ISL_448629, EPI_ISL_448630, EPI_ISL_448631, EPI_ISL_448632, EPI_ISL_448633, EPI_ISL_448634, EPI_ISL_448635, EPI_ISL_448636, EPI_ISL_448637, EPI_ISL_448638, EPI_ISL_448639, EPI_ISL_448640, EPI_ISL_448643, EPI_ISL_448644, EPI_ISL_448645, EPI_ISL_448646, EPI_ISL_448650, EPI_ISL_448651, EPI_ISL_448652, EPI_ISL_448654, EPI_ISL_448655, EPI_ISL_448656, EPI_ISL_448657, EPI_ISL_448658, EPI_ISL_448659, EPI_ISL_448660, EPI_ISL_448661, EPI_ISL_448662, EPI_ISL_448663, EPI_ISL_448664, EPI_ISL_448666, EPI_ISL_448667, EPI_ISL_448668, EPI_ISL_448669, EPI_ISL_448670, EPI_ISL_448671, EPI_ISL_448672, EPI_ISL_448673, EPI_ISL_448674, EPI_ISL_448675, EPI_ISL_448676, EPI_ISL_448678, EPI_ISL_448679, EPI_ISL_448680, EPI_ISL_448681, EPI_ISL_448682, EPI_ISL_448684, EPI_ISL_448685, EPI_ISL_448686, EPI_ISL_448687, EPI_ISL_448688, EPI_ISL_448689, EPI_ISL_448690, EPI_ISL_448691, EPI_ISL_448692, EPI_ISL_448694, EPI_ISL_448695, EPI_ISL_448696, EPI_ISL_448697, EPI_ISL_448698, EPI_ISL_448699, EPI_ISL_448700, EPI_ISL_448701, EPI_ISL_448702, EPI_ISL_448703, EPI_ISL_448704, EPI_ISL_448705, EPI_ISL_448706, EPI_ISL_448707, EPI_ISL_448708, EPI_ISL_448709, EPI_ISL_448710, EPI_ISL_448711, EPI_ISL_448712, EPI_ISL_448713, EPI_ISL_448714, EPI_ISL_448715, EPI_ISL_448716, EPI_ISL_448717, EPI_ISL_448718, EPI_ISL_448719, EPI_ISL_448720, EPI_ISL_448722, EPI_ISL_448723, EPI_ISL_448724, EPI_ISL_448725, EPI_ISL_448726, EPI_ISL_448727, EPI_ISL_448728, EPI_ISL_448729, EPI_ISL_448730, EPI_ISL_448731, EPI_ISL_448732, EPI_ISL_448733, EPI_ISL_448734, EPI_ISL_448735, EPI_ISL_448736, EPI_ISL_448737, EPI_ISL_448738, EPI_ISL_448739, EPI_ISL_448740, EPI_ISL_448743, EPI_ISL_448744, EPI_ISL_448745, EPI_ISL_448746, EPI_ISL_448747, EPI_ISL_448749, EPI_ISL_448750, EPI_ISL_448751, EPI_ISL_448752, EPI_ISL_448754, EPI_ISL_448755, EPI_ISL_448756, EPI_ISL_448757, EPI_ISL_448758, EPI_ISL_448759, EPI_ISL_448760, EPI_ISL_448762, EPI_ISL_448763, EPI_ISL_448764, EPI_ISL_448765, EPI_ISL_448766, EPI_ISL_448767, EPI_ISL_448768, EPI_ISL_448769, EPI_ISL_448770, EPI_ISL_448771, EPI_ISL_448772, EPI_ISL_448773, EPI_ISL_448774, EPI_ISL_448775, EPI_ISL_448776, EPI_ISL_448777, EPI_ISL_448778, EPI_ISL_448779, EPI_ISL_448780, EPI_ISL_448781, EPI_ISL_448782, EPI_ISL_448783, EPI_ISL_448784, EPI_ISL_448785, EPI_ISL_448786, EPI_ISL_448787, EPI_ISL_448788, EPI_ISL_448789, EPI_ISL_448790, EPI_ISL_448791, EPI_ISL_448792, EPI_ISL_448793, EPI_ISL_448794, EPI_ISL_448795, EPI_ISL_448796, EPI_ISL_448797, EPI_ISL_448799, EPI_ISL_448800, EPI_ISL_448801, EPI_ISL_448802, EPI_ISL_448803, EPI_ISL_448804, EPI_ISL_448806, EPI_ISL_448807, EPI_ISL_448808, EPI_ISL_448809, EPI_ISL_448810, EPI_ISL_448811, EPI_ISL_448812, EPI_ISL_448814, EPI_ISL_448816, EPI_ISL_448817, EPI_ISL_448818, EPI_ISL_448820, EPI_ISL_448821, EPI_ISL_448822, EPI_ISL_448824, EPI_ISL_448825 | | | |
| see above | Oxford Viromics, NDM, University of Oxford; Oxford University Hospitals; Basingstoke and North Hampshire Hospital | COVID-19 Genomics UK (COG-UK) Consortium | Tanya Golubchik, David Bonsall, George Macintyre, Amy Trebes, Mariateresa de Cesare, Catrin Moore, Alex Mobbs, Anita Justice, Robert Shaw, Monique Andersson, Emma Wise, Nathan Moore, Jessica Lynch, Nick Cortes, Stephen Kidd, David Buck, John Todd, Christophe Fraser |
| EPI_ISL_448827, EPI_ISL_448828, EPI_ISL_448834, EPI_ISL_448836, EPI_ISL_448841 | Virology Department, Sheffield Teaching Hospitals NHS Foundation Trust/Department of Infection, Immunity and Cardiovascular Disease, The Medical School, University of Sheffield | COVID-19 Genomics UK (COG-UK) Consortium | Thushan de Silva, Matthew Parker, Nikki Smith, Adri Angyal, Rebecca Brown, Luke Green, Rachel Tucker, Paul Parsons, Danielle Groves, Katie Johnson, Laura Carrilero, Alex Keeley, Dave Partridge, Matthew Wyles, Benjamin Lindsey, Mehmet Yavuz, Mohammad Raza, Cariad Evans |
| EPI_ISL_448844 | Virology Laboratory, Castle Hill Hospital, Hull University Teaching Hospitals NHS Trust/Department of Infection, Immunity and Cardiovascular Disease, The Medical School, University of Sheffield | COVID-19 Genomics UK (COG-UK) Consortium | Thushan de Silva, Matthew Parker, Nikki Smith, Adri Angyal, Rebecca Brown, Luke Green, Rachel Tucker, Paul Parsons, Danielle Groves, Katie Johnson, Laura Carrilero, Alex Keeley, Dave Partridge, Matthew Wyles, Benjamin Lindsey, Mehmet Yavuz, Mohammad Raza, Cariad Evans |
| EPI_ISL_448846, EPI_ISL_448860 | Virology Department, Sheffield Teaching Hospitals NHS Foundation Trust/Department of Infection, Immunity and Cardiovascular Disease, The Medical School, University of Sheffield | COVID-19 Genomics UK (COG-UK) Consortium | Thushan de Silva, Matthew Parker, Nikki Smith, Adri Angyal, Rebecca Brown, Luke Green, Rachel Tucker, Paul Parsons, Danielle Groves, Katie Johnson, Laura Carrilero, Alex Keeley, Dave Partridge, Matthew Wyles, Benjamin Lindsey, Mehmet Yavuz, Mohammad Raza, Cariad Evans |
| EPI_ISL_448864 | Virology Laboratory, Castle Hill Hospital, Hull University Teaching Hospitals NHS Trust/Department of Infection, Immunity and Cardiovascular Disease, The Medical School, University of Sheffield | COVID-19 Genomics UK (COG-UK) Consortium | Thushan de Silva, Matthew Parker, Nikki Smith, Adri Angyal, Rebecca Brown, Luke Green, Rachel Tucker, Paul Parsons, Danielle Groves, Katie Johnson, Laura Carrilero, Alex Keeley, Dave Partridge, Matthew Wyles, Benjamin Lindsey, Mehmet Yavuz, Mohammad Raza, Cariad Evans |
| EPI_ISL_448865 | Virology Department, Sheffield Teaching Hospitals NHS Foundation Trust/Department of Infection, Immunity and Cardiovascular Disease, The Medical School, University of Sheffield | COVID-19 Genomics UK (COG-UK) Consortium | Thushan de Silva, Matthew Parker, Nikki Smith, Adri Angyal, Rebecca Brown, Luke Green, Rachel Tucker, Paul Parsons, Danielle Groves, Katie Johnson, Laura Carrilero, Alex Keeley, Dave Partridge, Matthew Wyles, Benjamin Lindsey, Mehmet Yavuz, Mohammad Raza, Cariad Evans |
| EPI_ISL_448866 | Virology Laboratory, Castle Hill Hospital, Hull University Teaching Hospitals NHS Trust/Department of Infection, Immunity and Cardiovascular Disease, The Medical School, University of Sheffield | COVID-19 Genomics UK (COG-UK) Consortium | Thushan de Silva, Matthew Parker, Nikki Smith, Adri Angyal, Rebecca Brown, Luke Green, Rachel Tucker, Paul Parsons, Danielle Groves, Katie Johnson, Laura Carrilero, Alex Keeley, Dave Partridge, Matthew Wyles, Benjamin Lindsey, Mehmet Yavuz, Mohammad Raza, Cariad Evans |
| EPI_ISL_448867, EPI_ISL_448869, EPI_ISL_448870, EPI_ISL_448874, EPI_ISL_448877, EPI_ISL_448880, EPI_ISL_448881, EPI_ISL_448887 | Virology Department, Sheffield Teaching Hospitals NHS Foundation Trust/Department of Infection, Immunity and Cardiovascular Disease, The Medical School, University of Sheffield | COVID-19 Genomics UK (COG-UK) Consortium | Thushan de Silva, Matthew Parker, Nikki Smith, Adri Angyal, Rebecca Brown, Luke Green, Rachel Tucker, Paul Parsons, Danielle Groves, Katie Johnson, Laura Carrilero, Alex Keeley, Dave Partridge, Matthew Wyles, Benjamin Lindsey, Mehmet Yavuz, Mohammad Raza, Cariad Evans |
| EPI_ISL_448894, EPI_ISL_448896, EPI_ISL_448902 | Virology Laboratory, Castle Hill Hospital, Hull University Teaching Hospitals NHS Trust/Department of Infection, Immunity and Cardiovascular Disease, The Medical School, University of Sheffield | COVID-19 Genomics UK (COG-UK) Consortium | Thushan de Silva, Matthew Parker, Nikki Smith, Adri Angyal, Rebecca Brown, Luke Green, Rachel Tucker, Paul Parsons, Danielle Groves, Katie Johnson, Laura Carrilero, Alex Keeley, Dave Partridge, Matthew Wyles, Benjamin Lindsey, Mehmet Yavuz, Mohammad Raza, Cariad Evans |
| EPI_ISL_448903, EPI_ISL_448907, EPI_ISL_448908, EPI_ISL_448909, EPI_ISL_448913 | Virology Department, Sheffield Teaching Hospitals NHS Foundation Trust/Department of Infection, Immunity and Cardiovascular Disease, The Medical School, University of Sheffield | COVID-19 Genomics UK (COG-UK) Consortium | Thushan de Silva, Matthew Parker, Nikki Smith, Adri Angyal, Rebecca Brown, Luke Green, Rachel Tucker, Paul Parsons, Danielle Groves, Katie Johnson, Laura Carrilero, Alex Keeley, Dave Partridge, Matthew Wyles, Benjamin Lindsey, Mehmet Yavuz, Mohammad Raza, Cariad Evans |
| EPI_ISL_448918, EPI_ISL_448925, EPI_ISL_448928, EPI_ISL_448936, EPI_ISL_448939, EPI_ISL_448940, EPI_ISL_448942, EPI_ISL_448943, EPI_ISL_448945, EPI_ISL_448947, EPI_ISL_448950, EPI_ISL_448951, EPI_ISL_448955, EPI_ISL_448956, EPI_ISL_448958, EPI_ISL_448959, EPI_ISL_448961, EPI_ISL_448962, EPI_ISL_448965, EPI_ISL_448966, EPI_ISL_448967, EPI_ISL_448968, EPI_ISL_448970, EPI_ISL_448971, EPI_ISL_448972, EPI_ISL_448973, EPI_ISL_448974, EPI_ISL_448976, EPI_ISL_448977 | | | |

| | | | |
|---|---|---|---|
| see above | Regional Virus Laboratory, Belfast Health and Social Care Trust | COVID-19 Genomics UK (COG-UK) Consortium | Conall McCaughey, James McKenna, Tanya Curran, Susan Feeney, Alison Watt, Ciara Cox, Mairead Connor, Zoltan Molnar, David Simpson, Derek Fairley |
| EPI_ISL_448979, EPI_ISL_448980, EPI_ISL_448981, EPI_ISL_448982, EPI_ISL_448983, EPI_ISL_448984, EPI_ISL_448985, EPI_ISL_448986, EPI_ISL_448987, EPI_ISL_448988, EPI_ISL_448991, EPI_ISL_448992, EPI_ISL_448993, EPI_ISL_448994, EPI_ISL_448995, EPI_ISL_448996, EPI_ISL_448998, EPI_ISL_448999, EPI_ISL_449001, EPI_ISL_449002, EPI_ISL_449003, EPI_ISL_449004, EPI_ISL_449005, EPI_ISL_449006, EPI_ISL_449007, EPI_ISL_449008, EPI_ISL_449009, EPI_ISL_449010, EPI_ISL_449011, EPI_ISL_449012, EPI_ISL_449013, EPI_ISL_449015, EPI_ISL_449017, EPI_ISL_449019, EPI_ISL_449020, EPI_ISL_449021, EPI_ISL_449022, EPI_ISL_449023, EPI_ISL_449024, EPI_ISL_449025, EPI_ISL_449026, EPI_ISL_449027, EPI_ISL_449028, EPI_ISL_449029, EPI_ISL_449030, EPI_ISL_449031, EPI_ISL_449032, EPI_ISL_449034, EPI_ISL_449035, EPI_ISL_449036, EPI_ISL_449037, EPI_ISL_449038, EPI_ISL_449039, EPI_ISL_449040, EPI_ISL_449041, EPI_ISL_449042, EPI_ISL_449043, EPI_ISL_449044, EPI_ISL_449045, EPI_ISL_449046, EPI_ISL_449047, EPI_ISL_449048, EPI_ISL_449049, EPI_ISL_449050, EPI_ISL_449051, EPI_ISL_449053, EPI_ISL_449054, EPI_ISL_449055, EPI_ISL_449056, EPI_ISL_449057, EPI_ISL_449059, EPI_ISL_449060, EPI_ISL_449061, EPI_ISL_449062, EPI_ISL_449063, EPI_ISL_449065, EPI_ISL_449066, EPI_ISL_449068, EPI_ISL_449069, EPI_ISL_449070, EPI_ISL_449071, EPI_ISL_449072, EPI_ISL_449073, EPI_ISL_449074, EPI_ISL_449075, EPI_ISL_449077, EPI_ISL_449078, EPI_ISL_449079, EPI_ISL_449080, EPI_ISL_449081, EPI_ISL_449082, EPI_ISL_449083, EPI_ISL_449084, EPI_ISL_449085, EPI_ISL_449086, EPI_ISL_449087, EPI_ISL_449089, EPI_ISL_449090, EPI_ISL_449091, EPI_ISL_449092, EPI_ISL_449093, EPI_ISL_449094, EPI_ISL_449095, EPI_ISL_449096, EPI_ISL_449097, EPI_ISL_449099, EPI_ISL_449100, EPI_ISL_449101, EPI_ISL_449102, EPI_ISL_449103, EPI_ISL_449104, EPI_ISL_449105, EPI_ISL_449106, EPI_ISL_449107, EPI_ISL_449109, EPI_ISL_449110, EPI_ISL_449111, EPI_ISL_449112, EPI_ISL_449115, EPI_ISL_449116, EPI_ISL_449117, EPI_ISL_449118, EPI_ISL_449119, EPI_ISL_449121, EPI_ISL_449122, EPI_ISL_449124, EPI_ISL_449125, EPI_ISL_449126, EPI_ISL_449127, EPI_ISL_449128, EPI_ISL_449129, EPI_ISL_449130, EPI_ISL_449131, EPI_ISL_449132, EPI_ISL_449133, EPI_ISL_449135, EPI_ISL_449136, EPI_ISL_449137, EPI_ISL_449138, EPI_ISL_449139, EPI_ISL_449140, EPI_ISL_449141, EPI_ISL_449142, EPI_ISL_449143, EPI_ISL_449144, EPI_ISL_449145, EPI_ISL_449146, EPI_ISL_449148, EPI_ISL_449149, EPI_ISL_449150, EPI_ISL_449151, EPI_ISL_449152, EPI_ISL_449153, EPI_ISL_449154, EPI_ISL_449155, EPI_ISL_449156, EPI_ISL_449157, EPI_ISL_449158, EPI_ISL_449159, EPI_ISL_449160, EPI_ISL_449161, EPI_ISL_449162, EPI_ISL_449163, EPI_ISL_449164, EPI_ISL_449165, EPI_ISL_449166, EPI_ISL_449167, EPI_ISL_449168, EPI_ISL_449169, EPI_ISL_449170, EPI_ISL_449171, EPI_ISL_449172, EPI_ISL_449173, EPI_ISL_449174, EPI_ISL_449175 | | | |
| see above | Quadram Institute Bioscience | COVID-19 Genomics UK (COG-UK) Consortium | Dave J. Baker, Gemma L. Kay, Alp Aydin, Thanh Le-Viet, Steven Rudder, Ana P. Tedim, Anastasia Kolyva, Maria Diaz, Leonardo de Oliveira Martins, Nabil-Fareed Alikhan, Lizzie Meadows, Rachael Stanley, Ngozi Elumogo, Muhammed Yasir, Nicholas M. Thomson, Alexander J Trotter, Rachel Gilroy, Samuel Bloomfield, Claire Stuart, Andrew Bell, Reenesh Prakash, Samir Dervisevic, Alison E. Mather, John Wain, Mark Webber, Andrew J. Page, Justin O'Grady |
| EPI_ISL_449176, EPI_ISL_449178, EPI_ISL_449179, EPI_ISL_449187, EPI_ISL_449188, EPI_ISL_449189, EPI_ISL_449190, EPI_ISL_449191, EPI_ISL_449192, EPI_ISL_449193, EPI_ISL_449194, EPI_ISL_449195, EPI_ISL_449196, EPI_ISL_449197, EPI_ISL_449198, EPI_ISL_449199, EPI_ISL_449200, EPI_ISL_449201, EPI_ISL_449203, EPI_ISL_449204, EPI_ISL_449206, EPI_ISL_449207, EPI_ISL_449208, EPI_ISL_449209, EPI_ISL_449210, EPI_ISL_449211, EPI_ISL_449212, EPI_ISL_449213, EPI_ISL_449214, EPI_ISL_449215, EPI_ISL_449216, EPI_ISL_449217, EPI_ISL_449218, EPI_ISL_449219, EPI_ISL_449220, EPI_ISL_449221, EPI_ISL_449222, EPI_ISL_449223, EPI_ISL_449224, EPI_ISL_449225, EPI_ISL_449226, EPI_ISL_449227, EPI_ISL_449228, EPI_ISL_449229, EPI_ISL_449230, EPI_ISL_449231, EPI_ISL_449232, EPI_ISL_449233, EPI_ISL_449234, EPI_ISL_449235, EPI_ISL_449236, EPI_ISL_449237, EPI_ISL_449238, EPI_ISL_449240, EPI_ISL_449241, EPI_ISL_449242, EPI_ISL_449243, EPI_ISL_449244, EPI_ISL_449245, EPI_ISL_449246, EPI_ISL_449247, EPI_ISL_449248, EPI_ISL_449249, EPI_ISL_449250, EPI_ISL_449251, EPI_ISL_449252, EPI_ISL_449253, EPI_ISL_449254, EPI_ISL_449255, EPI_ISL_449256, EPI_ISL_449257, EPI_ISL_449258 | | | |
| see above | West of Scotland Specialist Virology Centre, NHSGGC / MRC-University of Glasgow Centre for Virus Research | COVID-19 Genomics UK (COG-UK) Consortium | Ana da Silva Filipe, Natasha Johnson, Kathy Smollett, Daniel Mair, Stephen Carmichael, Lily Tong, Jenna Nichols, Elihu Aranday-Cortes, Kirstyn Brunker, Yasmin Parr, Kyriaki Nomikou, Sarah McDonald, Marc Niebel, Patawee Asamaphan, Richard Orton, Joseph Hughes, Sreenu Vattipally, David L Robertson, Alasdair MacLean, Rory Gunson, Kathy Li, Natasha Jesudason, Rajiv Shah, James Shepherd, Antonia Ho, Emma Thomson |
| EPI_ISL_449260, EPI_ISL_449261, EPI_ISL_449264, EPI_ISL_449265, EPI_ISL_449271, EPI_ISL_449272, EPI_ISL_449274, EPI_ISL_449275, EPI_ISL_449276, EPI_ISL_449277, EPI_ISL_449278, EPI_ISL_449279, EPI_ISL_449280, EPI_ISL_449281, EPI_ISL_449282, EPI_ISL_449283, EPI_ISL_449284, EPI_ISL_449286, EPI_ISL_449288, EPI_ISL_449289, EPI_ISL_449290, EPI_ISL_449291, EPI_ISL_449292, EPI_ISL_449293, EPI_ISL_449294, EPI_ISL_449295, EPI_ISL_449296, EPI_ISL_449297, EPI_ISL_449298, EPI_ISL_449299, EPI_ISL_449300, EPI_ISL_449301, EPI_ISL_449302, EPI_ISL_449303, EPI_ISL_449306, EPI_ISL_449309, EPI_ISL_449311, EPI_ISL_449312, EPI_ISL_449313, EPI_ISL_449315, EPI_ISL_449316, EPI_ISL_449318, EPI_ISL_449319, EPI_ISL_449320, EPI_ISL_449321, EPI_ISL_449322, EPI_ISL_449323, EPI_ISL_449324, EPI_ISL_449325, EPI_ISL_449326, EPI_ISL_449327, EPI_ISL_449328, EPI_ISL_449329 | | | |
| see above | Virology Department, Royal Infirmary of Edinburgh, NHS Lothian / School of Biological Sciences, University of Edinburgh / Institute of Genetics and Molecular Medicine, University of Edinburgh | COVID-19 Genomics UK (COG-UK) Consortium | McHugh M, Dewar R, Rooke S, Gallagher M, Balcaza C, O'Toole Á, Scher E, Hill V, McCrone JT, Colquhoun R, Yu X, Jackson B, Rambaut A, Williams TC, Templeton K |
| EPI_ISL_449605, EPI_ISL_449608, EPI_ISL_449611, EPI_ISL_449614, EPI_ISL_449623 | Liverpool Clinical Laboratories | COVID-19 Genomics UK (COG-UK) Consortium | Sam Haldenby, Anita Lucaci, Steve Paterson, Julian Hiscox, Alistair Darby, M Almsaud, A Alrezaihi, Muhannad Alruwaili, Stuart D Armstrong, Jones Benjamin , Eleanor G Bentley, Anu Chawla, Jordan J Clark, Angela Cowell, Richard Eccles, Isabel García-Dorival, Matthew Gemmell, Alessandro Gerada, PKF Gilmore, Richard Gregory, Ximeng Han, Catherine Hartley, Margaret Hughes, Miren Iturriza-Gomara, James Johnson, L Luu, Jenifer Manson , Charlotte Nelson, Elaine O'Toole, Cassie Olateju, Rebekah Penrice-Randal , Lucille Rainbow, N.P Randle, Trevor Ian Robinson, Parul Sharma, Ghada T Shawli, James P Stewart , Neil Swainston, Ecaterina Vamos, Joanne Watts, Mark Whitehead |
| EPI_ISL_449626, EPI_ISL_449627, EPI_ISL_449629, EPI_ISL_449630, EPI_ISL_449631, EPI_ISL_449632, EPI_ISL_449633, EPI_ISL_449636, EPI_ISL_449637, EPI_ISL_449639, EPI_ISL_449640, EPI_ISL_449644, EPI_ISL_449645, EPI_ISL_449646, EPI_ISL_449647, EPI_ISL_449648, EPI_ISL_449649, EPI_ISL_449650, EPI_ISL_449651, EPI_ISL_449652, EPI_ISL_449653, EPI_ISL_449654, EPI_ISL_449655, EPI_ISL_449656, EPI_ISL_449657, EPI_ISL_449658, EPI_ISL_449659, EPI_ISL_449660, EPI_ISL_449661, EPI_ISL_449662, EPI_ISL_449663, EPI_ISL_449664, EPI_ISL_449665, EPI_ISL_449666, EPI_ISL_449667, EPI_ISL_449668, EPI_ISL_449669, EPI_ISL_449670, EPI_ISL_449672, EPI_ISL_449673, EPI_ISL_449674, EPI_ISL_449676, EPI_ISL_449678, EPI_ISL_449679, EPI_ISL_449680, EPI_ISL_449681, EPI_ISL_449682, EPI_ISL_449683, EPI_ISL_449685, EPI_ISL_449686, EPI_ISL_449687, EPI_ISL_449688, EPI_ISL_449689, EPI_ISL_449690, EPI_ISL_449691, EPI_ISL_449692, EPI_ISL_449694, EPI_ISL_449696, EPI_ISL_449698, EPI_ISL_449702, EPI_ISL_449703, EPI_ISL_449704, EPI_ISL_449705, EPI_ISL_449706, EPI_ISL_449707, EPI_ISL_449708, EPI_ISL_449709, EPI_ISL_449710, EPI_ISL_449711, EPI_ISL_449712, EPI_ISL_449713, EPI_ISL_449714, EPI_ISL_449715, EPI_ISL_449716, EPI_ISL_449718, EPI_ISL_449719, EPI_ISL_449720, EPI_ISL_449721, EPI_ISL_449722, EPI_ISL_449723, EPI_ISL_449724, EPI_ISL_449725, EPI_ISL_449726, EPI_ISL_449727, EPI_ISL_449728, EPI_ISL_449729 | | | |
| see above | University College London, Great Ormond Street Hospital for Children NHS Foundation Trust, Imperial College Healthcare NHS Trust | COVID-19 Genomics UK (COG-UK) Consortium | Sergi Castellano, Rachel Williams, Mark Kristiansen, Paola Resende Silva, Sunando Roy, Tony Brooks, Helena Tutill, Paola Niola, Patricia Dyal, Charlotte Williams, Leysa Forrest, Yasmin Panchbhaya, Jacqueline Findlay, Sam Weeks, Julianne Brown, Kathryn Harris, Paul Randell, James Price, Alison Holmes, Judith Breuer |
| EPI_ISL_449791, EPI_ISL_449792, EPI_ISL_449793, EPI_ISL_449794 | Dept. of Medical Microbiology, Stavanger University Hospital, Helse Stavanger HF | Norwegian Institute of Public Health, Department of Virology | Kathrine Stene-Johansen, Kamilla Heddeland Instefjord, Hilde Elshaug, Rasmus Riis Kopperud, Karoline Bragstad, Olav Hungnes |
| EPI_ISL_449799 | National Laboratory for Health, Environment and Food | National Laboratory for Health, Environment and Food | Mahnic A., Hedzet S., Janezic S., Duh D., Zavrsnik J., Blazun Vosner H., Rupnik M. |
| EPI_ISL_449801, EPI_ISL_449802, EPI_ISL_449804, EPI_ISL_449805, EPI_ISL_449806, EPI_ISL_449809, EPI_ISL_449810, EPI_ISL_449812, EPI_ISL_449815, EPI_ISL_449816, EPI_ISL_449817, EPI_ISL_449818, EPI_ISL_449819, EPI_ISL_449820, EPI_ISL_449821, EPI_ISL_449822, EPI_ISL_449823, EPI_ISL_449824, EPI_ISL_449825, EPI_ISL_449826, EPI_ISL_449827, EPI_ISL_449828, EPI_ISL_449829, EPI_ISL_449830, EPI_ISL_449831, EPI_ISL_449833, EPI_ISL_449834, EPI_ISL_449835, EPI_ISL_449836, EPI_ISL_449837, EPI_ISL_449838 | | | |
| see above | Utah Public Health Laboratory | Utah Public Health Laboratory | Erin Young, Kelly Oakeson |
| EPI_ISL_449839, EPI_ISL_449840, EPI_ISL_449841, EPI_ISL_449842, EPI_ISL_449843, EPI_ISL_449844, EPI_ISL_449845, EPI_ISL_449846, EPI_ISL_449847, EPI_ISL_449848, EPI_ISL_449849, EPI_ISL_449850, EPI_ISL_449851, EPI_ISL_449852, EPI_ISL_449853, EPI_ISL_449854, EPI_ISL_449855, EPI_ISL_449856, EPI_ISL_449857, EPI_ISL_449858, EPI_ISL_449859, EPI_ISL_449860, EPI_ISL_449861, EPI_ISL_449862, EPI_ISL_449863, EPI_ISL_449864, EPI_ISL_449865, EPI_ISL_449866, EPI_ISL_449867, EPI_ISL_449868, EPI_ISL_449869, EPI_ISL_449870, EPI_ISL_449871, EPI_ISL_449872, EPI_ISL_449873, EPI_ISL_449874, EPI_ISL_449875, EPI_ISL_449876, EPI_ISL_449877, EPI_ISL_449878, EPI_ISL_449879, EPI_ISL_449880, EPI_ISL_449881, EPI_ISL_449882, EPI_ISL_449883, EPI_ISL_449884, EPI_ISL_449885, EPI_ISL_449886, EPI_ISL_449887, EPI_ISL_449888, EPI_ISL_449889, EPI_ISL_449890, EPI_ISL_449891, EPI_ISL_449892, EPI_ISL_449894, EPI_ISL_449895, EPI_ISL_449896, EPI_ISL_449897, EPI_ISL_449898, EPI_ISL_449899, EPI_ISL_449900, EPI_ISL_449901, EPI_ISL_449902, EPI_ISL_449903, EPI_ISL_449904, EPI_ISL_449905, EPI_ISL_449906, EPI_ISL_449907, EPI_ISL_449908, EPI_ISL_449909, EPI_ISL_449910, EPI_ISL_449911, EPI_ISL_449912, EPI_ISL_449913, EPI_ISL_449914, EPI_ISL_449915, EPI_ISL_449917, EPI_ISL_449918, EPI_ISL_449919, EPI_ISL_449920, EPI_ISL_449921, EPI_ISL_449922, EPI_ISL_449923, EPI_ISL_449925, EPI_ISL_449926, EPI_ISL_449927, EPI_ISL_449928, EPI_ISL_449929, EPI_ISL_449930, EPI_ISL_449931, EPI_ISL_449932, EPI_ISL_449933, EPI_ISL_449934, EPI_ISL_449935, EPI_ISL_449936, EPI_ISL_449937, EPI_ISL_449938, EPI_ISL_449939, EPI_ISL_449940, EPI_ISL_449941, EPI_ISL_449943, EPI_ISL_449944, EPI_ISL_449945, EPI_ISL_449946, EPI_ISL_449947, EPI_ISL_449948, EPI_ISL_449949, EPI_ISL_449951, EPI_ISL_449952, EPI_ISL_449954, EPI_ISL_449955, EPI_ISL_449956, EPI_ISL_449957, EPI_ISL_449958, EPI_ISL_449959, EPI_ISL_449960, EPI_ISL_449962, EPI_ISL_449963, EPI_ISL_449964, EPI_ISL_449965, EPI_ISL_449966, EPI_ISL_449967, EPI_ISL_449968, EPI_ISL_449969, EPI_ISL_449970, EPI_ISL_449971, EPI_ISL_449972, EPI_ISL_449973, EPI_ISL_449974, EPI_ISL_449975, EPI_ISL_449977, EPI_ISL_449978, EPI_ISL_449979, EPI_ISL_449980, EPI_ISL_449981, EPI_ISL_449982, EPI_ISL_449983, EPI_ISL_449984, EPI_ISL_449985, EPI_ISL_449986, EPI_ISL_449987, EPI_ISL_449988, EPI_ISL_449989, EPI_ISL_449990, EPI_ISL_449991, EPI_ISL_449992, EPI_ISL_449994, EPI_ISL_449995, EPI_ISL_449996, EPI_ISL_449997, EPI_ISL_449998, EPI_ISL_449999, EPI_ISL_450000, EPI_ISL_450001, EPI_ISL_450002, EPI_ISL_450003 | | | |
| see above | Washington State Department of Health | Seattle Flu Study | Chu et al |
| EPI_ISL_450004, EPI_ISL_450005, EPI_ISL_450006, EPI_ISL_450007, EPI_ISL_450008, EPI_ISL_450009, EPI_ISL_450010, EPI_ISL_450011, EPI_ISL_450012, EPI_ISL_450013, EPI_ISL_450014, EPI_ISL_450015, EPI_ISL_450016, EPI_ISL_450017, EPI_ISL_450018, EPI_ISL_450019, EPI_ISL_450020, EPI_ISL_450021, EPI_ISL_450022, EPI_ISL_450023, EPI_ISL_450024, EPI_ISL_450025, EPI_ISL_450026, EPI_ISL_450027, EPI_ISL_450028, EPI_ISL_450029, EPI_ISL_450030, EPI_ISL_450031, EPI_ISL_450032, EPI_ISL_450033, EPI_ISL_450034, EPI_ISL_450035, EPI_ISL_450036, EPI_ISL_450037, EPI_ISL_450038, EPI_ISL_450039, EPI_ISL_450040, EPI_ISL_450041, EPI_ISL_450042, EPI_ISL_450043, EPI_ISL_450044, EPI_ISL_450045, EPI_ISL_450046, EPI_ISL_450047, EPI_ISL_450048, EPI_ISL_450049, EPI_ISL_450050, EPI_ISL_450051, EPI_ISL_450052, EPI_ISL_450053, EPI_ISL_450054, EPI_ISL_450055, EPI_ISL_450056, EPI_ISL_450058, EPI_ISL_450059, EPI_ISL_450060, EPI_ISL_450061, EPI_ISL_450062, EPI_ISL_450063, EPI_ISL_450064, EPI_ISL_450065, EPI_ISL_450066, EPI_ISL_450067, EPI_ISL_450068, EPI_ISL_450069, EPI_ISL_450070, EPI_ISL_450072, EPI_ISL_450074, EPI_ISL_450075, EPI_ISL_450076, EPI_ISL_450077, EPI_ISL_450078, EPI_ISL_450079, EPI_ISL_450080, EPI_ISL_450081, EPI_ISL_450082, EPI_ISL_450086, EPI_ISL_450087, EPI_ISL_450089, EPI_ISL_450090, EPI_ISL_450091, EPI_ISL_450092, EPI_ISL_450093, EPI_ISL_450094, EPI_ISL_450095, EPI_ISL_450096, EPI_ISL_450097, EPI_ISL_450098, EPI_ISL_450099, EPI_ISL_450100, EPI_ISL_450101, EPI_ISL_450102, EPI_ISL_450103, EPI_ISL_450104, EPI_ISL_450105, EPI_ISL_450106, EPI_ISL_450107, EPI_ISL_450108, EPI_ISL_450109, EPI_ISL_450110, EPI_ISL_450111, EPI_ISL_450112, EPI_ISL_450113, EPI_ISL_450114, EPI_ISL_450115, EPI_ISL_450116, EPI_ISL_450117, EPI_ISL_450118, EPI_ISL_450120, EPI_ISL_450121, EPI_ISL_450123, EPI_ISL_450124, EPI_ISL_450125, EPI_ISL_450126, EPI_ISL_450127, EPI_ISL_450128, EPI_ISL_450129, EPI_ISL_450130, EPI_ISL_450132, EPI_ISL_450133, EPI_ISL_450134, EPI_ISL_450135, EPI_ISL_450136, EPI_ISL_450137, EPI_ISL_450138, EPI_ISL_450139, EPI_ISL_450140, EPI_ISL_450141, EPI_ISL_450142, EPI_ISL_450143, EPI_ISL_450144, EPI_ISL_450145, EPI_ISL_450146, EPI_ISL_450147, EPI_ISL_450148, EPI_ISL_450149, EPI_ISL_450150, EPI_ISL_450151, EPI_ISL_450152, EPI_ISL_450153, EPI_ISL_450154, EPI_ISL_450155, EPI_ISL_450156, EPI_ISL_450157, EPI_ISL_450158, EPI_ISL_450159, EPI_ISL_450160, EPI_ISL_450161, EPI_ISL_450162, EPI_ISL_450163, EPI_ISL_450164, EPI_ISL_450165, EPI_ISL_450166, EPI_ISL_450167, EPI_ISL_450168, EPI_ISL_450169, EPI_ISL_450170 | | | |
| see above | MSHS Clinical Microbiology Laboratories | MSHS Pathogen Surveillance Program | Ana S. Gonzalez-Reiche, Mitchell Sullivan, Ajay Obla, Gopi Patel, Emilia Sordillo, Melissa Gitman, Alberto Paniz-mondolfi, Matthew Hernandez, Shelcie Fabre, Jose Polanco, Zenab Khan, Bremy Alburquerque, Jayeeta Dutta, Juan Soto, Shwetha Sridhar Hara, Ying-Chih Wang, Melissa Smith, Robert Sebra, Lisa Miorin, Wen-chun Liu, Randy Albrecht, Judith Aberg, Florian Krammer, Adolfo Garcia-Sastre, Viviana Simon, Harm van Bakel |
| EPI_ISL_450173, EPI_ISL_450174, EPI_ISL_450175, EPI_ISL_450176, EPI_ISL_450177, EPI_ISL_450178, EPI_ISL_450179, EPI_ISL_450180, EPI_ISL_450181, EPI_ISL_450182, EPI_ISL_450184, EPI_ISL_450185 | | | |
| see above | Robert Garry lab | Andersen lab at Scripps Research | Allison Smither, Gilberto Sabino-Santos, Patricia Snarski, Lilia Melnik, Antoinette Bell, Kaylynn Genemaras, Arnaud Drouin, Dahlene Fusco, Robert Garry with SEARCH Alliance San Diego |
| EPI_ISL_450186, EPI_ISL_450187, EPI_ISL_450188 | Biolab Diagnostic Laboratories | Andersen lab at Scripps Research | Issa Abu-Dayyeh, Ahmad Tibi, Lama Hussein, Lina Mohammad, Zein Naber, Amid Abdelnour with SEARCH Alliance San Diego |
| EPI_ISL_450190 | Rady's Childrens Hospital | Andersen lab at Scripps Research | SEARCH Alliance San Diego |
| EPI_ISL_450191 | Scripps Medical Laboratory | Andersen lab at Scripps Research | SEARCH Alliance San Diego with Michael Quigley, Ellen Stefanski, Ian Mchardy |
| EPI_ISL_450193, EPI_ISL_450194 | SA Pathology | SA Pathology | Lex Leong |
| EPI_ISL_450196 | bumrungrad international hospital | National Institute of Health. Department of medical Sciences, Ministry of Public Health, Thailand | Pilailuk,Okada; Siripaporn,Phuygun; Thanutsapa,Thanadachakul; Sittiporn,Parnmen;Warawan,Wongboot; Sunthareeya,Waicharoen; Malinee,Chittaganpitch |
| EPI_ISL_450197 | National Institute of Health. Department of medical Sciences, Ministry of Public Health, Thailand | National Institute of Health. Department of medical Sciences, Ministry of Public Health, Thailand | Pilailuk,Okada; Siripaporn,Phuygun; Thanutsapa,Thanadachakul; Sittiporn,Parnmen;Warawan,Wongboot; Sunthareeya,Waicharoen; Malinee,Chittaganpitch |
| EPI_ISL_450231 | Robert Garry lab | Andersen lab at Scripps Research | Allison Smither, Gilberto Sabino-Santos, Patricia Snarski, Lilia Melnik, Antoinette Bell, Kaylynn Genemaras, Arnaud Drouin, Dahlene Fusco, Robert Garry with SEARCH Alliance San Diego |
| EPI_ISL_450232, EPI_ISL_450233, EPI_ISL_450235, EPI_ISL_450236, EPI_ISL_450237, EPI_ISL_450238, EPI_ISL_450239, EPI_ISL_450240 | UCSF Clinical Microbiology Laboratory | Chiu Laboratory, University of California, San Francisco | Xianding Deng, Scot Federman, Wei Gu, and Charles Y. Chiu |
| EPI_ISL_450241, EPI_ISL_450242, EPI_ISL_450243, EPI_ISL_450244, EPI_ISL_450245, EPI_ISL_450246, EPI_ISL_450247, EPI_ISL_450248, EPI_ISL_450249, EPI_ISL_450250, EPI_ISL_450251, EPI_ISL_450252, EPI_ISL_450253, EPI_ISL_450254, EPI_ISL_450255, EPI_ISL_450256, EPI_ISL_450257, EPI_ISL_450258, EPI_ISL_450259, EPI_ISL_450260, EPI_ISL_450261, EPI_ISL_450262, EPI_ISL_450263, EPI_ISL_450264, EPI_ISL_450265, EPI_ISL_450266, EPI_ISL_450267, EPI_ISL_450268, EPI_ISL_450269, EPI_ISL_450270, EPI_ISL_450271, EPI_ISL_450272, EPI_ISL_450273, EPI_ISL_450274, EPI_ISL_450275, EPI_ISL_450276, EPI_ISL_450277, EPI_ISL_450278, EPI_ISL_450279, EPI_ISL_450280, EPI_ISL_450281, EPI_ISL_450282, EPI_ISL_450283, EPI_ISL_450284, EPI_ISL_450285, EPI_ISL_450286, EPI_ISL_450287, EPI_ISL_450288, EPI_ISL_450289, EPI_ISL_450290, EPI_ISL_450291, EPI_ISL_450292, EPI_ISL_450293 | | | |
| see above | WHO National Influenza Centre Russian Federation | WHO National Influenza Centre Russian Federation | Andrey Komissarov, Artem Fadeev, Mariia Sergeeva, Anna Ivanova, Tamila Musaeva, Ksenia Komissarova, Mariia Timofeeva, Veronica Eder, Mariia Pisareva, Daria Danilenko |
| EPI_ISL_450294 | Institute of Human Genetics, Polish Academy of Sciences Sanitary and Epidemiological Station in Poznań | Institute of Human Genetics, Polish Academy of Sciences | Szymon Hryhorowicz, Adam Ustaszewski, Emilia Lis, Marta Kaczmarek-Ryś, Michał Witt, Andrzej Pławski |
| EPI_ISL_450295 | Institute of Human Genetics, Polish Academy of Sciences; Sanitary and Epidemiological Station in Poznań | Institute of Human Genetics, Polish Academy of Sciences | Szymon Hryhorowicz, Adam Ustaszewski, Emilia Lis, Marta Kaczmarek-Ryś, Michał Witt, Andrzej Pławski |
| EPI_ISL_450302, EPI_ISL_450303 | Centre hospitalier Anna-Laberge | Laboratoire de santé publique du Québec | Sandrine Moreira, Ioannis Ragoussis, Guillaume Bourque, Jesse Shapiro, Mark Lathrop and Michel Roger on behalf of the CoVSeQ research group (http://covseq.ca/researchgroup) |
| EPI_ISL_450304, EPI_ISL_450305 | Hôpital Charles-LeMoyne | Laboratoire de santé publique du Québec | Sandrine Moreira, Ioannis Ragoussis, Guillaume Bourque, Jesse Shapiro, Mark Lathrop and Michel Roger on behalf of the CoVSeQ research group (http://covseq.ca/researchgroup) |
| EPI_ISL_450306 | CSSS Haut-Richelieu/Rouville (Hôpital) | Laboratoire de santé publique du Québec | Sandrine Moreira, Ioannis Ragoussis, Guillaume Bourque, Jesse Shapiro, Mark Lathrop and Michel Roger on behalf of the CoVSeQ research group (http://covseq.ca/researchgroup) |
| EPI_ISL_450307 | Hôpital du Suroît | Laboratoire de santé publique du Québec | Sandrine Moreira, Ioannis Ragoussis, Guillaume Bourque, Jesse Shapiro, Mark Lathrop and Michel Roger on behalf of the CoVSeQ research group (http://covseq.ca/researchgroup) |
| EPI_ISL_450308 | Hôpital Charles-LeMoyne | Laboratoire de santé publique du Québec | Sandrine Moreira, Ioannis Ragoussis, Guillaume Bourque, Jesse Shapiro, Mark Lathrop and Michel Roger on behalf of the CoVSeQ research group (http://covseq.ca/researchgroup) |
| EPI_ISL_450309 | Hôpital Pierre-Boucher | Laboratoire de santé publique du Québec | Sandrine Moreira, Ioannis Ragoussis, Guillaume Bourque, Jesse Shapiro, Mark Lathrop and Michel Roger on behalf of the CoVSeQ research group (http://covseq.ca/researchgroup) |
| EPI_ISL_450310 | Hôpital Charles-LeMoyne | Laboratoire de santé publique du Québec | Sandrine Moreira, Ioannis Ragoussis, Guillaume Bourque, Jesse Shapiro, Mark Lathrop and Michel Roger on behalf of the CoVSeQ research group (http://covseq.ca/researchgroup) |
| EPI_ISL_450311, EPI_ISL_450312, EPI_ISL_450313 | Hôpital du Suroît | Laboratoire de santé publique du Québec | Sandrine Moreira, Ioannis Ragoussis, Guillaume Bourque, Jesse Shapiro, Mark Lathrop and Michel Roger on behalf of the CoVSeQ research group (http://covseq.ca/researchgroup) |
| EPI_ISL_450314, EPI_ISL_450315 | Hôpital Pierre-Boucher | Laboratoire de santé publique du Québec | Sandrine Moreira, Ioannis Ragoussis, Guillaume Bourque, Jesse Shapiro, Mark Lathrop and Michel Roger on behalf of the CoVSeQ research group (http://covseq.ca/researchgroup) |

| | | | |
|---|---|---|---|
| EPI_ISL_450316 | Hôpital Charles-LeMoyne | Laboratoire de santé publique du Québec | Sandrine Moreira, Ioannis Ragoussis, Guillaume Bourque, Jesse Shapiro, Mark Lathrop and Michel Roger on behalf of the CoVSeQ research group (http://covseq.ca/researchgroup) |
| EPI_ISL_450317, EPI_ISL_450318 | Hôpital Pierre-Boucher | Laboratoire de santé publique du Québec | Sandrine Moreira, Ioannis Ragoussis, Guillaume Bourque, Jesse Shapiro, Mark Lathrop and Michel Roger on behalf of the CoVSeQ research group (http://covseq.ca/researchgroup) |
| EPI_ISL_450321 | NIV Pune | CSIR-Centre for Cellular and Molecular Biology | Dr V A Potdar, Dr ML Choudhary,Dr Priya Abraham,V. Vipat, S. Jadhav, U. Saha, H. Kengle, A. Awhale, A. Jagtap, A. Gondhalikar, V Malik, N Srivastava, S. Digraskar, P. Malsane, S. Hundekar, K. Patel, Yogesh Balakartik, M. Kakade, S. Jadhav, R. Gunjikar, V. Awtade, S. Bhorekar, P Shinde, S. Salve, B. Minhas S. Bharadwaj, H Kaushal Y. Gurav, S. Tomar,Payel Mukherjee, Sofia Banu, Priya Singh, Dhiviya Vedagiri, Divya Gupta, Vishal Sah, Santosh Kumar Kuncha, Krishnan Harinivas Harshan, Archana Bharadwaj Siva, Karthik Bharadwaj Tallapaka, Shagufta Khan, Lamuk Zaveri, Namami Gaur, Sakshi Shambhavi, Tulasi Nagabandi, Purushotham Vodnala,G. Aditya Kumar, Koushick Sivakumar, Pooja Ramesh Gupta, Rajan Kumar Jha, Shraddha Vijay Lahoti, Deepak Kumar, Devi Prasad Vijayashankara, Disha Nanda, Divya Das, Jotin Gogoi, Manish |
| EPI_ISL_450322 | NIV Pune | CSIR-Centre for Cellular and Molecular Biology | Dr V A Potdar, Dr ML Choudhary,Dr Priya Abraham,V. Vipat, S. Jadhav, U. Saha, H. Kengle, A. Awhale, A. Jagtap, A. Gondhalikar, V Malik, N Srivastava, S. Digraskar, P. Malsane, S. Hundekar, K. Patel, Yogesh Balakartik, M. Kakade, S. Jadhav, R. Gunjikar, V. Awtade, S. Bhorekar, P Shinde, S. Salve, B. Minhas S. Bharadwaj, H Kaushal Y. Gurav, S. Tomar,Sofia Banu, Payel Mukherjee, Priya Singh, Dhiviya Vedagiri, Divya Gupta, Vishal Sah, Santosh Kumar Kuncha, Krishnan Harinivas Harshan, Archana Bharadwaj Siva, Karthik Bharadwaj Tallapaka, Shagufta Khan, Lamuk Zaveri, Namami Gaur, Sakshi Shambhavi, Tulasi Nagabandi, Purushotham Vodnala, Disha Nanda, Divya Das, Jotin Gogoi, Manish Bhattacharjee, Ravi Prasad Mukku, Renu Sudhakar, Somesh Gorde, Gangumala Srinivas Reddy, Sujoy Deb, Swati Bayyana, Zeba Rizvi, Rakesh K Mishra |
| EPI_ISL_450323 | NIV Pune | CSIR-Centre for Cellular and Molecular Biology | Dr V A Potdar, Dr ML Choudhary,Dr Priya Abraham,V. Vipat, S. Jadhav, U. Saha, H. Kengle, A. Awhale, A. Jagtap, A. Gondhalikar, V Malik, N Srivastava, S. Digraskar, P. Malsane, S. Hundekar, K. Patel, Yogesh Balakartik, M. Kakade, S. Jadhav, R. Gunjikar, V. Awtade, S. Bhorekar, P Shinde, S. Salve, B. Minhas S. Bharadwaj, H Kaushal Y. Gurav, S. Tomar,Payel Mukherjee, Sofia Banu, Priya Singh, Dhiviya Vedagiri, Divya Gupta, Vishal Sah, Santosh Kumar Kuncha, Krishnan Harinivas Harshan, Archana Bharadwaj Siva, Karthik Bharadwaj Tallapaka, Shagufta Khan, Lamuk Zaveri, Namami Gaur, Sakshi Shambhavi, Tulasi Nagabandi, Purushotham Vodnala,G. Aditya Kumar, Koushick Sivakumar, Pooja Ramesh Gupta, Rajan Kumar Jha, Shraddha Vijay Lahoti, Deepak Kumar, Devi Prasad Vijayashankara, Disha Nanda, Divya Das, Jotin Gogoi, Manish |
| EPI_ISL_450324 | NIV Pune | CSIR-Centre for Cellular and Molecular Biology | Dr V A Potdar, Dr ML Choudhary,Dr Priya Abraham,V. Vipat, S. Jadhav, U. Saha, H. Kengle, A. Awhale, A. Jagtap, A. Gondhalikar, V Malik, N Srivastava, S. Digraskar, P. Malsane, S. Hundekar, K. Patel, Yogesh Balakartik, M. Kakade, S. Jadhav, R. Gunjikar, V. Awtade, S. Bhorekar, P Shinde, S. Salve, B. Minhas S. Bharadwaj, H Kaushal Y. Gurav, S. Tomar,Sofia Banu, Payel Mukherjee, Priya Singh, Dhiviya Vedagiri, Divya Gupta, Vishal Sah, Santosh Kumar Kuncha, Krishnan Harinivas Harshan, Archana Bharadwaj Siva, Karthik Bharadwaj Tallapaka, Shagufta Khan, Lamuk Zaveri, Namami Gaur, Sakshi Shambhavi, Tulasi Nagabandi, Purushotham Vodnala, Disha Nanda, Divya Das, Jotin Gogoi, Manish Bhattacharjee, Ravi Prasad Mukku, Renu Sudhakar, Somesh Gorde, Gangumala Srinivas Reddy, Sujoy Deb, Swati Bayyana, Zeba Rizvi, Rakesh K Mishra |
| EPI_ISL_450325 | NIV Pune | CSIR-Centre for Cellular and Molecular Biology | Dr V A Potdar, Dr ML Choudhary,Dr Priya Abraham,V. Vipat, S. Jadhav, U. Saha, H. Kengle, A. Awhale, A. Jagtap, A. Gondhalikar, V Malik, N Srivastava, S. Digraskar, P. Malsane, S. Hundekar, K. Patel, Yogesh Balakartik, M. Kakade, S. Jadhav, R. Gunjikar, V. Awtade, S. Bhorekar, P Shinde, S. Salve, B. Minhas S. Bharadwaj, H Kaushal Y. Gurav, S. Tomar,Payel Mukherjee, Sofia Banu, Priya Singh, Dhiviya Vedagiri, Divya Gupta, Vishal Sah, Santosh Kumar Kuncha, Krishnan Harinivas Harshan, Archana Bharadwaj Siva, Karthik Bharadwaj Tallapaka, Shagufta Khan, Lamuk Zaveri, Namami Gaur, Sakshi Shambhavi, Tulasi Nagabandi, Purushotham Vodnala,G. Aditya Kumar, Koushick Sivakumar, Pooja Ramesh Gupta, Rajan Kumar Jha, Shraddha Vijay Lahoti, Deepak Kumar, Devi Prasad Vijayashankara, Disha Nanda, Divya Das, Jotin Gogoi, Manish |
| EPI_ISL_450326 | CSIR-Centre for Cellular and Molecular Biology | CSIR-Centre for Cellular and Molecular Biology | Payel Mukherjee, Sofia Banu, Priya Singh, Dhiviya Vedagiri, Divya Gupta, Vishal Sah, Santosh Kumar Kuncha, Krishnan Harinivas Harshan, Archana Bharadwaj Siva, Karthik Bharadwaj Tallapaka, Shagufta Khan, Lamuk Zaveri, Namami Gaur, Sakshi Shambhavi, Tulasi Nagabandi, Purushotham Vodnala,G. Aditya Kumar, Koushick Sivakumar, Pooja Ramesh Gupta, Rajan Kumar Jha, Shraddha Vijay Lahoti, Deepak Kumar, Devi Prasad Vijayashankara, Disha Nanda, Divya Das, Jotin Gogoi, Manish Bhattacharjee, Rakesh K Mishra, Divya Tej Sowpati |
| EPI_ISL_450327 | CSIR-Centre for Cellular and Molecular Biology | CSIR-Centre for Cellular and Molecular Biology | Sofia Banu, Payel Mukherjee, Priya Singh, Dhiviya Vedagiri, Divya Gupta, Vishal Sah, Santosh Kumar Kuncha, Krishnan Harinivas Harshan, Archana Bharadwaj Siva, Karthik Bharadwaj Tallapaka, Shagufta Khan, Lamuk Zaveri, Namami Gaur, Sakshi Shambhavi, Tulasi Nagabandi, Purushotham Vodnala, Disha Nanda, Divya Das, Jotin Gogoi, Manish Bhattacharjee, Ravi Prasad Mukku, Renu Sudhakar, Somesh Gorde, Gangumala Srinivas Reddy, Sujoy Deb, Swati Bayyana, Zeba Rizvi, Rakesh K Mishra, Divya Tej Sowpati |
| EPI_ISL_450328 | CSIR-Centre for Cellular and Molecular Biology | CSIR-Centre for Cellular and Molecular Biology | Shagufta Khan, Lamuk Zaveri, Namami Gaur, Sakshi Shambhavi, Tulasi Nagabandi, Purushotham Vodnala, Payel Mukherjee, Sofia Banu, Priya Singh, Dhiviya Vedagiri, Divya Gupta, Vishal Sah, Santosh Kumar Kuncha, Krishnan Harinivas Harshan, Archana Bharadwaj Siva, Karthik Bharadwaj Tallapaka, Zeba Rizvi, Zuberwasim Sayyad, Kakade Aishwarya Arun, Amrutha H C, Ananga Ghosh, Kezia J Ann, Radhika Khandelwal, Roshan Maku Venkata, Shemin Mansuri, Sonu Uday, Sudipta Mondal, Rakesh K Mishra, Divya Tej Sowpati |
| EPI_ISL_450329 | CSIR-Centre for Cellular and Molecular Biology | CSIR-Centre for Cellular and Molecular Biology | Namami Gaur, Sakshi Shambhavi, Lamuk Zaveri, Shagufta Khan, Tulasi Nagabandi, Purushotham Vodnala, Payel Mukherjee, Sofia Banu, Priya Singh, Dhiviya Vedagiri, Divya Gupta, Vishal Sah, Santosh Kumar Kuncha, Krishnan Harinivas Harshan, Archana Bharadwaj Siva, Karthik Bharadwaj Tallapaka, Sonu Uday, Sudipta Mondal, Annapoorna P Karthyayani, Debabrata Jana, Debrya Saha, Gokulan C G, Gunjan Purohit, Hanuman Tulashiram Kale, Pankaj Kumar, Prachand Issarapu, Preethi Jampala Rakesh K Mishra, Divya Tej Sowpati |
| EPI_ISL_450330 | CSIR-Centre for Cellular and Molecular Biology | CSIR-Centre for Cellular and Molecular Biology | Sakshi Shambhavi, Lamuk Zaveri, Shagufta Khan, Namami Gaur, Tulasi Nagabandi, Purushotham Vodnala, Payel Mukherjee, Sofia Banu, Priya Singh, Dhiviya Vedagiri, Divya Gupta, Vishal Sah, Santosh Kumar Kuncha, Krishnan Harinivas Harshan, Archana Bharadwaj Siva, Karthik Bharadwaj Tallapaka,Preethi Jampala, Sharada Ravi Iyer, Sulagana Mukherjee, Swetha Sundar, Peddapuvala Sai Uday Kiran, Umesh Kumar, Unis Ahmad Bhat, Ajay Sarawagi, Priyanka Pant, Rajkanwar Nathawat, Nikhil Hajirnis, Pratheusa Maccha, M Soujanya Reddy Rakesh K Mishra, Divya Tej Sowpati |
| EPI_ISL_450331 | CSIR-Centre for Cellular and Molecular Biology | CSIR-Centre for Cellular and Molecular Biology | Tulasi Nagabandi, Namami Gaur, Sakshi Shambhavi, Lamuk Zaveri, Shagufta Khan, Purushotham Vodnala, Payel Mukherjee, Sofia Banu, Priya Singh, Dhiviya Vedagiri, Divya Gupta, Vishal Sah, Santosh Kumar Kuncha, Krishnan Harinivas Harshan, Archana Bharadwaj Siva, Karthik Bharadwaj Tallapaka,G. Aditya Kumar, Koushick Sivakumar, Pooja Ramesh Gupta, Rajan Kumar Jha, Shraddha Vijay Lahoti, Deepak Kumar, Devi Prasad Vijayashankara, Disha Nanda, Divya Das, Jotin Gogoi, Manish Bhattacharjee, Rakesh K Mishra, Divya Tej Sowpati |
| EPI_ISL_450332 | CSIR-Centre for Cellular and Molecular Biology | CSIR-Centre for Cellular and Molecular Biology | Payel Mukherjee, Sofia Banu, Priya Singh, Dhiviya Vedagiri, Divya Gupta, Vishal Sah, Santosh Kumar Kuncha, Krishnan Harinivas Harshan, Archana Bharadwaj Siva, Karthik Bharadwaj Tallapaka, Shagufta Khan, Lamuk Zaveri, Namami Gaur, Sakshi Shambhavi, Tulasi Nagabandi, Purushotham Vodnala,G. Aditya Kumar, Koushick Sivakumar, Pooja Ramesh Gupta, Rajan Kumar Jha, Shraddha Vijay Lahoti, Deepak Kumar, Devi Prasad Vijayashankara, Disha Nanda, Divya Das, Jotin Gogoi, Manish Bhattacharjee, Rakesh K Mishra, Divya Tej Sowpati |
| EPI_ISL_450337 | Hospital Universitari Vall d'Hebron - Vall d'Hebron Institut de Recerca | Hospital Universitari Vall d'Hebron | Cristina Andrés, Maria Piñana, Damir Garcia-Cehic, Mercedes Guerrero-Murillo, Ariadna Rando, Juliana Esperalba, Maria Gema Codina, Maria Carmen Martín, Tomàs Pumarola, Josep Quer, Andrés Antón |
| EPI_ISL_450338 | Institute of Human Genetics, Polish Academy of Sciences; Sanitary and Epidemiological Station in Poznań | Institute of Human Genetics, Polish Academy of Sciences | Szymon Hryhorowicz, Adam Ustaszewski, Emilia Lis, Marta Kaczmarek-Ryś, Michał Witt, Andrzej Pławski |
| EPI_ISL_450339 | Bangladesh Institute of Tropical & Infectious Diseases, COVID-19 Testing Laboratory | Basic and Applied Research on Jute Project | Rasel Ahmed, Md. Sabbir Hossain, Shah Md Tamim Kabir, Emdadul Mannan Emdad, Md. Nazmul Haq Rony, Eaftekhar Ahmed Rana, Paritous Kumar Biswas, M A Hassan Chowdhury, Md. Shakeel Ahmed, Md. Samiul Haque, Md. Monjurul Alam, Md. Sharifur Rahman, A S M Anwarul Huq, Md. Shahidul Islam, Goutam Buddha Das, AMAM Zonaed Siddiki |
| EPI_ISL_450341 | Bangladesh Institute of Tropical & Infectious Diseases, COVID-19 Testing Laboratory | Basic and Applied Research on Jute Project | Md. Sabbir Hossain, Rasel Ahmed, Shah Md Tamim Kabir, Emdadul Mannan Emdad, Md. Nazmul Haq Rony, Eaftekhar Ahmed Rana, Paritous Kumar Biswas, M A Hassan Chowdhury, Md. Shakeel Ahmed, Md. Samiul Haque, Md. Monjurul Alam, Md. Sharifur Rahman, A S M Anwarul Huq, Md. Shahidul Islam, Goutam Buddha Das, AMAM Zonaed Siddiki |
| EPI_ISL_450343 | Bangladesh Institute of Tropical & Infectious Diseases, COVID-19 Testing Laboratory | Basic and Applied Research on Jute Project | Md. Sabbir Hossain, Rasel Ahmed, Shah Md Tamim Kabir, Emdadul Mannan Emdad, Md. Nazmul Haq Rony, Eaftekhar Ahmed Rana, Paritous Kumar Biswas, M A Hassan Chowdhury, Md. Shakeel Ahmed, Md. Samiul Haque, Md. Monjurul Alam, Md. Sharifur Rahman, A S M Anwarul Huq, Md. Shahidul Islam, Goutam Buddha Das, AMAM Zonaed Siddiki |
| EPI_ISL_450344 | Bangladesh Institute of Tropical & Infectious Diseases, COVID-19 Testing Laboratory | Basic and Applied Research on Jute Project | Rasel Ahmed, Md. Sabbir Hossain, Shah Md Tamim Kabir, Emdadul Mannan Emdad, Md. Nazmul Haq Rony, Eaftekhar Ahmed Rana, Paritous Kumar Biswas, M A Hassan Chowdhury, Md. Shakeel Ahmed, Md. Samiul Haque, Md. Monjurul Alam, Md. Sharifur Rahman, A S M Anwarul Huq, Md. Shahidul Islam, Goutam Buddha Das, AMAM Zonaed Siddiki |
| EPI_ISL_450345 | Bangladesh Institute of Tropical & Infectious Diseases, COVID-19 Testing Laboratory | Basic and Applied Research on Jute Project | Md. Sabbir Hossain, Rasel Ahmed, Shah Md Tamim Kabir, Emdadul Mannan Emdad, Md. Nazmul Haq Rony, Eaftekhar Ahmed Rana, Paritous Kumar Biswas, M A Hassan Chowdhury, Md. Shakeel Ahmed, Md. Samiul Haque, Md. Monjurul Alam, Md. Sharifur Rahman, A S M Anwarul Huq, Md. Shahidul Islam, Goutam Buddha Das, AMAM Zonaed Siddiki |
| EPI_ISL_450346, EPI_ISL_450347, EPI_ISL_450348, EPI_ISL_450349, EPI_ISL_450350, EPI_ISL_450351, EPI_ISL_450352 | St.Olavs hospital/NTNU | Institute of Genomics Core Facility, University of Tartu | Aleksandr Ianevski, Tuuli Reisberg, Janne-Fossum Malmring, Svein Arne Nordbø, Denis Kainov |
| EPI_ISL_450393, EPI_ISL_450394, EPI_ISL_450396, EPI_ISL_450397, EPI_ISL_450398, EPI_ISL_450399, EPI_ISL_450400, EPI_ISL_450401, EPI_ISL_450402 | NYU Langone Health | Departments of Pathology and Medicine, New York University School of Medicine | Maria Aguero-Rosenfeld, Brendan Belovarac, Margaret Black, Ludovic Boytard, John Cadley, Paolo Cotzia, John Chen, Dacia Dimartino, Xiaojun Feng, Tatyana Gindin, Emily Guzman, Adriana Heguy, Megan Hogan, Emily Huang, George Jour, Alireza Khodadadi-Jamayran, Lawrence H. Lin, Raven Luther, Andrew Lytle, Christian Marier, Matthew T. Maurano, Mark J. Mulligan, Peter Meyn, Raquel Ordonez Ciriza, Iman Osman, Jared Pinnell, Vanessa Raabe, Sitharam Ramaswami, Amy Rapkiewicz, Andre M. Ribeiro-dos-Santos, Marie Samanovic-Golden, Antonio Serrano, Guomiao Shen, Matija Snuderl, Theodore Vougiouklakis, Nick Vulpescu, Gael Westby, Paul Zappile, Yutong Zhang |
| EPI_ISL_450445, EPI_ISL_450446, EPI_ISL_450447, EPI_ISL_450448, EPI_ISL_450449, EPI_ISL_450450, EPI_ISL_450451, EPI_ISL_450452, EPI_ISL_450453, EPI_ISL_450454, EPI_ISL_450455, EPI_ISL_450456, EPI_ISL_450457, EPI_ISL_450458, EPI_ISL_450459, EPI_ISL_450460, EPI_ISL_450461, EPI_ISL_450462, EPI_ISL_450463, EPI_ISL_450464, EPI_ISL_450465, EPI_ISL_450466, EPI_ISL_450467, EPI_ISL_450468, EPI_ISL_450469, EPI_ISL_450470, EPI_ISL_450471, EPI_ISL_450472, EPI_ISL_450473, EPI_ISL_450474, EPI_ISL_450475, EPI_ISL_450476, EPI_ISL_450477, EPI_ISL_450478, EPI_ISL_450479, EPI_ISL_450480, EPI_ISL_450481 | | | |
| see above | Stanford clinical virology lab | Chan-Zuckerberg Biohub | Benjamin Pinksy, Katharine Walter, Victoria N. Parikh, John Gorzynski, Hannah N. DeJong, Matthew T. Wheeler, Jason Andrews, Manuel Rivas, Carlos Bustamante, Euan Ashley, with CZB Cliahub Consortium |
| EPI_ISL_450496, EPI_ISL_450497 | National Public Health Surveillance Laboratory, Vilnius, Lithuania | Charite Universitaetsmedizin Berlin, Institute of Virology | Victor M Corman, Jorn Beheim-Schwarzbach, Talitha Veith, Barbara Muehlemann, Julia Schneider, Terry Jones, Ana Steponkiene, Christian Drosten |
| EPI_ISL_450498 | Health Board Laboratory of Communicable Diseases | Charite Universitatsmedizin Berlin, Institute of Virology | Victor M Corman, Jörn Beheim-Schwarzbach, Barbara Mühlemann, Talitha Veith, Julia Schneider, Liidia Dotsenko, Natalja Kuznetsova, Terry Jones, Christian Drosten |
| EPI_ISL_450508, EPI_ISL_450509, EPI_ISL_450511, EPI_ISL_450512, EPI_ISL_450515 | Rafik Hariri University Hospital | Rafik Hariri University Hospital | Rita Feghali |
| EPI_ISL_450518, EPI_ISL_450519 | E. Gulbja Laboratorija | Latvian Biomedical Research and Study Centre | Ivars Silamiķelis, Kaspars Megnis, Monta Ustinova, Ņikita Zrelovs, Vita Rovīte, Mikus Gavars, Dmitrijs Perminovs, Uga Dumpis, Jānis Kloviņš |
| EPI_ISL_450520, EPI_ISL_450521, EPI_ISL_450522, EPI_ISL_450523, EPI_ISL_450524 | Centrālā Laboratorija | Latvian Biomedical Research and Study Centre | Ivars Silamiķelis, Kaspars Megnis, Monta Ustinova, Ņikita Zrelovs, Vita Rovīte, Stella Lapiņa, Jana Osīte, Marta Priedīte, Uga Dumpis, Jānis Kloviņš |
| EPI_ISL_450525, EPI_ISL_450526, EPI_ISL_450527, EPI_ISL_450528, EPI_ISL_450529, EPI_ISL_450530 | Hematology Laboratory, Section of Molecular Diagnostics, University Clinical Centre, Medical University of Gdansk | Department of Virology, Faculty of Medicine, University of Helsinki, Helsinki, Finland | Maciej Grzybek, Marlena Robakowska, Aneta Szulc, Olli Vapalahti, Teemu Smura |
| EPI_ISL_450538, EPI_ISL_450539, EPI_ISL_450541, EPI_ISL_450542, EPI_ISL_450543, EPI_ISL_450544, EPI_ISL_450546, EPI_ISL_450547, EPI_ISL_450548, EPI_ISL_450551, EPI_ISL_450553, EPI_ISL_450554, EPI_ISL_450559, EPI_ISL_450560, EPI_ISL_450561, EPI_ISL_450563, EPI_ISL_450564, EPI_ISL_450566, EPI_ISL_450567 | | | |
| see above | Utah Public Health Laboratory | Utah Public Health Laboratory | Erin Young, Kelly Oakeson |
| EPI_ISL_450572, EPI_ISL_450574, EPI_ISL_450576, EPI_ISL_450577, EPI_ISL_450580, EPI_ISL_450581, EPI_ISL_450585, EPI_ISL_450588, EPI_ISL_450592 | Michigan Department of Health and Human Services, Bureau of Laboratories | Michigan Department of Health and Human Services, Bureau of Laboratories | Blankenship HM; Riner D; Soehnlen MK |
| EPI_ISL_450601, EPI_ISL_450604, EPI_ISL_450606, EPI_ISL_450607, EPI_ISL_450610, EPI_ISL_450612, EPI_ISL_450614, EPI_ISL_450616, EPI_ISL_450618, EPI_ISL_450620, EPI_ISL_450623, EPI_ISL_450631, EPI_ISL_450635, EPI_ISL_450636 | | | |
| see above | Michigan Department of Health and Human Services, Bureau of Laboratories | Michigan Department of Health and Human Services, Bureau of Laboratories | Blankenship HM, Riner D, Soehnlen MK |
| EPI_ISL_450639 | Laboratoire de microbiologie, Hopital de Verdun | Smith Laboratory, Centre de Recherche CHU Sainte-Justine | Martin Smith, Marieke Rozendaal, Ivan Pavlov |
| EPI_ISL_450700, EPI_ISL_450701, EPI_ISL_450702, EPI_ISL_450703, EPI_ISL_450704, EPI_ISL_450705, EPI_ISL_450706, EPI_ISL_450707, EPI_ISL_450708, EPI_ISL_450709, EPI_ISL_450710, EPI_ISL_450711, EPI_ISL_450712, EPI_ISL_450713, EPI_ISL_450714, EPI_ISL_450715, EPI_ISL_450716, EPI_ISL_450717, EPI_ISL_450718, EPI_ISL_450719, EPI_ISL_450720, EPI_ISL_450721, EPI_ISL_450722 | | | |

| | | | |
|---|---|---|---|
| see above | University of Wisconsin-Madison AIDS Vaccine Research Laboratories | University of Wisconsin-Madison AIDS Vaccine Research Laboratories | Gage Moreno, Katarina Braun, et al. AIDS Vaccine Research Laboratories |
| EPI_ISL_450723 | Ramathibodi Hospital | COVID-19 Network Investigations (CONI) Alliance | Elizabeth Batty, Wasun Chantratita, Thanat Chookajorn, Stefan Fernandez, Angkana Huang, Anthony R. Jones, Khajohn Joonsalak, Chonticha Klungtong, Theerarat Kochakarn, Namfon Kotanan, Krittikorn Kumpornsin, Wudtichai Manasatienkij, Bhakbhoom Panthan, Ekawat Pasomsub, Insee Sensorn, Arporn Wangwiwatsin |
| EPI_ISL_450724, EPI_ISL_450725, EPI_ISL_450726, EPI_ISL_450727, EPI_ISL_450728, EPI_ISL_450729, EPI_ISL_450730, EPI_ISL_450731, EPI_ISL_450732, EPI_ISL_450734, EPI_ISL_450736, EPI_ISL_450737 | | | |
| see above | Hospital AZ Rivierenland | Institute of Tropical Medicine | Philippe Selhorst, Colin Anthony |
| EPI_ISL_450738, EPI_ISL_450739, EPI_ISL_450740, EPI_ISL_450741, EPI_ISL_450742, EPI_ISL_450743, EPI_ISL_450744, EPI_ISL_450745 | OUCRU/HTD | OUCRU/HTD | Nguyen Van Vinh Chau, Nguyen Thi Thu Hong, Nguyen Thi Han Ny, Le Nguyen Truc Nhu, Nghiem My Ngoc, Vo Thanh Lam, Nguyen Thanh Dung, Lam Minh Yen, Ngo Ngoc Quang Minh, Le Manh Hung, Nguyen Tri Dung, Dinh Nguyen Huy Man, Lam Anh Nguyet, Tran Chanh Xuan, Tran Tinh Hien, Nguyen Thanh Phong, Tran Nguyen Hoang Tu, Tran Tan Thanh, Nguyen Thanh Truong, Nguyen Tan Binh, Tang Chi Thuong, Guy Thwaites, and Le Van Tan, for OUCRU COVID-19 research group* |
| EPI_ISL_450746 | Laboratory of Molecular Biology, Diagnostyka sp. z o.o. | Laboratory of Recombinant Vaccines | Lukasz Rabalski, Anna Piotrowska-Mietelska, Maciej Kosinski, Boguslaw Szewczyk, Krystyna Bienkowska-Szewczyk |
| EPI_ISL_450747 | Sunnybrook Health Sciences Centre | Department of Laboratory Medicine and Molecular Diagnostics, Sunnybrook Health Sciences Centre | Jalees A. Nasir, Robert A. Kozak, Patryk Aftanas, Amogelang R. Raphenya, Kendrick M. Smith, Finlay Maguire, Hassaan Maan, Muhannad Alruwaili, Arinjay Banerjee, Hamza Mbareche, Brian P. Alcock, Natalie C. Knox, Karen Mossman, Bo Wang, Julian A. Hiscox, Andrew G. McArthur, Samira Mubareka |
| EPI_ISL_450748, EPI_ISL_450749, EPI_ISL_450750, EPI_ISL_450751, EPI_ISL_450752, EPI_ISL_450753, EPI_ISL_450754, EPI_ISL_450755, EPI_ISL_450756, EPI_ISL_450757, EPI_ISL_450758, EPI_ISL_450759, EPI_ISL_450760, EPI_ISL_450761, EPI_ISL_450762, EPI_ISL_450763, EPI_ISL_450764, EPI_ISL_450765, EPI_ISL_450766, EPI_ISL_450767, EPI_ISL_450768, EPI_ISL_450769, EPI_ISL_450770, EPI_ISL_450771, EPI_ISL_450772, EPI_ISL_450773, EPI_ISL_450774, EPI_ISL_450775, EPI_ISL_450776, EPI_ISL_450777, EPI_ISL_450778, EPI_ISL_450779, EPI_ISL_450780 | | | |
| see above | Minnesota Department of Health, Public Health Laboratory | Minnesota Department of Health, Public Health Laboratory | Matt Plumb, Jacob Garfin, and Xiong Wang |
| EPI_ISL_450781 | Government Medical College-Bhavnagar | Gujarat Biotechnology Research Centre | Kairavi Desai, Saklain Malek, Shirish Patel, Ramesh Pandit, Tejas Shah, Ankit Hinsu, Pritesh Sabara, Apurvasinh Puvar, Janvi Raval, Zarna Patel, Monika Gandhi, Pinal Trivedi, Maharshi Pandya, Amit Kanani, Nidhi Patel, Nitin Savaliya, Raghawendra Kumar, Dinesh Kumar, Zuber Saiyed, Komal Patel, Labdhi Pandya, Snehal Bagatharia, Bhavesh Modi, Gaurishankar Shrimali, R D Dixit, A M Kadri, Akanksha Verma, Chaitanya Joshi, Madhvi Joshi |
| EPI_ISL_450782 | Government Medical College-Bhavnagar | Gujarat Biotechnology Research Centre | Saklain Malek, Shirish Patel, Kairavi Desai, Tejas Shah, Ankit Hinsu, Pritesh Sabara, Apurvasinh Puvar, Janvi Raval, Zarna Patel, Monika Gandhi, Pinal Trivedi, Maharshi Pandya, Amit Kanani, Nidhi Patel, Nitin Savaliya, Raghawendra Kumar, Dinesh Kumar, Zuber Saiyed, Komal Patel, Labdhi Pandya, Snehal Bagatharia, Ramesh Pandit, Bhavesh Modi, Gaurishankar Shrimali, R D Dixit, A M Kadri, Priti Pandita, Chaitanya Joshi, Madhvi Joshi |
| EPI_ISL_450783 | Government Medical College-Bhavnagar | Gujarat Biotechnology Research Centre | Shirish Patel, Kairavi Desai, Saklain Malek, Ankit Hinsu, Pritesh Sabara, Apurvasinh Puvar, Janvi Raval, Zarna Patel, Monika Gandhi, Pinal Trivedi, Maharshi Pandya, Amit Kanani, Nidhi Patel, Nitin Savaliya, Raghawendra Kumar, Dinesh Kumar, Zuber Saiyed, Komal Patel, Labdhi Pandya, Snehal Bagatharia, Ramesh Pandit, Tejas Shah, Bhavesh Modi, Gaurishankar Shrimali, R D Dixit, A M Kadri, Neha Rajpara, Chaitanya Joshi, Madhvi Joshi |
| EPI_ISL_450784 | Government Medical College-Bhavnagar | Gujarat Biotechnology Research Centre | Zarna Patel, Ramesh Pandit, Tejas Shah, Ankit Hinsu, Pritesh Sabara, Apurvasinh Puvar, Janvi Raval, Monika Gandhi, Pinal Trivedi, Maharshi Pandya, Amit Kanani, Nidhi Patel, Nitin Savaliya, Raghawendra Kumar, Dinesh Kumar, Zuber Saiyed, Komal Patel, Labdhi Pandya, Snehal Bagatharia, Kairavi Desai, Saklain Malek, Shirish Patel, Bhavesh Modi, Gaurishankar Shrimali, R D Dixit, A M Kadri, Afzal Ansari, Chaitanya Joshi, Madhvi Joshi |
| EPI_ISL_450785 | Pandit Deendayal Upadhyay Government Medical College, Rajkot | Gujarat Biotechnology Research Centre | Prakash Modi, Sejul Antala, Manish Pattani, Apurvasinh Puvar, Janvi Raval, Zarna Patel, Monika Gandhi, Pinal Trivedi, Maharshi Pandya, Amit Kanani, Nidhi Patel, Nitin Savaliya, Raghawendra Kumar, Dinesh Kumar, Zuber Saiyed, Komal Patel, Labdhi Pandya, Snehal Bagatharia, Ramesh Pandit, Tejas Shah, Ankit Hinsu, Pritesh Sabara, Bhavesh Modi, Gaurishankar Shrimali, R D Dixit, A M Kadri, Neelam Nathani, Chaitanya Joshi, Madhvi Joshi |
| EPI_ISL_450786 | Pandit Deendayal Upadhyay Government Medical College, Rajkot | Gujarat Biotechnology Research Centre | Sejul Antala, Manish Pattani, Prakash Modi, Janvi Raval, Zarna Patel, Monika Gandhi, Pinal Trivedi, Maharshi Pandya, Amit Kanani, Nidhi Patel, Nitin Savaliya, Raghawendra Kumar, Dinesh Kumar, Zuber Saiyed, Komal Patel, Labdhi Pandya, Snehal Bagatharia, Ramesh Pandit, Tejas Shah, Ankit Hinsu, Pritesh Sabara, Apurvasinh Puvar, Bhavesh Modi, Gaurishankar Shrimali, R D Dixit, A M Kadri, Armi Chaudhari, Chaitanya Joshi, Madhvi Joshi |
| EPI_ISL_450787 | Pandit Deendayal Upadhyay Government Medical College, Rajkot | Gujarat Biotechnology Research Centre | Manish Pattani, Prakash Modi, Sejul Antala, Zarna Patel, Monika Gandhi, Pinal Trivedi, Maharshi Pandya, Amit Kanani, Nidhi Patel, Nitin Savaliya, Raghawendra Kumar, Dinesh Kumar, Zuber Saiyed, Komal Patel, Labdhi Pandya, Snehal Bagatharia, Ramesh Pandit, Tejas Shah, Ankit Hinsu, Pritesh Sabara, Apurvasinh Puvar, Janvi Raval, Bhavesh Modi, Gaurishankar Shrimali, R D Dixit, A M Kadri, Bhavya Jindal, Chaitanya Joshi, Madhvi Joshi |
| EPI_ISL_450788 | Pandit Deendayal Upadhyay Government Medical College, Rajkot | Gujarat Biotechnology Research Centre | Zarna Patel, Tejas Shah, Ankit Hinsu, Pritesh Sabara, Apurvasinh Puvar, Janvi Raval, Monika Gandhi, Pinal Trivedi, Maharshi Pandya, Amit Kanani, Nidhi Patel, Nitin Savaliya, Raghawendra Kumar, Dinesh Kumar, Zuber Saiyed, Komal Patel, Labdhi Pandya, Snehal Bagatharia, Prakash Modi, Sejul Antala, Manish Pattani, Ramesh Pandit, Bhavesh Modi, Gaurishankar Shrimali, R D Dixit, A M Kadri, Camellia Chakraborty, Chaitanya Joshi, Madhvi Joshi |
| EPI_ISL_450789 | Pandit Deendayal Upadhyay Government Medical College, Rajkot | Gujarat Biotechnology Research Centre | Ankit Hinsu, Pritesh Sabara, Apurvasinh Puvar, Janvi Raval, Zarna Patel, Monika Gandhi, Pinal Trivedi, Maharshi Pandya, Amit Kanani, Nidhi Patel, Nitin Savaliya, Raghawendra Kumar, Dinesh Kumar, Zuber Saiyed, Komal Patel, Labdhi Pandya, Snehal Bagatharia, Prakash Modi, Sejul Antala, Manish Pattani, Ramesh Pandit, Tejas Shah, Bhavesh Modi, Gaurishankar Shrimali, R D Dixit, A M Kadri, Siddhant Kumar, Chaitanya Joshi, Madhvi Joshi |
| EPI_ISL_450790 | Pandit Deendayal Upadhyay Government Medical College, Rajkot | Gujarat Biotechnology Research Centre | Zarna Patel, Pritesh Sabara, Apurvasinh Puvar, Janvi Raval, Monika Gandhi, Pinal Trivedi, Maharshi Pandya, Amit Kanani, Nidhi Patel, Nitin Savaliya, Raghawendra Kumar, Dinesh Kumar, Zuber Saiyed, Komal Patel, Labdhi Pandya, Snehal Bagatharia, Prakash Modi, Sejul Antala, Manish Pattani, Ramesh Pandit, Tejas Shah, Ankit Hinsu, Bhavesh Modi, Gaurishankar Shrimali, R D Dixit, A M Kadri, Sharmistha Majumdar, Chaitanya Joshi, Madhvi Joshi |
| EPI_ISL_450791 | Pandit Deendayal Upadhyay Government Medical College, Rajkot | Gujarat Biotechnology Research Centre | Zarna Patel, Apurvasinh Puvar, Janvi Raval, Monika Gandhi, Pinal Trivedi, Maharshi Pandya, Amit Kanani, Nidhi Patel, Nitin Savaliya, Raghawendra Kumar, Dinesh Kumar, Zuber Saiyed, Komal Patel, Labdhi Pandya, Snehal Bagatharia, Prakash Modi, Sejul Antala, Manish Pattani, Ramesh Pandit, Tejas Shah, Ankit Hinsu, Pritesh Sabara, Bhavesh Modi, Gaurishankar Shrimali, R D Dixit, A M Kadri, Pooja P Doshi, Chaitanya Joshi, Madhvi Joshi |
| EPI_ISL_450792 | Jamaica Ministry of Health and Wellness | Pathogen Discovery, Respiratory Viruses Branch, Division of Viral Diseases, Centers for Disease Control and Prevention | Krista Queen, Yan Li, Anna Montmayeur, Ying Tao, Jing Zhang, Anna Uehara, Clinton R. Paden, Rachel Marine, Haibin Wang, Jasmine Padilla, Justin Lee, Zachary Weiner, Bettina Bankamp, Suxiang Tong |
| EPI_ISL_450793, EPI_ISL_450794, EPI_ISL_450795, EPI_ISL_450796, EPI_ISL_450797 | Jamaica Ministry of Health and Wellness | Pathogen Discovery, Respiratory Viruses Branch, Division of Viral Diseases, Centers for Disease Control and Prevention | Yan Li, Anna Montmayeur, Ying Tao, Krista Queen, Jing Zhang, Anna Uehara, Clinton R. Paden, Rachel Marine, Haibin Wang, Zachary Weiner, Bettina Bankamp, Suxiang Tong |
| EPI_ISL_450798 | Jamaica Ministry of Health and Wellness | Pathogen Discovery, Respiratory Viruses Branch, Division of Viral Diseases, Centers for Disease Control and Prevention | Krista Queen, Yan Li, Anna Montmayeur, Ying Tao, Jing Zhang, Anna Uehara, Clinton R. Paden, Rachel Marine, Haibin Wang, Jasmine Padilla, Justin Lee, Zachary Weiner, Bettina Bankamp, Suxiang Tong |
| EPI_ISL_450799 | Jamaica Ministry of Health and Wellness | Pathogen Discovery, Respiratory Viruses Branch, Division of Viral Diseases, Centers for Disease Control and Prevention | Yan Li, Anna Montmayeur, Ying Tao, Krista Queen, Jing Zhang, Anna Uehara, Clinton R. Paden, Rachel Marine, Haibin Wang, Zachary Weiner, Bettina Bankamp, Suxiang Tong |
| EPI_ISL_450800 | AR Dept. of Health-Public Health Lab | Pathogen Discovery, Respiratory Viruses Branch, Division of Viral Diseases, Centers for Disease Control and Prevention | Yan Li, Anna Montmayeur, Ying Tao, Krista Queen, Jing Zhang, Anna Uehara, Clinton R. Paden, Rachel Marine, Haibin Wang, Zachary Weiner, Bettina Bankamp, Suxiang Tong |
| EPI_ISL_450801 | Georgia Department of Health | Pathogen Discovery, Respiratory Viruses Branch, Division of Viral Diseases, Centers for Disease Control and Prevention | Yan Li, Anna Montmayeur, Ying Tao, Krista Queen, Jing Zhang, Anna Uehara, Clinton R. Paden, Rachel Marine, Haibin Wang, Zachary Weiner, Bettina Bankamp, Suxiang Tong |
| EPI_ISL_450802, EPI_ISL_450803 | PA Department of Health, Bureau of Laboratories | Pathogen Discovery, Respiratory Viruses Branch, Division of Viral Diseases, Centers for Disease Control and Prevention | Yan Li, Anna Montmayeur, Ying Tao, Krista Queen, Jing Zhang, Anna Uehara, Clinton R. Paden, Rachel Marine, Haibin Wang, Zachary Weiner, Bettina Bankamp, Suxiang Tong |
| EPI_ISL_450804, EPI_ISL_450806 | VI-US Virgin Islands Department of Health | Pathogen Discovery, Respiratory Viruses Branch, Division of Viral Diseases, Centers for Disease Control and Prevention | Yan Li, Anna Montmayeur, Ying Tao, Krista Queen, Jing Zhang, Anna Uehara, Clinton R. Paden, Rachel Marine, Haibin Wang, Zachary Weiner, Bettina Bankamp, Suxiang Tong |
| EPI_ISL_450807 | Victoria Vard och Halsa | The Public Health Agency of Sweden | Sarah Henriksson, Anna-Malin Linde, Maria Lind Karlberg, Oskar Karlsson Lindsjo, Olov Svartstrom, Anna Risberg, Theresa Enkirch, Mia Brytting, Karin Tegmark-Wisell |
| EPI_ISL_450808 | Saroledens Familjelakare | The Public Health Agency of Sweden | Katarina Jarbur, Anna-Malin Linde, Maria Lind Karlberg, Oskar Karlsson Lindsjo, Olov Svartstrom, Anna Risberg, Theresa Enkirch, Mia Brytting, Karin Tegmark-Wisell |
| EPI_ISL_450809 | Kungsholmsdoktorn | The Public Health Agency of Sweden | Linus Hammar, Anna-Malin Linde, Maria Lind Karlberg, Oskar Karlsson Lindsjo, Olov Svartstrom, Anna Risberg, Theresa Enkirch, Mia Brytting, Karin Tegmark-Wisell |
| EPI_ISL_450810 | Saroledens Familjelakare | The Public Health Agency of Sweden | Katarina Jarbur, Anna-Malin Linde, Maria Lind Karlberg, Oskar Karlsson Lindsjo, Olov Svartstrom, Anna Risberg, Theresa Enkirch, Mia Brytting, Karin Tegmark-Wisell |
| EPI_ISL_450811 | Knivsta VC | The Public Health Agency of Sweden | Johanna Carlson, Anna-Malin Linde, Maria Lind Karlberg, Oskar Karlsson Lindsjo, Olov Svartstrom, Anna Risberg, Theresa Enkirch, Mia Brytting, Karin Tegmark-Wisell |
| EPI_ISL_450812 | Uppsala Narakut Aleris | The Public Health Agency of Sweden | Annika Nilsson, Anna-Malin Linde, Maria Lind Karlberg, Oskar Karlsson Lindsjo, Olov Svartstrom, Anna Risberg, Theresa Enkirch, Mia Brytting, Karin Tegmark-Wisell |
| EPI_ISL_450813 | Bla Kustens halsocentral | The Public Health Agency of Sweden | Olof Norrby, Anna-Malin Linde, Maria Lind Karlberg, Oskar Karlsson Lindsjo, Olov Svartstrom, Anna Risberg, Theresa Enkirch, Mia Brytting, Karin Tegmark-Wisell |
| EPI_ISL_450814 | Huslakarna Varmbadhuset Varberg | The Public Health Agency of Sweden | Johanna Hilmersson, Anna-Malin Linde, Maria Lind Karlberg, Oskar Karlsson Lindsjo, Olov Svartstrom, Anna Risberg, Theresa Enkirch, Mia Brytting, Karin Tegmark-Wisell |
| EPI_ISL_450815 | Narhalsan Molnlycke, Barn och ungdomsmedicin | The Public Health Agency of Sweden | Mats Reimer, Anna-Malin Linde, Maria Lind Karlberg, Oskar Karlsson Lindsjo, Olov Svartstrom, Anna Risberg, Theresa Enkirch, Mia Brytting, Karin Tegmark-Wisell |
| EPI_ISL_450816 | orestadsklinikens VC | The Public Health Agency of Sweden | Lisa Kjellberg / Laura Plavitu, Anna-Malin Linde, Maria Lind Karlberg, Oskar Karlsson Lindsjo, Olov Svartstrom, Anna Risberg, Theresa Enkirch, Mia Brytting, Karin Tegmark-Wisell |
| EPI_ISL_450817 | Narhalsan Backa vardcentral | The Public Health Agency of Sweden | Mats Olsson, Anna-Malin Linde, Maria Lind Karlberg, Oskar Karlsson Lindsjo, Olov Svartstrom, Anna Risberg, Theresa Enkirch, Mia Brytting, Karin Tegmark-Wisell |
| EPI_ISL_450818 | Aneby VC | The Public Health Agency of Sweden | Ken Granath, Anna-Malin Linde, Maria Lind Karlberg, Oskar Karlsson Lindsjo, Olov Svartstrom, Anna Risberg, Theresa Enkirch, Mia Brytting, Karin Tegmark-Wisell |
| EPI_ISL_450819 | Hovas Askim Familjelakare och BVC | The Public Health Agency of Sweden | Anna Wendel, Anna-Malin Linde, Maria Lind Karlberg, Oskar Karlsson Lindsjo, Olov Svartstrom, Anna Risberg, Theresa Enkirch, Mia Brytting, Karin Tegmark-Wisell |
| EPI_ISL_450820 | Narhalsan Molnlycke, Barn och ungdomsmedicin | The Public Health Agency of Sweden | Mats Reimer, Anna-Malin Linde, Maria Lind Karlberg, Oskar Karlsson Lindsjo, Olov Svartstrom, Anna Risberg, Theresa Enkirch, Mia Brytting, Karin Tegmark-Wisell |
| EPI_ISL_450821, EPI_ISL_450822 | Jarpens HC | The Public Health Agency of Sweden | Gunilla Johansson, Anna-Malin Linde, Maria Lind Karlberg, Oskar Karlsson Lindsjo, Olov Svartstrom, Anna Risberg, Theresa Enkirch, Mia Brytting, Karin Tegmark-Wisell |
| EPI_ISL_450823 | Narhalsan Backa vardcentral | The Public Health Agency of Sweden | Mats Olsson, Anna-Malin Linde, Maria Lind Karlberg, Oskar Karlsson Lindsjo, Olov Svartstrom, Anna Risberg, Theresa Enkirch, Mia Brytting, Karin Tegmark-Wisell |
| EPI_ISL_450824 | Ulltuna Vardcentral | The Public Health Agency of Sweden | Heidi Lindback, Anna-Malin Linde, Maria Lind Karlberg, Oskar Karlsson Lindsjo, Olov Svartstrom, Anna Risberg, Theresa Enkirch, Mia Brytting, Karin Tegmark-Wisell |
| EPI_ISL_450825 | Narhalsan Backa vardcentral | The Public Health Agency of Sweden | Mats Olsson, Anna-Malin Linde, Maria Lind Karlberg, Oskar Karlsson Lindsjo, Olov Svartstrom, Anna Risberg, Theresa Enkirch, Mia Brytting, Karin Tegmark-Wisell |
| EPI_ISL_450826, EPI_ISL_450827, EPI_ISL_450828 | Uppsala Narakut Aleris | The Public Health Agency of Sweden | Annika Nilsson, Anna-Malin Linde, Maria Lind Karlberg, Oskar Karlsson Lindsjo, Olov Svartstrom, Anna Risberg, Theresa Enkirch, Mia Brytting, Karin Tegmark-Wisell |
| EPI_ISL_450829 | Narhalsan Sjobo vardcentral | The Public Health Agency of Sweden | Lovisa Hjerten, Anna-Malin Linde, Maria Lind Karlberg, Oskar Karlsson Lindsjo, Olov Svartstrom, Anna Risberg, Theresa Enkirch, Mia Brytting, Karin Tegmark-Wisell |
| EPI_ISL_450830 | Narhalsan Olskroken VC | The Public Health Agency of Sweden | Mahin Ghoroghi, Anna-Malin Linde, Maria Lind Karlberg, Oskar Karlsson Lindsjo, Olov Svartstrom, Anna Risberg, Theresa Enkirch, Mia Brytting, Karin Tegmark-Wisell |
| EPI_ISL_450831 | Wetterhalsan | The Public Health Agency of Sweden | Anders Tengblad, Anna-Malin Linde, Maria Lind Karlberg, Oskar Karlsson Lindsjo, Olov Svartstrom, Anna Risberg, Theresa Enkirch, Mia Brytting, Karin Tegmark-Wisell |

| | | | |
|---|---|---|---|
| EPI_ISL_450832 | Byjorden vardcentral | The Public Health Agency of Sweden | Pernilla Brunman, Anna-Malin Linde, Maria Lind Karlberg, Oskar Karlsson Lindsjo, Olov Svartstrom, Anna Risberg, Theresa Enkirch, Mia Brytting, Karin Tegmark-Wisell |
| EPI_ISL_450833 | Wetterhalsan | The Public Health Agency of Sweden | Anders Tengblad, Anna-Malin Linde, Maria Lind Karlberg, Oskar Karlsson Lindsjo, Olov Svartstrom, Anna Risberg, Theresa Enkirch, Mia Brytting, Karin Tegmark-Wisell |
| EPI_ISL_450834 | Klinisk mikrobiologi och vardhygien Halmstad | The Public Health Agency of Sweden | Arne Kotz, Anna-Malin Linde, Maria Lind Karlberg, Oskar Karlsson Lindsjo, Olov Svartstrom, Anna Risberg, Theresa Enkirch, Mia Brytting, Karin Tegmark-Wisell |
| EPI_ISL_450835 | Unilabs Skovde | The Public Health Agency of Sweden | Tobias Kollberg, Helena Enroth, Anna-Malin Linde, Maria Lind Karlberg, Oskar Karlsson Lindsjo, Olov Svartstrom, Anna Risberg, Theresa Enkirch, Mia Brytting, Karin Tegmark-Wisell |
| EPI_ISL_450836, EPI_ISL_450837 | Laboratoriemedicin | The Public Health Agency of Sweden | Anna-Malin Linde, Maria Lind Karlberg, Oskar Karlsson Lindsjo, Olov Svartstrom, Anna Risberg, Theresa Enkirch, Mia Brytting, Karin Tegmark-Wisell |
| EPI_ISL_450839 | COVID-19 Laboratory Centre for Advanced Research in Sciences (CARS), University of Dhaka, Dhaka-1000, Bangladesh | DNA Solution Ltd | Sharif Akhteruzzaman, Zeba Islam Seraj, Nazmul Ahsan, Md Imdadul Hoque, MA Malek, Shahryar Nabi, Sabrina Moriom Elius, ABM Khademul Islam, Richard Malo, Imran Khan, Abu Sufian, Sabita Rezwana Rahman, Habibul Bari Shozib, Mamun Ahmed, AHM Nurun Nabi, Mohammad Riazul Islam, Md Mizanur Rahman, Md Ismail Hosen, Latiful Bari, Gazi Nurun Nahar, Haseena Khan, M Anwar Hossain. |
| EPI_ISL_450840 | COVID-19 Laboratory | DNA Solution Ltd. L-5 | Sharif Akhteruzzaman, Zeba Islam Seraj, Nazmul Ahsan, Md Imdadul Hoque, MA Malek, Shahryar Nabi, Sabrina Moriom Elius, ABM Khademul Islam, Richard Malo, Imran Khan, Abu Sufian, Sabita Rezwana Rahman, Habibul Bari Shozib, Mamun Ahmed, AHM Nurun Nabi, Mohammad Riazul Islam, Md Mizanur Rahman, Md Ismail Hosen, Latiful Bari, Gazi Nurun Nahar, Haseena Khan, M Anwar Hossain. |
| EPI_ISL_450841 | COVID-19 Laboratory | DNA Solution Ltd | Sharif Akhteruzzaman, Zeba Islam Seraj, Nazmul Ahsan, Md Imdadul Hoque, MA Malek, Shahryar Nabi, Sabrina Moriom Elius, ABM Khademul Islam, Richard Malo, Imran Khan, Abu Sufian, Sabita Rezwana Rahman, Habibul Bari Shozib, Mamun Ahmed, AHM Nurun Nabi, Mohammad Riazul Islam, Md Mizanur Rahman, Md Ismail Hosen, Latiful Bari, Gazi Nurun Nahar, Haseena Khan, M Anwar Hossain. |
| EPI_ISL_450843 | COVID-19 Laboratory | DNA Solution Ltd. | Sharif Akhteruzzaman, Zeba Islam Seraj, Nazmul Ahsan, Md Imdadul Hoque, MA Malek, Shahryar Nabi, Sabrina Moriom Elius, ABM Khademul Islam, Richard Malo, Imran Khan, Abu Sufian, Sabita Rezwana Rahman, Habibul Bari Shozib, Mamun Ahmed, AHM Nurun Nabi, Mohammad Riazul Islam, Md Mizanur Rahman, Md Ismail Hosen, Latiful Bari, Gazi Nurun Nahar, Haseena Khan, M Anwar Hossain. |
| EPI_ISL_450849, EPI_ISL_450850 | Florida Bureau of Public Health Laboratories | Florida Bureau of Public Health Laboratories | Sarah Schmedes, Jason Blanton |
| EPI_ISL_450873 | Evandro Chagas Institute | Evandro Chagas Institute | Santos, M.C.; Silva, A.M.; Junior, W.D.C.; Barbagelata, L.S.; Ferreira, J.A.; Sousa, E.M.A.; da Silva, P.S.; Martins, L.C.; Sousa Junior, E.C.; Viana, G.M.R |
| EPI_ISL_450874 | Evandro Chagas Institute | Evandro Chagas Institute | Santos, M.C.; Silva, A.M.; Junior, W.D.C.; Barbagelata, L.S.; Ferreira, J.A.; Sousa, E.M.A.; da Silva, P.S.; Martins, L.C.;Sousa Junior, E.C.;Viana, G.M.R |
| EPI_ISL_451076 | West China Hospital of Sichuan University | State Key Laboratory of Biotherapy of Sichuan University | Baowen Du, Minjin Wang, Chao Tanga, Chuan Chena, Yongzhao Zhou, Mingxia Yu, Han-Cheng Wei, Weimin Li, Jing-wen Lin, Jia Geng, Binwu Ying, Lu Chen |
| EPI_ISL_451077, EPI_ISL_451078, EPI_ISL_451079, EPI_ISL_451080, EPI_ISL_451081, EPI_ISL_451082, EPI_ISL_451083, EPI_ISL_451084, EPI_ISL_451087, EPI_ISL_451088, EPI_ISL_451089, EPI_ISL_451090, EPI_ISL_451091, EPI_ISL_451093, EPI_ISL_451094, EPI_ISL_451095, EPI_ISL_451096, EPI_ISL_451097, EPI_ISL_451098, EPI_ISL_451099, EPI_ISL_451100, EPI_ISL_451101, EPI_ISL_451102, EPI_ISL_451103, EPI_ISL_451104, EPI_ISL_451105, EPI_ISL_451106, EPI_ISL_451107, EPI_ISL_451108, EPI_ISL_451109, EPI_ISL_451110, EPI_ISL_451111, EPI_ISL_451112, EPI_ISL_451113, EPI_ISL_451114, EPI_ISL_451115, EPI_ISL_451116, EPI_ISL_451117, EPI_ISL_451118, EPI_ISL_451121, EPI_ISL_451122, EPI_ISL_451123, EPI_ISL_451125, EPI_ISL_451126, EPI_ISL_451128, EPI_ISL_451129, EPI_ISL_451131, EPI_ISL_451132, EPI_ISL_451133, EPI_ISL_451134, EPI_ISL_451135, EPI_ISL_451136, EPI_ISL_451137, EPI_ISL_451138, EPI_ISL_451139, EPI_ISL_451140, EPI_ISL_451141, EPI_ISL_451144, EPI_ISL_451145, EPI_ISL_451147, EPI_ISL_451148 | | | |
| see above | SA Pathology | SA Pathology | Lex Leong, Chuan Kok Lim, Mark Turra, Ivan Bastian, Geoff Higgins |
| EPI_ISL_451149 | M.P Shah Government Medocal college Jamnagar | Gujarat Biotechnology Research Centre | Janvi Raval, Zarna Patel, Monika Gandhi, Pinal Trivedi, Maharshi Pandya, Amit Kanani, Nidhi Patel, Nitin Savaliya, Raghawendra Kumar, Dinesh Kumar, Zuber Saiyed, Komal Patel, Labdhi Pandya, Snehal Bagatharia, Ramesh Pandit, Tejas Shah, Ankit Hinsu, Pritesh Sabara, Apurvasinh Puvar, Binita Aring, Bhavesh Modi, Gaurishankar Shrimali, R D Dixit, A M Kadri, Priti Pandita, Chaitanya Joshi, Madhvi Joshi, |
| EPI_ISL_451150 | M.P Shah Government Medocal college Jamnagar | Gujarat Biotechnology Research Centre | Zarna Patel, Monika Gandhi, Pinal Trivedi, Maharshi Pandya, Amit Kanani, Nidhi Patel, Nitin Savaliya, Raghawendra Kumar, Dinesh Kumar, Zuber Saiyed, Komal Patel, Labdhi Pandya, Snehal Bagatharia, Ramesh Pandit, Tejas Shah, Ankit Hinsu, Pritesh Sabara, Apurvasinh Puvar, Binita Aring, Janvi Raval, Bhavesh Modi, Gaurishankar Shrimali, R D Dixit, A M Kadri, Pragya Sharma, Chaitanya Joshi, Madhvi Joshi, |
| EPI_ISL_451151 | M.P Shah Government Medocal college Jamnagar | Gujarat Biotechnology Research Centre | Monika Gandhi, Pinal Trivedi, Maharshi Pandya, Amit Kanani, Nidhi Patel, Nitin Savaliya, Raghawendra Kumar, Dinesh Kumar, Zuber Saiyed, Komal Patel, Labdhi Pandya, Snehal Bagatharia, Ramesh Pandit, Tejas Shah, Ankit Hinsu, Pritesh Sabara, Apurvasinh Puvar, Binita Aring, Janvi Raval, Zarna Patel, Bhavesh Modi, Gaurishankar Shrimali, R D Dixit, A M Kadri, Neha Rajpara, Chaitanya Joshi, Madhvi Joshi, |
| EPI_ISL_451152 | M.P Shah Government Medocal college Jamnagar | Gujarat Biotechnology Research Centre | Pinal Trivedi, Maharshi Pandya, Amit Kanani, Nidhi Patel, Nitin Savaliya, Raghawendra Kumar, Dinesh Kumar, Zuber Saiyed, Komal Patel, Labdhi Pandya, Snehal Bagatharia, Ramesh Pandit, Tejas Shah, Ankit Hinsu, Pritesh Sabara, Apurvasinh Puvar, Binita Aring, Janvi Raval, Zarna Patel, Monika Gandhi, Bhavesh Modi, Gaurishankar Shrimali, R D Dixit, A M Kadri, Afzal Ansari, Chaitanya Joshi, Madhvi Joshi, |
| EPI_ISL_451153 | M.P Shah Government Medocal college Jamnagar | Gujarat Biotechnology Research Centre | Maharshi Pandya, Amit Kanani, Nidhi Patel, Nitin Savaliya, Raghawendra Kumar, Dinesh Kumar, Zuber Saiyed, Komal Patel, Labdhi Pandya, Snehal Bagatharia, Ramesh Pandit, Tejas Shah, Ankit Hinsu, Pritesh Sabara, Apurvasinh Puvar, Binita Aring, Janvi Raval, Zarna Patel, Monika Gandhi, Pinal Trivedi, Bhavesh Modi, Gaurishankar Shrimali, R D Dixit, A M Kadri, Fenil Patel, Chaitanya Joshi, Madhvi Joshi, |
| EPI_ISL_451154 | Government Medical College, Vadodara | Gujarat Biotechnology Research Centre | Manish Pattani, Tanuja Javadekar , Maharshi Pandya, Amit Kanani, Nidhi Patel, Nitin Savaliya, Raghawendra Kumar, Dinesh Kumar, Zuber Saiyed, Komal Patel, Labdhi Pandya, Snehal Bagatharia, Ramesh Pandit, Tejas Shah, Ankit Hinsu, Pritesh Sabara, Apurvasinh Puvar, Janvi Raval, Zarna Patel, Monika Gandhi, Pinal Trivedi, Bhavesh Modi, Gaurishankar Shrimali, R D Dixit, A M Kadri, Neelam Nathani, Chaitanya Joshi, Madhvi Joshi |
| EPI_ISL_451155 | Government Medical College, Vadodara | Gujarat Biotechnology Research Centre | Tanuja Javadekar , Manish Pattani, Amit Kanani, Nidhi Patel, Nitin Savaliya, Raghawendra Kumar, Dinesh Kumar, Zuber Saiyed, Komal Patel, Labdhi Pandya, Snehal Bagatharia, Ramesh Pandit, Tejas Shah, Ankit Hinsu, Pritesh Sabara, Apurvasinh Puvar, Janvi Raval, Zarna Patel, Monika Gandhi, Pinal Trivedi, Maharshi Pandya, Bhavesh Modi, Gaurishankar Shrimali, R D Dixit, A M Kadri, Armi Chaudhari, Chaitanya Joshi, Madhvi Joshi |
| EPI_ISL_451156 | Government Medical College, Vadodara | Gujarat Biotechnology Research Centre | Amit Kanani, Nidhi Patel, Nitin Savaliya, Raghawendra Kumar, Dinesh Kumar, Zuber Saiyed, Komal Patel, Labdhi Pandya, Snehal Bagatharia, Ramesh Pandit, Tejas Shah, Ankit Hinsu, Pritesh Sabara, Apurvasinh Puvar, Janvi Raval, Zarna Patel, Monika Gandhi, Pinal Trivedi, Maharshi Pandya, Manish Pattani, Tanuja Javadekar , Bhavesh Modi, Gaurishankar Shrimali, R D Dixit, A M Kadri, Bhavya Jindal, Chaitanya Joshi, Madhvi Joshi |
| EPI_ISL_451157 | Government Medical College, Vadodara | Gujarat Biotechnology Research Centre | Nidhi Patel, Nitin Savaliya, Raghawendra Kumar, Dinesh Kumar, Zuber Saiyed, Komal Patel, Labdhi Pandya, Snehal Bagatharia, Ramesh Pandit, Tejas Shah, Ankit Hinsu, Pritesh Sabara, Apurvasinh Puvar, Janvi Raval, Zarna Patel, Monika Gandhi, Pinal Trivedi, Maharshi Pandya, Manish Pattani, Tanuja Javadekar , Amit Kanani, Bhavesh Modi, Gaurishankar Shrimali, R D Dixit, A M Kadri, Camellia Chakraborty, Chaitanya Joshi, Madhvi Joshi |
| EPI_ISL_451158 | Government Medical College, Vadodara | Gujarat Biotechnology Research Centre | Nitin Savaliya, Raghawendra Kumar, Dinesh Kumar, Zuber Saiyed, Komal Patel, Labdhi Pandya, Snehal Bagatharia, Ramesh Pandit, Tejas Shah, Ankit Hinsu, Pritesh Sabara, Apurvasinh Puvar, Janvi Raval, Zarna Patel, Monika Gandhi, Pinal Trivedi, Maharshi Pandya, Manish Pattani, Tanuja Javadekar , Amit Kanani, Nidhi Patel, Bhavesh Modi, Gaurishankar Shrimali, R D Dixit, A M Kadri, Siddhant Kumar, Chaitanya Joshi, Madhvi Joshi |
| EPI_ISL_451159 | Government Medical College, Vadodara | Gujarat Biotechnology Research Centre | Raghawendra Kumar, Dinesh Kumar, Zuber Saiyed, Komal Patel, Labdhi Pandya, Snehal Bagatharia, Ramesh Pandit, Tejas Shah, Ankit Hinsu, Pritesh Sabara, Apurvasinh Puvar, Janvi Raval, Zarna Patel, Monika Gandhi, Pinal Trivedi, Maharshi Pandya, Manish Pattani, Tanuja Javadekar , Amit Kanani, Nidhi Patel, Nitin Savaliya, Bhavesh Modi, Gaurishankar Shrimali, R D Dixit, A M Kadri, Sharmistha Majumdar, Chaitanya Joshi, Madhvi Joshi |
| EPI_ISL_451160 | Government Medical College, Vadodara | Gujarat Biotechnology Research Centre | Dinesh Kumar, Zuber Saiyed, Komal Patel, Labdhi Pandya, Snehal Bagatharia, Ramesh Pandit, Tejas Shah, Ankit Hinsu, Pritesh Sabara, Apurvasinh Puvar, Janvi Raval, Zarna Patel, Monika Gandhi, Pinal Trivedi, Maharshi Pandya, Manish Pattani, Tanuja Javadekar , Amit Kanani, Nidhi Patel, Nitin Savaliya, Raghawendra Kumar, Bhavesh Modi, Gaurishankar Shrimali, R D Dixit, A M Kadri, Pooja P Doshi, Chaitanya Joshi, Madhvi Joshi |
| EPI_ISL_451161 | Government Medical College, Vadodara | Gujarat Biotechnology Research Centre | Zuber Saiyed, Komal Patel, Labdhi Pandya, Snehal Bagatharia, Ramesh Pandit, Tejas Shah, Ankit Hinsu, Pritesh Sabara, Apurvasinh Puvar, Janvi Raval, Zarna Patel, Monika Gandhi, Pinal Trivedi, Maharshi Pandya, Manish Pattani, Tanuja Javadekar , Amit Kanani, Nidhi Patel, Nitin Savaliya, Raghawendra Kumar, Dinesh Kumar, Bhavesh Modi, Gaurishankar Shrimali, R D Dixit, A M Kadri, Akanksha Verma, Chaitanya Joshi, Madhvi Joshi |
| EPI_ISL_451162 | Government Medical College, Vadodara | Gujarat Biotechnology Research Centre | Komal Patel, Labdhi Pandya, Snehal Bagatharia, Ramesh Pandit, Tejas Shah, Ankit Hinsu, Pritesh Sabara, Apurvasinh Puvar, Janvi Raval, Zarna Patel, Monika Gandhi, Pinal Trivedi, Maharshi Pandya, Manish Pattani, Tanuja Javadekar , Amit Kanani, Nidhi Patel, Nitin Savaliya, Raghawendra Kumar, Dinesh Kumar, Zuber Saiyed, Bhavesh Modi, Gaurishankar Shrimali, R D Dixit, A M Kadri, Priti Pandita, Chaitanya Joshi, Madhvi Joshi |
| EPI_ISL_451163 | Government Medical College, Vadodara | Gujarat Biotechnology Research Centre | Labdhi Pandya, Snehal Bagatharia, Ramesh Pandit, Tejas Shah, Ankit Hinsu, Pritesh Sabara, Apurvasinh Puvar, Janvi Raval, Zarna Patel, Monika Gandhi, Pinal Trivedi, Maharshi Pandya, Manish Pattani, Tanuja Javadekar , Amit Kanani, Nidhi Patel, Nitin Savaliya, Raghawendra Kumar, Dinesh Kumar, Zuber Saiyed, Komal Patel, Bhavesh Modi, Gaurishankar Shrimali, R D Dixit, A M Kadri, Pragya Sharma, Chaitanya Joshi, Madhvi Joshi |
| EPI_ISL_451164, EPI_ISL_451166, EPI_ISL_451167, EPI_ISL_451168, EPI_ISL_451169, EPI_ISL_451170, EPI_ISL_451173 | Lab voor klinische biologie | Onderzoeksgroep Virologie | Laurens Lambrechts, Nick Vereecke, Marthe Pauwels, Jozefien De Clercq, Bruno Verhasselt, Linos Vandekerckhove, Hans Nauwynck, Sebastiaan Theuns |
| EPI_ISL_451175, EPI_ISL_451177, EPI_ISL_451178, EPI_ISL_451179, EPI_ISL_451180, EPI_ISL_451181 | Lab voor klinische biologie | Onderzoeksgroep Virologie | Nick Vereecke, Laurens Lambrechts, Marthe Pauwels, Jozefien De Clercq, Bruno Verhasselt, Linos Vandekerckhove, Hans Nauwynck, Sebastiaan Theuns |
| EPI_ISL_451183, EPI_ISL_451184, EPI_ISL_451186, EPI_ISL_451189, EPI_ISL_451190, EPI_ISL_451192, EPI_ISL_451193, EPI_ISL_451194, EPI_ISL_451195, EPI_ISL_451196, EPI_ISL_451197, EPI_ISL_451198, EPI_ISL_451199, EPI_ISL_451201, EPI_ISL_451202 | | | |
| see above | Uganda Virus Research Institute | MRC/UVRI & LSHTM Uganda Research Unit | Dan Lule Bugembe, John Kayiwa, My V.T Phan, Phionah Tushabe, Stephen Balinandi, Beatrice Dhaala, Deogratius Ssemwanga, Jonas Lexow, Henry Mwebesa, Jane Aceng, Henry Kyobe, Julius Lutwama, Pontiano Kaleebu, Matthew Cotten |
| EPI_ISL_451203, EPI_ISL_451204, EPI_ISL_451205, EPI_ISL_451206, EPI_ISL_451207, EPI_ISL_451208, EPI_ISL_451209, EPI_ISL_451210, EPI_ISL_451211, EPI_ISL_451212, EPI_ISL_451213, EPI_ISL_451214, EPI_ISL_451215, EPI_ISL_451216, EPI_ISL_451217, EPI_ISL_451218 | | | |
| see above | LSUHS Emerging Viral Threat Laboratory | Microbial Genome Sequencing Center | Jeremy P. Kamil, John A. Vanchiere, Rona S. Scott, Camille F. Abshire, Abida Siddiqa, Byeong-Jae Lee, Chan-ki Min, Md Maksudul Alam, Monica Gestal-Cartele, Edna Ondari, Adam Greer, Malgorzata Bienkowska-Haba, Katarzyna Zwolinska, Jason M. Bodily, Andrew D. Yurochko, Paul M. Weinberger, Christopher G. Kevil, Martin J. Sapp, Daniel J. Snyder, Vaughn S. Cooper |
| EPI_ISL_451219, EPI_ISL_451220, EPI_ISL_451221, EPI_ISL_451222, EPI_ISL_451223, EPI_ISL_451224, EPI_ISL_451225, EPI_ISL_451226, EPI_ISL_451227, EPI_ISL_451228, EPI_ISL_451229 | | | |
| see above | LSUHS Emerging Viral Threat Laboratory | Microbial Genome Sequencing Center | Rona S. Scott, Jeremy P. Kamil, John A. Vanchiere, Camille F. Abshire, Abida Siddiqa, Byeong-Jae Lee, Chan-ki Min, Md Maksudul Alam, Monica Gestal-Cartele, Edna Ondari, Adam Greer, Malgorzata Bienkowska-Haba, Katarzyna Zwolinska, Jason M. Bodily, Andrew D. Yurochko, Paul M. Weinberger, Christopher G. Kevil, Martin J. Sapp, Daniel J. Snyder, Vaughn S. Cooper |
| EPI_ISL_451230, EPI_ISL_451231, EPI_ISL_451232, EPI_ISL_451233, EPI_ISL_451234, EPI_ISL_451235, EPI_ISL_451236, EPI_ISL_451237, EPI_ISL_451238, EPI_ISL_451239, EPI_ISL_451240, EPI_ISL_451241, EPI_ISL_451242, EPI_ISL_451243, EPI_ISL_451244, EPI_ISL_451245, EPI_ISL_451246, EPI_ISL_451247 | | | |
| see above | LSUHS Emerging Viral Threat Laboratory | Microbial Genome Sequencing Center | John A. Vanchiere, Jeremy P. Kamil, Rona S. Scott, Camille F. Abshire, Abida Siddiqa, Byeong-Jae Lee, Chan-ki Min, Md Maksudul Alam, Monica Gestal-Cartele, Edna Ondari, Adam Greer, Malgorzata Bienkowska-Haba, Katarzyna Zwolinska, Jason M. Bodily, Andrew D. Yurochko, Paul M. Weinberger, Christopher G. Kevil, Martin J. Sapp, Daniel J. Snyder, Vaughn S. Cooper |
| EPI_ISL_451248, EPI_ISL_451249, EPI_ISL_451250, EPI_ISL_451251, EPI_ISL_451252, EPI_ISL_451253, EPI_ISL_451254, EPI_ISL_451255, EPI_ISL_451256, EPI_ISL_451261, EPI_ISL_451257, EPI_ISL_451258, EPI_ISL_451259, EPI_ISL_451260, EPI_ISL_451261, EPI_ISL_451262, EPI_ISL_451263, EPI_ISL_451264, EPI_ISL_451265, EPI_ISL_451266, EPI_ISL_451267, EPI_ISL_451268, EPI_ISL_451269, EPI_ISL_451270, EPI_ISL_451271, EPI_ISL_451272, EPI_ISL_451273, EPI_ISL_451274, EPI_ISL_451275, EPI_ISL_451276, EPI_ISL_451277, EPI_ISL_451278, EPI_ISL_451279, EPI_ISL_451280, EPI_ISL_451281, EPI_ISL_451282, EPI_ISL_451283, EPI_ISL_451284, EPI_ISL_451285, EPI_ISL_451286, EPI_ISL_451287, EPI_ISL_451288, EPI_ISL_451289, EPI_ISL_451290, EPI_ISL_451291, EPI_ISL_451292, EPI_ISL_451293, EPI_ISL_451294, EPI_ISL_451295, EPI_ISL_451296, EPI_ISL_451297 | | | |
| see above | LSUHS Emerging Viral Threat Laboratory | Microbial Genome Sequencing Center | Jeremy P. Kamil, John A. Vanchiere, Rona S. Scott, Camille F. Abshire, Abida Siddiqa, Byeong-Jae Lee, Chan-ki Min, Md Maksudul Alam, Monica Gestal-Cartele, Edna Ondari, Adam Greer, Malgorzata Bienkowska-Haba, Katarzyna Zwolinska, Jason M. Bodily, Andrew D. Yurochko, Paul M. Weinberger, Christopher G. Kevil, Martin J. Sapp, Daniel J. Snyder, Vaughn S. Cooper |
| EPI_ISL_451298 | Laboratory of Virology, INMI Lazzaro Spallanzani IRCCS | Laboratory of Virology, INMI Lazzaro Spallanzani IRCCS | Cesare E.M. Gruber, Martina Rueca, Barbara Bartolini, Francesco Messina, Antonino Di Caro, Maria R. Capobianchi, Giuseppe Ippolito |
| EPI_ISL_451299 | Laboratory of Virology, INMI Lazzaro Spallanzani IRCCS | Laboratory of Virology, INMI Lazzaro Spallanzani IRCCS | Martina Rueca, Cesare E.M. Gruber, Barbara Bartolini, Francesco Messina, Antonino Di Caro, Maria R. Capobianchi, Giuseppe Ippolito |
| EPI_ISL_451300 | Laboratory of Virology, INMI Lazzaro Spallanzani IRCCS | Laboratory of Virology, INMI Lazzaro Spallanzani IRCCS | Cesare E.M. Gruber, Martina Rueca, Barbara Bartolini, Francesco Messina, Antonino Di Caro, Maria R. Capobianchi, Giuseppe Ippolito |
| EPI_ISL_451301 | Laboratory of Virology, INMI Lazzaro Spallanzani IRCCS | Laboratory of Virology, INMI Lazzaro Spallanzani IRCCS | Martina Rueca, Cesare E.M. Gruber, Barbara Bartolini, Francesco Messina, Antonino Di Caro, Maria R. Capobianchi, Giuseppe Ippolito |
| EPI_ISL_451302 | Laboratory of Virology, INMI Lazzaro Spallanzani IRCCS | Laboratory of Virology, INMI Lazzaro Spallanzani IRCCS | Cesare E.M. Gruber, Martina Rueca, Barbara Bartolini, Francesco Messina, Antonino Di Caro, Maria R. Capobianchi, Giuseppe Ippolito |

| | | | |
|---|---|---|---|
| EPI_ISL_451303 | Laboratory of Virology, INMI Lazzaro Spallanzani IRCCS | Laboratory of Virology, INMI Lazzaro Spallanzani IRCCS | Martina Rueca, Cesare E.M. Gruber, Barbara Bartolini, Francesco Messina, Antonino Di Caro, Maria R. Capobianchi, Giuseppe Ippolito |
| EPI_ISL_451304 | Laboratory of Virology, INMI Lazzaro Spallanzani IRCCS | Laboratory of Virology, INMI Lazzaro Spallanzani IRCCS | Cesare E.M. Gruber, Martina Rueca, Barbara Bartolini, Francesco Messina, Antonino Di Caro, Maria R. Capobianchi, Giuseppe Ippolito |
| EPI_ISL_451306 | Molecular Virology Unit, Fondazione IRCCS Policlinico San Matteo , Pavia | Laboratory of Virology, INMI Lazzaro Spallanzani IRCCS | Antonio Piralla, Fausto Baldanti, Martina Rueca, Antonino Di Caro, Maria R. Capobianchi, Cesare E.M. Gruber, Barbara Bartolini |
| EPI_ISL_451307 | Molecular Virology Unit, Fondazione IRCCS Policlinico San Matteo , Pavia | Laboratory of Virology, INMI Lazzaro Spallanzani IRCCS | Fausto Baldanti, Antonio Piralla, Antonino Di Caro, Cesare E.M. Gruber, Martina Rueca, Barbara Bartolini, Maria R. Capobianchi |
| EPI_ISL_451308 | Molecular Virology Unit, Fondazione IRCCS Policlinico San Matteo , Pavia | Laboratory of Virology, INMI Lazzaro Spallanzani IRCCS | Antonio Piralla, Fausto Baldanti, Maria R. Capobianchi, Cesare E.M. Gruber, Martina Rueca, Barbara Bartolini, Antonino Di Caro |
| EPI_ISL_451309 | Molecular Virology Unit, Fondazione IRCCS Policlinico San Matteo , Pavia | Laboratory of Virology, INMI Lazzaro Spallanzani IRCCS | Fausto Baldanti, Antonio Piralla, Cesare E.M. Gruber, Maria R. Capobianchi, Antonino Di Caro, Martina Rueca, Barbara Bartolini |
| EPI_ISL_451310, EPI_ISL_451311 | Hellenic Pasteur Institute, National Influenza Reference laboratory of Southern Greece & Unit of Bioinformatics and Applied Genomics | Hellenic Pasteur Institute, National Influenza Reference laboratory of Southern Greece & Unit of Bioinformatics and Applied Genomics | Vasiliki Pogka, Timokratis Karamitros, Athanasios Kossyvakis, Antonios Kalliaropoulos, Horefti Elina, Evangelidou Maria, Androniki Voulgari-Kokota, Aspasia Kontou, Andreas Mentis |
| EPI_ISL_451313, EPI_ISL_451314, EPI_ISL_451315, EPI_ISL_451316, EPI_ISL_451318, EPI_ISL_451319, EPI_ISL_451320, EPI_ISL_451321, EPI_ISL_451322, EPI_ISL_451325, EPI_ISL_451326, EPI_ISL_451327, EPI_ISL_451328, EPI_ISL_451329, EPI_ISL_451330, EPI_ISL_451331, EPI_ISL_451334, EPI_ISL_451337, EPI_ISL_451338, EPI_ISL_451344, EPI_ISL_451345, EPI_ISL_451346, EPI_ISL_451348, EPI_ISL_451353, EPI_ISL_451354, EPI_ISL_451356, EPI_ISL_451357, EPI_ISL_451359, EPI_ISL_451360, EPI_ISL_451365, EPI_ISL_451369, EPI_ISL_451370, EPI_ISL_451371, EPI_ISL_451374, EPI_ISL_451376, EPI_ISL_451377, EPI_ISL_451378, EPI_ISL_451379, EPI_ISL_451380, EPI_ISL_451381, EPI_ISL_451382, EPI_ISL_451383, EPI_ISL_451384, EPI_ISL_451385, EPI_ISL_451386, EPI_ISL_451387, EPI_ISL_451388, EPI_ISL_451389, EPI_ISL_451390, EPI_ISL_451391, EPI_ISL_451392, EPI_ISL_451393, EPI_ISL_451394, EPI_ISL_451395, EPI_ISL_451398 | | | |
| see above | West China Hospital of Sichuan University | State Key Laboratory of Biotherapy of Sichuan University | Baowen Du, Minjin Wang, Chao Tang, Chuan Chen, Yongzhao Zhou, Mingxia Yu, Hancheng Wei, Weimin Li, Jing-wen Lin, Jia Geng, Binwu Ying, Lu Chen |
| EPI_ISL_451401, EPI_ISL_451402, EPI_ISL_451404, EPI_ISL_451405, EPI_ISL_451406, EPI_ISL_451407, EPI_ISL_451408, EPI_ISL_451409, EPI_ISL_451410, EPI_ISL_451411, EPI_ISL_451412, EPI_ISL_451413, EPI_ISL_451414, EPI_ISL_451415, EPI_ISL_451416, EPI_ISL_451417, EPI_ISL_451418, EPI_ISL_451419, EPI_ISL_451420, EPI_ISL_451421, EPI_ISL_451422, EPI_ISL_451423, EPI_ISL_451424, EPI_ISL_451426, EPI_ISL_451427, EPI_ISL_451428, EPI_ISL_451429, EPI_ISL_451430, EPI_ISL_451431, EPI_ISL_451432, EPI_ISL_451434, EPI_ISL_451435, EPI_ISL_451436, EPI_ISL_451437, EPI_ISL_451438, EPI_ISL_451439, EPI_ISL_451440, EPI_ISL_451441, EPI_ISL_451442, EPI_ISL_451443, EPI_ISL_451444, EPI_ISL_451445, EPI_ISL_451446, EPI_ISL_451447, EPI_ISL_451448, EPI_ISL_451449, EPI_ISL_451450, EPI_ISL_451451, EPI_ISL_451452, EPI_ISL_451453, EPI_ISL_451454, EPI_ISL_451455, EPI_ISL_451456, EPI_ISL_451457, EPI_ISL_451458, EPI_ISL_451459, EPI_ISL_451460, EPI_ISL_451461, EPI_ISL_451462, EPI_ISL_451463, EPI_ISL_451464, EPI_ISL_451465, EPI_ISL_451466, EPI_ISL_451467, EPI_ISL_451468, EPI_ISL_451469, EPI_ISL_451470, EPI_ISL_451471, EPI_ISL_451472, EPI_ISL_451474, EPI_ISL_451475, EPI_ISL_451476, EPI_ISL_451477, EPI_ISL_451478, EPI_ISL_451479 | | | |
| see above | NYU Langone Health | Departments of Pathology and Medicine, New York University School of Medicine | Maria Aguero-Rosenfeld, Brendan Belovarac, Margaret Black, Ludovic Boytard, John Cadley, Paolo Cotzia, John Chen, Dacia Dimartino, Xiaojun Feng, Tatyana Gindin, Emily Guzman, Adriana Heguy, Megan Hogan, Emily Huang, George Jour, Alireza Khodadadi-Jamayran, Lawrence H. Lin, Raven Luther, Andrew Lytle, Christian Marier, Matthew T. Maurano, Mark J. Mulligan, Peter Meyn, Raquel Ordonez Ciriza, Iman Osman, Jared Pinnell, Vanessa Raabe, Sitharam Ramaswami, Amy Rapkiewicz, Andre M. Ribeiro-dos-Santos, Marie Samanovic-Golden, Antonio Serrano, Guomiao Shen, Matija Snuderl, Theodore Vougiouklakis, Nick Vulpescu, Gael Westby, Paul Zappile, Yutong Zhang |
| EPI_ISL_451486 | Australian Clinical Labs | NSW Health Pathology - Institute of Clinical Pathology and Medical Research; Westmead Hospital; University of Sydney | CIDM-PH et al. |
| EPI_ISL_451487, EPI_ISL_451488 | Pathology North - NSW Health Pathology | NSW Health Pathology - Institute of Clinical Pathology and Medical Research; Westmead Hospital; University of Sydney | CIDM-PH et al. |
| EPI_ISL_451489 | Laverty Pathology | NSW Health Pathology - Institute of Clinical Pathology and Medical Research; Westmead Hospital; University of Sydney | CIDM-PH et al. |
| EPI_ISL_451490, EPI_ISL_451491, EPI_ISL_451492, EPI_ISL_451493, EPI_ISL_451494, EPI_ISL_451495, EPI_ISL_451496, EPI_ISL_451497, EPI_ISL_451498, EPI_ISL_451499, EPI_ISL_451500, EPI_ISL_451501, EPI_ISL_451502, EPI_ISL_451503, EPI_ISL_451504, EPI_ISL_451505, EPI_ISL_451506, EPI_ISL_451507, EPI_ISL_451508, EPI_ISL_451509, EPI_ISL_451510, EPI_ISL_451511, EPI_ISL_451512, EPI_ISL_451513, EPI_ISL_451514, EPI_ISL_451515, EPI_ISL_451516 | | | |
| see above | Pathology West - NSW Health Pathology | NSW Health Pathology - Institute of Clinical Pathology and Medical Research; Westmead Hospital; University of Sydney | CIDM-PH et al. |
| EPI_ISL_451517, EPI_ISL_451518, EPI_ISL_451519, EPI_ISL_451521 | South Eastern Area Laboratory Services | NSW Health Pathology - Institute of Clinical Pathology and Medical Research; Westmead Hospital; University of Sydney | CIDM-PH et al. |
| EPI_ISL_451522, EPI_ISL_451523, EPI_ISL_451524, EPI_ISL_451525, EPI_ISL_451526, EPI_ISL_451527, EPI_ISL_451528, EPI_ISL_451529 | Pathology West - NSW Health Pathology | NSW Health Pathology - Institute of Clinical Pathology and Medical Research; Westmead Hospital; University of Sydney | CIDM-PH et al. |
| EPI_ISL_451530, EPI_ISL_451531 | Pathology Sydney South West - NSW Health Pathology | NSW Health Pathology - Institute of Clinical Pathology and Medical Research; Westmead Hospital; University of Sydney | CIDM-PH et al. |
| EPI_ISL_451532 | Pathology West - NSW Health Pathology | NSW Health Pathology - Institute of Clinical Pathology and Medical Research; Westmead Hospital; University of Sydney | CIDM-PH et al. |
| EPI_ISL_451533 | Pathology Sydney South West - NSW Health Pathology | NSW Health Pathology - Institute of Clinical Pathology and Medical Research; Westmead Hospital; University of Sydney | CIDM-PH et al. |
| EPI_ISL_451534 | Medlab Pathology | NSW Health Pathology - Institute of Clinical Pathology and Medical Research; Westmead Hospital; University of Sydney | CIDM-PH et al. |
| EPI_ISL_451535 | Pathology West - NSW Health Pathology | NSW Health Pathology - Institute of Clinical Pathology and Medical Research; Westmead Hospital; University of Sydney | CIDM-PH et al. |
| EPI_ISL_451536 | Pathology Sydney South West - NSW Health Pathology | NSW Health Pathology - Institute of Clinical Pathology and Medical Research; Westmead Hospital; University of Sydney | CIDM-PH et al. |
| EPI_ISL_451537, EPI_ISL_451538, EPI_ISL_451539 | Pathology West - NSW Health Pathology | NSW Health Pathology - Institute of Clinical Pathology and Medical Research; Westmead Hospital; University of Sydney | CIDM-PH et al. |
| EPI_ISL_451540 | ACT pathology | NSW Health Pathology - Institute of Clinical Pathology and Medical Research; Westmead Hospital; University of Sydney | CIDM-PH et al. |
| EPI_ISL_451541 | Pathology West - NSW Health Pathology | NSW Health Pathology - Institute of Clinical Pathology and Medical Research; Westmead Hospital; University of Sydney | CIDM-PH et al. |
| EPI_ISL_451542, EPI_ISL_451543, EPI_ISL_451544 | Pathology Sydney South West - NSW Health Pathology | NSW Health Pathology - Institute of Clinical Pathology and Medical Research; Westmead Hospital; University of Sydney | CIDM-PH et al. |
| EPI_ISL_451545 | Laverty Pathology | NSW Health Pathology - Institute of Clinical Pathology and Medical Research; Westmead Hospital; University of Sydney | CIDM-PH et al. |
| EPI_ISL_451546, EPI_ISL_451547 | Pathology West - NSW Health Pathology | NSW Health Pathology - Institute of Clinical Pathology and Medical Research; Westmead Hospital; University of Sydney | CIDM-PH et al. |
| EPI_ISL_451548 | Childrens Hospital Westmead | NSW Health Pathology - Institute of Clinical Pathology and Medical Research; Westmead Hospital; University of Sydney | CIDM-PH et al. |
| EPI_ISL_451549 | Pathology Sydney South West - NSW Health Pathology | NSW Health Pathology - Institute of Clinical Pathology and Medical Research; Westmead Hospital; University of Sydney | CIDM-PH et al. |
| EPI_ISL_451550, EPI_ISL_451551 | Pathology West - NSW Health Pathology | NSW Health Pathology - Institute of Clinical Pathology and Medical Research; Westmead Hospital; University of Sydney | CIDM-PH et al. |
| EPI_ISL_451552 | Pathology North - NSW Health Pathology | NSW Health Pathology - Institute of Clinical Pathology and Medical Research; Westmead Hospital; University of Sydney | CIDM-PH et al. |
| EPI_ISL_451553, EPI_ISL_451554, EPI_ISL_451555 | Medlab Pathology | NSW Health Pathology - Institute of Clinical Pathology and Medical Research; Westmead Hospital; University of Sydney | CIDM-PH et al. |
| EPI_ISL_451556, EPI_ISL_451557 | Pathology West - NSW Health Pathology | NSW Health Pathology - Institute of Clinical Pathology and Medical Research; Westmead Hospital; University of Sydney | CIDM-PH et al. |
| EPI_ISL_451558 | Medlab Pathology | NSW Health Pathology - Institute of Clinical Pathology and Medical Research; Westmead Hospital; University of Sydney | CIDM-PH et al. |
| EPI_ISL_451559, EPI_ISL_451560, EPI_ISL_451561, EPI_ISL_451562, EPI_ISL_451563, EPI_ISL_451564, EPI_ISL_451565 | Pathology Sydney South West - NSW Health Pathology | NSW Health Pathology - Institute of Clinical Pathology and Medical Research; Westmead Hospital; University of Sydney | CIDM-PH et al. |
| EPI_ISL_451566 | Pathology West - NSW Health Pathology | NSW Health Pathology - Institute of Clinical Pathology and Medical Research; Westmead Hospital; University of Sydney | CIDM-PH et al. |
| EPI_ISL_451567 | Medlab Pathology | NSW Health Pathology - Institute of Clinical Pathology and Medical Research; Westmead Hospital; University of Sydney | CIDM-PH et al. |
| EPI_ISL_451568, EPI_ISL_451569, EPI_ISL_451570, EPI_ISL_451571 | Pathology West - NSW Health Pathology | NSW Health Pathology - Institute of Clinical Pathology and Medical Research; Westmead Hospital; University of Sydney | CIDM-PH et al. |
| EPI_ISL_451572 | Laverty Pathology | NSW Health Pathology - Institute of Clinical Pathology and Medical Research; Westmead Hospital; University of Sydney | CIDM-PH et al. |
| EPI_ISL_451573 | Pathology West - NSW Health Pathology | NSW Health Pathology - Institute of Clinical Pathology and Medical Research; Westmead Hospital; University of Sydney | CIDM-PH et al. |
| EPI_ISL_451574 | Australian Clinical Labs | NSW Health Pathology - Institute of Clinical Pathology and Medical Research; Westmead Hospital; University of Sydney | CIDM-PH et al. |
| EPI_ISL_451575, EPI_ISL_451576, | Pathology West - NSW Health Pathology | NSW Health Pathology - Institute of Clinical Pathology and Medical | CIDM-PH et al. |

| | | | |
|---|---|---|---|
| EPI_ISL_451577 | | Research; Westmead Hospital; University of Sydney | |
| EPI_ISL_451578 | Pathology Sydney South West - NSW Health Pathology | NSW Health Pathology - Institute of Clinical Pathology and Medical Research; Westmead Hospital; University of Sydney | CIDM-PH et al. |
| EPI_ISL_451579, EPI_ISL_451580 | Pathology West - NSW Health Pathology | NSW Health Pathology - Institute of Clinical Pathology and Medical Research; Westmead Hospital; University of Sydney | CIDM-PH et al. |
| EPI_ISL_451581 | Pathology Sydney South West - NSW Health Pathology | NSW Health Pathology - Institute of Clinical Pathology and Medical Research; Westmead Hospital; University of Sydney | CIDM-PH et al. |
| EPI_ISL_451582 | Pathology West - NSW Health Pathology | NSW Health Pathology - Institute of Clinical Pathology and Medical Research; Westmead Hospital; University of Sydney | CIDM-PH et al. |
| EPI_ISL_451583 | Medlab Pathology | NSW Health Pathology - Institute of Clinical Pathology and Medical Research; Westmead Hospital; University of Sydney | CIDM-PH et al. |
| EPI_ISL_451584, EPI_ISL_451586 | Pathology West - NSW Health Pathology | NSW Health Pathology - Institute of Clinical Pathology and Medical Research; Westmead Hospital; University of Sydney | CIDM-PH et al. |
| EPI_ISL_451587 | Pathology North Hunter- NSW Health Pathology | NSW Health Pathology - Institute of Clinical Pathology and Medical Research; Westmead Hospital; University of Sydney | CIDM-PH et al. |
| EPI_ISL_451588 | ACT pathology | NSW Health Pathology - Institute of Clinical Pathology and Medical Research; Westmead Hospital; University of Sydney | CIDM-PH et al. |
| EPI_ISL_451589 | Pathology North Hunter- NSW Health Pathology | NSW Health Pathology - Institute of Clinical Pathology and Medical Research; Westmead Hospital; University of Sydney | CIDM-PH et al. |
| EPI_ISL_451590, EPI_ISL_451591, EPI_ISL_451592, EPI_ISL_451593 | ACT pathology | NSW Health Pathology - Institute of Clinical Pathology and Medical Research; Westmead Hospital; University of Sydney | CIDM-PH et al. |
| EPI_ISL_451594 | Childrens Hospital Westmead | NSW Health Pathology - Institute of Clinical Pathology and Medical Research; Westmead Hospital; University of Sydney | CIDM-PH et al. |
| EPI_ISL_451595, EPI_ISL_451596 | ACT pathology | NSW Health Pathology - Institute of Clinical Pathology and Medical Research; Westmead Hospital; University of Sydney | CIDM-PH et al. |
| EPI_ISL_451597 | Medlab Pathology | NSW Health Pathology - Institute of Clinical Pathology and Medical Research; Westmead Hospital; University of Sydney | CIDM-PH et al. |
| EPI_ISL_451598 | ACT pathology | NSW Health Pathology - Institute of Clinical Pathology and Medical Research; Westmead Hospital; University of Sydney | CIDM-PH et al. |
| EPI_ISL_451599 | Australian Clinical Labs | NSW Health Pathology - Institute of Clinical Pathology and Medical Research; Westmead Hospital; University of Sydney | CIDM-PH et al. |
| EPI_ISL_451600 | Pathology North Hunter- NSW Health Pathology | NSW Health Pathology - Institute of Clinical Pathology and Medical Research; Westmead Hospital; University of Sydney | CIDM-PH et al. |
| EPI_ISL_451601 | Douglas Hanly Moir Pathology | NSW Health Pathology - Institute of Clinical Pathology and Medical Research; Westmead Hospital; University of Sydney | CIDM-PH et al. |
| EPI_ISL_451602 | Laverty Pathology | NSW Health Pathology - Institute of Clinical Pathology and Medical Research; Westmead Hospital; University of Sydney | CIDM-PH et al. |
| EPI_ISL_451603 | Pathology Sydney South West - NSW Health Pathology | NSW Health Pathology - Institute of Clinical Pathology and Medical Research; Westmead Hospital; University of Sydney | CIDM-PH et al. |
| EPI_ISL_451604 | Pathology North - NSW Health Pathology | NSW Health Pathology - Institute of Clinical Pathology and Medical Research; Westmead Hospital; University of Sydney | CIDM-PH et al. |
| EPI_ISL_451605 | Childrens Hospital Westmead | NSW Health Pathology - Institute of Clinical Pathology and Medical Research; Westmead Hospital; University of Sydney | CIDM-PH et al. |
| EPI_ISL_451606, EPI_ISL_451607 | Pathology West - NSW Health Pathology | NSW Health Pathology - Institute of Clinical Pathology and Medical Research; Westmead Hospital; University of Sydney | CIDM-PH et al. |
| EPI_ISL_451608 | Pathology Sydney South West - NSW Health Pathology | NSW Health Pathology - Institute of Clinical Pathology and Medical Research; Westmead Hospital; University of Sydney | CIDM-PH et al. |
| EPI_ISL_451609 | Laverty Pathology | NSW Health Pathology - Institute of Clinical Pathology and Medical Research; Westmead Hospital; University of Sydney | CIDM-PH et al. |
| EPI_ISL_451610 | Medlab Pathology | NSW Health Pathology - Institute of Clinical Pathology and Medical Research; Westmead Hospital; University of Sydney | CIDM-PH et al. |
| EPI_ISL_451611 | Laverty Pathology | NSW Health Pathology - Institute of Clinical Pathology and Medical Research; Westmead Hospital; University of Sydney | CIDM-PH et al. |
| EPI_ISL_451612 | ACT pathology | NSW Health Pathology - Institute of Clinical Pathology and Medical Research; Westmead Hospital; University of Sydney | CIDM-PH et al. |
| EPI_ISL_451613, EPI_ISL_451614, EPI_ISL_451615, EPI_ISL_451616, EPI_ISL_451617, EPI_ISL_451618, EPI_ISL_451619, EPI_ISL_451620, EPI_ISL_451621, EPI_ISL_451622, EPI_ISL_451623, EPI_ISL_451624, EPI_ISL_451625, EPI_ISL_451626, EPI_ISL_451627, EPI_ISL_451628, EPI_ISL_451629, EPI_ISL_451630 | | | |
| see above | Pathology West - NSW Health Pathology | NSW Health Pathology - Institute of Clinical Pathology and Medical Research; Westmead Hospital; University of Sydney | CIDM-PH et al. |
| EPI_ISL_451631 | Laverty Pathology | NSW Health Pathology - Institute of Clinical Pathology and Medical Research; Westmead Hospital; University of Sydney | CIDM-PH et al. |
| EPI_ISL_451632, EPI_ISL_451633, EPI_ISL_451634, EPI_ISL_451635, EPI_ISL_451636 | South Eastern Area Laboratory Services | NSW Health Pathology - Institute of Clinical Pathology and Medical Research; Westmead Hospital; University of Sydney | CIDM-PH et al. |
| EPI_ISL_451637, EPI_ISL_451638, EPI_ISL_451639, EPI_ISL_451640 | Pathology West - NSW Health Pathology | NSW Health Pathology - Institute of Clinical Pathology and Medical Research; Westmead Hospital; University of Sydney | CIDM-PH et al. |
| EPI_ISL_451641 | Laverty Pathology | NSW Health Pathology - Institute of Clinical Pathology and Medical Research; Westmead Hospital; University of Sydney | CIDM-PH et al. |
| EPI_ISL_451642, EPI_ISL_451643 | Pathology Sydney South West - NSW Health Pathology | NSW Health Pathology - Institute of Clinical Pathology and Medical Research; Westmead Hospital; University of Sydney | CIDM-PH et al. |
| EPI_ISL_451644, EPI_ISL_451645, EPI_ISL_451646, EPI_ISL_451647 | Laboratory of Molecular Biology, Diagnostyka sp. z o.o. | Laboratory of Recombinant Vaccines | Lukasz Rabalski, Anna Piotrowska-Mietelska, Maciej Kosinski, Boguslaw Szewczyk, Krystyna Bienkowska-Szewczyk |
| EPI_ISL_451648, EPI_ISL_451649, EPI_ISL_451650, EPI_ISL_451651, EPI_ISL_451652, EPI_ISL_451653, EPI_ISL_451654 | Hematology Laboratory, Section of Molecular Diagnostics, University Clinical Centre, Medical University of Gdansk | Laboratory of Recombinant Vaccines | Lukasz Rabalski, Adam Sodol, Aneta Szulc, Krzysztof Lewandowski, Ewa Milosz, Marlena Robakowska, Boguslaw Szewczyk, Krystyna Bienkowska-Szewczyk |
| EPI_ISL_451655, EPI_ISL_451656, EPI_ISL_451657, EPI_ISL_451658, EPI_ISL_451659, EPI_ISL_451660, EPI_ISL_451661, EPI_ISL_451662, EPI_ISL_451663, EPI_ISL_451664 | State Sanitary Inspectorate | Laboratory of Recombinant Vaccines | Lukasz Rabalski, Boguslaw Szewczyk, Krystyna Bienkowska-Szewczyk, Jaroslaw Pinkas |
| EPI_ISL_451665 | NYU Langone Health | Departments of Pathology and Medicine, New York University School of Medicine | Maria Aguero-Rosenfeld, Brendan Belovarac, Margaret Black, Ludovic Boytard, John Cadley, Paolo Cotzia, John Chen, Dacia Dimartino, Xiaojun Feng, Tatyana Gindin, Emily Guzman, Adriana Heguy, Megan Hogan, Emily Huang, George Jour, Alireza Khodadadi-Jamayran, Lawrence H. Lin, Raven Luther, Andrew Lytle, Christian Marier, Matthew T. Maurano, Mark J. Mulligan, Peter Meyn, Raquel Ordonez Ciriza, Iman Osman, Jared Pinnell, Vanessa Raabe, Sitharam Ramaswami, Amy Rapkiewicz, Andre M. Ribeiro-dos-Santos, Marie Samanovic-Golden, Antonio Serrano, Guomiao Shen, Matija Snuderl, Theodore Vougiouklakis, Nick Vulpescu, Gael Westby, Paul Zappile, Yutong Zhang |
| EPI_ISL_451666 | M.P Shah Government Medocal college Jamnagar | Gujarat Biotechnology Research Centre | Binita Aring, Janvi Raval, Zarna Patel, Monika Gandhi, Pinal Trivedi, Maharshi Pandya, Amit Kanani, Nidhi Patel, Nitin Savaliya, Raghawendra Kumar, Dinesh Kumar, Zuber Saiyed, Komal Patel, Labdhi Pandya, Snehal Bagatharia, Ramesh Pandit, Tejas Shah, Ankit Hinsu, Pritesh Sabara, Apurvasinh Puvar, Bhavesh Modi, Gaurishankar Shrimali, R D Dixit, A M Kadri, Akanksha Verma, Chaitanya Joshi, Madhvi Joshi, |
| EPI_ISL_451668, EPI_ISL_451669, EPI_ISL_451670, EPI_ISL_451672, EPI_ISL_451673, EPI_ISL_451674, EPI_ISL_451675, EPI_ISL_451676, EPI_ISL_451677, EPI_ISL_451678, EPI_ISL_451680, EPI_ISL_451681, EPI_ISL_451682, EPI_ISL_451683, EPI_ISL_451684, EPI_ISL_451685, EPI_ISL_451688, EPI_ISL_451689, EPI_ISL_451690, EPI_ISL_451691, EPI_ISL_451692, EPI_ISL_451693, EPI_ISL_451694, EPI_ISL_451695, EPI_ISL_451696, EPI_ISL_451697, EPI_ISL_451698, EPI_ISL_451699, EPI_ISL_451701, EPI_ISL_451702, EPI_ISL_451703, EPI_ISL_451704, EPI_ISL_451705, EPI_ISL_451706, EPI_ISL_451707, EPI_ISL_451708, EPI_ISL_451709, EPI_ISL_451711, EPI_ISL_451712, EPI_ISL_451713, EPI_ISL_451714, EPI_ISL_451716, EPI_ISL_451717, EPI_ISL_451718, EPI_ISL_451719, EPI_ISL_451720, EPI_ISL_451721, EPI_ISL_451722, EPI_ISL_451723, EPI_ISL_451724, EPI_ISL_451725, EPI_ISL_451727, EPI_ISL_451728, EPI_ISL_451730, EPI_ISL_451731, EPI_ISL_451732, EPI_ISL_451734, EPI_ISL_451736, EPI_ISL_451738, EPI_ISL_451740, EPI_ISL_451741, EPI_ISL_451742, EPI_ISL_451744, EPI_ISL_451745, EPI_ISL_451746, EPI_ISL_451747, EPI_ISL_451748, EPI_ISL_451749, EPI_ISL_451750, EPI_ISL_451751, EPI_ISL_451752, EPI_ISL_451753, EPI_ISL_451754, EPI_ISL_451755, EPI_ISL_451756, EPI_ISL_451758, EPI_ISL_451759, EPI_ISL_451760, EPI_ISL_451761, EPI_ISL_451763, EPI_ISL_451764, EPI_ISL_451765, EPI_ISL_451766, EPI_ISL_451767, EPI_ISL_451769, EPI_ISL_451770, EPI_ISL_451771, EPI_ISL_451772, EPI_ISL_451773, EPI_ISL_451776, EPI_ISL_451777, EPI_ISL_451778, EPI_ISL_451780, EPI_ISL_451781, EPI_ISL_451782, EPI_ISL_451784, EPI_ISL_451785, EPI_ISL_451786, EPI_ISL_451787, EPI_ISL_451788, EPI_ISL_451789, EPI_ISL_451790, EPI_ISL_451791, EPI_ISL_451792, EPI_ISL_451793, EPI_ISL_451794, EPI_ISL_451796, EPI_ISL_451797, EPI_ISL_451798, EPI_ISL_451799, EPI_ISL_451800, EPI_ISL_451801, EPI_ISL_451802, EPI_ISL_451803, EPI_ISL_451804, EPI_ISL_451805, EPI_ISL_451806, EPI_ISL_451807, EPI_ISL_451808, EPI_ISL_451809, EPI_ISL_451810, EPI_ISL_451811, EPI_ISL_451812, EPI_ISL_451813, EPI_ISL_451814, EPI_ISL_451815, EPI_ISL_451816, EPI_ISL_451817, EPI_ISL_451818, EPI_ISL_451819, EPI_ISL_451821, EPI_ISL_451822, EPI_ISL_451823, EPI_ISL_451824, EPI_ISL_451826, EPI_ISL_451827, EPI_ISL_451829, EPI_ISL_451830, EPI_ISL_451831, EPI_ISL_451832, EPI_ISL_451833, EPI_ISL_451834, EPI_ISL_451835, EPI_ISL_451837, EPI_ISL_451838, EPI_ISL_451839, EPI_ISL_451840, EPI_ISL_451842, EPI_ISL_451843, EPI_ISL_451844, EPI_ISL_451846, EPI_ISL_451848, EPI_ISL_451849, EPI_ISL_451850, EPI_ISL_451851, EPI_ISL_451852, EPI_ISL_451853, EPI_ISL_451854, EPI_ISL_451855, EPI_ISL_451859, EPI_ISL_451862, EPI_ISL_451863, EPI_ISL_451864, EPI_ISL_451867, EPI_ISL_451869, EPI_ISL_451870, EPI_ISL_451871, EPI_ISL_451872, | | | |

| | | | |
|---|---|---|---|
| EPI_ISL_451873, EPI_ISL_451874, EPI_ISL_451875, EPI_ISL_451877, EPI_ISL_451878, EPI_ISL_451879, EPI_ISL_451883, EPI_ISL_451884, EPI_ISL_451885, EPI_ISL_451886, EPI_ISL_451887, EPI_ISL_451888, EPI_ISL_451889, EPI_ISL_451891, EPI_ISL_451892, EPI_ISL_451894, EPI_ISL_451900, EPI_ISL_451902, EPI_ISL_451905, EPI_ISL_451906, EPI_ISL_451907, EPI_ISL_451908, EPI_ISL_451909, EPI_ISL_451911, EPI_ISL_451913, EPI_ISL_451914, EPI_ISL_451915, EPI_ISL_451916 | | | |
| see above | Viollier AG | Department of Biosystems Science and Engineering, ETH Zürich | Christian Beisel, Sarah Nadeau, Ivan Topolsky, Pedro Ferreira, Philipp Jablonski, Susana Posada-Céspedes, Tobias Schär, Ina Nissen, Natascha Santacroce, Elodie Burcklen, Christiane Beckmann, Maurice Redondo, Olivier Kobel, Christoph Noppen, Sophie Seidel, Noemie Santamaria de Souza, Niko Beerenwinkel, Tanja Stadler |
| EPI_ISL_451934 | Research Unit, University Hospital for Infectious Diseases "Dr. Fran Mihaljević" | Cicin Sain lab, Helmholtz Centre for Infection Research | Zeeshan Chaudhry, Kathrin Eschke, Željka Mačak Šafranko, Ivan-Christian Kurolt |
| EPI_ISL_451935 | CUB Hopital Erasme Laboratoire d'Anatomie Pathologique | CUB Hopital Erasme Laboratoire d'Anatomie Pathologique | Prof. Isabelle Salmon, Dr. Nicky D'Haene |
| EPI_ISL_451937, EPI_ISL_451940, EPI_ISL_451941, EPI_ISL_451944, EPI_ISL_451945 | Max von Pettenkofer Institute, Virology, National Reference Center for Retroviruses, LMU München | Laboratory for Functional Genome Analysis, Dept. Genomics, Gene Center of the LMU Munich | Max Muenchhoff, Stefan Krebs, Alexander Graf, Oliver Keppler, Helmut Blum |
| EPI_ISL_451948 | The Republican Research and Practical Center for Epidemiology and Microbiology | Charite Universitatsmedizin Berlin, Institute of Virology | Victor M Corman, Barbara Mühlemann, Talitha Veith, Jorn Beheim-Schwarzbach, Julia Schneider, Terry Jones, Natallia Shmialiova, Natallia Sivets, Christian Drosten |
| EPI_ISL_451958 | Jamil-ur-Rahman Center for Genome Research, Dr. Panjwani Center for Molecular Medicine and Drug Research, International Center for Chemical and Biological Sciences, University of Karachi | Jamil-ur-Rahman Center for Genome Research, Dr. Panjwani Center for Molecular Medicine and Drug Research, International Center for Chemical and Biological Sciences, University of Karachi | Shakeel,M., Raza,S.A., Khan,S., Khan,B.A., Zahid,M., Qureshi,M.A.and Khan,I.A |
| EPI_ISL_451961 | Istituto Zooprofilattico Sperimentale Puglia e Basilicata; Dipartimento di Bioscienze, Biotecnologie e Biofarmaceutica dell'Università degli Studi di Bari "A.Moro"; Istituto di Biomembrane. Bioenergetica e Biotecnologie Molecolari del Consiglio Nazionale delle Ricerche di Bari | Beaconlab (Bioinformatics Evolution and Comparative Genomics lab), Dept of Biosciences, University of Milan | Parisi A.,Pesole G., Manzari C., Chiara M. |
| EPI_ISL_451963 | Federal Budget Institution of Science, State Research Center for Applied Microbiology & Biotechnology | Federal Budget Institution of Science, State Research Center for Applied Microbiology & Biotechnology | Dyatlov I, Shemyakin I, Khramov M, Bogun A, Kislichkina A, Frolov V, Shishkina L, Sizova A, Chekan L, Blagodatskikh S, Podkopaev Y, Kosilova I, Koroleva-Ushakova A, Tyurin E, Galkina E, Slukina N, Shaikhutdinova R, Kalmantayev T, Kalmantayeva O, Fursova N, Silkina M, Gorbatov A, Titareva G, Firstova V, Makarova M, Gapelchenkova T, Solovieva A, Slukin P, Dentovskaya S, Detushev K, Vagayskaya A, Kartsev N, Detusheva E, Zeninskaya N, Ivanov S, Kartseva A, Platonov M, Hlyntseva A, Khomyakov A, Chernysh S, Krasilnikova E, Ryabko A, Solomentsev V, Teymurazov M, Bakhteeva I, Borzilov A, Skryabin Y, Kanashenko M, Abaimova A, Kolchanova A, Novikova T, Goncharova J, Timofeev V, Kuzina E, Fursov M, Zhumakaev R, Marin M, Denisenko E, Trunyakova A, Kuzin V |
| EPI_ISL_451966, EPI_ISL_451969, EPI_ISL_451970 | Federal Budget Institution of Science, State Research Center for Applied Microbiology & Biotechnology | Federal Budget Institution of Science, State Research Center for Applied Microbiology & Biotechnology | Dyatlov I, Shemyakin I, Khramov M, Bogun A, Kislichkina A, Frolov V, Shishkina L, Sizova A, Chekan L, Blagodatskikh S, Podkopaev Y, Kosilova I, Koroleva-Ushakova A, Tyurin E, Galkina E, Slukina N, Shaikhutdinova R, Kalmantayev T, Kalmantayeva O, Fursova N, Silkina M, Gorbatov A, Titareva G, Firstova V, Makarova M, Gapelchenkova T, Solovieva A, Slukin P, Dentovskaya S, Detushev K, Vagayskaya A, Kartsev N, Detusheva E, Zeninskaya N, Ivanov S, Kartseva A, Platonov M, Hlyntseva A, Khomyakov A, Chernysh S, Krasilnikova E, Ryabko A, Solomentsev V, Teymurazov M, Bakhteeva I, Borzilov A, Skryabin Y, Kanashenko M, Abaimova A, Kolchanova A, Novikova T, Goncharova J, Timofeev V, Kuzina E, Fursov M, Zhumakaev R, Marin M, Denisenko E, Trunyakova A, Kuzin V |
| EPI_ISL_451971 | 1. ViroGenetics - BSL3 Laboratory of Virology, Małopolska Centre of Biotechnology, Jagiellonian University; 2. II Department of Internal Medicine, Faculty of Medicine, Jagiellonian University Medical College; 3. DIAGNOSTYKA Ltd. | 1. ViroGenetics - BSL3 Laboratory of Virology, Małopolska Centre of Biotechnology, Jagiellonian University; 2. II Department of Internal Medicine, Faculty of Medicine, Jagiellonian University Medical College. | Marek Sanak, Marcin Surmiak, Monika Gąsecka-Czapla, Wojciech Branicki, Paweł P Łabaj, Marta Rogalska-Kupiec, Jakub Swadźba, Krzysztof Pyrć |
| EPI_ISL_451972 | 1. ViroGenetics - BSL3 Laboratory of Virology, Małopolska Centre of Biotechnology, Jagiellonian University; 2. II Department of Internal Medicine, Faculty of Medicine, Jagiellonian University Medical College; 3. DIAGNOSTYKA Ltd. | 1. ViroGenetics - BSL3 Laboratory of Virology, Małopolska Centre of Biotechnology, Jagiellonian University; 2. II Department of Internal Medicine, Faculty of Medicine, Jagiellonian University Medical College. | Marek Sanak, Marcin Surmiak, Monika Gąsecka-Czapla, Wojciech Branicki, Paweł P Łabaj, Marta Rogalska-Kupiec, Jakub Swadźba, Krzysztof Pyrć |
| EPI_ISL_451973, EPI_ISL_451974 | 1. ViroGenetics - BSL3 Laboratory of Virology, Małopolska Centre of Biotechnology, Jagiellonian University; 2. II Department of Internal Medicine, Faculty of Medicine, Jagiellonian University Medical College; 3. DIAGNOSTYKA Ltd. | 1. ViroGenetics - BSL3 Laboratory of Virology, Małopolska Centre of Biotechnology, Jagiellonian University; 2. II Department of Internal Medicine, Faculty of Medicine, Jagiellonian University Medical College. | Marek Sanak, Marcin Surmiak, Monika Gąsecka-Czapla, Wojciech Branicki, Paweł P Łabaj, Marta Rogalska-Kupiec, Jakub Swadźba, Krzysztof Pyrć |
| EPI_ISL_451975, EPI_ISL_451976, EPI_ISL_451977, EPI_ISL_451978, EPI_ISL_451979, EPI_ISL_451980, EPI_ISL_451981, EPI_ISL_451982, EPI_ISL_451983, EPI_ISL_451984, EPI_ISL_451985, EPI_ISL_451986, EPI_ISL_451987 | | | |
| see above | 1. ViroGenetics - BSL3 Laboratory of Virology, Małopolska Centre of Biotechnology, Jagiellonian University; 2. II Department of Internal Medicine, Faculty of Medicine, Jagiellonian University Medical College; 3. DIAGNOSTYKA Ltd. | 1. ViroGenetics - BSL3 Laboratory of Virology, Małopolska Centre of Biotechnology, Jagiellonian University; 2. II Department of Internal Medicine, Faculty of Medicine, Jagiellonian University Medical College. | Marek Sanak, Marcin Surmiak, Monika Gąsecka-Czapla, Wojciech Branicki, Paweł P Łabaj, Marta Rogalska-Kupiec, Jakub Swadźba, Krzysztof Pyrć |
| EPI_ISL_451988, EPI_ISL_451989, EPI_ISL_451990, EPI_ISL_451991, EPI_ISL_451992, EPI_ISL_451993, EPI_ISL_451994, EPI_ISL_451995, EPI_ISL_451996, EPI_ISL_451997, EPI_ISL_451998, EPI_ISL_451999, EPI_ISL_452000, EPI_ISL_452001, EPI_ISL_452002, EPI_ISL_452003, EPI_ISL_452004, EPI_ISL_452005, EPI_ISL_452007, EPI_ISL_452008, EPI_ISL_452009, EPI_ISL_452010, EPI_ISL_452012, EPI_ISL_452013, EPI_ISL_452014, EPI_ISL_452015, EPI_ISL_452016, EPI_ISL_452017, EPI_ISL_452018, EPI_ISL_452019, EPI_ISL_452020, EPI_ISL_452021, EPI_ISL_452022, EPI_ISL_452023, EPI_ISL_452024, EPI_ISL_452025, EPI_ISL_452026, EPI_ISL_452027, EPI_ISL_452028, EPI_ISL_452029, EPI_ISL_452030, EPI_ISL_452031, EPI_ISL_452032, EPI_ISL_452033, EPI_ISL_452034, EPI_ISL_452035, EPI_ISL_452037, EPI_ISL_452038, EPI_ISL_452039, EPI_ISL_452042, EPI_ISL_452043, EPI_ISL_452044, EPI_ISL_452045, EPI_ISL_452048, EPI_ISL_452049, EPI_ISL_452050, EPI_ISL_452052, EPI_ISL_452053, EPI_ISL_452054, EPI_ISL_452055, EPI_ISL_452056, EPI_ISL_452057, EPI_ISL_452058, EPI_ISL_452059, EPI_ISL_452060, EPI_ISL_452061, EPI_ISL_452062, EPI_ISL_452064, EPI_ISL_452065, EPI_ISL_452066, EPI_ISL_452067, EPI_ISL_452068, EPI_ISL_452069, EPI_ISL_452070, EPI_ISL_452074, EPI_ISL_452075, EPI_ISL_452076, EPI_ISL_452077, EPI_ISL_452078, EPI_ISL_452079, EPI_ISL_452080, EPI_ISL_452082, EPI_ISL_452083, EPI_ISL_452084, EPI_ISL_452085, EPI_ISL_452086, EPI_ISL_452087, EPI_ISL_452088, EPI_ISL_452089, EPI_ISL_452090, EPI_ISL_452091, EPI_ISL_452092, EPI_ISL_452093, EPI_ISL_452094, EPI_ISL_452098, EPI_ISL_452099, EPI_ISL_452100 | | | |
| see above | Department of Clinical Microbiology, Copenhagen University Hospital, Hvidovre, Kettegaard Alle 30, 2650 Hvidovre. | Albertsen lab, Department of Chemistry and Bioscience, Aalborg University, Denmark | Rasmus Kirkegaard |
| EPI_ISL_452101, EPI_ISL_452102 | Department of Virus and Microbiological Special Diagnostics, Statens Serum Institut, Copenhagen, Denmark, Artillerivej 5, 2300 Copenahgen S | Albertsen lab, Department of Chemistry and Bioscience, Aalborg University, Denmark | Rasmus Kirkegaard |
| EPI_ISL_452103, EPI_ISL_452104 | Max von Pettenkofer Institute, Virology, National Reference Center for Retroviruses, LMU München | Laboratory for Functional Genome Analysis, Dept. Genomics, Gene Center of the LMU Munich | Max Muenchhoff, Stefan Krebs, Alexander Graf, Oliver Keppler, Helmut Blum |
| EPI_ISL_452105 | Texas DSHS Lab Services | Pathogen Discovery, Respiratory Viruses Branch, Division of Viral Diseases, Centers for Disease Control and Prevention | Yan Li, Anna Montmayeur, Ying Tao, Krista Queen, Jing Zhang, Anna Uehara, Clinton R. Paden, Rachel Marine, Mary S. Keckler, Alison S. Laufer Halpin, Haibin Wang, Christopher A. Elkins, Zachary Weiner, Suxiang Tong |
| EPI_ISL_452106, EPI_ISL_452107 | LA Office of Public Health Laboratories | Pathogen Discovery, Respiratory Viruses Branch, Division of Viral Diseases, Centers for Disease Control and Prevention | Yan Li, Anna Montmayeur, Ying Tao, Krista Queen, Jing Zhang, Anna Uehara, Clinton R. Paden, Rachel Marine, Mary S. Keckler, Alison S. Laufer Halpin, Haibin Wang, Christopher A. Elkins, Zachary Weiner, Suxiang Tong |
| EPI_ISL_452108 | MN Department of Health | Pathogen Discovery, Respiratory Viruses Branch, Division of Viral Diseases, Centers for Disease Control and Prevention | Yan Li, Anna Montmayeur, Ying Tao, Krista Queen, Jing Zhang, Anna Uehara, Clinton R. Paden, Rachel Marine, Mary S. Keckler, Alison S. Laufer Halpin, Haibin Wang, Christopher A. Elkins, Zachary Weiner, Suxiang Tong |
| EPI_ISL_452109 | NM Department of Health | Pathogen Discovery, Respiratory Viruses Branch, Division of Viral Diseases, Centers for Disease Control and Prevention | Yan Li, Anna Montmayeur, Ying Tao, Krista Queen, Jing Zhang, Anna Uehara, Clinton R. Paden, Rachel Marine, Mary S. Keckler, Alison S. Laufer Halpin, Haibin Wang, Christopher A. Elkins, Zachary Weiner, Suxiang Tong |
| EPI_ISL_452110, EPI_ISL_452111 | FL Bureau of Public Health Laboratories | Pathogen Discovery, Respiratory Viruses Branch, Division of Viral Diseases, Centers for Disease Control and Prevention | Yan Li, Anna Montmayeur, Ying Tao, Krista Queen, Jing Zhang, Anna Uehara, Clinton R. Paden, Rachel Marine, Mary S. Keckler, Alison S. Laufer Halpin, Haibin Wang, Christopher A. Elkins, Zachary Weiner, Suxiang Tong |
| EPI_ISL_452112 | Texas DSHS Lab Services | Pathogen Discovery, Respiratory Viruses Branch, Division of Viral Diseases, Centers for Disease Control and Prevention | Yan Li, Anna Montmayeur, Ying Tao, Krista Queen, Jing Zhang, Anna Uehara, Clinton R. Paden, Rachel Marine, Mary S. Keckler, Alison S. Laufer Halpin, Haibin Wang, Christopher A. Elkins, Zachary Weiner, Suxiang Tong |
| EPI_ISL_452113 | MN Department of Health | Pathogen Discovery, Respiratory Viruses Branch, Division of Viral Diseases, Centers for Disease Control and Prevention | Yan Li, Anna Montmayeur, Ying Tao, Krista Queen, Jing Zhang, Anna Uehara, Clinton R. Paden, Rachel Marine, Mary S. Keckler, Alison S. Laufer Halpin, Haibin Wang, Christopher A. Elkins, Zachary Weiner, Suxiang Tong |
| EPI_ISL_452114 | KS Health and Environmental Laboratories | Pathogen Discovery, Respiratory Viruses Branch, Division of Viral Diseases, Centers for Disease Control and Prevention | Yan Li, Anna Montmayeur, Ying Tao, Krista Queen, Jing Zhang, Anna Uehara, Clinton R. Paden, Rachel Marine, Mary S. Keckler, Alison S. Laufer Halpin, Haibin Wang, Christopher A. Elkins, Zachary Weiner, Suxiang Tong |
| EPI_ISL_452115 | Texas DSHS Lab Services | Pathogen Discovery, Respiratory Viruses Branch, Division of Viral Diseases, Centers for Disease Control and Prevention | Yan Li, Anna Montmayeur, Ying Tao, Krista Queen, Jing Zhang, Anna Uehara, Clinton R. Paden, Rachel Marine, Mary S. Keckler, Alison S. Laufer Halpin, Haibin Wang, Christopher A. Elkins, Zachary Weiner, Suxiang Tong |
| EPI_ISL_452116, EPI_ISL_452117 | NJ Public Health and Environmental Laboratories | Pathogen Discovery, Respiratory Viruses Branch, Division of Viral Diseases, Centers for Disease Control and Prevention | Yan Li, Anna Montmayeur, Ying Tao, Krista Queen, Jing Zhang, Anna Uehara, Clinton R. Paden, Rachel Marine, Mary S. Keckler, Alison S. Laufer Halpin, Haibin Wang, Christopher A. Elkins, Zachary Weiner, Suxiang Tong |
| EPI_ISL_452118 | AZ SPHL, Arizona Department of Health Services | Pathogen Discovery, Respiratory Viruses Branch, Division of Viral Diseases, Centers for Disease Control and Prevention | Yan Li, Anna Montmayeur, Ying Tao, Krista Queen, Jing Zhang, Anna Uehara, Clinton R. Paden, Rachel Marine, Mary S. Keckler, Alison S. Laufer Halpin, Haibin Wang, Christopher A. Elkins, Zachary Weiner, Suxiang Tong |
| EPI_ISL_452119 | VI-US Virgin Islands Department of Health | Pathogen Discovery, Respiratory Viruses Branch, Division of Viral Diseases, Centers for Disease Control and Prevention | Yan Li, Anna Montmayeur, Ying Tao, Krista Queen, Jing Zhang, Anna Uehara, Clinton R. Paden, Rachel Marine, Mary S. Keckler, Alison S. Laufer Halpin, Haibin Wang, Christopher A. Elkins, Zachary Weiner, Suxiang Tong |
| EPI_ISL_452120 | Georgia Department of Health | Pathogen Discovery, Respiratory Viruses Branch, Division of Viral Diseases, Centers for Disease Control and Prevention | Jing Zhang, Anna Montmayeur, Yan Li, Ying Tao, Krista Queen, Anna Uehara, Clinton R. Paden, Rachel Marine, Mary S. Keckler, Alison S. Laufer Halpin, Haibin Wang, Christopher A. Elkins, Zachary Weiner, Suxiang Tong |
| EPI_ISL_452121, EPI_ISL_452122, EPI_ISL_452123 | VI-US Virgin Islands Department of Health | Pathogen Discovery, Respiratory Viruses Branch, Division of Viral Diseases, Centers for Disease Control and Prevention | Jing Zhang, Anna Montmayeur, Yan Li, Ying Tao, Krista Queen, Anna Uehara, Clinton R. Paden, Rachel Marine, Mary S. Keckler, Alison S. Laufer Halpin, Haibin Wang, Christopher A. Elkins, Zachary Weiner, Suxiang Tong |
| EPI_ISL_452124 | NC State Laboratory of Public Health | Pathogen Discovery, Respiratory Viruses Branch, Division of Viral Diseases, Centers for Disease Control and Prevention | Jing Zhang, Anna Montmayeur, Yan Li, Ying Tao, Krista Queen, Anna Uehara, Clinton R. Paden, Rachel Marine, Mary S. Keckler, Alison S. Laufer Halpin, Haibin Wang, Christopher A. Elkins, Zachary Weiner, Suxiang Tong |
| EPI_ISL_452125 | Georgia Department of Health | Pathogen Discovery, Respiratory Viruses Branch, Division of Viral Diseases, Centers for Disease Control and Prevention | Jing Zhang, Anna Montmayeur, Yan Li, Ying Tao, Krista Queen, Anna Uehara, Clinton R. Paden, Rachel Marine, Mary S. Keckler, Alison S. Laufer Halpin, Haibin Wang, Christopher A. Elkins, Zachary Weiner, Suxiang Tong |
| EPI_ISL_452126 | NC State Laboratory of Public Health | Pathogen Discovery, Respiratory Viruses Branch, Division of Viral Diseases, Centers for Disease Control and Prevention | Jing Zhang, Anna Montmayeur, Yan Li, Ying Tao, Krista Queen, Anna Uehara, Clinton R. Paden, Rachel Marine, Mary S. Keckler, Alison S. Laufer Halpin, Haibin Wang, Christopher A. Elkins, Zachary Weiner, Suxiang Tong |
| EPI_ISL_452127, EPI_ISL_452128, EPI_ISL_452129, EPI_ISL_452130 | CO Department of Public Health and Environment | Pathogen Discovery, Respiratory Viruses Branch, Division of Viral Diseases, Centers for Disease Control and Prevention | Jing Zhang, Anna Montmayeur, Yan Li, Ying Tao, Krista Queen, Anna Uehara, Clinton R. Paden, Rachel Marine, Mary S. Keckler, Alison S. Laufer Halpin, Haibin Wang, Christopher A. Elkins, Zachary Weiner, Suxiang Tong |
| EPI_ISL_452131 | IN State Department of Health Laboratory Services | Pathogen Discovery, Respiratory Viruses Branch, Division of Viral | Jing Zhang, Anna Montmayeur, Yan Li, Ying Tao, Krista Queen, Anna Uehara, Clinton R. Paden, Rachel Marine, Mary S. Keckler, Alison S. Laufer Halpin, Haibin Wang, Christopher A. Elkins, Zachary Weiner, Suxiang

| | | | |
|---|---|---|---|
| | | Diseases, Centers for Disease Control and Prevention | Tong |
| EPI_ISL_452132 | FL Bureau of Public Health Laboratories | Pathogen Discovery, Respiratory Viruses Branch, Division of Viral Diseases, Centers for Disease Control and Prevention | Krista Queen, Yan Li, Anna Montmayeur, Ying Tao, Jing Zhang, Anna Uehara, Clinton R. Paden, Rachel Marine, Mary S. Keckler, Alison S. Laufer Halpin, Haibin Wang, Christopher A. Elkins, Zachary Weiner, Suxiang Tong |
| EPI_ISL_452133 | MN Department of Health | Pathogen Discovery, Respiratory Viruses Branch, Division of Viral Diseases, Centers for Disease Control and Prevention | Krista Queen, Yan Li, Anna Montmayeur, Ying Tao, Jing Zhang, Anna Uehara, Clinton R. Paden, Rachel Marine, Mary S. Keckler, Alison S. Laufer Halpin, Haibin Wang, Christopher A. Elkins, Zachary Weiner, Suxiang Tong |
| EPI_ISL_452134 | IL Department of Public Health Chicago Laboratory | Pathogen Discovery, Respiratory Viruses Branch, Division of Viral Diseases, Centers for Disease Control and Prevention | Krista Queen, Yan Li, Anna Montmayeur, Ying Tao, Jing Zhang, Anna Uehara, Clinton R. Paden, Rachel Marine, Mary S. Keckler, Alison S. Laufer Halpin, Haibin Wang, Christopher A. Elkins, Zachary Weiner, Suxiang Tong |
| EPI_ISL_452137, EPI_ISL_452138 | VI-US Virgin Islands Department of Health | Pathogen Discovery, Respiratory Viruses Branch, Division of Viral Diseases, Centers for Disease Control and Prevention | Anna Uehara, Yan Li, Anna Montmayeur, Ying Tao, Krista Queen, Jing Zhang, Clinton R. Paden, Rachel Marine, Haibin Wang, Bettina Bankamp, Zachary Weiner, Suxiang Tong |
| EPI_ISL_452139 | Instituto de Diagnostico y Referencia Epidemiologicos (INDRE) | Instituto de diagnóstico y Referencia Epidemiologicos (INDRE) | Ramirez-Gonzalez Ernesto, Garces-Ayala Fabiola, Araiza-Rodriguez Adnan, Mendieta-Condado Edgar, Rodriguez-Maldonado Abril, Wong-Arambula Claudia, Barrera-Badillo Gisela, Hernandez-Rivas Lucia, Lopez-Martinez Irma |
| EPI_ISL_452140 | CUB Hopital Erasme Laboratoire d'Anatomie Pathologique | CUB Hopital Erasme Laboratoire d'Anatomie Pathologique | Isabelle Salmon, Nicky D'Haene |
| EPI_ISL_452142 | CUB Hopital Erasme Laboratoire d'Anatomie Pathologique | CUB Hopital Erasme Laboratoire d'Anatomie Pathologique | Isabelle Salmon, Nikcy D'Haene |
| EPI_ISL_452145 | Yale COVID-19 Biorepository | Grubaugh Lab - Yale School of Public Health | Joseph Fauver, Tara Alpert, Anderson Brito, Anne Wyllie, Chantal Vogels, Mary Petrone, Cole Jensen, Chaney Kalinich, Isabel Ott, Arnau Casanovas, Catherine Muenker, Adam Moore, Alice Lu, Maria Tokuyama, Patrick Wong, Peiwen Lu, Saad Omer, Richard Martinello, Allison Nelson, Shelli Farhadian, Akiko Iwasaki, Charlese Dela Cruz, Albert Ko, Nathan Grubaugh |
| EPI_ISL_452148, EPI_ISL_452149 | CUB Hopital Erasme Laboratoire d'Anatomie Pathologique | CUB Hopital Erasme Laboratoire d'Anatomie Pathologique | Isabelle Salmon, Nicky D'Haene |
| EPI_ISL_452150 | CUB Hopital Erasme Laboratoire d'Anatomie Pathologique | CUB Hopital Erasme Laboratoire d'Anatomie Pathologique | Prof. Isabelle Salmon, Dr Nicky D'Haene |
| EPI_ISL_452151 | CUB Hopital Erasme Laboratoire d'Anatomie Pathologique | CUB Hopital Erasme Laboratoire d'Anatomie Pathologique | Prof. Isabelle Salmon, Dr.Nikcy D'Haene |
| EPI_ISL_452152 | CUB Hopital Erasme Laboratoire d'Anatomie Pathologique | CUB Hopital Erasme Laboratoire d'Anatomie Pathologique | Prof. Isabelle Salmon, Dr Nicky D'Haene |
| EPI_ISL_452154, EPI_ISL_452157, EPI_ISL_452158, EPI_ISL_452161, EPI_ISL_452163, EPI_ISL_452166, EPI_ISL_452167, EPI_ISL_452168, EPI_ISL_452169, EPI_ISL_452171, EPI_ISL_452172, EPI_ISL_452173, EPI_ISL_452176 | | | |
| see above | Utah Public Health Laboratory | Utah Public Health Laboratory | Erin Young, Kelly Oakeson |
| EPI_ISL_452178, EPI_ISL_452179 | Laboratory Medicine | Department of Laboratory Medicine, Lin-Kou Chang Gung Memorial Hospital, Taoyuan, Taiwan | Kuo-Chien Tsao, Yu-Nong Gong, Shu-Li Yang, Yi-Chun Liu, Chung-Guei Huang, Mei-Jen Hsiao, Po-Wei Huang, Cheng-Ta Yang, Cheng-Hsun Chiu, Peng-Nien Huang, Kuo-Ming Lee, Guang-Wu Chen, Shin-Ru Shih |
| EPI_ISL_452181, EPI_ISL_452182, EPI_ISL_452183, EPI_ISL_452184, EPI_ISL_452185, EPI_ISL_452186, EPI_ISL_452187, EPI_ISL_452188, EPI_ISL_452189 | ULSS9 Distretto di Bussolengo | Istituto Zooprofilattico Sperimentale delle Venezie | Adelaide Milani, Alessia Schivo, Annalisa Salviato, Erika Giorgia Quaranta, Gianpiero Zamperin, Ambra Pastori, Bianca Zecchin, Alice Fusaro, Calogero Terregino, Antonia Ricci |
| EPI_ISL_452190, EPI_ISL_452191 | ULSS9 Distretto di San Bonifacio | Istituto Zooprofilattico Sperimentale delle Venezie | Adelaide Milani, Alessia Schivo, Annalisa Salviato, Erika Giorgia Quaranta, Gianpiero Zamperin, Ambra Pastori, Bianca Zecchin, Alice Fusaro, Calogero Terregino, Antonia Ricci |
| EPI_ISL_452192, EPI_ISL_452193, EPI_ISL_452194, EPI_ISL_452195, EPI_ISL_452196, EPI_ISL_452197, EPI_ISL_452198, EPI_ISL_452199, EPI_ISL_452200, EPI_ISL_452201, EPI_ISL_452202, EPI_ISL_452203, EPI_ISL_452204, EPI_ISL_452205, EPI_ISL_452206, EPI_ISL_452207, EPI_ISL_452208, EPI_ISL_452209, EPI_ISL_452210, EPI_ISL_452211, EPI_ISL_452212, EPI_ISL_452213, EPI_ISL_452214, EPI_ISL_452215, EPI_ISL_452216, EPI_ISL_452217 | | | |
| see above | NIV Influenza | NIV Influenza | Potdar V |
| EPI_ISL_452218, EPI_ISL_452219, EPI_ISL_452220, EPI_ISL_452221, EPI_ISL_452222, EPI_ISL_452223 | Goethe University Hospital Frankfurt | Institute for Medical Virology, Goethe University Hospital Frankfurt | Tuna Toptan, Sebastian Hoehl, Sandra Westhaus, Denisa Bojkova, Annemarie Berger, Björn Rotter, Klaus Hoffmeier, Jindrich Cinatl, Sandra Ciesek, and Marek Widera |
| EPI_ISL_452229, EPI_ISL_452230 | Narhalsan Backa vardcentral | The Public Health Agency of Sweden | Mats Olsson, Anna-Malin Linde, Maria Lind Karlberg, Oskar Karlsson Lindsjo, Olov Svartstrom, Anna Risberg, Theresa Enkirch, Mia Brytting, Karin Tegmark-Wisell |
| EPI_ISL_452231, EPI_ISL_452232 | Saroledens Familjelakare | The Public Health Agency of Sweden | Katarina Jarbur, Anna-Malin Linde, Maria Lind Karlberg, Oskar Karlsson Lindsjo, Olov Svartstrom, Anna Risberg, Theresa Enkirch, Mia Brytting, Karin Tegmark-Wisell |
| EPI_ISL_452233 | Victoria Vard och Halsa | The Public Health Agency of Sweden | Sarah Henriksson, Anna-Malin Linde, Maria Lind Karlberg, Oskar Karlsson Lindsjo, Olov Svartstrom, Anna Risberg, Theresa Enkirch, Mia Brytting, Karin Tegmark-Wisell |
| EPI_ISL_452234 | Din Klinik | The Public Health Agency of Sweden | Helene Warnborg, Anna-Malin Linde, Maria Lind Karlberg, Oskar Karlsson Lindsjo, Olov Svartstrom, Anna Risberg, Theresa Enkirch, Mia Brytting, Karin Tegmark-Wisell |
| EPI_ISL_452235 | Narhalsan Backa vardcentral | The Public Health Agency of Sweden | Mats Olsson, Anna-Malin Linde, Maria Lind Karlberg, Oskar Karlsson Lindsjo, Olov Svartstrom, Anna Risberg, Theresa Enkirch, Mia Brytting, Karin Tegmark-Wisell |
| EPI_ISL_452236 | VC Sorgenfrimottagningen | The Public Health Agency of Sweden | Lisa Esbjornsson Klemendz, Anna-Malin Linde, Maria Lind Karlberg, Oskar Karlsson Lindsjo, Olov Svartstrom, Anna Risberg, Theresa Enkirch, Mia Brytting, Karin Tegmark-Wisell |
| EPI_ISL_452237 | Huddinge VC | The Public Health Agency of Sweden | Anders Johansson, Anna-Malin Linde, Maria Lind Karlberg, Oskar Karlsson Lindsjo, Olov Svartstrom, Anna Risberg, Theresa Enkirch, Mia Brytting, Karin Tegmark-Wisell |
| EPI_ISL_452238 | Narhalsan Sjobo vardcentral | The Public Health Agency of Sweden | Lovisa Hjerten, Anna-Malin Linde, Maria Lind Karlberg, Oskar Karlsson Lindsjo, Olov Svartstrom, Anna Risberg, Theresa Enkirch, Mia Brytting, Karin Tegmark-Wisell |
| EPI_ISL_452239 | Narhalsan Backa vardcentral | The Public Health Agency of Sweden | Mats Olsson, Anna-Malin Linde, Maria Lind Karlberg, Oskar Karlsson Lindsjo, Olov Svartstrom, Anna Risberg, Theresa Enkirch, Mia Brytting, Karin Tegmark-Wisell |
| EPI_ISL_452240 | Huddinge VC | The Public Health Agency of Sweden | Anders Johansson, Anna-Malin Linde, Maria Lind Karlberg, Oskar Karlsson Lindsjo, Olov Svartstrom, Anna Risberg, Theresa Enkirch, Mia Brytting, Karin Tegmark-Wisell |
| EPI_ISL_452241 | Huslakarna Varmbadhuset Varberg | The Public Health Agency of Sweden | Johanna Hilmersson, Anna-Malin Linde, Maria Lind Karlberg, Oskar Karlsson Lindsjo, Olov Svartstrom, Anna Risberg, Theresa Enkirch, Mia Brytting, Karin Tegmark-Wisell |
| EPI_ISL_452242 | Wernstedt Medical AB | The Public Health Agency of Sweden | Eva Sandberg, Anna-Malin Linde, Maria Lind Karlberg, Oskar Karlsson Lindsjo, Olov Svartstrom, Anna Risberg, Theresa Enkirch, Mia Brytting, Karin Tegmark-Wisell |
| EPI_ISL_452251, EPI_ISL_452252, EPI_ISL_452254, EPI_ISL_452256, EPI_ISL_452257 | Respiratory Virus Unit, Microbiology Services Colindale, Public Health England | Respiratory Virus Unit, Microbiology Services Colindale, Public Health England | Steven Platt, Shahjahan Miah, Angie Lackenby, Omolola Akinbami, Tina Talts, Leena Bhaw, Richard Myers, Monica Galiano, Kirstin Edwards, Jonathan Hubb, Joanna Ellis, Maria Zambon |
| EPI_ISL_452262, EPI_ISL_452264, EPI_ISL_452266, EPI_ISL_452270, EPI_ISL_452272, EPI_ISL_452274, EPI_ISL_452278, EPI_ISL_452279, EPI_ISL_452280, EPI_ISL_452281, EPI_ISL_452282, EPI_ISL_452283, EPI_ISL_452284, EPI_ISL_452285, EPI_ISL_452287, EPI_ISL_452288, EPI_ISL_452289, EPI_ISL_452290, EPI_ISL_452291, EPI_ISL_452292, EPI_ISL_452294, EPI_ISL_452295, EPI_ISL_452297, EPI_ISL_452298, EPI_ISL_452300, EPI_ISL_452301, EPI_ISL_452302, EPI_ISL_452303, EPI_ISL_452305, EPI_ISL_452306, EPI_ISL_452309, EPI_ISL_452310, EPI_ISL_452311, EPI_ISL_452316, EPI_ISL_452317, EPI_ISL_452318, EPI_ISL_452319, EPI_ISL_452320, EPI_ISL_452321, EPI_ISL_452322, EPI_ISL_452323, EPI_ISL_452324, EPI_ISL_452325, EPI_ISL_452326 | | | |
| see above | Michigan Department of Health and Human Services, Bureau of Laboratories | Michigan Department of Health and Human Services, Bureau of Laboratories | Blankenship HM, Riner D, Soehnlen MK |
| EPI_ISL_452327, EPI_ISL_452328, EPI_ISL_452329, EPI_ISL_452330, EPI_ISL_452331, EPI_ISL_452332, EPI_ISL_452333, EPI_ISL_452334, EPI_ISL_452335, EPI_ISL_452336, EPI_ISL_452337, EPI_ISL_452338, EPI_ISL_452339, EPI_ISL_452340, EPI_ISL_452341, EPI_ISL_452342, EPI_ISL_452343, EPI_ISL_452344, EPI_ISL_452345, EPI_ISL_452346, EPI_ISL_452347, EPI_ISL_452348, EPI_ISL_452349, EPI_ISL_452350, EPI_ISL_452351, EPI_ISL_452352, EPI_ISL_452353, EPI_ISL_452354, EPI_ISL_452355, EPI_ISL_452356, EPI_ISL_452357, EPI_ISL_452358, EPI_ISL_452359, EPI_ISL_452360, EPI_ISL_452361, EPI_ISL_452362, EPI_ISL_452363, EPI_ISL_452364 | | | |
| see above | Laboratory of Infectious Diseases Center of Beijing Ditan Hospital | Laboratory of Infectious Diseases Center of Beijing Ditan Hospital | Siyuan Yang, Chengjie Jie, Fengting Yu, Yunxia Tang, Liting Yan, Linghang Wang |
| EPI_ISL_452367, EPI_ISL_452371, EPI_ISL_452372, EPI_ISL_452374, EPI_ISL_452379, EPI_ISL_452380, EPI_ISL_452381, EPI_ISL_452383, EPI_ISL_452385, EPI_ISL_452387, EPI_ISL_452392, EPI_ISL_452394, EPI_ISL_452396, EPI_ISL_452397, EPI_ISL_452398, EPI_ISL_452400, EPI_ISL_452402, EPI_ISL_452404, EPI_ISL_452405, EPI_ISL_452406, EPI_ISL_452407, EPI_ISL_452409, EPI_ISL_452410, EPI_ISL_452411, EPI_ISL_452413, EPI_ISL_452414, EPI_ISL_452426, EPI_ISL_452429, EPI_ISL_452430, EPI_ISL_452432, EPI_ISL_452433, EPI_ISL_452434, EPI_ISL_452436, EPI_ISL_452439, EPI_ISL_452441, EPI_ISL_452443, EPI_ISL_452444, EPI_ISL_452447, EPI_ISL_452448, EPI_ISL_452449, EPI_ISL_452452 | | | |
| see above | Servicio de Microbiología. HRU de Málaga. Servicio Andaluz de Salud | SeqCOVID-SPAIN consortium/IBV(CSIC) | Inmaculada de Toro Peinado, Maria Concepción Mediavilla Gradolph, Begoña Palop Borrás and SeqCOVID-SPAIN consortium |
| EPI_ISL_452453, EPI_ISL_452454, EPI_ISL_452457, EPI_ISL_452459, EPI_ISL_452460, EPI_ISL_452461, EPI_ISL_452462, EPI_ISL_452464, EPI_ISL_452465, EPI_ISL_452466, EPI_ISL_452467, EPI_ISL_452468, EPI_ISL_452469, EPI_ISL_452470, EPI_ISL_452471 | | | |
| see above | Hospital Universitario Puerta del Mar de Cádiz - INIBICA | SeqCOVID-SPAIN consortium/IBV(CSIC) | Salud Rodríguez-Pallares, Fátima-Galán-Sánchez, Manuel Rodríguez-Iglesias and SeqCOVID-SPAIN consortium |
| EPI_ISL_452473, EPI_ISL_452474, EPI_ISL_452476, EPI_ISL_452477, EPI_ISL_452479, EPI_ISL_452480, EPI_ISL_452481, EPI_ISL_452483, EPI_ISL_452484, EPI_ISL_452485, EPI_ISL_452486, EPI_ISL_452487, EPI_ISL_452488, EPI_ISL_452490, EPI_ISL_452491, EPI_ISL_452492, EPI_ISL_452493, EPI_ISL_452495, EPI_ISL_452496, EPI_ISL_452498, EPI_ISL_452500, EPI_ISL_452503, EPI_ISL_452505, EPI_ISL_452506, EPI_ISL_452507, EPI_ISL_452508, EPI_ISL_452509, EPI_ISL_452510, EPI_ISL_452513, EPI_ISL_452514, EPI_ISL_452516, EPI_ISL_452517, EPI_ISL_452518, EPI_ISL_452519, EPI_ISL_452520, EPI_ISL_452521, EPI_ISL_452523, EPI_ISL_452524, EPI_ISL_452526, EPI_ISL_452527, EPI_ISL_452528, EPI_ISL_452531, EPI_ISL_452532, EPI_ISL_452534, EPI_ISL_452535, EPI_ISL_452538, EPI_ISL_452539, EPI_ISL_452540, EPI_ISL_452541 | | | |
| see above | Clínica Universidad de Navarra. Servicio de Enfermedades Infecciosas y Microbiología clínica | SeqCOVID-SPAIN consortium/IBV(CSIC) | Mirian Fernández-Alonso, Jose Luis del Pozo and SeqCOVID-SPAIN consortium |
| EPI_ISL_452544, EPI_ISL_452546, EPI_ISL_452547, EPI_ISL_452549, EPI_ISL_452551, EPI_ISL_452552, EPI_ISL_452553, EPI_ISL_452554, EPI_ISL_452555, EPI_ISL_452556, EPI_ISL_452557, EPI_ISL_452558, EPI_ISL_452559, EPI_ISL_452560, EPI_ISL_452561, EPI_ISL_452562, EPI_ISL_452564, EPI_ISL_452566, EPI_ISL_452567, EPI_ISL_452570, EPI_ISL_452571, EPI_ISL_452572, EPI_ISL_452573, EPI_ISL_452574, EPI_ISL_452575, EPI_ISL_452576, EPI_ISL_452577, EPI_ISL_452578, EPI_ISL_452579, EPI_ISL_452580, EPI_ISL_452581, EPI_ISL_452582, EPI_ISL_452583, EPI_ISL_452584, EPI_ISL_452586, EPI_ISL_452588, EPI_ISL_452590, EPI_ISL_452592, EPI_ISL_452593, EPI_ISL_452594, EPI_ISL_452595, EPI_ISL_452596, EPI_ISL_452597, EPI_ISL_452598, EPI_ISL_452600, EPI_ISL_452601, EPI_ISL_452603, EPI_ISL_452604, EPI_ISL_452605, EPI_ISL_452606, EPI_ISL_452609, EPI_ISL_452610, EPI_ISL_452612, EPI_ISL_452613, EPI_ISL_452615 | | | |
| see above | Servicio de Microbiología y Parasitología clínica. UCEIMP. Hospital Universitario Virgen del Rocío/IBIS/CSIC/US. | SeqCOVID-SPAIN consortium/IBV(CSIC) | Guillermo Martín Gutiérrez, Ángel Rodríguez Villodres, Lidia Gálvez Benitez, Verónica González Galán, Javier Aznar Martín and SeqCOVID-SPAIN consortium |
| EPI_ISL_452618, EPI_ISL_452620, EPI_ISL_452621, EPI_ISL_452622, EPI_ISL_452623, EPI_ISL_452624, EPI_ISL_452625, EPI_ISL_452626, EPI_ISL_452627, EPI_ISL_452630, EPI_ISL_452631, EPI_ISL_452632, EPI_ISL_452633, EPI_ISL_452634, EPI_ISL_452635, EPI_ISL_452636, EPI_ISL_452637, EPI_ISL_452638, EPI_ISL_452639, EPI_ISL_452640, EPI_ISL_452641, EPI_ISL_452642, EPI_ISL_452643, EPI_ISL_452644, EPI_ISL_452645, EPI_ISL_452646, EPI_ISL_452647, EPI_ISL_452648, EPI_ISL_452649, EPI_ISL_452650, EPI_ISL_452651, EPI_ISL_452652, EPI_ISL_452653, EPI_ISL_452657, EPI_ISL_452658, EPI_ISL_452659, EPI_ISL_452661, EPI_ISL_452662, EPI_ISL_452663, EPI_ISL_452664, EPI_ISL_452665, EPI_ISL_452666, EPI_ISL_452669, EPI_ISL_452670, EPI_ISL_452671, EPI_ISL_452672, EPI_ISL_452675, EPI_ISL_452677, EPI_ISL_452678, EPI_ISL_452679, EPI_ISL_452680, EPI_ISL_452681, EPI_ISL_452682, EPI_ISL_452683, EPI_ISL_452684, EPI_ISL_452686, EPI_ISL_452688, EPI_ISL_452689, EPI_ISL_452691 | | | |
| see above | Servicio de Microbiología. Hospital Universitario Donostia. OSI Donostialdea. Área de Enfermedades Infecciosas, Grupo de Infección Respiratoria y Resistencia Antimicrobiana. Instituto de Investigación Sanitaria Biodonostia. | SeqCOVID-SPAIN consortium/IBV(CSIC) | Gustavo Cilla, Milagrosa Montes, Luis Piñeiro, Jose Maria Marimón and SeqCOVID-SPAIN consortium |
| EPI_ISL_452692, EPI_ISL_452693, EPI_ISL_452694, EPI_ISL_452695, EPI_ISL_452696, EPI_ISL_452698, EPI_ISL_452699, EPI_ISL_452700, EPI_ISL_452701, EPI_ISL_452703, EPI_ISL_452704, EPI_ISL_452707, EPI_ISL_452708, EPI_ISL_452709, EPI_ISL_452710, EPI_ISL_452711, EPI_ISL_452712, EPI_ISL_452713, EPI_ISL_452714, EPI_ISL_452715, EPI_ISL_452716, EPI_ISL_452717, EPI_ISL_452718, EPI_ISL_452719, EPI_ISL_452720, EPI_ISL_452721, EPI_ISL_452722, EPI_ISL_452723, EPI_ISL_452724, EPI_ISL_452725, EPI_ISL_452726, EPI_ISL_452727, EPI_ISL_452728, EPI_ISL_452729, EPI_ISL_452730, EPI_ISL_452731, EPI_ISL_452732, EPI_ISL_452733, EPI_ISL_452735, EPI_ISL_452736, EPI_ISL_452737, EPI_ISL_452738, EPI_ISL_452739, EPI_ISL_452740, EPI_ISL_452741, EPI_ISL_452742, EPI_ISL_452743, EPI_ISL_452744, EPI_ISL_452745, EPI_ISL_452746, EPI_ISL_452747, EPI_ISL_452748, EPI_ISL_452749, EPI_ISL_452750, EPI_ISL_452751, EPI_ISL_452753, EPI_ISL_452754, EPI_ISL_452755, EPI_ISL_452759, EPI_ISL_452761, EPI_ISL_452762, EPI_ISL_452763, EPI_ISL_452765, EPI_ISL_452766, EPI_ISL_452767, EPI_ISL_452768, EPI_ISL_452769, EPI_ISL_452770, EPI_ISL_452771, EPI_ISL_452772, EPI_ISL_452773, EPI_ISL_452774, EPI_ISL_452775, EPI_ISL_452777, EPI_ISL_452780, EPI_ISL_452782, EPI_ISL_452783, EPI_ISL_452784, EPI_ISL_452785, EPI_ISL_452786 | | | |
| see above | Hospital Universitario Araba. Vitoria-Gasteiz, | SeqCOVID-SPAIN consortium/IBV(CSIC) | Silvia Hernáez Crespo, Carmen Gómez González, Amaia Aguirre Quiñonero, Marina Fernández Torres, Maria Rosario Almela Ferrer, Maria Concepción Lecaroz Agara, Andrés Canut Blasco and SeqCOVID-SPAIN consortium |
| EPI_ISL_452790, EPI_ISL_452791, EPI_ISL_452792, EPI_ISL_452793, EPI_ISL_452794, EPI_ISL_452795 | ICAR-National Institute of High Security Animal Diseases | ICAR-National Institute of High Security Animal Diseases | Anamika Mishra, Ashutosh Aasdev, Sandeep Bhatia, Harshad Murugkar, Chakradhar Tosh, Niranjan Mishra, Shanmugasundaram Nagarajan, Katherukamem Rajukumar, Richa Sood, G Venkatesh, Atul Kumar Pateriya, Manoj Kumar, Shashi Bhushan Sudhakar, Fateh Singh, Sethil Kumar D, Senmannan Kalaiyarasu, Pradeep Gandhale, Naveen Kumar, Chandan Kumar Dubey, Sushil Tripathi, Sandeep Kumar Jhade, Meghna Tripathi, Suman Kumari Shah, Pushpendra Singh, Pushpendra Namdeo, Suman Mishra, Rupal Singh, Vishnupriya Patil, Dipesh Kumar Nayak, Vijendra Pal Singh, Ashwin Ashok Raut |

| | | | |
|---|---|---|---|
| EPI_ISL_452796, EPI_ISL_452797, EPI_ISL_452798, EPI_ISL_452799, EPI_ISL_452800, EPI_ISL_452801, EPI_ISL_452802, EPI_ISL_452803, EPI_ISL_452804, EPI_ISL_452805, EPI_ISL_452806, EPI_ISL_452807, EPI_ISL_452808, EPI_ISL_452809, EPI_ISL_452810, EPI_ISL_452811, EPI_ISL_452812, EPI_ISL_452813, EPI_ISL_452814, EPI_ISL_452815, EPI_ISL_452816, EPI_ISL_452817, EPI_ISL_452818, EPI_ISL_452819, EPI_ISL_452820, EPI_ISL_452821, EPI_ISL_452822, EPI_ISL_452823, EPI_ISL_452824, EPI_ISL_452825, EPI_ISL_452826, EPI_ISL_452827, EPI_ISL_452828, EPI_ISL_452829, EPI_ISL_452830, EPI_ISL_452831, EPI_ISL_452832, EPI_ISL_452833, EPI_ISL_452834, EPI_ISL_452835, EPI_ISL_452836, EPI_ISL_452837, EPI_ISL_452838, EPI_ISL_452839, EPI_ISL_452840, EPI_ISL_452841, EPI_ISL_452842, EPI_ISL_452843, EPI_ISL_452844, EPI_ISL_452845, EPI_ISL_452846, EPI_ISL_452847, EPI_ISL_452848, EPI_ISL_452849, EPI_ISL_452850, EPI_ISL_452851, EPI_ISL_452852, EPI_ISL_452853, EPI_ISL_452854, EPI_ISL_452855, EPI_ISL_452856 | | | |
| see above | Virginia DCLS | Virginia DCLS | Virginia DCLS |
| EPI_ISL_452857, EPI_ISL_452858, EPI_ISL_452859, EPI_ISL_452862, EPI_ISL_452863, EPI_ISL_452864, EPI_ISL_452865, EPI_ISL_452866, EPI_ISL_452867, EPI_ISL_452869, EPI_ISL_452871, EPI_ISL_452872, EPI_ISL_452876, EPI_ISL_452877, EPI_ISL_452878, EPI_ISL_452879, EPI_ISL_452880, EPI_ISL_452881, EPI_ISL_452882, EPI_ISL_452884, EPI_ISL_452885, EPI_ISL_452886, EPI_ISL_452888, EPI_ISL_452889, EPI_ISL_452890, EPI_ISL_452891, EPI_ISL_452892, EPI_ISL_452893, EPI_ISL_452894, EPI_ISL_452895, EPI_ISL_452896, EPI_ISL_452897, EPI_ISL_452898, EPI_ISL_452904, EPI_ISL_452906, EPI_ISL_452910, EPI_ISL_452912, EPI_ISL_452913, EPI_ISL_452914, EPI_ISL_452915, EPI_ISL_452916, EPI_ISL_452917, EPI_ISL_452918, EPI_ISL_452920, EPI_ISL_452926, EPI_ISL_452928, EPI_ISL_452929, EPI_ISL_452930, EPI_ISL_452932, EPI_ISL_452934, EPI_ISL_452936, EPI_ISL_452937, EPI_ISL_452938, EPI_ISL_452940, EPI_ISL_452942, EPI_ISL_452944, EPI_ISL_452946, EPI_ISL_452947, EPI_ISL_452948, EPI_ISL_452949, EPI_ISL_452950, EPI_ISL_452952, EPI_ISL_452953, EPI_ISL_452954, EPI_ISL_452955, EPI_ISL_452956, EPI_ISL_452957, EPI_ISL_452958, EPI_ISL_452959, EPI_ISL_452960, EPI_ISL_452961, EPI_ISL_452962, EPI_ISL_452963, EPI_ISL_452965, EPI_ISL_452968, EPI_ISL_452969, EPI_ISL_452971, EPI_ISL_452972, EPI_ISL_452973, EPI_ISL_452974, EPI_ISL_452977, EPI_ISL_452978, EPI_ISL_452979, EPI_ISL_452980, EPI_ISL_452982, EPI_ISL_452983, EPI_ISL_452984, EPI_ISL_452985, EPI_ISL_452986, EPI_ISL_452989, EPI_ISL_452993, EPI_ISL_452994, EPI_ISL_452996, EPI_ISL_452998, EPI_ISL_453000, EPI_ISL_453001, EPI_ISL_453003, EPI_ISL_453004, EPI_ISL_453005 | | | |
| see above | Department of Pathology, University of Cambridge | COVID-19 Genomics UK (COG-UK) Consortium | Luke W Meredith, M. Estée Török , Myra Hosmillo, William L. Hamilton, Martin D. Curran, Theresa Feltwell, Grant Hall, Anna Yakovleva, Fahad A Khokhar, Charlotte J. Houldcroft, Laura G Caller, Aminu S. Jahun, Sarah L. Caddy, Ian Goodfellow |
| EPI_ISL_453006, EPI_ISL_453013, EPI_ISL_453019, EPI_ISL_453030, EPI_ISL_453031, EPI_ISL_453034, EPI_ISL_453035, EPI_ISL_453037, EPI_ISL_453040, EPI_ISL_453041, EPI_ISL_453042, EPI_ISL_453044, EPI_ISL_453052, EPI_ISL_453056, EPI_ISL_453060, EPI_ISL_453061, EPI_ISL_453062, EPI_ISL_453063, EPI_ISL_453065, EPI_ISL_453069, EPI_ISL_453070, EPI_ISL_453072, EPI_ISL_453074, EPI_ISL_453075, EPI_ISL_453076, EPI_ISL_453078, EPI_ISL_453081, EPI_ISL_453083, EPI_ISL_453084, EPI_ISL_453085, EPI_ISL_453086, EPI_ISL_453088, EPI_ISL_453090, EPI_ISL_453097 | | | |
| see above | West of Scotland Specialist Virology Centre, NHSGGC / MRC-University of Glasgow Centre for Virus Research | COVID-19 Genomics UK (COG-UK) Consortium | Ana da Silva Filipe, Natasha Johnson, Kathy Smollett, Daniel Mair, Stephen Carmichael, Lily Tong, Jenna Nichols, Elihu Aranday-Cortes, Kirstyn Brunker, Yasmin Parr, Kyriaki Nomikou; Sarah McDonald, Marc Niebel, Patawee Asamaphan; Richard Orton, Joseph Hughes, Sreenu Vattipally, David L Robertson; Alasdair MacLean, Rory Gunson; Kathy Li, Natasha Jesudason, Rajiv Shah, James Shepherd, Antonia Ho, Emma Thomson |
| EPI_ISL_453099, EPI_ISL_453101, EPI_ISL_453102, EPI_ISL_453103, EPI_ISL_453104, EPI_ISL_453105, EPI_ISL_453106, EPI_ISL_453107, EPI_ISL_453108, EPI_ISL_453109, EPI_ISL_453111, EPI_ISL_453114, EPI_ISL_453115, EPI_ISL_453116, EPI_ISL_453117, EPI_ISL_453119, EPI_ISL_453121, EPI_ISL_453122, EPI_ISL_453125, EPI_ISL_453126, EPI_ISL_453127, EPI_ISL_453128, EPI_ISL_453129, EPI_ISL_453130, EPI_ISL_453132, EPI_ISL_453133, EPI_ISL_453141, EPI_ISL_453142, EPI_ISL_453143, EPI_ISL_453144, EPI_ISL_453145, EPI_ISL_453146, EPI_ISL_453147, EPI_ISL_453148, EPI_ISL_453149, EPI_ISL_453150, EPI_ISL_453152, EPI_ISL_453153, EPI_ISL_453154, EPI_ISL_453155, EPI_ISL_453156, EPI_ISL_453157, EPI_ISL_453158, EPI_ISL_453160, EPI_ISL_453161, EPI_ISL_453163, EPI_ISL_453164, EPI_ISL_453165, EPI_ISL_453166, EPI_ISL_453169, EPI_ISL_453170, EPI_ISL_453171, EPI_ISL_453172, EPI_ISL_453173, EPI_ISL_453174, EPI_ISL_453177, EPI_ISL_453178, EPI_ISL_453179, EPI_ISL_453180, EPI_ISL_453181, EPI_ISL_453182, EPI_ISL_453183, EPI_ISL_453184, EPI_ISL_453185, EPI_ISL_453187, EPI_ISL_453188, EPI_ISL_453190, EPI_ISL_453191, EPI_ISL_453192, EPI_ISL_453193, EPI_ISL_453194 | | | |
| see above | Virology Department, Royal Infirmary of Edinburgh, NHS Lothian / School of Biological Sciences, University of Edinburgh / Institute of Genetics and Molecular Medicine, University of Edinburgh | COVID-19 Genomics UK (COG-UK) Consortium | McHugh M, Dewar R, Rooke S, Gallagher M, Balcaza C, O'Toole Á, Scher E, Hill V, McCrone JT, Colquhoun R, Yu X, Jackson B, Rambaut A, Williams TC, Templeton K |
| EPI_ISL_453255, EPI_ISL_453327, EPI_ISL_453433, EPI_ISL_453435, EPI_ISL_453437, EPI_ISL_453441, EPI_ISL_453442, EPI_ISL_453446, EPI_ISL_453452, EPI_ISL_453453 | Liverpool Clinical Laboratories | COVID-19 Genomics UK (COG-UK) Consortium | Sam Haldenby, Anistair Lucaci, Steve Paterson, Julian Hiscox, Alistair Darby, M Almsaud, A Alrezaihi, Muhannad Alruwaili, Stuart D Armstrong, Jones Benjamin , Eleanor G Bentley, Anu Chawla, Jordan J Clark, Angela Cowell, Richard Eccles, Isabel García-Dorival, Matthew Gemmell, Alessandro Gerada, PKF Gilmore, Richard Gregory, Ximeng Han, Catherine Hartley, Margaret Hughes, Miren Iturriza-Gomara, James Johnson, L Luu, Jenifer Manson , Charlotte Nelson, Elaine O'Toole, Cassie Olateju, Rebekah Penrice-Randal , Lucille Rainbow, N.P Randle, Trevor Ian Robinson, Parul Sharma, Ghada T Shawli, James P Stewart , Neil Swainston, Ecaterina Vamos, Joanne Watts, Mark Whitehead |
| EPI_ISL_453460, EPI_ISL_453461, EPI_ISL_453462, EPI_ISL_453463, EPI_ISL_453464 | University College London, Great Ormond Street Hospital for Children NHS Foundation Trust, Imperial College Healthcare NHS Trust | COVID-19 Genomics UK (COG-UK) Consortium | Sergi Castellano, Rachel Williams, Mark Kristiansen, Paola Resende Silva, Sunando Roy, Tony Brooks, Helena Tutill, Paola Niola, Patricia Dyal, Charlotte Williams, Leysa Forrest, Yasmin Panchbhaya, Jacqueline Findlay, Sam Weeks, Julianne Brown, Kathryn Harris, Paul Randell, James Price, Alison Holmes, Judith Breuer |
| EPI_ISL_453465, EPI_ISL_453466, EPI_ISL_453467, EPI_ISL_453468, EPI_ISL_453469, EPI_ISL_453470, EPI_ISL_453471, EPI_ISL_453472, EPI_ISL_453473, EPI_ISL_453474, EPI_ISL_453475, EPI_ISL_453476, EPI_ISL_453477, EPI_ISL_453478, EPI_ISL_453479, EPI_ISL_453480, EPI_ISL_453481, EPI_ISL_453482, EPI_ISL_453483, EPI_ISL_453484, EPI_ISL_453485 | | | |
| see above | Regional Virus Laboratory, Belfast Health and Social Care Trust | COVID-19 Genomics UK (COG-UK) Consortium | Conall McCaughey, James McKenna, Tanya Curran, Susan Feeney, Alison Watt, Ciara Cox, Mairead Connor, Zoltan Molnar, David Simpson, Derek Fairley |
| EPI_ISL_453489, EPI_ISL_453492, EPI_ISL_453499, EPI_ISL_453507, EPI_ISL_453509, EPI_ISL_453534, EPI_ISL_453538, EPI_ISL_453542, EPI_ISL_453544 | Northumbria University / South Tees Hospitals NHS Foundation Trust / North Cumbria Integrated Care NHS Foundation Trust / North Tees and Hartlepool NHS Foundation Trust / Newcastle Hospitals NHS Foundation Trust | COVID-19 Genomics UK (COG-UK) Consortium | Darren L Smith,Andrew Nelson,Matthew Bashton,Greg R Young,Joshua Loh,John Allan,Mohammad A Tariq,Giles S Holt,Gary Black,Wen C Yew,Lynn Dover ,Paul Baker,Steve Liggett,Sarah Essex,Jane Greenaway ,Debra Padgett,Clive Graham,Garren Scott,Edward Barton ,Emma Swindells ,Brendan Payne,Jennifer Collins,Yusri Taha,Gary Eltringham |
| EPI_ISL_453545, EPI_ISL_453546, EPI_ISL_453547, EPI_ISL_453548, EPI_ISL_453550, EPI_ISL_453551, EPI_ISL_453552, EPI_ISL_453553, EPI_ISL_453554, EPI_ISL_453555, EPI_ISL_453556, EPI_ISL_453558, EPI_ISL_453559, EPI_ISL_453560, EPI_ISL_453561, EPI_ISL_453563, EPI_ISL_453565, EPI_ISL_453566, EPI_ISL_453567, EPI_ISL_453569, EPI_ISL_453570, EPI_ISL_453571, EPI_ISL_453572, EPI_ISL_453573, EPI_ISL_453574, EPI_ISL_453575, EPI_ISL_453577, EPI_ISL_453579, EPI_ISL_453580, EPI_ISL_453582, EPI_ISL_453583, EPI_ISL_453584, EPI_ISL_453585, EPI_ISL_453586, EPI_ISL_453587, EPI_ISL_453588, EPI_ISL_453591, EPI_ISL_453592, EPI_ISL_453593, EPI_ISL_453594, EPI_ISL_453595, EPI_ISL_453596, EPI_ISL_453597, EPI_ISL_453598, EPI_ISL_453599, EPI_ISL_453600, EPI_ISL_453601, EPI_ISL_453603, EPI_ISL_453604, EPI_ISL_453605, EPI_ISL_453606, EPI_ISL_453607, EPI_ISL_453608, EPI_ISL_453609, EPI_ISL_453610, EPI_ISL_453611, EPI_ISL_453612, EPI_ISL_453613 | | | |
| see above | Quadram Institute Bioscience | COVID-19 Genomics UK (COG-UK) Consortium | Dave J. Baker, Gemma L. Kay, Alp Aydin, Thanh Le-Viet, Steven Rudder, Ana P. Tedim, Anastasia Kolyva, Maria Diaz, Leonardo de Oliveira Martins, Nabil-Fareed Alikhan, Lizzie Meadows, Rachael Stanley, Ngozi Elumogo, Muhammed Yasir, Nicholas M. Thomson, Alexander J Trotter, Rachel Gilroy, Samuel Bloomfield, Claire Stuart, Andrew Bell, Reenesh Prakash, Samir Dervisevic, Alison E. Mather, John Wain, Mark Webber, Andrew J. Page, Justin O'Grady |
| EPI_ISL_453615, EPI_ISL_453616, EPI_ISL_453618, EPI_ISL_453619, EPI_ISL_453620, EPI_ISL_453621, EPI_ISL_453622, EPI_ISL_453623, EPI_ISL_453624, EPI_ISL_453625, EPI_ISL_453626, EPI_ISL_453627, EPI_ISL_453628, EPI_ISL_453629, EPI_ISL_453631, EPI_ISL_453632, EPI_ISL_453633, EPI_ISL_453634, EPI_ISL_453635, EPI_ISL_453637, EPI_ISL_453638, EPI_ISL_453639, EPI_ISL_453640, EPI_ISL_453641, EPI_ISL_453642, EPI_ISL_453645, EPI_ISL_453646, EPI_ISL_453647, EPI_ISL_453648, EPI_ISL_453649, EPI_ISL_453650, EPI_ISL_453651, EPI_ISL_453652, EPI_ISL_453655, EPI_ISL_453656, EPI_ISL_453657, EPI_ISL_453658, EPI_ISL_453659, EPI_ISL_453662, EPI_ISL_453663 | | | |
| see above | Queens Medical Centre, Clinical Microbiology Department / DeepSeq Nottingham | COVID-19 Genomics UK (COG-UK) Consortium | Gemma Clark, Wendy Smith, Manjinder Khakh, Hannah Howson-Wells, Jonathan Ball, Patrick McClure, Joseph Chappell, Theocharis Tsoleridis, Nadine Holmes, Matthew Carlisle, Christopher Moore, Fei Sang, Johnny Debebe, Victoria Wright, Matthew Loose |
| EPI_ISL_453670, EPI_ISL_453676, EPI_ISL_453677, EPI_ISL_453679, EPI_ISL_453684, EPI_ISL_453685, EPI_ISL_453687, EPI_ISL_453691 | Centre for Enzyme Innovation, University of Portsmouth / Translational Research Laboratory, Portsmouth Hospitals NHS Trust | COVID-19 Genomics UK (COG-UK) Consortium | Angela Beckett,,Yann Bourgeois,,Garry Scarlett,,Sharon Glaysher,,Scott Elliott,,Kelly Bicknell,,Robert Impey,,Allyson Lloyd,,Sarah Wyllie,,Ethan Butcher,,Anoop Chauhan,,Samuel Robson |
| EPI_ISL_453696, EPI_ISL_453698, EPI_ISL_453699, EPI_ISL_453701, EPI_ISL_453703, EPI_ISL_453704, EPI_ISL_453705, EPI_ISL_453706, EPI_ISL_453707, EPI_ISL_453708, EPI_ISL_453709, EPI_ISL_453710, EPI_ISL_453711, EPI_ISL_453712, EPI_ISL_453713, EPI_ISL_453714, EPI_ISL_453715, EPI_ISL_453717, EPI_ISL_453719, EPI_ISL_453721, EPI_ISL_453722, EPI_ISL_453724, EPI_ISL_453728, EPI_ISL_453729, EPI_ISL_453730, EPI_ISL_453732, EPI_ISL_453736, EPI_ISL_453737, EPI_ISL_453739, EPI_ISL_453740, EPI_ISL_453742, EPI_ISL_453744, EPI_ISL_453745, EPI_ISL_453746, EPI_ISL_453748, EPI_ISL_453749, EPI_ISL_453750, EPI_ISL_453751, EPI_ISL_453752, EPI_ISL_453753, EPI_ISL_453755, EPI_ISL_453756, EPI_ISL_453758, EPI_ISL_453759, EPI_ISL_453760, EPI_ISL_453761, EPI_ISL_453765, EPI_ISL_453766, EPI_ISL_453767, EPI_ISL_453768, EPI_ISL_453769, EPI_ISL_453770, EPI_ISL_453771, EPI_ISL_453772, EPI_ISL_453773, EPI_ISL_453774, EPI_ISL_453775, EPI_ISL_453776, EPI_ISL_453777 | | | |
| see above | Virology Department, Sheffield Teaching Hospitals NHS Foundation Trust/Department of Infection, Immunity and Cardiovascular Disease, The Medical School, University of Sheffield | COVID-19 Genomics UK (COG-UK) Consortium | Thushan de Silva, Matthew Parker, Nikki Smith, Adri Angyal, Rebecca Brown, Luke Green, Rachel Tucker, Paul Parsons, Danielle Groves, Katie Johnson, Laura Carrilero, Alex Keeley, Dave Partridge, Matthew Wyles, Benjamin Lindsey, Mehmet Yavuz, Mohammad Raza, Cariad Evans |
| EPI_ISL_453813, EPI_ISL_453814, EPI_ISL_453815, EPI_ISL_453816, EPI_ISL_453817, EPI_ISL_453818, EPI_ISL_453820, EPI_ISL_453821, EPI_ISL_453822, EPI_ISL_453823, EPI_ISL_453824, EPI_ISL_453825, EPI_ISL_453826, EPI_ISL_453827, EPI_ISL_453828, EPI_ISL_453829, EPI_ISL_453830, EPI_ISL_453832, EPI_ISL_453833, EPI_ISL_453834, EPI_ISL_453835, EPI_ISL_453836, EPI_ISL_453839, EPI_ISL_453840, EPI_ISL_453844, EPI_ISL_453845, EPI_ISL_453846, EPI_ISL_453847, EPI_ISL_453848, EPI_ISL_453850, EPI_ISL_453851, EPI_ISL_453852, EPI_ISL_453853, EPI_ISL_453854, EPI_ISL_453855, EPI_ISL_453856, EPI_ISL_453859, EPI_ISL_453860, EPI_ISL_453865, EPI_ISL_453866, EPI_ISL_453868, EPI_ISL_453869, EPI_ISL_453870, EPI_ISL_453872, EPI_ISL_453873, EPI_ISL_453875, EPI_ISL_453876, EPI_ISL_453877, EPI_ISL_453879, EPI_ISL_453881, EPI_ISL_453884, EPI_ISL_453885, EPI_ISL_453887, EPI_ISL_453889, EPI_ISL_453890, EPI_ISL_453891, EPI_ISL_453892, EPI_ISL_453893, EPI_ISL_453894, EPI_ISL_453895, EPI_ISL_453896, EPI_ISL_453898, EPI_ISL_453901, EPI_ISL_453906, EPI_ISL_453911, EPI_ISL_453912, EPI_ISL_453913, EPI_ISL_453914, EPI_ISL_453915, EPI_ISL_453916, EPI_ISL_453917, EPI_ISL_453922, EPI_ISL_453924, EPI_ISL_453925, EPI_ISL_453927, EPI_ISL_453929, EPI_ISL_453931, EPI_ISL_453933, EPI_ISL_453939, EPI_ISL_453942, EPI_ISL_453943, EPI_ISL_453944, EPI_ISL_453946, EPI_ISL_453947, EPI_ISL_453951, EPI_ISL_453958, EPI_ISL_453961, EPI_ISL_453962, EPI_ISL_453963, EPI_ISL_453964, EPI_ISL_453965, EPI_ISL_453966, EPI_ISL_453967, EPI_ISL_453969, EPI_ISL_453970, EPI_ISL_453971, EPI_ISL_453972, EPI_ISL_453973, EPI_ISL_453974, EPI_ISL_453975, EPI_ISL_453976, EPI_ISL_453977, EPI_ISL_453979, EPI_ISL_453981, EPI_ISL_453982, EPI_ISL_453984, EPI_ISL_453986, EPI_ISL_453987, EPI_ISL_453988, EPI_ISL_453989, EPI_ISL_453990, EPI_ISL_453992, EPI_ISL_453993, EPI_ISL_453996, EPI_ISL_453997, EPI_ISL_453998, EPI_ISL_453999, EPI_ISL_454000, EPI_ISL_454001, EPI_ISL_454003, EPI_ISL_454004, EPI_ISL_454005, EPI_ISL_454006, EPI_ISL_454007, EPI_ISL_454008, EPI_ISL_454009, EPI_ISL_454010, EPI_ISL_454011, EPI_ISL_454012, EPI_ISL_454013, EPI_ISL_454014, EPI_ISL_454015, EPI_ISL_454017, EPI_ISL_454018, EPI_ISL_454019, EPI_ISL_454020, EPI_ISL_454021, EPI_ISL_454022, EPI_ISL_454023, EPI_ISL_454025, EPI_ISL_454026, EPI_ISL_454029, EPI_ISL_454030, EPI_ISL_454031, EPI_ISL_454032, EPI_ISL_454033, EPI_ISL_454034, EPI_ISL_454035, EPI_ISL_454036, EPI_ISL_454037, EPI_ISL_454038, EPI_ISL_454039, EPI_ISL_454040, EPI_ISL_454041, EPI_ISL_454042, EPI_ISL_454043, EPI_ISL_454044, EPI_ISL_454045, EPI_ISL_454046, EPI_ISL_454047, EPI_ISL_454048, EPI_ISL_454049, EPI_ISL_454050, EPI_ISL_454051, EPI_ISL_454052, EPI_ISL_454053, EPI_ISL_454055, EPI_ISL_454057, EPI_ISL_454059, EPI_ISL_454060, EPI_ISL_454061, EPI_ISL_454062, EPI_ISL_454063, EPI_ISL_454064, EPI_ISL_454065, EPI_ISL_454066, EPI_ISL_454067, EPI_ISL_454068, EPI_ISL_454069, EPI_ISL_454070, EPI_ISL_454071, EPI_ISL_454072, EPI_ISL_454074, EPI_ISL_454076, EPI_ISL_454077, EPI_ISL_454078, EPI_ISL_454080, EPI_ISL_454081, EPI_ISL_454082, EPI_ISL_454083, EPI_ISL_454085, EPI_ISL_454086, EPI_ISL_454087, EPI_ISL_454089, EPI_ISL_454090, EPI_ISL_454091, EPI_ISL_454092, EPI_ISL_454093, EPI_ISL_454094, EPI_ISL_454095, EPI_ISL_454096, EPI_ISL_454097, EPI_ISL_454098, EPI_ISL_454099, EPI_ISL_454100, EPI_ISL_454101, EPI_ISL_454103, EPI_ISL_454104, EPI_ISL_454106, EPI_ISL_454111, EPI_ISL_454114, EPI_ISL_454115, EPI_ISL_454116, EPI_ISL_454118, EPI_ISL_454122, EPI_ISL_454123, EPI_ISL_454127, EPI_ISL_454128, EPI_ISL_454129, EPI_ISL_454130, EPI_ISL_454131, EPI_ISL_454132, EPI_ISL_454133, EPI_ISL_454134, EPI_ISL_454135, EPI_ISL_454137, EPI_ISL_454139, EPI_ISL_454141, EPI_ISL_454142, EPI_ISL_454143, EPI_ISL_454144, EPI_ISL_454145, EPI_ISL_454146, EPI_ISL_454147, EPI_ISL_454149, EPI_ISL_454150, EPI_ISL_454154, EPI_ISL_454158, EPI_ISL_454159, EPI_ISL_454162, EPI_ISL_454163, EPI_ISL_454167, EPI_ISL_454168, EPI_ISL_454169, EPI_ISL_454171, EPI_ISL_454172, EPI_ISL_454173, EPI_ISL_454174, EPI_ISL_454178, EPI_ISL_454181, EPI_ISL_454182, EPI_ISL_454183, EPI_ISL_454184, EPI_ISL_454185, EPI_ISL_454186, EPI_ISL_454188, EPI_ISL_454189, EPI_ISL_454190, EPI_ISL_454191, EPI_ISL_454192, EPI_ISL_454193, EPI_ISL_454194, EPI_ISL_454195, EPI_ISL_454196, EPI_ISL_454197, EPI_ISL_454198, EPI_ISL_454199, EPI_ISL_454200, EPI_ISL_454201, EPI_ISL_454202, EPI_ISL_454203, EPI_ISL_454204, EPI_ISL_454205, EPI_ISL_454206, EPI_ISL_454207, EPI_ISL_454208, EPI_ISL_454209, EPI_ISL_454210, EPI_ISL_454211, EPI_ISL_454213, EPI_ISL_454214, EPI_ISL_454215, EPI_ISL_454216, EPI_ISL_454217, EPI_ISL_454218, EPI_ISL_454219, EPI_ISL_454220, EPI_ISL_454221, EPI_ISL_454222, EPI_ISL_454223, EPI_ISL_454224, EPI_ISL_454225, EPI_ISL_454226, EPI_ISL_454227, EPI_ISL_454228, EPI_ISL_454229, EPI_ISL_454230, EPI_ISL_454232, EPI_ISL_454233, EPI_ISL_454234, EPI_ISL_454235, EPI_ISL_454236, EPI_ISL_454237, EPI_ISL_454238, EPI_ISL_454240, EPI_ISL_454241, EPI_ISL_454242, EPI_ISL_454243, EPI_ISL_454244, EPI_ISL_454245, EPI_ISL_454246, EPI_ISL_454247, EPI_ISL_454248, EPI_ISL_454249, EPI_ISL_454250, EPI_ISL_454251, EPI_ISL_454252, EPI_ISL_454253, EPI_ISL_454255, EPI_ISL_454257, EPI_ISL_454258, EPI_ISL_454260, EPI_ISL_454261, EPI_ISL_454262, EPI_ISL_454263, EPI_ISL_454264, EPI_ISL_454266, EPI_ISL_454267, EPI_ISL_454268, EPI_ISL_454269, EPI_ISL_454270, EPI_ISL_454271, EPI_ISL_454272, EPI_ISL_454274, EPI_ISL_454275, EPI_ISL_454276, EPI_ISL_454277, EPI_ISL_454278, EPI_ISL_454279, EPI_ISL_454280, EPI_ISL_454281, EPI_ISL_454282, EPI_ISL_454283, EPI_ISL_454284, EPI_ISL_454285, EPI_ISL_454287, EPI_ISL_454288, EPI_ISL_454289, EPI_ISL_454290, EPI_ISL_454291, EPI_ISL_454292, EPI_ISL_454293, EPI_ISL_454294, EPI_ISL_454295, EPI_ISL_454296, EPI_ISL_454297, EPI_ISL_454298, EPI_ISL_454300, EPI_ISL_454301, EPI_ISL_454302, EPI_ISL_454303, EPI_ISL_454304, EPI_ISL_454307, EPI_ISL_454311, EPI_ISL_454314, EPI_ISL_454315, EPI_ISL_454317, EPI_ISL_454318, EPI_ISL_454319, EPI_ISL_454320, EPI_ISL_454321, EPI_ISL_454322, EPI_ISL_454324, EPI_ISL_454326, EPI_ISL_454327, EPI_ISL_454328, EPI_ISL_454329, EPI_ISL_454330, EPI_ISL_454331, EPI_ISL_454332, EPI_ISL_454333, EPI_ISL_454334, EPI_ISL_454335, EPI_ISL_454337, EPI_ISL_454338, EPI_ISL_454339, EPI_ISL_454340, EPI_ISL_454341, EPI_ISL_454342, EPI_ISL_454343, EPI_ISL_454344, EPI_ISL_454345, EPI_ISL_454346, EPI_ISL_454347, EPI_ISL_454348, EPI_ISL_454349, EPI_ISL_454350, EPI_ISL_454351, EPI_ISL_454352 | | | |
| see above | unknown | Instituto Nacional de Saude (INSA) | Borges et al |
| EPI_ISL_454353 | Microbial Genome Sequencing Center; Microbial Genomic Epidemiology Laboratory | Microbial Genomic Epidemiology Laboratory, University of Pittsburgh | Mustapha M. Mustapha, Jane W. Marsh, Dan Snyder, Marissa P. Griffith, Stephanie L. Mitchell, Vatsala R. Srinivasa, Kady D. Waggle, Chinelo Ezeonwuku, Vaughn S. Cooper, Lee H. Harrison |
| EPI_ISL_454354 | UPMC Clinical Microbiology Laboratory | Microbial Genome Sequencing Center; Microbial Genomic Epidemiology Laboratory, University of Pittsburgh | Mustapha M. Mustapha, Jane W. Marsh, Dan Snyder, Marissa P. Griffith, Stephanie L. Mitchell, Vatsala R. Srinivasa, Kady D. Waggle, Chinelo Ezeonwuku, Vaughn S. Cooper, Lee H. Harrison |
| EPI_ISL_454355, EPI_ISL_454356, EPI_ISL_454357, EPI_ISL_454358, EPI_ISL_454359, EPI_ISL_454360, EPI_ISL_454361, EPI_ISL_454362, EPI_ISL_454363, EPI_ISL_454364, EPI_ISL_454365, EPI_ISL_454366, EPI_ISL_454367, EPI_ISL_454368, EPI_ISL_454369, EPI_ISL_454370, EPI_ISL_454371, EPI_ISL_454372, EPI_ISL_454373, EPI_ISL_454374, EPI_ISL_454375, EPI_ISL_454376, EPI_ISL_454377, EPI_ISL_454378, EPI_ISL_454379, EPI_ISL_454380, EPI_ISL_454381, EPI_ISL_454382, EPI_ISL_454383, EPI_ISL_454384, EPI_ISL_454385, EPI_ISL_454386, EPI_ISL_454387, EPI_ISL_454388, EPI_ISL_454389, EPI_ISL_454390, EPI_ISL_454391, EPI_ISL_454392, EPI_ISL_454393, EPI_ISL_454394, EPI_ISL_454395, EPI_ISL_454396, EPI_ISL_454397, EPI_ISL_454398, EPI_ISL_454399, EPI_ISL_454400, EPI_ISL_454401, EPI_ISL_454402, EPI_ISL_454403, EPI_ISL_454404, EPI_ISL_454405, EPI_ISL_454406, EPI_ISL_454407, EPI_ISL_454408, EPI_ISL_454409, EPI_ISL_454410, EPI_ISL_454411 | | | |
| see above | UPMC Clinical Microbiology Laboratory | Microbial Genome Sequencing Center, Microbial Genomic Epidemiological Laboratory | Mustapha M. Mustapha, Jane W. Marsh, Dan Snyder, Marissa P. Griffith, Stephanie L. Mitchell, Vatsala R. Srinivasa, Kady D. Waggle, Chinelo Ezeonwuku, Vaughn S. Cooper, Lee H. Harrison |
| EPI_ISL_454412, EPI_ISL_454413, EPI_ISL_454414, EPI_ISL_454415 | Dirk Dittmer | Dirk Dittmer | Bailey,A.G., Caro-Vegas,C.P., Dittmer,D., Eason,A.B., Juarez,A., Landis,J.T., McNamara,R.P., Miller,M.B., Moorad,R., Pluta,L.J., Seltzer,T.A., Thompson,C., Vahrson,W., Villamor,F. |
| EPI_ISL_454416 | Department of Medical Microbiology, Leiden University Medical Center | Department of Medical Microbiology, Leiden University Medical Center | Snijder,E.J., Ogando,N.S., Zevenhoven,J.C., Dalebout,T.J., de Vries,J.J. and Sidorov,I. |
| EPI_ISL_454417, EPI_ISL_454418 | Research and Experiment Center, Meizhou People Hospital | Research and Experiment Center, Meizhou People Hospital | Guo,X., Zeng,L. and Yu,Z. |
| EPI_ISL_454431, EPI_ISL_454435 | Maryland Department of Health Laboratories Administration | Maryland Department of Health Laboratories Administration | MDH Laboratories Administration |
| EPI_ISL_454436, EPI_ISL_454437, EPI_ISL_454438, EPI_ISL_454439, EPI_ISL_454440, EPI_ISL_454441, EPI_ISL_454442, EPI_ISL_454443, EPI_ISL_454444, EPI_ISL_454445, EPI_ISL_454446 | | | |
| see above | Halmstad klinisk mikrobiologi | The Public Health Agency of Sweden | Anna-Malin Linde, Maria Lind Karlberg, Mattias Haukland, Reza Advani, Olov Svartstrom, Oskar Karlsson Lindsjo, Petra Edquist, Shamam Muradrasoli, Anna Risberg, Karin Tegmark-Wisell |
| EPI_ISL_454447, EPI_ISL_454448, EPI_ISL_454449, EPI_ISL_454450, EPI_ISL_454451, EPI_ISL_454452, EPI_ISL_454453, EPI_ISL_454454, EPI_ISL_454455, EPI_ISL_454456, EPI_ISL_454457, EPI_ISL_454458, EPI_ISL_454459, EPI_ISL_454460, EPI_ISL_454461, EPI_ISL_454462, EPI_ISL_454463, EPI_ISL_454464, EPI_ISL_454465, EPI_ISL_454466, EPI_ISL_454467, EPI_ISL_454468, EPI_ISL_454469, EPI_ISL_454470, EPI_ISL_454471, EPI_ISL_454472, EPI_ISL_454473, EPI_ISL_454474, EPI_ISL_454475, EPI_ISL_454476, EPI_ISL_454477, EPI_ISL_454478, EPI_ISL_454479, EPI_ISL_454480, EPI_ISL_454481, EPI_ISL_454482, EPI_ISL_454483, EPI_ISL_454484, EPI_ISL_454485, EPI_ISL_454486, EPI_ISL_454487, EPI_ISL_454488, EPI_ISL_454489, EPI_ISL_454490, EPI_ISL_454491, EPI_ISL_454492, EPI_ISL_454493 | | | |
| see above | Karolinska Universitetslaboratoriet | The Public Health Agency of Sweden | Anna-Malin Linde, Maria Lind Karlberg, Mattias Haukland, Reza Advani, Olov Svartstrom, Oskar Karlsson Lindsjo, Petra Edquist, Shamam Muradrasoli, Anna Risberg, Karin Tegmark-Wisell |
| EPI_ISL_454494, EPI_ISL_454495, EPI_ISL_454496 | Skovde/Unilabs | The Public Health Agency of Sweden | Anna-Malin Linde, Maria Lind Karlberg, Mattias Haukland, Reza Advani, Olov Svartstrom, Oskar Karlsson Lindsjo, Petra Edquist, Shamam Muradrasoli, Anna Risberg, Karin Tegmark-Wisell |

| | | | |
|---|---|---|---|
| EPI_ISL_454497, EPI_ISL_454498 | RSE "National Center for Biotechnology" | RSE "National Center for Biotechnology" | Alexandr Shevtsov, Ilyas Akhmetollayev, Viktoriya Lutsay, Asylulan Amirgazin, Askar Abdaliyev, Akbota Rakhmetova, Zabira Aushakhmetova, Ruslan Kalendar, Yerlan Ramankulov |
| EPI_ISL_454499, EPI_ISL_454500, EPI_ISL_454501, EPI_ISL_454502, EPI_ISL_454503, EPI_ISL_454504, EPI_ISL_454505, EPI_ISL_454506, EPI_ISL_454507, EPI_ISL_454508, EPI_ISL_454509, EPI_ISL_454510, EPI_ISL_454511, EPI_ISL_454512, EPI_ISL_454513, EPI_ISL_454514, EPI_ISL_454515, EPI_ISL_454516, EPI_ISL_454517, EPI_ISL_454518, EPI_ISL_454519, EPI_ISL_454520 | | | |
| see above | RSE "National Center for Biotechnology" | RSE "National Center for Biotechnology" | Alexandr Shevtsov, Ilyas Akhmetollayev, Viktoriya Lutsay, Asylulan Amirgazin, Askar Abdaliyev, Akbota Rakhmetova, Zabira Aushakhmetova, Ruslan Kalendar, Yerlan Ramankulov |
| EPI_ISL_454521, EPI_ISL_454522, EPI_ISL_454524, EPI_ISL_454525, EPI_ISL_454526, EPI_ISL_454527, EPI_ISL_454528, EPI_ISL_454529, EPI_ISL_454530, EPI_ISL_454531, EPI_ISL_454532, EPI_ISL_454533, EPI_ISL_454534, EPI_ISL_454536, EPI_ISL_454537, EPI_ISL_454540, EPI_ISL_454542, EPI_ISL_454543, EPI_ISL_454544, EPI_ISL_454546, EPI_ISL_454547, EPI_ISL_454549, EPI_ISL_454551, EPI_ISL_454552, EPI_ISL_454556, EPI_ISL_454557, EPI_ISL_454558, EPI_ISL_454560, EPI_ISL_454561, EPI_ISL_454562, EPI_ISL_454563, EPI_ISL_454564, EPI_ISL_454565, EPI_ISL_454566, EPI_ISL_454567, EPI_ISL_454568, EPI_ISL_454569, EPI_ISL_454570 | | | |
| see above | NIV Influenza | NIV Influenza | Potdar V |
| EPI_ISL_454571 | National Center of Expertise | National Center for Expertise, National Center for Biotechnology, Kazakhstan | Abdaliyev Askar, Shevtsov Alexandr, Akhmetollayev Ilyas, Kalendar Ruslan, Rakhmetova Akbota, , Lutsay Viktoriya, Amirgazin Asylulan, Aushakhmetova Zabira, Ramankulov Yerlan |
| EPI_ISL_454572 | National Center of Expertise | National Center for Expertise, Kazakhstan National Center for Biotechnology, Kazakhstan | Abdaliyev Askar, Shevtsov Alexandr, Akhmetollayev Ilyas, Kalendar Ruslan, Rakhmetova Akbota, , Lutsay Viktoriya, Amirgazin Asylulan, Aushakhmetova Zabira, Ramankulov Yerlan |
| EPI_ISL_454574 | nstitute for Public Health | Laboratory for advanced genomics | Filip Rokić, Lovro Trgovec-Greif, Neven Sučić, Tomislav Rukavina, Igor Jurak, Oliver Vugrek |
| EPI_ISL_454575 | Laboratory of virology, National Center of Expertise | Laboratory of molecular-genetic research, National Center for Expertise, Kazakhstan National Center for Biotechnology, Kazakhstan | Abdaliyev Askar, Shevtsov Alexandr, Akhmetollayev Ilyas, Kalendar Ruslan, Rakhmetova Akbota, , Lutsay Viktoriya, Amirgazin Asylulan, Aushakhmetova Zabira, Ramankulov Yerlan |
| EPI_ISL_454576, EPI_ISL_454577 | Laboratory of virology, National Center of Expertise | Laboratory of molecular-genetic research, National Center for Expertise, Kazakhstan National Center for Biotechnology, Kazakhstan | Abdaliyev Askar, Shevtsov Alexandr, Akhmetollayev Ilyas, Kalendar Ruslan, Rakhmetova Akbota, , Lutsay Viktoriya, Amirgazin Asylulan, Aushakhmetova Zabira, Ramankulov Yerlan |
| EPI_ISL_454578 | University Hospital for Infectious Diseases "Dr. Fran Mihaljević", Research Unit | University of Zagreb, Centre for research and knowledge transfer in biotechnology | Ivan-Christian Kurolt, Jelena Ivancic Jelecki, Anamarija Slovic |
| EPI_ISL_454579, EPI_ISL_454580 | Laboratory of virology, National Center of Expertise | Laboratory of molecular-genetic research, National Center for Expertise, Kazakhstan National Center for Biotechnology, Kazakhstan | Abdaliyev Askar, Shevtsov Alexandr, Akhmetollayev Ilyas, Kalendar Ruslan, Rakhmetova Akbota, , Lutsay Viktoriya, Amirgazin Asylulan, Aushakhmetova Zabira, Ramankulov Yerlan |
| EPI_ISL_454581 | University Hospital for Infectious Diseases "Dr. Fran Mihaljević", Research Unit | University of Zagreb, Centre for research and knowledge transfer in biotechnology | Ivan-Christian Kurolt, Jelena Ivancic Jelecki, Anamarija Slovic |
| EPI_ISL_454582 | Laboratory of virology, National Center of Expertise | Laboratory of molecular-genetic research, National Center for Expertise, Kazakhstan National Center for Biotechnology, Kazakhstan | Abdaliyev Askar, Shevtsov Alexandr, Akhmetollayev Ilyas, Kalendar Ruslan, Rakhmetova Akbota, , Lutsay Viktoriya, Amirgazin Asylulan, Aushakhmetova Zabira, Ramankulov Yerlan |
| EPI_ISL_454583 | University Hospital for Infectious Diseases "Dr. Fran Mihaljević", Research Unit | University of Zagreb, Centre for research and knowledge transfer in biotechnology | Ivan-Christian Kurolt, Jelena Ivancic Jelecki, Anamarija Slovic |
| EPI_ISL_454584, EPI_ISL_454585, EPI_ISL_454586 | Laboratory of virology, National Center of Expertise | Laboratory of molecular-genetic research, National Center for Expertise, Kazakhstan National Center for Biotechnology, Kazakhstan | Abdaliyev Askar, Shevtsov Alexandr, Akhmetollayev Ilyas, Kalendar Ruslan, Rakhmetova Akbota, , Lutsay Viktoriya, Amirgazin Asylulan, Aushakhmetova Zabira, Ramankulov Yerlan |
| EPI_ISL_454587 | Laboratory of virology, National Center of Expertise | Laboratory of molecular-genetic research, National Center for Expertise, Kazakhstan National Center for Biotechnology, Kazakhstan | Abdaliyev Askar, Shevtsov Alexandr, Akhmetollayev Ilyas, Kalendar Ruslan, Rakhmetova Akbota, , Lutsay Viktoriya, Amirgazin Asylulan, Aushakhmetova Zabira, Ramankulov Yerlan |
| EPI_ISL_454588 | University Hospital for Infectious Diseases "Dr. Fran Mihaljević", Research Unit | University of Zagreb, Centre for research and knowledge transfer in biotechnology | Ivan-Christian Kurolt, Jelena Ivancic Jelecki, Anamarija Slovic |
| EPI_ISL_454589 | Laboratory of virology, National Center of Expertise | Laboratory of molecular-genetic research, National Center for Expertise, Kazakhstan National Center for Biotechnology, Kazakhstan | Abdaliyev Askar, Shevtsov Alexandr, Akhmetollayev Ilyas, Kalendar Ruslan, Rakhmetova Akbota, , Lutsay Viktoriya, Amirgazin Asylulan, Aushakhmetova Zabira, Ramankulov Yerlan |
| EPI_ISL_454590, EPI_ISL_454591 | Laboratory of virology, National Center of Expertise | Laboratory of molecular-genetic research, National Center for Expertise, Kazakhstan National Center for Biotechnology, Kazakhstan | Abdaliyev Askar, Shevtsov Alexandr, Akhmetollayev Ilyas, Kalendar Ruslan, Rakhmetova Akbota, , Lutsay Viktoriya, Amirgazin Asylulan, Aushakhmetova Zabira, Ramankulov Yerlan |
| EPI_ISL_454592 | University Hospital for Infectious Diseases "Dr. Fran Mihaljević", Research Unit | University of Zagreb, Centre for research and knowledge transfer in biotechnology | Ivan-Christian Kurolt, Jelena Ivancic Jelecki, Anamarija Slovic |
| EPI_ISL_454593, EPI_ISL_454594 | Laboratory of virology, National Center of Expertise | Laboratory of molecular-genetic research, National Center for Expertise, Kazakhstan National Center for Biotechnology, Kazakhstan | Abdaliyev Askar, Shevtsov Alexandr, Akhmetollayev Ilyas, Kalendar Ruslan, Rakhmetova Akbota, , Lutsay Viktoriya, Amirgazin Asylulan, Aushakhmetova Zabira, Ramankulov Yerlan |
| EPI_ISL_454596, EPI_ISL_454597 | Laboratory of virology, National Center of Expertise | Laboratory of molecular-genetic research, National Center for Expertise, Kazakhstan National Center for Biotechnology, Kazakhstan | Abdaliyev Askar, Shevtsov Alexandr, Akhmetollayev Ilyas, Kalendar Ruslan, Rakhmetova Akbota, , Lutsay Viktoriya, Amirgazin Asylulan, Aushakhmetova Zabira, Ramankulov Yerlan |
| EPI_ISL_454598 | Laboratory of virology, National Center of Expertise | Laboratory of molecular-genetic research, National Center for Expertise, Kazakhstan National Center for Biotechnology, Kazakhstan | Abdaliyev Askar, Shevtsov Alexandr, Akhmetollayev Ilyas, Kalendar Ruslan, Rakhmetova Akbota, , Lutsay Viktoriya, Amirgazin Asylulan, Aushakhmetova Zabira, Ramankulov Yerlan |
| EPI_ISL_454599, EPI_ISL_454600 | Laboratory of virology, National Center of Expertise | Laboratory of molecular-genetic research, National Center for Expertise, Kazakhstan National Center for Biotechnology, Kazakhstan | Abdaliyev Askar, Shevtsov Alexandr, Akhmetollayev Ilyas, Kalendar Ruslan, Rakhmetova Akbota, , Lutsay Viktoriya, Amirgazin Asylulan, Aushakhmetova Zabira, Ramankulov Yerlan |
| EPI_ISL_454601 | Laboratory of virology, National Center of Expertise | Laboratory of molecular-genetic research, National Center for Expertise, Kazakhstan National Center for Biotechnology, Kazakhstan | Abdaliyev Askar, Shevtsov Alexandr, Akhmetollayev Ilyas, Kalendar Ruslan, Rakhmetova Akbota, , Lutsay Viktoriya, Amirgazin Asylulan, Aushakhmetova Zabira, Ramankulov Yerlan |
| EPI_ISL_454602 | Croatian Institute of Public Health | University of Zagreb, Centre for research and knowledge transfer in biotechnology | Irena Tabain, Tatjana Vilibic-Cavlek, Jelena Ivancic Jelecki, Anamarija Slovic |
| EPI_ISL_454603 | Laboratory of virology, National Center of Expertise | Laboratory of molecular-genetic research, National Center for Expertise, Kazakhstan National Center for Biotechnology, Kazakhstan | Abdaliyev Askar, Shevtsov Alexandr, Akhmetollayev Ilyas, Kalendar Ruslan, Rakhmetova Akbota, , Lutsay Viktoriya, Amirgazin Asylulan, Aushakhmetova Zabira, Ramankulov Yerlan |
| EPI_ISL_454604 | Laboratory of virology, National Center of Expertise | Laboratory of molecular-genetic research, National Center for Expertise, Kazakhstan National Center for Biotechnology, Kazakhstan | Abdaliyev Askar, Shevtsov Alexandr, Akhmetollayev Ilyas, Kalendar Ruslan, Rakhmetova Akbota, , Lutsay Viktoriya, Amirgazin Asylulan, Aushakhmetova Zabira, Ramankulov Yerlan |
| EPI_ISL_454605, EPI_ISL_454606 | Institute for Public Health | Laboratory for advanced genomics | Filip Rokić, Lovro Trgovec-Greif, Neven Sučić, Tomislav Rukavina, Igor Jurak, Oliver Vugrek |
| EPI_ISL_454607, EPI_ISL_454608, EPI_ISL_454609, EPI_ISL_454610, EPI_ISL_454611, EPI_ISL_454612, EPI_ISL_454613 | Alameda County Public Health Lab | Chan-Zuckerberg Biohub | CZB Cliahub Consortium |
| EPI_ISL_454615, EPI_ISL_454617, EPI_ISL_454618, EPI_ISL_454619, EPI_ISL_454620, EPI_ISL_454621, EPI_ISL_454622, EPI_ISL_454624, EPI_ISL_454625, EPI_ISL_454626, EPI_ISL_454627, EPI_ISL_454628, EPI_ISL_454629, EPI_ISL_454630, EPI_ISL_454631, EPI_ISL_454632, EPI_ISL_454633, EPI_ISL_454634 | | | |
| see above | UCSF Clinical Microbiology Laboratory | Chan-Zuckerberg Biohub | CZB Cliahub Consortium |
| EPI_ISL_454635 | County Of San Luis Obispo Public Health Laboratory | Chan-Zuckerberg Biohub | CZB Cliahub Consortium |
| EPI_ISL_454636, EPI_ISL_454637, EPI_ISL_454638, EPI_ISL_454639, EPI_ISL_454640, EPI_ISL_454641 | Humboldt County Public Health Laboratory | Chan-Zuckerberg Biohub | CZB Cliahub Consortium |
| EPI_ISL_454642 | CT-Dr. Katherine A. Kelley State Public Health Lab | Pathogen Discovery, Respiratory Viruses Branch, Division of Viral Diseases, Centers for Disease Control and Prevention | Jing Zhang, Ying Tao, Clinton R. Paden, Anna Uehara, Krista Queen, Yan Li, Haibin Wang, Zachary Weiner, Bettina Bankamp, Suxiang Tong |
| EPI_ISL_454643, EPI_ISL_454644 | VI-US Virgin Islands Department of Health | Pathogen Discovery, Respiratory Viruses Branch, Division of Viral Diseases, Centers for Disease Control and Prevention | Jing Zhang, Ying Tao, Clinton R. Paden, Anna Uehara, Krista Queen, Yan Li, Haibin Wang, Zachary Weiner, Bettina Bankamp, Suxiang Tong |
| EPI_ISL_454645 | CT-Dr. Katherine A. Kelley State Public Health Lab | Pathogen Discovery, Respiratory Viruses Branch, Division of Viral Diseases, Centers for Disease Control and Prevention | Jing Zhang, Ying Tao, Clinton R. Paden, Anna Uehara, Krista Queen, Yan Li, Haibin Wang, Zachary Weiner, Bettina Bankamp, Suxiang Tong |
| EPI_ISL_454646 | City of El Paso Department of Public Health Laboratory | Pathogen Discovery, Respiratory Viruses Branch, Division of Viral Diseases, Centers for Disease Control and Prevention | Ying Tao, Clinton R. Paden, Jing Zhang, Anna Uehara, Krista Queen, Yan Li, Haibin Wang, Zachary Weiner, Michael Bowen, Suxiang Tong |
| EPI_ISL_454647, EPI_ISL_454648, EPI_ISL_454649, EPI_ISL_454650, EPI_ISL_454651, EPI_ISL_454652 | VI-US Virgin Islands Department of Health | Pathogen Discovery, Respiratory Viruses Branch, Division of Viral Diseases, Centers for Disease Control and Prevention | Ying Tao, Clinton R. Paden, Jing Zhang, Anna Uehara, Krista Queen, Yan Li, Haibin Wang, Zachary Weiner, Bettina Bankamp, Suxiang Tong |
| EPI_ISL_454654, EPI_ISL_454655, EPI_ISL_454656, EPI_ISL_454657, EPI_ISL_454658, EPI_ISL_454659, EPI_ISL_454661, EPI_ISL_454662, EPI_ISL_454663, EPI_ISL_454664, EPI_ISL_454665, EPI_ISL_454666, EPI_ISL_454667, EPI_ISL_454668, EPI_ISL_454670, EPI_ISL_454672, EPI_ISL_454673, EPI_ISL_454674, EPI_ISL_454675, EPI_ISL_454676, EPI_ISL_454677, EPI_ISL_454678, EPI_ISL_454679, EPI_ISL_454680, EPI_ISL_454681, EPI_ISL_454682, EPI_ISL_454683, EPI_ISL_454684, EPI_ISL_454685, EPI_ISL_454686, EPI_ISL_454687, EPI_ISL_454688, EPI_ISL_454689 | | | |
| see above | County of Santa Clara Public Health Department | Chan-Zuckerberg Biohub | CZB Cliahub Consortium |
| EPI_ISL_454690 | Emory Molecular Diagnostics Laboratory, Emory Healthcare | Piantadosi Lab, Emory Department of Pathology | Ahmed Babiker, Anne Piantadosi |
| EPI_ISL_454691 | Quest Diagnostics | Quest Diagnostics | Anderson,B.P., Rosenthal,S.H., Gerasimova,A., Kagan,R.M. and Owen, R. |
| EPI_ISL_454707, EPI_ISL_454708, EPI_ISL_454709, EPI_ISL_454710, EPI_ISL_454711, EPI_ISL_454714, EPI_ISL_454715, EPI_ISL_454717, EPI_ISL_454718, EPI_ISL_454721, EPI_ISL_454722, EPI_ISL_454723, EPI_ISL_454725, EPI_ISL_454727, EPI_ISL_454728, EPI_ISL_454729, EPI_ISL_454730, EPI_ISL_454731 | | | |
| see above | Utah Public Health Laboratory | Utah Public Health Laboratory | Erin Young, Kelly Oakeson |
| EPI_ISL_454732 | Russian State Collection of Viruses | Pathogenic Microorganisms Variability Laboratory | Denis Protsenko, Alexey Shchetinin, Maria Nikiforova, Elena Shidlovskaya, Nadezhda Kuznetsova, Vladimir Gushchin, Inna Dolzhikova, Daria Grousova, Andrey Botikov, Denis Logunov, Alexander Gintsburg, Alexey Mazus |
| EPI_ISL_454733 | Department of Medical, Biotechnologies University of Siena | Department of Medical, Biotechnologies University of Siena | Cusi,M.G., Pinzauti,D., Gandolfo,C., Anichini,G., Pozzi,G. and Santoro,F. |
| EPI_ISL_454749 | Japanese Quarantine Stations | Pathogen Genomics Center, National Institute of Infectious Diseases | Tsuyoshi Sekizuka, Kentaro Itokawa, Rina Tanaka, Masanori Hashino, Tsutomu Kageyama, Shinji Saito, Ikuyo Takayama, Hideki Hasegawa, Takuri Takahashi, Hajime Kamiya, Takuya Yamagishi, Motoi Suzuki, Takaji Wakita, Makoto Kuroda |
| EPI_ISL_454750, EPI_ISL_454751, EPI_ISL_454752, EPI_ISL_454753, EPI_ISL_454754, EPI_ISL_454755, EPI_ISL_454756, EPI_ISL_454757, EPI_ISL_454758, EPI_ISL_454759, EPI_ISL_454760, EPI_ISL_454761, EPI_ISL_454762, EPI_ISL_454763, EPI_ISL_454764, EPI_ISL_454765, EPI_ISL_454766, EPI_ISL_454767, EPI_ISL_454768, EPI_ISL_454769, EPI_ISL_454770, EPI_ISL_454771, EPI_ISL_454772, EPI_ISL_454773, | | | |

| | | | |
|---|---|---|---|
| EPI_ISL_454774, EPI_ISL_454775, EPI_ISL_454776, EPI_ISL_454777, EPI_ISL_454778, EPI_ISL_454779, EPI_ISL_454780, EPI_ISL_454781, EPI_ISL_454782, EPI_ISL_454783, EPI_ISL_454784, EPI_ISL_454785, EPI_ISL_454786, EPI_ISL_454787, EPI_ISL_454788, EPI_ISL_454789, EPI_ISL_454790, EPI_ISL_454791, EPI_ISL_454792, EPI_ISL_454793, EPI_ISL_454794 | | | |
| see above | Dutch COVID-19 response team | National Institute for Public Health and the Environment (RIVM) | Adam Meijer, Harry Vennema, Jeroen Cremer, Sharon van den Brink, Pieter Overduin, Florian Zwagemaker, Dennis Schmitz, Chantal Reusken, on behalf of the national COVID-19 response team |
| EPI_ISL_454795 | Veterinary Specialized Institute Kraljevo | Veterinary Specialized Institute Kraljevo | Vidanovic,D., Tesovic,B., Sekler,M., Dmitric,M., Debeljak,Z., Matovic,K., Vaskovic,N., Petrovic,T., Volkening,J. and Alfonso,C.L. |
| EPI_ISL_454796, EPI_ISL_454797, EPI_ISL_454798, EPI_ISL_454799, EPI_ISL_454800, EPI_ISL_454801, EPI_ISL_454802, EPI_ISL_454803, EPI_ISL_454804, EPI_ISL_454805, EPI_ISL_454806, EPI_ISL_454807, EPI_ISL_454808, EPI_ISL_454809, EPI_ISL_454810, EPI_ISL_454811, EPI_ISL_454812, EPI_ISL_454813, EPI_ISL_454814, EPI_ISL_454815, EPI_ISL_454816, EPI_ISL_454817, EPI_ISL_454818, EPI_ISL_454819, EPI_ISL_454820, EPI_ISL_454821, EPI_ISL_454822, EPI_ISL_454823, EPI_ISL_454824, EPI_ISL_454825, EPI_ISL_454826, EPI_ISL_454827, EPI_ISL_454828, EPI_ISL_454829 | | | |
| see above | Dirk Dittmer | Dirk Dittmer | Bailey,A.G., Caro-Vegas,C.P., Dittmer,D., Eason,A.B., Juarez,A., Landis,J.T., McNamara,R.P., Miller,M.B., Moorad,R., Pluta,L.J., Seltzer,T.A., Thompson,C., Vahrson,W., Villamor,F. |
| EPI_ISL_454830, EPI_ISL_454831, EPI_ISL_454832, EPI_ISL_454833 | SMS Medical College, Jaipur | CSIR Institute of Genomics and Integrative Biology | Sudhir Bhandari, Rahul Bhoyar, Mohammed Imran, Mohit Divakar, Disha Sharma, Anshul Kumar, Bani Jolly, Rahul Sahlot, Abhinav Jain, Paras Sehgal, Gyan Ranjan, Vinod Scaria, Sridhar Sivasubbu, Sandeep K Mathur |
| EPI_ISL_454858, EPI_ISL_454862, EPI_ISL_454865, EPI_ISL_454866, EPI_ISL_454867 | Translational Health Science and Technology Institute -ESIC medical college and hospital, Faridabad | THSTI Bioassay laboratory | Saurabh Kumar, Jigme Wangchuk, Anil Kumar Pandey, Asim Das, Guruprasad R. Medigeshi |
| EPI_ISL_454868, EPI_ISL_454869, EPI_ISL_454870, EPI_ISL_454871, EPI_ISL_454872, EPI_ISL_454873, EPI_ISL_454874, EPI_ISL_454875, EPI_ISL_454876, EPI_ISL_454877, EPI_ISL_454878, EPI_ISL_454879, EPI_ISL_454880, EPI_ISL_454881, EPI_ISL_454882, EPI_ISL_454883, EPI_ISL_454884, EPI_ISL_454885, EPI_ISL_454886, EPI_ISL_454887, EPI_ISL_454888, EPI_ISL_454889, EPI_ISL_454890, EPI_ISL_454891, EPI_ISL_454892, EPI_ISL_454893, EPI_ISL_454894, EPI_ISL_454895, EPI_ISL_454896, EPI_ISL_454897, EPI_ISL_454898, EPI_ISL_454899, EPI_ISL_454900, EPI_ISL_454901, EPI_ISL_454902 | | | |
| see above | Karolinska Universitetslaboratoriet | The Public Health Agency of Sweden | Anna-Malin Linde, Maria Lind Karlberg, Mattias Haukland, Reza Advani, Olov Svartstrom, Oskar Karlsson Lindsjo, Petra Edquist, Shamam Muradrasoli, Anna Risberg, Karin Tegmark-Wisell |
| EPI_ISL_454903 | Klinisk mikrobiologi, UAS | The Public Health Agency of Sweden | Anna-Malin Linde, Maria Lind Karlberg, Mattias Haukland, Reza Advani, Olov Svartstrom, Oskar Karlsson Lindsjo, Petra Edquist, Shamam Muradrasoli, Anna Risberg, Karin Tegmark-Wisell |
| EPI_ISL_454904, EPI_ISL_454905, EPI_ISL_454906, EPI_ISL_454907, EPI_ISL_454908, EPI_ISL_454909, EPI_ISL_454910, EPI_ISL_454911, EPI_ISL_454912, EPI_ISL_454913, EPI_ISL_454914, EPI_ISL_454915, EPI_ISL_454916, EPI_ISL_454917, EPI_ISL_454918, EPI_ISL_454919, EPI_ISL_454920, EPI_ISL_454921, EPI_ISL_454923, EPI_ISL_454924, EPI_ISL_454926, EPI_ISL_454927, EPI_ISL_454928, EPI_ISL_454929, EPI_ISL_454930, EPI_ISL_454931, EPI_ISL_454932, EPI_ISL_454933, EPI_ISL_454934, EPI_ISL_454935, EPI_ISL_454936, EPI_ISL_454937, EPI_ISL_454938, EPI_ISL_454939, EPI_ISL_454940, EPI_ISL_454941, EPI_ISL_454942, EPI_ISL_454943, EPI_ISL_454944, EPI_ISL_454946, EPI_ISL_454947, EPI_ISL_454948, EPI_ISL_454949, EPI_ISL_454950, EPI_ISL_454951, EPI_ISL_454952, EPI_ISL_454953, EPI_ISL_454954, EPI_ISL_454955, EPI_ISL_454956, EPI_ISL_454957, EPI_ISL_454958, EPI_ISL_454960, EPI_ISL_454961, EPI_ISL_454962, EPI_ISL_454963, EPI_ISL_454965, EPI_ISL_454967, EPI_ISL_454968, EPI_ISL_454969, EPI_ISL_454971, EPI_ISL_454972, EPI_ISL_454973, EPI_ISL_454974, EPI_ISL_454975, EPI_ISL_454976, EPI_ISL_454977, EPI_ISL_454978, EPI_ISL_454979, EPI_ISL_454980, EPI_ISL_454981, EPI_ISL_454982, EPI_ISL_454983, EPI_ISL_454984, EPI_ISL_454988, EPI_ISL_454989, EPI_ISL_454990, EPI_ISL_454991, EPI_ISL_454992, EPI_ISL_454993, EPI_ISL_454995, EPI_ISL_454996, EPI_ISL_454997 | | | |
| see above | Wuhan Chain Medical Labs (CMLabs) | State Key Laboratory of Biotherapy of Sichuan University | Baowen Du, Minjin Wang, Chao Tang, Chuan Chen, Yongzhao Zhou, Mingxia Yu, Hancheng Wei, Weimin Li, Jing-wen Lin, Jia Geng, Binwu Ying, Lu Chen |
| EPI_ISL_455015 | Pandit Deendayal Upadhyay Government Medical College, Rajkot | Gujarat Biotechnology Research Centre | Snehal Bagatharia, Prakash Modi, Sejul Antala, Manish Pattani, Ramesh Pandit, Tejas Shah, Ankit Hinsu, Pritesh Sabara, Apurvasinh Puvar, Janvi Raval, Zarna Patel, Monika Gandhi, Pinal Trivedi, Maharshi Pandya, Amit Kanani, Nidhi Patel, Nitin Savaliya, Raghawendra Kumar, Dinesh Kumar, Zuber Saiyed, Komal Patel, Labdhi Pandya, Neha Rajpara, Bhavesh Modi, Gaurishankar Shrimali, R D Dixit, A M Kadri, Umang Mishra, Chaitanya Joshi, Madhvi Joshi |
| EPI_ISL_455016 | Pandit Deendayal Upadhyay Government Medical College, Rajkot | Gujarat Biotechnology Research Centre | Prakash Modi, Sejul Antala, Manish Pattani, Ramesh Pandit, Tejas Shah, Ankit Hinsu, Pritesh Sabara, Apurvasinh Puvar, Janvi Raval, Zarna Patel, Monika Gandhi, Pinal Trivedi, Maharshi Pandya, Amit Kanani, Nidhi Patel, Nitin Savaliya, Raghawendra Kumar, Dinesh Kumar, Zuber Saiyed, Komal Patel, Labdhi Pandya, Snehal Bagatharia, Afzal Ansari, Bhavesh Modi, Gaurishankar Shrimali, R D Dixit, A M Kadri, Umang Mishra, Chaitanya Joshi, Madhvi Joshi |
| EPI_ISL_455017 | Government Medical College, Vadodara | Gujarat Biotechnology Research Centre | Tanuja Javadekar , R N Daveshwar, Ramesh Pandit, Tejas Shah, Ankit Hinsu, Pritesh Sabara, Apurvasinh Puvar, Janvi Raval, Zarna Patel, Monika Gandhi, Pinal Trivedi, Maharshi Pandya, Amit Kanani, Nidhi Patel, Nitin Savaliya, Raghawendra Kumar, Dinesh Kumar, Zuber Saiyed, Komal Patel, Labdhi Pandya, Snehal Bagatharia, Fenil Patel, Bhavesh Modi, Gaurishankar Shrimali, R D Dixit, A M Kadri, Umang Mishra, Chaitanya Joshi, Madhvi Joshi, |
| EPI_ISL_455018 | Government Medical College, Vadodara | Gujarat Biotechnology Research Centre | R N Daveshwar, Ramesh Pandit, Tejas Shah, Ankit Hinsu, Pritesh Sabara, Apurvasinh Puvar, Janvi Raval, Zarna Patel, Monika Gandhi, Pinal Trivedi, Maharshi Pandya, Amit Kanani, Nidhi Patel, Nitin Savaliya, Raghawendra Kumar, Dinesh Kumar, Zuber Saiyed, Komal Patel, Labdhi Pandya, Snehal Bagatharia, Tanuja Javadekar , Neelam Nathani, Bhavesh Modi, Gaurishankar Shrimali, R D Dixit, A M Kadri, Umang Mishra, Chaitanya Joshi, Madhvi Joshi, |
| EPI_ISL_455019 | Government Medical College, Vadodara | Gujarat Biotechnology Research Centre | Ramesh Pandit, Tejas Shah, Ankit Hinsu, Pritesh Sabara, Apurvasinh Puvar, Janvi Raval, Zarna Patel, Monika Gandhi, Pinal Trivedi, Maharshi Pandya, Amit Kanani, Nidhi Patel, Nitin Savaliya, Raghawendra Kumar, Dinesh Kumar, Zuber Saiyed, Komal Patel, Labdhi Pandya, Snehal Bagatharia, Tanuja Javadekar , R N Daveshwar, Armi Chaudhari, Bhavesh Modi, Gaurishankar Shrimali, R D Dixit, A M Kadri, Umang Mishra, Chaitanya Joshi, Madhvi Joshi, |
| EPI_ISL_455020 | Government Medical College, Vadodara | Gujarat Biotechnology Research Centre | Tejas Shah, Ankit Hinsu, Pritesh Sabara, Apurvasinh Puvar, Janvi Raval, Zarna Patel, Monika Gandhi, Pinal Trivedi, Maharshi Pandya, Amit Kanani, Nidhi Patel, Nitin Savaliya, Raghawendra Kumar, Dinesh Kumar, Zuber Saiyed, Komal Patel, Labdhi Pandya, Snehal Bagatharia, Tanuja Javadekar , R N Daveshwar, Ramesh Pandit, Bhavya Jindal, Bhavesh Modi, Gaurishankar Shrimali, R D Dixit, A M Kadri, Umang Mishra, Chaitanya Joshi, Madhvi Joshi, |
| EPI_ISL_455021 | Government Medical College, Vadodara | Gujarat Biotechnology Research Centre | Ankit Hinsu, Pritesh Sabara, Apurvasinh Puvar, Janvi Raval, Zarna Patel, Monika Gandhi, Pinal Trivedi, Maharshi Pandya, Amit Kanani, Nidhi Patel, Nitin Savaliya, Raghawendra Kumar, Dinesh Kumar, Zuber Saiyed, Komal Patel, Labdhi Pandya, Snehal Bagatharia, Tanuja Javadekar , R N Daveshwar, Ramesh Pandit, Tejas Shah, Camellia Chakraborty, Bhavesh Modi, Gaurishankar Shrimali, R D Dixit, A M Kadri, Umang Mishra, Chaitanya Joshi, Madhvi Joshi, |
| EPI_ISL_455022 | Government Medical College, Vadodara | Gujarat Biotechnology Research Centre | Pritesh Sabara, Apurvasinh Puvar, Janvi Raval, Zarna Patel, Monika Gandhi, Pinal Trivedi, Maharshi Pandya, Amit Kanani, Nidhi Patel, Nitin Savaliya, Raghawendra Kumar, Dinesh Kumar, Zuber Saiyed, Komal Patel, Labdhi Pandya, Snehal Bagatharia, Tanuja Javadekar , R N Daveshwar, Ramesh Pandit, Tejas Shah, Ankit Hinsu, Siddhant Kumar, Bhavesh Modi, Gaurishankar Shrimali, R D Dixit, A M Kadri, Umang Mishra, Chaitanya Joshi, Madhvi Joshi, |
| EPI_ISL_455023 | Government Medical College, Vadodara | Gujarat Biotechnology Research Centre | Apurvasinh Puvar, Janvi Raval, Zarna Patel, Monika Gandhi, Pinal Trivedi, Maharshi Pandya, Amit Kanani, Nidhi Patel, Nitin Savaliya, Raghawendra Kumar, Dinesh Kumar, Zuber Saiyed, Komal Patel, Labdhi Pandya, Snehal Bagatharia, Tanuja Javadekar , R N Daveshwar, Ramesh Pandit, Tejas Shah, Ankit Hinsu, Pritesh Sabara, Priyanka P Vatsa, Bhavesh Modi, Gaurishankar Shrimali, R D Dixit, A M Kadri, Umang Mishra, Chaitanya Joshi, Madhvi Joshi, |
| EPI_ISL_455024 | Government Medical College, Vadodara | Gujarat Biotechnology Research Centre | Janvi Raval, Zarna Patel, Monika Gandhi, Pinal Trivedi, Maharshi Pandya, Amit Kanani, Nidhi Patel, Nitin Savaliya, Raghawendra Kumar, Dinesh Kumar, Zuber Saiyed, Komal Patel, Labdhi Pandya, Snehal Bagatharia, Tanuja Javadekar , R N Daveshwar, Ramesh Pandit, Tejas Shah, Ankit Hinsu, Pritesh Sabara, Apurvasinh Puvar, Pooja P Doshi, Bhavesh Modi, Gaurishankar Shrimali, R D Dixit, A M Kadri, Umang Mishra, Chaitanya Joshi, Madhvi Joshi, |
| EPI_ISL_455025 | Government Medical College, Vadodara | Gujarat Biotechnology Research Centre | Zarna Patel, Monika Gandhi, Pinal Trivedi, Maharshi Pandya, Amit Kanani, Nidhi Patel, Nitin Savaliya, Raghawendra Kumar, Dinesh Kumar, Zuber Saiyed, Komal Patel, Labdhi Pandya, Snehal Bagatharia, Tanuja Javadekar , R N Daveshwar, Ramesh Pandit, Tejas Shah, Ankit Hinsu, Pritesh Sabara, Apurvasinh Puvar, Janvi Raval, Akanksha Verma, Bhavesh Modi, Gaurishankar Shrimali, R D Dixit, A M Kadri, Umang Mishra, Chaitanya Joshi, Madhvi Joshi, |
| EPI_ISL_455026 | Government Medical College, Vadodara | Gujarat Biotechnology Research Centre | Monika Gandhi, Pinal Trivedi, Maharshi Pandya, Amit Kanani, Nidhi Patel, Nitin Savaliya, Raghawendra Kumar, Dinesh Kumar, Zuber Saiyed, Komal Patel, Labdhi Pandya, Snehal Bagatharia, Tanuja Javadekar , R N Daveshwar, Ramesh Pandit, Tejas Shah, Ankit Hinsu, Pritesh Sabara, Apurvasinh Puvar, Janvi Raval, Zarna Patel, Priti Pandita, Bhavesh Modi, Gaurishankar Shrimali, R D Dixit, A M Kadri, Umang Mishra, Chaitanya Joshi, Madhvi Joshi, |
| EPI_ISL_455027 | Government Medical College, Vadodara | Gujarat Biotechnology Research Centre | Pinal Trivedi, Maharshi Pandya, Amit Kanani, Nidhi Patel, Nitin Savaliya, Raghawendra Kumar, Dinesh Kumar, Zuber Saiyed, Komal Patel, Labdhi Pandya, Snehal Bagatharia, Tanuja Javadekar , R N Daveshwar, Ramesh Pandit, Tejas Shah, Ankit Hinsu, Pritesh Sabara, Apurvasinh Puvar, Janvi Raval, Zarna Patel, Monika Gandhi, Pragya Sharma, Bhavesh Modi, Gaurishankar Shrimali, R D Dixit, A M Kadri, Umang Mishra, Chaitanya Joshi, Madhvi Joshi, |
| EPI_ISL_455028, EPI_ISL_455029, EPI_ISL_455030, EPI_ISL_455031, EPI_ISL_455032, EPI_ISL_455033, EPI_ISL_455034, EPI_ISL_455035 | Pathology West - NSW Health Pathology | NSW Health Pathology - Institute of Clinical Pathology and Medical Research; Westmead Hospital; University of Sydney | CIDM-PH et al. |
| EPI_ISL_455036 | South Eastern Area Laboratory Services | NSW Health Pathology - Institute of Clinical Pathology and Medical Research; Westmead Hospital; University of Sydney | CIDM-PH et al. |
| EPI_ISL_455037 | Pathology Sydney South West - NSW Health Pathology | NSW Health Pathology - Institute of Clinical Pathology and Medical Research; Westmead Hospital; University of Sydney | CIDM-PH et al. |
| EPI_ISL_455038, EPI_ISL_455039, EPI_ISL_455040 | Pathology West - NSW Health Pathology | NSW Health Pathology - Institute of Clinical Pathology and Medical Research; Westmead Hospital; University of Sydney | CIDM-PH et al. |
| EPI_ISL_455041 | Laverty Pathology | NSW Health Pathology - Institute of Clinical Pathology and Medical Research; Westmead Hospital; University of Sydney | CIDM-PH et al. |
| EPI_ISL_455042, EPI_ISL_455043 | ACT Pathology | NSW Health Pathology - Institute of Clinical Pathology and Medical Research; Westmead Hospital; University of Sydney | CIDM-PH et al. |
| EPI_ISL_455044, EPI_ISL_455045, EPI_ISL_455046, EPI_ISL_455047 | Pathology West - NSW Health Pathology | NSW Health Pathology - Institute of Clinical Pathology and Medical Research; Westmead Hospital; University of Sydney | CIDM-PH et al. |
| EPI_ISL_455048 | Laverty Pathology | NSW Health Pathology - Institute of Clinical Pathology and Medical Research; Westmead Hospital; University of Sydney | CIDM-PH et al. |
| EPI_ISL_455049 | Pathology West - NSW Health Pathology | NSW Health Pathology - Institute of Clinical Pathology and Medical Research; Westmead Hospital; University of Sydney | CIDM-PH et al. |
| EPI_ISL_455050 | ACT Pathology | NSW Health Pathology - Institute of Clinical Pathology and Medical Research; Westmead Hospital; University of Sydney | CIDM-PH et al. |
| EPI_ISL_455051, EPI_ISL_455052, EPI_ISL_455053 | Douglas Hanly Moir Pathology | NSW Health Pathology - Institute of Clinical Pathology and Medical Research; Westmead Hospital; University of Sydney | CIDM-PH et al. |
| EPI_ISL_455054 | South Eastern Area Laboratory Services | NSW Health Pathology - Institute of Clinical Pathology and Medical Research; Westmead Hospital; University of Sydney | CIDM-PH et al. |
| EPI_ISL_455055, EPI_ISL_455056, EPI_ISL_455057, EPI_ISL_455058, EPI_ISL_455059, EPI_ISL_455060, EPI_ISL_455061, EPI_ISL_455062 | Pathology West - NSW Health Pathology | NSW Health Pathology - Institute of Clinical Pathology and Medical Research; Westmead Hospital; University of Sydney | CIDM-PH et al. |
| EPI_ISL_455063 | Laverty Pathology | NSW Health Pathology - Institute of Clinical Pathology and Medical Research; Westmead Hospital; University of Sydney | CIDM-PH et al. |
| EPI_ISL_455064 | Pathology West - NSW Health Pathology | NSW Health Pathology - Institute of Clinical Pathology and Medical | CIDM-PH et al. |

| | | | |
|---|---|---|---|
| | | Research; Westmead Hospital; University of Sydney | |
| EPI_ISL_455065 | Douglas Hanly Moir Pathology | NSW Health Pathology - Institute of Clinical Pathology and Medical Research; Westmead Hospital; University of Sydney | CIDM-PH et al. |
| EPI_ISL_455066, EPI_ISL_455067 | Pathology West - NSW Health Pathology | NSW Health Pathology - Institute of Clinical Pathology and Medical Research; Westmead Hospital; University of Sydney | CIDM-PH et al. |
| EPI_ISL_455068 | Childrens Hospital Westmead | NSW Health Pathology - Institute of Clinical Pathology and Medical Research; Westmead Hospital; University of Sydney | CIDM-PH et al. |
| EPI_ISL_455069, EPI_ISL_455070 | Pathology West - NSW Health Pathology | NSW Health Pathology - Institute of Clinical Pathology and Medical Research; Westmead Hospital; University of Sydney | CIDM-PH et al. |
| EPI_ISL_455071 | Pathology Sydney South West - NSW Health Pathology | NSW Health Pathology - Institute of Clinical Pathology and Medical Research; Westmead Hospital; University of Sydney | CIDM-PH et al. |
| EPI_ISL_455072, EPI_ISL_455073 | Sullivan Nicolaides Pathology | NSW Health Pathology - Institute of Clinical Pathology and Medical Research; Westmead Hospital; University of Sydney | CIDM-PH et al. |
| EPI_ISL_455074 | ACT Pathology | NSW Health Pathology - Institute of Clinical Pathology and Medical Research; Westmead Hospital; University of Sydney | CIDM-PH et al. |
| EPI_ISL_455075, EPI_ISL_455076, EPI_ISL_455077, EPI_ISL_455078, EPI_ISL_455079, EPI_ISL_455080, EPI_ISL_455081, EPI_ISL_455082, EPI_ISL_455083, EPI_ISL_455084, EPI_ISL_455085, EPI_ISL_455086, EPI_ISL_455087, EPI_ISL_455088, EPI_ISL_455089, EPI_ISL_455090, EPI_ISL_455091, EPI_ISL_455092, EPI_ISL_455093 | | | |
| see above | South Eastern Area Laboratory Services | NSW Health Pathology - Institute of Clinical Pathology and Medical Research; Westmead Hospital; University of Sydney | CIDM-PH et al. |
| EPI_ISL_455094 | Pathology Sydney South West - NSW Health Pathology | NSW Health Pathology - Institute of Clinical Pathology and Medical Research; Westmead Hospital; University of Sydney | CIDM-PH et al. |
| EPI_ISL_455095 | Pathology West - NSW Health Pathology | NSW Health Pathology - Institute of Clinical Pathology and Medical Research; Westmead Hospital; University of Sydney | CIDM-PH et al. |
| EPI_ISL_455096, EPI_ISL_455097, EPI_ISL_455098, EPI_ISL_455099 | South Eastern Area Laboratory Services | NSW Health Pathology - Institute of Clinical Pathology and Medical Research; Westmead Hospital; University of Sydney | CIDM-PH et al. |
| EPI_ISL_455100 | Orsa VC | The Public Health Agency of Sweden | Anna-Karin Lundqvist, Anna-Malin Linde, Maria Lind Karlberg, Oskar Karlsson Lindsjo, Olov Svartstrom, Anna Risberg, Theresa Enkirch, Mia Brytting, Karin Tegmark-Wisell |
| EPI_ISL_455101 | Jourcentralen | The Public Health Agency of Sweden | Salvatore Ascione, Anna-Malin Linde, Maria Lind Karlberg, Oskar Karlsson Lindsjo, Olov Svartstrom, Anna Risberg, Theresa Enkirch, Mia Brytting, Karin Tegmark-Wisell |
| EPI_ISL_455102 | Kungsors VC | The Public Health Agency of Sweden | Jessica Karlsson, Anna-Malin Linde, Maria Lind Karlberg, Oskar Karlsson Lindsjo, Olov Svartstrom, Anna Risberg, Theresa Enkirch, Mia Brytting, Karin Tegmark-Wisell |
| EPI_ISL_455103 | Huslakarna Varmbadhuset Varberg | The Public Health Agency of Sweden | Johanna Hilmersson, Anna-Malin Linde, Maria Lind Karlberg, Oskar Karlsson Lindsjo, Olov Svartstrom, Anna Risberg, Theresa Enkirch, Mia Brytting, Karin Tegmark-Wisell |
| EPI_ISL_455104 | Ulltuna Vardcentral | The Public Health Agency of Sweden | Heidi Lindback, Anna-Malin Linde, Maria Lind Karlberg, Oskar Karlsson Lindsjo, Olov Svartstrom, Anna Risberg, Theresa Enkirch, Mia Brytting, Karin Tegmark-Wisell |
| EPI_ISL_455105 | Wetterhalsan | The Public Health Agency of Sweden | Anders Tengblad, Anna-Malin Linde, Maria Lind Karlberg, Oskar Karlsson Lindsjo, Olov Svartstrom, Anna Risberg, Theresa Enkirch, Mia Brytting, Karin Tegmark-Wisell |
| EPI_ISL_455106 | Surbrunns VC | The Public Health Agency of Sweden | Erik Embring, Anna-Malin Linde, Maria Lind Karlberg, Oskar Karlsson Lindsjo, Olov Svartstrom, Anna Risberg, Theresa Enkirch, Mia Brytting, Karin Tegmark-Wisell |
| EPI_ISL_455107 | Narhalsan Mellerud | The Public Health Agency of Sweden | Maria Nykvist, Anna-Malin Linde, Maria Lind Karlberg, Oskar Karlsson Lindsjo, Olov Svartstrom, Anna Risberg, Theresa Enkirch, Mia Brytting, Karin Tegmark-Wisell |
| EPI_ISL_455108 | Smedby HC | The Public Health Agency of Sweden | Susanne Brunby, Anna-Malin Linde, Maria Lind Karlberg, Oskar Karlsson Lindsjo, Olov Svartstrom, Anna Risberg, Theresa Enkirch, Mia Brytting, Karin Tegmark-Wisell |
| EPI_ISL_455109 | Ulltuna Vardcentral | The Public Health Agency of Sweden | Heidi Lindback, Anna-Malin Linde, Maria Lind Karlberg, Oskar Karlsson Lindsjo, Olov Svartstrom, Anna Risberg, Theresa Enkirch, Mia Brytting, Karin Tegmark-Wisell |
| EPI_ISL_455110 | Scania Halsocenter, B288 | The Public Health Agency of Sweden | Christina Lergin, Anna-Malin Linde, Maria Lind Karlberg, Oskar Karlsson Lindsjo, Olov Svartstrom, Anna Risberg, Theresa Enkirch, Mia Brytting, Karin Tegmark-Wisell |
| EPI_ISL_455111 | Olof Norrby | The Public Health Agency of Sweden | Bla Kustens halsocentral, Anna-Malin Linde, Maria Lind Karlberg, Oskar Karlsson Lindsjo, Olov Svartstrom, Anna Risberg, Theresa Enkirch, Mia Brytting, Karin Tegmark-Wisell |
| EPI_ISL_455112, EPI_ISL_455113, EPI_ISL_455114, EPI_ISL_455115, EPI_ISL_455116, EPI_ISL_455117, EPI_ISL_455118, EPI_ISL_455119, EPI_ISL_455120, EPI_ISL_455121, EPI_ISL_455122, EPI_ISL_455123, EPI_ISL_455125, EPI_ISL_455126, EPI_ISL_455129, EPI_ISL_455131, EPI_ISL_455132, EPI_ISL_455133, EPI_ISL_455134, EPI_ISL_455135, EPI_ISL_455141, EPI_ISL_455143, EPI_ISL_455144, EPI_ISL_455145, EPI_ISL_455147, EPI_ISL_455148, EPI_ISL_455149, EPI_ISL_455151, EPI_ISL_455152, EPI_ISL_455155, EPI_ISL_455156, EPI_ISL_455157, EPI_ISL_455159, EPI_ISL_455160, EPI_ISL_455161, EPI_ISL_455162, EPI_ISL_455163, EPI_ISL_455164, EPI_ISL_455165, EPI_ISL_455167, EPI_ISL_455168, EPI_ISL_455169, EPI_ISL_455170, EPI_ISL_455171, EPI_ISL_455172, EPI_ISL_455173, EPI_ISL_455174, EPI_ISL_455175, EPI_ISL_455176, EPI_ISL_455177, EPI_ISL_455178, EPI_ISL_455179, EPI_ISL_455180, EPI_ISL_455181, EPI_ISL_455184, EPI_ISL_455185, EPI_ISL_455186, EPI_ISL_455187, EPI_ISL_455188, EPI_ISL_455189, EPI_ISL_455190, EPI_ISL_455191, EPI_ISL_455192, EPI_ISL_455193, EPI_ISL_455195, EPI_ISL_455196, EPI_ISL_455198, EPI_ISL_455199, EPI_ISL_455200, EPI_ISL_455201, EPI_ISL_455202, EPI_ISL_455203, EPI_ISL_455205, EPI_ISL_455206, EPI_ISL_455208, EPI_ISL_455209, EPI_ISL_455210, EPI_ISL_455211, EPI_ISL_455212, EPI_ISL_455213, EPI_ISL_455214, EPI_ISL_455215, EPI_ISL_455216, EPI_ISL_455217, EPI_ISL_455219, EPI_ISL_455221, EPI_ISL_455222, EPI_ISL_455223, EPI_ISL_455224, EPI_ISL_455225, EPI_ISL_455226, EPI_ISL_455227, EPI_ISL_455228, EPI_ISL_455231, EPI_ISL_455232, EPI_ISL_455233, EPI_ISL_455235, EPI_ISL_455236, EPI_ISL_455237, EPI_ISL_455238, EPI_ISL_455241, EPI_ISL_455242, EPI_ISL_455244, EPI_ISL_455245, EPI_ISL_455246, EPI_ISL_455249, EPI_ISL_455251, EPI_ISL_455252, EPI_ISL_455253, EPI_ISL_455254, EPI_ISL_455255, EPI_ISL_455256, EPI_ISL_455257, EPI_ISL_455258, EPI_ISL_455259, EPI_ISL_455260, EPI_ISL_455261, EPI_ISL_455262, EPI_ISL_455263, EPI_ISL_455266, EPI_ISL_455269, EPI_ISL_455270, EPI_ISL_455274, EPI_ISL_455275, EPI_ISL_455276, EPI_ISL_455277, EPI_ISL_455278, EPI_ISL_455279, EPI_ISL_455280, EPI_ISL_455281, EPI_ISL_455283, EPI_ISL_455284, EPI_ISL_455285, EPI_ISL_455286, EPI_ISL_455287, EPI_ISL_455289, EPI_ISL_455290, EPI_ISL_455291, EPI_ISL_455292, EPI_ISL_455293, EPI_ISL_455296, EPI_ISL_455297, EPI_ISL_455298, EPI_ISL_455299, EPI_ISL_455300, EPI_ISL_455302, EPI_ISL_455303, EPI_ISL_455304, EPI_ISL_455305, EPI_ISL_455306, EPI_ISL_455307 | | | |
| see above | Dutch COVID-19 response team | Erasmus Medical Center | Bas Oude Munnink, David Nieuwenhuijse, Reina Sikkema, Claudia Schapendonk, Irina Chestakova, Anne van der Linden, Theo Bestebroer, Stefan van Nieuwkoop, Mark Pronk, Pascal Lexmond, Corien Swaan, Manon Haverkate, Madelief Mollers, Mart Stein, Sandra Kengne Kamga Mobou, Jeroen van Kampen, Jolanda Voermans, Aura Timen, Corine GeurtsvanKessel, Annemiek van der Eijk, Richard Molenkamp, Marion Koopmans, on behalf of the Dutch national COVID-19 response team. |
| EPI_ISL_455309, EPI_ISL_455311 | REGIONAL VRDL,ICMR-RMRC BBSR | Immunogenomics group, Institute of Life Sciences, Bhubaneswar | Sunil Raghav, Jyotirmayee Turuk, Arup Ghosh, Atimukta Jha, Viplov K. Biswas, Swati Madhulika, Manasi Priyadarshini, Shuchi Smita, Jaya Singh Khastri, Rupesh Dash, Soma Chattopadhyay, Ghulam Hussain Syed, Shanti Senapati, Tushar K. Beuria, Debdutta Bhattacharya, Rajeeb Swain, Punit Prasad, COVID-19 team of ILS & RMRC, Orissa COVID-19 study group, DBT's PAN-INDIA 1000 SARS-CoV2 RNA genome sequencing consortium, Sanghamitra Pati, Ajay Parida |
| EPI_ISL_455312 | Microbiology Unit, Department of Pathology & Laboratory Medicine, IIUM Medical Centre | SEA Microbiome Unit, Faculty of Industrial Sciences & Technology, Universiti Malaysia Pahang | Norhidayah Binti Kamarudin, Ahmad Hafiz Bin Zulkifly, Hajar Fauzan Ahmad, Muhammad Adam Lee Abdullah, Mohd Fazli Farida Asras, Ahmad Mahfuz Gazali, Mohd Nazli Bin Kamarulzaman, IIUM Medical Centre Covid19 Taskforce, UMP Covid19 Team |
| EPI_ISL_455313 | Microbiology Unit, Department of Pathology & Laboratory Medicine, IIUM Medical Centre | SEA Microbiome Unit, Faculty of Industrial Sciences & Technology, Universiti Malaysia Pahang | Hajar Fauzan Ahmad, Norhidayah Kamarudin, Ahmad Hafiz Zulkifly, IIUM Medical Centre Covid19 Taskforce, UMP Covid19 Team |
| EPI_ISL_455314 | Hospital Virgen del Rocio | Instituto de Salud Carlos III | Iglesias-Caballero, M. Molinero Calamita, M. González-Esguevillas, M. Camarero, S. Pozo, F. Casas, I. Jiménez, P. Jiménez, M. Zaballos, A. Monzón, S. Varona, S. Juliá, M. Cuesta, I, J. Lepe |
| EPI_ISL_455315, EPI_ISL_455316, EPI_ISL_455317, EPI_ISL_455318, EPI_ISL_455319, EPI_ISL_455320, EPI_ISL_455321, EPI_ISL_455322, EPI_ISL_455324 | Hospital Virgen de las Nieves | Instituto de Salud Carlos III | Iglesias-Caballero, M. Molinero Calamita, M. González-Esguevillas, M. Camarero, S. Pozo, F. Casas, I. Jiménez, P. Jiménez, M. Zaballos, A. Monzón, S. Varona, S. Juliá, M. Cuesta, I, S. Sanbonmatsu |
| EPI_ISL_455325 | Hospital Universitario de Canarias | Instituto de Salud Carlos III | Iglesias-Caballero, M. Molinero Calamita, M. González-Esguevillas, M. Camarero, S. Pozo, F. Casas, I. Jiménez, P. Jiménez, M. Zaballos, A. Monzón, S. Varona, S. Juliá, M. Cuesta, I, B. Castro |
| EPI_ISL_455326 | Hospital Universitario Insular de Gran Canaria | Instituto de Salud Carlos III | Iglesias-Caballero, M. Molinero Calamita, M. González-Esguevillas, M. Camarero, S. Pozo, F. Casas, I. Jiménez, P. Jiménez, M. Zaballos, A. Monzón, S. Varona, S. Juliá, M. Cuesta, I, A. Hernández |
| EPI_ISL_455327 | Consejeria de Sanidad y Asuntos Sociales | Instituto de Salud Carlos III | Iglesias-Caballero, M. Molinero Calamita, M. González-Esguevillas, M. Camarero, S. Pozo, F. Casas, I. Jiménez, P. Jiménez, M. Zaballos, A. Monzón, S. Varona, S. Juliá, M. Cuesta, I, G. Gutiérrez |
| EPI_ISL_455328, EPI_ISL_455329, EPI_ISL_455330, EPI_ISL_455331 | Complejo Hospitalario Universitario La Coruna | Instituto de Salud Carlos III | Iglesias-Caballero, M. Molinero Calamita, M. González-Esguevillas, M. Camarero, S. Pozo, F. Casas, I. Jiménez, P. Jiménez, M. Zaballos, A. Monzón, S. Varona, S. Juliá, M. Cuesta, I, M.A Canizares |
| EPI_ISL_455332 | Xerencia de Xestión Integrada de Pontevedra e o Salnés | Instituto de Salud Carlos III | Iglesias-Caballero, M. Molinero Calamita, M. González-Esguevillas, M. Camarero, S. Pozo, F. Casas, I. Jiménez, P. Jiménez, M. Zaballos, A. Monzón, S. Varona, S. Juliá, M. Cuesta, I, M. Garcia |
| EPI_ISL_455333 | Complejo Hospitalario Universitario de Santiago | Instituto de Salud Carlos III | Iglesias-Caballero, M. Molinero Calamita, M. González-Esguevillas, M. Camarero, S. Pozo, F. Casas, I. Jiménez, P. Jiménez, M. Zaballos, A. Monzón, S. Varona, S. Juliá, M. Cuesta, I, J. Llovo |
| EPI_ISL_455334, EPI_ISL_455335 | Complejo Hospitalario de Orense | Instituto de Salud Carlos III | Iglesias-Caballero, M. Molinero Calamita, M. González-Esguevillas, M. Camarero, S. Pozo, F. Casas, I. Jiménez, P. Jiménez, M. Zaballos, A. Monzón, S. Varona, S. Juliá, M. Cuesta, I, M. Paz |
| EPI_ISL_455336, EPI_ISL_455337, EPI_ISL_455338, EPI_ISL_455339, EPI_ISL_455340, EPI_ISL_455341, EPI_ISL_455342, EPI_ISL_455343 | Hospital San Pedro | Instituto de Salud Carlos III | Iglesias-Caballero, M. Molinero Calamita, M. González-Esguevillas, M. Camarero, S. Pozo, F. Casas, I. Jiménez, P. Jiménez, M. Zaballos, A. Monzón, S. Varona, S. Juliá, M. Cuesta, I, C. Alonso |
| EPI_ISL_455344, EPI_ISL_455345, EPI_ISL_455346, EPI_ISL_455347, EPI_ISL_455348, EPI_ISL_455349 | Hospital Comarcal de Melilla | Instituto de Salud Carlos III | Iglesias-Caballero, M. Molinero Calamita, M. González-Esguevillas, M. Camarero, S. Pozo, F. Casas, I. Jiménez, P. Jiménez, M. Zaballos, A. Monzón, S. Varona, S. Juliá, M. Cuesta, I, I. Pérez |
| EPI_ISL_455350, EPI_ISL_455351 | Hospital Txagorritxu | Instituto de Salud Carlos III | Iglesias-Caballero, M. Molinero Calamita, M. González-Esguevillas, M. Camarero, S. Pozo, F. Casas, I. Jiménez, P. Jiménez, M. Zaballos, A. Monzón, S. Varona, S. Juliá, M. Cuesta, I, C. Gómez |
| EPI_ISL_455352, EPI_ISL_455353, EPI_ISL_455354 | Hospital de Cruces | Instituto de Salud Carlos III | Iglesias-Caballero, M. Molinero Calamita, M. González-Esguevillas, M. Camarero, S. Pozo, F. Casas, I. Jiménez, P. Jiménez, M. Zaballos, A. Monzón, S. Varona, S. Juliá, M. Cuesta, I, M. Aranzamendi |
| EPI_ISL_455355, EPI_ISL_455356, EPI_ISL_455358, EPI_ISL_455359, EPI_ISL_455360, EPI_ISL_455361 | Emory Molecular Diagnostics Laboratory, Emory Healthcare | Piantadosi Lab, Emory Department of Pathology | Ahmed Babiker, Anne Piantadosi |
| EPI_ISL_455362 | Nigeria Centre for Disease Control (NCDC) | African Centre of Excellence for Genomics of Infectious Diseases (ACEGID), Redeemer's University, Ede, Osun State, Nigeria | Oluniyi P.E., Ajogbasile F.V., Kayode A., Olawoye I., Uwanibe J., Oguzie J., Olumade T., Folarin O.A., Ihekweazu C., Happi C.T. |
| EPI_ISL_455363, EPI_ISL_455364, EPI_ISL_455365, EPI_ISL_455366, EPI_ISL_455367, EPI_ISL_455368, EPI_ISL_455369, EPI_ISL_455370, EPI_ISL_455371, EPI_ISL_455372, EPI_ISL_455373, EPI_ISL_455374, EPI_ISL_455375, EPI_ISL_455376, EPI_ISL_455377, EPI_ISL_455378, EPI_ISL_455379, EPI_ISL_455380, EPI_ISL_455381, EPI_ISL_455382, EPI_ISL_455383, EPI_ISL_455384, EPI_ISL_455386, EPI_ISL_455388, EPI_ISL_455389, EPI_ISL_455390, EPI_ISL_455391, EPI_ISL_455392, EPI_ISL_455394, EPI_ISL_455395, EPI_ISL_455397, EPI_ISL_455398, EPI_ISL_455399, EPI_ISL_455406 | | | |
| see above | Wuhan Chain Medical Labs (CMLabs) | State Key Laboratory of Biotherapy of Sichuan University | Baowen Du, Minjin Wang, Chao Tang, Chuan Chen, Yongzhao Zhou, Mingxia Yu, Hancheng Wei, Weimin Li, Jing-wen Lin, Jia Geng, Binwu Ying, Lu Chen |
| EPI_ISL_455412, EPI_ISL_455413, EPI_ISL_455419 | Nigeria Centre for Disease Control (NCDC) | African Centre of Excellence for Genomics of Infectious Diseases (ACEGID), Redeemer's University, Ede, Osun State, Nigeria | Oluniyi P.E., Ajogbasile F.V., Kayode A., Oguzie J., Olawoye I., Uwanibe J., Olumade T., Folarin O.A., Ihekweazu C., Happi C.T. |
| EPI_ISL_455420 | National Institute of Laboratory Medicine and Referral Center | Genomic Research Lab, BCSIR | Abu Sayeed Mohammad Mahmud, Mohammad Samir Uzzaman, Eshrar Osman, Md. Ahasan Habib, Shahina Akhter, Tanjina Akhter Banu, Barna Goswami, Iffat Jahan, Tasnim Nafisa, Md. Maruf Ahmed Molla, |

| | | | |
|---|---|---|---|
| | | | MahmudaYeasmin, Sheikh Md. Selim Al Din, Utpal Chandra Ray, Md. Salim Khan |
| EPI_ISL_455422 | Nigeria Centre for Disease Control | African Centre of Excellence for Genomics of Infectious Diseases (ACEGID), Redeemer's University, Ede, Osun State, Nigeria | Oluniyi P.E., Ajogbasile F.V., Kayode A., Oguzie J., Olawoye I., Uwanibe J., Olumade T., Folarin O.A., Ihekweazu C., Happi C.T. |
| EPI_ISL_455423, EPI_ISL_455424 | Nigeria Centre for Disease Control (NCDC) | African Centre of Excellence for Genomics of Infectious Diseases (ACEGID), Redeemer's University, Ede, Osun State, Nigeria | Oluniyi P.E., Ajogbasile F.V., Kayode A., Oguzie J., Olawoye I., Uwanibe J., Olumade T., Folarin O.A., Ihekweazu C., Happi C.T. |
| EPI_ISL_455426 | Nigeria Centre for Disease Control | African Centre of Excellence for Genomics of Infectious Diseases (ACEGID), Redeemer's University, Ede, Osun State, Nigeria | Oluniyi P.E., Ajogbasile F.V., Kayode A., Oguzie J., Olawoye I., Uwanibe J., Olumade T., Folarin O.A., Ihekweazu C., Happi C.T. |
| EPI_ISL_455429, EPI_ISL_455431 | Nigeria Centre for Disease Control (NCDC) | African Centre of Excellence for Genomics of Infectious Diseases (ACEGID), Redeemer's University, Ede, Osun State, Nigeria | Oluniyi P.E., Ajogbasile F.V., Kayode A., Oguzie J., Olawoye I., Uwanibe J., Olumade T., Folarin O.A., Ihekweazu C., Happi C.T. |
| EPI_ISL_455434 | Instituto de Diagnostico y Referencia Epidemiologicos (INDRE) | Instituto de Diagnostico y Referencia Epidemiologicos (INDRE) | Taboada Ramírez Blanca. Ramirez-Gonzalez Ernesto, Garces-Ayala Fabiola, Araiza-Rodriguez Adnan, Mendieta-Condado Edgar, Rodriguez-Maldonado Abril, Wong-Arambula Claudia, Barrera-Badillo Gisela, Hernandez-Rivas Lucia, Lopez-Martinez Irma. |
| EPI_ISL_455435 | Instituto de Diagnostico y Referencia Epidemiologicos (INDRE) | Instituto de Diagnostico y Referencia Epidemiologicos (INDRE) | Garces-Ayala Fabiola. Taboada Ramírez Blanca. Ramirez-Gonzalez Ernesto, Araiza-Rodriguez Adnan , Mendieta-Condado Edgar, Rodriguez-Maldonado Abril, Wong-Arambula Claudia, Barrera-Badillo Gisela, Hernandez-Rivas Lucia, Lopez-Martinez Irma |
| EPI_ISL_455436 | Instituto de Diagnostico y Referencia Epidemiologicos (INDRE) | Instituto de Diagnostico y Referencia Epidemiologicos (INDRE) | Garces-Ayala Fabiola. Taboada Ramírez Blanca. Ramirez-Gonzalez Ernesto, Araiza-Rodriguez Adnan, Mendieta-Condado Edgar, Rodriguez-Maldonado Abril, Wong-Arambula Claudia, Barrera-Badillo Gisela, Hernandez-Rivas Lucia, Lopez-Martinez Irma. |
| EPI_ISL_455437, EPI_ISL_455438 | Instituto de Diagnostico y Referencia Epidemiologicos (INDRE) | Instituto de Diagnostico y Referencia Epidemiologicos (INDRE) | Araiza-Rodriguez Adnan, Garces-Ayala Fabiola. Ramirez-Gonzalez Ernesto, Mendieta-Condado Edgar, Rodriguez-Maldonado Abril, Wong-Arambula Claudia, Barrera-Badillo Gisela, Hernandez-Rivas Lucia, Lopez-Martinez Irma, Taboada Ramírez Blanca. |
| EPI_ISL_455439 | Instituto de Diagnostico y Referencia Epidemiologicos (INDRE) | Instituto de Diagnostico y Referencia Epidemiologicos (INDRE) | Mendieta-Condado Edgar, Araiza-Rodriguez Adnan, Garces-Ayala Fabiola, Rodriguez-Maldonado Abril, Wong-Arambula Claudia, Barrera-Badillo Gisela, Hernandez-Rivas Lucia, Lopez-Martinez Irma, Taboada Ramirez Blanca, Ramirez-Gonzalez Ernesto. |
| EPI_ISL_455440, EPI_ISL_455441, EPI_ISL_455442, EPI_ISL_455443, EPI_ISL_455444, EPI_ISL_455445, EPI_ISL_455446, EPI_ISL_455447, EPI_ISL_455448, EPI_ISL_455449, EPI_ISL_455450, EPI_ISL_455451, EPI_ISL_455452, EPI_ISL_455453 | | | |
| see above | 1. ViroGenetics - BSL3 Laboratory of Virology, Małopolska Centre of Biotechnology, Jagiellonian University; 2. II Department of Internal Medicine, Faculty of Medicine, Jagiellonian University Medical College; 3. Narodowy Instytut Zdrowia Publicznego – Państwowy Zakład Higieny (NIZP-PZH) | 1. ViroGenetics - BSL3 Laboratory of Virology, Małopolska Centre of Biotechnology, Jagiellonian University; 2. II Department of Internal Medicine, Faculty of Medicine, Jagiellonian University Medical College; 3. Narodowy Instytut Zdrowia Publicznego – Państwowy Zakład Higieny (NIZP-PZH). | Katarzyna Pancer, Marek Sanak, Aleksandra A. Zasada, Magdalena Rzeczkowska, Tomasz Wołkowicz, Katarzyna Zacharczuk, Agnieszka Kołakowska-Kulesza, Katarzyna Owczarek, Aleksandra Milewska, Natalia Wolaniuk, Ewelina Hallman-Szelińska, Paweł P Łabaj, Wojciech Branicki, Krzysztof Pyrć |
| EPI_ISL_455455 | Instituto de Diagnostico y Referencia Epidemiologicos (INDRE) | Instituto de Diagnostico y Referencia Epidemiologicos (INDRE) | Rodriguez-Maldonado Abril, Mendieta-Condado Edgar, Araiza-Rodriguez Adnan, Garces-Ayala Fabiola. Taboada Ramirez Blanca. Ramirez-Gonzalez Ernesto, Barrera-Badillo Gisela, Hernandez-Rivas Lucia, Lopez-Martinez Irma, Wong-Arambula Claudia. |
| EPI_ISL_455456 | Instituto de Diagnostico y Referencia Epidemiologicos (INDRE) | Instituto de Diagnostico y Referencia Epidemiologicos (INDRE) | Rodriguez-Maldonado Abril, Mendieta-Condado Edgar, Araiza-Rodriguez Adnan, Garces-Ayala Fabiola. Taboada Ramírez Blanca. Ramirez-Gonzalez Ernesto, , Barrera-Badillo Gisela, Hernandez-Rivas Lucia, Lopez-Martinez Irma, Wong-Arambula Claudia. |
| EPI_ISL_455458 | National Institute of Laboratory Medicine and Referral Center | Genomic Research Lab, BCSIR | Abu Sayeed Mohammad Mahmud, Mohammad Samir Uzzaman, Eshrar Osman, Md. Ahasan Habib, Shahina Akhter, Tanjina Akhter Banu, Barna Goswami, Iffat Jahan, Tasnim Nafisa, Md. Maruf Ahmed Molla, MahmudaYeasmin, Sheikh Md. Selim Al Din, Utpal Chandra Ray, Md. Salim Khan |
| EPI_ISL_455459 | National Institute of Laboratory Medicine and Referral Center | Genomic Research Lab, BCSIR | Abu Sayeed Mohammad Mahmud, Mohammad Samir Uzzaman, Eshrar Osman, Md. Ahasan Habib, Shahina Akhter, Tanjina Akhter Banu, Barna Goswami, Iffat Jahan, Tasnim Nafisa, Md. Maruf Ahmed Molla, MahmudaYeasmin, Sheikh Md. Selim Al Din, Utpal Chandra Ray, Md. Salim Khan |
| EPI_ISL_455460, EPI_ISL_455461, EPI_ISL_455462, EPI_ISL_455463, EPI_ISL_455464, EPI_ISL_455465, EPI_ISL_455466, EPI_ISL_455467 | Jiangxi Province Center for Disease Control and Prevention | Jiangxi Province Center for Disease Control and Prevention | JianXiong Li,Ying Xiong,Tian Gong,Yong Shi,Jun Zhou,Fang Xiao,ShiWen Liu,XiaoQing Liu,Gang Xu,DaJin Xiao,Xin Ran,YanNi Zhang |
| EPI_ISL_455468, EPI_ISL_455469, EPI_ISL_455470, EPI_ISL_455471, EPI_ISL_455472, EPI_ISL_455473, EPI_ISL_455474 | Laboratory for Respiratory Viruses, Cantacuzino National Military-Medical Institute for Research and Development | Cantacuzino Institute | M.Lazar, L.Ustea, A.Cretu, Tim Durfee |
| EPI_ISL_455475, EPI_ISL_455476, EPI_ISL_455477 | Laboratory for Respiratory Viruses, Cantacuzino National Military-Medical Institute for Research and Development | Cantacuzino Institute | M.Lazar, L.Ustea, A.Cretu, T.Durfee |
| EPI_ISL_455478 | REGIONAL VRDL,ICMR-RMRC BBSR | Immunogenomics group, Institute of Life Sciences, Bhubaneswar | Sunil Raghav, Jyotirmayee Turuk, Arup Ghosh, Atimukta Jha, Viplov K. Biswas, Swati Madhulika, Manasi Priyadarshini, Shuchi Smita, Jaya Singh Khastri, Rupesh Dash, Soma Chattopadhyay, Ghulam Hussain Syed, Shanti Senapati, Tushar K. Beuria, Debdutta Bhattacharya, Rajeeb Swain, Punit Prasad, COVID-19 team of ILS & RMRC, Orissa COVID-19 study group, DBT's PAN-INDIA 1000 SARS-CoV2 RNA genome sequencing consortium, Sanghamitra Pati, Ajay Parida |
| EPI_ISL_455480 | Veterinary Specialized Institute Kraljevo | Veterinary Specialized Institute Kraljevo | Vidanovic,D., Tesovic,B., Banovic Djeri,B., Sekler,M., Dmitric,M., Debeljak,Z., Matovic,K., Vaskovic,N., Petrovic,T., Volkening,J. and Alfonso,C.L. |
| EPI_ISL_455566, EPI_ISL_455567 | Institute for Public Health | Laboratory for advanced genomics | Filip Rokić, Lovro Trgovec-Greif, Neven Sučić, Tomislav Rukavina, Igor Jurak, Oliver Vugrek |
| EPI_ISL_455568, EPI_ISL_455569, EPI_ISL_455570, EPI_ISL_455571, EPI_ISL_455572 | Gundersen Molecular Diagnostics Laboratory | Kabara Cancer Research Institute | Craig S. Richmond, Paraic A. Kenny |
| EPI_ISL_455573, EPI_ISL_455574 | Gundersen Clinical Microbiology Laboratory | Kabara Cancer Research Institute | Craig S. Richmond, Paraic A. Kenny |
| EPI_ISL_455575, EPI_ISL_455576, EPI_ISL_455578 | Gundersen Molecular Diagnostics Laboratory | Kabara Cancer Research Institute | Craig S. Richmond, Paraic A. Kenny |
| EPI_ISL_455580, EPI_ISL_455581, EPI_ISL_455582 | Gundersen Clinical Microbiology Laboratory | Kabara Cancer Research Institute | Craig S. Richmond, Paraic A. Kenny |
| EPI_ISL_455583 | Central Chest Institute of Thailand | National Institute of Health. Department of medical Sciences, Ministry of Public Health, Thailand | Pilailuk,Okada; Siripaporn,Phuygun; Thanutsapa,Thanadachakul; Sittiporn,Parnmen;Warawan,Wongboot; Sunthareeya,Waicharoen; Malinee,Chittaganpitch |
| EPI_ISL_455584 | National Institute of Health. Department of medical Sciences, Ministry of Public Health, Thailand | National Institute of Health. Department of medical Sciences, Ministry of Public Health, Thailand | Pilailuk,Okada; Siripaporn,Phuygun; Thanutsapa,Thanadachakul; Sittiporn,Parnmen;Warawan,Wongboot; Sunthareeya,Waicharoen; Malinee,Chittaganpitch |
| EPI_ISL_455585 | Phramongkutklao Hospital | National Institute of Health. Department of medical Sciences, Ministry of Public Health, Thailand | Pilailuk,Okada; Siripaporn,Phuygun; Thanutsapa,Thanadachakul; Sittiporn,Parnmen;Warawan,Wongboot; Sunthareeya,Waicharoen; Malinee,Chittaganpitch |
| EPI_ISL_455586 | Siriraj hospital | National Institute of Health. Department of medical Sciences, Ministry of Public Health, Thailand | Pilailuk,Okada; Siripaporn,Phuygun; Thanutsapa,Thanadachakul; Sittiporn,Parnmen;Warawan,Wongboot; Sunthareeya,Waicharoen; Malinee,Chittaganpitch |
| EPI_ISL_455587 | H.R.H. Maha Chakri Sirindhorn Medical Center | National Institute of Health. Department of medical Sciences, Ministry of Public Health, Thailand | Pilailuk,Okada; Siripaporn,Phuygun; Thanutsapa,Thanadachakul; Sittiporn,Parnmen;Warawan,Wongboot; Sunthareeya,Waicharoen; Malinee,Chittaganpitch |
| EPI_ISL_455588 | Trang Hospital | National Institute of Health. Department of medical Sciences, Ministry of Public Health, Thailand | Pilailuk,Okada; Siripaporn,Phuygun; Thanutsapa,Thanadachakul; Sittiporn,Parnmen;Warawan,Wongboot; Sunthareeya,Waicharoen; Malinee,Chittaganpitch |
| EPI_ISL_455589, EPI_ISL_455590, EPI_ISL_455591, EPI_ISL_455592, EPI_ISL_455593 | National Institute of Health. Department of medical Sciences, Ministry of Public Health, Thailand | National Institute of Health. Department of medical Sciences, Ministry of Public Health, Thailand | Pilailuk,Okada; Siripaporn,Phuygun; Thanutsapa,Thanadachakul; Sittiporn,Parnmen;Warawan,Wongboot; Sunthareeya,Waicharoen; Malinee,Chittaganpitch |
| EPI_ISL_455594 | Central Chest Institute of Thailand | National Institute of Health. Department of medical Sciences, Ministry of Public Health, Thailand | Pilailuk,Okada; Siripaporn,Phuygun; Thanutsapa,Thanadachakul; Sittiporn,Parnmen;Warawan,Wongboot; Sunthareeya,Waicharoen; Malinee,Chittaganpitch |
| EPI_ISL_455595, EPI_ISL_455596, EPI_ISL_455597, EPI_ISL_455598, EPI_ISL_455599, EPI_ISL_455602, EPI_ISL_455603 | SA Pathology | VPRL | Beard, MR., Van Der Hoek, K., Lim, C.K., Leong, L.E.X., Coldbeck-Shackley, R., Shue, B., Kirby, E., Merrett, J., Llamas, B. |
| EPI_ISL_455604 | Ramkhamhaeng Hospital | National Institute of Health. Department of medical Sciences, Ministry of Public Health, Thailand | Pilailuk,Okada; Siripaporn,Phuygun; Thanutsapa,Thanadachakul; Sittiporn,Parnmen;Warawan,Wongboot; Sunthareeya,Waicharoen; Malinee,Chittaganpitch |
| EPI_ISL_455605 | Panyananthaphikkhu Chonprathan Medical Center | National Institute of Health. Department of medical Sciences, Ministry of Public Health, Thailand | Pilailuk,Okada; Siripaporn,Phuygun; Thanutsapa,Thanadachakul; Sittiporn,Parnmen;Warawan,Wongboot; Sunthareeya,Waicharoen; Malinee,Chittaganpitch |
| EPI_ISL_455606, EPI_ISL_455607 | Praram 9 Hospital | National Institute of Health. Department of medical Sciences, Ministry of Public Health, Thailand | Pilailuk,Okada; Siripaporn,Phuygun; Thanutsapa,Thanadachakul; Sittiporn,Parnmen;Warawan,Wongboot; Sunthareeya,Waicharoen; Malinee,Chittaganpitch |
| EPI_ISL_455608 | Phramongkutklao Hospital | National Institute of Health. Department of medical Sciences, Ministry of Public Health, Thailand | Pilailuk,Okada; Siripaporn,Phuygun; Thanutsapa,Thanadachakul; Sittiporn,Parnmen;Warawan,Wongboot; Sunthareeya,Waicharoen; Malinee,Chittaganpitch |
| EPI_ISL_455609, EPI_ISL_455610, EPI_ISL_455611, EPI_ISL_455612, EPI_ISL_455613, EPI_ISL_455614, EPI_ISL_455615, EPI_ISL_455616, EPI_ISL_455617, EPI_ISL_455618, EPI_ISL_455619, EPI_ISL_455620, EPI_ISL_455621, EPI_ISL_455622, EPI_ISL_455623 | | | |
| see above | Ochsner Health | Bioinfoexperts, LLC | Susanna L. Lamers, David J. Nolan, Rebecca Rose, Sissy Cross, David Moraga Amador, Tong Yang, Luke Caruso, Wayra Navia, Lydia Von Borstel, Xiao Hui Zhou, Amy Feehan, Julia-Garcia-Diaz |
| EPI_ISL_455624 | National Institute of Health. Department of medical Sciences, Ministry of Public Health, Thailand | National Institute of Health. Department of medical Sciences, Ministry of Public Health, Thailand | Pilailuk,Okada; Siripaporn,Phuygun; Thanutsapa,Thanadachakul; Sittiporn,Parnmen;Warawan,Wongboot; Sunthareeya,Waicharoen; Malinee,Chittaganpitch |
| EPI_ISL_455625, EPI_ISL_455626, EPI_ISL_455627 | unknown | Instituto Nacional de Saude (INSA) | Borges et al |

| | | | |
|---|---|---|---|
| EPI_ISL_455631, EPI_ISL_455632, EPI_ISL_455633, EPI_ISL_455635, EPI_ISL_455636, EPI_ISL_455639 | KRISP, KZN Research Innovation and Sequencing Platform | KRISP, KZN Research Innovation and Sequencing Platform | Giandhari J, Pillay S, Lessells R, Chimukangara B, Deforche K, Tegally H, Wilkinson E, de Oliveira T |
| EPI_ISL_455640, EPI_ISL_455641, EPI_ISL_455643, EPI_ISL_455644, EPI_ISL_455645, EPI_ISL_455646, EPI_ISL_455647, EPI_ISL_455648, EPI_ISL_455649, EPI_ISL_455650, EPI_ISL_455651, EPI_ISL_455652, EPI_ISL_455653, EPI_ISL_455654, EPI_ISL_455655, EPI_ISL_455656, EPI_ISL_455657, EPI_ISL_455658, EPI_ISL_455659, EPI_ISL_455660, EPI_ISL_455661, EPI_ISL_455662, EPI_ISL_455663, EPI_ISL_455664, EPI_ISL_455665, EPI_ISL_455666, EPI_ISL_455667, EPI_ISL_455668, EPI_ISL_455669, EPI_ISL_455670, EPI_ISL_455671, EPI_ISL_455672, EPI_ISL_455673, EPI_ISL_455674, EPI_ISL_455675, EPI_ISL_455676, EPI_ISL_455678, EPI_ISL_455679 | | | |
| see above | ICMR-National Institute of Cholera and Enteric Diseases | National Institute of Biomedical Genomics | Arindam Maitra, Mamta Chawla Sarkar, Sreedhar Chinnaswamy, Hasina Banu, Ananya Chatterjee, Shanta Dutta, Saumitra Das |
| EPI_ISL_455680 | Institute of pathogenic microbiology, Jiangsu Provincial Center for Disease Control and Prevention | Institute of pathogenic microbiology, Jiangsu Provincial Center for Disease Control and Prevention | Cui,L. |
| EPI_ISL_455683, EPI_ISL_455684, EPI_ISL_455685, EPI_ISL_455686, EPI_ISL_455687, EPI_ISL_455688, EPI_ISL_455689, EPI_ISL_455690, EPI_ISL_455691, EPI_ISL_455692, EPI_ISL_455693 | | | |
| see above | unknown | Department of Microbiology | Gao,Q., Bao,L., Mao,H., Wang,L., Xu,K., Yang,M., Li,Y., Zhu,L., Wang,N., Lv,Z., Gao,H., Ge,X., Kan,B., Hu,Y., Liu,J., Cai,F., Jiang,D., Yin,Y., Qin,C., Li,J., Gong,X., Lou,X., Shi,W., Wu,D., Zhang,H., Deng,W., Lu,J., Li,C., Wang,X., Yin,W., Zhang,Y., Sun,Y. |
| EPI_ISL_455694, EPI_ISL_455695, EPI_ISL_455696, EPI_ISL_455697, EPI_ISL_455698, EPI_ISL_455699, EPI_ISL_455700, EPI_ISL_455701, EPI_ISL_455702, EPI_ISL_455703, EPI_ISL_455704, EPI_ISL_455705, EPI_ISL_455706, EPI_ISL_455707, EPI_ISL_455708, EPI_ISL_455709, EPI_ISL_455710, EPI_ISL_455711, EPI_ISL_455712 | | | |
| see above | National Hospital of Tropical Diseases | Oxford University Clinical Research Unit, Hanoi, Vietnam | Nguyen Thi Tam, Van Dinh Trang, Nguyen Thu Trang, Nguyen Thi Ngoc Diep, Le Nguyen Minh Hoa, Pham Ngoc Thach, H. Rogier van Doorn, on behalf of the OUCRU COVID-19 research group |
| EPI_ISL_455714, EPI_ISL_455718 | National Hospital of Tropical Diseases | Oxford University Clinical Research Unit, Hanoi, Vietnam | Nguyen Thi Tam, Van Dinh Trang, Nguyen Thi Hong Thuong, Vu Thi Ngoc Bich, Nguyen Thu Trang, Nguyen Thi Ngoc Diep, Le Nguyen Minh Hoa, Pham Ngoc Thach, H. Rogier van Doorn, on behalf of the OUCRU COVID-19 research group |
| EPI_ISL_455727 | Servicio de Microbiologia. Hospital Clinico Universitario de Valencia | Sequencing and Bioinformatics Service and Molecular Epidemiology Research Group. FISABIO-Public Health, and SeqCOVID-Spain Consortium | David Navarro, Eliseo Albert, Maria Alma Bracho, Griselda De Marco, Lidia Ruiz Roldan, Neris Garcia-Gonzalez, Inma Galán Vendrell, Sandra Carbo, Loreto Ferrús Abad, Paula Ruiz-Hueso, Mariana Reyes-Prieto, Vicente Soriano Chirona, Ivan Ansari, Llúcia Martínez-Priego, Giuseppe 'Auria, Fernando Gonzalez-Candelas |
| EPI_ISL_455730 | Servicio de Microbiologia. Hospital Clinico Universitario de Valencia | Sequencing and Bioinformatics Service and Molecular Epidemiology Research Group. FISABIO-Public Health, and SeqCOVID-Spain Consortium | Mariana Reyes-Prieto, Vicente Soriano Chirona, Ivan Ansari, Llúcia Martínez-Priego, Giuseppe 'Auria, David Navarro, Eliseo Albert, Maria Alma Bracho, Lidia Ruiz Roldan, Neris Garcia-Gonzalez, Inma Galán Vendrell, Sandra Carbo, Loreto Ferrús Abad, Paula Ruiz-Hueso, Fernando Gonzalez-Candelas |
| EPI_ISL_455733 | Servicio de Microbiologia. Hospital Clinico Universitario de Valencia | Sequencing and Bioinformatics Service and Molecular Epidemiology Research Group. FISABIO-Public Health, and SeqCOVID-Spain Consortium | Loreto Ferrús Abad, Paula Ruiz-Hueso, Mariana Reyes-Prieto, Vicente Soriano Chirona, Ivan Ansari, Llúcia Martínez-Priego, Giuseppe 'Auria, David Navarro, Eliseo Albert, Maria Alma Bracho, Lidia Ruiz Roldan, Neris Garcia-Gonzalez, Inma Galán Vendrell, Sandra Carbo, Fernando Gonzalez-Candelas |
| EPI_ISL_455735, EPI_ISL_455736 | Servicio de Microbiologia. Hospital_Arnau_de_Vilanova | Sequencing and Bioinformatics Service and Molecular Epidemiology Research Group. FISABIO-Public Health, and SeqCOVID-Spain Consortium | Victoria Dominguez, Maria Alma Bracho, Griselda De Marco, Lidia Ruiz Roldan, Neris Garcia-Gonzalez, Inma Galán Vendrell, Sandra Carbo, Loreto Ferrús Abad, Paula Ruiz-Hueso, Mariana Reyes-Prieto, Vicente Soriano Chirona, Ivan Ansari, Llúcia Martínez-Priego, Giuseppe 'Auria, Fernando Gonzalez-Candelas |
| EPI_ISL_455741 | Servicio de Microbiologia. Hospital Clinico Universitario de Valencia | Sequencing and Bioinformatics Service and Molecular Epidemiology Research Group. FISABIO-Public Health, and SeqCOVID-Spain Consortium | David Navarro, Eliseo Albert, Maria Alma Bracho, Griselda De Marco, Lidia Ruiz Roldan, Neris Garcia-Gonzalez, Inma Galán Vendrell, Sandra Carbo, Loreto Ferrús Abad, Paula Ruiz-Hueso, Mariana Reyes-Prieto, Vicente Soriano Chirona, Ivan Ansari, Llúcia Martínez-Priego, Giuseppe 'Auria, Fernando Gonzalez-Candelas |
| EPI_ISL_455742 | Servicio de Microbiologia. Hospital Clinico Universitario de Valencia | Sequencing and Bioinformatics Service and Molecular Epidemiology Research Group. FISABIO-Public Health, and SeqCOVID-Spain Consortium | Neris Garcia-Gonzalez, Inma Galán Vendrell, Sandra Carbo, Loreto Ferrús Abad, Paula Ruiz-Hueso, Mariana Reyes-Prieto, Vicente Soriano Chirona, Ivan Ansari, Llúcia Martínez-Priego, Giuseppe 'Auria, David Navarro, Eliseo Albert, Maria Alma Bracho, Lidia Ruiz Roldan, Fernando Gonzalez-Candelas |
| EPI_ISL_455743 | Servicio de Microbiologia. Hospital Clinico Universitario de Valencia | Sequencing and Bioinformatics Service and Molecular Epidemiology Research Group. FISABIO-Public Health, and SeqCOVID-Spain Consortium | Eliseo Albert, Maria Alma Bracho, Griselda De Marco, Lidia Ruiz Roldan, Neris Garcia-Gonzalez, Inma Galán Vendrell, Sandra Carbo, Loreto Ferrús Abad, Paula Ruiz-Hueso, Mariana Reyes-Prieto, Vicente Soriano Chirona, Ivan Ansari, Llúcia Martínez-Priego, Giuseppe 'Auria, David Navarro, Fernando Gonzalez-Candelas |
| EPI_ISL_455751, EPI_ISL_455752, EPI_ISL_455754, EPI_ISL_455755, EPI_ISL_455757, EPI_ISL_455758, EPI_ISL_455760, EPI_ISL_455761, EPI_ISL_455763, EPI_ISL_455764, EPI_ISL_455765, EPI_ISL_455766, EPI_ISL_455767, EPI_ISL_455768, EPI_ISL_455770, EPI_ISL_455771, EPI_ISL_455775, EPI_ISL_455776, EPI_ISL_455777, EPI_ISL_455778, EPI_ISL_455779, EPI_ISL_455780, EPI_ISL_455782, EPI_ISL_455783, EPI_ISL_455784, EPI_ISL_455786, EPI_ISL_455787 | | | |
| see above | REGIONAL VRDL,ICMR-RMRC BBSR | Immunogenomics lab, Institute of Life Sciences, Bhubaneswar | Sunil Raghav, Jyotirmayee Turuk, Arup Ghosh, Atimukta Jha, Viplov K. Biswas, Swati Madhulika, Manasi Priyadarshini, Shuchi Smita, Jaya Singh Khastri, Rupesh Dash, Soma Chattopadhyay, Ghulam Hussain Syed, Shanti Senapati, Tushar K. Beuria, Debdutta Bhattacharya, Rajeeb Swain, Punit Prasad, COVID-19 team of ILS & RMRC, Orissa COVID-19 study group, DBT's PAN-INDIA 1000 SARS-CoV2 RNA genome sequencing consortium, Sanghamitra Pati, Ajay Parida |
| EPI_ISL_455790 | Institute for Medical Research, Infectious Disease Research Centre, National Institutes of Health, Ministry of Health Malaysia | Malaysia Genome Institute | Mohd Noor Mat Isa, Irni Suhayu Sapian, Yusuf Muhammad Noor, Jeyanthi Suppiah, Nurhezreen Md Iqbal, Enizza Kasim, Zarina Mohd Zawawi, Siti Noraini Othman, Mohd Faizal Abu Bakar, Shamsidar Sopie, Azrin Ahmad, Ravindran Thayan, Norazah Ahmad, Tahir Aris, Shahrul Hisham Zainal Ariffin |
| EPI_ISL_455791 | Institute for Medical Research, Infectious Disease Research Centre, National Institutes of Health, Ministry of Health Malaysia | Malaysia Genome Institute | Mohd Noor Mat Isa, Irni Suhayu Sapian, Yusuf Muhammad Noor, Jeyanthi Suppiah, Nurhezreen Md Iqbal, Enizza Kasim, Zarina Mohd Zawawi, Siti Noraini Othman, Mohd Faizal Abu Bakar, Shamsidar Sopie, Azrin Ahmad, Ravindran Thayan, Norazah Ahmad, Tahir Aris, Shahrul Hisham Zainal Ariffin |
| EPI_ISL_455792 | Institute for Medical Research, Infectious Disease Research Centre, National Institutes of Health, Ministry of Health Malaysia | Malaysia Genome Institute | Mohd Noor Mat Isa, Irni Suhayu Sapian, Yusuf Muhammad Noor, Jeyanthi Suppiah, Nurhezreen Md Iqbal, Enizza Kasim, Zarina Mohd Zawawi, Siti Noraini Othman, Mohd Faizal Abu Bakar, Shamsidar Sopie, Azrin Ahmad, Ravindran Thayan, Norazah Ahmad, Tahir Aris, Shahrul Hisham Zainal Ariffin |
| EPI_ISL_455793 | Institute for Medical Research, Infectious Disease Research Centre, National Institutes of Health, Ministry of Health Malaysia | Malaysia Genome Institute | Mohd Noor Mat Isa, Irni Suhayu Sapian, Yusuf Muhammad Noor, Jeyanthi Suppiah, Nurhezreen Md Iqbal, Enizza Kasim, Zarina Mohd Zawawi, Siti Noraini Othman, Mohd Faizal Abu Bakar, Shamsidar Sopie, Azrin Ahmad, Ravindran Thayan, Norazah Ahmad, Tahir Aris, Shahrul Hisham Zainal Ariffin |
| EPI_ISL_455840, EPI_ISL_455841 | Orebro klinisk mikrobiologi | The Public Health Agency of Sweden | Anna-Malin Linde, Maria Lind Karlberg, Mattias Haukland, Reza Advani, Olov Svartstrom, Oskar Karlsson Lindsjo, Petra Edquist, Shamam Muradrasoli, Anna Risberg, Karin Tegmark-Wisell |
| EPI_ISL_455842, EPI_ISL_455843, EPI_ISL_455844, EPI_ISL_455845 | Skovde/Unilabs | The Public Health Agency of Sweden | Anna-Malin Linde, Maria Lind Karlberg, Mattias Haukland, Reza Advani, Olov Svartstrom, Oskar Karlsson Lindsjo, Petra Edquist, Shamam Muradrasoli, Anna Risberg, Karin Tegmark-Wisell |
| EPI_ISL_455846, EPI_ISL_455847, EPI_ISL_455848, EPI_ISL_455849, EPI_ISL_455850, EPI_ISL_455851, EPI_ISL_455852, EPI_ISL_455853, EPI_ISL_455854, EPI_ISL_455855, EPI_ISL_455856, EPI_ISL_455857, EPI_ISL_455858, EPI_ISL_455859, EPI_ISL_455860, EPI_ISL_455861, EPI_ISL_455862, EPI_ISL_455863, EPI_ISL_455864, EPI_ISL_455865, EPI_ISL_455866, EPI_ISL_455867, EPI_ISL_455868, EPI_ISL_455869, EPI_ISL_455870, EPI_ISL_455871, EPI_ISL_455872, EPI_ISL_455873, EPI_ISL_455874, EPI_ISL_455875, EPI_ISL_455876, EPI_ISL_455877, EPI_ISL_455878, EPI_ISL_455879, EPI_ISL_455880, EPI_ISL_455881, EPI_ISL_455882, EPI_ISL_455883, EPI_ISL_455884, EPI_ISL_455885, EPI_ISL_455886, EPI_ISL_455887, EPI_ISL_455888, EPI_ISL_455889, EPI_ISL_455890, EPI_ISL_455891, EPI_ISL_455892, EPI_ISL_455893, EPI_ISL_455894, EPI_ISL_455895, EPI_ISL_455896, EPI_ISL_455897, EPI_ISL_455898, EPI_ISL_455899, EPI_ISL_455900, EPI_ISL_455901 | | | |
| see above | Karolinska Universitetslaboratoriet | The Public Health Agency of Sweden | Anna-Malin Linde, Maria Lind Karlberg, Mattias Haukland, Reza Advani, Olov Svartstrom, Oskar Karlsson Lindsjo, Petra Edquist, Shamam Muradrasoli, Anna Risberg, Karin Tegmark-Wisell |
| EPI_ISL_455902 | Halmstad klinisk mikrobiologi | The Public Health Agency of Sweden | Anna-Malin Linde, Maria Lind Karlberg, Mattias Haukland, Reza Advani, Olov Svartstrom, Oskar Karlsson Lindsjo, Petra Edquist, Shamam Muradrasoli, Anna Risberg, Karin Tegmark-Wisell |
| EPI_ISL_455903 | Folkhalsomyndigheten | The Public Health Agency of Sweden | Anna-Malin Linde, Maria Lind Karlberg, Mattias Haukland, Reza Advani, Olov Svartstrom, Oskar Karlsson Lindsjo, Petra Edquist, Shamam Muradrasoli, Anna Risberg, Karin Tegmark-Wisell |
| EPI_ISL_455904, EPI_ISL_455905, EPI_ISL_455906, EPI_ISL_455907 | Klinisk mikrobiologi, UAS | The Public Health Agency of Sweden | Anna-Malin Linde, Maria Lind Karlberg, Mattias Haukland, Reza Advani, Olov Svartstrom, Oskar Karlsson Lindsjo, Petra Edquist, Shamam Muradrasoli, Anna Risberg, Karin Tegmark-Wisell |
| EPI_ISL_455909, EPI_ISL_455910, EPI_ISL_455911, EPI_ISL_455912, EPI_ISL_455913, EPI_ISL_455915, EPI_ISL_455917, EPI_ISL_455922, EPI_ISL_455925, EPI_ISL_455926, EPI_ISL_455927, EPI_ISL_455928, EPI_ISL_455931, EPI_ISL_455932, EPI_ISL_455934, EPI_ISL_455935, EPI_ISL_455936, EPI_ISL_455938, EPI_ISL_455939, EPI_ISL_455940, EPI_ISL_455941, EPI_ISL_455942, EPI_ISL_455943, EPI_ISL_455947 | | | |
| see above | Ramathibodi Hospital | COVID-19 Network Investigations (CONI) Alliance | Elizabeth Batty, Wasun Chantratita, Thanat Chookajorn, Stefan Fernandez, Angkana Huang, Anthony R. Jones, Khajohn Joonsalak, Chonticha Klungtong, Theerarat Kochakarn, Namfon Kotanan, Krittikorn Kumpornsin, Wudtichai Manasatienkij, Bhakbhoom Panthan, Ekawat Pasomsub, Kingkan Rakmanee, Insee Sensorn, Janjira Thaipadungpanit, Arporn Wangwiwatsin,Treewat Watthanachockchai |
| EPI_ISL_455958, EPI_ISL_455961, EPI_ISL_455962, EPI_ISL_455963, EPI_ISL_455965, EPI_ISL_455966, EPI_ISL_455967, EPI_ISL_455968, EPI_ISL_455969, EPI_ISL_455970, EPI_ISL_455972, EPI_ISL_455973, EPI_ISL_455974, EPI_ISL_455975, EPI_ISL_455976, EPI_ISL_455977, EPI_ISL_455978, EPI_ISL_455979 | | | |
| see above | Department of Clinical Microbiology | GIGA Medical Genomics | Keith Durkin, Maria Artesi, Sébastien Bontems, Raphaël Boreux, Cécile Meex, Pierrette Melin, Marie-Pierre Hayette, Vincent Bours. |
| EPI_ISL_455980, EPI_ISL_455981, EPI_ISL_455982, EPI_ISL_455983, EPI_ISL_455984, EPI_ISL_455985, EPI_ISL_455986, EPI_ISL_455987, EPI_ISL_455988, EPI_ISL_455989 | LSUHS Emerging Viral Threat Laboratory | Microbial Genome Sequencing Center | Jeremy P. Kamil, John A. Vanchiere, Rona S. Scott, Camille F. Abshire, Abida Siddiqa, Byeong-Jae Lee, Chan-ki Min, Md Maksudul Alam, Monica Gestal-Cartele, Edna Ondari, Adam Greer, Malgorzata Bienkowska-Haba, Katarzyna Zwolinska, Michelle M. Arnold, Jason M. Bodily, Andrew D. Yurochko, Paul M. Weinberger, Christopher G. Kevil, Martin J. Sapp, Daniel J. Snyder, Vaughn S. Cooper |
| EPI_ISL_455990, EPI_ISL_455991, EPI_ISL_455992, EPI_ISL_455993, EPI_ISL_455994, EPI_ISL_455995, EPI_ISL_455996, EPI_ISL_455997, EPI_ISL_455998, EPI_ISL_455999 | LSUHS Emerging Viral Threat Laboratory | Microbial Genome Sequencing Center | John A. Vanchiere, Jeremy P. Kamil, Rona S. Scott, Camille F. Abshire, Abida Siddiqa, Byeong-Jae Lee, Chan-ki Min, Md Maksudul Alam, Monica Gestal-Cartele, Edna Ondari, Adam Greer, Malgorzata Bienkowska-Haba, Katarzyna Zwolinska, Michelle M. Arnold, Jason M. Bodily, Andrew D. Yurochko, Paul M. Weinberger, Christopher G. Kevil, Martin J. Sapp, Daniel J. Snyder, Vaughn S. Cooper |
| EPI_ISL_456000, EPI_ISL_456001, EPI_ISL_456002 | LSUHS Emerging Viral Threat Laboratory | Microbial Genome Sequencing Center | Rona S. Scott, Jeremy P. Kamil, John A. Vanchiere, Camille F. Abshire, Abida Siddiqa, Byeong-Jae Lee, Chan-ki Min, Md Maksudul Alam, Monica Gestal-Cartele, Edna Ondari, Adam Greer, Malgorzata Bienkowska-Haba, Katarzyna Zwolinska, Michelle M. Arnold, Jason M. Bodily, Andrew D. Yurochko, Paul M. Weinberger, Christopher G. Kevil, Martin J. Sapp, Daniel J. Snyder, Vaughn S. Cooper |
| EPI_ISL_456004, EPI_ISL_456006, EPI_ISL_456007, EPI_ISL_456008, EPI_ISL_456010, EPI_ISL_456011, EPI_ISL_456012, EPI_ISL_456013, EPI_ISL_456014, EPI_ISL_456015, EPI_ISL_456016, EPI_ISL_456017, EPI_ISL_456018, EPI_ISL_456019, EPI_ISL_456020, EPI_ISL_456021, EPI_ISL_456022, EPI_ISL_456023, EPI_ISL_456024, EPI_ISL_456025, EPI_ISL_456026, EPI_ISL_456027, EPI_ISL_456028, EPI_ISL_456029, EPI_ISL_456030, EPI_ISL_456031, EPI_ISL_456032, EPI_ISL_456033, EPI_ISL_456034, EPI_ISL_456035, EPI_ISL_456036, EPI_ISL_456037, EPI_ISL_456038, EPI_ISL_456039, EPI_ISL_456040, EPI_ISL_456041, EPI_ISL_456042, EPI_ISL_456043, EPI_ISL_456044, EPI_ISL_456045, EPI_ISL_456046, EPI_ISL_456047, EPI_ISL_456049, EPI_ISL_456050, EPI_ISL_456051, EPI_ISL_456052, EPI_ISL_456053, EPI_ISL_456054, EPI_ISL_456055, EPI_ISL_456056, EPI_ISL_456058, EPI_ISL_456060, EPI_ISL_456061, EPI_ISL_456062, EPI_ISL_456063, EPI_ISL_456065, EPI_ISL_456066 | | | |
| see above | NYU Langone Health | Departments of Pathology and Medicine, New York University School of Medicine | Maria Aguero-Rosenfeld, Brendan Belovarac, Margaret Black, Ludovic Boytard, John Cadley, Paolo Cotzia, John Chen, Dacia Dimartino, Xiaojun Feng, Tatyana Gindin, Emily Guzman, Adriana Heguy, Megan Hogan, Emily Huang, George Jour, Alireza Khodadadi-Jamayran, Lawrence H. Lin, Raven Luther, Andrew Lytle, Christian Marier, Matthew T. Maurano, Mark J. Mulligan, Peter Meyn, Raquel Ordonez Ciriza, Iman Osman, Jared Pinnell, Vanessa Raabe, Sitharam Ramaswami, Amy Rapkiewicz, Andre M. Ribeiro-dos-Santos, Marie Samanovic-Golden, Antonio Serrano, Guomiao Shen, Matija Snuderl, Theodore Vougiouklakis, Nick Vulpescu, Gael Westby, Paul Zappile, Yutong Zhang |
| EPI_ISL_456071, EPI_ISL_456072, EPI_ISL_456073, EPI_ISL_456074, EPI_ISL_456075 | Laboratory of Respiratory Viruses and Measles, Oswaldo Cruz Institute, FIOCRUZ | Laboratory of Respiratory Viruses and Measles, Oswaldo Cruz Institute, FIOCRUZ | Paola Resende, Luciana Appolinario, Fernando Motta, Aline Mattos, Milene Miranda, Cristiana Garcia, Braulia Caetano, Maria Ogrzewalska, Jonathan Lopes, Marilda Siqueira |
| EPI_ISL_456076, EPI_ISL_456077 | LACEN RJ - Laboratório Central de Saúde Pública Noel Nutels | Laboratory of Respiratory Viruses and Measles, Oswaldo Cruz Institute, FIOCRUZ | Paola Resende, Luciana Appolinario, Fernando Motta, Aline Mattos, Milene Miranda, Cristiana Garcia, Braulia Caetano, Maria Ogrzewalska, Jonathan Lopes, Marilda Siqueira |
| EPI_ISL_456079, EPI_ISL_456080, EPI_ISL_456081 | Laboratory of Respiratory Viruses and Measles, Oswaldo Cruz Institute, FIOCRUZ | Laboratory of Respiratory Viruses and Measles, Oswaldo Cruz Institute, FIOCRUZ | Paola Resende, Luciana Appolinario, Fernando Motta, Aline Mattos, Milene Miranda, Cristiana Garcia, Braulia Caetano, Maria Ogrzewalska, Jonathan Lopes, Marilda Siqueira |

| | | | |
|---|---|---|---|
| EPI_ISL_456082, EPI_ISL_456083 | LACEN RJ - Laboratório Central de Saúde Pública Noel Nutels | Laboratory of Respiratory Viruses and Measles, Oswaldo Cruz Institute, FIOCRUZ | Paola Resende, Luciana Appolinario, Fernando Motta, Aline Mattos, Milene Miranda, Cristiana Garcia, Braulia Caetano, Maria Ogrzewalska, Jonathan Lopes, Marilda Siqueira |
| EPI_ISL_456084, EPI_ISL_456085, EPI_ISL_456086, EPI_ISL_456087 | Laboratory of Respiratory Viruses and Measles, Oswaldo Cruz Institute, FIOCRUZ | Laboratory of Respiratory Viruses and Measles, Oswaldo Cruz Institute, FIOCRUZ | Paola Resende, Luciana Appolinario, Fernando Motta, Aline Mattos, Milene Miranda, Cristiana Garcia, Braulia Caetano, Maria Ogrzewalska, Jonathan Lopes, Marilda Siqueira |
| EPI_ISL_456088 | LACEN RJ - Laboratório Central de Saúde Pública Noel Nutels | Laboratory of Respiratory Viruses and Measles, Oswaldo Cruz Institute, FIOCRUZ | Paola Resende, Luciana Appolinario, Fernando Motta, Aline Mattos, Milene Miranda, Cristiana Garcia, Braulia Caetano, Maria Ogrzewalska, Jonathan Lopes, Marilda Siqueira |
| EPI_ISL_456089, EPI_ISL_456090, EPI_ISL_456091, EPI_ISL_456092, EPI_ISL_456093, EPI_ISL_456094, EPI_ISL_456095, EPI_ISL_456096, EPI_ISL_456097, EPI_ISL_456098, EPI_ISL_456099, EPI_ISL_456100, EPI_ISL_456101, EPI_ISL_456102, EPI_ISL_456103, EPI_ISL_456104, EPI_ISL_456105, EPI_ISL_456106 | | | |
| see above | Laboratory of Respiratory Viruses and Measles, Oswaldo Cruz Institute, FIOCRUZ | Laboratory of Respiratory Viruses and Measles, Oswaldo Cruz Institute, FIOCRUZ | Paola Resende, Luciana Appolinario, Fernando Motta, Aline Mattos, Milene Miranda, Cristiana Garcia, Braulia Caetano, Maria Ogrzewalska, Jonathan Lopes, Marilda Siqueira |
| EPI_ISL_456107, EPI_ISL_456108, EPI_ISL_456109, EPI_ISL_456111, EPI_ISL_456112, EPI_ISL_456113 | NYU Langone Health | Departments of Pathology and Medicine, New York University School of Medicine | Maria Aguero-Rosenfeld, Brendan Belovarac, Margaret Black, Ludovic Boytard, John Cadley, Paolo Cotzia, John Chen, Dacia Dimartino, Xiaojun Feng, Tatyana Gindin, Emily Guzman, Adriana Heguy, Megan Hogan, Emily Huang, George Jour, Alireza Khodadadi-Jamayran, Lawrence H. Lin, Raven Luther, Andrew Lytle, Christian Marier, Matthew T. Maurano, Mark J. Mulligan, Peter Meyn, Raquel Ordonez Ciriza, Iman Osman, Jared Pinnell, Vanessa Raabe, Sitharam Ramaswami, Amy Rapkiewicz, Andre M. Ribeiro-dos-Santos, Marie Samanovic-Golden, Antonio Serrano, Guomiao Shen, Matija Snuderl, Theodore Vougiouklakis, Nick Vulpescu, Gael Westby, Paul Zappile, Yutong Zhang |
| EPI_ISL_456117, EPI_ISL_456119, EPI_ISL_456120, EPI_ISL_456126, EPI_ISL_456127, EPI_ISL_456138, EPI_ISL_456144, EPI_ISL_456145, EPI_ISL_456146, EPI_ISL_456147, EPI_ISL_456148, EPI_ISL_456149, EPI_ISL_456150, EPI_ISL_456151, EPI_ISL_456152, EPI_ISL_456153, EPI_ISL_456154, EPI_ISL_456155 | | | |
| see above | Instituto Nacional de Salud - Unidad de Secuenciación y Análisis Genómico | Instituto Nacional de Salud, Universidad Cooperativa de Colombia, Instituto Alexander von Humboldt, Imperial College-London, London School of Hygiene & Tropical Medicine | Katherine Laiton-Donato, Diego A. Álvarez-Díaz, Carlos Franco-Muñoz, Jose A. Usme-Ciro, Gloria Puerto, Nicolas D. Franco-Sierra, Mailyn A.Gonzalez, Zulma M. Cucunubá, Christian Julian Villabona-Arenas, Liz Villabona-Arenas, Sussy Echeverria, Astrid C. Flórez, Sergio Gomez-Rangel, Luz Dary Rodriguez, Juliana Barbosa, Erika Ospitia, Diana Marcela Walteros-Acero, Martha Lucia Ospina Martinez, Marcela Mercado-Reyes. |
| EPI_ISL_456159 | Southern Community Labs Dunedin | Institute of Environmental Science and Research (ESR) | Matt Storey, Xiaoyun Ren, Anja Werno, Antje van der Linden, Arlo Upton, Chris Mansell, David Hammer, Dragana Drinkovic, Erasmus Smit, Gary McAuliffe, Hana Sofia Andersson, James Ussher, Jill Sherwood, Josh Freeman, Julia Howard, Juliet Elvy, Mary DeAlmeida, Matt Blakiston, Matthew Rogers, Max Bloomfield, Michael Addidle, Michelle Balm, Sally Roberts, Sarah Jefferies, Sharmini Muttaiyah, Susan Morpeth, Susan Taylor, Timothy Blackmore, Vani Sathyendran, Veronica Playle, Virginia Hope, Erasmus Smit, Lauren Jelly, Joep de Ligt |
| EPI_ISL_456160 | Waikato Hospital | Institute of Environmental Science and Research (ESR) | Matt Storey, Xiaoyun Ren, Anja Werno, Antje van der Linden, Arlo Upton, Chris Mansell, David Hammer, Dragana Drinkovic, Erasmus Smit, Gary McAuliffe, Hana Sofia Andersson, James Ussher, Jill Sherwood, Josh Freeman, Julia Howard, Juliet Elvy, Mary DeAlmeida, Matt Blakiston, Matthew Rogers, Max Bloomfield, Michael Addidle, Michelle Balm, Sally Roberts, Sarah Jefferies, Sharmini Muttaiyah, Susan Morpeth, Susan Taylor, Timothy Blackmore, Vani Sathyendran, Veronica Playle, Virginia Hope, Erasmus Smit, Lauren Jelly, Joep de Ligt |
| EPI_ISL_456161 | Wellington SCL | Institute of Environmental Science and Research (ESR) | Matt Storey, Xiaoyun Ren, Anja Werno, Antje van der Linden, Arlo Upton, Chris Mansell, David Hammer, Dragana Drinkovic, Erasmus Smit, Gary McAuliffe, Hana Sofia Andersson, James Ussher, Jill Sherwood, Josh Freeman, Julia Howard, Juliet Elvy, Mary DeAlmeida, Matt Blakiston, Matthew Rogers, Max Bloomfield, Michael Addidle, Michelle Balm, Sally Roberts, Sarah Jefferies, Sharmini Muttaiyah, Susan Morpeth, Susan Taylor, Timothy Blackmore, Vani Sathyendran, Veronica Playle, Virginia Hope, Erasmus Smit, Lauren Jelly, Joep de Ligt |
| EPI_ISL_456163 | PathLab Bay of Plenty | Institute of Environmental Science and Research (ESR) | Matt Storey, Xiaoyun Ren, Anja Werno, Antje van der Linden, Arlo Upton, Chris Mansell, David Hammer, Dragana Drinkovic, Erasmus Smit, Gary McAuliffe, Hana Sofia Andersson, James Ussher, Jill Sherwood, Josh Freeman, Julia Howard, Juliet Elvy, Mary DeAlmeida, Matt Blakiston, Matthew Rogers, Max Bloomfield, Michael Addidle, Michelle Balm, Sally Roberts, Sarah Jefferies, Sharmini Muttaiyah, Susan Morpeth, Susan Taylor, Timothy Blackmore, Vani Sathyendran, Veronica Playle, Virginia Hope, Erasmus Smit, Lauren Jelly, Joep de Ligt |
| EPI_ISL_456165 | Wellington SCL | Institute of Environmental Science and Research (ESR) | Matt Storey, Xiaoyun Ren, Anja Werno, Antje van der Linden, Arlo Upton, Chris Mansell, David Hammer, Dragana Drinkovic, Erasmus Smit, Gary McAuliffe, Hana Sofia Andersson, James Ussher, Jill Sherwood, Josh Freeman, Julia Howard, Juliet Elvy, Mary DeAlmeida, Matt Blakiston, Matthew Rogers, Max Bloomfield, Michael Addidle, Michelle Balm, Sally Roberts, Sarah Jefferies, Sharmini Muttaiyah, Susan Morpeth, Susan Taylor, Timothy Blackmore, Vani Sathyendran, Veronica Playle, Virginia Hope, Erasmus Smit, Lauren Jelly, Joep de Ligt |
| EPI_ISL_456167 | Southern Community Labs Dunedin | Institute of Environmental Science and Research (ESR) | Matt Storey, Xiaoyun Ren, Anja Werno, Antje van der Linden, Arlo Upton, Chris Mansell, David Hammer, Dragana Drinkovic, Erasmus Smit, Gary McAuliffe, Hana Sofia Andersson, James Ussher, Jill Sherwood, Josh Freeman, Julia Howard, Juliet Elvy, Mary DeAlmeida, Matt Blakiston, Matthew Rogers, Max Bloomfield, Michael Addidle, Michelle Balm, Sally Roberts, Sarah Jefferies, Sharmini Muttaiyah, Susan Morpeth, Susan Taylor, Timothy Blackmore, Vani Sathyendran, Veronica Playle, Virginia Hope, Erasmus Smit, Lauren Jelly, Joep de Ligt |
| EPI_ISL_456168 | Waikato Hospital | Institute of Environmental Science and Research (ESR) | Matt Storey, Xiaoyun Ren, Anja Werno, Antje van der Linden, Arlo Upton, Chris Mansell, David Hammer, Dragana Drinkovic, Erasmus Smit, Gary McAuliffe, Hana Sofia Andersson, James Ussher, Jill Sherwood, Josh Freeman, Julia Howard, Juliet Elvy, Mary DeAlmeida, Matt Blakiston, Matthew Rogers, Max Bloomfield, Michael Addidle, Michelle Balm, Sally Roberts, Sarah Jefferies, Sharmini Muttaiyah, Susan Morpeth, Susan Taylor, Timothy Blackmore, Vani Sathyendran, Veronica Playle, Virginia Hope, Erasmus Smit, Lauren Jelly, Joep de Ligt |
| EPI_ISL_456169, EPI_ISL_456170 | Wellington SCL | Institute of Environmental Science and Research (ESR) | Matt Storey, Xiaoyun Ren, Anja Werno, Antje van der Linden, Arlo Upton, Chris Mansell, David Hammer, Dragana Drinkovic, Erasmus Smit, Gary McAuliffe, Hana Sofia Andersson, James Ussher, Jill Sherwood, Josh Freeman, Julia Howard, Juliet Elvy, Mary DeAlmeida, Matt Blakiston, Matthew Rogers, Max Bloomfield, Michael Addidle, Michelle Balm, Sally Roberts, Sarah Jefferies, Sharmini Muttaiyah, Susan Morpeth, Susan Taylor, Timothy Blackmore, Vani Sathyendran, Veronica Playle, Virginia Hope, Erasmus Smit, Lauren Jelly, Joep de Ligt |
| EPI_ISL_456171 | PathLab Bay of Plenty | Institute of Environmental Science and Research (ESR) | Matt Storey, Xiaoyun Ren, Anja Werno, Antje van der Linden, Arlo Upton, Chris Mansell, David Hammer, Dragana Drinkovic, Erasmus Smit, Gary McAuliffe, Hana Sofia Andersson, James Ussher, Jill Sherwood, Josh Freeman, Julia Howard, Juliet Elvy, Mary DeAlmeida, Matt Blakiston, Matthew Rogers, Max Bloomfield, Michael Addidle, Michelle Balm, Sally Roberts, Sarah Jefferies, Sharmini Muttaiyah, Susan Morpeth, Susan Taylor, Timothy Blackmore, Vani Sathyendran, Veronica Playle, Virginia Hope, Erasmus Smit, Lauren Jelly, Joep de Ligt |
| EPI_ISL_456172, EPI_ISL_456173, EPI_ISL_456174, EPI_ISL_456175, EPI_ISL_456176, EPI_ISL_456177, EPI_ISL_456178, EPI_ISL_456179, EPI_ISL_456180 | LabPLUS | Institute of Environmental Science and Research (ESR) | Matt Storey, Xiaoyun Ren, Anja Werno, Antje van der Linden, Arlo Upton, Chris Mansell, David Hammer, Dragana Drinkovic, Erasmus Smit, Gary McAuliffe, Hana Sofia Andersson, James Ussher, Jill Sherwood, Josh Freeman, Julia Howard, Juliet Elvy, Mary DeAlmeida, Matt Blakiston, Matthew Rogers, Max Bloomfield, Michael Addidle, Michelle Balm, Sally Roberts, Sarah Jefferies, Sharmini Muttaiyah, Susan Morpeth, Susan Taylor, Timothy Blackmore, Vani Sathyendran, Veronica Playle, Virginia Hope, Erasmus Smit, Lauren Jelly, Joep de Ligt |
| EPI_ISL_456181 | Southern Community Labs Dunedin | Institute of Environmental Science and Research (ESR) | Matt Storey, Xiaoyun Ren, Anja Werno, Antje van der Linden, Arlo Upton, Chris Mansell, David Hammer, Dragana Drinkovic, Erasmus Smit, Gary McAuliffe, Hana Sofia Andersson, James Ussher, Jill Sherwood, Josh Freeman, Julia Howard, Juliet Elvy, Mary DeAlmeida, Matt Blakiston, Matthew Rogers, Max Bloomfield, Michael Addidle, Michelle Balm, Sally Roberts, Sarah Jefferies, Sharmini Muttaiyah, Susan Morpeth, Susan Taylor, Timothy Blackmore, Vani Sathyendran, Veronica Playle, Virginia Hope, Erasmus Smit, Lauren Jelly, Joep de Ligt |
| EPI_ISL_456182, EPI_ISL_456183, EPI_ISL_456184, EPI_ISL_456185, EPI_ISL_456186, EPI_ISL_456187, EPI_ISL_456188 | Wellington SCL | Institute of Environmental Science and Research (ESR) | Matt Storey, Xiaoyun Ren, Anja Werno, Antje van der Linden, Arlo Upton, Chris Mansell, David Hammer, Dragana Drinkovic, Erasmus Smit, Gary McAuliffe, Hana Sofia Andersson, James Ussher, Jill Sherwood, Josh Freeman, Julia Howard, Juliet Elvy, Mary DeAlmeida, Matt Blakiston, Matthew Rogers, Max Bloomfield, Michael Addidle, Michelle Balm, Sally Roberts, Sarah Jefferies, Sharmini Muttaiyah, Susan Morpeth, Susan Taylor, Timothy Blackmore, Vani Sathyendran, Veronica Playle, Virginia Hope, Erasmus Smit, Lauren Jelly, Joep de Ligt |
| EPI_ISL_456190, EPI_ISL_456191, EPI_ISL_456192, EPI_ISL_456193 | Waikato Hospital | Institute of Environmental Science and Research (ESR) | Matt Storey, Xiaoyun Ren, Anja Werno, Antje van der Linden, Arlo Upton, Chris Mansell, David Hammer, Dragana Drinkovic, Erasmus Smit, Gary McAuliffe, Hana Sofia Andersson, James Ussher, Jill Sherwood, Josh Freeman, Julia Howard, Juliet Elvy, Mary DeAlmeida, Matt Blakiston, Matthew Rogers, Max Bloomfield, Michael Addidle, Michelle Balm, Sally Roberts, Sarah Jefferies, Sharmini Muttaiyah, Susan Morpeth, Susan Taylor, Timothy Blackmore, Vani Sathyendran, Veronica Playle, Virginia Hope, Erasmus Smit, Lauren Jelly, Joep de Ligt |
| EPI_ISL_456194, EPI_ISL_456195 | PathLab Bay of Plenty | Institute of Environmental Science and Research (ESR) | Matt Storey, Xiaoyun Ren, Anja Werno, Antje van der Linden, Arlo Upton, Chris Mansell, David Hammer, Dragana Drinkovic, Erasmus Smit, Gary McAuliffe, Hana Sofia Andersson, James Ussher, Jill Sherwood, Josh Freeman, Julia Howard, Juliet Elvy, Mary DeAlmeida, Matt Blakiston, Matthew Rogers, Max Bloomfield, Michael Addidle, Michelle Balm, Sally Roberts, Sarah Jefferies, Sharmini Muttaiyah, Susan Morpeth, Susan Taylor, Timothy Blackmore, Vani Sathyendran, Veronica Playle, Virginia Hope, Erasmus Smit, Lauren Jelly, Joep de Ligt |
| EPI_ISL_456197, EPI_ISL_456198, EPI_ISL_456199, EPI_ISL_456200, EPI_ISL_456201, EPI_ISL_456202, EPI_ISL_456203, EPI_ISL_456205, EPI_ISL_456206, EPI_ISL_456210, EPI_ISL_456212 | | | |
| see above | LabPLUS | Institute of Environmental Science and Research (ESR) | Matt Storey, Xiaoyun Ren, Anja Werno, Antje van der Linden, Arlo Upton, Chris Mansell, David Hammer, Dragana Drinkovic, Erasmus Smit, Gary McAuliffe, Hana Sofia Andersson, James Ussher, Jill Sherwood, Josh Freeman, Julia Howard, Juliet Elvy, Mary DeAlmeida, Matt Blakiston, Matthew Rogers, Max Bloomfield, Michael Addidle, Michelle Balm, Sally Roberts, Sarah Jefferies, Sharmini Muttaiyah, Susan Morpeth, Susan Taylor, Timothy Blackmore, Vani Sathyendran, Veronica Playle, Virginia Hope, Erasmus Smit, Lauren Jelly, Joep de Ligt |
| EPI_ISL_456213, EPI_ISL_456214 | Middlemore Hospital | Institute of Environmental Science and Research (ESR) | Matt Storey, Xiaoyun Ren, Anja Werno, Antje van der Linden, Arlo Upton, Chris Mansell, David Hammer, Dragana Drinkovic, Erasmus Smit, Gary McAuliffe, Hana Sofia Andersson, James Ussher, Jill Sherwood, Josh Freeman, Julia Howard, Juliet Elvy, Mary DeAlmeida, Matt Blakiston, Matthew Rogers, Max Bloomfield, Michael Addidle, Michelle Balm, Sally Roberts, Sarah Jefferies, Sharmini Muttaiyah, Susan Morpeth, Susan Taylor, Timothy Blackmore, Vani Sathyendran, Veronica Playle, Virginia Hope, Erasmus Smit, Lauren Jelly, Joep de Ligt |
| EPI_ISL_456215 | PathLab Bay of Plenty | Institute of Environmental Science and Research (ESR) | Matt Storey, Xiaoyun Ren, Anja Werno, Antje van der Linden, Arlo Upton, Chris Mansell, David Hammer, Dragana Drinkovic, Erasmus Smit, Gary McAuliffe, Hana Sofia Andersson, James Ussher, Jill Sherwood, Josh Freeman, Julia Howard, Juliet Elvy, Mary DeAlmeida, Matt Blakiston, Matthew Rogers, Max Bloomfield, Michael Addidle, Michelle Balm, Sally Roberts, Sarah Jefferies, Sharmini Muttaiyah, Susan Morpeth, Susan Taylor, Timothy Blackmore, Vani Sathyendran, Veronica Playle, Virginia Hope, Erasmus Smit, Lauren Jelly, Joep de Ligt |
| EPI_ISL_456217, EPI_ISL_456218, EPI_ISL_456219, EPI_ISL_456220 | Southern Community Labs Dunedin | Institute of Environmental Science and Research (ESR) | Matt Storey, Xiaoyun Ren, Anja Werno, Antje van der Linden, Arlo Upton, Chris Mansell, David Hammer, Dragana Drinkovic, Erasmus Smit, Gary McAuliffe, Hana Sofia Andersson, James Ussher, Jill Sherwood, Josh Freeman, Julia Howard, Juliet Elvy, Mary DeAlmeida, Matt Blakiston, Matthew Rogers, Max Bloomfield, Michael Addidle, Michelle Balm, Sally Roberts, Sarah Jefferies, Sharmini Muttaiyah, Susan Morpeth, Susan Taylor, Timothy Blackmore, Vani Sathyendran, Veronica Playle, Virginia Hope, Erasmus Smit, Lauren Jelly, Joep de Ligt |
| EPI_ISL_456221, EPI_ISL_456222, EPI_ISL_456223, EPI_ISL_456224 | PathLab Bay of Plenty | Institute of Environmental Science and Research (ESR) | Matt Storey, Xiaoyun Ren, Anja Werno, Antje van der Linden, Arlo Upton, Chris Mansell, David Hammer, Dragana Drinkovic, Erasmus Smit, Gary McAuliffe, Hana Sofia Andersson, James Ussher, Jill Sherwood, Josh Freeman, Julia Howard, Juliet Elvy, Mary DeAlmeida, Matt Blakiston, Matthew Rogers, Max Bloomfield, Michael Addidle, Michelle Balm, Sally Roberts, Sarah Jefferies, Sharmini Muttaiyah, Susan Morpeth, Susan Taylor, Timothy Blackmore, Vani Sathyendran, Veronica Playle, Virginia Hope, Erasmus Smit, Lauren Jelly, Joep de Ligt |
| EPI_ISL_456225, EPI_ISL_456228, EPI_ISL_456230, EPI_ISL_456233, EPI_ISL_456234, EPI_ISL_456236, EPI_ISL_456237, EPI_ISL_456238, EPI_ISL_456242, EPI_ISL_456243, EPI_ISL_456244, EPI_ISL_456245, EPI_ISL_456246, EPI_ISL_456247, EPI_ISL_456248, EPI_ISL_456249, EPI_ISL_456250, EPI_ISL_456251, EPI_ISL_456252, EPI_ISL_456253, EPI_ISL_456254 | | | |
| see above | Southern Community Labs Dunedin | Institute of Environmental Science and Research (ESR) | Matt Storey, Xiaoyun Ren, Anja Werno, Antje van der Linden, Arlo Upton, Chris Mansell, David Hammer, Dragana Drinkovic, Erasmus Smit, Gary McAuliffe, Hana Sofia Andersson, James Ussher, Jill Sherwood, Josh Freeman, Julia Howard, Juliet Elvy, Mary DeAlmeida, Matt Blakiston, Matthew Rogers, Max Bloomfield, Michael Addidle, Michelle Balm, Sally Roberts, Sarah Jefferies, Sharmini Muttaiyah, Susan Morpeth, Susan Taylor, Timothy Blackmore, Vani Sathyendran, Veronica Playle, Virginia Hope, Erasmus Smit, Lauren Jelly, Joep de Ligt |
| EPI_ISL_456255, EPI_ISL_456256, EPI_ISL_456257, EPI_ISL_456258 | Middlemore Hospital | Institute of Environmental Science and Research (ESR) | Matt Storey, Xiaoyun Ren, Anja Werno, Antje van der Linden, Arlo Upton, Chris Mansell, David Hammer, Dragana Drinkovic, Erasmus Smit, Gary McAuliffe, Hana Sofia Andersson, James Ussher, Jill Sherwood, Josh Freeman, Julia Howard, Juliet Elvy, Mary DeAlmeida, Matt Blakiston, Matthew Rogers, Max Bloomfield, Michael Addidle, Michelle Balm, Sally Roberts, Sarah Jefferies, Sharmini Muttaiyah, Susan Morpeth, Susan Taylor, Timothy Blackmore, Vani Sathyendran, Veronica Playle, Virginia Hope, Erasmus Smit, Lauren Jelly, Joep de Ligt |
| EPI_ISL_456261, EPI_ISL_456262, EPI_ISL_456263, EPI_ISL_456264, EPI_ISL_456265, EPI_ISL_456266, EPI_ISL_456267, EPI_ISL_456268, EPI_ISL_456270, EPI_ISL_456271, EPI_ISL_456272, EPI_ISL_456273, EPI_ISL_456276, EPI_ISL_456278, EPI_ISL_456279, EPI_ISL_456280, EPI_ISL_456281, EPI_ISL_456282, EPI_ISL_456283, EPI_ISL_456284, EPI_ISL_456287, EPI_ISL_456288, EPI_ISL_456289, EPI_ISL_456290, EPI_ISL_456292 | | | |
| see above | Southern Community Labs Dunedin | Institute of Environmental Science and Research (ESR) | Matt Storey, Xiaoyun Ren, Anja Werno, Antje van der Linden, Arlo Upton, Chris Mansell, David Hammer, Dragana Drinkovic, Erasmus Smit, Gary McAuliffe, Hana Sofia Andersson, James Ussher, Jill Sherwood, Josh Freeman, Julia Howard, Juliet Elvy, Mary DeAlmeida, Matt Blakiston, Matthew Rogers, Max Bloomfield, Michael Addidle, Michelle Balm, Sally Roberts, Sarah Jefferies, Sharmini Muttaiyah, Susan Morpeth, Susan Taylor, Timothy Blackmore, Vani Sathyendran, Veronica Playle, Virginia Hope, Erasmus Smit, Lauren Jelly, Joep de Ligt |
| EPI_ISL_456294 | North Shore Hospital | Institute of Environmental Science and Research (ESR) | Matt Storey, Xiaoyun Ren, Anja Werno, Antje van der Linden, Arlo Upton, Chris Mansell, David Hammer, Dragana Drinkovic, Erasmus Smit, Gary McAuliffe, Hana Sofia Andersson, James Ussher, Jill Sherwood, Josh Freeman, Julia Howard, Juliet Elvy, Mary DeAlmeida, Matt Blakiston, Matthew Rogers, Max Bloomfield, Michael Addidle, Michelle Balm, Sally Roberts, Sarah Jefferies, Sharmini Muttaiyah, Susan Morpeth, Susan Taylor, Timothy Blackmore, Vani Sathyendran, Veronica Playle, Virginia Hope, Erasmus Smit, Lauren Jelly, Joep de Ligt |
| EPI_ISL_456295, EPI_ISL_456296, EPI_ISL_456297, EPI_ISL_456299 | Southern Community Labs Dunedin | Institute of Environmental Science and Research (ESR) | Matt Storey, Xiaoyun Ren, Anja Werno, Antje van der Linden, Arlo Upton, Chris Mansell, David Hammer, Dragana Drinkovic, Erasmus Smit, Gary McAuliffe, Hana Sofia Andersson, James Ussher, Jill Sherwood, Josh Freeman, Julia Howard, Juliet Elvy, Mary DeAlmeida, Matt Blakiston, Matthew Rogers, Max Bloomfield, Michael Addidle, Michelle Balm, Sally Roberts, Sarah Jefferies, Sharmini Muttaiyah, Susan Morpeth, Susan |

| | | | |
|---|---|---|---|
| | | | Taylor, Timothy Blackmore, Vani Sathyendran, Veronica Playle, Virginia Hope, Erasmus Smit, Lauren Jelly, Joep de Ligt |
| EPI_ISL_456300 | LabPLUS | Institute of Environmental Science and Research (ESR) | Matt Storey, Xiaoyun Ren, Anja Werno, Antje van der Linden, Arlo Upton, Chris Mansell, David Hammer, Dragana Drinkovic, Erasmus Smit, Gary McAuliffe, Hana Sofia Andersson, James Ussher, Jill Sherwood, Josh Freeman, Julia Howard, Juliet Elvy, Mary DeAlmeida, Matt Blakiston, Matthew Rogers, Max Bloomfield, Michael Addidle, Michelle Balm, Sally Roberts, Sarah Jefferies, Sharmini Muttaiyah, Susan Morpeth, Susan Taylor, Timothy Blackmore, Vani Sathyendran, Veronica Playle, Virginia Hope, Erasmus Smit, Lauren Jelly, Joep de Ligt |
| EPI_ISL_456302, EPI_ISL_456303, EPI_ISL_456304, EPI_ISL_456305, EPI_ISL_456308, EPI_ISL_456309, EPI_ISL_456310, EPI_ISL_456314, EPI_ISL_456315, EPI_ISL_456316, EPI_ISL_456317 | | | |
| see above | Southern Community Labs Dunedin | Institute of Environmental Science and Research (ESR) | Matt Storey, Xiaoyun Ren, Anja Werno, Antje van der Linden, Arlo Upton, Chris Mansell, David Hammer, Dragana Drinkovic, Erasmus Smit, Gary McAuliffe, Hana Sofia Andersson, James Ussher, Jill Sherwood, Josh Freeman, Julia Howard, Juliet Elvy, Mary DeAlmeida, Matt Blakiston, Matthew Rogers, Max Bloomfield, Michael Addidle, Michelle Balm, Sally Roberts, Sarah Jefferies, Sharmini Muttaiyah, Susan Morpeth, Susan Taylor, Timothy Blackmore, Vani Sathyendran, Veronica Playle, Virginia Hope, Erasmus Smit, Lauren Jelly, Joep de Ligt |
| EPI_ISL_456319, EPI_ISL_456320 | Wellington SCL | Institute of Environmental Science and Research (ESR) | Matt Storey, Xiaoyun Ren, Anja Werno, Antje van der Linden, Arlo Upton, Chris Mansell, David Hammer, Dragana Drinkovic, Erasmus Smit, Gary McAuliffe, Hana Sofia Andersson, James Ussher, Jill Sherwood, Josh Freeman, Julia Howard, Juliet Elvy, Mary DeAlmeida, Matt Blakiston, Matthew Rogers, Max Bloomfield, Michael Addidle, Michelle Balm, Sally Roberts, Sarah Jefferies, Sharmini Muttaiyah, Susan Morpeth, Susan Taylor, Timothy Blackmore, Vani Sathyendran, Veronica Playle, Virginia Hope, Erasmus Smit, Lauren Jelly, Joep de Ligt |
| EPI_ISL_456321, EPI_ISL_456322 | Canterbury Health Laboratories | Institute of Environmental Science and Research (ESR) | Matt Storey, Xiaoyun Ren, Anja Werno, Antje van der Linden, Arlo Upton, Chris Mansell, David Hammer, Dragana Drinkovic, Erasmus Smit, Gary McAuliffe, Hana Sofia Andersson, James Ussher, Jill Sherwood, Josh Freeman, Julia Howard, Juliet Elvy, Mary DeAlmeida, Matt Blakiston, Matthew Rogers, Max Bloomfield, Michael Addidle, Michelle Balm, Sally Roberts, Sarah Jefferies, Sharmini Muttaiyah, Susan Morpeth, Susan Taylor, Timothy Blackmore, Vani Sathyendran, Veronica Playle, Virginia Hope, Erasmus Smit, Lauren Jelly, Joep de Ligt |
| EPI_ISL_456324, EPI_ISL_456326, EPI_ISL_456327, EPI_ISL_456328, EPI_ISL_456329, EPI_ISL_456330, EPI_ISL_456331, EPI_ISL_456332, EPI_ISL_456334, EPI_ISL_456335, EPI_ISL_456336, EPI_ISL_456338, EPI_ISL_456339, EPI_ISL_456340, EPI_ISL_456342 | | | |
| see above | Wellington SCL | Institute of Environmental Science and Research (ESR) | Matt Storey, Xiaoyun Ren, Anja Werno, Antje van der Linden, Arlo Upton, Chris Mansell, David Hammer, Dragana Drinkovic, Erasmus Smit, Gary McAuliffe, Hana Sofia Andersson, James Ussher, Jill Sherwood, Josh Freeman, Julia Howard, Juliet Elvy, Mary DeAlmeida, Matt Blakiston, Matthew Rogers, Max Bloomfield, Michael Addidle, Michelle Balm, Sally Roberts, Sarah Jefferies, Sharmini Muttaiyah, Susan Morpeth, Susan Taylor, Timothy Blackmore, Vani Sathyendran, Veronica Playle, Virginia Hope, Erasmus Smit, Lauren Jelly, Joep de Ligt |
| EPI_ISL_456344, EPI_ISL_456347 | Southern Community Labs Dunedin | Institute of Environmental Science and Research (ESR) | Matt Storey, Xiaoyun Ren, Anja Werno, Antje van der Linden, Arlo Upton, Chris Mansell, David Hammer, Dragana Drinkovic, Erasmus Smit, Gary McAuliffe, Hana Sofia Andersson, James Ussher, Jill Sherwood, Josh Freeman, Julia Howard, Juliet Elvy, Mary DeAlmeida, Matt Blakiston, Matthew Rogers, Max Bloomfield, Michael Addidle, Michelle Balm, Sally Roberts, Sarah Jefferies, Sharmini Muttaiyah, Susan Morpeth, Susan Taylor, Timothy Blackmore, Vani Sathyendran, Veronica Playle, Virginia Hope, Erasmus Smit, Lauren Jelly, Joep de Ligt |
| EPI_ISL_456348, EPI_ISL_456352, EPI_ISL_456353, EPI_ISL_456355, EPI_ISL_456356, EPI_ISL_456358, EPI_ISL_456360, EPI_ISL_456362, EPI_ISL_456363, EPI_ISL_456364, EPI_ISL_456365, EPI_ISL_456366, EPI_ISL_456367, EPI_ISL_456369, EPI_ISL_456370, EPI_ISL_456371, EPI_ISL_456374, EPI_ISL_456375 | | | |
| see above | Canterbury Health Laboratories | Institute of Environmental Science and Research (ESR) | Matt Storey, Xiaoyun Ren, Anja Werno, Antje van der Linden, Arlo Upton, Chris Mansell, David Hammer, Dragana Drinkovic, Erasmus Smit, Gary McAuliffe, Hana Sofia Andersson, James Ussher, Jill Sherwood, Josh Freeman, Julia Howard, Juliet Elvy, Mary DeAlmeida, Matt Blakiston, Matthew Rogers, Max Bloomfield, Michael Addidle, Michelle Balm, Sally Roberts, Sarah Jefferies, Sharmini Muttaiyah, Susan Morpeth, Susan Taylor, Timothy Blackmore, Vani Sathyendran, Veronica Playle, Virginia Hope, Erasmus Smit, Lauren Jelly, Joep de Ligt |
| EPI_ISL_456376 | Middlemore Hospital | Institute of Environmental Science and Research (ESR) | Matt Storey, Xiaoyun Ren, Anja Werno, Antje van der Linden, Arlo Upton, Chris Mansell, David Hammer, Dragana Drinkovic, Erasmus Smit, Gary McAuliffe, Hana Sofia Andersson, James Ussher, Jill Sherwood, Josh Freeman, Julia Howard, Juliet Elvy, Mary DeAlmeida, Matt Blakiston, Matthew Rogers, Max Bloomfield, Michael Addidle, Michelle Balm, Sally Roberts, Sarah Jefferies, Sharmini Muttaiyah, Susan Morpeth, Susan Taylor, Timothy Blackmore, Vani Sathyendran, Veronica Playle, Virginia Hope, Erasmus Smit, Lauren Jelly, Joep de Ligt |
| EPI_ISL_456377, EPI_ISL_456378 | MedLab Central Ltd | Institute of Environmental Science and Research (ESR) | Matt Storey, Xiaoyun Ren, Anja Werno, Antje van der Linden, Arlo Upton, Chris Mansell, David Hammer, Dragana Drinkovic, Erasmus Smit, Gary McAuliffe, Hana Sofia Andersson, James Ussher, Jill Sherwood, Josh Freeman, Julia Howard, Juliet Elvy, Mary DeAlmeida, Matt Blakiston, Matthew Rogers, Max Bloomfield, Michael Addidle, Michelle Balm, Sally Roberts, Sarah Jefferies, Sharmini Muttaiyah, Susan Morpeth, Susan Taylor, Timothy Blackmore, Vani Sathyendran, Veronica Playle, Virginia Hope, Erasmus Smit, Lauren Jelly, Joep de Ligt |
| EPI_ISL_456379, EPI_ISL_456380, EPI_ISL_456381, EPI_ISL_456382, EPI_ISL_456383, EPI_ISL_456384 | LabPLUS | Institute of Environmental Science and Research (ESR) | Matt Storey, Xiaoyun Ren, Anja Werno, Antje van der Linden, Arlo Upton, Chris Mansell, David Hammer, Dragana Drinkovic, Erasmus Smit, Gary McAuliffe, Hana Sofia Andersson, James Ussher, Jill Sherwood, Josh Freeman, Julia Howard, Juliet Elvy, Mary DeAlmeida, Matt Blakiston, Matthew Rogers, Max Bloomfield, Michael Addidle, Michelle Balm, Sally Roberts, Sarah Jefferies, Sharmini Muttaiyah, Susan Morpeth, Susan Taylor, Timothy Blackmore, Vani Sathyendran, Veronica Playle, Virginia Hope, Erasmus Smit, Lauren Jelly, Joep de Ligt |
| EPI_ISL_456385, EPI_ISL_456386 | Middlemore Hospital | Institute of Environmental Science and Research (ESR) | Matt Storey, Xiaoyun Ren, Anja Werno, Antje van der Linden, Arlo Upton, Chris Mansell, David Hammer, Dragana Drinkovic, Erasmus Smit, Gary McAuliffe, Hana Sofia Andersson, James Ussher, Jill Sherwood, Josh Freeman, Julia Howard, Juliet Elvy, Mary DeAlmeida, Matt Blakiston, Matthew Rogers, Max Bloomfield, Michael Addidle, Michelle Balm, Sally Roberts, Sarah Jefferies, Sharmini Muttaiyah, Susan Morpeth, Susan Taylor, Timothy Blackmore, Vani Sathyendran, Veronica Playle, Virginia Hope, Erasmus Smit, Lauren Jelly, Joep de Ligt |
| EPI_ISL_456388, EPI_ISL_456390, EPI_ISL_456393, EPI_ISL_456394 | LabPLUS | Institute of Environmental Science and Research (ESR) | Matt Storey, Xiaoyun Ren, Anja Werno, Antje van der Linden, Arlo Upton, Chris Mansell, David Hammer, Dragana Drinkovic, Erasmus Smit, Gary McAuliffe, Hana Sofia Andersson, James Ussher, Jill Sherwood, Josh Freeman, Julia Howard, Juliet Elvy, Mary DeAlmeida, Matt Blakiston, Matthew Rogers, Max Bloomfield, Michael Addidle, Michelle Balm, Sally Roberts, Sarah Jefferies, Sharmini Muttaiyah, Susan Morpeth, Susan Taylor, Timothy Blackmore, Vani Sathyendran, Veronica Playle, Virginia Hope, Erasmus Smit, Lauren Jelly, Joep de Ligt |
| EPI_ISL_456395, EPI_ISL_456396, EPI_ISL_456397 | North Shore Hospital | Institute of Environmental Science and Research (ESR) | Matt Storey, Xiaoyun Ren, Anja Werno, Antje van der Linden, Arlo Upton, Chris Mansell, David Hammer, Dragana Drinkovic, Erasmus Smit, Gary McAuliffe, Hana Sofia Andersson, James Ussher, Jill Sherwood, Josh Freeman, Julia Howard, Juliet Elvy, Mary DeAlmeida, Matt Blakiston, Matthew Rogers, Max Bloomfield, Michael Addidle, Michelle Balm, Sally Roberts, Sarah Jefferies, Sharmini Muttaiyah, Susan Morpeth, Susan Taylor, Timothy Blackmore, Vani Sathyendran, Veronica Playle, Virginia Hope, Erasmus Smit, Lauren Jelly, Joep de Ligt |
| EPI_ISL_456402 | Wellington SCL | Institute of Environmental Science and Research (ESR) | Matt Storey, Xiaoyun Ren, Anja Werno, Antje van der Linden, Arlo Upton, Chris Mansell, David Hammer, Dragana Drinkovic, Erasmus Smit, Gary McAuliffe, Hana Sofia Andersson, James Ussher, Jill Sherwood, Josh Freeman, Julia Howard, Juliet Elvy, Mary DeAlmeida, Matt Blakiston, Matthew Rogers, Max Bloomfield, Michael Addidle, Michelle Balm, Sally Roberts, Sarah Jefferies, Sharmini Muttaiyah, Susan Morpeth, Susan Taylor, Timothy Blackmore, Vani Sathyendran, Veronica Playle, Virginia Hope, Erasmus Smit, Lauren Jelly, Joep de Ligt |
| EPI_ISL_456403 | Middlemore Hospital | Institute of Environmental Science and Research (ESR) | Matt Storey, Xiaoyun Ren, Anja Werno, Antje van der Linden, Arlo Upton, Chris Mansell, David Hammer, Dragana Drinkovic, Erasmus Smit, Gary McAuliffe, Hana Sofia Andersson, James Ussher, Jill Sherwood, Josh Freeman, Julia Howard, Juliet Elvy, Mary DeAlmeida, Matt Blakiston, Matthew Rogers, Max Bloomfield, Michael Addidle, Michelle Balm, Sally Roberts, Sarah Jefferies, Sharmini Muttaiyah, Susan Morpeth, Susan Taylor, Timothy Blackmore, Vani Sathyendran, Veronica Playle, Virginia Hope, Erasmus Smit, Lauren Jelly, Joep de Ligt |
| EPI_ISL_456410, EPI_ISL_456411, EPI_ISL_456414, EPI_ISL_456417, EPI_ISL_456419, EPI_ISL_456421, EPI_ISL_456426, EPI_ISL_456427, EPI_ISL_456429, EPI_ISL_456433, EPI_ISL_456435, EPI_ISL_456440, EPI_ISL_456441, EPI_ISL_456442, EPI_ISL_456443, EPI_ISL_456445, EPI_ISL_456446, EPI_ISL_456448, EPI_ISL_456451, EPI_ISL_456454, EPI_ISL_456456, EPI_ISL_456457, EPI_ISL_456460, EPI_ISL_456462, EPI_ISL_456463, EPI_ISL_456464, EPI_ISL_456468, EPI_ISL_456471, EPI_ISL_456472, EPI_ISL_456476, EPI_ISL_456477 | | | |
| see above | Victorian Infectious Diseases Reference Laboratory (VIDRL) | Microbiological Diagnostic Unit Public Health Laboratory and Victorian Infectious Diseases Reference Laboratory, Doherty Institute | Caly L., Seemann T., Sait, M., Schultz M., Druce J., Sherry, N. |
| EPI_ISL_456478, EPI_ISL_456479 | Microbiological Diagnostic Unit Public Health Laboratory | Microbiological Diagnostic Unit Public Health Laboratory, The Peter Doherty Institute for Infection and Immunity | Seemann T., Schultz M., Sait, M., Sherry, N. |
| EPI_ISL_456480, EPI_ISL_456482, EPI_ISL_456483, EPI_ISL_456484, EPI_ISL_456492, EPI_ISL_456493, EPI_ISL_456495, EPI_ISL_456499, EPI_ISL_456500, EPI_ISL_456501, EPI_ISL_456502, EPI_ISL_456503, EPI_ISL_456504, EPI_ISL_456506, EPI_ISL_456507, EPI_ISL_456508, EPI_ISL_456509, EPI_ISL_456510, EPI_ISL_456511, EPI_ISL_456512, EPI_ISL_456513, EPI_ISL_456516, EPI_ISL_456518, EPI_ISL_456519, EPI_ISL_456520, EPI_ISL_456521, EPI_ISL_456526, EPI_ISL_456529, EPI_ISL_456530, EPI_ISL_456531, EPI_ISL_456536, EPI_ISL_456538, EPI_ISL_456539, EPI_ISL_456540, EPI_ISL_456541, EPI_ISL_456542, EPI_ISL_456543, EPI_ISL_456548, EPI_ISL_456550, EPI_ISL_456553, EPI_ISL_456555, EPI_ISL_456557, EPI_ISL_456558, EPI_ISL_456559, EPI_ISL_456560, EPI_ISL_456562, EPI_ISL_456563, EPI_ISL_456565, EPI_ISL_456566, EPI_ISL_456568, EPI_ISL_456573, EPI_ISL_456574, EPI_ISL_456575, EPI_ISL_456580, EPI_ISL_456581, EPI_ISL_456589 | | | |
| see above | Victorian Infectious Diseases Reference Laboratory (VIDRL) | Microbiological Diagnostic Unit Public Health Laboratory and Victorian Infectious Diseases Reference Laboratory, Doherty Institute | Caly L., Seemann T., Sait, M., Schultz M., Druce J., Sherry, N. |
| EPI_ISL_456595 | Microbiological Diagnostic Unit Public Health Laboratory | Microbiological Diagnostic Unit Public Health Laboratory, The Peter Doherty Institute for Infection and Immunity | Seemann T., Schultz M., Sait, M., Sherry, N. |
| EPI_ISL_456596, EPI_ISL_456597, EPI_ISL_456600, EPI_ISL_456606, EPI_ISL_456607, EPI_ISL_456608, EPI_ISL_456611 | National Health Laboratory, Timor-Leste | Microbiological Diagnostic Unit Public Health Laboratory, The Peter Doherty Institute for Infection and Immunity | Soares da Silva, E., Dolores de Jesus da Costa, M., Salles de Sousa, A., Jayanti Pereira Tilman, A., Antonia da Costa, E., Barreto, I., Marr, I., Wapling, J., Francis, J., Ximenes, J., Canisia, D., Freeman, K., Dakh, F., Douglas, N., Baird, R., Caly, L., Seemann, T., Sait, M., Schultz, M., Sherry, N. |
| EPI_ISL_456616, EPI_ISL_456617, EPI_ISL_456619, EPI_ISL_456620, EPI_ISL_456630, EPI_ISL_456633, EPI_ISL_456643, EPI_ISL_456647, EPI_ISL_456649 | Victorian Infectious Diseases Reference Laboratory (VIDRL) | Microbiological Diagnostic Unit Public Health Laboratory and Victorian Infectious Diseases Reference Laboratory, Doherty Institute | Caly L., Seemann T., Sait, M., Schultz M., Druce J., Sherry, N. |
| EPI_ISL_456658, EPI_ISL_456659, EPI_ISL_456661, EPI_ISL_456663, EPI_ISL_456664, EPI_ISL_456665, EPI_ISL_456669, EPI_ISL_456671, EPI_ISL_456672, EPI_ISL_456673, EPI_ISL_456675, EPI_ISL_456676 | | | |
| see above | University of Birmingham | COVID-19 Genomics UK (COG-UK) Consortium | Loman Lab: Claire McMurray, Joanne Stockton, Samuel Nicholls, Radoslaw Poplawski, Will Rowe, Josh Quick, Nicholas Loman // UHB Lab: Celina M Whalley, Andrew Bosworth, Charlotte Poxon, Kasun Wanigasooriya, Oliver Pickles, Mike Kidd, Alex Richter, Andrew D Beggs // PHE Heartlands Lab: Husam Osman, Andrew Bosworth |
| EPI_ISL_456680, EPI_ISL_456688, EPI_ISL_456697, EPI_ISL_456699, EPI_ISL_456702, EPI_ISL_456704, EPI_ISL_456705, EPI_ISL_456707, EPI_ISL_456708, EPI_ISL_456709, EPI_ISL_456711, EPI_ISL_456713, EPI_ISL_456716, EPI_ISL_456717, EPI_ISL_456718, EPI_ISL_456719, EPI_ISL_456720, EPI_ISL_456721, EPI_ISL_456722, EPI_ISL_456723, EPI_ISL_456724, EPI_ISL_456727, EPI_ISL_456728, EPI_ISL_456729, EPI_ISL_456730, EPI_ISL_456731, EPI_ISL_456732, EPI_ISL_456733, EPI_ISL_456734, EPI_ISL_456736, EPI_ISL_456739, EPI_ISL_456740, EPI_ISL_456743, EPI_ISL_456745, EPI_ISL_456747, EPI_ISL_456749, EPI_ISL_456750, EPI_ISL_456751, EPI_ISL_456754, EPI_ISL_456755 | | | |
| see above | Department of Pathology, University of Cambridge | COVID-19 Genomics UK (COG-UK) Consortium | Luke W Meredith, M. Estée Török, Myra Hosmillo, William L. Hamilton, Martin D. Curran, Theresa Feltwell, Grant Hall, Anna Yakovleva, Fahad A Khokhar, Charlotte J. Houldcroft, Laura G Caller, Aminu S. Jahun, Sarah L. Caddy, Ian Goodfellow |
| EPI_ISL_456756, EPI_ISL_456757, EPI_ISL_456758, EPI_ISL_456759, EPI_ISL_456766, EPI_ISL_456767, EPI_ISL_456771, EPI_ISL_456773, EPI_ISL_456774, EPI_ISL_456776, EPI_ISL_456777, EPI_ISL_456778, EPI_ISL_456779, EPI_ISL_456780, EPI_ISL_456781, EPI_ISL_456782, EPI_ISL_456783, EPI_ISL_456784, EPI_ISL_456785, EPI_ISL_456786, EPI_ISL_456787, EPI_ISL_456788, EPI_ISL_456789, EPI_ISL_456790, EPI_ISL_456791, EPI_ISL_456792, EPI_ISL_456793, EPI_ISL_456794, EPI_ISL_456795, EPI_ISL_456796, EPI_ISL_456797, EPI_ISL_456798, EPI_ISL_456799, EPI_ISL_456801, EPI_ISL_456802, EPI_ISL_456803, EPI_ISL_456804, EPI_ISL_456805, EPI_ISL_456806, EPI_ISL_456807, EPI_ISL_456808, EPI_ISL_456809, EPI_ISL_456810, EPI_ISL_456811, EPI_ISL_456813, EPI_ISL_456815, EPI_ISL_456818, EPI_ISL_456819, EPI_ISL_456821, EPI_ISL_456824, EPI_ISL_456826, EPI_ISL_456827, EPI_ISL_456829, EPI_ISL_456830, EPI_ISL_456831, EPI_ISL_456833, EPI_ISL_456834, EPI_ISL_456835, EPI_ISL_456838, EPI_ISL_456839, EPI_ISL_456840, EPI_ISL_456842, EPI_ISL_456847, EPI_ISL_456848, EPI_ISL_456851, EPI_ISL_456852, EPI_ISL_456854, EPI_ISL_456855, EPI_ISL_456856, EPI_ISL_456863, EPI_ISL_456864, EPI_ISL_456865, EPI_ISL_456866, EPI_ISL_456868, EPI_ISL_456870, EPI_ISL_456871, EPI_ISL_456872, EPI_ISL_456874, EPI_ISL_456875, EPI_ISL_456877, EPI_ISL_456878, EPI_ISL_456880, EPI_ISL_456883, EPI_ISL_456884, EPI_ISL_456885, EPI_ISL_456886, EPI_ISL_456887, EPI_ISL_456889 | | | |
| see above | West of Scotland Specialist Virology Centre, NHSGGC / MRC-University of Glasgow Centre for Virus Research | COVID-19 Genomics UK (COG-UK) Consortium | Ana da Silva Filipe, Natasha Johnson, Kathy Smollett, Daniel Mair, Stephen Carmichael, Lily Tong, Jenna Nichols, Elihu Aranday-Cortes, Kirstyn Brunker, Yasmin Parr, Kyriaki Nomikou; Sarah McDonald, Marc Niebel, Patawee Asamaphan; Richard Orton, Joseph Hughes, Sreenu Vattipally, David L Robertson; Alasdair MacLean, Rory Gunson; Kathy Li, Natasha Jesudason, Rajiv Shah, James Shepherd, Antonia Ho, Emma Thomson |
| EPI_ISL_456890, EPI_ISL_456891, EPI_ISL_456893, EPI_ISL_456894, EPI_ISL_456896, EPI_ISL_456897, EPI_ISL_456898, EPI_ISL_456901, EPI_ISL_456902, EPI_ISL_456903, EPI_ISL_456904, EPI_ISL_456905, EPI_ISL_456906, EPI_ISL_456908, EPI_ISL_456909, EPI_ISL_456910, EPI_ISL_456911, EPI_ISL_456913, EPI_ISL_456914, EPI_ISL_456915, EPI_ISL_456916, EPI_ISL_456918, EPI_ISL_456920, EPI_ISL_456921, EPI_ISL_456922, EPI_ISL_456923, EPI_ISL_456924, EPI_ISL_456925, EPI_ISL_456926, EPI_ISL_456927, EPI_ISL_456928, EPI_ISL_456930, EPI_ISL_456931, EPI_ISL_456932, EPI_ISL_456933, EPI_ISL_456934, EPI_ISL_456938, EPI_ISL_456939, EPI_ISL_456941, EPI_ISL_456942, EPI_ISL_456943, EPI_ISL_456944, EPI_ISL_456945, EPI_ISL_456946, EPI_ISL_456947, EPI_ISL_456948, EPI_ISL_456949, EPI_ISL_456951, EPI_ISL_456952, EPI_ISL_456953, EPI_ISL_456954, EPI_ISL_456955, EPI_ISL_456956, EPI_ISL_456957, EPI_ISL_456958, EPI_ISL_456960, EPI_ISL_456961, EPI_ISL_456962, EPI_ISL_456964, EPI_ISL_456965, EPI_ISL_456966, EPI_ISL_456967, EPI_ISL_456973, EPI_ISL_456974, EPI_ISL_456975, EPI_ISL_456976, EPI_ISL_456977, EPI_ISL_456978, EPI_ISL_456979, EPI_ISL_456984, EPI_ISL_456985, EPI_ISL_456986, EPI_ISL_456987, EPI_ISL_456988, EPI_ISL_456989, EPI_ISL_456991, EPI_ISL_456992, EPI_ISL_456993, EPI_ISL_456994, EPI_ISL_456995, EPI_ISL_456996, EPI_ISL_456997, EPI_ISL_456998, EPI_ISL_456999, EPI_ISL_457000, EPI_ISL_457002, EPI_ISL_457003, EPI_ISL_457004, EPI_ISL_457005, EPI_ISL_457006, EPI_ISL_457009, EPI_ISL_457010, EPI_ISL_457011, EPI_ISL_457012, EPI_ISL_457013, EPI_ISL_457015, EPI_ISL_457018, EPI_ISL_457019, EPI_ISL_457020, EPI_ISL_457021, EPI_ISL_457023 | | | |
| see above | Virology Department, Royal Infirmary of Edinburgh, NHS Lothian / School of Biological Sciences, University of Edinburgh / Institute of Genetics and Molecular Medicine, University of Edinburgh | COVID-19 Genomics UK (COG-UK) Consortium | McHugh M, Dewar R, Rooke S, Gallagher M, Balcaza C, O'Toole Á, Scher E, Hill V, McCrone JT, Colquhoun R, Yu X, Jackson B, Rambaut A, Williams TC, Templeton K |
| EPI_ISL_457025, EPI_ISL_457026, EPI_ISL_457027, EPI_ISL_457028, EPI_ISL_457029, EPI_ISL_457030, EPI_ISL_457031, EPI_ISL_457035, EPI_ISL_457036, EPI_ISL_457038, EPI_ISL_457040, EPI_ISL_457041, EPI_ISL_457043, EPI_ISL_457044, EPI_ISL_457047, EPI_ISL_457049, EPI_ISL_457052, EPI_ISL_457053, EPI_ISL_457054, EPI_ISL_457055, EPI_ISL_457056, EPI_ISL_457057, EPI_ISL_457058, EPI_ISL_457059, EPI_ISL_457060, EPI_ISL_457061, EPI_ISL_457062, EPI_ISL_457063, EPI_ISL_457064, EPI_ISL_457065, EPI_ISL_457066, EPI_ISL_457067, EPI_ISL_457068, EPI_ISL_457069, EPI_ISL_457070, EPI_ISL_457071, EPI_ISL_457073, EPI_ISL_457074, EPI_ISL_457075, EPI_ISL_457076, EPI_ISL_457077, EPI_ISL_457078, EPI_ISL_457079, EPI_ISL_457081, EPI_ISL_457083, EPI_ISL_457086, EPI_ISL_457087, EPI_ISL_457088, EPI_ISL_457089, EPI_ISL_457090, EPI_ISL_457091, EPI_ISL_457092, EPI_ISL_457093, EPI_ISL_457094, EPI_ISL_457095, EPI_ISL_457096, EPI_ISL_457098, EPI_ISL_457099, EPI_ISL_457101, EPI_ISL_457103, EPI_ISL_457104, EPI_ISL_457105, EPI_ISL_457106, EPI_ISL_457108, EPI_ISL_457109, EPI_ISL_457111, EPI_ISL_457112, EPI_ISL_457114, EPI_ISL_457115, EPI_ISL_457116, EPI_ISL_457117, EPI_ISL_457119, | | | |

| | | | |
|---|---|---|---|
| EPI_ISL_457122, EPI_ISL_457123, EPI_ISL_457124, EPI_ISL_457125, EPI_ISL_457126, EPI_ISL_457127, EPI_ISL_457128, EPI_ISL_457129, EPI_ISL_457130, EPI_ISL_457131, EPI_ISL_457132, EPI_ISL_457133, EPI_ISL_457134, EPI_ISL_457135, EPI_ISL_457136, EPI_ISL_457137, EPI_ISL_457139, EPI_ISL_457140, EPI_ISL_457141, EPI_ISL_457142, EPI_ISL_457143, EPI_ISL_457144, EPI_ISL_457145, EPI_ISL_457146, EPI_ISL_457147, EPI_ISL_457148, EPI_ISL_457149, EPI_ISL_457150, EPI_ISL_457151, EPI_ISL_457152, EPI_ISL_457153, EPI_ISL_457154, EPI_ISL_457155, EPI_ISL_457156, EPI_ISL_457157, EPI_ISL_457159, EPI_ISL_457160, EPI_ISL_457161, EPI_ISL_457162, EPI_ISL_457163, EPI_ISL_457164, EPI_ISL_457165, EPI_ISL_457166, EPI_ISL_457167, EPI_ISL_457168, EPI_ISL_457169, EPI_ISL_457170, EPI_ISL_457171, EPI_ISL_457173, EPI_ISL_457176, EPI_ISL_457177, EPI_ISL_457178, EPI_ISL_457179, EPI_ISL_457180, EPI_ISL_457181, EPI_ISL_457182, EPI_ISL_457184, EPI_ISL_457185, EPI_ISL_457189, EPI_ISL_457191, EPI_ISL_457192, EPI_ISL_457194, EPI_ISL_457195, EPI_ISL_457196, EPI_ISL_457198, EPI_ISL_457200, EPI_ISL_457201, EPI_ISL_457204, EPI_ISL_457205, EPI_ISL_457206, EPI_ISL_457207, EPI_ISL_457208, EPI_ISL_457209, EPI_ISL_457210, EPI_ISL_457211, EPI_ISL_457212, EPI_ISL_457213, EPI_ISL_457214, EPI_ISL_457217, EPI_ISL_457218, EPI_ISL_457220, EPI_ISL_457221, EPI_ISL_457222, EPI_ISL_457223, EPI_ISL_457224, EPI_ISL_457225, EPI_ISL_457229, EPI_ISL_457232, EPI_ISL_457233, EPI_ISL_457234, EPI_ISL_457236, EPI_ISL_457238, EPI_ISL_457239, EPI_ISL_457240, EPI_ISL_457241, EPI_ISL_457243, EPI_ISL_457245, EPI_ISL_457246, EPI_ISL_457247, EPI_ISL_457248, EPI_ISL_457249, EPI_ISL_457250, EPI_ISL_457251, EPI_ISL_457252, EPI_ISL_457253, EPI_ISL_457254, EPI_ISL_457255, EPI_ISL_457256, EPI_ISL_457258 | | | |
| see above | University of Exeter | COVID-19 Genomics UK (COG-UK) Consortium | Ben Temperton,Aaron Jeffries,Michelle Michelsen,Joanna Warwick-Dugdale,Audrey Farbos,Robyn Manley,Stephen Michell,Jane Masoli |
| EPI_ISL_457270, EPI_ISL_457271, EPI_ISL_457272, EPI_ISL_457273, EPI_ISL_457274, EPI_ISL_457275, EPI_ISL_457277, EPI_ISL_457278, EPI_ISL_457280, EPI_ISL_457281, EPI_ISL_457283, EPI_ISL_457284, EPI_ISL_457285, EPI_ISL_457286, EPI_ISL_457287, EPI_ISL_457288, EPI_ISL_457289, EPI_ISL_457290, EPI_ISL_457291, EPI_ISL_457292, EPI_ISL_457293, EPI_ISL_457294, EPI_ISL_457295, EPI_ISL_457296, EPI_ISL_457297, EPI_ISL_457298, EPI_ISL_457299, EPI_ISL_457301 | | | |
| see above | University College London, Great Ormond Street Hospital for Children NHS Foundation Trust, Imperial College Healthcare NHS Trust | COVID-19 Genomics UK (COG-UK) Consortium | Sergi Castellano, Rachel Williams, Mark Kristiansen, Paola Resende Silva, Sunando Roy, Tony Brooks, Helena Tutill, Paola Niola, Patricia Dyal, Charlotte Williams, Leysa Forrest, Yasmin Panchbhaya, Jacqueline Findlay, Sam Weeks, Julianne Brown, Kathryn Harris, Paul Randell, James Price, Alison Holmes, Judith Breuer |
| EPI_ISL_457303, EPI_ISL_457305, EPI_ISL_457311, EPI_ISL_457312, EPI_ISL_457316, EPI_ISL_457319, EPI_ISL_457320, EPI_ISL_457325 | Northumbria University / South Tees Hospitals NHS Foundation Trust / North Cumbria Integrated Care NHS Foundation Trust / North Tees and Hartlepool NHS Foundation Trust / Newcastle Hospitals NHS Foundation Trust | COVID-19 Genomics UK (COG-UK) Consortium | Darren L Smith,Andrew Nelson,Matthew Bashton,Greg R Young,Joshua Loh,John Allan,Mohammad A Tariq,Giles S Holt,Gary Black,Wen C Yew,Lynn Dover,Paul Baker,Steve Liggett,Sarah Essex,Jane Greenaway,Debra Padgett,Clive Graham,Garren Scott,Edward Barton,Emma Swindells,Brendan Payne,Jennifer Collins,Yusri Taha,Gary Eltringham |
| EPI_ISL_457326, EPI_ISL_457328, EPI_ISL_457329, EPI_ISL_457330, EPI_ISL_457331, EPI_ISL_457332, EPI_ISL_457333, EPI_ISL_457334, EPI_ISL_457335, EPI_ISL_457336, EPI_ISL_457337, EPI_ISL_457338, EPI_ISL_457339, EPI_ISL_457340, EPI_ISL_457341, EPI_ISL_457342, EPI_ISL_457344, EPI_ISL_457345, EPI_ISL_457346, EPI_ISL_457347, EPI_ISL_457350, EPI_ISL_457351, EPI_ISL_457352, EPI_ISL_457353, EPI_ISL_457354, EPI_ISL_457355, EPI_ISL_457356, EPI_ISL_457357, EPI_ISL_457358, EPI_ISL_457360, EPI_ISL_457361, EPI_ISL_457362, EPI_ISL_457365, EPI_ISL_457366, EPI_ISL_457367, EPI_ISL_457368, EPI_ISL_457369, EPI_ISL_457370, EPI_ISL_457371, EPI_ISL_457373, EPI_ISL_457374, EPI_ISL_457375, EPI_ISL_457377, EPI_ISL_457378, EPI_ISL_457379, EPI_ISL_457380, EPI_ISL_457381, EPI_ISL_457382, EPI_ISL_457383, EPI_ISL_457385, EPI_ISL_457386, EPI_ISL_457387, EPI_ISL_457388, EPI_ISL_457389, EPI_ISL_457390, EPI_ISL_457391, EPI_ISL_457392, EPI_ISL_457394, EPI_ISL_457395, EPI_ISL_457397, EPI_ISL_457401, EPI_ISL_457402, EPI_ISL_457403, EPI_ISL_457405, EPI_ISL_457406, EPI_ISL_457407, EPI_ISL_457408, EPI_ISL_457409, EPI_ISL_457411, EPI_ISL_457412, EPI_ISL_457413, EPI_ISL_457414, EPI_ISL_457415, EPI_ISL_457416, EPI_ISL_457417, EPI_ISL_457418, EPI_ISL_457419, EPI_ISL_457420, EPI_ISL_457423, EPI_ISL_457425, EPI_ISL_457426, EPI_ISL_457427, EPI_ISL_457430, EPI_ISL_457431, EPI_ISL_457433, EPI_ISL_457434, EPI_ISL_457436, EPI_ISL_457437, EPI_ISL_457438, EPI_ISL_457440, EPI_ISL_457441, EPI_ISL_457442, EPI_ISL_457445, EPI_ISL_457446, EPI_ISL_457447, EPI_ISL_457448, EPI_ISL_457449, EPI_ISL_457451, EPI_ISL_457452, EPI_ISL_457455, EPI_ISL_457456, EPI_ISL_457457, EPI_ISL_457458, EPI_ISL_457459, EPI_ISL_457460, EPI_ISL_457461, EPI_ISL_457462, EPI_ISL_457463, EPI_ISL_457464, EPI_ISL_457465, EPI_ISL_457467, EPI_ISL_457468, EPI_ISL_457469, EPI_ISL_457470, EPI_ISL_457471, EPI_ISL_457472, EPI_ISL_457473, EPI_ISL_457474, EPI_ISL_457475, EPI_ISL_457476, EPI_ISL_457477, EPI_ISL_457478, EPI_ISL_457479, EPI_ISL_457480, EPI_ISL_457482, EPI_ISL_457483, EPI_ISL_457484, EPI_ISL_457485, EPI_ISL_457486, EPI_ISL_457487, EPI_ISL_457488, EPI_ISL_457489, EPI_ISL_457490, EPI_ISL_457491, EPI_ISL_457492, EPI_ISL_457493, EPI_ISL_457494, EPI_ISL_457495, EPI_ISL_457496, EPI_ISL_457497, EPI_ISL_457498, EPI_ISL_457499, EPI_ISL_457500, EPI_ISL_457501, EPI_ISL_457503, EPI_ISL_457504, EPI_ISL_457505, EPI_ISL_457506, EPI_ISL_457507, EPI_ISL_457509, EPI_ISL_457510, EPI_ISL_457511, EPI_ISL_457512, EPI_ISL_457513, EPI_ISL_457514, EPI_ISL_457515, EPI_ISL_457516, EPI_ISL_457517, EPI_ISL_457518, EPI_ISL_457520, EPI_ISL_457521, EPI_ISL_457523, EPI_ISL_457524, EPI_ISL_457525, EPI_ISL_457527, EPI_ISL_457528, EPI_ISL_457529, EPI_ISL_457530, EPI_ISL_457531, EPI_ISL_457532, EPI_ISL_457533, EPI_ISL_457534, EPI_ISL_457535, EPI_ISL_457538, EPI_ISL_457539, EPI_ISL_457540, EPI_ISL_457541, EPI_ISL_457542, EPI_ISL_457543, EPI_ISL_457544, EPI_ISL_457545, EPI_ISL_457546, EPI_ISL_457547, EPI_ISL_457548, EPI_ISL_457550, EPI_ISL_457551, EPI_ISL_457552, EPI_ISL_457553, EPI_ISL_457555, EPI_ISL_457556, EPI_ISL_457558, EPI_ISL_457559, EPI_ISL_457560, EPI_ISL_457561, EPI_ISL_457562, EPI_ISL_457563, EPI_ISL_457565, EPI_ISL_457566, EPI_ISL_457567, EPI_ISL_457568, EPI_ISL_457569, EPI_ISL_457570, EPI_ISL_457571, EPI_ISL_457572 | | | |
| see above | Quadram Institute Bioscience | COVID-19 Genomics UK (COG-UK) Consortium | Dave J. Baker, Gemma L. Kay, Alp Aydin, Thanh Le-Viet, Steven Rudder, Ana P. Tedim, Anastasia Kolyva, Maria Diaz, Leonardo de Oliveira Martins, Nabil-Fareed Alikhan, Lizzie Meadows, Rachael Stanley, Ngozi Elumogo, Muhammed Yasir, Nicholas M. Thomson, Alexander J Trotter, Rachel Gilroy, Samuel Bloomfield, Claire Stuart, Andrew Bell, Reenesh Prakash, Samir Dervisevic, Alison E. Mather, John Wain, Mark Webber, Andrew J. Page, Justin O’Grady |
| EPI_ISL_457574, EPI_ISL_457575, EPI_ISL_457577, EPI_ISL_457578, EPI_ISL_457579 | Queens Medical Centre, Clinical Microbiology Department / DeepSeq Nottingham | COVID-19 Genomics UK (COG-UK) Consortium | Gemma Clark, Wendy Smith, Manjinder Khakh, Hannah Howson-Wells, Jonathan Ball, Patrick McClure, Joseph Chappell, Theocharis Tsoleridis, Nadine Holmes, Matthew Carlisle, Christopher Moore, Fei Sang, Johnny Debebe, Victoria Wright, Matthew Loose |
| EPI_ISL_457581, EPI_ISL_457582, EPI_ISL_457583, EPI_ISL_457584, EPI_ISL_457586, EPI_ISL_457592, EPI_ISL_457593, EPI_ISL_457594, EPI_ISL_457596, EPI_ISL_457597, EPI_ISL_457598, EPI_ISL_457599, EPI_ISL_457602, EPI_ISL_457603, EPI_ISL_457604, EPI_ISL_457605, EPI_ISL_457606, EPI_ISL_457611, EPI_ISL_457617, EPI_ISL_457618, EPI_ISL_457620, EPI_ISL_457624, EPI_ISL_457627, EPI_ISL_457631, EPI_ISL_457633, EPI_ISL_457634, EPI_ISL_457636, EPI_ISL_457637, EPI_ISL_457640, EPI_ISL_457641, EPI_ISL_457643, EPI_ISL_457647, EPI_ISL_457648, EPI_ISL_457652, EPI_ISL_457654, EPI_ISL_457655, EPI_ISL_457657, EPI_ISL_457658, EPI_ISL_457659, EPI_ISL_457660, EPI_ISL_457661, EPI_ISL_457667, EPI_ISL_457670, EPI_ISL_457672, EPI_ISL_457675, EPI_ISL_457676, EPI_ISL_457679, EPI_ISL_457680, EPI_ISL_457683, EPI_ISL_457685 | | | |
| see above | Virology Department, Sheffield Teaching Hospitals NHS Foundation Trust/Department of Infection, Immunity and Cardiovascular Disease, The Medical School, University of Sheffield | COVID-19 Genomics UK (COG-UK) Consortium | Thushan de Silva, Matthew Parker, Nikki Smith, Adri Angyal, Rebecca Brown, Luke Green, Rachel Tucker, Paul Parsons, Danielle Groves, Katie Johnson, Laura Carrilero, Alex Keeley, Dave Partridge, Matthew Wyles, Benjamin Lindsey, Mehmet Yavuz, Mohammad Raza, Cariad Evans |
| EPI_ISL_457687, EPI_ISL_457688, EPI_ISL_457689, EPI_ISL_457690, EPI_ISL_457691, EPI_ISL_457692, EPI_ISL_457693, EPI_ISL_457694, EPI_ISL_457695, EPI_ISL_457696, EPI_ISL_457697, EPI_ISL_457698 | | | |
| see above | The First Affiliated Hospital of Guangzhou Medical University, Guangzhou, China | BGI-shenzhen & The First Affiliated Hospital of Guangzhou Medical University | Yanqun Wang, Daxi Wang, Lu Zhang, Wanying Sun, Zhaoyong Zhang et al. |
| EPI_ISL_457699, EPI_ISL_457700 | Department of Infectious Diseases, Istituto Superiore di Sanità, Roma , Italy | Army Medical and Veterinary Research Center | Paola Stefanelli, Alessandra Lo Presti, Stefano Fiore, Antonella Marchi, Eleonora Benedetti, Concetta Fabiani Silvia Fillo, Giovanni Faggioni, Riccardo De Sanctis, Antonella Fortunato, Anna Anselmo, Francesco Giordani, Vanessa Vera Fain, Nino D’Amore, Florigio Lista |
| EPI_ISL_457701 | Oman-NIC | Oman-NIC | Samira Al-Maruqi, Fahad Zadjali, Amina Al Jardani, Khulood Al-Mammary, Hanan Al-kindi, Fatma BaAlawi, Hamida AL Barwani, Zeyana AL-Dahmani, Intisar Al-Shukri, Aisha Al-Busaidi, Aisha Al-Amri, Ahlam Al-Amri, Mohammed Al-Tobi, Samiha Al Kharusi, Abdulla Balkhair |
| EPI_ISL_457702 | Oman-NIC | Microbiology laboratory- Sultan Qaboos University Hospital | Fahad Zadjali, Samira Al-Maruqi, Amina Al Jardani, Khulood Al-Mammary, Hanan Al-kindi, Fatma BaAlawi, Hamida AL Barwani, Zeyana AL-Dahmani, Intisar Al-Shukri, Aisha Al-Busaidi, Aisha Al-Amri, Ahlam Al-Amri, Mohammed Al-Tobi, Samiha Al Kharusi, Abdulla Balkhair |
| EPI_ISL_457703 | Oman-NIC | Department of Microbiology and Immunology- SQUH | Fahad Zadjali, Samira Al-Maruqi, Amina Al Jardani, Khulood Al-Mammary, Hanan Al-kindi, Fatma BaAlawi, Hamida AL Barwani, Zeyana AL-Dahmani, Intisar Al-Shukri, Aisha Al-Busaidi, Aisha Al-Amri, Ahlam Al-Amri, Mohammed Al-Tobi, Samiha Al Kharusi, Abdulla Balkhair |
| EPI_ISL_457704 | Oman-NIC | Oman-NIC | Samira Al-Maruqi, Fahad Zadjali, Amina Al Jardani, Khulood Al-Mammary, Hanan Al-kindi, Fatma BaAlawi, Hamida AL Barwani, Zeyana AL-Dahmani, Intisar Al-Shukri, Aisha Al-Busaidi, Aisha Al-Amri, Ahlam Al-Amri, Mohammed Al-Tobi, Samiha Al Kharusi, Abdulla Balkhair |
| EPI_ISL_457705 | OMAN-NIC | Department of Microbiology and Immunology- SQUH | Fahad Zadjali, Samira Al-Maruqi, Amina Al Jardani, Khulood Al-Mammary, Hanan Al-kindi, Fatma BaAlawi, Hamida AL Barwani, Zeyana AL-Dahmani, Intisar Al-Shukri, Aisha Al-Busaidi, Aisha Al-Amri, Ahlam Al-Amri, Mohammed Al-Tobi, Samiha Al Kharusi, Abdulla Balkhair |
| EPI_ISL_457706 | Oman-NIC | Oman-NIC | Samira Al-Maruqi, Fahad Zadjali, Amina Al Jardani, Khulood Al-Mammary, Hanan Al-kindi, Fatma BaAlawi, Hamida AL Barwani, Zeyana AL-Dahmani, Intisar Al-Shukri, Aisha Al-Busaidi, Aisha Al-Amri, Ahlam Al-Amri, Mohammed Al-Tobi, Samiha Al Kharusi, Abdulla Balkhair |
| EPI_ISL_457707 | Oman-NIC | Department of Microbiology and Immunology- SQUH | Fahad Zadjali, Samira Al-Maruqi, Amina Al Jardani, Khulood Al-Mammary, Hanan Al-kindi, Fatma BaAlawi, Hamida AL Barwani, Zeyana AL-Dahmani, Intisar Al-Shukri, Aisha Al-Busaidi, Aisha Al-Amri, Ahlam Al-Amri, Mohammed Al-Tobi, Samiha Al Kharusi, Abdulla Balkhair |
| EPI_ISL_457719 | SYNLAB Eesti OÜ | Charite Universitatsmedizin Berlin, Institute of Virology | Victor M Corman, Jorn Beheim-Schwarzbach, Barbara Mühlemann, Talitha Veith, Julia Schneider, Paul Naaber, Terry Jones, Christian Drosten |
| EPI_ISL_457721 | Department of Infectious Diseases, Istituto Superiore di Sanità, Roma , Italy | Army Medical and Veterinary Research Center | Paola Stefanelli, Alessandra Lo Presti, Stefano Fiore, Antonella Marchi, Eleonora Benedetti, Concetta Fabiani Silvia Fillo, Giovanni Faggioni, Riccardo De Sanctis, Antonella Fortunato, Anna Anselmo, Francesco Giordani, Vanessa Vera Fain, Nino D’Amore, Florigio Lista |
| EPI_ISL_457722, EPI_ISL_457723 | SYNLAB Eesti OÜ | Charite Universitatsmedizin Berlin, Institute of Virology | Victor M Corman, Jorn Beheim-Schwarzbach, Barbara Mühlemann, Talitha Veith, Julia Schneider, Paul Naaber, Terry Jones, Christian Drosten |
| EPI_ISL_457724 | Department of Infectious Diseases, Istituto Superiore di Sanità, Roma , Italy | Army Medical and Veterinary Research Center | Paola Stefanelli, Alessandra Lo Presti, Stefano Fiore, Antonella Marchi, Eleonora Benedetti, Concetta Fabiani Silvia Fillo, Giovanni Faggioni, Riccardo De Sanctis, Antonella Fortunato, Anna Anselmo, Francesco Giordani, Vanessa Vera Fain, Nino D’Amore, Florigio Lista |
| EPI_ISL_457725 | SYNLAB Eesti OÜ | Charite Universitatsmedizin Berlin, Institute of Virology | Victor M Corman, Jorn Beheim-Schwarzbach, Barbara Mühlemann, Talitha Veith, Julia Schneider, Paul Naaber, Terry Jones, Christian Drosten |
| EPI_ISL_457726 | TSGH-CP molecular lab | TSGH-CP molecular lab | Cherng-Lih Perng, Ming-Jr JIAN, Chih-Kai Chang, Jung-Chung Lin, Kuo-Ming Yeh, Chien-Wen Chen, Sheng-Kang Chiu, Hsing-Yi Chung, Shih-Hung Tsai, Kuo-Sheng Hung, Tien-Yao Chang, Feng-Yee Chang, Hung-Sheng Shang |
| EPI_ISL_457727 | SYNLAB Eesti OÜ | Charite Universitatsmedizin Berlin, Institute of Virology | Victor M Corman, Jorn Beheim-Schwarzbach, Barbara Mühlemann, Talitha Veith, Julia Schneider, Paul Naaber, Terry Jones, Christian Drosten |
| EPI_ISL_457728 | Department of Infectious Diseases, Istituto Superiore di Sanità, Roma , Italy | Army Medical and Veterinary Research Center | Paola Stefanelli, Alessandra Lo Presti, Stefano Fiore, Antonella Marchi, Eleonora Benedetti, Concetta Fabiani Silvia Fillo, Giovanni Faggioni, Riccardo De Sanctis, Antonella Fortunato, Anna Anselmo, Francesco Giordani, Vanessa Vera Fain, Nino D’Amore, Florigio Lista |
| EPI_ISL_457729 | SYNLAB Eesti OÜ | Charite Universitatsmedizin Berlin, Institute of Virology | Victor M Corman, Jorn Beheim-Schwarzbach, Barbara Mühlemann, Talitha Veith, Julia Schneider, Paul Naaber, Terry Jones, Christian Drosten |
| EPI_ISL_457730 | TSGH-CP molecular lab | TSGH-CP molecular lab | Cherng-Lih Perng, Ming-Jr JIAN, Chih-Kai Chang, Jung-Chung Lin, Kuo-Ming Yeh, Chien-Wen Chen, Sheng-Kang Chiu, Hsing-Yi Chung, Shih-Hung Tsai, Kuo-Sheng Hung, Tien-Yao Chang, Feng-Yee Chang, Hung-Sheng Shang |
| EPI_ISL_457732 | Department of Infectious Diseases, Istituto Superiore di Sanità, Roma , Italy | Army Medical and Veterinary Research Center | Paola Stefanelli, Alessandra Lo Presti, Stefano Fiore, Antonella Marchi, Eleonora Benedetti, Concetta Fabiani Silvia Fillo, Giovanni Faggioni, Riccardo De Sanctis, Antonella Fortunato, Anna Anselmo, Francesco Giordani, Vanessa Vera Fain, Nino D’Amore, Florigio Lista |
| EPI_ISL_457733 | TSGH-CP molecular lab | TSGH-CP molecular lab | Cherng-Lih Perng, Ming-Jr JIAN, Chih-Kai Chang, Jung-Chung Lin, Kuo-Ming Yeh, Chien-Wen Chen, Sheng-Kang Chiu, Hsing-Yi Chung, Shih-Hung Tsai, Kuo-Sheng Hung, Tien-Yao Chang, Feng-Yee Chang, Hung-Sheng Shang |
| EPI_ISL_457735 | SYNLAB Eesti OÜ | Charite Universitatsmedizin Berlin, Institute of Virology | Victor M Corman, Jorn Beheim-Schwarzbach, Barbara Mühlemann, Talitha Veith, Julia Schneider, Paul Naaber, Terry Jones, Christian Drosten |
| EPI_ISL_457736 | Department of Infectious Diseases, Istituto Superiore di Sanità, Roma , Italy | Army Medical and Veterinary Research Center | Paola Stefanelli, Alessandra Lo Presti, Stefano Fiore, Antonella Marchi, Eleonora Benedetti, Concetta Fabiani Silvia Fillo, Giovanni Faggioni, Riccardo De Sanctis, Antonella Fortunato, Anna Anselmo, Francesco Giordani, Vanessa Vera Fain, Nino D’Amore, Florigio Lista |
| EPI_ISL_457737, EPI_ISL_457740 | SYNLAB Eesti OÜ | Charite Universitatsmedizin Berlin, Institute of Virology | Victor M Corman, Jorn Beheim-Schwarzbach, Barbara Mühlemann, Talitha Veith, Julia Schneider, Paul Naaber, Terry Jones, Christian Drosten |
| EPI_ISL_457749 | Department of Infectious Diseases, Istituto Superiore di Sanità, Roma , Italy | Army Medical and Veterinary Research Center | Paola Stefanelli, Alessandra Lo Presti, Stefano Fiore, Antonella Marchi, Eleonora Benedetti, Concetta Fabiani Silvia Fillo, Giovanni Faggioni, Riccardo De Sanctis, Antonella Fortunato, Anna Anselmo, Francesco Giordani, Vanessa Vera Fain, Nino D’Amore, Florigio Lista |
| EPI_ISL_457750 | Centogene AG | Centogene AG | Prof. Dr. Peter Bauer, Dr. Krishna Kumar Kandaswamy |
| EPI_ISL_457752, EPI_ISL_457753, EPI_ISL_457754, EPI_ISL_457755, EPI_ISL_457756, EPI_ISL_457757, EPI_ISL_457758, EPI_ISL_457759, EPI_ISL_457760, EPI_ISL_457761, EPI_ISL_457764, EPI_ISL_457765, EPI_ISL_457766, EPI_ISL_457768, EPI_ISL_457769, EPI_ISL_457780, EPI_ISL_457781, EPI_ISL_457782, EPI_ISL_457783, EPI_ISL_457784, EPI_ISL_457785, EPI_ISL_457787, EPI_ISL_457788, EPI_ISL_457789, EPI_ISL_457790, EPI_ISL_457791, EPI_ISL_457792, EPI_ISL_457795, EPI_ISL_457796, EPI_ISL_457797, EPI_ISL_457798, EPI_ISL_457799, EPI_ISL_457800, EPI_ISL_457801, EPI_ISL_457802, EPI_ISL_457803, EPI_ISL_457804, EPI_ISL_457805, EPI_ISL_457806, EPI_ISL_457807, EPI_ISL_457808, EPI_ISL_457809, EPI_ISL_457810, EPI_ISL_457811, EPI_ISL_457812, EPI_ISL_457813, EPI_ISL_457814, EPI_ISL_457816, EPI_ISL_457817, EPI_ISL_457818, EPI_ISL_457819, EPI_ISL_457820, EPI_ISL_457821, EPI_ISL_457822, EPI_ISL_457823 | | | |
| see above | Johns Hopkins Hospital Department of Pathology | Johns Hopkins Hospital Department of Pathology | Peter M. Thielen, Thomas Mehoke, Shirlee Wohl, Srividya Ramakrishnan, Melanie Kirsche, Amanda Ernlund, Craig Howser, Kristina Zudock, Oluwaseun Falade-Nwulia, Norah Sadowski, Paul Morris, Mark Hopkins, Yunfan Fan, Nidia Trovao, Victoria Gniazdowski, Michael C. Schatz, Stuart C. Ray, Winston Timp, Heba H. Mostafa |
| EPI_ISL_457824 | Bezmialem Vakif University, Dept Microbiology, Medical School, Fatih, Istanbul, Turkey | Bezmialem Vakif University, Medical School & Beykoz Institute of Life Sciences & Biotechnology | Mehmet Z. Doymaz, Merve Kalkan, Nesibe Cetin, Elif Karaaslan, Bilge Sumbul, Filiz Guney |
| EPI_ISL_457825 | Army Medical Research Center - Scientific Department | Army Medical and Veterinary Research Center | Silvia Fillo, Giovanni Faggioni, Riccardo De Sanctis, Antonella Fortunato, Anna Anselmo, Francesco Giordani, Vanessa Vera Fain, Nino D’Amore, Florigio Lista |
| EPI_ISL_457826 | Army Medical Center - Scientific Department | Army Medical and Veterinary Research Center | Silvia Fillo, Giovanni Faggioni, Riccardo De Sanctis, Antonella Fortunato, Anna Anselmo, Francesco Giordani, Vanessa Vera Fain, Nino D’Amore, Florigio Lista |
| EPI_ISL_457827, EPI_ISL_457828, EPI_ISL_457829, EPI_ISL_457833, EPI_ISL_457843 | National Public Health Laboratory | KEMRI-Wellcome Trust Research Programme/KEMRI-CGMR-C Kilifi | Githinji G. et al 2020 |

| | | | |
|---|---|---|---|
| EPI_ISL_457854, EPI_ISL_457867, EPI_ISL_457868, EPI_ISL_457875, EPI_ISL_457884, EPI_ISL_457897, EPI_ISL_457906, EPI_ISL_457913, EPI_ISL_457915, EPI_ISL_457920, EPI_ISL_457921, EPI_ISL_457928 | | | |
| see above | KEMRI-CGMR-C | KEMRI-Wellcome Trust Research Programme/KEMRI-CGMR-C Kilifi | Githinji G. et al 2020 |
| EPI_ISL_457937, EPI_ISL_457938, EPI_ISL_457939 | Oman-NIC | Oman-NIC | Samira Al-Maruqi, Fahad Zadjali, Amina Al Jardani, Khulood Al-Mammary, Hanan Al-kindi, Fatma BaAlawi, Hamida AL Barwani, Zeyana AL-Dahmani, Intisar Al-Shukri, Aisha Al-Busaidi, Aisha Al-Amri, Ahlam Al-Amri, Mohammed Al-Tobi, Samiha Al Kharusi, Abdulla Balkhair |
| EPI_ISL_457946, EPI_ISL_457948, EPI_ISL_457952, EPI_ISL_457953, EPI_ISL_457956, EPI_ISL_457957, EPI_ISL_457958, EPI_ISL_457959, EPI_ISL_457961, EPI_ISL_457963, EPI_ISL_457965, EPI_ISL_457972 | | | |
| see above | Laboratorio de Biología Molecular Asociación Española Primera en Salud | Departments of Pathology and Medicine, New York University School of Medicine | Maria Victoria Elizondo, Maria Noel Zubillaga, Gonzalo Manrique, Paul Zappile, Gael Westby, Matthew T Maurano, Christian Marier, Adriana Heguy |
| EPI_ISL_457974, EPI_ISL_457975, EPI_ISL_457976, EPI_ISL_457977, EPI_ISL_457978, EPI_ISL_457979, EPI_ISL_457980 | Oman-NIC | Oman-NIC | Samira Al-Maruqi, Fahad Zadjali, Amina Al Jardani, Khulood Al-Mammary, Hanan Al-kindi, Fatma BaAlawi, Hamida AL Barwani, Zeyana AL-Dahmani, Intisar Al-Shukri, Aisha Al-Busaidi, Aisha Al-Amri, Ahlam Al-Amri, Mohammed Al-Tobi, Samiha Al Kharusi, Abdulla Balkhair |
| EPI_ISL_457981 | Oman-NIC | Department of Microbiology and Immunology-SQUH | Fahad Zadjali, Samira Al-Maruqi, Amina Al Jardani, Khulood Al-Mammary, Hanan Al-kindi, Fatma BaAlawi, Hamida AL Barwani, Zeyana AL-Dahmani, Intisar Al-Shukri, Aisha Al-Busaidi, Aisha Al-Amri, Ahlam Al-Amri, Mohammed Al-Tobi, Samiha Al Kharusi, Abdulla Balkhair |
| EPI_ISL_457985, EPI_ISL_457986, EPI_ISL_457987, EPI_ISL_457988, EPI_ISL_457989, EPI_ISL_457990, EPI_ISL_457991, EPI_ISL_457992, EPI_ISL_457993, EPI_ISL_457994, EPI_ISL_457995, EPI_ISL_457996, EPI_ISL_457997, EPI_ISL_457998 | | | |
| see above | Oman-NIC | Oman-NIC | Samira Al-Maruqi, Fahad Zadjali, Amina Al Jardani, Khulood Al-Mammary, Hanan Al-kindi, Fatma BaAlawi, Hamida AL Barwani, Zeyana AL-Dahmani, Intisar Al-Shukri, Aisha Al-Busaidi, Aisha Al-Amri, Ahlam Al-Amri, Mohammed Al-Tobi, Samiha Al Kharusi, Abdulla Balkhair |
| EPI_ISL_458002, EPI_ISL_458003, EPI_ISL_458004, EPI_ISL_458005, EPI_ISL_458006 | Dirk Dittmer | Dirk Dittmer | Bailey,A.G., Caro-Vegas,C.P., Dittmer,D., Eason,A.B., Juarez,A., Landis,J.T., McNamara,R.P., Miller,M.B., Moorad,R., Pluta,L.J., Seltzer,T.A., Thompson,C., Vahrson,W., Villamor,F. |
| EPI_ISL_458007 | Dirk Dittmer | Dirk Dittmer | Aubrey,B.G., Caro-Vegas,C.P., Dittmer,D., Eason,A.B., Juarez,A., Landis,J.T., McNamara,R.P., Miller,M.B., Moorad,R., Pluta,L.J., Seltzer,T.A., Thompson,C., Vahrson,W., Villamor,F. |
| EPI_ISL_458017, EPI_ISL_458018, EPI_ISL_458019, EPI_ISL_458020, EPI_ISL_458021 | NYU Langone Health | Departments of Pathology and Medicine, New York University School of Medicine | Maria Aguero-Rosenfeld, Brendan Belovarac, Margaret Black, Ludovic Boytard, John Cadley, Paolo Cotzia, John Chen, Dacia Dimartino, Xiaojun Feng, Tatyana Gindin, Emily Guzman, Adriana Heguy, Megan Hogan, Emily Huang, George Jour, Alireza Khodadadi-Jamayran, Lawrence H. Lin, Raven Luther, Andrew Lytle, Christian Marier, Matthew T. Maurano, Mark J. Mulligan, Peter Meyn, Raquel Ordonez Ciriza, Iman Osman, Jared Pinnell, Vanessa Raabe, Sitharam Ramaswami, Amy Rapkiewicz, Andre M. Ribeiro-dos-Santos, Marie Samanovic-Golden, Antonio Serrano, Guomiao Shen, Matija Snuderl, Theodore Vougiouklakis, Nick Vulpescu, Gael Westby, Paul Zappile, Yutong Zhang |
| EPI_ISL_458024 | Hospital for Tropical Diseases | COVID-19 Network Investigations (CONI) Alliance | Elizabeth Batty, Nantarat Chantawat, Wasun Chantratita, Thanat Chookajorn, Stefan Fernandez, Angkana Huang, Weena Janwitthayanan, Akanitt Jittmittraphap, Anthony R. Jones, Khajohn Joonsalak, Chonticha Klungtong, Theerarat Kochakarn, Namfon Kotanan, Krittikorn Kumpornsin, Pornsawan Leaungwutiwong, Wudtichai Manasatienkij, Bhakbhoom Panthan, Ekawat Pasomsub, Kingkan Rakmanee, Insee Sensorn, Janjira Thaipadungpanit, Arporn Wangwiwatsin,Treewat Watthanachockchai |
| EPI_ISL_458029 | TSGH-CP molecular lab | TSGH-CP molecular lab | Cherng-Lih Perng, Ming-Jr JIAN, Chih-Kai Chang, Jung-Chung Lin, Kuo-Ming Yeh, Chien-Wen Chen, Sheng-Kang Chiu, Hsing-Yi Chung, Shih-Hung Tsai, Kuo-Sheng Hung, Tien-Yao Chang, Feng-Yee Chang, Hung-Sheng Shang |
| EPI_ISL_458030 | King Institute of Preventive Medicine & Research | CSIR-Centre for Cellular and Molecular Biology | K.Kaveri,S.Sivasubramanian,S.Vennila,P.Padmapriya,R.Kiruba,S.Magesh,G. Dhinakar Raj, G. Ravikumar, P. Azhahianambi,K Thangaraj,Payel Mukherjee, Sofia Banu, Priya Singh, Dhiviya Vedagiri, Divya Gupta, Vishal Sah, Santosh Kumar Kuncha, Krishnan Harinivas Harshan, Archana Bharadwaj Siva, Karthik Bharadwaj Tallapaka, Shagufta Khan, Lamuk Zaveri, Namami Gaur, Sakshi Shambhavi, Tulasi Nagabandi, Purushotham Vodnala, Rakesh K Mishra, Divya Tej Sowpati |
| EPI_ISL_458031 | King Institute of Preventive Medicine & Research | CSIR-Centre for Cellular and Molecular Biology | K.Kaveri,S.Sivasubramanian,S.Vennila,P.Padmapriya,R.Kiruba,S.Magesh,G. Dhinakar Raj, G. Ravikumar, P. Azhahianambi, K Thangaraj,Sofia Banu, Payel Mukherjee, Priya Singh, Dhiviya Vedagiri, Divya Gupta, Vishal Sah, Santosh Kumar Kuncha, Krishnan Harinivas Harshan, Archana Bharadwaj Siva, Karthik Bharadwaj Tallapaka, Shagufta Khan, Lamuk Zaveri, Namami Gaur, Sakshi Shambhavi, Tulasi Nagabandi, Purushotham Vodnala, Rakesh K Mishra, Divya Tej Sowpati |
| EPI_ISL_458032 | King Institute of Preventive Medicine & Research | CSIR-Centre for Cellular and Molecular Biology | K.Kaveri,S.Sivasubramanian,S.Vennila,P.Padmapriya,R.Kiruba,S.Magesh,G. Dhinakar Raj, G. Ravikumar, P. Azhahianambi, K Thangaraj,Shagufta Khan, Lamuk Zaveri, Namami Gaur, Sakshi Shambhavi, Tulasi Nagabandi, Purushotham Vodnala, Payel Mukherjee, Sofia Banu, Priya Singh, Dhiviya Vedagiri, Divya Gupta, Vishal Sah, Santosh Kumar Kuncha, Krishnan Harinivas Harshan, Archana Bharadwaj Siva, Karthik Bharadwaj Tallapaka, Rakesh K Mishra, Divya Tej Sowpati |
| EPI_ISL_458033 | King Institute of Preventive Medicine & Research | CSIR-Centre for Cellular and Molecular Biology | K.Kaveri,S.Sivasubramanian,S.Vennila,P.Padmapriya,R.Kiruba,S.Magesh,G. Dhinakar Raj, G. Ravikumar, P. Azhahianambi, K Thangaraj,Lamuk Zaveri, Shagufta Khan, Namami Gaur, Sakshi Shambhavi, Tulasi Nagabandi, Purushotham Vodnala, Payel Mukherjee, Sofia Banu, Priya Singh, Dhiviya Vedagiri, Divya Gupta, Vishal Sah, Santosh Kumar Kuncha, Krishnan Harinivas Harshan, Archana Bharadwaj Siva, Karthik Bharadwaj Tallapaka, Rakesh K Mishra, Divya Tej Sowpati |
| EPI_ISL_458034 | King Institute of Preventive Medicine & Research | CSIR-Centre for Cellular and Molecular Biology | K.Kaveri,S.Sivasubramanian,S.Vennila,P.Padmapriya,R.Kiruba,S.Magesh,G. Dhinakar Raj, G. Ravikumar, P. Azhahianambi, K Thangaraj, Namami Gaur, Sakshi Shambhavi, Lamuk Zaveri, Shagufta Khan, Tulasi Nagabandi, Purushotham Vodnala, Payel Mukherjee, Sofia Banu, Priya Singh, Dhiviya Vedagiri, Divya Gupta, Vishal Sah, Santosh Kumar Kuncha, Krishnan Harinivas Harshan, Archana Bharadwaj Siva, Karthik Bharadwaj Tallapaka, Rakesh K Mishra, Divya Tej Sowpati |
| EPI_ISL_458035 | King Institute of Preventive Medicine & Research | CSIR-Centre for Cellular and Molecular Biology | K.Kaveri,S.Sivasubramanian,S.Vennila,P.Padmapriya,R.Kiruba,S.Magesh,G. Dhinakar Raj, G. Ravikumar, P. Azhahianambi, K Thangaraj, Tulasi Nagabandi, Namami Gaur, Sakshi Shambhavi, Lamuk Zaveri, Shagufta Khan, Purushotham Vodnala, Payel Mukherjee, Sofia Banu, Priya Singh, Dhiviya Vedagiri, Divya Gupta, Vishal Sah, Santosh Kumar Kuncha, Krishnan Harinivas Harshan, Archana Bharadwaj Siva, Karthik Bharadwaj Tallapaka, Rakesh K Mishra, Divya Tej Sowpati |
| EPI_ISL_458036 | King Institute of Preventive Medicine & Research | CSIR-Centre for Cellular and Molecular Biology | K.Kaveri,S.Sivasubramanian,S.Vennila,P.Padmapriya,R.Kiruba,S.Magesh,G. Dhinakar Raj, G. Ravikumar, R. P. Aravindh Babu, K Thangaraj, Payel Mukherjee, Sofia Banu, Priya Singh, Dhiviya Vedagiri, Divya Gupta, Vishal Sah, Santosh Kumar Kuncha, Krishnan Harinivas Harshan, Archana Bharadwaj Siva, Karthik Bharadwaj Tallapaka, Shagufta Khan, Lamuk Zaveri, Namami Gaur, Sakshi Shambhavi, Tulasi Nagabandi, Purushotham Vodnala, Rakesh K Mishra, Divya Tej Sowpati |
| EPI_ISL_458037 | King Institute of Preventive Medicine & Research | CSIR-Centre for Cellular and Molecular Biology | K.Kaveri,S.Sivasubramanian,S.Vennila,P.Padmapriya,R.Kiruba,S.Magesh,G. Dhinakar Raj, G. Ravikumar, R. P. Aravindh Babu, K Thangaraj, Sofia Banu, Payel Mukherjee, Priya Singh, Dhiviya Vedagiri, Divya Gupta, Vishal Sah, Santosh Kumar Kuncha, Krishnan Harinivas Harshan, Archana Bharadwaj Siva, Karthik Bharadwaj Tallapaka, Shagufta Khan, Lamuk Zaveri, Namami Gaur, Sakshi Shambhavi, Tulasi Nagabandi, Purushotham Vodnala, Rakesh K Mishra, Divya Tej Sowpati |
| EPI_ISL_458038 | King Institute of Preventive Medicine & Research | CSIR-Centre for Cellular and Molecular Biology | K.Kaveri,S.Sivasubramanian,S.Vennila,P.Padmapriya,R.Kiruba,S.Magesh,G. Dhinakar Raj, G. Ravikumar, R. P. Aravindh Babu, K Thangaraj, Shagufta Khan, Lamuk Zaveri, Namami Gaur, Sakshi Shambhavi, Tulasi Nagabandi, Purushotham Vodnala, Payel Mukherjee, Sofia Banu, Priya Singh, Dhiviya Vedagiri, Divya Gupta, Vishal Sah, Santosh Kumar Kuncha, Krishnan Harinivas Harshan, Archana Bharadwaj Siva, Karthik Bharadwaj Tallapaka, Rakesh K Mishra, Divya Tej Sowpati |
| EPI_ISL_458039 | King Institute of Preventive Medicine & Research | CSIR-Centre for Cellular and Molecular Biology | K.Kaveri,S.Sivasubramanian,S.Vennila,P.Padmapriya,R.Kiruba,S.Magesh,G. Dhinakar Raj, G. Ravikumar, R. P. Aravindh Babu, K Thangaraj, Lamuk Zaveri, Shagufta Khan, Namami Gaur, Sakshi Shambhavi, Tulasi Nagabandi, Purushotham Vodnala, Payel Mukherjee, Sofia Banu, Priya Singh, Dhiviya Vedagiri, Divya Gupta, Vishal Sah, Santosh Kumar Kuncha, Krishnan Harinivas Harshan, Archana Bharadwaj Siva, Karthik Bharadwaj Tallapaka, Rakesh K Mishra, Divya Tej Sowpati |
| EPI_ISL_458040 | King Institute of Preventive Medicine & Research | CSIR-Centre for Cellular and Molecular Biology | K.Kaveri,S.Sivasubramanian,S.Vennila,P.Padmapriya,R.Kiruba,S.Magesh,G. Dhinakar Raj, G. Ravikumar, R. P. Aravindh Babu, K Thangaraj, Namami Gaur, Sakshi Shambhavi, Lamuk Zaveri, Shagufta Khan, Tulasi Nagabandi, Purushotham Vodnala, Payel Mukherjee, Sofia Banu, Priya Singh, Dhiviya Vedagiri, Divya Gupta, Vishal Sah, Santosh Kumar Kuncha, Krishnan Harinivas Harshan, Archana Bharadwaj Siva, Karthik Bharadwaj Tallapaka, Rakesh K Mishra, Divya Tej Sowpati |
| EPI_ISL_458041 | King Institute of Preventive Medicine & Research | CSIR-Centre for Cellular and Molecular Biology | K.Kaveri,S.Sivasubramanian,S.Vennila,P.Padmapriya,R.Kiruba,S.Magesh,G. Dhinakar Raj, G. Ravikumar, R. P. Aravindh Babu, K Thangaraj, Tulasi Nagabandi, Namami Gaur, Sakshi Shambhavi, Lamuk Zaveri, Shagufta Khan, Purushotham Vodnala, Payel Mukherjee, Sofia Banu, Priya Singh, Dhiviya Vedagiri, Divya Gupta, Vishal Sah, Santosh Kumar Kuncha, Krishnan Harinivas Harshan, Archana Bharadwaj Siva, Karthik Bharadwaj Tallapaka, Rakesh K Mishra, Divya Tej Sowpati |
| EPI_ISL_458042 | King Institute of Preventive Medicine & Research | CSIR-Centre for Cellular and Molecular Biology | K.Kaveri,S.Sivasubramanian,S.Vennila,P.Padmapriya,R.Kiruba,S.Magesh,G. Dhinakar Raj, G. Ravikumar, M. Sekar, K Thangaraj, Payel Mukherjee, Sofia Banu, Priya Singh, Dhiviya Vedagiri, Divya Gupta, Vishal Sah, Santosh Kumar Kuncha, Krishnan Harinivas Harshan, Archana Bharadwaj Siva, Karthik Bharadwaj Tallapaka, Shagufta Khan, Lamuk Zaveri, Namami Gaur, Sakshi Shambhavi, Tulasi Nagabandi, Purushotham Vodnala, Rakesh K Mishra, Divya Tej Sowpati |
| EPI_ISL_458043 | King Institute of Preventive Medicine & Research | CSIR-Centre for Cellular and Molecular Biology | K.Kaveri,S.Sivasubramanian,S.Vennila,P.Padmapriya,R.Kiruba,S.Magesh,G. Dhinakar Raj, G. Ravikumar, M. Sekar, K Thangaraj,Sofia Banu, Payel Mukherjee, Priya Singh, Dhiviya Vedagiri, Divya Gupta, Vishal Sah, Santosh Kumar Kuncha, Krishnan Harinivas Harshan, Archana Bharadwaj Siva, Karthik Bharadwaj Tallapaka, Shagufta Khan, Lamuk Zaveri, Namami Gaur, Sakshi Shambhavi, Tulasi Nagabandi, Purushotham Vodnala, Rakesh K Mishra, Divya Tej Sowpati |
| EPI_ISL_458044 | King Institute of Preventive Medicine & Research | CSIR-Centre for Cellular and Molecular Biology | K.Kaveri,S.Sivasubramanian,S.Vennila,P.Padmapriya,R.Kiruba,S.Magesh,G. Dhinakar Raj, G. Ravikumar, M. Sekar, K Thangaraj,Shagufta Khan, Lamuk Zaveri, Namami Gaur, Sakshi Shambhavi, Tulasi Nagabandi, Purushotham Vodnala, Payel Mukherjee, Sofia Banu, Priya Singh, Dhiviya Vedagiri, Divya Gupta, Vishal Sah, Santosh Kumar Kuncha, Krishnan Harinivas Harshan, Archana Bharadwaj Siva, Karthik Bharadwaj Tallapaka, Rakesh K Mishra, Divya Tej Sowpati |
| EPI_ISL_458045 | CSIR-Centre for Cellular and Molecular Biology | CSIR-Centre for Cellular and Molecular Biology | Payel Mukherjee, Sofia Banu, Priya Singh, Dhiviya Vedagiri, Divya Gupta, Vishal Sah, Santosh Kumar Kuncha, Krishnan Harinivas Harshan, Archana Bharadwaj Siva, Karthik Bharadwaj Tallapaka, Shagufta Khan, Lamuk Zaveri, Namami Gaur, Sakshi Shambhavi, Tulasi Nagabandi, Purushotham Vodnala, G. Aditya Kumar, Koushick Sivakumar, Pooja Ramesh Gupta, Rajan Kumar Jha, Shraddha Vijay Lahoti, Rakesh K Mishra, Divya Tej Sowpati |
| EPI_ISL_458046 | CSIR-Centre for Cellular and Molecular Biology | CSIR-Centre for Cellular and Molecular Biology | Sofia Banu, Payel Mukherjee, Priya Singh, Dhiviya Vedagiri, Divya Gupta, Vishal Sah, Santosh Kumar Kuncha, Krishnan Harinivas Harshan, Archana Bharadwaj Siva, Karthik Bharadwaj Tallapaka, Shagufta Khan, Lamuk Zaveri, Namami Gaur, Sakshi Shambhavi, Tulasi Nagabandi, Purushotham Vodnala, Deepak Kumar, Devi Prasad Vijayashankar, Disha Nanda, Divya Das, Jotin Gogoi, Manish Bhattacharjee, Rakesh K Mishra, Divya Tej Sowpati |
| EPI_ISL_458047 | CSIR-Centre for Cellular and Molecular Biology | CSIR-Centre for Cellular and Molecular Biology | Shagufta Khan, Lamuk Zaveri, Namami Gaur, Sakshi Shambhavi, Tulasi Nagabandi, Purushotham Vodnala, Payel Mukherjee, Sofia Banu, Priya Singh, Dhiviya Vedagiri, Divya Gupta, Vishal Sah, Santosh Kumar Kuncha, Krishnan Harinivas Harshan, Archana Bharadwaj Siva, Karthik Bharadwaj Tallapaka, Disha Nanda, Divya Das, Jotin Gogoi, Manish Bhattacharjee, Ravi Prasad Mukku, Rakesh K Mishra, Divya Tej Sowpati |
| EPI_ISL_458048 | CSIR-Centre for Cellular and Molecular Biology | CSIR-Centre for Cellular and Molecular Biology | Lamuk Zaveri, Shagufta Khan, Namami Gaur, Sakshi Shambhavi, Tulasi Nagabandi, Purushotham Vodnala, Payel Mukherjee, Sofia Banu, Priya Singh, Dhiviya Vedagiri, Divya Gupta, Vishal Sah, Santosh Kumar Kuncha, Krishnan Harinivas Harshan, Archana Bharadwaj Siva, Karthik Bharadwaj Tallapaka, Renu Sudhakar, Somesh Gorde, Gangumala Srinivas Reddy, Sujoy Deb, Swati Bayyana, Rakesh K Mishra, Divya Tej Sowpati |
| EPI_ISL_458049 | CSIR-Centre for Cellular and Molecular Biology | CSIR-Centre for Cellular and Molecular Biology | Namami Gaur, Sakshi Shambhavi, Lamuk Zaveri, Shagufta Khan, Tulasi Nagabandi, Purushotham Vodnala, Payel Mukherjee, Sofia Banu, Priya Singh, Dhiviya Vedagiri, Divya Gupta, Vishal Sah, Santosh Kumar Kuncha, Krishnan Harinivas Harshan, Archana Bharadwaj Siva, Karthik Bharadwaj Tallapaka, Zeba Rizvi, Zuberwasim Sayyad, Kakade Aishwarya Arun, Amrutha H C, Ananga Ghosh, Rakesh K Mishra, Divya Tej Sowpati |
| EPI_ISL_458050 | CSIR-Centre for Cellular and Molecular Biology | CSIR-Centre for Cellular and Molecular Biology | Tulasi Nagabandi, Namami Gaur, Sakshi Shambhavi, Lamuk Zaveri, Shagufta Khan, Purushotham Vodnala, Payel Mukherjee, Sofia Banu, Priya Singh, Dhiviya Vedagiri, Divya Gupta, Vishal Sah, Santosh Kumar Kuncha, Krishnan Harinivas Harshan, Archana Bharadwaj Siva, Karthik Bharadwaj Tallapaka,Kezia J Ann, Radhika Khandelwal, Roshan Maku Venkata, Shemin Mansuri, Sonu Uday, Rakesh K Mishra, Divya Tej Sowpati |
| EPI_ISL_458051 | CSIR-Centre for Cellular and Molecular Biology | CSIR-Centre for Cellular and Molecular Biology | Payel Mukherjee, Sofia Banu, Priya Singh, Dhiviya Vedagiri, Divya Gupta, Vishal Sah, Santosh Kumar Kuncha, Krishnan Harinivas Harshan, Archana Bharadwaj Siva, Karthik Bharadwaj Tallapaka, Shagufta Khan, Lamuk Zaveri, Namami Gaur, Sakshi Shambhavi, Tulasi Nagabandi, Purushotham Vodnala, Gokulan C G, Gunjan Purohit, Hanuman Tulashiram Kale, Pankaj Kumar, Prachand Issarapu, Rakesh K Mishra, Divya Tej Sowpati |

| | | | |
|---|---|---|---|
| EPI_ISL_458052 | CSIR-Centre for Cellular and Molecular Biology | CSIR-Centre for Cellular and Molecular Biology | Sofia Banu, Payel Mukherjee, Priya Singh, Dhiviya Vedagiri, Divya Gupta, Vishal Sah, Santosh Kumar Kuncha, Krishnan Harinivas Harshan, Archana Bharadwaj Siva, Karthik Bharadwaj Tallapaka, Shagufta Khan, Lamuk Zaveri, Namami Gaur, Sakshi Shambhavi, Tulasi Nagabandi, Purushotham Vodnala,Preethi Jampala, Sharada Ravi Iyer, Sulagana Mukherjee, Swetha Sundar, Peddapuvala Sai Uday Kiran, Rakesh K Mishra, Divya Tej Sowpati |
| EPI_ISL_458053 | CSIR-Centre for Cellular and Molecular Biology | CSIR-Centre for Cellular and Molecular Biology | Shagufta Khan, Lamuk Zaveri, Namami Gaur, Sakshi Shambhavi, Tulasi Nagabandi, Purushotham Vodnala, Payel Mukherjee, Sofia Banu, Priya Singh, Dhiviya Vedagiri, Divya Gupta, Vishal Sah, Santosh Kumar Kuncha, Krishnan Harinivas Harshan, Archana Bharadwaj Siva, Karthik Bharadwaj Tallapaka,Umesh Kumar, Unis Ahmad Bhat, Ajay Sarawagi, Priyanka Pant, Rajkanwar Nathawat, Rakesh K Mishra, Divya Tej Sowpati |
| EPI_ISL_458054 | CSIR-Centre for Cellular and Molecular Biology | CSIR-Centre for Cellular and Molecular Biology | Lamuk Zaveri, Shagufta Khan, Namami Gaur, Sakshi Shambhavi, Tulasi Nagabandi, Purushotham Vodnala, Payel Mukherjee, Sofia Banu, Priya Singh, Dhiviya Vedagiri, Divya Gupta, Vishal Sah, Santosh Kumar Kuncha, Krishnan Harinivas Harshan, Archana Bharadwaj Siva, Karthik Bharadwaj Tallapaka,Umesh Kumar, Unis Ahmad Bhat, Ajay Sarawagi, Priyanka Pant, Rajkanwar Nathawat, Rakesh K Mishra, Divya Tej Sowpati |
| EPI_ISL_458055 | CSIR-Centre for Cellular and Molecular Biology | CSIR-Centre for Cellular and Molecular Biology | Namami Gaur, Sakshi Shambhavi, Lamuk Zaveri, Shagufta Khan, Tulasi Nagabandi, Purushotham Vodnala, Payel Mukherjee, Sofia Banu, Priya Singh, Dhiviya Vedagiri, Divya Gupta, Vishal Sah, Santosh Kumar Kuncha, Krishnan Harinivas Harshan, Archana Bharadwaj Siva, Karthik Bharadwaj Tallapaka, Nikhil Hajirnis, Pratheusa Maccha, M Soujanya Reddy,G. Aditya Kumar, Koushick Sivakumar, Rakesh K Mishra, Divya Tej Sowpati |
| EPI_ISL_458056 | CSIR-Centre for Cellular and Molecular Biology | CSIR-Centre for Cellular and Molecular Biology | Tulasi Nagabandi, Namami Gaur, Sakshi Shambhavi, Lamuk Zaveri, Shagufta Khan, Purushotham Vodnala, Payel Mukherjee, Sofia Banu, Priya Singh, Dhiviya Vedagiri, Divya Gupta, Vishal Sah, Santosh Kumar Kuncha, Krishnan Harinivas Harshan, Archana Bharadwaj Siva, Karthik Bharadwaj Tallapaka,G. Aditya Kumar, Koushick Sivakumar, Pooja Ramesh Gupta, Rajan Kumar Jha, Shraddha Vijay Lahoti, Rakesh K Mishra, Divya Tej Sowpati |
| EPI_ISL_458057 | CSIR-Centre for Cellular and Molecular Biology | CSIR-Centre for Cellular and Molecular Biology | Payel Mukherjee, Sofia Banu, Priya Singh, Dhiviya Vedagiri, Divya Gupta, Vishal Sah, Santosh Kumar Kuncha, Krishnan Harinivas Harshan, Archana Bharadwaj Siva, Karthik Bharadwaj Tallapaka, Shagufta Khan, Lamuk Zaveri, Namami Gaur, Sakshi Shambhavi, Tulasi Nagabandi, Purushotham Vodnala,Deepak Kumar, Devi Prasad Vijayashankar, Disha Nanda, Divya Das, Jotin Gogoi, Manish Bhattacharjee, Rakesh K Mishra, Divya Tej Sowpati |
| EPI_ISL_458058 | CSIR-Centre for Cellular and Molecular Biology | CSIR-Centre for Cellular and Molecular Biology | Sofia Banu, Payel Mukherjee, Priya Singh, Dhiviya Vedagiri, Divya Gupta, Vishal Sah, Santosh Kumar Kuncha, Krishnan Harinivas Harshan, Archana Bharadwaj Siva, Karthik Bharadwaj Tallapaka, Shagufta Khan, Lamuk Zaveri, Namami Gaur, Sakshi Shambhavi, Tulasi Nagabandi, Purushotham Vodnala, Disha Nanda, Divya Das, Jotin Gogoi, Manish Bhattacharjee, Ravi Prasad Mukku, Rakesh K Mishra, Divya Tej Sowpati |
| EPI_ISL_458059 | CSIR-Centre for Cellular and Molecular Biology | CSIR-Centre for Cellular and Molecular Biology | Shagufta Khan, Lamuk Zaveri, Namami Gaur, Sakshi Shambhavi, Tulasi Nagabandi, Purushotham Vodnala, Payel Mukherjee, Sofia Banu, Priya Singh, Dhiviya Vedagiri, Divya Gupta, Vishal Sah, Santosh Kumar Kuncha, Krishnan Harinivas Harshan, Archana Bharadwaj Siva, Karthik Bharadwaj Tallapaka, Renu Sudhakar, Somesh Gorde, Gangumala Srinivas Reddy, Sujoy Deb, Swati Bayyana, Rakesh K Mishra, Divya Tej Sowpati |
| EPI_ISL_458060 | CSIR-Centre for Cellular and Molecular Biology | CSIR-Centre for Cellular and Molecular Biology | Lamuk Zaveri, Shagufta Khan, Namami Gaur, Sakshi Shambhavi, Tulasi Nagabandi, Purushotham Vodnala, Payel Mukherjee, Sofia Banu, Priya Singh, Dhiviya Vedagiri, Divya Gupta, Vishal Sah, Santosh Kumar Kuncha, Krishnan Harinivas Harshan, Archana Bharadwaj Siva, Karthik Bharadwaj Tallapaka,Zeba Rizvi, Zuberwasim Sayyad, Kakade Aishwarya Arun, Amrutha H C, Ananga Ghosh, Rakesh K Mishra, Divya Tej Sowpati |
| EPI_ISL_458061 | CSIR-Centre for Cellular and Molecular Biology | CSIR-Centre for Cellular and Molecular Biology | Namami Gaur, Sakshi Shambhavi, Lamuk Zaveri, Shagufta Khan, Tulasi Nagabandi, Purushotham Vodnala, Payel Mukherjee, Sofia Banu, Priya Singh, Dhiviya Vedagiri, Divya Gupta, Vishal Sah, Santosh Kumar Kuncha, Krishnan Harinivas Harshan, Archana Bharadwaj Siva, Karthik Bharadwaj Tallapaka,Kezia J Ann, Radhika Khandelwal, Roshan Maku Venkata, Shemin Mansuri, Sonu Uday, Rakesh K Mishra, Divya Tej Sowpati |
| EPI_ISL_458062 | CSIR-Centre for Cellular and Molecular Biology | CSIR-Centre for Cellular and Molecular Biology | Payel Mukherjee, Sofia Banu, Priya Singh, Dhiviya Vedagiri, Divya Gupta, Vishal Sah, Santosh Kumar Kuncha, Krishnan Harinivas Harshan, Archana Bharadwaj Siva, Karthik Bharadwaj Tallapaka, Shagufta Khan, Lamuk Zaveri, Namami Gaur, Sakshi Shambhavi, Tulasi Nagabandi, Purushotham Vodnala, Rakesh K Mishra, Sonu Uday, Sudipta Mondal, Annapoorna P Karthyayani, Debabrata Jana, Debrya Saha, Divya Tej Sowpati |
| EPI_ISL_458063 | CSIR-Centre for Cellular and Molecular Biology | CSIR-Centre for Cellular and Molecular Biology | Sofia Banu, Payel Mukherjee, Priya Singh, Dhiviya Vedagiri, Divya Gupta, Vishal Sah, Santosh Kumar Kuncha, Krishnan Harinivas Harshan, Archana Bharadwaj Siva, Karthik Bharadwaj Tallapaka, Shagufta Khan, Lamuk Zaveri, Namami Gaur, Sakshi Shambhavi, Tulasi Nagabandi, Purushotham Vodnala, Gokulan C G, Gunjan Purohit, Hanuman Tulashiram Kale, Pankaj Kumar, Prachand Issarapu, Rakesh K Mishra, Divya Tej Sowpati |
| EPI_ISL_458064 | CSIR-Centre for Cellular and Molecular Biology | CSIR-Centre for Cellular and Molecular Biology | Shagufta Khan, Lamuk Zaveri, Namami Gaur, Sakshi Shambhavi, Tulasi Nagabandi, Purushotham Vodnala, Payel Mukherjee, Sofia Banu, Priya Singh, Dhiviya Vedagiri, Divya Gupta, Vishal Sah, Santosh Kumar Kuncha, Krishnan Harinivas Harshan, Archana Bharadwaj Siva, Karthik Bharadwaj Tallapaka,Preethi Jampala, Sharada Ravi Iyer, Sulagana Mukherjee, Swetha Sundar, Peddapuvala Sai Uday Kiran Rakesh K Mishra, Divya Tej Sowpati |
| EPI_ISL_458065 | CSIR-Centre for Cellular and Molecular Biology | CSIR-Centre for Cellular and Molecular Biology | Lamuk Zaveri, Shagufta Khan, Namami Gaur, Sakshi Shambhavi, Tulasi Nagabandi, Purushotham Vodnala, Payel Mukherjee, Sofia Banu, Priya Singh, Dhiviya Vedagiri, Divya Gupta, Vishal Sah, Santosh Kumar Kuncha, Krishnan Harinivas Harshan, Archana Bharadwaj Siva, Karthik Bharadwaj Tallapaka,Umesh Kumar, Unis Ahmad Bhat, Ajay Sarawagi, Priyanka Pant, Rajkanwar Nathawat, Rakesh K Mishra, Divya Tej Sowpati |
| EPI_ISL_458066, EPI_ISL_458067, EPI_ISL_458068, EPI_ISL_458069 | Osmania Medical College | CSIR-Centre for Cellular and Molecular Biology | Shashikala Reddy, Mahboob Khan,Payel Mukherjee, Sofia Banu, Priya Singh, Dhiviya Vedagiri, Divya Gupta, Vishal Sah, Santosh Kumar Kuncha, Krishnan Harinivas Harshan, Archana Bharadwaj Siva, Karthik Bharadwaj Tallapaka, Shagufta Khan, Lamuk Zaveri, Namami Gaur, Sakshi Shambhavi, Tulasi Nagabandi, Purushotham Vodnala, Rakesh K Mishra, Divya Tej Sowpati |
| EPI_ISL_458070 | CSIR-Centre for Cellular and Molecular Biology | CSIR-Centre for Cellular and Molecular Biology | Sakshi Shambhavi, Lamuk Zaveri, Shagufta Khan, Namami Gaur, Tulasi Nagabandi, Purushotham Vodnala, Payel Mukherjee, Sofia Banu, Priya Singh, Dhiviya Vedagiri, Divya Gupta, Vishal Sah, Santosh Kumar Kuncha, Krishnan Harinivas Harshan, Archana Bharadwaj Siva, Karthik Bharadwaj Tallapaka,Nikhil Hajirnis, Pratheusa Maccha, M Soujanya Reddy,G. Aditya Kumar, Koushick Sivakumar,Disha Nanda, Divya Das, Jotin Gogoi, Manish Bhattacharjee, Ravi Prasad Mukku, Rakesh K Mishra, Divya Tej Sowpati |
| EPI_ISL_458071 | CSIR-Centre for Cellular and Molecular Biology | CSIR-Centre for Cellular and Molecular Biology | Sakshi Shambhavi, Lamuk Zaveri, Shagufta Khan, Namami Gaur, Tulasi Nagabandi, Purushotham Vodnala, Payel Mukherjee, Sofia Banu, Priya Singh, Dhiviya Vedagiri, Divya Gupta, Vishal Sah, Santosh Kumar Kuncha, Krishnan Harinivas Harshan, Archana Bharadwaj Siva, Karthik Bharadwaj Tallapaka,Nikhil Hajirnis, Pratheusa Maccha, M Soujanya Reddy,G. Aditya Kumar, Koushick Sivakumar, Rakesh K Mishra, Divya Tej Sowpati |
| EPI_ISL_458072, EPI_ISL_458073, EPI_ISL_458074, EPI_ISL_458075, EPI_ISL_458076, EPI_ISL_458077 | CSIR-Centre for Cellular and Molecular Biology | CSIR-Centre for Cellular and Molecular Biology | Dhiviya Vedagiri, Divya Gupta, Vishal Sah, Payel Mukherjee, Sofia Banu, Priya Singh, Santosh Kumar Kuncha, Archana Bharadwaj Siva, Karthik Bharadwaj Tallapaka, Shagufta Khan, Lamuk Zaveri, Namami Gaur, Sakshi Shambhavi, Tulasi Nagabandi, Purushotham Vodnala, Rakesh K Mishra, Divya Tej Sowpati, Krishnan Harinivas Harshan |
| EPI_ISL_458079 | Mitra Keluarga Hospital Kenjeran | Institute of Tropical Disease, Universitas Airlangga | Aldise M Nastri, Jezzy R Dewantari, Rima R Prasetya, Krisnoadi Rahardjo, Anastasia W Jefuna, Gatot Soegiarto, Laksmi Wulandari, Retno A Setyoningrum, Resti Yudhawati, Yohko K Shimizu, Mitsuhiro Nishimura, Yasuko Mori, Soetjipto, Kazufumi Shimizu, Maria I Lusida |
| EPI_ISL_458080 | CSIR-Centre for Cellular and Molecular Biology | CSIR-Centre for Cellular and Molecular Biology | Sakshi Shambhavi, Lamuk Zaveri, Shagufta Khan, Namami Gaur, Tulasi Nagabandi, Purushotham Vodnala, Payel Mukherjee, Sofia Banu, Priya Singh, Dhiviya Vedagiri, Divya Gupta, Vishal Sah, Santosh Kumar Kuncha, Krishnan Harinivas Harshan, Archana Bharadwaj Siva, Karthik Bharadwaj Tallapaka, G. Aditya Kumar, Koushick Sivakumar, Pooja Ramesh Gupta, Rajan Kumar Jha, Shraddha Vijay Lahoti, Rakesh K Mishra, Divya Tej Sowpati |
| EPI_ISL_458081 | RSUD Bangil Pasuruan | Institute of Tropical Disease, Universitas Airlangga | Jezzy R Dewantari, Rima R Prasetya, Krisnoadi Rahardjo, Aldise M Nastri, Arma Roosalina, Gatot Soegiarto, Laksmi Wulandari, Retno A Setyoningrum, Resti Yudhawati, Yohko K Shimizu, Mitsuhiro Nishimura, Yasuko Mori, Soetjipto, Kazufumi Shimizu, Maria I Lusida |
| EPI_ISL_458083 | Adi Husada Undaan Hospital | Institute of Tropical Disease, Universitas Airlangga | Rima R Prasetya, Krisnoadi Rahardjo, Aldise M Nastri, Jezzy R Dewantari, Irawati Marga, Gatot Soegiarto, Laksmi Wulandari, Retno A Setyoningrum, Resti Yudhawati, Yohko K Shimizu, Mitsuhiro Nishimura, Yasuko Mori, Soetjipto, Kazufumi Shimizu, Maria I Lusida |
| EPI_ISL_458084 | Laboratorio Biologia Molecolare Sars Cov2 - UOC Laboratorio Analisi - Servizio Medicina di Laboratorio, Ospedale "San Francesco" - ATS-ASSL Nuoro | Laboratorio specialistico UOC Ematologia - Ospedale "San Francesco" - ATS-ASSL Nuoro | Piras Giovanna, Fancello Tatiana, Asproni Rosanna, Fiamma Maura, Monne Maria Itria, Toja Alessandro, Sanna Filomena, Floris Anna Rita, Sulis Vincenzo, Palmas Angelo Domenico, Casu Gavino, Lo Maglio Iana, Mameli Giuseppe. |
| EPI_ISL_458085 | Laboratorio Biologia Molecolare Sars Cov2 - UOC Laboratorio Analisi - Servizio Medicina di Laboratorio , Ospedale "San Francesco" - ATS- ASSL Nuoro | Laboratorio specialistico UOC Ematologia - Ospedale "San Francesco" - ATS-ASSL Nuoro | Piras Giovanna, Fancello Tatiana, Asproni Rosanna, Fiamma Maura, Monne Maria Itria, Toja Alessandro, Sanna Filomena, Floris Anna Rita, Sulis Vincenzo, Palmas Angelo Domenico, Casu Gavino, Lo Maglio Iana, Mameli Giuseppe. |
| EPI_ISL_458086 | B.J. Medical College and Civil hospital | Gujarat Biotechnology Research Centre | Dhaval Vaghela, Ramesh Patel, Pranay Shah, Kamlesh J Upadhyay, Ramesh Pandit, Tejas Shah, Ankit Hinsu, Pritesh Sabara, Apurvasinh Puvar, Janvi Raval, Zarna Patel, Monika Gandhi, Pinal Trivedi, Maharshi Pandya, Amit Kanani, Nidhi Patel, Nitin Savaliya, Raghawendra Kumar, Dinesh Kumar, Zuber Saiyed, Komal Patel, Labdhi Pandya, Snehal Bagatharia, Neha Rajpara, Bhavesh Modi, Gaurishankar Shrimali, R D Dixit, A M Kadri, Umang Mishra, Chaitanya Joshi, Madhvi Joshi |
| EPI_ISL_458087 | B.J. Medical College and Civil hospital | Gujarat Biotechnology Research Centre | Ramesh Patel, Pranay Shah, Kamlesh J Upadhyay, Ramesh Pandit, Tejas Shah, Ankit Hinsu, Pritesh Sabara, Apurvasinh Puvar, Janvi Raval, Zarna Patel, Monika Gandhi, Pinal Trivedi, Maharshi Pandya, Amit Kanani, Nidhi Patel, Nitin Savaliya, Raghawendra Kumar, Dinesh Kumar, Zuber Saiyed, Komal Patel, Labdhi Pandya, Snehal Bagatharia, Dhaval Vaghela, Afzal Ansari, Bhavesh Modi, Gaurishankar Shrimali, R D Dixit, A M Kadri, Umang Mishra, Chaitanya Joshi, Madhvi Joshi |
| EPI_ISL_458088 | B.J. Medical College and Civil hospital | Gujarat Biotechnology Research Centre | Pranay Shah, Kamlesh J Upadhyay, Ramesh Pandit, Tejas Shah, Ankit Hinsu, Pritesh Sabara, Apurvasinh Puvar, Janvi Raval, Zarna Patel, Monika Gandhi, Pinal Trivedi, Maharshi Pandya, Amit Kanani, Nidhi Patel, Nitin Savaliya, Raghawendra Kumar, Dinesh Kumar, Zuber Saiyed, Komal Patel, Labdhi Pandya, Snehal Bagatharia, Dhaval Vaghela, Ramesh Patel, Fenil Patel, Bhavesh Modi, Gaurishankar Shrimali, R D Dixit, A M Kadri, Umang Mishra, Chaitanya Joshi, Madhvi Joshi |
| EPI_ISL_458089 | B.J. Medical College and Civil hospital | Gujarat Biotechnology Research Centre | Pranay Shah, Kamlesh J Upadhyay, Ramesh Pandit, Tejas Shah, Ankit Hinsu, Pritesh Sabara, Apurvasinh Puvar, Janvi Raval, Zarna Patel, Monika Gandhi, Pinal Trivedi, Maharshi Pandya, Amit Kanani, Nidhi Patel, Nitin Savaliya, Raghawendra Kumar, Dinesh Kumar, Zuber Saiyed, Komal Patel, Labdhi Pandya, Snehal Bagatharia, Dhaval Vaghela, Ramesh Patel, Neelam Nathani, Bhavesh Modi, Gaurishankar Shrimali, R D Dixit, A M Kadri, Umang Mishra, Chaitanya Joshi, Madhvi Joshi |
| EPI_ISL_458090 | B.J. Medical College and Civil hospital | Gujarat Biotechnology Research Centre | Kamlesh J Upadhyay, Ramesh Pandit, Tejas Shah, Ankit Hinsu, Pritesh Sabara, Apurvasinh Puvar, Janvi Raval, Zarna Patel, Monika Gandhi, Pinal Trivedi, Maharshi Pandya, Amit Kanani, Nidhi Patel, Nitin Savaliya, Raghawendra Kumar, Dinesh Kumar, Zuber Saiyed, Komal Patel, Labdhi Pandya, Snehal Bagatharia, Dhaval Vaghela, Ramesh Patel, Pranay Shah, Armi Chaudhari, Bhavesh Modi, Gaurishankar Shrimali, R D Dixit, A M Kadri, Umang Mishra, Chaitanya Joshi, Madhvi Joshi |
| EPI_ISL_458091 | B.J. Medical College and Civil hospital | Gujarat Biotechnology Research Centre | Maharshi Pandya, Amit Kanani, Nidhi Patel, Nitin Savaliya, Raghawendra Kumar, Dinesh Kumar, Zuber Saiyed, Komal Patel, Labdhi Pandya, Snehal Bagatharia, Dhaval Vaghela, Ramesh Patel, Pranay Shah, Kamlesh J Upadhyay, Ramesh Pandit, Tejas Shah, Ankit Hinsu, Pritesh Sabara, Apurvasinh Puvar, Janvi Raval, Zarna Patel, Monika Gandhi, Pinal Trivedi, Bhavya Jindal, Bhavesh Modi, Gaurishankar Shrimali, R D Dixit, A M Kadri, Umang Mishra, Chaitanya Joshi, Madhvi Joshi |
| EPI_ISL_458092 | B.J. Medical College and Civil hospital | Gujarat Biotechnology Research Centre | Amit Kanani, Nidhi Patel, Nitin Savaliya, Raghawendra Kumar, Dinesh Kumar, Zuber Saiyed, Komal Patel, Labdhi Pandya, Snehal Bagatharia, Dhaval Vaghela, Ramesh Patel, Pranay Shah, Kamlesh J Upadhyay, Ramesh Pandit, Tejas Shah, Ankit Hinsu, Pritesh Sabara, Apurvasinh Puvar, Janvi Raval, Zarna Patel, Monika Gandhi, Pinal Trivedi, Maharshi Pandya, Camellia Chakraborty, Bhavesh Modi, Gaurishankar Shrimali, R D Dixit, A M Kadri, Umang Mishra, Chaitanya Joshi, Madhvi Joshi |
| EPI_ISL_458093 | B.J. Medical College and Civil hospital | Gujarat Biotechnology Research Centre | Nidhi Patel, Nitin Savaliya, Raghawendra Kumar, Dinesh Kumar, Zuber Saiyed, Komal Patel, Labdhi Pandya, Snehal Bagatharia, Dhaval Vaghela, Ramesh Patel, Pranay Shah, Kamlesh J Upadhyay, Ramesh Pandit, Tejas Shah, Ankit Hinsu, Pritesh Sabara, Apurvasinh Puvar, Janvi Raval, Zarna Patel, Monika Gandhi, Pinal Trivedi, Maharshi Pandya, Amit Kanani, Siddhant Kumar, Bhavesh Modi, Gaurishankar Shrimali, R D Dixit, A M Kadri, Umang Mishra, Chaitanya Joshi, Madhvi Joshi |
| EPI_ISL_458094 | B.J. Medical College and Civil hospital | Gujarat Biotechnology Research Centre | Nitin Savaliya, Raghawendra Kumar, Dinesh Kumar, Zuber Saiyed, Komal Patel, Labdhi Pandya, Snehal Bagatharia, Dhaval Vaghela, Ramesh Patel, Pranay Shah, Kamlesh J Upadhyay, Ramesh Pandit, Tejas Shah, Ankit Hinsu, Pritesh Sabara, Apurvasinh Puvar, Janvi Raval, Zarna Patel, Monika Gandhi, Pinal Trivedi, Maharshi Pandya, Amit Kanani, Nidhi Patel, Priyanka P Vatsa, Bhavesh Modi, Gaurishankar Shrimali, R D Dixit, A M Kadri, Umang Mishra, Chaitanya Joshi, Madhvi Joshi |
| EPI_ISL_458095 | B.J. Medical College and Civil hospital | Gujarat Biotechnology Research Centre | Raghawendra Kumar, Dinesh Kumar, Zuber Saiyed, Komal Patel, Labdhi Pandya, Snehal Bagatharia, Dhaval Vaghela, Ramesh Patel, Pranay Shah, Kamlesh J Upadhyay, Ramesh Pandit, Tejas Shah, Ankit Hinsu, Pritesh Sabara, Apurvasinh Puvar, Janvi Raval, Zarna Patel, Monika Gandhi, Pinal Trivedi, Maharshi Pandya, Amit Kanani, Nidhi Patel, Nitin Savaliya, Pooja P Doshi, Bhavesh Modi, Gaurishankar Shrimali, R D Dixit, A M Kadri, Umang Mishra, Chaitanya Joshi, Madhvi Joshi |

| | | | |
|---|---|---|---|
| EPI_ISL_458096 | B.J. Medical College and Civil hospital | Gujarat Biotechnology Research Centre | Dinesh Kumar, Zuber Saiyed, Komal Patel, Labdhi Pandya, Snehal Bagatharia, Dhaval Vaghela, Ramesh Patel, Pranay Shah, Kamlesh J Upadhyay, Ramesh Pandit, Tejas Shah, Ankit Hinsu, Pritesh Sabara, Apurvasinh Puvar, Janvi Raval, Zarna Patel, Monika Gandhi, Pinal Trivedi, Maharshi Pandya, Amit Kanani, Nidhi Patel, Nitin Savaliya, Raghawendra Kumar, Akanksha Verma, Bhavesh Modi, Gaurishankar Shrimali, R D Dixit, A M Kadri, Umang Mishra, Chaitanya Joshi, Madhvi Joshi |
| EPI_ISL_458097 | B.J. Medical College and Civil hospital | Gujarat Biotechnology Research Centre | Zuber Saiyed, Komal Patel, Labdhi Pandya, Snehal Bagatharia, Dhaval Vaghela, Ramesh Patel, Pranay Shah, Kamlesh J Upadhyay, Ramesh Pandit, Tejas Shah, Ankit Hinsu, Pritesh Sabara, Apurvasinh Puvar, Janvi Raval, Zarna Patel, Monika Gandhi, Pinal Trivedi, Maharshi Pandya, Amit Kanani, Nidhi Patel, Nitin Savaliya, Raghawendra Kumar, Dinesh Kumar, Priti Pandita, Bhavesh Modi, Gaurishankar Shrimali, R D Dixit, A M Kadri, Umang Mishra, Chaitanya Joshi, Madhvi Joshi |
| EPI_ISL_458098 | B.J. Medical College and Civil hospital | Gujarat Biotechnology Research Centre | Komal Patel, Labdhi Pandya, Snehal Bagatharia, Dhaval Vaghela, Ramesh Patel, Pranay Shah, Kamlesh J Upadhyay, Ramesh Pandit, Tejas Shah, Ankit Hinsu, Pritesh Sabara, Apurvasinh Puvar, Janvi Raval, Zarna Patel, Monika Gandhi, Pinal Trivedi, Maharshi Pandya, Amit Kanani, Nidhi Patel, Nitin Savaliya, Raghawendra Kumar, Dinesh Kumar, Zuber Saiyed, Pragya Sharma, Bhavesh Modi, Gaurishankar Shrimali, R D Dixit, A M Kadri, Umang Mishra, Chaitanya Joshi, Madhvi Joshi |
| EPI_ISL_458099 | B.J. Medical College and Civil hospital | Gujarat Biotechnology Research Centre | Labdhi Pandya, Snehal Bagatharia, Dhaval Vaghela, Ramesh Patel, Pranay Shah, Kamlesh J Upadhyay, Ramesh Pandit, Tejas Shah, Ankit Hinsu, Pritesh Sabara, Apurvasinh Puvar, Janvi Raval, Zarna Patel, Monika Gandhi, Pinal Trivedi, Maharshi Pandya, Amit Kanani, Nidhi Patel, Nitin Savaliya, Raghawendra Kumar, Dinesh Kumar, Zuber Saiyed, Komal Patel, Neha Rajpara, Bhavesh Modi, Gaurishankar Shrimali, R D Dixit, A M Kadri, Umang Mishra, Chaitanya Joshi, Madhvi Joshi |
| EPI_ISL_458100 | B.J. Medical College and Civil hospital | Gujarat Biotechnology Research Centre | Snehal Bagatharia, Dhaval Vaghela, Ramesh Patel, Pranay Shah, Kamlesh J Upadhyay, Ramesh Pandit, Tejas Shah, Ankit Hinsu, Pritesh Sabara, Apurvasinh Puvar, Janvi Raval, Zarna Patel, Monika Gandhi, Pinal Trivedi, Maharshi Pandya, Amit Kanani, Nidhi Patel, Nitin Savaliya, Raghawendra Kumar, Dinesh Kumar, Zuber Saiyed, Komal Patel, Labdhi Pandya, Afzal Ansari, Bhavesh Modi, Gaurishankar Shrimali, R D Dixit, A M Kadri, Umang Mishra, Chaitanya Joshi, Madhvi Joshi |
| EPI_ISL_458101 | B.J. Medical College and Civil hospital | Gujarat Biotechnology Research Centre | Dhaval Vaghela, Ramesh Patel, Pranay Shah, Kamlesh J Upadhyay, Ramesh Pandit, Tejas Shah, Ankit Hinsu, Pritesh Sabara, Apurvasinh Puvar, Janvi Raval, Zarna Patel, Monika Gandhi, Pinal Trivedi, Maharshi Pandya, Amit Kanani, Nidhi Patel, Nitin Savaliya, Raghawendra Kumar, Dinesh Kumar, Zuber Saiyed, Komal Patel, Labdhi Pandya, Snehal Bagatharia, Fenil Patel, Bhavesh Modi, Gaurishankar Shrimali, R D Dixit, A M Kadri, Umang Mishra, Chaitanya Joshi, Madhvi Joshi |
| EPI_ISL_458102 | B.J. Medical College and Civil hospital | Gujarat Biotechnology Research Centre | Ramesh Patel, Pranay Shah, Kamlesh J Upadhyay, Ramesh Pandit, Tejas Shah, Ankit Hinsu, Pritesh Sabara, Apurvasinh Puvar, Janvi Raval, Zarna Patel, Monika Gandhi, Pinal Trivedi, Maharshi Pandya, Amit Kanani, Nidhi Patel, Nitin Savaliya, Raghawendra Kumar, Dinesh Kumar, Zuber Saiyed, Komal Patel, Labdhi Pandya, Snehal Bagatharia, Dhaval Vaghela, Neelam Nathani, Bhavesh Modi, Gaurishankar Shrimali, R D Dixit, A M Kadri, Umang Mishra, Chaitanya Joshi, Madhvi Joshi |
| EPI_ISL_458103 | Gujarat Biotechnology Research Centre | Gujarat Biotechnology Research Centre | Ramesh Pandit, Tejas Shah, Ankit Hinsu, Pritesh Sabara, Apurvasinh Puvar, Janvi Raval, Zarna Patel, Monika Gandhi, Pinal Trivedi, Maharshi Pandya, Amit Kanani, Nidhi Patel, Nitin Savaliya, Raghawendra Kumar, Dinesh Kumar, Zuber Saiyed, Komal Patel, Labdhi Pandya, Snehal Bagatharia, Armi Chaudhari, Bhavesh Modi, Gaurishankar Shrimali, R D Dixit, A M Kadri, Umang Mishra, Chaitanya Joshi, Madhvi Joshi, , , , , |
| EPI_ISL_458104 | Gujarat Biotechnology Research Centre | Gujarat Biotechnology Research Centre | Tejas Shah, Ankit Hinsu, Pritesh Sabara, Apurvasinh Puvar, Janvi Raval, Zarna Patel, Monika Gandhi, Pinal Trivedi, Maharshi Pandya, Amit Kanani, Nidhi Patel, Nitin Savaliya, Raghawendra Kumar, Dinesh Kumar, Zuber Saiyed, Komal Patel, Labdhi Pandya, Snehal Bagatharia, Ramesh Pandit, Bhavya Jindal, Bhavesh Modi, Gaurishankar Shrimali, R D Dixit, A M Kadri, Umang Mishra, Chaitanya Joshi, Madhvi Joshi, , , , |
| EPI_ISL_458105 | Gujarat Biotechnology Research Centre | Gujarat Biotechnology Research Centre | Ankit Hinsu, Pritesh Sabara, Apurvasinh Puvar, Janvi Raval, Zarna Patel, Monika Gandhi, Pinal Trivedi, Maharshi Pandya, Amit Kanani, Nidhi Patel, Nitin Savaliya, Raghawendra Kumar, Dinesh Kumar, Zuber Saiyed, Komal Patel, Labdhi Pandya, Snehal Bagatharia, Ramesh Pandit, Tejas Shah, Camellia Chakraborty, Bhavesh Modi, Gaurishankar Shrimali, R D Dixit, A M Kadri, Umang Mishra, Chaitanya Joshi, Madhvi Joshi, , , , |
| EPI_ISL_458106 | Gujarat Biotechnology Research Centre | Gujarat Biotechnology Research Centre | Pritesh Sabara, Apurvasinh Puvar, Janvi Raval, Zarna Patel, Monika Gandhi, Pinal Trivedi, Maharshi Pandya, Amit Kanani, Nidhi Patel, Nitin Savaliya, Raghawendra Kumar, Dinesh Kumar, Zuber Saiyed, Komal Patel, Labdhi Pandya, Snehal Bagatharia, Ramesh Pandit, Tejas Shah, Ankit Hinsu, Siddhant Kumar, Bhavesh Modi, Gaurishankar Shrimali, R D Dixit, A M Kadri, Umang Mishra, Chaitanya Joshi, Madhvi Joshi, , , , |
| EPI_ISL_458107 | Gujarat Biotechnology Research Centre | Gujarat Biotechnology Research Centre | Apurvasinh Puvar, Janvi Raval, Zarna Patel, Monika Gandhi, Pinal Trivedi, Maharshi Pandya, Amit Kanani, Nidhi Patel, Nitin Savaliya, Raghawendra Kumar, Dinesh Kumar, Zuber Saiyed, Komal Patel, Labdhi Pandya, Snehal Bagatharia, Ramesh Pandit, Tejas Shah, Ankit Hinsu, Pritesh Sabara, Priyanka P Vatsa, Bhavesh Modi, Gaurishankar Shrimali, R D Dixit, A M Kadri, Umang Mishra, Chaitanya Joshi, Madhvi Joshi, , , , |
| EPI_ISL_458108 | Gujarat Biotechnology Research Centre | Gujarat Biotechnology Research Centre | Janvi Raval, Zarna Patel, Monika Gandhi, Pinal Trivedi, Maharshi Pandya, Amit Kanani, Nidhi Patel, Nitin Savaliya, Raghawendra Kumar, Dinesh Kumar, Zuber Saiyed, Komal Patel, Labdhi Pandya, Snehal Bagatharia, Ramesh Pandit, Tejas Shah, Ankit Hinsu, Pritesh Sabara, Apurvasinh Puvar, Pooja P Doshi, Bhavesh Modi, Gaurishankar Shrimali, R D Dixit, A M Kadri, Umang Mishra, Chaitanya Joshi, Madhvi Joshi, , , , |
| EPI_ISL_458109 | Gujarat Biotechnology Research Centre | Gujarat Biotechnology Research Centre | Zarna Patel, Monika Gandhi, Pinal Trivedi, Maharshi Pandya, Amit Kanani, Nidhi Patel, Nitin Savaliya, Raghawendra Kumar, Dinesh Kumar, Zuber Saiyed, Komal Patel, Labdhi Pandya, Snehal Bagatharia, Ramesh Pandit, Tejas Shah, Ankit Hinsu, Pritesh Sabara, Apurvasinh Puvar, Janvi Raval, Akanksha Verma, Bhavesh Modi, Gaurishankar Shrimali, R D Dixit, A M Kadri, Umang Mishra, Chaitanya Joshi, Madhvi Joshi, , , , |
| EPI_ISL_458110 | Gujarat Biotechnology Research Centre | Gujarat Biotechnology Research Centre | Monika Gandhi, Pinal Trivedi, Maharshi Pandya, Amit Kanani, Nidhi Patel, Nitin Savaliya, Raghawendra Kumar, Dinesh Kumar, Zuber Saiyed, Komal Patel, Labdhi Pandya, Snehal Bagatharia, Ramesh Pandit, Tejas Shah, Ankit Hinsu, Pritesh Sabara, Apurvasinh Puvar, Janvi Raval, Zarna Patel, Priti Pandita, Bhavesh Modi, Gaurishankar Shrimali, R D Dixit, A M Kadri, Umang Mishra, Chaitanya Joshi, Madhvi Joshi, , , , |
| EPI_ISL_458111 | Gujarat Biotechnology Research Centre | Gujarat Biotechnology Research Centre | Pinal Trivedi, Maharshi Pandya, Amit Kanani, Nidhi Patel, Nitin Savaliya, Raghawendra Kumar, Dinesh Kumar, Zuber Saiyed, Komal Patel, Labdhi Pandya, Snehal Bagatharia, Ramesh Pandit, Tejas Shah, Ankit Hinsu, Pritesh Sabara, Apurvasinh Puvar, Janvi Raval, Zarna Patel, Monika Gandhi, Pragya Sharma, Bhavesh Modi, Gaurishankar Shrimali, R D Dixit, A M Kadri, Umang Mishra, Chaitanya Joshi, Madhvi Joshi, , , , |
| EPI_ISL_458112 | Gujarat Biotechnology Research Centre | Gujarat Biotechnology Research Centre | Maharshi Pandya, Amit Kanani, Nidhi Patel, Nitin Savaliya, Raghawendra Kumar, Dinesh Kumar, Zuber Saiyed, Komal Patel, Labdhi Pandya, Snehal Bagatharia, Ramesh Pandit, Tejas Shah, Ankit Hinsu, Pritesh Sabara, Apurvasinh Puvar, Janvi Raval, Zarna Patel, Monika Gandhi, Pinal Trivedi, Neha Rajpara, Bhavesh Modi, Gaurishankar Shrimali, R D Dixit, A M Kadri, Umang Mishra, Chaitanya Joshi, Madhvi Joshi, , , , |
| EPI_ISL_458113 | Gujarat Biotechnology Research Centre | Gujarat Biotechnology Research Centre | Amit Kanani, Nidhi Patel, Nitin Savaliya, Raghawendra Kumar, Dinesh Kumar, Zuber Saiyed, Komal Patel, Labdhi Pandya, Snehal Bagatharia, Ramesh Pandit, Tejas Shah, Ankit Hinsu, Pritesh Sabara, Apurvasinh Puvar, Janvi Raval, Zarna Patel, Monika Gandhi, Pinal Trivedi, Maharshi Pandya, Afzal Ansari, Bhavesh Modi, Gaurishankar Shrimali, R D Dixit, A M Kadri, Umang Mishra, Chaitanya Joshi, Madhvi Joshi, , , , |
| EPI_ISL_458116, EPI_ISL_458117, EPI_ISL_458118, EPI_ISL_458119, EPI_ISL_458120, EPI_ISL_458121, EPI_ISL_458122, EPI_ISL_458123, EPI_ISL_458124 | Oman National Influenza Centre | Department of Microbiology and Immunology-SQUH | Fahad Zadjali, Samira Al-Maruqi, Amina Al Jardani, Khulood Al-Mammary, Hanan Al-kindi, Fatma BaAlawi, Hamida Al Barwani, Zeyana AL-Dahmani, Intisar Al-Shukri, Aisha Al-Busaidi, Aisha Al-Amri, Ahlam Al-Amri, Mohammed Al-Tobi, Samiha Al Kharusi, Abdulla Balkhair |
| EPI_ISL_458130, EPI_ISL_458131 | Hospital Universitari Vall d'Hebron - Vall d'Hebron Institut de Recerca | Hospital Universitari Vall d'Hebron | Cristina Andrés, Maria Piñana, Damir Garcia-Cehic, Mercedes Guerrero-Murillo, Ariadna Rando, Josep Gregori, Juliana Esperalba, Maria Gema Codina, Maria Carmen Martin, Tomàs Pumarola, Josep Quer, Andrés Antón |
| EPI_ISL_458132 | Hospital Universitari Vall d'Hebron - Vall d'Hebron Institut de Rercerca | Hospital Universitari Vall d'Hebron | Cristina Andrés, Maria Piñana, Damir Garcia-Cehic, Mercedes Guerrero-Murillo, Ariadna Rando, Josep Gregori, Juliana Esperalba, Maria Gema Codina, Maria Carmen Martin, Tomàs Pumarola, Josep Quer, Andrés Antón |
| EPI_ISL_458133 | National Institute of Biotechnology | Bioinformatics Division, National Institute of Biotechnology | Mohammad Uzzal Hossain, Md. Moniruzzaman, Md. Salim Khan, Md. Nazrul Islam, Md. Hadisur Rahman, Arittra Bhattacharjee, Md. Ruhul Amin, Asif Rashid, Chaman Ara Keya, Keshob Chandra Das, Md. Salimullah |
| EPI_ISL_458138, EPI_ISL_458139, EPI_ISL_458140, EPI_ISL_458141, EPI_ISL_458142, EPI_ISL_458143, EPI_ISL_458144, EPI_ISL_458145, EPI_ISL_458146, EPI_ISL_458147, EPI_ISL_458148, EPI_ISL_458149 | | | |
| see above | Evandro Chagas Institute | Evandro Chagas Institute | Santos, M.C.; Silva, A.M.; Junior, W.D.C.; Barbagelata, L.S.; Ferreira, J.A.; Sousa, E.M.A.; da Silva, P.S.; Resque, H.R; Martins, L.C.; Sousa Junior, E.C.;Viana, G.M.R |
| EPI_ISL_458150 | ANOUAL | ANOUAL | Jouali Farah, El Ansari Fatima Zahra, Marchoudi Nabila, Kasmi Yassine, Chenaoui Mohamed, El Aliani Aissam, Benhida Rachid, Azami Nawfel, Kitane Driss Lahlou, Loukman Salma, Fekkak Jamal |
| EPI_ISL_458156, EPI_ISL_458157, EPI_ISL_458158, EPI_ISL_458159, EPI_ISL_458160, EPI_ISL_458161, EPI_ISL_458162, EPI_ISL_458163, EPI_ISL_458164, EPI_ISL_458165, EPI_ISL_458166, EPI_ISL_458167, EPI_ISL_458168, EPI_ISL_458169, EPI_ISL_458170, EPI_ISL_458171, EPI_ISL_458172, EPI_ISL_458173, EPI_ISL_458174, EPI_ISL_458175, EPI_ISL_458176, EPI_ISL_458177, EPI_ISL_458178, EPI_ISL_458179, EPI_ISL_458180, EPI_ISL_458181, EPI_ISL_458182, EPI_ISL_458183, EPI_ISL_458184, EPI_ISL_458185, EPI_ISL_458186, EPI_ISL_458187, EPI_ISL_458188, EPI_ISL_458189, EPI_ISL_458190, EPI_ISL_458191, EPI_ISL_458192, EPI_ISL_458193, EPI_ISL_458194, EPI_ISL_458195, EPI_ISL_458196, EPI_ISL_458197, EPI_ISL_458198, EPI_ISL_458199, EPI_ISL_458200, EPI_ISL_458201, EPI_ISL_458202, EPI_ISL_458203, EPI_ISL_458204, EPI_ISL_458205, EPI_ISL_458206, EPI_ISL_458207, EPI_ISL_458208, EPI_ISL_458209, EPI_ISL_458210, EPI_ISL_458211, EPI_ISL_458212, EPI_ISL_458213, EPI_ISL_458214, EPI_ISL_458215, EPI_ISL_458216, EPI_ISL_458217, EPI_ISL_458218, EPI_ISL_458219, EPI_ISL_458220, EPI_ISL_458221, EPI_ISL_458222, EPI_ISL_458223, EPI_ISL_458224, EPI_ISL_458225, EPI_ISL_458226, EPI_ISL_458227, EPI_ISL_458228, EPI_ISL_458229, EPI_ISL_458230, EPI_ISL_458231, EPI_ISL_458232, EPI_ISL_458233, EPI_ISL_458234, EPI_ISL_458235 | | | |
| see above | KU Leuven, Rega Institute, Clinical and Epidemiological Virology | KU Leuven, Rega Institute, Clinical and Epidemiological Virology | Tony Wawina-Bokalanga, Bert Vanmechelen, Joan Marti-Carerras, Piet Maes |
| EPI_ISL_458237, EPI_ISL_458238, EPI_ISL_458239, EPI_ISL_458240, EPI_ISL_458241, EPI_ISL_458242, EPI_ISL_458243, EPI_ISL_458244, EPI_ISL_458245, EPI_ISL_458246, EPI_ISL_458247, EPI_ISL_458248, EPI_ISL_458249, EPI_ISL_458250, EPI_ISL_458251, EPI_ISL_458252, EPI_ISL_458253, EPI_ISL_458254, EPI_ISL_458255, EPI_ISL_458256, EPI_ISL_458257, EPI_ISL_458258, EPI_ISL_458260, EPI_ISL_458261, EPI_ISL_458262, EPI_ISL_458264, EPI_ISL_458265, EPI_ISL_458266, EPI_ISL_458267, EPI_ISL_458270, EPI_ISL_458271, EPI_ISL_458272, EPI_ISL_458274, EPI_ISL_458275, EPI_ISL_458276, EPI_ISL_458277, EPI_ISL_458278, EPI_ISL_458279, EPI_ISL_458280, EPI_ISL_458281, EPI_ISL_458282, EPI_ISL_458283, EPI_ISL_458284 | | | |
| see above | Scripps Medical Laboratory | Andersen lab at Scripps Research | SEARCH Alliance San Diego with Michael Quigley, Ellen Stefanski, Ian Mchardy |
| EPI_ISL_458286 | unknown | Bundeswehr Institute of Microbiology | Handrick,S., Bestehorn-Willmann,M.S., Eckstein,S., Walter,M.C., Antwerpen,M.H., Rehn,A., Naija,H., Stoecker,K., Woelfel,R. and Ben Moussa,M. |
| EPI_ISL_458287 | Biosafety Department PCL3 | Biosafety Department PCL3 | Lemriss,S., Souiri,A. and El Kabbaj,S. |
| EPI_ISL_458291 | Dirk Dittmer | Dirk Dittmer | Bailey,A.G., Caro-Vegas,C.P., Dittmer,D., Eason,A.B., Juarez,A., Landis,J.T., McNamara,R.P., Miller,M.B., Moorad,R., Pluta,L.J., Seltzer,T.A., Thompson,C., Vahrson,W., Villamor,F. |
| EPI_ISL_458292, EPI_ISL_458293 | Dirk Dittmer | Dirk Dittmer | Aubrey,B.G., Caro-Vegas,C.P., Dittmer,D., Eason,A.B., Juarez,A., Landis,J.T., Mcnamara,R.P., Miller,M.B., Moorad,R., Pluta,L.J., Seltzer,T.A., Thompson,C., Vahrson,W., Villamor,F. |
| EPI_ISL_458294, EPI_ISL_458295, EPI_ISL_458296, EPI_ISL_458297 | Dirk Dittmer | Dirk Dittmer | Bailey,A.G., Caro-Vegas,C.P., Dittmer,D., Eason,A.B., Juarez,A., Landis,J.T., McNamara,R.P., Miller,M.B., Moorad,R., Pluta,L.J., Seltzer,T.A., Thompson,C., Vahrson,W., Villamor,F. |
| EPI_ISL_458298 | CSIR-Centre for Cellular and Molecular Biology | CSIR-Centre for Cellular and Molecular Biology | Sakshi Shambhavi, Lamuk Zaveri, Shagufta Khan, Namami Gaur, Tulasi Nagabandi, Purushotham Vodnala, Payel Mukherjee, Sofia Banu, Priya Singh, Dhiviya Vedagiri, Divya Gupta, Vishal Sah, Santosh Kumar Kuncha, Krishnan Harinivas Harshan, Archana Bharadwaj Siva, Karthik Bharadwaj Tallapaka, Deepak Kumar, Devi Prasad Vijayashankar, Disha Nanda, Divya Das, Jotin Gogoi, Manish Bhattacharjee, Rakesh K Mishra, Divya Tej Sowpati |
| EPI_ISL_458300, EPI_ISL_458301, EPI_ISL_458302, EPI_ISL_458303, EPI_ISL_458304, EPI_ISL_458305, EPI_ISL_458306, EPI_ISL_458308, EPI_ISL_458311, EPI_ISL_458313, EPI_ISL_458316, EPI_ISL_458317, EPI_ISL_458319, EPI_ISL_458320, EPI_ISL_458321, EPI_ISL_458323, EPI_ISL_458324, EPI_ISL_458329, EPI_ISL_458330, EPI_ISL_458332, EPI_ISL_458333, EPI_ISL_458334, EPI_ISL_458337, EPI_ISL_458338, EPI_ISL_458339, EPI_ISL_458341, EPI_ISL_458343, EPI_ISL_458344, EPI_ISL_458345, EPI_ISL_458348, EPI_ISL_458349, EPI_ISL_458350, EPI_ISL_458351, EPI_ISL_458352, EPI_ISL_458353, EPI_ISL_458354, EPI_ISL_458355, EPI_ISL_458357, EPI_ISL_458358, EPI_ISL_458359, EPI_ISL_458360, EPI_ISL_458361, EPI_ISL_458362, EPI_ISL_458363, EPI_ISL_458365, EPI_ISL_458367, EPI_ISL_458368, EPI_ISL_458370, EPI_ISL_458371, EPI_ISL_458372, EPI_ISL_458375, EPI_ISL_458377, EPI_ISL_458379, EPI_ISL_458381, EPI_ISL_458382, EPI_ISL_458384, EPI_ISL_458385, EPI_ISL_458391, EPI_ISL_458394, EPI_ISL_458395, EPI_ISL_458396, EPI_ISL_458397, EPI_ISL_458400, EPI_ISL_458403, EPI_ISL_458405, EPI_ISL_458406, EPI_ISL_458407, EPI_ISL_458410, EPI_ISL_458412, EPI_ISL_458413, EPI_ISL_458414, EPI_ISL_458415, EPI_ISL_458416, EPI_ISL_458419, EPI_ISL_458420, EPI_ISL_458421, EPI_ISL_458422, EPI_ISL_458423, EPI_ISL_458425, EPI_ISL_458426, EPI_ISL_458427, EPI_ISL_458430, EPI_ISL_458431, EPI_ISL_458434, EPI_ISL_458435, EPI_ISL_458436, EPI_ISL_458437, EPI_ISL_458438, EPI_ISL_458439, EPI_ISL_458440, EPI_ISL_458441, EPI_ISL_458442, EPI_ISL_458443, EPI_ISL_458445, EPI_ISL_458446, EPI_ISL_458447, EPI_ISL_458448, EPI_ISL_458449, EPI_ISL_458451, EPI_ISL_458452, EPI_ISL_458453, EPI_ISL_458455, EPI_ISL_458456, EPI_ISL_458457, EPI_ISL_458459, EPI_ISL_458460, EPI_ISL_458461, EPI_ISL_458462, EPI_ISL_458465, EPI_ISL_458466, EPI_ISL_458468, EPI_ISL_458469, EPI_ISL_458470, EPI_ISL_458471, EPI_ISL_458472, EPI_ISL_458474, EPI_ISL_458476, EPI_ISL_458479, EPI_ISL_458480, EPI_ISL_458483, EPI_ISL_458484, EPI_ISL_458485, EPI_ISL_458486, EPI_ISL_458489, EPI_ISL_458490, EPI_ISL_458492, EPI_ISL_458495, EPI_ISL_458496, EPI_ISL_458497, EPI_ISL_458500, EPI_ISL_458501, EPI_ISL_458502, EPI_ISL_458503, EPI_ISL_458507, EPI_ISL_458509, EPI_ISL_458510, EPI_ISL_458511, EPI_ISL_458512, EPI_ISL_458513, EPI_ISL_458515 | | | |
| see above | PHE South West Regional Laboratory, National Infection Service | Wellcome Sanger Institute for the COVID-19 Genomics UK Consortium | Stephanie Hutchings, Hannah Pymont, Dr Peter Muir, Barry Vipond, Rich Hopes; and Alex Alderton, Roberto Amato, Sonia Goncalves, Ewan Harrison, David K. Jackson, Ian Johnston, Dominic Kwiatkowski, Cordelia Langford, John Sillitoe on behalf of the Wellcome Sanger Institute COVID-19 Surveillance Team (http://www.sanger.ac.uk/covid-team) |
| EPI_ISL_458529, EPI_ISL_458533, EPI_ISL_458552, EPI_ISL_458556, EPI_ISL_458559, EPI_ISL_458576 | Department of Pathology, University of Cambridge | Wellcome Sanger Institute for the COVID-19 Genomics UK Consortium | Luke W Meredith, M. Estée Török , Myra Hosmillo, William L. Hamilton, Martin D. Curran, Theresa Feltwell, Grant Hall, Anna Yakovleva, Fahad A Khokhar, Charlotte J. Houldcroft, Laura G Caller, Aminu S. Jahun, Sarah L. Caddy, Ian Goodfellow; and Alex Alderton, Roberto Amato, Sonia Goncalves, Ewan Harrison, David K. Jackson, Ian Johnston, Dominic Kwiatkowski, Cordelia Langford, John Sillitoe on behalf of the Wellcome Sanger Institute COVID-19 Surveillance Team (http://www.sanger.ac.uk/covid-team) |
| EPI_ISL_458579, EPI_ISL_458580, EPI_ISL_458581, EPI_ISL_458583, EPI_ISL_458584, EPI_ISL_458588, EPI_ISL_458589, EPI_ISL_458591, EPI_ISL_458593, EPI_ISL_458597, EPI_ISL_458600, EPI_ISL_458601, EPI_ISL_458602, EPI_ISL_458603, EPI_ISL_458604, EPI_ISL_458605, EPI_ISL_458606, EPI_ISL_458607, EPI_ISL_458609, EPI_ISL_458610, EPI_ISL_458611, EPI_ISL_458612, EPI_ISL_458613, EPI_ISL_458614, EPI_ISL_458615, EPI_ISL_458616, EPI_ISL_458617, EPI_ISL_458618, EPI_ISL_458619, EPI_ISL_458620, EPI_ISL_458621, EPI_ISL_458622, EPI_ISL_458623, EPI_ISL_458624, EPI_ISL_458625, EPI_ISL_458626, EPI_ISL_458628, EPI_ISL_458631, EPI_ISL_458632, EPI_ISL_458633, EPI_ISL_458635, EPI_ISL_458636, EPI_ISL_458637, EPI_ISL_458638, EPI_ISL_458640, EPI_ISL_458641, EPI_ISL_458642, EPI_ISL_458643, EPI_ISL_458644, EPI_ISL_458645, EPI_ISL_458646, EPI_ISL_458647, EPI_ISL_458648, EPI_ISL_458649, EPI_ISL_458650, EPI_ISL_458651, EPI_ISL_458652, EPI_ISL_458653, EPI_ISL_458654, EPI_ISL_458656, EPI_ISL_458657, EPI_ISL_458660, EPI_ISL_458661, EPI_ISL_458662, EPI_ISL_458664, EPI_ISL_458667, EPI_ISL_458668, EPI_ISL_458669, EPI_ISL_458670, EPI_ISL_458671, EPI_ISL_458672, EPI_ISL_458674, EPI_ISL_458675, EPI_ISL_458676, EPI_ISL_458677, EPI_ISL_458678, EPI_ISL_458679, EPI_ISL_458681, EPI_ISL_458684, EPI_ISL_458686, EPI_ISL_458687, EPI_ISL_458688, EPI_ISL_458689, EPI_ISL_458690, EPI_ISL_458691, EPI_ISL_458693, EPI_ISL_458694, EPI_ISL_458696, EPI_ISL_458697, EPI_ISL_458699, EPI_ISL_458700, EPI_ISL_458701, EPI_ISL_458702, EPI_ISL_458703, EPI_ISL_458706, EPI_ISL_458707, EPI_ISL_458708, EPI_ISL_458709, EPI_ISL_458710, EPI_ISL_458711, EPI_ISL_458713, EPI_ISL_458714, EPI_ISL_458715, EPI_ISL_458717, EPI_ISL_458718 | | | |
| see above | NU-OMICS DNA Sequencing research facility, Northumbria University | Wellcome Sanger Institute for the COVID-19 Genomics UK Consortium | Chris Duncan, Sheia Waugh, Shirelle Burton-Fanning, Gary Eltringham, Jennifer Collins, Brendan Payne, Yusri Taha, Emma Swindells, Jane Greenaway, Edward Barton, Garren Scott, Debra Padgett, Clive Graham, Sarah Essex, Steve Ligget, Paul Baker, Lynn Dover, Wen Yew, Gary Black, John Allan, Joshua Loh, Greg Young, Matthew Bashton, Andrew Nelson, Darren Smith and Alex Alderton, Roberto Amato, Sonia Goncalves, Ewan Harrison, David K. Jackson, Ian Johnston, Dominic Kwiatkowski, Cordelia Langford, John Sillitoe on behalf of the Wellcome Sanger Institute COVID-19 Surveillance Team (http://www.sanger.ac.uk/covid-team) |

| | | | |
|---|---|---|---|
| EPI_ISL_458721, EPI_ISL_458722, EPI_ISL_458723, EPI_ISL_458725, EPI_ISL_458726, EPI_ISL_458727, EPI_ISL_458728, EPI_ISL_458732, EPI_ISL_458733, EPI_ISL_458736, EPI_ISL_458739, EPI_ISL_458743, EPI_ISL_458745, EPI_ISL_458746, EPI_ISL_458748, EPI_ISL_458749, EPI_ISL_458750, EPI_ISL_458751, EPI_ISL_458752, EPI_ISL_458755, EPI_ISL_458756, EPI_ISL_458757, EPI_ISL_458758, EPI_ISL_458759, EPI_ISL_458760, EPI_ISL_458761, EPI_ISL_458762, EPI_ISL_458763, EPI_ISL_458764, EPI_ISL_458765, EPI_ISL_458766, EPI_ISL_458767, EPI_ISL_458768, EPI_ISL_458769, EPI_ISL_458770, EPI_ISL_458771, EPI_ISL_458772, EPI_ISL_458773, EPI_ISL_458774, EPI_ISL_458775, EPI_ISL_458777, EPI_ISL_458778, EPI_ISL_458779, EPI_ISL_458781, EPI_ISL_458782, EPI_ISL_458783, EPI_ISL_458784, EPI_ISL_458785, EPI_ISL_458787, EPI_ISL_458791, EPI_ISL_458792, EPI_ISL_458793, EPI_ISL_458794, EPI_ISL_458796, EPI_ISL_458797, EPI_ISL_458798, EPI_ISL_458799, EPI_ISL_458801, EPI_ISL_458803, EPI_ISL_458804, EPI_ISL_458808, EPI_ISL_458810, EPI_ISL_458811, EPI_ISL_458814, EPI_ISL_458815, EPI_ISL_458816, EPI_ISL_458817, EPI_ISL_458819, EPI_ISL_458820, EPI_ISL_458821, EPI_ISL_458822, EPI_ISL_458824, EPI_ISL_458825, EPI_ISL_458826, EPI_ISL_458827, EPI_ISL_458829, EPI_ISL_458830, EPI_ISL_458831, EPI_ISL_458832, EPI_ISL_458833, EPI_ISL_458835, EPI_ISL_458836, EPI_ISL_458838, EPI_ISL_458839, EPI_ISL_458840, EPI_ISL_458842, EPI_ISL_458844, EPI_ISL_458847, EPI_ISL_458848, EPI_ISL_458851, EPI_ISL_458854, EPI_ISL_458856, EPI_ISL_458858, EPI_ISL_458859, EPI_ISL_458861, EPI_ISL_458862, EPI_ISL_458863, EPI_ISL_458864, EPI_ISL_458866, EPI_ISL_458868, EPI_ISL_458871, EPI_ISL_458877, EPI_ISL_458878, EPI_ISL_458879, EPI_ISL_458880, EPI_ISL_458883, EPI_ISL_458885, EPI_ISL_458886, EPI_ISL_458888, EPI_ISL_458892, EPI_ISL_458894, EPI_ISL_458899, EPI_ISL_458903, EPI_ISL_458904, EPI_ISL_458908, EPI_ISL_458909, EPI_ISL_458911, EPI_ISL_458912, EPI_ISL_458913, EPI_ISL_458914, EPI_ISL_458915, EPI_ISL_458917, EPI_ISL_458918, EPI_ISL_458919, EPI_ISL_458922, EPI_ISL_458923, EPI_ISL_458924, EPI_ISL_458925, EPI_ISL_458926, EPI_ISL_458927, EPI_ISL_458928, EPI_ISL_458930, EPI_ISL_458931, EPI_ISL_458932, EPI_ISL_458934, EPI_ISL_458937, EPI_ISL_458938, EPI_ISL_458939, EPI_ISL_458941, EPI_ISL_458943, EPI_ISL_458945, EPI_ISL_458946, EPI_ISL_458947, EPI_ISL_458949, EPI_ISL_458950, EPI_ISL_458951, EPI_ISL_458952, EPI_ISL_458953, EPI_ISL_458955, EPI_ISL_458956, EPI_ISL_458957, EPI_ISL_458960, EPI_ISL_458961, EPI_ISL_458962, EPI_ISL_458963, EPI_ISL_458964, EPI_ISL_458965, EPI_ISL_458966, EPI_ISL_458967, EPI_ISL_458969, EPI_ISL_458971, EPI_ISL_458972, EPI_ISL_458974, EPI_ISL_458975, EPI_ISL_458976, EPI_ISL_458977, EPI_ISL_458978, EPI_ISL_458979, EPI_ISL_458981, EPI_ISL_458982, EPI_ISL_458983, EPI_ISL_458984, EPI_ISL_458985, EPI_ISL_458986, EPI_ISL_458987, EPI_ISL_458989, EPI_ISL_458990, EPI_ISL_458991, EPI_ISL_458992, EPI_ISL_458994, EPI_ISL_458995, EPI_ISL_458998, EPI_ISL_459001, EPI_ISL_459002, EPI_ISL_459004, EPI_ISL_459005, EPI_ISL_459006, EPI_ISL_459007, EPI_ISL_459008, EPI_ISL_459010, EPI_ISL_459012, EPI_ISL_459014, EPI_ISL_459015, EPI_ISL_459016, EPI_ISL_459017, EPI_ISL_459020, EPI_ISL_459021, EPI_ISL_459022, EPI_ISL_459024, EPI_ISL_459025, EPI_ISL_459026, EPI_ISL_459027, EPI_ISL_459028, EPI_ISL_459029, EPI_ISL_459030, EPI_ISL_459033, EPI_ISL_459034, EPI_ISL_459035, EPI_ISL_459036, EPI_ISL_459038, EPI_ISL_459040, EPI_ISL_459041, EPI_ISL_459042, EPI_ISL_459046, EPI_ISL_459047, EPI_ISL_459048, EPI_ISL_459050, EPI_ISL_459051, EPI_ISL_459052, EPI_ISL_459053, EPI_ISL_459055, EPI_ISL_459057, EPI_ISL_459058, EPI_ISL_459059, EPI_ISL_459060, EPI_ISL_459061, EPI_ISL_459064, EPI_ISL_459065, EPI_ISL_459066, EPI_ISL_459068, EPI_ISL_459069, EPI_ISL_459070, EPI_ISL_459071, EPI_ISL_459076, EPI_ISL_459077, EPI_ISL_459078, EPI_ISL_459079, EPI_ISL_459080, EPI_ISL_459081, EPI_ISL_459082, EPI_ISL_459083, EPI_ISL_459085, EPI_ISL_459086, EPI_ISL_459090, EPI_ISL_459091, EPI_ISL_459093 | | | |
| see above | PHE South West Regional Laboratory, National Infection Service | Wellcome Sanger Institute for the COVID-19 Genomics UK Consortium | Stephanie Hutchings, Hannah Pymont, Dr Peter Muir, Barry Vipond, Rich Hopes; and Alex Alderton, Roberto Amato, Sonia Goncalves, Ewan Harrison, David K. Jackson, Ian Johnston, Dominic Kwiatkowski, Cordelia Langford, John Sillitoe on behalf of the Wellcome Sanger Institute COVID-19 Surveillance Team (http://www.sanger.ac.uk/covid-team) |
| EPI_ISL_459096, EPI_ISL_459097, EPI_ISL_459098, EPI_ISL_459099, EPI_ISL_459100, EPI_ISL_459102, EPI_ISL_459103, EPI_ISL_459104, EPI_ISL_459105, EPI_ISL_459106, EPI_ISL_459107, EPI_ISL_459109, EPI_ISL_459111, EPI_ISL_459112, EPI_ISL_459113, EPI_ISL_459114, EPI_ISL_459115, EPI_ISL_459116, EPI_ISL_459118, EPI_ISL_459119, EPI_ISL_459120, EPI_ISL_459121, EPI_ISL_459122, EPI_ISL_459124, EPI_ISL_459125, EPI_ISL_459127, EPI_ISL_459130, EPI_ISL_459131, EPI_ISL_459132, EPI_ISL_459133, EPI_ISL_459134, EPI_ISL_459136, EPI_ISL_459137, EPI_ISL_459138, EPI_ISL_459140, EPI_ISL_459141, EPI_ISL_459142, EPI_ISL_459145, EPI_ISL_459146, EPI_ISL_459147, EPI_ISL_459149, EPI_ISL_459151, EPI_ISL_459153, EPI_ISL_459154, EPI_ISL_459155, EPI_ISL_459156, EPI_ISL_459157, EPI_ISL_459158, EPI_ISL_459161, EPI_ISL_459164, EPI_ISL_459165 | | | |
| see above | NHSGGC West of Scotland Specialist Virology Centre / MRC-University of Glasgow Centre for Virus Research | Wellcome Sanger Institute for the COVID-19 Genomics UK Consortium | Ana da Silva Filipe, Natasha Johnson, Kathy Smollett, Daniel Mair, Stephen Carmichael, Lily Tong, Jenna Nichols, Elihu Aranday-Cortes, Kirstyn Brunker, Yasmin Parr, Kyriaki Nomikou; Sarah McDonald, Marc Niebel, Patawee Asamaphan; Richard Orton, Joseph Hughes, Sreenu Vattipally, David L Robertson; Alasdair MacLean, Rory Gunson; Kathy Li, Natasha Jesudason, Rajiv Shah, James Shepherd, Antonia Ho, Alice Broos, Emma Thomson and Alex Alderton, Roberto Amato, Sonia Goncalves, Ewan Harrison, David K. Jackson, Ian Johnston, Dominic Kwiatkowski, Cordelia Langford, John Sillitoe on behalf of the Wellcome Sanger Institute COVID-19 Surveillance Team (http://www.sanger.ac.uk/covid-team) |
| EPI_ISL_459166, EPI_ISL_459167, EPI_ISL_459169, EPI_ISL_459171, EPI_ISL_459174, EPI_ISL_459178, EPI_ISL_459179, EPI_ISL_459194, EPI_ISL_459204, EPI_ISL_459207, EPI_ISL_459210, EPI_ISL_459211, EPI_ISL_459219, EPI_ISL_459227, EPI_ISL_459235, EPI_ISL_459240, EPI_ISL_459251, EPI_ISL_459252, EPI_ISL_459255, EPI_ISL_459257, EPI_ISL_459259, EPI_ISL_459262, EPI_ISL_459263, EPI_ISL_459264, EPI_ISL_459265, EPI_ISL_459266, EPI_ISL_459269, EPI_ISL_459271, EPI_ISL_459274, EPI_ISL_459284, EPI_ISL_459287, EPI_ISL_459289, EPI_ISL_459292, EPI_ISL_459295, EPI_ISL_459296, EPI_ISL_459309, EPI_ISL_459310, EPI_ISL_459313, EPI_ISL_459315, EPI_ISL_459316, EPI_ISL_459317, EPI_ISL_459320, EPI_ISL_459321 | | | |
| see above | Department of Pathology, University of Cambridge | Wellcome Sanger Institute for the COVID-19 Genomics UK Consortium | Luke W Meredith, M. Estée Török , Myra Hosmillo, William L. Hamilton, Martin D. Curran, Theresa Feltwell, Grant Hall, Anna Yakovleva, Fahad A Khokhar, Charlotte J. Houldcroft, Laura G Caller, Aminu S. Jahun, Sarah L. Caddy, Ian Goodfellow; and Alex Alderton, Roberto Amato, Sonia Goncalves, Ewan Harrison, David K. Jackson, Ian Johnston, Dominic Kwiatkowski, Cordelia Langford, John Sillitoe on behalf of the Wellcome Sanger Institute COVID-19 Surveillance Team (http://www.sanger.ac.uk/covid-team) |
| EPI_ISL_459324, EPI_ISL_459325, EPI_ISL_459326, EPI_ISL_459331, EPI_ISL_459332, EPI_ISL_459333, EPI_ISL_459334, EPI_ISL_459335, EPI_ISL_459336, EPI_ISL_459338, EPI_ISL_459341, EPI_ISL_459343, EPI_ISL_459344, EPI_ISL_459345, EPI_ISL_459347, EPI_ISL_459348, EPI_ISL_459349, EPI_ISL_459351, EPI_ISL_459353, EPI_ISL_459354, EPI_ISL_459360, EPI_ISL_459363, EPI_ISL_459365, EPI_ISL_459367, EPI_ISL_459368, EPI_ISL_459369, EPI_ISL_459370, EPI_ISL_459373, EPI_ISL_459375, EPI_ISL_459376, EPI_ISL_459378, EPI_ISL_459379, EPI_ISL_459380, EPI_ISL_459382, EPI_ISL_459387, EPI_ISL_459389, EPI_ISL_459391, EPI_ISL_459392, EPI_ISL_459394, EPI_ISL_459395, EPI_ISL_459399, EPI_ISL_459401, EPI_ISL_459402, EPI_ISL_459403, EPI_ISL_459404, EPI_ISL_459406, EPI_ISL_459407 | | | |
| see above | Regional Virus Laboratory, Belfast Health and Social Care Trust | Wellcome Sanger Institute for the COVID-19 Genomics UK Consortium | Conall McCaughey, James McKenna, Tanya Curran, Susan Feeney, Alison Watt, Ciara Cox, Mairead Connor, Zoltan Molnar, David Simpson, Derek Fairley; and Alex Alderton, Roberto Amato, Sonia Goncalves, Ewan Harrison, David K. Jackson, Ian Johnston, Dominic Kwiatkowski, Cordelia Langford, John Sillitoe on behalf of the Wellcome Sanger Institute COVID-19 Surveillance Team (http://www.sanger.ac.uk/covid-team) |
| EPI_ISL_459412, EPI_ISL_459413, EPI_ISL_459416, EPI_ISL_459421, EPI_ISL_459423, EPI_ISL_459424, EPI_ISL_459425, EPI_ISL_459434, EPI_ISL_459435, EPI_ISL_459437, EPI_ISL_459442, EPI_ISL_459448, EPI_ISL_459449, EPI_ISL_459453, EPI_ISL_459455, EPI_ISL_459456, EPI_ISL_459459, EPI_ISL_459461, EPI_ISL_459463, EPI_ISL_459466, EPI_ISL_459467, EPI_ISL_459468, EPI_ISL_459476, EPI_ISL_459477, EPI_ISL_459478, EPI_ISL_459479, EPI_ISL_459481, EPI_ISL_459493, EPI_ISL_459496, EPI_ISL_459499, EPI_ISL_459501, EPI_ISL_459504 | | | |
| see above | Department of Pathology, University of Cambridge | Wellcome Sanger Institute for the COVID-19 Genomics UK Consortium | Luke W Meredith, M. Estée Török , Myra Hosmillo, William L. Hamilton, Martin D. Curran, Theresa Feltwell, Grant Hall, Anna Yakovleva, Fahad A Khokhar, Charlotte J. Houldcroft, Laura G Caller, Aminu S. Jahun, Sarah L. Caddy, Ian Goodfellow; and Alex Alderton, Roberto Amato, Sonia Goncalves, Ewan Harrison, David K. Jackson, Ian Johnston, Dominic Kwiatkowski, Cordelia Langford, John Sillitoe on behalf of the Wellcome Sanger Institute COVID-19 Surveillance Team (http://www.sanger.ac.uk/covid-team) |
| EPI_ISL_459506, EPI_ISL_459508, EPI_ISL_459509, EPI_ISL_459510, EPI_ISL_459513, EPI_ISL_459514, EPI_ISL_459517, EPI_ISL_459518, EPI_ISL_459519, EPI_ISL_459520, EPI_ISL_459521, EPI_ISL_459522, EPI_ISL_459524, EPI_ISL_459525, EPI_ISL_459526, EPI_ISL_459527, EPI_ISL_459528, EPI_ISL_459529, EPI_ISL_459530, EPI_ISL_459532, EPI_ISL_459533, EPI_ISL_459534, EPI_ISL_459535, EPI_ISL_459536, EPI_ISL_459538, EPI_ISL_459539, EPI_ISL_459542, EPI_ISL_459544, EPI_ISL_459546, EPI_ISL_459547, EPI_ISL_459548, EPI_ISL_459550, EPI_ISL_459552, EPI_ISL_459555, EPI_ISL_459556, EPI_ISL_459557, EPI_ISL_459558, EPI_ISL_459559, EPI_ISL_459560, EPI_ISL_459561, EPI_ISL_459562, EPI_ISL_459563, EPI_ISL_459565, EPI_ISL_459566, EPI_ISL_459568, EPI_ISL_459569, EPI_ISL_459572, EPI_ISL_459574, EPI_ISL_459575, EPI_ISL_459577, EPI_ISL_459578, EPI_ISL_459580, EPI_ISL_459581, EPI_ISL_459583, EPI_ISL_459584, EPI_ISL_459586, EPI_ISL_459587, EPI_ISL_459589, EPI_ISL_459590, EPI_ISL_459591, EPI_ISL_459593, EPI_ISL_459594, EPI_ISL_459595, EPI_ISL_459596, EPI_ISL_459597, EPI_ISL_459598, EPI_ISL_459600, EPI_ISL_459602, EPI_ISL_459603, EPI_ISL_459605, EPI_ISL_459606, EPI_ISL_459607, EPI_ISL_459608, EPI_ISL_459609, EPI_ISL_459610, EPI_ISL_459613, EPI_ISL_459614, EPI_ISL_459616, EPI_ISL_459617, EPI_ISL_459619, EPI_ISL_459621, EPI_ISL_459623, EPI_ISL_459624, EPI_ISL_459625, EPI_ISL_459626, EPI_ISL_459628, EPI_ISL_459629, EPI_ISL_459630, EPI_ISL_459631, EPI_ISL_459632, EPI_ISL_459635, EPI_ISL_459636, EPI_ISL_459637, EPI_ISL_459638, EPI_ISL_459640, EPI_ISL_459641, EPI_ISL_459643, EPI_ISL_459644, EPI_ISL_459645, EPI_ISL_459646, EPI_ISL_459647, EPI_ISL_459648, EPI_ISL_459649, EPI_ISL_459651, EPI_ISL_459652, EPI_ISL_459654, EPI_ISL_459656, EPI_ISL_459657, EPI_ISL_459659, EPI_ISL_459660, EPI_ISL_459661, EPI_ISL_459662, EPI_ISL_459665, EPI_ISL_459666, EPI_ISL_459667, EPI_ISL_459668, EPI_ISL_459670, EPI_ISL_459672, EPI_ISL_459678, EPI_ISL_459679, EPI_ISL_459680, EPI_ISL_459681, EPI_ISL_459682, EPI_ISL_459684, EPI_ISL_459686, EPI_ISL_459688, EPI_ISL_459689, EPI_ISL_459690, EPI_ISL_459691, EPI_ISL_459692, EPI_ISL_459693, EPI_ISL_459694, EPI_ISL_459695, EPI_ISL_459696, EPI_ISL_459697, EPI_ISL_459699, EPI_ISL_459700, EPI_ISL_459701, EPI_ISL_459702, EPI_ISL_459703, EPI_ISL_459704, EPI_ISL_459705, EPI_ISL_459707, EPI_ISL_459708, EPI_ISL_459709, EPI_ISL_459710, EPI_ISL_459711, EPI_ISL_459712, EPI_ISL_459713, EPI_ISL_459714, EPI_ISL_459716, EPI_ISL_459718, EPI_ISL_459720, EPI_ISL_459721, EPI_ISL_459724 | | | |
| see above | NHSGGC West of Scotland Specialist Virology Centre / MRC-University of Glasgow Centre for Virus Research | Wellcome Sanger Institute for the COVID-19 Genomics UK Consortium | Ana da Silva Filipe, Natasha Johnson, Kathy Smollett, Daniel Mair, Stephen Carmichael, Lily Tong, Jenna Nichols, Elihu Aranday-Cortes, Kirstyn Brunker, Yasmin Parr, Kyriaki Nomikou; Sarah McDonald, Marc Niebel, Patawee Asamaphan; Richard Orton, Joseph Hughes, Sreenu Vattipally, David L Robertson; Alasdair MacLean, Rory Gunson; Kathy Li, Natasha Jesudason, Rajiv Shah, James Shepherd, Antonia Ho, Alice Broos, Emma Thomson and Alex Alderton, Roberto Amato, Sonia Goncalves, Ewan Harrison, David K. Jackson, Ian Johnston, Dominic Kwiatkowski, Cordelia Langford, John Sillitoe on behalf of the Wellcome Sanger Institute COVID-19 Surveillance Team (http://www.sanger.ac.uk/covid-team) |
| EPI_ISL_459725, EPI_ISL_459726 | PHE South West Regional Laboratory, National Infection Service | Wellcome Sanger Institute for the COVID-19 Genomics UK Consortium | Stephanie Hutchings, Hannah Pymont, Dr Peter Muir, Barry Vipond, Rich Hopes; and Alex Alderton, Roberto Amato, Sonia Goncalves, Ewan Harrison, David K. Jackson, Ian Johnston, Dominic Kwiatkowski, Cordelia Langford, John Sillitoe on behalf of the Wellcome Sanger Institute COVID-19 Surveillance Team (http://www.sanger.ac.uk/covid-team) |
| EPI_ISL_459856, EPI_ISL_459857, EPI_ISL_459858, EPI_ISL_459859, EPI_ISL_459860, EPI_ISL_459861, EPI_ISL_459862, EPI_ISL_459863 | Center for Genome Regulation (CRG) | Center for Mathematical Modeling and Center for Genome Regulation. Santiago, Chile | Gaete A, Travisany D, Palma R, Urra C, Varas M, Allende ML, Maass A, González M. |
| EPI_ISL_459866 | Kingston Health Sciences Center | Queen's Genomics Lab at Ongwanada (Q-GLO) | Sjaarda CP, Rustom N, Huang D, Perez-Patrigeon S, Hudson ML, Wong H,Guan H, Ayub M, Soares CN, Colautti R, Evans GA, Sheth P |
| EPI_ISL_459867 | Kingston Health Sciences Center | Queen's Genomics Lab at Ongwanada (Q-GLO) | Sjaarda CP, Rustom N, Huang D, Perez-Patrigeon S, Hudson ML, Wong H,Guan H, Ayub M, Soares CN, Colautti R, Evans GA, Sheth P |
| EPI_ISL_459868, EPI_ISL_459869, EPI_ISL_459871, EPI_ISL_459872, EPI_ISL_459873, EPI_ISL_459874, EPI_ISL_459875, EPI_ISL_459877, EPI_ISL_459879, EPI_ISL_459880, EPI_ISL_459881, EPI_ISL_459882, EPI_ISL_459883, EPI_ISL_459885, EPI_ISL_459886, EPI_ISL_459887, EPI_ISL_459888, EPI_ISL_459889, EPI_ISL_459890, EPI_ISL_459891, EPI_ISL_459892 | | | |
| see above | Kingston Health Sciences Center | Queen's Genomics Lab at Ongwanada (Q-GLO) | Sjaarda CP, Rustom N, Huang D, Perez-Patrigeon S, Hudson ML, Wong H,Guan H, Ayub M, Soares CN, Colautti R, Evans GA, Sheth P |
| EPI_ISL_459893, EPI_ISL_459894, EPI_ISL_459896, EPI_ISL_459898, EPI_ISL_459899, EPI_ISL_459900, EPI_ISL_459901, EPI_ISL_459903, EPI_ISL_459905, EPI_ISL_459906 | Laboratoire National de Sante, Microbiology, Virology | Laboratoire National de Sante, Microbiology, Epidemiology and Microbial Genomics | Anke Wienecke-Baldacchino, Jessica Tapp, Guillaume Fournier, Tamir Abdelrahman, Trung Nguyen Nguyen, Catherine Ragimbeau |
| EPI_ISL_459909 | Zoonotic and Exotic infection Diseases Division, Harbin Veterinary Research Institute, CAAS | Zoonotic and Exotic infection Diseases Division, Harbin Veterinary Research Institute, CAAS | Zhigao Bu, Jinliang Wang |
| EPI_ISL_459911 | Devki Devi Foundation, a unit of Max Healthcare | CSIR-IGIB/Max | Rajesh Pandey#, Samreen Siddiqui, Pooja Sharma, Bansidhar Tarai, Vivekanand A, Bharathram Uppili, Saruchi Wadhwa, Nishu Tyagi, Mitali Mukerji, Poonam Das, Sujeet Jha, Mohammed Faruq, Vinita Jha, Anurag Agrawal |
| EPI_ISL_459913, EPI_ISL_459914, EPI_ISL_459915, EPI_ISL_459916, EPI_ISL_459917, EPI_ISL_459918, EPI_ISL_459919, EPI_ISL_459920, EPI_ISL_459921, EPI_ISL_459922, EPI_ISL_459923, EPI_ISL_459924, EPI_ISL_459926, EPI_ISL_459927, EPI_ISL_459932, EPI_ISL_459933, EPI_ISL_459934, EPI_ISL_459935, EPI_ISL_459936, EPI_ISL_459937, EPI_ISL_459938, EPI_ISL_459939, EPI_ISL_459940, EPI_ISL_459941, EPI_ISL_459942, EPI_ISL_459943 | | | |
| see above | Devki Devi Foundation, a unit of Max Healthcare | CSIR-IGIB/Max | Rajesh Pandey#, Samreen Siddiqui, Pooja Sharma, Bansidhar Tarai, Vivekanand A, Bharathram Uppili, Saruchi Wadhwa, Nishu Tyagi, Mitali Mukerji, Bansidhar Tarai, Poonam Das, Sujeet Jha, Mohammed Faruq, Vinita Jha, Anurag Agrawal |
| EPI_ISL_459955 | Institute for Medical Research, Infectious Disease Research Centre, National Institutes of Health, Ministry of Health Malaysia | Institute for Medical Research, Infectious Disease Research Centre, National Institutes of Health, Ministry of Health Malaysia | Suppiah J, Mohd-Zawawi Z, Kamel KA, Ellan K, Kalyanasundram J, Mohd-Zain R, Thayan R |
| EPI_ISL_459957 | Institute for Medical Research, Infectious Disease Research Centre, National Institutes of Health, Minis | Institute for Medical Research, Infectious Disease Research Centre, National Institutes of Health, Minis | Suppiah J, Mohd-Zawawi Z, Kamel KA, Ellan K, Kalyanasundram J, Mohd-Zain R, Thayan R |
| EPI_ISL_459958, EPI_ISL_459960, EPI_ISL_459961 | Respiratory Virus Unit, Microbiology Services Colindale, Public Health England | Respiratory Virus Unit, Microbiology Services Colindale, Public Health England | Steven Platt, Shahjahan Miah, Angie Lackenby, Omolola Akinbami, Tina Talts, Leena Bhaw, Richard Myers, Monica Galiano, Kirstin Edwards, Jonathan Hubb, Joanna Ellis, Maria Zambon |
| EPI_ISL_459962, EPI_ISL_459963 | Centogene AG | Centogene AG | Prof. Dr. Peter Bauer, Dr. Krishna Kumar Kandaswamy |
| EPI_ISL_459964 | Centogene AG | Centogene AG | Prof. Dr. Peter Bauer, Dr. Krishna Kumar Kandaswamy |
| EPI_ISL_459965, EPI_ISL_459966, EPI_ISL_459967, EPI_ISL_459968, EPI_ISL_459972, EPI_ISL_459973, EPI_ISL_459974, EPI_ISL_459975, EPI_ISL_459976, EPI_ISL_459977, EPI_ISL_459978, EPI_ISL_459979, EPI_ISL_459980, EPI_ISL_459981, EPI_ISL_459982, EPI_ISL_459983, EPI_ISL_459984 | | | |
| see above | Institut Pasteur du Maroc | Institut Pasteur du Maroc | Marion Barbet, Sylvie Behillil, Méline Bizard, Angela Brisebarre, Camille Capel, Etienne Simon-Lorière, Vincent Enouf, Maud Vanpeene, Sylvie van der Werf, Latifa Anga, Abdellah Faouzi, Anass Abbad, Mjid Eloualid, Jalal Nourlil, Anderrahmane Maaroufi |
| EPI_ISL_459994, EPI_ISL_459995, EPI_ISL_459996, EPI_ISL_459997, EPI_ISL_459998, EPI_ISL_459999, EPI_ISL_460003, EPI_ISL_460004, EPI_ISL_460008, EPI_ISL_460009, EPI_ISL_460012, EPI_ISL_460015, EPI_ISL_460016, EPI_ISL_460017, EPI_ISL_460018, EPI_ISL_460020, EPI_ISL_460021, EPI_ISL_460023, EPI_ISL_460025, EPI_ISL_460026, EPI_ISL_460028, EPI_ISL_460031, EPI_ISL_460033, EPI_ISL_460034, EPI_ISL_460040, EPI_ISL_460042 | | | |
| see above | Michigan Department of Health and Human Services, Bureau of Laboratories | Michigan Department of Health and Human Services, Bureau of Laboratories | Blankenship HM, Riner D, Soehnlen MK |
| EPI_ISL_460045, EPI_ISL_460046, EPI_ISL_460047, EPI_ISL_460048, EPI_ISL_460049, EPI_ISL_460050, EPI_ISL_460051, EPI_ISL_460052, EPI_ISL_460053, EPI_ISL_460054, EPI_ISL_460055, EPI_ISL_460056, EPI_ISL_460057, EPI_ISL_460058, EPI_ISL_460059, EPI_ISL_460060, EPI_ISL_460061, EPI_ISL_460062, EPI_ISL_460063, EPI_ISL_460064, EPI_ISL_460065, EPI_ISL_460066, EPI_ISL_460067, EPI_ISL_460068, EPI_ISL_460069, EPI_ISL_460070, EPI_ISL_460071, EPI_ISL_460072, EPI_ISL_460073, EPI_ISL_460075, EPI_ISL_460076, EPI_ISL_460077, EPI_ISL_460078 | | | |
| see above | Minnesota Department of Health, Public Health Laboratory | Minnesota Department of Health, Public Health Laboratory | Matt Plumb, Jacob Garfin, and Xiong Wang |
| EPI_ISL_460079 | Molecular Virology Unit, Fondazione IRCCS Policlinico San Matteo , Pavia | Laboratory of Virology, INMI Lazzaro Spallanzani IRCCS | Barbara Bartolini, Cesare E.M. Gruber, Maria R. Capobianchi, Martina Rueca, Antonio Piralla, Fausto Baldanti, Antonino Di Caro |
| EPI_ISL_460080 | Molecular Virology Unit, Fondazione IRCCS Policlinico San Matteo , Pavia | Laboratory of Virology, INMI Lazzaro Spallanzani IRCCS | Antonio Piralla, Barbara Bartolini, Fausto Baldanti, Martina Rueca, Antonino Di Caro, Cesare E.M. Gruber, Maria R. Capobianchi |
| EPI_ISL_460081 | Molecular Virology Unit, Fondazione IRCCS Policlinico San Matteo , Pavia | Laboratory of Virology, INMI Lazzaro Spallanzani IRCCS | Fausto Baldanti, Martina Rueca, Antonio Piralla, Antonino Di Caro, Maria R. Capobianchi, Cesare E.M. Gruber, Barbara Bartolini |
| EPI_ISL_460082 | Molecular Virology Unit, Fondazione IRCCS Policlinico San Matteo , Pavia | Laboratory of Virology, INMI Lazzaro Spallanzani IRCCS | Martina Rueca, Cesare E.M. Gruber, Antonio Piralla, Antonino Di Caro, Barbara Bartolini, Maria R. Capobianchi, Fausto Baldanti |
| EPI_ISL_460083 | Molecular Virology Unit, Fondazione IRCCS Policlinico San Matteo , Pavia | Laboratory of Virology, INMI Lazzaro Spallanzani IRCCS | Martina Rueca, Antonino Di Caro, Cesare E.M. Gruber, Barbara Bartolini, Fausto Baldanti, Antonio Piralla, Maria R. Capobianchi |
| EPI_ISL_460084 | Molecular Virology Unit, Fondazione IRCCS Policlinico San Matteo , Pavia | Laboratory of Virology, INMI Lazzaro Spallanzani IRCCS | Fausto Baldanti, Antonio Piralla, Martina Rueca, Barbara Bartolini, Maria R. Capobianchi, Cesare E.M. Gruber, Antonino Di Caro |
| EPI_ISL_460085 | Molecular Virology Unit, Fondazione IRCCS Policlinico San Matteo , Pavia | Laboratory of Virology, INMI Lazzaro Spallanzani IRCCS | Cesare E.M. Gruber, Maria R. Capobianchi, Barbara Bartolini, Fausto Baldanti, Martina Rueca, Antonio Piralla, Antonino Di Caro |

| | | | |
|---|---|---|---|
| EPI_ISL_460086 | Molecular Virology Unit, Fondazione IRCCS Policlinico San Matteo , Pavia | Laboratory of Virology, INMI Lazzaro Spallanzani IRCCS | Maria R. Capobianchi, Fausto Baldanti, Antonio Piralla, Antonino Di Caro, Barbara Bartolini, Cesare E.M. Gruber, Martina Rueca |
| EPI_ISL_460087 | Molecular Virology Unit, Fondazione IRCCS Policlinico San Matteo , Pavia | Laboratory of Virology, INMI Lazzaro Spallanzani IRCCS | Cesare E.M. Gruber, Maria R. Capobianchi, Martina Rueca, Barbara Bartolini, Antonino Di Caro, Antonio Piralla, Fausto Baldanti |
| EPI_ISL_460088 | Molecular Virology Unit, Fondazione IRCCS Policlinico San Matteo , Pavia | Laboratory of Virology, INMI Lazzaro Spallanzani IRCCS | Martina Rueca, Barbara Bartolini, Fausto Baldanti, Maria R. Capobianchi, Cesare E.M. Gruber, Antonino Di Caro, Antonio Piralla |
| EPI_ISL_460089 | Molecular Virology Unit, Fondazione IRCCS Policlinico San Matteo , Pavia | Laboratory of Virology, INMI Lazzaro Spallanzani IRCCS | Antonino Di Caro, Barbara Bartolini, Martina Rueca, Cesare E.M. Gruber, Antonio Piralla, Fausto Baldanti, Maria R. Capobianchi |
| EPI_ISL_460090 | Molecular Virology Unit, Fondazione IRCCS Policlinico San Matteo , Pavia | Laboratory of Virology, INMI Lazzaro Spallanzani IRCCS | Antonio Piralla, Cesare E.M. Gruber, Antonino Di Caro, Maria R. Capobianchi, Martina Rueca, Barbara Bartolini, Fausto Baldanti |
| EPI_ISL_460091 | Molecular Virology Unit, Fondazione IRCCS Policlinico San Matteo , Pavia | Laboratory of Virology, INMI Lazzaro Spallanzani IRCCS | Antonino Di Caro, Antonio Piralla, Martina Rueca, Fausto Baldanti, Barbara Bartolini, Maria R. Capobianchi, Cesare E.M. Gruber |
| EPI_ISL_460092 | Molecular Virology Unit, Fondazione IRCCS Policlinico San Matteo , Pavia | Laboratory of Virology, INMI Lazzaro Spallanzani IRCCS | Cesare E.M. Gruber, Martina Rueca, Maria R. Capobianchi, Antonino Di Caro, Antonio Piralla, Barbara Bartolini, Fausto Baldanti |
| EPI_ISL_460093 | Molecular Virology Unit, Fondazione IRCCS Policlinico San Matteo , Pavia | Laboratory of Virology, INMI Lazzaro Spallanzani IRCCS | Maria R. Capobianchi, Antonio Piralla, Antonino Di Caro, Fausto Baldanti, Martina Rueca, Cesare E.M. Gruber, Barbara Bartolini |
| EPI_ISL_460094 | Molecular Virology Unit, Fondazione IRCCS Policlinico San Matteo , Pavia | Laboratory of Virology, INMI Lazzaro Spallanzani IRCCS | Barbara Bartolini, Maria R. Capobianchi, Antonino Di Caro, Antonio Piralla, Cesare E.M. Gruber, Martina Rueca, Fausto Baldanti |
| EPI_ISL_460095 | Molecular Virology Unit, Fondazione IRCCS Policlinico San Matteo , Pavia | Laboratory of Virology, INMI Lazzaro Spallanzani IRCCS | Barbara Bartolini, Antonino Di Caro, Fausto Baldanti, Cesare E.M. Gruber, Maria R. Capobianchi, Martina Rueca, Antonio Piralla |
| EPI_ISL_460096 | Molecular diagnostic laboratory of Federal Budget Institution of Science "Central Research Institute of Epidemiology" of The Federal Service on Customers' Rights Protection and Human Well-being Surveillance | Group of Genomics and Postgenomic Technologies of Central Research Institute of Epidemilology | Speranskaya AS, Kaptelova VV, Samoilov AE, Korneenko EV, Tivanova EV, Shipulina OY, Akimkin VG |
| EPI_ISL_460097, EPI_ISL_460098, EPI_ISL_460099, EPI_ISL_460101, EPI_ISL_460102, EPI_ISL_460103, EPI_ISL_460104, EPI_ISL_460105, EPI_ISL_460106, EPI_ISL_460107, EPI_ISL_460108, EPI_ISL_460109, EPI_ISL_460110, EPI_ISL_460111, EPI_ISL_460112, EPI_ISL_460114, EPI_ISL_460115, EPI_ISL_460116, EPI_ISL_460117, EPI_ISL_460118, EPI_ISL_460119, EPI_ISL_460120, EPI_ISL_460121, EPI_ISL_460122, EPI_ISL_460123, EPI_ISL_460124, EPI_ISL_460125, EPI_ISL_460126, EPI_ISL_460128, EPI_ISL_460129, EPI_ISL_460130, EPI_ISL_460131, EPI_ISL_460132, EPI_ISL_460133, EPI_ISL_460134, EPI_ISL_460135, EPI_ISL_460137, EPI_ISL_460138, EPI_ISL_460139, EPI_ISL_460140, EPI_ISL_460141, EPI_ISL_460142, EPI_ISL_460143, EPI_ISL_460144, EPI_ISL_460145, EPI_ISL_460146, EPI_ISL_460147, EPI_ISL_460148, EPI_ISL_460149, EPI_ISL_460150, EPI_ISL_460153, EPI_ISL_460154, EPI_ISL_460155, EPI_ISL_460156, EPI_ISL_460157, EPI_ISL_460158, EPI_ISL_460159, EPI_ISL_460160, EPI_ISL_460161, EPI_ISL_460162, EPI_ISL_460163, EPI_ISL_460164, EPI_ISL_460165, EPI_ISL_460166, EPI_ISL_460168, EPI_ISL_460169, EPI_ISL_460170, EPI_ISL_460171, EPI_ISL_460172, EPI_ISL_460173, EPI_ISL_460174, EPI_ISL_460175, EPI_ISL_460176, EPI_ISL_460177, EPI_ISL_460178, EPI_ISL_460179, EPI_ISL_460180, EPI_ISL_460181, EPI_ISL_460182, EPI_ISL_460183, EPI_ISL_460184, EPI_ISL_460185, EPI_ISL_460186, EPI_ISL_460187, EPI_ISL_460188, EPI_ISL_460189, EPI_ISL_460190, EPI_ISL_460191, EPI_ISL_460195, EPI_ISL_460196, EPI_ISL_460197, EPI_ISL_460198, EPI_ISL_460201, EPI_ISL_460202, EPI_ISL_460203, EPI_ISL_460204, EPI_ISL_460205, EPI_ISL_460206, EPI_ISL_460207, EPI_ISL_460208, EPI_ISL_460209, EPI_ISL_460210, EPI_ISL_460212, EPI_ISL_460213, EPI_ISL_460214, EPI_ISL_460215, EPI_ISL_460216, EPI_ISL_460217, EPI_ISL_460220, EPI_ISL_460221, EPI_ISL_460223, EPI_ISL_460224, EPI_ISL_460225, EPI_ISL_460226, EPI_ISL_460227, EPI_ISL_460229, EPI_ISL_460230, EPI_ISL_460231, EPI_ISL_460232, EPI_ISL_460233, EPI_ISL_460234, EPI_ISL_460235, EPI_ISL_460237, EPI_ISL_460238, EPI_ISL_460239, EPI_ISL_460240, EPI_ISL_460241, EPI_ISL_460242, EPI_ISL_460243, EPI_ISL_460244, EPI_ISL_460245, EPI_ISL_460246, EPI_ISL_460248, EPI_ISL_460249, EPI_ISL_460250, EPI_ISL_460252, EPI_ISL_460253, EPI_ISL_460254, EPI_ISL_460255, EPI_ISL_460256, EPI_ISL_460257, EPI_ISL_460258, EPI_ISL_460261, EPI_ISL_460263, EPI_ISL_460264, EPI_ISL_460265, EPI_ISL_460266, EPI_ISL_460267, EPI_ISL_460268, EPI_ISL_460269, EPI_ISL_460270, EPI_ISL_460271, EPI_ISL_460272, EPI_ISL_460273, EPI_ISL_460274, EPI_ISL_460275, EPI_ISL_460276, EPI_ISL_460277, EPI_ISL_460278, EPI_ISL_460279, EPI_ISL_460280, EPI_ISL_460281, EPI_ISL_460282, EPI_ISL_460283, EPI_ISL_460284, EPI_ISL_460285, EPI_ISL_460286, EPI_ISL_460288, EPI_ISL_460289, EPI_ISL_460290, EPI_ISL_460291, EPI_ISL_460292, EPI_ISL_460293, EPI_ISL_460294, EPI_ISL_460295, EPI_ISL_460296, EPI_ISL_460297, EPI_ISL_460298, EPI_ISL_460299, EPI_ISL_460300, EPI_ISL_460301, EPI_ISL_460302, EPI_ISL_460304, EPI_ISL_460305, EPI_ISL_460306, EPI_ISL_460307, EPI_ISL_460308, EPI_ISL_460309, EPI_ISL_460310, EPI_ISL_460311, EPI_ISL_460312, EPI_ISL_460313, EPI_ISL_460314, EPI_ISL_460315, EPI_ISL_460316, EPI_ISL_460317, EPI_ISL_460319, EPI_ISL_460321, EPI_ISL_460322, EPI_ISL_460323, EPI_ISL_460324, EPI_ISL_460325, EPI_ISL_460326, EPI_ISL_460327, EPI_ISL_460328, EPI_ISL_460329, EPI_ISL_460330, EPI_ISL_460331, EPI_ISL_460332, EPI_ISL_460333, EPI_ISL_460334, EPI_ISL_460335, EPI_ISL_460336, EPI_ISL_460337, EPI_ISL_460338, EPI_ISL_460339, EPI_ISL_460340, EPI_ISL_460341, EPI_ISL_460342, EPI_ISL_460343, EPI_ISL_460344, EPI_ISL_460345, EPI_ISL_460346, EPI_ISL_460347, EPI_ISL_460348, EPI_ISL_460349, EPI_ISL_460351, EPI_ISL_460352, EPI_ISL_460353, EPI_ISL_460354, EPI_ISL_460356, EPI_ISL_460357, EPI_ISL_460358, EPI_ISL_460359, EPI_ISL_460360, EPI_ISL_460361, EPI_ISL_460362, EPI_ISL_460364, EPI_ISL_460365, EPI_ISL_460366, EPI_ISL_460368, EPI_ISL_460369, EPI_ISL_460370, EPI_ISL_460371, EPI_ISL_460372, EPI_ISL_460373, EPI_ISL_460376, EPI_ISL_460377, EPI_ISL_460378, EPI_ISL_460379, EPI_ISL_460380, EPI_ISL_460381, EPI_ISL_460382, EPI_ISL_460383, EPI_ISL_460384, EPI_ISL_460385, EPI_ISL_460386, EPI_ISL_460388, EPI_ISL_460389, EPI_ISL_460390, EPI_ISL_460391, EPI_ISL_460392, EPI_ISL_460394, EPI_ISL_460396, EPI_ISL_460397, EPI_ISL_460398, EPI_ISL_460399, EPI_ISL_460400, EPI_ISL_460402, EPI_ISL_460403, EPI_ISL_460404, EPI_ISL_460406, EPI_ISL_460407, EPI_ISL_460408, EPI_ISL_460409, EPI_ISL_460410, EPI_ISL_460412, EPI_ISL_460413, EPI_ISL_460414, EPI_ISL_460415, EPI_ISL_460417, EPI_ISL_460418, EPI_ISL_460419, EPI_ISL_460420, EPI_ISL_460421, EPI_ISL_460422, EPI_ISL_460425, EPI_ISL_460426, EPI_ISL_460427, EPI_ISL_460428, EPI_ISL_460429, EPI_ISL_460430, EPI_ISL_460431, EPI_ISL_460432, EPI_ISL_460434, EPI_ISL_460435, EPI_ISL_460436, EPI_ISL_460437, EPI_ISL_460440, EPI_ISL_460441, EPI_ISL_460442, EPI_ISL_460443, EPI_ISL_460444, EPI_ISL_460445, EPI_ISL_460446, EPI_ISL_460447, EPI_ISL_460448, EPI_ISL_460450, EPI_ISL_460451, EPI_ISL_460453, EPI_ISL_460454, EPI_ISL_460455, EPI_ISL_460456, EPI_ISL_460457, EPI_ISL_460458, EPI_ISL_460459, EPI_ISL_460460, EPI_ISL_460461, EPI_ISL_460462, EPI_ISL_460463, EPI_ISL_460464, EPI_ISL_460465, EPI_ISL_460466, EPI_ISL_460467, EPI_ISL_460468, EPI_ISL_460469, EPI_ISL_460470, EPI_ISL_460471, EPI_ISL_460472 | | | |
| see above | Massachusetts General Hospital | Infectious Disease Program, Broad Institute of Harvard and MIT | Lemieux,J.E., Siddle,K.J., Shaw,B., Adams,G., Pierce,V., Turbett,S., Anahtar,M., Branda,J., Slater,D., Harris,J., Lin,A.E., Gladden-Young,A., Lagerborg,K., Rudy,M., DeRuff,K., Carter,A., Normandin,E., Bauer,M., Reilly,S., Tomkins-Tinch,C., Loreth,C., Chaluvadi,S., Neumann,A., Cusick,C., Chapman,S.B., Gnirke,A., Flowers,K., Cerrato,F., Birren,B.W., Gallagher,G., Smole,S., Park,D.J., MacInnis,B.L., Ryan,E., LaRocque,R., Rosenberg,E., Sabeti,P.C. |
| EPI_ISL_460556, EPI_ISL_460557, EPI_ISL_460558, EPI_ISL_460559, EPI_ISL_460560, EPI_ISL_460569, EPI_ISL_460571, EPI_ISL_460572, EPI_ISL_460573, EPI_ISL_460574, EPI_ISL_460581, EPI_ISL_460582, EPI_ISL_460583, EPI_ISL_460584, EPI_ISL_460585, EPI_ISL_460586, EPI_ISL_460587, EPI_ISL_460588, EPI_ISL_460599, EPI_ISL_460600, EPI_ISL_460601, EPI_ISL_460602 | | | |
| see above | Michigan Department of Health and Human Services, Bureau of Laboratories | Michigan Department of Health and Human Services, Bureau of Laboratories | Blankenship HM, Riner D, Soehnlen MK |
| EPI_ISL_460604 | Molecular diagnostic laboratory of Federal Budget Institution of Science "Central Research Institute of Epidemiology" of The Federal Service on Customers' Rights Protection and Human Well-being Surveillance | Group of Genomics and Postgenomic Technologies of Central Research Institute of Epidemilology | Speranskaya AS, Kaptelova VV, Samoilov AE, Korneenko EV, Tivanova EV, Shipulina OY, Akimkin VG |
| EPI_ISL_460605 | Molecular diagnostic laboratory of Federal Budget Institution of Science "Central Research Institute of Epidemiology" of The Federal Service on Customers' Rights Protection and Human Well-being Surveillance | Group of Genomics and Postgenomic Technologies of Central Research Institute of Epidemilology | Speranskaya AS, Kaptelova VV, Samoilov AE, Korneenko EV, Tivanova EV, Shipulina OY, Akimkin VG |
| EPI_ISL_460609, EPI_ISL_460610, EPI_ISL_460611, EPI_ISL_460612, EPI_ISL_460614, EPI_ISL_460615, EPI_ISL_460616 | BCCDC Public Health Laboratory | BCCDC Public Health Laboratory | Harrigan, Prystajecky, Krajden, Lee, Kamelian, Lapointe, Choi, Hoang, Sekirov, Levett, Tyson, Li, Gilmour |
| EPI_ISL_460617, EPI_ISL_460618, EPI_ISL_460619 | unknown | Physiology | Pence,S., Caykara,B., Pence,H.H., Tekin,S., Yiyit,N., Cevher Keskin,B., Kara,A. |
| EPI_ISL_460621, EPI_ISL_460622, EPI_ISL_460623, EPI_ISL_460624, EPI_ISL_460625, EPI_ISL_460626, EPI_ISL_460627, EPI_ISL_460628, EPI_ISL_460629, EPI_ISL_460630, EPI_ISL_460632, EPI_ISL_460633 | | | |
| see above | UW Virology Lab | UW Virology Lab | Pavitra Roychoudhury, Amin Addetia, Hong Xie, Lasata Shrestha, Truong Nguyen, Meei-Li Huang, Keith Jerome, Alexander Greninger |
| EPI_ISL_460635 | UHCW Pathology / University of Warwick | University of Warwick, for the COVID-19 Genomics (COG) UK Consortium | Richard Stark, Chrystala Constantinidou, Meera Unnikrishnan, Laura Baxter, Jeff Cheng, Grace Taylor-Joyce, Hannah Elizabeth Bridgewater, Lucy Frost, Sarojini Pandey, Paul Brown, Tauqeer Alam, Sascha Ott, Dimitris Grammatopoulos |
| EPI_ISL_460636, EPI_ISL_460637, EPI_ISL_460638, EPI_ISL_460639, EPI_ISL_460640, EPI_ISL_460641, EPI_ISL_460642, EPI_ISL_460643, EPI_ISL_460644, EPI_ISL_460646, EPI_ISL_460647, EPI_ISL_460648, EPI_ISL_460649, EPI_ISL_460650, EPI_ISL_460651, EPI_ISL_460652, EPI_ISL_460653, EPI_ISL_460654, EPI_ISL_460655, EPI_ISL_460656, EPI_ISL_460657, EPI_ISL_460658, EPI_ISL_460659, EPI_ISL_460660, EPI_ISL_460661, EPI_ISL_460662, EPI_ISL_460663, EPI_ISL_460664, EPI_ISL_460666, EPI_ISL_460667, EPI_ISL_460668, EPI_ISL_460669, EPI_ISL_460670, EPI_ISL_460671, EPI_ISL_460672, EPI_ISL_460673, EPI_ISL_460674, EPI_ISL_460675, EPI_ISL_460676, EPI_ISL_460677, EPI_ISL_460682, EPI_ISL_460683, EPI_ISL_460684, EPI_ISL_460685, EPI_ISL_460686, EPI_ISL_460687, EPI_ISL_460688, EPI_ISL_460690, EPI_ISL_460691, EPI_ISL_460692, EPI_ISL_460693, EPI_ISL_460694, EPI_ISL_460695, EPI_ISL_460696, EPI_ISL_460697, EPI_ISL_460698, EPI_ISL_460699, EPI_ISL_460700, EPI_ISL_460701, EPI_ISL_460702, EPI_ISL_460703, EPI_ISL_460704, EPI_ISL_460705, EPI_ISL_460706, EPI_ISL_460707, EPI_ISL_460708, EPI_ISL_460709, EPI_ISL_460710, EPI_ISL_460711, EPI_ISL_460712, EPI_ISL_460713, EPI_ISL_460716, EPI_ISL_460717, EPI_ISL_460718, EPI_ISL_460720, EPI_ISL_460722, EPI_ISL_460724, EPI_ISL_460725, EPI_ISL_460726, EPI_ISL_460727, EPI_ISL_460728, EPI_ISL_460729, EPI_ISL_460730, EPI_ISL_460731, EPI_ISL_460732, EPI_ISL_460733, EPI_ISL_460734, EPI_ISL_460735, EPI_ISL_460736, EPI_ISL_460737, EPI_ISL_460739, EPI_ISL_460740, EPI_ISL_460741, EPI_ISL_460742, EPI_ISL_460743, EPI_ISL_460746, EPI_ISL_460747, EPI_ISL_460748, EPI_ISL_460749, EPI_ISL_460750, EPI_ISL_460752, EPI_ISL_460753, EPI_ISL_460754, EPI_ISL_460755, EPI_ISL_460756, EPI_ISL_460757, EPI_ISL_460758, EPI_ISL_460760, EPI_ISL_460762, EPI_ISL_460763, EPI_ISL_460764, EPI_ISL_460766, EPI_ISL_460767, EPI_ISL_460768, EPI_ISL_460770, EPI_ISL_460771, EPI_ISL_460772, EPI_ISL_460773, EPI_ISL_460774, EPI_ISL_460775, EPI_ISL_460776, EPI_ISL_460777, EPI_ISL_460778, EPI_ISL_460779, EPI_ISL_460780, EPI_ISL_460781, EPI_ISL_460782, EPI_ISL_460785, EPI_ISL_460786, EPI_ISL_460787, EPI_ISL_460788, EPI_ISL_460789, EPI_ISL_460790, EPI_ISL_460791, EPI_ISL_460792, EPI_ISL_460793, EPI_ISL_460794, EPI_ISL_460795, EPI_ISL_460797, EPI_ISL_460798, EPI_ISL_460799, EPI_ISL_460800, EPI_ISL_460801, EPI_ISL_460802, EPI_ISL_460803, EPI_ISL_460804, EPI_ISL_460805, EPI_ISL_460806, EPI_ISL_460807, EPI_ISL_460808, EPI_ISL_460809, EPI_ISL_460810, EPI_ISL_460811, EPI_ISL_460813, EPI_ISL_460814, EPI_ISL_460815, EPI_ISL_460816, EPI_ISL_460817, EPI_ISL_460818, EPI_ISL_460819, EPI_ISL_460820, EPI_ISL_460821, EPI_ISL_460822, EPI_ISL_460823, EPI_ISL_460824, EPI_ISL_460825, EPI_ISL_460826, EPI_ISL_460827, EPI_ISL_460828, EPI_ISL_460829, EPI_ISL_460830, EPI_ISL_460831, EPI_ISL_460832, EPI_ISL_460833, EPI_ISL_460834, EPI_ISL_460835, EPI_ISL_460836, EPI_ISL_460837, EPI_ISL_460838, EPI_ISL_460839, EPI_ISL_460840, EPI_ISL_460841, EPI_ISL_460842, EPI_ISL_460843, EPI_ISL_460844, EPI_ISL_460845, EPI_ISL_460847, EPI_ISL_460849, EPI_ISL_460850, EPI_ISL_460851, EPI_ISL_460852, EPI_ISL_460853, EPI_ISL_460854, EPI_ISL_460855, EPI_ISL_460856, EPI_ISL_460859, EPI_ISL_460860, EPI_ISL_460861, EPI_ISL_460862, EPI_ISL_460863, EPI_ISL_460864, EPI_ISL_460865, EPI_ISL_460866, EPI_ISL_460867, EPI_ISL_460868, EPI_ISL_460869, EPI_ISL_460870, EPI_ISL_460871, EPI_ISL_460872, EPI_ISL_460873, EPI_ISL_460874, EPI_ISL_460875, EPI_ISL_460876, EPI_ISL_460877, EPI_ISL_460878, EPI_ISL_460879, EPI_ISL_460880, EPI_ISL_460881, EPI_ISL_460882, EPI_ISL_460883, EPI_ISL_460884, EPI_ISL_460885, EPI_ISL_460886, EPI_ISL_460887, EPI_ISL_460888, EPI_ISL_460889, EPI_ISL_460890, EPI_ISL_460891, EPI_ISL_460903, EPI_ISL_460904, EPI_ISL_460905, EPI_ISL_460906, EPI_ISL_460907, EPI_ISL_460908, EPI_ISL_460909, EPI_ISL_460910, EPI_ISL_460911, EPI_ISL_460912, EPI_ISL_460913, EPI_ISL_460914, EPI_ISL_460915, EPI_ISL_460917, EPI_ISL_460918, EPI_ISL_460919, EPI_ISL_460920, EPI_ISL_460921, EPI_ISL_460922, EPI_ISL_460923, EPI_ISL_460924, EPI_ISL_460926, EPI_ISL_460927, EPI_ISL_460928, EPI_ISL_460929, EPI_ISL_460930, EPI_ISL_460931, EPI_ISL_460932, EPI_ISL_460933, EPI_ISL_460934, EPI_ISL_460935, EPI_ISL_460936, EPI_ISL_460937, EPI_ISL_460938, EPI_ISL_460939, EPI_ISL_460940, EPI_ISL_460941, EPI_ISL_460942, EPI_ISL_460943, EPI_ISL_460944, EPI_ISL_460945, EPI_ISL_460946, EPI_ISL_460947, EPI_ISL_460948, EPI_ISL_460949, EPI_ISL_460950, EPI_ISL_460951, EPI_ISL_460952, EPI_ISL_460953, EPI_ISL_460955, EPI_ISL_460956, EPI_ISL_460959, EPI_ISL_460960, EPI_ISL_460961, EPI_ISL_460962, EPI_ISL_460963, EPI_ISL_460964, EPI_ISL_460965, EPI_ISL_460966, EPI_ISL_460967, EPI_ISL_460968, EPI_ISL_460969, EPI_ISL_460970, EPI_ISL_460971, EPI_ISL_460972, EPI_ISL_460973, EPI_ISL_460974, EPI_ISL_460975, EPI_ISL_460976, EPI_ISL_460977, EPI_ISL_460978, EPI_ISL_460979, EPI_ISL_460980, EPI_ISL_460981, EPI_ISL_460982, EPI_ISL_460983, EPI_ISL_460984, EPI_ISL_460985, EPI_ISL_460986, EPI_ISL_460987, EPI_ISL_460989, EPI_ISL_460990, EPI_ISL_460992, EPI_ISL_460993, EPI_ISL_460994, EPI_ISL_460995, EPI_ISL_460996, EPI_ISL_460997, EPI_ISL_460998, EPI_ISL_460999, EPI_ISL_461000, EPI_ISL_461001, EPI_ISL_461002, EPI_ISL_461003, EPI_ISL_461004, EPI_ISL_461006, EPI_ISL_461007, EPI_ISL_461008, EPI_ISL_461009, EPI_ISL_461010, EPI_ISL_461011, EPI_ISL_461012, EPI_ISL_461013, EPI_ISL_461015, EPI_ISL_461016, EPI_ISL_461017, EPI_ISL_461018, EPI_ISL_461019, EPI_ISL_461020, EPI_ISL_461021, EPI_ISL_461022, EPI_ISL_461025, EPI_ISL_461026, EPI_ISL_461027, EPI_ISL_461029, EPI_ISL_461030, EPI_ISL_461031, EPI_ISL_461032, EPI_ISL_461033, EPI_ISL_461034, EPI_ISL_461035, EPI_ISL_461036, EPI_ISL_461037, EPI_ISL_461038, EPI_ISL_461039, EPI_ISL_461040, EPI_ISL_461041, EPI_ISL_461042, EPI_ISL_461043, EPI_ISL_461044, EPI_ISL_461045, EPI_ISL_461046, EPI_ISL_461047, EPI_ISL_461048, EPI_ISL_461051, EPI_ISL_461052, EPI_ISL_461053, EPI_ISL_461054, EPI_ISL_461055, EPI_ISL_461056, EPI_ISL_461057, EPI_ISL_461058, EPI_ISL_461059, EPI_ISL_461060, EPI_ISL_461063, EPI_ISL_461064, EPI_ISL_461065, EPI_ISL_461066, EPI_ISL_461068, EPI_ISL_461069, EPI_ISL_461070, EPI_ISL_461071, EPI_ISL_461074, EPI_ISL_461075, EPI_ISL_461076, EPI_ISL_461077, EPI_ISL_461078, EPI_ISL_461079, EPI_ISL_461082, EPI_ISL_461083, EPI_ISL_461084, EPI_ISL_461085, EPI_ISL_461086, EPI_ISL_461087, EPI_ISL_461088, EPI_ISL_461089, EPI_ISL_461090, EPI_ISL_461091, EPI_ISL_461092, EPI_ISL_461093, EPI_ISL_461094, EPI_ISL_461095, EPI_ISL_461097, EPI_ISL_461098, EPI_ISL_461100, EPI_ISL_461101, EPI_ISL_461102, EPI_ISL_461103, EPI_ISL_461104, EPI_ISL_461105, EPI_ISL_461106, EPI_ISL_461108, EPI_ISL_461109, EPI_ISL_461111, EPI_ISL_461112, EPI_ISL_461113, EPI_ISL_461114, EPI_ISL_461115, EPI_ISL_461116, EPI_ISL_461118, EPI_ISL_461120, EPI_ISL_461121, EPI_ISL_461122, EPI_ISL_461123, EPI_ISL_461125, EPI_ISL_461126, EPI_ISL_461127, EPI_ISL_461128, EPI_ISL_461129, EPI_ISL_461130, EPI_ISL_461131, EPI_ISL_461132, EPI_ISL_461133, EPI_ISL_461134, EPI_ISL_461135, EPI_ISL_461136, EPI_ISL_461137, EPI_ISL_461139, EPI_ISL_461140, EPI_ISL_461141, EPI_ISL_461143, EPI_ISL_461144, EPI_ISL_461145, EPI_ISL_461146, EPI_ISL_461147, EPI_ISL_461148, EPI_ISL_461149, EPI_ISL_461150, EPI_ISL_461151, EPI_ISL_461152, EPI_ISL_461153, EPI_ISL_461155, EPI_ISL_461156, EPI_ISL_461157, EPI_ISL_461158, EPI_ISL_461159, EPI_ISL_461160, EPI_ISL_461161, EPI_ISL_461162, EPI_ISL_461163, EPI_ISL_461164, EPI_ISL_461165, EPI_ISL_461167, EPI_ISL_461168, EPI_ISL_461169, EPI_ISL_461171, EPI_ISL_461172, EPI_ISL_461173, EPI_ISL_461174, EPI_ISL_461175, EPI_ISL_461176, EPI_ISL_461177, EPI_ISL_461178, EPI_ISL_461179, EPI_ISL_461180, EPI_ISL_461181, EPI_ISL_461182, EPI_ISL_461183, EPI_ISL_461184, EPI_ISL_461185, EPI_ISL_461186, EPI_ISL_461188, EPI_ISL_461189, EPI_ISL_461191, EPI_ISL_461192, EPI_ISL_461193, EPI_ISL_461194, EPI_ISL_461195, EPI_ISL_461197, EPI_ISL_461198, EPI_ISL_461199, EPI_ISL_461200, EPI_ISL_461201, EPI_ISL_461202, EPI_ISL_461204, EPI_ISL_461206, EPI_ISL_461207, EPI_ISL_461208, EPI_ISL_461209, EPI_ISL_461210, EPI_ISL_461211, EPI_ISL_461212, EPI_ISL_461213, EPI_ISL_461214, EPI_ISL_461215, EPI_ISL_461216, EPI_ISL_461217, EPI_ISL_461218, EPI_ISL_461219, EPI_ISL_461221, EPI_ISL_461222, EPI_ISL_461223, EPI_ISL_461224, EPI_ISL_461226, EPI_ISL_461227, EPI_ISL_461228, EPI_ISL_461229, EPI_ISL_461230, EPI_ISL_461232, EPI_ISL_461233, EPI_ISL_461234, EPI_ISL_461235, EPI_ISL_461236, EPI_ISL_461237, EPI_ISL_461238, EPI_ISL_461239, EPI_ISL_461240, EPI_ISL_461241, EPI_ISL_461242, EPI_ISL_461243, EPI_ISL_461244, EPI_ISL_461245, EPI_ISL_461246, EPI_ISL_461247, EPI_ISL_461250, EPI_ISL_461251, EPI_ISL_461254, EPI_ISL_461255, EPI_ISL_461257, EPI_ISL_461258, EPI_ISL_461259, EPI_ISL_461260, EPI_ISL_461261, EPI_ISL_461264, EPI_ISL_461265, EPI_ISL_461266, EPI_ISL_461268, EPI_ISL_461270, EPI_ISL_461271, EPI_ISL_461272, EPI_ISL_461273, EPI_ISL_461274, EPI_ISL_461275, EPI_ISL_461276, EPI_ISL_461277, EPI_ISL_461280, EPI_ISL_461281, EPI_ISL_461282, EPI_ISL_461283, EPI_ISL_461284, EPI_ISL_461285, EPI_ISL_461286, EPI_ISL_461287, EPI_ISL_461288, EPI_ISL_461289, EPI_ISL_461291, EPI_ISL_461292, EPI_ISL_461293, EPI_ISL_461294, EPI_ISL_461295, EPI_ISL_461296, EPI_ISL_461297, EPI_ISL_461298, EPI_ISL_461299, EPI_ISL_461300, EPI_ISL_461302, EPI_ISL_461303, EPI_ISL_461305, EPI_ISL_461307, EPI_ISL_461308, EPI_ISL_461309, EPI_ISL_461310, EPI_ISL_461311, EPI_ISL_461312, EPI_ISL_461313, EPI_ISL_461314, EPI_ISL_461315, EPI_ISL_461316, EPI_ISL_461317, EPI_ISL_461319, EPI_ISL_461320, EPI_ISL_461321, EPI_ISL_461322, EPI_ISL_461323, EPI_ISL_461324, EPI_ISL_461325, EPI_ISL_461326, EPI_ISL_461328, EPI_ISL_461329, EPI_ISL_461331, EPI_ISL_461332, EPI_ISL_461333, EPI_ISL_461334, EPI_ISL_461335, EPI_ISL_461337, EPI_ISL_461338, EPI_ISL_461339, EPI_ISL_461340, EPI_ISL_461341, EPI_ISL_461342, EPI_ISL_461343, EPI_ISL_461344, EPI_ISL_461345, EPI_ISL_461346, EPI_ISL_461347, EPI_ISL_461348, EPI_ISL_461349, EPI_ISL_461350, EPI_ISL_461351, EPI_ISL_461352, EPI_ISL_461353, EPI_ISL_461355, EPI_ISL_461356, EPI_ISL_461357, EPI_ISL_461358, EPI_ISL_461359, EPI_ISL_461360, EPI_ISL_461361, EPI_ISL_461363, EPI_ISL_461364, EPI_ISL_461365, EPI_ISL_461366, EPI_ISL_461367, EPI_ISL_461368, EPI_ISL_461370, EPI_ISL_461373, EPI_ISL_461374, EPI_ISL_461375, EPI_ISL_461376, EPI_ISL_461377, EPI_ISL_461378, EPI_ISL_461379, EPI_ISL_461380, EPI_ISL_461381, EPI_ISL_461382, EPI_ISL_461383, EPI_ISL_461384, EPI_ISL_461385, EPI_ISL_461386, EPI_ISL_461387, EPI_ISL_461388, EPI_ISL_461389, EPI_ISL_461390, EPI_ISL_461391, EPI_ISL_461392, EPI_ISL_461393, EPI_ISL_461394, EPI_ISL_461395, EPI_ISL_461396, EPI_ISL_461397, EPI_ISL_461398 | | | |
| see above | Dutch COVID-19 response team | Erasmus Medical Center | Bas Oude Munnink, David Nieuwenhuijse, Reina Sikkema, Claudia Schapendonk, Irina Chestakova, Anne van der Linden, Theo Bestebroer, Stefan van Nieuwkoop, Mark Pronk, Pascal Lexmond, Corien Swaan, Manon Haverkate, Madelief Mollers, Mart Stein, Sandra Kengne Kamga Mobou, Jeroen van Kampen, Jolanda Voermans, Aura Timen, Corine GeurtsvanKessel, Annemiek van der Eijk, Richard Molenkamp, Marion Koopmans, on behalf of the Dutch national COVID-19 response team. |
| EPI_ISL_461399, EPI_ISL_461402, EPI_ISL_461403, EPI_ISL_461404, EPI_ISL_461405, EPI_ISL_461406, EPI_ISL_461407, EPI_ISL_461408, EPI_ISL_461409, EPI_ISL_461410, EPI_ISL_461411, EPI_ISL_461412, EPI_ISL_461413, EPI_ISL_461414, EPI_ISL_461415, EPI_ISL_461416, EPI_ISL_461417, EPI_ISL_461418, EPI_ISL_461419, EPI_ISL_461420, EPI_ISL_461421, EPI_ISL_461422, EPI_ISL_461423, EPI_ISL_461424, EPI_ISL_461425, EPI_ISL_461426, EPI_ISL_461427, EPI_ISL_461428, EPI_ISL_461429, EPI_ISL_461430, EPI_ISL_461432, EPI_ISL_461434, EPI_ISL_461435, EPI_ISL_461436, EPI_ISL_461437, EPI_ISL_461438, EPI_ISL_461440, EPI_ISL_461441, EPI_ISL_461442, EPI_ISL_461444, EPI_ISL_461445, EPI_ISL_461446, EPI_ISL_461447, EPI_ISL_461448, EPI_ISL_461449, EPI_ISL_461450, EPI_ISL_461451, EPI_ISL_461452, EPI_ISL_461453, EPI_ISL_461454, EPI_ISL_461455, EPI_ISL_461456, EPI_ISL_461457, EPI_ISL_461458, EPI_ISL_461459, EPI_ISL_461460, EPI_ISL_461461, EPI_ISL_461462, EPI_ISL_461463, EPI_ISL_461464, EPI_ISL_461465, EPI_ISL_461467, EPI_ISL_461468, EPI_ISL_461469, EPI_ISL_461470, EPI_ISL_461471, EPI_ISL_461472, EPI_ISL_461473, EPI_ISL_461474, EPI_ISL_461475, EPI_ISL_461476, EPI_ISL_461477 | | | |
| see above | UW Virology Lab | UW Virology Lab | Pavitra Roychoudhury, Amin Addetia, Hong Xie, Lasata Shrestha, Truong Nguyen, Meei-Li Huang, Keith Jerome, Alexander Greninger |
| EPI_ISL_461478 | Government Medical College, Vadodara | Gujarat Biotechnology Research Centre | Fenil Patel, Nidhi Patel, Nitin Savaliya, Raghawendra Kumar, Dinesh Kumar, Zuber Saiyed, Komal Patel, Labdhi Pandya, Snehal Bagatharia, Tanuja Javadekar , R N Daveshwar, Tejas Shah, Ankit Hinsu, Pritesh Sabara, Apurvasinh Puvar, Janvi Raval, Zarna Patel, Monika Gandhi, Pinal Trivedi, Maharshi Pandya, R D Dixit, A M Kadri, Harsh Bakshi, Chaitanya Joshi, Madhvi Joshi, |
| EPI_ISL_461479 | Government Medical College, Vadodara | Gujarat Biotechnology Research Centre | Neelam Nathani, Nitin Savaliya, Raghawendra Kumar, Dinesh Kumar, Zuber Saiyed, Komal Patel, Labdhi Pandya, Snehal Bagatharia, Tanuja Javadekar , R N Daveshwar, Tejas Shah, Ankit Hinsu, Pritesh Sabara, Apurvasinh Puvar, Janvi Raval, Zarna Patel, Monika Gandhi, Pinal Trivedi, Maharshi Pandya, Nidhi Patel, R D Dixit, A M Kadri, Harsh Bakshi, Chaitanya Joshi, Madhvi Joshi, |
| EPI_ISL_461480 | Government Medical College, Vadodara | Gujarat Biotechnology Research Centre | Armi Chaudhari, Raghawendra Kumar, Dinesh Kumar, Zuber Saiyed, Komal Patel, Labdhi Pandya, Snehal Bagatharia, Tanuja Javadekar , R N Daveshwar, Tejas Shah, Ankit Hinsu, Pritesh Sabara, Apurvasinh Puvar, Janvi Raval, Zarna Patel, Monika Gandhi, Pinal Trivedi, Maharshi Pandya, Nidhi Patel, Nitin Savaliya, R D Dixit, A M Kadri, Harsh Bakshi, Chaitanya Joshi, Madhvi Joshi, |
| EPI_ISL_461481 | Pandit Deendayal Upadhyay Government Medical College, Rajkot | Gujarat Biotechnology Research Centre | Bhavya Jindal, Dinesh Kumar, Zuber Saiyed, Komal Patel, Labdhi Pandya, Snehal Bagatharia, Prakash Modi, Sejul Antala, Manish Pattani, Tejas Shah, Ankit Hinsu, Pritesh Sabara, Apurvasinh Puvar, Janvi Raval, Zarna Patel, Monika Gandhi, Pinal Trivedi, Maharshi Pandya, Nidhi Patel, Nitin Savaliya, Raghawendra Kumar, R D Dixit, A M Kadri, Harsh Bakshi, Chaitanya Joshi, Madhvi Joshi |
| EPI_ISL_461482 | Pandit Deendayal Upadhyay Government Medical College, Rajkot | Gujarat Biotechnology Research Centre | Anjali Rajwar, Zuber Saiyed, Komal Patel, Labdhi Pandya, Snehal Bagatharia, Prakash Modi, Sejul Antala, Manish Pattani, Tejas Shah, Ankit Hinsu, Pritesh Sabara, Apurvasinh Puvar, Janvi Raval, Zarna Patel, Monika Gandhi, Pinal Trivedi, Maharshi Pandya, Nidhi Patel, Nitin Savaliya, Raghawendra Kumar, Dinesh Kumar, R D Dixit, A M Kadri, Harsh Bakshi, Chaitanya Joshi, Madhvi Joshi |
| EPI_ISL_461483 | B.J. Medical College and Civil hospital | Gujarat Biotechnology Research Centre | Dipeshwari Shewale, Komal Patel, Labdhi Pandya, Snehal Bagatharia, Pranay Shah, Kamlesh J Upadhyay, Tejas Shah, Ankit Hinsu, Pritesh Sabara, Apurvasinh Puvar, Janvi Raval, Zarna Patel, Monika Gandhi, Pinal Trivedi, Maharshi Pandya, Nidhi Patel, Nitin Savaliya, Raghawendra Kumar, Dinesh Kumar, Zuber Saiyed, R D Dixit, A M Kadri, Harsh Bakshi, Chaitanya Joshi, Madhvi Joshi, |

| | | | |
|---|---|---|---|
| EPI_ISL_461484 | B.J. Medical College and Civil hospital | Gujarat Biotechnology Research Centre | Priyanka P Vatsa, Labdhi Pandya, Snehal Bagatharia, Pranay Shah, Kamlesh J Upadhyay, Tejas Shah, Ankit Hinsu, Pritesh Sabara, Apurvasinh Puvar, Janvi Raval, Zarna Patel, Monika Gandhi, Pinal Trivedi, Maharshi Pandya, Nidhi Patel, Nitin Savaliya, Raghawendra Kumar, Dinesh Kumar, Zuber Saiyed, Komal Patel, R D Dixit, A M Kadri, Harsh Bakshi, Chaitanya Joshi, Madhvi Joshi, |
| EPI_ISL_461485 | B.J. Medical College and Civil hospital | Gujarat Biotechnology Research Centre | Pooja P Doshi, Snehal Bagatharia, Pranay Shah, Kamlesh J Upadhyay, Tejas Shah, Ankit Hinsu, Pritesh Sabara, Apurvasinh Puvar, Janvi Raval, Zarna Patel, Monika Gandhi, Pinal Trivedi, Maharshi Pandya, Nidhi Patel, Nitin Savaliya, Raghawendra Kumar, Dinesh Kumar, Zuber Saiyed, Komal Patel, Labdhi Pandya, R D Dixit, A M Kadri, Harsh Bakshi, Chaitanya Joshi, Madhvi Joshi, |
| EPI_ISL_461486 | B.J. Medical College and Civil hospital | Gujarat Biotechnology Research Centre | Akanksha Verma, Pranay Shah, Kamlesh J Upadhyay, Tejas Shah, Ankit Hinsu, Pritesh Sabara, Apurvasinh Puvar, Janvi Raval, Zarna Patel, Monika Gandhi, Pinal Trivedi, Maharshi Pandya, Nidhi Patel, Nitin Savaliya, Raghawendra Kumar, Dinesh Kumar, Zuber Saiyed, Komal Patel, Labdhi Pandya, Snehal Bagatharia, R D Dixit, A M Kadri, Harsh Bakshi, Chaitanya Joshi, Madhvi Joshi, |
| EPI_ISL_461487 | B.J. Medical College and Civil hospital | Gujarat Biotechnology Research Centre | Priti Pandita, Kamlesh J Upadhyay, Tejas Shah, Ankit Hinsu, Pritesh Sabara, Apurvasinh Puvar, Janvi Raval, Zarna Patel, Monika Gandhi, Pinal Trivedi, Maharshi Pandya, Nidhi Patel, Nitin Savaliya, Raghawendra Kumar, Dinesh Kumar, Zuber Saiyed, Komal Patel, Labdhi Pandya, Snehal Bagatharia, Pranay Shah, R D Dixit, A M Kadri, Harsh Bakshi, Chaitanya Joshi, Madhvi Joshi, |
| EPI_ISL_461488 | B.J. Medical College and Civil hospital | Gujarat Biotechnology Research Centre | Pragya Sharma, Tejas Shah, Ankit Hinsu, Pritesh Sabara, Apurvasinh Puvar, Janvi Raval, Zarna Patel, Monika Gandhi, Pinal Trivedi, Maharshi Pandya, Nidhi Patel, Nitin Savaliya, Raghawendra Kumar, Dinesh Kumar, Zuber Saiyed, Komal Patel, Labdhi Pandya, Snehal Bagatharia, Pranay Shah, Kamlesh J Upadhyay, R D Dixit, A M Kadri, Harsh Bakshi, Chaitanya Joshi, Madhvi Joshi, |
| EPI_ISL_461489 | B.J. Medical College and Civil hospital | Gujarat Biotechnology Research Centre | Neha Rajpara, Ankit Hinsu, Pritesh Sabara, Apurvasinh Puvar, Janvi Raval, Zarna Patel, Monika Gandhi, Pinal Trivedi, Maharshi Pandya, Nidhi Patel, Nitin Savaliya, Raghawendra Kumar, Dinesh Kumar, Zuber Saiyed, Komal Patel, Labdhi Pandya, Snehal Bagatharia, Pranay Shah, Kamlesh J Upadhyay, Tejas Shah, R D Dixit, A M Kadri, Harsh Bakshi, Chaitanya Joshi, Madhvi Joshi, |
| EPI_ISL_461490 | B.J. Medical College and Civil hospital | Gujarat Biotechnology Research Centre | Afzal Ansari, Pritesh Sabara, Apurvasinh Puvar, Janvi Raval, Zarna Patel, Monika Gandhi, Pinal Trivedi, Maharshi Pandya, Nidhi Patel, Nitin Savaliya, Raghawendra Kumar, Dinesh Kumar, Zuber Saiyed, Komal Patel, Labdhi Pandya, Snehal Bagatharia, Pranay Shah, Kamlesh J Upadhyay, Tejas Shah, Ankit Hinsu, R D Dixit, A M Kadri, Harsh Bakshi, Chaitanya Joshi, Madhvi Joshi, |
| EPI_ISL_461491 | B.J. Medical College and Civil hospital | Gujarat Biotechnology Research Centre | Fenil Patel, Apurvasinh Puvar, Janvi Raval, Zarna Patel, Monika Gandhi, Pinal Trivedi, Maharshi Pandya, Nidhi Patel, Nitin Savaliya, Raghawendra Kumar, Dinesh Kumar, Zuber Saiyed, Komal Patel, Labdhi Pandya, Snehal Bagatharia, Pranay Shah, Kamlesh J Upadhyay, Tejas Shah, Ankit Hinsu, Pritesh Sabara, R D Dixit, A M Kadri, Harsh Bakshi, Chaitanya Joshi, Madhvi Joshi, |
| EPI_ISL_461492 | B.J. Medical College and Civil hospital | Gujarat Biotechnology Research Centre | Neelam Nathani, Janvi Raval, Zarna Patel, Monika Gandhi, Pinal Trivedi, Maharshi Pandya, Nidhi Patel, Nitin Savaliya, Raghawendra Kumar, Dinesh Kumar, Zuber Saiyed, Komal Patel, Labdhi Pandya, Snehal Bagatharia, Pranay Shah, Kamlesh J Upadhyay, Tejas Shah, Ankit Hinsu, Pritesh Sabara, Apurvasinh Puvar, R D Dixit, A M Kadri, Harsh Bakshi, Chaitanya Joshi, Madhvi Joshi, |
| EPI_ISL_461493 | B.J. Medical College and Civil hospital | Gujarat Biotechnology Research Centre | Armi Chaudhari, Zarna Patel, Monika Gandhi, Pinal Trivedi, Maharshi Pandya, Nidhi Patel, Nitin Savaliya, Raghawendra Kumar, Dinesh Kumar, Zuber Saiyed, Komal Patel, Labdhi Pandya, Snehal Bagatharia, Pranay Shah, Kamlesh J Upadhyay, Tejas Shah, Ankit Hinsu, Pritesh Sabara, Apurvasinh Puvar, Janvi Raval, R D Dixit, A M Kadri, Harsh Bakshi, Chaitanya Joshi, Madhvi Joshi, |
| EPI_ISL_461494 | B.J. Medical College and Civil hospital | Gujarat Biotechnology Research Centre | Bhavya Jindal, Monika Gandhi, Pinal Trivedi, Maharshi Pandya, Nidhi Patel, Nitin Savaliya, Raghawendra Kumar, Dinesh Kumar, Zuber Saiyed, Komal Patel, Labdhi Pandya, Snehal Bagatharia, Pranay Shah, Kamlesh J Upadhyay, Tejas Shah, Ankit Hinsu, Pritesh Sabara, Apurvasinh Puvar, Janvi Raval, Zarna Patel, R D Dixit, A M Kadri, Harsh Bakshi, Chaitanya Joshi, Madhvi Joshi, |
| EPI_ISL_461495 | B.J. Medical College and Civil hospital | Gujarat Biotechnology Research Centre | Anjali Rajwar, Pinal Trivedi, Maharshi Pandya, Nidhi Patel, Nitin Savaliya, Raghawendra Kumar, Dinesh Kumar, Zuber Saiyed, Komal Patel, Labdhi Pandya, Snehal Bagatharia, Pranay Shah, Kamlesh J Upadhyay, Tejas Shah, Ankit Hinsu, Pritesh Sabara, Apurvasinh Puvar, Janvi Raval, Zarna Patel, Monika Gandhi, R D Dixit, A M Kadri, Harsh Bakshi, Chaitanya Joshi, Madhvi Joshi, |
| EPI_ISL_461496 | B.J. Medical College and Civil hospital | Gujarat Biotechnology Research Centre | Dipeshwari Shewale, Maharshi Pandya, Nidhi Patel, Nitin Savaliya, Raghawendra Kumar, Dinesh Kumar, Zuber Saiyed, Komal Patel, Labdhi Pandya, Snehal Bagatharia, Pranay Shah, Kamlesh J Upadhyay, Tejas Shah, Ankit Hinsu, Pritesh Sabara, Apurvasinh Puvar, Janvi Raval, Zarna Patel, Monika Gandhi, Pinal Trivedi, R D Dixit, A M Kadri, Harsh Bakshi, Chaitanya Joshi, Madhvi Joshi, |
| EPI_ISL_461497 | B.J. Medical College and Civil hospital | Gujarat Biotechnology Research Centre | Priyanka P Vatsa, Nidhi Patel, Nitin Savaliya, Raghawendra Kumar, Dinesh Kumar, Zuber Saiyed, Komal Patel, Labdhi Pandya, Snehal Bagatharia, Pranay Shah, Kamlesh J Upadhyay, Tejas Shah, Ankit Hinsu, Pritesh Sabara, Apurvasinh Puvar, Janvi Raval, Zarna Patel, Monika Gandhi, Pinal Trivedi, Maharshi Pandya, R D Dixit, A M Kadri, Harsh Bakshi, Chaitanya Joshi, Madhvi Joshi, |
| EPI_ISL_461498 | B.J. Medical College and Civil hospital | Gujarat Biotechnology Research Centre | Pooja P Doshi, Nitin Savaliya, Raghawendra Kumar, Dinesh Kumar, Zuber Saiyed, Komal Patel, Labdhi Pandya, Snehal Bagatharia, Pranay Shah, Kamlesh J Upadhyay, Tejas Shah, Ankit Hinsu, Pritesh Sabara, Apurvasinh Puvar, Janvi Raval, Zarna Patel, Monika Gandhi, Pinal Trivedi, Maharshi Pandya, Nidhi Patel, R D Dixit, A M Kadri, Harsh Bakshi, Chaitanya Joshi, Madhvi Joshi, |
| EPI_ISL_461499 | B.J. Medical College and Civil hospital | Gujarat Biotechnology Research Centre | Akanksha Verma, Raghawendra Kumar, Dinesh Kumar, Zuber Saiyed, Komal Patel, Labdhi Pandya, Snehal Bagatharia, Pranay Shah, Kamlesh J Upadhyay, Tejas Shah, Ankit Hinsu, Pritesh Sabara, Apurvasinh Puvar, Janvi Raval, Zarna Patel, Monika Gandhi, Pinal Trivedi, Maharshi Pandya, Nidhi Patel, Nitin Savaliya, R D Dixit, A M Kadri, Harsh Bakshi, Chaitanya Joshi, Madhvi Joshi, |
| EPI_ISL_461500 | B.J. Medical College and Civil hospital | Gujarat Biotechnology Research Centre | Priti Pandita, Dinesh Kumar, Zuber Saiyed, Komal Patel, Labdhi Pandya, Snehal Bagatharia, Pranay Shah, Kamlesh J Upadhyay, Tejas Shah, Ankit Hinsu, Pritesh Sabara, Apurvasinh Puvar, Janvi Raval, Zarna Patel, Monika Gandhi, Pinal Trivedi, Maharshi Pandya, Nidhi Patel, Nitin Savaliya, Raghawendra Kumar, R D Dixit, A M Kadri, Harsh Bakshi, Chaitanya Joshi, Madhvi Joshi, |
| EPI_ISL_461501 | B.J. Medical College and Civil hospital | Gujarat Biotechnology Research Centre | Pragya Sharma, Zuber Saiyed, Komal Patel, Labdhi Pandya, Snehal Bagatharia, Pranay Shah, Kamlesh J Upadhyay, Tejas Shah, Ankit Hinsu, Pritesh Sabara, Apurvasinh Puvar, Janvi Raval, Zarna Patel, Monika Gandhi, Pinal Trivedi, Maharshi Pandya, Nidhi Patel, Nitin Savaliya, Raghawendra Kumar, Dinesh Kumar, R D Dixit, A M Kadri, Harsh Bakshi, Chaitanya Joshi, Madhvi Joshi, |
| EPI_ISL_461502 | B.J. Medical College and Civil hospital | Gujarat Biotechnology Research Centre | Neha Rajpara, Komal Patel, Labdhi Pandya, Snehal Bagatharia, Pranay Shah, Kamlesh J Upadhyay, Tejas Shah, Ankit Hinsu, Pritesh Sabara, Apurvasinh Puvar, Janvi Raval, Zarna Patel, Monika Gandhi, Pinal Trivedi, Maharshi Pandya, Nidhi Patel, Nitin Savaliya, Raghawendra Kumar, Dinesh Kumar, Zuber Saiyed, R D Dixit, A M Kadri, Harsh Bakshi, Chaitanya Joshi, Madhvi Joshi, |
| EPI_ISL_461503 | B.J. Medical College and Civil hospital | Gujarat Biotechnology Research Centre | Afzal Ansari, Labdhi Pandya, Snehal Bagatharia, Pranay Shah, Kamlesh J Upadhyay, Tejas Shah, Ankit Hinsu, Pritesh Sabara, Apurvasinh Puvar, Janvi Raval, Zarna Patel, Monika Gandhi, Pinal Trivedi, Maharshi Pandya, Nidhi Patel, Nitin Savaliya, Raghawendra Kumar, Dinesh Kumar, Zuber Saiyed, Komal Patel, R D Dixit, A M Kadri, Harsh Bakshi, Chaitanya Joshi, Madhvi Joshi, |
| EPI_ISL_461504 | B.J. Medical College and Civil hospital | Gujarat Biotechnology Research Centre | Snehal Bagatharia, Pranay Shah, Kamlesh J Upadhyay, Tejas Shah, Ankit Hinsu, Pritesh Sabara, Apurvasinh Puvar, Janvi Raval, Zarna Patel, Monika Gandhi, Pinal Trivedi, Maharshi Pandya, Nidhi Patel, Nitin Savaliya, Raghawendra Kumar, Dinesh Kumar, Zuber Saiyed, Komal Patel, Labdhi Pandya, Fenil Patel, R D Dixit, A M Kadri, Harsh Bakshi, Chaitanya Joshi, Madhvi Joshi, |
| EPI_ISL_461505 | B.J. Medical College and Civil hospital | Gujarat Biotechnology Research Centre | Pranay Shah, Kamlesh J Upadhyay, Tejas Shah, Ankit Hinsu, Pritesh Sabara, Apurvasinh Puvar, Janvi Raval, Zarna Patel, Monika Gandhi, Pinal Trivedi, Maharshi Pandya, Nidhi Patel, Nitin Savaliya, Raghawendra Kumar, Dinesh Kumar, Zuber Saiyed, Komal Patel, Labdhi Pandya, Snehal Bagatharia, Neelam Nathani, R D Dixit, A M Kadri, Harsh Bakshi, Chaitanya Joshi, Madhvi Joshi, |
| EPI_ISL_461506 | B.J. Medical College and Civil hospital | Gujarat Biotechnology Research Centre | Kamlesh J Upadhyay, Tejas Shah, Ankit Hinsu, Pritesh Sabara, Apurvasinh Puvar, Janvi Raval, Zarna Patel, Monika Gandhi, Pinal Trivedi, Maharshi Pandya, Nidhi Patel, Nitin Savaliya, Raghawendra Kumar, Dinesh Kumar, Zuber Saiyed, Komal Patel, Labdhi Pandya, Snehal Bagatharia, Pranay Shah, Armi Chaudhari, R D Dixit, A M Kadri, Harsh Bakshi, Chaitanya Joshi, Madhvi Joshi, |
| EPI_ISL_461508, EPI_ISL_461509, EPI_ISL_461510, EPI_ISL_461513, EPI_ISL_461514, EPI_ISL_461515, EPI_ISL_461517, EPI_ISL_461518, EPI_ISL_461520, EPI_ISL_461521, EPI_ISL_461523, EPI_ISL_461524, EPI_ISL_461525, EPI_ISL_461526, EPI_ISL_461527, EPI_ISL_461528, EPI_ISL_461529, EPI_ISL_461530, EPI_ISL_461531, EPI_ISL_461532, EPI_ISL_461534, EPI_ISL_461535, EPI_ISL_461537, EPI_ISL_461538, EPI_ISL_461541, EPI_ISL_461542, EPI_ISL_461543, EPI_ISL_461544, EPI_ISL_461545 | | | |
| see above | University of Birmingham | COVID-19 Genomics UK (COG-UK) Consortium | Loman Lab: Claire McMurray, Joanne Stockton, Samuel Nicholls, Radoslaw Poplawski, Will Rowe, Josh Quick, Nicholas Loman // UHB Lab: Celina M Whalley, Andrew Bosworth, Charlotte Poxon, Kasun Wanigasooriya, Oliver Pickles, Mike Kidd, Alex Richter, Andrew D Beggs // PHE Heartlands Lab: Husam Osman, Andrew Bosworth |
| EPI_ISL_461546, EPI_ISL_461547, EPI_ISL_461548, EPI_ISL_461552, EPI_ISL_461553, EPI_ISL_461561, EPI_ISL_461562, EPI_ISL_461563, EPI_ISL_461571, EPI_ISL_461572, EPI_ISL_461573, EPI_ISL_461574, EPI_ISL_461575, EPI_ISL_461578, EPI_ISL_461579, EPI_ISL_461580, EPI_ISL_461581, EPI_ISL_461582, EPI_ISL_461583, EPI_ISL_461586, EPI_ISL_461587 | | | |
| see above | Department of Pathology, University of Cambridge | COVID-19 Genomics UK (COG-UK) Consortium | Luke W Meredith, M. Estée Török, Myra Hosmillo, William L. Hamilton, Martin D. Curran, Theresa Feltwell, Grant Hall, Anna Yakovleva, Fahad A Khokhar, Charlotte J. Houldcroft, Laura G Caller, Aminu S. Jahun, Sarah L. Caddy, Ian Goodfellow |
| EPI_ISL_461589, EPI_ISL_461592, EPI_ISL_461595, EPI_ISL_461596, EPI_ISL_461597, EPI_ISL_461598, EPI_ISL_461599, EPI_ISL_461600, EPI_ISL_461601, EPI_ISL_461603, EPI_ISL_461604, EPI_ISL_461605, EPI_ISL_461606, EPI_ISL_461607, EPI_ISL_461610, EPI_ISL_461611, EPI_ISL_461612, EPI_ISL_461613, EPI_ISL_461614, EPI_ISL_461615, EPI_ISL_461616, EPI_ISL_461617, EPI_ISL_461618, EPI_ISL_461619, EPI_ISL_461620, EPI_ISL_461621, EPI_ISL_461622, EPI_ISL_461623, EPI_ISL_461624, EPI_ISL_461625, EPI_ISL_461626, EPI_ISL_461628, EPI_ISL_461629, EPI_ISL_461630, EPI_ISL_461631, EPI_ISL_461633, EPI_ISL_461635, EPI_ISL_461637, EPI_ISL_461638, EPI_ISL_461639, EPI_ISL_461641, EPI_ISL_461642, EPI_ISL_461643, EPI_ISL_461644, EPI_ISL_461647, EPI_ISL_461649, EPI_ISL_461652, EPI_ISL_461655, EPI_ISL_461656, EPI_ISL_461657, EPI_ISL_461659, EPI_ISL_461660, EPI_ISL_461661, EPI_ISL_461662, EPI_ISL_461664, EPI_ISL_461665, EPI_ISL_461667, EPI_ISL_461668, EPI_ISL_461669, EPI_ISL_461670, EPI_ISL_461674, EPI_ISL_461675, EPI_ISL_461676, EPI_ISL_461678, EPI_ISL_461685, EPI_ISL_461688, EPI_ISL_461691, EPI_ISL_461698, EPI_ISL_461699, EPI_ISL_461705 | | | |
| see above | West of Scotland Specialist Virology Centre, NHSGGC / MRC-University of Glasgow Centre for Virus Research | COVID-19 Genomics UK (COG-UK) Consortium | Ana da Silva Filipe, Natasha Johnson, Kathy Smollett, Daniel Mair, Stephen Carmichael, Lily Tong, Jenna Nichols, Elihu Aranday-Cortes, Kirstyn Brunker, Yasmin Parr, Kyriaki Nomikou; Sarah McDonald, Marc Niebel, Patawee Asamaphan; Richard Orton, Joseph Hughes, Sreenu Vattipally, David L Robertson; Alasdair MacLean, Rory Gunson; Kathy Li, Natasha Jesudason, Rajiv Shah, James Shepherd, Antonia Ho, Emma Thomson |
| EPI_ISL_461706, EPI_ISL_461707, EPI_ISL_461708, EPI_ISL_461710, EPI_ISL_461711, EPI_ISL_461712, EPI_ISL_461713, EPI_ISL_461714, EPI_ISL_461716, EPI_ISL_461718, EPI_ISL_461719, EPI_ISL_461720, EPI_ISL_461721, EPI_ISL_461723, EPI_ISL_461725, EPI_ISL_461726, EPI_ISL_461727, EPI_ISL_461728, EPI_ISL_461729, EPI_ISL_461730, EPI_ISL_461731, EPI_ISL_461733, EPI_ISL_461735, EPI_ISL_461736, EPI_ISL_461737, EPI_ISL_461738, EPI_ISL_461739, EPI_ISL_461741, EPI_ISL_461742, EPI_ISL_461743, EPI_ISL_461744, EPI_ISL_461745, EPI_ISL_461746, EPI_ISL_461748, EPI_ISL_461749, EPI_ISL_461751, EPI_ISL_461753, EPI_ISL_461754, EPI_ISL_461755, EPI_ISL_461756, EPI_ISL_461757, EPI_ISL_461759, EPI_ISL_461761, EPI_ISL_461762 | | | |
| see above | Virology Department, Royal Infirmary of Edinburgh, NHS Lothian / School of Biological Sciences, University of Edinburgh / Institute of Genetics and Molecular Medicine, University of Edinburgh | COVID-19 Genomics UK (COG-UK) Consortium | McHugh M, Dewar R, Rooke S, Gallagher M, Balcaza C, O'Toole Á, Scher E, Hill V, McCrone JT, Colquhoun R, Yu X, Jackson B, Rambaut A, Williams TC, Templeton K |
| EPI_ISL_461763, EPI_ISL_461764, EPI_ISL_461765, EPI_ISL_461766, EPI_ISL_461767, EPI_ISL_461768, EPI_ISL_461769, EPI_ISL_461770, EPI_ISL_461771 | University College London, Great Ormond Street Hospital for Children NHS Foundation Trust, Imperial College Healthcare NHS Trust | COVID-19 Genomics UK (COG-UK) Consortium | Sergi Castellano, Rachel Williams, Mark Kristiansen, Paola Resende Silva, Sunando Roy, Tony Brooks, Helena Tutill, Paola Niola, Patricia Dyal, Charlotte Williams, Leysa Forrest, Yasmin Panchbhaya, Jacqueline Findlay, Sam Weeks, Julianne Brown, Kathryn Harris, Paul Randell, James Price, Alison Holmes, Judith Breuer |
| EPI_ISL_461791, EPI_ISL_461792, EPI_ISL_461793, EPI_ISL_461795, EPI_ISL_461797, EPI_ISL_461801 | Northumbria University / South Tees Hospitals NHS Foundation Trust / North Cumbria Integrated Care NHS Foundation Trust / North Tees and Hartlepool NHS Foundation Trust / Newcastle Hospitals NHS Foundation Trust | COVID-19 Genomics UK (COG-UK) Consortium | Darren L Smith,Andrew Nelson,Matthew Bashton,Greg R Young,Joshua Loh,John Allan,Mohammad A Tariq,Giles S Holt,Gary Black,Wen C Yew,Lynn Dover,Paul Baker,Steve Liggett,Sarah Essex,Jane Greenaway,Debra Padgett,Clive Graham,Garren Scott,Edward Barton,Emma Swindells,Brendan Payne,Jennifer Collins,Yusri Taha,Gary Eltringham |
| EPI_ISL_461805, EPI_ISL_461806, EPI_ISL_461809, EPI_ISL_461810, EPI_ISL_461811, EPI_ISL_461812, EPI_ISL_461813, EPI_ISL_461814, EPI_ISL_461816, EPI_ISL_461817, EPI_ISL_461820, EPI_ISL_461821, EPI_ISL_461822, EPI_ISL_461823, EPI_ISL_461825, EPI_ISL_461827, EPI_ISL_461828, EPI_ISL_461829, EPI_ISL_461836, EPI_ISL_461839, EPI_ISL_461844, EPI_ISL_461850, EPI_ISL_461857, EPI_ISL_461858, EPI_ISL_461860, EPI_ISL_461861, EPI_ISL_461862, EPI_ISL_461863, EPI_ISL_461864, EPI_ISL_461865, EPI_ISL_461866, EPI_ISL_461867, EPI_ISL_461868, EPI_ISL_461870, EPI_ISL_461871, EPI_ISL_461872, EPI_ISL_461876, EPI_ISL_461879, EPI_ISL_461880, EPI_ISL_461881, EPI_ISL_461883, EPI_ISL_461884, EPI_ISL_461885, EPI_ISL_461887, EPI_ISL_461888, EPI_ISL_461890 | | | |
| see above | Quadram Institute Bioscience | COVID-19 Genomics UK (COG-UK) Consortium | Dave J. Baker, Gemma L. Kay, Alp Aydin, Thanh Le-Viet, Steven Rudder, Ana P. Tedim, Anastasia Kolyva, Maria Diaz, Leonardo de Oliveira Martins, Nabil-Fareed Alikhan, Lizzie Meadows, Rachael Stanley, Ngozi Elumogo, Muhammed Yasir, Nicholas M. Thomson, Alexander J Trotter, Rachel Gilroy, Samuel Bloomfield, Claire Stuart, Andrew Bell, Reenesh Prakash, Samir Dervisevic, Alison E. Mather, John Wain, Mark Webber, Andrew J. Page, Justin O'Grady |
| EPI_ISL_461894, EPI_ISL_461895, EPI_ISL_461896, EPI_ISL_461897, EPI_ISL_461898, EPI_ISL_461899, EPI_ISL_461900, EPI_ISL_461901, EPI_ISL_461902, EPI_ISL_461905, EPI_ISL_461906, EPI_ISL_461907, EPI_ISL_461908, EPI_ISL_461909, EPI_ISL_461910, EPI_ISL_461911, EPI_ISL_461912, EPI_ISL_461915, EPI_ISL_461916, EPI_ISL_461917, EPI_ISL_461918, EPI_ISL_461919, EPI_ISL_461920, EPI_ISL_461921, EPI_ISL_461922, EPI_ISL_461923, EPI_ISL_461924, EPI_ISL_461925, EPI_ISL_461926, EPI_ISL_461927, EPI_ISL_461928, EPI_ISL_461929, EPI_ISL_461930, EPI_ISL_461931, EPI_ISL_461932, EPI_ISL_461934, EPI_ISL_461935, EPI_ISL_461936, EPI_ISL_461937, EPI_ISL_461940, EPI_ISL_461942, EPI_ISL_461943, EPI_ISL_461945, EPI_ISL_461946, EPI_ISL_461947, EPI_ISL_461948, EPI_ISL_461949, EPI_ISL_461950, EPI_ISL_461951, EPI_ISL_461952, EPI_ISL_461953, EPI_ISL_461954, EPI_ISL_461958, EPI_ISL_461960, EPI_ISL_461961, EPI_ISL_461962, EPI_ISL_461963, EPI_ISL_461964, EPI_ISL_461965, EPI_ISL_461967, EPI_ISL_461968, EPI_ISL_461969 | | | |
| see above | Queens Medical Centre, Clinical Microbiology Department / DeepSeq Nottingham | COVID-19 Genomics UK (COG-UK) Consortium | Gemma Clark, Wendy Smith, Manjinder Khakh, Hannah Howson-Wells, Jonathan Ball, Patrick McClure, Joseph Chappell, Theocharis Tsoleridis, Nadine Holmes, Matthew Carlisle, Christopher Moore, Fei Sang, Johnny Debebe, Victoria Wright, Matthew Loose |
| EPI_ISL_461971, EPI_ISL_461972, EPI_ISL_461973, EPI_ISL_461974, EPI_ISL_461975, EPI_ISL_461976, EPI_ISL_461977, EPI_ISL_461979, EPI_ISL_461981, EPI_ISL_461983, EPI_ISL_461986, EPI_ISL_461987, EPI_ISL_461989, EPI_ISL_461991, EPI_ISL_461998 | | | |
| see above | Centre for Enzyme Innovation, University of Portsmouth / Translational Research Laboratory, Portsmouth Hospitals NHS Trust | COVID-19 Genomics UK (COG-UK) Consortium | Angela Beckett,Yann Bourgeois,Garry Scarlett,Sharon Glaysher,Scott Elliott,Kelly Bicknell,Robert Impey,Allyson Lloyd,Sarah Wyllie,Ethan Butcher,Anoop Chauhan,Samuel Robson |
| EPI_ISL_461999, EPI_ISL_462000, EPI_ISL_462001, EPI_ISL_462002, EPI_ISL_462003, EPI_ISL_462004, EPI_ISL_462005, EPI_ISL_462007, EPI_ISL_462008, EPI_ISL_462009, EPI_ISL_462010, EPI_ISL_462011, EPI_ISL_462016, EPI_ISL_462017, EPI_ISL_462018, EPI_ISL_462020, EPI_ISL_462021, EPI_ISL_462023, EPI_ISL_462025, EPI_ISL_462029, EPI_ISL_462030, EPI_ISL_462032, EPI_ISL_462034, EPI_ISL_462036, EPI_ISL_462038, EPI_ISL_462039, EPI_ISL_462041, EPI_ISL_462043, EPI_ISL_462044, EPI_ISL_462045, EPI_ISL_462046, EPI_ISL_462047, EPI_ISL_462048, EPI_ISL_462049, EPI_ISL_462050, EPI_ISL_462051, EPI_ISL_462052, EPI_ISL_462053, EPI_ISL_462054, EPI_ISL_462056, EPI_ISL_462058, EPI_ISL_462059, EPI_ISL_462060, EPI_ISL_462061, EPI_ISL_462064, EPI_ISL_462065, EPI_ISL_462066, EPI_ISL_462067, EPI_ISL_462069, EPI_ISL_462070, EPI_ISL_462072, EPI_ISL_462074, EPI_ISL_462075, EPI_ISL_462076, EPI_ISL_462078, EPI_ISL_462079, EPI_ISL_462081 | | | |
| see above | Virology Department, Sheffield Teaching Hospitals NHS Foundation Trust/Department of Infection, Immunity and Cardiovascular Disease, The Medical School, University of Sheffield | COVID-19 Genomics UK (COG-UK) Consortium | Thushan de Silva, Matthew Parker, Nikki Smith, Adri Angyal, Rebecca Brown, Luke Green, Rachel Tucker, Paul Parsons, Danielle Groves, Katie Johnson, Laura Carrilero, Alex Keeley, Dave Partridge, Matthew Wyles, Benjamin Lindsey, Mehmet Yavuz, Mohammad Raza, Cariad Evans |
| EPI_ISL_462090 | National Institute of Laboratory Medicine and Referral Center | Genomic Research Lab, BCSIR | Barna Goswami, Abu Sayeed Mohammad Mahmud, Mohammad Samir Uzzaman, Eshrar Osman, Md. Ahasan Habib, Shahina Akter, Tanjina Akhter Banu, Iffat Jahan, Md. Saddam Hossain, Tasnim Nafisa, Md. Maruf |

| | | | |
|---|---|---|---|
| | | | Ahmed Molla, Mahmuda Yeasmin, Asish Kumar Ghos, Bayzid Bin Monir, Arifa Akram, Sheikh Md. Selim Al Din, Salek Ahmed Sajib, Utpal Chandra Ray, Md. Salim Khan |
| EPI_ISL_462091 | National Institute of Laboratory Medicine and Referral Center | Genomic Research Lab, BCSIR | Iffat Jahan, Abu Sayeed Mohammad Mahmud, Mohammad Samir Uzzaman, Eshrar Osman, Md. Ahasan Habib, Shahina Akter, Tanjina Akhter Banu, Barna Goswami, Md. Saddam Hossain, Tasnim Nafisa, Md. Maruf Ahmed Molla, Mahmuda Yeasmin, Asish Kumar Ghos, Bayzid Bin Monir, Arifa Akram, Sheikh Md. Selim Al Din, Salek Ahmed Sajib, Utpal Chandra Ray, Md. Salim Khan |
| EPI_ISL_462092 | National Institute of Laboratory Medicine and Referral Center | Genomic Research Lab, BCSIR | Shahina Akter, Abu Sayeed Mohammad Mahmud, Mohammad Samir Uzzaman, Eshrar Osman, Md. Ahasan Habib, Tanjina Akhter Banu, Barna Goswami, Iffat Jahan, Md. Saddam Hossain, Tasnim Nafisa, Md. Maruf Ahmed Molla, Mahmuda Yeasmin, Asish Kumar Ghos, Bayzid Bin Monir, Arifa Akram, Sheikh Md. Selim Al Din, Salek Ahmed Sajib, Utpal Chandra Ray, Md. Salim Khan |
| EPI_ISL_462093 | National Institute of Laboratory Medicine and Referral Center | Genomic Research Lab, BCSIR | Abu Sayeed Mohammad Mahmud, Mohammad Samir Uzzaman, Eshrar Osman, Md. Ahasan Habib, Tanjina Akhter Banu, Shahina Akter, Barna Goswami, Iffat Jahan, Md. Saddam Hossain, Tasnim Nafisa, Md. Maruf Ahmed Molla, Mahmuda Yeasmin, Asish Kumar Ghosh, Bayzid Bin Monir, Arifa Akram, Sheikh Md. Selim Al Din, Salek Ahmed Sajib, Utpal Chandra Ray, Md. Salim Khan |
| EPI_ISL_462094, EPI_ISL_462095 | National Institute of Laboratory Medicine and Referral Center | Genomic Research Lab, BCSIR | Abu Sayeed Mohammad Mahmud, Mohammad Samir Uzzaman, Eshrar Osman, Md. Ahasan Habib, Tanjina Akhter Banu, Shahina Akter, Barna Goswami, Iffat Jahan, Md. Saddam Hossain, Tasnim Nafisa, Md. Maruf Ahmed Molla, Mahmuda Yeasmin, Asish Kumar Ghosh, Bayzid Bin Monir, Arifa Akram, Sheikh Md. Selim Al Din, Salek Ahmed Sajib, Utpal Chandra Ray, Md. Salim Khan |
| EPI_ISL_462096 | National Institute of Laboratory Medicine and Referral Center | Genomic Research Lab, BCSIR | Abu Sayeed Mohammad Mahmud, Mohammad Samir Uzzaman, Eshrar Osman, Md. Ahasan Habib, Tanjina Akhter Banu, Shahina Akter, Barna Goswami, Iffat Jahan, Md. Saddam Hossain, Tasnim Nafisa, Md. Maruf Ahmed Molla, Mahmuda Yeasmin, Asish Kumar Ghosh, Bayzid Bin Monir, Arifa Akram, Sheikh Md. Selim Al Din, Salek Ahmed Sajib, Utpal Chandra Ray, Md. Salim Khan |
| EPI_ISL_462097 | National Institute of Laboratory Medicine and Referral Center | Genomic Research Lab, BCSIR | Abu Sayeed Mohammad Mahmud, Mohammad Samir Uzzaman, Eshrar Osman, Md. Ahasan Habib, Tanjina Akhter Banu, Shahina Akter, Barna Goswami, Iffat Jahan, Md. Saddam Hossain, Tasnim Nafisa, Md. Maruf Ahmed Molla, Mahmuda Yeasmin, Asish Kumar Ghosh, Bayzid Bin Monir, Arifa Akram, Sheikh Md. Selim Al Din, Salek Ahmed Sajib, Utpal Chandra Ray, Md. Salim Khan |
| EPI_ISL_462098 | National Institute of Laboratory Medicine and Referral Center | Genomic Research Lab, BCSIR | Abu Sayeed Mohammad Mahmud, Mohammad Samir Uzzaman, Eshrar Osman, Md. Ahasan Habib, Tanjina Akhter Banu, Shahina Akter, Barna Goswami, Iffat Jahan, Md. Saddam Hossain, Tasnim Nafisa, Md. Maruf Ahmed Molla, Mahmuda Yeasmin, Asish Kumar Ghosh, Bayzid Bin Monir, Arifa Akram, Sheikh Md. Selim Al Din, Salek Ahmed Sajib, Utpal Chandra Ray, Md. Salim Khan |
| EPI_ISL_462149, EPI_ISL_462150 | Molecular diagnostic laboratory of Federal Budget Institution of Science "Central Research Institute of Epidemiology" of The Federal Service on Customers' Rights Protection and Human Well-being Surveillance | Group of Genomics and Postgenomic Technologies of Central Research Institute of Epidemilology | Speranskaya AS, Kaptelova VV, Samoilov AE, Korneenko EV, Sizova TV, Tivanova EV, Shipulina OY, Akimkin VG |
| EPI_ISL_462434, EPI_ISL_462435, EPI_ISL_462436, EPI_ISL_462437, EPI_ISL_462438 | unknown | Laboratory Diagnostic | Vidanovic,D., Tesovic,B., Banovic Djeri,B., Knezevic,A., Jankovic,M., Sekler,M., Dmitric,M., Petrovic,T., Volkening,J., Afonso,C.L. |
| EPI_ISL_462439, EPI_ISL_462440, EPI_ISL_462441, EPI_ISL_462442, EPI_ISL_462443, EPI_ISL_462444, EPI_ISL_462445 | unknown | Ryota Kumagai Tokyo Metropolitan Institute of Public Health | Asakura,H., Kumagai,R., Yoshida,I., Nagashima,M., Chiba,T., Sadamasu,K. |
| EPI_ISL_462446 | unknown | Ryota Kumagai Tokyo Metropolitan Institute of Public Health | Kumagai,R., Yoshida,I., Asakura,H., Nagashima,M., Chiba,T., Sadamasu,K. |
| EPI_ISL_462995, EPI_ISL_462996, EPI_ISL_462997, EPI_ISL_462998, EPI_ISL_462999, EPI_ISL_463000 | Molecular Genetics | Molecular Genetics | Gomez,J., Coto,E. |
| EPI_ISL_463001, EPI_ISL_463002, EPI_ISL_463003, EPI_ISL_463004, EPI_ISL_463005, EPI_ISL_463006 | unknown | Clinical virology | Fares,W., Triki,H. |
| EPI_ISL_468044, EPI_ISL_468045, EPI_ISL_468046, EPI_ISL_468047, EPI_ISL_468048, EPI_ISL_468049, EPI_ISL_468050, EPI_ISL_468051, EPI_ISL_468052, EPI_ISL_468053, EPI_ISL_468054, EPI_ISL_468055 | | | |
| see above | Egyptian National Cancer Institute (ENCI) | Egyptian National Cancer Institute (ENCI) | Zekri, Abdel Rahman N, Amer,K.E., Ahmed,O.S., Soliman,H.K., Hafez,M.M., Bahnassy,A.A., Abdelhamid,W., Gad,A., Ali,M., Hassan,W., Samir,M., Raouf,A., Hamdy,M.S., Soliman,M.S., Elsissy,M.H., Elkhateeb,S.M., Ezzelarab,M.H., Abouelhoda, Mohamed |
| EPI_ISL_468056 | Egyptian National Cancer Institute (ENCI) | Egyptian National Cancer Institute (ENCI) | Zekri, Abdel Rahman N., Amer,K.E., Ahmed,O.S., Soliman,H.K., Ali,M.A., Hassan,W.A., Mahmoud,A.A., Khattab,A.A., Hafez,M.M., Abouelhoda, Mohamed |
| EPI_ISL_468057, EPI_ISL_468058, EPI_ISL_468059 | Egyptian National Cancer Institute (ENCI) | Egyptian National Cancer Institute (ENCI) | Zekri, Abdel Rahman N, Amer,K.E., Ahmed,O.S., Soliman,H.K., Hafez,M.M., Bahnassy,A.A., Abdelhamid,W., Gad,A., Ali,M., Hassan,W., Samir,M., Raouf,A., Hamdy,M.S., Soliman,M.S., Elsissy,M.H., Elkhateeb,S.M., Ezzelarab,M.H., Abouelhoda, Mohamed |
| EPI_ISL_468060, EPI_ISL_468061, EPI_ISL_468062 | Egyptian National Cancer Institute (ENCI) | Egyptian National Cancer Institute (ENCI) | Zekri, Abdel Rahman N., Amer,K.E., Ahmed,O.S., Soliman,H.K., Ali,M.A., Hassan,W.A., Mahmoud,A.A., Khattab,A.A., Hafez,M.M., Abouelhoda, Mohamed |
| EPI_ISL_468724, EPI_ISL_468725 | unknown | Contact:Ryota Kumagai Tokyo Metropolitan Institute of Public Health | Kumagai,R., Yoshida,I., Asakura,H., Nagashima,M., Chiba,T., Sadamasu,K. |
| EPI_ISL_475722 | Egyptian National Cancer Institute (ENCI) | Egyptian National Cancer Institute (ENCI) | Zekri, Abdel Rahman N, Amer,K.E., Ahmed,O.S., Soliman,H.K., Hafez,M.M., Bahnassy,A.A., Abdelhamid,W., Gad,A., Ali,M., Hassan,W., Samir,M., Raouf,A., Hamdy,M.S., Soliman,M.S., Elsissy,M.H., Elkhateeb,S.M., Ezzelarab,M.H., Abouelhoda, Mohamed |
| EPI_ISL_524450 | Microbiology & Immunology, University of North Carolina | Microbiology & Immunology, University of North Carolina | Bailey,A.G., Caro-Vegas,C.P., Dittmer,D., Eason,A.B., Juarez,A., Landis,J.T., McNamara,R.P., Miller,M.B., Moorad,R., Pluta,L.J., Seltzer,T.A., Thompson,C., Vahrson,W., Villamor,F. |
| EPI_ISL_529146 | Goethe University Hospital Frankfurt, Institute for Medical Virology | Goethe University Hospital Frankfurt, Institute for Medical Virology | Grabmair,T.T., Hoehl,S., Westhaus,S., Bojkova,D., Berger,A., Rotter,B., Hoffmeier,K., Cinatl,J., Ciesek,S., Widera,M. |
