## Supplementary material for "Analysis of the Dynamics and Distribution of SARS-CoV-2 Mutations and its Possible Structural and Functional Implications": GISAID_acknowledgements_part_4

We gratefully acknowledge the following Authors from the Originating laboratories responsible for obtaining the specimens, as well as the Submitting laboratories where the genome data were generated and shared via GISAID, on which this research is based.

All Submitters of data may be contacted directly via www.gisaid.org

| Accession ID | Originating Laboratory | Submitting Laboratory | Authors |
|---|---|---|---|
| EPI_ISL_462151, EPI_ISL_462152, EPI_ISL_462153, EPI_ISL_462154, EPI_ISL_462155, EPI_ISL_462156, EPI_ISL_462157, EPI_ISL_462158, EPI_ISL_462159, EPI_ISL_462160, EPI_ISL_462161, EPI_ISL_462162, EPI_ISL_462163, EPI_ISL_462164, EPI_ISL_462165, EPI_ISL_462166, EPI_ISL_462167, EPI_ISL_462168, EPI_ISL_462169, EPI_ISL_462170, EPI_ISL_462171, EPI_ISL_462172, EPI_ISL_462173, EPI_ISL_462174, EPI_ISL_462175, EPI_ISL_462176, EPI_ISL_462177, EPI_ISL_462178, EPI_ISL_462179, EPI_ISL_462180, EPI_ISL_462181, EPI_ISL_462182, EPI_ISL_462183, EPI_ISL_462184, EPI_ISL_462185, EPI_ISL_462186, EPI_ISL_462187, EPI_ISL_462188, EPI_ISL_462189, EPI_ISL_462190, EPI_ISL_462191, EPI_ISL_462192, EPI_ISL_462193, EPI_ISL_462194, EPI_ISL_462195, EPI_ISL_462196, EPI_ISL_462197, EPI_ISL_462198, EPI_ISL_462199, EPI_ISL_462200, EPI_ISL_462201, EPI_ISL_462202, EPI_ISL_462203, EPI_ISL_462204, EPI_ISL_462205, EPI_ISL_462206, EPI_ISL_462207, EPI_ISL_462208, EPI_ISL_462209, EPI_ISL_462210, EPI_ISL_462211, EPI_ISL_462212, EPI_ISL_462213, EPI_ISL_462214, EPI_ISL_462215, EPI_ISL_462216, EPI_ISL_462217, EPI_ISL_462218, EPI_ISL_462219, EPI_ISL_462220, EPI_ISL_462221, EPI_ISL_462222, EPI_ISL_462223, EPI_ISL_462224, EPI_ISL_462225, EPI_ISL_462226, EPI_ISL_462227, EPI_ISL_462228, EPI_ISL_462229, EPI_ISL_462230, EPI_ISL_462231, EPI_ISL_462232, EPI_ISL_462233, EPI_ISL_462234, EPI_ISL_462235, EPI_ISL_462236, EPI_ISL_462237, EPI_ISL_462238, EPI_ISL_462239, EPI_ISL_462240, EPI_ISL_462241, EPI_ISL_462242, EPI_ISL_462243, EPI_ISL_462244, EPI_ISL_462245, EPI_ISL_462246, EPI_ISL_462247, EPI_ISL_462248, EPI_ISL_462249, EPI_ISL_462250, EPI_ISL_462251, EPI_ISL_462252, EPI_ISL_462253, EPI_ISL_462254, EPI_ISL_462255, EPI_ISL_462256, EPI_ISL_462257, EPI_ISL_462258, EPI_ISL_462259, EPI_ISL_462260, EPI_ISL_462261, EPI_ISL_462262, EPI_ISL_462263, EPI_ISL_462264, EPI_ISL_462265, EPI_ISL_462266, EPI_ISL_462267, EPI_ISL_462268, EPI_ISL_462269, EPI_ISL_462270, EPI_ISL_462271, EPI_ISL_462272, EPI_ISL_462273, EPI_ISL_462274, EPI_ISL_462275 | | | |
| see above | KU Leuven, Rega Institute, Clinical and Epidemiological Virology | KU Leuven, Rega Institute, Clinical and Epidemiological Virology | Tony Wawina-Bokalanga, Bert Vanmechelen, Joan Marti-Carerras, Piet Maes |
| EPI_ISL_462276, EPI_ISL_462277, EPI_ISL_462278, EPI_ISL_462279, EPI_ISL_462280, EPI_ISL_462281, EPI_ISL_462282, EPI_ISL_462283, EPI_ISL_462284, EPI_ISL_462285, EPI_ISL_462286, EPI_ISL_462287, EPI_ISL_462288, EPI_ISL_462290, EPI_ISL_462291, EPI_ISL_462292, EPI_ISL_462293, EPI_ISL_462295, EPI_ISL_462296, EPI_ISL_462297, EPI_ISL_462298, EPI_ISL_462299, EPI_ISL_462300, EPI_ISL_462301, EPI_ISL_462302, EPI_ISL_462303, EPI_ISL_462305, EPI_ISL_462306, EPI_ISL_462307, EPI_ISL_462308, EPI_ISL_462309, EPI_ISL_462310, EPI_ISL_462311, EPI_ISL_462312, EPI_ISL_462313, EPI_ISL_462314, EPI_ISL_462315, EPI_ISL_462316, EPI_ISL_462317, EPI_ISL_462318, EPI_ISL_462319, EPI_ISL_462320, EPI_ISL_462321, EPI_ISL_462322, EPI_ISL_462323, EPI_ISL_462324, EPI_ISL_462325, EPI_ISL_462326, EPI_ISL_462328, EPI_ISL_462338, EPI_ISL_462340, EPI_ISL_462341, EPI_ISL_462342, EPI_ISL_462343, EPI_ISL_462344, EPI_ISL_462345, EPI_ISL_462346, EPI_ISL_462347, EPI_ISL_462348, EPI_ISL_462349, EPI_ISL_462350, EPI_ISL_462351, EPI_ISL_462352, EPI_ISL_462353, EPI_ISL_462354, EPI_ISL_462355, EPI_ISL_462356, EPI_ISL_462357, EPI_ISL_462358, EPI_ISL_462359, EPI_ISL_462360, EPI_ISL_462361, EPI_ISL_462369, EPI_ISL_462370, EPI_ISL_462371, EPI_ISL_462372, EPI_ISL_462373, EPI_ISL_462374, EPI_ISL_462375, EPI_ISL_462376, EPI_ISL_462377, EPI_ISL_462378, EPI_ISL_462379, EPI_ISL_462380, EPI_ISL_462381, EPI_ISL_462382, EPI_ISL_462383, EPI_ISL_462384, EPI_ISL_462385, EPI_ISL_462386, EPI_ISL_462387, EPI_ISL_462388, EPI_ISL_462389, EPI_ISL_462390, EPI_ISL_462391, EPI_ISL_462392, EPI_ISL_462393, EPI_ISL_462394, EPI_ISL_462395, EPI_ISL_462396, EPI_ISL_462397, EPI_ISL_462398, EPI_ISL_462399, EPI_ISL_462400, EPI_ISL_462401, EPI_ISL_462402, EPI_ISL_462403, EPI_ISL_462404, EPI_ISL_462405, EPI_ISL_462406, EPI_ISL_462407, EPI_ISL_462408, EPI_ISL_462409, EPI_ISL_462410, EPI_ISL_462411, EPI_ISL_462412, EPI_ISL_462413, EPI_ISL_462414, EPI_ISL_462415, EPI_ISL_462416, EPI_ISL_462417, EPI_ISL_462418, EPI_ISL_462419, EPI_ISL_462420, EPI_ISL_462421, EPI_ISL_462422, EPI_ISL_462423, EPI_ISL_462424, EPI_ISL_462425, EPI_ISL_462426, EPI_ISL_462427, EPI_ISL_462428, EPI_ISL_462429, EPI_ISL_462430, EPI_ISL_462431, EPI_ISL_462432, EPI_ISL_462433 | | | |
| see above | National Public Health Laboratory, National Centre for Infectious Diseases | National Public Health Laboratory, National Centre for Infectious Diseases | Mak TM, Octavia S, Chavatte JM, Cui L, Lin RTP |
| EPI_ISL_462447, EPI_ISL_462448 | Fundació Lluita contra la SIDA (FLSida)/Hospital Universitari Germans Trias i Pujol | IrsiCaixa AIDS Research Lab | Marc Noguera-Julian, Mariona Parera, Maria Pilar Armengol, Marc Corbacho, Maria Ubals, Oriol Mitjà, Lidia Ruiz, Nuria Izquierdo, Jorge Carrillo, Roger Paredes, Julia Blanco, Joaquim Segalés, Bonaventura Clotet |
| EPI_ISL_462449 | Fundació Lluita contra la SIDA (FLSida)/Hospital Universitari Germans Trias i Pujol | IrsiCaixa AIDS Research Lab | Marc Noguera-Julian, Mariona Parera, Maria Pilar Armengol, Marc Corbacho, Maria Ubals, Oriol Mitjà, Lidia Ruiz, Nuria Izquierdo, Jorge Carrillo, Roger Paredes, Julia Blanco, Bonaventura Clotet |
| EPI_ISL_462450, EPI_ISL_462451, EPI_ISL_462452, EPI_ISL_462453, EPI_ISL_462454, EPI_ISL_462455, EPI_ISL_462456, EPI_ISL_462457, EPI_ISL_462458, EPI_ISL_462459, EPI_ISL_462460, EPI_ISL_462461, EPI_ISL_462462, EPI_ISL_462463, EPI_ISL_462464, EPI_ISL_462465, EPI_ISL_462466, EPI_ISL_462467, EPI_ISL_462468, EPI_ISL_462469, EPI_ISL_462470, EPI_ISL_462471, EPI_ISL_462472, EPI_ISL_462473, EPI_ISL_462474, EPI_ISL_462475, EPI_ISL_462476 | | | |
| see above | Clinical Center, University of Sarajevo | Charite Universitatsmedizin Berlin, Institute of Virology | Victor M Corman, Jorn Beheim-Schwarzbach, Barbara Muehlemann, Talitha Veith, Julia Schneider, Terry Jones, Amela Dedeic-Ljubovic, Irma Salimovic-Besic, Suzana Arapcic, Almedina Hadzihasanovic-Moro, Selma Mutevelic, Christian Drosten |
| EPI_ISL_462478 | Fundación Jiménez Díaz | Instituto de Salud Carlos III | Iglesias-Caballero, M. Molinero Calamita, M. González-Esguevillas, M. Camarero, S. Pozo, F. Casas, I. Jiménez, P. Jiménez, M. Zaballos, A. Monzón, S. Varona, S. Juliá, M. Cuesta, I, R. Fernández |
| EPI_ISL_462479 | Hospital Clinic | Instituto de Salud Carlos III | Iglesias-Caballero, M. Molinero Calamita, M. González-Esguevillas, M. Camarero, S. Pozo, F. Casas, I. Jiménez, P. Jiménez, M. Zaballos, A. Monzón, S. Varona, S. Juliá, M. Cuesta, I, M.A Marcos |
| EPI_ISL_462480 | Institute of Human Genetics, Polish Academy of Sciences | Institute of Human Genetics, Polish Academy of Sciences | Szymon Hryhorowicz, Adam Ustaszewski, Emilia Lis, Marta Kaczmarek-Ryś, Michał Witt, Andrzej Pławski |
| EPI_ISL_462673, EPI_ISL_462674, EPI_ISL_462675, EPI_ISL_462678, EPI_ISL_462679, EPI_ISL_462680, EPI_ISL_462681, EPI_ISL_462683, EPI_ISL_462684, EPI_ISL_462685, EPI_ISL_462686, EPI_ISL_462687, EPI_ISL_462688, EPI_ISL_462689, EPI_ISL_462690, EPI_ISL_462691, EPI_ISL_462692, EPI_ISL_462693, EPI_ISL_462694, EPI_ISL_462695, EPI_ISL_462696, EPI_ISL_462697, EPI_ISL_462698, EPI_ISL_462699, EPI_ISL_462700, EPI_ISL_462701, EPI_ISL_462706, EPI_ISL_462707, EPI_ISL_462708, EPI_ISL_462719, EPI_ISL_462720, EPI_ISL_462725, EPI_ISL_462731, EPI_ISL_462732, EPI_ISL_462733, EPI_ISL_462734, EPI_ISL_462736 | | | |
| see above | Michigan Department of Health and Human Services, Bureau of Laboratories | Michigan Department of Health and Human Services, Bureau of Laboratories | Blankenship HM, Riner D, Soehnlen MK |
| EPI_ISL_462757, EPI_ISL_462758, EPI_ISL_462759, EPI_ISL_462760, EPI_ISL_462761, EPI_ISL_462765, EPI_ISL_462766, EPI_ISL_462767, EPI_ISL_462768, EPI_ISL_462769, EPI_ISL_462771, EPI_ISL_462772, EPI_ISL_462774, EPI_ISL_462775, EPI_ISL_462776, EPI_ISL_462777, EPI_ISL_462778, EPI_ISL_462779, EPI_ISL_462780, EPI_ISL_462781, EPI_ISL_462782, EPI_ISL_462783, EPI_ISL_462784, EPI_ISL_462786, EPI_ISL_462787, EPI_ISL_462789, EPI_ISL_462790, EPI_ISL_462791, EPI_ISL_462792, EPI_ISL_462793, EPI_ISL_462794, EPI_ISL_462798, EPI_ISL_462802, EPI_ISL_462805, EPI_ISL_462806, EPI_ISL_462807, EPI_ISL_462808, EPI_ISL_462809, EPI_ISL_462810, EPI_ISL_462811, EPI_ISL_462812, EPI_ISL_462813, EPI_ISL_462814, EPI_ISL_462815, EPI_ISL_462816, EPI_ISL_462818, EPI_ISL_462819, EPI_ISL_462820, EPI_ISL_462821, EPI_ISL_462822, EPI_ISL_462824, EPI_ISL_462825, EPI_ISL_462826, EPI_ISL_462827, EPI_ISL_462828, EPI_ISL_462829, EPI_ISL_462830, EPI_ISL_462831, EPI_ISL_462833, EPI_ISL_462834, EPI_ISL_462835, EPI_ISL_462836, EPI_ISL_462837, EPI_ISL_462838, EPI_ISL_462840, EPI_ISL_462841, EPI_ISL_462842, EPI_ISL_462843 | | | |
| see above | BCCDC Public Health Laboratory | BCCDC Public Health Laboratory | Harrigan, Prystajecky, Krajden, Lee, Kamelian, Lapointe, Choi, Hoang, Sekirov, Levett, Tyson, Li, Gilmour |
| EPI_ISL_462845, EPI_ISL_462846, EPI_ISL_462847, EPI_ISL_462848, EPI_ISL_462849, EPI_ISL_462850, EPI_ISL_462851, EPI_ISL_462852, EPI_ISL_462853, EPI_ISL_462854, EPI_ISL_462855, EPI_ISL_462856, EPI_ISL_462857, EPI_ISL_462858, EPI_ISL_462859, EPI_ISL_462860, EPI_ISL_462861, EPI_ISL_462862, EPI_ISL_462864, EPI_ISL_462865, EPI_ISL_462866, EPI_ISL_462867, EPI_ISL_462868, EPI_ISL_462869, EPI_ISL_462870, EPI_ISL_462871, EPI_ISL_462872, EPI_ISL_462873, EPI_ISL_462874, EPI_ISL_462875, EPI_ISL_462876, EPI_ISL_462877, EPI_ISL_462878, EPI_ISL_462879, EPI_ISL_462880, EPI_ISL_462881, EPI_ISL_462882, EPI_ISL_462883, EPI_ISL_462884, EPI_ISL_462885, EPI_ISL_462886, EPI_ISL_462887, EPI_ISL_462888, EPI_ISL_462889, EPI_ISL_462890, EPI_ISL_462891, EPI_ISL_462892, EPI_ISL_462893, EPI_ISL_462894, EPI_ISL_462895, EPI_ISL_462897, EPI_ISL_462898, EPI_ISL_462899, EPI_ISL_462900, EPI_ISL_462901, EPI_ISL_462902, EPI_ISL_462903, EPI_ISL_462904, EPI_ISL_462905, EPI_ISL_462906, EPI_ISL_462907, EPI_ISL_462908, EPI_ISL_462909, EPI_ISL_462910, EPI_ISL_462911 | | | |
| see above | Minnesota Department of Health, Public Health Laboratory | Minnesota Department of Health, Public Health Laboratory | Matt Plumb, Jacob Garfin, and Xiong Wang |
| EPI_ISL_462912, EPI_ISL_462915, EPI_ISL_462917, EPI_ISL_462918, EPI_ISL_462919, EPI_ISL_462924, EPI_ISL_462927, EPI_ISL_462929, EPI_ISL_462932, EPI_ISL_462933, EPI_ISL_462935, EPI_ISL_462936, EPI_ISL_462940, EPI_ISL_462952, EPI_ISL_462956, EPI_ISL_462957, EPI_ISL_462958, EPI_ISL_462965, EPI_ISL_462966, EPI_ISL_462969, EPI_ISL_462975, EPI_ISL_462976, EPI_ISL_462977, EPI_ISL_462979, EPI_ISL_462982 | | | |
| see above | Wyoming Public Health Laboratory | Center for Global Health, University of New Mexico Health Sciences Center | Daryl Domman, Kurt Schwalm, Rob Christensen, Wanda Manley, Cari Sloma, Noah Hull, Darrell Dinwiddie |
| EPI_ISL_462991 | Microbiology Division | Microbiology Division | Flores,H. |
| EPI_ISL_462992 | Nigerian Institute of Medical Research | Nigerian Institute of Medical Research | Saibu,J.O., Onwuamah,C.K., Okwuraiwe,A.P., Amoo,O.S., Salu,O.B., Ige,F.A., Liboro,G., Odewale,E., Adesegun,A., Abosede,O., Ahmed,R., Sokei,J., Oyefolu,A., Adegbola,R., Salako,B., Omilabu,S. and Audu,R. |
| EPI_ISL_463007 | Department of Laboratory Medicine, National Taiwan University Hospital | Microbial Genomics Core Lab, National Taiwan University Centers of Genomic and Precision Medicine | Shiou-Hwei Yeh, You-Yu Lin, Ya-Yun Lai, Chiao-Ling Li, Shan-Chwen Chang, Pei-Jer Chen, Sui-Yuan Chang |
| EPI_ISL_463008 | Institute of Molecular Virology, University Münster | Institute of Molecular Virology, University Münster | Angeles Mecate Zambrano, Linda Brunotte, Stephan Ludwig, Joachim Kühn, Alexander Mellmann |
| EPI_ISL_463010, EPI_ISL_463013, EPI_ISL_463014, EPI_ISL_463017, EPI_ISL_463018, EPI_ISL_463019, EPI_ISL_463020, EPI_ISL_463022, EPI_ISL_463023, EPI_ISL_463024, EPI_ISL_463025, EPI_ISL_463026, EPI_ISL_463027, EPI_ISL_463029 | | | |
| see above | Institute of Life Sciences, Bhubaneswar | Immunogenomics lab, Institute of Life Sciences, Bhubaneswar | Sunil Raghav, Arup Ghosh, Atimukta Jha, Viplov K. Biswas, Swati Madhulika, Manasi Priyadarshini, Shuchi Smita, Kaushik Sen, Hiren G. Dodia, Deepak Singh, Jeky Chawla, Shamima Ansari, Rupesh Dash, Soma Chattopadhyay, Ghulam Hussain Syed, Shanti Senapati, Tushar K. Beuria, Rajeeb Swain, Punit Prasad, ILS COVID-19 TEAM, Orissa COVID-19 Study Group, DBT's PAN-INDIA 1000 SARS-CoV2 RNA genome sequencing consortium, Ajay Parida |
| EPI_ISL_463031, EPI_ISL_463032, EPI_ISL_463033, EPI_ISL_463034, EPI_ISL_463036, EPI_ISL_463037, EPI_ISL_463038, EPI_ISL_463039, EPI_ISL_463041, EPI_ISL_463043, EPI_ISL_463049, EPI_ISL_463051 | | | |
| see above | Institute of Life Sciences, Bhubaneswar | Immunogenomics lab, Institute of Life Sciences, Bhubaneswar | Sunil Raghav, Arup Ghosh, Atimukta Jha, Viplov K. Biswas, Swati Madhulika, Manasi Priyadarshini, Shuchi Smita, O. P. Shriwas, Priyanka Mohapatra, Satya Ranjan Sahu, Aliva Minz, Debyashrita Barik, Rupesh Dash, Soma Chattopadhyay, Ghulam Hussain Syed, Shanti Senapati, Tushar K. Beuria, Rajeeb Swain, Punit Prasad, ILS COVID-19 TEAM, Orissa COVID-19 Study Group, DBT's PAN-INDIA 1000 SARS-CoV2 RNA genome sequencing consortium, Ajay Parida |
| EPI_ISL_463052, EPI_ISL_463054, EPI_ISL_463056, EPI_ISL_463058, EPI_ISL_463060, EPI_ISL_463061, EPI_ISL_463063, EPI_ISL_463064, EPI_ISL_463067 | Institute of Life Sciences, Bhubaneswar | Immunogenomics lab, Institute of Life Sciences, Bhubaneswar | Sunil Raghav, Arup Ghosh, Atimukta Jha, Viplov K. Biswas, Swati Madhulika, Manasi Priyadarshini, Shuchi Smita, Sifu Agarwal, Sanchari Chatterjee, Avula Kiran, Parej Nath, Supriya Suman, Rina Yadav, Rupesh Dash, Soma Chattopadhyay, Ghulam Hussain Syed, Shanti Senapati, Tushar K. Beuria, Rajeeb Swain, Punit Prasad, ILS COVID-19 TEAM, Orissa COVID-19 Study Group, DBT's PAN-INDIA 1000 SARS-CoV2 RNA genome sequencing consortium, Ajay Parida |
| EPI_ISL_463073, EPI_ISL_463074, EPI_ISL_463076, EPI_ISL_463078, EPI_ISL_463081, EPI_ISL_463085, EPI_ISL_463087, EPI_ISL_463088, EPI_ISL_463089 | Institute of Life Sciences, Bhubaneswar | Immunogenomics lab, Institute of Life Sciences, Bhubaneswar | Sunil Raghav, Arup Ghosh, Atimukta Jha, Viplov K. Biswas, Swati Madhulika, Manasi Priyadarshini, Shuchi Smita, Kautilya Kumar Jena, Sandhya Suranjika, Neha Singh, Eshna Laha, Saiket De, Rupesh Dash, Soma Chattopadhyay, Ghulam Hussain Syed, Shanti Senapati, Tushar K. Beuria, Rajeeb Swain, Punit Prasad, ILS COVID-19 TEAM, Orissa COVID-19 Study Group, DBT's PAN-INDIA 1000 SARS-CoV2 RNA genome sequencing consortium, Ajay Parida |
| EPI_ISL_463094, EPI_ISL_463095, EPI_ISL_463096, EPI_ISL_463098, EPI_ISL_463099, EPI_ISL_463100, EPI_ISL_463101, EPI_ISL_463103, EPI_ISL_463104, EPI_ISL_463105, EPI_ISL_463106, EPI_ISL_463107, EPI_ISL_463108, EPI_ISL_463109, EPI_ISL_463110, EPI_ISL_463111, EPI_ISL_463112, EPI_ISL_463113, EPI_ISL_463114, EPI_ISL_463115, EPI_ISL_463116, EPI_ISL_463117, EPI_ISL_463118, EPI_ISL_463119, EPI_ISL_463120, EPI_ISL_463121, EPI_ISL_463122, EPI_ISL_463123, EPI_ISL_463124, EPI_ISL_463125, EPI_ISL_463126, EPI_ISL_463127, EPI_ISL_463128, EPI_ISL_463129, EPI_ISL_463130, EPI_ISL_463131, EPI_ISL_463132, EPI_ISL_463133, EPI_ISL_463134, EPI_ISL_463135, EPI_ISL_463136, EPI_ISL_463137 | | | |
| see above | Virginia DCLS | Virginia DCLS | Virginia DCLS |
| EPI_ISL_463138, EPI_ISL_463141, EPI_ISL_463151, EPI_ISL_463156, EPI_ISL_463157, EPI_ISL_463158, EPI_ISL_463164 | Yale Clinical Virology Laboratory | Grubaugh Lab - Yale School of Public Health | Joseph Fauver, Tara Alpert, Anderson Brito, Anne Wyllie, Chantal Vogels, Mary Petrone, Cole Jensen, Chaney Kalinich, Isabel Ott, Arnau Casanovas, Catherine Muenker, Adam Moore, Alice Lu, Maria Tokuyama, Patrick Wong, Peiwen Lu, Saad Omer, Richard Martinello, Allison Nelson, Shelli Farhadian, Akiko Iwasaki, Charlese Dela Cruz, Albert Ko, Nathan Grubaugh |
| EPI_ISL_463186, EPI_ISL_463187, EPI_ISL_463189, EPI_ISL_463191, EPI_ISL_463193, EPI_ISL_463194, EPI_ISL_463195, EPI_ISL_463196, EPI_ISL_463198, EPI_ISL_463199, EPI_ISL_463200, EPI_ISL_463201, EPI_ISL_463202, EPI_ISL_463203, EPI_ISL_463204, EPI_ISL_463205, EPI_ISL_463206, EPI_ISL_463207, EPI_ISL_463208, EPI_ISL_463209, EPI_ISL_463210, EPI_ISL_463211, EPI_ISL_463212, EPI_ISL_463213, EPI_ISL_463214, EPI_ISL_463215, EPI_ISL_463216, EPI_ISL_463217, EPI_ISL_463219, EPI_ISL_463220, EPI_ISL_463221, EPI_ISL_463222, EPI_ISL_463223, EPI_ISL_463224, EPI_ISL_463225, EPI_ISL_463226, EPI_ISL_463227, EPI_ISL_463228, EPI_ISL_463229, EPI_ISL_463230, EPI_ISL_463231, EPI_ISL_463232, EPI_ISL_463235, EPI_ISL_463236, EPI_ISL_463237, EPI_ISL_463238, EPI_ISL_463239, EPI_ISL_463240, EPI_ISL_463241, EPI_ISL_463242, EPI_ISL_463243, EPI_ISL_463244, EPI_ISL_463245, EPI_ISL_463246, EPI_ISL_463247, EPI_ISL_463248, EPI_ISL_463249, EPI_ISL_463250, EPI_ISL_463251, EPI_ISL_463252, EPI_ISL_463253, EPI_ISL_463254, EPI_ISL_463255, EPI_ISL_463256, EPI_ISL_463257, EPI_ISL_463258, EPI_ISL_463259, EPI_ISL_463260, EPI_ISL_463261, EPI_ISL_463262, EPI_ISL_463263, EPI_ISL_463264, EPI_ISL_463265, EPI_ISL_463266, EPI_ISL_463267, EPI_ISL_463268, EPI_ISL_463270, EPI_ISL_463271, EPI_ISL_463272, EPI_ISL_463273, EPI_ISL_463274, EPI_ISL_463275, EPI_ISL_463276 | | | |
| see above | BCCDC Public Health Laboratory | BCCDC Public Health Laboratory | Richard Harrigan, Hope Lapointe, Jinny Choi, Kimia Kamelian, John Tyson,Terry Snutch, Linda Hoang, Inna Sekirov, Paul Levett, Mel Krajden, Natalie Prystajecky |
| EPI_ISL_463277, EPI_ISL_463278, EPI_ISL_463279, EPI_ISL_463280, EPI_ISL_463281, EPI_ISL_463282, EPI_ISL_463283, EPI_ISL_463284, EPI_ISL_463285, EPI_ISL_463286, EPI_ISL_463287, EPI_ISL_463288, EPI_ISL_463289, EPI_ISL_463290, EPI_ISL_463291, EPI_ISL_463292, EPI_ISL_463293, EPI_ISL_463294, EPI_ISL_463295, EPI_ISL_463296, EPI_ISL_463297, EPI_ISL_463298, EPI_ISL_463299, EPI_ISL_463300 | | | |
| see above | Ochsner Health | Bioinfoexperts, LLC | Susanna L. Lamers, David J. Nolan, Rebecca Rose, Sissy Cross, David Moraga Amador, Tong Yang, Luke Caruso, Wayra Navia, Lydia Von Borstel, Xiao Hui Zhou, Amy Feehan, Julia-Garcia-Diaz |
| EPI_ISL_463325, EPI_ISL_463326, EPI_ISL_463327, EPI_ISL_463328, EPI_ISL_463329, EPI_ISL_463330, EPI_ISL_463331, EPI_ISL_463332, EPI_ISL_463333, EPI_ISL_463334, EPI_ISL_463335, EPI_ISL_463337, EPI_ISL_463338, EPI_ISL_463339, EPI_ISL_463340, EPI_ISL_463341, EPI_ISL_463342, EPI_ISL_463343, EPI_ISL_463344, EPI_ISL_463346, EPI_ISL_463347, EPI_ISL_463348, EPI_ISL_463349, EPI_ISL_463350, EPI_ISL_463351, EPI_ISL_463352, EPI_ISL_463353, EPI_ISL_463354, EPI_ISL_463355, EPI_ISL_463356, EPI_ISL_463357, EPI_ISL_463358, EPI_ISL_463359, EPI_ISL_463360, EPI_ISL_463361, EPI_ISL_463362, EPI_ISL_463363, EPI_ISL_463364, EPI_ISL_463365, EPI_ISL_463366, EPI_ISL_463367, EPI_ISL_463369, EPI_ISL_463370, EPI_ISL_463371, EPI_ISL_463378, EPI_ISL_463379, EPI_ISL_463381, EPI_ISL_463382, | | | |

| | | | |
|---|---|---|---|
| EPI_ISL_463383, EPI_ISL_463385, EPI_ISL_463386, EPI_ISL_463388, EPI_ISL_463389, EPI_ISL_463390, EPI_ISL_463391, EPI_ISL_463392, EPI_ISL_463394, EPI_ISL_463395, EPI_ISL_463396, EPI_ISL_463397, EPI_ISL_463398, EPI_ISL_463399, EPI_ISL_463400, EPI_ISL_463401, EPI_ISL_463402, EPI_ISL_463403, EPI_ISL_463406, EPI_ISL_463407, EPI_ISL_463408, EPI_ISL_463409, EPI_ISL_463410, EPI_ISL_463411, EPI_ISL_463412, EPI_ISL_463413, EPI_ISL_463414, EPI_ISL_463415, EPI_ISL_463416, EPI_ISL_463417, EPI_ISL_463418, EPI_ISL_463419, EPI_ISL_463420, EPI_ISL_463421, EPI_ISL_463422, EPI_ISL_463423, EPI_ISL_463424, EPI_ISL_463425, EPI_ISL_463426, EPI_ISL_463427, EPI_ISL_463428, EPI_ISL_463429, EPI_ISL_463430, EPI_ISL_463431, EPI_ISL_463432, EPI_ISL_463433, EPI_ISL_463434, EPI_ISL_463435, EPI_ISL_463436, EPI_ISL_463437, EPI_ISL_463438, EPI_ISL_463439, EPI_ISL_463440, EPI_ISL_463441, EPI_ISL_463442, EPI_ISL_463443, EPI_ISL_463444, EPI_ISL_463446, EPI_ISL_463449, EPI_ISL_463450, EPI_ISL_463451, EPI_ISL_463452, EPI_ISL_463453, EPI_ISL_463454, EPI_ISL_463455, EPI_ISL_463456, EPI_ISL_463457, EPI_ISL_463458, EPI_ISL_463459, EPI_ISL_463460, EPI_ISL_463461, EPI_ISL_463462, EPI_ISL_463463, EPI_ISL_463464, EPI_ISL_463465, EPI_ISL_463466, EPI_ISL_463467, EPI_ISL_463472, EPI_ISL_463478, EPI_ISL_463479, EPI_ISL_463480, EPI_ISL_463481, EPI_ISL_463482, EPI_ISL_463483, EPI_ISL_463484, EPI_ISL_463485, EPI_ISL_463486, EPI_ISL_463487, EPI_ISL_463488, EPI_ISL_463489, EPI_ISL_463490, EPI_ISL_463491, EPI_ISL_463492, EPI_ISL_463493, EPI_ISL_463494, EPI_ISL_463495, EPI_ISL_463496, EPI_ISL_463497, EPI_ISL_463498, EPI_ISL_463499, EPI_ISL_463500, EPI_ISL_463501, EPI_ISL_463502, EPI_ISL_463503, EPI_ISL_463504, EPI_ISL_463505, EPI_ISL_463506, EPI_ISL_463507, EPI_ISL_463508, EPI_ISL_463509, EPI_ISL_463510, EPI_ISL_463511, EPI_ISL_463512, EPI_ISL_463513, EPI_ISL_463514, EPI_ISL_463515, EPI_ISL_463516, EPI_ISL_463517, EPI_ISL_463518, EPI_ISL_463519, EPI_ISL_463520, EPI_ISL_463521, EPI_ISL_463522, EPI_ISL_463523, EPI_ISL_463524, EPI_ISL_463525, EPI_ISL_463526, EPI_ISL_463527, EPI_ISL_463528, EPI_ISL_463529, EPI_ISL_463530, EPI_ISL_463531, EPI_ISL_463532, EPI_ISL_463533, EPI_ISL_463534, EPI_ISL_463535, EPI_ISL_463536, EPI_ISL_463538, EPI_ISL_463539, EPI_ISL_463540, EPI_ISL_463541, EPI_ISL_463542, EPI_ISL_463543, EPI_ISL_463544, EPI_ISL_463545, EPI_ISL_463546, EPI_ISL_463547, EPI_ISL_463548, EPI_ISL_463549, EPI_ISL_463550, EPI_ISL_463551, EPI_ISL_463552, EPI_ISL_463553, EPI_ISL_463554, EPI_ISL_463555, EPI_ISL_463556, EPI_ISL_463557, EPI_ISL_463558, EPI_ISL_463559, EPI_ISL_463560, EPI_ISL_463561, EPI_ISL_463562, EPI_ISL_463563, EPI_ISL_463564, EPI_ISL_463565, EPI_ISL_463566, EPI_ISL_463567, EPI_ISL_463568, EPI_ISL_463569, EPI_ISL_463570, EPI_ISL_463571, EPI_ISL_463572, EPI_ISL_463573, EPI_ISL_463574, EPI_ISL_463575, EPI_ISL_463576, EPI_ISL_463577, EPI_ISL_463579, EPI_ISL_463580, EPI_ISL_463581, EPI_ISL_463582, EPI_ISL_463583, EPI_ISL_463584, EPI_ISL_463585, EPI_ISL_463586, EPI_ISL_463587, EPI_ISL_463588, EPI_ISL_463589, EPI_ISL_463590, EPI_ISL_463591, EPI_ISL_463592, EPI_ISL_463593, EPI_ISL_463594, EPI_ISL_463595, EPI_ISL_463596, EPI_ISL_463597, EPI_ISL_463598, EPI_ISL_463599, EPI_ISL_463600, EPI_ISL_463601, EPI_ISL_463602, EPI_ISL_463603, EPI_ISL_463604, EPI_ISL_463605, EPI_ISL_463607, EPI_ISL_463608, EPI_ISL_463609, EPI_ISL_463610, EPI_ISL_463611, EPI_ISL_463612, EPI_ISL_463613, EPI_ISL_463614, EPI_ISL_463615, EPI_ISL_463616, EPI_ISL_463617, EPI_ISL_463618, EPI_ISL_463619, EPI_ISL_463620, EPI_ISL_463621, EPI_ISL_463622, EPI_ISL_463623, EPI_ISL_463624, EPI_ISL_463625, EPI_ISL_463626, EPI_ISL_463627, EPI_ISL_463628, EPI_ISL_463629, EPI_ISL_463630, EPI_ISL_463632, EPI_ISL_463633, EPI_ISL_463634, EPI_ISL_463635, EPI_ISL_463636, EPI_ISL_463637, EPI_ISL_463638, EPI_ISL_463640, EPI_ISL_463641, EPI_ISL_463642, EPI_ISL_463643, EPI_ISL_463644, EPI_ISL_463645, EPI_ISL_463646, EPI_ISL_463647, EPI_ISL_463648, EPI_ISL_463649, EPI_ISL_463650, EPI_ISL_463651, EPI_ISL_463654, EPI_ISL_463655, EPI_ISL_463656, EPI_ISL_463657, EPI_ISL_463658, EPI_ISL_463659, EPI_ISL_463660, EPI_ISL_463661, EPI_ISL_463662, EPI_ISL_463663, EPI_ISL_463664, EPI_ISL_463665, EPI_ISL_463666, EPI_ISL_463667, EPI_ISL_463668, EPI_ISL_463669, EPI_ISL_463670, EPI_ISL_463671, EPI_ISL_463672, EPI_ISL_463673, EPI_ISL_463674, EPI_ISL_463675, EPI_ISL_463676, EPI_ISL_463677, EPI_ISL_463678, EPI_ISL_463679, EPI_ISL_463680, EPI_ISL_463681, EPI_ISL_463682, EPI_ISL_463684, EPI_ISL_463686, EPI_ISL_463687, EPI_ISL_463688, EPI_ISL_463689, EPI_ISL_463690, EPI_ISL_463691, EPI_ISL_463692, EPI_ISL_463694, EPI_ISL_463696, EPI_ISL_463697, EPI_ISL_463698, EPI_ISL_463699, EPI_ISL_463700, EPI_ISL_463701, EPI_ISL_463702, EPI_ISL_463703, EPI_ISL_463704, EPI_ISL_463705 | | | |
| see above | Washington State Department of Health | Seattle Flu Study | Chu et al |
| EPI_ISL_463740 | Mohammed Bin Rashid University of Medicine and Health Sciences | Al Jalila Genomics Center | Ahmad Abou Tayoun, Tom Loney, Hamda Khansaheb, Sathishkumar Ramaswamy, Divinlal Harilal, Zulfa Omar Deesi, Rupa Murthy Varghese, Hanan Al Suwaidi, Abdulmajeed Alkhaja, Mohammed Uddin, Rifat Hamoudi, Rabih Halwani, Abiola Catherine Senok, Qutayba Hamid, Norbert Nowotny, Alawi Alsheikh-Ali |
| EPI_ISL_463741, EPI_ISL_463742, EPI_ISL_463743, EPI_ISL_463744, EPI_ISL_463745, EPI_ISL_463746, EPI_ISL_463747, EPI_ISL_463748 | Department of Molecular Virology, Cyprus Institute of Neurology and Genetics | Department of Molecular Virology, Cyprus Institute of Neurology and Genetics | Jan Richter, George Krashias, Christina Tryfonos, Stavros Bashiardes, Dana Koptides, Christina Christodoulou |
| EPI_ISL_463889, EPI_ISL_463894, EPI_ISL_463895, EPI_ISL_463896, EPI_ISL_463897, EPI_ISL_463901 | Shaoxing Center for Disease Control and Prevention | Department of Pathology and Laboratory Medicine, University of California Los Angeles | Jinkun Chen, Evann E. Hilt, Huan Wu, Zhuojing Jiang, QinChao Zhang, JiLing Wang, Yifang Wang, Fan Li, Ziqin Li, Jialiang Tang, Shangxin Yang |
| EPI_ISL_463904, EPI_ISL_463905, EPI_ISL_463906, EPI_ISL_463908, EPI_ISL_463909, EPI_ISL_463910, EPI_ISL_463912, EPI_ISL_463913, EPI_ISL_463914, EPI_ISL_463917, EPI_ISL_463918, EPI_ISL_463919, EPI_ISL_463920, EPI_ISL_463922, EPI_ISL_463923, EPI_ISL_463927, EPI_ISL_463928, EPI_ISL_463929, EPI_ISL_463930, EPI_ISL_463931, EPI_ISL_463933, EPI_ISL_463934, EPI_ISL_463936, EPI_ISL_463940, EPI_ISL_463941, EPI_ISL_463942, EPI_ISL_463943, EPI_ISL_463946, EPI_ISL_463947, EPI_ISL_463948, EPI_ISL_463953, EPI_ISL_463954, EPI_ISL_463956, EPI_ISL_463957, EPI_ISL_463961, EPI_ISL_463962, EPI_ISL_463963, EPI_ISL_463965, EPI_ISL_463968 | | | |
| see above | Laboratoire de microbiologie, Hopital de Verdun | Smith Laboratory, Centre de Recherche CHU Sainte-Justine | Martin Smith, Marieke Rozendaal, Ivan Pavlov |
| EPI_ISL_463970, EPI_ISL_463971, EPI_ISL_463975, EPI_ISL_463976, EPI_ISL_463977, EPI_ISL_463979, EPI_ISL_463980, EPI_ISL_463981, EPI_ISL_463982, EPI_ISL_463984, EPI_ISL_463988, EPI_ISL_463989, EPI_ISL_463991, EPI_ISL_463992, EPI_ISL_463993, EPI_ISL_463994 | | | |
| see above | Toronto Invasive Bacterial Diseases Network | McMaster University | Allison McGeer, Patryk Aftanas, Angel Li, Kuganya Nirmalarajah, Samira Mubareka, Andrew G. McArthur |
| EPI_ISL_463995, EPI_ISL_463996, EPI_ISL_463997, EPI_ISL_463998, EPI_ISL_463999, EPI_ISL_464001, EPI_ISL_464004, EPI_ISL_464007, EPI_ISL_464008, EPI_ISL_464011, EPI_ISL_464012, EPI_ISL_464013, EPI_ISL_464016, EPI_ISL_464018, EPI_ISL_464019, EPI_ISL_464020, EPI_ISL_464021, EPI_ISL_464022, EPI_ISL_464024, EPI_ISL_464026, EPI_ISL_464028, EPI_ISL_464029, EPI_ISL_464030, EPI_ISL_464033, EPI_ISL_464036, EPI_ISL_464037, EPI_ISL_464038, EPI_ISL_464040, EPI_ISL_464041, EPI_ISL_464042, EPI_ISL_464046, EPI_ISL_464047, EPI_ISL_464048, EPI_ISL_464050, EPI_ISL_464054, EPI_ISL_464056, EPI_ISL_464058, EPI_ISL_464059, EPI_ISL_464061, EPI_ISL_464062, EPI_ISL_464063, EPI_ISL_464064 | | | |
| see above | Unity Health Toronto | Ontario Institute for Cancer Research | Ramzi Fattouh,Larissa M. Matukas,Mark Downing,Annette Gower,Karel Boissinot,Samira Mubareka,TIBDN,Ilinca Lungu,Bernard Lam,Jeremy Johns,Paul Krzyzanowski,Richard de Borja,Philip Zuzarte,Jared Simpson |
| EPI_ISL_464065, EPI_ISL_464066, EPI_ISL_464067, EPI_ISL_464068, EPI_ISL_464069, EPI_ISL_464070, EPI_ISL_464071, EPI_ISL_464072, EPI_ISL_464073, EPI_ISL_464074, EPI_ISL_464075, EPI_ISL_464076, EPI_ISL_464077, EPI_ISL_464078, EPI_ISL_464079, EPI_ISL_464080, EPI_ISL_464081, EPI_ISL_464082, EPI_ISL_464083, EPI_ISL_464084, EPI_ISL_464085, EPI_ISL_464086, EPI_ISL_464087, EPI_ISL_464088, EPI_ISL_464089, EPI_ISL_464090 | | | |
| see above | KU Leuven, Rega Institute, Clinical and Epidemiological Virology | KU Leuven, Rega Institute, Clinical and Epidemiological Virology | Tony Wawina-Bokalanga, Bert Vanmechelen, Joan Marti-Carerras, Piet Maes |
| EPI_ISL_464092, EPI_ISL_464093, EPI_ISL_464094 | Laboratory Medicine | Department of Laboratory Medicine, Lin-Kou Chang Gung Memorial Hospital, Taoyuan, Taiwan | Kuo-Chien Tsao, Yu-Nong Gong, Shu-Li Yang, Yi-Chun Liu, Chung-Guei Huang, Mei-Jen Hsiao, Po-Wei Huang, Cheng-Ta Yang, Cheng-Hsun Chiu, Peng-Nien Huang, Kuo-Ming Lee, Guang-Wu Chen, Shin-Ru Shih |
| EPI_ISL_464112, EPI_ISL_464113, EPI_ISL_464114, EPI_ISL_464118, EPI_ISL_464119, EPI_ISL_464121, EPI_ISL_464123, EPI_ISL_464126, EPI_ISL_464127, EPI_ISL_464128, EPI_ISL_464129, EPI_ISL_464130, EPI_ISL_464131, EPI_ISL_464132, EPI_ISL_464133, EPI_ISL_464137, EPI_ISL_464138, EPI_ISL_464145, EPI_ISL_464153, EPI_ISL_464155, EPI_ISL_464157 | | | |
| see above | National Health Laboratory Service (NHLS), Tygerberg | Division of Medical Virology, Stellenbosch University and National Health Laboratory Service (NHLS) | Susan Engelbrecht, Kayla Delaney, Bronwyn Kleinhans, Houriiyah Tegally, Eduan Wilkindon, Gert van Zyl, Wolfgang Preiser, Tulio de Oliveira |
| EPI_ISL_464159, EPI_ISL_464160 | National Institute of Laboratory Medicine and Referral Center | Genomic Research Lab, BCSIR | Shahina Akter, Abu Sayeed Mohammad Mahmud, Mohammad Samir Uzzaman, Eshrar Osman, Md. Ahasan Habib, Tanjina Akhter Banu, Md. Murshed Hasan Sarker, Barna Goswami, Iffat Jahan, Md. Saddam Hossain, Tasnim Nafisa, Md. Maruf Ahmed Molla, Mahmuda Yeasmin, Asish Kumar Ghosh, Arifa Akram, A. K. M. Shamsuzzaman, Sheikh Md. Selim Al Din, Utpal Chandra Ray, Salek Ahmed Sajib, Md. Salim Khan |
| EPI_ISL_464161, EPI_ISL_464162 | National Institute of Laboratory Medicine and Referral Center | Genomic Research Lab, BCSIR | Md. Ahasan Habib, Abu Sayeed Mohammad Mahmud, Mohammad Samir Uzzaman, Eshrar Osman, Shahina Akter, Tanjina Akhter Banu, Md. Murshed Hasan Sarker, Barna Goswami, Iffat Jahan, Md. Saddam Hossain, Tasnim Nafisa, Md. Maruf Ahmed Molla, Mahmuda Yeasmin, Asish Kumar Ghosh, Arifa Akram, A. K. M. Shamsuzzaman, Sheikh Md. Selim Al Din, Utpal Chandra Ray, Salek Ahmed Sajib, Md. Salim Khan |
| EPI_ISL_464163, EPI_ISL_464164 | National Institute of Laboratory Medicine and Referral Center | Genomic Research Lab, BCSIR | Tanjina Akhter Banu, Abu Sayeed Mohammad Mahmud, Mohammad Samir Uzzaman, Eshrar Osman, Md. Ahasan Habib, Shahina Akter, Md. Murshed Hasan Sarker, Barna Goswami, Iffat Jahan, Md. Saddam Hossain, Tasnim Nafisa, Md. Maruf Ahmed Molla, Mahmuda Yeasmin, Asish Kumar Ghosh, Arifa Akram, A. K. M. Shamsuzzaman, Sheikh Md. Selim Al Din, Utpal Chandra Ray, Salek Ahmed Sajib, Md. Salim Khan |
| EPI_ISL_464165, EPI_ISL_464166 | National Institute of Laboratory Medicine and Referral Center | Genomic Research Lab, BCSIR | Barna Goswami, Abu Sayeed Mohammad Mahmud, Mohammad Samir Uzzaman, Eshrar Osman, Md. Ahasan Habib, Shahina Akter, Tanjina Akhter Banu, Md. Murshed Hasan Sarker, Iffat Jahan, Md. Saddam Hossain, Tasnim Nafisa, Md. Maruf Ahmed Molla, Mahmuda Yeasmin, Asish Kumar Ghosh, Arifa Akram, A. K. M. Shamsuzzaman, Sheikh Md. Selim Al Din, Utpal Chandra Ray, Salek Ahmed Sajib, Md. Salim Khan |
| EPI_ISL_464167 | VI-US Virgin Islands Department of Health | Pathogen Discovery, Respiratory Viruses Branch, Division of Viral Diseases, Centers for Disease Control and Prevention | Krista Queen, Ying Tao, Jing Zhang, Yan Li, Anna Uehara, Clinton R. Paden, Mary S. Keckler, Alison S. Laufer Halpin, Haibin Wang, Jasmine Padilla, Justin Lee, Christopher A. Elkins, Suxiang Tong |
| EPI_ISL_464168, EPI_ISL_464169, EPI_ISL_464175, EPI_ISL_464178, EPI_ISL_464180, EPI_ISL_464181, EPI_ISL_464182, EPI_ISL_464183, EPI_ISL_464184, EPI_ISL_464185, EPI_ISL_464186, EPI_ISL_464187, EPI_ISL_464188, EPI_ISL_464189, EPI_ISL_464190, EPI_ISL_464192, EPI_ISL_464193, EPI_ISL_464194, EPI_ISL_464195, EPI_ISL_464196, EPI_ISL_464197, EPI_ISL_464198, EPI_ISL_464199, EPI_ISL_464200, EPI_ISL_464201, EPI_ISL_464202, EPI_ISL_464204, EPI_ISL_464206, EPI_ISL_464207, EPI_ISL_464208, EPI_ISL_464209, EPI_ISL_464210, EPI_ISL_464212, EPI_ISL_464214, EPI_ISL_464215, EPI_ISL_464216, EPI_ISL_464217, EPI_ISL_464218, EPI_ISL_464219, EPI_ISL_464222, EPI_ISL_464223, EPI_ISL_464224, EPI_ISL_464227, EPI_ISL_464228, EPI_ISL_464229, EPI_ISL_464238, EPI_ISL_464241, EPI_ISL_464248, EPI_ISL_464252, EPI_ISL_464253, EPI_ISL_464258, EPI_ISL_464260, EPI_ISL_464262, EPI_ISL_464266, EPI_ISL_464270, EPI_ISL_464275, EPI_ISL_464278, EPI_ISL_464283, EPI_ISL_464288, EPI_ISL_464291, EPI_ISL_464292, EPI_ISL_464293, EPI_ISL_464294, EPI_ISL_464295, EPI_ISL_464296, EPI_ISL_464297, EPI_ISL_464298, EPI_ISL_464300, EPI_ISL_464302, EPI_ISL_464305, EPI_ISL_464307, EPI_ISL_464310, EPI_ISL_464311, EPI_ISL_464312, EPI_ISL_464314, EPI_ISL_464315, EPI_ISL_464318, EPI_ISL_464321, EPI_ISL_464323, EPI_ISL_464334, EPI_ISL_464335, EPI_ISL_464339, EPI_ISL_464342, EPI_ISL_464343, EPI_ISL_464344, EPI_ISL_464347, EPI_ISL_464348, EPI_ISL_464350, EPI_ISL_464355, EPI_ISL_464356, EPI_ISL_464357, EPI_ISL_464358, EPI_ISL_464359, EPI_ISL_464361, EPI_ISL_464362, EPI_ISL_464363, EPI_ISL_464364, EPI_ISL_464365, EPI_ISL_464367, EPI_ISL_464368, EPI_ISL_464369, EPI_ISL_464370, EPI_ISL_464371, EPI_ISL_464372, EPI_ISL_464374, EPI_ISL_464375, EPI_ISL_464377, EPI_ISL_464378, EPI_ISL_464380, EPI_ISL_464382, EPI_ISL_464383, EPI_ISL_464388, EPI_ISL_464389, EPI_ISL_464390, EPI_ISL_464392, EPI_ISL_464393, EPI_ISL_464395, EPI_ISL_464396, EPI_ISL_464397, EPI_ISL_464399, EPI_ISL_464401, EPI_ISL_464402, EPI_ISL_464403, EPI_ISL_464404, EPI_ISL_464408, EPI_ISL_464409, EPI_ISL_464410, EPI_ISL_464411, EPI_ISL_464413, EPI_ISL_464415, EPI_ISL_464416, EPI_ISL_464417, EPI_ISL_464419, EPI_ISL_464420, EPI_ISL_464422, EPI_ISL_464423, EPI_ISL_464424, EPI_ISL_464425, EPI_ISL_464426, EPI_ISL_464427, EPI_ISL_464428, EPI_ISL_464429, EPI_ISL_464430, EPI_ISL_464431, EPI_ISL_464432, EPI_ISL_464433, EPI_ISL_464434, EPI_ISL_464435, EPI_ISL_464437, EPI_ISL_464438, EPI_ISL_464439, EPI_ISL_464440, EPI_ISL_464441, EPI_ISL_464442, EPI_ISL_464443, EPI_ISL_464444, EPI_ISL_464446, EPI_ISL_464447, EPI_ISL_464448, EPI_ISL_464449, EPI_ISL_464450, EPI_ISL_464451, EPI_ISL_464452, EPI_ISL_464456, EPI_ISL_464457, EPI_ISL_464458, EPI_ISL_464459, EPI_ISL_464460, EPI_ISL_464461, EPI_ISL_464463, EPI_ISL_464464, EPI_ISL_464465, EPI_ISL_464466, EPI_ISL_464467, EPI_ISL_464468, EPI_ISL_464472, EPI_ISL_464473, EPI_ISL_464475, EPI_ISL_464476, EPI_ISL_464479, EPI_ISL_464480, EPI_ISL_464481, EPI_ISL_464482, EPI_ISL_464483, EPI_ISL_464485, EPI_ISL_464488, EPI_ISL_464489, EPI_ISL_464492, EPI_ISL_464497, EPI_ISL_464498, EPI_ISL_464505, EPI_ISL_464506, EPI_ISL_464507, EPI_ISL_464508, EPI_ISL_464517, EPI_ISL_464518, EPI_ISL_464520, EPI_ISL_464525, EPI_ISL_464527, EPI_ISL_464528, EPI_ISL_464529, EPI_ISL_464532, EPI_ISL_464534, EPI_ISL_464535, EPI_ISL_464541, EPI_ISL_464545, EPI_ISL_464546, EPI_ISL_464549, EPI_ISL_464554, EPI_ISL_464555, EPI_ISL_464557, EPI_ISL_464559, EPI_ISL_464561, EPI_ISL_464562, EPI_ISL_464563, EPI_ISL_464567, EPI_ISL_464583, EPI_ISL_464595, EPI_ISL_464614, EPI_ISL_464643, EPI_ISL_464649, EPI_ISL_464660, EPI_ISL_464664, EPI_ISL_464672, EPI_ISL_464682, EPI_ISL_464684, EPI_ISL_464711, EPI_ISL_464714, EPI_ISL_464715, EPI_ISL_464720, EPI_ISL_464721, EPI_ISL_464726, EPI_ISL_464728, EPI_ISL_464730, EPI_ISL_464732, EPI_ISL_464734, EPI_ISL_464735, EPI_ISL_464738, EPI_ISL_464741, EPI_ISL_464742, EPI_ISL_464743, EPI_ISL_464744, EPI_ISL_464745, EPI_ISL_464751, EPI_ISL_464755, EPI_ISL_464757, EPI_ISL_464758, EPI_ISL_464759, EPI_ISL_464761, EPI_ISL_464765, EPI_ISL_464767, EPI_ISL_464769, EPI_ISL_464770, EPI_ISL_464771, EPI_ISL_464772, EPI_ISL_464774, EPI_ISL_464776, EPI_ISL_464777, EPI_ISL_464778, EPI_ISL_464779, EPI_ISL_464780, EPI_ISL_464781, EPI_ISL_464782, EPI_ISL_464783, EPI_ISL_464784, EPI_ISL_464785, EPI_ISL_464786, EPI_ISL_464788, EPI_ISL_464789, EPI_ISL_464790, EPI_ISL_464791, EPI_ISL_464792, EPI_ISL_464793, EPI_ISL_464794, EPI_ISL_464795, EPI_ISL_464796, EPI_ISL_464797, EPI_ISL_464798, EPI_ISL_464799, EPI_ISL_464800, EPI_ISL_464801, EPI_ISL_464803, EPI_ISL_464806, EPI_ISL_464807, EPI_ISL_464808, EPI_ISL_464810, EPI_ISL_464813, EPI_ISL_464814, EPI_ISL_464816, EPI_ISL_464817, EPI_ISL_464818, EPI_ISL_464820, EPI_ISL_464821, EPI_ISL_464822, EPI_ISL_464824, EPI_ISL_464826, EPI_ISL_464830, EPI_ISL_464831, EPI_ISL_464833, EPI_ISL_464835, EPI_ISL_464836, EPI_ISL_464838, EPI_ISL_464839, EPI_ISL_464841, EPI_ISL_464842, EPI_ISL_464843, EPI_ISL_464844, EPI_ISL_464849, EPI_ISL_464850, EPI_ISL_464852, EPI_ISL_464855, EPI_ISL_464859, EPI_ISL_464861, EPI_ISL_464862, EPI_ISL_464863, EPI_ISL_464864, EPI_ISL_464866, EPI_ISL_464868, EPI_ISL_464869, EPI_ISL_464871, EPI_ISL_464873, EPI_ISL_464874, EPI_ISL_464876, EPI_ISL_464877, EPI_ISL_464879, EPI_ISL_464880, EPI_ISL_464882, EPI_ISL_464883, EPI_ISL_464894, EPI_ISL_464897, EPI_ISL_464898, EPI_ISL_464900, EPI_ISL_464908, EPI_ISL_464909, EPI_ISL_464912, EPI_ISL_464917, EPI_ISL_464920, EPI_ISL_464921, EPI_ISL_464923, EPI_ISL_464925, EPI_ISL_464926, EPI_ISL_464932, EPI_ISL_464936, EPI_ISL_464938, EPI_ISL_464939, EPI_ISL_464944, EPI_ISL_464946, EPI_ISL_464947, EPI_ISL_464948, EPI_ISL_464949, EPI_ISL_464950, EPI_ISL_464951, EPI_ISL_464954, EPI_ISL_464955, EPI_ISL_464956, EPI_ISL_464957, EPI_ISL_464958, EPI_ISL_464959, EPI_ISL_464961, EPI_ISL_464962, EPI_ISL_464963, EPI_ISL_464971, EPI_ISL_464972, EPI_ISL_464973, EPI_ISL_464974, EPI_ISL_464976, EPI_ISL_464978, EPI_ISL_464979, EPI_ISL_464980, EPI_ISL_464985, EPI_ISL_464989, EPI_ISL_464991, EPI_ISL_464996, EPI_ISL_464997, EPI_ISL_464999, EPI_ISL_465001, EPI_ISL_465002, EPI_ISL_465012, EPI_ISL_465013, EPI_ISL_465024, EPI_ISL_465028, EPI_ISL_465029, EPI_ISL_465030, EPI_ISL_465031, EPI_ISL_465033, EPI_ISL_465034, EPI_ISL_465035, EPI_ISL_465036, EPI_ISL_465037, EPI_ISL_465038, EPI_ISL_465039, EPI_ISL_465040, EPI_ISL_465041, EPI_ISL_465042, EPI_ISL_465043, EPI_ISL_465044, EPI_ISL_465045, EPI_ISL_465046, EPI_ISL_465049, EPI_ISL_465050, EPI_ISL_465052, EPI_ISL_465054, EPI_ISL_465057, EPI_ISL_465058, EPI_ISL_465059, EPI_ISL_465060, EPI_ISL_465063, EPI_ISL_465064, EPI_ISL_465065, EPI_ISL_465068, EPI_ISL_465070, EPI_ISL_465071, EPI_ISL_465073, EPI_ISL_465075, EPI_ISL_465078, EPI_ISL_465079, EPI_ISL_465081, EPI_ISL_465082, EPI_ISL_465083, EPI_ISL_465084, EPI_ISL_465085, EPI_ISL_465086, EPI_ISL_465087, EPI_ISL_465091, EPI_ISL_465092, EPI_ISL_465093, EPI_ISL_465096, EPI_ISL_465099, EPI_ISL_465100, EPI_ISL_465101, EPI_ISL_465104, EPI_ISL_465105, EPI_ISL_465108, EPI_ISL_465115, EPI_ISL_465120, EPI_ISL_465125, EPI_ISL_465130, EPI_ISL_465131, EPI_ISL_465135, EPI_ISL_465136, EPI_ISL_465143, EPI_ISL_465145, EPI_ISL_465155 | | | |
| see above | Respiratory Virus Unit, Microbiology Services Colindale, Public Health England | Respiratory Virus Unit, Microbiology Services Colindale, Public Health England | PHE Covid Sequencing Team |
| EPI_ISL_465163, EPI_ISL_465164 | National Institute of Laboratory Medicine and Referral Center | Genomic Research Lab, BCSIR | Iffat Jahan, Abu Sayeed Mohammad Mahmud, Mohammad Samir Uzzaman, Eshrar Osman, Md. Ahasan Habib, Shahina Akter, Tanjina Akhter Banu, Md. Murshed Hasan Sarker, Barna Goswami, Md. Saddam Hossain, Tasnim Nafisa, Md. Maruf Ahmed Molla, Mahmuda Yeasmin, Asish Kumar Ghosh, Arifa Akram, A. K. M. Shamsuzzaman, Sheikh Md. Selim Al Din, Utpal Chandra Ray, Salek Ahmed Sajib, Md. Salim Khan |
| EPI_ISL_465167, EPI_ISL_465168, EPI_ISL_465169, EPI_ISL_465170, EPI_ISL_465173, EPI_ISL_465174, EPI_ISL_465178, EPI_ISL_465180, EPI_ISL_465181, EPI_ISL_465182, EPI_ISL_465183, EPI_ISL_465184, EPI_ISL_465186, EPI_ISL_465192, EPI_ISL_465193, EPI_ISL_465194, EPI_ISL_465195, EPI_ISL_465196, EPI_ISL_465197, EPI_ISL_465198, EPI_ISL_465199, EPI_ISL_465202, EPI_ISL_465206, EPI_ISL_465208, EPI_ISL_465210, EPI_ISL_465211, EPI_ISL_465213, EPI_ISL_465218, EPI_ISL_465222, EPI_ISL_465224, EPI_ISL_465225, EPI_ISL_465227, EPI_ISL_465241, EPI_ISL_465243, EPI_ISL_465244, EPI_ISL_465245, EPI_ISL_465248, EPI_ISL_465253, EPI_ISL_465254, EPI_ISL_465259, EPI_ISL_465260, EPI_ISL_465261, EPI_ISL_465262, EPI_ISL_465263, EPI_ISL_465266, EPI_ISL_465270, EPI_ISL_465271, EPI_ISL_465272, EPI_ISL_465273, EPI_ISL_465274, EPI_ISL_465275, EPI_ISL_465276, EPI_ISL_465277, EPI_ISL_465278, EPI_ISL_465279, EPI_ISL_465280, EPI_ISL_465281, EPI_ISL_465282, EPI_ISL_465283, EPI_ISL_465285, EPI_ISL_465286, EPI_ISL_465287, EPI_ISL_465289, EPI_ISL_465290, EPI_ISL_465294, EPI_ISL_465297, EPI_ISL_465298, EPI_ISL_465300, EPI_ISL_465302, EPI_ISL_465304, EPI_ISL_465307, EPI_ISL_465309, EPI_ISL_465311, EPI_ISL_465312, EPI_ISL_465313, EPI_ISL_465315, EPI_ISL_465318, EPI_ISL_465319, EPI_ISL_465320, EPI_ISL_465322, EPI_ISL_465324, EPI_ISL_465332, EPI_ISL_465339, EPI_ISL_465343, EPI_ISL_465344, EPI_ISL_465347, EPI_ISL_465348, EPI_ISL_465349, EPI_ISL_465350, EPI_ISL_465351, EPI_ISL_465352, EPI_ISL_465353, EPI_ISL_465357, EPI_ISL_465359, EPI_ISL_465361, EPI_ISL_465370, EPI_ISL_465371, EPI_ISL_465381, EPI_ISL_465382, EPI_ISL_465383, EPI_ISL_465384, EPI_ISL_465386, EPI_ISL_465388, EPI_ISL_465389, EPI_ISL_465390, EPI_ISL_465396, EPI_ISL_465397, EPI_ISL_465398, EPI_ISL_465404, EPI_ISL_465406, EPI_ISL_465407, EPI_ISL_465411, EPI_ISL_465412, EPI_ISL_465417, EPI_ISL_465428, EPI_ISL_465430, EPI_ISL_465433, EPI_ISL_465436, EPI_ISL_465440, EPI_ISL_465441, EPI_ISL_465448, EPI_ISL_465450, EPI_ISL_465453, EPI_ISL_465454, EPI_ISL_465458, EPI_ISL_465464, EPI_ISL_465466, EPI_ISL_465471, EPI_ISL_465474, EPI_ISL_465475, EPI_ISL_465476, EPI_ISL_465477, EPI_ISL_465482, EPI_ISL_465485, EPI_ISL_465488, EPI_ISL_465491, EPI_ISL_465492, EPI_ISL_465493, EPI_ISL_465494, EPI_ISL_465495, EPI_ISL_465498, EPI_ISL_465499, EPI_ISL_465500, EPI_ISL_465501, EPI_ISL_465503, EPI_ISL_465505, EPI_ISL_465507, EPI_ISL_465508, EPI_ISL_465510, EPI_ISL_465513, EPI_ISL_465514, EPI_ISL_465517, EPI_ISL_465519, EPI_ISL_465520, EPI_ISL_465521, EPI_ISL_465523, EPI_ISL_465525, EPI_ISL_465526, EPI_ISL_465527, EPI_ISL_465528, EPI_ISL_465531, EPI_ISL_465532, EPI_ISL_465533, EPI_ISL_465534, EPI_ISL_465536, EPI_ISL_465537, EPI_ISL_465539, EPI_ISL_465543, EPI_ISL_465546, EPI_ISL_465549, EPI_ISL_465551, EPI_ISL_465552, EPI_ISL_465554, EPI_ISL_465555, EPI_ISL_465556, EPI_ISL_465557, EPI_ISL_465558, EPI_ISL_465559, EPI_ISL_465560, EPI_ISL_465561, EPI_ISL_465563, EPI_ISL_465564, EPI_ISL_465565, EPI_ISL_465566, EPI_ISL_465569, EPI_ISL_465570, EPI_ISL_465574, EPI_ISL_465576, EPI_ISL_465577, EPI_ISL_465579, EPI_ISL_465580, EPI_ISL_465581, EPI_ISL_465582, EPI_ISL_465583, EPI_ISL_465584, EPI_ISL_465586, EPI_ISL_465587, EPI_ISL_465589, EPI_ISL_465590, EPI_ISL_465591, EPI_ISL_465592, EPI_ISL_465593, EPI_ISL_465594, EPI_ISL_465595, EPI_ISL_465596, EPI_ISL_465597, EPI_ISL_465598, EPI_ISL_465600, EPI_ISL_465601, EPI_ISL_465602, EPI_ISL_465603, EPI_ISL_465605, EPI_ISL_465607, EPI_ISL_465608, EPI_ISL_465610, EPI_ISL_465611, EPI_ISL_465612, EPI_ISL_465613, EPI_ISL_465614, EPI_ISL_465615, EPI_ISL_465616, EPI_ISL_465617, EPI_ISL_465618, EPI_ISL_465620, EPI_ISL_465621, EPI_ISL_465623, EPI_ISL_465626, EPI_ISL_465628, EPI_ISL_465633, EPI_ISL_465640, EPI_ISL_465649, EPI_ISL_465652, EPI_ISL_465661, EPI_ISL_465662, EPI_ISL_465663, EPI_ISL_465664, EPI_ISL_465665, EPI_ISL_465666, EPI_ISL_465668, EPI_ISL_465673, EPI_ISL_465676 | | | |
| see above | Respiratory Virus Unit, Microbiology Services Colindale, Public Health England | Respiratory Virus Unit, Microbiology Services Colindale, Public Health England | PHE Covid Sequencing Team |
| EPI_ISL_465679, EPI_ISL_465680 | Hôpital du Suroît | Laboratoire de santé publique du Québec | Sandrine Moreira, Ioannis Ragoussis, Guillaume Bourque, Jesse Shapiro, Mark Lathrop and Michel Roger on behalf of the CoVSeQ research group (http://covseq.ca/researchgroup) |
| EPI_ISL_465681, EPI_ISL_465682 | Hôpital Charles-LeMoyne | Laboratoire de santé publique du Québec | Sandrine Moreira, Ioannis Ragoussis, Guillaume Bourque, Jesse Shapiro, Mark Lathrop and Michel Roger on behalf of the CoVSeQ research group (http://covseq.ca/researchgroup) |
| EPI_ISL_465683, EPI_ISL_465684 | Hôpital de Maria | Laboratoire de santé publique du Québec | Sandrine Moreira, Ioannis Ragoussis, Guillaume Bourque, Jesse Shapiro, Mark Lathrop and Michel Roger on behalf of the CoVSeQ research group (http://covseq.ca/researchgroup) |
| EPI_ISL_465685 | CSSS Haut-Richelieu/Rouville (Hôpital) | Laboratoire de santé publique du Québec | Sandrine Moreira, Ioannis Ragoussis, Guillaume Bourque, Jesse Shapiro, Mark Lathrop and Michel Roger on behalf of the CoVSeQ research group (http://covseq.ca/researchgroup) |
| EPI_ISL_465686 | Hôpital de Hull | Laboratoire de santé publique du Québec | Sandrine Moreira, Ioannis Ragoussis, Guillaume Bourque, Jesse Shapiro, Mark Lathrop and Michel Roger on behalf of the CoVSeQ research group (http://covseq.ca/researchgroup) |

| | | | |
|---|---|---|---|
| EPI_ISL_465687 | CSSS Haut-Richelieu/Rouville (Hôpital) | Laboratoire de santé publique du Québec | Sandrine Moreira, Ioannis Ragoussis, Guillaume Bourque, Jesse Shapiro, Mark Lathrop and Michel Roger on behalf of the CoVSeQ research group (http://covseq.ca/researchgroup) |
| EPI_ISL_465688, EPI_ISL_465689, EPI_ISL_465690 | Centre hospitalier Anna-Laberge | Laboratoire de santé publique du Québec | Sandrine Moreira, Ioannis Ragoussis, Guillaume Bourque, Jesse Shapiro, Mark Lathrop and Michel Roger on behalf of the CoVSeQ research group (http://covseq.ca/researchgroup) |
| EPI_ISL_465691 | Hôpital de Gatineau | Laboratoire de santé publique du Québec | Sandrine Moreira, Ioannis Ragoussis, Guillaume Bourque, Jesse Shapiro, Mark Lathrop and Michel Roger on behalf of the CoVSeQ research group (http://covseq.ca/researchgroup) |
| EPI_ISL_465692 | CSSS Haut-Richelieu/Rouville (Hôpital) | Laboratoire de santé publique du Québec | Sandrine Moreira, Ioannis Ragoussis, Guillaume Bourque, Jesse Shapiro, Mark Lathrop and Michel Roger on behalf of the CoVSeQ research group (http://covseq.ca/researchgroup) |
| EPI_ISL_465693 | Hôpital du Suroît | Laboratoire de santé publique du Québec | Sandrine Moreira, Ioannis Ragoussis, Guillaume Bourque, Jesse Shapiro, Mark Lathrop and Michel Roger on behalf of the CoVSeQ research group (http://covseq.ca/researchgroup) |
| EPI_ISL_465694 | Hôpital Charles-LeMoyne | Laboratoire de santé publique du Québec | Sandrine Moreira, Ioannis Ragoussis, Guillaume Bourque, Jesse Shapiro, Mark Lathrop and Michel Roger on behalf of the CoVSeQ research group (http://covseq.ca/researchgroup) |
| EPI_ISL_465695, EPI_ISL_465696 | Hôpital Pierre-Boucher | Laboratoire de santé publique du Québec | Sandrine Moreira, Ioannis Ragoussis, Guillaume Bourque, Jesse Shapiro, Mark Lathrop and Michel Roger on behalf of the CoVSeQ research group (http://covseq.ca/researchgroup) |
| EPI_ISL_465697 | Centre de santé Inuulitsivik | Laboratoire de santé publique du Québec | Sandrine Moreira, Ioannis Ragoussis, Guillaume Bourque, Jesse Shapiro, Mark Lathrop and Michel Roger on behalf of the CoVSeQ research group (http://covseq.ca/researchgroup) |
| EPI_ISL_465698, EPI_ISL_465699 | Hôpital Charles-LeMoyne | Laboratoire de santé publique du Québec | Sandrine Moreira, Ioannis Ragoussis, Guillaume Bourque, Jesse Shapiro, Mark Lathrop and Michel Roger on behalf of the CoVSeQ research group (http://covseq.ca/researchgroup) |
| EPI_ISL_465700 | Hôpital Pierre-Boucher | Laboratoire de santé publique du Québec | Sandrine Moreira, Ioannis Ragoussis, Guillaume Bourque, Jesse Shapiro, Mark Lathrop and Michel Roger on behalf of the CoVSeQ research group (http://covseq.ca/researchgroup) |
| EPI_ISL_465701 | Hôpital du Suroît | Laboratoire de santé publique du Québec | Sandrine Moreira, Ioannis Ragoussis, Guillaume Bourque, Jesse Shapiro, Mark Lathrop and Michel Roger on behalf of the CoVSeQ research group (http://covseq.ca/researchgroup) |
| EPI_ISL_465702, EPI_ISL_465703 | Hôpital Pierre-Boucher | Laboratoire de santé publique du Québec | Sandrine Moreira, Ioannis Ragoussis, Guillaume Bourque, Jesse Shapiro, Mark Lathrop and Michel Roger on behalf of the CoVSeQ research group (http://covseq.ca/researchgroup) |
| EPI_ISL_465704, EPI_ISL_465706, EPI_ISL_465708, EPI_ISL_465709, EPI_ISL_465710, EPI_ISL_465712, EPI_ISL_465713, EPI_ISL_465714, EPI_ISL_465717, EPI_ISL_465718, EPI_ISL_465725, EPI_ISL_465728, EPI_ISL_465732, EPI_ISL_465734, EPI_ISL_465739, EPI_ISL_465740, EPI_ISL_465742, EPI_ISL_465748, EPI_ISL_465753, EPI_ISL_465754, EPI_ISL_465757, EPI_ISL_465759, EPI_ISL_465766, EPI_ISL_465767, EPI_ISL_465768, EPI_ISL_465769, EPI_ISL_465774, EPI_ISL_465780, EPI_ISL_465788, EPI_ISL_465794, EPI_ISL_465796, EPI_ISL_465808, EPI_ISL_465811, EPI_ISL_465812, EPI_ISL_465818, EPI_ISL_465819, EPI_ISL_465820, EPI_ISL_465821, EPI_ISL_465822, EPI_ISL_465823, EPI_ISL_465824, EPI_ISL_465834, EPI_ISL_465854, EPI_ISL_465855, EPI_ISL_465856, EPI_ISL_465858, EPI_ISL_465859, EPI_ISL_465860, EPI_ISL_465861, EPI_ISL_465864, EPI_ISL_465865, EPI_ISL_465866, EPI_ISL_465867, EPI_ISL_465868, EPI_ISL_465869, EPI_ISL_465870, EPI_ISL_465871, EPI_ISL_465872, EPI_ISL_465873, EPI_ISL_465874, EPI_ISL_465875, EPI_ISL_465876, EPI_ISL_465877, EPI_ISL_465878, EPI_ISL_465879, EPI_ISL_465881, EPI_ISL_465883, EPI_ISL_465884, EPI_ISL_465885, EPI_ISL_465887, EPI_ISL_465888, EPI_ISL_465890, EPI_ISL_465895, EPI_ISL_465896, EPI_ISL_465904, EPI_ISL_465906, EPI_ISL_465914, EPI_ISL_465920, EPI_ISL_465921, EPI_ISL_465922, EPI_ISL_465923, EPI_ISL_465926, EPI_ISL_465932, EPI_ISL_465935, EPI_ISL_465936, EPI_ISL_465937, EPI_ISL_465939, EPI_ISL_465940, EPI_ISL_465941, EPI_ISL_465943, EPI_ISL_465944, EPI_ISL_465946, EPI_ISL_465960, EPI_ISL_465962, EPI_ISL_465966, EPI_ISL_465971, EPI_ISL_465973, EPI_ISL_465975, EPI_ISL_465976, EPI_ISL_465977, EPI_ISL_465978, EPI_ISL_465980, EPI_ISL_465984, EPI_ISL_465987, EPI_ISL_465989, EPI_ISL_465990, EPI_ISL_465991, EPI_ISL_465992, EPI_ISL_465993, EPI_ISL_465994, EPI_ISL_465995, EPI_ISL_465996, EPI_ISL_465997, EPI_ISL_465999, EPI_ISL_466000, EPI_ISL_466001, EPI_ISL_466002, EPI_ISL_466003, EPI_ISL_466004, EPI_ISL_466005, EPI_ISL_466006, EPI_ISL_466008, EPI_ISL_466014, EPI_ISL_466015, EPI_ISL_466016, EPI_ISL_466017, EPI_ISL_466018, EPI_ISL_466023, EPI_ISL_466025, EPI_ISL_466026, EPI_ISL_466028, EPI_ISL_466029, EPI_ISL_466030, EPI_ISL_466032, EPI_ISL_466033, EPI_ISL_466034, EPI_ISL_466036, EPI_ISL_466037, EPI_ISL_466038, EPI_ISL_466039, EPI_ISL_466040, EPI_ISL_466041, EPI_ISL_466043, EPI_ISL_466044, EPI_ISL_466045, EPI_ISL_466046, EPI_ISL_466047, EPI_ISL_466048, EPI_ISL_466050, EPI_ISL_466051, EPI_ISL_466052, EPI_ISL_466054, EPI_ISL_466057, EPI_ISL_466059, EPI_ISL_466061, EPI_ISL_466062, EPI_ISL_466063, EPI_ISL_466065, EPI_ISL_466066, EPI_ISL_466068, EPI_ISL_466069, EPI_ISL_466071, EPI_ISL_466074, EPI_ISL_466075, EPI_ISL_466076, EPI_ISL_466077, EPI_ISL_466081, EPI_ISL_466082, EPI_ISL_466084, EPI_ISL_466086, EPI_ISL_466088, EPI_ISL_466089, EPI_ISL_466090, EPI_ISL_466092, EPI_ISL_466094, EPI_ISL_466095, EPI_ISL_466096, EPI_ISL_466097, EPI_ISL_466098, EPI_ISL_466099, EPI_ISL_466101, EPI_ISL_466102, EPI_ISL_466103, EPI_ISL_466107, EPI_ISL_466109, EPI_ISL_466110, EPI_ISL_466113, EPI_ISL_466115, EPI_ISL_466120, EPI_ISL_466122, EPI_ISL_466124, EPI_ISL_466129, EPI_ISL_466131, EPI_ISL_466132, EPI_ISL_466135, EPI_ISL_466138, EPI_ISL_466139, EPI_ISL_466141, EPI_ISL_466142, EPI_ISL_466144, EPI_ISL_466145, EPI_ISL_466146, EPI_ISL_466147, EPI_ISL_466148, EPI_ISL_466149, EPI_ISL_466151, EPI_ISL_466152, EPI_ISL_466153, EPI_ISL_466155, EPI_ISL_466156, EPI_ISL_466157, EPI_ISL_466158, EPI_ISL_466159, EPI_ISL_466161, EPI_ISL_466162, EPI_ISL_466163, EPI_ISL_466164, EPI_ISL_466165, EPI_ISL_466166, EPI_ISL_466167, EPI_ISL_466168, EPI_ISL_466169, EPI_ISL_466171, EPI_ISL_466173, EPI_ISL_466174, EPI_ISL_466176, EPI_ISL_466178, EPI_ISL_466180, EPI_ISL_466181, EPI_ISL_466185, EPI_ISL_466186, EPI_ISL_466189, EPI_ISL_466191, EPI_ISL_466192, EPI_ISL_466195, EPI_ISL_466196, EPI_ISL_466197, EPI_ISL_466198, EPI_ISL_466199, EPI_ISL_466200, EPI_ISL_466201, EPI_ISL_466203, EPI_ISL_466204, EPI_ISL_466205, EPI_ISL_466206, EPI_ISL_466207, EPI_ISL_466208, EPI_ISL_466212, EPI_ISL_466213, EPI_ISL_466216, EPI_ISL_466217, EPI_ISL_466218, EPI_ISL_466219, EPI_ISL_466224, EPI_ISL_466225, EPI_ISL_466226, EPI_ISL_466228, EPI_ISL_466230, EPI_ISL_466231, EPI_ISL_466232, EPI_ISL_466233, EPI_ISL_466234, EPI_ISL_466235, EPI_ISL_466237, EPI_ISL_466238, EPI_ISL_466239, EPI_ISL_466240, EPI_ISL_466241, EPI_ISL_466242, EPI_ISL_466246, EPI_ISL_466248, EPI_ISL_466250, EPI_ISL_466252, EPI_ISL_466253, EPI_ISL_466254, EPI_ISL_466255, EPI_ISL_466257, EPI_ISL_466258, EPI_ISL_466259, EPI_ISL_466260, EPI_ISL_466261, EPI_ISL_466262, EPI_ISL_466267, EPI_ISL_466271, EPI_ISL_466272, EPI_ISL_466273, EPI_ISL_466275, EPI_ISL_466278, EPI_ISL_466280, EPI_ISL_466282, EPI_ISL_466286, EPI_ISL_466290, EPI_ISL_466291, EPI_ISL_466293, EPI_ISL_466298, EPI_ISL_466299, EPI_ISL_466308, EPI_ISL_466311, EPI_ISL_466317, EPI_ISL_466320, EPI_ISL_466322, EPI_ISL_466323, EPI_ISL_466324, EPI_ISL_466325, EPI_ISL_466326, EPI_ISL_466327, EPI_ISL_466328, EPI_ISL_466336, EPI_ISL_466338, EPI_ISL_466340, EPI_ISL_466342, EPI_ISL_466343, EPI_ISL_466346, EPI_ISL_466350, EPI_ISL_466351, EPI_ISL_466352, EPI_ISL_466355, EPI_ISL_466356, EPI_ISL_466359, EPI_ISL_466360, EPI_ISL_466361, EPI_ISL_466363, EPI_ISL_466365, EPI_ISL_466366, EPI_ISL_466367, EPI_ISL_466385, EPI_ISL_466389, EPI_ISL_466391, EPI_ISL_466392, EPI_ISL_466393, EPI_ISL_466395, EPI_ISL_466399, EPI_ISL_466400, EPI_ISL_466401, EPI_ISL_466408, EPI_ISL_466409, EPI_ISL_466410, EPI_ISL_466415, EPI_ISL_466432, EPI_ISL_466434, EPI_ISL_466436, EPI_ISL_466445, EPI_ISL_466447, EPI_ISL_466452, EPI_ISL_466457, EPI_ISL_466460, EPI_ISL_466462, EPI_ISL_466463, EPI_ISL_466472, EPI_ISL_466474, EPI_ISL_466485, EPI_ISL_466492, EPI_ISL_466493, EPI_ISL_466496, EPI_ISL_466498, EPI_ISL_466506, EPI_ISL_466507, EPI_ISL_466508, EPI_ISL_466511, EPI_ISL_466514, EPI_ISL_466516, EPI_ISL_466520, EPI_ISL_466525, EPI_ISL_466530, EPI_ISL_466532, EPI_ISL_466533, EPI_ISL_466535, EPI_ISL_466537, EPI_ISL_466538, EPI_ISL_466539, EPI_ISL_466543, EPI_ISL_466548, EPI_ISL_466550, EPI_ISL_466552, EPI_ISL_466554, EPI_ISL_466556, EPI_ISL_466557, EPI_ISL_466558, EPI_ISL_466559, EPI_ISL_466561, EPI_ISL_466562, EPI_ISL_466563, EPI_ISL_466564, EPI_ISL_466569, EPI_ISL_466570, EPI_ISL_466572, EPI_ISL_466573, EPI_ISL_466574, EPI_ISL_466575, EPI_ISL_466577, EPI_ISL_466580, EPI_ISL_466583, EPI_ISL_466584, EPI_ISL_466585, EPI_ISL_466586, EPI_ISL_466588, EPI_ISL_466589, EPI_ISL_466590, EPI_ISL_466591, EPI_ISL_466592, EPI_ISL_466593, EPI_ISL_466594, EPI_ISL_466596, EPI_ISL_466597, EPI_ISL_466598, EPI_ISL_466599, EPI_ISL_466600, EPI_ISL_466601, EPI_ISL_466603, EPI_ISL_466604, EPI_ISL_466605, EPI_ISL_466606, EPI_ISL_466608, EPI_ISL_466610, EPI_ISL_466611, EPI_ISL_466612, EPI_ISL_466613, EPI_ISL_466615, EPI_ISL_466616, EPI_ISL_466617, EPI_ISL_466618, EPI_ISL_466619, EPI_ISL_466620, EPI_ISL_466622, EPI_ISL_466623, EPI_ISL_466624, EPI_ISL_466625 | | | |
| see above | Respiratory Virus Unit, Microbiology Services Colindale, Public Health England | Respiratory Virus Unit, Microbiology Services Colindale, Public Health England | PHE Covid Sequencing Team |
| EPI_ISL_466626, EPI_ISL_466627, EPI_ISL_466628, EPI_ISL_466629, EPI_ISL_466630, EPI_ISL_466636, EPI_ISL_466637, EPI_ISL_466638, EPI_ISL_466639, EPI_ISL_466644, EPI_ISL_466645 | | | |
| see above | National Institute of Laboratory Medicine and Referral Center | Genomic Research Lab, BCSIR | Abu Sayeed Mohammad Mahmud, Mohammad Samir Uzzaman, Eshrar Osman, Md. Ahasan Habib, Shahina Akter, Tanjina Akhter Banu, Md. Murshed Hasan Sarker, Iffat Jahan, Barna Goswami, Md. Saddam Hossain, Tasnim Nafisa, Md. Maruf Ahmed Molla, Mahmuda Yeasmin, Asish Kumar Ghosh, Arifa Akram, A. K. M. Shamsuzzaman, Sheikh Md. Selim Al Din, Utpal Chandra Ray, Salek Ahmed Sajib, Md. Salim Khan |
| EPI_ISL_466648 | Innovative Genomics Institute, UCB | Innovative Genomics Institute, UCB | Stacia Wyman, Haridha Shivram, Liana Lareau, Shana McDevitt, Justin Choi |
| EPI_ISL_466649, EPI_ISL_466650 | National Institute of Laboratory Medicine and Referral Center | Genomic Research Lab, BCSIR | Abu Sayeed Mohammad Mahmud, Mohammad Samir Uzzaman, Eshrar Osman, Md. Ahasan Habib, Shahina Akter, Tanjina Akhter Banu, Md. Murshed Hasan Sarker, Iffat Jahan, Barna Goswami, Md. Saddam Hossain, Tasnim Nafisa, Md. Maruf Ahmed Molla, Mahmuda Yeasmin, Asish Kumar Ghosh, Arifa Akram, A. K. M. Shamsuzzaman, Sheikh Md. Selim Al Din, Utpal Chandra Ray, Salek Ahmed Sajib, Md. Salim Khan |
| EPI_ISL_466652, EPI_ISL_466653, EPI_ISL_466655, EPI_ISL_466658, EPI_ISL_466661, EPI_ISL_466665, EPI_ISL_466666, EPI_ISL_466667, EPI_ISL_466668, EPI_ISL_466669, EPI_ISL_466670, EPI_ISL_466671, EPI_ISL_466672, EPI_ISL_466675, EPI_ISL_466677, EPI_ISL_466678, EPI_ISL_466680 | | | |
| see above | Nebraska Public Health Laboratory | UNMC COVID-19 Response Team | UNMC COVID-19 Response Team |
| EPI_ISL_466686, EPI_ISL_466687, EPI_ISL_466688, EPI_ISL_466689, EPI_ISL_466690, EPI_ISL_466691, EPI_ISL_466693, EPI_ISL_466694 | National Institute of Laboratory Medicine and Referral Center | Genomic Research Lab, BCSIR | Abu Sayeed Mohammad Mahmud, Mohammad Samir Uzzaman, Eshrar Osman, Md. Ahasan Habib, Shahina Akter, Tanjina Akhter Banu, Md. Murshed Hasan Sarker, Iffat Jahan, Barna Goswami, Md. Saddam Hossain, Tasnim Nafisa, Md. Maruf Ahmed Molla, Mahmuda Yeasmin, Asish Kumar Ghosh, Arifa Akram, A. K. M. Shamsuzzaman, Sheikh Md. Selim Al Din, Utpal Chandra Ray, Salek Ahmed Sajib, Md. Salim Khan |
| EPI_ISL_466696, EPI_ISL_466697, EPI_ISL_466698, EPI_ISL_466699, EPI_ISL_466700, EPI_ISL_466701, EPI_ISL_466702, EPI_ISL_466703, EPI_ISL_466704, EPI_ISL_466705, EPI_ISL_466706, EPI_ISL_466707, EPI_ISL_466708, EPI_ISL_466709, EPI_ISL_466710, EPI_ISL_466711, EPI_ISL_466712, EPI_ISL_466713, EPI_ISL_466714, EPI_ISL_466715, EPI_ISL_466716, EPI_ISL_466717, EPI_ISL_466718, EPI_ISL_466719, EPI_ISL_466720, EPI_ISL_466721, EPI_ISL_466722, EPI_ISL_466723, EPI_ISL_466724, EPI_ISL_466725, EPI_ISL_466726, EPI_ISL_466727, EPI_ISL_466728, EPI_ISL_466729, EPI_ISL_466730, EPI_ISL_466731, EPI_ISL_466732, EPI_ISL_466733, EPI_ISL_466734, EPI_ISL_466735, EPI_ISL_466736, EPI_ISL_466737, EPI_ISL_466738, EPI_ISL_466739, EPI_ISL_466740, EPI_ISL_466741, EPI_ISL_466742, EPI_ISL_466743, EPI_ISL_466745, EPI_ISL_466747, EPI_ISL_466748, EPI_ISL_466749, EPI_ISL_466750, EPI_ISL_466751, EPI_ISL_466752, EPI_ISL_466753, EPI_ISL_466754, EPI_ISL_466755, EPI_ISL_466756, EPI_ISL_466757, EPI_ISL_466758, EPI_ISL_466759, EPI_ISL_466760, EPI_ISL_466761, EPI_ISL_466762, EPI_ISL_466763, EPI_ISL_466764, EPI_ISL_466765, EPI_ISL_466766, EPI_ISL_466767, EPI_ISL_466768, EPI_ISL_466769, EPI_ISL_466770, EPI_ISL_466772, EPI_ISL_466773, EPI_ISL_466774, EPI_ISL_466775, EPI_ISL_466776, EPI_ISL_466777, EPI_ISL_466779, EPI_ISL_466781, EPI_ISL_466782, EPI_ISL_466783, EPI_ISL_466784, EPI_ISL_466785, EPI_ISL_466787, EPI_ISL_466788, EPI_ISL_466789, EPI_ISL_466792, EPI_ISL_466793, EPI_ISL_466794, EPI_ISL_466795, EPI_ISL_466796, EPI_ISL_466798, EPI_ISL_466800, EPI_ISL_466802, EPI_ISL_466803, EPI_ISL_466806, EPI_ISL_466807, EPI_ISL_466808, EPI_ISL_466809, EPI_ISL_466810, EPI_ISL_466811, EPI_ISL_466812, EPI_ISL_466814, EPI_ISL_466815, EPI_ISL_466816, EPI_ISL_466819, EPI_ISL_466820, EPI_ISL_466821, EPI_ISL_466822, EPI_ISL_466824, EPI_ISL_466826, EPI_ISL_466829, EPI_ISL_466831, EPI_ISL_466832, EPI_ISL_466834, EPI_ISL_466838 | | | |
| see above | BCCDC Public Health Laboratory | BCCDC Public Health Laboratory | Richard Harrigan, Hope Lapointe, Jinny Choi, Kimia Kamelian, John Tyson,Terry Snutch, Linda Hoang, Inna Sekirov, Paul Levett, Mel Krajden, Natalie Prystajecky |
| EPI_ISL_466839 | National Genomics Core-Center for DNA Fingerprinting and Diagnostics | National Genomics Core- Center for DNA Fingerprinting and Diagnostics (NGC-CDFD)-DBT's PAN-INDIA-1000 Genome consortium | Bala Pratyusha, Vinay Donipadi, G Shashikanth, Amrita Bhattacherjee, Rajeshree Sanyal, Raju Kumar, Ajay Kumar Chaudhary, Akash Chinchole, Brahmaji Sontyana, C. Arun Kumar, R HARINARAYANAN, RASHNA BHANDARI, MURALI DHARAN BASHYAM, DEBASHIS MITRA, DIVYA VASHISHT, ASHWIN DALAL |
| EPI_ISL_466841 | National Genomics Core-Center for DNA Fingerprinting and Diagnostics | National Genomics Core- Center for DNA Fingerprinting and Diagnostics (NGC-CDFD)-DBT's PAN-INDIA-1000 Genome consortium | Bala Pratyusha, Vinay Donipadi, G Shashikanth, Amrita Bhattacherjee, Nalini Raghunathan, Rajeshree Sanyal, Raju Kumar, Ajay Kumar Chaudhary, Akash Chinchole, Brahmaji Sontyana, C. Arun Kumar, R Harinarayanan, Rashna Bhandari, Murali Dharan Bashyam, Debashis Mitra, Divya Vashisht, Ashwin Dalal |
| EPI_ISL_466850 | National Genomics Core-Center for DNA Fingerprinting and Diagnostics | National Genomics Core- Center for DNA Fingerprinting and Diagnostics (NGC-CDFD)-DBT's PAN-INDIA-1000 Genome consortium | Bala Pratyusha, Vinay Donipadi, G Shashikanth, Amrita Bhattacherjee, J. Mallikarjun, K. Viswakalyan, Kaisar Ahmad Lone, Kausika Kumar Malik, N. Sudheer, Neeraj Kumar, R HARINARAYANAN, RASHNA BHANDARI, MURALI DHARAN BASHYAM, DEBASHIS MITRA, DIVYA VASHISHT, ASHWIN DALAL |
| EPI_ISL_466853, EPI_ISL_466855, EPI_ISL_466857 | National Genomics Core-Center for DNA Fingerprinting and Diagnostics | National Genomics Core- Center for DNA Fingerprinting and Diagnostics (NGC-CDFD)-DBT's PAN-INDIA-1000 Genome consortium | Bala Pratyusha, Vinay Donipadi, G Shashikanth, Amrita Bhattacherjee, Niteen Pathak, Pradipta Hore, Rahul Baroi, Sayantan Goswami, Shaffiqu T S, Shalini Aricthota, R HARINARAYANAN, RASHNA BHANDARI, MURALI DHARAN BASHYAM, DEBASHIS MITRA, DIVYA VASHISHT, ASHWIN DALAL |
| EPI_ISL_466860 | National Genomics Core-Center for DNA Fingerprinting and Diagnostics | National Genomics Core- Center for DNA Fingerprinting and Diagnostics (NGC-CDFD)-DBT's PAN-INDIA-1000 Genome consortium | Bala Pratyusha, Vinay Donipadi, G Shashikanth, Amrita Bhattacherjee, Sobhan Babu, SPR Prasad, Yogesh Patidar, Arjita Jaiswal, Arpita Singh, Devanshi Gupta, R HARINARAYANAN, RASHNA BHANDARI, MURALI DHARAN BASHYAM, DEBASHIS MITRA, DIVYA VASHISHT, ASHWIN DALAL |
| EPI_ISL_466863, EPI_ISL_466864, EPI_ISL_466866, EPI_ISL_466867 | National Genomics Core-Center for DNA Fingerprinting and Diagnostics | National Genomics Core- Center for DNA Fingerprinting and Diagnostics (NGC-CDFD)-DBT's PAN-INDIA-1000 Genome consortium | Bala Pratyusha, Vinay Donipadi, G Shashikanth, Amrita Bhattacherjee, Romila Moirangthem, Sanjana Sarkar, Shivani Yadav, Shubhra Ganguli, Suchitra Upreti, Swathi Chodisetty , R HARINARAYANAN, RASHNA BHANDARI, MURALI DHARAN BASHYAM, DEBASHIS MITRA, DIVYA VASHISHT, ASHWIN DALAL |
| EPI_ISL_466870, EPI_ISL_466871, EPI_ISL_466872 | National Genomics Core-Center for DNA Fingerprinting and Diagnostics | National Genomics Core- Center for DNA Fingerprinting and Diagnostics (NGC-CDFD)-DBT's PAN-INDIA-1000 Genome consortium | Bala Pratyusha, Vinay Donipadi, G Shashikanth, Amrita Bhattacherjee, Vani Singh, Shubhra Ganguli, Suchitra Upreti, Swathi Chodisetty , Vani Singh , R HARINARAYANAN, RASHNA BHANDARI, MURALI DHARAN BASHYAM, DEBASHIS MITRA, DIVYA VASHISHT, ASHWIN DALAL |
| EPI_ISL_466873 | Centre for Clinical Infection and Diagnostics Research and Genomics Innovation Unit | Respiratory Virus Unit, Microbiology Services Colindale, Public Health England | PHE Covid Sequencing Team, Chloe Fisher, Luke Snell, Gaia Nebbia, Ali Awan |
| EPI_ISL_466875, EPI_ISL_466877, EPI_ISL_466878, EPI_ISL_466879, EPI_ISL_466880, EPI_ISL_466881, EPI_ISL_466882, EPI_ISL_466883, EPI_ISL_466884, EPI_ISL_466885, EPI_ISL_466888, EPI_ISL_466889, EPI_ISL_466891, EPI_ISL_466892, EPI_ISL_466893, EPI_ISL_466895, EPI_ISL_466896, EPI_ISL_466897, EPI_ISL_466899, EPI_ISL_466900, EPI_ISL_466901, EPI_ISL_466902, EPI_ISL_466903, EPI_ISL_466904, EPI_ISL_466905, EPI_ISL_466906, EPI_ISL_466907, EPI_ISL_466909, EPI_ISL_466910, EPI_ISL_466911, EPI_ISL_466913, EPI_ISL_466916, EPI_ISL_466917, EPI_ISL_466918, EPI_ISL_466919, EPI_ISL_466920, EPI_ISL_466921, EPI_ISL_466922, EPI_ISL_466923, EPI_ISL_466924, EPI_ISL_466925 | | | |
| see above | Max von Pettenkofer Institute, Virology, National Reference Center for Retroviruses, LMU München | Laboratory for Functional Genome Analysis, Dept. Genomics, Gene Center of the LMU Munich | Max Muenchhoff, Stefan Krebs, Alexander Graf, Oliver Keppler, Helmut Blum |
| EPI_ISL_466927, EPI_ISL_466928, EPI_ISL_466929, EPI_ISL_466930, EPI_ISL_466931, EPI_ISL_466932, EPI_ISL_466933, EPI_ISL_466935, EPI_ISL_466936, EPI_ISL_466937, EPI_ISL_466938, EPI_ISL_466939, EPI_ISL_466940, EPI_ISL_466941, EPI_ISL_466942, EPI_ISL_466943, EPI_ISL_466945, EPI_ISL_466946, EPI_ISL_466947, EPI_ISL_466948, EPI_ISL_466949, EPI_ISL_466950, EPI_ISL_466951, EPI_ISL_466952, EPI_ISL_466953, EPI_ISL_466954, EPI_ISL_466955, EPI_ISL_466956, EPI_ISL_466958, EPI_ISL_466959, EPI_ISL_466960, EPI_ISL_466961, EPI_ISL_466962, EPI_ISL_466964, EPI_ISL_466965, EPI_ISL_466966, EPI_ISL_466967, EPI_ISL_466968, EPI_ISL_466969, EPI_ISL_466970, EPI_ISL_466971, EPI_ISL_466972, EPI_ISL_466973, EPI_ISL_466974, EPI_ISL_466975, EPI_ISL_466976, EPI_ISL_466977, EPI_ISL_466979, EPI_ISL_466980, EPI_ISL_466981, EPI_ISL_466982, EPI_ISL_466983, EPI_ISL_466984, EPI_ISL_466985, EPI_ISL_466986, EPI_ISL_466987, EPI_ISL_466988, EPI_ISL_466989, EPI_ISL_466990, EPI_ISL_466991, EPI_ISL_466992, EPI_ISL_466995, EPI_ISL_466996, EPI_ISL_466997, EPI_ISL_466999, EPI_ISL_467002, EPI_ISL_467003, EPI_ISL_467004, EPI_ISL_467006, EPI_ISL_467009, EPI_ISL_467012, EPI_ISL_467014 | | | |
| see above | Viollier AG | Department of Biosystems Science and Engineering, ETH Zürich | Christian Beisel, Sarah Nadeau, Ivan Topolsky, Pedro Ferreira, Philipp Jablonski, Susana Posada-Céspedes, Tobias Schär, Ina Nissen, Natascha Santacroce, Elodie Burcklen, Christiane Beckmann, Maurice Redondo, Olivier Kobel, Christoph Noppen, Sophie Seidel, Noemie Santamaria de Souza, Niko Beerenwinkel, Tanja Stadler |
| EPI_ISL_467029 | GMERS Medical College and Hospital, Gandhinagar | Gujarat Biotechnology Research Centre | Seema Bhatt, Gaurishankar Shrimali, Bhavesh Modi, Bharti Rajani, Tejas Shah, Ankit Hinsu, Pritesh Sabara, Apurvasinh Puvar, Janvi Raval, Zarna Patel, Monika Gandhi, Pinal Trivedi, Maharshi Pandya, Nidhi Patel, Nitin Savaliya, Raghawendra Kumar, Dinesh Kumar, Zuber Saiyed, Komal Patel, Labdhi Pandya, Snehal Bagatharia, Bhavya Jindal, R D Dixit, A M Kadri, Harsh Bakshi, Chaitanya Joshi, Madhvi Joshi |
| EPI_ISL_467030 | GMERS Medical College and Hospital, Gandhinagar | Gujarat Biotechnology Research Centre | Gaurishankar Shrimali, Bhavesh Modi, Bharti Rajani, Tejas Shah, Ankit Hinsu, Pritesh Sabara, Apurvasinh Puvar, Janvi Raval, Zarna Patel, Monika Gandhi, Pinal Trivedi, Maharshi Pandya, Nidhi Patel, Nitin Savaliya, Raghawendra Kumar, Dinesh Kumar, Zuber Saiyed, Komal Patel, Labdhi Pandya, Snehal Bagatharia, Seema Bhatt, Priyanka P Vatsa, R D Dixit, A M Kadri, Harsh Bakshi, Chaitanya Joshi, Madhvi Joshi |
| EPI_ISL_467031 | GMERS Medical College and Hospital, Gandhinagar | Gujarat Biotechnology Research Centre | Bhavesh Modi, Bharti Rajani, Tejas Shah, Ankit Hinsu, Pritesh Sabara, Apurvasinh Puvar, Janvi Raval, Zarna Patel, Monika Gandhi, Pinal Trivedi, Maharshi Pandya, Nidhi Patel, Nitin Savaliya, Raghawendra Kumar, Dinesh Kumar, Zuber Saiyed, Komal Patel, Labdhi Pandya, Snehal Bagatharia, Seema Bhatt, Gaurishankar Shrimali, Pooja P Doshi, R D Dixit, A M Kadri, Harsh Bakshi, Chaitanya Joshi, Madhvi Joshi |
| EPI_ISL_467032 | GMERS Medical College and Hospital, Gandhinagar | Gujarat Biotechnology Research Centre | Bharti Rajani, Tejas Shah, Ankit Hinsu, Pritesh Sabara, Apurvasinh Puvar, Janvi Raval, Zarna Patel, Monika Gandhi, Pinal Trivedi, Maharshi Pandya, Nidhi Patel, Nitin Savaliya, Raghawendra Kumar, Dinesh Kumar, Zuber Saiyed, Komal Patel, Labdhi Pandya, Snehal Bagatharia, Seema Bhatt, Gaurishankar Shrimali, Bhavesh Modi, Akanksha Verma, R D Dixit, A M Kadri, Harsh Bakshi, Chaitanya Joshi, Madhvi Joshi |
| EPI_ISL_467033 | GMERS Medical College and Hospital, Gandhinagar | Gujarat Biotechnology Research Centre | Tejas Shah, Ankit Hinsu, Pritesh Sabara, Apurvasinh Puvar, Janvi Raval, Zarna Patel, Monika Gandhi, Pinal Trivedi, Maharshi Pandya, Nidhi Patel, Nitin Savaliya, Raghawendra Kumar, Dinesh Kumar, Zuber Saiyed, Komal Patel, Labdhi Pandya, Snehal Bagatharia, Seema Bhatt, Gaurishankar Shrimali, Bhavesh Modi, Bharti Rajani, Priti Pandita, R D Dixit, A M Kadri, Harsh Bakshi, Chaitanya Joshi, Madhvi Joshi |

| | | | |
|---|---|---|---|
| EPI_ISL_467034 | GMERS Medical College and Hospital, Gandhinagar | Gujarat Biotechnology Research Centre | Ankit Hinsu, Pritesh Sabara, Apurvasinh Puvar, Janvi Raval, Zarna Patel, Monika Gandhi, Pinal Trivedi, Maharshi Pandya, Nidhi Patel, Nitin Savaliya, Raghawendra Kumar, Dinesh Kumar, Zuber Saiyed, Komal Patel, Labdhi Pandya, Snehal Bagatharia, Seema Bhatt, Gaurishankar Shrimali, Bhavesh Modi, Bharti Rajani, Tejas Shah, Pragya Sharma, R D Dixit, A M Kadri, Harsh Bakshi, Chaitanya Joshi, Madhvi Joshi |
| EPI_ISL_467035 | GMERS Medical College and Hospital, Gandhinagar | Gujarat Biotechnology Research Centre | Pritesh Sabara, Apurvasinh Puvar, Janvi Raval, Zarna Patel, Monika Gandhi, Pinal Trivedi, Maharshi Pandya, Nidhi Patel, Nitin Savaliya, Raghawendra Kumar, Dinesh Kumar, Zuber Saiyed, Komal Patel, Labdhi Pandya, Snehal Bagatharia, Seema Bhatt, Gaurishankar Shrimali, Bhavesh Modi, Bharti Rajani, Tejas Shah, Ankit Hinsu, Neha Rajpara, R D Dixit, A M Kadri, Harsh Bakshi, Chaitanya Joshi, Madhvi Joshi |
| EPI_ISL_467036 | GMERS Medical College and Hospital, Gandhinagar | Gujarat Biotechnology Research Centre | Apurvasinh Puvar, Janvi Raval, Zarna Patel, Monika Gandhi, Pinal Trivedi, Maharshi Pandya, Nidhi Patel, Nitin Savaliya, Raghawendra Kumar, Dinesh Kumar, Zuber Saiyed, Komal Patel, Labdhi Pandya, Snehal Bagatharia, Seema Bhatt, Gaurishankar Shrimali, Bhavesh Modi, Bharti Rajani, Tejas Shah, Ankit Hinsu, Pritesh Sabara, Afzal Ansari, R D Dixit, A M Kadri, Harsh Bakshi, Chaitanya Joshi, Madhvi Joshi |
| EPI_ISL_467037 | GMERS Medical College and Hospital, Gandhinagar | Gujarat Biotechnology Research Centre | Janvi Raval, Zarna Patel, Monika Gandhi, Pinal Trivedi, Maharshi Pandya, Nidhi Patel, Nitin Savaliya, Raghawendra Kumar, Dinesh Kumar, Zuber Saiyed, Komal Patel, Labdhi Pandya, Snehal Bagatharia, Seema Bhatt, Gaurishankar Shrimali, Bhavesh Modi, Bharti Rajani, Tejas Shah, Ankit Hinsu, Pritesh Sabara, Apurvasinh Puvar, Fenil Patel, R D Dixit, A M Kadri, Harsh Bakshi, Chaitanya Joshi, Madhvi Joshi |
| EPI_ISL_467038 | GMERS Medical College and Hospital, Gandhinagar | Gujarat Biotechnology Research Centre | Zarna Patel, Monika Gandhi, Pinal Trivedi, Maharshi Pandya, Nidhi Patel, Nitin Savaliya, Raghawendra Kumar, Dinesh Kumar, Zuber Saiyed, Komal Patel, Labdhi Pandya, Snehal Bagatharia, Seema Bhatt, Gaurishankar Shrimali, Bhavesh Modi, Bharti Rajani, Tejas Shah, Ankit Hinsu, Pritesh Sabara, Apurvasinh Puvar, Janvi Raval, Neelam Nathani, R D Dixit, A M Kadri, Harsh Bakshi, Chaitanya Joshi, Madhvi Joshi |
| EPI_ISL_467039 | Government Medical College, Vadodara | Gujarat Biotechnology Research Centre | Meenakshi Shah, Neena Doshi, Varsha Godbole, Tejas Shah, Ankit Hinsu, Pritesh Sabara, Apurvasinh Puvar, Janvi Raval, Zarna Patel, Monika Gandhi, Pinal Trivedi, Maharshi Pandya, Nidhi Patel, Nitin Savaliya, Raghawendra Kumar, Dinesh Kumar, Zuber Saiyed, Komal Patel, Labdhi Pandya, Snehal Bagatharia, Armi Chaudhari, R D Dixit, A M Kadri, Harsh Bakshi, Chaitanya Joshi, Madhvi Joshi, |
| EPI_ISL_467040 | Government Medical College, Vadodara | Gujarat Biotechnology Research Centre | Neena Doshi, Varsha Godbole, Tejas Shah, Ankit Hinsu, Pritesh Sabara, Apurvasinh Puvar, Janvi Raval, Zarna Patel, Monika Gandhi, Pinal Trivedi, Maharshi Pandya, Nidhi Patel, Nitin Savaliya, Raghawendra Kumar, Dinesh Kumar, Zuber Saiyed, Komal Patel, Labdhi Pandya, Snehal Bagatharia, Meenakshi Shah, Bhavya Jindal, R D Dixit, A M Kadri, Harsh Bakshi, Chaitanya Joshi, Madhvi Joshi, |
| EPI_ISL_467041 | B.J. Medical College and Civil hospital | Gujarat Biotechnology Research Centre | Monika Gandhi, Pinal Trivedi, Maharshi Pandya, Nidhi Patel, Nitin Savaliya, Raghawendra Kumar, Dinesh Kumar, Zuber Saiyed, Komal Patel, Labdhi Pandya, Snehal Bagatharia, Pranay Shah, Kamlesh J Upadhyay, Nirav Mungalpara, Tejas Shah, Ankit Hinsu, Pritesh Sabara, Apurvasinh Puvar, Janvi Raval, Zarna Patel, Priyanka P Vatsa, R D Dixit, A M Kadri, Harsh Bakshi, Chaitanya Joshi, Madhvi Joshi, |
| EPI_ISL_467042 | B.J. Medical College and Civil hospital | Gujarat Biotechnology Research Centre | Pinal Trivedi, Maharshi Pandya, Nidhi Patel, Nitin Savaliya, Raghawendra Kumar, Dinesh Kumar, Zuber Saiyed, Komal Patel, Labdhi Pandya, Snehal Bagatharia, Pranay Shah, Kamlesh J Upadhyay, Nirav Mungalpara, Tejas Shah, Ankit Hinsu, Pritesh Sabara, Apurvasinh Puvar, Janvi Raval, Zarna Patel, Monika Gandhi, Pooja P Doshi, R D Dixit, A M Kadri, Harsh Bakshi, Chaitanya Joshi, Madhvi Joshi, |
| EPI_ISL_467043 | B.J. Medical College and Civil hospital | Gujarat Biotechnology Research Centre | Maharshi Pandya, Nidhi Patel, Nitin Savaliya, Raghawendra Kumar, Dinesh Kumar, Zuber Saiyed, Komal Patel, Labdhi Pandya, Snehal Bagatharia, Pranay Shah, Kamlesh J Upadhyay, Nirav Mungalpara, Tejas Shah, Ankit Hinsu, Pritesh Sabara, Apurvasinh Puvar, Janvi Raval, Zarna Patel, Monika Gandhi, Pinal Trivedi, Akanksha Verma, R D Dixit, A M Kadri, Harsh Bakshi, Chaitanya Joshi, Madhvi Joshi, |
| EPI_ISL_467044 | B.J. Medical College and Civil hospital | Gujarat Biotechnology Research Centre | Nidhi Patel, Nitin Savaliya, Raghawendra Kumar, Dinesh Kumar, Zuber Saiyed, Komal Patel, Labdhi Pandya, Snehal Bagatharia, Pranay Shah, Kamlesh J Upadhyay, Nirav Mungalpara, Tejas Shah, Ankit Hinsu, Pritesh Sabara, Apurvasinh Puvar, Janvi Raval, Zarna Patel, Monika Gandhi, Pinal Trivedi, Maharshi Pandya, Priti Pandita, R D Dixit, A M Kadri, Harsh Bakshi, Chaitanya Joshi, Madhvi Joshi, |
| EPI_ISL_467045 | B.J. Medical College and Civil hospital | Gujarat Biotechnology Research Centre | Nitin Savaliya, Raghawendra Kumar, Dinesh Kumar, Zuber Saiyed, Komal Patel, Labdhi Pandya, Snehal Bagatharia, Pranay Shah, Kamlesh J Upadhyay, Nirav Mungalpara, Tejas Shah, Ankit Hinsu, Pritesh Sabara, Apurvasinh Puvar, Janvi Raval, Zarna Patel, Monika Gandhi, Pinal Trivedi, Maharshi Pandya, Nidhi Patel, Pragya Sharma, R D Dixit, A M Kadri, Harsh Bakshi, Chaitanya Joshi, Madhvi Joshi, |
| EPI_ISL_467046 | B.J. Medical College and Civil hospital | Gujarat Biotechnology Research Centre | Raghawendra Kumar, Dinesh Kumar, Zuber Saiyed, Komal Patel, Labdhi Pandya, Snehal Bagatharia, Pranay Shah, Kamlesh J Upadhyay, Nirav Mungalpara, Tejas Shah, Ankit Hinsu, Pritesh Sabara, Apurvasinh Puvar, Janvi Raval, Zarna Patel, Monika Gandhi, Pinal Trivedi, Maharshi Pandya, Nidhi Patel, Nitin Savaliya, Neha Rajpara, R D Dixit, A M Kadri, Harsh Bakshi, Chaitanya Joshi, Madhvi Joshi, |
| EPI_ISL_467047 | B.J. Medical College and Civil hospital | Gujarat Biotechnology Research Centre | Dinesh Kumar, Zuber Saiyed, Komal Patel, Labdhi Pandya, Snehal Bagatharia, Pranay Shah, Kamlesh J Upadhyay, Nirav Mungalpara, Tejas Shah, Ankit Hinsu, Pritesh Sabara, Apurvasinh Puvar, Janvi Raval, Zarna Patel, Monika Gandhi, Pinal Trivedi, Maharshi Pandya, Nidhi Patel, Nitin Savaliya, Raghawendra Kumar, Afzal Ansari, R D Dixit, A M Kadri, Harsh Bakshi, Chaitanya Joshi, Madhvi Joshi, |
| EPI_ISL_467048 | B.J. Medical College and Civil hospital | Gujarat Biotechnology Research Centre | Zuber Saiyed, Komal Patel, Labdhi Pandya, Snehal Bagatharia, Pranay Shah, Kamlesh J Upadhyay, Nirav Mungalpara, Tejas Shah, Ankit Hinsu, Pritesh Sabara, Apurvasinh Puvar, Janvi Raval, Zarna Patel, Monika Gandhi, Pinal Trivedi, Maharshi Pandya, Nidhi Patel, Nitin Savaliya, Raghawendra Kumar, Dinesh Kumar, Fenil Patel, R D Dixit, A M Kadri, Harsh Bakshi, Chaitanya Joshi, Madhvi Joshi, |
| EPI_ISL_467049 | B.J. Medical College and Civil hospital | Gujarat Biotechnology Research Centre | Komal Patel, Labdhi Pandya, Snehal Bagatharia, Pranay Shah, Kamlesh J Upadhyay, Nirav Mungalpara, Tejas Shah, Ankit Hinsu, Pritesh Sabara, Apurvasinh Puvar, Janvi Raval, Zarna Patel, Monika Gandhi, Pinal Trivedi, Maharshi Pandya, Nidhi Patel, Nitin Savaliya, Raghawendra Kumar, Dinesh Kumar, Zuber Saiyed, Neelam Nathani, R D Dixit, A M Kadri, Harsh Bakshi, Chaitanya Joshi, Madhvi Joshi, |
| EPI_ISL_467050 | B.J. Medical College and Civil hospital | Gujarat Biotechnology Research Centre | Labdhi Pandya, Snehal Bagatharia, Pranay Shah, Kamlesh J Upadhyay, Nirav Mungalpara, Tejas Shah, Ankit Hinsu, Pritesh Sabara, Apurvasinh Puvar, Janvi Raval, Zarna Patel, Monika Gandhi, Pinal Trivedi, Maharshi Pandya, Nidhi Patel, Nitin Savaliya, Raghawendra Kumar, Dinesh Kumar, Zuber Saiyed, Komal Patel, Armi Chaudhari, R D Dixit, A M Kadri, Harsh Bakshi, Chaitanya Joshi, Madhvi Joshi, |
| EPI_ISL_467051 | B.J. Medical College and Civil hospital | Gujarat Biotechnology Research Centre | Snehal Bagatharia, Pranay Shah, Kamlesh J Upadhyay, Nirav Mungalpara, Tejas Shah, Ankit Hinsu, Pritesh Sabara, Apurvasinh Puvar, Janvi Raval, Zarna Patel, Monika Gandhi, Pinal Trivedi, Maharshi Pandya, Nidhi Patel, Nitin Savaliya, Raghawendra Kumar, Dinesh Kumar, Zuber Saiyed, Komal Patel, Labdhi Pandya, Bhavya Jindal, R D Dixit, A M Kadri, Harsh Bakshi, Chaitanya Joshi, Madhvi Joshi, |
| EPI_ISL_467052 | B.J. Medical College and Civil hospital | Gujarat Biotechnology Research Centre | Pranay Shah, Kamlesh J Upadhyay, Nirav Mungalpara, Tejas Shah, Ankit Hinsu, Pritesh Sabara, Apurvasinh Puvar, Janvi Raval, Zarna Patel, Monika Gandhi, Pinal Trivedi, Maharshi Pandya, Nidhi Patel, Nitin Savaliya, Raghawendra Kumar, Dinesh Kumar, Zuber Saiyed, Komal Patel, Labdhi Pandya, Snehal Bagatharia, Priyanka P Vatsa, R D Dixit, A M Kadri, Harsh Bakshi, Chaitanya Joshi, Madhvi Joshi, |
| EPI_ISL_467053 | B.J. Medical College and Civil hospital | Gujarat Biotechnology Research Centre | Kamlesh J Upadhyay, Nirav Mungalpara, Tejas Shah, Ankit Hinsu, Pritesh Sabara, Apurvasinh Puvar, Janvi Raval, Zarna Patel, Monika Gandhi, Pinal Trivedi, Maharshi Pandya, Nidhi Patel, Nitin Savaliya, Raghawendra Kumar, Dinesh Kumar, Zuber Saiyed, Komal Patel, Labdhi Pandya, Snehal Bagatharia, Pranay Shah, Pooja P Doshi, R D Dixit, A M Kadri, Harsh Bakshi, Chaitanya Joshi, Madhvi Joshi, |
| EPI_ISL_467054 | B.J. Medical College and Civil hospital | Gujarat Biotechnology Research Centre | Nirav Mungalpara, Tejas Shah, Ankit Hinsu, Pritesh Sabara, Apurvasinh Puvar, Janvi Raval, Zarna Patel, Monika Gandhi, Pinal Trivedi, Maharshi Pandya, Nidhi Patel, Nitin Savaliya, Raghawendra Kumar, Dinesh Kumar, Zuber Saiyed, Komal Patel, Labdhi Pandya, Snehal Bagatharia, Pranay Shah, Kamlesh J Upadhyay, Akanksha Verma, R D Dixit, A M Kadri, Harsh Bakshi, Chaitanya Joshi, Madhvi Joshi, |
| EPI_ISL_467055, EPI_ISL_467056, EPI_ISL_467058 | Servicio de Microbiología, Hospital Universitario Son Espases | SeqCOVID-SPAIN consortium/IBV(CSIC) | Carla López-Causapé, Jordi Reina y Antonio Oliver and SeqCOVID-SPAIN consortium |
| EPI_ISL_467059, EPI_ISL_467061, EPI_ISL_467062, EPI_ISL_467063 | Hospital Universitario Virgen de las Nieves de Granada-SAS | SeqCOVID-SPAIN consortium/IBV(CSIC) | Mercedes Pérez Ruiz, Sara Sanbonmatsu Gámez, Irene Pedrosa Corral, José M. Navarro-Marí and SeqCOVID-SPAIN consortium |
| EPI_ISL_467064, EPI_ISL_467066, EPI_ISL_467068, EPI_ISL_467069, EPI_ISL_467070, EPI_ISL_467071, EPI_ISL_467072, EPI_ISL_467073, EPI_ISL_467074, EPI_ISL_467075, EPI_ISL_467076, EPI_ISL_467077, EPI_ISL_467078, EPI_ISL_467079, EPI_ISL_467080, EPI_ISL_467081, EPI_ISL_467082, EPI_ISL_467083, EPI_ISL_467084, EPI_ISL_467085 | | | |
| see above | Hospital Universitario Puerta del Mar de Cádiz - INIBICA | SeqCOVID-SPAIN consortium/IBV(CSIC) | Salud Rodríguez-Pallares, Fátima-Galán-Sánchez, Manuel Rodríguez-Iglesias and SeqCOVID-SPAIN consortium |
| EPI_ISL_467086, EPI_ISL_467087, EPI_ISL_467088, EPI_ISL_467089, EPI_ISL_467090, EPI_ISL_467091 | Hospital Universitario de Gran Canaria Dr. Negrín | SeqCOVID-SPAIN consortium/IBV(CSIC) | M. Carmen Pérez González, Francisco J. Chamizo López, Ana Bordes Benítez and SeqCOVID-SPAIN consortium |
| EPI_ISL_467092, EPI_ISL_467093, EPI_ISL_467094, EPI_ISL_467095, EPI_ISL_467096, EPI_ISL_467097, EPI_ISL_467098, EPI_ISL_467099, EPI_ISL_467100, EPI_ISL_467101, EPI_ISL_467103, EPI_ISL_467104, EPI_ISL_467105, EPI_ISL_467106, EPI_ISL_467107, EPI_ISL_467108, EPI_ISL_467109, EPI_ISL_467110, EPI_ISL_467111, EPI_ISL_467113, EPI_ISL_467114, EPI_ISL_467115, EPI_ISL_467116, EPI_ISL_467117, EPI_ISL_467118, EPI_ISL_467120, EPI_ISL_467121, EPI_ISL_467122, EPI_ISL_467123, EPI_ISL_467124, EPI_ISL_467125, EPI_ISL_467126, EPI_ISL_467127, EPI_ISL_467128, EPI_ISL_467129, EPI_ISL_467130, EPI_ISL_467131, EPI_ISL_467132, EPI_ISL_467134, EPI_ISL_467136, EPI_ISL_467137, EPI_ISL_467138, EPI_ISL_467139, EPI_ISL_467140, EPI_ISL_467141, EPI_ISL_467142, EPI_ISL_467143, EPI_ISL_467144, EPI_ISL_467145, EPI_ISL_467146, EPI_ISL_467147, EPI_ISL_467148, EPI_ISL_467149, EPI_ISL_467150, EPI_ISL_467151, EPI_ISL_467152, EPI_ISL_467153, EPI_ISL_467154, EPI_ISL_467155, EPI_ISL_467156, EPI_ISL_467157, EPI_ISL_467158, EPI_ISL_467159, EPI_ISL_467160, EPI_ISL_467161, EPI_ISL_467162, EPI_ISL_467163, EPI_ISL_467164, EPI_ISL_467165, EPI_ISL_467166, EPI_ISL_467167, EPI_ISL_467168, EPI_ISL_467169, EPI_ISL_467170, EPI_ISL_467171, EPI_ISL_467172, EPI_ISL_467173, EPI_ISL_467174, EPI_ISL_467175, EPI_ISL_467176, EPI_ISL_467177, EPI_ISL_467178, EPI_ISL_467179, EPI_ISL_467180, EPI_ISL_467181, EPI_ISL_467182, EPI_ISL_467183 | | | |
| see above | Hospital Universitario Araba. Vitoria-Gasteiz | SeqCOVID-SPAIN consortium/IBV(CSIC) | Silvia Hernáez Crespo, Carmen Gómez González, Amaia Aguirre Quiñonero, Marina Fernández Torres, Mª Rosario Almela Ferrer, Mª Concepción Lecaroz Agara, Andrés Canut Blasco. and SeqCOVID-SPAIN consortium |
| EPI_ISL_467184, EPI_ISL_467185, EPI_ISL_467186, EPI_ISL_467188, EPI_ISL_467189, EPI_ISL_467190, EPI_ISL_467191, EPI_ISL_467192, EPI_ISL_467193, EPI_ISL_467194, EPI_ISL_467195, EPI_ISL_467196, EPI_ISL_467197, EPI_ISL_467198, EPI_ISL_467199, EPI_ISL_467200, EPI_ISL_467201, EPI_ISL_467202, EPI_ISL_467205, EPI_ISL_467206, EPI_ISL_467207, EPI_ISL_467208, EPI_ISL_467209, EPI_ISL_467211, EPI_ISL_467212, EPI_ISL_467213, EPI_ISL_467214, EPI_ISL_467215, EPI_ISL_467216, EPI_ISL_467217, EPI_ISL_467218, EPI_ISL_467219, EPI_ISL_467220, EPI_ISL_467221, EPI_ISL_467222, EPI_ISL_467223, EPI_ISL_467224, EPI_ISL_467225, EPI_ISL_467226, EPI_ISL_467227, EPI_ISL_467228, EPI_ISL_467229, EPI_ISL_467230, EPI_ISL_467231, EPI_ISL_467232, EPI_ISL_467233, EPI_ISL_467234, EPI_ISL_467235, EPI_ISL_467236, EPI_ISL_467237, EPI_ISL_467238, EPI_ISL_467239, EPI_ISL_467240, EPI_ISL_467241, EPI_ISL_467242, EPI_ISL_467243, EPI_ISL_467244, EPI_ISL_467245, EPI_ISL_467247, EPI_ISL_467248, EPI_ISL_467249, EPI_ISL_467250, EPI_ISL_467251, EPI_ISL_467252, EPI_ISL_467253, EPI_ISL_467254, EPI_ISL_467255, EPI_ISL_467256, EPI_ISL_467257, EPI_ISL_467259, EPI_ISL_467260, EPI_ISL_467261 | | | |
| see above | Hospital General Universitario Gregorio Marañón | SeqCOVID-SPAIN consortium/IBV(CSIC) | Laura Pérez-Lago, Marta Herranz, Jon Sicilia, Julia Suárez, Pilar Catalán, Patricia Muñoz, Darío García de Viedma and SeqCOVID-SPAIN consortium |
| EPI_ISL_467262, EPI_ISL_467263, EPI_ISL_467265, EPI_ISL_467266, EPI_ISL_467267, EPI_ISL_467269, EPI_ISL_467271, EPI_ISL_467272, EPI_ISL_467273, EPI_ISL_467274, EPI_ISL_467275, EPI_ISL_467276, EPI_ISL_467277, EPI_ISL_467278, EPI_ISL_467279, EPI_ISL_467280, EPI_ISL_467281, EPI_ISL_467283, EPI_ISL_467284, EPI_ISL_467285, EPI_ISL_467288, EPI_ISL_467289, EPI_ISL_467290, EPI_ISL_467291, EPI_ISL_467292, EPI_ISL_467293, EPI_ISL_467294, EPI_ISL_467295, EPI_ISL_467296, EPI_ISL_467297 | | | |
| see above | Hospital Clínico Universitario de Santiago de Compostela | SeqCOVID-SPAIN consortium/IBV(CSIC) | José Javier Costa Alcalde, Antonio Aguilera Guirao, Mª Luisa Pérez del Molino Bernal, Amparo Coira Nieto, Gema Barbeito Castiñeiras, Rocio Trastoy Pena and SeqCOVID-SPAIN consortium |
| EPI_ISL_467300 | General Hospital "Abdulah Nakas" | Alea Genetic Center | Rijad Konjhodzic; Lana Salihefendic; Teufik Goletic; Sead Jazic; Dino Pecar; Nihad Fejzic; Damir Marjanovic; Enis Kandic |
| EPI_ISL_467301 | Washington University in St. Louis | Washington University in St. Louis | David Wang, Carey-Ann Burnham, Scott Handley, Lindsay Droit, Stephen Tahan |
| EPI_ISL_467302 | Washington University in St. Louis | Washington University in St. Louis | David Wang, Carey-Ann Burnham, Scott Handley, Lindsay Droit, Stephen Tahan |
| EPI_ISL_467304 | Washington University in St. Louis | Washington University in St. Louis | David Wang, Carey-Ann Burnham, Scott Handley, Lindsay Droit, Stephen Tahan |
| EPI_ISL_467305, EPI_ISL_467306, EPI_ISL_467309, EPI_ISL_467310, EPI_ISL_467312, EPI_ISL_467313, EPI_ISL_467314, EPI_ISL_467315, EPI_ISL_467317, EPI_ISL_467318, EPI_ISL_467320, EPI_ISL_467321, EPI_ISL_467322, EPI_ISL_467324, EPI_ISL_467327, EPI_ISL_467329, EPI_ISL_467330, EPI_ISL_467331, EPI_ISL_467332, EPI_ISL_467333, EPI_ISL_467334, EPI_ISL_467335, EPI_ISL_467336, EPI_ISL_467337, EPI_ISL_467338, EPI_ISL_467339, EPI_ISL_467340, EPI_ISL_467341, EPI_ISL_467342, EPI_ISL_467343 | | | |
| see above | BCCDC Public Health Laboratory | BCCDC Public Health Laboratory | Richard Harrigan, Hope Lapointe, Jinny Choi, Kimia Kamelian, John Tyson,Terry Snutch, Linda Hoang, Inna Sekirov, Paul Levett, Mel Krajden, Natalie Prystajecky |

| | | | |
|---|---|---|---|
| EPI_ISL_467344, EPI_ISL_467345, EPI_ISL_467346, EPI_ISL_467347, EPI_ISL_467348, EPI_ISL_467349, EPI_ISL_467350, EPI_ISL_467351, EPI_ISL_467352, EPI_ISL_467353, EPI_ISL_467354, EPI_ISL_467355, EPI_ISL_467356, EPI_ISL_467357, EPI_ISL_467358, EPI_ISL_467359, EPI_ISL_467360, EPI_ISL_467361, EPI_ISL_467362, EPI_ISL_467363, EPI_ISL_467364, EPI_ISL_467365, EPI_ISL_467366, EPI_ISL_467367, EPI_ISL_467368, EPI_ISL_467369, EPI_ISL_467370, EPI_ISL_467371 | | | |
| see above | Laboratory of Respiratory Viruses and Measles, Oswaldo Cruz Institute, FIOCRUZ | Laboratory of Respiratory Viruses and Measles, Oswaldo Cruz Institute, FIOCRUZ | Paola Resende, Luciana Appolinario, Fernando Motta, Ana Carolina Paixão, Ana Carolina Mendonça, Aline Mattos, Milene Miranda, Cristiana Garcia, Braulia Caetano, Maria Ogrzewalska, Jonathan Lopes, Marilda Siqueira |
| EPI_ISL_467372 | Arizona State University Health Services | Arizona State University | Peter T. Skidmore, Rabia Maqsood, LaRinda A. Holland, Emily A. Kaelin, Lily I. Wu, Arvind Varsani, Rolf U. Halden, Brenda G. Hogue, Matthew Scotch, Efrem S. Lim |
| EPI_ISL_467373 | Arizona State University Health Services | Arizona State University | Peter T. Skidmore, Rabia Maqsood, LaRinda A. Holland, Emily A. Kaelin, Lily I. Wu, Arvind Varsani, Rolf U. Halden, Brenda G. Hogue, Matthew Scotch, Efrem S. Lim |
| EPI_ISL_467374 | Dinkes Samarinda | Eijkman Institute for Molecular Biology, Ministry of Research and Technology/National Agency for Research and Innovation | Edison Johar, Frilasita A Yudhaputri, Hidayat Trimarsanto, David H Muljono, Safarina G Malik, Khin Saw Myint, Amin Soebandrio |
| EPI_ISL_467375 | RSUP Prof. Dr. R. Kandou Manado | Eijkman Institute for Molecular Biology, Ministry of Research and Technology/National Agency for Research and Innovation | Edison Johar, Frilasita A Yudhaputri, Hidayat Trimarsanto, David H Muljono, Safarina G Malik, Khin Saw Myint, Amin Soebandrio |
| EPI_ISL_467376 | RSUP Fatmawati | Eijkman Institute for Molecular Biology, Ministry of Research and Technology/National Agency for Research and Innovation | Edison Johar, Frilasita A Yudhaputri, Hidayat Trimarsanto, David H Muljono, Safarina G Malik, Khin Saw Myint, Amin Soebandrio |
| EPI_ISL_467377, EPI_ISL_467378, EPI_ISL_467379, EPI_ISL_467380, EPI_ISL_467381, EPI_ISL_467384, EPI_ISL_467385, EPI_ISL_467387, EPI_ISL_467391, EPI_ISL_467394, EPI_ISL_467395, EPI_ISL_467396, EPI_ISL_467398, EPI_ISL_467399, EPI_ISL_467400, EPI_ISL_467405, EPI_ISL_467409, EPI_ISL_467411, EPI_ISL_467412, EPI_ISL_467413, EPI_ISL_467414, EPI_ISL_467416, EPI_ISL_467417, EPI_ISL_467418, EPI_ISL_467420, EPI_ISL_467421, EPI_ISL_467422 | | | |
| see above | NYU Langone Health | Departments of Pathology and Medicine, New York University School of Medicine | Maria Aguero-Rosenfeld, Brendan Belovarac, Margaret Black, Ludovic Boytard, John Cadley, Paolo Cotzia, John Chen, Dacia Dimartino, Xiaojun Feng, Tatyana Gindin, Emily Guzman, Adriana Heguy, Megan Hogan, Emily Huang, George Jour, Alireza Khodadadi-Jamayran, Lawrence H. Lin, Raven Luther, Andrew Lytle, Christian Marier, Matthew T. Maurano, Mark J. Mulligan, Peter Meyn, Raquel Ordonez Ciriza, Iman Osman, Jared Pinnell, Vanessa Raabe, Sitharam Ramaswami, Amy Rapkiewicz, Andre M. Ribeiro-dos-Santos, Marie Samanovic-Golden, Antonio Serrano, Guomiao Shen, Matija Snuderl, Theodore Vougiouklakis, Nick Vulpescu, Gael Westby, Paul Zappile, Yutong Zhang |
| EPI_ISL_467423, EPI_ISL_467424, EPI_ISL_467425, EPI_ISL_467426, EPI_ISL_467427, EPI_ISL_467428, EPI_ISL_467429 | BCCDC Public Health Laboratory | BCCDC Public Health Laboratory | Richard Harrigan, Hope Lapointe, Jinny Choi, Kimia Kamelian, John Tyson,Terry Snutch, Linda Hoang, Inna Sekirov, Paul Levett, Mel Krajden, Natalie Prystajecky |
| EPI_ISL_467430 | Zoonotic and Exotic infection Diseases Division | Zoontic and Exotic infection Diseases Division | Jinlang Wang, Zhigao Bu |
| EPI_ISL_467431 | Molecular Diagnostics Services (MDS) | KRISP, KZN Research Innovation and Sequencing Platform | Giandhari J, Pillay S, Lessells R, Chimukangara B, Mdlalose K, York D, Khan S, Tegally H, Wilkinson E, de Oliveira T |
| EPI_ISL_467432, EPI_ISL_467433, EPI_ISL_467434, EPI_ISL_467435 | AMPATH-DBN | KRISP, KZN Research Innovation and Sequencing Platform | Giandhari J, Pillay S, Lessells R, Chimukangara B, Mdlalose K, York D, Khan S, Tegally H, Wilkinson E, de Oliveira T |
| EPI_ISL_467437, EPI_ISL_467441, EPI_ISL_467442, EPI_ISL_467443 | NHLS-IALCH | KRISP, KZN Research Innovation and Sequencing Platform | Giandhari J, Pillay S, Lessells R, Chimukangara B, Mdlalose K, York D, Khan S, Tegally H, Wilkinson E, de Oliveira T |
| EPI_ISL_467444, EPI_ISL_467445, EPI_ISL_467446 | Molecular Diagnostics Services (MDS) | KRISP, KZN Research Innovation and Sequencing Platform | Giandhari J, Pillay S, Lessells R, Chimukangara B, Mdlalose K, York D, Khan S, Tegally H, Wilkinson E, de Oliveira T |
| EPI_ISL_467449, EPI_ISL_467450, EPI_ISL_467451, EPI_ISL_467453, EPI_ISL_467454, EPI_ISL_467455, EPI_ISL_467456, EPI_ISL_467457, EPI_ISL_467460, EPI_ISL_467461, EPI_ISL_467462, EPI_ISL_467465, EPI_ISL_467466, EPI_ISL_467467, EPI_ISL_467468, EPI_ISL_467469, EPI_ISL_467470, EPI_ISL_467471, EPI_ISL_467472, EPI_ISL_467473, EPI_ISL_467474 | | | |
| see above | AMPATH-DBN | KRISP, KZN Research Innovation and Sequencing Platform | Giandhari J, Pillay S, Lessells R, Chimukangara B, Mdlalose K, York D, Khan S, Tegally H, Wilkinson E, de Oliveira T |
| EPI_ISL_467475, EPI_ISL_467476, EPI_ISL_467477, EPI_ISL_467478, EPI_ISL_467479, EPI_ISL_467480, EPI_ISL_467481, EPI_ISL_467482, EPI_ISL_467483, EPI_ISL_467484, EPI_ISL_467485, EPI_ISL_467486, EPI_ISL_467488, EPI_ISL_467489, EPI_ISL_467490, EPI_ISL_467491 | | | |
| see above | Molecular Diagnostics Services (MDS) | KRISP, KZN Research Innovation and Sequencing Platform | Giandhari J, Pillay S, Lessells R, Chimukangara B, Mdlalose K, York D, Khan S, Tegally H, Wilkinson E, de Oliveira T |
| EPI_ISL_467493 | NHLS-IALCH | KRISP, KZN Research Innovation and Sequencing Platform | Giandhari J, Pillay S, Lessells R, Chimukangara B, Mdlalose K, York D, Khan S, Tegally H, Wilkinson E, de Oliveira T |
| EPI_ISL_467494, EPI_ISL_467495, EPI_ISL_467496, EPI_ISL_467497, EPI_ISL_467498, EPI_ISL_467499, EPI_ISL_467500, EPI_ISL_467501, EPI_ISL_467502, EPI_ISL_467503, EPI_ISL_467504, EPI_ISL_467506 | | | |
| see above | Molecular Diagnostics Services (MDS) | KRISP, KZN Research Innovation and Sequencing Platform | Giandhari J, Pillay S, Lessells R, Chimukangara B, Mdlalose K, York D, Khan S, Tegally H, Wilkinson E, de Oliveira T |
| EPI_ISL_467507, EPI_ISL_467508, EPI_ISL_467509, EPI_ISL_467511, EPI_ISL_467512, EPI_ISL_467513, EPI_ISL_467514, EPI_ISL_467515 | NHLS-IALCH | KRISP, KZN Research Innovation and Sequencing Platform | Giandhari J, Pillay S, Lessells R, Chimukangara B, Mdlalose K, York D, Khan S, Tegally H, Wilkinson E, de Oliveira T |
| EPI_ISL_467516 | CAPRISA | KRISP, KZN Research Innovation and Sequencing Platform | Giandhari J, Pillay S, Lessells R, Chimukangara B, Mdlalose K, York D, Khan S, Tegally H, Wilkinson E, de Oliveira T |
| EPI_ISL_467517, EPI_ISL_467518, EPI_ISL_467519, EPI_ISL_467520, EPI_ISL_467521, EPI_ISL_467522, EPI_ISL_467523, EPI_ISL_467524 | NHLS-IALCH | KRISP, KZN Research Innovation and Sequencing Platform | Giandhari J, Pillay S, Lessells R, Chimukangara B, Mdlalose K, York D, Khan S, Tegally H, Wilkinson E, de Oliveira T |
| EPI_ISL_467529, EPI_ISL_467536, EPI_ISL_467545, EPI_ISL_467551, EPI_ISL_467552, EPI_ISL_467563, EPI_ISL_467564, EPI_ISL_467572, EPI_ISL_467577, EPI_ISL_467583, EPI_ISL_467590, EPI_ISL_467591, EPI_ISL_467594, EPI_ISL_467596, EPI_ISL_467598, EPI_ISL_467599, EPI_ISL_467600, EPI_ISL_467604, EPI_ISL_467608, EPI_ISL_467610, EPI_ISL_467611, EPI_ISL_467612, EPI_ISL_467624, EPI_ISL_467630, EPI_ISL_467632, EPI_ISL_467633, EPI_ISL_467634, EPI_ISL_467644, EPI_ISL_467651, EPI_ISL_467652, EPI_ISL_467653, EPI_ISL_467654, EPI_ISL_467661, EPI_ISL_467663 | | | |
| see above | New Mexico Department of Health Scientific Laboratory Division | Center for Global Health, University of New Mexico Health Sciences Center | Daryl Domman, Kurt Schwalm, Twila Kunde, Joseph Hicks, Michael Edwards, Darrell Dinwiddie |
| EPI_ISL_467695, EPI_ISL_467699, EPI_ISL_467708, EPI_ISL_467713, EPI_ISL_467718 | PHE South West Regional Laboratory, National Infection Service | Wellcome Sanger Institute for the COVID-19 Genomics UK Consortium | Stephanie Hutchings, Hannah Pymont, Dr Peter Muir, Barry Vipond, Rich Hopes; and Alex Alderton, Roberto Amato, Sonia Goncalves, Ewan Harrison, David K. Jackson, Ian Johnston, Dominic Kwiatkowski, Cordelia Langford, John Sillitoe on behalf of the Wellcome Sanger Institute COVID-19 Surveillance Team (http://www.sanger.ac.uk/covid-team) |
| EPI_ISL_467774, EPI_ISL_467775 | Molecular diagnostic laboratory of Federal Budget Institution of Science "Central Research Institute of Epidemiology" of The Federal Service on Customers' Rights Protection and Human Well-being Surveillance | Group of Genomics and Postgenomic Technologies of Central Research Institute of Epidemiology | Speranskaya AS, Kaptelova VV, Samoilov AE, Korneenko EV, Sizova TV, Tivanova EV, Shipulina OY, Akimkin VG |
| EPI_ISL_467778, EPI_ISL_467779, EPI_ISL_467780, EPI_ISL_467781 | National Influenza Centre Romania | Charite Universitatsmedizin Berlin, Institute of Virology | Victor M Corman, Jorn Beheim-Schwarzbach, Barbara Muehlemann, Talitha Veith, Julia Schneider, Terry Jones, L. Ustea, N. Paraschiv, M. Lazar, Christian Drosten |
| EPI_ISL_467782, EPI_ISL_467783, EPI_ISL_467784, EPI_ISL_467785, EPI_ISL_467786, EPI_ISL_467787, EPI_ISL_467788, EPI_ISL_467789, EPI_ISL_467790, EPI_ISL_467791, EPI_ISL_467792, EPI_ISL_467793, EPI_ISL_467794, EPI_ISL_467795, EPI_ISL_467796, EPI_ISL_467797, EPI_ISL_467798, EPI_ISL_467799, EPI_ISL_467800, EPI_ISL_467801, EPI_ISL_467802, EPI_ISL_467803, EPI_ISL_467804, EPI_ISL_467805, EPI_ISL_467806, EPI_ISL_467807, EPI_ISL_467808 | | | |
| see above | Virginia DCLS | Virginia DCLS | Virginia DCLS |
| EPI_ISL_467809 | Cedars-Sinai Medical Center, Department of Pathology & Laboratory Medicine, Molecular Pathology Laboratory | Cedars-Sinai Medical Center, Molecular Pathology Laboratory of Department of Pathology & Laboratory Medicine and Genomic Core | Wenjuan Zhang, John Paul Govindavari, Brian Davis, Stephanie Chen, Jong Taek Kim, Jianbo Song, Jean Lopategui, Jasmine T Plummer, Eric Vail |
| EPI_ISL_467811, EPI_ISL_467812, EPI_ISL_467813, EPI_ISL_467814, EPI_ISL_467815, EPI_ISL_467816, EPI_ISL_467817, EPI_ISL_467818, EPI_ISL_467819, EPI_ISL_467820, EPI_ISL_467821, EPI_ISL_467822, EPI_ISL_467823, EPI_ISL_467824, EPI_ISL_467825, EPI_ISL_467827, EPI_ISL_467828, EPI_ISL_467829, EPI_ISL_467830, EPI_ISL_467831, EPI_ISL_467832, EPI_ISL_467833, EPI_ISL_467834, EPI_ISL_467835, EPI_ISL_467836, EPI_ISL_467837, EPI_ISL_467838, EPI_ISL_467839, EPI_ISL_467840, EPI_ISL_467841, EPI_ISL_467842, EPI_ISL_467843, EPI_ISL_467844, EPI_ISL_467845, EPI_ISL_467846, EPI_ISL_467847, EPI_ISL_467848, EPI_ISL_467849, EPI_ISL_467850, EPI_ISL_467851, EPI_ISL_467852, EPI_ISL_467853, EPI_ISL_467854, EPI_ISL_467855, EPI_ISL_467856, EPI_ISL_467857, EPI_ISL_467858, EPI_ISL_467859, EPI_ISL_467860, EPI_ISL_467861, EPI_ISL_467862, EPI_ISL_467863, EPI_ISL_467864, EPI_ISL_467865, EPI_ISL_467866, EPI_ISL_467867, EPI_ISL_467868, EPI_ISL_467869, EPI_ISL_467870, EPI_ISL_467871, EPI_ISL_467872, EPI_ISL_467873, EPI_ISL_467874, EPI_ISL_467875, EPI_ISL_467876, EPI_ISL_467877, EPI_ISL_467878, EPI_ISL_467879, EPI_ISL_467880, EPI_ISL_467881, EPI_ISL_467882, EPI_ISL_467883, EPI_ISL_467884, EPI_ISL_467885, EPI_ISL_467886, EPI_ISL_467887, EPI_ISL_467888, EPI_ISL_467889, EPI_ISL_467890, EPI_ISL_467891, EPI_ISL_467892, EPI_ISL_467893, EPI_ISL_467894, EPI_ISL_467895, EPI_ISL_467896, EPI_ISL_467897, EPI_ISL_467898, EPI_ISL_467899, EPI_ISL_467900, EPI_ISL_467901, EPI_ISL_467902, EPI_ISL_467903, EPI_ISL_467904, EPI_ISL_467905, EPI_ISL_467906, EPI_ISL_467907, EPI_ISL_467908, EPI_ISL_467909, EPI_ISL_467910, EPI_ISL_467911, EPI_ISL_467912, EPI_ISL_467913, EPI_ISL_467914, EPI_ISL_467915, EPI_ISL_467916, EPI_ISL_467917, EPI_ISL_467918, EPI_ISL_467919, EPI_ISL_467920, EPI_ISL_467921, EPI_ISL_467922, EPI_ISL_467923, EPI_ISL_467924, EPI_ISL_467925, EPI_ISL_467926, EPI_ISL_467927 | | | |
| see above | Quest Diagnostics | Quest Diagnostics | Anderson,B.P., Rosenthal,S.H., Gerasimova,A., Kagan,R.M. and Owen, R. |
| EPI_ISL_467928, EPI_ISL_467929, EPI_ISL_467930, EPI_ISL_467932, EPI_ISL_467933, EPI_ISL_467934, EPI_ISL_467936, EPI_ISL_467937, EPI_ISL_467938, EPI_ISL_467941, EPI_ISL_467942, EPI_ISL_467943, EPI_ISL_467944 | | | |
| see above | Virginia DCLS | Virginia DCLS | Virginia DCLS |
| EPI_ISL_467945 | Montefiore Medical Center, Dept. of Pathology, Clinical Virology | Albert Einstein College of Medicine, Dept. of Microbiology & Immunology, Chandran lab | J. Maximilian Fels, Saad Khan, Ryan Forster, Karin A. Skalina, Ariel S. Wirchnianski, Denise Haslwanter, Catalina Florez, Robert H. Bortz III, M. Eugenia Dieterle, Ethan Laudermilch, Rohit K. Jangra, Amanda Mengotto, Duncan Kimmel, Shahina B. Maqbool, John M. Greally, Wendy A. Szymczak, Amy S. Fox, Michael B. Prystowsky, D. Yitzchak Goldstein, Johanna P. Daily, Libusha Kelly, Kartik Chandran |
| EPI_ISL_467946, EPI_ISL_467948 | Innovative Genomics Institute, UC Berkeley | Innovative Genomics Institute, UC Berkeley | Stacia Wyman, Haridha Shivram, Liana Lareau, Shana McDevitt, Justin Choi |
| EPI_ISL_467950, EPI_ISL_467951, EPI_ISL_467952, EPI_ISL_467953, EPI_ISL_467954, EPI_ISL_467955, EPI_ISL_467956, EPI_ISL_467957, EPI_ISL_467958, EPI_ISL_467959, EPI_ISL_467960, EPI_ISL_467961, EPI_ISL_467962 | | | |
| see above | San Diego County Public Health Laboratory | Andersen lab at Scripps Research | SEARCH Alliance San Diego with Tracy Basler, Jovan Shephard, Brett Austin |
| EPI_ISL_467963 | San Diego County Public Health Laboratory | Andersen lab at Scripps Research | SEARCH Alliance San Diego with Michael Quigley, Ellen Stefanski, Ian Mchardy |
| EPI_ISL_467964, EPI_ISL_467965, EPI_ISL_467966 | San Diego County Public Health Laboratory | Andersen lab at Scripps Research | SEARCH Alliance San Diego with Tracy Basler, Jovan Shephard, Brett Austin |
| EPI_ISL_467967 | Scripps Medical Laboratory | Andersen lab at Scripps Research | SEARCH Alliance San Diego with Tracy Basler, Jovan Shephard, Brett Austin |
| EPI_ISL_467968, EPI_ISL_467969, EPI_ISL_467970, EPI_ISL_467971 | San Diego County Public Health Laboratory | Andersen lab at Scripps Research | SEARCH Alliance San Diego with Tracy Basler, Jovan Shephard, Brett Austin |
| EPI_ISL_467972 | Scripps Medical Laboratory | Andersen lab at Scripps Research | SEARCH Alliance San Diego with Michael Quigley, Ellen Stefanski, Ian Mchardy |
| EPI_ISL_467973, EPI_ISL_467974, EPI_ISL_467975 | San Diego County Public Health Laboratory | Andersen lab at Scripps Research | SEARCH Alliance San Diego with Tracy Basler, Jovan Shephard, Brett Austin |
| EPI_ISL_467976 | Rady's Childrens Hospital | Andersen lab at Scripps Research | SEARCH Alliance San Diego |
| EPI_ISL_467978, EPI_ISL_467979, EPI_ISL_467980 | San Diego County Public Health Laboratory | Andersen lab at Scripps Research | SEARCH Alliance San Diego with Tracy Basler, Jovan Shephard, Brett Austin |
| EPI_ISL_467981 | Rady's Childrens Hospital | Andersen lab at Scripps Research | SEARCH Alliance San Diego |

| | | | |
|---|---|---|---|
| EPI_ISL_467982, EPI_ISL_467983 | San Diego County Public Health Laboratory | Andersen lab at Scripps Research | SEARCH Alliance San Diego with Tracy Basler, Jovan Shephard, Brett Austin |
| EPI_ISL_467984 | Rady's Childrens Hospital | Andersen lab at Scripps Research | SEARCH Alliance San Diego |
| EPI_ISL_467990 | SA Pathology | SA Pathology | Leong, LEX, Lim, CK, Turra, M, Bastian, I, Higgins, G |
| EPI_ISL_467991, EPI_ISL_467994, EPI_ISL_467995, EPI_ISL_467996, EPI_ISL_467999, EPI_ISL_468000, EPI_ISL_468001, EPI_ISL_468002, EPI_ISL_468003, EPI_ISL_468004, EPI_ISL_468005, EPI_ISL_468006, EPI_ISL_468007, EPI_ISL_468008, EPI_ISL_468009, EPI_ISL_468010, EPI_ISL_468011, EPI_ISL_468012, EPI_ISL_468013, EPI_ISL_468014, EPI_ISL_468015, EPI_ISL_468016, EPI_ISL_468017, EPI_ISL_468018, EPI_ISL_468019, EPI_ISL_468020, EPI_ISL_468021, EPI_ISL_468022, EPI_ISL_468023, EPI_ISL_468024, EPI_ISL_468025, EPI_ISL_468026, EPI_ISL_468027, EPI_ISL_468028, EPI_ISL_468029, EPI_ISL_468030, EPI_ISL_468031, EPI_ISL_468032, EPI_ISL_468033, EPI_ISL_468034, EPI_ISL_468035, EPI_ISL_468036, EPI_ISL_468037, EPI_ISL_468038, EPI_ISL_468039, EPI_ISL_468040, EPI_ISL_468041, EPI_ISL_468042, EPI_ISL_468043 | | | |
| see above | SA Pathology | SA Pathology | Lex Leong, Chuan Kok Lim, Mark Turra, Ivan Bastian, Geoff Higgins |
| EPI_ISL_468066 | Physiology, Istanbul Medeniyet University | Physiology, Istanbul Medeniyet University | Pence,S., Caykara,B., Pence,H.H., Tekin,S., Yiyit,N., Cevher Keskin,B. and Kara,A. |
| EPI_ISL_468070, EPI_ISL_468071, EPI_ISL_468072, EPI_ISL_468073 | Child Health Research Foundation | Child Health Research Foundation | Senjuti Saha, Roly Malaker, Md Saiful Islam Sajib, Hafizur Rahman, Maksuda Islam, Samir K Saha |
| EPI_ISL_468075, EPI_ISL_468076, EPI_ISL_468077, EPI_ISL_468078 | Child Health Research Foundation | Child Health Research Foundation | Senjuti Saha, Roly Malaker, Md Saiful Islam Sajib, Hafizur Rahman, Afroza Akter Tanni, Syed Muktadir Al Sium, Maksuda Islam, Samir K Saha |
| EPI_ISL_468081, EPI_ISL_468082, EPI_ISL_468083, EPI_ISL_468084, EPI_ISL_468089, EPI_ISL_468095, EPI_ISL_468103, EPI_ISL_468105, EPI_ISL_468109, EPI_ISL_468110, EPI_ISL_468112, EPI_ISL_468113, EPI_ISL_468117, EPI_ISL_468120, EPI_ISL_468123, EPI_ISL_468125, EPI_ISL_468126 | | | |
| see above | OHSU Lab Services Molecular Microbiology Lab | Oregon SARS-CoV-2 Genome Sequencing Center | Brendan L. O'Connell, Ruth V. Nichols, Alec J. Hirsch, Guang Fan, Daniel N. Streblow, William B. Messer, Andrew C. Adey, Benjamin N. Bimber, Brian J. O'Roak |
| EPI_ISL_468134, EPI_ISL_468135, EPI_ISL_468136, EPI_ISL_468137, EPI_ISL_468138, EPI_ISL_468139, EPI_ISL_468140, EPI_ISL_468141, EPI_ISL_468143, EPI_ISL_468144, EPI_ISL_468145, EPI_ISL_468146, EPI_ISL_468147, EPI_ISL_468148, EPI_ISL_468149, EPI_ISL_468150, EPI_ISL_468153, EPI_ISL_468154, EPI_ISL_468155, EPI_ISL_468156, EPI_ISL_468158 | | | |
| see above | [Romania, Bucharest] National Institute for Infectious Diseases "Prof. Dr. Matei Balş" | [Romania, Bucharest] National Institute for Infectious Diseases "Prof. Dr. Matei Balş" | Leontina Banica, Marius Cotic, Corina Casangiu, Marius Surleac, Simona Paraschiv |
| EPI_ISL_468159 | unknown | Department of Virology, Public Health Laboratories Division, National Institute of Health | Massab Umair, Aamer Ikram, Muhammad Salman, Adnan Khurshid, Nazish Badar, Shannon Whitmer, John Klena |
| EPI_ISL_468160 | unknown | Department of Virology, Public Health Laboratories Division, National Institute of Health | Massab Umair, Aamer Ikram, Muhammad Salman, Adnan Khurshid, Nazish Badar, Shannon Whitmer, John Klena |
| EPI_ISL_468161 | Department of Virology, Public Health Laboratories Division, National Institute of Health | Department of Virology, Public Health Laboratories Division, National Institute of Health | Massab Umair, Aamer Ikram, Muhammad Salman, Adnan Khurshid, Nazish Badar, Shannon Whitmer, John Klena |
| EPI_ISL_468162 | unknown | Department of Virology, Public Health Laboratories Division, National Institute of Health | Massab Umair, Aamer Ikram, Muhammad Salman, Adnan Khurshid, Nazish Badar, Shannon Whitmer, John Klena |
| EPI_ISL_468163 | Department of Virology, Public Health Laboratories Division, National Institute of Health | Department of Virology, Public Health Laboratories Division, National Institute of Health | Massab Umair, Aamer Ikram, Muhammad Salman, Adnan Khurshid, Nazish Badar, Shannon Whitmer, John Klena |
| EPI_ISL_468203, EPI_ISL_468207, EPI_ISL_468208, EPI_ISL_468213, EPI_ISL_468221, EPI_ISL_468222, EPI_ISL_468223, EPI_ISL_468224, EPI_ISL_468226, EPI_ISL_468227, EPI_ISL_468228, EPI_ISL_468229, EPI_ISL_468232, EPI_ISL_468233, EPI_ISL_468235, EPI_ISL_468240, EPI_ISL_468241, EPI_ISL_468243, EPI_ISL_468244, EPI_ISL_468247, EPI_ISL_468248, EPI_ISL_468250, EPI_ISL_468254, EPI_ISL_468255, EPI_ISL_468257, EPI_ISL_468263, EPI_ISL_468266, EPI_ISL_468268, EPI_ISL_468275, EPI_ISL_468276, EPI_ISL_468279, EPI_ISL_468280, EPI_ISL_468282, EPI_ISL_468284, EPI_ISL_468285, EPI_ISL_468286, EPI_ISL_468293, EPI_ISL_468298, EPI_ISL_468299, EPI_ISL_468303 | | | |
| see above | Viollier AG | Department of Biosystems Science and Engineering, ETH Zürich | Christian Beisel, Sarah Nadeau, Ivan Topolsky, Pedro Ferreira, Philipp Jablonski, Susana Posada-Céspedes, Tobias Schär, Ina Nissen, Natascha Santacroce, Elodie Burcklen, Christiane Beckmann, Maurice Redondo, Olivier Kobel, Christoph Noppen, Sophie Seidel, Noemie Santamaria de Souza, Niko Beerenwinkel, Tanja Stadler |
| EPI_ISL_468305, EPI_ISL_468307 | Centro de Vigilancia a Saude de Diadema | Instituto Adolfo Lutz, Interdiciplinary Procedures Center, Strategic Laboratory | Claudio Tavares Sacchi, Claudia Regina Gonçalves, Erica Valessa Ramos Gomes |
| EPI_ISL_468308 | Hospital Municipal do Tatuape Carmino Caricchio | Instituto Adolfo Lutz, Interdiciplinary Procedures Center, Strategic Laboratory | Claudio Tavares Sacchi, Claudia Regina Gonçalves, Erica Valessa Ramos Gomes |
| EPI_ISL_468310 | Hospital Sao Paulo de Ensino da UNIFESP | Instituto Adolfo Lutz, Interdiciplinary Procedures Center, Strategic Laboratory | Claudio Tavares Sacchi, Claudia Regina Gonçalves, Erica Valessa Ramos Gomes |
| EPI_ISL_468311, EPI_ISL_468312 | Hospital Municipal Dr Ignacio Proenca de Gouvea | Instituto Adolfo Lutz, Interdiciplinary Procedures Center, Strategic Laboratory | Claudio Tavares Sacchi, Claudia Regina Gonçalves, Erica Valessa Ramos Gomes |
| EPI_ISL_468313 | Vigilancia Epidemiologica de São Bernardo do Campo | Instituto Adolfo Lutz, Interdiciplinary Procedures Center, Strategic Laboratory | Claudio Tavares Sacchi, Claudia Regina Gonçalves, Erica Valessa Ramos Gomes |
| EPI_ISL_468314 | CTA Centro de Testagem e Aconselhamento | Instituto Adolfo Lutz, Interdiciplinary Procedures Center, Strategic Laboratory | Claudio Tavares Sacchi, Claudia Regina Gonçalves, Erica Valessa Ramos Gomes |
| EPI_ISL_468315 | Hospital Municipal do Tatuape Carmino Caricchio | Instituto Adolfo Lutz, Interdiciplinary Procedures Center, Strategic Laboratory | Claudio Tavares Sacchi, Claudia Regina Gonçalves, Erica Valessa Ramos Gomes |
| EPI_ISL_468316 | UPA Vila Assis | Instituto Adolfo Lutz, Interdiciplinary Procedures Center, Strategic Laboratory | Claudio Tavares Sacchi, Claudia Regina Gonçalves, Erica Valessa Ramos Gomes |
| EPI_ISL_468318 | Hospital Universitario da USP | Instituto Adolfo Lutz, Interdiciplinary Procedures Center, Strategic Laboratory | Claudio Tavares Sacchi, Claudia Regina Gonçalves, Erica Valessa Ramos Gomes |
| EPI_ISL_468319 | Vigilancia Epidemiologica de São Bernardo do Campo | Instituto Adolfo Lutz, Interdiciplinary Procedures Center, Strategic Laboratory | Claudio Tavares Sacchi, Claudia Regina Gonçalves, Erica Valessa Ramos Gomes |
| EPI_ISL_468320 | Secretaria Municipal de Saude de Hortolandia | Instituto Adolfo Lutz, Interdiciplinary Procedures Center, Strategic Laboratory | Claudio Tavares Sacchi, Claudia Regina Gonçalves, Erica Valessa Ramos Gomes |
| EPI_ISL_468321 | Hospital Universitario da USP | Instituto Adolfo Lutz, Interdiciplinary Procedures Center, Strategic Laboratory | Claudio Tavares Sacchi, Claudia Regina Gonçalves, Erica Valessa Ramos Gomes |
| EPI_ISL_468339, EPI_ISL_468340 | Microbiology Service, University Hospital of A Coruna-Biomedical Research Institute | Genomes & Disease, Center for Research in Molecular Medicine and Chronic Diseases, University of Santiago de Compostela | Kelly Conde, Jorge Arca, Soraya Rumbo, Juan A. Vallejo, M Poza, G Bou, Ana Pequeno-Valtierra, Jorge Rodriguez-Castro, Javier Temes, Daniel Garcia-Souto, Martin Santamarina, Cristina Gomez, Jose M. C. Tubio |
| EPI_ISL_468345, EPI_ISL_468346, EPI_ISL_468347, EPI_ISL_468348, EPI_ISL_468349, EPI_ISL_468350, EPI_ISL_468351, EPI_ISL_468353, EPI_ISL_468354, EPI_ISL_468355, EPI_ISL_468356 | | | |
| see above | County of Santa Clara Public Health Department | Chan-Zuckerberg Biohub | CZB Cliahub Consortium |
| EPI_ISL_468358, EPI_ISL_468360, EPI_ISL_468361, EPI_ISL_468363, EPI_ISL_468364, EPI_ISL_468365, EPI_ISL_468367, EPI_ISL_468368, EPI_ISL_468370, EPI_ISL_468372, EPI_ISL_468373, EPI_ISL_468374, EPI_ISL_468377, EPI_ISL_468380, EPI_ISL_468382, EPI_ISL_468383, EPI_ISL_468385, EPI_ISL_468387 | | | |
| see above | Alameda County Public Health Lab | Chan-Zuckerberg Biohub | CZB Cliahub Consortium |
| EPI_ISL_468388, EPI_ISL_468389, EPI_ISL_468390, EPI_ISL_468391, EPI_ISL_468392, EPI_ISL_468393, EPI_ISL_468394, EPI_ISL_468395, EPI_ISL_468396, EPI_ISL_468397, EPI_ISL_468398, EPI_ISL_468399, EPI_ISL_468400, EPI_ISL_468401, EPI_ISL_468402, EPI_ISL_468403, EPI_ISL_468404, EPI_ISL_468405, EPI_ISL_468406, EPI_ISL_468407, EPI_ISL_468409, EPI_ISL_468410, EPI_ISL_468411, EPI_ISL_468413, EPI_ISL_468414, EPI_ISL_468415, EPI_ISL_468417, EPI_ISL_468418, EPI_ISL_468419, EPI_ISL_468420, EPI_ISL_468421, EPI_ISL_468422, EPI_ISL_468424, EPI_ISL_468425, EPI_ISL_468426, EPI_ISL_468427, EPI_ISL_468429, EPI_ISL_468430, EPI_ISL_468431, EPI_ISL_468432, EPI_ISL_468433, EPI_ISL_468435, EPI_ISL_468436, EPI_ISL_468437 | | | |
| see above | County of San Luis Obispo Public Health Laboratory | Chan-Zuckerberg Biohub | CZB Cliahub Consortium |
| EPI_ISL_468438, EPI_ISL_468440, EPI_ISL_468446, EPI_ISL_468447, EPI_ISL_468448, EPI_ISL_468449, EPI_ISL_468450, EPI_ISL_468452, EPI_ISL_468453, EPI_ISL_468456, EPI_ISL_468458, EPI_ISL_468459, EPI_ISL_468461 | | | |
| see above | Humboldt County Public Health Laboratory | Chan-Zuckerberg Biohub | CZB Cliahub Consortium |
| EPI_ISL_468462, EPI_ISL_468463, EPI_ISL_468464, EPI_ISL_468465, EPI_ISL_468466, EPI_ISL_468467, EPI_ISL_468468, EPI_ISL_468469, EPI_ISL_468470, EPI_ISL_468471, EPI_ISL_468472, EPI_ISL_468473, EPI_ISL_468475, EPI_ISL_468476, EPI_ISL_468477, EPI_ISL_468478, EPI_ISL_468479, EPI_ISL_468480, EPI_ISL_468481, EPI_ISL_468482, EPI_ISL_468483, EPI_ISL_468484, EPI_ISL_468485, EPI_ISL_468486, EPI_ISL_468487, EPI_ISL_468489, EPI_ISL_468490, EPI_ISL_468491, EPI_ISL_468492, EPI_ISL_468493, EPI_ISL_468494, EPI_ISL_468495, EPI_ISL_468496, EPI_ISL_468497, EPI_ISL_468498, EPI_ISL_468499, EPI_ISL_468500, EPI_ISL_468501, EPI_ISL_468502, EPI_ISL_468503, EPI_ISL_468504, EPI_ISL_468505 | | | |
| see above | Ventura County Public Health Lab | Chan-Zuckerberg Biohub | CZB Cliahub Consortium |
| EPI_ISL_468506, EPI_ISL_468507, EPI_ISL_468508, EPI_ISL_468509, EPI_ISL_468510, EPI_ISL_468511, EPI_ISL_468512, EPI_ISL_468513, EPI_ISL_468515, EPI_ISL_468517, EPI_ISL_468518, EPI_ISL_468519, EPI_ISL_468520, EPI_ISL_468521, EPI_ISL_468522, EPI_ISL_468523, EPI_ISL_468524, EPI_ISL_468525, EPI_ISL_468526, EPI_ISL_468527, EPI_ISL_468528, EPI_ISL_468529, EPI_ISL_468530, EPI_ISL_468531, EPI_ISL_468532, EPI_ISL_468533, EPI_ISL_468534, EPI_ISL_468535, EPI_ISL_468536, EPI_ISL_468537, EPI_ISL_468538, EPI_ISL_468539, EPI_ISL_468540, EPI_ISL_468541, EPI_ISL_468542, EPI_ISL_468543, EPI_ISL_468544, EPI_ISL_468545, EPI_ISL_468546, EPI_ISL_468547, EPI_ISL_468548, EPI_ISL_468549, EPI_ISL_468551, EPI_ISL_468552, EPI_ISL_468553, EPI_ISL_468554, EPI_ISL_468557, EPI_ISL_468558 | | | |
| see above | San Joaquin County Public Health Lab | Chan-Zuckerberg Biohub | CZB Cliahub Consortium |
| EPI_ISL_468560, EPI_ISL_468561, EPI_ISL_468562, EPI_ISL_468563, EPI_ISL_468564, EPI_ISL_468565, EPI_ISL_468566, EPI_ISL_468567, EPI_ISL_468568, EPI_ISL_468569, EPI_ISL_468570, EPI_ISL_468571, EPI_ISL_468572, EPI_ISL_468573, EPI_ISL_468574, EPI_ISL_468575, EPI_ISL_468576, EPI_ISL_468577, EPI_ISL_468578, EPI_ISL_468579, EPI_ISL_468580, EPI_ISL_468581, EPI_ISL_468582, EPI_ISL_468583, EPI_ISL_468584, EPI_ISL_468585, EPI_ISL_468586, EPI_ISL_468587, EPI_ISL_468588, EPI_ISL_468589, EPI_ISL_468590 | | | |
| see above | Quest Diagnostics | Quest Diagnostics | Anderson,B.P., Rosenthal,S.H., Gerasimova,A., Kagan,R.M. and Owen, R. |
| EPI_ISL_468591 | Institute for Public Health | Laboratory for advanced genomics | Filip Rokić, Lovro Trgovec-Greif, Neven Sučić, Tomislav Rukavina, Igor Jurak, Oliver Vugrek |
| EPI_ISL_468592, EPI_ISL_468593, EPI_ISL_468594, EPI_ISL_468595, EPI_ISL_468596, EPI_ISL_468597, EPI_ISL_468598, EPI_ISL_468599, EPI_ISL_468600, EPI_ISL_468601, EPI_ISL_468602, EPI_ISL_468603, EPI_ISL_468605, EPI_ISL_468607, EPI_ISL_468608, EPI_ISL_468611, EPI_ISL_468612, EPI_ISL_468614 | | | |
| see above | Orange County Public Health Lab | Chan-Zuckerberg Biohub | CZB Cliahub Consortium |
| EPI_ISL_468615, EPI_ISL_468616, EPI_ISL_468617, EPI_ISL_468618, EPI_ISL_468621, EPI_ISL_468622, EPI_ISL_468624, EPI_ISL_468625, EPI_ISL_468626, EPI_ISL_468627, EPI_ISL_468628, EPI_ISL_468629, EPI_ISL_468631, EPI_ISL_468632, EPI_ISL_468633, EPI_ISL_468634, EPI_ISL_468636, EPI_ISL_468637, EPI_ISL_468638, EPI_ISL_468639, EPI_ISL_468640, EPI_ISL_468642, EPI_ISL_468645, EPI_ISL_468646, EPI_ISL_468647, EPI_ISL_468648, EPI_ISL_468649, EPI_ISL_468650, EPI_ISL_468651, EPI_ISL_468652 | | | |
| see above | Contra Costa Public Health Lab | Chan-Zuckerberg Biohub | CZB Cliahub Consortium |
| EPI_ISL_468656 | Institute for Public Health | Laboratory for advanced genomics | Filip Rokić, Lovro Trgovec-Greif, Neven Sučić, Tomislav Rukavina, Igor Jurak, Oliver Vugrek |
| EPI_ISL_468658, EPI_ISL_468659, EPI_ISL_468660, EPI_ISL_468661, EPI_ISL_468662, EPI_ISL_468664, EPI_ISL_468665, EPI_ISL_468666, EPI_ISL_468668, EPI_ISL_468669, EPI_ISL_468670, EPI_ISL_468671, EPI_ISL_468672, EPI_ISL_468673, EPI_ISL_468674, EPI_ISL_468675, EPI_ISL_468676, EPI_ISL_468677, EPI_ISL_468678, EPI_ISL_468679, EPI_ISL_468680, EPI_ISL_468683, EPI_ISL_468684, EPI_ISL_468685, EPI_ISL_468686, EPI_ISL_468687, EPI_ISL_468688, EPI_ISL_468689, EPI_ISL_468691, EPI_ISL_468693, EPI_ISL_468694, EPI_ISL_468695, EPI_ISL_468696, EPI_ISL_468698, EPI_ISL_468699, EPI_ISL_468700 | | | |
| see above | BCCDC Public Health Laboratory | BCCDC Public Health Laboratory | Richard Harrigan, Hope Lapointe, Jinny Choi, Kimia Kamelian, John Tyson,Terry Snutch, Linda Hoang, Inna Sekirov, Paul Levett, Mel Krajden, Natalie Prystajecky |
| EPI_ISL_468701, EPI_ISL_468702, EPI_ISL_468703, EPI_ISL_468705, EPI_ISL_468706, EPI_ISL_468707, EPI_ISL_468708, EPI_ISL_468709, EPI_ISL_468710, EPI_ISL_468711, EPI_ISL_468712, EPI_ISL_468713, EPI_ISL_468714, EPI_ISL_468715, EPI_ISL_468716, EPI_ISL_468717 | | | |
| see above | Ochsner Health | Bioinfoexperts, LLC | Rebecca Rose, Amy Feehan, David J. Nolan, Sissy Cross, David Moraga Amador, Tong Yang, Luke Caruso, Wayra Navia, Lydia Von Borstel, Xiao Hui Zhou, Julia-Garcia-Diaz, Susanna L. Lamers |
| EPI_ISL_468718 | Environmental and Global Health | Environmental and Global Health | Stephenson,C.J., Subramaniam,K., Waltzek,T.B., Merck,L.H., Gibson,J.C., Morris,J.G. |
| EPI_ISL_468719 | University of Florida | University of Florida | Elbadry,M.A., Subramaniam,K., Waltzek,T.B., Stephenson,C.J., Gibson,J.C., Alam,M.M., Lauzardo,M., Morris,J.G., Lednicky,J.A. |
| EPI_ISL_468720 | University of Florida | University of Florida | Stephenson,C.J., Subramaniam,K., Waltzek,T.B., Lauzardo,M., Morris,J.G., Lednicky,J.A. |
| EPI_ISL_468721 | University of Florida | University of Florida | Stephenson,C.J., Subramaniam,K., Waltzek,T.B., Lauzardo,M., Gibson,J.C., Morris,J.G., Lednicky,J.A. |
| EPI_ISL_468722 | University of Florida | University of Florida | Elbadry,M.A., Subramaniam,K., Waltzek,T.B., Lauzardo,M., Morris,J.G., Lednicky,J.A. |
| EPI_ISL_468723 | University of Florida | University of Florida | Stephenson,C.J., Subramaniam,K., Waltzek,T.B., Lauzardo,M., Morris,J.G., Lednicky,J.A. |
| EPI_ISL_468726 | unknown | Department of Microbiology | Peng,H., Tang,H., Jiang,L., Qi,Z., Zhao,P. |

| | | | |
|---|---|---|---|
| EPI_ISL_468727, EPI_ISL_468728, EPI_ISL_468729, EPI_ISL_468730, EPI_ISL_468731, EPI_ISL_468732, EPI_ISL_468733, EPI_ISL_468734, EPI_ISL_468735 | Lab voor klinische biologie | Onderzoeksgroep Virologie | Laurens Lambrechts, Nick Vereecke, Marthe Pauwels, Bruno Verhasselt, Linos Vandekerckhove, Hans Nauwynck, Sebastiaan Theuns |
| EPI_ISL_468736, EPI_ISL_468737, EPI_ISL_468738, EPI_ISL_468739, EPI_ISL_468740, EPI_ISL_468741, EPI_ISL_468743, EPI_ISL_468744, EPI_ISL_468745, EPI_ISL_468746 | Lab voor klinische biologie | Onderzoeksgroep Virologie | Nick Vereecke, Laurens Lambrechts, Marthe Pauwels, Bruno Verhasselt, Linos Vandekerckhove, Hans Nauwynck, Sebastiaan Theuns |
| EPI_ISL_468747, EPI_ISL_468748, EPI_ISL_468749, EPI_ISL_468750, EPI_ISL_468751 | Facultad de Medicina UC | Center for Mathematical Modeling and Center for Genome Regulation. Santiago, Chile | Gaete A, Travisany D, Palma R, Urra C, Varas M, Allende ML, Maass A, González M, Ferres M. |
| EPI_ISL_468752 | Center for Genome Regulation (CRG) | Center for Mathematical Modeling and Center for Genome Regulation. Santiago, Chile | Gaete A, Travisany D, Palma R, Urra C, Varas M, Allende ML, Maass A, González M. |
| EPI_ISL_468757, EPI_ISL_468758, EPI_ISL_468759 | Laboratorio de Biologia Molecular, Facultad de Medicina, Universidad de Atacama | Center for Mathematical Modeling and Center for Genome Regulation. Santiago, Chile | Gaete A, Travisany D, Palma R, Urra C, Varas M, Allende ML, Maass A, González M, C Echeverria |
| EPI_ISL_468760 | Center for Genome Regulation (CRG) | Center for Mathematical Modeling and Center for Genome Regulation. Santiago, Chile | Gaete A, Travisany D, Palma R, Urra C, Varas M, Allende ML, Maass A, González M. |
| EPI_ISL_468761, EPI_ISL_468763, EPI_ISL_468764 | Centro de Investigación Biomédica de La Rioja - Hospital San Pedro Logroño | SeqCOVID-SPAIN consortium/IBV(CSIC) | María de Toro, José Manuel Azcona Gutiérrez, María Pilar Bea Escudero, Miriam Blasco Alberdi and SeqCOVID-SPAIN consortium |
| EPI_ISL_468765, EPI_ISL_468766, EPI_ISL_468768, EPI_ISL_468769, EPI_ISL_468770, EPI_ISL_468771, EPI_ISL_468772, EPI_ISL_468773, EPI_ISL_468774, EPI_ISL_468775, EPI_ISL_468776, EPI_ISL_468778, EPI_ISL_468779, EPI_ISL_468780, EPI_ISL_468781, EPI_ISL_468782, EPI_ISL_468783, EPI_ISL_468784, EPI_ISL_468785, EPI_ISL_468786, EPI_ISL_468788, EPI_ISL_468790, EPI_ISL_468791, EPI_ISL_468792, EPI_ISL_468793, EPI_ISL_468794, EPI_ISL_468795, EPI_ISL_468797, EPI_ISL_468798, EPI_ISL_468799, EPI_ISL_468800, EPI_ISL_468801, EPI_ISL_468802, EPI_ISL_468803, EPI_ISL_468804, EPI_ISL_468805, EPI_ISL_468806, EPI_ISL_468807, EPI_ISL_468808, EPI_ISL_468809, EPI_ISL_468810, EPI_ISL_468811, EPI_ISL_468812, EPI_ISL_468813, EPI_ISL_468814, EPI_ISL_468815, EPI_ISL_468816, EPI_ISL_468817, EPI_ISL_468818, EPI_ISL_468819, EPI_ISL_468820, EPI_ISL_468822, EPI_ISL_468823, EPI_ISL_468824, EPI_ISL_468825, EPI_ISL_468826, EPI_ISL_468828, EPI_ISL_468829, EPI_ISL_468830, EPI_ISL_468831, EPI_ISL_468832, EPI_ISL_468833, EPI_ISL_468834, EPI_ISL_468835, EPI_ISL_468836, EPI_ISL_468838, EPI_ISL_468840, EPI_ISL_468841, EPI_ISL_468842, EPI_ISL_468843, EPI_ISL_468845, EPI_ISL_468846, EPI_ISL_468847, EPI_ISL_468849, EPI_ISL_468850, EPI_ISL_468851, EPI_ISL_468852, EPI_ISL_468853, EPI_ISL_468854, EPI_ISL_468855, EPI_ISL_468856, EPI_ISL_468858, EPI_ISL_468859 | | | |
| see above | Servicio de Microbiología, Hospital Miguel Servet, Zaragoza | SeqCOVID-SPAIN consortium/IBV(CSIC) | Antonio Rezusta López, Alexander Tristancho Baró, Ana Milagro, Yolanda Gracia Grataloup, Nieves Martínez Cameo and SeqCOVID-SPAIN consortium |
| EPI_ISL_468860, EPI_ISL_468861, EPI_ISL_468862, EPI_ISL_468863, EPI_ISL_468864, EPI_ISL_468866, EPI_ISL_468867, EPI_ISL_468868, EPI_ISL_468869, EPI_ISL_468870, EPI_ISL_468871, EPI_ISL_468872, EPI_ISL_468873, EPI_ISL_468874, EPI_ISL_468875, EPI_ISL_468876, EPI_ISL_468877, EPI_ISL_468878, EPI_ISL_468879, EPI_ISL_468880, EPI_ISL_468881, EPI_ISL_468882, EPI_ISL_468883, EPI_ISL_468885, EPI_ISL_468887, EPI_ISL_468888, EPI_ISL_468890, EPI_ISL_468891, EPI_ISL_468892, EPI_ISL_468893, EPI_ISL_468894, EPI_ISL_468895, EPI_ISL_468896, EPI_ISL_468897, EPI_ISL_468898, EPI_ISL_468899, EPI_ISL_468900, EPI_ISL_468902, EPI_ISL_468903, EPI_ISL_468904, EPI_ISL_468905, EPI_ISL_468906, EPI_ISL_468907, EPI_ISL_468908, EPI_ISL_468909, EPI_ISL_468910, EPI_ISL_468911, EPI_ISL_468912, EPI_ISL_468913 | | | |
| see above | Servicio de Microbiología. Hospital Universitario Donostia. OSI Donostialdea. Área de Enfermedades Infecciosas, Grupo de Infección Respiratoria y Resistencia Antimicrobiana. Instituto de Investigación Sanitaria Biodonostia. | SeqCOVID-SPAIN consortium/IBV(CSIC) | Gustavo Cilla, Milagrosa Montes, Luis Piñeiro, Jose Maria Marimón and SeqCOVID-SPAIN consortium |
| EPI_ISL_468914 | Istituto Zooprofilattico Sperimentale Puglia e Basilicata; Dipartimento di Bioscienze, Biotecnologie e Biofarmaceutica dell'Università degli Studi di Bari "A.Moro"; Istituto di Biomembrane. Bioenergetica e Biotecnologie Molecolari del Consiglio Nazionale delle Ricerche di Bari | Beaconlab (Bioinformatics, Evolution and Comparative Genomics lab), Dept of Biosciences, University on Milan | Parisi A.,Pesole G., Manzari C., Chiara M. |
| EPI_ISL_468915, EPI_ISL_468916, EPI_ISL_468917, EPI_ISL_468918, EPI_ISL_468919, EPI_ISL_468920, EPI_ISL_468921, EPI_ISL_468922, EPI_ISL_468923, EPI_ISL_468924, EPI_ISL_468925, EPI_ISL_468926, EPI_ISL_468927, EPI_ISL_468928, EPI_ISL_468929, EPI_ISL_468930, EPI_ISL_468931, EPI_ISL_468932, EPI_ISL_468933, EPI_ISL_468934, EPI_ISL_468935, EPI_ISL_468936, EPI_ISL_468937, EPI_ISL_468938, EPI_ISL_468939, EPI_ISL_468940, EPI_ISL_468941, EPI_ISL_468942, EPI_ISL_468943, EPI_ISL_468944, EPI_ISL_468945, EPI_ISL_468946, EPI_ISL_468947, EPI_ISL_468948, EPI_ISL_468949, EPI_ISL_468950, EPI_ISL_468951 | | | |
| see above | Servicio de Microbiología. Hospital Universitario Donostia. OSI Donostialdea. Área de Enfermedades Infecciosas, Grupo de Infección Respiratoria y Resistencia Antimicrobiana. Instituto de Investigación Sanitaria Biodonostia. | SeqCOVID-SPAIN consortium/IBV(CSIC) | Gustavo Cilla, Milagrosa Montes, Luis Piñeiro, Jose Maria Marimón and SeqCOVID-SPAIN consortium |
| EPI_ISL_468952, EPI_ISL_468953, EPI_ISL_468954, EPI_ISL_468959, EPI_ISL_468960, EPI_ISL_468961, EPI_ISL_468962, EPI_ISL_468966, EPI_ISL_468968, EPI_ISL_468972, EPI_ISL_468974, EPI_ISL_468975, EPI_ISL_468976, EPI_ISL_468977, EPI_ISL_468979, EPI_ISL_468980, EPI_ISL_468982, EPI_ISL_468983, EPI_ISL_468984, EPI_ISL_468986, EPI_ISL_468987, EPI_ISL_468988, EPI_ISL_468989, EPI_ISL_468990, EPI_ISL_468991, EPI_ISL_468992, EPI_ISL_468993, EPI_ISL_468997, EPI_ISL_468998, EPI_ISL_468999, EPI_ISL_469000, EPI_ISL_469001, EPI_ISL_469003, EPI_ISL_469004, EPI_ISL_469006, EPI_ISL_469008, EPI_ISL_469009, EPI_ISL_469010, EPI_ISL_469012, EPI_ISL_469013, EPI_ISL_469014, EPI_ISL_469015 | | | |
| see above | Servicio de Microbiología, Hospital Universitario Son Espases | SeqCOVID-SPAIN consortium/IBV(CSIC) | Carla López-Causapé, Jordi Reina, Antonio Oliver and SeqCOVID-SPAIN consortium |
| EPI_ISL_469017 | LNR National Reference Laboratory, Mohammed VI University of Health Sciences | Medical Biotechnology Laboratory, Rabat Medical and Pharmacy School, Mohammed The Vth University in Rabat | Meriem LAAMARTI, Souad KARTTI, Rokaia LAAMRTI , M.W. CHEMAO-ELFIHRI, Loubna ALLAM, Mouna OUADGHIRI, Imane SMYEJ, Jalila RAHOUI, Houda BENRAHMA, Jalil El Atar, Idrissa Diawara, Rachid EL JAOUDI, Laila SBABOU, Chakib NEJJARI, Saaid AMZAZI, Rachid MENTAG, Lahcen BELYAMANI and Azeddine IBRAHIMI |
| EPI_ISL_469020, EPI_ISL_469022 | Istituto Zooprofilattico Sperimentale Puglia e Basilicata; Dipartimento di Bioscienze, Biotecnologie e Biofarmaceutica dell'Università degli Studi di Bari "A.Moro"; Istituto di Biomembrane. Bioenergetica e Biotecnologie Molecolari del Consiglio Nazionale delle Ricerche di Bari | Beaconlab (Bioinformatics, Evolution and Comparative Genomics lab), Dept of Biosciences, University on Milan | Parisi A.,Pesole G., Manzari C., Chiara M. |
| EPI_ISL_469023 | Istituto Zooprofilattico Sperimentale Puglia e Basilicata; Dipartimento di Bioscienze, Biotecnologie e Biofarmaceutica dell'Università degli Studi di Bari "A.Moro"; Istituto di Biomembrane. Bioenergetica e Biotecnologie Molecolari del Consiglio Nazionale delle Ricerche di Bari | Beaconlab (Bioinformatics, Evolution and Comparative Genomics lab), Dept of Biosciences, University on Milan | Parisi A.,Pesole G., Manzari C., Chiara M |
| EPI_ISL_469024 | B.J. Medical College and Civil hospital | Gujarat Biotechnology Research Centre | Tejas Shah, Ankit Hinsu, Pritesh Sabara, Apurvasinh Puvar, Janvi Raval, Zarna Patel, Monika Gandhi, Pinal Trivedi, Maharshi Pandya, Nidhi Patel, Nitin Savaliya, Raghawendra Kumar, Dinesh Kumar, Zuber Saiyed, Komal Patel, Labdhi Pandya, Snehal Bagatharia, Pranay Shah, Kamlesh J Upadhyay, Nirav Mungalpara, Priti Pandita, R D Dixit, A M Kadri, Harsh Bakshi, Chaitanya Joshi, Madhvi Joshi |
| EPI_ISL_469025 | B.J. Medical College and Civil hospital | Gujarat Biotechnology Research Centre | Ankit Hinsu, Pritesh Sabara, Apurvasinh Puvar, Janvi Raval, Zarna Patel, Monika Gandhi, Pinal Trivedi, Maharshi Pandya, Nidhi Patel, Nitin Savaliya, Raghawendra Kumar, Dinesh Kumar, Zuber Saiyed, Komal Patel, Labdhi Pandya, Snehal Bagatharia, Pranay Shah, Kamlesh J Upadhyay, Nirav Mungalpara, Tejas Shah, Pragya Sharma, R D Dixit, A M Kadri, Harsh Bakshi, Chaitanya Joshi, Madhvi Joshi |
| EPI_ISL_469026 | B.J. Medical College and Civil hospital | Gujarat Biotechnology Research Centre | Pritesh Sabara, Apurvasinh Puvar, Janvi Raval, Zarna Patel, Monika Gandhi, Pinal Trivedi, Maharshi Pandya, Nidhi Patel, Nitin Savaliya, Raghawendra Kumar, Dinesh Kumar, Zuber Saiyed, Komal Patel, Labdhi Pandya, Snehal Bagatharia, Pranay Shah, Kamlesh J Upadhyay, Nirav Mungalpara, Tejas Shah, Ankit Hinsu, Neha Rajpara, R D Dixit, A M Kadri, Harsh Bakshi, Chaitanya Joshi, Madhvi Joshi |
| EPI_ISL_469027 | B.J. Medical College and Civil hospital | Gujarat Biotechnology Research Centre | Apurvasinh Puvar, Janvi Raval, Zarna Patel, Monika Gandhi, Pinal Trivedi, Maharshi Pandya, Nidhi Patel, Nitin Savaliya, Raghawendra Kumar, Dinesh Kumar, Zuber Saiyed, Komal Patel, Labdhi Pandya, Snehal Bagatharia, Pranay Shah, Kamlesh J Upadhyay, Nirav Mungalpara, Tejas Shah, Ankit Hinsu, Pritesh Sabara, Afzal Ansari, R D Dixit, A M Kadri, Harsh Bakshi, Chaitanya Joshi, Madhvi Joshi |
| EPI_ISL_469028 | B.J. Medical College and Civil hospital | Gujarat Biotechnology Research Centre | Janvi Raval, Zarna Patel, Monika Gandhi, Pinal Trivedi, Maharshi Pandya, Nidhi Patel, Nitin Savaliya, Raghawendra Kumar, Dinesh Kumar, Zuber Saiyed, Komal Patel, Labdhi Pandya, Snehal Bagatharia, Pranay Shah, Kamlesh J Upadhyay, Nirav Mungalpara, Tejas Shah, Ankit Hinsu, Pritesh Sabara, Apurvasinh Puvar, Fenil Patel, R D Dixit, A M Kadri, Harsh Bakshi, Chaitanya Joshi, Madhvi Joshi |
| EPI_ISL_469029 | Government Medical College, Vadodara | Gujarat Biotechnology Research Centre | Zarna Patel, Monika Gandhi, Pinal Trivedi, Maharshi Pandya, Nidhi Patel, Nitin Savaliya, Raghawendra Kumar, Dinesh Kumar, Zuber Saiyed, Komal Patel, Labdhi Pandya, Snehal Bagatharia, Meenakshi Shah, Neena Doshi, Varsha Godbole, Tejas Shah, Ankit Hinsu, Pritesh Sabara, Apurvasinh Puvar, Janvi Raval, Neelam Nathani, R D Dixit, A M Kadri, Harsh Bakshi, Chaitanya Joshi, Madhvi Joshi |
| EPI_ISL_469030 | Government Medical College, Vadodara | Gujarat Biotechnology Research Centre | Monika Gandhi, Pinal Trivedi, Maharshi Pandya, Nidhi Patel, Nitin Savaliya, Raghawendra Kumar, Dinesh Kumar, Zuber Saiyed, Komal Patel, Labdhi Pandya, Snehal Bagatharia, Meenakshi Shah, Neena Doshi, Varsha Godbole, Tejas Shah, Ankit Hinsu, Pritesh Sabara, Apurvasinh Puvar, Janvi Raval, Zarna Patel, Armi Chaudhari, R D Dixit, A M Kadri, Harsh Bakshi, Chaitanya Joshi, Madhvi Joshi |
| EPI_ISL_469031 | Government Medical College, Vadodara | Gujarat Biotechnology Research Centre | Pinal Trivedi, Maharshi Pandya, Nidhi Patel, Nitin Savaliya, Raghawendra Kumar, Dinesh Kumar, Zuber Saiyed, Komal Patel, Labdhi Pandya, Snehal Bagatharia, Meenakshi Shah, Neena Doshi, Varsha Godbole, Tejas Shah, Ankit Hinsu, Pritesh Sabara, Apurvasinh Puvar, Janvi Raval, Zarna Patel, Monika Gandhi, Bhavya Jindal, R D Dixit, A M Kadri, Harsh Bakshi, Chaitanya Joshi, Madhvi Joshi |
| EPI_ISL_469032 | Government Medical College, Vadodara | Gujarat Biotechnology Research Centre | Maharshi Pandya, Nidhi Patel, Nitin Savaliya, Raghawendra Kumar, Dinesh Kumar, Zuber Saiyed, Komal Patel, Labdhi Pandya, Snehal Bagatharia, Meenakshi Shah, Neena Doshi, Varsha Godbole, Tejas Shah, Ankit Hinsu, Pritesh Sabara, Apurvasinh Puvar, Janvi Raval, Zarna Patel, Monika Gandhi, Pinal Trivedi, Pragya Sharma, R D Dixit, A M Kadri, Harsh Bakshi, Chaitanya Joshi, Madhvi Joshi |
| EPI_ISL_469033 | Government Medical College, Vadodara | Gujarat Biotechnology Research Centre | Nidhi Patel, Nitin Savaliya, Raghawendra Kumar, Dinesh Kumar, Zuber Saiyed, Komal Patel, Labdhi Pandya, Snehal Bagatharia, Meenakshi Shah, Neena Doshi, Varsha Godbole, Tejas Shah, Ankit Hinsu, Pritesh Sabara, Apurvasinh Puvar, Janvi Raval, Zarna Patel, Monika Gandhi, Pinal Trivedi, Maharshi Pandya, Priyanka P Vatsa, R D Dixit, A M Kadri, Harsh Bakshi, Chaitanya Joshi, Madhvi Joshi |
| EPI_ISL_469034 | Government Medical College, Vadodara | Gujarat Biotechnology Research Centre | Nitin Savaliya, Raghawendra Kumar, Dinesh Kumar, Zuber Saiyed, Komal Patel, Labdhi Pandya, Snehal Bagatharia, Meenakshi Shah, Neena Doshi, Varsha Godbole, Tejas Shah, Ankit Hinsu, Pritesh Sabara, Apurvasinh Puvar, Janvi Raval, Zarna Patel, Monika Gandhi, Pinal Trivedi, Maharshi Pandya, Nidhi Patel, Pooja P Doshi, R D Dixit, A M Kadri, Harsh Bakshi, Chaitanya Joshi, Madhvi Joshi |
| EPI_ISL_469035 | Government Medical College, Vadodara | Gujarat Biotechnology Research Centre | Raghawendra Kumar, Dinesh Kumar, Zuber Saiyed, Komal Patel, Labdhi Pandya, Snehal Bagatharia, Meenakshi Shah, Neena Doshi, Varsha Godbole, Tejas Shah, Ankit Hinsu, Pritesh Sabara, Apurvasinh Puvar, Janvi Raval, Zarna Patel, Monika Gandhi, Pinal Trivedi, Maharshi Pandya, Nidhi Patel, Nitin Savaliya, Akanksha Verma, R D Dixit, A M Kadri, Harsh Bakshi, Chaitanya Joshi, Madhvi Joshi |
| EPI_ISL_469036 | Government Medical College, Vadodara | Gujarat Biotechnology Research Centre | Dinesh Kumar, Zuber Saiyed, Komal Patel, Labdhi Pandya, Snehal Bagatharia, Meenakshi Shah, Neena Doshi, Varsha Godbole, Tejas Shah, Ankit Hinsu, Pritesh Sabara, Apurvasinh Puvar, Janvi Raval, Zarna Patel, Monika Gandhi, Pinal Trivedi, Maharshi Pandya, Nidhi Patel, Nitin Savaliya, Raghawendra Kumar, Priti Pandita, R D Dixit, A M Kadri, Harsh Bakshi, Chaitanya Joshi, Madhvi Joshi |
| EPI_ISL_469037 | GMERS Medical College & Hospital | Gujarat Biotechnology Research Centre | Zuber Saiyed, Komal Patel, Labdhi Pandya, Snehal Bagatharia, Meenakshi Shah, Neena Doshi, Varsha Godbole, Tejas Shah, Ankit Hinsu, Pritesh Sabara, Apurvasinh Puvar, Janvi Raval, Zarna Patel, Monika Gandhi, Pinal Trivedi, Maharshi Pandya, Nidhi Patel, Nitin Savaliya, Raghawendra Kumar, Dinesh Kumar, Pragya Sharma, R D Dixit, A M Kadri, Harsh Bakshi, Chaitanya Joshi, Madhvi Joshi |

| | | | |
|---|---|---|---|
| EPI_ISL_469038 | GMERS Medical College & Hospital | Gujarat Biotechnology Research Centre | Komal Patel, Labdhi Pandya, Snehal Bagatharia, Meenakshi Shah, Neena Doshi, Varsha Godbole, Tejas Shah, Ankit Hinsu, Pritesh Sabara, Apurvasinh Puvar, Janvi Raval, Zarna Patel, Monika Gandhi, Pinal Trivedi, Maharshi Pandya, Nidhi Patel, Nitin Savaliya, Raghawendra Kumar, Dinesh Kumar, Zuber Saiyed, Neha Rajpara, R D Dixit, A M Kadri, Harsh Bakshi, Chaitanya Joshi, Madhvi Joshi |
| EPI_ISL_469039 | GMERS Medical College & Hospital | Gujarat Biotechnology Research Centre | Labdhi Pandya, Snehal Bagatharia, Meenakshi Shah, Neena Doshi, Varsha Godbole, Tejas Shah, Ankit Hinsu, Pritesh Sabara, Apurvasinh Puvar, Janvi Raval, Zarna Patel, Monika Gandhi, Pinal Trivedi, Maharshi Pandya, Nidhi Patel, Nitin Savaliya, Raghawendra Kumar, Dinesh Kumar, Zuber Saiyed, Komal Patel, Afzal Ansari, R D Dixit, A M Kadri, Harsh Bakshi, Chaitanya Joshi, Madhvi Joshi |
| EPI_ISL_469040 | GMERS Medical College & Hospital | Gujarat Biotechnology Research Centre | Snehal Bagatharia, Meenakshi Shah, Neena Doshi, Varsha Godbole, Tejas Shah, Ankit Hinsu, Pritesh Sabara, Apurvasinh Puvar, Janvi Raval, Zarna Patel, Monika Gandhi, Pinal Trivedi, Maharshi Pandya, Nidhi Patel, Nitin Savaliya, Raghawendra Kumar, Dinesh Kumar, Zuber Saiyed, Komal Patel, Labdhi Pandya, Fenil Patel, R D Dixit, A M Kadri, Harsh Bakshi, Chaitanya Joshi, Madhvi Joshi |
| EPI_ISL_469041 | GMERS Medical College & Hospital | Gujarat Biotechnology Research Centre | Meenakshi Shah, Neena Doshi, Varsha Godbole, Tejas Shah, Ankit Hinsu, Pritesh Sabara, Apurvasinh Puvar, Janvi Raval, Zarna Patel, Monika Gandhi, Pinal Trivedi, Maharshi Pandya, Nidhi Patel, Nitin Savaliya, Raghawendra Kumar, Dinesh Kumar, Zuber Saiyed, Komal Patel, Labdhi Pandya, Snehal Bagatharia, Neelam Nathani, R D Dixit, A M Kadri, Harsh Bakshi, Chaitanya Joshi, Madhvi Joshi |
| EPI_ISL_469042 | GMERS Medical College & Hospital | Gujarat Biotechnology Research Centre | Neena Doshi, Varsha Godbole, Tejas Shah, Ankit Hinsu, Pritesh Sabara, Apurvasinh Puvar, Janvi Raval, Zarna Patel, Monika Gandhi, Pinal Trivedi, Maharshi Pandya, Nidhi Patel, Nitin Savaliya, Raghawendra Kumar, Dinesh Kumar, Zuber Saiyed, Komal Patel, Labdhi Pandya, Snehal Bagatharia, Meenakshi Shah, Armi Chaudhari, R D Dixit, A M Kadri, Harsh Bakshi, Chaitanya Joshi, Madhvi Joshi |
| EPI_ISL_469043 | Dr. N. D. Desai Medical College & Hospital | Gujarat Biotechnology Research Centre | J G Buch, Jigar Gusani, Supreet Prabhu, Tejas Shah, Ankit Hinsu, Pritesh Sabara, Apurvasinh Puvar, Janvi Raval, Zarna Patel, Monika Gandhi, Pinal Trivedi, Maharshi Pandya, Nidhi Patel, Nitin Savaliya, Raghawendra Kumar, Dinesh Kumar, Zuber Saiyed, Komal Patel, Labdhi Pandya, Snehal Bagatharia, Bhavya Jindal, R D Dixit, A M Kadri, Harsh Bakshi, Chaitanya Joshi, Madhvi Joshi |
| EPI_ISL_469044 | Dr. N. D. Desai Medical College & Hospital | Gujarat Biotechnology Research Centre | Jigar Gusani, Supreet Prabhu, Tejas Shah, Ankit Hinsu, Pritesh Sabara, Apurvasinh Puvar, Janvi Raval, Zarna Patel, Monika Gandhi, Pinal Trivedi, Maharshi Pandya, Nidhi Patel, Nitin Savaliya, Raghawendra Kumar, Dinesh Kumar, Zuber Saiyed, Komal Patel, Labdhi Pandya, Snehal Bagatharia, J G Buch, Neha Rajpara, R D Dixit, A M Kadri, Harsh Bakshi, Chaitanya Joshi, Madhvi Joshi |
| EPI_ISL_469045 | Dr. N. D. Desai Medical College & Hospital | Gujarat Biotechnology Research Centre | Supreet Prabhu, Tejas Shah, Ankit Hinsu, Pritesh Sabara, Apurvasinh Puvar, Janvi Raval, Zarna Patel, Monika Gandhi, Pinal Trivedi, Maharshi Pandya, Nidhi Patel, Nitin Savaliya, Raghawendra Kumar, Dinesh Kumar, Zuber Saiyed, Komal Patel, Labdhi Pandya, Snehal Bagatharia, J G Buch, Jigar Gusani, Priyanka P Vatsa, R D Dixit, A M Kadri, Harsh Bakshi, Chaitanya Joshi, Madhvi Joshi |
| EPI_ISL_469046 | Dr. N. D. Desai Medical College & Hospital | Gujarat Biotechnology Research Centre | Tejas Shah, Ankit Hinsu, Pritesh Sabara, Apurvasinh Puvar, Janvi Raval, Zarna Patel, Monika Gandhi, Pinal Trivedi, Maharshi Pandya, Nidhi Patel, Nitin Savaliya, Raghawendra Kumar, Dinesh Kumar, Zuber Saiyed, Komal Patel, Labdhi Pandya, Snehal Bagatharia, J G Buch, Jigar Gusani, Supreet Prabhu, Pooja P Doshi, R D Dixit, A M Kadri, Harsh Bakshi, Chaitanya Joshi, Madhvi Joshi |
| EPI_ISL_469047 | Dr. N. D. Desai Medical College & Hospital | Gujarat Biotechnology Research Centre | Ankit Hinsu, Pritesh Sabara, Apurvasinh Puvar, Janvi Raval, Zarna Patel, Monika Gandhi, Pinal Trivedi, Maharshi Pandya, Nidhi Patel, Nitin Savaliya, Raghawendra Kumar, Dinesh Kumar, Zuber Saiyed, Komal Patel, Labdhi Pandya, Snehal Bagatharia, J G Buch, Jigar Gusani, Supreet Prabhu, Tejas Shah, Akanksha Verma, R D Dixit, A M Kadri, Harsh Bakshi, Chaitanya Joshi, Madhvi Joshi |
| EPI_ISL_469048 | Banas Medical College and Research Institute | Gujarat Biotechnology Research Centre | Radhika Khara, Sunil R Joshi, Viren s Doshi, Zarna Patel, Monika Gandhi, Pinal Trivedi, Maharshi Pandya, Nidhi Patel, Nitin Savaliya, Raghawendra Kumar, Dinesh Kumar, Zuber Saiyed, Komal Patel, Labdhi Pandya, Snehal Bagatharia, Tejas Shah, Ankit Hinsu, Pritesh Sabara, Apurvasinh Puvar, Janvi Raval, Priti Pandita, R D Dixit, A M Kadri, Harsh Bakshi, Chaitanya Joshi, Madhvi Joshi |
| EPI_ISL_469049, EPI_ISL_469051, EPI_ISL_469052, EPI_ISL_469053, EPI_ISL_469054 | LNR National Reference Laboratory, Mohammed VI University of Health Sciences | Medical Biotechnology Laboratory, Rabat Medical and Pharmacy School, Mohammed The Vth University in Rabat | Meriem LAAMARTI, Souad KARTTI, Rokaia LAAMRTI , M.W. CHEMAO-ELFIHRI, Loubna ALLAM, Mouna OUADGHIRI, Imane SMYEJ, Jalila RAHOUI, Houda BENRAHMA, Jalil El Atar, Idrissa Diawara, Rachid EL JAOUDI, Laila SBABOU, Chakib NEJJARI, Saaid AMZAZI, Rachid MENTAG, Lahcen BELYAMANI and Azeddine IBRAHIMI |
| EPI_ISL_469055, EPI_ISL_469056 | Jourcentralen | The Public Health Agency of Sweden | Oskar Karlsson Lindsjo, Maria Lind Karlberg, Mattias Haukland, Reza Advani, Olov Svartstrom, Anna-Malin Linde, Sandra Broddesson, Petra Edquist, Shamam Muradrasoli, Anna Risberg, Karin Tegmark-Wisell |
| EPI_ISL_469057 | Inger Landgren | The Public Health Agency of Sweden | Oskar Karlsson Lindsjo, Maria Lind Karlberg, Mattias Haukland, Reza Advani, Olov Svartstrom, Anna-Malin Linde, Sandra Broddesson, Petra Edquist, Shamam Muradrasoli, Anna Risberg, Karin Tegmark-Wisell |
| EPI_ISL_469058 | Narhalsan Sjobo vardcentral | The Public Health Agency of Sweden | Oskar Karlsson Lindsjo, Maria Lind Karlberg, Mattias Haukland, Reza Advani, Olov Svartstrom, Anna-Malin Linde, Sandra Broddesson, Petra Edquist, Shamam Muradrasoli, Anna Risberg, Karin Tegmark-Wisell |
| EPI_ISL_469059 | Hovas Askim Familjelakare och BVC | The Public Health Agency of Sweden | Oskar Karlsson Lindsjo, Maria Lind Karlberg, Mattias Haukland, Reza Advani, Olov Svartstrom, Anna-Malin Linde, Sandra Broddesson, Petra Edquist, Shamam Muradrasoli, Anna Risberg, Karin Tegmark-Wisell |
| EPI_ISL_469060, EPI_ISL_469061 | Narhalsan Sjobo vardcentral | The Public Health Agency of Sweden | Oskar Karlsson Lindsjo, Maria Lind Karlberg, Mattias Haukland, Reza Advani, Olov Svartstrom, Anna-Malin Linde, Sandra Broddesson, Petra Edquist, Shamam Muradrasoli, Anna Risberg, Karin Tegmark-Wisell |
| EPI_ISL_469062 | Huddinge VC | The Public Health Agency of Sweden | Oskar Karlsson Lindsjo, Maria Lind Karlberg, Mattias Haukland, Reza Advani, Olov Svartstrom, Anna-Malin Linde, Sandra Broddesson, Petra Edquist, Shamam Muradrasoli, Anna Risberg, Karin Tegmark-Wisell |
| EPI_ISL_469063 | Ulltuna Vardcentral | The Public Health Agency of Sweden | Oskar Karlsson Lindsjo, Maria Lind Karlberg, Mattias Haukland, Reza Advani, Olov Svartstrom, Anna-Malin Linde, Sandra Broddesson, Petra Edquist, Shamam Muradrasoli, Anna Risberg, Karin Tegmark-Wisell |
| EPI_ISL_469064, EPI_ISL_469065 | Huddinge VC | The Public Health Agency of Sweden | Oskar Karlsson Lindsjo, Maria Lind Karlberg, Mattias Haukland, Reza Advani, Olov Svartstrom, Anna-Malin Linde, Sandra Broddesson, Petra Edquist, Shamam Muradrasoli, Anna Risberg, Karin Tegmark-Wisell |
| EPI_ISL_469066 | Surbrunns VC | The Public Health Agency of Sweden | Oskar Karlsson Lindsjo, Maria Lind Karlberg, Mattias Haukland, Reza Advani, Olov Svartstrom, Anna-Malin Linde, Sandra Broddesson, Petra Edquist, Shamam Muradrasoli, Anna Risberg, Karin Tegmark-Wisell |
| EPI_ISL_469067 | Kungsholmsdoktorn | The Public Health Agency of Sweden | Oskar Karlsson Lindsjo, Maria Lind Karlberg, Mattias Haukland, Reza Advani, Olov Svartstrom, Anna-Malin Linde, Sandra Broddesson, Petra Edquist, Shamam Muradrasoli, Anna Risberg, Karin Tegmark-Wisell |
| EPI_ISL_469068 | Hovas Askim Familjelakare och BVC | The Public Health Agency of Sweden | Oskar Karlsson Lindsjo, Maria Lind Karlberg, Mattias Haukland, Reza Advani, Olov Svartstrom, Anna-Malin Linde, Sandra Broddesson, Petra Edquist, Shamam Muradrasoli, Anna Risberg, Karin Tegmark-Wisell |
| EPI_ISL_469069 | Narhalsan Olskroken VC | The Public Health Agency of Sweden | Oskar Karlsson Lindsjo, Maria Lind Karlberg, Mattias Haukland, Reza Advani, Olov Svartstrom, Anna-Malin Linde, Sandra Broddesson, Petra Edquist, Shamam Muradrasoli, Anna Risberg, Karin Tegmark-Wisell |
| EPI_ISL_469070 | Surbrunns VC | The Public Health Agency of Sweden | Oskar Karlsson Lindsjo, Maria Lind Karlberg, Mattias Haukland, Reza Advani, Olov Svartstrom, Anna-Malin Linde, Sandra Broddesson, Petra Edquist, Shamam Muradrasoli, Anna Risberg, Karin Tegmark-Wisell |
| EPI_ISL_469071 | Wasterlakarna | The Public Health Agency of Sweden | Oskar Karlsson Lindsjo, Maria Lind Karlberg, Mattias Haukland, Reza Advani, Olov Svartstrom, Anna-Malin Linde, Sandra Broddesson, Petra Edquist, Shamam Muradrasoli, Anna Risberg, Karin Tegmark-Wisell |
| EPI_ISL_469072 | Ulltuna Vardcentral | The Public Health Agency of Sweden | Oskar Karlsson Lindsjo, Maria Lind Karlberg, Mattias Haukland, Reza Advani, Olov Svartstrom, Anna-Malin Linde, Sandra Broddesson, Petra Edquist, Shamam Muradrasoli, Anna Risberg, Karin Tegmark-Wisell |
| EPI_ISL_469073, EPI_ISL_469074 | Halmstad klinisk mikrobiologi | The Public Health Agency of Sweden | Oskar Karlsson Lindsjo, Maria Lind Karlberg, Mattias Haukland, Reza Advani, Olov Svartstrom, Anna-Malin Linde, Sandra Broddesson, Petra Edquist, Shamam Muradrasoli, Anna Risberg, Karin Tegmark-Wisell |
| EPI_ISL_469075 | Karolinska Universitetslaboratoriet | The Public Health Agency of Sweden | Oskar Karlsson Lindsjo, Maria Lind Karlberg, Mattias Haukland, Reza Advani, Olov Svartstrom, Anna-Malin Linde, Sandra Broddesson, Petra Edquist, Shamam Muradrasoli, Anna Risberg, Karin Tegmark-Wisell |
| EPI_ISL_469076 | Uppsala klinisk mikrobiologi | The Public Health Agency of Sweden | Oskar Karlsson Lindsjo, Maria Lind Karlberg, Mattias Haukland, Reza Advani, Olov Svartstrom, Anna-Malin Linde, Sandra Broddesson, Petra Edquist, Shamam Muradrasoli, Anna Risberg, Karin Tegmark-Wisell |
| EPI_ISL_469077, EPI_ISL_469078, EPI_ISL_469079 | Karolinska Universitetslaboratoriet | The Public Health Agency of Sweden | Oskar Karlsson Lindsjo, Maria Lind Karlberg, Mattias Haukland, Reza Advani, Olov Svartstrom, Anna-Malin Linde, Sandra Broddesson, Petra Edquist, Shamam Muradrasoli, Anna Risberg, Karin Tegmark-Wisell |
| EPI_ISL_469081, EPI_ISL_469082, EPI_ISL_469083, EPI_ISL_469084, EPI_ISL_469085, EPI_ISL_469086, EPI_ISL_469087, EPI_ISL_469088, EPI_ISL_469089, EPI_ISL_469090, EPI_ISL_469091, EPI_ISL_469092, EPI_ISL_469093, EPI_ISL_469094, EPI_ISL_469095, EPI_ISL_469096, EPI_ISL_469098, EPI_ISL_469099, EPI_ISL_469100, EPI_ISL_469101, EPI_ISL_469102, EPI_ISL_469103, EPI_ISL_469104, EPI_ISL_469105, EPI_ISL_469106, EPI_ISL_469107, EPI_ISL_469108, EPI_ISL_469109, EPI_ISL_469110, EPI_ISL_469111, EPI_ISL_469112, EPI_ISL_469113, EPI_ISL_469114, EPI_ISL_469115, EPI_ISL_469116, EPI_ISL_469117, EPI_ISL_469118, EPI_ISL_469119, EPI_ISL_469120, EPI_ISL_469121, EPI_ISL_469124, EPI_ISL_469125, EPI_ISL_469126, EPI_ISL_469127, EPI_ISL_469128, EPI_ISL_469129, EPI_ISL_469130, EPI_ISL_469131, EPI_ISL_469132, EPI_ISL_469133, EPI_ISL_469134, EPI_ISL_469135, EPI_ISL_469136, EPI_ISL_469137, EPI_ISL_469138, EPI_ISL_469139, EPI_ISL_469140, EPI_ISL_469141, EPI_ISL_469142, EPI_ISL_469143, EPI_ISL_469144, EPI_ISL_469145, EPI_ISL_469146, EPI_ISL_469147, EPI_ISL_469148, EPI_ISL_469149, EPI_ISL_469150, EPI_ISL_469151, EPI_ISL_469152, EPI_ISL_469153, EPI_ISL_469154, EPI_ISL_469155 | | | |
| see above | National Public Health Laboratory, National Centre for Infectious Diseases | National Public Health Laboratory, National Centre for Infectious Diseases | Mak TM, Octavia S, Chavatte JM, Cui L, Lin RTP |
| EPI_ISL_469173, EPI_ISL_469175, EPI_ISL_469177, EPI_ISL_469179, EPI_ISL_469181, EPI_ISL_469182, EPI_ISL_469183, EPI_ISL_469184, EPI_ISL_469186, EPI_ISL_469187, EPI_ISL_469189, EPI_ISL_469194, EPI_ISL_469195, EPI_ISL_469196, EPI_ISL_469197, EPI_ISL_469201, EPI_ISL_469203, EPI_ISL_469204, EPI_ISL_469208 | | | |
| see above | Yale Clinical Virology Laboratory | Grubaugh Lab - Yale School of Public Health | Joseph Fauver, Tara Alpert, Anderson Brito, Anne Wyllie, Chantal Vogels, Mary Petrone, Cole Jensen, Chaney Kalinich, Isabel Ott, Arnau Casanovas, Catherine Muenker, Adam Moore, Alice Lu, Maria Tokuyama, Patrick Wong, Peiwen Lu, Saad Omer, Richard Martinello, Allison Nelson, Shelli Farhadian, Akiko Iwasaki, Charlese Dela Cruz, Albert Ko, Nathan Grubaugh |
| EPI_ISL_469209, EPI_ISL_469210, EPI_ISL_469211, EPI_ISL_469212, EPI_ISL_469213, EPI_ISL_469214, EPI_ISL_469215, EPI_ISL_469216, EPI_ISL_469218, EPI_ISL_469222, EPI_ISL_469223 | | | |
| see above | BCCDC Public Health Laboratory | BCCDC Public Health Laboratory | Richard Harrigan, Hope Lapointe, Jinny Choi, Kimia Kamelian, John Tyson,Terry Snutch, Linda Hoang, Inna Sekirov, Paul Levett, Mel Krajden, Natalie Prystajecky |
| EPI_ISL_469224, EPI_ISL_469225, EPI_ISL_469226, EPI_ISL_469227, EPI_ISL_469228, EPI_ISL_469229, EPI_ISL_469230, EPI_ISL_469231, EPI_ISL_469232, EPI_ISL_469233, EPI_ISL_469234, EPI_ISL_469235, EPI_ISL_469236, EPI_ISL_469237, EPI_ISL_469238, EPI_ISL_469239, EPI_ISL_469240 | | | |
| see above | Public Health Laboratory | National Microbiology Laboratory | Anna Majer, Shari Tyson, Grace Seo, Kristyn Burak, Philip Mabon, Elsie Grudeski, Rhiannon Huzarewich, Russell Mandes, Jennifer Tanner, Natalie Knox, Morag Graham, Gary Van Domselaar, Robert Needle, Yang Yu, Adel Malek, Laura Gilbert, George Zahariadis, Nathalie Bastien, Yan Li, Timothy Booth, Matthew Gilmour |
| EPI_ISL_469241, EPI_ISL_469242, EPI_ISL_469243, EPI_ISL_469244, EPI_ISL_469245, EPI_ISL_469246, EPI_ISL_469247, EPI_ISL_469248, EPI_ISL_469249, EPI_ISL_469250, EPI_ISL_469251, EPI_ISL_469252 | | | |
| see above | Special Infectious Agents Unit | Special Infectious Agents Unit | Azhar,E.I., Hassan,A.M., Tolah,A.M., Uthman,N.A., Al-Sobahy,T.L., Farraj,S.A., El-Kafrawy,S.A. |
| EPI_ISL_469253 | Second Military Medical University, Department of Microbiology | Second Military Medical University, Department of Microbiology | Peng,H., Tang,H., Jiang,L., Qi,Z. and Zhao,P. |
| EPI_ISL_469254 | National Institute for Viral Disease Control and Prevention, China CDC | Institute of Viral Disease Control and Prevention, China CDC | Wenjie Tan, Lijuan Chen, Peihua NiuBaoying Huang, Li Zhao, Yubai Bi, Wenling Wang, Roujian Lu, Dayan Wang, Wenbo Xu, George Fu Gao, Chun Huang, Guizhen Wu |
| EPI_ISL_469255 | National Institute for Viral Disease Control and Prevention, China CDC | Institute of Viral Disease Control and Prevention, China CDC | Xiang ZhaoLijuan Chen, Dayan Wang, Yong Zhang, Yao MengZhixiao ChenYuchao Wu, Jun Han, Weifeng Shi, Yanhai Wang, William J. Liu, Shiwen Wang, George F. Gao, Wenbo Xu, Chun Huang, Guizhen Wu |
| EPI_ISL_469274 | National Public Health Laboratory, National Centre for Infectious Diseases | National Public Health Laboratory, National Centre for Infectious Diseases | Mak TM, Octavia S, Chavatte JM, Cui L, Lin RTP |
| EPI_ISL_469275 | Egyptian National Cancer Institute (ENCI) | Human Genome Center | Zekri, Abdel Rahman N, Amer,K.E., Ahmed,O.S., Soliman,H.K., Hafez,M.M., Bahnassy,A.A., Abdelhamid,W., Gad,A., Ali,M., Hassan,W., Samir,M., Raouf,A., Hamdy,M.S., Soliman,M.S., Elsissy,M.H., Elkhateeb,S.M., Ezzelarab,M.H., Abouelhoda, Mohamed |
| EPI_ISL_469276, EPI_ISL_469277, EPI_ISL_469278, EPI_ISL_469279, | Mohammed Bin Rashid University of Medicine and Health Sciences | Al Jalila Genomics Center | Ahmad Abou Tayoun, Tom Loney, Hamda Khansaheb, Sathishkumar Ramaswamy, Divinlal Harilal, Zulfa Omar Deesi, Rupa Murthy Varghese, Hanan Al Suwaidi, Abdulmajeed Alkhaja, Mohammed Uddin, Rifat Hamoudi, Rabih Halwani, Abiola Catherine Senok, Qutayba Hamid, Norbert Nowotny, Alawi Alsheikh-Ali |

| | | | |
|---|---|---|---|
| EPI_ISL_469280, EPI_ISL_469281 | | | |
| EPI_ISL_469282 | Service de Virologie Hôpital Saint-Louis | Laboratory of Cell Biology of viral infection, Unit INSERM-U944 | Laurent Meertens, Lucie Bonnet-Madin, Constance Delaugerre, Ali Amara |
| EPI_ISL_469283, EPI_ISL_469284 | Service de Virologie Hôpital Saint-Louis | Laboratory Cell Biology of Viral Infection-INSERM unit 944 | Laurent Meertens, Lucie Bonnet-Madin, Séverine Mercier-Delarue, Maud SALMONA, Constance Delaugerre, Ali Amara |
| EPI_ISL_469286 | National Institute of Laboratory Medicine and Referral Center | Genomic Research Lab, BCSIR | Tanjina Akhter Banu, Abu Sayeed Mohammad Mahmud, Mohammad Samir Uzzaman, Eshrar Osman, Md. Ahasan Habib, Shahina Akter, Md. Murshed Hasan Sarkar, Iffat Jahan, Barna Goswami, Md. Saddam Hossain, Tasnim Nafisa, Md. Maruf Ahmed Molla, Mahmuda Yeasmin, Asish Kumar Ghosh, Bayzid Bin Monir, A. K. M. Shamsuzzaman, Sheikh Md. Selim Al Din, Utpal Chandra Ray, Salek Ahmed Sajib, Md. Salim Khan |
| EPI_ISL_469287, EPI_ISL_469288, EPI_ISL_469289, EPI_ISL_469290, EPI_ISL_469291, EPI_ISL_469292, EPI_ISL_469293, EPI_ISL_469294, EPI_ISL_469295, EPI_ISL_469296 | Keio University Hospital | Keio University Hospital | Kenjiro Kosaki |
| EPI_ISL_469297 | National Institute of Laboratory Medicine and Referral Center | Genomic Research Lab, BCSIR | Barna Goswami, Abu Sayeed Mohammad Mahmud, Mohammad Samir Uzzaman, Eshrar Osman, Md. Ahasan Habib, Shahina Akter, Tanjina Akhter Banu, Md. Murshed Hasan Sarkar, Iffat Jahan, Md. Saddam Hossain, Tasnim Nafisa, Md. Maruf Ahmed Molla, Mahmuda Yeasmin, Asish Kumar Ghosh, Bayzid Bin Monir, A. K. M. Shamsuzzaman, Sheikh Md. Selim Al Din, Utpal Chandra Ray, Salek Ahmed Sajib, Md. Salim Khan |
| EPI_ISL_469298 | National Institute of Laboratory Medicine and Referral Center | Genomic Research Lab, BCSIR | Md. Murshed Hasan Sarkar, Abu Sayeed Mohammad Mahmud, Mohammad Samir Uzzaman, Eshrar Osman, Md. Ahasan Habib, Shahina Akter, Tanjina Akhter Banu, Barna Goswami, Iffat Jahan, Md. Saddam Hossain, Tasnim Nafisa, Md. Maruf Ahmed Molla, Mahmuda Yeasmin, Asish Kumar Ghosh, Bayzid Bin Monir, A. K. M. Shamsuzzaman, Sheikh Md. Selim Al Din, Utpal Chandra Ray, Salek Ahmed Sajib, Md. Salim Khan |
| EPI_ISL_469299 | National Institute of Laboratory Medicine and Referral Center | Genomic Research Lab, BCSIR | Iffat Jahan, Abu Sayeed Mohammad Mahmud, Mohammad Samir Uzzaman, Eshrar Osman, Md. Ahasan Habib, Shahina Akter, Tanjina Akhter Banu, Md. Murshed Hasan Sarkar, Barna Goswami, Md. Saddam Hossain, Tasnim Nafisa, Md. Maruf Ahmed Molla, Mahmuda Yeasmin, Asish Kumar Ghosh, Bayzid Bin Monir, A. K. M. Shamsuzzaman, Sheikh Md. Selim Al Din, Utpal Chandra Ray, Salek Ahmed Sajib, Md. Salim Khan |
| EPI_ISL_469300 | National Institute of Laboratory Medicine and Referral Center | Genomic Research Lab, BCSIR | Abu Sayeed Mohammad Mahmud, Mohammad Samir Uzzaman, Eshrar Osman, Md. Ahasan Habib, Shahina Akter, Tanjina Akhter Banu, Md. Murshed Hasan Sarkar, Barna Goswami, Iffat Jahan, Md. Saddam Hossain, Tasnim Nafisa, Md. Maruf Ahmed Molla, Mahmuda Yeasmin, Asish Kumar Ghosh, Bayzid Bin Monir, A. K. M. Shamsuzzaman, Sheikh Md. Selim Al Din, Utpal Chandra Ray, Salek Ahmed Sajib, Md. Salim Khan |
| EPI_ISL_469302, EPI_ISL_469303, EPI_ISL_469307, EPI_ISL_469309, EPI_ISL_469312, EPI_ISL_469313, EPI_ISL_469314, EPI_ISL_469315, EPI_ISL_469316, EPI_ISL_469319, EPI_ISL_469324, EPI_ISL_469325, EPI_ISL_469328, EPI_ISL_469329, EPI_ISL_469330, EPI_ISL_469335, EPI_ISL_469336, EPI_ISL_469340, EPI_ISL_469344 | | | |
| see above | NU-OMICS DNA Sequencing research facility, Northumbria University | Wellcome Sanger Institute for the COVID-19 Genomics UK Consortium | Chris Duncan, Sheia Waugh, Shirelle Burton-Fanning, Gary Eltringham, Jennifer Collins, Brendan Payne, Yusri Taha, Emma Swindells, Jane Greenaway, Edward Barton, Garren Scott, Debra Padgett, Clive Graham, Sarah Essex, Steve Ligget, Paul Baker, Lynn Dover, Wen Yew, Gary Black, John Allan, Joshua Loh, Greg Young, Matthew Bashton, Andrew Nelson, Darren Smith and Alex Alderton, Roberto Amato, Sonia Goncalves, Ewan Harrison, David K. Jackson, Ian Johnston, Dominic Kwiatkowski, Cordelia Langford, John Sillitoe on behalf of the Wellcome Sanger Institute COVID-19 Surveillance Team (http://www.sanger.ac.uk/covid-team) |
| EPI_ISL_469345, EPI_ISL_469346, EPI_ISL_469347, EPI_ISL_469348, EPI_ISL_469349, EPI_ISL_469350, EPI_ISL_469353, EPI_ISL_469354, EPI_ISL_469355, EPI_ISL_469356, EPI_ISL_469357, EPI_ISL_469358, EPI_ISL_469359, EPI_ISL_469360, EPI_ISL_469361, EPI_ISL_469362, EPI_ISL_469364, EPI_ISL_469365, EPI_ISL_469366, EPI_ISL_469368, EPI_ISL_469369, EPI_ISL_469370, EPI_ISL_469372, EPI_ISL_469373, EPI_ISL_469375, EPI_ISL_469376, EPI_ISL_469377, EPI_ISL_469379, EPI_ISL_469380, EPI_ISL_469381, EPI_ISL_469382, EPI_ISL_469383, EPI_ISL_469384, EPI_ISL_469385, EPI_ISL_469386, EPI_ISL_469387, EPI_ISL_469388, EPI_ISL_469389, EPI_ISL_469390, EPI_ISL_469391, EPI_ISL_469392, EPI_ISL_469394, EPI_ISL_469395, EPI_ISL_469396, EPI_ISL_469397, EPI_ISL_469398, EPI_ISL_469400, EPI_ISL_469401, EPI_ISL_469402, EPI_ISL_469403, EPI_ISL_469404, EPI_ISL_469405, EPI_ISL_469406, EPI_ISL_469407, EPI_ISL_469408, EPI_ISL_469409, EPI_ISL_469410, EPI_ISL_469412, EPI_ISL_469413, EPI_ISL_469414, EPI_ISL_469415, EPI_ISL_469416, EPI_ISL_469417, EPI_ISL_469418, EPI_ISL_469419, EPI_ISL_469420, EPI_ISL_469421, EPI_ISL_469422, EPI_ISL_469423, EPI_ISL_469424, EPI_ISL_469427, EPI_ISL_469428, EPI_ISL_469430, EPI_ISL_469431, EPI_ISL_469432, EPI_ISL_469434, EPI_ISL_469435, EPI_ISL_469436, EPI_ISL_469437, EPI_ISL_469438, EPI_ISL_469439, EPI_ISL_469440, EPI_ISL_469442, EPI_ISL_469443, EPI_ISL_469444, EPI_ISL_469445, EPI_ISL_469446, EPI_ISL_469448, EPI_ISL_469449, EPI_ISL_469451, EPI_ISL_469452, EPI_ISL_469453, EPI_ISL_469454, EPI_ISL_469455, EPI_ISL_469456, EPI_ISL_469457, EPI_ISL_469458, EPI_ISL_469459, EPI_ISL_469462, EPI_ISL_469463, EPI_ISL_469464, EPI_ISL_469465, EPI_ISL_469466, EPI_ISL_469468, EPI_ISL_469470, EPI_ISL_469471, EPI_ISL_469472, EPI_ISL_469474, EPI_ISL_469475, EPI_ISL_469476, EPI_ISL_469477, EPI_ISL_469478, EPI_ISL_469479, EPI_ISL_469480, EPI_ISL_469481, EPI_ISL_469483, EPI_ISL_469484, EPI_ISL_469485, EPI_ISL_469486, EPI_ISL_469488, EPI_ISL_469489, EPI_ISL_469490, EPI_ISL_469492, EPI_ISL_469493, EPI_ISL_469494, EPI_ISL_469495, EPI_ISL_469497, EPI_ISL_469498, EPI_ISL_469499, EPI_ISL_469500, EPI_ISL_469502, EPI_ISL_469503, EPI_ISL_469504, EPI_ISL_469505, EPI_ISL_469506, EPI_ISL_469507, EPI_ISL_469508, EPI_ISL_469509, EPI_ISL_469510, EPI_ISL_469513, EPI_ISL_469515, EPI_ISL_469516, EPI_ISL_469517, EPI_ISL_469518, EPI_ISL_469519, EPI_ISL_469520, EPI_ISL_469521, EPI_ISL_469523, EPI_ISL_469524, EPI_ISL_469525, EPI_ISL_469526, EPI_ISL_469529, EPI_ISL_469531, EPI_ISL_469532, EPI_ISL_469535, EPI_ISL_469536, EPI_ISL_469537, EPI_ISL_469538, EPI_ISL_469539, EPI_ISL_469543, EPI_ISL_469544, EPI_ISL_469546, EPI_ISL_469547, EPI_ISL_469548, EPI_ISL_469549, EPI_ISL_469550, EPI_ISL_469551, EPI_ISL_469553, EPI_ISL_469556, EPI_ISL_469557, EPI_ISL_469558, EPI_ISL_469559, EPI_ISL_469560, EPI_ISL_469562, EPI_ISL_469564, EPI_ISL_469566, EPI_ISL_469568, EPI_ISL_469570, EPI_ISL_469571, EPI_ISL_469573, EPI_ISL_469574, EPI_ISL_469577, EPI_ISL_469579, EPI_ISL_469580, EPI_ISL_469581, EPI_ISL_469582, EPI_ISL_469583, EPI_ISL_469584, EPI_ISL_469586, EPI_ISL_469588, EPI_ISL_469591, EPI_ISL_469600, EPI_ISL_469603, EPI_ISL_469604, EPI_ISL_469605, EPI_ISL_469607, EPI_ISL_469608, EPI_ISL_469609, EPI_ISL_469610, EPI_ISL_469612, EPI_ISL_469613, EPI_ISL_469614, EPI_ISL_469615, EPI_ISL_469616, EPI_ISL_469621, EPI_ISL_469624, EPI_ISL_469625, EPI_ISL_469626, EPI_ISL_469629, EPI_ISL_469630, EPI_ISL_469632, EPI_ISL_469634, EPI_ISL_469636, EPI_ISL_469637, EPI_ISL_469638, EPI_ISL_469639, EPI_ISL_469641, EPI_ISL_469643, EPI_ISL_469644, EPI_ISL_469645, EPI_ISL_469649, EPI_ISL_469651, EPI_ISL_469652, EPI_ISL_469654, EPI_ISL_469657, EPI_ISL_469659, EPI_ISL_469661, EPI_ISL_469663, EPI_ISL_469664, EPI_ISL_469665, EPI_ISL_469666, EPI_ISL_469667, EPI_ISL_469668, EPI_ISL_469670, EPI_ISL_469674, EPI_ISL_469675, EPI_ISL_469676, EPI_ISL_469677, EPI_ISL_469679, EPI_ISL_469680, EPI_ISL_469681, EPI_ISL_469682, EPI_ISL_469683, EPI_ISL_469684, EPI_ISL_469686, EPI_ISL_469687, EPI_ISL_469691, EPI_ISL_469692, EPI_ISL_469693, EPI_ISL_469694, EPI_ISL_469696, EPI_ISL_469697, EPI_ISL_469700, EPI_ISL_469701, EPI_ISL_469704, EPI_ISL_469705, EPI_ISL_469706, EPI_ISL_469708, EPI_ISL_469709, EPI_ISL_469710, EPI_ISL_469711, EPI_ISL_469712, EPI_ISL_469713, EPI_ISL_469714, EPI_ISL_469715, EPI_ISL_469720, EPI_ISL_469721, EPI_ISL_469723, EPI_ISL_469724, EPI_ISL_469725, EPI_ISL_469726, EPI_ISL_469727, EPI_ISL_469728, EPI_ISL_469730, EPI_ISL_469731, EPI_ISL_469733, EPI_ISL_469735, EPI_ISL_469737, EPI_ISL_469738, EPI_ISL_469739, EPI_ISL_469740, EPI_ISL_469741, EPI_ISL_469742, EPI_ISL_469743, EPI_ISL_469744, EPI_ISL_469745, EPI_ISL_469746, EPI_ISL_469747, EPI_ISL_469749, EPI_ISL_469750, EPI_ISL_469752, EPI_ISL_469753, EPI_ISL_469755, EPI_ISL_469758, EPI_ISL_469760, EPI_ISL_469763, EPI_ISL_469765, EPI_ISL_469766, EPI_ISL_469767, EPI_ISL_469768, EPI_ISL_469771, EPI_ISL_469772, EPI_ISL_469773, EPI_ISL_469774, EPI_ISL_469775, EPI_ISL_469777, EPI_ISL_469779, EPI_ISL_469781, EPI_ISL_469782, EPI_ISL_469785, EPI_ISL_469786, EPI_ISL_469787, EPI_ISL_469788, EPI_ISL_469789, EPI_ISL_469792, EPI_ISL_469793, EPI_ISL_469798, EPI_ISL_469799, EPI_ISL_469801, EPI_ISL_469802, EPI_ISL_469803, EPI_ISL_469804, EPI_ISL_469809, EPI_ISL_469810, EPI_ISL_469811, EPI_ISL_469814, EPI_ISL_469818, EPI_ISL_469819, EPI_ISL_469820, EPI_ISL_469822, EPI_ISL_469827, EPI_ISL_469829, EPI_ISL_469831, EPI_ISL_469833, EPI_ISL_469837, EPI_ISL_469840, EPI_ISL_469841 | | | |
| see above | PHE South West Regional Laboratory, National Infection Service | Wellcome Sanger Institute for the COVID-19 Genomics UK Consortium | Stephanie Hutchings, Hannah Pymont, Dr Peter Muir, Barry Vipond, Rich Hopes; and Alex Alderton, Roberto Amato, Sonia Goncalves, Ewan Harrison, David K. Jackson, Ian Johnston, Dominic Kwiatkowski, Cordelia Langford, John Sillitoe on behalf of the Wellcome Sanger Institute COVID-19 Surveillance Team (http://www.sanger.ac.uk/covid-team) |
| EPI_ISL_469845, EPI_ISL_469846, EPI_ISL_469848, EPI_ISL_469850, EPI_ISL_469853, EPI_ISL_469854, EPI_ISL_469855, EPI_ISL_469856, EPI_ISL_469857, EPI_ISL_469858, EPI_ISL_469859, EPI_ISL_469860, EPI_ISL_469861, EPI_ISL_469862, EPI_ISL_469864, EPI_ISL_469865, EPI_ISL_469866, EPI_ISL_469867, EPI_ISL_469868, EPI_ISL_469872, EPI_ISL_469874, EPI_ISL_469875, EPI_ISL_469878, EPI_ISL_469879, EPI_ISL_469880, EPI_ISL_469882, EPI_ISL_469883, EPI_ISL_469884, EPI_ISL_469887, EPI_ISL_469888, EPI_ISL_469889 | | | |
| see above | Regional Virus Laboratory, Belfast Health and Social Care Trust | Wellcome Sanger Institute for the COVID-19 Genomics UK Consortium | Conall McCaughey, James McKenna, Tanya Curran, Susan Feeney, Alison Watt, Ciara Cox, Mairead Connor, Zoltan Molnar, David Simpson, Derek Fairley; and Alex Alderton, Roberto Amato, Sonia Goncalves, Ewan Harrison, David K. Jackson, Ian Johnston, Dominic Kwiatkowski, Cordelia Langford, John Sillitoe on behalf of the Wellcome Sanger Institute COVID-19 Surveillance Team (http://www.sanger.ac.uk/covid-team) |
| EPI_ISL_469896, EPI_ISL_469899, EPI_ISL_469903, EPI_ISL_469906, EPI_ISL_469911 | Department of Pathology, University of Cambridge | Wellcome Sanger Institute for the COVID-19 Genomics UK Consortium | Luke W Meredith, M. Estée Török , Myra Hosmillo, William L. Hamilton, Martin D. Curran, Theresa Feltwell, Grant Hall, Anna Yakovleva, Fahad A Khokhar, Charlotte J. Houldcroft, Laura G Caller, Aminu S. Jahun, Sarah L. Caddy, Ian Goodfellow; and Alex Alderton, Roberto Amato, Sonia Goncalves, Ewan Harrison, David K. Jackson, Ian Johnston, Dominic Kwiatkowski, Cordelia Langford, John Sillitoe on behalf of the Wellcome Sanger Institute COVID-19 Surveillance Team (http://www.sanger.ac.uk/covid-team) |
| EPI_ISL_469914 | PHE South West Regional Laboratory, National Infection Service | Wellcome Sanger Institute for the COVID-19 Genomics UK Consortium | Stephanie Hutchings, Hannah Pymont, Dr Peter Muir, Barry Vipond, Rich Hopes; and Alex Alderton, Roberto Amato, Sonia Goncalves, Ewan Harrison, David K. Jackson, Ian Johnston, Dominic Kwiatkowski, Cordelia Langford, John Sillitoe on behalf of the Wellcome Sanger Institute COVID-19 Surveillance Team (http://www.sanger.ac.uk/covid-team) |
| EPI_ISL_469915 | NU-OMICS DNA Sequencing research facility, Northumbria University | Wellcome Sanger Institute for the COVID-19 Genomics UK Consortium | Chris Duncan, Sheia Waugh, Shirelle Burton-Fanning, Gary Eltringham, Jennifer Collins, Brendan Payne, Yusri Taha, Emma Swindells, Jane Greenaway, Edward Barton, Garren Scott, Debra Padgett, Clive Graham, Sarah Essex, Steve Ligget, Paul Baker, Lynn Dover, Wen Yew, Gary Black, John Allan, Joshua Loh, Greg Young, Matthew Bashton, Andrew Nelson, Darren Smith and Alex Alderton, Roberto Amato, Sonia Goncalves, Ewan Harrison, David K. Jackson, Ian Johnston, Dominic Kwiatkowski, Cordelia Langford, John Sillitoe on behalf of the Wellcome Sanger Institute COVID-19 Surveillance Team (http://www.sanger.ac.uk/covid-team) |
| EPI_ISL_469918, EPI_ISL_469925, EPI_ISL_469926, EPI_ISL_469928 | Virology Department, Sheffield Teaching Hospitals NHS Foundation Trust / Department of Infection, Immunity and Cardiovascular Disease, The Medical School, University of Sheffield | Wellcome Sanger Institute for the COVID-19 Genomics UK Consortium | Thushan de Silva, Matthew Parker,Adri Angyal, Rebecca Brown, Luke Green, Rachel Tucker, Paul Parsons, Danielle Groves, Alex Keeley, Dave Partridge, Matthew Wyles, Benjamin Lindsey, Mehmet Yavuz, Mohammad Raza, Cariad Evans and Alex Alderton, Roberto Amato, Sonia Goncalves, Ewan Harrison, David K. Jackson, Ian Johnston, Dominic Kwiatkowski, Cordelia Langford, John Sillitoe on behalf of the Wellcome Sanger Institute COVID-19 Surveillance Team (http://www.sanger.ac.uk/covid-team) |
| EPI_ISL_469930, EPI_ISL_469931, EPI_ISL_469932, EPI_ISL_469933, EPI_ISL_469934, EPI_ISL_469935, EPI_ISL_469940, EPI_ISL_469941, EPI_ISL_469943, EPI_ISL_469944, EPI_ISL_469945, EPI_ISL_469946, EPI_ISL_469947, EPI_ISL_469948, EPI_ISL_469949, EPI_ISL_469950, EPI_ISL_469952, EPI_ISL_469954, EPI_ISL_469956, EPI_ISL_469957, EPI_ISL_469959, EPI_ISL_469960, EPI_ISL_469961, EPI_ISL_469962, EPI_ISL_469963, EPI_ISL_469964, EPI_ISL_469965, EPI_ISL_469966, EPI_ISL_469967, EPI_ISL_469968, EPI_ISL_469969, EPI_ISL_469971, EPI_ISL_469972, EPI_ISL_469973, EPI_ISL_469974, EPI_ISL_469976, EPI_ISL_469977, EPI_ISL_469978, EPI_ISL_469980, EPI_ISL_469981, EPI_ISL_469982, EPI_ISL_469983, EPI_ISL_469984, EPI_ISL_469985, EPI_ISL_469986, EPI_ISL_469988, EPI_ISL_469989, EPI_ISL_469990, EPI_ISL_469991, EPI_ISL_469993, EPI_ISL_469994, EPI_ISL_469997, EPI_ISL_469999, EPI_ISL_470000, EPI_ISL_470001, EPI_ISL_470005, EPI_ISL_470006, EPI_ISL_470007, EPI_ISL_470009, EPI_ISL_470010 | | | |
| see above | NHSGGC West of Scotland Specialist Virology Centre / MRC-University of Glasgow Centre for Virus Research | Wellcome Sanger Institute for the COVID-19 Genomics UK Consortium | Ana da Silva Filipe, Natasha Johnson, Kathy Smollett, Daniel Mair, Stephen Carmichael, Lily Tong, Jenna Nichols, Elihu Aranday-Cortes, Kirstyn Brunker, Yasmin Parr, Kyriaki Nomikou; Sarah McDonald, Marc Niebel, Patawee Asamaphan; Richard Orton, Joseph Hughes, Sreenu Vattipally, David L Robertson; Alasdair MacLean, Rory Gunson; Kathy Li, Natasha Jesudason, Rajiv Shah, James Shepherd, Antonia Ho, Alice Broos, Emma Thomson and Alex Alderton, Roberto Amato, Sonia Goncalves, Ewan Harrison, David K. Jackson, Ian Johnston, Dominic Kwiatkowski, Cordelia Langford, John Sillitoe on behalf of the Wellcome Sanger Institute COVID-19 Surveillance Team (http://www.sanger.ac.uk/covid-team) |
| EPI_ISL_470014, EPI_ISL_470015, EPI_ISL_470016, EPI_ISL_470018, EPI_ISL_470021, EPI_ISL_470022, EPI_ISL_470024, EPI_ISL_470025, EPI_ISL_470026, EPI_ISL_470027, EPI_ISL_470029, EPI_ISL_470030, EPI_ISL_470031, EPI_ISL_470032, EPI_ISL_470033, EPI_ISL_470034, EPI_ISL_470036, EPI_ISL_470038, EPI_ISL_470041, EPI_ISL_470042, EPI_ISL_470044, EPI_ISL_470045, EPI_ISL_470047, EPI_ISL_470048, EPI_ISL_470050, EPI_ISL_470053, EPI_ISL_470055, EPI_ISL_470060, EPI_ISL_470063, EPI_ISL_470064, EPI_ISL_470065, EPI_ISL_470070, EPI_ISL_470071, EPI_ISL_470072, EPI_ISL_470075, EPI_ISL_470076, EPI_ISL_470077, EPI_ISL_470079, EPI_ISL_470080, EPI_ISL_470083, EPI_ISL_470084, EPI_ISL_470085, EPI_ISL_470086, EPI_ISL_470087, EPI_ISL_470088 | | | |
| see above | Regional Virus Laboratory, Belfast Health and Social Care Trust | Wellcome Sanger Institute for the COVID-19 Genomics UK Consortium | Conall McCaughey, James McKenna, Tanya Curran, Susan Feeney, Alison Watt, Ciara Cox, Mairead Connor, Zoltan Molnar, David Simpson, Derek Fairley; and Alex Alderton, Roberto Amato, Sonia Goncalves, Ewan Harrison, David K. Jackson, Ian Johnston, Dominic Kwiatkowski, Cordelia Langford, John Sillitoe on behalf of the Wellcome Sanger Institute COVID-19 Surveillance Team (http://www.sanger.ac.uk/covid-team) |
| EPI_ISL_470092, EPI_ISL_470096, EPI_ISL_470098, EPI_ISL_470099, EPI_ISL_470101, EPI_ISL_470102, EPI_ISL_470103, EPI_ISL_470104, EPI_ISL_470105, EPI_ISL_470107, EPI_ISL_470108, EPI_ISL_470109, EPI_ISL_470110, EPI_ISL_470114, EPI_ISL_470115, EPI_ISL_470116, EPI_ISL_470117, EPI_ISL_470118, EPI_ISL_470119, EPI_ISL_470120, EPI_ISL_470121, EPI_ISL_470122, EPI_ISL_470123, EPI_ISL_470124, EPI_ISL_470125, EPI_ISL_470127, EPI_ISL_470128, EPI_ISL_470130, EPI_ISL_470131, EPI_ISL_470132, EPI_ISL_470133, EPI_ISL_470136, EPI_ISL_470137, EPI_ISL_470140, EPI_ISL_470141, EPI_ISL_470142, EPI_ISL_470143, EPI_ISL_470144, EPI_ISL_470146, EPI_ISL_470147, EPI_ISL_470148, EPI_ISL_470150, EPI_ISL_470152, EPI_ISL_470153, EPI_ISL_470154, EPI_ISL_470155, EPI_ISL_470156, EPI_ISL_470158, EPI_ISL_470159, EPI_ISL_470160, EPI_ISL_470161, EPI_ISL_470163, EPI_ISL_470166, EPI_ISL_470167, EPI_ISL_470170, EPI_ISL_470171, EPI_ISL_470172, EPI_ISL_470173, EPI_ISL_470174, EPI_ISL_470181, EPI_ISL_470182, EPI_ISL_470183, EPI_ISL_470184, EPI_ISL_470185, EPI_ISL_470186, EPI_ISL_470187, EPI_ISL_470188, EPI_ISL_470190, EPI_ISL_470192, EPI_ISL_470194, EPI_ISL_470197, EPI_ISL_470198, EPI_ISL_470200, EPI_ISL_470201, EPI_ISL_470202, EPI_ISL_470203, EPI_ISL_470204, EPI_ISL_470205, EPI_ISL_470206, EPI_ISL_470208, EPI_ISL_470209, EPI_ISL_470212, EPI_ISL_470214, EPI_ISL_470215, EPI_ISL_470217, EPI_ISL_470218, EPI_ISL_470219, EPI_ISL_470221, EPI_ISL_470223, EPI_ISL_470225, EPI_ISL_470227, EPI_ISL_470229, EPI_ISL_470231, EPI_ISL_470232, EPI_ISL_470233, EPI_ISL_470235, EPI_ISL_470236, EPI_ISL_470237, EPI_ISL_470239, EPI_ISL_470241, EPI_ISL_470244, EPI_ISL_470245, EPI_ISL_470246, EPI_ISL_470248, EPI_ISL_470249, EPI_ISL_470250, EPI_ISL_470251, EPI_ISL_470252, EPI_ISL_470254, EPI_ISL_470255, EPI_ISL_470256, EPI_ISL_470258, EPI_ISL_470259, EPI_ISL_470260, EPI_ISL_470261, EPI_ISL_470265, EPI_ISL_470267, EPI_ISL_470268, EPI_ISL_470270, EPI_ISL_470271, EPI_ISL_470273, EPI_ISL_470274, EPI_ISL_470275, EPI_ISL_470276, EPI_ISL_470277, EPI_ISL_470278, EPI_ISL_470279, EPI_ISL_470280, EPI_ISL_470281, EPI_ISL_470282, EPI_ISL_470283, EPI_ISL_470284, EPI_ISL_470285, EPI_ISL_470286, EPI_ISL_470289, EPI_ISL_470290, EPI_ISL_470291, EPI_ISL_470292, EPI_ISL_470293, EPI_ISL_470297, EPI_ISL_470298, EPI_ISL_470299, EPI_ISL_470300, EPI_ISL_470302, EPI_ISL_470303, EPI_ISL_470307, EPI_ISL_470309, EPI_ISL_470310, EPI_ISL_470311, EPI_ISL_470312, EPI_ISL_470313, EPI_ISL_470314, EPI_ISL_470315, EPI_ISL_470316, EPI_ISL_470319, EPI_ISL_470320, EPI_ISL_470323, EPI_ISL_470324, EPI_ISL_470325, EPI_ISL_470328, EPI_ISL_470329, EPI_ISL_470330, EPI_ISL_470331, EPI_ISL_470332, EPI_ISL_470334, EPI_ISL_470335, EPI_ISL_470336, EPI_ISL_470340, EPI_ISL_470341, EPI_ISL_470342, EPI_ISL_470343, EPI_ISL_470344, EPI_ISL_470345, EPI_ISL_470347, EPI_ISL_470348, EPI_ISL_470349, EPI_ISL_470350, EPI_ISL_470351, EPI_ISL_470353 | | | |
| see above | Department of Pathology, University of Cambridge | Wellcome Sanger Institute for the COVID-19 Genomics UK Consortium | Luke W Meredith, M. Estée Török , Myra Hosmillo, William L. Hamilton, Martin D. Curran, Theresa Feltwell, Grant Hall, Anna Yakovleva, Fahad A Khokhar, Charlotte J. Houldcroft, Laura G Caller, Aminu S. Jahun, Sarah L. Caddy, Ian Goodfellow; and Alex Alderton, Roberto Amato, Sonia Goncalves, Ewan Harrison, David K. Jackson, Ian Johnston, Dominic Kwiatkowski, Cordelia Langford, John Sillitoe on behalf of the Wellcome Sanger Institute COVID-19 Surveillance Team (http://www.sanger.ac.uk/covid-team) |
| EPI_ISL_470356, EPI_ISL_470357 | PHE South West Regional Laboratory, National Infection Service | Wellcome Sanger Institute for the COVID-19 Genomics UK Consortium | Stephanie Hutchings, Hannah Pymont, Dr Peter Muir, Barry Vipond, Rich Hopes; and Alex Alderton, Roberto Amato, Sonia Goncalves, Ewan Harrison, David K. Jackson, Ian Johnston, Dominic Kwiatkowski, Cordelia Langford, John Sillitoe on behalf of the Wellcome Sanger Institute COVID-19 Surveillance Team (http://www.sanger.ac.uk/covid-team) |
| EPI_ISL_470359, EPI_ISL_470360, EPI_ISL_470361, EPI_ISL_470363, EPI_ISL_470364, EPI_ISL_470366, EPI_ISL_470367, EPI_ISL_470368, EPI_ISL_470370, EPI_ISL_470373, EPI_ISL_470374, EPI_ISL_470375, EPI_ISL_470379, EPI_ISL_470380, EPI_ISL_470381, EPI_ISL_470382, EPI_ISL_470383, EPI_ISL_470385, EPI_ISL_470386, EPI_ISL_470387, EPI_ISL_470388, EPI_ISL_470390, EPI_ISL_470392, EPI_ISL_470393, EPI_ISL_470394, EPI_ISL_470395, EPI_ISL_470397, EPI_ISL_470398, EPI_ISL_470400, EPI_ISL_470401, EPI_ISL_470402, EPI_ISL_470403, EPI_ISL_470404, EPI_ISL_470405, EPI_ISL_470406, EPI_ISL_470407, EPI_ISL_470408, EPI_ISL_470409, EPI_ISL_470411, EPI_ISL_470413, EPI_ISL_470416, EPI_ISL_470417, EPI_ISL_470418, EPI_ISL_470419, EPI_ISL_470424, EPI_ISL_470425, EPI_ISL_470426, EPI_ISL_470427, EPI_ISL_470430, EPI_ISL_470431, EPI_ISL_470432, EPI_ISL_470433, EPI_ISL_470434, EPI_ISL_470437, EPI_ISL_470438, EPI_ISL_470439, EPI_ISL_470440, EPI_ISL_470441, EPI_ISL_470442, EPI_ISL_470443, EPI_ISL_470445, EPI_ISL_470446, EPI_ISL_470447, EPI_ISL_470449, EPI_ISL_470451, EPI_ISL_470452, EPI_ISL_470453, EPI_ISL_470454, EPI_ISL_470455, EPI_ISL_470456, EPI_ISL_470457, EPI_ISL_470459, EPI_ISL_470460, EPI_ISL_470461, EPI_ISL_470462, EPI_ISL_470464, EPI_ISL_470465, EPI_ISL_470466, EPI_ISL_470468, EPI_ISL_470471, EPI_ISL_470472, EPI_ISL_470473, EPI_ISL_470474, EPI_ISL_470477, EPI_ISL_470478, EPI_ISL_470479, EPI_ISL_470480, EPI_ISL_470481, EPI_ISL_470482, EPI_ISL_470483, EPI_ISL_470485, EPI_ISL_470486, EPI_ISL_470487, EPI_ISL_470488, EPI_ISL_470490, EPI_ISL_470491, EPI_ISL_470495, EPI_ISL_470496, EPI_ISL_470497, EPI_ISL_470498, EPI_ISL_470499, EPI_ISL_470500, EPI_ISL_470501, EPI_ISL_470502, EPI_ISL_470503, EPI_ISL_470504, EPI_ISL_470505, EPI_ISL_470506, EPI_ISL_470507, EPI_ISL_470508, EPI_ISL_470510, EPI_ISL_470511, EPI_ISL_470512, EPI_ISL_470513, EPI_ISL_470514, EPI_ISL_470515, EPI_ISL_470516, EPI_ISL_470519, EPI_ISL_470520, EPI_ISL_470521, EPI_ISL_470522, EPI_ISL_470523, EPI_ISL_470524, EPI_ISL_470526, EPI_ISL_470527, EPI_ISL_470528 | | | |
| see above | Department of Pathology, University of Cambridge | Wellcome Sanger Institute for the COVID-19 Genomics UK Consortium | Luke W Meredith, M. Estée Török , Myra Hosmillo, William L. Hamilton, Martin D. Curran, Theresa Feltwell, Grant Hall, Anna Yakovleva, Fahad A Khokhar, Charlotte J. Houldcroft, Laura G Caller, Aminu S. Jahun, Sarah L. Caddy, Ian Goodfellow; and Alex Alderton, Roberto Amato, Sonia Goncalves, Ewan Harrison, David K. Jackson, Ian Johnston, Dominic Kwiatkowski, Cordelia Langford, John Sillitoe on behalf of the Wellcome Sanger Institute COVID-19 Surveillance Team (http://www.sanger.ac.uk/covid-team) |
| EPI_ISL_470529, EPI_ISL_470530 | PHE South West Regional Laboratory, National Infection Service | Wellcome Sanger Institute for the COVID-19 Genomics UK Consortium | Stephanie Hutchings, Hannah Pymont, Dr Peter Muir, Barry Vipond, Rich Hopes; and Alex Alderton, Roberto Amato, Sonia Goncalves, Ewan Harrison, David K. Jackson, Ian Johnston, Dominic |

| | | | |
|---|---|---|---|
| | | | Kwiatkowski, Cordelia Langford, John Sillitoe on behalf of the Wellcome Sanger Institute COVID-19 Surveillance Team (http://www.sanger.ac.uk/covid-team) |
| EPI_ISL_470531, EPI_ISL_470532, EPI_ISL_470533, EPI_ISL_470535, EPI_ISL_470536, EPI_ISL_470537 | Department of Pathology, University of Cambridge | Wellcome Sanger Institute for the COVID-19 Genomics UK Consortium | Luke W Meredith, M. Estée Török , Myra Hosmillo, William L. Hamilton, Martin D. Curran, Theresa Feltwell, Grant Hall, Anna Yakovleva, Fahad A Khokhar, Charlotte J. Houldcroft, Laura G Caller, Aminu S. Jahun, Sarah L. Caddy, Ian Goodfellow; and Alex Alderton, Roberto Amato, Sonia Goncalves, Ewan Harrison, David K. Jackson, Ian Johnston, Dominic Kwiatkowski, Cordelia Langford, John Sillitoe on behalf of the Wellcome Sanger Institute COVID-19 Surveillance Team (http://www.sanger.ac.uk/covid-team) |
| EPI_ISL_470539 | Molecular diagnostic laboratory of Federal Budget Institution of Science "Central Research Institute of Epidemiology" of The Federal Service on Customers' Rights Protection and Human Well-being Surveillance | Group of Genomics and Postgenomic Technologies of Central Research Institute of Epidemiology | Speranskaya AS, Kaptelova VV, Samoilov AE, Korneenko EV, Sizova TV, Tivanova EV, Shipulina OY, Akimkin VG |
| EPI_ISL_470540 | Wisconsin State Laboratory of Hygiene Communicable Disease Division | Wisconsin State Laboratory of Hygiene Communicable Disease Division | Kelsey R Florek |
| EPI_ISL_470544, EPI_ISL_470549, EPI_ISL_470550, EPI_ISL_470551, EPI_ISL_470552, EPI_ISL_470553, EPI_ISL_470554, EPI_ISL_470559, EPI_ISL_470565, EPI_ISL_470566, EPI_ISL_470656, EPI_ISL_470657, EPI_ISL_470659, EPI_ISL_470660, EPI_ISL_470662, EPI_ISL_470664, EPI_ISL_470666, EPI_ISL_470668, EPI_ISL_470669, EPI_ISL_470670, EPI_ISL_470673, EPI_ISL_470677, EPI_ISL_470678, EPI_ISL_470680, EPI_ISL_470681, EPI_ISL_470683, EPI_ISL_470684, EPI_ISL_470685, EPI_ISL_470686, EPI_ISL_470687, EPI_ISL_470688, EPI_ISL_470691, EPI_ISL_470694, EPI_ISL_470696, EPI_ISL_470697, EPI_ISL_470699, EPI_ISL_470700, EPI_ISL_470701, EPI_ISL_470703, EPI_ISL_470704, EPI_ISL_470705, EPI_ISL_470706, EPI_ISL_470708, EPI_ISL_470711, EPI_ISL_470712, EPI_ISL_470716, EPI_ISL_470717 | | | |
| see above | Utah Public Health Laboratory | Utah Public Health Laboratory | Erin Young, Kelly Oakeson |
| EPI_ISL_470721, EPI_ISL_470724, EPI_ISL_470725, EPI_ISL_470728, EPI_ISL_470732, EPI_ISL_470734, EPI_ISL_470738, EPI_ISL_470740, EPI_ISL_470741, EPI_ISL_470743, EPI_ISL_470745 | | | |
| see above | Utah Public Health Laboratory | Utah Public Health Laboratory | Heidi Butz, Erin Young, Kelly Oakeson |
| EPI_ISL_470747, EPI_ISL_470748, EPI_ISL_470749, EPI_ISL_470750, EPI_ISL_470751, EPI_ISL_470752, EPI_ISL_470753, EPI_ISL_470754, EPI_ISL_470755, EPI_ISL_470756, EPI_ISL_470757, EPI_ISL_470758, EPI_ISL_470759, EPI_ISL_470760, EPI_ISL_470761, EPI_ISL_470762, EPI_ISL_470763, EPI_ISL_470764, EPI_ISL_470765, EPI_ISL_470766, EPI_ISL_470767, EPI_ISL_470768, EPI_ISL_470769, EPI_ISL_470770, EPI_ISL_470771, EPI_ISL_470772, EPI_ISL_470773, EPI_ISL_470774, EPI_ISL_470775, EPI_ISL_470776, EPI_ISL_470777, EPI_ISL_470778, EPI_ISL_470779, EPI_ISL_470780, EPI_ISL_470781, EPI_ISL_470782, EPI_ISL_470783, EPI_ISL_470784, EPI_ISL_470785, EPI_ISL_470786, EPI_ISL_470787, EPI_ISL_470788, EPI_ISL_470789 | | | |
| see above | Minnesota Department of Health, Public Health Laboratory | Minnesota Department of Health, Public Health Laboratory | Matt Plumb, Jacob Garfin, and Xiong Wang |
| EPI_ISL_470790, EPI_ISL_470791, EPI_ISL_470792, EPI_ISL_470793, EPI_ISL_470794, EPI_ISL_470795, EPI_ISL_470796, EPI_ISL_470797, EPI_ISL_470798, EPI_ISL_470799, EPI_ISL_470800 | | | |
| see above | M Health Fairview | Minnesota Department of Health, Public Health Laboratory | Matt Plumb, Jacob Garfin, and Xiong Wang |
| EPI_ISL_470801 | Virology | Virology | Hossain,M.E., Hasan,R., Miah,M., Hasan,M.M., Sumaiya,M.K., Rahman,M.M., Alam,M.S., Clemens,J.D., Ahmed,T., Rahman,M.Z. and Rahman,M. |
| EPI_ISL_470830, EPI_ISL_470831, EPI_ISL_470833, EPI_ISL_470835, EPI_ISL_470839, EPI_ISL_470840, EPI_ISL_470841, EPI_ISL_470843, EPI_ISL_470845, EPI_ISL_470846, EPI_ISL_470847, EPI_ISL_470848, EPI_ISL_470849, EPI_ISL_470851, EPI_ISL_470853, EPI_ISL_470854, EPI_ISL_470855, EPI_ISL_470856, EPI_ISL_470858, EPI_ISL_470861, EPI_ISL_470862, EPI_ISL_470863, EPI_ISL_470864, EPI_ISL_470865, EPI_ISL_470867, EPI_ISL_470868, EPI_ISL_470869, EPI_ISL_470872, EPI_ISL_470873 | | | |
| see above | PathWest Laboratory Medicine WA | PathWest Laboratory Medicine WA | Chisha Sikazwe, Jurissa Lang, Avram Levy, David Smith and David Speers |
| EPI_ISL_470876 | Department for Virology, Molecular Biology and Genome Research, R. G. Lugar Center for Public Health Research, National Center for Disease Control and Public Health (NCDC) of Georgia. | Department for Virology, Molecular Biology and Genome Research, R. G. Lugar Center for Public Health Research, National Center for Disease Control and Public Health (NCDC) of Georgia. | Giorgi Tomashvili, Meri Pantsulaia, Gvantsa Brachveli, Gvantsa Chanturia, Ann Machablishvili, Nato Kotaria, Marine Murtskhvaladze, Lela Sabadze, Mari Gavashelidze, Ana Papkiauri, Tata Imnadze, Tamar Jashiashvili, Tea Tevdoradze, Ketevan Sidamonidze, Ekaterine Khmaladze, Ekaterine Zhghenti, Roena Sukhiashvili, Mariam Zakalashvili, Lela Urushadze, Magda Dgebuadze, Davit Tsaguria, Ekaterine Zangaladze, Nino Berishvili, Adam Kotorashvili, Maia Alkhazashvili, Irma Burjanadze, Anna Kasradze, Khatuna Zakhashvili, Paata Imnadze, Amiran Gamkrelidze. |
| EPI_ISL_470877 | Department for Virology, Molecular Biology and Genome Research, R. G. Lugar Center for Public Health Research, National Center for Disease Control and Public Health (NCDC) of Georgia. | Department for Virology, Molecular Biology and Genome Research, R. G. Lugar Center for Public Health Research, National Center for Disease Control and Public Health (NCDC) of Georgia. | Gvantsa Brachveli, Meri Pantsulaia, Giorgi Tomashvili, Gvantsa Chanturia, Ann Machablishvili, Nato Kotaria, Marine Murtskhvaladze, Lela Sabadze, Mari Gavashelidze, Ana Papkiauri, Tata Imnadze, Tamar Jashiashvili, Tea Tevdoradze, Ketevan Sidamonidze, Ekaterine Khmaladze, Ekaterine Zhghenti, Roena Sukhiashvili, Mariam Zakalashvili, Lela Urushadze, Magda Dgebuadze, Davit Tsaguria, Ekaterine Zangaladze, Nino Berishvili, Adam Kotorashvili, Maia Alkhazashvili, Irma Burjanadze, Anna Kasradze, Khatuna Zakhashvili, Paata Imnadze, Amiran Gamkrelidze. |
| EPI_ISL_470882 | Foerde Hospital, Department of Microbiology | Norwegian Institute of Public Health, Department of Virology | Kathrine Stene-Johansen, Kamilla Heddeland Instefjord, Hilde Elshaug, Rasmus Riis Kopperud, Karoline Bragstad, Olav Hungnes |
| EPI_ISL_470896 | Russian State Collection of Viruses | Pathogenic Microorganisms Variability Laboratory | Alexey Shchetinin, Maria Nikiforova, Elena Shidlovskaya, Nadezhda Kuznetsova, Inna Dolzhikova, Daria Grousova, Andrey Botikov, Denis Logunov, Alexander Gintsburg, Vladimir Gushchin |
| EPI_ISL_470897, EPI_ISL_470898, EPI_ISL_470899 | Pathogenic Microorganisms Variability Laboratory | Pathogenic Microorganisms Variability Laboratory | Alexey Shchetinin, Maria Nikiforova, Elena Shidlovskaya, Nadezhda Kuznetsova, Andrey Botikov, Alexander Gintsburg, Vladimir Gushchin |
| EPI_ISL_470900, EPI_ISL_470901, EPI_ISL_470902 | Influenza etiology and epidemiology laboratory | Pathogenic Microorganisms Variability Laboratory | Alexey Shchetinin, Maria Nikiforova, Elena Shidlovskaya, Nadezhda Kuznetsova, Vladimir Gushchin, Inna Dolzhikova, Daria Grousova, Andrey Botikov, Denis Logunov, Kirill Krasnoslobotsev, Svetlana Trushakova, Elena Burtseva, Ludmila Kolobukhina, Svetlana Smetanina, Alexander Gintsburg |
| EPI_ISL_470903, EPI_ISL_470904 | Influenza etiology and epidemiology laboratory | Pathogenic Microorganisms Variability Laboratory | Alexey Shchetinin, Maria Nikiforova, Elena Shidlovskaya, Nadezhda Kuznetsova, Vladimir Gushchin, Inna Dolzhikova, Daria Grousova, Andrey Botikov, Denis Logunov, Anna Ignatjeva, Evgeniya Mukasheva, Elena Burtseva, Ludmila Kolobukhina, Svetlana Smetanina, Alexander Gintsburg |
| EPI_ISL_471144, EPI_ISL_471145, EPI_ISL_471146, EPI_ISL_471147, EPI_ISL_471148, EPI_ISL_471149, EPI_ISL_471150, EPI_ISL_471151, EPI_ISL_471152, EPI_ISL_471153, EPI_ISL_471154, EPI_ISL_471155, EPI_ISL_471156 | | | |
| see above | Gundersen Molecular Diagnostics Laboratory | Kabara Cancer Research Institute | Craig S. Richmond, Paraic A. Kenny |
| EPI_ISL_471157 | Gundersen Clinical Microbiology Laboratory | Kabara Cancer Research Institute | Craig S. Richmond, Paraic A. Kenny |
| EPI_ISL_471158, EPI_ISL_471163, EPI_ISL_471164, EPI_ISL_471167, EPI_ISL_471171 | MRCG at LSHTM Genomics lab | MRCG at LSHTM Genomics lab | Sesay et al |
| EPI_ISL_471172 | Unilabs Laboratory Medicine | Norwegian Institute of Public Health, Department of Virology | Kathrine Stene-Johansen, Kamilla Heddeland Instefjord, Hilde Elshaug, Rasmus Riis Kopperud, Karoline Bragstad, Olav Hungnes |
| EPI_ISL_471173, EPI_ISL_471176 | Hospital of Southern Norway - Kristiansand, Department of Medical Microbiology | Norwegian Institute of Public Health, Department of Virology | Kathrine Stene-Johansen, Kamilla Heddeland Instefjord, Hilde Elshaug, Rasmus Riis Kopperud, Karoline Bragstad, Olav Hungnes |
| EPI_ISL_471177 | Oslo University Hospital, Department of Medical Microbiology | Norwegian Institute of Public Health, Department of Virology | Kathrine Stene-Johansen, Kamilla Heddeland Instefjord, Hilde Elshaug, Rasmus Riis Kopperud, Karoline Bragstad, Olav Hungnes |
| EPI_ISL_471178, EPI_ISL_471180, EPI_ISL_471181, EPI_ISL_471183, EPI_ISL_471185, EPI_ISL_471187, EPI_ISL_471188, EPI_ISL_471189, EPI_ISL_471190, EPI_ISL_471191, EPI_ISL_471192, EPI_ISL_471194, EPI_ISL_471196, EPI_ISL_471197, EPI_ISL_471199, EPI_ISL_471200, EPI_ISL_471201, EPI_ISL_471202, EPI_ISL_471203, EPI_ISL_471205, EPI_ISL_471206, EPI_ISL_471207, EPI_ISL_471208, EPI_ISL_471209, EPI_ISL_471211, EPI_ISL_471212, EPI_ISL_471213, EPI_ISL_471214, EPI_ISL_471215, EPI_ISL_471218, EPI_ISL_471219, EPI_ISL_471220, EPI_ISL_471221, EPI_ISL_471224, EPI_ISL_471225, EPI_ISL_471227, EPI_ISL_471229, EPI_ISL_471230, EPI_ISL_471231, EPI_ISL_471235, EPI_ISL_471240, EPI_ISL_471241, EPI_ISL_471242, EPI_ISL_471244, EPI_ISL_471245, EPI_ISL_471246, EPI_ISL_471247, EPI_ISL_471248, EPI_ISL_471249, EPI_ISL_471250, EPI_ISL_471251, EPI_ISL_471252, EPI_ISL_471253, EPI_ISL_471254, EPI_ISL_471255, EPI_ISL_471256, EPI_ISL_471258, EPI_ISL_471260, EPI_ISL_471261, EPI_ISL_471262, EPI_ISL_471264, EPI_ISL_471265, EPI_ISL_471266 | | | |
| see above | Wisconsin State Laboratory of Hygiene Communicable Disease Division | Wisconsin State Laboratory of Hygiene Communicable Disease Division | Kelsey R. Florek, Abigail C. Shockey |
| EPI_ISL_471267 | Hospital IESS Babahoyo | Institute of Microbiology, Universidad San Francisco de Quito | Sully Márquez, Belén Prado-Vivar, Juan José Guadalupe, Bernardo Gutiérrez, Francisco Cordova, Ninfa Henriquez, Killen Briones-Zamora, Killen Briones-Claudette, Verónica Barragán, Patricio Rojas-Silva, Gabriel Trueba, Michelle Grunauer, Paúl Cárdenas |
| EPI_ISL_471268 | Hospital IESS Babahoyo | Institute of Microbiology, Universidad San Francisco de Quito | Belén Prado-Vivar, Sully Márquez, Juan José Guadalupe, Bernardo Gutiérrez, Francisco Cordova, Ninfa Henriquez, Killen Briones-Zamora, Killen Briones-Claudette, Verónica Barragán, Patricio Rojas-Silva, Gabriel Trueba, Michelle Grunauer, Paúl Cárdenas |
| EPI_ISL_471269 | Hospital Oncológico Solca Núcleo de Quito | Institute of Microbiology, Universidad San Francisco de Quito | Sully Márquez, Belén Prado-Vivar, Juan José Guadalupe, Bernardo Gutiérrez, Marcos Di Stefano, Grace Salazar, Verónica Barragán, Patricio Rojas-Silva, Gabriel Trueba, Michelle Grunauer, Paúl Cárdenas |
| EPI_ISL_471270 | Hospital Oncológico Solca Núcleo de Quito | Institute of Microbiology, Universidad San Francisco de Quito | Sully Márquez, Belén Prado-Vivar, Juan José Guadalupe, Bernardo Gutiérrez, Marcos Di Stefano, Grace Salazar, Verónica Barragán, Patricio Rojas-Silva, Gabriel Trueba, Michelle Grunauer, Paúl Cárdenas |
| EPI_ISL_471271 | Hospital Oncológico Solca Núcleo de Quito | Institute of Microbiology, Universidad San Francisco de Quito | Sully Márquez, Belén Prado-Vivar, Juan José Guadalupe, Bernardo Gutiérrez, Marcos Di Stefano, Grace Salazar, Verónica Barragán, Patricio Rojas-Silva, Gabriel Trueba, Michelle Grunauer, Paúl Cárdenas |
| EPI_ISL_471396, EPI_ISL_471397, EPI_ISL_471398, EPI_ISL_471400, EPI_ISL_471401, EPI_ISL_471402, EPI_ISL_471403, EPI_ISL_471404, EPI_ISL_471405, EPI_ISL_471406, EPI_ISL_471407, EPI_ISL_471408, EPI_ISL_471409, EPI_ISL_471410, EPI_ISL_471411, EPI_ISL_471412, EPI_ISL_471414, EPI_ISL_471415 | | | |
| see above | Viral Respiratory Lab, National Institute for Biomedical Research (INRB) | Pathogen Sequencing Lab, National Institute for Biomedical Research (INRB) | Placide Mbala-Kingebeni, Edith Nkwembe, Eddy Kinganda-Lusamaki, Amuri Aziza, Francisca Muyembe Mawete, Catherine Pratt, Matthias Pauthner, Josh Quick, Allison Black, James Hadfield, Trevor Bedford, Ian Goodfellow, Andrew Rambaut, Nick Loman, Kristian Andersen, Michael Wiley, Steve Ahuka-Mundeke, Jean-Jacques Muyembe Tamfum |
| EPI_ISL_471416, EPI_ISL_471417, EPI_ISL_471419, EPI_ISL_471420, EPI_ISL_471422, EPI_ISL_471423, EPI_ISL_471424 | Laboratory for Respiratory Viruses, National Influenza Centre, Cantacuzino National Military-Medical Institute for Research and Development | Cantacuzino Institute | Luiza Ustea, Nicoleta Paraschiv, Tim Durfee, Mihaela Lazar |
| EPI_ISL_471425 | Division of Viral Diseases, Center for Laboratory Control of Infectious Diseases, Korea Centers for Diseases Control and Prevention | Division of Viral Diseases, Center for Laboratory Control of Infectious Diseases, Korea Centers for Diseases Control and Prevention | Jeong-Min Kim, Yoon-Seok Chung, Namjoo Lee, Mi-Seon Kim, Sang Hee Woo, Hye-Jun Jo, Sehee Park, Heui Man Kim, Jun-Sub Kim, Junhyeong Jang, Dong Hyun Song, Daesang Lee, Seong Tae Jeong, Myung Guk Han |
| EPI_ISL_471426 | Division of Viral Diseases, Center for Laboratory Control of Infectious Diseases, Korea Centers for Diseases Control and Prevention | Division of Viral Diseases, Center for Laboratory Control of Infectious Diseases, Korea Centers for Diseases Control and Prevention | Jeong-Min Kim, Yoon-Seok Chung, Namjoo Lee, Mi-Seon Kim, Sang Hee Woo, Hye-Jun Jo, Sehee Park, Heui Man Kim, Jun-Sub Kim, Junhyeong Jang, Dong Hyun Song, Daesang Lee, Seong Tae Jeong, Myung Guk Han |
| EPI_ISL_471427, EPI_ISL_471429, EPI_ISL_471430, EPI_ISL_471431, EPI_ISL_471435, EPI_ISL_471436 | Department of Clinical Microbiology | GIGA Medical Genomics | Keith Durkin, Maria Artesi, Sébastien Bontems, Raphaël Boreux, Cécile Meex, Axelle Chaslain, Céline Fombellida-Lopez, Pierrette Melin, Marie-Pierre Hayette, Vincent Bours. |
| EPI_ISL_471438, EPI_ISL_471439, EPI_ISL_471440, EPI_ISL_471441, EPI_ISL_471442, EPI_ISL_471443, EPI_ISL_471444 | Division of Viral Diseases, Center for Laboratory Control of Infectious Diseases, Korea Centers for Diseases Control and Prevention | Division of Viral Diseases, Center for Laboratory Control of Infectious Diseases, Korea Centers for Diseases Control and Prevention | Jeong-Min Kim, Yoon-Seok Chung, Namjoo Lee, Sang Hee Woo, Hye-Jun Jo, Heui Man Kim, Jun-Sub Kim, Dong Hyun Song, Daesang Lee, Seong Tae Jeong, Myung Guk Han |
| EPI_ISL_471445 | Division of Viral Diseases, Center for Laboratory Control of Infectious Diseases, Korea Centers for Diseases Control and Prevention | Division of Viral Diseases, Center for Laboratory Control of Infectious Diseases, Korea Centers for Diseases Control and Prevention | Jeong-Min Kim, Yoon-Seok Chung, Namjoo Lee, Sang Hee Woo, Hye-Jun Jo, Heui Man Kim, Jun-Sub Kim, Myung Guk Han |
| EPI_ISL_471446, EPI_ISL_471447, EPI_ISL_471448, EPI_ISL_471449, EPI_ISL_471450, EPI_ISL_471451, EPI_ISL_471452 | Division of Viral Diseases, Center for Laboratory Control of Infectious Diseases, Korea Centers for Diseases Control and Prevention | Division of Viral Diseases, Center for Laboratory Control of Infectious Diseases, Korea Centers for Diseases Control and Prevention | Jeong-Min Kim, Yoon-Seok Chung, Namjoo Lee, Sang Hee Woo, Hye-Jun Jo, Heui Man Kim, Jun-Sub Kim, Dong Hyun Song, Daesang Lee, Seong Tae Jeong, Myung Guk Han |
| EPI_ISL_471453 | Division of Viral Diseases, Center for Laboratory Control of Infectious Diseases, Korea Centers for Diseases Control and Prevention | Division of Viral Diseases, Center for Laboratory Control of Infectious Diseases, Korea Centers for Diseases Control and Prevention | Jeong-Min Kim, Yoon-Seok Chung, Namjoo Lee, Sang Hee Woo, Hye-Jun Jo, Heui Man Kim, Jun-Sub Kim, Myung Guk Han |
| EPI_ISL_471454, EPI_ISL_471455 | Division of Viral Diseases, Center for Laboratory Control of Infectious Diseases, Korea Centers for Diseases Control and Prevention | Division of Viral Diseases, Center for Laboratory Control of Infectious Diseases, Korea Centers for Diseases Control and Prevention | Jeong-Min Kim, Yoon-Seok Chung, Namjoo Lee, Sang Hee Woo, Hye-Jun Jo, Heui Man Kim, Jun-Sub Kim, Dong Hyun Song, Daesang Lee, Seong Tae Jeong, Myung Guk Han |
| EPI_ISL_471456, EPI_ISL_471457, | Centre de Virologie des Maladies Tropicales | Functional Genomic Platform/Service Analyses Biologique/UATRS/ Centre National | Hicham ANNAZ, Elmostafa EL FAHIME, Marouane MELLOUL, Yassine AKHOUAD, Mly Abdelaziz ELALAOUI, Ahmed REGGAD, Sanaa ALAOUI-Amine , Rachid ABI, Rida TAGAJDID, Zhor KASMY, Safaa |

| | | | |
|---|---|---|---|
| EPI_ISL_471458, EPI_ISL_471459, EPI_ISL_471460 | | Pour la Recherche Scientifique Et Technique (CNRST) | ELKORCHI, Nadia TOUIL, Farida HILALI, Abdelkader LAATIRIS , Abdelillah LARAQUI, Tahra BAJJOU , Yassine SEKHSOKH , Idriss-Amine LAHLOU, Mostafa ELOUENNASS, Khalid ENNIBI |
| EPI_ISL_471472 | Hospital Universitari Germans Trias i Pujol(HUGTiP)/Fundació Lluita contra la SIDA (FLSida)/IRTA-CReSA | IrsiCaixa AIDS Research Lab | Marc Noguera-Julian, Pilar Armengol, Jordi Rodón, Julia Vergara, Lidia Ruiz, Nuria Izquierdo, Jorge Carrillo, Roger Paredes, Albert Bensaid, Julia Blanco, Joaquim Segalés, Bonaventura Clotet |
| EPI_ISL_471510, EPI_ISL_471512, EPI_ISL_471515, EPI_ISL_471516, EPI_ISL_471517, EPI_ISL_471518, EPI_ISL_471520, EPI_ISL_471521, EPI_ISL_471524, EPI_ISL_471527 | Respiratory Virus Unit, Microbiology Services Colindale, Public Health England | Respiratory Virus Unit, Microbiology Services Colindale, Public Health England | PHE Covid Sequencing Team |
| EPI_ISL_471528, EPI_ISL_471530 | The National Institute of Public Health | State Veterinary Institute Prague and The National Institute of Public Health | Nagy,A;Jirincova,H;Novakova,L;Trnka,D;Vecerova,J |
| EPI_ISL_471539 | Hospital Universitario da USP Sao Paulo | Instituto Adolfo Lutz, Interdiciplinary Procedures Center, Strategic Laboratory | Claudio Tavares Sacchi, Claudia Regina Gonçalves, Erica Valessa Ramos Gomes |
| EPI_ISL_471540 | The National Institute of Public Health | State Veterinary Institute Prague and The National Institute of Public Health | Nagy,A;Jirincova,H;Novakova,L;Trnka,D;Vecerova,J |
| EPI_ISL_471541 | Hospital Geral Santa Marcelina | Instituto Adolfo Lutz, Interdiciplinary Procedures Center, Strategic Laboratory | Claudio Tavares Sacchi, Claudia Regina Gonçalves, Erica Valessa Ramos Gomes |
| EPI_ISL_471542 | Secretaria de Saude de Mogi das Cruzes | Instituto Adolfo Lutz, Interdiciplinary Procedures Center, Strategic Laboratory | Claudio Tavares Sacchi, Claudia Regina Gonçalves, Erica Valessa Ramos Gomes |
| EPI_ISL_471543 | Centro de Saude I Tacito Leite de Carvalho e Silva | Instituto Adolfo Lutz, Interdiciplinary Procedures Center, Strategic Laboratory | Claudio Tavares Sacchi, Claudia Regina Gonçalves, Erica Valessa Ramos Gomes |
| EPI_ISL_471544 | The National Institute of Public Health | State Veterinary Institute Prague and The National Institute of Public Health | Nagy,A;Jirincova,H;Novakova,L;Trnka,D;Vecerova,J |
| EPI_ISL_471545 | Hospital Sao Paulo de Ensino da Unifesp | Instituto Adolfo Lutz, Interdiciplinary Procedures Center, Strategic Laboratory | Claudio Tavares Sacchi, Claudia Regina Gonçalves, Erica Valessa Ramos Gomes |
| EPI_ISL_471546 | AMA DR Jose Soares Hungria | Instituto Adolfo Lutz, Interdiciplinary Procedures Center, Strategic Laboratory | Claudio Tavares Sacchi, Claudia Regina Gonçalves, Erica Valessa Ramos Gomes |
| EPI_ISL_471547 | The National Institute of Public Health | State Veterinary Institute Prague and The National Institute of Public Health | Nagy,A;Jirincova,H;Novakova,L;Trnka,D;Vecerova,J |
| EPI_ISL_471548 | Hospital do Servidor Público Estadual Francisco Morato de Oliveira | Instituto Adolfo Lutz, Interdiciplinary Procedures Center, Strategic Laboratory | Claudio Tavares Sacchi, Claudia Regina Gonçalves, Erica Valessa Ramos Gomes |
| EPI_ISL_471549 | Hospital Municipal Carmen Prudente | Instituto Adolfo Lutz, Interdiciplinary Procedures Center, Strategic Laboratory | Claudio Tavares Sacchi, Claudia Regina Gonçalves, Erica Valessa Ramos Gomes |
| EPI_ISL_471550 | The National Institute of Public Health | State Veterinary Institute Prague and The National Institute of Public Health | Nagy,A;Jirincova,H;Novakova,L;Trnka,D;Vecerova,J |
| EPI_ISL_471551 | Hospital Sao Paulo de Ensino da Unifesp | Instituto Adolfo Lutz, Interdiciplinary Procedures Center, Strategic Laboratory | Claudio Tavares Sacchi, Claudia Regina Gonçalves, Erica Valessa Ramos Gomes |
| EPI_ISL_471552 | Hospital Sancta Maggiore | Instituto Adolfo Lutz, Interdiciplinary Procedures Center, Strategic Laboratory | Claudio Tavares Sacchi, Claudia Regina Gonçalves, Erica Valessa Ramos Gomes |
| EPI_ISL_471553 | The National Institute of Public Health | State Veterinary Institute Prague and The National Institute of Public Health | Nagy,A;Jirincova,H;Novakova,L;Trnka,D;Vecerova,J |
| EPI_ISL_471554 | Hospital Bosque da Saúde | Instituto Adolfo Lutz, Interdiciplinary Procedures Center, Strategic Laboratory | Claudio Tavares Sacchi, Claudia Regina Gonçalves, Erica Valessa Ramos Gomes |
| EPI_ISL_471555 | The National Institute of Public Health | State Veterinary Institute Prague and The National Institute of Public Health | Nagy,A;Jirincova,H;Novakova,L;Trnka,D;Vecerova,J |
| EPI_ISL_471556 | Pronto Socorro Jose Ibrahin | Instituto Adolfo Lutz, Interdiciplinary Procedures Center, Strategic Laboratory | Claudio Tavares Sacchi, Claudia Regina Gonçalves, Erica Valessa Ramos Gomes |
| EPI_ISL_471562, EPI_ISL_471581, EPI_ISL_471582 | Hosp. Municipal Prof. Dr. Alípio Corrêa Netto | Instituto Adolfo Lutz, Interdiciplinary Procedures Center, Strategic Laboratory | Claudio Tavares Sacchi, Claudia Regina Gonçalves, Erica Valessa Ramos Gomes |
| EPI_ISL_471583, EPI_ISL_471584 | King Institute of Preventive Medicine & Research | CSIR-Centre for Cellular and Molecular Biology | K.Kaveri,S.Sivasubramanian,S.Vennila,P.Padmapriya,R.Kiruba,S.Magesh,G. Dhinakar Raj, G. Ravikumar, P. Azhahianambi,K Thangaraj,Payel Mukherjee, Sofia Banu, Priya Singh, Dhiviya Vedagiri, Divya Gupta, Vishal Sah, Santosh Kumar Kuncha, Krishnan Harinivas Harshan, Archana Bharadwaj Siva, Karthik Bharadwaj Tallapaka, Shagufta Khan, Lamuk Zaveri, Namami Gaur, Sakshi Shambhavi, Tulasi Nagabandi, Purushotham Vodnala, Rakesh K Mishra, Divya Tej Sowpati |
| EPI_ISL_471585 | CSIR-Centre for Cellular and Molecular Biology | CSIR-Centre for Cellular and Molecular Biology | Dhiviya Vedagiri, Divya Gupta, Vishal Sah, Payel Mukherjee, Sofia Banu, Priya Singh, Santosh Kumar Kuncha, Archana Bharadwaj Siva, Karthik Bharadwaj Tallapaka, Shagufta Khan, Lamuk Zaveri, Namami Gaur, Sakshi Shambhavi, Tulasi Nagabandi, Purushotham Vodnala, Rakesh K Mishra, Divya Tej Sowpati, Krishnan Harinivas Harshan |
| EPI_ISL_471586 | CSIR-Centre for Cellular and Molecular Biology | CSIR-Centre for Cellular and Molecular Biology | Lamuk Zaveri, Shagufta Khan, Namami Gaur, Sakshi Shambhavi, Tulasi Nagabandi, Purushotham Vodnala, Payel Mukherjee, Sofia Banu, Priya Singh, Dhiviya Vedagiri, Divya Gupta, Vishal Sah, Santosh Kumar Kuncha, Krishnan Harinivas Harshan, Archana Bharadwaj Siva, Karthik Bharadwaj Tallapaka,Zeba Rizvi, Zuberwasim Sayyad, Kakade Aishwarya Arun, Amrutha H C, Ananga Ghosh, Rakesh K Mishra, Divya Tej Sowpati |
| EPI_ISL_471587 | CSIR-Centre for Cellular and Molecular Biology | CSIR-Centre for Cellular and Molecular Biology | Dhiviya Vedagiri, Divya Gupta, Vishal Sah, Payel Mukherjee, Sofia Banu, Priya Singh, Santosh Kumar Kuncha, Archana Bharadwaj Siva, Karthik Bharadwaj Tallapaka, Shagufta Khan, Lamuk Zaveri, Namami Gaur, Sakshi Shambhavi, Tulasi Nagabandi, Purushotham Vodnala, Rakesh K Mishra, Divya Tej Sowpati, Krishnan Harinivas Harshan |
| EPI_ISL_471588 | CSIR-Centre for Cellular and Molecular Biology | CSIR-Centre for Cellular and Molecular Biology | Lamuk Zaveri, Shagufta Khan, Namami Gaur, Sakshi Shambhavi, Tulasi Nagabandi, Purushotham Vodnala, Payel Mukherjee, Sofia Banu, Priya Singh, Dhiviya Vedagiri, Divya Gupta, Vishal Sah, Santosh Kumar Kuncha, Krishnan Harinivas Harshan, Archana Bharadwaj Siva, Karthik Bharadwaj Tallapaka, Renu Sudhakar, Somesh Gorde, Gangumala Srinivas Reddy, Sujoy Deb, Swati Bayyana, Rakesh K Mishra, Divya Tej Sowpati |
| EPI_ISL_471589 | CSIR-Centre for Cellular and Molecular Biology | CSIR-Centre for Cellular and Molecular Biology | Lamuk Zaveri, Shagufta Khan, Namami Gaur, Sakshi Shambhavi, Tulasi Nagabandi, Purushotham Vodnala, Payel Mukherjee, Sofia Banu, Priya Singh, Dhiviya Vedagiri, Divya Gupta, Vishal Sah, Santosh Kumar Kuncha, Krishnan Harinivas Harshan, Archana Bharadwaj Siva, Karthik Bharadwaj Tallapaka,Umesh Kumar, Unis Ahmad Bhat, Ajay Sarawagi, Priyanka Pant, Rajkanwar Nathawat, Rakesh K Mishra, Divya Tej Sowpati |
| EPI_ISL_471590 | CSIR-Centre for Cellular and Molecular Biology | CSIR-Centre for Cellular and Molecular Biology | Lamuk Zaveri, Shagufta Khan, Namami Gaur, Sakshi Shambhavi, Tulasi Nagabandi, Purushotham Vodnala, Payel Mukherjee, Sofia Banu, Priya Singh, Dhiviya Vedagiri, Divya Gupta, Vishal Sah, Santosh Kumar Kuncha, Krishnan Harinivas Harshan, Archana Bharadwaj Siva, Karthik Bharadwaj Tallapaka,Zeba Rizvi, Zuberwasim Sayyad, Kakade Aishwarya Arun, Amrutha H C, Ananga Ghosh, Rakesh K Mishra, Divya Tej Sowpati |
| EPI_ISL_471591 | CSIR-Centre for Cellular and Molecular Biology | CSIR-Centre for Cellular and Molecular Biology | Namami Gaur, Sakshi Shambhavi, Lamuk Zaveri, Shagufta Khan, Tulasi Nagabandi, Purushotham Vodnala, Payel Mukherjee, Sofia Banu, Priya Singh, Dhiviya Vedagiri, Divya Gupta, Vishal Sah, Santosh Kumar Kuncha, Krishnan Harinivas Harshan, Archana Bharadwaj Siva, Karthik Bharadwaj Tallapaka, Zeba Rizvi, Zuberwasim Sayyad, Kakade Aishwarya Arun, Amrutha H C, Ananga Ghosh, Rakesh K Mishra, Divya Tej Sowpati |
| EPI_ISL_471592 | CSIR-Centre for Cellular and Molecular Biology | CSIR-Centre for Cellular and Molecular Biology | Namami Gaur, Sakshi Shambhavi, Lamuk Zaveri, Shagufta Khan, Tulasi Nagabandi, Purushotham Vodnala, Payel Mukherjee, Sofia Banu, Priya Singh, Dhiviya Vedagiri, Divya Gupta, Vishal Sah, Santosh Kumar Kuncha, Krishnan Harinivas Harshan, Archana Bharadwaj Siva, Karthik Bharadwaj Tallapaka, Nikhil Hajirnis, Pratheusa Maccha, M Soujanya Reddy,G. Aditya Kumar, Koushick Sivakumar, Rakesh K Mishra, Divya Tej Sowpati |
| EPI_ISL_471593 | CSIR-Centre for Cellular and Molecular Biology | CSIR-Centre for Cellular and Molecular Biology | Namami Gaur, Sakshi Shambhavi, Lamuk Zaveri, Shagufta Khan, Tulasi Nagabandi, Purushotham Vodnala, Payel Mukherjee, Sofia Banu, Priya Singh, Dhiviya Vedagiri, Divya Gupta, Vishal Sah, Santosh Kumar Kuncha, Krishnan Harinivas Harshan, Archana Bharadwaj Siva, Karthik Bharadwaj Tallapaka, Zeba Rizvi, Zuberwasim Sayyad, Kakade Aishwarya Arun, Amrutha H C, Ananga Ghosh, Rakesh K Mishra, Divya Tej Sowpati |
| EPI_ISL_471594 | CSIR-Centre for Cellular and Molecular Biology | CSIR-Centre for Cellular and Molecular Biology | Namami Gaur, Sakshi Shambhavi, Lamuk Zaveri, Shagufta Khan, Tulasi Nagabandi, Purushotham Vodnala, Payel Mukherjee, Sofia Banu, Priya Singh, Dhiviya Vedagiri, Divya Gupta, Vishal Sah, Santosh Kumar Kuncha, Krishnan Harinivas Harshan, Archana Bharadwaj Siva, Karthik Bharadwaj Tallapaka,Kezia J Ann, Radhika Khandelwal, Roshan Maku Venkata, Shemin Mansuri, Sonu Uday, Rakesh K Mishra, Divya Tej Sowpati |
| EPI_ISL_471595 | CSIR-Centre for Cellular and Molecular Biology | CSIR-Centre for Cellular and Molecular Biology | Payel Mukherjee, Sofia Banu, Priya Singh, Dhiviya Vedagiri, Divya Gupta, Vishal Sah, Santosh Kumar Kuncha, Krishnan Harinivas Harshan, Archana Bharadwaj Siva, Karthik Bharadwaj Tallapaka, Shagufta Khan, Lamuk Zaveri, Namami Gaur, Sakshi Shambhavi, Tulasi Nagabandi, Purushotham Vodnala, G. Aditya Kumar, Koushick Sivakumar, Pooja Ramesh Gupta, Rajan Kumar Jha, Shraddha Vijay Lahoti, Rakesh K Mishra, Divya Tej Sowpati |
| EPI_ISL_471596 | CSIR-Centre for Cellular and Molecular Biology | CSIR-Centre for Cellular and Molecular Biology | Payel Mukherjee, Sofia Banu, Priya Singh, Dhiviya Vedagiri, Divya Gupta, Vishal Sah, Santosh Kumar Kuncha, Krishnan Harinivas Harshan, Archana Bharadwaj Siva, Karthik Bharadwaj Tallapaka, Shagufta Khan, Lamuk Zaveri, Namami Gaur, Sakshi Shambhavi, Tulasi Nagabandi, Purushotham Vodnala, Gokulan C G, Gunjan Purohit, Hanuman Tulashiram Kale, Pankaj Kumar, Prachand Issarapu, Rakesh K Mishra, Divya Tej Sowpati |
| EPI_ISL_471597 | CSIR-Centre for Cellular and Molecular Biology | CSIR-Centre for Cellular and Molecular Biology | Payel Mukherjee, Sofia Banu, Priya Singh, Dhiviya Vedagiri, Divya Gupta, Vishal Sah, Santosh Kumar Kuncha, Krishnan Harinivas Harshan, Archana Bharadwaj Siva, Karthik Bharadwaj Tallapaka, Shagufta Khan, Lamuk Zaveri, Namami Gaur, Sakshi Shambhavi, Tulasi Nagabandi, Purushotham Vodnala, Rakesh K Mishra, Sonu Uday, Sudipta Mondal, Annapoorna P Karthyayani, Debabrata Jana, Debrya Saha, Divya Tej Sowpati |
| EPI_ISL_471598 | CSIR-Centre for Cellular and Molecular Biology | CSIR-Centre for Cellular and Molecular Biology | Payel Mukherjee, Sofia Banu, Priya Singh, Dhiviya Vedagiri, Divya Gupta, Vishal Sah, Santosh Kumar Kuncha, Krishnan Harinivas Harshan, Archana Bharadwaj Siva, Karthik Bharadwaj Tallapaka, Shagufta Khan, Lamuk Zaveri, Namami Gaur, Sakshi Shambhavi, Tulasi Nagabandi, Purushotham Vodnala,Deepak Kumar, Devi Prasad Vijayashankar, Disha Nanda, Divya Das, Jotin Gogoi, Manish Bhattacharjee, Rakesh K Mishra, Divya Tej Sowpati |
| EPI_ISL_471599 | CSIR-Centre for Cellular and Molecular Biology | CSIR-Centre for Cellular and Molecular Biology | Sakshi Shambhavi, Lamuk Zaveri, Shagufta Khan, Namami Gaur, Tulasi Nagabandi, Purushotham Vodnala, Payel Mukherjee, Sofia Banu, Priya Singh, Dhiviya Vedagiri, Divya Gupta, Vishal Sah, Santosh Kumar Kuncha, Krishnan Harinivas Harshan, Archana Bharadwaj Siva, Karthik Bharadwaj Tallapaka, Deepak Kumar, Devi Prasad Vijayashankar, Disha Nanda, Divya Das, Jotin Gogoi, Manish Bhattacharjee, Rakesh K Mishra, Divya Tej Sowpati |
| EPI_ISL_471600 | CSIR-Centre for Cellular and Molecular Biology | CSIR-Centre for Cellular and Molecular Biology | Sakshi Shambhavi, Lamuk Zaveri, Shagufta Khan, Namami Gaur, Tulasi Nagabandi, Purushotham Vodnala, Payel Mukherjee, Sofia Banu, Priya Singh, Dhiviya Vedagiri, Divya Gupta, Vishal Sah, Santosh Kumar Kuncha, Krishnan Harinivas Harshan, Archana Bharadwaj Siva, Karthik Bharadwaj Tallapaka, G. Aditya Kumar, Koushick Sivakumar, Pooja Ramesh Gupta, Rajan Kumar Jha, Shraddha Vijay Lahoti, Rakesh K Mishra, Divya Tej Sowpati |
| EPI_ISL_471601 | CSIR-Centre for Cellular and Molecular Biology | CSIR-Centre for Cellular and Molecular Biology | Sakshi Shambhavi, Lamuk Zaveri, Shagufta Khan, Namami Gaur, Tulasi Nagabandi, Purushotham Vodnala, Payel Mukherjee, Sofia Banu, Priya Singh, Dhiviya Vedagiri, Divya Gupta, Vishal Sah, Santosh Kumar Kuncha, Krishnan Harinivas Harshan, Archana Bharadwaj Siva, Karthik Bharadwaj Tallapaka,Nikhil Hajirnis, Pratheusa Maccha, M Soujanya Reddy,G. Aditya Kumar, Koushick Sivakumar, Rakesh K Mishra, Divya Tej Sowpati |
| EPI_ISL_471602 | CSIR-Centre for Cellular and Molecular Biology | CSIR-Centre for Cellular and Molecular Biology | Sakshi Shambhavi, Lamuk Zaveri, Shagufta Khan, Namami Gaur, Tulasi Nagabandi, Purushotham Vodnala, Payel Mukherjee, Sofia Banu, Priya Singh, Dhiviya Vedagiri, Divya Gupta, Vishal Sah, Santosh Kumar Kuncha, Krishnan Harinivas Harshan, Archana Bharadwaj Siva, Karthik Bharadwaj Tallapaka,Nikhil Hajirnis, Pratheusa Maccha, M Soujanya Reddy,G. Aditya Kumar, Koushick Sivakumar,Disha Nanda, Divya Das, Jotin Gogoi, Manish Bhattacharjee, Ravi Prasad Mukku, Rakesh K Mishra, Divya Tej Sowpati |
| EPI_ISL_471603 | CSIR-Centre for Cellular and Molecular Biology | CSIR-Centre for Cellular and Molecular Biology | Shagufta Khan, Lamuk Zaveri, Namami Gaur, Sakshi Shambhavi, Tulasi Nagabandi, Purushotham Vodnala, Payel Mukherjee, Sofia Banu, Priya Singh, Dhiviya Vedagiri, Divya Gupta, Vishal Sah, Santosh Kumar Kuncha, Krishnan Harinivas Harshan, Archana Bharadwaj Siva, Karthik Bharadwaj Tallapaka, Disha Nanda, Divya Das, Jotin Gogoi, Manish Bhattacharjee, Ravi Prasad Mukku, Rakesh K Mishra, Divya Tej Sowpati |
| EPI_ISL_471604 | CSIR-Centre for Cellular and Molecular Biology | CSIR-Centre for Cellular and Molecular Biology | Shagufta Khan, Lamuk Zaveri, Namami Gaur, Sakshi Shambhavi, Tulasi Nagabandi, Purushotham Vodnala, Payel Mukherjee, Sofia Banu, Priya Singh, Dhiviya Vedagiri, Divya Gupta, Vishal Sah, Santosh Kumar Kuncha, Krishnan Harinivas Harshan, Archana Bharadwaj Siva, Karthik Bharadwaj Tallapaka, Renu Sudhakar, Somesh Gorde, Gangumala Srinivas Reddy, Sujoy Deb, Swati Bayyana, Rakesh K Mishra, Divya Tej Sowpati |

| | | | |
|---|---|---|---|
| EPI_ISL_471605 | CSIR-Centre for Cellular and Molecular Biology | CSIR-Centre for Cellular and Molecular Biology | Shagufta Khan, Lamuk Zaveri, Namami Gaur, Sakshi Shambhavi, Tulasi Nagabandi, Purushotham Vodnala, Payel Mukherjee, Sofia Banu, Priya Singh, Dhiviya Vedagiri, Divya Gupta, Vishal Sah, Santosh Kumar Kuncha, Krishnan Harinivas Harshan, Archana Bharadwaj Siva, Karthik Bharadwaj Tallapaka,Preethi Jampala, Sharada Ravi Iyer, Sulagana Mukherjee, Swetha Sundar, Peddapuvala Sai Uday Kiran Rakesh K Mishra, Divya Tej Sowpati |
| EPI_ISL_471606 | CSIR-Centre for Cellular and Molecular Biology | CSIR-Centre for Cellular and Molecular Biology | Shagufta Khan, Lamuk Zaveri, Namami Gaur, Sakshi Shambhavi, Tulasi Nagabandi, Purushotham Vodnala, Payel Mukherjee, Sofia Banu, Priya Singh, Dhiviya Vedagiri, Divya Gupta, Vishal Sah, Santosh Kumar Kuncha, Krishnan Harinivas Harshan, Archana Bharadwaj Siva, Karthik Bharadwaj Tallapaka,Umesh Kumar, Unis Ahmad Bhat, Ajay Sarawagi, Priyanka Pant, Rajkanwar Nathawat, Rakesh K Mishra, Divya Tej Sowpati |
| EPI_ISL_471607 | CSIR-Centre for Cellular and Molecular Biology | CSIR-Centre for Cellular and Molecular Biology | Sofia Banu, Payel Mukherjee, Priya Singh, Dhiviya Vedagiri, Divya Gupta, Vishal Sah, Santosh Kumar Kuncha, Krishnan Harinivas Harshan, Archana Bharadwaj Siva, Karthik Bharadwaj Tallapaka, Shagufta Khan, Lamuk Zaveri, Namami Gaur, Sakshi Shambhavi, Tulasi Nagabandi, Purushotham Vodnala, Deepak Kumar, Devi Prasad Vijayashankar, Disha Nanda, Divya Das, Jotin Gogoi, Manish Bhattacharjee, Rakesh K Mishra, Divya Tej Sowpati |
| EPI_ISL_471608 | CSIR-Centre for Cellular and Molecular Biology | CSIR-Centre for Cellular and Molecular Biology | Sofia Banu, Payel Mukherjee, Priya Singh, Dhiviya Vedagiri, Divya Gupta, Vishal Sah, Santosh Kumar Kuncha, Krishnan Harinivas Harshan, Archana Bharadwaj Siva, Karthik Bharadwaj Tallapaka, Shagufta Khan, Lamuk Zaveri, Namami Gaur, Sakshi Shambhavi, Tulasi Nagabandi, Purushotham Vodnala, Disha Nanda, Divya Das, Jotin Gogoi, Manish Bhattacharjee, Ravi Prasad Mukku, Rakesh K Mishra, Divya Tej Sowpati |
| EPI_ISL_471609 | CSIR-Centre for Cellular and Molecular Biology | CSIR-Centre for Cellular and Molecular Biology | Sofia Banu, Payel Mukherjee, Priya Singh, Dhiviya Vedagiri, Divya Gupta, Vishal Sah, Santosh Kumar Kuncha, Krishnan Harinivas Harshan, Archana Bharadwaj Siva, Karthik Bharadwaj Tallapaka, Shagufta Khan, Lamuk Zaveri, Namami Gaur, Sakshi Shambhavi, Tulasi Nagabandi, Purushotham Vodnala, Gokulan C G, Gunjan Purohit, Hanuman Tulashiram Kale, Pankaj Kumar, Prachand Issarapu, Rakesh K Mishra, Divya Tej Sowpati |
| EPI_ISL_471610 | CSIR-Centre for Cellular and Molecular Biology | CSIR-Centre for Cellular and Molecular Biology | Sofia Banu, Payel Mukherjee, Priya Singh, Dhiviya Vedagiri, Divya Gupta, Vishal Sah, Santosh Kumar Kuncha, Krishnan Harinivas Harshan, Archana Bharadwaj Siva, Karthik Bharadwaj Tallapaka, Shagufta Khan, Lamuk Zaveri, Namami Gaur, Sakshi Shambhavi, Tulasi Nagabandi, Purushotham Vodnala,Preethi Jampala, Sharada Ravi Iyer, Sulagana Mukherjee, Swetha Sundar, Peddapuvala Sai Uday Kiran, Rakesh K Mishra, Divya Tej Sowpati |
| EPI_ISL_471611 | CSIR-Centre for Cellular and Molecular Biology | CSIR-Centre for Cellular and Molecular Biology | Tulasi Nagabandi, Namami Gaur, Sakshi Shambhavi, Lamuk Zaveri, Shagufta Khan, Purushotham Vodnala, Payel Mukherjee, Sofia Banu, Priya Singh, Dhiviya Vedagiri, Divya Gupta, Vishal Sah, Santosh Kumar Kuncha, Krishnan Harinivas Harshan, Archana Bharadwaj Siva, Karthik Bharadwaj Tallapaka,G. Aditya Kumar, Koushick Sivakumar, Pooja Ramesh Gupta, Rajan Kumar Jha, Shraddha Vijay Lahoti, Rakesh K Mishra, Divya Tej Sowpati |
| EPI_ISL_471612 | CSIR-Centre for Cellular and Molecular Biology | CSIR-Centre for Cellular and Molecular Biology | Tulasi Nagabandi, Namami Gaur, Sakshi Shambhavi, Lamuk Zaveri, Shagufta Khan, Purushotham Vodnala, Payel Mukherjee, Sofia Banu, Priya Singh, Dhiviya Vedagiri, Divya Gupta, Vishal Sah, Santosh Kumar Kuncha, Krishnan Harinivas Harshan, Archana Bharadwaj Siva, Karthik Bharadwaj Tallapaka,Kezia J Ann, Radhika Khandelwal, Roshan Maku Venkata, Shemin Mansuri, Sonu Uday, Rakesh K Mishra, Divya Tej Sowpati |
| EPI_ISL_471613 | CSIR-Centre for Cellular and Molecular Biology | CSIR-Centre for Cellular and Molecular Biology | Tulasi Nagabandi, Namami Gaur, Sakshi Shambhavi, Lamuk Zaveri, Shagufta Khan, Purushotham Vodnala, Payel Mukherjee, Sofia Banu, Priya Singh, Dhiviya Vedagiri, Divya Gupta, Vishal Sah, Santosh Kumar Kuncha, Krishnan Harinivas Harshan, Archana Bharadwaj Siva, Karthik Bharadwaj Tallapaka,G. Aditya Kumar, Koushick Sivakumar, Pooja Ramesh Gupta, Rajan Kumar Jha, Shraddha Vijay Lahoti, Rakesh K Mishra, Divya Tej Sowpati |
| EPI_ISL_471614 | CSIR-Centre for Cellular and Molecular Biology | CSIR-Centre for Cellular and Molecular Biology | Tulasi Nagabandi, Namami Gaur, Sakshi Shambhavi, Lamuk Zaveri, Shagufta Khan, Purushotham Vodnala, Payel Mukherjee, Sofia Banu, Priya Singh, Dhiviya Vedagiri, Divya Gupta, Vishal Sah, Santosh Kumar Kuncha, Krishnan Harinivas Harshan, Archana Bharadwaj Siva, Karthik Bharadwaj Tallapaka,Kezia J Ann, Radhika Khandelwal, Roshan Maku Venkata, Shemin Mansuri, Sonu Uday, Rakesh K Mishra, Divya Tej Sowpati |
| EPI_ISL_471615 | CSIR-Centre for Cellular and Molecular Biology | CSIR-Centre for Cellular and Molecular Biology | Lamuk Zaveri, Shagufta Khan, Namami Gaur, Sakshi Shambhavi, Tulasi Nagabandi, Purushotham Vodnala, Payel Mukherjee, Sofia Banu, Priya Singh, Dhiviya Vedagiri, Divya Gupta, Vishal Sah, Santosh Kumar Kuncha, Krishnan Harinivas Harshan, Archana Bharadwaj Siva, Karthik Bharadwaj Tallapaka,Zeba Rizvi, Zuberwasim Sayyad, Kakade Aishwarya Arun, Amrutha H C, Ananga Ghosh, Rakesh K Mishra, Divya Tej Sowpati |
| EPI_ISL_471616 | CSIR-Centre for Cellular and Molecular Biology | CSIR-Centre for Cellular and Molecular Biology | Lamuk Zaveri, Shagufta Khan, Namami Gaur, Sakshi Shambhavi, Tulasi Nagabandi, Purushotham Vodnala, Payel Mukherjee, Sofia Banu, Priya Singh, Dhiviya Vedagiri, Divya Gupta, Vishal Sah, Santosh Kumar Kuncha, Krishnan Harinivas Harshan, Archana Bharadwaj Siva, Karthik Bharadwaj Tallapaka, Renu Sudhakar, Somesh Gorde, Gangumala Srinivas Reddy, Sujoy Deb, Swati Bayyana, Rakesh K Mishra, Divya Tej Sowpati |
| EPI_ISL_471617 | CSIR-Centre for Cellular and Molecular Biology | CSIR-Centre for Cellular and Molecular Biology | Lamuk Zaveri, Shagufta Khan, Namami Gaur, Sakshi Shambhavi, Tulasi Nagabandi, Purushotham Vodnala, Payel Mukherjee, Sofia Banu, Priya Singh, Dhiviya Vedagiri, Divya Gupta, Vishal Sah, Santosh Kumar Kuncha, Krishnan Harinivas Harshan, Archana Bharadwaj Siva, Karthik Bharadwaj Tallapaka,Umesh Kumar, Unis Ahmad Bhat, Ajay Sarawagi, Priyanka Pant, Rajkanwar Nathawat, Rakesh K Mishra, Divya Tej Sowpati |
| EPI_ISL_471618 | CSIR-Centre for Cellular and Molecular Biology | CSIR-Centre for Cellular and Molecular Biology | Lamuk Zaveri, Shagufta Khan, Namami Gaur, Sakshi Shambhavi, Tulasi Nagabandi, Purushotham Vodnala, Payel Mukherjee, Sofia Banu, Priya Singh, Dhiviya Vedagiri, Divya Gupta, Vishal Sah, Santosh Kumar Kuncha, Krishnan Harinivas Harshan, Archana Bharadwaj Siva, Karthik Bharadwaj Tallapaka,Zeba Rizvi, Zuberwasim Sayyad, Kakade Aishwarya Arun, Amrutha H C, Ananga Ghosh, Rakesh K Mishra, Divya Tej Sowpati |
| EPI_ISL_471619 | CSIR-Centre for Cellular and Molecular Biology | CSIR-Centre for Cellular and Molecular Biology | Namami Gaur, Sakshi Shambhavi, Lamuk Zaveri, Shagufta Khan, Tulasi Nagabandi, Purushotham Vodnala, Payel Mukherjee, Sofia Banu, Priya Singh, Dhiviya Vedagiri, Divya Gupta, Vishal Sah, Santosh Kumar Kuncha, Krishnan Harinivas Harshan, Archana Bharadwaj Siva, Karthik Bharadwaj Tallapaka, Zeba Rizvi, Zuberwasim Sayyad, Kakade Aishwarya Arun, Amrutha H C, Ananga Ghosh, Rakesh K Mishra, Divya Tej Sowpati |
| EPI_ISL_471620 | CSIR-Centre for Cellular and Molecular Biology | CSIR-Centre for Cellular and Molecular Biology | Namami Gaur, Sakshi Shambhavi, Lamuk Zaveri, Shagufta Khan, Tulasi Nagabandi, Purushotham Vodnala, Payel Mukherjee, Sofia Banu, Priya Singh, Dhiviya Vedagiri, Divya Gupta, Vishal Sah, Santosh Kumar Kuncha, Krishnan Harinivas Harshan, Archana Bharadwaj Siva, Karthik Bharadwaj Tallapaka, Nikhil Hajirnis, Pratheusa Maccha, M Soujanya Reddy,G. Aditya Kumar, Koushick Sivakumar, Rakesh K Mishra, Divya Tej Sowpati |
| EPI_ISL_471621 | CSIR-Centre for Cellular and Molecular Biology | CSIR-Centre for Cellular and Molecular Biology | Namami Gaur, Sakshi Shambhavi, Lamuk Zaveri, Shagufta Khan, Tulasi Nagabandi, Purushotham Vodnala, Payel Mukherjee, Sofia Banu, Priya Singh, Dhiviya Vedagiri, Divya Gupta, Vishal Sah, Santosh Kumar Kuncha, Krishnan Harinivas Harshan, Archana Bharadwaj Siva, Karthik Bharadwaj Tallapaka, Zeba Rizvi, Zuberwasim Sayyad, Kakade Aishwarya Arun, Amrutha H C, Ananga Ghosh, Rakesh K Mishra, Divya Tej Sowpati |
| EPI_ISL_471622 | CSIR-Centre for Cellular and Molecular Biology | CSIR-Centre for Cellular and Molecular Biology | Namami Gaur, Sakshi Shambhavi, Lamuk Zaveri, Shagufta Khan, Tulasi Nagabandi, Purushotham Vodnala, Payel Mukherjee, Sofia Banu, Priya Singh, Dhiviya Vedagiri, Divya Gupta, Vishal Sah, Santosh Kumar Kuncha, Krishnan Harinivas Harshan, Archana Bharadwaj Siva, Karthik Bharadwaj Tallapaka,Kezia J Ann, Radhika Khandelwal, Roshan Maku Venkata, Shemin Mansuri, Sonu Uday, Rakesh K Mishra, Divya Tej Sowpati |
| EPI_ISL_471623 | CSIR-Centre for Cellular and Molecular Biology | CSIR-Centre for Cellular and Molecular Biology | Payel Mukherjee, Sofia Banu, Priya Singh, Dhiviya Vedagiri, Divya Gupta, Vishal Sah, Santosh Kumar Kuncha, Krishnan Harinivas Harshan, Archana Bharadwaj Siva, Karthik Bharadwaj Tallapaka, Shagufta Khan, Lamuk Zaveri, Namami Gaur, Sakshi Shambhavi, Tulasi Nagabandi, Purushotham Vodnala, G. Aditya Kumar, Koushick Sivakumar, Pooja Ramesh Gupta, Rajan Kumar Jha, Shraddha Vijay Lahoti, Rakesh K Mishra, Divya Tej Sowpati |
| EPI_ISL_471624 | CSIR-Centre for Cellular and Molecular Biology | CSIR-Centre for Cellular and Molecular Biology | Payel Mukherjee, Sofia Banu, Priya Singh, Dhiviya Vedagiri, Divya Gupta, Vishal Sah, Santosh Kumar Kuncha, Krishnan Harinivas Harshan, Archana Bharadwaj Siva, Karthik Bharadwaj Tallapaka, Shagufta Khan, Lamuk Zaveri, Namami Gaur, Sakshi Shambhavi, Tulasi Nagabandi, Purushotham Vodnala, Gokulan C G, Gunjan Purohit, Hanuman Tulashiram Kale, Pankaj Kumar, Prachand Issarapu, Rakesh K Mishra, Divya Tej Sowpati |
| EPI_ISL_471625 | CSIR-Centre for Cellular and Molecular Biology | CSIR-Centre for Cellular and Molecular Biology | Payel Mukherjee, Sofia Banu, Priya Singh, Dhiviya Vedagiri, Divya Gupta, Vishal Sah, Santosh Kumar Kuncha, Krishnan Harinivas Harshan, Archana Bharadwaj Siva, Karthik Bharadwaj Tallapaka, Shagufta Khan, Lamuk Zaveri, Namami Gaur, Sakshi Shambhavi, Tulasi Nagabandi, Purushotham Vodnala, Rakesh K Mishra, Sonu Uday, Sudipta Mondal, Annapoorna P Karthyayani, Debabrata Jana, Debrya Saha, Divya Tej Sowpati |
| EPI_ISL_471626 | CSIR-Centre for Cellular and Molecular Biology | CSIR-Centre for Cellular and Molecular Biology | Payel Mukherjee, Sofia Banu, Priya Singh, Dhiviya Vedagiri, Divya Gupta, Vishal Sah, Santosh Kumar Kuncha, Krishnan Harinivas Harshan, Archana Bharadwaj Siva, Karthik Bharadwaj Tallapaka, Shagufta Khan, Lamuk Zaveri, Namami Gaur, Sakshi Shambhavi, Tulasi Nagabandi, Purushotham Vodnala,Deepak Kumar, Devi Prasad Vijayashankar, Disha Nanda, Divya Das, Jotin Gogoi, Manish Bhattacharjee, Rakesh K Mishra, Divya Tej Sowpati |
| EPI_ISL_471627 | CSIR-Centre for Cellular and Molecular Biology | CSIR-Centre for Cellular and Molecular Biology | Sakshi Shambhavi, Lamuk Zaveri, Shagufta Khan, Namami Gaur, Tulasi Nagabandi, Purushotham Vodnala, Payel Mukherjee, Sofia Banu, Priya Singh, Dhiviya Vedagiri, Divya Gupta, Vishal Sah, Santosh Kumar Kuncha, Krishnan Harinivas Harshan, Archana Bharadwaj Siva, Karthik Bharadwaj Tallapaka, Deepak Kumar, Devi Prasad Vijayashankar, Disha Nanda, Divya Das, Jotin Gogoi, Manish Bhattacharjee, Rakesh K Mishra, Divya Tej Sowpati |
| EPI_ISL_471628 | CSIR-Centre for Cellular and Molecular Biology | CSIR-Centre for Cellular and Molecular Biology | Sakshi Shambhavi, Lamuk Zaveri, Shagufta Khan, Namami Gaur, Tulasi Nagabandi, Purushotham Vodnala, Payel Mukherjee, Sofia Banu, Priya Singh, Dhiviya Vedagiri, Divya Gupta, Vishal Sah, Santosh Kumar Kuncha, Krishnan Harinivas Harshan, Archana Bharadwaj Siva, Karthik Bharadwaj Tallapaka, G. Aditya Kumar, Koushick Sivakumar, Pooja Ramesh Gupta, Rajan Kumar Jha, Shraddha Vijay Lahoti, Rakesh K Mishra, Divya Tej Sowpati |
| EPI_ISL_471629 | CSIR-Centre for Cellular and Molecular Biology | CSIR-Centre for Cellular and Molecular Biology | Sakshi Shambhavi, Lamuk Zaveri, Shagufta Khan, Namami Gaur, Tulasi Nagabandi, Purushotham Vodnala, Payel Mukherjee, Sofia Banu, Priya Singh, Dhiviya Vedagiri, Divya Gupta, Vishal Sah, Santosh Kumar Kuncha, Krishnan Harinivas Harshan, Archana Bharadwaj Siva, Karthik Bharadwaj Tallapaka,Nikhil Hajirnis, Pratheusa Maccha, M Soujanya Reddy,G. Aditya Kumar, Koushick Sivakumar, Rakesh K Mishra, Divya Tej Sowpati |
| EPI_ISL_471630 | CSIR-Centre for Cellular and Molecular Biology | CSIR-Centre for Cellular and Molecular Biology | Sakshi Shambhavi, Lamuk Zaveri, Shagufta Khan, Namami Gaur, Tulasi Nagabandi, Purushotham Vodnala, Payel Mukherjee, Sofia Banu, Priya Singh, Dhiviya Vedagiri, Divya Gupta, Vishal Sah, Santosh Kumar Kuncha, Krishnan Harinivas Harshan, Archana Bharadwaj Siva, Karthik Bharadwaj Tallapaka,Nikhil Hajirnis, Pratheusa Maccha, M Soujanya Reddy,G. Aditya Kumar, Koushick Sivakumar,Disha Nanda, Divya Das, Jotin Gogoi, Manish Bhattacharjee, Ravi Prasad Mukku, Rakesh K Mishra, Divya Tej Sowpati |
| EPI_ISL_471631 | CSIR-Centre for Cellular and Molecular Biology | CSIR-Centre for Cellular and Molecular Biology | Shagufta Khan, Lamuk Zaveri, Namami Gaur, Sakshi Shambhavi, Tulasi Nagabandi, Purushotham Vodnala, Payel Mukherjee, Sofia Banu, Priya Singh, Dhiviya Vedagiri, Divya Gupta, Vishal Sah, Santosh Kumar Kuncha, Krishnan Harinivas Harshan, Archana Bharadwaj Siva, Karthik Bharadwaj Tallapaka, Disha Nanda, Divya Das, Jotin Gogoi, Manish Bhattacharjee, Ravi Prasad Mukku, Rakesh K Mishra, Divya Tej Sowpati |
| EPI_ISL_471632 | CSIR-Centre for Cellular and Molecular Biology | CSIR-Centre for Cellular and Molecular Biology | Shagufta Khan, Lamuk Zaveri, Namami Gaur, Sakshi Shambhavi, Tulasi Nagabandi, Purushotham Vodnala, Payel Mukherjee, Sofia Banu, Priya Singh, Dhiviya Vedagiri, Divya Gupta, Vishal Sah, Santosh Kumar Kuncha, Krishnan Harinivas Harshan, Archana Bharadwaj Siva, Karthik Bharadwaj Tallapaka, Renu Sudhakar, Somesh Gorde, Gangumala Srinivas Reddy, Sujoy Deb, Swati Bayyana, Rakesh K Mishra, Divya Tej Sowpati |
| EPI_ISL_471633 | CSIR-Centre for Cellular and Molecular Biology | CSIR-Centre for Cellular and Molecular Biology | Shagufta Khan, Lamuk Zaveri, Namami Gaur, Sakshi Shambhavi, Tulasi Nagabandi, Purushotham Vodnala, Payel Mukherjee, Sofia Banu, Priya Singh, Dhiviya Vedagiri, Divya Gupta, Vishal Sah, Santosh Kumar Kuncha, Krishnan Harinivas Harshan, Archana Bharadwaj Siva, Karthik Bharadwaj Tallapaka,Preethi Jampala, Sharada Ravi Iyer, Sulagana Mukherjee, Swetha Sundar, Peddapuvala Sai Uday Kiran Rakesh K Mishra, Divya Tej Sowpati |
| EPI_ISL_471634 | CSIR-Centre for Cellular and Molecular Biology | CSIR-Centre for Cellular and Molecular Biology | Shagufta Khan, Lamuk Zaveri, Namami Gaur, Sakshi Shambhavi, Tulasi Nagabandi, Purushotham Vodnala, Payel Mukherjee, Sofia Banu, Priya Singh, Dhiviya Vedagiri, Divya Gupta, Vishal Sah, Santosh Kumar Kuncha, Krishnan Harinivas Harshan, Archana Bharadwaj Siva, Karthik Bharadwaj Tallapaka,Umesh Kumar, Unis Ahmad Bhat, Ajay Sarawagi, Priyanka Pant, Rajkanwar Nathawat, Rakesh K Mishra, Divya Tej Sowpati |
| EPI_ISL_471635 | CSIR-Centre for Cellular and Molecular Biology | CSIR-Centre for Cellular and Molecular Biology | Sofia Banu, Payel Mukherjee, Priya Singh, Dhiviya Vedagiri, Divya Gupta, Vishal Sah, Santosh Kumar Kuncha, Krishnan Harinivas Harshan, Archana Bharadwaj Siva, Karthik Bharadwaj Tallapaka, Shagufta Khan, Lamuk Zaveri, Namami Gaur, Sakshi Shambhavi, Tulasi Nagabandi, Purushotham Vodnala, Deepak Kumar, Devi Prasad Vijayashankar, Disha Nanda, Divya Das, Jotin Gogoi, Manish Bhattacharjee, Rakesh K Mishra, Divya Tej Sowpati |
| EPI_ISL_471636 | CSIR-Centre for Cellular and Molecular Biology | CSIR-Centre for Cellular and Molecular Biology | Sofia Banu, Payel Mukherjee, Priya Singh, Dhiviya Vedagiri, Divya Gupta, Vishal Sah, Santosh Kumar Kuncha, Krishnan Harinivas Harshan, Archana Bharadwaj Siva, Karthik Bharadwaj Tallapaka, Shagufta Khan, Lamuk Zaveri, Namami Gaur, Sakshi Shambhavi, Tulasi Nagabandi, Purushotham Vodnala, Disha Nanda, Divya Das, Jotin Gogoi, Manish Bhattacharjee, Ravi Prasad Mukku, Rakesh K Mishra, Divya Tej Sowpati |

| | | | |
|---|---|---|---|
| EPI_ISL_471637 | CSIR-Centre for Cellular and Molecular Biology | CSIR-Centre for Cellular and Molecular Biology | Sofia Banu, Payel Mukherjee, Priya Singh, Dhiviya Vedagiri, Divya Gupta, Vishal Sah, Santosh Kumar Kuncha, Krishnan Harinivas Harshan, Archana Bharadwaj Siva, Karthik Bharadwaj Tallapaka, Shagufta Khan, Lamuk Zaveri, Namami Gaur, Sakshi Shambhavi, Tulasi Nagabandi, Purushotham Vodnala, Gokulan C G, Gunjan Purohit, Hanuman Tulashiram Kale, Pankaj Kumar, Prachand Issarapu, Rakesh K Mishra, Divya Tej Sowpati |
| EPI_ISL_471638 | CSIR-Centre for Cellular and Molecular Biology | CSIR-Centre for Cellular and Molecular Biology | Sofia Banu, Payel Mukherjee, Priya Singh, Dhiviya Vedagiri, Divya Gupta, Vishal Sah, Santosh Kumar Kuncha, Krishnan Harinivas Harshan, Archana Bharadwaj Siva, Karthik Bharadwaj Tallapaka, Shagufta Khan, Lamuk Zaveri, Namami Gaur, Sakshi Shambhavi, Tulasi Nagabandi, Purushotham Vodnala,Preethi Jampala, Sharada Ravi Iyer, Sulagana Mukherjee, Swetha Sundar, Peddapuvala Sai Uday Kiran, Rakesh K Mishra, Divya Tej Sowpati |
| EPI_ISL_471639 | CSIR-Centre for Cellular and Molecular Biology | CSIR-Centre for Cellular and Molecular Biology | Tulasi Nagabandi, Namami Gaur, Sakshi Shambhavi, Lamuk Zaveri, Shagufta Khan, Purushotham Vodnala, Payel Mukherjee, Sofia Banu, Priya Singh, Dhiviya Vedagiri, Divya Gupta, Vishal Sah, Santosh Kumar Kuncha, Krishnan Harinivas Harshan, Archana Bharadwaj Siva, Karthik Bharadwaj Tallapaka,G. Aditya Kumar, Koushick Sivakumar, Pooja Ramesh Gupta, Rajan Kumar Jha, Shraddha Vijay Lahoti, Rakesh K Mishra, Divya Tej Sowpati |
| EPI_ISL_471640 | CSIR-Centre for Cellular and Molecular Biology | CSIR-Centre for Cellular and Molecular Biology | Tulasi Nagabandi, Namami Gaur, Sakshi Shambhavi, Lamuk Zaveri, Shagufta Khan, Purushotham Vodnala, Payel Mukherjee, Sofia Banu, Priya Singh, Dhiviya Vedagiri, Divya Gupta, Vishal Sah, Santosh Kumar Kuncha, Krishnan Harinivas Harshan, Archana Bharadwaj Siva, Karthik Bharadwaj Tallapaka,Kezia J Ann, Radhika Khandelwal, Roshan Maku Venkata, Shemin Mansuri, Sonu Uday, Rakesh K Mishra, Divya Tej Sowpati |
| EPI_ISL_471641, EPI_ISL_471642 | CSIR-Centre for Cellular and Molecular Biology | CSIR-Centre for Cellular and Molecular Biology | Dhiviya Vedagiri, Divya Gupta, Vishal Sah, Payel Mukherjee, Sofia Banu, Priya Singh, Santosh Kumar Kuncha, Archana Bharadwaj Siva, Karthik Bharadwaj Tallapaka, Shagufta Khan, Lamuk Zaveri, Namami Gaur, Sakshi Shambhavi, Tulasi Nagabandi, Purushotham Vodnala, Rakesh K Mishra, Divya Tej Sowpati, Krishnan Harinivas Harshan |
| EPI_ISL_471643 | CSIR-Centre for Cellular and Molecular Biology | CSIR-Centre for Cellular and Molecular Biology | Tulasi Nagabandi, Namami Gaur, Sakshi Shambhavi, Lamuk Zaveri, Shagufta Khan, Purushotham Vodnala, Payel Mukherjee, Sofia Banu, Priya Singh, Dhiviya Vedagiri, Divya Gupta, Vishal Sah, Santosh Kumar Kuncha, Krishnan Harinivas Harshan, Archana Bharadwaj Siva, Karthik Bharadwaj Tallapaka,G. Aditya Kumar, Koushick Sivakumar, Pooja Ramesh Gupta, Rajan Kumar Jha, Shraddha Vijay Lahoti, Rakesh K Mishra, Divya Tej Sowpati |
| EPI_ISL_471644 | CSIR-Centre for Cellular and Molecular Biology | CSIR-Centre for Cellular and Molecular Biology | Tulasi Nagabandi, Namami Gaur, Sakshi Shambhavi, Lamuk Zaveri, Shagufta Khan, Purushotham Vodnala, Payel Mukherjee, Sofia Banu, Priya Singh, Dhiviya Vedagiri, Divya Gupta, Vishal Sah, Santosh Kumar Kuncha, Krishnan Harinivas Harshan, Archana Bharadwaj Siva, Karthik Bharadwaj Tallapaka,Kezia J Ann, Radhika Khandelwal, Roshan Maku Venkata, Shemin Mansuri, Sonu Uday, Rakesh K Mishra, Divya Tej Sowpati |
| EPI_ISL_471645, EPI_ISL_471646 | CSIR-Centre for Cellular and Molecular Biology | CSIR-Centre for Cellular and Molecular Biology | Dhiviya Vedagiri, Divya Gupta, Vishal Sah, Payel Mukherjee, Sofia Banu, Priya Singh, Santosh Kumar Kuncha, Archana Bharadwaj Siva, Karthik Bharadwaj Tallapaka, Shagufta Khan, Lamuk Zaveri, Namami Gaur, Sakshi Shambhavi, Tulasi Nagabandi, Purushotham Vodnala, Rakesh K Mishra, Divya Tej Sowpati, Krishnan Harinivas Harshan |
| EPI_ISL_471647 | Hospital Municipal de Barueri Dr. Francisco Moran | Instituto Adolfo Lutz, Interdiciplinary Procedures Center, Strategic Laboratory | Claudio Tavares Sacchi, Claudia Regina Gonçalves, Erica Valessa Ramos Gomes |
| EPI_ISL_471648 | UBS e Pronto Socorro Jd. Jacira | Instituto Adolfo Lutz, Interdiciplinary Procedures Center, Strategic Laboratory | Claudio Tavares Sacchi, Claudia Regina Gonçalves, Erica Valessa Ramos Gomes |
| EPI_ISL_471679, EPI_ISL_471685, EPI_ISL_471686, EPI_ISL_471687, EPI_ISL_471689, EPI_ISL_471691, EPI_ISL_471692, EPI_ISL_471693, EPI_ISL_471695, EPI_ISL_471696, EPI_ISL_471701, EPI_ISL_471702, EPI_ISL_471704, EPI_ISL_471707, EPI_ISL_471708, EPI_ISL_471709, EPI_ISL_471712, EPI_ISL_471713, EPI_ISL_471714, EPI_ISL_471715, EPI_ISL_471716, EPI_ISL_471717, EPI_ISL_471720, EPI_ISL_471721, EPI_ISL_471722, EPI_ISL_471724, EPI_ISL_471725, EPI_ISL_471728, EPI_ISL_471729, EPI_ISL_471731, EPI_ISL_471732, EPI_ISL_471739, EPI_ISL_471740, EPI_ISL_471742, EPI_ISL_471743, EPI_ISL_471745, EPI_ISL_471749, EPI_ISL_471750, EPI_ISL_471754, EPI_ISL_471767, EPI_ISL_471768, EPI_ISL_471769, EPI_ISL_471770, EPI_ISL_471771, EPI_ISL_471772, EPI_ISL_471787, EPI_ISL_471788, EPI_ISL_471789, EPI_ISL_471790, EPI_ISL_471791, EPI_ISL_471792, EPI_ISL_471793, EPI_ISL_471794, EPI_ISL_471795, EPI_ISL_471796, EPI_ISL_471797, EPI_ISL_471807, EPI_ISL_471808, EPI_ISL_471809, EPI_ISL_471810, EPI_ISL_471812, EPI_ISL_471813, EPI_ISL_471814, EPI_ISL_471815, EPI_ISL_471817, EPI_ISL_471818, EPI_ISL_471819, EPI_ISL_471820, EPI_ISL_471821, EPI_ISL_471822, EPI_ISL_471823, EPI_ISL_471825, EPI_ISL_471826, EPI_ISL_471830, EPI_ISL_471831, EPI_ISL_471832, EPI_ISL_471833, EPI_ISL_471834, EPI_ISL_471835, EPI_ISL_471837, EPI_ISL_471838, EPI_ISL_471839, EPI_ISL_471840, EPI_ISL_471842, EPI_ISL_471843, EPI_ISL_471844, EPI_ISL_471847, EPI_ISL_471848, EPI_ISL_471850, EPI_ISL_471851, EPI_ISL_471852, EPI_ISL_471853, EPI_ISL_471854, EPI_ISL_471855, EPI_ISL_471856, EPI_ISL_471857, EPI_ISL_471858, EPI_ISL_471859, EPI_ISL_471871, EPI_ISL_471872, EPI_ISL_471873, EPI_ISL_471874, EPI_ISL_471875, EPI_ISL_471876, EPI_ISL_471877, EPI_ISL_471878, EPI_ISL_471879, EPI_ISL_471880, EPI_ISL_471881, EPI_ISL_471882, EPI_ISL_471883, EPI_ISL_471884, EPI_ISL_471885, EPI_ISL_471886, EPI_ISL_471887, EPI_ISL_471888, EPI_ISL_471889, EPI_ISL_471890, EPI_ISL_471891, EPI_ISL_471896, EPI_ISL_471897, EPI_ISL_471898, EPI_ISL_471899, EPI_ISL_471900, EPI_ISL_471901, EPI_ISL_471902, EPI_ISL_471903, EPI_ISL_471904, EPI_ISL_471905, EPI_ISL_471906, EPI_ISL_471907, EPI_ISL_471908, EPI_ISL_471909, EPI_ISL_471910 | | | |
| see above | Michigan Department of Health and Human Services, Bureau of Laboratories | Michigan Department of Health and Human Services, Bureau of Laboratories | Blankenship HM, Riner D, Soehnlen MK |
| EPI_ISL_471911, EPI_ISL_471912, EPI_ISL_471913, EPI_ISL_471914, EPI_ISL_471915, EPI_ISL_471916, EPI_ISL_471917, EPI_ISL_471918, EPI_ISL_471919, EPI_ISL_471921, EPI_ISL_471922, EPI_ISL_471924, EPI_ISL_471925, EPI_ISL_471926, EPI_ISL_471927, EPI_ISL_471929, EPI_ISL_471930, EPI_ISL_471931, EPI_ISL_471932, EPI_ISL_471933, EPI_ISL_471934, EPI_ISL_471935, EPI_ISL_471936, EPI_ISL_471937, EPI_ISL_471938, EPI_ISL_471941, EPI_ISL_471943, EPI_ISL_471944, EPI_ISL_471946, EPI_ISL_471947, EPI_ISL_471949, EPI_ISL_471950, EPI_ISL_471951, EPI_ISL_471955, EPI_ISL_471956, EPI_ISL_471957, EPI_ISL_471959, EPI_ISL_471960, EPI_ISL_471963, EPI_ISL_471964, EPI_ISL_471966, EPI_ISL_471967, EPI_ISL_471971, EPI_ISL_471972, EPI_ISL_471973, EPI_ISL_471975, EPI_ISL_471976, EPI_ISL_471977, EPI_ISL_471978, EPI_ISL_471979, EPI_ISL_471981, EPI_ISL_471982, EPI_ISL_471987, EPI_ISL_471989, EPI_ISL_471990, EPI_ISL_471991, EPI_ISL_471992 | | | |
| see above | University of Exeter | COVID-19 Genomics UK (COG-UK) Consortium | Ben Temperton,Aaron Jeffries,Michelle Michelsen,Joanna Warwick-Dugdale,Audrey Farbos,Robyn Manley,Stephen Michell,Jane Masoli |
| EPI_ISL_471993, EPI_ISL_471994, EPI_ISL_471995, EPI_ISL_471996, EPI_ISL_471997, EPI_ISL_471998, EPI_ISL_471999, EPI_ISL_472000, EPI_ISL_472001, EPI_ISL_472003, EPI_ISL_472004, EPI_ISL_472005, EPI_ISL_472006, EPI_ISL_472007, EPI_ISL_472008, EPI_ISL_472009, EPI_ISL_472010, EPI_ISL_472011, EPI_ISL_472012, EPI_ISL_472013, EPI_ISL_472014, EPI_ISL_472015, EPI_ISL_472016, EPI_ISL_472018, EPI_ISL_472019, EPI_ISL_472020, EPI_ISL_472021, EPI_ISL_472022, EPI_ISL_472057, EPI_ISL_472090, EPI_ISL_472091, EPI_ISL_472092, EPI_ISL_472093, EPI_ISL_472096, EPI_ISL_472097, EPI_ISL_472098, EPI_ISL_472099, EPI_ISL_472100, EPI_ISL_472102, EPI_ISL_472104, EPI_ISL_472105, EPI_ISL_472106, EPI_ISL_472107, EPI_ISL_472109, EPI_ISL_472111, EPI_ISL_472113, EPI_ISL_472114, EPI_ISL_472115, EPI_ISL_472116, EPI_ISL_472118, EPI_ISL_472119, EPI_ISL_472121, EPI_ISL_472122, EPI_ISL_472124, EPI_ISL_472126, EPI_ISL_472127, EPI_ISL_472129, EPI_ISL_472131, EPI_ISL_472132, EPI_ISL_472133, EPI_ISL_472134, EPI_ISL_472135 | | | |
| see above | Liverpool Clinical Laboratories | COVID-19 Genomics UK (COG-UK) Consortium | Sam Haldenby, Anita Lucaci, Steve Paterson, Julian Hiscox, Alistair Darby, M Almsaud, A Alrezaihi, Muhannad Alruwaili, Stuart D Armstrong, Jones Benjamin, Eleanor G Bentley, Anu Chawla, Jordan J Clark, Angela Cowell, Richard Eccles, Isabel García-Dorival, Matthew Gemmell, Alessandro Gerada, PKF Gilmore, Richard Gregory, Ximeng Han, Catherine Hartley, Margaret Hughes, Miren Iturriza-Gomara, James Johnson, L Luu, Jenifer Manson, Charlotte Nelson, Elaine O'Toole, Cassie Olateju, Rebekah Penrice-Randal , Lucille Rainbow, N.P Randle, Trevor Ian Robinson, Parul Sharma, Ghada T Shawli, James P Stewart, Neil Swainston, Ecaterina Vamos, Joanne Watts, Mark Whitehead |
| EPI_ISL_472228, EPI_ISL_472249, EPI_ISL_472255, EPI_ISL_472256 | Northumbria University / South Tees Hospitals NHS Foundation Trust / North Cumbria Integrated Care NHS Foundation Trust / North Tees and Hartlepool NHS Foundation Trust / Newcastle Hospitals NHS Foundation Trust | COVID-19 Genomics UK (COG-UK) Consortium | Darren L Smith,Andrew Nelson,Matthew Bashton,Greg R Young,Joshua Loh,John Allan,Mohammad A Tariq,Giles S Holt,Gary Black,Wen C Yew,Lynn Dover,Paul Baker,Steve Liggett,Sarah Essex,Jane Greenaway,Debra Padgett,Clive Graham,Garren Scott,Edward Barton,Emma Swindells,Brendan Payne,Jennifer Collins,Yusri Taha,Gary Eltringham |
| EPI_ISL_472291, EPI_ISL_472296, EPI_ISL_472300, EPI_ISL_472301, EPI_ISL_472302, EPI_ISL_472303, EPI_ISL_472305, EPI_ISL_472306, EPI_ISL_472307, EPI_ISL_472310, EPI_ISL_472311, EPI_ISL_472313, EPI_ISL_472314, EPI_ISL_472316, EPI_ISL_472317, EPI_ISL_472320, EPI_ISL_472322, EPI_ISL_472326, EPI_ISL_472328, EPI_ISL_472330, EPI_ISL_472334, EPI_ISL_472336, EPI_ISL_472342, EPI_ISL_472343, EPI_ISL_472347, EPI_ISL_472349, EPI_ISL_472354, EPI_ISL_472356, EPI_ISL_472361, EPI_ISL_472363, EPI_ISL_472365, EPI_ISL_472366, EPI_ISL_472367, EPI_ISL_472368, EPI_ISL_472371, EPI_ISL_472375, EPI_ISL_472376, EPI_ISL_472381 | | | |
| see above | Quadram Institute Bioscience | COVID-19 Genomics UK (COG-UK) Consortium | Dave J. Baker, Gemma L. Kay, Alp Aydin, Thanh Le-Viet, Steven Rudder, Ana P. Tedim, Anastasia Kolyva, Maria Diaz, Leonardo de Oliveira Martins, Nabil-Fareed Alikhan, Lizzie Meadows, Rachael Stanley, Ngozi Elumogo, Muhammed Yasir, Nicholas M. Thomson, Alexander J Trotter, Rachel Gilroy, Samuel Bloomfield, Claire Stuart, Andrew Bell, Reenesh Prakash, Samir Dervisevic, Alison E. Mather, John Wain, Mark Webber, Andrew J. Page, Justin O'Grady |
| EPI_ISL_472384, EPI_ISL_472385, EPI_ISL_472386, EPI_ISL_472387, EPI_ISL_472388, EPI_ISL_472390, EPI_ISL_472392, EPI_ISL_472393, EPI_ISL_472394, EPI_ISL_472395, EPI_ISL_472396, EPI_ISL_472398, EPI_ISL_472399, EPI_ISL_472400, EPI_ISL_472401, EPI_ISL_472402, EPI_ISL_472403, EPI_ISL_472404, EPI_ISL_472405, EPI_ISL_472406, EPI_ISL_472407, EPI_ISL_472408, EPI_ISL_472409, EPI_ISL_472410, EPI_ISL_472411, EPI_ISL_472412, EPI_ISL_472413, EPI_ISL_472414, EPI_ISL_472415, EPI_ISL_472416, EPI_ISL_472418, EPI_ISL_472419, EPI_ISL_472420, EPI_ISL_472422, EPI_ISL_472423, EPI_ISL_472424, EPI_ISL_472425, EPI_ISL_472426, EPI_ISL_472427, EPI_ISL_472428, EPI_ISL_472429 | | | |
| see above | Queens Medical Centre, Clinical Microbiology Department / DeepSeq Nottingham | COVID-19 Genomics UK (COG-UK) Consortium | Gemma Clark, Wendy Smith, Manjinder Khakh, Vicki M Fleming, Michelle M Lister, Hannah Howson-Wells, Jonathan Ball, Patrick McClure, Joseph Chappell, Theocharis Tsoleridis, Nadine Holmes, Matthew Carlisle, Christopher Moore, Fei Sang, Johnny Debebe, Victoria Wright, Matthew Loose |
| EPI_ISL_472437, EPI_ISL_472439, EPI_ISL_472445, EPI_ISL_472450, EPI_ISL_472452, EPI_ISL_472454, EPI_ISL_472459, EPI_ISL_472461, EPI_ISL_472479, EPI_ISL_472480, EPI_ISL_472482, EPI_ISL_472489, EPI_ISL_472498, EPI_ISL_472509, EPI_ISL_472513, EPI_ISL_472516, EPI_ISL_472522, EPI_ISL_472523, EPI_ISL_472530, EPI_ISL_472538, EPI_ISL_472541, EPI_ISL_472542, EPI_ISL_472545, EPI_ISL_472547, EPI_ISL_472553, EPI_ISL_472561, EPI_ISL_472565, EPI_ISL_472579, EPI_ISL_472580, EPI_ISL_472589, EPI_ISL_472598, EPI_ISL_472602, EPI_ISL_472603, EPI_ISL_472605, EPI_ISL_472606, EPI_ISL_472615, EPI_ISL_472618, EPI_ISL_472622, EPI_ISL_472623, EPI_ISL_472625, EPI_ISL_472626, EPI_ISL_472636, EPI_ISL_472637, EPI_ISL_472639, EPI_ISL_472653, EPI_ISL_472655, EPI_ISL_472664, EPI_ISL_472666, EPI_ISL_472669, EPI_ISL_472670, EPI_ISL_472676, EPI_ISL_472684, EPI_ISL_472692, EPI_ISL_472693, EPI_ISL_472699, EPI_ISL_472701, EPI_ISL_472716, EPI_ISL_472729, EPI_ISL_472734, EPI_ISL_472735, EPI_ISL_472737, EPI_ISL_472740, EPI_ISL_472742, EPI_ISL_472746, EPI_ISL_472747, EPI_ISL_472749, EPI_ISL_472750, EPI_ISL_472751, EPI_ISL_472753, EPI_ISL_472754, EPI_ISL_472757, EPI_ISL_472758, EPI_ISL_472760, EPI_ISL_472761, EPI_ISL_472762, EPI_ISL_472763, EPI_ISL_472764, EPI_ISL_472766, EPI_ISL_472768, EPI_ISL_472769, EPI_ISL_472770, EPI_ISL_472771, EPI_ISL_472773, EPI_ISL_472775, EPI_ISL_472776, EPI_ISL_472777, EPI_ISL_472778, EPI_ISL_472779, EPI_ISL_472780, EPI_ISL_472782, EPI_ISL_472786, EPI_ISL_472789, EPI_ISL_472790, EPI_ISL_472791, EPI_ISL_472792, EPI_ISL_472793, EPI_ISL_472795, EPI_ISL_472798, EPI_ISL_472799, EPI_ISL_472800, EPI_ISL_472801, EPI_ISL_472805, EPI_ISL_472808, EPI_ISL_472809, EPI_ISL_472810, EPI_ISL_472812, EPI_ISL_472813, EPI_ISL_472814, EPI_ISL_472815, EPI_ISL_472818, EPI_ISL_472820, EPI_ISL_472821, EPI_ISL_472823, EPI_ISL_472824, EPI_ISL_472825, EPI_ISL_472830, EPI_ISL_472831, EPI_ISL_472833, EPI_ISL_472834, EPI_ISL_472835, EPI_ISL_472838, EPI_ISL_472839, EPI_ISL_472843, EPI_ISL_472844, EPI_ISL_472845, EPI_ISL_472846, EPI_ISL_472848, EPI_ISL_472850, EPI_ISL_472851, EPI_ISL_472854, EPI_ISL_472856, EPI_ISL_472857, EPI_ISL_472858, EPI_ISL_472861, EPI_ISL_472870, EPI_ISL_472873, EPI_ISL_472874, EPI_ISL_472875, EPI_ISL_472878, EPI_ISL_472879, EPI_ISL_472880, EPI_ISL_472882, EPI_ISL_472883, EPI_ISL_472884, EPI_ISL_472886, EPI_ISL_472889, EPI_ISL_472890, EPI_ISL_472892, EPI_ISL_472895, EPI_ISL_472896, EPI_ISL_472898, EPI_ISL_472899, EPI_ISL_472901, EPI_ISL_472902, EPI_ISL_472904, EPI_ISL_472905, EPI_ISL_472906, EPI_ISL_472913, EPI_ISL_472914, EPI_ISL_472916, EPI_ISL_472922, EPI_ISL_472926, EPI_ISL_472927, EPI_ISL_472934, EPI_ISL_472935, EPI_ISL_472937, EPI_ISL_472938, EPI_ISL_472939, EPI_ISL_472940, EPI_ISL_472942, EPI_ISL_472947, EPI_ISL_472948, EPI_ISL_472949, EPI_ISL_472952, EPI_ISL_472955, EPI_ISL_472956, EPI_ISL_472957, EPI_ISL_472960, EPI_ISL_472961, EPI_ISL_472963, EPI_ISL_472965, EPI_ISL_472966, EPI_ISL_472967, EPI_ISL_472968, EPI_ISL_472972, EPI_ISL_472974, EPI_ISL_472975, EPI_ISL_472976, EPI_ISL_472979, EPI_ISL_472980, EPI_ISL_472983, EPI_ISL_472988, EPI_ISL_472990, EPI_ISL_472991, EPI_ISL_472994, EPI_ISL_472995, EPI_ISL_472997, EPI_ISL_472998, EPI_ISL_473002, EPI_ISL_473004, EPI_ISL_473005, EPI_ISL_473009, EPI_ISL_473010, EPI_ISL_473014, EPI_ISL_473015, EPI_ISL_473019, EPI_ISL_473020, EPI_ISL_473021, EPI_ISL_473024, EPI_ISL_473026, EPI_ISL_473027, EPI_ISL_473029, EPI_ISL_473031, EPI_ISL_473032, EPI_ISL_473033, EPI_ISL_473034, EPI_ISL_473035, EPI_ISL_473036, EPI_ISL_473037, EPI_ISL_473038, EPI_ISL_473039, EPI_ISL_473044, EPI_ISL_473045, EPI_ISL_473046, EPI_ISL_473048, EPI_ISL_473051, EPI_ISL_473052, EPI_ISL_473053, EPI_ISL_473055, EPI_ISL_473056, EPI_ISL_473059, EPI_ISL_473074, EPI_ISL_473079, EPI_ISL_473081, EPI_ISL_473085, EPI_ISL_473088, EPI_ISL_473090, EPI_ISL_473092, EPI_ISL_473095, EPI_ISL_473097, EPI_ISL_473102, EPI_ISL_473104, EPI_ISL_473105, EPI_ISL_473107, EPI_ISL_473115, EPI_ISL_473119, EPI_ISL_473122, EPI_ISL_473125, EPI_ISL_473126, EPI_ISL_473127, EPI_ISL_473132, EPI_ISL_473134, EPI_ISL_473139, EPI_ISL_473141, EPI_ISL_473146, EPI_ISL_473147, EPI_ISL_473149, EPI_ISL_473155, EPI_ISL_473158, EPI_ISL_473162, EPI_ISL_473163, EPI_ISL_473166, EPI_ISL_473168, EPI_ISL_473170, EPI_ISL_473173, EPI_ISL_473178, EPI_ISL_473179, EPI_ISL_473183, EPI_ISL_473186, EPI_ISL_473188, EPI_ISL_473189, EPI_ISL_473194, EPI_ISL_473198, EPI_ISL_473199, EPI_ISL_473200, EPI_ISL_473202, EPI_ISL_473206, EPI_ISL_473210, EPI_ISL_473211, EPI_ISL_473223, EPI_ISL_473230, EPI_ISL_473231, EPI_ISL_473232, EPI_ISL_473236, EPI_ISL_473238, EPI_ISL_473239, EPI_ISL_473241, EPI_ISL_473246, EPI_ISL_473249, EPI_ISL_473250, EPI_ISL_473254, EPI_ISL_473259, EPI_ISL_473261, EPI_ISL_473266, EPI_ISL_473283, EPI_ISL_473285, EPI_ISL_473286, EPI_ISL_473287, EPI_ISL_473289, EPI_ISL_473291, EPI_ISL_473292, EPI_ISL_473294, EPI_ISL_473296, EPI_ISL_473297, EPI_ISL_473300, EPI_ISL_473303, EPI_ISL_473305 | | | |
| see above | Wales Specialist Virology Centre Sequencing lab: Pathogen Genomics Unit | COVID-19 Genomics UK (COG-UK) Consortium | Catherine Moore, Johnathan Evans, Laura Gifford, Malorie Perry, Simon Cottrell, Angela Marchbank, Alec Birchley, Alexander Adams, Amy Gaskin, Bree Gatica-Wilcox, Jason Coombes, Joel Southgate, Lauren Gilbert, Lee Graham, Nicole Pacchiarini, Sara Kumziene-Summerhayes, Sarah Taylor, Sophie Jones, Sara Rey, Matthew Bull, Joanne Watkins, Sally Corden, Tom Connor |
| EPI_ISL_473306, EPI_ISL_473308, EPI_ISL_473310, EPI_ISL_473311, EPI_ISL_473312, EPI_ISL_473313, EPI_ISL_473314, EPI_ISL_473315, EPI_ISL_473316, EPI_ISL_473317, EPI_ISL_473318, EPI_ISL_473319, EPI_ISL_473321, EPI_ISL_473322, EPI_ISL_473323, EPI_ISL_473324, EPI_ISL_473328, EPI_ISL_473329, EPI_ISL_473330, EPI_ISL_473331, EPI_ISL_473332, EPI_ISL_473333, EPI_ISL_473336, EPI_ISL_473338, EPI_ISL_473340, EPI_ISL_473343, EPI_ISL_473344, EPI_ISL_473345, EPI_ISL_473348, EPI_ISL_473349, EPI_ISL_473350, EPI_ISL_473352, EPI_ISL_473354, EPI_ISL_473356, EPI_ISL_473357, EPI_ISL_473358, EPI_ISL_473360, EPI_ISL_473361, EPI_ISL_473362, EPI_ISL_473363, EPI_ISL_473365, EPI_ISL_473371, EPI_ISL_473374, EPI_ISL_473376, EPI_ISL_473377, EPI_ISL_473379, EPI_ISL_473381, EPI_ISL_473382, EPI_ISL_473383, EPI_ISL_473384, EPI_ISL_473385, EPI_ISL_473386, EPI_ISL_473388, EPI_ISL_473389, EPI_ISL_473390, EPI_ISL_473392, EPI_ISL_473394, EPI_ISL_473400, EPI_ISL_473403, EPI_ISL_473406, EPI_ISL_473407, EPI_ISL_473408, EPI_ISL_473410, EPI_ISL_473412, EPI_ISL_473414, EPI_ISL_473415, EPI_ISL_473416, EPI_ISL_473418, EPI_ISL_473419, EPI_ISL_473422, EPI_ISL_473423, EPI_ISL_473424, EPI_ISL_473427, EPI_ISL_473429, EPI_ISL_473434, EPI_ISL_473435, EPI_ISL_473436, EPI_ISL_473438, EPI_ISL_473441, EPI_ISL_473442, EPI_ISL_473443, EPI_ISL_473444, EPI_ISL_473445, EPI_ISL_473446, EPI_ISL_473447, EPI_ISL_473448, EPI_ISL_473450 | | | |
| see above | University of Birmingham | COVID-19 Genomics UK (COG-UK) Consortium | Institute of Microbiology, University of Birmingham: Claire McMurray, Joanne Stockton, Samuel Nicholls, Radoslaw Poplawski, Will Rowe, Josh Quick, Nicholas Loman, University of Birmingham Testing Laboratory: Celina M Whalley, Andrew Bosworth, Charlotte Poxon, Kasun Wanigasooriya, Oliver Pickles, Mike Kidd, Alex Richter, Andrew D Beggs PHE Heartlands Lab: Husam Osman, Andrew Bosworth. Queen Elizabeth Hospital: Anna Casey |
| EPI_ISL_473454, EPI_ISL_473455, EPI_ISL_473459, EPI_ISL_473463, EPI_ISL_473464, EPI_ISL_473466, EPI_ISL_473471, EPI_ISL_473472, EPI_ISL_473474, EPI_ISL_473477, EPI_ISL_473479, EPI_ISL_473481, EPI_ISL_473482, EPI_ISL_473483, EPI_ISL_473484, EPI_ISL_473487, EPI_ISL_473488, EPI_ISL_473489, EPI_ISL_473490, EPI_ISL_473491, EPI_ISL_473492, EPI_ISL_473493, EPI_ISL_473496, EPI_ISL_473498, EPI_ISL_473499, EPI_ISL_473500, EPI_ISL_473501, EPI_ISL_473503, EPI_ISL_473504, EPI_ISL_473507 | | | |
| see above | Department of Pathology, University of Cambridge | COVID-19 Genomics UK (COG-UK) Consortium | Luke W Meredith, M. Estée Török, Myra Hosmillo, William L. Hamilton, Martin D. Curran, Theresa Feltwell, Grant Hall, Anna Yakovleva, Fahad A Khokhar, Charlotte J. Houldcroft, Laura G Caller, Aminu S. Jahun, Sarah L. Caddy, Yasmin Chaudhry, Malte Pinckert, Ian Goodfellow |
| EPI_ISL_473509, EPI_ISL_473510, EPI_ISL_473511, EPI_ISL_473512, EPI_ISL_473513, EPI_ISL_473514, EPI_ISL_473515, EPI_ISL_473516, EPI_ISL_473517, EPI_ISL_473518, EPI_ISL_473519, EPI_ISL_473520, EPI_ISL_473521, EPI_ISL_473522, EPI_ISL_473524, EPI_ISL_473525, EPI_ISL_473526, EPI_ISL_473527, EPI_ISL_473528, EPI_ISL_473529, EPI_ISL_473532, EPI_ISL_473533, EPI_ISL_473534, EPI_ISL_473535, EPI_ISL_473536, EPI_ISL_473537, EPI_ISL_473540, EPI_ISL_473541, EPI_ISL_473543, EPI_ISL_473544, EPI_ISL_473546, EPI_ISL_473547, EPI_ISL_473548, EPI_ISL_473549, EPI_ISL_473550, EPI_ISL_473551, EPI_ISL_473552, EPI_ISL_473553, EPI_ISL_473554, EPI_ISL_473555, EPI_ISL_473557, EPI_ISL_473558, EPI_ISL_473559, EPI_ISL_473560, EPI_ISL_473561, EPI_ISL_473562, EPI_ISL_473563, EPI_ISL_473565, EPI_ISL_473566, EPI_ISL_473567, EPI_ISL_473568, EPI_ISL_473569, EPI_ISL_473570, EPI_ISL_473572, EPI_ISL_473573, EPI_ISL_473574, EPI_ISL_473575, EPI_ISL_473576, EPI_ISL_473577, EPI_ISL_473578, EPI_ISL_473579, EPI_ISL_473580, EPI_ISL_473581, EPI_ISL_473582, EPI_ISL_473583, EPI_ISL_473584, EPI_ISL_473585, EPI_ISL_473586, EPI_ISL_473587, EPI_ISL_473588, EPI_ISL_473589, EPI_ISL_473590, EPI_ISL_473591, EPI_ISL_473592, EPI_ISL_473593, EPI_ISL_473594, EPI_ISL_473595, EPI_ISL_473596, EPI_ISL_473597, EPI_ISL_473598, EPI_ISL_473599, EPI_ISL_473600, EPI_ISL_473601, EPI_ISL_473602, EPI_ISL_473603, EPI_ISL_473604, EPI_ISL_473605, EPI_ISL_473606, EPI_ISL_473608, EPI_ISL_473609, EPI_ISL_473610, EPI_ISL_473612, EPI_ISL_473613, EPI_ISL_473614, EPI_ISL_473615, EPI_ISL_473616, EPI_ISL_473617, EPI_ISL_473618, EPI_ISL_473619, EPI_ISL_473621, EPI_ISL_473623, EPI_ISL_473624, EPI_ISL_473628, EPI_ISL_473629, EPI_ISL_473630, EPI_ISL_473631, EPI_ISL_473632, EPI_ISL_473633, EPI_ISL_473635, EPI_ISL_473636, EPI_ISL_473637, EPI_ISL_473642, EPI_ISL_473643, EPI_ISL_473645, EPI_ISL_473646, EPI_ISL_473647, EPI_ISL_473648, EPI_ISL_473649, EPI_ISL_473651, EPI_ISL_473653, EPI_ISL_473654, EPI_ISL_473655, EPI_ISL_473656, EPI_ISL_473659, EPI_ISL_473660, EPI_ISL_473662, EPI_ISL_473663, EPI_ISL_473666, EPI_ISL_473667, EPI_ISL_473670, EPI_ISL_473672, EPI_ISL_473673, EPI_ISL_473674, EPI_ISL_473676, EPI_ISL_473679, EPI_ISL_473680, EPI_ISL_473681, EPI_ISL_473684, EPI_ISL_473686, EPI_ISL_473689, EPI_ISL_473694, EPI_ISL_473697, EPI_ISL_473702, EPI_ISL_473705, EPI_ISL_473706, EPI_ISL_473707, EPI_ISL_473709, EPI_ISL_473712, EPI_ISL_473713, EPI_ISL_473715, EPI_ISL_473717, EPI_ISL_473719, EPI_ISL_473721, EPI_ISL_473722, EPI_ISL_473723, EPI_ISL_473724, EPI_ISL_473725, EPI_ISL_473731, EPI_ISL_473732, EPI_ISL_473734, EPI_ISL_473745, EPI_ISL_473746, EPI_ISL_473748, EPI_ISL_473749, EPI_ISL_473750, EPI_ISL_473753, EPI_ISL_473754, EPI_ISL_473761, EPI_ISL_473762, EPI_ISL_473764, EPI_ISL_473766, EPI_ISL_473768, EPI_ISL_473769, EPI_ISL_473770, EPI_ISL_473771, EPI_ISL_473772, EPI_ISL_473773, EPI_ISL_473774, EPI_ISL_473776, EPI_ISL_473777, EPI_ISL_473778, EPI_ISL_473779, EPI_ISL_473780, EPI_ISL_473781, EPI_ISL_473782 | | | |
| see above | West of Scotland Specialist Virology Centre, NHSGGC / MRC-University of Glasgow Centre for Virus Research | COVID-19 Genomics UK (COG-UK) Consortium | Ana da Silva Filipe, Natasha Johnson, Kathy Smollett, Daniel Mair, Stephen Carmichael, Lily Tong, Jenna Nichols, Elihu Aranday-Cortes, Kirstyn Brunker, Yasmin Parr, Alice Broos, Kyriaki Nomikou; Sarah McDonald, Marc Niebel, Patawee Asamaphan; Richard Orton, Joseph Hughes, Sreenu Vattipally, David L Robertson; Alasdair MacLean, Rory Gunson; Kathy Li, Natasha Jesudason, Rajiv Shah, James Shepherd, Antonia Ho, Emma Thomson |
| EPI_ISL_473783, EPI_ISL_473784, EPI_ISL_473785, EPI_ISL_473786, EPI_ISL_473787, EPI_ISL_473788, EPI_ISL_473789, EPI_ISL_473790, EPI_ISL_473791, EPI_ISL_473792, EPI_ISL_473793, EPI_ISL_473794, EPI_ISL_473795, EPI_ISL_473796, EPI_ISL_473797, EPI_ISL_473798, EPI_ISL_473799, EPI_ISL_473800, EPI_ISL_473801, EPI_ISL_473803, EPI_ISL_473804, EPI_ISL_473805, EPI_ISL_473806, EPI_ISL_473807, EPI_ISL_473808, EPI_ISL_473809, EPI_ISL_473810, EPI_ISL_473811, EPI_ISL_473812, EPI_ISL_473813, EPI_ISL_473814, EPI_ISL_473815, EPI_ISL_473817, EPI_ISL_473818, EPI_ISL_473819, EPI_ISL_473820, EPI_ISL_473821, EPI_ISL_473822, EPI_ISL_473823, EPI_ISL_473824, EPI_ISL_473825, EPI_ISL_473826, EPI_ISL_473827, EPI_ISL_473828, EPI_ISL_473829, EPI_ISL_473830, EPI_ISL_473831, EPI_ISL_473832, EPI_ISL_473833, EPI_ISL_473834, EPI_ISL_473836, EPI_ISL_473838, EPI_ISL_473839, EPI_ISL_473840, EPI_ISL_473841, EPI_ISL_473842, EPI_ISL_473843, EPI_ISL_473844, EPI_ISL_473845, EPI_ISL_473846, EPI_ISL_473847, EPI_ISL_473848, EPI_ISL_473849, EPI_ISL_473850, EPI_ISL_473851, EPI_ISL_473852, EPI_ISL_473854, EPI_ISL_473855, EPI_ISL_473856, EPI_ISL_473857, EPI_ISL_473858, EPI_ISL_473859, EPI_ISL_473860, EPI_ISL_473861, EPI_ISL_473862, EPI_ISL_473863, EPI_ISL_473864, EPI_ISL_473865, EPI_ISL_473866, EPI_ISL_473867, EPI_ISL_473868, EPI_ISL_473869, EPI_ISL_473870, EPI_ISL_473871, EPI_ISL_473872, EPI_ISL_473873, EPI_ISL_473874, EPI_ISL_473875, EPI_ISL_473876, EPI_ISL_473877, EPI_ISL_473878, EPI_ISL_473880, EPI_ISL_473881, EPI_ISL_473882, EPI_ISL_473883, EPI_ISL_473884, | | | |

| | | | |
|---|---|---|---|
| EPI_ISL_473885, EPI_ISL_473886, EPI_ISL_473887, EPI_ISL_473889, EPI_ISL_473890, EPI_ISL_473891, EPI_ISL_473892, EPI_ISL_473893, EPI_ISL_473894, EPI_ISL_473895, EPI_ISL_473896, EPI_ISL_473897, EPI_ISL_473898, EPI_ISL_473899, EPI_ISL_473900, EPI_ISL_473901, EPI_ISL_473902, EPI_ISL_473903, EPI_ISL_473904, EPI_ISL_473905, EPI_ISL_473906, EPI_ISL_473907, EPI_ISL_473908, EPI_ISL_473909, EPI_ISL_473910, EPI_ISL_473911, EPI_ISL_473912, EPI_ISL_473913, EPI_ISL_473914, EPI_ISL_473915, EPI_ISL_473916, EPI_ISL_473917, EPI_ISL_473918, EPI_ISL_473919, EPI_ISL_473920, EPI_ISL_473921, EPI_ISL_473922, EPI_ISL_473924, EPI_ISL_473925, EPI_ISL_473926, EPI_ISL_473927, EPI_ISL_473928, EPI_ISL_473929, EPI_ISL_473930, EPI_ISL_473931, EPI_ISL_473932, EPI_ISL_473933, EPI_ISL_473934, EPI_ISL_473935, EPI_ISL_473936, EPI_ISL_473938, EPI_ISL_473940, EPI_ISL_473946, EPI_ISL_473947, EPI_ISL_473948, EPI_ISL_473949, EPI_ISL_473952, EPI_ISL_473953, EPI_ISL_473954 | | | |
| see above | Virology Department, Royal Infirmary of Edinburgh, NHS Lothian / School of Biological Sciences, University of Edinburgh / Institute of Genetics and Molecular Medicine, University of Edinburgh | COVID-19 Genomics UK (COG-UK) Consortium | McHugh M, Dewar R, Rooke S, Gallagher M, Balcaza C, O'Toole Á, Scher E, Hill V, McCrone JT, Colquhoun R, Yu X, Jackson B, Rambaut A, Williams TC, Templeton K |
| EPI_ISL_473958, EPI_ISL_473959, EPI_ISL_473960, EPI_ISL_473961, EPI_ISL_473962, EPI_ISL_473965, EPI_ISL_473966, EPI_ISL_473967, EPI_ISL_473969, EPI_ISL_473970, EPI_ISL_473971, EPI_ISL_473975, EPI_ISL_473976, EPI_ISL_473977, EPI_ISL_473978, EPI_ISL_473981, EPI_ISL_473983, EPI_ISL_473984, EPI_ISL_473986, EPI_ISL_473988, EPI_ISL_473990, EPI_ISL_473991, EPI_ISL_473992, EPI_ISL_473993, EPI_ISL_473994, EPI_ISL_473998, EPI_ISL_474000, EPI_ISL_474001, EPI_ISL_474006, EPI_ISL_474007, EPI_ISL_474009, EPI_ISL_474010, EPI_ISL_474011, EPI_ISL_474012, EPI_ISL_474013, EPI_ISL_474020, EPI_ISL_474021, EPI_ISL_474022, EPI_ISL_474026, EPI_ISL_474030, EPI_ISL_474034, EPI_ISL_474036, EPI_ISL_474040, EPI_ISL_474043, EPI_ISL_474044, EPI_ISL_474046, EPI_ISL_474047, EPI_ISL_474048, EPI_ISL_474050, EPI_ISL_474052, EPI_ISL_474055, EPI_ISL_474056, EPI_ISL_474058, EPI_ISL_474059, EPI_ISL_474060, EPI_ISL_474061, EPI_ISL_474062, EPI_ISL_474064, EPI_ISL_474065, EPI_ISL_474067, EPI_ISL_474069, EPI_ISL_474071, EPI_ISL_474073, EPI_ISL_474075, EPI_ISL_474077, EPI_ISL_474078, EPI_ISL_474080, EPI_ISL_474081, EPI_ISL_474084, EPI_ISL_474085, EPI_ISL_474095, EPI_ISL_474096, EPI_ISL_474098, EPI_ISL_474099, EPI_ISL_474100, EPI_ISL_474101, EPI_ISL_474103, EPI_ISL_474104, EPI_ISL_474105, EPI_ISL_474110, EPI_ISL_474112, EPI_ISL_474113, EPI_ISL_474115, EPI_ISL_474116, EPI_ISL_474117, EPI_ISL_474118, EPI_ISL_474122, EPI_ISL_474124, EPI_ISL_474125, EPI_ISL_474126, EPI_ISL_474131, EPI_ISL_474135, EPI_ISL_474136, EPI_ISL_474137, EPI_ISL_474140, EPI_ISL_474141, EPI_ISL_474147, EPI_ISL_474150, EPI_ISL_474152, EPI_ISL_474155, EPI_ISL_474156, EPI_ISL_474157, EPI_ISL_474159, EPI_ISL_474160, EPI_ISL_474161, EPI_ISL_474162, EPI_ISL_474164, EPI_ISL_474165, EPI_ISL_474167, EPI_ISL_474168, EPI_ISL_474169, EPI_ISL_474171, EPI_ISL_474173, EPI_ISL_474175, EPI_ISL_474177, EPI_ISL_474178, EPI_ISL_474179, EPI_ISL_474180, EPI_ISL_474181, EPI_ISL_474182, EPI_ISL_474183, EPI_ISL_474184, EPI_ISL_474187, EPI_ISL_474189, EPI_ISL_474190, EPI_ISL_474193, EPI_ISL_474195, EPI_ISL_474198, EPI_ISL_474201, EPI_ISL_474202, EPI_ISL_474204, EPI_ISL_474205, EPI_ISL_474206, EPI_ISL_474208, EPI_ISL_474209, EPI_ISL_474210, EPI_ISL_474211, EPI_ISL_474212, EPI_ISL_474214, EPI_ISL_474215, EPI_ISL_474216, EPI_ISL_474217, EPI_ISL_474220, EPI_ISL_474225, EPI_ISL_474239, EPI_ISL_474240, EPI_ISL_474248, EPI_ISL_474250, EPI_ISL_474253, EPI_ISL_474254, EPI_ISL_474259, EPI_ISL_474260, EPI_ISL_474265, EPI_ISL_474275, EPI_ISL_474291, EPI_ISL_474293, EPI_ISL_474296, EPI_ISL_474301, EPI_ISL_474302, EPI_ISL_474303, EPI_ISL_474316, EPI_ISL_474318, EPI_ISL_474322, EPI_ISL_474323, EPI_ISL_474333, EPI_ISL_474336, EPI_ISL_474340, EPI_ISL_474343, EPI_ISL_474349, EPI_ISL_474352, EPI_ISL_474353, EPI_ISL_474355, EPI_ISL_474356, EPI_ISL_474358, EPI_ISL_474359, EPI_ISL_474360, EPI_ISL_474361, EPI_ISL_474362, EPI_ISL_474364, EPI_ISL_474367, EPI_ISL_474368, EPI_ISL_474369, EPI_ISL_474370, EPI_ISL_474375, EPI_ISL_474376, EPI_ISL_474377, EPI_ISL_474378, EPI_ISL_474382, EPI_ISL_474384, EPI_ISL_474387, EPI_ISL_474388, EPI_ISL_474391, EPI_ISL_474395, EPI_ISL_474397, EPI_ISL_474398, EPI_ISL_474403, EPI_ISL_474421, EPI_ISL_474464, EPI_ISL_474466, EPI_ISL_474467, EPI_ISL_474468, EPI_ISL_474471, EPI_ISL_474477, EPI_ISL_474478, EPI_ISL_474479, EPI_ISL_474480, EPI_ISL_474486, EPI_ISL_474490, EPI_ISL_474494, EPI_ISL_474497, EPI_ISL_474498, EPI_ISL_474501, EPI_ISL_474504, EPI_ISL_474509, EPI_ISL_474517, EPI_ISL_474522, EPI_ISL_474525, EPI_ISL_474526, EPI_ISL_474537, EPI_ISL_474540, EPI_ISL_474542, EPI_ISL_474543, EPI_ISL_474545, EPI_ISL_474551, EPI_ISL_474553, EPI_ISL_474556, EPI_ISL_474557, EPI_ISL_474566, EPI_ISL_474576, EPI_ISL_474580, EPI_ISL_474588, EPI_ISL_474592, EPI_ISL_474595, EPI_ISL_474597, EPI_ISL_474600, EPI_ISL_474607, EPI_ISL_474608, EPI_ISL_474609, EPI_ISL_474615, EPI_ISL_474616, EPI_ISL_474622, EPI_ISL_474624, EPI_ISL_474626, EPI_ISL_474631, EPI_ISL_474634, EPI_ISL_474637, EPI_ISL_474639, EPI_ISL_474650, EPI_ISL_474651, EPI_ISL_474656, EPI_ISL_474657, EPI_ISL_474658, EPI_ISL_474660, EPI_ISL_474664, EPI_ISL_474671, EPI_ISL_474675, EPI_ISL_474676, EPI_ISL_474677, EPI_ISL_474680, EPI_ISL_474682, EPI_ISL_474684, EPI_ISL_474686, EPI_ISL_474689, EPI_ISL_474692, EPI_ISL_474693, EPI_ISL_474699, EPI_ISL_474704, EPI_ISL_474740, EPI_ISL_474748, EPI_ISL_474755, EPI_ISL_474768, EPI_ISL_474779, EPI_ISL_474780, EPI_ISL_474785, EPI_ISL_474794, EPI_ISL_474795 | | | |
| see above | Originating lab: Wales Specialist Virology Centre Sequencing lab: Pathogen Genomics Unit | COVID-19 Genomics UK (COG-UK) Consortium | Catherine Moore, Johnathan Evans, Laura Gifford, Malorie Perry, Simon Cottrell, Angela Marchbank, Alec Birchley, Alexander Adams, Amy Gaskin, Bree Gatica-Wilcox, Jason Coombes, Joel Southgate, Lauren Gilbert, Lee Graham, Nicole Pacchiarini, Sara Kumziene-Summerhayes, Sarah Taylor, Sophie Jones, Sara Rey, Matthew Bull, Joanne Watkins, Sally Corden, Tom Connor |
| EPI_ISL_474797 | Hospital Universitario Virgen de las Nieves de Granada-SAS | SeqCOVID-SPAIN consortium/IBV(CSIC) | Mercedes Pérez Ruiz, Sara Sanbonmatsu Gámez, Irene Pedrosa Corral, José M. Navarro-Marí and SeqCOVID-SPAIN consortium |
| EPI_ISL_474798, EPI_ISL_474801, EPI_ISL_474802, EPI_ISL_474803, EPI_ISL_474804, EPI_ISL_474805, EPI_ISL_474806, EPI_ISL_474807, EPI_ISL_474808, EPI_ISL_474809, EPI_ISL_474810, EPI_ISL_474811, EPI_ISL_474812, EPI_ISL_474813, EPI_ISL_474815, EPI_ISL_474816, EPI_ISL_474818, EPI_ISL_474819, EPI_ISL_474820, EPI_ISL_474822, EPI_ISL_474823, EPI_ISL_474824, EPI_ISL_474825, EPI_ISL_474826, EPI_ISL_474827, EPI_ISL_474828, EPI_ISL_474830, EPI_ISL_474831 | | | |
| see above | Complejo Hospitalario Universitario de Albacete | SeqCOVID-SPAIN consortium/IBV(CSIC) | Encarnacion Simarro Córdoba, Julia Lozano Serra, Lorena Robles Fonseca , Monica Parra Grandes, Caridad Sainz de Baranda Camino and SeqCOVID-SPAIN consortium |
| EPI_ISL_474832, EPI_ISL_474833, EPI_ISL_474834, EPI_ISL_474836 | Hospital Universitario Virgen de las Nieves de Granada-SAS | SeqCOVID-SPAIN consortium/IBV(CSIC) | Mercedes Pérez Ruiz, Sara Sanbonmatsu Gámez, Irene Pedrosa Corral, José M. Navarro-Marí and SeqCOVID-SPAIN consortium |
| EPI_ISL_474837, EPI_ISL_474838 | Complejo Hospitalario Universitario de Albacete | SeqCOVID-SPAIN consortium/IBV(CSIC) | Encarnacion Simarro Córdoba, Julia Lozano Serra, Lorena Robles Fonseca , Monica Parra Grandes, Caridad Sainz de Baranda Camino and SeqCOVID-SPAIN consortium |
| EPI_ISL_474841, EPI_ISL_474842, EPI_ISL_474843, EPI_ISL_474845 | Hospital Universitario Virgen de las Nieves de Granada-SAS | SeqCOVID-SPAIN consortium/IBV(CSIC) | Mercedes Pérez Ruiz, Sara Sanbonmatsu Gámez, Irene Pedrosa Corral, José M. Navarro-Marí and SeqCOVID-SPAIN consortium |
| EPI_ISL_474847 | Complejo Hospitalario Universitario de Albacete | SeqCOVID-SPAIN consortium/IBV(CSIC) | Encarnacion Simarro Córdoba, Julia Lozano Serra, Lorena Robles Fonseca , Monica Parra Grandes, Caridad Sainz de Baranda Camino and SeqCOVID-SPAIN consortium |
| EPI_ISL_474850, EPI_ISL_474851, EPI_ISL_474852 | Hospital Universitario Virgen de las Nieves de Granada-SAS | SeqCOVID-SPAIN consortium/IBV(CSIC) | Mercedes Pérez Ruiz, Sara Sanbonmatsu Gámez, Irene Pedrosa Corral, José M. Navarro-Marí and SeqCOVID-SPAIN consortium |
| EPI_ISL_474853 | Complejo Hospitalario Universitario de Albacete | SeqCOVID-SPAIN consortium/IBV(CSIC) | Encarnacion Simarro Córdoba, Julia Lozano Serra, Lorena Robles Fonseca , Monica Parra Grandes, Caridad Sainz de Baranda Camino and SeqCOVID-SPAIN consortium |
| EPI_ISL_474854, EPI_ISL_474855, EPI_ISL_474856, EPI_ISL_474857, EPI_ISL_474859, EPI_ISL_474860, EPI_ISL_474862, EPI_ISL_474864, EPI_ISL_474867, EPI_ISL_474868, EPI_ISL_474869, EPI_ISL_474870, EPI_ISL_474871, EPI_ISL_474872, EPI_ISL_474873, EPI_ISL_474874, EPI_ISL_474875, EPI_ISL_474876, EPI_ISL_474879, EPI_ISL_474880, EPI_ISL_474881, EPI_ISL_474882, EPI_ISL_474883, EPI_ISL_474886, EPI_ISL_474887, EPI_ISL_474888, EPI_ISL_474889, EPI_ISL_474890, EPI_ISL_474891, EPI_ISL_474892, EPI_ISL_474894, EPI_ISL_474897, EPI_ISL_474898, EPI_ISL_474900 | | | |
| see above | Hospital Universitario Virgen de las Nieves de Granada-SAS | SeqCOVID-SPAIN consortium/IBV(CSIC) | Mercedes Pérez Ruiz, Sara Sanbonmatsu Gámez, Irene Pedrosa Corral, José M. Navarro-Marí and SeqCOVID-SPAIN consortium |
| EPI_ISL_474901, EPI_ISL_474902, EPI_ISL_474903, EPI_ISL_474904 | Complejo Hospitalario Universitario de Albacete | SeqCOVID-SPAIN consortium/IBV(CSIC) | Encarnacion Simarro Córdoba, Julia Lozano Serra, Lorena Robles Fonseca , Monica Parra Grandes, Caridad Sainz de Baranda Camino and SeqCOVID-SPAIN consortium |
| EPI_ISL_474908, EPI_ISL_474909 | Hospital Universitario Virgen de las Nieves de Granada-SAS | SeqCOVID-SPAIN consortium/IBV(CSIC) | Mercedes Pérez Ruiz, Sara Sanbonmatsu Gámez, Irene Pedrosa Corral, José M. Navarro-Marí and SeqCOVID-SPAIN consortium |
| EPI_ISL_474910, EPI_ISL_474911, EPI_ISL_474912, EPI_ISL_474913, EPI_ISL_474914, EPI_ISL_474915, EPI_ISL_474916, EPI_ISL_474917 | Hospital Universitario de Gran Canaria Dr. Negrín | SeqCOVID-SPAIN consortium/IBV(CSIC) | M. Carmen Pérez González, Francisco J. Chamizo López, Ana Bordes Benítez and SeqCOVID-SPAIN consortium |
| EPI_ISL_474919 | Complejo Hospitalario Universitario de Albacete | SeqCOVID-SPAIN consortium/IBV(CSIC) | Encarnacion Simarro Córdoba, Julia Lozano Serra, Lorena Robles Fonseca , Monica Parra Grandes, Caridad Sainz de Baranda Camino and SeqCOVID-SPAIN consortium |
| EPI_ISL_474920 | Hospital Universitario Virgen de las Nieves de Granada-SAS | SeqCOVID-SPAIN consortium/IBV(CSIC) | Mercedes Pérez Ruiz, Sara Sanbonmatsu Gámez, Irene Pedrosa Corral, José M. Navarro-Marí and SeqCOVID-SPAIN consortium |
| EPI_ISL_474921 | Complejo Hospitalario Universitario de Albacete | SeqCOVID-SPAIN consortium/IBV(CSIC) | Encarnacion Simarro Córdoba, Julia Lozano Serra, Lorena Robles Fonseca , Monica Parra Grandes, Caridad Sainz de Baranda Camino and SeqCOVID-SPAIN consortium |
| EPI_ISL_474922, EPI_ISL_474923, EPI_ISL_474926, EPI_ISL_474927, EPI_ISL_474928, EPI_ISL_474930 | Hospital Universitario Virgen de las Nieves de Granada-SAS | SeqCOVID-SPAIN consortium/IBV(CSIC) | Mercedes Pérez Ruiz, Sara Sanbonmatsu Gámez, Irene Pedrosa Corral, José M. Navarro-Marí and SeqCOVID-SPAIN consortium |
| EPI_ISL_474933 | Complejo Hospitalario Universitario de Albacete | SeqCOVID-SPAIN consortium/IBV(CSIC) | Encarnacion Simarro Córdoba, Julia Lozano Serra, Lorena Robles Fonseca , Monica Parra Grandes, Caridad Sainz de Baranda Camino and SeqCOVID-SPAIN consortium |
| EPI_ISL_474934, EPI_ISL_474935, EPI_ISL_474936, EPI_ISL_474939 | Hospital Universitario Virgen de las Nieves de Granada-SAS | SeqCOVID-SPAIN consortium/IBV(CSIC) | Mercedes Pérez Ruiz, Sara Sanbonmatsu Gámez, Irene Pedrosa Corral, José M. Navarro-Marí and SeqCOVID-SPAIN consortium |
| EPI_ISL_474940 | Complejo Hospitalario Universitario de Albacete | SeqCOVID-SPAIN consortium/IBV(CSIC) | Encarnacion Simarro Córdoba, Julia Lozano Serra, Lorena Robles Fonseca , Monica Parra Grandes, Caridad Sainz de Baranda Camino and SeqCOVID-SPAIN consortium |
| EPI_ISL_474943, EPI_ISL_474944 | Hospital Universitario Virgen de las Nieves de Granada-SAS | SeqCOVID-SPAIN consortium/IBV(CSIC) | Mercedes Pérez Ruiz, Sara Sanbonmatsu Gámez, Irene Pedrosa Corral, José M. Navarro-Marí and SeqCOVID-SPAIN consortium |
| EPI_ISL_474946, EPI_ISL_474947 | Complejo Hospitalario Universitario de Albacete | SeqCOVID-SPAIN consortium/IBV(CSIC) | Encarnacion Simarro Córdoba, Julia Lozano Serra, Lorena Robles Fonseca , Monica Parra Grandes, Caridad Sainz de Baranda Camino and SeqCOVID-SPAIN consortium |
| EPI_ISL_474948, EPI_ISL_474949, EPI_ISL_474950 | Hospital Universitario Virgen de las Nieves de Granada-SAS | SeqCOVID-SPAIN consortium/IBV(CSIC) | Mercedes Pérez Ruiz, Sara Sanbonmatsu Gámez, Irene Pedrosa Corral, José M. Navarro-Marí and SeqCOVID-SPAIN consortium |
| EPI_ISL_474951, EPI_ISL_474952, EPI_ISL_474953, EPI_ISL_474954, EPI_ISL_474955, EPI_ISL_474956 | Complejo Hospitalario Universitario de Albacete | SeqCOVID-SPAIN consortium/IBV(CSIC) | Encarnacion Simarro Córdoba, Julia Lozano Serra, Lorena Robles Fonseca , Monica Parra Grandes, Caridad Sainz de Baranda Camino and SeqCOVID-SPAIN consortium |
| EPI_ISL_474958 | Israeli Central Virology laboratory | Israel Central Virology laboratory | Neta Zuckerman, Efrat Dahan Bucris, Oran Erster, Ella Mendelson, Michal Mandelboim |
| EPI_ISL_474959, EPI_ISL_474960, EPI_ISL_474961, EPI_ISL_474962, EPI_ISL_474963, EPI_ISL_474964, EPI_ISL_474965, EPI_ISL_474966, EPI_ISL_474967, EPI_ISL_474968, EPI_ISL_474969, EPI_ISL_474970, EPI_ISL_474971, EPI_ISL_474972, EPI_ISL_474973, EPI_ISL_474974, EPI_ISL_474975, EPI_ISL_474977, EPI_ISL_474978, EPI_ISL_474979, EPI_ISL_474980, EPI_ISL_474981, EPI_ISL_474982, EPI_ISL_474983, EPI_ISL_474984, EPI_ISL_474985, EPI_ISL_474986, EPI_ISL_474987, EPI_ISL_474988, EPI_ISL_474989, EPI_ISL_474990, EPI_ISL_474991, EPI_ISL_474992, EPI_ISL_474993, EPI_ISL_474994, EPI_ISL_474995, EPI_ISL_474996, EPI_ISL_474997, EPI_ISL_474999, EPI_ISL_475000, EPI_ISL_475001, EPI_ISL_475002, EPI_ISL_475003, EPI_ISL_475004, EPI_ISL_475005, EPI_ISL_475006, EPI_ISL_475007, EPI_ISL_475008, EPI_ISL_475009, EPI_ISL_475010, EPI_ISL_475011, EPI_ISL_475012, EPI_ISL_475013, EPI_ISL_475014, EPI_ISL_475015, EPI_ISL_475016, EPI_ISL_475017, EPI_ISL_475018, EPI_ISL_475019, EPI_ISL_475020, EPI_ISL_475021, EPI_ISL_475022, EPI_ISL_475023, EPI_ISL_475024, EPI_ISL_475025 | | | |
| see above | Israel Central Virology laboratory | Israel Central Virology laboratory | Neta Zuckerman, Efrat Dahan Bucris, Oran Erster, Ella Mendelson, Michal Mandelboim |
| EPI_ISL_475026 | Banas Medical College and Research Institute | Gujarat Biotechnology Research Centre | Sunil R Joshi, Viren s Doshi, Pritesh Sabara, Apurvasinh Puvar, Janvi Raval, Zarna Patel, Monika Gandhi, Pinal Trivedi, Maharshi Pandya, Nidhi Patel, Nitin Savaliya, Raghawendra Kumar, Dinesh Kumar, Zuber Saiyed, Komal Patel, Labdhi Pandya, Snehal Bagatharia, Radhika Khara, Neha Rajpara, R D Dixit, A M Kadri, Harsh Bakshi, Chaitanya Joshi, Madhvi Joshi |
| EPI_ISL_475027 | Banas Medical College and Research Institute | Gujarat Biotechnology Research Centre | Viren s Doshi, Pritesh Sabara, Apurvasinh Puvar, Janvi Raval, Zarna Patel, Monika Gandhi, Pinal Trivedi, Maharshi Pandya, Nidhi Patel, Nitin Savaliya, Raghawendra Kumar, Dinesh Kumar, Zuber Saiyed, Komal Patel, Labdhi Pandya, Snehal Bagatharia, Radhika Khara, Sunil R Joshi, Afzal Ansari, R D Dixit, A M Kadri, Harsh Bakshi, Chaitanya Joshi, Madhvi Joshi |
| EPI_ISL_475028 | Banas Medical College and Research Institute | Gujarat Biotechnology Research Centre | Pritesh Sabara, Apurvasinh Puvar, Janvi Raval, Zarna Patel, Monika Gandhi, Pinal Trivedi, Maharshi Pandya, Nidhi Patel, Nitin Savaliya, Raghawendra Kumar, Dinesh Kumar, Zuber Saiyed, Komal Patel, Labdhi Pandya, Snehal Bagatharia, Radhika Khara, Sunil R Joshi, Viren s Doshi, Fenil Patel, R D Dixit, A M Kadri, Harsh Bakshi, Chaitanya Joshi, Madhvi Joshi |
| EPI_ISL_475029 | Banas Medical College and Research Institute | Gujarat Biotechnology Research Centre | Apurvasinh Puvar, Janvi Raval, Zarna Patel, Monika Gandhi, Pinal Trivedi, Maharshi Pandya, Nidhi Patel, Nitin Savaliya, Raghawendra Kumar, Dinesh Kumar, Zuber Saiyed, Komal Patel, Labdhi Pandya, Snehal Bagatharia, Radhika Khara, Sunil R Joshi, Viren s Doshi, Pritesh Sabara, Neelam Nathani, R D Dixit, A M Kadri, Harsh Bakshi, Chaitanya Joshi, Madhvi Joshi |
| EPI_ISL_475030 | Department of MicroBiology, Government Medical College, Surat | Gujarat Biotechnology Research Centre | Janvi Raval, Zarna Patel, Monika Gandhi, Pinal Trivedi, Maharshi Pandya, Nidhi Patel, Nitin Savaliya, Raghawendra Kumar, Dinesh Kumar, Zuber Saiyed, Komal Patel, Labdhi Pandya, Snehal Bagatharia, Naresh Chauhan, Summaiya Mullan, Amit gamit, Pritesh Sabara, Apurvasinh Puvar, Armi Chaudhari, R D Dixit, A M Kadri, Harsh Bakshi, Chaitanya Joshi, Madhvi Joshi |
| EPI_ISL_475031 | Department of MicroBiology, Government Medical College, Surat | Gujarat Biotechnology Research Centre | Zarna Patel, Monika Gandhi, Pinal Trivedi, Maharshi Pandya, Nidhi Patel, Nitin Savaliya, Raghawendra Kumar, Dinesh Kumar, Zuber Saiyed, Komal Patel, Labdhi Pandya, Snehal Bagatharia, Naresh Chauhan, Summaiya Mullan, Amit gamit, Pritesh Sabara, Apurvasinh Puvar, Janvi Raval, Bhavya Jindal, R D Dixit, A M Kadri, Harsh Bakshi, Chaitanya Joshi, Madhvi Joshi |
| EPI_ISL_475032 | Department of MicroBiology, Government Medical College, Surat | Gujarat Biotechnology Research Centre | Monika Gandhi, Pinal Trivedi, Maharshi Pandya, Nidhi Patel, Nitin Savaliya, Raghawendra Kumar, Dinesh Kumar, Zuber Saiyed, Komal Patel, Labdhi Pandya, Snehal Bagatharia, Naresh Chauhan, Summaiya Mullan, Amit gamit, Pritesh Sabara, Apurvasinh Puvar, Janvi Raval, Zarna Patel, Priyanka P Vatsa, R D Dixit, A M Kadri, Harsh Bakshi, Chaitanya Joshi, Madhvi Joshi |
| EPI_ISL_475033 | Department of MicroBiology, Government Medical College, Surat | Gujarat Biotechnology Research Centre | Pinal Trivedi, Maharshi Pandya, Nidhi Patel, Nitin Savaliya, Raghawendra Kumar, Dinesh Kumar, Zuber Saiyed, Komal Patel, Labdhi Pandya, Snehal Bagatharia, Naresh Chauhan, Summaiya Mullan, Amit gamit, Pritesh Sabara, Apurvasinh Puvar, Janvi Raval, Zarna Patel, Monika Gandhi, Pooja P Doshi, R D Dixit, A M Kadri, Harsh Bakshi, Chaitanya Joshi, Madhvi Joshi |
| EPI_ISL_475034 | Department of MicroBiology, Government Medical College, Surat | Gujarat Biotechnology Research Centre | Maharshi Pandya, Nidhi Patel, Nitin Savaliya, Raghawendra Kumar, Dinesh Kumar, Zuber Saiyed, Komal Patel, Labdhi Pandya, Snehal Bagatharia, Naresh Chauhan, Summaiya Mullan, Amit gamit, Pritesh Sabara, Apurvasinh Puvar, Janvi Raval, Zarna Patel, Monika Gandhi, Pinal Trivedi, Akanksha Verma, R D Dixit, A M Kadri, Harsh Bakshi, Chaitanya Joshi, Madhvi Joshi |
| EPI_ISL_475035 | Department of MicroBiology, Government Medical College, Surat | Gujarat Biotechnology Research Centre | Nidhi Patel, Nitin Savaliya, Raghawendra Kumar, Dinesh Kumar, Zuber Saiyed, Komal Patel, Labdhi Pandya, Snehal Bagatharia, Naresh Chauhan, Summaiya Mullan, Amit gamit, Pritesh Sabara, Apurvasinh Puvar, Janvi Raval, Zarna Patel, Monika Gandhi, Pinal Trivedi, Maharshi Pandya, Priti Pandita, R D Dixit, A M Kadri, Harsh Bakshi, Chaitanya Joshi, Madhvi Joshi |
| EPI_ISL_475036 | Department of MicroBiology, Government Medical College, Surat | Gujarat Biotechnology Research Centre | Nitin Savaliya, Raghawendra Kumar, Dinesh Kumar, Zuber Saiyed, Komal Patel, Labdhi Pandya, Snehal Bagatharia, Naresh Chauhan, Summaiya Mullan, Amit gamit, Pritesh Sabara, Apurvasinh Puvar, Janvi Raval, Zarna Patel, Monika Gandhi, Pinal Trivedi, Maharshi Pandya, Nidhi Patel, Pragya Sharma, R D Dixit, A M Kadri, Harsh Bakshi, Chaitanya Joshi, Madhvi Joshi |
| EPI_ISL_475037 | Department of MicroBiology, Government Medical College, Surat | Gujarat Biotechnology Research Centre | Raghawendra Kumar, Dinesh Kumar, Zuber Saiyed, Komal Patel, Labdhi Pandya, Snehal Bagatharia, Naresh Chauhan, Summaiya Mullan, Amit gamit, Pritesh Sabara, Apurvasinh Puvar, Janvi Raval, Zarna Patel, Monika Gandhi, Pinal Trivedi, Maharshi Pandya, Nidhi Patel, Nitin Savaliya, Neha Rajpara, R D Dixit, A M Kadri, Harsh Bakshi, Chaitanya Joshi, Madhvi Joshi |
| EPI_ISL_475038 | Department of MicroBiology, Government Medical College, Surat | Gujarat Biotechnology Research Centre | Dinesh Kumar, Zuber Saiyed, Komal Patel, Labdhi Pandya, Snehal Bagatharia, Naresh Chauhan, Summaiya Mullan, Amit gamit, Pritesh Sabara, Apurvasinh Puvar, Janvi Raval, Zarna Patel, Monika |

| | | | |
|---|---|---|---|
| | | | Gandhi, Pinal Trivedi, Maharshi Pandya, Nidhi Patel, Nitin Savaliya, Raghawendra Kumar, Afzal Ansari, R D Dixit, A M Kadri, Harsh Bakshi, Chaitanya Joshi, Madhvi Joshi |
| EPI_ISL_475039 | Department of MicroBiology, Government Medical College, Surat | Gujarat Biotechnology Research Centre | Zuber Saiyed, Komal Patel, Labdhi Pandya, Snehal Bagatharia, Naresh Chauhan, Summaiya Mullan, Amit gamit, Pritesh Sabara, Apurvasinh Puvar, Janvi Raval, Zarna Patel, Monika Gandhi, Pinal Trivedi, Maharshi Pandya, Nidhi Patel, Nitin Savaliya, Raghawendra Kumar, Dinesh Kumar, Fenil Patel, R D Dixit, A M Kadri, Harsh Bakshi, Chaitanya Joshi, Madhvi Joshi |
| EPI_ISL_475040 | Department of MicroBiology, Government Medical College, Surat | Gujarat Biotechnology Research Centre | Komal Patel, Labdhi Pandya, Snehal Bagatharia, Naresh Chauhan, Summaiya Mullan, Amit gamit, Pritesh Sabara, Apurvasinh Puvar, Janvi Raval, Zarna Patel, Monika Gandhi, Pinal Trivedi, Maharshi Pandya, Nidhi Patel, Nitin Savaliya, Raghawendra Kumar, Dinesh Kumar, Zuber Saiyed, Neelam Nathani, R D Dixit, A M Kadri, Harsh Bakshi, Chaitanya Joshi, Madhvi Joshi |
| EPI_ISL_475043 | Department of MicroBiology, Government Medical College, Surat | Gujarat Biotechnology Research Centre | Naresh Chauhan, Summaiya Mullan, Amit gamit, Pritesh Sabara, Apurvasinh Puvar, Janvi Raval, Zarna Patel, Monika Gandhi, Pinal Trivedi, Maharshi Pandya, Nidhi Patel, Nitin Savaliya, Raghawendra Kumar, Dinesh Kumar, Zuber Saiyed, Komal Patel, Labdhi Pandya, Snehal Bagatharia, Priyanka P Vatsa, R D Dixit, A M Kadri, Harsh Bakshi, Chaitanya Joshi, Madhvi Joshi |
| EPI_ISL_475044 | Department of MicroBiology, Government Medical College, Surat | Gujarat Biotechnology Research Centre | Summaiya Mullan, Amit gamit, Pritesh Sabara, Apurvasinh Puvar, Janvi Raval, Zarna Patel, Monika Gandhi, Pinal Trivedi, Maharshi Pandya, Nidhi Patel, Nitin Savaliya, Raghawendra Kumar, Dinesh Kumar, Zuber Saiyed, Komal Patel, Labdhi Pandya, Snehal Bagatharia, Naresh Chauhan, Pooja P Doshi, R D Dixit, A M Kadri, Harsh Bakshi, Chaitanya Joshi, Madhvi Joshi |
| EPI_ISL_475045 | Department of MicroBiology, Government Medical College, Surat | Gujarat Biotechnology Research Centre | Amit gamit, Pritesh Sabara, Apurvasinh Puvar, Janvi Raval, Zarna Patel, Monika Gandhi, Pinal Trivedi, Maharshi Pandya, Nidhi Patel, Nitin Savaliya, Raghawendra Kumar, Dinesh Kumar, Zuber Saiyed, Komal Patel, Labdhi Pandya, Snehal Bagatharia, Naresh Chauhan, Summaiya Mullan, Akanksha Verma, R D Dixit, A M Kadri, Harsh Bakshi, Chaitanya Joshi, Madhvi Joshi |
| EPI_ISL_475046 | Department of MicroBiology, Government Medical College, Surat | Gujarat Biotechnology Research Centre | Pritesh Sabara, Apurvasinh Puvar, Janvi Raval, Zarna Patel, Monika Gandhi, Pinal Trivedi, Maharshi Pandya, Nidhi Patel, Nitin Savaliya, Raghawendra Kumar, Dinesh Kumar, Zuber Saiyed, Komal Patel, Labdhi Pandya, Snehal Bagatharia, Naresh Chauhan, Summaiya Mullan, Amit gamit, Priti Pandita, R D Dixit, A M Kadri, Harsh Bakshi, Chaitanya Joshi, Madhvi Joshi |
| EPI_ISL_475047 | GMERS Medical College & Hospital | Gujarat Biotechnology Research Centre | Apurvasinh Puvar, Janvi Raval, Zarna Patel, Monika Gandhi, Pinal Trivedi, Maharshi Pandya, Nidhi Patel, Nitin Savaliya, Raghawendra Kumar, Dinesh Kumar, Zuber Saiyed, Komal Patel, Labdhi Pandya, Snehal Bagatharia, Meenakshi Shah, Neena Doshi, Varsha Godbole, Pritesh Sabara, Pragya Sharma, R D Dixit, A M Kadri, Harsh Bakshi, Chaitanya Joshi, Madhvi Joshi |
| EPI_ISL_475048 | GMERS Medical College & Hospital | Gujarat Biotechnology Research Centre | Janvi Raval, Zarna Patel, Monika Gandhi, Pinal Trivedi, Maharshi Pandya, Nidhi Patel, Nitin Savaliya, Raghawendra Kumar, Dinesh Kumar, Zuber Saiyed, Komal Patel, Labdhi Pandya, Snehal Bagatharia, Meenakshi Shah, Neena Doshi, Varsha Godbole, Pritesh Sabara, Apurvasinh Puvar, Neha Rajpara, R D Dixit, A M Kadri, Harsh Bakshi, Chaitanya Joshi, Madhvi Joshi |
| EPI_ISL_475049 | GMERS Medical College & Hospital | Gujarat Biotechnology Research Centre | Zarna Patel, Monika Gandhi, Pinal Trivedi, Maharshi Pandya, Nidhi Patel, Nitin Savaliya, Raghawendra Kumar, Dinesh Kumar, Zuber Saiyed, Komal Patel, Labdhi Pandya, Snehal Bagatharia, Meenakshi Shah, Neena Doshi, Varsha Godbole, Pritesh Sabara, Apurvasinh Puvar, Janvi Raval, Afzal Ansari, R D Dixit, A M Kadri, Harsh Bakshi, Chaitanya Joshi, Madhvi Joshi |
| EPI_ISL_475050 | GMERS Medical College & Hospital | Gujarat Biotechnology Research Centre | Monika Gandhi, Pinal Trivedi, Maharshi Pandya, Nidhi Patel, Nitin Savaliya, Raghawendra Kumar, Dinesh Kumar, Zuber Saiyed, Komal Patel, Labdhi Pandya, Snehal Bagatharia, Meenakshi Shah, Neena Doshi, Varsha Godbole, Pritesh Sabara, Apurvasinh Puvar, Janvi Raval, Zarna Patel, Fenil Patel, R D Dixit, A M Kadri, Harsh Bakshi, Chaitanya Joshi, Madhvi Joshi |
| EPI_ISL_475051 | GMERS Medical College & Hospital | Gujarat Biotechnology Research Centre | Pinal Trivedi, Maharshi Pandya, Nidhi Patel, Nitin Savaliya, Raghawendra Kumar, Dinesh Kumar, Zuber Saiyed, Komal Patel, Labdhi Pandya, Snehal Bagatharia, Meenakshi Shah, Neena Doshi, Varsha Godbole, Pritesh Sabara, Apurvasinh Puvar, Janvi Raval, Zarna Patel, Monika Gandhi, Neelam Nathani, R D Dixit, A M Kadri, Harsh Bakshi, Chaitanya Joshi, Madhvi Joshi |
| EPI_ISL_475052 | GMERS Medical College & Hospital | Gujarat Biotechnology Research Centre | Maharshi Pandya, Nidhi Patel, Nitin Savaliya, Raghawendra Kumar, Dinesh Kumar, Zuber Saiyed, Komal Patel, Labdhi Pandya, Snehal Bagatharia, Meenakshi Shah, Neena Doshi, Varsha Godbole, Pritesh Sabara, Apurvasinh Puvar, Janvi Raval, Zarna Patel, Monika Gandhi, Pinal Trivedi, Armi Chaudhari, R D Dixit, A M Kadri, Harsh Bakshi, Chaitanya Joshi, Madhvi Joshi |
| EPI_ISL_475053 | GMERS Medical College & Hospital | Gujarat Biotechnology Research Centre | Nidhi Patel, Nitin Savaliya, Raghawendra Kumar, Dinesh Kumar, Zuber Saiyed, Komal Patel, Labdhi Pandya, Snehal Bagatharia, Meenakshi Shah, Neena Doshi, Varsha Godbole, Pritesh Sabara, Apurvasinh Puvar, Janvi Raval, Zarna Patel, Monika Gandhi, Pinal Trivedi, Maharshi Pandya, Bhavya Jindal, R D Dixit, A M Kadri, Harsh Bakshi, Chaitanya Joshi, Madhvi Joshi |
| EPI_ISL_475054 | GMERS Medical College & Hospital | Gujarat Biotechnology Research Centre | Nitin Savaliya, Raghawendra Kumar, Dinesh Kumar, Zuber Saiyed, Komal Patel, Labdhi Pandya, Snehal Bagatharia, Meenakshi Shah, Neena Doshi, Varsha Godbole, Pritesh Sabara, Apurvasinh Puvar, Janvi Raval, Zarna Patel, Monika Gandhi, Pinal Trivedi, Maharshi Pandya, Nidhi Patel, Priyanka P Vatsa, R D Dixit, A M Kadri, Harsh Bakshi, Chaitanya Joshi, Madhvi Joshi |
| EPI_ISL_475055 | GMERS Medical College & Hospital | Gujarat Biotechnology Research Centre | Raghawendra Kumar, Dinesh Kumar, Zuber Saiyed, Komal Patel, Labdhi Pandya, Snehal Bagatharia, Meenakshi Shah, Neena Doshi, Varsha Godbole, Pritesh Sabara, Apurvasinh Puvar, Janvi Raval, Zarna Patel, Monika Gandhi, Pinal Trivedi, Maharshi Pandya, Nidhi Patel, Nitin Savaliya, Pooja P Doshi, R D Dixit, A M Kadri, Harsh Bakshi, Chaitanya Joshi, Madhvi Joshi |
| EPI_ISL_475056 | Dr. N. D. Desai Medical College & Hospital | Gujarat Biotechnology Research Centre | Dinesh Kumar, Zuber Saiyed, Komal Patel, Labdhi Pandya, Snehal Bagatharia, J G Buch, Jigar Gusani, Supreet Prabhu, Pritesh Sabara, Apurvasinh Puvar, Janvi Raval, Zarna Patel, Monika Gandhi, Pinal Trivedi, Maharshi Pandya, Nidhi Patel, Nitin Savaliya, Raghawendra Kumar, Akanksha Verma, R D Dixit, A M Kadri, Harsh Bakshi, Chaitanya Joshi, Madhvi Joshi |
| EPI_ISL_475057 | Dr. N. D. Desai Medical College & Hospital | Gujarat Biotechnology Research Centre | Zuber Saiyed, Komal Patel, Labdhi Pandya, Supreet Prabhu, Snehal Bagatharia, Jigar Gusani, J G Buch, Pritesh Sabara, Apurvasinh Puvar, Janvi Raval, Zarna Patel, Monika Gandhi, Pinal Trivedi, Maharshi Pandya, Nidhi Patel, Nitin Savaliya, Raghawendra Kumar, Dinesh Kumar, Priti Pandita, R D Dixit, A M Kadri, Harsh Bakshi, Chaitanya Joshi, Madhvi Joshi |
| EPI_ISL_475058 | GAIMS & G K General Hospital | Gujarat Biotechnology Research Centre | Babulal Babhoria, Hitesh Assudani, Komal Patel, Labdhi Pandya, Snehal Bagatharia, Pritesh Sabara, Apurvasinh Puvar, Janvi Raval, Zarna Patel, Monika Gandhi, Pinal Trivedi, Maharshi Pandya, Nidhi Patel, Nitin Savaliya, Raghawendra Kumar, Dinesh Kumar, Zuber Saiyed, Pragya Sharma, R D Dixit, A M Kadri, Harsh Bakshi, Chaitanya Joshi, Madhvi Joshi |
| EPI_ISL_475059 | GAIMS & G K General Hospital | Gujarat Biotechnology Research Centre | Hitesh Assudani, Babulal Babhoria, Labdhi Pandya, Snehal Bagatharia, Pritesh Sabara, Apurvasinh Puvar, Janvi Raval, Zarna Patel, Monika Gandhi, Pinal Trivedi, Maharshi Pandya, Nidhi Patel, Nitin Savaliya, Raghawendra Kumar, Dinesh Kumar, Zuber Saiyed, Komal Patel, Neha Rajpara, R D Dixit, A M Kadri, Harsh Bakshi, Chaitanya Joshi, Madhvi Joshi |
| EPI_ISL_475060, EPI_ISL_475061, EPI_ISL_475062, EPI_ISL_475063, EPI_ISL_475064, EPI_ISL_475065, EPI_ISL_475066, EPI_ISL_475067, EPI_ISL_475068, EPI_ISL_475069, EPI_ISL_475070 | | | |
| see above | Lab voor klinische biologie | Onderzoeksgroep Virologie | Laurens Lambrechts, Nick Vereecke, Marthe Pauwels, Bruno Verhasselt, Linos Vandekerckhove, Hans Nauwynck, Sebastiaan Theuns |
| EPI_ISL_475071, EPI_ISL_475072, EPI_ISL_475073, EPI_ISL_475074, EPI_ISL_475075, EPI_ISL_475076, EPI_ISL_475077, EPI_ISL_475078, EPI_ISL_475079, EPI_ISL_475080, EPI_ISL_475081, EPI_ISL_475082 | | | |
| see above | Lab voor klinische biologie | Onderzoeksgroep Virologie | Nick Vereecke, Laurens Lambrechts, Marthe Pauwels, Bruno Verhasselt, Linos Vandekerckhove, Hans Nauwynck, Sebastiaan Theuns |
| EPI_ISL_475083, EPI_ISL_475084 | National Institute of Laboratory Medicine and Referral Center | Genomic Research Lab, BCSIR | Md. Murshed Hasan Sarkar, Abu Sayeed Mohammad Mahmud, Mohammad Samir Uzzaman, Eshrar Osman, Md. Ahasan Habib, Shahina Akter, Tanjina Akhter Banu, Barna Goswami, Iffat Jahan, Md. Saddam Hossain, Tasnim Nafisa, Md. Maruf Ahmed Molla, Mahmuda Yeasmin, Asish Kumar Ghosh, Bayzid Bin Monir, A. K. M. Shamsuzzaman, Sheikh Md. Selim Al Din, Utpal Chandra Ray, Salek Ahmed Sajib, Md. Salim Khan |
| EPI_ISL_475085 | Skovde/Unilabs | The Public Health Agency of Sweden | Oskar Karlsson Lindsjo, Maria Lind Karlberg, Mattias Haukland, Reza Advani, Olov Svartstrom, Anna-Malin Linde, Sandra Broddesson, Petra Edquist, Shamam Muradrasoli, Anna Risberg, Karin Tegmark-Wisell |
| EPI_ISL_475086, EPI_ISL_475087, EPI_ISL_475088, EPI_ISL_475089, EPI_ISL_475090, EPI_ISL_475091 | Karolinska Universitetslaboratoriet | The Public Health Agency of Sweden | Oskar Karlsson Lindsjo, Maria Lind Karlberg, Mattias Haukland, Reza Advani, Olov Svartstrom, Anna-Malin Linde, Sandra Broddesson, Petra Edquist, Shamam Muradrasoli, Anna Risberg, Karin Tegmark-Wisell |
| EPI_ISL_475092 | Skovde/Unilabs | The Public Health Agency of Sweden | Oskar Karlsson Lindsjo, Maria Lind Karlberg, Mattias Haukland, Reza Advani, Olov Svartstrom, Anna-Malin Linde, Sandra Broddesson, Petra Edquist, Shamam Muradrasoli, Anna Risberg, Karin Tegmark-Wisell |
| EPI_ISL_475093, EPI_ISL_475094 | Halmstad klinisk mikrobiologi | The Public Health Agency of Sweden | Oskar Karlsson Lindsjo, Maria Lind Karlberg, Mattias Haukland, Reza Advani, Olov Svartstrom, Anna-Malin Linde, Sandra Broddesson, Petra Edquist, Shamam Muradrasoli, Anna Risberg, Karin Tegmark-Wisell |
| EPI_ISL_475095 | Karolinska Universitetslaboratoriet | The Public Health Agency of Sweden | Oskar Karlsson Lindsjo, Maria Lind Karlberg, Mattias Haukland, Reza Advani, Olov Svartstrom, Anna-Malin Linde, Sandra Broddesson, Petra Edquist, Shamam Muradrasoli, Anna Risberg, Karin Tegmark-Wisell |
| EPI_ISL_475096, EPI_ISL_475097, EPI_ISL_475098 | Halmstad klinisk mikrobiologi | The Public Health Agency of Sweden | Oskar Karlsson Lindsjo, Maria Lind Karlberg, Mattias Haukland, Reza Advani, Olov Svartstrom, Anna-Malin Linde, Sandra Broddesson, Petra Edquist, Shamam Muradrasoli, Anna Risberg, Karin Tegmark-Wisell |
| EPI_ISL_475099 | Skovde/Unilabs | The Public Health Agency of Sweden | Oskar Karlsson Lindsjo, Maria Lind Karlberg, Mattias Haukland, Reza Advani, Olov Svartstrom, Anna-Malin Linde, Sandra Broddesson, Petra Edquist, Shamam Muradrasoli, Anna Risberg, Karin Tegmark-Wisell |
| EPI_ISL_475100 | Halmstad klinisk mikrobiologi | The Public Health Agency of Sweden | Oskar Karlsson Lindsjo, Maria Lind Karlberg, Mattias Haukland, Reza Advani, Olov Svartstrom, Anna-Malin Linde, Sandra Broddesson, Petra Edquist, Shamam Muradrasoli, Anna Risberg, Karin Tegmark-Wisell |
| EPI_ISL_475101, EPI_ISL_475102, EPI_ISL_475103, EPI_ISL_475104 | Klinisk Mikrobiologi | The Public Health Agency of Sweden | Oskar Karlsson Lindsjo, Maria Lind Karlberg, Mattias Haukland, Reza Advani, Olov Svartstrom, Anna-Malin Linde, Sandra Broddesson, Petra Edquist, Shamam Muradrasoli, Anna Risberg, Karin Tegmark-Wisell |
| EPI_ISL_475105, EPI_ISL_475106, EPI_ISL_475107, EPI_ISL_475108, EPI_ISL_475109 | Skovde/Unilabs | The Public Health Agency of Sweden | Oskar Karlsson Lindsjo, Maria Lind Karlberg, Mattias Haukland, Reza Advani, Olov Svartstrom, Anna-Malin Linde, Sandra Broddesson, Petra Edquist, Shamam Muradrasoli, Anna Risberg, Karin Tegmark-Wisell |
| EPI_ISL_475110 | Gavle klinisk mikrobiologi | The Public Health Agency of Sweden | Oskar Karlsson Lindsjo, Maria Lind Karlberg, Mattias Haukland, Reza Advani, Olov Svartstrom, Anna-Malin Linde, Sandra Broddesson, Petra Edquist, Shamam Muradrasoli, Anna Risberg, Karin Tegmark-Wisell |
| EPI_ISL_475111, EPI_ISL_475112, EPI_ISL_475113 | Skovde/Unilabs | The Public Health Agency of Sweden | Oskar Karlsson Lindsjo, Maria Lind Karlberg, Mattias Haukland, Reza Advani, Olov Svartstrom, Anna-Malin Linde, Sandra Broddesson, Petra Edquist, Shamam Muradrasoli, Anna Risberg, Karin Tegmark-Wisell |
| EPI_ISL_475114 | Halmstad klinisk mikrobiologi | The Public Health Agency of Sweden | Oskar Karlsson Lindsjo, Maria Lind Karlberg, Mattias Haukland, Reza Advani, Olov Svartstrom, Anna-Malin Linde, Sandra Broddesson, Petra Edquist, Shamam Muradrasoli, Anna Risberg, Karin Tegmark-Wisell |
| EPI_ISL_475115 | Gavle klinisk mikrobiologi | The Public Health Agency of Sweden | Oskar Karlsson Lindsjo, Maria Lind Karlberg, Mattias Haukland, Reza Advani, Olov Svartstrom, Anna-Malin Linde, Sandra Broddesson, Petra Edquist, Shamam Muradrasoli, Anna Risberg, Karin Tegmark-Wisell |
| EPI_ISL_475116, EPI_ISL_475117 | Halmstad klinisk mikrobiologi | The Public Health Agency of Sweden | Oskar Karlsson Lindsjo, Maria Lind Karlberg, Mattias Haukland, Reza Advani, Olov Svartstrom, Anna-Malin Linde, Sandra Broddesson, Petra Edquist, Shamam Muradrasoli, Anna Risberg, Karin Tegmark-Wisell |
| EPI_ISL_475118, EPI_ISL_475119 | Uppsala klinisk mikrobiologi | The Public Health Agency of Sweden | Oskar Karlsson Lindsjo, Maria Lind Karlberg, Mattias Haukland, Reza Advani, Olov Svartstrom, Anna-Malin Linde, Sandra Broddesson, Petra Edquist, Shamam Muradrasoli, Anna Risberg, Karin Tegmark-Wisell |
| EPI_ISL_475120, EPI_ISL_475121 | Halmstad klinisk mikrobiologi | The Public Health Agency of Sweden | Oskar Karlsson Lindsjo, Maria Lind Karlberg, Mattias Haukland, Reza Advani, Olov Svartstrom, Anna-Malin Linde, Sandra Broddesson, Petra Edquist, Shamam Muradrasoli, Anna Risberg, Karin Tegmark-Wisell |
| EPI_ISL_475122 | Umea klinisk mikrobiologi | The Public Health Agency of Sweden | Oskar Karlsson Lindsjo, Maria Lind Karlberg, Mattias Haukland, Reza Advani, Olov Svartstrom, Anna-Malin Linde, Sandra Broddesson, Petra Edquist, Shamam Muradrasoli, Anna Risberg, Karin Tegmark-Wisell |
| EPI_ISL_475123 | Halmstad klinisk mikrobiologi | The Public Health Agency of Sweden | Oskar Karlsson Lindsjo, Maria Lind Karlberg, Mattias Haukland, Reza Advani, Olov Svartstrom, Anna-Malin Linde, Sandra Broddesson, Petra Edquist, Shamam Muradrasoli, Anna Risberg, Karin Tegmark-Wisell |
| EPI_ISL_475124 | Ostersund klinisk mikrobiologi | The Public Health Agency of Sweden | Oskar Karlsson Lindsjo, Maria Lind Karlberg, Mattias Haukland, Reza Advani, Olov Svartstrom, Anna-Malin Linde, Sandra Broddesson, Petra Edquist, Shamam Muradrasoli, Anna Risberg, Karin Tegmark-Wisell |
| EPI_ISL_475125 | Halmstad klinisk mikrobiologi | The Public Health Agency of Sweden | Oskar Karlsson Lindsjo, Maria Lind Karlberg, Mattias Haukland, Reza Advani, Olov Svartstrom, Anna-Malin Linde, Sandra Broddesson, Petra Edquist, Shamam Muradrasoli, Anna Risberg, Karin Tegmark-Wisell |

| | | | |
|---|---|---|---|
| EPI_ISL_475126, EPI_ISL_475127 | Umea klinisk mikrobiologi | The Public Health Agency of Sweden | Oskar Karlsson Lindsjo, Maria Lind Karlberg, Mattias Haukland, Reza Advani, Olov Svartstrom, Anna-Malin Linde, Sandra Broddesson, Petra Edquist, Shamam Muradrasoli, Anna Risberg, Karin Tegmark-Wisell |
| EPI_ISL_475128, EPI_ISL_475129, EPI_ISL_475130, EPI_ISL_475131, EPI_ISL_475132, EPI_ISL_475133, EPI_ISL_475134, EPI_ISL_475135 | Orebro klinisk mikrobiologi | The Public Health Agency of Sweden | Oskar Karlsson Lindsjo, Maria Lind Karlberg, Mattias Haukland, Reza Advani, Olov Svartstrom, Anna-Malin Linde, Sandra Broddesson, Petra Edquist, Shamam Muradrasoli, Anna Risberg, Karin Tegmark-Wisell |
| EPI_ISL_475136 | Karolinska Universitetslaboratoriet | The Public Health Agency of Sweden | Oskar Karlsson Lindsjo, Maria Lind Karlberg, Mattias Haukland, Reza Advani, Olov Svartstrom, Anna-Malin Linde, Sandra Broddesson, Petra Edquist, Shamam Muradrasoli, Anna Risberg, Karin Tegmark-Wisell |
| EPI_ISL_475137, EPI_ISL_475138 | Kalmar klinisk mikrobiologi | The Public Health Agency of Sweden | Oskar Karlsson Lindsjo, Maria Lind Karlberg, Mattias Haukland, Reza Advani, Olov Svartstrom, Anna-Malin Linde, Sandra Broddesson, Petra Edquist, Shamam Muradrasoli, Anna Risberg, Karin Tegmark-Wisell |
| EPI_ISL_475139 | Orebro klinisk mikrobiologi | The Public Health Agency of Sweden | Oskar Karlsson Lindsjo, Maria Lind Karlberg, Mattias Haukland, Reza Advani, Olov Svartstrom, Anna-Malin Linde, Sandra Broddesson, Petra Edquist, Shamam Muradrasoli, Anna Risberg, Karin Tegmark-Wisell |
| EPI_ISL_475141, EPI_ISL_475142 | Karolinska Universitetslaboratoriet | The Public Health Agency of Sweden | Oskar Karlsson Lindsjo, Maria Lind Karlberg, Mattias Haukland, Reza Advani, Olov Svartstrom, Anna-Malin Linde, Sandra Broddesson, Petra Edquist, Shamam Muradrasoli, Anna Risberg, Karin Tegmark-Wisell |
| EPI_ISL_475143 | Umea klinisk mikrobiologi | The Public Health Agency of Sweden | Oskar Karlsson Lindsjo, Maria Lind Karlberg, Mattias Haukland, Reza Advani, Olov Svartstrom, Anna-Malin Linde, Sandra Broddesson, Petra Edquist, Shamam Muradrasoli, Anna Risberg, Karin Tegmark-Wisell |
| EPI_ISL_475144, EPI_ISL_475145, EPI_ISL_475146, EPI_ISL_475147, EPI_ISL_475148 | Klinisk mikrobiologi Vasternorrland | The Public Health Agency of Sweden | Oskar Karlsson Lindsjo, Maria Lind Karlberg, Mattias Haukland, Reza Advani, Olov Svartstrom, Anna-Malin Linde, Sandra Broddesson, Petra Edquist, Shamam Muradrasoli, Anna Risberg, Karin Tegmark-Wisell |
| EPI_ISL_475149, EPI_ISL_475150 | Karolinska Universitetslaboratoriet | The Public Health Agency of Sweden | Oskar Karlsson Lindsjo, Maria Lind Karlberg, Mattias Haukland, Reza Advani, Olov Svartstrom, Anna-Malin Linde, Sandra Broddesson, Petra Edquist, Shamam Muradrasoli, Anna Risberg, Karin Tegmark-Wisell |
| EPI_ISL_475151 | Uppsala klinisk mikrobiologi | The Public Health Agency of Sweden | Oskar Karlsson Lindsjo, Maria Lind Karlberg, Mattias Haukland, Reza Advani, Olov Svartstrom, Anna-Malin Linde, Sandra Broddesson, Petra Edquist, Shamam Muradrasoli, Anna Risberg, Karin Tegmark-Wisell |
| EPI_ISL_475152 | Folkhalsomyndigheten | The Public Health Agency of Sweden | Oskar Karlsson Lindsjo, Maria Lind Karlberg, Mattias Haukland, Reza Advani, Olov Svartstrom, Anna-Malin Linde, Sandra Broddesson, Petra Edquist, Shamam Muradrasoli, Anna Risberg, Karin Tegmark-Wisell |
| EPI_ISL_475153, EPI_ISL_475154, EPI_ISL_475155 | Klinisk mikrobiologi Vasternorrland | The Public Health Agency of Sweden | Oskar Karlsson Lindsjo, Maria Lind Karlberg, Mattias Haukland, Reza Advani, Olov Svartstrom, Anna-Malin Linde, Sandra Broddesson, Petra Edquist, Shamam Muradrasoli, Anna Risberg, Karin Tegmark-Wisell |
| EPI_ISL_475156, EPI_ISL_475157, EPI_ISL_475158, EPI_ISL_475159 | Halmstad klinisk mikrobiologi | The Public Health Agency of Sweden | Oskar Karlsson Lindsjo, Maria Lind Karlberg, Mattias Haukland, Reza Advani, Olov Svartstrom, Anna-Malin Linde, Sandra Broddesson, Petra Edquist, Shamam Muradrasoli, Anna Risberg, Karin Tegmark-Wisell |
| EPI_ISL_475160, EPI_ISL_475161, EPI_ISL_475162, EPI_ISL_475163 | Klinisk mikrobiologi Vasternorrland | The Public Health Agency of Sweden | Oskar Karlsson Lindsjo, Maria Lind Karlberg, Mattias Haukland, Reza Advani, Olov Svartstrom, Anna-Malin Linde, Sandra Broddesson, Petra Edquist, Shamam Muradrasoli, Anna Risberg, Karin Tegmark-Wisell |
| EPI_ISL_475164 | Halmstad klinisk mikrobiologi | The Public Health Agency of Sweden | Oskar Karlsson Lindsjo, Maria Lind Karlberg, Mattias Haukland, Reza Advani, Olov Svartstrom, Anna-Malin Linde, Sandra Broddesson, Petra Edquist, Shamam Muradrasoli, Anna Risberg, Karin Tegmark-Wisell |
| EPI_ISL_475165 | National Institute of Laboratory Medicine and Referral Center | Genomic Research Lab, BCSIR | Shahina Akter, Abu Sayeed Mohammad Mahmud, Mohammad Samir Uzzaman, Eshrar Osman, Md. Ahasan Habib, Tanjina Akhter Banu, Md. Murshed Hasan Sarkar, Barna Goswami, Iffat Jahan, Md. Saddam Hossain, Tasnim Nafisa, Md. Maruf Ahmed Molla, Mahmuda Yeasmin, Asish Kumar Ghosh, Bayzid Bin Monir, A. K. M. Shamsuzzaman, Sheikh Md. Selim Al Din, Utpal Chandra Ray, Salek Ahmed Sajib, Md. Salim Khan |
| EPI_ISL_475166 | National Institute of Laboratory Medicine and Referral Center | Genomic Research Lab, BCSIR | Tanjina Akhter Banu, Abu Sayeed Mohammad Mahmud, Mohammad Samir Uzzaman, Eshrar Osman, Md. Ahasan Habib, Shahina Akter, Md. Murshed Hasan Sarkar, Barna Goswami, Iffat Jahan, Md. Saddam Hossain, Tasnim Nafisa, Md. Maruf Ahmed Molla, Mahmuda Yeasmin, Asish Kumar Ghosh, Bayzid Bin Monir, A. K. M. Shamsuzzaman, Sheikh Md. Selim Al Din, Utpal Chandra Ray, Salek Ahmed Sajib, Md. Salim Khan |
| EPI_ISL_475167 | National Institute of Laboratory Medicine and Referral Center | Genomic Research Lab, BCSIR | Barna Goswami, Abu Sayeed Mohammad Mahmud, Mohammad Samir Uzzaman, Eshrar Osman, Md. Ahasan Habib, Shahina Akter, Tanjina Akhter Banu, Md. Murshed Hasan Sarkar, Iffat Jahan, Md. Saddam Hossain, Tasnim Nafisa, Md. Maruf Ahmed Molla, Mahmuda Yeasmin, Asish Kumar Ghosh, Bayzid Bin Monir, A. K. M. Shamsuzzaman, Sheikh Md. Selim Al Din, Utpal Chandra Ray, Salek Ahmed Sajib, Md. Salim Khan |
| EPI_ISL_475168 | National Institute of Laboratory Medicine and Referral Center | Genomic Research Lab, BCSIR | Iffat Jahan, Abu Sayeed Mohammad Mahmud, Mohammad Samir Uzzaman, Eshrar Osman, Md. Ahasan Habib, Shahina Akter, Tanjina Akhter Banu, Md. Murshed Hasan Sarkar, Barna Goswami, Md. Saddam Hossain, Tasnim Nafisa, Md. Maruf Ahmed Molla, Mahmuda Yeasmin, Asish Kumar Ghosh, Bayzid Bin Monir, A. K. M. Shamsuzzaman, Sheikh Md. Selim Al Din, Utpal Chandra Ray, Salek Ahmed Sajib, Md. Salim Khan |
| EPI_ISL_475169 | National Institute of Laboratory Medicine and Referral Center | Genomic Research Lab, BCSIR | Md. Saddam Hossain, Abu Sayeed Mohammad Mahmud, Mohammad Samir Uzzaman, Eshrar Osman, Md. Ahasan Habib, Shahina Akter, Tanjina Akhter Banu, Md. Murshed Hasan Sarkar, Barna Goswami, Iffat Jahan, Tasnim Nafisa, Md. Maruf Ahmed Molla, Mahmuda Yeasmin, Asish Kumar Ghosh, Bayzid Bin Monir, A. K. M. Shamsuzzaman, Sheikh Md. Selim Al Din, Utpal Chandra Ray, Salek Ahmed Sajib, Md. Salim Khan |
| EPI_ISL_475170, EPI_ISL_475171, EPI_ISL_475172, EPI_ISL_475173 | National Institute of Laboratory Medicine and Referral Center | Genomic Research Lab, BCSIR | Abu Sayeed Mohammad Mahmud, Mohammad Samir Uzzaman, Eshrar Osman, Md. Ahasan Habib, Shahina Akter, Tanjina Akhter Banu, Md. Murshed Hasan Sarkar, Barna Goswami, Iffat Jahan, Md. Saddam Hossain, Tasnim Nafisa, Md. Maruf Ahmed Molla, Mahmuda Yeasmin, Asish Kumar Ghosh, Bayzid Bin Monir, A. K. M. Shamsuzzaman, Sheikh Md. Selim Al Din, Utpal Chandra Ray, Salek Ahmed Sajib, Md. Salim Khan |
| EPI_ISL_475176, EPI_ISL_475178, EPI_ISL_475180, EPI_ISL_475184, EPI_ISL_475185, EPI_ISL_475186, EPI_ISL_475187, EPI_ISL_475188, EPI_ISL_475189, EPI_ISL_475190, EPI_ISL_475191, EPI_ISL_475192, EPI_ISL_475194, EPI_ISL_475195, EPI_ISL_475196, EPI_ISL_475197, EPI_ISL_475198, EPI_ISL_475200, EPI_ISL_475201, EPI_ISL_475203, EPI_ISL_475204, EPI_ISL_475205, EPI_ISL_475208, EPI_ISL_475209, EPI_ISL_475210, EPI_ISL_475211, EPI_ISL_475213, EPI_ISL_475214, EPI_ISL_475215, EPI_ISL_475216, EPI_ISL_475217, EPI_ISL_475219, EPI_ISL_475220, EPI_ISL_475221, EPI_ISL_475222, EPI_ISL_475225, EPI_ISL_475227, EPI_ISL_475228, EPI_ISL_475229, EPI_ISL_475231, EPI_ISL_475232, EPI_ISL_475233, EPI_ISL_475236, EPI_ISL_475237 | | | |
| see above | Nebraska Public Health Laboratory | UNMC COVID-19 Response Team | UNMC COVID-19 Response Team |
| EPI_ISL_475238 | National Institute of Laboratory Medicine and Referral Center | Genomic Research Lab, BCSIR | Abu Sayeed Mohammad Mahmud, Mohammad Samir Uzzaman, Eshrar Osman, Md. Ahasan Habib, Shahina Akter, Tanjina Akhter Banu, Md. Murshed Hasan Sarkar, Barna Goswami, Iffat Jahan, Md. Saddam Hossain, Tasnim Nafisa, Md. Maruf Ahmed Molla, Mahmuda Yeasmin, Asish Kumar Ghosh, Bayzid Bin Monir, A. K. M. Shamsuzzaman, Sheikh Md. Selim Al Din, Utpal Chandra Ray, Salek Ahmed Sajib, Md. Salim Khan |
| EPI_ISL_475240, EPI_ISL_475241, EPI_ISL_475242, EPI_ISL_475244, EPI_ISL_475247, EPI_ISL_475263, EPI_ISL_475266, EPI_ISL_475267, EPI_ISL_475268, EPI_ISL_475269, EPI_ISL_475271, EPI_ISL_475272, EPI_ISL_475273, EPI_ISL_475274, EPI_ISL_475276, EPI_ISL_475279, EPI_ISL_475280, EPI_ISL_475282, EPI_ISL_475283, EPI_ISL_475284, EPI_ISL_475285, EPI_ISL_475286, EPI_ISL_475287, EPI_ISL_475288, EPI_ISL_475289, EPI_ISL_475291, EPI_ISL_475292, EPI_ISL_475293, EPI_ISL_475296, EPI_ISL_475297, EPI_ISL_475298, EPI_ISL_475299, EPI_ISL_475300, EPI_ISL_475302, EPI_ISL_475303, EPI_ISL_475304, EPI_ISL_475306, EPI_ISL_475307, EPI_ISL_475310, EPI_ISL_475311, EPI_ISL_475312, EPI_ISL_475315, EPI_ISL_475316, EPI_ISL_475318, EPI_ISL_475319, EPI_ISL_475320, EPI_ISL_475321, EPI_ISL_475322, EPI_ISL_475323, EPI_ISL_475325, EPI_ISL_475326, EPI_ISL_475327, EPI_ISL_475328, EPI_ISL_475329, EPI_ISL_475330, EPI_ISL_475331, EPI_ISL_475333, EPI_ISL_475334, EPI_ISL_475335, EPI_ISL_475337, EPI_ISL_475339, EPI_ISL_475341 | | | |
| see above | Centre for Enzyme Innovation, University of Portsmouth / Translational Research Laboratory, Portsmouth Hospitals NHS Trust | COVID-19 Genomics UK (COG-UK) Consortium | Angela Beckett,Yann Bourgeois,Garry Scarlett,Sharon Glaysher,Scott Elliott,Kelly Bicknell,Robert Impey,Allyson Lloyd,Sarah Wyllie,Ethan Butcher,Anoop Chauhan,Samuel Robson |
| EPI_ISL_475343, EPI_ISL_475344, EPI_ISL_475347, EPI_ISL_475348, EPI_ISL_475356, EPI_ISL_475360, EPI_ISL_475367, EPI_ISL_475368, EPI_ISL_475370, EPI_ISL_475371, EPI_ISL_475372, EPI_ISL_475375, EPI_ISL_475378, EPI_ISL_475379, EPI_ISL_475384, EPI_ISL_475389, EPI_ISL_475391, EPI_ISL_475393, EPI_ISL_475401, EPI_ISL_475402, EPI_ISL_475403, EPI_ISL_475407, EPI_ISL_475409, EPI_ISL_475413, EPI_ISL_475415, EPI_ISL_475417, EPI_ISL_475418, EPI_ISL_475419, EPI_ISL_475420, EPI_ISL_475421, EPI_ISL_475423, EPI_ISL_475424, EPI_ISL_475425, EPI_ISL_475429, EPI_ISL_475434, EPI_ISL_475443, EPI_ISL_475445, EPI_ISL_475447, EPI_ISL_475448, EPI_ISL_475453, EPI_ISL_475454, EPI_ISL_475456, EPI_ISL_475457, EPI_ISL_475461, EPI_ISL_475466, EPI_ISL_475469, EPI_ISL_475471, EPI_ISL_475473, EPI_ISL_475476, EPI_ISL_475477, EPI_ISL_475478, EPI_ISL_475484, EPI_ISL_475485, EPI_ISL_475492, EPI_ISL_475497, EPI_ISL_475504, EPI_ISL_475507, EPI_ISL_475508, EPI_ISL_475509 | | | |
| see above | Virology Department, Sheffield Teaching Hospitals NHS Foundation Trust/Department of Infection, Immunity and Cardiovascular Disease, The Medical School, University of Sheffield | COVID-19 Genomics UK (COG-UK) Consortium | Thushan de Silva, Matthew Parker, Nikki Smith, Adri Angyal, Rebecca Brown, Luke Green, Rachel Tucker, Paul Parsons, Danielle Groves, Katie Johnson, Laura Carrilero, Alex Keeley, Dave Partridge, Matthew Wyles, Benjamin Lindsey, Mehmet Yavuz, Mohammad Raza, Cariad Evans |
| EPI_ISL_475511 | Orestadsklinikens VC | The Public Health Agency of Sweden | Oskar Karlsson Lindsjo, Maria Lind Karlberg, Mattias Haukland, Reza Advani, Olov Svartstrom, Anna-Malin Linde, Sandra Broddesson, Mia Brytting, Anna Risberg, Karin Tegmark-Wisell |
| EPI_ISL_475512 | Din Klinik | The Public Health Agency of Sweden | Oskar Karlsson Lindsjo, Maria Lind Karlberg, Mattias Haukland, Reza Advani, Olov Svartstrom, Anna-Malin Linde, Sandra Broddesson, Mia Brytting, Anna Risberg, Karin Tegmark-Wisell |
| EPI_ISL_475513 | Huddinge VC | The Public Health Agency of Sweden | Oskar Karlsson Lindsjo, Maria Lind Karlberg, Mattias Haukland, Reza Advani, Olov Svartstrom, Anna-Malin Linde, Sandra Broddesson, Mia Brytting, Anna Risberg, Karin Tegmark-Wisell |
| EPI_ISL_475514 | Uppsala Narakut Aleris | The Public Health Agency of Sweden | Oskar Karlsson Lindsjo, Maria Lind Karlberg, Mattias Haukland, Reza Advani, Olov Svartstrom, Anna-Malin Linde, Sandra Broddesson, Mia Brytting, Anna Risberg, Karin Tegmark-Wisell |
| EPI_ISL_475515 | Lakargruppen | The Public Health Agency of Sweden | Oskar Karlsson Lindsjo, Maria Lind Karlberg, Mattias Haukland, Reza Advani, Olov Svartstrom, Anna-Malin Linde, Sandra Broddesson, Mia Brytting, Anna Risberg, Karin Tegmark-Wisell |
| EPI_ISL_475516, EPI_ISL_475517 | Uppsala Narakut Aleris | The Public Health Agency of Sweden | Oskar Karlsson Lindsjo, Maria Lind Karlberg, Mattias Haukland, Reza Advani, Olov Svartstrom, Anna-Malin Linde, Sandra Broddesson, Mia Brytting, Anna Risberg, Karin Tegmark-Wisell |
| EPI_ISL_475518 | Trollbackens VC | The Public Health Agency of Sweden | Oskar Karlsson Lindsjo, Maria Lind Karlberg, Mattias Haukland, Reza Advani, Olov Svartstrom, Anna-Malin Linde, Sandra Broddesson, Mia Brytting, Anna Risberg, Karin Tegmark-Wisell |
| EPI_ISL_475519 | Orsa VC | The Public Health Agency of Sweden | Oskar Karlsson Lindsjo, Maria Lind Karlberg, Mattias Haukland, Reza Advani, Olov Svartstrom, Anna-Malin Linde, Sandra Broddesson, Mia Brytting, Anna Risberg, Karin Tegmark-Wisell |
| EPI_ISL_475520 | Vardcentralen Brinken | The Public Health Agency of Sweden | Oskar Karlsson Lindsjo, Maria Lind Karlberg, Mattias Haukland, Reza Advani, Olov Svartstrom, Anna-Malin Linde, Sandra Broddesson, Mia Brytting, Anna Risberg, Karin Tegmark-Wisell |
| EPI_ISL_475521 | Ulltuna Vardcentral | The Public Health Agency of Sweden | Oskar Karlsson Lindsjo, Maria Lind Karlberg, Mattias Haukland, Reza Advani, Olov Svartstrom, Anna-Malin Linde, Sandra Broddesson, Mia Brytting, Anna Risberg, Karin Tegmark-Wisell |
| EPI_ISL_475522, EPI_ISL_475523 | Huddinge VC | The Public Health Agency of Sweden | Oskar Karlsson Lindsjo, Maria Lind Karlberg, Mattias Haukland, Reza Advani, Olov Svartstrom, Anna-Malin Linde, Sandra Broddesson, Mia Brytting, Anna Risberg, Karin Tegmark-Wisell |
| EPI_ISL_475524 | Narhalsan Sjobo vardcentral | The Public Health Agency of Sweden | Oskar Karlsson Lindsjo, Maria Lind Karlberg, Mattias Haukland, Reza Advani, Olov Svartstrom, Anna-Malin Linde, Sandra Broddesson, Mia Brytting, Anna Risberg, Karin Tegmark-Wisell |
| EPI_ISL_475525 | Huddinge VC | The Public Health Agency of Sweden | Oskar Karlsson Lindsjo, Maria Lind Karlberg, Mattias Haukland, Reza Advani, Olov Svartstrom, Anna-Malin Linde, Sandra Broddesson, Mia Brytting, Anna Risberg, Karin Tegmark-Wisell |
| EPI_ISL_475526, EPI_ISL_475527 | Uppsala Narakut Aleris | The Public Health Agency of Sweden | Oskar Karlsson Lindsjo, Maria Lind Karlberg, Mattias Haukland, Reza Advani, Olov Svartstrom, Anna-Malin Linde, Sandra Broddesson, Mia Brytting, Anna Risberg, Karin Tegmark-Wisell |
| EPI_ISL_475528 | Omtanken Grimmered | The Public Health Agency of Sweden | Oskar Karlsson Lindsjo, Maria Lind Karlberg, Mattias Haukland, Reza Advani, Olov Svartstrom, Anna-Malin Linde, Sandra Broddesson, Mia Brytting, Anna Risberg, Karin Tegmark-Wisell |
| EPI_ISL_475529, EPI_ISL_475530, EPI_ISL_475531, EPI_ISL_475532 | Kungsors VC | The Public Health Agency of Sweden | Oskar Karlsson Lindsjo, Maria Lind Karlberg, Mattias Haukland, Reza Advani, Olov Svartstrom, Anna-Malin Linde, Sandra Broddesson, Mia Brytting, Anna Risberg, Karin Tegmark-Wisell |
| EPI_ISL_475533, EPI_ISL_475534, EPI_ISL_475535 | Omtanken Grimmered | The Public Health Agency of Sweden | Oskar Karlsson Lindsjo, Maria Lind Karlberg, Mattias Haukland, Reza Advani, Olov Svartstrom, Anna-Malin Linde, Sandra Broddesson, Mia Brytting, Anna Risberg, Karin Tegmark-Wisell |
| EPI_ISL_475536 | Follinge Halsocentral | The Public Health Agency of Sweden | Oskar Karlsson Lindsjo, Maria Lind Karlberg, Mattias Haukland, Reza Advani, Olov Svartstrom, Anna-Malin Linde, Sandra Broddesson, Mia Brytting, Anna Risberg, Karin Tegmark-Wisell |
| EPI_ISL_475537 | Narhalsan Oden VC | The Public Health Agency of Sweden | Oskar Karlsson Lindsjo, Maria Lind Karlberg, Mattias Haukland, Reza Advani, Olov Svartstrom, Anna-Malin Linde, Sandra Broddesson, Mia Brytting, Anna Risberg, Karin Tegmark-Wisell |

| | | | |
|---|---|---|---|
| EPI_ISL_475538 | Omtanken Grimmered | The Public Health Agency of Sweden | Oskar Karlsson Lindsjo, Maria Lind Karlberg, Mattias Haukland, Reza Advani, Olov Svartstrom, Anna-Malin Linde, Sandra Broddesson, Mia Brytting, Anna Risberg, Karin Tegmark-Wisell |
| EPI_ISL_475539 | Narhalsan Sjobo vardcentral | The Public Health Agency of Sweden | Oskar Karlsson Lindsjo, Maria Lind Karlberg, Mattias Haukland, Reza Advani, Olov Svartstrom, Anna-Malin Linde, Sandra Broddesson, Mia Brytting, Anna Risberg, Karin Tegmark-Wisell |
| EPI_ISL_475540 | Bla Kustens halsocentral | The Public Health Agency of Sweden | Oskar Karlsson Lindsjo, Maria Lind Karlberg, Mattias Haukland, Reza Advani, Olov Svartstrom, Anna-Malin Linde, Sandra Broddesson, Mia Brytting, Anna Risberg, Karin Tegmark-Wisell |
| EPI_ISL_475541 | Follinge Halsocentral | The Public Health Agency of Sweden | Oskar Karlsson Lindsjo, Maria Lind Karlberg, Mattias Haukland, Reza Advani, Olov Svartstrom, Anna-Malin Linde, Sandra Broddesson, Mia Brytting, Anna Risberg, Karin Tegmark-Wisell |
| EPI_ISL_475542 | Kungsholmsdoktorn | The Public Health Agency of Sweden | Oskar Karlsson Lindsjo, Maria Lind Karlberg, Mattias Haukland, Reza Advani, Olov Svartstrom, Anna-Malin Linde, Sandra Broddesson, Mia Brytting, Anna Risberg, Karin Tegmark-Wisell |
| EPI_ISL_475543 | Surbrunns VC | The Public Health Agency of Sweden | Oskar Karlsson Lindsjo, Maria Lind Karlberg, Mattias Haukland, Reza Advani, Olov Svartstrom, Anna-Malin Linde, Sandra Broddesson, Mia Brytting, Anna Risberg, Karin Tegmark-Wisell |
| EPI_ISL_475544, EPI_ISL_475545, EPI_ISL_475546, EPI_ISL_475547 | Karolinska Universitetslaboratoriet | The Public Health Agency of Sweden | Oskar Karlsson Lindsjo, Maria Lind Karlberg, Mattias Haukland, Reza Advani, Olov Svartstrom, Anna-Malin Linde, Sandra Broddesson, Shaman Muradrasoli, Anna Risberg, Karin Tegmark-Wisell |
| EPI_ISL_475548 | Halmstad klinisk mikrobiologi | The Public Health Agency of Sweden | Oskar Karlsson Lindsjo, Maria Lind Karlberg, Mattias Haukland, Reza Advani, Olov Svartstrom, Anna-Malin Linde, Sandra Broddesson, Shaman Muradrasoli, Anna Risberg, Karin Tegmark-Wisell |
| EPI_ISL_475549, EPI_ISL_475550 | Skovde/Unilabs | The Public Health Agency of Sweden | Oskar Karlsson Lindsjo, Maria Lind Karlberg, Mattias Haukland, Reza Advani, Olov Svartstrom, Anna-Malin Linde, Sandra Broddesson, Shaman Muradrasoli, Anna Risberg, Karin Tegmark-Wisell |
| EPI_ISL_475551, EPI_ISL_475552 | Karolinska Universitetslaboratoriet | The Public Health Agency of Sweden | Oskar Karlsson Lindsjo, Maria Lind Karlberg, Mattias Haukland, Reza Advani, Olov Svartstrom, Anna-Malin Linde, Sandra Broddesson, Shaman Muradrasoli, Anna Risberg, Karin Tegmark-Wisell |
| EPI_ISL_475553 | Halmstad klinisk mikrobiologi | The Public Health Agency of Sweden | Oskar Karlsson Lindsjo, Maria Lind Karlberg, Mattias Haukland, Reza Advani, Olov Svartstrom, Anna-Malin Linde, Sandra Broddesson, Shaman Muradrasoli, Anna Risberg, Karin Tegmark-Wisell |
| EPI_ISL_475554, EPI_ISL_475555 | Skovde/Unilabs | The Public Health Agency of Sweden | Oskar Karlsson Lindsjo, Maria Lind Karlberg, Mattias Haukland, Reza Advani, Olov Svartstrom, Anna-Malin Linde, Sandra Broddesson, Shaman Muradrasoli, Anna Risberg, Karin Tegmark-Wisell |
| EPI_ISL_475556, EPI_ISL_475557 | Halmstad klinisk mikrobiologi | The Public Health Agency of Sweden | Oskar Karlsson Lindsjo, Maria Lind Karlberg, Mattias Haukland, Reza Advani, Olov Svartstrom, Anna-Malin Linde, Sandra Broddesson, Shaman Muradrasoli, Anna Risberg, Karin Tegmark-Wisell |
| EPI_ISL_475558, EPI_ISL_475560, EPI_ISL_475561 | Karolinska Universitetslaboratoriet | The Public Health Agency of Sweden | Oskar Karlsson Lindsjo, Maria Lind Karlberg, Mattias Haukland, Reza Advani, Olov Svartstrom, Anna-Malin Linde, Sandra Broddesson, Shaman Muradrasoli, Anna Risberg, Karin Tegmark-Wisell |
| EPI_ISL_475562, EPI_ISL_475563 | Din Klinik | The Public Health Agency of Sweden | Oskar Karlsson Lindsjo, Maria Lind Karlberg, Mattias Haukland, Reza Advani, Olov Svartstrom, Anna-Malin Linde, Sandra Broddesson, Mia Brytting, Anna Risberg, Karin Tegmark-Wisell |
| EPI_ISL_475564 | Surbrunns VC | The Public Health Agency of Sweden | Oskar Karlsson Lindsjo, Maria Lind Karlberg, Mattias Haukland, Reza Advani, Olov Svartstrom, Anna-Malin Linde, Sandra Broddesson, Mia Brytting, Anna Risberg, Karin Tegmark-Wisell |
| EPI_ISL_475566 | Vardcentralen Brinken | The Public Health Agency of Sweden | Oskar Karlsson Lindsjo, Maria Lind Karlberg, Mattias Haukland, Reza Advani, Olov Svartstrom, Anna-Malin Linde, Sandra Broddesson, Mia Brytting, Anna Risberg, Karin Tegmark-Wisell |
| EPI_ISL_475567 | Huddinge VC | The Public Health Agency of Sweden | Oskar Karlsson Lindsjo, Maria Lind Karlberg, Mattias Haukland, Reza Advani, Olov Svartstrom, Anna-Malin Linde, Sandra Broddesson, Mia Brytting, Anna Risberg, Karin Tegmark-Wisell |
| EPI_ISL_475568 | Kungsors VC | The Public Health Agency of Sweden | Oskar Karlsson Lindsjo, Maria Lind Karlberg, Mattias Haukland, Reza Advani, Olov Svartstrom, Anna-Malin Linde, Sandra Broddesson, Mia Brytting, Anna Risberg, Karin Tegmark-Wisell |
| EPI_ISL_475569 | Kungsholmsdoktorn | The Public Health Agency of Sweden | Oskar Karlsson Lindsjo, Maria Lind Karlberg, Mattias Haukland, Reza Advani, Olov Svartstrom, Anna-Malin Linde, Sandra Broddesson, Mia Brytting, Anna Risberg, Karin Tegmark-Wisell |
| EPI_ISL_475570 | Genome Center | Genome Center | A. S. M. Rubayet- Ul- Alam, Ovinu Kibria Islam, Md. Shazid Hasan, Hassan M. Al-Emran, Shireen Nigar, Selina Akter, Pravas Chandra Roy, Md. Tanvir Islam, Shovon Lal Sarkar, M. Shaminur Rahman, M. Rafiul Islam, Habiba Ibnat, Md Nur Kabidul Azam, Chakrabortty Atonu, Proshanto Kumar Das, Md. Hasan al Pramanik, Md. Zannat Ali, Shohanur Rahaman, Md. Aminul Islam, Ashok Kumar, Md. Nazmul Hasan, Md. Iqbal Kabir Jahid, Md. Anwar Hossain |
| EPI_ISL_475571 | Genome Center | Genome Center | Hassan M. Al-Emran, Md. Shazid Hasan, Ovinu Kibria Islam, A. S. M. Rubayet- Ul- Alam, Pravas Chandra Roy, Selina Akter, Shireen Nigar, Shovon Lal Sarkar, Md. Tanvir Islam, Mithun Talukder Md. Tawyabur, Md. Taijul Islam, Provakar Mondol, Md. Muzahidul Islam, Md. Iqbal Kabir Jahid Md. Anwar Hossain |
| EPI_ISL_475572 | Imperial College London | Imperial College London | Jie Zhou, Wendy Barclay |
| EPI_ISL_475573 | Genome Center | Genome Center | Md. Shazid Hasan, Hassan M. Al-Emran, Ovinu Kibria Islam, A. S. M. Rubayet- Ul- Alam, Selina Akter, Shireen Nigar, Md. Tanvir Islam, Pravas Chandra Roy, Shovon Lal Sarkar, Md. Nazmul Hasan, Tanay Chakrovarty, Md. Ali Ahasan Setu, Sourav Dutta, Ruhul Amin, Md. Iqbal Kabir Jahid, Md. Anwar Hossain |
| EPI_ISL_475574, EPI_ISL_475576, EPI_ISL_475577, EPI_ISL_475578, EPI_ISL_475579, EPI_ISL_475580, EPI_ISL_475582, EPI_ISL_475583, EPI_ISL_475585, EPI_ISL_475588, EPI_ISL_475590, EPI_ISL_475593, EPI_ISL_475594, EPI_ISL_475595, EPI_ISL_475596, EPI_ISL_475598, EPI_ISL_475599, EPI_ISL_475600, EPI_ISL_475601, EPI_ISL_475602, EPI_ISL_475603, EPI_ISL_475604, EPI_ISL_475605, EPI_ISL_475606, EPI_ISL_475607, EPI_ISL_475608, EPI_ISL_475609, EPI_ISL_475610, EPI_ISL_475611, EPI_ISL_475612, EPI_ISL_475613, EPI_ISL_475614, EPI_ISL_475615, EPI_ISL_475616, EPI_ISL_475617, EPI_ISL_475618, EPI_ISL_475619, EPI_ISL_475621, EPI_ISL_475622, EPI_ISL_475625, EPI_ISL_475627, EPI_ISL_475628, EPI_ISL_475629, EPI_ISL_475630, EPI_ISL_475631, EPI_ISL_475632, EPI_ISL_475633, EPI_ISL_475634, EPI_ISL_475635, EPI_ISL_475636, EPI_ISL_475637, EPI_ISL_475638, EPI_ISL_475639, EPI_ISL_475640, EPI_ISL_475641, EPI_ISL_475643, EPI_ISL_475644, EPI_ISL_475645, EPI_ISL_475646, EPI_ISL_475647, EPI_ISL_475648, EPI_ISL_475650, EPI_ISL_475651, EPI_ISL_475652, EPI_ISL_475653, EPI_ISL_475654, EPI_ISL_475655, EPI_ISL_475656, EPI_ISL_475657, EPI_ISL_475658, EPI_ISL_475659, EPI_ISL_475660, EPI_ISL_475661, EPI_ISL_475662, EPI_ISL_475663, EPI_ISL_475664, EPI_ISL_475666, EPI_ISL_475667, EPI_ISL_475668, EPI_ISL_475669, EPI_ISL_475670, EPI_ISL_475671, EPI_ISL_475673, EPI_ISL_475674, EPI_ISL_475675, EPI_ISL_475677, EPI_ISL_475678, EPI_ISL_475679, EPI_ISL_475680, EPI_ISL_475681, EPI_ISL_475682, EPI_ISL_475683, EPI_ISL_475684, EPI_ISL_475685, EPI_ISL_475686, EPI_ISL_475687, EPI_ISL_475688, EPI_ISL_475689, EPI_ISL_475690, EPI_ISL_475691, EPI_ISL_475692, EPI_ISL_475694, EPI_ISL_475695, EPI_ISL_475697, EPI_ISL_475698, EPI_ISL_475699, EPI_ISL_475700, EPI_ISL_475701, EPI_ISL_475702, EPI_ISL_475703, EPI_ISL_475705, EPI_ISL_475706, EPI_ISL_475708, EPI_ISL_475709, EPI_ISL_475710, EPI_ISL_475711, EPI_ISL_475712, EPI_ISL_475713, EPI_ISL_475714, EPI_ISL_475715, EPI_ISL_475716 | | | |
| see above | Cedars-Sinai Medical Center, Department of Pathology & Laboratory Medicine, Molecular Pathology Laboratory | Cedars-Sinai Medical Center, Molecular Pathology Laboratory of Department of Pathology & Laboratory Medicine and Genomic Core | Wenjuan Zhang, John Paul Govindavari, Brian Davis, Stephanie Chen, Jong Taek Kim, Jianbo Song, Jean Lopategui, Jasmine T Plummer, Eric Vail |
| EPI_ISL_475717, EPI_ISL_475718, EPI_ISL_475719, EPI_ISL_475720, EPI_ISL_475721 | Microbiology, University Hospital Donostia | Microbiology, University Hospital Donostia | Cilla,G., Montes,M., Pineiro,L., Marimon,J.M. |
| EPI_ISL_475723, EPI_ISL_475724 | Egyptian National Cancer Institute (ENCI) | Egyptian National Cancer Institute (ENCI) | Zekri, Abdel Rahman N, Amer,K.E., Ahmed,O.S., Soliman,H.K., Hafez,M.M., Bahnassy,A.A., Abdelhamid,W., Gad,A., Ali,M., Hassan,W., Samir,M., Raouf,A., Hamdy,M.S., Soliman,M.S., Elsissy,M.H., Elkhateeb,S.M., Ezzelarab,M.H., Abouelhoda, Mohamed |
| EPI_ISL_475726, EPI_ISL_475727, EPI_ISL_475728, EPI_ISL_475730, EPI_ISL_475743 | Utah Public Health Laboratory | Utah Public Health Laboratory | Erin Young, Kelly Oakeson |
| EPI_ISL_475745, EPI_ISL_475746, EPI_ISL_475747, EPI_ISL_475748, EPI_ISL_475749, EPI_ISL_475751, EPI_ISL_475752, EPI_ISL_475753 | Medical Ain Shams Research Institute (MASRI), Ain Shams University | Medical Ain Shams Research Institute (MASRI), Ain Shams University | Hesham Elghazaly , Sara Hassan Agwa, Mahmoud Elmeteini , Ahmad Moustafa , Ashraf Omar, Osama Mansour, Samia Abdo, Hala Hafez, Ghada Ismael , Shaimaa Moustafa , Aya Mohamed, Reham Mamdouh , Hoda Abd Elsatar, Manal Hamdy Elsaid, Fatma Ebied |
| EPI_ISL_475754 | National Institute of Laboratory Medicine and Referral Center | Genomic Research Lab, BCSIR | Shahina Akter, Abu Sayeed Mohammad Mahmud, Mohammad Samir Uzzaman, Eshrar Osman, Md. Ahasan Habib, Tanjina Akhter Banu, Md. Murshed Hasan Sarkar, Barna Goswami, Iffat Jahan, Md. Saddam Hossain, Tasnim Nafisa, Md. Maruf Ahmed Molla, Mahmuda Yeasmin, Asish Kumar Ghosh, Arifa Akram, A. K. M. Shamsuzzaman, Sheikh Md. Selim Al Din, Utpal Chandra Ray, Salek Ahmed Sajib, Md. Salim Khan |
| EPI_ISL_475755 | National Institute of Laboratory Medicine and Referral Center | Genomic Research Lab, BCSIR | Md. Murshed Hasan Sarkar, Abu Sayeed Mohammad Mahmud, Mohammad Samir Uzzaman, Eshrar Osman, Md. Ahasan Habib, Shahina Akter, Tanjina Akhter Banu, Barna Goswami, Iffat Jahan, Md. Saddam Hossain, Tasnim Nafisa, Md. Maruf Ahmed Molla, Mahmuda Yeasmin, Asish Kumar Ghosh, Arifa Akram, A. K. M. Shamsuzzaman, Sheikh Md. Selim Al Din, Utpal Chandra Ray, Salek Ahmed Sajib, Md. Salim Khan |
| EPI_ISL_475756 | National Institute of Laboratory Medicine and Referral Center | Genomic Research Lab, BCSIR | Tanjina Akhter Banu, Abu Sayeed Mohammad Mahmud, Mohammad Samir Uzzaman, Eshrar Osman, Md. Ahasan Habib, Shahina Akter, Md. Murshed Hasan Sarkar, Barna Goswami, Iffat Jahan, Md. Saddam Hossain, Tasnim Nafisa, Md. Maruf Ahmed Molla, Mahmuda Yeasmin, Asish Kumar Ghosh, Arifa Akram, A. K. M. Shamsuzzaman, Sheikh Md. Selim Al Din, Utpal Chandra Ray, Salek Ahmed Sajib, Md. Salim Khan |
| EPI_ISL_475757 | National Institute of Laboratory Medicine and Referral Center | Genomic Research Lab, BCSIR | Barna Goswami, Abu Sayeed Mohammad Mahmud, Mohammad Samir Uzzaman, Eshrar Osman, Md. Ahasan Habib, Shahina Akter, Tanjina Akhter Banu, Md. Murshed Hasan Sarkar, Iffat Jahan, Md. Saddam Hossain, Tasnim Nafisa, Md. Maruf Ahmed Molla, Mahmuda Yeasmin, Asish Kumar Ghosh, Arifa Akram, A. K. M. Shamsuzzaman, Sheikh Md. Selim Al Din, Utpal Chandra Ray, Salek Ahmed Sajib, Md. Salim Khan |
| EPI_ISL_475758 | National Institute of Laboratory Medicine and Referral Center | Genomic Research Lab, BCSIR | Iffat Jahan, Abu Sayeed Mohammad Mahmud, Mohammad Samir Uzzaman, Eshrar Osman, Md. Ahasan Habib, Shahina Akter, Tanjina Akhter Banu, Md. Murshed Hasan Sarkar, Barna Goswami, Md. Saddam Hossain, Tasnim Nafisa, Md. Maruf Ahmed Molla, Mahmuda Yeasmin, Asish Kumar Ghosh, Arifa Akram, A. K. M. Shamsuzzaman, Sheikh Md. Selim Al Din, Utpal Chandra Ray, Salek Ahmed Sajib, Md. Salim Khan |
| EPI_ISL_475759 | National Institute of Laboratory Medicine and Referral Center | Genomic Research Lab, BCSIR | Md. Saddam Hossain, Abu Sayeed Mohammad Mahmud, Mohammad Samir Uzzaman, Eshrar Osman, Md. Ahasan Habib, Shahina Akter, Tanjina Akhter Banu, Md. Murshed Hasan Sarkar, Barna Goswami, Iffat Jahan, Tasnim Nafisa, Md. Maruf Ahmed Molla, Mahmuda Yeasmin, Asish Kumar Ghosh, Arifa Akram, A. K. M. Shamsuzzaman, Sheikh Md. Selim Al Din, Utpal Chandra Ray, Salek Ahmed Sajib, Md. Salim Khan |
| EPI_ISL_475760, EPI_ISL_475761 | National Institute of Laboratory Medicine and Referral Center | Genomic Research Lab, BCSIR | Abu Sayeed Mohammad Mahmud, Mohammad Samir Uzzaman, Eshrar Osman, Md. Ahasan Habib, Shahina Akter, Tanjina Akhter Banu, Md. Murshed Hasan Sarkar, Barna Goswami, Iffat Jahan, Md. Saddam Hossain, Tasnim Nafisa, Md. Maruf Ahmed Molla, Mahmuda Yeasmin, Asish Kumar Ghosh, Arifa Akram, A. K. M. Shamsuzzaman, Sheikh Md. Selim Al Din, Utpal Chandra Ray, Salek Ahmed Sajib, Md. Salim Khan |
| EPI_ISL_475762 | Oklahoma State Department of Health | França Lab | Caio Martinelle B. de França, Graham Wiley, Samuel T. Dunn, and Matthew J. Miller. |
| EPI_ISL_475766 | Universitaetsklinik für Innere Medizin II Innsbruck | Bergthaler laboratory, CeMM Research Center for Molecular Medicine of the Austrian Academy of Sciences | Alexandra Popa, Benedikt Agerer, Henrique Colaco, Lukas Endler, Jakob-Wendelin Genger, Alexander Lercher, Mark Smyth, Thomas Penz, Michael Schuster, Jan Laine, Martin Senekowitsch, Judith Aberle, Stephan Aberle, Peter Hufnagl, Daniela Schmid, Franz Allerberger, Elisabeth Puchhammer-Stoeckl, Manfred Nairz, Guenter Weiss, Gregor Hörmann, Kinga Rigler-Hohenwarter, Rainer Gattringer, Wegene Borena, Dorothee von Laer, Christoph Bock, Andreas Bergthaler |
| EPI_ISL_475769 | Institut für Virologie am Department für Hygiene, Mikrobiologie und Public Health | Bergthaler laboratory, CeMM Research Center for Molecular Medicine of the Austrian Academy of Sciences | Alexandra Popa, Benedikt Agerer, Henrique Colaco, Lukas Endler, Jakob-Wendelin Genger, Alexander Lercher, Mark Smyth, Thomas Penz, Michael Schuster, Jan Laine, Martin Senekowitsch, Judith Aberle, Stephan Aberle, Peter Hufnagl, Daniela Schmid, Franz Allerberger, Elisabeth Puchhammer-Stoeckl, Manfred Nairz, Guenter Weiss, Gregor Hörmann, Kinga Rigler-Hohenwarter, Rainer Gattringer, Wegene Borena, Dorothee von Laer, Christoph Bock, Andreas Bergthaler |
| EPI_ISL_475770, EPI_ISL_475771, EPI_ISL_475772, EPI_ISL_475773, EPI_ISL_475774, EPI_ISL_475775, EPI_ISL_475776, EPI_ISL_475779, EPI_ISL_475780, EPI_ISL_475781, EPI_ISL_475782, EPI_ISL_475783, EPI_ISL_475784, EPI_ISL_475785, EPI_ISL_475786, EPI_ISL_475787, EPI_ISL_475790, EPI_ISL_475791, EPI_ISL_475792, EPI_ISL_475793, EPI_ISL_475794, EPI_ISL_475795, EPI_ISL_475797, EPI_ISL_475798, EPI_ISL_475799, EPI_ISL_475800, EPI_ISL_475801, EPI_ISL_475802, EPI_ISL_475803, EPI_ISL_475804, EPI_ISL_475805, EPI_ISL_475806, EPI_ISL_475808, EPI_ISL_475809, EPI_ISL_475810, EPI_ISL_475811, EPI_ISL_475812 | | | |
| see above | Center for Virology, Medical University of Vienna | Bergthaler laboratory, CeMM Research Center for Molecular Medicine of the Austrian Academy of Sciences | Alexandra Popa, Benedikt Agerer, Henrique Colaco, Lukas Endler, Jakob-Wendelin Genger, Alexander Lercher, Mark Smyth, Thomas Penz, Michael Schuster, Jan Laine, Martin Senekowitsch, Judith Aberle, Stephan Aberle, Peter Hufnagl, Daniela Schmid, Franz Allerberger, Elisabeth Puchhammer-Stoeckl, Manfred Nairz, Guenter Weiss, Gregor Hörmann, Kinga Rigler-Hohenwarter, Rainer Gattringer, Wegene Borena, Dorothee von Laer, Christoph Bock, Andreas Bergthaler |
| EPI_ISL_475813, EPI_ISL_475814, EPI_ISL_475815, EPI_ISL_475816, EPI_ISL_475817, EPI_ISL_475818, EPI_ISL_475819, EPI_ISL_475820, EPI_ISL_475821, EPI_ISL_475822, EPI_ISL_475823, EPI_ISL_475824, EPI_ISL_475825, EPI_ISL_475826, EPI_ISL_475827, EPI_ISL_475828, EPI_ISL_475829 | | | |
| see above | Institut für Virologie am Department für Hygiene, Mikrobiologie und Public Health | Bergthaler laboratory, CeMM Research Center for Molecular Medicine of the Austrian Academy of Sciences | Alexandra Popa, Benedikt Agerer, Henrique Colaco, Lukas Endler, Jakob-Wendelin Genger, Alexander Lercher, Mark Smyth, Thomas Penz, Michael Schuster, Jan Laine, Martin Senekowitsch, Judith Aberle, Stephan Aberle, Peter Hufnagl, Daniela Schmid, Franz Allerberger, Elisabeth Puchhammer-Stoeckl, Manfred Nairz, Guenter Weiss, Gregor Hörmann, Kinga Rigler-Hohenwarter, Rainer Gattringer, Wegene Borena, Dorothee von Laer, Christoph Bock, Andreas Bergthaler |
| EPI_ISL_475830, EPI_ISL_475831, EPI_ISL_475832, EPI_ISL_475833, EPI_ISL_475834, EPI_ISL_475835, EPI_ISL_475836, EPI_ISL_475837, EPI_ISL_475838, EPI_ISL_475839, EPI_ISL_475840, EPI_ISL_475841, EPI_ISL_475842, EPI_ISL_475843, EPI_ISL_475844, EPI_ISL_475845, EPI_ISL_475848, EPI_ISL_475849, EPI_ISL_475850, EPI_ISL_475851, EPI_ISL_475852, EPI_ISL_475853, EPI_ISL_475854, EPI_ISL_475855, EPI_ISL_475859, EPI_ISL_475860, EPI_ISL_475861, EPI_ISL_475862, EPI_ISL_475863, EPI_ISL_475864, EPI_ISL_475865, EPI_ISL_475866, EPI_ISL_475869, EPI_ISL_475870, EPI_ISL_475871, EPI_ISL_475872, EPI_ISL_475873, EPI_ISL_475874, EPI_ISL_475875, EPI_ISL_475876, EPI_ISL_475878, EPI_ISL_475879, EPI_ISL_475880, EPI_ISL_475882, EPI_ISL_475883, EPI_ISL_475884, EPI_ISL_475885, EPI_ISL_475886 | | | |
| see above | Austrian Agency for Health and Food Safety (AGES) | Bergthaler laboratory, CeMM Research Center for Molecular Medicine of the Austrian | Alexandra Popa, Benedikt Agerer, Henrique Colaco, Lukas Endler, Jakob-Wendelin Genger, Alexander Lercher, Mark Smyth, Thomas Penz, Michael Schuster, Jan Laine, Martin Senekowitsch, Judith |

| | | | |
|---|---|---|---|
| | | Academy of Sciences | Aberle, Stephan Aberle, Peter Hufnagl, Daniela Schmid, Franz Allerberger, Elisabeth Puchhammer-Stoeckl, Manfred Nairz, Guenter Weiss, Gregor Hörmann, Kinga Rigler-Hohenwarter, Rainer Gattringer, Wegene Borena, Dorothee von Laer, Christoph Bock, Andreas Bergthaler |
| EPI_ISL_475889, EPI_ISL_475890, EPI_ISL_475891, EPI_ISL_475893, EPI_ISL_475894, EPI_ISL_475895, EPI_ISL_475896, EPI_ISL_475897, EPI_ISL_475899, EPI_ISL_475900, EPI_ISL_475901, EPI_ISL_475902, EPI_ISL_475903, EPI_ISL_475904, EPI_ISL_475905, EPI_ISL_475906, EPI_ISL_475907, EPI_ISL_475908, EPI_ISL_475909 | | | |
| see above | Zentralinstitut für medizinische und chemische Labordiagnostik, Universitätskliniken Insbruck | Bergthaler laboratory, CeMM Research Center for Molecular Medicine of the Austrian Academy of Sciences | Alexandra Popa, Benedikt Agerer, Henrique Colaco, Lukas Endler, Jakob-Wendelin Genger, Alexander Lercher, Mark Smyth, Thomas Penz, Michael Schuster, Jan Laine, Martin Senekowitsch, Judith Aberle, Stephan Aberle, Peter Hufnagl, Daniela Schmid, Franz Allerberger, Elisabeth Puchhammer-Stoeckl, Manfred Nairz, Guenter Weiss, Gregor Hörmann, Kinga Rigler-Hohenwarter, Rainer Gattringer, Wegene Borena, Dorothee von Laer, Christoph Bock, Andreas Bergthaler |
| EPI_ISL_475910, EPI_ISL_475911, EPI_ISL_475912, EPI_ISL_475913, EPI_ISL_475914, EPI_ISL_475915 | Klinikum Wels-Grieskirchen | Bergthaler laboratory, CeMM Research Center for Molecular Medicine of the Austrian Academy of Sciences | Alexandra Popa, Benedikt Agerer, Henrique Colaco, Lukas Endler, Jakob-Wendelin Genger, Alexander Lercher, Mark Smyth, Thomas Penz, Michael Schuster, Jan Laine, Martin Senekowitsch, Judith Aberle, Stephan Aberle, Peter Hufnagl, Daniela Schmid, Franz Allerberger, Elisabeth Puchhammer-Stoeckl, Manfred Nairz, Guenter Weiss, Gregor Hörmann, Kinga Rigler-Hohenwarter, Rainer Gattringer, Wegene Borena, Dorothee von Laer, Christoph Bock, Andreas Bergthaler |
| EPI_ISL_475919, EPI_ISL_475923, EPI_ISL_475924, EPI_ISL_475926, EPI_ISL_475927, EPI_ISL_475928 | Institut für Virologie am Department für Hygiene, Mikrobiologie und Public Health | Bergthaler laboratory, CeMM Research Center for Molecular Medicine of the Austrian Academy of Sciences | Alexandra Popa, Benedikt Agerer, Henrique Colaco, Lukas Endler, Jakob-Wendelin Genger, Alexander Lercher, Mark Smyth, Thomas Penz, Michael Schuster, Jan Laine, Martin Senekowitsch, Judith Aberle, Stephan Aberle, Peter Hufnagl, Daniela Schmid, Franz Allerberger, Elisabeth Puchhammer-Stoeckl, Manfred Nairz, Guenter Weiss, Gregor Hörmann, Kinga Rigler-Hohenwarter, Rainer Gattringer, Wegene Borena, Dorothee von Laer, Christoph Bock, Andreas Bergthaler |
| EPI_ISL_475929, EPI_ISL_475930, EPI_ISL_475931, EPI_ISL_475932, EPI_ISL_475934, EPI_ISL_475935, EPI_ISL_475936 | Universitaetsklinik für Innere Medizin II Innsbruck | Bergthaler laboratory, CeMM Research Center for Molecular Medicine of the Austrian Academy of Sciences | Alexandra Popa, Benedikt Agerer, Henrique Colaco, Lukas Endler, Jakob-Wendelin Genger, Alexander Lercher, Mark Smyth, Thomas Penz, Michael Schuster, Jan Laine, Martin Senekowitsch, Judith Aberle, Stephan Aberle, Peter Hufnagl, Daniela Schmid, Franz Allerberger, Elisabeth Puchhammer-Stoeckl, Manfred Nairz, Guenter Weiss, Gregor Hörmann, Kinga Rigler-Hohenwarter, Rainer Gattringer, Wegene Borena, Dorothee von Laer, Christoph Bock, Andreas Bergthaler |
| EPI_ISL_475937, EPI_ISL_475938, EPI_ISL_475939, EPI_ISL_475940, EPI_ISL_475941, EPI_ISL_475942, EPI_ISL_475943, EPI_ISL_475944, EPI_ISL_475945, EPI_ISL_475946, EPI_ISL_475947, EPI_ISL_475948, EPI_ISL_475949, EPI_ISL_475950, EPI_ISL_475951, EPI_ISL_475953, EPI_ISL_475954, EPI_ISL_475955, EPI_ISL_475957, EPI_ISL_475958, EPI_ISL_475959, EPI_ISL_475960, EPI_ISL_475962, EPI_ISL_475963, EPI_ISL_475966, EPI_ISL_475967, EPI_ISL_475968, EPI_ISL_475969, EPI_ISL_475970, EPI_ISL_475971, EPI_ISL_475972, EPI_ISL_475973, EPI_ISL_475979, EPI_ISL_475981, EPI_ISL_475983, EPI_ISL_475985, EPI_ISL_475986, EPI_ISL_475990, EPI_ISL_475991, EPI_ISL_475993, EPI_ISL_475995, EPI_ISL_475997 | | | |
| see above | National Public Health Laboratory, National Centre for Infectious Diseases | National Public Health Laboratory, National Centre for Infectious Diseases | Mak TM, Octavia S, Chavatte JM, Cui L, Lin RTP |
| EPI_ISL_476018, EPI_ISL_476019, EPI_ISL_476020, EPI_ISL_476021 | Washington University in St. Louis | Washington University in St. Louis | David Wang, Carey-Ann Burnham, Scott Handley, Lindsay Droit, Stephen Tahan |
| EPI_ISL_476022 | Defence Research & Development Establishment | Defence Research & Development Establishment | Shashi Sharma, Paban Kumar Dash, Jyoti S Kumar, Sushil Kumar Sharma, Ambuj Shrivastava |
| EPI_ISL_476023 | Defence Research & Development Establishment (DRDE) | Defence Research & Development Establishment (DRDE) | Shashi Sharma, Paban Kumar Dash, Sushil Kumar Sharma, Ambuj Shrivastava, Jyoti S. Kumar |
| EPI_ISL_476027, EPI_ISL_476028, EPI_ISL_476029, EPI_ISL_476030, EPI_ISL_476032, EPI_ISL_476033, EPI_ISL_476034, EPI_ISL_476038, EPI_ISL_476039, EPI_ISL_476040, EPI_ISL_476041, EPI_ISL_476042, EPI_ISL_476043, EPI_ISL_476044, EPI_ISL_476045, EPI_ISL_476046, EPI_ISL_476048, EPI_ISL_476049, EPI_ISL_476050, EPI_ISL_476051, EPI_ISL_476053, EPI_ISL_476054, EPI_ISL_476055, EPI_ISL_476056, EPI_ISL_476058, EPI_ISL_476059, EPI_ISL_476060, EPI_ISL_476062, EPI_ISL_476063, EPI_ISL_476064 | | | |
| see above | Michigan Department of Health and Human Services, Bureau of Laboratories | Michigan Department of Health and Human Services, Bureau of Laboratories | Blankenship HM, Riner D, Soehnlen MK |
| EPI_ISL_476067 | The National Institute of Public Health | State Veterinary Institute Prague and The National Institute of Public Health | Nagy,A;Jirincova,H;Novakova,L;Trnka,D;Vecerova,J |
| EPI_ISL_476068, EPI_ISL_476069, EPI_ISL_476070, EPI_ISL_476071, EPI_ISL_476072, EPI_ISL_476073, EPI_ISL_476074, EPI_ISL_476075, EPI_ISL_476076, EPI_ISL_476077 | University of Debrecen, Department of Medical Microbiology | National Laboratory of Virology, Szentágothai Research Centre | Endre Gábor Tóth, Balázs Somogyi, Brigitta Zana, Eszter Csoma, Ferenc Jakab, Gábor Kemenesi |
| EPI_ISL_476078 | University of Szeged, Institute of Clinical Microbiology | National Laboratory of Virology, Szentágothai Research Centre | Endre Gábor Tóth, Balázs Somogyi, Brigitta Zana, Terhes Gabriella, Ferenc Jakab, Gábor Kemenesi |
| EPI_ISL_476079, EPI_ISL_476080, EPI_ISL_476081, EPI_ISL_476082, EPI_ISL_476083, EPI_ISL_476084, EPI_ISL_476085, EPI_ISL_476086, EPI_ISL_476087, EPI_ISL_476088, EPI_ISL_476089, EPI_ISL_476090, EPI_ISL_476091, EPI_ISL_476093, EPI_ISL_476094, EPI_ISL_476095, EPI_ISL_476096, EPI_ISL_476097, EPI_ISL_476098, EPI_ISL_476099, EPI_ISL_476100, EPI_ISL_476101, EPI_ISL_476102, EPI_ISL_476103, EPI_ISL_476104, EPI_ISL_476105, EPI_ISL_476106, EPI_ISL_476107, EPI_ISL_476108, EPI_ISL_476110, EPI_ISL_476112, EPI_ISL_476113, EPI_ISL_476114, EPI_ISL_476115, EPI_ISL_476116, EPI_ISL_476117, EPI_ISL_476118, EPI_ISL_476119, EPI_ISL_476120, EPI_ISL_476121, EPI_ISL_476122, EPI_ISL_476123, EPI_ISL_476124, EPI_ISL_476126, EPI_ISL_476127, EPI_ISL_476128, EPI_ISL_476129, EPI_ISL_476130, EPI_ISL_476131, EPI_ISL_476132, EPI_ISL_476133, EPI_ISL_476134 | | | |
| see above | Viollier AG | Department of Biosystems Science and Engineering, ETH Zürich | Christian Beisel, Sarah Nadeau, Ivan Topolsky, Pedro Ferreira, Philipp Jablonski, Susana Posada-Céspedes, Tobias Schär, Ina Nissen, Natascha Santacroce, Elodie Burcklen, Christiane Beckmann, Maurice Redondo, Olivier Kobel, Christoph Noppen, Sophie Seidel, Noemie Santamaria de Souza, Niko Beerenwinkel, Tanja Stadler |
| EPI_ISL_476135 | Achima Care Fristadens VC | The Public Health Agency of Sweden | Oskar Karlsson Lindsjo, Maria Lind Karlberg, Mattias Haukland, Reza Advani, Olov Svartstrom, Anna-Malin Linde, Sandra Broddesson, Petra Edquist, Mia Brytting, Anna Risberg, Karin Tegmark-Wisell |
| EPI_ISL_476136 | Surbrunns VC | The Public Health Agency of Sweden | Oskar Karlsson Lindsjo, Maria Lind Karlberg, Mattias Haukland, Reza Advani, Olov Svartstrom, Anna-Malin Linde, Sandra Broddesson, Petra Edquist, Mia Brytting, Anna Risberg, Karin Tegmark-Wisell |
| EPI_ISL_476137 | Wasterlakarna | The Public Health Agency of Sweden | Oskar Karlsson Lindsjo, Maria Lind Karlberg, Mattias Haukland, Reza Advani, Olov Svartstrom, Anna-Malin Linde, Sandra Broddesson, Petra Edquist, Mia Brytting, Anna Risberg, Karin Tegmark-Wisell |
| EPI_ISL_476138 | Ulltuna Vardcentral | The Public Health Agency of Sweden | Oskar Karlsson Lindsjo, Maria Lind Karlberg, Mattias Haukland, Reza Advani, Olov Svartstrom, Anna-Malin Linde, Sandra Broddesson, Petra Edquist, Mia Brytting, Anna Risberg, Karin Tegmark-Wisell |
| EPI_ISL_476139 | Folkhalsomyndigheten | The Public Health Agency of Sweden | Oskar Karlsson Lindsjo, Maria Lind Karlberg, Mattias Haukland, Reza Advani, Olov Svartstrom, Anna-Malin Linde, Sandra Broddesson, Petra Edquist, Shamam Muradrasoli, Anna Risberg, Karin Tegmark-Wisell |
| EPI_ISL_476140, EPI_ISL_476141, EPI_ISL_476142 | Klinisk Mikrobiologi | The Public Health Agency of Sweden | Oskar Karlsson Lindsjo, Maria Lind Karlberg, Mattias Haukland, Reza Advani, Olov Svartstrom, Anna-Malin Linde, Sandra Broddesson, Petra Edquist, Shamam Muradrasoli, Anna Risberg, Karin Tegmark-Wisell |
| EPI_ISL_476143, EPI_ISL_476144 | Skovde/Unilabs | The Public Health Agency of Sweden | Oskar Karlsson Lindsjo, Maria Lind Karlberg, Mattias Haukland, Reza Advani, Olov Svartstrom, Anna-Malin Linde, Sandra Broddesson, Petra Edquist, Shamam Muradrasoli, Anna Risberg, Karin Tegmark-Wisell |
| EPI_ISL_476145, EPI_ISL_476146, EPI_ISL_476147 | Ostersund klinisk mikrobiologi | The Public Health Agency of Sweden | Oskar Karlsson Lindsjo, Maria Lind Karlberg, Mattias Haukland, Reza Advani, Olov Svartstrom, Anna-Malin Linde, Sandra Broddesson, Petra Edquist, Shamam Muradrasoli, Anna Risberg, Karin Tegmark-Wisell |
| EPI_ISL_476148, EPI_ISL_476149 | Institut Pasteur Dakar | Institut Pasteur de Dakar | Ndongo Dia, Moussa Moise Diagne, Mamadou Diop, Ousmane Faye, Amadou Alpha Sall |
| EPI_ISL_476221 | Laboratory Fleury | Instituto de Medicina Tropical da Univesidade de São Paulo | Samples: Celso Granato; Sequencing: Ingra Morales Claro, Jaqueline Goes de Jesus, Erika Regina Manuli, Flavia Cristina da Silva Sales, Thais de Moura Coletti, Camila Alves Maia da Silva, Mariana Severo Ramundo, Giula Magalhaes Ferreira, Darlan da Silva Candido, Julien Theze, Nuno Faria, Ester Sabino |
| EPI_ISL_476282, EPI_ISL_476288, EPI_ISL_476289, EPI_ISL_476297 | DB Diagnósticos do Brasil | Instituto de Medicina Tropical da Univesidade de São Paulo | Samples: Nelson Gaburo Jr; Sequencing: Ingra Morales Claro, Jaqueline Goes de Jesus, Erika Regina Manuli, Flavia Cristina da Silva Sales, Thais de Moura Coletti, Camila Alves Maia da Silva, Mariana Severo Ramundo, Giula Magalhaes Ferreira, Darlan da Silva Candido, Julien Theze, Nuno Faria, Ester Sabino |
| EPI_ISL_476341 | Laboratório de Patologia Clínica - UNICAMP | Laboratório de Estudos de Vírus Emergentes - UNICAMP | José Luiz Proença-Modena, Magnun Nueldo Nunes dos Santos, Angelica Schreiber, Julia Forato,Camila Simeoni, Marcilio Jorge Fumagalli, Mariene Ribeiro Amorim, Darlan da Silva Candido, Nuno Rodrigues Faria, Julien Theze, Luiz Gonzaga,Jaqueline Goes Jesus e William Marciel de Souza |
| EPI_ISL_476373 | Hospital da Clínicas da Faculdade de Medicina da Universidade de São Paulo | Instituto de Medicina Tropical da Univesidade de São Paulo | Samples: Ingra Morales Claro, Erika Regina Manuli, Cecilia Salete Alencar, Carolina S. Lazar, Silvia F. Costa; Sequencing: Ingra Morales Claro, Jaqueline Goes de Jesus, Erika Regina Manuli, Flavia Cristina da Silva Sales, Thais de Moura Coletti, Camila Alves Maia da Silva, Mariana Severo Ramundo, Giula Magalhaes Ferreira, Darlan da Silva Candido, Julien Theze, Nuno Faria, Ester Sabino |
| EPI_ISL_476395, EPI_ISL_476398 | Laboratório de Patologia Clínica - UNICAMP | Laboratório de Estudos de Vírus Emergentes - UNICAMP | José Luiz Proença-Modena, Magnun Nueldo Nunes dos Santos, Angelica Schreiber, Julia Forato,Camila Simeoni, Marcilio Jorge Fumagalli, Mariene Ribeiro Amorim, Darlan da Silva Candido, Nuno Rodrigues Faria, Julien Theze, Luiz Gonzaga,Jaqueline Goes Jesus e William Marciel de Souza |
| EPI_ISL_476435, EPI_ISL_476439, EPI_ISL_476445, EPI_ISL_476446, EPI_ISL_476469, EPI_ISL_476490 | Hospital da Clínicas da Faculdade de Medicina da Universidade de São Paulo | Instituto de Medicina Tropical da Univesidade de São Paulo | Samples: Ingra Morales Claro, Erika Regina Manuli, Cecilia Salete Alencar, Carolina S. Lazar, Silvia F. Costa; Sequencing: Ingra Morales Claro, Jaqueline Goes de Jesus, Erika Regina Manuli, Flavia Cristina da Silva Sales, Thais de Moura Coletti, Camila Alves Maia da Silva, Mariana Severo Ramundo, Giula Magalhaes Ferreira, Darlan da Silva Candido, Julien Theze, Nuno Faria, Ester Sabino |
| EPI_ISL_476492 | Institut Pasteur Dakar | Institut Pasteur de Dakar | Ndongo Dia, Moussa Moise Diagne, Mamadou Diop, Ousmane Faye, Amadou Alpha Sall |
| EPI_ISL_476493 | Institut Pasteur Dakar | Institut Pasteur de Dakar | Ndongo Dia, Moussa Moise Diagne, Mamadou Diop, Ousmane Faye, Amadou alpha Sall |
| EPI_ISL_476494 | Institut Pasteur Dakar | Institut Pasteur de Dakar | Ndongo Dia, Moussa Moise Diagne, Mamadou Diop, Ousmane Faye, Amadou Alpha Sall |
| EPI_ISL_476495 | Institut Pasteur Dakar | Institut Pasteur de Dakar | Ndongo Dia, Moussa Moise Diagne, Mamadou Diop, Ousmane Faye, Amadou alpha Sall |
| EPI_ISL_476496 | Hospital Garrahan | Héritas | Dalmacio Pereyra, Roberta Crespo, Mauricio Grisolia, Cristian Rohr, Andrea Mangano, Maria Florencia Fernandez, Fabian Fay, Martin Vazquez |
| EPI_ISL_476498, EPI_ISL_476499, EPI_ISL_476500, EPI_ISL_476501, EPI_ISL_476502, EPI_ISL_476503, EPI_ISL_476505, EPI_ISL_476506, EPI_ISL_476507, EPI_ISL_476508, EPI_ISL_476509, EPI_ISL_476510, EPI_ISL_476511, EPI_ISL_476512, EPI_ISL_476513 | | | |
| see above | Laboratoire de microbiologie, Hopital de Verdun | Smith Laboratory, Centre de Recherche CHU Sainte-Justine | Martin Smith, Marieke Rozendaal, Ivan Pavlov |
| EPI_ISL_476514 | Institut Pasteur Dakar | Institut Pasteur de Dakar | Ndongo Dia, Moussa Moise Diagne, Mamadou Diop, Ousmane Faye, Amadou Alpha Sall |
| EPI_ISL_476516 | Institut Pasteur Dakar | Institut Pasteur de Dakar | Ndongo Dia, Moussa Moise Diagne, mamadou Diop, Ousmane Faye, Amadou Alpha Sall |
| EPI_ISL_476517, EPI_ISL_476518, EPI_ISL_476521, EPI_ISL_476522, EPI_ISL_476523, EPI_ISL_476525, EPI_ISL_476526, EPI_ISL_476527, EPI_ISL_476528, EPI_ISL_476531, EPI_ISL_476542, EPI_ISL_476545, EPI_ISL_476547, EPI_ISL_476548, EPI_ISL_476549, EPI_ISL_476550, EPI_ISL_476552, EPI_ISL_476553, EPI_ISL_476554, EPI_ISL_476556 | | | |
| see above | Yale Clinical Virology Laboratory | Grubaugh Lab - Yale School of Public Health | Joseph Fauver, Tara Alpert, Anderson Brito, Anne Wyllie, Chantal Vogels, Mary Petrone, Cole Jensen, Chaney Kalinich, Isabel Ott, Arnau Casanovas, Catherine Muenker, Adam Moore, Alice Lu, Maria Tokuyama, Patrick Wong, Peiwen Lu, Saad Omer, Richard Martinello, Allison Nelson, Shelli Farhadian, Akiko Iwasaki, Charlese Dela Cruz, Albert Ko, Nathan Grubaugh |
| EPI_ISL_476558 | Institut Pasteur Dakar | Institut Pasteur de Dakar | Ndongo Dia, Moussa Moise Diagne, Mamadou Diop, Ousmane Faye, Amadou Alpha Sall |
| EPI_ISL_476559 | unknown | Laboratoire Sciences et Technologies de la Santé (STS) Institut Supérieur des Sciences de la Santé Université Hassan 1er, Settat, Morocco | Hajar Lemriss, Sanaâ Lemriss, Amal Souiri, Narjis Amar, Mustapha Mouallif, Touria Essayagh, Jawad Bouzid, Saâd EL Kabbaj, Abderraouf Hilali |
| EPI_ISL_476561 | Hospital Garrahan | Héritas | Roberta Crespo, Dalmacio Pereyra, Mauricio Grisolia, Cristian Rohr, Andrea Mangano, Maria Florencia Fernandez, Fabian Fay, Martin Vazquez |
| EPI_ISL_476562 | Institut Pasteur Dakar | Institut Pasteur de Dakar | Ndongo Dia, Moussa Moise Diagne, Mamadou Diop, Ousmane Faye, Amadou Alpha Sall |
| EPI_ISL_476563 | Hospital de Pediatría "Prof. Dr. Juan P Garrahan" | Héritas | Dalmacio Pereyra, Roberta Crespo, Mauricio Grisolia, Cristian Rohr, Andrea Mangano, Maria Florencia Fernandez, Fabian Fay, Martin Vazquez |
| EPI_ISL_476564 | Institut Pasteur Dakar | Institut Pasteur de Dakar | Ndongo Dia, Moussa Moise Diagne, Mamadou diop, Ousmane Faye, Amadou alpha Sall |

| | | | |
|---|---|---|---|
| EPI_ISL_476565 | Hospital de Pediatría "Prof. Dr. Juan P Garrahan" | Héritas | Andrea Mangano, Maria Florencia Fernandez, Dalmacio Pereyra, Roberta Crespo, Mauricio Grisolia, Cristian Rohr, Fabian Fay, Martin Vazquez |
| EPI_ISL_476567 | Hospital de Pediatría "Prof. Dr. Juan P Garrahan" | Héritas | Dalmacio Pereyra, Roberta Crespo, Mauricio Grisolia, Cristian Rohr, Andrea Mangano, Maria Florencia Fernandez, Fabian Fay, Martin Vazquez |
| EPI_ISL_476568 | Hospital de Pediatría "Prof. Dr. Juan P Garrahan" | Héritas | Cristian Rohr, Andrea Mangano, Maria Florencia Fernandez, Dalmacio Pereyra, Roberta Crespo, Mauricio Grisolia, Fabian Fay, Martin Vazquez |
| EPI_ISL_476569 | Institut Pasteur Dakar | Institut Pasteur de Dakar | Ndongo Dia, Moussa Moise, Mamadou Diop, Ousmane Faye, Amadou Alpha Sall |
| EPI_ISL_476571 | Hospital de Pediatría "Prof. Dr. Juan P Garrahan" | Héritas | Dalmacio Pereyra, Roberta Crespo, Mauricio Grisolia, Cristian Rohr, Andrea Mangano, Maria Florencia Fernandez, Fabian Fay, Martin Vazquez |
| EPI_ISL_476572 | Institut Pasteur Dakar | Institut Pasteur de Dakar | Ndongo Dia, Moussa Moise Diagne, Mamadou Diop, Ousmane Faye, Amadou Alpha Sall |
| EPI_ISL_476573 | Hospital de Pediatría "Prof. Dr. Juan P Garrahan" | Héritas | Dalmacio Pereyra, Roberta Crespo, Mauricio Grisolia, Cristian Rohr, Andrea Mangano, Maria Florencia Fernandez, Fabian Fay, Martin Vazquez |
| EPI_ISL_476574 | Institut Pasteur Dakar | Institut Pasteur de Dakar | Ndongo Dia, Moussa Moise Diagne, Mamadou Diop, Ousmane Faye, Amadou Alpha Sall |
| EPI_ISL_476702 | Incubadora Venezolana de Ciencia, Venezuela | Incubadora Venezolana de Ciencia, Venezuela / Instituto Nacional de Salud, Bogotá, Colombia / Grupo de Investigaciones Microbiológicas-UR (GIMUR), Departamento de Biología, Facultad de Ciencias Naturales, Universidad del Rosario, Bogotá, Colombia / Icahn School of Medicine at Mount Sinai, New York, USA | Alberto Paniz-Mondolfi, Marina Muñoz, Luis Perez-Garcia, Lourdes Delgado, Carolina Florez, Sergio Gomez, Angelica Rico, Lisseth Pardo, Esther C. Barros, Carolina Hernández, Jesús E. Jaimes, Aníbal A. Teherán, Ana S. Gonzalez-Reiche, Matthew M. Hernandez, Emilia Mia Sordillo, Viviana Simon, Harm van Bakel, Juan David Ramírez |
| EPI_ISL_476704 | Incubadora Venezolana de Ciencia, Venezuela | Incubadora Venezolana de Ciencia, Venezuela / Instituto Nacional de Salud, Bogotá, Colombia / Grupo de Investigaciones Microbiológicas-UR (GIMUR), Departamento de Biología, Facultad de Ciencias Naturales, Universidad del Rosario, Bogotá, Colombia / Icahn School of Medicine at Mount Sinai, New York, USA | Alberto Paniz-Mondolfi, Marina Muñoz, Luis Perez-Garcia, Lourdes Delgado, Carolina Florez, Sergio Gomez, Angelica Rico, Lisseth Pardo, Esther C. Barros, Carolina Hernández, Jesús E. Jaimes, Aníbal A. Teherán, Ana S. Gonzalez-Reiche, Matthew M. Hernandez, Emilia Mia Sordillo, Viviana Simon, Harm van Bakel, Juan David Ramírez |
| EPI_ISL_476705 | Labor Kneißler GmbH & Co. KG | Heinrich Pette Institute, Leibniz Institute for Experimental Virology | Thomas Günther, Adam Grundhoff, Manja Czech-Sioli, Nicole Fischer, Matthias Ottinger, Melanie M. Brinkmann |
| EPI_ISL_476706, EPI_ISL_476707, EPI_ISL_476708, EPI_ISL_476709, EPI_ISL_476710, EPI_ISL_476711, EPI_ISL_476712, EPI_ISL_476713, EPI_ISL_476714, EPI_ISL_476715, EPI_ISL_476716, EPI_ISL_476717, EPI_ISL_476718, EPI_ISL_476719, EPI_ISL_476720, EPI_ISL_476721, EPI_ISL_476722, EPI_ISL_476723, EPI_ISL_476724, EPI_ISL_476725, EPI_ISL_476726, EPI_ISL_476727, EPI_ISL_476728, EPI_ISL_476729, EPI_ISL_476730, EPI_ISL_476731, EPI_ISL_476732, EPI_ISL_476733, EPI_ISL_476734, EPI_ISL_476735, EPI_ISL_476736, EPI_ISL_476737, EPI_ISL_476738, EPI_ISL_476739, EPI_ISL_476740, EPI_ISL_476741, EPI_ISL_476742, EPI_ISL_476743, EPI_ISL_476744, EPI_ISL_476745, EPI_ISL_476746, EPI_ISL_476747, EPI_ISL_476748, EPI_ISL_476749, EPI_ISL_476750, EPI_ISL_476751, EPI_ISL_476752, EPI_ISL_476753, EPI_ISL_476754, EPI_ISL_476755, EPI_ISL_476756, EPI_ISL_476757, EPI_ISL_476758, EPI_ISL_476759, EPI_ISL_476760, EPI_ISL_476761, EPI_ISL_476762, EPI_ISL_476763, EPI_ISL_476764, EPI_ISL_476765, EPI_ISL_476766 | | | |
| see above | Minnesota Department of Health, Public Health Laboratory | Minnesota Department of Health, Public Health Laboratory | Matt Plumb, Jacob Garfin, and Xiong Wang |
| EPI_ISL_476768, EPI_ISL_476772, EPI_ISL_476773, EPI_ISL_476774, EPI_ISL_476775, EPI_ISL_476778, EPI_ISL_476780, EPI_ISL_476781, EPI_ISL_476782, EPI_ISL_476783, EPI_ISL_476784, EPI_ISL_476785, EPI_ISL_476786, EPI_ISL_476787, EPI_ISL_476788, EPI_ISL_476791, EPI_ISL_476792 | | | |
| see above | Stanford clinical virology lab | Chan-Zuckerberg Biohub | Benjamin Pinksy, Katharine Walter, Victoria N. Parikh, John Gorzynski, Hannah N. DeJong, Matthew T. Wheeler, Jason Andrews, Manuel Rivas, Carlos Bustamante, Euan Ashley, with CZB Cliahub Consortium |
| EPI_ISL_476801, EPI_ISL_476802, EPI_ISL_476803, EPI_ISL_476804 | Hong Kong Department of Health | School of Public Health, The University of Hong Kong | Dominic N.C. Tsang, Daniel K.W. Chu, Leo L.M. Poon, Malik Peiris |
| EPI_ISL_476814, EPI_ISL_476818, EPI_ISL_476819 | Department of Laboratory Medicine, Tan Tock Seng Hospital | Department of Laboratory Medicine, Tan Tock Seng Hospital | Chen YYC, Zair X, Li C, Tang WY, Maurer-Stroh S, Barkham TMS, Nagarajan N, Sessions OM |
| EPI_ISL_476822, EPI_ISL_476823, EPI_ISL_476824, EPI_ISL_476825, EPI_ISL_476826, EPI_ISL_476827, EPI_ISL_476828, EPI_ISL_476829, EPI_ISL_476830, EPI_ISL_476831 | Laboratoire des Fièvres Hémorragiques Virales du Benin | Charité-Universitätsmedizin Berlin | Yadouleton, Anges; Sander Anna-Lena; Moreira-Soto Andres; Drexler, Jan Felix |
| EPI_ISL_476832 | Medical Biology Department, Kocaeli University | Medical Genetics Department, Kocaeli University | Savli H, Cine N, Sunnetci-Akkoyunlu D, Eren-Keskin S, Ilgazli A, Akhan S, Karadenizli A, Kasap M, Sayan M, Akpinar G, Canturk NZ. |
| EPI_ISL_476833, EPI_ISL_476834 | Laboratoire des Fièvres Hémorragiques Virales du Benin | Charité-Universitätsmedizin Berlin | Yadouleton, Anges; Sander Anna-Lena; Moreira-Soto Andres; Drexler, Jan Felix |
| EPI_ISL_476840, EPI_ISL_476842, EPI_ISL_476844, EPI_ISL_476846, EPI_ISL_476848, EPI_ISL_476849, EPI_ISL_476850, EPI_ISL_476852, EPI_ISL_476853, EPI_ISL_476854 | Defence Research & Development Establishment (DRDE) | Defence Research & Development Establishment (DRDE) | Shashi Sharma, Paban Kumar Dash, Sushil Kumar Sharma, Ambuj Shrivastava, Jyoti S. Kumar |
| EPI_ISL_476855 | GMERS Medical College & Hospital, Gotri, Vadodara | Gujarat Biotechnology Research Centre | Apurvasinh Puvar, Janvi Raval, Zarna Patel, Monika Gandhi, Pinal Trivedi, Maharshi Pandya, Nidhi Patel, Nitin Savaliya, Raghawendra Kumar, Dinesh Kumar, Zuber Saiyed, Komal Patel, Labdhi Pandya, Afzal Ansari, Nikha Trivedi, Meenakshi Shah, Neena Doshi, Varsha Godbole, R D Dixit, A M Kadri, Harsh Bakshi, Chaitanya Joshi, Madhvi Joshi |
| EPI_ISL_476856 | GMERS Medical College & Hospital, Gotri, Vadodara | Gujarat Biotechnology Research Centre | Janvi Raval, Zarna Patel, Monika Gandhi, Pinal Trivedi, Maharshi Pandya, Nidhi Patel, Nitin Savaliya, Raghawendra Kumar, Dinesh Kumar, Zuber Saiyed, Komal Patel, Labdhi Pandya, Afzal Ansari, Nikha Trivedi, Meenakshi Shah, Neena Doshi, Varsha Godbole, Apurvasinh Puvar, R D Dixit, A M Kadri, Harsh Bakshi, Chaitanya Joshi, Madhvi Joshi |
| EPI_ISL_476857 | GMERS Medical College & Hospital, Gotri, Vadodara | Gujarat Biotechnology Research Centre | Zarna Patel, Monika Gandhi, Pinal Trivedi, Maharshi Pandya, Nidhi Patel, Nitin Savaliya, Raghawendra Kumar, Dinesh Kumar, Zuber Saiyed, Komal Patel, Labdhi Pandya, Afzal Ansari, Nikha Trivedi, Meenakshi Shah, Neena Doshi, Varsha Godbole, Apurvasinh Puvar, Janvi Raval, R D Dixit, A M Kadri, Harsh Bakshi, Chaitanya Joshi, Madhvi Joshi |
| EPI_ISL_476858 | GMERS Medical College & Hospital, Gotri, Vadodara | Gujarat Biotechnology Research Centre | Monika Gandhi, Pinal Trivedi, Maharshi Pandya, Nidhi Patel, Nitin Savaliya, Raghawendra Kumar, Dinesh Kumar, Zuber Saiyed, Komal Patel, Labdhi Pandya, Afzal Ansari, Nikha Trivedi, Meenakshi Shah, Neena Doshi, Varsha Godbole, Apurvasinh Puvar, Janvi Raval, Zarna Patel, R D Dixit, A M Kadri, Harsh Bakshi, Chaitanya Joshi, Madhvi Joshi |
| EPI_ISL_476859 | GMERS Medical College & Hospital, Gotri, Vadodara | Gujarat Biotechnology Research Centre | Pinal Trivedi, Maharshi Pandya, Nidhi Patel, Nitin Savaliya, Raghawendra Kumar, Dinesh Kumar, Zuber Saiyed, Komal Patel, Labdhi Pandya, Afzal Ansari, Nikha Trivedi, Meenakshi Shah, Neena Doshi, Varsha Godbole, Apurvasinh Puvar, Janvi Raval, Zarna Patel, Monika Gandhi, R D Dixit, A M Kadri, Harsh Bakshi, Chaitanya Joshi, Madhvi Joshi |
| EPI_ISL_476860 | GMERS Medical College & Hospital, Gotri, Vadodara | Gujarat Biotechnology Research Centre | Maharshi Pandya, Nidhi Patel, Nitin Savaliya, Raghawendra Kumar, Dinesh Kumar, Zuber Saiyed, Komal Patel, Labdhi Pandya, Afzal Ansari, Nikha Trivedi, Meenakshi Shah, Neena Doshi, Varsha Godbole, Apurvasinh Puvar, Janvi Raval, Zarna Patel, Monika Gandhi, Pinal Trivedi, R D Dixit, A M Kadri, Harsh Bakshi, Chaitanya Joshi, Madhvi Joshi |
| EPI_ISL_476861 | GMERS Medical College & Hospital, Gotri, Vadodara | Gujarat Biotechnology Research Centre | Nidhi Patel, Nitin Savaliya, Raghawendra Kumar, Dinesh Kumar, Zuber Saiyed, Komal Patel, Labdhi Pandya, Afzal Ansari, Nikha Trivedi, Meenakshi Shah, Neena Doshi, Varsha Godbole, Apurvasinh Puvar, Janvi Raval, Zarna Patel, Monika Gandhi, Pinal Trivedi, Maharshi Pandya, R D Dixit, A M Kadri, Harsh Bakshi, Chaitanya Joshi, Madhvi Joshi |
| EPI_ISL_476862 | GMERS Medical College & Hospital, Gotri, Vadodara | Gujarat Biotechnology Research Centre | Nitin Savaliya, Raghawendra Kumar, Dinesh Kumar, Zuber Saiyed, Komal Patel, Labdhi Pandya, Afzal Ansari, Nikha Trivedi, Meenakshi Shah, Neena Doshi, Varsha Godbole, Apurvasinh Puvar, Janvi Raval, Zarna Patel, Monika Gandhi, Pinal Trivedi, Maharshi Pandya, Nidhi Patel, R D Dixit, A M Kadri, Harsh Bakshi, Chaitanya Joshi, Madhvi Joshi |
| EPI_ISL_476863 | GMERS Medical College and Hospital, Gandhinagar | Gujarat Biotechnology Research Centre | Raghawendra Kumar, Dinesh Kumar, Zuber Saiyed, Komal Patel, Labdhi Pandya, Afzal Ansari, Nikha Trivedi, Seema Bhatt, Gaurishankar Shrimali, Bhavesh Modi, Bharti Rajani, Apurvasinh Puvar, Janvi Raval, Zarna Patel, Monika Gandhi, Pinal Trivedi, Maharshi Pandya, Nidhi Patel, Nitin Savaliya, R D Dixit, A M Kadri, Harsh Bakshi, Chaitanya Joshi, Madhvi Joshi |
| EPI_ISL_476864 | GMERS Medical College and Hospital, Gandhinagar | Gujarat Biotechnology Research Centre | Dinesh Kumar, Zuber Saiyed, Komal Patel, Labdhi Pandya, Afzal Ansari, Nikha Trivedi, Seema Bhatt, Gaurishankar Shrimali, Bhavesh Modi, Bharti Rajani, Apurvasinh Puvar, Janvi Raval, Zarna Patel, Monika Gandhi, Pinal Trivedi, Maharshi Pandya, Nidhi Patel, Nitin Savaliya, Raghawendra Kumar R D Dixit, A M Kadri, Harsh Bakshi, Chaitanya Joshi, Madhvi Joshi |
| EPI_ISL_476865 | GMERS Medical College and Hospital, Gandhinagar | Gujarat Biotechnology Research Centre | Zuber Saiyed, Komal Patel, Labdhi Pandya, Afzal Ansari, Nikha Trivedi, Seema Bhatt, Gaurishankar Shrimali, Bhavesh Modi, Bharti Rajani, Apurvasinh Puvar, Janvi Raval, Zarna Patel, Monika Gandhi, Pinal Trivedi, Maharshi Pandya, Nidhi Patel, Nitin Savaliya, Raghawendra Kumar, Dinesh Kumar, R D Dixit, A M Kadri, Harsh Bakshi, Chaitanya Joshi, Madhvi Joshi |
| EPI_ISL_476866 | GMERS Medical College and Hospital, Gandhinagar | Gujarat Biotechnology Research Centre | Komal Patel, Labdhi Pandya, Afzal Ansari, Nikha Trivedi, Seema Bhatt, Gaurishankar Shrimali, Bhavesh Modi, Bharti Rajani, Apurvasinh Puvar, Janvi Raval, Zarna Patel, Monika Gandhi, Pinal Trivedi, Maharshi Pandya, Nidhi Patel, Nitin Savaliya, Raghawendra Kumar, Dinesh Kumar, Zuber Saiyed, R D Dixit, A M Kadri, Harsh Bakshi, Chaitanya Joshi, Madhvi Joshi |
| EPI_ISL_476867 | Banas Medical College and Research Institute | Gujarat Biotechnology Research Centre | Labdhi Pandya, Afzal Ansari, Nikha Trivedi, Radhika Khara, Sunil R Joshi, Viren s Doshi, Apurvasinh Puvar, Janvi Raval, Zarna Patel, Monika Gandhi, Pinal Trivedi, Maharshi Pandya, Nidhi Patel, Nitin Savaliya, Raghawendra Kumar, Dinesh Kumar, Zuber Saiyed, Komal Patel, R D Dixit, A M Kadri, Harsh Bakshi, Chaitanya Joshi, Madhvi Joshi |
| EPI_ISL_476868 | Banas Medical College and Research Institute | Gujarat Biotechnology Research Centre | Afzal Ansari, Nikha Trivedi, Radhika Khara, Sunil R Joshi, Viren s Doshi, Apurvasinh Puvar, Janvi Raval, Zarna Patel, Monika Gandhi, Pinal Trivedi, Maharshi Pandya, Nidhi Patel, Nitin Savaliya, Raghawendra Kumar, Dinesh Kumar, Zuber Saiyed, Komal Patel, Labdhi Pandya, R D Dixit, A M Kadri, Harsh Bakshi, Chaitanya Joshi, Madhvi Joshi |
| EPI_ISL_476869 | Department of MicroBiology, Government Medical College, Surat | Gujarat Biotechnology Research Centre | Nikha Trivedi, Naresh Chauhan, Summaiya Mullan, Amit gamit, Apurvasinh Puvar, Janvi Raval, Zarna Patel, Monika Gandhi, Pinal Trivedi, Maharshi Pandya, Nidhi Patel, Nitin Savaliya, Raghawendra Kumar, Dinesh Kumar, Zuber Saiyed, Komal Patel, Labdhi Pandya, Afzal Ansari, R D Dixit, A M Kadri, Harsh Bakshi, Chaitanya Joshi, Madhvi Joshi |
| EPI_ISL_476870 | Department of MicroBiology, Government Medical College, Surat | Gujarat Biotechnology Research Centre | Naresh Chauhan, Summaiya Mullan, Amit gamit, Apurvasinh Puvar, Janvi Raval, Zarna Patel, Monika Gandhi, Pinal Trivedi, Maharshi Pandya, Nidhi Patel, Nitin Savaliya, Raghawendra Kumar, Dinesh Kumar, Zuber Saiyed, Komal Patel, Labdhi Pandya, Afzal Ansari, Nikha Trivedi, R D Dixit, A M Kadri, Harsh Bakshi, Chaitanya Joshi, Madhvi Joshi |
| EPI_ISL_476871 | Department of MicroBiology, Government Medical College, Surat | Gujarat Biotechnology Research Centre | Summaiya Mullan, Amit gamit, Apurvasinh Puvar, Janvi Raval, Zarna Patel, Monika Gandhi, Pinal Trivedi, Maharshi Pandya, Nidhi Patel, Nitin Savaliya, Raghawendra Kumar, Dinesh Kumar, Zuber Saiyed, Komal Patel, Labdhi Pandya, Afzal Ansari, Nikha Trivedi, Naresh Chauhan, R D Dixit, A M Kadri, Harsh Bakshi, Chaitanya Joshi, Madhvi Joshi |
| EPI_ISL_476872 | Department of MicroBiology, Government Medical College, Surat | Gujarat Biotechnology Research Centre | Amit gamit, Apurvasinh Puvar, Janvi Raval, Zarna Patel, Monika Gandhi, Pinal Trivedi, Maharshi Pandya, Nidhi Patel, Nitin Savaliya, Raghawendra Kumar, Dinesh Kumar, Zuber Saiyed, Komal Patel, Labdhi Pandya, Afzal Ansari, Nikha Trivedi, Naresh Chauhan, Summaiya Mullan, R D Dixit, A M Kadri, Harsh Bakshi, Chaitanya Joshi, Madhvi Joshi |
| EPI_ISL_476873 | Department of MicroBiology, Government Medical College, Surat | Gujarat Biotechnology Research Centre | Apurvasinh Puvar, Janvi Raval, Zarna Patel, Monika Gandhi, Pinal Trivedi, Maharshi Pandya, Nidhi Patel, Nitin Savaliya, Raghawendra Kumar, Dinesh Kumar, Zuber Saiyed, Komal Patel, Labdhi Pandya, Afzal Ansari, Nikha Trivedi, Naresh Chauhan, Summaiya Mullan, Amit gamit, R D Dixit, A M Kadri, Harsh Bakshi, Chaitanya Joshi, Madhvi Joshi |
| EPI_ISL_476874 | Department of MicroBiology, Government Medical College, Surat | Gujarat Biotechnology Research Centre | Janvi Raval, Zarna Patel, Monika Gandhi, Pinal Trivedi, Maharshi Pandya, Nidhi Patel, Nitin Savaliya, Raghawendra Kumar, Dinesh Kumar, Zuber Saiyed, Komal Patel, Labdhi Pandya, Afzal Ansari, Nikha Trivedi, Naresh Chauhan, Summaiya Mullan, Amit gamit, Apurvasinh Puvar, R D Dixit, A M Kadri, Harsh Bakshi, Chaitanya Joshi, Madhvi Joshi |
| EPI_ISL_476875 | Department of MicroBiology, Government Medical College, Surat | Gujarat Biotechnology Research Centre | Zarna Patel, Monika Gandhi, Pinal Trivedi, Maharshi Pandya, Nidhi Patel, Nitin Savaliya, Raghawendra Kumar, Dinesh Kumar, Zuber Saiyed, Komal Patel, Labdhi Pandya, Afzal Ansari, Nikha Trivedi, Naresh Chauhan, Summaiya Mullan, Amit gamit, Apurvasinh Puvar, Janvi Raval, R D Dixit, A M Kadri, Harsh Bakshi, Chaitanya Joshi, Madhvi Joshi |
| EPI_ISL_476876 | Department of MicroBiology, Government Medical College, Surat | Gujarat Biotechnology Research Centre | Pinal Trivedi, Maharshi Pandya, Nidhi Patel, Nitin Savaliya, Raghawendra Kumar, Dinesh Kumar, Zuber Saiyed, Komal Patel, Labdhi Pandya, Afzal Ansari, Nikha Trivedi, Naresh Chauhan, Summaiya Mullan, Amit gamit, Apurvasinh Puvar, Janvi Raval, Zarna Patel, Monika Gandhi, R D Dixit, A M Kadri, Harsh Bakshi, Chaitanya Joshi, Madhvi Joshi |
| EPI_ISL_476877 | Department of MicroBiology, Government Medical College, Surat | Gujarat Biotechnology Research Centre | Maharshi Pandya, Nidhi Patel, Nitin Savaliya, Raghawendra Kumar, Dinesh Kumar, Zuber Saiyed, Komal Patel, Labdhi Pandya, Afzal Ansari, Nikha Trivedi, Naresh Chauhan, Summaiya Mullan, Amit gamit, Apurvasinh Puvar, Janvi Raval, Zarna Patel, Monika Gandhi, Pinal Trivedi, R D Dixit, A M Kadri, Harsh Bakshi, Chaitanya Joshi, Madhvi Joshi |
| EPI_ISL_476878 | Department of MicroBiology, Government Medical College, Surat | Gujarat Biotechnology Research Centre | Nidhi Patel, Nitin Savaliya, Raghawendra Kumar, Dinesh Kumar, Zuber Saiyed, Komal Patel, Labdhi Pandya, Afzal Ansari, Nikha Trivedi, Naresh Chauhan, Summaiya Mullan, Amit gamit, Apurvasinh Puvar, Janvi Raval, Zarna Patel, Monika Gandhi, Pinal Trivedi, Maharshi Pandya, R D Dixit, A M Kadri, Harsh Bakshi, Chaitanya Joshi, Madhvi Joshi |
| EPI_ISL_476879 | Department of MicroBiology, Government Medical College, Surat | Gujarat Biotechnology Research Centre | Nitin Savaliya, Raghawendra Kumar, Dinesh Kumar, Zuber Saiyed, Komal Patel, Labdhi Pandya, Afzal Ansari, Nikha Trivedi, Naresh Chauhan, Summaiya Mullan, Amit gamit, Apurvasinh Puvar, Janvi Raval, Zarna Patel, Monika Gandhi, Pinal Trivedi, Maharshi Pandya, Nidhi Patel, R D Dixit, A M Kadri, Harsh Bakshi, Chaitanya Joshi, Madhvi Joshi |

| | | | |
|---|---|---|---|
| EPI_ISL_476880 | Department of MicroBiology, Government Medical College, Surat | Gujarat Biotechnology Research Centre | Raghawendra Kumar, Dinesh Kumar, Zuber Saiyed, Komal Patel, Labdhi Pandya, Afzal Ansari, Nikha Trivedi, Naresh Chauhan, Summaiya Mullan, Amit gamit, Apurvasinh Puvar, Janvi Raval, Zarna Patel, Monika Gandhi, Pinal Trivedi, Maharshi Pandya, Nidhi Patel, Nitin Savaliya, R D Dixit, A M Kadri, Harsh Bakshi, Chaitanya Joshi, Madhvi Joshi |
| EPI_ISL_476881 | Department of MicroBiology, Government Medical College, Surat | Gujarat Biotechnology Research Centre | Dinesh Kumar, Zuber Saiyed, Komal Patel, Labdhi Pandya, Afzal Ansari, Nikha Trivedi, Naresh Chauhan, Summaiya Mullan, Amit gamit, Apurvasinh Puvar, Janvi Raval, Zarna Patel, Monika Gandhi, Pinal Trivedi, Maharshi Pandya, Nidhi Patel, Nitin Savaliya, Raghawendra Kumar, R D Dixit, A M Kadri, Harsh Bakshi, Chaitanya Joshi, Madhvi Joshi |
| EPI_ISL_476882 | Department of MicroBiology, Government Medical College, Surat | Gujarat Biotechnology Research Centre | Zuber Saiyed, Komal Patel, Labdhi Pandya, Afzal Ansari, Nikha Trivedi, Naresh Chauhan, Summaiya Mullan, Amit gamit, Apurvasinh Puvar, Janvi Raval, Zarna Patel, Monika Gandhi, Pinal Trivedi, Maharshi Pandya, Nidhi Patel, Nitin Savaliya, Raghawendra Kumar, Dinesh Kumar, R D Dixit, A M Kadri, Harsh Bakshi, Chaitanya Joshi, Madhvi Joshi |
| EPI_ISL_476883, EPI_ISL_476884, EPI_ISL_476885, EPI_ISL_476886, EPI_ISL_476887, EPI_ISL_476888, EPI_ISL_476889, EPI_ISL_476890, EPI_ISL_476891, EPI_ISL_476892, EPI_ISL_476893, EPI_ISL_476894, EPI_ISL_476895, EPI_ISL_476896 | | | |
| see above | Defence Research & Development Establishment (DRDE) | Defence Research & Development Establishment (DRDE) | Shashi Sharma, Paban Kumar Dash, Sushil Kumar Sharma, Ambuj Shrivastava, Jyoti S. Kumar |
| EPI_ISL_476897 | University of South Carolina Functional Genomics Core | University of South Carolina Functional Genomics Core | Michael Shtutman |
| EPI_ISL_476898, EPI_ISL_476899 | Alaska State Virology Laboratory | Alaska State Virology Laboratory | Jack Chen, Ph.D. |
| EPI_ISL_476900, EPI_ISL_476902, EPI_ISL_476903, EPI_ISL_476905, EPI_ISL_476906, EPI_ISL_476907, EPI_ISL_476908, EPI_ISL_476909, EPI_ISL_476910, EPI_ISL_476911, EPI_ISL_476912, EPI_ISL_476914, EPI_ISL_476915, EPI_ISL_476916, EPI_ISL_476917, EPI_ISL_476918, EPI_ISL_476919, EPI_ISL_476920, EPI_ISL_476921, EPI_ISL_476922, EPI_ISL_476923, EPI_ISL_476924, EPI_ISL_476925, EPI_ISL_476926, EPI_ISL_476927, EPI_ISL_476928, EPI_ISL_476929, EPI_ISL_476930, EPI_ISL_476931, EPI_ISL_476932, EPI_ISL_476933, EPI_ISL_476935, EPI_ISL_476936, EPI_ISL_476937, EPI_ISL_476938, EPI_ISL_476939, EPI_ISL_476940 | | | |
| see above | UW Virology Lab | UW Virology Lab | Pavitra Roychoudhury, Hong Xie, Lasata Shrestha, Amin Addetia, Truong Nguyen, Victoria M Rachleff, Meei-Li Huang, Keith R Jerome, Alexander Greninger |
| EPI_ISL_476941, EPI_ISL_476942, EPI_ISL_476943, EPI_ISL_476944, EPI_ISL_476945, EPI_ISL_476946, EPI_ISL_476947, EPI_ISL_476948, EPI_ISL_476949, EPI_ISL_476950, EPI_ISL_476951, EPI_ISL_476952, EPI_ISL_476953, EPI_ISL_476954, EPI_ISL_476955, EPI_ISL_476956, EPI_ISL_476957, EPI_ISL_476958, EPI_ISL_476959, EPI_ISL_476960, EPI_ISL_476961, EPI_ISL_476962, EPI_ISL_476963, EPI_ISL_476964, EPI_ISL_476965, EPI_ISL_476966, EPI_ISL_476967, EPI_ISL_476968, EPI_ISL_476969, EPI_ISL_476970, EPI_ISL_476971, EPI_ISL_476972, EPI_ISL_476973, EPI_ISL_476974, EPI_ISL_476975, EPI_ISL_476976, EPI_ISL_476977, EPI_ISL_476978, EPI_ISL_476979, EPI_ISL_476980, EPI_ISL_476981, EPI_ISL_476982, EPI_ISL_476983, EPI_ISL_476984, EPI_ISL_476985, EPI_ISL_476986, EPI_ISL_476987, EPI_ISL_476988, EPI_ISL_476989, EPI_ISL_476990, EPI_ISL_476991, EPI_ISL_476992, EPI_ISL_476993, EPI_ISL_476994, EPI_ISL_476995, EPI_ISL_476996, EPI_ISL_476997, EPI_ISL_476998, EPI_ISL_476999, EPI_ISL_477000, EPI_ISL_477001, EPI_ISL_477002, EPI_ISL_477003, EPI_ISL_477004, EPI_ISL_477005, EPI_ISL_477006, EPI_ISL_477007 | | | |
| see above | KU Leuven, Rega Institute, Clinical and Epidemiological Virology | KU Leuven, Rega Institute, Clinical and Epidemiological Virology | Tony Wawina-Bokalanga, Joan Marti-Carerras, Bert Vanmechelen, Piet Maes |
| EPI_ISL_477008, EPI_ISL_477009, EPI_ISL_477010, EPI_ISL_477011, EPI_ISL_477012, EPI_ISL_477013 | University of Debrecen, Department of Medical Microbiology | National Laboratory of Virology, Szentágothai Research Centre | Endre Gábor Tóth, Balázs Somogyi, Brigitta Zana, Eszter Csoma, Ferenc Jakab, Gábor Kemenesi |
| EPI_ISL_477014 | Institute of Microbiology, Universidad San Francisco de Quito | Institute of Microbiology, Universidad San Francisco de Quito | Belen Prado-Vivar, Sully Marquez, Juan Jose Guadalupe, Monica Becerra-Wong, Carla Torres, Bernardo Gutierrez, Francisco Mora, Juan Gaviria, Alejandra Ramones, Franklin Espinoza, Edison Ligña, Jorge Reyes, Patricio Rojas-Silva, Veronica Barragan, Gabriel Trueba, Michelle Grunauer, Paul Cardenas |
| EPI_ISL_477015 | Institute of Microbiology, Universidad San Francisco de Quito | Institute of Microbiology, Universidad San Francisco de Quito | Sully Márquez, Belén Prado-Vivar, Juan José Guadalupe, Monica Becerra-Wong, Carla Torres, Bernardo Gutiérrez, Jorge Luis Velez, Verónica Barragán, Patricio Rojas-Silva, Gabriel Trueba, Michelle Grunauer, Paúl Cárdenas |
| EPI_ISL_477016 | Institute of Microbiology, Universidad San Francisco de Quito | Institute of Microbiology, Universidad San Francisco de Quito | Juan José Guadalupe, Sully Márquez, Belén Prado-Vivar, Monica Becerra-Wong, Carla Torres, Bernardo Gutiérrez, Jorge Luis Velez, Verónica Barragán, Patricio Rojas-Silva, Gabriel Trueba, Michelle Grunauer, Paúl Cárdenas |
| EPI_ISL_477020, EPI_ISL_477021, EPI_ISL_477022, EPI_ISL_477023, EPI_ISL_477024, EPI_ISL_477029, EPI_ISL_477031, EPI_ISL_477032, EPI_ISL_477033, EPI_ISL_477034, EPI_ISL_477035, EPI_ISL_477036, EPI_ISL_477037, EPI_ISL_477038, EPI_ISL_477039, EPI_ISL_477040, EPI_ISL_477041, EPI_ISL_477045, EPI_ISL_477046, EPI_ISL_477047, EPI_ISL_477048, EPI_ISL_477049, EPI_ISL_477050, EPI_ISL_477051, EPI_ISL_477052, EPI_ISL_477054, EPI_ISL_477056, EPI_ISL_477057, EPI_ISL_477058, EPI_ISL_477059, EPI_ISL_477060, EPI_ISL_477061, EPI_ISL_477062, EPI_ISL_477063, EPI_ISL_477064, EPI_ISL_477065, EPI_ISL_477066, EPI_ISL_477067, EPI_ISL_477068, EPI_ISL_477069, EPI_ISL_477070, EPI_ISL_477071, EPI_ISL_477072, EPI_ISL_477073, EPI_ISL_477074, EPI_ISL_477075, EPI_ISL_477076, EPI_ISL_477079, EPI_ISL_477080, EPI_ISL_477081, EPI_ISL_477085, EPI_ISL_477086, EPI_ISL_477087, EPI_ISL_477088, EPI_ISL_477090, EPI_ISL_477091, EPI_ISL_477092, EPI_ISL_477093, EPI_ISL_477094, EPI_ISL_477095, EPI_ISL_477096, EPI_ISL_477098, EPI_ISL_477099, EPI_ISL_477100, EPI_ISL_477101, EPI_ISL_477102, EPI_ISL_477103, EPI_ISL_477105, EPI_ISL_477106, EPI_ISL_477107, EPI_ISL_477108, EPI_ISL_477109, EPI_ISL_477110, EPI_ISL_477111, EPI_ISL_477112, EPI_ISL_477113, EPI_ISL_477114, EPI_ISL_477115, EPI_ISL_477117, EPI_ISL_477118, EPI_ISL_477119 | | | |
| see above | BCCDC Public Health Laboratory | BCCDC Public Health Laboratory | Richard Harrigan, Hope Lapointe, Jinny Choi, Kimia Kamelian, John Tyson,Terry Snutch, Linda Hoang, Inna Sekirov, Paul Levett, Mel Krajden, Natalie Prystajecky |
| EPI_ISL_477125, EPI_ISL_477126, EPI_ISL_477127, EPI_ISL_477128, EPI_ISL_477129, EPI_ISL_477130, EPI_ISL_477131, EPI_ISL_477132, EPI_ISL_477133, EPI_ISL_477134, EPI_ISL_477135, EPI_ISL_477136, EPI_ISL_477138, EPI_ISL_477139, EPI_ISL_477140 | | | |
| see above | Child Health Research Foundation | Child Health Research Foundation | Senjuti Saha, Md Saiful Islam Sajib, Roly Malaker, Md Hafizur Rahman, Afroza Akter Tanni, Syed Muktadir Al Sium, Maksuda Islam, Samir K Saha |
| EPI_ISL_477143, EPI_ISL_477150, EPI_ISL_477155, EPI_ISL_477156 | Institut Pasteur Dakar | Institut Pasteur de Dakar | Ndongo Dia, Moussa Moise Diagne, Mamadou Diop, Mamadou Malado Jallow, Marie Henriette Dior Ndione, Safietou Sankhe, Ousmane Faye, Amadou Alpha Sall. |
| EPI_ISL_477161 | Egyptian National Cancer Institute (ENCI) | Egyptian National Cancer Institute (ENCI) | Zekri, Abdel Rahman N, Amer,K.E., Ahmed,O.S., Soliman,H.K., Hafez,M.M., Bahnassy,A.A., Abdelhamid,W., Gad,A., Ali,M., Hassan,W., Samir,M., Raouf,A., Hamdy,M.S., Soliman,M.S., Elsissy,M.H., Elkhateeb,S.M., Ezzelarab,M.H., Abouelhoda, Mohamed |
| EPI_ISL_477163 | Laboratory of Dr. John Lednicky | University of Florida | John A. Lednicky, Maha A. Elbadry, Kuttichantran Subramaniam, Thomas B. Waltzek, John Glenn Morris, Jr. |
| EPI_ISL_477169 | Department for Virology, Molecular Biology and Genome Research, R. G. Lugar Center for Public Health Research, National Center for Disease Control and Public Health (NCDC) of Georgia. | Department for Virology, Molecular Biology and Genome Research, R. G. Lugar Center for Public Health Research, National Center for Disease Control and Public Health (NCDC) of Georgia. | Tata Imnadze, Giorgi Tomashvili, Meri Pantsulaia, Gvantsa Brachveli, Gvantsa Chanturia, Ann Machablishvili, Nato Kotaria, Marine Murtskhvaladze, Lela Sabadze, Meri Gavashelidze, Ana Papkiauri, Tamar Jashiashvili, Tea Tevdoradze, Ketevan Sidamonidze, Ekaterine Khmaladze, Ekaterine Zhghenti, Roena Sukhiashvili, Mariam Zakalashvili, Lela Urushadze, Magda Dgebuadze, Davit Tsaguria, Ekaterine Zangaladze, Nino Berishvili, Adam Kotorashvili, Maia Alkhazashvili, Irma Burjanadze, Anna Kasradze, Khatuna Zakhashvili, Paata Imnadze, Amiran Gamkrelidze. |
| EPI_ISL_477170 | Department of Laboratory, Medicine Tan Tock Seng Hospital | Department of Laboratory Medicine Tan Tock Seng Hospital | Chen YYC, Zair X, Li C, Tang WY, Maurer-Stroh S, Barkham TMS, Nagarajan N, Sessions OM |
| EPI_ISL_477171 | Department of Laboratory, Medicine Tan Tock Seng Hospital | Department of Laboratory Medicine Tan Tock Seng Hospital | Chen YYC, Zair X, Li C, Tang WY, Maurer-Stroh S, Barkham TMS, Nagarajan N, Sessions OM |
| EPI_ISL_477172, EPI_ISL_477174, EPI_ISL_477175, EPI_ISL_477177, EPI_ISL_477178, EPI_ISL_477180, EPI_ISL_477182 | Department of Laboratory Medicine Tan Tock Seng Hospital | Department of Laboratory Medicine Tan Tock Seng Hospital | Chen YYC, Zair X, Li C, Tang WY, Maurer-Stroh S, Barkham TMS, Nagarajan N, Sessions OM |
| EPI_ISL_477183 | Department of MicroBiology, Government Medical College, Surat | Gujarat Biotechnology Research Centre | Monika Gandhi, Pinal Trivedi, Maharshi Pandya, Nidhi Patel, Nitin Savaliya, Raghawendra Kumar, Dinesh Kumar, Zuber Saiyed, Komal Patel, Labdhi Pandya, Afzal Ansari, Nikha Trivedi, Naresh Chauhan, Summaiya Mullan, Amit gamit, Apurvasinh Puvar, Janvi Raval, Zarna Patel, R D Dixit, A M Kadri, Harsh Bakshi, Chaitanya Joshi, Madhvi Joshi |
| EPI_ISL_477184, EPI_ISL_477187, EPI_ISL_477188, EPI_ISL_477189, EPI_ISL_477190, EPI_ISL_477191, EPI_ISL_477192 | Department of Laboratory Medicine Tan Tock Seng Hospital | Department of Laboratory Medicine Tan Tock Seng Hospital | Chen YYC, Zair X, Li C, Tang WY, Maurer-Stroh S, Barkham TMS, Nagarajan N, Sessions OM |
| EPI_ISL_477193 | Istituto Zooprofilattico Sperimentale Puglia e Basilicata; | Beaconlab (Bioinformatics, Evolution and Comparative Genomics lab), Dept of Biosciences, University on Mila | Parisi A.,Pesole G., Manzari C., Chiara M. |
| EPI_ISL_477199, EPI_ISL_477200 | Istituto Zooprofilattico Sperimentale Puglia e Basilicata; | Beaconlab (Bioinformatics, Evolution and Comparative Genomics lab), Dept of Biosciences, University on Milan | Parisi A.,Pesole G., Manzari C., Chiara M. |
| EPI_ISL_477203 | Istituto Zooprofilattico Sperimentale Puglia e Basilicata; | Beaconlab (Bioinformatics, Evolution and Comparative Genomics lab), Dept of Biosciences, University on Mila | Parisi A.,Pesole G., Manzari C., Chiara M. |
| EPI_ISL_477205, EPI_ISL_477207, EPI_ISL_477210, EPI_ISL_477211, EPI_ISL_477239, EPI_ISL_477241, EPI_ISL_477242, EPI_ISL_477246, EPI_ISL_477249, EPI_ISL_477250, EPI_ISL_477255, EPI_ISL_477261 | | | |
| see above | Institute for Stem Cell Science and Regenerative Medicine | National Centre for Biological Sciences | Farhan Ali, Vanessa Molin Paynter, Srikar Krishna, Mohak Sharda, Shah-e-Jahan Gulzar, Awadhesh Pandit, Varadha Sundarmurthy, Uma Ramakrishnan, Dasaradhi Palakodeti, Aswin Seshasayee |
| EPI_ISL_477272, EPI_ISL_477273, EPI_ISL_477274, EPI_ISL_477275, EPI_ISL_477276 | Mayo Clinic & Mayo Clinic Laboratories | Minnesota Department of Health, Public Health Laboratory | Matt Plumb, Jacob Garfin, Kelly Pung, and Xiong Wang |
| EPI_ISL_477277, EPI_ISL_477278, EPI_ISL_477279, EPI_ISL_477280, EPI_ISL_477281, EPI_ISL_477282, EPI_ISL_477283, EPI_ISL_477284, EPI_ISL_477285, EPI_ISL_477286, EPI_ISL_477287, EPI_ISL_477288, EPI_ISL_477289, EPI_ISL_477290 | | | |
| see above | M Health Fairview | Minnesota Department of Health, Public Health Laboratory | Matt Plumb, Jacob Garfin, Kelly Pung, and Xiong Wang |
| EPI_ISL_477291, EPI_ISL_477292, EPI_ISL_477293, EPI_ISL_477294, EPI_ISL_477295, EPI_ISL_477296, EPI_ISL_477297, EPI_ISL_477298, EPI_ISL_477299, EPI_ISL_477300, EPI_ISL_477301, EPI_ISL_477302, EPI_ISL_477303, EPI_ISL_477304, EPI_ISL_477305, EPI_ISL_477306, EPI_ISL_477307, EPI_ISL_477308, EPI_ISL_477309, EPI_ISL_477310, EPI_ISL_477311, EPI_ISL_477312 | | | |
| see above | Mayo Clinic & Mayo Clinic Laboratories | Minnesota Department of Health, Public Health Laboratory | Matt Plumb, Jacob Garfin, Kelly Pung, and Xiong Wang |
| EPI_ISL_477615, EPI_ISL_477616, EPI_ISL_477617, EPI_ISL_477618, EPI_ISL_477619, EPI_ISL_477620, EPI_ISL_477621, EPI_ISL_477622, EPI_ISL_477623, EPI_ISL_477624, EPI_ISL_477625 | | | |
| see above | University of Szeged, Institute of Clinical Microbiology | National Laboratory of Virology, Szentágothai Research Centre | Endre Gábor Tóth, Balázs Somogyi, Brigitta Zana, Terhes Gabriella, Ferenc Jakab, Gábor Kemenesi |
| EPI_ISL_477631, EPI_ISL_477632, EPI_ISL_477633, EPI_ISL_477634, EPI_ISL_477635, EPI_ISL_477636, EPI_ISL_477637, EPI_ISL_477638, EPI_ISL_477639, EPI_ISL_477640, EPI_ISL_477641, EPI_ISL_477642, EPI_ISL_477643, EPI_ISL_477644, EPI_ISL_477645, EPI_ISL_477646, EPI_ISL_477647, EPI_ISL_477648, EPI_ISL_477649, EPI_ISL_477650, EPI_ISL_477651, EPI_ISL_477652, EPI_ISL_477653, EPI_ISL_477654, EPI_ISL_477655, EPI_ISL_477656, EPI_ISL_477657, EPI_ISL_477658, EPI_ISL_477659, EPI_ISL_477660, EPI_ISL_477661, EPI_ISL_477662, EPI_ISL_477663, EPI_ISL_477664, EPI_ISL_477665, EPI_ISL_477666, EPI_ISL_477667, EPI_ISL_477668, EPI_ISL_477669, EPI_ISL_477670, EPI_ISL_477671, EPI_ISL_477672 | | | |
| see above | Virginia DCLS | Virginia DCLS | Virginia DCLS |
| EPI_ISL_477673, EPI_ISL_477674, EPI_ISL_477675, EPI_ISL_477676, EPI_ISL_477677, EPI_ISL_477678, EPI_ISL_477679, EPI_ISL_477680, EPI_ISL_477681, EPI_ISL_477682, EPI_ISL_477683, EPI_ISL_477684, EPI_ISL_477685, EPI_ISL_477686, EPI_ISL_477687, EPI_ISL_477689, EPI_ISL_477690, EPI_ISL_477691, EPI_ISL_477692, EPI_ISL_477693, EPI_ISL_477694, EPI_ISL_477695, EPI_ISL_477696, EPI_ISL_477697, EPI_ISL_477698, EPI_ISL_477699, EPI_ISL_477700, EPI_ISL_477701, EPI_ISL_477702, EPI_ISL_477703, EPI_ISL_477704, EPI_ISL_477705, EPI_ISL_477706, EPI_ISL_477707, EPI_ISL_477708, EPI_ISL_477709, EPI_ISL_477710, EPI_ISL_477711, EPI_ISL_477712, EPI_ISL_477713, EPI_ISL_477714, EPI_ISL_477715, EPI_ISL_477716, EPI_ISL_477717, EPI_ISL_477718, EPI_ISL_477719, EPI_ISL_477720, EPI_ISL_477721, EPI_ISL_477723, EPI_ISL_477724 | | | |
| see above | UW Virology Lab | UW Virology Lab | Pavitra Roychoudhury, Hong Xie, Lasata Shrestha, Amin Addetia, Truong Nguyen, Victoria M Rachleff, Meei-Li Huang, Keith R Jerome, Alexander Greninger |
| EPI_ISL_477730, EPI_ISL_477735, EPI_ISL_477741, EPI_ISL_477742, EPI_ISL_477744, EPI_ISL_477746, EPI_ISL_477747, EPI_ISL_477749, EPI_ISL_477751, EPI_ISL_477752, EPI_ISL_477756, EPI_ISL_477758, EPI_ISL_477762, EPI_ISL_477763, EPI_ISL_477768, EPI_ISL_477769, EPI_ISL_477774, EPI_ISL_477776, EPI_ISL_477779 | | | |
| see above | University of Birmingham | COVID-19 Genomics UK (COG-UK) Consortium | Institute of Microbiology, University of Birmingham: Claire McMurray, Joanne Stockton, Samuel Nicholls, Radoslaw Poplawski, Will Rowe, Josh Quick, Nicholas Loman. University of Birmingham Testing Laboratory: Celina M Whalley, Andrew Bosworth, Charlotte Poxon, Kasun Wanigasooriya, Oliver Pickles, Mike Kidd, Alex Richter, Andrew D Beggs PHE Heartlands Lab: Husam Osman, Andrew Bosworth. Queen Elizabeth Hospital: Anna Casey |
| EPI_ISL_477796, EPI_ISL_477798, EPI_ISL_477800, EPI_ISL_477802, EPI_ISL_477805, EPI_ISL_477806, EPI_ISL_477807, EPI_ISL_477808, EPI_ISL_477809, EPI_ISL_477810, EPI_ISL_477811, EPI_ISL_477812, EPI_ISL_477813, EPI_ISL_477814 | | | |
| see above | Department of Pathology, University of Cambridge | COVID-19 Genomics UK (COG-UK) Consortium | Luke W Meredith, M. Estée Török, Myra Hosmillo, William L. Hamilton, Martin D. Curran, Theresa Feltwell, Grant Hall, Anna Yakovleva, Fahad A Khokhar, Charlotte J. Houldcroft, Laura G Caller, Aminu S. Jahun, Sarah L. Caddy, Yasmin Chaudhry, Malte Pinckert, Ian Goodfellow |
| EPI_ISL_477822, EPI_ISL_477823, EPI_ISL_477825, EPI_ISL_477827, EPI_ISL_477828, EPI_ISL_477829, EPI_ISL_477830, EPI_ISL_477831, EPI_ISL_477833, EPI_ISL_477834, EPI_ISL_477838, EPI_ISL_477839, EPI_ISL_477841, EPI_ISL_477842, EPI_ISL_477843, EPI_ISL_477845, EPI_ISL_477849, EPI_ISL_477852, EPI_ISL_477853, EPI_ISL_477855, EPI_ISL_477856, EPI_ISL_477857, EPI_ISL_477858, EPI_ISL_477859, EPI_ISL_477861, EPI_ISL_477862, EPI_ISL_477866, EPI_ISL_477867, EPI_ISL_477869, EPI_ISL_477870, EPI_ISL_477871, EPI_ISL_477872, EPI_ISL_477874, EPI_ISL_477875, EPI_ISL_477876, EPI_ISL_477877, EPI_ISL_477879, EPI_ISL_477881, EPI_ISL_477883, EPI_ISL_477884, EPI_ISL_477885, EPI_ISL_477886, EPI_ISL_477887, EPI_ISL_477889, EPI_ISL_477890, EPI_ISL_477894, EPI_ISL_477895, EPI_ISL_477896, EPI_ISL_477897, EPI_ISL_477898, EPI_ISL_477899, EPI_ISL_477901, EPI_ISL_477902, EPI_ISL_477903, EPI_ISL_477904, EPI_ISL_477905, EPI_ISL_477906, EPI_ISL_477907, EPI_ISL_477909, EPI_ISL_477910, EPI_ISL_477911, EPI_ISL_477912, EPI_ISL_477914, EPI_ISL_477915, EPI_ISL_477916, EPI_ISL_477918, EPI_ISL_477919, EPI_ISL_477920, EPI_ISL_477922, EPI_ISL_477924, EPI_ISL_477925, EPI_ISL_477926, EPI_ISL_477928, EPI_ISL_477929, EPI_ISL_477930, EPI_ISL_477932, EPI_ISL_477933, EPI_ISL_477934, EPI_ISL_477935, EPI_ISL_477936, EPI_ISL_477938, EPI_ISL_477939, EPI_ISL_477940, EPI_ISL_477941, EPI_ISL_477942, EPI_ISL_477943, EPI_ISL_477944, EPI_ISL_477946, EPI_ISL_477947, EPI_ISL_477949, EPI_ISL_477951, EPI_ISL_477952, EPI_ISL_477953, EPI_ISL_477954, EPI_ISL_477955, EPI_ISL_477956, EPI_ISL_477957, EPI_ISL_477958, EPI_ISL_477960, EPI_ISL_477961, EPI_ISL_477962, EPI_ISL_477964, EPI_ISL_477965, EPI_ISL_477967, EPI_ISL_477969, EPI_ISL_477970, EPI_ISL_477971, EPI_ISL_477972, EPI_ISL_477973, EPI_ISL_477974, EPI_ISL_477975, EPI_ISL_477976, EPI_ISL_477977, EPI_ISL_477978, EPI_ISL_477979, EPI_ISL_477980, EPI_ISL_477981, EPI_ISL_477983, EPI_ISL_477984, EPI_ISL_477985, | | | |

| | | | |
|---|---|---|---|
| EPI_ISL_477986, EPI_ISL_477987, EPI_ISL_477988, EPI_ISL_477989, EPI_ISL_477990, EPI_ISL_477991, EPI_ISL_477992, EPI_ISL_477993, EPI_ISL_477994, EPI_ISL_477995, EPI_ISL_477996, EPI_ISL_477997, EPI_ISL_477998, EPI_ISL_477999, EPI_ISL_478000, EPI_ISL_478001, EPI_ISL_478002, EPI_ISL_478003, EPI_ISL_478004, EPI_ISL_478005, EPI_ISL_478006, EPI_ISL_478008, EPI_ISL_478009, EPI_ISL_478010, EPI_ISL_478011, EPI_ISL_478012, EPI_ISL_478013, EPI_ISL_478014, EPI_ISL_478015, EPI_ISL_478016, EPI_ISL_478017, EPI_ISL_478018, EPI_ISL_478019, EPI_ISL_478020, EPI_ISL_478022, EPI_ISL_478023, EPI_ISL_478024, EPI_ISL_478025, EPI_ISL_478027, EPI_ISL_478028, EPI_ISL_478029, EPI_ISL_478030, EPI_ISL_478031, EPI_ISL_478032, EPI_ISL_478035, EPI_ISL_478036, EPI_ISL_478039, EPI_ISL_478045, EPI_ISL_478047, EPI_ISL_478048, EPI_ISL_478049, EPI_ISL_478050, EPI_ISL_478051, EPI_ISL_478053, EPI_ISL_478058, EPI_ISL_478059, EPI_ISL_478060, EPI_ISL_478062, EPI_ISL_478064, EPI_ISL_478065, EPI_ISL_478066, EPI_ISL_478068, EPI_ISL_478070, EPI_ISL_478071, EPI_ISL_478079, EPI_ISL_478080, EPI_ISL_478084, EPI_ISL_478085, EPI_ISL_478086, EPI_ISL_478089, EPI_ISL_478090, EPI_ISL_478091, EPI_ISL_478093, EPI_ISL_478094, EPI_ISL_478095, EPI_ISL_478097, EPI_ISL_478098, EPI_ISL_478105, EPI_ISL_478107, EPI_ISL_478108, EPI_ISL_478109, EPI_ISL_478112, EPI_ISL_478114, EPI_ISL_478115, EPI_ISL_478116, EPI_ISL_478117, EPI_ISL_478118, EPI_ISL_478119, EPI_ISL_478121, EPI_ISL_478122, EPI_ISL_478124, EPI_ISL_478125, EPI_ISL_478126, EPI_ISL_478127, EPI_ISL_478128, EPI_ISL_478129, EPI_ISL_478132, EPI_ISL_478133, EPI_ISL_478134, EPI_ISL_478135, EPI_ISL_478136, EPI_ISL_478137, EPI_ISL_478139, EPI_ISL_478140, EPI_ISL_478142, EPI_ISL_478143, EPI_ISL_478144, EPI_ISL_478146, EPI_ISL_478147, EPI_ISL_478148, EPI_ISL_478149, EPI_ISL_478150, EPI_ISL_478151, EPI_ISL_478153, EPI_ISL_478154, EPI_ISL_478155, EPI_ISL_478156, EPI_ISL_478157, EPI_ISL_478158, EPI_ISL_478159, EPI_ISL_478160 | | | |
| see above | West of Scotland Specialist Virology Centre, NHSGGC / MRC-University of Glasgow Centre for Virus Research | COVID-19 Genomics UK (COG-UK) Consortium | Ana da Silva Filipe, Natasha Johnson, Kathy Smollett, Daniel Mair, Stephen Carmichael, Lily Tong, Jenna Nichols, Elihu Aranday-Cortes, Kirstyn Brunker, Yasmin Parr, Alice Broos, Kyriaki Nomikou; Sarah McDonald, Marc Niebel, Patawee Asamaphan; Richard Orton, Joseph Hughes, Sreenu Vattipally, David L Robertson; Alasdair MacLean, Rory Gunson; Kathy Li, Natasha Jesudason, Rajiv Shah, James Shepherd, Antonia Ho, Emma Thomson |
| EPI_ISL_478165, EPI_ISL_478166, EPI_ISL_478167, EPI_ISL_478171, EPI_ISL_478173, EPI_ISL_478174, EPI_ISL_478176, EPI_ISL_478177, EPI_ISL_478178, EPI_ISL_478179, EPI_ISL_478180, EPI_ISL_478182, EPI_ISL_478183, EPI_ISL_478184, EPI_ISL_478186, EPI_ISL_478188, EPI_ISL_478190, EPI_ISL_478192, EPI_ISL_478193, EPI_ISL_478194, EPI_ISL_478195, EPI_ISL_478199, EPI_ISL_478200, EPI_ISL_478201, EPI_ISL_478203, EPI_ISL_478204, EPI_ISL_478205, EPI_ISL_478206, EPI_ISL_478207, EPI_ISL_478208, EPI_ISL_478209, EPI_ISL_478210, EPI_ISL_478211, EPI_ISL_478212, EPI_ISL_478213, EPI_ISL_478214, EPI_ISL_478215, EPI_ISL_478216, EPI_ISL_478217, EPI_ISL_478218, EPI_ISL_478221, EPI_ISL_478224, EPI_ISL_478225, EPI_ISL_478226, EPI_ISL_478227, EPI_ISL_478230, EPI_ISL_478232, EPI_ISL_478233, EPI_ISL_478249, EPI_ISL_478250, EPI_ISL_478251, EPI_ISL_478252, EPI_ISL_478254, EPI_ISL_478255, EPI_ISL_478256, EPI_ISL_478257, EPI_ISL_478258, EPI_ISL_478259, EPI_ISL_478260, EPI_ISL_478261, EPI_ISL_478262, EPI_ISL_478263, EPI_ISL_478265, EPI_ISL_478266, EPI_ISL_478267, EPI_ISL_478268, EPI_ISL_478269, EPI_ISL_478270 | | | |
| see above | Virology Department, Royal Infirmary of Edinburgh, NHS Lothian / School of Biological Sciences, University of Edinburgh / Institute of Genetics and Molecular Medicine, University of Edinburgh | COVID-19 Genomics UK (COG-UK) Consortium | McHugh M, Dewar R, Rooke S, Gallagher M, Balcaza C, O'Toole Á, Scher E, Hill V, McCrone JT, Colquhoun R, Yu X, Jackson B, Rambaut A, Williams TC, Templeton K |
| EPI_ISL_478289, EPI_ISL_478290, EPI_ISL_478292, EPI_ISL_478293, EPI_ISL_478294, EPI_ISL_478295, EPI_ISL_478296, EPI_ISL_478297, EPI_ISL_478298, EPI_ISL_478299, EPI_ISL_478301, EPI_ISL_478304, EPI_ISL_478305, EPI_ISL_478306, EPI_ISL_478307, EPI_ISL_478311, EPI_ISL_478312, EPI_ISL_478315, EPI_ISL_478316, EPI_ISL_478317, EPI_ISL_478320, EPI_ISL_478321, EPI_ISL_478324, EPI_ISL_478325, EPI_ISL_478327, EPI_ISL_478328, EPI_ISL_478329, EPI_ISL_478330, EPI_ISL_478331, EPI_ISL_478334, EPI_ISL_478335, EPI_ISL_478336, EPI_ISL_478338, EPI_ISL_478343, EPI_ISL_478345, EPI_ISL_478347, EPI_ISL_478349, EPI_ISL_478350, EPI_ISL_478351, EPI_ISL_478353, EPI_ISL_478355, EPI_ISL_478359, EPI_ISL_478360, EPI_ISL_478361, EPI_ISL_478362, EPI_ISL_478370, EPI_ISL_478371 | | | |
| see above | University Hospitals Of Leicester NHS Trust and DeepSeq Nottingham | COVID-19 Genomics UK (COG-UK) Consortium | Christopher Holmes, Paul Bird, Thomas Helmer, Karlie Fallon, Julian Tang, Jonathan Ball, Patrick McClure, Joeseph Chappell, Nadine Holmes, Matthew Carlisle, Christopher Moore, Fei Sang, Johnny Debebe, Victoria Wright, Matthew Loose |
| EPI_ISL_478392, EPI_ISL_478394, EPI_ISL_478395, EPI_ISL_478396 | Liverpool Clinical Laboratories | COVID-19 Genomics UK (COG-UK) Consortium | Sam Haldenby, Anita Lucaci, Steve Paterson, Julian Hiscox, Alistair Darby, M Almsaud, A Alrezaihi, Muhannad Alruwaili, Stuart D Armstrong, Jones Benjamin, Eleanor G Bentley, Anu Chawla, Jordan J Clark, Angela Cowell, Richard Eccles, Isabel García-Dorival, Matthew Gemmell, Alessandro Gerada, PKF Gilmore, Richard Gregory, Ximeng Han, Catherine Hartley, Margaret Hughes, Miren Iturriza-Gomara, James Johnson, L Luu, Jenifer Manson, Charlotte Nelson, Elaine O'Toole, Cassie Olateju, Rebekah Penrice-Randal , Lucille Rainbow, N.P Randle, Trevor Ian Robinson, Parul Sharma, Ghada T Shawli, James P Stewart, Neil Swainston, Ecaterina Vamos, Joanne Watts, Mark Whitehead |
| EPI_ISL_478404, EPI_ISL_478405, EPI_ISL_478406, EPI_ISL_478410, EPI_ISL_478411, EPI_ISL_478412, EPI_ISL_478413, EPI_ISL_478414, EPI_ISL_478415, EPI_ISL_478417, EPI_ISL_478418, EPI_ISL_478419, EPI_ISL_478420, EPI_ISL_478421, EPI_ISL_478422, EPI_ISL_478424, EPI_ISL_478426, EPI_ISL_478427, EPI_ISL_478428, EPI_ISL_478429, EPI_ISL_478430, EPI_ISL_478433, EPI_ISL_478434, EPI_ISL_478435, EPI_ISL_478438, EPI_ISL_478440, EPI_ISL_478441, EPI_ISL_478443, EPI_ISL_478444, EPI_ISL_478445, EPI_ISL_478450, EPI_ISL_478451, EPI_ISL_478452, EPI_ISL_478453, EPI_ISL_478454, EPI_ISL_478455, EPI_ISL_478456, EPI_ISL_478457, EPI_ISL_478458, EPI_ISL_478461, EPI_ISL_478462, EPI_ISL_478463, EPI_ISL_478464, EPI_ISL_478465, EPI_ISL_478466, EPI_ISL_478468, EPI_ISL_478469, EPI_ISL_478470 | | | |
| see above | University College London, Great Ormond Street Hospital for Children NHS Foundation Trust, Imperial College Healthcare NHS Trust | COVID-19 Genomics UK (COG-UK) Consortium | Sergi Castellano, Rachel Williams, Mark Kristiansen, Paola Resende Silva, Sunando Roy, Tony Brooks, Helena Tutill, Paola Niola, Patricia Dyal, Charlotte Williams, Leysa Forrest, Yasmin Panchbhaya, Jacqueline Findlay, Samuel Weeks, Julianne Brown, Kathryn Harris, Paul Randell, James Price, Alison Holmes, Judith Breuer |
| EPI_ISL_478513, EPI_ISL_478515, EPI_ISL_478516, EPI_ISL_478517, EPI_ISL_478519, EPI_ISL_478520, EPI_ISL_478521, EPI_ISL_478522, EPI_ISL_478524, EPI_ISL_478525, EPI_ISL_478527, EPI_ISL_478528, EPI_ISL_478529, EPI_ISL_478530, EPI_ISL_478531, EPI_ISL_478532, EPI_ISL_478536, EPI_ISL_478537, EPI_ISL_478539, EPI_ISL_478541, EPI_ISL_478543, EPI_ISL_478544, EPI_ISL_478546, EPI_ISL_478547, EPI_ISL_478548, EPI_ISL_478549, EPI_ISL_478550, EPI_ISL_478551, EPI_ISL_478553, EPI_ISL_478555, EPI_ISL_478556, EPI_ISL_478558, EPI_ISL_478559, EPI_ISL_478561, EPI_ISL_478564, EPI_ISL_478565, EPI_ISL_478566, EPI_ISL_478567, EPI_ISL_478568, EPI_ISL_478569, EPI_ISL_478571, EPI_ISL_478573, EPI_ISL_478575, EPI_ISL_478576, EPI_ISL_478577, EPI_ISL_478578, EPI_ISL_478579, EPI_ISL_478580, EPI_ISL_478581, EPI_ISL_478582, EPI_ISL_478583, EPI_ISL_478584, EPI_ISL_478585, EPI_ISL_478586, EPI_ISL_478587, EPI_ISL_478588, EPI_ISL_478591, EPI_ISL_478592, EPI_ISL_478593, EPI_ISL_478594, EPI_ISL_478596, EPI_ISL_478608, EPI_ISL_478614, EPI_ISL_478616, EPI_ISL_478619, EPI_ISL_478621, EPI_ISL_478622, EPI_ISL_478623, EPI_ISL_478624, EPI_ISL_478626, EPI_ISL_478627, EPI_ISL_478628, EPI_ISL_478630, EPI_ISL_478632, EPI_ISL_478636, EPI_ISL_478637, EPI_ISL_478638, EPI_ISL_478645, EPI_ISL_478648, EPI_ISL_478652, EPI_ISL_478655, EPI_ISL_478657, EPI_ISL_478658, EPI_ISL_478661, EPI_ISL_478662 | | | |
| see above | Northumbria University / South Tees Hospitals NHS Foundation Trust / North Cumbria Integrated Care NHS Foundation Trust / North Tees and Hartlepool NHS Foundation Trust / Newcastle Hospitals NHS Foundation Trust | COVID-19 Genomics UK (COG-UK) Consortium | Darren L Smith,Andrew Nelson,Matthew Bashton,Greg R Young,Joshua Loh,John Allan,Mohammad A Tariq,Giles S Holt,Gary Black,Wen C Yew,Lynn Dover,Paul Baker,Steve Liggett,Sarah Essex,Jane Greenaway,Debra Padgett,Clive Graham,Garren Scott,Edward Barton,Emma Swindells,Brendan Payne,Jennifer Collins,Yusri Taha,Gary Eltringham |
| EPI_ISL_478669, EPI_ISL_478670 | unknown | Microbiology, Koc University | Can,F., Ozer,B., Nurtop,E., Dogan,O. |
| EPI_ISL_478672 | Egyptian National Cancer Institute (ENCI) | Egyptian National Cancer Institute (ENCI) | Zekri, Abdel Rahman N, Amer,K.E., Ahmed,O.S., Soliman,H.K., Hafez,M.M., Bahnassy,A.A., Abdelhamid,W., Gad,A., Ali,M., Hassan,W., Samir,M., Raouf,A., Hamdy,M.S., Soliman,M.S., Elsissy,M.H., Elkhateeb,S.M., Ezzelarab,M.H., Abouelhoda, Mohamed |
| EPI_ISL_478673, EPI_ISL_478674 | Pathology North - Royal North Shore Hospital - NSW Health Pathology | NSW Health Pathology - Institute of Clinical Pathology and Medical Research; Westmead Hospital; University of Sydney | CIDM-PH et al. |
| EPI_ISL_478675, EPI_ISL_478676, EPI_ISL_478677, EPI_ISL_478678, EPI_ISL_478679, EPI_ISL_478680, EPI_ISL_478681, EPI_ISL_478682 | Sydney South West Pathology Service (SSWPS) - Liverpool Hospital - NSW Health Pathology | NSW Health Pathology - Institute of Clinical Pathology and Medical Research; Westmead Hospital; University of Sydney | CIDM-PH et al. |
| EPI_ISL_478683, EPI_ISL_478684, EPI_ISL_478685, EPI_ISL_478686, EPI_ISL_478687, EPI_ISL_478688, EPI_ISL_478689, EPI_ISL_478690, EPI_ISL_478691, EPI_ISL_478692, EPI_ISL_478693, EPI_ISL_478694, EPI_ISL_478695, EPI_ISL_478696, EPI_ISL_478697, EPI_ISL_478698, EPI_ISL_478699, EPI_ISL_478700, EPI_ISL_478701, EPI_ISL_478702, EPI_ISL_478703, EPI_ISL_478704, EPI_ISL_478705 | | | |
| see above | South Eastern Area Laboratory Services (SEALS) | NSW Health Pathology - Institute of Clinical Pathology and Medical Research; Westmead Hospital; University of Sydney | CIDM-PH et al. |
| EPI_ISL_478706, EPI_ISL_478707 | Sydney South West Pathology Service (SSWPS) - Liverpool Hospital - NSW Health Pathology | NSW Health Pathology - Institute of Clinical Pathology and Medical Research; Westmead Hospital; University of Sydney | CIDM-PH et al. |
| EPI_ISL_478708, EPI_ISL_478709, EPI_ISL_478710, EPI_ISL_478711 | South Eastern Area Laboratory Services (SEALS) | NSW Health Pathology - Institute of Clinical Pathology and Medical Research; Westmead Hospital; University of Sydney | CIDM-PH et al. |
| EPI_ISL_478712, EPI_ISL_478713, EPI_ISL_478714, EPI_ISL_478715, EPI_ISL_478716 | Sydney South West Pathology Service (SSWPS) - Liverpool Hospital - NSW Health Pathology | NSW Health Pathology - Institute of Clinical Pathology and Medical Research; Westmead Hospital; University of Sydney | CIDM-PH et al. |
| EPI_ISL_478717, EPI_ISL_478718 | Sydney South West Pathology Service (SSWPS) - Concord Repatriation General Hospital - NSW Health Pathology | NSW Health Pathology - Institute of Clinical Pathology and Medical Research; Westmead Hospital; University of Sydney | CIDM-PH et al. |
| EPI_ISL_478719, EPI_ISL_478720 | Quadram Institute Bioscience | COVID-19 Genomics UK (COG-UK) Consortium | Dave J. Baker, Gemma L. Kay, Alp Aydin, Thanh Le-Viet, Steven Rudder, Ana P. Tedim, Anastasia Kolyva, Maria Diaz, Leonardo de Oliveira Martins, Nabil-Fareed Alikhan, Lizzie Meadows, Rachael Stanley, Ngozi Elumogo, Muhammed Yasir, Nicholas M. Thomson, Alexander J Trotter, Rachel Gilroy, Samuel Bloomfield, Claire Stuart, Andrew Bell, Reenesh Prakash, Samir Dervisevic, Alison E. Mather, John Wain, Mark Webber, Andrew J. Page, Justin O'Grady |
| EPI_ISL_478721, EPI_ISL_478722, EPI_ISL_478724, EPI_ISL_478725 | Queens Medical Centre, Clinical Microbiology Department / DeepSeq Nottingham | COVID-19 Genomics UK (COG-UK) Consortium | Gemma Clark, Wendy Smith, Manjinder Khakh, Vicki M Fleming, Michelle M Lister, Hannah Howson-Wells, Jonathan Ball, Patrick McClure, Joseph Chappell, Theocharis Tsoleridis, Nadine Holmes, Matthew Carlisle, Christopher Moore, Fei Sang, Johnny Debebe, Victoria Wright, Matthew Loose |
| EPI_ISL_478729, EPI_ISL_478730, EPI_ISL_478731, EPI_ISL_478732, EPI_ISL_478733, EPI_ISL_478734, EPI_ISL_478735, EPI_ISL_478736, EPI_ISL_478737, EPI_ISL_478738, EPI_ISL_478739, EPI_ISL_478740, EPI_ISL_478742, EPI_ISL_478743, EPI_ISL_478744, EPI_ISL_478745, EPI_ISL_478746, EPI_ISL_478747, EPI_ISL_478748, EPI_ISL_478752, EPI_ISL_478753, EPI_ISL_478754, EPI_ISL_478755, EPI_ISL_478756, EPI_ISL_478757, EPI_ISL_478758, EPI_ISL_478759, EPI_ISL_478760, EPI_ISL_478761, EPI_ISL_478762, EPI_ISL_478763, EPI_ISL_478764, EPI_ISL_478765, EPI_ISL_478768, EPI_ISL_478769, EPI_ISL_478771, EPI_ISL_478772, EPI_ISL_478773, EPI_ISL_478774, EPI_ISL_478775, EPI_ISL_478776, EPI_ISL_478777, EPI_ISL_478778, EPI_ISL_478779, EPI_ISL_478780, EPI_ISL_478781, EPI_ISL_478782, EPI_ISL_478783, EPI_ISL_478784, EPI_ISL_478786, EPI_ISL_478787, EPI_ISL_478790, EPI_ISL_478791, EPI_ISL_478792, EPI_ISL_478793, EPI_ISL_478794, EPI_ISL_478795, EPI_ISL_478796, EPI_ISL_478797, EPI_ISL_478798, EPI_ISL_478799, EPI_ISL_478801, EPI_ISL_478802, EPI_ISL_478803, EPI_ISL_478804, EPI_ISL_478805, EPI_ISL_478806, EPI_ISL_478807, EPI_ISL_478808, EPI_ISL_478809, EPI_ISL_478810, EPI_ISL_478811, EPI_ISL_478812, EPI_ISL_478813, EPI_ISL_478814, EPI_ISL_478815, EPI_ISL_478816, EPI_ISL_478817, EPI_ISL_478818, EPI_ISL_478819, EPI_ISL_478820, EPI_ISL_478821, EPI_ISL_478822, EPI_ISL_478823, EPI_ISL_478824, EPI_ISL_478825, EPI_ISL_478826, EPI_ISL_478827, EPI_ISL_478828, EPI_ISL_478829, EPI_ISL_478830, EPI_ISL_478831, EPI_ISL_478832, EPI_ISL_478833, EPI_ISL_478834, EPI_ISL_478835, EPI_ISL_478836, EPI_ISL_478837, EPI_ISL_478838, EPI_ISL_478839, EPI_ISL_478840, EPI_ISL_478841, EPI_ISL_478842, EPI_ISL_478843, EPI_ISL_478844, EPI_ISL_478845, EPI_ISL_478846, EPI_ISL_478847, EPI_ISL_478848, EPI_ISL_478849, EPI_ISL_478850, EPI_ISL_478851, EPI_ISL_478852, EPI_ISL_478853, EPI_ISL_478854, EPI_ISL_478855, EPI_ISL_478856, EPI_ISL_478857, EPI_ISL_478859, EPI_ISL_478860, EPI_ISL_478861, EPI_ISL_478862, EPI_ISL_478863, EPI_ISL_478864, EPI_ISL_478865, EPI_ISL_478866, EPI_ISL_478868, EPI_ISL_478869, EPI_ISL_478870, EPI_ISL_478872, EPI_ISL_478873, EPI_ISL_478874, EPI_ISL_478875, EPI_ISL_478876, EPI_ISL_478877, EPI_ISL_478878, EPI_ISL_478879, EPI_ISL_478880, EPI_ISL_478881, EPI_ISL_478882, EPI_ISL_478883, EPI_ISL_478885, EPI_ISL_478886, EPI_ISL_478887, EPI_ISL_478888, EPI_ISL_478889, EPI_ISL_478891, EPI_ISL_478892, EPI_ISL_478893, EPI_ISL_478894, EPI_ISL_478895, EPI_ISL_478896, EPI_ISL_478898, EPI_ISL_478899, EPI_ISL_478902, EPI_ISL_478903, EPI_ISL_478905, EPI_ISL_478906, EPI_ISL_478907, EPI_ISL_478908, EPI_ISL_478909, EPI_ISL_478910, EPI_ISL_478911, EPI_ISL_478913, EPI_ISL_478914, EPI_ISL_478917, EPI_ISL_478918, EPI_ISL_478919, EPI_ISL_478920, EPI_ISL_478921, EPI_ISL_478922, EPI_ISL_478923, EPI_ISL_478924, EPI_ISL_478925, EPI_ISL_478926, EPI_ISL_478927, EPI_ISL_478928, EPI_ISL_478929, EPI_ISL_478931, EPI_ISL_478932, EPI_ISL_478933, EPI_ISL_478934, EPI_ISL_478936, EPI_ISL_478937, EPI_ISL_478939, EPI_ISL_478940, EPI_ISL_478941, EPI_ISL_478943, EPI_ISL_478945, EPI_ISL_478948, EPI_ISL_478949, EPI_ISL_478950, EPI_ISL_478952, EPI_ISL_478954, EPI_ISL_478955, EPI_ISL_478956, EPI_ISL_478957, EPI_ISL_478959, EPI_ISL_478960, EPI_ISL_478961, EPI_ISL_478962, EPI_ISL_478963, EPI_ISL_478965, EPI_ISL_478967, EPI_ISL_478968, EPI_ISL_478969, EPI_ISL_478970, EPI_ISL_478971, EPI_ISL_478972, EPI_ISL_478973, EPI_ISL_478974, EPI_ISL_478975, EPI_ISL_478976, EPI_ISL_478977, EPI_ISL_478979, EPI_ISL_478980, EPI_ISL_478981, EPI_ISL_478982, EPI_ISL_478983, EPI_ISL_478984, EPI_ISL_478985, EPI_ISL_478986, EPI_ISL_478987, EPI_ISL_478988, EPI_ISL_478989, EPI_ISL_478990, EPI_ISL_478991, EPI_ISL_478992, EPI_ISL_478993, EPI_ISL_478994, EPI_ISL_478995, EPI_ISL_478996, EPI_ISL_478997, EPI_ISL_478998, EPI_ISL_478999, EPI_ISL_479001, EPI_ISL_479003, EPI_ISL_479005, EPI_ISL_479006, EPI_ISL_479007, EPI_ISL_479011, EPI_ISL_479012, EPI_ISL_479013, EPI_ISL_479014, EPI_ISL_479015, EPI_ISL_479016, EPI_ISL_479017, EPI_ISL_479018, EPI_ISL_479019, EPI_ISL_479020, EPI_ISL_479022, EPI_ISL_479023, EPI_ISL_479024, EPI_ISL_479025, EPI_ISL_479026, EPI_ISL_479027, EPI_ISL_479030, EPI_ISL_479031, EPI_ISL_479032, EPI_ISL_479033, EPI_ISL_479035, EPI_ISL_479036, EPI_ISL_479037, EPI_ISL_479038, EPI_ISL_479040, EPI_ISL_479041, EPI_ISL_479042, EPI_ISL_479043, EPI_ISL_479044, EPI_ISL_479045, EPI_ISL_479046, EPI_ISL_479047, EPI_ISL_479048, EPI_ISL_479050, EPI_ISL_479051, EPI_ISL_479052, EPI_ISL_479054, EPI_ISL_479055, EPI_ISL_479056, EPI_ISL_479057, EPI_ISL_479058, EPI_ISL_479059, EPI_ISL_479060, EPI_ISL_479061, EPI_ISL_479062, EPI_ISL_479063, EPI_ISL_479064, EPI_ISL_479066, EPI_ISL_479067, EPI_ISL_479068, EPI_ISL_479069, EPI_ISL_479070, EPI_ISL_479071, EPI_ISL_479072, EPI_ISL_479073, EPI_ISL_479074, EPI_ISL_479076, EPI_ISL_479077, EPI_ISL_479079, EPI_ISL_479080, EPI_ISL_479081, EPI_ISL_479082, EPI_ISL_479083, EPI_ISL_479084, EPI_ISL_479085, EPI_ISL_479086, EPI_ISL_479088, EPI_ISL_479089, EPI_ISL_479090, EPI_ISL_479091, EPI_ISL_479092, EPI_ISL_479093, EPI_ISL_479094, EPI_ISL_479095, EPI_ISL_479096, EPI_ISL_479097, EPI_ISL_479098, EPI_ISL_479099, EPI_ISL_479100, EPI_ISL_479101, EPI_ISL_479102, EPI_ISL_479103, EPI_ISL_479104, EPI_ISL_479105, EPI_ISL_479106, EPI_ISL_479107, EPI_ISL_479108, EPI_ISL_479110, EPI_ISL_479111, EPI_ISL_479113, EPI_ISL_479114, EPI_ISL_479115, EPI_ISL_479116, EPI_ISL_479117, EPI_ISL_479119, EPI_ISL_479120, EPI_ISL_479121, EPI_ISL_479122, EPI_ISL_479123, EPI_ISL_479124, EPI_ISL_479126, EPI_ISL_479127, EPI_ISL_479130, EPI_ISL_479131, EPI_ISL_479132, EPI_ISL_479133, EPI_ISL_479134, EPI_ISL_479135, EPI_ISL_479136, EPI_ISL_479137, EPI_ISL_479138, EPI_ISL_479140, EPI_ISL_479141, EPI_ISL_479142, EPI_ISL_479143, EPI_ISL_479144, EPI_ISL_479146, EPI_ISL_479147, EPI_ISL_479148, EPI_ISL_479150, EPI_ISL_479152, EPI_ISL_479153, EPI_ISL_479154, EPI_ISL_479155, EPI_ISL_479156, EPI_ISL_479157, EPI_ISL_479158, EPI_ISL_479160, EPI_ISL_479162, EPI_ISL_479165, EPI_ISL_479166, EPI_ISL_479167, EPI_ISL_479168, EPI_ISL_479169, EPI_ISL_479170, EPI_ISL_479171, EPI_ISL_479172, EPI_ISL_479173, EPI_ISL_479174 | | | |
| see above | Oxford Viromics, NDM, University of Oxford; Oxford University Hospitals; Basingstoke and North Hampshire Hospital | COVID-19 Genomics UK (COG-UK) Consortium | Tanya Golubchik, David Bonsall, George Macintyre, Amy Trebes, Mariateresa de Cesare, Catrin Moore, Alex Mobbs, Anita Justice, Robert Shaw, Monique Andersson, Timothy Peto, Emma Wise, Nathan Moore, Jessica Lynch, Nick Cortes, Matilde Mori, Stephen Kidd, David Buck, John Todd, Christophe Fraser |
| EPI_ISL_479175, EPI_ISL_479176, EPI_ISL_479177, EPI_ISL_479178, EPI_ISL_479179, EPI_ISL_479180, EPI_ISL_479181, EPI_ISL_479182, EPI_ISL_479183, EPI_ISL_479184, EPI_ISL_479185, EPI_ISL_479186, EPI_ISL_479187, EPI_ISL_479188, EPI_ISL_479189, EPI_ISL_479190, EPI_ISL_479191, EPI_ISL_479192, EPI_ISL_479193, EPI_ISL_479194 | | | |
| see above | Centre for Enzyme Innovation, University of Portsmouth / Translational Research Laboratory, Portsmouth Hospitals NHS Trust | COVID-19 Genomics UK (COG-UK) Consortium | Angela Beckett,Yann Bourgeois,Garry Scarlett,Sharon Glaysher,Scott Elliott,Kelly Bicknell,Robert Impey,Allyson Lloyd,Sarah Wyllie,Ethan Butcher,Anoop Chauhan,Samuel Robson |
| EPI_ISL_479198, EPI_ISL_479201, EPI_ISL_479214, EPI_ISL_479216, EPI_ISL_479218, EPI_ISL_479221, EPI_ISL_479223, EPI_ISL_479224, EPI_ISL_479225, EPI_ISL_479226, EPI_ISL_479236, EPI_ISL_479242, EPI_ISL_479248, EPI_ISL_479258, EPI_ISL_479266, EPI_ISL_479267, EPI_ISL_479270, EPI_ISL_479275, EPI_ISL_479276, EPI_ISL_479279, EPI_ISL_479280, EPI_ISL_479283 | | | |
| see above | Virology Department, Sheffield Teaching Hospitals NHS Foundation Trust/Department of Infection, Immunity and Cardiovascular Disease, The Medical School, University of Sheffield | COVID-19 Genomics UK (COG-UK) Consortium | Thushan de Silva, Matthew Parker, Nikki Smith, Adri Angyal, Rebecca Brown, Luke Green, Rachel Tucker, Paul Parsons, Danielle Groves, Katie Johnson, Laura Carrilero, Alex Keeley, Dave Partridge, Matthew Wyles, Benjamin Lindsey, Mehmet Yavuz, Mohammad Raza, Cariad Evans |
| EPI_ISL_479294, EPI_ISL_479297, EPI_ISL_479344, EPI_ISL_479346, EPI_ISL_479349, EPI_ISL_479355, EPI_ISL_479361, EPI_ISL_479362, EPI_ISL_479365, EPI_ISL_479397, EPI_ISL_479403, EPI_ISL_479410, EPI_ISL_479412, EPI_ISL_479420, EPI_ISL_479431, EPI_ISL_479436, EPI_ISL_479439, EPI_ISL_479441, EPI_ISL_479442, EPI_ISL_479443, EPI_ISL_479445, EPI_ISL_479451, EPI_ISL_479455, EPI_ISL_479463, EPI_ISL_479480 | | | |
| see above | Wales Specialist Virology Centre Sequencing lab: Pathogen Genomics Unit | COVID-19 Genomics UK (COG-UK) Consortium | Catherine Moore, Johnathan Evans, Laura Gifford, Malorie Perry, Simon Cottrell, Angela Marchbank, Alec Birchley, Alexander Adams, Amy Gaskin, Bree Gatica-Wilcox, Jason Coombes, Joel Southgate, Lauren Gilbert, Lee Graham, Nicole Pacchiarini, Sara Kumziene-Summerhayes, Sarah Taylor, Sophie Jones, Sara Rey, Matthew Bull, Joanne Watkins, Sally Corden, Tom Connor |
| EPI_ISL_479482, EPI_ISL_479483, EPI_ISL_479484, EPI_ISL_479485, EPI_ISL_479486, EPI_ISL_479487, EPI_ISL_479488, EPI_ISL_479489, EPI_ISL_479490, EPI_ISL_479491, EPI_ISL_479492 | | | |
| see above | Department of Laboratory Medicine Tan Tock Seng Hospital | Department of Laboratory Medicine Tan Tock Seng Hospital | Chen YYC, Zair X, Li C, Tang WY, Maurer-Stroh S, Barkham TMS, Nagarajan N, Sessions OM |
| EPI_ISL_479493, EPI_ISL_479494, EPI_ISL_479495, EPI_ISL_479496, EPI_ISL_479497, EPI_ISL_479498, EPI_ISL_479499, EPI_ISL_479500, EPI_ISL_479501, EPI_ISL_479502, EPI_ISL_479503, EPI_ISL_479504, EPI_ISL_479505, EPI_ISL_479506, EPI_ISL_479507, EPI_ISL_479508, EPI_ISL_479509, EPI_ISL_479510, EPI_ISL_479511, EPI_ISL_479512, EPI_ISL_479513, EPI_ISL_479514, EPI_ISL_479515, EPI_ISL_479516, EPI_ISL_479517, EPI_ISL_479518, EPI_ISL_479519, EPI_ISL_479520, EPI_ISL_479521, EPI_ISL_479522, EPI_ISL_479523, EPI_ISL_479524, EPI_ISL_479526, EPI_ISL_479527, EPI_ISL_479528, EPI_ISL_479529, EPI_ISL_479530, EPI_ISL_479531, EPI_ISL_479532, EPI_ISL_479533, EPI_ISL_479534, EPI_ISL_479535, EPI_ISL_479536, EPI_ISL_479537, EPI_ISL_479538, EPI_ISL_479539, EPI_ISL_479540, EPI_ISL_479541, EPI_ISL_479542, EPI_ISL_479543, EPI_ISL_479544, EPI_ISL_479545, EPI_ISL_479546, EPI_ISL_479547, EPI_ISL_479548, EPI_ISL_479550, EPI_ISL_479551, EPI_ISL_479552, EPI_ISL_479553, EPI_ISL_479554, EPI_ISL_479555, EPI_ISL_479556, EPI_ISL_479557, EPI_ISL_479558, EPI_ISL_479559, EPI_ISL_479560, EPI_ISL_479562, EPI_ISL_479563, EPI_ISL_479564, EPI_ISL_479565, EPI_ISL_479566, EPI_ISL_479567, | | | |

| | | | |
|---|---|---|---|
| EPI_ISL_479568, EPI_ISL_479570, EPI_ISL_479571, EPI_ISL_479572 | | | |
| see above | NIV Influenza | NIV Influenza | Potdar V |
| EPI_ISL_479574, EPI_ISL_479575, EPI_ISL_479576, EPI_ISL_479577, EPI_ISL_479578, EPI_ISL_479580, EPI_ISL_479581, EPI_ISL_479582, EPI_ISL_479583, EPI_ISL_479584, EPI_ISL_479585, EPI_ISL_479586, EPI_ISL_479587, EPI_ISL_479588, EPI_ISL_479589, EPI_ISL_479590, EPI_ISL_479591, EPI_ISL_479592, EPI_ISL_479593, EPI_ISL_479594, EPI_ISL_479595, EPI_ISL_479596, EPI_ISL_479597, EPI_ISL_479598, EPI_ISL_479599, EPI_ISL_479600, EPI_ISL_479601, EPI_ISL_479602, EPI_ISL_479603 | | | |
| see above | National Public Health Laboratory, National Centre for Infectious Diseases | National Public Health Laboratory, National Centre for Infectious Diseases | Mak TM, Octavia S, Zhou Z, Chavatte JM, Cui L, Lin RTP |
| EPI_ISL_479616, EPI_ISL_479617 | Laboratory of Molecular Virology of the International Centre for Genetic Engineering and Biotechnology (ICGEB) | ARGO Open Lab Platform for Genome Sequencing | Licastro, D, Rajasekharan S, Dal Monego S, Segat L, D'Agaro P, Salton F, Confalonieri P, Confalonieri M Marcello A |
| EPI_ISL_479618 | Laboratory of Molecular Virology of the International Centre for Genetic Engineering and Biotechnology (ICGEB) | ARGO Open Lab Platform for Genome Sequencing | Licastro, D, Rajasekharan S, Dal Monego S, Segat L, D'Agaro P, Salton F, Confalonieri P, Confalonieri M, Marcello A |
| EPI_ISL_479619 | Laboratory of Molecular Virology of the International Centre for Genetic Engineering and Biotechnology (ICGEB) | ARGO Open Lab Platform for Genome Sequencing | Licastro, D, Rajasekharan S, Dal Monego S, Segat L, D'Agaro P, Salton F, Confalonieri P, Confalonieri M, Marcello A |
| EPI_ISL_479620, EPI_ISL_479621, EPI_ISL_479622, EPI_ISL_479623, EPI_ISL_479624 | Molecular diagnostic laboratory of Federal Budget Institution of Science "Central Research Institute of Epidemiology" of The Federal Service on Customers' Rights Protection and Human Well-being Surveillance | Group of Genomics and Postgenomic Technologies of Central Research Institute of Epidemilology | Speranskaya AS, Kaptelova VV,Valdokhina AV, Bulanenko VP, Samoilov AE, Korneenko EV, Sizova TV, Tivanova EV, Shipulina OY, Akimkin VG |
| EPI_ISL_479664, EPI_ISL_479665, EPI_ISL_479667, EPI_ISL_479668, EPI_ISL_479671, EPI_ISL_479672, EPI_ISL_479674 | Center for Genomics and System Biology, New York University | Center for Genomics and System Biology, New York University | Roder,A., Banakis,S., Johnson,K., Khalfan,M., Borenstein,E.S., Samanovic,M., Cornelius,A., Herati,R., Ulrich,R., Fleming,A., Kottkamp,A., Raabe,V., Mulligan,M.J., Gresham,D., Ghedin,E. |
| EPI_ISL_479676, EPI_ISL_479677, EPI_ISL_479678, EPI_ISL_479679, EPI_ISL_479680, EPI_ISL_479681, EPI_ISL_479682, EPI_ISL_479683, EPI_ISL_479684, EPI_ISL_479685 | unknown | Contact:Hiroyuki Asakura Tokyo Metropolitan Institute of Public Health, Department of Microbiology | Asakura,H., Yoshida,I., Kumagai,R., Nagashima,M., Chiba,T., Sadamasu,K. |
| EPI_ISL_479686, EPI_ISL_479687, EPI_ISL_479688, EPI_ISL_479689, EPI_ISL_479690, EPI_ISL_479691, EPI_ISL_479692, EPI_ISL_479693, EPI_ISL_479694, EPI_ISL_479695, EPI_ISL_479696, EPI_ISL_479697, EPI_ISL_479698, EPI_ISL_479699, EPI_ISL_479700, EPI_ISL_479701, EPI_ISL_479703, EPI_ISL_479704, EPI_ISL_479705, EPI_ISL_479706, EPI_ISL_479707, EPI_ISL_479708, EPI_ISL_479710, EPI_ISL_479711, EPI_ISL_479712, EPI_ISL_479713, EPI_ISL_479714, EPI_ISL_479715, EPI_ISL_479716, EPI_ISL_479717, EPI_ISL_479718, EPI_ISL_479719, EPI_ISL_479720, EPI_ISL_479721, EPI_ISL_479722, EPI_ISL_479723, EPI_ISL_479724, EPI_ISL_479725, EPI_ISL_479726, EPI_ISL_479727 | | | |
| see above | Egyptian National Cancer Institute (ENCI) | Egyptian National Cancer Institute (ENCI) | Zekri, Abdel Rahman N, Amer,K.E., Ahmed,O.S., Soliman,H.K., Hafez,M.M., Bahnassy,A.A., Abdelhamid,W., Gad,A., Ali,M., Hassan,W., Samir,M., Raouf,A., Hamdy,M.S., Soliman,M.S., Elsissy,M.H., Elkhateeb,S.M., Ezzelarab,M.H., Abouelhoda, Mohamed |
| EPI_ISL_479728 | Egyptian National Cancer Institute (ENCI) | Egyptian National Cancer Institute (ENCI) | Zekri,A.N., Amer,K.E., Ahmed,O.S., Soliman,H.K., Bahnassy,A.A., Ali,M., Abdelhamid,W., Gad,A., Hassan,W., Samir,M., Raouf,A., Hamdy,M.S., Soliman,M.S., Elsissy,M.H., Elkhateeb,S.M., Ezzelarab,M.H., Abouelhoda,M. |
| EPI_ISL_479729, EPI_ISL_479730, EPI_ISL_479731, EPI_ISL_479733, EPI_ISL_479734, EPI_ISL_479735 | Egyptian National Cancer Institute (ENCI) | Egyptian National Cancer Institute (ENCI) | Zekri, Abdel Rahman N, Amer,K.E., Ahmed,O.S., Soliman,H.K., Hafez,M.M., Bahnassy,A.A., Abdelhamid,W., Gad,A., Ali,M., Hassan,W., Samir,M., Raouf,A., Hamdy,M.S., Soliman,M.S., Elsissy,M.H., Elkhateeb,S.M., Ezzelarab,M.H., Abouelhoda, Mohamed |
| EPI_ISL_486890, EPI_ISL_486891, EPI_ISL_486892, EPI_ISL_486893, EPI_ISL_486894, EPI_ISL_486895, EPI_ISL_486896 | Tokyo Metoropolitan Institute of Public Health | Tokyo Metoropolitan Institute of Public Health | Asakura,H., Yoshida,I., Kumagai,R., Nagashima,M., Chiba,T., Sadamasu,K. |
| EPI_ISL_605882, EPI_ISL_605883, EPI_ISL_605884, EPI_ISL_605885, EPI_ISL_605886, EPI_ISL_605887, EPI_ISL_605888, EPI_ISL_605889, EPI_ISL_605890, EPI_ISL_605891, EPI_ISL_605892, EPI_ISL_605893, EPI_ISL_605894, EPI_ISL_605895, EPI_ISL_605896, EPI_ISL_605897, EPI_ISL_605898, EPI_ISL_605899, EPI_ISL_605900, EPI_ISL_605901, EPI_ISL_605902, EPI_ISL_605903, EPI_ISL_605904, EPI_ISL_605905, EPI_ISL_605906, EPI_ISL_605907 | | | |
| see above | NGS Lab, DNA SOLUTION LTD. | NGS Lab, DNA SOLUTION LTD. | Khan,M.I., Hasan,K.N., Sufian,A., Polol,M.N.I., Khaleque,A., Rahman,M., Chowdhury,M., Haider,H.U., Razu,M.H., Khan,M., Rabbi,M.F.A. |
| EPI_ISL_605909, EPI_ISL_605910, EPI_ISL_605911, EPI_ISL_605912, EPI_ISL_605913 | NGS Lab, DNA SOLUTION LTD. | NGS Lab, DNA SOLUTION LTD. | Khan,M.I., Hasan,K.N., Sufian,A., Hosen,M.B., Polol,M.N.I., Khaleque,A., Rahman,M., Chowdhury,M., Haider,H.U., Razu,M.H., Khan,M., Rabbi,M.F.A. |
| EPI_ISL_605914, EPI_ISL_605915, EPI_ISL_605916, EPI_ISL_605917, EPI_ISL_605918, EPI_ISL_605919, EPI_ISL_605920, EPI_ISL_605921, EPI_ISL_605922, EPI_ISL_605923 | NGS Lab, DNA SOLUTION LTD. | NGS Lab, DNA SOLUTION LTD. | Khan,M.I., Hasan,K.N., Sufian,A., Hosen,M.B., Khaleque,A., Rahman,M., Chowdhury,M., Haider,H.U., Razu,M.H., Khan,M., Rabbi,M.F.A. |
