## Supplementary material for "Analysis of the Dynamics and Distribution of SARS-CoV-2 Mutations and its Possible Structural and Functional Implications": GISAID_acknowledgements_part_6

We gratefully acknowledge the following Authors from the Originating laboratories responsible for obtaining the specimens, as well as the Submitting laboratories where the genome data were generated and shared via GISAID, on which this research is based.

All Submitters of data may be contacted directly via www.gisaid.org

| Accession ID | Originating Laboratory | Submitting Laboratory | Authors |
|---|---|---|---|
| EPI_ISL_495918, EPI_ISL_495919, EPI_ISL_495920, EPI_ISL_495921, EPI_ISL_495922, EPI_ISL_495923, EPI_ISL_495924, EPI_ISL_495926, EPI_ISL_495927, EPI_ISL_495928, EPI_ISL_495929, EPI_ISL_495930, EPI_ISL_495931, EPI_ISL_495932, EPI_ISL_495934, EPI_ISL_495935, EPI_ISL_495936, EPI_ISL_495937, EPI_ISL_495938, EPI_ISL_495940, EPI_ISL_495943, EPI_ISL_495944, EPI_ISL_495946, EPI_ISL_495947, EPI_ISL_495948, EPI_ISL_495949, EPI_ISL_495950, EPI_ISL_495951, EPI_ISL_495952, EPI_ISL_495954, EPI_ISL_495955, EPI_ISL_495956, EPI_ISL_495959, EPI_ISL_495962, EPI_ISL_495963, EPI_ISL_495964, EPI_ISL_495965, EPI_ISL_495966, EPI_ISL_495967, EPI_ISL_495968, EPI_ISL_495970, EPI_ISL_495972, EPI_ISL_495973, EPI_ISL_495974, EPI_ISL_495975, EPI_ISL_495977, EPI_ISL_495978, EPI_ISL_495979, EPI_ISL_495980, EPI_ISL_495981, EPI_ISL_495982, EPI_ISL_495983, EPI_ISL_495984, EPI_ISL_495985, EPI_ISL_495986, EPI_ISL_495987, EPI_ISL_495988, EPI_ISL_495989, EPI_ISL_495991, EPI_ISL_495992, EPI_ISL_495993, EPI_ISL_495995, EPI_ISL_495996, EPI_ISL_495997, EPI_ISL_495998, EPI_ISL_495999, EPI_ISL_496000, EPI_ISL_496001, EPI_ISL_496002, EPI_ISL_496003, EPI_ISL_496004, EPI_ISL_496005, EPI_ISL_496006, EPI_ISL_496007, EPI_ISL_496008, EPI_ISL_496009, EPI_ISL_496011, EPI_ISL_496012, EPI_ISL_496013, EPI_ISL_496014, EPI_ISL_496016, EPI_ISL_496018, EPI_ISL_496019, EPI_ISL_496021, EPI_ISL_496022, EPI_ISL_496023, EPI_ISL_496024, EPI_ISL_496025, EPI_ISL_496026, EPI_ISL_496027, EPI_ISL_496028, EPI_ISL_496029, EPI_ISL_496030, EPI_ISL_496031, EPI_ISL_496032, EPI_ISL_496034, EPI_ISL_496035, EPI_ISL_496036, EPI_ISL_496037, EPI_ISL_496038, EPI_ISL_496039, EPI_ISL_496040, EPI_ISL_496042, EPI_ISL_496043, EPI_ISL_496044, EPI_ISL_496045, EPI_ISL_496046, EPI_ISL_496047, EPI_ISL_496048, EPI_ISL_496049, EPI_ISL_496050, EPI_ISL_496051, EPI_ISL_496052, EPI_ISL_496053, EPI_ISL_496054, EPI_ISL_496055, EPI_ISL_496056, EPI_ISL_496057, EPI_ISL_496058, EPI_ISL_496059, EPI_ISL_496060, EPI_ISL_496061, EPI_ISL_496062, EPI_ISL_496063, EPI_ISL_496064, EPI_ISL_496065, EPI_ISL_496066, EPI_ISL_496067, EPI_ISL_496068, EPI_ISL_496069, EPI_ISL_496070, EPI_ISL_496071, EPI_ISL_496073, EPI_ISL_496074, EPI_ISL_496076, EPI_ISL_496077, EPI_ISL_496078, EPI_ISL_496079, EPI_ISL_496080, EPI_ISL_496081, EPI_ISL_496082, EPI_ISL_496083, EPI_ISL_496084, EPI_ISL_496085, EPI_ISL_496086, EPI_ISL_496087, EPI_ISL_496088, EPI_ISL_496089, EPI_ISL_496090, EPI_ISL_496091, EPI_ISL_496092, EPI_ISL_496094, EPI_ISL_496096, EPI_ISL_496097, EPI_ISL_496098, EPI_ISL_496099, EPI_ISL_496100, EPI_ISL_496101, EPI_ISL_496102, EPI_ISL_496103, EPI_ISL_496104, EPI_ISL_496105, EPI_ISL_496106, EPI_ISL_496107, EPI_ISL_496108, EPI_ISL_496110, EPI_ISL_496111, EPI_ISL_496112, EPI_ISL_496113, EPI_ISL_496114, EPI_ISL_496115, EPI_ISL_496116, EPI_ISL_496117, EPI_ISL_496118, EPI_ISL_496119, EPI_ISL_496120, EPI_ISL_496121, EPI_ISL_496122, EPI_ISL_496123, EPI_ISL_496124, EPI_ISL_496125, EPI_ISL_496126, EPI_ISL_496127, EPI_ISL_496128, EPI_ISL_496129, EPI_ISL_496130, EPI_ISL_496132, EPI_ISL_496133, EPI_ISL_496134, EPI_ISL_496135, EPI_ISL_496136, EPI_ISL_496137, EPI_ISL_496138, EPI_ISL_496139, EPI_ISL_496140, EPI_ISL_496141, EPI_ISL_496142, EPI_ISL_496143, EPI_ISL_496144, EPI_ISL_496145, EPI_ISL_496146, EPI_ISL_496147, EPI_ISL_496148, EPI_ISL_496149, EPI_ISL_496150, EPI_ISL_496151, EPI_ISL_496152, EPI_ISL_496153, EPI_ISL_496154, EPI_ISL_496155, EPI_ISL_496156, EPI_ISL_496157, EPI_ISL_496158, EPI_ISL_496159, EPI_ISL_496160, EPI_ISL_496161, EPI_ISL_496162, EPI_ISL_496163, EPI_ISL_496164, EPI_ISL_496165, EPI_ISL_496166, EPI_ISL_496167, EPI_ISL_496168, EPI_ISL_496169, EPI_ISL_496171, EPI_ISL_496173, EPI_ISL_496174, EPI_ISL_496175, EPI_ISL_496176, EPI_ISL_496177, EPI_ISL_496178, EPI_ISL_496179, EPI_ISL_496180, EPI_ISL_496181, EPI_ISL_496182, EPI_ISL_496183, EPI_ISL_496184, EPI_ISL_496186, EPI_ISL_496187, EPI_ISL_496188, EPI_ISL_496190, EPI_ISL_496191, EPI_ISL_496192, EPI_ISL_496193, EPI_ISL_496194, EPI_ISL_496195, EPI_ISL_496196, EPI_ISL_496197, EPI_ISL_496199, EPI_ISL_496200, EPI_ISL_496201, EPI_ISL_496202, EPI_ISL_496203, EPI_ISL_496204, EPI_ISL_496205, EPI_ISL_496206, EPI_ISL_496207, EPI_ISL_496208, EPI_ISL_496209, EPI_ISL_496210, EPI_ISL_496211, EPI_ISL_496212, EPI_ISL_496213, EPI_ISL_496214, EPI_ISL_496215, EPI_ISL_496216, EPI_ISL_496217, EPI_ISL_496218, EPI_ISL_496219, EPI_ISL_496220, EPI_ISL_496221, EPI_ISL_496222, EPI_ISL_496223, EPI_ISL_496224, EPI_ISL_496225, EPI_ISL_496226, EPI_ISL_496227, EPI_ISL_496228, EPI_ISL_496229, EPI_ISL_496230, EPI_ISL_496231, EPI_ISL_496232, EPI_ISL_496233, EPI_ISL_496234, EPI_ISL_496235, EPI_ISL_496236, EPI_ISL_496237, EPI_ISL_496238, EPI_ISL_496239, EPI_ISL_496240, EPI_ISL_496241, EPI_ISL_496242, EPI_ISL_496243, EPI_ISL_496244, EPI_ISL_496245, EPI_ISL_496246, EPI_ISL_496248, EPI_ISL_496249, EPI_ISL_496250, EPI_ISL_496252, EPI_ISL_496253, EPI_ISL_496255, EPI_ISL_496256, EPI_ISL_496257, EPI_ISL_496258, EPI_ISL_496259, EPI_ISL_496260, EPI_ISL_496261, EPI_ISL_496263, EPI_ISL_496264, EPI_ISL_496265, EPI_ISL_496266, EPI_ISL_496267, EPI_ISL_496268, EPI_ISL_496269, EPI_ISL_496270, EPI_ISL_496271, EPI_ISL_496272, EPI_ISL_496273, EPI_ISL_496274, EPI_ISL_496275, EPI_ISL_496276, EPI_ISL_496277, EPI_ISL_496278, EPI_ISL_496280, EPI_ISL_496281, EPI_ISL_496282, EPI_ISL_496283, EPI_ISL_496285, EPI_ISL_496286, EPI_ISL_496287, EPI_ISL_496288, EPI_ISL_496289, EPI_ISL_496290, EPI_ISL_496291, EPI_ISL_496292, EPI_ISL_496294, EPI_ISL_496295, EPI_ISL_496296, EPI_ISL_496297, EPI_ISL_496298, EPI_ISL_496299, EPI_ISL_496300, EPI_ISL_496301, EPI_ISL_496302, EPI_ISL_496303, EPI_ISL_496304, EPI_ISL_496305, EPI_ISL_496306, EPI_ISL_496307, EPI_ISL_496308, EPI_ISL_496309, EPI_ISL_496310, EPI_ISL_496311, EPI_ISL_496312, EPI_ISL_496313, EPI_ISL_496314, EPI_ISL_496315, EPI_ISL_496316, EPI_ISL_496317, EPI_ISL_496318, EPI_ISL_496319, EPI_ISL_496320, EPI_ISL_496321, EPI_ISL_496322, EPI_ISL_496323, EPI_ISL_496324, EPI_ISL_496325, EPI_ISL_496326, EPI_ISL_496327, EPI_ISL_496328, EPI_ISL_496329, EPI_ISL_496330, EPI_ISL_496331, EPI_ISL_496332, EPI_ISL_496333, EPI_ISL_496334, EPI_ISL_496335, EPI_ISL_496336, EPI_ISL_496337, EPI_ISL_496338 | | | |
| see above | Washington State Department of Health | Seattle Flu Study | Deborah A. Nickerson, Chris D. Frazar, Jover Lee, Benjamin Pelle, Matthew Richardson, Amanda Adler, Elisabeth Brandstetter, Peter D. Han, Kairsten Fay, Misja Ilcisin, Kirsten Lacombe, Thomas R. Sibley, Melissa Truong, Caitlin R. Wolf, Romesh Gautom, Geoff Melly, Brian Hiatt, Philip Dykema, Scott Lindquist, Michael Boeckh, Janet A. Englund, Michael Famulare, Barry R. Lutz, Mark J. Rieder, Lea M. Starita, Matthew Thompson, Helen Y. Chu, Jay Shendure, Trevor Bedford |
| EPI_ISL_496350, EPI_ISL_496369, EPI_ISL_496374 | Infectolab | Andersen lab at Scripps Research | SEARCH Alliance San Diego with Samuel Navarro Alvarez, Carlos A. Cota Haros, Octavio Renteria Pacheco |
| EPI_ISL_496492, EPI_ISL_496497, EPI_ISL_496498, EPI_ISL_496499, EPI_ISL_496504, EPI_ISL_496506, EPI_ISL_496509, EPI_ISL_496512, EPI_ISL_496513, EPI_ISL_496514 | Viral Respiratory Lab, National Institute for Biomedical Research (INRB) | Pathogen Sequencing Lab, National Institute for Biomedical Research (INRB) | Placide Mbala-Kingebeni, Edith Nkwembe, Eddy Kinganda-Lusamaki, Amuri Aziza, Francisca Muyembe Mawete, Emmanuel Lokilo Lofiko, Catherine Pratt, Matthias Pauthner, Josh Quick, Allison Black, James Hadfield, Trevor Bedford, Ian Goodfellow, Andrew Rambaut, Nick Loman, Kristian Andersen, Michael Wiley, Steve Ahuka-Mundeke, Jean-Jacques Muyembe Tamfum |
| EPI_ISL_496518, EPI_ISL_496519, EPI_ISL_496520 | Armed Forces Medical College | National Centre For Cell Science | Dhiraj Paul, Kunal Jani, Radha Chauhan, Janesh Kumar, Vasudevan Seshadri, Girdhari Lal, Rajesh Karyakarte, Suvarna Joshi, Murlidhar Tambe, Sourav Sen, Santosh Karade, Kavita Bala Anand, Shelinder Pal Singh Shergill, Rajiv Mohan Gupta, Manoj Kumar Bhat, Arvind Sahu, Maharashtra COVID-19 Study Group, DBT's PAN-INDIA 1000 SARS-CoV2 RNA genome sequencing consortium, Yogesh S Shouche |
| EPI_ISL_496521, EPI_ISL_496522, EPI_ISL_496523 | B.J. Govt. Medical College | National Centre For Cell Science | Dhiraj Paul, Kunal Jani, Radha Chauhan, Janesh Kumar, Vasudevan Seshadri, Girdhari Lal, Rajesh Karyakarte, Suvarna Joshi, Murlidhar Tambe, Sourav Sen, Santosh Karade, Kavita Bala Anand, Shelinder Pal Singh Shergill, Rajiv Mohan Gupta, Manoj Kumar Bhat, Arvind Sahu, Maharashtra COVID-19 Study Group, DBT's PAN-INDIA 1000 SARS-CoV2 RNA genome sequencing consortium, Yogesh S Shouche |
| EPI_ISL_496524, EPI_ISL_496525, EPI_ISL_496526 | National Centre For Cell Science | National Centre For Cell Science | Dhiraj Paul, Kunal Jani, Radha Chauhan, Janesh Kumar, Vasudevan Seshadri, Girdhari Lal, Rajesh Karyakarte, Suvarna Joshi, Murlidhar Tambe, Sourav Sen, Santosh Karade, Kavita Bala Anand, Shelinder Pal Singh Shergill, Rajiv Mohan Gupta, Manoj Kumar Bhat, Arvind Sahu, Maharashtra COVID-19 Study Group, DBT's PAN-INDIA 1000 SARS-CoV2 RNA genome sequencing consortium, Yogesh S Shouche |
| EPI_ISL_496529 | Armed Forces Medical College | National Centre For Cell Science | Dhiraj Paul, Kunal Jani, Radha Chauhan, Janesh Kumar, Vasudevan Seshadri, Girdhari Lal, Rajesh Karyakarte, Suvarna Joshi, Murlidhar Tambe, Sourav Sen, Santosh Karade, Kavita Bala Anand, Shelinder Pal Singh Shergill, Rajiv Mohan Gupta, Manoj Kumar Bhat, Arvind Sahu, Maharashtra COVID-19 Study Group, DBT's PAN-INDIA 1000 SARS-CoV2 RNA genome sequencing consortium, Yogesh S Shouche |
| EPI_ISL_496530 | B.J. Govt. Medical College | National Centre For Cell Science | Dhiraj Paul, Kunal Jani, Radha Chauhan, Janesh Kumar, Vasudevan Seshadri, Girdhari Lal, Rajesh Karyakarte, Suvarna Joshi, Murlidhar Tambe, Sourav Sen, Santosh Karade, Kavita Bala Anand, Shelinder Pal Singh Shergill, Rajiv Mohan Gupta, Manoj Kumar Bhat, Arvind Sahu, Maharashtra COVID-19 Study Group, DBT's PAN-INDIA 1000 SARS-CoV2 RNA genome sequencing consortium, Yogesh S Shouche |
| EPI_ISL_496531, EPI_ISL_496532 | National Centre For Cell Science | National Centre For Cell Science | Dhiraj Paul, Kunal Jani, Radha Chauhan, Janesh Kumar, Vasudevan Seshadri, Girdhari Lal, Rajesh Karyakarte, Suvarna Joshi, Murlidhar Tambe, Sourav Sen, Santosh Karade, Kavita Bala Anand, Shelinder Pal Singh Shergill, Rajiv Mohan Gupta, Manoj Kumar Bhat, Arvind Sahu, Maharashtra COVID-19 Study Group, DBT's PAN-INDIA 1000 SARS-CoV2 RNA genome sequencing consortium, Yogesh S Shouche |
| EPI_ISL_496533 | Armed Forces Medical College | National Centre For Cell Science | Dhiraj Paul, Kunal Jani, Radha Chauhan, Janesh Kumar, Vasudevan Seshadri, Girdhari Lal, Rajesh Karyakarte, Suvarna Joshi, Murlidhar Tambe, Sourav Sen, Santosh Karade, Kavita Bala Anand, Shelinder Pal Singh Shergill, Rajiv Mohan Gupta, Manoj Kumar Bhat, Arvind Sahu, Maharashtra COVID-19 Study Group, DBT's PAN-INDIA 1000 SARS-CoV2 RNA genome sequencing consortium, Yogesh S Shouche |
| EPI_ISL_496534 | B.J. Govt. Medical College | National Centre For Cell Science | Dhiraj Paul, Kunal Jani, Radha Chauhan, Janesh Kumar, Vasudevan Seshadri, Girdhari Lal, Rajesh Karyakarte, Suvarna Joshi, Murlidhar Tambe, Sourav Sen, Santosh Karade, Kavita Bala Anand, Shelinder Pal Singh Shergill, Rajiv Mohan Gupta, Manoj Kumar Bhat, Arvind Sahu, Maharashtra COVID-19 Study Group, DBT's PAN-INDIA 1000 SARS-CoV2 RNA genome sequencing consortium, Yogesh S Shouche |
| EPI_ISL_496535 | National Centre For Cell Science | National Centre For Cell Science | Dhiraj Paul, Kunal Jani, Radha Chauhan, Janesh Kumar, Vasudevan Seshadri, Girdhari Lal, Rajesh Karyakarte, Suvarna Joshi, Murlidhar Tambe, Sourav Sen, Santosh Karade, Kavita Bala Anand, Shelinder Pal Singh Shergill, Rajiv Mohan Gupta, Manoj Kumar Bhat, Arvind Sahu, Maharashtra COVID-19 Study Group, DBT's PAN-INDIA 1000 SARS-CoV2 RNA genome sequencing consortium, Yogesh S Shouche |
| EPI_ISL_496537, EPI_ISL_496538, EPI_ISL_496539, EPI_ISL_496540, EPI_ISL_496541, EPI_ISL_496542, EPI_ISL_496543, EPI_ISL_496544, EPI_ISL_496545 | Armed Forces Medical College | National Centre For Cell Science | Dhiraj Paul, Kunal Jani, Radha Chauhan, Janesh Kumar, Vasudevan Seshadri, Girdhari Lal, Rajesh Karyakarte, Suvarna Joshi, Murlidhar Tambe, Sourav Sen, Santosh Karade, Kavita Bala Anand, Shelinder Pal Singh Shergill, Rajiv Mohan Gupta, Manoj Kumar Bhat, Arvind Sahu, Maharashtra COVID-19 Study Group, DBT's PAN-INDIA 1000 SARS-CoV2 RNA genome sequencing consortium, Yogesh S Shouche |
| EPI_ISL_496546, EPI_ISL_496547, EPI_ISL_496548, EPI_ISL_496549, EPI_ISL_496550, EPI_ISL_496551, EPI_ISL_496552, EPI_ISL_496553, EPI_ISL_496554 | B.J. Govt. Medical College | National Centre For Cell Science | Dhiraj Paul, Kunal Jani, Radha Chauhan, Janesh Kumar, Vasudevan Seshadri, Girdhari Lal, Rajesh Karyakarte, Suvarna Joshi, Murlidhar Tambe, Sourav Sen, Santosh Karade, Kavita Bala Anand, Shelinder Pal Singh Shergill, Rajiv Mohan Gupta, Manoj Kumar Bhat, Arvind Sahu, Maharashtra COVID-19 Study Group, DBT's PAN-INDIA 1000 SARS-CoV2 RNA genome sequencing consortium, Yogesh S Shouche |
| EPI_ISL_496555, EPI_ISL_496556, EPI_ISL_496557, EPI_ISL_496558, EPI_ISL_496559, EPI_ISL_496560, EPI_ISL_496561, EPI_ISL_496562, EPI_ISL_496563, EPI_ISL_496564, EPI_ISL_496565, EPI_ISL_496566, EPI_ISL_496567, EPI_ISL_496568, EPI_ISL_496569, EPI_ISL_496570, EPI_ISL_496571, EPI_ISL_496572, EPI_ISL_496573, EPI_ISL_496574, EPI_ISL_496575, EPI_ISL_496576, EPI_ISL_496577, EPI_ISL_496578, EPI_ISL_496579, EPI_ISL_496580, EPI_ISL_496581, EPI_ISL_496582, EPI_ISL_496583, EPI_ISL_496584, EPI_ISL_496585, EPI_ISL_496586, EPI_ISL_496587 | | | |
| see above | National Centre For Cell Science | National Centre For Cell Science | Dhiraj Paul, Kunal Jani, Radha Chauhan, Janesh Kumar, Vasudevan Seshadri, Girdhari Lal, Rajesh Karyakarte, Suvarna Joshi, Murlidhar Tambe, Sourav Sen, Santosh Karade, Kavita Bala Anand, Shelinder Pal Singh Shergill, Rajiv Mohan Gupta, Manoj Kumar Bhat, Arvind Sahu, Maharashtra COVID-19 Study Group, DBT's PAN-INDIA 1000 SARS-CoV2 RNA genome sequencing consortium, Yogesh S Shouche |
| EPI_ISL_496602 | Armed Forces Medical College | National Centre For Cell Science | Dhiraj Paul, Kunal Jani, Radha Chauhan, Janesh Kumar, Vasudevan Seshadri, Girdhari Lal, Rajesh Karyakarte, Suvarna Joshi, Murlidhar Tambe, Sourav Sen, Santosh Karade, Kavita Bala Anand, Shelinder Pal Singh Shergill, Rajiv Mohan Gupta, Manoj Kumar Bhat, Arvind Sahu, Maharashtra COVID-19 Study Group, DBT's PAN-INDIA 1000 SARS-CoV2 RNA genome sequencing consortium, Yogesh S Shouche |
| EPI_ISL_496916, EPI_ISL_496917 | Minnesota Department of Health, Public Health Laboratory | Minnesota Department of Health, Public Health Laboratory | Matt Plumb, Jacob Garfin, and Xiong Wang |
| EPI_ISL_496918 | Mayo Clinic & Mayo Clinic Laboratories | Minnesota Department of Health, Public Health Laboratory | Matt Plumb, Jacob Garfin, and Xiong Wang |
| EPI_ISL_496919, EPI_ISL_496920, EPI_ISL_496921, EPI_ISL_496922, EPI_ISL_496923, EPI_ISL_496924, EPI_ISL_496925, EPI_ISL_496926, EPI_ISL_496927, EPI_ISL_496928, EPI_ISL_496929, EPI_ISL_496930 | | | |
| see above | Minnesota Department of Health, Public Health Laboratory | Minnesota Department of Health, Public Health Laboratory | Matt Plumb, Jacob Garfin, and Xiong Wang |
| EPI_ISL_496931, EPI_ISL_496932, EPI_ISL_496933, EPI_ISL_496934, EPI_ISL_496935, EPI_ISL_496936, EPI_ISL_496937, EPI_ISL_496938, EPI_ISL_496939, EPI_ISL_496940, EPI_ISL_496941, EPI_ISL_496942, EPI_ISL_496943, EPI_ISL_496945, EPI_ISL_496947, EPI_ISL_496948, EPI_ISL_496949, EPI_ISL_496951, EPI_ISL_496952, EPI_ISL_496953, EPI_ISL_496954, EPI_ISL_496955, EPI_ISL_496956, EPI_ISL_496957, EPI_ISL_496958, EPI_ISL_496960, EPI_ISL_496961, EPI_ISL_496963, EPI_ISL_496964, EPI_ISL_496965, EPI_ISL_496966, EPI_ISL_496967, EPI_ISL_496968, EPI_ISL_496969, EPI_ISL_496970, EPI_ISL_496972, EPI_ISL_496974, EPI_ISL_496975, EPI_ISL_496976, EPI_ISL_496977, EPI_ISL_496978, EPI_ISL_496979, EPI_ISL_496980, EPI_ISL_496981, EPI_ISL_496982, EPI_ISL_496983, EPI_ISL_496984, EPI_ISL_496985, EPI_ISL_496987, EPI_ISL_496988, EPI_ISL_496989, EPI_ISL_496990, EPI_ISL_496991, EPI_ISL_496992, EPI_ISL_496993, EPI_ISL_496994, EPI_ISL_496995, EPI_ISL_496996, EPI_ISL_496997, EPI_ISL_496998, EPI_ISL_496999, EPI_ISL_497000, EPI_ISL_497003, EPI_ISL_497004, EPI_ISL_497005, EPI_ISL_497006, EPI_ISL_497007, EPI_ISL_497008, EPI_ISL_497009, EPI_ISL_497010, EPI_ISL_497011, EPI_ISL_497012, EPI_ISL_497013, EPI_ISL_497014, EPI_ISL_497015, EPI_ISL_497016, EPI_ISL_497017, EPI_ISL_497018, EPI_ISL_497019, EPI_ISL_497020, EPI_ISL_497021, EPI_ISL_497022, EPI_ISL_497023, EPI_ISL_497024, EPI_ISL_497025, EPI_ISL_497026, EPI_ISL_497027, EPI_ISL_497028, EPI_ISL_497029, EPI_ISL_497030, EPI_ISL_497031, EPI_ISL_497032, EPI_ISL_497033, EPI_ISL_497034, EPI_ISL_497035, EPI_ISL_497036, EPI_ISL_497037, EPI_ISL_497038, EPI_ISL_497039, EPI_ISL_497040, EPI_ISL_497041, EPI_ISL_497042, EPI_ISL_497043, EPI_ISL_497044, EPI_ISL_497045, EPI_ISL_497046, EPI_ISL_497048, EPI_ISL_497049, EPI_ISL_497050, EPI_ISL_497051, EPI_ISL_497052, EPI_ISL_497054, EPI_ISL_497055, EPI_ISL_497056, EPI_ISL_497057, EPI_ISL_497058, EPI_ISL_497059, EPI_ISL_497060, EPI_ISL_497061, EPI_ISL_497062, EPI_ISL_497063, EPI_ISL_497064, EPI_ISL_497065, EPI_ISL_497066, EPI_ISL_497067, EPI_ISL_497068, EPI_ISL_497069, EPI_ISL_497070, EPI_ISL_497071, EPI_ISL_497072, EPI_ISL_497074, EPI_ISL_497075, EPI_ISL_497076, EPI_ISL_497077, EPI_ISL_497078, EPI_ISL_497079, EPI_ISL_497080, EPI_ISL_497081, EPI_ISL_497082, EPI_ISL_497083, EPI_ISL_497084, EPI_ISL_497085, EPI_ISL_497086, EPI_ISL_497087, EPI_ISL_497088, EPI_ISL_497089, EPI_ISL_497090, EPI_ISL_497091, EPI_ISL_497092, EPI_ISL_497093, EPI_ISL_497095, EPI_ISL_497096, EPI_ISL_497097, EPI_ISL_497098, EPI_ISL_497099, EPI_ISL_497100, EPI_ISL_497101, EPI_ISL_497102, EPI_ISL_497103, EPI_ISL_497104, EPI_ISL_497105, EPI_ISL_497106, EPI_ISL_497107, EPI_ISL_497109, EPI_ISL_497110, EPI_ISL_497111, EPI_ISL_497112, EPI_ISL_497113, EPI_ISL_497114, EPI_ISL_497115, EPI_ISL_497116, EPI_ISL_497117, EPI_ISL_497118, EPI_ISL_497119, EPI_ISL_497120, EPI_ISL_497121, EPI_ISL_497122, EPI_ISL_497123, EPI_ISL_497124, EPI_ISL_497125, EPI_ISL_497126, EPI_ISL_497127, EPI_ISL_497128, EPI_ISL_497129, EPI_ISL_497130, EPI_ISL_497131, EPI_ISL_497132, EPI_ISL_497133, EPI_ISL_497134, EPI_ISL_497135, EPI_ISL_497136, EPI_ISL_497137, EPI_ISL_497138, EPI_ISL_497139, EPI_ISL_497140, EPI_ISL_497141, EPI_ISL_497142, EPI_ISL_497143, EPI_ISL_497144, EPI_ISL_497145, EPI_ISL_497146, EPI_ISL_497147, EPI_ISL_497148, EPI_ISL_497149, EPI_ISL_497150, EPI_ISL_497151, EPI_ISL_497152, EPI_ISL_497154, EPI_ISL_497155, EPI_ISL_497156, EPI_ISL_497157, EPI_ISL_497158, EPI_ISL_497159, EPI_ISL_497160, EPI_ISL_497161, EPI_ISL_497162, EPI_ISL_497163, EPI_ISL_497164, EPI_ISL_497165, EPI_ISL_497166, EPI_ISL_497168, EPI_ISL_497169, EPI_ISL_497170, EPI_ISL_497171, EPI_ISL_497172, EPI_ISL_497173, EPI_ISL_497174, EPI_ISL_497175, EPI_ISL_497176, EPI_ISL_497177, EPI_ISL_497178, EPI_ISL_497179, EPI_ISL_497180, EPI_ISL_497181, EPI_ISL_497182, EPI_ISL_497183, EPI_ISL_497184, EPI_ISL_497185, EPI_ISL_497186, EPI_ISL_497187, | | | |

| | | | |
|---|---|---|---|
| EPI_ISL_497189, EPI_ISL_497190, EPI_ISL_497191, EPI_ISL_497192, EPI_ISL_497193, EPI_ISL_497194, EPI_ISL_497195, EPI_ISL_497196, EPI_ISL_497197, EPI_ISL_497198, EPI_ISL_497199, EPI_ISL_497200, EPI_ISL_497201, EPI_ISL_497202, EPI_ISL_497203, EPI_ISL_497205, EPI_ISL_497206, EPI_ISL_497207, EPI_ISL_497208, EPI_ISL_497209, EPI_ISL_497210, EPI_ISL_497211, EPI_ISL_497212, EPI_ISL_497213, EPI_ISL_497214, EPI_ISL_497215, EPI_ISL_497216, EPI_ISL_497217, EPI_ISL_497218, EPI_ISL_497219, EPI_ISL_497220, EPI_ISL_497221, EPI_ISL_497222, EPI_ISL_497223, EPI_ISL_497224, EPI_ISL_497225, EPI_ISL_497226, EPI_ISL_497228, EPI_ISL_497230, EPI_ISL_497231, EPI_ISL_497232, EPI_ISL_497233, EPI_ISL_497234, EPI_ISL_497235, EPI_ISL_497236, EPI_ISL_497237, EPI_ISL_497238, EPI_ISL_497240, EPI_ISL_497241, EPI_ISL_497242, EPI_ISL_497243, EPI_ISL_497244, EPI_ISL_497246, EPI_ISL_497247, EPI_ISL_497248, EPI_ISL_497249, EPI_ISL_497250, EPI_ISL_497251, EPI_ISL_497252, EPI_ISL_497253, EPI_ISL_497254, EPI_ISL_497255, EPI_ISL_497256, EPI_ISL_497257, EPI_ISL_497258, EPI_ISL_497259, EPI_ISL_497260, EPI_ISL_497261, EPI_ISL_497262, EPI_ISL_497263, EPI_ISL_497265, EPI_ISL_497266, EPI_ISL_497267, EPI_ISL_497268, EPI_ISL_497269, EPI_ISL_497270, EPI_ISL_497271, EPI_ISL_497272, EPI_ISL_497273, EPI_ISL_497274, EPI_ISL_497275, EPI_ISL_497276, EPI_ISL_497277, EPI_ISL_497278, EPI_ISL_497279, EPI_ISL_497280, EPI_ISL_497281, EPI_ISL_497282, EPI_ISL_497283, EPI_ISL_497284, EPI_ISL_497285, EPI_ISL_497286, EPI_ISL_497287, EPI_ISL_497288, EPI_ISL_497289, EPI_ISL_497290, EPI_ISL_497291, EPI_ISL_497292, EPI_ISL_497293, EPI_ISL_497294, EPI_ISL_497295, EPI_ISL_497296, EPI_ISL_497297, EPI_ISL_497298, EPI_ISL_497299, EPI_ISL_497300, EPI_ISL_497301, EPI_ISL_497302, EPI_ISL_497303, EPI_ISL_497304, EPI_ISL_497305, EPI_ISL_497306, EPI_ISL_497307, EPI_ISL_497308, EPI_ISL_497310, EPI_ISL_497311, EPI_ISL_497312, EPI_ISL_497313, EPI_ISL_497315, EPI_ISL_497316, EPI_ISL_497317, EPI_ISL_497318, EPI_ISL_497319, EPI_ISL_497321, EPI_ISL_497322, EPI_ISL_497323, EPI_ISL_497324, EPI_ISL_497325, EPI_ISL_497326, EPI_ISL_497327, EPI_ISL_497329, EPI_ISL_497330, EPI_ISL_497331, EPI_ISL_497332, EPI_ISL_497333, EPI_ISL_497335, EPI_ISL_497336, EPI_ISL_497337, EPI_ISL_497338, EPI_ISL_497339, EPI_ISL_497340, EPI_ISL_497341, EPI_ISL_497342, EPI_ISL_497343, EPI_ISL_497344, EPI_ISL_497345, EPI_ISL_497346, EPI_ISL_497347, EPI_ISL_497348, EPI_ISL_497349, EPI_ISL_497350, EPI_ISL_497351, EPI_ISL_497352, EPI_ISL_497353, EPI_ISL_497354, EPI_ISL_497356, EPI_ISL_497357, EPI_ISL_497358, EPI_ISL_497359, EPI_ISL_497360, EPI_ISL_497361, EPI_ISL_497362, EPI_ISL_497363, EPI_ISL_497364, EPI_ISL_497365, EPI_ISL_497366, EPI_ISL_497367, EPI_ISL_497368, EPI_ISL_497369, EPI_ISL_497370, EPI_ISL_497371, EPI_ISL_497372, EPI_ISL_497373, EPI_ISL_497374, EPI_ISL_497375, EPI_ISL_497376, EPI_ISL_497377, EPI_ISL_497378, EPI_ISL_497379, EPI_ISL_497380, EPI_ISL_497381, EPI_ISL_497382, EPI_ISL_497383, EPI_ISL_497384, EPI_ISL_497385, EPI_ISL_497386, EPI_ISL_497387, EPI_ISL_497388, EPI_ISL_497389, EPI_ISL_497390, EPI_ISL_497391, EPI_ISL_497392, EPI_ISL_497393, EPI_ISL_497394, EPI_ISL_497395, EPI_ISL_497396, EPI_ISL_497397, EPI_ISL_497398, EPI_ISL_497399, EPI_ISL_497400, EPI_ISL_497401, EPI_ISL_497402, EPI_ISL_497403, EPI_ISL_497404, EPI_ISL_497405, EPI_ISL_497406, EPI_ISL_497407, EPI_ISL_497408, EPI_ISL_497409, EPI_ISL_497410, EPI_ISL_497411, EPI_ISL_497412, EPI_ISL_497413, EPI_ISL_497414, EPI_ISL_497415, EPI_ISL_497416, EPI_ISL_497417, EPI_ISL_497418, EPI_ISL_497420, EPI_ISL_497421, EPI_ISL_497422, EPI_ISL_497423, EPI_ISL_497424, EPI_ISL_497425, EPI_ISL_497426, EPI_ISL_497427, EPI_ISL_497428, EPI_ISL_497429, EPI_ISL_497430, EPI_ISL_497431, EPI_ISL_497432, EPI_ISL_497433, EPI_ISL_497434, EPI_ISL_497435, EPI_ISL_497436, EPI_ISL_497437, EPI_ISL_497438, EPI_ISL_497440, EPI_ISL_497441, EPI_ISL_497442, EPI_ISL_497443, EPI_ISL_497444, EPI_ISL_497445, EPI_ISL_497446, EPI_ISL_497447, EPI_ISL_497448, EPI_ISL_497449, EPI_ISL_497450, EPI_ISL_497451, EPI_ISL_497453, EPI_ISL_497454, EPI_ISL_497455, EPI_ISL_497456, EPI_ISL_497457, EPI_ISL_497458, EPI_ISL_497459, EPI_ISL_497460, EPI_ISL_497461, EPI_ISL_497462, EPI_ISL_497463, EPI_ISL_497464, EPI_ISL_497465, EPI_ISL_497466, EPI_ISL_497467, EPI_ISL_497468, EPI_ISL_497469, EPI_ISL_497470, EPI_ISL_497471, EPI_ISL_497472, EPI_ISL_497473, EPI_ISL_497474, EPI_ISL_497475, EPI_ISL_497476, EPI_ISL_497477, EPI_ISL_497478, EPI_ISL_497479, EPI_ISL_497480, EPI_ISL_497481, EPI_ISL_497482, EPI_ISL_497483, EPI_ISL_497485, EPI_ISL_497486, EPI_ISL_497487, EPI_ISL_497488, EPI_ISL_497489, EPI_ISL_497490, EPI_ISL_497491, EPI_ISL_497492, EPI_ISL_497493, EPI_ISL_497494, EPI_ISL_497495, EPI_ISL_497496, EPI_ISL_497497, EPI_ISL_497498, EPI_ISL_497499, EPI_ISL_497500, EPI_ISL_497501, EPI_ISL_497502, EPI_ISL_497503, EPI_ISL_497504, EPI_ISL_497505, EPI_ISL_497506, EPI_ISL_497507, EPI_ISL_497508, EPI_ISL_497509, EPI_ISL_497510, EPI_ISL_497511, EPI_ISL_497512, EPI_ISL_497513, EPI_ISL_497514, EPI_ISL_497515, EPI_ISL_497516, EPI_ISL_497517, EPI_ISL_497518, EPI_ISL_497519, EPI_ISL_497520, EPI_ISL_497521, EPI_ISL_497522, EPI_ISL_497523, EPI_ISL_497524, EPI_ISL_497525, EPI_ISL_497526, EPI_ISL_497527, EPI_ISL_497528, EPI_ISL_497529, EPI_ISL_497530, EPI_ISL_497531, EPI_ISL_497532, EPI_ISL_497533, EPI_ISL_497534, EPI_ISL_497535, EPI_ISL_497536, EPI_ISL_497537, EPI_ISL_497538, EPI_ISL_497539, EPI_ISL_497540, EPI_ISL_497541, EPI_ISL_497542, EPI_ISL_497544, EPI_ISL_497545, EPI_ISL_497546, EPI_ISL_497547, EPI_ISL_497548, EPI_ISL_497549, EPI_ISL_497550, EPI_ISL_497551, EPI_ISL_497552, EPI_ISL_497553, EPI_ISL_497554, EPI_ISL_497555, EPI_ISL_497556, EPI_ISL_497557, EPI_ISL_497558, EPI_ISL_497559, EPI_ISL_497561, EPI_ISL_497562, EPI_ISL_497564, EPI_ISL_497565, EPI_ISL_497567, EPI_ISL_497569, EPI_ISL_497570, EPI_ISL_497571, EPI_ISL_497573, EPI_ISL_497574, EPI_ISL_497575, EPI_ISL_497576, EPI_ISL_497577, EPI_ISL_497578, EPI_ISL_497579, EPI_ISL_497580, EPI_ISL_497581, EPI_ISL_497582, EPI_ISL_497583, EPI_ISL_497584, EPI_ISL_497585, EPI_ISL_497586, EPI_ISL_497587, EPI_ISL_497588, EPI_ISL_497589, EPI_ISL_497590, EPI_ISL_497591, EPI_ISL_497592, EPI_ISL_497593, EPI_ISL_497594, EPI_ISL_497595, EPI_ISL_497596, EPI_ISL_497598, EPI_ISL_497599, EPI_ISL_497600, EPI_ISL_497601, EPI_ISL_497603, EPI_ISL_497604, EPI_ISL_497605, EPI_ISL_497607, EPI_ISL_497608, EPI_ISL_497609, EPI_ISL_497610, EPI_ISL_497611, EPI_ISL_497612, EPI_ISL_497613, EPI_ISL_497614, EPI_ISL_497615, EPI_ISL_497616, EPI_ISL_497617, EPI_ISL_497618, EPI_ISL_497619, EPI_ISL_497620, EPI_ISL_497623, EPI_ISL_497624, EPI_ISL_497626, EPI_ISL_497628, EPI_ISL_497629, EPI_ISL_497630, EPI_ISL_497631, EPI_ISL_497632, EPI_ISL_497633, EPI_ISL_497634, EPI_ISL_497636, EPI_ISL_497637, EPI_ISL_497638, EPI_ISL_497639, EPI_ISL_497640, EPI_ISL_497641, EPI_ISL_497642, EPI_ISL_497643, EPI_ISL_497644, EPI_ISL_497645, EPI_ISL_497646, EPI_ISL_497647, EPI_ISL_497648, EPI_ISL_497649, EPI_ISL_497650, EPI_ISL_497651, EPI_ISL_497652, EPI_ISL_497654, EPI_ISL_497655, EPI_ISL_497656, EPI_ISL_497657, EPI_ISL_497658, EPI_ISL_497659, EPI_ISL_497660, EPI_ISL_497661, EPI_ISL_497662, EPI_ISL_497663, EPI_ISL_497664, EPI_ISL_497665, EPI_ISL_497667, EPI_ISL_497668, EPI_ISL_497669, EPI_ISL_497670, EPI_ISL_497671, EPI_ISL_497672, EPI_ISL_497673, EPI_ISL_497674, EPI_ISL_497675, EPI_ISL_497676, EPI_ISL_497677, EPI_ISL_497679, EPI_ISL_497680, EPI_ISL_497681, EPI_ISL_497682, EPI_ISL_497683, EPI_ISL_497684, EPI_ISL_497685, EPI_ISL_497686, EPI_ISL_497687, EPI_ISL_497688, EPI_ISL_497689, EPI_ISL_497690, EPI_ISL_497691, EPI_ISL_497692, EPI_ISL_497693, EPI_ISL_497694, EPI_ISL_497695, EPI_ISL_497696, EPI_ISL_497697, EPI_ISL_497698, EPI_ISL_497699, EPI_ISL_497700, EPI_ISL_497701, EPI_ISL_497702, EPI_ISL_497703, EPI_ISL_497704, EPI_ISL_497705, EPI_ISL_497706, EPI_ISL_497707, EPI_ISL_497708, EPI_ISL_497709, EPI_ISL_497710, EPI_ISL_497711, EPI_ISL_497712, EPI_ISL_497713, EPI_ISL_497714, EPI_ISL_497715, EPI_ISL_497716, EPI_ISL_497717, EPI_ISL_497718, EPI_ISL_497719, EPI_ISL_497720, EPI_ISL_497721, EPI_ISL_497722, EPI_ISL_497723, EPI_ISL_497724, EPI_ISL_497725, EPI_ISL_497726 | | | |
| see above | Washington State Department of Health | Seattle Flu Study | Deborah A. Nickerson, Chris D. Frazar, Jover Lee, Benjamin Pelle, Matthew Richardson, Amanda Adler, Elisabeth Brandstetter, Peter D. Han, Kairsten Fay, Misja Ilcisin, Kirsten Lacombe, Thomas R. Sibley, Melissa Truong, Caitlin R. Wolf, Romesh Gautom, Geoff Melly, Brian Hiatt, Philip Dykema, Scott Lindquist, Michael Boeckh, Janet A. Englund, Michael Famulare, Barry R. Lutz, Mark J. Rieder, Lea M. Starita, Matthew Thompson, Helen Y. Chu, Jay Shendure, Trevor Bedford |
| EPI_ISL_497736, EPI_ISL_497738, EPI_ISL_497744, EPI_ISL_497745 | Instituto Nacional de Salud, Bogotá, Colombia | Instituto Nacional de Salud, Bogotá, Colombia | Katherine Laiton-Donato, Diego A. Álvarez-Díaz, Carlos Franco-Muñoz, Jonathan Reales, Diego Andrés Prada, Jose A. Usme-Ciro, Nicolas D. Franco-Sierra, Zulma M. Cucunubá, Christian Julian Villabona-Arenas, Liz Villabona-Arenas, Sussy Echeverria, Astrid C. Flórez, Carolina Ferro, Diana Marcela Walteros-Acero, Franklin Prieto, Carlos Andrés Durán, Martha Lucia Ospina Martinez, Marcela Mercado-Reyes |
| EPI_ISL_497758, EPI_ISL_497760 | CSIR-CDRI/SGPGI, Lucknow | CSIr-CDRI/SGPGI, Lucknow | Saumya Sarkar, Dharam Veer Singh, Rahul Vishvkarma, Ujjala Ghoshal, Uday Ghoshal, Ravishankar Ramachandran, Tapas Kumar Kundu, Rajender Singh |
| EPI_ISL_497762, EPI_ISL_497763 | CSIR-CDRI/SGPGI, Lucknow | CSIR-CDRI/SGPGI, Lucknow | Saumya Sarkar, Dharam Veer Singh, Rahul Vishvkarma, Ujjala Ghoshal, Uday Ghoshal, Ravishankar Ramachandran, Tapas Kumar Kundu, Rajender Singh |
| EPI_ISL_497764 | CSIR-CDRi/SGPGI, Lucknow | CSIR-CDRI/SGPGI, Lucknow | Saumya Sarkar, Dharam Veer Singh, Rahul Vishvkarma, Ujjala Ghoshal, Uday Ghoshal, Ravishankar Ramachandran, Tapas Kumar Kundu, Rajender Singh |
| EPI_ISL_497765 | CSIR-CDRI/SGPGI, Lucknow | CSIR-CDRI/SGPGI, Lucknow | Saumya Sarkar, Dharam Veer Singh, Rahul Vishvkarma, Ujjala Ghoshal, Uday Ghoshal, Ravishankar Ramachandran, Tapas Kumar Kundu, Rajender Singh |
| EPI_ISL_497766 | CSIR-CDRI/SGPGI, Lucknow | CSIR-CDRi/SGPGI, Lucknow | Saumya Sarkar, Dharam Veer Singh, Rahul Vishvkarma, Ujjala Ghoshal, Uday Ghoshal, Ravishankar Ramachandran, Tapas Kumar Kundu, Rajender Singh |
| EPI_ISL_497767 | CSIR-CDRI/SGPGI, Lucknow | CSIR-CDRI/SGPGI, Lucknow | Saumya Sarkar, Dharam Veer Singh, Rahul Vishvkarma, Ujjala Ghoshal, Uday Ghoshal, Ravishankar Ramachandran, Tapas Kumar Kundu, Rajender Singh |
| EPI_ISL_497769, EPI_ISL_497770, EPI_ISL_497771, EPI_ISL_497773, EPI_ISL_497775, EPI_ISL_497776, EPI_ISL_497777, EPI_ISL_497778, EPI_ISL_497780, EPI_ISL_497782, EPI_ISL_497783, EPI_ISL_497784, EPI_ISL_497785, EPI_ISL_497786, EPI_ISL_497790, EPI_ISL_497791, EPI_ISL_497793, EPI_ISL_497794, EPI_ISL_497795, EPI_ISL_497796, EPI_ISL_497797, EPI_ISL_497798, EPI_ISL_497799, EPI_ISL_497800, EPI_ISL_497801, EPI_ISL_497803, EPI_ISL_497806, EPI_ISL_497807, EPI_ISL_497808, EPI_ISL_497809, EPI_ISL_497810, EPI_ISL_497811, EPI_ISL_497812, EPI_ISL_497813, EPI_ISL_497814, EPI_ISL_497815, EPI_ISL_497816, EPI_ISL_497817, EPI_ISL_497818, EPI_ISL_497819, EPI_ISL_497820, EPI_ISL_497821, EPI_ISL_497822, EPI_ISL_497823, EPI_ISL_497824, EPI_ISL_497825, EPI_ISL_497826, EPI_ISL_497827, EPI_ISL_497829, EPI_ISL_497830, EPI_ISL_497831, EPI_ISL_497832, EPI_ISL_497833, EPI_ISL_497835, EPI_ISL_497836, EPI_ISL_497840, EPI_ISL_497841, EPI_ISL_497842, EPI_ISL_497844, EPI_ISL_497845, EPI_ISL_497846, EPI_ISL_497847, EPI_ISL_497848, EPI_ISL_497849, EPI_ISL_497850, EPI_ISL_497851, EPI_ISL_497853, EPI_ISL_497854, EPI_ISL_497855, EPI_ISL_497856, EPI_ISL_497857, EPI_ISL_497860, EPI_ISL_497861, EPI_ISL_497862, EPI_ISL_497863, EPI_ISL_497864, EPI_ISL_497865, EPI_ISL_497866, EPI_ISL_497867, EPI_ISL_497868, EPI_ISL_497869, EPI_ISL_497870 | | | |
| see above | Department of Microbiology, The University of Hong Kong | Department of Microbiology, The University of Hong Kong | Kelvin K.W. To, Kwok-Yung Yuen |
| EPI_ISL_497950 | Shaoxing CDC | Zhejiang Provincial Center for Disease Control and Prevention | Yin Chen, Yanjun Zhang, Haiyan Mao, Junhang Pan, Xiuyu Lou, Yi Sun, Hao Yan, Zhen Li, Wen Shi |
| EPI_ISL_497951 | Division of Viral Diseases, Center for Laboratory Control of Infectious Diseases, Korea Centers for Diseases Control and Prevention | Division of Viral Diseases, Center for Laboratory Control of Infectious Diseases, Korea Centers for Diseases Control and Prevention | Jeong-Min Kim, Yoon-Seok Chung, Namjoo Lee, Sang Hee Woo, Hye-Jun Jo, Heui Man Kim, Jun-Sub Kim, Myung Guk Han |
| EPI_ISL_497953, EPI_ISL_497954, EPI_ISL_497955, EPI_ISL_497956, EPI_ISL_497957, EPI_ISL_497958, EPI_ISL_497959, EPI_ISL_497960 | Division of Viral Diseases, Center for Laboratory Control of Infectious Diseases, Korea Centers for Diseases Control and Prevention | Division of Viral Diseases, Center for Laboratory Control of Infectious Diseases, Korea Centers for Diseases Control and Prevention | Jeong-Min Kim, Yoon-Seok Chung, Namjoo Lee, Sang Hee Woo, Hye-Jun Jo, Heui Man Kim, Jun-Sub Kim, Dong Hyun Song, Daesang Lee, Seong Tae Jeong, Myung Guk Han |
| EPI_ISL_497961, EPI_ISL_497962, EPI_ISL_497963, EPI_ISL_497964, EPI_ISL_497965, EPI_ISL_497966, EPI_ISL_497967 | Division of Viral Diseases, Center for Laboratory Control of Infectious Diseases, Korea Centers for Diseases Control and Prevention | Division of Viral Diseases, Center for Laboratory Control of Infectious Diseases, Korea Centers for Diseases Control and Prevention | Jeong-Min Kim, Yoon-Seok Chung, Namjoo Lee, Sang Hee Woo, Hye-Jun Jo, Heui Man Kim, Jun-Sub Kim, Myung Guk Han |
| EPI_ISL_497968, EPI_ISL_497969, EPI_ISL_497971 | Division of Viral Diseases, Center for Laboratory Control of Infectious Diseases, Korea Centers for Diseases Control and Prevention | Division of Viral Diseases, Center for Laboratory Control of Infectious Diseases, Korea Centers for Diseases Control and Prevention | Jeong-Min Kim, Yoon-Seok Chung, Namjoo Lee, Sang Hee Woo, Hye-Jun Jo, Heui Man Kim, Jun-Sub Kim, Dong Hyun Song, Daesang Lee, Seong Tae Jeong, Myung Guk Han |
| EPI_ISL_497972, EPI_ISL_497973, EPI_ISL_497975, EPI_ISL_497976, EPI_ISL_497977, EPI_ISL_497978, EPI_ISL_497979, EPI_ISL_497980, EPI_ISL_497981, EPI_ISL_497982, EPI_ISL_497983, EPI_ISL_497984 | | | |
| see above | Division of Viral Diseases, Center for Laboratory Control of Infectious Diseases, Korea Centers for Diseases Control and Prevention | Division of Viral Diseases, Center for Laboratory Control of Infectious Diseases, Korea Centers for Diseases Control and Prevention | Jeong-Min Kim, Yoon-Seok Chung, Namjoo Lee, Sang Hee Woo, Hye-Jun Jo, Heui Man Kim, Jun-Sub Kim, Myung Guk Han |
| EPI_ISL_497985 | Division of Viral Diseases, Center for Laboratory Control of Infectious Diseases, Korea Centers for Diseases Control and Prevention | Division of Viral Diseases, Center for Laboratory Control of Infectious Diseases, Korea Centers for Diseases Control and Prevention | Jeong-Min Kim, Yoon-Seok Chung, Namjoo Lee, Sang Hee Woo, Hye-Jun Jo, Heui Man Kim, Jun-Sub Kim, Dong Hyun Song, Daesang Lee, Seong Tae Jeong, Myung Guk Han |
| EPI_ISL_497986 | Division of Viral Diseases, Center for Laboratory Control of Infectious Diseases, Korea Centers for Diseases Control and Prevention | Division of Viral Diseases, Center for Laboratory Control of Infectious Diseases, Korea Centers for Diseases Control and Prevention | Jeong-Min Kim, Yoon-Seok Chung, Namjoo Lee, Sang Hee Woo, Hye-Jun Jo, Heui Man Kim, Jun-Sub Kim, Myung Guk Han |
| EPI_ISL_497987, EPI_ISL_497988, EPI_ISL_497989, EPI_ISL_497990 | Division of Viral Diseases, Center for Laboratory Control of Infectious Diseases, Korea Centers for Diseases Control and Prevention | Division of Viral Diseases, Center for Laboratory Control of Infectious Diseases, Korea Centers for Diseases Control and Prevention | Jeong-Min Kim, Yoon-Seok Chung, Namjoo Lee, Sang Hee Woo, Hye-Jun Jo, Heui Man Kim, Jun-Sub Kim, Dong Hyun Song, Daesang Lee, Seong Tae Jeong, Myung Guk Han |
| EPI_ISL_497991, EPI_ISL_497992, EPI_ISL_497993 | Division of Viral Diseases, Center for Laboratory Control of Infectious Diseases, Korea Centers for Diseases Control and Prevention | Division of Viral Diseases, Center for Laboratory Control of Infectious Diseases, Korea Centers for Diseases Control and Prevention | Jeong-Min Kim, Yoon-Seok Chung, Namjoo Lee, Sang Hee Woo, Hye-Jun Jo, Heui Man Kim, Jun-Sub Kim, Myung Guk Han |
| EPI_ISL_497994, EPI_ISL_497995 | Division of Viral Diseases, Center for Laboratory Control of Infectious Diseases, Korea Centers for Diseases Control and Prevention | Division of Viral Diseases, Center for Laboratory Control of Infectious Diseases, Korea Centers for Diseases Control and Prevention | Jeong-Min Kim, Yoon-Seok Chung, Namjoo Lee, Sang Hee Woo, Hye-Jun Jo, Heui Man Kim, Jun-Sub Kim, Dong Hyun Song, Daesang Lee, Seong Tae Jeong, Myung Guk Han |
| EPI_ISL_497996 | Division of Viral Diseases, Center for Laboratory Control of Infectious Diseases, Korea Centers for Diseases Control and Prevention | Division of Viral Diseases, Center for Laboratory Control of Infectious Diseases, Korea Centers for Diseases Control and Prevention | Jeong-Min Kim, Yoon-Seok Chung, Namjoo Lee, Sang Hee Woo, Hye-Jun Jo, Heui Man Kim, Jun-Sub Kim, Myung Guk Han |
| EPI_ISL_497997 | Division of Viral Diseases, Center for Laboratory Control of Infectious Diseases, Korea Centers for Diseases Control and Prevention | Division of Viral Diseases, Center for Laboratory Control of Infectious Diseases, Korea Centers for Diseases Control and Prevention | Jeong-Min Kim, Yoon-Seok Chung, Namjoo Lee, Sang Hee Woo, Hye-Jun Jo, Heui Man Kim, Jun-Sub Kim, Dong Hyun Song, Daesang Lee, Seong Tae Jeong, Myung Guk Han |
| EPI_ISL_497998, EPI_ISL_497999, EPI_ISL_498000, EPI_ISL_498001, EPI_ISL_498002, EPI_ISL_498003 | Division of Viral Diseases, Center for Laboratory Control of Infectious Diseases, Korea Centers for Diseases Control and Prevention | Division of Viral Diseases, Center for Laboratory Control of Infectious Diseases, Korea Centers for Diseases Control and Prevention | Jeong-Min Kim, Yoon-Seok Chung, Namjoo Lee, Sang Hee Woo, Hye-Jun Jo, Heui Man Kim, Jun-Sub Kim, Myung Guk Han |
| EPI_ISL_498004 | Division of Viral Diseases, Center for Laboratory Control of Infectious Diseases, Korea Centers for Diseases Control and Prevention | Division of Viral Diseases, Center for Laboratory Control of Infectious Diseases, Korea Centers for Diseases Control and Prevention | Jeong-Min Kim, Yoon-Seok Chung, Namjoo Lee, Sang Hee Woo, Hye-Jun Jo, Heui Man Kim, Jun-Sub Kim, Dong Hyun Song, Daesang Lee, Seong Tae Jeong, Myung Guk Han |
| EPI_ISL_498005 | Division of Viral Diseases, Center for Laboratory Control of Infectious Diseases, Korea Centers for Diseases Control and Prevention | Division of Viral Diseases, Center for Laboratory Control of Infectious Diseases, Korea Centers for Diseases Control and Prevention | Jeong-Min Kim, Yoon-Seok Chung, Namjoo Lee, Sang Hee Woo, Hye-Jun Jo, Heui Man Kim, Jun-Sub Kim, Myung Guk Han |
| EPI_ISL_498006, EPI_ISL_498007, EPI_ISL_498008, EPI_ISL_498009, EPI_ISL_498010, EPI_ISL_498011, EPI_ISL_498012, EPI_ISL_498013, EPI_ISL_498014, EPI_ISL_498015, EPI_ISL_498016 | | | |
| see above | Division of Viral Diseases, Center for Laboratory Control of Infectious Diseases, Korea Centers for Diseases Control and Prevention | Division of Viral Diseases, Center for Laboratory Control of Infectious Diseases, Korea Centers for Diseases Control and Prevention | Jeong-Min Kim, Yoon-Seok Chung, Namjoo Lee, Sang Hee Woo, Hye-Jun Jo, Heui Man Kim, Jun-Sub Kim, Dong Hyun Song, Daesang Lee, Seong Tae Jeong, Myung Guk Han |
| EPI_ISL_498017 | Division of Viral Diseases, Center for Laboratory Control of Infectious Diseases, Korea Centers for Diseases Control and Prevention | Division of Viral Diseases, Center for Laboratory Control of Infectious Diseases, Korea Centers for Diseases Control and Prevention | Jeong-Min Kim, Yoon-Seok Chung, Namjoo Lee, Sang Hee Woo, Hye-Jun Jo, Heui Man Kim, Jun-Sub Kim, Myung Guk Han |

| | | | |
|---|---|---|---|
| EPI_ISL_498018, EPI_ISL_498019, EPI_ISL_498020, EPI_ISL_498021, EPI_ISL_498022, EPI_ISL_498023, EPI_ISL_498024, EPI_ISL_498025, EPI_ISL_498026, EPI_ISL_498027, EPI_ISL_498028, EPI_ISL_498029, EPI_ISL_498030, EPI_ISL_498031, EPI_ISL_498032, EPI_ISL_498033, EPI_ISL_498034, EPI_ISL_498035 | | | |
| see above | Division of Viral Diseases, Center for Laboratory Control of Infectious Diseases, Korea Centers for Diseases Control and Prevention | Division of Viral Diseases, Center for Laboratory Control of Infectious Diseases, Korea Centers for Diseases Control and Prevention | Jeong-Min Kim, Yoon-Seok Chung, Namjoo Lee, Sang Hee Woo, Hye-Jun Jo, Heui Man Kim, Jun-Sub Kim, Dong Hyun Song, Daesang Lee, Seong Tae Jeong, Myung Guk Han |
| EPI_ISL_498036 | Division of Viral Diseases, Center for Laboratory Control of Infectious Diseases, Korea Centers for Diseases Control and Prevention | Division of Viral Diseases, Center for Laboratory Control of Infectious Diseases, Korea Centers for Diseases Control and Prevention | Jeong-Min Kim, Yoon-Seok Chung, Namjoo Lee, Sang Hee Woo, Hye-Jun Jo, Heui Man Kim, Jun-Sub Kim, Myung Guk Han |
| EPI_ISL_498037, EPI_ISL_498038, EPI_ISL_498039, EPI_ISL_498040, EPI_ISL_498041, EPI_ISL_498042, EPI_ISL_498043, EPI_ISL_498044, EPI_ISL_498045, EPI_ISL_498046, EPI_ISL_498047, EPI_ISL_498048, EPI_ISL_498049, EPI_ISL_498050, EPI_ISL_498051, EPI_ISL_498052, EPI_ISL_498053 | | | |
| see above | Division of Viral Diseases, Center for Laboratory Control of Infectious Diseases, Korea Centers for Diseases Control and Prevention | Division of Viral Diseases, Center for Laboratory Control of Infectious Diseases, Korea Centers for Diseases Control and Prevention | Jeong-Min Kim, Yoon-Seok Chung, Namjoo Lee, Sang Hee Woo, Hye-Jun Jo, Heui Man Kim, Jun-Sub Kim, Dong Hyun Song, Daesang Lee, Seong Tae Jeong, Myung Guk Han |
| EPI_ISL_498055, EPI_ISL_498060, EPI_ISL_498061, EPI_ISL_498062, EPI_ISL_498064, EPI_ISL_498065, EPI_ISL_498066, EPI_ISL_498067, EPI_ISL_498069, EPI_ISL_498070, EPI_ISL_498071, EPI_ISL_498072, EPI_ISL_498073, EPI_ISL_498074, EPI_ISL_498075, EPI_ISL_498077, EPI_ISL_498078, EPI_ISL_498079, EPI_ISL_498081, EPI_ISL_498082, EPI_ISL_498084, EPI_ISL_498085, EPI_ISL_498086, EPI_ISL_498087, EPI_ISL_498088, EPI_ISL_498089, EPI_ISL_498090, EPI_ISL_498091, EPI_ISL_498092, EPI_ISL_498093, EPI_ISL_498094, EPI_ISL_498095, EPI_ISL_498096, EPI_ISL_498097, EPI_ISL_498098, EPI_ISL_498099, EPI_ISL_498100, EPI_ISL_498101, EPI_ISL_498102, EPI_ISL_498103, EPI_ISL_498104, EPI_ISL_498105, EPI_ISL_498106, EPI_ISL_498107, EPI_ISL_498108, EPI_ISL_498109, EPI_ISL_498110, EPI_ISL_498111, EPI_ISL_498112, EPI_ISL_498113, EPI_ISL_498114, EPI_ISL_498115, EPI_ISL_498116, EPI_ISL_498117, EPI_ISL_498118, EPI_ISL_498119, EPI_ISL_498120, EPI_ISL_498121, EPI_ISL_498122, EPI_ISL_498123, EPI_ISL_498124, EPI_ISL_498125, EPI_ISL_498126 | | | |
| see above | NHLS-IALCH | KRISP, KZN Research Innovation and Sequencing Platform | Giandhari J, Pillay S, Lessells R, Chimukangara B, Mdlalose K, York D, Khan S, Tegally H, Wilkinson E, de Oliveira T |
| EPI_ISL_498127, EPI_ISL_498128, EPI_ISL_498129, EPI_ISL_498130, EPI_ISL_498131, EPI_ISL_498132, EPI_ISL_498133, EPI_ISL_498134, EPI_ISL_498135, EPI_ISL_498136, EPI_ISL_498137, EPI_ISL_498138, EPI_ISL_498139, EPI_ISL_498141, EPI_ISL_498143, EPI_ISL_498144, EPI_ISL_498146, EPI_ISL_498149, EPI_ISL_498150, EPI_ISL_498151 | | | |
| see above | Department of Clinical Microbiology | GIGA Medical Genomics | Keith Durkin, Maria Artesi, Sébastien Bontems, Raphaël Boreux, Cécile Meex, Axelle Chaslain, Céline Fombellida-Lopez, Pierrette Melin, Marie-Pierre Hayette, Vincent Bours. |
| EPI_ISL_498152, EPI_ISL_498153, EPI_ISL_498154, EPI_ISL_498155, EPI_ISL_498156, EPI_ISL_498157, EPI_ISL_498158, EPI_ISL_498159, EPI_ISL_498160, EPI_ISL_498161, EPI_ISL_498162, EPI_ISL_498163, EPI_ISL_498164, EPI_ISL_498165, EPI_ISL_498166, EPI_ISL_498167, EPI_ISL_498168, EPI_ISL_498169, EPI_ISL_498170 | | | |
| see above | Instituto Nacional de Salud, Bogotá, Colombia | Instituto Nacional de Salud, Bogotá, Colombia | Katherine Laiton-Donato, Diego A. Álvarez-Díaz, Carlos Franco-Muñoz, Jonathan Reales, Diego Andrés Prada, Jose A. Usme-Ciro, Nicolas D. Franco-Sierra, Zulma M. Cucunubá, Christian Julian Villabona-Arenas, Liz Villabona-Arenas, Sussy Echeverria, Astrid C. Flórez, Carolina Ferro, Diana Marcela Walteros-Acero, Franklin Prieto, Carlos Andrés Durán, Martha Lucia Ospina Martinez, Marcela Mercado-Reyes |
| EPI_ISL_498171, EPI_ISL_498172, EPI_ISL_498173, EPI_ISL_498174, EPI_ISL_498175, EPI_ISL_498176, EPI_ISL_498177, EPI_ISL_498178, EPI_ISL_498179, EPI_ISL_498180, EPI_ISL_498181, EPI_ISL_498182, EPI_ISL_498183, EPI_ISL_498184, EPI_ISL_498185, EPI_ISL_498186, EPI_ISL_498187, EPI_ISL_498188, EPI_ISL_498189, EPI_ISL_498190, EPI_ISL_498191, EPI_ISL_498192 | | | |
| see above | OUCRU | OUCRU | Nguyen Van Vinh Chau, Nguyen Thi Thu Hong, Nguyen Thi Han Ny, Le Nguyen Truc Nhu, Nghiem My Ngoc, Vo Thanh Lam, Nguyen Thanh Dung, Lam Minh Yen, Ngo Ngoc Quang Minh, Le Manh Hung, Nguyen Tri Dung, Dinh Nguyen Huy Man, Lam Anh Nguyet, Tran Chanh Xuan, Tran Tinh Hien, Nguyen Thanh Phong, Tran Nguyen Hoang Tu, Tran Tan Thanh, Nguyen Thanh Truong, Nguyen Tan Binh, Tang Chi Thuong, Guy Thwaites, and Le Van Tan, for OUCRU COVID-19 research group* |
| EPI_ISL_498226 | LIC | LIC | LIC |
| EPI_ISL_498227, EPI_ISL_498228 | National Institute of Laboratory Medicine and Referral Center | Genomic Research Lab, BCSIR | Shahina Akter, Abu Sayeed Mohammad Mahmud, Mohammad Samir Uzzaman, Eshrar Osman, Md. Ahasan Habib, Tanjina Akhter Banu, Md. Murshed Hasan Sarkar, Barna Goswami, Iffat Jahan, Md. Saddam Hossain, Tasnim Nafisa, Md. Maruf Ahmed Molla, Mahmuda Yeasmin, Asish Kumar Ghosh, A. K. M. Shamsuzzaman, Sheikh Md. Selim Al Din, Utpal Chandra Ray, Salek Ahmed Sajib, Md. Salim Khan |
| EPI_ISL_498229, EPI_ISL_498230, EPI_ISL_498236, EPI_ISL_498238, EPI_ISL_498242, EPI_ISL_498243, EPI_ISL_498249, EPI_ISL_498250 | Institut Pasteur de Dakar | Institut Pasteur de Dakar | Ndongo Dia, Moussa Moise Diagne, Mamadou Diop, Marie Henriette Dior Ndione, Mamadou Malado Jallow, Safietou Sankhe Mbengue, Ousmane Faye, Amadou Alpha Sall. |
| EPI_ISL_498253, EPI_ISL_498254 | National Institute of Laboratory Medicine and Referral Center | Genomic Research Lab, BCSIR | Tanjina Akhter Banu, Abu Sayeed Mohammad Mahmud, Mohammad Samir Uzzaman, Eshrar Osman, Md. Ahasan Habib, Shahina Akter, Md. Murshed Hasan Sarkar, Barna Goswami, Iffat Jahan, Md. Saddam Hossain, Tasnim Nafisa, Md. Maruf Ahmed Molla, Mahmuda Yeasmin, Asish Kumar Ghosh, A. K. M. Shamsuzzaman, Sheikh Md. Selim Al Din, Utpal Chandra Ray, Salek Ahmed Sajib, Md. Salim Khan |
| EPI_ISL_498257 | Hospital for Tropical Diseases | COVID-19 Network Investigations (CONI) Alliance | Elizabeth Batty, Nantarat Chantawat, Wasun Chantratita, Thanat Chookajorn, Stefan Fernandez, Angkana Huang, Weena Janwitthayanan, Akanitt Jittmittraphap, Anthony R. Jones, Khajohn Joonsalak, Chonticha Klungtong, Theerarat Kochakarn, Namfon Kotanan, Krittikorn Kumpornsin, Pornsawan Leaungwutiwong, Wudtichai Manasatienkij, Bhakbhoom Panthan, Ekawat Pasomsub, Kingkan Rakmanee, Insee Sensorn, Janjira Thaipadungpanit, Arporn Wangwiwatsin,Treewat Watthanachockchai |
| EPI_ISL_498264, EPI_ISL_498265 | Ramathibodi Hospital | COVID-19 Network Investigations (CONI) Alliance | Elizabeth Batty, Wasun Chantratita, Thanat Chookajorn, Stefan Fernandez, Angkana Huang, Anthony R. Jones, Khajohn Joonsalak, Chonticha Klungtong, Theerarat Kochakarn, Namfon Kotanan, Krittikorn Kumpornsin, Wudtichai Manasatienkij, Bhakbhoom Panthan, Ekawat Pasomsub, Kingkan Rakmanee, Insee Sensorn, Janjira Thaipadungpanit, Arporn Wangwiwatsin,Treewat Watthanachockchai |
| EPI_ISL_498267, EPI_ISL_498268 | National Institute of Laboratory Medicine and Referral Center | Genomic Research Lab, BCSIR | Barna Goswami, Abu Sayeed Mohammad Mahmud, Mohammad Samir Uzzaman, Eshrar Osman, Md. Ahasan Habib, Shahina Akter, Tanjina Akhter Banu, Md. Murshed Hasan Sarkar, Iffat Jahan, Md. Saddam Hossain, Tasnim Nafisa, Md. Maruf Ahmed Molla, Mahmuda Yeasmin, Asish Kumar Ghosh, A. K. M. Shamsuzzaman, Sheikh Md. Selim Al Din, Utpal Chandra Ray, Salek Ahmed Sajib, Md. Salim Khan |
| EPI_ISL_498270, EPI_ISL_498271 | Department of Microbiology, The University of Hong Kong | Department of Microbiology, The University of Hong Kong | Kelvin K.W. To, Kwok-Yung Yuen |
| EPI_ISL_498273 | National Institute of Laboratory Medicine and Referral Center | Genomic Research Lab, BCSIR | Iffat Jahan, Abu Sayeed Mohammad Mahmud, Mohammad Samir Uzzaman, Eshrar Osman, Md. Ahasan Habib, Shahina Akter, Tanjina Akhter Banu, Md. Murshed Hasan Sarkar, Barna Goswami, Md. Saddam Hossain, Tasnim Nafisa, Md. Maruf Ahmed Molla, Mahmuda Yeasmin, Asish Kumar Ghosh, A. K. M. Shamsuzzaman, Sheikh Md. Selim Al Din, Utpal Chandra Ray, Salek Ahmed Sajib, Md. Salim Khan |
| EPI_ISL_498274, EPI_ISL_498417 | National Institute of Laboratory Medicine and Referral Center | Genomic Research Lab, BCSIR | Tasnim Nafisa, Abu Sayeed Mohammad Mahmud, Mohammad Samir Uzzaman, Eshrar Osman, Md. Ahasan Habib, Shahina Akter, Tanjina Akhter Banu, Md. Murshed Hasan Sarkar, Barna Goswami, Iffat Jahan, Md. Saddam Hossain, Md. Maruf Ahmed Molla, Mahmuda Yeasmin, Asish Kumar Ghosh, A. K. M. Shamsuzzaman, Sheikh Md. Selim Al Din, Utpal Chandra Ray, Salek Ahmed Sajib, Md. Salim Khan |
| EPI_ISL_498419 | National Institute of Laboratory Medicine and Referral Center | Genomic Research Lab, BCSIR | Md. Maruf Ahmed Molla, Abu Sayeed Mohammad Mahmud, Mohammad Samir Uzzaman, Eshrar Osman, Md. Ahasan Habib, Shahina Akter, Tanjina Akhter Banu, Md. Murshed Hasan Sarkar, Barna Goswami, Iffat Jahan, Md. Saddam Hossain, Tasnim Nafisa, Mahmuda Yeasmin, Asish Kumar Ghosh, A. K. M. Shamsuzzaman, Sheikh Md. Selim Al Din, Utpal Chandra Ray, Salek Ahmed Sajib, Md. Salim Khan |
| EPI_ISL_498466, EPI_ISL_498467 | National Institute of Laboratory Medicine and Referral Center | Genomic Research Lab, BCSIR | Mahmuda Yeasmin, Abu Sayeed Mohammad Mahmud, Mohammad Samir Uzzaman, Eshrar Osman, Md. Ahasan Habib, Shahina Akter, Tanjina Akhter Banu, Md. Murshed Hasan Sarkar, Barna Goswami, Iffat Jahan, Md. Saddam Hossain, Tasnim Nafisa, Md. Maruf Ahmed Molla, Asish Kumar Ghosh, A. K. M. Shamsuzzaman, Sheikh Md. Selim Al Din, Utpal Chandra Ray, Salek Ahmed Sajib, Md. Salim Khan |
| EPI_ISL_498468, EPI_ISL_498472, EPI_ISL_498473, EPI_ISL_498477, EPI_ISL_498478, EPI_ISL_498480, EPI_ISL_498483, EPI_ISL_498485, EPI_ISL_498486, EPI_ISL_498489, EPI_ISL_498490, EPI_ISL_498491, EPI_ISL_498492, EPI_ISL_498493, EPI_ISL_498494, EPI_ISL_498495, EPI_ISL_498496, EPI_ISL_498497, EPI_ISL_498498, EPI_ISL_498499, EPI_ISL_498500, EPI_ISL_498501, EPI_ISL_498502, EPI_ISL_498503, EPI_ISL_498505, EPI_ISL_498506, EPI_ISL_498510, EPI_ISL_498511, EPI_ISL_498512, EPI_ISL_498513, EPI_ISL_498514, EPI_ISL_498515, EPI_ISL_498518, EPI_ISL_498520, EPI_ISL_498523, EPI_ISL_498525, EPI_ISL_498526, EPI_ISL_498529, EPI_ISL_498531, EPI_ISL_498537, EPI_ISL_498540, EPI_ISL_498542, EPI_ISL_498543, EPI_ISL_498544, EPI_ISL_498545, EPI_ISL_498546, EPI_ISL_498547, EPI_ISL_498548 | | | |
| see above | ACT Pathology | Schwessinger Lab | Ashley Jones, Benjamin Schwessinger, Robert Lanfear, Robyn N Hall, Megan McDonald, Ming-Dao Chia, Kevin Murray, Craig Kennedy, Karina Kennedy |
| EPI_ISL_498549, EPI_ISL_498550 | National Institute of Laboratory Medicine and Referral Center | Genomic Research Lab, BCSIR | Asish Kumar Ghosh, Abu Sayeed Mohammad Mahmud, Mohammad Samir Uzzaman, Eshrar Osman, Md. Ahasan Habib, Shahina Akter, Tanjina Akhter Banu, Md. Murshed Hasan Sarkar, Barna Goswami, Iffat Jahan, Md. Saddam Hossain, Tasnim Nafisa, Md. Maruf Ahmed Molla, Mahmuda Yeasmin, A. K. M. Shamsuzzaman, Sheikh Md. Selim Al Din, Utpal Chandra Ray, Salek Ahmed Sajib, Md. Salim Khan |
| EPI_ISL_498551, EPI_ISL_498552, EPI_ISL_498554 | Lebanese American University | Lebanese American University | Abi Habib,W., Abdallah,J., El Shesheny,R., Mokhbat,J., Webby,R.J., Goldstein,J. and Kayali,G. |
| EPI_ISL_498558, EPI_ISL_498559, EPI_ISL_498560, EPI_ISL_498561 | Laboratory of Molecular Virology International Center for Genetic Engineering and Biotechnology (ICGEB) | ARGO Open Lab Platform for Genome Sequencing | Licastro D, Rajasekharan S, Dal Monego S, Segat L, D'Agaro P, Marcello A |
| EPI_ISL_498562 | Laboratory of Molecular Virology International Center for Genetic Engineering and Biotechnology (ICGEB) | ARGO Open Lab Platform for Genome Sequencing | Licastro D, Rajasekharan S, Dal Monego S, Segat L, D'Agaro P, Marcello A |
| EPI_ISL_498563 | Laboratory of Molecular Virology International Center for Genetic Engineering and Biotechnology (ICGEB) | ARGO Open Lab Platform for Genome Sequencing | Licastro D, Rajasekharan S, Dal Monego S, Segat L, D'Agaro P, Marcello A |
| EPI_ISL_498564, EPI_ISL_498565, EPI_ISL_498566, EPI_ISL_498567, EPI_ISL_498568, EPI_ISL_498569, EPI_ISL_498570, EPI_ISL_498571, EPI_ISL_498581, EPI_ISL_498582, EPI_ISL_498583, EPI_ISL_498584, EPI_ISL_498585, EPI_ISL_498586, EPI_ISL_498587, EPI_ISL_498588, EPI_ISL_498589, EPI_ISL_498590, EPI_ISL_498591, EPI_ISL_498592, EPI_ISL_498593, EPI_ISL_498594, EPI_ISL_498595, EPI_ISL_498601, EPI_ISL_498602, EPI_ISL_498603, EPI_ISL_498604, EPI_ISL_498605, EPI_ISL_498608, EPI_ISL_498609, EPI_ISL_498610, EPI_ISL_498611, EPI_ISL_498613, EPI_ISL_498615, EPI_ISL_498616, EPI_ISL_498618 | | | |
| see above | National Public Health Laboratory, National Centre for Infectious Diseases | National Public Health Laboratory, National Centre for Infectious Diseases | Mak TM, Octavia S, Zhou Z, Chavatte JM, Cui L, Lin RTP |
| EPI_ISL_498620, EPI_ISL_498621, EPI_ISL_498622, EPI_ISL_498623, EPI_ISL_498624, EPI_ISL_498626, EPI_ISL_498627 | Viollier AG | Department of Biosystems Science and Engineering, ETH Zürich | Christian Beisel, Sarah Nadeau, Ivan Topolsky, Pedro Ferreira, Philipp Jablonski, Susana Posada-Céspedes, Tobias Schär, Ina Nissen, Natascha Santacroce, Elodie Burcklen, Christiane Beckmann, Maurice Redondo, Olivier Kobel, Christoph Noppen, Sophie Seidel, Noemie Santamaria de Souza, Niko Beerenwinkel, Tanja Stadler |
| EPI_ISL_498628, EPI_ISL_498629 | Department of Clinical Microbiology | GIGA Medical Genomics | Keith Durkin, Maria Artesi, Sébastien Bontems, Raphaël Boreux, Cécile Meex, Axelle Chaslain, Céline Fombellida-Lopez, Pierrette Melin, Marie-Pierre Hayette, Vincent Bours. |
| EPI_ISL_498630, EPI_ISL_498631, EPI_ISL_498632, EPI_ISL_498633, EPI_ISL_498634, EPI_ISL_498635, EPI_ISL_498637, EPI_ISL_498638, EPI_ISL_498639, EPI_ISL_498640, EPI_ISL_498641, EPI_ISL_498642, EPI_ISL_498643, EPI_ISL_498644, EPI_ISL_498645, EPI_ISL_498646, EPI_ISL_498647, EPI_ISL_498648, EPI_ISL_498649, EPI_ISL_498650, EPI_ISL_498651, EPI_ISL_498652, EPI_ISL_498653, EPI_ISL_498654, EPI_ISL_498655, EPI_ISL_498656, EPI_ISL_498657, EPI_ISL_498658, EPI_ISL_498659, EPI_ISL_498660, EPI_ISL_498661, EPI_ISL_498662, EPI_ISL_498663, EPI_ISL_498664, EPI_ISL_498665, EPI_ISL_498666, EPI_ISL_498667, EPI_ISL_498668, EPI_ISL_498670, EPI_ISL_498671, EPI_ISL_498687 | | | |
| see above | Utah Public Health Laboratory | Utah Public Health Laboratory | Heidi Butz, Erin Young, Kelly Oakeson |
| EPI_ISL_498691, EPI_ISL_498692, EPI_ISL_498693, EPI_ISL_498694 | National Institute for Viral Disease Control and Prevention, China CDC | National Institute for Viral Disease Control and Prevention, China CDC | Xiang Zhao,LingLing Mao,Yao Meng,Zhixiao Chen,Yuchao Wu,Yong ZhangBo ZhijianJianqun Zhang,Yang Song,Dayan Wang,WenQing YaoWenbo Xu |
| EPI_ISL_498695, EPI_ISL_498696, EPI_ISL_498697, EPI_ISL_498698, EPI_ISL_498699, EPI_ISL_498700, EPI_ISL_498701, EPI_ISL_498702, EPI_ISL_498703, EPI_ISL_498704, EPI_ISL_498705, EPI_ISL_498706, EPI_ISL_498707, EPI_ISL_498708, EPI_ISL_498709, EPI_ISL_498710, EPI_ISL_498711, EPI_ISL_498712, EPI_ISL_498713, EPI_ISL_498714, EPI_ISL_498715, EPI_ISL_498716, EPI_ISL_498717, EPI_ISL_498718, EPI_ISL_498719, EPI_ISL_498720, EPI_ISL_498721, EPI_ISL_498722, EPI_ISL_498723, EPI_ISL_498724, EPI_ISL_498725, EPI_ISL_498726, EPI_ISL_498727, EPI_ISL_498728, EPI_ISL_498729, EPI_ISL_498730, EPI_ISL_498731, EPI_ISL_498732, EPI_ISL_498733, EPI_ISL_498734, EPI_ISL_498735, EPI_ISL_498736, EPI_ISL_498737, EPI_ISL_498738, EPI_ISL_498739, EPI_ISL_498740, EPI_ISL_498741, EPI_ISL_498742, EPI_ISL_498743, EPI_ISL_498744, EPI_ISL_498745, EPI_ISL_498746, EPI_ISL_498747 | | | |

| | | | |
|---|---|---|---|
| see above | Quest Diagnostics | Quest Diagnostics | Rosenthal,S.H., Gerasimova,A., Kagan,R.M. and Owen, R. |
| EPI_ISL_498748, EPI_ISL_498749, EPI_ISL_498750 | Pathology West - NSW Health Pathology | NSW Health Pathology - Institute of Clinical Pathology and Medical Research; Westmead Hospital; University of Sydney | CIDM-PH et al. |
| EPI_ISL_498751 | South Eastern Area Laboratory Services (SEALS) | NSW Health Pathology - Institute of Clinical Pathology and Medical Research; Westmead Hospital; University of Sydney | CIDM-PH et al. |
| EPI_ISL_498761, EPI_ISL_498762 | Pathology West - NSW Health Pathology | NSW Health Pathology - Institute of Clinical Pathology and Medical Research; Westmead Hospital; University of Sydney | CIDM-PH et al. |
| EPI_ISL_498763, EPI_ISL_498764 | Sydney South West Pathology Service (SSWPS) - Liverpool Hospital - NSW Health Pathology | NSW Health Pathology - Institute of Clinical Pathology and Medical Research; Westmead Hospital; University of Sydney | CIDM-PH et al. |
| EPI_ISL_498765 | Sydney South West Pathology Service (SSWPS) - Concord Repatriation General Hospital - NSW Health Pathology | NSW Health Pathology - Institute of Clinical Pathology and Medical Research; Westmead Hospital; University of Sydney | CIDM-PH et al. |
| EPI_ISL_498766, EPI_ISL_498767 | Pathology West - NSW Health Pathology | NSW Health Pathology - Institute of Clinical Pathology and Medical Research; Westmead Hospital; University of Sydney | CIDM-PH et al. |
| EPI_ISL_498768 | South Eastern Area Laboratory Services (SEALS) | NSW Health Pathology - Institute of Clinical Pathology and Medical Research; Westmead Hospital; University of Sydney | CIDM-PH et al. |
| EPI_ISL_498769, EPI_ISL_498770, EPI_ISL_498771, EPI_ISL_498772 | Laverty Pathology | NSW Health Pathology - Institute of Clinical Pathology and Medical Research; Westmead Hospital; University of Sydney | CIDM-PH et al. |
| EPI_ISL_498773 | Douglass Hanly Moir Pathology | NSW Health Pathology - Institute of Clinical Pathology and Medical Research; Westmead Hospital; University of Sydney | CIDM-PH et al. |
| EPI_ISL_498774, EPI_ISL_498775 | Histopath | NSW Health Pathology - Institute of Clinical Pathology and Medical Research; Westmead Hospital; University of Sydney | CIDM-PH et al. |
| EPI_ISL_498776, EPI_ISL_498777 | Pathology West - NSW Health Pathology | NSW Health Pathology - Institute of Clinical Pathology and Medical Research; Westmead Hospital; University of Sydney | CIDM-PH et al. |
| EPI_ISL_498778 | Sydney South West Pathology Service (SSWPS) - Liverpool Hospital - NSW Health Pathology | NSW Health Pathology - Institute of Clinical Pathology and Medical Research; Westmead Hospital; University of Sydney | CIDM-PH et al. |
| EPI_ISL_498779, EPI_ISL_498780 | Pathology West - NSW Health Pathology | NSW Health Pathology - Institute of Clinical Pathology and Medical Research; Westmead Hospital; University of Sydney | CIDM-PH et al. |
| EPI_ISL_498781, EPI_ISL_498782 | Sydney South West Pathology Service (SSWPS) - Liverpool Hospital - NSW Health Pathology | NSW Health Pathology - Institute of Clinical Pathology and Medical Research; Westmead Hospital; University of Sydney | CIDM-PH et al. |
| EPI_ISL_498783, EPI_ISL_498784, EPI_ISL_498785, EPI_ISL_498786, EPI_ISL_498787, EPI_ISL_498788, EPI_ISL_498789, EPI_ISL_498790, EPI_ISL_498791, EPI_ISL_498792 | National Institute of Laboratory Medicine and Referral Center | Genomic Research Lab, BCSIR | Md. Saddam Hossain, Abu Sayeed Mohammad Mahmud, Mohammad Samir Uzzaman, Eshrar Osman, Md. Ahasan Habib, Shahina Akter, Tanjina Akhter Banu, Md. Murshed Hasan Sarkar, Barna Goswami, Iffat Jahan, Tasnim Nafisa, Md. Maruf Ahmed Molla, Mahmuda Yeasmin, Asish Kumar Ghosh, A. K. M. Shamsuzzaman, Sheikh Md. Selim Al Din, Utpal Chandra Ray, Salek Ahmed Sajib, Md. Salim Khan |
| EPI_ISL_498793, EPI_ISL_498794, EPI_ISL_498795, EPI_ISL_498796, EPI_ISL_498797, EPI_ISL_498800, EPI_ISL_498801, EPI_ISL_498802, EPI_ISL_498804, EPI_ISL_498805, EPI_ISL_498806 | | | |
| see above | National Institute of Laboratory Medicine and Referral Center | Genomic Research Lab, BCSIR | Md. Murshed Hasan Sarkar, Abu Sayeed Mohammad Mahmud, Mohammad Samir Uzzaman, Eshrar Osman, Md. Ahasan Habib, Shahina Akter, Tanjina Akhter Banu, Barna Goswami, Iffat Jahan, Md. Saddam Hossain, Tasnim Nafisa, Md. Maruf Ahmed Molla, Mahmuda Yeasmin, Asish Kumar Ghosh, A. K. M. Shamsuzzaman, Sheikh Md. Selim Al Din, Utpal Chandra Ray, Salek Ahmed Sajib, Md. Salim Khan |
| EPI_ISL_498808, EPI_ISL_498811, EPI_ISL_498814, EPI_ISL_498815, EPI_ISL_498816, EPI_ISL_498817, EPI_ISL_498818, EPI_ISL_498830, EPI_ISL_498892, EPI_ISL_498932, EPI_ISL_498933 | | | |
| see above | National Institute of Laboratory Medicine and Referral Center | Genomic Research Lab, BCSIR | Abu Sayeed Mohammad Mahmud, Mohammad Samir Uzzaman, Eshrar Osman, Md. Ahasan Habib, Shahina Akter, Tanjina Akhter Banu, Md. Murshed Hasan Sarkar, Barna Goswami, Iffat Jahan, Md. Saddam Hossain, Tasnim Nafisa, Md. Maruf Ahmed Molla, Mahmuda Yeasmin, Asish Kumar Ghosh, A. K. M. Shamsuzzaman, Sheikh Md. Selim Al Din, Utpal Chandra Ray, Salek Ahmed Sajib, Md. Salim Khan |
| EPI_ISL_499083 | Instituto de Virologia "Dr. J. M. Vanella", Facultad de Ciencias Medicas, Universidad Nacional de Cordoba. Laboratorio Central de la Provincia de Cordoba, Argentina. Ministerio de Salud de la provincia de Cordoba, Argentina. | Laboratorio de Virología, Hospital de Niños Ricardo Gutiérrez, CABA, Argentina. | Sandra Gallego, Brenda Konigheim, Sebastian Blanco, Lorena Spinsanti, Javier Aguilar, Adrian Diaz, Gonzalo Castro, Gabriela Barbas, Mercedes Nabaes, Stephanie Goya, Monica Natale, Silvina Lusso, Mariana Viegas. |
| EPI_ISL_499266, EPI_ISL_499267, EPI_ISL_499269 | Queens Medical Centre, Clinical Microbiology Department / DeepSeq Nottingham | COVID-19 Genomics UK (COG-UK) Consortium | Gemma Clark, Wendy Smith, Manjinder Khakh, Vicki M Fleming, Michelle M Lister, Hannah Howson-Wells, Jonathan Ball, Patrick McClure, Joseph Chappell, Theocharis Tsoleridis, Nadine Holmes, Matthew Carlisle, Christopher Moore, Fei Sang, Johnny Debebe, Victoria Wright, Matthew Loose |
| EPI_ISL_499272, EPI_ISL_499278, EPI_ISL_499281, EPI_ISL_499282, EPI_ISL_499283, EPI_ISL_499285, EPI_ISL_499288, EPI_ISL_499289, EPI_ISL_499294, EPI_ISL_499295, EPI_ISL_499296, EPI_ISL_499297, EPI_ISL_499298, EPI_ISL_499300, EPI_ISL_499302, EPI_ISL_499303, EPI_ISL_499304, EPI_ISL_499306, EPI_ISL_499307, EPI_ISL_499308, EPI_ISL_499309, EPI_ISL_499310, EPI_ISL_499311, EPI_ISL_499312, EPI_ISL_499314, EPI_ISL_499315, EPI_ISL_499316, EPI_ISL_499318, EPI_ISL_499322, EPI_ISL_499323, EPI_ISL_499324, EPI_ISL_499325, EPI_ISL_499327, EPI_ISL_499328, EPI_ISL_499329 | | | |
| see above | Centre for Enzyme Innovation, University of Portsmouth / Translational Research Laboratory, Portsmouth Hospitals NHS Trust | COVID-19 Genomics UK (COG-UK) Consortium | Angela Beckett,Yann Bourgeois,Garry Scarlett,Sharon Glaysher,Scott Elliott,Kelly Bicknell,Robert Impey,Allyson Lloyd,Sarah Wyllie,Ethan Butcher,Anoop Chauhan,Samuel Robson |
| EPI_ISL_499330, EPI_ISL_499332, EPI_ISL_499336, EPI_ISL_499344, EPI_ISL_499348 | Virology Department, Sheffield Teaching Hospitals NHS Foundation Trust/Department of Infection, Immunity and Cardiovascular Disease, The Medical School, University of Sheffield | COVID-19 Genomics UK (COG-UK) Consortium | Thushan de Silva, Matthew Parker, Nikki Smith, Adri Angyal, Rebecca Brown, Luke Green, Rachel Tucker, Paul Parsons, Danielle Groves, Katie Johnson, Laura Carrilero, Alex Keeley, Dave Partridge, Matthew Wyles, Benjamin Lindsey, Mehmet Yavuz, Mohammad Raza, Cariad Evans |
| EPI_ISL_499354 | West of Scotland Specialist Virology Centre, NHSGGC / MRC-University of Glasgow Centre for Virus Research | COVID-19 Genomics UK (COG-UK) Consortium | Ana da Silva Filipe, Natasha Johnson, Kathy Smollett, Daniel Mair, Stephen Carmichael, Lily Tong, Jenna Nichols, Elihu Aranday-Cortes, Kirstyn Brunker, Yasmin Parr, Alice Broos, Kyriaki Nomikou; Sarah McDonald, Marc Niebel, Patawee Asamaphan; Richard Orton, Joseph Hughes, Sreenu Vattipally, David L Robertson; Alasdair MacLean, Rory Gunson; Kathy Li, Natasha Jesudason, Rajiv Shah, James Shepherd, Antonia Ho, Emma Thomson |
| EPI_ISL_499355, EPI_ISL_499356, EPI_ISL_499357, EPI_ISL_499358, EPI_ISL_499360, EPI_ISL_499361, EPI_ISL_499362, EPI_ISL_499363, EPI_ISL_499364, EPI_ISL_499366, EPI_ISL_499367, EPI_ISL_499368, EPI_ISL_499369, EPI_ISL_499370, EPI_ISL_499372, EPI_ISL_499374, EPI_ISL_499375, EPI_ISL_499376, EPI_ISL_499377, EPI_ISL_499378, EPI_ISL_499379, EPI_ISL_499380, EPI_ISL_499381, EPI_ISL_499382, EPI_ISL_499384, EPI_ISL_499385, EPI_ISL_499386, EPI_ISL_499387, EPI_ISL_499388, EPI_ISL_499389, EPI_ISL_499390, EPI_ISL_499393, EPI_ISL_499395, EPI_ISL_499396 | | | |
| see above | Originating lab: Wales Specialist Virology Centre Sequencing lab: Pathogen Genomics Unit | COVID-19 Genomics UK (COG-UK) Consortium | Catherine Moore, Johnathan Evans, Laura Gifford, Malorie Perry, Simon Cottrell, Angela Marchbank, Alec Birchley, Alexander Adams, Amy Gaskin, Bree Gatica-Wilcox, Jason Coombes, Joel Southgate, Lauren Gilbert, Lee Graham, Nicole Pacchiarini, Sara Kumziene-Summerhayes, Sarah Taylor, Sophie Jones, Sara Rey, Matthew Bull, Joanne Watkins, Sally Corden, Tom Connor |
| EPI_ISL_499398, EPI_ISL_499399, EPI_ISL_499400, EPI_ISL_499401, EPI_ISL_499415, EPI_ISL_499416, EPI_ISL_499417, EPI_ISL_499418, EPI_ISL_499419, EPI_ISL_499420, EPI_ISL_499421, EPI_ISL_499422, EPI_ISL_499423, EPI_ISL_499424, EPI_ISL_499425, EPI_ISL_499426, EPI_ISL_499427, EPI_ISL_499428, EPI_ISL_499429, EPI_ISL_499430, EPI_ISL_499431, EPI_ISL_499432, EPI_ISL_499433, EPI_ISL_499434, EPI_ISL_499435, EPI_ISL_499436, EPI_ISL_499437, EPI_ISL_499438, EPI_ISL_499439, EPI_ISL_499440, EPI_ISL_499441, EPI_ISL_499442, EPI_ISL_499443, EPI_ISL_499444, EPI_ISL_499445, EPI_ISL_499446, EPI_ISL_499447, EPI_ISL_499448, EPI_ISL_499449, EPI_ISL_499450, EPI_ISL_499451, EPI_ISL_499452, EPI_ISL_499453, EPI_ISL_499454, EPI_ISL_499455, EPI_ISL_499456, EPI_ISL_499457, EPI_ISL_499458, EPI_ISL_499459 | | | |
| see above | Wales Specialist Virology Centre Sequencing lab: Pathogen Genomics Unit | COVID-19 Genomics UK (COG-UK) Consortium | Catherine Moore, Johnathan Evans, Laura Gifford, Malorie Perry, Simon Cottrell, Angela Marchbank, Alec Birchley, Alexander Adams, Amy Gaskin, Bree Gatica-Wilcox, Jason Coombes, Joel Southgate, Lauren Gilbert, Lee Graham, Nicole Pacchiarini, Sara Kumziene-Summerhayes, Sarah Taylor, Sophie Jones, Sara Rey, Matthew Bull, Joanne Watkins, Sally Corden, Tom Connor |
| EPI_ISL_499461, EPI_ISL_499462, EPI_ISL_499463, EPI_ISL_499464, EPI_ISL_499465, EPI_ISL_499466, EPI_ISL_499467, EPI_ISL_499468, EPI_ISL_499469, EPI_ISL_499640, EPI_ISL_499641, EPI_ISL_499642, EPI_ISL_499643, EPI_ISL_499644, EPI_ISL_499645, EPI_ISL_499646, EPI_ISL_499647, EPI_ISL_499648, EPI_ISL_499649, EPI_ISL_499651, EPI_ISL_499653, EPI_ISL_499654, EPI_ISL_499655, EPI_ISL_499656, EPI_ISL_499657, EPI_ISL_499658, EPI_ISL_499659, EPI_ISL_499660, EPI_ISL_499661, EPI_ISL_499662, EPI_ISL_499663, EPI_ISL_499664, EPI_ISL_499665, EPI_ISL_499666, EPI_ISL_499667, EPI_ISL_499668, EPI_ISL_499669, EPI_ISL_499670, EPI_ISL_499671, EPI_ISL_499672, EPI_ISL_499673, EPI_ISL_499674, EPI_ISL_499675, EPI_ISL_499676, EPI_ISL_499677, EPI_ISL_499678, EPI_ISL_499679, EPI_ISL_499680, EPI_ISL_499681, EPI_ISL_499682, EPI_ISL_499683, EPI_ISL_499684, EPI_ISL_499685, EPI_ISL_499686, EPI_ISL_499687, EPI_ISL_499688, EPI_ISL_499689, EPI_ISL_499690, EPI_ISL_499691, EPI_ISL_499692, EPI_ISL_499693, EPI_ISL_499694, EPI_ISL_499695, EPI_ISL_499696, EPI_ISL_499697, EPI_ISL_499698, EPI_ISL_499699, EPI_ISL_499700, EPI_ISL_499701, EPI_ISL_499702, EPI_ISL_499703, EPI_ISL_499704, EPI_ISL_499705, EPI_ISL_499706, EPI_ISL_499707, EPI_ISL_499708, EPI_ISL_499709, EPI_ISL_499710, EPI_ISL_499711, EPI_ISL_499712, EPI_ISL_499713, EPI_ISL_499714, EPI_ISL_499715, EPI_ISL_499716, EPI_ISL_499717, EPI_ISL_499718, EPI_ISL_499719, EPI_ISL_499720, EPI_ISL_499721, EPI_ISL_499722, EPI_ISL_499723, EPI_ISL_499724, EPI_ISL_499725, EPI_ISL_499726, EPI_ISL_499727, EPI_ISL_499728, EPI_ISL_499729, EPI_ISL_499730, EPI_ISL_499731, EPI_ISL_499732, EPI_ISL_499733, EPI_ISL_499734, EPI_ISL_499735, EPI_ISL_499736, EPI_ISL_499737, EPI_ISL_499738, EPI_ISL_499739, EPI_ISL_499740, EPI_ISL_499741, EPI_ISL_499742, EPI_ISL_499743, EPI_ISL_499744, EPI_ISL_499745, EPI_ISL_499746, EPI_ISL_499747, EPI_ISL_499748, EPI_ISL_499749, EPI_ISL_499750, EPI_ISL_499751, EPI_ISL_499752, EPI_ISL_499753, EPI_ISL_499754, EPI_ISL_499755, EPI_ISL_499756, EPI_ISL_499757, EPI_ISL_499758, EPI_ISL_499759, EPI_ISL_499760, EPI_ISL_499761, EPI_ISL_499762, EPI_ISL_499763, EPI_ISL_499764, EPI_ISL_499765, EPI_ISL_499766, EPI_ISL_499767, EPI_ISL_499768, EPI_ISL_499769, EPI_ISL_499770, EPI_ISL_499771, EPI_ISL_499772, EPI_ISL_499773, EPI_ISL_499774, EPI_ISL_499775, EPI_ISL_499776 | | | |
| see above | Liverpool Clinical Laboratories | COVID-19 Genomics UK (COG-UK) Consortium | Sam Haldenby, Anita Lucaci, Steve Paterson, Julian Hiscox, Alistair Darby, M Almsaud, A Alrezaihi, Muhannad Alruwaili, Stuart D Armstrong, Jones Benjamin, Eleanor G Bentley, Anu Chawla, Jordan J Clark, Angela Cowell, Richard Eccles, Isabel García-Dorival, Matthew Gemmell, Alessandro Gerada, PKF Gilmore, Richard Gregory, Ximeng Han, Catherine Hartley, Margaret Hughes, Miren Iturriza-Gomara, James Johnson, L Luu, Jenifer Manson, Charlotte Nelson, Elaine O'Toole, Cassie Olateju, Rebekah Penrice-Randal , Lucille Rainbow, N.P Randle, Trevor Ian Robinson, Parul Sharma, Ghada T Shawli, James P Stewart, Neil Swainston, Ecaterina Vamos, Joanne Watts, Mark Whitehead |
| EPI_ISL_499777, EPI_ISL_499778, EPI_ISL_499779, EPI_ISL_499780, EPI_ISL_499781, EPI_ISL_499782, EPI_ISL_499783, EPI_ISL_499784, EPI_ISL_499785, EPI_ISL_499786, EPI_ISL_499787, EPI_ISL_499788, EPI_ISL_499789, EPI_ISL_499790, EPI_ISL_499791, EPI_ISL_499792, EPI_ISL_499793, EPI_ISL_499794, EPI_ISL_499795, EPI_ISL_499796, EPI_ISL_499797, EPI_ISL_499798, EPI_ISL_499799, EPI_ISL_499800, EPI_ISL_499801, EPI_ISL_499802, EPI_ISL_499803, EPI_ISL_499804, EPI_ISL_499805, EPI_ISL_499806, EPI_ISL_499807 | | | |
| see above | Northumbria University / South Tees Hospitals NHS Foundation Trust / North Cumbria Integrated Care NHS Foundation Trust / North Tees and Hartlepool NHS Foundation Trust / Newcastle Hospitals NHS Foundation Trust | COVID-19 Genomics UK (COG-UK) Consortium | Darren L Smith,Andrew Nelson,Matthew Bashton,Greg R Young,Joshua Loh,John Allan,Mohammad A Tariq,Giles S Holt,Gary Black,Wen C Yew,Lynn Dover,Paul Baker,Steve Liggett,Sarah Essex,Jane Greenaway,Debra Padgett,Clive Graham,Garren Scott,Edward Barton,Emma Swindells,Brendan Payne,Jennifer Collins,Yusri Taha,Gary Eltringham |
| EPI_ISL_499808, EPI_ISL_499809 | Queens Medical Centre, Clinical Microbiology Department / DeepSeq Nottingham | COVID-19 Genomics UK (COG-UK) Consortium | Gemma Clark, Wendy Smith, Manjinder Khakh, Vicki M Fleming, Michelle M Lister, Hannah Howson-Wells, Jonathan Ball, Patrick McClure, Joseph Chappell, Theocharis Tsoleridis, Nadine Holmes, Matthew Carlisle, Christopher Moore, Fei Sang, Johnny Debebe, Victoria Wright, Matthew Loose |
| EPI_ISL_499810 | University Hospitals Of Leicester NHS Trust and DeepSeq Nottingham | COVID-19 Genomics UK (COG-UK) Consortium | Christopher Holmes, Paul Bird, Thomas Helmer, Karlie Fallon, Julian Tang, Jonathan Ball, Patrick McClure, Joeseph Chappell, Nadine Holmes, Matthew Carlisle, Christopher Moore, Fei Sang, Johnny Debebe, Victoria Wright, Matthew Loose |
| EPI_ISL_499811, EPI_ISL_499812, EPI_ISL_499813, EPI_ISL_499814, EPI_ISL_499815, EPI_ISL_499816, EPI_ISL_499817, EPI_ISL_499818, EPI_ISL_499819, EPI_ISL_499820, EPI_ISL_499821, EPI_ISL_499822, EPI_ISL_499823, EPI_ISL_499824 | | | |
| see above | Liverpool Clinical Laboratories | COVID-19 Genomics UK (COG-UK) Consortium | Sam Haldenby, Anita Lucaci, Steve Paterson, Julian Hiscox, Alistair Darby, M Almsaud, A Alrezaihi, Muhannad Alruwaili, Stuart D Armstrong, Jones Benjamin, Eleanor G Bentley, Anu Chawla, Jordan J |

| | | | |
|---|---|---|---|
| | | | Clark, Angela Cowell, Richard Eccles, Isabel García-Dorival, Matthew Gemmell, Alessandro Gerada, PKF Gilmore, Richard Gregory, Ximeng Han, Catherine Hartley, Margaret Hughes, Miren Iturriza-Gomara, James Johnson, L Luu, Jenifer Manson, Charlotte Nelson, Elaine O'Toole, Cassie Olateju, Rebekah Penrice-Randal , Lucille Rainbow, N.P Randle, Trevor Ian Robinson, Parul Sharma, Ghada T Shawli, James P Stewart, Neil Swainston, Ecaterina Vamos, Joanne Watts, Mark Whitehead |
| EPI_ISL_499968, EPI_ISL_499969, EPI_ISL_499970, EPI_ISL_499971, EPI_ISL_499972, EPI_ISL_499973 | Department of Pathology, University of Cambridge | COVID-19 Genomics UK (COG-UK) Consortium | Luke W Meredith, M. Estée Török, Myra Hosmillo, William L. Hamilton, Martin D. Curran, Theresa Feltwell, Grant Hall, Anna Yakovleva, Fahad A Khokhar, Charlotte J. Houldcroft, Laura G Caller, Aminu S. Jahun, Sarah L. Caddy, Yasmin Chaudhry, Malte Pinckert, Ian Goodfellow |
| EPI_ISL_499974, EPI_ISL_499975, EPI_ISL_499976, EPI_ISL_499977, EPI_ISL_499978, EPI_ISL_499979, EPI_ISL_499980, EPI_ISL_499981, EPI_ISL_499982, EPI_ISL_499983 | University Hospitals Of Leicester NHS Trust and DeepSeq Nottingham | COVID-19 Genomics UK (COG-UK) Consortium | Christopher Holmes, Paul Bird, Thomas Helmer, Karlie Fallon, Julian Tang, Jonathan Ball, Patrick McClure, Joeseph Chappell, Nadine Holmes, Matthew Carlisle, Christopher Moore, Fei Sang, Johnny Debebe, Victoria Wright, Matthew Loose |
| EPI_ISL_499984, EPI_ISL_499985, EPI_ISL_499986, EPI_ISL_499987, EPI_ISL_499988, EPI_ISL_499989, EPI_ISL_499990, EPI_ISL_499991, EPI_ISL_499992, EPI_ISL_499993, EPI_ISL_499994, EPI_ISL_499995, EPI_ISL_499996, EPI_ISL_499997, EPI_ISL_499998, EPI_ISL_499999, EPI_ISL_500000, EPI_ISL_500001, EPI_ISL_500002, EPI_ISL_500003, EPI_ISL_500004, EPI_ISL_500005, EPI_ISL_500006, EPI_ISL_500007, EPI_ISL_500008, EPI_ISL_500009, EPI_ISL_500010, EPI_ISL_500011, EPI_ISL_500012, EPI_ISL_500013, EPI_ISL_500014, EPI_ISL_500015, EPI_ISL_500016, EPI_ISL_500017, EPI_ISL_500018, EPI_ISL_500019, EPI_ISL_500020, EPI_ISL_500021, EPI_ISL_500022, EPI_ISL_500023, EPI_ISL_500024, EPI_ISL_500025, EPI_ISL_500026, EPI_ISL_500027, EPI_ISL_500028, EPI_ISL_500029, EPI_ISL_500030, EPI_ISL_500031, EPI_ISL_500032, EPI_ISL_500033, EPI_ISL_500034, EPI_ISL_500035, EPI_ISL_500036, EPI_ISL_500037, EPI_ISL_500038, EPI_ISL_500039, EPI_ISL_500040, EPI_ISL_500041, EPI_ISL_500042, EPI_ISL_500043, EPI_ISL_500044, EPI_ISL_500045, EPI_ISL_500046, EPI_ISL_500047, EPI_ISL_500048, EPI_ISL_500049, EPI_ISL_500050, EPI_ISL_500051, EPI_ISL_500052, EPI_ISL_500053, EPI_ISL_500054, EPI_ISL_500055, EPI_ISL_500056, EPI_ISL_500057, EPI_ISL_500058, EPI_ISL_500059, EPI_ISL_500060, EPI_ISL_500061, EPI_ISL_500062, EPI_ISL_500063, EPI_ISL_500064, EPI_ISL_500065, EPI_ISL_500066, EPI_ISL_500067, EPI_ISL_500068, EPI_ISL_500069, EPI_ISL_500070, EPI_ISL_500071, EPI_ISL_500072, EPI_ISL_500073, EPI_ISL_500074, EPI_ISL_500075, EPI_ISL_500076, EPI_ISL_500077, EPI_ISL_500078, EPI_ISL_500079, EPI_ISL_500080, EPI_ISL_500081, EPI_ISL_500082, EPI_ISL_500083, EPI_ISL_500084, EPI_ISL_500085, EPI_ISL_500086, EPI_ISL_500087, EPI_ISL_500088, EPI_ISL_500089, EPI_ISL_500090, EPI_ISL_500091, EPI_ISL_500092, EPI_ISL_500093, EPI_ISL_500094, EPI_ISL_500095, EPI_ISL_500096, EPI_ISL_500097, EPI_ISL_500098, EPI_ISL_500099, EPI_ISL_500100, EPI_ISL_500101, EPI_ISL_500102, EPI_ISL_500103, EPI_ISL_500104, EPI_ISL_500105, EPI_ISL_500106, EPI_ISL_500107, EPI_ISL_500108, EPI_ISL_500109, EPI_ISL_500110, EPI_ISL_500111, EPI_ISL_500112, EPI_ISL_500113, EPI_ISL_500114, EPI_ISL_500115, EPI_ISL_500116, EPI_ISL_500117, EPI_ISL_500118, EPI_ISL_500119, EPI_ISL_500120, EPI_ISL_500121, EPI_ISL_500122, EPI_ISL_500123, EPI_ISL_500124, EPI_ISL_500125, EPI_ISL_500126, EPI_ISL_500127, EPI_ISL_500128, EPI_ISL_500129, EPI_ISL_500130, EPI_ISL_500131, EPI_ISL_500132, EPI_ISL_500133, EPI_ISL_500134, EPI_ISL_500135, EPI_ISL_500136, EPI_ISL_500137, EPI_ISL_500138, EPI_ISL_500139, EPI_ISL_500140, EPI_ISL_500141, EPI_ISL_500142, EPI_ISL_500143, EPI_ISL_500144, EPI_ISL_500145, EPI_ISL_500146, EPI_ISL_500147, EPI_ISL_500148, EPI_ISL_500149, EPI_ISL_500150, EPI_ISL_500151, EPI_ISL_500152, EPI_ISL_500153, EPI_ISL_500154, EPI_ISL_500155, EPI_ISL_500156 | | | |
| see above | Liverpool Clinical Laboratories | COVID-19 Genomics UK (COG-UK) Consortium | Sam Haldenby, Anita Lucaci, Steve Paterson, Julian Hiscox, Alistair Darby, M Almsaud, A Alrezaihi, Muhannad Alruwaili, Stuart D Armstrong, Jones Benjamin, Eleanor G Bentley, Anu Chawla, Jordan J Clark, Angela Cowell, Richard Eccles, Isabel García-Dorival, Matthew Gemmell, Alessandro Gerada, PKF Gilmore, Richard Gregory, Ximeng Han, Catherine Hartley, Margaret Hughes, Miren Iturriza-Gomara, James Johnson, L Luu, Jenifer Manson, Charlotte Nelson, Elaine O'Toole, Cassie Olateju, Rebekah Penrice-Randal , Lucille Rainbow, N.P Randle, Trevor Ian Robinson, Parul Sharma, Ghada T Shawli, James P Stewart, Neil Swainston, Ecaterina Vamos, Joanne Watts, Mark Whitehead |
| EPI_ISL_500287 | Servicio de Microbiología, Hospital Miguel Servet, Zaragoza | SeqCOVID-SPAIN consortium/IBV(CSIC) | Antonio Rezusta López, Alexander Tristancho Baró, Ana Milagro, Yolanda Gracia Grataloup, Nieves Martínez Cameo and SeqCOVID-SPAIN consortium |
| EPI_ISL_500292, EPI_ISL_500294, EPI_ISL_500296, EPI_ISL_500297, EPI_ISL_500298, EPI_ISL_500300, EPI_ISL_500301, EPI_ISL_500302, EPI_ISL_500303, EPI_ISL_500304, EPI_ISL_500305, EPI_ISL_500306, EPI_ISL_500307, EPI_ISL_500308, EPI_ISL_500309, EPI_ISL_500311, EPI_ISL_500315, EPI_ISL_500319, EPI_ISL_500320, EPI_ISL_500321, EPI_ISL_500322, EPI_ISL_500323, EPI_ISL_500324, EPI_ISL_500325, EPI_ISL_500326, EPI_ISL_500328, EPI_ISL_500329, EPI_ISL_500331, EPI_ISL_500332, EPI_ISL_500333, EPI_ISL_500334, EPI_ISL_500335 | | | |
| see above | Servicio de Microbiología. Hospital Universitario Donostia. OSI Donostialdea. Área de Enfermedades Infecciosas, Grupo de Infección Respiratoria y Resistencia Antimicrobiana. Instituto de Investigación Sanitaria Biodonostia | SeqCOVID-SPAIN consortium/IBV(CSIC) | Gustavo Cilla, Milagrosa Montes, Luis Piñeiro, Jose Maria Marimón and SeqCOVID-SPAIN consortium |
| EPI_ISL_500337, EPI_ISL_500344, EPI_ISL_500347, EPI_ISL_500349, EPI_ISL_500354 | Servicio de Microbiología, Hospital Miguel Servet, Zaragoza | SeqCOVID-SPAIN consortium/IBV(CSIC) | Antonio Rezusta López, Alexander Tristancho Baró, Ana Milagro, Yolanda Gracia Grataloup, Nieves Martínez Cameo and SeqCOVID-SPAIN consortium |
| EPI_ISL_500369, EPI_ISL_500372, EPI_ISL_500375, EPI_ISL_500376, EPI_ISL_500379, EPI_ISL_500386, EPI_ISL_500391, EPI_ISL_500396, EPI_ISL_500398, EPI_ISL_500399, EPI_ISL_500400, EPI_ISL_500401, EPI_ISL_500402, EPI_ISL_500407, EPI_ISL_500408, EPI_ISL_500409, EPI_ISL_500410, EPI_ISL_500411, EPI_ISL_500412, EPI_ISL_500413, EPI_ISL_500416, EPI_ISL_500417, EPI_ISL_500418, EPI_ISL_500421, EPI_ISL_500423, EPI_ISL_500425, EPI_ISL_500427, EPI_ISL_500428, EPI_ISL_500429, EPI_ISL_500430, EPI_ISL_500431, EPI_ISL_500433, EPI_ISL_500439, EPI_ISL_500445, EPI_ISL_500449, EPI_ISL_500450, EPI_ISL_500451, EPI_ISL_500454, EPI_ISL_500455, EPI_ISL_500456, EPI_ISL_500458 | | | |
| see above | Centro de Investigación Biomédica de La Rioja - Hospital San Pedro Logroño | SeqCOVID-SPAIN consortium/IBV(CSIC) | María de Toro, José Manuel Azcona Gutiérrez, María Pilar Bea Escudero, Miriam Blasco Alberdi and SeqCOVID-SPAIN consortium |
| EPI_ISL_500459 | Quest Diagnostics | Quest Diagnostics | Rosenthal,S.H., Gerasimova,A., Kagan,R.M., Owen, R. AND Lacbawan, F. |
| EPI_ISL_500460, EPI_ISL_500461, EPI_ISL_500462, EPI_ISL_500463, EPI_ISL_500471, EPI_ISL_500473, EPI_ISL_500475, EPI_ISL_500477, EPI_ISL_500480, EPI_ISL_500481, EPI_ISL_500482, EPI_ISL_500483, EPI_ISL_500484, EPI_ISL_500485, EPI_ISL_500486 | | | |
| see above | LACEN/PE | WallauLab, Aggeu Magalhaes Institute | Marcelo Henrique Santos Paiva, Duschinka Ribeiro Duarte Guedes, Cássia Docena, Matheus Filgueira Bezerra, Filipe Zimmer Dezordi, Laís Ceschini Machado, Larissa Krokovsky, Elisama Helvecio, Alexandre Freitas da Silva, Luydson Richardson Silva Vasconcelos, Antonio Mauro Rezende, Severino Jefferson Ribeiro da Silva, Kamila Gaudêncio da Silva Sales, Bruna Santos Lima Figueiredo de Sá, Derciliano Lopes da Cruz, Claudio Eduardo Cavalcanti, Armando de Menezes Neto, Caroline Targino Alves da Silva, Renata Pessôa Germano Mendes, Maria Almerice Lopes da Silva, Tiago Gräf, Paola Cristina Resende, Gonzalo Bello, Michelle da Silva Barros, Wheverton Ricardo Correia do Nascimento, Rodrigo Moraes Loyo Arcoverde, Luciane Caroline Albuquerque Bezerra, Sinval Pinto Brandão Filho, Constância Flávia Junqueira Ayres, Gabriel Luz Wallau |
| EPI_ISL_500500, EPI_ISL_500501 | University of Washington Virology Lab | University of Washington Virology Lab | Pavitra Roychoudhury, Hong Xie, Lasata Shrestha, Amin Addetia, Truong Nguyen, Victoria M Rachleff, Meei-Li Huang, Keith R Jerome, Alexander Greninger |
| EPI_ISL_500503, EPI_ISL_500504, EPI_ISL_500505, EPI_ISL_500506, EPI_ISL_500507, EPI_ISL_500508, EPI_ISL_500509, EPI_ISL_500510, EPI_ISL_500511, EPI_ISL_500512, EPI_ISL_500513, EPI_ISL_500514, EPI_ISL_500515, EPI_ISL_500516, EPI_ISL_500517, EPI_ISL_500518, EPI_ISL_500519, EPI_ISL_500520, EPI_ISL_500521, EPI_ISL_500522, EPI_ISL_500523, EPI_ISL_500524, EPI_ISL_500525, EPI_ISL_500526, EPI_ISL_500527, EPI_ISL_500528, EPI_ISL_500529, EPI_ISL_500530, EPI_ISL_500531, EPI_ISL_500533, EPI_ISL_500534, EPI_ISL_500535, EPI_ISL_500536, EPI_ISL_500537, EPI_ISL_500538 | | | |
| see above | Mayo Clinic Laboratories | University of Washington Virology Lab | Pavitra Roychoudhury, Hong Xie, Lasata Shrestha, Amin Addetia, Truong Nguyen, Victoria M Rachleff, Meei-Li Huang, Keith R Jerome, Alexander Greninger |
| EPI_ISL_500539, EPI_ISL_500540, EPI_ISL_500541, EPI_ISL_500542, EPI_ISL_500543, EPI_ISL_500544, EPI_ISL_500545, EPI_ISL_500546, EPI_ISL_500547, EPI_ISL_500548, EPI_ISL_500550, EPI_ISL_500551, EPI_ISL_500552, EPI_ISL_500553, EPI_ISL_500554, EPI_ISL_500555, EPI_ISL_500556, EPI_ISL_500557, EPI_ISL_500558, EPI_ISL_500559, EPI_ISL_500560, EPI_ISL_500561, EPI_ISL_500562, EPI_ISL_500563, EPI_ISL_500564, EPI_ISL_500565, EPI_ISL_500566, EPI_ISL_500567, EPI_ISL_500568, EPI_ISL_500569, EPI_ISL_500570, EPI_ISL_500571, EPI_ISL_500572 | | | |
| see above | Singapore General Hospital | Department of Microbiology | Nurdyana Abdul Rahman, Kun Lee Lim, Chenhao Li, Kian Sing Chan, Lynette Oon, Kern Rei Chng, Niranjan Nagarajan, Karrie Ko |
| EPI_ISL_500573, EPI_ISL_500574, EPI_ISL_500575, EPI_ISL_500576, EPI_ISL_500577, EPI_ISL_500578, EPI_ISL_500580, EPI_ISL_500581, EPI_ISL_500582, EPI_ISL_500583, EPI_ISL_500584, EPI_ISL_500585, EPI_ISL_500586, EPI_ISL_500587, EPI_ISL_500588, EPI_ISL_500589, EPI_ISL_500591, EPI_ISL_500593, EPI_ISL_500594, EPI_ISL_500595 | | | |
| see above | National Virus Reference Laboratory | National Virus Reference Laboratory | Michael Carr, Gabriel Gonzalez, Jonathan Dean, Suzie Coughlan, Cillian F De Gascun |
| EPI_ISL_500596, EPI_ISL_500597, EPI_ISL_500598, EPI_ISL_500599, EPI_ISL_500600, EPI_ISL_500601, EPI_ISL_500602, EPI_ISL_500603, EPI_ISL_500604, EPI_ISL_500605, EPI_ISL_500606, EPI_ISL_500607, EPI_ISL_500608, EPI_ISL_500609, EPI_ISL_500610, EPI_ISL_500611, EPI_ISL_500612, EPI_ISL_500613, EPI_ISL_500614, EPI_ISL_500615, EPI_ISL_500616, EPI_ISL_500617, EPI_ISL_500618, EPI_ISL_500619, EPI_ISL_500620, EPI_ISL_500621, EPI_ISL_500622, EPI_ISL_500623, EPI_ISL_500624, EPI_ISL_500625, EPI_ISL_500626, EPI_ISL_500627, EPI_ISL_500628, EPI_ISL_500629, EPI_ISL_500630, EPI_ISL_500631, EPI_ISL_500632, EPI_ISL_500633, EPI_ISL_500634, EPI_ISL_500635, EPI_ISL_500636, EPI_ISL_500637, EPI_ISL_500638, EPI_ISL_500639, EPI_ISL_500640, EPI_ISL_500641, EPI_ISL_500642, EPI_ISL_500643, EPI_ISL_500644, EPI_ISL_500645, EPI_ISL_500646, EPI_ISL_500647, EPI_ISL_500648, EPI_ISL_500649, EPI_ISL_500650, EPI_ISL_500651, EPI_ISL_500652, EPI_ISL_500653, EPI_ISL_500654, EPI_ISL_500655, EPI_ISL_500656, EPI_ISL_500657, EPI_ISL_500658, EPI_ISL_500659, EPI_ISL_500660, EPI_ISL_500661, EPI_ISL_500662, EPI_ISL_500663, EPI_ISL_500664, EPI_ISL_500665, EPI_ISL_500666, EPI_ISL_500667, EPI_ISL_500668, EPI_ISL_500669, EPI_ISL_500670, EPI_ISL_500671, EPI_ISL_500672, EPI_ISL_500673, EPI_ISL_500674, EPI_ISL_500675, EPI_ISL_500676, EPI_ISL_500677, EPI_ISL_500678, EPI_ISL_500679, EPI_ISL_500680, EPI_ISL_500681, EPI_ISL_500682, EPI_ISL_500683, EPI_ISL_500684, EPI_ISL_500685, EPI_ISL_500686, EPI_ISL_500687, EPI_ISL_500688, EPI_ISL_500689, EPI_ISL_500690, EPI_ISL_500691, EPI_ISL_500692, EPI_ISL_500693, EPI_ISL_500694, EPI_ISL_500695, EPI_ISL_500696, EPI_ISL_500697, EPI_ISL_500698, EPI_ISL_500699, EPI_ISL_500700, EPI_ISL_500701, EPI_ISL_500702, EPI_ISL_500703, EPI_ISL_500704, EPI_ISL_500705, EPI_ISL_500706 | | | |
| see above | Area of Virology, Serology and Virology Division (SAViD), New South Wales Health Pathology Randwick | Area of Virology, Serology and Virology Division (SAViD), New South Wales Health Pathology Randwick | Rawlinson, W. |
| EPI_ISL_500709, EPI_ISL_500710, EPI_ISL_500711, EPI_ISL_500715 | Respiratory Virus Unit, Microbiology Services Colindale, Public Health England | Respiratory Virus Unit, Microbiology Services Colindale, Public Health England | PHE Covid Sequencing Team |
| EPI_ISL_500717 | Area of Virology, Serology and Virology Division (SAViD), New South Wales Health Pathology Randwick | Area of Virology, Serology and Virology Division (SAViD), New South Wales Health Pathology Randwick | Rawlinson, W |
| EPI_ISL_500769, EPI_ISL_500770, EPI_ISL_500771, EPI_ISL_500773, EPI_ISL_500774 | Furst Medical Laboratory | Norwegian Institute of Public Health, Department of Virology | Kathrine Stene-Johansen, Kamilla Heddeland Instefjord, Hilde Elshaug, Rasmus Riis Kopperud, Karoline Bragstad, Olav Hungnes |
| EPI_ISL_500775 | Hospital of Southern Norway - Kristiansand, Department of Medical Microbiology | Norwegian Institute of Public Health, Department of Virology | Kathrine Stene-Johansen, Kamilla Heddeland Instefjord, Hilde Elshaug, Rasmus Riis Kopperud, Karoline Bragstad, Olav Hungnes |
| EPI_ISL_500776, EPI_ISL_500777, EPI_ISL_500778 | Foerde Hospital, Department of Microbiology | Norwegian Institute of Public Health, Department of Virology | Kathrine Stene-Johansen, Kamilla Heddeland Instefjord, Hilde Elshaug, Rasmus Riis Kopperud, Karoline Bragstad, Olav Hungnes |
| EPI_ISL_500779, EPI_ISL_500780, EPI_ISL_500782, EPI_ISL_500783 | Akershus University Hospital, Department for Microbiology and Infectious Disease Control | Norwegian Institute of Public Health, Department of Virology | Kathrine Stene-Johansen, Kamilla Heddeland Instefjord, Hilde Elshaug, Rasmus Riis Kopperud, Karoline Bragstad, Olav Hungnes |
| EPI_ISL_500785, EPI_ISL_500786, EPI_ISL_500787, EPI_ISL_500789, EPI_ISL_500790 | Furst Medical Laboratory | Norwegian Institute of Public Health, Department of Virology | Kathrine Stene-Johansen, Kamilla Heddeland Instefjord, Hilde Elshaug, Rasmus Riis Kopperud, Karoline Bragstad, Olav Hungnes |
| EPI_ISL_500791, EPI_ISL_500792 | Hospital of Southern Norway - Kristiansand, Department of Medical Microbiology | Norwegian Institute of Public Health, Department of Virology | Kathrine Stene-Johansen, Kamilla Heddeland Instefjord, Hilde Elshaug, Rasmus Riis Kopperud, Karoline Bragstad, Olav Hungnes |
| EPI_ISL_500794, EPI_ISL_500795, EPI_ISL_500796 | Akershus University Hospital, Department for Microbiology and Infectious Disease Control | Norwegian Institute of Public Health, Department of Virology | Kathrine Stene-Johansen, Kamilla Heddeland Instefjord, Hilde Elshaug, Rasmus Riis Kopperud, Karoline Bragstad, Olav Hungnes |
| EPI_ISL_500797, EPI_ISL_500799 | Hospital of Southern Norway - Kristiansand, Department of Medical Microbiology | Norwegian Institute of Public Health, Department of Virology | Kathrine Stene-Johansen, Kamilla Heddeland Instefjord, Hilde Elshaug, Rasmus Riis Kopperud, Karoline Bragstad, Olav Hungnes |
| EPI_ISL_500831, EPI_ISL_500832, EPI_ISL_500833, EPI_ISL_500834, EPI_ISL_500835, EPI_ISL_500836, EPI_ISL_500838, EPI_ISL_500839, EPI_ISL_500840, EPI_ISL_500841, EPI_ISL_500842, EPI_ISL_500843, EPI_ISL_500844, EPI_ISL_500846, EPI_ISL_500847, EPI_ISL_500848, EPI_ISL_500849, EPI_ISL_500850, EPI_ISL_500851, EPI_ISL_500852, EPI_ISL_500853, EPI_ISL_500854, EPI_ISL_500855, EPI_ISL_500856, EPI_ISL_500857, EPI_ISL_500858, EPI_ISL_500859, EPI_ISL_500860, EPI_ISL_500861, EPI_ISL_500862, EPI_ISL_500863 | | | |
| see above | Virginia DCLS | Virginia DCLS | Virginia DCLS |
| EPI_ISL_500872 | LACEN/PE | WallauLab, Aggeu Magalhaes Institute | Marcelo Henrique Santos Paiva, Duschinka Ribeiro Duarte Guedes, Cássia Docena, Matheus Filgueira Bezerra, Filipe Zimmer Dezordi, Laís Ceschini Machado, Larissa Krokovsky, Elisama Helvecio, Alexandre Freitas da Silva, Luydson Richardson Silva Vasconcelos, Antonio Mauro Rezende, Severino Jefferson Ribeiro da Silva, Kamila Gaudêncio da Silva Sales, Bruna Santos Lima Figueiredo de Sá, Derciliano Lopes da Cruz, Claudio Eduardo Cavalcanti, Armando de Menezes Neto, Caroline Targino Alves da Silva, Renata Pessôa Germano Mendes, Maria Almerice Lopes da Silva, Tiago Gräf, Paola Cristina Resende, Gonzalo Bello, Michelle da Silva Barros, Wheverton Ricardo Correia do Nascimento, Rodrigo Moraes Loyo Arcoverde, Luciane Caroline Albuquerque Bezerra, Sinval Pinto Brandão Filho, Constância Flávia Junqueira Ayres, Gabriel Luz Wallau |

| | | | |
|---|---|---|---|
| EPI_ISL_500878, EPI_ISL_500879, EPI_ISL_500880, EPI_ISL_500881, EPI_ISL_500882, EPI_ISL_500883, EPI_ISL_500884, EPI_ISL_500885, EPI_ISL_500886, EPI_ISL_500887, EPI_ISL_500889, EPI_ISL_500890, EPI_ISL_500891, EPI_ISL_500892, EPI_ISL_500893, EPI_ISL_500894, EPI_ISL_500895, EPI_ISL_500896, EPI_ISL_500897, EPI_ISL_500898, EPI_ISL_500899, EPI_ISL_500901, EPI_ISL_500902, EPI_ISL_500903, EPI_ISL_500904, EPI_ISL_500905, EPI_ISL_500906, EPI_ISL_500908, EPI_ISL_500909, EPI_ISL_500910, EPI_ISL_500911, EPI_ISL_500912, EPI_ISL_500913, EPI_ISL_500914, EPI_ISL_500916, EPI_ISL_500917, EPI_ISL_500919, EPI_ISL_500920, EPI_ISL_500921, EPI_ISL_500922, EPI_ISL_500923, EPI_ISL_500924, EPI_ISL_500925, EPI_ISL_500927, EPI_ISL_500929, EPI_ISL_500932, EPI_ISL_500933, EPI_ISL_500934, EPI_ISL_500935, EPI_ISL_500936, EPI_ISL_500937, EPI_ISL_500938, EPI_ISL_500939, EPI_ISL_500940, EPI_ISL_500941, EPI_ISL_500942, EPI_ISL_500943, EPI_ISL_500944, EPI_ISL_500945 | | | |
| see above | Viollier AG | Department of Biosystems Science and Engineering, ETH Zürich | Christian Beisel, Sarah Nadeau, Ivan Topolsky, Pedro Ferreira, Philipp Jablonski, Susana Posada-Céspedes, Tobias Schär, Ina Nissen, Natascha Santacroce, Elodie Burcklen, Christiane Beckmann, Maurice Redondo, Olivier Kobel, Christoph Noppen, Sophie Seidel, Noemie Santamaria de Souza, Niko Beerenwinkel, Tanja Stadler |
| EPI_ISL_500946 | GMERS Medical College & Hospital, Gotri, Vadodara | Gujarat Biotechnology Research Centre | Zuber Saiyed, Komal Patel, Labdhi Pandya, Afzal Ansari, Nikha Trivedi, Meenakshi Shah, Neena Doshi, Varsha Godbole, Apurvasinh Puvar, Janvi Raval, Zarna Patel, Monika Gandhi, Pinal Trivedi, Maharshi Pandya, Nidhi Patel, Nitin Savaliya, Raghawendra Kumar, Dinesh Kumar, R D Dixit, A M Kadri, Harsh Bakshi, Chaitanya Joshi, Madhvi Joshi |
| EPI_ISL_500947 | Department of MicroBiology, Government Medical College, Surat | Gujarat Biotechnology Research Centre | Maharshi Pandya, Nidhi Patel, Nitin Savaliya, Raghawendra Kumar, Dinesh Kumar, Zuber Saiyed, Komal Patel, Labdhi Pandya, Afzal Ansari, Nikha Trivedi, Naresh Chauhan, Summaiya Mullan, Amit gamit, Apurvasinh Puvar, Janvi Raval, Zarna Patel, Monika Gandhi, Pinal Trivedi, R D Dixit, A M Kadri, Harsh Bakshi, Chaitanya Joshi, Madhvi Joshi |
| EPI_ISL_500948 | Department of MicroBiology, Government Medical College, Surat | Gujarat Biotechnology Research Centre | Zuber Saiyed, Komal Patel, Labdhi Pandya, Afzal Ansari, Nikha Trivedi, Naresh Chauhan, Summaiya Mullan, Amit gamit, Apurvasinh Puvar, Janvi Raval, Zarna Patel, Monika Gandhi, Pinal Trivedi, Maharshi Pandya, Nidhi Patel, Nitin Savaliya, Raghawendra Kumar, Dinesh Kumar, R D Dixit, A M Kadri, Harsh Bakshi, Chaitanya Joshi, Madhvi Joshi |
| EPI_ISL_500949 | Department of MicroBiology, Government Medical College, Surat | Gujarat Biotechnology Research Centre | Komal Patel, Labdhi Pandya, Afzal Ansari, Nikha Trivedi, Naresh Chauhan, Summaiya Mullan, Amit gamit, Apurvasinh Puvar, Janvi Raval, Zarna Patel, Monika Gandhi, Pinal Trivedi, Maharshi Pandya, Nidhi Patel, Nitin Savaliya, Raghawendra Kumar, Dinesh Kumar, Zuber Saiyed, R D Dixit, A M Kadri, Harsh Bakshi, Chaitanya Joshi, Madhvi Joshi |
| EPI_ISL_500950 | Department of MicroBiology, Government Medical College, Surat | Gujarat Biotechnology Research Centre | Labdhi Pandya, Afzal Ansari, Nikha Trivedi, Naresh Chauhan, Summaiya Mullan, Amit gamit, Apurvasinh Puvar, Janvi Raval, Zarna Patel, Monika Gandhi, Pinal Trivedi, Maharshi Pandya, Nidhi Patel, Nitin Savaliya, Raghawendra Kumar, Dinesh Kumar, Zuber Saiyed, Komal Patel, R D Dixit, A M Kadri, Harsh Bakshi, Chaitanya Joshi, Madhvi Joshi |
| EPI_ISL_500952, EPI_ISL_500953 | Respiratory Virus Unit, Microbiology Services Colindale, Public Health England | Respiratory Virus Unit, Microbiology Services Colindale, Public Health England | PHE Covid Sequencing Team |
| EPI_ISL_500954, EPI_ISL_500955, EPI_ISL_500957, EPI_ISL_500958, EPI_ISL_500959, EPI_ISL_500960, EPI_ISL_500961, EPI_ISL_500962, EPI_ISL_500963, EPI_ISL_500964, EPI_ISL_500965, EPI_ISL_500966, EPI_ISL_500967, EPI_ISL_500968, EPI_ISL_500970, EPI_ISL_500971, EPI_ISL_500972, EPI_ISL_500973, EPI_ISL_500974, EPI_ISL_500975, EPI_ISL_500976, EPI_ISL_500977, EPI_ISL_500978, EPI_ISL_500979, EPI_ISL_500980, EPI_ISL_500981, EPI_ISL_500982, EPI_ISL_500983, EPI_ISL_500984, EPI_ISL_500985, EPI_ISL_500986, EPI_ISL_500987, EPI_ISL_500988, EPI_ISL_500989, EPI_ISL_500990, EPI_ISL_500991, EPI_ISL_500992, EPI_ISL_500993, EPI_ISL_500994, EPI_ISL_500996, EPI_ISL_500997, EPI_ISL_500998, EPI_ISL_501001, EPI_ISL_501002, EPI_ISL_501003, EPI_ISL_501004, EPI_ISL_501006, EPI_ISL_501007, EPI_ISL_501008, EPI_ISL_501010, EPI_ISL_501011, EPI_ISL_501012, EPI_ISL_501014, EPI_ISL_501015, EPI_ISL_501016, EPI_ISL_501017, EPI_ISL_501018, EPI_ISL_501019, EPI_ISL_501020, EPI_ISL_501021, EPI_ISL_501022, EPI_ISL_501023, EPI_ISL_501024, EPI_ISL_501025, EPI_ISL_501026, EPI_ISL_501027, EPI_ISL_501028, EPI_ISL_501029, EPI_ISL_501030, EPI_ISL_501031, EPI_ISL_501032, EPI_ISL_501033, EPI_ISL_501034, EPI_ISL_501035, EPI_ISL_501036, EPI_ISL_501038, EPI_ISL_501039, EPI_ISL_501040, EPI_ISL_501041, EPI_ISL_501042, EPI_ISL_501043, EPI_ISL_501044, EPI_ISL_501045, EPI_ISL_501046, EPI_ISL_501047, EPI_ISL_501048, EPI_ISL_501049, EPI_ISL_501050, EPI_ISL_501051, EPI_ISL_501052, EPI_ISL_501053, EPI_ISL_501054, EPI_ISL_501055, EPI_ISL_501056, EPI_ISL_501057, EPI_ISL_501058, EPI_ISL_501059, EPI_ISL_501060, EPI_ISL_501061, EPI_ISL_501062, EPI_ISL_501063, EPI_ISL_501064, EPI_ISL_501065, EPI_ISL_501066, EPI_ISL_501067, EPI_ISL_501068, EPI_ISL_501069, EPI_ISL_501071 | | | |
| see above | Regional Virus Laboratory, Belfast Health and Social Care Trust | Wellcome Sanger Institute for the COVID-19 Genomics UK Consortium | Conall McCaughey, James McKenna, Tanya Curran, Susan Feeney, Alison Watt, Ciara Cox, Mairead Connor, Zoltan Molnar, David Simpson, Derek Fairley; and Alex Alderton, Roberto Amato, Sonia Goncalves, Ewan Harrison, David K. Jackson, Ian Johnston, Dominic Kwiatkowski, Cordelia Langford, John Sillitoe on behalf of the Wellcome Sanger Institute COVID-19 Surveillance Team (http://www.sanger.ac.uk/covid-team) |
| EPI_ISL_501072, EPI_ISL_501073, EPI_ISL_501074, EPI_ISL_501075, EPI_ISL_501076, EPI_ISL_501077, EPI_ISL_501078, EPI_ISL_501079, EPI_ISL_501080, EPI_ISL_501081, EPI_ISL_501082 | | | |
| see above | Mayo Clinic Laboratories | University of Washington Virology Lab | Pavitra Roychoudhury, Hong Xie, Lasata Shrestha, Amin Addetia, Truong Nguyen, Victoria M Rachleff, Meei-Li Huang, Keith R Jerome, Alexander Greninger |
| EPI_ISL_501083, EPI_ISL_501084, EPI_ISL_501085, EPI_ISL_501086, EPI_ISL_501087, EPI_ISL_501088, EPI_ISL_501089, EPI_ISL_501090, EPI_ISL_501091, EPI_ISL_501092, EPI_ISL_501093, EPI_ISL_501095, EPI_ISL_501096, EPI_ISL_501097, EPI_ISL_501098, EPI_ISL_501099, EPI_ISL_501100, EPI_ISL_501101, EPI_ISL_501102, EPI_ISL_501103, EPI_ISL_501104, EPI_ISL_501105, EPI_ISL_501106, EPI_ISL_501107, EPI_ISL_501108, EPI_ISL_501109, EPI_ISL_501110, EPI_ISL_501111, EPI_ISL_501112, EPI_ISL_501113, EPI_ISL_501114, EPI_ISL_501115, EPI_ISL_501116, EPI_ISL_501117, EPI_ISL_501118, EPI_ISL_501120, EPI_ISL_501121, EPI_ISL_501122, EPI_ISL_501123, EPI_ISL_501124, EPI_ISL_501125, EPI_ISL_501126, EPI_ISL_501127, EPI_ISL_501128, EPI_ISL_501131, EPI_ISL_501132, EPI_ISL_501133, EPI_ISL_501134, EPI_ISL_501135, EPI_ISL_501136, EPI_ISL_501137, EPI_ISL_501138, EPI_ISL_501140, EPI_ISL_501141, EPI_ISL_501142, EPI_ISL_501143, EPI_ISL_501144, EPI_ISL_501145, EPI_ISL_501146, EPI_ISL_501147, EPI_ISL_501148, EPI_ISL_501149, EPI_ISL_501150, EPI_ISL_501151, EPI_ISL_501152, EPI_ISL_501153, EPI_ISL_501154, EPI_ISL_501155, EPI_ISL_501156, EPI_ISL_501157, EPI_ISL_501158, EPI_ISL_501159, EPI_ISL_501160, EPI_ISL_501161, EPI_ISL_501162, EPI_ISL_501163, EPI_ISL_501164 | | | |
| see above | University of Washington Virology Lab | University of Washington Virology Lab | Pavitra Roychoudhury, Hong Xie, Lasata Shrestha, Amin Addetia, Truong Nguyen, Victoria M Rachleff, Meei-Li Huang, Keith R Jerome, Alexander Greninger |
| EPI_ISL_501165 | Mayo Clinic Laboratories | University of Washington Virology Lab | Pavitra Roychoudhury, Hong Xie, Lasata Shrestha, Amin Addetia, Truong Nguyen, Victoria M Rachleff, Meei-Li Huang, Keith R Jerome, Alexander Greninger |
| EPI_ISL_501167, EPI_ISL_501169, EPI_ISL_501171, EPI_ISL_501172, EPI_ISL_501173 | Baylor College of Medicine | Baylor College of Medicine: HGSC | Vasanthi Avadhanula, Erin Nicholson, David Henke, Pedro Piedra, Harsha Doddapaneni, Donna Muzny, Qingchang Meng, Hsu Chao, Zeineen Momin, Hua Shen, George Weissenberger, Kavya Kottapalli, Yimiti Meiheerguli, Sejal Salvi, Ginger Metcalf, Vipin Menon, Sara J.J. Cregeen, Matthew C. Ross, Tulin Ayvaz, Richard Sucgang, Kristi L. Hoffman, Matthew Wong, Joseph F. Petrosino |
| EPI_ISL_501176, EPI_ISL_501177, EPI_ISL_501178, EPI_ISL_501179, EPI_ISL_501180, EPI_ISL_501181, EPI_ISL_501182, EPI_ISL_501183, EPI_ISL_501184, EPI_ISL_501185, EPI_ISL_501186, EPI_ISL_501187, EPI_ISL_501188, EPI_ISL_501189, EPI_ISL_501190, EPI_ISL_501191, EPI_ISL_501192, EPI_ISL_501193, EPI_ISL_501194, EPI_ISL_501195, EPI_ISL_501196, EPI_ISL_501197, EPI_ISL_501198, EPI_ISL_501199, EPI_ISL_501200, EPI_ISL_501201, EPI_ISL_501202, EPI_ISL_501203, EPI_ISL_501204, EPI_ISL_501205, EPI_ISL_501206, EPI_ISL_501207, EPI_ISL_501208, EPI_ISL_501209, EPI_ISL_501210, EPI_ISL_501211, EPI_ISL_501212, EPI_ISL_501213, EPI_ISL_501214, EPI_ISL_501215, EPI_ISL_501216, EPI_ISL_501217, EPI_ISL_501218, EPI_ISL_501219, EPI_ISL_501220, EPI_ISL_501221, EPI_ISL_501222, EPI_ISL_501223, EPI_ISL_501224, EPI_ISL_501225, EPI_ISL_501226, EPI_ISL_501227, EPI_ISL_501228 | | | |
| see above | Department of Medical Microbiology, University Malaya Medical Centre | Department of Medical Microbiology, Faculty of Medicine, University of Malaya | Yoong Min CHONG, Jennifer Chong, I-Ching SAM, Yoke Fun CHAN, University Malaya Medical Centre COVID Team |
| EPI_ISL_501230 | Hellenic Pasteur Institute, Public Health Laboratories | Hellenic Pasteur Institute, National Influenza Reference laboratory of Southern Greece & Unit of Bioinformatics and Applied Genomics | Vasiliki Pogka, Timokratis Karamitros, Athanasios Kossyvakis, Antonios Kalliaropoulos, Horefti Elina, Evangelidou Maria, Androniki Voulgari-Kokota, Aspasia Kontou, Andreas Mentis |
| EPI_ISL_501233, EPI_ISL_501236, EPI_ISL_501248, EPI_ISL_501249, EPI_ISL_501250, EPI_ISL_501251 | Hellenic Pasteur Institute, National Influenza Reference laboratory of Southern Greece & Unit of Bioinformatics and Applied Genomics | Hellenic Pasteur Institute, National Influenza Reference laboratory of Southern Greece & Unit of Bioinformatics and Applied Genomics | Vasiliki Pogka, Timokratis Karamitros, Athanasios Kossyvakis, Antonios Kalliaropoulos, Horefti Elina, Evangelidou Maria, Androniki Voulgari-Kokota, Aspasia Kontou, Andreas Mentis |
| EPI_ISL_501259, EPI_ISL_501268, EPI_ISL_501273, EPI_ISL_501274 | National Virus Reference Laboratory | National Virus Reference Laboratory | Michael Carr, Gabriel Gonzalez, Jonathan Dean, Suzie Coughlan, Cillian F De Gascun |
| EPI_ISL_501275 | E. Gulbja Laboratorija | Latvian Biomedical Research and Study Centre | Ivars Silamiķelis, Kaspars Megnis, Monta Ustinova, Ņikita Zrelovs, Vita Rovīte, Mikus Gavars, Dmitrijs Perminovs, Uga Dumpis, Jānis Kloviņš |
| EPI_ISL_501284, EPI_ISL_501285 | E. Gulbja Laboratorija | Latvian Biomedical Research and Study Centre | Ivars Silamiķelis, Kaspars Megnis, Monta Ustinova, Ņikita Zrelovs, Vita Rovīte, Mikus Gavars, Dmitrijs Perminovs, Uga Dumpis, Jānis Kloviņš |
| EPI_ISL_501286, EPI_ISL_501287, EPI_ISL_501288, EPI_ISL_501289 | Centrālā laboratorija | Latvian Biomedical Research and Study Centre | Ivars Silamiķelis, Kaspars Megnis, Monta Ustinova, Ņikita Zrelovs, Vita Rovīte, Stella Lapiņa, Jana Osīte, Marta Priedīte, Uga Dumpis, Jānis Kloviņš |
| EPI_ISL_501556, EPI_ISL_501559, EPI_ISL_501560, EPI_ISL_501561, EPI_ISL_501563, EPI_ISL_501564, EPI_ISL_501565, EPI_ISL_501566, EPI_ISL_501567, EPI_ISL_501568, EPI_ISL_501570, EPI_ISL_501572, EPI_ISL_501573, EPI_ISL_501574, EPI_ISL_501575, EPI_ISL_501578, EPI_ISL_501580, EPI_ISL_501581, EPI_ISL_501583, EPI_ISL_501586, EPI_ISL_501588, EPI_ISL_501589, EPI_ISL_501590, EPI_ISL_501591, EPI_ISL_501592, EPI_ISL_501593, EPI_ISL_501595, EPI_ISL_501596, EPI_ISL_501597, EPI_ISL_501598, EPI_ISL_501599, EPI_ISL_501601 | | | |
| see above | PHE South West Regional Laboratory, National Infection Service | Wellcome Sanger Institute for the COVID-19 Genomics UK Consortium | Stephanie Hutchings, Hannah Pymont, Dr Peter Muir, Barry Vipond, Rich Hopes; and Alex Alderton, Roberto Amato, Sonia Goncalves, Ewan Harrison, David K. Jackson, Ian Johnston, Dominic Kwiatkowski, Cordelia Langford, John Sillitoe on behalf of the Wellcome Sanger Institute COVID-19 Surveillance Team (http://www.sanger.ac.uk/covid-team) |
| EPI_ISL_501602 | Department of Medical Microbiology, Western Sussex Hospitals NHS Foundation Trust, St Richard's Hospital | Wellcome Sanger Institute for the COVID-19 Genomics UK Consortium | Manasa Mutingwende, Sarah Lowdon, Olga Podplomyk, Michelle Erkiert, Jonathan Lewis, Paul Randell and Alex Alderton, Roberto Amato, Sonia Goncalves, Ewan Harrison, David K. Jackson, Ian Johnston, Dominic Kwiatkowski, Cordelia Langford, John Sillitoe on behalf of the Wellcome Sanger Institute COVID-19 Surveillance Team (http://www.sanger.ac.uk/covid-team) |
| EPI_ISL_501606, EPI_ISL_501607, EPI_ISL_501609, EPI_ISL_501611 | PHE South West Regional Laboratory, National Infection Service | Wellcome Sanger Institute for the COVID-19 Genomics UK Consortium | Stephanie Hutchings, Hannah Pymont, Dr Peter Muir, Barry Vipond, Rich Hopes; and Alex Alderton, Roberto Amato, Sonia Goncalves, Ewan Harrison, David K. Jackson, Ian Johnston, Dominic Kwiatkowski, Cordelia Langford, John Sillitoe on behalf of the Wellcome Sanger Institute COVID-19 Surveillance Team (http://www.sanger.ac.uk/covid-team) |
| EPI_ISL_501613 | Department of Medical Microbiology, Western Sussex Hospitals NHS Foundation Trust, St Richard's Hospital | Wellcome Sanger Institute for the COVID-19 Genomics UK Consortium | Manasa Mutingwende, Sarah Lowdon, Olga Podplomyk, Michelle Erkiert, Jonathan Lewis, Paul Randell and Alex Alderton, Roberto Amato, Sonia Goncalves, Ewan Harrison, David K. Jackson, Ian Johnston, Dominic Kwiatkowski, Cordelia Langford, John Sillitoe on behalf of the Wellcome Sanger Institute COVID-19 Surveillance Team (http://www.sanger.ac.uk/covid-team) |
| EPI_ISL_501614, EPI_ISL_501615, EPI_ISL_501616, EPI_ISL_501617, EPI_ISL_501618, EPI_ISL_501620, EPI_ISL_501621, EPI_ISL_501622, EPI_ISL_501623, EPI_ISL_501624, EPI_ISL_501625, EPI_ISL_501626, EPI_ISL_501627, EPI_ISL_501628, EPI_ISL_501629 | | | |
| see above | Lab Microbiology, Pathology Department, William Harvey Hospital | Wellcome Sanger Institute for the COVID-19 Genomics UK Consortium | Samuel Moses, Hannah Lowe, Felicity Ryan and Alex Alderton, Roberto Amato, Sonia Goncalves, Ewan Harrison, David K. Jackson, Ian Johnston, Dominic Kwiatkowski, Cordelia Langford, John Sillitoe on behalf of the Wellcome Sanger Institute COVID-19 Surveillance Team (http://www.sanger.ac.uk/covid-team) |
| EPI_ISL_501632 | Virology Department, Royal Infirmary of Edinburgh, NHS Lothian / School of Biological Sciences, University of Edinburgh | Wellcome Sanger Institute for the COVID-19 Genomics UK Consortium | McHugh M, Dewar R, Rooke S, O'Toole Á, Scher E, Hill V, McCrone JT, Colquhoun R, Yu X, Jackson B, Rambaut A, Templeton K and Alex Alderton, Roberto Amato, Sonia Goncalves, Ewan Harrison, David K. Jackson, Ian Johnston, Dominic Kwiatkowski, Cordelia Langford, John Sillitoe on behalf of the Wellcome Sanger Institute COVID-19 Surveillance Team (http://www.sanger.ac.uk/covid-team) |
| EPI_ISL_501633, EPI_ISL_501634, EPI_ISL_501635 | NHSGGC West of Scotland Specialist Virology Centre / MRC-University of Glasgow Centre for Virus Research | Wellcome Sanger Institute for the COVID-19 Genomics UK Consortium | Ana da Silva Filipe, Natasha Johnson, Kathy Smollett, Daniel Mair, Stephen Carmichael, Lily Tong, Jenna Nichols, Elihu Aranday-Cortes, Kirstyn Brunker, Yasmin Parr, Kyriaki Nomikou; Sarah McDonald, Marc Niebel, Patawee Asamaphan; Richard Orton, Joseph Hughes, Sreenu Vattipally, David L Robertson; Alasdair MacLean, Rory Gunson; Kathy Li, Natasha Jesudason, Rajiv Shah, James Shepherd, Antonia Ho, Alice Broos, Emma Thomson and Alex Alderton, Roberto Amato, Sonia Goncalves, Ewan Harrison, David K. Jackson, Ian Johnston, Dominic Kwiatkowski, Cordelia Langford, John Sillitoe on behalf of the Wellcome Sanger Institute COVID-19 Surveillance Team (http://www.sanger.ac.uk/covid-team) |
| EPI_ISL_501808, EPI_ISL_501817 | Centrālā laboratorija | Latvian Biomedical Research and Study Centre | Ivars Silamiķelis, Kaspars Megnis, Monta Ustinova, Ņikita Zrelovs, Vita Rovīte, Stella Lapiņa, Jana Osīte, Marta Priedīte, Uga Dumpis, Jānis Kloviņš |
| EPI_ISL_501823, EPI_ISL_501829, EPI_ISL_501833, EPI_ISL_501839, EPI_ISL_501849, EPI_ISL_501894 | E. Gulbja Laboratorija | Latvian Biomedical Research and Study Centre | Ivars Silamiķelis, Kaspars Megnis, Monta Ustinova, Ņikita Zrelovs, Vita Rovīte, Mikus Gavars, Dmitrijs Perminovs, Uga Dumpis, Jānis Kloviņš |
| EPI_ISL_501895, EPI_ISL_501896, EPI_ISL_501915, EPI_ISL_501922 | Centrālā laboratorija | Latvian Biomedical Research and Study Centre | Ivars Silamiķelis, Kaspars Megnis, Monta Ustinova, Ņikita Zrelovs, Vita Rovīte, Stella Lapiņa, Jana Osīte, Marta Priedīte, Uga Dumpis, Jānis Kloviņš |
| EPI_ISL_501929, EPI_ISL_501936 | E. Gulbja Laboratorija | Latvian Biomedical Research and Study Centre | Ivars Silamiķelis, Kaspars Megnis, Monta Ustinova, Ņikita Zrelovs, Vita Rovīte, Mikus Gavars, Dmitrijs Perminovs, Uga Dumpis, Jānis Kloviņš |
| EPI_ISL_502779 | LACEN/PE | LABBE, Federal University of Pernambuco | WILSON JOSE DA SILVA JUNIOR, HEIDI LACERDA ALVES DA CRUZ, MARCOS DA SILVEIRA REGUEIRA NETO, BRUNO SAMPAIO, SERGIO DE SA LEITAO PAIVA JUNIOR, ZILDENE DE SOUSA SILVEIRA, MAIRA GALDINO DA ROCHA PITTA, MICHELLY CRISTINY PEREIRA, MARCOS ANTONIO DE MORAIS JUNIOR, ANTONIO CARLOS DE FREITAS, VALDIR DE QUEIROZ BALBINO. |
| EPI_ISL_502875 | LACEN/PE | LABBE, Federal University of Pernambuco | WILSON JOSE DA SILVA JUNIOR, HEIDI LACERDA ALVES DA CRUZ, MARCOS DA SILVEIRA REGUEIRA NETO, BRUNO SAMPAIO, SERGIO DE SA LEITAO PAIVA JUNIOR, ZILDENE DE SOUSA SILVEIRA, MAIRA GALDINO DA ROCHA PITTA, MICHELLY CRISTINY PEREIRA, REGINALDO GONCALVES DE LIMA NETO, MARCOS ANTONIO DE MORAIS JUNIOR, ANTONIO CARLOS DE FREITAS, VALDIR DE QUEIROZ BALBINO. |
| EPI_ISL_503958, EPI_ISL_504040 | National Institute of Laboratory Medicine and Referral Center | Genomic Research Lab, BCSIR | Md. Ahasan Habib, Abu Sayeed Mohammad Mahmud, Mohammad Samir Uzzaman, Eshrar Osman, Shahina Akter, Tanjina Akhter Banu, Md. Murshed Hasan Sarkar, Barna Goswami, Iffat Jahan, Md. Saddam Hossain, Tarannum Taznin, Tasnim Nafisa, Md. Maruf Ahmed Molla, Mahmuda Yeasmin, Asish Kumar Ghosh, A. K. M. Shamsuzzaman, Sheikh Md. Selim Al Din, Utpal Chandra Ray, Salek Ahmed Sajib, Md. Salim Khan |
| EPI_ISL_504137, EPI_ISL_504176, EPI_ISL_504177, EPI_ISL_504178, EPI_ISL_504179, EPI_ISL_504180, | National Institute of Laboratory Medicine and Referral Center | Genomic Research Lab, BCSIR | Abu Sayeed Mohammad Mahmud, Mohammad Samir Uzzaman, Eshrar Osman, Md. Ahasan Habib, Shahina Akter, Tanjina Akhter Banu, Md. Murshed Hasan Sarkar, Barna Goswami, Iffat Jahan, Md. Saddam Hossain, Tarannum Taznin, Tasnim Nafisa, Md. Maruf Ahmed Molla, Mahmuda Yeasmin, Asish Kumar Ghosh, A. K. M. Shamsuzzaman, Sheikh Md. Selim Al Din, Utpal Chandra Ray, Salek Ahmed Sajib, Md. Salim Khan |

| | | | |
|---|---|---|---|
| EPI_ISL_504181, EPI_ISL_504182, EPI_ISL_504183, EPI_ISL_504184 | | | |
| EPI_ISL_504185 | Discovery DNA | Discovery DNA | Dustin Hittel, Marina Kerr, Leo Dimnik, Desmond Koo, Alice Li, Aneal Khan |
| EPI_ISL_505003 | Biology Dpt | Microbiology and Infections Diseases | Emmanuelle Billon-Denis, Audrey Ferrier-Rembert, Annabelle Garnier, Laurence Cheutin, Clarisse Vigne, Emilie Tessier, Jessica Denis, Olivier Gorgé, Flora Nolent, Isabelle Drouet, Noémie Verguet, Olivier Ferraris, Jean-Nicolas Tournier |
| EPI_ISL_506947, EPI_ISL_506948, EPI_ISL_506949, EPI_ISL_506950, EPI_ISL_506951, EPI_ISL_506952, EPI_ISL_506953, EPI_ISL_506954, EPI_ISL_506955, EPI_ISL_506956 | Division of Viral Diseases, Center for Laboratory Control of Infectious Diseases, Korea Centers for Diseases Control and Prevention | Division of Viral Diseases, Center for Laboratory Control of Infectious Diseases, Korea Centers for Diseases Control and Prevention | Jeong-Min Kim, Yoon-Seok Chung, Namjoo Lee, Sang Hee Woo, Hye-Jun Jo, Heui Man Kim, Jun-Sub Kim, Dong Hyun Song, Daesang Lee, Seong Tae Jeong, Myung Guk Han |
| EPI_ISL_506957, EPI_ISL_506958, EPI_ISL_506959, EPI_ISL_506960, EPI_ISL_506961, EPI_ISL_506962, EPI_ISL_506963, EPI_ISL_506964 | Division of Viral Diseases, Center for Laboratory Control of Infectious Diseases, Korea Centers for Diseases Control and Prevention | Division of Viral Diseases, Center for Laboratory Control of Infectious Diseases, Korea Centers for Diseases Control and Prevention | Jeong-Min Kim, Yoon-Seok Chung, Namjoo Lee, Sang Hee Woo, Hye-Jun Jo, Heui Man Kim, Jun-Sub Kim, Myung Guk Han |
| EPI_ISL_506965, EPI_ISL_506966, EPI_ISL_506967, EPI_ISL_506968, EPI_ISL_506969, EPI_ISL_506970, EPI_ISL_506971, EPI_ISL_506972, EPI_ISL_506973, EPI_ISL_506974 | Division of Viral Diseases, Center for Laboratory Control of Infectious Diseases, Korea Centers for Diseases Control and Prevention | Division of Viral Diseases, Center for Laboratory Control of Infectious Diseases, Korea Centers for Diseases Control and Prevention | Jeong-Min Kim, Yoon-Seok Chung, Namjoo Lee, Sang Hee Woo, Hye-Jun Jo, Heui Man Kim, Jun-Sub Kim, Dong Hyun Song, Daesang Lee, Seong Tae Jeong, Myung Guk Han |
| EPI_ISL_506975 | Division of Viral Diseases, Center for Laboratory Control of Infectious Diseases, Korea Centers for Diseases Control and Prevention | Division of Viral Diseases, Center for Laboratory Control of Infectious Diseases, Korea Centers for Diseases Control and Prevention | Jeong-Min Kim, Yoon-Seok Chung, Namjoo Lee, Sang Hee Woo, Hye-Jun Jo, Heui Man Kim, Jun-Sub Kim, Myung Guk Han |
| EPI_ISL_506976, EPI_ISL_506977, EPI_ISL_506978, EPI_ISL_506979, EPI_ISL_506980, EPI_ISL_506981, EPI_ISL_506982, EPI_ISL_506983, EPI_ISL_506984, EPI_ISL_506985, EPI_ISL_506986, EPI_ISL_506987, EPI_ISL_506988, EPI_ISL_506989, EPI_ISL_506990, EPI_ISL_506991, EPI_ISL_506992, EPI_ISL_506993 | | | |
| see above | Division of Viral Diseases, Center for Laboratory Control of Infectious Diseases, Korea Centers for Diseases Control and Prevention | Division of Viral Diseases, Center for Laboratory Control of Infectious Diseases, Korea Centers for Diseases Control and Prevention | Jeong-Min Kim, Yoon-Seok Chung, Namjoo Lee, Sang Hee Woo, Hye-Jun Jo, Heui Man Kim, Jun-Sub Kim, Dong Hyun Song, Daesang Lee, Seong Tae Jeong, Myung Guk Han |
| EPI_ISL_506994, EPI_ISL_506995 | Division of Viral Diseases, Center for Laboratory Control of Infectious Diseases, Korea Centers for Diseases Control and Prevention | Division of Viral Diseases, Center for Laboratory Control of Infectious Diseases, Korea Centers for Diseases Control and Prevention | Jeong-Min Kim, Yoon-Seok Chung, Namjoo Lee, Sang Hee Woo, Hye-Jun Jo, Heui Man Kim, Jun-Sub Kim, Myung Guk Han |
| EPI_ISL_506996 | Department of Medical Microbiology, University Malaya Medical Centre | Department of Medical Microbiology, Faculty of Medicine, University of Malaya | Yoong Min CHONG, Jennifer Chong, I-Ching SAM, Yoke Fun CHAN, University Malaya Medical Centre COVID Team |
| EPI_ISL_507002, EPI_ISL_507003, EPI_ISL_507004, EPI_ISL_507005, EPI_ISL_507006 | Department of Laboratory Medicine, Tan Tock Seng Hospital | Department of Laboratory Medicine, Tan Tock Seng Hospital | Chen YYC, Zair X, Li C, Tang WY, Maurer-Stroh S, Barkham TMS, Nagarajan N, Sessions OM |
| EPI_ISL_507008 | National Veterinary Institute | National Veterinary Institute | Siamak Zohari |
| EPI_ISL_507010, EPI_ISL_507011, EPI_ISL_507012, EPI_ISL_507013, EPI_ISL_507014, EPI_ISL_507015, EPI_ISL_507016, EPI_ISL_507017, EPI_ISL_507018, EPI_ISL_507019, EPI_ISL_507020, EPI_ISL_507021, EPI_ISL_507022, EPI_ISL_507023, EPI_ISL_507024, EPI_ISL_507025, EPI_ISL_507026, EPI_ISL_507027, EPI_ISL_507028, EPI_ISL_507029, EPI_ISL_507030, EPI_ISL_507031, EPI_ISL_507032, EPI_ISL_507033, EPI_ISL_507034, EPI_ISL_507035, EPI_ISL_507036, EPI_ISL_507037, EPI_ISL_507038 | | | |
| see above | unknown | Infectious Diseases Research, King Abdullah International Medical Research Center (KAIMRC) | Alghoribi,M.F. |
| EPI_ISL_507039 | Department of Microbiology, College of Medicine and Medical Research Institute Chungbuk National University | Department of Microbiology, College of Medicine and Medical Research Institute Chungbuk National University | Young-Il Kim, Mark Anthony B. Casel, Se-Mi Kim, Seong-Gyu Kim, Su-Jin Park, Eun-Ha Kim, Hye Won Jeong, Young Ki Choi |
| EPI_ISL_507040, EPI_ISL_507041, EPI_ISL_507042, EPI_ISL_507043, EPI_ISL_507047, EPI_ISL_507048, EPI_ISL_507049, EPI_ISL_507050, EPI_ISL_507052, EPI_ISL_507053, EPI_ISL_507054, EPI_ISL_507055, EPI_ISL_507056, EPI_ISL_507057, EPI_ISL_507058, EPI_ISL_507059, EPI_ISL_507060, EPI_ISL_507062, EPI_ISL_507063, EPI_ISL_507064, EPI_ISL_507065, EPI_ISL_507066, EPI_ISL_507067, EPI_ISL_507068, EPI_ISL_507069, EPI_ISL_507070, EPI_ISL_507071, EPI_ISL_507072, EPI_ISL_507073, EPI_ISL_507074, EPI_ISL_507075, EPI_ISL_507076, EPI_ISL_507077, EPI_ISL_507078, EPI_ISL_507079, EPI_ISL_507080, EPI_ISL_507081, EPI_ISL_507082, EPI_ISL_507083, EPI_ISL_507084, EPI_ISL_507086, EPI_ISL_507087, EPI_ISL_507089, EPI_ISL_507090, EPI_ISL_507091, EPI_ISL_507092, EPI_ISL_507093, EPI_ISL_507094, EPI_ISL_507096, EPI_ISL_507097, EPI_ISL_507098, EPI_ISL_507103, EPI_ISL_507104, EPI_ISL_507105, EPI_ISL_507107, EPI_ISL_507108 | | | |
| see above | University College London Hospital | COVID-19 Genomics UK (COG-UK) Consortium | Judith Heaney, Matthew Byott, Catherine Houlihan, Dan Frampton, Stuart Kirk, Moira Spyer and Eleni Nastouli |
| EPI_ISL_507110, EPI_ISL_507111, EPI_ISL_507112, EPI_ISL_507113, EPI_ISL_507114, EPI_ISL_507116, EPI_ISL_507117, EPI_ISL_507118, EPI_ISL_507120, EPI_ISL_507121, EPI_ISL_507123, EPI_ISL_507124, EPI_ISL_507125, EPI_ISL_507126, EPI_ISL_507127, EPI_ISL_507129 | | | |
| see above | Northumbria University / South Tees Hospitals NHS Foundation Trust / North Cumbria Integrated Care NHS Foundation Trust / North Tees and Hartlepool NHS Foundation Trust / Newcastle Hospitals NHS Foundation Trust | COVID-19 Genomics UK (COG-UK) Consortium | Darren L Smith,Andrew Nelson,Matthew Bashton,Greg R Young,Joshua Loh,John Allan,Mohammad A Tariq,Giles S Holt,Gary Black,Wen C Yew,Lynn Dover,Paul Baker,Steve Liggett,Sarah Essex,Jane Greenaway,Debra Padgett,Clive Graham,Garren Scott,Edward Barton,Emma Swindells,Brendan Payne,Jennifer Collins,Yusri Taha,Gary Eltringham |
| EPI_ISL_507131, EPI_ISL_507132, EPI_ISL_507133, EPI_ISL_507134, EPI_ISL_507135, EPI_ISL_507137, EPI_ISL_507139, EPI_ISL_507140, EPI_ISL_507141, EPI_ISL_507142, EPI_ISL_507143, EPI_ISL_507146, EPI_ISL_507147, EPI_ISL_507148, EPI_ISL_507149, EPI_ISL_507152, EPI_ISL_507153, EPI_ISL_507155, EPI_ISL_507159 | | | |
| see above | Centre for Enzyme Innovation, University of Portsmouth / Translational Research Laboratory, Portsmouth Hospitals NHS Trust | COVID-19 Genomics UK (COG-UK) Consortium | Angela Beckett,Yann Bourgeois,Garry Scarlett,Sharon Glaysher,Scott Elliott,Kelly Bicknell,Robert Impey,Allyson Lloyd,Sarah Wyllie,Ethan Butcher,Anoop Chauhan,Samuel Robson |
| EPI_ISL_507161, EPI_ISL_507163, EPI_ISL_507165, EPI_ISL_507178, EPI_ISL_507179 | Virology Department, Sheffield Teaching Hospitals NHS Foundation Trust/Department of Infection, Immunity and Cardiovascular Disease, The Medical School, University of Sheffield | COVID-19 Genomics UK (COG-UK) Consortium | Thushan de Silva, Matthew Parker, Nikki Smith, Adri Angyal, Rebecca Brown, Luke Green, Rachel Tucker, Paul Parsons, Danielle Groves, Katie Johnson, Laura Carrilero, Alex Keeley, Dave Partridge, Matthew Wyles, Benjamin Lindsey, Mehmet Yavuz, Mohammad Raza, Cariad Evans |
| EPI_ISL_507180, EPI_ISL_507181, EPI_ISL_507184, EPI_ISL_507185, EPI_ISL_507188, EPI_ISL_507192, EPI_ISL_507196, EPI_ISL_507200, EPI_ISL_507201 | Virology Department, Royal Infirmary of Edinburgh, NHS Lothian / School of Biological Sciences, University of Edinburgh / Institute of Genetics and Molecular Medicine, University of Edinburgh | COVID-19 Genomics UK (COG-UK) Consortium | McHugh M, Dewar R, Rooke S, Gallagher M, Balcaza C, O'Toole Á, Scher E, Hill V, McCrone JT, Colquhoun R, Yu X, Jackson B, Rambaut A, Williams TC, Templeton K |
| EPI_ISL_507205 | WHO National Influenza Centre Russian Federation | WHO National Influenza Centre Russian Federation | Andrey Komissarov, Artem Fadeev, Mariia Sergeeva, Anna Ivanova, Daria Danilenko |
| EPI_ISL_507206, EPI_ISL_507207, EPI_ISL_507208, EPI_ISL_507209, EPI_ISL_507210, EPI_ISL_507211, EPI_ISL_507212, EPI_ISL_507213, EPI_ISL_507214, EPI_ISL_507215 | Department of Experimental Modeling and Pathogenesis of Infectious Diseases | WHO National Influenza Centre Russian Federation | Andrey Komissarov, Artem Fadeev, Mariia Sergeeva, Anna Ivanova, Daria Danilenko |
| EPI_ISL_507216, EPI_ISL_507217, EPI_ISL_507218, EPI_ISL_507219, EPI_ISL_507220, EPI_ISL_507221, EPI_ISL_507222, EPI_ISL_507223, EPI_ISL_507224, EPI_ISL_507225, EPI_ISL_507226, EPI_ISL_507227, EPI_ISL_507228, EPI_ISL_507229, EPI_ISL_507230, EPI_ISL_507231, EPI_ISL_507232, EPI_ISL_507233, EPI_ISL_507234, EPI_ISL_507235, EPI_ISL_507236, EPI_ISL_507237, EPI_ISL_507238, EPI_ISL_507239, EPI_ISL_507240, EPI_ISL_507241, EPI_ISL_507242, EPI_ISL_507243, EPI_ISL_507244, EPI_ISL_507245, EPI_ISL_507246, EPI_ISL_507247, EPI_ISL_507248, EPI_ISL_507249, EPI_ISL_507250, EPI_ISL_507251, EPI_ISL_507252, EPI_ISL_507253, EPI_ISL_507254, EPI_ISL_507255, EPI_ISL_507256, EPI_ISL_507257, EPI_ISL_507258, EPI_ISL_507259, EPI_ISL_507260, EPI_ISL_507261, EPI_ISL_507262, EPI_ISL_507263, EPI_ISL_507264, EPI_ISL_507265, EPI_ISL_507266, EPI_ISL_507267, EPI_ISL_507268, EPI_ISL_507269, EPI_ISL_507270, EPI_ISL_507271, EPI_ISL_507272, EPI_ISL_507273, EPI_ISL_507274, EPI_ISL_507275, EPI_ISL_507276, EPI_ISL_507277, EPI_ISL_507278, EPI_ISL_507279, EPI_ISL_507280, EPI_ISL_507281, EPI_ISL_507282, EPI_ISL_507283, EPI_ISL_507284, EPI_ISL_507285, EPI_ISL_507286, EPI_ISL_507287, EPI_ISL_507288, EPI_ISL_507289, EPI_ISL_507290, EPI_ISL_507291, EPI_ISL_507292, EPI_ISL_507293, EPI_ISL_507294, EPI_ISL_507295, EPI_ISL_507296 | | | |
| see above | WHO National Influenza Centre Russian Federation | WHO National Influenza Centre Russian Federation | Andrey Komissarov, Artem Fadeev, Mariia Sergeeva, Anna Ivanova, Daria Danilenko |
| EPI_ISL_507297, EPI_ISL_507298, EPI_ISL_507299 | Republican Medical Genetics Center | WHO National Influenza Centre Russian Federation | Andrey Komissarov, Artem Fadeev, Mariia Sergeeva, Anna Ivanova, Ildar Minniakhmetov, Rita Khusainova, Daria Danilenko |
| EPI_ISL_507300 | WHO National Influenza Centre Russian Federation | WHO National Influenza Centre Russian Federation | Andrey Komissarov, Artem Fadeev, Mariia Sergeeva, Anna Ivanova, Daria Danilenko |
| EPI_ISL_507431, EPI_ISL_507434, EPI_ISL_507435, EPI_ISL_507437, EPI_ISL_507438, EPI_ISL_507441, EPI_ISL_507442, EPI_ISL_507444, EPI_ISL_507445, EPI_ISL_507446, EPI_ISL_507459, EPI_ISL_507460, EPI_ISL_507462, EPI_ISL_507464, EPI_ISL_507465, EPI_ISL_507466, EPI_ISL_507467, EPI_ISL_507468, EPI_ISL_507469, EPI_ISL_507470, EPI_ISL_507471, EPI_ISL_507528, EPI_ISL_507602, EPI_ISL_507603, EPI_ISL_507604, EPI_ISL_507605, EPI_ISL_507606, EPI_ISL_507607, EPI_ISL_507608, EPI_ISL_507609, EPI_ISL_507610, EPI_ISL_507611, EPI_ISL_507612, EPI_ISL_507613, EPI_ISL_507614, EPI_ISL_507615, EPI_ISL_507616, EPI_ISL_507617, EPI_ISL_507618, EPI_ISL_507619, EPI_ISL_507620, EPI_ISL_507621, EPI_ISL_507622, EPI_ISL_507623, EPI_ISL_507682, EPI_ISL_507684, EPI_ISL_507685, EPI_ISL_507686, EPI_ISL_507688, EPI_ISL_507689, EPI_ISL_507690, EPI_ISL_507691, EPI_ISL_507692, EPI_ISL_507693, EPI_ISL_507694, EPI_ISL_507695, EPI_ISL_507696, EPI_ISL_507697, EPI_ISL_507698, EPI_ISL_507699, EPI_ISL_507700, EPI_ISL_507701, EPI_ISL_507702, EPI_ISL_507703, EPI_ISL_507704, EPI_ISL_507705, EPI_ISL_507706, EPI_ISL_507707, EPI_ISL_507708, EPI_ISL_507709, EPI_ISL_507710, EPI_ISL_507711, EPI_ISL_507712, EPI_ISL_507713, EPI_ISL_507714, EPI_ISL_507715, EPI_ISL_507716, EPI_ISL_507717, EPI_ISL_507718, EPI_ISL_507719, EPI_ISL_507720, EPI_ISL_507739, EPI_ISL_507740, EPI_ISL_507743, EPI_ISL_507744, EPI_ISL_507745, EPI_ISL_507746, EPI_ISL_507747, EPI_ISL_507748, EPI_ISL_507760, EPI_ISL_507761, EPI_ISL_507763, EPI_ISL_507764, EPI_ISL_507777, EPI_ISL_507778, EPI_ISL_507779, EPI_ISL_507780, EPI_ISL_507781, EPI_ISL_507782, EPI_ISL_507783, EPI_ISL_507784, EPI_ISL_507785, EPI_ISL_507786, EPI_ISL_507787, EPI_ISL_507788, EPI_ISL_507789, EPI_ISL_507870, EPI_ISL_507871, EPI_ISL_507874, EPI_ISL_507879, EPI_ISL_507881, EPI_ISL_507882, EPI_ISL_507883, EPI_ISL_507884, EPI_ISL_507885, EPI_ISL_507886, EPI_ISL_507887, EPI_ISL_507888, EPI_ISL_507889, EPI_ISL_507890, EPI_ISL_507891, EPI_ISL_507892, EPI_ISL_507893, EPI_ISL_507894, EPI_ISL_507895, EPI_ISL_507896, EPI_ISL_507897, EPI_ISL_507898, EPI_ISL_507899, EPI_ISL_507900, EPI_ISL_507901, EPI_ISL_507902, EPI_ISL_507903, EPI_ISL_507904, EPI_ISL_507905, EPI_ISL_507906, EPI_ISL_507907, EPI_ISL_507908, EPI_ISL_507909, EPI_ISL_507910, EPI_ISL_507911, EPI_ISL_507912, EPI_ISL_507913, EPI_ISL_507914, EPI_ISL_507915, EPI_ISL_507916, EPI_ISL_507917, EPI_ISL_507919, EPI_ISL_507920, EPI_ISL_507921, EPI_ISL_507922, EPI_ISL_507923, EPI_ISL_507924, EPI_ISL_507925, EPI_ISL_507926, EPI_ISL_507927, EPI_ISL_507928, EPI_ISL_507930, EPI_ISL_507931, EPI_ISL_507932, EPI_ISL_507933 | | | |
| see above | Michigan Department of Health and Human Services, Bureau of Laboratories | Michigan Department of Health and Human Services, Bureau of Laboratories | Blankenship HM, Riner D, Soehnlen MK |
| EPI_ISL_507934, EPI_ISL_507935, EPI_ISL_507936, EPI_ISL_507937, EPI_ISL_507938, EPI_ISL_507939, EPI_ISL_507940, EPI_ISL_507941, EPI_ISL_507942, EPI_ISL_507943, EPI_ISL_507944, EPI_ISL_507945, EPI_ISL_507946, EPI_ISL_507947, EPI_ISL_507948, EPI_ISL_507949, EPI_ISL_507950, EPI_ISL_507951, EPI_ISL_507952, EPI_ISL_507953, EPI_ISL_507954, EPI_ISL_507955, EPI_ISL_507956 | | | |
| see above | Minnesota Department of Health, Public Health Laboratory | Minnesota Department of Health, Public Health Laboratory | Matt Plumb, Jacob Garfin, and Xiong Wang |
| EPI_ISL_507957, EPI_ISL_507958, EPI_ISL_507959, EPI_ISL_507960, EPI_ISL_507961 | Mayo Clinic & Mayo Clinic Laboratories | Minnesota Department of Health, Public Health Laboratory | Matt Plumb, Jacob Garfin, and Xiong Wang |
| EPI_ISL_507962 | Children's Hospitals and Clinics of Minnesota | Minnesota Department of Health, Public Health Laboratory | Matt Plumb, Jacob Garfin, and Xiong Wang |

| | | | |
|---|---|---|---|
| EPI_ISL_507963, EPI_ISL_507964, EPI_ISL_507965, EPI_ISL_507966, EPI_ISL_507967, EPI_ISL_507968, EPI_ISL_507969, EPI_ISL_507970, EPI_ISL_507971, EPI_ISL_507972 | Avera Mckennan Laboratory | Minnesota Department of Health, Public Health Laboratory | Matt Plumb, Jacob Garfin, and Xiong Wang |
| EPI_ISL_507973, EPI_ISL_507974, EPI_ISL_507975, EPI_ISL_507976, EPI_ISL_507977, EPI_ISL_507978, EPI_ISL_507979, EPI_ISL_507980, EPI_ISL_507981, EPI_ISL_507982, EPI_ISL_507983 | | | |
| see above | Minnesota Department of Health, Public Health Laboratory | Minnesota Department of Health, Public Health Laboratory | Matt Plumb, Jacob Garfin, and Xiong Wang |
| EPI_ISL_507986, EPI_ISL_507987, EPI_ISL_507991, EPI_ISL_507995, EPI_ISL_508002, EPI_ISL_508013, EPI_ISL_508014, EPI_ISL_508018, EPI_ISL_508026, EPI_ISL_508029, EPI_ISL_508057, EPI_ISL_508063, EPI_ISL_508066, EPI_ISL_508091, EPI_ISL_508093, EPI_ISL_508097, EPI_ISL_508099, EPI_ISL_508100, EPI_ISL_508101 | | | |
| see above | New Mexico Department of Health Scientific Laboratory Division | Center for Global Health, University of New Mexico Health Sciences Center | Daryl Domman, Kurt Schwalm, Twila Kunde, Joseph Hicks, Michael Edwards, Darrell Dinwiddie |
| EPI_ISL_508122, EPI_ISL_508123, EPI_ISL_508124, EPI_ISL_508125, EPI_ISL_508126, EPI_ISL_508127, EPI_ISL_508128, EPI_ISL_508129, EPI_ISL_508130, EPI_ISL_508131, EPI_ISL_508132, EPI_ISL_508133, EPI_ISL_508134, EPI_ISL_508135, EPI_ISL_508136, EPI_ISL_508137, EPI_ISL_508138, EPI_ISL_508139, EPI_ISL_508140, EPI_ISL_508141, EPI_ISL_508142, EPI_ISL_508143 | | | |
| see above | SA Pathology | SA Pathology | Lex Leong, Chuan Kok Lim, Mark Turra, Ivan Bastian, Geoff Higgins |
| EPI_ISL_508156, EPI_ISL_508157, EPI_ISL_508158, EPI_ISL_508159, EPI_ISL_508160, EPI_ISL_508161, EPI_ISL_508162, EPI_ISL_508163, EPI_ISL_508164, EPI_ISL_508165, EPI_ISL_508166, EPI_ISL_508167, EPI_ISL_508168, EPI_ISL_508169, EPI_ISL_508170, EPI_ISL_508171, EPI_ISL_508172, EPI_ISL_508173, EPI_ISL_508174, EPI_ISL_508175, EPI_ISL_508176, EPI_ISL_508177, EPI_ISL_508178, EPI_ISL_508179, EPI_ISL_508180, EPI_ISL_508181, EPI_ISL_508182, EPI_ISL_508183, EPI_ISL_508184, EPI_ISL_508185, EPI_ISL_508186, EPI_ISL_508187, EPI_ISL_508188, EPI_ISL_508189, EPI_ISL_508190, EPI_ISL_508191, EPI_ISL_508193, EPI_ISL_508194, EPI_ISL_508195, EPI_ISL_508196, EPI_ISL_508197, EPI_ISL_508198, EPI_ISL_508199, EPI_ISL_508200, EPI_ISL_508201, EPI_ISL_508202, EPI_ISL_508203, EPI_ISL_508204, EPI_ISL_508205, EPI_ISL_508206 | | | |
| see above | All india institute of Medical Sciences Rishikesh | National Institute of Biomedical Genomics | Arindam Maitra, Deepjyoti Kalita, Amit Mangla, Ravi Kant, Saumitra Das |
| EPI_ISL_508207, EPI_ISL_508208, EPI_ISL_508209, EPI_ISL_508210, EPI_ISL_508211, EPI_ISL_508212, EPI_ISL_508213, EPI_ISL_508214, EPI_ISL_508215, EPI_ISL_508216, EPI_ISL_508217, EPI_ISL_508218, EPI_ISL_508219, EPI_ISL_508220, EPI_ISL_508221, EPI_ISL_508222, EPI_ISL_508223, EPI_ISL_508224, EPI_ISL_508225, EPI_ISL_508226, EPI_ISL_508227, EPI_ISL_508228, EPI_ISL_508229, EPI_ISL_508230, EPI_ISL_508231, EPI_ISL_508232, EPI_ISL_508233, EPI_ISL_508234, EPI_ISL_508235, EPI_ISL_508236, EPI_ISL_508237, EPI_ISL_508238, EPI_ISL_508239, EPI_ISL_508240, EPI_ISL_508241, EPI_ISL_508242, EPI_ISL_508243, EPI_ISL_508244, EPI_ISL_508245, EPI_ISL_508246, EPI_ISL_508247, EPI_ISL_508248, EPI_ISL_508249, EPI_ISL_508250, EPI_ISL_508251, EPI_ISL_508252, EPI_ISL_508253, EPI_ISL_508254, EPI_ISL_508255, EPI_ISL_508256, EPI_ISL_508257, EPI_ISL_508258, EPI_ISL_508259, EPI_ISL_508260, EPI_ISL_508261, EPI_ISL_508262, EPI_ISL_508263, EPI_ISL_508264, EPI_ISL_508265, EPI_ISL_508266, EPI_ISL_508267, EPI_ISL_508268, EPI_ISL_508269, EPI_ISL_508270, EPI_ISL_508271, EPI_ISL_508272, EPI_ISL_508273, EPI_ISL_508274, EPI_ISL_508275, EPI_ISL_508276, EPI_ISL_508277, EPI_ISL_508278, EPI_ISL_508279, EPI_ISL_508280, EPI_ISL_508281, EPI_ISL_508282, EPI_ISL_508283, EPI_ISL_508284, EPI_ISL_508285, EPI_ISL_508286 | | | |
| see above | Government Medical College | National Institute of Biomedical Genomics | Arindam Maitra, Jyoti Iravane, Dhaval Khatri, Maitrik Dave, Saumitra Das |
| EPI_ISL_508287, EPI_ISL_508288, EPI_ISL_508289, EPI_ISL_508290, EPI_ISL_508291, EPI_ISL_508292, EPI_ISL_508293, EPI_ISL_508294, EPI_ISL_508295, EPI_ISL_508296, EPI_ISL_508297, EPI_ISL_508298, EPI_ISL_508299, EPI_ISL_508300, EPI_ISL_508301, EPI_ISL_508302, EPI_ISL_508303, EPI_ISL_508304, EPI_ISL_508305, EPI_ISL_508306, EPI_ISL_508307, EPI_ISL_508308, EPI_ISL_508309, EPI_ISL_508310, EPI_ISL_508311, EPI_ISL_508312, EPI_ISL_508313, EPI_ISL_508315, EPI_ISL_508316, EPI_ISL_508317, EPI_ISL_508318, EPI_ISL_508319, EPI_ISL_508320, EPI_ISL_508321, EPI_ISL_508322, EPI_ISL_508323, EPI_ISL_508324, EPI_ISL_508325, EPI_ISL_508326, EPI_ISL_508327, EPI_ISL_508328, EPI_ISL_508329, EPI_ISL_508330, EPI_ISL_508331, EPI_ISL_508332, EPI_ISL_508333, EPI_ISL_508334, EPI_ISL_508335, EPI_ISL_508336, EPI_ISL_508337 | | | |
| see above | Indian Institute of Science | National Institute of Biomedical Genomics | Arindam Maitra, Bharath K Sundararaj, Harsha Raheja, N. Srinivasan, Deepak K Saini, Amit Singh, Saumitra Das |
| EPI_ISL_508338, EPI_ISL_508339, EPI_ISL_508340, EPI_ISL_508341, EPI_ISL_508342, EPI_ISL_508343, EPI_ISL_508344, EPI_ISL_508345, EPI_ISL_508346, EPI_ISL_508347, EPI_ISL_508348, EPI_ISL_508349, EPI_ISL_508350, EPI_ISL_508351, EPI_ISL_508352, EPI_ISL_508353, EPI_ISL_508354, EPI_ISL_508355, EPI_ISL_508356, EPI_ISL_508357, EPI_ISL_508358, EPI_ISL_508359, EPI_ISL_508360, EPI_ISL_508361, EPI_ISL_508362, EPI_ISL_508363, EPI_ISL_508364, EPI_ISL_508365, EPI_ISL_508366, EPI_ISL_508367, EPI_ISL_508368, EPI_ISL_508369, EPI_ISL_508370, EPI_ISL_508371, EPI_ISL_508372, EPI_ISL_508373, EPI_ISL_508374, EPI_ISL_508375, EPI_ISL_508376, EPI_ISL_508377, EPI_ISL_508378, EPI_ISL_508379, EPI_ISL_508380, EPI_ISL_508381, EPI_ISL_508382, EPI_ISL_508383, EPI_ISL_508384, EPI_ISL_508385, EPI_ISL_508386, EPI_ISL_508387, EPI_ISL_508388, EPI_ISL_508389, EPI_ISL_508391, EPI_ISL_508392, EPI_ISL_508393, EPI_ISL_508394, EPI_ISL_508395, EPI_ISL_508396, EPI_ISL_508397, EPI_ISL_508398, EPI_ISL_508399, EPI_ISL_508400, EPI_ISL_508401, EPI_ISL_508402, EPI_ISL_508403, EPI_ISL_508404, EPI_ISL_508405, EPI_ISL_508406, EPI_ISL_508407, EPI_ISL_508408, EPI_ISL_508409, EPI_ISL_508410, EPI_ISL_508411, EPI_ISL_508412, EPI_ISL_508413, EPI_ISL_508414 | | | |
| see above | Institute of Post Graduate Medical Education & Research | National Institute of Biomedical Genomics | Arindam Maitra, Aritra Biswas, Jayeeta Haldar, Raja Ray, Monimoy Banerjee, Saumitra Das |
| EPI_ISL_508415, EPI_ISL_508416, EPI_ISL_508417, EPI_ISL_508418, EPI_ISL_508419, EPI_ISL_508420, EPI_ISL_508421, EPI_ISL_508422 | Maulana Azad Medical College | National Institute of Biomedical Genomics | Arindam Maitra, Sonal Saxena, Vikas Manchanda, Oves Siddiqui, Saumitra Das |
| EPI_ISL_508423, EPI_ISL_508424, EPI_ISL_508425, EPI_ISL_508426, EPI_ISL_508427, EPI_ISL_508428, EPI_ISL_508429, EPI_ISL_508430, EPI_ISL_508431, EPI_ISL_508432, EPI_ISL_508433, EPI_ISL_508434, EPI_ISL_508435, EPI_ISL_508436, EPI_ISL_508437, EPI_ISL_508438, EPI_ISL_508439, EPI_ISL_508440 | | | |
| see above | Mahatma Gandhi Institute of Medical Sciences | National Institute of Biomedical Genomics | Arindam Maitra, Vijayshri Deotale, Rahul Narang, Deepashri Maraskolhe, Saumitra Das |
| EPI_ISL_508444, EPI_ISL_508445, EPI_ISL_508446, EPI_ISL_508447, EPI_ISL_508448, EPI_ISL_508449, EPI_ISL_508450, EPI_ISL_508451, EPI_ISL_508452, EPI_ISL_508453, EPI_ISL_508454, EPI_ISL_508456, EPI_ISL_508457, EPI_ISL_508460, EPI_ISL_508463, EPI_ISL_508464, EPI_ISL_508465, EPI_ISL_508468, EPI_ISL_508469, EPI_ISL_508471, EPI_ISL_508475, EPI_ISL_508481, EPI_ISL_508482, EPI_ISL_508483, EPI_ISL_508487 | | | |
| see above | ICMR-National Institute of Cholera and Enteric Diseases | National Institute of Biomedical Genomics | Arindam Maitra, Mamta Chawla Sarkar, Sreedhar Chinnaswamy, Hasina Banu, Ananya Chatterjee, Shanta Dutta, Saumitra Das |
| EPI_ISL_508489, EPI_ISL_508490, EPI_ISL_508491, EPI_ISL_508492, EPI_ISL_508493, EPI_ISL_508494, EPI_ISL_508495, EPI_ISL_508496, EPI_ISL_508497, EPI_ISL_508498, EPI_ISL_508499, EPI_ISL_508500, EPI_ISL_508501, EPI_ISL_508502, EPI_ISL_508503, EPI_ISL_508504, EPI_ISL_508505, EPI_ISL_508506, EPI_ISL_508507, EPI_ISL_508508, EPI_ISL_508509 | | | |
| see above | Translational Health Science and Technology Institute | National Institute of Biomedical Genomics | Arindam Maitra, Guruprasad Medigeshi, Sharanabasava Patil, Anbalagan Ananthraj, Madhu Pareek, Imran Khan, Gagandeep Kang, Saumitra Das |
| EPI_ISL_508605, EPI_ISL_508606, EPI_ISL_508607 | SA Pathology | SA Pathology | Lex Leong, Chuan Kok Lim, Mark Turra, Ivan Bastian, Geoff Higgins |
| EPI_ISL_508627, EPI_ISL_508639, EPI_ISL_508640, EPI_ISL_508641, EPI_ISL_508642, EPI_ISL_508643, EPI_ISL_508646, EPI_ISL_508647, EPI_ISL_508653, EPI_ISL_508661, EPI_ISL_508663, EPI_ISL_508667, EPI_ISL_508676 | | | |
| see above | Departamento de Microbiología, CDB, Hospital Clínic, Barcelona | SeqCOVID-SPAIN consortium/IBV(CSIC) | Andrea Vergara, Mikel Martínez, Elisa Rubio, Jéssica Navero, Aida Peiró and SeqCOVID-SPAIN consortium |
| EPI_ISL_508688, EPI_ISL_508690, EPI_ISL_508691, EPI_ISL_508693, EPI_ISL_508694, EPI_ISL_508696, EPI_ISL_508697, EPI_ISL_508699, EPI_ISL_508700, EPI_ISL_508701, EPI_ISL_508702, EPI_ISL_508703, EPI_ISL_508704 | | | |
| see above | Institut für Virologie und Epidemiologie der Viruskrankheiten, Universitätsklinikum Tübingen | NGS Competence Center Tübingen, Institut für Medizinische Mikrobiologie und Hygiene, Universitätsklinikum Tübingen | Angel Angelov |
| EPI_ISL_508708, EPI_ISL_508709, EPI_ISL_508711, EPI_ISL_508712, EPI_ISL_508713, EPI_ISL_508716, EPI_ISL_508717, EPI_ISL_508718, EPI_ISL_508719, EPI_ISL_508720, EPI_ISL_508721, EPI_ISL_508723, EPI_ISL_508724, EPI_ISL_508725, EPI_ISL_508726, EPI_ISL_508727, EPI_ISL_508729, EPI_ISL_508730, EPI_ISL_508731, EPI_ISL_508732, EPI_ISL_508733, EPI_ISL_508734, EPI_ISL_508735, EPI_ISL_508736, EPI_ISL_508737, EPI_ISL_508743, EPI_ISL_508744, EPI_ISL_508745, EPI_ISL_508746, EPI_ISL_508748, EPI_ISL_508749, EPI_ISL_508750, EPI_ISL_508752, EPI_ISL_508753, EPI_ISL_508754, EPI_ISL_508755, EPI_ISL_508756, EPI_ISL_508757, EPI_ISL_508758, EPI_ISL_508759, EPI_ISL_508760, EPI_ISL_508762, EPI_ISL_508765, EPI_ISL_508766, EPI_ISL_508767, EPI_ISL_508769, EPI_ISL_508771, EPI_ISL_508772, EPI_ISL_508773, EPI_ISL_508774, EPI_ISL_508775, EPI_ISL_508776, EPI_ISL_508777, EPI_ISL_508778, EPI_ISL_508779, EPI_ISL_508780, EPI_ISL_508781, EPI_ISL_508782, EPI_ISL_508784, EPI_ISL_508786, EPI_ISL_508787, EPI_ISL_508788, EPI_ISL_508790, EPI_ISL_508791, EPI_ISL_508792, EPI_ISL_508793, EPI_ISL_508794, EPI_ISL_508795, EPI_ISL_508796, EPI_ISL_508797, EPI_ISL_508798, EPI_ISL_508799, EPI_ISL_508800, EPI_ISL_508801, EPI_ISL_508802, EPI_ISL_508803, EPI_ISL_508804, EPI_ISL_508805, EPI_ISL_508807, EPI_ISL_508821, EPI_ISL_508836, EPI_ISL_508837, EPI_ISL_508838, EPI_ISL_508839, EPI_ISL_508840, EPI_ISL_508841 | | | |
| see above | Florida Bureau of Public Health Laboratories | Florida Bureau of Public Health Laboratories | Sarah Schmedes, Jason Blanton |
| EPI_ISL_508862, EPI_ISL_508863 | Virology Unit, Institut Pasteur de Madagascar | Virology Unit, Institut Pasteur de Madagascar | Christian Ranaivoson, Cara Brook, Norosoa Razanajatovo, Vida Ahyong, Tsiry Randriambolamanantsoa, Michelle Tan, Vololoniaina Raharinosy, Helisoa Razafimanjato, Cristina M. Tato, Joseph L. DeRisi, Soa Fy Andriamandimby, Jean-Michel Heraud |
| EPI_ISL_508864 | Division of Infectious Diseases and Hospital Epidemiology, University Hospital Zürich | Institute of Medical Virology, University of Zurich | Verena Kufner, Maryam Zaheri, Dana Weissberg, Jürg Böni, Silvana K. Rampini, Peter W. Schreiber, Irene A. Abela, Hugo Sax, Aline Wolfensberger, Michael Huber |
| EPI_ISL_508865 | Division of Infectious Diseases and Hospital Epidemiology, University Hospital Zürich | Institute of Medical Virology, University of Zurich | Maryam Zaheri, Verena Kufner, Dana Weissberg, Jürg Böni, Silvana K. Rampini, Peter W. Schreiber, Irene A. Abela, Hugo Sax, Aline Wolfensberger, Michael Huber |
| EPI_ISL_508866 | Division of Infectious Diseases and Hospital Epidemiology, University Hospital Zürich | Institute of Medical Virology, University of Zurich | Verena Kufner, Maryam Zaheri, Dana Weissberg, Jürg Böni, Silvana K. Rampini, Peter W. Schreiber, Irene A. Abela, Hugo Sax, Aline Wolfensberger, Michael Huber |
| EPI_ISL_508867 | Division of Infectious Diseases and Hospital Epidemiology, University Hospital Zürich | Institute of Medical Virology, University of Zurich | Maryam Zaheri, Verena Kufner, Dana Weissberg, Jürg Böni, Silvana K. Rampini, Peter W. Schreiber, Irene A. Abela, Hugo Sax, Aline Wolfensberger, Michael Huber |
| EPI_ISL_508868 | Division of Infectious Diseases and Hospital Epidemiology, University Hospital Zürich | Institute of Medical Virology, University of Zurich | Verena Kufner, Maryam Zaheri, Dana Weissberg, Jürg Böni, Silvana K. Rampini, Peter W. Schreiber, Irene A. Abela, Hugo Sax, Aline Wolfensberger, Michael Huber |
| EPI_ISL_508869 | Division of Infectious Diseases and Hospital Epidemiology, University Hospital Zürich | Institute of Medical Virology, University of Zurich | Maryam Zaheri, Verena Kufner, Dana Weissberg, Jürg Böni, Silvana K. Rampini, Peter W. Schreiber, Irene A. Abela, Hugo Sax, Aline Wolfensberger, Michael Huber |
| EPI_ISL_508870 | Division of Infectious Diseases and Hospital Epidemiology, University Hospital Zürich | Institute of Medical Virology, University of Zurich | Verena Kufner, Maryam Zaheri, Dana Weissberg, Jürg Böni, Silvana K. Rampini, Peter W. Schreiber, Irene A. Abela, Hugo Sax, Aline Wolfensberger, Michael Huber |
| EPI_ISL_508871, EPI_ISL_508872, EPI_ISL_508873, EPI_ISL_508874 | Institut des Agents Infectieux (IAI), Hospices Civils de Lyon | CNR Virus des Infections Respiratoires - France SUD | Antonin Bal, Gregory Destras, Gwendolyne Burfin, Solenne Brun, Carine Moustaud, Raphaelle Lamy, Alexandre Gaymard, Maude Bouscambert-Duchamp, Florence Morfin-Sherpa, Martine Valette, Bruno Lina, Laurence Josset |
| EPI_ISL_508875, EPI_ISL_508876 | Centre Hospitalier de Macon | CNR Virus des Infections Respiratoires - France SUD | Antonin Bal, Gregory Destras, Gwendolyne Burfin, Solenne Brun, Carine Moustaud, Raphaelle Lamy, Alexandre Gaymard, Maude Bouscambert-Duchamp, Florence Morfin-Sherpa, Martine Valette, Bruno Lina, Laurence Josset |
| EPI_ISL_508877 | Institut des Agents Infectieux (IAI), Hospices Civils de Lyon | CNR Virus des Infections Respiratoires - France SUD | Antonin Bal, Gregory Destras, Gwendolyne Burfin, Solenne Brun, Carine Moustaud, Raphaelle Lamy, Alexandre Gaymard, Maude Bouscambert-Duchamp, Florence Morfin-Sherpa, Martine Valette, Bruno Lina, Laurence Josset |
| EPI_ISL_508878 | GH Les Portes du Sud | CNR Virus des Infections Respiratoires - France SUD | Antonin Bal, Gregory Destras, Gwendolyne Burfin, Solenne Brun, Carine Moustaud, Raphaelle Lamy, Alexandre Gaymard, Maude Bouscambert-Duchamp, Florence Morfin-Sherpa, Martine Valette, Bruno Lina, Laurence Josset |
| EPI_ISL_508879, EPI_ISL_508880 | Centre Hospitalier Saint Joseph Saint Luc | CNR Virus des Infections Respiratoires - France SUD | Antonin Bal, Gregory Destras, Gwendolyne Burfin, Solenne Brun, Carine Moustaud, Raphaelle Lamy, Alexandre Gaymard, Maude Bouscambert-Duchamp, Florence Morfin-Sherpa, Martine Valette, Bruno Lina, Laurence Josset |
| EPI_ISL_508881 | Centre Hospitalier de Valence | CNR Virus des Infections Respiratoires - France SUD | Antonin Bal, Gregory Destras, Gwendolyne Burfin, Solenne Brun, Carine Moustaud, Raphaelle Lamy, Alexandre Gaymard, Maude Bouscambert-Duchamp, Florence Morfin-Sherpa, Martine Valette, Bruno Lina, Laurence Josset |
| EPI_ISL_508882, EPI_ISL_508883, EPI_ISL_508884, EPI_ISL_508885, EPI_ISL_508886, EPI_ISL_508887, EPI_ISL_508888, EPI_ISL_508889, EPI_ISL_508890, EPI_ISL_508891, EPI_ISL_508892, EPI_ISL_508893, EPI_ISL_508894, EPI_ISL_508895, EPI_ISL_508896, EPI_ISL_508897, EPI_ISL_508898, EPI_ISL_508899, EPI_ISL_508900, EPI_ISL_508901, EPI_ISL_508902, EPI_ISL_508903, EPI_ISL_508904, EPI_ISL_508905, EPI_ISL_508906, EPI_ISL_508907, EPI_ISL_508908, EPI_ISL_508909, EPI_ISL_508910, EPI_ISL_508911 | | | |
| see above | Institut des Agents Infectieux (IAI), Hospices Civils de Lyon | CNR Virus des Infections Respiratoires - France SUD | Antonin Bal, Gregory Destras, Gwendolyne Burfin, Solenne Brun, Carine Moustaud, Raphaelle Lamy, Alexandre Gaymard, Maude Bouscambert-Duchamp, Florence Morfin-Sherpa, Martine Valette, Bruno Lina, Laurence Josset |

EPI_ISL_508912, EPI_ISL_508913, EPI_ISL_508914, EPI_ISL_508915, EPI_ISL_508916, EPI_ISL_508917, EPI_ISL_508918, EPI_ISL_508921, EPI_ISL_508922, EPI_ISL_508923, EPI_ISL_508924, EPI_ISL_508925, EPI_ISL_508926, EPI_ISL_508927, EPI_ISL_508928, EPI_ISL_508929, EPI_ISL_508930

| | | | |
|---|---|---|---|
| see above | CNR Virus des Infections Respiratoires - France SUD | CNR Virus des Infections Respiratoires - France SUD | Antonin Bal, Gregory Destras, Gwendolyne Burfin, Solenne Brun, Carine Moustaud, Raphaelle Lamy, Alexandre Gaymard, Maude Bouscambert-Duchamp, Florence Morfin-Sherpa, Martine Valette, Bruno Lina, Laurence Josset |
| EPI_ISL_508931 | L'Hôpital Nord-Ouest Tarare-Grandris | CNR Virus des Infections Respiratoires - France SUD | Antonin Bal, Gregory Destras, Gwendolyne Burfin, Solenne Brun, Carine Moustaud, Raphaelle Lamy, Alexandre Gaymard, Maude Bouscambert-Duchamp, Florence Morfin-Sherpa, Martine Valette, Bruno Lina, Laurence Josset |
| EPI_ISL_508932 | Centre Hospitalier de Villefranche | CNR Virus des Infections Respiratoires - France SUD | Antonin Bal, Gregory Destras, Gwendolyne Burfin, Solenne Brun, Carine Moustaud, Raphaelle Lamy, Alexandre Gaymard, Maude Bouscambert-Duchamp, Florence Morfin-Sherpa, Martine Valette, Bruno Lina, Laurence Josset |
| EPI_ISL_508933 | CNR Virus des Infections Respiratoires - France SUD | CNR Virus des Infections Respiratoires - France SUD | Antonin Bal, Gregory Destras, Gwendolyne Burfin, Solenne Brun, Carine Moustaud, Raphaelle Lamy, Alexandre Gaymard, Maude Bouscambert-Duchamp, Florence Morfin-Sherpa, Martine Valette, Bruno Lina, Laurence Josset |
| EPI_ISL_508934, EPI_ISL_508935, EPI_ISL_508936 | Centre hospitalier Métropole Savoie | CNR Virus des Infections Respiratoires - France SUD | Antonin Bal, Gregory Destras, Gwendolyne Burfin, Solenne Brun, Carine Moustaud, Raphaelle Lamy, Alexandre Gaymard, Maude Bouscambert-Duchamp, Florence Morfin-Sherpa, Martine Valette, Bruno Lina, Laurence Josset |
| EPI_ISL_508937 | CNR Virus des Infections Respiratoires - France SUD | CNR Virus des Infections Respiratoires - France SUD | Antonin Bal, Gregory Destras, Gwendolyne Burfin, Solenne Brun, Carine Moustaud, Raphaelle Lamy, Alexandre Gaymard, Maude Bouscambert-Duchamp, Florence Morfin-Sherpa, Martine Valette, Bruno Lina, Laurence Josset |
| EPI_ISL_508938 | Centre Hospitalier de Bourg en Bresse | CNR Virus des Infections Respiratoires - France SUD | Antonin Bal, Gregory Destras, Gwendolyne Burfin, Solenne Brun, Carine Moustaud, Raphaelle Lamy, Alexandre Gaymard, Maude Bouscambert-Duchamp, Florence Morfin-Sherpa, Martine Valette, Bruno Lina, Laurence Josset |
| EPI_ISL_508939, EPI_ISL_508940, EPI_ISL_508942 | CNR Virus des Infections Respiratoires - France SUD | CNR Virus des Infections Respiratoires - France SUD | Antonin Bal, Gregory Destras, Gwendolyne Burfin, Solenne Brun, Carine Moustaud, Raphaelle Lamy, Alexandre Gaymard, Maude Bouscambert-Duchamp, Florence Morfin-Sherpa, Martine Valette, Bruno Lina, Laurence Josset |
| EPI_ISL_508943 | Centre Hospitalier de Macon | CNR Virus des Infections Respiratoires - France SUD | Antonin Bal, Gregory Destras, Gwendolyne Burfin, Solenne Brun, Carine Moustaud, Raphaelle Lamy, Alexandre Gaymard, Maude Bouscambert-Duchamp, Florence Morfin-Sherpa, Martine Valette, Bruno Lina, Laurence Josset |
| EPI_ISL_508944 | Centre Hospitalier de Bourg en Bresse | CNR Virus des Infections Respiratoires - France SUD | Antonin Bal, Gregory Destras, Gwendolyne Burfin, Solenne Brun, Carine Moustaud, Raphaelle Lamy, Alexandre Gaymard, Maude Bouscambert-Duchamp, Florence Morfin-Sherpa, Martine Valette, Bruno Lina, Laurence Josset |
| EPI_ISL_508945 | CNR Virus des Infections Respiratoires - France SUD | CNR Virus des Infections Respiratoires - France SUD | Antonin Bal, Gregory Destras, Gwendolyne Burfin, Solenne Brun, Carine Moustaud, Raphaelle Lamy, Alexandre Gaymard, Maude Bouscambert-Duchamp, Florence Morfin-Sherpa, Martine Valette, Bruno Lina, Laurence Josset |
| EPI_ISL_508946 | Centre Hospitalier de Macon | CNR Virus des Infections Respiratoires - France SUD | Antonin Bal, Gregory Destras, Gwendolyne Burfin, Solenne Brun, Carine Moustaud, Raphaelle Lamy, Alexandre Gaymard, Maude Bouscambert-Duchamp, Florence Morfin-Sherpa, Martine Valette, Bruno Lina, Laurence Josset |
| EPI_ISL_508947 | Centre Hospitalier Lucien Hussel | CNR Virus des Infections Respiratoires - France SUD | Antonin Bal, Gregory Destras, Gwendolyne Burfin, Solenne Brun, Carine Moustaud, Raphaelle Lamy, Alexandre Gaymard, Maude Bouscambert-Duchamp, Florence Morfin-Sherpa, Martine Valette, Bruno Lina, Laurence Josset |
| EPI_ISL_508948 | CNR Virus des Infections Respiratoires - France SUD | CNR Virus des Infections Respiratoires - France SUD | Antonin Bal, Gregory Destras, Gwendolyne Burfin, Solenne Brun, Carine Moustaud, Raphaelle Lamy, Alexandre Gaymard, Maude Bouscambert-Duchamp, Florence Morfin-Sherpa, Martine Valette, Bruno Lina, Laurence Josset |
| EPI_ISL_508949, EPI_ISL_508950 | Centre Hospitalier de Macon | CNR Virus des Infections Respiratoires - France SUD | Antonin Bal, Gregory Destras, Gwendolyne Burfin, Solenne Brun, Carine Moustaud, Raphaelle Lamy, Alexandre Gaymard, Maude Bouscambert-Duchamp, Florence Morfin-Sherpa, Martine Valette, Bruno Lina, Laurence Josset |
| EPI_ISL_508951 | CNR Virus des Infections Respiratoires - France SUD | CNR Virus des Infections Respiratoires - France SUD | Antonin Bal, Gregory Destras, Gwendolyne Burfin, Solenne Brun, Carine Moustaud, Raphaelle Lamy, Alexandre Gaymard, Maude Bouscambert-Duchamp, Florence Morfin-Sherpa, Martine Valette, Bruno Lina, Laurence Josset |
| EPI_ISL_508952 | Centre Hospitalier de Macon | CNR Virus des Infections Respiratoires - France SUD | Antonin Bal, Gregory Destras, Gwendolyne Burfin, Solenne Brun, Carine Moustaud, Raphaelle Lamy, Alexandre Gaymard, Maude Bouscambert-Duchamp, Florence Morfin-Sherpa, Martine Valette, Bruno Lina, Laurence Josset |
| EPI_ISL_508953, EPI_ISL_508954, EPI_ISL_508955, EPI_ISL_508956, EPI_ISL_508957 | CNR Virus des Infections Respiratoires - France SUD | CNR Virus des Infections Respiratoires - France SUD | Antonin Bal, Gregory Destras, Gwendolyne Burfin, Solenne Brun, Carine Moustaud, Raphaelle Lamy, Alexandre Gaymard, Maude Bouscambert-Duchamp, Florence Morfin-Sherpa, Martine Valette, Bruno Lina, Laurence Josset |
| EPI_ISL_508958 | Centre Hospitalier Alpes Leman | CNR Virus des Infections Respiratoires - France SUD | Antonin Bal, Gregory Destras, Gwendolyne Burfin, Solenne Brun, Carine Moustaud, Raphaelle Lamy, Alexandre Gaymard, Maude Bouscambert-Duchamp, Florence Morfin-Sherpa, Martine Valette, Bruno Lina, Laurence Josset |
| EPI_ISL_508959, EPI_ISL_508960 | Centre Hospitalier Saint Joseph Saint Luc | CNR Virus des Infections Respiratoires - France SUD | Antonin Bal, Gregory Destras, Gwendolyne Burfin, Solenne Brun, Carine Moustaud, Raphaelle Lamy, Alexandre Gaymard, Maude Bouscambert-Duchamp, Florence Morfin-Sherpa, Martine Valette, Bruno Lina, Laurence Josset |
| EPI_ISL_508961, EPI_ISL_508962, EPI_ISL_508964, EPI_ISL_508965 | CNR Virus des Infections Respiratoires - France SUD | CNR Virus des Infections Respiratoires - France SUD | Antonin Bal, Gregory Destras, Gwendolyne Burfin, Solenne Brun, Carine Moustaud, Raphaelle Lamy, Alexandre Gaymard, Maude Bouscambert-Duchamp, Florence Morfin-Sherpa, Martine Valette, Bruno Lina, Laurence Josset |
| EPI_ISL_508966 | Centre Hospitalier Pierre Oudot | CNR Virus des Infections Respiratoires - France SUD | Antonin Bal, Gregory Destras, Gwendolyne Burfin, Solenne Brun, Carine Moustaud, Raphaelle Lamy, Alexandre Gaymard, Maude Bouscambert-Duchamp, Florence Morfin-Sherpa, Martine Valette, Bruno Lina, Laurence Josset |
| EPI_ISL_508967 | CNR Virus des Infections Respiratoires - France SUD | CNR Virus des Infections Respiratoires - France SUD | Antonin Bal, Gregory Destras, Gwendolyne Burfin, Solenne Brun, Carine Moustaud, Raphaelle Lamy, Alexandre Gaymard, Maude Bouscambert-Duchamp, Florence Morfin-Sherpa, Martine Valette, Bruno Lina, Laurence Josset |
| EPI_ISL_508968 | Centre Hospitalier de Bourg en Bresse | CNR Virus des Infections Respiratoires - France SUD | Antonin Bal, Gregory Destras, Gwendolyne Burfin, Solenne Brun, Carine Moustaud, Raphaelle Lamy, Alexandre Gaymard, Maude Bouscambert-Duchamp, Florence Morfin-Sherpa, Martine Valette, Bruno Lina, Laurence Josset |
| EPI_ISL_508969, EPI_ISL_508970, EPI_ISL_508972, EPI_ISL_508973, EPI_ISL_508974 | CNR Virus des Infections Respiratoires - France SUD | CNR Virus des Infections Respiratoires - France SUD | Antonin Bal, Gregory Destras, Gwendolyne Burfin, Solenne Brun, Carine Moustaud, Raphaelle Lamy, Alexandre Gaymard, Maude Bouscambert-Duchamp, Florence Morfin-Sherpa, Martine Valette, Bruno Lina, Laurence Josset |
| EPI_ISL_508975 | Centre Hospitalier Pierre Oudot | CNR Virus des Infections Respiratoires - France SUD | Antonin Bal, Gregory Destras, Gwendolyne Burfin, Solenne Brun, Carine Moustaud, Raphaelle Lamy, Alexandre Gaymard, Maude Bouscambert-Duchamp, Florence Morfin-Sherpa, Martine Valette, Bruno Lina, Laurence Josset |
| EPI_ISL_508976, EPI_ISL_508977 | CNR Virus des Infections Respiratoires - France SUD | CNR Virus des Infections Respiratoires - France SUD | Antonin Bal, Gregory Destras, Gwendolyne Burfin, Solenne Brun, Carine Moustaud, Raphaelle Lamy, Alexandre Gaymard, Maude Bouscambert-Duchamp, Florence Morfin-Sherpa, Martine Valette, Bruno Lina, Laurence Josset |
| EPI_ISL_508978 | Centre Hospitalier de Villefranche | CNR Virus des Infections Respiratoires - France SUD | Antonin Bal, Gregory Destras, Gwendolyne Burfin, Solenne Brun, Carine Moustaud, Raphaelle Lamy, Alexandre Gaymard, Maude Bouscambert-Duchamp, Florence Morfin-Sherpa, Martine Valette, Bruno Lina, Laurence Josset |
| EPI_ISL_508979, EPI_ISL_508980, EPI_ISL_508981, EPI_ISL_508982, EPI_ISL_508983, EPI_ISL_508984, EPI_ISL_508985 | CNR Virus des Infections Respiratoires - France SUD | CNR Virus des Infections Respiratoires - France SUD | Antonin Bal, Gregory Destras, Gwendolyne Burfin, Solenne Brun, Carine Moustaud, Raphaelle Lamy, Alexandre Gaymard, Maude Bouscambert-Duchamp, Florence Morfin-Sherpa, Martine Valette, Bruno Lina, Laurence Josset |
| EPI_ISL_508986, EPI_ISL_508987 | CNR Virus des Infections Respiratoires - France SUD | CNR Virus des Infections Respiratoires - France SUD | Antonin Bal, Gregory Destras, Gwendolyne Burfin, Solenne Brun, Alexandre Gaymard, Maude Bouscambert-Duchamp, Florence Morfin-Sherpa, Martine Valette, Bruno Lina, Laurence Josset |
| EPI_ISL_508990, EPI_ISL_508991, EPI_ISL_508993, EPI_ISL_508994, EPI_ISL_508995, EPI_ISL_508996, EPI_ISL_508997 | CNR Virus des Infections Respiratoires - France SUD | CNR Virus des Infections Respiratoires - France SUD | Antonin Bal, Gregory Destras, Gwendolyne Burfin, Solenne Brun, Carine Moustaud, Raphaelle Lamy, Alexandre Gaymard, Maude Bouscambert-Duchamp, Florence Morfin-Sherpa, Martine Valette, Bruno Lina, Laurence Josset |
| EPI_ISL_508998 | Centre Hospitalier Pierre Oudot | CNR Virus des Infections Respiratoires - France SUD | Antonin Bal, Gregory Destras, Gwendolyne Burfin, Solenne Brun, Carine Moustaud, Raphaelle Lamy, Alexandre Gaymard, Maude Bouscambert-Duchamp, Florence Morfin-Sherpa, Martine Valette, Bruno Lina, Laurence Josset |
| EPI_ISL_508999, EPI_ISL_509000, EPI_ISL_509001, EPI_ISL_509002 | CNR Virus des Infections Respiratoires - France SUD | CNR Virus des Infections Respiratoires - France SUD | Antonin Bal, Gregory Destras, Gwendolyne Burfin, Solenne Brun, Carine Moustaud, Raphaelle Lamy, Alexandre Gaymard, Maude Bouscambert-Duchamp, Florence Morfin-Sherpa, Martine Valette, Bruno Lina, Laurence Josset |
| EPI_ISL_509003, EPI_ISL_509004 | GH Les Portes du Sud | CNR Virus des Infections Respiratoires - France SUD | Antonin Bal, Gregory Destras, Gwendolyne Burfin, Solenne Brun, Carine Moustaud, Raphaelle Lamy, Alexandre Gaymard, Maude Bouscambert-Duchamp, Florence Morfin-Sherpa, Martine Valette, Bruno Lina, Laurence Josset |
| EPI_ISL_509005 | Centre Hospitalier de Macon | CNR Virus des Infections Respiratoires - France SUD | Antonin Bal, Gregory Destras, Gwendolyne Burfin, Solenne Brun, Carine Moustaud, Raphaelle Lamy, Alexandre Gaymard, Maude Bouscambert-Duchamp, Florence Morfin-Sherpa, Martine Valette, Bruno Lina, Laurence Josset |
| EPI_ISL_509006 | Centre Hospitalier de Villefranche | CNR Virus des Infections Respiratoires - France SUD | Antonin Bal, Gregory Destras, Gwendolyne Burfin, Solenne Brun, Carine Moustaud, Raphaelle Lamy, Alexandre Gaymard, Maude Bouscambert-Duchamp, Florence Morfin-Sherpa, Martine Valette, Bruno Lina, Laurence Josset |
| EPI_ISL_509007, EPI_ISL_509008, EPI_ISL_509009, EPI_ISL_509010, EPI_ISL_509011 | Institut des Agents Infectieux (IAI), Hospices Civils de Lyon | CNR Virus des Infections Respiratoires - France SUD | Antonin Bal, Gregory Destras, Gwendolyne Burfin, Solenne Brun, Carine Moustaud, Raphaelle Lamy, Alexandre Gaymard, Maude Bouscambert-Duchamp, Florence Morfin-Sherpa, Martine Valette, Bruno Lina, Laurence Josset |
| EPI_ISL_509012 | Centre Hospitalier Alpes Leman | CNR Virus des Infections Respiratoires - France SUD | Antonin Bal, Gregory Destras, Gwendolyne Burfin, Solenne Brun, Carine Moustaud, Raphaelle Lamy, Alexandre Gaymard, Maude Bouscambert-Duchamp, Florence Morfin-Sherpa, Martine Valette, Bruno Lina, Laurence Josset |
| EPI_ISL_509013, EPI_ISL_509014 | Institut des Agents Infectieux (IAI), Hospices Civils de Lyon | CNR Virus des Infections Respiratoires - France SUD | Antonin Bal, Gregory Destras, Gwendolyne Burfin, Solenne Brun, Carine Moustaud, Raphaelle Lamy, Alexandre Gaymard, Maude Bouscambert-Duchamp, Florence Morfin-Sherpa, Martine Valette, Bruno Lina, Laurence Josset |
| EPI_ISL_509015 | Centre Hospitalier de Villefranche | CNR Virus des Infections Respiratoires - France SUD | Antonin Bal, Gregory Destras, Gwendolyne Burfin, Solenne Brun, Carine Moustaud, Raphaelle Lamy, Alexandre Gaymard, Maude Bouscambert-Duchamp, Florence Morfin-Sherpa, Martine Valette, Bruno Lina, Laurence Josset |
| EPI_ISL_509016 | Centre Hospitalier du Haut-Bugey | CNR Virus des Infections Respiratoires - France SUD | Antonin Bal, Gregory Destras, Gwendolyne Burfin, Solenne Brun, Carine Moustaud, Raphaelle Lamy, Alexandre Gaymard, Maude Bouscambert-Duchamp, Florence Morfin-Sherpa, Martine Valette, Bruno Lina, Laurence Josset |

EPI_ISL_509017, EPI_ISL_509018, EPI_ISL_509019, EPI_ISL_509020, EPI_ISL_509021, EPI_ISL_509022, EPI_ISL_509023, EPI_ISL_509024, EPI_ISL_509026, EPI_ISL_509028, EPI_ISL_509029, EPI_ISL_509031, EPI_ISL_509033, EPI_ISL_509035, EPI_ISL_509037, EPI_ISL_509038, EPI_ISL_509051

| | | | |
|---|---|---|---|
| see above | genXone SA, Molecular Diagnostics Laboratory / NZOZ | genXone SA, Research & Development Laboratory | Maciej Sykulski, Grzegorz Nowicki, Monika Mańkowska-Woźniak, Jakub Grabowski, Natalia Drwęska-Matelska, Łukasz Krych, Michał Kaszuba |
| EPI_ISL_509059, EPI_ISL_509060, EPI_ISL_509061, EPI_ISL_509062, EPI_ISL_509064, EPI_ISL_509065, EPI_ISL_509066, EPI_ISL_509067, EPI_ISL_509069, EPI_ISL_509070, EPI_ISL_509071, EPI_ISL_509072, EPI_ISL_509073, EPI_ISL_509074, EPI_ISL_509075, EPI_ISL_509076, EPI_ISL_509079, EPI_ISL_509080, EPI_ISL_509081, EPI_ISL_509082, EPI_ISL_509083, EPI_ISL_509084, EPI_ISL_509085, EPI_ISL_509086, EPI_ISL_509087, EPI_ISL_509088, EPI_ISL_509089, EPI_ISL_509090, EPI_ISL_509091, EPI_ISL_509092, EPI_ISL_509093, EPI_ISL_509094, EPI_ISL_509095, EPI_ISL_509097, EPI_ISL_509099, EPI_ISL_509101, EPI_ISL_509102, EPI_ISL_509103, EPI_ISL_509104, EPI_ISL_509106, EPI_ISL_509107, EPI_ISL_509108, EPI_ISL_509109, EPI_ISL_509110, EPI_ISL_509111, EPI_ISL_509112, EPI_ISL_509113, EPI_ISL_509114, EPI_ISL_509115, EPI_ISL_509116, EPI_ISL_509117, EPI_ISL_509118, EPI_ISL_509119, EPI_ISL_509120, EPI_ISL_509121, EPI_ISL_509122, EPI_ISL_509123, EPI_ISL_509124, EPI_ISL_509125, EPI_ISL_509126, EPI_ISL_509127, EPI_ISL_509128, EPI_ISL_509129, EPI_ISL_509130, EPI_ISL_509131, EPI_ISL_509132, EPI_ISL_509133, EPI_ISL_509134, EPI_ISL_509135, EPI_ISL_509136, EPI_ISL_509137, EPI_ISL_509138, EPI_ISL_509139, EPI_ISL_509140, EPI_ISL_509141, EPI_ISL_509142, EPI_ISL_509143, EPI_ISL_509144, EPI_ISL_509145, EPI_ISL_509146, EPI_ISL_509147, EPI_ISL_509148, EPI_ISL_509149, EPI_ISL_509150, EPI_ISL_509151, EPI_ISL_509153, EPI_ISL_509154, EPI_ISL_509155, EPI_ISL_509156, EPI_ISL_509157, EPI_ISL_509159, EPI_ISL_509160, EPI_ISL_509161, EPI_ISL_509162, EPI_ISL_509163, EPI_ISL_509164, EPI_ISL_509165, EPI_ISL_509166, EPI_ISL_509167, EPI_ISL_509168, EPI_ISL_509169, EPI_ISL_509170, EPI_ISL_509171, EPI_ISL_509174, EPI_ISL_509175, EPI_ISL_509176, EPI_ISL_509181, EPI_ISL_509182, EPI_ISL_509183, EPI_ISL_509184, EPI_ISL_509185, EPI_ISL_509186, EPI_ISL_509187, EPI_ISL_509188, EPI_ISL_509189, EPI_ISL_509191, EPI_ISL_509192, EPI_ISL_509193, EPI_ISL_509194, EPI_ISL_509195, EPI_ISL_509197, EPI_ISL_509198, EPI_ISL_509199, EPI_ISL_509201, EPI_ISL_509202, EPI_ISL_509203, EPI_ISL_509204, EPI_ISL_509205, EPI_ISL_509206, EPI_ISL_509208, EPI_ISL_509209, EPI_ISL_509210, EPI_ISL_509211, EPI_ISL_509212, EPI_ISL_509213, EPI_ISL_509214 | | | |
| see above | OHSU Lab Services Molecular Microbiology Lab | Oregon SARS-CoV-2 Genome Sequencing Center | Brendan L. O'Connell, Ruth V. Nichols, Sally B. Grindstaff, Alec J. Hirsch, Guang Fan, Daniel N. Streblow, William B. Messer, Andrew C. Adey, Benjamin N. Bimber, Brian J. O'Roak |
| EPI_ISL_509222 | Division of Infectious Diseases and Hospital Epidemiology, University Hospital Zürich | Institute of Medical Virology, University of Zurich | Maryam Zaheri, Verena Kufner, Dana Weissberg, Jürg Böni, Silvana K. Rampini, Peter W. Schreiber, Irene A. Abela, Hugo Sax, Aline Wolfensberger, Michael Huber |
| EPI_ISL_509223, EPI_ISL_509224, EPI_ISL_509225, EPI_ISL_509226, EPI_ISL_509228, EPI_ISL_509229, EPI_ISL_509230, EPI_ISL_509231, EPI_ISL_509232, EPI_ISL_509233, EPI_ISL_509234, EPI_ISL_509235, EPI_ISL_509236, EPI_ISL_509237, EPI_ISL_509238, EPI_ISL_509239, EPI_ISL_509240, EPI_ISL_509241, EPI_ISL_509242, EPI_ISL_509243, EPI_ISL_509244, EPI_ISL_509245, EPI_ISL_509246, EPI_ISL_509247, EPI_ISL_509248, EPI_ISL_509249, EPI_ISL_509250, EPI_ISL_509251, EPI_ISL_509252, EPI_ISL_509253, EPI_ISL_509254, EPI_ISL_509255, EPI_ISL_509256, EPI_ISL_509257, EPI_ISL_509258, EPI_ISL_509260, EPI_ISL_509261, EPI_ISL_509262, EPI_ISL_509263, EPI_ISL_509264, EPI_ISL_509265, EPI_ISL_509266, EPI_ISL_509267, EPI_ISL_509269, EPI_ISL_509270, EPI_ISL_509271, EPI_ISL_509274, EPI_ISL_509275, EPI_ISL_509276, EPI_ISL_509279, EPI_ISL_509281, EPI_ISL_509283, EPI_ISL_509284, EPI_ISL_509285, EPI_ISL_509286, EPI_ISL_509287, EPI_ISL_509288, EPI_ISL_509289, EPI_ISL_509290, EPI_ISL_509292, EPI_ISL_509293, EPI_ISL_509294, EPI_ISL_509295, EPI_ISL_509296, EPI_ISL_509297, EPI_ISL_509299, EPI_ISL_509300, EPI_ISL_509301, EPI_ISL_509302, EPI_ISL_509303, EPI_ISL_509304, EPI_ISL_509305, EPI_ISL_509306, EPI_ISL_509307, EPI_ISL_509308, EPI_ISL_509309, EPI_ISL_509310, EPI_ISL_509311, EPI_ISL_509312, EPI_ISL_509313, EPI_ISL_509314, EPI_ISL_509315, EPI_ISL_509316, EPI_ISL_509317, EPI_ISL_509318, EPI_ISL_509319, EPI_ISL_509320, EPI_ISL_509321, EPI_ISL_509322, EPI_ISL_509323, EPI_ISL_509324, EPI_ISL_509325, EPI_ISL_509326, EPI_ISL_509327, EPI_ISL_509328, EPI_ISL_509329, EPI_ISL_509330, EPI_ISL_509331, EPI_ISL_509332, EPI_ISL_509333, EPI_ISL_509334, EPI_ISL_509335, EPI_ISL_509336, EPI_ISL_509337, EPI_ISL_509338, EPI_ISL_509339, EPI_ISL_509340, EPI_ISL_509341, EPI_ISL_509342, EPI_ISL_509343, EPI_ISL_509344, EPI_ISL_509345, EPI_ISL_509346, EPI_ISL_509347, EPI_ISL_509348, EPI_ISL_509349, EPI_ISL_509350, EPI_ISL_509351, EPI_ISL_509352, EPI_ISL_509353, EPI_ISL_509354, EPI_ISL_509355, EPI_ISL_509356, EPI_ISL_509357, EPI_ISL_509358, EPI_ISL_509360, EPI_ISL_509361, EPI_ISL_509362, EPI_ISL_509363, EPI_ISL_509364, EPI_ISL_509365, EPI_ISL_509366, EPI_ISL_509367, EPI_ISL_509368, EPI_ISL_509369 | | | |
| see above | NHLS-IALCH | KRISP, KZN Research Innovation and Sequencing Platform | Giandhari J, Pillay S, Lessells R, Mdlalose K, York D, Tegally H, Wilkinson E, de Oliveira T |
| EPI_ISL_509373, EPI_ISL_509374, EPI_ISL_509375, EPI_ISL_509377, EPI_ISL_509378, EPI_ISL_509379, EPI_ISL_509380, EPI_ISL_509381, EPI_ISL_509382, EPI_ISL_509383, EPI_ISL_509384, EPI_ISL_509385, EPI_ISL_509386, EPI_ISL_509387, EPI_ISL_509388, EPI_ISL_509389, EPI_ISL_509390, EPI_ISL_509391, EPI_ISL_509392, EPI_ISL_509393, EPI_ISL_509394 | | | |
| see above | Singapore General Hospital | Department of Microbiology | Nurdyana Abdul Rahman, Kun Lee Lim, Chenhao Li, Kian Sing Chan, Lynette Oon, Kern Rei Chng, Niranjan Nagarajan, Karrie Ko |
| EPI_ISL_509416 | Acibadem Labcell Cellular Therapy Laboratory | Acibadem Mehmet Ali Aydinlar University School of Medicine, Medical Genetics Department | Ozden Hatirnaz Ng, Sezer Akyoney, Ilayda Sahin, Gunseli Bayram Akcapinar, Ozkan Ozdemir, Derya Dilek Kancagi, Gozde Sir Karakus, Bulut Yurtsever, Cihan Tastan, Ercument Ovali, Ugur Ozbek |
| EPI_ISL_509424, EPI_ISL_509425, EPI_ISL_509426, EPI_ISL_509427, EPI_ISL_509428, EPI_ISL_509429 | Microbiology and Immunology, University of South Alabama | Microbiology and Immunology, University of South Alabama | Wood,R.R., Roberts,R.A., Houserova,D., Borchert,G.M., Fouty,B., Rayner,J.O. |
| EPI_ISL_509430, EPI_ISL_509431, EPI_ISL_509432, EPI_ISL_509433, EPI_ISL_509434, EPI_ISL_509435 | Centro de Desenvolvimento Tecnologico em Saude, Fundacao Oswaldo Cruz | Centro de Desenvolvimento Tecnologico em Saude, Fundacao Oswaldo Cruz | Souza,T.M., Fintelman-Rodrigues,N., De Paula,A.D., Saraiva,F.B., Ferreira,M.A., Sacramento,C.Q., Medeiros,M.A. |
| EPI_ISL_509436, EPI_ISL_509439, EPI_ISL_509440, EPI_ISL_509442, EPI_ISL_509443 | The Princess Alexandra Hospital | Wellcome Sanger Institute for the COVID-19 Genomics UK Consortium | Nick Levene, Louise Lopez, Lynn Monaghan, Jessica Scott, Claudia McCrea and Alex Alderton, Roberto Amato, Sonia Goncalves, Ewan Harrison, David K. Jackson, Ian Johnston, Dominic Kwiatkowski, Cordelia Langford, John Sillitoe on behalf of the Wellcome Sanger Institute COVID-19 Surveillance Team (http://www.sanger.ac.uk/covid-team) |
| EPI_ISL_509444, EPI_ISL_509445, EPI_ISL_509446, EPI_ISL_509447, EPI_ISL_509448, EPI_ISL_509449, EPI_ISL_509450, EPI_ISL_509451, EPI_ISL_509452, EPI_ISL_509453, EPI_ISL_509454, EPI_ISL_509455, EPI_ISL_509456, EPI_ISL_509457, EPI_ISL_509458, EPI_ISL_509459, EPI_ISL_509460, EPI_ISL_509461, EPI_ISL_509462, EPI_ISL_509463, EPI_ISL_509464, EPI_ISL_509465, EPI_ISL_509466, EPI_ISL_509467, EPI_ISL_509468, EPI_ISL_509469, EPI_ISL_509470, EPI_ISL_509471, EPI_ISL_509472, EPI_ISL_509473, EPI_ISL_509474, EPI_ISL_509475, EPI_ISL_509476, EPI_ISL_509477, EPI_ISL_509478, EPI_ISL_509479, EPI_ISL_509480, EPI_ISL_509481, EPI_ISL_509482, EPI_ISL_509483, EPI_ISL_509484, EPI_ISL_509485, EPI_ISL_509486, EPI_ISL_509487, EPI_ISL_509488, EPI_ISL_509489, EPI_ISL_509490, EPI_ISL_509491 | | | |
| see above | Maryland Department of Health | Maryland Department of Health | Keller,E. |
| EPI_ISL_509492, EPI_ISL_509493, EPI_ISL_509494, EPI_ISL_509495, EPI_ISL_509496, EPI_ISL_509497, EPI_ISL_509498, EPI_ISL_509499, EPI_ISL_509500, EPI_ISL_509501, EPI_ISL_509502, EPI_ISL_509503, EPI_ISL_509504, EPI_ISL_509505, EPI_ISL_509506, EPI_ISL_509507, EPI_ISL_509508, EPI_ISL_509509, EPI_ISL_509510, EPI_ISL_509511, EPI_ISL_509512, EPI_ISL_509513, EPI_ISL_509514, EPI_ISL_509515, EPI_ISL_509516, EPI_ISL_509517, EPI_ISL_509518, EPI_ISL_509519, EPI_ISL_509520, EPI_ISL_509521, EPI_ISL_509522, EPI_ISL_509523 | | | |
| see above | Area of Virology, Serology and Virology Division (SAViD), New South Wales Health Pathology Randwick | Area of Virology, Serology and Virology Division (SAViD), New South Wales Health Pathology Randwick | Rawlinson, W. |
| EPI_ISL_509528, EPI_ISL_509529, EPI_ISL_509530, EPI_ISL_509531, EPI_ISL_509534, EPI_ISL_509535, EPI_ISL_509536, EPI_ISL_509537, EPI_ISL_509540, EPI_ISL_509541, EPI_ISL_509547, EPI_ISL_509549, EPI_ISL_509550, EPI_ISL_509553, EPI_ISL_509554, EPI_ISL_509558, EPI_ISL_509559, EPI_ISL_509560, EPI_ISL_509563, EPI_ISL_509568, EPI_ISL_509571, EPI_ISL_509572, EPI_ISL_509576, EPI_ISL_509577, EPI_ISL_509580, EPI_ISL_509585, EPI_ISL_509588, EPI_ISL_509589, EPI_ISL_509590, EPI_ISL_509592 | | | |
| see above | Utah Public Health Laboratory | Utah Public Health Laboratory | Heidi Butz, Erin Young, Kelly Oakeson |
| EPI_ISL_509616, EPI_ISL_509617, EPI_ISL_509618 | Hospital Universitario Araba. Vitoria-Gasteiz | SeqCOVID-SPAIN consortium/IBV(CSIC) | Silvia Hernáez Crespo, Carmen Gómez González, Amaia Aguirre Quiñonero, Marina Fernández Torres, Mª Rosario Almela Ferrer, Mª Concepción Lecaroz Agara, Andrés Canut Blasco and SeqCOVID-SPAIN consortium |
| EPI_ISL_509619, EPI_ISL_509620, EPI_ISL_509621, EPI_ISL_509622, EPI_ISL_509623, EPI_ISL_509624, EPI_ISL_509625, EPI_ISL_509626, EPI_ISL_509627, EPI_ISL_509628, EPI_ISL_509629, EPI_ISL_509630, EPI_ISL_509631, EPI_ISL_509632 | | | |
| see above | Servicio de Microbiología. HRU de Málaga. Servicio Andaluz de Salud | SeqCOVID-SPAIN consortium/IBV(CSIC) | Inmaculada de Toro Peinado. MªConcepción Mediavilla Gradolph. Begoña Palop Borrás and SeqCOVID-SPAIN consortium |
| EPI_ISL_509633, EPI_ISL_509634, EPI_ISL_509635, EPI_ISL_509636, EPI_ISL_509637, EPI_ISL_509638, EPI_ISL_509639, EPI_ISL_509640, EPI_ISL_509641, EPI_ISL_509642, EPI_ISL_509643, EPI_ISL_509644, EPI_ISL_509645, EPI_ISL_509646, EPI_ISL_509647, EPI_ISL_509648, EPI_ISL_509649, EPI_ISL_509650, EPI_ISL_509651, EPI_ISL_509652, EPI_ISL_509653, EPI_ISL_509654 | | | |
| see above | Servicio de Microbiología. Hospital Universitario Donostia. OSI Donostialdea. Área de Enfermedades Infecciosas, Grupo de Infección Respiratoria y Resistencia Antimicrobiana. Instituto de Investigación Sanitaria Biodonostia | SeqCOVID-SPAIN consortium/IBV(CSIC) | Gustavo Cilla, Milagrosa Montes, Luis Piñeiro, Jose Maria Marimón and SeqCOVID-SPAIN consortium |
| EPI_ISL_509662 | Georgia Department of Health | Pathogen Discovery, Respiratory Viruses Branch, Division of Viral Diseases, Centers for Disease Control and Prevention | Yan Li, Anna Montmayeur, Jing Zhang, Krista Queen, Anna Uehara, Ying Tao, Rachel Marine, Clinton R. Paden, Haibin Wang, Suxiang Tong |
| EPI_ISL_509663, EPI_ISL_509664, EPI_ISL_509665, EPI_ISL_509666, EPI_ISL_509667, EPI_ISL_509668, EPI_ISL_509670, EPI_ISL_509671, EPI_ISL_509672, EPI_ISL_509673, EPI_ISL_509674, EPI_ISL_509675, EPI_ISL_509676, EPI_ISL_509677, EPI_ISL_509678, EPI_ISL_509679, EPI_ISL_509680, EPI_ISL_509681, EPI_ISL_509682 | | | |
| see above | AR Dept. of Health-Public Health Lab | Pathogen Discovery, Respiratory Viruses Branch, Division of Viral Diseases, Centers for Disease Control and Prevention | Yan Li, Anna Montmayeur, Jing Zhang, Krista Queen, Anna Uehara, Ying Tao, Rachel Marine, Clinton R. Paden, Haibin Wang, Suxiang Tong |
| EPI_ISL_509683, EPI_ISL_509684, EPI_ISL_509685 | AR Dept. of Health-Public Health Lab | Pathogen Discovery, Respiratory Viruses Branch, Division of Viral Diseases, Centers for Disease Control and Prevention | Jing Zhang, Yan Li, Anna Montmayeur, Krista Queen, Anna Uehara, Ying Tao, Rachel Marine, Clinton R. Paden, Haibin Wang, Suxiang Tong |
| EPI_ISL_509686 | M Health Fairview | Minnesota Department of Health, Public Health Laboratory | Matt Plumb, Jacob Garfin, and Xiong Wang |
| EPI_ISL_509688 | Alabama Department of Public Health Bureau of Clinical Laboratories | Pathogen Discovery, Respiratory Viruses Branch, Division of Viral Diseases, Centers for Disease Control and Prevention | Ying Tao, Jing Zhang, Krista Queen, Anna Uehara, Yan Li, Clinton Paden, Haibin Wang, Suxiang Tong |
| EPI_ISL_509689, EPI_ISL_509690, EPI_ISL_509691, EPI_ISL_509692, EPI_ISL_509693, EPI_ISL_509694 | Utah Public Health Laboratory | Pathogen Discovery, Respiratory Viruses Branch, Division of Viral Diseases, Centers for Disease Control and Prevention | Ying Tao, Jing Zhang, Krista Queen, Anna Uehara, Yan Li, Clinton Paden, Haibin Wang, Suxiang Tong |
| EPI_ISL_509695, EPI_ISL_509696, EPI_ISL_509697, EPI_ISL_509698, EPI_ISL_509699, EPI_ISL_509700, EPI_ISL_509701, EPI_ISL_509702, EPI_ISL_509703 | Guatemala Ministry of Public Health | Pathogen Discovery, Respiratory Viruses Branch, Division of Viral Diseases, Centers for Disease Control and Prevention | Ying Tao, Jing Zhang, Krista Queen, Anna Uehara, Yan Li, Clinton Paden, Haibin Wang, Suxiang Tong |
| EPI_ISL_509704, EPI_ISL_509705, EPI_ISL_509706 | Wisconsin Department of Health Services | Pathogen Discovery, Respiratory Viruses Branch, Division of Viral Diseases, Centers for Disease Control and Prevention | Ying Tao, Jing Zhang, Krista Queen, Anna Uehara, Yan Li, Clinton Paden, Haibin Wang, Suxiang Tong |
| EPI_ISL_509707, EPI_ISL_509708, EPI_ISL_509709 | Utah Public Health Laboratory | Pathogen Discovery, Respiratory Viruses Branch, Division of Viral Diseases, Centers for Disease Control and Prevention | Jing Zhang, Ying Tao, Krista Queen, Anna Uehara, Yan Li, Clinton Paden, Haibin Wang, Suxiang Tong |
| EPI_ISL_509710 | Guatemala Ministry of Public Health | Pathogen Discovery, Respiratory Viruses Branch, Division of Viral Diseases, Centers for Disease Control and Prevention | Jing Zhang, Ying Tao, Krista Queen, Anna Uehara, Yan Li, Clinton Paden, Haibin Wang, Suxiang Tong |
| EPI_ISL_509711, EPI_ISL_509712, EPI_ISL_509713, EPI_ISL_509714 | Belize Ministry of Health | Pathogen Discovery, Respiratory Viruses Branch, Division of Viral Diseases, Centers for Disease Control and Prevention | Jing Zhang, Ying Tao, Krista Queen, Anna Uehara, Yan Li, Clinton Paden, Haibin Wang, Suxiang Tong |
| EPI_ISL_509715 | Wisconsin Department of Health Services | Pathogen Discovery, Respiratory Viruses Branch, Division of Viral Diseases, Centers for Disease Control and Prevention | Jing Zhang, Ying Tao, Krista Queen, Anna Uehara, Yan Li, Clinton Paden, Haibin Wang, Suxiang Tong |
| EPI_ISL_509716, EPI_ISL_509717, EPI_ISL_509718, EPI_ISL_509719, EPI_ISL_509721, EPI_ISL_509722, EPI_ISL_509723, EPI_ISL_509724, EPI_ISL_509725, EPI_ISL_509726, EPI_ISL_509727, EPI_ISL_509728, EPI_ISL_509729, EPI_ISL_509730, EPI_ISL_509731, EPI_ISL_509732, EPI_ISL_509733, EPI_ISL_509734, EPI_ISL_509735, EPI_ISL_509736, EPI_ISL_509737, EPI_ISL_509738, EPI_ISL_509739, EPI_ISL_509740, EPI_ISL_509741, EPI_ISL_509742, EPI_ISL_509743, EPI_ISL_509745, EPI_ISL_509746, EPI_ISL_509747, EPI_ISL_509748, EPI_ISL_509749, EPI_ISL_509750, EPI_ISL_509751, EPI_ISL_509752, EPI_ISL_509753, EPI_ISL_509754, EPI_ISL_509755, EPI_ISL_509756, EPI_ISL_509757, EPI_ISL_509758, EPI_ISL_509759, EPI_ISL_509760, EPI_ISL_509761, EPI_ISL_509762, EPI_ISL_509763, EPI_ISL_509764, EPI_ISL_509765, EPI_ISL_509766, EPI_ISL_509767, EPI_ISL_509769, EPI_ISL_509771, EPI_ISL_509772, EPI_ISL_509773, EPI_ISL_509774, EPI_ISL_509776, EPI_ISL_509778, EPI_ISL_509780, EPI_ISL_509781, EPI_ISL_509782, EPI_ISL_509783, EPI_ISL_509786, EPI_ISL_509787, EPI_ISL_509788, EPI_ISL_509789, EPI_ISL_509791, EPI_ISL_509792, EPI_ISL_509793, EPI_ISL_509794, EPI_ISL_509795, EPI_ISL_509796, EPI_ISL_509797 | | | |
| see above | Florida Bureau of Public Health Laboratories | Florida Bureau of Public Health Laboratories | Sarah Schmedes, Jason Blanton |
| EPI_ISL_509799, EPI_ISL_509800, EPI_ISL_509802, EPI_ISL_509803, EPI_ISL_509804, EPI_ISL_509805, EPI_ISL_509806, EPI_ISL_509807, EPI_ISL_509809, EPI_ISL_509810, EPI_ISL_509811, EPI_ISL_509812, EPI_ISL_509813, EPI_ISL_509814, EPI_ISL_509815, EPI_ISL_509816, EPI_ISL_509817, EPI_ISL_509818, EPI_ISL_509819, EPI_ISL_509820, EPI_ISL_509821, EPI_ISL_509822, EPI_ISL_509823, EPI_ISL_509824, EPI_ISL_509825, EPI_ISL_509826, EPI_ISL_509827, EPI_ISL_509828, EPI_ISL_509829, EPI_ISL_509830, EPI_ISL_509831, EPI_ISL_509832, EPI_ISL_509833, EPI_ISL_509834, EPI_ISL_509835, EPI_ISL_509836, EPI_ISL_509837, EPI_ISL_509838, EPI_ISL_509839, EPI_ISL_509840, EPI_ISL_509841, EPI_ISL_509842, EPI_ISL_509843, EPI_ISL_509844, EPI_ISL_509845, EPI_ISL_509847, EPI_ISL_509848, EPI_ISL_509849, EPI_ISL_509850, EPI_ISL_509851, EPI_ISL_509852, EPI_ISL_509853, EPI_ISL_509854, EPI_ISL_509855, EPI_ISL_509856, EPI_ISL_509857, EPI_ISL_509859, EPI_ISL_509860, EPI_ISL_509861, EPI_ISL_509863, EPI_ISL_509864, EPI_ISL_509865, EPI_ISL_509866, EPI_ISL_509867, EPI_ISL_509868, EPI_ISL_509869, EPI_ISL_509870, EPI_ISL_509871, EPI_ISL_509872, EPI_ISL_509874, EPI_ISL_509875, EPI_ISL_509876, EPI_ISL_509877, EPI_ISL_509878, EPI_ISL_509879, EPI_ISL_509880, EPI_ISL_509881, EPI_ISL_509882, EPI_ISL_509883, EPI_ISL_509884, EPI_ISL_509885, EPI_ISL_509886, EPI_ISL_509888, EPI_ISL_509889, EPI_ISL_509890, EPI_ISL_509891, EPI_ISL_509892, EPI_ISL_509893, EPI_ISL_509894, EPI_ISL_509895, EPI_ISL_509896, EPI_ISL_509897, EPI_ISL_509898, EPI_ISL_509899, EPI_ISL_509900, EPI_ISL_509901, EPI_ISL_509902, EPI_ISL_509903, EPI_ISL_509904, EPI_ISL_509905, EPI_ISL_509906, EPI_ISL_509907, EPI_ISL_509908, EPI_ISL_509909, EPI_ISL_509910, EPI_ISL_509911, EPI_ISL_509912, EPI_ISL_509914, EPI_ISL_509915, EPI_ISL_509916, EPI_ISL_509917, EPI_ISL_509918, EPI_ISL_509919, EPI_ISL_509920, EPI_ISL_509921, EPI_ISL_509922, EPI_ISL_509923, EPI_ISL_509924, EPI_ISL_509925, EPI_ISL_509926, EPI_ISL_509927, EPI_ISL_509928, EPI_ISL_509929, EPI_ISL_509930, EPI_ISL_509931, EPI_ISL_509932, EPI_ISL_509933, EPI_ISL_509934, EPI_ISL_509935, EPI_ISL_509936, EPI_ISL_509939, EPI_ISL_509940, EPI_ISL_509941, EPI_ISL_509942, EPI_ISL_509943, EPI_ISL_509944, EPI_ISL_509945, EPI_ISL_509946, EPI_ISL_509947, EPI_ISL_509948, EPI_ISL_509949, EPI_ISL_509950, EPI_ISL_509951, EPI_ISL_509952, EPI_ISL_509953, EPI_ISL_509954, EPI_ISL_509955, EPI_ISL_509956, EPI_ISL_509957, EPI_ISL_509958, EPI_ISL_509959, EPI_ISL_509960, EPI_ISL_509961, EPI_ISL_509962, EPI_ISL_509963, EPI_ISL_509964, EPI_ISL_509965, EPI_ISL_509966, EPI_ISL_509967, EPI_ISL_509969, EPI_ISL_509970, EPI_ISL_509971, EPI_ISL_509972, EPI_ISL_509973, EPI_ISL_509974, EPI_ISL_509975, EPI_ISL_509976, EPI_ISL_509977, EPI_ISL_509978, EPI_ISL_509979, EPI_ISL_509980, EPI_ISL_509981, EPI_ISL_509982, EPI_ISL_509983, EPI_ISL_509984, EPI_ISL_509985, EPI_ISL_509986, EPI_ISL_509987, EPI_ISL_509988, EPI_ISL_509990, EPI_ISL_509991, EPI_ISL_509992, EPI_ISL_509994, EPI_ISL_509995, EPI_ISL_509996, EPI_ISL_509997, EPI_ISL_509998 | | | |
| see above | University of Wisconsin-Madison AIDS Vaccine | University of Wisconsin-Madison AIDS Vaccine Research Laboratories | Gage Moreno, Katarina Braun, et al. AIDS Vaccine Research Laboratories |

| | | | |
|---|---|---|---|
| | Research Laboratories | | |
| EPI_ISL_510052, EPI_ISL_510053, EPI_ISL_510055, EPI_ISL_510058, EPI_ISL_510059 | Servicio de Microbiología. HRU de Málaga. Servicio Andaluz de Salud | SeqCOVID-SPAIN consortium/IBV(CSIC) | Inmaculada de Toro Peinado. MªConcepción Mediavilla Gradolph. Begoña Palop Borrás and SeqCOVID-SPAIN consortium |
| EPI_ISL_510066, EPI_ISL_510067, EPI_ISL_510068, EPI_ISL_510069, EPI_ISL_510072, EPI_ISL_510075, EPI_ISL_510077 | Instituto de Investigaciones Biomédicas de Barcelona (CSIC), Hospital Clinic i Provincial de Barcelona, Instituto de Biomedicina de Valencia (CSIC), Hospital de Sant Pau | SeqCOVID-SPAIN consortium/IBV(CSIC) | Anna M. Planas, Mª Angeles Marcos, Miguel J. Martínez, Andrea Vergara, Alex Soriano, Jordi Pérez Tur, Israel Fernández Cadenas and SeqCOVID-SPAIN consortium |
| EPI_ISL_510106, EPI_ISL_510110, EPI_ISL_510111, EPI_ISL_510112, EPI_ISL_510113, EPI_ISL_510117, EPI_ISL_510122, EPI_ISL_510128, EPI_ISL_510129, EPI_ISL_510131, EPI_ISL_510133, EPI_ISL_510135, EPI_ISL_510145, EPI_ISL_510146, EPI_ISL_510149, EPI_ISL_510165, EPI_ISL_510166, EPI_ISL_510167, EPI_ISL_510168, EPI_ISL_510169, EPI_ISL_510170, EPI_ISL_510171, EPI_ISL_510172, EPI_ISL_510173, EPI_ISL_510174, EPI_ISL_510176, EPI_ISL_510178, EPI_ISL_510180, EPI_ISL_510182, EPI_ISL_510183, EPI_ISL_510185, EPI_ISL_510186, EPI_ISL_510190, EPI_ISL_510191, EPI_ISL_510192, EPI_ISL_510193, EPI_ISL_510194, EPI_ISL_510195, EPI_ISL_510196, EPI_ISL_510199, EPI_ISL_510201, EPI_ISL_510202, EPI_ISL_510203, EPI_ISL_510204, EPI_ISL_510205, EPI_ISL_510207, EPI_ISL_510211, EPI_ISL_510212, EPI_ISL_510213, EPI_ISL_510216, EPI_ISL_510217, EPI_ISL_510218, EPI_ISL_510220, EPI_ISL_510221, EPI_ISL_510222, EPI_ISL_510223, EPI_ISL_510224, EPI_ISL_510225, EPI_ISL_510226, EPI_ISL_510227, EPI_ISL_510228, EPI_ISL_510231, EPI_ISL_510232, EPI_ISL_510233, EPI_ISL_510236, EPI_ISL_510237, EPI_ISL_510238, EPI_ISL_510240, EPI_ISL_510243, EPI_ISL_510244 | | | |
| see above | Hospital General Universitario Gregorio Marañón | SeqCOVID-SPAIN consortium/IBV(CSIC) | Laura Pérez-Lago, Marta Herranz, Jon Sicilia, Julia Suárez, Pilar Catalán, Patricia Muñoz, Darío García de Viedma and SeqCOVID-SPAIN consortium |
| EPI_ISL_510247, EPI_ISL_510248, EPI_ISL_510249, EPI_ISL_510250, EPI_ISL_510251, EPI_ISL_510252, EPI_ISL_510253, EPI_ISL_510256, EPI_ISL_510257, EPI_ISL_510258, EPI_ISL_510259, EPI_ISL_510260, EPI_ISL_510261, EPI_ISL_510262, EPI_ISL_510263, EPI_ISL_510264, EPI_ISL_510265, EPI_ISL_510266 | | | |
| see above | Hospital de la Santa Creu i Sant Pau. Servicio de Microbiología | SeqCOVID-SPAIN consortium/IBV(CSIC) | Ferran Navarro, Núria Rabella, Elisenda Miró and SeqCOVID-SPAIN consortium |
| EPI_ISL_510268, EPI_ISL_510269, EPI_ISL_510271, EPI_ISL_510272, EPI_ISL_510273, EPI_ISL_510274, EPI_ISL_510275, EPI_ISL_510277, EPI_ISL_510279, EPI_ISL_510280, EPI_ISL_510281, EPI_ISL_510283, EPI_ISL_510284, EPI_ISL_510285, EPI_ISL_510286, EPI_ISL_510288, EPI_ISL_510290, EPI_ISL_510291, EPI_ISL_510292, EPI_ISL_510293, EPI_ISL_510294, EPI_ISL_510296, EPI_ISL_510297, EPI_ISL_510298, EPI_ISL_510300, EPI_ISL_510301, EPI_ISL_510302, EPI_ISL_510303 | | | |
| see above | Hospital Clínico Universitario de Santiago de Compostela | SeqCOVID-SPAIN consortium/IBV(CSIC) | José Javier Costa Alcalde, Antonio Aguilera Guirao, Mª Luisa Pérez del Molino Bernal, Amparo Coira Nieto, Gema Barbeito Castiñeiras, Rocio Trastoy Pena and SeqCOVID-SPAIN consortium |
| EPI_ISL_510305, EPI_ISL_510306, EPI_ISL_510307, EPI_ISL_510309, EPI_ISL_510310, EPI_ISL_510311, EPI_ISL_510312, EPI_ISL_510313, EPI_ISL_510314, EPI_ISL_510315, EPI_ISL_510316, EPI_ISL_510317, EPI_ISL_510318, EPI_ISL_510319, EPI_ISL_510320, EPI_ISL_510321, EPI_ISL_510322, EPI_ISL_510323, EPI_ISL_510324, EPI_ISL_510325, EPI_ISL_510326, EPI_ISL_510327, EPI_ISL_510328, EPI_ISL_510329, EPI_ISL_510330, EPI_ISL_510331 | | | |
| see above | Hospital San Pedro de Alcántara (Cáceres) | SeqCOVID-SPAIN consortium/IBV(CSIC) | Cristina Muñoz Cuevas, Guadalupe Rodriguez Rodriguez and SeqCOVID-SPAIN consortium |
| EPI_ISL_510334, EPI_ISL_510338, EPI_ISL_510340, EPI_ISL_510343, EPI_ISL_510344, EPI_ISL_510346, EPI_ISL_510349, EPI_ISL_510351, EPI_ISL_510352, EPI_ISL_510354, EPI_ISL_510356, EPI_ISL_510357, EPI_ISL_510358, EPI_ISL_510359, EPI_ISL_510360, EPI_ISL_510361, EPI_ISL_510365, EPI_ISL_510366, EPI_ISL_510367, EPI_ISL_510371, EPI_ISL_510372, EPI_ISL_510373, EPI_ISL_510374, EPI_ISL_510375, EPI_ISL_510376, EPI_ISL_510377, EPI_ISL_510378, EPI_ISL_510379, EPI_ISL_510382, EPI_ISL_510383, EPI_ISL_510385, EPI_ISL_510386, EPI_ISL_510387, EPI_ISL_510388, EPI_ISL_510390 | | | |
| see above | Servicio de Microbiología, Hospital Miguel Servet, Zaragoza | SeqCOVID-SPAIN consortium/IBV(CSIC) | Antonio Rezusta López, Alexander Tristancho Baró, Ana Milagro, Yolanda Gracia Grataloup, Nieves Martínez Cameo and SeqCOVID-SPAIN consortium |
| EPI_ISL_510391, EPI_ISL_510392, EPI_ISL_510393, EPI_ISL_510394, EPI_ISL_510395, EPI_ISL_510396, EPI_ISL_510397, EPI_ISL_510398, EPI_ISL_510399, EPI_ISL_510400, EPI_ISL_510401, EPI_ISL_510402, EPI_ISL_510403, EPI_ISL_510404, EPI_ISL_510405, EPI_ISL_510406, EPI_ISL_510407, EPI_ISL_510409, EPI_ISL_510410, EPI_ISL_510411, EPI_ISL_510413, EPI_ISL_510415, EPI_ISL_510416, EPI_ISL_510418, EPI_ISL_510419, EPI_ISL_510420, EPI_ISL_510421, EPI_ISL_510422, EPI_ISL_510423 | | | |
| see above | Servicio de Microbiología. Hospital Universitario Donostia. OSI Donostialdea. Área de Enfermedades Infecciosas, Grupo de Infección Respiratoria y Resistencia Antimicrobiana. Instituto de Investigación Sanitaria Biodonostia | SeqCOVID-SPAIN consortium/IBV(CSIC) | Gustavo Cilla, Milagrosa Montes, Luis Piñeiro, Jose Maria Marimón and SeqCOVID-SPAIN consortium |
| EPI_ISL_510424, EPI_ISL_510427, EPI_ISL_510428, EPI_ISL_510431, EPI_ISL_510433, EPI_ISL_510434, EPI_ISL_510435, EPI_ISL_510436, EPI_ISL_510437, EPI_ISL_510439, EPI_ISL_510440, EPI_ISL_510441, EPI_ISL_510444, EPI_ISL_510445, EPI_ISL_510446, EPI_ISL_510447, EPI_ISL_510448, EPI_ISL_510449, EPI_ISL_510450, EPI_ISL_510451, EPI_ISL_510452, EPI_ISL_510454, EPI_ISL_510455, EPI_ISL_510456, EPI_ISL_510458, EPI_ISL_510459, EPI_ISL_510460 | | | |
| see above | Hospital Universitario Virgen de las Nieves de Granada-SAS | SeqCOVID-SPAIN consortium/IBV(CSIC) | Mercedes Pérez Ruiz, Sara Sanbonmatsu Gámez, Irene Pedrosa Corral, José M. Navarro-Marí and SeqCOVID-SPAIN consortium |
| EPI_ISL_510463, EPI_ISL_510464, EPI_ISL_510465 | Instituto de Investigaciones Biomédicas de Barcelona (CSIC), Hospital Clinic i Provincial de Barcelona, Instituto de Biomedicina de Valencia (CSIC), Hospital de Sant Pau | SeqCOVID-SPAIN consortium/IBV(CSIC) | Anna M. Planas, Mª Angeles Marcos, Miguel J. Martínez, Andrea Vergara, Alex Soriano, Jordi Pérez Tur, Israel Fernández Cadenas and SeqCOVID-SPAIN consortium |
| EPI_ISL_510467, EPI_ISL_510468, EPI_ISL_510469, EPI_ISL_510470, EPI_ISL_510471, EPI_ISL_510473, EPI_ISL_510474, EPI_ISL_510475, EPI_ISL_510476, EPI_ISL_510477, EPI_ISL_510478, EPI_ISL_510479, EPI_ISL_510480, EPI_ISL_510481, EPI_ISL_510482, EPI_ISL_510483, EPI_ISL_510484, EPI_ISL_510485, EPI_ISL_510486, EPI_ISL_510487, EPI_ISL_510488, EPI_ISL_510489, EPI_ISL_510490, EPI_ISL_510491, EPI_ISL_510492, EPI_ISL_510493, EPI_ISL_510495, EPI_ISL_510496, EPI_ISL_510497, EPI_ISL_510498, EPI_ISL_510499, EPI_ISL_510500, EPI_ISL_510501, EPI_ISL_510502, EPI_ISL_510503, EPI_ISL_510504, EPI_ISL_510505, EPI_ISL_510506, EPI_ISL_510507, EPI_ISL_510508, EPI_ISL_510509 | | | |
| see above | Servicio de Microbiología. Hospital Universitario Donostia. OSI Donostialdea. Área de Enfermedades Infecciosas, Grupo de Infección Respiratoria y Resistencia Antimicrobiana. Instituto de Investigación Sanitaria Biodonostia | SeqCOVID-SPAIN consortium/IBV(CSIC) | Gustavo Cilla, Milagrosa Montes, Luis Piñeiro, Jose Maria Marimón and SeqCOVID-SPAIN consortium |
| EPI_ISL_510510, EPI_ISL_510511, EPI_ISL_510512, EPI_ISL_510513, EPI_ISL_510514, EPI_ISL_510515, EPI_ISL_510516, EPI_ISL_510517, EPI_ISL_510518, EPI_ISL_510520, EPI_ISL_510522, EPI_ISL_510523 | | | |
| see above | Servicio de Microbiología, Laboratori Clínic Metropolitana Nord. Hospital Universitari Germans Trias i Pujol. Institut d'Investigació en Ciències de la Salut Germans Trias i Pujol (IGTP) | SeqCOVID-SPAIN consortium/IBV(CSIC) | Elisa Martró, Antoni E. Bordoy, Anna Not, Adrián Antuori, Anabel Fernández, Nona Romaní and SeqCOVID-SPAIN consortium |
| EPI_ISL_510532 | Biological prevention, army | Biological prevention, army | Seadawy,M.G., ELnabrawy,H.A., Shamel,M.D., Elhoseiny,M.F., Gad,A.F., Hassan,W.A., Raouf,A.A., Harty,B.E., ElGohary,A.A., Karam,M.A., Amer,k.E., Elnakeeb,M.A., Elnagdy,T.A., Ali,M.A., Kandeil,A.M. and Soliman,Y.A. |
| EPI_ISL_510535 | Molecular Virology, Instituto Carlos Chagas / Fiocruz Paraná | Universidade Federal do Parana (UFPR) | Suzukawa,A., Tscha,M., Zanluca,C., Raboni,S., Duarte dos Santos,C. |
| EPI_ISL_510536 | Centro de Desenvolvimento Tecnologico em Saude, Fundacao Oswaldo Cruz | Centro de Desenvolvimento Tecnologico em Saude, Fundacao Oswaldo Cruz | Souza,T.M., Fintelman-Rodrigues,N., De Paula,A.D., Saraiva,F.B., Ferreira,M.A., Sacramento,C.Q. and Medeiros,M.A. |
| EPI_ISL_510538, EPI_ISL_510539 | Department of Microbiology, The University of Hong Kong | Department of Microbiology, The University of Hong Kong | Kelvin K.W. To, Kwok-Yung Yuen |
| EPI_ISL_510541 | Centro de Desenvolvimento Tecnologico em Saude, Fundacao Oswaldo Cruz | Centro de Desenvolvimento Tecnologico em Saude, Fundacao Oswaldo Cruz | Souza,T.M., Fintelman-Rodrigues,N., De Paula,A.D., Saraiva,F.B., Ferreira,M.A., Sacramento,C.Q. and Medeiros,M.A. |
| EPI_ISL_510542, EPI_ISL_510544 | SA Pathology | SA Pathology | Lex Leong, Chuan Kok Lim, Mark Turra, Ivan Bastian, Geoff Higgins |
| EPI_ISL_510547, EPI_ISL_510548, EPI_ISL_510549, EPI_ISL_510550, EPI_ISL_510551, EPI_ISL_510552, EPI_ISL_510553, EPI_ISL_510554, EPI_ISL_510555, EPI_ISL_510556, EPI_ISL_510557, EPI_ISL_510558, EPI_ISL_510559, EPI_ISL_510560, EPI_ISL_510561, EPI_ISL_510562, EPI_ISL_510563, EPI_ISL_510564, EPI_ISL_510565, EPI_ISL_510566, EPI_ISL_510567, EPI_ISL_510568, EPI_ISL_510569, EPI_ISL_510570, EPI_ISL_510571, EPI_ISL_510572, EPI_ISL_510573, EPI_ISL_510574, EPI_ISL_510575, EPI_ISL_510576, EPI_ISL_510577, EPI_ISL_510578, EPI_ISL_510579, EPI_ISL_510580, EPI_ISL_510581, EPI_ISL_510582, EPI_ISL_510583, EPI_ISL_510584, EPI_ISL_510585, EPI_ISL_510586, EPI_ISL_510587, EPI_ISL_510588, EPI_ISL_510589, EPI_ISL_510590, EPI_ISL_510591, EPI_ISL_510592, EPI_ISL_510593, EPI_ISL_510594, EPI_ISL_510595, EPI_ISL_510596, EPI_ISL_510597, EPI_ISL_510598, EPI_ISL_510599, EPI_ISL_510600, EPI_ISL_510601, EPI_ISL_510602, EPI_ISL_510603, EPI_ISL_510604, EPI_ISL_510605, EPI_ISL_510606, EPI_ISL_510607, EPI_ISL_510608, EPI_ISL_510609, EPI_ISL_510610, EPI_ISL_510611, EPI_ISL_510612, EPI_ISL_510613, EPI_ISL_510614, EPI_ISL_510615, EPI_ISL_510616, EPI_ISL_510617, EPI_ISL_510618, EPI_ISL_510619, EPI_ISL_510620, EPI_ISL_510621, EPI_ISL_510622, EPI_ISL_510623, EPI_ISL_510624, EPI_ISL_510625, EPI_ISL_510626, EPI_ISL_510627, EPI_ISL_510628, EPI_ISL_510629, EPI_ISL_510630, EPI_ISL_510631, EPI_ISL_510632, EPI_ISL_510633, EPI_ISL_510634, EPI_ISL_510635, EPI_ISL_510636, EPI_ISL_510637, EPI_ISL_510638, EPI_ISL_510639, EPI_ISL_510640, EPI_ISL_510641, EPI_ISL_510642, EPI_ISL_510643, EPI_ISL_510644, EPI_ISL_510645, EPI_ISL_510646, EPI_ISL_510647, EPI_ISL_510648, EPI_ISL_510649, EPI_ISL_510650, EPI_ISL_510651, EPI_ISL_510652, EPI_ISL_510653, EPI_ISL_510654, EPI_ISL_510655, EPI_ISL_510656, EPI_ISL_510657, EPI_ISL_510658, EPI_ISL_510659, EPI_ISL_510660, EPI_ISL_510661, EPI_ISL_510662, EPI_ISL_510663, EPI_ISL_510664, EPI_ISL_510665, EPI_ISL_510666, EPI_ISL_510667, EPI_ISL_510668, EPI_ISL_510669, EPI_ISL_510670, EPI_ISL_510671, EPI_ISL_510672, EPI_ISL_510673, EPI_ISL_510674, EPI_ISL_510675, EPI_ISL_510676, EPI_ISL_510677, EPI_ISL_510678, EPI_ISL_510679, EPI_ISL_510680, EPI_ISL_510681, EPI_ISL_510682, EPI_ISL_510683, EPI_ISL_510684, EPI_ISL_510685, EPI_ISL_510686, EPI_ISL_510687, EPI_ISL_510688 | | | |
| see above | Division of Viral Diseases, Center for Laboratory Control of Infectious Diseases, Korea Centers for Diseases Control and Prevention | Division of Viral Diseases, Center for Laboratory Control of Infectious Diseases, Korea Centers for Diseases Control and Prevention | Jeong-Min Kim, Yoon-Seok Chung, Namjoo Lee, Sang Hee Woo, Hye-Jun Jo, Heui Man Kim, Jun-Sub Kim, Myung Guk Han |
| EPI_ISL_510689 | Hospital Universitari Germans Trias i Pujol(HUGTiP)/Fundació Lluita contra la SIDA (FLSida)/IRTA-CReSA | IrsiCaixa AIDS Research Lab | Pilar Armengol, Marc Noguera-Julian, Jordi Rodón, Julia Vergara, Lidia Ruiz, Nuria Izquierdo, Jorge Carrillo, Roger Paredes, Albert Bensaid, Julia Blanco, Joaquim Segalés, Bonaventura Clotet |
| EPI_ISL_510690, EPI_ISL_510691, EPI_ISL_510692, EPI_ISL_510693, EPI_ISL_510694, EPI_ISL_510695, EPI_ISL_510696, EPI_ISL_510698, EPI_ISL_510699, EPI_ISL_510701, EPI_ISL_510702, EPI_ISL_510703, EPI_ISL_510704, EPI_ISL_510705, EPI_ISL_510706, EPI_ISL_510707, EPI_ISL_510708, EPI_ISL_510709, EPI_ISL_510710, EPI_ISL_510711, EPI_ISL_510712, EPI_ISL_510713, EPI_ISL_510714, EPI_ISL_510715, EPI_ISL_510716, EPI_ISL_510717, EPI_ISL_510718, EPI_ISL_510719, EPI_ISL_510720, EPI_ISL_510721, EPI_ISL_510722, EPI_ISL_510723, EPI_ISL_510724, EPI_ISL_510725, EPI_ISL_510726, EPI_ISL_510727, EPI_ISL_510728, EPI_ISL_510729, EPI_ISL_510730, EPI_ISL_510731, EPI_ISL_510732, EPI_ISL_510733, EPI_ISL_510734, EPI_ISL_510735, EPI_ISL_510736, EPI_ISL_510737, EPI_ISL_510738, EPI_ISL_510739, EPI_ISL_510740, EPI_ISL_510741, EPI_ISL_510742, EPI_ISL_510743, EPI_ISL_510744, EPI_ISL_510745, EPI_ISL_510746, EPI_ISL_510747, EPI_ISL_510748, EPI_ISL_510749, EPI_ISL_510750, EPI_ISL_510751, EPI_ISL_510753, EPI_ISL_510754, EPI_ISL_510755, EPI_ISL_510756, EPI_ISL_510757, EPI_ISL_510758, EPI_ISL_510759, EPI_ISL_510760, EPI_ISL_510761, EPI_ISL_510762, EPI_ISL_510764, EPI_ISL_510765, EPI_ISL_510766, EPI_ISL_510767, EPI_ISL_510768, EPI_ISL_510769, EPI_ISL_510770, EPI_ISL_510771, EPI_ISL_510772, EPI_ISL_510773, EPI_ISL_510774, EPI_ISL_510775, EPI_ISL_510776, EPI_ISL_510777, EPI_ISL_510778, EPI_ISL_510779, EPI_ISL_510780, EPI_ISL_510781, EPI_ISL_510784, EPI_ISL_510785, EPI_ISL_510786, EPI_ISL_510787, EPI_ISL_510788, EPI_ISL_510789, EPI_ISL_510793, EPI_ISL_510794, EPI_ISL_510795, EPI_ISL_510796, EPI_ISL_510798, EPI_ISL_510800, EPI_ISL_510804, EPI_ISL_510805, EPI_ISL_510806, EPI_ISL_510807, EPI_ISL_510808, EPI_ISL_510809 | | | |
| see above | Viollier AG | Department of Biosystems Science and Engineering, ETH Zürich | Christian Beisel, Sarah Nadeau, Ivan Topolsky, Pedro Ferreira, Philipp Jablonski, Susana Posada-Céspedes, Tobias Schär, Ina Nissen, Natascha Santacroce, Elodie Burcklen, Christiane Beckmann, Maurice Redondo, Olivier Kobel, Christoph Noppen, Sophie Seidel, Noemie Santamaria de Souza, Niko Beerenwinkel, Tanja Stadler |
| EPI_ISL_510810, EPI_ISL_510811, EPI_ISL_510812, EPI_ISL_510813, EPI_ISL_510814, EPI_ISL_510815, EPI_ISL_510816 | NA | The Public Health Agency of Sweden | Oskar Karlsson Lindsjo, Maria Lind Karlberg, Mattias Haukland, Reza Advani, Olov Svartstrom, Anna-Malin Linde, Sandra Broddesson, Petra Edquist, Mia Brytting, Anna Risberg, Karin Tegmark-Wisell |
| EPI_ISL_510817 | Orebro klinisk mikrobiologi | The Public Health Agency of Sweden | Oskar Karlsson Lindsjo, Maria Lind Karlberg, Mattias Haukland, Reza Advani, Olov Svartstrom, Anna-Malin Linde, Sandra Broddesson, Petra Edquist, Mia Brytting, Anna Risberg, Karin Tegmark-Wisell |
| EPI_ISL_510818, EPI_ISL_510819 | Klinisk mikrobiologi Vasternorrland | The Public Health Agency of Sweden | Oskar Karlsson Lindsjo, Maria Lind Karlberg, Mattias Haukland, Reza Advani, Olov Svartstrom, Anna-Malin Linde, Sandra Broddesson, Petra Edquist, Mia Brytting, Anna Risberg, Karin Tegmark-Wisell |
| EPI_ISL_510820, EPI_ISL_510821, EPI_ISL_510822, EPI_ISL_510823, EPI_ISL_510824 | Klinisk mikrobiologi centralsjukhuset Karlstad | The Public Health Agency of Sweden | Oskar Karlsson Lindsjo, Maria Lind Karlberg, Mattias Haukland, Reza Advani, Olov Svartstrom, Anna-Malin Linde, Sandra Broddesson, Petra Edquist, Mia Brytting, Anna Risberg, Karin Tegmark-Wisell |
| EPI_ISL_510825, EPI_ISL_510826 | Kalmar klinisk mikrobiologi | The Public Health Agency of Sweden | Oskar Karlsson Lindsjo, Maria Lind Karlberg, Mattias Haukland, Reza Advani, Olov Svartstrom, Anna-Malin Linde, Sandra Broddesson, Petra Edquist, Mia Brytting, Anna Risberg, Karin Tegmark-Wisell |

| | | | |
|---|---|---|---|
| EPI_ISL_510827, EPI_ISL_510828, EPI_ISL_510829, EPI_ISL_510830 | Unilabs Eskilstuna | The Public Health Agency of Sweden | Oskar Karlsson Lindsjo, Maria Lind Karlberg, Mattias Haukland, Reza Advani, Olov Svartstrom, Anna-Malin Linde, Sandra Broddesson, Petra Edquist, Mia Brytting, Anna Risberg, Karin Tegmark-Wisell |
| EPI_ISL_510831 | Halmstad klinisk mikrobiologi | The Public Health Agency of Sweden | Oskar Karlsson Lindsjo, Maria Lind Karlberg, Mattias Haukland, Reza Advani, Olov Svartstrom, Anna-Malin Linde, Sandra Broddesson, Petra Edquist, Mia Brytting, Anna Risberg, Karin Tegmark-Wisell |
| EPI_ISL_510832, EPI_ISL_510833 | Kalmar klinisk mikrobiologi | The Public Health Agency of Sweden | Oskar Karlsson Lindsjo, Maria Lind Karlberg, Mattias Haukland, Reza Advani, Olov Svartstrom, Anna-Malin Linde, Sandra Broddesson, Petra Edquist, Mia Brytting, Anna Risberg, Karin Tegmark-Wisell |
| EPI_ISL_510834, EPI_ISL_510835 | Karolinska Universitetslaboratoriet | The Public Health Agency of Sweden | Oskar Karlsson Lindsjo, Maria Lind Karlberg, Mattias Haukland, Reza Advani, Olov Svartstrom, Anna-Malin Linde, Sandra Broddesson, Petra Edquist, Mia Brytting, Anna Risberg, Karin Tegmark-Wisell |
| EPI_ISL_510836, EPI_ISL_510837 | Kalmar klinisk mikrobiologi | The Public Health Agency of Sweden | Oskar Karlsson Lindsjo, Maria Lind Karlberg, Mattias Haukland, Reza Advani, Olov Svartstrom, Anna-Malin Linde, Sandra Broddesson, Petra Edquist, Mia Brytting, Anna Risberg, Karin Tegmark-Wisell |
| EPI_ISL_510838, EPI_ISL_510839, EPI_ISL_510840 | Karolinska Universitetslaboratoriet | The Public Health Agency of Sweden | Oskar Karlsson Lindsjo, Maria Lind Karlberg, Mattias Haukland, Reza Advani, Olov Svartstrom, Anna-Malin Linde, Sandra Broddesson, Petra Edquist, Mia Brytting, Anna Risberg, Karin Tegmark-Wisell |
| EPI_ISL_510841, EPI_ISL_510842, EPI_ISL_510843, EPI_ISL_510844 | Klinisk mikrobiologi NAL Trollhattan | The Public Health Agency of Sweden | Oskar Karlsson Lindsjo, Maria Lind Karlberg, Mattias Haukland, Reza Advani, Olov Svartstrom, Anna-Malin Linde, Sandra Broddesson, Petra Edquist, Mia Brytting, Anna Risberg, Karin Tegmark-Wisell |
| EPI_ISL_510845, EPI_ISL_510846, EPI_ISL_510847, EPI_ISL_510848, EPI_ISL_510849, EPI_ISL_510850, EPI_ISL_510851 | Skovde/Unilabs | The Public Health Agency of Sweden | Oskar Karlsson Lindsjo, Maria Lind Karlberg, Mattias Haukland, Reza Advani, Olov Svartstrom, Anna-Malin Linde, Sandra Broddesson, Petra Edquist, Mia Brytting, Anna Risberg, Karin Tegmark-Wisell |
| EPI_ISL_510852 | Karolinska Universitetslaboratoriet | The Public Health Agency of Sweden | Oskar Karlsson Lindsjo, Maria Lind Karlberg, Mattias Haukland, Reza Advani, Olov Svartstrom, Anna-Malin Linde, Sandra Broddesson, Petra Edquist, Mia Brytting, Anna Risberg, Karin Tegmark-Wisell |
| EPI_ISL_510853, EPI_ISL_510854, EPI_ISL_510855, EPI_ISL_510856, EPI_ISL_510857, EPI_ISL_510858 | Klinisk mikrobiologi Vasternorrland | The Public Health Agency of Sweden | Oskar Karlsson Lindsjo, Maria Lind Karlberg, Mattias Haukland, Reza Advani, Olov Svartstrom, Anna-Malin Linde, Sandra Broddesson, Petra Edquist, Mia Brytting, Anna Risberg, Karin Tegmark-Wisell |
| EPI_ISL_510859 | Kalmar klinisk mikrobiologi | The Public Health Agency of Sweden | Oskar Karlsson Lindsjo, Maria Lind Karlberg, Mattias Haukland, Reza Advani, Olov Svartstrom, Anna-Malin Linde, Sandra Broddesson, Petra Edquist, Mia Brytting, Anna Risberg, Karin Tegmark-Wisell |
| EPI_ISL_510860, EPI_ISL_510861, EPI_ISL_510862, EPI_ISL_510863 | Karolinska Universitetslaboratoriet | The Public Health Agency of Sweden | Oskar Karlsson Lindsjo, Maria Lind Karlberg, Mattias Haukland, Reza Advani, Olov Svartstrom, Anna-Malin Linde, Sandra Broddesson, Petra Edquist, Mia Brytting, Anna Risberg, Karin Tegmark-Wisell |
| EPI_ISL_510864, EPI_ISL_510865 | Kalmar klinisk mikrobiologi | The Public Health Agency of Sweden | Oskar Karlsson Lindsjo, Maria Lind Karlberg, Mattias Haukland, Reza Advani, Olov Svartstrom, Anna-Malin Linde, Sandra Broddesson, Petra Edquist, Mia Brytting, Anna Risberg, Karin Tegmark-Wisell |
| EPI_ISL_510866 | Halmstad klinisk mikrobiologi | The Public Health Agency of Sweden | Oskar Karlsson Lindsjo, Maria Lind Karlberg, Mattias Haukland, Reza Advani, Olov Svartstrom, Anna-Malin Linde, Sandra Broddesson, Petra Edquist, Mia Brytting, Anna Risberg, Karin Tegmark-Wisell |
| EPI_ISL_510867 | Klinisk mikrobiologi centralsjukhuset Karlstad | The Public Health Agency of Sweden | Oskar Karlsson Lindsjo, Maria Lind Karlberg, Mattias Haukland, Reza Advani, Olov Svartstrom, Anna-Malin Linde, Sandra Broddesson, Petra Edquist, Mia Brytting, Anna Risberg, Karin Tegmark-Wisell |
| EPI_ISL_510868, EPI_ISL_510869, EPI_ISL_510870 | Klinisk mikrobiologi NAL Trollhattan | The Public Health Agency of Sweden | Oskar Karlsson Lindsjo, Maria Lind Karlberg, Mattias Haukland, Reza Advani, Olov Svartstrom, Anna-Malin Linde, Sandra Broddesson, Petra Edquist, Mia Brytting, Anna Risberg, Karin Tegmark-Wisell |
| EPI_ISL_510871, EPI_ISL_510872, EPI_ISL_510873, EPI_ISL_510874, EPI_ISL_510875, EPI_ISL_510876, EPI_ISL_510877 | Klinsisk mikrobiologi Linkoping | The Public Health Agency of Sweden | Oskar Karlsson Lindsjo, Maria Lind Karlberg, Mattias Haukland, Reza Advani, Olov Svartstrom, Anna-Malin Linde, Sandra Broddesson, Petra Edquist, Mia Brytting, Anna Risberg, Karin Tegmark-Wisell |
| EPI_ISL_510882, EPI_ISL_510883, EPI_ISL_510884, EPI_ISL_510885, EPI_ISL_510886, EPI_ISL_510887, EPI_ISL_510888 | New Mexico Department of Health Scientific Laboratory | New Mexico Department of Health Scientific Laboratory | Ellie Johnson, Anastacia Griego-Fisher |
| EPI_ISL_510889, EPI_ISL_510890, EPI_ISL_510891, EPI_ISL_510892, EPI_ISL_510893, EPI_ISL_510896, EPI_ISL_510898, EPI_ISL_510900, EPI_ISL_510901, EPI_ISL_510902, EPI_ISL_510903, EPI_ISL_510904, EPI_ISL_510905, EPI_ISL_510906, EPI_ISL_510907, EPI_ISL_510909, EPI_ISL_510912, EPI_ISL_510915, EPI_ISL_510916, EPI_ISL_510921, EPI_ISL_510923, EPI_ISL_510924, EPI_ISL_510925, EPI_ISL_510926, EPI_ISL_510927, EPI_ISL_510928, EPI_ISL_510929, EPI_ISL_510931, EPI_ISL_510932, EPI_ISL_510933, EPI_ISL_510934, EPI_ISL_510935, EPI_ISL_510936, EPI_ISL_510937, EPI_ISL_510938, EPI_ISL_510939, EPI_ISL_510940, EPI_ISL_510941, EPI_ISL_510942, EPI_ISL_510943, EPI_ISL_510944, EPI_ISL_510945, EPI_ISL_510946, EPI_ISL_510947, EPI_ISL_510948, EPI_ISL_510949, EPI_ISL_510950, EPI_ISL_510951, EPI_ISL_510952, EPI_ISL_510953, EPI_ISL_510954, EPI_ISL_510956, EPI_ISL_510958, EPI_ISL_510959, EPI_ISL_510960, EPI_ISL_510961, EPI_ISL_510962, EPI_ISL_510963, EPI_ISL_510964, EPI_ISL_510965, EPI_ISL_510966, EPI_ISL_510967, EPI_ISL_510968, EPI_ISL_510969, EPI_ISL_510970, EPI_ISL_510971, EPI_ISL_510972, EPI_ISL_510973, EPI_ISL_510974, EPI_ISL_510975, EPI_ISL_510976, EPI_ISL_510978, EPI_ISL_510981, EPI_ISL_510982, EPI_ISL_510983, EPI_ISL_510984, EPI_ISL_510987, EPI_ISL_510991, EPI_ISL_510992, EPI_ISL_510994, EPI_ISL_510995, EPI_ISL_510997, EPI_ISL_510999, EPI_ISL_511000, EPI_ISL_511001, EPI_ISL_511003, EPI_ISL_511004, EPI_ISL_511006, EPI_ISL_511007, EPI_ISL_511008, EPI_ISL_511009, EPI_ISL_511010, EPI_ISL_511011, EPI_ISL_511012, EPI_ISL_511014, EPI_ISL_511016, EPI_ISL_511017, EPI_ISL_511019, EPI_ISL_511021, EPI_ISL_511022, EPI_ISL_511025, EPI_ISL_511027, EPI_ISL_511028, EPI_ISL_511029, EPI_ISL_511030, EPI_ISL_511031, EPI_ISL_511032, EPI_ISL_511033, EPI_ISL_511034, EPI_ISL_511035, EPI_ISL_511036, EPI_ISL_511037, EPI_ISL_511039, EPI_ISL_511040, EPI_ISL_511041, EPI_ISL_511042, EPI_ISL_511043, EPI_ISL_511045, EPI_ISL_511046, EPI_ISL_511047, EPI_ISL_511050, EPI_ISL_511052, EPI_ISL_511053, EPI_ISL_511054, EPI_ISL_511055, EPI_ISL_511056, EPI_ISL_511057, EPI_ISL_511058, EPI_ISL_511059, EPI_ISL_511060, EPI_ISL_511061, EPI_ISL_511062, EPI_ISL_511063, EPI_ISL_511064, EPI_ISL_511065, EPI_ISL_511066, EPI_ISL_511067, EPI_ISL_511068, EPI_ISL_511069, EPI_ISL_511070, EPI_ISL_511071, EPI_ISL_511072, EPI_ISL_511073, EPI_ISL_511074, EPI_ISL_511076, EPI_ISL_511077, EPI_ISL_511079, EPI_ISL_511080, EPI_ISL_511082, EPI_ISL_511083, EPI_ISL_511085, EPI_ISL_511086, EPI_ISL_511087, EPI_ISL_511089, EPI_ISL_511090, EPI_ISL_511091, EPI_ISL_511092, EPI_ISL_511093, EPI_ISL_511094, EPI_ISL_511095, EPI_ISL_511096, EPI_ISL_511097, EPI_ISL_511098, EPI_ISL_511099, EPI_ISL_511100, EPI_ISL_511101, EPI_ISL_511102, EPI_ISL_511103, EPI_ISL_511104, EPI_ISL_511107, EPI_ISL_511108, EPI_ISL_511109, EPI_ISL_511110, EPI_ISL_511111, EPI_ISL_511112, EPI_ISL_511113, EPI_ISL_511114, EPI_ISL_511115, EPI_ISL_511116, EPI_ISL_511117, EPI_ISL_511118, EPI_ISL_511119, EPI_ISL_511120, EPI_ISL_511121, EPI_ISL_511122, EPI_ISL_511123, EPI_ISL_511124, EPI_ISL_511125, EPI_ISL_511126, EPI_ISL_511127, EPI_ISL_511128, EPI_ISL_511129, EPI_ISL_511130, EPI_ISL_511131, EPI_ISL_511132, EPI_ISL_511133, EPI_ISL_511134, EPI_ISL_511135, EPI_ISL_511136, EPI_ISL_511137, EPI_ISL_511138, EPI_ISL_511139, EPI_ISL_511140, EPI_ISL_511141, EPI_ISL_511144, EPI_ISL_511145, EPI_ISL_511146, EPI_ISL_511147, EPI_ISL_511148, EPI_ISL_511149, EPI_ISL_511150, EPI_ISL_511151, EPI_ISL_511152, EPI_ISL_511154, EPI_ISL_511155, EPI_ISL_511156, EPI_ISL_511157, EPI_ISL_511158, EPI_ISL_511159, EPI_ISL_511160, EPI_ISL_511161, EPI_ISL_511162, EPI_ISL_511163, EPI_ISL_511164, EPI_ISL_511165, EPI_ISL_511166, EPI_ISL_511167, EPI_ISL_511168, EPI_ISL_511169, EPI_ISL_511170, EPI_ISL_511172, EPI_ISL_511173, EPI_ISL_511174, EPI_ISL_511175, EPI_ISL_511176, EPI_ISL_511177, EPI_ISL_511180, EPI_ISL_511181 | | | |
| see above | Instituto Nacional de Saude (INSA) | Instituto Nacional de Saude (INSA) | Borges et al |
| EPI_ISL_511182, EPI_ISL_511183, EPI_ISL_511184, EPI_ISL_511185, EPI_ISL_511186, EPI_ISL_511187, EPI_ISL_511188, EPI_ISL_511190, EPI_ISL_511191, EPI_ISL_511192, EPI_ISL_511193, EPI_ISL_511194, EPI_ISL_511195, EPI_ISL_511196, EPI_ISL_511197, EPI_ISL_511198, EPI_ISL_511199, EPI_ISL_511200, EPI_ISL_511201, EPI_ISL_511202, EPI_ISL_511203, EPI_ISL_511204, EPI_ISL_511205, EPI_ISL_511206, EPI_ISL_511207, EPI_ISL_511208, EPI_ISL_511209, EPI_ISL_511211, EPI_ISL_511212, EPI_ISL_511213, EPI_ISL_511214, EPI_ISL_511215, EPI_ISL_511216, EPI_ISL_511217, EPI_ISL_511218, EPI_ISL_511219, EPI_ISL_511220, EPI_ISL_511221, EPI_ISL_511222, EPI_ISL_511223, EPI_ISL_511224, EPI_ISL_511225, EPI_ISL_511226, EPI_ISL_511228, EPI_ISL_511229, EPI_ISL_511230, EPI_ISL_511231, EPI_ISL_511232, EPI_ISL_511233, EPI_ISL_511234, EPI_ISL_511235, EPI_ISL_511236, EPI_ISL_511237, EPI_ISL_511238, EPI_ISL_511239, EPI_ISL_511240, EPI_ISL_511241, EPI_ISL_511242, EPI_ISL_511243, EPI_ISL_511244, EPI_ISL_511245, EPI_ISL_511246, EPI_ISL_511247, EPI_ISL_511248, EPI_ISL_511249, EPI_ISL_511250, EPI_ISL_511251, EPI_ISL_511252, EPI_ISL_511253, EPI_ISL_511254, EPI_ISL_511255, EPI_ISL_511256, EPI_ISL_511259, EPI_ISL_511261, EPI_ISL_511264, EPI_ISL_511267, EPI_ISL_511268, EPI_ISL_511269, EPI_ISL_511271, EPI_ISL_511272, EPI_ISL_511274, EPI_ISL_511275, EPI_ISL_511276, EPI_ISL_511277, EPI_ISL_511281, EPI_ISL_511283, EPI_ISL_511284, EPI_ISL_511285, EPI_ISL_511286, EPI_ISL_511287, EPI_ISL_511288, EPI_ISL_511289, EPI_ISL_511290, EPI_ISL_511294, EPI_ISL_511295, EPI_ISL_511296, EPI_ISL_511297, EPI_ISL_511298, EPI_ISL_511299, EPI_ISL_511301, EPI_ISL_511302, EPI_ISL_511307, EPI_ISL_511310, EPI_ISL_511311, EPI_ISL_511312, EPI_ISL_511313, EPI_ISL_511314 | | | |
| see above | Instituto Nacional de Saude (INSA) and Instituto Gulbenkian de Ciencia (IGC) | Instituto Nacional de Saude (INSA) and Instituto Gulbenkian de Ciencia (IGC) | Borges et al |
| EPI_ISL_511316, EPI_ISL_511324, EPI_ISL_511326, EPI_ISL_511327, EPI_ISL_511328, EPI_ISL_511331, EPI_ISL_511332, EPI_ISL_511333, EPI_ISL_511334, EPI_ISL_511336, EPI_ISL_511337, EPI_ISL_511338, EPI_ISL_511339, EPI_ISL_511341, EPI_ISL_511343, EPI_ISL_511345, EPI_ISL_511346, EPI_ISL_511347, EPI_ISL_511348, EPI_ISL_511350, EPI_ISL_511351, EPI_ISL_511352, EPI_ISL_511353, EPI_ISL_511355, EPI_ISL_511357, EPI_ISL_511358, EPI_ISL_511359, EPI_ISL_511360, EPI_ISL_511361, EPI_ISL_511362, EPI_ISL_511363, EPI_ISL_511364, EPI_ISL_511366, EPI_ISL_511367, EPI_ISL_511368, EPI_ISL_511369, EPI_ISL_511370, EPI_ISL_511371, EPI_ISL_511372, EPI_ISL_511373, EPI_ISL_511374, EPI_ISL_511375, EPI_ISL_511376, EPI_ISL_511377, EPI_ISL_511378, EPI_ISL_511379, EPI_ISL_511381, EPI_ISL_511382, EPI_ISL_511383, EPI_ISL_511384, EPI_ISL_511385, EPI_ISL_511386, EPI_ISL_511387, EPI_ISL_511388, EPI_ISL_511389, EPI_ISL_511390, EPI_ISL_511391, EPI_ISL_511392, EPI_ISL_511393, EPI_ISL_511394, EPI_ISL_511395, EPI_ISL_511396, EPI_ISL_511397, EPI_ISL_511398, EPI_ISL_511399, EPI_ISL_511400, EPI_ISL_511401, EPI_ISL_511402, EPI_ISL_511403, EPI_ISL_511404, EPI_ISL_511405, EPI_ISL_511406, EPI_ISL_511407, EPI_ISL_511408, EPI_ISL_511409, EPI_ISL_511410, EPI_ISL_511411, EPI_ISL_511412, EPI_ISL_511413, EPI_ISL_511414, EPI_ISL_511415, EPI_ISL_511416, EPI_ISL_511417, EPI_ISL_511418, EPI_ISL_511419, EPI_ISL_511420, EPI_ISL_511422, EPI_ISL_511423, EPI_ISL_511424, EPI_ISL_511425, EPI_ISL_511426, EPI_ISL_511427, EPI_ISL_511428, EPI_ISL_511429, EPI_ISL_511430, EPI_ISL_511431, EPI_ISL_511432, EPI_ISL_511433, EPI_ISL_511434, EPI_ISL_511435, EPI_ISL_511436, EPI_ISL_511437, EPI_ISL_511438, EPI_ISL_511439, EPI_ISL_511441, EPI_ISL_511442, EPI_ISL_511444, EPI_ISL_511446, EPI_ISL_511447, EPI_ISL_511448, EPI_ISL_511449, EPI_ISL_511450, EPI_ISL_511451, EPI_ISL_511452, EPI_ISL_511453, EPI_ISL_511455, EPI_ISL_511456, EPI_ISL_511457, EPI_ISL_511458, EPI_ISL_511459, EPI_ISL_511460, EPI_ISL_511461, EPI_ISL_511462, EPI_ISL_511463, EPI_ISL_511465, EPI_ISL_511466, EPI_ISL_511467, EPI_ISL_511469, EPI_ISL_511470, EPI_ISL_511471, EPI_ISL_511472, EPI_ISL_511473, EPI_ISL_511474, EPI_ISL_511477, EPI_ISL_511479, EPI_ISL_511481, EPI_ISL_511483 | | | |
| see above | Instituto Nacional de Saude (INSA) | Instituto Nacional de Saude (INSA) | Borges et al |
| EPI_ISL_511484, EPI_ISL_511485, EPI_ISL_511486, EPI_ISL_511487, EPI_ISL_511488, EPI_ISL_511489, EPI_ISL_511490, EPI_ISL_511491, EPI_ISL_511492, EPI_ISL_511493, EPI_ISL_511494, EPI_ISL_511495, EPI_ISL_511496, EPI_ISL_511497, EPI_ISL_511499, EPI_ISL_511500, EPI_ISL_511501, EPI_ISL_511502, EPI_ISL_511503, EPI_ISL_511504, EPI_ISL_511505, EPI_ISL_511506, EPI_ISL_511507, EPI_ISL_511508 | | | |
| see above | Instituto Nacional de Saude (INSA) | Instituto Nacional de Saude (INSA) and Instituto Gulbenkian de Ciencia (IGC) | Borges et al |
| EPI_ISL_511509, EPI_ISL_511510, EPI_ISL_511512, EPI_ISL_511513, EPI_ISL_511514, EPI_ISL_511515, EPI_ISL_511516, EPI_ISL_511517, EPI_ISL_511518, EPI_ISL_511520, EPI_ISL_511521, EPI_ISL_511522, EPI_ISL_511523, EPI_ISL_511524, EPI_ISL_511525, EPI_ISL_511526, EPI_ISL_511527, EPI_ISL_511528, EPI_ISL_511529, EPI_ISL_511530, EPI_ISL_511531, EPI_ISL_511532, EPI_ISL_511533, EPI_ISL_511534, EPI_ISL_511535, EPI_ISL_511536, EPI_ISL_511537, EPI_ISL_511539, EPI_ISL_511541, EPI_ISL_511542, EPI_ISL_511543, EPI_ISL_511544, EPI_ISL_511545, EPI_ISL_511546, EPI_ISL_511549, EPI_ISL_511551, EPI_ISL_511553, EPI_ISL_511556, EPI_ISL_511557, EPI_ISL_511558, EPI_ISL_511559, EPI_ISL_511560, EPI_ISL_511561, EPI_ISL_511563, EPI_ISL_511565, EPI_ISL_511566, EPI_ISL_511567, EPI_ISL_511569, EPI_ISL_511570, EPI_ISL_511571, EPI_ISL_511576, EPI_ISL_511584, EPI_ISL_511594, EPI_ISL_511598, EPI_ISL_511600, EPI_ISL_511603, EPI_ISL_511611, EPI_ISL_511612, EPI_ISL_511613, EPI_ISL_511614, EPI_ISL_511615, EPI_ISL_511616, EPI_ISL_511617, EPI_ISL_511619, EPI_ISL_511620, EPI_ISL_511621, EPI_ISL_511622, EPI_ISL_511623, EPI_ISL_511624, EPI_ISL_511625, EPI_ISL_511626, EPI_ISL_511627, EPI_ISL_511628, EPI_ISL_511629, EPI_ISL_511630, EPI_ISL_511631, EPI_ISL_511632, EPI_ISL_511633, EPI_ISL_511634, EPI_ISL_511635, EPI_ISL_511636, EPI_ISL_511637, EPI_ISL_511638, EPI_ISL_511639, EPI_ISL_511640, EPI_ISL_511642, EPI_ISL_511643, EPI_ISL_511645, EPI_ISL_511646, EPI_ISL_511647, EPI_ISL_511648, EPI_ISL_511649, EPI_ISL_511650, EPI_ISL_511651, EPI_ISL_511652, EPI_ISL_511653, EPI_ISL_511654, EPI_ISL_511655, EPI_ISL_511656, EPI_ISL_511657, EPI_ISL_511658, EPI_ISL_511659, EPI_ISL_511660, EPI_ISL_511661, EPI_ISL_511663, EPI_ISL_511668, EPI_ISL_511669, EPI_ISL_511670, EPI_ISL_511671, EPI_ISL_511672, EPI_ISL_511673, EPI_ISL_511674, EPI_ISL_511675, EPI_ISL_511676, EPI_ISL_511677, EPI_ISL_511678, EPI_ISL_511679, EPI_ISL_511680, EPI_ISL_511683, EPI_ISL_511684, EPI_ISL_511685, EPI_ISL_511687, EPI_ISL_511688, EPI_ISL_511689, EPI_ISL_511691, EPI_ISL_511692, EPI_ISL_511693, EPI_ISL_511695, EPI_ISL_511696, EPI_ISL_511702, EPI_ISL_511706, EPI_ISL_511708, EPI_ISL_511709, EPI_ISL_511710, EPI_ISL_511711, EPI_ISL_511713, EPI_ISL_511714, EPI_ISL_511715, EPI_ISL_511716, EPI_ISL_511717, EPI_ISL_511719, EPI_ISL_511720, EPI_ISL_511721, EPI_ISL_511722, EPI_ISL_511723, EPI_ISL_511724, EPI_ISL_511725, EPI_ISL_511726, EPI_ISL_511727, EPI_ISL_511728, EPI_ISL_511729, EPI_ISL_511730, EPI_ISL_511733, EPI_ISL_511741, EPI_ISL_511748 | | | |
| see above | Instituto Nacional de Saude (INSA) | Instituto Nacional de Saude (INSA) | Borges et al |
| EPI_ISL_511752, EPI_ISL_511753, EPI_ISL_511754, EPI_ISL_511755, EPI_ISL_511756, EPI_ISL_511757, EPI_ISL_511759, EPI_ISL_511760, EPI_ISL_511761, EPI_ISL_511762, EPI_ISL_511763, EPI_ISL_511764, EPI_ISL_511765, EPI_ISL_511766, EPI_ISL_511767, EPI_ISL_511768, EPI_ISL_511769, EPI_ISL_511770, EPI_ISL_511771 | | | |
| see above | Instituto Nacional de Saude (INSA) | Instituto Nacional de Saude (INSA) and Instituto Gulbenkian de Ciencia (IGC) | Borges et al |
| EPI_ISL_511805, EPI_ISL_511806, EPI_ISL_511807, EPI_ISL_511808, EPI_ISL_511809, EPI_ISL_511810, EPI_ISL_511812, EPI_ISL_511813, EPI_ISL_511814, EPI_ISL_511815, EPI_ISL_511816, EPI_ISL_511817, EPI_ISL_511819, EPI_ISL_511821, EPI_ISL_511822, EPI_ISL_511823, EPI_ISL_511824, EPI_ISL_511825, EPI_ISL_511828, EPI_ISL_511829, EPI_ISL_511830, EPI_ISL_511831, EPI_ISL_511832, EPI_ISL_511833, EPI_ISL_511834, EPI_ISL_511835, EPI_ISL_511837, EPI_ISL_511839, EPI_ISL_511840, EPI_ISL_511842, EPI_ISL_511843, EPI_ISL_511845, EPI_ISL_511848, EPI_ISL_511849, EPI_ISL_511850, EPI_ISL_511851 | | | |
| see above | Innovative Genomics Institute, UC Berkeley | Innovative Genomics Institute, UC Berkeley | Stacia Wyman, Haridha Shivram, Liana Lareau, Shana McDevitt, Justin Choi |
| EPI_ISL_511853, EPI_ISL_511854, EPI_ISL_511855, EPI_ISL_511856, EPI_ISL_511860, EPI_ISL_511861 | UW Virology Lab | UW Virology Lab | Pavitra Roychoudhury, Amin Addetia, Hong Xie, Lasata Shrestha, Truong Nguyen, Meei-Li Huang, Keith Jerome, Alexander Greninger |
| EPI_ISL_511862 | Innovative Genomics Institute, UC Berkeley | Innovative Genomics Institute, UC Berkeley | Stacia Wyman, Haridha Shivram, Liana Lareau, Shana McDevitt, Justin Choi |
| EPI_ISL_511869, EPI_ISL_511871, EPI_ISL_511874, EPI_ISL_511875 | Johns Hopkins Hospital Department of Pathology | Johns Hopkins Hospital Department of Pathology | Peter M. Thielen, Thomas Mehoke, Shirlee Wohl, Srividya Ramakrishnan, Melanie Kirsche, Amanda Ernlund, Craig Howser, Kristina Zudock, Oluwaseun Falade-Nwulia, Norah Sadowski, Paul Morris, Mark Hopkins, Yunfan Fan, Nidia Trovao, Victoria Gniazdowski, Michael C. Schatz, Stuart C. Ray, Winston Timp, Heba H. Mostafa |
| EPI_ISL_511879 | Laboratorium Kesehatan Provinsi Jawa Barat | Molecular Genetics Laboratory-Faculty of Medicine-Universitas Padjadjaran; School of Life Sciences and Technology & School of Pharmacy-Institut Teknologi Bandung; Laboratorium Kesehatan Provinsi Jawa Barat | Marselina Irasonia Tan, Yunia Sribudiani, Catur Riani, Azzania Fibriani, Husna Nugrahapraja, Tarwadi, Ema Rahmawati, Hesti Lina Wiraswati, Lia Faridah, Savira Ekawardhani, Ryan Bayusantika Ristandi, Rifky Waluyajati Rachman, Cut Nur Cinthia Alamanda, Hammam Riza, Soni Solistia Wirawan, Agung Eru Wibowo |
| EPI_ISL_511891, EPI_ISL_511892, EPI_ISL_511893, EPI_ISL_511894, EPI_ISL_511895, EPI_ISL_511896, EPI_ISL_511897, EPI_ISL_511898 | National Hospital of Tropical Diseases | Oxford University Clinical Research Unit, Hanoi, Vietnam | Nguyen Thi Tam, Van Dinh Trang, Nguyen Thi Hong Thuong, Vu Thi Ngoc Bich, Nguyen Thu Trang, Nguyen Thi Ngoc Diep, Le Nguyen Minh Hoa, Pham Ngoc Thach, H. Rogier van Doorn, on behalf of the OUCRU COVID-19 research group |
| EPI_ISL_511899 | ICMR-National Institute of Cholera and Enteric Diseases | National Institute of Biomedical Genomics - DBT's PAN-INDIA 1000 SARS--CoV-2 RNA Genome Sequencing Consortium | Arindam Maitra, Mamta Chawla Sarkar, Sreedhar Chinnaswamy, Hasina Banu, Ananya Chatterjee, Shanta Dutta, Saumitra Das |
| EPI_ISL_511900, EPI_ISL_511901, EPI_ISL_511902, EPI_ISL_511903, EPI_ISL_511904, EPI_ISL_511905, EPI_ISL_511906, EPI_ISL_511907 | Institute of Post Graduate Medical Education & Research | National Institute of Biomedical Genomics - DBT's PAN-INDIA 1000 SARS--CoV-2 RNA Genome Sequencing Consortium | Arindam Maitra, Aritra Biswas, Jayeeta Haldar, Raja Ray, Monimoy Banerjee, Saumitra Das |
| EPI_ISL_511908, EPI_ISL_511909, | All india institute of Medical Sciences Rishikesh | National Institute of Biomedical Genomics - DBT's PAN-INDIA 1000 SARS--CoV-2 RNA Genome Sequencing | Arindam Maitra, Deepjyoti Kalita, Amit Mangla, Ravi Kant, Saumitra Das |

| | | | |
|---|---|---|---|
| EPI_ISL_511910, EPI_ISL_511912, EPI_ISL_511921, EPI_ISL_511922 | | Consortium | |
| EPI_ISL_511924, EPI_ISL_511926, EPI_ISL_511927, EPI_ISL_511928, EPI_ISL_511929 | Mahatma Gandhi Institute of Medical Sciences | National Institute of Biomedical Genomics - DBT's PAN-INDIA 1000 SARS--CoV-2 RNA Genome Sequencing Consortium | Arindam Maitra, Vijayshri Deotale, Rahul Narang, Deepashri Maraskolhe, Saumitra Das |
| EPI_ISL_511930, EPI_ISL_511931, EPI_ISL_511932, EPI_ISL_511933, EPI_ISL_511934, EPI_ISL_511935, EPI_ISL_511939 | Government Medical College | National Institute of Biomedical Genomics - DBT's PAN-INDIA 1000 SARS--CoV-2 RNA Genome Sequencing Consortium | Arindam Maitra, Jyoti Iravane, Dhaval Khatri, Maitrik Dave, Saumitra Das |
| EPI_ISL_511943, EPI_ISL_511944 | Translational Health Science and Technology Institute | National Institute of Biomedical Genomics - DBT's PAN-INDIA 1000 SARS--CoV-2 RNA Genome Sequencing Consortium | Arindam Maitra, Guruprasad Medigeshi, Sharanabasava Patil, Anbalagan Ananthraj, Madhu Pareek, Imran Khan, Gagandeep Kang, Saumitra Das |
| EPI_ISL_511955, EPI_ISL_511956, EPI_ISL_511957, EPI_ISL_511958, EPI_ISL_511960, EPI_ISL_511964, EPI_ISL_511966, EPI_ISL_511967, EPI_ISL_511968 | PHE South West Regional Laboratory, National Infection Service | Wellcome Sanger Institute for the COVID-19 Genomics UK Consortium | Stephanie Hutchings, Hannah Pymont, Dr Peter Muir, Barry Vipond, Rich Hopes; and Alex Alderton, Roberto Amato, Sonia Goncalves, Ewan Harrison, David K. Jackson, Ian Johnston, Dominic Kwiatkowski, Cordelia Langford, John Sillitoe on behalf of the Wellcome Sanger Institute COVID-19 Surveillance Team (http://www.sanger.ac.uk/covid-team) |
| EPI_ISL_511970 | Department of Pathology, University of Cambridge | Wellcome Sanger Institute for the COVID-19 Genomics UK Consortium | Luke W Meredith, M. Estée Török , Myra Hosmillo, William L. Hamilton, Martin D. Curran, Theresa Feltwell, Grant Hall, Anna Yakovleva, Fahad A Khokhar, Charlotte J. Houldcroft, Laura G Caller, Aminu S. Jahun, Sarah L. Caddy, Ian Goodfellow; and Alex Alderton, Roberto Amato, Sonia Goncalves, Ewan Harrison, David K. Jackson, Ian Johnston, Dominic Kwiatkowski, Cordelia Langford, John Sillitoe on behalf of the Wellcome Sanger Institute COVID-19 Surveillance Team (http://www.sanger.ac.uk/covid-team) |
| EPI_ISL_511971, EPI_ISL_511972, EPI_ISL_511973, EPI_ISL_511974, EPI_ISL_511976, EPI_ISL_511978, EPI_ISL_511979, EPI_ISL_511980, EPI_ISL_511981 | PHE South West Regional Laboratory, National Infection Service | Wellcome Sanger Institute for the COVID-19 Genomics UK Consortium | Stephanie Hutchings, Hannah Pymont, Dr Peter Muir, Barry Vipond, Rich Hopes; and Alex Alderton, Roberto Amato, Sonia Goncalves, Ewan Harrison, David K. Jackson, Ian Johnston, Dominic Kwiatkowski, Cordelia Langford, John Sillitoe on behalf of the Wellcome Sanger Institute COVID-19 Surveillance Team (http://www.sanger.ac.uk/covid-team) |
| EPI_ISL_511985, EPI_ISL_511986, EPI_ISL_511987, EPI_ISL_511988, EPI_ISL_511989, EPI_ISL_511990, EPI_ISL_511991, EPI_ISL_511992, EPI_ISL_511993, EPI_ISL_511994, EPI_ISL_511997, EPI_ISL_511998, EPI_ISL_511999, EPI_ISL_512000, EPI_ISL_512001, EPI_ISL_512002, EPI_ISL_512003, EPI_ISL_512004, EPI_ISL_512005, EPI_ISL_512006, EPI_ISL_512009, EPI_ISL_512010, EPI_ISL_512011, EPI_ISL_512012, EPI_ISL_512013, EPI_ISL_512014, EPI_ISL_512015, EPI_ISL_512016, EPI_ISL_512017, EPI_ISL_512018, EPI_ISL_512020, EPI_ISL_512021, EPI_ISL_512023, EPI_ISL_512024, EPI_ISL_512025, EPI_ISL_512027, EPI_ISL_512028, EPI_ISL_512029, EPI_ISL_512030, EPI_ISL_512031, EPI_ISL_512032, EPI_ISL_512033, EPI_ISL_512034, EPI_ISL_512035, EPI_ISL_512036, EPI_ISL_512037, EPI_ISL_512038, EPI_ISL_512039, EPI_ISL_512040, EPI_ISL_512042, EPI_ISL_512043, EPI_ISL_512044, EPI_ISL_512045, EPI_ISL_512046, EPI_ISL_512047, EPI_ISL_512048, EPI_ISL_512049, EPI_ISL_512050, EPI_ISL_512051, EPI_ISL_512052, EPI_ISL_512053, EPI_ISL_512054 | | | |
| see above | Viollier AG | Department of Biosystems Science and Engineering, ETH Zürich | Christian Beisel, Sarah Nadeau, Ivan Topolsky, Pedro Ferreira, Philipp Jablonski, Susana Posada-Céspedes, Tobias Schär, Ina Nissen, Natascha Santacroce, Elodie Burcklen, Christiane Beckmann, Maurice Redondo, Olivier Kobel, Christoph Noppen, Sophie Seidel, Noemie Santamaria de Souza, Niko Beerenwinkel, Tanja Stadler |
| EPI_ISL_512058 | B.J. Medical College and Civil hospital, Ahmedabad | Gujarat Biotechnology Research Centre | Monika Gandhi, Pinal Trivedi, Maharshi Pandya, Nidhi Patel, Nitin Savaliya, Raghawendra Kumar, Dinesh Kumar, Zuber Saiyed, Komal Patel, Labdhi Pandya, Afzal Ansari, Nikha Trivedi, Pranay Shah, Kamlesh J Upadhyay, Sanjay Kapadia, Apurvasinh Puvar, Janvi Raval, Zarna Patel, R D Dixit, A M Kadri, Harsh Bakshi, Chaitanya Joshi, Madhvi Joshi |
| EPI_ISL_512059 | B.J. Medical College and Civil hospital, Ahmedabad | Gujarat Biotechnology Research Centre | Pinal Trivedi, Maharshi Pandya, Nidhi Patel, Nitin Savaliya, Raghawendra Kumar, Dinesh Kumar, Zuber Saiyed, Komal Patel, Labdhi Pandya, Afzal Ansari, Nikha Trivedi, Pranay Shah, Kamlesh J Upadhyay, Sanjay Kapadia, Apurvasinh Puvar, Janvi Raval, Zarna Patel, Monika Gandhi, R D Dixit, A M Kadri, Harsh Bakshi, Chaitanya Joshi, Madhvi Joshi |
| EPI_ISL_512060 | B.J. Medical College and Civil hospital, Ahmedabad | Gujarat Biotechnology Research Centre | Maharshi Pandya, Nidhi Patel, Nitin Savaliya, Raghawendra Kumar, Dinesh Kumar, Zuber Saiyed, Komal Patel, Labdhi Pandya, Afzal Ansari, Nikha Trivedi, Pranay Shah, Kamlesh J Upadhyay, Sanjay Kapadia, Apurvasinh Puvar, Janvi Raval, Zarna Patel, Monika Gandhi, Pinal Trivedi, R D Dixit, A M Kadri, Harsh Bakshi, Chaitanya Joshi, Madhvi Joshi |
| EPI_ISL_512063 | B.J. Medical College and Civil hospital, Ahmedabad | Gujarat Biotechnology Research Centre | Raghawendra Kumar, Dinesh Kumar, Zuber Saiyed, Komal Patel, Labdhi Pandya, Afzal Ansari, Nikha Trivedi, Pranay Shah, Kamlesh J Upadhyay, Sanjay Kapadia, Apurvasinh Puvar, Janvi Raval, Zarna Patel, Monika Gandhi, Pinal Trivedi, Maharshi Pandya, Nidhi Patel, Nitin Savaliya, R D Dixit, A M Kadri, Harsh Bakshi, Chaitanya Joshi, Madhvi Joshi |
| EPI_ISL_512064 | B.J. Medical College and Civil hospital, Ahmedabad | Gujarat Biotechnology Research Centre | Dinesh Kumar, Zuber Saiyed, Komal Patel, Labdhi Pandya, Afzal Ansari, Nikha Trivedi, Pranay Shah, Kamlesh J Upadhyay, Sanjay Kapadia, Apurvasinh Puvar, Janvi Raval, Zarna Patel, Monika Gandhi, Pinal Trivedi, Maharshi Pandya, Nidhi Patel, Nitin Savaliya, Raghawendra Kumar, R D Dixit, A M Kadri, Harsh Bakshi, Chaitanya Joshi, Madhvi Joshi |
| EPI_ISL_512065 | B.J. Medical College and Civil hospital, Ahmedabad | Gujarat Biotechnology Research Centre | Zuber Saiyed, Komal Patel, Labdhi Pandya, Afzal Ansari, Nikha Trivedi, Pranay Shah, Kamlesh J Upadhyay, Sanjay Kapadia, Apurvasinh Puvar, Janvi Raval, Zarna Patel, Monika Gandhi, Pinal Trivedi, Maharshi Pandya, Nidhi Patel, Nitin Savaliya, Raghawendra Kumar, Dinesh Kumar, R D Dixit, A M Kadri, Harsh Bakshi, Chaitanya Joshi, Madhvi Joshi |
| EPI_ISL_512066 | Sardar Vallabhbhai Patel Institute of Medical Sciences & Research | Gujarat Biotechnology Research Centre | Komal Patel, Labdhi Pandya, Afzal Ansari, Nikha Trivedi, Pranay Shah, Kamlesh J Upadhyay, Sanjay Kapadia, Apurvasinh Puvar, Janvi Raval, Zarna Patel, Monika Gandhi, Pinal Trivedi, Maharshi Pandya, Nidhi Patel, Nitin Savaliya, Raghawendra Kumar, Dinesh Kumar, Zuber Saiyed, R D Dixit, A M Kadri, Harsh Bakshi, Chaitanya Joshi, Madhvi Joshi |
| EPI_ISL_512067 | Sardar Vallabhbhai Patel Institute of Medical Sciences & Research | Gujarat Biotechnology Research Centre | Labdhi Pandya, Afzal Ansari, Nikha Trivedi, Pranay Shah, Kamlesh J Upadhyay, Sanjay Kapadia, Apurvasinh Puvar, Janvi Raval, Zarna Patel, Monika Gandhi, Pinal Trivedi, Maharshi Pandya, Nidhi Patel, Nitin Savaliya, Raghawendra Kumar, Dinesh Kumar, Zuber Saiyed, Komal Patel, R D Dixit, A M Kadri, Harsh Bakshi, Chaitanya Joshi, Madhvi Joshi |
| EPI_ISL_512068 | Sardar Vallabhbhai Patel Institute of Medical Sciences & Research | Gujarat Biotechnology Research Centre | Afzal Ansari, Nikha Trivedi, Pranay Shah, Kamlesh J Upadhyay, Sanjay Kapadia, Apurvasinh Puvar, Janvi Raval, Zarna Patel, Monika Gandhi, Pinal Trivedi, Maharshi Pandya, Nidhi Patel, Nitin Savaliya, Raghawendra Kumar, Dinesh Kumar, Zuber Saiyed, Komal Patel, Labdhi Pandya, R D Dixit, A M Kadri, Harsh Bakshi, Chaitanya Joshi, Madhvi Joshi |
| EPI_ISL_512069 | Sardar Vallabhbhai Patel Institute of Medical Sciences & Research | Gujarat Biotechnology Research Centre | Nikha Trivedi, Pranay Shah, Kamlesh J Upadhyay, Sanjay Kapadia, Apurvasinh Puvar, Janvi Raval, Zarna Patel, Monika Gandhi, Pinal Trivedi, Maharshi Pandya, Nidhi Patel, Nitin Savaliya, Raghawendra Kumar, Dinesh Kumar, Zuber Saiyed, Komal Patel, Labdhi Pandya, Afzal Ansari, R D Dixit, A M Kadri, Harsh Bakshi, Chaitanya Joshi, Madhvi Joshi |
| EPI_ISL_512070 | Department of MicroBiology, Government Medical College, Surat | Gujarat Biotechnology Research Centre | Naresh Chauhan, Summaiya Mullan, Amit gamit, Apurvasinh Puvar, Janvi Raval, Zarna Patel, Monika Gandhi, Pinal Trivedi, Maharshi Pandya, Nidhi Patel, Nitin Savaliya, Raghawendra Kumar, Dinesh Kumar, Zuber Saiyed, Komal Patel, Labdhi Pandya, Afzal Ansari, Nikha Trivedi, R D Dixit, A M Kadri, Harsh Bakshi, Chaitanya Joshi, Madhvi Joshi |
| EPI_ISL_512071 | Department of MicroBiology, Government Medical College, Surat | Gujarat Biotechnology Research Centre | Summaiya Mullan, Amit gamit, Apurvasinh Puvar, Janvi Raval, Zarna Patel, Monika Gandhi, Pinal Trivedi, Maharshi Pandya, Nidhi Patel, Nitin Savaliya, Raghawendra Kumar, Dinesh Kumar, Zuber Saiyed, Komal Patel, Labdhi Pandya, Afzal Ansari, Nikha Trivedi, Naresh Chauhan, R D Dixit, A M Kadri, Harsh Bakshi, Chaitanya Joshi, Madhvi Joshi |
| EPI_ISL_512072 | Saikrishna Hospital, Mehsana | Gujarat Biotechnology Research Centre | Harshadbhai Parmar, Apurvasinh Puvar, Janvi Raval, Zarna Patel, Monika Gandhi, Pinal Trivedi, Maharshi Pandya, Nidhi Patel, Nitin Savaliya, Raghawendra Kumar, Dinesh Kumar, Zuber Saiyed, Komal Patel, Labdhi Pandya, Afzal Ansari, Nikha Trivedi, R D Dixit, A M Kadri, Harsh Bakshi, Chaitanya Joshi, Madhvi Joshi |
| EPI_ISL_512073 | Saikrishna Hospital, Mehsana | Gujarat Biotechnology Research Centre | Apurvasinh Puvar, Janvi Raval, Zarna Patel, Monika Gandhi, Pinal Trivedi, Maharshi Pandya, Nidhi Patel, Nitin Savaliya, Raghawendra Kumar, Dinesh Kumar, Zuber Saiyed, Komal Patel, Labdhi Pandya, Afzal Ansari, Nikha Trivedi, Harshadbhai Parmar, R D Dixit, A M Kadri, Harsh Bakshi, Chaitanya Joshi, Madhvi Joshi |
| EPI_ISL_512074 | Saikrishna Hospital, Mehsana | Gujarat Biotechnology Research Centre | Janvi Raval, Zarna Patel, Monika Gandhi, Pinal Trivedi, Maharshi Pandya, Nidhi Patel, Nitin Savaliya, Raghawendra Kumar, Dinesh Kumar, Zuber Saiyed, Komal Patel, Labdhi Pandya, Afzal Ansari, Nikha Trivedi, Harshadbhai Parmar, Apurvasinh Puvar, R D Dixit, A M Kadri, Harsh Bakshi, Chaitanya Joshi, Madhvi Joshi |
| EPI_ISL_512075 | Dr. RSS Hospital, Modasa | Gujarat Biotechnology Research Centre | Monika Gandhi, Pinal Trivedi, Maharshi Pandya, Nidhi Patel, Nitin Savaliya, Raghawendra Kumar, Dinesh Kumar, Zuber Saiyed, Komal Patel, Labdhi Pandya, Afzal Ansari, Nikha Trivedi, Harsh Chaudhari, Apurvasinh Puvar, Janvi Raval, Zarna Patel, R D Dixit, A M Kadri, Harsh Bakshi, Chaitanya Joshi, Madhvi Joshi |
| EPI_ISL_512076 | Dr. RSS Hospital, Modasa | Gujarat Biotechnology Research Centre | Pinal Trivedi, Maharshi Pandya, Nidhi Patel, Nitin Savaliya, Raghawendra Kumar, Dinesh Kumar, Zuber Saiyed, Komal Patel, Labdhi Pandya, Afzal Ansari, Nikha Trivedi, Harsh Chaudhari, Apurvasinh Puvar, Janvi Raval, Zarna Patel, Monika Gandhi, R D Dixit, A M Kadri, Harsh Bakshi, Chaitanya Joshi, Madhvi Joshi |
| EPI_ISL_512077 | Dr. RSS Hospital, Modasa | Gujarat Biotechnology Research Centre | Maharshi Pandya, Nidhi Patel, Nitin Savaliya, Raghawendra Kumar, Dinesh Kumar, Zuber Saiyed, Komal Patel, Labdhi Pandya, Afzal Ansari, Nikha Trivedi, Harsh Chaudhari, Apurvasinh Puvar, Janvi Raval, Zarna Patel, Monika Gandhi, Pinal Trivedi, R D Dixit, A M Kadri, Harsh Bakshi, Chaitanya Joshi, Madhvi Joshi |
| EPI_ISL_512086 | UW Virology lab | UW Virology lab | Pavitra Roychoudhury, Amin Addetia, Hong Xie, Lasata Shrestha, Truong Nguyen, Meei-Li Huang, Keith Jerome, Alexander Greninger |
| EPI_ISL_512087, EPI_ISL_512088, EPI_ISL_512089, EPI_ISL_512091, EPI_ISL_512092, EPI_ISL_512093, EPI_ISL_512094, EPI_ISL_512097, EPI_ISL_512098, EPI_ISL_512099, EPI_ISL_512100, EPI_ISL_512101, EPI_ISL_512103, EPI_ISL_512104, EPI_ISL_512105, EPI_ISL_512106 | | | |
| see above | National Virus Reference Laboratory | National Virus Reference Laboratory | Michael Carr, Gabriel Gonzalez, Jonathan Dean, Aditi Chaturvedi, Suzie Coughlan, Cillian F De Gascun |
| EPI_ISL_512144, EPI_ISL_512148, EPI_ISL_512156, EPI_ISL_512157 | Alaska State Virology Laboratory | Alaska State Virology Laboratory | Chen J et al with Pathogenomics group Dagdag R, Redlinger M, Milton E, George W, Kovalenko A, Drown DM, Bortz E |
| EPI_ISL_512158, EPI_ISL_512160, EPI_ISL_512161, EPI_ISL_512162, EPI_ISL_512163, EPI_ISL_512164, EPI_ISL_512165, EPI_ISL_512166, EPI_ISL_512167, EPI_ISL_512168, EPI_ISL_512169, EPI_ISL_512170, EPI_ISL_512171, EPI_ISL_512172, EPI_ISL_512173, EPI_ISL_512174, EPI_ISL_512175, EPI_ISL_512176, EPI_ISL_512178, EPI_ISL_512179, EPI_ISL_512180, EPI_ISL_512181, EPI_ISL_512182, EPI_ISL_512183, EPI_ISL_512184, EPI_ISL_512185, EPI_ISL_512186, EPI_ISL_512187, EPI_ISL_512188, EPI_ISL_512189, EPI_ISL_512191, EPI_ISL_512192, EPI_ISL_512193, EPI_ISL_512194, EPI_ISL_512195, EPI_ISL_512196, EPI_ISL_512197, EPI_ISL_512198, EPI_ISL_512199, EPI_ISL_512200, EPI_ISL_512201, EPI_ISL_512203, EPI_ISL_512204, EPI_ISL_512206, EPI_ISL_512207, EPI_ISL_512208, EPI_ISL_512209, EPI_ISL_512211, EPI_ISL_512212, EPI_ISL_512213, EPI_ISL_512214, EPI_ISL_512215, EPI_ISL_512220, EPI_ISL_512221, EPI_ISL_512222, EPI_ISL_512223, EPI_ISL_512224, EPI_ISL_512225, EPI_ISL_512226, EPI_ISL_512227, EPI_ISL_512228, EPI_ISL_512229, EPI_ISL_512230, EPI_ISL_512231, EPI_ISL_512232, EPI_ISL_512233, EPI_ISL_512235, EPI_ISL_512236, EPI_ISL_512237, EPI_ISL_512238, EPI_ISL_512239, EPI_ISL_512240, EPI_ISL_512241, EPI_ISL_512242, EPI_ISL_512243, EPI_ISL_512244, EPI_ISL_512245, EPI_ISL_512246, EPI_ISL_512247, EPI_ISL_512248, EPI_ISL_512250, EPI_ISL_512251, EPI_ISL_512252, EPI_ISL_512253, EPI_ISL_512254, EPI_ISL_512256, EPI_ISL_512257, EPI_ISL_512259, EPI_ISL_512260, EPI_ISL_512261, EPI_ISL_512262, EPI_ISL_512264, EPI_ISL_512265, EPI_ISL_512266, EPI_ISL_512267, EPI_ISL_512268, EPI_ISL_512269, EPI_ISL_512270, EPI_ISL_512271, EPI_ISL_512273 | | | |
| see above | San Diego County Public Health Laboratory | Andersen lab at Scripps Research | SEARCH Alliance San Diego with Tracy Basler, Jovan Shephard, Brett Austin |
| EPI_ISL_512296, EPI_ISL_512297, EPI_ISL_512298, EPI_ISL_512299, EPI_ISL_512300, EPI_ISL_512301, EPI_ISL_512302, EPI_ISL_512303 | Hematology Laboratory, Section of Molecular Diagnostics, University Clinical Centre, Medical University of Gdansk | Department of Virology, Faculty of Medicine, University of Helsinki, Helsinki, Finland | Maciej Grzybek, Marlena Robakowska, Aneta Szulc, Ewa Miłosz, Olli Vapalahti, Teemu Smura |
| EPI_ISL_512313, EPI_ISL_512314 | E. Gulbja Laboratorija | Latvian Biomedical Research and Study Centre | Ivars Silamiķelis, Kaspars Megnis, Monta Ustinova, Ņikita Zrelovs, Vita Rovīte, Mikus Gavars, Dmitrijs Perminovs, Uga Dumpis, Jānis Kloviņš |
| EPI_ISL_512332, EPI_ISL_512333 | Department of Pathology, University of Cambridge | COVID-19 Genomics UK (COG-UK) Consortium | Luke W Meredith, M. Estée Török, Myra Hosmillo, William L. Hamilton, Martin D. Curran, Theresa Feltwell, Grant Hall, Anna Yakovleva, Fahad A Khokhar, Charlotte J. Houldcroft, Laura G Caller, Aminu S. Jahun, Sarah L. Caddy, Yasmin Chaudhry, Malte Pinckert, Ian Goodfellow |
| EPI_ISL_512339 | University of Exeter | COVID-19 Genomics UK (COG-UK) Consortium | Ben Temperton,Aaron Jeffries,Michelle Michelsen,Joanna Warwick-Dugdale,Audrey Farbos,Robyn Manley,Stephen Michell,Jane Masoli |
| EPI_ISL_512342, EPI_ISL_512344 | Liverpool Clinical Laboratories | COVID-19 Genomics UK (COG-UK) Consortium | Sam Haldenby, Anita Lucaci, Steve Paterson, Julian Hiscox, Alistair Darby, M Almsaud, A Alrezaihi, Muhannad Alruwaili, Stuart D Armstrong, Jones Benjamin, Eleanor G Bentley, Anu Chawla, Jordan J Clark, Angela Cowell, Richard Eccles, Isabel García-Dorival, Matthew Gemmell, Alessandro Gerada, PKF Gilmore, Richard Gregory, Ximeng Han, Catherine Hartley, Margaret Hughes, Miren Iturriza-Gomara, James Johnson, L Luu, Jenifer Manson, Charlotte Nelson, Elaine O'Toole, Cassie Olateju, Rebekah Penrice-Randal , Lucille Rainbow, N.P Randle, Trevor Ian Robinson, Parul Sharma, Ghada T Shawli, James P Stewart, Neil Swainston, Ecaterina Vamos, Joanne Watts, Mark Whitehead |
| EPI_ISL_512348, EPI_ISL_512349, EPI_ISL_512350, EPI_ISL_512351, EPI_ISL_512353, EPI_ISL_512354, EPI_ISL_512356, EPI_ISL_512358, EPI_ISL_512359, EPI_ISL_512360, EPI_ISL_512361 | | | |
| see above | Northumbria University / South Tees Hospitals NHS Foundation Trust / North Cumbria Integrated Care NHS Foundation Trust / North Tees and Hartlepool | COVID-19 Genomics UK (COG-UK) Consortium | Darren L Smith,Andrew Nelson,Matthew Bashton,Greg R Young,Joshua Loh,John Allan,Mohammad A Tariq,Giles S Holt,Gary Black,Wen C Yew,Lynn Dover,Paul Baker,Steve Liggett,Sarah Essex,Jane Greenaway,Debra Padgett,Clive Graham,Garren Scott,Edward Barton,Emma Swindells,Brendan Payne,Jennifer Collins,Yusri Taha,Gary Eltringham |

| | | | |
|---|---|---|---|
| | NHS Foundation Trust / Newcastle Hospitals NHS Foundation Trust | | |
| EPI_ISL_512380, EPI_ISL_512381, EPI_ISL_512382 | Queens Medical Centre, Clinical Microbiology Department / DeepSeq Nottingham | COVID-19 Genomics UK (COG-UK) Consortium | Gemma Clark, Wendy Smith, Manjinder Khakh, Vicki M Fleming, Michelle M Lister, Hannah Howson-Wells, Jonathan Ball, Patrick McClure, Joseph Chappell, Theocharis Tsoleridis, Nadine Holmes, Matthew Carlisle, Christopher Moore, Fei Sang, Johnny Debebe, Victoria Wright, Matthew Loose |
| EPI_ISL_512389, EPI_ISL_512395, EPI_ISL_512400, EPI_ISL_512407, EPI_ISL_512409, EPI_ISL_512410, EPI_ISL_512411, EPI_ISL_512412, EPI_ISL_512415, EPI_ISL_512416, EPI_ISL_512417, EPI_ISL_512426, EPI_ISL_512430, EPI_ISL_512432, EPI_ISL_512433 | | | |
| see above | Centre for Enzyme Innovation, University of Portsmouth / Translational Research Laboratory, Portsmouth Hospitals NHS Trust | COVID-19 Genomics UK (COG-UK) Consortium | Angela Beckett,Yann Bourgeois,Garry Scarlett,Sharon Glaysher,Scott Elliott,Kelly Bicknell,Robert Impey,Allyson Lloyd,Sarah Wyllie,Ethan Butcher,Anoop Chauhan,Samuel Robson |
| EPI_ISL_512436, EPI_ISL_512438, EPI_ISL_512439, EPI_ISL_512440, EPI_ISL_512443, EPI_ISL_512446, EPI_ISL_512447, EPI_ISL_512448, EPI_ISL_512449, EPI_ISL_512452, EPI_ISL_512453, EPI_ISL_512454, EPI_ISL_512455, EPI_ISL_512456, EPI_ISL_512457, EPI_ISL_512458, EPI_ISL_512459, EPI_ISL_512460, EPI_ISL_512461, EPI_ISL_512462, EPI_ISL_512463, EPI_ISL_512467, EPI_ISL_512468, EPI_ISL_512469, EPI_ISL_512470, EPI_ISL_512471, EPI_ISL_512472, EPI_ISL_512474, EPI_ISL_512475, EPI_ISL_512476, EPI_ISL_512477, EPI_ISL_512478, EPI_ISL_512479, EPI_ISL_512480 | | | |
| see above | West of Scotland Specialist Virology Centre, NHSGGC / MRC-University of Glasgow Centre for Virus Research | COVID-19 Genomics UK (COG-UK) Consortium | Ana da Silva Filipe, Natasha Johnson, Kathy Smollett, Daniel Mair, Stephen Carmichael, Lily Tong, Jenna Nichols, Elihu Aranday-Cortes, Kirstyn Brunker, Yasmin Parr, Alice Broos, Kyriaki Nomikou; Sarah McDonald, Marc Niebel, Patawee Asamaphan; Richard Orton, Joseph Hughes, Sreenu Vattipally, David L Robertson; Alasdair MacLean, Rory Gunson; Kathy Li, Natasha Jesudason, Rajiv Shah, James Shepherd, Antonia Ho, Emma Thomson |
| EPI_ISL_512481, EPI_ISL_512482, EPI_ISL_512483, EPI_ISL_512484, EPI_ISL_512485, EPI_ISL_512486, EPI_ISL_512487, EPI_ISL_512488, EPI_ISL_512489, EPI_ISL_512490, EPI_ISL_512491, EPI_ISL_512492, EPI_ISL_512493, EPI_ISL_512494, EPI_ISL_512495, EPI_ISL_512496, EPI_ISL_512497, EPI_ISL_512498, EPI_ISL_512499, EPI_ISL_512500, EPI_ISL_512501, EPI_ISL_512502, EPI_ISL_512503, EPI_ISL_512504, EPI_ISL_512505, EPI_ISL_512506, EPI_ISL_512507, EPI_ISL_512508, EPI_ISL_512509, EPI_ISL_512510, EPI_ISL_512511, EPI_ISL_512513, EPI_ISL_512515, EPI_ISL_512517, EPI_ISL_512518, EPI_ISL_512519, EPI_ISL_512520, EPI_ISL_512521, EPI_ISL_512522, EPI_ISL_512523, EPI_ISL_512524, EPI_ISL_512527, EPI_ISL_512528, EPI_ISL_512531, EPI_ISL_512532, EPI_ISL_512533, EPI_ISL_512535, EPI_ISL_512536, EPI_ISL_512537, EPI_ISL_512538, EPI_ISL_512539, EPI_ISL_512540, EPI_ISL_512541, EPI_ISL_512542, EPI_ISL_512543, EPI_ISL_512544, EPI_ISL_512545 | | | |
| see above | Wales Specialist Virology Centre Sequencing lab: Pathogen Genomics Unit | COVID-19 Genomics UK (COG-UK) Consortium | Catherine Moore, Johnathan Evans, Laura Gifford, Malorie Perry, Simon Cottrell, Angela Marchbank, Alec Birchley, Alexander Adams, Amy Gaskin, Bree Gatica-Wilcox, Jason Coombes, Joel Southgate, Lauren Gilbert, Lee Graham, Nicole Pacchiarini, Sara Kumziene-Summerhayes, Sarah Taylor, Sophie Jones, Sara Rey, Matthew Bull, Joanne Watkins, Sally Corden, Tom Connor |
| EPI_ISL_512547, EPI_ISL_512549, EPI_ISL_512550, EPI_ISL_512552, EPI_ISL_512554, EPI_ISL_512555, EPI_ISL_512556, EPI_ISL_512557, EPI_ISL_512558, EPI_ISL_512559, EPI_ISL_512560, EPI_ISL_512561, EPI_ISL_512562, EPI_ISL_512563, EPI_ISL_512564, EPI_ISL_512565, EPI_ISL_512566, EPI_ISL_512567, EPI_ISL_512568, EPI_ISL_512570, EPI_ISL_512571, EPI_ISL_512572, EPI_ISL_512573, EPI_ISL_512575, EPI_ISL_512578, EPI_ISL_512579 | | | |
| see above | Florida Bureau of Public Health Laboratories | Florida Bureau of Public Health Laboratories | Sarah Schmedes, Jason Blanton |
| EPI_ISL_512597, EPI_ISL_512598, EPI_ISL_512599, EPI_ISL_512600, EPI_ISL_512603, EPI_ISL_512604, EPI_ISL_512605, EPI_ISL_512606, EPI_ISL_512607, EPI_ISL_512608, EPI_ISL_512609, EPI_ISL_512610, EPI_ISL_512611, EPI_ISL_512612, EPI_ISL_512614, EPI_ISL_512615, EPI_ISL_512616, EPI_ISL_512617, EPI_ISL_512618, EPI_ISL_512620, EPI_ISL_512621, EPI_ISL_512622, EPI_ISL_512623, EPI_ISL_512624, EPI_ISL_512625, EPI_ISL_512626, EPI_ISL_512628, EPI_ISL_512629, EPI_ISL_512631, EPI_ISL_512632, EPI_ISL_512634, EPI_ISL_512636, EPI_ISL_512638, EPI_ISL_512640, EPI_ISL_512641, EPI_ISL_512642, EPI_ISL_512643 | | | |
| see above | National Laboratory for Influenza/Virology reference laboratory, Public Health Center of the Ministry of Health of Ukraine | Respiratory Virus Unit, Microbiology Services Colindale, Public Health England | PHE Covid Sequencing Team, Dr. Iryna Demchyshyna |
| EPI_ISL_512645, EPI_ISL_512646 | E. Gulbja Laboratorija | Latvian Biomedical Research and Study Centre | Ivars Silamiķelis, Kaspars Megnis, Monta Ustinova, Ņikita Zrelovs, Vita Rovīte, Mikus Gavars, Dmitrijs Perminovs, Uga Dumpis, Jānis Kloviņš |
| EPI_ISL_512647, EPI_ISL_512648 | Latvijas Infektoloģijas centrs | Latvian Biomedical Research and Study Centre | Ivars Silamiķelis, Kaspars Megnis, Monta Ustinova, Ņikita Zrelovs, Vita Rovīte, Jeļena Storoženko, Tatjana Kolupajeva, Oksana Savicka, Uga Dumpis, Jānis Kloviņš |
| EPI_ISL_512649 | Latvijas Infektoloģijas centrs | Latvian Biomedical Research and Study Centre | Ivars Silamiķelis, Kaspars Megnis, Monta Ustinova, Ņikita Zrelovs, Vita Rovīte, Jeļena Storoženko, Tatjana Kolupajeva, Oksana Savicka, Uga Dumpis, Jānis Kloviņš |
| EPI_ISL_512650, EPI_ISL_512651, EPI_ISL_512652 | Latvijas Infektoloģijas centrs | Latvian Biomedical Research and Study Centre | Ivars Silamiķelis, Kaspars Megnis, Monta Ustinova, Ņikita Zrelovs, Vita Rovīte, Jeļena Storoženko, Tatjana Kolupajeva, Oksana Savicka, Uga Dumpis, Jānis Kloviņš |
| EPI_ISL_512653 | Area De Salud Desamparados 1 - Clinica Dr. Marcial Fallas [Grifo Alto/Desampara | Inciensa, Instituto Costarricense de Investigación y Enseñanza en Nutrición y Salud | Francisco Duarte, Hebleen Porras, Claudio Soto-Garita, Estela Cordero, Adriana Godinez & Melany Calderon |
| EPI_ISL_512654 | Hospital Dr. Rafael A. Calderon Guardia [San Jose/San Jose] | Inciensa, Instituto Costarricense de Investigación y Enseñanza en Nutrición y Salud | Francisco Duarte, Hebleen Porras, Claudio Soto-Garita, Estela Cordero, Adriana Godinez & Melany Calderon |
| EPI_ISL_512655 | Hospital De Niños Dr. Carlos Saenz Herrera [San Jose/San Jose] | Inciensa, Instituto Costarricense de Investigación y Enseñanza en Nutrición y Salud | Francisco Duarte, Hebleen Porras, Claudio Soto-Garita, Estela Cordero, Adriana Godinez & Melany Calderon |
| EPI_ISL_512656 | Area De Salud Pavas (Coopesalud) [Pavas/San Jose] | Inciensa, Instituto Costarricense de Investigación y Enseñanza en Nutrición y Salud | Francisco Duarte, Hebleen Porras, Claudio Soto-Garita, Estela Cordero, Adriana Godinez & Melany Calderon |
| EPI_ISL_512657 | Area De Salud Tibas-Uruca-Merced - Clinica Dr. Clorito Picado [Tibas/San Jose] | Inciensa, Instituto Costarricense de Investigación y Enseñanza en Nutrición y Salud | Francisco Duarte, Hebleen Porras, Claudio Soto-Garita, Estela Cordero, Adriana Godinez & Melany Calderon |
| EPI_ISL_512658 | Area De Salud Orotina-San Mateo [Orotina/Alajuela] | Inciensa, Instituto Costarricense de Investigación y Enseñanza en Nutrición y Salud | Francisco Duarte, Hebleen Porras, Claudio Soto-Garita, Estela Cordero, Adriana Godinez & Melany Calderon |
| EPI_ISL_512659 | Area De Salud Fortuna | Inciensa, Instituto Costarricense de Investigación y Enseñanza en Nutrición y Salud | Francisco Duarte, Hebleen Porras, Claudio Soto-Garita, Estela Cordero, Adriana Godinez & Melany Calderon |
| EPI_ISL_512660, EPI_ISL_512661 | Area De Salud Los Chiles | Inciensa, Instituto Costarricense de Investigación y Enseñanza en Nutrición y Salud | Francisco Duarte, Hebleen Porras, Claudio Soto-Garita, Estela Cordero, Adriana Godinez & Melany Calderon |
| EPI_ISL_512662 | Area De Salud La Cruz | Inciensa, Instituto Costarricense de Investigación y Enseñanza en Nutrición y Salud | Francisco Duarte, Hebleen Porras, Claudio Soto-Garita, Estela Cordero, Adriana Godinez & Melany Calderon |
| EPI_ISL_512663, EPI_ISL_512664 | Area De Salud Alajuela Norte - Clinica Dr. Marcial Rodriguez | Inciensa, Instituto Costarricense de Investigación y Enseñanza en Nutrición y Salud | Francisco Duarte, Hebleen Porras, Claudio Soto-Garita, Estela Cordero, Adriana Godinez & Melany Calderon |
| EPI_ISL_512665 | Hospital De Las Mujeres Dr. Adolfo Carit | Inciensa, Instituto Costarricense de Investigación y Enseñanza en Nutrición y Salud | Francisco Duarte, Hebleen Porras, Claudio Soto-Garita, Estela Cordero, Adriana Godinez & Melany Calderon |
| EPI_ISL_512666, EPI_ISL_512667 | Area De Salud La Cruz | Inciensa, Instituto Costarricense de Investigación y Enseñanza en Nutrición y Salud | Francisco Duarte, Hebleen Porras, Claudio Soto-Garita, Estela Cordero, Adriana Godinez & Melany Calderon |
| EPI_ISL_512668 | Area De Salud Corredores | Inciensa, Instituto Costarricense de Investigación y Enseñanza en Nutrición y Salud | Francisco Duarte, Hebleen Porras, Claudio Soto-Garita, Estela Cordero, Adriana Godinez & Melany Calderon |
| EPI_ISL_512669 | Area De Salud Alajuela Norte - Clinica Dr. Marcial Rodriguez | Inciensa, Instituto Costarricense de Investigación y Enseñanza en Nutrición y Salud | Francisco Duarte, Hebleen Porras, Claudio Soto-Garita, Estela Cordero, Adriana Godinez & Melany Calderon |
| EPI_ISL_512670 | Centro Nacional De Rehabilitacion Humberto Araya Rojas (Cenare) | Inciensa, Instituto Costarricense de Investigación y Enseñanza en Nutrición y Salud | Francisco Duarte, Hebleen Porras, Claudio Soto-Garita, Estela Cordero, Adriana Godinez & Melany Calderon |
| EPI_ISL_512671 | Area De Salud La Cruz | Inciensa, Instituto Costarricense de Investigación y Enseñanza en Nutrición y Salud | Francisco Duarte, Hebleen Porras, Claudio Soto-Garita, Estela Cordero, Adriana Godinez & Melany Calderon |
| EPI_ISL_512672, EPI_ISL_512673, EPI_ISL_512674, EPI_ISL_512675 | Hospital De Niños Dr. Carlos Saenz Herrera [San Jose/San Jose] | Inciensa, Instituto Costarricense de Investigación y Enseñanza en Nutrición y Salud | Francisco Duarte, Hebleen Porras, Claudio Soto-Garita, Estela Cordero, Adriana Godinez & Melany Calderon |
| EPI_ISL_512676 | Florida Bureau of Public Health Laboratories | Florida Bureau of Public Health Laboratories | Sarah Schmedes, Jason Blanton |
| EPI_ISL_512712, EPI_ISL_512713, EPI_ISL_512714, EPI_ISL_512715, EPI_ISL_512716, EPI_ISL_512717, EPI_ISL_512719, EPI_ISL_512720, EPI_ISL_512722, EPI_ISL_512723, EPI_ISL_512724, EPI_ISL_512725, EPI_ISL_512727, EPI_ISL_512728, EPI_ISL_512730, EPI_ISL_512732, EPI_ISL_512734, EPI_ISL_512735, EPI_ISL_512737, EPI_ISL_512738, EPI_ISL_512739, EPI_ISL_512741, EPI_ISL_512742, EPI_ISL_512743, EPI_ISL_512744, EPI_ISL_512747, EPI_ISL_512748, EPI_ISL_512749, EPI_ISL_512750, EPI_ISL_512752, EPI_ISL_512754, EPI_ISL_512755, EPI_ISL_512756, EPI_ISL_512757, EPI_ISL_512758, EPI_ISL_512759, EPI_ISL_512760, EPI_ISL_512761, EPI_ISL_512762, EPI_ISL_512763, EPI_ISL_512764, EPI_ISL_512765 | | | |
| see above | PathWest Laboratory Medicine WA | PathWest Laboratory Medicine WA Microbial Surveillance Unit | PathWest Laboratory Medicine WA Microbial Surveillance Unit |
| EPI_ISL_512785, EPI_ISL_512787, EPI_ISL_512789, EPI_ISL_512790, EPI_ISL_512795, EPI_ISL_512797, EPI_ISL_512798, EPI_ISL_512799, EPI_ISL_512800, EPI_ISL_512801, EPI_ISL_512802, EPI_ISL_512803, EPI_ISL_512805, EPI_ISL_512806, EPI_ISL_512807, EPI_ISL_512808, EPI_ISL_512809 | | | |
| see above | Public Health, United States Air Force School of Aerospace Medicine | Public Health, United States Air Force School of Aerospace Medicine | Fries,A.C., Purves,S.M., Meyer,J.R., Javorina,A.K., Connors,B.C., Macias,E.A., Lambert,A.W., Chapleau,R.R., Starr,C.R. |
| EPI_ISL_512810 | National Laboratory for Influenza/Virology reference laboratory, Public Health Center of the Ministry of Health of Ukraine | Respiratory Virus Unit, Microbiology Services Colindale, Public Health England | PHE Covid Sequencing Team, Dr. Iryna Demchyshyna |
| EPI_ISL_512811, EPI_ISL_512812, EPI_ISL_512813, EPI_ISL_512814, EPI_ISL_512815, EPI_ISL_512816, EPI_ISL_512817, EPI_ISL_512818, EPI_ISL_512820, EPI_ISL_512821 | Kenema Government Hospital, Ministry of Health and Sanitation | Kenema Government Hospital, Ministry of Health and Sanitation | Goba,A., Momoh,M., Sandi,J., Tomkins-Tinch,C., Siddle,K., Mehta,S., Oluniyi,P., Jalloh,S., Park,D., Andersen,K., Garry,R., Happi,C., Grant,D., Olawoye,I. |
| EPI_ISL_512822, EPI_ISL_512823, EPI_ISL_512824, EPI_ISL_512825, EPI_ISL_512826, EPI_ISL_512827, EPI_ISL_512828, EPI_ISL_512829, EPI_ISL_512830, EPI_ISL_512832, EPI_ISL_512833, EPI_ISL_512834, EPI_ISL_512835, EPI_ISL_512836, EPI_ISL_512837, EPI_ISL_512838, EPI_ISL_512839, EPI_ISL_512840, EPI_ISL_512841, EPI_ISL_512842 | | | |
| see above | National Public Health Laboratory, National Centre for Infectious Diseases | National Public Health Laboratory, National Centre for Infectious Diseases | Mak TM, Octavia S, Zhou Z, Chavatte JM, Cui L, Lin RTP |
| EPI_ISL_512844 | Department of Medical Research | DMR_Myanmar | Myat Htut Nyunt, Hnin Ohnmar Soe, Kay Thi Aye, Wah Wah Aung,Yi Yi Kyaw, Aung Kyaw Kyaw, Theingi Win Myat, Phyu Win Ei, Aung Zaw Latt, Nan Aye Thida Oo, Lai Lai San, Su Mon Win, Ni Ni Zaw, Htin Lin, Hlaing Myat Thu, Zaw Than Htun |
| EPI_ISL_512846 | O.I.J. MORGUE JUDICIAL | Inciensa, Instituto Costarricense de Investigación y Enseñanza en Nutrición y Salud | Francisco Duarte, Hebleen Porras, Claudio Soto-Garita, Estela Cordero, Adriana Godinez & Melany Calderon |
| EPI_ISL_512849, EPI_ISL_512851, EPI_ISL_512852, EPI_ISL_512856, EPI_ISL_512857, EPI_ISL_512858, EPI_ISL_512859, EPI_ISL_512860, EPI_ISL_512861, EPI_ISL_512862, EPI_ISL_512863, EPI_ISL_512864, EPI_ISL_512866, EPI_ISL_512869 | | | |
| see above | Ramathibodi Hospital | COVID-19 Network Investigations (CONI) Alliance | Elizabeth Batty, Wasun Chantratita, Thanat Chookajorn, Stefan Fernandez, Angkana Huang, Anthony R. Jones, Khajohn Joonsalak, Chonticha Klungtong, Theerarat Kochakarn, Namfon Kotanan, Krittikorn Kumpornsin, Wudtichai Manasatienkij, Bhakbhoom Panthan, Ekawat Pasomsub, Kingkan Rakmanee, Insee Sensorn, Janjira Thaipadungpanit, Arporn Wangwiwatsin,Treewat Watthanachockchai |
| EPI_ISL_512874, EPI_ISL_512875, EPI_ISL_512876, EPI_ISL_512877, EPI_ISL_512878, EPI_ISL_512879, EPI_ISL_512880, EPI_ISL_512881, EPI_ISL_512882, EPI_ISL_512883, EPI_ISL_512884, EPI_ISL_512885, EPI_ISL_512886, EPI_ISL_512887, EPI_ISL_512888, EPI_ISL_512889, EPI_ISL_512890, EPI_ISL_512891, EPI_ISL_512892, EPI_ISL_512893, EPI_ISL_512894, EPI_ISL_512895, EPI_ISL_512896, EPI_ISL_512898, EPI_ISL_512899, EPI_ISL_512900, EPI_ISL_512901, EPI_ISL_512902, EPI_ISL_512903 | | | |
| see above | Pathogen Genomics Lab King Abdullah University of Science and Technology(KAUST) | Pathogen Genomics Lab King Abdullah University of Science and Technology(KAUST) | Raeece Naeem, Rahul P Salunke, Sharif Hala, Sara Mfarrej, Amit Kumar Subudhi, Fadwa Alofi, Fathia Ben Rached, Afrah Alsomali, Asim Khogeer, Ahmad Bakur Mahmoud, Anwar Hashem, Naif Almontashiri, Arnab Pain |
| EPI_ISL_512904, EPI_ISL_512905, EPI_ISL_512906, EPI_ISL_512907 | Pathogen Genomics Lab King Abdullah University of Science and Technology(KAUST) | Pathogen Genomics Lab King Abdullah University of Science and Technology(KAUST) | Fathia Ben Rached, Raeece Naeem, Sharif Hala, Fadwa Alofi, Rahul P Salunke, Sara Mfarrej, Amit Kumar Subudhi, Afrah Alsomali, Asim Khogeer, Ahmad Bakur Mahmoud, Anwar Hashem, Naif Almontashiri, Arnab Pain |
| EPI_ISL_512908, EPI_ISL_512909, EPI_ISL_512911, EPI_ISL_512912, | Pathogen Genomics Lab King Abdullah University of Science and Technology(KAUST) | Pathogen Genomics Lab King Abdullah University of Science and Technology(KAUST) | Sharif Hala, Fadwa Alofi, Sara Mfarrej, Amit Kumar Subudhi, Rahul P Salunke, Fathia Ben Rached, Amanda Ooi, Luke Esau, Afrah Alsomali, Asim Khogeer, Jumana Taha, Abdulaziz Alahmadi, Kahled Algithami, Raeece Naeem, Anwar Hashem, Naif Almontashiri, Arnab Pain |

| | | | |
|---|---|---|---|
| EPI_ISL_512913, EPI_ISL_512914, EPI_ISL_512915, EPI_ISL_512916, EPI_ISL_512917 | | | |
| EPI_ISL_512918, EPI_ISL_512919, EPI_ISL_512920, EPI_ISL_512921, EPI_ISL_512922, EPI_ISL_512923, EPI_ISL_512924, EPI_ISL_512926, EPI_ISL_512927, EPI_ISL_512928, EPI_ISL_512929, EPI_ISL_512930, EPI_ISL_512931, EPI_ISL_512932, EPI_ISL_512933, EPI_ISL_512934, EPI_ISL_512935, EPI_ISL_512936, EPI_ISL_512937, EPI_ISL_512938, EPI_ISL_512939, EPI_ISL_512941, EPI_ISL_512942, EPI_ISL_512943, EPI_ISL_512944, EPI_ISL_512945 | | | |
| see above | Pathogen Genomics Lab King Abdullah University of Science and Technology(KAUST) | Pathogen Genomics Lab King Abdullah University of Science and Technology(KAUST) | Fadwa Alofi, Sharif Hala, Rahul P Salunke, Sara Mfarrej, Amit Kumar Subudhi, Fathia Ben Rached, Amanda, Luke, Afrah Alsomali, Asim Khogeer, Jumana Taha, Abdulaziz Alahmadi, Kahled Algithami, Raeece Naeem, Anwar Hashem, Naif Almontashiri, Arnab Pain |
| EPI_ISL_512946, EPI_ISL_512947, EPI_ISL_512948, EPI_ISL_512949, EPI_ISL_512950, EPI_ISL_512951, EPI_ISL_512952, EPI_ISL_512953, EPI_ISL_512954, EPI_ISL_512955, EPI_ISL_512956, EPI_ISL_512957, EPI_ISL_512959, EPI_ISL_512963, EPI_ISL_512964, EPI_ISL_512966, EPI_ISL_512967, EPI_ISL_512969, EPI_ISL_512970, EPI_ISL_512971, EPI_ISL_512973, EPI_ISL_512974, EPI_ISL_512975, EPI_ISL_512976, EPI_ISL_512977, EPI_ISL_512979, EPI_ISL_512980, EPI_ISL_512981, EPI_ISL_512982, EPI_ISL_512983, EPI_ISL_512984, EPI_ISL_512986, EPI_ISL_512987, EPI_ISL_512988 | | | |
| see above | Pathogen Genomics Lab King Abdullah University of Science and Technology(KAUST) | Pathogen Genomics Lab King Abdullah University of Science and Technology(KAUST) | Sara Mfarrej, Raeece Naeem, Rahul P Salunke, Sharif Hala, Fadwa Alofi, Amit Kumar Subudhi, Fathia Ben Rached, Afrah Alsomali, Jumana Taha, Abdulaziz Alahmadi, Asim Khogeer, Nashwa Al-khotani, Anwar Hashem, Naif Almontashiri, Arnab Pain |
| EPI_ISL_512989, EPI_ISL_512990, EPI_ISL_512991, EPI_ISL_512992, EPI_ISL_512993, EPI_ISL_512994, EPI_ISL_512995, EPI_ISL_512996, EPI_ISL_512997, EPI_ISL_512998, EPI_ISL_512999, EPI_ISL_513000, EPI_ISL_513001 | | | |
| see above | Pathogen Genomics Lab King Abdullah University of Science and Technology(KAUST) | Pathogen Genomics Lab King Abdullah University of Science and Technology(KAUST) | Amit Kumar Subudhi, Rahul P Salunke, Sara Mfarrej, Sharif Hala, Fadwa Alofi, Fathia Ben Rached, Afrah Alsomali, Asim Khogeer, Nashwa Al-khotani, Raeece Naeem, Anwar Hashem, Naif Almontashiri, Arnab Pain |
| EPI_ISL_513002, EPI_ISL_513003, EPI_ISL_513004, EPI_ISL_513005, EPI_ISL_513006, EPI_ISL_513008, EPI_ISL_513009, EPI_ISL_513010, EPI_ISL_513012, EPI_ISL_513013, EPI_ISL_513014, EPI_ISL_513015, EPI_ISL_513016, EPI_ISL_513017, EPI_ISL_513018, EPI_ISL_513019, EPI_ISL_513020, EPI_ISL_513021, EPI_ISL_513022, EPI_ISL_513023, EPI_ISL_513024, EPI_ISL_513025, EPI_ISL_513026 | | | |
| see above | Pathogen Genomics Lab King Abdullah University of Science and Technology(KAUST) | Pathogen Genomics Lab King Abdullah University of Science and Technology(KAUST) | Afrah Alsomali, Fathia Ben Rached, Raeece Naeem, Sharif Hala,Rahul P Salunke, Amanda Ooi, Luke Esau, Sara Mfarrej, Amit Kumar Subudhi, Fadwa Alofi, Asim Khogeer, Kahled Algithami, Anwar Hashem, Naif Almontashiri, Arnab Pain |
| EPI_ISL_513027, EPI_ISL_513028, EPI_ISL_513029, EPI_ISL_513032, EPI_ISL_513033, EPI_ISL_513034, EPI_ISL_513035, EPI_ISL_513036, EPI_ISL_513037, EPI_ISL_513038, EPI_ISL_513039, EPI_ISL_513040, EPI_ISL_513041, EPI_ISL_513042, EPI_ISL_513043, EPI_ISL_513045, EPI_ISL_513046, EPI_ISL_513047, EPI_ISL_513048, EPI_ISL_513049, EPI_ISL_513050, EPI_ISL_513051, EPI_ISL_513052, EPI_ISL_513053, EPI_ISL_513054, EPI_ISL_513055, EPI_ISL_513056, EPI_ISL_513057, EPI_ISL_513058, EPI_ISL_513059, EPI_ISL_513060, EPI_ISL_513061, EPI_ISL_513062, EPI_ISL_513063 | | | |
| see above | Pathogen Genomics Lab King Abdullah University of Science and Technology(KAUST) | Pathogen Genomics Lab King Abdullah University of Science and Technology(KAUST) | Rahul P Salunke, Sharif Hala, Raeece Naeem, Sara Mfarrej, Amit Kumar Subudhi, Amanda Ooi, Luke Esau, Fadwa Alofi, Fathia Ben Rached, Afrah Alsomali, Asim Khogeer, Ahmad Bakur Mahmoud, Anwar Hashem, Naif Almontashiri, Arnab Pain |
| EPI_ISL_513064, EPI_ISL_513065, EPI_ISL_513066, EPI_ISL_513067, EPI_ISL_513069, EPI_ISL_513070, EPI_ISL_513071, EPI_ISL_513072, EPI_ISL_513073, EPI_ISL_513074, EPI_ISL_513075, EPI_ISL_513076 | | | |
| see above | Pathogen Genomics Lab King Abdullah University of Science and Technology(KAUST) | Pathogen Genomics Lab King Abdullah University of Science and Technology(KAUST) | Raeece Naeem, Rahul P Salunke, Sharif Hala, Sara Mfarrej, Amit Kumar Subudhi, Fadwa Alofi, Fathia Ben Rached, Afrah Alsomali, Asim Khogeer, Ahmad Bakur Mahmoud, Anwar Hashem, Naif Almontashiri, Arnab Pain |
| EPI_ISL_513077, EPI_ISL_513078, EPI_ISL_513079, EPI_ISL_513080, EPI_ISL_513081, EPI_ISL_513082, EPI_ISL_513083, EPI_ISL_513085, EPI_ISL_513086, EPI_ISL_513087, EPI_ISL_513089, EPI_ISL_513090, EPI_ISL_513091, EPI_ISL_513092, EPI_ISL_513093, EPI_ISL_513094, EPI_ISL_513095, EPI_ISL_513096, EPI_ISL_513097, EPI_ISL_513098, EPI_ISL_513099, EPI_ISL_513100, EPI_ISL_513101, EPI_ISL_513102, EPI_ISL_513103, EPI_ISL_513104, EPI_ISL_513105, EPI_ISL_513106, EPI_ISL_513107, EPI_ISL_513108, EPI_ISL_513109, EPI_ISL_513111, EPI_ISL_513112, EPI_ISL_513113, EPI_ISL_513114, EPI_ISL_513115, EPI_ISL_513116, EPI_ISL_513117, EPI_ISL_513118, EPI_ISL_513119, EPI_ISL_513120 | | | |
| see above | Pathogen Genomics Lab King Abdullah University of Science and Technology(KAUST) | Pathogen Genomics Lab King Abdullah University of Science and Technology(KAUST) | Fathia Ben Rached, Raeece Naeem, Sharif Hala, Fadwa Alofi, Rahul P Salunke, Sara Mfarrej, Amit Kumar Subudhi, Afrah Alsomali, Asim Khogeer, Ahmad Bakur Mahmoud, Anwar Hashem, Naif Almontashiri, Arnab Pain |
| EPI_ISL_513121, EPI_ISL_513122, EPI_ISL_513123, EPI_ISL_513124, EPI_ISL_513125, EPI_ISL_513126, EPI_ISL_513127, EPI_ISL_513128, EPI_ISL_513129, EPI_ISL_513131, EPI_ISL_513132, EPI_ISL_513133, EPI_ISL_513134, EPI_ISL_513135, EPI_ISL_513136, EPI_ISL_513137, EPI_ISL_513138, EPI_ISL_513139, EPI_ISL_513141, EPI_ISL_513142, EPI_ISL_513143, EPI_ISL_513144, EPI_ISL_513145, EPI_ISL_513146 | | | |
| see above | Pathogen Genomics Lab King Abdullah University of Science and Technology(KAUST) | Pathogen Genomics Lab King Abdullah University of Science and Technology(KAUST) | Sharif Hala, Fadwa Alofi, Sara Mfarrej, Amit Kumar Subudhi, Rahul P Salunke, Fathia Ben Rached, Amanda Ooi, Luke Esau, Afrah Alsomali, Asim Khogeer, Jumana Taha, Abdulaziz Alahmadi, Kahled Algithami, Raeece Naeem, Anwar Hashem, Naif Almontashiri, Arnab Pain |
| EPI_ISL_513147, EPI_ISL_513148, EPI_ISL_513149, EPI_ISL_513150, EPI_ISL_513151, EPI_ISL_513152, EPI_ISL_513153, EPI_ISL_513154, EPI_ISL_513156, EPI_ISL_513157, EPI_ISL_513158, EPI_ISL_513159, EPI_ISL_513160, EPI_ISL_513161, EPI_ISL_513162, EPI_ISL_513163, EPI_ISL_513164, EPI_ISL_513166, EPI_ISL_513167 | | | |
| see above | Pathogen Genomics Lab King Abdullah University of Science and Technology(KAUST) | Pathogen Genomics Lab King Abdullah University of Science and Technology(KAUST) | Fadwa Alofi, Sharif Hala, Rahul P Salunke, Sara Mfarrej, Amit Kumar Subudhi, Fathia Ben Rached, Amanda, Luke, Afrah Alsomali, Asim Khogeer, Jumana Taha, Abdulaziz Alahmadi, Kahled Algithami, Raeece Naeem, Anwar Hashem, Naif Almontashiri, Arnab Pain |
| EPI_ISL_513168, EPI_ISL_513169, EPI_ISL_513170, EPI_ISL_513171, EPI_ISL_513172, EPI_ISL_513173, EPI_ISL_513174 | Pathogen Genomics Lab King Abdullah University of Science and Technology(KAUST) | Pathogen Genomics Lab King Abdullah University of Science and Technology(KAUST) | Sara Mfarrej, Raeece Naeem, Rahul P Salunke, Sharif Hala, Fadwa Alofi, Amit Kumar Subudhi, Fathia Ben Rached, Afrah Alsomali, Jumana Taha, Abdulaziz Alahmadi, Asim Khogeer, Nashwa Al-khotani, Anwar Hashem, Naif Almontashiri, Arnab Pain |
| EPI_ISL_513175, EPI_ISL_513176, EPI_ISL_513177, EPI_ISL_513178, EPI_ISL_513179, EPI_ISL_513180, EPI_ISL_513181, EPI_ISL_513182, EPI_ISL_513183, EPI_ISL_513184, EPI_ISL_513185, EPI_ISL_513186, EPI_ISL_513187, EPI_ISL_513188, EPI_ISL_513189 | | | |
| see above | Pathogen Genomics Lab King Abdullah University of Science and Technology(KAUST) | Pathogen Genomics Lab King Abdullah University of Science and Technology(KAUST) | Amit Kumar Subudhi, Rahul P Salunke, Sara Mfarrej, Sharif Hala, Fadwa Alofi, Fathia Ben Rached, Afrah Alsomali, Asim Khogeer, Nashwa Al-khotani, Raeece Naeem, Anwar Hashem, Naif Almontashiri, Arnab Pain |
| EPI_ISL_513190, EPI_ISL_513191, EPI_ISL_513192, EPI_ISL_513193, EPI_ISL_513194, EPI_ISL_513195, EPI_ISL_513196, EPI_ISL_513197 | Pathogen Genomics Lab King Abdullah University of Science and Technology(KAUST) | Pathogen Genomics Lab King Abdullah University of Science and Technology(KAUST) | Afrah Alsomali, Fathia Ben Rached, Raeece Naeem, Sharif Hala,Rahul P Salunke, Amanda Ooi, Luke Esau, Sara Mfarrej, Amit Kumar Subudhi, Fadwa Alofi, Asim Khogeer, Kahled Algithami, Anwar Hashem, Naif Almontashiri, Arnab Pain |
| EPI_ISL_513198, EPI_ISL_513199, EPI_ISL_513200, EPI_ISL_513201, EPI_ISL_513202, EPI_ISL_513203, EPI_ISL_513204, EPI_ISL_513205, EPI_ISL_513206, EPI_ISL_513207 | Pathogen Genomics Lab King Abdullah University of Science and Technology(KAUST) | Pathogen Genomics Lab King Abdullah University of Science and Technology(KAUST) | Rahul P Salunke, Sharif Hala, Raeece Naeem, Sara Mfarrej, Amit Kumar Subudhi, Amanda Ooi, Luke Esau, Fadwa Alofi, Fathia Ben Rached, Afrah Alsomali, Asim Khogeer, Ahmad Bakur Mahmoud, Anwar Hashem, Naif Almontashiri, Arnab Pain |
| EPI_ISL_513208, EPI_ISL_513209, EPI_ISL_513210, EPI_ISL_513211, EPI_ISL_513212 | Pathogen Genomics Lab King Abdullah University of Science and Technology(KAUST) | Pathogen Genomics Lab King Abdullah University of Science and Technology(KAUST) | Raeece Naeem, Rahul P Salunke, Sharif Hala, Sara Mfarrej, Amit Kumar Subudhi, Fadwa Alofi, Fathia Ben Rached, Afrah Alsomali, Asim Khogeer, Ahmad Bakur Mahmoud, Anwar Hashem, Naif Almontashiri, Arnab Pain |
| EPI_ISL_513214, EPI_ISL_513215, EPI_ISL_513216, EPI_ISL_513217, EPI_ISL_513218, EPI_ISL_513219, EPI_ISL_513221, EPI_ISL_513222, EPI_ISL_513224, EPI_ISL_513226 | Pathogen Genomics Lab King Abdullah University of Science and Technology(KAUST) | Pathogen Genomics Lab King Abdullah University of Science and Technology(KAUST) | Sharif Hala, Fadwa Alofi, Sara Mfarrej, Amit Kumar Subudhi, Rahul P Salunke, Fathia Ben Rached, Amanda Ooi, Luke Esau, Afrah Alsomali, Asim Khogeer, Jumana Taha, Abdulaziz Alahmadi, Kahled Algithami, Raeece Naeem, Anwar Hashem, Naif Almontashiri, Arnab Pain |
| EPI_ISL_513228, EPI_ISL_513229, EPI_ISL_513230, EPI_ISL_513231, EPI_ISL_513233, EPI_ISL_513234, EPI_ISL_513235, EPI_ISL_513236, EPI_ISL_513238, EPI_ISL_513240, EPI_ISL_513241, EPI_ISL_513242, EPI_ISL_513243, EPI_ISL_513244, EPI_ISL_513245, EPI_ISL_513246, EPI_ISL_513247, EPI_ISL_513248 | | | |
| see above | Pathogen Genomics Lab King Abdullah University of Science and Technology(KAUST) | Pathogen Genomics Lab King Abdullah University of Science and Technology(KAUST) | Amit Kumar Subudhi, Rahul P Salunke, Sara Mfarrej, Sharif Hala, Fadwa Alofi, Fathia Ben Rached, Afrah Alsomali, Asim Khogeer, Nashwa Al-khotani, Raeece Naeem, Anwar Hashem, Naif Almontashiri, Arnab Pain |
| EPI_ISL_513249, EPI_ISL_513250, EPI_ISL_513251, EPI_ISL_513252, EPI_ISL_513253, EPI_ISL_513254, EPI_ISL_513257, EPI_ISL_513258, EPI_ISL_513259, EPI_ISL_513260, EPI_ISL_513261, EPI_ISL_513262, EPI_ISL_513263, EPI_ISL_513264 | | | |
| see above | Pathogen Genomics Lab King Abdullah University of Science and Technology(KAUST) | Pathogen Genomics Lab King Abdullah University of Science and Technology(KAUST) | Afrah Alsomali, Fathia Ben Rached, Raeece Naeem, Sharif Hala,Rahul P Salunke, Amanda Ooi, Luke Esau, Sara Mfarrej, Amit Kumar Subudhi, Fadwa Alofi, Asim Khogeer, Kahled Algithami, Anwar Hashem, Naif Almontashiri, Arnab Pain |
| EPI_ISL_513265, EPI_ISL_513266, EPI_ISL_513267, EPI_ISL_513268, EPI_ISL_513269, EPI_ISL_513270, EPI_ISL_513271, EPI_ISL_513272, EPI_ISL_513273, EPI_ISL_513274, EPI_ISL_513275, EPI_ISL_513276, EPI_ISL_513277, EPI_ISL_513278, EPI_ISL_513279, EPI_ISL_513281, EPI_ISL_513282, EPI_ISL_513283, EPI_ISL_513284, EPI_ISL_513285, EPI_ISL_513286, EPI_ISL_513287, EPI_ISL_513288, EPI_ISL_513289, EPI_ISL_513290, EPI_ISL_513291, EPI_ISL_513292, EPI_ISL_513293 | | | |
| see above | University of Miami Immunology and Histocompatibility Laboratory | University of Miami Immunology and Histocompatibility Laboratory | Emilio Margolles-Clark, PhD and Phillip Ruiz, MD, PhD |
| EPI_ISL_513310 | Public Health, United States Air Force School of Aerospace Medicine | Public Health, United States Air Force School of Aerospace Medicine | Fries,A.C., Purves,S.M., Meyer,J.R., Javorina,A.K., Connors,B.C., Macias,E.A., Lambert,A.W., Chapleau,R.R. and Starr,C.R. |
| EPI_ISL_513312 | Public Health, United States Air Force School of Aerospace Medicine | Public Health, United States Air Force School of Aerospace Medicine | Fries,A.C., Purves,S.M., Meyer,J.R., Javorina,A.K., Connors,B.C., Macias,E.A., Lambert,A.W., Chapleau,R.R. and Starr,C.R. |
| EPI_ISL_513314 | The National Institute of Public Health | The National Institute of Public Health and State Veterinary Institute Prague | Nagy,A;Jirincova,H;Novakova,L;Trnka,D;Vecerova,J |
| EPI_ISL_513315, EPI_ISL_513316, EPI_ISL_513317 | South Eastern Area Laboratory Services (SEALS) | NSW Health Pathology - Institute of Clinical Pathology and Medical Research; Westmead Hospital; University of Sydney | CIDM-PH et al. |
| EPI_ISL_513318, EPI_ISL_513319, EPI_ISL_513320 | Histopath | NSW Health Pathology - Institute of Clinical Pathology and Medical Research; Westmead Hospital; University of Sydney | CIDM-PH et al. |
| EPI_ISL_513321, EPI_ISL_513322 | Pathology West - NSW Health Pathology | NSW Health Pathology - Institute of Clinical Pathology and Medical Research; Westmead Hospital; University of Sydney | CIDM-PH et al. |
| EPI_ISL_513323 | Sydney South West Pathology Service (SSWPS) - Liverpool Hospital - NSW Health Pathology | NSW Health Pathology - Institute of Clinical Pathology and Medical Research; Westmead Hospital; University of Sydney | CIDM-PH et al. |
| EPI_ISL_513324, EPI_ISL_513325, EPI_ISL_513326 | Pathology West - NSW Health Pathology | NSW Health Pathology - Institute of Clinical Pathology and Medical Research; Westmead Hospital; University of Sydney | CIDM-PH et al. |
| EPI_ISL_513327, EPI_ISL_513328, EPI_ISL_513329 | Sydney South West Pathology Service (SSWPS) - Liverpool Hospital - NSW Health Pathology | NSW Health Pathology - Institute of Clinical Pathology and Medical Research; Westmead Hospital; University of Sydney | CIDM-PH et al. |
| EPI_ISL_513330 | St Vincent's Pathology (SydPath) | NSW Health Pathology - Institute of Clinical Pathology and Medical Research; Westmead Hospital; University of Sydney | CIDM-PH et al. |
| EPI_ISL_513331 | Australian Clinical Labs | NSW Health Pathology - Institute of Clinical Pathology and Medical Research; Westmead Hospital; University of Sydney | CIDM-PH et al. |
| EPI_ISL_513332 | Pathology West - NSW Health Pathology | NSW Health Pathology - Institute of Clinical Pathology and Medical Research; Westmead Hospital; University of | CIDM-PH et al. |

| | | | |
|---|---|---|---|
| | | Sydney | |
| EPI_ISL_513333, EPI_ISL_513334 | Sydney South West Pathology Service (SSWPS) - Liverpool Hospital - NSW Health Pathology | NSW Health Pathology - Institute of Clinical Pathology and Medical Research; Westmead Hospital; University of Sydney | CIDM-PH et al. |
| EPI_ISL_513335 | Pathology North - Hunter - NSW Health Pathology | NSW Health Pathology - Institute of Clinical Pathology and Medical Research; Westmead Hospital; University of Sydney | CIDM-PH et al. |
| EPI_ISL_513336, EPI_ISL_513337 | Sydney South West Pathology Service (SSWPS) - Liverpool Hospital - NSW Health Pathology | NSW Health Pathology - Institute of Clinical Pathology and Medical Research; Westmead Hospital; University of Sydney | CIDM-PH et al. |
| EPI_ISL_513338, EPI_ISL_513339 | Pathology West - NSW Health Pathology | NSW Health Pathology - Institute of Clinical Pathology and Medical Research; Westmead Hospital; University of Sydney | CIDM-PH et al. |
| EPI_ISL_513340 | Austech Medical Laboratories | NSW Health Pathology - Institute of Clinical Pathology and Medical Research; Westmead Hospital; University of Sydney | CIDM-PH et al. |
| EPI_ISL_513341 | Medlab Pathology | NSW Health Pathology - Institute of Clinical Pathology and Medical Research; Westmead Hospital; University of Sydney | CIDM-PH et al. |
| EPI_ISL_513342 | Pathology West - NSW Health Pathology | NSW Health Pathology - Institute of Clinical Pathology and Medical Research; Westmead Hospital; University of Sydney | CIDM-PH et al. |
| EPI_ISL_513343, EPI_ISL_513344 | Children Westmead Hospital | NSW Health Pathology - Institute of Clinical Pathology and Medical Research; Westmead Hospital; University of Sydney | CIDM-PH et al. |
| EPI_ISL_513345, EPI_ISL_513346 | Pathology West - NSW Health Pathology | NSW Health Pathology - Institute of Clinical Pathology and Medical Research; Westmead Hospital; University of Sydney | CIDM-PH et al. |
| EPI_ISL_513347 | St Vincent's Pathology (SydPath) | NSW Health Pathology - Institute of Clinical Pathology and Medical Research; Westmead Hospital; University of Sydney | CIDM-PH et al. |
| EPI_ISL_513348, EPI_ISL_513349 | 4Cyte Pathology | NSW Health Pathology - Institute of Clinical Pathology and Medical Research; Westmead Hospital; University of Sydney | CIDM-PH et al. |
| EPI_ISL_513350, EPI_ISL_513351 | Pathology West - NSW Health Pathology | NSW Health Pathology - Institute of Clinical Pathology and Medical Research; Westmead Hospital; University of Sydney | CIDM-PH et al. |
| EPI_ISL_513352 | South Eastern Area Laboratory Services (SEALS) | NSW Health Pathology - Institute of Clinical Pathology and Medical Research; Westmead Hospital; University of Sydney | CIDM-PH et al. |
| EPI_ISL_513353, EPI_ISL_513354, EPI_ISL_513355, EPI_ISL_513356 | St Vincent's Pathology (SydPath) | NSW Health Pathology - Institute of Clinical Pathology and Medical Research; Westmead Hospital; University of Sydney | CIDM-PH et al. |
| EPI_ISL_513357 | Douglas Hanly Moir | NSW Health Pathology - Institute of Clinical Pathology and Medical Research; Westmead Hospital; University of Sydney | CIDM-PH et al. |
| EPI_ISL_513358, EPI_ISL_513359 | Pathology West - NSW Health Pathology | NSW Health Pathology - Institute of Clinical Pathology and Medical Research; Westmead Hospital; University of Sydney | CIDM-PH et al. |
| EPI_ISL_513360 | Pathology North - Hunter - NSW Health Pathology | NSW Health Pathology - Institute of Clinical Pathology and Medical Research; Westmead Hospital; University of Sydney | CIDM-PH et al. |
| EPI_ISL_513361, EPI_ISL_513362, EPI_ISL_513363, EPI_ISL_513364, EPI_ISL_513365 | St Vincent's Pathology (SydPath) | NSW Health Pathology - Institute of Clinical Pathology and Medical Research; Westmead Hospital; University of Sydney | CIDM-PH et al. |
| EPI_ISL_513366, EPI_ISL_513367 | Sydney South West Pathology Service (SSWPS) - Liverpool Hospital - NSW Health Pathology | NSW Health Pathology - Institute of Clinical Pathology and Medical Research; Westmead Hospital; University of Sydney | CIDM-PH et al. |
| EPI_ISL_513368 | Pathology North - Hunter - NSW Health Pathology | NSW Health Pathology - Institute of Clinical Pathology and Medical Research; Westmead Hospital; University of Sydney | CIDM-PH et al. |
| EPI_ISL_513369, EPI_ISL_513370, EPI_ISL_513371 | Pathology West - NSW Health Pathology | NSW Health Pathology - Institute of Clinical Pathology and Medical Research; Westmead Hospital; University of Sydney | CIDM-PH et al. |
| EPI_ISL_513372 | St Vincent's Pathology (SydPath) | NSW Health Pathology - Institute of Clinical Pathology and Medical Research; Westmead Hospital; University of Sydney | CIDM-PH et al. |
| EPI_ISL_513373, EPI_ISL_513374 | Sydney South West Pathology Service (SSWPS) - Liverpool Hospital - NSW Health Pathology | NSW Health Pathology - Institute of Clinical Pathology and Medical Research; Westmead Hospital; University of Sydney | CIDM-PH et al. |
| EPI_ISL_513375, EPI_ISL_513376 | Pathology North - Royal North Shore Hospital - NSW Health Pathology | NSW Health Pathology - Institute of Clinical Pathology and Medical Research; Westmead Hospital; University of Sydney | CIDM-PH et al. |
| EPI_ISL_513377 | South Eastern Area Laboratory Services (SEALS) | NSW Health Pathology - Institute of Clinical Pathology and Medical Research; Westmead Hospital; University of Sydney | CIDM-PH et al. |
| EPI_ISL_513378 | Pathology West - NSW Health Pathology | NSW Health Pathology - Institute of Clinical Pathology and Medical Research; Westmead Hospital; University of Sydney | CIDM-PH et al. |
| EPI_ISL_513379 | St Vincent's Pathology (SydPath) | NSW Health Pathology - Institute of Clinical Pathology and Medical Research; Westmead Hospital; University of Sydney | CIDM-PH et al. |
| EPI_ISL_513380 | Pathology West - NSW Health Pathology | NSW Health Pathology - Institute of Clinical Pathology and Medical Research; Westmead Hospital; University of Sydney | CIDM-PH et al. |
| EPI_ISL_513381, EPI_ISL_513382, EPI_ISL_513383 | Sydney South West Pathology Service (SSWPS) - Liverpool Hospital - NSW Health Pathology | NSW Health Pathology - Institute of Clinical Pathology and Medical Research; Westmead Hospital; University of Sydney | CIDM-PH et al. |
| EPI_ISL_513384, EPI_ISL_513385 | St Vincent's Pathology (SydPath) | NSW Health Pathology - Institute of Clinical Pathology and Medical Research; Westmead Hospital; University of Sydney | CIDM-PH et al. |
| EPI_ISL_513386, EPI_ISL_513387, EPI_ISL_513388 | Pathology North - NSW Health Pathology | NSW Health Pathology - Institute of Clinical Pathology and Medical Research; Westmead Hospital; University of Sydney | CIDM-PH et al. |
| EPI_ISL_513389 | Douglas Hanly Moir | NSW Health Pathology - Institute of Clinical Pathology and Medical Research; Westmead Hospital; University of Sydney | CIDM-PH et al. |
| EPI_ISL_513390, EPI_ISL_513391, EPI_ISL_513392, EPI_ISL_513393, EPI_ISL_513394, EPI_ISL_513395 | Pathology West - NSW Health Pathology | NSW Health Pathology - Institute of Clinical Pathology and Medical Research; Westmead Hospital; University of Sydney | CIDM-PH et al. |
| EPI_ISL_513396, EPI_ISL_513397, EPI_ISL_513398, EPI_ISL_513399 | St Vincent's Pathology (SydPath) | NSW Health Pathology - Institute of Clinical Pathology and Medical Research; Westmead Hospital; University of Sydney | CIDM-PH et al. |
| EPI_ISL_513400 | Sydney South West Pathology Service (SSWPS) - Royal Prince Alfred Hospital - NSW Health Pathology | NSW Health Pathology - Institute of Clinical Pathology and Medical Research; Westmead Hospital; University of Sydney | CIDM-PH et al. |
| EPI_ISL_513401 | Sydney South West Pathology Service (SSWPS) - Liverpool Hospital - NSW Health Pathology | NSW Health Pathology - Institute of Clinical Pathology and Medical Research; Westmead Hospital; University of Sydney | CIDM-PH et al. |
| EPI_ISL_513402 | 4Cyte Pathology | NSW Health Pathology - Institute of Clinical Pathology and Medical Research; Westmead Hospital; University of Sydney | CIDM-PH et al. |
| EPI_ISL_513403, EPI_ISL_513404, EPI_ISL_513405, EPI_ISL_513406, EPI_ISL_513407, EPI_ISL_513408 | Laverty Pathology | NSW Health Pathology - Institute of Clinical Pathology and Medical Research; Westmead Hospital; University of Sydney | CIDM-PH et al. |
| EPI_ISL_513409, EPI_ISL_513410, EPI_ISL_513411, EPI_ISL_513412 | Sydney South West Pathology Service (SSWPS) - Liverpool Hospital - NSW Health Pathology | NSW Health Pathology - Institute of Clinical Pathology and Medical Research; Westmead Hospital; University of Sydney | CIDM-PH et al. |
| EPI_ISL_513413 | 4Cyte Pathology | NSW Health Pathology - Institute of Clinical Pathology and Medical Research; Westmead Hospital; University of Sydney | CIDM-PH et al. |
| EPI_ISL_513414, EPI_ISL_513415, EPI_ISL_513416, EPI_ISL_513418, EPI_ISL_513419, EPI_ISL_513420, EPI_ISL_513421, EPI_ISL_513422, EPI_ISL_513423, EPI_ISL_513424, EPI_ISL_513425, EPI_ISL_513426, EPI_ISL_513427, EPI_ISL_513428, EPI_ISL_513429, EPI_ISL_513430, EPI_ISL_513431, EPI_ISL_513432, EPI_ISL_513433, EPI_ISL_513434, EPI_ISL_513435, EPI_ISL_513436, EPI_ISL_513437, EPI_ISL_513438, EPI_ISL_513439, EPI_ISL_513440, EPI_ISL_513441, EPI_ISL_513442, EPI_ISL_513443, EPI_ISL_513444, EPI_ISL_513447, EPI_ISL_513448, EPI_ISL_513449, EPI_ISL_513450, EPI_ISL_513451, EPI_ISL_513452, EPI_ISL_513453, EPI_ISL_513454, EPI_ISL_513455, EPI_ISL_513456, EPI_ISL_513457, EPI_ISL_513458, EPI_ISL_513459, EPI_ISL_513460, EPI_ISL_513461, EPI_ISL_513462, EPI_ISL_513463, EPI_ISL_513464, EPI_ISL_513465, EPI_ISL_513466, EPI_ISL_513467, EPI_ISL_513468, EPI_ISL_513469, EPI_ISL_513470, EPI_ISL_513471, EPI_ISL_513472, EPI_ISL_513473, EPI_ISL_513474, EPI_ISL_513475, EPI_ISL_513476, EPI_ISL_513477, EPI_ISL_513478, EPI_ISL_513479, EPI_ISL_513480, EPI_ISL_513481, EPI_ISL_513482, EPI_ISL_513483, EPI_ISL_513484, EPI_ISL_513485, EPI_ISL_513486, EPI_ISL_513488, EPI_ISL_513489, EPI_ISL_513490, EPI_ISL_513491, EPI_ISL_513492, EPI_ISL_513493, EPI_ISL_513494, EPI_ISL_513495, EPI_ISL_513496, EPI_ISL_513497, EPI_ISL_513498, EPI_ISL_513499, EPI_ISL_513500, EPI_ISL_513501, EPI_ISL_513502, EPI_ISL_513503, EPI_ISL_513504, EPI_ISL_513505, EPI_ISL_513506, EPI_ISL_513507, EPI_ISL_513508, EPI_ISL_513509, EPI_ISL_513510 | | | |
| see above | Maine HETL | Tewhey Lab, The Jackson Laboratory | Matluk,N., Dewey,H., Barter,M., Lynch,R., Munger,H. and Tewhey,R. |
| EPI_ISL_513511, EPI_ISL_513512 | The National Institute of Public Health | The National Institute of Public Health and State Veterinary Institute Prague | Nagy,A;Jirincova,H;Novakova,L;Trnka,D;Vecerova,J |

| | | | |
|---|---|---|---|
| EPI_ISL_513513, EPI_ISL_513514, EPI_ISL_513515, EPI_ISL_513518, EPI_ISL_513519, EPI_ISL_513520, EPI_ISL_513521, EPI_ISL_513522, EPI_ISL_513524, EPI_ISL_513525, EPI_ISL_513526, EPI_ISL_513527, EPI_ISL_513529, EPI_ISL_513530, EPI_ISL_513531, EPI_ISL_513532, EPI_ISL_513533, EPI_ISL_513534, EPI_ISL_513535, EPI_ISL_513536, EPI_ISL_513537, EPI_ISL_513538, EPI_ISL_513539, EPI_ISL_513540, EPI_ISL_513541, EPI_ISL_513542, EPI_ISL_513543, EPI_ISL_513544, EPI_ISL_513545, EPI_ISL_513546, EPI_ISL_513547, EPI_ISL_513548, EPI_ISL_513549, EPI_ISL_513550, EPI_ISL_513551, EPI_ISL_513553, EPI_ISL_513556, EPI_ISL_513557, EPI_ISL_513558, EPI_ISL_513559, EPI_ISL_513560, EPI_ISL_513561, EPI_ISL_513562, EPI_ISL_513563, EPI_ISL_513564, EPI_ISL_513567, EPI_ISL_513568, EPI_ISL_513571, EPI_ISL_513572, EPI_ISL_513573, EPI_ISL_513574, EPI_ISL_513575, EPI_ISL_513576, EPI_ISL_513577, EPI_ISL_513578, EPI_ISL_513579, EPI_ISL_513580, EPI_ISL_513581, EPI_ISL_513582, EPI_ISL_513583 | | | |
| see above | Programa de Oncovirologia, Instituto Nacional de Câncer | Programa de Oncovirologia, Instituto Nacional de Câncer | Juliana D. Siqueira, Livia R. Goes, Brunna M. Alves, Claudia Cicala,James Arthos, João P.B. Viola, Andreia C. de Melo, Marcelo A. Soares |
| EPI_ISL_513584 | PHE South West Regional Laboratory, National Infection Service | Wellcome Sanger Institute for the COVID-19 Genomics UK Consortium | Stephanie Hutchings, Hannah Pymont, Dr Peter Muir, Barry Vipond, Rich Hopes; and Alex Alderton, Roberto Amato, Sonia Goncalves, Ewan Harrison, David K. Jackson, Ian Johnston, Dominic Kwiatkowski, Cordelia Langford, John Sillitoe on behalf of the Wellcome Sanger Institute COVID-19 Surveillance Team (http://www.sanger.ac.uk/covid-team) |
| EPI_ISL_513585, EPI_ISL_513587, EPI_ISL_513588 | Microbiology, Pathology, Kettering General Hospital | Wellcome Sanger Institute for the COVID-19 Genomics UK Consortium | Sahar El-dirdiri, Anita Kenyon, Thomas Davis and Alex Alderton, Roberto Amato, Sonia Goncalves, Ewan Harrison, David K. Jackson, Ian Johnston, Dominic Kwiatkowski, Cordelia Langford, John Sillitoe on behalf of the Wellcome Sanger Institute COVID-19 Surveillance Team (http://www.sanger.ac.uk/covid-team) |
| EPI_ISL_513592, EPI_ISL_513597, EPI_ISL_513598, EPI_ISL_513600, EPI_ISL_513601, EPI_ISL_513602, EPI_ISL_513604, EPI_ISL_513605, EPI_ISL_513606, EPI_ISL_513607, EPI_ISL_513608, EPI_ISL_513609, EPI_ISL_513610, EPI_ISL_513613, EPI_ISL_513615, EPI_ISL_513616, EPI_ISL_513617, EPI_ISL_513618, EPI_ISL_513630 | | | |
| see above | Viral Respiratory Lab, National Institute for Biomedical Research (INRB) | Pathogen Sequencing Lab, National Institute for Biomedical Research (INRB) | Placide Mbala-Kingebeni, Edith Nkwembe, Eddy Kinganda-Lusamaki, Amuri Aziza, Francisca Muyembe Mawete, Emmanuel Lokilo Lofiko, Catherine Pratt, Matthias Pauthner, Josh Quick, Allison Black, James Hadfield, Trevor Bedford, Ian Goodfellow, Andrew Rambaut, Nick Loman, Kristian Andersen, Michael Wiley, Steve Ahuka-Mundeke, Jean-Jacques Muyembe Tamfum |
| EPI_ISL_513632, EPI_ISL_513634, EPI_ISL_513635, EPI_ISL_513637 | University of Washington Virology Lab | University of Washington Virology Lab | Pavitra Roychoudhury, Hong Xie, Lasata Shrestha, Amin Addetia, Truong Nguyen, Victoria M Rachleff, Meei-Li Huang, Keith R Jerome, Alexander Greninger |
| EPI_ISL_513645, EPI_ISL_513647, EPI_ISL_513648, EPI_ISL_513652, EPI_ISL_513657, EPI_ISL_513671, EPI_ISL_513675, EPI_ISL_513677, EPI_ISL_513678, EPI_ISL_513680, EPI_ISL_513683, EPI_ISL_513684, EPI_ISL_513685, EPI_ISL_513686, EPI_ISL_513687, EPI_ISL_513688, EPI_ISL_513690, EPI_ISL_513691, EPI_ISL_513692, EPI_ISL_513696, EPI_ISL_513697, EPI_ISL_513700, EPI_ISL_513701, EPI_ISL_513704 | | | |
| see above | Utah Public Health Laboratory | Utah Public Health Laboratory | Heidi Butz, Erin Young, Kelly Oakeson |
| EPI_ISL_513707, EPI_ISL_513708, EPI_ISL_513709, EPI_ISL_513710, EPI_ISL_513711, EPI_ISL_513712, EPI_ISL_513713, EPI_ISL_513714, EPI_ISL_513715, EPI_ISL_513716, EPI_ISL_513717, EPI_ISL_513718, EPI_ISL_513719, EPI_ISL_513720, EPI_ISL_513721, EPI_ISL_513723, EPI_ISL_513724, EPI_ISL_513725, EPI_ISL_513726, EPI_ISL_513727, EPI_ISL_513728, EPI_ISL_513729, EPI_ISL_513734, EPI_ISL_513735, EPI_ISL_513737, EPI_ISL_513738, EPI_ISL_513739, EPI_ISL_513740, EPI_ISL_513741, EPI_ISL_513742, EPI_ISL_513743, EPI_ISL_513744, EPI_ISL_513745, EPI_ISL_513746, EPI_ISL_513747, EPI_ISL_513748, EPI_ISL_513749, EPI_ISL_513750, EPI_ISL_513751, EPI_ISL_513752, EPI_ISL_513753, EPI_ISL_513754, EPI_ISL_513755, EPI_ISL_513756, EPI_ISL_513757, EPI_ISL_513758, EPI_ISL_513760, EPI_ISL_513761, EPI_ISL_513762, EPI_ISL_513766, EPI_ISL_513767, EPI_ISL_513768, EPI_ISL_513769, EPI_ISL_513770, EPI_ISL_513771, EPI_ISL_513772 | | | |
| see above | Orange County Public Health Lab | Chan-Zuckerberg Biohub | CZB Cliahub Consortium |
| EPI_ISL_513773, EPI_ISL_513774, EPI_ISL_513775, EPI_ISL_513776, EPI_ISL_513777, EPI_ISL_513778, EPI_ISL_513779, EPI_ISL_513780, EPI_ISL_513781, EPI_ISL_513782, EPI_ISL_513783, EPI_ISL_513784, EPI_ISL_513786, EPI_ISL_513788 | | | |
| see above | County of Santa Clara Public Health Department | Chan-Zuckerberg Biohub | CZB Cliahub Consortium |
| EPI_ISL_513789, EPI_ISL_513790, EPI_ISL_513791, EPI_ISL_513792, EPI_ISL_513793, EPI_ISL_513794, EPI_ISL_513795, EPI_ISL_513796, EPI_ISL_513797, EPI_ISL_513798, EPI_ISL_513799, EPI_ISL_513800, EPI_ISL_513801, EPI_ISL_513802, EPI_ISL_513803, EPI_ISL_513804, EPI_ISL_513805, EPI_ISL_513806, EPI_ISL_513814, EPI_ISL_513815, EPI_ISL_513816, EPI_ISL_513817, EPI_ISL_513820, EPI_ISL_513822, EPI_ISL_513824, EPI_ISL_513826, EPI_ISL_513828, EPI_ISL_513829, EPI_ISL_513830, EPI_ISL_513832, EPI_ISL_513833, EPI_ISL_513835, EPI_ISL_513836, EPI_ISL_513837, EPI_ISL_513838, EPI_ISL_513839 | | | |
| see above | Orange County Public Health Laboratory | Chan-Zuckerberg Biohub | CZB Cliahub Consortium |
| EPI_ISL_513840, EPI_ISL_513841, EPI_ISL_513842, EPI_ISL_513843, EPI_ISL_513844, EPI_ISL_513845, EPI_ISL_513846, EPI_ISL_513847, EPI_ISL_513848, EPI_ISL_513850, EPI_ISL_513852, EPI_ISL_513853, EPI_ISL_513854, EPI_ISL_513855 | | | |
| see above | Humboldt County Public Health Laboratory | Chan-Zuckerberg Biohub | CZB Cliahub Consortium |
| EPI_ISL_513856, EPI_ISL_513857, EPI_ISL_513858, EPI_ISL_513859, EPI_ISL_513860, EPI_ISL_513861, EPI_ISL_513862, EPI_ISL_513863, EPI_ISL_513864, EPI_ISL_513865, EPI_ISL_513866, EPI_ISL_513867, EPI_ISL_513869, EPI_ISL_513870, EPI_ISL_513872, EPI_ISL_513873, EPI_ISL_513874, EPI_ISL_513875, EPI_ISL_513876, EPI_ISL_513878, EPI_ISL_513879, EPI_ISL_513880, EPI_ISL_513882 | | | |
| see above | San Francisco Public Health Laboratory | Chan-Zuckerberg Biohub | CZB Cliahub Consortium |
| EPI_ISL_513883, EPI_ISL_513884, EPI_ISL_513885, EPI_ISL_513889, EPI_ISL_513891, EPI_ISL_513892, EPI_ISL_513893, EPI_ISL_513894, EPI_ISL_513895, EPI_ISL_513896, EPI_ISL_513897, EPI_ISL_513898, EPI_ISL_513899, EPI_ISL_513900, EPI_ISL_513901, EPI_ISL_513902, EPI_ISL_513903, EPI_ISL_513905, EPI_ISL_513907, EPI_ISL_513908, EPI_ISL_513909, EPI_ISL_513910 | | | |
| see above | UCSF Clinical Microbiology Laboratory | Chan-Zuckerberg Biohub | CZB Cliahub Consortium |
| EPI_ISL_513911 | SA Pathology | SA Pathology | Lex Leong, Chuan Kok Lim, Mark Turra, Ivan Bastian, Geoff Higgins |
| EPI_ISL_513925 | Microbiology & Bioinformatics and Biostatistics, Kohat University of Science and Technology (Pakistan) & Shanghai Jiao Tong University (China) | Microbiology & Bioinformatics and Biostatistics, Kohat University of Science and Technology (Pakistan) & Shanghai Jiao Tong University (China) | Khan,M.T., Khan,T.A., Ali,S., Khan,A.S., Muhammad,N. and Wei,D.Q. |
| EPI_ISL_514062, EPI_ISL_514063, EPI_ISL_514064 | Navy and Marine Corps Public Health Center | Pathogen Discovery, Respiratory Viruses Branch, Division of Viral Diseases, Centers for Disease Control and Prevention | Yan Li, Anna Montmayeur, Krista Queen, Jing Zhang, Ying Tao, Anna Uehara, Rachel Marine, Clinton R. Paden, Haibin Wang, Suxiang Tong |
| EPI_ISL_514065 | Navy and Marine Corps Public Health Center | Pathogen Discovery, Respiratory Viruses Branch, Division of Viral Diseases, Centers for Disease Control and Prevention | Krista Queen, Yan Li, Anna Montmayeur, Jing Zhang, Ying Tao, Anna Uehara, Rachel Marine, Clinton R. Paden, Haibin Wang, Suxiang Tong |
| EPI_ISL_514066 | Navy and Marine Corps Public Health Center | Pathogen Discovery, Respiratory Viruses Branch, Division of Viral Diseases, Centers for Disease Control and Prevention | Yan Li, Anna Montmayeur, Krista Queen, Jing Zhang, Ying Tao, Anna Uehara, Rachel Marine, Clinton R. Paden, Haibin Wang, Suxiang Tong |
| EPI_ISL_514067, EPI_ISL_514068 | Navy and Marine Corps Public Health Center | Pathogen Discovery, Respiratory Viruses Branch, Division of Viral Diseases, Centers for Disease Control and Prevention | Krista Queen, Yan Li, Anna Montmayeur, Jing Zhang, Ying Tao, Anna Uehara, Rachel Marine, Clinton R. Paden, Haibin Wang, Suxiang Tong |
| EPI_ISL_514069, EPI_ISL_514070 | Navy and Marine Corps Public Health Center | Pathogen Discovery, Respiratory Viruses Branch, Division of Viral Diseases, Centers for Disease Control and Prevention | Yan Li, Anna Montmayeur, Krista Queen, Jing Zhang, Ying Tao, Anna Uehara, Rachel Marine, Clinton R. Paden, Haibin Wang, Suxiang Tong |
| EPI_ISL_514071, EPI_ISL_514072 | Navy and Marine Corps Public Health Center | Pathogen Discovery, Respiratory Viruses Branch, Division of Viral Diseases, Centers for Disease Control and Prevention | Krista Queen, Yan Li, Anna Montmayeur, Jing Zhang, Ying Tao, Anna Uehara, Rachel Marine, Clinton R. Paden, Haibin Wang, Suxiang Tong |
| EPI_ISL_514073, EPI_ISL_514074 | Navy and Marine Corps Public Health Center | Pathogen Discovery, Respiratory Viruses Branch, Division of Viral Diseases, Centers for Disease Control and Prevention | Yan Li, Anna Montmayeur, Krista Queen, Jing Zhang, Ying Tao, Anna Uehara, Rachel Marine, Clinton R. Paden, Haibin Wang, Suxiang Tong |
| EPI_ISL_514075 | Navy and Marine Corps Public Health Center | Pathogen Discovery, Respiratory Viruses Branch, Division of Viral Diseases, Centers for Disease Control and Prevention | Krista Queen, Yan Li, Anna Montmayeur, Jing Zhang, Ying Tao, Anna Uehara, Rachel Marine, Clinton R. Paden, Haibin Wang, Suxiang Tong |
| EPI_ISL_514077, EPI_ISL_514078, EPI_ISL_514079 | Navy and Marine Corps Public Health Center | Pathogen Discovery, Respiratory Viruses Branch, Division of Viral Diseases, Centers for Disease Control and Prevention | Yan Li, Anna Montmayeur, Krista Queen, Jing Zhang, Ying Tao, Anna Uehara, Rachel Marine, Clinton R. Paden, Haibin Wang, Suxiang Tong |
| EPI_ISL_514080, EPI_ISL_514081, EPI_ISL_514082, EPI_ISL_514083 | Navy and Marine Corps Public Health Center | Pathogen Discovery, Respiratory Viruses Branch, Division of Viral Diseases, Centers for Disease Control and Prevention | Krista Queen, Yan Li, Anna Montmayeur, Jing Zhang, Ying Tao, Anna Uehara, Rachel Marine, Clinton R. Paden, Haibin Wang, Suxiang Tong |
| EPI_ISL_514084 | Navy and Marine Corps Public Health Center | Pathogen Discovery, Respiratory Viruses Branch, Division of Viral Diseases, Centers for Disease Control and Prevention | Yan Li, Anna Montmayeur, Krista Queen, Jing Zhang, Ying Tao, Anna Uehara, Rachel Marine, Clinton R. Paden, Haibin Wang, Suxiang Tong |
| EPI_ISL_514085, EPI_ISL_514086, EPI_ISL_514087 | Navy and Marine Corps Public Health Center | Pathogen Discovery, Respiratory Viruses Branch, Division of Viral Diseases, Centers for Disease Control and Prevention | Krista Queen, Yan Li, Anna Montmayeur, Jing Zhang, Ying Tao, Anna Uehara, Rachel Marine, Clinton R. Paden, Haibin Wang, Suxiang Tong |
| EPI_ISL_514088 | Navy and Marine Corps Public Health Center | Pathogen Discovery, Respiratory Viruses Branch, Division of Viral Diseases, Centers for Disease Control and Prevention | Yan Li, Anna Montmayeur, Krista Queen, Jing Zhang, Ying Tao, Anna Uehara, Rachel Marine, Clinton R. Paden, Haibin Wang, Suxiang Tong |
| EPI_ISL_514089 | Navy and Marine Corps Public Health Center | Pathogen Discovery, Respiratory Viruses Branch, Division of Viral Diseases, Centers for Disease Control and Prevention | Krista Queen, Yan Li, Anna Montmayeur, Jing Zhang, Ying Tao, Anna Uehara, Rachel Marine, Clinton R. Paden, Haibin Wang, Suxiang Tong |
| EPI_ISL_514090, EPI_ISL_514091, EPI_ISL_514092 | Navy and Marine Corps Public Health Center | Pathogen Discovery, Respiratory Viruses Branch, Division of Viral Diseases, Centers for Disease Control and Prevention | Yan Li, Anna Montmayeur, Krista Queen, Jing Zhang, Ying Tao, Anna Uehara, Rachel Marine, Clinton R. Paden, Haibin Wang, Suxiang Tong |
| EPI_ISL_514114, EPI_ISL_514115, EPI_ISL_514117, EPI_ISL_514118, EPI_ISL_514119, EPI_ISL_514122 | Viral Respiratory Lab, National Institute for Biomedical Research (INRB) | Pathogen Sequencing Lab, National Institute for Biomedical Research (INRB) | Placide Mbala-Kingebeni, Edith Nkwembe, Eddy Kinganda-Lusamaki, Amuri Aziza, Francisca Muyembe Mawete, Emmanuel Lokilo Lofiko, Catherine Pratt, Matthias Pauthner, Josh Quick, Allison Black, James Hadfield, Trevor Bedford, Ian Goodfellow, Andrew Rambaut, Nick Loman, Kristian Andersen, Michael Wiley, Steve Ahuka-Mundeke, Jean-Jacques Muyembe Tamfum |
| EPI_ISL_514128 | Navy and Marine Corps Public Health Center | Centers for Disease Control and Prevention | Krista Queen, Yan Li, Anna Montmayeur, Jing Zhang, Ying Tao, Anna Uehara, Rachel Marine, Clinton R. Paden, Haibin Wang, Suxiang Tong |
| EPI_ISL_514129, EPI_ISL_514130 | National Institute of Laboratory Medicine and Referral Center | Genomic Research Lab, BCSIR | Md. Murshed Hasan Sarkar, Abu Sayeed Mohammad Mahmud, Mohammad Samir Uzzaman, Eshrar Osman, Md. Ahashan Habib, Shahina Akter, Tanjina Akhter Banu, Barna Goswami, Iffat Jahan, Md. Saddam Hossain, Tasnim Nafisa, Md. Maruf Ahmed Molla, Mahmuda Yeasmin, Asish Kumar Ghosh, A. K. M. Shamsuzzaman, Sheikh Md. Selim Al Din, Utpal Chandra Ray, Salek Ahmed Sajib, Md. Salim Khan |
| EPI_ISL_514131 | Rondônia Central Public Health Laboratory (LACEN/RO), vinculated to State Health Secretariat of Rondônia (SESAU/RO) | Molecular Virology Laboratory of Oswaldo Cruz Foundation of Rondônia | Luan Felipo Botelho-Souza, Felipe Souza Nogueira-Lima, Tárcio Peixoto Roca, Alcione de Oliveira dos Santos, Felipe Gomes Naveca, Adriana Cristina Salvador Maia, Cicileia Correia da Silva, Aline Linhares Ferreira de Melo Mendonça, Celina Aparecida Bertoni Lugtenburg, Camila Flávia Gomes Azzi, Juliana Loca Furtado, Suelen Cavalcante, Rita de Cássia Pontello Rampazzo, Caio Henrique Nemeth Santos, Alice Paula Di Sabatino Guimarães, Jansen Fernandes de Medeiros, Fernando Rodrigues Máximo, Juan Miguel Vilallobos-Salcedo and Deusilene Souza Vieira1 |
| EPI_ISL_514132 | Rondônia Central Public Health Laboratory (LACEN/RO), vinculated to State Health Secretariat of Rondônia (SESAU/RO) | Molecular Virology Laboratory of Oswaldo Cruz Foundation of Rondônia | Luan Felipo Botelho-Souza, Felipe Souza Nogueira-Lima, Tárcio Peixoto Roca, Alcione de Oliveira dos Santos, Felipe Gomes Naveca, Adriana Cristina Salvador Maia, Cicileia Correia da Silva, Aline Linhares Ferreira de Melo Mendonça, Celina Aparecida Bertoni Lugtenburg, Camila Flávia Gomes Azzi, Juliana Loca Furtado, Suelen Cavalcante, Rita de Cássia Pontello Rampazzo, Caio Henrique Nemeth Santos, Alice Paula Di Sabatino Guimarães, Jansen Fernandes de Medeiros, Fernando Rodrigues Máximo, Juan Miguel Vilallobos-Salcedo and Deusilene Souza Vieira. |
| EPI_ISL_514133, EPI_ISL_514134 | Rondônia Central Public Health Laboratory (LACEN/RO), vinculated to State Health Secretariat of Rondônia (SESAU/RO) | Molecular Virology Laboratory of Oswaldo Cruz Foundation of Rondônia | Luan Felipo Botelho-Souza, Felipe Souza Nogueira-Lima, Tárcio Peixoto Roca, Alcione de Oliveira dos Santos, Felipe Gomes Naveca, Adriana Cristina Salvador Maia, Cicileia Correia da Silva, Aline Linhares Ferreira de Melo Mendonça, Celina Aparecida Bertoni Lugtenburg, Camila Flávia Gomes Azzi, Juliana Loca Furtado, Suelen Cavalcante, Rita de Cássia Pontello Rampazzo, Caio Henrique Nemeth Santos, Alice Paula Di Sabatino Guimarães, Jansen Fernandes de Medeiros, Fernando Rodrigues Máximo, Juan Miguel Vilallobos-Salcedo and Deusilene Souza Vieira |
| EPI_ISL_514135 | Rondônia Central Public Health Laboratory (LACEN/RO), vinculated to State Health Secretariat of Rondônia (SESAU/RO) | Molecular Virology Laboratory of Oswaldo Cruz Foundation of Rondônia | Luan Felipo Botelho-Souza, Felipe Souza Nogueira-Lima, Tárcio Peixoto Roca, Alcione de Oliveira dos Santos, Felipe Gomes Naveca, Adriana Cristina Salvador Maia, Cicileia Correia da Silva, Aline Linhares Ferreira de Melo Mendonça, Celina Aparecida Bertoni Lugtenburg, Camila Flávia Gomes Azzi, Juliana Loca Furtado, Suelen Cavalcante, Rita de Cássia Pontello Rampazzo, Caio Henrique Nemeth Santos, Alice Paula Di Sabatino Guimarães, Jansen Fernandes de Medeiros, Fernando Rodrigues Máximo, Juan Miguel Vilallobos-Salcedo and Deusilene Souza Vieira |
| EPI_ISL_514136 | Rondônia Central Public Health Laboratory (LACEN/RO), vinculated to State Health Secretariat of Rondônia (SESAU/RO) | Molecular Virology Laboratory of Oswaldo Cruz Foundation of Rondônia | Luan Felipo Botelho-Souza, Felipe Souza Nogueira-Lima, Tárcio Peixoto Roca, Alcione de Oliveira dos Santos, Felipe Gomes Naveca, Adriana Cristina Salvador Maia, Cicileia Correia da Silva, Aline Linhares Ferreira de Melo Mendonça, Celina Aparecida Bertoni Lugtenburg, Camila Flávia Gomes Azzi, Juliana Loca Furtado, Suelen Cavalcante, Rita de Cássia Pontello Rampazzo, Caio Henrique Nemeth Santos, Alice Paula Di Sabatino Guimarães, Jansen Fernandes de Medeiros, Fernando Rodrigues Máximo, Juan Miguel Vilallobos-Salcedo and Deusilene Souza Vieira |
| EPI_ISL_514137 | Rondônia Central Public Health Laboratory (LACEN/RO), vinculated to State Health Secretariat | Molecular Virology Laboratory of Oswaldo Cruz Foundation of Rondônia | Luan Felipo Botelho-Souza, Felipe Souza Nogueira-Lima, Tárcio Peixoto Roca, Alcione de Oliveira dos Santos, Felipe Gomes Naveca, Adriana Cristina Salvador Maia, Cicileia Correia da Silva, Aline Linhares Ferreira de Melo Mendonça, Celina Aparecida Bertoni Lugtenburg, Camila Flávia Gomes Azzi, Juliana Loca Furtado, Suelen Cavalcante, Rita de Cássia Pontello Rampazzo, Caio Henrique |

| | | | |
|---|---|---|---|
| | of Rondônia (SESAU/RO) | | Nemeth Santos, Alice Paula Di Sabatino Guimarães, Jansen Fernandes de Medeiros, Fernando Rodrigues Máximo, Juan Miguel Villallobos-Salcedo and Deusilene Souza Vieira |
| EPI_ISL_514138 | Rondônia Central Public Health Laboratory (LACEN/RO), vinculated to State Health Secretariat of Rondônia (SESAU/RO) | Molecular Virology Laboratory of Oswaldo Cruz Foundation of Rondônia | Luan Felipo Botelho-Souza, Felipe Souza Nogueira-Lima, Tárcio Peixoto Roca, Alcione de Oliveira dos Santos, Felipe Gomes Naveca, Adriana Cristina Salvador Maia, Cicileia Correia da Silva, Aline Linhares Ferreira de Melo Mendonça, Celina Aparecida Bertoni Lugtenburg, Camila Flávia Gomes Azzi, Juliana Loca Furtado, Suelen Cavalcante, Rita de Cássia Pontello Rampazzo, Caio Henrique Nemeth Santos, Alice Paula Di Sabatino Guimarães, Jansen Fernandes de Medeiros, Fernando Rodrigues Máximo, Juan Miguel Villallobos-Salcedo and Deusilene Souza Vieira |
| EPI_ISL_514140, EPI_ISL_514141, EPI_ISL_514142, EPI_ISL_514143, EPI_ISL_514144, EPI_ISL_514145, EPI_ISL_514146, EPI_ISL_514147, EPI_ISL_514148, EPI_ISL_514151, EPI_ISL_514152, EPI_ISL_514153, EPI_ISL_514155, EPI_ISL_514156, EPI_ISL_514157, EPI_ISL_514158, EPI_ISL_514159, EPI_ISL_514160, EPI_ISL_514161, EPI_ISL_514162, EPI_ISL_514163, EPI_ISL_514168, EPI_ISL_514169, EPI_ISL_514171, EPI_ISL_514176, EPI_ISL_514177, EPI_ISL_514178, EPI_ISL_514180, EPI_ISL_514183, EPI_ISL_514184, EPI_ISL_514186, EPI_ISL_514187, EPI_ISL_514188, EPI_ISL_514189, EPI_ISL_514197, EPI_ISL_514201, EPI_ISL_514202, EPI_ISL_514203, EPI_ISL_514204, EPI_ISL_514205, EPI_ISL_514207, EPI_ISL_514211, EPI_ISL_514212, EPI_ISL_514213, EPI_ISL_514214, EPI_ISL_514215, EPI_ISL_514216, EPI_ISL_514219, EPI_ISL_514220, EPI_ISL_514222, EPI_ISL_514223, EPI_ISL_514224, EPI_ISL_514225 | | | |
| see above | Florida Bureau of Public Health Laboratories | Florida Bureau of Public Health Laboratories | Sarah Schmedes, Jason Blanton |
| EPI_ISL_514226 | Laboratorio de Referencia Nacional de Virus Respiratorio. Instituto Nacional de Salud Perú | Laboratorio de Referencia Nacional de Biotecnología y Biología Molecular. Instituto Nacional de Salud Perú | Carlos Padilla Rojas, Karolyn Vega Chozo, Priscila Lope Pari, Omar Caceres Rey, Marco Galarza Perez, Maribel Huaringa Nuñez, Johanna Balbuena Torrez, Henri Bailon Calderon, Nancy Rojas Serrano |
| EPI_ISL_514227 | Laboratorio de Referencia Nacional de Virus Respiratorio. Instituto Nacional de Salud. Peru | Laboratorio de Referencia Nacional de Biotecnologia y Biologia Molecular.Instituto Nacional de Salud.Peru | Carlos Padilla Rojas, Karolyn Vega Chozo, Priscila Lope Pari, Omar Caceres Rey, Marco Galarza Perez, Maribel Huaringa Nuñez, Johanna Balbuena Torrez, Henri Bailon Calderon, Nancy Rojas Serrano |
| EPI_ISL_514228, EPI_ISL_514229 | National Institute of Laboratory Medicine and Referral Center | Genomic Research Lab, BCSIR | Barna Goswami, Abu Sayeed Mohammad Mahmud, Mohammad Samir Uzzaman, Eshrar Osman, Md. Ahashan Habib, Shahina Akter, Tanjina Akhter Banu, Md. Murshed Hasan Sarkar, Iffat Jahan, Md. Saddam Hossain, Tasnim Nafisa, Md. Maruf Ahmed Molla, Mahmuda Yeasmin, Asish Kumar Ghosh, A. K. M. Shamsuzzaman, Sheikh Md. Selim Al Din, Utpal Chandra Ray, Salek Ahmed Sajib, Md. Salim Khan |
| EPI_ISL_514231, EPI_ISL_514232 | National Institute of Laboratory Medicine and Referral Center | Genomic Research Lab, BCSIR | Shahina Akter, Abu Sayeed Mohammad Mahmud, Mohammad Samir Uzzaman, Eshrar Osman, Md. Ahashan Habib, Tanjina Akhter Banu, Md. Murshed Hasan Sarkar, Barna Goswami, Iffat Jahan, Md. Saddam Hossain, Tasnim Nafisa, Md. Maruf Ahmed Molla, Mahmuda Yeasmin, Asish Kumar Ghosh, A. K. M. Shamsuzzaman, Sheikh Md. Selim Al Din, Utpal Chandra Ray, Salek Ahmed Sajib, Md. Salim Khan |
| EPI_ISL_514233, EPI_ISL_514234, EPI_ISL_514235 | National Institute of Laboratory Medicine and Referral Center | Genomic Research Lab, BCSIR | Md. Murshed Hasan Sarkar, Abu Sayeed Mohammad Mahmud, Mohammad Samir Uzzaman, Eshrar Osman, Md. Ahashan Habib, Shahina Akter, Tanjina Akhter Banu, Barna Goswami, Iffat Jahan, Md. Saddam Hossain, Tasnim Nafisa, Md. Maruf Ahmed Molla, Mahmuda Yeasmin, Asish Kumar Ghosh, A. K. M. Shamsuzzaman, Sheikh Md. Selim Al Din, Utpal Chandra Ray, Salek Ahmed Sajib, Md. Salim Khan |
| EPI_ISL_514236 | National Institute of Laboratory Medicine and Referral Center | Genomic Research Lab, BCSIR | Md. Murshed Hasan Sarkar, Abu Sayeed Mohammad Mahmud, Mohammad Samir Uzzaman, Eshrar Osman, Md. Ahashan Habib, Shahina Akter, Tanjina Akhter Banu, Barna Goswami, Iffat Jahan, Md. Saddam Hossain, Tasnim Nafisa, Md. Maruf Ahmed Molla, Mahmuda Yeasmin, Asish Kumar Ghosh, A. K. M. Shamsuzzaman, Sheikh Md. Selim Al Din, Utpal Chandra Ray, Salek Ahmed Sajib, Md. Salim Khan |
| EPI_ISL_514237, EPI_ISL_514238, EPI_ISL_514239, EPI_ISL_514240, EPI_ISL_514241 | National Institute of Laboratory Medicine and Referral Center | Genomic Research Lab, BCSIR | Md. Saddam Hossain, Abu Sayeed Mohammad Mahmud, Mohammad Samir Uzzaman, Eshrar Osman, Md. Ahashan Habib, Shahina Akter, Tanjina Akhter Banu, Md. Murshed Hasan Sarkar, Barna Goswami, Iffat Jahan, Tasnim Nafisa, Md. Maruf Ahmed Molla, Mahmuda Yeasmin, Asish Kumar Ghosh, A. K. M. Shamsuzzaman, Sheikh Md. Selim Al Din, Utpal Chandra Ray, Salek Ahmed Sajib, Md. Salim Khan |
| EPI_ISL_514242, EPI_ISL_514243, EPI_ISL_514244 | National Institute of Laboratory Medicine and Referral Center | Genomic Research Lab, BCSIR | Tanjina Akhter Banu, Abu Sayeed Mohammad Mahmud, Mohammad Samir Uzzaman, Eshrar Osman, Md. Ahashan Habib, Shahina Akter, Md. Murshed Hasan Sarkar, Barna Goswami, Iffat Jahan, Md. Saddam Hossain, Tasnim Nafisa, Md. Maruf Ahmed Molla, Mahmuda Yeasmin, Asish Kumar Ghosh, A. K. M. Shamsuzzaman, Sheikh Md. Selim Al Din, Utpal Chandra Ray, Salek Ahmed Sajib, Md. Salim Khan |
| EPI_ISL_514245, EPI_ISL_514246, EPI_ISL_514247 | National Institute of Laboratory Medicine and Referral Center | Genomic Research Lab, BCSIR | Iffat Jahan, Abu Sayeed Mohammad Mahmud, Mohammad Samir Uzzaman, Eshrar Osman, Md. Ahashan Habib, Shahina Akter, Tanjina Akhter Banu, Md. Murshed Hasan Sarkar, Barna Goswami, Md. Saddam Hossain, Tasnim Nafisa, Md. Maruf Ahmed Molla, Mahmuda Yeasmin, Asish Kumar Ghosh, A. K. M. Shamsuzzaman, Sheikh Md. Selim Al Din, Utpal Chandra Ray, Salek Ahmed Sajib, Md. Salim Khan |
| EPI_ISL_514248, EPI_ISL_514249, EPI_ISL_514250, EPI_ISL_514251, EPI_ISL_514252 | National Institute of Laboratory Medicine and Referral Center | Genomic Research Lab, BCSIR | Abu Sayeed Mohammad Mahmud, Mohammad Samir Uzzaman, Eshrar Osman, Md. Ahashan Habib, Shahina Akter, Tanjina Akhter Banu, Md. Murshed Hasan Sarkar, Barna Goswami, Iffat Jahan, Md. Saddam Hossain, Tasnim Nafisa, Md. Maruf Ahmed Molla, Mahmuda Yeasmin, Asish Kumar Ghosh, A. K. M. Shamsuzzaman, Sheikh Md. Selim Al Din, Utpal Chandra Ray, Salek Ahmed Sajib, Md. Salim Khan |
| EPI_ISL_514253 | Advanced Biotechnology Laboratory | Genomic Research Lab, BCSIR | Abu Sayeed Mohammad Mahmud, Mohammad Samir Uzzaman, Eshrar Osman, Hossain Uddin Shekhar, M. Aftab Uddin, Md. Bayejid Hosen, Eunus Ali, Md. Ahashan Habib, Shahina Akter, Tanjina Akhter Banu, Md. Murshed Hasan Sarkar, Barna Goswami, Iffat Jahan, Md. Saddam Hossain, Utpal Chandra Ray, Salek Ahmed Sajib, Md. Salim Khan |
| EPI_ISL_514264, EPI_ISL_514265 | Laboratorio de Referencia Nacional de Virus Respiratorio. Instituto Nacional de Salud. Perú | Laboratorio de Referencia Nacional de Biotecnologia y Biologia Molecular.Instituto Nacional de Salud.Perú | Carlos Padilla Rojas, Karolyn Vega Chozo, Priscila Lope Pari, Omar Caceres Rey, Marco Galarza Perez, Maribel Huaringa Nuñez, Johanna Balbuena Torrez, Henri Bailon Calderon, Nancy Rojas Serrano |
| EPI_ISL_514266, EPI_ISL_514268, EPI_ISL_514269, EPI_ISL_514270, EPI_ISL_514271, EPI_ISL_514272, EPI_ISL_514274 | Israel Central Virology laboratory | Israel Central Virology laboratory | Neta Zuckerman, Efrat Dahan Bucris, Oran Erster, Ella Mendelson, Michal Mandelboim |
| EPI_ISL_514275 | Israeli Central Virology laboratory | Israel Central Virology laboratory | Neta Zuckerman, Efrat Dahan Bucris, Oran Erster, Ella Mendelson, Michal Mandelboim |
| EPI_ISL_514278, EPI_ISL_514279, EPI_ISL_514280, EPI_ISL_514281, EPI_ISL_514283, EPI_ISL_514284, EPI_ISL_514286, EPI_ISL_514287, EPI_ISL_514288, EPI_ISL_514289, EPI_ISL_514290, EPI_ISL_514291, EPI_ISL_514292, EPI_ISL_514293, EPI_ISL_514294, EPI_ISL_514295, EPI_ISL_514296, EPI_ISL_514299, EPI_ISL_514300, EPI_ISL_514301, EPI_ISL_514302, EPI_ISL_514303, EPI_ISL_514305, EPI_ISL_514306, EPI_ISL_514308, EPI_ISL_514309, EPI_ISL_514310, EPI_ISL_514311, EPI_ISL_514313, EPI_ISL_514314, EPI_ISL_514315, EPI_ISL_514316 | | | |
| see above | Israel Central Virology laboratory | Israel Central Virology laboratory | Neta Zuckerman, Efrat Dahan Bucris, Oran Erster, Ella Mendelson, Michal Mandelboim |
| EPI_ISL_514317 | Laboratorio de Referencia Nacional de Virus Respiratorio. Instituto Nacional de Salud Perú | Laboratorio de Referencia Nacional de Biotecnología y Biología Molecular. Instituto Nacional de Salud Perú | Carlos Padilla Rojas, Karolyn Vega Chozo, Priscila Lope Pari, Omar Caceres Rey, Marco Galarza Perez, Maribel Huaringa Nuñez, Johanna Balbuena Torres, Henri Bailon Calderon, Nancy Rojas Serrano. |
| EPI_ISL_514338, EPI_ISL_514339, EPI_ISL_514340, EPI_ISL_514341, EPI_ISL_514342 | Laboratorio de Referencia Nacional de Virus Respiratorio. Instituto Nacional de Salud Perú | Laboratorio de Referencia Nacional de Biotecnología y Biología Molecular. Instituto Nacional de Salud Perú | Carlos Padilla Rojas, Karolyn Vega Chozo, Priscila Lope Pari, Omar Caceres Rey, Marco Galarza Perez, Maribel Huaringa Nuñez, Johanna Balbuena Torres, Henri Bailon Calderon, Nancy Rojas Serrano. |
| EPI_ISL_514343, EPI_ISL_514344, EPI_ISL_514345, EPI_ISL_514346, EPI_ISL_514347, EPI_ISL_514348, EPI_ISL_514349, EPI_ISL_514350, EPI_ISL_514351, EPI_ISL_514352 | Respiratory Virus Unit, Microbiology Services Colindale, Public Health England | Respiratory Virus Unit, Microbiology Services Colindale, Public Health England | PHE Covid Sequencing Team |
| EPI_ISL_514354 | General Hospital - Prilep | Research Center for Genetic Engineering and Biotechnology "Georgi D. Efremov" , Macedonian Academy of Sciences and Arts | RCGEB - MASA |
| EPI_ISL_514355, EPI_ISL_514360, EPI_ISL_514367, EPI_ISL_514375, EPI_ISL_514382, EPI_ISL_514424 | National Institute for Communicable Diseases of the National Health Laboratory Service | National Institute for Communicable Diseases of the National Health Laboratory Service | Allam M, Ismail A, Khumalo Z, Kwenda S, Mtshali P, Mnyameni F, Mohale T, Bhiman JN |
| EPI_ISL_514425, EPI_ISL_514426, EPI_ISL_514427, EPI_ISL_514428, EPI_ISL_514429, EPI_ISL_514430, EPI_ISL_514431 | Laboratory Diagnostic, Veterinary Specialized Institute Kraljevo | Laboratory Diagnostic, Veterinary Specialized Institute Kraljevo | Vidanovic,D., Tesovic,B., Knezevic,A., Jankovic,M., Sekler,M., Banovic Djeri,B., Volkening,J., Afonso,C., Petrovic,T. |
| EPI_ISL_514432 | Prof. Massimo Zollo CEINGE TASK-FORCE COVID19 - Regione Campania | Prof. Massimo Zollo CEINGE TASK-FORCE COVID19 - Regione Campania | Veronica Ferrucci, Dae young Kong, Fatemeh asadzadeh, Laura Marrone, Roberto Siciliano, Rino Cerino, Giovanna Fusco, Marika Comegna, Angelo Boccia, Maurizio Viscardi, Giorgia Borriello, Sergio Brandi, Claudia Tiberio, Luigi Atripaldi, Giovanni Paolella, Giuseppe Castaldo, Stefano Pascarella, Martina Bianchi, Lorenzo Chiariotti, Jae Myun Lee, Jae Ho Jung, Kyong Seop Yun, Hong Yeoul Kim and Massimo Zollo |
| EPI_ISL_514435 | Dr. RSS Hospital, Modasa | Gujarat Biotechnology Research Centre | Harsh Chaudhari, Zarna Patel, Monika Gandhi, Pinal Trivedi, Maharshi Pandya, Nidhi Patel, Nitin Savaliya, Raghawendra Kumar, Dinesh Kumar, Zuber Saiyed, Komal Patel, Labdhi Pandya, Afzal Ansari, Nikha Trivedi, Apurvasinh Puvar, Janvi Raval, R D Dixit, A M Kadri, Harsh Bakshi, Chaitanya Joshi, Madhvi Joshi |
| EPI_ISL_514436 | Dr. RSS Hospital, Modasa | Gujarat Biotechnology Research Centre | Zarna Patel, Monika Gandhi, Pinal Trivedi, Maharshi Pandya, Nidhi Patel, Nitin Savaliya, Raghawendra Kumar, Dinesh Kumar, Zuber Saiyed, Komal Patel, Labdhi Pandya, Afzal Ansari, Nikha Trivedi, Harsh Chaudhari, Apurvasinh Puvar, Janvi Raval, R D Dixit, A M Kadri, Harsh Bakshi, Chaitanya Joshi, Madhvi Joshi |
| EPI_ISL_514437 | Dr. RSS Hospital, Modasa | Gujarat Biotechnology Research Centre | Nidhi Patel, Nitin Savaliya, Raghawendra Kumar, Dinesh Kumar, Zuber Saiyed, Komal Patel, Labdhi Pandya, Afzal Ansari, Nikha Trivedi, Harsh Chaudhari, Apurvasinh Puvar, Janvi Raval, Zarna Patel, Monika Gandhi, Pinal Trivedi, Maharshi Pandya, R D Dixit, A M Kadri, Harsh Bakshi, Chaitanya Joshi, Madhvi Joshi |
| EPI_ISL_514440 | NSTU COVID-19 Diagnostic Center | NSU Genome Research Institute (NGRI), North South University | Dr. Muhammad Maqsud Hossain, Aura Rahman, Prof. Firoz Ahmed, Tahrima Huq, Abdus Sadique, Tamanna Afroze, Jahidul Alam, Md Aminul Islam, Prof. Md. Didar-Ul-Alam, Prof. Kazi Nadim Hasan, Prof. Abdul Khaleque, Prof, Hasan Mahmud Reza |
| EPI_ISL_514443, EPI_ISL_514445, EPI_ISL_514448, EPI_ISL_514449, EPI_ISL_514450, EPI_ISL_514451 | Department of Pathology, University of Cambridge | COVID-19 Genomics UK (COG-UK) Consortium | Luke W Meredith, M. Estée Török, Myra Hosmillo, William L. Hamilton, Martin D. Curran, Theresa Feltwell, Grant Hall, Anna Yakovleva, Fahad A Khokhar, Charlotte J. Houldcroft, Laura G Caller, Aminu S. Jahun, Sarah L. Caddy, Yasmin Chaudhry, Malte Pinckert, Ian Goodfellow |
| EPI_ISL_514452, EPI_ISL_514453, EPI_ISL_514454 | Queens Medical Centre, Clinical Microbiology Department / DeepSeq Nottingham | COVID-19 Genomics UK (COG-UK) Consortium | Gemma Clark, Wendy Smith, Manjinder Khakh, Vicki M Fleming, Michelle M Lister, Hannah Howson-Wells, Jonathan Ball, Patrick McClure, Joseph Chappell, Theocharis Tsoleridis, Nadine Holmes, Matthew Carlisle, Christopher Moore, Fei Sang, Johnny Debebe, Victoria Wright, Matthew Loose |
| EPI_ISL_514457, EPI_ISL_514458, EPI_ISL_514459, EPI_ISL_514460, EPI_ISL_514461, EPI_ISL_514462, EPI_ISL_514463, EPI_ISL_514465, EPI_ISL_514466, EPI_ISL_514467, EPI_ISL_514468, EPI_ISL_514469, EPI_ISL_514472, EPI_ISL_514473, EPI_ISL_514474, EPI_ISL_514475, EPI_ISL_514476, EPI_ISL_514477, EPI_ISL_514478, EPI_ISL_514479, EPI_ISL_514480, EPI_ISL_514482, EPI_ISL_514484, EPI_ISL_514485, EPI_ISL_514486, EPI_ISL_514490, EPI_ISL_514505, EPI_ISL_514506, EPI_ISL_514508, EPI_ISL_514510 | | | |
| see above | Centre for Enzyme Innovation, University of Portsmouth / Translational Research Laboratory, Portsmouth Hospitals NHS Trust | COVID-19 Genomics UK (COG-UK) Consortium | Angela Beckett,Yann Bourgeois,Garry Scarlett,Sharon Glaysher,Scott Elliott,Kelly Bicknell,Robert Impey,Allyson Lloyd,Sarah Wyllie,Ethan Butcher,Anoop Chauhan,Samuel Robson |
| EPI_ISL_514514, EPI_ISL_514515 | West of Scotland Specialist Virology Centre, NHSGGC / MRC-University of Glasgow Centre for Virus Research | COVID-19 Genomics UK (COG-UK) Consortium | Ana da Silva Filipe, Natasha Johnson, Kathy Smollett, Daniel Mair, Stephen Carmichael, Lily Tong, Jenna Nichols, Elihu Aranday-Cortes, Kirstyn Brunker, Yasmin Parr, Alice Broos, Kyriaki Nomikou; Sarah McDonald, Marc Niebel, Patawee Asamaphan; Richard Orton, Joseph Hughes, Sreenu Vattipally, David L Robertson; Alasdair MacLean, Rory Gunson; Kathy Li, Natasha Jesudason, Rajiv Shah, James Shepherd, Antonia Ho, Emma Thomson |
| EPI_ISL_514516, EPI_ISL_514517, EPI_ISL_514518, EPI_ISL_514519, EPI_ISL_514520, EPI_ISL_514521, EPI_ISL_514522, EPI_ISL_514523, EPI_ISL_514524, EPI_ISL_514526, EPI_ISL_514527, EPI_ISL_514528, EPI_ISL_514529, EPI_ISL_514530, EPI_ISL_514531, EPI_ISL_514533, EPI_ISL_514534, EPI_ISL_514535, EPI_ISL_514536, EPI_ISL_514537, EPI_ISL_514539, EPI_ISL_514540, EPI_ISL_514541, EPI_ISL_514542, EPI_ISL_514543, EPI_ISL_514544, EPI_ISL_514545, EPI_ISL_514546, EPI_ISL_514547, EPI_ISL_514548, EPI_ISL_514549, EPI_ISL_514550, EPI_ISL_514551, EPI_ISL_514552, EPI_ISL_514553, EPI_ISL_514554, EPI_ISL_514555, EPI_ISL_514557, EPI_ISL_514558, EPI_ISL_514559, EPI_ISL_514560, EPI_ISL_514561, EPI_ISL_514562 | | | |
| see above | Virology Department, Royal Infirmary of Edinburgh, | COVID-19 Genomics UK (COG-UK) Consortium | McHugh M, Dewar R, Rooke S, Gallagher M, Balcaza C, O'Toole Á, Scher E, Hill V, McCrone JT, Colquhoun R, Yu X, Jackson B, Rambaut A, Williams TC, Templeton K |

| | | | |
|---|---|---|---|
| | NHS Lothian / School of Biological Sciences, University of Edinburgh / Institute of Genetics and Molecular Medicine, University of Edinburgh | | |
| EPI_ISL_514563, EPI_ISL_514564, EPI_ISL_514565, EPI_ISL_514566, EPI_ISL_514567, EPI_ISL_514568, EPI_ISL_514569, EPI_ISL_514570, EPI_ISL_514571, EPI_ISL_514572, EPI_ISL_514573, EPI_ISL_514574, EPI_ISL_514575, EPI_ISL_514576, EPI_ISL_514577, EPI_ISL_514578, EPI_ISL_514579 | | | |
| see above | Wales Specialist Virology Centre Sequencing lab: Pathogen Genomics Unit | COVID-19 Genomics UK (COG-UK) Consortium | Catherine Moore, Johnathan Evans, Laura Gifford, Malorie Perry, Simon Cottrell, Angela Marchbank, Alec Birchley, Alexander Adams, Amy Gaskin, Bree Gatica-Wilcox, Jason Coombes, Joel Southgate, Lauren Gilbert, Lee Graham, Nicole Pacchiarini, Sara Kumziene-Summerhayes, Sarah Taylor, Sophie Jones, Sara Rey, Matthew Bull, Joanne Watkins, Sally Corden, Tom Connor |
| EPI_ISL_514580 | NSTU COVID-19 Diagnostic Center | NSU Genome Research Institute (NGRI), North South University | Dr. Muhammad Maqsud Hossain, Aura Rahman, Prof. Firoz Ahmed, Tahrima Huq, Abdus Sadique, Jahidul Alam, Tamanna Afroze, Md Aminul Islam, Prof. Md. Didar-Ul-Alam, Prof. Kazi Nadim Hasan, Prof. Abdul Khaleque, Prof, Hasan Mahmud Reza |
| EPI_ISL_514581 | B.J. Medical College and Civil hospital, Ahmedabad | Gujarat Biotechnology Research Centre | Nitin Savaliya, Raghawendra Kumar, Dinesh Kumar, Zuber Saiyed, Komal Patel, Labdhi Pandya, Afzal Ansari, Nikha Trivedi, Pranay Shah, Kamlesh J Upadhyay, Sanjay Kapadia, Apurvasinh Puvar, Janvi Raval, Zarna Patel, Monika Gandhi, Pinal Trivedi, Maharshi Pandya, Nidhi Patel, R D Dixit, A M Kadri, Harsh Bakshi, Chaitanya Joshi, Madhvi Joshi |
| EPI_ISL_514582 | B.J. Medical College and Civil hospital, Ahmedabad | Gujarat Biotechnology Research Centre | Raghawendra Kumar, Dinesh Kumar, Zuber Saiyed, Komal Patel, Labdhi Pandya, Afzal Ansari, Nikha Trivedi, Pranay Shah, Kamlesh J Upadhyay, Sanjay Kapadia, Apurvasinh Puvar, Janvi Raval, Zarna Patel, Monika Gandhi, Pinal Trivedi, Maharshi Pandya, Nidhi Patel, Nitin Savaliya, R D Dixit, A M Kadri, Harsh Bakshi, Chaitanya Joshi, Madhvi Joshi |
| EPI_ISL_514583 | B.J. Medical College and Civil hospital, Ahmedabad | Gujarat Biotechnology Research Centre | Dinesh Kumar, Zuber Saiyed, Komal Patel, Labdhi Pandya, Afzal Ansari, Nikha Trivedi, Pranay Shah, Kamlesh J Upadhyay, Sanjay Kapadia, Apurvasinh Puvar, Janvi Raval, Zarna Patel, Monika Gandhi, Pinal Trivedi, Maharshi Pandya, Nidhi Patel, Nitin Savaliya, Raghawendra Kumar, R D Dixit, A M Kadri, Harsh Bakshi, Chaitanya Joshi, Madhvi Joshi |
| EPI_ISL_514584 | B.J. Medical College and Civil hospital, Ahmedabad | Gujarat Biotechnology Research Centre | Zuber Saiyed, Komal Patel, Labdhi Pandya, Afzal Ansari, Nikha Trivedi, Pranay Shah, Kamlesh J Upadhyay, Sanjay Kapadia, Apurvasinh Puvar, Janvi Raval, Zarna Patel, Monika Gandhi, Pinal Trivedi, Maharshi Pandya, Nidhi Patel, Nitin Savaliya, Raghawendra Kumar, Dinesh Kumar, R D Dixit, A M Kadri, Harsh Bakshi, Chaitanya Joshi, Madhvi Joshi |
| EPI_ISL_514585 | B.J. Medical College and Civil hospital, Ahmedabad | Gujarat Biotechnology Research Centre | Komal Patel, Labdhi Pandya, Afzal Ansari, Nikha Trivedi, Pranay Shah, Kamlesh J Upadhyay, Sanjay Kapadia, Apurvasinh Puvar, Janvi Raval, Zarna Patel, Monika Gandhi, Pinal Trivedi, Maharshi Pandya, Nidhi Patel, Nitin Savaliya, Raghawendra Kumar, Dinesh Kumar, Zuber Saiyed, R D Dixit, A M Kadri, Harsh Bakshi, Chaitanya Joshi, Madhvi Joshi |
| EPI_ISL_514586 | B.J. Medical College and Civil hospital, Ahmedabad | Gujarat Biotechnology Research Centre | Labdhi Pandya, Afzal Ansari, Nikha Trivedi, Pranay Shah, Kamlesh J Upadhyay, Sanjay Kapadia, Apurvasinh Puvar, Janvi Raval, Zarna Patel, Monika Gandhi, Pinal Trivedi, Maharshi Pandya, Nidhi Patel, Nitin Savaliya, Raghawendra Kumar, Dinesh Kumar, Zuber Saiyed, Komal Patel, R D Dixit, A M Kadri, Harsh Bakshi, Chaitanya Joshi, Madhvi Joshi |
| EPI_ISL_514587 | B.J. Medical College and Civil hospital, Ahmedabad | Gujarat Biotechnology Research Centre | Afzal Ansari, Nikha Trivedi, Pranay Shah, Kamlesh J Upadhyay, Sanjay Kapadia, Apurvasinh Puvar, Janvi Raval, Zarna Patel, Monika Gandhi, Pinal Trivedi, Maharshi Pandya, Nidhi Patel, Nitin Savaliya, Raghawendra Kumar, Dinesh Kumar, Zuber Saiyed, Komal Patel, Labdhi Pandya, R D Dixit, A M Kadri, Harsh Bakshi, Chaitanya Joshi, Madhvi Joshi |
| EPI_ISL_514588 | B.J. Medical College and Civil hospital, Ahmedabad | Gujarat Biotechnology Research Centre | Nikha Trivedi, Pranay Shah, Kamlesh J Upadhyay, Sanjay Kapadia, Apurvasinh Puvar, Janvi Raval, Zarna Patel, Monika Gandhi, Pinal Trivedi, Maharshi Pandya, Nidhi Patel, Nitin Savaliya, Raghawendra Kumar, Dinesh Kumar, Zuber Saiyed, Komal Patel, Labdhi Pandya, Afzal Ansari, R D Dixit, A M Kadri, Harsh Bakshi, Chaitanya Joshi, Madhvi Joshi |
| EPI_ISL_514589 | B.J. Medical College and Civil hospital, Ahmedabad | Gujarat Biotechnology Research Centre | Pranay Shah, Kamlesh J Upadhyay, Sanjay Kapadia, Apurvasinh Puvar, Janvi Raval, Zarna Patel, Monika Gandhi, Pinal Trivedi, Maharshi Pandya, Nidhi Patel, Nitin Savaliya, Raghawendra Kumar, Dinesh Kumar, Zuber Saiyed, Komal Patel, Labdhi Pandya, Afzal Ansari, Nikha Trivedi, R D Dixit, A M Kadri, Harsh Bakshi, Chaitanya Joshi, Madhvi Joshi |
| EPI_ISL_514590 | B.J. Medical College and Civil hospital, Ahmedabad | Gujarat Biotechnology Research Centre | Kamlesh J Upadhyay, Sanjay Kapadia, Apurvasinh Puvar, Janvi Raval, Zarna Patel, Monika Gandhi, Pinal Trivedi, Maharshi Pandya, Nidhi Patel, Nitin Savaliya, Raghawendra Kumar, Dinesh Kumar, Zuber Saiyed, Komal Patel, Labdhi Pandya, Afzal Ansari, Nikha Trivedi, Pranay Shah, R D Dixit, A M Kadri, Harsh Bakshi, Chaitanya Joshi, Madhvi Joshi |
| EPI_ISL_514591 | B.J. Medical College and Civil hospital, Ahmedabad | Gujarat Biotechnology Research Centre | Sanjay Kapadia,Apurvasinh Puvar,Janvi Raval,Zarna Patel, Monika Gandhi, Pinal Trivedi, Maharshi Pandya, Nidhi Patel, Nitin Savaliya, Raghawendra Kumar, Dinesh Kumar, Zuber Saiyed, Komal Patel, Labdhi Pandya, Afzal Ansari, Nikha Trivedi, Pranay Shah, Kamlesh J Upadhyay, R D Dixit, A M Kadri, Harsh Bakshi, Chaitanya Joshi, Madhvi Joshi |
| EPI_ISL_514592 | B.J. Medical College and Civil hospital, Ahmedabad | Gujarat Biotechnology Research Centre | Apurvasinh Puvar, Janvi Raval, Zarna Patel, Monika Gandhi, Pinal Trivedi, Maharshi Pandya, Nidhi Patel, Nitin Savaliya, Raghawendra Kumar, Dinesh Kumar, Zuber Saiyed, Komal Patel, Labdhi Pandya, Afzal Ansari, Nikha Trivedi, Pranay Shah, Kamlesh J Upadhyay, Sanjay Kapadia, R D Dixit, A M Kadri, Harsh Bakshi, Chaitanya Joshi, Madhvi Joshi |
| EPI_ISL_514593 | B.J. Medical College and Civil hospital, Ahmedabad | Gujarat Biotechnology Research Centre | Janvi Raval, Zarna Patel, Monika Gandhi, Pinal Trivedi, Maharshi Pandya, Nidhi Patel, Nitin Savaliya, Raghawendra Kumar, Dinesh Kumar, Zuber Saiyed, Komal Patel, Labdhi Pandya, Afzal Ansari, Nikha Trivedi, Pranay Shah, Kamlesh J Upadhyay, Sanjay Kapadia, Apurvasinh Puvar, R D Dixit, A M Kadri, Harsh Bakshi, Chaitanya Joshi, Madhvi Joshi |
| EPI_ISL_514594 | B.J. Medical College and Civil hospital, Ahmedabad | Gujarat Biotechnology Research Centre | Zarna Patel, Monika Gandhi, Pinal Trivedi, Maharshi Pandya, Nidhi Patel, Nitin Savaliya, Raghawendra Kumar, Dinesh Kumar, Zuber Saiyed, Komal Patel, Labdhi Pandya, Afzal Ansari, Nikha Trivedi, Pranay Shah, Kamlesh J Upadhyay, Sanjay Kapadia, Apurvasinh Puvar, Janvi Raval, R D Dixit, A M Kadri, Harsh Bakshi, Chaitanya Joshi, Madhvi Joshi |
| EPI_ISL_514595 | B.J. Medical College and Civil hospital, Ahmedabad | Gujarat Biotechnology Research Centre | Monika Gandhi, Pinal Trivedi, Maharshi Pandya, Nidhi Patel, Nitin Savaliya, Raghawendra Kumar, Dinesh Kumar, Zuber Saiyed, Komal Patel, Labdhi Pandya, Afzal Ansari, Nikha Trivedi, Pranay Shah, Kamlesh J Upadhyay, Sanjay Kapadia, Apurvasinh Puvar, Janvi Raval, Zarna Patel, R D Dixit, A M Kadri, Harsh Bakshi, Chaitanya Joshi, Madhvi Joshi |
| EPI_ISL_514596 | B.J. Medical College and Civil hospital, Ahmedabad | Gujarat Biotechnology Research Centre | Pinal Trivedi, Maharshi Pandya, Nidhi Patel, Nitin Savaliya, Raghawendra Kumar, Dinesh Kumar, Zuber Saiyed, Komal Patel, Labdhi Pandya, Afzal Ansari, Nikha Trivedi, Pranay Shah, Kamlesh J Upadhyay, Sanjay Kapadia, Apurvasinh Puvar, Janvi Raval, Zarna Patel, Monika Gandhi, R D Dixit, A M Kadri, Harsh Bakshi, Chaitanya Joshi, Madhvi Joshi |
| EPI_ISL_514597 | B.J. Medical College and Civil hospital, Ahmedabad | Gujarat Biotechnology Research Centre | Maharshi Pandya, Nidhi Patel, Nitin Savaliya, Raghawendra Kumar, Dinesh Kumar, Zuber Saiyed, Komal Patel, Labdhi Pandya, Afzal Ansari, Nikha Trivedi, Pranay Shah, Kamlesh J Upadhyay, Sanjay Kapadia, Apurvasinh Puvar, Janvi Raval, Zarna Patel, Monika Gandhi, Pinal Trivedi, R D Dixit, A M Kadri, Harsh Bakshi, Chaitanya Joshi, Madhvi Joshi |
| EPI_ISL_514598 | B.J. Medical College and Civil hospital, Ahmedabad | Gujarat Biotechnology Research Centre | Nidhi Patel, Nitin Savaliya, Raghawendra Kumar, Dinesh Kumar, Zuber Saiyed, Komal Patel, Labdhi Pandya, Afzal Ansari, Nikha Trivedi, Pranay Shah, Kamlesh J Upadhyay, Sanjay Kapadia, Apurvasinh Puvar, Janvi Raval, Zarna Patel, Monika Gandhi, Pinal Trivedi, Maharshi Pandya, R D Dixit, A M Kadri, Harsh Bakshi, Chaitanya Joshi, Madhvi Joshi |
| EPI_ISL_514599 | B.J. Medical College and Civil hospital, Ahmedabad | Gujarat Biotechnology Research Centre | Nitin Savaliya, Raghawendra Kumar, Dinesh Kumar, Zuber Saiyed, Komal Patel, Labdhi Pandya, Afzal Ansari, Nikha Trivedi, Pranay Shah, Kamlesh J Upadhyay, Sanjay Kapadia, Apurvasinh Puvar, Janvi Raval, Zarna Patel, Monika Gandhi, Pinal Trivedi, Maharshi Pandya, Nidhi Patel, R D Dixit, A M Kadri, Harsh Bakshi, Chaitanya Joshi, Madhvi Joshi |
| EPI_ISL_514600 | B.J. Medical College and Civil hospital, Ahmedabad | Gujarat Biotechnology Research Centre | Raghawendra Kumar, Dinesh Kumar, Zuber Saiyed, Komal Patel, Labdhi Pandya, Afzal Ansari, Nikha Trivedi, Pranay Shah, Kamlesh J Upadhyay, Sanjay Kapadia, Apurvasinh Puvar, Janvi Raval, Zarna Patel, Monika Gandhi, Pinal Trivedi, Maharshi Pandya, Nidhi Patel, Nitin Savaliya, R D Dixit, A M Kadri, Harsh Bakshi, Chaitanya Joshi, Madhvi Joshi |
| EPI_ISL_514601 | B.J. Medical College and Civil hospital, Ahmedabad | Gujarat Biotechnology Research Centre | Dinesh Kumar, Zuber Saiyed, Komal Patel, Labdhi Pandya, Afzal Ansari, Nikha Trivedi, Pranay Shah, Kamlesh J Upadhyay, Sanjay Kapadia, Apurvasinh Puvar, Janvi Raval, Zarna Patel, Monika Gandhi, Pinal Trivedi, Maharshi Pandya, Nidhi Patel, Nitin Savaliya, Raghawendra Kumar, R D Dixit, A M Kadri, Harsh Bakshi, Chaitanya Joshi, Madhvi Joshi |
| EPI_ISL_514602 | B.J. Medical College and Civil hospital, Ahmedabad | Gujarat Biotechnology Research Centre | Zuber Saiyed, Komal Patel, Labdhi Pandya, Afzal Ansari, Nikha Trivedi, Pranay Shah, Kamlesh J Upadhyay, Sanjay Kapadia, Apurvasinh Puvar, Janvi Raval, Zarna Patel, Monika Gandhi, Pinal Trivedi, Maharshi Pandya, Nidhi Patel, Nitin Savaliya, Raghawendra Kumar, Dinesh Kumar, R D Dixit, A M Kadri, Harsh Bakshi, Chaitanya Joshi, Madhvi Joshi |
| EPI_ISL_514603 | B.J. Medical College and Civil hospital, Ahmedabad | Gujarat Biotechnology Research Centre | Komal Patel, Labdhi Pandya, Afzal Ansari, Nikha Trivedi, Pranay Shah, Kamlesh J Upadhyay, Sanjay Kapadia, Apurvasinh Puvar, Janvi Raval, Zarna Patel, Monika Gandhi, Pinal Trivedi, Maharshi Pandya, Nidhi Patel, Nitin Savaliya, Raghawendra Kumar, Dinesh Kumar, Zuber Saiyed, R D Dixit, A M Kadri, Harsh Bakshi, Chaitanya Joshi, Madhvi Joshi |
| EPI_ISL_514604 | B.J. Medical College and Civil hospital, Ahmedabad | Gujarat Biotechnology Research Centre | Labdhi Pandya, Afzal Ansari, Nikha Trivedi, Pranay Shah, Kamlesh J Upadhyay, Sanjay Kapadia, Apurvasinh Puvar, Janvi Raval, Zarna Patel, Monika Gandhi, Pinal Trivedi, Maharshi Pandya, Nidhi Patel, Nitin Savaliya, Raghawendra Kumar, Dinesh Kumar, Zuber Saiyed, Komal Patel, R D Dixit, A M Kadri, Harsh Bakshi, Chaitanya Joshi, Madhvi Joshi |
| EPI_ISL_514605 | B.J. Medical College and Civil hospital, Ahmedabad | Gujarat Biotechnology Research Centre | Afzal Ansari, Nikha Trivedi, Pranay Shah, Kamlesh J Upadhyay, Sanjay Kapadia, Apurvasinh Puvar, Janvi Raval, Zarna Patel, Monika Gandhi, Pinal Trivedi, Maharshi Pandya, Nidhi Patel, Nitin Savaliya, Raghawendra Kumar, Dinesh Kumar, Zuber Saiyed, Komal Patel, Labdhi Pandya, R D Dixit, A M Kadri, Harsh Bakshi, Chaitanya Joshi, Madhvi Joshi |
| EPI_ISL_514606 | B.J. Medical College and Civil hospital, Ahmedabad | Gujarat Biotechnology Research Centre | Nikha Trivedi, Pranay Shah, Kamlesh J Upadhyay, Sanjay Kapadia, Apurvasinh Puvar, Janvi Raval, Zarna Patel, Monika Gandhi, Pinal Trivedi, Maharshi Pandya, Nidhi Patel, Nitin Savaliya, Raghawendra Kumar, Dinesh Kumar, Zuber Saiyed, Komal Patel, Labdhi Pandya, Afzal Ansari, R D Dixit, A M Kadri, Harsh Bakshi, Chaitanya Joshi, Madhvi Joshi |
| EPI_ISL_514607 | B.J. Medical College and Civil hospital, Ahmedabad | Gujarat Biotechnology Research Centre | Pranay Shah, Kamlesh J Upadhyay, Sanjay Kapadia, Apurvasinh Puvar, Janvi Raval, Zarna Patel, Monika Gandhi, Pinal Trivedi, Maharshi Pandya, Nidhi Patel, Nitin Savaliya, Raghawendra Kumar, Dinesh Kumar, Zuber Saiyed, Komal Patel, Labdhi Pandya, Afzal Ansari, Nikha Trivedi, R D Dixit, A M Kadri, Harsh Bakshi, Chaitanya Joshi, Madhvi Joshi |
| EPI_ISL_514608 | B.J. Medical College and Civil hospital, Ahmedabad | Gujarat Biotechnology Research Centre | Kamlesh J Upadhyay, Sanjay Kapadia, Apurvasinh Puvar, Janvi Raval, Zarna Patel, Monika Gandhi, Pinal Trivedi, Maharshi Pandya, Nidhi Patel, Nitin Savaliya, Raghawendra Kumar, Dinesh Kumar, Zuber Saiyed, Komal Patel, Labdhi Pandya, Afzal Ansari, Nikha Trivedi, Pranay Shah, R D Dixit, A M Kadri, Harsh Bakshi, Chaitanya Joshi, Madhvi Joshi |
| EPI_ISL_514609 | B.J. Medical College and Civil hospital, Ahmedabad | Gujarat Biotechnology Research Centre | Sanjay Kapadia, Apurvasinh Puvar, Janvi Raval, Zarna Patel, Monika Gandhi, Pinal Trivedi, Maharshi Pandya, Nidhi Patel, Nitin Savaliya, Raghawendra Kumar, Dinesh Kumar, Zuber Saiyed, Komal Patel, Labdhi Pandya, Afzal Ansari, Nikha Trivedi, Pranay Shah, Kamlesh J Upadhyay, R D Dixit, A M Kadri, Harsh Bakshi, Chaitanya Joshi, Madhvi Joshi |
| EPI_ISL_514610 | GMERS Medical College & Hospital, Gotri, Vadodara | Gujarat Biotechnology Research Centre | Meenakshi Shah, Neena Doshi, Varsha Godbole, Apurvasinh Puvar, Janvi Raval, Zarna Patel, Monika Gandhi, Pinal Trivedi, Maharshi Pandya, Nidhi Patel, Nitin Savaliya, Raghawendra Kumar, Dinesh Kumar, Zuber Saiyed, Komal Patel, Labdhi Pandya, Afzal Ansari, Nikha Trivedi, Pranay Shah, Kamlesh J Upadhyay, Sanjay Kapadia, R D Dixit, A M Kadri, Harsh Bakshi, Chaitanya Joshi, Madhvi Joshi, |
| EPI_ISL_514611 | GMERS Medical College and Hospital, Gandhinagar | Gujarat Biotechnology Research Centre | Seema Bhatt, Gaurishankar Shrimali, Bhavesh Modi, Bharti Rajani, Apurvasinh Puvar, Janvi Raval, Zarna Patel, Monika Gandhi, Pinal Trivedi, Maharshi Pandya, Nidhi Patel, Nitin Savaliya, Raghawendra Kumar, Dinesh Kumar, Zuber Saiyed, Komal Patel, Labdhi Pandya, Afzal Ansari, Nikha Trivedi, Pranay Shah, Kamlesh J Upadhyay, Sanjay Kapadia, R D Dixit, A M Kadri, Harsh Bakshi, Chaitanya Joshi, Madhvi Joshi |
| EPI_ISL_514612 | GMERS Medical College and Hospital, Gandhinagar | Gujarat Biotechnology Research Centre | Gaurishankar Shrimali, Bhavesh Modi, Bharti Rajani, Apurvasinh Puvar, Janvi Raval, Zarna Patel, Monika Gandhi, Pinal Trivedi, Maharshi Pandya, Nidhi Patel, Nitin Savaliya, Raghawendra Kumar, Dinesh Kumar, Zuber Saiyed, Komal Patel, Labdhi Pandya, Afzal Ansari, Nikha Trivedi, Pranay Shah, Kamlesh J Upadhyay, Sanjay Kapadia, Seema Bhatt, R D Dixit, A M Kadri, Harsh Bakshi, Chaitanya Joshi, Madhvi Joshi |
| EPI_ISL_514613 | NSTU COVID-19 Diagnostic Center | NSU Genome Research Institute (NGRI), North South University | Dr. Muhammad Maqsud Hossain, Aura Rahman, Prof. Firoz Ahmed, Tahrima Huq, Abdus Sadique, Jahidul Alam, Tamanna Afroze, Md Aminul Islam, Prof. Md. Didar-Ul-Alam, Prof. Kazi Nadim Hasan, Prof. Abdul Khaleque, Prof, Hasan Mahmud Reza |
| EPI_ISL_514615 | NSTU COVID-19 Diagnostic Center | NSU Genome Research Institute (NGRI), North South University | Dr. Muhammad Maqsud Hossain, Aura Rahman, Prof. Firoz Ahmed, Tahrima Huq, Abdus Sadique, Jahidul Alam, Tamanna Afroze, Md Aminul Islam, Prof. Md. Didar-Ul-Alam, Prof. Kazi Nadim Hasan, Prof. Abdul Khaleque, Prof, Hasan Mahmud Reza |
| EPI_ISL_514616, EPI_ISL_514617, EPI_ISL_514618 | Minnesota Department of Health, Public Health Laboratory | Minnesota Department of Health, Public Health Laboratory | Matt Plumb, Jacob Garfin, and Xiong Wang |
| EPI_ISL_514619, EPI_ISL_514620, EPI_ISL_514621, EPI_ISL_514622, EPI_ISL_514623, EPI_ISL_514624, EPI_ISL_514625, EPI_ISL_514626, EPI_ISL_514627, EPI_ISL_514628 | Mayo Clinic & Mayo Clinic Laboratories | Minnesota Department of Health, Public Health Laboratory | Matt Plumb, Jacob Garfin, and Xiong Wang |
| EPI_ISL_514629, EPI_ISL_514630, EPI_ISL_514631, EPI_ISL_514632, EPI_ISL_514633, EPI_ISL_514634, EPI_ISL_514635 | M Health Fairview | Minnesota Department of Health, Public Health Laboratory | Matt Plumb, Jacob Garfin, and Xiong Wang |
| EPI_ISL_514636 | Mayo Clinic & Mayo Clinic Laboratories | Minnesota Department of Health, Public Health Laboratory | Matt Plumb, Jacob Garfin, and Xiong Wang |
| EPI_ISL_514637 | M Health Fairview St. Joseph's Hospital | Minnesota Department of Health, Public Health Laboratory | Matt Plumb, Jacob Garfin, and Xiong Wang |

| | | | |
|---|---|---|---|
| EPI_ISL_514638, EPI_ISL_514639, EPI_ISL_514640 | Mayo Clinic & Mayo Clinic Laboratories | Minnesota Department of Health, Public Health Laboratory | Matt Plumb, Jacob Garfin, and Xiong Wang |
| EPI_ISL_514641 | St. Luke's Hospital | Minnesota Department of Health, Public Health Laboratory | Matt Plumb, Jacob Garfin, and Xiong Wang |
| EPI_ISL_514642 | Essentia Health-St. Mary's Medical Center | Minnesota Department of Health, Public Health Laboratory | Matt Plumb, Jacob Garfin, and Xiong Wang |
| EPI_ISL_514643 | M Health Fairview St. Joseph's Hospital | Minnesota Department of Health, Public Health Laboratory | Matt Plumb, Jacob Garfin, and Xiong Wang |
| EPI_ISL_514644 | Mayo Clinic & Mayo Clinic Laboratories | Minnesota Department of Health, Public Health Laboratory | Matt Plumb, Jacob Garfin, and Xiong Wang |
| EPI_ISL_514645 | Avera Mckennan Laboratory | Minnesota Department of Health, Public Health Laboratory | Matt Plumb, Jacob Garfin, and Xiong Wang |
| EPI_ISL_514646 | St. Luke's Hospital | Minnesota Department of Health, Public Health Laboratory | Matt Plumb, Jacob Garfin, and Xiong Wang |
| EPI_ISL_514647 | Allina Health Laboratory | Minnesota Department of Health, Public Health Laboratory | Matt Plumb, Jacob Garfin, and Xiong Wang |
| EPI_ISL_514648 | Essentia Health-St. Joseph's Medical Center | Minnesota Department of Health, Public Health Laboratory | Matt Plumb, Jacob Garfin, and Xiong Wang |
| EPI_ISL_514649 | Allina Health Laboratory | Minnesota Department of Health, Public Health Laboratory | Matt Plumb, Jacob Garfin, and Xiong Wang |
| EPI_ISL_514650 | M Health Fairview St. Joseph's Hospital | Minnesota Department of Health, Public Health Laboratory | Matt Plumb, Jacob Garfin, and Xiong Wang |
| EPI_ISL_514651 | Allina Health Laboratory | Minnesota Department of Health, Public Health Laboratory | Matt Plumb, Jacob Garfin, and Xiong Wang |
| EPI_ISL_514652 | M Health Fairview St. Joseph's Hospital | Minnesota Department of Health, Public Health Laboratory | Matt Plumb, Jacob Garfin, and Xiong Wang |
| EPI_ISL_514653, EPI_ISL_514654, EPI_ISL_514655, EPI_ISL_514656 | Mayo Clinic & Mayo Clinic Laboratories | Minnesota Department of Health, Public Health Laboratory | Matt Plumb, Jacob Garfin, and Xiong Wang |
| EPI_ISL_514657 | M Health Fairview St. Joseph's Hospital | Minnesota Department of Health, Public Health Laboratory | Matt Plumb, Jacob Garfin, and Xiong Wang |
| EPI_ISL_514658, EPI_ISL_514659, EPI_ISL_514660, EPI_ISL_514661, EPI_ISL_514662, EPI_ISL_514663, EPI_ISL_514664, EPI_ISL_514665, EPI_ISL_514666, EPI_ISL_514667, EPI_ISL_514668, EPI_ISL_514669, EPI_ISL_514670, EPI_ISL_514671, EPI_ISL_514672 | | | |
| see above | Minnesota Department of Health, Public Health Laboratory | Minnesota Department of Health, Public Health Laboratory | Matt Plumb, Jacob Garfin, and Xiong Wang |
| EPI_ISL_514673 | Nevada State Public Health Laboratory | Nevada State Public Health Laboratory | Richard Tillet, Joel R. Sevinsky, Paul Hartley, Heather Kerwin, David Jackson, Subhash C. Verma, Cyprian Rosetto, Andrew Gorzalski, Chris Laverdure, Natalie Crawford, Stephanie Van Hooser, and Mark Pandori |
| EPI_ISL_514750 | Pirogov Russian National Research Medical University | Pirogov Russian National Research Medical University, Research and Development | Blagodatskikh, K.A. |
| EPI_ISL_514751 | CoronaNet Lab- TaskForce Regione Campania, CEINGE Biotecnologie Avanzate, Via G. Salvatore | CoronaNet Lab- TaskForce Regione Campania, CEINGE Biotecnologie Avanzate, Via G. Salvatore | Zollo,M., Ferrucci,V., Kong,Dy., Asadzadeh,F., Marrone,L.,Siciliano,R., Cerino,R., Fusco,G., Comegna,M., Boccia,A.,Viscardi,M., Borriello,G., Brandi,S., Tiberio,C., Atripaldi,L.,Paolella,G., Castaldo,G., Pascarella,S., Bianchi,M., Chiariotti,L.,Lee,J.M., Jung,J.H., Yun,K.S. and Kim,H.Y. |
| EPI_ISL_514753 | Yaftabad Hospital, COVID Lab Center | University of Tabriz | Shahabzadeh,Z., Hosseinzadeh Gharajeh,N., Hashemian,S.M. and Barati,O. |
| EPI_ISL_514756, EPI_ISL_514757, EPI_ISL_514758, EPI_ISL_514759, EPI_ISL_514760, EPI_ISL_514761, EPI_ISL_514762, EPI_ISL_514763, EPI_ISL_514764, EPI_ISL_514765, EPI_ISL_514766, EPI_ISL_514767, EPI_ISL_514768, EPI_ISL_514769, EPI_ISL_514770, EPI_ISL_514771, EPI_ISL_514772, EPI_ISL_514773, EPI_ISL_514774, EPI_ISL_514775, EPI_ISL_514776, EPI_ISL_514777, EPI_ISL_514778, EPI_ISL_514779, EPI_ISL_514780, EPI_ISL_514781, EPI_ISL_514782, EPI_ISL_514783, EPI_ISL_514784, EPI_ISL_514785, EPI_ISL_514786, EPI_ISL_514787, EPI_ISL_514788, EPI_ISL_514789, EPI_ISL_514790, EPI_ISL_514791, EPI_ISL_514792, EPI_ISL_514793, EPI_ISL_514794, EPI_ISL_514795, EPI_ISL_514796, EPI_ISL_514797, EPI_ISL_514798, EPI_ISL_514799, EPI_ISL_514800, EPI_ISL_514801, EPI_ISL_514802, EPI_ISL_514803, EPI_ISL_514804, EPI_ISL_514805, EPI_ISL_514806, EPI_ISL_514807, EPI_ISL_514808, EPI_ISL_514809, EPI_ISL_514810, EPI_ISL_514811, EPI_ISL_514812, EPI_ISL_514813, EPI_ISL_514814, EPI_ISL_514815, EPI_ISL_514816, EPI_ISL_514817, EPI_ISL_514818, EPI_ISL_514819, EPI_ISL_514820, EPI_ISL_514821, EPI_ISL_514822, EPI_ISL_514823, EPI_ISL_514824, EPI_ISL_514825, EPI_ISL_514826, EPI_ISL_514827, EPI_ISL_514828, EPI_ISL_514829, EPI_ISL_514830, EPI_ISL_514831, EPI_ISL_514832, EPI_ISL_514833, EPI_ISL_514834, EPI_ISL_514835, EPI_ISL_514836, EPI_ISL_514837, EPI_ISL_514838, EPI_ISL_514839, EPI_ISL_514840, EPI_ISL_514841, EPI_ISL_514842, EPI_ISL_514843, EPI_ISL_514844, EPI_ISL_514845, EPI_ISL_514846, EPI_ISL_514847, EPI_ISL_514848, EPI_ISL_514849, EPI_ISL_514850, EPI_ISL_514851, EPI_ISL_514852, EPI_ISL_514853, EPI_ISL_514854, EPI_ISL_514855, EPI_ISL_514856, EPI_ISL_514857, EPI_ISL_514858, EPI_ISL_514859, EPI_ISL_514860, EPI_ISL_514861, EPI_ISL_514862, EPI_ISL_514863, EPI_ISL_514864, EPI_ISL_514865, EPI_ISL_514866, EPI_ISL_514867, EPI_ISL_514868, EPI_ISL_514869, EPI_ISL_514870, EPI_ISL_514871, EPI_ISL_514872, EPI_ISL_514873, EPI_ISL_514874, EPI_ISL_514875, EPI_ISL_514876, EPI_ISL_514877, EPI_ISL_514878, EPI_ISL_514879, EPI_ISL_514880, EPI_ISL_514881, EPI_ISL_514882, EPI_ISL_514883, EPI_ISL_514884, EPI_ISL_514885, EPI_ISL_514886, EPI_ISL_514887, EPI_ISL_514888, EPI_ISL_514889, EPI_ISL_514890, EPI_ISL_514891, EPI_ISL_514892, EPI_ISL_514893, EPI_ISL_514894, EPI_ISL_514895, EPI_ISL_514896, EPI_ISL_514897, EPI_ISL_514898, EPI_ISL_514899, EPI_ISL_514900, EPI_ISL_514901, EPI_ISL_514902, EPI_ISL_514903, EPI_ISL_514904, EPI_ISL_514905, EPI_ISL_514906, EPI_ISL_514907, EPI_ISL_514908, EPI_ISL_514909, EPI_ISL_514910, EPI_ISL_514911, EPI_ISL_514912, EPI_ISL_514913, EPI_ISL_514914, EPI_ISL_514915, EPI_ISL_514916, EPI_ISL_514917, EPI_ISL_514918, EPI_ISL_514919, EPI_ISL_514920, EPI_ISL_514921, EPI_ISL_514922, EPI_ISL_514923, EPI_ISL_514924, EPI_ISL_514925, EPI_ISL_514926, EPI_ISL_514927, EPI_ISL_514928, EPI_ISL_514929, EPI_ISL_514930, EPI_ISL_514931, EPI_ISL_514932, EPI_ISL_514933, EPI_ISL_514934, EPI_ISL_514935, EPI_ISL_514936, EPI_ISL_514937, EPI_ISL_514938, EPI_ISL_514939, EPI_ISL_514940, EPI_ISL_514941, EPI_ISL_514942, EPI_ISL_514943, EPI_ISL_514944, EPI_ISL_514945, EPI_ISL_514946, EPI_ISL_514947, EPI_ISL_514948, EPI_ISL_514949, EPI_ISL_514950, EPI_ISL_514951, EPI_ISL_514952, EPI_ISL_514953, EPI_ISL_514954, EPI_ISL_514955, EPI_ISL_514956, EPI_ISL_514957, EPI_ISL_514958, EPI_ISL_514959, EPI_ISL_514960, EPI_ISL_514961, EPI_ISL_514962, EPI_ISL_514963, EPI_ISL_514964, EPI_ISL_514965, EPI_ISL_514966, EPI_ISL_514967, EPI_ISL_514968, EPI_ISL_514969, EPI_ISL_514970, EPI_ISL_514971, EPI_ISL_514972, EPI_ISL_514973, EPI_ISL_514974, EPI_ISL_514975, EPI_ISL_514976, EPI_ISL_514977, EPI_ISL_514978, EPI_ISL_514979, EPI_ISL_514980, EPI_ISL_514981, EPI_ISL_514982, EPI_ISL_514983, EPI_ISL_514984, EPI_ISL_514985, EPI_ISL_514986, EPI_ISL_514987, EPI_ISL_514988, EPI_ISL_514989, EPI_ISL_514990, EPI_ISL_514991, EPI_ISL_514992, EPI_ISL_514993, EPI_ISL_514994, EPI_ISL_514995, EPI_ISL_514996, EPI_ISL_514997, EPI_ISL_514998, EPI_ISL_514999, EPI_ISL_515000, EPI_ISL_515001, EPI_ISL_515002, EPI_ISL_515003, EPI_ISL_515004, EPI_ISL_515005, EPI_ISL_515006, EPI_ISL_515007, EPI_ISL_515008, EPI_ISL_515009, EPI_ISL_515010, EPI_ISL_515011, EPI_ISL_515012, EPI_ISL_515013, EPI_ISL_515014, EPI_ISL_515015, EPI_ISL_515016, EPI_ISL_515017, EPI_ISL_515018, EPI_ISL_515019, EPI_ISL_515020, EPI_ISL_515021, EPI_ISL_515022, EPI_ISL_515023, EPI_ISL_515024, EPI_ISL_515025, EPI_ISL_515026, EPI_ISL_515027, EPI_ISL_515028, EPI_ISL_515029, EPI_ISL_515030, EPI_ISL_515031, EPI_ISL_515032, EPI_ISL_515033, EPI_ISL_515034, EPI_ISL_515035, EPI_ISL_515036, EPI_ISL_515037, EPI_ISL_515038, EPI_ISL_515039, EPI_ISL_515040, EPI_ISL_515041, EPI_ISL_515042, EPI_ISL_515043, EPI_ISL_515044, EPI_ISL_515045, EPI_ISL_515046, EPI_ISL_515047, EPI_ISL_515048, EPI_ISL_515049, EPI_ISL_515050, EPI_ISL_515052, EPI_ISL_515053, EPI_ISL_515054 | | | |
| see above | Division of Viral Diseases, Center for Laboratory Control of Infectious Diseases, Korea Centers for Diseases Control and Prevention | Division of Viral Diseases, Center for Laboratory Control of Infectious Diseases, Korea Centers for Diseases Control and Prevention | Jeong-Min Kim, Yoon-Seok Chung, Namjoo Lee, Sang Hee Woo, Hye-Jun Jo, Heui Man Kim, Jun-Sub Kim, Myung Guk Han |
| EPI_ISL_515055, EPI_ISL_515056, EPI_ISL_515057, EPI_ISL_515058, EPI_ISL_515059, EPI_ISL_515060, EPI_ISL_515061, EPI_ISL_515062, EPI_ISL_515063, EPI_ISL_515064, EPI_ISL_515065, EPI_ISL_515066, EPI_ISL_515067, EPI_ISL_515068, EPI_ISL_515069, EPI_ISL_515070, EPI_ISL_515071, EPI_ISL_515073, EPI_ISL_515074, EPI_ISL_515075, EPI_ISL_515076, EPI_ISL_515077, EPI_ISL_515078, EPI_ISL_515079, EPI_ISL_515080, EPI_ISL_515081 | | | |
| see above | Department of Clinical Microbiology | GIGA Medical Genomics | Keith Durkin, Maria Artesi, Sebastien Bontems, Raphael Boreux, Cecile Meex, Axelle Chaslain, Celine Fombellida-Lopez, Pierrette Melin, Marie-Pierre Hayette, Vincent Bours. |
| EPI_ISL_515082, EPI_ISL_515084, EPI_ISL_515090, EPI_ISL_515096, EPI_ISL_515098, EPI_ISL_515100, EPI_ISL_515101, EPI_ISL_515102, EPI_ISL_515107, EPI_ISL_515108, EPI_ISL_515110, EPI_ISL_515111 | | | |
| see above | Department of Biochemistry, Cell and Molecular Biology | WACCBIP, University of Ghana | Ngoi,J.M., Quashie,P., Morang'a,C.M., Amuzu,D.S., Adu,B., Kumordjie,S., Eshun,M., Boatemaa,L., Magnussen,V., Kotey,E., Tei-Maya,F., Arjarquah,A., Mutungi,J.K., Bediako,Y., Asante,I., Bonney,E., Kyei,G.B., Bonney,K., Amenga-Etego,L.N., Anang,A.K., Awandare,G.A., Ampofo,W. |
| EPI_ISL_515156, EPI_ISL_515166, EPI_ISL_515172 | National Institute for Communicable Diseases of the National Health Laboratory Service | National Institute for Communicable Diseases of the National Health Laboratory Service | Allam M, Ismail A, Khumalo Z, Kwenda S, Mtshali P, Mnyameni F, Mohale T, Bhiman JN |
| EPI_ISL_515181, EPI_ISL_515182, EPI_ISL_515183 | Kumasi Centre for Collaborative Research in Tropical Medicine, Kumasi. | Institute of Virology, Charité – Universitätsmedizin Berlin | Augustina Sylverken, Philip El-Duah, Michael Owusu, Julia Schneider, Richmond Yeboah, Richmond Gorman, Eric Adu, Sherihane Aryeetey, Jesse Addo Asamoah,Jörn Beheim-Schwarzbach, Victor Max Corman, Christian Drosten, Richard Phillips. |
| EPI_ISL_515185, EPI_ISL_515186 | Latvijas Infektoloģijas centrs | Latvian Biomedical Research and Study Centre | Ivars Silamiķelis, Kaspars Megnis, Monta Ustinova, Ņikita Zrelovs, Vita Rovīte, Jeļena Storoženko, Tatjana Kolupajeva, Oksana Savicka, Uga Dumpis, Jānis Kloviņš |
| EPI_ISL_515187, EPI_ISL_515188, EPI_ISL_515189, EPI_ISL_515190, EPI_ISL_515191, EPI_ISL_515192, EPI_ISL_515193, EPI_ISL_515194, EPI_ISL_515195 | E. Gulbja Laboratorija | Latvian Biomedical Research and Study Centre | Ivars Silamiķelis, Kaspars Megnis, Monta Ustinova, Ņikita Zrelovs, Vita Rovīte, Mikus Gavars, Dmitrijs Perminovs, Uga Dumpis, Jānis Kloviņš |
| EPI_ISL_515196 | Centrālā laboratorija | Latvian Biomedical Research and Study Centre | Ivars Silamiķelis, Kaspars Megnis, Monta Ustinova, Ņikita Zrelovs, Vita Rovīte, Stella Lapiņa, Jana Osīte, Marta Priedīte, Uga Dumpis, Jānis Kloviņš |
| EPI_ISL_515197, EPI_ISL_515202, EPI_ISL_515203, EPI_ISL_515209, EPI_ISL_515210, EPI_ISL_515211, EPI_ISL_515212, EPI_ISL_515214, EPI_ISL_515216, EPI_ISL_515220, EPI_ISL_515222, EPI_ISL_515224, EPI_ISL_515225, EPI_ISL_515226, EPI_ISL_515227, EPI_ISL_515228, EPI_ISL_515229, EPI_ISL_515230, EPI_ISL_515231, EPI_ISL_515233, EPI_ISL_515234, EPI_ISL_515235, EPI_ISL_515237, EPI_ISL_515238, EPI_ISL_515241, EPI_ISL_515243, EPI_ISL_515244, EPI_ISL_515245 | | | |
| see above | Laboratoire de microbiologie, Hopital de Verdun | Smith Laboratory, Centre de Recherche CHU Sainte-Justine | Martin Smith, Marieke Rozendaal, Ivan Pavlov |
| EPI_ISL_515250, EPI_ISL_515251, EPI_ISL_515252, EPI_ISL_515253, EPI_ISL_515254, EPI_ISL_515259 | Texas Department of State Health Services | Texas Department of State Health Services | Rashmi Tuladhar, Bonnie Oh, Cara Akrout, Jenny Zhang, Maliha Rahman, Anita Pokharel, Myong Koag, Chun Wang, Rachel Lee, Grace Kubin |
| EPI_ISL_515262 | Minnesota Department of Health, Public Health Laboratory | Minnesota Department of Health, Public Health Laboratory | Matt Plumb, Jacob Garfin, and Xiong Wang |
| EPI_ISL_515263, EPI_ISL_515264, EPI_ISL_515265, EPI_ISL_515266 | Mayo Clinic & Mayo Clinic Laboratories | Minnesota Department of Health, Public Health Laboratory | Matt Plumb, Jacob Garfin, and Xiong Wang |
| EPI_ISL_515267, EPI_ISL_515268 | M Health Fairview | Minnesota Department of Health, Public Health Laboratory | Matt Plumb, Jacob Garfin, and Xiong Wang |
| EPI_ISL_515269 | Minnesota Department of Health, Public Health Laboratory | Minnesota Department of Health, Public Health Laboratory | Matt Plumb, Jacob Garfin, and Xiong Wang |
| EPI_ISL_515270, EPI_ISL_515271, EPI_ISL_515272, EPI_ISL_515273, EPI_ISL_515274, EPI_ISL_515275, EPI_ISL_515276, EPI_ISL_515277, EPI_ISL_515278, EPI_ISL_515279, EPI_ISL_515280, EPI_ISL_515281, EPI_ISL_515282, EPI_ISL_515283, EPI_ISL_515284, EPI_ISL_515285 | | | |
| see above | University of Washington Virology Lab | University of Washington Virology Lab | Pavitra Roychoudhury, Hong Xie, Lasata Shrestha, Amin Addetia, Truong Nguyen, Victoria M Rachleff, Meei-Li Huang, Keith R Jerome, Alexander Greninger |
| EPI_ISL_515287, EPI_ISL_515288, EPI_ISL_515289, EPI_ISL_515290, EPI_ISL_515291 | National Institute of Health. Department of medical Sciences, Ministry of Public Health, Thailand | National Institute of Health. Department of medical Sciences, Ministry of Public Health, Thailand | Pilailuk,Okada; Siripaporn,Phuygun; Thanutsapa,Thanadachakul; Sittiporn,Parnmen;Warawan,Wongboot; Sunthareeya,Waicharoen; Malinee,Chittaganpitch |
| EPI_ISL_515292 | Ramkhamhaeng Hospital | National Institute of Health. Department of medical Sciences, Ministry of Public Health, Thailand | Pilailuk,Okada; Siripaporn,Phuygun; Thanutsapa,Thanadachakul; Sittiporn,Parnmen;Warawan,Wongboot; Sunthareeya,Waicharoen; Malinee,Chittaganpitch |
| EPI_ISL_515293, EPI_ISL_515294, EPI_ISL_515295, EPI_ISL_515296, EPI_ISL_515297, EPI_ISL_515299, EPI_ISL_515300, EPI_ISL_515301, EPI_ISL_515302, EPI_ISL_515303, EPI_ISL_515304, EPI_ISL_515305, EPI_ISL_515306, EPI_ISL_515307, EPI_ISL_515308, EPI_ISL_515309, EPI_ISL_515311, EPI_ISL_515312, EPI_ISL_515313, EPI_ISL_515314, EPI_ISL_515315, EPI_ISL_515316, EPI_ISL_515317, EPI_ISL_515318, EPI_ISL_515319, EPI_ISL_515320, EPI_ISL_515321, EPI_ISL_515322, EPI_ISL_515324, EPI_ISL_515325, EPI_ISL_515327, EPI_ISL_515328, EPI_ISL_515329, EPI_ISL_515330, EPI_ISL_515331, EPI_ISL_515332, EPI_ISL_515333, EPI_ISL_515334, EPI_ISL_515336, EPI_ISL_515337, EPI_ISL_515338, EPI_ISL_515339, EPI_ISL_515340, EPI_ISL_515341, EPI_ISL_515342, EPI_ISL_515344, EPI_ISL_515345, EPI_ISL_515346, EPI_ISL_515347, EPI_ISL_515348, EPI_ISL_515349, EPI_ISL_515350, EPI_ISL_515351, EPI_ISL_515352, EPI_ISL_515353, EPI_ISL_515354, EPI_ISL_515355, EPI_ISL_515356, EPI_ISL_515357, EPI_ISL_515358, EPI_ISL_515359, EPI_ISL_515361, EPI_ISL_515362, EPI_ISL_515363, EPI_ISL_515364, EPI_ISL_515365, EPI_ISL_515366, EPI_ISL_515367, EPI_ISL_515368, EPI_ISL_515369, EPI_ISL_515370, EPI_ISL_515371, EPI_ISL_515372, EPI_ISL_515373, EPI_ISL_515374, EPI_ISL_515375, EPI_ISL_515376, EPI_ISL_515377, EPI_ISL_515378, EPI_ISL_515379, EPI_ISL_515381, EPI_ISL_515382, EPI_ISL_515384, EPI_ISL_515385, EPI_ISL_515386, EPI_ISL_515387, EPI_ISL_515388, EPI_ISL_515389, EPI_ISL_515390, EPI_ISL_515391, EPI_ISL_515392, EPI_ISL_515393, EPI_ISL_515394, EPI_ISL_515397, EPI_ISL_515399, EPI_ISL_515401, EPI_ISL_515403, EPI_ISL_515404, EPI_ISL_515405, EPI_ISL_515406, EPI_ISL_515407, EPI_ISL_515409, EPI_ISL_515410, EPI_ISL_515411, EPI_ISL_515412, EPI_ISL_515413, EPI_ISL_515414, EPI_ISL_515415, EPI_ISL_515416, EPI_ISL_515417, EPI_ISL_515418, EPI_ISL_515419, EPI_ISL_515420, EPI_ISL_515421, EPI_ISL_515422, EPI_ISL_515423, EPI_ISL_515424, EPI_ISL_515425, EPI_ISL_515426, EPI_ISL_515427, EPI_ISL_515428, EPI_ISL_515429, EPI_ISL_515430, EPI_ISL_515431, EPI_ISL_515432, EPI_ISL_515433, EPI_ISL_515434, EPI_ISL_515436, EPI_ISL_515437, EPI_ISL_515438, EPI_ISL_515439, EPI_ISL_515440, EPI_ISL_515441, EPI_ISL_515442, EPI_ISL_515443, EPI_ISL_515444, EPI_ISL_515445, EPI_ISL_515446, EPI_ISL_515447, EPI_ISL_515448, EPI_ISL_515449, EPI_ISL_515450, EPI_ISL_515451, EPI_ISL_515452, EPI_ISL_515453, EPI_ISL_515454, EPI_ISL_515455, EPI_ISL_515456, EPI_ISL_515457, EPI_ISL_515459, EPI_ISL_515460, EPI_ISL_515461, EPI_ISL_515462 | | | |
| see above | Nevada State Public Health Laboratory | Nevada State Public Health Laboratory | Richard Tillett, Joel R. Sevinsky, Paul Hartley, Heather Kerwin, David Jackson, Subhash C. Verma, Cyprian Rossetto, Andrew Gorzalski, Chris Laverdure, Natalie Crawford, Stephanie Van Hooser, and Mark Pandori |
| EPI_ISL_515463 | National Institute of Health. Department of medical Sciences, Ministry of Public Health, Thailand | National Institute of Health. Department of medical Sciences, Ministry of Public Health, Thailand | Pilailuk,Okada; Siripaporn,Phuygun; Thanutsapa,Thanadachakul; Sittiporn,Parnmen;Warawan,Wongboot; Sunthareeya,Waicharoen; Malinee,Chittaganpitch |

| | | | |
|---|---|---|---|
| EPI_ISL_515464 | Siriraj Hospital | National Institute of Health. Department of medical Sciences, Ministry of Public Health, Thailand | Pilailuk,Okada; Siripaporn,Phuygun; Thanutsapa,Thanadachakul; Sittiporn,Parnmen;Warawan,Wongboot; Sunthareeya,Waicharoen; Malinee,Chittaganpitch |
| EPI_ISL_515465 | Ramathibodi Hospital | National Institute of Health. Department of medical Sciences, Ministry of Public Health, Thailand | Pilailuk,Okada; Siripaporn,Phuygun; Thanutsapa,Thanadachakul; Sittiporn,Parnmen;Warawan,Wongboot; Sunthareeya,Waicharoen; Malinee,Chittaganpitch |
| EPI_ISL_515466 | Discovery DNA | Discovery DNA | Dustin Hittel, Marina Kerr, Leo Dimnik, Desmond Koo, Alice Li, Aneal Khan |
| EPI_ISL_515468 | Ramathibodi Hospital | National Institute of Health. Department of medical Sciences, Ministry of Public Health, Thailand | Pilailuk,Okada; Siripaporn,Phuygun; Thanutsapa,Thanadachakul; Sittiporn,Parnmen;Warawan,Wongboot; Sunthareeya,Waicharoen; Malinee,Chittaganpitch |
| EPI_ISL_515469 | Bamrasnaradura hospital | National Institute of Health. Department of medical Sciences, Ministry of Public Health, Thailand | Pilailuk,Okada; Siripaporn,Phuygun; Thanutsapa,Thanadachakul; Sittiporn,Parnmen;Warawan,Wongboot; Sunthareeya,Waicharoen; Malinee,Chittaganpitch |
| EPI_ISL_515470 | National Institute of Health. Department of medical Sciences, Ministry of Public Health, Thailand | National Institute of Health. Department of medical Sciences, Ministry of Public Health, Thailand | Pilailuk,Okada; Siripaporn,Phuygun; Thanutsapa,Thanadachakul; Sittiporn,Parnmen;Warawan,Wongboot; Sunthareeya,Waicharoen; Malinee,Chittaganpitch |
| EPI_ISL_515472, EPI_ISL_515473, EPI_ISL_515475, EPI_ISL_515476, EPI_ISL_515477 | National Institute of Health, Department of Medical Sciences, Ministry of Public Health, Thailand | National Institute of Health, Department of Medical Sciences, Ministry of Public Health, Thailand | Pilailuk Okada; Siripaporn Phuygun; Thanutsapa Thanadachakul; Sittiporn Parnmen; Warawan Wongboot; Sunthareeya Waicharoen; Malinee Chittaganpitch |
| EPI_ISL_515520 | Hospital Municipal do Tatuape Carmino Caricchio | Instituto Adolfo Lutz, Interdiciplinary Procedures Center, Strategic Laboratory | Claudio Tavares Sacchi, Claudia Regina Gonçalves, Erica Valessa Ramos Gomes |
| EPI_ISL_515521 | Hospital Municipal Dr Waldemar Tebaldi | Instituto Adolfo Lutz, Interdiciplinary Procedures Center, Strategic Laboratory | Claudio Tavares Sacchi, Claudia Regina Gonçalves, Erica Valessa Ramos Gomes |
| EPI_ISL_515522 | UPA 24HS de Itatiba | Instituto Adolfo Lutz, Interdiciplinary Procedures Center, Strategic Laboratory | Claudio Tavares Sacchi, Claudia Regina Gonçalves, Erica Valessa Ramos Gomes |
| EPI_ISL_515523, EPI_ISL_515524 | PS Municipal Dr Lauro Ribas Braga | Instituto Adolfo Lutz, Interdiciplinary Procedures Center, Strategic Laboratory | Claudio Tavares Sacchi, Claudia Regina Gonçalves, Erica Valessa Ramos Gomes |
| EPI_ISL_515525 | National Influenza Center - Instituto Adolfo Lutz | Instituto Adolfo Lutz, Interdiciplinary Procedures Center, Strategic Laboratory | Claudio Tavares Sacchi, Claudia Regina Gonçalves, Erica Valessa Ramos Gomes |
| EPI_ISL_515526 | Hospital Municipal do Tatuape Carmino Caricchio | Instituto Adolfo Lutz, Interdiciplinary Procedures Center, Strategic Laboratory | Claudio Tavares Sacchi, Claudia Regina Gonçalves, Erica Valessa Ramos Gomes |
| EPI_ISL_515527 | Hospital Santa Clara | Instituto Adolfo Lutz, Interdiciplinary Procedures Center, Strategic Laboratory | Claudio Tavares Sacchi, Claudia Regina Gonçalves, Erica Valessa Ramos Gomes |
| EPI_ISL_515528 | Hospital Sao Paulo de Ensino da Unifesp | Instituto Adolfo Lutz, Interdiciplinary Procedures Center, Strategic Laboratory | Claudio Tavares Sacchi, Claudia Regina Gonçalves, Erica Valessa Ramos Gomes |
| EPI_ISL_515529 | Pronto Socorro Municipal Julio Tupy | Instituto Adolfo Lutz, Interdiciplinary Procedures Center, Strategic Laboratory | Claudio Tavares Sacchi, Claudia Regina Gonçalves, Erica Valessa Ramos Gomes |
| EPI_ISL_515541 | Hospital Montemagno | Instituto Adolfo Lutz, Interdiciplinary Procedures Center, Strategic Laboratory | Claudio Tavares Sacchi, Claudia Regina Gonçalves, Erica Valessa Ramos Gomes |
| EPI_ISL_515542 | Vigilância Epidemiológica de Leme | Instituto Adolfo Lutz, Interdiciplinary Procedures Center, Strategic Laboratory | Claudio Tavares Sacchi, Claudia Regina Gonçalves, Erica Valessa Ramos Gomes |
| EPI_ISL_515543 | Serviço de Vigilância Sanitária e Epidemiológica | Instituto Adolfo Lutz, Interdiciplinary Procedures Center, Strategic Laboratory | Claudio Tavares Sacchi, Claudia Regina Gonçalves, Erica Valessa Ramos Gomes |
| EPI_ISL_515544 | Ama Dr Jose Soares Hungria | Instituto Adolfo Lutz, Interdiciplinary Procedures Center, Strategic Laboratory | Claudio Tavares Sacchi, Claudia Regina Gonçalves, Erica Valessa Ramos Gomes |
| EPI_ISL_515545 | Hospital Sao Paulo de Ensino da Unifesp | Instituto Adolfo Lutz, Interdiciplinary Procedures Center, Strategic Laboratory | Claudio Tavares Sacchi, Claudia Regina Gonçalves, Erica Valessa Ramos Gomes |
| EPI_ISL_515546 | Hospital Municipal do Tatuape Carmino Caricchio | Instituto Adolfo Lutz, Interdiciplinary Procedures Center, Strategic Laboratory | Claudio Tavares Sacchi, Claudia Regina Gonçalves, Erica Valessa Ramos Gomes |
| EPI_ISL_515547 | Centro Medico da Policia Militar do Estado de Sao Paulo | Instituto Adolfo Lutz, Interdiciplinary Procedures Center, Strategic Laboratory | Claudio Tavares Sacchi, Claudia Regina Gonçalves, Erica Valessa Ramos Gomes |
| EPI_ISL_515548 | Hospital Municipal Dr. Jose Soares Hungria | Instituto Adolfo Lutz, Interdiciplinary Procedures Center, Strategic Laboratory | Claudio Tavares Sacchi, Claudia Regina Gonçalves, Erica Valessa Ramos Gomes |
| EPI_ISL_515550 | UPA Vila Santa Catarina | Instituto Adolfo Lutz, Interdiciplinary Procedures Center, Strategic Laboratory | Claudio Tavares Sacchi, Claudia Regina Gonçalves, Erica Valessa Ramos Gomes |
| EPI_ISL_515551, EPI_ISL_515552 | Hospital Municipal do Tatuape Carmino Caricchio | Instituto Adolfo Lutz, Interdiciplinary Procedures Center, Strategic Laboratory | Claudio Tavares Sacchi, Claudia Regina Gonçalves, Erica Valessa Ramos Gomes |
| EPI_ISL_515553 | Hospital Municipal Dr. Ignacio Proença de Gouvea | Instituto Adolfo Lutz, Interdiciplinary Procedures Center, Strategic Laboratory | Claudio Tavares Sacchi, Claudia Regina Gonçalves, Erica Valessa Ramos Gomes |
| EPI_ISL_515554 | Pronto Socorro Municipal de Perus | Instituto Adolfo Lutz, Interdiciplinary Procedures Center, Strategic Laboratory | Claudio Tavares Sacchi, Claudia Regina Gonçalves, Erica Valessa Ramos Gomes |
| EPI_ISL_515555 | Hospital Geral de Vila Nova Cachoeirinha | Instituto Adolfo Lutz, Interdiciplinary Procedures Center, Strategic Laboratory | Claudio Tavares Sacchi, Claudia Regina Gonçalves, Erica Valessa Ramos Gomes |
| EPI_ISL_515557 | Hospital Municipal Dr. Moysés Deutsch | Instituto Adolfo Lutz, Interdiciplinary Procedures Center, Strategic Laboratory | Claudio Tavares Sacchi, Claudia Regina Gonçalves, Erica Valessa Ramos Gomes |
| EPI_ISL_515559, EPI_ISL_515560 | Hospital Sao Paulo de Ensino da Unifesp | Instituto Adolfo Lutz, Interdiciplinary Procedures Center, Strategic Laboratory | Claudio Tavares Sacchi, Claudia Regina Gonçalves, Erica Valessa Ramos Gomes |
| EPI_ISL_515561 | Hospital Montemagno | Instituto Adolfo Lutz, Interdiciplinary Procedures Center, Strategic Laboratory | Claudio Tavares Sacchi, Claudia Regina Gonçalves, Erica Valessa Ramos Gomes |
| EPI_ISL_515562 | Hospital Municipal Doutor Alexandre Zaio | Instituto Adolfo Lutz, Interdiciplinary Procedures Center, Strategic Laboratory | Claudio Tavares Sacchi, Claudia Regina Gonçalves, Erica Valessa Ramos Gomes |
| EPI_ISL_515563 | Hospital Municipal Dr. Jose Soares Hungria | Instituto Adolfo Lutz, Interdiciplinary Procedures Center, Strategic Laboratory | Claudio Tavares Sacchi, Claudia Regina Gonçalves, Erica Valessa Ramos Gomes |
| EPI_ISL_515564 | Hosp. Municipal Prof. Dr. Alípio Corrêa Netto | Instituto Adolfo Lutz, Interdiciplinary Procedures Center, Strategic Laboratory | Claudio Tavares Sacchi, Claudia Regina Gonçalves, Erica Valessa Ramos Gomes |
| EPI_ISL_515565 | Hospital do Servidor Público Estadual Francisco Morato de Oliveira | Instituto Adolfo Lutz, Interdiciplinary Procedures Center, Strategic Laboratory | Claudio Tavares Sacchi, Claudia Regina Gonçalves, Erica Valessa Ramos Gomes |
| EPI_ISL_515566 | PS Municipal Dr Lauro Ribas Braga | Instituto Adolfo Lutz, Interdiciplinary Procedures Center, Strategic Laboratory | Claudio Tavares Sacchi, Claudia Regina Gonçalves, Erica Valessa Ramos Gomes |
| EPI_ISL_515568, EPI_ISL_515569, EPI_ISL_515570, EPI_ISL_515573, EPI_ISL_515574, EPI_ISL_515575, EPI_ISL_515579, EPI_ISL_515580, EPI_ISL_515582, EPI_ISL_515584, EPI_ISL_515585, EPI_ISL_515586, EPI_ISL_515589, EPI_ISL_515592, EPI_ISL_515593, EPI_ISL_515594, EPI_ISL_515596, EPI_ISL_515597, EPI_ISL_515600, EPI_ISL_515601, EPI_ISL_515602, EPI_ISL_515603, EPI_ISL_515604, EPI_ISL_515606, EPI_ISL_515607, EPI_ISL_515610, EPI_ISL_515612, EPI_ISL_515613, EPI_ISL_515615, EPI_ISL_515616, EPI_ISL_515619, EPI_ISL_515622, EPI_ISL_515623, EPI_ISL_515627, EPI_ISL_515628, EPI_ISL_515631, EPI_ISL_515633, EPI_ISL_515634, EPI_ISL_515635, EPI_ISL_515639, EPI_ISL_515640, EPI_ISL_515641, EPI_ISL_515642, EPI_ISL_515643, EPI_ISL_515644, EPI_ISL_515645, EPI_ISL_515647, EPI_ISL_515649, EPI_ISL_515650, EPI_ISL_515651, EPI_ISL_515652, EPI_ISL_515653, EPI_ISL_515654, EPI_ISL_515655, EPI_ISL_515656, EPI_ISL_515657, EPI_ISL_515658, EPI_ISL_515659, EPI_ISL_515660, EPI_ISL_515661, EPI_ISL_515662, EPI_ISL_515665, EPI_ISL_515666, EPI_ISL_515668, EPI_ISL_515669, EPI_ISL_515671, EPI_ISL_515672, EPI_ISL_515674, EPI_ISL_515675, EPI_ISL_515676, EPI_ISL_515678, EPI_ISL_515679, EPI_ISL_515680, EPI_ISL_515681, EPI_ISL_515682, EPI_ISL_515683, EPI_ISL_515684, EPI_ISL_515685, EPI_ISL_515686, EPI_ISL_515687, EPI_ISL_515688, EPI_ISL_515689, EPI_ISL_515690, EPI_ISL_515691, EPI_ISL_515692, EPI_ISL_515693, EPI_ISL_515694, EPI_ISL_515696, EPI_ISL_515697, EPI_ISL_515698, EPI_ISL_515699, EPI_ISL_515700, EPI_ISL_515701, EPI_ISL_515702, EPI_ISL_515703, EPI_ISL_515704, EPI_ISL_515705, EPI_ISL_515706, EPI_ISL_515707, EPI_ISL_515708, EPI_ISL_515709, EPI_ISL_515710, EPI_ISL_515711, EPI_ISL_515712, EPI_ISL_515714, EPI_ISL_515715, EPI_ISL_515716, EPI_ISL_515718, EPI_ISL_515719, EPI_ISL_515720, EPI_ISL_515721, EPI_ISL_515722, EPI_ISL_515723, EPI_ISL_515724, EPI_ISL_515725, EPI_ISL_515728, EPI_ISL_515729, EPI_ISL_515730, EPI_ISL_515731, EPI_ISL_515732, EPI_ISL_515733, EPI_ISL_515734, EPI_ISL_515736, EPI_ISL_515737, EPI_ISL_515738, EPI_ISL_515739, EPI_ISL_515740, EPI_ISL_515741, EPI_ISL_515743, EPI_ISL_515744, EPI_ISL_515745, EPI_ISL_515747, EPI_ISL_515748, EPI_ISL_515749, EPI_ISL_515751, EPI_ISL_515752, EPI_ISL_515753, EPI_ISL_515754, EPI_ISL_515756, EPI_ISL_515757, EPI_ISL_515758, EPI_ISL_515759, EPI_ISL_515760, EPI_ISL_515762, EPI_ISL_515763, EPI_ISL_515764, EPI_ISL_515767, EPI_ISL_515769, EPI_ISL_515771, EPI_ISL_515773, EPI_ISL_515775, EPI_ISL_515776, EPI_ISL_515777, EPI_ISL_515778, EPI_ISL_515780, EPI_ISL_515781, EPI_ISL_515782, EPI_ISL_515783, EPI_ISL_515784, EPI_ISL_515787, EPI_ISL_515789, EPI_ISL_515790, EPI_ISL_515791, EPI_ISL_515792, EPI_ISL_515794, EPI_ISL_515796, EPI_ISL_515797, EPI_ISL_515799 | | | |
| see above | NHLS-IALCH | KRISP, KZN Research Innovation and Sequencing Platform | Giandhari J, Pillay S, Lessells R, Mdlalose K, York D, Khan S, Tegally H, Wilkinson E, de Oliveira T |
| EPI_ISL_515801, EPI_ISL_515802, EPI_ISL_515803, EPI_ISL_515804, EPI_ISL_515805, EPI_ISL_515806, EPI_ISL_515807, EPI_ISL_515808, EPI_ISL_515809, EPI_ISL_515810, EPI_ISL_515811, EPI_ISL_515812, EPI_ISL_515813, EPI_ISL_515814, EPI_ISL_515815, EPI_ISL_515816, EPI_ISL_515817, EPI_ISL_515818, EPI_ISL_515819, EPI_ISL_515820, EPI_ISL_515821, EPI_ISL_515822, EPI_ISL_515823, EPI_ISL_515824, EPI_ISL_515825, EPI_ISL_515826, EPI_ISL_515827, EPI_ISL_515828, EPI_ISL_515829, EPI_ISL_515830, EPI_ISL_515831, EPI_ISL_515832, EPI_ISL_515833, EPI_ISL_515835, EPI_ISL_515836, EPI_ISL_515837, EPI_ISL_515838, EPI_ISL_515839, EPI_ISL_515840, EPI_ISL_515841, EPI_ISL_515842, EPI_ISL_515843, EPI_ISL_515844, EPI_ISL_515845, EPI_ISL_515846, EPI_ISL_515847, EPI_ISL_515848, EPI_ISL_515849, EPI_ISL_515850, EPI_ISL_515851, EPI_ISL_515852, EPI_ISL_515853, EPI_ISL_515854, EPI_ISL_515855, EPI_ISL_515856, EPI_ISL_515857, EPI_ISL_515858, EPI_ISL_515859, EPI_ISL_515860, EPI_ISL_515861, EPI_ISL_515862, EPI_ISL_515863, EPI_ISL_515864, EPI_ISL_515865, EPI_ISL_515866, EPI_ISL_515867, EPI_ISL_515868, EPI_ISL_515869, EPI_ISL_515870, EPI_ISL_515871, EPI_ISL_515872, EPI_ISL_515873, EPI_ISL_515874, EPI_ISL_515875, EPI_ISL_515876, EPI_ISL_515877, EPI_ISL_515878, EPI_ISL_515880, EPI_ISL_515881, EPI_ISL_515882, EPI_ISL_515883, EPI_ISL_515884, EPI_ISL_515885, EPI_ISL_515886, EPI_ISL_515887, EPI_ISL_515888, EPI_ISL_515889, EPI_ISL_515890, EPI_ISL_515891, EPI_ISL_515892, EPI_ISL_515893 | | | |
| see above | Medical Disagnostics Services (MDS) | KRISP, KZN Research Innovation and Sequencing Platform | Giandhari J, Pillay S, Lessells R, ChimukangaraB, Mdlalose K, York D, Khan S, Tegally H, Wilkinson E, de Oliveira T |
| EPI_ISL_515894, EPI_ISL_515896, EPI_ISL_515897, EPI_ISL_515898, EPI_ISL_515899, EPI_ISL_515900, EPI_ISL_515901, EPI_ISL_515902, EPI_ISL_515903, EPI_ISL_515905, EPI_ISL_515906, EPI_ISL_515907, EPI_ISL_515908, EPI_ISL_515909, EPI_ISL_515910, EPI_ISL_515911, EPI_ISL_515913, EPI_ISL_515917, EPI_ISL_515918, EPI_ISL_515919, EPI_ISL_515920, EPI_ISL_515922, EPI_ISL_515923, EPI_ISL_515924, EPI_ISL_515925, EPI_ISL_515928, EPI_ISL_515929 | | | |
| see above | California Department of Public Health | California Department of Public Health | CDPH IDLB COVIDNet |
| EPI_ISL_515938, EPI_ISL_515941, EPI_ISL_515942 | CV RAMAN HOSPITAL | Department of Neurovirology, National Institute of Mental Health and Neuroscience (NIMHANS) | Chitra Pattabiraman,Vijayalakshmi Reddy, Harsha PK, Risha Rasheed, Pramada Prasad, Shafeeq S Hameed, Manjunatha Venkataswamy, Anita Desai, Ravi Vasanthapuram |
| EPI_ISL_515945, EPI_ISL_515947 | DH | Department of Neurovirology, National Institute of Mental Health and Neuroscience (NIMHANS) | Chitra Pattabiraman,Vijayalakshmi Reddy, Harsha PK, Risha Rasheed, Pramada Prasad, Shafeeq S Hameed, Manjunatha Venkataswamy, Anita Desai, Ravi Vasanthapuram |
| EPI_ISL_515952 | VICTORIA HOSPITAL | Department of Neurovirology, National Institute of Mental Health and Neuroscience (NIMHANS) | Chitra Pattabiraman,Vijayalakshmi Reddy, Harsha PK, Risha Rasheed, Pramada Prasad, Shafeeq S Hameed, Manjunatha Venkataswamy, Anita Desai, Ravi Vasanthapuram |
| EPI_ISL_515954, EPI_ISL_515957, EPI_ISL_515958, EPI_ISL_515959, EPI_ISL_515961 | DH | Department of Neurovirology, National Institute of Mental Health and Neuroscience (NIMHANS) | Chitra Pattabiraman,Vijayalakshmi Reddy, Harsha PK, Risha Rasheed, Pramada Prasad, Shafeeq S Hameed, Manjunatha Venkataswamy, Anita Desai, Ravi Vasanthapuram |
| EPI_ISL_515963, EPI_ISL_515964, EPI_ISL_515966, EPI_ISL_515967 | VICTORIA HOSPITAL | Department of Neurovirology, National Institute of Mental Health and Neuroscience (NIMHANS) | Chitra Pattabiraman,Vijayalakshmi Reddy, Harsha PK, Risha Rasheed, Pramada Prasad, Shafeeq S Hameed, Manjunatha Venkataswamy, Anita Desai, Ravi Vasanthapuram |
| EPI_ISL_515968, EPI_ISL_515969, EPI_ISL_515970, EPI_ISL_515971, EPI_ISL_515972, EPI_ISL_515973 | MIMS | Department of Neurovirology, National Institute of Mental Health and Neuroscience (NIMHANS) | Chitra Pattabiraman,Vijayalakshmi Reddy, Harsha PK, Risha Rasheed, Pramada Prasad, Shafeeq S Hameed, Manjunatha Venkataswamy, Anita Desai, Ravi Vasanthapuram |
| EPI_ISL_515974 | Department of Virus and Microbiological Special Diagnostics, Statens Serum Institut | Albertsen lab, Department of Chemistry and Bioscience, Aalborg University, Denmark | Danish Corona Danica Consortia |
| EPI_ISL_516079, EPI_ISL_516080, EPI_ISL_516081, EPI_ISL_516082, EPI_ISL_516083, EPI_ISL_516084, EPI_ISL_516085, EPI_ISL_516086, EPI_ISL_516087, EPI_ISL_516088 | Biomedical Sciences and Public Health, Polytechnic University of Marche | Biomedical Sciences and Public Health, Polytechnic University of Marche | Bagnarelli,P., Caucci,S., Di Sante,L., Menzo,S., Alessandrini,F., Onofri,V., Turchi,C., Melchionda,F., Tagliabracci,A. |
| EPI_ISL_516267, EPI_ISL_516268, EPI_ISL_516269, EPI_ISL_516270, EPI_ISL_516272, EPI_ISL_516273, EPI_ISL_516274, EPI_ISL_516275, EPI_ISL_516277, EPI_ISL_516278, EPI_ISL_516281, EPI_ISL_516282, EPI_ISL_516283, EPI_ISL_516285, EPI_ISL_516287, EPI_ISL_516294, EPI_ISL_516295, EPI_ISL_516296, EPI_ISL_516297, EPI_ISL_516302, EPI_ISL_516306, EPI_ISL_516308, EPI_ISL_516311, EPI_ISL_516314, EPI_ISL_516316, EPI_ISL_516317, EPI_ISL_516321, EPI_ISL_516322, EPI_ISL_516324, EPI_ISL_516329, EPI_ISL_516330, EPI_ISL_516331, EPI_ISL_516332, EPI_ISL_516334, EPI_ISL_516335, EPI_ISL_516348, EPI_ISL_516351, EPI_ISL_516359, EPI_ISL_516360, EPI_ISL_516361, EPI_ISL_516362, EPI_ISL_516363, EPI_ISL_516365, EPI_ISL_516367, EPI_ISL_516371, EPI_ISL_516372, EPI_ISL_516373, EPI_ISL_516376, EPI_ISL_516380, EPI_ISL_516412 | | | |
| see above | Michigan Department of Health and Human Services, Bureau of Laboratories | Michigan Department of Health and Human Services, Bureau of Laboratories | Blankenship HM, Riner D, Soehnlen MK |
| EPI_ISL_516413 | General Hospital - Kumanovo | Research Center for Genetic Engineering and Biotechnology "Georgi D. Efremov” , Macedonian Academy of Sciences and Arts | RCGEB - MASA |
| EPI_ISL_516414, EPI_ISL_516415, EPI_ISL_516416, EPI_ISL_516417, EPI_ISL_516418, EPI_ISL_516419, EPI_ISL_516420, EPI_ISL_516421, EPI_ISL_516422, EPI_ISL_516423, EPI_ISL_516424, EPI_ISL_516425 | | | |
| see above | Center for public health - Skopje | Research Center for Genetic Engineering and Biotechnology "Georgi D. Efremov” , Macedonian Academy of | RCGEB - MASA |

| | | | |
|---|---|---|---|
| | | Sciences and Arts | |
| EPI_ISL_516426, EPI_ISL_516427, EPI_ISL_516428, EPI_ISL_516429, EPI_ISL_516430, EPI_ISL_516431 | Clinical Hospital - Shtip | Research Center for Genetic Engineering and Biotechnology "Georgi D. Efremov" , Macedonian Academy of Sciences and Arts | RCGEB - MASA |
| EPI_ISL_516432, EPI_ISL_516433, EPI_ISL_516434, EPI_ISL_516435, EPI_ISL_516436, EPI_ISL_516437, EPI_ISL_516438, EPI_ISL_516439, EPI_ISL_516440, EPI_ISL_516441, EPI_ISL_516442, EPI_ISL_516443, EPI_ISL_516444, EPI_ISL_516445, EPI_ISL_516446, EPI_ISL_516447, EPI_ISL_516448, EPI_ISL_516449, EPI_ISL_516451, EPI_ISL_516452, EPI_ISL_516453, EPI_ISL_516454, EPI_ISL_516455, EPI_ISL_516456, EPI_ISL_516457, EPI_ISL_516458, EPI_ISL_516459, EPI_ISL_516460, EPI_ISL_516461, EPI_ISL_516462, EPI_ISL_516463, EPI_ISL_516464, EPI_ISL_516465, EPI_ISL_516466, EPI_ISL_516467, EPI_ISL_516468, EPI_ISL_516469, EPI_ISL_516470, EPI_ISL_516471, EPI_ISL_516472, EPI_ISL_516473, EPI_ISL_516474, EPI_ISL_516475, EPI_ISL_516476, EPI_ISL_516477, EPI_ISL_516478, EPI_ISL_516479, EPI_ISL_516480, EPI_ISL_516481, EPI_ISL_516482, EPI_ISL_516483, EPI_ISL_516484, EPI_ISL_516485, EPI_ISL_516486, EPI_ISL_516487, EPI_ISL_516488, EPI_ISL_516489, EPI_ISL_516490, EPI_ISL_516491, EPI_ISL_516492, EPI_ISL_516493, EPI_ISL_516494, EPI_ISL_516495, EPI_ISL_516496, EPI_ISL_516497, EPI_ISL_516498, EPI_ISL_516499, EPI_ISL_516500, EPI_ISL_516501, EPI_ISL_516502, EPI_ISL_516503, EPI_ISL_516504, EPI_ISL_516505, EPI_ISL_516506, EPI_ISL_516507, EPI_ISL_516509, EPI_ISL_516510, EPI_ISL_516511, EPI_ISL_516512, EPI_ISL_516513, EPI_ISL_516514, EPI_ISL_516515, EPI_ISL_516516, EPI_ISL_516517, EPI_ISL_516518, EPI_ISL_516519, EPI_ISL_516520, EPI_ISL_516521, EPI_ISL_516522, EPI_ISL_516523, EPI_ISL_516524, EPI_ISL_516525, EPI_ISL_516526, EPI_ISL_516527, EPI_ISL_516528, EPI_ISL_516529, EPI_ISL_516530, EPI_ISL_516531, EPI_ISL_516532, EPI_ISL_516533, EPI_ISL_516534, EPI_ISL_516535, EPI_ISL_516536, EPI_ISL_516537, EPI_ISL_516538, EPI_ISL_516539, EPI_ISL_516540, EPI_ISL_516541, EPI_ISL_516542, EPI_ISL_516543, EPI_ISL_516544, EPI_ISL_516545, EPI_ISL_516546, EPI_ISL_516547, EPI_ISL_516548, EPI_ISL_516549 | | | |
| see above | University of Wisconsin-Madison AIDS Vaccine Research Laboratories | University of Wisconsin-Madison AIDS Vaccine Research Laboratories | Gage Moreno, Katarina Braun, et al. AIDS Vaccine Research Laboratories |
| EPI_ISL_516551 | SA Pathology | SA Pathology | Lex Leong, Julien Soubrier, Luke Walters, Chuan Kok Lim, Song Gao, Mark Turra, Karin Kassahn, Ivan Bastian, Geoff Higgins |
| EPI_ISL_516552, EPI_ISL_516553, EPI_ISL_516554, EPI_ISL_516555, EPI_ISL_516556, EPI_ISL_516559, EPI_ISL_516560, EPI_ISL_516561, EPI_ISL_516562, EPI_ISL_516563, EPI_ISL_516564, EPI_ISL_516565, EPI_ISL_516566, EPI_ISL_516568, EPI_ISL_516570, EPI_ISL_516571, EPI_ISL_516572, EPI_ISL_516573, EPI_ISL_516574, EPI_ISL_516575, EPI_ISL_516576, EPI_ISL_516577, EPI_ISL_516578, EPI_ISL_516581, EPI_ISL_516582, EPI_ISL_516583, EPI_ISL_516584, EPI_ISL_516585, EPI_ISL_516586, EPI_ISL_516587, EPI_ISL_516588, EPI_ISL_516589, EPI_ISL_516590, EPI_ISL_516591, EPI_ISL_516592, EPI_ISL_516593, EPI_ISL_516594, EPI_ISL_516595, EPI_ISL_516596, EPI_ISL_516597, EPI_ISL_516598, EPI_ISL_516599, EPI_ISL_516600, EPI_ISL_516601, EPI_ISL_516604, EPI_ISL_516605, EPI_ISL_516606 | | | |
| see above | Viollier AG | Department of Biosystems Science and Engineering, ETH Zürich | Christian Beisel, Sarah Nadeau, Ivan Topolsky, Pedro Ferreira, Philipp Jablonski, Susana Posada-Céspedes, Tobias Schär, Ina Nissen, Natascha Santacroce, Elodie Burcklen, Christiane Beckmann, Maurice Redondo, Olivier Kobel, Christoph Noppen, Sophie Seidel, Noemie Santamaria de Souza, Niko Beerenwinkel, Tanja Stadler |
| EPI_ISL_516609 | Instituto de Diagnostico y Referencia Epidemiologicos (INDRE) | Instituto de Diagnostico y Referencia Epidemiologicos (INDRE) | Ernesto Ramirez-Gonzalez, Abril Rodriguez-Maldonado, Claudia Wong-Arambula , Natividad Cruz-Ortiz, Tatiana Nunez-Garcia, Dayanira Arellano-Suarez, Adnan Araiza-Rodriguez, Fabiola Garces-Ayala, Lucia Hernandez-Rivas, Irma Lopez-Martinez, Gisela Barrera-Badillo. |
| EPI_ISL_516611 | Instituto de Diagnostico y Referencia Epidemiologicos (INDRE) | Instituto de Diagnostico y Referencia Epidemiologicos (INDRE) | Gisela Barrera-Badillo , Abril Rodriguez-Maldonado, Claudia Wong-Arambula , Natividad Cruz-Ortiz, Tatiana Nunez-Garcia, Dayanira Arellano-Suarez, Fabiola Garces-Ayala, Edgar Mendieta-Condado, Lucia Hernandez-Rivas, Irma Lopez-Martinez, Ernesto Ramirez-Gonzalez. |
| EPI_ISL_516613 | Instituto de Diagnostico y Referencia Epidemiologicos (INDRE) | Instituto de Diagnostico y Referencia Epidemiologicos (INDRE) | Ernesto Ramirez-Gonzalez, Abril Rodriguez-Maldonado, Claudia Wong-Arambula , Natividad Cruz-Ortiz, Tatiana Nunez-Garcia, Dayanira Arellano-Suarez, Adnan Araiza-Rodriguez, Edgar Mendieta-Condado, Lucia Hernandez-Rivas, Irma Lopez-Martinez, Gisela Barrera-Badillo. |
| EPI_ISL_516618 | Instituto de Diagnostico y Referencia Epidemiologicos (INDRE) | Instituto de Diagnostico y Referencia Epidemiologicos (INDRE) | Gisela Barrera-Badillo , Abril Rodriguez-Maldonado, Claudia Wong-Arambula , Natividad Cruz-Ortiz, Tatiana Nunez-Garcia, Dayanira Arellano-Suarez, Fabiola Garces-Ayala, Edgar Mendieta-Condado, Lucia Hernandez-Rivas, Irma Lopez-Martinez, Ernesto Ramirez-Gonzalez. |
| EPI_ISL_516620 | Instituto de Diagnostico y Referencia Epidemiologicos (INDRE) | Instituto de Diagnostico y Referencia Epidemiologicos (INDRE) | Gisela Barrera-Badillo , Abril Rodriguez-Maldonado, Claudia Wong-Arambula , Natividad Cruz-Ortiz, Tatiana Nunez-Garcia, Dayanira Arellano-Suarez, Adnan Araiza-Rodriguez, Edgar Mendieta-Condado, Lucia Hernandez-Rivas, Irma Lopez-Martinez, Ernesto Ramirez-Gonzalez. |
| EPI_ISL_516622, EPI_ISL_516624, EPI_ISL_516625 | Instituto de Diagnostico y Referencia Epidemiologicos (INDRE) | Instituto de Diagnostico y Referencia Epidemiologicos (INDRE) | Gisela Barrera-Badillo , Abril Rodriguez-Maldonado, Claudia Wong-Arambula , Natividad Cruz-Ortiz, Tatiana Nunez-Garcia, Dayanira Arellano-Suarez, Fabiola Garces-Ayala, Edgar Mendieta-Condado, Lucia Hernandez-Rivas, Irma Lopez-Martinez, Ernesto Ramirez-Gonzalez. |
| EPI_ISL_516626 | Laboratorio de Referencia Nacional de Virus Respiratorio. Centro Nacional de Salud Publica. Instituto Nacional de Salud Peru. | Laboratorio de Referencia Nacional de Biotecnología y Biología Molecular. Centro Nacional de Salud Publica. Instituto Nacional de Salud Peru. | Carlos Padilla Rojas, Karolyn Vega Chozo, Priscila Lope Pari, Omar Caceres Rey, Marco Galarza Perez, Maribel Huaringa Nuñez, Johanna Balbuena Torres, Henri Bailon Calderon, Nancy Rojas Serrano. |
| EPI_ISL_516627 | SYNLAB Eesti OÜ | Charite Universitatsmedizin Berlin, Institute of Virology | Victor M Corman, Jorn Beheim-Schwarzbach, Barbara Mühlemann, Talitha Veith, Julia Schneider, Paul Naaber, Terry Jones, Christian Drosten |
| EPI_ISL_516628 | Laboratorio de Referencia Nacional de Virus Respiratorio. Centro Nacional de Salud Publica. Instituto Nacional de Salud Peru. | Laboratorio de Referencia Nacional de Biotecnología y Biología Molecular. Centro Nacional de Salud Publica. Instituto Nacional de Salud Peru. | Carlos Padilla Rojas, Karolyn Vega Chozo, Priscila Lope Pari, Omar Caceres Rey, Marco Galarza Perez, Maribel Huaringa Nuñez, Johanna Balbuena Torres, Henri Bailon Calderon, Nancy Rojas Serrano. |
| EPI_ISL_516629, EPI_ISL_516630, EPI_ISL_516631, EPI_ISL_516632, EPI_ISL_516633, EPI_ISL_516634, EPI_ISL_516635, EPI_ISL_516636, EPI_ISL_516637, EPI_ISL_516638, EPI_ISL_516639, EPI_ISL_516640, EPI_ISL_516641, EPI_ISL_516642, EPI_ISL_516644, EPI_ISL_516645 | | | |
| see above | Charité Universitätsmedizin Berlin, Institut für Virologie/Labor Berlin | Charité Universitätsmedizin Berlin, Institut für Virologie/Labor Berlin | Victor M Corman, Barbara Mühlemann, Jörn Beheim-Schwarzbach, Julia Schneider, Talitha Veith, Terry Jones, Christian Drosten |
| EPI_ISL_516646, EPI_ISL_516647 | Laboratorio de Referencia Nacional de Virus Respiratorio. Centro Nacional de Salud Publica. Instituto Nacional de Salud Peru. | Laboratorio de Referencia Nacional de Biotecnología y Biología Molecular. Centro Nacional de Salud Publica. Instituto Nacional de Salud Peru. | Carlos Padilla Rojas, Karolyn Vega Chozo, Priscila Lope Pari, Omar Caceres Rey, Marco Galarza Perez, Maribel Huaringa Nuñez, Johanna Balbuena Torres, Henri Bailon Calderon, Nancy Rojas Serrano. |
| EPI_ISL_516648 | Institute of Microbiology, Universidad San Francisco de Quito | Institute of Microbiology, Universidad San Francisco de Quito | Juan José Guadalupe, Monica Becerra-Wong, Prado-Vivar, Sully Márquez, Bernardo Gutiérrez, Nabih Dahik, Carlos Mena, Ligia Briceño, Verónica Barragán, Patricio Rojas-Silva, Gabriel Trueba, Michelle Grunauer, Paúl Cárdenas |
| EPI_ISL_516649 | Institute of Microbiology, Universidad San Francisco de Quito | Institute of Microbiology, Universidad San Francisco de Quito | Juan José Guadalupe, Monica Becerra-Wong, Bernardo Gutiérrez, Nabih Dahik, Carlos Mena, Ligia Briceño, Verónica Barragán, Patricio Rojas-Silva, Gabriel Trueba, Michelle Grunauer, Paúl Cárdenas |
| EPI_ISL_516650 | Institute of Microbiology, Universidad San Francisco de Quito | Institute of Microbiology, Universidad San Francisco de Quito | Prado-Vivar, Sully Márquez, Juan José Guadalupe, Monica Becerra-Wong, Bernardo Gutiérrez, Verónica Barragán, Patricio Rojas-Silva, Gabriel Trueba, Michelle Grunauer, Paúl Cárdenas |
| EPI_ISL_516651 | Institute of Microbiology, Universidad San Francisco de Quito | Institute of Microbiology, Universidad San Francisco de Quito | Prado-Vivar, Sully Márquez, Juan José Guadalupe, Monica Becerra-Wong, Bernardo Gutiérrez, Nabih Dahik, Carlos Mena, Eulalia Pazmiño, Carolina Pacheco, DamarisSandoya, Verónica Barragán, Patricio Rojas-Silva, Gabriel Trueba, Michelle Grunauer, Paúl Cárdenas |
| EPI_ISL_516652 | Institute of Microbiology, Universidad San Francisco de Quito | Institute of Microbiology, Universidad San Francisco de Quito | Prado-Vivar, Sully Márquez, Juan José Guadalupe, Monica Becerra-Wong, Bernardo Gutiérrez, Nabih Dahik, Carlos Mena, Edy Quizhpe, Yomara Napa, Verónica Barragán, Patricio Rojas-Silva, Gabriel Trueba, Michelle Grunauer, Paúl Cárdenas |
| EPI_ISL_516653, EPI_ISL_516654 | Laboratorio de Referencia Nacional de Virus Respiratorio. Centro Nacional de Salud Publica. Instituto Nacional de Salud Peru. | Laboratorio de Referencia Nacional de Biotecnología y Biología Molecular. Centro Nacional de Salud Publica. Instituto Nacional de Salud Peru. | Carlos Padilla Rojas, Karolyn Vega Chozo, Priscila Lope Pari, Omar Caceres Rey, Marco Galarza Perez, Maribel Huaringa Nuñez, Johanna Balbuena Torres, Henri Bailon Calderon, Nancy Rojas Serrano. |
| EPI_ISL_516655, EPI_ISL_516656, EPI_ISL_516657, EPI_ISL_516659, EPI_ISL_516660, EPI_ISL_516661, EPI_ISL_516662, EPI_ISL_516663, EPI_ISL_516664, EPI_ISL_516665, EPI_ISL_516666, EPI_ISL_516667, EPI_ISL_516668, EPI_ISL_516669, EPI_ISL_516670, EPI_ISL_516671, EPI_ISL_516672, EPI_ISL_516673, EPI_ISL_516674, EPI_ISL_516675, EPI_ISL_516676, EPI_ISL_516677, EPI_ISL_516678, EPI_ISL_516679, EPI_ISL_516680, EPI_ISL_516681, EPI_ISL_516682, EPI_ISL_516683, EPI_ISL_516684 | | | |
| see above | Virginia DCLS | Virginia DCLS | Virginia DCLS |
| EPI_ISL_516685, EPI_ISL_516686 | Laboratorio de Referencia Nacional de Virus Respiratorio. Centro Nacional de Salud Publica. Instituto Nacional de Salud Peru. | Laboratorio de Referencia Nacional de Biotecnología y Biología Molecular. Centro Nacional de Salud Publica. Instituto Nacional de Salud Peru. | Carlos Padilla Rojas, Karolyn Vega Chozo, Priscila Lope Pari, Omar Caceres Rey, Marco Galarza Perez, Maribel Huaringa Nuñez, Johanna Balbuena Torres, Henri Bailon Calderon, Nancy Rojas Serrano. |
| EPI_ISL_516687, EPI_ISL_516688, EPI_ISL_516689, EPI_ISL_516690, EPI_ISL_516691, EPI_ISL_516692, EPI_ISL_516693, EPI_ISL_516694, EPI_ISL_516695, EPI_ISL_516696, EPI_ISL_516698, EPI_ISL_516699, EPI_ISL_516700, EPI_ISL_516701, EPI_ISL_516702, EPI_ISL_516703, EPI_ISL_516704, EPI_ISL_516705, EPI_ISL_516706, EPI_ISL_516707, EPI_ISL_516708, EPI_ISL_516709, EPI_ISL_516710, EPI_ISL_516711, EPI_ISL_516712, EPI_ISL_516713, EPI_ISL_516714, EPI_ISL_516715, EPI_ISL_516716, EPI_ISL_516717, EPI_ISL_516718, EPI_ISL_516719, EPI_ISL_516720 | | | |
| see above | Virginia DCLS | Virginia DCLS | Virginia DCLS |
| EPI_ISL_516721, EPI_ISL_516722 | Laboratorio de Referencia Nacional de Virus Respiratorio. Centro Nacional de Salud Publica. Instituto Nacional de Salud Peru. | Laboratorio de Referencia Nacional de Biotecnología y Biología Molecular. Centro Nacional de Salud Publica. Instituto Nacional de Salud Peru. | Carlos Padilla Rojas, Karolyn Vega Chozo, Priscila Lope Pari, Omar Caceres Rey, Marco Galarza Perez, Maribel Huaringa Nuñez, Johanna Balbuena Torres, Henri Bailon Calderon, Nancy Rojas Serrano. |
| EPI_ISL_516723, EPI_ISL_516724, EPI_ISL_516725, EPI_ISL_516726, EPI_ISL_516727, EPI_ISL_516728, EPI_ISL_516729, EPI_ISL_516730, EPI_ISL_516731, EPI_ISL_516732, EPI_ISL_516733, EPI_ISL_516734, EPI_ISL_516735, EPI_ISL_516736, EPI_ISL_516738, EPI_ISL_516739, EPI_ISL_516740, EPI_ISL_516741, EPI_ISL_516742, EPI_ISL_516743, EPI_ISL_516744, EPI_ISL_516745, EPI_ISL_516746, EPI_ISL_516747 | | | |
| see above | UCSF Clinical Microbiology Laboratory | Chan-Zuckerberg Biohub | CZB Cliahub Consortium |
| EPI_ISL_516748, EPI_ISL_516749, EPI_ISL_516750, EPI_ISL_516752, EPI_ISL_516753, EPI_ISL_516755, EPI_ISL_516756, EPI_ISL_516757, EPI_ISL_516758, EPI_ISL_516762, EPI_ISL_516763, EPI_ISL_516764, EPI_ISL_516765, EPI_ISL_516766, EPI_ISL_516767, EPI_ISL_516768, EPI_ISL_516770, EPI_ISL_516771, EPI_ISL_516772, EPI_ISL_516773, EPI_ISL_516774, EPI_ISL_516775, EPI_ISL_516776, EPI_ISL_516777, EPI_ISL_516778, EPI_ISL_516779 | | | |
| see above | van Bakel Laboratory, Genetics and Genomics Sciences, Icahn School of Medicine at Mount Sinai | van Bakel Laboratory, Genetics and Genomics Sciences, Icahn School of Medicine at Mount Sinai | Andrew G. Letizia, Irene Ramos, Ajay Obla, Carl Goforth, Dawn Weir, Yongchao Ge, Marcas M. Bamman, Jayeeta Dutta, Ethan Ellis, Luis Estrella, Mary-Catherine George, Ana S. Gonzalez-Reiche, Darnell Graham, Adriana van de Guchte, Ramiro Gutierrez, Franca Jones, Aspasia Kalomoiri, Rhonda Lizewski, Stephen Lizewski, Jan Marayag, Nada Marjanovic, Eugene V. Millar, Venugopalan Nair, German Nudelman, Edgar Nunez, Brian Pike, James Regeimbal, Stas Rirak , Ernesto Santa Ana, Rachel S. Gelernter Sealfon, Robert Sebra, Mark Simons, Alessandra Soares-Schanoski, Michael Termini, Sindhu Vangeti, Carlos Williams, Harm van Bakel, Stuart C. Sealfon |
| EPI_ISL_516783, EPI_ISL_516784, EPI_ISL_516785 | Center for Laboratory Control of Infectious Diseases, Korea Centers for Diseases Control and Prevention | Center for Laboratory Control of Infectious Diseases, Korea Centers for Diseases Control and Prevention | Junyoung Kim, Ae Kyung Park, Eunkyung Shin, Jin Sun No, Jeong-Min Kim, Yoon-Seok Chung, Heui Man Kim, Myung Guk Han |
| EPI_ISL_516798, EPI_ISL_516799 | Department of Microbiology, The University of Hong Kong | Department of Microbiology, The University of Hong Kong | Kelvin K.W. To, Kwok-Yung Yuen |
| EPI_ISL_516800 | Rumah Sakit Akademik Universitas Gadjah Mada | Genetics Working Group (Pokja Genetik) Faculty of Medicine, Public Health and Nursing Universitas Gadjah Mada (FK-KMK UGM); Disease Investigation Center Wates Ministry of Agriculture Indonesia; Department of Microbiology FK-KMK UGM; Laboratorium Diagnostik Yayasan Tahija World Mosquito Program (WMP) Yogyakarta Center for Tropical Medicine FK-KMK UGM; Integrated Research center FK-KMK UGM; Department of Computer Science and Electronics FMIPA UGM | Gunadi, Hendra Wibawa, . Marcellus, Mohamad S. Hakim, Edwin W. Daniwijaya, Ludhang P. Rizki, Endah Supriyati, Eggi Arguni, Titik Nuryastuti, Tri Wibawa, Dwi AA Nugrahaningsih, Afiahayati, Siswanto, Alvin S. Kalim, Desyifa Mursalin |
| EPI_ISL_522407 | Department of Infection Prevention and Infectious Diseases, University Hospital Regensburg | Department of Infection Prevention and Infectious Diseases, University Hospital Regensburg | Fritsch,J., Holzmann,T., Schneider-Brachert,W. |
| EPI_ISL_524426 | Egyptian National Cancer Institute (ENCI) | Egyptian National Cancer Institute (ENCI) | Zekri, Abdel Rahman N., Amer,K.E., Ahmed,O.S., Soliman,H.K., Ali,M.A., Hassan,W.A., Mahmoud,A.A., Khattab,A.A., Hafez,M.M., Abouelhoda, Mohamed |
| EPI_ISL_524427 | Egyptian National Cancer Institute (ENCI) | Egyptian National Cancer Institute (ENCI) | Zekri, Abdel Rahman N, Amer,K.E., Ahmed,O.S., Soliman,H.K., Hafez,M.M., Bahnassy,A.A., Abdelhamid,W., Gad,A., Ali,M., Hassan,W., Samir,M., Raouf,A., Hamdy,M.S., Soliman,M.S., Elsissy,M.H., Elkhateeb,S.M., Ezzelarab,M.H., Abouelhoda, Mohamed |
| EPI_ISL_524435 | Department of Immunology, The Scripps Research Institute | Department of Immunology, The Scripps Research Institute | SEARCH Alliance,S.D., Pride,D., Shin,J.H. |

| | | | |
|---|---|---|---|
| EPI_ISL_525472, EPI_ISL_525473 | Institute of Clinical Microbiology and Hygiene, University Hospital Regensburg | Institute of Clinical Microbiology and Hygiene, University Hospital Regensburg | Hiergeist, A. |
| EPI_ISL_529139 | Centro de Desenvolvimento Tecnologico em Saude, Fundacao Oswaldo Cruz | Centro de Desenvolvimento Tecnologico em Saude, Fundacao Oswaldo Cruz | Souza,T.M., Fintelman-Rodrigues,N., De Paula,A.D., Saraiva,F.B., Ferreira,M.A., Sacramento,C.Q., Medeiros,M.A. |
| EPI_ISL_529140 | Centro de Desenvolvimento Tecnologico em Saude, Fundacao Oswaldo Cruz | Centro de Desenvolvimento Tecnologico em Saude, Fundacao Oswaldo Cruz | Souza,T.M., Fintelman-Rodrigues,N., De Paula,A.D., Saraiva,F.B., Ferreira,M.A., Sacramento,C.Q., Medeiros,M.A. |
| EPI_ISL_529141, EPI_ISL_529142, EPI_ISL_529143, EPI_ISL_529144, EPI_ISL_529145 | Egyptian National Cancer Institute (ENCI) | Egyptian National Cancer Institute (ENCI) | Zekri, Abdel Rahman N., Amer,K.E., Ahmed,O.S., Soliman,H.K., Ali,M.A., Hassan,W.A., Mahmoud,A.A., Khattab,A.A., Hafez,M.M., Abouelhoda, Mohamed |
| EPI_ISL_529151, EPI_ISL_529152, EPI_ISL_529153 | Department of Immunology, The Scripps Research Institute | Andersen lab at Scripps Research | Quigley, M., Stefanski, E., Mchardy, I. with SEARCH Alliance San Diego |
| EPI_ISL_529154 | Department of Immunology, The Scripps Research Institute | Andersen lab at Scripps Research | Basler, T., Shephard, J., Austin, B. with SEARCH Alliance San Diego |
| EPI_ISL_529162, EPI_ISL_529164, EPI_ISL_529165, EPI_ISL_529166 | Department of Immunology, The Scripps Research Institute | Andersen lab at Scripps Research | Allison Smither, Gilberto Sabino-Santos, Patricia Snarski, Lilia Melnik, Antoinette Bell, Kaylynn Genemaras, Arnaud Drouin, Dahlene Fusco, Robert Garry with SEARCH Alliance San Diego |
| EPI_ISL_529167, EPI_ISL_529168 | Maryland Department of Health Laboratory | Maryland Department of Health Laboratory | Eric Keller, Maryland Department of Health Laboratories Administration |
| EPI_ISL_529171, EPI_ISL_529173, EPI_ISL_529174, EPI_ISL_529175, EPI_ISL_529176, EPI_ISL_529178, EPI_ISL_529180, EPI_ISL_529181, EPI_ISL_529182, EPI_ISL_529183, EPI_ISL_529184, EPI_ISL_529185, EPI_ISL_529186, EPI_ISL_529187, EPI_ISL_529188, EPI_ISL_529190, EPI_ISL_529191, EPI_ISL_529192, EPI_ISL_529193, EPI_ISL_529194, EPI_ISL_529196, EPI_ISL_529197, EPI_ISL_529198 | | | |
| see above | South Carolina Department of Health and Environmental Control | South Carolina Department of Health and Environmental Control | Haley V. Flores |
