## Supplementary material for "Analysis of the Dynamics and Distribution of SARS-CoV-2 Mutations and its Possible Structural and Functional Implications": GISAID_acknowledgements_part_7

We gratefully acknowledge the following Authors from the Originating laboratories responsible for obtaining the specimens, as well as the Submitting laboratories where the genome data were generated and shared via GISAID, on which this research is based.

All Submitters of data may be contacted directly via www.gisaid.org

| Accession ID | Originating Laboratory | Submitting Laboratory | Authors |
|---|---|---|---|
| EPI_ISL_516801 | Department of Laboratory Medicine, Tan Tock Seng Hospital | Department of Laboratory Medicine, Tan Tock Seng Hospital | Chen YYC, Zair X, Li C, Tang WY, Maurer-Stroh S, Barkham TMS, Nagarajan N, Sessions OM |
| EPI_ISL_516802, EPI_ISL_516803 | Department of Laboratory Medicine, Tan Tock Seng Hospital | Department of Laboratory Medicine, Tan Tock Seng Hospital | Chen YYC, Zair X, Li C, Tang WY, Maurer-Stroh S, Barkham TMS, Nagarajan N, Sessions OM |
| EPI_ISL_516804 | Department of Laboratory Medicine, Tan Tock Seng Hospital | Department of Laboratory Medicine, Tan Tock Seng Hospital | Chen YYC, Zair X, Li C, Tang WY, Maurer-Stroh S, Barkham TMS, Nagarajan N, Sessions OM |
| EPI_ISL_516805 | Department of Laboratory Medicine, Tan Tock Seng Hospital | Department of Laboratory Medicine, Tan Tock Seng Hospital | Chen YYC, Zair X, Li C, Tang WY, Maurer-Stroh S, Barkham TMS, Nagarajan N, Sessions OM |
| EPI_ISL_516806 | Rumah Sakit PKU Gamping | Genetics Working Group (Pokja Genetik) Faculty of Medicine, Public Health and Nursing Universitas Gadjah Mada (FK-KMK UGM); Disease Investigation Center Wates Ministry of Agriculture Indonesia; Department of Microbiology FK-KMK UGM; Laboratorium Diagnostik Yayasan Tahija World Mosquito Program (WMP) Yogyakarta Center for Tropical Medicine FK-KMK UGM; Integrated Research center FK-KMK UGM; Department of Computer Science and Electronics FMIPA UGM | Gunadi, Hendra Wibawa, . Marcellus, Mohamad S. Hakim, Edwin W. Daniwijaya, Ludhang P. Rizki, Endah Supriyati, Eggi Arguni, Titik Nuryastuti, Tri Wibawa, Dwi AA Nugrahaningsih, . Afiahayati , . Siswanto, Ardorisye Saptaty Fornia, Kemala Athollah |
| EPI_ISL_516807, EPI_ISL_516808, EPI_ISL_516809, EPI_ISL_516810, EPI_ISL_516811, EPI_ISL_516812, EPI_ISL_516813, EPI_ISL_516814, EPI_ISL_516815, EPI_ISL_516816, EPI_ISL_516817, EPI_ISL_516818, EPI_ISL_516819, EPI_ISL_516820, EPI_ISL_516821, EPI_ISL_516822, EPI_ISL_516823, EPI_ISL_516824, EPI_ISL_516825, EPI_ISL_516826 | | | |
| see above | National Public Health Laboratory, National Centre for Infectious Diseases | National Public Health Laboratory, National Centre for Infectious Diseases | Mak TM, Octavia S, Zhou Z, Cui L, Lin RTP |
| EPI_ISL_516829 | RSUD Nyi Ageng Serang | Genetics Working Group (Pokja Genetik) Faculty of Medicine, Public Health and Nursing Universitas Gadjah Mada (FK-KMK UGM), Disease Investigation Center Wates Ministry of Agriculture Indonesia, Department of Microbiology FK-KMK UGM, Laboratorium Diagnostik Yayasan Tahija World Mosquito Program (WMP) Yogyakarta Center for Tropical Medicine FK-KMK UGM, Integrated Research Center FK-KMK UGM, Department of Computer Science and Electronics FMIPA UGM | Gunadi, Hendra Wibawa, . Marcellus, Mohamad S. Hakim, Edwin W. Daniwijaya, Ludhang P. Rizki, Endah Supriyati, Eggi Arguni, Titik Nuryastuti, Tri Wibawa, Dwi AA Nugrahaningsih, Afiahayati , . Siswanto, Beby Dewi Sartika, Dyah Ayu Puspitarani |
| EPI_ISL_516831, EPI_ISL_516832, EPI_ISL_516833, EPI_ISL_516834, EPI_ISL_516835, EPI_ISL_516836, EPI_ISL_516841, EPI_ISL_516842, EPI_ISL_516843, EPI_ISL_516844, EPI_ISL_516846, EPI_ISL_516848, EPI_ISL_516852, EPI_ISL_516857, EPI_ISL_516858, EPI_ISL_516863, EPI_ISL_516864, EPI_ISL_516865, EPI_ISL_516866, EPI_ISL_516868, EPI_ISL_516870, EPI_ISL_516873, EPI_ISL_516874, EPI_ISL_516876, EPI_ISL_516877, EPI_ISL_516878, EPI_ISL_516880, EPI_ISL_516883 | | | |
| see above | North West London Pathology, Imperial College Healthcare NHS Trust | Wellcome Sanger Institute for the COVID-19 Genomics UK Consortium | Ling Li, Paul Randell, David Muir, Frankie Bolt, Alison Holmes, James Price, Aileen Rowan, Graham Taylor, Anjna Badhan, Carolina Herrera and Alex Alderton, Roberto Amato, Sonia Goncalves, Ewan Harrison, David K. Jackson, Ian Johnston, Dominic Kwiatkowski, Cordelia Langford, John Sillitoe on behalf of the Wellcome Sanger Institute COVID-19 Surveillance Team (http://www.sanger.ac.uk/covid-team) |
| EPI_ISL_516884, EPI_ISL_516885, EPI_ISL_516886, EPI_ISL_516887, EPI_ISL_516888, EPI_ISL_516889, EPI_ISL_516890, EPI_ISL_516891 | Israel Central Virology laboratory | Israel Central Virology laboratory | Neta Zuckerman, Efrat Dahan Bucris, Oran Erster, Ella Mendelson, Michal Mandelboim |
| EPI_ISL_516892 | Israeli Central Virology laboratory | Israel Central Virology laboratory | Neta Zuckerman, Efrat Dahan Bucris, Oran Erster, Ella Mendelson, Michal Mandelboim |
| EPI_ISL_516893, EPI_ISL_516894, EPI_ISL_516895, EPI_ISL_516896, EPI_ISL_516897, EPI_ISL_516898, EPI_ISL_516899, EPI_ISL_516900, EPI_ISL_516901, EPI_ISL_516902, EPI_ISL_516903, EPI_ISL_516905, EPI_ISL_516906, EPI_ISL_516913, EPI_ISL_516915, EPI_ISL_516917, EPI_ISL_516919, EPI_ISL_516920, EPI_ISL_516921 | | | |
| see above | Israel Central Virology laboratory | Israel Central Virology laboratory | Neta Zuckerman, Efrat Dahan Bucris, Oran Erster, Ella Mendelson, Michal Mandelboim |
| EPI_ISL_516922, EPI_ISL_516923, EPI_ISL_516924, EPI_ISL_516925, EPI_ISL_516926, EPI_ISL_516927, EPI_ISL_516928, EPI_ISL_516929, EPI_ISL_516930, EPI_ISL_516931, EPI_ISL_516932, EPI_ISL_516933 | | | |
| see above | Department for Molecular Diagnostics, Centre for Medical Microbiology, Institute of Public Health of Montenegro | Charité Universitätsmedizin Berlin, Institut für Virologie | Victor M Corman, Terry Jones, Jörn Beheim-Schwarzbach, Barbara Muehlemann, Talitha Veith, Julia Schneider, Marija Govedarica and Danijela Vujošević, Christian Drosten |
| EPI_ISL_516934, EPI_ISL_516938 | Nicolae Testemitanu State University of Medicine and Pharmacy | International Centre for Genetic Engineering and Biotechnology (ICGEB) and ARGO Open Lab Platform for Genome Sequencing | Ulinici M, Licastro D, Dal Monego S, Rajasekharan S, Marcello A |
| EPI_ISL_516940, EPI_ISL_516942, EPI_ISL_516943, EPI_ISL_516946, EPI_ISL_516948, EPI_ISL_516949, EPI_ISL_516969, EPI_ISL_516974, EPI_ISL_516976, EPI_ISL_516977, EPI_ISL_516980, EPI_ISL_516981, EPI_ISL_516982, EPI_ISL_516983, EPI_ISL_516984, EPI_ISL_516986 | | | |
| see above | King Georges Medical University | CSIR-National Botanical Research Institute | Priti Prasad, Shantanu Prakash, Kishan Sahu, Babita Singh, Suruchi Shukla, Hricha Mishra, Danish Nasar Khan , Om Prakash, MLB Bhatt, SK Barik, Mehar H.Asif,Samir V. Sawant,Amita Jain, Sumit Kr. Bag |
| EPI_ISL_516987 | Laboratorio de Referencia Nacional de Virus Respiratorio. Centro Nacional de Salud Publica. Instituto Nacional de Salud Peru. | Laboratorio de Referencia Nacional de Biotecnología y Biología Molecular. Centro Nacional de Salud Publica. Instituto Nacional de Salud Peru. | Carlos Padilla Rojas, Karolyn Vega Chozo, Priscila Lope Pari, Omar Caceres Rey, Marco Galarza Perez, Maribel Huaringa Nuñez, Johanna Balbuena Torres, Henri Bailon Calderon, Nancy Rojas Serrano. |
| EPI_ISL_516988, EPI_ISL_516989 | Public Health Authority of the Slovak Republic, Department of Medical Microbiology | Charite Universitatsmedizin Berlin, Institute of Virology | Victor M Corman, Terry Jones, Jörn Beheim-Schwarzbach, Barbara Muehlemann, Talitha Veith, Julia Schneider, Mgr. Edita Staronova, Christian Drosten |
| EPI_ISL_516993, EPI_ISL_516995, EPI_ISL_516996, EPI_ISL_516999, EPI_ISL_517000 | Department of Pathology, University of Cambridge | COVID-19 Genomics UK (COG-UK) Consortium | Luke W Meredith, M. Estée Török, Myra Hosmillo, William L. Hamilton, Martin D. Curran, Theresa Feltwell, Grant Hall, Anna Yakovleva, Fahad A Khokhar, Charlotte J. Houldcroft, Laura G Caller, Aminu S. Jahun, Sarah L. Caddy, Yasmin Chaudhry, Malte Pinckert, Ian Goodfellow |
| EPI_ISL_517001, EPI_ISL_517002, EPI_ISL_517003, EPI_ISL_517004, EPI_ISL_517005, EPI_ISL_517006, EPI_ISL_517007, EPI_ISL_517008, EPI_ISL_517010, EPI_ISL_517011, EPI_ISL_517012, EPI_ISL_517013, EPI_ISL_517014, EPI_ISL_517015, EPI_ISL_517016, EPI_ISL_517017, EPI_ISL_517018, EPI_ISL_517020, EPI_ISL_517021, EPI_ISL_517022, EPI_ISL_517023, EPI_ISL_517024, EPI_ISL_517025, EPI_ISL_517027, EPI_ISL_517029, EPI_ISL_517031, EPI_ISL_517033, EPI_ISL_517034, EPI_ISL_517036, EPI_ISL_517037, EPI_ISL_517039, EPI_ISL_517040, EPI_ISL_517041, EPI_ISL_517044, EPI_ISL_517045, EPI_ISL_517047, EPI_ISL_517048, EPI_ISL_517049, EPI_ISL_517050, EPI_ISL_517051, EPI_ISL_517052, EPI_ISL_517055, EPI_ISL_517057, EPI_ISL_517058, EPI_ISL_517059, EPI_ISL_517060, EPI_ISL_517061, EPI_ISL_517063, EPI_ISL_517064, EPI_ISL_517066, EPI_ISL_517068, EPI_ISL_517070, EPI_ISL_517071, EPI_ISL_517072, EPI_ISL_517073, EPI_ISL_517074, EPI_ISL_517075, EPI_ISL_517076, EPI_ISL_517077, EPI_ISL_517078, EPI_ISL_517079, EPI_ISL_517080, EPI_ISL_517081, EPI_ISL_517082, EPI_ISL_517083, EPI_ISL_517084, EPI_ISL_517085, EPI_ISL_517087, EPI_ISL_517093, EPI_ISL_517095, EPI_ISL_517108, EPI_ISL_517116, EPI_ISL_517121, EPI_ISL_517123, EPI_ISL_517129, EPI_ISL_517139, EPI_ISL_517144, EPI_ISL_517146, EPI_ISL_517151, EPI_ISL_517155, EPI_ISL_517159, EPI_ISL_517160, EPI_ISL_517165, EPI_ISL_517166, EPI_ISL_517170, EPI_ISL_517171, EPI_ISL_517172, EPI_ISL_517173, EPI_ISL_517174, EPI_ISL_517175, EPI_ISL_517176, EPI_ISL_517177, EPI_ISL_517178, EPI_ISL_517180, EPI_ISL_517182, EPI_ISL_517183, EPI_ISL_517184, EPI_ISL_517185, EPI_ISL_517186, EPI_ISL_517187, EPI_ISL_517188, EPI_ISL_517189, EPI_ISL_517190, EPI_ISL_517191, EPI_ISL_517192, EPI_ISL_517193, EPI_ISL_517195, EPI_ISL_517196, EPI_ISL_517198, EPI_ISL_517199, EPI_ISL_517200, EPI_ISL_517201, EPI_ISL_517203, EPI_ISL_517204, EPI_ISL_517205, EPI_ISL_517206, EPI_ISL_517207, EPI_ISL_517208, EPI_ISL_517209, EPI_ISL_517210, EPI_ISL_517211, EPI_ISL_517212, EPI_ISL_517213, EPI_ISL_517214, EPI_ISL_517215, EPI_ISL_517216, EPI_ISL_517217, EPI_ISL_517218, EPI_ISL_517219, EPI_ISL_517220, EPI_ISL_517221, EPI_ISL_517222, EPI_ISL_517223, EPI_ISL_517224, EPI_ISL_517225, EPI_ISL_517226, EPI_ISL_517227, EPI_ISL_517231, EPI_ISL_517232, EPI_ISL_517233, EPI_ISL_517234, EPI_ISL_517235, EPI_ISL_517236, EPI_ISL_517237, EPI_ISL_517238, EPI_ISL_517240, EPI_ISL_517241, EPI_ISL_517242, EPI_ISL_517243, EPI_ISL_517244, EPI_ISL_517246, EPI_ISL_517247, EPI_ISL_517248, EPI_ISL_517249, EPI_ISL_517250, EPI_ISL_517251, EPI_ISL_517252, EPI_ISL_517253, EPI_ISL_517254, EPI_ISL_517255, EPI_ISL_517256, EPI_ISL_517257, EPI_ISL_517258, EPI_ISL_517259, EPI_ISL_517260, EPI_ISL_517261, EPI_ISL_517262, EPI_ISL_517263, EPI_ISL_517264, EPI_ISL_517265, EPI_ISL_517266, EPI_ISL_517267, EPI_ISL_517268, EPI_ISL_517269, EPI_ISL_517270, EPI_ISL_517271, EPI_ISL_517272, EPI_ISL_517273, EPI_ISL_517275, EPI_ISL_517276, EPI_ISL_517278, EPI_ISL_517279, EPI_ISL_517281, EPI_ISL_517282, EPI_ISL_517284, EPI_ISL_517286, EPI_ISL_517287, EPI_ISL_517288, EPI_ISL_517289, EPI_ISL_517291, EPI_ISL_517292, EPI_ISL_517293, EPI_ISL_517294, EPI_ISL_517296, EPI_ISL_517297, EPI_ISL_517300, EPI_ISL_517301, EPI_ISL_517302, EPI_ISL_517303, EPI_ISL_517304, EPI_ISL_517305, EPI_ISL_517306, EPI_ISL_517307, EPI_ISL_517308, EPI_ISL_517309, EPI_ISL_517310, EPI_ISL_517311, EPI_ISL_517312, EPI_ISL_517313, EPI_ISL_517314, EPI_ISL_517315, EPI_ISL_517316, EPI_ISL_517317, EPI_ISL_517318, EPI_ISL_517319, EPI_ISL_517320, EPI_ISL_517322, EPI_ISL_517323, EPI_ISL_517324, EPI_ISL_517325, EPI_ISL_517326, EPI_ISL_517327, EPI_ISL_517328, EPI_ISL_517329, EPI_ISL_517330, EPI_ISL_517331, EPI_ISL_517332, EPI_ISL_517333, EPI_ISL_517334, EPI_ISL_517335, EPI_ISL_517336, EPI_ISL_517337, EPI_ISL_517338, EPI_ISL_517339, EPI_ISL_517340, EPI_ISL_517341, EPI_ISL_517343, EPI_ISL_517344, EPI_ISL_517345, EPI_ISL_517346, EPI_ISL_517347, EPI_ISL_517348, EPI_ISL_517349, EPI_ISL_517350, EPI_ISL_517351, EPI_ISL_517352, EPI_ISL_517353, EPI_ISL_517354, EPI_ISL_517355, EPI_ISL_517356, EPI_ISL_517357, EPI_ISL_517358, EPI_ISL_517359, EPI_ISL_517360, EPI_ISL_517361, EPI_ISL_517362, EPI_ISL_517363, EPI_ISL_517364, EPI_ISL_517365, EPI_ISL_517366, EPI_ISL_517367, EPI_ISL_517368, EPI_ISL_517369, EPI_ISL_517370, EPI_ISL_517371, EPI_ISL_517373, EPI_ISL_517380, EPI_ISL_517385, EPI_ISL_517396, EPI_ISL_517397, EPI_ISL_517399, EPI_ISL_517410, EPI_ISL_517416, EPI_ISL_517422, EPI_ISL_517432, EPI_ISL_517433, EPI_ISL_517438, EPI_ISL_517444, EPI_ISL_517448, EPI_ISL_517449, EPI_ISL_517452, EPI_ISL_517464, EPI_ISL_517465, EPI_ISL_517466, EPI_ISL_517469, EPI_ISL_517470, EPI_ISL_517471, EPI_ISL_517472, EPI_ISL_517473, EPI_ISL_517475, EPI_ISL_517476, EPI_ISL_517477, EPI_ISL_517479, EPI_ISL_517480, EPI_ISL_517481, EPI_ISL_517482, EPI_ISL_517483, EPI_ISL_517484, EPI_ISL_517485, EPI_ISL_517486, EPI_ISL_517487, EPI_ISL_517488, EPI_ISL_517489, EPI_ISL_517490, EPI_ISL_517491, EPI_ISL_517492, EPI_ISL_517493, EPI_ISL_517494, EPI_ISL_517495, EPI_ISL_517496, EPI_ISL_517497, EPI_ISL_517498, EPI_ISL_517500, EPI_ISL_517501, EPI_ISL_517502, EPI_ISL_517503, EPI_ISL_517504, EPI_ISL_517505, EPI_ISL_517506, EPI_ISL_517507, EPI_ISL_517508, EPI_ISL_517509 | | | |
| see above | Liverpool Clinical Laboratories | COVID-19 Genomics UK (COG-UK) Consortium | Sam Haldenby, Anita Lucaci, Steve Paterson, Julian Hiscox, Alistair Darby, M Almsaud, A Alrezaihi, Muhannad Alruwaili, Stuart D Armstrong, Jones Benjamin, Eleanor G Bentley, Anu Chawla, Jordan J Clark, Angela Cowell, Richard Eccles, Isabel García-Dorival, Matthew Gemmell, Alessandro Gerada, PKF Gilmore, Richard Gregory, Ximeng Han, Catherine Hartley, Margaret Hughes, Miren Iturriza-Gomara, James Johnson, L Luu, Jenifer Manson, Charlotte Nelson, Elaine O'Toole, Cassie Olateju, Rebekah Penrice-Randal , Lucille Rainbow, N.P Randle, Trevor Ian Robinson, Parul Sharma, Ghada T Shawli, James P Stewart, Neil Swainston, Ecaterina Vamos, Joanne Watts, Mark Whitehead |
| EPI_ISL_517512, EPI_ISL_517513, EPI_ISL_517517, EPI_ISL_517519, EPI_ISL_517522, EPI_ISL_517526, EPI_ISL_517529 | Centre for Enzyme Innovation, University of Portsmouth / Translational Research Laboratory, Portsmouth Hospitals NHS Trust | COVID-19 Genomics UK (COG-UK) Consortium | Angela Beckett,Yann Bourgeois,Garry Scarlett,Sharon Glaysher,Scott Elliott,Kelly Bicknell,Robert Impey,Allyson Lloyd,Sarah Wyllie,Ethan Butcher,Anoop Chauhan,Samuel Robson |
| EPI_ISL_517531 | Laboratorio de Referencia Nacional de Virus Respiratorio. Centro Nacional de Salud Publica. Instituto Nacional de Salud Peru. | Laboratorio de Referencia Nacional de Biotecnología y Biología Molecular. Centro Nacional de Salud Publica. Instituto Nacional de Salud Peru. | Carlos Padilla Rojas, Karolyn Vega Chozo, Priscila Lope Pari, Omar Caceres Rey, Marco Galarza Perez, Maribel Huaringa Nuñez, Johanna Balbuena Torres, Henri Bailon Calderon, Nancy Rojas Serrano. |
| EPI_ISL_517532, EPI_ISL_517535, EPI_ISL_517536, EPI_ISL_517537, EPI_ISL_517540, EPI_ISL_517542 | Centre for Enzyme Innovation, University of Portsmouth / Translational Research Laboratory, Portsmouth Hospitals NHS Trust | COVID-19 Genomics UK (COG-UK) Consortium | Angela Beckett,Yann Bourgeois,Garry Scarlett,Sharon Glaysher,Scott Elliott,Kelly Bicknell,Robert Impey,Allyson Lloyd,Sarah Wyllie,Ethan Butcher,Anoop Chauhan,Samuel Robson |
| EPI_ISL_517543, EPI_ISL_517544, EPI_ISL_517546, EPI_ISL_517548, EPI_ISL_517550, EPI_ISL_517552, EPI_ISL_517556, EPI_ISL_517560, EPI_ISL_517561, EPI_ISL_517570, EPI_ISL_517572, EPI_ISL_517575, EPI_ISL_517576 | | | |
| see above | Virology Department, Sheffield Teaching Hospitals NHS Foundation Trust/Department of Infection, Immunity and Cardiovascular Disease, The Medical School, University of Sheffield | COVID-19 Genomics UK (COG-UK) Consortium | Thushan de Silva, Matthew Parker, Nikki Smith, Adri Angyal, Rebecca Brown, Luke Green, Rachel Tucker, Paul Parsons, Danielle Groves, Katie Johnson, Laura Carrilero, Alex Keeley, Dave Partridge, Matthew Wyles, Benjamin Lindsey, Mehmet Yavuz, Mohammad Raza, Cariad Evans |
| EPI_ISL_517580, EPI_ISL_517581, EPI_ISL_517582 | West of Scotland Specialist Virology Centre, NHSGGC / MRC-University of Glasgow Centre for Virus Research | COVID-19 Genomics UK (COG-UK) Consortium | Ana da Silva Filipe, Natasha Johnson, Kathy Smollett, Daniel Mair, Stephen Carmichael, Lily Tong, Jenna Nichols, Elihu Aranday-Cortes, Kirstyn Brunker, Yasmin Parr, Alice Broos, Kyriaki Nomikou; Sarah McDonald, Marc Niebel, Patawee Asamaphan; Richard Orton, Joseph Hughes, Sreenu Vattipally, David L Robertson; Alasdair MacLean, Rory Gunson; Kathy Li, Natasha Jesudason, Rajiv Shah, James Shepherd, Antonia Ho, Emma Thomson |
| EPI_ISL_517583, EPI_ISL_517584, EPI_ISL_517585, EPI_ISL_517586, EPI_ISL_517587, EPI_ISL_517588, EPI_ISL_517589 | Virology Department, Royal Infirmary of Edinburgh, NHS Lothian / School of Biological Sciences, University of Edinburgh / Institute of Genetics and Molecular Medicine, University of Edinburgh | COVID-19 Genomics UK (COG-UK) Consortium | McHugh M, Dewar R, Rooke S, Gallagher M, Balcaza C, O'Toole Á, Scher E, Hill V, McCrone JT, Colquhoun R, Yu X, Jackson B, Rambaut A, Williams TC, Templeton K |
| EPI_ISL_517590, EPI_ISL_517591, EPI_ISL_517592, EPI_ISL_517593, EPI_ISL_517594, EPI_ISL_517595, EPI_ISL_517596, EPI_ISL_517597, EPI_ISL_517598, EPI_ISL_517599, EPI_ISL_517600, EPI_ISL_517601, EPI_ISL_517605, EPI_ISL_517607, EPI_ISL_517609 | | | |

| | | | |
|---|---|---|---|
| see above | Wales Specialist Virology Centre Sequencing lab: Pathogen Genomics Unit | COVID-19 Genomics UK (COG-UK) Consortium | Catherine Moore, Johnathan Evans, Laura Gifford, Malorie Perry, Simon Cottrell, Angela Marchbank, Alec Birchley, Alexander Adams, Amy Gaskin, Bree Gatica-Wilcox, Jason Coombes, Joel Southgate, Lauren Gilbert, Lee Graham, Nicole Pacchiarini, Sara Kumziene-Summerhayes, Sarah Taylor, Sophie Jones, Sara Rey, Matthew Bull, Joanne Watkins, Sally Corden, Tom Connor |
| EPI_ISL_517611 | Institute of Medical Microbiology, University Medical Center Goettingen | Institute of Human Genetics, University Medical Center Goettingen | Arne Zibat, Gabriela Salinas-Riester, Andreas E Zautner, Maren Sitte, Antje Dickmanns, Maren Stegmann, Uwe Groß, Matthias Dobbelstein, Bernd Wollnik |
| EPI_ISL_517613, EPI_ISL_517615, EPI_ISL_517616, EPI_ISL_517620, EPI_ISL_517621, EPI_ISL_517622, EPI_ISL_517623, EPI_ISL_517624, EPI_ISL_517625, EPI_ISL_517626, EPI_ISL_517627, EPI_ISL_517628, EPI_ISL_517629, EPI_ISL_517630, EPI_ISL_517631, EPI_ISL_517633, EPI_ISL_517636, EPI_ISL_517637, EPI_ISL_517638, EPI_ISL_517639, EPI_ISL_517640, EPI_ISL_517641, EPI_ISL_517644, EPI_ISL_517645, EPI_ISL_517646, EPI_ISL_517647, EPI_ISL_517648, EPI_ISL_517650, EPI_ISL_517651, EPI_ISL_517652, EPI_ISL_517653, EPI_ISL_517654, EPI_ISL_517657, EPI_ISL_517658, EPI_ISL_517659 | | | |
| see above | Academic Hospital Paramaribo | Erasmus Medical Center | Bas Oude Munnink, Dion Gajadin, Ed Ijzerman, Emmanuelle Munger, Gary Gummels, Ingrid Krishnadath, Lycke Woittiez, Marion Koopmans, Mireille Van de Veer, Princes Wongsowidjojo, Radjesh Ori, Rohma Banwari, Stephen Vreden |
| EPI_ISL_517686, EPI_ISL_517687, EPI_ISL_517713 | Laboratorio de Referencia Nacional de Virus Respiratorio. Centro Nacional de Salud Publica. Instituto Nacional de Salud Peru. | Laboratorio de Referencia Nacional de Biotecnología y Biología Molecular. Centro Nacional de Salud Publica. Instituto Nacional de Salud Peru. | Carlos Padilla Rojas, Karolyn Vega Chozo, Priscila Lope Pari, Omar Caceres Rey, Marco Galarza Perez, Maribel Huaringa Nuñez, Johanna Balbuena Torres, Henri Bailon Calderon, Nancy Rojas Serrano. |
| EPI_ISL_517770 | Laboratorio de Referencia Nacional de Virus Respiratorio. Centro Nacional de Salud Publica. Instituto Nacional de Salud Peru | Laboratorio de Referencia Nacional de Biotecnología y Biología Molecular. Centro Nacional de Salud Publica. Instituto Nacional de Salud Peru. | Carlos Padilla Rojas, Karolyn Vega Chozo, Priscila Lope Pari, Omar Caceres Rey, Marco Galarza Perez, Maribel Huaringa Nuñez, Johanna Balbuena Torres, Henri Bailon Calderon, Nancy Rojas Serrano. |
| EPI_ISL_517772, EPI_ISL_517778, EPI_ISL_517790, EPI_ISL_517791, EPI_ISL_517792, EPI_ISL_517794, EPI_ISL_517795, EPI_ISL_517797, EPI_ISL_517798, EPI_ISL_517800, EPI_ISL_517801, EPI_ISL_517802, EPI_ISL_517803, EPI_ISL_517804, EPI_ISL_517805, EPI_ISL_517806, EPI_ISL_517807, EPI_ISL_517808, EPI_ISL_517809, EPI_ISL_517810, EPI_ISL_517812, EPI_ISL_517813, EPI_ISL_517814, EPI_ISL_517815, EPI_ISL_517816, EPI_ISL_517817, EPI_ISL_517819, EPI_ISL_517821, EPI_ISL_517823, EPI_ISL_517825, EPI_ISL_517827, EPI_ISL_517828, EPI_ISL_517829, EPI_ISL_517830, EPI_ISL_517831, EPI_ISL_517832, EPI_ISL_517833, EPI_ISL_517834, EPI_ISL_517835, EPI_ISL_517836, EPI_ISL_517839, EPI_ISL_517840, EPI_ISL_517842, EPI_ISL_517843, EPI_ISL_517844, EPI_ISL_517846, EPI_ISL_517847, EPI_ISL_517849, EPI_ISL_517850, EPI_ISL_517852, EPI_ISL_517854, EPI_ISL_517857, EPI_ISL_517858, EPI_ISL_517859, EPI_ISL_517860, EPI_ISL_517861, EPI_ISL_517862, EPI_ISL_517863, EPI_ISL_517864, EPI_ISL_517865, EPI_ISL_517866, EPI_ISL_517867, EPI_ISL_517868, EPI_ISL_517869, EPI_ISL_517870, EPI_ISL_517871, EPI_ISL_517873, EPI_ISL_517875, EPI_ISL_517877, EPI_ISL_517878, EPI_ISL_517879, EPI_ISL_517880, EPI_ISL_517882, EPI_ISL_517883, EPI_ISL_517885, EPI_ISL_517888, EPI_ISL_517889, EPI_ISL_517890, EPI_ISL_517892, EPI_ISL_517893, EPI_ISL_517894, EPI_ISL_517895, EPI_ISL_517896, EPI_ISL_517897, EPI_ISL_517899, EPI_ISL_517900, EPI_ISL_517902, EPI_ISL_517903, EPI_ISL_517904, EPI_ISL_517905, EPI_ISL_517906, EPI_ISL_517907, EPI_ISL_517908, EPI_ISL_517909, EPI_ISL_517910, EPI_ISL_517911, EPI_ISL_517912, EPI_ISL_517914, EPI_ISL_517915, EPI_ISL_517916, EPI_ISL_517918, EPI_ISL_517920, EPI_ISL_517921, EPI_ISL_517922, EPI_ISL_517923, EPI_ISL_517924, EPI_ISL_517925, EPI_ISL_517926, EPI_ISL_517928, EPI_ISL_517929, EPI_ISL_517930, EPI_ISL_517931, EPI_ISL_517932, EPI_ISL_517933, EPI_ISL_517934, EPI_ISL_517936, EPI_ISL_517937, EPI_ISL_517939, EPI_ISL_517940, EPI_ISL_517941, EPI_ISL_517944, EPI_ISL_517945, EPI_ISL_517946, EPI_ISL_517947, EPI_ISL_517949, EPI_ISL_517950, EPI_ISL_517951, EPI_ISL_517952, EPI_ISL_517953, EPI_ISL_517954, EPI_ISL_517956 | | | |
| see above | Florida Bureau of Public Health Laboratories | Florida Bureau of Public Health Laboratories | Sarah Schmedes, Jason Blanton |
| EPI_ISL_517958 | Laboratorio de Referencia Nacional de Virus Respiratorio. Centro Nacional de Salud Publica. Instituto Nacional de Salud Peru. | Laboratorio de Referencia Nacional de Biotecnología y Biología Molecular. Centro Nacional de Salud Publica. Instituto Nacional de Salud Peru. | Carlos Padilla Rojas, Karolyn Vega Chozo, Priscila Lope Pari, Omar Caceres Rey, Marco Galarza Perez, Maribel Huaringa Nuñez, Johanna Balbuena Torres, Henri Bailon Calderon, Nancy Rojas Serrano. |
| EPI_ISL_517959 | Laboratorio de Referencia Nacional de Virus Respiratorio. Centro Nacional de Salud Publica. Instituto Nacional de Salud Peru | Laboratorio de Referencia Nacional de Biotecnología y Biología Molecular. Centro Nacional de Salud Publica. Instituto Nacional de Salud Peru. | Carlos Padilla Rojas, Karolyn Vega Chozo, Priscila Lope Pari, Omar Caceres Rey, Marco Galarza Perez, Maribel Huaringa Nuñez, Johanna Balbuena Torres, Henri Bailon Calderon, Nancy Rojas Serrano. |
| EPI_ISL_517969 | Texas Department of State Health Services | Texas Department of State Health Services | Rashmi Tuladhar, Bonnie Oh, Cara Akrout, Jenny Zhang, Maliha Rahman, Anita Pokharel, Myong Koag, Chun Wang, Rachel Lee, Grace Kubin |
| EPI_ISL_517996, EPI_ISL_517997, EPI_ISL_517998, EPI_ISL_517999, EPI_ISL_518001, EPI_ISL_518002, EPI_ISL_518003, EPI_ISL_518004, EPI_ISL_518005, EPI_ISL_518006, EPI_ISL_518007, EPI_ISL_518008, EPI_ISL_518009, EPI_ISL_518010, EPI_ISL_518011, EPI_ISL_518013, EPI_ISL_518014, EPI_ISL_518015, EPI_ISL_518016 | | | |
| see above | Singapore General Hospital | Department of Microbiology | Nurdyana Abdul Rahman, Kun Lee Lim, Chenhao Li, Kian Sing Chan, Lynette Oon, Kern Rei Chng, Niranjan Nagarajan, Karrie Ko |
| EPI_ISL_518034, EPI_ISL_518035, EPI_ISL_518037, EPI_ISL_518038, EPI_ISL_518039, EPI_ISL_518040, EPI_ISL_518041, EPI_ISL_518042, EPI_ISL_518043, EPI_ISL_518044, EPI_ISL_518045, EPI_ISL_518046, EPI_ISL_518047, EPI_ISL_518048, EPI_ISL_518049, EPI_ISL_518050, EPI_ISL_518051, EPI_ISL_518052 | | | |
| see above | NHLS-IALCH | KRISP, KZN Research Innovation and Sequencing Platform | Giandhari J, Pillay S, Lessells R, Mdlalose K, York D, Khan S, Tegally H, Wilkinson E, de Oliveira T |
| EPI_ISL_518063, EPI_ISL_518064, EPI_ISL_518065, EPI_ISL_518066, EPI_ISL_518067, EPI_ISL_518068, EPI_ISL_518069, EPI_ISL_518070, EPI_ISL_518071, EPI_ISL_518072, EPI_ISL_518073, EPI_ISL_518074, EPI_ISL_518075, EPI_ISL_518076, EPI_ISL_518077, EPI_ISL_518078, EPI_ISL_518079, EPI_ISL_518080 | | | |
| see above | Microbiological Diagnostic Unit - Public Health Laboratory (MDU-PHL) | MDU-PHL | Seemann T., Schultz M., Sait, M., Sherry, N. |
| EPI_ISL_518081, EPI_ISL_518082, EPI_ISL_518083, EPI_ISL_518084, EPI_ISL_518086, EPI_ISL_518088, EPI_ISL_518089, EPI_ISL_518090, EPI_ISL_518091, EPI_ISL_518092 | Victorian Infectious Diseases Reference Laboratory (VIDRL) | VIDRL and MDU-PHL | Caly L., Seemann T., Sait, M., Schultz M., Druce J., Sherry, N. |
| EPI_ISL_518093, EPI_ISL_518094, EPI_ISL_518096, EPI_ISL_518097 | Microbiological Diagnostic Unit - Public Health Laboratory (MDU-PHL) | MDU-PHL | Seemann T., Schultz M., Sait, M., Sherry, N. |
| EPI_ISL_518099, EPI_ISL_518100, EPI_ISL_518101, EPI_ISL_518102, EPI_ISL_518104, EPI_ISL_518105, EPI_ISL_518106, EPI_ISL_518107, EPI_ISL_518108, EPI_ISL_518109, EPI_ISL_518110, EPI_ISL_518111, EPI_ISL_518112, EPI_ISL_518113, EPI_ISL_518114, EPI_ISL_518115, EPI_ISL_518116, EPI_ISL_518118, EPI_ISL_518119, EPI_ISL_518120, EPI_ISL_518121, EPI_ISL_518122, EPI_ISL_518123, EPI_ISL_518124, EPI_ISL_518125, EPI_ISL_518126, EPI_ISL_518127, EPI_ISL_518128, EPI_ISL_518129, EPI_ISL_518130, EPI_ISL_518131, EPI_ISL_518132, EPI_ISL_518133, EPI_ISL_518134, EPI_ISL_518135, EPI_ISL_518136, EPI_ISL_518138, EPI_ISL_518139, EPI_ISL_518140, EPI_ISL_518141, EPI_ISL_518142, EPI_ISL_518143, EPI_ISL_518144, EPI_ISL_518145, EPI_ISL_518146, EPI_ISL_518147, EPI_ISL_518148, EPI_ISL_518149, EPI_ISL_518150, EPI_ISL_518151, EPI_ISL_518152, EPI_ISL_518153, EPI_ISL_518154, EPI_ISL_518155, EPI_ISL_518156, EPI_ISL_518157, EPI_ISL_518158, EPI_ISL_518159, EPI_ISL_518160, EPI_ISL_518161, EPI_ISL_518162, EPI_ISL_518163, EPI_ISL_518165, EPI_ISL_518166, EPI_ISL_518167, EPI_ISL_518169, EPI_ISL_518170, EPI_ISL_518171, EPI_ISL_518174, EPI_ISL_518175, EPI_ISL_518176, EPI_ISL_518177, EPI_ISL_518179, EPI_ISL_518180, EPI_ISL_518181, EPI_ISL_518182, EPI_ISL_518183, EPI_ISL_518184, EPI_ISL_518185, EPI_ISL_518186, EPI_ISL_518188, EPI_ISL_518191, EPI_ISL_518194, EPI_ISL_518195, EPI_ISL_518196, EPI_ISL_518198, EPI_ISL_518199, EPI_ISL_518200, EPI_ISL_518201, EPI_ISL_518202, EPI_ISL_518203, EPI_ISL_518204, EPI_ISL_518205, EPI_ISL_518206, EPI_ISL_518207, EPI_ISL_518208, EPI_ISL_518209, EPI_ISL_518210, EPI_ISL_518211, EPI_ISL_518212, EPI_ISL_518213, EPI_ISL_518214, EPI_ISL_518215, EPI_ISL_518217, EPI_ISL_518218, EPI_ISL_518219, EPI_ISL_518220, EPI_ISL_518221, EPI_ISL_518222, EPI_ISL_518223, EPI_ISL_518224, EPI_ISL_518226, EPI_ISL_518227, EPI_ISL_518228, EPI_ISL_518229, EPI_ISL_518230 | | | |
| see above | Victorian Infectious Diseases Reference Laboratory (VIDRL) | VIDRL and MDU-PHL | Caly L., Seemann T., Sait, M., Schultz M., Druce J., Sherry, N. |
| EPI_ISL_518231, EPI_ISL_518232, EPI_ISL_518234, EPI_ISL_518235, EPI_ISL_518236, EPI_ISL_518238 | Microbiological Diagnostic Unit - Public Health Laboratory (MDU-PHL) | MDU-PHL | Seemann T., Schultz M., Sait, M., Sherry, N. |
| EPI_ISL_518239, EPI_ISL_518240 | Victorian Infectious Diseases Reference Laboratory (VIDRL) | VIDRL and MDU-PHL | Caly L., Seemann T., Sait, M., Schultz M., Druce J., Sherry, N. |
| EPI_ISL_518241, EPI_ISL_518242 | Microbiological Diagnostic Unit - Public Health Laboratory (MDU-PHL) | MDU-PHL | Seemann T., Schultz M., Sait, M., Sherry, N. |
| EPI_ISL_518244, EPI_ISL_518246, EPI_ISL_518247, EPI_ISL_518248, EPI_ISL_518249 | Victorian Infectious Diseases Reference Laboratory (VIDRL) | VIDRL and MDU-PHL | Caly L., Seemann T., Sait, M., Schultz M., Druce J., Sherry, N. |
| EPI_ISL_518250, EPI_ISL_518251, EPI_ISL_518252, EPI_ISL_518253, EPI_ISL_518254, EPI_ISL_518255, EPI_ISL_518256, EPI_ISL_518257, EPI_ISL_518258, EPI_ISL_518259, EPI_ISL_518260, EPI_ISL_518261, EPI_ISL_518262, EPI_ISL_518263, EPI_ISL_518264, EPI_ISL_518265, EPI_ISL_518266, EPI_ISL_518267, EPI_ISL_518268, EPI_ISL_518269, EPI_ISL_518270, EPI_ISL_518271, EPI_ISL_518272, EPI_ISL_518273, EPI_ISL_518274, EPI_ISL_518275, EPI_ISL_518276, EPI_ISL_518277, EPI_ISL_518278, EPI_ISL_518282, EPI_ISL_518283, EPI_ISL_518284, EPI_ISL_518285, EPI_ISL_518287, EPI_ISL_518288, EPI_ISL_518289, EPI_ISL_518290, EPI_ISL_518292, EPI_ISL_518293, EPI_ISL_518294, EPI_ISL_518295, EPI_ISL_518296, EPI_ISL_518297, EPI_ISL_518298, EPI_ISL_518299, EPI_ISL_518300, EPI_ISL_518301, EPI_ISL_518302, EPI_ISL_518304, EPI_ISL_518305, EPI_ISL_518306, EPI_ISL_518307, EPI_ISL_518308, EPI_ISL_518309, EPI_ISL_518311, EPI_ISL_518312, EPI_ISL_518313, EPI_ISL_518314, EPI_ISL_518316, EPI_ISL_518318, EPI_ISL_518319, EPI_ISL_518321, EPI_ISL_518322, EPI_ISL_518323, EPI_ISL_518324, EPI_ISL_518325, EPI_ISL_518326, EPI_ISL_518328, EPI_ISL_518329, EPI_ISL_518330, EPI_ISL_518331, EPI_ISL_518332, EPI_ISL_518333, EPI_ISL_518334, EPI_ISL_518335, EPI_ISL_518336, EPI_ISL_518337, EPI_ISL_518339, EPI_ISL_518340, EPI_ISL_518341, EPI_ISL_518342, EPI_ISL_518343, EPI_ISL_518344, EPI_ISL_518345, EPI_ISL_518346, EPI_ISL_518347, EPI_ISL_518349, EPI_ISL_518350, EPI_ISL_518351, EPI_ISL_518352, EPI_ISL_518353, EPI_ISL_518354, EPI_ISL_518355, EPI_ISL_518356, EPI_ISL_518357, EPI_ISL_518358, EPI_ISL_518359, EPI_ISL_518360, EPI_ISL_518361, EPI_ISL_518362, EPI_ISL_518363, EPI_ISL_518364, EPI_ISL_518365, EPI_ISL_518367, EPI_ISL_518368, EPI_ISL_518369, EPI_ISL_518370, EPI_ISL_518371, EPI_ISL_518374, EPI_ISL_518375, EPI_ISL_518376, EPI_ISL_518378, EPI_ISL_518380, EPI_ISL_518381, EPI_ISL_518382, EPI_ISL_518383, EPI_ISL_518384, EPI_ISL_518386, EPI_ISL_518387 | | | |
| see above | Microbiological Diagnostic Unit - Public Health Laboratory (MDU-PHL) | MDU-PHL | Seemann T., Schultz M., Sait, M., Sherry, N. |
| EPI_ISL_518389, EPI_ISL_518391, EPI_ISL_518393, EPI_ISL_518394, EPI_ISL_518395, EPI_ISL_518396, EPI_ISL_518397, EPI_ISL_518398, EPI_ISL_518399, EPI_ISL_518401 | Victorian Infectious Diseases Reference Laboratory (VIDRL) | VIDRL and MDU-PHL | Caly L., Seemann T., Sait, M., Schultz M., Druce J., Sherry, N. |
| EPI_ISL_518402, EPI_ISL_518403, EPI_ISL_518404, EPI_ISL_518405, EPI_ISL_518406, EPI_ISL_518407, EPI_ISL_518410, EPI_ISL_518411, EPI_ISL_518412, EPI_ISL_518415, EPI_ISL_518416, EPI_ISL_518422 | | | |
| see above | Microbiological Diagnostic Unit - Public Health Laboratory (MDU-PHL) | MDU-PHL | Seemann T., Schultz M., Sait, M., Sherry, N. |
| EPI_ISL_518424, EPI_ISL_518426, EPI_ISL_518428, EPI_ISL_518429, EPI_ISL_518431, EPI_ISL_518433, EPI_ISL_518434, EPI_ISL_518435, EPI_ISL_518436, EPI_ISL_518438, EPI_ISL_518439, EPI_ISL_518440, EPI_ISL_518442, EPI_ISL_518443, EPI_ISL_518444, EPI_ISL_518445, EPI_ISL_518446, EPI_ISL_518448, EPI_ISL_518449, EPI_ISL_518452, EPI_ISL_518453, EPI_ISL_518454, EPI_ISL_518456, EPI_ISL_518457, EPI_ISL_518459, EPI_ISL_518460, EPI_ISL_518461, EPI_ISL_518462, EPI_ISL_518463, EPI_ISL_518465, EPI_ISL_518466, EPI_ISL_518467, EPI_ISL_518469, EPI_ISL_518470, EPI_ISL_518471 | | | |
| see above | Victorian Infectious Diseases Reference Laboratory (VIDRL) | VIDRL and MDU-PHL | Caly L., Seemann T., Sait, M., Schultz M., Druce J., Sherry, N. |
| EPI_ISL_518472, EPI_ISL_518473, EPI_ISL_518474, EPI_ISL_518475, EPI_ISL_518476, EPI_ISL_518477, EPI_ISL_518479, EPI_ISL_518480, EPI_ISL_518482, EPI_ISL_518483, EPI_ISL_518484, EPI_ISL_518485, EPI_ISL_518486, EPI_ISL_518487, EPI_ISL_518488, EPI_ISL_518490, EPI_ISL_518491, EPI_ISL_518492, EPI_ISL_518493, EPI_ISL_518494, EPI_ISL_518495, EPI_ISL_518496, EPI_ISL_518497, EPI_ISL_518498, EPI_ISL_518499, EPI_ISL_518500, EPI_ISL_518501, EPI_ISL_518502, EPI_ISL_518503, EPI_ISL_518504, EPI_ISL_518505, EPI_ISL_518506, EPI_ISL_518507, EPI_ISL_518508, EPI_ISL_518509, EPI_ISL_518510, EPI_ISL_518511, EPI_ISL_518514, EPI_ISL_518515, EPI_ISL_518516, EPI_ISL_518517, EPI_ISL_518518, EPI_ISL_518519, EPI_ISL_518520, EPI_ISL_518522, EPI_ISL_518523, EPI_ISL_518524, EPI_ISL_518525, EPI_ISL_518526, EPI_ISL_518527, EPI_ISL_518528, EPI_ISL_518529, EPI_ISL_518530, EPI_ISL_518531, EPI_ISL_518533, EPI_ISL_518534, EPI_ISL_518535, EPI_ISL_518536, EPI_ISL_518537, EPI_ISL_518538, EPI_ISL_518539, EPI_ISL_518540, EPI_ISL_518541, EPI_ISL_518542, EPI_ISL_518544, EPI_ISL_518545, EPI_ISL_518546, EPI_ISL_518547, EPI_ISL_518548, EPI_ISL_518549, EPI_ISL_518550, EPI_ISL_518551, EPI_ISL_518552, EPI_ISL_518553, EPI_ISL_518554, EPI_ISL_518555, EPI_ISL_518556, EPI_ISL_518557, EPI_ISL_518558, EPI_ISL_518559, EPI_ISL_518560, EPI_ISL_518561, EPI_ISL_518562, EPI_ISL_518563, EPI_ISL_518564, EPI_ISL_518565, EPI_ISL_518567, EPI_ISL_518568, EPI_ISL_518569, EPI_ISL_518570, EPI_ISL_518571, EPI_ISL_518572, EPI_ISL_518573, EPI_ISL_518574, EPI_ISL_518576, EPI_ISL_518577, EPI_ISL_518578, EPI_ISL_518579, EPI_ISL_518580, EPI_ISL_518581, EPI_ISL_518582, EPI_ISL_518584, EPI_ISL_518586, EPI_ISL_518587, EPI_ISL_518588, EPI_ISL_518589, EPI_ISL_518590, EPI_ISL_518591, EPI_ISL_518592, EPI_ISL_518594, EPI_ISL_518595, EPI_ISL_518596, EPI_ISL_518597, EPI_ISL_518599, EPI_ISL_518600, EPI_ISL_518601, EPI_ISL_518602, EPI_ISL_518603, EPI_ISL_518604, EPI_ISL_518605, EPI_ISL_518606, EPI_ISL_518607, EPI_ISL_518608, EPI_ISL_518609, EPI_ISL_518610, EPI_ISL_518611, EPI_ISL_518612, EPI_ISL_518613, EPI_ISL_518614, EPI_ISL_518615, EPI_ISL_518616, EPI_ISL_518617, EPI_ISL_518618, EPI_ISL_518619, EPI_ISL_518620, EPI_ISL_518621, EPI_ISL_518622, EPI_ISL_518623, EPI_ISL_518624, EPI_ISL_518625, EPI_ISL_518626, EPI_ISL_518627, EPI_ISL_518628, EPI_ISL_518629, EPI_ISL_518630, EPI_ISL_518631, EPI_ISL_518632, EPI_ISL_518633, EPI_ISL_518634, EPI_ISL_518635, EPI_ISL_518636, EPI_ISL_518637, EPI_ISL_518638, EPI_ISL_518639, EPI_ISL_518640, EPI_ISL_518641, EPI_ISL_518642, EPI_ISL_518643, EPI_ISL_518644, EPI_ISL_518645, EPI_ISL_518646, EPI_ISL_518647, EPI_ISL_518649 | | | |
| see above | Microbiological Diagnostic Unit - Public Health Laboratory (MDU-PHL) | MDU-PHL | Seemann T., Schultz M., Sait, M., Sherry, N. |
| EPI_ISL_518651 | Microbiological Diagnostic Unit - Public Health Laboratory (MDU-PHL) | Microbiological Diagnostic Unit - Public Health Laboratory (MDU-PHL) | Seemann T., Schultz M., Sait, M., Sherry, N. |
| EPI_ISL_518652, EPI_ISL_518653, EPI_ISL_518654, EPI_ISL_518655, EPI_ISL_518656, EPI_ISL_518657, EPI_ISL_518658, EPI_ISL_518659, EPI_ISL_518660, EPI_ISL_518662, EPI_ISL_518664, EPI_ISL_518665, EPI_ISL_518666, EPI_ISL_518667, EPI_ISL_518669, EPI_ISL_518670, EPI_ISL_518671, EPI_ISL_518672, EPI_ISL_518673, EPI_ISL_518674, EPI_ISL_518675, EPI_ISL_518676, EPI_ISL_518677, EPI_ISL_518678, EPI_ISL_518679, EPI_ISL_518680, EPI_ISL_518681, EPI_ISL_518682, EPI_ISL_518683, EPI_ISL_518684, EPI_ISL_518685, EPI_ISL_518686, EPI_ISL_518687, EPI_ISL_518688, EPI_ISL_518689, EPI_ISL_518690, EPI_ISL_518691, EPI_ISL_518692, EPI_ISL_518693, EPI_ISL_518694, EPI_ISL_518695, EPI_ISL_518696, EPI_ISL_518697, EPI_ISL_518698, EPI_ISL_518699, EPI_ISL_518700, EPI_ISL_518701, EPI_ISL_518702, EPI_ISL_518703, EPI_ISL_518704, EPI_ISL_518705, EPI_ISL_518706, EPI_ISL_518707, EPI_ISL_518708, EPI_ISL_518709, EPI_ISL_518711, EPI_ISL_518712, EPI_ISL_518713, EPI_ISL_518714, EPI_ISL_518715, EPI_ISL_518716, EPI_ISL_518717, EPI_ISL_518718, EPI_ISL_518719, EPI_ISL_518720, EPI_ISL_518721, EPI_ISL_518722 | | | |
| see above | Microbiological Diagnostic Unit - Public Health Laboratory (MDU-PHL) | MDU-PHL | Seemann T., Schultz M., Sait, M., Sherry, N. |
| EPI_ISL_518723 | Victorian Infectious Diseases Reference Laboratory (VIDRL) | VIDRL and MDU-PHL | Caly L., Seemann T., Sait, M., Schultz M., Druce J., Sherry, N. |
| EPI_ISL_518724, EPI_ISL_518725, EPI_ISL_518726, EPI_ISL_518727, EPI_ISL_518728, EPI_ISL_518729, EPI_ISL_518730, EPI_ISL_518731, EPI_ISL_518732, EPI_ISL_518733, EPI_ISL_518734, EPI_ISL_518735, EPI_ISL_518736, EPI_ISL_518737, EPI_ISL_518738, EPI_ISL_518739, EPI_ISL_518740, EPI_ISL_518741, EPI_ISL_518742, EPI_ISL_518743, EPI_ISL_518744, EPI_ISL_518745, EPI_ISL_518746, EPI_ISL_518747, EPI_ISL_518748, EPI_ISL_518749, EPI_ISL_518750, EPI_ISL_518751, EPI_ISL_518752, EPI_ISL_518753, EPI_ISL_518754, EPI_ISL_518755, EPI_ISL_518756, EPI_ISL_518757, EPI_ISL_518758, EPI_ISL_518759, EPI_ISL_518760, EPI_ISL_518761, EPI_ISL_518762, EPI_ISL_518763, EPI_ISL_518765, EPI_ISL_518766, EPI_ISL_518767, EPI_ISL_518768, EPI_ISL_518769, EPI_ISL_518770, EPI_ISL_518771, EPI_ISL_518773, EPI_ISL_518774, EPI_ISL_518775, EPI_ISL_518776, EPI_ISL_518777, EPI_ISL_518778, EPI_ISL_518779, EPI_ISL_518780, EPI_ISL_518781, EPI_ISL_518782, EPI_ISL_518783, EPI_ISL_518784, EPI_ISL_518785, EPI_ISL_518786, EPI_ISL_518787, EPI_ISL_518788, EPI_ISL_518789, EPI_ISL_518790, EPI_ISL_518791, EPI_ISL_518792, EPI_ISL_518795, EPI_ISL_518796 | | | |

| | | | |
|---|---|---|---|
| see above | Microbiological Diagnostic Unit - Public Health Laboratory (MDU-PHL) | MDU-PHL | Seemann T., Schultz M., Sait, M., Sherry, N. |
| EPI_ISL_518799, EPI_ISL_518801, EPI_ISL_518802, EPI_ISL_518803, EPI_ISL_518804, EPI_ISL_518805, EPI_ISL_518806, EPI_ISL_518807, EPI_ISL_518808, EPI_ISL_518809, EPI_ISL_518810, EPI_ISL_518812, EPI_ISL_518814, EPI_ISL_518816, EPI_ISL_518817 | | | |
| see above | Academic Hospital Paramaribo | Erasmus Medical Center | Bas Oude Munnink, Dion Gajadin, Ed Ijzerman, Emmanuelle Munger, Gary Gummels, Ingrid Krishnadath, Lycke Woittiez, Marion Koopmans, Mireille Van de Veer, Princes Wongsowidjojo, Radjesh Ori, Rohma Banwari, Stephen Vreden |
| EPI_ISL_518819 | Qadr Hospital, Tangerang, Banten | Biosafety Level-3 Laboratory, Indonesian Institute of Sciences (LIPI) | Anik Budhi Dharmayanthi, Syam Budi Iryanto, Andri Wardiana, Anggia Prasetyoputri, Isa Nuryana, Ade Andriani, Ario Betha Juanssilfero, Asep M Ridwanuloh, Ahmad Fathoni, Rifki Sadikin, Ratih Asmana Ningrum, Wien Kusharyoto, Puspita Lisdiyanti |
| EPI_ISL_518821, EPI_ISL_518822, EPI_ISL_518823 | Oman-National Influenza Center | Biotechnology & OMICs Laboratory, Natural & Medical Sciences Research Center, University of Nizwa | Samira Al-Mahruqi, Abdul Latif Khan, Samiha Al-Kharusi, Adil Khan , Ahmed Al-Rawahi, Sajjad Asaf, Amina Al-Jardani, Hanan Al-Kindi, Intisar Al-Shukri, Adil Al-Wahaibi, Seif Al-Abri, Ahmed Al-Harrasi |
| EPI_ISL_518824 | Oman-National Influenza Center | Biotechnology & OMICs Laboratory, Natural & Medical Sciences Research Center, University of Nizwa | Abdul Latif Khan, Samira Al-Mahruqi, Ahmed Al-Harrasi, Samiha Al-Kharusi, Adil Khan, Ahmed Al-Rawahi, Sajjad Asaf, Amina Al-Jardani, Hanan Al-Kindi, Intisar Al-Shukri, Ahlam Al-Amri, Aisha Al-Amri, Aisha Al-Busaidi, Adil Al-Wahaibi, Seif Al-Abri. |
| EPI_ISL_518825 | Oman-National Influenza Center | Biotechnology & OMICs Laboratory, Natural & Medical Sciences Research Center, University of Nizwa | Samira Al-Mahruqi, Abdul Latif Khan, Samiha Al-Kharusi, Adil Khan , Ahmed Al-Rawahi, Sajjad Asaf, Amina Al-Jardani, Hanan Al-Kindi, Intisar Al-Shukri, Adil Al-Wahaibi, Seif Al-Abri, Ahmed Al-Harrasi |
| EPI_ISL_518826, EPI_ISL_518827, EPI_ISL_518828, EPI_ISL_518829, EPI_ISL_518831 | Oman-National Influenza Center | Biotechnology & OMICs Laboratory, Natural & Medical Sciences Research Center, University of Nizwa | Sajjad Asaf, Samiha Al-Kharusi, Ahmed Al-Harrasi, Samira Al-Mahruqi, Adil Khan, Ahmed Al-Rawahi, Abdul Latif Khan, Amina Al-Jardani, Hanan Al-Kindi, Intisar Al-Shukri, Ahlam Al-Amri, Aisha Al-Amri, Aisha Al-Busaidi, Adil Al-Wahaibi, Seif Al-Abri. |
| EPI_ISL_518833, EPI_ISL_518834, EPI_ISL_518835, EPI_ISL_518836 | Oman-National Influenza Center | Biotechnology & OMICs Laboratory, Natural & Medical Sciences Research Center, University of Nizwa | Samira Al-Mahruqi, Abdul Latif Khan, Samiha Al-Kharusi, Adil Khan , Ahmed Al-Rawahi, Sajjad Asaf, Amina Al-Jardani, Hanan Al-Kindi, Intisar Al-Shukri, Adil Al-Wahaibi, Seif Al-Abri, Ahmed Al-Harrasi |
| EPI_ISL_518837, EPI_ISL_518838, EPI_ISL_518839, EPI_ISL_518840, EPI_ISL_518841, EPI_ISL_518842, EPI_ISL_518843, EPI_ISL_518844, EPI_ISL_518845 | Oman-National Influenza Center | Biotechnology & OMICs Laboratory, Natural & Medical Sciences Research Center, University of Nizwa | Samiha Al-Kharusi, Sajjad Asaf, Abdul Latif Khan, Samira Al-Mahruqi, Adil Khan, Ahmed Al-Rawahi, Amina Al-Jardani, Hanan Al-Kindi, Intisar Al-Shukri, Ahlam Al-Amri, Aisha Al-Amri, Aisha Al-Busaidi, Adil Al-Wahaibi, Seif Al-Abri, Ahmed Al-Harrasi |
| EPI_ISL_518847, EPI_ISL_518849, EPI_ISL_518850, EPI_ISL_518851, EPI_ISL_518852, EPI_ISL_518853, EPI_ISL_518854 | Oman-National Influenza Center | Biotechnology & OMICs Laboratory, Natural & Medical Sciences Research Center, University of Nizwa | Abdul Latif Khan, Samira Al-Mahruqi, Ahmed Al-Harrasi, Samiha Al-Kharusi, Adil Khan, Ahmed Al-Rawahi, Sajjad Asaf, Amina Al-Jardani, Hanan Al-Kindi, Intisar Al-Shukri, Ahlam Al-Amri, Aisha Al-Amri, Aisha Al-Busaidi, Adil Al-Wahaibi, Seif Al-Abri. |
| EPI_ISL_518855 | Minnesota Department of Health, Public Health Laboratory | Minnesota Department of Health, Public Health Laboratory | Matt Plumb, Jacob Garfin, and Xiong Wang |
| EPI_ISL_518856, EPI_ISL_518857, EPI_ISL_518858, EPI_ISL_518859, EPI_ISL_518860, EPI_ISL_518861, EPI_ISL_518862, EPI_ISL_518863, EPI_ISL_518864, EPI_ISL_518865, EPI_ISL_518866, EPI_ISL_518867, EPI_ISL_518868, EPI_ISL_518869, EPI_ISL_518870, EPI_ISL_518871, EPI_ISL_518872, EPI_ISL_518873, EPI_ISL_518874, EPI_ISL_518875, EPI_ISL_518876, EPI_ISL_518877, EPI_ISL_518878, EPI_ISL_518879, EPI_ISL_518880, EPI_ISL_518881, EPI_ISL_518882, EPI_ISL_518883, EPI_ISL_518884, EPI_ISL_518885, EPI_ISL_518886, EPI_ISL_518887, EPI_ISL_518888, EPI_ISL_518889, EPI_ISL_518890, EPI_ISL_518891, EPI_ISL_518892, EPI_ISL_518893, EPI_ISL_518894, EPI_ISL_518895, EPI_ISL_518896, EPI_ISL_518897, EPI_ISL_518898, EPI_ISL_518899, EPI_ISL_518900 | | | |
| see above | Mayo Clinic & Mayo Clinic Laboratories | Minnesota Department of Health, Public Health Laboratory | Matt Plumb, Jacob Garfin, and Xiong Wang |
| EPI_ISL_518901, EPI_ISL_518902, EPI_ISL_518903, EPI_ISL_518904, EPI_ISL_518905, EPI_ISL_518906, EPI_ISL_518907, EPI_ISL_518908, EPI_ISL_518909, EPI_ISL_518910, EPI_ISL_518911, EPI_ISL_518912, EPI_ISL_518913, EPI_ISL_518914, EPI_ISL_518915, EPI_ISL_518917, EPI_ISL_518919, EPI_ISL_518920, EPI_ISL_518921, EPI_ISL_518922, EPI_ISL_518923, EPI_ISL_518924, EPI_ISL_518925, EPI_ISL_518926, EPI_ISL_518927, EPI_ISL_518928, EPI_ISL_518929, EPI_ISL_518930, EPI_ISL_518931, EPI_ISL_518932, EPI_ISL_518933, EPI_ISL_518934, EPI_ISL_518937, EPI_ISL_518938, EPI_ISL_518939, EPI_ISL_518940, EPI_ISL_518941, EPI_ISL_518943, EPI_ISL_518944, EPI_ISL_518945, EPI_ISL_518946, EPI_ISL_518947, EPI_ISL_518948, EPI_ISL_518949, EPI_ISL_518950, EPI_ISL_518951, EPI_ISL_518952, EPI_ISL_518954, EPI_ISL_518955, EPI_ISL_518956, EPI_ISL_518957, EPI_ISL_518958, EPI_ISL_518959, EPI_ISL_518960, EPI_ISL_518961, EPI_ISL_518962, EPI_ISL_518963, EPI_ISL_518964, EPI_ISL_518965, EPI_ISL_518966, EPI_ISL_518967, EPI_ISL_518968, EPI_ISL_518969, EPI_ISL_518971, EPI_ISL_518972, EPI_ISL_518973, EPI_ISL_518975 | | | |
| see above | Microbiological Diagnostic Unit - Public Health Laboratory (MDU-PHL) | MDU-PHL | Seemann T., Schultz M., Sait, M., Sherry, N. |
| EPI_ISL_518976 | Victorian Infectious Diseases Reference Laboratory (VIDRL) | VIDRL and MDU-PHL | Caly L., Seemann T., Sait, M., Schultz M., Druce J., Sherry, N. |
| EPI_ISL_518977, EPI_ISL_518978, EPI_ISL_518979, EPI_ISL_518980 | Microbiological Diagnostic Unit - Public Health Laboratory (MDU-PHL) | MDU-PHL | Seemann T., Schultz M., Sait, M., Sherry, N. |
| EPI_ISL_518981, EPI_ISL_518982, EPI_ISL_518983, EPI_ISL_518984, EPI_ISL_518985, EPI_ISL_518986, EPI_ISL_518987, EPI_ISL_518988, EPI_ISL_518989, EPI_ISL_518990, EPI_ISL_518991, EPI_ISL_518992, EPI_ISL_518993, EPI_ISL_518994, EPI_ISL_518995, EPI_ISL_518996, EPI_ISL_518997, EPI_ISL_518998, EPI_ISL_518999, EPI_ISL_519000, EPI_ISL_519001, EPI_ISL_519002, EPI_ISL_519003, EPI_ISL_519004, EPI_ISL_519006, EPI_ISL_519008, EPI_ISL_519009, EPI_ISL_519010, EPI_ISL_519011, EPI_ISL_519013, EPI_ISL_519014, EPI_ISL_519015, EPI_ISL_519016, EPI_ISL_519017, EPI_ISL_519018, EPI_ISL_519019, EPI_ISL_519020, EPI_ISL_519022, EPI_ISL_519023, EPI_ISL_519025, EPI_ISL_519026, EPI_ISL_519027, EPI_ISL_519028, EPI_ISL_519030, EPI_ISL_519031, EPI_ISL_519032, EPI_ISL_519033, EPI_ISL_519034, EPI_ISL_519035, EPI_ISL_519036, EPI_ISL_519037, EPI_ISL_519038, EPI_ISL_519039, EPI_ISL_519040, EPI_ISL_519041, EPI_ISL_519042, EPI_ISL_519043, EPI_ISL_519044, EPI_ISL_519045, EPI_ISL_519046, EPI_ISL_519047, EPI_ISL_519048, EPI_ISL_519049, EPI_ISL_519050, EPI_ISL_519051 | | | |
| see above | Victorian Infectious Diseases Reference Laboratory (VIDRL) | VIDRL and MDU-PHL | Caly L., Seemann T., Sait, M., Schultz M., Druce J., Sherry, N. |
| EPI_ISL_519052, EPI_ISL_519053, EPI_ISL_519054, EPI_ISL_519055, EPI_ISL_519056, EPI_ISL_519057, EPI_ISL_519058, EPI_ISL_519059, EPI_ISL_519060, EPI_ISL_519061, EPI_ISL_519062, EPI_ISL_519063, EPI_ISL_519064, EPI_ISL_519065, EPI_ISL_519066, EPI_ISL_519067, EPI_ISL_519068, EPI_ISL_519069, EPI_ISL_519070, EPI_ISL_519071, EPI_ISL_519072, EPI_ISL_519073, EPI_ISL_519074, EPI_ISL_519075, EPI_ISL_519076, EPI_ISL_519077, EPI_ISL_519078, EPI_ISL_519079, EPI_ISL_519080, EPI_ISL_519081, EPI_ISL_519082, EPI_ISL_519083, EPI_ISL_519084, EPI_ISL_519085, EPI_ISL_519086, EPI_ISL_519087, EPI_ISL_519088, EPI_ISL_519089, EPI_ISL_519090, EPI_ISL_519091, EPI_ISL_519092, EPI_ISL_519093, EPI_ISL_519094, EPI_ISL_519095, EPI_ISL_519096, EPI_ISL_519097, EPI_ISL_519098, EPI_ISL_519099, EPI_ISL_519100, EPI_ISL_519101, EPI_ISL_519102, EPI_ISL_519103, EPI_ISL_519104, EPI_ISL_519105, EPI_ISL_519106, EPI_ISL_519107, EPI_ISL_519108, EPI_ISL_519109, EPI_ISL_519110, EPI_ISL_519111, EPI_ISL_519113, EPI_ISL_519114, EPI_ISL_519115, EPI_ISL_519116, EPI_ISL_519117, EPI_ISL_519118, EPI_ISL_519119, EPI_ISL_519120, EPI_ISL_519121, EPI_ISL_519122, EPI_ISL_519123, EPI_ISL_519124, EPI_ISL_519125, EPI_ISL_519126, EPI_ISL_519127, EPI_ISL_519128, EPI_ISL_519130, EPI_ISL_519131, EPI_ISL_519132, EPI_ISL_519133, EPI_ISL_519134, EPI_ISL_519135, EPI_ISL_519136, EPI_ISL_519137, EPI_ISL_519138, EPI_ISL_519139, EPI_ISL_519140, EPI_ISL_519141, EPI_ISL_519142, EPI_ISL_519143, EPI_ISL_519144, EPI_ISL_519145, EPI_ISL_519146, EPI_ISL_519147, EPI_ISL_519148, EPI_ISL_519149, EPI_ISL_519150, EPI_ISL_519151, EPI_ISL_519152, EPI_ISL_519153, EPI_ISL_519154, EPI_ISL_519155, EPI_ISL_519156, EPI_ISL_519157, EPI_ISL_519158, EPI_ISL_519159, EPI_ISL_519160, EPI_ISL_519161, EPI_ISL_519163, EPI_ISL_519164, EPI_ISL_519166, EPI_ISL_519167, EPI_ISL_519168, EPI_ISL_519169, EPI_ISL_519170, EPI_ISL_519171, EPI_ISL_519172, EPI_ISL_519173, EPI_ISL_519174, EPI_ISL_519176, EPI_ISL_519177, EPI_ISL_519178, EPI_ISL_519179, EPI_ISL_519180, EPI_ISL_519181, EPI_ISL_519182, EPI_ISL_519183, EPI_ISL_519184, EPI_ISL_519185, EPI_ISL_519187, EPI_ISL_519189, EPI_ISL_519190, EPI_ISL_519191, EPI_ISL_519193, EPI_ISL_519194, EPI_ISL_519195, EPI_ISL_519196, EPI_ISL_519198, EPI_ISL_519199, EPI_ISL_519200, EPI_ISL_519201, EPI_ISL_519203, EPI_ISL_519204, EPI_ISL_519205, EPI_ISL_519206, EPI_ISL_519207, EPI_ISL_519208, EPI_ISL_519209, EPI_ISL_519210, EPI_ISL_519211, EPI_ISL_519212, EPI_ISL_519213, EPI_ISL_519214, EPI_ISL_519215, EPI_ISL_519216, EPI_ISL_519217, EPI_ISL_519219, EPI_ISL_519220, EPI_ISL_519221, EPI_ISL_519222, EPI_ISL_519223, EPI_ISL_519224, EPI_ISL_519225, EPI_ISL_519226, EPI_ISL_519228, EPI_ISL_519229, EPI_ISL_519231, EPI_ISL_519232, EPI_ISL_519233, EPI_ISL_519234, EPI_ISL_519235, EPI_ISL_519236, EPI_ISL_519237, EPI_ISL_519238, EPI_ISL_519239, EPI_ISL_519240, EPI_ISL_519241, EPI_ISL_519242, EPI_ISL_519243, EPI_ISL_519244, EPI_ISL_519245, EPI_ISL_519246, EPI_ISL_519247, EPI_ISL_519248, EPI_ISL_519249, EPI_ISL_519250, EPI_ISL_519251, EPI_ISL_519252, EPI_ISL_519253, EPI_ISL_519254, EPI_ISL_519255 | | | |
| see above | Microbiological Diagnostic Unit - Public Health Laboratory (MDU-PHL) | MDU-PHL | Seemann T., Schultz M., Sait, M., Sherry, N. |
| EPI_ISL_519257, EPI_ISL_519258, EPI_ISL_519259, EPI_ISL_519260, EPI_ISL_519261, EPI_ISL_519262, EPI_ISL_519263, EPI_ISL_519264, EPI_ISL_519265, EPI_ISL_519266, EPI_ISL_519267, EPI_ISL_519269, EPI_ISL_519270, EPI_ISL_519274, EPI_ISL_519275, EPI_ISL_519277, EPI_ISL_519279, EPI_ISL_519280, EPI_ISL_519281, EPI_ISL_519282, EPI_ISL_519283, EPI_ISL_519284, EPI_ISL_519285, EPI_ISL_519286, EPI_ISL_519287, EPI_ISL_519288, EPI_ISL_519289, EPI_ISL_519290, EPI_ISL_519291, EPI_ISL_519292, EPI_ISL_519293, EPI_ISL_519295, EPI_ISL_519296, EPI_ISL_519297, EPI_ISL_519298, EPI_ISL_519300, EPI_ISL_519302, EPI_ISL_519303, EPI_ISL_519304, EPI_ISL_519305, EPI_ISL_519306, EPI_ISL_519307, EPI_ISL_519308, EPI_ISL_519309, EPI_ISL_519310, EPI_ISL_519311, EPI_ISL_519312, EPI_ISL_519313, EPI_ISL_519314, EPI_ISL_519315, EPI_ISL_519317, EPI_ISL_519318, EPI_ISL_519319, EPI_ISL_519320, EPI_ISL_519321, EPI_ISL_519322, EPI_ISL_519323, EPI_ISL_519324, EPI_ISL_519325, EPI_ISL_519326 | | | |
| see above | Victorian Infectious Diseases Reference Laboratory (VIDRL) | VIDRL and MDU-PHL | Caly L., Seemann T., Sait, M., Schultz M., Druce J., Sherry, N. |
| EPI_ISL_519329, EPI_ISL_519330 | Microbiological Diagnostic Unit - Public Health Laboratory (MDU-PHL) | MDU-PHL | Seemann T., Schultz M., Sait, M., Sherry, N. |
| EPI_ISL_519331, EPI_ISL_519332 | Victorian Infectious Diseases Reference Laboratory (VIDRL) | VIDRL and MDU-PHL | Caly L., Seemann T., Sait, M., Schultz M., Druce J., Sherry, N. |
| EPI_ISL_519333, EPI_ISL_519334, EPI_ISL_519336, EPI_ISL_519337, EPI_ISL_519338, EPI_ISL_519339, EPI_ISL_519340, EPI_ISL_519341, EPI_ISL_519342, EPI_ISL_519343, EPI_ISL_519344, EPI_ISL_519345, EPI_ISL_519346, EPI_ISL_519347, EPI_ISL_519348, EPI_ISL_519349, EPI_ISL_519350, EPI_ISL_519352, EPI_ISL_519353, EPI_ISL_519354, EPI_ISL_519355, EPI_ISL_519357, EPI_ISL_519358, EPI_ISL_519360, EPI_ISL_519361, EPI_ISL_519362, EPI_ISL_519363, EPI_ISL_519364, EPI_ISL_519365, EPI_ISL_519366, EPI_ISL_519367, EPI_ISL_519368, EPI_ISL_519369, EPI_ISL_519371, EPI_ISL_519372, EPI_ISL_519373, EPI_ISL_519374, EPI_ISL_519375, EPI_ISL_519376, EPI_ISL_519377, EPI_ISL_519378, EPI_ISL_519379, EPI_ISL_519380, EPI_ISL_519381, EPI_ISL_519382, EPI_ISL_519383, EPI_ISL_519384 | | | |
| see above | Microbiological Diagnostic Unit - Public Health Laboratory (MDU-PHL) | MDU-PHL | Seemann T., Schultz M., Sait, M., Sherry, N. |
| EPI_ISL_519385 | Victorian Infectious Diseases Reference Laboratory (VIDRL) | VIDRL and MDU-PHL | Caly L., Seemann T., Sait, M., Schultz M., Druce J., Sherry, N. |
| EPI_ISL_519386, EPI_ISL_519387, EPI_ISL_519389, EPI_ISL_519390, EPI_ISL_519392, EPI_ISL_519393, EPI_ISL_519394, EPI_ISL_519395, EPI_ISL_519396, EPI_ISL_519397, EPI_ISL_519398, EPI_ISL_519399, EPI_ISL_519400, EPI_ISL_519401, EPI_ISL_519402, EPI_ISL_519403, EPI_ISL_519405, EPI_ISL_519406, EPI_ISL_519408, EPI_ISL_519409, EPI_ISL_519410, EPI_ISL_519416, EPI_ISL_519418, EPI_ISL_519422, EPI_ISL_519423, EPI_ISL_519424, EPI_ISL_519426, EPI_ISL_519428, EPI_ISL_519433, EPI_ISL_519436, EPI_ISL_519438, EPI_ISL_519439, EPI_ISL_519440, EPI_ISL_519442, EPI_ISL_519443, EPI_ISL_519446, EPI_ISL_519447, EPI_ISL_519449, EPI_ISL_519452, EPI_ISL_519453, EPI_ISL_519454, EPI_ISL_519458, EPI_ISL_519459, EPI_ISL_519465, EPI_ISL_519470, EPI_ISL_519471, EPI_ISL_519472, EPI_ISL_519473, EPI_ISL_519474, EPI_ISL_519476, EPI_ISL_519478, EPI_ISL_519481, EPI_ISL_519482, EPI_ISL_519484, EPI_ISL_519485, EPI_ISL_519503, EPI_ISL_519504, EPI_ISL_519505, EPI_ISL_519506, EPI_ISL_519507, EPI_ISL_519508, EPI_ISL_519518 | | | |
| see above | Microbiological Diagnostic Unit - Public Health Laboratory (MDU-PHL) | MDU-PHL | Seemann T., Schultz M., Sait, M., Sherry, N. |
| EPI_ISL_519520 | Victorian Infectious Diseases Reference Laboratory (VIDRL) | VIDRL and MDU-PHL | Caly L., Seemann T., Sait, M., Schultz M., Druce J., Sherry, N. |
| EPI_ISL_519522, EPI_ISL_519523, EPI_ISL_519524, EPI_ISL_519525, EPI_ISL_519526, EPI_ISL_519527, EPI_ISL_519528, EPI_ISL_519529, EPI_ISL_519530, EPI_ISL_519531, EPI_ISL_519532, EPI_ISL_519533, EPI_ISL_519534, EPI_ISL_519535, EPI_ISL_519536, EPI_ISL_519537, EPI_ISL_519538, EPI_ISL_519539, EPI_ISL_519540, EPI_ISL_519541, EPI_ISL_519542, EPI_ISL_519543, EPI_ISL_519544, EPI_ISL_519545, EPI_ISL_519546, EPI_ISL_519547, EPI_ISL_519548, EPI_ISL_519549, EPI_ISL_519551, EPI_ISL_519552, EPI_ISL_519553, EPI_ISL_519554, EPI_ISL_519555, EPI_ISL_519556, EPI_ISL_519557, EPI_ISL_519558, EPI_ISL_519559, EPI_ISL_519560, EPI_ISL_519561, EPI_ISL_519562, EPI_ISL_519563, EPI_ISL_519564, EPI_ISL_519565, EPI_ISL_519566, EPI_ISL_519567, EPI_ISL_519568, EPI_ISL_519569, EPI_ISL_519570, EPI_ISL_519571, EPI_ISL_519572, EPI_ISL_519573, EPI_ISL_519574, EPI_ISL_519575, EPI_ISL_519577, EPI_ISL_519578, EPI_ISL_519579, EPI_ISL_519580, EPI_ISL_519581, EPI_ISL_519582, EPI_ISL_519583, EPI_ISL_519584, EPI_ISL_519585, EPI_ISL_519586, EPI_ISL_519587, EPI_ISL_519588, EPI_ISL_519589, EPI_ISL_519590, EPI_ISL_519591, EPI_ISL_519592, EPI_ISL_519593, EPI_ISL_519594, EPI_ISL_519595, EPI_ISL_519596, EPI_ISL_519597, EPI_ISL_519598, EPI_ISL_519599, EPI_ISL_519600, EPI_ISL_519601, EPI_ISL_519602, EPI_ISL_519603, EPI_ISL_519604, EPI_ISL_519605, EPI_ISL_519606, EPI_ISL_519607, EPI_ISL_519608, EPI_ISL_519609, EPI_ISL_519610, EPI_ISL_519611, EPI_ISL_519612, EPI_ISL_519613, EPI_ISL_519614, EPI_ISL_519615, EPI_ISL_519616, EPI_ISL_519617, EPI_ISL_519618, EPI_ISL_519619, EPI_ISL_519620, EPI_ISL_519621, EPI_ISL_519622, EPI_ISL_519623, EPI_ISL_519625, EPI_ISL_519627, EPI_ISL_519628, EPI_ISL_519629, EPI_ISL_519631, EPI_ISL_519632, EPI_ISL_519633, EPI_ISL_519634, EPI_ISL_519635, EPI_ISL_519636, EPI_ISL_519637, EPI_ISL_519638, EPI_ISL_519639, EPI_ISL_519640, EPI_ISL_519641, EPI_ISL_519642, EPI_ISL_519643, EPI_ISL_519645, EPI_ISL_519646, EPI_ISL_519647, EPI_ISL_519648, EPI_ISL_519649, EPI_ISL_519650, EPI_ISL_519651, EPI_ISL_519652, EPI_ISL_519653, EPI_ISL_519654, EPI_ISL_519655, EPI_ISL_519656, EPI_ISL_519657, EPI_ISL_519658, EPI_ISL_519659, EPI_ISL_519660, EPI_ISL_519661, EPI_ISL_519662, EPI_ISL_519663, EPI_ISL_519664, EPI_ISL_519665, EPI_ISL_519666, EPI_ISL_519667, EPI_ISL_519668, EPI_ISL_519669, EPI_ISL_519670, EPI_ISL_519671, EPI_ISL_519672, EPI_ISL_519674, EPI_ISL_519675, EPI_ISL_519676, EPI_ISL_519677, EPI_ISL_519678, EPI_ISL_519679, EPI_ISL_519680, EPI_ISL_519681, EPI_ISL_519682, EPI_ISL_519683, EPI_ISL_519684, EPI_ISL_519685, EPI_ISL_519686, EPI_ISL_519687, EPI_ISL_519688, EPI_ISL_519689, EPI_ISL_519691, EPI_ISL_519692, EPI_ISL_519693, EPI_ISL_519694, EPI_ISL_519695, EPI_ISL_519696, EPI_ISL_519697, EPI_ISL_519698, EPI_ISL_519699, EPI_ISL_519700, EPI_ISL_519701, EPI_ISL_519702, EPI_ISL_519703, EPI_ISL_519704, EPI_ISL_519705, EPI_ISL_519706, EPI_ISL_519707, EPI_ISL_519708, EPI_ISL_519709, EPI_ISL_519710, EPI_ISL_519711, EPI_ISL_519712, EPI_ISL_519713, EPI_ISL_519714, EPI_ISL_519715, EPI_ISL_519716, EPI_ISL_519717, EPI_ISL_519718, EPI_ISL_519719, EPI_ISL_519720, EPI_ISL_519721, EPI_ISL_519722, EPI_ISL_519723, EPI_ISL_519724, EPI_ISL_519725, EPI_ISL_519726, EPI_ISL_519727, EPI_ISL_519728, EPI_ISL_519729, EPI_ISL_519730, EPI_ISL_519731, EPI_ISL_519732, EPI_ISL_519733, EPI_ISL_519734, EPI_ISL_519735, EPI_ISL_519736, EPI_ISL_519737, EPI_ISL_519738, EPI_ISL_519739, EPI_ISL_519740, EPI_ISL_519741, EPI_ISL_519742, EPI_ISL_519743, EPI_ISL_519744, EPI_ISL_519745, EPI_ISL_519746, EPI_ISL_519747, EPI_ISL_519748, EPI_ISL_519749, EPI_ISL_519750, EPI_ISL_519751, EPI_ISL_519752, EPI_ISL_519753, EPI_ISL_519754, EPI_ISL_519755, EPI_ISL_519757, EPI_ISL_519758, EPI_ISL_519759, EPI_ISL_519760, EPI_ISL_519761, EPI_ISL_519762, EPI_ISL_519763, EPI_ISL_519764, EPI_ISL_519765, EPI_ISL_519766, EPI_ISL_519767, EPI_ISL_519768, EPI_ISL_519769, EPI_ISL_519770, EPI_ISL_519771, EPI_ISL_519772, EPI_ISL_519773, EPI_ISL_519774, EPI_ISL_519775, EPI_ISL_519776, EPI_ISL_519777, EPI_ISL_519778, EPI_ISL_519779, EPI_ISL_519780, EPI_ISL_519781, EPI_ISL_519783, EPI_ISL_519784, EPI_ISL_519786, EPI_ISL_519787, EPI_ISL_519788, EPI_ISL_519789, EPI_ISL_519790, EPI_ISL_519791, EPI_ISL_519792, EPI_ISL_519793, EPI_ISL_519794, EPI_ISL_519795, EPI_ISL_519796, EPI_ISL_519797, EPI_ISL_519798, EPI_ISL_519799, EPI_ISL_519800, EPI_ISL_519801, EPI_ISL_519802, EPI_ISL_519803, EPI_ISL_519804, EPI_ISL_519805, EPI_ISL_519806, EPI_ISL_519807, EPI_ISL_519808, EPI_ISL_519809, EPI_ISL_519810, EPI_ISL_519811, EPI_ISL_519812, EPI_ISL_519813, EPI_ISL_519814, EPI_ISL_519815, EPI_ISL_519816, EPI_ISL_519818, EPI_ISL_519819, EPI_ISL_519820, EPI_ISL_519821, EPI_ISL_519822, EPI_ISL_519823, EPI_ISL_519824, EPI_ISL_519825, EPI_ISL_519826, EPI_ISL_519827, EPI_ISL_519828, EPI_ISL_519829, EPI_ISL_519830, EPI_ISL_519831, EPI_ISL_519832, EPI_ISL_519833, EPI_ISL_519834, EPI_ISL_519835, EPI_ISL_519836, EPI_ISL_519837, EPI_ISL_519838, EPI_ISL_519839, EPI_ISL_519840, EPI_ISL_519841, EPI_ISL_519842, EPI_ISL_519843, EPI_ISL_519844, EPI_ISL_519845, EPI_ISL_519846, EPI_ISL_519847, EPI_ISL_519848, EPI_ISL_519849, EPI_ISL_519850, EPI_ISL_519851, EPI_ISL_519852, EPI_ISL_519853, EPI_ISL_519854, EPI_ISL_519855, EPI_ISL_519856, EPI_ISL_519857, EPI_ISL_519858, EPI_ISL_519859, EPI_ISL_519860, EPI_ISL_519861, EPI_ISL_519862, EPI_ISL_519863, EPI_ISL_519864, EPI_ISL_519865, EPI_ISL_519866, EPI_ISL_519867, EPI_ISL_519868, EPI_ISL_519869, EPI_ISL_519870, EPI_ISL_519871, EPI_ISL_519872, EPI_ISL_519873, EPI_ISL_519874, EPI_ISL_519875, EPI_ISL_519876, EPI_ISL_519877, EPI_ISL_519878, EPI_ISL_519880, EPI_ISL_519881, EPI_ISL_519882, EPI_ISL_519883, EPI_ISL_519884, EPI_ISL_519885, EPI_ISL_519886, EPI_ISL_519887, EPI_ISL_519888, EPI_ISL_519889, EPI_ISL_519890, EPI_ISL_519891, EPI_ISL_519892, EPI_ISL_519893, EPI_ISL_519894, EPI_ISL_519896, EPI_ISL_519897, EPI_ISL_519898, EPI_ISL_519899, EPI_ISL_519900, EPI_ISL_519901, EPI_ISL_519902, EPI_ISL_519903, EPI_ISL_519904, EPI_ISL_519905, EPI_ISL_519906, EPI_ISL_519907, EPI_ISL_519908, EPI_ISL_519909, EPI_ISL_519910, EPI_ISL_519911, EPI_ISL_519912, EPI_ISL_519913, EPI_ISL_519914, EPI_ISL_519915, EPI_ISL_519916, EPI_ISL_519917, EPI_ISL_519918, EPI_ISL_519919, EPI_ISL_519920, EPI_ISL_519921, EPI_ISL_519922, EPI_ISL_519923, EPI_ISL_519924, EPI_ISL_519926, EPI_ISL_519927, EPI_ISL_519928, EPI_ISL_519929, EPI_ISL_519930, EPI_ISL_519931, EPI_ISL_519932, EPI_ISL_519933, EPI_ISL_519934, EPI_ISL_519935, EPI_ISL_519936, EPI_ISL_519937, EPI_ISL_519938, EPI_ISL_519939, EPI_ISL_519940, EPI_ISL_519941, EPI_ISL_519942, EPI_ISL_519943, EPI_ISL_519944, EPI_ISL_519945, EPI_ISL_519946, EPI_ISL_519947 | | | |
| see above | Microbiological Diagnostic Unit - Public Health Laboratory (MDU-PHL) | MDU-PHL | Seemann T., Schultz M., Sait, M., Sherry, N. |
| EPI_ISL_519948, EPI_ISL_519950 | Victorian Infectious Diseases Reference Laboratory (VIDRL) | VIDRL and MDU-PHL | Caly L., Seemann T., Sait, M., Schultz M., Druce J., Sherry, N. |
| EPI_ISL_519951, EPI_ISL_519952, EPI_ISL_519953, EPI_ISL_519954, EPI_ISL_519955, EPI_ISL_519956, EPI_ISL_519957, EPI_ISL_519958, EPI_ISL_519959, EPI_ISL_519960, EPI_ISL_519961, EPI_ISL_519962, EPI_ISL_519963, EPI_ISL_519964, EPI_ISL_519965, EPI_ISL_519966, EPI_ISL_519967, EPI_ISL_519968, EPI_ISL_519969, EPI_ISL_519971, EPI_ISL_519972, EPI_ISL_519973, EPI_ISL_519974, EPI_ISL_519975, EPI_ISL_519976, EPI_ISL_519977, EPI_ISL_519978, EPI_ISL_519979, EPI_ISL_519980, EPI_ISL_519981, EPI_ISL_519982, EPI_ISL_519983, EPI_ISL_519984, EPI_ISL_519985, EPI_ISL_519986, EPI_ISL_519987, EPI_ISL_519988, EPI_ISL_519989, EPI_ISL_519990, EPI_ISL_519991, EPI_ISL_519992, EPI_ISL_519993, EPI_ISL_519994, EPI_ISL_519995, EPI_ISL_519996, EPI_ISL_519997, EPI_ISL_519998, EPI_ISL_519999, EPI_ISL_520000, EPI_ISL_520001, EPI_ISL_520002, EPI_ISL_520003, EPI_ISL_520004, EPI_ISL_520005, EPI_ISL_520006, EPI_ISL_520007, EPI_ISL_520008, EPI_ISL_520009, EPI_ISL_520010, EPI_ISL_520011, EPI_ISL_520012, EPI_ISL_520013, EPI_ISL_520014, EPI_ISL_520015, EPI_ISL_520016, EPI_ISL_520017, EPI_ISL_520018, EPI_ISL_520019, EPI_ISL_520020, EPI_ISL_520021, EPI_ISL_520022, EPI_ISL_520023, | | | |

| | | | |
|---|---|---|---|
| EPI_ISL_520024, EPI_ISL_520025, EPI_ISL_520026, EPI_ISL_520027, EPI_ISL_520028, EPI_ISL_520029, EPI_ISL_520030, EPI_ISL_520031, EPI_ISL_520032, EPI_ISL_520033, EPI_ISL_520034, EPI_ISL_520035, EPI_ISL_520036, EPI_ISL_520037, EPI_ISL_520038, EPI_ISL_520039, EPI_ISL_520040, EPI_ISL_520041, EPI_ISL_520042, EPI_ISL_520043, EPI_ISL_520044, EPI_ISL_520045, EPI_ISL_520046, EPI_ISL_520047, EPI_ISL_520048, EPI_ISL_520049, EPI_ISL_520050, EPI_ISL_520052, EPI_ISL_520053, EPI_ISL_520054, EPI_ISL_520055, EPI_ISL_520056, EPI_ISL_520057, EPI_ISL_520058, EPI_ISL_520059, EPI_ISL_520060, EPI_ISL_520061, EPI_ISL_520062, EPI_ISL_520063, EPI_ISL_520065, EPI_ISL_520066, EPI_ISL_520067, EPI_ISL_520068, EPI_ISL_520069, EPI_ISL_520070, EPI_ISL_520071, EPI_ISL_520072, EPI_ISL_520073, EPI_ISL_520074, EPI_ISL_520076, EPI_ISL_520077, EPI_ISL_520078, EPI_ISL_520080, EPI_ISL_520081, EPI_ISL_520082, EPI_ISL_520083, EPI_ISL_520084, EPI_ISL_520085, EPI_ISL_520086, EPI_ISL_520087, EPI_ISL_520088, EPI_ISL_520089, EPI_ISL_520090, EPI_ISL_520091, EPI_ISL_520092, EPI_ISL_520093, EPI_ISL_520094, EPI_ISL_520095, EPI_ISL_520096, EPI_ISL_520097, EPI_ISL_520098, EPI_ISL_520099, EPI_ISL_520100, EPI_ISL_520101, EPI_ISL_520102, EPI_ISL_520103, EPI_ISL_520104, EPI_ISL_520105, EPI_ISL_520106, EPI_ISL_520107, EPI_ISL_520108, EPI_ISL_520109, EPI_ISL_520110, EPI_ISL_520111, EPI_ISL_520112, EPI_ISL_520113, EPI_ISL_520114, EPI_ISL_520115, EPI_ISL_520116, EPI_ISL_520117, EPI_ISL_520118, EPI_ISL_520119, EPI_ISL_520121, EPI_ISL_520122, EPI_ISL_520123, EPI_ISL_520124, EPI_ISL_520125, EPI_ISL_520126, EPI_ISL_520127, EPI_ISL_520128, EPI_ISL_520129, EPI_ISL_520130, EPI_ISL_520131, EPI_ISL_520132, EPI_ISL_520133, EPI_ISL_520134, EPI_ISL_520135, EPI_ISL_520136, EPI_ISL_520137, EPI_ISL_520138, EPI_ISL_520139, EPI_ISL_520140, EPI_ISL_520141, EPI_ISL_520142, EPI_ISL_520143, EPI_ISL_520144, EPI_ISL_520145, EPI_ISL_520146, EPI_ISL_520147, EPI_ISL_520148, EPI_ISL_520149, EPI_ISL_520150, EPI_ISL_520151, EPI_ISL_520152, EPI_ISL_520153, EPI_ISL_520154, EPI_ISL_520155, EPI_ISL_520156, EPI_ISL_520157, EPI_ISL_520158, EPI_ISL_520159, EPI_ISL_520160, EPI_ISL_520161, EPI_ISL_520162, EPI_ISL_520163, EPI_ISL_520164, EPI_ISL_520165, EPI_ISL_520166, EPI_ISL_520167, EPI_ISL_520168, EPI_ISL_520169, EPI_ISL_520170, EPI_ISL_520171, EPI_ISL_520172, EPI_ISL_520173, EPI_ISL_520174, EPI_ISL_520175, EPI_ISL_520176, EPI_ISL_520177, EPI_ISL_520178, EPI_ISL_520179, EPI_ISL_520180 | | | |
| see above | Microbiological Diagnostic Unit - Public Health Laboratory (MDU-PHL) | MDU-PHL | Seemann T., Schultz M., Sait, M., Sherry, N. |
| EPI_ISL_520183, EPI_ISL_520184, EPI_ISL_520185, EPI_ISL_520186, EPI_ISL_520188, EPI_ISL_520189, EPI_ISL_520190, EPI_ISL_520192, EPI_ISL_520193, EPI_ISL_520194, EPI_ISL_520195, EPI_ISL_520196, EPI_ISL_520197, EPI_ISL_520198, EPI_ISL_520199, EPI_ISL_520200, EPI_ISL_520202, EPI_ISL_520203, EPI_ISL_520204, EPI_ISL_520205, EPI_ISL_520206, EPI_ISL_520207, EPI_ISL_520208, EPI_ISL_520209, EPI_ISL_520210, EPI_ISL_520211, EPI_ISL_520212, EPI_ISL_520213, EPI_ISL_520214 | | | |
| see above | Victorian Infectious Diseases Reference Laboratory (VIDRL) | VIDRL and MDU-PHL | Caly L., Seemann T., Sait, M., Schultz M., Druce J., Sherry, N. |
| EPI_ISL_520215, EPI_ISL_520216, EPI_ISL_520217, EPI_ISL_520218, EPI_ISL_520219, EPI_ISL_520220, EPI_ISL_520221, EPI_ISL_520222, EPI_ISL_520223, EPI_ISL_520224, EPI_ISL_520225, EPI_ISL_520226, EPI_ISL_520227, EPI_ISL_520228, EPI_ISL_520229, EPI_ISL_520230, EPI_ISL_520233, EPI_ISL_520234, EPI_ISL_520235, EPI_ISL_520237, EPI_ISL_520238, EPI_ISL_520240, EPI_ISL_520241, EPI_ISL_520243, EPI_ISL_520244, EPI_ISL_520245, EPI_ISL_520246, EPI_ISL_520247, EPI_ISL_520248 | | | |
| see above | Microbiological Diagnostic Unit - Public Health Laboratory (MDU-PHL) | MDU-PHL | Seemann T., Schultz M., Sait, M., Sherry, N. |
| EPI_ISL_520249, EPI_ISL_520250, EPI_ISL_520251, EPI_ISL_520252, EPI_ISL_520254, EPI_ISL_520255, EPI_ISL_520256, EPI_ISL_520257, EPI_ISL_520259, EPI_ISL_520260, EPI_ISL_520261, EPI_ISL_520262, EPI_ISL_520263, EPI_ISL_520264, EPI_ISL_520265, EPI_ISL_520267, EPI_ISL_520268, EPI_ISL_520272, EPI_ISL_520273, EPI_ISL_520274, EPI_ISL_520275, EPI_ISL_520276, EPI_ISL_520277, EPI_ISL_520278, EPI_ISL_520279, EPI_ISL_520280, EPI_ISL_520281, EPI_ISL_520282, EPI_ISL_520283, EPI_ISL_520284, EPI_ISL_520287, EPI_ISL_520289, EPI_ISL_520290, EPI_ISL_520291, EPI_ISL_520292, EPI_ISL_520293, EPI_ISL_520294, EPI_ISL_520295, EPI_ISL_520297, EPI_ISL_520300, EPI_ISL_520302, EPI_ISL_520303, EPI_ISL_520304, EPI_ISL_520305, EPI_ISL_520306, EPI_ISL_520308, EPI_ISL_520309, EPI_ISL_520311, EPI_ISL_520312, EPI_ISL_520313, EPI_ISL_520314, EPI_ISL_520315, EPI_ISL_520316, EPI_ISL_520317, EPI_ISL_520320, EPI_ISL_520322, EPI_ISL_520323, EPI_ISL_520325, EPI_ISL_520326, EPI_ISL_520327, EPI_ISL_520328, EPI_ISL_520329, EPI_ISL_520330, EPI_ISL_520331, EPI_ISL_520332, EPI_ISL_520333, EPI_ISL_520334, EPI_ISL_520335, EPI_ISL_520336, EPI_ISL_520337, EPI_ISL_520338, EPI_ISL_520339, EPI_ISL_520340, EPI_ISL_520341, EPI_ISL_520342, EPI_ISL_520343, EPI_ISL_520344, EPI_ISL_520345, EPI_ISL_520346, EPI_ISL_520347, EPI_ISL_520349, EPI_ISL_520350, EPI_ISL_520351, EPI_ISL_520352, EPI_ISL_520353, EPI_ISL_520354, EPI_ISL_520356, EPI_ISL_520357, EPI_ISL_520358, EPI_ISL_520360, EPI_ISL_520361, EPI_ISL_520362, EPI_ISL_520363, EPI_ISL_520364, EPI_ISL_520365, EPI_ISL_520366, EPI_ISL_520367, EPI_ISL_520368, EPI_ISL_520369, EPI_ISL_520370, EPI_ISL_520371, EPI_ISL_520372, EPI_ISL_520373, EPI_ISL_520374, EPI_ISL_520375, EPI_ISL_520377, EPI_ISL_520378, EPI_ISL_520379, EPI_ISL_520381, EPI_ISL_520383, EPI_ISL_520385, EPI_ISL_520386, EPI_ISL_520387, EPI_ISL_520388, EPI_ISL_520389, EPI_ISL_520390 | | | |
| see above | Victorian Infectious Diseases Reference Laboratory (VIDRL) | VIDRL and MDU-PHL | Caly L., Seemann T., Sait, M., Schultz M., Druce J., Sherry, N. |
| EPI_ISL_520421, EPI_ISL_520449, EPI_ISL_520507, EPI_ISL_520508, EPI_ISL_520509, EPI_ISL_520510, EPI_ISL_520511, EPI_ISL_520512, EPI_ISL_520513, EPI_ISL_520516, EPI_ISL_520517, EPI_ISL_520518, EPI_ISL_520519, EPI_ISL_520520, EPI_ISL_520521, EPI_ISL_520522, EPI_ISL_520523, EPI_ISL_520524, EPI_ISL_520525, EPI_ISL_520526, EPI_ISL_520528, EPI_ISL_520529, EPI_ISL_520530, EPI_ISL_520531, EPI_ISL_520532, EPI_ISL_520533, EPI_ISL_520534, EPI_ISL_520535, EPI_ISL_520536, EPI_ISL_520537, EPI_ISL_520538, EPI_ISL_520539, EPI_ISL_520540, EPI_ISL_520541, EPI_ISL_520542, EPI_ISL_520543, EPI_ISL_520544, EPI_ISL_520545, EPI_ISL_520546, EPI_ISL_520547, EPI_ISL_520548, EPI_ISL_520549, EPI_ISL_520550, EPI_ISL_520551, EPI_ISL_520552, EPI_ISL_520554, EPI_ISL_520555, EPI_ISL_520556, EPI_ISL_520557, EPI_ISL_520558, EPI_ISL_520559, EPI_ISL_520560, EPI_ISL_520561, EPI_ISL_520562, EPI_ISL_520563, EPI_ISL_520564, EPI_ISL_520565, EPI_ISL_520566, EPI_ISL_520567, EPI_ISL_520568, EPI_ISL_520569, EPI_ISL_520570, EPI_ISL_520571, EPI_ISL_520572, EPI_ISL_520573, EPI_ISL_520574, EPI_ISL_520575, EPI_ISL_520576, EPI_ISL_520577, EPI_ISL_520578, EPI_ISL_520579, EPI_ISL_520580, EPI_ISL_520581, EPI_ISL_520582, EPI_ISL_520583, EPI_ISL_520584, EPI_ISL_520585, EPI_ISL_520586, EPI_ISL_520587, EPI_ISL_520588, EPI_ISL_520589, EPI_ISL_520590, EPI_ISL_520591, EPI_ISL_520592, EPI_ISL_520593, EPI_ISL_520594, EPI_ISL_520596, EPI_ISL_520597, EPI_ISL_520598, EPI_ISL_520599, EPI_ISL_520600, EPI_ISL_520601, EPI_ISL_520602, EPI_ISL_520603, EPI_ISL_520606, EPI_ISL_520607, EPI_ISL_520608, EPI_ISL_520609, EPI_ISL_520610, EPI_ISL_520611, EPI_ISL_520612, EPI_ISL_520613, EPI_ISL_520614, EPI_ISL_520615, EPI_ISL_520616, EPI_ISL_520617, EPI_ISL_520618, EPI_ISL_520619, EPI_ISL_520620, EPI_ISL_520621, EPI_ISL_520622, EPI_ISL_520623, EPI_ISL_520624, EPI_ISL_520625, EPI_ISL_520626, EPI_ISL_520627, EPI_ISL_520628, EPI_ISL_520629, EPI_ISL_520630, EPI_ISL_520631, EPI_ISL_520632, EPI_ISL_520633, EPI_ISL_520634, EPI_ISL_520635, EPI_ISL_520636, EPI_ISL_520637, EPI_ISL_520638, EPI_ISL_520639, EPI_ISL_520640, EPI_ISL_520641, EPI_ISL_520642, EPI_ISL_520643, EPI_ISL_520645, EPI_ISL_520646, EPI_ISL_520647, EPI_ISL_520648, EPI_ISL_520649, EPI_ISL_520650, EPI_ISL_520651, EPI_ISL_520652, EPI_ISL_520653, EPI_ISL_520654, EPI_ISL_520656, EPI_ISL_520657, EPI_ISL_520658, EPI_ISL_520659, EPI_ISL_520660, EPI_ISL_520661, EPI_ISL_520662 | | | |
| see above | Microbiological Diagnostic Unit - Public Health Laboratory (MDU-PHL) | MDU-PHL | Seemann T., Schultz M., Sait, M., Sherry, N. |
| EPI_ISL_520663, EPI_ISL_520664, EPI_ISL_520666, EPI_ISL_520667, EPI_ISL_520668, EPI_ISL_520669, EPI_ISL_520670, EPI_ISL_520671, EPI_ISL_520672, EPI_ISL_520673, EPI_ISL_520674, EPI_ISL_520675, EPI_ISL_520676, EPI_ISL_520677, EPI_ISL_520678, EPI_ISL_520679, EPI_ISL_520680, EPI_ISL_520681, EPI_ISL_520682, EPI_ISL_520683, EPI_ISL_520684, EPI_ISL_520685, EPI_ISL_520686, EPI_ISL_520687, EPI_ISL_520688, EPI_ISL_520689, EPI_ISL_520690, EPI_ISL_520691, EPI_ISL_520692, EPI_ISL_520693, EPI_ISL_520694, EPI_ISL_520695, EPI_ISL_520696, EPI_ISL_520697, EPI_ISL_520698, EPI_ISL_520700, EPI_ISL_520701, EPI_ISL_520702, EPI_ISL_520703, EPI_ISL_520704, EPI_ISL_520705, EPI_ISL_520706, EPI_ISL_520707, EPI_ISL_520708, EPI_ISL_520709, EPI_ISL_520710, EPI_ISL_520711, EPI_ISL_520712, EPI_ISL_520713, EPI_ISL_520714, EPI_ISL_520715, EPI_ISL_520716, EPI_ISL_520717, EPI_ISL_520718, EPI_ISL_520719, EPI_ISL_520720, EPI_ISL_520721, EPI_ISL_520722, EPI_ISL_520723, EPI_ISL_520724, EPI_ISL_520725, EPI_ISL_520726, EPI_ISL_520727, EPI_ISL_520728, EPI_ISL_520729, EPI_ISL_520730, EPI_ISL_520731, EPI_ISL_520732, EPI_ISL_520733, EPI_ISL_520734, EPI_ISL_520735, EPI_ISL_520736, EPI_ISL_520737, EPI_ISL_520738, EPI_ISL_520739, EPI_ISL_520740, EPI_ISL_520741, EPI_ISL_520742, EPI_ISL_520743, EPI_ISL_520744 | | | |
| see above | Mohammed Bin Rashid University of Medicine and Health Sciences | Al Jalila Genomics Center | Ahmad Abou Tayoun, Tom Loney, Hamda Khansaheb, Sathishkumar Ramaswamy, Divinlal Harilal, Zulfa Omar Deesi, Rupa Murthy Varghese, Hanan Al Suwaidi, Abdulmajeed Alkhaja, Mohammed Uddin, Rifat Hamoudi, Rabih Halwani, Abiola Catherine Senok, Qutayba Hamid, Norbert Nowotny, Alawi Alsheikh-Ali |
| EPI_ISL_520745, EPI_ISL_520746, EPI_ISL_520747, EPI_ISL_520748, EPI_ISL_520749, EPI_ISL_520750, EPI_ISL_520752, EPI_ISL_520753, EPI_ISL_520754, EPI_ISL_520755, EPI_ISL_520756, EPI_ISL_520757, EPI_ISL_520758, EPI_ISL_520759, EPI_ISL_520760, EPI_ISL_520761, EPI_ISL_520762 | | | |
| see above | Victorian Infectious Diseases Reference Laboratory (VIDRL) | VIDRL and MDU-PHL | Caly L., Seemann T., Sait, M., Schultz M., Druce J., Sherry, N. |
| EPI_ISL_520765, EPI_ISL_520766, EPI_ISL_520767, EPI_ISL_520768, EPI_ISL_520769, EPI_ISL_520770, EPI_ISL_520771, EPI_ISL_520772, EPI_ISL_520773, EPI_ISL_520774, EPI_ISL_520776, EPI_ISL_520777, EPI_ISL_520778, EPI_ISL_520779, EPI_ISL_520780, EPI_ISL_520781, EPI_ISL_520782, EPI_ISL_520783, EPI_ISL_520784, EPI_ISL_520785, EPI_ISL_520786, EPI_ISL_520787, EPI_ISL_520788, EPI_ISL_520789, EPI_ISL_520790 | | | |
| see above | Microbiological Diagnostic Unit - Public Health Laboratory (MDU-PHL) | MDU-PHL | Seemann T., Schultz M., Sait, M., Sherry, N. |
| EPI_ISL_520791, EPI_ISL_520792, EPI_ISL_520793, EPI_ISL_520794, EPI_ISL_520795, EPI_ISL_520796, EPI_ISL_520797, EPI_ISL_520798, EPI_ISL_520799, EPI_ISL_520800, EPI_ISL_520801, EPI_ISL_520802, EPI_ISL_520803, EPI_ISL_520804, EPI_ISL_520805, EPI_ISL_520806, EPI_ISL_520807, EPI_ISL_520808, EPI_ISL_520809, EPI_ISL_520811, EPI_ISL_520812, EPI_ISL_520813, EPI_ISL_520814, EPI_ISL_520815, EPI_ISL_520816, EPI_ISL_520818, EPI_ISL_520819, EPI_ISL_520820, EPI_ISL_520821, EPI_ISL_520822, EPI_ISL_520823, EPI_ISL_520824, EPI_ISL_520825, EPI_ISL_520826, EPI_ISL_520827, EPI_ISL_520828, EPI_ISL_520829, EPI_ISL_520830, EPI_ISL_520831, EPI_ISL_520832, EPI_ISL_520833, EPI_ISL_520834, EPI_ISL_520835, EPI_ISL_520836, EPI_ISL_520837, EPI_ISL_520838, EPI_ISL_520840, EPI_ISL_520841, EPI_ISL_520842, EPI_ISL_520843 | | | |
| see above | Victorian Infectious Diseases Reference Laboratory (VIDRL) | VIDRL and MDU-PHL | Caly L., Seemann T., Sait, M., Schultz M., Druce J., Sherry, N. |
| EPI_ISL_520844, EPI_ISL_520845, EPI_ISL_520846, EPI_ISL_520847, EPI_ISL_520848, EPI_ISL_520849, EPI_ISL_520850, EPI_ISL_520851, EPI_ISL_520852, EPI_ISL_520853, EPI_ISL_520854, EPI_ISL_520855, EPI_ISL_520856, EPI_ISL_520857, EPI_ISL_520858, EPI_ISL_520859, EPI_ISL_520860, EPI_ISL_520861, EPI_ISL_520862, EPI_ISL_520863 | | | |
| see above | Microbiological Diagnostic Unit - Public Health Laboratory (MDU-PHL) | MDU-PHL | Seemann T., Schultz M., Sait, M., Sherry, N. |
| EPI_ISL_520864, EPI_ISL_520865, EPI_ISL_520866, EPI_ISL_520867, EPI_ISL_520868, EPI_ISL_520869, EPI_ISL_520870, EPI_ISL_520871, EPI_ISL_520872, EPI_ISL_520873, EPI_ISL_520874, EPI_ISL_520875, EPI_ISL_520876, EPI_ISL_520877, EPI_ISL_520878, EPI_ISL_520879, EPI_ISL_520880, EPI_ISL_520881, EPI_ISL_520882, EPI_ISL_520883, EPI_ISL_520884, EPI_ISL_520885, EPI_ISL_520886, EPI_ISL_520887, EPI_ISL_520888, EPI_ISL_520889, EPI_ISL_520890, EPI_ISL_520891, EPI_ISL_520892, EPI_ISL_520893, EPI_ISL_520894, EPI_ISL_520895, EPI_ISL_520896, EPI_ISL_520897, EPI_ISL_520898, EPI_ISL_520899, EPI_ISL_520900, EPI_ISL_520901, EPI_ISL_520902, EPI_ISL_520903, EPI_ISL_520904, EPI_ISL_520905, EPI_ISL_520906, EPI_ISL_520907, EPI_ISL_520908, EPI_ISL_520909, EPI_ISL_520910, EPI_ISL_520911, EPI_ISL_520913, EPI_ISL_520914, EPI_ISL_520915, EPI_ISL_520916, EPI_ISL_520917, EPI_ISL_520918, EPI_ISL_520919, EPI_ISL_520920, EPI_ISL_520921, EPI_ISL_520922, EPI_ISL_520923, EPI_ISL_520924, EPI_ISL_520925, EPI_ISL_520926, EPI_ISL_520927, EPI_ISL_520928 | | | |
| see above | Victorian Infectious Diseases Reference Laboratory (VIDRL) | VIDRL and MDU-PHL | Caly L., Seemann T., Sait, M., Schultz M., Druce J., Sherry, N. |
| EPI_ISL_520929, EPI_ISL_520930, EPI_ISL_520931, EPI_ISL_520932, EPI_ISL_520933, EPI_ISL_520934, EPI_ISL_520935, EPI_ISL_520936, EPI_ISL_520937, EPI_ISL_520938, EPI_ISL_520939, EPI_ISL_520940, EPI_ISL_520941, EPI_ISL_520942, EPI_ISL_520943, EPI_ISL_520944, EPI_ISL_520945, EPI_ISL_520946, EPI_ISL_520947, EPI_ISL_520949, EPI_ISL_520950, EPI_ISL_520951, EPI_ISL_520952, EPI_ISL_520954, EPI_ISL_520955, EPI_ISL_520956, EPI_ISL_520957, EPI_ISL_520958, EPI_ISL_520959, EPI_ISL_520960, EPI_ISL_520961, EPI_ISL_520962, EPI_ISL_520964, EPI_ISL_520965, EPI_ISL_520966, EPI_ISL_520967, EPI_ISL_520968, EPI_ISL_520969, EPI_ISL_520970, EPI_ISL_520971, EPI_ISL_520972, EPI_ISL_520973, EPI_ISL_520974, EPI_ISL_520975, EPI_ISL_520976, EPI_ISL_520977, EPI_ISL_520978, EPI_ISL_520979, EPI_ISL_520980, EPI_ISL_520981, EPI_ISL_520982, EPI_ISL_520983, EPI_ISL_520984, EPI_ISL_520985, EPI_ISL_520986, EPI_ISL_520987, EPI_ISL_520988, EPI_ISL_520989, EPI_ISL_520990, EPI_ISL_520991, EPI_ISL_520992, EPI_ISL_520993, EPI_ISL_520994, EPI_ISL_520995, EPI_ISL_520996, EPI_ISL_520997, EPI_ISL_520998, EPI_ISL_520999, EPI_ISL_521000, EPI_ISL_521001, EPI_ISL_521002, EPI_ISL_521003, EPI_ISL_521004, EPI_ISL_521005, EPI_ISL_521006, EPI_ISL_521007, EPI_ISL_521008, EPI_ISL_521009, EPI_ISL_521010, EPI_ISL_521011, EPI_ISL_521012, EPI_ISL_521013, EPI_ISL_521014, EPI_ISL_521015, EPI_ISL_521016, EPI_ISL_521017, EPI_ISL_521018, EPI_ISL_521019, EPI_ISL_521020, EPI_ISL_521021, EPI_ISL_521022, EPI_ISL_521023, EPI_ISL_521024, EPI_ISL_521025, EPI_ISL_521026, EPI_ISL_521027, EPI_ISL_521028, EPI_ISL_521029, EPI_ISL_521030, EPI_ISL_521031, EPI_ISL_521032, EPI_ISL_521033, EPI_ISL_521034, EPI_ISL_521035, EPI_ISL_521036, EPI_ISL_521037, EPI_ISL_521038, EPI_ISL_521039, EPI_ISL_521040, EPI_ISL_521041, EPI_ISL_521042, EPI_ISL_521043, EPI_ISL_521044, EPI_ISL_521045, EPI_ISL_521046, EPI_ISL_521047, EPI_ISL_521048, EPI_ISL_521049, EPI_ISL_521050, EPI_ISL_521051, EPI_ISL_521052, EPI_ISL_521053, EPI_ISL_521054, EPI_ISL_521055, EPI_ISL_521056, EPI_ISL_521057, EPI_ISL_521058, EPI_ISL_521059, EPI_ISL_521060, EPI_ISL_521061, EPI_ISL_521062, EPI_ISL_521063, EPI_ISL_521064, EPI_ISL_521065, EPI_ISL_521066, EPI_ISL_521067, EPI_ISL_521068, EPI_ISL_521069, EPI_ISL_521070, EPI_ISL_521071, EPI_ISL_521072, EPI_ISL_521073, EPI_ISL_521074, EPI_ISL_521076, EPI_ISL_521077, EPI_ISL_521078, EPI_ISL_521079, EPI_ISL_521080, EPI_ISL_521081, EPI_ISL_521082, EPI_ISL_521083, EPI_ISL_521084, EPI_ISL_521085, EPI_ISL_521086, EPI_ISL_521087, EPI_ISL_521088, EPI_ISL_521089, EPI_ISL_521091, EPI_ISL_521092, EPI_ISL_521093, EPI_ISL_521094, EPI_ISL_521095, EPI_ISL_521096, EPI_ISL_521097, EPI_ISL_521098, EPI_ISL_521099, EPI_ISL_521100, EPI_ISL_521101 | | | |
| see above | Microbiological Diagnostic Unit - Public Health Laboratory (MDU-PHL) | MDU-PHL | Seemann T., Schultz M., Sait, M., Sherry, N. |
| EPI_ISL_521102, EPI_ISL_521104, EPI_ISL_521105, EPI_ISL_521106, EPI_ISL_521107, EPI_ISL_521108, EPI_ISL_521110, EPI_ISL_521111, EPI_ISL_521112, EPI_ISL_521114, EPI_ISL_521115, EPI_ISL_521116, EPI_ISL_521117, EPI_ISL_521118, EPI_ISL_521119, EPI_ISL_521120, EPI_ISL_521121, EPI_ISL_521123, EPI_ISL_521124, EPI_ISL_521125, EPI_ISL_521128, EPI_ISL_521131, EPI_ISL_521133, EPI_ISL_521134, EPI_ISL_521136, EPI_ISL_521137, EPI_ISL_521138, EPI_ISL_521139, EPI_ISL_521140, EPI_ISL_521141, EPI_ISL_521142, EPI_ISL_521143, EPI_ISL_521144, EPI_ISL_521145, EPI_ISL_521146, EPI_ISL_521147, EPI_ISL_521148, EPI_ISL_521150, EPI_ISL_521151, EPI_ISL_521152 | | | |
| see above | Victorian Infectious Diseases Reference Laboratory (VIDRL) | VIDRL and MDU-PHL | Caly L., Seemann T., Sait, M., Schultz M., Druce J., Sherry, N. |
| EPI_ISL_521153 | Mayo Clinic & Mayo Clinic Laboratories | Minnesota Department of Health, Public Health Laboratory | Matt Plumb, Jacob Garfin, and Xiong Wang |
| EPI_ISL_521154, EPI_ISL_521156, EPI_ISL_521157, EPI_ISL_521158, EPI_ISL_521159, EPI_ISL_521160, EPI_ISL_521161, EPI_ISL_521162, EPI_ISL_521163, EPI_ISL_521165, EPI_ISL_521166, EPI_ISL_521167, EPI_ISL_521168, EPI_ISL_521170, EPI_ISL_521171, EPI_ISL_521172, EPI_ISL_521173, EPI_ISL_521174, EPI_ISL_521175, EPI_ISL_521176, EPI_ISL_521177, EPI_ISL_521178, EPI_ISL_521179, EPI_ISL_521180, EPI_ISL_521181, EPI_ISL_521182, EPI_ISL_521183, EPI_ISL_521184, EPI_ISL_521185, EPI_ISL_521186, EPI_ISL_521187, EPI_ISL_521188, EPI_ISL_521189, EPI_ISL_521190, EPI_ISL_521191, EPI_ISL_521192, EPI_ISL_521193, EPI_ISL_521194, EPI_ISL_521195, EPI_ISL_521196, EPI_ISL_521197, EPI_ISL_521198, EPI_ISL_521199, EPI_ISL_521200, EPI_ISL_521201, EPI_ISL_521202, EPI_ISL_521203, EPI_ISL_521204, EPI_ISL_521206, EPI_ISL_521207, EPI_ISL_521208, EPI_ISL_521210, EPI_ISL_521211, EPI_ISL_521212, EPI_ISL_521213, EPI_ISL_521214, EPI_ISL_521215, EPI_ISL_521217, EPI_ISL_521218, EPI_ISL_521219, EPI_ISL_521220, EPI_ISL_521221, EPI_ISL_521222, EPI_ISL_521227, EPI_ISL_521228, EPI_ISL_521229, EPI_ISL_521230, EPI_ISL_521231, EPI_ISL_521234, EPI_ISL_521235, EPI_ISL_521236, EPI_ISL_521237, EPI_ISL_521238, EPI_ISL_521239, EPI_ISL_521240, EPI_ISL_521241, EPI_ISL_521242, EPI_ISL_521243, EPI_ISL_521244, EPI_ISL_521245, EPI_ISL_521246, EPI_ISL_521247, EPI_ISL_521248, EPI_ISL_521249, EPI_ISL_521250, EPI_ISL_521251, EPI_ISL_521252, EPI_ISL_521253, EPI_ISL_521254, EPI_ISL_521255, EPI_ISL_521256, EPI_ISL_521257, EPI_ISL_521258, EPI_ISL_521259, EPI_ISL_521260, EPI_ISL_521261, EPI_ISL_521262, EPI_ISL_521263, EPI_ISL_521264, EPI_ISL_521265, EPI_ISL_521266, EPI_ISL_521267, EPI_ISL_521268, EPI_ISL_521269 | | | |
| see above | Microbiological Diagnostic Unit - Public Health Laboratory (MDU-PHL) | MDU-PHL | Seemann T., Schultz M., Sait, M., Sherry, N. |
| EPI_ISL_521270, EPI_ISL_521271, EPI_ISL_521272, EPI_ISL_521274, EPI_ISL_521275, EPI_ISL_521276, EPI_ISL_521277 | Victorian Infectious Diseases Reference Laboratory (VIDRL) | VIDRL and MDU-PHL | Caly L., Seemann T., Sait, M., Schultz M., Druce J., Sherry, N. |
| EPI_ISL_521278, EPI_ISL_521279, EPI_ISL_521280, EPI_ISL_521281, EPI_ISL_521282, EPI_ISL_521283, EPI_ISL_521284, EPI_ISL_521285, EPI_ISL_521287, EPI_ISL_521288, EPI_ISL_521289, EPI_ISL_521290, EPI_ISL_521291, EPI_ISL_521292, EPI_ISL_521293, EPI_ISL_521294, EPI_ISL_521295, EPI_ISL_521296, EPI_ISL_521297, EPI_ISL_521298, EPI_ISL_521299, EPI_ISL_521300, EPI_ISL_521301, EPI_ISL_521302, EPI_ISL_521303, EPI_ISL_521304, EPI_ISL_521305, EPI_ISL_521306, EPI_ISL_521307, EPI_ISL_521308, EPI_ISL_521309, EPI_ISL_521310, EPI_ISL_521311, EPI_ISL_521312, EPI_ISL_521313, EPI_ISL_521314 | | | |
| see above | Microbiological Diagnostic Unit - Public Health Laboratory (MDU-PHL) | MDU-PHL | Seemann T., Schultz M., Sait, M., Sherry, N. |
| EPI_ISL_521315, EPI_ISL_521316, EPI_ISL_521317, EPI_ISL_521318, EPI_ISL_521320, EPI_ISL_521321, EPI_ISL_521325, EPI_ISL_521326, EPI_ISL_521327, EPI_ISL_521328, EPI_ISL_521329, EPI_ISL_521330, EPI_ISL_521331, EPI_ISL_521332, EPI_ISL_521334, EPI_ISL_521335, EPI_ISL_521338, EPI_ISL_521339, EPI_ISL_521340, EPI_ISL_521341, EPI_ISL_521342, EPI_ISL_521343, EPI_ISL_521345, EPI_ISL_521346, EPI_ISL_521347, EPI_ISL_521348, EPI_ISL_521349, EPI_ISL_521350, EPI_ISL_521352, EPI_ISL_521353, EPI_ISL_521354, EPI_ISL_521355, EPI_ISL_521356, EPI_ISL_521357, EPI_ISL_521358, EPI_ISL_521359, EPI_ISL_521360, EPI_ISL_521361, EPI_ISL_521362, EPI_ISL_521363, EPI_ISL_521364, EPI_ISL_521365, EPI_ISL_521366, EPI_ISL_521367, EPI_ISL_521368, EPI_ISL_521369, EPI_ISL_521370, EPI_ISL_521371, EPI_ISL_521372, EPI_ISL_521374, EPI_ISL_521376, EPI_ISL_521377, EPI_ISL_521379, EPI_ISL_521380, EPI_ISL_521381, EPI_ISL_521382, EPI_ISL_521383, EPI_ISL_521384, EPI_ISL_521385, EPI_ISL_521386, EPI_ISL_521387, EPI_ISL_521388, EPI_ISL_521389, EPI_ISL_521391, EPI_ISL_521394, EPI_ISL_521395, EPI_ISL_521396, EPI_ISL_521398, EPI_ISL_521399, EPI_ISL_521400, EPI_ISL_521402, EPI_ISL_521403, EPI_ISL_521404, EPI_ISL_521405, EPI_ISL_521406, EPI_ISL_521407, EPI_ISL_521409, EPI_ISL_521410, EPI_ISL_521411, EPI_ISL_521412, EPI_ISL_521415, EPI_ISL_521416, EPI_ISL_521418, EPI_ISL_521419, EPI_ISL_521420, EPI_ISL_521421, EPI_ISL_521422, EPI_ISL_521423, EPI_ISL_521424, EPI_ISL_521425, EPI_ISL_521426, EPI_ISL_521427, EPI_ISL_521428, EPI_ISL_521430, EPI_ISL_521431, EPI_ISL_521432, EPI_ISL_521433, EPI_ISL_521434, EPI_ISL_521435, EPI_ISL_521436, EPI_ISL_521438, EPI_ISL_521440, EPI_ISL_521441, EPI_ISL_521442, EPI_ISL_521443, EPI_ISL_521444, EPI_ISL_521445, EPI_ISL_521447, EPI_ISL_521448, EPI_ISL_521449 | | | |
| see above | Victorian Infectious Diseases Reference Laboratory (VIDRL) | VIDRL and MDU-PHL | Caly L., Seemann T., Sait, M., Schultz M., Druce J., Sherry, N. |
| EPI_ISL_521450, EPI_ISL_521452, EPI_ISL_521453, EPI_ISL_521454, EPI_ISL_521455, EPI_ISL_521456, EPI_ISL_521457, EPI_ISL_521458, EPI_ISL_521459, EPI_ISL_521460, EPI_ISL_521461 | | | |
| see above | Microbiological Diagnostic Unit - Public Health Laboratory (MDU-PHL) | MDU-PHL | Seemann T., Schultz M., Sait, M., Sherry, N. |
| EPI_ISL_521463, EPI_ISL_521464, EPI_ISL_521465, EPI_ISL_521467, EPI_ISL_521468, EPI_ISL_521469, EPI_ISL_521470, EPI_ISL_521471, EPI_ISL_521472, EPI_ISL_521473, EPI_ISL_521474, EPI_ISL_521475, EPI_ISL_521476, EPI_ISL_521477, EPI_ISL_521478, EPI_ISL_521480, EPI_ISL_521481, EPI_ISL_521482, EPI_ISL_521483, EPI_ISL_521485, EPI_ISL_521486, EPI_ISL_521487, EPI_ISL_521488, EPI_ISL_521489, EPI_ISL_521490, EPI_ISL_521491, EPI_ISL_521492, EPI_ISL_521493, EPI_ISL_521494, EPI_ISL_521495, EPI_ISL_521496, EPI_ISL_521497, EPI_ISL_521498, EPI_ISL_521500, EPI_ISL_521502, EPI_ISL_521503, EPI_ISL_521504, EPI_ISL_521505, EPI_ISL_521506, EPI_ISL_521507, EPI_ISL_521508, EPI_ISL_521509, EPI_ISL_521510, EPI_ISL_521511, EPI_ISL_521512, EPI_ISL_521514, EPI_ISL_521515, EPI_ISL_521516, EPI_ISL_521517, EPI_ISL_521518, EPI_ISL_521519, EPI_ISL_521520, EPI_ISL_521521, EPI_ISL_521522, EPI_ISL_521523, EPI_ISL_521524, EPI_ISL_521525, EPI_ISL_521526, EPI_ISL_521527, EPI_ISL_521528, EPI_ISL_521529, EPI_ISL_521530, EPI_ISL_521531, EPI_ISL_521532, EPI_ISL_521533, EPI_ISL_521534, EPI_ISL_521535, EPI_ISL_521536, EPI_ISL_521537, EPI_ISL_521538, EPI_ISL_521539, EPI_ISL_521540, | | | |

| | | | |
|---|---|---|---|
| EPI_ISL_521541, EPI_ISL_521543, EPI_ISL_521545, EPI_ISL_521546, EPI_ISL_521548, EPI_ISL_521549, EPI_ISL_521550, EPI_ISL_521551, EPI_ISL_521552, EPI_ISL_521553, EPI_ISL_521554, EPI_ISL_521555, EPI_ISL_521556, EPI_ISL_521557, EPI_ISL_521558, EPI_ISL_521560, EPI_ISL_521561 | | | |
| see above | Victorian Infectious Diseases Reference Laboratory (VIDRL) | VIDRL and MDU-PHL | Caly L., Seemann T., Sait, M., Schultz M., Druce J., Sherry, N. |
| EPI_ISL_521562, EPI_ISL_521563, EPI_ISL_521564, EPI_ISL_521565, EPI_ISL_521566, EPI_ISL_521567, EPI_ISL_521568, EPI_ISL_521569, EPI_ISL_521570, EPI_ISL_521571, EPI_ISL_521573, EPI_ISL_521574, EPI_ISL_521575, EPI_ISL_521576, EPI_ISL_521577, EPI_ISL_521578, EPI_ISL_521579, EPI_ISL_521580, EPI_ISL_521581, EPI_ISL_521582, EPI_ISL_521583, EPI_ISL_521584, EPI_ISL_521585, EPI_ISL_521586, EPI_ISL_521587, EPI_ISL_521588, EPI_ISL_521589, EPI_ISL_521590, EPI_ISL_521591, EPI_ISL_521592, EPI_ISL_521593, EPI_ISL_521594, EPI_ISL_521596, EPI_ISL_521597, EPI_ISL_521599, EPI_ISL_521600, EPI_ISL_521601, EPI_ISL_521602, EPI_ISL_521603, EPI_ISL_521604, EPI_ISL_521606, EPI_ISL_521607, EPI_ISL_521608, EPI_ISL_521609, EPI_ISL_521610, EPI_ISL_521611, EPI_ISL_521612, EPI_ISL_521613, EPI_ISL_521614, EPI_ISL_521615, EPI_ISL_521616, EPI_ISL_521617, EPI_ISL_521618, EPI_ISL_521619, EPI_ISL_521622, EPI_ISL_521624, EPI_ISL_521625, EPI_ISL_521626, EPI_ISL_521627, EPI_ISL_521628, EPI_ISL_521629, EPI_ISL_521630, EPI_ISL_521631, EPI_ISL_521632, EPI_ISL_521633, EPI_ISL_521634, EPI_ISL_521635, EPI_ISL_521636, EPI_ISL_521637, EPI_ISL_521638, EPI_ISL_521639, EPI_ISL_521640, EPI_ISL_521641, EPI_ISL_521642, EPI_ISL_521643, EPI_ISL_521644, EPI_ISL_521645, EPI_ISL_521646, EPI_ISL_521647, EPI_ISL_521648, EPI_ISL_521649, EPI_ISL_521650, EPI_ISL_521651, EPI_ISL_521652, EPI_ISL_521653, EPI_ISL_521654, EPI_ISL_521655, EPI_ISL_521656, EPI_ISL_521657, EPI_ISL_521658, EPI_ISL_521659, EPI_ISL_521660, EPI_ISL_521661, EPI_ISL_521662, EPI_ISL_521663, EPI_ISL_521664, EPI_ISL_521665, EPI_ISL_521666, EPI_ISL_521667, EPI_ISL_521668, EPI_ISL_521669, EPI_ISL_521670, EPI_ISL_521671, EPI_ISL_521672, EPI_ISL_521673, EPI_ISL_521674, EPI_ISL_521675, EPI_ISL_521676, EPI_ISL_521677, EPI_ISL_521678, EPI_ISL_521679, EPI_ISL_521680, EPI_ISL_521681, EPI_ISL_521683, EPI_ISL_521684, EPI_ISL_521685, EPI_ISL_521686, EPI_ISL_521687, EPI_ISL_521688, EPI_ISL_521689, EPI_ISL_521690, EPI_ISL_521691, EPI_ISL_521692, EPI_ISL_521693, EPI_ISL_521694, EPI_ISL_521695, EPI_ISL_521696, EPI_ISL_521697, EPI_ISL_521698, EPI_ISL_521699, EPI_ISL_521700, EPI_ISL_521701, EPI_ISL_521702, EPI_ISL_521703, EPI_ISL_521704, EPI_ISL_521705, EPI_ISL_521706, EPI_ISL_521707, EPI_ISL_521708, EPI_ISL_521709, EPI_ISL_521710, EPI_ISL_521711, EPI_ISL_521712, EPI_ISL_521713, EPI_ISL_521714, EPI_ISL_521715, EPI_ISL_521716, EPI_ISL_521717, EPI_ISL_521718, EPI_ISL_521719, EPI_ISL_521720, EPI_ISL_521721, EPI_ISL_521722, EPI_ISL_521723, EPI_ISL_521724, EPI_ISL_521725, EPI_ISL_521726, EPI_ISL_521727, EPI_ISL_521729, EPI_ISL_521730, EPI_ISL_521731, EPI_ISL_521732, EPI_ISL_521733, EPI_ISL_521734, EPI_ISL_521736, EPI_ISL_521737, EPI_ISL_521738, EPI_ISL_521739, EPI_ISL_521740, EPI_ISL_521741, EPI_ISL_521742, EPI_ISL_521743, EPI_ISL_521744, EPI_ISL_521745, EPI_ISL_521746, EPI_ISL_521747, EPI_ISL_521748, EPI_ISL_521749, EPI_ISL_521750, EPI_ISL_521751, EPI_ISL_521752, EPI_ISL_521753, EPI_ISL_521754, EPI_ISL_521755, EPI_ISL_521756, EPI_ISL_521757, EPI_ISL_521758, EPI_ISL_521760, EPI_ISL_521761, EPI_ISL_521762, EPI_ISL_521763, EPI_ISL_521764, EPI_ISL_521765, EPI_ISL_521766, EPI_ISL_521767, EPI_ISL_521768, EPI_ISL_521769, EPI_ISL_521770, EPI_ISL_521771, EPI_ISL_521772, EPI_ISL_521773, EPI_ISL_521774, EPI_ISL_521776, EPI_ISL_521777, EPI_ISL_521778, EPI_ISL_521779, EPI_ISL_521780, EPI_ISL_521781, EPI_ISL_521782, EPI_ISL_521783, EPI_ISL_521784, EPI_ISL_521785, EPI_ISL_521786, EPI_ISL_521787, EPI_ISL_521788, EPI_ISL_521789, EPI_ISL_521790, EPI_ISL_521791, EPI_ISL_521792, EPI_ISL_521793, EPI_ISL_521795, EPI_ISL_521796, EPI_ISL_521797, EPI_ISL_521798, EPI_ISL_521799, EPI_ISL_521800, EPI_ISL_521801, EPI_ISL_521802, EPI_ISL_521803, EPI_ISL_521804, EPI_ISL_521805, EPI_ISL_521806, EPI_ISL_521807, EPI_ISL_521808, EPI_ISL_521809, EPI_ISL_521810, EPI_ISL_521811, EPI_ISL_521812, EPI_ISL_521813, EPI_ISL_521814, EPI_ISL_521815, EPI_ISL_521816, EPI_ISL_521817, EPI_ISL_521818, EPI_ISL_521819, EPI_ISL_521820, EPI_ISL_521821, EPI_ISL_521822, EPI_ISL_521823, EPI_ISL_521824, EPI_ISL_521826, EPI_ISL_521827, EPI_ISL_521828, EPI_ISL_521829, EPI_ISL_521830, EPI_ISL_521831, EPI_ISL_521832, EPI_ISL_521833, EPI_ISL_521834, EPI_ISL_521835, EPI_ISL_521836, EPI_ISL_521837, EPI_ISL_521838, EPI_ISL_521839 | | | |
| see above | Microbiological Diagnostic Unit - Public Health Laboratory (MDU-PHL) | MDU-PHL | Seemann T., Schultz M., Sait, M., Sherry, N. |
| EPI_ISL_521842, EPI_ISL_521843, EPI_ISL_521844, EPI_ISL_521845, EPI_ISL_521847, EPI_ISL_521849, EPI_ISL_521850, EPI_ISL_521851, EPI_ISL_521852, EPI_ISL_521853, EPI_ISL_521854, EPI_ISL_521856, EPI_ISL_521857, EPI_ISL_521858, EPI_ISL_521859 | | | |
| see above | Victorian Infectious Diseases Reference Laboratory (VIDRL) | VIDRL and MDU-PHL | Caly L., Seemann T., Sait, M., Schultz M., Druce J., Sherry, N. |
| EPI_ISL_521860, EPI_ISL_521861, EPI_ISL_521862 | Royal Darwin Hospital Pathology | MDU-PHL | Meumann, E., Caly L., Seemann T., Sait, M., Schultz M., Druce J., Sherry, N. |
| EPI_ISL_521871, EPI_ISL_521872, EPI_ISL_521873, EPI_ISL_521874, EPI_ISL_521875, EPI_ISL_521876, EPI_ISL_521877, EPI_ISL_521878, EPI_ISL_521879, EPI_ISL_521880, EPI_ISL_521881, EPI_ISL_521882, EPI_ISL_521883, EPI_ISL_521884 | | | |
| see above | Victorian Infectious Diseases Reference Laboratory (VIDRL) | VIDRL and MDU-PHL | Caly L., Seemann T., Sait, M., Schultz M., Druce J., Sherry, N. |
| EPI_ISL_521887 | Microbiological Diagnostic Unit - Public Health Laboratory (MDU-PHL) | MDU-PHL | Seemann T., Schultz M., Sait, M., Sherry, N. |
| EPI_ISL_521892, EPI_ISL_521893, EPI_ISL_521894, EPI_ISL_521895, EPI_ISL_521897, EPI_ISL_521898, EPI_ISL_521900, EPI_ISL_521901, EPI_ISL_521902, EPI_ISL_521903, EPI_ISL_521904 | | | |
| see above | Victorian Infectious Diseases Reference Laboratory (VIDRL) | VIDRL and MDU-PHL | Caly L., Seemann T., Sait, M., Schultz M., Druce J., Sherry, N. |
| EPI_ISL_521906, EPI_ISL_521907, EPI_ISL_521909, EPI_ISL_521910 | Microbiological Diagnostic Unit - Public Health Laboratory (MDU-PHL) | MDU-PHL | Seemann T., Schultz M., Sait, M., Sherry, N. |
| EPI_ISL_521912, EPI_ISL_521913, EPI_ISL_521914, EPI_ISL_521916, EPI_ISL_521917, EPI_ISL_521918, EPI_ISL_521919, EPI_ISL_521920, EPI_ISL_521922, EPI_ISL_521924, EPI_ISL_521925 | | | |
| see above | Victorian Infectious Diseases Reference Laboratory (VIDRL) | VIDRL and MDU-PHL | Caly L., Seemann T., Sait, M., Schultz M., Druce J., Sherry, N. |
| EPI_ISL_521927, EPI_ISL_521928, EPI_ISL_521929, EPI_ISL_521930, EPI_ISL_521931, EPI_ISL_521932, EPI_ISL_521933, EPI_ISL_521934, EPI_ISL_521935, EPI_ISL_521936, EPI_ISL_521937, EPI_ISL_521938, EPI_ISL_521939, EPI_ISL_521940, EPI_ISL_521941, EPI_ISL_521942, EPI_ISL_521943, EPI_ISL_521944 | | | |
| see above | Microbiological Diagnostic Unit - Public Health Laboratory (MDU-PHL) | MDU-PHL | Seemann T., Schultz M., Sait, M., Sherry, N. |
| EPI_ISL_521945, EPI_ISL_521946, EPI_ISL_521947, EPI_ISL_521948, EPI_ISL_521950, EPI_ISL_521951, EPI_ISL_521952, EPI_ISL_521953, EPI_ISL_521954, EPI_ISL_521956, EPI_ISL_521957, EPI_ISL_521959, EPI_ISL_521961, EPI_ISL_521963, EPI_ISL_521964, EPI_ISL_521965, EPI_ISL_521966, EPI_ISL_521967, EPI_ISL_521968, EPI_ISL_521969, EPI_ISL_521971 | | | |
| see above | Victorian Infectious Diseases Reference Laboratory (VIDRL) | VIDRL and MDU-PHL | Caly L., Seemann T., Sait, M., Schultz M., Druce J., Sherry, N. |
| EPI_ISL_521972, EPI_ISL_521973, EPI_ISL_521974 | Microbiological Diagnostic Unit - Public Health Laboratory (MDU-PHL) | MDU-PHL | Seemann T., Schultz M., Sait, M., Sherry, N. |
| EPI_ISL_521975, EPI_ISL_521976, EPI_ISL_521977, EPI_ISL_521978, EPI_ISL_521981, EPI_ISL_521982, EPI_ISL_521984, EPI_ISL_521985, EPI_ISL_521986, EPI_ISL_521988 | Victorian Infectious Diseases Reference Laboratory (VIDRL) | VIDRL and MDU-PHL | Caly L., Seemann T., Sait, M., Schultz M., Druce J., Sherry, N. |
| EPI_ISL_521989, EPI_ISL_521991, EPI_ISL_521992, EPI_ISL_521993, EPI_ISL_521994, EPI_ISL_521995, EPI_ISL_521996, EPI_ISL_521997, EPI_ISL_521998, EPI_ISL_521999, EPI_ISL_522000, EPI_ISL_522001, EPI_ISL_522002, EPI_ISL_522003, EPI_ISL_522004, EPI_ISL_522005, EPI_ISL_522006, EPI_ISL_522007, EPI_ISL_522008, EPI_ISL_522009, EPI_ISL_522010, EPI_ISL_522011, EPI_ISL_522012, EPI_ISL_522013, EPI_ISL_522014, EPI_ISL_522015, EPI_ISL_522016, EPI_ISL_522017, EPI_ISL_522018, EPI_ISL_522019, EPI_ISL_522020, EPI_ISL_522021, EPI_ISL_522022, EPI_ISL_522023, EPI_ISL_522024, EPI_ISL_522025, EPI_ISL_522026, EPI_ISL_522027, EPI_ISL_522028, EPI_ISL_522029, EPI_ISL_522030, EPI_ISL_522031, EPI_ISL_522032, EPI_ISL_522033, EPI_ISL_522034, EPI_ISL_522035, EPI_ISL_522036, EPI_ISL_522037, EPI_ISL_522038, EPI_ISL_522039, EPI_ISL_522040, EPI_ISL_522041, EPI_ISL_522042, EPI_ISL_522043, EPI_ISL_522044, EPI_ISL_522045, EPI_ISL_522046, EPI_ISL_522047, EPI_ISL_522048, EPI_ISL_522049, EPI_ISL_522050, EPI_ISL_522051, EPI_ISL_522052, EPI_ISL_522053, EPI_ISL_522054 | | | |
| see above | Microbiological Diagnostic Unit - Public Health Laboratory (MDU-PHL) | MDU-PHL | Seemann T., Schultz M., Sait, M., Sherry, N. |
| EPI_ISL_522055, EPI_ISL_522056, EPI_ISL_522057, EPI_ISL_522058, EPI_ISL_522059, EPI_ISL_522061, EPI_ISL_522062, EPI_ISL_522063, EPI_ISL_522064, EPI_ISL_522065, EPI_ISL_522066, EPI_ISL_522067, EPI_ISL_522068, EPI_ISL_522069, EPI_ISL_522070, EPI_ISL_522071, EPI_ISL_522072, EPI_ISL_522073, EPI_ISL_522074, EPI_ISL_522075, EPI_ISL_522077, EPI_ISL_522078, EPI_ISL_522079, EPI_ISL_522080, EPI_ISL_522081, EPI_ISL_522082, EPI_ISL_522083, EPI_ISL_522084, EPI_ISL_522085, EPI_ISL_522086, EPI_ISL_522087, EPI_ISL_522088, EPI_ISL_522089, EPI_ISL_522091, EPI_ISL_522093, EPI_ISL_522094, EPI_ISL_522096, EPI_ISL_522097, EPI_ISL_522098, EPI_ISL_522099, EPI_ISL_522101, EPI_ISL_522102, EPI_ISL_522103, EPI_ISL_522104, EPI_ISL_522106, EPI_ISL_522107, EPI_ISL_522108, EPI_ISL_522109 | | | |
| see above | Victorian Infectious Diseases Reference Laboratory (VIDRL) | VIDRL and MDU-PHL | Caly L., Seemann T., Sait, M., Schultz M., Druce J., Sherry, N. |
| EPI_ISL_522111, EPI_ISL_522112, EPI_ISL_522113, EPI_ISL_522114, EPI_ISL_522115, EPI_ISL_522116, EPI_ISL_522117, EPI_ISL_522118 | Microbiological Diagnostic Unit - Public Health Laboratory (MDU-PHL) | MDU-PHL | Seemann T., Schultz M., Sait, M., Sherry, N. |
| EPI_ISL_522120, EPI_ISL_522121, EPI_ISL_522122, EPI_ISL_522123, EPI_ISL_522124, EPI_ISL_522125, EPI_ISL_522126, EPI_ISL_522127, EPI_ISL_522128, EPI_ISL_522129, EPI_ISL_522130, EPI_ISL_522131, EPI_ISL_522132, EPI_ISL_522133, EPI_ISL_522134, EPI_ISL_522135, EPI_ISL_522136, EPI_ISL_522137, EPI_ISL_522138, EPI_ISL_522139, EPI_ISL_522140, EPI_ISL_522141, EPI_ISL_522142, EPI_ISL_522143, EPI_ISL_522144, EPI_ISL_522145, EPI_ISL_522146, EPI_ISL_522147, EPI_ISL_522148, EPI_ISL_522149, EPI_ISL_522150, EPI_ISL_522151, EPI_ISL_522152, EPI_ISL_522153, EPI_ISL_522154, EPI_ISL_522155, EPI_ISL_522157, EPI_ISL_522158, EPI_ISL_522159, EPI_ISL_522160, EPI_ISL_522161, EPI_ISL_522162, EPI_ISL_522163, EPI_ISL_522164, EPI_ISL_522165, EPI_ISL_522166, EPI_ISL_522167, EPI_ISL_522168, EPI_ISL_522169, EPI_ISL_522170, EPI_ISL_522171, EPI_ISL_522172, EPI_ISL_522173, EPI_ISL_522174, EPI_ISL_522175, EPI_ISL_522176, EPI_ISL_522177, EPI_ISL_522178, EPI_ISL_522179, EPI_ISL_522180, EPI_ISL_522181, EPI_ISL_522182, EPI_ISL_522183, EPI_ISL_522184, EPI_ISL_522185, EPI_ISL_522186, EPI_ISL_522187 | | | |
| see above | Victorian Infectious Diseases Reference Laboratory (VIDRL) | VIDRL and MDU-PHL | Caly L., Seemann T., Sait, M., Schultz M., Druce J., Sherry, N. |
| EPI_ISL_522188, EPI_ISL_522189, EPI_ISL_522192, EPI_ISL_522193, EPI_ISL_522194, EPI_ISL_522195, EPI_ISL_522196, EPI_ISL_522197, EPI_ISL_522198, EPI_ISL_522199, EPI_ISL_522200, EPI_ISL_522201, EPI_ISL_522202, EPI_ISL_522203, EPI_ISL_522204, EPI_ISL_522205, EPI_ISL_522206, EPI_ISL_522207, EPI_ISL_522208, EPI_ISL_522209, EPI_ISL_522210, EPI_ISL_522211, EPI_ISL_522212, EPI_ISL_522213, EPI_ISL_522214, EPI_ISL_522215, EPI_ISL_522216, EPI_ISL_522217, EPI_ISL_522218, EPI_ISL_522219, EPI_ISL_522220, EPI_ISL_522221, EPI_ISL_522222, EPI_ISL_522223, EPI_ISL_522224, EPI_ISL_522225, EPI_ISL_522226, EPI_ISL_522227, EPI_ISL_522228, EPI_ISL_522229, EPI_ISL_522230, EPI_ISL_522231 | | | |
| see above | Microbiological Diagnostic Unit - Public Health Laboratory (MDU-PHL) | MDU-PHL | Seemann T., Schultz M., Sait, M., Sherry, N. |
| EPI_ISL_522233, EPI_ISL_522234, EPI_ISL_522235, EPI_ISL_522236, EPI_ISL_522237, EPI_ISL_522238, EPI_ISL_522239, EPI_ISL_522240, EPI_ISL_522241, EPI_ISL_522242 | Victorian Infectious Diseases Reference Laboratory (VIDRL) | VIDRL and MDU-PHL | Caly L., Seemann T., Sait, M., Schultz M., Druce J., Sherry, N. |
| EPI_ISL_522243, EPI_ISL_522244, EPI_ISL_522245, EPI_ISL_522247, EPI_ISL_522249, EPI_ISL_522250, EPI_ISL_522251, EPI_ISL_522252, EPI_ISL_522253 | Microbiological Diagnostic Unit - Public Health Laboratory (MDU-PHL) | MDU-PHL | Seemann T., Schultz M., Sait, M., Sherry, N. |
| EPI_ISL_522254, EPI_ISL_522255, EPI_ISL_522258, EPI_ISL_522260, EPI_ISL_522261, EPI_ISL_522262, EPI_ISL_522264, EPI_ISL_522270, EPI_ISL_522272, EPI_ISL_522274, EPI_ISL_522276, EPI_ISL_522279, EPI_ISL_522281, EPI_ISL_522282, EPI_ISL_522285, EPI_ISL_522287, EPI_ISL_522288, EPI_ISL_522290, EPI_ISL_522291, EPI_ISL_522294, EPI_ISL_522295, EPI_ISL_522296, EPI_ISL_522300, EPI_ISL_522302, EPI_ISL_522303, EPI_ISL_522304, EPI_ISL_522305, EPI_ISL_522307, EPI_ISL_522311, EPI_ISL_522312, EPI_ISL_522315, EPI_ISL_522322, EPI_ISL_522323, EPI_ISL_522331, EPI_ISL_522336, EPI_ISL_522337, EPI_ISL_522340, EPI_ISL_522341, EPI_ISL_522343 | | | |
| see above | Utah Public Health Laboratory | Utah Public Health Laboratory | Erin Young, Kelly Oakeson |
| EPI_ISL_522349, EPI_ISL_522350 | KU Leuven, Rega Institute, Clinical and Epidemiological Virology | KU Leuven, Rega Institute, Clinical and Epidemiological Virology | Tony Wawina-Bokalanga, Joan Marti-Carerras, Bert Vanmechelen, Piet Maes |
| EPI_ISL_522378, EPI_ISL_522393, EPI_ISL_522394 | Texas Department of State Health Services | Texas Department of State Health Services | Rashmi Tuladhar, Bonnie Oh, Cara Akrout, Jenny Zhang, Maliha Rahman, Anita Pokharel, Myong Koag, Chun Wang, Rachel Lee, Grace Kubin |
| EPI_ISL_522396, EPI_ISL_522397, EPI_ISL_522398, EPI_ISL_522399, EPI_ISL_522400, EPI_ISL_522401, EPI_ISL_522402, EPI_ISL_522403, EPI_ISL_522404, EPI_ISL_522405 | Alaska State Virology Laboratory | Alaska State Virology Laboratory | Jack Chen, Ph.D. |
| EPI_ISL_522437 | Mahatma Gandhi Institute of Medical Sciences | National Institute of Biomedical Genomics - DBT's PAN-INDIA 1000 SARS--CoV-2 RNA Genome Sequencing Consortium | Arindam Maitra, Vijayshri Deotale, Rahul Narang, Deepashri Maraskolhe, Saumitra Das |
| EPI_ISL_522440, EPI_ISL_522441, EPI_ISL_522442, EPI_ISL_522443, EPI_ISL_522444, EPI_ISL_522445, EPI_ISL_522446, EPI_ISL_522447, EPI_ISL_522448, EPI_ISL_522449, EPI_ISL_522450, EPI_ISL_522451, EPI_ISL_522452, EPI_ISL_522453, EPI_ISL_522455, EPI_ISL_522456, EPI_ISL_522458, EPI_ISL_522459, EPI_ISL_522460, EPI_ISL_522461, EPI_ISL_522462, EPI_ISL_522463 | | | |
| see above | Center for Laboratory Control of Infectious Diseases, Korea Centers for Diseases Control and Prevention | Center for Laboratory Control of Infectious Diseases, Korea Centers for Diseases Control and Prevention | Junyoung Kim, Ae Kyung Park, Eunkyung Shin, Jin Sun No, Jeong-Min Kim, Yoon-Seok Chung, Heui Man Kim, Myung Guk Han |
| EPI_ISL_522465, EPI_ISL_522466 | Division of Viral Diseases, Center for Laboratory Control of Infectious Diseases, Korea Centers for Diseases Control and Prevention | Division of Viral Diseases, Center for Laboratory Control of Infectious Diseases, Korea Centers for Diseases Control and Prevention | Jeong-Min Kim, Yoon-Seok Chung, Namjoo Lee, Sang Hee Woo, Hye-Jun Jo, Heui Man Kim, Jun-Sub Kim, Myung Guk Han |
| EPI_ISL_522467, EPI_ISL_522468, EPI_ISL_522469, EPI_ISL_522470, | Center for Laboratory Control of Infectious Diseases, Korea Centers for Diseases Control and Prevention | Center for Laboratory Control of Infectious Diseases, Korea Centers for Diseases Control and Prevention | Junyoung Kim, Ae Kyung Park, Eunkyung Shin, Jin Sun No, Jeong-Min Kim, Yoon-Seok Chung, Heui Man Kim, Myung Guk Han |

| | | | |
|---|---|---|---|
| EPI_ISL_522472, EPI_ISL_522473 | | | |
| EPI_ISL_522474, EPI_ISL_522475 | Division of Viral Diseases, Center for Laboratory Control of Infectious Diseases, Korea Centers for Diseases Control and Prevention | Division of Viral Diseases, Center for Laboratory Control of Infectious Diseases, Korea Centers for Diseases Control and Prevention | Jeong-Min Kim, Yoon-Seok Chung, Namjoo Lee, Sang Hee Woo, Hye-Jun Jo, Heui Man Kim, Jun-Sub Kim, Myung Guk Han |
| EPI_ISL_522476 | Center for Laboratory Control of Infectious Diseases, Korea Centers for Diseases Control and Prevention | Center for Laboratory Control of Infectious Diseases, Korea Centers for Diseases Control and Prevention | Junyoung Kim, Ae Kyung Park, Eunkyung Shin, Jin Sun No, Jeong-Min Kim, Yoon-Seok Chung, Heui Man Kim, Myung Guk Han |
| EPI_ISL_522477, EPI_ISL_522478 | Division of Viral Diseases, Center for Laboratory Control of Infectious Diseases, Korea Centers for Diseases Control and Prevention | Division of Viral Diseases, Center for Laboratory Control of Infectious Diseases, Korea Centers for Diseases Control and Prevention | Jeong-Min Kim, Yoon-Seok Chung, Namjoo Lee, Sang Hee Woo, Hye-Jun Jo, Heui Man Kim, Jun-Sub Kim, Myung Guk Han |
| EPI_ISL_522479, EPI_ISL_522480, EPI_ISL_522481, EPI_ISL_522482, EPI_ISL_522483 | Center for Laboratory Control of Infectious Diseases, Korea Centers for Diseases Control and Prevention | Center for Laboratory Control of Infectious Diseases, Korea Centers for Diseases Control and Prevention | Junyoung Kim, Ae Kyung Park, Eunkyung Shin, Jin Sun No, Jeong-Min Kim, Yoon-Seok Chung, Heui Man Kim, Myung Guk Han |
| EPI_ISL_522484 | Division of Viral Diseases, Center for Laboratory Control of Infectious Diseases, Korea Centers for Diseases Control and Prevention | Division of Viral Diseases, Center for Laboratory Control of Infectious Diseases, Korea Centers for Diseases Control and Prevention | Jeong-Min Kim, Yoon-Seok Chung, Namjoo Lee, Sang Hee Woo, Hye-Jun Jo, Heui Man Kim, Jun-Sub Kim, Myung Guk Han |
| EPI_ISL_522485 | Center for Laboratory Control of Infectious Diseases, Korea Centers for Diseases Control and Prevention | Center for Laboratory Control of Infectious Diseases, Korea Centers for Diseases Control and Prevention | Junyoung Kim, Ae Kyung Park, Eunkyung Shin, Jin Sun No, Jeong-Min Kim, Yoon-Seok Chung, Heui Man Kim, Myung Guk Han |
| EPI_ISL_522486, EPI_ISL_522487, EPI_ISL_522488, EPI_ISL_522489, EPI_ISL_522490 | Division of Viral Diseases, Center for Laboratory Control of Infectious Diseases, Korea Centers for Diseases Control and Prevention | Division of Viral Diseases, Center for Laboratory Control of Infectious Diseases, Korea Centers for Diseases Control and Prevention | Jeong-Min Kim, Yoon-Seok Chung, Namjoo Lee, Sang Hee Woo, Hye-Jun Jo, Heui Man Kim, Jun-Sub Kim, Myung Guk Han |
| EPI_ISL_522491, EPI_ISL_522492, EPI_ISL_522493, EPI_ISL_522495, EPI_ISL_522496 | Center for Laboratory Control of Infectious Diseases, Korea Centers for Diseases Control and Prevention | Center for Laboratory Control of Infectious Diseases, Korea Centers for Diseases Control and Prevention | Junyoung Kim, Ae Kyung Park, Eunkyung Shin, Jin Sun No, Jeong-Min Kim, Yoon-Seok Chung, Heui Man Kim, Myung Guk Han |
| EPI_ISL_522498, EPI_ISL_522499, EPI_ISL_522500, EPI_ISL_522501 | Division of Viral Diseases, Center for Laboratory Control of Infectious Diseases, Korea Centers for Diseases Control and Prevention | Division of Viral Diseases, Center for Laboratory Control of Infectious Diseases, Korea Centers for Diseases Control and Prevention | Jeong-Min Kim, Yoon-Seok Chung, Namjoo Lee, Sang Hee Woo, Hye-Jun Jo, Heui Man Kim, Jun-Sub Kim, Myung Guk Han |
| EPI_ISL_522502 | Center for Laboratory Control of Infectious Diseases, Korea Centers for Diseases Control and Prevention | Center for Laboratory Control of Infectious Diseases, Korea Centers for Diseases Control and Prevention | Junyoung Kim, Ae Kyung Park, Eunkyung Shin, Jin Sun No, Jeong-Min Kim, Yoon-Seok Chung, Heui Man Kim, Myung Guk Han |
| EPI_ISL_522503, EPI_ISL_522504 | Division of Viral Diseases, Center for Laboratory Control of Infectious Diseases, Korea Centers for Diseases Control and Prevention | Division of Viral Diseases, Center for Laboratory Control of Infectious Diseases, Korea Centers for Diseases Control and Prevention | Jeong-Min Kim, Yoon-Seok Chung, Namjoo Lee, Sang Hee Woo, Hye-Jun Jo, Heui Man Kim, Jun-Sub Kim, Myung Guk Han |
| EPI_ISL_522505 | Center for Laboratory Control of Infectious Diseases, Korea Centers for Diseases Control and Prevention | Center for Laboratory Control of Infectious Diseases, Korea Centers for Diseases Control and Prevention | Junyoung Kim, Ae Kyung Park, Eunkyung Shin, Jin Sun No, Jeong-Min Kim, Yoon-Seok Chung, Heui Man Kim, Myung Guk Han |
| EPI_ISL_522506, EPI_ISL_522507, EPI_ISL_522509 | Division of Viral Diseases, Center for Laboratory Control of Infectious Diseases, Korea Centers for Diseases Control and Prevention | Division of Viral Diseases, Center for Laboratory Control of Infectious Diseases, Korea Centers for Diseases Control and Prevention | Jeong-Min Kim, Yoon-Seok Chung, Namjoo Lee, Sang Hee Woo, Hye-Jun Jo, Heui Man Kim, Jun-Sub Kim, Myung Guk Han |
| EPI_ISL_522510 | Center for Laboratory Control of Infectious Diseases, Korea Centers for Diseases Control and Prevention | Center for Laboratory Control of Infectious Diseases, Korea Centers for Diseases Control and Prevention | Junyoung Kim, Ae Kyung Park, Eunkyung Shin, Jin Sun No, Jeong-Min Kim, Yoon-Seok Chung, Heui Man Kim, Myung Guk Han |
| EPI_ISL_522511, EPI_ISL_522512, EPI_ISL_522513, EPI_ISL_522514 | Division of Viral Diseases, Center for Laboratory Control of Infectious Diseases, Korea Centers for Diseases Control and Prevention | Division of Viral Diseases, Center for Laboratory Control of Infectious Diseases, Korea Centers for Diseases Control and Prevention | Jeong-Min Kim, Yoon-Seok Chung, Namjoo Lee, Sang Hee Woo, Hye-Jun Jo, Heui Man Kim, Jun-Sub Kim, Myung Guk Han |
| EPI_ISL_522515, EPI_ISL_522516 | Center for Laboratory Control of Infectious Diseases, Korea Centers for Diseases Control and Prevention | Center for Laboratory Control of Infectious Diseases, Korea Centers for Diseases Control and Prevention | Junyoung Kim, Ae Kyung Park, Eunkyung Shin, Jin Sun No, Jeong-Min Kim, Yoon-Seok Chung, Heui Man Kim, Myung Guk Han |
| EPI_ISL_522517 | Division of Viral Diseases, Center for Laboratory Control of Infectious Diseases, Korea Centers for Diseases Control and Prevention | Division of Viral Diseases, Center for Laboratory Control of Infectious Diseases, Korea Centers for Diseases Control and Prevention | Jeong-Min Kim, Yoon-Seok Chung, Namjoo Lee, Sang Hee Woo, Hye-Jun Jo, Heui Man Kim, Jun-Sub Kim, Myung Guk Han |
| EPI_ISL_522518 | Center for Laboratory Control of Infectious Diseases, Korea Centers for Diseases Control and Prevention | Center for Laboratory Control of Infectious Diseases, Korea Centers for Diseases Control and Prevention | Junyoung Kim, Ae Kyung Park, Eunkyung Shin, Jin Sun No, Jeong-Min Kim, Yoon-Seok Chung, Heui Man Kim, Myung Guk Han |
| EPI_ISL_522519, EPI_ISL_522520, EPI_ISL_522521, EPI_ISL_522522, EPI_ISL_522523, EPI_ISL_522524, EPI_ISL_522525, EPI_ISL_522526, EPI_ISL_522527, EPI_ISL_522528, EPI_ISL_522529, EPI_ISL_522530, EPI_ISL_522531, EPI_ISL_522532, EPI_ISL_522533, EPI_ISL_522534, EPI_ISL_522535, EPI_ISL_522536, EPI_ISL_522537, EPI_ISL_522538, EPI_ISL_522539, EPI_ISL_522540, EPI_ISL_522541, EPI_ISL_522542, EPI_ISL_522543, EPI_ISL_522544 | | | |
| see above | Division of Viral Diseases, Center for Laboratory Control of Infectious Diseases, Korea Centers for Diseases Control and Prevention | Division of Viral Diseases, Center for Laboratory Control of Infectious Diseases, Korea Centers for Diseases Control and Prevention | Jeong-Min Kim, Yoon-Seok Chung, Namjoo Lee, Sang Hee Woo, Hye-Jun Jo, Heui Man Kim, Jun-Sub Kim, Myung Guk Han |
| EPI_ISL_522547, EPI_ISL_522548 | Platforme CYROI | UMR PIMIT Université de La Réunion | David Wilkinson, Camille Lebarbenchon, Patrick Mavingui |
| EPI_ISL_522549, EPI_ISL_522550 | Félix Guyon Hospital | UMR PIMIT Université de La Réunion | David Wilkinson, Camille Lebarbenchon, Patrick Mavingui |
| EPI_ISL_522556, EPI_ISL_522558, EPI_ISL_522559, EPI_ISL_522563, EPI_ISL_522564, EPI_ISL_522565, EPI_ISL_522568, EPI_ISL_522574, EPI_ISL_522575, EPI_ISL_522576, EPI_ISL_522599, EPI_ISL_522610, EPI_ISL_522630, EPI_ISL_522638, EPI_ISL_522646, EPI_ISL_522650, EPI_ISL_522652, EPI_ISL_522653, EPI_ISL_522654, EPI_ISL_522655, EPI_ISL_522656, EPI_ISL_522657, EPI_ISL_522659, EPI_ISL_522662, EPI_ISL_522665, EPI_ISL_522666, EPI_ISL_522668, EPI_ISL_522669, EPI_ISL_522671, EPI_ISL_522672, EPI_ISL_522673, EPI_ISL_522674, EPI_ISL_522675, EPI_ISL_522676, EPI_ISL_522678, EPI_ISL_522679, EPI_ISL_522680, EPI_ISL_522681, EPI_ISL_522683, EPI_ISL_522685, EPI_ISL_522689, EPI_ISL_522690, EPI_ISL_522691, EPI_ISL_522692, EPI_ISL_522693, EPI_ISL_522694, EPI_ISL_522695, EPI_ISL_522696, EPI_ISL_522697, EPI_ISL_522699, EPI_ISL_522700, EPI_ISL_522701, EPI_ISL_522702, EPI_ISL_522704, EPI_ISL_522705, EPI_ISL_522714, EPI_ISL_522715, EPI_ISL_522716, EPI_ISL_522728, EPI_ISL_522731, EPI_ISL_522734, EPI_ISL_522737, EPI_ISL_522740, EPI_ISL_522748, EPI_ISL_522767 | | | |
| see above | Royal Hobart Hospital Microbiology Department | MDU-PHL | Cooley L., van Haeften R., Seemann T., Sait M., Schultz, M.B., Sherry N. |
| EPI_ISL_522777, EPI_ISL_522778, EPI_ISL_522779, EPI_ISL_522780, EPI_ISL_522781, EPI_ISL_522782, EPI_ISL_522783, EPI_ISL_522784, EPI_ISL_522785, EPI_ISL_522786, EPI_ISL_522787, EPI_ISL_522788, EPI_ISL_522789, EPI_ISL_522790, EPI_ISL_522791, EPI_ISL_522792, EPI_ISL_522793, EPI_ISL_522794, EPI_ISL_522795, EPI_ISL_522796, EPI_ISL_522797, EPI_ISL_522798, EPI_ISL_522799, EPI_ISL_522800, EPI_ISL_522801, EPI_ISL_522802, EPI_ISL_522803, EPI_ISL_522804, EPI_ISL_522805, EPI_ISL_522806, EPI_ISL_522807, EPI_ISL_522808, EPI_ISL_522809, EPI_ISL_522810, EPI_ISL_522811, EPI_ISL_522812, EPI_ISL_522813, EPI_ISL_522814, EPI_ISL_522815, EPI_ISL_522816, EPI_ISL_522817, EPI_ISL_522818, EPI_ISL_522819, EPI_ISL_522820, EPI_ISL_522821, EPI_ISL_522822, EPI_ISL_522823 | | | |
| see above | Virginia DCLS | Virginia DCLS | Virginia DCLS |
| EPI_ISL_522824, EPI_ISL_522825, EPI_ISL_522826, EPI_ISL_522827, EPI_ISL_522828, EPI_ISL_522829, EPI_ISL_522830, EPI_ISL_522831, EPI_ISL_522832, EPI_ISL_522833, EPI_ISL_522834, EPI_ISL_522836, EPI_ISL_522837, EPI_ISL_522838, EPI_ISL_522839, EPI_ISL_522840, EPI_ISL_522842, EPI_ISL_522843, EPI_ISL_522844, EPI_ISL_522845, EPI_ISL_522846, EPI_ISL_522848, EPI_ISL_522849, EPI_ISL_522850, EPI_ISL_522851, EPI_ISL_522852, EPI_ISL_522853, EPI_ISL_522854 | | | |
| see above | Maryland Department of Health | Maryland Department of Health | Maryland Department of Health Laboratories Administration |
| EPI_ISL_522855 | ULSS9 Distretto di Bussolengo | Istituto Zooprofilattico Sperimentale delle Venezie | Adelaide Milani, Alessia Schivo, Annalisa Salviato, Erika Giorgia Quaranta, Gianpiero Zamperin, Ambra Pastori, Bianca Zecchin, Alice Fusaro, Calogero Terregino, Antonia Ricci |
| EPI_ISL_522856 | ULSS9 Distretto di San Bonifacio | Istituto Zooprofilattico Sperimentale delle Venezie | Adelaide Milani, Alessia Schivo, Annalisa Salviato, Erika Giorgia Quaranta, Gianpiero Zamperin, Ambra Pastori, Bianca Zecchin, Alice Fusaro, Calogero Terregino, Antonia Ricci |
| EPI_ISL_522857 | ULSS9 Scaligera | Istituto Zooprofilattico Sperimentale delle Venezie | Adelaide Milani, Alessia Schivo, Annalisa Salviato, Erika Giorgia Quaranta, Gianpiero Zamperin, Ambra Pastori, Bianca Zecchin, Alice Fusaro, Calogero Terregino, Antonia Ricci |
| EPI_ISL_522858 | ULSS9 Distretto di San Bonifacio | Istituto Zooprofilattico Sperimentale delle Venezie | Adelaide Milani, Alessia Schivo, Annalisa Salviato, Erika Giorgia Quaranta, Gianpiero Zamperin, Ambra Pastori, Bianca Zecchin, Alice Fusaro, Calogero Terregino, Antonia Ricci |
| EPI_ISL_522859 | ULSS9 Scaligera | Istituto Zooprofilattico Sperimentale delle Venezie | Adelaide Milani, Alessia Schivo, Annalisa Salviato, Erika Giorgia Quaranta, Gianpiero Zamperin, Ambra Pastori, Bianca Zecchin, Alice Fusaro, Calogero Terregino, Antonia Ricci |
| EPI_ISL_522860, EPI_ISL_522861, EPI_ISL_522862, EPI_ISL_522863, EPI_ISL_522864, EPI_ISL_522865, EPI_ISL_522866, EPI_ISL_522867, EPI_ISL_522868 | ULSS9 Distretto di Bussolengo | Istituto Zooprofilattico Sperimentale delle Venezie | Adelaide Milani, Alessia Schivo, Annalisa Salviato, Erika Giorgia Quaranta, Gianpiero Zamperin, Ambra Pastori, Bianca Zecchin, Alice Fusaro, Calogero Terregino, Antonia Ricci |
| EPI_ISL_522869, EPI_ISL_522870, EPI_ISL_522871 | Maryland Department of Health | Maryland Department of Health | Maryland Department of Health Laboratories Administration |
| EPI_ISL_522874 | Instituto Nacional de Medicina Genomica | Instituto Nacional de Medicina Genomica | Hidalgo-Miranda A, Mendoza-Vargas A, Reyes-Grajeda JP, Cisneros-Villanueva M, Cedro-Tanda A, Hurtado-Cordova E, Peñaloza-Figueroa F, Herrera-Montalvo LA |
| EPI_ISL_522877 | Instituto Nacional de Medicina Genomica | Instituto Nacional de Medicina Genomica | Hidalgo-Miranda A, Mendoza-Vargas A, Reyes-Grajeda JP, Cisneros-Villanueva M, Cedro-Tanda A, Hurtado-Cordova E, Peñaloza-Figueroa F, Herrera-Montalvo LA |
| EPI_ISL_522878 | Instituto Nacional de Medicina Genomica | Instituto Nacional de Medicina Genomica | Hidalgo-Miranda A, Mendoza-Vargas A, Reyes-Grajeda JP, Cisneros-Villanueva M, Hurtado-Cordova E, Cedro-Tanda A, Peñaloza-Figueroa F, Herrera-Montalvo LA |
| EPI_ISL_522880 | Instituto Nacional de Medicina Genómica | Instituto Nacional de Medicina Genómica | Hidalgo-Miranda A, Mendoza-Vargas A, Reyes-Grajeda JP, Cisneros-Villanueva M, Cedro-Tanda A, Hurtado-Cordova E, Peñaloza-Figueroa F, Herrera-Montalvo LA |
| EPI_ISL_522881, EPI_ISL_522882, EPI_ISL_522883, EPI_ISL_522884, EPI_ISL_522885, EPI_ISL_522886, EPI_ISL_522887, EPI_ISL_522888, EPI_ISL_522889, EPI_ISL_522890, EPI_ISL_522891, EPI_ISL_522892, EPI_ISL_522893, EPI_ISL_522895, EPI_ISL_522896, EPI_ISL_522898, EPI_ISL_522899, EPI_ISL_522900, EPI_ISL_522901, EPI_ISL_522902, EPI_ISL_522903, EPI_ISL_522904, EPI_ISL_522905, EPI_ISL_522906, EPI_ISL_522907, EPI_ISL_522908, EPI_ISL_522909, EPI_ISL_522910, EPI_ISL_522911, EPI_ISL_522912, EPI_ISL_522913, EPI_ISL_522914, EPI_ISL_522915, EPI_ISL_522917, EPI_ISL_522918, EPI_ISL_522924, EPI_ISL_522925, EPI_ISL_522926, EPI_ISL_522927, EPI_ISL_522928, EPI_ISL_522929, EPI_ISL_522931, EPI_ISL_522932, EPI_ISL_522933, EPI_ISL_522935, EPI_ISL_522936, EPI_ISL_522937, EPI_ISL_522938, EPI_ISL_522939 | | | |
| see above | University of Wisconsin-Madison AIDS Vaccine Research Laboratories | University of Wisconsin-Madison AIDS Vaccine Research Laboratories | Gage Moreno, Katarina Braun, et al. AIDS Vaccine Research Laboratories |
| EPI_ISL_522940 | Instituto Nacional de Medicina Genómica | Instituto Nacional de Medicina Genómica | Hidalgo-Miranda A, Mendoza-Vargas A, Reyes-Grajeda JP, Cisneros-Villanueva M, Cedro-Tanda A, Hurtado-Cordova E, Peñaloza-Figueroa F, Herrera-Montalvo LA |
| EPI_ISL_522941 | Instituto Nacional de Medicina Genómica | Instituto Nacional de Medicina Genómica | Hidalgo-Miranda A, Mendoza-Vargas A, Reyes-Grajeda JP, Cisneros-Villanueva M, Cedro-Tanda A, Hurtado-Cordova E, Peñaloza-Figueroa F, Herrera-Montalvo LA |
| EPI_ISL_522942 | Instituto Nacional de Medicina Genómica | Instituto Nacional de Medicina Genómica | Hidalgo-Miranda A, Mendoza-Vargas A, Reyes-Grajeda JP, Cisneros-Villanueva M, Cedro-Tanda A, Hurtado-Cordova E, Peñaloza-Figueroa F, Herrera-Montalvo LA |
| EPI_ISL_522949, EPI_ISL_522953, | Texas Department of State Health Services | Texas Department of State Health Services | Rashmi Tuladhar, Bonnie Oh, Cara Akrout, Jenny Zhang, Maliha Rahman, Anita Pokharel, Myong Koag, Chun Wang, Rachel Lee, Grace Kubin |

| | | | |
|---|---|---|---|
| EPI_ISL_522960, EPI_ISL_522966, EPI_ISL_522969, EPI_ISL_522971 | | | |
| EPI_ISL_522978, EPI_ISL_522980, EPI_ISL_522981 | Instituto Nacional de Medicina Genómica | Instituto Nacional de Medicina Genómica | Hidalgo-Miranda A, Mendoza-Vargas A, Reyes-Grajeda JP, Cisneros-Villanueva M, Cedro-Tanda A, Hurtado-Cordova E, Peñaloza-Figueroa F, Herrera-Montalvo LA |
| EPI_ISL_522982 | Instituto Nacional de Medicina Genómica | Instituto Nacional de Medicina Genómica | Hidalgo-Miranda A, Mendoza-Vargas A, Reyes-Grajeda JP, Cisneros-Villanueva M, Cedro-Tanda A, Hurtado-Cordova E, Peñaloza-Figueroa F, Herrera-Montalvo LA |
| EPI_ISL_522984 | Instituto Nacional de Medicina Genómica | Instituto Nacional de Medicina Genómica | Hidalgo-Miranda A, Mendoza-Vargas A, Reyes-Grajeda JP, Cisneros-Villanueva M, Cedro-Tanda A, Hurtado-Cordova E, Peñaloza-Figueroa F, Herrera-Montalvo LA |
| EPI_ISL_522986 | Instituto Nacional de Medicina Genómica | Instituto Nacional de Medicina Genómica | Hidalgo-Miranda A, Mendoza-Vargas A, Reyes-Grajeda JP, Cisneros-Villanueva M, Cedro-Tanda A, Hurtado-Cordova E, Peñaloza-Figueroa F, Herrera-Montalvo LA |
| EPI_ISL_523121, EPI_ISL_523122, EPI_ISL_523123, EPI_ISL_523124, EPI_ISL_523125, EPI_ISL_523126, EPI_ISL_523127, EPI_ISL_523128, EPI_ISL_523129, EPI_ISL_523130, EPI_ISL_523132, EPI_ISL_523133, EPI_ISL_523134, EPI_ISL_523138, EPI_ISL_523139, EPI_ISL_523140, EPI_ISL_523141, EPI_ISL_523142, EPI_ISL_523143, EPI_ISL_523144, EPI_ISL_523145, EPI_ISL_523146, EPI_ISL_523147, EPI_ISL_523148, EPI_ISL_523149, EPI_ISL_523150, EPI_ISL_523151, EPI_ISL_523152, EPI_ISL_523153, EPI_ISL_523154, EPI_ISL_523156, EPI_ISL_523157, EPI_ISL_523158, EPI_ISL_523159, EPI_ISL_523160, EPI_ISL_523161, EPI_ISL_523167, EPI_ISL_523168, EPI_ISL_523170, EPI_ISL_523171, EPI_ISL_523172, EPI_ISL_523173, EPI_ISL_523174, EPI_ISL_523175, EPI_ISL_523176, EPI_ISL_523177, EPI_ISL_523178, EPI_ISL_523179, EPI_ISL_523180, EPI_ISL_523181, EPI_ISL_523182, EPI_ISL_523183, EPI_ISL_523184, EPI_ISL_523185, EPI_ISL_523186, EPI_ISL_523187, EPI_ISL_523188, EPI_ISL_523189, EPI_ISL_523191, EPI_ISL_523192, EPI_ISL_523193, EPI_ISL_523195, EPI_ISL_523196, EPI_ISL_523197, EPI_ISL_523199, EPI_ISL_523200, EPI_ISL_523202, EPI_ISL_523203, EPI_ISL_523204, EPI_ISL_523205, EPI_ISL_523207, EPI_ISL_523208, EPI_ISL_523209, EPI_ISL_523210, EPI_ISL_523211, EPI_ISL_523213, EPI_ISL_523214, EPI_ISL_523215, EPI_ISL_523216, EPI_ISL_523217, EPI_ISL_523218, EPI_ISL_523220, EPI_ISL_523221, EPI_ISL_523222, EPI_ISL_523223, EPI_ISL_523225, EPI_ISL_523226, EPI_ISL_523227, EPI_ISL_523228, EPI_ISL_523229, EPI_ISL_523230, EPI_ISL_523231, EPI_ISL_523233, EPI_ISL_523235, EPI_ISL_523236, EPI_ISL_523238, EPI_ISL_523239, EPI_ISL_523240, EPI_ISL_523241, EPI_ISL_523245, EPI_ISL_523246, EPI_ISL_523247, EPI_ISL_523248, EPI_ISL_523249, EPI_ISL_523250, EPI_ISL_523251, EPI_ISL_523252, EPI_ISL_523254, EPI_ISL_523257, EPI_ISL_523258, EPI_ISL_523259, EPI_ISL_523260, EPI_ISL_523261, EPI_ISL_523262, EPI_ISL_523263, EPI_ISL_523264, EPI_ISL_523265, EPI_ISL_523267, EPI_ISL_523268, EPI_ISL_523269, EPI_ISL_523270, EPI_ISL_523272, EPI_ISL_523273, EPI_ISL_523274, EPI_ISL_523275, EPI_ISL_523276, EPI_ISL_523277, EPI_ISL_523278, EPI_ISL_523279, EPI_ISL_523280, EPI_ISL_523281, EPI_ISL_523282, EPI_ISL_523283, EPI_ISL_523284, EPI_ISL_523285, EPI_ISL_523286, EPI_ISL_523288, EPI_ISL_523289, EPI_ISL_523290, EPI_ISL_523291, EPI_ISL_523292, EPI_ISL_523293, EPI_ISL_523294, EPI_ISL_523295, EPI_ISL_523296, EPI_ISL_523297, EPI_ISL_523298, EPI_ISL_523299, EPI_ISL_523301, EPI_ISL_523303, EPI_ISL_523304, EPI_ISL_523305, EPI_ISL_523306, EPI_ISL_523307, EPI_ISL_523308, EPI_ISL_523309, EPI_ISL_523310, EPI_ISL_523311, EPI_ISL_523312, EPI_ISL_523313, EPI_ISL_523314, EPI_ISL_523315, EPI_ISL_523316, EPI_ISL_523317, EPI_ISL_523318, EPI_ISL_523319, EPI_ISL_523320, EPI_ISL_523321, EPI_ISL_523322, EPI_ISL_523323, EPI_ISL_523324, EPI_ISL_523325, EPI_ISL_523326, EPI_ISL_523327, EPI_ISL_523328, EPI_ISL_523329, EPI_ISL_523330, EPI_ISL_523331, EPI_ISL_523332, EPI_ISL_523333, EPI_ISL_523334, EPI_ISL_523335, EPI_ISL_523336, EPI_ISL_523337, EPI_ISL_523340, EPI_ISL_523341, EPI_ISL_523342, EPI_ISL_523343, EPI_ISL_523345, EPI_ISL_523347, EPI_ISL_523348, EPI_ISL_523349, EPI_ISL_523350, EPI_ISL_523351, EPI_ISL_523352, EPI_ISL_523353, EPI_ISL_523354, EPI_ISL_523355, EPI_ISL_523356, EPI_ISL_523357, EPI_ISL_523358, EPI_ISL_523359, EPI_ISL_523361, EPI_ISL_523364, EPI_ISL_523366, EPI_ISL_523367, EPI_ISL_523368, EPI_ISL_523369, EPI_ISL_523370, EPI_ISL_523371, EPI_ISL_523372, EPI_ISL_523373, EPI_ISL_523374, EPI_ISL_523375, EPI_ISL_523376, EPI_ISL_523377, EPI_ISL_523378, EPI_ISL_523380, EPI_ISL_523381, EPI_ISL_523382, EPI_ISL_523383, EPI_ISL_523384, EPI_ISL_523385, EPI_ISL_523386, EPI_ISL_523388, EPI_ISL_523389, EPI_ISL_523394, EPI_ISL_523395, EPI_ISL_523396, EPI_ISL_523397, EPI_ISL_523401, EPI_ISL_523402, EPI_ISL_523403, EPI_ISL_523405, EPI_ISL_523407, EPI_ISL_523408, EPI_ISL_523409, EPI_ISL_523410, EPI_ISL_523412, EPI_ISL_523413, EPI_ISL_523414, EPI_ISL_523415, EPI_ISL_523416, EPI_ISL_523417, EPI_ISL_523420, EPI_ISL_523421, EPI_ISL_523422, EPI_ISL_523423, EPI_ISL_523424, EPI_ISL_523425, EPI_ISL_523426, EPI_ISL_523427, EPI_ISL_523428, EPI_ISL_523429, EPI_ISL_523430, EPI_ISL_523431, EPI_ISL_523432, EPI_ISL_523433, EPI_ISL_523434, EPI_ISL_523435, EPI_ISL_523436, EPI_ISL_523437, EPI_ISL_523439, EPI_ISL_523441, EPI_ISL_523442, EPI_ISL_523443, EPI_ISL_523444, EPI_ISL_523445, EPI_ISL_523446, EPI_ISL_523447, EPI_ISL_523448, EPI_ISL_523449, EPI_ISL_523450, EPI_ISL_523451, EPI_ISL_523452, EPI_ISL_523453, EPI_ISL_523454, EPI_ISL_523455, EPI_ISL_523456, EPI_ISL_523457, EPI_ISL_523458, EPI_ISL_523459, EPI_ISL_523460, EPI_ISL_523461, EPI_ISL_523462, EPI_ISL_523463, EPI_ISL_523464, EPI_ISL_523465, EPI_ISL_523466, EPI_ISL_523467, EPI_ISL_523468, EPI_ISL_523469, EPI_ISL_523470, EPI_ISL_523471, EPI_ISL_523472, EPI_ISL_523473, EPI_ISL_523474, EPI_ISL_523475, EPI_ISL_523476, EPI_ISL_523479, EPI_ISL_523483, EPI_ISL_523484, EPI_ISL_523485, EPI_ISL_523486, EPI_ISL_523487, EPI_ISL_523490, EPI_ISL_523491, EPI_ISL_523493, EPI_ISL_523494, EPI_ISL_523495, EPI_ISL_523497, EPI_ISL_523498, EPI_ISL_523499 | | | |
| see above | Dutch COVID-19 response team | Erasmus Medical Center | Bas Oude Munnink, David Nieuwenhuijse, Reina Sikkema, Claudia Schapendonk, Irina Chestakova, Anne van der Linden, Theo Bestebroer, Stefan van Nieuwkoop, Mark Pronk, Pascal Lexmond, Corien Swaan, Manon Haverkate, Madelief Mollers, Mart Stein, Sandra Kengne Kamga Mobou, Jeroen van Kampen, Jolanda Voermans, Aura Timen, Corine GeurtsvanKessel, Annemiek van der Eijk, Richard Molenkamp, Marion Koopmans, on behalf of the Dutch national COVID-19 response team. |
| EPI_ISL_523500 | Instituto Nacional de Medicina Genómica | Instituto Nacional de Medicina Genómica | Hidalgo-Miranda A, Mendoza-Vargas A, Reyes-Grajeda JP, Cisneros-Villanueva M, Cedro-Tanda A, Hurtado-Cordova E, Peñaloza-Figueroa F, Herrera-Montalvo LA |
| EPI_ISL_523501, EPI_ISL_523502, EPI_ISL_523503, EPI_ISL_523504, EPI_ISL_523505, EPI_ISL_523506, EPI_ISL_523507, EPI_ISL_523508, EPI_ISL_523509, EPI_ISL_523513, EPI_ISL_523514, EPI_ISL_523515, EPI_ISL_523516, EPI_ISL_523517, EPI_ISL_523518, EPI_ISL_523520, EPI_ISL_523521, EPI_ISL_523523, EPI_ISL_523524, EPI_ISL_523525, EPI_ISL_523526, EPI_ISL_523527, EPI_ISL_523528, EPI_ISL_523529, EPI_ISL_523530, EPI_ISL_523531, EPI_ISL_523532, EPI_ISL_523533, EPI_ISL_523534, EPI_ISL_523536, EPI_ISL_523537, EPI_ISL_523539, EPI_ISL_523540, EPI_ISL_523541, EPI_ISL_523542, EPI_ISL_523543, EPI_ISL_523544, EPI_ISL_523545, EPI_ISL_523546, EPI_ISL_523547, EPI_ISL_523549, EPI_ISL_523550, EPI_ISL_523551, EPI_ISL_523552, EPI_ISL_523553, EPI_ISL_523554, EPI_ISL_523555, EPI_ISL_523556, EPI_ISL_523557, EPI_ISL_523558, EPI_ISL_523559, EPI_ISL_523560, EPI_ISL_523563, EPI_ISL_523564, EPI_ISL_523565, EPI_ISL_523566, EPI_ISL_523568, EPI_ISL_523569, EPI_ISL_523570, EPI_ISL_523571, EPI_ISL_523572, EPI_ISL_523573, EPI_ISL_523575, EPI_ISL_523576, EPI_ISL_523577, EPI_ISL_523578, EPI_ISL_523579, EPI_ISL_523580, EPI_ISL_523583, EPI_ISL_523584, EPI_ISL_523585, EPI_ISL_523586, EPI_ISL_523587, EPI_ISL_523588, EPI_ISL_523591, EPI_ISL_523592, EPI_ISL_523593, EPI_ISL_523596, EPI_ISL_523597, EPI_ISL_523598, EPI_ISL_523599, EPI_ISL_523600, EPI_ISL_523601, EPI_ISL_523602, EPI_ISL_523603, EPI_ISL_523604, EPI_ISL_523605, EPI_ISL_523606, EPI_ISL_523607, EPI_ISL_523608, EPI_ISL_523609, EPI_ISL_523610, EPI_ISL_523611, EPI_ISL_523612, EPI_ISL_523613, EPI_ISL_523614, EPI_ISL_523616, EPI_ISL_523618, EPI_ISL_523619, EPI_ISL_523620, EPI_ISL_523621, EPI_ISL_523622, EPI_ISL_523623, EPI_ISL_523625, EPI_ISL_523626, EPI_ISL_523627, EPI_ISL_523628, EPI_ISL_523629, EPI_ISL_523630, EPI_ISL_523631, EPI_ISL_523632, EPI_ISL_523633, EPI_ISL_523634, EPI_ISL_523635, EPI_ISL_523636, EPI_ISL_523637, EPI_ISL_523638, EPI_ISL_523639, EPI_ISL_523640, EPI_ISL_523641, EPI_ISL_523643, EPI_ISL_523644, EPI_ISL_523645, EPI_ISL_523646, EPI_ISL_523647, EPI_ISL_523648, EPI_ISL_523649, EPI_ISL_523650, EPI_ISL_523651, EPI_ISL_523652, EPI_ISL_523654, EPI_ISL_523657, EPI_ISL_523658, EPI_ISL_523659, EPI_ISL_523660, EPI_ISL_523661, EPI_ISL_523662, EPI_ISL_523663, EPI_ISL_523664, EPI_ISL_523665, EPI_ISL_523666, EPI_ISL_523667, EPI_ISL_523668, EPI_ISL_523669, EPI_ISL_523670, EPI_ISL_523671, EPI_ISL_523672, EPI_ISL_523673, EPI_ISL_523674, EPI_ISL_523675, EPI_ISL_523676, EPI_ISL_523677, EPI_ISL_523678, EPI_ISL_523679, EPI_ISL_523680, EPI_ISL_523681, EPI_ISL_523682, EPI_ISL_523683, EPI_ISL_523684, EPI_ISL_523685, EPI_ISL_523686, EPI_ISL_523687, EPI_ISL_523689, EPI_ISL_523690, EPI_ISL_523692, EPI_ISL_523693, EPI_ISL_523694, EPI_ISL_523695, EPI_ISL_523696, EPI_ISL_523697, EPI_ISL_523698, EPI_ISL_523699, EPI_ISL_523700, EPI_ISL_523701, EPI_ISL_523702, EPI_ISL_523703, EPI_ISL_523704, EPI_ISL_523705, EPI_ISL_523706, EPI_ISL_523707, EPI_ISL_523708, EPI_ISL_523709, EPI_ISL_523710, EPI_ISL_523711, EPI_ISL_523712, EPI_ISL_523713, EPI_ISL_523714, EPI_ISL_523715, EPI_ISL_523716, EPI_ISL_523717, EPI_ISL_523718, EPI_ISL_523719, EPI_ISL_523720, EPI_ISL_523721, EPI_ISL_523722, EPI_ISL_523723, EPI_ISL_523724, EPI_ISL_523725, EPI_ISL_523726, EPI_ISL_523727, EPI_ISL_523728, EPI_ISL_523729, EPI_ISL_523730, EPI_ISL_523731, EPI_ISL_523732, EPI_ISL_523733, EPI_ISL_523734, EPI_ISL_523735, EPI_ISL_523736, EPI_ISL_523737, EPI_ISL_523738, EPI_ISL_523739, EPI_ISL_523740, EPI_ISL_523741, EPI_ISL_523742, EPI_ISL_523744, EPI_ISL_523745, EPI_ISL_523746, EPI_ISL_523747, EPI_ISL_523748, EPI_ISL_523749, EPI_ISL_523750, EPI_ISL_523751, EPI_ISL_523752, EPI_ISL_523753, EPI_ISL_523754, EPI_ISL_523755, EPI_ISL_523756, EPI_ISL_523757, EPI_ISL_523758, EPI_ISL_523759, EPI_ISL_523760, EPI_ISL_523761, EPI_ISL_523762, EPI_ISL_523763, EPI_ISL_523764, EPI_ISL_523765, EPI_ISL_523766, EPI_ISL_523767, EPI_ISL_523768, EPI_ISL_523769, EPI_ISL_523770, EPI_ISL_523771, EPI_ISL_523772, EPI_ISL_523773, EPI_ISL_523774, EPI_ISL_523775, EPI_ISL_523776, EPI_ISL_523777, EPI_ISL_523778, EPI_ISL_523779, EPI_ISL_523780, EPI_ISL_523782, EPI_ISL_523783, EPI_ISL_523784, EPI_ISL_523785, EPI_ISL_523786, EPI_ISL_523787, EPI_ISL_523788, EPI_ISL_523789, EPI_ISL_523790, EPI_ISL_523791, EPI_ISL_523792, EPI_ISL_523793, EPI_ISL_523794, EPI_ISL_523795, EPI_ISL_523796, EPI_ISL_523797, EPI_ISL_523798, EPI_ISL_523799, EPI_ISL_523800, EPI_ISL_523802, EPI_ISL_523803, EPI_ISL_523804, EPI_ISL_523805, EPI_ISL_523806, EPI_ISL_523807, EPI_ISL_523808, EPI_ISL_523809 | | | |
| see above | Dutch COVID-19 response team | Erasmus Medical Center | Bas Oude Munnink, David Nieuwenhuijse, Reina Sikkema, Claudia Schapendonk, Irina Chestakova, Anne van der Linden, Theo Bestebroer, Stefan van Nieuwkoop, Mark Pronk, Pascal Lexmond, Corien Swaan, Manon Haverkate, Madelief Mollers, Mart Stein, Sandra Kengne Kamga Mobou, Jeroen van Kampen, Jolanda Voermans, Aura Timen, Corine GeurtsvanKessel, Annemiek van der Eijk, Richard Molenkamp, Marion Koopmans, on behalf of the Dutch national COVID-19 response team. |
| EPI_ISL_523810 | Laboratorio de Referencia Nacional de Virus Respiratorio. Centro Nacional de Salud Publica. Instituto Nacional de Salud Peru. | Laboratorio de Referencia Nacional de Biotecnología y Biología Molecular. Centro Nacional de Salud Publica. Instituto Nacional de Salud Peru. | Carlos Padilla Rojas, Karolyn Vega Chozo, Priscila Lope Pari, Omar Caceres Rey, Marco Galarza Perez, Maribel Huaringa Nuñez, Johanna Balbuena Torres, Henri Bailon Calderon, Nancy Rojas Serrano. |
| EPI_ISL_523811 | Universidad Iberoamericana, Instituto de Medicina Tropical & Salud Global | International Centre for Genetic Engineering and Biotechnology (ICGEB) and ARGO Open Lab Platform | Robert Paulino-Ramirez, Eileen Riego, Alejandro Vallejo Degaudenzi, Victor Virgilio Calderon, Leandro Tapia, Danilo Licastro, Simeone Dal Monego, Sreejith Rajasekharan and Alessandro Marcello. |
| EPI_ISL_523856, EPI_ISL_523857, EPI_ISL_523858, EPI_ISL_523859, EPI_ISL_523860, EPI_ISL_523861, EPI_ISL_523862, EPI_ISL_523863, EPI_ISL_523864, EPI_ISL_523865, EPI_ISL_523866, EPI_ISL_523867, EPI_ISL_523868, EPI_ISL_523869, EPI_ISL_523871, EPI_ISL_523872, EPI_ISL_523873, EPI_ISL_523874, EPI_ISL_523875, EPI_ISL_523876, EPI_ISL_523879, EPI_ISL_523880, EPI_ISL_523881, EPI_ISL_523882, EPI_ISL_523883, EPI_ISL_523884, EPI_ISL_523885, EPI_ISL_523887, EPI_ISL_523888, EPI_ISL_523889, EPI_ISL_523891, EPI_ISL_523892, EPI_ISL_523894, EPI_ISL_523895, EPI_ISL_523896, EPI_ISL_523898, EPI_ISL_523899, EPI_ISL_523900, EPI_ISL_523901, EPI_ISL_523902, EPI_ISL_523903, EPI_ISL_523905, EPI_ISL_523906, EPI_ISL_523907, EPI_ISL_523908, EPI_ISL_523909, EPI_ISL_523910, EPI_ISL_523911, EPI_ISL_523913, EPI_ISL_523915, EPI_ISL_523916, EPI_ISL_523917, EPI_ISL_523918, EPI_ISL_523919, EPI_ISL_523920, EPI_ISL_523921, EPI_ISL_523922, EPI_ISL_523923, EPI_ISL_523924, EPI_ISL_523925, EPI_ISL_523926 | | | |
| see above | Viollier AG | Department of Biosystems Science and Engineering, ETH Zürich | Christian Beisel, Sarah Nadeau, Ivan Topolsky, Pedro Ferreira, Philipp Jablonski, Susana Posada-Céspedes, Tobias Schär, Ina Nissen, Natascha Santacroce, Elodie Burcklen, Christiane Beckmann, Maurice Redondo, Olivier Kobel, Christoph Noppen, Sophie Seidel, Noemie Santamaria de Souza, Niko Beerenwinkel, Tanja Stadler |
| EPI_ISL_523927, EPI_ISL_523929, EPI_ISL_523931, EPI_ISL_523932, EPI_ISL_523933, EPI_ISL_523934, EPI_ISL_523935, EPI_ISL_523937, EPI_ISL_523938, EPI_ISL_523939, EPI_ISL_523940, EPI_ISL_523941, EPI_ISL_523942, EPI_ISL_523943, EPI_ISL_523944, EPI_ISL_523945, EPI_ISL_523947, EPI_ISL_523948, EPI_ISL_523949 | | | |
| see above | Center of Medical Microbiology, Virology, and Hospital Hygiene, University of Duesseldorf | Center of Medical Microbiology, Virology, and Hospital Hygiene, University of Duesseldorf | Maximilian Damagnez, Alexander Dilthey, Torsten Houwaart, Malte Kohns Vasconcelos, Marek Korencak, Jessica Nicolai, Klaus Pfeffer, Hendrik Streeck, Daniel Strelow, Jörg Timm, Andreas Walker, Tobias Wienemann |
| EPI_ISL_523950 | Center of Medical Microbiology, Virology, and Hospital Hygiene, University of Duesseldorf | Center of Medical Microbiology, Virology, and Hospital Hygiene, Heinrich Heine University Düsseldorf | Maximilian Damagnez, Alexander Dilthey, Torsten Houwaart, Malte Kohns Vasconcelos, Marek Korencak, Jessica Nicolai, Klaus Pfeffer, Hendrik Streeck, Daniel Strelow, Jörg Timm, Andreas Walker, Tobias Wienemann |
| EPI_ISL_523951 | Respiratory Virus Unit, Microbiology Services Colindale, Public Health England | Respiratory Virus Unit, Microbiology Services Colindale, Public Health England | PHE Covid Sequencing Team |
| EPI_ISL_523952 | Mohammed Bin Rashid University of Medicine and Health Sciences | Al Jalila Genomics Center | Ahmad Abou Tayoun, Tom Loney, Hamda Khansaheb, Sathishkumar Ramaswamy, Divinlal Harilal, Zulfa Omar Deesi, Rupa Murthy Varghese, Hanan Al Suwaidi, Abdulmajeed Alkhaja, Mohammed Uddin, Rifat Hamoudi, Rabih Halwani, Abiola Catherine Senok, Qutayba Hamid, Norbert Nowotny, Alawi Alsheikh-Ali |
| EPI_ISL_523954 | Laboratorio de Referencia Nacional de Virus Respiratorio. Centro Nacional de Salud Publica. Instituto Nacional de Salud Peru. | Laboratorio de Referencia Nacional de Biotecnología y Biología Molecular. Centro Nacional de Salud Publica. Instituto Nacional de Salud Peru. | Carlos Padilla Rojas, Karolyn Vega Chozo, Priscila Lope Pari, Omar Caceres Rey, Marco Galarza Perez, Maribel Huaringa Nuñez, Johanna Balbuena Torres, Henri Bailon Calderon, Nancy Rojas Serrano. |
| EPI_ISL_523955 | Hospital Municipal do Tatuape Carmino Caricchio | Instituto Adolfo Lutz, Interdiciplinary Procedures Center, Strategic Laboratory | Claudio Tavares Sacchi, Claudia Regina Gonçalves, Erica Valessa Ramos Gomes |
| EPI_ISL_523956 | Hospital Regional de Assis | Instituto Adolfo Lutz, Interdiciplinary Procedures Center, Strategic Laboratory | Claudio Tavares Sacchi, Claudia Regina Gonçalves, Erica Valessa Ramos Gomes |
| EPI_ISL_523957 | Hospital Itamaraty | Instituto Adolfo Lutz, Interdiciplinary Procedures Center, Strategic Laboratory | Claudio Tavares Sacchi, Claudia Regina Gonçalves, Erica Valessa Ramos Gomes |
| EPI_ISL_523958, EPI_ISL_523959 | Pronto Socorro Municipal de Perus | Instituto Adolfo Lutz, Interdiciplinary Procedures Center, Strategic Laboratory | Claudio Tavares Sacchi, Claudia Regina Gonçalves, Erica Valessa Ramos Gomes |
| EPI_ISL_523960 | National Agency for Public Health, Republic of Moldova | Charite Universitatsmedizin Berlin, Institute of Virology | Victor M Corman, Jörn Beheim-Schwarzbach, Barbara Mühlemann, Talitha Veith, Julia Schneider, Ala Halacu, Mariana Apostol, Terry Jones, Christian Drosten |
| EPI_ISL_523961 | Pronto Socorro Municipal 21 de Junho | Instituto Adolfo Lutz, Interdiciplinary Procedures Center, Strategic Laboratory | Claudio Tavares Sacchi, Claudia Regina Gonçalves, Erica Valessa Ramos Gomes |
| EPI_ISL_523962 | National Agency for Public Health, Republic of Moldova | Charite Universitatsmedizin Berlin, Institute of Virology | Victor M Corman, Joern Beheim-Schwarzbach, Barbara Mühlemann, Talitha Veith, Julia Schneider, Elizabeta Jancheska, Maja Kuzmanovska, Golubinka Bosevska, Terry Jones, Christian Drosten |
| EPI_ISL_523963 | UBS Vila Silvia | Instituto Adolfo Lutz, Interdiciplinary Procedures Center, Strategic Laboratory | Claudio Tavares Sacchi, Claudia Regina Gonçalves, Erica Valessa Ramos Gomes |
| EPI_ISL_523964 | National Agency for Public Health, Republic of Moldova | Charite Universitatsmedizin Berlin, Institute of Virology | Victor M Corman, Jörn Beheim-Schwarzbach, Barbara Mühlemann, Talitha Veith, Julia Schneider, Ala Halacu, Mariana Apostol, Terry Jones, Christian Drosten |
| EPI_ISL_523965 | Hospital do Servidor Público Estadual Francisco Morato de Oliveira | Instituto Adolfo Lutz, Interdiciplinary Procedures Center, Strategic Laboratory | Claudio Tavares Sacchi, Claudia Regina Gonçalves, Erica Valessa Ramos Gomes |
| EPI_ISL_523966 | National Agency for Public Health, Republic of Moldova | Charite Universitatsmedizin Berlin, Institute of Virology | Victor M Corman, Jörn Beheim-Schwarzbach, Barbara Mühlemann, Talitha Veith, Julia Schneider, Ala Halacu, Mariana Apostol, Terry Jones, Christian Drosten |
| EPI_ISL_523967 | Hospital Sancta Maggiore | Instituto Adolfo Lutz, Interdiciplinary Procedures Center, Strategic Laboratory | Claudio Tavares Sacchi, Claudia Regina Gonçalves, Erica Valessa Ramos Gomes |
| EPI_ISL_523968 | National Agency for Public Health, Republic of Moldova | Charite Universitatsmedizin Berlin, Institute of Virology | Victor M Corman, Jörn Beheim-Schwarzbach, Barbara Mühlemann, Talitha Veith, Julia Schneider, Ala Halacu, Mariana Apostol, Terry Jones, Christian Drosten |
| EPI_ISL_523969 | Hospital Sao Paulo de Ensino da Unifesp | Instituto Adolfo Lutz, Interdiciplinary Procedures Center, Strategic Laboratory | Claudio Tavares Sacchi, Claudia Regina Gonçalves, Erica Valessa Ramos Gomes |
| EPI_ISL_523970 | Conjunto Hospitalar do Mandaqui | Instituto Adolfo Lutz, Interdiciplinary Procedures Center, Strategic Laboratory | Claudio Tavares Sacchi, Claudia Regina Gonçalves, Erica Valessa Ramos Gomes |
| EPI_ISL_523971 | Hospital Geral Santa Marcelina | Instituto Adolfo Lutz, Interdiciplinary Procedures Center, Strategic Laboratory | Claudio Tavares Sacchi, Claudia Regina Gonçalves, Erica Valessa Ramos Gomes |
| EPI_ISL_523972 | Hospital do Servidor Público Estadual Francisco Morato de Oliveira | Instituto Adolfo Lutz, Interdiciplinary Procedures Center, Strategic Laboratory | Claudio Tavares Sacchi, Claudia Regina Gonçalves, Erica Valessa Ramos Gomes |
| EPI_ISL_523973 | PS Municipal Dona Maria Antonieta Ferreira de Barros | Instituto Adolfo Lutz, Interdiciplinary Procedures Center, Strategic Laboratory | Claudio Tavares Sacchi, Claudia Regina Gonçalves, Erica Valessa Ramos Gomes |
| EPI_ISL_523974 | Hospital Municipal do Tatuape Carmino Caricchio | Instituto Adolfo Lutz, Interdiciplinary Procedures Center, Strategic Laboratory | Claudio Tavares Sacchi, Claudia Regina Gonçalves, Erica Valessa Ramos Gomes |
| EPI_ISL_523975 | UPA Tito Lopes | Instituto Adolfo Lutz, Interdiciplinary Procedures Center, Strategic Laboratory | Claudio Tavares Sacchi, Claudia Regina Gonçalves, Erica Valessa Ramos Gomes |
| EPI_ISL_523976 | Hospital Municipal do Tatuape Carmino Caricchio | Instituto Adolfo Lutz, Interdiciplinary Procedures Center, Strategic Laboratory | Claudio Tavares Sacchi, Claudia Regina Gonçalves, Erica Valessa Ramos Gomes |
| EPI_ISL_523977 | Hosp. Municipal Prof. Dr. Alípio Corrêa Netto | Instituto Adolfo Lutz, Interdiciplinary Procedures Center, Strategic Laboratory | Claudio Tavares Sacchi, Claudia Regina Gonçalves, Erica Valessa Ramos Gomes |
| EPI_ISL_523978 | Hospital do Servidor Público Estadual Francisco Morato de Oliveira | Instituto Adolfo Lutz, Interdiciplinary Procedures Center, Strategic Laboratory | Claudio Tavares Sacchi, Claudia Regina Gonçalves, Erica Valessa Ramos Gomes |
| EPI_ISL_523979 | Laboratorio de Referencia Nacional de Virus Respiratorio. Centro Nacional de Salud Publica. Instituto Nacional de Salud Peru. | Laboratorio de Referencia Nacional de Biotecnología y Biología Molecular. Centro Nacional de Salud Publica. Instituto Nacional de Salud Peru. | Carlos Padilla Rojas, Karolyn Vega Chozo, Priscila Lope Pari, Omar Caceres Rey, Marco Galarza Perez, Maribel Huaringa Nuñez, Johanna Balbuena Torres, Henri Bailon Calderon, Nancy Rojas Serrano. |
| EPI_ISL_523980 | UPA Tito Lopes | Instituto Adolfo Lutz, Interdiciplinary Procedures Center, Strategic Laboratory | Claudio Tavares Sacchi, Claudia Regina Gonçalves, Erica Valessa Ramos Gomes |
| EPI_ISL_523981 | Hospital Sao Paulo de Ensino da Unifesp | Instituto Adolfo Lutz, Interdiciplinary Procedures Center, Strategic Laboratory | Claudio Tavares Sacchi, Claudia Regina Gonçalves, Erica Valessa Ramos Gomes |
| EPI_ISL_523982 | Hospital do Servidor Público Estadual Francisco Morato de Oliveira | Instituto Adolfo Lutz, Interdiciplinary Procedures Center, Strategic Laboratory | Claudio Tavares Sacchi, Claudia Regina Gonçalves, Erica Valessa Ramos Gomes |

| | | | |
|---|---|---|---|
| EPI_ISL_523983 | UPA Campo Limpo | Instituto Adolfo Lutz, Interdiciplinary Procedures Center, Strategic Laboratory | Claudio Tavares Sacchi, Claudia Regina Gonçalves, Erica Valessa Ramos Gomes |
| EPI_ISL_523984 | Ama Dr Jose Soares Hungria | Instituto Adolfo Lutz, Interdiciplinary Procedures Center, Strategic Laboratory | Claudio Tavares Sacchi, Claudia Regina Gonçalves, Erica Valessa Ramos Gomes |
| EPI_ISL_523985 | Hospital Municipal Dr. Benedicto Montenegro | Instituto Adolfo Lutz, Interdiciplinary Procedures Center, Strategic Laboratory | Claudio Tavares Sacchi, Claudia Regina Gonçalves, Erica Valessa Ramos Gomes |
| EPI_ISL_523986 | Ama Dr Jose Soares Hungria | Instituto Adolfo Lutz, Interdiciplinary Procedures Center, Strategic Laboratory | Claudio Tavares Sacchi, Claudia Regina Gonçalves, Erica Valessa Ramos Gomes |
| EPI_ISL_523988 | Hospital Sao Paulo de Ensino da Unifesp | Instituto Adolfo Lutz, Interdiciplinary Procedures Center, Strategic Laboratory | Claudio Tavares Sacchi, Claudia Regina Gonçalves, Erica Valessa Ramos Gomes |
| EPI_ISL_523989 | AMA Jardim Joamar | Instituto Adolfo Lutz, Interdiciplinary Procedures Center, Strategic Laboratory | Claudio Tavares Sacchi, Claudia Regina Gonçalves, Erica Valessa Ramos Gomes |
| EPI_ISL_523990 | AMA Jardim Peri | Instituto Adolfo Lutz, Interdiciplinary Procedures Center, Strategic Laboratory | Claudio Tavares Sacchi, Claudia Regina Gonçalves, Erica Valessa Ramos Gomes |
| EPI_ISL_523991, EPI_ISL_523992 | Hospital Municipal Carmen Prudente | Instituto Adolfo Lutz, Interdiciplinary Procedures Center, Strategic Laboratory | Claudio Tavares Sacchi, Claudia Regina Gonçalves, Erica Valessa Ramos Gomes |
| EPI_ISL_523993 | UPA Campo Limpo | Instituto Adolfo Lutz, Interdiciplinary Procedures Center, Strategic Laboratory | Claudio Tavares Sacchi, Claudia Regina Gonçalves, Erica Valessa Ramos Gomes |
| EPI_ISL_523994 | Laboratorio de Referencia Nacional de Virus Respiratorio. Centro Nacional de Salud Publica. Instituto Nacional de Salud Peru. | Laboratorio de Referencia Nacional de Biotecnología y Biología Molecular. Centro Nacional de Salud Publica. Instituto Nacional de Salud Peru. | Carlos Padilla Rojas, Karolyn Vega Chozo, Priscila Lope Pari, Omar Caceres Rey, Marco Galarza Perez, Maribel Huaringa Nuñez, Johanna Balbuena Torres, Henri Bailon Calderon, Nancy Rojas Serrano. |
| EPI_ISL_523995, EPI_ISL_523996, EPI_ISL_523997, EPI_ISL_523998, EPI_ISL_523999, EPI_ISL_524000, EPI_ISL_524001, EPI_ISL_524002, EPI_ISL_524003, EPI_ISL_524004, EPI_ISL_524005, EPI_ISL_524006, EPI_ISL_524007, EPI_ISL_524008, EPI_ISL_524009, EPI_ISL_524010, EPI_ISL_524011, EPI_ISL_524012, EPI_ISL_524013, EPI_ISL_524014, EPI_ISL_524015, EPI_ISL_524016, EPI_ISL_524017, EPI_ISL_524018, EPI_ISL_524019, EPI_ISL_524020, EPI_ISL_524021, EPI_ISL_524022, EPI_ISL_524023, EPI_ISL_524024, EPI_ISL_524025, EPI_ISL_524026, EPI_ISL_524027, EPI_ISL_524028, EPI_ISL_524029, EPI_ISL_524030, EPI_ISL_524031, EPI_ISL_524032, EPI_ISL_524033, EPI_ISL_524034, EPI_ISL_524035, EPI_ISL_524036, EPI_ISL_524037, EPI_ISL_524038, EPI_ISL_524039, EPI_ISL_524040, EPI_ISL_524041, EPI_ISL_524042, EPI_ISL_524043, EPI_ISL_524044, EPI_ISL_524045, EPI_ISL_524046, EPI_ISL_524047, EPI_ISL_524048, EPI_ISL_524049, EPI_ISL_524050, EPI_ISL_524051, EPI_ISL_524052 | | | |
| see above | WHO National Influenza Centre Russian Federation | WHO National Influenza Centre Russian Federation | Andrey Komissarov, Artem Fadeev, Mariia Sergeeva, Anna Ivanova, Daria Danilenko |
| EPI_ISL_524074, EPI_ISL_524075, EPI_ISL_524076, EPI_ISL_524077, EPI_ISL_524078, EPI_ISL_524080, EPI_ISL_524081, EPI_ISL_524082, EPI_ISL_524084, EPI_ISL_524086, EPI_ISL_524087, EPI_ISL_524089, EPI_ISL_524090, EPI_ISL_524093, EPI_ISL_524096, EPI_ISL_524098, EPI_ISL_524100, EPI_ISL_524101, EPI_ISL_524102, EPI_ISL_524104, EPI_ISL_524105, EPI_ISL_524107, EPI_ISL_524108, EPI_ISL_524109, EPI_ISL_524111, EPI_ISL_524113, EPI_ISL_524114, EPI_ISL_524115, EPI_ISL_524117, EPI_ISL_524118, EPI_ISL_524121, EPI_ISL_524122, EPI_ISL_524123, EPI_ISL_524124, EPI_ISL_524125, EPI_ISL_524126, EPI_ISL_524127, EPI_ISL_524128, EPI_ISL_524129, EPI_ISL_524130, EPI_ISL_524133, EPI_ISL_524134, EPI_ISL_524135, EPI_ISL_524136, EPI_ISL_524138, EPI_ISL_524139, EPI_ISL_524141, EPI_ISL_524142, EPI_ISL_524143, EPI_ISL_524145, EPI_ISL_524147, EPI_ISL_524148, EPI_ISL_524149, EPI_ISL_524151, EPI_ISL_524152, EPI_ISL_524154, EPI_ISL_524155, EPI_ISL_524157, EPI_ISL_524161, EPI_ISL_524162, EPI_ISL_524163, EPI_ISL_524164, EPI_ISL_524165, EPI_ISL_524166, EPI_ISL_524167, EPI_ISL_524170, EPI_ISL_524171, EPI_ISL_524172, EPI_ISL_524173, EPI_ISL_524174, EPI_ISL_524175, EPI_ISL_524176, EPI_ISL_524178, EPI_ISL_524179, EPI_ISL_524180, EPI_ISL_524181, EPI_ISL_524182, EPI_ISL_524183, EPI_ISL_524184, EPI_ISL_524186, EPI_ISL_524187, EPI_ISL_524188, EPI_ISL_524190, EPI_ISL_524191, EPI_ISL_524192, EPI_ISL_524193, EPI_ISL_524194, EPI_ISL_524196, EPI_ISL_524200, EPI_ISL_524201, EPI_ISL_524202, EPI_ISL_524203, EPI_ISL_524204, EPI_ISL_524206, EPI_ISL_524209, EPI_ISL_524210, EPI_ISL_524214, EPI_ISL_524215, EPI_ISL_524216, EPI_ISL_524217, EPI_ISL_524220, EPI_ISL_524224, EPI_ISL_524225, EPI_ISL_524226, EPI_ISL_524227, EPI_ISL_524228, EPI_ISL_524231, EPI_ISL_524232, EPI_ISL_524236, EPI_ISL_524237, EPI_ISL_524239, EPI_ISL_524240, EPI_ISL_524242, EPI_ISL_524245, EPI_ISL_524247, EPI_ISL_524249, EPI_ISL_524250, EPI_ISL_524251, EPI_ISL_524253, EPI_ISL_524255, EPI_ISL_524256, EPI_ISL_524257, EPI_ISL_524260, EPI_ISL_524261, EPI_ISL_524262, EPI_ISL_524263, EPI_ISL_524265, EPI_ISL_524266, EPI_ISL_524269, EPI_ISL_524271, EPI_ISL_524272, EPI_ISL_524274, EPI_ISL_524275, EPI_ISL_524277, EPI_ISL_524279, EPI_ISL_524280, EPI_ISL_524281, EPI_ISL_524283, EPI_ISL_524284, EPI_ISL_524285, EPI_ISL_524288, EPI_ISL_524289, EPI_ISL_524290, EPI_ISL_524291, EPI_ISL_524292, EPI_ISL_524293, EPI_ISL_524295, EPI_ISL_524296, EPI_ISL_524297, EPI_ISL_524299, EPI_ISL_524303, EPI_ISL_524304, EPI_ISL_524305, EPI_ISL_524307, EPI_ISL_524309, EPI_ISL_524311, EPI_ISL_524313, EPI_ISL_524314, EPI_ISL_524316, EPI_ISL_524320, EPI_ISL_524321, EPI_ISL_524322, EPI_ISL_524323, EPI_ISL_524324, EPI_ISL_524325, EPI_ISL_524327, EPI_ISL_524333, EPI_ISL_524335, EPI_ISL_524337, EPI_ISL_524341, EPI_ISL_524342, EPI_ISL_524343, EPI_ISL_524344, EPI_ISL_524345, EPI_ISL_524346, EPI_ISL_524348, EPI_ISL_524350, EPI_ISL_524351, EPI_ISL_524352, EPI_ISL_524353, EPI_ISL_524354, EPI_ISL_524355, EPI_ISL_524357, EPI_ISL_524358, EPI_ISL_524359, EPI_ISL_524360, EPI_ISL_524362, EPI_ISL_524365, EPI_ISL_524367, EPI_ISL_524368, EPI_ISL_524370, EPI_ISL_524371, EPI_ISL_524372, EPI_ISL_524373, EPI_ISL_524374, EPI_ISL_524375, EPI_ISL_524376, EPI_ISL_524377, EPI_ISL_524378, EPI_ISL_524381, EPI_ISL_524382, EPI_ISL_524384, EPI_ISL_524385, EPI_ISL_524387, EPI_ISL_524388, EPI_ISL_524389, EPI_ISL_524392, EPI_ISL_524393, EPI_ISL_524394, EPI_ISL_524396, EPI_ISL_524398, EPI_ISL_524400, EPI_ISL_524401, EPI_ISL_524402, EPI_ISL_524403, EPI_ISL_524404, EPI_ISL_524405, EPI_ISL_524408, EPI_ISL_524412, EPI_ISL_524413, EPI_ISL_524414, EPI_ISL_524415, EPI_ISL_524416, EPI_ISL_524417, EPI_ISL_524418, EPI_ISL_524419, EPI_ISL_524420, EPI_ISL_524421, EPI_ISL_524424, EPI_ISL_524425 | | | |
| see above | Utah Public Health Laboratory | Utah Public Health Laboratory | Erin L. Young, Kelly Oakeson, Tara Gallagher, Michael T. Pyne, E. Susan Slechta, Melanie A. Mallory, Jeffrey B. Stevenson, Salika M. Shakir, David R. Hillyard |
| EPI_ISL_524430 | Dirk Dittmer | Dirk Dittmer | Bailey,A.G., Caro-Vegas,C.P., Dittmer,D., Eason,A.B., Juarez,A., Landis,J.T., McNamara,R.P., Miller,M.B., Moorad,R., Pluta,L.J.,Seltzer,T.A., Thompson,C., Vahrson,W. and Villamor,F. |
| EPI_ISL_524445, EPI_ISL_524447, EPI_ISL_524448, EPI_ISL_524449 | Singapore General Hospital | Department of Microbiology | Nurdyana Abdul Rahman, Kun Lee Lim, Chenhao Li, Kian Sing Chan, Lynette Oon, Kern Rei Chng, Niranjan Nagarajan, Karrie Ko |
| EPI_ISL_524462 | Hospital Metropolitano | Instituto Adolfo Lutz, Interdiciplinary Procedures Center, Strategic Laboratory | Claudio Tavares Sacchi, Claudia Regina Gonçalves, Erica Valessa Ramos Gomes |
| EPI_ISL_524463 | Hospital Regional de Cotia | Instituto Adolfo Lutz, Interdiciplinary Procedures Center, Strategic Laboratory | Claudio Tavares Sacchi, Claudia Regina Gonçalves, Erica Valessa Ramos Gomes |
| EPI_ISL_524464 | Santa Casa de Santa Isabel | Instituto Adolfo Lutz, Interdiciplinary Procedures Center, Strategic Laboratory | Claudio Tavares Sacchi, Claudia Regina Gonçalves, Erica Valessa Ramos Gomes |
| EPI_ISL_524465 | PS Municipal Dr. Caetano Virgílio Neto | Instituto Adolfo Lutz, Interdiciplinary Procedures Center, Strategic Laboratory | Claudio Tavares Sacchi, Claudia Regina Gonçalves, Erica Valessa Ramos Gomes |
| EPI_ISL_524466 | PS Municipal Dr Lauro Ribas Braga | Instituto Adolfo Lutz, Interdiciplinary Procedures Center, Strategic Laboratory | Claudio Tavares Sacchi, Claudia Regina Gonçalves, Erica Valessa Ramos Gomes |
| EPI_ISL_524467 | Hospital Municipal Dr. Moysés Deutsch | Instituto Adolfo Lutz, Interdiciplinary Procedures Center, Strategic Laboratory | Claudio Tavares Sacchi, Claudia Regina Gonçalves, Erica Valessa Ramos Gomes |
| EPI_ISL_524468 | Hospital Municipal Vereador Jose Storopolli | Instituto Adolfo Lutz, Interdiciplinary Procedures Center, Strategic Laboratory | Claudio Tavares Sacchi, Claudia Regina Gonçalves, Erica Valessa Ramos Gomes |
| EPI_ISL_524469 | Santa Casa de Misericordia de Sao Paulo | Instituto Adolfo Lutz, Interdiciplinary Procedures Center, Strategic Laboratory | Claudio Tavares Sacchi, Claudia Regina Gonçalves, Erica Valessa Ramos Gomes |
| EPI_ISL_524470 | Hospital do Servidor Público Estadual Francisco Morato de Oliveira | Instituto Adolfo Lutz, Interdiciplinary Procedures Center, Strategic Laboratory | Claudio Tavares Sacchi, Claudia Regina Gonçalves, Erica Valessa Ramos Gomes |
| EPI_ISL_524471, EPI_ISL_524473 | Laboratorio de Referencia Nacional de Virus Respiratorio. Centro Nacional de Salud Publica. Instituto Nacional de Salud Peru. | Laboratorio de Referencia Nacional de Biotecnología y Biología Molecular. Centro Nacional de Salud Publica. Instituto Nacional de Salud Peru. | Carlos Padilla Rojas, Karolyn Vega Chozo, Priscila Lope Pari, Omar Caceres Rey, Marco Galarza Perez, Maribel Huaringa Nuñez, Johanna Balbuena Torres, Henri Bailon Calderon, Nancy Rojas Serrano. |
| EPI_ISL_524474 | Department of Infectious Diseases, Cantonal Hospital Baden | Institute of Medical Virology, University of Zurich | Stefan Schmutz, Maryam Zaheri, Verena Kufner, Gabriela Ziltener, Patrick Redli, Fiona Steiner, Jon Huder, Riccarda Capaul, Andrea Zbinden, Jürg Böni, Michael Huber, Alexandra Trkola |
| EPI_ISL_524475 | Division of Infectious Diseases, University Hospital Zürich | Institute of Medical Virology, University of Zurich | Stefan Schmutz, Maryam Zaheri, Verena Kufner, Gabriela Ziltener, Patrick Redli, Fiona Steiner, Jon Huder, Riccarda Capaul, Andrea Zbinden, Jürg Böni, Michael Huber, Alexandra Trkola |
| EPI_ISL_524476 | Bülach Hospital | Institute of Medical Virology, University of Zurich | Stefan Schmutz, Maryam Zaheri, Verena Kufner, Gabriela Ziltener, Patrick Redli, Fiona Steiner, Jon Huder, Riccarda Capaul, Andrea Zbinden, Jürg Böni, Michael Huber, Alexandra Trkola |
| EPI_ISL_524477 | Division of Infectious Diseases, University Hospital Zürich | Institute of Medical Virology, University of Zurich | Stefan Schmutz, Maryam Zaheri, Verena Kufner, Gabriela Ziltener, Patrick Redli, Fiona Steiner, Jon Huder, Riccarda Capaul, Andrea Zbinden, Jürg Böni, Michael Huber, Alexandra Trkola |
| EPI_ISL_524478 | Cantonal Hospital Winterthur | Institute of Medical Virology, University of Zurich | Stefan Schmutz, Maryam Zaheri, Verena Kufner, Gabriela Ziltener, Patrick Redli, Fiona Steiner, Jon Huder, Riccarda Capaul, Andrea Zbinden, Jürg Böni, Michael Huber, Alexandra Trkola |
| EPI_ISL_524479 | Klinik Hirslanden Zürich | Institute of Medical Virology, University of Zurich | Stefan Schmutz, Maryam Zaheri, Verena Kufner, Gabriela Ziltener, Patrick Redli, Fiona Steiner, Jon Huder, Riccarda Capaul, Andrea Zbinden, Jürg Böni, Michael Huber, Alexandra Trkola |
| EPI_ISL_524480, EPI_ISL_524481 | Department of Infectious Diseases, Cantonal Hospital Baden | Institute of Medical Virology, University of Zurich | Stefan Schmutz, Maryam Zaheri, Verena Kufner, Gabriela Ziltener, Patrick Redli, Fiona Steiner, Jon Huder, Riccarda Capaul, Andrea Zbinden, Jürg Böni, Michael Huber, Alexandra Trkola |
| EPI_ISL_524482 | University Hospital Zürich | Institute of Medical Virology, University of Zurich | Verena Kufner, Maryam Zaheri, Gabriela Ziltener, Stefan Schmutz, Patrick Redli, Fiona Steiner, Jon Huder, Riccarda Capaul, Andrea Zbinden, Jürg Böni, Michael Huber, Alexandra Trkola |
| EPI_ISL_524483 | Division of Infectious Diseases, University Hospital Zürich | Institute of Medical Virology, University of Zurich | Verena Kufner, Maryam Zaheri, Gabriela Ziltener, Stefan Schmutz, Patrick Redli, Fiona Steiner, Jon Huder, Riccarda Capaul, Andrea Zbinden, Jürg Böni, Michael Huber, Alexandra Trkola |
| EPI_ISL_524484 | Hospital Schwyz | Institute of Medical Virology, University of Zurich | Verena Kufner, Maryam Zaheri, Gabriela Ziltener, Stefan Schmutz, Patrick Redli, Fiona Steiner, Jon Huder, Riccarda Capaul, Andrea Zbinden, Jürg Böni, Michael Huber, Alexandra Trkola |
| EPI_ISL_524485 | Division of Infectious Diseases, University Hospital Zürich | Institute of Medical Virology, University of Zurich | Verena Kufner, Maryam Zaheri, Gabriela Ziltener, Stefan Schmutz, Patrick Redli, Fiona Steiner, Jon Huder, Riccarda Capaul, Andrea Zbinden, Jürg Böni, Michael Huber, Alexandra Trkola |
| EPI_ISL_524486, EPI_ISL_524487, EPI_ISL_524488, EPI_ISL_524489, EPI_ISL_524490, EPI_ISL_524491, EPI_ISL_524492 | PHE South West Regional Laboratory, National Infection Service | Wellcome Sanger Institute for the COVID-19 Genomics UK Consortium | Stephanie Hutchings, Hannah Pymont, Dr Peter Muir, Barry Vipond, Rich Hopes; and Alex Alderton, Roberto Amato, Sonia Goncalves, Ewan Harrison, David K. Jackson, Ian Johnston, Dominic Kwiatkowski, Cordelia Langford, John Sillitoe on behalf of the Wellcome Sanger Institute COVID-19 Surveillance Team (http://www.sanger.ac.uk/covid-team) |
| EPI_ISL_524494 | NHSGGC West of Scotland Specialist Virology Centre / MRC-University of Glasgow Centre for Virus Research | Wellcome Sanger Institute for the COVID-19 Genomics UK Consortium | Ana da Silva Filipe, Natasha Johnson, Kathy Smollett, Daniel Mair, Stephen Carmichael, Lily Tong, Jenna Nichols, Elihu Aranday-Cortes, Kirstyn Brunker, Yasmin Parr, Kyriaki Nomikou; Sarah McDonald, Marc Niebel, Patawee Asamaphan; Richard Orton, Joseph Hughes, Sreenu Vattipally, David L Robertson; Alasdair MacLean, Rory Gunson; Kathy Li, Natasha Jesudason, Rajiv Shah, James Shepherd, Antonia Ho, Alice Broos, Emma Thomson and Alex Alderton, Roberto Amato, Sonia Goncalves, Ewan Harrison, David K. Jackson, Ian Johnston, Dominic Kwiatkowski, Cordelia Langford, John Sillitoe on behalf of the Wellcome Sanger Institute COVID-19 Surveillance Team (http://www.sanger.ac.uk/covid-team) |
| EPI_ISL_524495, EPI_ISL_524497, EPI_ISL_524498 | Department of Pathology, University of Cambridge | Wellcome Sanger Institute for the COVID-19 Genomics UK Consortium | Luke W Meredith, M. Estée Török , Myra Hosmillo, William L. Hamilton, Martin D. Curran, Theresa Feltwell, Grant Hall, Anna Yakovleva, Fahad A Khokhar, Charlotte J. Houldcroft, Laura G Caller, Aminu S. Jahun, Sarah L. Caddy, Ian Goodfellow; and Alex Alderton, Roberto Amato, Sonia Goncalves, Ewan Harrison, David K. Jackson, Ian Johnston, Dominic Kwiatkowski, Cordelia Langford, John Sillitoe on behalf of the Wellcome Sanger Institute COVID-19 Surveillance Team (http://www.sanger.ac.uk/covid-team) |
| EPI_ISL_524500 | NHSGGC West of Scotland Specialist Virology Centre / MRC-University of Glasgow Centre for Virus Research | Wellcome Sanger Institute for the COVID-19 Genomics UK Consortium | Ana da Silva Filipe, Natasha Johnson, Kathy Smollett, Daniel Mair, Stephen Carmichael, Lily Tong, Jenna Nichols, Elihu Aranday-Cortes, Kirstyn Brunker, Yasmin Parr, Kyriaki Nomikou; Sarah McDonald, Marc Niebel, Patawee Asamaphan; Richard Orton, Joseph Hughes, Sreenu Vattipally, David L Robertson; Alasdair MacLean, Rory Gunson; Kathy Li, Natasha Jesudason, Rajiv Shah, James Shepherd, Antonia Ho, Alice Broos, Emma Thomson and Alex Alderton, Roberto Amato, Sonia Goncalves, Ewan Harrison, David K. Jackson, Ian Johnston, Dominic Kwiatkowski, Cordelia Langford, John Sillitoe on behalf of the Wellcome Sanger Institute COVID-19 Surveillance Team (http://www.sanger.ac.uk/covid-team) |
| EPI_ISL_524501, EPI_ISL_524509 | Department of Pathology, University of Cambridge | Wellcome Sanger Institute for the COVID-19 Genomics UK Consortium | Luke W Meredith, M. Estée Török , Myra Hosmillo, William L. Hamilton, Martin D. Curran, Theresa Feltwell, Grant Hall, Anna Yakovleva, Fahad A Khokhar, Charlotte J. Houldcroft, Laura G Caller, Aminu S. Jahun, Sarah L. Caddy, Ian Goodfellow; and Alex Alderton, Roberto Amato, Sonia Goncalves, Ewan Harrison, David K. Jackson, Ian Johnston, Dominic Kwiatkowski, Cordelia Langford, John Sillitoe on behalf of the Wellcome Sanger Institute COVID-19 Surveillance Team (http://www.sanger.ac.uk/covid-team) |
| EPI_ISL_524513, EPI_ISL_524514, EPI_ISL_524516, EPI_ISL_524518, EPI_ISL_524519, EPI_ISL_524520, EPI_ISL_524526, EPI_ISL_524527 | NHSGGC West of Scotland Specialist Virology Centre / MRC-University of Glasgow Centre for Virus Research | Wellcome Sanger Institute for the COVID-19 Genomics UK Consortium | Ana da Silva Filipe, Natasha Johnson, Kathy Smollett, Daniel Mair, Stephen Carmichael, Lily Tong, Jenna Nichols, Elihu Aranday-Cortes, Kirstyn Brunker, Yasmin Parr, Kyriaki Nomikou; Sarah McDonald, Marc Niebel, Patawee Asamaphan; Richard Orton, Joseph Hughes, Sreenu Vattipally, David L Robertson; Alasdair MacLean, Rory Gunson; Kathy Li, Natasha Jesudason, Rajiv Shah, James Shepherd, Antonia Ho, Alice Broos, Emma Thomson and Alex Alderton, Roberto Amato, Sonia Goncalves, Ewan Harrison, David K. Jackson, Ian Johnston, Dominic Kwiatkowski, Cordelia Langford, John Sillitoe on behalf of the Wellcome Sanger Institute COVID-19 Surveillance Team (http://www.sanger.ac.uk/covid-team) |
| EPI_ISL_524530, EPI_ISL_524531 | Department of Pathology, University of Cambridge | Wellcome Sanger Institute for the COVID-19 Genomics UK Consortium | Luke W Meredith, M. Estée Török , Myra Hosmillo, William L. Hamilton, Martin D. Curran, Theresa Feltwell, Grant Hall, Anna Yakovleva, Fahad A Khokhar, Charlotte J. Houldcroft, Laura G Caller, Aminu S. Jahun, Sarah L. Caddy, Ian Goodfellow; and Alex Alderton, Roberto Amato, Sonia Goncalves, Ewan Harrison, David K. Jackson, Ian |

| | | | |
|---|---|---|---|
| | | | Johnston, Dominic Kwiatkowski, Cordelia Langford, John Sillitoe on behalf of the Wellcome Sanger Institute COVID-19 Surveillance Team (http://www.sanger.ac.uk/covid-team) |
| EPI_ISL_524534, EPI_ISL_524535, EPI_ISL_524536 | NHSGGC West of Scotland Specialist Virology Centre / MRC-University of Glasgow Centre for Virus Research | Wellcome Sanger Institute for the COVID-19 Genomics UK Consortium | Ana da Silva Filipe, Natasha Johnson, Kathy Smollett, Daniel Mair, Stephen Carmichael, Lily Tong, Jenna Nichols, Elihu Aranday-Cortes, Kirstyn Brunker, Yasmin Parr, Kyriaki Nomikou; Sarah McDonald, Marc Niebel, Patawee Asamaphan; Richard Orton, Joseph Hughes, Sreenu Vattipally, David L Robertson; Alasdair MacLean, Rory Gunson; Kathy Li, Natasha Jesudason, Rajiv Shah, James Shepherd, Antonia Ho, Alice Broos, Emma Thomson and Alex Alderton, Roberto Amato, Sonia Goncalves, Ewan Harrison, David K. Jackson, Ian Johnston, Dominic Kwiatkowski, Cordelia Langford, John Sillitoe on behalf of the Wellcome Sanger Institute COVID-19 Surveillance Team (http://www.sanger.ac.uk/covid-team) |
| EPI_ISL_524537 | Department of Pathology, University of Cambridge | Wellcome Sanger Institute for the COVID-19 Genomics UK Consortium | Luke W Meredith, M. Estée Török , Myra Hosmillo, William L. Hamilton, Martin D. Curran, Theresa Feltwell, Grant Hall, Anna Yakovleva, Fahad A Khokhar, Charlotte J. Houldcroft, Laura G Caller, Aminu S. Jahun, Sarah L. Caddy, Ian Goodfellow; and Alex Alderton, Roberto Amato, Sonia Goncalves, Ewan Harrison, David K. Jackson, Ian Johnston, Dominic Kwiatkowski, Cordelia Langford, John Sillitoe on behalf of the Wellcome Sanger Institute COVID-19 Surveillance Team (http://www.sanger.ac.uk/covid-team) |
| EPI_ISL_524539, EPI_ISL_524545, EPI_ISL_524550 | NHSGGC West of Scotland Specialist Virology Centre / MRC-University of Glasgow Centre for Virus Research | Wellcome Sanger Institute for the COVID-19 Genomics UK Consortium | Ana da Silva Filipe, Natasha Johnson, Kathy Smollett, Daniel Mair, Stephen Carmichael, Lily Tong, Jenna Nichols, Elihu Aranday-Cortes, Kirstyn Brunker, Yasmin Parr, Kyriaki Nomikou; Sarah McDonald, Marc Niebel, Patawee Asamaphan; Richard Orton, Joseph Hughes, Sreenu Vattipally, David L Robertson; Alasdair MacLean, Rory Gunson; Kathy Li, Natasha Jesudason, Rajiv Shah, James Shepherd, Antonia Ho, Alice Broos, Emma Thomson and Alex Alderton, Roberto Amato, Sonia Goncalves, Ewan Harrison, David K. Jackson, Ian Johnston, Dominic Kwiatkowski, Cordelia Langford, John Sillitoe on behalf of the Wellcome Sanger Institute COVID-19 Surveillance Team (http://www.sanger.ac.uk/covid-team) |
| EPI_ISL_524551 | Department of Pathology, University of Cambridge | Wellcome Sanger Institute for the COVID-19 Genomics UK Consortium | Luke W Meredith, M. Estée Török , Myra Hosmillo, William L. Hamilton, Martin D. Curran, Theresa Feltwell, Grant Hall, Anna Yakovleva, Fahad A Khokhar, Charlotte J. Houldcroft, Laura G Caller, Aminu S. Jahun, Sarah L. Caddy, Ian Goodfellow; and Alex Alderton, Roberto Amato, Sonia Goncalves, Ewan Harrison, David K. Jackson, Ian Johnston, Dominic Kwiatkowski, Cordelia Langford, John Sillitoe on behalf of the Wellcome Sanger Institute COVID-19 Surveillance Team (http://www.sanger.ac.uk/covid-team) |
| EPI_ISL_524552, EPI_ISL_524556 | NHSGGC West of Scotland Specialist Virology Centre / MRC-University of Glasgow Centre for Virus Research | Wellcome Sanger Institute for the COVID-19 Genomics UK Consortium | Ana da Silva Filipe, Natasha Johnson, Kathy Smollett, Daniel Mair, Stephen Carmichael, Lily Tong, Jenna Nichols, Elihu Aranday-Cortes, Kirstyn Brunker, Yasmin Parr, Kyriaki Nomikou; Sarah McDonald, Marc Niebel, Patawee Asamaphan; Richard Orton, Joseph Hughes, Sreenu Vattipally, David L Robertson; Alasdair MacLean, Rory Gunson; Kathy Li, Natasha Jesudason, Rajiv Shah, James Shepherd, Antonia Ho, Alice Broos, Emma Thomson and Alex Alderton, Roberto Amato, Sonia Goncalves, Ewan Harrison, David K. Jackson, Ian Johnston, Dominic Kwiatkowski, Cordelia Langford, John Sillitoe on behalf of the Wellcome Sanger Institute COVID-19 Surveillance Team (http://www.sanger.ac.uk/covid-team) |
| EPI_ISL_524560 | Department of Pathology, University of Cambridge | Wellcome Sanger Institute for the COVID-19 Genomics UK Consortium | Luke W Meredith, M. Estée Török , Myra Hosmillo, William L. Hamilton, Martin D. Curran, Theresa Feltwell, Grant Hall, Anna Yakovleva, Fahad A Khokhar, Charlotte J. Houldcroft, Laura G Caller, Aminu S. Jahun, Sarah L. Caddy, Ian Goodfellow; and Alex Alderton, Roberto Amato, Sonia Goncalves, Ewan Harrison, David K. Jackson, Ian Johnston, Dominic Kwiatkowski, Cordelia Langford, John Sillitoe on behalf of the Wellcome Sanger Institute COVID-19 Surveillance Team (http://www.sanger.ac.uk/covid-team) |
| EPI_ISL_524562, EPI_ISL_524563 | NHSGGC West of Scotland Specialist Virology Centre / MRC-University of Glasgow Centre for Virus Research | Wellcome Sanger Institute for the COVID-19 Genomics UK Consortium | Ana da Silva Filipe, Natasha Johnson, Kathy Smollett, Daniel Mair, Stephen Carmichael, Lily Tong, Jenna Nichols, Elihu Aranday-Cortes, Kirstyn Brunker, Yasmin Parr, Kyriaki Nomikou; Sarah McDonald, Marc Niebel, Patawee Asamaphan; Richard Orton, Joseph Hughes, Sreenu Vattipally, David L Robertson; Alasdair MacLean, Rory Gunson; Kathy Li, Natasha Jesudason, Rajiv Shah, James Shepherd, Antonia Ho, Alice Broos, Emma Thomson and Alex Alderton, Roberto Amato, Sonia Goncalves, Ewan Harrison, David K. Jackson, Ian Johnston, Dominic Kwiatkowski, Cordelia Langford, John Sillitoe on behalf of the Wellcome Sanger Institute COVID-19 Surveillance Team (http://www.sanger.ac.uk/covid-team) |
| EPI_ISL_524564 | Department of Pathology, University of Cambridge | Wellcome Sanger Institute for the COVID-19 Genomics UK Consortium | Luke W Meredith, M. Estée Török , Myra Hosmillo, William L. Hamilton, Martin D. Curran, Theresa Feltwell, Grant Hall, Anna Yakovleva, Fahad A Khokhar, Charlotte J. Houldcroft, Laura G Caller, Aminu S. Jahun, Sarah L. Caddy, Ian Goodfellow; and Alex Alderton, Roberto Amato, Sonia Goncalves, Ewan Harrison, David K. Jackson, Ian Johnston, Dominic Kwiatkowski, Cordelia Langford, John Sillitoe on behalf of the Wellcome Sanger Institute COVID-19 Surveillance Team (http://www.sanger.ac.uk/covid-team) |
| EPI_ISL_524566 | NHSGGC West of Scotland Specialist Virology Centre / MRC-University of Glasgow Centre for Virus Research | Wellcome Sanger Institute for the COVID-19 Genomics UK Consortium | Ana da Silva Filipe, Natasha Johnson, Kathy Smollett, Daniel Mair, Stephen Carmichael, Lily Tong, Jenna Nichols, Elihu Aranday-Cortes, Kirstyn Brunker, Yasmin Parr, Kyriaki Nomikou; Sarah McDonald, Marc Niebel, Patawee Asamaphan; Richard Orton, Joseph Hughes, Sreenu Vattipally, David L Robertson; Alasdair MacLean, Rory Gunson; Kathy Li, Natasha Jesudason, Rajiv Shah, James Shepherd, Antonia Ho, Alice Broos, Emma Thomson and Alex Alderton, Roberto Amato, Sonia Goncalves, Ewan Harrison, David K. Jackson, Ian Johnston, Dominic Kwiatkowski, Cordelia Langford, John Sillitoe on behalf of the Wellcome Sanger Institute COVID-19 Surveillance Team (http://www.sanger.ac.uk/covid-team) |
| EPI_ISL_524568, EPI_ISL_524570 | Department of Pathology, University of Cambridge | Wellcome Sanger Institute for the COVID-19 Genomics UK Consortium | Luke W Meredith, M. Estée Török , Myra Hosmillo, William L. Hamilton, Martin D. Curran, Theresa Feltwell, Grant Hall, Anna Yakovleva, Fahad A Khokhar, Charlotte J. Houldcroft, Laura G Caller, Aminu S. Jahun, Sarah L. Caddy, Ian Goodfellow; and Alex Alderton, Roberto Amato, Sonia Goncalves, Ewan Harrison, David K. Jackson, Ian Johnston, Dominic Kwiatkowski, Cordelia Langford, John Sillitoe on behalf of the Wellcome Sanger Institute COVID-19 Surveillance Team (http://www.sanger.ac.uk/covid-team) |
| EPI_ISL_524572 | NHSGGC West of Scotland Specialist Virology Centre / MRC-University of Glasgow Centre for Virus Research | Wellcome Sanger Institute for the COVID-19 Genomics UK Consortium | Ana da Silva Filipe, Natasha Johnson, Kathy Smollett, Daniel Mair, Stephen Carmichael, Lily Tong, Jenna Nichols, Elihu Aranday-Cortes, Kirstyn Brunker, Yasmin Parr, Kyriaki Nomikou; Sarah McDonald, Marc Niebel, Patawee Asamaphan; Richard Orton, Joseph Hughes, Sreenu Vattipally, David L Robertson; Alasdair MacLean, Rory Gunson; Kathy Li, Natasha Jesudason, Rajiv Shah, James Shepherd, Antonia Ho, Alice Broos, Emma Thomson and Alex Alderton, Roberto Amato, Sonia Goncalves, Ewan Harrison, David K. Jackson, Ian Johnston, Dominic Kwiatkowski, Cordelia Langford, John Sillitoe on behalf of the Wellcome Sanger Institute COVID-19 Surveillance Team (http://www.sanger.ac.uk/covid-team) |
| EPI_ISL_524573, EPI_ISL_524575, EPI_ISL_524576, EPI_ISL_524577, EPI_ISL_524578, EPI_ISL_524583, EPI_ISL_524584, EPI_ISL_524586, EPI_ISL_524587, EPI_ISL_524589, EPI_ISL_524591, EPI_ISL_524593, EPI_ISL_524594, EPI_ISL_524596, EPI_ISL_524597, EPI_ISL_524598, EPI_ISL_524599, EPI_ISL_524600, EPI_ISL_524601, EPI_ISL_524603, EPI_ISL_524604, EPI_ISL_524606, EPI_ISL_524608, EPI_ISL_524609, EPI_ISL_524610, EPI_ISL_524612, EPI_ISL_524615, EPI_ISL_524616, EPI_ISL_524617, EPI_ISL_524618 | | | |
| see above | Department of Pathology, University of Cambridge | Wellcome Sanger Institute for the COVID-19 Genomics UK Consortium | Luke W Meredith, M. Estée Török , Myra Hosmillo, William L. Hamilton, Martin D. Curran, Theresa Feltwell, Grant Hall, Anna Yakovleva, Fahad A Khokhar, Charlotte J. Houldcroft, Laura G Caller, Aminu S. Jahun, Sarah L. Caddy, Ian Goodfellow; and Alex Alderton, Roberto Amato, Sonia Goncalves, Ewan Harrison, David K. Jackson, Ian Johnston, Dominic Kwiatkowski, Cordelia Langford, John Sillitoe on behalf of the Wellcome Sanger Institute COVID-19 Surveillance Team (http://www.sanger.ac.uk/covid-team) |
| EPI_ISL_524619, EPI_ISL_524620, EPI_ISL_524621, EPI_ISL_524622, EPI_ISL_524623, EPI_ISL_524624, EPI_ISL_524625, EPI_ISL_524628, EPI_ISL_524629, EPI_ISL_524630, EPI_ISL_524631, EPI_ISL_524632, EPI_ISL_524633, EPI_ISL_524634, EPI_ISL_524636, EPI_ISL_524638, EPI_ISL_524640, EPI_ISL_524641, EPI_ISL_524642, EPI_ISL_524643, EPI_ISL_524645, EPI_ISL_524646, EPI_ISL_524647, EPI_ISL_524648, EPI_ISL_524649, EPI_ISL_524651, EPI_ISL_524652, EPI_ISL_524653, EPI_ISL_524654, EPI_ISL_524656, EPI_ISL_524658, EPI_ISL_524659, EPI_ISL_524663, EPI_ISL_524665, EPI_ISL_524668, EPI_ISL_524669, EPI_ISL_524670, EPI_ISL_524671, EPI_ISL_524673, EPI_ISL_524674, EPI_ISL_524676, EPI_ISL_524680, EPI_ISL_524682, EPI_ISL_524683, EPI_ISL_524684, EPI_ISL_524686, EPI_ISL_524688, EPI_ISL_524689, EPI_ISL_524690, EPI_ISL_524691, EPI_ISL_524692, EPI_ISL_524693, EPI_ISL_524694, EPI_ISL_524696, EPI_ISL_524698, EPI_ISL_524699, EPI_ISL_524700, EPI_ISL_524702, EPI_ISL_524703, EPI_ISL_524704, EPI_ISL_524705, EPI_ISL_524706, EPI_ISL_524707, EPI_ISL_524708, EPI_ISL_524709, EPI_ISL_524710, EPI_ISL_524711, EPI_ISL_524712 | | | |
| see above | North West London Pathology, Imperial College Healthcare NHS Trust | Wellcome Sanger Institute for the COVID-19 Genomics UK Consortium | Ling Li, Paul Randell, David Muir, Frankie Bolt, Alison Holmes, James Price, Aileen Rowan, Graham Taylor, Anjna Badhan, Carolina Herrera and Alex Alderton, Roberto Amato, Sonia Goncalves, Ewan Harrison, David K. Jackson, Ian Johnston, Dominic Kwiatkowski, Cordelia Langford, John Sillitoe on behalf of the Wellcome Sanger Institute COVID-19 Surveillance Team (http://www.sanger.ac.uk/covid-team) |
| EPI_ISL_524713 | B.J. Medical College and Civil hospital, Ahmedabad | Gujarat Biotechnology Research Centre | Apurvasinh Puvar, Janvi Raval, Zarna Patel, Monika Gandhi, Pinal Trivedi, Maharshi Pandya, Nidhi Patel, Nitin Savaliya, Raghawendra Kumar, Dinesh Kumar, Zuber Saiyed, Komal Patel, Labdhi Pandya, Afzal Ansari, Nikha Trivedi, Pranay Shah, Kamlesh J Upadhyay, Sanjay Kapadia, R D Dixit, A M Kadri, Harsh Bakshi, Chaitanya Joshi, Madhvi Joshi |
| EPI_ISL_524714 | B.J. Medical College and Civil hospital, Ahmedabad | Gujarat Biotechnology Research Centre | Janvi Raval, Zarna Patel, Monika Gandhi, Pinal Trivedi, Maharshi Pandya, Nidhi Patel, Nitin Savaliya, Raghawendra Kumar, Dinesh Kumar, Zuber Saiyed, Komal Patel, Labdhi Pandya, Afzal Ansari, Nikha Trivedi, Pranay Shah, Kamlesh J Upadhyay, Sanjay Kapadia, Apurvasinh Puvar, R D Dixit, A M Kadri, Harsh Bakshi, Chaitanya Joshi, Madhvi Joshi |
| EPI_ISL_524715 | B.J. Medical College and Civil hospital, Ahmedabad | Gujarat Biotechnology Research Centre | Zarna Patel, Monika Gandhi, Pinal Trivedi, Maharshi Pandya, Nidhi Patel, Nitin Savaliya, Raghawendra Kumar, Dinesh Kumar, Zuber Saiyed, Komal Patel, Labdhi Pandya, Afzal Ansari, Nikha Trivedi, Pranay Shah, Kamlesh J Upadhyay, Sanjay Kapadia, Apurvasinh Puvar, Janvi Raval, R D Dixit, A M Kadri, Harsh Bakshi, Chaitanya Joshi, Madhvi Joshi |
| EPI_ISL_524716 | B.J. Medical College and Civil hospital, Ahmedabad | Gujarat Biotechnology Research Centre | Monika Gandhi, Pinal Trivedi, Maharshi Pandya, Nidhi Patel, Nitin Savaliya, Raghawendra Kumar, Dinesh Kumar, Zuber Saiyed, Komal Patel, Labdhi Pandya, Afzal Ansari, Nikha Trivedi, Pranay Shah, Kamlesh J Upadhyay, Sanjay Kapadia, Apurvasinh Puvar, Janvi Raval, Zarna Patel, R D Dixit, A M Kadri, Harsh Bakshi, Chaitanya Joshi, Madhvi Joshi |
| EPI_ISL_524717 | B.J. Medical College and Civil hospital, Ahmedabad | Gujarat Biotechnology Research Centre | Pinal Trivedi, Maharshi Pandya, Nidhi Patel, Nitin Savaliya, Raghawendra Kumar, Dinesh Kumar, Zuber Saiyed, Komal Patel, Labdhi Pandya, Afzal Ansari, Nikha Trivedi, Pranay Shah, Kamlesh J Upadhyay, Sanjay Kapadia, Apurvasinh Puvar, Janvi Raval, Zarna Patel, Monika Gandhi, R D Dixit, A M Kadri, Harsh Bakshi, Chaitanya Joshi, Madhvi Joshi |
| EPI_ISL_524718 | B.J. Medical College and Civil hospital, Ahmedabad | Gujarat Biotechnology Research Centre | Maharshi Pandya, Nidhi Patel, Nitin Savaliya, Raghawendra Kumar, Dinesh Kumar, Zuber Saiyed, Komal Patel, Labdhi Pandya, Afzal Ansari, Nikha Trivedi, Pranay Shah, Kamlesh J Upadhyay, Sanjay Kapadia, Apurvasinh Puvar, Janvi Raval, Zarna Patel, Monika Gandhi, Pinal Trivedi, R D Dixit, A M Kadri, Harsh Bakshi, Chaitanya Joshi, Madhvi Joshi |
| EPI_ISL_524719 | B.J. Medical College and Civil hospital, Ahmedabad | Gujarat Biotechnology Research Centre | Nidhi Patel, Nitin Savaliya, Raghawendra Kumar, Dinesh Kumar, Zuber Saiyed, Komal Patel, Labdhi Pandya, Afzal Ansari, Nikha Trivedi, Pranay Shah, Kamlesh J Upadhyay, Sanjay Kapadia, Apurvasinh Puvar, Janvi Raval, Zarna Patel, Monika Gandhi, Pinal Trivedi, Maharshi Pandya, R D Dixit, A M Kadri, Harsh Bakshi, Chaitanya Joshi, Madhvi Joshi |
| EPI_ISL_524720 | B.J. Medical College and Civil hospital, Ahmedabad | Gujarat Biotechnology Research Centre | Nitin Savaliya, Raghawendra Kumar, Dinesh Kumar, Zuber Saiyed, Komal Patel, Labdhi Pandya, Afzal Ansari, Nikha Trivedi, Pranay Shah, Kamlesh J Upadhyay, Sanjay Kapadia, Apurvasinh Puvar, Janvi Raval, Zarna Patel, Monika Gandhi, Pinal Trivedi, Maharshi Pandya, Nidhi Patel, R D Dixit, A M Kadri, Harsh Bakshi, Chaitanya Joshi, Madhvi Joshi |
| EPI_ISL_524721 | B.J. Medical College and Civil hospital, Ahmedabad | Gujarat Biotechnology Research Centre | Raghawendra Kumar, Dinesh Kumar, Zuber Saiyed, Komal Patel, Labdhi Pandya, Afzal Ansari, Nikha Trivedi, Pranay Shah, Kamlesh J Upadhyay, Sanjay Kapadia, Apurvasinh Puvar, Janvi Raval, Zarna Patel, Monika Gandhi, Pinal Trivedi, Maharshi Pandya, Nidhi Patel, Nitin Savaliya, R D Dixit, A M Kadri, Harsh Bakshi, Chaitanya Joshi, Madhvi Joshi |
| EPI_ISL_524722 | B.J. Medical College and Civil hospital, Ahmedabad | Gujarat Biotechnology Research Centre | Dinesh Kumar, Zuber Saiyed, Komal Patel, Labdhi Pandya, Afzal Ansari, Nikha Trivedi, Pranay Shah, Kamlesh J Upadhyay, Sanjay Kapadia, Apurvasinh Puvar, Janvi Raval, Zarna Patel, Monika Gandhi, Pinal Trivedi, Maharshi Pandya, Nidhi Patel, Nitin Savaliya, Raghawendra Kumar, R D Dixit, A M Kadri, Harsh Bakshi, Chaitanya Joshi, Madhvi Joshi |
| EPI_ISL_524723 | B.J. Medical College and Civil hospital, Ahmedabad | Gujarat Biotechnology Research Centre | Zuber Saiyed, Komal Patel, Labdhi Pandya, Afzal Ansari, Nikha Trivedi, Pranay Shah, Kamlesh J Upadhyay, Sanjay Kapadia, Apurvasinh Puvar, Janvi Raval, Zarna Patel, Monika Gandhi, Pinal Trivedi, Maharshi Pandya, Nidhi Patel, Nitin Savaliya, Raghawendra Kumar, Dinesh Kumar, R D Dixit, A M Kadri, Harsh Bakshi, Chaitanya Joshi, Madhvi Joshi |
| EPI_ISL_524724 | B.J. Medical College and Civil hospital, Ahmedabad | Gujarat Biotechnology Research Centre | Komal Patel, Labdhi Pandya, Afzal Ansari, Nikha Trivedi, Pranay Shah, Kamlesh J Upadhyay, Sanjay Kapadia, Apurvasinh Puvar, Janvi Raval, Zarna Patel, Monika Gandhi, Pinal Trivedi, Maharshi Pandya, Nidhi Patel, Nitin Savaliya, Raghawendra Kumar, Dinesh Kumar, Zuber Saiyed, R D Dixit, A M Kadri, Harsh Bakshi, Chaitanya Joshi, Madhvi Joshi |
| EPI_ISL_524725 | B.J. Medical College and Civil hospital, Ahmedabad | Gujarat Biotechnology Research Centre | Labdhi Pandya, Afzal Ansari, Nikha Trivedi, Pranay Shah, Kamlesh J Upadhyay, Sanjay Kapadia, Apurvasinh Puvar, Janvi Raval, Zarna Patel, Monika Gandhi, Pinal Trivedi, Maharshi Pandya, Nidhi Patel, Nitin Savaliya, Raghawendra Kumar, Dinesh Kumar, Zuber Saiyed, Komal Patel, R D Dixit, A M Kadri, Harsh Bakshi, Chaitanya Joshi, Madhvi Joshi |
| EPI_ISL_524726 | B.J. Medical College and Civil hospital, Ahmedabad | Gujarat Biotechnology Research Centre | Afzal Ansari, Nikha Trivedi, Pranay Shah, Kamlesh J Upadhyay, Sanjay Kapadia, Apurvasinh Puvar, Janvi Raval, Zarna Patel, Monika Gandhi, Pinal Trivedi, Maharshi Pandya, Nidhi Patel, Nitin Savaliya, Raghawendra Kumar, Dinesh Kumar, Zuber Saiyed, Komal Patel, Labdhi Pandya, R D Dixit, A M Kadri, Harsh Bakshi, Chaitanya Joshi, Madhvi Joshi |
| EPI_ISL_524727 | B.J. Medical College and Civil hospital, Ahmedabad | Gujarat Biotechnology Research Centre | Nikha Trivedi, Pranay Shah, Kamlesh J Upadhyay, Sanjay Kapadia, Apurvasinh Puvar, Janvi Raval, Zarna Patel, Monika Gandhi, Pinal Trivedi, Maharshi Pandya, Nidhi Patel, Nitin Savaliya, Raghawendra Kumar, Dinesh Kumar, Zuber Saiyed, Komal Patel, Labdhi Pandya, Afzal Ansari, R D Dixit, A M Kadri, Harsh Bakshi, Chaitanya Joshi, Madhvi Joshi |
| EPI_ISL_524728 | B.J. Medical College and Civil hospital, Ahmedabad | Gujarat Biotechnology Research Centre | Pranay Shah, Kamlesh J Upadhyay, Sanjay Kapadia, Apurvasinh Puvar, Janvi Raval, Zarna Patel, Monika Gandhi, Pinal Trivedi, Maharshi Pandya, Nidhi Patel, Nitin Savaliya, Raghawendra Kumar, Dinesh Kumar, Zuber Saiyed, Komal Patel, Labdhi Pandya, Afzal Ansari, Nikha Trivedi, R D Dixit, A M Kadri, Harsh Bakshi, Chaitanya Joshi, Madhvi Joshi |
| EPI_ISL_524729 | B.J. Medical College and Civil hospital, Ahmedabad | Gujarat Biotechnology Research Centre | Kamlesh J Upadhyay, Sanjay Kapadia, Apurvasinh Puvar, Janvi Raval, Zarna Patel, Monika Gandhi, Pinal Trivedi, Maharshi Pandya, Nidhi Patel, Nitin Savaliya, Raghawendra Kumar, Dinesh Kumar, Zuber Saiyed, Komal Patel, Labdhi Pandya, Afzal Ansari, Nikha Trivedi, Pranay Shah, R D Dixit, A M Kadri, Harsh Bakshi, Chaitanya Joshi, Madhvi Joshi |
| EPI_ISL_524730 | GMERS Medical College & Hospital, Gotri, Vadodara | Gujarat Biotechnology Research Centre | Meenakshi Shah, Neena Doshi, Varsha Godbole, Apurvasinh Puvar, Janvi Raval, Zarna Patel, Monika Gandhi, Pinal Trivedi, Maharshi Pandya, Nidhi Patel, Nitin Savaliya, Raghawendra Kumar, Dinesh Kumar, Zuber Saiyed, Komal Patel, Labdhi Pandya, Afzal Ansari, Nikha Trivedi, R D Dixit, A M Kadri, Harsh Bakshi, Chaitanya Joshi, Madhvi Joshi |

| | | | |
|---|---|---|---|
| EPI_ISL_524731 | GMERS Medical College & Hospital, Gotri, Vadodara | Gujarat Biotechnology Research Centre | Neena Doshi, Varsha Godbole, Apurvasinh Puvar, Janvi Raval, Zarna Patel, Monika Gandhi, Pinal Trivedi, Maharshi Pandya, Nidhi Patel, Nitin Savaliya, Raghawendra Kumar, Dinesh Kumar, Zuber Saiyed, Komal Patel, Labdhi Pandya, Afzal Ansari, Nikha Trivedi, Meenakshi Shah, R D Dixit, A M Kadri, Harsh Bakshi, Chaitanya Joshi, Madhvi Joshi |
| EPI_ISL_524732 | Dr. RSS Hospital, Modasa | Gujarat Biotechnology Research Centre | Harsh Chaudhari, Apurvasinh Puvar, Janvi Raval, Zarna Patel, Monika Gandhi, Pinal Trivedi, Maharshi Pandya, Nidhi Patel, Nitin Savaliya, Raghawendra Kumar, Dinesh Kumar, Zuber Saiyed, Komal Patel, Labdhi Pandya, Afzal Ansari, Nikha Trivedi, R D Dixit, A M Kadri, Harsh Bakshi, Chaitanya Joshi, Madhvi Joshi, Neena Doshi, Varsha Godbole |
| EPI_ISL_524733 | GMERS Medical College and Hospital, Dharpur, Patan | Gujarat Biotechnology Research Centre | A N Parmar, Apurvasinh Puvar, Janvi Raval, Zarna Patel, Monika Gandhi, Pinal Trivedi, Maharshi Pandya, Nidhi Patel, Nitin Savaliya, Raghawendra Kumar, Dinesh Kumar, Zuber Saiyed, Komal Patel, Labdhi Pandya, Afzal Ansari, Nikha Trivedi, R D Dixit, A M Kadri, Harsh Bakshi, Chaitanya Joshi, Madhvi Joshi |
| EPI_ISL_524734 | GMERS Medical College and Hospital, Dharpur, Patan | Gujarat Biotechnology Research Centre | Apurvasinh Puvar, Janvi Raval, Zarna Patel, Monika Gandhi, Pinal Trivedi, Maharshi Pandya, Nidhi Patel, Nitin Savaliya, Raghawendra Kumar, Dinesh Kumar, Zuber Saiyed, Komal Patel, Labdhi Pandya, Afzal Ansari, Nikha Trivedi, A N Parmar, R D Dixit, A M Kadri, Harsh Bakshi, Chaitanya Joshi, Madhvi Joshi |
| EPI_ISL_524735 | Government General Hospital, Jam Khambhaliya, Devbhoomi Dwarka | Gujarat Biotechnology Research Centre | Harish Matani, Janvi Raval, Zarna Patel, Monika Gandhi, Pinal Trivedi, Maharshi Pandya, Nidhi Patel, Nitin Savaliya, Raghawendra Kumar, Dinesh Kumar, Zuber Saiyed, Komal Patel, Labdhi Pandya, Afzal Ansari, Nikha Trivedi, Apurvasinh Puvar, R D Dixit, A M Kadri, Harsh Bakshi, Chaitanya Joshi, Madhvi Joshi |
| EPI_ISL_524736 | Government General Hospital, Jam Khambhaliya, Devbhoomi Dwarka | Gujarat Biotechnology Research Centre | Janvi Raval, Zarna Patel, Monika Gandhi, Pinal Trivedi, Maharshi Pandya, Nidhi Patel, Nitin Savaliya, Raghawendra Kumar, Dinesh Kumar, Zuber Saiyed, Komal Patel, Labdhi Pandya, Afzal Ansari, Nikha Trivedi, Harish Matani, Apurvasinh Puvar, R D Dixit, A M Kadri, Harsh Bakshi, Chaitanya Joshi, Madhvi Joshi |
| EPI_ISL_524737 | GMERS Medical College and Hospital, Gandhinagar | Gujarat Biotechnology Research Centre | Zarna Patel, Monika Gandhi, Pinal Trivedi, Maharshi Pandya, Nidhi Patel, Nitin Savaliya, Raghawendra Kumar, Dinesh Kumar, Zuber Saiyed, Komal Patel, Labdhi Pandya, Afzal Ansari, Nikha Trivedi, Seema Bhatt, Gaurishankar Shrimali, Bhavesh Modi, Bharti Rajani, Apurvasinh Puvar, Janvi Raval, R D Dixit, A M Kadri, Harsh Bakshi, Chaitanya Joshi, Madhvi Joshi |
| EPI_ISL_524738 | GMERS Medical College and Hospital, Gandhinagar | Gujarat Biotechnology Research Centre | Monika Gandhi, Pinal Trivedi, Maharshi Pandya, Nidhi Patel, Nitin Savaliya, Raghawendra Kumar, Dinesh Kumar, Zuber Saiyed, Komal Patel, Labdhi Pandya, Afzal Ansari, Nikha Trivedi, Seema Bhatt, Gaurishankar Shrimali, Bhavesh Modi, Bharti Rajani, Apurvasinh Puvar, Janvi Raval, Zarna Patel, R D Dixit, A M Kadri, Harsh Bakshi, Chaitanya Joshi, Madhvi Joshi |
| EPI_ISL_524739 | GMERS Medical College and Hospital, Gandhinagar | Gujarat Biotechnology Research Centre | Pinal Trivedi, Maharshi Pandya, Nidhi Patel, Nitin Savaliya, Raghawendra Kumar, Dinesh Kumar, Zuber Saiyed, Komal Patel, Labdhi Pandya, Afzal Ansari, Nikha Trivedi, Seema Bhatt, Gaurishankar Shrimali, Bhavesh Modi, Bharti Rajani, Apurvasinh Puvar, Janvi Raval, Zarna Patel, Monika Gandhi, R D Dixit, A M Kadri, Harsh Bakshi, Chaitanya Joshi, Madhvi Joshi |
| EPI_ISL_524740 | GMERS Medical College and Hospital, Gandhinagar | Gujarat Biotechnology Research Centre | Maharshi Pandya, Nidhi Patel, Nitin Savaliya, Raghawendra Kumar, Dinesh Kumar, Zuber Saiyed, Komal Patel, Labdhi Pandya, Afzal Ansari, Nikha Trivedi, Seema Bhatt, Gaurishankar Shrimali, Bhavesh Modi, Bharti Rajani, Apurvasinh Puvar, Janvi Raval, Zarna Patel, Monika Gandhi, Pinal Trivedi, R D Dixit, A M Kadri, Harsh Bakshi, Chaitanya Joshi, Madhvi Joshi |
| EPI_ISL_524741 | GMERS Medical College and Hospital, Gandhinagar | Gujarat Biotechnology Research Centre | Nidhi Patel, Nitin Savaliya, Raghawendra Kumar, Dinesh Kumar, Zuber Saiyed, Komal Patel, Labdhi Pandya, Afzal Ansari, Nikha Trivedi, Seema Bhatt, Gaurishankar Shrimali, Bhavesh Modi, Bharti Rajani, Apurvasinh Puvar, Janvi Raval, Zarna Patel, Monika Gandhi, Pinal Trivedi, Maharshi Pandya, R D Dixit, A M Kadri, Harsh Bakshi, Chaitanya Joshi, Madhvi Joshi |
| EPI_ISL_524742 | GMERS Medical College and Hospital, Gandhinagar | Gujarat Biotechnology Research Centre | Dinesh Kumar, Zuber Saiyed, Komal Patel, Labdhi Pandya, Afzal Ansari, Nikha Trivedi, Seema Bhatt, Gaurishankar Shrimali, Bhavesh Modi, Bharti Rajani, Apurvasinh Puvar, Janvi Raval, Zarna Patel, Monika Gandhi, Pinal Trivedi, Maharshi Pandya, Nidhi Patel, Nitin Savaliya, Raghawendra Kumar, R D Dixit, A M Kadri, Harsh Bakshi, Chaitanya Joshi, Madhvi Joshi |
| EPI_ISL_524743 | GMERS Medical College and Hospital, Gandhinagar | Gujarat Biotechnology Research Centre | Zuber Saiyed, Komal Patel, Labdhi Pandya, Afzal Ansari, Nikha Trivedi, Seema Bhatt, Gaurishankar Shrimali, Bhavesh Modi, Bharti Rajani, Apurvasinh Puvar, Janvi Raval, Zarna Patel, Monika Gandhi, Pinal Trivedi, Maharshi Pandya, Nidhi Patel, Nitin Savaliya, Raghawendra Kumar, Dinesh Kumar, R D Dixit, A M Kadri, Harsh Bakshi, Chaitanya Joshi, Madhvi Joshi |
| EPI_ISL_524744 | GMERS Medical College and Hospital, Gandhinagar | Gujarat Biotechnology Research Centre | Komal Patel, Labdhi Pandya, Afzal Ansari, Nikha Trivedi, Seema Bhatt, Gaurishankar Shrimali, Bhavesh Modi, Bharti Rajani, Apurvasinh Puvar, Janvi Raval, Zarna Patel, Monika Gandhi, Pinal Trivedi, Maharshi Pandya, Nidhi Patel, Nitin Savaliya, Raghawendra Kumar, Dinesh Kumar, Zuber Saiyed, R D Dixit, A M Kadri, Harsh Bakshi, Chaitanya Joshi, Madhvi Joshi |
| EPI_ISL_524745 | GMERS Medical College and Hospital, Gandhinagar | Gujarat Biotechnology Research Centre | Labdhi Pandya, Afzal Ansari, Nikha Trivedi, Seema Bhatt, Gaurishankar Shrimali, Bhavesh Modi, Bharti Rajani, Apurvasinh Puvar, Janvi Raval, Zarna Patel, Monika Gandhi, Pinal Trivedi, Maharshi Pandya, Nidhi Patel, Nitin Savaliya, Raghawendra Kumar, Dinesh Kumar, Zuber Saiyed, Komal Patel, R D Dixit, A M Kadri, Harsh Bakshi, Chaitanya Joshi, Madhvi Joshi |
| EPI_ISL_524746 | Sir M P Shah Government Medical College, Jamnagar | Gujarat Biotechnology Research Centre | Binita Aring, Afzal Ansari, Nikha Trivedi, Apurvasinh Puvar, Janvi Raval, Zarna Patel, Monika Gandhi, Pinal Trivedi, Maharshi Pandya, Nidhi Patel, Nitin Savaliya, Raghawendra Kumar, Dinesh Kumar, Zuber Saiyed, Komal Patel, Labdhi Pandya, R D Dixit, A M Kadri, Harsh Bakshi, Chaitanya Joshi, Madhvi Joshi |
| EPI_ISL_524747 | Sir M P Shah Government Medical College, Jamnagar | Gujarat Biotechnology Research Centre | Afzal Ansari, Nikha Trivedi, Binita Aring, Apurvasinh Puvar, Janvi Raval, Zarna Patel, Monika Gandhi, Pinal Trivedi, Maharshi Pandya, Nidhi Patel, Nitin Savaliya, Raghawendra Kumar, Dinesh Kumar, Zuber Saiyed, Komal Patel, Labdhi Pandya, R D Dixit, A M Kadri, Harsh Bakshi, Chaitanya Joshi, Madhvi Joshi |
| EPI_ISL_524748 | Sir M P Shah Government Medical College, Jamnagar | Gujarat Biotechnology Research Centre | Nikha Trivedi, Apurvasinh Puvar, Janvi Raval, Binita Aring, Zarna Patel, Monika Gandhi, Pinal Trivedi, Maharshi Pandya, Nidhi Patel, Nitin Savaliya, Raghawendra Kumar, Dinesh Kumar, Zuber Saiyed, Komal Patel, Labdhi Pandya, Afzal Ansari, R D Dixit, A M Kadri, Harsh Bakshi, Chaitanya Joshi, Madhvi Joshi |
| EPI_ISL_524749 | Sir M P Shah Government Medical College, Jamnagar | Gujarat Biotechnology Research Centre | Apurvasinh Puvar, Janvi Raval, Zarna Patel, Monika Gandhi, Pinal Trivedi, Maharshi Pandya, Nidhi Patel, Nitin Savaliya, Raghawendra Kumar, Dinesh Kumar, Zuber Saiyed, Komal Patel, Labdhi Pandya, Afzal Ansari, Nikha Trivedi, Binita Aring, R D Dixit, A M Kadri, Harsh Bakshi, Chaitanya Joshi, Madhvi Joshi |
| EPI_ISL_524750 | Sir M P Shah Government Medical College, Jamnagar | Gujarat Biotechnology Research Centre | Janvi Raval, Zarna Patel, Monika Gandhi, Pinal Trivedi, Maharshi Pandya, Nidhi Patel, Nitin Savaliya, Raghawendra Kumar, Dinesh Kumar, Zuber Saiyed, Komal Patel, Labdhi Pandya, Afzal Ansari, Nikha Trivedi, Binita Aring, Apurvasinh Puvar, R D Dixit, A M Kadri, Harsh Bakshi, Chaitanya Joshi, Madhvi Joshi |
| EPI_ISL_524751 | Sir M P Shah Government Medical College, Jamnagar | Gujarat Biotechnology Research Centre | Zarna Patel, Monika Gandhi, Pinal Trivedi, Maharshi Pandya, Nidhi Patel, Nitin Savaliya, Raghawendra Kumar, Dinesh Kumar, Zuber Saiyed, Komal Patel, Labdhi Pandya, Afzal Ansari, Nikha Trivedi, Binita Aring, Apurvasinh Puvar, Janvi Raval, R D Dixit, A M Kadri, Harsh Bakshi, Chaitanya Joshi, Madhvi Joshi |
| EPI_ISL_524752 | GMERS Medical College and Hospital, Dharpur, Patan | Gujarat Biotechnology Research Centre | Monika Gandhi, Pinal Trivedi, Maharshi Pandya, Nidhi Patel, Nitin Savaliya, Raghawendra Kumar, Dinesh Kumar, Zuber Saiyed, Komal Patel, Labdhi Pandya, Afzal Ansari, Nikha Trivedi, A N Parmar, Apurvasinh Puvar, Janvi Raval, Zarna Patel, R D Dixit, A M Kadri, Harsh Bakshi, Chaitanya Joshi, Madhvi Joshi |
| EPI_ISL_524753 | GMERS Medical College and Hospital, Dharpur, Patan | Gujarat Biotechnology Research Centre | Pinal Trivedi, Maharshi Pandya, Nidhi Patel, Nitin Savaliya, Raghawendra Kumar, Dinesh Kumar, Zuber Saiyed, Komal Patel, Labdhi Pandya, Afzal Ansari, Nikha Trivedi, A N Parmar, Apurvasinh Puvar, Janvi Raval, Zarna Patel, Monika Gandhi, R D Dixit, A M Kadri, Harsh Bakshi, Chaitanya Joshi, Madhvi Joshi |
| EPI_ISL_524754 | GMERS Medical College and Hospital, Dharpur, Patan | Gujarat Biotechnology Research Centre | Maharshi Pandya, Nidhi Patel, Nitin Savaliya, Raghawendra Kumar, Dinesh Kumar, Zuber Saiyed, Komal Patel, Labdhi Pandya, Afzal Ansari, Nikha Trivedi, A N Parmar, Apurvasinh Puvar, Janvi Raval, Zarna Patel, Monika Gandhi, Pinal Trivedi, R D Dixit, A M Kadri, Harsh Bakshi, Chaitanya Joshi, Madhvi Joshi |
| EPI_ISL_524755 | GMERS Medical College and Hospital, Dharpur, Patan | Gujarat Biotechnology Research Centre | Nidhi Patel, Nitin Savaliya, Raghawendra Kumar, Dinesh Kumar, Zuber Saiyed, Komal Patel, Labdhi Pandya, Afzal Ansari, Nikha Trivedi, A N Parmar, Apurvasinh Puvar, Janvi Raval, Zarna Patel, Monika Gandhi, Pinal Trivedi, Maharshi Pandya, R D Dixit, A M Kadri, Harsh Bakshi, Chaitanya Joshi, Madhvi Joshi |
| EPI_ISL_524756 | GMERS Medical College and Hospital, Dharpur, Patan | Gujarat Biotechnology Research Centre | Nitin Savaliya, Raghawendra Kumar, Dinesh Kumar, Zuber Saiyed, Komal Patel, Labdhi Pandya, Afzal Ansari, Nikha Trivedi, A N Parmar, Apurvasinh Puvar, Janvi Raval, Zarna Patel, Monika Gandhi, Pinal Trivedi, Maharshi Pandya, Nidhi Patel, R D Dixit, A M Kadri, Harsh Bakshi, Chaitanya Joshi, Madhvi Joshi |
| EPI_ISL_524757 | GMERS Medical College and Hospital, Dharpur, Patan | Gujarat Biotechnology Research Centre | Raghawendra Kumar, Dinesh Kumar, Zuber Saiyed, Komal Patel, Labdhi Pandya, Afzal Ansari, Nikha Trivedi, A N Parmar, Apurvasinh Puvar, Janvi Raval, Zarna Patel, Monika Gandhi, Pinal Trivedi, Maharshi Pandya, Nidhi Patel, Nitin Savaliya, R D Dixit, A M Kadri, Harsh Bakshi, Chaitanya Joshi, Madhvi Joshi |
| EPI_ISL_524758 | GMERS Medical College and Hospital, Dharpur, Patan | Gujarat Biotechnology Research Centre | Dinesh Kumar, Zuber Saiyed, Komal Patel, Labdhi Pandya, Afzal Ansari, Nikha Trivedi, A N Parmar, Apurvasinh Puvar, Janvi Raval, Zarna Patel, Monika Gandhi, Pinal Trivedi, Maharshi Pandya, Nidhi Patel, Nitin Savaliya, Raghawendra Kumar, R D Dixit, A M Kadri, Harsh Bakshi, Chaitanya Joshi, Madhvi Joshi |
| EPI_ISL_524759 | GMERS Medical College and Hospital, Dharpur, Patan | Gujarat Biotechnology Research Centre | Zuber Saiyed, Komal Patel, Labdhi Pandya, Afzal Ansari, Nikha Trivedi, A N Parmar, Apurvasinh Puvar, Janvi Raval, Zarna Patel, Monika Gandhi, Pinal Trivedi, Maharshi Pandya, Nidhi Patel, Nitin Savaliya, Raghawendra Kumar, Dinesh Kumar, R D Dixit, A M Kadri, Harsh Bakshi, Chaitanya Joshi, Madhvi Joshi |
| EPI_ISL_524760 | GMERS Medical College and Hospital, Dharpur, Patan | Gujarat Biotechnology Research Centre | Komal Patel, Labdhi Pandya, Afzal Ansari, Nikha Trivedi, A N Parmar, Apurvasinh Puvar, Janvi Raval, Zarna Patel, Monika Gandhi, Pinal Trivedi, Maharshi Pandya, Nidhi Patel, Nitin Savaliya, Raghawendra Kumar, Dinesh Kumar, Zuber Saiyed, R D Dixit, A M Kadri, Harsh Bakshi, Chaitanya Joshi, Madhvi Joshi |
| EPI_ISL_524761 | GMERS Medical College and Hospital, Dharpur, Patan | Gujarat Biotechnology Research Centre | Labdhi Pandya, Afzal Ansari, Nikha Trivedi, A N Parmar, Apurvasinh Puvar, Janvi Raval, Zarna Patel, Monika Gandhi, Pinal Trivedi, Maharshi Pandya, Nidhi Patel, Nitin Savaliya, Raghawendra Kumar, Dinesh Kumar, Zuber Saiyed, Komal Patel, R D Dixit, A M Kadri, Harsh Bakshi, Chaitanya Joshi, Madhvi Joshi |
| EPI_ISL_524762 | GMERS Medical College and Hospital, Dharpur, Patan | Gujarat Biotechnology Research Centre | Afzal Ansari, Nikha Trivedi, A N Parmar, Apurvasinh Puvar, Janvi Raval, Zarna Patel, Monika Gandhi, Pinal Trivedi, Maharshi Pandya, Nidhi Patel, Nitin Savaliya, Raghawendra Kumar, Dinesh Kumar, Zuber Saiyed, Komal Patel, Labdhi Pandya, R D Dixit, A M Kadri, Harsh Bakshi, Chaitanya Joshi, Madhvi Joshi |
| EPI_ISL_524763 | GMERS Medical College and Hospital, Dharpur, Patan | Gujarat Biotechnology Research Centre | Nikha Trivedi, A N Parmar, Apurvasinh Puvar, Janvi Raval, Zarna Patel, Monika Gandhi, Pinal Trivedi, Maharshi Pandya, Nidhi Patel, Nitin Savaliya, Raghawendra Kumar, Dinesh Kumar, Zuber Saiyed, Komal Patel, Labdhi Pandya, Afzal Ansari, R D Dixit, A M Kadri, Harsh Bakshi, Chaitanya Joshi, Madhvi Joshi |
| EPI_ISL_524764 | GMERS Medical College and Hospital, Dharpur, Patan | Gujarat Biotechnology Research Centre | A N Parmar, Apurvasinh Puvar, Janvi Raval, Zarna Patel, Monika Gandhi, Pinal Trivedi, Maharshi Pandya, Nidhi Patel, Nitin Savaliya, Raghawendra Kumar, Dinesh Kumar, Zuber Saiyed, Komal Patel, Labdhi Pandya, Afzal Ansari, Nikha Trivedi, R D Dixit, A M Kadri, Harsh Bakshi, Chaitanya Joshi, Madhvi Joshi |
| EPI_ISL_524765 | GMERS Medical College and Hospital, Dharpur, Patan | Gujarat Biotechnology Research Centre | Apurvasinh Puvar, Janvi Raval, Zarna Patel, Monika Gandhi, Pinal Trivedi, Maharshi Pandya, Nidhi Patel, Nitin Savaliya, Raghawendra Kumar, Dinesh Kumar, Zuber Saiyed, Komal Patel, Labdhi Pandya, Afzal Ansari, Nikha Trivedi, A N Parmar, R D Dixit, A M Kadri, Harsh Bakshi, Chaitanya Joshi, Madhvi Joshi |
| EPI_ISL_524766 | GMERS Medical College and Hospital, Dharpur, Patan | Gujarat Biotechnology Research Centre | Janvi Raval, Zarna Patel, Monika Gandhi, Pinal Trivedi, Maharshi Pandya, Nidhi Patel, Nitin Savaliya, Raghawendra Kumar, Dinesh Kumar, Zuber Saiyed, Komal Patel, Labdhi Pandya, Afzal Ansari, Nikha Trivedi, Apurvasinh Puvar, A N Parmar, R D Dixit, A M Kadri, Harsh Bakshi, Chaitanya Joshi, Madhvi Joshi |
| EPI_ISL_524767, EPI_ISL_524768, EPI_ISL_524769, EPI_ISL_524770, EPI_ISL_524771, EPI_ISL_524772, EPI_ISL_524773, EPI_ISL_524774, EPI_ISL_524775, EPI_ISL_524776, EPI_ISL_524778, EPI_ISL_524779, EPI_ISL_524780 | | | |
| see above | Utah Public Health Laboratory | Utah Public Health Laboratory | Erin L. Young, Kelly Oakeson, Tara Gallagher, Michael T. Pyne, E. Susan Slechta, Melanie A. Mallory, Jeffrey B. Stevenson, Salika M. Shakir, David R. Hillyard |
| EPI_ISL_524782 | University Hospital Zürich | Institute of Medical Virology, University of Zurich | Stefan Schmutz, Maryam Zaheri, Verena Kufner, Gabriela Ziltener, Patrick Redli, Fiona Steiner, Jon Huder, Riccarda Capaul, Andrea Zbinden, Jürg Böni, Michael Huber, Alexandra Trkola |
| EPI_ISL_524783, EPI_ISL_524784, EPI_ISL_524785, EPI_ISL_524786, EPI_ISL_524787, EPI_ISL_524788, EPI_ISL_524789, EPI_ISL_524790, EPI_ISL_524791, EPI_ISL_524792, EPI_ISL_524793, EPI_ISL_524794, EPI_ISL_524795, EPI_ISL_524796, EPI_ISL_524797, EPI_ISL_524798, EPI_ISL_524799, EPI_ISL_524800, EPI_ISL_524801 | | | |
| see above | Evandro Chagas Institute | Evandro Chagas Institute | Santos, M.C.; Silva, A.M.; Junior, W.D.C.; Barbagelata, L.S.; Ferreira, J.A.; Sousa, E.M.A.; da Silva, P.S.; Resque, H.R; Martins, L.C.; Sousa Junior, E.C.;Viana, G.M.R |
| EPI_ISL_524802, EPI_ISL_524804, EPI_ISL_524805, EPI_ISL_524806, EPI_ISL_524807, EPI_ISL_524808, EPI_ISL_524809, EPI_ISL_524810, EPI_ISL_524811, EPI_ISL_524812, EPI_ISL_524813, EPI_ISL_524814, EPI_ISL_524815, EPI_ISL_524817, EPI_ISL_524818, EPI_ISL_524819, EPI_ISL_524821, EPI_ISL_524822, EPI_ISL_524824, EPI_ISL_524825, EPI_ISL_524826, EPI_ISL_524828, EPI_ISL_524829, EPI_ISL_524830, EPI_ISL_524831, EPI_ISL_524832, EPI_ISL_524833, EPI_ISL_524834, EPI_ISL_524835, EPI_ISL_524836, EPI_ISL_524837, EPI_ISL_524838, EPI_ISL_524840, EPI_ISL_524841, EPI_ISL_524842, EPI_ISL_524843, EPI_ISL_524844, EPI_ISL_524845, EPI_ISL_524846, EPI_ISL_524848, EPI_ISL_524849, EPI_ISL_524850, EPI_ISL_524851, EPI_ISL_524852, EPI_ISL_524853, EPI_ISL_524855, EPI_ISL_524856, EPI_ISL_524857, EPI_ISL_524858, EPI_ISL_524859, EPI_ISL_524861, EPI_ISL_524862, EPI_ISL_524863, EPI_ISL_524864, EPI_ISL_524867, EPI_ISL_524868, EPI_ISL_524869, EPI_ISL_524870, EPI_ISL_524871, EPI_ISL_524872 | | | |
| see above | Utah Public Health Laboratory | Utah Public Health Laboratory | Erin L. Young, Kelly Oakeson, Tara Gallagher, Michael T. Pyne, E. Susan Slechta, Melanie A. Mallory, Jeffrey B. Stevenson, Salika M. Shakir, David R. Hillyard |

| | | | |
|---|---|---|---|
| EPI_ISL_524873, EPI_ISL_524875, EPI_ISL_524876, EPI_ISL_524877, EPI_ISL_524878, EPI_ISL_524879, EPI_ISL_524880, EPI_ISL_524881, EPI_ISL_524882, EPI_ISL_524884, EPI_ISL_524885, EPI_ISL_524886, EPI_ISL_524887, EPI_ISL_524888, EPI_ISL_524889, EPI_ISL_524890, EPI_ISL_524891, EPI_ISL_524892, EPI_ISL_524893, EPI_ISL_524894, EPI_ISL_524895, EPI_ISL_524896, EPI_ISL_524897, EPI_ISL_524898, EPI_ISL_524899, EPI_ISL_524900, EPI_ISL_524901 | | | |
| see above | MD PHL | MD PHL | Maryland Department of Health Laboratories Administration |
| EPI_ISL_524905, EPI_ISL_524908, EPI_ISL_524910, EPI_ISL_524911, EPI_ISL_524915, EPI_ISL_524917, EPI_ISL_524919, EPI_ISL_524923, EPI_ISL_524925, EPI_ISL_524926, EPI_ISL_524927, EPI_ISL_524928, EPI_ISL_524931, EPI_ISL_524932, EPI_ISL_524933, EPI_ISL_524934, EPI_ISL_524935, EPI_ISL_524938, EPI_ISL_524939, EPI_ISL_524940, EPI_ISL_524942, EPI_ISL_524943, EPI_ISL_524944, EPI_ISL_524945, EPI_ISL_524946, EPI_ISL_524948, EPI_ISL_524949, EPI_ISL_524950, EPI_ISL_524952, EPI_ISL_524954, EPI_ISL_524955, EPI_ISL_524957, EPI_ISL_524958, EPI_ISL_524959, EPI_ISL_524960, EPI_ISL_524963, EPI_ISL_524964, EPI_ISL_524965, EPI_ISL_524968, EPI_ISL_524969, EPI_ISL_524971, EPI_ISL_524976, EPI_ISL_524979, EPI_ISL_524980, EPI_ISL_524984, EPI_ISL_524985, EPI_ISL_524986, EPI_ISL_524988, EPI_ISL_524990, EPI_ISL_524991, EPI_ISL_524992, EPI_ISL_524993, EPI_ISL_524994, EPI_ISL_524995, EPI_ISL_524996, EPI_ISL_524997, EPI_ISL_524999, EPI_ISL_525002, EPI_ISL_525010, EPI_ISL_525011, EPI_ISL_525014, EPI_ISL_525016, EPI_ISL_525020, EPI_ISL_525021, EPI_ISL_525022, EPI_ISL_525023, EPI_ISL_525025, EPI_ISL_525027, EPI_ISL_525028, EPI_ISL_525029, EPI_ISL_525030, EPI_ISL_525032, EPI_ISL_525033, EPI_ISL_525034, EPI_ISL_525036, EPI_ISL_525037, EPI_ISL_525041, EPI_ISL_525042, EPI_ISL_525043, EPI_ISL_525044, EPI_ISL_525045, EPI_ISL_525048, EPI_ISL_525049, EPI_ISL_525050, EPI_ISL_525052, EPI_ISL_525058, EPI_ISL_525060, EPI_ISL_525062, EPI_ISL_525065, EPI_ISL_525068, EPI_ISL_525070, EPI_ISL_525072, EPI_ISL_525074, EPI_ISL_525076, EPI_ISL_525077, EPI_ISL_525079, EPI_ISL_525080, EPI_ISL_525082, EPI_ISL_525083, EPI_ISL_525092, EPI_ISL_525094, EPI_ISL_525102, EPI_ISL_525103, EPI_ISL_525104, EPI_ISL_525106, EPI_ISL_525108, EPI_ISL_525109, EPI_ISL_525110, EPI_ISL_525124, EPI_ISL_525128, EPI_ISL_525129, EPI_ISL_525131, EPI_ISL_525133, EPI_ISL_525136, EPI_ISL_525138, EPI_ISL_525143, EPI_ISL_525148, EPI_ISL_525149, EPI_ISL_525150, EPI_ISL_525151, EPI_ISL_525152, EPI_ISL_525154, EPI_ISL_525155, EPI_ISL_525158, EPI_ISL_525159, EPI_ISL_525160, EPI_ISL_525162, EPI_ISL_525166, EPI_ISL_525173, EPI_ISL_525177, EPI_ISL_525180, EPI_ISL_525183, EPI_ISL_525186, EPI_ISL_525187 | | | |
| see above | Utah Public Health Laboratory | Utah Public Health Laboratory | Erin L. Young, Kelly Oakeson, Tara Gallagher, Michael T. Pyne, E. Susan Slechta, Melanie A. Mallory, Jeffrey B. Stevenson, Salika M. Shakir, David R. Hillyard |
| EPI_ISL_525189, EPI_ISL_525190, EPI_ISL_525191, EPI_ISL_525192, EPI_ISL_525193, EPI_ISL_525194, EPI_ISL_525195, EPI_ISL_525196, EPI_ISL_525197, EPI_ISL_525198, EPI_ISL_525199, EPI_ISL_525200, EPI_ISL_525201, EPI_ISL_525202, EPI_ISL_525203, EPI_ISL_525204, EPI_ISL_525205 | | | |
| see above | Virginia DCLS | Virginia DCLS | Virginia DCLS |
| EPI_ISL_525206 | Laboratorio de Referencia Nacional de Virus Respiratorio. Centro Nacional de Salud Publica. Instituto Nacional de Salud Peru. | Laboratorio de Referencia Nacional de Biotecnología y Biología Molecular. Centro Nacional de Salud Publica. Instituto Nacional de Salud Peru. | Carlos Padilla Rojas, Karolyn Vega Chozo, Priscila Lope Pari, Omar Caceres Rey, Marco Galarza Perez, Maribel Huaringa Nuñez, Johanna Balbuena Torres, Henri Bailon Calderon, Nancy Rojas Serrano. |
| EPI_ISL_525208, EPI_ISL_525209, EPI_ISL_525211, EPI_ISL_525213, EPI_ISL_525215, EPI_ISL_525216, EPI_ISL_525217, EPI_ISL_525219, EPI_ISL_525221, EPI_ISL_525223, EPI_ISL_525224, EPI_ISL_525225, EPI_ISL_525227, EPI_ISL_525228, EPI_ISL_525229, EPI_ISL_525231, EPI_ISL_525233, EPI_ISL_525236, EPI_ISL_525237, EPI_ISL_525239, EPI_ISL_525241, EPI_ISL_525242, EPI_ISL_525244, EPI_ISL_525246, EPI_ISL_525247, EPI_ISL_525248, EPI_ISL_525250, EPI_ISL_525251, EPI_ISL_525252, EPI_ISL_525255, EPI_ISL_525256, EPI_ISL_525258, EPI_ISL_525259, EPI_ISL_525262, EPI_ISL_525263, EPI_ISL_525265, EPI_ISL_525266, EPI_ISL_525268, EPI_ISL_525270, EPI_ISL_525271, EPI_ISL_525273, EPI_ISL_525275, EPI_ISL_525277, EPI_ISL_525280, EPI_ISL_525281, EPI_ISL_525282, EPI_ISL_525283, EPI_ISL_525284, EPI_ISL_525286, EPI_ISL_525287, EPI_ISL_525288, EPI_ISL_525290, EPI_ISL_525291, EPI_ISL_525292, EPI_ISL_525293, EPI_ISL_525295, EPI_ISL_525297, EPI_ISL_525299, EPI_ISL_525300, EPI_ISL_525302, EPI_ISL_525304, EPI_ISL_525305, EPI_ISL_525306, EPI_ISL_525307, EPI_ISL_525309, EPI_ISL_525310, EPI_ISL_525311, EPI_ISL_525312, EPI_ISL_525313, EPI_ISL_525316, EPI_ISL_525317, EPI_ISL_525319, EPI_ISL_525320, EPI_ISL_525321, EPI_ISL_525322, EPI_ISL_525325, EPI_ISL_525326, EPI_ISL_525328, EPI_ISL_525330, EPI_ISL_525334 | | | |
| see above | Utah Public Health Laboratory | Utah Public Health Laboratory | Erin L. Young, Kelly Oakeson, Tara Gallagher, Michael T. Pyne, E. Susan Slechta, Melanie A. Mallory, Jeffrey B. Stevenson, Salika M. Shakir, David R. Hillyard |
| EPI_ISL_525338, EPI_ISL_525339, EPI_ISL_525340, EPI_ISL_525343, EPI_ISL_525344, EPI_ISL_525345, EPI_ISL_525346, EPI_ISL_525347, EPI_ISL_525348, EPI_ISL_525350, EPI_ISL_525351 | | | |
| see above | Utah Public Health Laboratory | Utah Public Health Laboratory | Erin Young, Kelly Oakeson |
| EPI_ISL_525352, EPI_ISL_525353, EPI_ISL_525356, EPI_ISL_525357, EPI_ISL_525358, EPI_ISL_525359, EPI_ISL_525360, EPI_ISL_525361, EPI_ISL_525362, EPI_ISL_525364, EPI_ISL_525365, EPI_ISL_525366, EPI_ISL_525367, EPI_ISL_525368, EPI_ISL_525369, EPI_ISL_525370, EPI_ISL_525371, EPI_ISL_525372, EPI_ISL_525373, EPI_ISL_525374, EPI_ISL_525375, EPI_ISL_525376, EPI_ISL_525382, EPI_ISL_525383, EPI_ISL_525384, EPI_ISL_525385, EPI_ISL_525386, EPI_ISL_525387, EPI_ISL_525388, EPI_ISL_525390, EPI_ISL_525394, EPI_ISL_525396, EPI_ISL_525397, EPI_ISL_525398, EPI_ISL_525399, EPI_ISL_525400, EPI_ISL_525401, EPI_ISL_525402, EPI_ISL_525403, EPI_ISL_525404, EPI_ISL_525405, EPI_ISL_525406, EPI_ISL_525407, EPI_ISL_525408, EPI_ISL_525409, EPI_ISL_525410, EPI_ISL_525411, EPI_ISL_525412, EPI_ISL_525413, EPI_ISL_525414, EPI_ISL_525415, EPI_ISL_525416, EPI_ISL_525417, EPI_ISL_525418 | | | |
| see above | National Virus Reference Laboratory | National Virus Reference Laboratory | Michael Carr, Gabriel Gonzalez, Jonathan Dean, Aditi Chaturvedi, Suzie Coughlan, Cillian F De Gascun |
| EPI_ISL_525419 | GMERS Medical College and Hospital, Gandhinagar | Gujarat Biotechnology Research Centre | Nitin Savaliya, Raghawendra Kumar, Dinesh Kumar, Zuber Saiyed, Komal Patel, Labdhi Pandya, Afzal Ansari, Nikha Trivedi, Seema Bhatt, Gaurishankar Shrimali, Bhavesh Modi, Bharti Rajani, Apurvasinh Puvar, Janvi Raval, Zarna Patel, Monika Gandhi, Pinal Trivedi, Maharshi Pandya, Nidhi Patel, R D Dixit, A M Kadri, Harsh Bakshi, Chaitanya Joshi, Madhvi Joshi |
| EPI_ISL_525420 | GMERS Medical College and Hospital, Gandhinagar | Gujarat Biotechnology Research Centre | Raghawendra Kumar, Dinesh Kumar, Zuber Saiyed, Komal Patel, Labdhi Pandya, Afzal Ansari, Nikha Trivedi, Seema Bhatt, Gaurishankar Shrimali, Bhavesh Modi, Bharti Rajani, Apurvasinh Puvar, Janvi Raval, Zarna Patel, Monika Gandhi, Pinal Trivedi, Maharshi Pandya, Nidhi Patel, Nitin Savaliya, R D Dixit, A M Kadri, Harsh Bakshi, Chaitanya Joshi, Madhvi Joshi |
| EPI_ISL_525421 | B.J. Medical College and Civil hospital, Ahmedabad | Gujarat Biotechnology Research Centre | Zarna Patel, Monika Gandhi, Pinal Trivedi, Maharshi Pandya, Nidhi Patel, Nitin Savaliya, Raghawendra Kumar, Dinesh Kumar, Zuber Saiyed, Komal Patel, Labdhi Pandya, Afzal Ansari, Nikha Trivedi, Pranay Shah, Kamlesh J Upadhyay, Sanjay Kapadia, Apurvasinh Puvar, Janvi Raval, R D Dixit, A M Kadri, Harsh Bakshi, Chaitanya Joshi, Madhvi Joshi |
| EPI_ISL_525422 | B.J. Medical College and Civil hospital, Ahmedabad | Gujarat Biotechnology Research Centre | Monika Gandhi, Pinal Trivedi, Maharshi Pandya, Nidhi Patel, Nitin Savaliya, Raghawendra Kumar, Dinesh Kumar, Zuber Saiyed, Komal Patel, Labdhi Pandya, Afzal Ansari, Nikha Trivedi, Pranay Shah, Kamlesh J Upadhyay, Sanjay Kapadia, Apurvasinh Puvar, Janvi Raval, Zarna Patel, R D Dixit, A M Kadri, Harsh Bakshi, Chaitanya Joshi, Madhvi Joshi |
| EPI_ISL_525423 | Oman-National Influenza Center | Biotechnology & OMICs Laboratory | Sajjad Asaf, Samiha Al-Kharusi, Ahmed Al-Harrasi, Samira Al-Mahruqi, Adil Khan, Ahmed Al-Rawahi, Abdul Latif Khan, Amina Al-Jardani, Hanan Al-Kindi, Intisar Al-Shukri, Ahlam Al-Amri, Aisha Al-Amri, Aisha Al-Busaidi, Adil Al-Wahaibi, Seif Al-Abri. |
| EPI_ISL_525424 | Oman-National Influenza Center | Biotechnology & OMICs Laboratory | Samira Al-Mahruqi, Abdul Latif Khan, Samiha Al-Kharusi, Adil Khan , Ahmed Al-Rawahi, Sajjad Asaf, Amina Al-Jardani, Hanan Al-Kindi, Intisar Al-Shukri, Ahlam Al-Amri, Aisha Al-Amri, Aisha Al-Busaidi, Adil Al-Wahaibi, Seif Al-Abri, Ahmed Al-Harrasi |
| EPI_ISL_525425 | Oman-National Influenza Center | Biotechnology & OMICs Laboratory | Samira Al-Mahruqi, Abdul Latif Khan, Samiha Al-Kharusi, Adil Khan , Ahmed Al-Rawahi, Sajjad Asaf, Amina Al-Jardani, Hanan Al-Kindi, Intisar Al-Shukri, Adil Al-Wahaibi, Seif Al-Abri, Ahmed Al-Harrasi |
| EPI_ISL_525426, EPI_ISL_525427, EPI_ISL_525428 | Oman-National Influenza Center | Biotechnology & OMICs Laboratory | Sajjad Asaf, Samiha Al-Kharusi, Ahmed Al-Harrasi, Samira Al-Mahruqi, Adil Khan, Ahmed Al-Rawahi, Abdul Latif Khan, Amina Al-Jardani, Hanan Al-Kindi, Intisar Al-Shukri, Ahlam Al-Amri, Aisha Al-Amri, Aisha Al-Busaidi, Adil Al-Wahaibi, Seif Al-Abri. |
| EPI_ISL_525429 | Oman-National Influenza Center | Biotechnology & OMICs Laboratory | Samira Al-Mahruqi, Abdul Latif Khan, Samiha Al-Kharusi, Adil Khan , Ahmed Al-Rawahi, Sajjad Asaf, Amina Al-Jardani, Hanan Al-Kindi, Intisar Al-Shukri, Adil Al-Wahaibi, Seif Al-Abri, Ahmed Al-Harrasi |
| EPI_ISL_525430 | Institute of Microbiology, Universidad San Francisco de Quito | Institute of Microbiology, Universidad San Francisco de Quito | Juan José Guadalupe, Monica Becerra-Wong, Belén Prado-Vivar, Sully Márquez, Bernardo Gutiérrez, Verónica Barragán, Patricio Rojas-Silva, Gabriel Trueba, Michelle Grunauer, Paúl Cárdenas |
| EPI_ISL_525431, EPI_ISL_525432 | Institute of Microbiology, Universidad San Francisco de Quito | Institute of Microbiology, Universidad San Francisco de Quito | Juan José Guadalupe, Monica Becerra-Wong, Belén Prado-Vivar, Sully Márquez, Ligia Briceño, Carlos Mena, Nabih Dahik, Bernardo Gutiérrez, Verónica Barragán, Patricio Rojas-Silva, Gabriel Trueba, Michelle Grunauer, Paúl Cárdenas |
| EPI_ISL_525433, EPI_ISL_525434 | Institute of Microbiology, Universidad San Francisco de Quito | Institute of Microbiology, Universidad San Francisco de Quito | Juan José Guadalupe, Monica Becerra-Wong, Belén Prado-Vivar, Sully Márquez, Bernardo Gutiérrez, Eulalia Pazmiño, Carolina Pacheco, Damaris Zandoya, Carlos Mena, Nabih Dahik, Verónica Barragán, Patricio Rojas-Silva, Gabriel Trueba, Michelle Grunauer, Paúl Cárdenas |
| EPI_ISL_525435 | Institute of Microbiology, Universidad San Francisco de Quito | Institute of Microbiology, Universidad San Francisco de Quito | Juan José Guadalupe, Monica Becerra-Wong, Belén Prado-Vivar, Sully Márquez, Bernardo Gutiérrez, Yomara Napa, Edy Quizhpe, Carlos Mena, Nabih Dahik, Verónica Barragán, Patricio Rojas-Silva, Gabriel Trueba, Michelle Grunauer, Paúl Cárdenas |
| EPI_ISL_525436, EPI_ISL_525437 | Institute of Microbiology, Universidad San Francisco de Quito | Institute of Microbiology, Universidad San Francisco de Quito | Juan José Guadalupe, Monica Becerra-Wong, Belén Prado-Vivar, Sully Márquez, Bernardo Gutiérrez, Dayron Brossard, Carlos Mena, Nabih Dahik, Verónica Barragán, Patricio Rojas-Silva, Gabriel Trueba, Michelle Grunauer, Paúl Cárdenas |
| EPI_ISL_525438 | Institute of Microbiology, Universidad San Francisco de Quito | Institute of Microbiology, Universidad San Francisco de Quito | Diego Egas, Juan José Guadalupe, Monica Becerra-Wong, Belén Prado-Vivar, Sully Márquez, Bernardo Gutiérrez, Stalin Castillo, Eddy Chavez, Francisco Rodriguez, Verónica Barragán, Patricio Rojas-Silva, Gabriel Trueba, Michelle Grunauer, Paúl Cárdenas |
| EPI_ISL_525466 | Oman-National Influenza Center | Biotechnology & OMICs Laboratory | Samiha Al-Kharusi, Sajjad Asaf, Abdul Latif Khan, Samira Al-Mahruqi, Adil Khan, Ahmed Al-Rawahi, Amina Al-Jardani, Hanan Al-Kindi, Intisar Al-Shukri, Aisha Al-Busaidi, Adil Al-Wahaibi, Seif Al-Abri, Ahmed Al-Harrasi |
| EPI_ISL_525467 | Universidad Iberoamericana | International Centre for Genetic Engineering and Biotechnology (ICGEB) and ARGO Open Lab Platform | Robert Paulino-Ramirez, Eileen Riego, Alejandro Vallejo Degaudenzi, Victor Virgilio Calderon, Leandro Tapia, Patricia Leon, Danilo Licastro, Simeone Dal Monego, Sreejith Rajasekharan and Alessandro Marcello. |
| EPI_ISL_525468 | Universidad Iberoamericana | International Centre for Genetic Engineering and Biotechnology (ICGEB) and ARGO Open Lab Platform | Robert Paulino-Ramirez, Eileen Riego, Alejandro Vallejo Degaudenzi, Victor Virgilio Calderon, Leandro Tapia,Patricia Leon,Danilo Licastro, Simeone Dal Monego, Sreejith Rajasekharan and Alessandro Marcello. |
| EPI_ISL_525469 | Universidad Iberoamericana | International Centre for Genetic Engineering and Biotechnology (ICGEB) and ARGO Open Lab Platform | Robert Paulino-Ramirez, Eileen Riego, Alejandro Vallejo Degaudenzi, Victor Virgilio Calderon, Leandro Tapia, Patricia Leon, Danilo Licastro, Simeone Dal Monego, Sreejith Rajasekharan and Alessandro Marcello. |
| EPI_ISL_525474 | Centre for Dengue Research | Centre for Dengue Research, USJ, SL | Chandima Jeewandara, Deshni Jayathilaka, Dinuka Ariyaratne, Laksiri Gomes, Diyanath Ranasinghe, Dinuka Guruge, Ruwan Wijayamuni, Gathsaurie Neelika Malavige |
| EPI_ISL_525476 | Centre for Dengue Research | Centre for Dengue Research | Chandima Jeewandara, Deshni Jayathilaka, Dinuka Ariyaratne, Laksiri Gomes, Diyanath Ranasinghe, Dinuka Guruge, Ruwan Wijayamuni, Gathsaurie Neelika Malavige |
| EPI_ISL_525478, EPI_ISL_525479, EPI_ISL_525481 | Centre for Dengue Research | Centre for Dengue Research | Chandima Jeewandara, Deshni Jayathilaka, Dinuka Ariyaratne, Laksiri Gomes, Diyanath Ranasinghe, Ananda Wijewickrama, Eranga Narangoda, Damayanthi Idampitiya, Gathsaurie Neelika Malavige |
| EPI_ISL_525486, EPI_ISL_525489 | Centre for Dengue Research | Centre for Dengue Research | Chandima Jeewandara, Deshni Jayathilaka, Dinuka Ariyaratne, Laksiri Gomes, Diyanath Ranasinghe, Ananda Wijewickrama, Malika Karunaratne, Eranga Narangoda, Damayanthi Idampitiya, Gathsaurie Neelika Malavige |
| EPI_ISL_525492 | RSUP dr. SOERADJI TIRTONEGORO | Genetics Working Group (Pokja Genetik) Faculty of Medicine, Public Health and Nursing Universitas Gadjah Mada (FK-KMK UGM), Disease Investigation Center Wates Ministry of Agriculture Indonesia, Department of Microbiology FK-KMK UGM, Laboratorium Diagnostik Yayasan Tahija World Mosquito Program (WMP) Yogyakarta Center for Tropical Medicine FK-KMK UGM, Integrated Research Center FK-KMK UGM, Department of Computer Science and Electronics FMIPA UGM | Gunadi, Hendra Wibawa, . Marcellus, Mohamad S. Hakim, Edwin W. Daniwijaya, Ludhang P. Rizki, Endah Supriyati, Eggi Arguni, Titik Nuryastuti, Tri Wibawa, Dwi AA Nugrahaningsih, Afiahayati , . Siswanto, Kurniyanto, Indah Juliana, Alvin S. Kalim, Dwiki Afandy |
| EPI_ISL_525495, EPI_ISL_525496 | Laboratory of Molecular Virology of the International Centre for Genetic Engineering and Biotechnology (ICGEB) | ARGO Open Lab Platform for Genome Sequencing | Licastro D, Rajasekharan S, Dal Monego S, Segat L, D'Agaro P, Marcello A |
| EPI_ISL_525536, EPI_ISL_525537 | CNR Virus des Infections Respiratoires - France SUD | CNR Virus des Infections Respiratoires - France SUD | Antonin Bal, Gregory Destras, Gwendolyne Burfin, Solenne Brun, Alexandre Gaymard, Maude Bouscambert-Duchamp, Florence Morfin-Sherpa, Martine Valette, Bruno Lina, Laurence Josset |
| EPI_ISL_525539 | Centre Hospitalier de Bourg en Bresse | CNR Virus des Infections Respiratoires - France SUD | Antonin Bal, Gregory Destras, Gwendolyne Burfin, Solenne Brun, Alexandre Gaymard, Maude Bouscambert-Duchamp, Florence Morfin-Sherpa, Martine Valette, Bruno Lina, Laurence Josset |
| EPI_ISL_525540, EPI_ISL_525541, EPI_ISL_525542, EPI_ISL_525543 | CNR Virus des Infections Respiratoires - France SUD | CNR Virus des Infections Respiratoires - France SUD | Antonin Bal, Gregory Destras, Gwendolyne Burfin, Solenne Brun, Alexandre Gaymard, Maude Bouscambert-Duchamp, Florence Morfin-Sherpa, Martine Valette, Bruno Lina, Laurence Josset |
| EPI_ISL_525553, EPI_ISL_525557, EPI_ISL_525560, EPI_ISL_525561, EPI_ISL_525569, EPI_ISL_525570, EPI_ISL_525573, EPI_ISL_525574 | Istituto Zooprofilattico Sperimentale Puglia e Basilicata; Dipartimento di Bioscienze, Biotecnologie e Biofarmaceutica dell'Università degli Studi di Bari "A.Moro"; Istituto di Biomembrane. Bioenergetica e Biotecnologie Molecolari del Consiglio Nazionale delle Ricerche di Bari | Beaconlab (Bioinformatics, Evolution and Comparative Genomics lab), Dept of Biosciences, University on Milan | Parisi A.,Pesole G., Manzari C., Chiara M |
| EPI_ISL_525575, EPI_ISL_525576, EPI_ISL_525577, EPI_ISL_525578, EPI_ISL_525579, EPI_ISL_525580, EPI_ISL_525582, EPI_ISL_525583, EPI_ISL_525584, EPI_ISL_525585, EPI_ISL_525587, EPI_ISL_525588, EPI_ISL_525589, EPI_ISL_525590, EPI_ISL_525591, EPI_ISL_525592, EPI_ISL_525593, EPI_ISL_525594, EPI_ISL_525595, EPI_ISL_525596, EPI_ISL_525597, EPI_ISL_525598, EPI_ISL_525599, EPI_ISL_525600, EPI_ISL_525601, EPI_ISL_525602, EPI_ISL_525605, EPI_ISL_525606, EPI_ISL_525607, EPI_ISL_525608, EPI_ISL_525609, EPI_ISL_525610, EPI_ISL_525611, EPI_ISL_525612, EPI_ISL_525613, EPI_ISL_525614, EPI_ISL_525615, EPI_ISL_525616, EPI_ISL_525617, EPI_ISL_525618, EPI_ISL_525619, EPI_ISL_525620, EPI_ISL_525621, EPI_ISL_525622, EPI_ISL_525623, EPI_ISL_525624, EPI_ISL_525625, EPI_ISL_525626, | | | |

| | | | |
|---|---|---|---|
| EPI_ISL_525627, EPI_ISL_525628, EPI_ISL_525629, EPI_ISL_525630, EPI_ISL_525631, EPI_ISL_525632, EPI_ISL_525633, EPI_ISL_525634, EPI_ISL_525635, EPI_ISL_525636, EPI_ISL_525637, EPI_ISL_525638, EPI_ISL_525639, EPI_ISL_525640, EPI_ISL_525641, EPI_ISL_525642, EPI_ISL_525643, EPI_ISL_525644, EPI_ISL_525645, EPI_ISL_525646, EPI_ISL_525647, EPI_ISL_525648, EPI_ISL_525649, EPI_ISL_525650, EPI_ISL_525651, EPI_ISL_525652, EPI_ISL_525653, EPI_ISL_525654, EPI_ISL_525655, EPI_ISL_525656, EPI_ISL_525657, EPI_ISL_525658, EPI_ISL_525659, EPI_ISL_525660, EPI_ISL_525663, EPI_ISL_525664, EPI_ISL_525665, EPI_ISL_525666, EPI_ISL_525667, EPI_ISL_525668, EPI_ISL_525669, EPI_ISL_525670, EPI_ISL_525671, EPI_ISL_525672, EPI_ISL_525674, EPI_ISL_525675, EPI_ISL_525676, EPI_ISL_525677, EPI_ISL_525678, EPI_ISL_525679, EPI_ISL_525680, EPI_ISL_525681, EPI_ISL_525682, EPI_ISL_525683, EPI_ISL_525684, EPI_ISL_525685, EPI_ISL_525686 | | | |
| see above | Wadsworth Center, New York State Department of Health | Wadsworth Center, New York State Department of Health | Kirsten St. George, Daryl M. Lamson, Sara Griesemer, Jonathan Plitnick, Navjot Singh, Matthew D. Shudt, Erica Lasek-Nesselquist |
| EPI_ISL_525700, EPI_ISL_525701 | Seattle Flu Study | Seattle Flu Study | Deborah A. Nickerson, Chris D. Frazar, Jover Lee, Benjamin Pelle, Matthew Richardson, Amanda Adler, Elisabeth Brandstetter, Peter D. Han, Kairsten Fay, Misja Ilcisin, Kirsten Lacombe, Thomas R. Sibley, Melissa Truong, Caitlin R. Wolf, Michael Boeckh, Janet A. Englund, Michael Famulare, Barry R. Lutz, Mark J. Rieder, Lea M. Starita, Matthew Thompson, Jay Shendure, Trevor Bedford, Helen Y. Chu |
| EPI_ISL_525702, EPI_ISL_525703, EPI_ISL_525704, EPI_ISL_525705 | Seattle Flu Study | Seattle Flu Study | Deborah A. Nickerson, Chris D. Frazar, Jover Lee, Benjamin Pelle, Matthew Richardson, Amanda Adler, Elisabeth Brandstetter, Peter D. Han, Kairsten Fay, Misja Ilcisin, Kirsten Lacombe, Thomas R. Sibley, Melissa Truong, Caitlin R. Wolf, Karen Cowgill, Stephanie Schrag, Jeff Duchin, Michael Boeckh, Janet A. Englund, Michael Famulare, Barry R. Lutz, Mark J. Rieder, Lea M. Starita, Matthew Thompson, Helen Y. Chu, Trevor Bedford, Jay Shendure |
| EPI_ISL_525706 | Seattle Flu Study | Seattle Flu Study | Deborah A. Nickerson, Chris D. Frazar, Jover Lee, Benjamin Pelle, Matthew Richardson, Amanda Adler, Elisabeth Brandstetter, Peter D. Han, Kairsten Fay, Misja Ilcisin, Kirsten Lacombe, Thomas R. Sibley, Melissa Truong, Caitlin R. Wolf, Michael Boeckh, Janet A. Englund, Michael Famulare, Barry R. Lutz, Mark J. Rieder, Lea M. Starita, Matthew Thompson, Jay Shendure, Trevor Bedford, Helen Y. Chu |
| EPI_ISL_525708, EPI_ISL_525709, EPI_ISL_525710, EPI_ISL_525711, EPI_ISL_525712, EPI_ISL_525713, EPI_ISL_525714, EPI_ISL_525715, EPI_ISL_525716, EPI_ISL_525717, EPI_ISL_525718, EPI_ISL_525719, EPI_ISL_525720, EPI_ISL_525721, EPI_ISL_525722, EPI_ISL_525723, EPI_ISL_525724, EPI_ISL_525725, EPI_ISL_525726 | | | |
| see above | Seattle Flu Study | Seattle Flu Study | Deborah A. Nickerson, Chris D. Frazar, Jover Lee, Benjamin Pelle, Matthew Richardson, Amanda Adler, Elisabeth Brandstetter, Peter D. Han, Kairsten Fay, Misja Ilcisin, Kirsten Lacombe, Thomas R. Sibley, Melissa Truong, Caitlin R. Wolf, Karen Cowgill, Stephanie Schrag, Jeff Duchin, Michael Boeckh, Janet A. Englund, Michael Famulare, Barry R. Lutz, Mark J. Rieder, Lea M. Starita, Matthew Thompson, Helen Y. Chu, Trevor Bedford, Jay Shendure |
| EPI_ISL_525728 | Seattle Flu Study | Seattle Flu Study | Deborah A. Nickerson, Chris D. Frazar, Jover Lee, Benjamin Pelle, Matthew Richardson, Amanda Adler, Elisabeth Brandstetter, Peter D. Han, Kairsten Fay, Misja Ilcisin, Kirsten Lacombe, Thomas R. Sibley, Melissa Truong, Caitlin R. Wolf, Michael Boeckh, Janet A. Englund, Michael Famulare, Barry R. Lutz, Mark J. Rieder, Lea M. Starita, Matthew Thompson, Jay Shendure, Trevor Bedford, Helen Y. Chu |
| EPI_ISL_525729, EPI_ISL_525730, EPI_ISL_525731, EPI_ISL_525732, EPI_ISL_525733, EPI_ISL_525734, EPI_ISL_525735, EPI_ISL_525736, EPI_ISL_525737, EPI_ISL_525738, EPI_ISL_525739, EPI_ISL_525741 | | | |
| see above | Seattle Flu Study | Seattle Flu Study | Deborah A. Nickerson, Chris D. Frazar, Jover Lee, Benjamin Pelle, Matthew Richardson, Amanda Adler, Elisabeth Brandstetter, Peter D. Han, Kairsten Fay, Misja Ilcisin, Kirsten Lacombe, Thomas R. Sibley, Melissa Truong, Caitlin R. Wolf, Karen Cowgill, Stephanie Schrag, Jeff Duchin, Michael Boeckh, Janet A. Englund, Michael Famulare, Barry R. Lutz, Mark J. Rieder, Lea M. Starita, Matthew Thompson, Helen Y. Chu, Trevor Bedford, Jay Shendure |
| EPI_ISL_525742 | Seattle Flu Study | Seattle Flu Study | Deborah A. Nickerson, Chris D. Frazar, Jover Lee, Benjamin Pelle, Matthew Richardson, Amanda Adler, Elisabeth Brandstetter, Peter D. Han, Kairsten Fay, Misja Ilcisin, Kirsten Lacombe, Thomas R. Sibley, Melissa Truong, Caitlin R. Wolf, Michael Boeckh, Janet A. Englund, Michael Famulare, Barry R. Lutz, Mark J. Rieder, Lea M. Starita, Matthew Thompson, Jay Shendure, Trevor Bedford, Helen Y. Chu |
| EPI_ISL_525743, EPI_ISL_525744, EPI_ISL_525746, EPI_ISL_525747, EPI_ISL_525748, EPI_ISL_525749, EPI_ISL_525751, EPI_ISL_525753, EPI_ISL_525754, EPI_ISL_525755, EPI_ISL_525756 | | | |
| see above | Seattle Flu Study | Seattle Flu Study | Deborah A. Nickerson, Chris D. Frazar, Jover Lee, Benjamin Pelle, Matthew Richardson, Amanda Adler, Elisabeth Brandstetter, Peter D. Han, Kairsten Fay, Misja Ilcisin, Kirsten Lacombe, Thomas R. Sibley, Melissa Truong, Caitlin R. Wolf, Karen Cowgill, Stephanie Schrag, Jeff Duchin, Michael Boeckh, Janet A. Englund, Michael Famulare, Barry R. Lutz, Mark J. Rieder, Lea M. Starita, Matthew Thompson, Helen Y. Chu, Trevor Bedford, Jay Shendure |
| EPI_ISL_525758, EPI_ISL_525759, EPI_ISL_525760 | Alaska State Virology Laboratory | Alaska State Virology Laboratory | Jack Chen, Ph.D. |
| EPI_ISL_525763, EPI_ISL_525788 | Texas Department of State Health Services | Texas Department of State Health Services | Jenny Zhang, Rashmi Tuladhar, Bonnie Oh, Maliha Rahman, Anita Pokharel, Myong Koag, Chun Wang, Rachel Lee, Grace Kubin |
| EPI_ISL_525802 | OHSU Lab Services Molecular Microbiology Lab | Oregon SARS-CoV-2 Genome Sequencing Center | Brendan L. O'Connell, Ruth V. Nichols, Alec J. Hirsch, Guang Fan, Daniel N. Streblow, William B. Messer, Andrew C. Adey, Benjamin N. Bimber, Brian J. O’Roak |
| EPI_ISL_525803 | OHSU Lab Services Molecular Microbiology Lab | Ginkgo Bioworks Clinical Laboratory | Brendan L. O'Connell, Ruth V. Nichols, Alec J. Hirsch, Guang Fan, Daniel N. Streblow, Malaika Mckenzie-Bennett, James McGann, Jim Griffin, Keith Robison, Alex Plocik, Becky Schilling, Rebecca Littlefield, Michelle Spencer, Birgitte Simen, William B. Messer, Andrew C. Adey, Benjamin N. Bimber, Brian J. O’Roak |
| EPI_ISL_525804, EPI_ISL_525805, EPI_ISL_525806, EPI_ISL_525807 | OHSU Lab Services Molecular Microbiology Lab | Oregon SARS-CoV-2 Genome Sequencing Center | Brendan L. O'Connell, Ruth V. Nichols, Alec J. Hirsch, Guang Fan, Daniel N. Streblow, William B. Messer, Andrew C. Adey, Benjamin N. Bimber, Brian J. O’Roak |
| EPI_ISL_525808, EPI_ISL_525809, EPI_ISL_525813, EPI_ISL_525815, EPI_ISL_525816, EPI_ISL_525817, EPI_ISL_525818, EPI_ISL_525823 | OHSU Lab Services Molecular Microbiology Lab | Ginkgo Bioworks Clinical Laboratory | Brendan L. O'Connell, Ruth V. Nichols, Alec J. Hirsch, Guang Fan, Daniel N. Streblow, Malaika Mckenzie-Bennett, James McGann, Jim Griffin, Keith Robison, Alex Plocik, Becky Schilling, Rebecca Littlefield, Michelle Spencer, Birgitte Simen, William B. Messer, Andrew C. Adey, Benjamin N. Bimber, Brian J. O’Roak |
| EPI_ISL_525825 | OHSU Lab Services Molecular Microbiology Lab | Oregon SARS-CoV-2 Genome Sequencing Center | Brendan L. O'Connell, Ruth V. Nichols, Alec J. Hirsch, Guang Fan, Daniel N. Streblow, William B. Messer, Andrew C. Adey, Benjamin N. Bimber, Brian J. O’Roak |
| EPI_ISL_525826 | OHSU Lab Services Molecular Microbiology Lab | Ginkgo Bioworks Clinical Laboratory | Brendan L. O'Connell, Ruth V. Nichols, Alec J. Hirsch, Guang Fan, Daniel N. Streblow, Malaika Mckenzie-Bennett, James McGann, Jim Griffin, Keith Robison, Alex Plocik, Becky Schilling, Rebecca Littlefield, Michelle Spencer, Birgitte Simen, William B. Messer, Andrew C. Adey, Benjamin N. Bimber, Brian J. O’Roak |
| EPI_ISL_525827 | OHSU Lab Services Molecular Microbiology Lab | Oregon SARS-CoV-2 Genome Sequencing Center | Brendan L. O'Connell, Ruth V. Nichols, Alec J. Hirsch, Guang Fan, Daniel N. Streblow, William B. Messer, Andrew C. Adey, Benjamin N. Bimber, Brian J. O’Roak |
| EPI_ISL_525828, EPI_ISL_525829, EPI_ISL_525830, EPI_ISL_525831, EPI_ISL_525832 | OHSU Lab Services Molecular Microbiology Lab | Ginkgo Bioworks Clinical Laboratory | Brendan L. O'Connell, Ruth V. Nichols, Alec J. Hirsch, Guang Fan, Daniel N. Streblow, Malaika Mckenzie-Bennett, James McGann, Jim Griffin, Keith Robison, Alex Plocik, Becky Schilling, Rebecca Littlefield, Michelle Spencer, Birgitte Simen, William B. Messer, Andrew C. Adey, Benjamin N. Bimber, Brian J. O’Roak |
| EPI_ISL_525833, EPI_ISL_525841 | OHSU Lab Services Molecular Microbiology Lab | Oregon SARS-CoV-2 Genome Sequencing Center | Brendan L. O'Connell, Ruth V. Nichols, Alec J. Hirsch, Guang Fan, Daniel N. Streblow, William B. Messer, Andrew C. Adey, Benjamin N. Bimber, Brian J. O’Roak |
| EPI_ISL_525842, EPI_ISL_525843, EPI_ISL_525845, EPI_ISL_525846 | OHSU Lab Services Molecular Microbiology Lab | Ginkgo Bioworks Clinical Laboratory | Brendan L. O'Connell, Ruth V. Nichols, Alec J. Hirsch, Guang Fan, Daniel N. Streblow, Malaika Mckenzie-Bennett, James McGann, Jim Griffin, Keith Robison, Alex Plocik, Becky Schilling, Rebecca Littlefield, Michelle Spencer, Birgitte Simen, William B. Messer, Andrew C. Adey, Benjamin N. Bimber, Brian J. O’Roak |
| EPI_ISL_525847 | OHSU Lab Services Molecular Microbiology Lab | Oregon SARS-CoV-2 Genome Sequencing Center | Brendan L. O'Connell, Ruth V. Nichols, Alec J. Hirsch, Guang Fan, Daniel N. Streblow, William B. Messer, Andrew C. Adey, Benjamin N. Bimber, Brian J. O’Roak |
| EPI_ISL_525848 | OHSU Lab Services Molecular Microbiology Lab | Ginkgo Bioworks Clinical Laboratory | Brendan L. O'Connell, Ruth V. Nichols, Alec J. Hirsch, Guang Fan, Daniel N. Streblow, Malaika Mckenzie-Bennett, James McGann, Jim Griffin, Keith Robison, Alex Plocik, Becky Schilling, Rebecca Littlefield, Michelle Spencer, Birgitte Simen, William B. Messer, Andrew C. Adey, Benjamin N. Bimber, Brian J. O’Roak |
| EPI_ISL_525849, EPI_ISL_525851 | OHSU Lab Services Molecular Microbiology Lab | Oregon SARS-CoV-2 Genome Sequencing Center | Brendan L. O'Connell, Ruth V. Nichols, Alec J. Hirsch, Guang Fan, Daniel N. Streblow, William B. Messer, Andrew C. Adey, Benjamin N. Bimber, Brian J. O’Roak |
| EPI_ISL_525852, EPI_ISL_525854 | OHSU Lab Services Molecular Microbiology Lab | Ginkgo Bioworks Clinical Laboratory | Brendan L. O'Connell, Ruth V. Nichols, Alec J. Hirsch, Guang Fan, Daniel N. Streblow, Malaika Mckenzie-Bennett, James McGann, Jim Griffin, Keith Robison, Alex Plocik, Becky Schilling, Rebecca Littlefield, Michelle Spencer, Birgitte Simen, William B. Messer, Andrew C. Adey, Benjamin N. Bimber, Brian J. O’Roak |
| EPI_ISL_525856, EPI_ISL_525857, EPI_ISL_525858, EPI_ISL_525860, EPI_ISL_525862 | OHSU Lab Services Molecular Microbiology Lab | Oregon SARS-CoV-2 Genome Sequencing Center | Brendan L. O'Connell, Ruth V. Nichols, Alec J. Hirsch, Guang Fan, Daniel N. Streblow, William B. Messer, Andrew C. Adey, Benjamin N. Bimber, Brian J. O’Roak |
| EPI_ISL_525863, EPI_ISL_525864, EPI_ISL_525867 | OHSU Lab Services Molecular Microbiology Lab | Ginkgo Bioworks Clinical Laboratory | Brendan L. O'Connell, Ruth V. Nichols, Alec J. Hirsch, Guang Fan, Daniel N. Streblow, Malaika Mckenzie-Bennett, James McGann, Jim Griffin, Keith Robison, Alex Plocik, Becky Schilling, Rebecca Littlefield, Michelle Spencer, Birgitte Simen, William B. Messer, Andrew C. Adey, Benjamin N. Bimber, Brian J. O’Roak |
| EPI_ISL_525868 | OHSU Lab Services Molecular Microbiology Lab | Oregon SARS-CoV-2 Genome Sequencing Center | Brendan L. O'Connell, Ruth V. Nichols, Alec J. Hirsch, Guang Fan, Daniel N. Streblow, William B. Messer, Andrew C. Adey, Benjamin N. Bimber, Brian J. O’Roak |
| EPI_ISL_525869 | OHSU Lab Services Molecular Microbiology Lab | Ginkgo Bioworks Clinical Laboratory | Brendan L. O'Connell, Ruth V. Nichols, Alec J. Hirsch, Guang Fan, Daniel N. Streblow, Malaika Mckenzie-Bennett, James McGann, Jim Griffin, Keith Robison, Alex Plocik, Becky Schilling, Rebecca Littlefield, Michelle Spencer, Birgitte Simen, William B. Messer, Andrew C. Adey, Benjamin N. Bimber, Brian J. O’Roak |
| EPI_ISL_525870 | OHSU Lab Services Molecular Microbiology Lab | Oregon SARS-CoV-2 Genome Sequencing Center | Brendan L. O'Connell, Ruth V. Nichols, Alec J. Hirsch, Guang Fan, Daniel N. Streblow, William B. Messer, Andrew C. Adey, Benjamin N. Bimber, Brian J. O’Roak |
| EPI_ISL_525873 | OHSU Lab Services Molecular Microbiology Lab | Ginkgo Bioworks Clinical Laboratory | Brendan L. O'Connell, Ruth V. Nichols, Alec J. Hirsch, Guang Fan, Daniel N. Streblow, Malaika Mckenzie-Bennett, James McGann, Jim Griffin, Keith Robison, Alex Plocik, Becky Schilling, Rebecca Littlefield, Michelle Spencer, Birgitte Simen, William B. Messer, Andrew C. Adey, Benjamin N. Bimber, Brian J. O’Roak |
| EPI_ISL_525874, EPI_ISL_525876 | OHSU Lab Services Molecular Microbiology Lab | Oregon SARS-CoV-2 Genome Sequencing Center | Brendan L. O'Connell, Ruth V. Nichols, Alec J. Hirsch, Guang Fan, Daniel N. Streblow, William B. Messer, Andrew C. Adey, Benjamin N. Bimber, Brian J. O’Roak |
| EPI_ISL_525877 | OHSU Lab Services Molecular Microbiology Lab | Ginkgo Bioworks Clinical Laboratory | Brendan L. O'Connell, Ruth V. Nichols, Alec J. Hirsch, Guang Fan, Daniel N. Streblow, Malaika Mckenzie-Bennett, James McGann, Jim Griffin, Keith Robison, Alex Plocik, Becky Schilling, Rebecca Littlefield, Michelle Spencer, Birgitte Simen, William B. Messer, Andrew C. Adey, Benjamin N. Bimber, Brian J. O’Roak |
| EPI_ISL_525878, EPI_ISL_525880, EPI_ISL_525881, EPI_ISL_525883, EPI_ISL_525885, EPI_ISL_525891, EPI_ISL_525892, EPI_ISL_525893, EPI_ISL_525894, EPI_ISL_525895, EPI_ISL_525896, EPI_ISL_525897, EPI_ISL_525898, EPI_ISL_525899, EPI_ISL_525900, EPI_ISL_525901, EPI_ISL_525902, EPI_ISL_525903, EPI_ISL_525904, EPI_ISL_525905, EPI_ISL_525906, EPI_ISL_525907, EPI_ISL_525908, EPI_ISL_525909, EPI_ISL_525910, EPI_ISL_525911, EPI_ISL_525912, EPI_ISL_525913, EPI_ISL_525914, EPI_ISL_525915, EPI_ISL_525916, EPI_ISL_525917, EPI_ISL_525918, EPI_ISL_525919, EPI_ISL_525920, EPI_ISL_525921, EPI_ISL_525922, EPI_ISL_525923, EPI_ISL_525924, EPI_ISL_525925, EPI_ISL_525927, EPI_ISL_525928, EPI_ISL_525929, EPI_ISL_525930, EPI_ISL_525931, EPI_ISL_525932, EPI_ISL_525933, EPI_ISL_525934, EPI_ISL_525935, EPI_ISL_525936, EPI_ISL_525937, EPI_ISL_525938, EPI_ISL_525939, EPI_ISL_525940, EPI_ISL_525941, EPI_ISL_525942, EPI_ISL_525943, EPI_ISL_525944, EPI_ISL_525945, EPI_ISL_525947, EPI_ISL_525948, EPI_ISL_525949, EPI_ISL_525950, EPI_ISL_525951, EPI_ISL_525952, EPI_ISL_525953, EPI_ISL_525954, EPI_ISL_525955, EPI_ISL_525956, EPI_ISL_525957, EPI_ISL_525958, EPI_ISL_525959, EPI_ISL_525960, EPI_ISL_525961, EPI_ISL_525962, EPI_ISL_525963, EPI_ISL_525964, EPI_ISL_525965, EPI_ISL_525966, EPI_ISL_525967, EPI_ISL_525969, EPI_ISL_525970, EPI_ISL_525971, EPI_ISL_525972, EPI_ISL_525973, EPI_ISL_525974, EPI_ISL_525975, EPI_ISL_525976, EPI_ISL_525977, EPI_ISL_525979, EPI_ISL_525980, EPI_ISL_525981, EPI_ISL_525982, EPI_ISL_525983, EPI_ISL_525984, EPI_ISL_525985, EPI_ISL_525986, EPI_ISL_525987, EPI_ISL_525988, EPI_ISL_525989, EPI_ISL_525990, EPI_ISL_525991, EPI_ISL_525992, EPI_ISL_525993, EPI_ISL_525994, EPI_ISL_525995, EPI_ISL_525996, EPI_ISL_525997, EPI_ISL_525998, EPI_ISL_525999, EPI_ISL_526000, EPI_ISL_526001, EPI_ISL_526002, EPI_ISL_526003, EPI_ISL_526004, EPI_ISL_526005, EPI_ISL_526006, EPI_ISL_526007, EPI_ISL_526008, EPI_ISL_526009, EPI_ISL_526010, EPI_ISL_526011, EPI_ISL_526012, EPI_ISL_526013, EPI_ISL_526014, EPI_ISL_526015, EPI_ISL_526016, EPI_ISL_526017, EPI_ISL_526018, EPI_ISL_526019, EPI_ISL_526020, EPI_ISL_526021, EPI_ISL_526022, EPI_ISL_526024, EPI_ISL_526025, EPI_ISL_526026, EPI_ISL_526027, EPI_ISL_526028, EPI_ISL_526029, EPI_ISL_526030, EPI_ISL_526031, EPI_ISL_526032, EPI_ISL_526033, EPI_ISL_526034, EPI_ISL_526035, EPI_ISL_526036, EPI_ISL_526037, EPI_ISL_526038, EPI_ISL_526039, EPI_ISL_526040, EPI_ISL_526041, EPI_ISL_526043, EPI_ISL_526044, EPI_ISL_526045, EPI_ISL_526046, EPI_ISL_526047, EPI_ISL_526048, EPI_ISL_526049, EPI_ISL_526050, EPI_ISL_526051, EPI_ISL_526052, EPI_ISL_526053, EPI_ISL_526054, EPI_ISL_526055, EPI_ISL_526056, EPI_ISL_526057, EPI_ISL_526058, EPI_ISL_526059, EPI_ISL_526060, EPI_ISL_526061, EPI_ISL_526062, EPI_ISL_526063, EPI_ISL_526064, EPI_ISL_526065, EPI_ISL_526066, EPI_ISL_526067, EPI_ISL_526068, EPI_ISL_526069, EPI_ISL_526071, EPI_ISL_526072, EPI_ISL_526073, EPI_ISL_526074, EPI_ISL_526075, EPI_ISL_526076, EPI_ISL_526078, EPI_ISL_526079, EPI_ISL_526080, EPI_ISL_526082, EPI_ISL_526083, EPI_ISL_526084, EPI_ISL_526085, EPI_ISL_526086 | | | |
| see above | OHSU Lab Services Molecular Microbiology Lab | Oregon SARS-CoV-2 Genome Sequencing Center | Brendan L. O'Connell, Ruth V. Nichols, Alec J. Hirsch, Guang Fan, Daniel N. Streblow, William B. Messer, Andrew C. Adey, Benjamin N. Bimber, Brian J. O’Roak |
| EPI_ISL_526087, EPI_ISL_526088, EPI_ISL_526089 | OHSU Lab Services Molecular Microbiology Lab | Ginkgo Bioworks Clinical Laboratory | Brendan L. O'Connell, Ruth V. Nichols, Alec J. Hirsch, Guang Fan, Daniel N. Streblow, Malaika Mckenzie-Bennett, James McGann, Jim Griffin, Keith Robison, Alex Plocik, Becky Schilling, Rebecca Littlefield, Michelle Spencer, Birgitte Simen, William B. Messer, Andrew C. Adey, Benjamin N. Bimber, Brian J. O’Roak |
| EPI_ISL_526091 | OHSU Lab Services Molecular Microbiology Lab | Oregon SARS-CoV-2 Genome Sequencing Center | Brendan L. O'Connell, Ruth V. Nichols, Alec J. Hirsch, Guang Fan, Daniel N. Streblow, William B. Messer, Andrew C. Adey, Benjamin N. Bimber, Brian J. O’Roak |
| EPI_ISL_526092, EPI_ISL_526093 | OHSU Lab Services Molecular Microbiology Lab | Ginkgo Bioworks Clinical Laboratory | Brendan L. O'Connell, Ruth V. Nichols, Alec J. Hirsch, Guang Fan, Daniel N. Streblow, Malaika Mckenzie-Bennett, James McGann, Jim Griffin, Keith Robison, Alex Plocik, Becky Schilling, Rebecca Littlefield, Michelle Spencer, Birgitte Simen, William B. Messer, Andrew C. Adey, Benjamin N. Bimber, Brian J. O’Roak |
| EPI_ISL_526094, EPI_ISL_526095, EPI_ISL_526096 | OHSU Lab Services Molecular Microbiology Lab | Oregon SARS-CoV-2 Genome Sequencing Center | Brendan L. O'Connell, Ruth V. Nichols, Alec J. Hirsch, Guang Fan, Daniel N. Streblow, William B. Messer, Andrew C. Adey, Benjamin N. Bimber, Brian J. O’Roak |
| EPI_ISL_526097 | OHSU Lab Services Molecular Microbiology Lab | Ginkgo Bioworks Clinical Laboratory | Brendan L. O'Connell, Ruth V. Nichols, Alec J. Hirsch, Guang Fan, Daniel N. Streblow, Malaika Mckenzie-Bennett, James McGann, Jim Griffin, Keith Robison, Alex Plocik, Becky Schilling, Rebecca Littlefield, Michelle Spencer, Birgitte Simen, William B. Messer, Andrew C. Adey, Benjamin N. Bimber, Brian J. O’Roak |

| | | | |
|---|---|---|---|
| EPI_ISL_526098, EPI_ISL_526099, EPI_ISL_526100, EPI_ISL_526101, EPI_ISL_526102, EPI_ISL_526103, EPI_ISL_526104, EPI_ISL_526105, EPI_ISL_526106, EPI_ISL_526107, EPI_ISL_526108, EPI_ISL_526109, EPI_ISL_526110, EPI_ISL_526111, EPI_ISL_526112, EPI_ISL_526113, EPI_ISL_526114 | | | |
| see above | OHSU Lab Services Molecular Microbiology Lab | Oregon SARS-CoV-2 Genome Sequencing Center | Brendan L. O'Connell, Ruth V. Nichols, Alec J. Hirsch, Guang Fan, Daniel N. Streblow, William B. Messer, Andrew C. Adey, Benjamin N. Bimber, Brian J. O'Roak |
| EPI_ISL_526115, EPI_ISL_526116, EPI_ISL_526117, EPI_ISL_526118 | Pathology West - NSW Health Pathology | NSW Health Pathology - Institute of Clinical Pathology and Medical Research; Westmead Hospital; University of Sydney | CIDM-PH et al. |
| EPI_ISL_526119, EPI_ISL_526120 | Sydney South West Pathology Service (SSWPS) - Liverpool Hospital - NSW Health Pathology | NSW Health Pathology - Institute of Clinical Pathology and Medical Research; Westmead Hospital; University of Sydney | CIDM-PH et al. |
| EPI_ISL_526121, EPI_ISL_526122, EPI_ISL_526123 | St Vincent's Pathology (SydPath) | NSW Health Pathology - Institute of Clinical Pathology and Medical Research; Westmead Hospital; University of Sydney | CIDM-PH et al. |
| EPI_ISL_526124 | Sydney South West Pathology Service (SSWPS) - Liverpool Hospital - NSW Health Pathology | NSW Health Pathology - Institute of Clinical Pathology and Medical Research; Westmead Hospital; University of Sydney | CIDM-PH et al. |
| EPI_ISL_526125 | Sydney South West Pathology Service (SSWPS) - Concord Repatriation General Hospital - NSW Health Pathology | NSW Health Pathology - Institute of Clinical Pathology and Medical Research; Westmead Hospital; University of Sydney | CIDM-PH et al. |
| EPI_ISL_526126, EPI_ISL_526127 | Sydney South West Pathology Service (SSWPS) - Liverpool Hospital - NSW Health Pathology | NSW Health Pathology - Institute of Clinical Pathology and Medical Research; Westmead Hospital; University of Sydney | CIDM-PH et al. |
| EPI_ISL_526128 | Pathology North - Royal North Shore Hospital - NSW Health Pathology | NSW Health Pathology - Institute of Clinical Pathology and Medical Research; Westmead Hospital; University of Sydney | CIDM-PH et al. |
| EPI_ISL_526129 | The Children's Hospital at Westmead | NSW Health Pathology - Institute of Clinical Pathology and Medical Research; Westmead Hospital; University of Sydney | CIDM-PH et al. |
| EPI_ISL_526130 | Pathology North - Royal North Shore Hospital - NSW Health Pathology | NSW Health Pathology - Institute of Clinical Pathology and Medical Research; Westmead Hospital; University of Sydney | CIDM-PH et al. |
| EPI_ISL_526131 | Sydney South West Pathology Service (SSWPS) - Liverpool Hospital - NSW Health Pathology | NSW Health Pathology - Institute of Clinical Pathology and Medical Research; Westmead Hospital; University of Sydney | CIDM-PH et al. |
| EPI_ISL_526132, EPI_ISL_526133 | Pathology North - Royal North Shore Hospital - NSW Health Pathology | NSW Health Pathology - Institute of Clinical Pathology and Medical Research; Westmead Hospital; University of Sydney | CIDM-PH et al. |
| EPI_ISL_526134, EPI_ISL_526135, EPI_ISL_526136, EPI_ISL_526137 | Sydney South West Pathology Service (SSWPS) - Royal Prince Alfred Hospital - NSW Health Pathology | NSW Health Pathology - Institute of Clinical Pathology and Medical Research; Westmead Hospital; University of Sydney | CIDM-PH et al. |
| EPI_ISL_526138, EPI_ISL_526139, EPI_ISL_526140, EPI_ISL_526141 | Pathology West - NSW Health Pathology | NSW Health Pathology - Institute of Clinical Pathology and Medical Research; Westmead Hospital; University of Sydney | CIDM-PH et al. |
| EPI_ISL_526142 | South Eastern Area Laboratory Services (SEALS) | NSW Health Pathology - Institute of Clinical Pathology and Medical Research; Westmead Hospital; University of Sydney | CIDM-PH et al. |
| EPI_ISL_526143 | Laverty Pathology | NSW Health Pathology - Institute of Clinical Pathology and Medical Research; Westmead Hospital; University of Sydney | CIDM-PH et al. |
| EPI_ISL_526144, EPI_ISL_526145, EPI_ISL_526146, EPI_ISL_526147, EPI_ISL_526148, EPI_ISL_526149 | South Eastern Area Laboratory Services (SEALS) | NSW Health Pathology - Institute of Clinical Pathology and Medical Research; Westmead Hospital; University of Sydney | CIDM-PH et al. |
| EPI_ISL_526150, EPI_ISL_526151 | Pathology West - NSW Health Pathology | NSW Health Pathology - Institute of Clinical Pathology and Medical Research; Westmead Hospital; University of Sydney | CIDM-PH et al. |
| EPI_ISL_526152 | Pathology North - Hunter - NSW Health Pathology | NSW Health Pathology - Institute of Clinical Pathology and Medical Research; Westmead Hospital; University of Sydney | CIDM-PH et al. |
| EPI_ISL_526153 | Douglass Hanly Moir Pathology | NSW Health Pathology - Institute of Clinical Pathology and Medical Research; Westmead Hospital; University of Sydney | CIDM-PH et al. |
| EPI_ISL_526154, EPI_ISL_526155 | Pathology North - Royal North Shore Hospital - NSW Health Pathology | NSW Health Pathology - Institute of Clinical Pathology and Medical Research; Westmead Hospital; University of Sydney | CIDM-PH et al. |
| EPI_ISL_526156 | Pathology West - NSW Health Pathology | NSW Health Pathology - Institute of Clinical Pathology and Medical Research; Westmead Hospital; University of Sydney | CIDM-PH et al. |
| EPI_ISL_526157, EPI_ISL_526158, EPI_ISL_526159, EPI_ISL_526160 | Sydney South West Pathology Service (SSWPS) - Liverpool Hospital - NSW Health Pathology | NSW Health Pathology - Institute of Clinical Pathology and Medical Research; Westmead Hospital; University of Sydney | CIDM-PH et al. |
| EPI_ISL_526161, EPI_ISL_526162, EPI_ISL_526163, EPI_ISL_526164, EPI_ISL_526165 | Pathology West - NSW Health Pathology | NSW Health Pathology - Institute of Clinical Pathology and Medical Research; Westmead Hospital; University of Sydney | CIDM-PH et al. |
| EPI_ISL_526166, EPI_ISL_526167, EPI_ISL_526168, EPI_ISL_526169 | Austech Medical Laboratories | NSW Health Pathology - Institute of Clinical Pathology and Medical Research; Westmead Hospital; University of Sydney | CIDM-PH et al. |
| EPI_ISL_526170 | Pathology North - Royal North Shore Hospital - NSW Health Pathology | NSW Health Pathology - Institute of Clinical Pathology and Medical Research; Westmead Hospital; University of Sydney | CIDM-PH et al. |
| EPI_ISL_526171 | St Vincent's Pathology (SydPath) | NSW Health Pathology - Institute of Clinical Pathology and Medical Research; Westmead Hospital; University of Sydney | CIDM-PH et al. |
| EPI_ISL_526172, EPI_ISL_526173, EPI_ISL_526174, EPI_ISL_526175, EPI_ISL_526176 | Sydney South West Pathology Service (SSWPS) - Liverpool Hospital - NSW Health Pathology | NSW Health Pathology - Institute of Clinical Pathology and Medical Research; Westmead Hospital; University of Sydney | CIDM-PH et al. |
| EPI_ISL_526177, EPI_ISL_526178, EPI_ISL_526179, EPI_ISL_526180, EPI_ISL_526181, EPI_ISL_526182, EPI_ISL_526183, EPI_ISL_526184, EPI_ISL_526185 | St Vincent's Pathology (SydPath) | NSW Health Pathology - Institute of Clinical Pathology and Medical Research; Westmead Hospital; University of Sydney | CIDM-PH et al. |
| EPI_ISL_526186 | 4Cyte Pathology | NSW Health Pathology - Institute of Clinical Pathology and Medical Research; Westmead Hospital; University of Sydney | CIDM-PH et al. |
| EPI_ISL_526187, EPI_ISL_526188, EPI_ISL_526189, EPI_ISL_526190 | St Vincent's Pathology (SydPath) | NSW Health Pathology - Institute of Clinical Pathology and Medical Research; Westmead Hospital; University of Sydney | CIDM-PH et al. |
| EPI_ISL_526191 | South Eastern Area Laboratory Services (SEALS) | NSW Health Pathology - Institute of Clinical Pathology and Medical Research; Westmead Hospital; University of Sydney | CIDM-PH et al. |
| EPI_ISL_526192 | Medlab Pathology | NSW Health Pathology - Institute of Clinical Pathology and Medical Research; Westmead Hospital; University of Sydney | CIDM-PH et al. |
| EPI_ISL_526193 | Sydney South West Pathology Service (SSWPS) - Liverpool Hospital - NSW Health Pathology | NSW Health Pathology - Institute of Clinical Pathology and Medical Research; Westmead Hospital; University of Sydney | CIDM-PH et al. |
| EPI_ISL_526194, EPI_ISL_526195 | St Vincent's Pathology (SydPath) | NSW Health Pathology - Institute of Clinical Pathology and Medical Research; Westmead Hospital; University of Sydney | CIDM-PH et al. |
| EPI_ISL_526196 | Sydney South West Pathology Service (SSWPS) - Royal Prince Alfred Hospital - NSW Health Pathology | NSW Health Pathology - Institute of Clinical Pathology and Medical Research; Westmead Hospital; University of Sydney | CIDM-PH et al. |
| EPI_ISL_526197, EPI_ISL_526198 | Sydney South West Pathology Service (SSWPS) - Liverpool Hospital - NSW Health Pathology | NSW Health Pathology - Institute of Clinical Pathology and Medical Research; Westmead Hospital; University of Sydney | CIDM-PH et al. |
| EPI_ISL_526199 | Pathology West - NSW Health Pathology | NSW Health Pathology - Institute of Clinical Pathology and Medical Research; Westmead Hospital; University of Sydney | CIDM-PH et al. |
| EPI_ISL_526200 | St Vincent's Pathology (SydPath) | NSW Health Pathology - Institute of Clinical Pathology and Medical Research; Westmead Hospital; University of Sydney | CIDM-PH et al. |
| EPI_ISL_526201 | Sydney South West Pathology Service (SSWPS) - Royal Prince Alfred Hospital - NSW Health Pathology | NSW Health Pathology - Institute of Clinical Pathology and Medical Research; Westmead Hospital; University of Sydney | CIDM-PH et al. |
| EPI_ISL_526202, EPI_ISL_526203, EPI_ISL_526204 | Laverty Pathology | NSW Health Pathology - Institute of Clinical Pathology and Medical Research; Westmead Hospital; University of Sydney | CIDM-PH et al. |
| EPI_ISL_526205 | The Children's Hospital at Westmead | NSW Health Pathology - Institute of Clinical Pathology and Medical Research; Westmead Hospital; University of Sydney | CIDM-PH et al. |
| EPI_ISL_526206 | Sydney South West Pathology Service (SSWPS) - Liverpool Hospital | NSW Health Pathology - Institute of Clinical Pathology and Medical Research; Westmead Hospital; University of | CIDM-PH et al. |

| | | | |
|---|---|---|---|
| | - NSW Health Pathology | Sydney | |
| EPI_ISL_526207, EPI_ISL_526208, EPI_ISL_526209 | Sydney South West Pathology Service (SSWPS) - Royal Prince Alfred Hospital - NSW Health Pathology | NSW Health Pathology - Institute of Clinical Pathology and Medical Research; Westmead Hospital; University of Sydney | CIDM-PH et al. |
| EPI_ISL_526210, EPI_ISL_526211 | Sydney South West Pathology Service (SSWPS) - Liverpool Hospital - NSW Health Pathology | NSW Health Pathology - Institute of Clinical Pathology and Medical Research; Westmead Hospital; University of Sydney | CIDM-PH et al. |
| EPI_ISL_526212 | Australian Clinical Labs (formerly Healthscope Pathology) | NSW Health Pathology - Institute of Clinical Pathology and Medical Research; Westmead Hospital; University of Sydney | CIDM-PH et al. |
| EPI_ISL_526213, EPI_ISL_526214 | Douglass Hanly Moir Pathology | NSW Health Pathology - Institute of Clinical Pathology and Medical Research; Westmead Hospital; University of Sydney | CIDM-PH et al. |
| EPI_ISL_526215, EPI_ISL_526216, EPI_ISL_526217, EPI_ISL_526218, EPI_ISL_526219, EPI_ISL_526220, EPI_ISL_526221, EPI_ISL_526222, EPI_ISL_526223, EPI_ISL_526224, EPI_ISL_526225, EPI_ISL_526226, EPI_ISL_526227, EPI_ISL_526228, EPI_ISL_526229, EPI_ISL_526230, EPI_ISL_526231, EPI_ISL_526232, EPI_ISL_526233, EPI_ISL_526234, EPI_ISL_526235, EPI_ISL_526236, EPI_ISL_526237, EPI_ISL_526238 | | | |
| see above | Hungarian Defence Forces Military Medical Centre | National Laboratory of Virology, Szentágothai Research Centre | Endre Gábor Tóth, Balázs Somogyi, Bálint Eszenyi, Ferenc Jakab, Gábor Kemenesi |
| EPI_ISL_526259, EPI_ISL_526268, EPI_ISL_526275, EPI_ISL_526284 | Unity Health Toronto | Ontario Institute for Cancer Research | Ramzi Fattouh, Larissa M. Matukas, Mark Downing, Annette Gower, Karel Boissinot, Samira Mubareka, TIBDN, Ilinca Lungu, Bernard Lam, Jeremy Johns, Paul Krzyzanowski, Richard de Borja, Felicia Vincelli, Philip Zuzarte, Jared Simpson |
| EPI_ISL_526288, EPI_ISL_526289, EPI_ISL_526290, EPI_ISL_526291, EPI_ISL_526294, EPI_ISL_526295, EPI_ISL_526296, EPI_ISL_526298, EPI_ISL_526300, EPI_ISL_526301, EPI_ISL_526304, EPI_ISL_526305, EPI_ISL_526309, EPI_ISL_526310, EPI_ISL_526312, EPI_ISL_526313, EPI_ISL_526316, EPI_ISL_526317, EPI_ISL_526318, EPI_ISL_526319, EPI_ISL_526323, EPI_ISL_526324, EPI_ISL_526325, EPI_ISL_526327, EPI_ISL_526330, EPI_ISL_526331, EPI_ISL_526332, EPI_ISL_526333, EPI_ISL_526334, EPI_ISL_526336 | | | |
| see above | University of Birmingham | COVID-19 Genomics UK (COG-UK) Consortium | Institute of Microbiology, University of Birmingham: Claire McMurray, Joanne Stockton, Samuel Nicholls, Radoslaw Poplawski, Will Rowe, Josh Quick, Nicholas Loman. University of Birmingham Testing Laboratory: Celina M Whalley, Andrew Bosworth, Charlotte Poxon, Kasun Wanigasooriya, Oliver Pickles, Mike Kidd, Alex Richter, Andrew D Beggs PHE Heartlands Lab: Husam Osman, Andrew Bosworth. Queen Elizabeth Hospital: Anna Casey |
| EPI_ISL_526337, EPI_ISL_526338, EPI_ISL_526339, EPI_ISL_526340, EPI_ISL_526341, EPI_ISL_526343, EPI_ISL_526346, EPI_ISL_526348, EPI_ISL_526350, EPI_ISL_526351, EPI_ISL_526352, EPI_ISL_526353, EPI_ISL_526355, EPI_ISL_526356, EPI_ISL_526358, EPI_ISL_526360, EPI_ISL_526361, EPI_ISL_526362, EPI_ISL_526366 | | | |
| see above | Queens Medical Centre, Clinical Microbiology Department / DeepSeq Nottingham | COVID-19 Genomics UK (COG-UK) Consortium | Gemma Clark, Wendy Smith, Manjinder Khakh, Vicki M Fleming, Michelle M Lister, Hannah Howson-Wells, Jonathan Ball, Patrick McClure, Joseph Chappell, Theocharis Tsoleridis, Nadine Holmes, Matthew Carlisle, Christopher Moore, Fei Sang, Johnny Debebe, Victoria Wright, Matthew Loose |
| EPI_ISL_526368, EPI_ISL_526369, EPI_ISL_526370, EPI_ISL_526371, EPI_ISL_526372, EPI_ISL_526373, EPI_ISL_526374, EPI_ISL_526375, EPI_ISL_526376, EPI_ISL_526377, EPI_ISL_526378, EPI_ISL_526379, EPI_ISL_526380, EPI_ISL_526381, EPI_ISL_526382, EPI_ISL_526383, EPI_ISL_526384, EPI_ISL_526385, EPI_ISL_526386, EPI_ISL_526387, EPI_ISL_526388, EPI_ISL_526389, EPI_ISL_526390, EPI_ISL_526391, EPI_ISL_526393, EPI_ISL_526394 | | | |
| see above | Liverpool Clinical Laboratories | COVID-19 Genomics UK (COG-UK) Consortium | Sam Haldenby, Anita Lucaci, Steve Paterson, Julian Hiscox, Alistair Darby, M Almsaud, A Alrezaihi, Muhannad Alruwaili, Stuart D Armstrong, Jones Benjamin, Eleanor G Bentley, Anu Chawla, Jordan J Clark, Angela Cowell, Richard Eccles, Isabel García-Dorival, Matthew Gemmell, Alessandro Gerada, PKF Gilmore, Richard Gregory, Ximeng Han, Catherine Hartley, Margaret Hughes, Miren Iturriza-Gomara, James Johnson, L Luu, Jenifer Manson, Charlotte Nelson, Elaine O'Toole, Cassie Olateju, Rebekah Penrice-Randal , Lucille Rainbow, N.P Randle, Trevor Ian Robinson, Parul Sharma, Ghada T Shawli, James P Stewart, Neil Swainston, Ecaterina Vamos, Joanne Watts, Mark Whitehead |
| EPI_ISL_526395, EPI_ISL_526396, EPI_ISL_526397, EPI_ISL_526398, EPI_ISL_526399, EPI_ISL_526401, EPI_ISL_526402, EPI_ISL_526403, EPI_ISL_526404, EPI_ISL_526405, EPI_ISL_526406, EPI_ISL_526407, EPI_ISL_526408, EPI_ISL_526410, EPI_ISL_526411, EPI_ISL_526412, EPI_ISL_526413, EPI_ISL_526414, EPI_ISL_526415, EPI_ISL_526418, EPI_ISL_526419, EPI_ISL_526420, EPI_ISL_526421, EPI_ISL_526422, EPI_ISL_526423, EPI_ISL_526424, EPI_ISL_526425, EPI_ISL_526427, EPI_ISL_526430 | | | |
| see above | Queens Medical Centre, Clinical Microbiology Department / DeepSeq Nottingham | COVID-19 Genomics UK (COG-UK) Consortium | Gemma Clark, Wendy Smith, Manjinder Khakh, Vicki M Fleming, Michelle M Lister, Hannah Howson-Wells, Jonathan Ball, Patrick McClure, Joseph Chappell, Theocharis Tsoleridis, Nadine Holmes, Matthew Carlisle, Christopher Moore, Fei Sang, Johnny Debebe, Victoria Wright, Matthew Loose |
| EPI_ISL_526434, EPI_ISL_526435, EPI_ISL_526436 | Centre for Enzyme Innovation, University of Portsmouth / Translational Research Laboratory, Portsmouth Hospitals NHS Trust | COVID-19 Genomics UK (COG-UK) Consortium | Angela Beckett,Yann Bourgeois,Garry Scarlett,Sharon Glaysher,Scott Elliott,Kelly Bicknell,Robert Impey,Allyson Lloyd,Sarah Wyllie,Ethan Butcher,Anoop Chauhan,Samuel Robson |
| EPI_ISL_526458 | West of Scotland Specialist Virology Centre, NHSGGC / MRC-University of Glasgow Centre for Virus Research | COVID-19 Genomics UK (COG-UK) Consortium | Ana da Silva Filipe, Natasha Johnson, Kathy Smollett, Daniel Mair, Stephen Carmichael, Lily Tong, Jenna Nichols, Elihu Aranday-Cortes, Kirstyn Brunker, Yasmin Parr, Alice Broos, Kyriaki Nomikou; Sarah McDonald, Marc Niebel, Patawee Asamaphan; Richard Orton, Joseph Hughes, Sreenu Vattipally, David L Robertson; Alasdair MacLean, Rory Gunson; Kathy Li, Natasha Jesudason, Rajiv Shah, James Shepherd, Antonia Ho, Emma Thomson |
| EPI_ISL_526459, EPI_ISL_526460, EPI_ISL_526465, EPI_ISL_526466, EPI_ISL_526467, EPI_ISL_526469, EPI_ISL_526470, EPI_ISL_526471, EPI_ISL_526472, EPI_ISL_526473, EPI_ISL_526474, EPI_ISL_526475, EPI_ISL_526476, EPI_ISL_526477, EPI_ISL_526478, EPI_ISL_526479, EPI_ISL_526480, EPI_ISL_526481, EPI_ISL_526482, EPI_ISL_526483, EPI_ISL_526484, EPI_ISL_526485, EPI_ISL_526486, EPI_ISL_526487, EPI_ISL_526488, EPI_ISL_526489, EPI_ISL_526494, EPI_ISL_526495, EPI_ISL_526496, EPI_ISL_526497, EPI_ISL_526498, EPI_ISL_526499, EPI_ISL_526500, EPI_ISL_526501, EPI_ISL_526502, EPI_ISL_526503, EPI_ISL_526504, EPI_ISL_526505, EPI_ISL_526506, EPI_ISL_526507, EPI_ISL_526508, EPI_ISL_526509, EPI_ISL_526510, EPI_ISL_526511, EPI_ISL_526512, EPI_ISL_526513, EPI_ISL_526514, EPI_ISL_526515, EPI_ISL_526521, EPI_ISL_526526, EPI_ISL_526532, EPI_ISL_526533, EPI_ISL_526534, EPI_ISL_526536 | | | |
| see above | Virology Department, Royal Infirmary of Edinburgh, NHS Lothian / School of Biological Sciences, University of Edinburgh / Institute of Genetics and Molecular Medicine, University of Edinburgh | COVID-19 Genomics UK (COG-UK) Consortium | McHugh M, Dewar R, Rooke S, Gallagher M, Balcaza C, O'Toole Á, Scher E, Hill V, McCrone JT, Colquhoun R, Yu X, Jackson B, Rambaut A, Williams TC, Templeton K |
| EPI_ISL_526537, EPI_ISL_526538, EPI_ISL_526539, EPI_ISL_526545, EPI_ISL_526547, EPI_ISL_526550 | Respiratory Virus Unit, Microbiology Services Colindale, Public Health England | Respiratory Virus Unit, Microbiology Services Colindale, Public Health England | PHE Covid Sequencing Team |
| EPI_ISL_526551 | OHSU Lab Services Molecular Microbiology Lab | Oregon SARS-CoV-2 Genome Sequencing Center | Brendan L. O'Connell, Ruth V. Nichols, Alec J. Hirsch, Guang Fan, Daniel N. Streblow, William B. Messer, Andrew C. Adey, Benjamin N. Bimber, Brian J. O'Roak |
| EPI_ISL_526553, EPI_ISL_526555, EPI_ISL_526563, EPI_ISL_526567, EPI_ISL_526568, EPI_ISL_526569, EPI_ISL_526570, EPI_ISL_526571, EPI_ISL_526572, EPI_ISL_526573, EPI_ISL_526575, EPI_ISL_526580, EPI_ISL_526581, EPI_ISL_526583, EPI_ISL_526585, EPI_ISL_526586, EPI_ISL_526588, EPI_ISL_526589, EPI_ISL_526590, EPI_ISL_526591, EPI_ISL_526592, EPI_ISL_526593, EPI_ISL_526594, EPI_ISL_526595, EPI_ISL_526596, EPI_ISL_526597, EPI_ISL_526598, EPI_ISL_526599, EPI_ISL_526600, EPI_ISL_526602, EPI_ISL_526603, EPI_ISL_526605, EPI_ISL_526606, EPI_ISL_526607, EPI_ISL_526608, EPI_ISL_526610, EPI_ISL_526611, EPI_ISL_526612, EPI_ISL_526613, EPI_ISL_526614, EPI_ISL_526616, EPI_ISL_526617, EPI_ISL_526618, EPI_ISL_526619, EPI_ISL_526620, EPI_ISL_526621, EPI_ISL_526622, EPI_ISL_526623, EPI_ISL_526624, EPI_ISL_526625, EPI_ISL_526626, EPI_ISL_526627, EPI_ISL_526628, EPI_ISL_526629, EPI_ISL_526630, EPI_ISL_526631, EPI_ISL_526632, EPI_ISL_526633, EPI_ISL_526634, EPI_ISL_526635, EPI_ISL_526636, EPI_ISL_526637, EPI_ISL_526638, EPI_ISL_526643, EPI_ISL_526644, EPI_ISL_526645, EPI_ISL_526646, EPI_ISL_526648, EPI_ISL_526649, EPI_ISL_526651, EPI_ISL_526652, EPI_ISL_526653, EPI_ISL_526656, EPI_ISL_526659, EPI_ISL_526660, EPI_ISL_526661, EPI_ISL_526662, EPI_ISL_526663, EPI_ISL_526664, EPI_ISL_526665, EPI_ISL_526666, EPI_ISL_526667, EPI_ISL_526669, EPI_ISL_526670, EPI_ISL_526671, EPI_ISL_526672, EPI_ISL_526673, EPI_ISL_526674, EPI_ISL_526675, EPI_ISL_526676, EPI_ISL_526677, EPI_ISL_526678, EPI_ISL_526679, EPI_ISL_526680, EPI_ISL_526681, EPI_ISL_526682 | | | |
| see above | Florida Bureau of Public Health Laboratories | Florida Bureau of Public Health Laboratories | Sarah Schmedes, Jason Blanton |
| EPI_ISL_526686, EPI_ISL_526687, EPI_ISL_526688 | Faith Laboratory, Immunology Institute, Icahn School of Medicine at Mount Sinai | van Bakel Laboratory, Genetics and Genomics Sciences, Icahn School of Medicine at Mount Sinai | Graham J. Britton, Alice Chen-Liaw, Francesca Cossarini, Alexandra Livanos, Matthew P. Spindler, Tamar Plitt, Joseph Eggers, Ilaria Mogno, Ana S. Gonzalez-Reiche, Sophia Sui, Michael Tankelevich, Lauren Tal Grinspan, Rebekah E. Dixon, Divya Jha, Gustavo Martinez-Delgado, Fatima Amanat, Daisy Hoagland, Benjamin R. tenOever, Marla C. Dubinsky, Miriam Merad, Harm Van Bakel, Florian Krammer, Gerold Bongers, Saurabh Mehandru and Jeremiah J. Faith |
| EPI_ISL_526689 | Pathology West - NSW Health Pathology | NSW Health Pathology - Institute of Clinical Pathology and Medical Research; Westmead Hospital; University of Sydney | CIDM-PH et al. |
| EPI_ISL_526697 | South Eastern Area Laboratory Services (SEALS) | NSW Health Pathology - Institute of Clinical Pathology and Medical Research; Westmead Hospital; University of Sydney | CIDM-PH et al. |
| EPI_ISL_526700, EPI_ISL_526701, EPI_ISL_526702, EPI_ISL_526703, EPI_ISL_526704, EPI_ISL_526705, EPI_ISL_526706, EPI_ISL_526707, EPI_ISL_526708, EPI_ISL_526709, EPI_ISL_526710, EPI_ISL_526711, EPI_ISL_526712, EPI_ISL_526713, EPI_ISL_526714, EPI_ISL_526715, EPI_ISL_526716, EPI_ISL_526717, EPI_ISL_526718, EPI_ISL_526719, EPI_ISL_526720, EPI_ISL_526721, EPI_ISL_526722, EPI_ISL_526723, EPI_ISL_526724, EPI_ISL_526725, EPI_ISL_526726, EPI_ISL_526727, EPI_ISL_526728, EPI_ISL_526729 | | | |
| see above | Division of Viral Diseases, Center for Laboratory Control of Infectious Diseases, Korea Centers for Diseases Control and Prevention | Division of Viral Diseases, Center for Laboratory Control of Infectious Diseases, Korea Centers for Diseases Control and Prevention | Jeong-Min Kim, Yoon-Seok Chung, Namjoo Lee, Sang Hee Woo, Hye-Jun Jo, Heui Man Kim, Jun-Sub Kim, Myung Guk Han |
| EPI_ISL_526730, EPI_ISL_526731 | Center for Laboratory Control of Infectious Diseases, Korea Centers for Diseases Control and Prevention | Center for Laboratory Control of Infectious Diseases, Korea Centers for Diseases Control and Prevention | Junyoung Kim, Ae Kyung Park, Eunkyung Shin, Jin Sun No, Jeong-Min Kim, Yoon-Seok Chung, Heui Man Kim, Myung Guk Han |
| EPI_ISL_526732, EPI_ISL_526733 | Division of Viral Diseases, Center for Laboratory Control of Infectious Diseases, Korea Centers for Diseases Control and Prevention | Division of Viral Diseases, Center for Laboratory Control of Infectious Diseases, Korea Centers for Diseases Control and Prevention | Jeong-Min Kim, Yoon-Seok Chung, Namjoo Lee, Sang Hee Woo, Hye-Jun Jo, Heui Man Kim, Jun-Sub Kim, Myung Guk Han |
| EPI_ISL_526734, EPI_ISL_526735, EPI_ISL_526736, EPI_ISL_526737, EPI_ISL_526738, EPI_ISL_526739, EPI_ISL_526740, EPI_ISL_526741, EPI_ISL_526742, EPI_ISL_526743, EPI_ISL_526744, EPI_ISL_526745, EPI_ISL_526746 | | | |
| see above | Center for Laboratory Control of Infectious Diseases, Korea Centers for Diseases Control and Prevention | Center for Laboratory Control of Infectious Diseases, Korea Centers for Diseases Control and Prevention | Junyoung Kim, Ae Kyung Park, Eunkyung Shin, Jin Sun No, Jeong-Min Kim, Yoon-Seok Chung, Heui Man Kim, Myung Guk Han |
| EPI_ISL_526754, EPI_ISL_526755, EPI_ISL_526756, EPI_ISL_526757, EPI_ISL_526758, EPI_ISL_526759, EPI_ISL_526762, EPI_ISL_526763, EPI_ISL_526764, EPI_ISL_526765, EPI_ISL_526766, EPI_ISL_526767, EPI_ISL_526769, EPI_ISL_526770, EPI_ISL_526772, EPI_ISL_526773 | | | |
| see above | Respiratory Virus Unit, Microbiology Services Colindale, Public Health England | Respiratory Virus Unit, Microbiology Services Colindale, Public Health England | PHE Covid Sequencing Team |
| EPI_ISL_526775, EPI_ISL_526776, EPI_ISL_526777, EPI_ISL_526778, EPI_ISL_526779, EPI_ISL_526780, EPI_ISL_526781, EPI_ISL_526782, EPI_ISL_526783, EPI_ISL_526784, EPI_ISL_526785, EPI_ISL_526786, EPI_ISL_526787, EPI_ISL_526788, EPI_ISL_526789, EPI_ISL_526790, EPI_ISL_526791, EPI_ISL_526792, EPI_ISL_526793, EPI_ISL_526794, EPI_ISL_526795, EPI_ISL_526796, EPI_ISL_526797, EPI_ISL_526798, EPI_ISL_526799, EPI_ISL_526800, EPI_ISL_526801, EPI_ISL_526802, EPI_ISL_526803, EPI_ISL_526804, EPI_ISL_526805, EPI_ISL_526806, EPI_ISL_526807, EPI_ISL_526808, EPI_ISL_526809, EPI_ISL_526810, EPI_ISL_526811, EPI_ISL_526812, EPI_ISL_526813, EPI_ISL_526814, EPI_ISL_526815, EPI_ISL_526816, EPI_ISL_526817, EPI_ISL_526818, EPI_ISL_526820, EPI_ISL_526821, EPI_ISL_526822, EPI_ISL_526823, EPI_ISL_526824, EPI_ISL_526825, EPI_ISL_526826, EPI_ISL_526827, EPI_ISL_526828, EPI_ISL_526829, EPI_ISL_526830, EPI_ISL_526831, EPI_ISL_526832, EPI_ISL_526833, EPI_ISL_526834, EPI_ISL_526835, EPI_ISL_526836, EPI_ISL_526837, EPI_ISL_526838, EPI_ISL_526839, EPI_ISL_526840, EPI_ISL_526841, EPI_ISL_526842, EPI_ISL_526843, EPI_ISL_526844, EPI_ISL_526845, EPI_ISL_526846, EPI_ISL_526847, EPI_ISL_526848, EPI_ISL_526849, EPI_ISL_526850, EPI_ISL_526851, EPI_ISL_526852, EPI_ISL_526853, EPI_ISL_526854, EPI_ISL_526855, EPI_ISL_526856, EPI_ISL_526857, EPI_ISL_526858, EPI_ISL_526859, EPI_ISL_526860, EPI_ISL_526861, EPI_ISL_526862, EPI_ISL_526863, EPI_ISL_526864, EPI_ISL_526865, EPI_ISL_526866, EPI_ISL_526867, EPI_ISL_526868, EPI_ISL_526869, EPI_ISL_526870, EPI_ISL_526871, EPI_ISL_526872, EPI_ISL_526873, EPI_ISL_526874, EPI_ISL_526875, EPI_ISL_526876, EPI_ISL_526877, EPI_ISL_526878, EPI_ISL_526879, EPI_ISL_526880, EPI_ISL_526881, EPI_ISL_526882, EPI_ISL_526883, EPI_ISL_526884, EPI_ISL_526885, EPI_ISL_526886, EPI_ISL_526887, EPI_ISL_526888, EPI_ISL_526889, EPI_ISL_526890, EPI_ISL_526891, EPI_ISL_526892, EPI_ISL_526893, EPI_ISL_526894, EPI_ISL_526895, EPI_ISL_526896, EPI_ISL_526897, EPI_ISL_526898, EPI_ISL_526899, EPI_ISL_526900, EPI_ISL_526901, EPI_ISL_526902, EPI_ISL_526903, EPI_ISL_526904, EPI_ISL_526905, EPI_ISL_526906, EPI_ISL_526907, EPI_ISL_526908, EPI_ISL_526909, EPI_ISL_526910, EPI_ISL_526911, EPI_ISL_526912, EPI_ISL_526913, EPI_ISL_526914, EPI_ISL_526915, EPI_ISL_526916, EPI_ISL_526917, EPI_ISL_526918, EPI_ISL_526919, EPI_ISL_526920, EPI_ISL_526921, EPI_ISL_526922, EPI_ISL_526923, EPI_ISL_526924, EPI_ISL_526925, EPI_ISL_526926, EPI_ISL_526927, EPI_ISL_526928, EPI_ISL_526929, EPI_ISL_526930, EPI_ISL_526931 | | | |
| see above | Virginia DCLS | Virginia DCLS | Virginia DCLS |
| EPI_ISL_526933, EPI_ISL_526934 | Instituto Nacional de Salud, Bogotá, Colombia | Instituto Nacional de Salud, Bogotá, Colombia | Katherine Laiton-Donato, Diego A. Álvarez-Díaz, Carlos Franco-Muñoz, Mauricio Pacheco-Montealegre, Jonathan Reales, Diego Andrés Prada, Jose A. Usme-Ciro, Zulma M. Cucunubá, Christian Julian Villabona-Arenas, Liz Villabona-Arenas, Sussy Echeverria, Astrid C. Flórez, Carolina Ferro, Diana Marcela Walteros-Acero, Franklin Prieto, Carlos Andrés Durán, Martha Lucia Ospina Martinez, Marcela Mercado-Reyes |
| EPI_ISL_526937, EPI_ISL_526938, EPI_ISL_526939, EPI_ISL_526940, EPI_ISL_526941, EPI_ISL_526943, EPI_ISL_526946 | Faroese National Reference Laboratory for Fish and Animal Diseases | Faroese National Reference Laboratory for Fish and Animal Diseases | Maria Marjunardóttir Dahl, Petra Elisabeth Petersen, Debes Hammershaimb Christiansen |
| EPI_ISL_526949, EPI_ISL_526950, EPI_ISL_526951, EPI_ISL_526955, EPI_ISL_526956, EPI_ISL_526958 | Instituto Nacional de Salud, Bogotá, Colombia | Instituto Nacional de Salud, Bogotá, Colombia | Katherine Laiton-Donato, Diego A. Álvarez-Díaz, Carlos Franco-Muñoz, Mauricio Pacheco-Montealegre, Jonathan Reales, Diego Andrés Prada, Jose A. Usme-Ciro, Zulma M. Cucunubá, Christian Julian Villabona-Arenas, Liz Villabona-Arenas, Sussy Echeverria, Astrid C. Flórez, Carolina Ferro, Diana Marcela Walteros-Acero, Franklin Prieto, Carlos Andrés Durán, Martha Lucia Ospina Martinez, Marcela Mercado-Reyes |
| EPI_ISL_526963, EPI_ISL_526964, | Instituto Nacional de Salud, Bogotá, Colombia | Instituto Nacional de Salud, Bogotá, Colombia | Katherine Laiton-Donato, Diego A. Álvarez-Díaz, Carlos Franco-Muñoz, Jonathan Reales, Diego Andrés Prada, Jeadran Malagón-Rojas, Felix Betzler, Wendy K. Jo, Edmilson F. de |

| | | | |
|---|---|---|---|
| EPI_ISL_526965 | | | Oliveira-Filho, Carolina Ferro, Diana Marcela Walteros-Acero, Franklin Prieto, Carlos Andrés Durán, Martha Lucia Ospina Martinez, Marcela Mercado-Reyes |
| EPI_ISL_526967 | Instituto Nacional de Salud, Bogotá, Colombia | Instituto Nacional de Salud, Bogotá, Colombia | Katherine Laiton-Donato, Diego A. Álvarez-Díaz, Carlos Franco-Muñoz, Mauricio Pacheco-Montealegre, Jonathan Reales, Diego Andrés Prada, Jose A. Usme-Ciro, Zulma M. Cucunubá, Christian Julian Villabona-Arenas, Liz Villabona-Arenas, Sussy Echeverria, Astrid C. Flórez, Carolina Ferro, Diana Marcela Walteros-Acero, Franklin Prieto, Carlos Andrés Durán, Martha Lucia Ospina Martinez, Marcela Mercado-Reyes |
| EPI_ISL_526971 | Instituto Nacional de Salud, Bogotá, Colombia | Instituto Nacional de Salud, Bogotá, Colombia | Katherine Laiton-Donato, Diego A. Álvarez-Díaz, Carlos Franco-Muñoz, Mauricio Pacheco-Montealegre, Jonathan Reales, Diego Andrés Prada, Jose A. Usme-Ciro, Zulma M. Cucunubá, Christian Julian Villabona-Arenas, Liz Villabona-Arenas, Sussy Echeverria, Astrid C. Flórez, Carolina Ferro, Diana Marcela Walteros-Acero, Franklin Prieto, Carlos Andrés Durán, Martha Lucia Ospina Martinez, Marcela Mercado-Reyes |
| EPI_ISL_526975, EPI_ISL_526976, EPI_ISL_526977, EPI_ISL_526978, EPI_ISL_526979, EPI_ISL_526980, EPI_ISL_526981, EPI_ISL_526982, EPI_ISL_526983, EPI_ISL_526984, EPI_ISL_526985, EPI_ISL_526986, EPI_ISL_526987, EPI_ISL_526988, EPI_ISL_526989, EPI_ISL_526990, EPI_ISL_526991, EPI_ISL_526992, EPI_ISL_526993, EPI_ISL_526994, EPI_ISL_526995, EPI_ISL_526996 | | | |
| see above | Biological prevention, army | Biological prevention, army | Seadawy, M.G., Gad, A.F., Harty, B.E., Elhosieny, M.F., Shamel, M.D. |
| EPI_ISL_526997, EPI_ISL_526998 | Biological prevention, army | Biological prevention, army | Seadawy, M.G., Harty,B.E., Gad,A.F., Elhoseiny,M.F., Shamel,M.D., Shabaan,A.E., Ageez,A.M. |
| EPI_ISL_526999, EPI_ISL_527000, EPI_ISL_527001 | Biological prevention, army | Biological prevention, army | Seadawy,M.G., Harty,B.E., Gad,A.F., Elhoseiny,M.F., Shamel,M.D., Shabaan,A.E., Ageez,A.M. |
| EPI_ISL_527002 | Biological prevention, army | Biological prevention, army | Seadawy, M.G., Harty,B.E., Gad,A.F., Elhoseiny,M.F., Shamel,M.D., Shabaan,A.E., Ageez,A.M. |
| EPI_ISL_527003, EPI_ISL_527004, EPI_ISL_527005, EPI_ISL_527006 | Biological prevention, army | Biological prevention, army | Seadawy,M.G., Harty,B.E., Gad,A.F., Elhoseiny,M.F., Shamel,M.D., Shabaan,A.E., Ageez,A.M. |
| EPI_ISL_527007 | Biological Prevention, Army | Biological Prevention, Army | Seadawy, M.G., Harty,B.E., Gad,A.F., Elhoseiny,M.F., Shamel,M.D., Shabaan,A.E., Ageez,A.M. |
| EPI_ISL_527008, EPI_ISL_527009, EPI_ISL_527010, EPI_ISL_527011, EPI_ISL_527012, EPI_ISL_527013, EPI_ISL_527014, EPI_ISL_527015, EPI_ISL_527016, EPI_ISL_527017, EPI_ISL_527018, EPI_ISL_527019, EPI_ISL_527020, EPI_ISL_527021, EPI_ISL_527022, EPI_ISL_527023, EPI_ISL_527024, EPI_ISL_527025, EPI_ISL_527026, EPI_ISL_527027, EPI_ISL_527028, EPI_ISL_527029, EPI_ISL_527030, EPI_ISL_527031, EPI_ISL_527032, EPI_ISL_527033, EPI_ISL_527034, EPI_ISL_527035, EPI_ISL_527036, EPI_ISL_527037, EPI_ISL_527038, EPI_ISL_527039, EPI_ISL_527040, EPI_ISL_527041, EPI_ISL_527042, EPI_ISL_527043, EPI_ISL_527044, EPI_ISL_527045, EPI_ISL_527046, EPI_ISL_527047, EPI_ISL_527048, EPI_ISL_527049, EPI_ISL_527050, EPI_ISL_527051, EPI_ISL_527052, EPI_ISL_527053, EPI_ISL_527054, EPI_ISL_527055, EPI_ISL_527056, EPI_ISL_527057, EPI_ISL_527058, EPI_ISL_527059, EPI_ISL_527060, EPI_ISL_527061, EPI_ISL_527062, EPI_ISL_527063, EPI_ISL_527064, EPI_ISL_527179, EPI_ISL_527180 | | | |
| see above | Area of Virology, Serology and Virology Division (SAViD), New South Wales Health Pathology Randwick | Area of Virology, Serology and Virology Division (SAViD), New South Wales Health Pathology Randwick | Rawlinson, W. |
| EPI_ISL_527329, EPI_ISL_527331, EPI_ISL_527332, EPI_ISL_527333, EPI_ISL_527334, EPI_ISL_527335, EPI_ISL_527336, EPI_ISL_527337, EPI_ISL_527339, EPI_ISL_527343, EPI_ISL_527344, EPI_ISL_527345, EPI_ISL_527346, EPI_ISL_527347, EPI_ISL_527348, EPI_ISL_527349, EPI_ISL_527351, EPI_ISL_527352, EPI_ISL_527353, EPI_ISL_527354, EPI_ISL_527355, EPI_ISL_527356, EPI_ISL_527357, EPI_ISL_527358 | | | |
| see above | Respiratory Virus Unit, Microbiology Services Colindale, Public Health England | Respiratory Virus Unit, Microbiology Services Colindale, Public Health England | PHE Covid Sequencing Team |
| EPI_ISL_527359, EPI_ISL_527360, EPI_ISL_527361, EPI_ISL_527362, EPI_ISL_527363, EPI_ISL_527364, EPI_ISL_527365, EPI_ISL_527366, EPI_ISL_527367, EPI_ISL_527368, EPI_ISL_527370, EPI_ISL_527371, EPI_ISL_527372, EPI_ISL_527373, EPI_ISL_527374, EPI_ISL_527375, EPI_ISL_527376, EPI_ISL_527377, EPI_ISL_527379 | | | |
| see above | National Public Health Laboratory, National Centre for Infectious Diseases | National Public Health Laboratory, National Centre for Infectious Diseases | Mak TM, Octavia S, Zhou Z, Cui L, Lin RTP |
| EPI_ISL_527380 | Istituto Zooprofilattico Sperimentale Puglia e Basilicata; Dipartimento di Bioscienze, Biotecnologie e Biofarmaceutica dell'Università degli Studi di Bari "A.Moro"; Istituto di Biomembrane. Bioenergetica e Biotecnologie Molecolari del Consiglio Nazionale delle Ricerche di Bari | Beaconlab (Bioinformatics, Evolution and Comparative Genomics lab), Dept of Biosciences, University on Milan | Parisi A.,Pesole G., Manzari C., Chiara M |
| EPI_ISL_527382, EPI_ISL_527383, EPI_ISL_527384, EPI_ISL_527385, EPI_ISL_527386, EPI_ISL_527387, EPI_ISL_527388, EPI_ISL_527389, EPI_ISL_527390, EPI_ISL_527391, EPI_ISL_527392, EPI_ISL_527393, EPI_ISL_527394, EPI_ISL_527395, EPI_ISL_527396, EPI_ISL_527397, EPI_ISL_527398, EPI_ISL_527399 | | | |
| see above | University of Miami Immunology and Histocompatibility Laboratory | University of Miami Immunology and Histocompatibility Laboratory | Emilio Margolles-Clark, PhD and Phillip Ruiz, MD, PhD |
| EPI_ISL_527400 | Area of Virology, Serology and Virology Division (SAViD), New South Wales Health Pathology Randwick | Area of Virology, Serology and Virology Division (SAViD), New South Wales Health Pathology Randwick | Rawlinson, W., Deveson, I., Bull, R. |
| EPI_ISL_527401, EPI_ISL_527402, EPI_ISL_527403, EPI_ISL_527404, EPI_ISL_527405, EPI_ISL_527406, EPI_ISL_527407, EPI_ISL_527408, EPI_ISL_527409, EPI_ISL_527410, EPI_ISL_527411, EPI_ISL_527412, EPI_ISL_527413, EPI_ISL_527414, EPI_ISL_527415, EPI_ISL_527416, EPI_ISL_527417, EPI_ISL_527418, EPI_ISL_527419, EPI_ISL_527420, EPI_ISL_527421, EPI_ISL_527422, EPI_ISL_527423, EPI_ISL_527424, EPI_ISL_527425, EPI_ISL_527426, EPI_ISL_527427, EPI_ISL_527428, EPI_ISL_527429, EPI_ISL_527430, EPI_ISL_527431, EPI_ISL_527432, EPI_ISL_527433, EPI_ISL_527434, EPI_ISL_527435, EPI_ISL_527436, EPI_ISL_527437, EPI_ISL_527438, EPI_ISL_527439, EPI_ISL_527440, EPI_ISL_527441, EPI_ISL_527442, EPI_ISL_527443, EPI_ISL_527444, EPI_ISL_527445, EPI_ISL_527446, EPI_ISL_527447, EPI_ISL_527448, EPI_ISL_527449, EPI_ISL_527450, EPI_ISL_527451, EPI_ISL_527452, EPI_ISL_527453, EPI_ISL_527454, EPI_ISL_527455, EPI_ISL_527456, EPI_ISL_527457, EPI_ISL_527458, EPI_ISL_527459, EPI_ISL_527460, EPI_ISL_527462, EPI_ISL_527463, EPI_ISL_527464, EPI_ISL_527465, EPI_ISL_527466, EPI_ISL_527468, EPI_ISL_527469, EPI_ISL_527470, EPI_ISL_527471, EPI_ISL_527472, EPI_ISL_527473, EPI_ISL_527474, EPI_ISL_527475, EPI_ISL_527476, EPI_ISL_527477, EPI_ISL_527478, EPI_ISL_527479, EPI_ISL_527480, EPI_ISL_527481, EPI_ISL_527482, EPI_ISL_527483, EPI_ISL_527484, EPI_ISL_527485, EPI_ISL_527486, EPI_ISL_527487, EPI_ISL_527488 | | | |
| see above | Colorado State University - Ebel Lab | Colorado State University - Ebel Lab | Greg Ebel et al. |
| EPI_ISL_527489, EPI_ISL_527543, EPI_ISL_527544, EPI_ISL_527545, EPI_ISL_527546, EPI_ISL_527547, EPI_ISL_527548, EPI_ISL_527549, EPI_ISL_527550, EPI_ISL_527551, EPI_ISL_527552, EPI_ISL_527553, EPI_ISL_527554, EPI_ISL_527555, EPI_ISL_527556, EPI_ISL_527557, EPI_ISL_527558, EPI_ISL_527559, EPI_ISL_527560, EPI_ISL_527561, EPI_ISL_527562, EPI_ISL_527563, EPI_ISL_527564, EPI_ISL_527565, EPI_ISL_527566, EPI_ISL_527567, EPI_ISL_527568, EPI_ISL_527569, EPI_ISL_527570, EPI_ISL_527571, EPI_ISL_527572 | | | |
| see above | Viral Respiratory Lab, National Institute for Biomedical Research (INRB) | Pathogen Sequencing Lab, National Institute for Biomedical Research (INRB) | Placide Mbala-Kingebeni, Edith Nkwembe, Eddy Kinganda-Lusamaki, Amuri Aziza, Francisca Muyembe Mawete, Emmanuel Lokilo Lofiko, Catherine Pratt, Matthias Pauthner, Josh Quick, Allison Black, James Hadfield, Trevor Bedford, Ian Goodfellow, Andrew Rambaut, Nick Loman, Kristian Andersen, Michael Wiley, Steve Ahuka-Mundeke, Jean-Jacques Muyembe Tamfum |
| EPI_ISL_527573, EPI_ISL_527574, EPI_ISL_527575, EPI_ISL_527576, EPI_ISL_527577, EPI_ISL_527578, EPI_ISL_527579, EPI_ISL_527580, EPI_ISL_527581, EPI_ISL_527582, EPI_ISL_527583, EPI_ISL_527584, EPI_ISL_527585, EPI_ISL_527586, EPI_ISL_527587, EPI_ISL_527588, EPI_ISL_527589, EPI_ISL_527590, EPI_ISL_527591, EPI_ISL_527592, EPI_ISL_527593, EPI_ISL_527594, EPI_ISL_527595, EPI_ISL_527596, EPI_ISL_527597, EPI_ISL_527598, EPI_ISL_527599, EPI_ISL_527600, EPI_ISL_527601, EPI_ISL_527602, EPI_ISL_527603, EPI_ISL_527604, EPI_ISL_527605, EPI_ISL_527606, EPI_ISL_527607, EPI_ISL_527608, EPI_ISL_527609, EPI_ISL_527610, EPI_ISL_527611, EPI_ISL_527612, EPI_ISL_527613, EPI_ISL_527614, EPI_ISL_527615, EPI_ISL_527616, EPI_ISL_527617, EPI_ISL_527618, EPI_ISL_527619, EPI_ISL_527620, EPI_ISL_527621, EPI_ISL_527622, EPI_ISL_527623, EPI_ISL_527624, EPI_ISL_527625, EPI_ISL_527626, EPI_ISL_527627, EPI_ISL_527628, EPI_ISL_527629 | | | |
| see above | Minnesota Department of Health, Public Health Laboratory | Minnesota Department of Health, Public Health Laboratory | Matt Plumb, Jacob Garfin, and Xiong Wang |
| EPI_ISL_527630, EPI_ISL_527631 | Mayo Clinic & Mayo Clinic Laboratories | Minnesota Department of Health, Public Health Laboratory | Matt Plumb, Jacob Garfin, and Xiong Wang |
| EPI_ISL_527632, EPI_ISL_527633, EPI_ISL_527634, EPI_ISL_527635, EPI_ISL_527636, EPI_ISL_527637, EPI_ISL_527638, EPI_ISL_527639, EPI_ISL_527640, EPI_ISL_527641, EPI_ISL_527642, EPI_ISL_527643, EPI_ISL_527644, EPI_ISL_527645, EPI_ISL_527646, EPI_ISL_527647, EPI_ISL_527648, EPI_ISL_527649, EPI_ISL_527650, EPI_ISL_527651, EPI_ISL_527652, EPI_ISL_527653, EPI_ISL_527654, EPI_ISL_527655, EPI_ISL_527656, EPI_ISL_527657 | | | |
| see above | AR Dept. of Health-Public Health Lab | Pathogen Discovery, Respiratory Viruses Branch, Division of Viral Diseases, Centers for Disease Control and Prevention | Ying Tao, Jing Zhang, Yan Li, Krista Queen, Anna Uehara, Clinton Paden, Haibin Wang, Suxiang Tong |
| EPI_ISL_527658, EPI_ISL_527659, EPI_ISL_527660, EPI_ISL_527661, EPI_ISL_527662, EPI_ISL_527663, EPI_ISL_527664, EPI_ISL_527665 | GA Department of Public Health Laboratory | Pathogen Discovery, Respiratory Viruses Branch, Division of Viral Diseases, Centers for Disease Control and Prevention | Yan Li, Anna Montmayeur, Jing Zhang, Krista Queen, Ying Tao, Anna Uehara, Rachel Marine, Clinton R. Paden, Haibin Wang, Suxiang Tong |
| EPI_ISL_527666 | IN State Department of Health Laboratory Services | Pathogen Discovery, Respiratory Viruses Branch, Division of Viral Diseases, Centers for Disease Control and Prevention | Krista Queen, Brian Lynch, Yan Li, Anna Montmayeur, Jing Zhang, Ying Tao, Anna Uehara, Rachel Marine, Clinton R. Paden, Haibin Wang, Suxiang Tong |
| EPI_ISL_527667, EPI_ISL_527668, EPI_ISL_527669, EPI_ISL_527670, EPI_ISL_527671, EPI_ISL_527672, EPI_ISL_527673, EPI_ISL_527674, EPI_ISL_527675, EPI_ISL_527676, EPI_ISL_527677, EPI_ISL_527678, EPI_ISL_527679, EPI_ISL_527680, EPI_ISL_527681, EPI_ISL_527682, EPI_ISL_527683, EPI_ISL_527684, EPI_ISL_527685, EPI_ISL_527686, EPI_ISL_527687, EPI_ISL_527688, EPI_ISL_527689, EPI_ISL_527690, EPI_ISL_527691 | | | |
| see above | MN PHL Division, Minnesota Department of Health | Pathogen Discovery, Respiratory Viruses Branch, Division of Viral Diseases, Centers for Disease Control and Prevention | Yan Li, Anna Montmayeur, Jing Zhang, Krista Queen, Ying Tao, Anna Uehara, Rachel Marine, Clinton R. Paden, Haibin Wang, Suxiang Tong |
| EPI_ISL_527692 | MN PHL Division, Minnesota Department of Health | Pathogen Discovery, Respiratory Viruses Branch, Division of Viral Diseases, Centers for Disease Control and Prevention | Krista Queen, Brian Lynch, Yan Li, Anna Montmayeur, Jing Zhang, Ying Tao, Anna Uehara, Rachel Marine, Clinton R. Paden, Haibin Wang, Suxiang Tong |
| EPI_ISL_527693, EPI_ISL_527694, EPI_ISL_527695, EPI_ISL_527696, EPI_ISL_527697, EPI_ISL_527698 | MN PHL Division, Minnesota Department of Health | Pathogen Discovery, Respiratory Viruses Branch, Division of Viral Diseases, Centers for Disease Control and Prevention | Yan Li, Anna Montmayeur, Jing Zhang, Krista Queen, Ying Tao, Anna Uehara, Rachel Marine, Clinton R. Paden, Haibin Wang, Suxiang Tong |
| EPI_ISL_527699 | MN PHL Division, Minnesota Department of Health | Pathogen Discovery, Respiratory Viruses Branch, Division of Viral Diseases, Centers for Disease Control and Prevention | Krista Queen, Brian Lynch, Yan Li, Anna Montmayeur, Jing Zhang, Ying Tao, Anna Uehara, Rachel Marine, Clinton R. Paden, Haibin Wang, Suxiang Tong |
| EPI_ISL_527700 | MN PHL Division, Minnesota Department of Health | Pathogen Discovery, Respiratory Viruses Branch, Division of Viral Diseases, Centers for Disease Control and Prevention | Yan Li, Anna Montmayeur, Jing Zhang, Krista Queen, Ying Tao, Anna Uehara, Rachel Marine, Clinton R. Paden, Haibin Wang, Suxiang Tong |
| EPI_ISL_527701 | MN PHL Division, Minnesota Department of Health | Pathogen Discovery, Respiratory Viruses Branch, Division of Viral Diseases, Centers for Disease Control and Prevention | Krista Queen, Brian Lynch, Yan Li, Anna Montmayeur, Jing Zhang, Ying Tao, Anna Uehara, Rachel Marine, Clinton R. Paden, Haibin Wang, Suxiang Tong |
| EPI_ISL_527702, EPI_ISL_527703, EPI_ISL_527704, EPI_ISL_527705, EPI_ISL_527706, EPI_ISL_527707, EPI_ISL_527708, EPI_ISL_527709, EPI_ISL_527710, EPI_ISL_527711, EPI_ISL_527712, EPI_ISL_527713 | | | |
| see above | MN PHL Division, Minnesota Department of Health | Pathogen Discovery, Respiratory Viruses Branch, Division of Viral Diseases, Centers for Disease Control and Prevention | Yan Li, Anna Montmayeur, Jing Zhang, Krista Queen, Ying Tao, Anna Uehara, Rachel Marine, Clinton R. Paden, Haibin Wang, Suxiang Tong |
| EPI_ISL_527714 | MN PHL Division, Minnesota Department of Health | Pathogen Discovery, Respiratory Viruses Branch, Division of Viral Diseases, Centers for Disease Control and Prevention | Krista Queen, Brian Lynch, Yan Li, Anna Montmayeur, Jing Zhang, Ying Tao, Anna Uehara, Rachel Marine, Clinton R. Paden, Haibin Wang, Suxiang Tong |
| EPI_ISL_527715 | MN PHL Division, Minnesota Department of Health | Pathogen Discovery, Respiratory Viruses Branch, Division of Viral Diseases, Centers for Disease Control and Prevention | Yan Li, Anna Montmayeur, Jing Zhang, Krista Queen, Ying Tao, Anna Uehara, Rachel Marine, Clinton R. Paden, Haibin Wang, Suxiang Tong |
| EPI_ISL_527716, EPI_ISL_527717 | MN PHL Division, Minnesota Department of Health | Pathogen Discovery, Respiratory Viruses Branch, Division of Viral Diseases, Centers for Disease Control and Prevention | Krista Queen, Brian Lynch, Yan Li, Anna Montmayeur, Jing Zhang, Ying Tao, Anna Uehara, Rachel Marine, Clinton R. Paden, Haibin Wang, Suxiang Tong |
| EPI_ISL_527718, EPI_ISL_527719 | MN PHL Division, Minnesota Department of Health | Pathogen Discovery, Respiratory Viruses Branch, Division of Viral Diseases, Centers for Disease Control and Prevention | Yan Li, Anna Montmayeur, Jing Zhang, Krista Queen, Ying Tao, Anna Uehara, Rachel Marine, Clinton R. Paden, Haibin Wang, Suxiang Tong |
| EPI_ISL_527720 | MN PHL Division, Minnesota Department of Health | Pathogen Discovery, Respiratory Viruses Branch, Division of Viral Diseases, Centers for Disease Control and Prevention | Krista Queen, Brian Lynch, Yan Li, Anna Montmayeur, Jing Zhang, Ying Tao, Anna Uehara, Rachel Marine, Clinton R. Paden, Haibin Wang, Suxiang Tong |

| | | | |
|---|---|---|---|
| EPI_ISL_527721, EPI_ISL_527722, EPI_ISL_527723 | MN PHL Division, Minnesota Department of Health | Pathogen Discovery, Respiratory Viruses Branch, Division of Viral Diseases, Centers for Disease Control and Prevention | Yan Li, Anna Montmayeur, Jing Zhang, Krista Queen, Ying Tao, Anna Uehara, Rachel Marine, Clinton R. Paden, Haibin Wang, Suxiang Tong |
| EPI_ISL_527724 | MN PHL Division, Minnesota Department of Health | Pathogen Discovery, Respiratory Viruses Branch, Division of Viral Diseases, Centers for Disease Control and Prevention | Krista Queen, Brian Lynch, Yan Li, Anna Montmayeur, Jing Zhang, Ying Tao, Anna Uehara, Rachel Marine, Clinton R. Paden, Haibin Wang, Suxiang Tong |
| EPI_ISL_527725, EPI_ISL_527726, EPI_ISL_527727, EPI_ISL_527728, EPI_ISL_527729, EPI_ISL_527730 | MN PHL Division, Minnesota Department of Health | Pathogen Discovery, Respiratory Viruses Branch, Division of Viral Diseases, Centers for Disease Control and Prevention | Yan Li, Anna Montmayeur, Jing Zhang, Krista Queen, Ying Tao, Anna Uehara, Rachel Marine, Clinton R. Paden, Haibin Wang, Suxiang Tong |
| EPI_ISL_527731 | MN PHL Division, Minnesota Department of Health | Pathogen Discovery, Respiratory Viruses Branch, Division of Viral Diseases, Centers for Disease Control and Prevention | Krista Queen, Brian Lynch, Yan Li, Anna Montmayeur, Jing Zhang, Ying Tao, Anna Uehara, Rachel Marine, Clinton R. Paden, Haibin Wang, Suxiang Tong |
| EPI_ISL_527732, EPI_ISL_527733 | MN PHL Division, Minnesota Department of Health | Pathogen Discovery, Respiratory Viruses Branch, Division of Viral Diseases, Centers for Disease Control and Prevention | Yan Li, Anna Montmayeur, Jing Zhang, Krista Queen, Ying Tao, Anna Uehara, Rachel Marine, Clinton R. Paden, Haibin Wang, Suxiang Tong |
| EPI_ISL_527734, EPI_ISL_527735 | MN PHL Division, Minnesota Department of Health | Pathogen Discovery, Respiratory Viruses Branch, Division of Viral Diseases, Centers for Disease Control and Prevention | Krista Queen, Brian Lynch, Yan Li, Anna Montmayeur, Jing Zhang, Ying Tao, Anna Uehara, Rachel Marine, Clinton R. Paden, Haibin Wang, Suxiang Tong |
| EPI_ISL_527736 | MN PHL Division, Minnesota Department of Health | Pathogen Discovery, Respiratory Viruses Branch, Division of Viral Diseases, Centers for Disease Control and Prevention | Yan Li, Anna Montmayeur, Jing Zhang, Krista Queen, Ying Tao, Anna Uehara, Rachel Marine, Clinton R. Paden, Haibin Wang, Suxiang Tong |
| EPI_ISL_527737 | WI State Laboratory of Hygiene | Pathogen Discovery, Respiratory Viruses Branch, Division of Viral Diseases, Centers for Disease Control and Prevention | Ying Tao, Jing Zhang, Yan Li, Krista Queen, Anna Uehara, Clinton Paden, Haibin Wang, Suxiang Tong |
| EPI_ISL_527739 | Hospital Mexico [San Jose/San Jose] | Inciensa, Instituto Costarricense de Investigación y Enseñanza en Nutrición y Salud | Francisco Duarte, Hebleen Porras, Claudio Soto-Garita, Estela Cordero, Adriana Godinez & Melany Calderon |
| EPI_ISL_527740 | Area De Salud Alajuela Norte - Clinica Dr. Marcial Rodriguez | Inciensa, Instituto Costarricense de Investigación y Enseñanza en Nutrición y Salud | Francisco Duarte, Hebleen Porras, Claudio Soto-Garita, Estela Cordero, Adriana Godinez & Melany Calderon |
| EPI_ISL_527741 | Hospital Metropolitano | Inciensa, Instituto Costarricense de Investigación y Enseñanza en Nutrición y Salud | Francisco Duarte, Hebleen Porras, Claudio Soto-Garita, Estela Cordero, Adriana Godinez & Melany Calderon |
| EPI_ISL_527742 | Centro Nacional De Rehabilitacion Humberto Araya Rojas (Cenare) | Inciensa, Instituto Costarricense de Investigación y Enseñanza en Nutrición y Salud | Francisco Duarte, Hebleen Porras, Claudio Soto-Garita, Estela Cordero, Adriana Godinez & Melany Calderon |
| EPI_ISL_527743, EPI_ISL_527744 | Hospital Cima | Inciensa, Instituto Costarricense de Investigación y Enseñanza en Nutrición y Salud | Francisco Duarte, Hebleen Porras, Claudio Soto-Garita, Estela Cordero, Adriana Godinez & Melany Calderon |
| EPI_ISL_527746, EPI_ISL_527747 | Area De Salud La Cruz | Inciensa, Instituto Costarricense de Investigación y Enseñanza en Nutrición y Salud | Francisco Duarte, Hebleen Porras, Claudio Soto-Garita, Estela Cordero, Adriana Godinez & Melany Calderon |
| EPI_ISL_527748 | Area De Salud Corredores | Inciensa, Instituto Costarricense de Investigación y Enseñanza en Nutrición y Salud | Francisco Duarte, Hebleen Porras, Claudio Soto-Garita, Estela Cordero, Adriana Godinez & Melany Calderon |
| EPI_ISL_527749 | Area De Salud Alajuela Norte - Clinica Dr. Marcial Rodriguez | Inciensa, Instituto Costarricense de Investigación y Enseñanza en Nutrición y Salud | Francisco Duarte, Hebleen Porras, Claudio Soto-Garita, Estela Cordero, Adriana Godinez & Melany Calderon |
| EPI_ISL_527750 | Hospital De Niños Dr. Carlos Saenz Herrera [San Jose/San Jose] | Inciensa, Instituto Costarricense de Investigación y Enseñanza en Nutrición y Salud | Francisco Duarte, Hebleen Porras, Claudio Soto-Garita, Estela Cordero, Adriana Godinez & Melany Calderon |
| EPI_ISL_527751 | Area De Salud Corredores | Inciensa, Instituto Costarricense de Investigación y Enseñanza en Nutrición y Salud | Francisco Duarte, Hebleen Porras, Claudio Soto-Garita, Estela Cordero, Adriana Godinez & Melany Calderon |
| EPI_ISL_527752 | Hospital Dr. Rafael A. Calderon Guardia | Inciensa, Instituto Costarricense de Investigación y Enseñanza en Nutrición y Salud | Francisco Duarte, Hebleen Porras, Claudio Soto-Garita, Estela Cordero, Adriana Godinez & Melany Calderon |
| EPI_ISL_527753, EPI_ISL_527754 | Hospital San Vicente De Paul | Inciensa, Instituto Costarricense de Investigación y Enseñanza en Nutrición y Salud | Francisco Duarte, Hebleen Porras, Claudio Soto-Garita, Estela Cordero, Adriana Godinez & Melany Calderon |
| EPI_ISL_527755 | Centro Nacional De Rehabilitacion Humberto Araya Rojas (Cenare) | Inciensa, Instituto Costarricense de Investigación y Enseñanza en Nutrición y Salud | Francisco Duarte, Hebleen Porras, Claudio Soto-Garita, Estela Cordero, Adriana Godinez & Melany Calderon |
| EPI_ISL_527756 | Area De Salud Aserri | Inciensa, Instituto Costarricense de Investigación y Enseñanza en Nutrición y Salud | Francisco Duarte, Hebleen Porras, Claudio Soto-Garita, Estela Cordero, Adriana Godinez & Melany Calderon |
| EPI_ISL_527757 | Area De Salud Goicoechea 1 | Inciensa, Instituto Costarricense de Investigación y Enseñanza en Nutrición y Salud | Francisco Duarte, Hebleen Porras, Claudio Soto-Garita, Estela Cordero, Adriana Godinez & Melany Calderon |
| EPI_ISL_527787, EPI_ISL_527789 | Laboratorio de Referencia Nacional de Virus Respiratorio. Centro Nacional de Salud Publica. Instituto Nacional de Salud Peru. | Laboratorio de Referencia Nacional de Biotecnología y Biología Molecular. Centro Nacional de Salud Publica. Instituto Nacional de Salud Peru. | Carlos Padilla Rojas, Karolyn Vega Chozo, Priscila Lope Pari, Omar Caceres Rey, Marco Galarza Perez, Maribel Huaringa Nuñez, Johanna Balbuena Torres, Henri Bailon Calderon, Nancy Rojas Serrano. |
| EPI_ISL_527794, EPI_ISL_527795, EPI_ISL_527796, EPI_ISL_527797, EPI_ISL_527799, EPI_ISL_527801, EPI_ISL_527802, EPI_ISL_527803, EPI_ISL_527804, EPI_ISL_527806, EPI_ISL_527808 | | | |
| see above | University of Wisconsin-Madison AIDS Vaccine Research Laboratories | University of Wisconsin-Madison AIDS Vaccine Research Laboratories | Gage Moreno, Katarina Braun, et al. AIDS Vaccine Research Laboratories |
| EPI_ISL_527809, EPI_ISL_527810, EPI_ISL_527811 | Institute of Microbiology, Universidad San Francisco de Quito | Institute of Microbiology, Universidad San Francisco de Quito | Belén Prado-Vivar, Sully Márquez, Juan José Guadalupe, Monica Becerra-Wong, Bernardo Gutiérrez, Stephanie Arregui, Rene Bracho, Karina Barragan, Anita Garcia, Carlos Tobar, Verónica Barragán, Patricio Rojas-Silva, Gabriel Trueba, Michelle Grunauer, Paúl Cárdenas |
| EPI_ISL_527812, EPI_ISL_527813, EPI_ISL_527814, EPI_ISL_527815, EPI_ISL_527816, EPI_ISL_527817 | Institute of Microbiology, Universidad San Francisco de Quito | Institute of Microbiology, Universidad San Francisco de Quito | Belén Prado-Vivar, Sully Márquez, Juan José Guadalupe, Monica Becerra-Wong, Bernardo Gutiérrez, Khurram Mahbbob, Verónica Barragán, Patricio Rojas-Silva, Gabriel Trueba, Michelle Grunauer, Paúl Cárdenas |
| EPI_ISL_527818, EPI_ISL_527819 | Centro de Investigaciones, Universidad de Especialidades Espíritu Santo | Institute of Microbiology, Universidad San Francisco de Quito | Derly Andrade, Juan Carlos Fernandez, Belén Prado-Vivar, Sully Márquez, Juan José Guadalupe, Monica Becerra-Wong, Bernardo Gutiérrez, Gabriel Morey, Ruben Armas, Jose Pedro Barberan, Fernando Espinoza, Edith Lopez, Verónica Barragán, Patricio Rojas-Silva, Gabriel Trueba, Michelle Grunauer, Paúl Cárdenas |
| EPI_ISL_527838 | Texas Department of State Health Services | Texas Department of State Health Services | Bonnie Oh, Rashmi Tuladhar, Jenny Zhang, Maliha Rahman, Anita Pokharel, Myong Koag, Chun Wang, Rachel Lee, Grace Kubin |
| EPI_ISL_527856 | Hospital Municipal Prof. Waldomiro de Paula | Instituto Adolfo Lutz, Interdiciplinary Procedures Center, Strategic Laboratory | Claudio Tavares Sacchi, Claudia Regina Gonçalves, Erica Valessa Ramos Gomes |
| EPI_ISL_527857 | Hospital Regional Vale do Ribeira | Instituto Adolfo Lutz, Interdiciplinary Procedures Center, Strategic Laboratory | Claudio Tavares Sacchi, Claudia Regina Gonçalves, Erica Valessa Ramos Gomes |
| EPI_ISL_527858 | Pronto Atendimento Sancta Maggiore Jardim Paulista | Instituto Adolfo Lutz, Interdiciplinary Procedures Center, Strategic Laboratory | Claudio Tavares Sacchi, Claudia Regina Gonçalves, Erica Valessa Ramos Gomes |
| EPI_ISL_527859 | Hospital Municipal Vereador Jose Storopolli | Instituto Adolfo Lutz, Interdiciplinary Procedures Center, Strategic Laboratory | Claudio Tavares Sacchi, Claudia Regina Gonçalves, Erica Valessa Ramos Gomes |
| EPI_ISL_527860 | Hospital Municipal de Parelheiros Josanias Castanha Braga | Instituto Adolfo Lutz, Interdiciplinary Procedures Center, Strategic Laboratory | Claudio Tavares Sacchi, Claudia Regina Gonçalves, Erica Valessa Ramos Gomes |
| EPI_ISL_527861 | Hospital e Maternidade Celso Pierro | Instituto Adolfo Lutz, Interdiciplinary Procedures Center, Strategic Laboratory | Av. Dr. Arnaldo, 355 - Brazil, Cerqueira Cesar, São Paulo - SP, 01246-1301 |
| EPI_ISL_527862 | Hospital Municipal de Urgência | Instituto Adolfo Lutz, Interdiciplinary Procedures Center, Strategic Laboratory | Claudio Tavares Sacchi, Claudia Regina Gonçalves, Erica Valessa Ramos Gomes |
| EPI_ISL_527863 | Hospital Municipal do Tatuape Carmino Caricchio | Instituto Adolfo Lutz, Interdiciplinary Procedures Center, Strategic Laboratory | Claudio Tavares Sacchi, Claudia Regina Gonçalves, Erica Valessa Ramos Gomes |
| EPI_ISL_527864 | Hospital e Pronto Socorro Comunitário Vila Iolanda | Instituto Adolfo Lutz, Interdiciplinary Procedures Center, Strategic Laboratory | Claudio Tavares Sacchi, Claudia Regina Gonçalves, Erica Valessa Ramos Gomes |
| EPI_ISL_527865 | Hospital e Maternidade São Cristóvão | Instituto Adolfo Lutz, Interdiciplinary Procedures Center, Strategic Laboratory | Claudio Tavares Sacchi, Claudia Regina Gonçalves, Erica Valessa Ramos Gomes |
| EPI_ISL_527866 | PS Municipal Dr Lauro Ribas Braga | Instituto Adolfo Lutz, Interdiciplinary Procedures Center, Strategic Laboratory | Av. Dr. Arnaldo, 355 - Brazil, Cerqueira Cesar, São Paulo - SP, 01246-1301 |
| EPI_ISL_527867 | Pronto Socorro Municipal - Balneario São José | Instituto Adolfo Lutz, Interdiciplinary Procedures Center, Strategic Laboratory | Claudio Tavares Sacchi, Claudia Regina Gonçalves, Erica Valessa Ramos Gomes |
| EPI_ISL_527868 | Hospital e Maternidade do Braz | Instituto Adolfo Lutz, Interdiciplinary Procedures Center, Strategic Laboratory | Claudio Tavares Sacchi, Claudia Regina Gonçalves, Erica Valessa Ramos Gomes |
| EPI_ISL_527869 | Hospital Municipal Carmen Prudente | Instituto Adolfo Lutz, Interdiciplinary Procedures Center, Strategic Laboratory | Claudio Tavares Sacchi, Claudia Regina Gonçalves, Erica Valessa Ramos Gomes |
| EPI_ISL_527870 | Hospital Municipal Mário Gatti | Instituto Adolfo Lutz, Interdiciplinary Procedures Center, Strategic Laboratory | Claudio Tavares Sacchi, Claudia Regina Gonçalves, Erica Valessa Ramos Gomes |
| EPI_ISL_527873, EPI_ISL_527874, EPI_ISL_527876, EPI_ISL_527878, EPI_ISL_527879, EPI_ISL_527880, EPI_ISL_527881, EPI_ISL_527882, EPI_ISL_527883, EPI_ISL_527884, EPI_ISL_527887, EPI_ISL_527888, EPI_ISL_527889, EPI_ISL_527890, EPI_ISL_527891, EPI_ISL_527892, EPI_ISL_527893, EPI_ISL_527898, EPI_ISL_527901, EPI_ISL_527903, EPI_ISL_527904, EPI_ISL_527905, EPI_ISL_527910, EPI_ISL_527911, EPI_ISL_527912, EPI_ISL_527914, EPI_ISL_527915 | | | |
| see above | Nigeria Centre for Disease Control (NCDC) | African Centre of Excellence for Genomics of Infectious Diseases (ACEGID), Redeemer's University, Ede, Osun State, Nigeria | Oluniyi P.E. et al |
| EPI_ISL_527916, EPI_ISL_527917, EPI_ISL_527926, EPI_ISL_527928, EPI_ISL_527929, EPI_ISL_527930, EPI_ISL_527931, EPI_ISL_527932, EPI_ISL_527933, EPI_ISL_527934, EPI_ISL_527941, EPI_ISL_527947, EPI_ISL_527954, EPI_ISL_527955, EPI_ISL_527956, EPI_ISL_527957, EPI_ISL_527958, EPI_ISL_527963, EPI_ISL_527964, EPI_ISL_527966, EPI_ISL_527967, EPI_ISL_527968, EPI_ISL_527970, EPI_ISL_527972, EPI_ISL_527973, EPI_ISL_527974, EPI_ISL_527983, EPI_ISL_527984, EPI_ISL_527986, EPI_ISL_527987, EPI_ISL_527990, EPI_ISL_527991, EPI_ISL_527992, EPI_ISL_527994, EPI_ISL_527996, EPI_ISL_527998, EPI_ISL_528001, EPI_ISL_528002, EPI_ISL_528003, EPI_ISL_528004, EPI_ISL_528005, EPI_ISL_528006, EPI_ISL_528007, EPI_ISL_528008, EPI_ISL_528009, EPI_ISL_528010, EPI_ISL_528011, EPI_ISL_528012, EPI_ISL_528013, EPI_ISL_528014, EPI_ISL_528015, EPI_ISL_528016, EPI_ISL_528017, EPI_ISL_528020, EPI_ISL_528021, EPI_ISL_528022, EPI_ISL_528023, EPI_ISL_528024, EPI_ISL_528025, EPI_ISL_528027, EPI_ISL_528028, EPI_ISL_528029, EPI_ISL_528030, EPI_ISL_528031, EPI_ISL_528032, EPI_ISL_528033, EPI_ISL_528034, EPI_ISL_528037, EPI_ISL_528039, EPI_ISL_528040, EPI_ISL_528041, EPI_ISL_528042, EPI_ISL_528043, EPI_ISL_528044, EPI_ISL_528045, EPI_ISL_528046, EPI_ISL_528047, EPI_ISL_528048, EPI_ISL_528049, EPI_ISL_528050, EPI_ISL_528051, EPI_ISL_528052, EPI_ISL_528053, EPI_ISL_528054, EPI_ISL_528055, EPI_ISL_528056, EPI_ISL_528057, EPI_ISL_528058, EPI_ISL_528059, EPI_ISL_528060, EPI_ISL_528061, EPI_ISL_528063, EPI_ISL_528064, EPI_ISL_528065, EPI_ISL_528066, EPI_ISL_528067, EPI_ISL_528068, EPI_ISL_528069, EPI_ISL_528070, EPI_ISL_528072, EPI_ISL_528073, EPI_ISL_528074, EPI_ISL_528075, EPI_ISL_528076, EPI_ISL_528077, EPI_ISL_528079, EPI_ISL_528080, EPI_ISL_528081, EPI_ISL_528082, EPI_ISL_528083, EPI_ISL_528084, EPI_ISL_528085, EPI_ISL_528086, EPI_ISL_528087, EPI_ISL_528088, EPI_ISL_528089, EPI_ISL_528090, EPI_ISL_528091, EPI_ISL_528092, EPI_ISL_528093, EPI_ISL_528095, EPI_ISL_528096, EPI_ISL_528097, EPI_ISL_528099, EPI_ISL_528100, EPI_ISL_528103, EPI_ISL_528104, EPI_ISL_528105, EPI_ISL_528106, EPI_ISL_528107, EPI_ISL_528109, EPI_ISL_528110, EPI_ISL_528111, EPI_ISL_528112, EPI_ISL_528113, EPI_ISL_528114, EPI_ISL_528115, EPI_ISL_528117, EPI_ISL_528118, EPI_ISL_528119, EPI_ISL_528120, EPI_ISL_528121, EPI_ISL_528122, EPI_ISL_528123, EPI_ISL_528124, EPI_ISL_528125, EPI_ISL_528126, EPI_ISL_528127, EPI_ISL_528128, EPI_ISL_528129, EPI_ISL_528130, EPI_ISL_528131, EPI_ISL_528132, EPI_ISL_528133, EPI_ISL_528134, EPI_ISL_528135, EPI_ISL_528137, EPI_ISL_528138, EPI_ISL_528139, EPI_ISL_528140, EPI_ISL_528141, EPI_ISL_528142, EPI_ISL_528143, EPI_ISL_528146, EPI_ISL_528147, EPI_ISL_528148, EPI_ISL_528149, EPI_ISL_528151, EPI_ISL_528154, EPI_ISL_528155, EPI_ISL_528157, EPI_ISL_528158, EPI_ISL_528162, EPI_ISL_528163, EPI_ISL_528164, EPI_ISL_528165, EPI_ISL_528166, EPI_ISL_528167, EPI_ISL_528168, EPI_ISL_528169, EPI_ISL_528171, EPI_ISL_528172, EPI_ISL_528173, EPI_ISL_528174, EPI_ISL_528175, EPI_ISL_528176, EPI_ISL_528178, EPI_ISL_528179, EPI_ISL_528180, EPI_ISL_528181, EPI_ISL_528183, EPI_ISL_528185, EPI_ISL_528186, EPI_ISL_528187, EPI_ISL_528188, EPI_ISL_528190, EPI_ISL_528191, EPI_ISL_528193, EPI_ISL_528194, EPI_ISL_528196, EPI_ISL_528198, EPI_ISL_528199, EPI_ISL_528200, EPI_ISL_528201, EPI_ISL_528202, EPI_ISL_528204, EPI_ISL_528206, EPI_ISL_528208, EPI_ISL_528212, EPI_ISL_528213, EPI_ISL_528214, EPI_ISL_528215, EPI_ISL_528216, EPI_ISL_528217, EPI_ISL_528218, EPI_ISL_528222, EPI_ISL_528223, EPI_ISL_528224, EPI_ISL_528225, EPI_ISL_528227, EPI_ISL_528229, EPI_ISL_528231, EPI_ISL_528232, EPI_ISL_528233, EPI_ISL_528235, EPI_ISL_528236, EPI_ISL_528237, EPI_ISL_528239, EPI_ISL_528242, EPI_ISL_528243, EPI_ISL_528244, EPI_ISL_528245, EPI_ISL_528246, EPI_ISL_528247, EPI_ISL_528249, EPI_ISL_528251, EPI_ISL_528252, EPI_ISL_528254, EPI_ISL_528255, EPI_ISL_528257, EPI_ISL_528258, EPI_ISL_528259, EPI_ISL_528261, EPI_ISL_528262, EPI_ISL_528263, EPI_ISL_528264, EPI_ISL_528265, EPI_ISL_528267, EPI_ISL_528270, EPI_ISL_528271, EPI_ISL_528272, EPI_ISL_528273, EPI_ISL_528274, EPI_ISL_528277, EPI_ISL_528279, EPI_ISL_528280, EPI_ISL_528281, EPI_ISL_528283, EPI_ISL_528284, EPI_ISL_528285, EPI_ISL_528286, EPI_ISL_528287, EPI_ISL_528288, EPI_ISL_528292, EPI_ISL_528293, EPI_ISL_528294, EPI_ISL_528295, EPI_ISL_528296, EPI_ISL_528297, EPI_ISL_528299, EPI_ISL_528301, EPI_ISL_528302, EPI_ISL_528304, EPI_ISL_528309, EPI_ISL_528310, EPI_ISL_528311, EPI_ISL_528313, EPI_ISL_528315, EPI_ISL_528316, EPI_ISL_528317, EPI_ISL_528318, EPI_ISL_528319, EPI_ISL_528320, EPI_ISL_528323, EPI_ISL_528325, EPI_ISL_528326, EPI_ISL_528327, EPI_ISL_528328, EPI_ISL_528330, EPI_ISL_528331, EPI_ISL_528332, EPI_ISL_528337, EPI_ISL_528340, EPI_ISL_528341, EPI_ISL_528343, EPI_ISL_528344, EPI_ISL_528345, EPI_ISL_528347, EPI_ISL_528348, EPI_ISL_528349, EPI_ISL_528350, EPI_ISL_528351, EPI_ISL_528352, EPI_ISL_528354, EPI_ISL_528355, EPI_ISL_528359, EPI_ISL_528360, EPI_ISL_528362, EPI_ISL_528363, EPI_ISL_528364, EPI_ISL_528365, EPI_ISL_528366, EPI_ISL_528367, EPI_ISL_528368, EPI_ISL_528369, EPI_ISL_528370, EPI_ISL_528371, EPI_ISL_528373, EPI_ISL_528374, EPI_ISL_528377, EPI_ISL_528381 | | | |
| see above | University Hospital Basel, Clinical Virology | University Hospital Basel, Clinical Bacteriology | Madlen Stange, Alfredo Mari, Tim Roloff, Helena MB Seth-Smith, Michael Schweitzer, Myrta Brunner, Karoline Leuzinger, Kirstine K. Soegaard, Alexander Gensch, Sarah Tschudin-Sutter, Simon Fuchs, Julia Bielicki, Hans Pargger, Martin Siegemund, Christian Nickel, Roland Bingisser, Michael Osthoff, Stefano Bassetti, Rita Schneider-Sliwa, Manuel Battegay, Hans Hirsch, Adrian Egli |
| EPI_ISL_528382, EPI_ISL_528383, EPI_ISL_528384, EPI_ISL_528385 | Translational Health Science and Technology Institute -ESIC medical college and hospital, Faridabad | THSTI Bioassay laboratory | Saurabh Kumar, Jigme Wangchuk, Anil Kumar Pandey, Asim Das, Guruprasad R. Medigeshi |
| EPI_ISL_528386 | Viral vaccines, VSVRI- Veterinary serum and vaccine research institute | Viral vaccines, VSVRI- Veterinary serum and vaccine research institute | Saleh,A.A., Saad,M.A. |
| EPI_ISL_528388, EPI_ISL_528389, EPI_ISL_528391, EPI_ISL_528392, EPI_ISL_528398, EPI_ISL_528399, EPI_ISL_528400, EPI_ISL_528401, EPI_ISL_528403, EPI_ISL_528404, EPI_ISL_528405, EPI_ISL_528406, EPI_ISL_528407, EPI_ISL_528409, EPI_ISL_528411, EPI_ISL_528413, EPI_ISL_528414, EPI_ISL_528415, EPI_ISL_528430, EPI_ISL_528432, EPI_ISL_528433, EPI_ISL_528434, EPI_ISL_528435, EPI_ISL_528438 | | | |

| | | | |
|---|---|---|---|
| see above | Respiratory Virus Unit, Microbiology Services Colindale, Public Health England | Respiratory Virus Unit, Microbiology Services Colindale, Public Health England | PHE Covid Sequencing Team |
| EPI_ISL_528440, EPI_ISL_528441, EPI_ISL_528442, EPI_ISL_528443, EPI_ISL_528444, EPI_ISL_528445, EPI_ISL_528446, EPI_ISL_528447, EPI_ISL_528449, EPI_ISL_528450, EPI_ISL_528451, EPI_ISL_528452, EPI_ISL_528456, EPI_ISL_528457, EPI_ISL_528459, EPI_ISL_528460, EPI_ISL_528461, EPI_ISL_528462, EPI_ISL_528463, EPI_ISL_528464, EPI_ISL_528467, EPI_ISL_528468, EPI_ISL_528471, EPI_ISL_528472, EPI_ISL_528475, EPI_ISL_528477, EPI_ISL_528479, EPI_ISL_528480, EPI_ISL_528481, EPI_ISL_528482, EPI_ISL_528483, EPI_ISL_528484 | | | |
| see above | National Virus Reference Laboratory | National Virus Reference Laboratory | Michael Carr, Gabriel Gonzalez, Jonathan Dean, Suzie Coughlan, Cillian F De Gascun |
| EPI_ISL_528486, EPI_ISL_528493, EPI_ISL_528494, EPI_ISL_528495, EPI_ISL_528498, EPI_ISL_528513, EPI_ISL_528526 | Alaska State Virology Laboratory | Alaska State Virology Laboratory | Chen J et al with Pathogenomics group Dagdag R, Redlinger M, Milton E, George W, Kovalenko A, Drown DM, Bortz E |
| EPI_ISL_528538 | Alsafar | Alsafar | Andreas Henschel, Gihan Elsir Ahmed Daw Elbait, Samuel Feng, Rifat, Ernesto Damiani, Guan Tay, Habiba Alsafar |
| EPI_ISL_528539 | LVM/UFRJ | LNCC | Gustavo M. Romário M. de Souza; Bruno B. Bezerra; Lucio A. Caldas; Fabio Limonte; Elena Cobos; Sharton V. A. Coelho; Luiz Almeida; Luiza Higga; Isadora A. Correa; Diana Marianni; Luciana B. Arruda; Marcelo Bozza; Orlando Ferreira; Wanderley de Souza; Ana Teresa R. Vasconcelos; Terezinha M. Castineiras; Amilcar Tanuri; Luciana J. Costa. |
| EPI_ISL_528548, EPI_ISL_528549 | National Genomics Core-Center for DNA Fingerprinting and Diagnostics | National Genomics Core- Center for DNA Fingerprinting and Diagnostics (NGC-CDFD)- DBT's PAN-INDIA-1000 Genome consortium | G Shashikanth, Heena Shah, Bala Pratyusha, Vinay Donipadi, S Vasantha Rani, M Sri Lalitha, R. Angalena, Usha Rani Dutta, Nimmala Naresh, Nanci Rani K, Ch Venkateshwar Goud, Devinder Singh Negi, R Harinarayanan, Rashna Bhandari, Murali Dharan Bashyam, Debashish Mitra, Divya Vashisht, Ashwin Dalal |
| EPI_ISL_528555, EPI_ISL_528556 | National Genomics Core-Center for DNA Fingerprinting and Diagnostics | National Genomics Core- Center for DNA Fingerprinting and Diagnostics (NGC-CDFD)- DBT's PAN-INDIA-1000 Genome consortium | Heena Shah, G Shashikanth, Bala Pratyusha, Vinay Donipadi, Shruti Dasgupta, Kandali Sreethi Sreenivasulu Reddy, Chandra Shekhar Singh, Sunke Vijayakumar, R Lakshmi Vaishna, Jenige Aravindh Kumar, Muthulakshmi, V Naga Sailaja, R Harinarayanan, Rashna Bhandari, Murali Dharan Bashyam, Debashish Mitra, Divya Vashisht, Ashwin Dalal |
| EPI_ISL_528560, EPI_ISL_528562 | National Genomics Core-Center for DNA Fingerprinting and Diagnostics | National Genomics Core- Center for DNA Fingerprinting and Diagnostics (NGC-CDFD)- DBT's PAN-INDIA-1000 Genome consortium | Heena Shah, G Shashikanth, Bala Pratyusha, Vinay Donipadi, Binod Bihari Pradhan, Jamal Md Nurul Jain, Srinivas G, C Bala Maddeleti, R. Manorama, T. Navaneetha, Surya Vamshi, Chendra Shekar P, R Harinarayanan, Rashna Bhandari, Murali Dharan Bashyam, Debashish Mitra, Divya Vashisht, Ashwin Dalal |
| EPI_ISL_528576 | National Genomics Core-Center for DNA Fingerprinting and Diagnostics | National Genomics Core- Center for DNA Fingerprinting and Diagnostics (NGC-CDFD)- DBT's PAN-INDIA-1000 Genome consortium | Vinay Donipadi, G Shashikanth, Heena Shah, Bala Pratyusha, Parveen Kumar, Sandip Patra, Mugdha Singh, Reelina Basu, Dhanraj Adey, Bharath Kumar, Bhavani Sontam, Shaik Nasar Vali, R Harinarayanan, Rashna Bhandari, Murali Dharan Bashyam, Debashish Mitra, Divya Vashisht, Ashwin Dalal |
| EPI_ISL_528581, EPI_ISL_528586 | National Genomics Core-Center for DNA Fingerprinting and Diagnostics | National Genomics Core- Center for DNA Fingerprinting and Diagnostics (NGC-CDFD)- DBT's PAN-INDIA-1000 Genome consortium | G Shashikanth, Heena Shah, Bala Pratyusha, Vinay Donipadi, Bathula Siddardha, Vineesha Oddi, Lavanya Banda, Surya Chodisetty, Abhijeeth Singh Thakur, Mohammad Mudassir, Nalini Raghunathan, Rajeshree Sanyal, R Harinarayanan, Rashna Bhandari, Murali Dharan Bashyam, Debashish Mitra, Divya Vashisht, Ashwin Dalal |
| EPI_ISL_528637, EPI_ISL_528638 | LVM/UFRJ | Bioinformatics Laboratory / LNCC | Gustavo D. P. Silva; M. Romário M. de Souza; Bruno B. Bezerra; Lucio A. Caldas; Fabio Limonte; Elena Cobos; Sharton V. A. Coelho; Luiz Almeida; Luiza Higga; Isadora A. Correa; Diana Marianni; Luciana B. Arruda; Marcelo Bozza; Orlando Ferreira; Wanderley de Souza; Ana Teresa R. Vasconcelos; Terezinha M. Castineiras; Amilcar Tanuri; Luciana J. Costa |
| EPI_ISL_528639, EPI_ISL_528640, EPI_ISL_528641, EPI_ISL_528642, EPI_ISL_528643, EPI_ISL_528644, EPI_ISL_528645, EPI_ISL_528646, EPI_ISL_528647, EPI_ISL_528648, EPI_ISL_528649, EPI_ISL_528650, EPI_ISL_528651, EPI_ISL_528652, EPI_ISL_528654, EPI_ISL_528655, EPI_ISL_528656, EPI_ISL_528657, EPI_ISL_528658, EPI_ISL_528659, EPI_ISL_528660, EPI_ISL_528661, EPI_ISL_528662 | | | |
| see above | Virginia DCLS | Virginia DCLS | Virginia DCLS |
| EPI_ISL_528663, EPI_ISL_528664, EPI_ISL_528665, EPI_ISL_528666, EPI_ISL_528667, EPI_ISL_528668, EPI_ISL_528669, EPI_ISL_528670, EPI_ISL_528671, EPI_ISL_528672, EPI_ISL_528673, EPI_ISL_528674, EPI_ISL_528675, EPI_ISL_528676, EPI_ISL_528677, EPI_ISL_528678, EPI_ISL_528679, EPI_ISL_528680, EPI_ISL_528681, EPI_ISL_528682 | | | |
| see above | Virginia Division of Consolidated Laboratory Services (DCLS) | Virginia Division of Consolidated Laboratory Services (DCLS) | Virginia DCLS |
| EPI_ISL_528683 | NCDC Institute of Genomics and Integrative Biology | NCDC Institute of Genomics and Integrative Biology | Vivekanand A, Mahesh S. Dhar, Bharathram Uppili, Akshay Kanakan, Simmi Tiwari, RadhaKrishnan VS, Robin Marwal, Azka Khan, Ajit Shewale, Pooja Sharma, Tushar Nale, Rajesh Pandey, Sandhya Kabra, Mohammed Faruq, Sujeet Singh, Anurag Agrawal, Partha Rakshit |
| EPI_ISL_528686, EPI_ISL_528687, EPI_ISL_528688, EPI_ISL_528689, EPI_ISL_528690, EPI_ISL_528691, EPI_ISL_528692, EPI_ISL_528693, EPI_ISL_528694, EPI_ISL_528695, EPI_ISL_528696, EPI_ISL_528697, EPI_ISL_528698, EPI_ISL_528699, EPI_ISL_528700, EPI_ISL_528701, EPI_ISL_528702, EPI_ISL_528703, EPI_ISL_528704, EPI_ISL_528705, EPI_ISL_528706, EPI_ISL_528707, EPI_ISL_528708, EPI_ISL_528709, EPI_ISL_528710, EPI_ISL_528711, EPI_ISL_528712, EPI_ISL_528713, EPI_ISL_528714, EPI_ISL_528715, EPI_ISL_528716, EPI_ISL_528717, EPI_ISL_528718, EPI_ISL_528719, EPI_ISL_528720, EPI_ISL_528721 | | | |
| see above | Alsafar - Khalifa University Abu Dhabi | Alsafar - Khalifa University Abu Dhabi | Andreas Henschel, Gihan Daw Elbait, Samuel Feng, Rifat Hamoudi, Ernesto Damiani, Guan Tay, Habiba Alsafar |
| EPI_ISL_528738, EPI_ISL_528739, EPI_ISL_528740, EPI_ISL_528741 | Malaysia Genome Institute | Malaysia Genome Institute | Mohd Noor Mat Isa, Irni Suhayu Sapian, Yusuf Muhammad Noor, Nurhezreen Md Iqbal, Mohd Faizal Abu Bakar, Enizza Kasim, Shamsidar Sopie, Siti Noraini Othman, Azrin Ahmad, Nor Azfa Johari, Shahrul Hisham Zainal Ariffin |
| EPI_ISL_528742 | Malaysia Genome Institute | Malaysia Genome Institute | Mohd Noor Mat Isa, Irni Suhayu Sapian, Gan Han Ming, Yusuf Muhammad Noor, Tan Ju Lin, Nurhezreen Md Iqbal, Mohd Faizal Abu Bakar, Enizza Kasim, Shamsidar Sopie, Siti Noraini Othman, Azrin Ahmad, Nor Azfa Johari, Shahrul Hisham Zainal Ariffin |
| EPI_ISL_528743, EPI_ISL_528744 | Malaysia Genome Institute | Malaysia Genome Institute | Mohd Noor Mat Isa, Irni Suhayu Sapian, Yusuf Muhammad Noor, Nurhezreen Md Iqbal, Mohd Faizal Abu Bakar, Enizza Kasim, Shamsidar Sopie, Siti Noraini Othman, Azrin Ahmad, Nor Azfa Johari, Shahrul Hisham Zainal Ariffin |
| EPI_ISL_528745 | Laboratorium Kesehatan Provinsi Jawa Barat | School of Life Sciences and Technology & School of Pharmacy-Institut Teknologi Bandung; Molecular Genetics Laboratory-Faculty of Medicine-Universitas Padjadjaran; Laboratorium Kesehatan Provinsi Jawa Barat | Marselina Irasonia Tan, Yunia Sribudiani, Catur Riani, Azzania Fibriani, Husna Nugrahapraja, Tarwadi, Ema Rahmawati, Savira Ekawardhani, Hesti Lina Wiraswati, Ryan Bayusantika Ristandi, Rifky Waluyajati Rachman, Cut Nur Cinthia Alamanda, Lia Faridah, Tri Hanggono Achmad, Mas Rizky A.A. Syamsunarno, Fensi Amalina, Hammam Riza, Sony Solistia Wirawan, Agung Eru Wibowo, Irvan Faizal |
| EPI_ISL_528746 | Immanuel Hospital | Molecular Genetics Laboratory-Faculty of Medicine-Universitas Padjadjaran; School of Life Sciences and Technology & School of Pharmacy-Institut Teknologi Bandung; Laboratorium Kesehatan Provinsi Jawa Barat | Yunia Sribudiani,Tri Hanggono Achmad, Mas Rizky A.A. Syamsunarno, Fensi Amalina, Catur Riani, Azzania Fibriani, Husna Nugrahapraja, Marselina Irasonia Tan, Tarwadi, Ema Rahmawati, Savira Ekawardhani, Hesti Lina Wiraswati, Ryan Bayusantika Ristandi, Rifky Waluyajati Rachman, Cut Nur Cinthia Alamanda, Lia Faridah, Hammam Riza, Sony Solistia Wirawan, Agung Eru Wibowo, Irvan Faizal |
| EPI_ISL_528747 | Santo Borromeus Hospital | School of Pharmacy & School of Life Sciences and Technology - Institut Teknologi Bandung; Molecular Genetics Laboratory-Faculty of Medicine-Universitas Padjadjaran; Laboratorium Kesehatan Provinsi Jawa Barat | Catur Riani, Marselina Irasonia Tan, Yunia Sribudiani, Azzania Fibriani, Husna Nugrahapraja, Tarwadi, Ema Rahmawati, Savira Ekawardhani, Hesti Lina Wiraswati, Ryan Bayusantika Ristandi, Rifky Waluyajati Rachman, Cut Nur Cinthia Alamanda, Lia Faridah, Miftahul Faridl, Karimatu Khoirunnisa, Hammam Riza, Sony Solistia Wirawan, Agung Eru Wibowo, Irvan Faizal |
| EPI_ISL_528748 | Dinkes Provinsi Jawa Barat | School of Life Sciences and Technology & School of Pharmacy-Institut Teknologi Bandung; Molecular Genetics Laboratory-Faculty of Medicine-Universitas Padjadjaran; Laboratorium Kesehatan Provinsi Jawa Barat | Azzania Fibriani, Catur Riani, Marselina Irasonia Tan, Yunia Sribudiani, Husna Nugrahapraja, Tarwadi, Ema Rahmawati, Savira Ekawardhani, Hesti Lina Wiraswati, Ryan Bayusantika Ristandi, Rifky Waluyajati Rachman, Cut Nur Cinthia Alamanda, Lia Faridah, Gusti Ayu Prani Pradani, Adelina Khristiani Rahayu, Hammam Riza, Sony Solistia Wirawan, Agung Eru Wibowo, Irvan Faizal |
| EPI_ISL_528749 | Santosa Hospital Bandung Central | School of Life Sciences and Technology & School of Pharmacy-Institut Teknologi Bandung; Molecular Genetics Laboratory-Faculty of Medicine-Universitas Padjadjaran; Laboratorium Kesehatan Provinsi Jawa Barat | Husna Nugrahapraja, Azzania Fibriani, Catur Riani, Marselina Irasonia Tan, Yunia Sribudiani, Tarwadi, Ema Rahmawati, Savira Ekawardhani, Hesti Lina Wiraswati, Ryan Bayusantika Ristandi, Rifky Waluyajati Rachman, Cut Nur Cinthia Alamanda, Lia Faridah, Davin H. E. Setiamarga, Rizki Mardian, Hammam Riza, Sony Solistia Wirawan, Agung Eru Wibowo, Irvan Faizal |
| EPI_ISL_528750 | Santo Borromeus Hospital | School of Life Sciences and Technology & School of Pharmacy-Institut Teknologi Bandung; Molecular Genetics Laboratory-Faculty of Medicine-Universitas Padjadjaran; Laboratorium Kesehatan Provinsi Jawa Barat | Marselina Irasonia Tan, Yunia Sribudiani, Catur Riani, Azzania Fibriani, Husna Nugrahapraja, Tarwadi, Ema Rahmawati, Savira Ekawardhani, Hesti Lina Wiraswati, Ryan Bayusantika Ristandi, Rifky Waluyajati Rachman, Cut Nur Cinthia Alamanda, Lia Faridah, Miftahul Faridl, Karimatu Khoirunnisa, Hammam Riza, Sony Solistia Wirawan, Agung Eru Wibowo, Irvan Faizal |
| EPI_ISL_528751 | Santo Borromeus Hospital | Molecular Genetics Laboratory-Faculty of Medicine-Universitas Padjadjaran; School of Life Sciences and Technology & School of Pharmacy-Institut Teknologi Bandung; Laboratorium Kesehatan Provinsi Jawa Barat | Yunia Sribudiani, Tri Hanggono Achmad, Mas Rizky A.A. Syamsunarno, Fensi Amalina, Catur Riani, Azzania Fibriani, Husna Nugrahapraja, Marselina Irasonia Tan, Tarwadi, Ema Rahmawati, Savira Ekawardhani, Hesti Lina Wiraswati, Ryan Bayusantika Ristandi, Rifky Waluyajati Rachman, Cut Nur Cinthia Alamanda, Lia Faridah, Hammam Riza, Sony Solistia Wirawan, Agung Eru Wibowo, Irvan Faizal |
| EPI_ISL_528752 | Dr. H. A. Rotinsulu Lung Hospital | School of Pharmacy & School of Life Sciences and Technology - Institut Teknologi Bandung; Molecular Genetics Laboratory-Faculty of Medicine-Universitas Padjadjaran; Laboratorium Kesehatan Provinsi Jawa Barat | Catur Riani, Marselina Irasonia Tan, Yunia Sribudiani, Azzania Fibriani, Husna Nugrahapraja, Tarwadi, Ema Rahmawati, Savira Ekawardhani, Hesti Lina Wiraswati, Ryan Bayusantika Ristandi, Rifky Waluyajati Rachman, Cut Nur Cinthia Alamanda, Lia Faridah, Gusti Ayu Prani Pradani, Adelina Khristiani Rahayu, Hammam Riza, Sony Solistia Wirawan, Agung Eru Wibowo, Irvan Faizal |
| EPI_ISL_528753 | Dinkes Kota Bogor | School of Life Sciences and Technology & School of Pharmacy-Institut Teknologi Bandung; Molecular Genetics Laboratory-Faculty of Medicine-Universitas Padjadjaran; Laboratorium Kesehatan Provinsi Jawa Barat | Azzania Fibriani, Catur Riani, Marselina Irasonia Tan, Yunia Sribudiani, Husna Nugrahapraja, Tarwadi, Ema Rahmawati, Savira Ekawardhani, Hesti Lina Wiraswati, Ryan Bayusantika Ristandi, Rifky Waluyajati Rachman, Cut Nur Cinthia Alamanda, Lia Faridah, Davin H. E. Setiamarga, Rizki Mardian , Hammam Riza, Sony Solistia Wirawan, Agung Eru Wibowo, Irvan Faizal |
| EPI_ISL_528759 | Santo Borromeus Hospital | School of Life Sciences and Technology & School of Pharmacy-Institut Teknologi Bandung; Molecular Genetics Laboratory-Faculty of Medicine-Universitas Padjadjaran; Laboratorium Kesehatan Provinsi Jawa Barat | Husna Nugrahapraja, Azzania Fibriani, Catur Riani, Marselina Irasonia Tan, Yunia Sribudiani, Tarwadi, Ema Rahmawati, Savira Ekawardhani, Hesti Lina Wiraswati, Ryan Bayusantika Ristandi, Rifky Waluyajati Rachman, Cut Nur Cinthia Alamanda, Lia Faridah, Tri Hanggono Achmad, Mas Rizky A.A. Syamsunarno, Fensi Amalina, Hammam Riza, Sony Solistia Wirawan, Agung Eru Wibowo, Irvan Faizal |
| EPI_ISL_528788, EPI_ISL_528790, EPI_ISL_528791, EPI_ISL_528793, EPI_ISL_528796, EPI_ISL_528799, EPI_ISL_528801, EPI_ISL_528802, EPI_ISL_528803, EPI_ISL_528805, EPI_ISL_528806, EPI_ISL_528807, EPI_ISL_528808 | | | |
| see above | Microbiology Department, Barking Havering and Redbridge University Hospitals NHS trust | Wellcome Sanger Institute for the COVID-19 Genomics UK Consortium | Amy Ash, Fatima Ali, Cherian Koshy and Alex Alderton, Roberto Amato, Sonia Goncalves, Ewan Harrison, David K. Jackson, Ian Johnston, Dominic Kwiatkowski, Cordelia Langford, John Sillitoe on behalf of the Wellcome Sanger Institute COVID-19 Surveillance Team (http://www.sanger.ac.uk/covid-team) |
| EPI_ISL_528809 | Department of Medicine, Gandhi hospital, Hyderabad | CSIR-Centre for Cellular and Molecular Biology | Rajarao Mesipogu , Thrilok Chander Bingi ,Vinayasekhar Aedula,Tulasi Nagabandi, Namami Gaur, Sakshi Shambhavi, Lamuk Zaveri, Shagufta Khan, Nikhil Hajirnis, M Soujanya Reddy, Pratheusa Maccha, Purushotham Vodnala, Payel Mukherjee, Sofia Banu, Priya Singh, Onkar Kulkarni, Dhiviya Vedagiri, Divya Gupta, Vishal Sah, Santosh Kumar Kuncha, Krishnan Harinivas Harshan, Archana Bharadwaj Siva, Karthik Bharadwaj Tallapaka,G. Aditya Kumar, Koushick Sivakumar, Pooja Ramesh Gupta, Rajan Kumar Jha, Shraddha Vijay Lahoti, Rakesh K Mishra, Divya Tej Sowpati |
| EPI_ISL_528810, EPI_ISL_528811, EPI_ISL_528812 | Department of Medicine, Gandhi hospital, Hyderabad | CSIR-Centre for Cellular and Molecular Biology | Thrilok Chander Bingi,Rajarao Mesipogu ,Vinayasekhar Aedula,Lamuk Zaveri, Shagufta Khan, Namami Gaur, Sakshi Shambhavi, Nikhil Hajirnis, M Soujanya Reddy, Pratheusa Maccha, Tulasi Nagabandi, Purushotham Vodnala, Payel Mukherjee, Sofia Banu, Priya Singh, Onkar Kulkarni, Dhiviya Vedagiri, Divya Gupta, Vishal Sah, Santosh Kumar Kuncha, Krishnan Harinivas Harshan, Archana Bharadwaj Siva, Karthik Bharadwaj Tallapaka, Renu Sudhakar, Somesh Gorde, Gangumala Srinivas Reddy, Sujoy Deb, Swati Bayyana, Rakesh K Mishra, Divya Tej Sowpati |
| EPI_ISL_528813 | Department of Medicine, Gandhi hospital, Hyderabad | CSIR-Centre for Cellular and Molecular Biology | Vinayasekhar Aedula,Thrilok Chander Bingi, Rajarao Mesipogu, Shagufta Khan, Lamuk Zaveri, Namami Gaur, Sakshi Shambhavi, Nikhil Hajirnis, M Soujanya Reddy, Pratheusa Maccha,Tulasi Nagabandi, Purushotham Vodnala, Payel Mukherjee, Sofia Banu, Priya Singh, Onkar Kulkarni, Dhiviya Vedagiri, Divya Gupta, Vishal Sah, Santosh Kumar Kuncha, Krishnan Harinivas Harshan, Archana Bharadwaj Siva, Karthik Bharadwaj Tallapaka,Umesh Kumar, Unis Ahmad Bhat, Ajay Sarawagi, Priyanka Pant, Rajkanwar Nathawat, Rakesh K Mishra, Divya Tej Sowpati |
| EPI_ISL_528814 | Department of Medicine, Gandhi hospital, Hyderabad | CSIR-Centre for Cellular and Molecular Biology | Rajarao Mesipogu , Thrilok Chander Bingi ,Vinayasekhar Aedula,Tulasi Nagabandi, Namami Gaur, Sakshi Shambhavi, Lamuk Zaveri, Shagufta Khan, Nikhil Hajirnis, M Soujanya Reddy, Pratheusa Maccha, Purushotham Vodnala, Payel Mukherjee, Sofia Banu, Priya Singh, Onkar Kulkarni, Dhiviya Vedagiri, Divya Gupta, Vishal Sah, Santosh Kumar Kuncha, Krishnan Harinivas Harshan, Archana Bharadwaj Siva, Karthik Bharadwaj Tallapaka,G. Aditya Kumar, Koushick Sivakumar, Pooja Ramesh Gupta, Rajan Kumar Jha, Shraddha Vijay Lahoti, Rakesh K Mishra, Divya Tej Sowpati |
| EPI_ISL_528815 | Department of Medicine, Gandhi hospital, Hyderabad | CSIR-Centre for Cellular and Molecular Biology | Vinayasekhar Aedula,Thrilok Chander Bingi, Rajarao Mesipogu, Shagufta Khan, Lamuk Zaveri, Namami Gaur, Sakshi Shambhavi, Nikhil Hajirnis, M Soujanya Reddy, Pratheusa Maccha,Tulasi Nagabandi, Purushotham Vodnala, Payel Mukherjee, Sofia Banu, Priya Singh, Onkar Kulkarni, Dhiviya Vedagiri, Divya Gupta, Vishal Sah, Santosh Kumar Kuncha, Krishnan Harinivas Harshan, Archana Bharadwaj Siva, Karthik Bharadwaj Tallapaka,Umesh Kumar, Unis Ahmad Bhat, Ajay Sarawagi, Priyanka Pant, Rajkanwar Nathawat, Rakesh K Mishra, Divya Tej Sowpati |

| | | | |
|---|---|---|---|
| EPI_ISL_528816 | Department of Medicine, Gandhi hospital, Hyderabad | CSIR-Centre for Cellular and Molecular Biology | Rajarao Mesipogu , Thrilok Chander Bingi ,Vinayasekhar Aedula,Tulasi Nagabandi, Namami Gaur, Sakshi Shambhavi, Lamuk Zaveri, Shagufta Khan, Nikhil Hajirnis, M Soujanya Reddy, Pratheusa Maccha, Purushotham Vodnala, Payel Mukherjee, Sofia Banu, Priya Singh, Onkar Kulkarni, Dhiviya Vedagiri, Divya Gupta, Vishal Sah, Santosh Kumar Kuncha, Krishnan Harinivas Harshan, Archana Bharadwaj Siva, Karthik Bharadwaj Tallapaka,G. Aditya Kumar, Koushick Sivakumar, Pooja Ramesh Gupta, Rajan Kumar Jha, Shraddha Vijay Lahoti, Rakesh K Mishra, Divya Tej Sowpati |
| EPI_ISL_528817 | Department of Medicine, Gandhi hospital, Hyderabad | CSIR-Centre for Cellular and Molecular Biology | Thrilok Chander Bingi,Rajarao Mesipogu ,Vinayasekhar Aedula,Lamuk Zaveri, Shagufta Khan, Namami Gaur, Sakshi Shambhavi, Nikhil Hajirnis, M Soujanya Reddy, Pratheusa Maccha, Tulasi Nagabandi, Purushotham Vodnala, Payel Mukherjee, Sofia Banu, Priya Singh, Onkar Kulkarni, Dhiviya Vedagiri, Divya Gupta, Vishal Sah, Santosh Kumar Kuncha, Krishnan Harinivas Harshan, Archana Bharadwaj Siva, Karthik Bharadwaj Tallapaka, Renu Sudhakar, Somesh Gorde, Gangumala Srinivas Reddy, Sujoy Deb, Swati Bayyana, Rakesh K Mishra, Divya Tej Sowpati |
| EPI_ISL_528818 | Department of Medicine, Gandhi hospital, Hyderabad | CSIR-Centre for Cellular and Molecular Biology | Vinayasekhar Aedula,Thrilok Chander Bingi, Rajarao Mesipogu, Shagufta Khan, Lamuk Zaveri, Namami Gaur, Sakshi Shambhavi, Nikhil Hajirnis, M Soujanya Reddy, Pratheusa Maccha,Tulasi Nagabandi, Purushotham Vodnala, Payel Mukherjee, Sofia Banu, Priya Singh, Onkar Kulkarni, Dhiviya Vedagiri, Divya Gupta, Vishal Sah, Santosh Kumar Kuncha, Krishnan Harinivas Harshan, Archana Bharadwaj Siva, Karthik Bharadwaj Tallapaka,Umesh Kumar, Unis Ahmad Bhat, Ajay Sarawagi, Priyanka Pant, Rajkanwar Nathawat, Rakesh K Mishra, Divya Tej Sowpati |
| EPI_ISL_528819, EPI_ISL_528820 | Department of Medicine, Gandhi hospital, Hyderabad | CSIR-Centre for Cellular and Molecular Biology | Rajarao Mesipogu , Thrilok Chander Bingi ,Vinayasekhar Aedula,Tulasi Nagabandi, Namami Gaur, Sakshi Shambhavi, Lamuk Zaveri, Shagufta Khan, Nikhil Hajirnis, M Soujanya Reddy, Pratheusa Maccha, Purushotham Vodnala, Payel Mukherjee, Sofia Banu, Priya Singh, Onkar Kulkarni, Dhiviya Vedagiri, Divya Gupta, Vishal Sah, Santosh Kumar Kuncha, Krishnan Harinivas Harshan, Archana Bharadwaj Siva, Karthik Bharadwaj Tallapaka,G. Aditya Kumar, Koushick Sivakumar, Pooja Ramesh Gupta, Rajan Kumar Jha, Shraddha Vijay Lahoti, Rakesh K Mishra, Divya Tej Sowpati |
| EPI_ISL_528821 | Department of Medicine, Gandhi hospital, Hyderabad | CSIR-Centre for Cellular and Molecular Biology | Thrilok Chander Bingi,Rajarao Mesipogu ,Vinayasekhar Aedula,Lamuk Zaveri, Shagufta Khan, Namami Gaur, Sakshi Shambhavi, Nikhil Hajirnis, M Soujanya Reddy, Pratheusa Maccha, Tulasi Nagabandi, Purushotham Vodnala, Payel Mukherjee, Sofia Banu, Priya Singh, Onkar Kulkarni, Dhiviya Vedagiri, Divya Gupta, Vishal Sah, Santosh Kumar Kuncha, Krishnan Harinivas Harshan, Archana Bharadwaj Siva, Karthik Bharadwaj Tallapaka, Renu Sudhakar, Somesh Gorde, Gangumala Srinivas Reddy, Sujoy Deb, Swati Bayyana, Rakesh K Mishra, Divya Tej Sowpati |
| EPI_ISL_528822 | Department of Medicine, Gandhi hospital, Hyderabad | CSIR-Centre for Cellular and Molecular Biology | Vinayasekhar Aedula,Thrilok Chander Bingi, Rajarao Mesipogu, Shagufta Khan, Lamuk Zaveri, Namami Gaur, Sakshi Shambhavi, Nikhil Hajirnis, M Soujanya Reddy, Pratheusa Maccha,Tulasi Nagabandi, Purushotham Vodnala, Payel Mukherjee, Sofia Banu, Priya Singh, Onkar Kulkarni, Dhiviya Vedagiri, Divya Gupta, Vishal Sah, Santosh Kumar Kuncha, Krishnan Harinivas Harshan, Archana Bharadwaj Siva, Karthik Bharadwaj Tallapaka,Umesh Kumar, Unis Ahmad Bhat, Ajay Sarawagi, Priyanka Pant, Rajkanwar Nathawat, Rakesh K Mishra, Divya Tej Sowpati |
| EPI_ISL_528823 | CSIR-Centre for Cellular and Molecular Biology | CSIR-Centre for Cellular and Molecular Biology | Shagufta Khan, Lamuk Zaveri, Namami Gaur, Sakshi Shambhavi, Nikhil Hajirnis, M Soujanya Reddy, Pratheusa Maccha, Tulasi Nagabandi, Purushotham Vodnala, Payel Mukherjee, Sofia Banu, Priya Singh, Onkar Kulkarni, Dhiviya Vedagiri, Divya Gupta, Vishal Sah, Santosh Kumar Kuncha, Krishnan Harinivas Harshan, Archana Bharadwaj Siva, Karthik Bharadwaj Tallapaka, Renu Sudhakar, Somesh Gorde, Gangumala Srinivas Reddy, Sujoy Deb, Swati Bayyana, Rakesh K Mishra, Divya Tej Sowpati |
| EPI_ISL_528824 | CSIR-Centre for Cellular and Molecular Biology | CSIR-Centre for Cellular and Molecular Biology | Tulasi Nagabandi, Namami Gaur, Sakshi Shambhavi, Lamuk Zaveri, Shagufta Khan, Nikhil Hajirnis, M Soujanya Reddy, Pratheusa Maccha, Purushotham Vodnala, Payel Mukherjee, Sofia Banu, Priya Singh, Onkar Kulkarni, Dhiviya Vedagiri, Divya Gupta, Vishal Sah, Santosh Kumar Kuncha, Krishnan Harinivas Harshan, Archana Bharadwaj Siva, Karthik Bharadwaj Tallapaka,G. Aditya Kumar, Koushick Sivakumar, Pooja Ramesh Gupta, Rajan Kumar Jha, Shraddha Vijay Lahoti, Rakesh K Mishra, Divya Tej Sowpati |
| EPI_ISL_528825, EPI_ISL_528826 | CSIR-Centre for Cellular and Molecular Biology | CSIR-Centre for Cellular and Molecular Biology | Shagufta Khan, Lamuk Zaveri, Namami Gaur, Sakshi Shambhavi, Nikhil Hajirnis, M Soujanya Reddy, Pratheusa Maccha, Tulasi Nagabandi, Purushotham Vodnala, Payel Mukherjee, Sofia Banu, Priya Singh, Onkar Kulkarni, Dhiviya Vedagiri, Divya Gupta, Vishal Sah, Santosh Kumar Kuncha, Krishnan Harinivas Harshan, Archana Bharadwaj Siva, Karthik Bharadwaj Tallapaka, Renu Sudhakar, Somesh Gorde, Gangumala Srinivas Reddy, Sujoy Deb, Swati Bayyana, Rakesh K Mishra, Divya Tej Sowpati |
| EPI_ISL_528827 | CSIR-Centre for Cellular and Molecular Biology | CSIR-Centre for Cellular and Molecular Biology | Namami Gaur, Sakshi Shambhavi, Lamuk Zaveri, Shagufta Khan, Nikhil Hajirnis, M Soujanya Reddy, Pratheusa Maccha, Tulasi Nagabandi, Purushotham Vodnala, Payel Mukherjee, Sofia Banu, Priya Singh, Onkar Kulkarni, Dhiviya Vedagiri, Divya Gupta, Vishal Sah, Santosh Kumar Kuncha, Krishnan Harinivas Harshan, Archana Bharadwaj Siva, Karthik Bharadwaj Tallapaka, Zeba Rizvi, Zuberwasim Sayyad, Kakade Aishwarya Arun, Amrutha H C, Ananga Ghosh, Rakesh K Mishra, Divya Tej Sowpati |
| EPI_ISL_528828 | CSIR-Centre for Cellular and Molecular Biology | CSIR-Centre for Cellular and Molecular Biology | Sofia Banu, Payel Mukherjee, Priya Singh,Onkar Kulkarni, Dhiviya Vedagiri, Divya Gupta, Vishal Sah, Santosh Kumar Kuncha, Krishnan Harinivas Harshan, Archana Bharadwaj Siva, Karthik Bharadwaj Tallapaka, Shagufta Khan, Lamuk Zaveri, Namami Gaur, Sakshi Shambhavi, Nikhil Hajirnis, M Soujanya Reddy, Pratheusa Maccha,Tulasi Nagabandi, Purushotham Vodnala, Deepak Kumar, Devi Prasad Vijayashankar, Disha Nanda, Divya Das, Jotin Gogoi, Manish Bhattacharjee, Rakesh K Mishra, Divya Tej Sowpati |
| EPI_ISL_528829 | CSIR-Centre for Cellular and Molecular Biology | CSIR-Centre for Cellular and Molecular Biology | Sakshi Shambhavi, Lamuk Zaveri, Shagufta Khan, Namami Gaur, Nikhil Hajirnis, M Soujanya Reddy, Pratheusa Maccha, Tulasi Nagabandi, Purushotham Vodnala, Payel Mukherjee, Sofia Banu, Priya Singh, Onkar Kulkarni, Dhiviya Vedagiri, Divya Gupta, Vishal Sah, Santosh Kumar Kuncha, Krishnan Harinivas Harshan, Archana Bharadwaj Siva, Karthik Bharadwaj Tallapaka, Deepak Kumar, Devi Prasad Vijayashankar, Disha Nanda, Divya Das, Jotin Gogoi, Manish Bhattacharjee, Rakesh K Mishra, Divya Tej Sowpati |
| EPI_ISL_528830 | CSIR-Centre for Cellular and Molecular Biology | CSIR-Centre for Cellular and Molecular Biology | Tulasi Nagabandi, Namami Gaur, Sakshi Shambhavi, Lamuk Zaveri, Shagufta Khan, Nikhil Hajirnis, M Soujanya Reddy, Pratheusa Maccha, Purushotham Vodnala, Payel Mukherjee, Sofia Banu, Priya Singh, Onkar Kulkarni, Dhiviya Vedagiri, Divya Gupta, Vishal Sah, Santosh Kumar Kuncha, Krishnan Harinivas Harshan, Archana Bharadwaj Siva, Karthik Bharadwaj Tallapaka,G. Aditya Kumar, Koushick Sivakumar, Pooja Ramesh Gupta, Rajan Kumar Jha, Shraddha Vijay Lahoti, Rakesh K Mishra, Divya Tej Sowpati |
| EPI_ISL_528831 | CSIR-Centre for Cellular and Molecular Biology | CSIR-Centre for Cellular and Molecular Biology | Lamuk Zaveri, Shagufta Khan, Namami Gaur, Sakshi Shambhavi, Nikhil Hajirnis, M Soujanya Reddy, Pratheusa Maccha, Tulasi Nagabandi, Purushotham Vodnala, Payel Mukherjee, Sofia Banu, Priya Singh, Onkar Kulkarni, Dhiviya Vedagiri, Divya Gupta, Vishal Sah, Santosh Kumar Kuncha, Krishnan Harinivas Harshan, Archana Bharadwaj Siva, Karthik Bharadwaj Tallapaka, Renu Sudhakar, Somesh Gorde, Gangumala Srinivas Reddy, Sujoy Deb, Swati Bayyana, Rakesh K Mishra, Divya Tej Sowpati |
| EPI_ISL_528832 | CSIR-Centre for Cellular and Molecular Biology | CSIR-Centre for Cellular and Molecular Biology | Sofia Banu, Payel Mukherjee, Priya Singh,Onkar Kulkarni, Dhiviya Vedagiri, Divya Gupta, Vishal Sah, Santosh Kumar Kuncha, Krishnan Harinivas Harshan, Archana Bharadwaj Siva, Karthik Bharadwaj Tallapaka, Shagufta Khan, Lamuk Zaveri, Namami Gaur, Sakshi Shambhavi, Nikhil Hajirnis, M Soujanya Reddy, Pratheusa Maccha,Tulasi Nagabandi, Purushotham Vodnala, Deepak Kumar, Devi Prasad Vijayashankar, Disha Nanda, Divya Das, Jotin Gogoi, Manish Bhattacharjee, Rakesh K Mishra, Divya Tej Sowpati |
| EPI_ISL_528833 | CSIR-Centre for Cellular and Molecular Biology | CSIR-Centre for Cellular and Molecular Biology | Lamuk Zaveri, Shagufta Khan, Namami Gaur, Sakshi Shambhavi, Nikhil Hajirnis, M Soujanya Reddy, Pratheusa Maccha, Tulasi Nagabandi, Purushotham Vodnala, Payel Mukherjee, Sofia Banu, Priya Singh, Onkar Kulkarni, Dhiviya Vedagiri, Divya Gupta, Vishal Sah, Santosh Kumar Kuncha, Krishnan Harinivas Harshan, Archana Bharadwaj Siva, Karthik Bharadwaj Tallapaka, Renu Sudhakar, Somesh Gorde, Gangumala Srinivas Reddy, Sujoy Deb, Swati Bayyana, Rakesh K Mishra, Divya Tej Sowpati |
| EPI_ISL_528834 | CSIR-Centre for Cellular and Molecular Biology | CSIR-Centre for Cellular and Molecular Biology | Pratheusa Maccha, Sakshi Shambhavi, Lamuk Zaveri, Shagufta Khan, Namami Gaur, Nikhil Hajirnis, M Soujanya Reddy, Tulasi Nagabandi, Purushotham Vodnala, Payel Mukherjee, Sofia Banu, Priya Singh,Onkar Kulkarni, Dhiviya Vedagiri, Divya Gupta, Vishal Sah, Santosh Kumar Kuncha, Krishnan Harinivas Harshan, Archana Bharadwaj Siva, Karthik Bharadwaj Tallapaka, G. Aditya Kumar, Koushick Sivakumar,Disha Nanda, Divya Das, Jotin Gogoi, Manish Bhattacharjee, Ravi Prasad Mukku, Rakesh K Mishra, Divya Tej Sowpati |
| EPI_ISL_528835 | CSIR-Centre for Cellular and Molecular Biology | CSIR-Centre for Cellular and Molecular Biology | M Soujanya Reddy, Nikhil Hajirnis, Pratheusa Maccha, Namami Gaur, Sakshi Shambhavi, Lamuk Zaveri, Shagufta Khan, Tulasi Nagabandi, Purushotham Vodnala, Payel Mukherjee, Sofia Banu, Priya Singh, Onkar Kulkarni, Dhiviya Vedagiri, Divya Gupta, Vishal Sah, Santosh Kumar Kuncha, Krishnan Harinivas Harshan, Archana Bharadwaj Siva, Karthik Bharadwaj Tallapaka, Zeba Rizvi, Zuberwasim Sayyad, Kakade Aishwarya Arun, Amrutha H C, Ananga Ghosh, Rakesh K Mishra, Divya Tej Sowpati |
| EPI_ISL_528836 | CSIR-Centre for Cellular and Molecular Biology | CSIR-Centre for Cellular and Molecular Biology | Tulasi Nagabandi, Namami Gaur, Sakshi Shambhavi, Lamuk Zaveri, Shagufta Khan, Nikhil Hajirnis, M Soujanya Reddy, Pratheusa Maccha, Purushotham Vodnala, Payel Mukherjee, Sofia Banu, Priya Singh, Onkar Kulkarni, Dhiviya Vedagiri, Divya Gupta, Vishal Sah, Santosh Kumar Kuncha, Krishnan Harinivas Harshan, Archana Bharadwaj Siva, Karthik Bharadwaj Tallapaka,G. Aditya Kumar, Koushick Sivakumar, Pooja Ramesh Gupta, Rajan Kumar Jha, Shraddha Vijay Lahoti, Rakesh K Mishra, Divya Tej Sowpati |
| EPI_ISL_528837 | CSIR-Centre for Cellular and Molecular Biology | CSIR-Centre for Cellular and Molecular Biology | Sakshi Shambhavi, Lamuk Zaveri, Shagufta Khan, Namami Gaur, Nikhil Hajirnis, M Soujanya Reddy, Pratheusa Maccha, Tulasi Nagabandi, Purushotham Vodnala, Payel Mukherjee, Sofia Banu, Priya Singh, Onkar Kulkarni, Dhiviya Vedagiri, Divya Gupta, Vishal Sah, Santosh Kumar Kuncha, Krishnan Harinivas Harshan, Archana Bharadwaj Siva, Karthik Bharadwaj Tallapaka, Deepak Kumar, Devi Prasad Vijayashankar, Disha Nanda, Divya Das, Jotin Gogoi, Manish Bhattacharjee, Rakesh K Mishra, Divya Tej Sowpati |
| EPI_ISL_528838 | CSIR-Centre for Cellular and Molecular Biology | CSIR-Centre for Cellular and Molecular Biology | Payel Mukherjee, Sofia Banu, Priya Singh, Onkar Kulkarni, Dhiviya Vedagiri, Divya Gupta, Vishal Sah, Santosh Kumar Kuncha, Krishnan Harinivas Harshan, Archana Bharadwaj Siva, Karthik Bharadwaj Tallapaka, Shagufta Khan, Lamuk Zaveri, Nikhil Hajirnis, M Soujanya Reddy, Pratheusa Maccha, Namami Gaur, Sakshi Shambhavi, Tulasi Nagabandi, Purushotham Vodnala, Rakesh K Mishra, Sonu Uday, Sudipta Mondal, Annapoorna P Karthyayani, Debabrata Jana, Debrya Saha, Divya Tej Sowpati |
| EPI_ISL_528839 | CSIR-Centre for Cellular and Molecular Biology | CSIR-Centre for Cellular and Molecular Biology | Pratheusa Maccha, Sakshi Shambhavi, Lamuk Zaveri, Shagufta Khan, Namami Gaur, Nikhil Hajirnis, M Soujanya Reddy, Tulasi Nagabandi, Purushotham Vodnala, Payel Mukherjee, Sofia Banu, Priya Singh,Onkar Kulkarni, Dhiviya Vedagiri, Divya Gupta, Vishal Sah, Santosh Kumar Kuncha, Krishnan Harinivas Harshan, Archana Bharadwaj Siva, Karthik Bharadwaj Tallapaka, G. Aditya Kumar, Koushick Sivakumar,Disha Nanda, Divya Das, Jotin Gogoi, Manish Bhattacharjee, Ravi Prasad Mukku, Rakesh K Mishra, Divya Tej Sowpati |
| EPI_ISL_528840 | CSIR-Centre for Cellular and Molecular Biology | CSIR-Centre for Cellular and Molecular Biology | Tulasi Nagabandi, Namami Gaur, Sakshi Shambhavi, Lamuk Zaveri, Shagufta Khan, Nikhil Hajirnis, M Soujanya Reddy, Pratheusa Maccha, Purushotham Vodnala, Payel Mukherjee, Sofia Banu, Priya Singh, Onkar Kulkarni, Dhiviya Vedagiri, Divya Gupta, Vishal Sah, Santosh Kumar Kuncha, Krishnan Harinivas Harshan, Archana Bharadwaj Siva, Karthik Bharadwaj Tallapaka,G. Aditya Kumar, Koushick Sivakumar, Pooja Ramesh Gupta, Rajan Kumar Jha, Shraddha Vijay Lahoti, Rakesh K Mishra, Divya Tej Sowpati |
| EPI_ISL_528841, EPI_ISL_528842 | CSIR-Centre for Cellular and Molecular Biology | CSIR-Centre for Cellular and Molecular Biology | Payel Mukherjee, Sofia Banu, Priya Singh, Onkar Kulkarni, Dhiviya Vedagiri, Divya Gupta, Vishal Sah, Santosh Kumar Kuncha, Krishnan Harinivas Harshan, Archana Bharadwaj Siva, Karthik Bharadwaj Tallapaka, Shagufta Khan, Lamuk Zaveri, Nikhil Hajirnis, M Soujanya Reddy, Pratheusa Maccha, Namami Gaur, Sakshi Shambhavi, Tulasi Nagabandi, Purushotham Vodnala, Rakesh K Mishra, Sonu Uday, Sudipta Mondal, Annapoorna P Karthyayani, Debabrata Jana, Debrya Saha, Divya Tej Sowpati |
| EPI_ISL_528843, EPI_ISL_528844 | CSIR-Centre for Cellular and Molecular Biology | CSIR-Centre for Cellular and Molecular Biology | Namami Gaur, Sakshi Shambhavi, Lamuk Zaveri, Shagufta Khan, Nikhil Hajirnis, M Soujanya Reddy, Pratheusa Maccha, Tulasi Nagabandi, Purushotham Vodnala, Payel Mukherjee, Sofia Banu, Priya Singh, Onkar Kulkarni, Dhiviya Vedagiri, Divya Gupta, Vishal Sah, Santosh Kumar Kuncha, Krishnan Harinivas Harshan, Archana Bharadwaj Siva, Karthik Bharadwaj Tallapaka, Zeba Rizvi, Zuberwasim Sayyad, Kakade Aishwarya Arun, Amrutha H C, Ananga Ghosh, Rakesh K Mishra, Divya Tej Sowpati |
| EPI_ISL_528845 | CSIR-Centre for Cellular and Molecular Biology | CSIR-Centre for Cellular and Molecular Biology | Payel Mukherjee, Sofia Banu, Priya Singh, Onkar Kulkarni, Dhiviya Vedagiri, Divya Gupta, Vishal Sah, Santosh Kumar Kuncha, Krishnan Harinivas Harshan, Archana Bharadwaj Siva, Karthik Bharadwaj Tallapaka, Shagufta Khan, Lamuk Zaveri, Nikhil Hajirnis, M Soujanya Reddy, Pratheusa Maccha, Namami Gaur, Sakshi Shambhavi, Tulasi Nagabandi, Purushotham Vodnala, Rakesh K Mishra, Sonu Uday, Sudipta Mondal, Annapoorna P Karthyayani, Debabrata Jana, Debrya Saha, Divya Tej Sowpati |
| EPI_ISL_528846 | CSIR-Centre for Cellular and Molecular Biology | CSIR-Centre for Cellular and Molecular Biology | Namami Gaur, Sakshi Shambhavi, Lamuk Zaveri, Shagufta Khan, Nikhil Hajirnis, M Soujanya Reddy, Pratheusa Maccha, Tulasi Nagabandi, Purushotham Vodnala, Payel Mukherjee, Sofia Banu, Priya Singh, Onkar Kulkarni, Dhiviya Vedagiri, Divya Gupta, Vishal Sah, Santosh Kumar Kuncha, Krishnan Harinivas Harshan, Archana Bharadwaj Siva, Karthik Bharadwaj Tallapaka, Zeba Rizvi, Zuberwasim Sayyad, Kakade Aishwarya Arun, Amrutha H C, Ananga Ghosh, Rakesh K Mishra, Divya Tej Sowpati |
| EPI_ISL_528847 | CSIR-Centre for Cellular and Molecular Biology | CSIR-Centre for Cellular and Molecular Biology | Shagufta Khan, Lamuk Zaveri, Namami Gaur, Sakshi Shambhavi, Nikhil Hajirnis, M Soujanya Reddy, Pratheusa Maccha, Tulasi Nagabandi, Purushotham Vodnala, Payel Mukherjee, Sofia Banu, Priya Singh, Onkar Kulkarni, Dhiviya Vedagiri, Divya Gupta, Vishal Sah, Santosh Kumar Kuncha, Krishnan Harinivas Harshan, Archana Bharadwaj Siva, Karthik Bharadwaj Tallapaka, Renu Sudhakar, Somesh Gorde, Gangumala Srinivas Reddy, Sujoy Deb, Swati Bayyana, Rakesh K Mishra, Divya Tej Sowpati |
| EPI_ISL_528848 | CSIR-Centre for Cellular and Molecular Biology | CSIR-Centre for Cellular and Molecular Biology | Sofia Banu, Payel Mukherjee, Priya Singh,Onkar Kulkarni, Dhiviya Vedagiri, Divya Gupta, Vishal Sah, Santosh Kumar Kuncha, Krishnan Harinivas Harshan, Archana Bharadwaj Siva, Karthik Bharadwaj Tallapaka, Shagufta Khan, Lamuk Zaveri, Namami Gaur, Sakshi Shambhavi, Nikhil Hajirnis, M Soujanya Reddy, Pratheusa Maccha,Tulasi Nagabandi, Purushotham Vodnala, Deepak Kumar, Devi Prasad Vijayashankar, Disha Nanda, Divya Das, Jotin Gogoi, Manish Bhattacharjee, Rakesh K Mishra, Divya Tej Sowpati |
| EPI_ISL_528849 | CSIR-Centre for Cellular and Molecular Biology | CSIR-Centre for Cellular and Molecular Biology | Nikhil Hajirnis, M Soujanya Reddy, Pratheusa Maccha, Lamuk Zaveri, Shagufta Khan, Namami Gaur, Sakshi Shambhavi, Tulasi Nagabandi, Purushotham Vodnala, Payel Mukherjee, Sofia Banu, Priya Singh, Onkar Kulkarni, Dhiviya Vedagiri, Divya Gupta, Vishal Sah, Santosh Kumar Kuncha, Krishnan Harinivas Harshan, Archana Bharadwaj Siva, |

| | | | |
|---|---|---|---|
| | | | Karthik Bharadwaj Tallapaka,Zeba Rizvi, Zuberwasim Sayyad, Kakade Aishwarya Arun, Amrutha H C, Ananga Ghosh, Rakesh K Mishra, Divya Tej Sowpati |
| EPI_ISL_528850 | CSIR-Centre for Cellular and Molecular Biology | CSIR-Centre for Cellular and Molecular Biology | Lamuk Zaveri, Shagufta Khan, Namami Gaur, Sakshi Shambhavi, Nikhil Hajirnis, M Soujanya Reddy, Pratheusa Maccha, Tulasi Nagabandi, Purushotham Vodnala, Payel Mukherjee, Sofia Banu, Priya Singh, Onkar Kulkarni, Dhiviya Vedagiri, Divya Gupta, Vishal Sah, Santosh Kumar Kuncha, Krishnan Harinivas Harshan, Archana Bharadwaj Siva, Karthik Bharadwaj Tallapaka, Renu Sudhakar, Somesh Gorde, Gangumala Srinivas Reddy, Sujoy Deb, Swati Bayyana, Rakesh K Mishra, Divya Tej Sowpati |
| EPI_ISL_528851 | CSIR-Centre for Cellular and Molecular Biology | CSIR-Centre for Cellular and Molecular Biology | Payel Mukherjee, Sofia Banu, Priya Singh, Onkar Kulkarni, Dhiviya Vedagiri, Divya Gupta, Vishal Sah, Santosh Kumar Kuncha, Krishnan Harinivas Harshan, Archana Bharadwaj Siva, Karthik Bharadwaj Tallapaka, Shagufta Khan, Lamuk Zaveri, Nikhil Hajirnis, M Soujanya Reddy, Pratheusa Maccha, Namami Gaur, Sakshi Shambhavi, Tulasi Nagabandi, Purushotham Vodnala, Rakesh K Mishra, Sonu Uday, Sudipta Mondal, Annapoorna P Karthyayani, Debabrata Jana, Debrya Saha, Divya Tej Sowpati |
| EPI_ISL_528852 | CSIR-Centre for Cellular and Molecular Biology | CSIR-Centre for Cellular and Molecular Biology | Namami Gaur, Sakshi Shambhavi, Lamuk Zaveri, Shagufta Khan, Nikhil Hajirnis, M Soujanya Reddy, Pratheusa Maccha, Tulasi Nagabandi, Purushotham Vodnala, Payel Mukherjee, Sofia Banu, Priya Singh, Onkar Kulkarni, Dhiviya Vedagiri, Divya Gupta, Vishal Sah, Santosh Kumar Kuncha, Krishnan Harinivas Harshan, Archana Bharadwaj Siva, Karthik Bharadwaj Tallapaka, Zeba Rizvi, Zuberwasim Sayyad, Kakade Aishwarya Arun, Amrutha H C, Ananga Ghosh, Rakesh K Mishra, Divya Tej Sowpati |
| EPI_ISL_528853 | CSIR-Centre for Cellular and Molecular Biology | CSIR-Centre for Cellular and Molecular Biology | Tulasi Nagabandi, Namami Gaur, Sakshi Shambhavi, Lamuk Zaveri, Shagufta Khan, Nikhil Hajirnis, M Soujanya Reddy, Pratheusa Maccha, Purushotham Vodnala, Payel Mukherjee, Sofia Banu, Priya Singh, Onkar Kulkarni, Dhiviya Vedagiri, Divya Gupta, Vishal Sah, Santosh Kumar Kuncha, Krishnan Harinivas Harshan, Archana Bharadwaj Siva, Karthik Bharadwaj Tallapaka,G. Aditya Kumar, Koushick Sivakumar, Pooja Ramesh Gupta, Rajan Kumar Jha, Shraddha Vijay Lahoti, Rakesh K Mishra, Divya Tej Sowpati |
| EPI_ISL_528854 | CSIR-Centre for Cellular and Molecular Biology | CSIR-Centre for Cellular and Molecular Biology | Payel Mukherjee, Sofia Banu, Priya Singh, Onkar Kulkarni, Dhiviya Vedagiri, Divya Gupta, Vishal Sah, Santosh Kumar Kuncha, Krishnan Harinivas Harshan, Archana Bharadwaj Siva, Karthik Bharadwaj Tallapaka, Shagufta Khan, Lamuk Zaveri, Nikhil Hajirnis, M Soujanya Reddy, Pratheusa Maccha, Namami Gaur, Sakshi Shambhavi, Tulasi Nagabandi, Purushotham Vodnala, Rakesh K Mishra, Sonu Uday, Sudipta Mondal, Annapoorna P Karthyayani, Debabrata Jana, Debrya Saha, Divya Tej Sowpati |
| EPI_ISL_528855 | CSIR-Centre for Cellular and Molecular Biology | CSIR-Centre for Cellular and Molecular Biology | Nikhil Hajirnis, M Soujanya Reddy, Pratheusa Maccha, Lamuk Zaveri, Shagufta Khan, Namami Gaur, Sakshi Shambhavi, Tulasi Nagabandi, Purushotham Vodnala, Payel Mukherjee, Sofia Banu, Priya Singh, Onkar Kulkarni, Dhiviya Vedagiri, Divya Gupta, Vishal Sah, Santosh Kumar Kuncha, Krishnan Harinivas Harshan, Archana Bharadwaj Siva, Karthik Bharadwaj Tallapaka,Zeba Rizvi, Zuberwasim Sayyad, Kakade Aishwarya Arun, Amrutha H C, Ananga Ghosh, Rakesh K Mishra, Divya Tej Sowpati |
| EPI_ISL_528856 | CSIR-Centre for Cellular and Molecular Biology | CSIR-Centre for Cellular and Molecular Biology | Pratheusa Maccha, Sakshi Shambhavi, Lamuk Zaveri, Shagufta Khan, Namami Gaur, Nikhil Hajirnis, M Soujanya Reddy, Tulasi Nagabandi, Purushotham Vodnala, Payel Mukherjee, Sofia Banu, Priya Singh,Onkar Kulkarni, Dhiviya Vedagiri, Divya Gupta, Vishal Sah, Santosh Kumar Kuncha, Krishnan Harinivas Harshan, Archana Bharadwaj Siva, Karthik Bharadwaj Tallapaka, G. Aditya Kumar, Koushick Sivakumar,Disha Nanda, Divya Das, Jotin Gogoi, Manish Bhattacharjee, Ravi Prasad Mukku, Rakesh K Mishra, Divya Tej Sowpati |
| EPI_ISL_528857 | CSIR-Centre for Cellular and Molecular Biology | CSIR-Centre for Cellular and Molecular Biology | Nikhil Hajirnis, M Soujanya Reddy, Pratheusa Maccha, Lamuk Zaveri, Shagufta Khan, Namami Gaur, Sakshi Shambhavi, Tulasi Nagabandi, Purushotham Vodnala, Payel Mukherjee, Sofia Banu, Priya Singh, Onkar Kulkarni, Dhiviya Vedagiri, Divya Gupta, Vishal Sah, Santosh Kumar Kuncha, Krishnan Harinivas Harshan, Archana Bharadwaj Siva, Karthik Bharadwaj Tallapaka,Zeba Rizvi, Zuberwasim Sayyad, Kakade Aishwarya Arun, Amrutha H C, Ananga Ghosh, Rakesh K Mishra, Divya Tej Sowpati |
| EPI_ISL_528858, EPI_ISL_528859 | CSIR-Centre for Cellular and Molecular Biology | CSIR-Centre for Cellular and Molecular Biology | Pratheusa Maccha, Sakshi Shambhavi, Lamuk Zaveri, Shagufta Khan, Namami Gaur, Nikhil Hajirnis, M Soujanya Reddy, Tulasi Nagabandi, Purushotham Vodnala, Payel Mukherjee, Sofia Banu, Priya Singh,Onkar Kulkarni, Dhiviya Vedagiri, Divya Gupta, Vishal Sah, Santosh Kumar Kuncha, Krishnan Harinivas Harshan, Archana Bharadwaj Siva, Karthik Bharadwaj Tallapaka, G. Aditya Kumar, Koushick Sivakumar,Disha Nanda, Divya Das, Jotin Gogoi, Manish Bhattacharjee, Ravi Prasad Mukku, Rakesh K Mishra, Divya Tej Sowpati |
| EPI_ISL_528860 | CSIR-Centre for Cellular and Molecular Biology | CSIR-Centre for Cellular and Molecular Biology | Tulasi Nagabandi, Namami Gaur, Sakshi Shambhavi, Lamuk Zaveri, Shagufta Khan, Nikhil Hajirnis, M Soujanya Reddy, Pratheusa Maccha, Purushotham Vodnala, Payel Mukherjee, Sofia Banu, Priya Singh, Onkar Kulkarni, Dhiviya Vedagiri, Divya Gupta, Vishal Sah, Santosh Kumar Kuncha, Krishnan Harinivas Harshan, Archana Bharadwaj Siva, Karthik Bharadwaj Tallapaka,G. Aditya Kumar, Koushick Sivakumar, Pooja Ramesh Gupta, Rajan Kumar Jha, Shraddha Vijay Lahoti, Rakesh K Mishra, Divya Tej Sowpati |
| EPI_ISL_528861 | CSIR-Centre for Cellular and Molecular Biology | CSIR-Centre for Cellular and Molecular Biology | Sakshi Shambhavi, Lamuk Zaveri, Shagufta Khan, Namami Gaur, Nikhil Hajirnis, M Soujanya Reddy, Pratheusa Maccha, Tulasi Nagabandi, Purushotham Vodnala, Payel Mukherjee, Sofia Banu, Priya Singh, Onkar Kulkarni, Dhiviya Vedagiri, Divya Gupta, Vishal Sah, Santosh Kumar Kuncha, Krishnan Harinivas Harshan, Archana Bharadwaj Siva, Karthik Bharadwaj Tallapaka, Deepak Kumar, Devi Prasad Vijayashankar, Disha Nanda, Divya Das, Jotin Gogoi, Manish Bhattacharjee, Rakesh K Mishra, Divya Tej Sowpati |
| EPI_ISL_528862 | CSIR-Centre for Cellular and Molecular Biology | CSIR-Centre for Cellular and Molecular Biology | Lamuk Zaveri, Shagufta Khan, Namami Gaur, Sakshi Shambhavi, Nikhil Hajirnis, M Soujanya Reddy, Pratheusa Maccha, Tulasi Nagabandi, Purushotham Vodnala, Payel Mukherjee, Sofia Banu, Priya Singh, Onkar Kulkarni, Dhiviya Vedagiri, Divya Gupta, Vishal Sah, Santosh Kumar Kuncha, Krishnan Harinivas Harshan, Archana Bharadwaj Siva, Karthik Bharadwaj Tallapaka, Renu Sudhakar, Somesh Gorde, Gangumala Srinivas Reddy, Sujoy Deb, Swati Bayyana, Rakesh K Mishra, Divya Tej Sowpati |
| EPI_ISL_528863 | CSIR-Centre for Cellular and Molecular Biology | CSIR-Centre for Cellular and Molecular Biology | Payel Mukherjee, Sofia Banu, Priya Singh, Onkar Kulkarni, Dhiviya Vedagiri, Divya Gupta, Vishal Sah, Santosh Kumar Kuncha, Krishnan Harinivas Harshan, Archana Bharadwaj Siva, Karthik Bharadwaj Tallapaka, Shagufta Khan, Lamuk Zaveri, Nikhil Hajirnis, M Soujanya Reddy, Pratheusa Maccha, Namami Gaur, Sakshi Shambhavi, Tulasi Nagabandi, Purushotham Vodnala, Rakesh K Mishra, Sonu Uday, Sudipta Mondal, Annapoorna P Karthyayani, Debabrata Jana, Debrya Saha, Divya Tej Sowpati |
| EPI_ISL_528864 | CSIR-Centre for Cellular and Molecular Biology | CSIR-Centre for Cellular and Molecular Biology | Sofia Banu, Payel Mukherjee, Priya Singh,Onkar Kulkarni, Dhiviya Vedagiri, Divya Gupta, Vishal Sah, Santosh Kumar Kuncha, Krishnan Harinivas Harshan, Archana Bharadwaj Siva, Karthik Bharadwaj Tallapaka, Shagufta Khan, Lamuk Zaveri, Namami Gaur, Sakshi Shambhavi, Nikhil Hajirnis, M Soujanya Reddy, Pratheusa Maccha,Tulasi Nagabandi, Purushotham Vodnala, Deepak Kumar, Devi Prasad Vijayashankar, Disha Nanda, Divya Das, Jotin Gogoi, Manish Bhattacharjee, Rakesh K Mishra, Divya Tej Sowpati |
| EPI_ISL_528865 | CSIR-Centre for Cellular and Molecular Biology | CSIR-Centre for Cellular and Molecular Biology | Shagufta Khan, Lamuk Zaveri, Namami Gaur, Sakshi Shambhavi, Nikhil Hajirnis, M Soujanya Reddy, Pratheusa Maccha, Tulasi Nagabandi, Purushotham Vodnala, Payel Mukherjee, Sofia Banu, Priya Singh, Onkar Kulkarni, Dhiviya Vedagiri, Divya Gupta, Vishal Sah, Santosh Kumar Kuncha, Krishnan Harinivas Harshan, Archana Bharadwaj Siva, Karthik Bharadwaj Tallapaka, Renu Sudhakar, Somesh Gorde, Gangumala Srinivas Reddy, Sujoy Deb, Swati Bayyana, Rakesh K Mishra, Divya Tej Sowpati |
| EPI_ISL_528866 | CSIR-Centre for Cellular and Molecular Biology | CSIR-Centre for Cellular and Molecular Biology | Sakshi Shambhavi, Lamuk Zaveri, Shagufta Khan, Namami Gaur, Nikhil Hajirnis, M Soujanya Reddy, Pratheusa Maccha, Tulasi Nagabandi, Purushotham Vodnala, Payel Mukherjee, Sofia Banu, Priya Singh, Onkar Kulkarni, Dhiviya Vedagiri, Divya Gupta, Vishal Sah, Santosh Kumar Kuncha, Krishnan Harinivas Harshan, Archana Bharadwaj Siva, Karthik Bharadwaj Tallapaka, Deepak Kumar, Devi Prasad Vijayashankar, Disha Nanda, Divya Das, Jotin Gogoi, Manish Bhattacharjee, Rakesh K Mishra, Divya Tej Sowpati |
| EPI_ISL_528867 | CSIR-Centre for Cellular and Molecular Biology | CSIR-Centre for Cellular and Molecular Biology | M Soujanya Reddy, Nikhil Hajirnis, Pratheusa Maccha, Namami Gaur, Sakshi Shambhavi, Lamuk Zaveri, Shagufta Khan, Tulasi Nagabandi, Purushotham Vodnala, Payel Mukherjee, Sofia Banu, Priya Singh, Onkar Kulkarni, Dhiviya Vedagiri, Divya Gupta, Vishal Sah, Santosh Kumar Kuncha, Krishnan Harinivas Harshan, Archana Bharadwaj Siva, Karthik Bharadwaj Tallapaka, Zeba Rizvi, Zuberwasim Sayyad, Kakade Aishwarya Arun, Amrutha H C, Ananga Ghosh, Rakesh K Mishra, Divya Tej Sowpati |
| EPI_ISL_528868, EPI_ISL_528869 | CSIR-Centre for Cellular and Molecular Biology | CSIR-Centre for Cellular and Molecular Biology | Shagufta Khan, Lamuk Zaveri, Namami Gaur, Sakshi Shambhavi, Nikhil Hajirnis, M Soujanya Reddy, Pratheusa Maccha, Tulasi Nagabandi, Purushotham Vodnala, Payel Mukherjee, Sofia Banu, Priya Singh, Onkar Kulkarni, Dhiviya Vedagiri, Divya Gupta, Vishal Sah, Santosh Kumar Kuncha, Krishnan Harinivas Harshan, Archana Bharadwaj Siva, Karthik Bharadwaj Tallapaka, Renu Sudhakar, Somesh Gorde, Gangumala Srinivas Reddy, Sujoy Deb, Swati Bayyana, Rakesh K Mishra, Divya Tej Sowpati |
| EPI_ISL_528919 | Ospedale Civile S. Liberatore-Atri | Istituto Zooprofilattico Sperimentale dell'Abruzzo e Molise "G.Caporale" | Lorusso A, Marcacci M, Di Domenico M, Curini V, Ancora M, Cammà C, Rinaldi A, Mangone I, Di Pasquale A, Puglia I, Savini G. |
| EPI_ISL_528920, EPI_ISL_528921 | Presidio Ospedaliero "Santo Spirito"-Pescara | Istituto Zooprofilattico Sperimentale dell'Abruzzo e Molise "G.Caporale" | Lorusso A, Marcacci M, Di Domenico M, Curini V, Ancora M, Cammà C, Rinaldi A, Mangone I, Di Pasquale A, Puglia I, Savini G. |
| EPI_ISL_528922, EPI_ISL_528924 | Ospedale "Giuseppe Mazzini"-Teramo | Istituto Zooprofilattico Sperimentale dell'Abruzzo e Molise "G.Caporale" | Lorusso A, Marcacci M, Di Domenico M, Curini V, Ancora M, Cammà C, Rinaldi A, Mangone I, Di Pasquale A, Puglia I, Savini G. |
| EPI_ISL_528925 | Ospedale Regionale San Salvatore-L'Aquila | Istituto Zooprofilattico Sperimentale dell'Abruzzo e Molise "G.Caporale" | Lorusso A, Marcacci M, Di Domenico M, Curini V, Ancora M, Cammà C, Rinaldi A, Mangone I, Di Pasquale A, Puglia I, Savini G. |
| EPI_ISL_528926, EPI_ISL_528927, EPI_ISL_528928 | Ospedale "Giuseppe Mazzini"-Teramo | Istituto Zooprofilattico Sperimentale dell'Abruzzo e Molise "G.Caporale" | Lorusso A, Marcacci M, Di Domenico M, Curini V, Ancora M, Cammà C, Rinaldi A, Mangone I, Di Pasquale A, Puglia I, Savini G. |
| EPI_ISL_528929 | Ospedale Civile S. Liberatore-Atri | Istituto Zooprofilattico Sperimentale dell'Abruzzo e Molise "G.Caporale" | Lorusso A, Marcacci M, Di Domenico M, Curini V, Ancora M, Cammà C, Rinaldi A, Mangone I, Di Pasquale A, Puglia I, Savini G. |
| EPI_ISL_528930 | Respiratory Virus Unit, Microbiology Services Colindale, Public Health England | Respiratory Virus Unit, Microbiology Services Colindale, Public Health England | PHE Covid Sequencing Team |
| EPI_ISL_528934, EPI_ISL_528935, EPI_ISL_528936, EPI_ISL_528937, EPI_ISL_528938, EPI_ISL_528939, EPI_ISL_528940, EPI_ISL_528941, EPI_ISL_528942, EPI_ISL_528943, EPI_ISL_528944, EPI_ISL_528945, EPI_ISL_528946, EPI_ISL_528947, EPI_ISL_528948, EPI_ISL_528949 | | | |
| see above | Agenzia di Tutela della Salute di Bergamo | Istituto Zooprofilattico Sperimentale dell'Abruzzo e Molise "G.Caporale" | Lorusso A, Marcacci M, Di Domenico M, Curini V, Ancora M, Cammà C, Rinaldi A, Mangone I, Di Pasquale A, Puglia I, Savini G. |
| EPI_ISL_528950 | Respiratory Virus Unit, Microbiology Services Colindale, Public Health England | Respiratory Virus Unit, Microbiology Services Colindale, Public Health England | PHE Covid Sequencing Team |
| EPI_ISL_528990 | Ospedale Civile Maria SS. dello Splendore | Istituto Zooprofilattico Sperimentale dell'Abruzzo e Molise "G.Caporale" | Lorusso A, Marcacci M, Di Domenico M, Curini V, Ancora M, Cammà C, Rinaldi A, Mangone I, Di Pasquale A, Puglia I, Savini G. |
| EPI_ISL_528991 | Ospedale SS Annunziata-Sulmona | Istituto Zooprofilattico Sperimentale dell'Abruzzo e Molise "G.Caporale" | Lorusso A, Marcacci M, Di Domenico M, Curini V, Ancora M, Cammà C, Rinaldi A, Mangone I, Di Pasquale A, Puglia I, Savini G. |
| EPI_ISL_528993 | Ospedale Civile S. Liberatore-Atri | Istituto Zooprofilattico Sperimentale dell'Abruzzo e Molise "G.Caporale" | Lorusso A, Marcacci M, Di Domenico M, Curini V, Ancora M, Cammà C, Rinaldi A, Mangone I, Di Pasquale A, Puglia I, Savini G. |
| EPI_ISL_528994, EPI_ISL_528995, EPI_ISL_528996, EPI_ISL_528997, EPI_ISL_528998, EPI_ISL_528999, EPI_ISL_529000, EPI_ISL_529001, EPI_ISL_529003, EPI_ISL_529004, EPI_ISL_529005 | | | |
| see above | Servizio di igiene epidemiologia e sanità pubblica (SIESP)-Chieti | Istituto Zooprofilattico Sperimentale dell'Abruzzo e Molise "G.Caporale" | Lorusso A, Marcacci M, Di Domenico M, Curini V, Ancora M, Cammà C, Rinaldi A, Mangone I, Di Pasquale A, Puglia I, Savini G. |
| EPI_ISL_529006 | Servizio di igiene e sanità pubblica (SIESP)-Teramo | Istituto Zooprofilattico Sperimentale dell'Abruzzo e Molise "G.Caporale" | Lorusso A, Marcacci M, Di Domenico M, Curini V, Ancora M, Cammà C, Rinaldi A, Mangone I, Di Pasquale A, Puglia I, Savini G. |
| EPI_ISL_529007, EPI_ISL_529009 | Ospedale Civile S. Liberatore-Atri | Istituto Zooprofilattico Sperimentale dell'Abruzzo e Molise "G.Caporale" | Lorusso A, Marcacci M, Di Domenico M, Curini V, Ancora M, Cammà C, Rinaldi A, Mangone I, Di Pasquale A, Puglia I, Savini G. |
| EPI_ISL_529010, EPI_ISL_529011, EPI_ISL_529012 | Servizio Igiene Epidemiologia e Sanità Pubblica (SIESP)-L'Aquila | Istituto Zooprofilattico Sperimentale dell'Abruzzo e Molise "G.Caporale" | Lorusso A, Marcacci M, Di Domenico M, Curini V, Ancora M, Cammà C, Rinaldi A, Mangone I, Di Pasquale A, Puglia I, Savini G. |
| EPI_ISL_529013 | Presidio Ospedaliero "S.Filippo e Nicola"-Avezzano | Istituto Zooprofilattico Sperimentale dell'Abruzzo e Molise "G.Caporale" | Lorusso A, Marcacci M, Di Domenico M, Curini V, Ancora M, Cammà C, Rinaldi A, Mangone I, Di Pasquale A, Puglia I, Savini G. |
| EPI_ISL_529014, EPI_ISL_529015 | Ospedale "Ss. Annunziata" | Istituto Zooprofilattico Sperimentale dell'Abruzzo e Molise "G.Caporale" | Lorusso A, Marcacci M, Di Domenico M, Curini V, Ancora M, Cammà C, Rinaldi A, Mangone I, Di Pasquale A, Puglia I, Savini G. |
| EPI_ISL_529016 | Ospedale SS Annunziata-Sulmona | Istituto Zooprofilattico Sperimentale dell'Abruzzo e Molise "G.Caporale" | Lorusso A, Marcacci M, Di Domenico M, Curini V, Ancora M, Cammà C, Rinaldi A, Mangone I, Di Pasquale A, Puglia I, Savini G. |
| EPI_ISL_529018 | Ospedale "Giuseppe Mazzini"-Teramo | Istituto Zooprofilattico Sperimentale dell'Abruzzo e Molise "G.Caporale" | Lorusso A, Marcacci M, Di Domenico M, Curini V, Ancora M, Cammà C, Rinaldi A, Mangone I, Di Pasquale A, Puglia I, Savini G. |
| EPI_ISL_529019 | RSA/RP Villa San Giovanni - Gruppo Edos | Istituto Zooprofilattico Sperimentale dell'Abruzzo e Molise "G.Caporale" | Lorusso A, Marcacci M, Di Domenico M, Curini V, Ancora M, Cammà C, Rinaldi A, Mangone I, Di Pasquale A, Puglia I, Savini G. |
| EPI_ISL_529020, EPI_ISL_529021 | Ospedale Civile S. Liberatore-Atri | Istituto Zooprofilattico Sperimentale dell'Abruzzo e Molise "G.Caporale" | Lorusso A, Marcacci M, Di Domenico M, Curini V, Ancora M, Cammà C, Rinaldi A, Mangone I, Di Pasquale A, Puglia I, Savini G. |
| EPI_ISL_529022 | Ospedale "Ss. Annunziata" | Istituto Zooprofilattico Sperimentale dell'Abruzzo e Molise "G.Caporale" | Lorusso A, Marcacci M, Di Domenico M, Curini V, Ancora M, Cammà C, Rinaldi A, Mangone I, Di Pasquale A, Puglia I, Savini G. |
| EPI_ISL_529026 | Ospedale "Giuseppe Mazzini"-Teramo | Istituto Zooprofilattico Sperimentale dell'Abruzzo e Molise "G.Caporale" | Lorusso A, Marcacci M, Di Domenico M, Curini V, Ancora M, Cammà C, Rinaldi A, Mangone I, Di Pasquale A, Puglia I, Savini G. |
| EPI_ISL_529028 | Respiratory Virus Unit, Microbiology Services Colindale, Public Health England | Respiratory Virus Unit, Microbiology Services Colindale, Public Health England | PHE Covid Sequencing Team |

| | | | |
|---|---|---|---|
| EPI_ISL_529031 | Central Molecular Microbiology Laboratory, Clinical and Chemical Pathology Department, Faculty of Medicine, CAIRO UNIVERSITY | Next Generation Sequencing Reference Laboratory, Faculty of Medicine, Cairo University and The Center for Genome and Microbiome Research, Faculty of Pharmacy, CAIRO UNIVERSITY | May Sherif Soliman, May Abdelfattah, Ramy Karam Aziz |
| EPI_ISL_529032 | Central Molecular Microbiology Laboratory and Next Generation Sequencing Reference Laboratory, Clinical and Chemical Pathology Department, Faculty of Medicine, CAIRO UNIVERSITY | Next Generation Sequencing Reference Laboratory, Faculty of Medicine, CAIRO UNIVERSITY and The Center for Genome and Microbiome Research, Faculty of Pharmacy, CAIRO UNIVERSITY | May Sherif Soliman, May Abdelfattah, Ramy Karam Aziz |
| EPI_ISL_529033, EPI_ISL_529034, EPI_ISL_529035, EPI_ISL_529036 | Wadsworth Center, New York State Department of Health | Wadsworth Center, New York State Department of Health | Kirsten St. George, Daryl M. Lamson, Sara Griesemer, Jonathan Plitnick, Navjot Singh, Matthew D. Shudt, Erica Lasek-Nesselquist |
| EPI_ISL_529065, EPI_ISL_529066, EPI_ISL_529067, EPI_ISL_529068, EPI_ISL_529069, EPI_ISL_529070, EPI_ISL_529071, EPI_ISL_529072, EPI_ISL_529073, EPI_ISL_529074, EPI_ISL_529075, EPI_ISL_529076, EPI_ISL_529077, EPI_ISL_529078, EPI_ISL_529079 | | | |
| see above | Laboratorio de Referencia Nacional de Virus Respiratorios, Instituto Nacional de Salud Peru | Laboratorio de Genómica Microbiana, Universidad Peruana Cayetano Heredia | Pablo Tsukayama, Alejandra Dávila-Barclay, Luis González, Pedro E. Romero, Brenda Ayzanoa, Janet Huancachoque, Pool Marcos, Maribel Huaringa |
| EPI_ISL_529080, EPI_ISL_529081, EPI_ISL_529082, EPI_ISL_529083, EPI_ISL_529084, EPI_ISL_529085, EPI_ISL_529086, EPI_ISL_529087, EPI_ISL_529088, EPI_ISL_529089, EPI_ISL_529090, EPI_ISL_529091, EPI_ISL_529092, EPI_ISL_529093, EPI_ISL_529094, EPI_ISL_529095, EPI_ISL_529096, EPI_ISL_529097, EPI_ISL_529098, EPI_ISL_529099, EPI_ISL_529100, EPI_ISL_529101, EPI_ISL_529103, EPI_ISL_529105, EPI_ISL_529106, EPI_ISL_529107, EPI_ISL_529108 | | | |
| see above | Microbiology Division, SC DHEC | Microbiology Division, SC DHEC | Flores,H. |
| EPI_ISL_529135 | Department of Virology III, National Institute of Infectious Diseases | Pathogen Genomics Center, National Institute of Infectious Diseases | Tsuyoshi Sekizuka, Shutoku Matsuyama, Kentaro Itokawa, Rina Tanaka, Masanori Hashino, Makoto Takeda, Takaji Wakita, Makoto Kuroda |
| EPI_ISL_529138 | RSAL Dr. Ramelan Surabaya | Institute of Tropical Disease, Universitas Airlangga | Jezzy R Dewantari, Rima R Prasetya, Krisnoadi Rahardjo, Aldise M Nastri, Radito Soesanto, Gatot Soegiarto, Laksmi Wulandari, Retno A Setyoningrum, Resti Yudhawati, Yohko K Shimizu, Mitsuhiro Nishimura, Yasuko Mori, Soetjipto, Kazufumi Shimizu, Maria I Lusida |
| EPI_ISL_529218, EPI_ISL_529219, EPI_ISL_529220, EPI_ISL_529221, EPI_ISL_529222, EPI_ISL_529223 | University of Birmingham | COVID-19 Genomics UK (COG-UK) Consortium | Institute of Microbiology, University of Birmingham: Claire McMurray, Joanne Stockton, Samuel Nicholls, Radoslaw Poplawski, Will Rowe, Josh Quick, Nicholas Loman. University of Birmingham Testing Laboratory: Celina M Whalley, Andrew Bosworth, Charlotte Poxon, Kasun Wanigasooriya, Oliver Pickles, Mike Kidd, Alex Richter, Andrew D Beggs PHE Heartlands Lab: Husam Osman, Andrew Bosworth. Queen Elizabeth Hospital: Anna Casey |
| EPI_ISL_529224, EPI_ISL_529225 | Department of Pathology, University of Cambridge | COVID-19 Genomics UK (COG-UK) Consortium | Luke W Meredith, M. Estée Török, Myra Hosmillo, William L. Hamilton, Martin D. Curran, Theresa Feltwell, Grant Hall, Anna Yakovleva, Fahad A Khokhar, Charlotte J. Houldcroft, Laura G Caller, Aminu S. Jahun, Sarah L. Caddy, Yasmin Chaudhry, Malte Pinckert, Ian Goodfellow |
| EPI_ISL_529226, EPI_ISL_529227, EPI_ISL_529228, EPI_ISL_529229, EPI_ISL_529230, EPI_ISL_529231, EPI_ISL_529232, EPI_ISL_529233 | Wales Specialist Virology Centre Sequencing lab: Pathogen Genomics Unit | COVID-19 Genomics UK (COG-UK) Consortium | Catherine Moore, Johnathan Evans, Laura Gifford, Malorie Perry, Simon Cottrell, Angela Marchbank, Alec Birchley, Alexander Adams, Amy Gaskin, Bree Gatica-Wilcox, Jason Coombes, Joel Southgate, Lauren Gilbert, Lee Graham, Nicole Pacchiarini, Sara Kumziene-Summerhayes, Sarah Taylor, Sophie Jones, Sara Rey, Matthew Bull, Joanne Watkins, Sally Corden, Tom Connor |
| EPI_ISL_529234, EPI_ISL_529235 | University of Birmingham | COVID-19 Genomics UK (COG-UK) Consortium | Institute of Microbiology, University of Birmingham: Claire McMurray, Joanne Stockton, Samuel Nicholls, Radoslaw Poplawski, Will Rowe, Josh Quick, Nicholas Loman. University of Birmingham Testing Laboratory: Celina M Whalley, Andrew Bosworth, Charlotte Poxon, Kasun Wanigasooriya, Oliver Pickles, Mike Kidd, Alex Richter, Andrew D Beggs PHE Heartlands Lab: Husam Osman, Andrew Bosworth. Queen Elizabeth Hospital: Anna Casey |
| EPI_ISL_529236 | Department of Pathology, University of Cambridge | COVID-19 Genomics UK (COG-UK) Consortium | Luke W Meredith, M. Estée Török, Myra Hosmillo, William L. Hamilton, Martin D. Curran, Theresa Feltwell, Grant Hall, Anna Yakovleva, Fahad A Khokhar, Charlotte J. Houldcroft, Laura G Caller, Aminu S. Jahun, Sarah L. Caddy, Yasmin Chaudhry, Malte Pinckert, Ian Goodfellow |
| EPI_ISL_529237 | University of Birmingham | COVID-19 Genomics UK (COG-UK) Consortium | Institute of Microbiology, University of Birmingham: Claire McMurray, Joanne Stockton, Samuel Nicholls, Radoslaw Poplawski, Will Rowe, Josh Quick, Nicholas Loman. University of Birmingham Testing Laboratory: Celina M Whalley, Andrew Bosworth, Charlotte Poxon, Kasun Wanigasooriya, Oliver Pickles, Mike Kidd, Alex Richter, Andrew D Beggs PHE Heartlands Lab: Husam Osman, Andrew Bosworth. Queen Elizabeth Hospital: Anna Casey |
| EPI_ISL_529238 | Quadram Institute Bioscience | COVID-19 Genomics UK (COG-UK) Consortium | Dave J. Baker, Gemma L. Kay, Alp Aydin, Thanh Le-Viet, Steven Rudder, Ana P. Tedim, Anastasia Kolyva, Maria Diaz, Leonardo de Oliveira Martins, Nabil-Fareed Alikhan, Lizzie Meadows, Rachael Stanley, Ngozi Elumogo, Muhammed Yasir, Nicholas M. Thomson, Alexander J Trotter, Rachel Gilroy, Samuel Bloomfield, Claire Stuart, Andrew Bell, Reenesh Prakash, Samir Dervisevic, Alison E. Mather, John Wain, Mark Webber, Andrew J. Page, Justin O'Grady |
| EPI_ISL_529239 | University of Birmingham | COVID-19 Genomics UK (COG-UK) Consortium | Institute of Microbiology, University of Birmingham: Claire McMurray, Joanne Stockton, Samuel Nicholls, Radoslaw Poplawski, Will Rowe, Josh Quick, Nicholas Loman. University of Birmingham Testing Laboratory: Celina M Whalley, Andrew Bosworth, Charlotte Poxon, Kasun Wanigasooriya, Oliver Pickles, Mike Kidd, Alex Richter, Andrew D Beggs PHE Heartlands Lab: Husam Osman, Andrew Bosworth. Queen Elizabeth Hospital: Anna Casey |
| EPI_ISL_529240, EPI_ISL_529241, EPI_ISL_529242, EPI_ISL_529243, EPI_ISL_529244, EPI_ISL_529245, EPI_ISL_529246, EPI_ISL_529247, EPI_ISL_529248, EPI_ISL_529249, EPI_ISL_529250 | | | |
| see above | Quadram Institute Bioscience | COVID-19 Genomics UK (COG-UK) Consortium | Dave J. Baker, Gemma L. Kay, Alp Aydin, Thanh Le-Viet, Steven Rudder, Ana P. Tedim, Anastasia Kolyva, Maria Diaz, Leonardo de Oliveira Martins, Nabil-Fareed Alikhan, Lizzie Meadows, Rachael Stanley, Ngozi Elumogo, Muhammed Yasir, Nicholas M. Thomson, Alexander J Trotter, Rachel Gilroy, Samuel Bloomfield, Claire Stuart, Andrew Bell, Reenesh Prakash, Samir Dervisevic, Alison E. Mather, John Wain, Mark Webber, Andrew J. Page, Justin O'Grady |
| EPI_ISL_529251, EPI_ISL_529252 | University of Birmingham | COVID-19 Genomics UK (COG-UK) Consortium | Institute of Microbiology, University of Birmingham: Claire McMurray, Joanne Stockton, Samuel Nicholls, Radoslaw Poplawski, Will Rowe, Josh Quick, Nicholas Loman. University of Birmingham Testing Laboratory: Celina M Whalley, Andrew Bosworth, Charlotte Poxon, Kasun Wanigasooriya, Oliver Pickles, Mike Kidd, Alex Richter, Andrew D Beggs PHE Heartlands Lab: Husam Osman, Andrew Bosworth. Queen Elizabeth Hospital: Anna Casey |
| EPI_ISL_529253 | Wales Specialist Virology Centre Sequencing lab: Pathogen Genomics Unit | COVID-19 Genomics UK (COG-UK) Consortium | Catherine Moore, Johnathan Evans, Laura Gifford, Malorie Perry, Simon Cottrell, Angela Marchbank, Alec Birchley, Alexander Adams, Amy Gaskin, Bree Gatica-Wilcox, Jason Coombes, Joel Southgate, Lauren Gilbert, Lee Graham, Nicole Pacchiarini, Sara Kumziene-Summerhayes, Sarah Taylor, Sophie Jones, Sara Rey, Matthew Bull, Joanne Watkins, Sally Corden, Tom Connor |
| EPI_ISL_529528, EPI_ISL_529529, EPI_ISL_529530, EPI_ISL_529531, EPI_ISL_529532, EPI_ISL_529533, EPI_ISL_529534, EPI_ISL_529535, EPI_ISL_529536, EPI_ISL_529537, EPI_ISL_529538, EPI_ISL_529539, EPI_ISL_529540, EPI_ISL_529541, EPI_ISL_529542, EPI_ISL_529543, EPI_ISL_529544, EPI_ISL_529545, EPI_ISL_529546, EPI_ISL_529547, EPI_ISL_529548, EPI_ISL_529549, EPI_ISL_529550, EPI_ISL_529551, EPI_ISL_529552, EPI_ISL_529553, EPI_ISL_529554, EPI_ISL_529555, EPI_ISL_529556 | | | |
| see above | Queens Medical Centre, Clinical Microbiology Department / DeepSeq Nottingham | COVID-19 Genomics UK (COG-UK) Consortium | Gemma Clark, Wendy Smith, Manjinder Khakh, Vicki M Fleming, Michelle M Lister, Hannah Howson-Wells, Jonathan Ball, Patrick McClure, Joseph Chappell, Theocharis Tsoleridis, Nadine Holmes, Matthew Carlisle, Christopher Moore, Fei Sang, Johnny Debebe, Victoria Wright, Matthew Loose |
| EPI_ISL_529557, EPI_ISL_529558, EPI_ISL_529559, EPI_ISL_529560, EPI_ISL_529561, EPI_ISL_529562, EPI_ISL_529563, EPI_ISL_529564, EPI_ISL_529565, EPI_ISL_529566, EPI_ISL_529567, EPI_ISL_529568, EPI_ISL_529569, EPI_ISL_529570, EPI_ISL_529571, EPI_ISL_529572, EPI_ISL_529573, EPI_ISL_529574, EPI_ISL_529575, EPI_ISL_529576, EPI_ISL_529577, EPI_ISL_529578, EPI_ISL_529579, EPI_ISL_529580, EPI_ISL_529581, EPI_ISL_529582, EPI_ISL_529583, EPI_ISL_529584, EPI_ISL_529585, EPI_ISL_529586, EPI_ISL_529587, EPI_ISL_529588, EPI_ISL_529589 | | | |
| see above | Quadram Institute Bioscience | COVID-19 Genomics UK (COG-UK) Consortium | Dave J. Baker, Gemma L. Kay, Alp Aydin, Thanh Le-Viet, Steven Rudder, Ana P. Tedim, Anastasia Kolyva, Maria Diaz, Leonardo de Oliveira Martins, Nabil-Fareed Alikhan, Lizzie Meadows, Rachael Stanley, Ngozi Elumogo, Muhammed Yasir, Nicholas M. Thomson, Alexander J Trotter, Rachel Gilroy, Samuel Bloomfield, Claire Stuart, Andrew Bell, Reenesh Prakash, Samir Dervisevic, Alison E. Mather, John Wain, Mark Webber, Andrew J. Page, Justin O'Grady |
| EPI_ISL_529590, EPI_ISL_529591, EPI_ISL_529592, EPI_ISL_529593, EPI_ISL_529594, EPI_ISL_529595, EPI_ISL_529596, EPI_ISL_529597, EPI_ISL_529598, EPI_ISL_529599, EPI_ISL_529600, EPI_ISL_529601, EPI_ISL_529602, EPI_ISL_529603, EPI_ISL_529604, EPI_ISL_529605, EPI_ISL_529606, EPI_ISL_529607, EPI_ISL_529608, EPI_ISL_529609, EPI_ISL_529610, EPI_ISL_529611, EPI_ISL_529612, EPI_ISL_529613, EPI_ISL_529614, EPI_ISL_529615, EPI_ISL_529616, EPI_ISL_529617, EPI_ISL_529618, EPI_ISL_529619, EPI_ISL_529620, EPI_ISL_529621, EPI_ISL_529622, EPI_ISL_529623, EPI_ISL_529624, EPI_ISL_529625, EPI_ISL_529626, EPI_ISL_529627, EPI_ISL_529628, EPI_ISL_529629, EPI_ISL_529630, EPI_ISL_529631, EPI_ISL_529632, EPI_ISL_529633, EPI_ISL_529634, EPI_ISL_529635, EPI_ISL_529636, EPI_ISL_529637, EPI_ISL_529638, EPI_ISL_529639, EPI_ISL_529640, EPI_ISL_529641, EPI_ISL_529642, EPI_ISL_529643, EPI_ISL_529644, EPI_ISL_529645, EPI_ISL_529646, EPI_ISL_529647, EPI_ISL_529648, EPI_ISL_529649, EPI_ISL_529650, EPI_ISL_529651, EPI_ISL_529652, EPI_ISL_529653, EPI_ISL_529654, EPI_ISL_529655 | | | |
| see above | University of Birmingham | COVID-19 Genomics UK (COG-UK) Consortium | Institute of Microbiology, University of Birmingham: Claire McMurray, Joanne Stockton, Samuel Nicholls, Radoslaw Poplawski, Will Rowe, Josh Quick, Nicholas Loman. University of Birmingham Testing Laboratory: Celina M Whalley, Andrew Bosworth, Charlotte Poxon, Kasun Wanigasooriya, Oliver Pickles, Mike Kidd, Alex Richter, Andrew D Beggs PHE Heartlands Lab: Husam Osman, Andrew Bosworth. Queen Elizabeth Hospital: Anna Casey |
| EPI_ISL_529656, EPI_ISL_529657, EPI_ISL_529658, EPI_ISL_529659, EPI_ISL_529660 | Department of Pathology, University of Cambridge | COVID-19 Genomics UK (COG-UK) Consortium | Luke W Meredith, M. Estée Török, Myra Hosmillo, William L. Hamilton, Martin D. Curran, Theresa Feltwell, Grant Hall, Anna Yakovleva, Fahad A Khokhar, Charlotte J. Houldcroft, Laura G Caller, Aminu S. Jahun, Sarah L. Caddy, Yasmin Chaudhry, Malte Pinckert, Ian Goodfellow |
| EPI_ISL_529661, EPI_ISL_529662, EPI_ISL_529663, EPI_ISL_529664, EPI_ISL_529665, EPI_ISL_529666 | University of Birmingham | COVID-19 Genomics UK (COG-UK) Consortium | Institute of Microbiology, University of Birmingham: Claire McMurray, Joanne Stockton, Samuel Nicholls, Radoslaw Poplawski, Will Rowe, Josh Quick, Nicholas Loman. University of Birmingham Testing Laboratory: Celina M Whalley, Andrew Bosworth, Charlotte Poxon, Kasun Wanigasooriya, Oliver Pickles, Mike Kidd, Alex Richter, Andrew D Beggs PHE Heartlands Lab: Husam Osman, Andrew Bosworth. Queen Elizabeth Hospital: Anna Casey |
| EPI_ISL_529667 | Queens Medical Centre, Clinical Microbiology Department / DeepSeq Nottingham | COVID-19 Genomics UK (COG-UK) Consortium | Gemma Clark, Wendy Smith, Manjinder Khakh, Vicki M Fleming, Michelle M Lister, Hannah Howson-Wells, Jonathan Ball, Patrick McClure, Joseph Chappell, Theocharis Tsoleridis, Nadine Holmes, Matthew Carlisle, Christopher Moore, Fei Sang, Johnny Debebe, Victoria Wright, Matthew Loose |
| EPI_ISL_529668, EPI_ISL_529669, EPI_ISL_529670, EPI_ISL_529671, EPI_ISL_529672, EPI_ISL_529673, EPI_ISL_529674 | Centre for Enzyme Innovation, University of Portsmouth / Translational Research Laboratory, Portsmouth Hospitals NHS Trust | COVID-19 Genomics UK (COG-UK) Consortium | Angela Beckett,Yann Bourgeois,Garry Scarlett,Sharon Glaysher,Scott Elliott,Kelly Bicknell,Robert Impey,Allyson Lloyd,Sarah Wyllie,Ethan Butcher,Anoop Chauhan,Samuel Robson |
| EPI_ISL_529675, EPI_ISL_529676, EPI_ISL_529677, EPI_ISL_529678, EPI_ISL_529679, EPI_ISL_529680, EPI_ISL_529681, EPI_ISL_529682, EPI_ISL_529683 | Virology Department, Sheffield Teaching Hospitals NHS Foundation Trust/Department of Infection, Immunity and Cardiovascular Disease, The Medical School, University of Sheffield | COVID-19 Genomics UK (COG-UK) Consortium | Thushan de Silva, Matthew Parker, Nikki Smith, Adri Angyal, Rebecca Brown, Luke Green, Rachel Tucker, Paul Parsons, Danielle Groves, Katie Johnson, Laura Carrilero, Alex Keeley, Dave Partridge, Matthew Wyles, Benjamin Lindsey, Mehmet Yavuz, Mohammad Raza, Cariad Evans |
| EPI_ISL_529684, EPI_ISL_529685, EPI_ISL_529686, EPI_ISL_529687, EPI_ISL_529688 | West of Scotland Specialist Virology Centre, NHSGGC / MRC-University of Glasgow Centre for Virus Research | COVID-19 Genomics UK (COG-UK) Consortium | Ana da Silva Filipe, Natasha Johnson, Kathy Smollett, Daniel Mair, Stephen Carmichael, Lily Tong, Jenna Nichols, Elihu Aranday-Cortes, Kirstyn Brunker, Yasmin Parr, Alice Broos, Kyriaki Nomikou; Sarah McDonald, Marc Niebel, Patawee Asamaphan; Richard Orton, Joseph Hughes, Sreenu Vattipally, David L Robertson; Alasdair MacLean, Rory Gunson; Kathy Li, Natasha Jesudason, Rajiv Shah, James Shepherd, Antonia Ho, Emma Thomson |
| EPI_ISL_529689, EPI_ISL_529690, EPI_ISL_529691, EPI_ISL_529692, EPI_ISL_529693, EPI_ISL_529694, EPI_ISL_529695, EPI_ISL_529696 | Virology Department, Royal Infirmary of Edinburgh, NHS Lothian / School of Biological Sciences, University of Edinburgh / Institute of Genetics and Molecular Medicine, University of Edinburgh | COVID-19 Genomics UK (COG-UK) Consortium | McHugh M, Dewar R, Rooke S, Gallagher M, Balcaza C, O'Toole Á, Scher E, Hill V, McCrone JT, Colquhoun R, Yu X, Jackson B, Rambaut A, Williams TC, Templeton K |
| EPI_ISL_529697, EPI_ISL_529698, EPI_ISL_529699, EPI_ISL_529700, EPI_ISL_529701, EPI_ISL_529702, EPI_ISL_529703, EPI_ISL_529704, EPI_ISL_529705, EPI_ISL_529706, EPI_ISL_529707, EPI_ISL_529708, EPI_ISL_529709, EPI_ISL_529710, EPI_ISL_529711, EPI_ISL_529712, EPI_ISL_529713, EPI_ISL_529714, EPI_ISL_529715, EPI_ISL_529716, EPI_ISL_529717 | | | |
| see above | Wales Specialist Virology Centre Sequencing lab: Pathogen Genomics Unit | COVID-19 Genomics UK (COG-UK) Consortium | Catherine Moore, Johnathan Evans, Laura Gifford, Malorie Perry, Simon Cottrell, Angela Marchbank, Alec Birchley, Alexander Adams, Amy Gaskin, Bree Gatica-Wilcox, Jason Coombes, Joel Southgate, Lauren Gilbert, Lee Graham, Nicole Pacchiarini, Sara Kumziene-Summerhayes, Sarah Taylor, Sophie Jones, Sara Rey, Matthew Bull, Joanne Watkins, Sally Corden, Tom Connor |
| EPI_ISL_529720, EPI_ISL_529721, EPI_ISL_529723, EPI_ISL_529724, EPI_ISL_529726, EPI_ISL_529728, EPI_ISL_529729, EPI_ISL_529730, EPI_ISL_529731, EPI_ISL_529732, EPI_ISL_529733, EPI_ISL_529734, EPI_ISL_529735, EPI_ISL_529736, EPI_ISL_529737, EPI_ISL_529738, EPI_ISL_529739, EPI_ISL_529740, EPI_ISL_529741, EPI_ISL_529742, EPI_ISL_529743, EPI_ISL_529744, EPI_ISL_529745, EPI_ISL_529747, EPI_ISL_529748, EPI_ISL_529749, EPI_ISL_529750, EPI_ISL_529751, EPI_ISL_529752, EPI_ISL_529753, EPI_ISL_529755, EPI_ISL_529756, EPI_ISL_529757, EPI_ISL_529758, EPI_ISL_529759, EPI_ISL_529760, EPI_ISL_529761, EPI_ISL_529762, EPI_ISL_529763, EPI_ISL_529764, EPI_ISL_529765, EPI_ISL_529766, EPI_ISL_529767, EPI_ISL_529769, EPI_ISL_529770, EPI_ISL_529771, EPI_ISL_529772, EPI_ISL_529773, | | | |

| | | | |
|---|---|---|---|
| EPI_ISL_529774, EPI_ISL_529775, EPI_ISL_529776, EPI_ISL_529777, EPI_ISL_529778, EPI_ISL_529779, EPI_ISL_529780, EPI_ISL_529782, EPI_ISL_529783, EPI_ISL_529784, EPI_ISL_529785, EPI_ISL_529786, EPI_ISL_529787, EPI_ISL_529788, EPI_ISL_529789, EPI_ISL_529790, EPI_ISL_529791, EPI_ISL_529792, EPI_ISL_529793, EPI_ISL_529794, EPI_ISL_529796, EPI_ISL_529797, EPI_ISL_529798, EPI_ISL_529799, EPI_ISL_529800, EPI_ISL_529801, EPI_ISL_529802 | | | |
| see above | NHLS-IALCH | KRISP, KZN Research Innovation and Sequencing Platform | Giandhari J, Pillay S, Lessells R, Mdlalose K, York D, Khan S, Tegally H, Wilkinson E, de Oliveira T |
| EPI_ISL_529818, EPI_ISL_529830, EPI_ISL_529832, EPI_ISL_529850, EPI_ISL_529874, EPI_ISL_529888 | Michigan Department of Health and Human Services, Bureau of Laboratories | Michigan Department of Health and Human Services, Bureau of Laboratories | Blankenship HM, Riner D, Soehnlen MK |
| EPI_ISL_529908, EPI_ISL_529909, EPI_ISL_529910, EPI_ISL_529912, EPI_ISL_529913, EPI_ISL_529914, EPI_ISL_529915, EPI_ISL_529916, EPI_ISL_529917, EPI_ISL_529918, EPI_ISL_529919, EPI_ISL_529920, EPI_ISL_529921, EPI_ISL_529923, EPI_ISL_529924, EPI_ISL_529925, EPI_ISL_529926, EPI_ISL_529927, EPI_ISL_529928, EPI_ISL_529929, EPI_ISL_529930, EPI_ISL_529931, EPI_ISL_529932, EPI_ISL_529933, EPI_ISL_529934, EPI_ISL_529935, EPI_ISL_529936 | | | |
| see above | Virginia Division of Consolidated Laboratory Services | Virginia Division of Consolidated Laboratory Services | Virginia DCLS |
| EPI_ISL_529937, EPI_ISL_529938, EPI_ISL_529939, EPI_ISL_529940, EPI_ISL_529941, EPI_ISL_529942, EPI_ISL_529943, EPI_ISL_529944, EPI_ISL_529945, EPI_ISL_529946, EPI_ISL_529947, EPI_ISL_529948, EPI_ISL_529949, EPI_ISL_529950, EPI_ISL_529951, EPI_ISL_529952, EPI_ISL_529953, EPI_ISL_529954, EPI_ISL_529955, EPI_ISL_529956, EPI_ISL_529957, EPI_ISL_529958, EPI_ISL_529959, EPI_ISL_529960 | | | |
| see above | Virginia DCLS | Virginia DCLS | Virginia DCLS |
| EPI_ISL_529961 | Husada Utama Hospital | Institute of Tropical Disease, Universitas Airlangga | Krisnoadi Rahardjo, Aldise M Nastri, Jezzy R Dewantari, Rima R Prasetya, Didi Dewanto, Gatot Soegiarto, Laksmi Wulandari, Retno A Setyoningrum, Resti Yudhawati, Yohko K Shimizu, Mitsuhiro Nishimura, Yasuko Mori, Soetjipto, Kazufumi Shimizu, Maria I Lusida |
| EPI_ISL_529962 | Universitas Airlangga Hospital | Institute of Tropical Disease, Universitas Airlangga | Jezzy R Dewantari, Rima R Prasetya, Krisnoadi Rahardjo, Aldise M Nastri, Nasronudin, Gatot Soegiarto, Laksmi Wulandari, Retno A Setyoningrum, Resti Yudhawati, Yohko K Shimizu, Mitsuhiro Nishimura, Yasuko Mori, Soetjipto, Kazufumi Shimizu, Maria I Lusida |
| EPI_ISL_529963 | Universitas Airlangga Hospital | Institute of Tropical Disease, Universitas Airlangga | Rima R Prasetya, Krisnoadi Rahardjo, Aldise M Nastri, Jezzy R Dewantari, Nasronudin, Gatot Soegiarto, Laksmi Wulandari, Retno A Setyoningrum, Resti Yudhawati, Yohko K Shimizu, Mitsuhiro Nishimura, Yasuko Mori, Soetjipto, Kazufumi Shimizu, Maria I Lusida |
| EPI_ISL_529964 | RSUD Bangil Pasuruan | Institute of Tropical Disease, Universitas Airlangga | Aldise M Nastri, Jezzy R Dewantari, Rima R Prasetya, Krisnoadi Rahardjo, Arma Roosalina, Gatot Soegiarto, Laksmi Wulandari, Retno A Setyoningrum, Resti Yudhawati, Yohko K Shimizu, Mitsuhiro Nishimura, Yasuko Mori, Soetjipto, Kazufumi Shimizu, Maria I Lusida |
| EPI_ISL_529965 | RSUD Dr. Soetomo | Institute of Tropical Disease, Universitas Airlangga | Krisnoadi Rahardjo, Aldise M Nastri, Jezzy R Dewantari, Rima R Prasetya, Joni Wahyuhadi, Gatot Soegiarto, Laksmi Wulandari, Retno A Setyoningrum, Resti Yudhawati, Yohko K Shimizu, Mitsuhiro Nishimura, Yasuko Mori, Soetjipto, Kazufumi Shimizu, Maria I Lusida |
| EPI_ISL_529966 | RSUD Sidoarjo | Institute of Tropical Disease, Universitas Airlangga | Kazufumi Shimizu, Krisnoadi Rahardjo, Aldise M Nastri, Jezzy R Dewantari, Rima R Prasetya, Atok Irawan, Gatot Soegiarto, Laksmi Wulandari, Retno A Setyoningrum, Resti Yudhawati, Yohko K Shimizu, Mitsuhiro Nishimura, Yasuko Mori, Soetjipto, Maria I Lusida |
| EPI_ISL_529967 | RSUD Sidoarjo | Institute of Tropical Disease, Universitas Airlangga | Aldise M Nastri, Jezzy R Dewantari, Rima R Prasetya, Krisnoadi Rahardjo, Atok Irawan, Gatot Soegiarto, Laksmi Wulandari, Retno A Setyoningrum, Resti Yudhawati, Yohko K Shimizu, Mitsuhiro Nishimura, Yasuko Mori, Soetjipto, Kazufumi Shimizu, Maria I Lusida |
| EPI_ISL_529972, EPI_ISL_529973, EPI_ISL_529974, EPI_ISL_529975, EPI_ISL_529976, EPI_ISL_529977, EPI_ISL_529978, EPI_ISL_529979, EPI_ISL_529980, EPI_ISL_529981, EPI_ISL_529982, EPI_ISL_529983, EPI_ISL_529984, EPI_ISL_529985, EPI_ISL_529986, EPI_ISL_529987, EPI_ISL_529988 | | | |
| see above | Hospital Universitario 12 de Octubre | Hospital Universitario 12 de Octubre | Raúl Recio, Sara González, Esther Viedma, Elias Dahdouh, Fernando Lázaro, Natalia Stella, Julio García, Juan Carlos Galán, Rafael Cantón, Mª Dolores Folgueira, Rafael Delgado, Jesús Mingorance |
| EPI_ISL_529989, EPI_ISL_529990, EPI_ISL_529991, EPI_ISL_529992, EPI_ISL_529993, EPI_ISL_529994, EPI_ISL_529995, EPI_ISL_529996, EPI_ISL_529997, EPI_ISL_529998, EPI_ISL_529999, EPI_ISL_530000, EPI_ISL_530001, EPI_ISL_530002, EPI_ISL_530003, EPI_ISL_530004, EPI_ISL_530005, EPI_ISL_530006 | | | |
| see above | Hospital Universitario 12 de Octubre | Hospital Universitario 12 de Octubre | Sara González, Esther Viedma, Raúl Recio, Elias Dahdouh, Fernando Lázaro, Natalia Stella, Julio García, Juan Carlos Galán, Rafael Cantón, Mª Dolores Folgueira, Rafael Delgado, Jesús Mingorance |
| EPI_ISL_530007, EPI_ISL_530008, EPI_ISL_530009, EPI_ISL_530010, EPI_ISL_530011, EPI_ISL_530012, EPI_ISL_530013, EPI_ISL_530014, EPI_ISL_530015, EPI_ISL_530016, EPI_ISL_530017, EPI_ISL_530018, EPI_ISL_530019, EPI_ISL_530020, EPI_ISL_530021, EPI_ISL_530022, EPI_ISL_530023, EPI_ISL_530024 | | | |
| see above | Hospital Universitario 12 de Octubre | Hospital Universitario 12 de Octubre | Esther Viedma, Raúl Recio, Sara González, Elias Dahdouh, Fernando Lázaro, Natalia Stella, Julio García, Juan Carlos Galán, Rafael Cantón, Mª Dolores Folgueira, Rafael Delgado, Jesús Mingorance |
| EPI_ISL_530025, EPI_ISL_530027, EPI_ISL_530028, EPI_ISL_530029, EPI_ISL_530030, EPI_ISL_530031, EPI_ISL_530032, EPI_ISL_530033, EPI_ISL_530034, EPI_ISL_530035, EPI_ISL_530036, EPI_ISL_530037, EPI_ISL_530038, EPI_ISL_530039, EPI_ISL_530040, EPI_ISL_530041, EPI_ISL_530042, EPI_ISL_530043 | | | |
| see above | Hospital Universitario La Paz | Hospital Universitario La Paz | María Rodríguez, Elias Dahdouh, Sara González, Raúl Recio, Fernando Lázaro, Esther Viedma, Natalia Stella, Julio García, Juan Carlos Galán, Rafael Cantón, Mª Dolores Folgueira, Rafael Delgado, Jesús Mingorance |
| EPI_ISL_530044, EPI_ISL_530045, EPI_ISL_530046, EPI_ISL_530047, EPI_ISL_530048, EPI_ISL_530049, EPI_ISL_530050, EPI_ISL_530051, EPI_ISL_530052, EPI_ISL_530053, EPI_ISL_530054, EPI_ISL_530055, EPI_ISL_530056, EPI_ISL_530057, EPI_ISL_530058, EPI_ISL_530059, EPI_ISL_530060, EPI_ISL_530061, EPI_ISL_530062, EPI_ISL_530063, EPI_ISL_530064, EPI_ISL_530065, EPI_ISL_530066, EPI_ISL_530067, EPI_ISL_530068, EPI_ISL_530069 | | | |
| see above | Hospital Universitario La Paz | Hospital Universitario La Paz | Elias Dahdouh, Sara González, Raúl Recio, Fernando Lázaro, Esther Viedma, Natalia Stella, Julio García, Juan Carlos Galán, Rafael Cantón, Mª Dolores Folgueira, Rafael Delgado, Jesús Mingorance |
| EPI_ISL_530070, EPI_ISL_530071, EPI_ISL_530072, EPI_ISL_530073, EPI_ISL_530074, EPI_ISL_530075, EPI_ISL_530077, EPI_ISL_530078, EPI_ISL_530079, EPI_ISL_530080, EPI_ISL_530081, EPI_ISL_530082, EPI_ISL_530083, EPI_ISL_530084, EPI_ISL_530085, EPI_ISL_530086, EPI_ISL_530087, EPI_ISL_530088, EPI_ISL_530089, EPI_ISL_530090, EPI_ISL_530091, EPI_ISL_530092, EPI_ISL_530093 | | | |
| see above | Hospital Universitario La Paz | Hospital Universitario La Paz | María Rodríguez, Elias Dahdouh, Sara González, Raúl Recio, Fernando Lázaro, Esther Viedma, Natalia Stella, Julio García, Juan Carlos Galán, Rafael Cantón, Mª Dolores Folgueira, Rafael Delgado, Jesús Mingorance |
| EPI_ISL_530094, EPI_ISL_530095, EPI_ISL_530096, EPI_ISL_530097, EPI_ISL_530098, EPI_ISL_530099 | Hospital Universitario Ramón y Cajal | Hospital Universitario 12 de Octubre | Raúl Recio, Sara González, Elias Dahdouh, Fernando Lázaro, Esther Viedma, Natalia Stella, Julio García, Juan Carlos Galán, Rafael Cantón, Mª Dolores Folgueira, Rafael Delgado, Jesús Mingorance |
| EPI_ISL_530100, EPI_ISL_530101, EPI_ISL_530102, EPI_ISL_530103, EPI_ISL_530104, EPI_ISL_530105, EPI_ISL_530106, EPI_ISL_530107, EPI_ISL_530108, EPI_ISL_530109, EPI_ISL_530110, EPI_ISL_530111, EPI_ISL_530112, EPI_ISL_530113 | | | |
| see above | Hospital Universitario Ramón y Cajal | Hospital Universitario La Paz | Raúl Recio, Sara González, Elias Dahdouh, Fernando Lázaro, Esther Viedma, Natalia Stella, Julio García, Juan Carlos Galán, Rafael Cantón, Mª Dolores Folgueira, Rafael Delgado, Jesús Mingorance |
| EPI_ISL_530114, EPI_ISL_530115, EPI_ISL_530116, EPI_ISL_530117, EPI_ISL_530118 | Hospital Universitario Ramón y Cajal | Hospital Universitario La Paz | María Rodríguez, Elias Dahdouh, Sara González, Raúl Recio, Fernando Lázaro, Esther Viedma, Natalia Stella, Julio García, Juan Carlos Galán, Rafael Cantón, Mª Dolores Folgueira, Rafael Delgado, Jesús Mingorance |
| EPI_ISL_530122 | University Hospital Basel, Clinical Virology | University Hospital Basel, Clinical Bacteriology | Madlen Stange, Alfredo Mari, Tim Roloff, Helena MB Seth-Smith, Michael Schweitzer, Myrta Brunner, Karoline Leuzinger, Kirstine K. Soegaard, Alexander Gensch, Sarah Tschudin-Sutter, Simon Fuchs, Julia Bielicki, Hans Pargger, Martin Siegemund, Christian Nickel, Roland Bingisser, Michael Osthoff, Stefano Bassetti, Rita Schneider-Sliwa, Manuel Battegay, Hans Hirsch, Adrian Egli |
| EPI_ISL_530124, EPI_ISL_530125 | Seattle Flu Study | Seattle Flu Study | Deborah A. Nickerson, Chris D. Frazar, Jover Lee, Benjamin Pelle, Matthew Richardson, Amanda Adler, Elisabeth Brandstetter, Peter D. Han, Kairsten Fay, Misja Ilcisin, Kirsten Lacombe, Thomas R. Sibley, Melissa Truong, Caitlin R. Wolf, Michael Boeckh, Janet A. Englund, Michael Famulare, Barry R. Lutz, Mark J. Rieder, Lea M. Starita, Matthew Thompson, Jay Shendure, Trevor Bedford, Helen Y. Chu |
| EPI_ISL_530126, EPI_ISL_530127, EPI_ISL_530128, EPI_ISL_530129, EPI_ISL_530131 | Seattle Flu Study | Seattle Flu Study | Deborah A. Nickerson, Chris D. Frazar, Jover Lee, Benjamin Pelle, Matthew Richardson, Amanda Adler, Elisabeth Brandstetter, Peter D. Han, Kairsten Fay, Misja Ilcisin, Kirsten Lacombe, Thomas R. Sibley, Melissa Truong, Caitlin R. Wolf, Karen Cowgill, Stephanie Schrag, Jeff Duchin, Michael Boeckh, Janet A. Englund, Michael Famulare, Barry R. Lutz, Mark J. Rieder, Lea M. Starita, Matthew Thompson, Helen Y. Chu, Trevor Bedford, Jay Shendure |
| EPI_ISL_530132 | Seattle Flu Study | Seattle Flu Study | Deborah A. Nickerson, Chris D. Frazar, Jover Lee, Benjamin Pelle, Matthew Richardson, Amanda Adler, Elisabeth Brandstetter, Peter D. Han, Kairsten Fay, Misja Ilcisin, Kirsten Lacombe, Thomas R. Sibley, Melissa Truong, Caitlin R. Wolf, Michael Boeckh, Janet A. Englund, Michael Famulare, Barry R. Lutz, Mark J. Rieder, Lea M. Starita, Matthew Thompson, Jay Shendure, Trevor Bedford, Helen Y. Chu |
| EPI_ISL_530136, EPI_ISL_530137, EPI_ISL_530138, EPI_ISL_530139, EPI_ISL_530140, EPI_ISL_530141, EPI_ISL_530142, EPI_ISL_530143, EPI_ISL_530144, EPI_ISL_530145, EPI_ISL_530146, EPI_ISL_530147, EPI_ISL_530148, EPI_ISL_530149, EPI_ISL_530150, EPI_ISL_530151, EPI_ISL_530152, EPI_ISL_530154, EPI_ISL_530155, EPI_ISL_530156, EPI_ISL_530157, EPI_ISL_530158, EPI_ISL_530159, EPI_ISL_530161, EPI_ISL_530162 | | | |
| see above | Seattle Flu Study | Seattle Flu Study | Deborah A. Nickerson, Chris D. Frazar, Jover Lee, Benjamin Pelle, Matthew Richardson, Amanda Adler, Elisabeth Brandstetter, Peter D. Han, Kairsten Fay, Misja Ilcisin, Kirsten Lacombe, Thomas R. Sibley, Melissa Truong, Caitlin R. Wolf, Karen Cowgill, Stephanie Schrag, Jeff Duchin, Michael Boeckh, Janet A. Englund, Michael Famulare, Barry R. Lutz, Mark J. Rieder, Lea M. Starita, Matthew Thompson, Helen Y. Chu, Trevor Bedford, Jay Shendure |
| EPI_ISL_530163, EPI_ISL_530164, EPI_ISL_530165, EPI_ISL_530166, EPI_ISL_530167, EPI_ISL_530168 | M Health Fairview | Minnesota Department of Health, Public Health Laboratory | Matt Plumb, Jacob Garfin, and Xiong Wang |
| EPI_ISL_530169, EPI_ISL_530170, EPI_ISL_530171 | Hennepin County Medical Center | Minnesota Department of Health, Public Health Laboratory | Matt Plumb, Jacob Garfin, and Xiong Wang |
| EPI_ISL_530172, EPI_ISL_530173, EPI_ISL_530174, EPI_ISL_530175, EPI_ISL_530176, EPI_ISL_530177, EPI_ISL_530178, EPI_ISL_530179, EPI_ISL_530180, EPI_ISL_530181, EPI_ISL_530182, EPI_ISL_530183, EPI_ISL_530184, EPI_ISL_530185, EPI_ISL_530186, EPI_ISL_530187, EPI_ISL_530188, EPI_ISL_530189, EPI_ISL_530190, EPI_ISL_530191, EPI_ISL_530192, EPI_ISL_530193, EPI_ISL_530194, EPI_ISL_530195, EPI_ISL_530196, EPI_ISL_530197, EPI_ISL_530198, EPI_ISL_530199, EPI_ISL_530200, EPI_ISL_530201, EPI_ISL_530202, EPI_ISL_530203, EPI_ISL_530204, EPI_ISL_530205, EPI_ISL_530206, EPI_ISL_530207, EPI_ISL_530208, EPI_ISL_530209, EPI_ISL_530210, EPI_ISL_530211, EPI_ISL_530212, EPI_ISL_530213, EPI_ISL_530215, EPI_ISL_530216, EPI_ISL_530217, EPI_ISL_530218, EPI_ISL_530219, EPI_ISL_530220, EPI_ISL_530221, EPI_ISL_530222, EPI_ISL_530223, EPI_ISL_530224 | | | |
| see above | Minnesota Department of Health, Public Health Laboratory | Minnesota Department of Health, Public Health Laboratory | Matt Plumb, Jacob Garfin, and Xiong Wang |
| EPI_ISL_582135, EPI_ISL_582136, EPI_ISL_582137, EPI_ISL_582138, EPI_ISL_582139, EPI_ISL_582140, EPI_ISL_582141, EPI_ISL_582142, EPI_ISL_582143, EPI_ISL_582144, EPI_ISL_582145, EPI_ISL_582146, EPI_ISL_582147 | | | |
| see above | Tokyo Metropolitan Institute of Public Health, Department of Microbiology | Tokyo Metropolitan Institute of Public Health, Department of Microbiology | Asakura,H., Yoshida,I., Kumagai,R., Nagashima,M., Chiba,T. and Sadamasu,K. |
