## Supplementary material for "Analysis of the Dynamics and Distribution of SARS-CoV-2 Mutations and its Possible Structural and Functional Implications": GISAID_acknowledgements_part_8

We gratefully acknowledge the following Authors from the Originating laboratories responsible for obtaining the specimens, as well as the Submitting laboratories where the genome data were generated and shared via GISAID, on which this research is based.

All Submitters of data may be contacted directly via www.gisaid.org

| Accession ID | Originating Laboratory | Submitting Laboratory | Authors |
|---|---|---|---|
| EPI_ISL_530225, EPI_ISL_530226, EPI_ISL_530227, EPI_ISL_530228, EPI_ISL_530229, EPI_ISL_530230, EPI_ISL_530231, EPI_ISL_530232, EPI_ISL_530233, EPI_ISL_530234, EPI_ISL_530235, EPI_ISL_530236, EPI_ISL_530237, EPI_ISL_530238, EPI_ISL_530239, EPI_ISL_530240, EPI_ISL_530241, EPI_ISL_530242, EPI_ISL_530243, EPI_ISL_530244, EPI_ISL_530245, EPI_ISL_530246, EPI_ISL_530247, EPI_ISL_530248, EPI_ISL_530249, EPI_ISL_530250, EPI_ISL_530251, EPI_ISL_530252, EPI_ISL_530253, EPI_ISL_530254, EPI_ISL_530255, EPI_ISL_530256, EPI_ISL_530257, EPI_ISL_530258, EPI_ISL_530259, EPI_ISL_530260, EPI_ISL_530261, EPI_ISL_530262, EPI_ISL_530263, EPI_ISL_530264, EPI_ISL_530265, EPI_ISL_530266, EPI_ISL_530267, EPI_ISL_530268, EPI_ISL_530269, EPI_ISL_530270, EPI_ISL_530271, EPI_ISL_530272, EPI_ISL_530273, EPI_ISL_530274, EPI_ISL_530275, EPI_ISL_530276, EPI_ISL_530277, EPI_ISL_530278, EPI_ISL_530279, EPI_ISL_530280, EPI_ISL_530281, EPI_ISL_530282, EPI_ISL_530283 | | | |
| see above | Queensland Health Forensic and Scientific Services, Public Health Virology | Public Health Virology Laboratory, Forensic and Scientific Services, Queensland Health | Son Nguyen et al |
| EPI_ISL_530334, EPI_ISL_530335, EPI_ISL_530336, EPI_ISL_530337, EPI_ISL_530338, EPI_ISL_530339, EPI_ISL_530340 | Area of Virology, Serology and Virology Division (SAViD), New South Wales Health Pathology Randwick | Area of Virology, Serology and Virology Division (SAViD), New South Wales Health Pathology Randwick | Rawlinson, W., Deveson, I., Bull, R., Van Hal, S. |
| EPI_ISL_530347 | The National Institute of Public Health | State Veterinary Institute Prague | Nagy,A;Jirincova,H;Novakova,L;Trnka,D;Vecerova,J |
| EPI_ISL_530348, EPI_ISL_530349, EPI_ISL_530350 | The National Institute of Public Health | State Veterinary Institute Prague | Nagy,A;Jirincova,H;Novakova,L;Trnka,D;Vecerova,J |
| EPI_ISL_530352, EPI_ISL_530353, EPI_ISL_530358, EPI_ISL_530360 | Lighthouse Lab in Glasgow | Wellcome Sanger Institute for the COVID-19 Genomics UK Consortium | Harper VanSteenhouse, Yumi Kasai, David Gray, Carol Clugston, Anna Dominiczak and Alex Alderton, Roberto Amato, Sonia Goncalves, Ewan Harrison, David K. Jackson, Ian Johnston, Dominic Kwiatkowski, Cordelia Langford, John Sillitoe |
| EPI_ISL_530361 | NHSGGC West of Scotland Specialist Virology Centre / MRC-University of Glasgow Centre for Virus Research | Wellcome Sanger Institute for the COVID-19 Genomics UK Consortium | Ana da Silva Filipe, Natasha Johnson, Kathy Smollett, Daniel Mair, Stephen Carmichael, Lily Tong, Jenna Nichols, Elihu Aranday-Cortes, Kirstyn Brunker, Yasmin Parr, Kyriaki Nomikou; Sarah McDonald, Marc Niebel, Patawee Asamaphan; Richard Orton, Joseph Hughes, Sreenu Vattipally, David L Robertson; Alasdair MacLean, Rory Gunson; Kathy Li, Natasha Jesudason, Rajiv Shah, James Shepherd, Antonia Ho, Alice Broos, Emma Thomson and Alex Alderton, Roberto Amato, Sonia Goncalves, Ewan Harrison, David K. Jackson, Ian Johnston, Dominic Kwiatkowski, Cordelia Langford, John Sillitoe |
| EPI_ISL_530362, EPI_ISL_530363 | Lighthouse Lab in Glasgow | Wellcome Sanger Institute for the COVID-19 Genomics UK Consortium | Harper VanSteenhouse, Yumi Kasai, David Gray, Carol Clugston, Anna Dominiczak and Alex Alderton, Roberto Amato, Sonia Goncalves, Ewan Harrison, David K. Jackson, Ian Johnston, Dominic Kwiatkowski, Cordelia Langford, John Sillitoe |
| EPI_ISL_530366 | NHSGGC West of Scotland Specialist Virology Centre / MRC-University of Glasgow Centre for Virus Research | Wellcome Sanger Institute for the COVID-19 Genomics UK Consortium | Ana da Silva Filipe, Natasha Johnson, Kathy Smollett, Daniel Mair, Stephen Carmichael, Lily Tong, Jenna Nichols, Elihu Aranday-Cortes, Kirstyn Brunker, Yasmin Parr, Kyriaki Nomikou; Sarah McDonald, Marc Niebel, Patawee Asamaphan; Richard Orton, Joseph Hughes, Sreenu Vattipally, David L Robertson; Alasdair MacLean, Rory Gunson; Kathy Li, Natasha Jesudason, Rajiv Shah, James Shepherd, Antonia Ho, Alice Broos, Emma Thomson and Alex Alderton, Roberto Amato, Sonia Goncalves, Ewan Harrison, David K. Jackson, Ian Johnston, Dominic Kwiatkowski, Cordelia Langford, John Sillitoe |
| EPI_ISL_530374, EPI_ISL_530375, EPI_ISL_530378, EPI_ISL_530380, EPI_ISL_530382, EPI_ISL_530383, EPI_ISL_530384 | Lighthouse Lab in Glasgow | Wellcome Sanger Institute for the COVID-19 Genomics UK Consortium | Harper VanSteenhouse, Yumi Kasai, David Gray, Carol Clugston, Anna Dominiczak and Alex Alderton, Roberto Amato, Sonia Goncalves, Ewan Harrison, David K. Jackson, Ian Johnston, Dominic Kwiatkowski, Cordelia Langford, John Sillitoe |
| EPI_ISL_530386 | NHSGGC West of Scotland Specialist Virology Centre / MRC-University of Glasgow Centre for Virus Research | Wellcome Sanger Institute for the COVID-19 Genomics UK Consortium | Ana da Silva Filipe, Natasha Johnson, Kathy Smollett, Daniel Mair, Stephen Carmichael, Lily Tong, Jenna Nichols, Elihu Aranday-Cortes, Kirstyn Brunker, Yasmin Parr, Kyriaki Nomikou; Sarah McDonald, Marc Niebel, Patawee Asamaphan; Richard Orton, Joseph Hughes, Sreenu Vattipally, David L Robertson; Alasdair MacLean, Rory Gunson; Kathy Li, Natasha Jesudason, Rajiv Shah, James Shepherd, Antonia Ho, Alice Broos, Emma Thomson and Alex Alderton, Roberto Amato, Sonia Goncalves, Ewan Harrison, David K. Jackson, Ian Johnston, Dominic Kwiatkowski, Cordelia Langford, John Sillitoe |
| EPI_ISL_530387, EPI_ISL_530390, EPI_ISL_530394, EPI_ISL_530395, EPI_ISL_530396, EPI_ISL_530397, EPI_ISL_530399, EPI_ISL_530400, EPI_ISL_530401, EPI_ISL_530402, EPI_ISL_530403, EPI_ISL_530404, EPI_ISL_530406, EPI_ISL_530407, EPI_ISL_530408, EPI_ISL_530409 | | | |
| see above | Lighthouse Lab in Glasgow | Wellcome Sanger Institute for the COVID-19 Genomics UK Consortium | Harper VanSteenhouse, Yumi Kasai, David Gray, Carol Clugston, Anna Dominiczak and Alex Alderton, Roberto Amato, Sonia Goncalves, Ewan Harrison, David K. Jackson, Ian Johnston, Dominic Kwiatkowski, Cordelia Langford, John Sillitoe |
| EPI_ISL_530410 | NHSGGC West of Scotland Specialist Virology Centre / MRC-University of Glasgow Centre for Virus Research | Wellcome Sanger Institute for the COVID-19 Genomics UK Consortium | Ana da Silva Filipe, Natasha Johnson, Kathy Smollett, Daniel Mair, Stephen Carmichael, Lily Tong, Jenna Nichols, Elihu Aranday-Cortes, Kirstyn Brunker, Yasmin Parr, Kyriaki Nomikou; Sarah McDonald, Marc Niebel, Patawee Asamaphan; Richard Orton, Joseph Hughes, Sreenu Vattipally, David L Robertson; Alasdair MacLean, Rory Gunson; Kathy Li, Natasha Jesudason, Rajiv Shah, James Shepherd, Antonia Ho, Alice Broos, Emma Thomson and Alex Alderton, Roberto Amato, Sonia Goncalves, Ewan Harrison, David K. Jackson, Ian Johnston, Dominic Kwiatkowski, Cordelia Langford, John Sillitoe |
| EPI_ISL_530411, EPI_ISL_530412, EPI_ISL_530414, EPI_ISL_530415, EPI_ISL_530416, EPI_ISL_530417, EPI_ISL_530418, EPI_ISL_530419, EPI_ISL_530421, EPI_ISL_530422, EPI_ISL_530423, EPI_ISL_530424, EPI_ISL_530425, EPI_ISL_530427, EPI_ISL_530428 | | | |
| see above | Lighthouse Lab in Glasgow | Wellcome Sanger Institute for the COVID-19 Genomics UK Consortium | Harper VanSteenhouse, Yumi Kasai, David Gray, Carol Clugston, Anna Dominiczak and Alex Alderton, Roberto Amato, Sonia Goncalves, Ewan Harrison, David K. Jackson, Ian Johnston, Dominic Kwiatkowski, Cordelia Langford, John Sillitoe |
| EPI_ISL_530429 | NHSGGC West of Scotland Specialist Virology Centre / MRC-University of Glasgow Centre for Virus Research | Wellcome Sanger Institute for the COVID-19 Genomics UK Consortium | Ana da Silva Filipe, Natasha Johnson, Kathy Smollett, Daniel Mair, Stephen Carmichael, Lily Tong, Jenna Nichols, Elihu Aranday-Cortes, Kirstyn Brunker, Yasmin Parr, Kyriaki Nomikou; Sarah McDonald, Marc Niebel, Patawee Asamaphan; Richard Orton, Joseph Hughes, Sreenu Vattipally, David L Robertson; Alasdair MacLean, Rory Gunson; Kathy Li, Natasha Jesudason, Rajiv Shah, James Shepherd, Antonia Ho, Alice Broos, Emma Thomson and Alex Alderton, Roberto Amato, Sonia Goncalves, Ewan Harrison, David K. Jackson, Ian Johnston, Dominic Kwiatkowski, Cordelia Langford, John Sillitoe |
| EPI_ISL_530430, EPI_ISL_530431, EPI_ISL_530433, EPI_ISL_530434, EPI_ISL_530436, EPI_ISL_530438, EPI_ISL_530439, EPI_ISL_530441, EPI_ISL_530442, EPI_ISL_530443, EPI_ISL_530444, EPI_ISL_530445, EPI_ISL_530447, EPI_ISL_530449, EPI_ISL_530450, EPI_ISL_530451, EPI_ISL_530452, EPI_ISL_530454, EPI_ISL_530455, EPI_ISL_530457, EPI_ISL_530458, EPI_ISL_530459, EPI_ISL_530460, EPI_ISL_530461, EPI_ISL_530462, EPI_ISL_530463, EPI_ISL_530464, EPI_ISL_530465, EPI_ISL_530466, EPI_ISL_530467 | | | |
| see above | Lighthouse Lab in Glasgow | Wellcome Sanger Institute for the COVID-19 Genomics UK Consortium | Harper VanSteenhouse, Yumi Kasai, David Gray, Carol Clugston, Anna Dominiczak and Alex Alderton, Roberto Amato, Sonia Goncalves, Ewan Harrison, David K. Jackson, Ian Johnston, Dominic Kwiatkowski, Cordelia Langford, John Sillitoe |
| EPI_ISL_530468 | NHSGGC West of Scotland Specialist Virology Centre / MRC-University of Glasgow Centre for Virus Research | Wellcome Sanger Institute for the COVID-19 Genomics UK Consortium | Ana da Silva Filipe, Natasha Johnson, Kathy Smollett, Daniel Mair, Stephen Carmichael, Lily Tong, Jenna Nichols, Elihu Aranday-Cortes, Kirstyn Brunker, Yasmin Parr, Kyriaki Nomikou; Sarah McDonald, Marc Niebel, Patawee Asamaphan; Richard Orton, Joseph Hughes, Sreenu Vattipally, David L Robertson; Alasdair MacLean, Rory Gunson; Kathy Li, Natasha Jesudason, Rajiv Shah, James Shepherd, Antonia Ho, Alice Broos, Emma Thomson and Alex Alderton, Roberto Amato, Sonia Goncalves, Ewan Harrison, David K. Jackson, Ian Johnston, Dominic Kwiatkowski, Cordelia Langford, John Sillitoe |
| EPI_ISL_530470, EPI_ISL_530471, EPI_ISL_530473, EPI_ISL_530474, EPI_ISL_530476, EPI_ISL_530477, EPI_ISL_530478 | Lighthouse Lab in Glasgow | Wellcome Sanger Institute for the COVID-19 Genomics UK Consortium | Harper VanSteenhouse, Yumi Kasai, David Gray, Carol Clugston, Anna Dominiczak and Alex Alderton, Roberto Amato, Sonia Goncalves, Ewan Harrison, David K. Jackson, Ian Johnston, Dominic Kwiatkowski, Cordelia Langford, John Sillitoe |
| EPI_ISL_530480 | NHSGGC West of Scotland Specialist Virology Centre / MRC-University of Glasgow Centre for Virus Research | Wellcome Sanger Institute for the COVID-19 Genomics UK Consortium | Ana da Silva Filipe, Natasha Johnson, Kathy Smollett, Daniel Mair, Stephen Carmichael, Lily Tong, Jenna Nichols, Elihu Aranday-Cortes, Kirstyn Brunker, Yasmin Parr, Kyriaki Nomikou; Sarah McDonald, Marc Niebel, Patawee Asamaphan; Richard Orton, Joseph Hughes, Sreenu Vattipally, David L Robertson; Alasdair MacLean, Rory Gunson; Kathy Li, Natasha Jesudason, Rajiv Shah, James Shepherd, Antonia Ho, Alice Broos, Emma Thomson and Alex Alderton, Roberto Amato, Sonia Goncalves, Ewan Harrison, David K. Jackson, Ian Johnston, Dominic Kwiatkowski, Cordelia Langford, John Sillitoe |
| EPI_ISL_530483, EPI_ISL_530484, EPI_ISL_530486, EPI_ISL_530487 | Lighthouse Lab in Glasgow | Wellcome Sanger Institute for the COVID-19 Genomics UK Consortium | Harper VanSteenhouse, Yumi Kasai, David Gray, Carol Clugston, Anna Dominiczak and Alex Alderton, Roberto Amato, Sonia Goncalves, Ewan Harrison, David K. Jackson, Ian Johnston, Dominic Kwiatkowski, Cordelia Langford, John Sillitoe |
| EPI_ISL_530488, EPI_ISL_530489 | NHSGGC West of Scotland Specialist Virology Centre / MRC-University of Glasgow Centre for Virus Research | Wellcome Sanger Institute for the COVID-19 Genomics UK Consortium | Ana da Silva Filipe, Natasha Johnson, Kathy Smollett, Daniel Mair, Stephen Carmichael, Lily Tong, Jenna Nichols, Elihu Aranday-Cortes, Kirstyn Brunker, Yasmin Parr, Kyriaki Nomikou; Sarah McDonald, Marc Niebel, Patawee Asamaphan; Richard Orton, Joseph Hughes, Sreenu Vattipally, David L Robertson; Alasdair MacLean, Rory Gunson; Kathy Li, Natasha Jesudason, Rajiv Shah, James Shepherd, Antonia Ho, Alice Broos, Emma Thomson and Alex Alderton, Roberto Amato, Sonia Goncalves, Ewan Harrison, David K. Jackson, Ian Johnston, Dominic Kwiatkowski, Cordelia Langford, John Sillitoe |
| EPI_ISL_530490, EPI_ISL_530491, EPI_ISL_530493, EPI_ISL_530494, EPI_ISL_530495, EPI_ISL_530496, EPI_ISL_530498, EPI_ISL_530499, EPI_ISL_530501, EPI_ISL_530503, EPI_ISL_530504, EPI_ISL_530505, EPI_ISL_530506, EPI_ISL_530507, EPI_ISL_530508, EPI_ISL_530511, EPI_ISL_530513, EPI_ISL_530514, EPI_ISL_530515, EPI_ISL_530516, EPI_ISL_530517, EPI_ISL_530518, EPI_ISL_530519, EPI_ISL_530520, EPI_ISL_530521 | | | |
| see above | Lighthouse Lab in Glasgow | Wellcome Sanger Institute for the COVID-19 Genomics UK Consortium | Harper VanSteenhouse, Yumi Kasai, David Gray, Carol Clugston, Anna Dominiczak and Alex Alderton, Roberto Amato, Sonia Goncalves, Ewan Harrison, David K. Jackson, Ian Johnston, Dominic Kwiatkowski, Cordelia Langford, John Sillitoe |
| EPI_ISL_530523, EPI_ISL_530528 | NHSGGC West of Scotland Specialist Virology Centre / MRC-University of Glasgow Centre for Virus Research | Wellcome Sanger Institute for the COVID-19 Genomics UK Consortium | Ana da Silva Filipe, Natasha Johnson, Kathy Smollett, Daniel Mair, Stephen Carmichael, Lily Tong, Jenna Nichols, Elihu Aranday-Cortes, Kirstyn Brunker, Yasmin Parr, Kyriaki Nomikou; Sarah McDonald, Marc Niebel, Patawee Asamaphan; Richard Orton, Joseph Hughes, Sreenu Vattipally, David L Robertson; Alasdair MacLean, Rory Gunson; Kathy Li, Natasha Jesudason, Rajiv Shah, James Shepherd, Antonia Ho, Alice Broos, Emma Thomson and Alex Alderton, Roberto Amato, Sonia Goncalves, Ewan Harrison, David K. Jackson, Ian Johnston, Dominic Kwiatkowski, Cordelia Langford, John Sillitoe |
| EPI_ISL_530530, EPI_ISL_530531, EPI_ISL_530532 | Lighthouse Lab in Glasgow | Wellcome Sanger Institute for the COVID-19 Genomics UK Consortium | Harper VanSteenhouse, Yumi Kasai, David Gray, Carol Clugston, Anna Dominiczak and Alex Alderton, Roberto Amato, Sonia Goncalves, Ewan Harrison, David K. Jackson, Ian Johnston, Dominic Kwiatkowski, Cordelia Langford, John Sillitoe |
| EPI_ISL_530533 | NHSGGC West of Scotland Specialist Virology Centre / MRC-University of Glasgow Centre for Virus Research | Wellcome Sanger Institute for the COVID-19 Genomics UK Consortium | Ana da Silva Filipe, Natasha Johnson, Kathy Smollett, Daniel Mair, Stephen Carmichael, Lily Tong, Jenna Nichols, Elihu Aranday-Cortes, Kirstyn Brunker, Yasmin Parr, Kyriaki Nomikou; Sarah McDonald, Marc Niebel, Patawee Asamaphan; Richard Orton, Joseph Hughes, Sreenu Vattipally, David L Robertson; Alasdair MacLean, Rory Gunson; Kathy Li, Natasha Jesudason, Rajiv Shah, James Shepherd, Antonia Ho, Alice Broos, Emma Thomson and Alex Alderton, Roberto Amato, Sonia Goncalves, Ewan Harrison, David K. Jackson, Ian Johnston, Dominic Kwiatkowski, Cordelia Langford, John Sillitoe |
| EPI_ISL_530534, EPI_ISL_530536, EPI_ISL_530537, EPI_ISL_530538, EPI_ISL_530540, EPI_ISL_530541 | Lighthouse Lab in Glasgow | Wellcome Sanger Institute for the COVID-19 Genomics UK Consortium | Harper VanSteenhouse, Yumi Kasai, David Gray, Carol Clugston, Anna Dominiczak and Alex Alderton, Roberto Amato, Sonia Goncalves, Ewan Harrison, David K. Jackson, Ian Johnston, Dominic Kwiatkowski, Cordelia Langford, John Sillitoe |
| EPI_ISL_530542 | NHSGGC West of Scotland Specialist Virology Centre / MRC-University of Glasgow Centre for Virus Research | Wellcome Sanger Institute for the COVID-19 Genomics UK Consortium | Ana da Silva Filipe, Natasha Johnson, Kathy Smollett, Daniel Mair, Stephen Carmichael, Lily Tong, Jenna Nichols, Elihu Aranday-Cortes, Kirstyn Brunker, Yasmin Parr, Kyriaki Nomikou; Sarah McDonald, Marc Niebel, Patawee Asamaphan; Richard Orton, Joseph Hughes, Sreenu Vattipally, David L Robertson; Alasdair MacLean, Rory Gunson; Kathy Li, Natasha Jesudason, Rajiv Shah, James Shepherd, Antonia Ho, Alice Broos, Emma Thomson and Alex Alderton, Roberto Amato, Sonia Goncalves, Ewan Harrison, David K. Jackson, Ian Johnston, Dominic Kwiatkowski, Cordelia Langford, John Sillitoe |
| EPI_ISL_530543, EPI_ISL_530544, EPI_ISL_530545, EPI_ISL_530546, EPI_ISL_530547, EPI_ISL_530548, EPI_ISL_530549, EPI_ISL_530550, EPI_ISL_530551, EPI_ISL_530552 | Lighthouse Lab in Glasgow | Wellcome Sanger Institute for the COVID-19 Genomics UK Consortium | Harper VanSteenhouse, Yumi Kasai, David Gray, Carol Clugston, Anna Dominiczak and Alex Alderton, Roberto Amato, Sonia Goncalves, Ewan Harrison, David K. Jackson, Ian Johnston, Dominic Kwiatkowski, Cordelia Langford, John Sillitoe |
| EPI_ISL_530553 | NHSGGC West of Scotland Specialist Virology Centre / MRC-University of Glasgow Centre for Virus Research | Wellcome Sanger Institute for the COVID-19 Genomics UK Consortium | Ana da Silva Filipe, Natasha Johnson, Kathy Smollett, Daniel Mair, Stephen Carmichael, Lily Tong, Jenna Nichols, Elihu Aranday-Cortes, Kirstyn Brunker, Yasmin Parr, Kyriaki Nomikou; Sarah McDonald, Marc Niebel, Patawee Asamaphan; Richard Orton, Joseph Hughes, Sreenu Vattipally, David L Robertson; Alasdair MacLean, Rory Gunson; Kathy Li, Natasha Jesudason, Rajiv Shah, James Shepherd, Antonia Ho, Alice Broos, Emma Thomson and Alex Alderton, Roberto Amato, Sonia Goncalves, Ewan Harrison, David K. Jackson, Ian Johnston, Dominic Kwiatkowski, Cordelia Langford, John Sillitoe |
| EPI_ISL_530554, EPI_ISL_530555, EPI_ISL_530556, | Lighthouse Lab in Glasgow | Wellcome Sanger Institute for the | Harper VanSteenhouse, Yumi Kasai, David Gray, Carol Clugston, Anna Dominiczak and Alex Alderton, Roberto Amato, Sonia Goncalves, Ewan Harrison, David K. Jackson, Ian Johnston, Dominic Kwiatkowski, Cordelia Langford, John Sillitoe |

| | | | |
|---|---|---|---|
| EPI_ISL_530557, EPI_ISL_530559, EPI_ISL_530560, EPI_ISL_530561 | | COVID-19 Genomics UK Consortium | |
| EPI_ISL_530562 | NHSGGC West of Scotland Specialist Virology Centre / MRC-University of Glasgow Centre for Virus Research | Wellcome Sanger Institute for the COVID-19 Genomics UK Consortium | Ana da Silva Filipe, Natasha Johnson, Kathy Smollett, Daniel Mair, Stephen Carmichael, Lily Tong, Jenna Nichols, Elihu Aranday-Cortes, Kirstyn Brunker, Yasmin Parr, Kyriaki Nomikou; Sarah McDonald, Marc Niebel, Patawee Asamaphan; Richard Orton, Joseph Hughes, Sreenu Vattipally, David L Robertson; Alasdair MacLean, Rory Gunson; Kathy Li, Natasha Jesudason, Rajiv Shah, James Shepherd, Antonia Ho, Alice Broos, Emma Thomson and Alex Alderton, Roberto Amato, Sonia Goncalves, Ewan Harrison, David K. Jackson, Ian Johnston, Dominic Kwiatkowski, Cordelia Langford, John Sillitoe |
| EPI_ISL_530563, EPI_ISL_530564, EPI_ISL_530565, EPI_ISL_530566, EPI_ISL_530567, EPI_ISL_530568, EPI_ISL_530570, EPI_ISL_530571, EPI_ISL_530572, EPI_ISL_530573, EPI_ISL_530574, EPI_ISL_530575, EPI_ISL_530576, EPI_ISL_530578, EPI_ISL_530580, EPI_ISL_530581, EPI_ISL_530582, EPI_ISL_530583, EPI_ISL_530584, EPI_ISL_530585, EPI_ISL_530586, EPI_ISL_530587, EPI_ISL_530589, EPI_ISL_530590, EPI_ISL_530591, EPI_ISL_530593, EPI_ISL_530594, EPI_ISL_530595 | | | |
| see above | Lighthouse Lab in Glasgow | Wellcome Sanger Institute for the COVID-19 Genomics UK Consortium | Harper VanSteenhouse, Yumi Kasai, David Gray, Carol Clugston, Anna Dominiczak and Alex Alderton, Roberto Amato, Sonia Goncalves, Ewan Harrison, David K. Jackson, Ian Johnston, Dominic Kwiatkowski, Cordelia Langford, John Sillitoe |
| EPI_ISL_530596 | NHSGGC West of Scotland Specialist Virology Centre / MRC-University of Glasgow Centre for Virus Research | Wellcome Sanger Institute for the COVID-19 Genomics UK Consortium | Ana da Silva Filipe, Natasha Johnson, Kathy Smollett, Daniel Mair, Stephen Carmichael, Lily Tong, Jenna Nichols, Elihu Aranday-Cortes, Kirstyn Brunker, Yasmin Parr, Kyriaki Nomikou; Sarah McDonald, Marc Niebel, Patawee Asamaphan; Richard Orton, Joseph Hughes, Sreenu Vattipally, David L Robertson; Alasdair MacLean, Rory Gunson; Kathy Li, Natasha Jesudason, Rajiv Shah, James Shepherd, Antonia Ho, Alice Broos, Emma Thomson and Alex Alderton, Roberto Amato, Sonia Goncalves, Ewan Harrison, David K. Jackson, Ian Johnston, Dominic Kwiatkowski, Cordelia Langford, John Sillitoe |
| EPI_ISL_530597, EPI_ISL_530598, EPI_ISL_530599, EPI_ISL_530601, EPI_ISL_530602, EPI_ISL_530603, EPI_ISL_530606, EPI_ISL_530607, EPI_ISL_530608, EPI_ISL_530609, EPI_ISL_530610, EPI_ISL_530611, EPI_ISL_530612, EPI_ISL_530613, EPI_ISL_530615, EPI_ISL_530616, EPI_ISL_530617, EPI_ISL_530618, EPI_ISL_530619, EPI_ISL_530620, EPI_ISL_530621, EPI_ISL_530623, EPI_ISL_530624, EPI_ISL_530626, EPI_ISL_530627, EPI_ISL_530628, EPI_ISL_530629, EPI_ISL_530631, EPI_ISL_530632, EPI_ISL_530633, EPI_ISL_530634, EPI_ISL_530635, EPI_ISL_530637, EPI_ISL_530638, EPI_ISL_530639, EPI_ISL_530641, EPI_ISL_530643, EPI_ISL_530644, EPI_ISL_530646, EPI_ISL_530648, EPI_ISL_530650, EPI_ISL_530652, EPI_ISL_530653, EPI_ISL_530654, EPI_ISL_530656 | | | |
| see above | Lighthouse Lab in Glasgow | Wellcome Sanger Institute for the COVID-19 Genomics UK Consortium | Harper VanSteenhouse, Yumi Kasai, David Gray, Carol Clugston, Anna Dominiczak and Alex Alderton, Roberto Amato, Sonia Goncalves, Ewan Harrison, David K. Jackson, Ian Johnston, Dominic Kwiatkowski, Cordelia Langford, John Sillitoe |
| EPI_ISL_530659 | NHSGGC West of Scotland Specialist Virology Centre / MRC-University of Glasgow Centre for Virus Research | Wellcome Sanger Institute for the COVID-19 Genomics UK Consortium | Ana da Silva Filipe, Natasha Johnson, Kathy Smollett, Daniel Mair, Stephen Carmichael, Lily Tong, Jenna Nichols, Elihu Aranday-Cortes, Kirstyn Brunker, Yasmin Parr, Kyriaki Nomikou; Sarah McDonald, Marc Niebel, Patawee Asamaphan; Richard Orton, Joseph Hughes, Sreenu Vattipally, David L Robertson; Alasdair MacLean, Rory Gunson; Kathy Li, Natasha Jesudason, Rajiv Shah, James Shepherd, Antonia Ho, Alice Broos, Emma Thomson and Alex Alderton, Roberto Amato, Sonia Goncalves, Ewan Harrison, David K. Jackson, Ian Johnston, Dominic Kwiatkowski, Cordelia Langford, John Sillitoe |
| EPI_ISL_530660, EPI_ISL_530661, EPI_ISL_530662, EPI_ISL_530664, EPI_ISL_530665, EPI_ISL_530667, EPI_ISL_530668, EPI_ISL_530669, EPI_ISL_530670, EPI_ISL_530671, EPI_ISL_530672 | | | |
| see above | Lighthouse Lab in Glasgow | Wellcome Sanger Institute for the COVID-19 Genomics UK Consortium | Harper VanSteenhouse, Yumi Kasai, David Gray, Carol Clugston, Anna Dominiczak and Alex Alderton, Roberto Amato, Sonia Goncalves, Ewan Harrison, David K. Jackson, Ian Johnston, Dominic Kwiatkowski, Cordelia Langford, John Sillitoe |
| EPI_ISL_530673 | NHSGGC West of Scotland Specialist Virology Centre / MRC-University of Glasgow Centre for Virus Research | Wellcome Sanger Institute for the COVID-19 Genomics UK Consortium | Ana da Silva Filipe, Natasha Johnson, Kathy Smollett, Daniel Mair, Stephen Carmichael, Lily Tong, Jenna Nichols, Elihu Aranday-Cortes, Kirstyn Brunker, Yasmin Parr, Kyriaki Nomikou; Sarah McDonald, Marc Niebel, Patawee Asamaphan; Richard Orton, Joseph Hughes, Sreenu Vattipally, David L Robertson; Alasdair MacLean, Rory Gunson; Kathy Li, Natasha Jesudason, Rajiv Shah, James Shepherd, Antonia Ho, Alice Broos, Emma Thomson and Alex Alderton, Roberto Amato, Sonia Goncalves, Ewan Harrison, David K. Jackson, Ian Johnston, Dominic Kwiatkowski, Cordelia Langford, John Sillitoe |
| EPI_ISL_530674, EPI_ISL_530677, EPI_ISL_530678, EPI_ISL_530679, EPI_ISL_530680 | Lighthouse Lab in Glasgow | Wellcome Sanger Institute for the COVID-19 Genomics UK Consortium | Harper VanSteenhouse, Yumi Kasai, David Gray, Carol Clugston, Anna Dominiczak and Alex Alderton, Roberto Amato, Sonia Goncalves, Ewan Harrison, David K. Jackson, Ian Johnston, Dominic Kwiatkowski, Cordelia Langford, John Sillitoe |
| EPI_ISL_530682 | NHSGGC West of Scotland Specialist Virology Centre / MRC-University of Glasgow Centre for Virus Research | Wellcome Sanger Institute for the COVID-19 Genomics UK Consortium | Ana da Silva Filipe, Natasha Johnson, Kathy Smollett, Daniel Mair, Stephen Carmichael, Lily Tong, Jenna Nichols, Elihu Aranday-Cortes, Kirstyn Brunker, Yasmin Parr, Kyriaki Nomikou; Sarah McDonald, Marc Niebel, Patawee Asamaphan; Richard Orton, Joseph Hughes, Sreenu Vattipally, David L Robertson; Alasdair MacLean, Rory Gunson; Kathy Li, Natasha Jesudason, Rajiv Shah, James Shepherd, Antonia Ho, Alice Broos, Emma Thomson and Alex Alderton, Roberto Amato, Sonia Goncalves, Ewan Harrison, David K. Jackson, Ian Johnston, Dominic Kwiatkowski, Cordelia Langford, John Sillitoe |
| EPI_ISL_530683, EPI_ISL_530684, EPI_ISL_530685, EPI_ISL_530687, EPI_ISL_530688, EPI_ISL_530690, EPI_ISL_530691, EPI_ISL_530692, EPI_ISL_530693, EPI_ISL_530694, EPI_ISL_530696, EPI_ISL_530697, EPI_ISL_530698, EPI_ISL_530699, EPI_ISL_530700, EPI_ISL_530701, EPI_ISL_530702, EPI_ISL_530703, EPI_ISL_530704, EPI_ISL_530705, EPI_ISL_530706, EPI_ISL_530707 | | | |
| see above | Lighthouse Lab in Glasgow | Wellcome Sanger Institute for the COVID-19 Genomics UK Consortium | Harper VanSteenhouse, Yumi Kasai, David Gray, Carol Clugston, Anna Dominiczak and Alex Alderton, Roberto Amato, Sonia Goncalves, Ewan Harrison, David K. Jackson, Ian Johnston, Dominic Kwiatkowski, Cordelia Langford, John Sillitoe |
| EPI_ISL_530708 | NHSGGC West of Scotland Specialist Virology Centre / MRC-University of Glasgow Centre for Virus Research | Wellcome Sanger Institute for the COVID-19 Genomics UK Consortium | Ana da Silva Filipe, Natasha Johnson, Kathy Smollett, Daniel Mair, Stephen Carmichael, Lily Tong, Jenna Nichols, Elihu Aranday-Cortes, Kirstyn Brunker, Yasmin Parr, Kyriaki Nomikou; Sarah McDonald, Marc Niebel, Patawee Asamaphan; Richard Orton, Joseph Hughes, Sreenu Vattipally, David L Robertson; Alasdair MacLean, Rory Gunson; Kathy Li, Natasha Jesudason, Rajiv Shah, James Shepherd, Antonia Ho, Alice Broos, Emma Thomson and Alex Alderton, Roberto Amato, Sonia Goncalves, Ewan Harrison, David K. Jackson, Ian Johnston, Dominic Kwiatkowski, Cordelia Langford, John Sillitoe |
| EPI_ISL_530709, EPI_ISL_530710, EPI_ISL_530711, EPI_ISL_530712, EPI_ISL_530713, EPI_ISL_530715, EPI_ISL_530716, EPI_ISL_530717, EPI_ISL_530718, EPI_ISL_530719, EPI_ISL_530720, EPI_ISL_530721 | | | |
| see above | Lighthouse Lab in Glasgow | Wellcome Sanger Institute for the COVID-19 Genomics UK Consortium | Harper VanSteenhouse, Yumi Kasai, David Gray, Carol Clugston, Anna Dominiczak and Alex Alderton, Roberto Amato, Sonia Goncalves, Ewan Harrison, David K. Jackson, Ian Johnston, Dominic Kwiatkowski, Cordelia Langford, John Sillitoe |
| EPI_ISL_530722 | NHSGGC West of Scotland Specialist Virology Centre / MRC-University of Glasgow Centre for Virus Research | Wellcome Sanger Institute for the COVID-19 Genomics UK Consortium | Ana da Silva Filipe, Natasha Johnson, Kathy Smollett, Daniel Mair, Stephen Carmichael, Lily Tong, Jenna Nichols, Elihu Aranday-Cortes, Kirstyn Brunker, Yasmin Parr, Kyriaki Nomikou; Sarah McDonald, Marc Niebel, Patawee Asamaphan; Richard Orton, Joseph Hughes, Sreenu Vattipally, David L Robertson; Alasdair MacLean, Rory Gunson; Kathy Li, Natasha Jesudason, Rajiv Shah, James Shepherd, Antonia Ho, Alice Broos, Emma Thomson and Alex Alderton, Roberto Amato, Sonia Goncalves, Ewan Harrison, David K. Jackson, Ian Johnston, Dominic Kwiatkowski, Cordelia Langford, John Sillitoe |
| EPI_ISL_530723, EPI_ISL_530724, EPI_ISL_530725, EPI_ISL_530726, EPI_ISL_530727, EPI_ISL_530729, EPI_ISL_530730, EPI_ISL_530731, EPI_ISL_530732, EPI_ISL_530733, EPI_ISL_530734, EPI_ISL_530736, EPI_ISL_530737, EPI_ISL_530738, EPI_ISL_530741, EPI_ISL_530742 | | | |
| see above | Lighthouse Lab in Glasgow | Wellcome Sanger Institute for the COVID-19 Genomics UK Consortium | Harper VanSteenhouse, Yumi Kasai, David Gray, Carol Clugston, Anna Dominiczak and Alex Alderton, Roberto Amato, Sonia Goncalves, Ewan Harrison, David K. Jackson, Ian Johnston, Dominic Kwiatkowski, Cordelia Langford, John Sillitoe |
| EPI_ISL_530743 | NHSGGC West of Scotland Specialist Virology Centre / MRC-University of Glasgow Centre for Virus Research | Wellcome Sanger Institute for the COVID-19 Genomics UK Consortium | Ana da Silva Filipe, Natasha Johnson, Kathy Smollett, Daniel Mair, Stephen Carmichael, Lily Tong, Jenna Nichols, Elihu Aranday-Cortes, Kirstyn Brunker, Yasmin Parr, Kyriaki Nomikou; Sarah McDonald, Marc Niebel, Patawee Asamaphan; Richard Orton, Joseph Hughes, Sreenu Vattipally, David L Robertson; Alasdair MacLean, Rory Gunson; Kathy Li, Natasha Jesudason, Rajiv Shah, James Shepherd, Antonia Ho, Alice Broos, Emma Thomson and Alex Alderton, Roberto Amato, Sonia Goncalves, Ewan Harrison, David K. Jackson, Ian Johnston, Dominic Kwiatkowski, Cordelia Langford, John Sillitoe |
| EPI_ISL_530745, EPI_ISL_530747, EPI_ISL_530748, EPI_ISL_530749, EPI_ISL_530750, EPI_ISL_530751, EPI_ISL_530752, EPI_ISL_530753 | Lighthouse Lab in Glasgow | Wellcome Sanger Institute for the COVID-19 Genomics UK Consortium | Harper VanSteenhouse, Yumi Kasai, David Gray, Carol Clugston, Anna Dominiczak and Alex Alderton, Roberto Amato, Sonia Goncalves, Ewan Harrison, David K. Jackson, Ian Johnston, Dominic Kwiatkowski, Cordelia Langford, John Sillitoe |
| EPI_ISL_530754 | NHSGGC West of Scotland Specialist Virology Centre / MRC-University of Glasgow Centre for Virus Research | Wellcome Sanger Institute for the COVID-19 Genomics UK Consortium | Ana da Silva Filipe, Natasha Johnson, Kathy Smollett, Daniel Mair, Stephen Carmichael, Lily Tong, Jenna Nichols, Elihu Aranday-Cortes, Kirstyn Brunker, Yasmin Parr, Kyriaki Nomikou; Sarah McDonald, Marc Niebel, Patawee Asamaphan; Richard Orton, Joseph Hughes, Sreenu Vattipally, David L Robertson; Alasdair MacLean, Rory Gunson; Kathy Li, Natasha Jesudason, Rajiv Shah, James Shepherd, Antonia Ho, Alice Broos, Emma Thomson and Alex Alderton, Roberto Amato, Sonia Goncalves, Ewan Harrison, David K. Jackson, Ian Johnston, Dominic Kwiatkowski, Cordelia Langford, John Sillitoe |
| EPI_ISL_530755, EPI_ISL_530756, EPI_ISL_530758, EPI_ISL_530759, EPI_ISL_530760, EPI_ISL_530763 | Lighthouse Lab in Glasgow | Wellcome Sanger Institute for the COVID-19 Genomics UK Consortium | Harper VanSteenhouse, Yumi Kasai, David Gray, Carol Clugston, Anna Dominiczak and Alex Alderton, Roberto Amato, Sonia Goncalves, Ewan Harrison, David K. Jackson, Ian Johnston, Dominic Kwiatkowski, Cordelia Langford, John Sillitoe |
| EPI_ISL_530764 | NHSGGC West of Scotland Specialist Virology Centre / MRC-University of Glasgow Centre for Virus Research | Wellcome Sanger Institute for the COVID-19 Genomics UK Consortium | Ana da Silva Filipe, Natasha Johnson, Kathy Smollett, Daniel Mair, Stephen Carmichael, Lily Tong, Jenna Nichols, Elihu Aranday-Cortes, Kirstyn Brunker, Yasmin Parr, Kyriaki Nomikou; Sarah McDonald, Marc Niebel, Patawee Asamaphan; Richard Orton, Joseph Hughes, Sreenu Vattipally, David L Robertson; Alasdair MacLean, Rory Gunson; Kathy Li, Natasha Jesudason, Rajiv Shah, James Shepherd, Antonia Ho, Alice Broos, Emma Thomson and Alex Alderton, Roberto Amato, Sonia Goncalves, Ewan Harrison, David K. Jackson, Ian Johnston, Dominic Kwiatkowski, Cordelia Langford, John Sillitoe |
| EPI_ISL_530765 | Lighthouse Lab in Glasgow | Wellcome Sanger Institute for the COVID-19 Genomics UK Consortium | Harper VanSteenhouse, Yumi Kasai, David Gray, Carol Clugston, Anna Dominiczak and Alex Alderton, Roberto Amato, Sonia Goncalves, Ewan Harrison, David K. Jackson, Ian Johnston, Dominic Kwiatkowski, Cordelia Langford, John Sillitoe |
| EPI_ISL_530766, EPI_ISL_530767, EPI_ISL_530768 | NHSGGC West of Scotland Specialist Virology Centre / MRC-University of Glasgow Centre for Virus Research | Wellcome Sanger Institute for the COVID-19 Genomics UK Consortium | Ana da Silva Filipe, Natasha Johnson, Kathy Smollett, Daniel Mair, Stephen Carmichael, Lily Tong, Jenna Nichols, Elihu Aranday-Cortes, Kirstyn Brunker, Yasmin Parr, Kyriaki Nomikou; Sarah McDonald, Marc Niebel, Patawee Asamaphan; Richard Orton, Joseph Hughes, Sreenu Vattipally, David L Robertson; Alasdair MacLean, Rory Gunson; Kathy Li, Natasha Jesudason, Rajiv Shah, James Shepherd, Antonia Ho, Alice Broos, Emma Thomson and Alex Alderton, Roberto Amato, Sonia Goncalves, Ewan Harrison, David K. Jackson, Ian Johnston, Dominic Kwiatkowski, Cordelia Langford, John Sillitoe |
| EPI_ISL_530770, EPI_ISL_530772, EPI_ISL_530773, EPI_ISL_530774, EPI_ISL_530775, EPI_ISL_530776, EPI_ISL_530777, EPI_ISL_530778, EPI_ISL_530779, EPI_ISL_530780, EPI_ISL_530782, EPI_ISL_530783, EPI_ISL_530784, EPI_ISL_530785, EPI_ISL_530786, EPI_ISL_530787, EPI_ISL_530788, EPI_ISL_530789, EPI_ISL_530791, EPI_ISL_530792, EPI_ISL_530793, EPI_ISL_530794, EPI_ISL_530796, EPI_ISL_530797, EPI_ISL_530798, EPI_ISL_530801, EPI_ISL_530802, EPI_ISL_530803, EPI_ISL_530804, EPI_ISL_530805, EPI_ISL_530806, EPI_ISL_530807, EPI_ISL_530808, EPI_ISL_530809, EPI_ISL_530810, EPI_ISL_530811, EPI_ISL_530812, EPI_ISL_530813, EPI_ISL_530814, EPI_ISL_530815, EPI_ISL_530816, EPI_ISL_530817, EPI_ISL_530818, EPI_ISL_530819, EPI_ISL_530820, EPI_ISL_530821, EPI_ISL_530822, EPI_ISL_530823, EPI_ISL_530824, EPI_ISL_530825 | | | |
| see above | Lighthouse Lab in Glasgow | Wellcome Sanger Institute for the COVID-19 Genomics UK Consortium | Harper VanSteenhouse, Yumi Kasai, David Gray, Carol Clugston, Anna Dominiczak and Alex Alderton, Roberto Amato, Sonia Goncalves, Ewan Harrison, David K. Jackson, Ian Johnston, Dominic Kwiatkowski, Cordelia Langford, John Sillitoe |
| EPI_ISL_530826 | NHSGGC West of Scotland Specialist Virology Centre / MRC-University of Glasgow Centre for Virus Research | Wellcome Sanger Institute for the COVID-19 Genomics UK Consortium | Ana da Silva Filipe, Natasha Johnson, Kathy Smollett, Daniel Mair, Stephen Carmichael, Lily Tong, Jenna Nichols, Elihu Aranday-Cortes, Kirstyn Brunker, Yasmin Parr, Kyriaki Nomikou; Sarah McDonald, Marc Niebel, Patawee Asamaphan; Richard Orton, Joseph Hughes, Sreenu Vattipally, David L Robertson; Alasdair MacLean, Rory Gunson; Kathy Li, Natasha Jesudason, Rajiv Shah, James Shepherd, Antonia Ho, Alice Broos, Emma Thomson and Alex Alderton, Roberto Amato, Sonia Goncalves, Ewan Harrison, David K. Jackson, Ian Johnston, Dominic Kwiatkowski, Cordelia Langford, John Sillitoe |
| EPI_ISL_530827, EPI_ISL_530828, EPI_ISL_530829, EPI_ISL_530830, EPI_ISL_530831, EPI_ISL_530832, EPI_ISL_530833, EPI_ISL_530834, EPI_ISL_530835 | Lighthouse Lab in Glasgow | Wellcome Sanger Institute for the COVID-19 Genomics UK Consortium | Harper VanSteenhouse, Yumi Kasai, David Gray, Carol Clugston, Anna Dominiczak and Alex Alderton, Roberto Amato, Sonia Goncalves, Ewan Harrison, David K. Jackson, Ian Johnston, Dominic Kwiatkowski, Cordelia Langford, John Sillitoe |
| EPI_ISL_530836 | NHSGGC West of Scotland Specialist Virology Centre / MRC-University of Glasgow Centre for Virus Research | Wellcome Sanger Institute for the COVID-19 Genomics UK Consortium | Ana da Silva Filipe, Natasha Johnson, Kathy Smollett, Daniel Mair, Stephen Carmichael, Lily Tong, Jenna Nichols, Elihu Aranday-Cortes, Kirstyn Brunker, Yasmin Parr, Kyriaki Nomikou; Sarah McDonald, Marc Niebel, Patawee Asamaphan; Richard Orton, Joseph Hughes, Sreenu Vattipally, David L Robertson; Alasdair MacLean, Rory Gunson; Kathy Li, Natasha Jesudason, Rajiv Shah, James Shepherd, Antonia Ho, Alice Broos, Emma Thomson and Alex Alderton, Roberto Amato, Sonia Goncalves, Ewan Harrison, David K. Jackson, Ian Johnston, Dominic Kwiatkowski, Cordelia Langford, John Sillitoe |
| EPI_ISL_530837, EPI_ISL_530838, EPI_ISL_530839, EPI_ISL_530840, EPI_ISL_530841, EPI_ISL_530842, EPI_ISL_530843, EPI_ISL_530844 | Lighthouse Lab in Glasgow | Wellcome Sanger Institute for the COVID-19 Genomics UK Consortium | Harper VanSteenhouse, Yumi Kasai, David Gray, Carol Clugston, Anna Dominiczak and Alex Alderton, Roberto Amato, Sonia Goncalves, Ewan Harrison, David K. Jackson, Ian Johnston, Dominic Kwiatkowski, Cordelia Langford, John Sillitoe |
| EPI_ISL_530845 | NHSGGC West of Scotland Specialist Virology Centre / MRC-University of Glasgow Centre for Virus Research | Wellcome Sanger Institute for the COVID-19 Genomics UK Consortium | Ana da Silva Filipe, Natasha Johnson, Kathy Smollett, Daniel Mair, Stephen Carmichael, Lily Tong, Jenna Nichols, Elihu Aranday-Cortes, Kirstyn Brunker, Yasmin Parr, Kyriaki Nomikou; Sarah McDonald, Marc Niebel, Patawee Asamaphan; Richard Orton, Joseph Hughes, Sreenu Vattipally, David L Robertson; Alasdair MacLean, Rory Gunson; Kathy Li, Natasha Jesudason, Rajiv Shah, James Shepherd, Antonia Ho, Alice Broos, Emma Thomson and Alex Alderton, Roberto Amato, Sonia Goncalves, Ewan Harrison, David K. Jackson, Ian Johnston, Dominic Kwiatkowski, Cordelia Langford, John Sillitoe |
| EPI_ISL_530848, EPI_ISL_530849, EPI_ISL_530850, EPI_ISL_530851, EPI_ISL_530852, EPI_ISL_530854, EPI_ISL_530857, EPI_ISL_530858, EPI_ISL_530859, EPI_ISL_530860, EPI_ISL_530861 | | | |
| see above | Lighthouse Lab in Glasgow | Wellcome Sanger Institute for the COVID-19 Genomics UK Consortium | Harper VanSteenhouse, Yumi Kasai, David Gray, Carol Clugston, Anna Dominiczak and Alex Alderton, Roberto Amato, Sonia Goncalves, Ewan Harrison, David K. Jackson, Ian Johnston, Dominic Kwiatkowski, Cordelia Langford, John Sillitoe |
| EPI_ISL_530862 | NHSGGC West of Scotland Specialist Virology Centre / MRC-University of Glasgow Centre for Virus Research | Wellcome Sanger Institute for the COVID-19 Genomics UK Consortium | Ana da Silva Filipe, Natasha Johnson, Kathy Smollett, Daniel Mair, Stephen Carmichael, Lily Tong, Jenna Nichols, Elihu Aranday-Cortes, Kirstyn Brunker, Yasmin Parr, Kyriaki Nomikou; Sarah McDonald, Marc Niebel, Patawee Asamaphan; Richard Orton, Joseph Hughes, Sreenu Vattipally, David L Robertson; Alasdair MacLean, Rory Gunson; Kathy Li, Natasha Jesudason, Rajiv Shah, James Shepherd, Antonia Ho, Alice Broos, Emma Thomson and Alex Alderton, Roberto Amato, Sonia Goncalves, Ewan Harrison, David K. Jackson, Ian Johnston, Dominic Kwiatkowski, Cordelia Langford, John Sillitoe |
| EPI_ISL_530863, EPI_ISL_530864, EPI_ISL_530865, EPI_ISL_530866, EPI_ISL_530867, EPI_ISL_530869, | Lighthouse Lab in Glasgow | Wellcome Sanger Institute for the COVID-19 Genomics UK Consortium | Harper VanSteenhouse, Yumi Kasai, David Gray, Carol Clugston, Anna Dominiczak and Alex Alderton, Roberto Amato, Sonia Goncalves, Ewan Harrison, David K. Jackson, Ian Johnston, Dominic Kwiatkowski, Cordelia Langford, John Sillitoe |

| | | | |
|---|---|---|---|
| EPI_ISL_530870 | | | |
| EPI_ISL_530871 | NHSGGC West of Scotland Specialist Virology Centre / MRC-University of Glasgow Centre for Virus Research | Wellcome Sanger Institute for the COVID-19 Genomics UK Consortium | Ana da Silva Filipe, Natasha Johnson, Kathy Smollett, Daniel Mair, Stephen Carmichael, Lily Tong, Jenna Nichols, Elihu Aranday-Cortes, Kirstyn Brunker, Yasmin Parr, Kyriaki Nomikou; Sarah McDonald, Marc Niebel, Patawee Asamaphan; Richard Orton, Joseph Hughes, Sreenu Vattipally, David L Robertson; Alasdair MacLean, Rory Gunson; Kathy Li, Natasha Jesudason, Rajiv Shah, James Shepherd, Antonia Ho, Alice Broos, Emma Thomson and Alex Alderton, Roberto Amato, Sonia Goncalves, Ewan Harrison, David K. Jackson, Ian Johnston, Dominic Kwiatkowski, Cordelia Langford, John Sillitoe |
| EPI_ISL_530872, EPI_ISL_530876, EPI_ISL_530877, EPI_ISL_530878, EPI_ISL_530879, EPI_ISL_530880, EPI_ISL_530881, EPI_ISL_530882, EPI_ISL_530883, EPI_ISL_530884, EPI_ISL_530885, EPI_ISL_530886, EPI_ISL_530887, EPI_ISL_530888, EPI_ISL_530890, EPI_ISL_530891, EPI_ISL_530892, EPI_ISL_530893, EPI_ISL_530894, EPI_ISL_530895, EPI_ISL_530896, EPI_ISL_530897, EPI_ISL_530900, EPI_ISL_530901, EPI_ISL_530902, EPI_ISL_530903, EPI_ISL_530905 | | | |
| see above | Lighthouse Lab in Glasgow | Wellcome Sanger Institute for the COVID-19 Genomics UK Consortium | Harper VanSteenhouse, Yumi Kasai, David Gray, Carol Clugston, Anna Dominiczak and Alex Alderton, Roberto Amato, Sonia Goncalves, Ewan Harrison, David K. Jackson, Ian Johnston, Dominic Kwiatkowski, Cordelia Langford, John Sillitoe |
| EPI_ISL_530906 | NHSGGC West of Scotland Specialist Virology Centre / MRC-University of Glasgow Centre for Virus Research | Wellcome Sanger Institute for the COVID-19 Genomics UK Consortium | Ana da Silva Filipe, Natasha Johnson, Kathy Smollett, Daniel Mair, Stephen Carmichael, Lily Tong, Jenna Nichols, Elihu Aranday-Cortes, Kirstyn Brunker, Yasmin Parr, Kyriaki Nomikou; Sarah McDonald, Marc Niebel, Patawee Asamaphan; Richard Orton, Joseph Hughes, Sreenu Vattipally, David L Robertson; Alasdair MacLean, Rory Gunson; Kathy Li, Natasha Jesudason, Rajiv Shah, James Shepherd, Antonia Ho, Alice Broos, Emma Thomson and Alex Alderton, Roberto Amato, Sonia Goncalves, Ewan Harrison, David K. Jackson, Ian Johnston, Dominic Kwiatkowski, Cordelia Langford, John Sillitoe |
| EPI_ISL_530908, EPI_ISL_530909, EPI_ISL_530910, EPI_ISL_530911, EPI_ISL_530912, EPI_ISL_530913, EPI_ISL_530914, EPI_ISL_530915, EPI_ISL_530916, EPI_ISL_530917, EPI_ISL_530919, EPI_ISL_530920, EPI_ISL_530922 | | | |
| see above | Lighthouse Lab in Glasgow | Wellcome Sanger Institute for the COVID-19 Genomics UK Consortium | Harper VanSteenhouse, Yumi Kasai, David Gray, Carol Clugston, Anna Dominiczak and Alex Alderton, Roberto Amato, Sonia Goncalves, Ewan Harrison, David K. Jackson, Ian Johnston, Dominic Kwiatkowski, Cordelia Langford, John Sillitoe |
| EPI_ISL_530923 | NHSGGC West of Scotland Specialist Virology Centre / MRC-University of Glasgow Centre for Virus Research | Wellcome Sanger Institute for the COVID-19 Genomics UK Consortium | Ana da Silva Filipe, Natasha Johnson, Kathy Smollett, Daniel Mair, Stephen Carmichael, Lily Tong, Jenna Nichols, Elihu Aranday-Cortes, Kirstyn Brunker, Yasmin Parr, Kyriaki Nomikou; Sarah McDonald, Marc Niebel, Patawee Asamaphan; Richard Orton, Joseph Hughes, Sreenu Vattipally, David L Robertson; Alasdair MacLean, Rory Gunson; Kathy Li, Natasha Jesudason, Rajiv Shah, James Shepherd, Antonia Ho, Alice Broos, Emma Thomson and Alex Alderton, Roberto Amato, Sonia Goncalves, Ewan Harrison, David K. Jackson, Ian Johnston, Dominic Kwiatkowski, Cordelia Langford, John Sillitoe |
| EPI_ISL_530924, EPI_ISL_530926, EPI_ISL_530927, EPI_ISL_530928 | Lighthouse Lab in Glasgow | Wellcome Sanger Institute for the COVID-19 Genomics UK Consortium | Harper VanSteenhouse, Yumi Kasai, David Gray, Carol Clugston, Anna Dominiczak and Alex Alderton, Roberto Amato, Sonia Goncalves, Ewan Harrison, David K. Jackson, Ian Johnston, Dominic Kwiatkowski, Cordelia Langford, John Sillitoe |
| EPI_ISL_530929 | NHSGGC West of Scotland Specialist Virology Centre / MRC-University of Glasgow Centre for Virus Research | Wellcome Sanger Institute for the COVID-19 Genomics UK Consortium | Ana da Silva Filipe, Natasha Johnson, Kathy Smollett, Daniel Mair, Stephen Carmichael, Lily Tong, Jenna Nichols, Elihu Aranday-Cortes, Kirstyn Brunker, Yasmin Parr, Kyriaki Nomikou; Sarah McDonald, Marc Niebel, Patawee Asamaphan; Richard Orton, Joseph Hughes, Sreenu Vattipally, David L Robertson; Alasdair MacLean, Rory Gunson; Kathy Li, Natasha Jesudason, Rajiv Shah, James Shepherd, Antonia Ho, Alice Broos, Emma Thomson and Alex Alderton, Roberto Amato, Sonia Goncalves, Ewan Harrison, David K. Jackson, Ian Johnston, Dominic Kwiatkowski, Cordelia Langford, John Sillitoe |
| EPI_ISL_530930, EPI_ISL_530932, EPI_ISL_530933, EPI_ISL_530934, EPI_ISL_530935, EPI_ISL_530937, EPI_ISL_530938, EPI_ISL_530939, EPI_ISL_530940, EPI_ISL_530941, EPI_ISL_530942, EPI_ISL_530944, EPI_ISL_530945, EPI_ISL_530946, EPI_ISL_530947, EPI_ISL_530948, EPI_ISL_530949, EPI_ISL_530950, EPI_ISL_530951, EPI_ISL_530952 | | | |
| see above | Lighthouse Lab in Glasgow | Wellcome Sanger Institute for the COVID-19 Genomics UK Consortium | Harper VanSteenhouse, Yumi Kasai, David Gray, Carol Clugston, Anna Dominiczak and Alex Alderton, Roberto Amato, Sonia Goncalves, Ewan Harrison, David K. Jackson, Ian Johnston, Dominic Kwiatkowski, Cordelia Langford, John Sillitoe |
| EPI_ISL_530953 | NHSGGC West of Scotland Specialist Virology Centre / MRC-University of Glasgow Centre for Virus Research | Wellcome Sanger Institute for the COVID-19 Genomics UK Consortium | Ana da Silva Filipe, Natasha Johnson, Kathy Smollett, Daniel Mair, Stephen Carmichael, Lily Tong, Jenna Nichols, Elihu Aranday-Cortes, Kirstyn Brunker, Yasmin Parr, Kyriaki Nomikou; Sarah McDonald, Marc Niebel, Patawee Asamaphan; Richard Orton, Joseph Hughes, Sreenu Vattipally, David L Robertson; Alasdair MacLean, Rory Gunson; Kathy Li, Natasha Jesudason, Rajiv Shah, James Shepherd, Antonia Ho, Alice Broos, Emma Thomson and Alex Alderton, Roberto Amato, Sonia Goncalves, Ewan Harrison, David K. Jackson, Ian Johnston, Dominic Kwiatkowski, Cordelia Langford, John Sillitoe |
| EPI_ISL_530954, EPI_ISL_530955, EPI_ISL_530956, EPI_ISL_530957 | Lighthouse Lab in Glasgow | Wellcome Sanger Institute for the COVID-19 Genomics UK Consortium | Harper VanSteenhouse, Yumi Kasai, David Gray, Carol Clugston, Anna Dominiczak and Alex Alderton, Roberto Amato, Sonia Goncalves, Ewan Harrison, David K. Jackson, Ian Johnston, Dominic Kwiatkowski, Cordelia Langford, John Sillitoe |
| EPI_ISL_530959 | NHSGGC West of Scotland Specialist Virology Centre / MRC-University of Glasgow Centre for Virus Research | Wellcome Sanger Institute for the COVID-19 Genomics UK Consortium | Ana da Silva Filipe, Natasha Johnson, Kathy Smollett, Daniel Mair, Stephen Carmichael, Lily Tong, Jenna Nichols, Elihu Aranday-Cortes, Kirstyn Brunker, Yasmin Parr, Kyriaki Nomikou; Sarah McDonald, Marc Niebel, Patawee Asamaphan; Richard Orton, Joseph Hughes, Sreenu Vattipally, David L Robertson; Alasdair MacLean, Rory Gunson; Kathy Li, Natasha Jesudason, Rajiv Shah, James Shepherd, Antonia Ho, Alice Broos, Emma Thomson and Alex Alderton, Roberto Amato, Sonia Goncalves, Ewan Harrison, David K. Jackson, Ian Johnston, Dominic Kwiatkowski, Cordelia Langford, John Sillitoe |
| EPI_ISL_530960 | Lighthouse Lab in Glasgow | Wellcome Sanger Institute for the COVID-19 Genomics UK Consortium | Harper VanSteenhouse, Yumi Kasai, David Gray, Carol Clugston, Anna Dominiczak and Alex Alderton, Roberto Amato, Sonia Goncalves, Ewan Harrison, David K. Jackson, Ian Johnston, Dominic Kwiatkowski, Cordelia Langford, John Sillitoe |
| EPI_ISL_530961, EPI_ISL_530962 | NHSGGC West of Scotland Specialist Virology Centre / MRC-University of Glasgow Centre for Virus Research | Wellcome Sanger Institute for the COVID-19 Genomics UK Consortium | Ana da Silva Filipe, Natasha Johnson, Kathy Smollett, Daniel Mair, Stephen Carmichael, Lily Tong, Jenna Nichols, Elihu Aranday-Cortes, Kirstyn Brunker, Yasmin Parr, Kyriaki Nomikou; Sarah McDonald, Marc Niebel, Patawee Asamaphan; Richard Orton, Joseph Hughes, Sreenu Vattipally, David L Robertson; Alasdair MacLean, Rory Gunson; Kathy Li, Natasha Jesudason, Rajiv Shah, James Shepherd, Antonia Ho, Alice Broos, Emma Thomson and Alex Alderton, Roberto Amato, Sonia Goncalves, Ewan Harrison, David K. Jackson, Ian Johnston, Dominic Kwiatkowski, Cordelia Langford, John Sillitoe |
| EPI_ISL_530964, EPI_ISL_530966, EPI_ISL_530967, EPI_ISL_530968, EPI_ISL_530969, EPI_ISL_530970 | Lighthouse Lab in Glasgow | Wellcome Sanger Institute for the COVID-19 Genomics UK Consortium | Harper VanSteenhouse, Yumi Kasai, David Gray, Carol Clugston, Anna Dominiczak and Alex Alderton, Roberto Amato, Sonia Goncalves, Ewan Harrison, David K. Jackson, Ian Johnston, Dominic Kwiatkowski, Cordelia Langford, John Sillitoe |
| EPI_ISL_530973 | NHSGGC West of Scotland Specialist Virology Centre / MRC-University of Glasgow Centre for Virus Research | Wellcome Sanger Institute for the COVID-19 Genomics UK Consortium | Ana da Silva Filipe, Natasha Johnson, Kathy Smollett, Daniel Mair, Stephen Carmichael, Lily Tong, Jenna Nichols, Elihu Aranday-Cortes, Kirstyn Brunker, Yasmin Parr, Kyriaki Nomikou; Sarah McDonald, Marc Niebel, Patawee Asamaphan; Richard Orton, Joseph Hughes, Sreenu Vattipally, David L Robertson; Alasdair MacLean, Rory Gunson; Kathy Li, Natasha Jesudason, Rajiv Shah, James Shepherd, Antonia Ho, Alice Broos, Emma Thomson and Alex Alderton, Roberto Amato, Sonia Goncalves, Ewan Harrison, David K. Jackson, Ian Johnston, Dominic Kwiatkowski, Cordelia Langford, John Sillitoe |
| EPI_ISL_530974 | Lighthouse Lab in Glasgow | Wellcome Sanger Institute for the COVID-19 Genomics UK Consortium | Harper VanSteenhouse, Yumi Kasai, David Gray, Carol Clugston, Anna Dominiczak and Alex Alderton, Roberto Amato, Sonia Goncalves, Ewan Harrison, David K. Jackson, Ian Johnston, Dominic Kwiatkowski, Cordelia Langford, John Sillitoe |
| EPI_ISL_530976 | NHSGGC West of Scotland Specialist Virology Centre / MRC-University of Glasgow Centre for Virus Research | Wellcome Sanger Institute for the COVID-19 Genomics UK Consortium | Ana da Silva Filipe, Natasha Johnson, Kathy Smollett, Daniel Mair, Stephen Carmichael, Lily Tong, Jenna Nichols, Elihu Aranday-Cortes, Kirstyn Brunker, Yasmin Parr, Kyriaki Nomikou; Sarah McDonald, Marc Niebel, Patawee Asamaphan; Richard Orton, Joseph Hughes, Sreenu Vattipally, David L Robertson; Alasdair MacLean, Rory Gunson; Kathy Li, Natasha Jesudason, Rajiv Shah, James Shepherd, Antonia Ho, Alice Broos, Emma Thomson and Alex Alderton, Roberto Amato, Sonia Goncalves, Ewan Harrison, David K. Jackson, Ian Johnston, Dominic Kwiatkowski, Cordelia Langford, John Sillitoe |
| EPI_ISL_530977, EPI_ISL_530978 | Lighthouse Lab in Glasgow | Wellcome Sanger Institute for the COVID-19 Genomics UK Consortium | Harper VanSteenhouse, Yumi Kasai, David Gray, Carol Clugston, Anna Dominiczak and Alex Alderton, Roberto Amato, Sonia Goncalves, Ewan Harrison, David K. Jackson, Ian Johnston, Dominic Kwiatkowski, Cordelia Langford, John Sillitoe |
| EPI_ISL_530979 | NHSGGC West of Scotland Specialist Virology Centre / MRC-University of Glasgow Centre for Virus Research | Wellcome Sanger Institute for the COVID-19 Genomics UK Consortium | Ana da Silva Filipe, Natasha Johnson, Kathy Smollett, Daniel Mair, Stephen Carmichael, Lily Tong, Jenna Nichols, Elihu Aranday-Cortes, Kirstyn Brunker, Yasmin Parr, Kyriaki Nomikou; Sarah McDonald, Marc Niebel, Patawee Asamaphan; Richard Orton, Joseph Hughes, Sreenu Vattipally, David L Robertson; Alasdair MacLean, Rory Gunson; Kathy Li, Natasha Jesudason, Rajiv Shah, James Shepherd, Antonia Ho, Alice Broos, Emma Thomson and Alex Alderton, Roberto Amato, Sonia Goncalves, Ewan Harrison, David K. Jackson, Ian Johnston, Dominic Kwiatkowski, Cordelia Langford, John Sillitoe |
| EPI_ISL_530980, EPI_ISL_530981, EPI_ISL_530982 | Lighthouse Lab in Glasgow | Wellcome Sanger Institute for the COVID-19 Genomics UK Consortium | Harper VanSteenhouse, Yumi Kasai, David Gray, Carol Clugston, Anna Dominiczak and Alex Alderton, Roberto Amato, Sonia Goncalves, Ewan Harrison, David K. Jackson, Ian Johnston, Dominic Kwiatkowski, Cordelia Langford, John Sillitoe |
| EPI_ISL_530983 | NHSGGC West of Scotland Specialist Virology Centre / MRC-University of Glasgow Centre for Virus Research | Wellcome Sanger Institute for the COVID-19 Genomics UK Consortium | Ana da Silva Filipe, Natasha Johnson, Kathy Smollett, Daniel Mair, Stephen Carmichael, Lily Tong, Jenna Nichols, Elihu Aranday-Cortes, Kirstyn Brunker, Yasmin Parr, Kyriaki Nomikou; Sarah McDonald, Marc Niebel, Patawee Asamaphan; Richard Orton, Joseph Hughes, Sreenu Vattipally, David L Robertson; Alasdair MacLean, Rory Gunson; Kathy Li, Natasha Jesudason, Rajiv Shah, James Shepherd, Antonia Ho, Alice Broos, Emma Thomson and Alex Alderton, Roberto Amato, Sonia Goncalves, Ewan Harrison, David K. Jackson, Ian Johnston, Dominic Kwiatkowski, Cordelia Langford, John Sillitoe |
| EPI_ISL_530984 | Lighthouse Lab in Glasgow | Wellcome Sanger Institute for the COVID-19 Genomics UK Consortium | Harper VanSteenhouse, Yumi Kasai, David Gray, Carol Clugston, Anna Dominiczak and Alex Alderton, Roberto Amato, Sonia Goncalves, Ewan Harrison, David K. Jackson, Ian Johnston, Dominic Kwiatkowski, Cordelia Langford, John Sillitoe |
| EPI_ISL_530985, EPI_ISL_530986 | NHSGGC West of Scotland Specialist Virology Centre / MRC-University of Glasgow Centre for Virus Research | Wellcome Sanger Institute for the COVID-19 Genomics UK Consortium | Ana da Silva Filipe, Natasha Johnson, Kathy Smollett, Daniel Mair, Stephen Carmichael, Lily Tong, Jenna Nichols, Elihu Aranday-Cortes, Kirstyn Brunker, Yasmin Parr, Kyriaki Nomikou; Sarah McDonald, Marc Niebel, Patawee Asamaphan; Richard Orton, Joseph Hughes, Sreenu Vattipally, David L Robertson; Alasdair MacLean, Rory Gunson; Kathy Li, Natasha Jesudason, Rajiv Shah, James Shepherd, Antonia Ho, Alice Broos, Emma Thomson and Alex Alderton, Roberto Amato, Sonia Goncalves, Ewan Harrison, David K. Jackson, Ian Johnston, Dominic Kwiatkowski, Cordelia Langford, John Sillitoe |
| EPI_ISL_530988, EPI_ISL_530989, EPI_ISL_530991, EPI_ISL_530992, EPI_ISL_530993, EPI_ISL_530994, EPI_ISL_530995 | Lighthouse Lab in Glasgow | Wellcome Sanger Institute for the COVID-19 Genomics UK Consortium | Harper VanSteenhouse, Yumi Kasai, David Gray, Carol Clugston, Anna Dominiczak and Alex Alderton, Roberto Amato, Sonia Goncalves, Ewan Harrison, David K. Jackson, Ian Johnston, Dominic Kwiatkowski, Cordelia Langford, John Sillitoe |
| EPI_ISL_530997 | NHSGGC West of Scotland Specialist Virology Centre / MRC-University of Glasgow Centre for Virus Research | Wellcome Sanger Institute for the COVID-19 Genomics UK Consortium | Ana da Silva Filipe, Natasha Johnson, Kathy Smollett, Daniel Mair, Stephen Carmichael, Lily Tong, Jenna Nichols, Elihu Aranday-Cortes, Kirstyn Brunker, Yasmin Parr, Kyriaki Nomikou; Sarah McDonald, Marc Niebel, Patawee Asamaphan; Richard Orton, Joseph Hughes, Sreenu Vattipally, David L Robertson; Alasdair MacLean, Rory Gunson; Kathy Li, Natasha Jesudason, Rajiv Shah, James Shepherd, Antonia Ho, Alice Broos, Emma Thomson and Alex Alderton, Roberto Amato, Sonia Goncalves, Ewan Harrison, David K. Jackson, Ian Johnston, Dominic Kwiatkowski, Cordelia Langford, John Sillitoe |
| EPI_ISL_530998, EPI_ISL_530999, EPI_ISL_531000, EPI_ISL_531001 | Lighthouse Lab in Glasgow | Wellcome Sanger Institute for the COVID-19 Genomics UK Consortium | Harper VanSteenhouse, Yumi Kasai, David Gray, Carol Clugston, Anna Dominiczak and Alex Alderton, Roberto Amato, Sonia Goncalves, Ewan Harrison, David K. Jackson, Ian Johnston, Dominic Kwiatkowski, Cordelia Langford, John Sillitoe |
| EPI_ISL_531002 | NHSGGC West of Scotland Specialist Virology Centre / MRC-University of Glasgow Centre for Virus Research | Wellcome Sanger Institute for the COVID-19 Genomics UK Consortium | Ana da Silva Filipe, Natasha Johnson, Kathy Smollett, Daniel Mair, Stephen Carmichael, Lily Tong, Jenna Nichols, Elihu Aranday-Cortes, Kirstyn Brunker, Yasmin Parr, Kyriaki Nomikou; Sarah McDonald, Marc Niebel, Patawee Asamaphan; Richard Orton, Joseph Hughes, Sreenu Vattipally, David L Robertson; Alasdair MacLean, Rory Gunson; Kathy Li, Natasha Jesudason, Rajiv Shah, James Shepherd, Antonia Ho, Alice Broos, Emma Thomson and Alex Alderton, Roberto Amato, Sonia Goncalves, Ewan Harrison, David K. Jackson, Ian Johnston, Dominic Kwiatkowski, Cordelia Langford, John Sillitoe |
| EPI_ISL_531003 | Lighthouse Lab in Glasgow | Wellcome Sanger Institute for the COVID-19 Genomics UK Consortium | Harper VanSteenhouse, Yumi Kasai, David Gray, Carol Clugston, Anna Dominiczak and Alex Alderton, Roberto Amato, Sonia Goncalves, Ewan Harrison, David K. Jackson, Ian Johnston, Dominic Kwiatkowski, Cordelia Langford, John Sillitoe |
| EPI_ISL_531004 | NHSGGC West of Scotland Specialist Virology Centre / MRC-University of Glasgow Centre for Virus Research | Wellcome Sanger Institute for the COVID-19 Genomics UK Consortium | Ana da Silva Filipe, Natasha Johnson, Kathy Smollett, Daniel Mair, Stephen Carmichael, Lily Tong, Jenna Nichols, Elihu Aranday-Cortes, Kirstyn Brunker, Yasmin Parr, Kyriaki Nomikou; Sarah McDonald, Marc Niebel, Patawee Asamaphan; Richard Orton, Joseph Hughes, Sreenu Vattipally, David L Robertson; Alasdair MacLean, Rory Gunson; Kathy Li, Natasha Jesudason, Rajiv Shah, James Shepherd, Antonia Ho, Alice Broos, Emma Thomson and Alex Alderton, Roberto Amato, Sonia Goncalves, Ewan Harrison, David K. Jackson, Ian Johnston, Dominic Kwiatkowski, Cordelia Langford, John Sillitoe |
| EPI_ISL_531005 | Lighthouse Lab in Glasgow | Wellcome Sanger Institute for the COVID-19 Genomics UK Consortium | Harper VanSteenhouse, Yumi Kasai, David Gray, Carol Clugston, Anna Dominiczak and Alex Alderton, Roberto Amato, Sonia Goncalves, Ewan Harrison, David K. Jackson, Ian Johnston, Dominic Kwiatkowski, Cordelia Langford, John Sillitoe |
| EPI_ISL_531006 | NHSGGC West of Scotland Specialist Virology Centre / MRC-University of Glasgow Centre for Virus Research | Wellcome Sanger Institute for the COVID-19 Genomics UK Consortium | Ana da Silva Filipe, Natasha Johnson, Kathy Smollett, Daniel Mair, Stephen Carmichael, Lily Tong, Jenna Nichols, Elihu Aranday-Cortes, Kirstyn Brunker, Yasmin Parr, Kyriaki Nomikou; Sarah McDonald, Marc Niebel, Patawee Asamaphan; Richard Orton, Joseph Hughes, Sreenu Vattipally, David L Robertson; Alasdair MacLean, Rory Gunson; Kathy Li, Natasha Jesudason, Rajiv Shah, James Shepherd, Antonia Ho, Alice Broos, Emma Thomson and Alex Alderton, Roberto Amato, Sonia Goncalves, Ewan Harrison, David K. Jackson, Ian Johnston, Dominic Kwiatkowski, Cordelia Langford, John Sillitoe |
| EPI_ISL_531007, EPI_ISL_531009, EPI_ISL_531010, EPI_ISL_531011, EPI_ISL_531013, EPI_ISL_531014, EPI_ISL_531015, EPI_ISL_531016 | Lighthouse Lab in Glasgow | Wellcome Sanger Institute for the COVID-19 Genomics UK Consortium | Harper VanSteenhouse, Yumi Kasai, David Gray, Carol Clugston, Anna Dominiczak and Alex Alderton, Roberto Amato, Sonia Goncalves, Ewan Harrison, David K. Jackson, Ian Johnston, Dominic Kwiatkowski, Cordelia Langford, John Sillitoe |
| EPI_ISL_531017 | NHSGGC West of Scotland Specialist Virology Centre / MRC-University of Glasgow Centre for Virus Research | Wellcome Sanger Institute for the COVID-19 Genomics UK Consortium | Ana da Silva Filipe, Natasha Johnson, Kathy Smollett, Daniel Mair, Stephen Carmichael, Lily Tong, Jenna Nichols, Elihu Aranday-Cortes, Kirstyn Brunker, Yasmin Parr, Kyriaki Nomikou; Sarah McDonald, Marc Niebel, Patawee Asamaphan; Richard Orton, Joseph Hughes, Sreenu Vattipally, David L Robertson; Alasdair MacLean, Rory Gunson; Kathy Li, Natasha Jesudason, Rajiv Shah, James Shepherd, Antonia Ho, Alice Broos, Emma Thomson and Alex Alderton, Roberto Amato, Sonia Goncalves, Ewan Harrison, David K. Jackson, Ian Johnston, Dominic Kwiatkowski, Cordelia Langford, John Sillitoe |
| EPI_ISL_531018 | Lighthouse Lab in Glasgow | Wellcome Sanger Institute for the COVID-19 Genomics UK Consortium | Harper VanSteenhouse, Yumi Kasai, David Gray, Carol Clugston, Anna Dominiczak and Alex Alderton, Roberto Amato, Sonia Goncalves, Ewan Harrison, David K. Jackson, Ian Johnston, Dominic Kwiatkowski, Cordelia Langford, John Sillitoe |
| EPI_ISL_531020 | NHSGGC West of Scotland Specialist Virology Centre / MRC-University | Wellcome Sanger Institute for the | Ana da Silva Filipe, Natasha Johnson, Kathy Smollett, Daniel Mair, Stephen Carmichael, Lily Tong, Jenna Nichols, Elihu Aranday-Cortes, Kirstyn Brunker, Yasmin Parr, Kyriaki Nomikou; Sarah McDonald, Marc Niebel, Patawee Asamaphan; |

| | | | |
|---|---|---|---|
| | of Glasgow Centre for Virus Research | COVID-19 Genomics UK Consortium | Richard Orton, Joseph Hughes, Sreenu Vattipally, David L Robertson; Alasdair MacLean, Rory Gunson; Kathy Li, Natasha Jesudason, Rajiv Shah, James Shepherd, Antonia Ho, Alice Broos, Emma Thomson and Alex Alderton, Roberto Amato, Sonia Goncalves, Ewan Harrison, David K. Jackson, Ian Johnston, Dominic Kwiatkowski, Cordelia Langford, John Sillitoe |
| EPI_ISL_531021, EPI_ISL_531022, EPI_ISL_531023, EPI_ISL_531024, EPI_ISL_531026, EPI_ISL_531027, EPI_ISL_531028, EPI_ISL_531030 | Lighthouse Lab in Glasgow | Wellcome Sanger Institute for the COVID-19 Genomics UK Consortium | Harper VanSteenhouse, Yumi Kasai, David Gray, Carol Clugston, Anna Dominiczak and Alex Alderton, Roberto Amato, Sonia Goncalves, Ewan Harrison, David K. Jackson, Ian Johnston, Dominic Kwiatkowski, Cordelia Langford, John Sillitoe |
| EPI_ISL_531032, EPI_ISL_531033 | NHSGGC West of Scotland Specialist Virology Centre / MRC-University of Glasgow Centre for Virus Research | Wellcome Sanger Institute for the COVID-19 Genomics UK Consortium | Ana da Silva Filipe, Natasha Johnson, Kathy Smollett, Daniel Mair, Stephen Carmichael, Lily Tong, Jenna Nichols, Elihu Aranday-Cortes, Kirstyn Brunker, Yasmin Parr, Kyriaki Nomikou; Sarah McDonald, Marc Niebel, Patawee Asamaphan; Richard Orton, Joseph Hughes, Sreenu Vattipally, David L Robertson; Alasdair MacLean, Rory Gunson; Kathy Li, Natasha Jesudason, Rajiv Shah, James Shepherd, Antonia Ho, Alice Broos, Emma Thomson and Alex Alderton, Roberto Amato, Sonia Goncalves, Ewan Harrison, David K. Jackson, Ian Johnston, Dominic Kwiatkowski, Cordelia Langford, John Sillitoe |
| EPI_ISL_531034, EPI_ISL_531035, EPI_ISL_531036, EPI_ISL_531037 | Lighthouse Lab in Glasgow | Wellcome Sanger Institute for the COVID-19 Genomics UK Consortium | Harper VanSteenhouse, Yumi Kasai, David Gray, Carol Clugston, Anna Dominiczak and Alex Alderton, Roberto Amato, Sonia Goncalves, Ewan Harrison, David K. Jackson, Ian Johnston, Dominic Kwiatkowski, Cordelia Langford, John Sillitoe |
| EPI_ISL_531038 | NHSGGC West of Scotland Specialist Virology Centre / MRC-University of Glasgow Centre for Virus Research | Wellcome Sanger Institute for the COVID-19 Genomics UK Consortium | Ana da Silva Filipe, Natasha Johnson, Kathy Smollett, Daniel Mair, Stephen Carmichael, Lily Tong, Jenna Nichols, Elihu Aranday-Cortes, Kirstyn Brunker, Yasmin Parr, Kyriaki Nomikou; Sarah McDonald, Marc Niebel, Patawee Asamaphan; Richard Orton, Joseph Hughes, Sreenu Vattipally, David L Robertson; Alasdair MacLean, Rory Gunson; Kathy Li, Natasha Jesudason, Rajiv Shah, James Shepherd, Antonia Ho, Alice Broos, Emma Thomson and Alex Alderton, Roberto Amato, Sonia Goncalves, Ewan Harrison, David K. Jackson, Ian Johnston, Dominic Kwiatkowski, Cordelia Langford, John Sillitoe |
| EPI_ISL_531039 | Lighthouse Lab in Glasgow | Wellcome Sanger Institute for the COVID-19 Genomics UK Consortium | Harper VanSteenhouse, Yumi Kasai, David Gray, Carol Clugston, Anna Dominiczak and Alex Alderton, Roberto Amato, Sonia Goncalves, Ewan Harrison, David K. Jackson, Ian Johnston, Dominic Kwiatkowski, Cordelia Langford, John Sillitoe |
| EPI_ISL_531040 | NHSGGC West of Scotland Specialist Virology Centre / MRC-University of Glasgow Centre for Virus Research | Wellcome Sanger Institute for the COVID-19 Genomics UK Consortium | Ana da Silva Filipe, Natasha Johnson, Kathy Smollett, Daniel Mair, Stephen Carmichael, Lily Tong, Jenna Nichols, Elihu Aranday-Cortes, Kirstyn Brunker, Yasmin Parr, Kyriaki Nomikou; Sarah McDonald, Marc Niebel, Patawee Asamaphan; Richard Orton, Joseph Hughes, Sreenu Vattipally, David L Robertson; Alasdair MacLean, Rory Gunson; Kathy Li, Natasha Jesudason, Rajiv Shah, James Shepherd, Antonia Ho, Alice Broos, Emma Thomson and Alex Alderton, Roberto Amato, Sonia Goncalves, Ewan Harrison, David K. Jackson, Ian Johnston, Dominic Kwiatkowski, Cordelia Langford, John Sillitoe |
| EPI_ISL_531041, EPI_ISL_531042, EPI_ISL_531043, EPI_ISL_531044 | Lighthouse Lab in Glasgow | Wellcome Sanger Institute for the COVID-19 Genomics UK Consortium | Harper VanSteenhouse, Yumi Kasai, David Gray, Carol Clugston, Anna Dominiczak and Alex Alderton, Roberto Amato, Sonia Goncalves, Ewan Harrison, David K. Jackson, Ian Johnston, Dominic Kwiatkowski, Cordelia Langford, John Sillitoe |
| EPI_ISL_531045 | NHSGGC West of Scotland Specialist Virology Centre / MRC-University of Glasgow Centre for Virus Research | Wellcome Sanger Institute for the COVID-19 Genomics UK Consortium | Ana da Silva Filipe, Natasha Johnson, Kathy Smollett, Daniel Mair, Stephen Carmichael, Lily Tong, Jenna Nichols, Elihu Aranday-Cortes, Kirstyn Brunker, Yasmin Parr, Kyriaki Nomikou; Sarah McDonald, Marc Niebel, Patawee Asamaphan; Richard Orton, Joseph Hughes, Sreenu Vattipally, David L Robertson; Alasdair MacLean, Rory Gunson; Kathy Li, Natasha Jesudason, Rajiv Shah, James Shepherd, Antonia Ho, Alice Broos, Emma Thomson and Alex Alderton, Roberto Amato, Sonia Goncalves, Ewan Harrison, David K. Jackson, Ian Johnston, Dominic Kwiatkowski, Cordelia Langford, John Sillitoe |
| EPI_ISL_531046, EPI_ISL_531047 | Lighthouse Lab in Glasgow | Wellcome Sanger Institute for the COVID-19 Genomics UK Consortium | Harper VanSteenhouse, Yumi Kasai, David Gray, Carol Clugston, Anna Dominiczak and Alex Alderton, Roberto Amato, Sonia Goncalves, Ewan Harrison, David K. Jackson, Ian Johnston, Dominic Kwiatkowski, Cordelia Langford, John Sillitoe |
| EPI_ISL_531048, EPI_ISL_531049 | NHSGGC West of Scotland Specialist Virology Centre / MRC-University of Glasgow Centre for Virus Research | Wellcome Sanger Institute for the COVID-19 Genomics UK Consortium | Ana da Silva Filipe, Natasha Johnson, Kathy Smollett, Daniel Mair, Stephen Carmichael, Lily Tong, Jenna Nichols, Elihu Aranday-Cortes, Kirstyn Brunker, Yasmin Parr, Kyriaki Nomikou; Sarah McDonald, Marc Niebel, Patawee Asamaphan; Richard Orton, Joseph Hughes, Sreenu Vattipally, David L Robertson; Alasdair MacLean, Rory Gunson; Kathy Li, Natasha Jesudason, Rajiv Shah, James Shepherd, Antonia Ho, Alice Broos, Emma Thomson and Alex Alderton, Roberto Amato, Sonia Goncalves, Ewan Harrison, David K. Jackson, Ian Johnston, Dominic Kwiatkowski, Cordelia Langford, John Sillitoe |
| EPI_ISL_531050 | Lighthouse Lab in Glasgow | Wellcome Sanger Institute for the COVID-19 Genomics UK Consortium | Harper VanSteenhouse, Yumi Kasai, David Gray, Carol Clugston, Anna Dominiczak and Alex Alderton, Roberto Amato, Sonia Goncalves, Ewan Harrison, David K. Jackson, Ian Johnston, Dominic Kwiatkowski, Cordelia Langford, John Sillitoe |
| EPI_ISL_531052 | NHSGGC West of Scotland Specialist Virology Centre / MRC-University of Glasgow Centre for Virus Research | Wellcome Sanger Institute for the COVID-19 Genomics UK Consortium | Ana da Silva Filipe, Natasha Johnson, Kathy Smollett, Daniel Mair, Stephen Carmichael, Lily Tong, Jenna Nichols, Elihu Aranday-Cortes, Kirstyn Brunker, Yasmin Parr, Kyriaki Nomikou; Sarah McDonald, Marc Niebel, Patawee Asamaphan; Richard Orton, Joseph Hughes, Sreenu Vattipally, David L Robertson; Alasdair MacLean, Rory Gunson; Kathy Li, Natasha Jesudason, Rajiv Shah, James Shepherd, Antonia Ho, Alice Broos, Emma Thomson and Alex Alderton, Roberto Amato, Sonia Goncalves, Ewan Harrison, David K. Jackson, Ian Johnston, Dominic Kwiatkowski, Cordelia Langford, John Sillitoe |
| EPI_ISL_531053 | Lighthouse Lab in Glasgow | Wellcome Sanger Institute for the COVID-19 Genomics UK Consortium | Harper VanSteenhouse, Yumi Kasai, David Gray, Carol Clugston, Anna Dominiczak and Alex Alderton, Roberto Amato, Sonia Goncalves, Ewan Harrison, David K. Jackson, Ian Johnston, Dominic Kwiatkowski, Cordelia Langford, John Sillitoe |
| EPI_ISL_531054 | NHSGGC West of Scotland Specialist Virology Centre / MRC-University of Glasgow Centre for Virus Research | Wellcome Sanger Institute for the COVID-19 Genomics UK Consortium | Ana da Silva Filipe, Natasha Johnson, Kathy Smollett, Daniel Mair, Stephen Carmichael, Lily Tong, Jenna Nichols, Elihu Aranday-Cortes, Kirstyn Brunker, Yasmin Parr, Kyriaki Nomikou; Sarah McDonald, Marc Niebel, Patawee Asamaphan; Richard Orton, Joseph Hughes, Sreenu Vattipally, David L Robertson; Alasdair MacLean, Rory Gunson; Kathy Li, Natasha Jesudason, Rajiv Shah, James Shepherd, Antonia Ho, Alice Broos, Emma Thomson and Alex Alderton, Roberto Amato, Sonia Goncalves, Ewan Harrison, David K. Jackson, Ian Johnston, Dominic Kwiatkowski, Cordelia Langford, John Sillitoe |
| EPI_ISL_531055 | Lighthouse Lab in Glasgow | Wellcome Sanger Institute for the COVID-19 Genomics UK Consortium | Harper VanSteenhouse, Yumi Kasai, David Gray, Carol Clugston, Anna Dominiczak and Alex Alderton, Roberto Amato, Sonia Goncalves, Ewan Harrison, David K. Jackson, Ian Johnston, Dominic Kwiatkowski, Cordelia Langford, John Sillitoe |
| EPI_ISL_531057 | NHSGGC West of Scotland Specialist Virology Centre / MRC-University of Glasgow Centre for Virus Research | Wellcome Sanger Institute for the COVID-19 Genomics UK Consortium | Ana da Silva Filipe, Natasha Johnson, Kathy Smollett, Daniel Mair, Stephen Carmichael, Lily Tong, Jenna Nichols, Elihu Aranday-Cortes, Kirstyn Brunker, Yasmin Parr, Kyriaki Nomikou; Sarah McDonald, Marc Niebel, Patawee Asamaphan; Richard Orton, Joseph Hughes, Sreenu Vattipally, David L Robertson; Alasdair MacLean, Rory Gunson; Kathy Li, Natasha Jesudason, Rajiv Shah, James Shepherd, Antonia Ho, Alice Broos, Emma Thomson and Alex Alderton, Roberto Amato, Sonia Goncalves, Ewan Harrison, David K. Jackson, Ian Johnston, Dominic Kwiatkowski, Cordelia Langford, John Sillitoe |
| EPI_ISL_531058, EPI_ISL_531059, EPI_ISL_531060, EPI_ISL_531061, EPI_ISL_531062, EPI_ISL_531063, EPI_ISL_531064 | Lighthouse Lab in Glasgow | Wellcome Sanger Institute for the COVID-19 Genomics UK Consortium | Harper VanSteenhouse, Yumi Kasai, David Gray, Carol Clugston, Anna Dominiczak and Alex Alderton, Roberto Amato, Sonia Goncalves, Ewan Harrison, David K. Jackson, Ian Johnston, Dominic Kwiatkowski, Cordelia Langford, John Sillitoe |
| EPI_ISL_531065 | NHSGGC West of Scotland Specialist Virology Centre / MRC-University of Glasgow Centre for Virus Research | Wellcome Sanger Institute for the COVID-19 Genomics UK Consortium | Ana da Silva Filipe, Natasha Johnson, Kathy Smollett, Daniel Mair, Stephen Carmichael, Lily Tong, Jenna Nichols, Elihu Aranday-Cortes, Kirstyn Brunker, Yasmin Parr, Kyriaki Nomikou; Sarah McDonald, Marc Niebel, Patawee Asamaphan; Richard Orton, Joseph Hughes, Sreenu Vattipally, David L Robertson; Alasdair MacLean, Rory Gunson; Kathy Li, Natasha Jesudason, Rajiv Shah, James Shepherd, Antonia Ho, Alice Broos, Emma Thomson and Alex Alderton, Roberto Amato, Sonia Goncalves, Ewan Harrison, David K. Jackson, Ian Johnston, Dominic Kwiatkowski, Cordelia Langford, John Sillitoe |
| EPI_ISL_531067, EPI_ISL_531068, EPI_ISL_531069, EPI_ISL_531070, EPI_ISL_531071, EPI_ISL_531072, EPI_ISL_531073, EPI_ISL_531074, EPI_ISL_531075, EPI_ISL_531076, EPI_ISL_531077, EPI_ISL_531078 | | | |
| see above | Lighthouse Lab in Glasgow | Wellcome Sanger Institute for the COVID-19 Genomics UK Consortium | Harper VanSteenhouse, Yumi Kasai, David Gray, Carol Clugston, Anna Dominiczak and Alex Alderton, Roberto Amato, Sonia Goncalves, Ewan Harrison, David K. Jackson, Ian Johnston, Dominic Kwiatkowski, Cordelia Langford, John Sillitoe |
| EPI_ISL_531079 | NHSGGC West of Scotland Specialist Virology Centre / MRC-University of Glasgow Centre for Virus Research | Wellcome Sanger Institute for the COVID-19 Genomics UK Consortium | Ana da Silva Filipe, Natasha Johnson, Kathy Smollett, Daniel Mair, Stephen Carmichael, Lily Tong, Jenna Nichols, Elihu Aranday-Cortes, Kirstyn Brunker, Yasmin Parr, Kyriaki Nomikou; Sarah McDonald, Marc Niebel, Patawee Asamaphan; Richard Orton, Joseph Hughes, Sreenu Vattipally, David L Robertson; Alasdair MacLean, Rory Gunson; Kathy Li, Natasha Jesudason, Rajiv Shah, James Shepherd, Antonia Ho, Alice Broos, Emma Thomson and Alex Alderton, Roberto Amato, Sonia Goncalves, Ewan Harrison, David K. Jackson, Ian Johnston, Dominic Kwiatkowski, Cordelia Langford, John Sillitoe |
| EPI_ISL_531080, EPI_ISL_531081, EPI_ISL_531082, EPI_ISL_531084, EPI_ISL_531085, EPI_ISL_531086 | Lighthouse Lab in Glasgow | Wellcome Sanger Institute for the COVID-19 Genomics UK Consortium | Harper VanSteenhouse, Yumi Kasai, David Gray, Carol Clugston, Anna Dominiczak and Alex Alderton, Roberto Amato, Sonia Goncalves, Ewan Harrison, David K. Jackson, Ian Johnston, Dominic Kwiatkowski, Cordelia Langford, John Sillitoe |
| EPI_ISL_531087 | NHSGGC West of Scotland Specialist Virology Centre / MRC-University of Glasgow Centre for Virus Research | Wellcome Sanger Institute for the COVID-19 Genomics UK Consortium | Ana da Silva Filipe, Natasha Johnson, Kathy Smollett, Daniel Mair, Stephen Carmichael, Lily Tong, Jenna Nichols, Elihu Aranday-Cortes, Kirstyn Brunker, Yasmin Parr, Kyriaki Nomikou; Sarah McDonald, Marc Niebel, Patawee Asamaphan; Richard Orton, Joseph Hughes, Sreenu Vattipally, David L Robertson; Alasdair MacLean, Rory Gunson; Kathy Li, Natasha Jesudason, Rajiv Shah, James Shepherd, Antonia Ho, Alice Broos, Emma Thomson and Alex Alderton, Roberto Amato, Sonia Goncalves, Ewan Harrison, David K. Jackson, Ian Johnston, Dominic Kwiatkowski, Cordelia Langford, John Sillitoe |
| EPI_ISL_531088 | Lighthouse Lab in Glasgow | Wellcome Sanger Institute for the COVID-19 Genomics UK Consortium | Harper VanSteenhouse, Yumi Kasai, David Gray, Carol Clugston, Anna Dominiczak and Alex Alderton, Roberto Amato, Sonia Goncalves, Ewan Harrison, David K. Jackson, Ian Johnston, Dominic Kwiatkowski, Cordelia Langford, John Sillitoe |
| EPI_ISL_531091 | NHSGGC West of Scotland Specialist Virology Centre / MRC-University of Glasgow Centre for Virus Research | Wellcome Sanger Institute for the COVID-19 Genomics UK Consortium | Ana da Silva Filipe, Natasha Johnson, Kathy Smollett, Daniel Mair, Stephen Carmichael, Lily Tong, Jenna Nichols, Elihu Aranday-Cortes, Kirstyn Brunker, Yasmin Parr, Kyriaki Nomikou; Sarah McDonald, Marc Niebel, Patawee Asamaphan; Richard Orton, Joseph Hughes, Sreenu Vattipally, David L Robertson; Alasdair MacLean, Rory Gunson; Kathy Li, Natasha Jesudason, Rajiv Shah, James Shepherd, Antonia Ho, Alice Broos, Emma Thomson and Alex Alderton, Roberto Amato, Sonia Goncalves, Ewan Harrison, David K. Jackson, Ian Johnston, Dominic Kwiatkowski, Cordelia Langford, John Sillitoe |
| EPI_ISL_531094, EPI_ISL_531095 | Lighthouse Lab in Glasgow | Wellcome Sanger Institute for the COVID-19 Genomics UK Consortium | Harper VanSteenhouse, Yumi Kasai, David Gray, Carol Clugston, Anna Dominiczak and Alex Alderton, Roberto Amato, Sonia Goncalves, Ewan Harrison, David K. Jackson, Ian Johnston, Dominic Kwiatkowski, Cordelia Langford, John Sillitoe |
| EPI_ISL_531096 | NHSGGC West of Scotland Specialist Virology Centre / MRC-University of Glasgow Centre for Virus Research | Wellcome Sanger Institute for the COVID-19 Genomics UK Consortium | Ana da Silva Filipe, Natasha Johnson, Kathy Smollett, Daniel Mair, Stephen Carmichael, Lily Tong, Jenna Nichols, Elihu Aranday-Cortes, Kirstyn Brunker, Yasmin Parr, Kyriaki Nomikou; Sarah McDonald, Marc Niebel, Patawee Asamaphan; Richard Orton, Joseph Hughes, Sreenu Vattipally, David L Robertson; Alasdair MacLean, Rory Gunson; Kathy Li, Natasha Jesudason, Rajiv Shah, James Shepherd, Antonia Ho, Alice Broos, Emma Thomson and Alex Alderton, Roberto Amato, Sonia Goncalves, Ewan Harrison, David K. Jackson, Ian Johnston, Dominic Kwiatkowski, Cordelia Langford, John Sillitoe |
| EPI_ISL_531097, EPI_ISL_531098 | Lighthouse Lab in Glasgow | Wellcome Sanger Institute for the COVID-19 Genomics UK Consortium | Harper VanSteenhouse, Yumi Kasai, David Gray, Carol Clugston, Anna Dominiczak and Alex Alderton, Roberto Amato, Sonia Goncalves, Ewan Harrison, David K. Jackson, Ian Johnston, Dominic Kwiatkowski, Cordelia Langford, John Sillitoe |
| EPI_ISL_531099, EPI_ISL_531100 | NHSGGC West of Scotland Specialist Virology Centre / MRC-University of Glasgow Centre for Virus Research | Wellcome Sanger Institute for the COVID-19 Genomics UK Consortium | Ana da Silva Filipe, Natasha Johnson, Kathy Smollett, Daniel Mair, Stephen Carmichael, Lily Tong, Jenna Nichols, Elihu Aranday-Cortes, Kirstyn Brunker, Yasmin Parr, Kyriaki Nomikou; Sarah McDonald, Marc Niebel, Patawee Asamaphan; Richard Orton, Joseph Hughes, Sreenu Vattipally, David L Robertson; Alasdair MacLean, Rory Gunson; Kathy Li, Natasha Jesudason, Rajiv Shah, James Shepherd, Antonia Ho, Alice Broos, Emma Thomson and Alex Alderton, Roberto Amato, Sonia Goncalves, Ewan Harrison, David K. Jackson, Ian Johnston, Dominic Kwiatkowski, Cordelia Langford, John Sillitoe |
| EPI_ISL_531101, EPI_ISL_531102 | Lighthouse Lab in Glasgow | Wellcome Sanger Institute for the COVID-19 Genomics UK Consortium | Harper VanSteenhouse, Yumi Kasai, David Gray, Carol Clugston, Anna Dominiczak and Alex Alderton, Roberto Amato, Sonia Goncalves, Ewan Harrison, David K. Jackson, Ian Johnston, Dominic Kwiatkowski, Cordelia Langford, John Sillitoe |
| EPI_ISL_531103, EPI_ISL_531104 | NHSGGC West of Scotland Specialist Virology Centre / MRC-University of Glasgow Centre for Virus Research | Wellcome Sanger Institute for the COVID-19 Genomics UK Consortium | Ana da Silva Filipe, Natasha Johnson, Kathy Smollett, Daniel Mair, Stephen Carmichael, Lily Tong, Jenna Nichols, Elihu Aranday-Cortes, Kirstyn Brunker, Yasmin Parr, Kyriaki Nomikou; Sarah McDonald, Marc Niebel, Patawee Asamaphan; Richard Orton, Joseph Hughes, Sreenu Vattipally, David L Robertson; Alasdair MacLean, Rory Gunson; Kathy Li, Natasha Jesudason, Rajiv Shah, James Shepherd, Antonia Ho, Alice Broos, Emma Thomson and Alex Alderton, Roberto Amato, Sonia Goncalves, Ewan Harrison, David K. Jackson, Ian Johnston, Dominic Kwiatkowski, Cordelia Langford, John Sillitoe |
| EPI_ISL_531105, EPI_ISL_531106, EPI_ISL_531107, EPI_ISL_531108, EPI_ISL_531109, EPI_ISL_531111, EPI_ISL_531112, EPI_ISL_531113 | Lighthouse Lab in Glasgow | Wellcome Sanger Institute for the COVID-19 Genomics UK Consortium | Harper VanSteenhouse, Yumi Kasai, David Gray, Carol Clugston, Anna Dominiczak and Alex Alderton, Roberto Amato, Sonia Goncalves, Ewan Harrison, David K. Jackson, Ian Johnston, Dominic Kwiatkowski, Cordelia Langford, John Sillitoe |
| EPI_ISL_531116 | NHSGGC West of Scotland Specialist Virology Centre / MRC-University of Glasgow Centre for Virus Research | Wellcome Sanger Institute for the COVID-19 Genomics UK Consortium | Ana da Silva Filipe, Natasha Johnson, Kathy Smollett, Daniel Mair, Stephen Carmichael, Lily Tong, Jenna Nichols, Elihu Aranday-Cortes, Kirstyn Brunker, Yasmin Parr, Kyriaki Nomikou; Sarah McDonald, Marc Niebel, Patawee Asamaphan; Richard Orton, Joseph Hughes, Sreenu Vattipally, David L Robertson; Alasdair MacLean, Rory Gunson; Kathy Li, Natasha Jesudason, Rajiv Shah, James Shepherd, Antonia Ho, Alice Broos, Emma Thomson and Alex Alderton, Roberto Amato, Sonia Goncalves, Ewan Harrison, David K. Jackson, Ian Johnston, Dominic Kwiatkowski, Cordelia Langford, John Sillitoe |
| EPI_ISL_531117 | Lighthouse Lab in Glasgow | Wellcome Sanger Institute for the COVID-19 Genomics UK Consortium | Harper VanSteenhouse, Yumi Kasai, David Gray, Carol Clugston, Anna Dominiczak and Alex Alderton, Roberto Amato, Sonia Goncalves, Ewan Harrison, David K. Jackson, Ian Johnston, Dominic Kwiatkowski, Cordelia Langford, John Sillitoe |
| EPI_ISL_531118 | NHSGGC West of Scotland Specialist Virology Centre / MRC-University of Glasgow Centre for Virus Research | Wellcome Sanger Institute for the COVID-19 Genomics UK Consortium | Ana da Silva Filipe, Natasha Johnson, Kathy Smollett, Daniel Mair, Stephen Carmichael, Lily Tong, Jenna Nichols, Elihu Aranday-Cortes, Kirstyn Brunker, Yasmin Parr, Kyriaki Nomikou; Sarah McDonald, Marc Niebel, Patawee Asamaphan; Richard Orton, Joseph Hughes, Sreenu Vattipally, David L Robertson; Alasdair MacLean, Rory Gunson; Kathy Li, Natasha Jesudason, Rajiv Shah, James Shepherd, Antonia Ho, Alice Broos, Emma Thomson and Alex Alderton, Roberto Amato, Sonia Goncalves, Ewan Harrison, David K. Jackson, Ian Johnston, Dominic Kwiatkowski, Cordelia Langford, John Sillitoe |
| EPI_ISL_531119, EPI_ISL_531120, EPI_ISL_531122 | Lighthouse Lab in Glasgow | Wellcome Sanger Institute for the COVID-19 Genomics UK Consortium | Harper VanSteenhouse, Yumi Kasai, David Gray, Carol Clugston, Anna Dominiczak and Alex Alderton, Roberto Amato, Sonia Goncalves, Ewan Harrison, David K. Jackson, Ian Johnston, Dominic Kwiatkowski, Cordelia Langford, John Sillitoe |

| | | | |
|---|---|---|---|
| EPI_ISL_531123 | NHSGGC West of Scotland Specialist Virology Centre / MRC-University of Glasgow Centre for Virus Research | Wellcome Sanger Institute for the COVID-19 Genomics UK Consortium | Ana da Silva Filipe, Natasha Johnson, Kathy Smollett, Daniel Mair, Stephen Carmichael, Lily Tong, Jenna Nichols, Elihu Aranday-Cortes, Kirstyn Brunker, Yasmin Parr, Kyriaki Nomikou; Sarah McDonald, Marc Niebel, Patawee Asamaphan; Richard Orton, Joseph Hughes, Sreenu Vattipally, David L Robertson; Alasdair MacLean, Rory Gunson; Kathy Li, Natasha Jesudason, Rajiv Shah, James Shepherd, Antonia Ho, Alice Broos, Emma Thomson and Alex Alderton, Roberto Amato, Sonia Goncalves, Ewan Harrison, David K. Jackson, Ian Johnston, Dominic Kwiatkowski, Cordelia Langford, John Sillitoe |
| EPI_ISL_531124, EPI_ISL_531125 | Lighthouse Lab in Glasgow | Wellcome Sanger Institute for the COVID-19 Genomics UK Consortium | Harper VanSteenhouse, Yumi Kasai, David Gray, Carol Clugston, Anna Dominiczak and Alex Alderton, Roberto Amato, Sonia Goncalves, Ewan Harrison, David K. Jackson, Ian Johnston, Dominic Kwiatkowski, Cordelia Langford, John Sillitoe |
| EPI_ISL_531126 | NHSGGC West of Scotland Specialist Virology Centre / MRC-University of Glasgow Centre for Virus Research | Wellcome Sanger Institute for the COVID-19 Genomics UK Consortium | Ana da Silva Filipe, Natasha Johnson, Kathy Smollett, Daniel Mair, Stephen Carmichael, Lily Tong, Jenna Nichols, Elihu Aranday-Cortes, Kirstyn Brunker, Yasmin Parr, Kyriaki Nomikou; Sarah McDonald, Marc Niebel, Patawee Asamaphan; Richard Orton, Joseph Hughes, Sreenu Vattipally, David L Robertson; Alasdair MacLean, Rory Gunson; Kathy Li, Natasha Jesudason, Rajiv Shah, James Shepherd, Antonia Ho, Alice Broos, Emma Thomson and Alex Alderton, Roberto Amato, Sonia Goncalves, Ewan Harrison, David K. Jackson, Ian Johnston, Dominic Kwiatkowski, Cordelia Langford, John Sillitoe |
| EPI_ISL_531127, EPI_ISL_531128, EPI_ISL_531129 | Lighthouse Lab in Glasgow | Wellcome Sanger Institute for the COVID-19 Genomics UK Consortium | Harper VanSteenhouse, Yumi Kasai, David Gray, Carol Clugston, Anna Dominiczak and Alex Alderton, Roberto Amato, Sonia Goncalves, Ewan Harrison, David K. Jackson, Ian Johnston, Dominic Kwiatkowski, Cordelia Langford, John Sillitoe |
| EPI_ISL_531130 | NHSGGC West of Scotland Specialist Virology Centre / MRC-University of Glasgow Centre for Virus Research | Wellcome Sanger Institute for the COVID-19 Genomics UK Consortium | Ana da Silva Filipe, Natasha Johnson, Kathy Smollett, Daniel Mair, Stephen Carmichael, Lily Tong, Jenna Nichols, Elihu Aranday-Cortes, Kirstyn Brunker, Yasmin Parr, Kyriaki Nomikou; Sarah McDonald, Marc Niebel, Patawee Asamaphan; Richard Orton, Joseph Hughes, Sreenu Vattipally, David L Robertson; Alasdair MacLean, Rory Gunson; Kathy Li, Natasha Jesudason, Rajiv Shah, James Shepherd, Antonia Ho, Alice Broos, Emma Thomson and Alex Alderton, Roberto Amato, Sonia Goncalves, Ewan Harrison, David K. Jackson, Ian Johnston, Dominic Kwiatkowski, Cordelia Langford, John Sillitoe |
| EPI_ISL_531133, EPI_ISL_531134, EPI_ISL_531135, EPI_ISL_531136, EPI_ISL_531137, EPI_ISL_531138, EPI_ISL_531139, EPI_ISL_531140, EPI_ISL_531141 | Lighthouse Lab in Glasgow | Wellcome Sanger Institute for the COVID-19 Genomics UK Consortium | Harper VanSteenhouse, Yumi Kasai, David Gray, Carol Clugston, Anna Dominiczak and Alex Alderton, Roberto Amato, Sonia Goncalves, Ewan Harrison, David K. Jackson, Ian Johnston, Dominic Kwiatkowski, Cordelia Langford, John Sillitoe |
| EPI_ISL_531142 | NHSGGC West of Scotland Specialist Virology Centre / MRC-University of Glasgow Centre for Virus Research | Wellcome Sanger Institute for the COVID-19 Genomics UK Consortium | Ana da Silva Filipe, Natasha Johnson, Kathy Smollett, Daniel Mair, Stephen Carmichael, Lily Tong, Jenna Nichols, Elihu Aranday-Cortes, Kirstyn Brunker, Yasmin Parr, Kyriaki Nomikou; Sarah McDonald, Marc Niebel, Patawee Asamaphan; Richard Orton, Joseph Hughes, Sreenu Vattipally, David L Robertson; Alasdair MacLean, Rory Gunson; Kathy Li, Natasha Jesudason, Rajiv Shah, James Shepherd, Antonia Ho, Alice Broos, Emma Thomson and Alex Alderton, Roberto Amato, Sonia Goncalves, Ewan Harrison, David K. Jackson, Ian Johnston, Dominic Kwiatkowski, Cordelia Langford, John Sillitoe |
| EPI_ISL_531145, EPI_ISL_531146 | Lighthouse Lab in Glasgow | Wellcome Sanger Institute for the COVID-19 Genomics UK Consortium | Harper VanSteenhouse, Yumi Kasai, David Gray, Carol Clugston, Anna Dominiczak and Alex Alderton, Roberto Amato, Sonia Goncalves, Ewan Harrison, David K. Jackson, Ian Johnston, Dominic Kwiatkowski, Cordelia Langford, John Sillitoe |
| EPI_ISL_531147 | NHSGGC West of Scotland Specialist Virology Centre / MRC-University of Glasgow Centre for Virus Research | Wellcome Sanger Institute for the COVID-19 Genomics UK Consortium | Ana da Silva Filipe, Natasha Johnson, Kathy Smollett, Daniel Mair, Stephen Carmichael, Lily Tong, Jenna Nichols, Elihu Aranday-Cortes, Kirstyn Brunker, Yasmin Parr, Kyriaki Nomikou; Sarah McDonald, Marc Niebel, Patawee Asamaphan; Richard Orton, Joseph Hughes, Sreenu Vattipally, David L Robertson; Alasdair MacLean, Rory Gunson; Kathy Li, Natasha Jesudason, Rajiv Shah, James Shepherd, Antonia Ho, Alice Broos, Emma Thomson and Alex Alderton, Roberto Amato, Sonia Goncalves, Ewan Harrison, David K. Jackson, Ian Johnston, Dominic Kwiatkowski, Cordelia Langford, John Sillitoe |
| EPI_ISL_531150, EPI_ISL_531151, EPI_ISL_531152 | Lighthouse Lab in Glasgow | Wellcome Sanger Institute for the COVID-19 Genomics UK Consortium | Harper VanSteenhouse, Yumi Kasai, David Gray, Carol Clugston, Anna Dominiczak and Alex Alderton, Roberto Amato, Sonia Goncalves, Ewan Harrison, David K. Jackson, Ian Johnston, Dominic Kwiatkowski, Cordelia Langford, John Sillitoe |
| EPI_ISL_531154, EPI_ISL_531155, EPI_ISL_531156, EPI_ISL_531157 | NHSGGC West of Scotland Specialist Virology Centre / MRC-University of Glasgow Centre for Virus Research | Wellcome Sanger Institute for the COVID-19 Genomics UK Consortium | Ana da Silva Filipe, Natasha Johnson, Kathy Smollett, Daniel Mair, Stephen Carmichael, Lily Tong, Jenna Nichols, Elihu Aranday-Cortes, Kirstyn Brunker, Yasmin Parr, Kyriaki Nomikou; Sarah McDonald, Marc Niebel, Patawee Asamaphan; Richard Orton, Joseph Hughes, Sreenu Vattipally, David L Robertson; Alasdair MacLean, Rory Gunson; Kathy Li, Natasha Jesudason, Rajiv Shah, James Shepherd, Antonia Ho, Alice Broos, Emma Thomson and Alex Alderton, Roberto Amato, Sonia Goncalves, Ewan Harrison, David K. Jackson, Ian Johnston, Dominic Kwiatkowski, Cordelia Langford, John Sillitoe |
| EPI_ISL_531158, EPI_ISL_531161, EPI_ISL_531163, EPI_ISL_531164, EPI_ISL_531165 | Lighthouse Lab in Glasgow | Wellcome Sanger Institute for the COVID-19 Genomics UK Consortium | Harper VanSteenhouse, Yumi Kasai, David Gray, Carol Clugston, Anna Dominiczak and Alex Alderton, Roberto Amato, Sonia Goncalves, Ewan Harrison, David K. Jackson, Ian Johnston, Dominic Kwiatkowski, Cordelia Langford, John Sillitoe |
| EPI_ISL_531166 | NHSGGC West of Scotland Specialist Virology Centre / MRC-University of Glasgow Centre for Virus Research | Wellcome Sanger Institute for the COVID-19 Genomics UK Consortium | Ana da Silva Filipe, Natasha Johnson, Kathy Smollett, Daniel Mair, Stephen Carmichael, Lily Tong, Jenna Nichols, Elihu Aranday-Cortes, Kirstyn Brunker, Yasmin Parr, Kyriaki Nomikou; Sarah McDonald, Marc Niebel, Patawee Asamaphan; Richard Orton, Joseph Hughes, Sreenu Vattipally, David L Robertson; Alasdair MacLean, Rory Gunson; Kathy Li, Natasha Jesudason, Rajiv Shah, James Shepherd, Antonia Ho, Alice Broos, Emma Thomson and Alex Alderton, Roberto Amato, Sonia Goncalves, Ewan Harrison, David K. Jackson, Ian Johnston, Dominic Kwiatkowski, Cordelia Langford, John Sillitoe |
| EPI_ISL_531167, EPI_ISL_531168, EPI_ISL_531169, EPI_ISL_531170, EPI_ISL_531171, EPI_ISL_531173 | Lighthouse Lab in Glasgow | Wellcome Sanger Institute for the COVID-19 Genomics UK Consortium | Harper VanSteenhouse, Yumi Kasai, David Gray, Carol Clugston, Anna Dominiczak and Alex Alderton, Roberto Amato, Sonia Goncalves, Ewan Harrison, David K. Jackson, Ian Johnston, Dominic Kwiatkowski, Cordelia Langford, John Sillitoe |
| EPI_ISL_531174, EPI_ISL_531175 | NHSGGC West of Scotland Specialist Virology Centre / MRC-University of Glasgow Centre for Virus Research | Wellcome Sanger Institute for the COVID-19 Genomics UK Consortium | Ana da Silva Filipe, Natasha Johnson, Kathy Smollett, Daniel Mair, Stephen Carmichael, Lily Tong, Jenna Nichols, Elihu Aranday-Cortes, Kirstyn Brunker, Yasmin Parr, Kyriaki Nomikou; Sarah McDonald, Marc Niebel, Patawee Asamaphan; Richard Orton, Joseph Hughes, Sreenu Vattipally, David L Robertson; Alasdair MacLean, Rory Gunson; Kathy Li, Natasha Jesudason, Rajiv Shah, James Shepherd, Antonia Ho, Alice Broos, Emma Thomson and Alex Alderton, Roberto Amato, Sonia Goncalves, Ewan Harrison, David K. Jackson, Ian Johnston, Dominic Kwiatkowski, Cordelia Langford, John Sillitoe |
| EPI_ISL_531176, EPI_ISL_531177, EPI_ISL_531178, EPI_ISL_531179, EPI_ISL_531180, EPI_ISL_531181, EPI_ISL_531183, EPI_ISL_531184, EPI_ISL_531185, EPI_ISL_531186 | Lighthouse Lab in Glasgow | Wellcome Sanger Institute for the COVID-19 Genomics UK Consortium | Harper VanSteenhouse, Yumi Kasai, David Gray, Carol Clugston, Anna Dominiczak and Alex Alderton, Roberto Amato, Sonia Goncalves, Ewan Harrison, David K. Jackson, Ian Johnston, Dominic Kwiatkowski, Cordelia Langford, John Sillitoe |
| EPI_ISL_531189 | NHSGGC West of Scotland Specialist Virology Centre / MRC-University of Glasgow Centre for Virus Research | Wellcome Sanger Institute for the COVID-19 Genomics UK Consortium | Ana da Silva Filipe, Natasha Johnson, Kathy Smollett, Daniel Mair, Stephen Carmichael, Lily Tong, Jenna Nichols, Elihu Aranday-Cortes, Kirstyn Brunker, Yasmin Parr, Kyriaki Nomikou; Sarah McDonald, Marc Niebel, Patawee Asamaphan; Richard Orton, Joseph Hughes, Sreenu Vattipally, David L Robertson; Alasdair MacLean, Rory Gunson; Kathy Li, Natasha Jesudason, Rajiv Shah, James Shepherd, Antonia Ho, Alice Broos, Emma Thomson and Alex Alderton, Roberto Amato, Sonia Goncalves, Ewan Harrison, David K. Jackson, Ian Johnston, Dominic Kwiatkowski, Cordelia Langford, John Sillitoe |
| EPI_ISL_531192, EPI_ISL_531193, EPI_ISL_531194, EPI_ISL_531195, EPI_ISL_531196, EPI_ISL_531197 | Lighthouse Lab in Glasgow | Wellcome Sanger Institute for the COVID-19 Genomics UK Consortium | Harper VanSteenhouse, Yumi Kasai, David Gray, Carol Clugston, Anna Dominiczak and Alex Alderton, Roberto Amato, Sonia Goncalves, Ewan Harrison, David K. Jackson, Ian Johnston, Dominic Kwiatkowski, Cordelia Langford, John Sillitoe |
| EPI_ISL_531198 | NHSGGC West of Scotland Specialist Virology Centre / MRC-University of Glasgow Centre for Virus Research | Wellcome Sanger Institute for the COVID-19 Genomics UK Consortium | Ana da Silva Filipe, Natasha Johnson, Kathy Smollett, Daniel Mair, Stephen Carmichael, Lily Tong, Jenna Nichols, Elihu Aranday-Cortes, Kirstyn Brunker, Yasmin Parr, Kyriaki Nomikou; Sarah McDonald, Marc Niebel, Patawee Asamaphan; Richard Orton, Joseph Hughes, Sreenu Vattipally, David L Robertson; Alasdair MacLean, Rory Gunson; Kathy Li, Natasha Jesudason, Rajiv Shah, James Shepherd, Antonia Ho, Alice Broos, Emma Thomson and Alex Alderton, Roberto Amato, Sonia Goncalves, Ewan Harrison, David K. Jackson, Ian Johnston, Dominic Kwiatkowski, Cordelia Langford, John Sillitoe |
| EPI_ISL_531199, EPI_ISL_531200, EPI_ISL_531201, EPI_ISL_531203, EPI_ISL_531206 | Lighthouse Lab in Glasgow | Wellcome Sanger Institute for the COVID-19 Genomics UK Consortium | Harper VanSteenhouse, Yumi Kasai, David Gray, Carol Clugston, Anna Dominiczak and Alex Alderton, Roberto Amato, Sonia Goncalves, Ewan Harrison, David K. Jackson, Ian Johnston, Dominic Kwiatkowski, Cordelia Langford, John Sillitoe |
| EPI_ISL_531207 | NHSGGC West of Scotland Specialist Virology Centre / MRC-University of Glasgow Centre for Virus Research | Wellcome Sanger Institute for the COVID-19 Genomics UK Consortium | Ana da Silva Filipe, Natasha Johnson, Kathy Smollett, Daniel Mair, Stephen Carmichael, Lily Tong, Jenna Nichols, Elihu Aranday-Cortes, Kirstyn Brunker, Yasmin Parr, Kyriaki Nomikou; Sarah McDonald, Marc Niebel, Patawee Asamaphan; Richard Orton, Joseph Hughes, Sreenu Vattipally, David L Robertson; Alasdair MacLean, Rory Gunson; Kathy Li, Natasha Jesudason, Rajiv Shah, James Shepherd, Antonia Ho, Alice Broos, Emma Thomson and Alex Alderton, Roberto Amato, Sonia Goncalves, Ewan Harrison, David K. Jackson, Ian Johnston, Dominic Kwiatkowski, Cordelia Langford, John Sillitoe |
| EPI_ISL_531209, EPI_ISL_531210 | Lighthouse Lab in Glasgow | Wellcome Sanger Institute for the COVID-19 Genomics UK Consortium | Harper VanSteenhouse, Yumi Kasai, David Gray, Carol Clugston, Anna Dominiczak and Alex Alderton, Roberto Amato, Sonia Goncalves, Ewan Harrison, David K. Jackson, Ian Johnston, Dominic Kwiatkowski, Cordelia Langford, John Sillitoe |
| EPI_ISL_531212 | NHSGGC West of Scotland Specialist Virology Centre / MRC-University of Glasgow Centre for Virus Research | Wellcome Sanger Institute for the COVID-19 Genomics UK Consortium | Ana da Silva Filipe, Natasha Johnson, Kathy Smollett, Daniel Mair, Stephen Carmichael, Lily Tong, Jenna Nichols, Elihu Aranday-Cortes, Kirstyn Brunker, Yasmin Parr, Kyriaki Nomikou; Sarah McDonald, Marc Niebel, Patawee Asamaphan; Richard Orton, Joseph Hughes, Sreenu Vattipally, David L Robertson; Alasdair MacLean, Rory Gunson; Kathy Li, Natasha Jesudason, Rajiv Shah, James Shepherd, Antonia Ho, Alice Broos, Emma Thomson and Alex Alderton, Roberto Amato, Sonia Goncalves, Ewan Harrison, David K. Jackson, Ian Johnston, Dominic Kwiatkowski, Cordelia Langford, John Sillitoe |
| EPI_ISL_531213 | Lighthouse Lab in Glasgow | Wellcome Sanger Institute for the COVID-19 Genomics UK Consortium | Harper VanSteenhouse, Yumi Kasai, David Gray, Carol Clugston, Anna Dominiczak and Alex Alderton, Roberto Amato, Sonia Goncalves, Ewan Harrison, David K. Jackson, Ian Johnston, Dominic Kwiatkowski, Cordelia Langford, John Sillitoe |
| EPI_ISL_531214 | NHSGGC West of Scotland Specialist Virology Centre / MRC-University of Glasgow Centre for Virus Research | Wellcome Sanger Institute for the COVID-19 Genomics UK Consortium | Ana da Silva Filipe, Natasha Johnson, Kathy Smollett, Daniel Mair, Stephen Carmichael, Lily Tong, Jenna Nichols, Elihu Aranday-Cortes, Kirstyn Brunker, Yasmin Parr, Kyriaki Nomikou; Sarah McDonald, Marc Niebel, Patawee Asamaphan; Richard Orton, Joseph Hughes, Sreenu Vattipally, David L Robertson; Alasdair MacLean, Rory Gunson; Kathy Li, Natasha Jesudason, Rajiv Shah, James Shepherd, Antonia Ho, Alice Broos, Emma Thomson and Alex Alderton, Roberto Amato, Sonia Goncalves, Ewan Harrison, David K. Jackson, Ian Johnston, Dominic Kwiatkowski, Cordelia Langford, John Sillitoe |
| EPI_ISL_531215, EPI_ISL_531216, EPI_ISL_531217, EPI_ISL_531218, EPI_ISL_531219, EPI_ISL_531220, EPI_ISL_531222 | Lighthouse Lab in Glasgow | Wellcome Sanger Institute for the COVID-19 Genomics UK Consortium | Harper VanSteenhouse, Yumi Kasai, David Gray, Carol Clugston, Anna Dominiczak and Alex Alderton, Roberto Amato, Sonia Goncalves, Ewan Harrison, David K. Jackson, Ian Johnston, Dominic Kwiatkowski, Cordelia Langford, John Sillitoe |
| EPI_ISL_531223 | NHSGGC West of Scotland Specialist Virology Centre / MRC-University of Glasgow Centre for Virus Research | Wellcome Sanger Institute for the COVID-19 Genomics UK Consortium | Ana da Silva Filipe, Natasha Johnson, Kathy Smollett, Daniel Mair, Stephen Carmichael, Lily Tong, Jenna Nichols, Elihu Aranday-Cortes, Kirstyn Brunker, Yasmin Parr, Kyriaki Nomikou; Sarah McDonald, Marc Niebel, Patawee Asamaphan; Richard Orton, Joseph Hughes, Sreenu Vattipally, David L Robertson; Alasdair MacLean, Rory Gunson; Kathy Li, Natasha Jesudason, Rajiv Shah, James Shepherd, Antonia Ho, Alice Broos, Emma Thomson and Alex Alderton, Roberto Amato, Sonia Goncalves, Ewan Harrison, David K. Jackson, Ian Johnston, Dominic Kwiatkowski, Cordelia Langford, John Sillitoe |
| EPI_ISL_531226, EPI_ISL_531227, EPI_ISL_531229, EPI_ISL_531230, EPI_ISL_531232, EPI_ISL_531233, EPI_ISL_531235, EPI_ISL_531236, EPI_ISL_531237, EPI_ISL_531239, EPI_ISL_531240, EPI_ISL_531241, EPI_ISL_531242, EPI_ISL_531244, EPI_ISL_531245, EPI_ISL_531247 | | | |
| see above | Lighthouse Lab in Glasgow | Wellcome Sanger Institute for the COVID-19 Genomics UK Consortium | Harper VanSteenhouse, Yumi Kasai, David Gray, Carol Clugston, Anna Dominiczak and Alex Alderton, Roberto Amato, Sonia Goncalves, Ewan Harrison, David K. Jackson, Ian Johnston, Dominic Kwiatkowski, Cordelia Langford, John Sillitoe |
| EPI_ISL_531249 | NHSGGC West of Scotland Specialist Virology Centre / MRC-University of Glasgow Centre for Virus Research | Wellcome Sanger Institute for the COVID-19 Genomics UK Consortium | Ana da Silva Filipe, Natasha Johnson, Kathy Smollett, Daniel Mair, Stephen Carmichael, Lily Tong, Jenna Nichols, Elihu Aranday-Cortes, Kirstyn Brunker, Yasmin Parr, Kyriaki Nomikou; Sarah McDonald, Marc Niebel, Patawee Asamaphan; Richard Orton, Joseph Hughes, Sreenu Vattipally, David L Robertson; Alasdair MacLean, Rory Gunson; Kathy Li, Natasha Jesudason, Rajiv Shah, James Shepherd, Antonia Ho, Alice Broos, Emma Thomson and Alex Alderton, Roberto Amato, Sonia Goncalves, Ewan Harrison, David K. Jackson, Ian Johnston, Dominic Kwiatkowski, Cordelia Langford, John Sillitoe |
| EPI_ISL_531250, EPI_ISL_531251, EPI_ISL_531253, EPI_ISL_531254, EPI_ISL_531255, EPI_ISL_531256, EPI_ISL_531257, EPI_ISL_531258, EPI_ISL_531259, EPI_ISL_531260, EPI_ISL_531261, EPI_ISL_531262, EPI_ISL_531263, EPI_ISL_531264, EPI_ISL_531265 | | | |
| see above | Lighthouse Lab in Glasgow | Wellcome Sanger Institute for the COVID-19 Genomics UK Consortium | Harper VanSteenhouse, Yumi Kasai, David Gray, Carol Clugston, Anna Dominiczak and Alex Alderton, Roberto Amato, Sonia Goncalves, Ewan Harrison, David K. Jackson, Ian Johnston, Dominic Kwiatkowski, Cordelia Langford, John Sillitoe |
| EPI_ISL_531266 | NHSGGC West of Scotland Specialist Virology Centre / MRC-University of Glasgow Centre for Virus Research | Wellcome Sanger Institute for the COVID-19 Genomics UK Consortium | Ana da Silva Filipe, Natasha Johnson, Kathy Smollett, Daniel Mair, Stephen Carmichael, Lily Tong, Jenna Nichols, Elihu Aranday-Cortes, Kirstyn Brunker, Yasmin Parr, Kyriaki Nomikou; Sarah McDonald, Marc Niebel, Patawee Asamaphan; Richard Orton, Joseph Hughes, Sreenu Vattipally, David L Robertson; Alasdair MacLean, Rory Gunson; Kathy Li, Natasha Jesudason, Rajiv Shah, James Shepherd, Antonia Ho, Alice Broos, Emma Thomson and Alex Alderton, Roberto Amato, Sonia Goncalves, Ewan Harrison, David K. Jackson, Ian Johnston, Dominic Kwiatkowski, Cordelia Langford, John Sillitoe |
| EPI_ISL_531267, EPI_ISL_531268, EPI_ISL_531269, EPI_ISL_531270, EPI_ISL_531271, EPI_ISL_531272, EPI_ISL_531273, EPI_ISL_531274, EPI_ISL_531275, EPI_ISL_531276, EPI_ISL_531277, EPI_ISL_531278, EPI_ISL_531280, EPI_ISL_531281, EPI_ISL_531282, EPI_ISL_531283, EPI_ISL_531284, EPI_ISL_531285, EPI_ISL_531286, EPI_ISL_531287, EPI_ISL_531288, EPI_ISL_531290, EPI_ISL_531292, EPI_ISL_531293, EPI_ISL_531295, EPI_ISL_531296, EPI_ISL_531297, EPI_ISL_531298, EPI_ISL_531299, EPI_ISL_531300, EPI_ISL_531301, EPI_ISL_531303, EPI_ISL_531304, EPI_ISL_531305, EPI_ISL_531306, EPI_ISL_531307, EPI_ISL_531308, EPI_ISL_531309, EPI_ISL_531310, EPI_ISL_531311, EPI_ISL_531312, EPI_ISL_531313, EPI_ISL_531314, EPI_ISL_531315, EPI_ISL_531316, EPI_ISL_531317, EPI_ISL_531318, EPI_ISL_531319, EPI_ISL_531320 | | | |
| see above | Lighthouse Lab in Glasgow | Wellcome Sanger Institute for the COVID-19 Genomics UK Consortium | Harper VanSteenhouse, Yumi Kasai, David Gray, Carol Clugston, Anna Dominiczak and Alex Alderton, Roberto Amato, Sonia Goncalves, Ewan Harrison, David K. Jackson, Ian Johnston, Dominic Kwiatkowski, Cordelia Langford, John Sillitoe |
| EPI_ISL_531321 | NHSGGC West of Scotland Specialist Virology Centre / MRC-University of Glasgow Centre for Virus Research | Wellcome Sanger Institute for the COVID-19 Genomics UK Consortium | Ana da Silva Filipe, Natasha Johnson, Kathy Smollett, Daniel Mair, Stephen Carmichael, Lily Tong, Jenna Nichols, Elihu Aranday-Cortes, Kirstyn Brunker, Yasmin Parr, Kyriaki Nomikou; Sarah McDonald, Marc Niebel, Patawee Asamaphan; Richard Orton, Joseph Hughes, Sreenu Vattipally, David L Robertson; Alasdair MacLean, Rory Gunson; Kathy Li, Natasha Jesudason, Rajiv Shah, James Shepherd, Antonia Ho, Alice Broos, Emma Thomson and Alex Alderton, Roberto Amato, Sonia Goncalves, Ewan Harrison, David K. Jackson, Ian Johnston, Dominic Kwiatkowski, Cordelia Langford, John Sillitoe |
| EPI_ISL_531322, EPI_ISL_531323, EPI_ISL_531324, EPI_ISL_531325, EPI_ISL_531326, EPI_ISL_531327, | Lighthouse Lab in Glasgow | Wellcome Sanger Institute for the COVID-19 Genomics UK Consortium | Harper VanSteenhouse, Yumi Kasai, David Gray, Carol Clugston, Anna Dominiczak and Alex Alderton, Roberto Amato, Sonia Goncalves, Ewan Harrison, David K. Jackson, Ian Johnston, Dominic Kwiatkowski, Cordelia Langford, John Sillitoe |

| | | | |
|---|---|---|---|
| EPI_ISL_531329 | | | |
| EPI_ISL_531330 | NHSGGC West of Scotland Specialist Virology Centre / MRC-University of Glasgow Centre for Virus Research | Wellcome Sanger Institute for the COVID-19 Genomics UK Consortium | Ana da Silva Filipe, Natasha Johnson, Kathy Smollett, Daniel Mair, Stephen Carmichael, Lily Tong, Jenna Nichols, Elihu Aranday-Cortes, Kirstyn Brunker, Yasmin Parr, Kyriaki Nomikou; Sarah McDonald, Marc Niebel, Patawee Asamaphan; Richard Orton, Joseph Hughes, Sreenu Vattipally, David L Robertson; Alasdair MacLean, Rory Gunson; Kathy Li, Natasha Jesudason, Rajiv Shah, James Shepherd, Antonia Ho, Alice Broos, Emma Thomson and Alex Alderton, Roberto Amato, Sonia Goncalves, Ewan Harrison, David K. Jackson, Ian Johnston, Dominic Kwiatkowski, Cordelia Langford, John Sillitoe |
| EPI_ISL_531331, EPI_ISL_531332, EPI_ISL_531334, EPI_ISL_531338, EPI_ISL_531339, EPI_ISL_531340, EPI_ISL_531341, EPI_ISL_531342, EPI_ISL_531343, EPI_ISL_531344, EPI_ISL_531345, EPI_ISL_531346, EPI_ISL_531347, EPI_ISL_531348, EPI_ISL_531349, EPI_ISL_531350, EPI_ISL_531351, EPI_ISL_531353, EPI_ISL_531354, EPI_ISL_531356, EPI_ISL_531357, EPI_ISL_531358, EPI_ISL_531359, EPI_ISL_531360, EPI_ISL_531362, EPI_ISL_531363, EPI_ISL_531364 | | | |
| see above | Lighthouse Lab in Glasgow | Wellcome Sanger Institute for the COVID-19 Genomics UK Consortium | Harper VanSteenhouse, Yumi Kasai, David Gray, Carol Clugston, Anna Dominiczak and Alex Alderton, Roberto Amato, Sonia Goncalves, Ewan Harrison, David K. Jackson, Ian Johnston, Dominic Kwiatkowski, Cordelia Langford, John Sillitoe |
| EPI_ISL_531365 | NHSGGC West of Scotland Specialist Virology Centre / MRC-University of Glasgow Centre for Virus Research | Wellcome Sanger Institute for the COVID-19 Genomics UK Consortium | Ana da Silva Filipe, Natasha Johnson, Kathy Smollett, Daniel Mair, Stephen Carmichael, Lily Tong, Jenna Nichols, Elihu Aranday-Cortes, Kirstyn Brunker, Yasmin Parr, Kyriaki Nomikou; Sarah McDonald, Marc Niebel, Patawee Asamaphan; Richard Orton, Joseph Hughes, Sreenu Vattipally, David L Robertson; Alasdair MacLean, Rory Gunson; Kathy Li, Natasha Jesudason, Rajiv Shah, James Shepherd, Antonia Ho, Alice Broos, Emma Thomson and Alex Alderton, Roberto Amato, Sonia Goncalves, Ewan Harrison, David K. Jackson, Ian Johnston, Dominic Kwiatkowski, Cordelia Langford, John Sillitoe |
| EPI_ISL_531366, EPI_ISL_531367, EPI_ISL_531368, EPI_ISL_531369, EPI_ISL_531370, EPI_ISL_531371, EPI_ISL_531372, EPI_ISL_531373 | Lighthouse Lab in Glasgow | Wellcome Sanger Institute for the COVID-19 Genomics UK Consortium | Harper VanSteenhouse, Yumi Kasai, David Gray, Carol Clugston, Anna Dominiczak and Alex Alderton, Roberto Amato, Sonia Goncalves, Ewan Harrison, David K. Jackson, Ian Johnston, Dominic Kwiatkowski, Cordelia Langford, John Sillitoe |
| EPI_ISL_531374 | NHSGGC West of Scotland Specialist Virology Centre / MRC-University of Glasgow Centre for Virus Research | Wellcome Sanger Institute for the COVID-19 Genomics UK Consortium | Ana da Silva Filipe, Natasha Johnson, Kathy Smollett, Daniel Mair, Stephen Carmichael, Lily Tong, Jenna Nichols, Elihu Aranday-Cortes, Kirstyn Brunker, Yasmin Parr, Kyriaki Nomikou; Sarah McDonald, Marc Niebel, Patawee Asamaphan; Richard Orton, Joseph Hughes, Sreenu Vattipally, David L Robertson; Alasdair MacLean, Rory Gunson; Kathy Li, Natasha Jesudason, Rajiv Shah, James Shepherd, Antonia Ho, Alice Broos, Emma Thomson and Alex Alderton, Roberto Amato, Sonia Goncalves, Ewan Harrison, David K. Jackson, Ian Johnston, Dominic Kwiatkowski, Cordelia Langford, John Sillitoe |
| EPI_ISL_531375, EPI_ISL_531376, EPI_ISL_531377, EPI_ISL_531378, EPI_ISL_531380 | Lighthouse Lab in Glasgow | Wellcome Sanger Institute for the COVID-19 Genomics UK Consortium | Harper VanSteenhouse, Yumi Kasai, David Gray, Carol Clugston, Anna Dominiczak and Alex Alderton, Roberto Amato, Sonia Goncalves, Ewan Harrison, David K. Jackson, Ian Johnston, Dominic Kwiatkowski, Cordelia Langford, John Sillitoe |
| EPI_ISL_531382 | NHSGGC West of Scotland Specialist Virology Centre / MRC-University of Glasgow Centre for Virus Research | Wellcome Sanger Institute for the COVID-19 Genomics UK Consortium | Ana da Silva Filipe, Natasha Johnson, Kathy Smollett, Daniel Mair, Stephen Carmichael, Lily Tong, Jenna Nichols, Elihu Aranday-Cortes, Kirstyn Brunker, Yasmin Parr, Kyriaki Nomikou; Sarah McDonald, Marc Niebel, Patawee Asamaphan; Richard Orton, Joseph Hughes, Sreenu Vattipally, David L Robertson; Alasdair MacLean, Rory Gunson; Kathy Li, Natasha Jesudason, Rajiv Shah, James Shepherd, Antonia Ho, Alice Broos, Emma Thomson and Alex Alderton, Roberto Amato, Sonia Goncalves, Ewan Harrison, David K. Jackson, Ian Johnston, Dominic Kwiatkowski, Cordelia Langford, John Sillitoe |
| EPI_ISL_531383, EPI_ISL_531385, EPI_ISL_531387, EPI_ISL_531389, EPI_ISL_531390, EPI_ISL_531392, EPI_ISL_531393, EPI_ISL_531394, EPI_ISL_531396, EPI_ISL_531397, EPI_ISL_531398, EPI_ISL_531399, EPI_ISL_531400, EPI_ISL_531401, EPI_ISL_531402, EPI_ISL_531403, EPI_ISL_531404, EPI_ISL_531406, EPI_ISL_531407, EPI_ISL_531408, EPI_ISL_531409, EPI_ISL_531410, EPI_ISL_531411 | | | |
| see above | Lighthouse Lab in Glasgow | Wellcome Sanger Institute for the COVID-19 Genomics UK Consortium | Harper VanSteenhouse, Yumi Kasai, David Gray, Carol Clugston, Anna Dominiczak and Alex Alderton, Roberto Amato, Sonia Goncalves, Ewan Harrison, David K. Jackson, Ian Johnston, Dominic Kwiatkowski, Cordelia Langford, John Sillitoe |
| EPI_ISL_531412 | NHSGGC West of Scotland Specialist Virology Centre / MRC-University of Glasgow Centre for Virus Research | Wellcome Sanger Institute for the COVID-19 Genomics UK Consortium | Ana da Silva Filipe, Natasha Johnson, Kathy Smollett, Daniel Mair, Stephen Carmichael, Lily Tong, Jenna Nichols, Elihu Aranday-Cortes, Kirstyn Brunker, Yasmin Parr, Kyriaki Nomikou; Sarah McDonald, Marc Niebel, Patawee Asamaphan; Richard Orton, Joseph Hughes, Sreenu Vattipally, David L Robertson; Alasdair MacLean, Rory Gunson; Kathy Li, Natasha Jesudason, Rajiv Shah, James Shepherd, Antonia Ho, Alice Broos, Emma Thomson and Alex Alderton, Roberto Amato, Sonia Goncalves, Ewan Harrison, David K. Jackson, Ian Johnston, Dominic Kwiatkowski, Cordelia Langford, John Sillitoe |
| EPI_ISL_531413, EPI_ISL_531414, EPI_ISL_531415 | Lighthouse Lab in Glasgow | Wellcome Sanger Institute for the COVID-19 Genomics UK Consortium | Harper VanSteenhouse, Yumi Kasai, David Gray, Carol Clugston, Anna Dominiczak and Alex Alderton, Roberto Amato, Sonia Goncalves, Ewan Harrison, David K. Jackson, Ian Johnston, Dominic Kwiatkowski, Cordelia Langford, John Sillitoe |
| EPI_ISL_531416 | NHSGGC West of Scotland Specialist Virology Centre / MRC-University of Glasgow Centre for Virus Research | Wellcome Sanger Institute for the COVID-19 Genomics UK Consortium | Ana da Silva Filipe, Natasha Johnson, Kathy Smollett, Daniel Mair, Stephen Carmichael, Lily Tong, Jenna Nichols, Elihu Aranday-Cortes, Kirstyn Brunker, Yasmin Parr, Kyriaki Nomikou; Sarah McDonald, Marc Niebel, Patawee Asamaphan; Richard Orton, Joseph Hughes, Sreenu Vattipally, David L Robertson; Alasdair MacLean, Rory Gunson; Kathy Li, Natasha Jesudason, Rajiv Shah, James Shepherd, Antonia Ho, Alice Broos, Emma Thomson and Alex Alderton, Roberto Amato, Sonia Goncalves, Ewan Harrison, David K. Jackson, Ian Johnston, Dominic Kwiatkowski, Cordelia Langford, John Sillitoe |
| EPI_ISL_531417, EPI_ISL_531418, EPI_ISL_531419 | Lighthouse Lab in Glasgow | Wellcome Sanger Institute for the COVID-19 Genomics UK Consortium | Harper VanSteenhouse, Yumi Kasai, David Gray, Carol Clugston, Anna Dominiczak and Alex Alderton, Roberto Amato, Sonia Goncalves, Ewan Harrison, David K. Jackson, Ian Johnston, Dominic Kwiatkowski, Cordelia Langford, John Sillitoe |
| EPI_ISL_531420, EPI_ISL_531422 | NHSGGC West of Scotland Specialist Virology Centre / MRC-University of Glasgow Centre for Virus Research | Wellcome Sanger Institute for the COVID-19 Genomics UK Consortium | Ana da Silva Filipe, Natasha Johnson, Kathy Smollett, Daniel Mair, Stephen Carmichael, Lily Tong, Jenna Nichols, Elihu Aranday-Cortes, Kirstyn Brunker, Yasmin Parr, Kyriaki Nomikou; Sarah McDonald, Marc Niebel, Patawee Asamaphan; Richard Orton, Joseph Hughes, Sreenu Vattipally, David L Robertson; Alasdair MacLean, Rory Gunson; Kathy Li, Natasha Jesudason, Rajiv Shah, James Shepherd, Antonia Ho, Alice Broos, Emma Thomson and Alex Alderton, Roberto Amato, Sonia Goncalves, Ewan Harrison, David K. Jackson, Ian Johnston, Dominic Kwiatkowski, Cordelia Langford, John Sillitoe |
| EPI_ISL_531424, EPI_ISL_531425, EPI_ISL_531426, EPI_ISL_531427, EPI_ISL_531428, EPI_ISL_531429, EPI_ISL_531431 | Lighthouse Lab in Glasgow | Wellcome Sanger Institute for the COVID-19 Genomics UK Consortium | Harper VanSteenhouse, Yumi Kasai, David Gray, Carol Clugston, Anna Dominiczak and Alex Alderton, Roberto Amato, Sonia Goncalves, Ewan Harrison, David K. Jackson, Ian Johnston, Dominic Kwiatkowski, Cordelia Langford, John Sillitoe |
| EPI_ISL_531432 | NHSGGC West of Scotland Specialist Virology Centre / MRC-University of Glasgow Centre for Virus Research | Wellcome Sanger Institute for the COVID-19 Genomics UK Consortium | Ana da Silva Filipe, Natasha Johnson, Kathy Smollett, Daniel Mair, Stephen Carmichael, Lily Tong, Jenna Nichols, Elihu Aranday-Cortes, Kirstyn Brunker, Yasmin Parr, Kyriaki Nomikou; Sarah McDonald, Marc Niebel, Patawee Asamaphan; Richard Orton, Joseph Hughes, Sreenu Vattipally, David L Robertson; Alasdair MacLean, Rory Gunson; Kathy Li, Natasha Jesudason, Rajiv Shah, James Shepherd, Antonia Ho, Alice Broos, Emma Thomson and Alex Alderton, Roberto Amato, Sonia Goncalves, Ewan Harrison, David K. Jackson, Ian Johnston, Dominic Kwiatkowski, Cordelia Langford, John Sillitoe |
| EPI_ISL_531433, EPI_ISL_531434, EPI_ISL_531435, EPI_ISL_531436, EPI_ISL_531437, EPI_ISL_531439, EPI_ISL_531440, EPI_ISL_531441 | Lighthouse Lab in Glasgow | Wellcome Sanger Institute for the COVID-19 Genomics UK Consortium | Harper VanSteenhouse, Yumi Kasai, David Gray, Carol Clugston, Anna Dominiczak and Alex Alderton, Roberto Amato, Sonia Goncalves, Ewan Harrison, David K. Jackson, Ian Johnston, Dominic Kwiatkowski, Cordelia Langford, John Sillitoe |
| EPI_ISL_531442 | NHSGGC West of Scotland Specialist Virology Centre / MRC-University of Glasgow Centre for Virus Research | Wellcome Sanger Institute for the COVID-19 Genomics UK Consortium | Ana da Silva Filipe, Natasha Johnson, Kathy Smollett, Daniel Mair, Stephen Carmichael, Lily Tong, Jenna Nichols, Elihu Aranday-Cortes, Kirstyn Brunker, Yasmin Parr, Kyriaki Nomikou; Sarah McDonald, Marc Niebel, Patawee Asamaphan; Richard Orton, Joseph Hughes, Sreenu Vattipally, David L Robertson; Alasdair MacLean, Rory Gunson; Kathy Li, Natasha Jesudason, Rajiv Shah, James Shepherd, Antonia Ho, Alice Broos, Emma Thomson and Alex Alderton, Roberto Amato, Sonia Goncalves, Ewan Harrison, David K. Jackson, Ian Johnston, Dominic Kwiatkowski, Cordelia Langford, John Sillitoe |
| EPI_ISL_531443, EPI_ISL_531444, EPI_ISL_531446, EPI_ISL_531447, EPI_ISL_531448, EPI_ISL_531449, EPI_ISL_531450, EPI_ISL_531451, EPI_ISL_531452, EPI_ISL_531453, EPI_ISL_531454, EPI_ISL_531455, EPI_ISL_531456, EPI_ISL_531457, EPI_ISL_531458, EPI_ISL_531459, EPI_ISL_531461, EPI_ISL_531462, EPI_ISL_531463 | | | |
| see above | Lighthouse Lab in Glasgow | Wellcome Sanger Institute for the COVID-19 Genomics UK Consortium | Harper VanSteenhouse, Yumi Kasai, David Gray, Carol Clugston, Anna Dominiczak and Alex Alderton, Roberto Amato, Sonia Goncalves, Ewan Harrison, David K. Jackson, Ian Johnston, Dominic Kwiatkowski, Cordelia Langford, John Sillitoe |
| EPI_ISL_531464 | NHSGGC West of Scotland Specialist Virology Centre / MRC-University of Glasgow Centre for Virus Research | Wellcome Sanger Institute for the COVID-19 Genomics UK Consortium | Ana da Silva Filipe, Natasha Johnson, Kathy Smollett, Daniel Mair, Stephen Carmichael, Lily Tong, Jenna Nichols, Elihu Aranday-Cortes, Kirstyn Brunker, Yasmin Parr, Kyriaki Nomikou; Sarah McDonald, Marc Niebel, Patawee Asamaphan; Richard Orton, Joseph Hughes, Sreenu Vattipally, David L Robertson; Alasdair MacLean, Rory Gunson; Kathy Li, Natasha Jesudason, Rajiv Shah, James Shepherd, Antonia Ho, Alice Broos, Emma Thomson and Alex Alderton, Roberto Amato, Sonia Goncalves, Ewan Harrison, David K. Jackson, Ian Johnston, Dominic Kwiatkowski, Cordelia Langford, John Sillitoe |
| EPI_ISL_531465, EPI_ISL_531466, EPI_ISL_531467, EPI_ISL_531468, EPI_ISL_531469 | Lighthouse Lab in Glasgow | Wellcome Sanger Institute for the COVID-19 Genomics UK Consortium | Harper VanSteenhouse, Yumi Kasai, David Gray, Carol Clugston, Anna Dominiczak and Alex Alderton, Roberto Amato, Sonia Goncalves, Ewan Harrison, David K. Jackson, Ian Johnston, Dominic Kwiatkowski, Cordelia Langford, John Sillitoe |
| EPI_ISL_531470 | NHSGGC West of Scotland Specialist Virology Centre / MRC-University of Glasgow Centre for Virus Research | Wellcome Sanger Institute for the COVID-19 Genomics UK Consortium | Ana da Silva Filipe, Natasha Johnson, Kathy Smollett, Daniel Mair, Stephen Carmichael, Lily Tong, Jenna Nichols, Elihu Aranday-Cortes, Kirstyn Brunker, Yasmin Parr, Kyriaki Nomikou; Sarah McDonald, Marc Niebel, Patawee Asamaphan; Richard Orton, Joseph Hughes, Sreenu Vattipally, David L Robertson; Alasdair MacLean, Rory Gunson; Kathy Li, Natasha Jesudason, Rajiv Shah, James Shepherd, Antonia Ho, Alice Broos, Emma Thomson and Alex Alderton, Roberto Amato, Sonia Goncalves, Ewan Harrison, David K. Jackson, Ian Johnston, Dominic Kwiatkowski, Cordelia Langford, John Sillitoe |
| EPI_ISL_531471, EPI_ISL_531472 | Lighthouse Lab in Glasgow | Wellcome Sanger Institute for the COVID-19 Genomics UK Consortium | Harper VanSteenhouse, Yumi Kasai, David Gray, Carol Clugston, Anna Dominiczak and Alex Alderton, Roberto Amato, Sonia Goncalves, Ewan Harrison, David K. Jackson, Ian Johnston, Dominic Kwiatkowski, Cordelia Langford, John Sillitoe |
| EPI_ISL_531473 | NHSGGC West of Scotland Specialist Virology Centre / MRC-University of Glasgow Centre for Virus Research | Wellcome Sanger Institute for the COVID-19 Genomics UK Consortium | Ana da Silva Filipe, Natasha Johnson, Kathy Smollett, Daniel Mair, Stephen Carmichael, Lily Tong, Jenna Nichols, Elihu Aranday-Cortes, Kirstyn Brunker, Yasmin Parr, Kyriaki Nomikou; Sarah McDonald, Marc Niebel, Patawee Asamaphan; Richard Orton, Joseph Hughes, Sreenu Vattipally, David L Robertson; Alasdair MacLean, Rory Gunson; Kathy Li, Natasha Jesudason, Rajiv Shah, James Shepherd, Antonia Ho, Alice Broos, Emma Thomson and Alex Alderton, Roberto Amato, Sonia Goncalves, Ewan Harrison, David K. Jackson, Ian Johnston, Dominic Kwiatkowski, Cordelia Langford, John Sillitoe |
| EPI_ISL_531475, EPI_ISL_531479 | Lighthouse Lab in Glasgow | Wellcome Sanger Institute for the COVID-19 Genomics UK Consortium | Harper VanSteenhouse, Yumi Kasai, David Gray, Carol Clugston, Anna Dominiczak and Alex Alderton, Roberto Amato, Sonia Goncalves, Ewan Harrison, David K. Jackson, Ian Johnston, Dominic Kwiatkowski, Cordelia Langford, John Sillitoe |
| EPI_ISL_531480 | NHSGGC West of Scotland Specialist Virology Centre / MRC-University of Glasgow Centre for Virus Research | Wellcome Sanger Institute for the COVID-19 Genomics UK Consortium | Ana da Silva Filipe, Natasha Johnson, Kathy Smollett, Daniel Mair, Stephen Carmichael, Lily Tong, Jenna Nichols, Elihu Aranday-Cortes, Kirstyn Brunker, Yasmin Parr, Kyriaki Nomikou; Sarah McDonald, Marc Niebel, Patawee Asamaphan; Richard Orton, Joseph Hughes, Sreenu Vattipally, David L Robertson; Alasdair MacLean, Rory Gunson; Kathy Li, Natasha Jesudason, Rajiv Shah, James Shepherd, Antonia Ho, Alice Broos, Emma Thomson and Alex Alderton, Roberto Amato, Sonia Goncalves, Ewan Harrison, David K. Jackson, Ian Johnston, Dominic Kwiatkowski, Cordelia Langford, John Sillitoe |
| EPI_ISL_531482 | Lighthouse Lab in Glasgow | Wellcome Sanger Institute for the COVID-19 Genomics UK Consortium | Harper VanSteenhouse, Yumi Kasai, David Gray, Carol Clugston, Anna Dominiczak and Alex Alderton, Roberto Amato, Sonia Goncalves, Ewan Harrison, David K. Jackson, Ian Johnston, Dominic Kwiatkowski, Cordelia Langford, John Sillitoe |
| EPI_ISL_531483, EPI_ISL_531484, EPI_ISL_531485 | NHSGGC West of Scotland Specialist Virology Centre / MRC-University of Glasgow Centre for Virus Research | Wellcome Sanger Institute for the COVID-19 Genomics UK Consortium | Ana da Silva Filipe, Natasha Johnson, Kathy Smollett, Daniel Mair, Stephen Carmichael, Lily Tong, Jenna Nichols, Elihu Aranday-Cortes, Kirstyn Brunker, Yasmin Parr, Kyriaki Nomikou; Sarah McDonald, Marc Niebel, Patawee Asamaphan; Richard Orton, Joseph Hughes, Sreenu Vattipally, David L Robertson; Alasdair MacLean, Rory Gunson; Kathy Li, Natasha Jesudason, Rajiv Shah, James Shepherd, Antonia Ho, Alice Broos, Emma Thomson and Alex Alderton, Roberto Amato, Sonia Goncalves, Ewan Harrison, David K. Jackson, Ian Johnston, Dominic Kwiatkowski, Cordelia Langford, John Sillitoe |
| EPI_ISL_531486, EPI_ISL_531487, EPI_ISL_531489, EPI_ISL_531490, EPI_ISL_531491 | Lighthouse Lab in Glasgow | Wellcome Sanger Institute for the COVID-19 Genomics UK Consortium | Harper VanSteenhouse, Yumi Kasai, David Gray, Carol Clugston, Anna Dominiczak and Alex Alderton, Roberto Amato, Sonia Goncalves, Ewan Harrison, David K. Jackson, Ian Johnston, Dominic Kwiatkowski, Cordelia Langford, John Sillitoe |
| EPI_ISL_531492 | NHSGGC West of Scotland Specialist Virology Centre / MRC-University of Glasgow Centre for Virus Research | Wellcome Sanger Institute for the COVID-19 Genomics UK Consortium | Ana da Silva Filipe, Natasha Johnson, Kathy Smollett, Daniel Mair, Stephen Carmichael, Lily Tong, Jenna Nichols, Elihu Aranday-Cortes, Kirstyn Brunker, Yasmin Parr, Kyriaki Nomikou; Sarah McDonald, Marc Niebel, Patawee Asamaphan; Richard Orton, Joseph Hughes, Sreenu Vattipally, David L Robertson; Alasdair MacLean, Rory Gunson; Kathy Li, Natasha Jesudason, Rajiv Shah, James Shepherd, Antonia Ho, Alice Broos, Emma Thomson and Alex Alderton, Roberto Amato, Sonia Goncalves, Ewan Harrison, David K. Jackson, Ian Johnston, Dominic Kwiatkowski, Cordelia Langford, John Sillitoe |
| EPI_ISL_531494 | Lighthouse Lab in Glasgow | Wellcome Sanger Institute for the COVID-19 Genomics UK Consortium | Harper VanSteenhouse, Yumi Kasai, David Gray, Carol Clugston, Anna Dominiczak and Alex Alderton, Roberto Amato, Sonia Goncalves, Ewan Harrison, David K. Jackson, Ian Johnston, Dominic Kwiatkowski, Cordelia Langford, John Sillitoe |
| EPI_ISL_531495 | NHSGGC West of Scotland Specialist Virology Centre / MRC-University of Glasgow Centre for Virus Research | Wellcome Sanger Institute for the COVID-19 Genomics UK Consortium | Ana da Silva Filipe, Natasha Johnson, Kathy Smollett, Daniel Mair, Stephen Carmichael, Lily Tong, Jenna Nichols, Elihu Aranday-Cortes, Kirstyn Brunker, Yasmin Parr, Kyriaki Nomikou; Sarah McDonald, Marc Niebel, Patawee Asamaphan; Richard Orton, Joseph Hughes, Sreenu Vattipally, David L Robertson; Alasdair MacLean, Rory Gunson; Kathy Li, Natasha Jesudason, Rajiv Shah, James Shepherd, Antonia Ho, Alice Broos, Emma Thomson and Alex Alderton, Roberto Amato, Sonia Goncalves, Ewan Harrison, David K. Jackson, Ian Johnston, Dominic Kwiatkowski, Cordelia Langford, John Sillitoe |
| EPI_ISL_531496, EPI_ISL_531497, EPI_ISL_531498, EPI_ISL_531499 | Lighthouse Lab in Glasgow | Wellcome Sanger Institute for the COVID-19 Genomics UK Consortium | Harper VanSteenhouse, Yumi Kasai, David Gray, Carol Clugston, Anna Dominiczak and Alex Alderton, Roberto Amato, Sonia Goncalves, Ewan Harrison, David K. Jackson, Ian Johnston, Dominic Kwiatkowski, Cordelia Langford, John Sillitoe |
| EPI_ISL_531500 | NHSGGC West of Scotland Specialist Virology Centre / MRC-University of Glasgow Centre for Virus Research | Wellcome Sanger Institute for the COVID-19 Genomics UK Consortium | Ana da Silva Filipe, Natasha Johnson, Kathy Smollett, Daniel Mair, Stephen Carmichael, Lily Tong, Jenna Nichols, Elihu Aranday-Cortes, Kirstyn Brunker, Yasmin Parr, Kyriaki Nomikou; Sarah McDonald, Marc Niebel, Patawee Asamaphan; Richard Orton, Joseph Hughes, Sreenu Vattipally, David L Robertson; Alasdair MacLean, Rory Gunson; Kathy Li, Natasha Jesudason, Rajiv Shah, James Shepherd, Antonia Ho, Alice Broos, Emma Thomson and Alex Alderton, Roberto Amato, Sonia Goncalves, Ewan Harrison, David K. Jackson, Ian Johnston, Dominic Kwiatkowski, Cordelia Langford, John Sillitoe |
| EPI_ISL_531501, EPI_ISL_531502, EPI_ISL_531503 | Lighthouse Lab in Glasgow | Wellcome Sanger Institute for the COVID-19 Genomics UK Consortium | Harper VanSteenhouse, Yumi Kasai, David Gray, Carol Clugston, Anna Dominiczak and Alex Alderton, Roberto Amato, Sonia Goncalves, Ewan Harrison, David K. Jackson, Ian Johnston, Dominic Kwiatkowski, Cordelia Langford, John Sillitoe |

| | | | |
|---|---|---|---|
| EPI_ISL_531504 | NHSGGC West of Scotland Specialist Virology Centre / MRC-University of Glasgow Centre for Virus Research | Wellcome Sanger Institute for the COVID-19 Genomics UK Consortium | Ana da Silva Filipe, Natasha Johnson, Kathy Smollett, Daniel Mair, Stephen Carmichael, Lily Tong, Jenna Nichols, Elihu Aranday-Cortes, Kirstyn Brunker, Yasmin Parr, Kyriaki Nomikou; Sarah McDonald, Marc Niebel, Patawee Asamaphan; Richard Orton, Joseph Hughes, Sreenu Vattipally, David L Robertson; Alasdair MacLean, Rory Gunson; Kathy Li, Natasha Jesudason, Rajiv Shah, James Shepherd, Antonia Ho, Alice Broos, Emma Thomson and Alex Alderton, Roberto Amato, Sonia Goncalves, Ewan Harrison, David K. Jackson, Ian Johnston, Dominic Kwiatkowski, Cordelia Langford, John Sillitoe |
| EPI_ISL_531505, EPI_ISL_531506, EPI_ISL_531507, EPI_ISL_531508, EPI_ISL_531509, EPI_ISL_531510, EPI_ISL_531511, EPI_ISL_531512, EPI_ISL_531513, EPI_ISL_531514, EPI_ISL_531515, EPI_ISL_531516, EPI_ISL_531517, EPI_ISL_531518, EPI_ISL_531519, EPI_ISL_531520, EPI_ISL_531521, EPI_ISL_531522 | | | |
| see above | Lighthouse Lab in Glasgow | Wellcome Sanger Institute for the COVID-19 Genomics UK Consortium | Harper VanSteenhouse, Yumi Kasai, David Gray, Carol Clugston, Anna Dominiczak and Alex Alderton, Roberto Amato, Sonia Goncalves, Ewan Harrison, David K. Jackson, Ian Johnston, Dominic Kwiatkowski, Cordelia Langford, John Sillitoe |
| EPI_ISL_531524 | NHSGGC West of Scotland Specialist Virology Centre / MRC-University of Glasgow Centre for Virus Research | Wellcome Sanger Institute for the COVID-19 Genomics UK Consortium | Ana da Silva Filipe, Natasha Johnson, Kathy Smollett, Daniel Mair, Stephen Carmichael, Lily Tong, Jenna Nichols, Elihu Aranday-Cortes, Kirstyn Brunker, Yasmin Parr, Kyriaki Nomikou; Sarah McDonald, Marc Niebel, Patawee Asamaphan; Richard Orton, Joseph Hughes, Sreenu Vattipally, David L Robertson; Alasdair MacLean, Rory Gunson; Kathy Li, Natasha Jesudason, Rajiv Shah, James Shepherd, Antonia Ho, Alice Broos, Emma Thomson and Alex Alderton, Roberto Amato, Sonia Goncalves, Ewan Harrison, David K. Jackson, Ian Johnston, Dominic Kwiatkowski, Cordelia Langford, John Sillitoe |
| EPI_ISL_531525, EPI_ISL_531526, EPI_ISL_531528, EPI_ISL_531529, EPI_ISL_531530, EPI_ISL_531531, EPI_ISL_531532, EPI_ISL_531533, EPI_ISL_531534, EPI_ISL_531535, EPI_ISL_531537, EPI_ISL_531538, EPI_ISL_531539, EPI_ISL_531540, EPI_ISL_531541, EPI_ISL_531542, EPI_ISL_531543, EPI_ISL_531544, EPI_ISL_531545 | | | |
| see above | Lighthouse Lab in Glasgow | Wellcome Sanger Institute for the COVID-19 Genomics UK Consortium | Harper VanSteenhouse, Yumi Kasai, David Gray, Carol Clugston, Anna Dominiczak and Alex Alderton, Roberto Amato, Sonia Goncalves, Ewan Harrison, David K. Jackson, Ian Johnston, Dominic Kwiatkowski, Cordelia Langford, John Sillitoe |
| EPI_ISL_531546, EPI_ISL_531548 | NHSGGC West of Scotland Specialist Virology Centre / MRC-University of Glasgow Centre for Virus Research | Wellcome Sanger Institute for the COVID-19 Genomics UK Consortium | Ana da Silva Filipe, Natasha Johnson, Kathy Smollett, Daniel Mair, Stephen Carmichael, Lily Tong, Jenna Nichols, Elihu Aranday-Cortes, Kirstyn Brunker, Yasmin Parr, Kyriaki Nomikou; Sarah McDonald, Marc Niebel, Patawee Asamaphan; Richard Orton, Joseph Hughes, Sreenu Vattipally, David L Robertson; Alasdair MacLean, Rory Gunson; Kathy Li, Natasha Jesudason, Rajiv Shah, James Shepherd, Antonia Ho, Alice Broos, Emma Thomson and Alex Alderton, Roberto Amato, Sonia Goncalves, Ewan Harrison, David K. Jackson, Ian Johnston, Dominic Kwiatkowski, Cordelia Langford, John Sillitoe |
| EPI_ISL_531550, EPI_ISL_531551, EPI_ISL_531552, EPI_ISL_531553, EPI_ISL_531554, EPI_ISL_531555 | Lighthouse Lab in Glasgow | Wellcome Sanger Institute for the COVID-19 Genomics UK Consortium | Harper VanSteenhouse, Yumi Kasai, David Gray, Carol Clugston, Anna Dominiczak and Alex Alderton, Roberto Amato, Sonia Goncalves, Ewan Harrison, David K. Jackson, Ian Johnston, Dominic Kwiatkowski, Cordelia Langford, John Sillitoe |
| EPI_ISL_531557 | NHSGGC West of Scotland Specialist Virology Centre / MRC-University of Glasgow Centre for Virus Research | Wellcome Sanger Institute for the COVID-19 Genomics UK Consortium | Ana da Silva Filipe, Natasha Johnson, Kathy Smollett, Daniel Mair, Stephen Carmichael, Lily Tong, Jenna Nichols, Elihu Aranday-Cortes, Kirstyn Brunker, Yasmin Parr, Kyriaki Nomikou; Sarah McDonald, Marc Niebel, Patawee Asamaphan; Richard Orton, Joseph Hughes, Sreenu Vattipally, David L Robertson; Alasdair MacLean, Rory Gunson; Kathy Li, Natasha Jesudason, Rajiv Shah, James Shepherd, Antonia Ho, Alice Broos, Emma Thomson and Alex Alderton, Roberto Amato, Sonia Goncalves, Ewan Harrison, David K. Jackson, Ian Johnston, Dominic Kwiatkowski, Cordelia Langford, John Sillitoe |
| EPI_ISL_531558, EPI_ISL_531559, EPI_ISL_531560, EPI_ISL_531561, EPI_ISL_531562 | Lighthouse Lab in Glasgow | Wellcome Sanger Institute for the COVID-19 Genomics UK Consortium | Harper VanSteenhouse, Yumi Kasai, David Gray, Carol Clugston, Anna Dominiczak and Alex Alderton, Roberto Amato, Sonia Goncalves, Ewan Harrison, David K. Jackson, Ian Johnston, Dominic Kwiatkowski, Cordelia Langford, John Sillitoe |
| EPI_ISL_531563, EPI_ISL_531564, EPI_ISL_531565 | NHSGGC West of Scotland Specialist Virology Centre / MRC-University of Glasgow Centre for Virus Research | Wellcome Sanger Institute for the COVID-19 Genomics UK Consortium | Ana da Silva Filipe, Natasha Johnson, Kathy Smollett, Daniel Mair, Stephen Carmichael, Lily Tong, Jenna Nichols, Elihu Aranday-Cortes, Kirstyn Brunker, Yasmin Parr, Kyriaki Nomikou; Sarah McDonald, Marc Niebel, Patawee Asamaphan; Richard Orton, Joseph Hughes, Sreenu Vattipally, David L Robertson; Alasdair MacLean, Rory Gunson; Kathy Li, Natasha Jesudason, Rajiv Shah, James Shepherd, Antonia Ho, Alice Broos, Emma Thomson and Alex Alderton, Roberto Amato, Sonia Goncalves, Ewan Harrison, David K. Jackson, Ian Johnston, Dominic Kwiatkowski, Cordelia Langford, John Sillitoe |
| EPI_ISL_531568 | Lighthouse Lab in Glasgow | Wellcome Sanger Institute for the COVID-19 Genomics UK Consortium | Harper VanSteenhouse, Yumi Kasai, David Gray, Carol Clugston, Anna Dominiczak and Alex Alderton, Roberto Amato, Sonia Goncalves, Ewan Harrison, David K. Jackson, Ian Johnston, Dominic Kwiatkowski, Cordelia Langford, John Sillitoe |
| EPI_ISL_531569, EPI_ISL_531570 | NHSGGC West of Scotland Specialist Virology Centre / MRC-University of Glasgow Centre for Virus Research | Wellcome Sanger Institute for the COVID-19 Genomics UK Consortium | Ana da Silva Filipe, Natasha Johnson, Kathy Smollett, Daniel Mair, Stephen Carmichael, Lily Tong, Jenna Nichols, Elihu Aranday-Cortes, Kirstyn Brunker, Yasmin Parr, Kyriaki Nomikou; Sarah McDonald, Marc Niebel, Patawee Asamaphan; Richard Orton, Joseph Hughes, Sreenu Vattipally, David L Robertson; Alasdair MacLean, Rory Gunson; Kathy Li, Natasha Jesudason, Rajiv Shah, James Shepherd, Antonia Ho, Alice Broos, Emma Thomson and Alex Alderton, Roberto Amato, Sonia Goncalves, Ewan Harrison, David K. Jackson, Ian Johnston, Dominic Kwiatkowski, Cordelia Langford, John Sillitoe |
| EPI_ISL_531571, EPI_ISL_531573, EPI_ISL_531574 | Lighthouse Lab in Glasgow | Wellcome Sanger Institute for the COVID-19 Genomics UK Consortium | Harper VanSteenhouse, Yumi Kasai, David Gray, Carol Clugston, Anna Dominiczak and Alex Alderton, Roberto Amato, Sonia Goncalves, Ewan Harrison, David K. Jackson, Ian Johnston, Dominic Kwiatkowski, Cordelia Langford, John Sillitoe |
| EPI_ISL_531575 | NHSGGC West of Scotland Specialist Virology Centre / MRC-University of Glasgow Centre for Virus Research | Wellcome Sanger Institute for the COVID-19 Genomics UK Consortium | Ana da Silva Filipe, Natasha Johnson, Kathy Smollett, Daniel Mair, Stephen Carmichael, Lily Tong, Jenna Nichols, Elihu Aranday-Cortes, Kirstyn Brunker, Yasmin Parr, Kyriaki Nomikou; Sarah McDonald, Marc Niebel, Patawee Asamaphan; Richard Orton, Joseph Hughes, Sreenu Vattipally, David L Robertson; Alasdair MacLean, Rory Gunson; Kathy Li, Natasha Jesudason, Rajiv Shah, James Shepherd, Antonia Ho, Alice Broos, Emma Thomson and Alex Alderton, Roberto Amato, Sonia Goncalves, Ewan Harrison, David K. Jackson, Ian Johnston, Dominic Kwiatkowski, Cordelia Langford, John Sillitoe |
| EPI_ISL_531576 | Lighthouse Lab in Glasgow | Wellcome Sanger Institute for the COVID-19 Genomics UK Consortium | Harper VanSteenhouse, Yumi Kasai, David Gray, Carol Clugston, Anna Dominiczak and Alex Alderton, Roberto Amato, Sonia Goncalves, Ewan Harrison, David K. Jackson, Ian Johnston, Dominic Kwiatkowski, Cordelia Langford, John Sillitoe |
| EPI_ISL_531577 | NHSGGC West of Scotland Specialist Virology Centre / MRC-University of Glasgow Centre for Virus Research | Wellcome Sanger Institute for the COVID-19 Genomics UK Consortium | Ana da Silva Filipe, Natasha Johnson, Kathy Smollett, Daniel Mair, Stephen Carmichael, Lily Tong, Jenna Nichols, Elihu Aranday-Cortes, Kirstyn Brunker, Yasmin Parr, Kyriaki Nomikou; Sarah McDonald, Marc Niebel, Patawee Asamaphan; Richard Orton, Joseph Hughes, Sreenu Vattipally, David L Robertson; Alasdair MacLean, Rory Gunson; Kathy Li, Natasha Jesudason, Rajiv Shah, James Shepherd, Antonia Ho, Alice Broos, Emma Thomson and Alex Alderton, Roberto Amato, Sonia Goncalves, Ewan Harrison, David K. Jackson, Ian Johnston, Dominic Kwiatkowski, Cordelia Langford, John Sillitoe |
| EPI_ISL_531578, EPI_ISL_531580, EPI_ISL_531581, EPI_ISL_531582 | Lighthouse Lab in Glasgow | Wellcome Sanger Institute for the COVID-19 Genomics UK Consortium | Harper VanSteenhouse, Yumi Kasai, David Gray, Carol Clugston, Anna Dominiczak and Alex Alderton, Roberto Amato, Sonia Goncalves, Ewan Harrison, David K. Jackson, Ian Johnston, Dominic Kwiatkowski, Cordelia Langford, John Sillitoe |
| EPI_ISL_531583 | NHSGGC West of Scotland Specialist Virology Centre / MRC-University of Glasgow Centre for Virus Research | Wellcome Sanger Institute for the COVID-19 Genomics UK Consortium | Ana da Silva Filipe, Natasha Johnson, Kathy Smollett, Daniel Mair, Stephen Carmichael, Lily Tong, Jenna Nichols, Elihu Aranday-Cortes, Kirstyn Brunker, Yasmin Parr, Kyriaki Nomikou; Sarah McDonald, Marc Niebel, Patawee Asamaphan; Richard Orton, Joseph Hughes, Sreenu Vattipally, David L Robertson; Alasdair MacLean, Rory Gunson; Kathy Li, Natasha Jesudason, Rajiv Shah, James Shepherd, Antonia Ho, Alice Broos, Emma Thomson and Alex Alderton, Roberto Amato, Sonia Goncalves, Ewan Harrison, David K. Jackson, Ian Johnston, Dominic Kwiatkowski, Cordelia Langford, John Sillitoe |
| EPI_ISL_531584, EPI_ISL_531586, EPI_ISL_531587, EPI_ISL_531588 | Lighthouse Lab in Glasgow | Wellcome Sanger Institute for the COVID-19 Genomics UK Consortium | Harper VanSteenhouse, Yumi Kasai, David Gray, Carol Clugston, Anna Dominiczak and Alex Alderton, Roberto Amato, Sonia Goncalves, Ewan Harrison, David K. Jackson, Ian Johnston, Dominic Kwiatkowski, Cordelia Langford, John Sillitoe |
| EPI_ISL_531590 | NHSGGC West of Scotland Specialist Virology Centre / MRC-University of Glasgow Centre for Virus Research | Wellcome Sanger Institute for the COVID-19 Genomics UK Consortium | Ana da Silva Filipe, Natasha Johnson, Kathy Smollett, Daniel Mair, Stephen Carmichael, Lily Tong, Jenna Nichols, Elihu Aranday-Cortes, Kirstyn Brunker, Yasmin Parr, Kyriaki Nomikou; Sarah McDonald, Marc Niebel, Patawee Asamaphan; Richard Orton, Joseph Hughes, Sreenu Vattipally, David L Robertson; Alasdair MacLean, Rory Gunson; Kathy Li, Natasha Jesudason, Rajiv Shah, James Shepherd, Antonia Ho, Alice Broos, Emma Thomson and Alex Alderton, Roberto Amato, Sonia Goncalves, Ewan Harrison, David K. Jackson, Ian Johnston, Dominic Kwiatkowski, Cordelia Langford, John Sillitoe |
| EPI_ISL_531592, EPI_ISL_531593, EPI_ISL_531594 | Lighthouse Lab in Glasgow | Wellcome Sanger Institute for the COVID-19 Genomics UK Consortium | Harper VanSteenhouse, Yumi Kasai, David Gray, Carol Clugston, Anna Dominiczak and Alex Alderton, Roberto Amato, Sonia Goncalves, Ewan Harrison, David K. Jackson, Ian Johnston, Dominic Kwiatkowski, Cordelia Langford, John Sillitoe |
| EPI_ISL_531595 | NHSGGC West of Scotland Specialist Virology Centre / MRC-University of Glasgow Centre for Virus Research | Wellcome Sanger Institute for the COVID-19 Genomics UK Consortium | Ana da Silva Filipe, Natasha Johnson, Kathy Smollett, Daniel Mair, Stephen Carmichael, Lily Tong, Jenna Nichols, Elihu Aranday-Cortes, Kirstyn Brunker, Yasmin Parr, Kyriaki Nomikou; Sarah McDonald, Marc Niebel, Patawee Asamaphan; Richard Orton, Joseph Hughes, Sreenu Vattipally, David L Robertson; Alasdair MacLean, Rory Gunson; Kathy Li, Natasha Jesudason, Rajiv Shah, James Shepherd, Antonia Ho, Alice Broos, Emma Thomson and Alex Alderton, Roberto Amato, Sonia Goncalves, Ewan Harrison, David K. Jackson, Ian Johnston, Dominic Kwiatkowski, Cordelia Langford, John Sillitoe |
| EPI_ISL_531596, EPI_ISL_531597 | Lighthouse Lab in Glasgow | Wellcome Sanger Institute for the COVID-19 Genomics UK Consortium | Harper VanSteenhouse, Yumi Kasai, David Gray, Carol Clugston, Anna Dominiczak and Alex Alderton, Roberto Amato, Sonia Goncalves, Ewan Harrison, David K. Jackson, Ian Johnston, Dominic Kwiatkowski, Cordelia Langford, John Sillitoe |
| EPI_ISL_531598, EPI_ISL_531599 | NHSGGC West of Scotland Specialist Virology Centre / MRC-University of Glasgow Centre for Virus Research | Wellcome Sanger Institute for the COVID-19 Genomics UK Consortium | Ana da Silva Filipe, Natasha Johnson, Kathy Smollett, Daniel Mair, Stephen Carmichael, Lily Tong, Jenna Nichols, Elihu Aranday-Cortes, Kirstyn Brunker, Yasmin Parr, Kyriaki Nomikou; Sarah McDonald, Marc Niebel, Patawee Asamaphan; Richard Orton, Joseph Hughes, Sreenu Vattipally, David L Robertson; Alasdair MacLean, Rory Gunson; Kathy Li, Natasha Jesudason, Rajiv Shah, James Shepherd, Antonia Ho, Alice Broos, Emma Thomson and Alex Alderton, Roberto Amato, Sonia Goncalves, Ewan Harrison, David K. Jackson, Ian Johnston, Dominic Kwiatkowski, Cordelia Langford, John Sillitoe |
| EPI_ISL_531600, EPI_ISL_531601, EPI_ISL_531602, EPI_ISL_531603, EPI_ISL_531604, EPI_ISL_531605, EPI_ISL_531606 | Lighthouse Lab in Glasgow | Wellcome Sanger Institute for the COVID-19 Genomics UK Consortium | Harper VanSteenhouse, Yumi Kasai, David Gray, Carol Clugston, Anna Dominiczak and Alex Alderton, Roberto Amato, Sonia Goncalves, Ewan Harrison, David K. Jackson, Ian Johnston, Dominic Kwiatkowski, Cordelia Langford, John Sillitoe |
| EPI_ISL_531607 | NHSGGC West of Scotland Specialist Virology Centre / MRC-University of Glasgow Centre for Virus Research | Wellcome Sanger Institute for the COVID-19 Genomics UK Consortium | Ana da Silva Filipe, Natasha Johnson, Kathy Smollett, Daniel Mair, Stephen Carmichael, Lily Tong, Jenna Nichols, Elihu Aranday-Cortes, Kirstyn Brunker, Yasmin Parr, Kyriaki Nomikou; Sarah McDonald, Marc Niebel, Patawee Asamaphan; Richard Orton, Joseph Hughes, Sreenu Vattipally, David L Robertson; Alasdair MacLean, Rory Gunson; Kathy Li, Natasha Jesudason, Rajiv Shah, James Shepherd, Antonia Ho, Alice Broos, Emma Thomson and Alex Alderton, Roberto Amato, Sonia Goncalves, Ewan Harrison, David K. Jackson, Ian Johnston, Dominic Kwiatkowski, Cordelia Langford, John Sillitoe |
| EPI_ISL_531608, EPI_ISL_531610, EPI_ISL_531611, EPI_ISL_531612, EPI_ISL_531613, EPI_ISL_531614, EPI_ISL_531616 | Lighthouse Lab in Glasgow | Wellcome Sanger Institute for the COVID-19 Genomics UK Consortium | Harper VanSteenhouse, Yumi Kasai, David Gray, Carol Clugston, Anna Dominiczak and Alex Alderton, Roberto Amato, Sonia Goncalves, Ewan Harrison, David K. Jackson, Ian Johnston, Dominic Kwiatkowski, Cordelia Langford, John Sillitoe |
| EPI_ISL_531617 | NHSGGC West of Scotland Specialist Virology Centre / MRC-University of Glasgow Centre for Virus Research | Wellcome Sanger Institute for the COVID-19 Genomics UK Consortium | Ana da Silva Filipe, Natasha Johnson, Kathy Smollett, Daniel Mair, Stephen Carmichael, Lily Tong, Jenna Nichols, Elihu Aranday-Cortes, Kirstyn Brunker, Yasmin Parr, Kyriaki Nomikou; Sarah McDonald, Marc Niebel, Patawee Asamaphan; Richard Orton, Joseph Hughes, Sreenu Vattipally, David L Robertson; Alasdair MacLean, Rory Gunson; Kathy Li, Natasha Jesudason, Rajiv Shah, James Shepherd, Antonia Ho, Alice Broos, Emma Thomson and Alex Alderton, Roberto Amato, Sonia Goncalves, Ewan Harrison, David K. Jackson, Ian Johnston, Dominic Kwiatkowski, Cordelia Langford, John Sillitoe |
| EPI_ISL_531619, EPI_ISL_531621, EPI_ISL_531622, EPI_ISL_531623, EPI_ISL_531624 | Lighthouse Lab in Glasgow | Wellcome Sanger Institute for the COVID-19 Genomics UK Consortium | Harper VanSteenhouse, Yumi Kasai, David Gray, Carol Clugston, Anna Dominiczak and Alex Alderton, Roberto Amato, Sonia Goncalves, Ewan Harrison, David K. Jackson, Ian Johnston, Dominic Kwiatkowski, Cordelia Langford, John Sillitoe |
| EPI_ISL_531625 | NHSGGC West of Scotland Specialist Virology Centre / MRC-University of Glasgow Centre for Virus Research | Wellcome Sanger Institute for the COVID-19 Genomics UK Consortium | Ana da Silva Filipe, Natasha Johnson, Kathy Smollett, Daniel Mair, Stephen Carmichael, Lily Tong, Jenna Nichols, Elihu Aranday-Cortes, Kirstyn Brunker, Yasmin Parr, Kyriaki Nomikou; Sarah McDonald, Marc Niebel, Patawee Asamaphan; Richard Orton, Joseph Hughes, Sreenu Vattipally, David L Robertson; Alasdair MacLean, Rory Gunson; Kathy Li, Natasha Jesudason, Rajiv Shah, James Shepherd, Antonia Ho, Alice Broos, Emma Thomson and Alex Alderton, Roberto Amato, Sonia Goncalves, Ewan Harrison, David K. Jackson, Ian Johnston, Dominic Kwiatkowski, Cordelia Langford, John Sillitoe |
| EPI_ISL_531627, EPI_ISL_531628, EPI_ISL_531629, EPI_ISL_531630 | Lighthouse Lab in Glasgow | Wellcome Sanger Institute for the COVID-19 Genomics UK Consortium | Harper VanSteenhouse, Yumi Kasai, David Gray, Carol Clugston, Anna Dominiczak and Alex Alderton, Roberto Amato, Sonia Goncalves, Ewan Harrison, David K. Jackson, Ian Johnston, Dominic Kwiatkowski, Cordelia Langford, John Sillitoe |
| EPI_ISL_531631 | NHSGGC West of Scotland Specialist Virology Centre / MRC-University of Glasgow Centre for Virus Research | Wellcome Sanger Institute for the COVID-19 Genomics UK Consortium | Ana da Silva Filipe, Natasha Johnson, Kathy Smollett, Daniel Mair, Stephen Carmichael, Lily Tong, Jenna Nichols, Elihu Aranday-Cortes, Kirstyn Brunker, Yasmin Parr, Kyriaki Nomikou; Sarah McDonald, Marc Niebel, Patawee Asamaphan; Richard Orton, Joseph Hughes, Sreenu Vattipally, David L Robertson; Alasdair MacLean, Rory Gunson; Kathy Li, Natasha Jesudason, Rajiv Shah, James Shepherd, Antonia Ho, Alice Broos, Emma Thomson and Alex Alderton, Roberto Amato, Sonia Goncalves, Ewan Harrison, David K. Jackson, Ian Johnston, Dominic Kwiatkowski, Cordelia Langford, John Sillitoe |
| EPI_ISL_531632, EPI_ISL_531633 | Lighthouse Lab in Glasgow | Wellcome Sanger Institute for the COVID-19 Genomics UK Consortium | Harper VanSteenhouse, Yumi Kasai, David Gray, Carol Clugston, Anna Dominiczak and Alex Alderton, Roberto Amato, Sonia Goncalves, Ewan Harrison, David K. Jackson, Ian Johnston, Dominic Kwiatkowski, Cordelia Langford, John Sillitoe |
| EPI_ISL_531634 | NHSGGC West of Scotland Specialist Virology Centre / MRC-University of Glasgow Centre for Virus Research | Wellcome Sanger Institute for the COVID-19 Genomics UK Consortium | Ana da Silva Filipe, Natasha Johnson, Kathy Smollett, Daniel Mair, Stephen Carmichael, Lily Tong, Jenna Nichols, Elihu Aranday-Cortes, Kirstyn Brunker, Yasmin Parr, Kyriaki Nomikou; Sarah McDonald, Marc Niebel, Patawee Asamaphan; Richard Orton, Joseph Hughes, Sreenu Vattipally, David L Robertson; Alasdair MacLean, Rory Gunson; Kathy Li, Natasha Jesudason, Rajiv Shah, James Shepherd, Antonia Ho, Alice Broos, Emma Thomson and Alex Alderton, Roberto Amato, Sonia Goncalves, Ewan Harrison, David K. Jackson, Ian Johnston, Dominic Kwiatkowski, Cordelia Langford, John Sillitoe |
| EPI_ISL_531635, EPI_ISL_531636 | Lighthouse Lab in Glasgow | Wellcome Sanger Institute for the COVID-19 Genomics UK Consortium | Harper VanSteenhouse, Yumi Kasai, David Gray, Carol Clugston, Anna Dominiczak and Alex Alderton, Roberto Amato, Sonia Goncalves, Ewan Harrison, David K. Jackson, Ian Johnston, Dominic Kwiatkowski, Cordelia Langford, John Sillitoe |
| EPI_ISL_531638, EPI_ISL_531639 | NHSGGC West of Scotland Specialist Virology Centre / MRC-University of Glasgow Centre for Virus Research | Wellcome Sanger Institute for the COVID-19 Genomics UK Consortium | Ana da Silva Filipe, Natasha Johnson, Kathy Smollett, Daniel Mair, Stephen Carmichael, Lily Tong, Jenna Nichols, Elihu Aranday-Cortes, Kirstyn Brunker, Yasmin Parr, Kyriaki Nomikou; Sarah McDonald, Marc Niebel, Patawee Asamaphan; Richard Orton, Joseph Hughes, Sreenu Vattipally, David L Robertson; Alasdair MacLean, Rory Gunson; Kathy Li, Natasha Jesudason, Rajiv Shah, James Shepherd, Antonia Ho, Alice Broos, Emma Thomson and Alex Alderton, Roberto Amato, Sonia Goncalves, Ewan Harrison, David K. Jackson, Ian Johnston, Dominic Kwiatkowski, Cordelia Langford, John Sillitoe |
| EPI_ISL_531642, EPI_ISL_531643, EPI_ISL_531644, | Lighthouse Lab in Glasgow | Wellcome Sanger Institute for the | Harper VanSteenhouse, Yumi Kasai, David Gray, Carol Clugston, Anna Dominiczak and Alex Alderton, Roberto Amato, Sonia Goncalves, Ewan Harrison, David K. Jackson, Ian Johnston, Dominic Kwiatkowski, Cordelia Langford, John Sillitoe |

| | | | |
|---|---|---|---|
| EPI_ISL_531645, EPI_ISL_531647 | | COVID-19 Genomics UK Consortium | |
| EPI_ISL_531649 | NHSGGC West of Scotland Specialist Virology Centre / MRC-University of Glasgow Centre for Virus Research | Wellcome Sanger Institute for the COVID-19 Genomics UK Consortium | Ana da Silva Filipe, Natasha Johnson, Kathy Smollett, Daniel Mair, Stephen Carmichael, Lily Tong, Jenna Nichols, Elihu Aranday-Cortes, Kirstyn Brunker, Yasmin Parr, Kyriaki Nomikou; Sarah McDonald, Marc Niebel, Patawee Asamaphan; Richard Orton, Joseph Hughes, Sreenu Vattipally, David L Robertson; Alasdair MacLean, Rory Gunson; Kathy Li, Natasha Jesudason, Rajiv Shah, James Shepherd, Antonia Ho, Alice Broos, Emma Thomson and Alex Alderton, Roberto Amato, Sonia Goncalves, Ewan Harrison, David K. Jackson, Ian Johnston, Dominic Kwiatkowski, Cordelia Langford, John Sillitoe |
| EPI_ISL_531650, EPI_ISL_531651, EPI_ISL_531652, EPI_ISL_531654, EPI_ISL_531656, EPI_ISL_531658 | Lighthouse Lab in Glasgow | Wellcome Sanger Institute for the COVID-19 Genomics UK Consortium | Harper VanSteenhouse, Yumi Kasai, David Gray, Carol Clugston, Anna Dominiczak and Alex Alderton, Roberto Amato, Sonia Goncalves, Ewan Harrison, David K. Jackson, Ian Johnston, Dominic Kwiatkowski, Cordelia Langford, John Sillitoe |
| EPI_ISL_531660 | NHSGGC West of Scotland Specialist Virology Centre / MRC-University of Glasgow Centre for Virus Research | Wellcome Sanger Institute for the COVID-19 Genomics UK Consortium | Ana da Silva Filipe, Natasha Johnson, Kathy Smollett, Daniel Mair, Stephen Carmichael, Lily Tong, Jenna Nichols, Elihu Aranday-Cortes, Kirstyn Brunker, Yasmin Parr, Kyriaki Nomikou; Sarah McDonald, Marc Niebel, Patawee Asamaphan; Richard Orton, Joseph Hughes, Sreenu Vattipally, David L Robertson; Alasdair MacLean, Rory Gunson; Kathy Li, Natasha Jesudason, Rajiv Shah, James Shepherd, Antonia Ho, Alice Broos, Emma Thomson and Alex Alderton, Roberto Amato, Sonia Goncalves, Ewan Harrison, David K. Jackson, Ian Johnston, Dominic Kwiatkowski, Cordelia Langford, John Sillitoe |
| EPI_ISL_531661, EPI_ISL_531662, EPI_ISL_531663, EPI_ISL_531664, EPI_ISL_531665, EPI_ISL_531666, EPI_ISL_531667 | Lighthouse Lab in Glasgow | Wellcome Sanger Institute for the COVID-19 Genomics UK Consortium | Harper VanSteenhouse, Yumi Kasai, David Gray, Carol Clugston, Anna Dominiczak and Alex Alderton, Roberto Amato, Sonia Goncalves, Ewan Harrison, David K. Jackson, Ian Johnston, Dominic Kwiatkowski, Cordelia Langford, John Sillitoe |
| EPI_ISL_531668 | NHSGGC West of Scotland Specialist Virology Centre / MRC-University of Glasgow Centre for Virus Research | Wellcome Sanger Institute for the COVID-19 Genomics UK Consortium | Ana da Silva Filipe, Natasha Johnson, Kathy Smollett, Daniel Mair, Stephen Carmichael, Lily Tong, Jenna Nichols, Elihu Aranday-Cortes, Kirstyn Brunker, Yasmin Parr, Kyriaki Nomikou; Sarah McDonald, Marc Niebel, Patawee Asamaphan; Richard Orton, Joseph Hughes, Sreenu Vattipally, David L Robertson; Alasdair MacLean, Rory Gunson; Kathy Li, Natasha Jesudason, Rajiv Shah, James Shepherd, Antonia Ho, Alice Broos, Emma Thomson and Alex Alderton, Roberto Amato, Sonia Goncalves, Ewan Harrison, David K. Jackson, Ian Johnston, Dominic Kwiatkowski, Cordelia Langford, John Sillitoe |
| EPI_ISL_531669, EPI_ISL_531671 | Lighthouse Lab in Glasgow | Wellcome Sanger Institute for the COVID-19 Genomics UK Consortium | Harper VanSteenhouse, Yumi Kasai, David Gray, Carol Clugston, Anna Dominiczak and Alex Alderton, Roberto Amato, Sonia Goncalves, Ewan Harrison, David K. Jackson, Ian Johnston, Dominic Kwiatkowski, Cordelia Langford, John Sillitoe |
| EPI_ISL_531672 | NHSGGC West of Scotland Specialist Virology Centre / MRC-University of Glasgow Centre for Virus Research | Wellcome Sanger Institute for the COVID-19 Genomics UK Consortium | Ana da Silva Filipe, Natasha Johnson, Kathy Smollett, Daniel Mair, Stephen Carmichael, Lily Tong, Jenna Nichols, Elihu Aranday-Cortes, Kirstyn Brunker, Yasmin Parr, Kyriaki Nomikou; Sarah McDonald, Marc Niebel, Patawee Asamaphan; Richard Orton, Joseph Hughes, Sreenu Vattipally, David L Robertson; Alasdair MacLean, Rory Gunson; Kathy Li, Natasha Jesudason, Rajiv Shah, James Shepherd, Antonia Ho, Alice Broos, Emma Thomson and Alex Alderton, Roberto Amato, Sonia Goncalves, Ewan Harrison, David K. Jackson, Ian Johnston, Dominic Kwiatkowski, Cordelia Langford, John Sillitoe |
| EPI_ISL_531673, EPI_ISL_531675, EPI_ISL_531676 | Lighthouse Lab in Glasgow | Wellcome Sanger Institute for the COVID-19 Genomics UK Consortium | Harper VanSteenhouse, Yumi Kasai, David Gray, Carol Clugston, Anna Dominiczak and Alex Alderton, Roberto Amato, Sonia Goncalves, Ewan Harrison, David K. Jackson, Ian Johnston, Dominic Kwiatkowski, Cordelia Langford, John Sillitoe |
| EPI_ISL_531677, EPI_ISL_531678 | NHSGGC West of Scotland Specialist Virology Centre / MRC-University of Glasgow Centre for Virus Research | Wellcome Sanger Institute for the COVID-19 Genomics UK Consortium | Ana da Silva Filipe, Natasha Johnson, Kathy Smollett, Daniel Mair, Stephen Carmichael, Lily Tong, Jenna Nichols, Elihu Aranday-Cortes, Kirstyn Brunker, Yasmin Parr, Kyriaki Nomikou; Sarah McDonald, Marc Niebel, Patawee Asamaphan; Richard Orton, Joseph Hughes, Sreenu Vattipally, David L Robertson; Alasdair MacLean, Rory Gunson; Kathy Li, Natasha Jesudason, Rajiv Shah, James Shepherd, Antonia Ho, Alice Broos, Emma Thomson and Alex Alderton, Roberto Amato, Sonia Goncalves, Ewan Harrison, David K. Jackson, Ian Johnston, Dominic Kwiatkowski, Cordelia Langford, John Sillitoe |
| EPI_ISL_531679, EPI_ISL_531680, EPI_ISL_531682, EPI_ISL_531683, EPI_ISL_531684, EPI_ISL_531685 | Lighthouse Lab in Glasgow | Wellcome Sanger Institute for the COVID-19 Genomics UK Consortium | Harper VanSteenhouse, Yumi Kasai, David Gray, Carol Clugston, Anna Dominiczak and Alex Alderton, Roberto Amato, Sonia Goncalves, Ewan Harrison, David K. Jackson, Ian Johnston, Dominic Kwiatkowski, Cordelia Langford, John Sillitoe |
| EPI_ISL_531689, EPI_ISL_531690, EPI_ISL_531691 | NHSGGC West of Scotland Specialist Virology Centre / MRC-University of Glasgow Centre for Virus Research | Wellcome Sanger Institute for the COVID-19 Genomics UK Consortium | Ana da Silva Filipe, Natasha Johnson, Kathy Smollett, Daniel Mair, Stephen Carmichael, Lily Tong, Jenna Nichols, Elihu Aranday-Cortes, Kirstyn Brunker, Yasmin Parr, Kyriaki Nomikou; Sarah McDonald, Marc Niebel, Patawee Asamaphan; Richard Orton, Joseph Hughes, Sreenu Vattipally, David L Robertson; Alasdair MacLean, Rory Gunson; Kathy Li, Natasha Jesudason, Rajiv Shah, James Shepherd, Antonia Ho, Alice Broos, Emma Thomson and Alex Alderton, Roberto Amato, Sonia Goncalves, Ewan Harrison, David K. Jackson, Ian Johnston, Dominic Kwiatkowski, Cordelia Langford, John Sillitoe |
| EPI_ISL_531692, EPI_ISL_531693, EPI_ISL_531694, EPI_ISL_531695, EPI_ISL_531696, EPI_ISL_531697, EPI_ISL_531698, EPI_ISL_531699, EPI_ISL_531700, EPI_ISL_531701 | Lighthouse Lab in Glasgow | Wellcome Sanger Institute for the COVID-19 Genomics UK Consortium | Harper VanSteenhouse, Yumi Kasai, David Gray, Carol Clugston, Anna Dominiczak and Alex Alderton, Roberto Amato, Sonia Goncalves, Ewan Harrison, David K. Jackson, Ian Johnston, Dominic Kwiatkowski, Cordelia Langford, John Sillitoe |
| EPI_ISL_531702, EPI_ISL_531703 | NHSGGC West of Scotland Specialist Virology Centre / MRC-University of Glasgow Centre for Virus Research | Wellcome Sanger Institute for the COVID-19 Genomics UK Consortium | Ana da Silva Filipe, Natasha Johnson, Kathy Smollett, Daniel Mair, Stephen Carmichael, Lily Tong, Jenna Nichols, Elihu Aranday-Cortes, Kirstyn Brunker, Yasmin Parr, Kyriaki Nomikou; Sarah McDonald, Marc Niebel, Patawee Asamaphan; Richard Orton, Joseph Hughes, Sreenu Vattipally, David L Robertson; Alasdair MacLean, Rory Gunson; Kathy Li, Natasha Jesudason, Rajiv Shah, James Shepherd, Antonia Ho, Alice Broos, Emma Thomson and Alex Alderton, Roberto Amato, Sonia Goncalves, Ewan Harrison, David K. Jackson, Ian Johnston, Dominic Kwiatkowski, Cordelia Langford, John Sillitoe |
| EPI_ISL_531704, EPI_ISL_531705, EPI_ISL_531707, EPI_ISL_531708, EPI_ISL_531709, EPI_ISL_531710, EPI_ISL_531711, EPI_ISL_531712, EPI_ISL_531713 | Lighthouse Lab in Glasgow | Wellcome Sanger Institute for the COVID-19 Genomics UK Consortium | Harper VanSteenhouse, Yumi Kasai, David Gray, Carol Clugston, Anna Dominiczak and Alex Alderton, Roberto Amato, Sonia Goncalves, Ewan Harrison, David K. Jackson, Ian Johnston, Dominic Kwiatkowski, Cordelia Langford, John Sillitoe |
| EPI_ISL_531714 | NHSGGC West of Scotland Specialist Virology Centre / MRC-University of Glasgow Centre for Virus Research | Wellcome Sanger Institute for the COVID-19 Genomics UK Consortium | Ana da Silva Filipe, Natasha Johnson, Kathy Smollett, Daniel Mair, Stephen Carmichael, Lily Tong, Jenna Nichols, Elihu Aranday-Cortes, Kirstyn Brunker, Yasmin Parr, Kyriaki Nomikou; Sarah McDonald, Marc Niebel, Patawee Asamaphan; Richard Orton, Joseph Hughes, Sreenu Vattipally, David L Robertson; Alasdair MacLean, Rory Gunson; Kathy Li, Natasha Jesudason, Rajiv Shah, James Shepherd, Antonia Ho, Alice Broos, Emma Thomson and Alex Alderton, Roberto Amato, Sonia Goncalves, Ewan Harrison, David K. Jackson, Ian Johnston, Dominic Kwiatkowski, Cordelia Langford, John Sillitoe |
| EPI_ISL_531715, EPI_ISL_531716, EPI_ISL_531718, EPI_ISL_531719, EPI_ISL_531720, EPI_ISL_531722, EPI_ISL_531723, EPI_ISL_531724, EPI_ISL_531725, EPI_ISL_531728 | Lighthouse Lab in Glasgow | Wellcome Sanger Institute for the COVID-19 Genomics UK Consortium | Harper VanSteenhouse, Yumi Kasai, David Gray, Carol Clugston, Anna Dominiczak and Alex Alderton, Roberto Amato, Sonia Goncalves, Ewan Harrison, David K. Jackson, Ian Johnston, Dominic Kwiatkowski, Cordelia Langford, John Sillitoe |
| EPI_ISL_531730 | NHSGGC West of Scotland Specialist Virology Centre / MRC-University of Glasgow Centre for Virus Research | Wellcome Sanger Institute for the COVID-19 Genomics UK Consortium | Ana da Silva Filipe, Natasha Johnson, Kathy Smollett, Daniel Mair, Stephen Carmichael, Lily Tong, Jenna Nichols, Elihu Aranday-Cortes, Kirstyn Brunker, Yasmin Parr, Kyriaki Nomikou; Sarah McDonald, Marc Niebel, Patawee Asamaphan; Richard Orton, Joseph Hughes, Sreenu Vattipally, David L Robertson; Alasdair MacLean, Rory Gunson; Kathy Li, Natasha Jesudason, Rajiv Shah, James Shepherd, Antonia Ho, Alice Broos, Emma Thomson and Alex Alderton, Roberto Amato, Sonia Goncalves, Ewan Harrison, David K. Jackson, Ian Johnston, Dominic Kwiatkowski, Cordelia Langford, John Sillitoe |
| EPI_ISL_531731 | Lighthouse Lab in Glasgow | Wellcome Sanger Institute for the COVID-19 Genomics UK Consortium | Harper VanSteenhouse, Yumi Kasai, David Gray, Carol Clugston, Anna Dominiczak and Alex Alderton, Roberto Amato, Sonia Goncalves, Ewan Harrison, David K. Jackson, Ian Johnston, Dominic Kwiatkowski, Cordelia Langford, John Sillitoe |
| EPI_ISL_531732 | NHSGGC West of Scotland Specialist Virology Centre / MRC-University of Glasgow Centre for Virus Research | Wellcome Sanger Institute for the COVID-19 Genomics UK Consortium | Ana da Silva Filipe, Natasha Johnson, Kathy Smollett, Daniel Mair, Stephen Carmichael, Lily Tong, Jenna Nichols, Elihu Aranday-Cortes, Kirstyn Brunker, Yasmin Parr, Kyriaki Nomikou; Sarah McDonald, Marc Niebel, Patawee Asamaphan; Richard Orton, Joseph Hughes, Sreenu Vattipally, David L Robertson; Alasdair MacLean, Rory Gunson; Kathy Li, Natasha Jesudason, Rajiv Shah, James Shepherd, Antonia Ho, Alice Broos, Emma Thomson and Alex Alderton, Roberto Amato, Sonia Goncalves, Ewan Harrison, David K. Jackson, Ian Johnston, Dominic Kwiatkowski, Cordelia Langford, John Sillitoe |
| EPI_ISL_531733, EPI_ISL_531739, EPI_ISL_531741 | Lighthouse Lab in Glasgow | Wellcome Sanger Institute for the COVID-19 Genomics UK Consortium | Harper VanSteenhouse, Yumi Kasai, David Gray, Carol Clugston, Anna Dominiczak and Alex Alderton, Roberto Amato, Sonia Goncalves, Ewan Harrison, David K. Jackson, Ian Johnston, Dominic Kwiatkowski, Cordelia Langford, John Sillitoe |
| EPI_ISL_531743 | NHSGGC West of Scotland Specialist Virology Centre / MRC-University of Glasgow Centre for Virus Research | Wellcome Sanger Institute for the COVID-19 Genomics UK Consortium | Ana da Silva Filipe, Natasha Johnson, Kathy Smollett, Daniel Mair, Stephen Carmichael, Lily Tong, Jenna Nichols, Elihu Aranday-Cortes, Kirstyn Brunker, Yasmin Parr, Kyriaki Nomikou; Sarah McDonald, Marc Niebel, Patawee Asamaphan; Richard Orton, Joseph Hughes, Sreenu Vattipally, David L Robertson; Alasdair MacLean, Rory Gunson; Kathy Li, Natasha Jesudason, Rajiv Shah, James Shepherd, Antonia Ho, Alice Broos, Emma Thomson and Alex Alderton, Roberto Amato, Sonia Goncalves, Ewan Harrison, David K. Jackson, Ian Johnston, Dominic Kwiatkowski, Cordelia Langford, John Sillitoe |
| EPI_ISL_531744, EPI_ISL_531745, EPI_ISL_531746, EPI_ISL_531748, EPI_ISL_531750, EPI_ISL_531751, EPI_ISL_531752 | Lighthouse Lab in Glasgow | Wellcome Sanger Institute for the COVID-19 Genomics UK Consortium | Harper VanSteenhouse, Yumi Kasai, David Gray, Carol Clugston, Anna Dominiczak and Alex Alderton, Roberto Amato, Sonia Goncalves, Ewan Harrison, David K. Jackson, Ian Johnston, Dominic Kwiatkowski, Cordelia Langford, John Sillitoe |
| EPI_ISL_531754, EPI_ISL_531755 | NHSGGC West of Scotland Specialist Virology Centre / MRC-University of Glasgow Centre for Virus Research | Wellcome Sanger Institute for the COVID-19 Genomics UK Consortium | Ana da Silva Filipe, Natasha Johnson, Kathy Smollett, Daniel Mair, Stephen Carmichael, Lily Tong, Jenna Nichols, Elihu Aranday-Cortes, Kirstyn Brunker, Yasmin Parr, Kyriaki Nomikou; Sarah McDonald, Marc Niebel, Patawee Asamaphan; Richard Orton, Joseph Hughes, Sreenu Vattipally, David L Robertson; Alasdair MacLean, Rory Gunson; Kathy Li, Natasha Jesudason, Rajiv Shah, James Shepherd, Antonia Ho, Alice Broos, Emma Thomson and Alex Alderton, Roberto Amato, Sonia Goncalves, Ewan Harrison, David K. Jackson, Ian Johnston, Dominic Kwiatkowski, Cordelia Langford, John Sillitoe |
| EPI_ISL_531756, EPI_ISL_531757, EPI_ISL_531759, EPI_ISL_531760, EPI_ISL_531761, EPI_ISL_531762, EPI_ISL_531763, EPI_ISL_531764, EPI_ISL_531766, EPI_ISL_531767 | Lighthouse Lab in Glasgow | Wellcome Sanger Institute for the COVID-19 Genomics UK Consortium | Harper VanSteenhouse, Yumi Kasai, David Gray, Carol Clugston, Anna Dominiczak and Alex Alderton, Roberto Amato, Sonia Goncalves, Ewan Harrison, David K. Jackson, Ian Johnston, Dominic Kwiatkowski, Cordelia Langford, John Sillitoe |
| EPI_ISL_531768 | NHSGGC West of Scotland Specialist Virology Centre / MRC-University of Glasgow Centre for Virus Research | Wellcome Sanger Institute for the COVID-19 Genomics UK Consortium | Ana da Silva Filipe, Natasha Johnson, Kathy Smollett, Daniel Mair, Stephen Carmichael, Lily Tong, Jenna Nichols, Elihu Aranday-Cortes, Kirstyn Brunker, Yasmin Parr, Kyriaki Nomikou; Sarah McDonald, Marc Niebel, Patawee Asamaphan; Richard Orton, Joseph Hughes, Sreenu Vattipally, David L Robertson; Alasdair MacLean, Rory Gunson; Kathy Li, Natasha Jesudason, Rajiv Shah, James Shepherd, Antonia Ho, Alice Broos, Emma Thomson and Alex Alderton, Roberto Amato, Sonia Goncalves, Ewan Harrison, David K. Jackson, Ian Johnston, Dominic Kwiatkowski, Cordelia Langford, John Sillitoe |
| EPI_ISL_531769, EPI_ISL_531770, EPI_ISL_531771, EPI_ISL_531772, EPI_ISL_531773, EPI_ISL_531774, EPI_ISL_531776, EPI_ISL_531777, EPI_ISL_531778, EPI_ISL_531780, EPI_ISL_531781, EPI_ISL_531782, EPI_ISL_531783, EPI_ISL_531785, EPI_ISL_531786, EPI_ISL_531787, EPI_ISL_531790, EPI_ISL_531791, EPI_ISL_531793, EPI_ISL_531794, EPI_ISL_531795, EPI_ISL_531796, EPI_ISL_531797, EPI_ISL_531798, EPI_ISL_531799, EPI_ISL_531800 | | | |
| see above | Lighthouse Lab in Glasgow | Wellcome Sanger Institute for the COVID-19 Genomics UK Consortium | Harper VanSteenhouse, Yumi Kasai, David Gray, Carol Clugston, Anna Dominiczak and Alex Alderton, Roberto Amato, Sonia Goncalves, Ewan Harrison, David K. Jackson, Ian Johnston, Dominic Kwiatkowski, Cordelia Langford, John Sillitoe |
| EPI_ISL_531801, EPI_ISL_531802 | NHSGGC West of Scotland Specialist Virology Centre / MRC-University of Glasgow Centre for Virus Research | Wellcome Sanger Institute for the COVID-19 Genomics UK Consortium | Ana da Silva Filipe, Natasha Johnson, Kathy Smollett, Daniel Mair, Stephen Carmichael, Lily Tong, Jenna Nichols, Elihu Aranday-Cortes, Kirstyn Brunker, Yasmin Parr, Kyriaki Nomikou; Sarah McDonald, Marc Niebel, Patawee Asamaphan; Richard Orton, Joseph Hughes, Sreenu Vattipally, David L Robertson; Alasdair MacLean, Rory Gunson; Kathy Li, Natasha Jesudason, Rajiv Shah, James Shepherd, Antonia Ho, Alice Broos, Emma Thomson and Alex Alderton, Roberto Amato, Sonia Goncalves, Ewan Harrison, David K. Jackson, Ian Johnston, Dominic Kwiatkowski, Cordelia Langford, John Sillitoe |
| EPI_ISL_531803, EPI_ISL_531804 | Lighthouse Lab in Glasgow | Wellcome Sanger Institute for the COVID-19 Genomics UK Consortium | Harper VanSteenhouse, Yumi Kasai, David Gray, Carol Clugston, Anna Dominiczak and Alex Alderton, Roberto Amato, Sonia Goncalves, Ewan Harrison, David K. Jackson, Ian Johnston, Dominic Kwiatkowski, Cordelia Langford, John Sillitoe |
| EPI_ISL_531805 | NHSGGC West of Scotland Specialist Virology Centre / MRC-University of Glasgow Centre for Virus Research | Wellcome Sanger Institute for the COVID-19 Genomics UK Consortium | Ana da Silva Filipe, Natasha Johnson, Kathy Smollett, Daniel Mair, Stephen Carmichael, Lily Tong, Jenna Nichols, Elihu Aranday-Cortes, Kirstyn Brunker, Yasmin Parr, Kyriaki Nomikou; Sarah McDonald, Marc Niebel, Patawee Asamaphan; Richard Orton, Joseph Hughes, Sreenu Vattipally, David L Robertson; Alasdair MacLean, Rory Gunson; Kathy Li, Natasha Jesudason, Rajiv Shah, James Shepherd, Antonia Ho, Alice Broos, Emma Thomson and Alex Alderton, Roberto Amato, Sonia Goncalves, Ewan Harrison, David K. Jackson, Ian Johnston, Dominic Kwiatkowski, Cordelia Langford, John Sillitoe |
| EPI_ISL_531806, EPI_ISL_531807, EPI_ISL_531808, EPI_ISL_531809, EPI_ISL_531810, EPI_ISL_531811 | Lighthouse Lab in Glasgow | Wellcome Sanger Institute for the COVID-19 Genomics UK Consortium | Harper VanSteenhouse, Yumi Kasai, David Gray, Carol Clugston, Anna Dominiczak and Alex Alderton, Roberto Amato, Sonia Goncalves, Ewan Harrison, David K. Jackson, Ian Johnston, Dominic Kwiatkowski, Cordelia Langford, John Sillitoe |
| EPI_ISL_531814, EPI_ISL_531815 | NHSGGC West of Scotland Specialist Virology Centre / MRC-University of Glasgow Centre for Virus Research | Wellcome Sanger Institute for the COVID-19 Genomics UK Consortium | Ana da Silva Filipe, Natasha Johnson, Kathy Smollett, Daniel Mair, Stephen Carmichael, Lily Tong, Jenna Nichols, Elihu Aranday-Cortes, Kirstyn Brunker, Yasmin Parr, Kyriaki Nomikou; Sarah McDonald, Marc Niebel, Patawee Asamaphan; Richard Orton, Joseph Hughes, Sreenu Vattipally, David L Robertson; Alasdair MacLean, Rory Gunson; Kathy Li, Natasha Jesudason, Rajiv Shah, James Shepherd, Antonia Ho, Alice Broos, Emma Thomson and Alex Alderton, Roberto Amato, Sonia Goncalves, Ewan Harrison, David K. Jackson, Ian Johnston, Dominic Kwiatkowski, Cordelia Langford, John Sillitoe |
| EPI_ISL_531816, EPI_ISL_531818, EPI_ISL_531819, EPI_ISL_531820 | Lighthouse Lab in Glasgow | Wellcome Sanger Institute for the COVID-19 Genomics UK Consortium | Harper VanSteenhouse, Yumi Kasai, David Gray, Carol Clugston, Anna Dominiczak and Alex Alderton, Roberto Amato, Sonia Goncalves, Ewan Harrison, David K. Jackson, Ian Johnston, Dominic Kwiatkowski, Cordelia Langford, John Sillitoe |
| EPI_ISL_531821 | NHSGGC West of Scotland Specialist Virology Centre / MRC-University of Glasgow Centre for Virus Research | Wellcome Sanger Institute for the COVID-19 Genomics UK Consortium | Ana da Silva Filipe, Natasha Johnson, Kathy Smollett, Daniel Mair, Stephen Carmichael, Lily Tong, Jenna Nichols, Elihu Aranday-Cortes, Kirstyn Brunker, Yasmin Parr, Kyriaki Nomikou; Sarah McDonald, Marc Niebel, Patawee Asamaphan; Richard Orton, Joseph Hughes, Sreenu Vattipally, David L Robertson; Alasdair MacLean, Rory Gunson; Kathy Li, Natasha Jesudason, Rajiv Shah, James Shepherd, Antonia Ho, Alice Broos, Emma Thomson and Alex Alderton, Roberto |

| | | | |
|---|---|---|---|
| | | | Amato, Sonia Goncalves, Ewan Harrison, David K. Jackson, Ian Johnston, Dominic Kwiatkowski, Cordelia Langford, John Sillitoe |
| EPI_ISL_531822 | Lighthouse Lab in Glasgow | Wellcome Sanger Institute for the COVID-19 Genomics UK Consortium | Harper VanSteenhouse, Yumi Kasai, David Gray, Carol Clugston, Anna Dominiczak and Alex Alderton, Roberto Amato, Sonia Goncalves, Ewan Harrison, David K. Jackson, Ian Johnston, Dominic Kwiatkowski, Cordelia Langford, John Sillitoe |
| EPI_ISL_531824 | NHSGGC West of Scotland Specialist Virology Centre / MRC-University of Glasgow Centre for Virus Research | Wellcome Sanger Institute for the COVID-19 Genomics UK Consortium | Ana da Silva Filipe, Natasha Johnson, Kathy Smollett, Daniel Mair, Stephen Carmichael, Lily Tong, Jenna Nichols, Elihu Aranday-Cortes, Kirstyn Brunker, Yasmin Parr, Kyriaki Nomikou; Sarah McDonald, Marc Niebel, Patawee Asamaphan; Richard Orton, Joseph Hughes, Sreenu Vattipally, David L Robertson; Alasdair MacLean, Rory Gunson; Kathy Li, Natasha Jesudason, Rajiv Shah, James Shepherd, Antonia Ho, Alice Broos, Emma Thomson and Alex Alderton, Roberto Amato, Sonia Goncalves, Ewan Harrison, David K. Jackson, Ian Johnston, Dominic Kwiatkowski, Cordelia Langford, John Sillitoe |
| EPI_ISL_531825, EPI_ISL_531827, EPI_ISL_531829, EPI_ISL_531830, EPI_ISL_531831 | Lighthouse Lab in Glasgow | Wellcome Sanger Institute for the COVID-19 Genomics UK Consortium | Harper VanSteenhouse, Yumi Kasai, David Gray, Carol Clugston, Anna Dominiczak and Alex Alderton, Roberto Amato, Sonia Goncalves, Ewan Harrison, David K. Jackson, Ian Johnston, Dominic Kwiatkowski, Cordelia Langford, John Sillitoe |
| EPI_ISL_531832 | NHSGGC West of Scotland Specialist Virology Centre / MRC-University of Glasgow Centre for Virus Research | Wellcome Sanger Institute for the COVID-19 Genomics UK Consortium | Ana da Silva Filipe, Natasha Johnson, Kathy Smollett, Daniel Mair, Stephen Carmichael, Lily Tong, Jenna Nichols, Elihu Aranday-Cortes, Kirstyn Brunker, Yasmin Parr, Kyriaki Nomikou; Sarah McDonald, Marc Niebel, Patawee Asamaphan; Richard Orton, Joseph Hughes, Sreenu Vattipally, David L Robertson; Alasdair MacLean, Rory Gunson; Kathy Li, Natasha Jesudason, Rajiv Shah, James Shepherd, Antonia Ho, Alice Broos, Emma Thomson and Alex Alderton, Roberto Amato, Sonia Goncalves, Ewan Harrison, David K. Jackson, Ian Johnston, Dominic Kwiatkowski, Cordelia Langford, John Sillitoe |
| EPI_ISL_531833, EPI_ISL_531834, EPI_ISL_531836, EPI_ISL_531837, EPI_ISL_531838, EPI_ISL_531839, EPI_ISL_531840, EPI_ISL_531841 | Lighthouse Lab in Glasgow | Wellcome Sanger Institute for the COVID-19 Genomics UK Consortium | Harper VanSteenhouse, Yumi Kasai, David Gray, Carol Clugston, Anna Dominiczak and Alex Alderton, Roberto Amato, Sonia Goncalves, Ewan Harrison, David K. Jackson, Ian Johnston, Dominic Kwiatkowski, Cordelia Langford, John Sillitoe |
| EPI_ISL_531842 | NHSGGC West of Scotland Specialist Virology Centre / MRC-University of Glasgow Centre for Virus Research | Wellcome Sanger Institute for the COVID-19 Genomics UK Consortium | Ana da Silva Filipe, Natasha Johnson, Kathy Smollett, Daniel Mair, Stephen Carmichael, Lily Tong, Jenna Nichols, Elihu Aranday-Cortes, Kirstyn Brunker, Yasmin Parr, Kyriaki Nomikou; Sarah McDonald, Marc Niebel, Patawee Asamaphan; Richard Orton, Joseph Hughes, Sreenu Vattipally, David L Robertson; Alasdair MacLean, Rory Gunson; Kathy Li, Natasha Jesudason, Rajiv Shah, James Shepherd, Antonia Ho, Alice Broos, Emma Thomson and Alex Alderton, Roberto Amato, Sonia Goncalves, Ewan Harrison, David K. Jackson, Ian Johnston, Dominic Kwiatkowski, Cordelia Langford, John Sillitoe |
| EPI_ISL_531843, EPI_ISL_531844, EPI_ISL_531845, EPI_ISL_531846, EPI_ISL_531848, EPI_ISL_531849, EPI_ISL_531852, EPI_ISL_531853, EPI_ISL_531854 | Lighthouse Lab in Glasgow | Wellcome Sanger Institute for the COVID-19 Genomics UK Consortium | Harper VanSteenhouse, Yumi Kasai, David Gray, Carol Clugston, Anna Dominiczak and Alex Alderton, Roberto Amato, Sonia Goncalves, Ewan Harrison, David K. Jackson, Ian Johnston, Dominic Kwiatkowski, Cordelia Langford, John Sillitoe |
| EPI_ISL_531856, EPI_ISL_531857, EPI_ISL_531858 | NHSGGC West of Scotland Specialist Virology Centre / MRC-University of Glasgow Centre for Virus Research | Wellcome Sanger Institute for the COVID-19 Genomics UK Consortium | Ana da Silva Filipe, Natasha Johnson, Kathy Smollett, Daniel Mair, Stephen Carmichael, Lily Tong, Jenna Nichols, Elihu Aranday-Cortes, Kirstyn Brunker, Yasmin Parr, Kyriaki Nomikou; Sarah McDonald, Marc Niebel, Patawee Asamaphan; Richard Orton, Joseph Hughes, Sreenu Vattipally, David L Robertson; Alasdair MacLean, Rory Gunson; Kathy Li, Natasha Jesudason, Rajiv Shah, James Shepherd, Antonia Ho, Alice Broos, Emma Thomson and Alex Alderton, Roberto Amato, Sonia Goncalves, Ewan Harrison, David K. Jackson, Ian Johnston, Dominic Kwiatkowski, Cordelia Langford, John Sillitoe |
| EPI_ISL_531859, EPI_ISL_531860, EPI_ISL_531861 | Lighthouse Lab in Glasgow | Wellcome Sanger Institute for the COVID-19 Genomics UK Consortium | Harper VanSteenhouse, Yumi Kasai, David Gray, Carol Clugston, Anna Dominiczak and Alex Alderton, Roberto Amato, Sonia Goncalves, Ewan Harrison, David K. Jackson, Ian Johnston, Dominic Kwiatkowski, Cordelia Langford, John Sillitoe |
| EPI_ISL_531862, EPI_ISL_531863 | NHSGGC West of Scotland Specialist Virology Centre / MRC-University of Glasgow Centre for Virus Research | Wellcome Sanger Institute for the COVID-19 Genomics UK Consortium | Ana da Silva Filipe, Natasha Johnson, Kathy Smollett, Daniel Mair, Stephen Carmichael, Lily Tong, Jenna Nichols, Elihu Aranday-Cortes, Kirstyn Brunker, Yasmin Parr, Kyriaki Nomikou; Sarah McDonald, Marc Niebel, Patawee Asamaphan; Richard Orton, Joseph Hughes, Sreenu Vattipally, David L Robertson; Alasdair MacLean, Rory Gunson; Kathy Li, Natasha Jesudason, Rajiv Shah, James Shepherd, Antonia Ho, Alice Broos, Emma Thomson and Alex Alderton, Roberto Amato, Sonia Goncalves, Ewan Harrison, David K. Jackson, Ian Johnston, Dominic Kwiatkowski, Cordelia Langford, John Sillitoe |
| EPI_ISL_531864 | Lighthouse Lab in Glasgow | Wellcome Sanger Institute for the COVID-19 Genomics UK Consortium | Harper VanSteenhouse, Yumi Kasai, David Gray, Carol Clugston, Anna Dominiczak and Alex Alderton, Roberto Amato, Sonia Goncalves, Ewan Harrison, David K. Jackson, Ian Johnston, Dominic Kwiatkowski, Cordelia Langford, John Sillitoe |
| EPI_ISL_531865 | NHSGGC West of Scotland Specialist Virology Centre / MRC-University of Glasgow Centre for Virus Research | Wellcome Sanger Institute for the COVID-19 Genomics UK Consortium | Ana da Silva Filipe, Natasha Johnson, Kathy Smollett, Daniel Mair, Stephen Carmichael, Lily Tong, Jenna Nichols, Elihu Aranday-Cortes, Kirstyn Brunker, Yasmin Parr, Kyriaki Nomikou; Sarah McDonald, Marc Niebel, Patawee Asamaphan; Richard Orton, Joseph Hughes, Sreenu Vattipally, David L Robertson; Alasdair MacLean, Rory Gunson; Kathy Li, Natasha Jesudason, Rajiv Shah, James Shepherd, Antonia Ho, Alice Broos, Emma Thomson and Alex Alderton, Roberto Amato, Sonia Goncalves, Ewan Harrison, David K. Jackson, Ian Johnston, Dominic Kwiatkowski, Cordelia Langford, John Sillitoe |
| EPI_ISL_531866, EPI_ISL_531867, EPI_ISL_531868, EPI_ISL_531869, EPI_ISL_531870, EPI_ISL_531871 | Lighthouse Lab in Glasgow | Wellcome Sanger Institute for the COVID-19 Genomics UK Consortium | Harper VanSteenhouse, Yumi Kasai, David Gray, Carol Clugston, Anna Dominiczak and Alex Alderton, Roberto Amato, Sonia Goncalves, Ewan Harrison, David K. Jackson, Ian Johnston, Dominic Kwiatkowski, Cordelia Langford, John Sillitoe |
| EPI_ISL_531873 | NHSGGC West of Scotland Specialist Virology Centre / MRC-University of Glasgow Centre for Virus Research | Wellcome Sanger Institute for the COVID-19 Genomics UK Consortium | Ana da Silva Filipe, Natasha Johnson, Kathy Smollett, Daniel Mair, Stephen Carmichael, Lily Tong, Jenna Nichols, Elihu Aranday-Cortes, Kirstyn Brunker, Yasmin Parr, Kyriaki Nomikou; Sarah McDonald, Marc Niebel, Patawee Asamaphan; Richard Orton, Joseph Hughes, Sreenu Vattipally, David L Robertson; Alasdair MacLean, Rory Gunson; Kathy Li, Natasha Jesudason, Rajiv Shah, James Shepherd, Antonia Ho, Alice Broos, Emma Thomson and Alex Alderton, Roberto Amato, Sonia Goncalves, Ewan Harrison, David K. Jackson, Ian Johnston, Dominic Kwiatkowski, Cordelia Langford, John Sillitoe |
| EPI_ISL_531874, EPI_ISL_531875, EPI_ISL_531876 | Lighthouse Lab in Glasgow | Wellcome Sanger Institute for the COVID-19 Genomics UK Consortium | Harper VanSteenhouse, Yumi Kasai, David Gray, Carol Clugston, Anna Dominiczak and Alex Alderton, Roberto Amato, Sonia Goncalves, Ewan Harrison, David K. Jackson, Ian Johnston, Dominic Kwiatkowski, Cordelia Langford, John Sillitoe |
| EPI_ISL_531877 | NHSGGC West of Scotland Specialist Virology Centre / MRC-University of Glasgow Centre for Virus Research | Wellcome Sanger Institute for the COVID-19 Genomics UK Consortium | Ana da Silva Filipe, Natasha Johnson, Kathy Smollett, Daniel Mair, Stephen Carmichael, Lily Tong, Jenna Nichols, Elihu Aranday-Cortes, Kirstyn Brunker, Yasmin Parr, Kyriaki Nomikou; Sarah McDonald, Marc Niebel, Patawee Asamaphan; Richard Orton, Joseph Hughes, Sreenu Vattipally, David L Robertson; Alasdair MacLean, Rory Gunson; Kathy Li, Natasha Jesudason, Rajiv Shah, James Shepherd, Antonia Ho, Alice Broos, Emma Thomson and Alex Alderton, Roberto Amato, Sonia Goncalves, Ewan Harrison, David K. Jackson, Ian Johnston, Dominic Kwiatkowski, Cordelia Langford, John Sillitoe |
| EPI_ISL_531879 | Lighthouse Lab in Glasgow | Wellcome Sanger Institute for the COVID-19 Genomics UK Consortium | Harper VanSteenhouse, Yumi Kasai, David Gray, Carol Clugston, Anna Dominiczak and Alex Alderton, Roberto Amato, Sonia Goncalves, Ewan Harrison, David K. Jackson, Ian Johnston, Dominic Kwiatkowski, Cordelia Langford, John Sillitoe |
| EPI_ISL_531880, EPI_ISL_531881 | NHSGGC West of Scotland Specialist Virology Centre / MRC-University of Glasgow Centre for Virus Research | Wellcome Sanger Institute for the COVID-19 Genomics UK Consortium | Ana da Silva Filipe, Natasha Johnson, Kathy Smollett, Daniel Mair, Stephen Carmichael, Lily Tong, Jenna Nichols, Elihu Aranday-Cortes, Kirstyn Brunker, Yasmin Parr, Kyriaki Nomikou; Sarah McDonald, Marc Niebel, Patawee Asamaphan; Richard Orton, Joseph Hughes, Sreenu Vattipally, David L Robertson; Alasdair MacLean, Rory Gunson; Kathy Li, Natasha Jesudason, Rajiv Shah, James Shepherd, Antonia Ho, Alice Broos, Emma Thomson and Alex Alderton, Roberto Amato, Sonia Goncalves, Ewan Harrison, David K. Jackson, Ian Johnston, Dominic Kwiatkowski, Cordelia Langford, John Sillitoe |
| EPI_ISL_531882, EPI_ISL_531883 | Lighthouse Lab in Glasgow | Wellcome Sanger Institute for the COVID-19 Genomics UK Consortium | Harper VanSteenhouse, Yumi Kasai, David Gray, Carol Clugston, Anna Dominiczak and Alex Alderton, Roberto Amato, Sonia Goncalves, Ewan Harrison, David K. Jackson, Ian Johnston, Dominic Kwiatkowski, Cordelia Langford, John Sillitoe |
| EPI_ISL_531884 | NHSGGC West of Scotland Specialist Virology Centre / MRC-University of Glasgow Centre for Virus Research | Wellcome Sanger Institute for the COVID-19 Genomics UK Consortium | Ana da Silva Filipe, Natasha Johnson, Kathy Smollett, Daniel Mair, Stephen Carmichael, Lily Tong, Jenna Nichols, Elihu Aranday-Cortes, Kirstyn Brunker, Yasmin Parr, Kyriaki Nomikou; Sarah McDonald, Marc Niebel, Patawee Asamaphan; Richard Orton, Joseph Hughes, Sreenu Vattipally, David L Robertson; Alasdair MacLean, Rory Gunson; Kathy Li, Natasha Jesudason, Rajiv Shah, James Shepherd, Antonia Ho, Alice Broos, Emma Thomson and Alex Alderton, Roberto Amato, Sonia Goncalves, Ewan Harrison, David K. Jackson, Ian Johnston, Dominic Kwiatkowski, Cordelia Langford, John Sillitoe |
| EPI_ISL_531885, EPI_ISL_531886 | Lighthouse Lab in Glasgow | Wellcome Sanger Institute for the COVID-19 Genomics UK Consortium | Harper VanSteenhouse, Yumi Kasai, David Gray, Carol Clugston, Anna Dominiczak and Alex Alderton, Roberto Amato, Sonia Goncalves, Ewan Harrison, David K. Jackson, Ian Johnston, Dominic Kwiatkowski, Cordelia Langford, John Sillitoe |
| EPI_ISL_531887 | NHSGGC West of Scotland Specialist Virology Centre / MRC-University of Glasgow Centre for Virus Research | Wellcome Sanger Institute for the COVID-19 Genomics UK Consortium | Ana da Silva Filipe, Natasha Johnson, Kathy Smollett, Daniel Mair, Stephen Carmichael, Lily Tong, Jenna Nichols, Elihu Aranday-Cortes, Kirstyn Brunker, Yasmin Parr, Kyriaki Nomikou; Sarah McDonald, Marc Niebel, Patawee Asamaphan; Richard Orton, Joseph Hughes, Sreenu Vattipally, David L Robertson; Alasdair MacLean, Rory Gunson; Kathy Li, Natasha Jesudason, Rajiv Shah, James Shepherd, Antonia Ho, Alice Broos, Emma Thomson and Alex Alderton, Roberto Amato, Sonia Goncalves, Ewan Harrison, David K. Jackson, Ian Johnston, Dominic Kwiatkowski, Cordelia Langford, John Sillitoe |
| EPI_ISL_531888 | Lighthouse Lab in Glasgow | Wellcome Sanger Institute for the COVID-19 Genomics UK Consortium | Harper VanSteenhouse, Yumi Kasai, David Gray, Carol Clugston, Anna Dominiczak and Alex Alderton, Roberto Amato, Sonia Goncalves, Ewan Harrison, David K. Jackson, Ian Johnston, Dominic Kwiatkowski, Cordelia Langford, John Sillitoe |
| EPI_ISL_531889, EPI_ISL_531890 | NHSGGC West of Scotland Specialist Virology Centre / MRC-University of Glasgow Centre for Virus Research | Wellcome Sanger Institute for the COVID-19 Genomics UK Consortium | Ana da Silva Filipe, Natasha Johnson, Kathy Smollett, Daniel Mair, Stephen Carmichael, Lily Tong, Jenna Nichols, Elihu Aranday-Cortes, Kirstyn Brunker, Yasmin Parr, Kyriaki Nomikou; Sarah McDonald, Marc Niebel, Patawee Asamaphan; Richard Orton, Joseph Hughes, Sreenu Vattipally, David L Robertson; Alasdair MacLean, Rory Gunson; Kathy Li, Natasha Jesudason, Rajiv Shah, James Shepherd, Antonia Ho, Alice Broos, Emma Thomson and Alex Alderton, Roberto Amato, Sonia Goncalves, Ewan Harrison, David K. Jackson, Ian Johnston, Dominic Kwiatkowski, Cordelia Langford, John Sillitoe |
| EPI_ISL_531891 | Lighthouse Lab in Glasgow | Wellcome Sanger Institute for the COVID-19 Genomics UK Consortium | Harper VanSteenhouse, Yumi Kasai, David Gray, Carol Clugston, Anna Dominiczak and Alex Alderton, Roberto Amato, Sonia Goncalves, Ewan Harrison, David K. Jackson, Ian Johnston, Dominic Kwiatkowski, Cordelia Langford, John Sillitoe |
| EPI_ISL_531892 | NHSGGC West of Scotland Specialist Virology Centre / MRC-University of Glasgow Centre for Virus Research | Wellcome Sanger Institute for the COVID-19 Genomics UK Consortium | Ana da Silva Filipe, Natasha Johnson, Kathy Smollett, Daniel Mair, Stephen Carmichael, Lily Tong, Jenna Nichols, Elihu Aranday-Cortes, Kirstyn Brunker, Yasmin Parr, Kyriaki Nomikou; Sarah McDonald, Marc Niebel, Patawee Asamaphan; Richard Orton, Joseph Hughes, Sreenu Vattipally, David L Robertson; Alasdair MacLean, Rory Gunson; Kathy Li, Natasha Jesudason, Rajiv Shah, James Shepherd, Antonia Ho, Alice Broos, Emma Thomson and Alex Alderton, Roberto Amato, Sonia Goncalves, Ewan Harrison, David K. Jackson, Ian Johnston, Dominic Kwiatkowski, Cordelia Langford, John Sillitoe |
| EPI_ISL_531893 | Lighthouse Lab in Glasgow | Wellcome Sanger Institute for the COVID-19 Genomics UK Consortium | Harper VanSteenhouse, Yumi Kasai, David Gray, Carol Clugston, Anna Dominiczak and Alex Alderton, Roberto Amato, Sonia Goncalves, Ewan Harrison, David K. Jackson, Ian Johnston, Dominic Kwiatkowski, Cordelia Langford, John Sillitoe |
| EPI_ISL_531894 | NHSGGC West of Scotland Specialist Virology Centre / MRC-University of Glasgow Centre for Virus Research | Wellcome Sanger Institute for the COVID-19 Genomics UK Consortium | Ana da Silva Filipe, Natasha Johnson, Kathy Smollett, Daniel Mair, Stephen Carmichael, Lily Tong, Jenna Nichols, Elihu Aranday-Cortes, Kirstyn Brunker, Yasmin Parr, Kyriaki Nomikou; Sarah McDonald, Marc Niebel, Patawee Asamaphan; Richard Orton, Joseph Hughes, Sreenu Vattipally, David L Robertson; Alasdair MacLean, Rory Gunson; Kathy Li, Natasha Jesudason, Rajiv Shah, James Shepherd, Antonia Ho, Alice Broos, Emma Thomson and Alex Alderton, Roberto Amato, Sonia Goncalves, Ewan Harrison, David K. Jackson, Ian Johnston, Dominic Kwiatkowski, Cordelia Langford, John Sillitoe |
| EPI_ISL_531895, EPI_ISL_531897, EPI_ISL_531898, EPI_ISL_531899, EPI_ISL_531900, EPI_ISL_531901, EPI_ISL_531903 | Lighthouse Lab in Glasgow | Wellcome Sanger Institute for the COVID-19 Genomics UK Consortium | Harper VanSteenhouse, Yumi Kasai, David Gray, Carol Clugston, Anna Dominiczak and Alex Alderton, Roberto Amato, Sonia Goncalves, Ewan Harrison, David K. Jackson, Ian Johnston, Dominic Kwiatkowski, Cordelia Langford, John Sillitoe |
| EPI_ISL_531905 | NHSGGC West of Scotland Specialist Virology Centre / MRC-University of Glasgow Centre for Virus Research | Wellcome Sanger Institute for the COVID-19 Genomics UK Consortium | Ana da Silva Filipe, Natasha Johnson, Kathy Smollett, Daniel Mair, Stephen Carmichael, Lily Tong, Jenna Nichols, Elihu Aranday-Cortes, Kirstyn Brunker, Yasmin Parr, Kyriaki Nomikou; Sarah McDonald, Marc Niebel, Patawee Asamaphan; Richard Orton, Joseph Hughes, Sreenu Vattipally, David L Robertson; Alasdair MacLean, Rory Gunson; Kathy Li, Natasha Jesudason, Rajiv Shah, James Shepherd, Antonia Ho, Alice Broos, Emma Thomson and Alex Alderton, Roberto Amato, Sonia Goncalves, Ewan Harrison, David K. Jackson, Ian Johnston, Dominic Kwiatkowski, Cordelia Langford, John Sillitoe |
| EPI_ISL_531906, EPI_ISL_531907, EPI_ISL_531908, EPI_ISL_531909, EPI_ISL_531910 | Lighthouse Lab in Glasgow | Wellcome Sanger Institute for the COVID-19 Genomics UK Consortium | Harper VanSteenhouse, Yumi Kasai, David Gray, Carol Clugston, Anna Dominiczak and Alex Alderton, Roberto Amato, Sonia Goncalves, Ewan Harrison, David K. Jackson, Ian Johnston, Dominic Kwiatkowski, Cordelia Langford, John Sillitoe |
| EPI_ISL_531912, EPI_ISL_531913 | NHSGGC West of Scotland Specialist Virology Centre / MRC-University of Glasgow Centre for Virus Research | Wellcome Sanger Institute for the COVID-19 Genomics UK Consortium | Ana da Silva Filipe, Natasha Johnson, Kathy Smollett, Daniel Mair, Stephen Carmichael, Lily Tong, Jenna Nichols, Elihu Aranday-Cortes, Kirstyn Brunker, Yasmin Parr, Kyriaki Nomikou; Sarah McDonald, Marc Niebel, Patawee Asamaphan; Richard Orton, Joseph Hughes, Sreenu Vattipally, David L Robertson; Alasdair MacLean, Rory Gunson; Kathy Li, Natasha Jesudason, Rajiv Shah, James Shepherd, Antonia Ho, Alice Broos, Emma Thomson and Alex Alderton, Roberto Amato, Sonia Goncalves, Ewan Harrison, David K. Jackson, Ian Johnston, Dominic Kwiatkowski, Cordelia Langford, John Sillitoe |
| EPI_ISL_531916, EPI_ISL_531917 | Lighthouse Lab in Glasgow | Wellcome Sanger Institute for the COVID-19 Genomics UK Consortium | Harper VanSteenhouse, Yumi Kasai, David Gray, Carol Clugston, Anna Dominiczak and Alex Alderton, Roberto Amato, Sonia Goncalves, Ewan Harrison, David K. Jackson, Ian Johnston, Dominic Kwiatkowski, Cordelia Langford, John Sillitoe |
| EPI_ISL_531918, EPI_ISL_531919 | NHSGGC West of Scotland Specialist Virology Centre / MRC-University of Glasgow Centre for Virus Research | Wellcome Sanger Institute for the COVID-19 Genomics UK Consortium | Ana da Silva Filipe, Natasha Johnson, Kathy Smollett, Daniel Mair, Stephen Carmichael, Lily Tong, Jenna Nichols, Elihu Aranday-Cortes, Kirstyn Brunker, Yasmin Parr, Kyriaki Nomikou; Sarah McDonald, Marc Niebel, Patawee Asamaphan; Richard Orton, Joseph Hughes, Sreenu Vattipally, David L Robertson; Alasdair MacLean, Rory Gunson; Kathy Li, Natasha Jesudason, Rajiv Shah, James Shepherd, Antonia Ho, Alice Broos, Emma Thomson and Alex Alderton, Roberto Amato, Sonia Goncalves, Ewan Harrison, David K. Jackson, Ian Johnston, Dominic Kwiatkowski, Cordelia Langford, John Sillitoe |
| EPI_ISL_531920, EPI_ISL_531921, EPI_ISL_531922, EPI_ISL_531923, EPI_ISL_531924 | Lighthouse Lab in Glasgow | Wellcome Sanger Institute for the COVID-19 Genomics UK Consortium | Harper VanSteenhouse, Yumi Kasai, David Gray, Carol Clugston, Anna Dominiczak and Alex Alderton, Roberto Amato, Sonia Goncalves, Ewan Harrison, David K. Jackson, Ian Johnston, Dominic Kwiatkowski, Cordelia Langford, John Sillitoe |
| EPI_ISL_531925 | NHSGGC West of Scotland Specialist Virology Centre / MRC-University of Glasgow Centre for Virus Research | Wellcome Sanger Institute for the COVID-19 Genomics UK Consortium | Ana da Silva Filipe, Natasha Johnson, Kathy Smollett, Daniel Mair, Stephen Carmichael, Lily Tong, Jenna Nichols, Elihu Aranday-Cortes, Kirstyn Brunker, Yasmin Parr, Kyriaki Nomikou; Sarah McDonald, Marc Niebel, Patawee Asamaphan; Richard Orton, Joseph Hughes, Sreenu Vattipally, David L Robertson; Alasdair MacLean, Rory Gunson; Kathy Li, Natasha Jesudason, Rajiv Shah, James Shepherd, Antonia Ho, Alice Broos, Emma Thomson and Alex Alderton, Roberto Amato, Sonia Goncalves, Ewan Harrison, David K. Jackson, Ian Johnston, Dominic Kwiatkowski, Cordelia Langford, John Sillitoe |

| | | | |
|---|---|---|---|
| EPI_ISL_531926, EPI_ISL_531927 | Lighthouse Lab in Glasgow | Wellcome Sanger Institute for the COVID-19 Genomics UK Consortium | Harper VanSteenhouse, Yumi Kasai, David Gray, Carol Clugston, Anna Dominiczak and Alex Alderton, Roberto Amato, Sonia Goncalves, Ewan Harrison, David K. Jackson, Ian Johnston, Dominic Kwiatkowski, Cordelia Langford, John Sillitoe |
| EPI_ISL_531929 | NHSGGC West of Scotland Specialist Virology Centre / MRC-University of Glasgow Centre for Virus Research | Wellcome Sanger Institute for the COVID-19 Genomics UK Consortium | Ana da Silva Filipe, Natasha Johnson, Kathy Smollett, Daniel Mair, Stephen Carmichael, Lily Tong, Jenna Nichols, Elihu Aranday-Cortes, Kirstyn Brunker, Yasmin Parr, Kyriaki Nomikou; Sarah McDonald, Marc Niebel, Patawee Asamaphan; Richard Orton, Joseph Hughes, Sreenu Vattipally, David L Robertson; Alasdair MacLean, Rory Gunson; Kathy Li, Natasha Jesudason, Rajiv Shah, James Shepherd, Antonia Ho, Alice Broos, Emma Thomson and Alex Alderton, Roberto Amato, Sonia Goncalves, Ewan Harrison, David K. Jackson, Ian Johnston, Dominic Kwiatkowski, Cordelia Langford, John Sillitoe |
| EPI_ISL_531930, EPI_ISL_531931 | Lighthouse Lab in Glasgow | Wellcome Sanger Institute for the COVID-19 Genomics UK Consortium | Harper VanSteenhouse, Yumi Kasai, David Gray, Carol Clugston, Anna Dominiczak and Alex Alderton, Roberto Amato, Sonia Goncalves, Ewan Harrison, David K. Jackson, Ian Johnston, Dominic Kwiatkowski, Cordelia Langford, John Sillitoe |
| EPI_ISL_531932, EPI_ISL_531933 | NHSGGC West of Scotland Specialist Virology Centre / MRC-University of Glasgow Centre for Virus Research | Wellcome Sanger Institute for the COVID-19 Genomics UK Consortium | Ana da Silva Filipe, Natasha Johnson, Kathy Smollett, Daniel Mair, Stephen Carmichael, Lily Tong, Jenna Nichols, Elihu Aranday-Cortes, Kirstyn Brunker, Yasmin Parr, Kyriaki Nomikou; Sarah McDonald, Marc Niebel, Patawee Asamaphan; Richard Orton, Joseph Hughes, Sreenu Vattipally, David L Robertson; Alasdair MacLean, Rory Gunson; Kathy Li, Natasha Jesudason, Rajiv Shah, James Shepherd, Antonia Ho, Alice Broos, Emma Thomson and Alex Alderton, Roberto Amato, Sonia Goncalves, Ewan Harrison, David K. Jackson, Ian Johnston, Dominic Kwiatkowski, Cordelia Langford, John Sillitoe |
| EPI_ISL_531934 | Lighthouse Lab in Glasgow | Wellcome Sanger Institute for the COVID-19 Genomics UK Consortium | Harper VanSteenhouse, Yumi Kasai, David Gray, Carol Clugston, Anna Dominiczak and Alex Alderton, Roberto Amato, Sonia Goncalves, Ewan Harrison, David K. Jackson, Ian Johnston, Dominic Kwiatkowski, Cordelia Langford, John Sillitoe |
| EPI_ISL_531935, EPI_ISL_531936 | NHSGGC West of Scotland Specialist Virology Centre / MRC-University of Glasgow Centre for Virus Research | Wellcome Sanger Institute for the COVID-19 Genomics UK Consortium | Ana da Silva Filipe, Natasha Johnson, Kathy Smollett, Daniel Mair, Stephen Carmichael, Lily Tong, Jenna Nichols, Elihu Aranday-Cortes, Kirstyn Brunker, Yasmin Parr, Kyriaki Nomikou; Sarah McDonald, Marc Niebel, Patawee Asamaphan; Richard Orton, Joseph Hughes, Sreenu Vattipally, David L Robertson; Alasdair MacLean, Rory Gunson; Kathy Li, Natasha Jesudason, Rajiv Shah, James Shepherd, Antonia Ho, Alice Broos, Emma Thomson and Alex Alderton, Roberto Amato, Sonia Goncalves, Ewan Harrison, David K. Jackson, Ian Johnston, Dominic Kwiatkowski, Cordelia Langford, John Sillitoe |
| EPI_ISL_531937, EPI_ISL_531938, EPI_ISL_531939, EPI_ISL_531940, EPI_ISL_531942 | Lighthouse Lab in Glasgow | Wellcome Sanger Institute for the COVID-19 Genomics UK Consortium | Harper VanSteenhouse, Yumi Kasai, David Gray, Carol Clugston, Anna Dominiczak and Alex Alderton, Roberto Amato, Sonia Goncalves, Ewan Harrison, David K. Jackson, Ian Johnston, Dominic Kwiatkowski, Cordelia Langford, John Sillitoe |
| EPI_ISL_531943 | NHSGGC West of Scotland Specialist Virology Centre / MRC-University of Glasgow Centre for Virus Research | Wellcome Sanger Institute for the COVID-19 Genomics UK Consortium | Ana da Silva Filipe, Natasha Johnson, Kathy Smollett, Daniel Mair, Stephen Carmichael, Lily Tong, Jenna Nichols, Elihu Aranday-Cortes, Kirstyn Brunker, Yasmin Parr, Kyriaki Nomikou; Sarah McDonald, Marc Niebel, Patawee Asamaphan; Richard Orton, Joseph Hughes, Sreenu Vattipally, David L Robertson; Alasdair MacLean, Rory Gunson; Kathy Li, Natasha Jesudason, Rajiv Shah, James Shepherd, Antonia Ho, Alice Broos, Emma Thomson and Alex Alderton, Roberto Amato, Sonia Goncalves, Ewan Harrison, David K. Jackson, Ian Johnston, Dominic Kwiatkowski, Cordelia Langford, John Sillitoe |
| EPI_ISL_531944 | Lighthouse Lab in Glasgow | Wellcome Sanger Institute for the COVID-19 Genomics UK Consortium | Harper VanSteenhouse, Yumi Kasai, David Gray, Carol Clugston, Anna Dominiczak and Alex Alderton, Roberto Amato, Sonia Goncalves, Ewan Harrison, David K. Jackson, Ian Johnston, Dominic Kwiatkowski, Cordelia Langford, John Sillitoe |
| EPI_ISL_531945 | NHSGGC West of Scotland Specialist Virology Centre / MRC-University of Glasgow Centre for Virus Research | Wellcome Sanger Institute for the COVID-19 Genomics UK Consortium | Ana da Silva Filipe, Natasha Johnson, Kathy Smollett, Daniel Mair, Stephen Carmichael, Lily Tong, Jenna Nichols, Elihu Aranday-Cortes, Kirstyn Brunker, Yasmin Parr, Kyriaki Nomikou; Sarah McDonald, Marc Niebel, Patawee Asamaphan; Richard Orton, Joseph Hughes, Sreenu Vattipally, David L Robertson; Alasdair MacLean, Rory Gunson; Kathy Li, Natasha Jesudason, Rajiv Shah, James Shepherd, Antonia Ho, Alice Broos, Emma Thomson and Alex Alderton, Roberto Amato, Sonia Goncalves, Ewan Harrison, David K. Jackson, Ian Johnston, Dominic Kwiatkowski, Cordelia Langford, John Sillitoe |
| EPI_ISL_531947, EPI_ISL_531948, EPI_ISL_531949, EPI_ISL_531950 | Lighthouse Lab in Glasgow | Wellcome Sanger Institute for the COVID-19 Genomics UK Consortium | Harper VanSteenhouse, Yumi Kasai, David Gray, Carol Clugston, Anna Dominiczak and Alex Alderton, Roberto Amato, Sonia Goncalves, Ewan Harrison, David K. Jackson, Ian Johnston, Dominic Kwiatkowski, Cordelia Langford, John Sillitoe |
| EPI_ISL_531951 | NHSGGC West of Scotland Specialist Virology Centre / MRC-University of Glasgow Centre for Virus Research | Wellcome Sanger Institute for the COVID-19 Genomics UK Consortium | Ana da Silva Filipe, Natasha Johnson, Kathy Smollett, Daniel Mair, Stephen Carmichael, Lily Tong, Jenna Nichols, Elihu Aranday-Cortes, Kirstyn Brunker, Yasmin Parr, Kyriaki Nomikou; Sarah McDonald, Marc Niebel, Patawee Asamaphan; Richard Orton, Joseph Hughes, Sreenu Vattipally, David L Robertson; Alasdair MacLean, Rory Gunson; Kathy Li, Natasha Jesudason, Rajiv Shah, James Shepherd, Antonia Ho, Alice Broos, Emma Thomson and Alex Alderton, Roberto Amato, Sonia Goncalves, Ewan Harrison, David K. Jackson, Ian Johnston, Dominic Kwiatkowski, Cordelia Langford, John Sillitoe |
| EPI_ISL_531952, EPI_ISL_531953 | Lighthouse Lab in Glasgow | Wellcome Sanger Institute for the COVID-19 Genomics UK Consortium | Harper VanSteenhouse, Yumi Kasai, David Gray, Carol Clugston, Anna Dominiczak and Alex Alderton, Roberto Amato, Sonia Goncalves, Ewan Harrison, David K. Jackson, Ian Johnston, Dominic Kwiatkowski, Cordelia Langford, John Sillitoe |
| EPI_ISL_531954 | NHSGGC West of Scotland Specialist Virology Centre / MRC-University of Glasgow Centre for Virus Research | Wellcome Sanger Institute for the COVID-19 Genomics UK Consortium | Ana da Silva Filipe, Natasha Johnson, Kathy Smollett, Daniel Mair, Stephen Carmichael, Lily Tong, Jenna Nichols, Elihu Aranday-Cortes, Kirstyn Brunker, Yasmin Parr, Kyriaki Nomikou; Sarah McDonald, Marc Niebel, Patawee Asamaphan; Richard Orton, Joseph Hughes, Sreenu Vattipally, David L Robertson; Alasdair MacLean, Rory Gunson; Kathy Li, Natasha Jesudason, Rajiv Shah, James Shepherd, Antonia Ho, Alice Broos, Emma Thomson and Alex Alderton, Roberto Amato, Sonia Goncalves, Ewan Harrison, David K. Jackson, Ian Johnston, Dominic Kwiatkowski, Cordelia Langford, John Sillitoe |
| EPI_ISL_531955, EPI_ISL_531956, EPI_ISL_531957, EPI_ISL_531958, EPI_ISL_531959, EPI_ISL_531960, EPI_ISL_531961, EPI_ISL_531962 | Lighthouse Lab in Glasgow | Wellcome Sanger Institute for the COVID-19 Genomics UK Consortium | Harper VanSteenhouse, Yumi Kasai, David Gray, Carol Clugston, Anna Dominiczak and Alex Alderton, Roberto Amato, Sonia Goncalves, Ewan Harrison, David K. Jackson, Ian Johnston, Dominic Kwiatkowski, Cordelia Langford, John Sillitoe |
| EPI_ISL_531963 | NHSGGC West of Scotland Specialist Virology Centre / MRC-University of Glasgow Centre for Virus Research | Wellcome Sanger Institute for the COVID-19 Genomics UK Consortium | Ana da Silva Filipe, Natasha Johnson, Kathy Smollett, Daniel Mair, Stephen Carmichael, Lily Tong, Jenna Nichols, Elihu Aranday-Cortes, Kirstyn Brunker, Yasmin Parr, Kyriaki Nomikou; Sarah McDonald, Marc Niebel, Patawee Asamaphan; Richard Orton, Joseph Hughes, Sreenu Vattipally, David L Robertson; Alasdair MacLean, Rory Gunson; Kathy Li, Natasha Jesudason, Rajiv Shah, James Shepherd, Antonia Ho, Alice Broos, Emma Thomson and Alex Alderton, Roberto Amato, Sonia Goncalves, Ewan Harrison, David K. Jackson, Ian Johnston, Dominic Kwiatkowski, Cordelia Langford, John Sillitoe |
| EPI_ISL_531964, EPI_ISL_531965, EPI_ISL_531966, EPI_ISL_531967, EPI_ISL_531968, EPI_ISL_531969, EPI_ISL_531970, EPI_ISL_531971, EPI_ISL_531972 | Lighthouse Lab in Glasgow | Wellcome Sanger Institute for the COVID-19 Genomics UK Consortium | Harper VanSteenhouse, Yumi Kasai, David Gray, Carol Clugston, Anna Dominiczak and Alex Alderton, Roberto Amato, Sonia Goncalves, Ewan Harrison, David K. Jackson, Ian Johnston, Dominic Kwiatkowski, Cordelia Langford, John Sillitoe |
| EPI_ISL_531973 | NHSGGC West of Scotland Specialist Virology Centre / MRC-University of Glasgow Centre for Virus Research | Wellcome Sanger Institute for the COVID-19 Genomics UK Consortium | Ana da Silva Filipe, Natasha Johnson, Kathy Smollett, Daniel Mair, Stephen Carmichael, Lily Tong, Jenna Nichols, Elihu Aranday-Cortes, Kirstyn Brunker, Yasmin Parr, Kyriaki Nomikou; Sarah McDonald, Marc Niebel, Patawee Asamaphan; Richard Orton, Joseph Hughes, Sreenu Vattipally, David L Robertson; Alasdair MacLean, Rory Gunson; Kathy Li, Natasha Jesudason, Rajiv Shah, James Shepherd, Antonia Ho, Alice Broos, Emma Thomson and Alex Alderton, Roberto Amato, Sonia Goncalves, Ewan Harrison, David K. Jackson, Ian Johnston, Dominic Kwiatkowski, Cordelia Langford, John Sillitoe |
| EPI_ISL_531974, EPI_ISL_531975, EPI_ISL_531976 | Lighthouse Lab in Glasgow | Wellcome Sanger Institute for the COVID-19 Genomics UK Consortium | Harper VanSteenhouse, Yumi Kasai, David Gray, Carol Clugston, Anna Dominiczak and Alex Alderton, Roberto Amato, Sonia Goncalves, Ewan Harrison, David K. Jackson, Ian Johnston, Dominic Kwiatkowski, Cordelia Langford, John Sillitoe |
| EPI_ISL_531978, EPI_ISL_531979 | NHSGGC West of Scotland Specialist Virology Centre / MRC-University of Glasgow Centre for Virus Research | Wellcome Sanger Institute for the COVID-19 Genomics UK Consortium | Ana da Silva Filipe, Natasha Johnson, Kathy Smollett, Daniel Mair, Stephen Carmichael, Lily Tong, Jenna Nichols, Elihu Aranday-Cortes, Kirstyn Brunker, Yasmin Parr, Kyriaki Nomikou; Sarah McDonald, Marc Niebel, Patawee Asamaphan; Richard Orton, Joseph Hughes, Sreenu Vattipally, David L Robertson; Alasdair MacLean, Rory Gunson; Kathy Li, Natasha Jesudason, Rajiv Shah, James Shepherd, Antonia Ho, Alice Broos, Emma Thomson and Alex Alderton, Roberto Amato, Sonia Goncalves, Ewan Harrison, David K. Jackson, Ian Johnston, Dominic Kwiatkowski, Cordelia Langford, John Sillitoe |
| EPI_ISL_531980, EPI_ISL_531981 | Lighthouse Lab in Glasgow | Wellcome Sanger Institute for the COVID-19 Genomics UK Consortium | Harper VanSteenhouse, Yumi Kasai, David Gray, Carol Clugston, Anna Dominiczak and Alex Alderton, Roberto Amato, Sonia Goncalves, Ewan Harrison, David K. Jackson, Ian Johnston, Dominic Kwiatkowski, Cordelia Langford, John Sillitoe |
| EPI_ISL_531982 | NHSGGC West of Scotland Specialist Virology Centre / MRC-University of Glasgow Centre for Virus Research | Wellcome Sanger Institute for the COVID-19 Genomics UK Consortium | Ana da Silva Filipe, Natasha Johnson, Kathy Smollett, Daniel Mair, Stephen Carmichael, Lily Tong, Jenna Nichols, Elihu Aranday-Cortes, Kirstyn Brunker, Yasmin Parr, Kyriaki Nomikou; Sarah McDonald, Marc Niebel, Patawee Asamaphan; Richard Orton, Joseph Hughes, Sreenu Vattipally, David L Robertson; Alasdair MacLean, Rory Gunson; Kathy Li, Natasha Jesudason, Rajiv Shah, James Shepherd, Antonia Ho, Alice Broos, Emma Thomson and Alex Alderton, Roberto Amato, Sonia Goncalves, Ewan Harrison, David K. Jackson, Ian Johnston, Dominic Kwiatkowski, Cordelia Langford, John Sillitoe |
| EPI_ISL_531983 | Lighthouse Lab in Glasgow | Wellcome Sanger Institute for the COVID-19 Genomics UK Consortium | Harper VanSteenhouse, Yumi Kasai, David Gray, Carol Clugston, Anna Dominiczak and Alex Alderton, Roberto Amato, Sonia Goncalves, Ewan Harrison, David K. Jackson, Ian Johnston, Dominic Kwiatkowski, Cordelia Langford, John Sillitoe |
| EPI_ISL_531984 | NHSGGC West of Scotland Specialist Virology Centre / MRC-University of Glasgow Centre for Virus Research | Wellcome Sanger Institute for the COVID-19 Genomics UK Consortium | Ana da Silva Filipe, Natasha Johnson, Kathy Smollett, Daniel Mair, Stephen Carmichael, Lily Tong, Jenna Nichols, Elihu Aranday-Cortes, Kirstyn Brunker, Yasmin Parr, Kyriaki Nomikou; Sarah McDonald, Marc Niebel, Patawee Asamaphan; Richard Orton, Joseph Hughes, Sreenu Vattipally, David L Robertson; Alasdair MacLean, Rory Gunson; Kathy Li, Natasha Jesudason, Rajiv Shah, James Shepherd, Antonia Ho, Alice Broos, Emma Thomson and Alex Alderton, Roberto Amato, Sonia Goncalves, Ewan Harrison, David K. Jackson, Ian Johnston, Dominic Kwiatkowski, Cordelia Langford, John Sillitoe |
| EPI_ISL_531985, EPI_ISL_531986, EPI_ISL_531988, EPI_ISL_531989 | Lighthouse Lab in Glasgow | Wellcome Sanger Institute for the COVID-19 Genomics UK Consortium | Harper VanSteenhouse, Yumi Kasai, David Gray, Carol Clugston, Anna Dominiczak and Alex Alderton, Roberto Amato, Sonia Goncalves, Ewan Harrison, David K. Jackson, Ian Johnston, Dominic Kwiatkowski, Cordelia Langford, John Sillitoe |
| EPI_ISL_531990 | NHSGGC West of Scotland Specialist Virology Centre / MRC-University of Glasgow Centre for Virus Research | Wellcome Sanger Institute for the COVID-19 Genomics UK Consortium | Ana da Silva Filipe, Natasha Johnson, Kathy Smollett, Daniel Mair, Stephen Carmichael, Lily Tong, Jenna Nichols, Elihu Aranday-Cortes, Kirstyn Brunker, Yasmin Parr, Kyriaki Nomikou; Sarah McDonald, Marc Niebel, Patawee Asamaphan; Richard Orton, Joseph Hughes, Sreenu Vattipally, David L Robertson; Alasdair MacLean, Rory Gunson; Kathy Li, Natasha Jesudason, Rajiv Shah, James Shepherd, Antonia Ho, Alice Broos, Emma Thomson and Alex Alderton, Roberto Amato, Sonia Goncalves, Ewan Harrison, David K. Jackson, Ian Johnston, Dominic Kwiatkowski, Cordelia Langford, John Sillitoe |
| EPI_ISL_531991, EPI_ISL_531993, EPI_ISL_531994 | Lighthouse Lab in Glasgow | Wellcome Sanger Institute for the COVID-19 Genomics UK Consortium | Harper VanSteenhouse, Yumi Kasai, David Gray, Carol Clugston, Anna Dominiczak and Alex Alderton, Roberto Amato, Sonia Goncalves, Ewan Harrison, David K. Jackson, Ian Johnston, Dominic Kwiatkowski, Cordelia Langford, John Sillitoe |
| EPI_ISL_531995 | NHSGGC West of Scotland Specialist Virology Centre / MRC-University of Glasgow Centre for Virus Research | Wellcome Sanger Institute for the COVID-19 Genomics UK Consortium | Ana da Silva Filipe, Natasha Johnson, Kathy Smollett, Daniel Mair, Stephen Carmichael, Lily Tong, Jenna Nichols, Elihu Aranday-Cortes, Kirstyn Brunker, Yasmin Parr, Kyriaki Nomikou; Sarah McDonald, Marc Niebel, Patawee Asamaphan; Richard Orton, Joseph Hughes, Sreenu Vattipally, David L Robertson; Alasdair MacLean, Rory Gunson; Kathy Li, Natasha Jesudason, Rajiv Shah, James Shepherd, Antonia Ho, Alice Broos, Emma Thomson and Alex Alderton, Roberto Amato, Sonia Goncalves, Ewan Harrison, David K. Jackson, Ian Johnston, Dominic Kwiatkowski, Cordelia Langford, John Sillitoe |
| EPI_ISL_531996, EPI_ISL_531997, EPI_ISL_531998, EPI_ISL_531999, EPI_ISL_532001 | Lighthouse Lab in Glasgow | Wellcome Sanger Institute for the COVID-19 Genomics UK Consortium | Harper VanSteenhouse, Yumi Kasai, David Gray, Carol Clugston, Anna Dominiczak and Alex Alderton, Roberto Amato, Sonia Goncalves, Ewan Harrison, David K. Jackson, Ian Johnston, Dominic Kwiatkowski, Cordelia Langford, John Sillitoe |
| EPI_ISL_532003, EPI_ISL_532004, EPI_ISL_532005 | NHSGGC West of Scotland Specialist Virology Centre / MRC-University of Glasgow Centre for Virus Research | Wellcome Sanger Institute for the COVID-19 Genomics UK Consortium | Ana da Silva Filipe, Natasha Johnson, Kathy Smollett, Daniel Mair, Stephen Carmichael, Lily Tong, Jenna Nichols, Elihu Aranday-Cortes, Kirstyn Brunker, Yasmin Parr, Kyriaki Nomikou; Sarah McDonald, Marc Niebel, Patawee Asamaphan; Richard Orton, Joseph Hughes, Sreenu Vattipally, David L Robertson; Alasdair MacLean, Rory Gunson; Kathy Li, Natasha Jesudason, Rajiv Shah, James Shepherd, Antonia Ho, Alice Broos, Emma Thomson and Alex Alderton, Roberto Amato, Sonia Goncalves, Ewan Harrison, David K. Jackson, Ian Johnston, Dominic Kwiatkowski, Cordelia Langford, John Sillitoe |
| EPI_ISL_532006, EPI_ISL_532008, EPI_ISL_532009, EPI_ISL_532010, EPI_ISL_532011, EPI_ISL_532014, EPI_ISL_532015, EPI_ISL_532017 | Lighthouse Lab in Glasgow | Wellcome Sanger Institute for the COVID-19 Genomics UK Consortium | Harper VanSteenhouse, Yumi Kasai, David Gray, Carol Clugston, Anna Dominiczak and Alex Alderton, Roberto Amato, Sonia Goncalves, Ewan Harrison, David K. Jackson, Ian Johnston, Dominic Kwiatkowski, Cordelia Langford, John Sillitoe |
| EPI_ISL_532018 | NHSGGC West of Scotland Specialist Virology Centre / MRC-University of Glasgow Centre for Virus Research | Wellcome Sanger Institute for the COVID-19 Genomics UK Consortium | Ana da Silva Filipe, Natasha Johnson, Kathy Smollett, Daniel Mair, Stephen Carmichael, Lily Tong, Jenna Nichols, Elihu Aranday-Cortes, Kirstyn Brunker, Yasmin Parr, Kyriaki Nomikou; Sarah McDonald, Marc Niebel, Patawee Asamaphan; Richard Orton, Joseph Hughes, Sreenu Vattipally, David L Robertson; Alasdair MacLean, Rory Gunson; Kathy Li, Natasha Jesudason, Rajiv Shah, James Shepherd, Antonia Ho, Alice Broos, Emma Thomson and Alex Alderton, Roberto Amato, Sonia Goncalves, Ewan Harrison, David K. Jackson, Ian Johnston, Dominic Kwiatkowski, Cordelia Langford, John Sillitoe |
| EPI_ISL_532019, EPI_ISL_532020, EPI_ISL_532021, EPI_ISL_532022, EPI_ISL_532023 | Lighthouse Lab in Glasgow | Wellcome Sanger Institute for the COVID-19 Genomics UK Consortium | Harper VanSteenhouse, Yumi Kasai, David Gray, Carol Clugston, Anna Dominiczak and Alex Alderton, Roberto Amato, Sonia Goncalves, Ewan Harrison, David K. Jackson, Ian Johnston, Dominic Kwiatkowski, Cordelia Langford, John Sillitoe |
| EPI_ISL_532024 | NHSGGC West of Scotland Specialist Virology Centre / MRC-University of Glasgow Centre for Virus Research | Wellcome Sanger Institute for the COVID-19 Genomics UK Consortium | Ana da Silva Filipe, Natasha Johnson, Kathy Smollett, Daniel Mair, Stephen Carmichael, Lily Tong, Jenna Nichols, Elihu Aranday-Cortes, Kirstyn Brunker, Yasmin Parr, Kyriaki Nomikou; Sarah McDonald, Marc Niebel, Patawee Asamaphan; Richard Orton, Joseph Hughes, Sreenu Vattipally, David L Robertson; Alasdair MacLean, Rory Gunson; Kathy Li, Natasha Jesudason, Rajiv Shah, James Shepherd, Antonia Ho, Alice Broos, Emma Thomson and Alex Alderton, Roberto Amato, Sonia Goncalves, Ewan Harrison, David K. Jackson, Ian Johnston, Dominic Kwiatkowski, Cordelia Langford, John Sillitoe |
| EPI_ISL_532025, EPI_ISL_532026, EPI_ISL_532027, EPI_ISL_532028, EPI_ISL_532029, EPI_ISL_532030, EPI_ISL_532031, EPI_ISL_532033, EPI_ISL_532034, EPI_ISL_532035, EPI_ISL_532036, EPI_ISL_532037 | | | |
| see above | Lighthouse Lab in Glasgow | Wellcome Sanger Institute for the COVID-19 Genomics UK Consortium | Harper VanSteenhouse, Yumi Kasai, David Gray, Carol Clugston, Anna Dominiczak and Alex Alderton, Roberto Amato, Sonia Goncalves, Ewan Harrison, David K. Jackson, Ian Johnston, Dominic Kwiatkowski, Cordelia Langford, John Sillitoe |
| EPI_ISL_532038 | NHSGGC West of Scotland Specialist Virology Centre / MRC-University of Glasgow Centre for Virus Research | Wellcome Sanger Institute for the COVID-19 Genomics UK Consortium | Ana da Silva Filipe, Natasha Johnson, Kathy Smollett, Daniel Mair, Stephen Carmichael, Lily Tong, Jenna Nichols, Elihu Aranday-Cortes, Kirstyn Brunker, Yasmin Parr, Kyriaki Nomikou; Sarah McDonald, Marc Niebel, Patawee Asamaphan; Richard Orton, Joseph Hughes, Sreenu Vattipally, David L Robertson; Alasdair MacLean, Rory Gunson; Kathy Li, Natasha Jesudason, Rajiv Shah, James Shepherd, Antonia Ho, Alice Broos, Emma Thomson and Alex Alderton, Roberto |

| | | | |
|---|---|---|---|
| | | | Amato, Sonia Goncalves, Ewan Harrison, David K. Jackson, Ian Johnston, Dominic Kwiatkowski, Cordelia Langford, John Sillitoe |
| EPI_ISL_532039, EPI_ISL_532040, EPI_ISL_532041, EPI_ISL_532042, EPI_ISL_532043, EPI_ISL_532044, EPI_ISL_532045, EPI_ISL_532046, EPI_ISL_532047 | Lighthouse Lab in Glasgow | Wellcome Sanger Institute for the COVID-19 Genomics UK Consortium | Harper VanSteenhouse, Yumi Kasai, David Gray, Carol Clugston, Anna Dominiczak and Alex Alderton, Roberto Amato, Sonia Goncalves, Ewan Harrison, David K. Jackson, Ian Johnston, Dominic Kwiatkowski, Cordelia Langford, John Sillitoe |
| EPI_ISL_532049 | NHSGGC West of Scotland Specialist Virology Centre / MRC-University of Glasgow Centre for Virus Research | Wellcome Sanger Institute for the COVID-19 Genomics UK Consortium | Ana da Silva Filipe, Natasha Johnson, Kathy Smollett, Daniel Mair, Stephen Carmichael, Lily Tong, Jenna Nichols, Elihu Aranday-Cortes, Kirstyn Brunker, Yasmin Parr, Kyriaki Nomikou; Sarah McDonald, Marc Niebel, Patawee Asamaphan; Richard Orton, Joseph Hughes, Sreenu Vattipally, David L Robertson; Alasdair MacLean, Rory Gunson; Kathy Li, Natasha Jesudason, Rajiv Shah, James Shepherd, Antonia Ho, Alice Broos, Emma Thomson and Alex Alderton, Roberto Amato, Sonia Goncalves, Ewan Harrison, David K. Jackson, Ian Johnston, Dominic Kwiatkowski, Cordelia Langford, John Sillitoe |
| EPI_ISL_532050, EPI_ISL_532052 | Lighthouse Lab in Glasgow | Wellcome Sanger Institute for the COVID-19 Genomics UK Consortium | Harper VanSteenhouse, Yumi Kasai, David Gray, Carol Clugston, Anna Dominiczak and Alex Alderton, Roberto Amato, Sonia Goncalves, Ewan Harrison, David K. Jackson, Ian Johnston, Dominic Kwiatkowski, Cordelia Langford, John Sillitoe |
| EPI_ISL_532053 | NHSGGC West of Scotland Specialist Virology Centre / MRC-University of Glasgow Centre for Virus Research | Wellcome Sanger Institute for the COVID-19 Genomics UK Consortium | Ana da Silva Filipe, Natasha Johnson, Kathy Smollett, Daniel Mair, Stephen Carmichael, Lily Tong, Jenna Nichols, Elihu Aranday-Cortes, Kirstyn Brunker, Yasmin Parr, Kyriaki Nomikou; Sarah McDonald, Marc Niebel, Patawee Asamaphan; Richard Orton, Joseph Hughes, Sreenu Vattipally, David L Robertson; Alasdair MacLean, Rory Gunson; Kathy Li, Natasha Jesudason, Rajiv Shah, James Shepherd, Antonia Ho, Alice Broos, Emma Thomson and Alex Alderton, Roberto Amato, Sonia Goncalves, Ewan Harrison, David K. Jackson, Ian Johnston, Dominic Kwiatkowski, Cordelia Langford, John Sillitoe |
| EPI_ISL_532054 | Lighthouse Lab in Glasgow | Wellcome Sanger Institute for the COVID-19 Genomics UK Consortium | Harper VanSteenhouse, Yumi Kasai, David Gray, Carol Clugston, Anna Dominiczak and Alex Alderton, Roberto Amato, Sonia Goncalves, Ewan Harrison, David K. Jackson, Ian Johnston, Dominic Kwiatkowski, Cordelia Langford, John Sillitoe |
| EPI_ISL_532055 | NHSGGC West of Scotland Specialist Virology Centre / MRC-University of Glasgow Centre for Virus Research | Wellcome Sanger Institute for the COVID-19 Genomics UK Consortium | Ana da Silva Filipe, Natasha Johnson, Kathy Smollett, Daniel Mair, Stephen Carmichael, Lily Tong, Jenna Nichols, Elihu Aranday-Cortes, Kirstyn Brunker, Yasmin Parr, Kyriaki Nomikou; Sarah McDonald, Marc Niebel, Patawee Asamaphan; Richard Orton, Joseph Hughes, Sreenu Vattipally, David L Robertson; Alasdair MacLean, Rory Gunson; Kathy Li, Natasha Jesudason, Rajiv Shah, James Shepherd, Antonia Ho, Alice Broos, Emma Thomson and Alex Alderton, Roberto Amato, Sonia Goncalves, Ewan Harrison, David K. Jackson, Ian Johnston, Dominic Kwiatkowski, Cordelia Langford, John Sillitoe |
| EPI_ISL_532056 | Lighthouse Lab in Glasgow | Wellcome Sanger Institute for the COVID-19 Genomics UK Consortium | Harper VanSteenhouse, Yumi Kasai, David Gray, Carol Clugston, Anna Dominiczak and Alex Alderton, Roberto Amato, Sonia Goncalves, Ewan Harrison, David K. Jackson, Ian Johnston, Dominic Kwiatkowski, Cordelia Langford, John Sillitoe |
| EPI_ISL_532057 | NHSGGC West of Scotland Specialist Virology Centre / MRC-University of Glasgow Centre for Virus Research | Wellcome Sanger Institute for the COVID-19 Genomics UK Consortium | Ana da Silva Filipe, Natasha Johnson, Kathy Smollett, Daniel Mair, Stephen Carmichael, Lily Tong, Jenna Nichols, Elihu Aranday-Cortes, Kirstyn Brunker, Yasmin Parr, Kyriaki Nomikou; Sarah McDonald, Marc Niebel, Patawee Asamaphan; Richard Orton, Joseph Hughes, Sreenu Vattipally, David L Robertson; Alasdair MacLean, Rory Gunson; Kathy Li, Natasha Jesudason, Rajiv Shah, James Shepherd, Antonia Ho, Alice Broos, Emma Thomson and Alex Alderton, Roberto Amato, Sonia Goncalves, Ewan Harrison, David K. Jackson, Ian Johnston, Dominic Kwiatkowski, Cordelia Langford, John Sillitoe |
| EPI_ISL_532058, EPI_ISL_532059, EPI_ISL_532060 | Lighthouse Lab in Glasgow | Wellcome Sanger Institute for the COVID-19 Genomics UK Consortium | Harper VanSteenhouse, Yumi Kasai, David Gray, Carol Clugston, Anna Dominiczak and Alex Alderton, Roberto Amato, Sonia Goncalves, Ewan Harrison, David K. Jackson, Ian Johnston, Dominic Kwiatkowski, Cordelia Langford, John Sillitoe |
| EPI_ISL_532062 | NHSGGC West of Scotland Specialist Virology Centre / MRC-University of Glasgow Centre for Virus Research | Wellcome Sanger Institute for the COVID-19 Genomics UK Consortium | Ana da Silva Filipe, Natasha Johnson, Kathy Smollett, Daniel Mair, Stephen Carmichael, Lily Tong, Jenna Nichols, Elihu Aranday-Cortes, Kirstyn Brunker, Yasmin Parr, Kyriaki Nomikou; Sarah McDonald, Marc Niebel, Patawee Asamaphan; Richard Orton, Joseph Hughes, Sreenu Vattipally, David L Robertson; Alasdair MacLean, Rory Gunson; Kathy Li, Natasha Jesudason, Rajiv Shah, James Shepherd, Antonia Ho, Alice Broos, Emma Thomson and Alex Alderton, Roberto Amato, Sonia Goncalves, Ewan Harrison, David K. Jackson, Ian Johnston, Dominic Kwiatkowski, Cordelia Langford, John Sillitoe |
| EPI_ISL_532063, EPI_ISL_532064, EPI_ISL_532065, EPI_ISL_532066, EPI_ISL_532067, EPI_ISL_532068, EPI_ISL_532069 | Lighthouse Lab in Glasgow | Wellcome Sanger Institute for the COVID-19 Genomics UK Consortium | Harper VanSteenhouse, Yumi Kasai, David Gray, Carol Clugston, Anna Dominiczak and Alex Alderton, Roberto Amato, Sonia Goncalves, Ewan Harrison, David K. Jackson, Ian Johnston, Dominic Kwiatkowski, Cordelia Langford, John Sillitoe |
| EPI_ISL_532070 | NHSGGC West of Scotland Specialist Virology Centre / MRC-University of Glasgow Centre for Virus Research | Wellcome Sanger Institute for the COVID-19 Genomics UK Consortium | Ana da Silva Filipe, Natasha Johnson, Kathy Smollett, Daniel Mair, Stephen Carmichael, Lily Tong, Jenna Nichols, Elihu Aranday-Cortes, Kirstyn Brunker, Yasmin Parr, Kyriaki Nomikou; Sarah McDonald, Marc Niebel, Patawee Asamaphan; Richard Orton, Joseph Hughes, Sreenu Vattipally, David L Robertson; Alasdair MacLean, Rory Gunson; Kathy Li, Natasha Jesudason, Rajiv Shah, James Shepherd, Antonia Ho, Alice Broos, Emma Thomson and Alex Alderton, Roberto Amato, Sonia Goncalves, Ewan Harrison, David K. Jackson, Ian Johnston, Dominic Kwiatkowski, Cordelia Langford, John Sillitoe |
| EPI_ISL_532072, EPI_ISL_532073, EPI_ISL_532074, EPI_ISL_532075 | Lighthouse Lab in Glasgow | Wellcome Sanger Institute for the COVID-19 Genomics UK Consortium | Harper VanSteenhouse, Yumi Kasai, David Gray, Carol Clugston, Anna Dominiczak and Alex Alderton, Roberto Amato, Sonia Goncalves, Ewan Harrison, David K. Jackson, Ian Johnston, Dominic Kwiatkowski, Cordelia Langford, John Sillitoe |
| EPI_ISL_532076, EPI_ISL_532077, EPI_ISL_532078 | NHSGGC West of Scotland Specialist Virology Centre / MRC-University of Glasgow Centre for Virus Research | Wellcome Sanger Institute for the COVID-19 Genomics UK Consortium | Ana da Silva Filipe, Natasha Johnson, Kathy Smollett, Daniel Mair, Stephen Carmichael, Lily Tong, Jenna Nichols, Elihu Aranday-Cortes, Kirstyn Brunker, Yasmin Parr, Kyriaki Nomikou; Sarah McDonald, Marc Niebel, Patawee Asamaphan; Richard Orton, Joseph Hughes, Sreenu Vattipally, David L Robertson; Alasdair MacLean, Rory Gunson; Kathy Li, Natasha Jesudason, Rajiv Shah, James Shepherd, Antonia Ho, Alice Broos, Emma Thomson and Alex Alderton, Roberto Amato, Sonia Goncalves, Ewan Harrison, David K. Jackson, Ian Johnston, Dominic Kwiatkowski, Cordelia Langford, John Sillitoe |
| EPI_ISL_532079 | Lighthouse Lab in Glasgow | Wellcome Sanger Institute for the COVID-19 Genomics UK Consortium | Harper VanSteenhouse, Yumi Kasai, David Gray, Carol Clugston, Anna Dominiczak and Alex Alderton, Roberto Amato, Sonia Goncalves, Ewan Harrison, David K. Jackson, Ian Johnston, Dominic Kwiatkowski, Cordelia Langford, John Sillitoe |
| EPI_ISL_532080, EPI_ISL_532081 | NHSGGC West of Scotland Specialist Virology Centre / MRC-University of Glasgow Centre for Virus Research | Wellcome Sanger Institute for the COVID-19 Genomics UK Consortium | Ana da Silva Filipe, Natasha Johnson, Kathy Smollett, Daniel Mair, Stephen Carmichael, Lily Tong, Jenna Nichols, Elihu Aranday-Cortes, Kirstyn Brunker, Yasmin Parr, Kyriaki Nomikou; Sarah McDonald, Marc Niebel, Patawee Asamaphan; Richard Orton, Joseph Hughes, Sreenu Vattipally, David L Robertson; Alasdair MacLean, Rory Gunson; Kathy Li, Natasha Jesudason, Rajiv Shah, James Shepherd, Antonia Ho, Alice Broos, Emma Thomson and Alex Alderton, Roberto Amato, Sonia Goncalves, Ewan Harrison, David K. Jackson, Ian Johnston, Dominic Kwiatkowski, Cordelia Langford, John Sillitoe |
| EPI_ISL_532082, EPI_ISL_532083, EPI_ISL_532084, EPI_ISL_532086, EPI_ISL_532087, EPI_ISL_532088, EPI_ISL_532089, EPI_ISL_532090, EPI_ISL_532091, EPI_ISL_532092 | Lighthouse Lab in Glasgow | Wellcome Sanger Institute for the COVID-19 Genomics UK Consortium | Harper VanSteenhouse, Yumi Kasai, David Gray, Carol Clugston, Anna Dominiczak and Alex Alderton, Roberto Amato, Sonia Goncalves, Ewan Harrison, David K. Jackson, Ian Johnston, Dominic Kwiatkowski, Cordelia Langford, John Sillitoe |
| EPI_ISL_532093, EPI_ISL_532094 | NHSGGC West of Scotland Specialist Virology Centre / MRC-University of Glasgow Centre for Virus Research | Wellcome Sanger Institute for the COVID-19 Genomics UK Consortium | Ana da Silva Filipe, Natasha Johnson, Kathy Smollett, Daniel Mair, Stephen Carmichael, Lily Tong, Jenna Nichols, Elihu Aranday-Cortes, Kirstyn Brunker, Yasmin Parr, Kyriaki Nomikou; Sarah McDonald, Marc Niebel, Patawee Asamaphan; Richard Orton, Joseph Hughes, Sreenu Vattipally, David L Robertson; Alasdair MacLean, Rory Gunson; Kathy Li, Natasha Jesudason, Rajiv Shah, James Shepherd, Antonia Ho, Alice Broos, Emma Thomson and Alex Alderton, Roberto Amato, Sonia Goncalves, Ewan Harrison, David K. Jackson, Ian Johnston, Dominic Kwiatkowski, Cordelia Langford, John Sillitoe |
| EPI_ISL_532095, EPI_ISL_532096, EPI_ISL_532097 | Lighthouse Lab in Glasgow | Wellcome Sanger Institute for the COVID-19 Genomics UK Consortium | Harper VanSteenhouse, Yumi Kasai, David Gray, Carol Clugston, Anna Dominiczak and Alex Alderton, Roberto Amato, Sonia Goncalves, Ewan Harrison, David K. Jackson, Ian Johnston, Dominic Kwiatkowski, Cordelia Langford, John Sillitoe |
| EPI_ISL_532098 | NHSGGC West of Scotland Specialist Virology Centre / MRC-University of Glasgow Centre for Virus Research | Wellcome Sanger Institute for the COVID-19 Genomics UK Consortium | Ana da Silva Filipe, Natasha Johnson, Kathy Smollett, Daniel Mair, Stephen Carmichael, Lily Tong, Jenna Nichols, Elihu Aranday-Cortes, Kirstyn Brunker, Yasmin Parr, Kyriaki Nomikou; Sarah McDonald, Marc Niebel, Patawee Asamaphan; Richard Orton, Joseph Hughes, Sreenu Vattipally, David L Robertson; Alasdair MacLean, Rory Gunson; Kathy Li, Natasha Jesudason, Rajiv Shah, James Shepherd, Antonia Ho, Alice Broos, Emma Thomson and Alex Alderton, Roberto Amato, Sonia Goncalves, Ewan Harrison, David K. Jackson, Ian Johnston, Dominic Kwiatkowski, Cordelia Langford, John Sillitoe |
| EPI_ISL_532099, EPI_ISL_532101 | Lighthouse Lab in Glasgow | Wellcome Sanger Institute for the COVID-19 Genomics UK Consortium | Harper VanSteenhouse, Yumi Kasai, David Gray, Carol Clugston, Anna Dominiczak and Alex Alderton, Roberto Amato, Sonia Goncalves, Ewan Harrison, David K. Jackson, Ian Johnston, Dominic Kwiatkowski, Cordelia Langford, John Sillitoe |
| EPI_ISL_532102 | NHSGGC West of Scotland Specialist Virology Centre / MRC-University of Glasgow Centre for Virus Research | Wellcome Sanger Institute for the COVID-19 Genomics UK Consortium | Ana da Silva Filipe, Natasha Johnson, Kathy Smollett, Daniel Mair, Stephen Carmichael, Lily Tong, Jenna Nichols, Elihu Aranday-Cortes, Kirstyn Brunker, Yasmin Parr, Kyriaki Nomikou; Sarah McDonald, Marc Niebel, Patawee Asamaphan; Richard Orton, Joseph Hughes, Sreenu Vattipally, David L Robertson; Alasdair MacLean, Rory Gunson; Kathy Li, Natasha Jesudason, Rajiv Shah, James Shepherd, Antonia Ho, Alice Broos, Emma Thomson and Alex Alderton, Roberto Amato, Sonia Goncalves, Ewan Harrison, David K. Jackson, Ian Johnston, Dominic Kwiatkowski, Cordelia Langford, John Sillitoe |
| EPI_ISL_532103, EPI_ISL_532104 | Lighthouse Lab in Glasgow | Wellcome Sanger Institute for the COVID-19 Genomics UK Consortium | Harper VanSteenhouse, Yumi Kasai, David Gray, Carol Clugston, Anna Dominiczak and Alex Alderton, Roberto Amato, Sonia Goncalves, Ewan Harrison, David K. Jackson, Ian Johnston, Dominic Kwiatkowski, Cordelia Langford, John Sillitoe |
| EPI_ISL_532105 | NHSGGC West of Scotland Specialist Virology Centre / MRC-University of Glasgow Centre for Virus Research | Wellcome Sanger Institute for the COVID-19 Genomics UK Consortium | Ana da Silva Filipe, Natasha Johnson, Kathy Smollett, Daniel Mair, Stephen Carmichael, Lily Tong, Jenna Nichols, Elihu Aranday-Cortes, Kirstyn Brunker, Yasmin Parr, Kyriaki Nomikou; Sarah McDonald, Marc Niebel, Patawee Asamaphan; Richard Orton, Joseph Hughes, Sreenu Vattipally, David L Robertson; Alasdair MacLean, Rory Gunson; Kathy Li, Natasha Jesudason, Rajiv Shah, James Shepherd, Antonia Ho, Alice Broos, Emma Thomson and Alex Alderton, Roberto Amato, Sonia Goncalves, Ewan Harrison, David K. Jackson, Ian Johnston, Dominic Kwiatkowski, Cordelia Langford, John Sillitoe |
| EPI_ISL_532106, EPI_ISL_532107, EPI_ISL_532109, EPI_ISL_532110, EPI_ISL_532111, EPI_ISL_532112, EPI_ISL_532113, EPI_ISL_532114, EPI_ISL_532115, EPI_ISL_532116, EPI_ISL_532117 | | | |
| see above | Lighthouse Lab in Glasgow | Wellcome Sanger Institute for the COVID-19 Genomics UK Consortium | Harper VanSteenhouse, Yumi Kasai, David Gray, Carol Clugston, Anna Dominiczak and Alex Alderton, Roberto Amato, Sonia Goncalves, Ewan Harrison, David K. Jackson, Ian Johnston, Dominic Kwiatkowski, Cordelia Langford, John Sillitoe |
| EPI_ISL_532118 | NHSGGC West of Scotland Specialist Virology Centre / MRC-University of Glasgow Centre for Virus Research | Wellcome Sanger Institute for the COVID-19 Genomics UK Consortium | Ana da Silva Filipe, Natasha Johnson, Kathy Smollett, Daniel Mair, Stephen Carmichael, Lily Tong, Jenna Nichols, Elihu Aranday-Cortes, Kirstyn Brunker, Yasmin Parr, Kyriaki Nomikou; Sarah McDonald, Marc Niebel, Patawee Asamaphan; Richard Orton, Joseph Hughes, Sreenu Vattipally, David L Robertson; Alasdair MacLean, Rory Gunson; Kathy Li, Natasha Jesudason, Rajiv Shah, James Shepherd, Antonia Ho, Alice Broos, Emma Thomson and Alex Alderton, Roberto Amato, Sonia Goncalves, Ewan Harrison, David K. Jackson, Ian Johnston, Dominic Kwiatkowski, Cordelia Langford, John Sillitoe |
| EPI_ISL_532119, EPI_ISL_532120, EPI_ISL_532121, EPI_ISL_532122 | Lighthouse Lab in Glasgow | Wellcome Sanger Institute for the COVID-19 Genomics UK Consortium | Harper VanSteenhouse, Yumi Kasai, David Gray, Carol Clugston, Anna Dominiczak and Alex Alderton, Roberto Amato, Sonia Goncalves, Ewan Harrison, David K. Jackson, Ian Johnston, Dominic Kwiatkowski, Cordelia Langford, John Sillitoe |
| EPI_ISL_532123 | NHSGGC West of Scotland Specialist Virology Centre / MRC-University of Glasgow Centre for Virus Research | Wellcome Sanger Institute for the COVID-19 Genomics UK Consortium | Ana da Silva Filipe, Natasha Johnson, Kathy Smollett, Daniel Mair, Stephen Carmichael, Lily Tong, Jenna Nichols, Elihu Aranday-Cortes, Kirstyn Brunker, Yasmin Parr, Kyriaki Nomikou; Sarah McDonald, Marc Niebel, Patawee Asamaphan; Richard Orton, Joseph Hughes, Sreenu Vattipally, David L Robertson; Alasdair MacLean, Rory Gunson; Kathy Li, Natasha Jesudason, Rajiv Shah, James Shepherd, Antonia Ho, Alice Broos, Emma Thomson and Alex Alderton, Roberto Amato, Sonia Goncalves, Ewan Harrison, David K. Jackson, Ian Johnston, Dominic Kwiatkowski, Cordelia Langford, John Sillitoe |
| EPI_ISL_532124 | Lighthouse Lab in Glasgow | Wellcome Sanger Institute for the COVID-19 Genomics UK Consortium | Harper VanSteenhouse, Yumi Kasai, David Gray, Carol Clugston, Anna Dominiczak and Alex Alderton, Roberto Amato, Sonia Goncalves, Ewan Harrison, David K. Jackson, Ian Johnston, Dominic Kwiatkowski, Cordelia Langford, John Sillitoe |
| EPI_ISL_532125 | NHSGGC West of Scotland Specialist Virology Centre / MRC-University of Glasgow Centre for Virus Research | Wellcome Sanger Institute for the COVID-19 Genomics UK Consortium | Ana da Silva Filipe, Natasha Johnson, Kathy Smollett, Daniel Mair, Stephen Carmichael, Lily Tong, Jenna Nichols, Elihu Aranday-Cortes, Kirstyn Brunker, Yasmin Parr, Kyriaki Nomikou; Sarah McDonald, Marc Niebel, Patawee Asamaphan; Richard Orton, Joseph Hughes, Sreenu Vattipally, David L Robertson; Alasdair MacLean, Rory Gunson; Kathy Li, Natasha Jesudason, Rajiv Shah, James Shepherd, Antonia Ho, Alice Broos, Emma Thomson and Alex Alderton, Roberto Amato, Sonia Goncalves, Ewan Harrison, David K. Jackson, Ian Johnston, Dominic Kwiatkowski, Cordelia Langford, John Sillitoe |
| EPI_ISL_532126, EPI_ISL_532127, EPI_ISL_532128 | Lighthouse Lab in Glasgow | Wellcome Sanger Institute for the COVID-19 Genomics UK Consortium | Harper VanSteenhouse, Yumi Kasai, David Gray, Carol Clugston, Anna Dominiczak and Alex Alderton, Roberto Amato, Sonia Goncalves, Ewan Harrison, David K. Jackson, Ian Johnston, Dominic Kwiatkowski, Cordelia Langford, John Sillitoe |
| EPI_ISL_532129, EPI_ISL_532130, EPI_ISL_532131 | NHSGGC West of Scotland Specialist Virology Centre / MRC-University of Glasgow Centre for Virus Research | Wellcome Sanger Institute for the COVID-19 Genomics UK Consortium | Ana da Silva Filipe, Natasha Johnson, Kathy Smollett, Daniel Mair, Stephen Carmichael, Lily Tong, Jenna Nichols, Elihu Aranday-Cortes, Kirstyn Brunker, Yasmin Parr, Kyriaki Nomikou; Sarah McDonald, Marc Niebel, Patawee Asamaphan; Richard Orton, Joseph Hughes, Sreenu Vattipally, David L Robertson; Alasdair MacLean, Rory Gunson; Kathy Li, Natasha Jesudason, Rajiv Shah, James Shepherd, Antonia Ho, Alice Broos, Emma Thomson and Alex Alderton, Roberto Amato, Sonia Goncalves, Ewan Harrison, David K. Jackson, Ian Johnston, Dominic Kwiatkowski, Cordelia Langford, John Sillitoe |
| EPI_ISL_532132, EPI_ISL_532133, EPI_ISL_532134, EPI_ISL_532135, EPI_ISL_532136, EPI_ISL_532137 | Lighthouse Lab in Glasgow | Wellcome Sanger Institute for the COVID-19 Genomics UK Consortium | Harper VanSteenhouse, Yumi Kasai, David Gray, Carol Clugston, Anna Dominiczak and Alex Alderton, Roberto Amato, Sonia Goncalves, Ewan Harrison, David K. Jackson, Ian Johnston, Dominic Kwiatkowski, Cordelia Langford, John Sillitoe |
| EPI_ISL_532138 | NHSGGC West of Scotland Specialist Virology Centre / MRC-University of Glasgow Centre for Virus Research | Wellcome Sanger Institute for the COVID-19 Genomics UK Consortium | Ana da Silva Filipe, Natasha Johnson, Kathy Smollett, Daniel Mair, Stephen Carmichael, Lily Tong, Jenna Nichols, Elihu Aranday-Cortes, Kirstyn Brunker, Yasmin Parr, Kyriaki Nomikou; Sarah McDonald, Marc Niebel, Patawee Asamaphan; Richard Orton, Joseph Hughes, Sreenu Vattipally, David L Robertson; Alasdair MacLean, Rory Gunson; Kathy Li, Natasha Jesudason, Rajiv Shah, James Shepherd, Antonia Ho, Alice Broos, Emma Thomson and Alex Alderton, Roberto Amato, Sonia Goncalves, Ewan Harrison, David K. Jackson, Ian Johnston, Dominic Kwiatkowski, Cordelia Langford, John Sillitoe |
| EPI_ISL_532140, EPI_ISL_532141, EPI_ISL_532143, EPI_ISL_532144, EPI_ISL_532146, EPI_ISL_532147, EPI_ISL_532148, EPI_ISL_532150, EPI_ISL_532151, EPI_ISL_532152, EPI_ISL_532156, EPI_ISL_532157, EPI_ISL_532158, EPI_ISL_532159, EPI_ISL_532160, EPI_ISL_532162, EPI_ISL_532164, EPI_ISL_532165, EPI_ISL_532166, EPI_ISL_532170, EPI_ISL_532171, EPI_ISL_532172, EPI_ISL_532173, EPI_ISL_532176 | | | |
| see above | Lighthouse Lab in Glasgow | Wellcome Sanger Institute for the | Harper VanSteenhouse, Yumi Kasai, David Gray, Carol Clugston, Anna Dominiczak and Alex Alderton, Roberto Amato, Sonia Goncalves, Ewan Harrison, David K. Jackson, Ian Johnston, Dominic Kwiatkowski, Cordelia Langford, John Sillitoe |

| | | | |
|---|---|---|---|
| | | COVID-19 Genomics UK Consortium | |
| EPI_ISL_532178 | NHSGGC West of Scotland Specialist Virology Centre / MRC-University of Glasgow Centre for Virus Research | Wellcome Sanger Institute for the COVID-19 Genomics UK Consortium | Ana da Silva Filipe, Natasha Johnson, Kathy Smollett, Daniel Mair, Stephen Carmichael, Lily Tong, Jenna Nichols, Elihu Aranday-Cortes, Kirstyn Brunker, Yasmin Parr, Kyriaki Nomikou; Sarah McDonald, Marc Niebel, Patawee Asamaphan; Richard Orton, Joseph Hughes, Sreenu Vattipally, David L Robertson; Alasdair MacLean, Rory Gunson; Kathy Li, Natasha Jesudason, Rajiv Shah, James Shepherd, Antonia Ho, Alice Broos, Emma Thomson and Alex Alderton, Roberto Amato, Sonia Goncalves, Ewan Harrison, David K. Jackson, Ian Johnston, Dominic Kwiatkowski, Cordelia Langford, John Sillitoe |
| EPI_ISL_532180, EPI_ISL_532181, EPI_ISL_532183, EPI_ISL_532184, EPI_ISL_532185, EPI_ISL_532188, EPI_ISL_532190, EPI_ISL_532191 | Lighthouse Lab in Glasgow | Wellcome Sanger Institute for the COVID-19 Genomics UK Consortium | Harper VanSteenhouse, Yumi Kasai, David Gray, Carol Clugston, Anna Dominiczak and Alex Alderton, Roberto Amato, Sonia Goncalves, Ewan Harrison, David K. Jackson, Ian Johnston, Dominic Kwiatkowski, Cordelia Langford, John Sillitoe |
| EPI_ISL_532192 | NHSGGC West of Scotland Specialist Virology Centre / MRC-University of Glasgow Centre for Virus Research | Wellcome Sanger Institute for the COVID-19 Genomics UK Consortium | Ana da Silva Filipe, Natasha Johnson, Kathy Smollett, Daniel Mair, Stephen Carmichael, Lily Tong, Jenna Nichols, Elihu Aranday-Cortes, Kirstyn Brunker, Yasmin Parr, Kyriaki Nomikou; Sarah McDonald, Marc Niebel, Patawee Asamaphan; Richard Orton, Joseph Hughes, Sreenu Vattipally, David L Robertson; Alasdair MacLean, Rory Gunson; Kathy Li, Natasha Jesudason, Rajiv Shah, James Shepherd, Antonia Ho, Alice Broos, Emma Thomson and Alex Alderton, Roberto Amato, Sonia Goncalves, Ewan Harrison, David K. Jackson, Ian Johnston, Dominic Kwiatkowski, Cordelia Langford, John Sillitoe |
| EPI_ISL_532193 | Lighthouse Lab in Glasgow | Wellcome Sanger Institute for the COVID-19 Genomics UK Consortium | Harper VanSteenhouse, Yumi Kasai, David Gray, Carol Clugston, Anna Dominiczak and Alex Alderton, Roberto Amato, Sonia Goncalves, Ewan Harrison, David K. Jackson, Ian Johnston, Dominic Kwiatkowski, Cordelia Langford, John Sillitoe |
| EPI_ISL_532195, EPI_ISL_532196 | NHSGGC West of Scotland Specialist Virology Centre / MRC-University of Glasgow Centre for Virus Research | Wellcome Sanger Institute for the COVID-19 Genomics UK Consortium | Ana da Silva Filipe, Natasha Johnson, Kathy Smollett, Daniel Mair, Stephen Carmichael, Lily Tong, Jenna Nichols, Elihu Aranday-Cortes, Kirstyn Brunker, Yasmin Parr, Kyriaki Nomikou; Sarah McDonald, Marc Niebel, Patawee Asamaphan; Richard Orton, Joseph Hughes, Sreenu Vattipally, David L Robertson; Alasdair MacLean, Rory Gunson; Kathy Li, Natasha Jesudason, Rajiv Shah, James Shepherd, Antonia Ho, Alice Broos, Emma Thomson and Alex Alderton, Roberto Amato, Sonia Goncalves, Ewan Harrison, David K. Jackson, Ian Johnston, Dominic Kwiatkowski, Cordelia Langford, John Sillitoe |
| EPI_ISL_532198, EPI_ISL_532199 | Lighthouse Lab in Glasgow | Wellcome Sanger Institute for the COVID-19 Genomics UK Consortium | Harper VanSteenhouse, Yumi Kasai, David Gray, Carol Clugston, Anna Dominiczak and Alex Alderton, Roberto Amato, Sonia Goncalves, Ewan Harrison, David K. Jackson, Ian Johnston, Dominic Kwiatkowski, Cordelia Langford, John Sillitoe |
| EPI_ISL_532200 | NHSGGC West of Scotland Specialist Virology Centre / MRC-University of Glasgow Centre for Virus Research | Wellcome Sanger Institute for the COVID-19 Genomics UK Consortium | Ana da Silva Filipe, Natasha Johnson, Kathy Smollett, Daniel Mair, Stephen Carmichael, Lily Tong, Jenna Nichols, Elihu Aranday-Cortes, Kirstyn Brunker, Yasmin Parr, Kyriaki Nomikou; Sarah McDonald, Marc Niebel, Patawee Asamaphan; Richard Orton, Joseph Hughes, Sreenu Vattipally, David L Robertson; Alasdair MacLean, Rory Gunson; Kathy Li, Natasha Jesudason, Rajiv Shah, James Shepherd, Antonia Ho, Alice Broos, Emma Thomson and Alex Alderton, Roberto Amato, Sonia Goncalves, Ewan Harrison, David K. Jackson, Ian Johnston, Dominic Kwiatkowski, Cordelia Langford, John Sillitoe |
| EPI_ISL_532201 | Lighthouse Lab in Glasgow | Wellcome Sanger Institute for the COVID-19 Genomics UK Consortium | Harper VanSteenhouse, Yumi Kasai, David Gray, Carol Clugston, Anna Dominiczak and Alex Alderton, Roberto Amato, Sonia Goncalves, Ewan Harrison, David K. Jackson, Ian Johnston, Dominic Kwiatkowski, Cordelia Langford, John Sillitoe |
| EPI_ISL_532202, EPI_ISL_532203 | NHSGGC West of Scotland Specialist Virology Centre / MRC-University of Glasgow Centre for Virus Research | Wellcome Sanger Institute for the COVID-19 Genomics UK Consortium | Ana da Silva Filipe, Natasha Johnson, Kathy Smollett, Daniel Mair, Stephen Carmichael, Lily Tong, Jenna Nichols, Elihu Aranday-Cortes, Kirstyn Brunker, Yasmin Parr, Kyriaki Nomikou; Sarah McDonald, Marc Niebel, Patawee Asamaphan; Richard Orton, Joseph Hughes, Sreenu Vattipally, David L Robertson; Alasdair MacLean, Rory Gunson; Kathy Li, Natasha Jesudason, Rajiv Shah, James Shepherd, Antonia Ho, Alice Broos, Emma Thomson and Alex Alderton, Roberto Amato, Sonia Goncalves, Ewan Harrison, David K. Jackson, Ian Johnston, Dominic Kwiatkowski, Cordelia Langford, John Sillitoe |
| EPI_ISL_532204, EPI_ISL_532205, EPI_ISL_532206, EPI_ISL_532209 | Lighthouse Lab in Glasgow | Wellcome Sanger Institute for the COVID-19 Genomics UK Consortium | Harper VanSteenhouse, Yumi Kasai, David Gray, Carol Clugston, Anna Dominiczak and Alex Alderton, Roberto Amato, Sonia Goncalves, Ewan Harrison, David K. Jackson, Ian Johnston, Dominic Kwiatkowski, Cordelia Langford, John Sillitoe |
| EPI_ISL_532210 | NHSGGC West of Scotland Specialist Virology Centre / MRC-University of Glasgow Centre for Virus Research | Wellcome Sanger Institute for the COVID-19 Genomics UK Consortium | Ana da Silva Filipe, Natasha Johnson, Kathy Smollett, Daniel Mair, Stephen Carmichael, Lily Tong, Jenna Nichols, Elihu Aranday-Cortes, Kirstyn Brunker, Yasmin Parr, Kyriaki Nomikou; Sarah McDonald, Marc Niebel, Patawee Asamaphan; Richard Orton, Joseph Hughes, Sreenu Vattipally, David L Robertson; Alasdair MacLean, Rory Gunson; Kathy Li, Natasha Jesudason, Rajiv Shah, James Shepherd, Antonia Ho, Alice Broos, Emma Thomson and Alex Alderton, Roberto Amato, Sonia Goncalves, Ewan Harrison, David K. Jackson, Ian Johnston, Dominic Kwiatkowski, Cordelia Langford, John Sillitoe |
| EPI_ISL_532212 | Lighthouse Lab in Glasgow | Wellcome Sanger Institute for the COVID-19 Genomics UK Consortium | Harper VanSteenhouse, Yumi Kasai, David Gray, Carol Clugston, Anna Dominiczak and Alex Alderton, Roberto Amato, Sonia Goncalves, Ewan Harrison, David K. Jackson, Ian Johnston, Dominic Kwiatkowski, Cordelia Langford, John Sillitoe |
| EPI_ISL_532214 | NHSGGC West of Scotland Specialist Virology Centre / MRC-University of Glasgow Centre for Virus Research | Wellcome Sanger Institute for the COVID-19 Genomics UK Consortium | Ana da Silva Filipe, Natasha Johnson, Kathy Smollett, Daniel Mair, Stephen Carmichael, Lily Tong, Jenna Nichols, Elihu Aranday-Cortes, Kirstyn Brunker, Yasmin Parr, Kyriaki Nomikou; Sarah McDonald, Marc Niebel, Patawee Asamaphan; Richard Orton, Joseph Hughes, Sreenu Vattipally, David L Robertson; Alasdair MacLean, Rory Gunson; Kathy Li, Natasha Jesudason, Rajiv Shah, James Shepherd, Antonia Ho, Alice Broos, Emma Thomson and Alex Alderton, Roberto Amato, Sonia Goncalves, Ewan Harrison, David K. Jackson, Ian Johnston, Dominic Kwiatkowski, Cordelia Langford, John Sillitoe |
| EPI_ISL_532216, EPI_ISL_532217, EPI_ISL_532220, EPI_ISL_532222, EPI_ISL_532223, EPI_ISL_532224, EPI_ISL_532226, EPI_ISL_532227, EPI_ISL_532228, EPI_ISL_532230, EPI_ISL_532231, EPI_ISL_532233, EPI_ISL_532234, EPI_ISL_532236, EPI_ISL_532237, EPI_ISL_532238, EPI_ISL_532239 | | | |
| see above | Lighthouse Lab in Glasgow | Wellcome Sanger Institute for the COVID-19 Genomics UK Consortium | Harper VanSteenhouse, Yumi Kasai, David Gray, Carol Clugston, Anna Dominiczak and Alex Alderton, Roberto Amato, Sonia Goncalves, Ewan Harrison, David K. Jackson, Ian Johnston, Dominic Kwiatkowski, Cordelia Langford, John Sillitoe |
| EPI_ISL_532241 | NHSGGC West of Scotland Specialist Virology Centre / MRC-University of Glasgow Centre for Virus Research | Wellcome Sanger Institute for the COVID-19 Genomics UK Consortium | Ana da Silva Filipe, Natasha Johnson, Kathy Smollett, Daniel Mair, Stephen Carmichael, Lily Tong, Jenna Nichols, Elihu Aranday-Cortes, Kirstyn Brunker, Yasmin Parr, Kyriaki Nomikou; Sarah McDonald, Marc Niebel, Patawee Asamaphan; Richard Orton, Joseph Hughes, Sreenu Vattipally, David L Robertson; Alasdair MacLean, Rory Gunson; Kathy Li, Natasha Jesudason, Rajiv Shah, James Shepherd, Antonia Ho, Alice Broos, Emma Thomson and Alex Alderton, Roberto Amato, Sonia Goncalves, Ewan Harrison, David K. Jackson, Ian Johnston, Dominic Kwiatkowski, Cordelia Langford, John Sillitoe |
| EPI_ISL_532243, EPI_ISL_532245, EPI_ISL_532247, EPI_ISL_532248, EPI_ISL_532251, EPI_ISL_532252, EPI_ISL_532253, EPI_ISL_532254, EPI_ISL_532255, EPI_ISL_532256, EPI_ISL_532257 | | | |
| see above | Lighthouse Lab in Glasgow | Wellcome Sanger Institute for the COVID-19 Genomics UK Consortium | Harper VanSteenhouse, Yumi Kasai, David Gray, Carol Clugston, Anna Dominiczak and Alex Alderton, Roberto Amato, Sonia Goncalves, Ewan Harrison, David K. Jackson, Ian Johnston, Dominic Kwiatkowski, Cordelia Langford, John Sillitoe |
| EPI_ISL_532258 | NHSGGC West of Scotland Specialist Virology Centre / MRC-University of Glasgow Centre for Virus Research | Wellcome Sanger Institute for the COVID-19 Genomics UK Consortium | Ana da Silva Filipe, Natasha Johnson, Kathy Smollett, Daniel Mair, Stephen Carmichael, Lily Tong, Jenna Nichols, Elihu Aranday-Cortes, Kirstyn Brunker, Yasmin Parr, Kyriaki Nomikou; Sarah McDonald, Marc Niebel, Patawee Asamaphan; Richard Orton, Joseph Hughes, Sreenu Vattipally, David L Robertson; Alasdair MacLean, Rory Gunson; Kathy Li, Natasha Jesudason, Rajiv Shah, James Shepherd, Antonia Ho, Alice Broos, Emma Thomson and Alex Alderton, Roberto Amato, Sonia Goncalves, Ewan Harrison, David K. Jackson, Ian Johnston, Dominic Kwiatkowski, Cordelia Langford, John Sillitoe |
| EPI_ISL_532260, EPI_ISL_532261, EPI_ISL_532262 | Lighthouse Lab in Glasgow | Wellcome Sanger Institute for the COVID-19 Genomics UK Consortium | Harper VanSteenhouse, Yumi Kasai, David Gray, Carol Clugston, Anna Dominiczak and Alex Alderton, Roberto Amato, Sonia Goncalves, Ewan Harrison, David K. Jackson, Ian Johnston, Dominic Kwiatkowski, Cordelia Langford, John Sillitoe |
| EPI_ISL_532263 | NHSGGC West of Scotland Specialist Virology Centre / MRC-University of Glasgow Centre for Virus Research | Wellcome Sanger Institute for the COVID-19 Genomics UK Consortium | Ana da Silva Filipe, Natasha Johnson, Kathy Smollett, Daniel Mair, Stephen Carmichael, Lily Tong, Jenna Nichols, Elihu Aranday-Cortes, Kirstyn Brunker, Yasmin Parr, Kyriaki Nomikou; Sarah McDonald, Marc Niebel, Patawee Asamaphan; Richard Orton, Joseph Hughes, Sreenu Vattipally, David L Robertson; Alasdair MacLean, Rory Gunson; Kathy Li, Natasha Jesudason, Rajiv Shah, James Shepherd, Antonia Ho, Alice Broos, Emma Thomson and Alex Alderton, Roberto Amato, Sonia Goncalves, Ewan Harrison, David K. Jackson, Ian Johnston, Dominic Kwiatkowski, Cordelia Langford, John Sillitoe |
| EPI_ISL_532265, EPI_ISL_532266, EPI_ISL_532267, EPI_ISL_532268, EPI_ISL_532269, EPI_ISL_532270 | Lighthouse Lab in Glasgow | Wellcome Sanger Institute for the COVID-19 Genomics UK Consortium | Harper VanSteenhouse, Yumi Kasai, David Gray, Carol Clugston, Anna Dominiczak and Alex Alderton, Roberto Amato, Sonia Goncalves, Ewan Harrison, David K. Jackson, Ian Johnston, Dominic Kwiatkowski, Cordelia Langford, John Sillitoe |
| EPI_ISL_532271, EPI_ISL_532272 | NHSGGC West of Scotland Specialist Virology Centre / MRC-University of Glasgow Centre for Virus Research | Wellcome Sanger Institute for the COVID-19 Genomics UK Consortium | Ana da Silva Filipe, Natasha Johnson, Kathy Smollett, Daniel Mair, Stephen Carmichael, Lily Tong, Jenna Nichols, Elihu Aranday-Cortes, Kirstyn Brunker, Yasmin Parr, Kyriaki Nomikou; Sarah McDonald, Marc Niebel, Patawee Asamaphan; Richard Orton, Joseph Hughes, Sreenu Vattipally, David L Robertson; Alasdair MacLean, Rory Gunson; Kathy Li, Natasha Jesudason, Rajiv Shah, James Shepherd, Antonia Ho, Alice Broos, Emma Thomson and Alex Alderton, Roberto Amato, Sonia Goncalves, Ewan Harrison, David K. Jackson, Ian Johnston, Dominic Kwiatkowski, Cordelia Langford, John Sillitoe |
| EPI_ISL_532273 | Lighthouse Lab in Glasgow | Wellcome Sanger Institute for the COVID-19 Genomics UK Consortium | Harper VanSteenhouse, Yumi Kasai, David Gray, Carol Clugston, Anna Dominiczak and Alex Alderton, Roberto Amato, Sonia Goncalves, Ewan Harrison, David K. Jackson, Ian Johnston, Dominic Kwiatkowski, Cordelia Langford, John Sillitoe |
| EPI_ISL_532275, EPI_ISL_532276, EPI_ISL_532278 | NHSGGC West of Scotland Specialist Virology Centre / MRC-University of Glasgow Centre for Virus Research | Wellcome Sanger Institute for the COVID-19 Genomics UK Consortium | Ana da Silva Filipe, Natasha Johnson, Kathy Smollett, Daniel Mair, Stephen Carmichael, Lily Tong, Jenna Nichols, Elihu Aranday-Cortes, Kirstyn Brunker, Yasmin Parr, Kyriaki Nomikou; Sarah McDonald, Marc Niebel, Patawee Asamaphan; Richard Orton, Joseph Hughes, Sreenu Vattipally, David L Robertson; Alasdair MacLean, Rory Gunson; Kathy Li, Natasha Jesudason, Rajiv Shah, James Shepherd, Antonia Ho, Alice Broos, Emma Thomson and Alex Alderton, Roberto Amato, Sonia Goncalves, Ewan Harrison, David K. Jackson, Ian Johnston, Dominic Kwiatkowski, Cordelia Langford, John Sillitoe |
| EPI_ISL_532279, EPI_ISL_532280, EPI_ISL_532282 | Lighthouse Lab in Glasgow | Wellcome Sanger Institute for the COVID-19 Genomics UK Consortium | Harper VanSteenhouse, Yumi Kasai, David Gray, Carol Clugston, Anna Dominiczak and Alex Alderton, Roberto Amato, Sonia Goncalves, Ewan Harrison, David K. Jackson, Ian Johnston, Dominic Kwiatkowski, Cordelia Langford, John Sillitoe |
| EPI_ISL_532283 | NHSGGC West of Scotland Specialist Virology Centre / MRC-University of Glasgow Centre for Virus Research | Wellcome Sanger Institute for the COVID-19 Genomics UK Consortium | Ana da Silva Filipe, Natasha Johnson, Kathy Smollett, Daniel Mair, Stephen Carmichael, Lily Tong, Jenna Nichols, Elihu Aranday-Cortes, Kirstyn Brunker, Yasmin Parr, Kyriaki Nomikou; Sarah McDonald, Marc Niebel, Patawee Asamaphan; Richard Orton, Joseph Hughes, Sreenu Vattipally, David L Robertson; Alasdair MacLean, Rory Gunson; Kathy Li, Natasha Jesudason, Rajiv Shah, James Shepherd, Antonia Ho, Alice Broos, Emma Thomson and Alex Alderton, Roberto Amato, Sonia Goncalves, Ewan Harrison, David K. Jackson, Ian Johnston, Dominic Kwiatkowski, Cordelia Langford, John Sillitoe |
| EPI_ISL_532284, EPI_ISL_532285, EPI_ISL_532286, EPI_ISL_532287 | Lighthouse Lab in Glasgow | Wellcome Sanger Institute for the COVID-19 Genomics UK Consortium | Harper VanSteenhouse, Yumi Kasai, David Gray, Carol Clugston, Anna Dominiczak and Alex Alderton, Roberto Amato, Sonia Goncalves, Ewan Harrison, David K. Jackson, Ian Johnston, Dominic Kwiatkowski, Cordelia Langford, John Sillitoe |
| EPI_ISL_532288 | NHSGGC West of Scotland Specialist Virology Centre / MRC-University of Glasgow Centre for Virus Research | Wellcome Sanger Institute for the COVID-19 Genomics UK Consortium | Ana da Silva Filipe, Natasha Johnson, Kathy Smollett, Daniel Mair, Stephen Carmichael, Lily Tong, Jenna Nichols, Elihu Aranday-Cortes, Kirstyn Brunker, Yasmin Parr, Kyriaki Nomikou; Sarah McDonald, Marc Niebel, Patawee Asamaphan; Richard Orton, Joseph Hughes, Sreenu Vattipally, David L Robertson; Alasdair MacLean, Rory Gunson; Kathy Li, Natasha Jesudason, Rajiv Shah, James Shepherd, Antonia Ho, Alice Broos, Emma Thomson and Alex Alderton, Roberto Amato, Sonia Goncalves, Ewan Harrison, David K. Jackson, Ian Johnston, Dominic Kwiatkowski, Cordelia Langford, John Sillitoe |
| EPI_ISL_532289, EPI_ISL_532290 | Lighthouse Lab in Glasgow | Wellcome Sanger Institute for the COVID-19 Genomics UK Consortium | Harper VanSteenhouse, Yumi Kasai, David Gray, Carol Clugston, Anna Dominiczak and Alex Alderton, Roberto Amato, Sonia Goncalves, Ewan Harrison, David K. Jackson, Ian Johnston, Dominic Kwiatkowski, Cordelia Langford, John Sillitoe |
| EPI_ISL_532293 | NHSGGC West of Scotland Specialist Virology Centre / MRC-University of Glasgow Centre for Virus Research | Wellcome Sanger Institute for the COVID-19 Genomics UK Consortium | Ana da Silva Filipe, Natasha Johnson, Kathy Smollett, Daniel Mair, Stephen Carmichael, Lily Tong, Jenna Nichols, Elihu Aranday-Cortes, Kirstyn Brunker, Yasmin Parr, Kyriaki Nomikou; Sarah McDonald, Marc Niebel, Patawee Asamaphan; Richard Orton, Joseph Hughes, Sreenu Vattipally, David L Robertson; Alasdair MacLean, Rory Gunson; Kathy Li, Natasha Jesudason, Rajiv Shah, James Shepherd, Antonia Ho, Alice Broos, Emma Thomson and Alex Alderton, Roberto Amato, Sonia Goncalves, Ewan Harrison, David K. Jackson, Ian Johnston, Dominic Kwiatkowski, Cordelia Langford, John Sillitoe |
| EPI_ISL_532295, EPI_ISL_532296, EPI_ISL_532297, EPI_ISL_532298, EPI_ISL_532299, EPI_ISL_532300 | Lighthouse Lab in Glasgow | Wellcome Sanger Institute for the COVID-19 Genomics UK Consortium | Harper VanSteenhouse, Yumi Kasai, David Gray, Carol Clugston, Anna Dominiczak and Alex Alderton, Roberto Amato, Sonia Goncalves, Ewan Harrison, David K. Jackson, Ian Johnston, Dominic Kwiatkowski, Cordelia Langford, John Sillitoe |
| EPI_ISL_532301 | NHSGGC West of Scotland Specialist Virology Centre / MRC-University of Glasgow Centre for Virus Research | Wellcome Sanger Institute for the COVID-19 Genomics UK Consortium | Ana da Silva Filipe, Natasha Johnson, Kathy Smollett, Daniel Mair, Stephen Carmichael, Lily Tong, Jenna Nichols, Elihu Aranday-Cortes, Kirstyn Brunker, Yasmin Parr, Kyriaki Nomikou; Sarah McDonald, Marc Niebel, Patawee Asamaphan; Richard Orton, Joseph Hughes, Sreenu Vattipally, David L Robertson; Alasdair MacLean, Rory Gunson; Kathy Li, Natasha Jesudason, Rajiv Shah, James Shepherd, Antonia Ho, Alice Broos, Emma Thomson and Alex Alderton, Roberto Amato, Sonia Goncalves, Ewan Harrison, David K. Jackson, Ian Johnston, Dominic Kwiatkowski, Cordelia Langford, John Sillitoe |
| EPI_ISL_532302, EPI_ISL_532303, EPI_ISL_532305, EPI_ISL_532308, EPI_ISL_532309, EPI_ISL_532310 | Lighthouse Lab in Glasgow | Wellcome Sanger Institute for the COVID-19 Genomics UK Consortium | Harper VanSteenhouse, Yumi Kasai, David Gray, Carol Clugston, Anna Dominiczak and Alex Alderton, Roberto Amato, Sonia Goncalves, Ewan Harrison, David K. Jackson, Ian Johnston, Dominic Kwiatkowski, Cordelia Langford, John Sillitoe |
| EPI_ISL_532311 | NHSGGC West of Scotland Specialist Virology Centre / MRC-University of Glasgow Centre for Virus Research | Wellcome Sanger Institute for the COVID-19 Genomics UK Consortium | Ana da Silva Filipe, Natasha Johnson, Kathy Smollett, Daniel Mair, Stephen Carmichael, Lily Tong, Jenna Nichols, Elihu Aranday-Cortes, Kirstyn Brunker, Yasmin Parr, Kyriaki Nomikou; Sarah McDonald, Marc Niebel, Patawee Asamaphan; Richard Orton, Joseph Hughes, Sreenu Vattipally, David L Robertson; Alasdair MacLean, Rory Gunson; Kathy Li, Natasha Jesudason, Rajiv Shah, James Shepherd, Antonia Ho, Alice Broos, Emma Thomson and Alex Alderton, Roberto Amato, Sonia Goncalves, Ewan Harrison, David K. Jackson, Ian Johnston, Dominic Kwiatkowski, Cordelia Langford, John Sillitoe |
| EPI_ISL_532312, EPI_ISL_532313, EPI_ISL_532314 | Lighthouse Lab in Glasgow | Wellcome Sanger Institute for the COVID-19 Genomics UK Consortium | Harper VanSteenhouse, Yumi Kasai, David Gray, Carol Clugston, Anna Dominiczak and Alex Alderton, Roberto Amato, Sonia Goncalves, Ewan Harrison, David K. Jackson, Ian Johnston, Dominic Kwiatkowski, Cordelia Langford, John Sillitoe |
| EPI_ISL_532316, EPI_ISL_532317 | NHSGGC West of Scotland Specialist Virology Centre / MRC-University of Glasgow Centre for Virus Research | Wellcome Sanger Institute for the COVID-19 Genomics UK Consortium | Ana da Silva Filipe, Natasha Johnson, Kathy Smollett, Daniel Mair, Stephen Carmichael, Lily Tong, Jenna Nichols, Elihu Aranday-Cortes, Kirstyn Brunker, Yasmin Parr, Kyriaki Nomikou; Sarah McDonald, Marc Niebel, Patawee Asamaphan; Richard Orton, Joseph Hughes, Sreenu Vattipally, David L Robertson; Alasdair MacLean, Rory Gunson; Kathy Li, Natasha Jesudason, Rajiv Shah, James Shepherd, Antonia Ho, Alice Broos, Emma Thomson and Alex Alderton, Roberto Amato, Sonia Goncalves, Ewan Harrison, David K. Jackson, Ian Johnston, Dominic Kwiatkowski, Cordelia Langford, John Sillitoe |
| EPI_ISL_532318, EPI_ISL_532319, EPI_ISL_532320 | Lighthouse Lab in Glasgow | Wellcome Sanger Institute for the | Harper VanSteenhouse, Yumi Kasai, David Gray, Carol Clugston, Anna Dominiczak and Alex Alderton, Roberto Amato, Sonia Goncalves, Ewan Harrison, David K. Jackson, Ian Johnston, Dominic Kwiatkowski, Cordelia Langford, John Sillitoe |

| | | | |
|---|---|---|---|
| | | COVID-19 Genomics UK Consortium | |
| EPI_ISL_532321 | NHSGGC West of Scotland Specialist Virology Centre / MRC-University of Glasgow Centre for Virus Research | Wellcome Sanger Institute for the COVID-19 Genomics UK Consortium | Ana da Silva Filipe, Natasha Johnson, Kathy Smollett, Daniel Mair, Stephen Carmichael, Lily Tong, Jenna Nichols, Elihu Aranday-Cortes, Kirstyn Brunker, Yasmin Parr, Kyriaki Nomikou; Sarah McDonald, Marc Niebel, Patawee Asamaphan; Richard Orton, Joseph Hughes, Sreenu Vattipally, David L Robertson; Alasdair MacLean, Rory Gunson; Kathy Li, Natasha Jesudason, Rajiv Shah, James Shepherd, Antonia Ho, Alice Broos, Emma Thomson and Alex Alderton, Roberto Amato, Sonia Goncalves, Ewan Harrison, David K. Jackson, Ian Johnston, Dominic Kwiatkowski, Cordelia Langford, John Sillitoe |
| EPI_ISL_532322, EPI_ISL_532323, EPI_ISL_532324, EPI_ISL_532325, EPI_ISL_532326 | Lighthouse Lab in Glasgow | Wellcome Sanger Institute for the COVID-19 Genomics UK Consortium | Harper VanSteenhouse, Yumi Kasai, David Gray, Carol Clugston, Anna Dominiczak and Alex Alderton, Roberto Amato, Sonia Goncalves, Ewan Harrison, David K. Jackson, Ian Johnston, Dominic Kwiatkowski, Cordelia Langford, John Sillitoe |
| EPI_ISL_532328 | NHSGGC West of Scotland Specialist Virology Centre / MRC-University of Glasgow Centre for Virus Research | Wellcome Sanger Institute for the COVID-19 Genomics UK Consortium | Ana da Silva Filipe, Natasha Johnson, Kathy Smollett, Daniel Mair, Stephen Carmichael, Lily Tong, Jenna Nichols, Elihu Aranday-Cortes, Kirstyn Brunker, Yasmin Parr, Kyriaki Nomikou; Sarah McDonald, Marc Niebel, Patawee Asamaphan; Richard Orton, Joseph Hughes, Sreenu Vattipally, David L Robertson; Alasdair MacLean, Rory Gunson; Kathy Li, Natasha Jesudason, Rajiv Shah, James Shepherd, Antonia Ho, Alice Broos, Emma Thomson and Alex Alderton, Roberto Amato, Sonia Goncalves, Ewan Harrison, David K. Jackson, Ian Johnston, Dominic Kwiatkowski, Cordelia Langford, John Sillitoe |
| EPI_ISL_532329, EPI_ISL_532330, EPI_ISL_532331, EPI_ISL_532333 | Lighthouse Lab in Glasgow | Wellcome Sanger Institute for the COVID-19 Genomics UK Consortium | Harper VanSteenhouse, Yumi Kasai, David Gray, Carol Clugston, Anna Dominiczak and Alex Alderton, Roberto Amato, Sonia Goncalves, Ewan Harrison, David K. Jackson, Ian Johnston, Dominic Kwiatkowski, Cordelia Langford, John Sillitoe |
| EPI_ISL_532334, EPI_ISL_532335 | NHSGGC West of Scotland Specialist Virology Centre / MRC-University of Glasgow Centre for Virus Research | Wellcome Sanger Institute for the COVID-19 Genomics UK Consortium | Ana da Silva Filipe, Natasha Johnson, Kathy Smollett, Daniel Mair, Stephen Carmichael, Lily Tong, Jenna Nichols, Elihu Aranday-Cortes, Kirstyn Brunker, Yasmin Parr, Kyriaki Nomikou; Sarah McDonald, Marc Niebel, Patawee Asamaphan; Richard Orton, Joseph Hughes, Sreenu Vattipally, David L Robertson; Alasdair MacLean, Rory Gunson; Kathy Li, Natasha Jesudason, Rajiv Shah, James Shepherd, Antonia Ho, Alice Broos, Emma Thomson and Alex Alderton, Roberto Amato, Sonia Goncalves, Ewan Harrison, David K. Jackson, Ian Johnston, Dominic Kwiatkowski, Cordelia Langford, John Sillitoe |
| EPI_ISL_532336, EPI_ISL_532337 | Lighthouse Lab in Glasgow | Wellcome Sanger Institute for the COVID-19 Genomics UK Consortium | Harper VanSteenhouse, Yumi Kasai, David Gray, Carol Clugston, Anna Dominiczak and Alex Alderton, Roberto Amato, Sonia Goncalves, Ewan Harrison, David K. Jackson, Ian Johnston, Dominic Kwiatkowski, Cordelia Langford, John Sillitoe |
| EPI_ISL_532338 | NHSGGC West of Scotland Specialist Virology Centre / MRC-University of Glasgow Centre for Virus Research | Wellcome Sanger Institute for the COVID-19 Genomics UK Consortium | Ana da Silva Filipe, Natasha Johnson, Kathy Smollett, Daniel Mair, Stephen Carmichael, Lily Tong, Jenna Nichols, Elihu Aranday-Cortes, Kirstyn Brunker, Yasmin Parr, Kyriaki Nomikou; Sarah McDonald, Marc Niebel, Patawee Asamaphan; Richard Orton, Joseph Hughes, Sreenu Vattipally, David L Robertson; Alasdair MacLean, Rory Gunson; Kathy Li, Natasha Jesudason, Rajiv Shah, James Shepherd, Antonia Ho, Alice Broos, Emma Thomson and Alex Alderton, Roberto Amato, Sonia Goncalves, Ewan Harrison, David K. Jackson, Ian Johnston, Dominic Kwiatkowski, Cordelia Langford, John Sillitoe |
| EPI_ISL_532339 | Lighthouse Lab in Glasgow | Wellcome Sanger Institute for the COVID-19 Genomics UK Consortium | Harper VanSteenhouse, Yumi Kasai, David Gray, Carol Clugston, Anna Dominiczak and Alex Alderton, Roberto Amato, Sonia Goncalves, Ewan Harrison, David K. Jackson, Ian Johnston, Dominic Kwiatkowski, Cordelia Langford, John Sillitoe |
| EPI_ISL_532340 | NHSGGC West of Scotland Specialist Virology Centre / MRC-University of Glasgow Centre for Virus Research | Wellcome Sanger Institute for the COVID-19 Genomics UK Consortium | Ana da Silva Filipe, Natasha Johnson, Kathy Smollett, Daniel Mair, Stephen Carmichael, Lily Tong, Jenna Nichols, Elihu Aranday-Cortes, Kirstyn Brunker, Yasmin Parr, Kyriaki Nomikou; Sarah McDonald, Marc Niebel, Patawee Asamaphan; Richard Orton, Joseph Hughes, Sreenu Vattipally, David L Robertson; Alasdair MacLean, Rory Gunson; Kathy Li, Natasha Jesudason, Rajiv Shah, James Shepherd, Antonia Ho, Alice Broos, Emma Thomson and Alex Alderton, Roberto Amato, Sonia Goncalves, Ewan Harrison, David K. Jackson, Ian Johnston, Dominic Kwiatkowski, Cordelia Langford, John Sillitoe |
| EPI_ISL_532341, EPI_ISL_532343, EPI_ISL_532344 | Lighthouse Lab in Glasgow | Wellcome Sanger Institute for the COVID-19 Genomics UK Consortium | Harper VanSteenhouse, Yumi Kasai, David Gray, Carol Clugston, Anna Dominiczak and Alex Alderton, Roberto Amato, Sonia Goncalves, Ewan Harrison, David K. Jackson, Ian Johnston, Dominic Kwiatkowski, Cordelia Langford, John Sillitoe |
| EPI_ISL_532347 | NHSGGC West of Scotland Specialist Virology Centre / MRC-University of Glasgow Centre for Virus Research | Wellcome Sanger Institute for the COVID-19 Genomics UK Consortium | Ana da Silva Filipe, Natasha Johnson, Kathy Smollett, Daniel Mair, Stephen Carmichael, Lily Tong, Jenna Nichols, Elihu Aranday-Cortes, Kirstyn Brunker, Yasmin Parr, Kyriaki Nomikou; Sarah McDonald, Marc Niebel, Patawee Asamaphan; Richard Orton, Joseph Hughes, Sreenu Vattipally, David L Robertson; Alasdair MacLean, Rory Gunson; Kathy Li, Natasha Jesudason, Rajiv Shah, James Shepherd, Antonia Ho, Alice Broos, Emma Thomson and Alex Alderton, Roberto Amato, Sonia Goncalves, Ewan Harrison, David K. Jackson, Ian Johnston, Dominic Kwiatkowski, Cordelia Langford, John Sillitoe |
| EPI_ISL_532348 | Lighthouse Lab in Glasgow | Wellcome Sanger Institute for the COVID-19 Genomics UK Consortium | Harper VanSteenhouse, Yumi Kasai, David Gray, Carol Clugston, Anna Dominiczak and Alex Alderton, Roberto Amato, Sonia Goncalves, Ewan Harrison, David K. Jackson, Ian Johnston, Dominic Kwiatkowski, Cordelia Langford, John Sillitoe |
| EPI_ISL_532349 | NHSGGC West of Scotland Specialist Virology Centre / MRC-University of Glasgow Centre for Virus Research | Wellcome Sanger Institute for the COVID-19 Genomics UK Consortium | Ana da Silva Filipe, Natasha Johnson, Kathy Smollett, Daniel Mair, Stephen Carmichael, Lily Tong, Jenna Nichols, Elihu Aranday-Cortes, Kirstyn Brunker, Yasmin Parr, Kyriaki Nomikou; Sarah McDonald, Marc Niebel, Patawee Asamaphan; Richard Orton, Joseph Hughes, Sreenu Vattipally, David L Robertson; Alasdair MacLean, Rory Gunson; Kathy Li, Natasha Jesudason, Rajiv Shah, James Shepherd, Antonia Ho, Alice Broos, Emma Thomson and Alex Alderton, Roberto Amato, Sonia Goncalves, Ewan Harrison, David K. Jackson, Ian Johnston, Dominic Kwiatkowski, Cordelia Langford, John Sillitoe |
| EPI_ISL_532350 | Lighthouse Lab in Glasgow | Wellcome Sanger Institute for the COVID-19 Genomics UK Consortium | Harper VanSteenhouse, Yumi Kasai, David Gray, Carol Clugston, Anna Dominiczak and Alex Alderton, Roberto Amato, Sonia Goncalves, Ewan Harrison, David K. Jackson, Ian Johnston, Dominic Kwiatkowski, Cordelia Langford, John Sillitoe |
| EPI_ISL_532351 | NHSGGC West of Scotland Specialist Virology Centre / MRC-University of Glasgow Centre for Virus Research | Wellcome Sanger Institute for the COVID-19 Genomics UK Consortium | Ana da Silva Filipe, Natasha Johnson, Kathy Smollett, Daniel Mair, Stephen Carmichael, Lily Tong, Jenna Nichols, Elihu Aranday-Cortes, Kirstyn Brunker, Yasmin Parr, Kyriaki Nomikou; Sarah McDonald, Marc Niebel, Patawee Asamaphan; Richard Orton, Joseph Hughes, Sreenu Vattipally, David L Robertson; Alasdair MacLean, Rory Gunson; Kathy Li, Natasha Jesudason, Rajiv Shah, James Shepherd, Antonia Ho, Alice Broos, Emma Thomson and Alex Alderton, Roberto Amato, Sonia Goncalves, Ewan Harrison, David K. Jackson, Ian Johnston, Dominic Kwiatkowski, Cordelia Langford, John Sillitoe |
| EPI_ISL_532352 | Lighthouse Lab in Glasgow | Wellcome Sanger Institute for the COVID-19 Genomics UK Consortium | Harper VanSteenhouse, Yumi Kasai, David Gray, Carol Clugston, Anna Dominiczak and Alex Alderton, Roberto Amato, Sonia Goncalves, Ewan Harrison, David K. Jackson, Ian Johnston, Dominic Kwiatkowski, Cordelia Langford, John Sillitoe |
| EPI_ISL_532353 | NHSGGC West of Scotland Specialist Virology Centre / MRC-University of Glasgow Centre for Virus Research | Wellcome Sanger Institute for the COVID-19 Genomics UK Consortium | Ana da Silva Filipe, Natasha Johnson, Kathy Smollett, Daniel Mair, Stephen Carmichael, Lily Tong, Jenna Nichols, Elihu Aranday-Cortes, Kirstyn Brunker, Yasmin Parr, Kyriaki Nomikou; Sarah McDonald, Marc Niebel, Patawee Asamaphan; Richard Orton, Joseph Hughes, Sreenu Vattipally, David L Robertson; Alasdair MacLean, Rory Gunson; Kathy Li, Natasha Jesudason, Rajiv Shah, James Shepherd, Antonia Ho, Alice Broos, Emma Thomson and Alex Alderton, Roberto Amato, Sonia Goncalves, Ewan Harrison, David K. Jackson, Ian Johnston, Dominic Kwiatkowski, Cordelia Langford, John Sillitoe |
| EPI_ISL_532354 | Lighthouse Lab in Glasgow | Wellcome Sanger Institute for the COVID-19 Genomics UK Consortium | Harper VanSteenhouse, Yumi Kasai, David Gray, Carol Clugston, Anna Dominiczak and Alex Alderton, Roberto Amato, Sonia Goncalves, Ewan Harrison, David K. Jackson, Ian Johnston, Dominic Kwiatkowski, Cordelia Langford, John Sillitoe |
| EPI_ISL_532355, EPI_ISL_532356 | NHSGGC West of Scotland Specialist Virology Centre / MRC-University of Glasgow Centre for Virus Research | Wellcome Sanger Institute for the COVID-19 Genomics UK Consortium | Ana da Silva Filipe, Natasha Johnson, Kathy Smollett, Daniel Mair, Stephen Carmichael, Lily Tong, Jenna Nichols, Elihu Aranday-Cortes, Kirstyn Brunker, Yasmin Parr, Kyriaki Nomikou; Sarah McDonald, Marc Niebel, Patawee Asamaphan; Richard Orton, Joseph Hughes, Sreenu Vattipally, David L Robertson; Alasdair MacLean, Rory Gunson; Kathy Li, Natasha Jesudason, Rajiv Shah, James Shepherd, Antonia Ho, Alice Broos, Emma Thomson and Alex Alderton, Roberto Amato, Sonia Goncalves, Ewan Harrison, David K. Jackson, Ian Johnston, Dominic Kwiatkowski, Cordelia Langford, John Sillitoe |
| EPI_ISL_532357, EPI_ISL_532359, EPI_ISL_532360, EPI_ISL_532362 | Lighthouse Lab in Glasgow | Wellcome Sanger Institute for the COVID-19 Genomics UK Consortium | Harper VanSteenhouse, Yumi Kasai, David Gray, Carol Clugston, Anna Dominiczak and Alex Alderton, Roberto Amato, Sonia Goncalves, Ewan Harrison, David K. Jackson, Ian Johnston, Dominic Kwiatkowski, Cordelia Langford, John Sillitoe |
| EPI_ISL_532363, EPI_ISL_532364 | NHSGGC West of Scotland Specialist Virology Centre / MRC-University of Glasgow Centre for Virus Research | Wellcome Sanger Institute for the COVID-19 Genomics UK Consortium | Ana da Silva Filipe, Natasha Johnson, Kathy Smollett, Daniel Mair, Stephen Carmichael, Lily Tong, Jenna Nichols, Elihu Aranday-Cortes, Kirstyn Brunker, Yasmin Parr, Kyriaki Nomikou; Sarah McDonald, Marc Niebel, Patawee Asamaphan; Richard Orton, Joseph Hughes, Sreenu Vattipally, David L Robertson; Alasdair MacLean, Rory Gunson; Kathy Li, Natasha Jesudason, Rajiv Shah, James Shepherd, Antonia Ho, Alice Broos, Emma Thomson and Alex Alderton, Roberto Amato, Sonia Goncalves, Ewan Harrison, David K. Jackson, Ian Johnston, Dominic Kwiatkowski, Cordelia Langford, John Sillitoe |
| EPI_ISL_532366, EPI_ISL_532367, EPI_ISL_532368, EPI_ISL_532369, EPI_ISL_532371, EPI_ISL_532372, EPI_ISL_532373, EPI_ISL_532374, EPI_ISL_532375, EPI_ISL_532377, EPI_ISL_532378, EPI_ISL_532380, EPI_ISL_532381, EPI_ISL_532382, EPI_ISL_532383 | | | |
| see above | Lighthouse Lab in Glasgow | Wellcome Sanger Institute for the COVID-19 Genomics UK Consortium | Harper VanSteenhouse, Yumi Kasai, David Gray, Carol Clugston, Anna Dominiczak and Alex Alderton, Roberto Amato, Sonia Goncalves, Ewan Harrison, David K. Jackson, Ian Johnston, Dominic Kwiatkowski, Cordelia Langford, John Sillitoe |
| EPI_ISL_532384 | NHSGGC West of Scotland Specialist Virology Centre / MRC-University of Glasgow Centre for Virus Research | Wellcome Sanger Institute for the COVID-19 Genomics UK Consortium | Ana da Silva Filipe, Natasha Johnson, Kathy Smollett, Daniel Mair, Stephen Carmichael, Lily Tong, Jenna Nichols, Elihu Aranday-Cortes, Kirstyn Brunker, Yasmin Parr, Kyriaki Nomikou; Sarah McDonald, Marc Niebel, Patawee Asamaphan; Richard Orton, Joseph Hughes, Sreenu Vattipally, David L Robertson; Alasdair MacLean, Rory Gunson; Kathy Li, Natasha Jesudason, Rajiv Shah, James Shepherd, Antonia Ho, Alice Broos, Emma Thomson and Alex Alderton, Roberto Amato, Sonia Goncalves, Ewan Harrison, David K. Jackson, Ian Johnston, Dominic Kwiatkowski, Cordelia Langford, John Sillitoe |
| EPI_ISL_532386, EPI_ISL_532387 | Lighthouse Lab in Glasgow | Wellcome Sanger Institute for the COVID-19 Genomics UK Consortium | Harper VanSteenhouse, Yumi Kasai, David Gray, Carol Clugston, Anna Dominiczak and Alex Alderton, Roberto Amato, Sonia Goncalves, Ewan Harrison, David K. Jackson, Ian Johnston, Dominic Kwiatkowski, Cordelia Langford, John Sillitoe |
| EPI_ISL_532388 | NHSGGC West of Scotland Specialist Virology Centre / MRC-University of Glasgow Centre for Virus Research | Wellcome Sanger Institute for the COVID-19 Genomics UK Consortium | Ana da Silva Filipe, Natasha Johnson, Kathy Smollett, Daniel Mair, Stephen Carmichael, Lily Tong, Jenna Nichols, Elihu Aranday-Cortes, Kirstyn Brunker, Yasmin Parr, Kyriaki Nomikou; Sarah McDonald, Marc Niebel, Patawee Asamaphan; Richard Orton, Joseph Hughes, Sreenu Vattipally, David L Robertson; Alasdair MacLean, Rory Gunson; Kathy Li, Natasha Jesudason, Rajiv Shah, James Shepherd, Antonia Ho, Alice Broos, Emma Thomson and Alex Alderton, Roberto Amato, Sonia Goncalves, Ewan Harrison, David K. Jackson, Ian Johnston, Dominic Kwiatkowski, Cordelia Langford, John Sillitoe |
| EPI_ISL_532389 | Lighthouse Lab in Glasgow | Wellcome Sanger Institute for the COVID-19 Genomics UK Consortium | Harper VanSteenhouse, Yumi Kasai, David Gray, Carol Clugston, Anna Dominiczak and Alex Alderton, Roberto Amato, Sonia Goncalves, Ewan Harrison, David K. Jackson, Ian Johnston, Dominic Kwiatkowski, Cordelia Langford, John Sillitoe |
| EPI_ISL_532390 | NHSGGC West of Scotland Specialist Virology Centre / MRC-University of Glasgow Centre for Virus Research | Wellcome Sanger Institute for the COVID-19 Genomics UK Consortium | Ana da Silva Filipe, Natasha Johnson, Kathy Smollett, Daniel Mair, Stephen Carmichael, Lily Tong, Jenna Nichols, Elihu Aranday-Cortes, Kirstyn Brunker, Yasmin Parr, Kyriaki Nomikou; Sarah McDonald, Marc Niebel, Patawee Asamaphan; Richard Orton, Joseph Hughes, Sreenu Vattipally, David L Robertson; Alasdair MacLean, Rory Gunson; Kathy Li, Natasha Jesudason, Rajiv Shah, James Shepherd, Antonia Ho, Alice Broos, Emma Thomson and Alex Alderton, Roberto Amato, Sonia Goncalves, Ewan Harrison, David K. Jackson, Ian Johnston, Dominic Kwiatkowski, Cordelia Langford, John Sillitoe |
| EPI_ISL_532391, EPI_ISL_532392, EPI_ISL_532393 | Lighthouse Lab in Glasgow | Wellcome Sanger Institute for the COVID-19 Genomics UK Consortium | Harper VanSteenhouse, Yumi Kasai, David Gray, Carol Clugston, Anna Dominiczak and Alex Alderton, Roberto Amato, Sonia Goncalves, Ewan Harrison, David K. Jackson, Ian Johnston, Dominic Kwiatkowski, Cordelia Langford, John Sillitoe |
| EPI_ISL_532394 | NHSGGC West of Scotland Specialist Virology Centre / MRC-University of Glasgow Centre for Virus Research | Wellcome Sanger Institute for the COVID-19 Genomics UK Consortium | Ana da Silva Filipe, Natasha Johnson, Kathy Smollett, Daniel Mair, Stephen Carmichael, Lily Tong, Jenna Nichols, Elihu Aranday-Cortes, Kirstyn Brunker, Yasmin Parr, Kyriaki Nomikou; Sarah McDonald, Marc Niebel, Patawee Asamaphan; Richard Orton, Joseph Hughes, Sreenu Vattipally, David L Robertson; Alasdair MacLean, Rory Gunson; Kathy Li, Natasha Jesudason, Rajiv Shah, James Shepherd, Antonia Ho, Alice Broos, Emma Thomson and Alex Alderton, Roberto Amato, Sonia Goncalves, Ewan Harrison, David K. Jackson, Ian Johnston, Dominic Kwiatkowski, Cordelia Langford, John Sillitoe |
| EPI_ISL_532395, EPI_ISL_532396, EPI_ISL_532397, EPI_ISL_532398 | Lighthouse Lab in Glasgow | Wellcome Sanger Institute for the COVID-19 Genomics UK Consortium | Harper VanSteenhouse, Yumi Kasai, David Gray, Carol Clugston, Anna Dominiczak and Alex Alderton, Roberto Amato, Sonia Goncalves, Ewan Harrison, David K. Jackson, Ian Johnston, Dominic Kwiatkowski, Cordelia Langford, John Sillitoe |
| EPI_ISL_532399 | NHSGGC West of Scotland Specialist Virology Centre / MRC-University of Glasgow Centre for Virus Research | Wellcome Sanger Institute for the COVID-19 Genomics UK Consortium | Ana da Silva Filipe, Natasha Johnson, Kathy Smollett, Daniel Mair, Stephen Carmichael, Lily Tong, Jenna Nichols, Elihu Aranday-Cortes, Kirstyn Brunker, Yasmin Parr, Kyriaki Nomikou; Sarah McDonald, Marc Niebel, Patawee Asamaphan; Richard Orton, Joseph Hughes, Sreenu Vattipally, David L Robertson; Alasdair MacLean, Rory Gunson; Kathy Li, Natasha Jesudason, Rajiv Shah, James Shepherd, Antonia Ho, Alice Broos, Emma Thomson and Alex Alderton, Roberto Amato, Sonia Goncalves, Ewan Harrison, David K. Jackson, Ian Johnston, Dominic Kwiatkowski, Cordelia Langford, John Sillitoe |
| EPI_ISL_532400, EPI_ISL_532401 | Lighthouse Lab in Glasgow | Wellcome Sanger Institute for the COVID-19 Genomics UK Consortium | Harper VanSteenhouse, Yumi Kasai, David Gray, Carol Clugston, Anna Dominiczak and Alex Alderton, Roberto Amato, Sonia Goncalves, Ewan Harrison, David K. Jackson, Ian Johnston, Dominic Kwiatkowski, Cordelia Langford, John Sillitoe |
| EPI_ISL_532402 | NHSGGC West of Scotland Specialist Virology Centre / MRC-University of Glasgow Centre for Virus Research | Wellcome Sanger Institute for the COVID-19 Genomics UK Consortium | Ana da Silva Filipe, Natasha Johnson, Kathy Smollett, Daniel Mair, Stephen Carmichael, Lily Tong, Jenna Nichols, Elihu Aranday-Cortes, Kirstyn Brunker, Yasmin Parr, Kyriaki Nomikou; Sarah McDonald, Marc Niebel, Patawee Asamaphan; Richard Orton, Joseph Hughes, Sreenu Vattipally, David L Robertson; Alasdair MacLean, Rory Gunson; Kathy Li, Natasha Jesudason, Rajiv Shah, James Shepherd, Antonia Ho, Alice Broos, Emma Thomson and Alex Alderton, Roberto Amato, Sonia Goncalves, Ewan Harrison, David K. Jackson, Ian Johnston, Dominic Kwiatkowski, Cordelia Langford, John Sillitoe |
| EPI_ISL_532403 | Lighthouse Lab in Glasgow | Wellcome Sanger Institute for the COVID-19 Genomics UK Consortium | Harper VanSteenhouse, Yumi Kasai, David Gray, Carol Clugston, Anna Dominiczak and Alex Alderton, Roberto Amato, Sonia Goncalves, Ewan Harrison, David K. Jackson, Ian Johnston, Dominic Kwiatkowski, Cordelia Langford, John Sillitoe |
| EPI_ISL_532404 | NHSGGC West of Scotland Specialist Virology Centre / MRC-University of Glasgow Centre for Virus Research | Wellcome Sanger Institute for the COVID-19 Genomics UK Consortium | Ana da Silva Filipe, Natasha Johnson, Kathy Smollett, Daniel Mair, Stephen Carmichael, Lily Tong, Jenna Nichols, Elihu Aranday-Cortes, Kirstyn Brunker, Yasmin Parr, Kyriaki Nomikou; Sarah McDonald, Marc Niebel, Patawee Asamaphan; Richard Orton, Joseph Hughes, Sreenu Vattipally, David L Robertson; Alasdair MacLean, Rory Gunson; Kathy Li, Natasha Jesudason, Rajiv Shah, James Shepherd, Antonia Ho, Alice Broos, Emma Thomson and Alex Alderton, Roberto Amato, Sonia Goncalves, Ewan Harrison, David K. Jackson, Ian Johnston, Dominic Kwiatkowski, Cordelia Langford, John Sillitoe |
| EPI_ISL_532406, EPI_ISL_532407, EPI_ISL_532408, EPI_ISL_532409, EPI_ISL_532410, EPI_ISL_532411, EPI_ISL_532412, EPI_ISL_532413 | Lighthouse Lab in Glasgow | Wellcome Sanger Institute for the COVID-19 Genomics UK Consortium | Harper VanSteenhouse, Yumi Kasai, David Gray, Carol Clugston, Anna Dominiczak and Alex Alderton, Roberto Amato, Sonia Goncalves, Ewan Harrison, David K. Jackson, Ian Johnston, Dominic Kwiatkowski, Cordelia Langford, John Sillitoe |

| | | | |
|---|---|---|---|
| EPI_ISL_532414 | NHSGGC West of Scotland Specialist Virology Centre / MRC-University of Glasgow Centre for Virus Research | Wellcome Sanger Institute for the COVID-19 Genomics UK Consortium | Ana da Silva Filipe, Natasha Johnson, Kathy Smollett, Daniel Mair, Stephen Carmichael, Lily Tong, Jenna Nichols, Elihu Aranday-Cortes, Kirstyn Brunker, Yasmin Parr, Kyriaki Nomikou; Sarah McDonald, Marc Niebel, Patawee Asamaphan; Richard Orton, Joseph Hughes, Sreenu Vattipally, David L Robertson; Alasdair MacLean, Rory Gunson; Kathy Li, Natasha Jesudason, Rajiv Shah, James Shepherd, Antonia Ho, Alice Broos, Emma Thomson and Alex Alderton, Roberto Amato, Sonia Goncalves, Ewan Harrison, David K. Jackson, Ian Johnston, Dominic Kwiatkowski, Cordelia Langford, John Sillitoe |
| EPI_ISL_532415, EPI_ISL_532416, EPI_ISL_532417, EPI_ISL_532418, EPI_ISL_532419, EPI_ISL_532420, EPI_ISL_532421, EPI_ISL_532424, EPI_ISL_532425, EPI_ISL_532426, EPI_ISL_532428, EPI_ISL_532429 | | | |
| see above | Lighthouse Lab in Glasgow | Wellcome Sanger Institute for the COVID-19 Genomics UK Consortium | Harper VanSteenhouse, Yumi Kasai, David Gray, Carol Clugston, Anna Dominiczak and Alex Alderton, Roberto Amato, Sonia Goncalves, Ewan Harrison, David K. Jackson, Ian Johnston, Dominic Kwiatkowski, Cordelia Langford, John Sillitoe |
| EPI_ISL_532430, EPI_ISL_532432 | NHSGGC West of Scotland Specialist Virology Centre / MRC-University of Glasgow Centre for Virus Research | Wellcome Sanger Institute for the COVID-19 Genomics UK Consortium | Ana da Silva Filipe, Natasha Johnson, Kathy Smollett, Daniel Mair, Stephen Carmichael, Lily Tong, Jenna Nichols, Elihu Aranday-Cortes, Kirstyn Brunker, Yasmin Parr, Kyriaki Nomikou; Sarah McDonald, Marc Niebel, Patawee Asamaphan; Richard Orton, Joseph Hughes, Sreenu Vattipally, David L Robertson; Alasdair MacLean, Rory Gunson; Kathy Li, Natasha Jesudason, Rajiv Shah, James Shepherd, Antonia Ho, Alice Broos, Emma Thomson and Alex Alderton, Roberto Amato, Sonia Goncalves, Ewan Harrison, David K. Jackson, Ian Johnston, Dominic Kwiatkowski, Cordelia Langford, John Sillitoe |
| EPI_ISL_532433, EPI_ISL_532436, EPI_ISL_532437, EPI_ISL_532439 | Lighthouse Lab in Glasgow | Wellcome Sanger Institute for the COVID-19 Genomics UK Consortium | Harper VanSteenhouse, Yumi Kasai, David Gray, Carol Clugston, Anna Dominiczak and Alex Alderton, Roberto Amato, Sonia Goncalves, Ewan Harrison, David K. Jackson, Ian Johnston, Dominic Kwiatkowski, Cordelia Langford, John Sillitoe |
| EPI_ISL_532440 | NHSGGC West of Scotland Specialist Virology Centre / MRC-University of Glasgow Centre for Virus Research | Wellcome Sanger Institute for the COVID-19 Genomics UK Consortium | Ana da Silva Filipe, Natasha Johnson, Kathy Smollett, Daniel Mair, Stephen Carmichael, Lily Tong, Jenna Nichols, Elihu Aranday-Cortes, Kirstyn Brunker, Yasmin Parr, Kyriaki Nomikou; Sarah McDonald, Marc Niebel, Patawee Asamaphan; Richard Orton, Joseph Hughes, Sreenu Vattipally, David L Robertson; Alasdair MacLean, Rory Gunson; Kathy Li, Natasha Jesudason, Rajiv Shah, James Shepherd, Antonia Ho, Alice Broos, Emma Thomson and Alex Alderton, Roberto Amato, Sonia Goncalves, Ewan Harrison, David K. Jackson, Ian Johnston, Dominic Kwiatkowski, Cordelia Langford, John Sillitoe |
| EPI_ISL_532442 | Lighthouse Lab in Glasgow | Wellcome Sanger Institute for the COVID-19 Genomics UK Consortium | Harper VanSteenhouse, Yumi Kasai, David Gray, Carol Clugston, Anna Dominiczak and Alex Alderton, Roberto Amato, Sonia Goncalves, Ewan Harrison, David K. Jackson, Ian Johnston, Dominic Kwiatkowski, Cordelia Langford, John Sillitoe |
| EPI_ISL_532444, EPI_ISL_532446 | NHSGGC West of Scotland Specialist Virology Centre / MRC-University of Glasgow Centre for Virus Research | Wellcome Sanger Institute for the COVID-19 Genomics UK Consortium | Ana da Silva Filipe, Natasha Johnson, Kathy Smollett, Daniel Mair, Stephen Carmichael, Lily Tong, Jenna Nichols, Elihu Aranday-Cortes, Kirstyn Brunker, Yasmin Parr, Kyriaki Nomikou; Sarah McDonald, Marc Niebel, Patawee Asamaphan; Richard Orton, Joseph Hughes, Sreenu Vattipally, David L Robertson; Alasdair MacLean, Rory Gunson; Kathy Li, Natasha Jesudason, Rajiv Shah, James Shepherd, Antonia Ho, Alice Broos, Emma Thomson and Alex Alderton, Roberto Amato, Sonia Goncalves, Ewan Harrison, David K. Jackson, Ian Johnston, Dominic Kwiatkowski, Cordelia Langford, John Sillitoe |
| EPI_ISL_532447, EPI_ISL_532449, EPI_ISL_532450, EPI_ISL_532451 | Lighthouse Lab in Glasgow | Wellcome Sanger Institute for the COVID-19 Genomics UK Consortium | Harper VanSteenhouse, Yumi Kasai, David Gray, Carol Clugston, Anna Dominiczak and Alex Alderton, Roberto Amato, Sonia Goncalves, Ewan Harrison, David K. Jackson, Ian Johnston, Dominic Kwiatkowski, Cordelia Langford, John Sillitoe |
| EPI_ISL_532452 | NHSGGC West of Scotland Specialist Virology Centre / MRC-University of Glasgow Centre for Virus Research | Wellcome Sanger Institute for the COVID-19 Genomics UK Consortium | Ana da Silva Filipe, Natasha Johnson, Kathy Smollett, Daniel Mair, Stephen Carmichael, Lily Tong, Jenna Nichols, Elihu Aranday-Cortes, Kirstyn Brunker, Yasmin Parr, Kyriaki Nomikou; Sarah McDonald, Marc Niebel, Patawee Asamaphan; Richard Orton, Joseph Hughes, Sreenu Vattipally, David L Robertson; Alasdair MacLean, Rory Gunson; Kathy Li, Natasha Jesudason, Rajiv Shah, James Shepherd, Antonia Ho, Alice Broos, Emma Thomson and Alex Alderton, Roberto Amato, Sonia Goncalves, Ewan Harrison, David K. Jackson, Ian Johnston, Dominic Kwiatkowski, Cordelia Langford, John Sillitoe |
| EPI_ISL_532453 | Lighthouse Lab in Glasgow | Wellcome Sanger Institute for the COVID-19 Genomics UK Consortium | Harper VanSteenhouse, Yumi Kasai, David Gray, Carol Clugston, Anna Dominiczak and Alex Alderton, Roberto Amato, Sonia Goncalves, Ewan Harrison, David K. Jackson, Ian Johnston, Dominic Kwiatkowski, Cordelia Langford, John Sillitoe |
| EPI_ISL_532454 | NHSGGC West of Scotland Specialist Virology Centre / MRC-University of Glasgow Centre for Virus Research | Wellcome Sanger Institute for the COVID-19 Genomics UK Consortium | Ana da Silva Filipe, Natasha Johnson, Kathy Smollett, Daniel Mair, Stephen Carmichael, Lily Tong, Jenna Nichols, Elihu Aranday-Cortes, Kirstyn Brunker, Yasmin Parr, Kyriaki Nomikou; Sarah McDonald, Marc Niebel, Patawee Asamaphan; Richard Orton, Joseph Hughes, Sreenu Vattipally, David L Robertson; Alasdair MacLean, Rory Gunson; Kathy Li, Natasha Jesudason, Rajiv Shah, James Shepherd, Antonia Ho, Alice Broos, Emma Thomson and Alex Alderton, Roberto Amato, Sonia Goncalves, Ewan Harrison, David K. Jackson, Ian Johnston, Dominic Kwiatkowski, Cordelia Langford, John Sillitoe |
| EPI_ISL_532455 | Lighthouse Lab in Glasgow | Wellcome Sanger Institute for the COVID-19 Genomics UK Consortium | Harper VanSteenhouse, Yumi Kasai, David Gray, Carol Clugston, Anna Dominiczak and Alex Alderton, Roberto Amato, Sonia Goncalves, Ewan Harrison, David K. Jackson, Ian Johnston, Dominic Kwiatkowski, Cordelia Langford, John Sillitoe |
| EPI_ISL_532456 | NHSGGC West of Scotland Specialist Virology Centre / MRC-University of Glasgow Centre for Virus Research | Wellcome Sanger Institute for the COVID-19 Genomics UK Consortium | Ana da Silva Filipe, Natasha Johnson, Kathy Smollett, Daniel Mair, Stephen Carmichael, Lily Tong, Jenna Nichols, Elihu Aranday-Cortes, Kirstyn Brunker, Yasmin Parr, Kyriaki Nomikou; Sarah McDonald, Marc Niebel, Patawee Asamaphan; Richard Orton, Joseph Hughes, Sreenu Vattipally, David L Robertson; Alasdair MacLean, Rory Gunson; Kathy Li, Natasha Jesudason, Rajiv Shah, James Shepherd, Antonia Ho, Alice Broos, Emma Thomson and Alex Alderton, Roberto Amato, Sonia Goncalves, Ewan Harrison, David K. Jackson, Ian Johnston, Dominic Kwiatkowski, Cordelia Langford, John Sillitoe |
| EPI_ISL_532457, EPI_ISL_532458, EPI_ISL_532459, EPI_ISL_532461, EPI_ISL_532462 | Lighthouse Lab in Glasgow | Wellcome Sanger Institute for the COVID-19 Genomics UK Consortium | Harper VanSteenhouse, Yumi Kasai, David Gray, Carol Clugston, Anna Dominiczak and Alex Alderton, Roberto Amato, Sonia Goncalves, Ewan Harrison, David K. Jackson, Ian Johnston, Dominic Kwiatkowski, Cordelia Langford, John Sillitoe |
| EPI_ISL_532463, EPI_ISL_532464 | NHSGGC West of Scotland Specialist Virology Centre / MRC-University of Glasgow Centre for Virus Research | Wellcome Sanger Institute for the COVID-19 Genomics UK Consortium | Ana da Silva Filipe, Natasha Johnson, Kathy Smollett, Daniel Mair, Stephen Carmichael, Lily Tong, Jenna Nichols, Elihu Aranday-Cortes, Kirstyn Brunker, Yasmin Parr, Kyriaki Nomikou; Sarah McDonald, Marc Niebel, Patawee Asamaphan; Richard Orton, Joseph Hughes, Sreenu Vattipally, David L Robertson; Alasdair MacLean, Rory Gunson; Kathy Li, Natasha Jesudason, Rajiv Shah, James Shepherd, Antonia Ho, Alice Broos, Emma Thomson and Alex Alderton, Roberto Amato, Sonia Goncalves, Ewan Harrison, David K. Jackson, Ian Johnston, Dominic Kwiatkowski, Cordelia Langford, John Sillitoe |
| EPI_ISL_532465, EPI_ISL_532466, EPI_ISL_532467, EPI_ISL_532468, EPI_ISL_532470, EPI_ISL_532471 | Lighthouse Lab in Glasgow | Wellcome Sanger Institute for the COVID-19 Genomics UK Consortium | Harper VanSteenhouse, Yumi Kasai, David Gray, Carol Clugston, Anna Dominiczak and Alex Alderton, Roberto Amato, Sonia Goncalves, Ewan Harrison, David K. Jackson, Ian Johnston, Dominic Kwiatkowski, Cordelia Langford, John Sillitoe |
| EPI_ISL_532472, EPI_ISL_532473 | NHSGGC West of Scotland Specialist Virology Centre / MRC-University of Glasgow Centre for Virus Research | Wellcome Sanger Institute for the COVID-19 Genomics UK Consortium | Ana da Silva Filipe, Natasha Johnson, Kathy Smollett, Daniel Mair, Stephen Carmichael, Lily Tong, Jenna Nichols, Elihu Aranday-Cortes, Kirstyn Brunker, Yasmin Parr, Kyriaki Nomikou; Sarah McDonald, Marc Niebel, Patawee Asamaphan; Richard Orton, Joseph Hughes, Sreenu Vattipally, David L Robertson; Alasdair MacLean, Rory Gunson; Kathy Li, Natasha Jesudason, Rajiv Shah, James Shepherd, Antonia Ho, Alice Broos, Emma Thomson and Alex Alderton, Roberto Amato, Sonia Goncalves, Ewan Harrison, David K. Jackson, Ian Johnston, Dominic Kwiatkowski, Cordelia Langford, John Sillitoe |
| EPI_ISL_532474, EPI_ISL_532476, EPI_ISL_532477, EPI_ISL_532478 | Lighthouse Lab in Glasgow | Wellcome Sanger Institute for the COVID-19 Genomics UK Consortium | Harper VanSteenhouse, Yumi Kasai, David Gray, Carol Clugston, Anna Dominiczak and Alex Alderton, Roberto Amato, Sonia Goncalves, Ewan Harrison, David K. Jackson, Ian Johnston, Dominic Kwiatkowski, Cordelia Langford, John Sillitoe |
| EPI_ISL_532480 | NHSGGC West of Scotland Specialist Virology Centre / MRC-University of Glasgow Centre for Virus Research | Wellcome Sanger Institute for the COVID-19 Genomics UK Consortium | Ana da Silva Filipe, Natasha Johnson, Kathy Smollett, Daniel Mair, Stephen Carmichael, Lily Tong, Jenna Nichols, Elihu Aranday-Cortes, Kirstyn Brunker, Yasmin Parr, Kyriaki Nomikou; Sarah McDonald, Marc Niebel, Patawee Asamaphan; Richard Orton, Joseph Hughes, Sreenu Vattipally, David L Robertson; Alasdair MacLean, Rory Gunson; Kathy Li, Natasha Jesudason, Rajiv Shah, James Shepherd, Antonia Ho, Alice Broos, Emma Thomson and Alex Alderton, Roberto Amato, Sonia Goncalves, Ewan Harrison, David K. Jackson, Ian Johnston, Dominic Kwiatkowski, Cordelia Langford, John Sillitoe |
| EPI_ISL_532481, EPI_ISL_532482, EPI_ISL_532483 | Lighthouse Lab in Glasgow | Wellcome Sanger Institute for the COVID-19 Genomics UK Consortium | Harper VanSteenhouse, Yumi Kasai, David Gray, Carol Clugston, Anna Dominiczak and Alex Alderton, Roberto Amato, Sonia Goncalves, Ewan Harrison, David K. Jackson, Ian Johnston, Dominic Kwiatkowski, Cordelia Langford, John Sillitoe |
| EPI_ISL_532484 | NHSGGC West of Scotland Specialist Virology Centre / MRC-University of Glasgow Centre for Virus Research | Wellcome Sanger Institute for the COVID-19 Genomics UK Consortium | Ana da Silva Filipe, Natasha Johnson, Kathy Smollett, Daniel Mair, Stephen Carmichael, Lily Tong, Jenna Nichols, Elihu Aranday-Cortes, Kirstyn Brunker, Yasmin Parr, Kyriaki Nomikou; Sarah McDonald, Marc Niebel, Patawee Asamaphan; Richard Orton, Joseph Hughes, Sreenu Vattipally, David L Robertson; Alasdair MacLean, Rory Gunson; Kathy Li, Natasha Jesudason, Rajiv Shah, James Shepherd, Antonia Ho, Alice Broos, Emma Thomson and Alex Alderton, Roberto Amato, Sonia Goncalves, Ewan Harrison, David K. Jackson, Ian Johnston, Dominic Kwiatkowski, Cordelia Langford, John Sillitoe |
| EPI_ISL_532485, EPI_ISL_532487, EPI_ISL_532488, EPI_ISL_532489, EPI_ISL_532491 | Lighthouse Lab in Glasgow | Wellcome Sanger Institute for the COVID-19 Genomics UK Consortium | Harper VanSteenhouse, Yumi Kasai, David Gray, Carol Clugston, Anna Dominiczak and Alex Alderton, Roberto Amato, Sonia Goncalves, Ewan Harrison, David K. Jackson, Ian Johnston, Dominic Kwiatkowski, Cordelia Langford, John Sillitoe |
| EPI_ISL_532492, EPI_ISL_532493 | NHSGGC West of Scotland Specialist Virology Centre / MRC-University of Glasgow Centre for Virus Research | Wellcome Sanger Institute for the COVID-19 Genomics UK Consortium | Ana da Silva Filipe, Natasha Johnson, Kathy Smollett, Daniel Mair, Stephen Carmichael, Lily Tong, Jenna Nichols, Elihu Aranday-Cortes, Kirstyn Brunker, Yasmin Parr, Kyriaki Nomikou; Sarah McDonald, Marc Niebel, Patawee Asamaphan; Richard Orton, Joseph Hughes, Sreenu Vattipally, David L Robertson; Alasdair MacLean, Rory Gunson; Kathy Li, Natasha Jesudason, Rajiv Shah, James Shepherd, Antonia Ho, Alice Broos, Emma Thomson and Alex Alderton, Roberto Amato, Sonia Goncalves, Ewan Harrison, David K. Jackson, Ian Johnston, Dominic Kwiatkowski, Cordelia Langford, John Sillitoe |
| EPI_ISL_532494, EPI_ISL_532495, EPI_ISL_532496, EPI_ISL_532497, EPI_ISL_532498, EPI_ISL_532499, EPI_ISL_532500, EPI_ISL_532501, EPI_ISL_532502, EPI_ISL_532503, EPI_ISL_532504, EPI_ISL_532505 | | | |
| see above | Lighthouse Lab in Glasgow | Wellcome Sanger Institute for the COVID-19 Genomics UK Consortium | Harper VanSteenhouse, Yumi Kasai, David Gray, Carol Clugston, Anna Dominiczak and Alex Alderton, Roberto Amato, Sonia Goncalves, Ewan Harrison, David K. Jackson, Ian Johnston, Dominic Kwiatkowski, Cordelia Langford, John Sillitoe |
| EPI_ISL_532506 | NHSGGC West of Scotland Specialist Virology Centre / MRC-University of Glasgow Centre for Virus Research | Wellcome Sanger Institute for the COVID-19 Genomics UK Consortium | Ana da Silva Filipe, Natasha Johnson, Kathy Smollett, Daniel Mair, Stephen Carmichael, Lily Tong, Jenna Nichols, Elihu Aranday-Cortes, Kirstyn Brunker, Yasmin Parr, Kyriaki Nomikou; Sarah McDonald, Marc Niebel, Patawee Asamaphan; Richard Orton, Joseph Hughes, Sreenu Vattipally, David L Robertson; Alasdair MacLean, Rory Gunson; Kathy Li, Natasha Jesudason, Rajiv Shah, James Shepherd, Antonia Ho, Alice Broos, Emma Thomson and Alex Alderton, Roberto Amato, Sonia Goncalves, Ewan Harrison, David K. Jackson, Ian Johnston, Dominic Kwiatkowski, Cordelia Langford, John Sillitoe |
| EPI_ISL_532507, EPI_ISL_532508, EPI_ISL_532509 | Lighthouse Lab in Glasgow | Wellcome Sanger Institute for the COVID-19 Genomics UK Consortium | Harper VanSteenhouse, Yumi Kasai, David Gray, Carol Clugston, Anna Dominiczak and Alex Alderton, Roberto Amato, Sonia Goncalves, Ewan Harrison, David K. Jackson, Ian Johnston, Dominic Kwiatkowski, Cordelia Langford, John Sillitoe |
| EPI_ISL_532510, EPI_ISL_532512 | NHSGGC West of Scotland Specialist Virology Centre / MRC-University of Glasgow Centre for Virus Research | Wellcome Sanger Institute for the COVID-19 Genomics UK Consortium | Ana da Silva Filipe, Natasha Johnson, Kathy Smollett, Daniel Mair, Stephen Carmichael, Lily Tong, Jenna Nichols, Elihu Aranday-Cortes, Kirstyn Brunker, Yasmin Parr, Kyriaki Nomikou; Sarah McDonald, Marc Niebel, Patawee Asamaphan; Richard Orton, Joseph Hughes, Sreenu Vattipally, David L Robertson; Alasdair MacLean, Rory Gunson; Kathy Li, Natasha Jesudason, Rajiv Shah, James Shepherd, Antonia Ho, Alice Broos, Emma Thomson and Alex Alderton, Roberto Amato, Sonia Goncalves, Ewan Harrison, David K. Jackson, Ian Johnston, Dominic Kwiatkowski, Cordelia Langford, John Sillitoe |
| EPI_ISL_532513 | Lighthouse Lab in Glasgow | Wellcome Sanger Institute for the COVID-19 Genomics UK Consortium | Harper VanSteenhouse, Yumi Kasai, David Gray, Carol Clugston, Anna Dominiczak and Alex Alderton, Roberto Amato, Sonia Goncalves, Ewan Harrison, David K. Jackson, Ian Johnston, Dominic Kwiatkowski, Cordelia Langford, John Sillitoe |
| EPI_ISL_532514, EPI_ISL_532515 | NHSGGC West of Scotland Specialist Virology Centre / MRC-University of Glasgow Centre for Virus Research | Wellcome Sanger Institute for the COVID-19 Genomics UK Consortium | Ana da Silva Filipe, Natasha Johnson, Kathy Smollett, Daniel Mair, Stephen Carmichael, Lily Tong, Jenna Nichols, Elihu Aranday-Cortes, Kirstyn Brunker, Yasmin Parr, Kyriaki Nomikou; Sarah McDonald, Marc Niebel, Patawee Asamaphan; Richard Orton, Joseph Hughes, Sreenu Vattipally, David L Robertson; Alasdair MacLean, Rory Gunson; Kathy Li, Natasha Jesudason, Rajiv Shah, James Shepherd, Antonia Ho, Alice Broos, Emma Thomson and Alex Alderton, Roberto Amato, Sonia Goncalves, Ewan Harrison, David K. Jackson, Ian Johnston, Dominic Kwiatkowski, Cordelia Langford, John Sillitoe |
| EPI_ISL_532516, EPI_ISL_532517 | Lighthouse Lab in Glasgow | Wellcome Sanger Institute for the COVID-19 Genomics UK Consortium | Harper VanSteenhouse, Yumi Kasai, David Gray, Carol Clugston, Anna Dominiczak and Alex Alderton, Roberto Amato, Sonia Goncalves, Ewan Harrison, David K. Jackson, Ian Johnston, Dominic Kwiatkowski, Cordelia Langford, John Sillitoe |
| EPI_ISL_532518 | NHSGGC West of Scotland Specialist Virology Centre / MRC-University of Glasgow Centre for Virus Research | Wellcome Sanger Institute for the COVID-19 Genomics UK Consortium | Ana da Silva Filipe, Natasha Johnson, Kathy Smollett, Daniel Mair, Stephen Carmichael, Lily Tong, Jenna Nichols, Elihu Aranday-Cortes, Kirstyn Brunker, Yasmin Parr, Kyriaki Nomikou; Sarah McDonald, Marc Niebel, Patawee Asamaphan; Richard Orton, Joseph Hughes, Sreenu Vattipally, David L Robertson; Alasdair MacLean, Rory Gunson; Kathy Li, Natasha Jesudason, Rajiv Shah, James Shepherd, Antonia Ho, Alice Broos, Emma Thomson and Alex Alderton, Roberto Amato, Sonia Goncalves, Ewan Harrison, David K. Jackson, Ian Johnston, Dominic Kwiatkowski, Cordelia Langford, John Sillitoe |
| EPI_ISL_532520, EPI_ISL_532521, EPI_ISL_532522 | Lighthouse Lab in Glasgow | Wellcome Sanger Institute for the COVID-19 Genomics UK Consortium | Harper VanSteenhouse, Yumi Kasai, David Gray, Carol Clugston, Anna Dominiczak and Alex Alderton, Roberto Amato, Sonia Goncalves, Ewan Harrison, David K. Jackson, Ian Johnston, Dominic Kwiatkowski, Cordelia Langford, John Sillitoe |
| EPI_ISL_532523 | NHSGGC West of Scotland Specialist Virology Centre / MRC-University of Glasgow Centre for Virus Research | Wellcome Sanger Institute for the COVID-19 Genomics UK Consortium | Ana da Silva Filipe, Natasha Johnson, Kathy Smollett, Daniel Mair, Stephen Carmichael, Lily Tong, Jenna Nichols, Elihu Aranday-Cortes, Kirstyn Brunker, Yasmin Parr, Kyriaki Nomikou; Sarah McDonald, Marc Niebel, Patawee Asamaphan; Richard Orton, Joseph Hughes, Sreenu Vattipally, David L Robertson; Alasdair MacLean, Rory Gunson; Kathy Li, Natasha Jesudason, Rajiv Shah, James Shepherd, Antonia Ho, Alice Broos, Emma Thomson and Alex Alderton, Roberto Amato, Sonia Goncalves, Ewan Harrison, David K. Jackson, Ian Johnston, Dominic Kwiatkowski, Cordelia Langford, John Sillitoe |
| EPI_ISL_532524, EPI_ISL_532525, EPI_ISL_532526, EPI_ISL_532527, EPI_ISL_532528, EPI_ISL_532529, EPI_ISL_532530 | Lighthouse Lab in Glasgow | Wellcome Sanger Institute for the COVID-19 Genomics UK Consortium | Harper VanSteenhouse, Yumi Kasai, David Gray, Carol Clugston, Anna Dominiczak and Alex Alderton, Roberto Amato, Sonia Goncalves, Ewan Harrison, David K. Jackson, Ian Johnston, Dominic Kwiatkowski, Cordelia Langford, John Sillitoe |
| EPI_ISL_532531 | NHSGGC West of Scotland Specialist Virology Centre / MRC-University of Glasgow Centre for Virus Research | Wellcome Sanger Institute for the COVID-19 Genomics UK Consortium | Ana da Silva Filipe, Natasha Johnson, Kathy Smollett, Daniel Mair, Stephen Carmichael, Lily Tong, Jenna Nichols, Elihu Aranday-Cortes, Kirstyn Brunker, Yasmin Parr, Kyriaki Nomikou; Sarah McDonald, Marc Niebel, Patawee Asamaphan; Richard Orton, Joseph Hughes, Sreenu Vattipally, David L Robertson; Alasdair MacLean, Rory Gunson; Kathy Li, Natasha Jesudason, Rajiv Shah, James Shepherd, Antonia Ho, Alice Broos, Emma Thomson and Alex Alderton, Roberto Amato, Sonia Goncalves, Ewan Harrison, David K. Jackson, Ian Johnston, Dominic Kwiatkowski, Cordelia Langford, John Sillitoe |
| EPI_ISL_532532, EPI_ISL_532533, EPI_ISL_532534, EPI_ISL_532535, EPI_ISL_532536, EPI_ISL_532537 | Lighthouse Lab in Glasgow | Wellcome Sanger Institute for the COVID-19 Genomics UK Consortium | Harper VanSteenhouse, Yumi Kasai, David Gray, Carol Clugston, Anna Dominiczak and Alex Alderton, Roberto Amato, Sonia Goncalves, Ewan Harrison, David K. Jackson, Ian Johnston, Dominic Kwiatkowski, Cordelia Langford, John Sillitoe |

| | | | |
|---|---|---|---|
| EPI_ISL_532538 | NHSGGC West of Scotland Specialist Virology Centre / MRC-University of Glasgow Centre for Virus Research | Wellcome Sanger Institute for the COVID-19 Genomics UK Consortium | Ana da Silva Filipe, Natasha Johnson, Kathy Smollett, Daniel Mair, Stephen Carmichael, Lily Tong, Jenna Nichols, Elihu Aranday-Cortes, Kirstyn Brunker, Yasmin Parr, Kyriaki Nomikou; Sarah McDonald, Marc Niebel, Patawee Asamaphan; Richard Orton, Joseph Hughes, Sreenu Vattipally, David L Robertson; Alasdair MacLean, Rory Gunson; Kathy Li, Natasha Jesudason, Rajiv Shah, James Shepherd, Antonia Ho, Alice Broos, Emma Thomson and Alex Alderton, Roberto Amato, Sonia Goncalves, Ewan Harrison, David K. Jackson, Ian Johnston, Dominic Kwiatkowski, Cordelia Langford, John Sillitoe |
| EPI_ISL_532539 | Lighthouse Lab in Glasgow | Wellcome Sanger Institute for the COVID-19 Genomics UK Consortium | Harper VanSteenhouse, Yumi Kasai, David Gray, Carol Clugston, Anna Dominiczak and Alex Alderton, Roberto Amato, Sonia Goncalves, Ewan Harrison, David K. Jackson, Ian Johnston, Dominic Kwiatkowski, Cordelia Langford, John Sillitoe |
| EPI_ISL_532540 | NHSGGC West of Scotland Specialist Virology Centre / MRC-University of Glasgow Centre for Virus Research | Wellcome Sanger Institute for the COVID-19 Genomics UK Consortium | Ana da Silva Filipe, Natasha Johnson, Kathy Smollett, Daniel Mair, Stephen Carmichael, Lily Tong, Jenna Nichols, Elihu Aranday-Cortes, Kirstyn Brunker, Yasmin Parr, Kyriaki Nomikou; Sarah McDonald, Marc Niebel, Patawee Asamaphan; Richard Orton, Joseph Hughes, Sreenu Vattipally, David L Robertson; Alasdair MacLean, Rory Gunson; Kathy Li, Natasha Jesudason, Rajiv Shah, James Shepherd, Antonia Ho, Alice Broos, Emma Thomson and Alex Alderton, Roberto Amato, Sonia Goncalves, Ewan Harrison, David K. Jackson, Ian Johnston, Dominic Kwiatkowski, Cordelia Langford, John Sillitoe |
| EPI_ISL_532541, EPI_ISL_532542, EPI_ISL_532543, EPI_ISL_532544, EPI_ISL_532545, EPI_ISL_532548, EPI_ISL_532550, EPI_ISL_532553, EPI_ISL_532554, EPI_ISL_532555, EPI_ISL_532556, EPI_ISL_532557, EPI_ISL_532558, EPI_ISL_532559, EPI_ISL_532561, EPI_ISL_532562 | | | |
| see above | Lighthouse Lab in Glasgow | Wellcome Sanger Institute for the COVID-19 Genomics UK Consortium | Harper VanSteenhouse, Yumi Kasai, David Gray, Carol Clugston, Anna Dominiczak and Alex Alderton, Roberto Amato, Sonia Goncalves, Ewan Harrison, David K. Jackson, Ian Johnston, Dominic Kwiatkowski, Cordelia Langford, John Sillitoe |
| EPI_ISL_532563, EPI_ISL_532564 | NHSGGC West of Scotland Specialist Virology Centre / MRC-University of Glasgow Centre for Virus Research | Wellcome Sanger Institute for the COVID-19 Genomics UK Consortium | Ana da Silva Filipe, Natasha Johnson, Kathy Smollett, Daniel Mair, Stephen Carmichael, Lily Tong, Jenna Nichols, Elihu Aranday-Cortes, Kirstyn Brunker, Yasmin Parr, Kyriaki Nomikou; Sarah McDonald, Marc Niebel, Patawee Asamaphan; Richard Orton, Joseph Hughes, Sreenu Vattipally, David L Robertson; Alasdair MacLean, Rory Gunson; Kathy Li, Natasha Jesudason, Rajiv Shah, James Shepherd, Antonia Ho, Alice Broos, Emma Thomson and Alex Alderton, Roberto Amato, Sonia Goncalves, Ewan Harrison, David K. Jackson, Ian Johnston, Dominic Kwiatkowski, Cordelia Langford, John Sillitoe |
| EPI_ISL_532565, EPI_ISL_532567, EPI_ISL_532569, EPI_ISL_532570, EPI_ISL_532571 | Lighthouse Lab in Glasgow | Wellcome Sanger Institute for the COVID-19 Genomics UK Consortium | Harper VanSteenhouse, Yumi Kasai, David Gray, Carol Clugston, Anna Dominiczak and Alex Alderton, Roberto Amato, Sonia Goncalves, Ewan Harrison, David K. Jackson, Ian Johnston, Dominic Kwiatkowski, Cordelia Langford, John Sillitoe |
| EPI_ISL_532572 | NHSGGC West of Scotland Specialist Virology Centre / MRC-University of Glasgow Centre for Virus Research | Wellcome Sanger Institute for the COVID-19 Genomics UK Consortium | Ana da Silva Filipe, Natasha Johnson, Kathy Smollett, Daniel Mair, Stephen Carmichael, Lily Tong, Jenna Nichols, Elihu Aranday-Cortes, Kirstyn Brunker, Yasmin Parr, Kyriaki Nomikou; Sarah McDonald, Marc Niebel, Patawee Asamaphan; Richard Orton, Joseph Hughes, Sreenu Vattipally, David L Robertson; Alasdair MacLean, Rory Gunson; Kathy Li, Natasha Jesudason, Rajiv Shah, James Shepherd, Antonia Ho, Alice Broos, Emma Thomson and Alex Alderton, Roberto Amato, Sonia Goncalves, Ewan Harrison, David K. Jackson, Ian Johnston, Dominic Kwiatkowski, Cordelia Langford, John Sillitoe |
| EPI_ISL_532573, EPI_ISL_532574, EPI_ISL_532575, EPI_ISL_532576, EPI_ISL_532577, EPI_ISL_532578, EPI_ISL_532579, EPI_ISL_532581 | Lighthouse Lab in Glasgow | Wellcome Sanger Institute for the COVID-19 Genomics UK Consortium | Harper VanSteenhouse, Yumi Kasai, David Gray, Carol Clugston, Anna Dominiczak and Alex Alderton, Roberto Amato, Sonia Goncalves, Ewan Harrison, David K. Jackson, Ian Johnston, Dominic Kwiatkowski, Cordelia Langford, John Sillitoe |
| EPI_ISL_532582 | NHSGGC West of Scotland Specialist Virology Centre / MRC-University of Glasgow Centre for Virus Research | Wellcome Sanger Institute for the COVID-19 Genomics UK Consortium | Ana da Silva Filipe, Natasha Johnson, Kathy Smollett, Daniel Mair, Stephen Carmichael, Lily Tong, Jenna Nichols, Elihu Aranday-Cortes, Kirstyn Brunker, Yasmin Parr, Kyriaki Nomikou; Sarah McDonald, Marc Niebel, Patawee Asamaphan; Richard Orton, Joseph Hughes, Sreenu Vattipally, David L Robertson; Alasdair MacLean, Rory Gunson; Kathy Li, Natasha Jesudason, Rajiv Shah, James Shepherd, Antonia Ho, Alice Broos, Emma Thomson and Alex Alderton, Roberto Amato, Sonia Goncalves, Ewan Harrison, David K. Jackson, Ian Johnston, Dominic Kwiatkowski, Cordelia Langford, John Sillitoe |
| EPI_ISL_532584, EPI_ISL_532585, EPI_ISL_532586 | Lighthouse Lab in Glasgow | Wellcome Sanger Institute for the COVID-19 Genomics UK Consortium | Harper VanSteenhouse, Yumi Kasai, David Gray, Carol Clugston, Anna Dominiczak and Alex Alderton, Roberto Amato, Sonia Goncalves, Ewan Harrison, David K. Jackson, Ian Johnston, Dominic Kwiatkowski, Cordelia Langford, John Sillitoe |
| EPI_ISL_532587 | NHSGGC West of Scotland Specialist Virology Centre / MRC-University of Glasgow Centre for Virus Research | Wellcome Sanger Institute for the COVID-19 Genomics UK Consortium | Ana da Silva Filipe, Natasha Johnson, Kathy Smollett, Daniel Mair, Stephen Carmichael, Lily Tong, Jenna Nichols, Elihu Aranday-Cortes, Kirstyn Brunker, Yasmin Parr, Kyriaki Nomikou; Sarah McDonald, Marc Niebel, Patawee Asamaphan; Richard Orton, Joseph Hughes, Sreenu Vattipally, David L Robertson; Alasdair MacLean, Rory Gunson; Kathy Li, Natasha Jesudason, Rajiv Shah, James Shepherd, Antonia Ho, Alice Broos, Emma Thomson and Alex Alderton, Roberto Amato, Sonia Goncalves, Ewan Harrison, David K. Jackson, Ian Johnston, Dominic Kwiatkowski, Cordelia Langford, John Sillitoe |
| EPI_ISL_532588, EPI_ISL_532589, EPI_ISL_532590, EPI_ISL_532591, EPI_ISL_532592, EPI_ISL_532593 | Lighthouse Lab in Glasgow | Wellcome Sanger Institute for the COVID-19 Genomics UK Consortium | Harper VanSteenhouse, Yumi Kasai, David Gray, Carol Clugston, Anna Dominiczak and Alex Alderton, Roberto Amato, Sonia Goncalves, Ewan Harrison, David K. Jackson, Ian Johnston, Dominic Kwiatkowski, Cordelia Langford, John Sillitoe |
| EPI_ISL_532594 | NHSGGC West of Scotland Specialist Virology Centre / MRC-University of Glasgow Centre for Virus Research | Wellcome Sanger Institute for the COVID-19 Genomics UK Consortium | Ana da Silva Filipe, Natasha Johnson, Kathy Smollett, Daniel Mair, Stephen Carmichael, Lily Tong, Jenna Nichols, Elihu Aranday-Cortes, Kirstyn Brunker, Yasmin Parr, Kyriaki Nomikou; Sarah McDonald, Marc Niebel, Patawee Asamaphan; Richard Orton, Joseph Hughes, Sreenu Vattipally, David L Robertson; Alasdair MacLean, Rory Gunson; Kathy Li, Natasha Jesudason, Rajiv Shah, James Shepherd, Antonia Ho, Alice Broos, Emma Thomson and Alex Alderton, Roberto Amato, Sonia Goncalves, Ewan Harrison, David K. Jackson, Ian Johnston, Dominic Kwiatkowski, Cordelia Langford, John Sillitoe |
| EPI_ISL_532595, EPI_ISL_532597 | Lighthouse Lab in Glasgow | Wellcome Sanger Institute for the COVID-19 Genomics UK Consortium | Harper VanSteenhouse, Yumi Kasai, David Gray, Carol Clugston, Anna Dominiczak and Alex Alderton, Roberto Amato, Sonia Goncalves, Ewan Harrison, David K. Jackson, Ian Johnston, Dominic Kwiatkowski, Cordelia Langford, John Sillitoe |
| EPI_ISL_532598 | NHSGGC West of Scotland Specialist Virology Centre / MRC-University of Glasgow Centre for Virus Research | Wellcome Sanger Institute for the COVID-19 Genomics UK Consortium | Ana da Silva Filipe, Natasha Johnson, Kathy Smollett, Daniel Mair, Stephen Carmichael, Lily Tong, Jenna Nichols, Elihu Aranday-Cortes, Kirstyn Brunker, Yasmin Parr, Kyriaki Nomikou; Sarah McDonald, Marc Niebel, Patawee Asamaphan; Richard Orton, Joseph Hughes, Sreenu Vattipally, David L Robertson; Alasdair MacLean, Rory Gunson; Kathy Li, Natasha Jesudason, Rajiv Shah, James Shepherd, Antonia Ho, Alice Broos, Emma Thomson and Alex Alderton, Roberto Amato, Sonia Goncalves, Ewan Harrison, David K. Jackson, Ian Johnston, Dominic Kwiatkowski, Cordelia Langford, John Sillitoe |
| EPI_ISL_532599, EPI_ISL_532600, EPI_ISL_532601, EPI_ISL_532602, EPI_ISL_532603, EPI_ISL_532604, EPI_ISL_532605, EPI_ISL_532607, EPI_ISL_532608, EPI_ISL_532609, EPI_ISL_532610, EPI_ISL_532613, EPI_ISL_532614, EPI_ISL_532615, EPI_ISL_532616, EPI_ISL_532617, EPI_ISL_532618, EPI_ISL_532619 | | | |
| see above | Lighthouse Lab in Glasgow | Wellcome Sanger Institute for the COVID-19 Genomics UK Consortium | Harper VanSteenhouse, Yumi Kasai, David Gray, Carol Clugston, Anna Dominiczak and Alex Alderton, Roberto Amato, Sonia Goncalves, Ewan Harrison, David K. Jackson, Ian Johnston, Dominic Kwiatkowski, Cordelia Langford, John Sillitoe |
| EPI_ISL_532620, EPI_ISL_532621 | NHSGGC West of Scotland Specialist Virology Centre / MRC-University of Glasgow Centre for Virus Research | Wellcome Sanger Institute for the COVID-19 Genomics UK Consortium | Ana da Silva Filipe, Natasha Johnson, Kathy Smollett, Daniel Mair, Stephen Carmichael, Lily Tong, Jenna Nichols, Elihu Aranday-Cortes, Kirstyn Brunker, Yasmin Parr, Kyriaki Nomikou; Sarah McDonald, Marc Niebel, Patawee Asamaphan; Richard Orton, Joseph Hughes, Sreenu Vattipally, David L Robertson; Alasdair MacLean, Rory Gunson; Kathy Li, Natasha Jesudason, Rajiv Shah, James Shepherd, Antonia Ho, Alice Broos, Emma Thomson and Alex Alderton, Roberto Amato, Sonia Goncalves, Ewan Harrison, David K. Jackson, Ian Johnston, Dominic Kwiatkowski, Cordelia Langford, John Sillitoe |
| EPI_ISL_532622, EPI_ISL_532623, EPI_ISL_532624 | Lighthouse Lab in Glasgow | Wellcome Sanger Institute for the COVID-19 Genomics UK Consortium | Harper VanSteenhouse, Yumi Kasai, David Gray, Carol Clugston, Anna Dominiczak and Alex Alderton, Roberto Amato, Sonia Goncalves, Ewan Harrison, David K. Jackson, Ian Johnston, Dominic Kwiatkowski, Cordelia Langford, John Sillitoe |
| EPI_ISL_532625, EPI_ISL_532626 | NHSGGC West of Scotland Specialist Virology Centre / MRC-University of Glasgow Centre for Virus Research | Wellcome Sanger Institute for the COVID-19 Genomics UK Consortium | Ana da Silva Filipe, Natasha Johnson, Kathy Smollett, Daniel Mair, Stephen Carmichael, Lily Tong, Jenna Nichols, Elihu Aranday-Cortes, Kirstyn Brunker, Yasmin Parr, Kyriaki Nomikou; Sarah McDonald, Marc Niebel, Patawee Asamaphan; Richard Orton, Joseph Hughes, Sreenu Vattipally, David L Robertson; Alasdair MacLean, Rory Gunson; Kathy Li, Natasha Jesudason, Rajiv Shah, James Shepherd, Antonia Ho, Alice Broos, Emma Thomson and Alex Alderton, Roberto Amato, Sonia Goncalves, Ewan Harrison, David K. Jackson, Ian Johnston, Dominic Kwiatkowski, Cordelia Langford, John Sillitoe |
| EPI_ISL_532627, EPI_ISL_532628, EPI_ISL_532629, EPI_ISL_532630, EPI_ISL_532631, EPI_ISL_532632, EPI_ISL_532634, EPI_ISL_532635, EPI_ISL_532637, EPI_ISL_532641, EPI_ISL_532642, EPI_ISL_532643, EPI_ISL_532644, EPI_ISL_532647 | | | |
| see above | Lighthouse Lab in Glasgow | Wellcome Sanger Institute for the COVID-19 Genomics UK Consortium | Harper VanSteenhouse, Yumi Kasai, David Gray, Carol Clugston, Anna Dominiczak and Alex Alderton, Roberto Amato, Sonia Goncalves, Ewan Harrison, David K. Jackson, Ian Johnston, Dominic Kwiatkowski, Cordelia Langford, John Sillitoe |
| EPI_ISL_532648 | NHSGGC West of Scotland Specialist Virology Centre / MRC-University of Glasgow Centre for Virus Research | Wellcome Sanger Institute for the COVID-19 Genomics UK Consortium | Ana da Silva Filipe, Natasha Johnson, Kathy Smollett, Daniel Mair, Stephen Carmichael, Lily Tong, Jenna Nichols, Elihu Aranday-Cortes, Kirstyn Brunker, Yasmin Parr, Kyriaki Nomikou; Sarah McDonald, Marc Niebel, Patawee Asamaphan; Richard Orton, Joseph Hughes, Sreenu Vattipally, David L Robertson; Alasdair MacLean, Rory Gunson; Kathy Li, Natasha Jesudason, Rajiv Shah, James Shepherd, Antonia Ho, Alice Broos, Emma Thomson and Alex Alderton, Roberto Amato, Sonia Goncalves, Ewan Harrison, David K. Jackson, Ian Johnston, Dominic Kwiatkowski, Cordelia Langford, John Sillitoe |
| EPI_ISL_532649, EPI_ISL_532650, EPI_ISL_532651, EPI_ISL_532652, EPI_ISL_532654, EPI_ISL_532656, EPI_ISL_532657 | Lighthouse Lab in Glasgow | Wellcome Sanger Institute for the COVID-19 Genomics UK Consortium | Harper VanSteenhouse, Yumi Kasai, David Gray, Carol Clugston, Anna Dominiczak and Alex Alderton, Roberto Amato, Sonia Goncalves, Ewan Harrison, David K. Jackson, Ian Johnston, Dominic Kwiatkowski, Cordelia Langford, John Sillitoe |
| EPI_ISL_532659 | Virology Department, Royal Infirmary of Edinburgh, NHS Lothian / School of Biological Sciences, University of Edinburgh | Wellcome Sanger Institute for the COVID-19 Genomics UK Consortium | McHugh M, Dewar R, Rooke S, O'Toole Á, Scher E, Hill V, McCrone JT, Colquhoun R, Yu X, Jackson B, Rambaut A, Templeton K and Alex Alderton, Roberto Amato, Sonia Goncalves, Ewan Harrison, David K. Jackson, Ian Johnston, Dominic Kwiatkowski, Cordelia Langford, John Sillitoe |
| EPI_ISL_532660, EPI_ISL_532662, EPI_ISL_532663, EPI_ISL_532665, EPI_ISL_532666, EPI_ISL_532667, EPI_ISL_532668 | Lighthouse Lab in Glasgow | Wellcome Sanger Institute for the COVID-19 Genomics UK Consortium | Harper VanSteenhouse, Yumi Kasai, David Gray, Carol Clugston, Anna Dominiczak and Alex Alderton, Roberto Amato, Sonia Goncalves, Ewan Harrison, David K. Jackson, Ian Johnston, Dominic Kwiatkowski, Cordelia Langford, John Sillitoe |
| EPI_ISL_532670 | NHSGGC West of Scotland Specialist Virology Centre / MRC-University of Glasgow Centre for Virus Research | Wellcome Sanger Institute for the COVID-19 Genomics UK Consortium | Ana da Silva Filipe, Natasha Johnson, Kathy Smollett, Daniel Mair, Stephen Carmichael, Lily Tong, Jenna Nichols, Elihu Aranday-Cortes, Kirstyn Brunker, Yasmin Parr, Kyriaki Nomikou; Sarah McDonald, Marc Niebel, Patawee Asamaphan; Richard Orton, Joseph Hughes, Sreenu Vattipally, David L Robertson; Alasdair MacLean, Rory Gunson; Kathy Li, Natasha Jesudason, Rajiv Shah, James Shepherd, Antonia Ho, Alice Broos, Emma Thomson and Alex Alderton, Roberto Amato, Sonia Goncalves, Ewan Harrison, David K. Jackson, Ian Johnston, Dominic Kwiatkowski, Cordelia Langford, John Sillitoe |
| EPI_ISL_532671, EPI_ISL_532674, EPI_ISL_532675, EPI_ISL_532676, EPI_ISL_532677, EPI_ISL_532678, EPI_ISL_532679, EPI_ISL_532680, EPI_ISL_532681, EPI_ISL_532682 | Lighthouse Lab in Glasgow | Wellcome Sanger Institute for the COVID-19 Genomics UK Consortium | Harper VanSteenhouse, Yumi Kasai, David Gray, Carol Clugston, Anna Dominiczak and Alex Alderton, Roberto Amato, Sonia Goncalves, Ewan Harrison, David K. Jackson, Ian Johnston, Dominic Kwiatkowski, Cordelia Langford, John Sillitoe |
| EPI_ISL_532683 | Virology Department, Royal Infirmary of Edinburgh, NHS Lothian / School of Biological Sciences, University of Edinburgh | Wellcome Sanger Institute for the COVID-19 Genomics UK Consortium | McHugh M, Dewar R, Rooke S, O'Toole Á, Scher E, Hill V, McCrone JT, Colquhoun R, Yu X, Jackson B, Rambaut A, Templeton K and Alex Alderton, Roberto Amato, Sonia Goncalves, Ewan Harrison, David K. Jackson, Ian Johnston, Dominic Kwiatkowski, Cordelia Langford, John Sillitoe |
| EPI_ISL_532685, EPI_ISL_532686, EPI_ISL_532687, EPI_ISL_532688 | Lighthouse Lab in Glasgow | Wellcome Sanger Institute for the COVID-19 Genomics UK Consortium | Harper VanSteenhouse, Yumi Kasai, David Gray, Carol Clugston, Anna Dominiczak and Alex Alderton, Roberto Amato, Sonia Goncalves, Ewan Harrison, David K. Jackson, Ian Johnston, Dominic Kwiatkowski, Cordelia Langford, John Sillitoe |
| EPI_ISL_532689 | Virology Department, Royal Infirmary of Edinburgh, NHS Lothian / School of Biological Sciences, University of Edinburgh | Wellcome Sanger Institute for the COVID-19 Genomics UK Consortium | McHugh M, Dewar R, Rooke S, O'Toole Á, Scher E, Hill V, McCrone JT, Colquhoun R, Yu X, Jackson B, Rambaut A, Templeton K and Alex Alderton, Roberto Amato, Sonia Goncalves, Ewan Harrison, David K. Jackson, Ian Johnston, Dominic Kwiatkowski, Cordelia Langford, John Sillitoe |
| EPI_ISL_532690, EPI_ISL_532691, EPI_ISL_532692, EPI_ISL_532693, EPI_ISL_532694, EPI_ISL_532695, EPI_ISL_532696 | Lighthouse Lab in Glasgow | Wellcome Sanger Institute for the COVID-19 Genomics UK Consortium | Harper VanSteenhouse, Yumi Kasai, David Gray, Carol Clugston, Anna Dominiczak and Alex Alderton, Roberto Amato, Sonia Goncalves, Ewan Harrison, David K. Jackson, Ian Johnston, Dominic Kwiatkowski, Cordelia Langford, John Sillitoe |
| EPI_ISL_532697 | NHSGGC West of Scotland Specialist Virology Centre / MRC-University of Glasgow Centre for Virus Research | Wellcome Sanger Institute for the COVID-19 Genomics UK Consortium | Ana da Silva Filipe, Natasha Johnson, Kathy Smollett, Daniel Mair, Stephen Carmichael, Lily Tong, Jenna Nichols, Elihu Aranday-Cortes, Kirstyn Brunker, Yasmin Parr, Kyriaki Nomikou; Sarah McDonald, Marc Niebel, Patawee Asamaphan; Richard Orton, Joseph Hughes, Sreenu Vattipally, David L Robertson; Alasdair MacLean, Rory Gunson; Kathy Li, Natasha Jesudason, Rajiv Shah, James Shepherd, Antonia Ho, Alice Broos, Emma Thomson and Alex Alderton, Roberto Amato, Sonia Goncalves, Ewan Harrison, David K. Jackson, Ian Johnston, Dominic Kwiatkowski, Cordelia Langford, John Sillitoe |
| EPI_ISL_532698, EPI_ISL_532699, EPI_ISL_532700, EPI_ISL_532701, EPI_ISL_532702, EPI_ISL_532703, EPI_ISL_532704 | Lighthouse Lab in Glasgow | Wellcome Sanger Institute for the COVID-19 Genomics UK Consortium | Harper VanSteenhouse, Yumi Kasai, David Gray, Carol Clugston, Anna Dominiczak and Alex Alderton, Roberto Amato, Sonia Goncalves, Ewan Harrison, David K. Jackson, Ian Johnston, Dominic Kwiatkowski, Cordelia Langford, John Sillitoe |
| EPI_ISL_532705 | Virology Department, Royal Infirmary of Edinburgh, NHS Lothian / School of Biological Sciences, University of Edinburgh | Wellcome Sanger Institute for the COVID-19 Genomics UK Consortium | McHugh M, Dewar R, Rooke S, O'Toole Á, Scher E, Hill V, McCrone JT, Colquhoun R, Yu X, Jackson B, Rambaut A, Templeton K and Alex Alderton, Roberto Amato, Sonia Goncalves, Ewan Harrison, David K. Jackson, Ian Johnston, Dominic Kwiatkowski, Cordelia Langford, John Sillitoe |
| EPI_ISL_532706, EPI_ISL_532708, EPI_ISL_532711, EPI_ISL_532712, EPI_ISL_532713, EPI_ISL_532714 | Lighthouse Lab in Glasgow | Wellcome Sanger Institute for the COVID-19 Genomics UK Consortium | Harper VanSteenhouse, Yumi Kasai, David Gray, Carol Clugston, Anna Dominiczak and Alex Alderton, Roberto Amato, Sonia Goncalves, Ewan Harrison, David K. Jackson, Ian Johnston, Dominic Kwiatkowski, Cordelia Langford, John Sillitoe |
| EPI_ISL_532715 | NHSGGC West of Scotland Specialist Virology Centre / MRC-University of Glasgow Centre for Virus Research | Wellcome Sanger Institute for the COVID-19 Genomics UK Consortium | Ana da Silva Filipe, Natasha Johnson, Kathy Smollett, Daniel Mair, Stephen Carmichael, Lily Tong, Jenna Nichols, Elihu Aranday-Cortes, Kirstyn Brunker, Yasmin Parr, Kyriaki Nomikou; Sarah McDonald, Marc Niebel, Patawee Asamaphan; Richard Orton, Joseph Hughes, Sreenu Vattipally, David L Robertson; Alasdair MacLean, Rory Gunson; Kathy Li, Natasha Jesudason, Rajiv Shah, James Shepherd, Antonia Ho, Alice Broos, Emma Thomson and Alex Alderton, Roberto Amato, Sonia Goncalves, Ewan Harrison, David K. Jackson, Ian Johnston, Dominic Kwiatkowski, Cordelia Langford, John Sillitoe |

| | | | |
|---|---|---|---|
| EPI_ISL_532716, EPI_ISL_532717, EPI_ISL_532718, EPI_ISL_532719, EPI_ISL_532721, EPI_ISL_532722, EPI_ISL_532724, EPI_ISL_532725, EPI_ISL_532726, EPI_ISL_532727, EPI_ISL_532729, EPI_ISL_532730, EPI_ISL_532731, EPI_ISL_532732 | | | |
| see above | Lighthouse Lab in Glasgow | Wellcome Sanger Institute for the COVID-19 Genomics UK Consortium | Harper VanSteenhouse, Yumi Kasai, David Gray, Carol Clugston, Anna Dominiczak and Alex Alderton, Roberto Amato, Sonia Goncalves, Ewan Harrison, David K. Jackson, Ian Johnston, Dominic Kwiatkowski, Cordelia Langford, John Sillitoe |
| EPI_ISL_532733 | NHSGGC West of Scotland Specialist Virology Centre / MRC-University of Glasgow Centre for Virus Research | Wellcome Sanger Institute for the COVID-19 Genomics UK Consortium | Ana da Silva Filipe, Natasha Johnson, Kathy Smollett, Daniel Mair, Stephen Carmichael, Lily Tong, Jenna Nichols, Elihu Aranday-Cortes, Kirstyn Brunker, Yasmin Parr, Kyriaki Nomikou; Sarah McDonald, Marc Niebel, Patawee Asamaphan; Richard Orton, Joseph Hughes, Sreenu Vattipally, David L Robertson; Alasdair MacLean, Rory Gunson; Kathy Li, Natasha Jesudason, Rajiv Shah, James Shepherd, Antonia Ho, Alice Broos, Emma Thomson and Alex Alderton, Roberto Amato, Sonia Goncalves, Ewan Harrison, David K. Jackson, Ian Johnston, Dominic Kwiatkowski, Cordelia Langford, John Sillitoe |
| EPI_ISL_532734, EPI_ISL_532735, EPI_ISL_532736, EPI_ISL_532737 | Lighthouse Lab in Glasgow | Wellcome Sanger Institute for the COVID-19 Genomics UK Consortium | Harper VanSteenhouse, Yumi Kasai, David Gray, Carol Clugston, Anna Dominiczak and Alex Alderton, Roberto Amato, Sonia Goncalves, Ewan Harrison, David K. Jackson, Ian Johnston, Dominic Kwiatkowski, Cordelia Langford, John Sillitoe |
| EPI_ISL_532738 | NHSGGC West of Scotland Specialist Virology Centre / MRC-University of Glasgow Centre for Virus Research | Wellcome Sanger Institute for the COVID-19 Genomics UK Consortium | Ana da Silva Filipe, Natasha Johnson, Kathy Smollett, Daniel Mair, Stephen Carmichael, Lily Tong, Jenna Nichols, Elihu Aranday-Cortes, Kirstyn Brunker, Yasmin Parr, Kyriaki Nomikou; Sarah McDonald, Marc Niebel, Patawee Asamaphan; Richard Orton, Joseph Hughes, Sreenu Vattipally, David L Robertson; Alasdair MacLean, Rory Gunson; Kathy Li, Natasha Jesudason, Rajiv Shah, James Shepherd, Antonia Ho, Alice Broos, Emma Thomson and Alex Alderton, Roberto Amato, Sonia Goncalves, Ewan Harrison, David K. Jackson, Ian Johnston, Dominic Kwiatkowski, Cordelia Langford, John Sillitoe |
| EPI_ISL_532740, EPI_ISL_532741, EPI_ISL_532742, EPI_ISL_532743 | Lighthouse Lab in Glasgow | Wellcome Sanger Institute for the COVID-19 Genomics UK Consortium | Harper VanSteenhouse, Yumi Kasai, David Gray, Carol Clugston, Anna Dominiczak and Alex Alderton, Roberto Amato, Sonia Goncalves, Ewan Harrison, David K. Jackson, Ian Johnston, Dominic Kwiatkowski, Cordelia Langford, John Sillitoe |
| EPI_ISL_532744 | NHSGGC West of Scotland Specialist Virology Centre / MRC-University of Glasgow Centre for Virus Research | Wellcome Sanger Institute for the COVID-19 Genomics UK Consortium | Ana da Silva Filipe, Natasha Johnson, Kathy Smollett, Daniel Mair, Stephen Carmichael, Lily Tong, Jenna Nichols, Elihu Aranday-Cortes, Kirstyn Brunker, Yasmin Parr, Kyriaki Nomikou; Sarah McDonald, Marc Niebel, Patawee Asamaphan; Richard Orton, Joseph Hughes, Sreenu Vattipally, David L Robertson; Alasdair MacLean, Rory Gunson; Kathy Li, Natasha Jesudason, Rajiv Shah, James Shepherd, Antonia Ho, Alice Broos, Emma Thomson and Alex Alderton, Roberto Amato, Sonia Goncalves, Ewan Harrison, David K. Jackson, Ian Johnston, Dominic Kwiatkowski, Cordelia Langford, John Sillitoe |
| EPI_ISL_532745 | Lighthouse Lab in Glasgow | Wellcome Sanger Institute for the COVID-19 Genomics UK Consortium | Harper VanSteenhouse, Yumi Kasai, David Gray, Carol Clugston, Anna Dominiczak and Alex Alderton, Roberto Amato, Sonia Goncalves, Ewan Harrison, David K. Jackson, Ian Johnston, Dominic Kwiatkowski, Cordelia Langford, John Sillitoe |
| EPI_ISL_532746 | NHSGGC West of Scotland Specialist Virology Centre / MRC-University of Glasgow Centre for Virus Research | Wellcome Sanger Institute for the COVID-19 Genomics UK Consortium | Ana da Silva Filipe, Natasha Johnson, Kathy Smollett, Daniel Mair, Stephen Carmichael, Lily Tong, Jenna Nichols, Elihu Aranday-Cortes, Kirstyn Brunker, Yasmin Parr, Kyriaki Nomikou; Sarah McDonald, Marc Niebel, Patawee Asamaphan; Richard Orton, Joseph Hughes, Sreenu Vattipally, David L Robertson; Alasdair MacLean, Rory Gunson; Kathy Li, Natasha Jesudason, Rajiv Shah, James Shepherd, Antonia Ho, Alice Broos, Emma Thomson and Alex Alderton, Roberto Amato, Sonia Goncalves, Ewan Harrison, David K. Jackson, Ian Johnston, Dominic Kwiatkowski, Cordelia Langford, John Sillitoe |
| EPI_ISL_532747, EPI_ISL_532748, EPI_ISL_532749 | Lighthouse Lab in Glasgow | Wellcome Sanger Institute for the COVID-19 Genomics UK Consortium | Harper VanSteenhouse, Yumi Kasai, David Gray, Carol Clugston, Anna Dominiczak and Alex Alderton, Roberto Amato, Sonia Goncalves, Ewan Harrison, David K. Jackson, Ian Johnston, Dominic Kwiatkowski, Cordelia Langford, John Sillitoe |
| EPI_ISL_532750 | NHSGGC West of Scotland Specialist Virology Centre / MRC-University of Glasgow Centre for Virus Research | Wellcome Sanger Institute for the COVID-19 Genomics UK Consortium | Ana da Silva Filipe, Natasha Johnson, Kathy Smollett, Daniel Mair, Stephen Carmichael, Lily Tong, Jenna Nichols, Elihu Aranday-Cortes, Kirstyn Brunker, Yasmin Parr, Kyriaki Nomikou; Sarah McDonald, Marc Niebel, Patawee Asamaphan; Richard Orton, Joseph Hughes, Sreenu Vattipally, David L Robertson; Alasdair MacLean, Rory Gunson; Kathy Li, Natasha Jesudason, Rajiv Shah, James Shepherd, Antonia Ho, Alice Broos, Emma Thomson and Alex Alderton, Roberto Amato, Sonia Goncalves, Ewan Harrison, David K. Jackson, Ian Johnston, Dominic Kwiatkowski, Cordelia Langford, John Sillitoe |
| EPI_ISL_532751, EPI_ISL_532752 | Lighthouse Lab in Glasgow | Wellcome Sanger Institute for the COVID-19 Genomics UK Consortium | Harper VanSteenhouse, Yumi Kasai, David Gray, Carol Clugston, Anna Dominiczak and Alex Alderton, Roberto Amato, Sonia Goncalves, Ewan Harrison, David K. Jackson, Ian Johnston, Dominic Kwiatkowski, Cordelia Langford, John Sillitoe |
| EPI_ISL_532753 | NHSGGC West of Scotland Specialist Virology Centre / MRC-University of Glasgow Centre for Virus Research | Wellcome Sanger Institute for the COVID-19 Genomics UK Consortium | Ana da Silva Filipe, Natasha Johnson, Kathy Smollett, Daniel Mair, Stephen Carmichael, Lily Tong, Jenna Nichols, Elihu Aranday-Cortes, Kirstyn Brunker, Yasmin Parr, Kyriaki Nomikou; Sarah McDonald, Marc Niebel, Patawee Asamaphan; Richard Orton, Joseph Hughes, Sreenu Vattipally, David L Robertson; Alasdair MacLean, Rory Gunson; Kathy Li, Natasha Jesudason, Rajiv Shah, James Shepherd, Antonia Ho, Alice Broos, Emma Thomson and Alex Alderton, Roberto Amato, Sonia Goncalves, Ewan Harrison, David K. Jackson, Ian Johnston, Dominic Kwiatkowski, Cordelia Langford, John Sillitoe |
| EPI_ISL_532754 | Lighthouse Lab in Glasgow | Wellcome Sanger Institute for the COVID-19 Genomics UK Consortium | Harper VanSteenhouse, Yumi Kasai, David Gray, Carol Clugston, Anna Dominiczak and Alex Alderton, Roberto Amato, Sonia Goncalves, Ewan Harrison, David K. Jackson, Ian Johnston, Dominic Kwiatkowski, Cordelia Langford, John Sillitoe |
| EPI_ISL_532755 | NHSGGC West of Scotland Specialist Virology Centre / MRC-University of Glasgow Centre for Virus Research | Wellcome Sanger Institute for the COVID-19 Genomics UK Consortium | Ana da Silva Filipe, Natasha Johnson, Kathy Smollett, Daniel Mair, Stephen Carmichael, Lily Tong, Jenna Nichols, Elihu Aranday-Cortes, Kirstyn Brunker, Yasmin Parr, Kyriaki Nomikou; Sarah McDonald, Marc Niebel, Patawee Asamaphan; Richard Orton, Joseph Hughes, Sreenu Vattipally, David L Robertson; Alasdair MacLean, Rory Gunson; Kathy Li, Natasha Jesudason, Rajiv Shah, James Shepherd, Antonia Ho, Alice Broos, Emma Thomson and Alex Alderton, Roberto Amato, Sonia Goncalves, Ewan Harrison, David K. Jackson, Ian Johnston, Dominic Kwiatkowski, Cordelia Langford, John Sillitoe |
| EPI_ISL_532756, EPI_ISL_532757, EPI_ISL_532758, EPI_ISL_532760, EPI_ISL_532761, EPI_ISL_532762, EPI_ISL_532763, EPI_ISL_532764, EPI_ISL_532766 | Lighthouse Lab in Glasgow | Wellcome Sanger Institute for the COVID-19 Genomics UK Consortium | Harper VanSteenhouse, Yumi Kasai, David Gray, Carol Clugston, Anna Dominiczak and Alex Alderton, Roberto Amato, Sonia Goncalves, Ewan Harrison, David K. Jackson, Ian Johnston, Dominic Kwiatkowski, Cordelia Langford, John Sillitoe |
| EPI_ISL_532767 | NHSGGC West of Scotland Specialist Virology Centre / MRC-University of Glasgow Centre for Virus Research | Wellcome Sanger Institute for the COVID-19 Genomics UK Consortium | Ana da Silva Filipe, Natasha Johnson, Kathy Smollett, Daniel Mair, Stephen Carmichael, Lily Tong, Jenna Nichols, Elihu Aranday-Cortes, Kirstyn Brunker, Yasmin Parr, Kyriaki Nomikou; Sarah McDonald, Marc Niebel, Patawee Asamaphan; Richard Orton, Joseph Hughes, Sreenu Vattipally, David L Robertson; Alasdair MacLean, Rory Gunson; Kathy Li, Natasha Jesudason, Rajiv Shah, James Shepherd, Antonia Ho, Alice Broos, Emma Thomson and Alex Alderton, Roberto Amato, Sonia Goncalves, Ewan Harrison, David K. Jackson, Ian Johnston, Dominic Kwiatkowski, Cordelia Langford, John Sillitoe |
| EPI_ISL_532768, EPI_ISL_532769, EPI_ISL_532771, EPI_ISL_532772, EPI_ISL_532773, EPI_ISL_532774, EPI_ISL_532775, EPI_ISL_532776, EPI_ISL_532777, EPI_ISL_532778, EPI_ISL_532779, EPI_ISL_532780, EPI_ISL_532783 | | | |
| see above | Lighthouse Lab in Glasgow | Wellcome Sanger Institute for the COVID-19 Genomics UK Consortium | Harper VanSteenhouse, Yumi Kasai, David Gray, Carol Clugston, Anna Dominiczak and Alex Alderton, Roberto Amato, Sonia Goncalves, Ewan Harrison, David K. Jackson, Ian Johnston, Dominic Kwiatkowski, Cordelia Langford, John Sillitoe |
| EPI_ISL_532784 | NHSGGC West of Scotland Specialist Virology Centre / MRC-University of Glasgow Centre for Virus Research | Wellcome Sanger Institute for the COVID-19 Genomics UK Consortium | Ana da Silva Filipe, Natasha Johnson, Kathy Smollett, Daniel Mair, Stephen Carmichael, Lily Tong, Jenna Nichols, Elihu Aranday-Cortes, Kirstyn Brunker, Yasmin Parr, Kyriaki Nomikou; Sarah McDonald, Marc Niebel, Patawee Asamaphan; Richard Orton, Joseph Hughes, Sreenu Vattipally, David L Robertson; Alasdair MacLean, Rory Gunson; Kathy Li, Natasha Jesudason, Rajiv Shah, James Shepherd, Antonia Ho, Alice Broos, Emma Thomson and Alex Alderton, Roberto Amato, Sonia Goncalves, Ewan Harrison, David K. Jackson, Ian Johnston, Dominic Kwiatkowski, Cordelia Langford, John Sillitoe |
| EPI_ISL_532785 | Lighthouse Lab in Glasgow | Wellcome Sanger Institute for the COVID-19 Genomics UK Consortium | Harper VanSteenhouse, Yumi Kasai, David Gray, Carol Clugston, Anna Dominiczak and Alex Alderton, Roberto Amato, Sonia Goncalves, Ewan Harrison, David K. Jackson, Ian Johnston, Dominic Kwiatkowski, Cordelia Langford, John Sillitoe |
| EPI_ISL_532786 | NHSGGC West of Scotland Specialist Virology Centre / MRC-University of Glasgow Centre for Virus Research | Wellcome Sanger Institute for the COVID-19 Genomics UK Consortium | Ana da Silva Filipe, Natasha Johnson, Kathy Smollett, Daniel Mair, Stephen Carmichael, Lily Tong, Jenna Nichols, Elihu Aranday-Cortes, Kirstyn Brunker, Yasmin Parr, Kyriaki Nomikou; Sarah McDonald, Marc Niebel, Patawee Asamaphan; Richard Orton, Joseph Hughes, Sreenu Vattipally, David L Robertson; Alasdair MacLean, Rory Gunson; Kathy Li, Natasha Jesudason, Rajiv Shah, James Shepherd, Antonia Ho, Alice Broos, Emma Thomson and Alex Alderton, Roberto Amato, Sonia Goncalves, Ewan Harrison, David K. Jackson, Ian Johnston, Dominic Kwiatkowski, Cordelia Langford, John Sillitoe |
| EPI_ISL_532787, EPI_ISL_532788, EPI_ISL_532790 | Lighthouse Lab in Glasgow | Wellcome Sanger Institute for the COVID-19 Genomics UK Consortium | Harper VanSteenhouse, Yumi Kasai, David Gray, Carol Clugston, Anna Dominiczak and Alex Alderton, Roberto Amato, Sonia Goncalves, Ewan Harrison, David K. Jackson, Ian Johnston, Dominic Kwiatkowski, Cordelia Langford, John Sillitoe |
| EPI_ISL_532791 | NHSGGC West of Scotland Specialist Virology Centre / MRC-University of Glasgow Centre for Virus Research | Wellcome Sanger Institute for the COVID-19 Genomics UK Consortium | Ana da Silva Filipe, Natasha Johnson, Kathy Smollett, Daniel Mair, Stephen Carmichael, Lily Tong, Jenna Nichols, Elihu Aranday-Cortes, Kirstyn Brunker, Yasmin Parr, Kyriaki Nomikou; Sarah McDonald, Marc Niebel, Patawee Asamaphan; Richard Orton, Joseph Hughes, Sreenu Vattipally, David L Robertson; Alasdair MacLean, Rory Gunson; Kathy Li, Natasha Jesudason, Rajiv Shah, James Shepherd, Antonia Ho, Alice Broos, Emma Thomson and Alex Alderton, Roberto Amato, Sonia Goncalves, Ewan Harrison, David K. Jackson, Ian Johnston, Dominic Kwiatkowski, Cordelia Langford, John Sillitoe |
| EPI_ISL_532792, EPI_ISL_532793, EPI_ISL_532794, EPI_ISL_532795, EPI_ISL_532796, EPI_ISL_532797, EPI_ISL_532798, EPI_ISL_532799, EPI_ISL_532800, EPI_ISL_532802, EPI_ISL_532803, EPI_ISL_532804, EPI_ISL_532805, EPI_ISL_532806, EPI_ISL_532807, EPI_ISL_532808, EPI_ISL_532809, EPI_ISL_532810 | | | |
| see above | Lighthouse Lab in Glasgow | Wellcome Sanger Institute for the COVID-19 Genomics UK Consortium | Harper VanSteenhouse, Yumi Kasai, David Gray, Carol Clugston, Anna Dominiczak and Alex Alderton, Roberto Amato, Sonia Goncalves, Ewan Harrison, David K. Jackson, Ian Johnston, Dominic Kwiatkowski, Cordelia Langford, John Sillitoe |
| EPI_ISL_532811 | NHSGGC West of Scotland Specialist Virology Centre / MRC-University of Glasgow Centre for Virus Research | Wellcome Sanger Institute for the COVID-19 Genomics UK Consortium | Ana da Silva Filipe, Natasha Johnson, Kathy Smollett, Daniel Mair, Stephen Carmichael, Lily Tong, Jenna Nichols, Elihu Aranday-Cortes, Kirstyn Brunker, Yasmin Parr, Kyriaki Nomikou; Sarah McDonald, Marc Niebel, Patawee Asamaphan; Richard Orton, Joseph Hughes, Sreenu Vattipally, David L Robertson; Alasdair MacLean, Rory Gunson; Kathy Li, Natasha Jesudason, Rajiv Shah, James Shepherd, Antonia Ho, Alice Broos, Emma Thomson and Alex Alderton, Roberto Amato, Sonia Goncalves, Ewan Harrison, David K. Jackson, Ian Johnston, Dominic Kwiatkowski, Cordelia Langford, John Sillitoe |
| EPI_ISL_532812, EPI_ISL_532813, EPI_ISL_532815, EPI_ISL_532816, EPI_ISL_532817, EPI_ISL_532819, EPI_ISL_532820, EPI_ISL_532822, EPI_ISL_532823, EPI_ISL_532825, EPI_ISL_532826, EPI_ISL_532827, EPI_ISL_532828, EPI_ISL_532829, EPI_ISL_532830, EPI_ISL_532831, EPI_ISL_532832, EPI_ISL_532833, EPI_ISL_532834, EPI_ISL_532836, EPI_ISL_532837 | | | |
| see above | Lighthouse Lab in Glasgow | Wellcome Sanger Institute for the COVID-19 Genomics UK Consortium | Harper VanSteenhouse, Yumi Kasai, David Gray, Carol Clugston, Anna Dominiczak and Alex Alderton, Roberto Amato, Sonia Goncalves, Ewan Harrison, David K. Jackson, Ian Johnston, Dominic Kwiatkowski, Cordelia Langford, John Sillitoe |
| EPI_ISL_532838 | NHSGGC West of Scotland Specialist Virology Centre / MRC-University of Glasgow Centre for Virus Research | Wellcome Sanger Institute for the COVID-19 Genomics UK Consortium | Ana da Silva Filipe, Natasha Johnson, Kathy Smollett, Daniel Mair, Stephen Carmichael, Lily Tong, Jenna Nichols, Elihu Aranday-Cortes, Kirstyn Brunker, Yasmin Parr, Kyriaki Nomikou; Sarah McDonald, Marc Niebel, Patawee Asamaphan; Richard Orton, Joseph Hughes, Sreenu Vattipally, David L Robertson; Alasdair MacLean, Rory Gunson; Kathy Li, Natasha Jesudason, Rajiv Shah, James Shepherd, Antonia Ho, Alice Broos, Emma Thomson and Alex Alderton, Roberto Amato, Sonia Goncalves, Ewan Harrison, David K. Jackson, Ian Johnston, Dominic Kwiatkowski, Cordelia Langford, John Sillitoe |
| EPI_ISL_532840, EPI_ISL_532842, EPI_ISL_532845, EPI_ISL_532847, EPI_ISL_532848, EPI_ISL_532849, EPI_ISL_532853, EPI_ISL_532854, EPI_ISL_532855, EPI_ISL_532856, EPI_ISL_532860, EPI_ISL_532861, EPI_ISL_532863 | | | |
| see above | Lighthouse Lab in Glasgow | Wellcome Sanger Institute for the COVID-19 Genomics UK Consortium | Harper VanSteenhouse, Yumi Kasai, David Gray, Carol Clugston, Anna Dominiczak and Alex Alderton, Roberto Amato, Sonia Goncalves, Ewan Harrison, David K. Jackson, Ian Johnston, Dominic Kwiatkowski, Cordelia Langford, John Sillitoe |
| EPI_ISL_532865 | NHSGGC West of Scotland Specialist Virology Centre / MRC-University of Glasgow Centre for Virus Research | Wellcome Sanger Institute for the COVID-19 Genomics UK Consortium | Ana da Silva Filipe, Natasha Johnson, Kathy Smollett, Daniel Mair, Stephen Carmichael, Lily Tong, Jenna Nichols, Elihu Aranday-Cortes, Kirstyn Brunker, Yasmin Parr, Kyriaki Nomikou; Sarah McDonald, Marc Niebel, Patawee Asamaphan; Richard Orton, Joseph Hughes, Sreenu Vattipally, David L Robertson; Alasdair MacLean, Rory Gunson; Kathy Li, Natasha Jesudason, Rajiv Shah, James Shepherd, Antonia Ho, Alice Broos, Emma Thomson and Alex Alderton, Roberto Amato, Sonia Goncalves, Ewan Harrison, David K. Jackson, Ian Johnston, Dominic Kwiatkowski, Cordelia Langford, John Sillitoe |
| EPI_ISL_532866, EPI_ISL_532867, EPI_ISL_532868, EPI_ISL_532869, EPI_ISL_532870, EPI_ISL_532871, EPI_ISL_532872 | Lighthouse Lab in Glasgow | Wellcome Sanger Institute for the COVID-19 Genomics UK Consortium | Harper VanSteenhouse, Yumi Kasai, David Gray, Carol Clugston, Anna Dominiczak and Alex Alderton, Roberto Amato, Sonia Goncalves, Ewan Harrison, David K. Jackson, Ian Johnston, Dominic Kwiatkowski, Cordelia Langford, John Sillitoe |
| EPI_ISL_532874, EPI_ISL_532876 | NHSGGC West of Scotland Specialist Virology Centre / MRC-University of Glasgow Centre for Virus Research | Wellcome Sanger Institute for the COVID-19 Genomics UK Consortium | Ana da Silva Filipe, Natasha Johnson, Kathy Smollett, Daniel Mair, Stephen Carmichael, Lily Tong, Jenna Nichols, Elihu Aranday-Cortes, Kirstyn Brunker, Yasmin Parr, Kyriaki Nomikou; Sarah McDonald, Marc Niebel, Patawee Asamaphan; Richard Orton, Joseph Hughes, Sreenu Vattipally, David L Robertson; Alasdair MacLean, Rory Gunson; Kathy Li, Natasha Jesudason, Rajiv Shah, James Shepherd, Antonia Ho, Alice Broos, Emma Thomson and Alex Alderton, Roberto Amato, Sonia Goncalves, Ewan Harrison, David K. Jackson, Ian Johnston, Dominic Kwiatkowski, Cordelia Langford, John Sillitoe |
| EPI_ISL_532877 | Lighthouse Lab in Glasgow | Wellcome Sanger Institute for the COVID-19 Genomics UK Consortium | Harper VanSteenhouse, Yumi Kasai, David Gray, Carol Clugston, Anna Dominiczak and Alex Alderton, Roberto Amato, Sonia Goncalves, Ewan Harrison, David K. Jackson, Ian Johnston, Dominic Kwiatkowski, Cordelia Langford, John Sillitoe |
| EPI_ISL_532878 | NHSGGC West of Scotland Specialist Virology Centre / MRC-University of Glasgow Centre for Virus Research | Wellcome Sanger Institute for the COVID-19 Genomics UK Consortium | Ana da Silva Filipe, Natasha Johnson, Kathy Smollett, Daniel Mair, Stephen Carmichael, Lily Tong, Jenna Nichols, Elihu Aranday-Cortes, Kirstyn Brunker, Yasmin Parr, Kyriaki Nomikou; Sarah McDonald, Marc Niebel, Patawee Asamaphan; Richard Orton, Joseph Hughes, Sreenu Vattipally, David L Robertson; Alasdair MacLean, Rory Gunson; Kathy Li, Natasha Jesudason, Rajiv Shah, James Shepherd, Antonia Ho, Alice Broos, Emma Thomson and Alex Alderton, Roberto Amato, Sonia Goncalves, Ewan Harrison, David K. Jackson, Ian Johnston, Dominic Kwiatkowski, Cordelia Langford, John Sillitoe |
| EPI_ISL_532879, EPI_ISL_532882, EPI_ISL_532883, EPI_ISL_532884 | Lighthouse Lab in Glasgow | Wellcome Sanger Institute for the COVID-19 Genomics UK Consortium | Harper VanSteenhouse, Yumi Kasai, David Gray, Carol Clugston, Anna Dominiczak and Alex Alderton, Roberto Amato, Sonia Goncalves, Ewan Harrison, David K. Jackson, Ian Johnston, Dominic Kwiatkowski, Cordelia Langford, John Sillitoe |
| EPI_ISL_532885 | NHSGGC West of Scotland Specialist Virology Centre / MRC-University of Glasgow Centre for Virus Research | Wellcome Sanger Institute for the COVID-19 Genomics UK Consortium | Ana da Silva Filipe, Natasha Johnson, Kathy Smollett, Daniel Mair, Stephen Carmichael, Lily Tong, Jenna Nichols, Elihu Aranday-Cortes, Kirstyn Brunker, Yasmin Parr, Kyriaki Nomikou; Sarah McDonald, Marc Niebel, Patawee Asamaphan; Richard Orton, Joseph Hughes, Sreenu Vattipally, David L Robertson; Alasdair MacLean, Rory Gunson; Kathy Li, Natasha Jesudason, Rajiv Shah, James Shepherd, Antonia Ho, Alice Broos, Emma Thomson and Alex Alderton, Roberto Amato, Sonia Goncalves, Ewan Harrison, David K. Jackson, Ian Johnston, Dominic Kwiatkowski, Cordelia Langford, John Sillitoe |

| | | | |
|---|---|---|---|
| EPI_ISL_532887, EPI_ISL_532888, EPI_ISL_532889, EPI_ISL_532890 | Lighthouse Lab in Glasgow | Wellcome Sanger Institute for the COVID-19 Genomics UK Consortium | Harper VanSteenhouse, Yumi Kasai, David Gray, Carol Clugston, Anna Dominiczak and Alex Alderton, Roberto Amato, Sonia Goncalves, Ewan Harrison, David K. Jackson, Ian Johnston, Dominic Kwiatkowski, Cordelia Langford, John Sillitoe |
| EPI_ISL_532891 | NHSGGC West of Scotland Specialist Virology Centre / MRC-University of Glasgow Centre for Virus Research | Wellcome Sanger Institute for the COVID-19 Genomics UK Consortium | Ana da Silva Filipe, Natasha Johnson, Kathy Smollett, Daniel Mair, Stephen Carmichael, Lily Tong, Jenna Nichols, Elihu Aranday-Cortes, Kirstyn Brunker, Yasmin Parr, Kyriaki Nomikou; Sarah McDonald, Marc Niebel, Patawee Asamaphan; Richard Orton, Joseph Hughes, Sreenu Vattipally, David L Robertson; Alasdair MacLean, Rory Gunson; Kathy Li, Natasha Jesudason, Rajiv Shah, James Shepherd, Antonia Ho, Alice Broos, Emma Thomson and Alex Alderton, Roberto Amato, Sonia Goncalves, Ewan Harrison, David K. Jackson, Ian Johnston, Dominic Kwiatkowski, Cordelia Langford, John Sillitoe |
| EPI_ISL_532892, EPI_ISL_532893 | Lighthouse Lab in Glasgow | Wellcome Sanger Institute for the COVID-19 Genomics UK Consortium | Harper VanSteenhouse, Yumi Kasai, David Gray, Carol Clugston, Anna Dominiczak and Alex Alderton, Roberto Amato, Sonia Goncalves, Ewan Harrison, David K. Jackson, Ian Johnston, Dominic Kwiatkowski, Cordelia Langford, John Sillitoe |
| EPI_ISL_532894 | NHSGGC West of Scotland Specialist Virology Centre / MRC-University of Glasgow Centre for Virus Research | Wellcome Sanger Institute for the COVID-19 Genomics UK Consortium | Ana da Silva Filipe, Natasha Johnson, Kathy Smollett, Daniel Mair, Stephen Carmichael, Lily Tong, Jenna Nichols, Elihu Aranday-Cortes, Kirstyn Brunker, Yasmin Parr, Kyriaki Nomikou; Sarah McDonald, Marc Niebel, Patawee Asamaphan; Richard Orton, Joseph Hughes, Sreenu Vattipally, David L Robertson; Alasdair MacLean, Rory Gunson; Kathy Li, Natasha Jesudason, Rajiv Shah, James Shepherd, Antonia Ho, Alice Broos, Emma Thomson and Alex Alderton, Roberto Amato, Sonia Goncalves, Ewan Harrison, David K. Jackson, Ian Johnston, Dominic Kwiatkowski, Cordelia Langford, John Sillitoe |
| EPI_ISL_532895 | Lighthouse Lab in Glasgow | Wellcome Sanger Institute for the COVID-19 Genomics UK Consortium | Harper VanSteenhouse, Yumi Kasai, David Gray, Carol Clugston, Anna Dominiczak and Alex Alderton, Roberto Amato, Sonia Goncalves, Ewan Harrison, David K. Jackson, Ian Johnston, Dominic Kwiatkowski, Cordelia Langford, John Sillitoe |
| EPI_ISL_532896 | NHSGGC West of Scotland Specialist Virology Centre / MRC-University of Glasgow Centre for Virus Research | Wellcome Sanger Institute for the COVID-19 Genomics UK Consortium | Ana da Silva Filipe, Natasha Johnson, Kathy Smollett, Daniel Mair, Stephen Carmichael, Lily Tong, Jenna Nichols, Elihu Aranday-Cortes, Kirstyn Brunker, Yasmin Parr, Kyriaki Nomikou; Sarah McDonald, Marc Niebel, Patawee Asamaphan; Richard Orton, Joseph Hughes, Sreenu Vattipally, David L Robertson; Alasdair MacLean, Rory Gunson; Kathy Li, Natasha Jesudason, Rajiv Shah, James Shepherd, Antonia Ho, Alice Broos, Emma Thomson and Alex Alderton, Roberto Amato, Sonia Goncalves, Ewan Harrison, David K. Jackson, Ian Johnston, Dominic Kwiatkowski, Cordelia Langford, John Sillitoe |
| EPI_ISL_532897, EPI_ISL_532898, EPI_ISL_532899, EPI_ISL_532900, EPI_ISL_532901, EPI_ISL_532903, EPI_ISL_532904 | Lighthouse Lab in Glasgow | Wellcome Sanger Institute for the COVID-19 Genomics UK Consortium | Harper VanSteenhouse, Yumi Kasai, David Gray, Carol Clugston, Anna Dominiczak and Alex Alderton, Roberto Amato, Sonia Goncalves, Ewan Harrison, David K. Jackson, Ian Johnston, Dominic Kwiatkowski, Cordelia Langford, John Sillitoe |
| EPI_ISL_532905, EPI_ISL_532906 | NHSGGC West of Scotland Specialist Virology Centre / MRC-University of Glasgow Centre for Virus Research | Wellcome Sanger Institute for the COVID-19 Genomics UK Consortium | Ana da Silva Filipe, Natasha Johnson, Kathy Smollett, Daniel Mair, Stephen Carmichael, Lily Tong, Jenna Nichols, Elihu Aranday-Cortes, Kirstyn Brunker, Yasmin Parr, Kyriaki Nomikou; Sarah McDonald, Marc Niebel, Patawee Asamaphan; Richard Orton, Joseph Hughes, Sreenu Vattipally, David L Robertson; Alasdair MacLean, Rory Gunson; Kathy Li, Natasha Jesudason, Rajiv Shah, James Shepherd, Antonia Ho, Alice Broos, Emma Thomson and Alex Alderton, Roberto Amato, Sonia Goncalves, Ewan Harrison, David K. Jackson, Ian Johnston, Dominic Kwiatkowski, Cordelia Langford, John Sillitoe |
| EPI_ISL_532907, EPI_ISL_532908, EPI_ISL_532911, EPI_ISL_532912 | Lighthouse Lab in Glasgow | Wellcome Sanger Institute for the COVID-19 Genomics UK Consortium | Harper VanSteenhouse, Yumi Kasai, David Gray, Carol Clugston, Anna Dominiczak and Alex Alderton, Roberto Amato, Sonia Goncalves, Ewan Harrison, David K. Jackson, Ian Johnston, Dominic Kwiatkowski, Cordelia Langford, John Sillitoe |
| EPI_ISL_532915 | NHSGGC West of Scotland Specialist Virology Centre / MRC-University of Glasgow Centre for Virus Research | Wellcome Sanger Institute for the COVID-19 Genomics UK Consortium | Ana da Silva Filipe, Natasha Johnson, Kathy Smollett, Daniel Mair, Stephen Carmichael, Lily Tong, Jenna Nichols, Elihu Aranday-Cortes, Kirstyn Brunker, Yasmin Parr, Kyriaki Nomikou; Sarah McDonald, Marc Niebel, Patawee Asamaphan; Richard Orton, Joseph Hughes, Sreenu Vattipally, David L Robertson; Alasdair MacLean, Rory Gunson; Kathy Li, Natasha Jesudason, Rajiv Shah, James Shepherd, Antonia Ho, Alice Broos, Emma Thomson and Alex Alderton, Roberto Amato, Sonia Goncalves, Ewan Harrison, David K. Jackson, Ian Johnston, Dominic Kwiatkowski, Cordelia Langford, John Sillitoe |
| EPI_ISL_532917, EPI_ISL_532918, EPI_ISL_532922, EPI_ISL_532923, EPI_ISL_532924, EPI_ISL_532927, EPI_ISL_532928, EPI_ISL_532930, EPI_ISL_532931 | Lighthouse Lab in Glasgow | Wellcome Sanger Institute for the COVID-19 Genomics UK Consortium | Harper VanSteenhouse, Yumi Kasai, David Gray, Carol Clugston, Anna Dominiczak and Alex Alderton, Roberto Amato, Sonia Goncalves, Ewan Harrison, David K. Jackson, Ian Johnston, Dominic Kwiatkowski, Cordelia Langford, John Sillitoe |
| EPI_ISL_532932, EPI_ISL_532933 | NHSGGC West of Scotland Specialist Virology Centre / MRC-University of Glasgow Centre for Virus Research | Wellcome Sanger Institute for the COVID-19 Genomics UK Consortium | Ana da Silva Filipe, Natasha Johnson, Kathy Smollett, Daniel Mair, Stephen Carmichael, Lily Tong, Jenna Nichols, Elihu Aranday-Cortes, Kirstyn Brunker, Yasmin Parr, Kyriaki Nomikou; Sarah McDonald, Marc Niebel, Patawee Asamaphan; Richard Orton, Joseph Hughes, Sreenu Vattipally, David L Robertson; Alasdair MacLean, Rory Gunson; Kathy Li, Natasha Jesudason, Rajiv Shah, James Shepherd, Antonia Ho, Alice Broos, Emma Thomson and Alex Alderton, Roberto Amato, Sonia Goncalves, Ewan Harrison, David K. Jackson, Ian Johnston, Dominic Kwiatkowski, Cordelia Langford, John Sillitoe |
| EPI_ISL_532934, EPI_ISL_532935, EPI_ISL_532938, EPI_ISL_532939, EPI_ISL_532941, EPI_ISL_532942, EPI_ISL_532944, EPI_ISL_532945, EPI_ISL_532947, EPI_ISL_532949, EPI_ISL_532950, EPI_ISL_532952, EPI_ISL_532953, EPI_ISL_532954, EPI_ISL_532955, EPI_ISL_532956, EPI_ISL_532957, EPI_ISL_532958, EPI_ISL_532959, EPI_ISL_532960, EPI_ISL_532961 | | | |
| see above | Lighthouse Lab in Glasgow | Wellcome Sanger Institute for the COVID-19 Genomics UK Consortium | Harper VanSteenhouse, Yumi Kasai, David Gray, Carol Clugston, Anna Dominiczak and Alex Alderton, Roberto Amato, Sonia Goncalves, Ewan Harrison, David K. Jackson, Ian Johnston, Dominic Kwiatkowski, Cordelia Langford, John Sillitoe |
| EPI_ISL_532962 | NHSGGC West of Scotland Specialist Virology Centre / MRC-University of Glasgow Centre for Virus Research | Wellcome Sanger Institute for the COVID-19 Genomics UK Consortium | Ana da Silva Filipe, Natasha Johnson, Kathy Smollett, Daniel Mair, Stephen Carmichael, Lily Tong, Jenna Nichols, Elihu Aranday-Cortes, Kirstyn Brunker, Yasmin Parr, Kyriaki Nomikou; Sarah McDonald, Marc Niebel, Patawee Asamaphan; Richard Orton, Joseph Hughes, Sreenu Vattipally, David L Robertson; Alasdair MacLean, Rory Gunson; Kathy Li, Natasha Jesudason, Rajiv Shah, James Shepherd, Antonia Ho, Alice Broos, Emma Thomson and Alex Alderton, Roberto Amato, Sonia Goncalves, Ewan Harrison, David K. Jackson, Ian Johnston, Dominic Kwiatkowski, Cordelia Langford, John Sillitoe |
| EPI_ISL_532963, EPI_ISL_532964, EPI_ISL_532965, EPI_ISL_532966, EPI_ISL_532967, EPI_ISL_532968, EPI_ISL_532970, EPI_ISL_532971, EPI_ISL_532972, EPI_ISL_532973, EPI_ISL_532974, EPI_ISL_532975, EPI_ISL_532977 | | | |
| see above | Lighthouse Lab in Glasgow | Wellcome Sanger Institute for the COVID-19 Genomics UK Consortium | Harper VanSteenhouse, Yumi Kasai, David Gray, Carol Clugston, Anna Dominiczak and Alex Alderton, Roberto Amato, Sonia Goncalves, Ewan Harrison, David K. Jackson, Ian Johnston, Dominic Kwiatkowski, Cordelia Langford, John Sillitoe |
| EPI_ISL_532978 | Virology Department, Royal Infirmary of Edinburgh, NHS Lothian / School of Biological Sciences, University of Edinburgh | Wellcome Sanger Institute for the COVID-19 Genomics UK Consortium | McHugh M, Dewar R, Rooke S, O'Toole Á, Scher E, Hill V, McCrone JT, Colquhoun R, Yu X, Jackson B, Rambaut A, Templeton K and Alex Alderton, Roberto Amato, Sonia Goncalves, Ewan Harrison, David K. Jackson, Ian Johnston, Dominic Kwiatkowski, Cordelia Langford, John Sillitoe |
| EPI_ISL_532979, EPI_ISL_532980, EPI_ISL_532981, EPI_ISL_532982 | Lighthouse Lab in Glasgow | Wellcome Sanger Institute for the COVID-19 Genomics UK Consortium | Harper VanSteenhouse, Yumi Kasai, David Gray, Carol Clugston, Anna Dominiczak and Alex Alderton, Roberto Amato, Sonia Goncalves, Ewan Harrison, David K. Jackson, Ian Johnston, Dominic Kwiatkowski, Cordelia Langford, John Sillitoe |
| EPI_ISL_532983 | Virology Department, Royal Infirmary of Edinburgh, NHS Lothian / School of Biological Sciences, University of Edinburgh | Wellcome Sanger Institute for the COVID-19 Genomics UK Consortium | McHugh M, Dewar R, Rooke S, O'Toole Á, Scher E, Hill V, McCrone JT, Colquhoun R, Yu X, Jackson B, Rambaut A, Templeton K and Alex Alderton, Roberto Amato, Sonia Goncalves, Ewan Harrison, David K. Jackson, Ian Johnston, Dominic Kwiatkowski, Cordelia Langford, John Sillitoe |
| EPI_ISL_532984, EPI_ISL_532985, EPI_ISL_532986, EPI_ISL_532987 | Lighthouse Lab in Glasgow | Wellcome Sanger Institute for the COVID-19 Genomics UK Consortium | Harper VanSteenhouse, Yumi Kasai, David Gray, Carol Clugston, Anna Dominiczak and Alex Alderton, Roberto Amato, Sonia Goncalves, Ewan Harrison, David K. Jackson, Ian Johnston, Dominic Kwiatkowski, Cordelia Langford, John Sillitoe |
| EPI_ISL_532988 | NHSGGC West of Scotland Specialist Virology Centre / MRC-University of Glasgow Centre for Virus Research | Wellcome Sanger Institute for the COVID-19 Genomics UK Consortium | Ana da Silva Filipe, Natasha Johnson, Kathy Smollett, Daniel Mair, Stephen Carmichael, Lily Tong, Jenna Nichols, Elihu Aranday-Cortes, Kirstyn Brunker, Yasmin Parr, Kyriaki Nomikou; Sarah McDonald, Marc Niebel, Patawee Asamaphan; Richard Orton, Joseph Hughes, Sreenu Vattipally, David L Robertson; Alasdair MacLean, Rory Gunson; Kathy Li, Natasha Jesudason, Rajiv Shah, James Shepherd, Antonia Ho, Alice Broos, Emma Thomson and Alex Alderton, Roberto Amato, Sonia Goncalves, Ewan Harrison, David K. Jackson, Ian Johnston, Dominic Kwiatkowski, Cordelia Langford, John Sillitoe |
| EPI_ISL_532989 | Lighthouse Lab in Glasgow | Wellcome Sanger Institute for the COVID-19 Genomics UK Consortium | Harper VanSteenhouse, Yumi Kasai, David Gray, Carol Clugston, Anna Dominiczak and Alex Alderton, Roberto Amato, Sonia Goncalves, Ewan Harrison, David K. Jackson, Ian Johnston, Dominic Kwiatkowski, Cordelia Langford, John Sillitoe |
| EPI_ISL_532990 | Virology Department, Royal Infirmary of Edinburgh, NHS Lothian / School of Biological Sciences, University of Edinburgh | Wellcome Sanger Institute for the COVID-19 Genomics UK Consortium | McHugh M, Dewar R, Rooke S, O'Toole Á, Scher E, Hill V, McCrone JT, Colquhoun R, Yu X, Jackson B, Rambaut A, Templeton K and Alex Alderton, Roberto Amato, Sonia Goncalves, Ewan Harrison, David K. Jackson, Ian Johnston, Dominic Kwiatkowski, Cordelia Langford, John Sillitoe |
| EPI_ISL_532991 | Lighthouse Lab in Glasgow | Wellcome Sanger Institute for the COVID-19 Genomics UK Consortium | Harper VanSteenhouse, Yumi Kasai, David Gray, Carol Clugston, Anna Dominiczak and Alex Alderton, Roberto Amato, Sonia Goncalves, Ewan Harrison, David K. Jackson, Ian Johnston, Dominic Kwiatkowski, Cordelia Langford, John Sillitoe |
| EPI_ISL_532992 | Virology Department, Royal Infirmary of Edinburgh, NHS Lothian / School of Biological Sciences, University of Edinburgh | Wellcome Sanger Institute for the COVID-19 Genomics UK Consortium | McHugh M, Dewar R, Rooke S, O'Toole Á, Scher E, Hill V, McCrone JT, Colquhoun R, Yu X, Jackson B, Rambaut A, Templeton K and Alex Alderton, Roberto Amato, Sonia Goncalves, Ewan Harrison, David K. Jackson, Ian Johnston, Dominic Kwiatkowski, Cordelia Langford, John Sillitoe |
| EPI_ISL_532993 | Lighthouse Lab in Glasgow | Wellcome Sanger Institute for the COVID-19 Genomics UK Consortium | Harper VanSteenhouse, Yumi Kasai, David Gray, Carol Clugston, Anna Dominiczak and Alex Alderton, Roberto Amato, Sonia Goncalves, Ewan Harrison, David K. Jackson, Ian Johnston, Dominic Kwiatkowski, Cordelia Langford, John Sillitoe |
| EPI_ISL_532994 | Virology Department, Royal Infirmary of Edinburgh, NHS Lothian / School of Biological Sciences, University of Edinburgh | Wellcome Sanger Institute for the COVID-19 Genomics UK Consortium | McHugh M, Dewar R, Rooke S, O'Toole Á, Scher E, Hill V, McCrone JT, Colquhoun R, Yu X, Jackson B, Rambaut A, Templeton K and Alex Alderton, Roberto Amato, Sonia Goncalves, Ewan Harrison, David K. Jackson, Ian Johnston, Dominic Kwiatkowski, Cordelia Langford, John Sillitoe |
| EPI_ISL_532995, EPI_ISL_532996, EPI_ISL_532997, EPI_ISL_532998, EPI_ISL_532999, EPI_ISL_533001, EPI_ISL_533002, EPI_ISL_533003, EPI_ISL_533004, EPI_ISL_533005, EPI_ISL_533007, EPI_ISL_533008, EPI_ISL_533009, EPI_ISL_533010, EPI_ISL_533011, EPI_ISL_533013, EPI_ISL_533014, EPI_ISL_533015 | | | |
| see above | Lighthouse Lab in Glasgow | Wellcome Sanger Institute for the COVID-19 Genomics UK Consortium | Harper VanSteenhouse, Yumi Kasai, David Gray, Carol Clugston, Anna Dominiczak and Alex Alderton, Roberto Amato, Sonia Goncalves, Ewan Harrison, David K. Jackson, Ian Johnston, Dominic Kwiatkowski, Cordelia Langford, John Sillitoe |
| EPI_ISL_533016 | NHSGGC West of Scotland Specialist Virology Centre / MRC-University of Glasgow Centre for Virus Research | Wellcome Sanger Institute for the COVID-19 Genomics UK Consortium | Ana da Silva Filipe, Natasha Johnson, Kathy Smollett, Daniel Mair, Stephen Carmichael, Lily Tong, Jenna Nichols, Elihu Aranday-Cortes, Kirstyn Brunker, Yasmin Parr, Kyriaki Nomikou; Sarah McDonald, Marc Niebel, Patawee Asamaphan; Richard Orton, Joseph Hughes, Sreenu Vattipally, David L Robertson; Alasdair MacLean, Rory Gunson; Kathy Li, Natasha Jesudason, Rajiv Shah, James Shepherd, Antonia Ho, Alice Broos, Emma Thomson and Alex Alderton, Roberto Amato, Sonia Goncalves, Ewan Harrison, David K. Jackson, Ian Johnston, Dominic Kwiatkowski, Cordelia Langford, John Sillitoe |
| EPI_ISL_533018, EPI_ISL_533019, EPI_ISL_533020, EPI_ISL_533021, EPI_ISL_533022, EPI_ISL_533023, EPI_ISL_533025, EPI_ISL_533026 | Lighthouse Lab in Glasgow | Wellcome Sanger Institute for the COVID-19 Genomics UK Consortium | Harper VanSteenhouse, Yumi Kasai, David Gray, Carol Clugston, Anna Dominiczak and Alex Alderton, Roberto Amato, Sonia Goncalves, Ewan Harrison, David K. Jackson, Ian Johnston, Dominic Kwiatkowski, Cordelia Langford, John Sillitoe |
| EPI_ISL_533027 | NHSGGC West of Scotland Specialist Virology Centre / MRC-University of Glasgow Centre for Virus Research | Wellcome Sanger Institute for the COVID-19 Genomics UK Consortium | Ana da Silva Filipe, Natasha Johnson, Kathy Smollett, Daniel Mair, Stephen Carmichael, Lily Tong, Jenna Nichols, Elihu Aranday-Cortes, Kirstyn Brunker, Yasmin Parr, Kyriaki Nomikou; Sarah McDonald, Marc Niebel, Patawee Asamaphan; Richard Orton, Joseph Hughes, Sreenu Vattipally, David L Robertson; Alasdair MacLean, Rory Gunson; Kathy Li, Natasha Jesudason, Rajiv Shah, James Shepherd, Antonia Ho, Alice Broos, Emma Thomson and Alex Alderton, Roberto Amato, Sonia Goncalves, Ewan Harrison, David K. Jackson, Ian Johnston, Dominic Kwiatkowski, Cordelia Langford, John Sillitoe |
| EPI_ISL_533028, EPI_ISL_533031, EPI_ISL_533032, EPI_ISL_533033, EPI_ISL_533034, EPI_ISL_533035, EPI_ISL_533036, EPI_ISL_533037, EPI_ISL_533038, EPI_ISL_533039, EPI_ISL_533040, EPI_ISL_533041, EPI_ISL_533042, EPI_ISL_533043, EPI_ISL_533044, EPI_ISL_533045, EPI_ISL_533047, EPI_ISL_533048, EPI_ISL_533050, EPI_ISL_533051, EPI_ISL_533052, EPI_ISL_533053, EPI_ISL_533056, EPI_ISL_533057, EPI_ISL_533058, EPI_ISL_533060, EPI_ISL_533061, EPI_ISL_533062, EPI_ISL_533063, EPI_ISL_533064, EPI_ISL_533065, EPI_ISL_533067, EPI_ISL_533068, EPI_ISL_533069, EPI_ISL_533070, EPI_ISL_533071 | | | |
| see above | Lighthouse Lab in Glasgow | Wellcome Sanger Institute for the COVID-19 Genomics UK Consortium | Harper VanSteenhouse, Yumi Kasai, David Gray, Carol Clugston, Anna Dominiczak and Alex Alderton, Roberto Amato, Sonia Goncalves, Ewan Harrison, David K. Jackson, Ian Johnston, Dominic Kwiatkowski, Cordelia Langford, John Sillitoe |
| EPI_ISL_533072 | NHSGGC West of Scotland Specialist Virology Centre / MRC-University of Glasgow Centre for Virus Research | Wellcome Sanger Institute for the COVID-19 Genomics UK Consortium | Ana da Silva Filipe, Natasha Johnson, Kathy Smollett, Daniel Mair, Stephen Carmichael, Lily Tong, Jenna Nichols, Elihu Aranday-Cortes, Kirstyn Brunker, Yasmin Parr, Kyriaki Nomikou; Sarah McDonald, Marc Niebel, Patawee Asamaphan; Richard Orton, Joseph Hughes, Sreenu Vattipally, David L Robertson; Alasdair MacLean, Rory Gunson; Kathy Li, Natasha Jesudason, Rajiv Shah, James Shepherd, Antonia Ho, Alice Broos, Emma Thomson and Alex Alderton, Roberto Amato, Sonia Goncalves, Ewan Harrison, David K. Jackson, Ian Johnston, Dominic Kwiatkowski, Cordelia Langford, John Sillitoe |
| EPI_ISL_533073, EPI_ISL_533074, EPI_ISL_533075, EPI_ISL_533076, EPI_ISL_533077, EPI_ISL_533078, EPI_ISL_533080, EPI_ISL_533081, EPI_ISL_533082, EPI_ISL_533083, EPI_ISL_533085, EPI_ISL_533088, EPI_ISL_533090, EPI_ISL_533092, EPI_ISL_533093, EPI_ISL_533095, EPI_ISL_533096, EPI_ISL_533097, EPI_ISL_533099, EPI_ISL_533100, EPI_ISL_533101, EPI_ISL_533102, EPI_ISL_533103, EPI_ISL_533104, EPI_ISL_533105, EPI_ISL_533106, EPI_ISL_533108, EPI_ISL_533109, EPI_ISL_533110, EPI_ISL_533111, EPI_ISL_533112, EPI_ISL_533113, EPI_ISL_533114, EPI_ISL_533115, EPI_ISL_533116, EPI_ISL_533118, EPI_ISL_533120, EPI_ISL_533121, EPI_ISL_533122, EPI_ISL_533123, EPI_ISL_533124, EPI_ISL_533125, EPI_ISL_533126, EPI_ISL_533127, EPI_ISL_533128, EPI_ISL_533129, EPI_ISL_533131, EPI_ISL_533132, EPI_ISL_533134, EPI_ISL_533137, EPI_ISL_533142, EPI_ISL_533143, EPI_ISL_533145, EPI_ISL_533146, EPI_ISL_533148, EPI_ISL_533149, EPI_ISL_533150, EPI_ISL_533151, EPI_ISL_533152, EPI_ISL_533153, EPI_ISL_533155, EPI_ISL_533156, EPI_ISL_533157, EPI_ISL_533158, EPI_ISL_533159, EPI_ISL_533160, EPI_ISL_533161, EPI_ISL_533162, EPI_ISL_533163, EPI_ISL_533164, EPI_ISL_533165, EPI_ISL_533167, EPI_ISL_533168, EPI_ISL_533169, EPI_ISL_533170, EPI_ISL_533171, EPI_ISL_533172, EPI_ISL_533173, EPI_ISL_533174, EPI_ISL_533176, EPI_ISL_533177, EPI_ISL_533178, EPI_ISL_533179, EPI_ISL_533181, EPI_ISL_533182, EPI_ISL_533183, EPI_ISL_533184, EPI_ISL_533185, EPI_ISL_533186, EPI_ISL_533187, EPI_ISL_533189, EPI_ISL_533190, EPI_ISL_533192, EPI_ISL_533193, EPI_ISL_533194, EPI_ISL_533195, EPI_ISL_533196, EPI_ISL_533197, EPI_ISL_533198, EPI_ISL_533200, EPI_ISL_533201, EPI_ISL_533202, EPI_ISL_533203, EPI_ISL_533205, EPI_ISL_533206, EPI_ISL_533208, EPI_ISL_533209, EPI_ISL_533210, EPI_ISL_533211, EPI_ISL_533212, EPI_ISL_533213, EPI_ISL_533214, EPI_ISL_533215, EPI_ISL_533216, EPI_ISL_533217, EPI_ISL_533218, EPI_ISL_533219, EPI_ISL_533220, EPI_ISL_533221, EPI_ISL_533223, EPI_ISL_533224, EPI_ISL_533225, EPI_ISL_533226, EPI_ISL_533227 | | | |
| see above | Lighthouse Lab in Glasgow | Wellcome Sanger Institute for the COVID-19 Genomics UK Consortium | Harper VanSteenhouse, Yumi Kasai, David Gray, Carol Clugston, Anna Dominiczak and Alex Alderton, Roberto Amato, Sonia Goncalves, Ewan Harrison, David K. Jackson, Ian Johnston, Dominic Kwiatkowski, Cordelia Langford, John Sillitoe |
| EPI_ISL_533228, EPI_ISL_533231, EPI_ISL_533232 | NHSGGC West of Scotland Specialist Virology Centre / MRC-University of Glasgow Centre for Virus Research | Wellcome Sanger Institute for the COVID-19 Genomics UK Consortium | Ana da Silva Filipe, Natasha Johnson, Kathy Smollett, Daniel Mair, Stephen Carmichael, Lily Tong, Jenna Nichols, Elihu Aranday-Cortes, Kirstyn Brunker, Yasmin Parr, Kyriaki Nomikou; Sarah McDonald, Marc Niebel, Patawee Asamaphan; Richard Orton, Joseph Hughes, Sreenu Vattipally, David L Robertson; Alasdair MacLean, Rory Gunson; Kathy Li, Natasha Jesudason, Rajiv Shah, James Shepherd, Antonia Ho, Alice Broos, Emma Thomson and Alex Alderton, Roberto |

| | | | |
|---|---|---|---|
| | | | Amato, Sonia Goncalves, Ewan Harrison, David K. Jackson, Ian Johnston, Dominic Kwiatkowski, Cordelia Langford, John Sillitoe |
| EPI_ISL_533233, EPI_ISL_533234, EPI_ISL_533235 | Lighthouse Lab in Glasgow | Wellcome Sanger Institute for the COVID-19 Genomics UK Consortium | Harper VanSteenhouse, Yumi Kasai, David Gray, Carol Clugston, Anna Dominiczak and Alex Alderton, Roberto Amato, Sonia Goncalves, Ewan Harrison, David K. Jackson, Ian Johnston, Dominic Kwiatkowski, Cordelia Langford, John Sillitoe |
| EPI_ISL_533236, EPI_ISL_533237 | NHSGGC West of Scotland Specialist Virology Centre / MRC-University of Glasgow Centre for Virus Research | Wellcome Sanger Institute for the COVID-19 Genomics UK Consortium | Ana da Silva Filipe, Natasha Johnson, Kathy Smollett, Daniel Mair, Stephen Carmichael, Lily Tong, Jenna Nichols, Elihu Aranday-Cortes, Kirstyn Brunker, Yasmin Parr, Kyriaki Nomikou; Sarah McDonald, Marc Niebel, Patawee Asamaphan; Richard Orton, Joseph Hughes, Sreenu Vattipally, David L Robertson; Alasdair MacLean, Rory Gunson; Kathy Li, Natasha Jesudason, Rajiv Shah, James Shepherd, Antonia Ho, Alice Broos, Emma Thomson and Alex Alderton, Roberto Amato, Sonia Goncalves, Ewan Harrison, David K. Jackson, Ian Johnston, Dominic Kwiatkowski, Cordelia Langford, John Sillitoe |
| EPI_ISL_533238, EPI_ISL_533239, EPI_ISL_533240 | Lighthouse Lab in Glasgow | Wellcome Sanger Institute for the COVID-19 Genomics UK Consortium | Harper VanSteenhouse, Yumi Kasai, David Gray, Carol Clugston, Anna Dominiczak and Alex Alderton, Roberto Amato, Sonia Goncalves, Ewan Harrison, David K. Jackson, Ian Johnston, Dominic Kwiatkowski, Cordelia Langford, John Sillitoe |
| EPI_ISL_533241 | NHSGGC West of Scotland Specialist Virology Centre / MRC-University of Glasgow Centre for Virus Research | Wellcome Sanger Institute for the COVID-19 Genomics UK Consortium | Ana da Silva Filipe, Natasha Johnson, Kathy Smollett, Daniel Mair, Stephen Carmichael, Lily Tong, Jenna Nichols, Elihu Aranday-Cortes, Kirstyn Brunker, Yasmin Parr, Kyriaki Nomikou; Sarah McDonald, Marc Niebel, Patawee Asamaphan; Richard Orton, Joseph Hughes, Sreenu Vattipally, David L Robertson; Alasdair MacLean, Rory Gunson; Kathy Li, Natasha Jesudason, Rajiv Shah, James Shepherd, Antonia Ho, Alice Broos, Emma Thomson and Alex Alderton, Roberto Amato, Sonia Goncalves, Ewan Harrison, David K. Jackson, Ian Johnston, Dominic Kwiatkowski, Cordelia Langford, John Sillitoe |
| EPI_ISL_533243, EPI_ISL_533244, EPI_ISL_533245, EPI_ISL_533246, EPI_ISL_533247, EPI_ISL_533248, EPI_ISL_533249 | Lighthouse Lab in Glasgow | Wellcome Sanger Institute for the COVID-19 Genomics UK Consortium | Harper VanSteenhouse, Yumi Kasai, David Gray, Carol Clugston, Anna Dominiczak and Alex Alderton, Roberto Amato, Sonia Goncalves, Ewan Harrison, David K. Jackson, Ian Johnston, Dominic Kwiatkowski, Cordelia Langford, John Sillitoe |
| EPI_ISL_533252 | NHSGGC West of Scotland Specialist Virology Centre / MRC-University of Glasgow Centre for Virus Research | Wellcome Sanger Institute for the COVID-19 Genomics UK Consortium | Ana da Silva Filipe, Natasha Johnson, Kathy Smollett, Daniel Mair, Stephen Carmichael, Lily Tong, Jenna Nichols, Elihu Aranday-Cortes, Kirstyn Brunker, Yasmin Parr, Kyriaki Nomikou; Sarah McDonald, Marc Niebel, Patawee Asamaphan; Richard Orton, Joseph Hughes, Sreenu Vattipally, David L Robertson; Alasdair MacLean, Rory Gunson; Kathy Li, Natasha Jesudason, Rajiv Shah, James Shepherd, Antonia Ho, Alice Broos, Emma Thomson and Alex Alderton, Roberto Amato, Sonia Goncalves, Ewan Harrison, David K. Jackson, Ian Johnston, Dominic Kwiatkowski, Cordelia Langford, John Sillitoe |
| EPI_ISL_533253, EPI_ISL_533254, EPI_ISL_533255 | Lighthouse Lab in Glasgow | Wellcome Sanger Institute for the COVID-19 Genomics UK Consortium | Harper VanSteenhouse, Yumi Kasai, David Gray, Carol Clugston, Anna Dominiczak and Alex Alderton, Roberto Amato, Sonia Goncalves, Ewan Harrison, David K. Jackson, Ian Johnston, Dominic Kwiatkowski, Cordelia Langford, John Sillitoe |
| EPI_ISL_533256 | NHSGGC West of Scotland Specialist Virology Centre / MRC-University of Glasgow Centre for Virus Research | Wellcome Sanger Institute for the COVID-19 Genomics UK Consortium | Ana da Silva Filipe, Natasha Johnson, Kathy Smollett, Daniel Mair, Stephen Carmichael, Lily Tong, Jenna Nichols, Elihu Aranday-Cortes, Kirstyn Brunker, Yasmin Parr, Kyriaki Nomikou; Sarah McDonald, Marc Niebel, Patawee Asamaphan; Richard Orton, Joseph Hughes, Sreenu Vattipally, David L Robertson; Alasdair MacLean, Rory Gunson; Kathy Li, Natasha Jesudason, Rajiv Shah, James Shepherd, Antonia Ho, Alice Broos, Emma Thomson and Alex Alderton, Roberto Amato, Sonia Goncalves, Ewan Harrison, David K. Jackson, Ian Johnston, Dominic Kwiatkowski, Cordelia Langford, John Sillitoe |
| EPI_ISL_533257, EPI_ISL_533258, EPI_ISL_533259, EPI_ISL_533260, EPI_ISL_533261 | Lighthouse Lab in Glasgow | Wellcome Sanger Institute for the COVID-19 Genomics UK Consortium | Harper VanSteenhouse, Yumi Kasai, David Gray, Carol Clugston, Anna Dominiczak and Alex Alderton, Roberto Amato, Sonia Goncalves, Ewan Harrison, David K. Jackson, Ian Johnston, Dominic Kwiatkowski, Cordelia Langford, John Sillitoe |
| EPI_ISL_533262 | NHSGGC West of Scotland Specialist Virology Centre / MRC-University of Glasgow Centre for Virus Research | Wellcome Sanger Institute for the COVID-19 Genomics UK Consortium | Ana da Silva Filipe, Natasha Johnson, Kathy Smollett, Daniel Mair, Stephen Carmichael, Lily Tong, Jenna Nichols, Elihu Aranday-Cortes, Kirstyn Brunker, Yasmin Parr, Kyriaki Nomikou; Sarah McDonald, Marc Niebel, Patawee Asamaphan; Richard Orton, Joseph Hughes, Sreenu Vattipally, David L Robertson; Alasdair MacLean, Rory Gunson; Kathy Li, Natasha Jesudason, Rajiv Shah, James Shepherd, Antonia Ho, Alice Broos, Emma Thomson and Alex Alderton, Roberto Amato, Sonia Goncalves, Ewan Harrison, David K. Jackson, Ian Johnston, Dominic Kwiatkowski, Cordelia Langford, John Sillitoe |
| EPI_ISL_533263, EPI_ISL_533264, EPI_ISL_533265, EPI_ISL_533266, EPI_ISL_533268, EPI_ISL_533269 | Lighthouse Lab in Glasgow | Wellcome Sanger Institute for the COVID-19 Genomics UK Consortium | Harper VanSteenhouse, Yumi Kasai, David Gray, Carol Clugston, Anna Dominiczak and Alex Alderton, Roberto Amato, Sonia Goncalves, Ewan Harrison, David K. Jackson, Ian Johnston, Dominic Kwiatkowski, Cordelia Langford, John Sillitoe |
| EPI_ISL_533271 | NHSGGC West of Scotland Specialist Virology Centre / MRC-University of Glasgow Centre for Virus Research | Wellcome Sanger Institute for the COVID-19 Genomics UK Consortium | Ana da Silva Filipe, Natasha Johnson, Kathy Smollett, Daniel Mair, Stephen Carmichael, Lily Tong, Jenna Nichols, Elihu Aranday-Cortes, Kirstyn Brunker, Yasmin Parr, Kyriaki Nomikou; Sarah McDonald, Marc Niebel, Patawee Asamaphan; Richard Orton, Joseph Hughes, Sreenu Vattipally, David L Robertson; Alasdair MacLean, Rory Gunson; Kathy Li, Natasha Jesudason, Rajiv Shah, James Shepherd, Antonia Ho, Alice Broos, Emma Thomson and Alex Alderton, Roberto Amato, Sonia Goncalves, Ewan Harrison, David K. Jackson, Ian Johnston, Dominic Kwiatkowski, Cordelia Langford, John Sillitoe |
| EPI_ISL_533273, EPI_ISL_533274, EPI_ISL_533275, EPI_ISL_533277, EPI_ISL_533278, EPI_ISL_533279, EPI_ISL_533280, EPI_ISL_533281, EPI_ISL_533282 | Lighthouse Lab in Glasgow | Wellcome Sanger Institute for the COVID-19 Genomics UK Consortium | Harper VanSteenhouse, Yumi Kasai, David Gray, Carol Clugston, Anna Dominiczak and Alex Alderton, Roberto Amato, Sonia Goncalves, Ewan Harrison, David K. Jackson, Ian Johnston, Dominic Kwiatkowski, Cordelia Langford, John Sillitoe |
| EPI_ISL_533283 | NHSGGC West of Scotland Specialist Virology Centre / MRC-University of Glasgow Centre for Virus Research | Wellcome Sanger Institute for the COVID-19 Genomics UK Consortium | Ana da Silva Filipe, Natasha Johnson, Kathy Smollett, Daniel Mair, Stephen Carmichael, Lily Tong, Jenna Nichols, Elihu Aranday-Cortes, Kirstyn Brunker, Yasmin Parr, Kyriaki Nomikou; Sarah McDonald, Marc Niebel, Patawee Asamaphan; Richard Orton, Joseph Hughes, Sreenu Vattipally, David L Robertson; Alasdair MacLean, Rory Gunson; Kathy Li, Natasha Jesudason, Rajiv Shah, James Shepherd, Antonia Ho, Alice Broos, Emma Thomson and Alex Alderton, Roberto Amato, Sonia Goncalves, Ewan Harrison, David K. Jackson, Ian Johnston, Dominic Kwiatkowski, Cordelia Langford, John Sillitoe |
| EPI_ISL_533284, EPI_ISL_533286, EPI_ISL_533287, EPI_ISL_533288, EPI_ISL_533290, EPI_ISL_533291, EPI_ISL_533293, EPI_ISL_533294, EPI_ISL_533295, EPI_ISL_533296, EPI_ISL_533297, EPI_ISL_533299, EPI_ISL_533300, EPI_ISL_533301, EPI_ISL_533302, EPI_ISL_533303, EPI_ISL_533304, EPI_ISL_533305, EPI_ISL_533306, EPI_ISL_533308, EPI_ISL_533309, EPI_ISL_533310 | | | |
| see above | Lighthouse Lab in Glasgow | Wellcome Sanger Institute for the COVID-19 Genomics UK Consortium | Harper VanSteenhouse, Yumi Kasai, David Gray, Carol Clugston, Anna Dominiczak and Alex Alderton, Roberto Amato, Sonia Goncalves, Ewan Harrison, David K. Jackson, Ian Johnston, Dominic Kwiatkowski, Cordelia Langford, John Sillitoe |
| EPI_ISL_533311 | NHSGGC West of Scotland Specialist Virology Centre / MRC-University of Glasgow Centre for Virus Research | Wellcome Sanger Institute for the COVID-19 Genomics UK Consortium | Ana da Silva Filipe, Natasha Johnson, Kathy Smollett, Daniel Mair, Stephen Carmichael, Lily Tong, Jenna Nichols, Elihu Aranday-Cortes, Kirstyn Brunker, Yasmin Parr, Kyriaki Nomikou; Sarah McDonald, Marc Niebel, Patawee Asamaphan; Richard Orton, Joseph Hughes, Sreenu Vattipally, David L Robertson; Alasdair MacLean, Rory Gunson; Kathy Li, Natasha Jesudason, Rajiv Shah, James Shepherd, Antonia Ho, Alice Broos, Emma Thomson and Alex Alderton, Roberto Amato, Sonia Goncalves, Ewan Harrison, David K. Jackson, Ian Johnston, Dominic Kwiatkowski, Cordelia Langford, John Sillitoe |
| EPI_ISL_533313, EPI_ISL_533314, EPI_ISL_533315, EPI_ISL_533316, EPI_ISL_533318, EPI_ISL_533319, EPI_ISL_533320, EPI_ISL_533321, EPI_ISL_533322, EPI_ISL_533323, EPI_ISL_533324, EPI_ISL_533325, EPI_ISL_533326, EPI_ISL_533329, EPI_ISL_533331, EPI_ISL_533332, EPI_ISL_533333, EPI_ISL_533335, EPI_ISL_533336, EPI_ISL_533337, EPI_ISL_533338, EPI_ISL_533339, EPI_ISL_533341, EPI_ISL_533342 | | | |
| see above | Lighthouse Lab in Glasgow | Wellcome Sanger Institute for the COVID-19 Genomics UK Consortium | Harper VanSteenhouse, Yumi Kasai, David Gray, Carol Clugston, Anna Dominiczak and Alex Alderton, Roberto Amato, Sonia Goncalves, Ewan Harrison, David K. Jackson, Ian Johnston, Dominic Kwiatkowski, Cordelia Langford, John Sillitoe |
| EPI_ISL_533343 | NHSGGC West of Scotland Specialist Virology Centre / MRC-University of Glasgow Centre for Virus Research | Wellcome Sanger Institute for the COVID-19 Genomics UK Consortium | Ana da Silva Filipe, Natasha Johnson, Kathy Smollett, Daniel Mair, Stephen Carmichael, Lily Tong, Jenna Nichols, Elihu Aranday-Cortes, Kirstyn Brunker, Yasmin Parr, Kyriaki Nomikou; Sarah McDonald, Marc Niebel, Patawee Asamaphan; Richard Orton, Joseph Hughes, Sreenu Vattipally, David L Robertson; Alasdair MacLean, Rory Gunson; Kathy Li, Natasha Jesudason, Rajiv Shah, James Shepherd, Antonia Ho, Alice Broos, Emma Thomson and Alex Alderton, Roberto Amato, Sonia Goncalves, Ewan Harrison, David K. Jackson, Ian Johnston, Dominic Kwiatkowski, Cordelia Langford, John Sillitoe |
| EPI_ISL_533344, EPI_ISL_533345 | Lighthouse Lab in Glasgow | Wellcome Sanger Institute for the COVID-19 Genomics UK Consortium | Harper VanSteenhouse, Yumi Kasai, David Gray, Carol Clugston, Anna Dominiczak and Alex Alderton, Roberto Amato, Sonia Goncalves, Ewan Harrison, David K. Jackson, Ian Johnston, Dominic Kwiatkowski, Cordelia Langford, John Sillitoe |
| EPI_ISL_533346 | NHSGGC West of Scotland Specialist Virology Centre / MRC-University of Glasgow Centre for Virus Research | Wellcome Sanger Institute for the COVID-19 Genomics UK Consortium | Ana da Silva Filipe, Natasha Johnson, Kathy Smollett, Daniel Mair, Stephen Carmichael, Lily Tong, Jenna Nichols, Elihu Aranday-Cortes, Kirstyn Brunker, Yasmin Parr, Kyriaki Nomikou; Sarah McDonald, Marc Niebel, Patawee Asamaphan; Richard Orton, Joseph Hughes, Sreenu Vattipally, David L Robertson; Alasdair MacLean, Rory Gunson; Kathy Li, Natasha Jesudason, Rajiv Shah, James Shepherd, Antonia Ho, Alice Broos, Emma Thomson and Alex Alderton, Roberto Amato, Sonia Goncalves, Ewan Harrison, David K. Jackson, Ian Johnston, Dominic Kwiatkowski, Cordelia Langford, John Sillitoe |
| EPI_ISL_533347, EPI_ISL_533348, EPI_ISL_533349 | Lighthouse Lab in Glasgow | Wellcome Sanger Institute for the COVID-19 Genomics UK Consortium | Harper VanSteenhouse, Yumi Kasai, David Gray, Carol Clugston, Anna Dominiczak and Alex Alderton, Roberto Amato, Sonia Goncalves, Ewan Harrison, David K. Jackson, Ian Johnston, Dominic Kwiatkowski, Cordelia Langford, John Sillitoe |
| EPI_ISL_533350, EPI_ISL_533351 | NHSGGC West of Scotland Specialist Virology Centre / MRC-University of Glasgow Centre for Virus Research | Wellcome Sanger Institute for the COVID-19 Genomics UK Consortium | Ana da Silva Filipe, Natasha Johnson, Kathy Smollett, Daniel Mair, Stephen Carmichael, Lily Tong, Jenna Nichols, Elihu Aranday-Cortes, Kirstyn Brunker, Yasmin Parr, Kyriaki Nomikou; Sarah McDonald, Marc Niebel, Patawee Asamaphan; Richard Orton, Joseph Hughes, Sreenu Vattipally, David L Robertson; Alasdair MacLean, Rory Gunson; Kathy Li, Natasha Jesudason, Rajiv Shah, James Shepherd, Antonia Ho, Alice Broos, Emma Thomson and Alex Alderton, Roberto Amato, Sonia Goncalves, Ewan Harrison, David K. Jackson, Ian Johnston, Dominic Kwiatkowski, Cordelia Langford, John Sillitoe |
| EPI_ISL_533352, EPI_ISL_533353, EPI_ISL_533356, EPI_ISL_533357, EPI_ISL_533360, EPI_ISL_533361, EPI_ISL_533362, EPI_ISL_533363, EPI_ISL_533364, EPI_ISL_533366, EPI_ISL_533368, EPI_ISL_533369, EPI_ISL_533370, EPI_ISL_533371, EPI_ISL_533372, EPI_ISL_533374, EPI_ISL_533375 | | | |
| see above | Lighthouse Lab in Glasgow | Wellcome Sanger Institute for the COVID-19 Genomics UK Consortium | Harper VanSteenhouse, Yumi Kasai, David Gray, Carol Clugston, Anna Dominiczak and Alex Alderton, Roberto Amato, Sonia Goncalves, Ewan Harrison, David K. Jackson, Ian Johnston, Dominic Kwiatkowski, Cordelia Langford, John Sillitoe |
| EPI_ISL_533376 | NHSGGC West of Scotland Specialist Virology Centre / MRC-University of Glasgow Centre for Virus Research | Wellcome Sanger Institute for the COVID-19 Genomics UK Consortium | Ana da Silva Filipe, Natasha Johnson, Kathy Smollett, Daniel Mair, Stephen Carmichael, Lily Tong, Jenna Nichols, Elihu Aranday-Cortes, Kirstyn Brunker, Yasmin Parr, Kyriaki Nomikou; Sarah McDonald, Marc Niebel, Patawee Asamaphan; Richard Orton, Joseph Hughes, Sreenu Vattipally, David L Robertson; Alasdair MacLean, Rory Gunson; Kathy Li, Natasha Jesudason, Rajiv Shah, James Shepherd, Antonia Ho, Alice Broos, Emma Thomson and Alex Alderton, Roberto Amato, Sonia Goncalves, Ewan Harrison, David K. Jackson, Ian Johnston, Dominic Kwiatkowski, Cordelia Langford, John Sillitoe |
| EPI_ISL_533377, EPI_ISL_533378, EPI_ISL_533379, EPI_ISL_533380, EPI_ISL_533381, EPI_ISL_533382, EPI_ISL_533383, EPI_ISL_533384, EPI_ISL_533385, EPI_ISL_533386, EPI_ISL_533388, EPI_ISL_533389, EPI_ISL_533390, EPI_ISL_533391, EPI_ISL_533392, EPI_ISL_533393 | | | |
| see above | Lighthouse Lab in Glasgow | Wellcome Sanger Institute for the COVID-19 Genomics UK Consortium | Harper VanSteenhouse, Yumi Kasai, David Gray, Carol Clugston, Anna Dominiczak and Alex Alderton, Roberto Amato, Sonia Goncalves, Ewan Harrison, David K. Jackson, Ian Johnston, Dominic Kwiatkowski, Cordelia Langford, John Sillitoe |
| EPI_ISL_533394 | NHSGGC West of Scotland Specialist Virology Centre / MRC-University of Glasgow Centre for Virus Research | Wellcome Sanger Institute for the COVID-19 Genomics UK Consortium | Ana da Silva Filipe, Natasha Johnson, Kathy Smollett, Daniel Mair, Stephen Carmichael, Lily Tong, Jenna Nichols, Elihu Aranday-Cortes, Kirstyn Brunker, Yasmin Parr, Kyriaki Nomikou; Sarah McDonald, Marc Niebel, Patawee Asamaphan; Richard Orton, Joseph Hughes, Sreenu Vattipally, David L Robertson; Alasdair MacLean, Rory Gunson; Kathy Li, Natasha Jesudason, Rajiv Shah, James Shepherd, Antonia Ho, Alice Broos, Emma Thomson and Alex Alderton, Roberto Amato, Sonia Goncalves, Ewan Harrison, David K. Jackson, Ian Johnston, Dominic Kwiatkowski, Cordelia Langford, John Sillitoe |
| EPI_ISL_533395, EPI_ISL_533397, EPI_ISL_533398 | Lighthouse Lab in Glasgow | Wellcome Sanger Institute for the COVID-19 Genomics UK Consortium | Harper VanSteenhouse, Yumi Kasai, David Gray, Carol Clugston, Anna Dominiczak and Alex Alderton, Roberto Amato, Sonia Goncalves, Ewan Harrison, David K. Jackson, Ian Johnston, Dominic Kwiatkowski, Cordelia Langford, John Sillitoe |
| EPI_ISL_533400 | Lighthouse Lab in Glasgow | Wellcome Sanger Institute for the COVID-19 Genomics UK Consortium | Harper VanSteenhouse, Yumi Kasai, David Gray, Carol Clugston, Anna Dominiczak and Alex Alderton, Roberto Amato, Sonia Goncalves, Ewan Harrison, David K. Jackson, Ian Johnston, Dominic Kwiatkowski, Cordelia Langford, John Sillitoe on behalf of the Wellcome Sanger Institute COVID-19 Surveillance Team |
| EPI_ISL_533401, EPI_ISL_533402, EPI_ISL_533404, EPI_ISL_533405 | Lighthouse Lab in Glasgow | Wellcome Sanger Institute for the COVID-19 Genomics UK Consortium | Harper VanSteenhouse, Yumi Kasai, David Gray, Carol Clugston, Anna Dominiczak and Alex Alderton, Roberto Amato, Sonia Goncalves, Ewan Harrison, David K. Jackson, Ian Johnston, Dominic Kwiatkowski, Cordelia Langford, John Sillitoe |
| EPI_ISL_533406 | NHSGGC West of Scotland Specialist Virology Centre / MRC-University of Glasgow Centre for Virus Research | Wellcome Sanger Institute for the COVID-19 Genomics UK Consortium | Ana da Silva Filipe, Natasha Johnson, Kathy Smollett, Daniel Mair, Stephen Carmichael, Lily Tong, Jenna Nichols, Elihu Aranday-Cortes, Kirstyn Brunker, Yasmin Parr, Kyriaki Nomikou; Sarah McDonald, Marc Niebel, Patawee Asamaphan; Richard Orton, Joseph Hughes, Sreenu Vattipally, David L Robertson; Alasdair MacLean, Rory Gunson; Kathy Li, Natasha Jesudason, Rajiv Shah, James Shepherd, Antonia Ho, Alice Broos, Emma Thomson and Alex Alderton, Roberto Amato, Sonia Goncalves, Ewan Harrison, David K. Jackson, Ian Johnston, Dominic Kwiatkowski, Cordelia Langford, John Sillitoe |
| EPI_ISL_533407, EPI_ISL_533408, EPI_ISL_533409, EPI_ISL_533410, EPI_ISL_533412, EPI_ISL_533413, EPI_ISL_533414, EPI_ISL_533415, EPI_ISL_533416, EPI_ISL_533417, EPI_ISL_533418 | | | |
| see above | Lighthouse Lab in Glasgow | Wellcome Sanger Institute for the COVID-19 Genomics UK Consortium | Harper VanSteenhouse, Yumi Kasai, David Gray, Carol Clugston, Anna Dominiczak and Alex Alderton, Roberto Amato, Sonia Goncalves, Ewan Harrison, David K. Jackson, Ian Johnston, Dominic Kwiatkowski, Cordelia Langford, John Sillitoe |
| EPI_ISL_533419 | NHSGGC West of Scotland Specialist Virology Centre / MRC-University of Glasgow Centre for Virus Research | Wellcome Sanger Institute for the COVID-19 Genomics UK Consortium | Ana da Silva Filipe, Natasha Johnson, Kathy Smollett, Daniel Mair, Stephen Carmichael, Lily Tong, Jenna Nichols, Elihu Aranday-Cortes, Kirstyn Brunker, Yasmin Parr, Kyriaki Nomikou; Sarah McDonald, Marc Niebel, Patawee Asamaphan; Richard Orton, Joseph Hughes, Sreenu Vattipally, David L Robertson; Alasdair MacLean, Rory Gunson; Kathy Li, Natasha Jesudason, Rajiv Shah, James Shepherd, Antonia Ho, Alice Broos, Emma Thomson and Alex Alderton, Roberto Amato, Sonia Goncalves, Ewan Harrison, David K. Jackson, Ian Johnston, Dominic Kwiatkowski, Cordelia Langford, John Sillitoe |
| EPI_ISL_533420 | Lighthouse Lab in Glasgow | Wellcome Sanger Institute for the COVID-19 Genomics UK Consortium | Harper VanSteenhouse, Yumi Kasai, David Gray, Carol Clugston, Anna Dominiczak and Alex Alderton, Roberto Amato, Sonia Goncalves, Ewan Harrison, David K. Jackson, Ian Johnston, Dominic Kwiatkowski, Cordelia Langford, John Sillitoe |
| EPI_ISL_533421 | NHSGGC West of Scotland Specialist Virology Centre / MRC-University of Glasgow Centre for Virus Research | Wellcome Sanger Institute for the COVID-19 Genomics UK Consortium | Ana da Silva Filipe, Natasha Johnson, Kathy Smollett, Daniel Mair, Stephen Carmichael, Lily Tong, Jenna Nichols, Elihu Aranday-Cortes, Kirstyn Brunker, Yasmin Parr, Kyriaki Nomikou; Sarah McDonald, Marc Niebel, Patawee Asamaphan; Richard Orton, Joseph Hughes, Sreenu Vattipally, David L Robertson; Alasdair MacLean, Rory Gunson; Kathy Li, Natasha Jesudason, Rajiv Shah, James Shepherd, Antonia Ho, Alice Broos, Emma Thomson and Alex Alderton, Roberto Amato, Sonia Goncalves, Ewan Harrison, David K. Jackson, Ian Johnston, Dominic Kwiatkowski, Cordelia Langford, John Sillitoe |

| | | | |
|---|---|---|---|
| EPI_ISL_533422, EPI_ISL_533424, EPI_ISL_533425, EPI_ISL_533426, EPI_ISL_533428, EPI_ISL_533429, EPI_ISL_533430, EPI_ISL_533431 | Lighthouse Lab in Glasgow | Wellcome Sanger Institute for the COVID-19 Genomics UK Consortium | Harper VanSteenhouse, Yumi Kasai, David Gray, Carol Clugston, Anna Dominiczak and Alex Alderton, Roberto Amato, Sonia Goncalves, Ewan Harrison, David K. Jackson, Ian Johnston, Dominic Kwiatkowski, Cordelia Langford, John Sillitoe |
| EPI_ISL_533433 | Lighthouse Lab in Glasgow | Wellcome Sanger Institute for the COVID-19 Genomics UK Consortium | Harper VanSteenhouse, Yumi Kasai, David Gray, Carol Clugston, Anna Dominiczak and Alex Alderton, Roberto Amato, Sonia Goncalves, Ewan Harrison, David K. Jackson, Ian Johnston, Dominic Kwiatkowski, Cordelia Langford, John Sillitoe on behalf of the Wellcome Sanger Institute COVID-19 Surveillance Team |
| EPI_ISL_534199 | E. Gulbja Laboratorija | Latvian Biomedical Research and Study Centre | Ivars Silamiķelis, Jānis Pjalkovskis, Kaspars Megnis, Monta Ustinova, Ņikita Zrelovs, Vita Rovīte, Mikus Gavars, Dmitrijs Perminovs, Uga Dumpis, Jānis Kloviņš |
| EPI_ISL_534200, EPI_ISL_534201, EPI_ISL_534202 | Centrālā laboratorija | Latvian Biomedical Research and Study Centre | Ivars Silamiķelis, Jānis Pjalkovskis, Kaspars Megnis, Monta Ustinova, Ņikita Zrelovs, Vita Rovīte, Stella Lapiņa, Jana Osīte, Marta Priedīte, Uga Dumpis, Jānis Kloviņš |
| EPI_ISL_534203 | E. Gulbja Laboratorija | Latvian Biomedical Research and Study Centre | Ivars Silamiķelis, Jānis Pjalkovskis, Kaspars Megnis, Monta Ustinova, Ņikita Zrelovs, Vita Rovīte, Mikus Gavars, Dmitrijs Perminovs, Uga Dumpis, Jānis Kloviņš |
| EPI_ISL_534204, EPI_ISL_534205 | Latvijas Infektoloģijas centrs | Latvian Biomedical Research and Study Centre | Ivars Silamiķelis, Jānis Pjalkovskis, Kaspars Megnis, Monta Ustinova, Ņikita Zrelovs, Vita Rovīte, Jeļena Storoženko, Tatjana Kolupajeva, Oksana Savicka, Uga Dumpis, Jānis Kloviņš |
| EPI_ISL_534206, EPI_ISL_534207, EPI_ISL_534208 | Centrālā laboratorija | Latvian Biomedical Research and Study Centre | Ivars Silamiķelis, Jānis Pjalkovskis, Kaspars Megnis, Monta Ustinova, Ņikita Zrelovs, Vita Rovīte, Stella Lapiņa, Jana Osīte, Marta Priedīte, Uga Dumpis, Jānis Kloviņš |
| EPI_ISL_534209 | Latvijas Infektoloģijas centrs | Latvian Biomedical Research and Study Centre | Ivars Silamiķelis, Jānis Pjalkovskis, Kaspars Megnis, Monta Ustinova, Ņikita Zrelovs, Vita Rovīte, Jeļena Storoženko, Tatjana Kolupajeva, Oksana Savicka, Uga Dumpis, Jānis Kloviņš |
| EPI_ISL_534210, EPI_ISL_534211 | Centrālā laboratorija | Latvian Biomedical Research and Study Centre | Ivars Silamiķelis, Jānis Pjalkovskis, Kaspars Megnis, Monta Ustinova, Ņikita Zrelovs, Vita Rovīte, Stella Lapiņa, Jana Osīte, Marta Priedīte, Uga Dumpis, Jānis Kloviņš |
| EPI_ISL_534212, EPI_ISL_534213, EPI_ISL_534214, EPI_ISL_534215, EPI_ISL_534216, EPI_ISL_534218, EPI_ISL_534219 | E. Gulbja Laboratorija | Latvian Biomedical Research and Study Centre | Ivars Silamiķelis, Jānis Pjalkovskis, Kaspars Megnis, Monta Ustinova, Ņikita Zrelovs, Vita Rovīte, Mikus Gavars, Dmitrijs Perminovs, Uga Dumpis, Jānis Kloviņš |
| EPI_ISL_534220 | Centrālā laboratorija | Latvian Biomedical Research and Study Centre | Ivars Silamiķelis, Jānis Pjalkovskis, Kaspars Megnis, Monta Ustinova, Ņikita Zrelovs, Vita Rovīte, Stella Lapiņa, Jana Osīte, Marta Priedīte, Uga Dumpis, Jānis Kloviņš |
| EPI_ISL_534221 | E. Gulbja Laboratorija | Latvian Biomedical Research and Study Centre | Ivars Silamiķelis, Jānis Pjalkovskis, Kaspars Megnis, Monta Ustinova, Ņikita Zrelovs, Vita Rovīte, Mikus Gavars, Dmitrijs Perminovs, Uga Dumpis, Jānis Kloviņš |
| EPI_ISL_534222, EPI_ISL_534223 | Centrālā laboratorija | Latvian Biomedical Research and Study Centre | Ivars Silamiķelis, Jānis Pjalkovskis, Kaspars Megnis, Monta Ustinova, Ņikita Zrelovs, Vita Rovīte, Stella Lapiņa, Jana Osīte, Marta Priedīte, Uga Dumpis, Jānis Kloviņš |
| EPI_ISL_534224 | Universitetssjukhuset i Linkoping | The Public Health Agency of Sweden | Anna-Malin Linde, Maria Lind Karlberg, Mattias Haukland, Reza Advani, Olov Svartstrom, Oskar Karlsson Lindsjo, Sandra Broddesson, Petra Edquist, Mia Brytting, Anna Risberg, Karin Tegmark-Wisell |
| EPI_ISL_534225, EPI_ISL_534226, EPI_ISL_534227, EPI_ISL_534228, EPI_ISL_534229 | Skanes universitetssjukhus Lund | The Public Health Agency of Sweden | Anna-Malin Linde, Maria Lind Karlberg, Mattias Haukland, Reza Advani, Olov Svartstrom, Oskar Karlsson Lindsjo, Sandra Broddesson, Petra Edquist, Mia Brytting, Anna Risberg, Karin Tegmark-Wisell |
| EPI_ISL_534230, EPI_ISL_534231, EPI_ISL_534232, EPI_ISL_534233 | Capio S:t Gorans sjukhus | The Public Health Agency of Sweden | Anna-Malin Linde, Maria Lind Karlberg, Mattias Haukland, Reza Advani, Olov Svartstrom, Oskar Karlsson Lindsjo, Sandra Broddesson, Petra Edquist, Mia Brytting, Anna Risberg, Karin Tegmark-Wisell |
| EPI_ISL_534234, EPI_ISL_534235, EPI_ISL_534236 | Karolinska universitetslaboratoriet SOLNA | The Public Health Agency of Sweden | Anna-Malin Linde, Maria Lind Karlberg, Mattias Haukland, Reza Advani, Olov Svartstrom, Oskar Karlsson Lindsjo, Sandra Broddesson, Petra Edquist, Mia Brytting, Anna Risberg, Karin Tegmark-Wisell |
| EPI_ISL_534237, EPI_ISL_534238 | Lanssjukhuset Kalmar | The Public Health Agency of Sweden | Anna-Malin Linde, Maria Lind Karlberg, Mattias Haukland, Reza Advani, Olov Svartstrom, Oskar Karlsson Lindsjo, Sandra Broddesson, Petra Edquist, Mia Brytting, Anna Risberg, Karin Tegmark-Wisell |
| EPI_ISL_534239, EPI_ISL_534240, EPI_ISL_534241, EPI_ISL_534242, EPI_ISL_534243 | Norra Alvsborgs lanssjukhus | The Public Health Agency of Sweden | Anna-Malin Linde, Maria Lind Karlberg, Mattias Haukland, Reza Advani, Olov Svartstrom, Oskar Karlsson Lindsjo, Sandra Broddesson, Petra Edquist, Mia Brytting, Anna Risberg, Karin Tegmark-Wisell |
| EPI_ISL_534244 | Sundsvalls sjukhus | The Public Health Agency of Sweden | Anna-Malin Linde, Maria Lind Karlberg, Mattias Haukland, Reza Advani, Olov Svartstrom, Oskar Karlsson Lindsjo, Sandra Broddesson, Petra Edquist, Mia Brytting, Anna Risberg, Karin Tegmark-Wisell |
| EPI_ISL_534245 | Kliniskt mikrobiologiska laboratoriet | The Public Health Agency of Sweden | Anna-Malin Linde, Maria Lind Karlberg, Mattias Haukland, Reza Advani, Olov Svartstrom, Oskar Karlsson Lindsjo, Sandra Broddesson, Petra Edquist, Mia Brytting, Anna Risberg, Karin Tegmark-Wisell |
| EPI_ISL_534246, EPI_ISL_534247 | Universitetssjukhuset i Linkoping | The Public Health Agency of Sweden | Anna-Malin Linde, Maria Lind Karlberg, Mattias Haukland, Reza Advani, Olov Svartstrom, Oskar Karlsson Lindsjo, Sandra Broddesson, Petra Edquist, Mia Brytting, Anna Risberg, Karin Tegmark-Wisell |
| EPI_ISL_534248 | Laboratoriemedicin Vasternorrland | The Public Health Agency of Sweden | Anna-Malin Linde, Maria Lind Karlberg, Mattias Haukland, Reza Advani, Olov Svartstrom, Oskar Karlsson Lindsjo, Sandra Broddesson, Petra Edquist, Mia Brytting, Anna Risberg, Karin Tegmark-Wisell |
| EPI_ISL_534249, EPI_ISL_534250, EPI_ISL_534251 | Gavle Sjukhus | The Public Health Agency of Sweden | Anna-Malin Linde, Maria Lind Karlberg, Mattias Haukland, Reza Advani, Olov Svartstrom, Oskar Karlsson Lindsjo, Sandra Broddesson, Petra Edquist, Mia Brytting, Anna Risberg, Karin Tegmark-Wisell |
| EPI_ISL_534252, EPI_ISL_534253, EPI_ISL_534254, EPI_ISL_534255 | Laboratoriemedicin Vasternorrland | The Public Health Agency of Sweden | Anna-Malin Linde, Maria Lind Karlberg, Mattias Haukland, Reza Advani, Olov Svartstrom, Oskar Karlsson Lindsjo, Sandra Broddesson, Petra Edquist, Mia Brytting, Anna Risberg, Karin Tegmark-Wisell |
| EPI_ISL_534256 | Klinisk mikrobiologi, Laboratoriemedicin Gavleborg | The Public Health Agency of Sweden | Anna-Malin Linde, Maria Lind Karlberg, Mattias Haukland, Reza Advani, Olov Svartstrom, Oskar Karlsson Lindsjo, Sandra Broddesson, Petra Edquist, Mia Brytting, Anna Risberg, Karin Tegmark-Wisell |
| EPI_ISL_534257, EPI_ISL_534258 | Laboratoriemedicin Vasternorrland | The Public Health Agency of Sweden | Anna-Malin Linde, Maria Lind Karlberg, Mattias Haukland, Reza Advani, Olov Svartstrom, Oskar Karlsson Lindsjo, Sandra Broddesson, Petra Edquist, Mia Brytting, Anna Risberg, Karin Tegmark-Wisell |
| EPI_ISL_534259 | Karolinska universitetslaboratoriet | The Public Health Agency of Sweden | Anna-Malin Linde, Maria Lind Karlberg, Mattias Haukland, Reza Advani, Olov Svartstrom, Oskar Karlsson Lindsjo, Sandra Broddesson, Petra Edquist, Mia Brytting, Anna Risberg, Karin Tegmark-Wisell |
| EPI_ISL_534311 | UPA III 26 de Agosto | Instituto Adolfo Lutz, Interdiciplinary Procedures Center, Strategic Laboratory | Claudio Tavares Sacchi, Claudia Regina Gonçalves, Erica Valessa Ramos Gomes |
| EPI_ISL_534312 | Distrito Sanitario Sul | Instituto Adolfo Lutz, Interdiciplinary Procedures Center, Strategic Laboratory | Claudio Tavares Sacchi, Claudia Regina Gonçalves, Erica Valessa Ramos Gomes |
| EPI_ISL_534313 | Hospital da Sta Casa de Sto Amaro | Instituto Adolfo Lutz, Interdiciplinary Procedures Center, Strategic Laboratory | Claudio Tavares Sacchi, Claudia Regina Gonçalves, Erica Valessa Ramos Gomes |
| EPI_ISL_534314 | Hospital Universitario da USP de SP | Instituto Adolfo Lutz, Interdiciplinary Procedures Center, Strategic Laboratory | Claudio Tavares Sacchi, Claudia Regina Gonçalves, Erica Valessa Ramos Gomes |
| EPI_ISL_534315 | Serviço de Verificação de Óbitos SVO Guaruulhos | Instituto Adolfo Lutz, Interdiciplinary Procedures Center, Strategic Laboratory | Claudio Tavares Sacchi, Claudia Regina Gonçalves, Erica Valessa Ramos Gomes |
| EPI_ISL_534316 | OS Mun Santana Lauro Ribas Braga | Instituto Adolfo Lutz, Interdiciplinary Procedures Center, Strategic Laboratory | Claudio Tavares Sacchi, Claudia Regina Gonçalves, Erica Valessa Ramos Gomes |
| EPI_ISL_534317 | Hospital Geral de Itapevi | Instituto Adolfo Lutz, Interdiciplinary Procedures Center, Strategic Laboratory | Claudio Tavares Sacchi, Claudia Regina Gonçalves, Erica Valessa Ramos Gomes |
| EPI_ISL_534318 | Hospital Municipal Antonio Giglio | Instituto Adolfo Lutz, Interdiciplinary Procedures Center, Strategic Laboratory | Claudio Tavares Sacchi, Claudia Regina Gonçalves, Erica Valessa Ramos Gomes |
| EPI_ISL_534319, EPI_ISL_534320 | Hospital do Serv Pub ESTAFCO Morato de Oliveira | Instituto Adolfo Lutz, Interdiciplinary Procedures Center, Strategic Laboratory | Claudio Tavares Sacchi, Claudia Regina Gonçalves, Erica Valessa Ramos Gomes |
| EPI_ISL_534321 | PS e Maternidade Nair Fonseca Leitao Arantes | Instituto Adolfo Lutz, Interdiciplinary Procedures Center, Strategic Laboratory | Claudio Tavares Sacchi, Claudia Regina Gonçalves, Erica Valessa Ramos Gomes |
| EPI_ISL_534322 | PS Mun Julio Tupy | Instituto Adolfo Lutz, Interdiciplinary Procedures Center, Strategic Laboratory | Claudio Tavares Sacchi, Claudia Regina Gonçalves, Erica Valessa Ramos Gomes |
| EPI_ISL_534323 | Hospital e Pronto Socorro Comunitario Vila Yolanda | Instituto Adolfo Lutz, Interdiciplinary Procedures Center, Strategic Laboratory | Claudio Tavares Sacchi, Claudia Regina Gonçalves, Erica Valessa Ramos Gomes |
| EPI_ISL_534324 | Hospital Mun Ver Jose Storopolli | Instituto Adolfo Lutz, Interdiciplinary Procedures Center, Strategic Laboratory | Claudio Tavares Sacchi, Claudia Regina Gonçalves, Erica Valessa Ramos Gomes |
| EPI_ISL_534325 | Unidade de Vigilancia em Saude de Guarulhos | Instituto Adolfo Lutz, Interdiciplinary Procedures Center, Strategic Laboratory | Claudio Tavares Sacchi, Claudia Regina Gonçalves, Erica Valessa Ramos Gomes |
| EPI_ISL_534326 | Notre Dame Intermedica Saude AS | Instituto Adolfo Lutz, Interdiciplinary Procedures Center, Strategic | Claudio Tavares Sacchi, Claudia Regina Gonçalves, Erica Valessa Ramos Gomes |

| | | | |
|---|---|---|---|
| | | Laboratory | |
| EPI_ISL_534327, EPI_ISL_534328 | Hospital Universitario 12 de Octubre | Hospital Universitario 12 de Octubre | Raúl Recio, Sara González, Esther Viedma, Elias Dahdouh, Fernando Lázaro, Natalia Stella, Julio García, Juan Carlos Galán, Rafael Cantón, Ma Dolores Folgueira, Rafael Delgado, Jesús Mingorance |
| EPI_ISL_534329 | Hospital Universitario 12 de Octubre | Hospital Universitario 12 de Octubre | Esther Viedma, Raúl Recio, Sara González, Elias Dahdouh, Fernando Lázaro, Natalia Stella, Julio García, Juan Carlos Galán, Rafael Cantón, Ma Dolores Folgueira, Rafael Delgado, Jesús Mingorance |
| EPI_ISL_534330, EPI_ISL_534331, EPI_ISL_534332, EPI_ISL_534333 | Hospital Universitario La Paz | Hospital Universitario La Paz | María Rodríguez, Elias Dahdouh, Sara González, Raúl Recio, Fernando Lázaro, Esther Viedma, Natalia Stella, Julio García, Juan Carlos Galán, Rafael Cantón, Ma Dolores Folgueira, Rafael Delgado, Jesús Mingorance |
| EPI_ISL_534334 | Hospital Universitario Ramón y Cajal | Hospital Universitario La Paz | María Rodríguez, Elias Dahdouh, Sara González, Raúl Recio, Fernando Lázaro, Esther Viedma, Natalia Stella, Julio García, Juan Carlos Galán, Rafael Cantón, Ma Dolores Folgueira, Rafael Delgado, Jesús Mingorance |
| EPI_ISL_534336 | Department of Laboratory Medicine, National Taiwan University Hospital | Microbial Genomics Core Lab, National Taiwan University Centers of Genomic and Precision Medicine | Shiou-Hwei Yeh, You-Yu Lin, Ya-Yun Lai, Chiao-Ling Li, Shan-Chwen Chang, Pei-Jer Chen, Sui-Yuan Chang |
| EPI_ISL_534337, EPI_ISL_534338, EPI_ISL_534339, EPI_ISL_534340, EPI_ISL_534341, EPI_ISL_534342, EPI_ISL_534343, EPI_ISL_534344, EPI_ISL_534345 | Molecular diagnostic laboratory of Federal Budget Institution of Science "Central Research Institute of Epidemiology" of The Federal Service on Customers' Rights Protection and Human Well-being Surveillance | Group of Genomics and Postgenomic Technologies of Central Research Institute of Epidemilology | Speranskaya AS, Kaptelova VV, Samoilov AE, Valdokhina AV, Bulanenko VP, Bukharina A.Y., Tivanova EV, Shipulina OY, Akimkin VG |
| EPI_ISL_534346, EPI_ISL_534347, EPI_ISL_534348, EPI_ISL_534349 | Molecular diagnostic laboratory of Federal Budget Institution of Science "Central Research Institute of Epidemiology" of The Federal Service on Customers' Rights Protection and Human Well-being Surveillance | Group of Genomics and Postgenomic Technologies of Central Research Institute of Epidemilology | Speranskaya AS, Kaptelova VV, Valdokhina AV, Bulanenko VP, Samoilov AE, Korneenko EV, Tivanova EV, Shipulina OY, Akimkin VG |
| EPI_ISL_534365, EPI_ISL_534366, EPI_ISL_534367, EPI_ISL_534368, EPI_ISL_534369, EPI_ISL_534371, EPI_ISL_534372, EPI_ISL_534373, EPI_ISL_534374, EPI_ISL_534375, EPI_ISL_534376, EPI_ISL_534377, EPI_ISL_534378, EPI_ISL_534380, EPI_ISL_534382, EPI_ISL_534383, EPI_ISL_534384, EPI_ISL_534385, EPI_ISL_534386, EPI_ISL_534387, EPI_ISL_534389, EPI_ISL_534390, EPI_ISL_534391, EPI_ISL_534392, EPI_ISL_534393, EPI_ISL_534394, EPI_ISL_534396, EPI_ISL_534397, EPI_ISL_534398, EPI_ISL_534399, EPI_ISL_534401, EPI_ISL_534402, EPI_ISL_534404, EPI_ISL_534405, EPI_ISL_534406, EPI_ISL_534407, EPI_ISL_534408, EPI_ISL_534409, EPI_ISL_534410, EPI_ISL_534411, EPI_ISL_534412, EPI_ISL_534413, EPI_ISL_534414, EPI_ISL_534415, EPI_ISL_534418, EPI_ISL_534419, EPI_ISL_534420, EPI_ISL_534421, EPI_ISL_534422, EPI_ISL_534423, EPI_ISL_534424, EPI_ISL_534425, EPI_ISL_534426, EPI_ISL_534427, EPI_ISL_534428, EPI_ISL_534429, EPI_ISL_534430, EPI_ISL_534431, EPI_ISL_534433, EPI_ISL_534434, EPI_ISL_534435, EPI_ISL_534436, EPI_ISL_534437, EPI_ISL_534438, EPI_ISL_534439, EPI_ISL_534440, EPI_ISL_534441, EPI_ISL_534442, EPI_ISL_534443, EPI_ISL_534444, EPI_ISL_534445, EPI_ISL_534446, EPI_ISL_534447, EPI_ISL_534449, EPI_ISL_534452, EPI_ISL_534453, EPI_ISL_534455, EPI_ISL_534456, EPI_ISL_534457, EPI_ISL_534458, EPI_ISL_534459, EPI_ISL_534461, EPI_ISL_534462, EPI_ISL_534463, EPI_ISL_534465, EPI_ISL_534467, EPI_ISL_534468, EPI_ISL_534469, EPI_ISL_534470, EPI_ISL_534471, EPI_ISL_534472, EPI_ISL_534473, EPI_ISL_534474, EPI_ISL_534476, EPI_ISL_534477, EPI_ISL_534478, EPI_ISL_534480, EPI_ISL_534481, EPI_ISL_534482, EPI_ISL_534483, EPI_ISL_534484, EPI_ISL_534486, EPI_ISL_534487, EPI_ISL_534488, EPI_ISL_534489, EPI_ISL_534490, EPI_ISL_534491, EPI_ISL_534493, EPI_ISL_534495, EPI_ISL_534496, EPI_ISL_534497, EPI_ISL_534498, EPI_ISL_534499, EPI_ISL_534500, EPI_ISL_534501, EPI_ISL_534502, EPI_ISL_534503, EPI_ISL_534505, EPI_ISL_534506, EPI_ISL_534507, EPI_ISL_534508, EPI_ISL_534509, EPI_ISL_534511, EPI_ISL_534515, EPI_ISL_534516, EPI_ISL_534517, EPI_ISL_534520, EPI_ISL_534521, EPI_ISL_534522, EPI_ISL_534524, EPI_ISL_534526, EPI_ISL_534527, EPI_ISL_534528, EPI_ISL_534529, EPI_ISL_534530, EPI_ISL_534531, EPI_ISL_534534, EPI_ISL_534536, EPI_ISL_534537, EPI_ISL_534538, EPI_ISL_534539, EPI_ISL_534541, EPI_ISL_534542, EPI_ISL_534544, EPI_ISL_534545, EPI_ISL_534546, EPI_ISL_534548, EPI_ISL_534549, EPI_ISL_534551, EPI_ISL_534552, EPI_ISL_534553, EPI_ISL_534554, EPI_ISL_534555, EPI_ISL_534556, EPI_ISL_534557, EPI_ISL_534558, EPI_ISL_534559, EPI_ISL_534560, EPI_ISL_534562, EPI_ISL_534563, EPI_ISL_534564, EPI_ISL_534565, EPI_ISL_534567, EPI_ISL_534568, EPI_ISL_534570, EPI_ISL_534572, EPI_ISL_534573, EPI_ISL_534574, EPI_ISL_534575, EPI_ISL_534576, EPI_ISL_534577, EPI_ISL_534578, EPI_ISL_534579, EPI_ISL_534580, EPI_ISL_534581, EPI_ISL_534582, EPI_ISL_534583, EPI_ISL_534584, EPI_ISL_534585, EPI_ISL_534586, EPI_ISL_534587, EPI_ISL_534588, EPI_ISL_534589, EPI_ISL_534590, EPI_ISL_534591, EPI_ISL_534592, EPI_ISL_534593, EPI_ISL_534594, EPI_ISL_534595, EPI_ISL_534596, EPI_ISL_534597, EPI_ISL_534598, EPI_ISL_534599, EPI_ISL_534601, EPI_ISL_534602, EPI_ISL_534603, EPI_ISL_534605, EPI_ISL_534606, EPI_ISL_534607, EPI_ISL_534608, EPI_ISL_534609, EPI_ISL_534610, EPI_ISL_534611, EPI_ISL_534613, EPI_ISL_534614, EPI_ISL_534615, EPI_ISL_534616, EPI_ISL_534617, EPI_ISL_534619, EPI_ISL_534620, EPI_ISL_534621, EPI_ISL_534623, EPI_ISL_534624, EPI_ISL_534625, EPI_ISL_534627, EPI_ISL_534628, EPI_ISL_534629, EPI_ISL_534630, EPI_ISL_534631, EPI_ISL_534632, EPI_ISL_534633, EPI_ISL_534636, EPI_ISL_534637, EPI_ISL_534638, EPI_ISL_534639, EPI_ISL_534640, EPI_ISL_534641, EPI_ISL_534642, EPI_ISL_534643, EPI_ISL_534645, EPI_ISL_534646, EPI_ISL_534647, EPI_ISL_534648, EPI_ISL_534650, EPI_ISL_534651, EPI_ISL_534652, EPI_ISL_534653, EPI_ISL_534656, EPI_ISL_534657, EPI_ISL_534658, EPI_ISL_534659, EPI_ISL_534661, EPI_ISL_534662, EPI_ISL_534663, EPI_ISL_534664, EPI_ISL_534665, EPI_ISL_534666, EPI_ISL_534667, EPI_ISL_534668, EPI_ISL_534669, EPI_ISL_534670, EPI_ISL_534671, EPI_ISL_534672, EPI_ISL_534673, EPI_ISL_534674, EPI_ISL_534675, EPI_ISL_534676, EPI_ISL_534677, EPI_ISL_534678, EPI_ISL_534679, EPI_ISL_534680, EPI_ISL_534681, EPI_ISL_534683, EPI_ISL_534684, EPI_ISL_534685, EPI_ISL_534686, EPI_ISL_534687, EPI_ISL_534688, EPI_ISL_534690 | | | |
| see above | NHSGGC West of Scotland Specialist Virology Centre / MRC-University of Glasgow Centre for Virus Research | Wellcome Sanger Institute for the COVID-19 Genomics UK Consortium | Ana da Silva Filipe, Natasha Johnson, Kathy Smollett, Daniel Mair, Stephen Carmichael, Lily Tong, Jenna Nichols, Elihu Aranday-Cortes, Kirstyn Brunker, Yasmin Parr, Kyriaki Nomikou; Sarah McDonald, Marc Niebel, Patawee Asamaphan; Richard Orton, Joseph Hughes, Sreenu Vattipally, David L Robertson; Alasdair MacLean, Rory Gunson; Kathy Li, Natasha Jesudason, Rajiv Shah, James Shepherd, Antonia Ho, Alice Broos, Emma Thomson and Alex Alderton, Roberto Amato, Sonia Goncalves, Ewan Harrison, David K. Jackson, Ian Johnston, Dominic Kwiatkowski, Cordelia Langford, John Sillitoe on behalf of the Wellcome Sanger Institute COVID-19 Surveillance Team (http://www.sanger.ac.uk/covid-team) |
| EPI_ISL_534693, EPI_ISL_534694, EPI_ISL_534695, EPI_ISL_534696, EPI_ISL_534697, EPI_ISL_534698 | University of Miami Immunology and Histocompatibility Laboratory | University of Miami Immunology and Histocompatibility Laboratory | Emilio Margolles-Clark, PhD and Phillip Ruiz, MD, PhD |
| EPI_ISL_534699, EPI_ISL_534700, EPI_ISL_534701, EPI_ISL_534702, EPI_ISL_534703, EPI_ISL_534704, EPI_ISL_534705, EPI_ISL_534706, EPI_ISL_534708, EPI_ISL_534709, EPI_ISL_534710, EPI_ISL_534711, EPI_ISL_534713, EPI_ISL_534716 | | | |
| see above | MD PHL | MD PHL | Maryland Department of Health Laboratories Administration |
| EPI_ISL_534717, EPI_ISL_534723, EPI_ISL_534724, EPI_ISL_534725, EPI_ISL_534726, EPI_ISL_534727, EPI_ISL_534728, EPI_ISL_534729 | Respiratory Virus Unit, Microbiology Services Colindale, Public Health England | Respiratory Virus Unit, Microbiology Services Colindale, Public Health England | PHE Covid Sequencing Team |
| EPI_ISL_534731 | Wadsworth Center, New York State Department of Health | Wadsworth Center, New York State Department of Health | Kirsten St. George, Daryl M. Lamson, Sara Griesemer, Jonathan Plitnick, Navjot Singh, Matthew D. Shudt, Erica Lasek-Nesselquist |
| EPI_ISL_534732, EPI_ISL_534748, EPI_ISL_534749, EPI_ISL_534750, EPI_ISL_534751, EPI_ISL_534752, EPI_ISL_534753, EPI_ISL_534754, EPI_ISL_534755 | Liverpool Clinical Laboratories | COVID-19 Genomics UK (COG-UK) Consortium | Sam Haldenby, Anita Lucaci, Steve Paterson, Julian Hiscox, Alistair Darby, M Almsaud, A Alrezaihi, Muhannad Alruwaili, Stuart D Armstrong, Jones Benjamin, Eleanor G Bentley, Anu Chawla, Jordan J Clark, Angela Cowell, Richard Eccles, Isabel García-Dorival, Matthew Gemmell, Alessandro Gerada, PKF Gilmore, Richard Gregory, Ximeng Han, Catherine Hartley, Margaret Hughes, Miren Iturriza-Gomara, James Johnson, L Luu, Jenifer Manson, Charlotte Nelson, Elaine O'Toole, Cassie Olateju, Rebekah Penrice-Randal , Lucille Rainbow, N.P Randle, Trevor Ian Robinson, Parul Sharma, Ghada T Shawli, James P Stewart, Neil Swainston, Ecaterina Vamos, Joanne Watts, Mark Whitehead |
| EPI_ISL_534763, EPI_ISL_534764, EPI_ISL_534765, EPI_ISL_534766, EPI_ISL_534767, EPI_ISL_534768, EPI_ISL_534769, EPI_ISL_534770, EPI_ISL_534771, EPI_ISL_534772, EPI_ISL_534774, EPI_ISL_534775, EPI_ISL_534776, EPI_ISL_534777, EPI_ISL_534778, EPI_ISL_534779, EPI_ISL_534780, EPI_ISL_534781, EPI_ISL_534782, EPI_ISL_534783, EPI_ISL_534784, EPI_ISL_534785, EPI_ISL_534786, EPI_ISL_534787, EPI_ISL_534789, EPI_ISL_534790, EPI_ISL_534791, EPI_ISL_534792, EPI_ISL_534793, EPI_ISL_534795, EPI_ISL_534796, EPI_ISL_534797, EPI_ISL_534798, EPI_ISL_534799, EPI_ISL_534800, EPI_ISL_534801, EPI_ISL_534802, EPI_ISL_534803, EPI_ISL_534804, EPI_ISL_534805, EPI_ISL_534806, EPI_ISL_534807, EPI_ISL_534808, EPI_ISL_534809, EPI_ISL_534810, EPI_ISL_534811, EPI_ISL_534813, EPI_ISL_534814, EPI_ISL_534815, EPI_ISL_534816, EPI_ISL_534817, EPI_ISL_534819, EPI_ISL_534820, EPI_ISL_534821, EPI_ISL_534822, EPI_ISL_534823, EPI_ISL_534825, EPI_ISL_534826, EPI_ISL_534827, EPI_ISL_534828, EPI_ISL_534829, EPI_ISL_534830, EPI_ISL_534831, EPI_ISL_534833, EPI_ISL_534834, EPI_ISL_534835, EPI_ISL_534836, EPI_ISL_534837, EPI_ISL_534838, EPI_ISL_534839, EPI_ISL_534840, EPI_ISL_534841, EPI_ISL_534842, EPI_ISL_534843, EPI_ISL_534844, EPI_ISL_534845, EPI_ISL_534846, EPI_ISL_534847, EPI_ISL_534848, EPI_ISL_534849, EPI_ISL_534850, EPI_ISL_534851, EPI_ISL_534852, EPI_ISL_534853, EPI_ISL_534854, EPI_ISL_534855, EPI_ISL_534856, EPI_ISL_534857, EPI_ISL_534858, EPI_ISL_534859, EPI_ISL_534860, EPI_ISL_534861, EPI_ISL_534862, EPI_ISL_534863, EPI_ISL_534865, EPI_ISL_534866, EPI_ISL_534867, EPI_ISL_534868, EPI_ISL_534870, EPI_ISL_534871, EPI_ISL_534872, EPI_ISL_534873, EPI_ISL_534874, EPI_ISL_534875, EPI_ISL_534876, EPI_ISL_534878, EPI_ISL_534879, EPI_ISL_534880, EPI_ISL_534881, EPI_ISL_534882, EPI_ISL_534883, EPI_ISL_534884, EPI_ISL_534885, EPI_ISL_534886, EPI_ISL_534887, EPI_ISL_534888, EPI_ISL_534889, EPI_ISL_534890, EPI_ISL_534891, EPI_ISL_534892, EPI_ISL_534893, EPI_ISL_534894, EPI_ISL_534895, EPI_ISL_534896, EPI_ISL_534897, EPI_ISL_534898, EPI_ISL_534899, EPI_ISL_534900, EPI_ISL_534901, EPI_ISL_534902, EPI_ISL_534903, EPI_ISL_534904, EPI_ISL_534905, EPI_ISL_534906, EPI_ISL_534907, EPI_ISL_534909, EPI_ISL_534910, EPI_ISL_534911, EPI_ISL_534912, EPI_ISL_534913, EPI_ISL_534914, EPI_ISL_534915, EPI_ISL_534917, EPI_ISL_534918, EPI_ISL_534920, EPI_ISL_534921, EPI_ISL_534922, EPI_ISL_534924, EPI_ISL_534925, EPI_ISL_534926, EPI_ISL_534927, EPI_ISL_534928, EPI_ISL_534929, EPI_ISL_534930, EPI_ISL_534931, EPI_ISL_534932, EPI_ISL_534933, EPI_ISL_534934, EPI_ISL_534935, EPI_ISL_534938, EPI_ISL_534939, EPI_ISL_534941, EPI_ISL_534942, EPI_ISL_534943, EPI_ISL_534944, EPI_ISL_534945, EPI_ISL_534947, EPI_ISL_534948, EPI_ISL_534950, EPI_ISL_534952, EPI_ISL_534953, EPI_ISL_534954, EPI_ISL_534955, EPI_ISL_534958, EPI_ISL_534959, EPI_ISL_534960, EPI_ISL_534961, EPI_ISL_534963, EPI_ISL_534964, EPI_ISL_534966, EPI_ISL_534967, EPI_ISL_534969, EPI_ISL_534970, EPI_ISL_534971, EPI_ISL_534972, EPI_ISL_534973, EPI_ISL_534975, EPI_ISL_534976, EPI_ISL_534978, EPI_ISL_534980, EPI_ISL_534981, EPI_ISL_534982, EPI_ISL_534983, EPI_ISL_534984, EPI_ISL_534985, EPI_ISL_534986, EPI_ISL_534987, EPI_ISL_534988, EPI_ISL_534989, EPI_ISL_534990, EPI_ISL_534993, EPI_ISL_534994, EPI_ISL_534995, EPI_ISL_534996, EPI_ISL_534997, EPI_ISL_534998, EPI_ISL_535000, EPI_ISL_535001, EPI_ISL_535002, EPI_ISL_535003, EPI_ISL_535004, EPI_ISL_535005, EPI_ISL_535006, EPI_ISL_535007, EPI_ISL_535008, EPI_ISL_535009, EPI_ISL_535010, EPI_ISL_535011, EPI_ISL_535012, EPI_ISL_535013, EPI_ISL_535014, EPI_ISL_535015, EPI_ISL_535017, EPI_ISL_535018, EPI_ISL_535019, EPI_ISL_535020, EPI_ISL_535021, EPI_ISL_535022, EPI_ISL_535023 | | | |
| see above | Oxford Viromics, NDM, University of Oxford; Oxford University Hospitals; Basingstoke and North Hampshire Hospital | COVID-19 Genomics UK (COG-UK) Consortium | Tanya Golubchik, David Bonsall, George Macintyre, Amy Trebes, Mariateresa de Cesare, Catrin Moore, Alex Mobbs, Anita Justice, Robert Shaw, Monique Andersson, Timothy Peto, Emma Wise, Nathan Moore, Jessica Lynch, Nick Cortes, Matilde Mori, Stephen Kidd, David Buck, John Todd, Christophe Fraser |
| EPI_ISL_535024, EPI_ISL_535025, EPI_ISL_535027, EPI_ISL_535028, EPI_ISL_535029, EPI_ISL_535031, EPI_ISL_535032, EPI_ISL_535036, EPI_ISL_535038 | Centre for Enzyme Innovation, University of Portsmouth / Translational Research Laboratory, Portsmouth Hospitals NHS Trust | COVID-19 Genomics UK (COG-UK) Consortium | Angela Beckett,Yann Bourgeois,Garry Scarlett,Sharon Glaysher,Scott Elliott,Kelly Bicknell,Robert Impey,Allyson Lloyd,Sarah Wyllie,Ethan Butcher,Anoop Chauhan,Samuel Robson |
| EPI_ISL_535043, EPI_ISL_535044, EPI_ISL_535047, EPI_ISL_535051, EPI_ISL_535052, EPI_ISL_535056, EPI_ISL_535060, EPI_ISL_535061, EPI_ISL_535062, EPI_ISL_535063, EPI_ISL_535064, EPI_ISL_535066, EPI_ISL_535069, EPI_ISL_535071, EPI_ISL_535073, EPI_ISL_535075, EPI_ISL_535077, EPI_ISL_535079, EPI_ISL_535080, EPI_ISL_535081, EPI_ISL_535082, EPI_ISL_535084, EPI_ISL_535085, EPI_ISL_535087, EPI_ISL_535088, EPI_ISL_535089, EPI_ISL_535091, EPI_ISL_535095, EPI_ISL_535096, EPI_ISL_535097, EPI_ISL_535099, EPI_ISL_535100 | | | |
| see above | Virology Department, Sheffield Teaching Hospitals NHS Foundation Trust/Department of Infection, Immunity and Cardiovascular Disease, The Medical School, University of Sheffield | COVID-19 Genomics UK (COG-UK) Consortium | Thushan de Silva, Matthew Parker, Nikki Smith, Adri Angyal, Rebecca Brown, Luke Green, Rachel Tucker, Paul Parsons, Danielle Groves, Katie Johnson, Laura Carrilero, Alex Keeley, Dave Partridge, Matthew Wyles, Benjamin Lindsey, Mehmet Yavuz, Mohammad Raza, Cariad Evans |
| EPI_ISL_535102, EPI_ISL_535103, EPI_ISL_535105, EPI_ISL_535106, EPI_ISL_535107, EPI_ISL_535108, EPI_ISL_535110, EPI_ISL_535111, EPI_ISL_535112, EPI_ISL_535113, EPI_ISL_535114, EPI_ISL_535116, EPI_ISL_535117, EPI_ISL_535118, EPI_ISL_535119, EPI_ISL_535122, EPI_ISL_535124, EPI_ISL_535127, EPI_ISL_535128, EPI_ISL_535129, EPI_ISL_535130, EPI_ISL_535132, EPI_ISL_535134, EPI_ISL_535135, EPI_ISL_535136, EPI_ISL_535137, EPI_ISL_535138, EPI_ISL_535141, EPI_ISL_535144, EPI_ISL_535145, EPI_ISL_535146, EPI_ISL_535147, EPI_ISL_535148, EPI_ISL_535149, EPI_ISL_535150, EPI_ISL_535151, EPI_ISL_535154, EPI_ISL_535156, EPI_ISL_535157, EPI_ISL_535159, EPI_ISL_535160, EPI_ISL_535162, EPI_ISL_535164, EPI_ISL_535166, EPI_ISL_535168, EPI_ISL_535169, EPI_ISL_535171, EPI_ISL_535172, EPI_ISL_535173, EPI_ISL_535175, EPI_ISL_535176, EPI_ISL_535177, EPI_ISL_535178, EPI_ISL_535180, EPI_ISL_535182 | | | |
| see above | West of Scotland Specialist Virology Centre, NHSGGC / MRC-University of Glasgow Centre for Virus Research | COVID-19 Genomics UK (COG-UK) Consortium | Ana da Silva Filipe, Natasha Johnson, Kathy Smollett, Daniel Mair, Stephen Carmichael, Lily Tong, Jenna Nichols, Elihu Aranday-Cortes, Yasmin Parr, Alice Broos, Kyriaki Nomikou; Sarah McDonald, Marc Niebel, Patawee Asamaphan; Richard Orton, Joseph Hughes, Sreenu Vattipally, David L Robertson; Alasdair MacLean, Rory Gunson; Kathy Li, Natasha Jesudason, Rajiv Shah, James Shepherd, Antonia Ho, Emma Thomson |
| EPI_ISL_535184, EPI_ISL_535185, EPI_ISL_535186, EPI_ISL_535188, EPI_ISL_535189, EPI_ISL_535190, EPI_ISL_535191, EPI_ISL_535192, EPI_ISL_535193, EPI_ISL_535194, EPI_ISL_535195, EPI_ISL_535196, EPI_ISL_535197, EPI_ISL_535198, EPI_ISL_535199, EPI_ISL_535201, EPI_ISL_535203, EPI_ISL_535204, EPI_ISL_535206, EPI_ISL_535207, EPI_ISL_535208, EPI_ISL_535209, EPI_ISL_535210, EPI_ISL_535212, EPI_ISL_535213, EPI_ISL_535214, EPI_ISL_535215, EPI_ISL_535217, EPI_ISL_535218, EPI_ISL_535219, EPI_ISL_535220, EPI_ISL_535225, EPI_ISL_535226, EPI_ISL_535227, EPI_ISL_535228, EPI_ISL_535230, EPI_ISL_535233, EPI_ISL_535234, EPI_ISL_535236, EPI_ISL_535238, EPI_ISL_535239, EPI_ISL_535242, EPI_ISL_535245, EPI_ISL_535246, EPI_ISL_535247, EPI_ISL_535248, EPI_ISL_535249, EPI_ISL_535250, EPI_ISL_535251, EPI_ISL_535254, EPI_ISL_535257, EPI_ISL_535260, EPI_ISL_535261 | | | |
| see above | Wales Specialist Virology Centre Sequencing lab: Pathogen Genomics Unit | COVID-19 Genomics UK (COG-UK) Consortium | Catherine Moore, Johnathan Evans, Laura Gifford, Malorie Perry, Simon Cottrell, Angela Marchbank, Alec Birchley, Alexander Adams, Amy Gaskin, Bree Gatica-Wilcox, Jason Coombes, Joel Southgate, Lauren Gilbert, Lee Graham, Nicole Pacchiarini, Sara Kumziene-Summerhayes, Sarah Taylor, Sophie Jones, Sara Rey, Matthew Bull, Joanne Watkins, Sally Corden, Tom Connor |
| EPI_ISL_535264, EPI_ISL_535265, EPI_ISL_535266, EPI_ISL_535268, EPI_ISL_535269, EPI_ISL_535270, EPI_ISL_535271, EPI_ISL_535272, EPI_ISL_535273, EPI_ISL_535274, EPI_ISL_535276, EPI_ISL_535278, EPI_ISL_535279, EPI_ISL_535280, EPI_ISL_535281, EPI_ISL_535282, EPI_ISL_535283, EPI_ISL_535284, EPI_ISL_535285, EPI_ISL_535286, EPI_ISL_535287, EPI_ISL_535288, EPI_ISL_535291, EPI_ISL_535292, EPI_ISL_535293, EPI_ISL_535295, EPI_ISL_535297, EPI_ISL_535298, EPI_ISL_535299, EPI_ISL_535300, EPI_ISL_535301, EPI_ISL_535302, EPI_ISL_535303 | | | |
| see above | New Mexico Department of Health Scientific Laboratory | New Mexico Department of Health Scientific Laboratory | Ellie Johnson, Anastacia Griego-Fisher, D'Eldra Malone |
| EPI_ISL_535305, EPI_ISL_535306, EPI_ISL_535307, EPI_ISL_535308, EPI_ISL_535309, EPI_ISL_535310, EPI_ISL_535311, EPI_ISL_535312, EPI_ISL_535313, EPI_ISL_535314, EPI_ISL_535315, EPI_ISL_535316, EPI_ISL_535317, EPI_ISL_535318, EPI_ISL_535319, EPI_ISL_535320, EPI_ISL_535321, EPI_ISL_535322, EPI_ISL_535323, EPI_ISL_535324, EPI_ISL_535325, EPI_ISL_535326, EPI_ISL_535327, EPI_ISL_535328, EPI_ISL_535329, EPI_ISL_535330, EPI_ISL_535331, EPI_ISL_535332, EPI_ISL_535333, EPI_ISL_535334, EPI_ISL_535335, EPI_ISL_535336, EPI_ISL_535337, EPI_ISL_535338, EPI_ISL_535339, EPI_ISL_535340 | | | |
| see above | LA Office of Public Health Laboratories | Pathogen Discovery, Respiratory Viruses Branch, Division of Viral Diseases, Centers for Disease Control and Prevention | Ying Tao, Jing Zhang, Yan Li, Krista Queen, Anna Uehara, Clinton Paden, Haibin Wang, Suxiang Tong |
| EPI_ISL_535341, EPI_ISL_535342 | LA Office of Public Health Laboratories | Pathogen Discovery, Respiratory Viruses Branch, Division of Viral Diseases, Centers for Disease Control and Prevention | Jing Zhang, Ying Tao, Yan Li, Krista Queen, Anna Uehara, Clinton Paden, Haibin Wang, Suxiang Tong |
| EPI_ISL_535343 | LA Office of Public Health Laboratories | Pathogen Discovery, Respiratory Viruses Branch, Division of Viral Diseases, Centers for Disease Control and Prevention | Ying Tao, Jing Zhang, Yan Li, Krista Queen, Anna Uehara, Clinton Paden, Haibin Wang, Suxiang Tong |
| EPI_ISL_535344, EPI_ISL_535345, EPI_ISL_535346 | LA Office of Public Health Laboratories | Pathogen Discovery, Respiratory Viruses Branch, Division of Viral Diseases, Centers for Disease Control | Jing Zhang, Ying Tao, Yan Li, Krista Queen, Anna Uehara, Clinton Paden, Haibin Wang, Suxiang Tong |

| | | | |
|---|---|---|---|
| | | and Prevention | |
| EPI_ISL_535347 | LA Office of Public Health Laboratories | Pathogen Discovery, Respiratory Viruses Branch, Division of Viral Diseases, Centers for Disease Control and Prevention | Ying Tao, Jing Zhang, Yan Li, Krista Queen, Anna Uehara, Clinton Paden, Haibin Wang, Suxiang Tong |
| EPI_ISL_535348 | LA Office of Public Health Laboratories | Pathogen Discovery, Respiratory Viruses Branch, Division of Viral Diseases, Centers for Disease Control and Prevention | Jing Zhang, Ying Tao, Yan Li, Krista Queen, Anna Uehara, Clinton Paden, Haibin Wang, Suxiang Tong |
| EPI_ISL_535349, EPI_ISL_535350 | LA Office of Public Health Laboratories | Pathogen Discovery, Respiratory Viruses Branch, Division of Viral Diseases, Centers for Disease Control and Prevention | Ying Tao, Jing Zhang, Yan Li, Krista Queen, Anna Uehara, Clinton Paden, Haibin Wang, Suxiang Tong |
| EPI_ISL_535351 | LA Office of Public Health Laboratories | Pathogen Discovery, Respiratory Viruses Branch, Division of Viral Diseases, Centers for Disease Control and Prevention | Jing Zhang, Ying Tao, Yan Li, Krista Queen, Anna Uehara, Clinton Paden, Haibin Wang, Suxiang Tong |
| EPI_ISL_535352, EPI_ISL_535353, EPI_ISL_535354, EPI_ISL_535355, EPI_ISL_535356 | LA Office of Public Health Laboratories | Pathogen Discovery, Respiratory Viruses Branch, Division of Viral Diseases, Centers for Disease Control and Prevention | Ying Tao, Jing Zhang, Yan Li, Krista Queen, Anna Uehara, Clinton Paden, Haibin Wang, Suxiang Tong |
| EPI_ISL_535357 | LA Office of Public Health Laboratories | Pathogen Discovery, Respiratory Viruses Branch, Division of Viral Diseases, Centers for Disease Control and Prevention | Jing Zhang, Ying Tao, Yan Li, Krista Queen, Anna Uehara, Clinton Paden, Haibin Wang, Suxiang Tong |
| EPI_ISL_535358 | LA Office of Public Health Laboratories | Pathogen Discovery, Respiratory Viruses Branch, Division of Viral Diseases, Centers for Disease Control and Prevention | Ying Tao, Jing Zhang, Yan Li, Krista Queen, Anna Uehara, Clinton Paden, Haibin Wang, Suxiang Tong |
| EPI_ISL_535359, EPI_ISL_535360 | RI State Health Laboratories | Pathogen Discovery, Respiratory Viruses Branch, Division of Viral Diseases, Centers for Disease Control and Prevention | Jing Zhang, Ying Tao, Yan Li, Krista Queen, Anna Uehara, Clinton Paden, Haibin Wang, Suxiang Tong |
| EPI_ISL_535361, EPI_ISL_535362, EPI_ISL_535363, EPI_ISL_535364 | Oklahoma Animal Disease Diagnostic Laboratory | Oklahoma Animal Disease Diagnostic Laboratory | Sai Narayanan, John C Ritchey, Girish Patil, Teluguakula Narasaraju, Sunil More, Jerry Malayer, Jeremiah Saliki, Anil Kaul, Akhilesh Ramachandran |
| EPI_ISL_535390, EPI_ISL_535392, EPI_ISL_535393, EPI_ISL_535394, EPI_ISL_535395, EPI_ISL_535396, EPI_ISL_535397, EPI_ISL_535398, EPI_ISL_535399, EPI_ISL_535400, EPI_ISL_535403, EPI_ISL_535404, EPI_ISL_535405, EPI_ISL_535406, EPI_ISL_535408, EPI_ISL_535410, EPI_ISL_535412, EPI_ISL_535413, EPI_ISL_535414, EPI_ISL_535415, EPI_ISL_535416, EPI_ISL_535417, EPI_ISL_535418, EPI_ISL_535420, EPI_ISL_535421, EPI_ISL_535422, EPI_ISL_535424, EPI_ISL_535425, EPI_ISL_535426, EPI_ISL_535427, EPI_ISL_535428, EPI_ISL_535430, EPI_ISL_535431, EPI_ISL_535432, EPI_ISL_535433, EPI_ISL_535434, EPI_ISL_535435, EPI_ISL_535436, EPI_ISL_535438, EPI_ISL_535439, EPI_ISL_535440, EPI_ISL_535441, EPI_ISL_535442, EPI_ISL_535443, EPI_ISL_535444, EPI_ISL_535445, EPI_ISL_535446, EPI_ISL_535447, EPI_ISL_535448, EPI_ISL_535449, EPI_ISL_535450, EPI_ISL_535451, EPI_ISL_535452, EPI_ISL_535454, EPI_ISL_535455, EPI_ISL_535456, EPI_ISL_535457, EPI_ISL_535458, EPI_ISL_535459, EPI_ISL_535460, EPI_ISL_535461, EPI_ISL_535462, EPI_ISL_535463, EPI_ISL_535464, EPI_ISL_535465, EPI_ISL_535466, EPI_ISL_535467, EPI_ISL_535468, EPI_ISL_535469, EPI_ISL_535470, EPI_ISL_535471, EPI_ISL_535472, EPI_ISL_535473, EPI_ISL_535475, EPI_ISL_535476, EPI_ISL_535477, EPI_ISL_535478, EPI_ISL_535479, EPI_ISL_535480, EPI_ISL_535481, EPI_ISL_535483, EPI_ISL_535484, EPI_ISL_535485, EPI_ISL_535487, EPI_ISL_535488, EPI_ISL_535496, EPI_ISL_535497, EPI_ISL_535498, EPI_ISL_535499, EPI_ISL_535500, EPI_ISL_535501, EPI_ISL_535502, EPI_ISL_535503, EPI_ISL_535504, EPI_ISL_535505, EPI_ISL_535506, EPI_ISL_535507, EPI_ISL_535508, EPI_ISL_535509, EPI_ISL_535510, EPI_ISL_535511, EPI_ISL_535512, EPI_ISL_535513, EPI_ISL_535514, EPI_ISL_535515, EPI_ISL_535516, EPI_ISL_535518, EPI_ISL_535519, EPI_ISL_535520, EPI_ISL_535522, EPI_ISL_535523, EPI_ISL_535526, EPI_ISL_535527, EPI_ISL_535529, EPI_ISL_535530, EPI_ISL_535531, EPI_ISL_535532, EPI_ISL_535533, EPI_ISL_535534, EPI_ISL_535536, EPI_ISL_535537, EPI_ISL_535539, EPI_ISL_535540, EPI_ISL_535541, EPI_ISL_535542, EPI_ISL_535543, EPI_ISL_535544, EPI_ISL_535545, EPI_ISL_535546, EPI_ISL_535547, EPI_ISL_535551, EPI_ISL_535552, EPI_ISL_535553, EPI_ISL_535554, EPI_ISL_535555, EPI_ISL_535556, EPI_ISL_535557, EPI_ISL_535558, EPI_ISL_535559, EPI_ISL_535560, EPI_ISL_535561, EPI_ISL_535562, EPI_ISL_535563, EPI_ISL_535564, EPI_ISL_535565, EPI_ISL_535566, EPI_ISL_535567, EPI_ISL_535568, EPI_ISL_535569, EPI_ISL_535570, EPI_ISL_535571, EPI_ISL_535572 | | | |
| see above | NHLS-IALCH | KRISP, KZN Research Innovation and Sequencing Platform | Giandhari J, Pillay S, Lessells R, Mdlalose K, York D, Khan S, Tegally H, Wilkinson E, de Oliveira T |
| EPI_ISL_535575 | Hospital Universitari Germans Trias i Pujol(HUGTiP)/Fundació Lluita contra la SIDA (FLSida) | IrsiCaixa AIDS Research Lab | Marc Noguera-Julian, Mariona Parera, Maria Pilar Armengol, Marta Massanella, Ester Ballana, Lidia Ruiz, Nuria Izquierdo, Jorge Carrillo, Roger Paredes, Julia Blanco, Joaquim Segalés, Bonaventura Clotet |
| EPI_ISL_535585, EPI_ISL_535586, EPI_ISL_535587, EPI_ISL_535588, EPI_ISL_535589, EPI_ISL_535590, EPI_ISL_535593, EPI_ISL_535595, EPI_ISL_535596, EPI_ISL_535597, EPI_ISL_535598, EPI_ISL_535599, EPI_ISL_535600, EPI_ISL_535602, EPI_ISL_535604, EPI_ISL_535606, EPI_ISL_535607, EPI_ISL_535608, EPI_ISL_535610, EPI_ISL_535612, EPI_ISL_535613, EPI_ISL_535614, EPI_ISL_535615, EPI_ISL_535618, EPI_ISL_535621, EPI_ISL_535623, EPI_ISL_535624, EPI_ISL_535625, EPI_ISL_535627, EPI_ISL_535629, EPI_ISL_535631, EPI_ISL_535632, EPI_ISL_535633, EPI_ISL_535634, EPI_ISL_535640, EPI_ISL_535641, EPI_ISL_535643, EPI_ISL_535644, EPI_ISL_535645, EPI_ISL_535646, EPI_ISL_535647 | | | |
| see above | Viollier AG | Department of Biosystems Science and Engineering, ETH Zürich | Christian Beisel, Sarah Nadeau, Ivan Topolsky, Pedro Ferreira, Philipp Jablonski, Susana Posada-Céspedes, Tobias Schär, Ina Nissen, Natascha Santacroce, Elodie Burcklen, Christiane Beckmann, Maurice Redondo, Olivier Kobel, Christoph Noppen, Sophie Seidel, Noemie Santamaria de Souza, Niko Beerenwinkel, Tanja Stadler |
| EPI_ISL_535650 | AR Dept. of Health-Public Health Lab | Pathogen Discovery, Respiratory Viruses Branch, Division of Viral Diseases, Centers for Disease Control and Prevention | Brian Lynch, Yan Li, Jing Zhang, Ying Tao, Krista Queen, Anna Uehara, Clinton R. Paden, Rachel Marine, Haibin Wang, Suxiang Tong |
| EPI_ISL_535651 | AR Dept. of Health-Public Health Lab | Pathogen Discovery, Respiratory Viruses Branch, Division of Viral Diseases, Centers for Disease Control and Prevention | Yan Li, Jing Zhang, Ying Tao, Krista Queen, Brian Lynch, Anna Uehara, Clinton R. Paden, Rachel Marine, Haibin Wang, Suxiang Tong |
| EPI_ISL_535652 | AR Dept. of Health-Public Health Lab | Pathogen Discovery, Respiratory Viruses Branch, Division of Viral Diseases, Centers for Disease Control and Prevention | Brian Lynch, Yan Li, Jing Zhang, Ying Tao, Krista Queen, Anna Uehara, Clinton R. Paden, Rachel Marine, Haibin Wang, Suxiang Tong |
| EPI_ISL_535653 | AR Dept. of Health-Public Health Lab | Pathogen Discovery, Respiratory Viruses Branch, Division of Viral Diseases, Centers for Disease Control and Prevention | Yan Li, Jing Zhang, Ying Tao, Krista Queen, Brian Lynch, Anna Uehara, Clinton R. Paden, Rachel Marine, Haibin Wang, Suxiang Tong |
| EPI_ISL_535654 | AR Dept. of Health-Public Health Lab | Pathogen Discovery, Respiratory Viruses Branch, Division of Viral Diseases, Centers for Disease Control and Prevention | Brian Lynch, Yan Li, Jing Zhang, Ying Tao, Krista Queen, Anna Uehara, Clinton R. Paden, Rachel Marine, Haibin Wang, Suxiang Tong |
| EPI_ISL_535655, EPI_ISL_535656, EPI_ISL_535657 | AR Dept. of Health-Public Health Lab | Pathogen Discovery, Respiratory Viruses Branch, Division of Viral Diseases, Centers for Disease Control and Prevention | Yan Li, Jing Zhang, Ying Tao, Krista Queen, Brian Lynch, Anna Uehara, Clinton R. Paden, Rachel Marine, Haibin Wang, Suxiang Tong |
| EPI_ISL_535658, EPI_ISL_535659 | AR Dept. of Health-Public Health Lab | Pathogen Discovery, Respiratory Viruses Branch, Division of Viral Diseases, Centers for Disease Control and Prevention | Brian Lynch, Yan Li, Jing Zhang, Ying Tao, Krista Queen, Anna Uehara, Clinton R. Paden, Rachel Marine, Haibin Wang, Suxiang Tong |
| EPI_ISL_535660 | AR Dept. of Health-Public Health Lab | Pathogen Discovery, Respiratory Viruses Branch, Division of Viral Diseases, Centers for Disease Control and Prevention | Yan Li, Jing Zhang, Ying Tao, Krista Queen, Brian Lynch, Anna Uehara, Clinton R. Paden, Rachel Marine, Haibin Wang, Suxiang Tong |
| EPI_ISL_535661 | AR Dept. of Health-Public Health Lab | Pathogen Discovery, Respiratory Viruses Branch, Division of Viral Diseases, Centers for Disease Control and Prevention | Brian Lynch, Yan Li, Jing Zhang, Ying Tao, Krista Queen, Anna Uehara, Clinton R. Paden, Rachel Marine, Haibin Wang, Suxiang Tong |
| EPI_ISL_535662, EPI_ISL_535663, EPI_ISL_535664, EPI_ISL_535665, EPI_ISL_535666, EPI_ISL_535667, EPI_ISL_535668, EPI_ISL_535669, EPI_ISL_535670, EPI_ISL_535671, EPI_ISL_535672, EPI_ISL_535673, EPI_ISL_535674 | | | |
| see above | CDPH, Microbial Diseases Laboratory | Pathogen Discovery, Respiratory Viruses Branch, Division of Viral Diseases, Centers for Disease Control and Prevention | Yan Li, Jing Zhang, Ying Tao, Krista Queen, Brian Lynch, Anna Uehara, Clinton R. Paden, Rachel Marine, Haibin Wang, Suxiang Tong |
| EPI_ISL_535675 | CDPH, Microbial Diseases Laboratory | Pathogen Discovery, Respiratory Viruses Branch, Division of Viral Diseases, Centers for Disease Control and Prevention | Brian Lynch, Yan Li, Jing Zhang, Ying Tao, Krista Queen, Anna Uehara, Clinton R. Paden, Rachel Marine, Haibin Wang, Suxiang Tong |
| EPI_ISL_535676, EPI_ISL_535677, EPI_ISL_535678, EPI_ISL_535679, EPI_ISL_535680, EPI_ISL_535681, EPI_ISL_535682, EPI_ISL_535683, EPI_ISL_535684, EPI_ISL_535685, EPI_ISL_535686, EPI_ISL_535687, EPI_ISL_535688, EPI_ISL_535689, EPI_ISL_535690, EPI_ISL_535691, EPI_ISL_535692, EPI_ISL_535693, EPI_ISL_535694, EPI_ISL_535695, EPI_ISL_535696, EPI_ISL_535697, EPI_ISL_535698, EPI_ISL_535699, EPI_ISL_535700 | | | |
| see above | CDPH, Microbial Diseases Laboratory | Pathogen Discovery, Respiratory | Yan Li, Jing Zhang, Ying Tao, Krista Queen, Brian Lynch, Anna Uehara, Clinton R. Paden, Rachel Marine, Haibin Wang, Suxiang Tong |

| | | | |
|---|---|---|---|
| | | Viruses Branch, Division of Viral Diseases, Centers for Disease Control and Prevention | |
| EPI_ISL_535701 | CDPH, Microbial Diseases Laboratory | Pathogen Discovery, Respiratory Viruses Branch, Division of Viral Diseases, Centers for Disease Control and Prevention | Brian Lynch, Yan Li, Jing Zhang, Ying Tao, Krista Queen, Anna Uehara, Clinton R. Paden, Rachel Marine, Haibin Wang, Suxiang Tong |
| EPI_ISL_535702 | CDPH, Microbial Diseases Laboratory | Pathogen Discovery, Respiratory Viruses Branch, Division of Viral Diseases, Centers for Disease Control and Prevention | Yan Li, Jing Zhang, Ying Tao, Krista Queen, Brian Lynch, Anna Uehara, Clinton R. Paden, Rachel Marine, Haibin Wang, Suxiang Tong |
| EPI_ISL_535703 | CDPH, Microbial Diseases Laboratory | Pathogen Discovery, Respiratory Viruses Branch, Division of Viral Diseases, Centers for Disease Control and Prevention | Ying Tao, Jing Zhang, Yan Li, Krista Queen, Anna Uehara, Clinton R. Paden, Haibin Wang, Suxiang Tong |
| EPI_ISL_535704, EPI_ISL_535705 | CDPH, Microbial Diseases Laboratory | Pathogen Discovery, Respiratory Viruses Branch, Division of Viral Diseases, Centers for Disease Control and Prevention | Yan Li, Jing Zhang, Ying Tao, Krista Queen, Brian Lynch, Anna Uehara, Clinton R. Paden, Rachel Marine, Haibin Wang, Suxiang Tong |
| EPI_ISL_535706 | CDPH, Microbial Diseases Laboratory | Pathogen Discovery, Respiratory Viruses Branch, Division of Viral Diseases, Centers for Disease Control and Prevention | Ying Tao, Jing Zhang, Yan Li, Krista Queen, Anna Uehara, Clinton R. Paden, Haibin Wang, Suxiang Tong |
| EPI_ISL_535707 | CDPH, Microbial Diseases Laboratory | Pathogen Discovery, Respiratory Viruses Branch, Division of Viral Diseases, Centers for Disease Control and Prevention | Yan Li, Jing Zhang, Ying Tao, Krista Queen, Brian Lynch, Anna Uehara, Clinton R. Paden, Rachel Marine, Haibin Wang, Suxiang Tong |
| EPI_ISL_535708 | CDPH, Microbial Diseases Laboratory | Pathogen Discovery, Respiratory Viruses Branch, Division of Viral Diseases, Centers for Disease Control and Prevention | Brian Lynch, Yan Li, Jing Zhang, Ying Tao, Krista Queen, Anna Uehara, Clinton R. Paden, Rachel Marine, Haibin Wang, Suxiang Tong |
| EPI_ISL_535709, EPI_ISL_535710, EPI_ISL_535711, EPI_ISL_535712, EPI_ISL_535713, EPI_ISL_535714, EPI_ISL_535715 | CDPH, Microbial Diseases Laboratory | Pathogen Discovery, Respiratory Viruses Branch, Division of Viral Diseases, Centers for Disease Control and Prevention | Yan Li, Jing Zhang, Ying Tao, Krista Queen, Brian Lynch, Anna Uehara, Clinton R. Paden, Rachel Marine, Haibin Wang, Suxiang Tong |
| EPI_ISL_535716 | Hôpital de Verdun | Laboratoire de santé publique du Québec | Sandrine Moreira, Ioannis Ragoussis, Guillaume Bourque, Jesse Shapiro, Mark Lathrop and Michel Roger |
| EPI_ISL_535717 | Hôpital Honoré-Mercier | Laboratoire de santé publique du Québec | Sandrine Moreira, Ioannis Ragoussis, Guillaume Bourque, Jesse Shapiro, Mark Lathrop and Michel Roger |
| EPI_ISL_535718 | Hôpital Pierre-Boucher | Laboratoire de santé publique du Québec | Sandrine Moreira, Ioannis Ragoussis, Guillaume Bourque, Jesse Shapiro, Mark Lathrop and Michel Roger |
| EPI_ISL_535719, EPI_ISL_535720 | Hôpital général Juif | Laboratoire de santé publique du Québec | Sandrine Moreira, Ioannis Ragoussis, Guillaume Bourque, Jesse Shapiro, Mark Lathrop and Michel Roger |
| EPI_ISL_535721 | Hôpital Notre-Dame | Laboratoire de santé publique du Québec | Sandrine Moreira, Ioannis Ragoussis, Guillaume Bourque, Jesse Shapiro, Mark Lathrop and Michel Roger |
| EPI_ISL_535722 | Centre de SSS de Trois-Rivières | Laboratoire de santé publique du Québec | Sandrine Moreira, Ioannis Ragoussis, Guillaume Bourque, Jesse Shapiro, Mark Lathrop and Michel Roger |
| EPI_ISL_535723 | Hôpital de Verdun | Laboratoire de santé publique du Québec | Sandrine Moreira, Ioannis Ragoussis, Guillaume Bourque, Jesse Shapiro, Mark Lathrop and Michel Roger |
| EPI_ISL_535724 | Centre de SSS La Pommeraie | Laboratoire de santé publique du Québec | Sandrine Moreira, Ioannis Ragoussis, Guillaume Bourque, Jesse Shapiro, Mark Lathrop and Michel Roger |
| EPI_ISL_535725 | Hôtel-Dieu de Lévis | Laboratoire de santé publique du Québec | Sandrine Moreira, Ioannis Ragoussis, Guillaume Bourque, Jesse Shapiro, Mark Lathrop and Michel Roger |
| EPI_ISL_535726 | CHUM - Microbiologie - Hôpital Saint-Luc | Laboratoire de santé publique du Québec | Sandrine Moreira, Ioannis Ragoussis, Guillaume Bourque, Jesse Shapiro, Mark Lathrop and Michel Roger |
| EPI_ISL_535727 | Centre de SSS de la Haute-Yamaska | Laboratoire de santé publique du Québec | Sandrine Moreira, Ioannis Ragoussis, Guillaume Bourque, Jesse Shapiro, Mark Lathrop and Michel Roger |
| EPI_ISL_535728 | Hôpital de Verdun | Laboratoire de santé publique du Québec | Sandrine Moreira, Ioannis Ragoussis, Guillaume Bourque, Jesse Shapiro, Mark Lathrop and Michel Roger |
| EPI_ISL_535729, EPI_ISL_535730 | CHUM - Microbiologie - Hôpital Saint-Luc | Laboratoire de santé publique du Québec | Sandrine Moreira, Ioannis Ragoussis, Guillaume Bourque, Jesse Shapiro, Mark Lathrop and Michel Roger |
| EPI_ISL_535731 | Hôpital Pierre-Le Gardeur | Laboratoire de santé publique du Québec | Sandrine Moreira, Ioannis Ragoussis, Guillaume Bourque, Jesse Shapiro, Mark Lathrop and Michel Roger |
| EPI_ISL_535732 | CHU Sainte-Justine | Laboratoire de santé publique du Québec | Sandrine Moreira, Ioannis Ragoussis, Guillaume Bourque, Jesse Shapiro, Mark Lathrop and Michel Roger |
| EPI_ISL_535734 | Hôpital Pierre-Boucher | Laboratoire de santé publique du Québec | Sandrine Moreira, Ioannis Ragoussis, Guillaume Bourque, Jesse Shapiro, Mark Lathrop and Michel Roger |
| EPI_ISL_535735 | CHUM - Microbiologie - Hôpital Saint-Luc | Laboratoire de santé publique du Québec | Sandrine Moreira, Ioannis Ragoussis, Guillaume Bourque, Jesse Shapiro, Mark Lathrop and Michel Roger |
| EPI_ISL_535736, EPI_ISL_535737 | CHUL-LABO MULTI / MICRO | Laboratoire de santé publique du Québec | Sandrine Moreira, Ioannis Ragoussis, Guillaume Bourque, Jesse Shapiro, Mark Lathrop and Michel Roger |
| EPI_ISL_535738 | Hôpital Charles-LeMoyne | Laboratoire de santé publique du Québec | Sandrine Moreira, Ioannis Ragoussis, Guillaume Bourque, Jesse Shapiro, Mark Lathrop and Michel Roger |
| EPI_ISL_535739 | Hôpital de l'Enfant-Jésus | Laboratoire de santé publique du Québec | Sandrine Moreira, Ioannis Ragoussis, Guillaume Bourque, Jesse Shapiro, Mark Lathrop and Michel Roger |
| EPI_ISL_535740 | CHUM - Microbiologie - Hôpital Saint-Luc | Laboratoire de santé publique du Québec | Sandrine Moreira, Ioannis Ragoussis, Guillaume Bourque, Jesse Shapiro, Mark Lathrop and Michel Roger |
| EPI_ISL_535741 | CHUL-LABO MULTI / MICRO | Laboratoire de santé publique du Québec | Sandrine Moreira, Ioannis Ragoussis, Guillaume Bourque, Jesse Shapiro, Mark Lathrop and Michel Roger |
| EPI_ISL_535742, EPI_ISL_535743 | Centre de SSS La Pommeraie | Laboratoire de santé publique du Québec | Sandrine Moreira, Ioannis Ragoussis, Guillaume Bourque, Jesse Shapiro, Mark Lathrop and Michel Roger |
| EPI_ISL_535744 | Hôpital de Lasalle | Laboratoire de santé publique du Québec | Sandrine Moreira, Ioannis Ragoussis, Guillaume Bourque, Jesse Shapiro, Mark Lathrop and Michel Roger |
| EPI_ISL_535745 | Hôpital Laurentien | Laboratoire de santé publique du Québec | Sandrine Moreira, Ioannis Ragoussis, Guillaume Bourque, Jesse Shapiro, Mark Lathrop and Michel Roger |
| EPI_ISL_535746 | Centre hospitalier de St-Mary | Laboratoire de santé publique du Québec | Sandrine Moreira, Ioannis Ragoussis, Guillaume Bourque, Jesse Shapiro, Mark Lathrop and Michel Roger |
| EPI_ISL_535748 | Centre de SSS de Trois-Rivières | Laboratoire de santé publique du Québec | Sandrine Moreira, Ioannis Ragoussis, Guillaume Bourque, Jesse Shapiro, Mark Lathrop and Michel Roger |
| EPI_ISL_535749 | CUSM-Site Glen-LAB Microbiologie | Laboratoire de santé publique du Québec | Sandrine Moreira, Ioannis Ragoussis, Guillaume Bourque, Jesse Shapiro, Mark Lathrop and Michel Roger |
| EPI_ISL_535750, EPI_ISL_535751 | Hôtel-Dieu de Lévis | Laboratoire de santé publique du Québec | Sandrine Moreira, Ioannis Ragoussis, Guillaume Bourque, Jesse Shapiro, Mark Lathrop and Michel Roger |

| | | | |
|---|---|---|---|
| EPI_ISL_535752 | Hôpital Pierre-Le Gardeur | Laboratoire de santé publique du Québec | Sandrine Moreira, Ioannis Ragoussis, Guillaume Bourque, Jesse Shapiro, Mark Lathrop and Michel Roger |
| EPI_ISL_535753, EPI_ISL_535754 | CHUM - Microbiologie - Hôpital Saint-Luc | Laboratoire de santé publique du Québec | Sandrine Moreira, Ioannis Ragoussis, Guillaume Bourque, Jesse Shapiro, Mark Lathrop and Michel Roger |
| EPI_ISL_535755 | Hôpital Maisonneuve-Rosemont | Laboratoire de santé publique du Québec | Sandrine Moreira, Ioannis Ragoussis, Guillaume Bourque, Jesse Shapiro, Mark Lathrop and Michel Roger |
| EPI_ISL_535756, EPI_ISL_535757 | Hôpital général Juif | Laboratoire de santé publique du Québec | Sandrine Moreira, Ioannis Ragoussis, Guillaume Bourque, Jesse Shapiro, Mark Lathrop and Michel Roger |
| EPI_ISL_535758, EPI_ISL_535759 | Hôtel-Dieu de Lévis | Laboratoire de santé publique du Québec | Sandrine Moreira, Ioannis Ragoussis, Guillaume Bourque, Jesse Shapiro, Mark Lathrop and Michel Roger |
| EPI_ISL_535761 | CUSM-Site Glen-LAB Microbiologie | Laboratoire de santé publique du Québec | Sandrine Moreira, Ioannis Ragoussis, Guillaume Bourque, Jesse Shapiro, Mark Lathrop and Michel Roger |
| EPI_ISL_535762 | Hôpital Pierre-Le Gardeur | Laboratoire de santé publique du Québec | Sandrine Moreira, Ioannis Ragoussis, Guillaume Bourque, Jesse Shapiro, Mark Lathrop and Michel Roger |
| EPI_ISL_535763 | Hôpital Pierre-Boucher | Laboratoire de santé publique du Québec | Sandrine Moreira, Ioannis Ragoussis, Guillaume Bourque, Jesse Shapiro, Mark Lathrop and Michel Roger |
| EPI_ISL_535764, EPI_ISL_535765 | CUSM-Site Glen-LAB Microbiologie | Laboratoire de santé publique du Québec | Sandrine Moreira, Ioannis Ragoussis, Guillaume Bourque, Jesse Shapiro, Mark Lathrop and Michel Roger |
| EPI_ISL_535766 | Hôpital Pierre-Le Gardeur | Laboratoire de santé publique du Québec | Sandrine Moreira, Ioannis Ragoussis, Guillaume Bourque, Jesse Shapiro, Mark Lathrop and Michel Roger |
| EPI_ISL_535767, EPI_ISL_535768 | Centre hospitalier de St-Mary | Laboratoire de santé publique du Québec | Sandrine Moreira, Ioannis Ragoussis, Guillaume Bourque, Jesse Shapiro, Mark Lathrop and Michel Roger |
| EPI_ISL_535769 | Hôpital général Juif | Laboratoire de santé publique du Québec | Sandrine Moreira, Ioannis Ragoussis, Guillaume Bourque, Jesse Shapiro, Mark Lathrop and Michel Roger |
| EPI_ISL_535770, EPI_ISL_535771 | Hôpital Pierre-Le Gardeur | Laboratoire de santé publique du Québec | Sandrine Moreira, Ioannis Ragoussis, Guillaume Bourque, Jesse Shapiro, Mark Lathrop and Michel Roger |
| EPI_ISL_535772, EPI_ISL_535773, EPI_ISL_535774, EPI_ISL_535775 | CHUL-LABO MULTI / MICRO | Laboratoire de santé publique du Québec | Sandrine Moreira, Ioannis Ragoussis, Guillaume Bourque, Jesse Shapiro, Mark Lathrop and Michel Roger |
| EPI_ISL_535776, EPI_ISL_535777, EPI_ISL_535778, EPI_ISL_535779, EPI_ISL_535780 | Centre de SSS de la Haute-Yamaska | Laboratoire de santé publique du Québec | Sandrine Moreira, Ioannis Ragoussis, Guillaume Bourque, Jesse Shapiro, Mark Lathrop and Michel Roger |
| EPI_ISL_535781, EPI_ISL_535782 | CHUL-LABO MULTI / MICRO | Laboratoire de santé publique du Québec | Sandrine Moreira, Ioannis Ragoussis, Guillaume Bourque, Jesse Shapiro, Mark Lathrop and Michel Roger |
| EPI_ISL_535783 | CHUM - Microbiologie - Hôpital Saint-Luc | Laboratoire de santé publique du Québec | Sandrine Moreira, Ioannis Ragoussis, Guillaume Bourque, Jesse Shapiro, Mark Lathrop and Michel Roger |
| EPI_ISL_535784, EPI_ISL_535785 | CHUL-LABO MULTI / MICRO | Laboratoire de santé publique du Québec | Sandrine Moreira, Ioannis Ragoussis, Guillaume Bourque, Jesse Shapiro, Mark Lathrop and Michel Roger |
| EPI_ISL_535786 | Hôpital Pierre-Boucher | Laboratoire de santé publique du Québec | Sandrine Moreira, Ioannis Ragoussis, Guillaume Bourque, Jesse Shapiro, Mark Lathrop and Michel Roger |
| EPI_ISL_535787, EPI_ISL_535788 | Centre Hospitalier Régional de Lanaudière | Laboratoire de santé publique du Québec | Sandrine Moreira, Ioannis Ragoussis, Guillaume Bourque, Jesse Shapiro, Mark Lathrop and Michel Roger |
| EPI_ISL_535789 | CHUM - Microbiologie - Hôpital Saint-Luc | Laboratoire de santé publique du Québec | Sandrine Moreira, Ioannis Ragoussis, Guillaume Bourque, Jesse Shapiro, Mark Lathrop and Michel Roger |
| EPI_ISL_535790 | Centre Hospitalier Régional de Lanaudière | Laboratoire de santé publique du Québec | Sandrine Moreira, Ioannis Ragoussis, Guillaume Bourque, Jesse Shapiro, Mark Lathrop and Michel Roger |
| EPI_ISL_535791 | Hôpital régional de Saint-Jérôme | Laboratoire de santé publique du Québec | Sandrine Moreira, Ioannis Ragoussis, Guillaume Bourque, Jesse Shapiro, Mark Lathrop and Michel Roger |
| EPI_ISL_535792 | Hôpital Fleurimont | Laboratoire de santé publique du Québec | Sandrine Moreira, Ioannis Ragoussis, Guillaume Bourque, Jesse Shapiro, Mark Lathrop and Michel Roger |
| EPI_ISL_535793 | Hôpital de Saint-Eustache | Laboratoire de santé publique du Québec | Sandrine Moreira, Ioannis Ragoussis, Guillaume Bourque, Jesse Shapiro, Mark Lathrop and Michel Roger |
| EPI_ISL_535794 | Hôpital de Hull | Laboratoire de santé publique du Québec | Sandrine Moreira, Ioannis Ragoussis, Guillaume Bourque, Jesse Shapiro, Mark Lathrop and Michel Roger |
| EPI_ISL_535795 | Hôpital Notre-Dame | Laboratoire de santé publique du Québec | Sandrine Moreira, Ioannis Ragoussis, Guillaume Bourque, Jesse Shapiro, Mark Lathrop and Michel Roger |
| EPI_ISL_535796 | Hôpital Charles-LeMoyne | Laboratoire de santé publique du Québec | Sandrine Moreira, Ioannis Ragoussis, Guillaume Bourque, Jesse Shapiro, Mark Lathrop and Michel Roger |
| EPI_ISL_535797 | Hôpital Honoré-Mercier | Laboratoire de santé publique du Québec | Sandrine Moreira, Ioannis Ragoussis, Guillaume Bourque, Jesse Shapiro, Mark Lathrop and Michel Roger |
| EPI_ISL_535798 | Hôpital régional de Saint-Jérôme | Laboratoire de santé publique du Québec | Sandrine Moreira, Ioannis Ragoussis, Guillaume Bourque, Jesse Shapiro, Mark Lathrop and Michel Roger |
| EPI_ISL_535799, EPI_ISL_535800 | CSSS Haut-Richelieu/Rouville (Hôpital) | Laboratoire de santé publique du Québec | Sandrine Moreira, Ioannis Ragoussis, Guillaume Bourque, Jesse Shapiro, Mark Lathrop and Michel Roger |
| EPI_ISL_535801 | CHU Sainte-Justine | Laboratoire de santé publique du Québec | Sandrine Moreira, Ioannis Ragoussis, Guillaume Bourque, Jesse Shapiro, Mark Lathrop and Michel Roger |
| EPI_ISL_535802 | CHUM - Microbiologie - Hôpital Saint-Luc | Laboratoire de santé publique du Québec | Sandrine Moreira, Ioannis Ragoussis, Guillaume Bourque, Jesse Shapiro, Mark Lathrop and Michel Roger |
| EPI_ISL_535803, EPI_ISL_535804, EPI_ISL_535805 | Centre de SSS de la Haute-Yamaska | Laboratoire de santé publique du Québec | Sandrine Moreira, Ioannis Ragoussis, Guillaume Bourque, Jesse Shapiro, Mark Lathrop and Michel Roger |
| EPI_ISL_535806 | CHUM - Microbiologie - Hôpital Saint-Luc | Laboratoire de santé publique du Québec | Sandrine Moreira, Ioannis Ragoussis, Guillaume Bourque, Jesse Shapiro, Mark Lathrop and Michel Roger |
| EPI_ISL_535807 | Centre hospitalier régional du Grand Portage | Laboratoire de santé publique du Québec | Sandrine Moreira, Ioannis Ragoussis, Guillaume Bourque, Jesse Shapiro, Mark Lathrop and Michel Roger |
| EPI_ISL_535808 | Centre de santé Chibougamau | Laboratoire de santé publique du Québec | Sandrine Moreira, Ioannis Ragoussis, Guillaume Bourque, Jesse Shapiro, Mark Lathrop and Michel Roger |
| EPI_ISL_535809 | Hôpital de Hull | Laboratoire de santé publique du Québec | Sandrine Moreira, Ioannis Ragoussis, Guillaume Bourque, Jesse Shapiro, Mark Lathrop and Michel Roger |
| EPI_ISL_535810 | Hôpital Sainte-Croix | Laboratoire de santé publique du Québec | Sandrine Moreira, Ioannis Ragoussis, Guillaume Bourque, Jesse Shapiro, Mark Lathrop and Michel Roger |
| EPI_ISL_535811 | CUSM-Site Glen-LAB Microbiologie | Laboratoire de santé publique du Québec | Sandrine Moreira, Ioannis Ragoussis, Guillaume Bourque, Jesse Shapiro, Mark Lathrop and Michel Roger |
| EPI_ISL_535812 | Hôpital du Sacré-Coeur de Montréal | Laboratoire de santé publique du Québec | Sandrine Moreira, Ioannis Ragoussis, Guillaume Bourque, Jesse Shapiro, Mark Lathrop and Michel Roger |
| EPI_ISL_535813, EPI_ISL_535815 | CSSS Haut-Richelieu/Rouville (Hôpital) | Laboratoire de santé publique du Québec | Sandrine Moreira, Ioannis Ragoussis, Guillaume Bourque, Jesse Shapiro, Mark Lathrop and Michel Roger |
| EPI_ISL_535816 | Centre de SSS de la Haute-Yamaska | Laboratoire de santé publique du Québec | Sandrine Moreira, Ioannis Ragoussis, Guillaume Bourque, Jesse Shapiro, Mark Lathrop and Michel Roger |
| EPI_ISL_535817 | Hôpital Maisonneuve-Rosemont | Laboratoire de santé publique du Québec | Sandrine Moreira, Ioannis Ragoussis, Guillaume Bourque, Jesse Shapiro, Mark Lathrop and Michel Roger |
| EPI_ISL_535818 | Hôpital Pierre-Boucher | Laboratoire de santé publique du Québec | Sandrine Moreira, Ioannis Ragoussis, Guillaume Bourque, Jesse Shapiro, Mark Lathrop and Michel Roger |

| | | | |
|---|---|---|---|
| EPI_ISL_535819 | Hôpital Charles-LeMoyne | Laboratoire de santé publique du Québec | Sandrine Moreira, Ioannis Ragoussis, Guillaume Bourque, Jesse Shapiro, Mark Lathrop and Michel Roger |
| EPI_ISL_535820, EPI_ISL_535821 | Hôtel-Dieu de Lévis | Laboratoire de santé publique du Québec | Sandrine Moreira, Ioannis Ragoussis, Guillaume Bourque, Jesse Shapiro, Mark Lathrop and Michel Roger |
| EPI_ISL_535822, EPI_ISL_535823, EPI_ISL_535824, EPI_ISL_535825 | CHUL-LABO MULTI / MICRO | Laboratoire de santé publique du Québec | Sandrine Moreira, Ioannis Ragoussis, Guillaume Bourque, Jesse Shapiro, Mark Lathrop and Michel Roger |
| EPI_ISL_535826 | CHUM - Microbiologie - Hôpital Saint-Luc | Laboratoire de santé publique du Québec | Sandrine Moreira, Ioannis Ragoussis, Guillaume Bourque, Jesse Shapiro, Mark Lathrop and Michel Roger |
| EPI_ISL_535828 | CHUL-LABO MULTI / MICRO | Laboratoire de santé publique du Québec | Sandrine Moreira, Ioannis Ragoussis, Guillaume Bourque, Jesse Shapiro, Mark Lathrop and Michel Roger |
| EPI_ISL_535829 | Hôpital de Chicoutimi | Laboratoire de santé publique du Québec | Sandrine Moreira, Ioannis Ragoussis, Guillaume Bourque, Jesse Shapiro, Mark Lathrop and Michel Roger |
| EPI_ISL_535830 | CHUM - Microbiologie - Hôpital Saint-Luc | Laboratoire de santé publique du Québec | Sandrine Moreira, Ioannis Ragoussis, Guillaume Bourque, Jesse Shapiro, Mark Lathrop and Michel Roger |
| EPI_ISL_535831 | Hôpital Maisonneuve-Rosemont | Laboratoire de santé publique du Québec | Sandrine Moreira, Ioannis Ragoussis, Guillaume Bourque, Jesse Shapiro, Mark Lathrop and Michel Roger |
| EPI_ISL_535832 | CHUM - Microbiologie - Hôpital Saint-Luc | Laboratoire de santé publique du Québec | Sandrine Moreira, Ioannis Ragoussis, Guillaume Bourque, Jesse Shapiro, Mark Lathrop and Michel Roger |
| EPI_ISL_535833, EPI_ISL_535834, EPI_ISL_535835, EPI_ISL_535836 | Hôtel-Dieu de Lévis | Laboratoire de santé publique du Québec | Sandrine Moreira, Ioannis Ragoussis, Guillaume Bourque, Jesse Shapiro, Mark Lathrop and Michel Roger |
| EPI_ISL_535838 | Hôpital régional de Saint-Jérôme | Laboratoire de santé publique du Québec | Sandrine Moreira, Ioannis Ragoussis, Guillaume Bourque, Jesse Shapiro, Mark Lathrop and Michel Roger |
| EPI_ISL_535839 | Hôpital Cité de la Santé | Laboratoire de santé publique du Québec | Sandrine Moreira, Ioannis Ragoussis, Guillaume Bourque, Jesse Shapiro, Mark Lathrop and Michel Roger |
| EPI_ISL_535840 | Hôpital Pierre-Boucher | Laboratoire de santé publique du Québec | Sandrine Moreira, Ioannis Ragoussis, Guillaume Bourque, Jesse Shapiro, Mark Lathrop and Michel Roger |
| EPI_ISL_535842, EPI_ISL_535844 | Centre Hospitalier Régional de Lanaudière | Laboratoire de santé publique du Québec | Sandrine Moreira, Ioannis Ragoussis, Guillaume Bourque, Jesse Shapiro, Mark Lathrop and Michel Roger |
| EPI_ISL_535850 | Hôpital Pierre-Le Gardeur | Laboratoire de santé publique du Québec | Sandrine Moreira, Ioannis Ragoussis, Guillaume Bourque, Jesse Shapiro, Mark Lathrop and Michel Roger |
| EPI_ISL_535851 | Hôpital Charles-LeMoyne | Laboratoire de santé publique du Québec | Sandrine Moreira, Ioannis Ragoussis, Guillaume Bourque, Jesse Shapiro, Mark Lathrop and Michel Roger |
| EPI_ISL_535854 | Centre de SSS de la Haute-Yamaska | Laboratoire de santé publique du Québec | Sandrine Moreira, Ioannis Ragoussis, Guillaume Bourque, Jesse Shapiro, Mark Lathrop and Michel Roger |
| EPI_ISL_535857 | Hôpital Pierre-Boucher | Laboratoire de santé publique du Québec | Sandrine Moreira, Ioannis Ragoussis, Guillaume Bourque, Jesse Shapiro, Mark Lathrop and Michel Roger |
| EPI_ISL_535858 | Hôpital de Gatineau | Laboratoire de santé publique du Québec | Sandrine Moreira, Ioannis Ragoussis, Guillaume Bourque, Jesse Shapiro, Mark Lathrop and Michel Roger |
| EPI_ISL_535862, EPI_ISL_535863 | Hôpital Pierre-Le Gardeur | Laboratoire de santé publique du Québec | Sandrine Moreira, Ioannis Ragoussis, Guillaume Bourque, Jesse Shapiro, Mark Lathrop and Michel Roger |
| EPI_ISL_535864 | CSSS Haut-Richelieu/Rouville (Hôpital) | Laboratoire de santé publique du Québec | Sandrine Moreira, Ioannis Ragoussis, Guillaume Bourque, Jesse Shapiro, Mark Lathrop and Michel Roger |
| EPI_ISL_535865 | Hôpital de Saint-Eustache | Laboratoire de santé publique du Québec | Sandrine Moreira, Ioannis Ragoussis, Guillaume Bourque, Jesse Shapiro, Mark Lathrop and Michel Roger |
| EPI_ISL_535866 | Hôpital Charles-LeMoyne | Laboratoire de santé publique du Québec | Sandrine Moreira, Ioannis Ragoussis, Guillaume Bourque, Jesse Shapiro, Mark Lathrop and Michel Roger |
| EPI_ISL_535868 | Hôpital du Suroît | Laboratoire de santé publique du Québec | Sandrine Moreira, Ioannis Ragoussis, Guillaume Bourque, Jesse Shapiro, Mark Lathrop and Michel Roger |
| EPI_ISL_535869 | Hôpital Charles-LeMoyne | Laboratoire de santé publique du Québec | Sandrine Moreira, Ioannis Ragoussis, Guillaume Bourque, Jesse Shapiro, Mark Lathrop and Michel Roger |
| EPI_ISL_535871 | Centre Hospitalier Régional de Lanaudière | Laboratoire de santé publique du Québec | Sandrine Moreira, Ioannis Ragoussis, Guillaume Bourque, Jesse Shapiro, Mark Lathrop and Michel Roger |
| EPI_ISL_535874 | Hôpital Charles-LeMoyne | Laboratoire de santé publique du Québec | Sandrine Moreira, Ioannis Ragoussis, Guillaume Bourque, Jesse Shapiro, Mark Lathrop and Michel Roger |
| EPI_ISL_535875 | Hôpital général du Lakeshore | Laboratoire de santé publique du Québec | Sandrine Moreira, Ioannis Ragoussis, Guillaume Bourque, Jesse Shapiro, Mark Lathrop and Michel Roger |
| EPI_ISL_535876, EPI_ISL_535877 | CSSS Haut-Richelieu/Rouville (Hôpital) | Laboratoire de santé publique du Québec | Sandrine Moreira, Ioannis Ragoussis, Guillaume Bourque, Jesse Shapiro, Mark Lathrop and Michel Roger |
| EPI_ISL_535879 | Hôpital Pierre-Boucher | Laboratoire de santé publique du Québec | Sandrine Moreira, Ioannis Ragoussis, Guillaume Bourque, Jesse Shapiro, Mark Lathrop and Michel Roger |
| EPI_ISL_535881, EPI_ISL_535882, EPI_ISL_535883 | Hôpital Pierre-Le Gardeur | Laboratoire de santé publique du Québec | Sandrine Moreira, Ioannis Ragoussis, Guillaume Bourque, Jesse Shapiro, Mark Lathrop and Michel Roger |
| EPI_ISL_535884 | CSSS Haut-Richelieu/Rouville (Hôpital) | Laboratoire de santé publique du Québec | Sandrine Moreira, Ioannis Ragoussis, Guillaume Bourque, Jesse Shapiro, Mark Lathrop and Michel Roger |
| EPI_ISL_535885 | Hôpital de Maria | Laboratoire de santé publique du Québec | Sandrine Moreira, Ioannis Ragoussis, Guillaume Bourque, Jesse Shapiro, Mark Lathrop and Michel Roger |
| EPI_ISL_535886, EPI_ISL_535887 | Hôpital de Hull | Laboratoire de santé publique du Québec | Sandrine Moreira, Ioannis Ragoussis, Guillaume Bourque, Jesse Shapiro, Mark Lathrop and Michel Roger |
| EPI_ISL_535888 | Hôpital Charles-LeMoyne | Laboratoire de santé publique du Québec | Sandrine Moreira, Ioannis Ragoussis, Guillaume Bourque, Jesse Shapiro, Mark Lathrop and Michel Roger |
| EPI_ISL_535889 | Hôpital de Saint-Eustache | Laboratoire de santé publique du Québec | Sandrine Moreira, Ioannis Ragoussis, Guillaume Bourque, Jesse Shapiro, Mark Lathrop and Michel Roger |
| EPI_ISL_535890, EPI_ISL_535891 | Hôpital Charles-LeMoyne | Laboratoire de santé publique du Québec | Sandrine Moreira, Ioannis Ragoussis, Guillaume Bourque, Jesse Shapiro, Mark Lathrop and Michel Roger |
| EPI_ISL_535892, EPI_ISL_535893 | Hôpital Pierre-Le Gardeur | Laboratoire de santé publique du Québec | Sandrine Moreira, Ioannis Ragoussis, Guillaume Bourque, Jesse Shapiro, Mark Lathrop and Michel Roger |
| EPI_ISL_535896 | Centre Hospitalier Régional de Lanaudière | Laboratoire de santé publique du Québec | Sandrine Moreira, Ioannis Ragoussis, Guillaume Bourque, Jesse Shapiro, Mark Lathrop and Michel Roger |
| EPI_ISL_535899, EPI_ISL_535901 | Hôpital Sainte-Croix | Laboratoire de santé publique du Québec | Sandrine Moreira, Ioannis Ragoussis, Guillaume Bourque, Jesse Shapiro, Mark Lathrop and Michel Roger |
| EPI_ISL_535902 | CSSS Haut-Richelieu/Rouville (Hôpital) | Laboratoire de santé publique du Québec | Sandrine Moreira, Ioannis Ragoussis, Guillaume Bourque, Jesse Shapiro, Mark Lathrop and Michel Roger |
| EPI_ISL_535903, EPI_ISL_535904 | Hôtel-Dieu de Sorel | Laboratoire de santé publique du Québec | Sandrine Moreira, Ioannis Ragoussis, Guillaume Bourque, Jesse Shapiro, Mark Lathrop and Michel Roger |
| EPI_ISL_535905 | Hôpital de Chicoutimi | Laboratoire de santé publique du Québec | Sandrine Moreira, Ioannis Ragoussis, Guillaume Bourque, Jesse Shapiro, Mark Lathrop and Michel Roger |
| EPI_ISL_535907 | Centre Hospitalier Régional de Lanaudière | Laboratoire de santé publique du Québec | Sandrine Moreira, Ioannis Ragoussis, Guillaume Bourque, Jesse Shapiro, Mark Lathrop and Michel Roger |
| EPI_ISL_535909 | Hôpital Pierre-Boucher | Laboratoire de santé publique du Québec | Sandrine Moreira, Ioannis Ragoussis, Guillaume Bourque, Jesse Shapiro, Mark Lathrop and Michel Roger |

| | | | |
|---|---|---|---|
| EPI_ISL_535910 | CSSS Haut-Richelieu/Rouville (Hôpital) | Laboratoire de santé publique du Québec | Sandrine Moreira, Ioannis Ragoussis, Guillaume Bourque, Jesse Shapiro, Mark Lathrop and Michel Roger |
| EPI_ISL_535911 | Hôpital de Hull | Laboratoire de santé publique du Québec | Sandrine Moreira, Ioannis Ragoussis, Guillaume Bourque, Jesse Shapiro, Mark Lathrop and Michel Roger |
| EPI_ISL_535912 | Hôpital Honoré-Mercier | Laboratoire de santé publique du Québec | Sandrine Moreira, Ioannis Ragoussis, Guillaume Bourque, Jesse Shapiro, Mark Lathrop and Michel Roger |
| EPI_ISL_535913 | Hôpital du Suroît | Laboratoire de santé publique du Québec | Sandrine Moreira, Ioannis Ragoussis, Guillaume Bourque, Jesse Shapiro, Mark Lathrop and Michel Roger |
| EPI_ISL_535914 | Hôpital du Centre-de-la-Mauricie | Laboratoire de santé publique du Québec | Sandrine Moreira, Ioannis Ragoussis, Guillaume Bourque, Jesse Shapiro, Mark Lathrop and Michel Roger |
| EPI_ISL_535915 | Hôpital Honoré-Mercier | Laboratoire de santé publique du Québec | Sandrine Moreira, Ioannis Ragoussis, Guillaume Bourque, Jesse Shapiro, Mark Lathrop and Michel Roger |
| EPI_ISL_535916 | Centre de SSS de Trois-Rivières | Laboratoire de santé publique du Québec | Sandrine Moreira, Ioannis Ragoussis, Guillaume Bourque, Jesse Shapiro, Mark Lathrop and Michel Roger |
| EPI_ISL_535917, EPI_ISL_535918, EPI_ISL_535919 | Hôpital du Suroît | Laboratoire de santé publique du Québec | Sandrine Moreira, Ioannis Ragoussis, Guillaume Bourque, Jesse Shapiro, Mark Lathrop and Michel Roger |
| EPI_ISL_535920, EPI_ISL_535921, EPI_ISL_535922, EPI_ISL_535923 | Hôpital Charles-LeMoyne | Laboratoire de santé publique du Québec | Sandrine Moreira, Ioannis Ragoussis, Guillaume Bourque, Jesse Shapiro, Mark Lathrop and Michel Roger |
| EPI_ISL_535925 | CSSS Haut-Richelieu/Rouville (Hôpital) | Laboratoire de santé publique du Québec | Sandrine Moreira, Ioannis Ragoussis, Guillaume Bourque, Jesse Shapiro, Mark Lathrop and Michel Roger |
| EPI_ISL_535926 | Hôpital de Saint-Eustache | Laboratoire de santé publique du Québec | Sandrine Moreira, Ioannis Ragoussis, Guillaume Bourque, Jesse Shapiro, Mark Lathrop and Michel Roger |
| EPI_ISL_535927 | Hôpital Charles-LeMoyne | Laboratoire de santé publique du Québec | Sandrine Moreira, Ioannis Ragoussis, Guillaume Bourque, Jesse Shapiro, Mark Lathrop and Michel Roger |
| EPI_ISL_535928 | CSSS Haut-Richelieu/Rouville (Hôpital) | Laboratoire de santé publique du Québec | Sandrine Moreira, Ioannis Ragoussis, Guillaume Bourque, Jesse Shapiro, Mark Lathrop and Michel Roger |
| EPI_ISL_535929, EPI_ISL_535930 | Hôpital de Saint-Eustache | Laboratoire de santé publique du Québec | Sandrine Moreira, Ioannis Ragoussis, Guillaume Bourque, Jesse Shapiro, Mark Lathrop and Michel Roger |
| EPI_ISL_535931, EPI_ISL_535932, EPI_ISL_535933, EPI_ISL_535934, EPI_ISL_535935 | Centre Hospitalier Régional de Lanaudière | Laboratoire de santé publique du Québec | Sandrine Moreira, Ioannis Ragoussis, Guillaume Bourque, Jesse Shapiro, Mark Lathrop and Michel Roger |
| EPI_ISL_535936 | Hôpital Pierre-Le Gardeur | Laboratoire de santé publique du Québec | Sandrine Moreira, Ioannis Ragoussis, Guillaume Bourque, Jesse Shapiro, Mark Lathrop and Michel Roger |
| EPI_ISL_535937 | Hôpital du Centre-de-la-Mauricie | Laboratoire de santé publique du Québec | Sandrine Moreira, Ioannis Ragoussis, Guillaume Bourque, Jesse Shapiro, Mark Lathrop and Michel Roger |
| EPI_ISL_535938 | Hôpital Pierre-Le Gardeur | Laboratoire de santé publique du Québec | Sandrine Moreira, Ioannis Ragoussis, Guillaume Bourque, Jesse Shapiro, Mark Lathrop and Michel Roger |
| EPI_ISL_535940 | Centre de SSS D'Arthabaska-et-de-l'érable - Hôtel-Dieu | Laboratoire de santé publique du Québec | Sandrine Moreira, Ioannis Ragoussis, Guillaume Bourque, Jesse Shapiro, Mark Lathrop and Michel Roger |
| EPI_ISL_535941, EPI_ISL_535942, EPI_ISL_535943 | Hôpital de Saint-Eustache | Laboratoire de santé publique du Québec | Sandrine Moreira, Ioannis Ragoussis, Guillaume Bourque, Jesse Shapiro, Mark Lathrop and Michel Roger |
| EPI_ISL_535944 | Hôpital Charles-LeMoyne | Laboratoire de santé publique du Québec | Sandrine Moreira, Ioannis Ragoussis, Guillaume Bourque, Jesse Shapiro, Mark Lathrop and Michel Roger |
| EPI_ISL_535945 | Hôpital de Mont-Laurier | Laboratoire de santé publique du Québec | Sandrine Moreira, Ioannis Ragoussis, Guillaume Bourque, Jesse Shapiro, Mark Lathrop and Michel Roger |
| EPI_ISL_535946 | CSSS Haut-Richelieu/Rouville (Hôpital) | Laboratoire de santé publique du Québec | Sandrine Moreira, Ioannis Ragoussis, Guillaume Bourque, Jesse Shapiro, Mark Lathrop and Michel Roger |
| EPI_ISL_535947 | Hôpital de Saint-Eustache | Laboratoire de santé publique du Québec | Sandrine Moreira, Ioannis Ragoussis, Guillaume Bourque, Jesse Shapiro, Mark Lathrop and Michel Roger |
| EPI_ISL_535948 | Hôpital Pierre-Boucher | Laboratoire de santé publique du Québec | Sandrine Moreira, Ioannis Ragoussis, Guillaume Bourque, Jesse Shapiro, Mark Lathrop and Michel Roger |
| EPI_ISL_535949, EPI_ISL_535950, EPI_ISL_535952, EPI_ISL_535953 | Hôpital Charles-LeMoyne | Laboratoire de santé publique du Québec | Sandrine Moreira, Ioannis Ragoussis, Guillaume Bourque, Jesse Shapiro, Mark Lathrop and Michel Roger |
| EPI_ISL_535954 | Hôpital Pierre-Le Gardeur | Laboratoire de santé publique du Québec | Sandrine Moreira, Ioannis Ragoussis, Guillaume Bourque, Jesse Shapiro, Mark Lathrop and Michel Roger |
| EPI_ISL_535955 | Centre Hospitalier Régional de Lanaudière | Laboratoire de santé publique du Québec | Sandrine Moreira, Ioannis Ragoussis, Guillaume Bourque, Jesse Shapiro, Mark Lathrop and Michel Roger |
| EPI_ISL_535956 | Hôpital Pierre-Le Gardeur | Laboratoire de santé publique du Québec | Sandrine Moreira, Ioannis Ragoussis, Guillaume Bourque, Jesse Shapiro, Mark Lathrop and Michel Roger |
| EPI_ISL_535957 | Hôpital Charles-LeMoyne | Laboratoire de santé publique du Québec | Sandrine Moreira, Ioannis Ragoussis, Guillaume Bourque, Jesse Shapiro, Mark Lathrop and Michel Roger |
| EPI_ISL_535959 | Centre hospitalier régional du Grand Portage | Laboratoire de santé publique du Québec | Sandrine Moreira, Ioannis Ragoussis, Guillaume Bourque, Jesse Shapiro, Mark Lathrop and Michel Roger |
| EPI_ISL_535962 | Hôpital Charles-LeMoyne | Laboratoire de santé publique du Québec | Sandrine Moreira, Ioannis Ragoussis, Guillaume Bourque, Jesse Shapiro, Mark Lathrop and Michel Roger |
| EPI_ISL_535963 | Hôpital d'Alma | Laboratoire de santé publique du Québec | Sandrine Moreira, Ioannis Ragoussis, Guillaume Bourque, Jesse Shapiro, Mark Lathrop and Michel Roger |
| EPI_ISL_535964, EPI_ISL_535965 | Hôpital de Chicoutimi | Laboratoire de santé publique du Québec | Sandrine Moreira, Ioannis Ragoussis, Guillaume Bourque, Jesse Shapiro, Mark Lathrop and Michel Roger |
| EPI_ISL_535966 | Hôpital Charles-LeMoyne | Laboratoire de santé publique du Québec | Sandrine Moreira, Ioannis Ragoussis, Guillaume Bourque, Jesse Shapiro, Mark Lathrop and Michel Roger |
| EPI_ISL_535967 | CSSS Haut-Richelieu/Rouville (Hôpital) | Laboratoire de santé publique du Québec | Sandrine Moreira, Ioannis Ragoussis, Guillaume Bourque, Jesse Shapiro, Mark Lathrop and Michel Roger |
| EPI_ISL_535969 | Hôpital Sainte-Croix | Laboratoire de santé publique du Québec | Sandrine Moreira, Ioannis Ragoussis, Guillaume Bourque, Jesse Shapiro, Mark Lathrop and Michel Roger |
| EPI_ISL_535971 | Hôpital du Centre-de-la-Mauricie | Laboratoire de santé publique du Québec | Sandrine Moreira, Ioannis Ragoussis, Guillaume Bourque, Jesse Shapiro, Mark Lathrop and Michel Roger |
| EPI_ISL_535972, EPI_ISL_535973 | Hôpital Charles-LeMoyne | Laboratoire de santé publique du Québec | Sandrine Moreira, Ioannis Ragoussis, Guillaume Bourque, Jesse Shapiro, Mark Lathrop and Michel Roger |
| EPI_ISL_535974, EPI_ISL_535975 | Hôpital du Centre-de-la-Mauricie | Laboratoire de santé publique du Québec | Sandrine Moreira, Ioannis Ragoussis, Guillaume Bourque, Jesse Shapiro, Mark Lathrop and Michel Roger |
| EPI_ISL_535976, EPI_ISL_535977 | Centre de SSS de Trois-Rivières | Laboratoire de santé publique du Québec | Sandrine Moreira, Ioannis Ragoussis, Guillaume Bourque, Jesse Shapiro, Mark Lathrop and Michel Roger |
| EPI_ISL_535978 | Hôpital du Centre-de-la-Mauricie | Laboratoire de santé publique du Québec | Sandrine Moreira, Ioannis Ragoussis, Guillaume Bourque, Jesse Shapiro, Mark Lathrop and Michel Roger |
| EPI_ISL_535979 | Centre de SSS de Trois-Rivières | Laboratoire de santé publique du Québec | Sandrine Moreira, Ioannis Ragoussis, Guillaume Bourque, Jesse Shapiro, Mark Lathrop and Michel Roger |
| EPI_ISL_535980, EPI_ISL_535981 | Centre hospitalier Anna-Laberge | Laboratoire de santé publique du Québec | Sandrine Moreira, Ioannis Ragoussis, Guillaume Bourque, Jesse Shapiro, Mark Lathrop and Michel Roger |
| EPI_ISL_535982 | Hôpital de Saint-Eustache | Laboratoire de santé publique du Québec | Sandrine Moreira, Ioannis Ragoussis, Guillaume Bourque, Jesse Shapiro, Mark Lathrop and Michel Roger |

| | | | |
|---|---|---|---|
| EPI_ISL_535983 | Hôpital Charles-LeMoyne | Laboratoire de santé publique du Québec | Sandrine Moreira, Ioannis Ragoussis, Guillaume Bourque, Jesse Shapiro, Mark Lathrop and Michel Roger |
| EPI_ISL_535984 | CSSS Haut-Richelieu/Rouville (Hôpital) | Laboratoire de santé publique du Québec | Sandrine Moreira, Ioannis Ragoussis, Guillaume Bourque, Jesse Shapiro, Mark Lathrop and Michel Roger |
| EPI_ISL_535985, EPI_ISL_535986, EPI_ISL_535987 | Hôpital de Rouyn-Noranda | Laboratoire de santé publique du Québec | Sandrine Moreira, Ioannis Ragoussis, Guillaume Bourque, Jesse Shapiro, Mark Lathrop and Michel Roger |
| EPI_ISL_535988 | Hôpital du Suroît | Laboratoire de santé publique du Québec | Sandrine Moreira, Ioannis Ragoussis, Guillaume Bourque, Jesse Shapiro, Mark Lathrop and Michel Roger |
| EPI_ISL_535989 | Hôpital de Hull | Laboratoire de santé publique du Québec | Sandrine Moreira, Ioannis Ragoussis, Guillaume Bourque, Jesse Shapiro, Mark Lathrop and Michel Roger |
| EPI_ISL_535990 | Hôtel-Dieu de Sorel | Laboratoire de santé publique du Québec | Sandrine Moreira, Ioannis Ragoussis, Guillaume Bourque, Jesse Shapiro, Mark Lathrop and Michel Roger |
| EPI_ISL_535991 | Hôpital et Centre d'Hébergement de Sept-Iles | Laboratoire de santé publique du Québec | Sandrine Moreira, Ioannis Ragoussis, Guillaume Bourque, Jesse Shapiro, Mark Lathrop and Michel Roger |
| EPI_ISL_535992 | Hôtel-Dieu de Sorel | Laboratoire de santé publique du Québec | Sandrine Moreira, Ioannis Ragoussis, Guillaume Bourque, Jesse Shapiro, Mark Lathrop and Michel Roger |
| EPI_ISL_535994 | Hôpital Laurentien | Laboratoire de santé publique du Québec | Sandrine Moreira, Ioannis Ragoussis, Guillaume Bourque, Jesse Shapiro, Mark Lathrop and Michel Roger |
| EPI_ISL_535995 | Hôpital de Hull | Laboratoire de santé publique du Québec | Sandrine Moreira, Ioannis Ragoussis, Guillaume Bourque, Jesse Shapiro, Mark Lathrop and Michel Roger |
| EPI_ISL_535996, EPI_ISL_535997, EPI_ISL_535998, EPI_ISL_535999 | Hôpital du Centre-de-la-Mauricie | Laboratoire de santé publique du Québec | Sandrine Moreira, Ioannis Ragoussis, Guillaume Bourque, Jesse Shapiro, Mark Lathrop and Michel Roger |
| EPI_ISL_536000 | Centre hospitalier Anna-Laberge | Laboratoire de santé publique du Québec | Sandrine Moreira, Ioannis Ragoussis, Guillaume Bourque, Jesse Shapiro, Mark Lathrop and Michel Roger |
| EPI_ISL_536001 | Hôpital du Centre-de-la-Mauricie | Laboratoire de santé publique du Québec | Sandrine Moreira, Ioannis Ragoussis, Guillaume Bourque, Jesse Shapiro, Mark Lathrop and Michel Roger |
| EPI_ISL_536002 | Centre Hospitalier Régional de Lanaudière | Laboratoire de santé publique du Québec | Sandrine Moreira, Ioannis Ragoussis, Guillaume Bourque, Jesse Shapiro, Mark Lathrop and Michel Roger |
| EPI_ISL_536003 | Hôpital Sainte-Croix | Laboratoire de santé publique du Québec | Sandrine Moreira, Ioannis Ragoussis, Guillaume Bourque, Jesse Shapiro, Mark Lathrop and Michel Roger |
| EPI_ISL_536004 | Centre hospitalier Anna-Laberge | Laboratoire de santé publique du Québec | Sandrine Moreira, Ioannis Ragoussis, Guillaume Bourque, Jesse Shapiro, Mark Lathrop and Michel Roger |
| EPI_ISL_536005, EPI_ISL_536006 | Centre Hospitalier Régional de Lanaudière | Laboratoire de santé publique du Québec | Sandrine Moreira, Ioannis Ragoussis, Guillaume Bourque, Jesse Shapiro, Mark Lathrop and Michel Roger |
| EPI_ISL_536007 | Centre de SSS de Trois-Rivières | Laboratoire de santé publique du Québec | Sandrine Moreira, Ioannis Ragoussis, Guillaume Bourque, Jesse Shapiro, Mark Lathrop and Michel Roger |
| EPI_ISL_536008 | Hôpital de Saint-Eustache | Laboratoire de santé publique du Québec | Sandrine Moreira, Ioannis Ragoussis, Guillaume Bourque, Jesse Shapiro, Mark Lathrop and Michel Roger |
| EPI_ISL_536009 | Centre Hospitalier Régional de Lanaudière | Laboratoire de santé publique du Québec | Sandrine Moreira, Ioannis Ragoussis, Guillaume Bourque, Jesse Shapiro, Mark Lathrop and Michel Roger |
| EPI_ISL_536010, EPI_ISL_536011 | Hôpital Sainte-Croix | Laboratoire de santé publique du Québec | Sandrine Moreira, Ioannis Ragoussis, Guillaume Bourque, Jesse Shapiro, Mark Lathrop and Michel Roger |
| EPI_ISL_536013 | Hôpital Honoré-Mercier | Laboratoire de santé publique du Québec | Sandrine Moreira, Ioannis Ragoussis, Guillaume Bourque, Jesse Shapiro, Mark Lathrop and Michel Roger |
| EPI_ISL_536014, EPI_ISL_536015, EPI_ISL_536016 | Centre Hospitalier Régional de Lanaudière | Laboratoire de santé publique du Québec | Sandrine Moreira, Ioannis Ragoussis, Guillaume Bourque, Jesse Shapiro, Mark Lathrop and Michel Roger |
| EPI_ISL_536017, EPI_ISL_536018, EPI_ISL_536020 | Hôpital Charles-LeMoyne | Laboratoire de santé publique du Québec | Sandrine Moreira, Ioannis Ragoussis, Guillaume Bourque, Jesse Shapiro, Mark Lathrop and Michel Roger |
| EPI_ISL_536021 | Hôpital Pierre-Boucher | Laboratoire de santé publique du Québec | Sandrine Moreira, Ioannis Ragoussis, Guillaume Bourque, Jesse Shapiro, Mark Lathrop and Michel Roger |
| EPI_ISL_536022 | Hôpital régional de Saint-Jérôme | Laboratoire de santé publique du Québec | Sandrine Moreira, Ioannis Ragoussis, Guillaume Bourque, Jesse Shapiro, Mark Lathrop and Michel Roger |
| EPI_ISL_536023 | Centre hospitalier Anna-Laberge | Laboratoire de santé publique du Québec | Sandrine Moreira, Ioannis Ragoussis, Guillaume Bourque, Jesse Shapiro, Mark Lathrop and Michel Roger |
| EPI_ISL_536024, EPI_ISL_536025 | Hôpital Charles-LeMoyne | Laboratoire de santé publique du Québec | Sandrine Moreira, Ioannis Ragoussis, Guillaume Bourque, Jesse Shapiro, Mark Lathrop and Michel Roger |
| EPI_ISL_536026 | Hôpital du Suroît | Laboratoire de santé publique du Québec | Sandrine Moreira, Ioannis Ragoussis, Guillaume Bourque, Jesse Shapiro, Mark Lathrop and Michel Roger |
| EPI_ISL_536028 | CSSS Haut-Richelieu/Rouville (Hôpital) | Laboratoire de santé publique du Québec | Sandrine Moreira, Ioannis Ragoussis, Guillaume Bourque, Jesse Shapiro, Mark Lathrop and Michel Roger |
| EPI_ISL_536029 | Hôpital Honoré-Mercier | Laboratoire de santé publique du Québec | Sandrine Moreira, Ioannis Ragoussis, Guillaume Bourque, Jesse Shapiro, Mark Lathrop and Michel Roger |
| EPI_ISL_536030 | Hôpital du Centre-de-la-Mauricie | Laboratoire de santé publique du Québec | Sandrine Moreira, Ioannis Ragoussis, Guillaume Bourque, Jesse Shapiro, Mark Lathrop and Michel Roger |
| EPI_ISL_536031 | Centre de SSS de Trois-Rivières | Laboratoire de santé publique du Québec | Sandrine Moreira, Ioannis Ragoussis, Guillaume Bourque, Jesse Shapiro, Mark Lathrop and Michel Roger |
| EPI_ISL_536032, EPI_ISL_536033 | Hôpital du Centre-de-la-Mauricie | Laboratoire de santé publique du Québec | Sandrine Moreira, Ioannis Ragoussis, Guillaume Bourque, Jesse Shapiro, Mark Lathrop and Michel Roger |
| EPI_ISL_536034, EPI_ISL_536035, EPI_ISL_536036 | Centre de SSS de Trois-Rivières | Laboratoire de santé publique du Québec | Sandrine Moreira, Ioannis Ragoussis, Guillaume Bourque, Jesse Shapiro, Mark Lathrop and Michel Roger |
| EPI_ISL_536037 | Hôpital du Centre-de-la-Mauricie | Laboratoire de santé publique du Québec | Sandrine Moreira, Ioannis Ragoussis, Guillaume Bourque, Jesse Shapiro, Mark Lathrop and Michel Roger |
| EPI_ISL_536038 | Centre de SSS de Trois-Rivières | Laboratoire de santé publique du Québec | Sandrine Moreira, Ioannis Ragoussis, Guillaume Bourque, Jesse Shapiro, Mark Lathrop and Michel Roger |
| EPI_ISL_536039 | Hôpital Pierre-Boucher | Laboratoire de santé publique du Québec | Sandrine Moreira, Ioannis Ragoussis, Guillaume Bourque, Jesse Shapiro, Mark Lathrop and Michel Roger |
| EPI_ISL_536040 | CSSS Haut-Richelieu/Rouville (Hôpital) | Laboratoire de santé publique du Québec | Sandrine Moreira, Ioannis Ragoussis, Guillaume Bourque, Jesse Shapiro, Mark Lathrop and Michel Roger |
| EPI_ISL_536041 | Hôpital Pierre-Boucher | Laboratoire de santé publique du Québec | Sandrine Moreira, Ioannis Ragoussis, Guillaume Bourque, Jesse Shapiro, Mark Lathrop and Michel Roger |
| EPI_ISL_536042, EPI_ISL_536044 | CSSS Haut-Richelieu/Rouville (Hôpital) | Laboratoire de santé publique du Québec | Sandrine Moreira, Ioannis Ragoussis, Guillaume Bourque, Jesse Shapiro, Mark Lathrop and Michel Roger |
| EPI_ISL_536045 | Centre hospitalier Anna-Laberge | Laboratoire de santé publique du Québec | Sandrine Moreira, Ioannis Ragoussis, Guillaume Bourque, Jesse Shapiro, Mark Lathrop and Michel Roger |
| EPI_ISL_536046 | Hôpital d'Alma | Laboratoire de santé publique du Québec | Sandrine Moreira, Ioannis Ragoussis, Guillaume Bourque, Jesse Shapiro, Mark Lathrop and Michel Roger |
| EPI_ISL_536047 | Centre Hospitalier Régional de Lanaudière | Laboratoire de santé publique du Québec | Sandrine Moreira, Ioannis Ragoussis, Guillaume Bourque, Jesse Shapiro, Mark Lathrop and Michel Roger |
| EPI_ISL_536049 | Hôpital de Saint-Eustache | Laboratoire de santé publique du Québec | Sandrine Moreira, Ioannis Ragoussis, Guillaume Bourque, Jesse Shapiro, Mark Lathrop and Michel Roger |

| | | | |
|---|---|---|---|
| EPI_ISL_536050 | Centre Hospitalier Régional de Lanaudière | Laboratoire de santé publique du Québec | Sandrine Moreira, Ioannis Ragoussis, Guillaume Bourque, Jesse Shapiro, Mark Lathrop and Michel Roger |
| EPI_ISL_536051 | Hôpital de Hull | Laboratoire de santé publique du Québec | Sandrine Moreira, Ioannis Ragoussis, Guillaume Bourque, Jesse Shapiro, Mark Lathrop and Michel Roger |
| EPI_ISL_536052, EPI_ISL_536053, EPI_ISL_536054 | Centre Hospitalier Régional de Lanaudière | Laboratoire de santé publique du Québec | Sandrine Moreira, Ioannis Ragoussis, Guillaume Bourque, Jesse Shapiro, Mark Lathrop and Michel Roger |
| EPI_ISL_536055 | Hôpital Honoré-Mercier | Laboratoire de santé publique du Québec | Sandrine Moreira, Ioannis Ragoussis, Guillaume Bourque, Jesse Shapiro, Mark Lathrop and Michel Roger |
| EPI_ISL_536056, EPI_ISL_536057 | Hôpital Sainte-Croix | Laboratoire de santé publique du Québec | Sandrine Moreira, Ioannis Ragoussis, Guillaume Bourque, Jesse Shapiro, Mark Lathrop and Michel Roger |
| EPI_ISL_536058 | Hôpital de Saint-Eustache | Laboratoire de santé publique du Québec | Sandrine Moreira, Ioannis Ragoussis, Guillaume Bourque, Jesse Shapiro, Mark Lathrop and Michel Roger |
| EPI_ISL_536059, EPI_ISL_536060 | Centre Hospitalier Régional de Lanaudière | Laboratoire de santé publique du Québec | Sandrine Moreira, Ioannis Ragoussis, Guillaume Bourque, Jesse Shapiro, Mark Lathrop and Michel Roger |
| EPI_ISL_536061, EPI_ISL_536062 | Hôpital Pierre-Le Gardeur | Laboratoire de santé publique du Québec | Sandrine Moreira, Ioannis Ragoussis, Guillaume Bourque, Jesse Shapiro, Mark Lathrop and Michel Roger |
| EPI_ISL_536063 | Centre Hospitalier Régional de Lanaudière | Laboratoire de santé publique du Québec | Sandrine Moreira, Ioannis Ragoussis, Guillaume Bourque, Jesse Shapiro, Mark Lathrop and Michel Roger |
| EPI_ISL_536064, EPI_ISL_536065, EPI_ISL_536066, EPI_ISL_536067, EPI_ISL_536068 | Hôpital Pierre-Le Gardeur | Laboratoire de santé publique du Québec | Sandrine Moreira, Ioannis Ragoussis, Guillaume Bourque, Jesse Shapiro, Mark Lathrop and Michel Roger |
| EPI_ISL_536069 | Hôpital de Saint-Eustache | Laboratoire de santé publique du Québec | Sandrine Moreira, Ioannis Ragoussis, Guillaume Bourque, Jesse Shapiro, Mark Lathrop and Michel Roger |
| EPI_ISL_536070 | Hôpital Pierre-Le Gardeur | Laboratoire de santé publique du Québec | Sandrine Moreira, Ioannis Ragoussis, Guillaume Bourque, Jesse Shapiro, Mark Lathrop and Michel Roger |
| EPI_ISL_536071, EPI_ISL_536072 | Hôpital de Saint-Eustache | Laboratoire de santé publique du Québec | Sandrine Moreira, Ioannis Ragoussis, Guillaume Bourque, Jesse Shapiro, Mark Lathrop and Michel Roger |
| EPI_ISL_536073, EPI_ISL_536074 | Centre Hospitalier Régional de Lanaudière | Laboratoire de santé publique du Québec | Sandrine Moreira, Ioannis Ragoussis, Guillaume Bourque, Jesse Shapiro, Mark Lathrop and Michel Roger |
| EPI_ISL_536075 | Hôpital Charles-LeMoyne | Laboratoire de santé publique du Québec | Sandrine Moreira, Ioannis Ragoussis, Guillaume Bourque, Jesse Shapiro, Mark Lathrop and Michel Roger |
| EPI_ISL_536076 | Centre Hospitalier Régional de Lanaudière | Laboratoire de santé publique du Québec | Sandrine Moreira, Ioannis Ragoussis, Guillaume Bourque, Jesse Shapiro, Mark Lathrop and Michel Roger |
| EPI_ISL_536077 | Hôpital Charles-LeMoyne | Laboratoire de santé publique du Québec | Sandrine Moreira, Ioannis Ragoussis, Guillaume Bourque, Jesse Shapiro, Mark Lathrop and Michel Roger |
| EPI_ISL_536079 | Centre Hospitalier Régional de Lanaudière | Laboratoire de santé publique du Québec | Sandrine Moreira, Ioannis Ragoussis, Guillaume Bourque, Jesse Shapiro, Mark Lathrop and Michel Roger |
| EPI_ISL_536080 | Centre hospitalier Anna-Laberge | Laboratoire de santé publique du Québec | Sandrine Moreira, Ioannis Ragoussis, Guillaume Bourque, Jesse Shapiro, Mark Lathrop and Michel Roger |
| EPI_ISL_536082 | Hôpital Charles-LeMoyne | Laboratoire de santé publique du Québec | Sandrine Moreira, Ioannis Ragoussis, Guillaume Bourque, Jesse Shapiro, Mark Lathrop and Michel Roger |
| EPI_ISL_536083, EPI_ISL_536084, EPI_ISL_536086 | Hôpital Pierre-Le Gardeur | Laboratoire de santé publique du Québec | Sandrine Moreira, Ioannis Ragoussis, Guillaume Bourque, Jesse Shapiro, Mark Lathrop and Michel Roger |
| EPI_ISL_536087, EPI_ISL_536088, EPI_ISL_536090 | Hôpital Charles-LeMoyne | Laboratoire de santé publique du Québec | Sandrine Moreira, Ioannis Ragoussis, Guillaume Bourque, Jesse Shapiro, Mark Lathrop and Michel Roger |
| EPI_ISL_536092 | Hôpital Pierre-Boucher | Laboratoire de santé publique du Québec | Sandrine Moreira, Ioannis Ragoussis, Guillaume Bourque, Jesse Shapiro, Mark Lathrop and Michel Roger |
| EPI_ISL_536094 | Hôpital Charles-LeMoyne | Laboratoire de santé publique du Québec | Sandrine Moreira, Ioannis Ragoussis, Guillaume Bourque, Jesse Shapiro, Mark Lathrop and Michel Roger |
| EPI_ISL_536095 | Hôpital du Suroît | Laboratoire de santé publique du Québec | Sandrine Moreira, Ioannis Ragoussis, Guillaume Bourque, Jesse Shapiro, Mark Lathrop and Michel Roger |
| EPI_ISL_536096 | Hôpital Pierre-Boucher | Laboratoire de santé publique du Québec | Sandrine Moreira, Ioannis Ragoussis, Guillaume Bourque, Jesse Shapiro, Mark Lathrop and Michel Roger |
| EPI_ISL_536097, EPI_ISL_536098 | Hôpital du Suroît | Laboratoire de santé publique du Québec | Sandrine Moreira, Ioannis Ragoussis, Guillaume Bourque, Jesse Shapiro, Mark Lathrop and Michel Roger |
| EPI_ISL_536099 | Hôpital Pierre-Boucher | Laboratoire de santé publique du Québec | Sandrine Moreira, Ioannis Ragoussis, Guillaume Bourque, Jesse Shapiro, Mark Lathrop and Michel Roger |
| EPI_ISL_536101 | Hôpital, CLSC et Centre d'Hébergement de Roberval | Laboratoire de santé publique du Québec | Sandrine Moreira, Ioannis Ragoussis, Guillaume Bourque, Jesse Shapiro, Mark Lathrop and Michel Roger |
| EPI_ISL_536102 | Hôpital du Suroît | Laboratoire de santé publique du Québec | Sandrine Moreira, Ioannis Ragoussis, Guillaume Bourque, Jesse Shapiro, Mark Lathrop and Michel Roger |
| EPI_ISL_536104 | Centre hospitalier Anna-Laberge | Laboratoire de santé publique du Québec | Sandrine Moreira, Ioannis Ragoussis, Guillaume Bourque, Jesse Shapiro, Mark Lathrop and Michel Roger |
| EPI_ISL_536105 | Hôpital du Suroît | Laboratoire de santé publique du Québec | Sandrine Moreira, Ioannis Ragoussis, Guillaume Bourque, Jesse Shapiro, Mark Lathrop and Michel Roger |
| EPI_ISL_536107 | Hôpital de Chicoutimi | Laboratoire de santé publique du Québec | Sandrine Moreira, Ioannis Ragoussis, Guillaume Bourque, Jesse Shapiro, Mark Lathrop and Michel Roger |
| EPI_ISL_536108 | Hôpital Le Royer | Laboratoire de santé publique du Québec | Sandrine Moreira, Ioannis Ragoussis, Guillaume Bourque, Jesse Shapiro, Mark Lathrop and Michel Roger |
| EPI_ISL_536109 | CSSS de Port-Cartier | Laboratoire de santé publique du Québec | Sandrine Moreira, Ioannis Ragoussis, Guillaume Bourque, Jesse Shapiro, Mark Lathrop and Michel Roger |
| EPI_ISL_536110 | CSSS Haut-Richelieu/Rouville (Hôpital) | Laboratoire de santé publique du Québec | Sandrine Moreira, Ioannis Ragoussis, Guillaume Bourque, Jesse Shapiro, Mark Lathrop and Michel Roger |
| EPI_ISL_536111, EPI_ISL_536112 | Hôpital de Maria | Laboratoire de santé publique du Québec | Sandrine Moreira, Ioannis Ragoussis, Guillaume Bourque, Jesse Shapiro, Mark Lathrop and Michel Roger |
| EPI_ISL_536113, EPI_ISL_536114 | Hôpital et Centre d'Hébergement de Sept-Iles | Laboratoire de santé publique du Québec | Sandrine Moreira, Ioannis Ragoussis, Guillaume Bourque, Jesse Shapiro, Mark Lathrop and Michel Roger |
| EPI_ISL_536115 | Centre hospitalier régional du Grand Portage | Laboratoire de santé publique du Québec | Sandrine Moreira, Ioannis Ragoussis, Guillaume Bourque, Jesse Shapiro, Mark Lathrop and Michel Roger |
| EPI_ISL_536117 | Hôpital de Hull | Laboratoire de santé publique du Québec | Sandrine Moreira, Ioannis Ragoussis, Guillaume Bourque, Jesse Shapiro, Mark Lathrop and Michel Roger |
| EPI_ISL_536120 | Centre Hospitalier Régional de Lanaudière | Laboratoire de santé publique du Québec | Sandrine Moreira, Ioannis Ragoussis, Guillaume Bourque, Jesse Shapiro, Mark Lathrop and Michel Roger |
| EPI_ISL_536121 | Hôtel-Dieu de Sorel | Laboratoire de santé publique du Québec | Sandrine Moreira, Ioannis Ragoussis, Guillaume Bourque, Jesse Shapiro, Mark Lathrop and Michel Roger |
| EPI_ISL_536122, EPI_ISL_536124 | Hôpital de Hull | Laboratoire de santé publique du Québec | Sandrine Moreira, Ioannis Ragoussis, Guillaume Bourque, Jesse Shapiro, Mark Lathrop and Michel Roger |
| EPI_ISL_536126 | Centre Hospitalier Régional de Lanaudière | Laboratoire de santé publique du Québec | Sandrine Moreira, Ioannis Ragoussis, Guillaume Bourque, Jesse Shapiro, Mark Lathrop and Michel Roger |
| EPI_ISL_536127 | Hôpital de Hull | Laboratoire de santé publique du Québec | Sandrine Moreira, Ioannis Ragoussis, Guillaume Bourque, Jesse Shapiro, Mark Lathrop and Michel Roger |

| | | | |
|---|---|---|---|
| EPI_ISL_536128 | Hôpital Pierre-Le Gardeur | Laboratoire de santé publique du Québec | Sandrine Moreira, Ioannis Ragoussis, Guillaume Bourque, Jesse Shapiro, Mark Lathrop and Michel Roger |
| EPI_ISL_536129, EPI_ISL_536130 | Centre Hospitalier Régional de Lanaudière | Laboratoire de santé publique du Québec | Sandrine Moreira, Ioannis Ragoussis, Guillaume Bourque, Jesse Shapiro, Mark Lathrop and Michel Roger |
| EPI_ISL_536131 | Hôpital Pierre-Le Gardeur | Laboratoire de santé publique du Québec | Sandrine Moreira, Ioannis Ragoussis, Guillaume Bourque, Jesse Shapiro, Mark Lathrop and Michel Roger |
| EPI_ISL_536132 | Hôpital de Hull | Laboratoire de santé publique du Québec | Sandrine Moreira, Ioannis Ragoussis, Guillaume Bourque, Jesse Shapiro, Mark Lathrop and Michel Roger |
| EPI_ISL_536134 | Centre Hospitalier Régional de Lanaudière | Laboratoire de santé publique du Québec | Sandrine Moreira, Ioannis Ragoussis, Guillaume Bourque, Jesse Shapiro, Mark Lathrop and Michel Roger |
| EPI_ISL_536137 | Hôpital Pierre-Le Gardeur | Laboratoire de santé publique du Québec | Sandrine Moreira, Ioannis Ragoussis, Guillaume Bourque, Jesse Shapiro, Mark Lathrop and Michel Roger |
| EPI_ISL_536138 | Centre Hospitalier Régional de Lanaudière | Laboratoire de santé publique du Québec | Sandrine Moreira, Ioannis Ragoussis, Guillaume Bourque, Jesse Shapiro, Mark Lathrop and Michel Roger |
| EPI_ISL_536139 | Hôpital Pierre-Le Gardeur | Laboratoire de santé publique du Québec | Sandrine Moreira, Ioannis Ragoussis, Guillaume Bourque, Jesse Shapiro, Mark Lathrop and Michel Roger |
| EPI_ISL_536141, EPI_ISL_536142 | Centre hospitalier Anna-Laberge | Laboratoire de santé publique du Québec | Sandrine Moreira, Ioannis Ragoussis, Guillaume Bourque, Jesse Shapiro, Mark Lathrop and Michel Roger |
| EPI_ISL_536143, EPI_ISL_536144 | Hôpital Pierre-Le Gardeur | Laboratoire de santé publique du Québec | Sandrine Moreira, Ioannis Ragoussis, Guillaume Bourque, Jesse Shapiro, Mark Lathrop and Michel Roger |
| EPI_ISL_536145 | Centre hospitalier Anna-Laberge | Laboratoire de santé publique du Québec | Sandrine Moreira, Ioannis Ragoussis, Guillaume Bourque, Jesse Shapiro, Mark Lathrop and Michel Roger |
| EPI_ISL_536146, EPI_ISL_536147, EPI_ISL_536148 | Hôpital Charles-LeMoyne | Laboratoire de santé publique du Québec | Sandrine Moreira, Ioannis Ragoussis, Guillaume Bourque, Jesse Shapiro, Mark Lathrop and Michel Roger |
| EPI_ISL_536150 | CSSS Haut-Richelieu/Rouville (Hôpital) | Laboratoire de santé publique du Québec | Sandrine Moreira, Ioannis Ragoussis, Guillaume Bourque, Jesse Shapiro, Mark Lathrop and Michel Roger |
| EPI_ISL_536151 | Centre Hospitalier Régional de Lanaudière | Laboratoire de santé publique du Québec | Sandrine Moreira, Ioannis Ragoussis, Guillaume Bourque, Jesse Shapiro, Mark Lathrop and Michel Roger |
| EPI_ISL_536152 | Hôpital Pierre-Le Gardeur | Laboratoire de santé publique du Québec | Sandrine Moreira, Ioannis Ragoussis, Guillaume Bourque, Jesse Shapiro, Mark Lathrop and Michel Roger |
| EPI_ISL_536153, EPI_ISL_536154 | Centre Hospitalier Régional de Lanaudière | Laboratoire de santé publique du Québec | Sandrine Moreira, Ioannis Ragoussis, Guillaume Bourque, Jesse Shapiro, Mark Lathrop and Michel Roger |
| EPI_ISL_536156 | Centre hospitalier Anna-Laberge | Laboratoire de santé publique du Québec | Sandrine Moreira, Ioannis Ragoussis, Guillaume Bourque, Jesse Shapiro, Mark Lathrop and Michel Roger |
| EPI_ISL_536157 | Hôpital Pierre-Le Gardeur | Laboratoire de santé publique du Québec | Sandrine Moreira, Ioannis Ragoussis, Guillaume Bourque, Jesse Shapiro, Mark Lathrop and Michel Roger |
| EPI_ISL_536158 | CSSS Haut-Richelieu/Rouville (Hôpital) | Laboratoire de santé publique du Québec | Sandrine Moreira, Ioannis Ragoussis, Guillaume Bourque, Jesse Shapiro, Mark Lathrop and Michel Roger |
| EPI_ISL_536159 | Centre hospitalier Anna-Laberge | Laboratoire de santé publique du Québec | Sandrine Moreira, Ioannis Ragoussis, Guillaume Bourque, Jesse Shapiro, Mark Lathrop and Michel Roger |
| EPI_ISL_536160 | CSSS Haut-Richelieu/Rouville (Hôpital) | Laboratoire de santé publique du Québec | Sandrine Moreira, Ioannis Ragoussis, Guillaume Bourque, Jesse Shapiro, Mark Lathrop and Michel Roger |
| EPI_ISL_536162 | Centre hospitalier Anna-Laberge | Laboratoire de santé publique du Québec | Sandrine Moreira, Ioannis Ragoussis, Guillaume Bourque, Jesse Shapiro, Mark Lathrop and Michel Roger |
| EPI_ISL_536163 | Hôpital du Suroît | Laboratoire de santé publique du Québec | Sandrine Moreira, Ioannis Ragoussis, Guillaume Bourque, Jesse Shapiro, Mark Lathrop and Michel Roger |
| EPI_ISL_536166 | CSSS Haut-Richelieu/Rouville (Hôpital) | Laboratoire de santé publique du Québec | Sandrine Moreira, Ioannis Ragoussis, Guillaume Bourque, Jesse Shapiro, Mark Lathrop and Michel Roger |
| EPI_ISL_536168, EPI_ISL_536169 | Hôpital de Hull | Laboratoire de santé publique du Québec | Sandrine Moreira, Ioannis Ragoussis, Guillaume Bourque, Jesse Shapiro, Mark Lathrop and Michel Roger |
| EPI_ISL_536171 | Hôpital de Papineau | Laboratoire de santé publique du Québec | Sandrine Moreira, Ioannis Ragoussis, Guillaume Bourque, Jesse Shapiro, Mark Lathrop and Michel Roger |
| EPI_ISL_536173, EPI_ISL_536174, EPI_ISL_536176, EPI_ISL_536178, EPI_ISL_536179 | Hôpital Pierre-Boucher | Laboratoire de santé publique du Québec | Sandrine Moreira, Ioannis Ragoussis, Guillaume Bourque, Jesse Shapiro, Mark Lathrop and Michel Roger |
| EPI_ISL_536180 | Hôpital Honoré-Mercier | Laboratoire de santé publique du Québec | Sandrine Moreira, Ioannis Ragoussis, Guillaume Bourque, Jesse Shapiro, Mark Lathrop and Michel Roger |
| EPI_ISL_536181 | CSSS Haut-Richelieu/Rouville (Hôpital) | Laboratoire de santé publique du Québec | Sandrine Moreira, Ioannis Ragoussis, Guillaume Bourque, Jesse Shapiro, Mark Lathrop and Michel Roger |
| EPI_ISL_536182, EPI_ISL_536183 | Centre hospitalier Anna-Laberge | Laboratoire de santé publique du Québec | Sandrine Moreira, Ioannis Ragoussis, Guillaume Bourque, Jesse Shapiro, Mark Lathrop and Michel Roger |
| EPI_ISL_536186, EPI_ISL_536187 | Hôpital Charles-LeMoyne | Laboratoire de santé publique du Québec | Sandrine Moreira, Ioannis Ragoussis, Guillaume Bourque, Jesse Shapiro, Mark Lathrop and Michel Roger |
| EPI_ISL_536195, EPI_ISL_536196 | Hôpital Pierre-Boucher | Laboratoire de santé publique du Québec | Sandrine Moreira, Ioannis Ragoussis, Guillaume Bourque, Jesse Shapiro, Mark Lathrop and Michel Roger |
| EPI_ISL_536198 | Hôpital du Suroît | Laboratoire de santé publique du Québec | Sandrine Moreira, Ioannis Ragoussis, Guillaume Bourque, Jesse Shapiro, Mark Lathrop and Michel Roger |
| EPI_ISL_536199 | Centre hospitalier régional du Grand Portage | Laboratoire de santé publique du Québec | Sandrine Moreira, Ioannis Ragoussis, Guillaume Bourque, Jesse Shapiro, Mark Lathrop and Michel Roger |
| EPI_ISL_536200, EPI_ISL_536201, EPI_ISL_536202 | Hôpital du Suroît | Laboratoire de santé publique du Québec | Sandrine Moreira, Ioannis Ragoussis, Guillaume Bourque, Jesse Shapiro, Mark Lathrop and Michel Roger |
| EPI_ISL_536205 | Hôpital régional de Rimouski | Laboratoire de santé publique du Québec | Sandrine Moreira, Ioannis Ragoussis, Guillaume Bourque, Jesse Shapiro, Mark Lathrop and Michel Roger |
| EPI_ISL_536206 | CSSS de la Minganie | Laboratoire de santé publique du Québec | Sandrine Moreira, Ioannis Ragoussis, Guillaume Bourque, Jesse Shapiro, Mark Lathrop and Michel Roger |
| EPI_ISL_536207, EPI_ISL_536208, EPI_ISL_536209 | Hôpital de Hull | Laboratoire de santé publique du Québec | Sandrine Moreira, Ioannis Ragoussis, Guillaume Bourque, Jesse Shapiro, Mark Lathrop and Michel Roger |
| EPI_ISL_536212 | Centre hospitalier Anna-Laberge | Laboratoire de santé publique du Québec | Sandrine Moreira, Ioannis Ragoussis, Guillaume Bourque, Jesse Shapiro, Mark Lathrop and Michel Roger |
| EPI_ISL_536213, EPI_ISL_536215, EPI_ISL_536216 | CSSS Haut-Richelieu/Rouville (Hôpital) | Laboratoire de santé publique du Québec | Sandrine Moreira, Ioannis Ragoussis, Guillaume Bourque, Jesse Shapiro, Mark Lathrop and Michel Roger |
| EPI_ISL_536218, EPI_ISL_536219, EPI_ISL_536220 | Centre hospitalier Anna-Laberge | Laboratoire de santé publique du Québec | Sandrine Moreira, Ioannis Ragoussis, Guillaume Bourque, Jesse Shapiro, Mark Lathrop and Michel Roger |
| EPI_ISL_536223 | Hôpital Charles-LeMoyne | Laboratoire de santé publique du Québec | Sandrine Moreira, Ioannis Ragoussis, Guillaume Bourque, Jesse Shapiro, Mark Lathrop and Michel Roger |
| EPI_ISL_536224 | Centre hospitalier Anna-Laberge | Laboratoire de santé publique du Québec | Sandrine Moreira, Ioannis Ragoussis, Guillaume Bourque, Jesse Shapiro, Mark Lathrop and Michel Roger |
| EPI_ISL_536225 | Hôpital Charles-LeMoyne | Laboratoire de santé publique du Québec | Sandrine Moreira, Ioannis Ragoussis, Guillaume Bourque, Jesse Shapiro, Mark Lathrop and Michel Roger |
| EPI_ISL_536228 | Hôpital du Suroît | Laboratoire de santé publique du Québec | Sandrine Moreira, Ioannis Ragoussis, Guillaume Bourque, Jesse Shapiro, Mark Lathrop and Michel Roger |

| | | | |
|---|---|---|---|
| EPI_ISL_536229 | Hôpital Pierre-Boucher | Laboratoire de santé publique du Québec | Sandrine Moreira, Ioannis Ragoussis, Guillaume Bourque, Jesse Shapiro, Mark Lathrop and Michel Roger |
| EPI_ISL_536230 | Hôpital de Hull | Laboratoire de santé publique du Québec | Sandrine Moreira, Ioannis Ragoussis, Guillaume Bourque, Jesse Shapiro, Mark Lathrop and Michel Roger |
| EPI_ISL_536232 | Centre hospitalier Anna-Laberge | Laboratoire de santé publique du Québec | Sandrine Moreira, Ioannis Ragoussis, Guillaume Bourque, Jesse Shapiro, Mark Lathrop and Michel Roger |
| EPI_ISL_536234, EPI_ISL_536236 | Hôpital Pierre-Boucher | Laboratoire de santé publique du Québec | Sandrine Moreira, Ioannis Ragoussis, Guillaume Bourque, Jesse Shapiro, Mark Lathrop and Michel Roger |
| EPI_ISL_536237 | CSSS Haut-Richelieu/Rouville (Hôpital) | Laboratoire de santé publique du Québec | Sandrine Moreira, Ioannis Ragoussis, Guillaume Bourque, Jesse Shapiro, Mark Lathrop and Michel Roger |
| EPI_ISL_536238, EPI_ISL_536239 | Hôpital de Hull | Laboratoire de santé publique du Québec | Sandrine Moreira, Ioannis Ragoussis, Guillaume Bourque, Jesse Shapiro, Mark Lathrop and Michel Roger |
| EPI_ISL_536240, EPI_ISL_536241 | Centre hospitalier Anna-Laberge | Laboratoire de santé publique du Québec | Sandrine Moreira, Ioannis Ragoussis, Guillaume Bourque, Jesse Shapiro, Mark Lathrop and Michel Roger |
| EPI_ISL_536242 | Hôpital de Hull | Laboratoire de santé publique du Québec | Sandrine Moreira, Ioannis Ragoussis, Guillaume Bourque, Jesse Shapiro, Mark Lathrop and Michel Roger |
| EPI_ISL_536243 | CSSS Haut-Richelieu/Rouville (Hôpital) | Laboratoire de santé publique du Québec | Sandrine Moreira, Ioannis Ragoussis, Guillaume Bourque, Jesse Shapiro, Mark Lathrop and Michel Roger |
| EPI_ISL_536244 | Hôpital de Hull | Laboratoire de santé publique du Québec | Sandrine Moreira, Ioannis Ragoussis, Guillaume Bourque, Jesse Shapiro, Mark Lathrop and Michel Roger |
| EPI_ISL_536246 | Hôpital du Centre-de-la-Mauricie | Laboratoire de santé publique du Québec | Sandrine Moreira, Ioannis Ragoussis, Guillaume Bourque, Jesse Shapiro, Mark Lathrop and Michel Roger |
| EPI_ISL_536247 | Hôpital Charles-LeMoyne | Laboratoire de santé publique du Québec | Sandrine Moreira, Ioannis Ragoussis, Guillaume Bourque, Jesse Shapiro, Mark Lathrop and Michel Roger |
| EPI_ISL_536248, EPI_ISL_536249, EPI_ISL_536250, EPI_ISL_536251 | Hôpital du Centre-de-la-Mauricie | Laboratoire de santé publique du Québec | Sandrine Moreira, Ioannis Ragoussis, Guillaume Bourque, Jesse Shapiro, Mark Lathrop and Michel Roger |
| EPI_ISL_536252 | Hôpital Sainte-Croix | Laboratoire de santé publique du Québec | Sandrine Moreira, Ioannis Ragoussis, Guillaume Bourque, Jesse Shapiro, Mark Lathrop and Michel Roger |
| EPI_ISL_536257 | Hôpital du Suroît | Laboratoire de santé publique du Québec | Sandrine Moreira, Ioannis Ragoussis, Guillaume Bourque, Jesse Shapiro, Mark Lathrop and Michel Roger |
| EPI_ISL_536260 | CSSS de la Minganie | Laboratoire de santé publique du Québec | Sandrine Moreira, Ioannis Ragoussis, Guillaume Bourque, Jesse Shapiro, Mark Lathrop and Michel Roger |
| EPI_ISL_536266 | Hôpital de Gatineau | Laboratoire de santé publique du Québec | Sandrine Moreira, Ioannis Ragoussis, Guillaume Bourque, Jesse Shapiro, Mark Lathrop and Michel Roger |
| EPI_ISL_536268, EPI_ISL_536269, EPI_ISL_536270 | Hôpital Charles-LeMoyne | Laboratoire de santé publique du Québec | Sandrine Moreira, Ioannis Ragoussis, Guillaume Bourque, Jesse Shapiro, Mark Lathrop and Michel Roger |
| EPI_ISL_536272 | CSSS Haut-Richelieu/Rouville (Hôpital) | Laboratoire de santé publique du Québec | Sandrine Moreira, Ioannis Ragoussis, Guillaume Bourque, Jesse Shapiro, Mark Lathrop and Michel Roger |
| EPI_ISL_536273, EPI_ISL_536274 | Hôpital du Suroît | Laboratoire de santé publique du Québec | Sandrine Moreira, Ioannis Ragoussis, Guillaume Bourque, Jesse Shapiro, Mark Lathrop and Michel Roger |
| EPI_ISL_536276 | Hôpital Pierre-Boucher | Laboratoire de santé publique du Québec | Sandrine Moreira, Ioannis Ragoussis, Guillaume Bourque, Jesse Shapiro, Mark Lathrop and Michel Roger |
| EPI_ISL_536279, EPI_ISL_536280, EPI_ISL_536281, EPI_ISL_536282 | Hôpital Charles-LeMoyne | Laboratoire de santé publique du Québec | Sandrine Moreira, Ioannis Ragoussis, Guillaume Bourque, Jesse Shapiro, Mark Lathrop and Michel Roger |
| EPI_ISL_536283 | Centre hospitalier Anna-Laberge | Laboratoire de santé publique du Québec | Sandrine Moreira, Ioannis Ragoussis, Guillaume Bourque, Jesse Shapiro, Mark Lathrop and Michel Roger |
| EPI_ISL_536284, EPI_ISL_536285 | Hôpital Charles-LeMoyne | Laboratoire de santé publique du Québec | Sandrine Moreira, Ioannis Ragoussis, Guillaume Bourque, Jesse Shapiro, Mark Lathrop and Michel Roger |
| EPI_ISL_536286 | CSSS Haut-Richelieu/Rouville (Hôpital) | Laboratoire de santé publique du Québec | Sandrine Moreira, Ioannis Ragoussis, Guillaume Bourque, Jesse Shapiro, Mark Lathrop and Michel Roger |
| EPI_ISL_536288 | Conseil Cri de la SSS de la Baie-James | Laboratoire de santé publique du Québec | Sandrine Moreira, Ioannis Ragoussis, Guillaume Bourque, Jesse Shapiro, Mark Lathrop and Michel Roger |
| EPI_ISL_536289 | Hôpital de Maria | Laboratoire de santé publique du Québec | Sandrine Moreira, Ioannis Ragoussis, Guillaume Bourque, Jesse Shapiro, Mark Lathrop and Michel Roger |
| EPI_ISL_536290 | Hôpital Charles-LeMoyne | Laboratoire de santé publique du Québec | Sandrine Moreira, Ioannis Ragoussis, Guillaume Bourque, Jesse Shapiro, Mark Lathrop and Michel Roger |
| EPI_ISL_536291 | Hôpital de Hull | Laboratoire de santé publique du Québec | Sandrine Moreira, Ioannis Ragoussis, Guillaume Bourque, Jesse Shapiro, Mark Lathrop and Michel Roger |
| EPI_ISL_536292, EPI_ISL_536293, EPI_ISL_536294 | Hôpital Charles-LeMoyne | Laboratoire de santé publique du Québec | Sandrine Moreira, Ioannis Ragoussis, Guillaume Bourque, Jesse Shapiro, Mark Lathrop and Michel Roger |
| EPI_ISL_536295 | Hôpital Pierre-Boucher | Laboratoire de santé publique du Québec | Sandrine Moreira, Ioannis Ragoussis, Guillaume Bourque, Jesse Shapiro, Mark Lathrop and Michel Roger |
| EPI_ISL_536296, EPI_ISL_536297, EPI_ISL_536298 | Hôpital Charles-LeMoyne | Laboratoire de santé publique du Québec | Sandrine Moreira, Ioannis Ragoussis, Guillaume Bourque, Jesse Shapiro, Mark Lathrop and Michel Roger |
| EPI_ISL_536299 | CSSS Haut-Richelieu/Rouville (Hôpital) | Laboratoire de santé publique du Québec | Sandrine Moreira, Ioannis Ragoussis, Guillaume Bourque, Jesse Shapiro, Mark Lathrop and Michel Roger |
| EPI_ISL_536300 | Hôpital Pierre-Boucher | Laboratoire de santé publique du Québec | Sandrine Moreira, Ioannis Ragoussis, Guillaume Bourque, Jesse Shapiro, Mark Lathrop and Michel Roger |
| EPI_ISL_536301, EPI_ISL_536302 | CSSS Haut-Richelieu/Rouville (Hôpital) | Laboratoire de santé publique du Québec | Sandrine Moreira, Ioannis Ragoussis, Guillaume Bourque, Jesse Shapiro, Mark Lathrop and Michel Roger |
| EPI_ISL_536303 | Hôpital Pierre-Boucher | Laboratoire de santé publique du Québec | Sandrine Moreira, Ioannis Ragoussis, Guillaume Bourque, Jesse Shapiro, Mark Lathrop and Michel Roger |
| EPI_ISL_536304 | CSSS Haut-Richelieu/Rouville (Hôpital) | Laboratoire de santé publique du Québec | Sandrine Moreira, Ioannis Ragoussis, Guillaume Bourque, Jesse Shapiro, Mark Lathrop and Michel Roger |
| EPI_ISL_536305, EPI_ISL_536306, EPI_ISL_536308, EPI_ISL_536309 | Hôpital Pierre-Boucher | Laboratoire de santé publique du Québec | Sandrine Moreira, Ioannis Ragoussis, Guillaume Bourque, Jesse Shapiro, Mark Lathrop and Michel Roger |
| EPI_ISL_536312 | Hôpital de Hull | Laboratoire de santé publique du Québec | Sandrine Moreira, Ioannis Ragoussis, Guillaume Bourque, Jesse Shapiro, Mark Lathrop and Michel Roger |
| EPI_ISL_536313, EPI_ISL_536314 | Hôpital Charles-LeMoyne | Laboratoire de santé publique du Québec | Sandrine Moreira, Ioannis Ragoussis, Guillaume Bourque, Jesse Shapiro, Mark Lathrop and Michel Roger |
| EPI_ISL_536319 | Hôpital du Suroît | Laboratoire de santé publique du Québec | Sandrine Moreira, Ioannis Ragoussis, Guillaume Bourque, Jesse Shapiro, Mark Lathrop and Michel Roger |
| EPI_ISL_536322, EPI_ISL_536323 | Hôpital Pierre-Boucher | Laboratoire de santé publique du Québec | Sandrine Moreira, Ioannis Ragoussis, Guillaume Bourque, Jesse Shapiro, Mark Lathrop and Michel Roger |
| EPI_ISL_536325 | Centre hospitalier Anna-Laberge | Laboratoire de santé publique du Québec | Sandrine Moreira, Ioannis Ragoussis, Guillaume Bourque, Jesse Shapiro, Mark Lathrop and Michel Roger |
| EPI_ISL_536328, EPI_ISL_536331 | Hôpital Charles-LeMoyne | Laboratoire de santé publique du Québec | Sandrine Moreira, Ioannis Ragoussis, Guillaume Bourque, Jesse Shapiro, Mark Lathrop and Michel Roger |
| EPI_ISL_536332, EPI_ISL_536335 | Hôpital Pierre-Boucher | Laboratoire de santé publique du Québec | Sandrine Moreira, Ioannis Ragoussis, Guillaume Bourque, Jesse Shapiro, Mark Lathrop and Michel Roger |

| | | | |
|---|---|---|---|
| EPI_ISL_536336 | Hôpital Charles-LeMoyne | Laboratoire de santé publique du Québec | Sandrine Moreira, Ioannis Ragoussis, Guillaume Bourque, Jesse Shapiro, Mark Lathrop and Michel Roger |
| EPI_ISL_536337 | CUSM-Site Glen-LAB Microbiologie | Laboratoire de santé publique du Québec | Sandrine Moreira, Ioannis Ragoussis, Guillaume Bourque, Jesse Shapiro, Mark Lathrop and Michel Roger |
| EPI_ISL_536338, EPI_ISL_536339, EPI_ISL_536340, EPI_ISL_536342 | Hôpital Charles-LeMoyne | Laboratoire de santé publique du Québec | Sandrine Moreira, Ioannis Ragoussis, Guillaume Bourque, Jesse Shapiro, Mark Lathrop and Michel Roger |
| EPI_ISL_536344 | CSSS Haut-Richelieu/Rouville (Hôpital) | Laboratoire de santé publique du Québec | Sandrine Moreira, Ioannis Ragoussis, Guillaume Bourque, Jesse Shapiro, Mark Lathrop and Michel Roger |
| EPI_ISL_536347 | Hôpital Pierre-Boucher | Laboratoire de santé publique du Québec | Sandrine Moreira, Ioannis Ragoussis, Guillaume Bourque, Jesse Shapiro, Mark Lathrop and Michel Roger |
| EPI_ISL_536348 | Centre hospitalier Anna-Laberge | Laboratoire de santé publique du Québec | Sandrine Moreira, Ioannis Ragoussis, Guillaume Bourque, Jesse Shapiro, Mark Lathrop and Michel Roger |
| EPI_ISL_536349 | Hôpital du Suroît | Laboratoire de santé publique du Québec | Sandrine Moreira, Ioannis Ragoussis, Guillaume Bourque, Jesse Shapiro, Mark Lathrop and Michel Roger |
| EPI_ISL_536350 | Hôpital Charles-LeMoyne | Laboratoire de santé publique du Québec | Sandrine Moreira, Ioannis Ragoussis, Guillaume Bourque, Jesse Shapiro, Mark Lathrop and Michel Roger |
| EPI_ISL_536351, EPI_ISL_536352 | Hôpital Pierre-Boucher | Laboratoire de santé publique du Québec | Sandrine Moreira, Ioannis Ragoussis, Guillaume Bourque, Jesse Shapiro, Mark Lathrop and Michel Roger |
| EPI_ISL_536353 | Centre de santé Inuulitsivik | Laboratoire de santé publique du Québec | Sandrine Moreira, Ioannis Ragoussis, Guillaume Bourque, Jesse Shapiro, Mark Lathrop and Michel Roger |
| EPI_ISL_536354, EPI_ISL_536355, EPI_ISL_536356, EPI_ISL_536357, EPI_ISL_536358, EPI_ISL_536359 | Hôpital du Suroît | Laboratoire de santé publique du Québec | Sandrine Moreira, Ioannis Ragoussis, Guillaume Bourque, Jesse Shapiro, Mark Lathrop and Michel Roger |
| EPI_ISL_536360 | Hôpital Pierre-Boucher | Laboratoire de santé publique du Québec | Sandrine Moreira, Ioannis Ragoussis, Guillaume Bourque, Jesse Shapiro, Mark Lathrop and Michel Roger |
| EPI_ISL_536361 | CSSS Haut-Richelieu/Rouville (Hôpital) | Laboratoire de santé publique du Québec | Sandrine Moreira, Ioannis Ragoussis, Guillaume Bourque, Jesse Shapiro, Mark Lathrop and Michel Roger |
| EPI_ISL_536362 | Hôpital du Suroît | Laboratoire de santé publique du Québec | Sandrine Moreira, Ioannis Ragoussis, Guillaume Bourque, Jesse Shapiro, Mark Lathrop and Michel Roger |
| EPI_ISL_536363 | Hôpital de Hull | Laboratoire de santé publique du Québec | Sandrine Moreira, Ioannis Ragoussis, Guillaume Bourque, Jesse Shapiro, Mark Lathrop and Michel Roger |
| EPI_ISL_536365, EPI_ISL_536366 | CUSM-Site Glen-LAB Microbiologie | Laboratoire de santé publique du Québec | Sandrine Moreira, Ioannis Ragoussis, Guillaume Bourque, Jesse Shapiro, Mark Lathrop and Michel Roger |
| EPI_ISL_536367 | Hôpital de Hull | Laboratoire de santé publique du Québec | Sandrine Moreira, Ioannis Ragoussis, Guillaume Bourque, Jesse Shapiro, Mark Lathrop and Michel Roger |
| EPI_ISL_536369, EPI_ISL_536370 | Centre hospitalier Anna-Laberge | Laboratoire de santé publique du Québec | Sandrine Moreira, Ioannis Ragoussis, Guillaume Bourque, Jesse Shapiro, Mark Lathrop and Michel Roger |
| EPI_ISL_536372, EPI_ISL_536373 | Hôpital Charles-LeMoyne | Laboratoire de santé publique du Québec | Sandrine Moreira, Ioannis Ragoussis, Guillaume Bourque, Jesse Shapiro, Mark Lathrop and Michel Roger |
| EPI_ISL_536375 | Hôpital Pierre-Boucher | Laboratoire de santé publique du Québec | Sandrine Moreira, Ioannis Ragoussis, Guillaume Bourque, Jesse Shapiro, Mark Lathrop and Michel Roger |
| EPI_ISL_536377, EPI_ISL_536378 | Centre hospitalier Anna-Laberge | Laboratoire de santé publique du Québec | Sandrine Moreira, Ioannis Ragoussis, Guillaume Bourque, Jesse Shapiro, Mark Lathrop and Michel Roger |
| EPI_ISL_536379 | Hôpital Honoré-Mercier | Laboratoire de santé publique du Québec | Sandrine Moreira, Ioannis Ragoussis, Guillaume Bourque, Jesse Shapiro, Mark Lathrop and Michel Roger |
| EPI_ISL_536381 | Centre Hospitalier Régional de Lanaudière | Laboratoire de santé publique du Québec | Sandrine Moreira, Ioannis Ragoussis, Guillaume Bourque, Jesse Shapiro, Mark Lathrop and Michel Roger |
| EPI_ISL_536383 | CSSS Haut-Richelieu/Rouville (Hôpital) | Laboratoire de santé publique du Québec | Sandrine Moreira, Ioannis Ragoussis, Guillaume Bourque, Jesse Shapiro, Mark Lathrop and Michel Roger |
| EPI_ISL_536385, EPI_ISL_536386, EPI_ISL_536387 | Hôpital de Hull | Laboratoire de santé publique du Québec | Sandrine Moreira, Ioannis Ragoussis, Guillaume Bourque, Jesse Shapiro, Mark Lathrop and Michel Roger |
| EPI_ISL_536388, EPI_ISL_536389 | Hôpital Pierre-Boucher | Laboratoire de santé publique du Québec | Sandrine Moreira, Ioannis Ragoussis, Guillaume Bourque, Jesse Shapiro, Mark Lathrop and Michel Roger |
| EPI_ISL_536390 | Hôpital du Suroît | Laboratoire de santé publique du Québec | Sandrine Moreira, Ioannis Ragoussis, Guillaume Bourque, Jesse Shapiro, Mark Lathrop and Michel Roger |
| EPI_ISL_536391 | Hôpital Pierre-Boucher | Laboratoire de santé publique du Québec | Sandrine Moreira, Ioannis Ragoussis, Guillaume Bourque, Jesse Shapiro, Mark Lathrop and Michel Roger |
| EPI_ISL_536392, EPI_ISL_536393 | Hôpital de Hull | Laboratoire de santé publique du Québec | Sandrine Moreira, Ioannis Ragoussis, Guillaume Bourque, Jesse Shapiro, Mark Lathrop and Michel Roger |
| EPI_ISL_536395, EPI_ISL_536397 | Hôpital Pierre-Boucher | Laboratoire de santé publique du Québec | Sandrine Moreira, Ioannis Ragoussis, Guillaume Bourque, Jesse Shapiro, Mark Lathrop and Michel Roger |
| EPI_ISL_536398 | Lithuanian University of Health Sciences Hospital, Department of Laboratory Medicine | Lithuanian University of Health Sciences, Molecular cardiology lab. | Lukas Zemaitis, Arnoldas Pautienius, Kamile Tamusauskaite, Dovydas Gecys, Vaiva Lesauskaite, Astra Vitkauskiene |
| EPI_ISL_536399 | Laboratory of Immunovirology. Universidad de Antioquia | Instituto Nacional de Salud - Unidad de Secuenciación y Genómica | Wbeimar Aguilar-Jimenez, Lizdany Flórez, Francisco J. Díaz, Katherine Laiton-Donato, Carlos Franco-Muñoz, Diego Álvarez-Diaz and Marcela Mercado-Reyes |
| EPI_ISL_536411 | Medtimes Molecular Laboratory | Medtimes Molecular Laboratory | Eric Chan, Winsome Wong, Jacqueline Tam, Isaac Chow |
| EPI_ISL_536412, EPI_ISL_536413, EPI_ISL_536414, EPI_ISL_536415, EPI_ISL_536416, EPI_ISL_536417, EPI_ISL_536418, EPI_ISL_536419, EPI_ISL_536420, EPI_ISL_536421, EPI_ISL_536422, EPI_ISL_536423, EPI_ISL_536424, EPI_ISL_536425, EPI_ISL_536426, EPI_ISL_536427, EPI_ISL_536428, EPI_ISL_536429, EPI_ISL_536430, EPI_ISL_536431, EPI_ISL_536432, EPI_ISL_536433, EPI_ISL_536434, EPI_ISL_536435, EPI_ISL_536436, EPI_ISL_536437, EPI_ISL_536438, EPI_ISL_536439, EPI_ISL_536440, EPI_ISL_536441, EPI_ISL_536442, EPI_ISL_536443, EPI_ISL_536444, EPI_ISL_536445, EPI_ISL_536446, EPI_ISL_536447, EPI_ISL_536448, EPI_ISL_536449, EPI_ISL_536450 | | | |
| see above | National Public Health Laboratory, National Centre for Infectious Diseases | National Public Health Laboratory, National Centre for Infectious Diseases | Mak TM, Octavia S, Zhou Z, Cui L, Lin RTP |
| EPI_ISL_536477, EPI_ISL_536478, EPI_ISL_536479, EPI_ISL_536480, EPI_ISL_536481, EPI_ISL_536482, EPI_ISL_536483, EPI_ISL_536484, EPI_ISL_536485, EPI_ISL_536486, EPI_ISL_536487, EPI_ISL_536488, EPI_ISL_536489, EPI_ISL_536490, EPI_ISL_536491, EPI_ISL_536492, EPI_ISL_536493, EPI_ISL_536494, EPI_ISL_536495, EPI_ISL_536496, EPI_ISL_536498, EPI_ISL_536499, EPI_ISL_536500, EPI_ISL_536502, EPI_ISL_536503, EPI_ISL_536504, EPI_ISL_536505, EPI_ISL_536506, EPI_ISL_536508, EPI_ISL_536509, EPI_ISL_536510, EPI_ISL_536511, EPI_ISL_536512, EPI_ISL_536513, EPI_ISL_536514, EPI_ISL_536515, EPI_ISL_536516, EPI_ISL_536517, EPI_ISL_536518, EPI_ISL_536519, EPI_ISL_536520, EPI_ISL_536521, EPI_ISL_536522, EPI_ISL_536523, EPI_ISL_536524, EPI_ISL_536527, EPI_ISL_536528, EPI_ISL_536529, EPI_ISL_536531, EPI_ISL_536533, EPI_ISL_536534, EPI_ISL_536537, EPI_ISL_536538, EPI_ISL_536539, EPI_ISL_536540, EPI_ISL_536541, EPI_ISL_536542, EPI_ISL_536543, EPI_ISL_536545, EPI_ISL_536548, EPI_ISL_536549, EPI_ISL_536550, EPI_ISL_536551, EPI_ISL_536552, EPI_ISL_536553, EPI_ISL_536554, EPI_ISL_536555, EPI_ISL_536556, EPI_ISL_536557, EPI_ISL_536558, EPI_ISL_536559, EPI_ISL_536560, EPI_ISL_536561, EPI_ISL_536562 | | | |
| see above | Instituto Nacional de Salud | Laboratorio de Infecciones Respiratorias Agudas | Eduardo Juscamayta Lopez, David Tarazona, Faviola Valdivia Guerrero, Nancy Rojas Serrano, Dennis Carhuaricra, Lenin Maturrano Hernandez, Ronnie Gavilan Chavez |
| EPI_ISL_536572, EPI_ISL_536573, EPI_ISL_536574, EPI_ISL_536575, EPI_ISL_536576, EPI_ISL_536577, EPI_ISL_536578, EPI_ISL_536579, EPI_ISL_536580, EPI_ISL_536581, EPI_ISL_536582, EPI_ISL_536583, EPI_ISL_536584, EPI_ISL_536585, EPI_ISL_536586, EPI_ISL_536587, EPI_ISL_536588, EPI_ISL_536589, EPI_ISL_536590, EPI_ISL_536591, EPI_ISL_536592, EPI_ISL_536593, EPI_ISL_536594, EPI_ISL_536595, EPI_ISL_536596, EPI_ISL_536597, EPI_ISL_536598, EPI_ISL_536599, EPI_ISL_536601, EPI_ISL_536602, EPI_ISL_536603, EPI_ISL_536604, EPI_ISL_536605, EPI_ISL_536606, EPI_ISL_536607, EPI_ISL_536608, EPI_ISL_536609, EPI_ISL_536610, EPI_ISL_536611, EPI_ISL_536612, EPI_ISL_536613, EPI_ISL_536614, EPI_ISL_536615, EPI_ISL_536616, EPI_ISL_536617, EPI_ISL_536618, EPI_ISL_536619, EPI_ISL_536620, EPI_ISL_536621, EPI_ISL_536622, EPI_ISL_536624, EPI_ISL_536625, EPI_ISL_536626, EPI_ISL_536627, EPI_ISL_536628, EPI_ISL_536629, EPI_ISL_536630, EPI_ISL_536631, EPI_ISL_536632, EPI_ISL_536633, EPI_ISL_536634, EPI_ISL_536635, EPI_ISL_536636, EPI_ISL_536637, EPI_ISL_536638, EPI_ISL_536639, EPI_ISL_536640, EPI_ISL_536641, EPI_ISL_536642, EPI_ISL_536643, EPI_ISL_536644, EPI_ISL_536645, EPI_ISL_536646, EPI_ISL_536647, EPI_ISL_536648, EPI_ISL_536649, EPI_ISL_536650, EPI_ISL_536651, EPI_ISL_536652, EPI_ISL_536653, EPI_ISL_536654, EPI_ISL_536655 | | | |
| see above | University of Wisconsin-Madison AIDS Vaccine Research Laboratories | University of Wisconsin-Madison AIDS Vaccine Research Laboratories | Gage Moreno, Katarina Braun, et al. AIDS Vaccine Research Laboratories |
| EPI_ISL_536656, EPI_ISL_536657 | University of Wisconsin-Madison Campus AIDS Vaccine Research Laboratories | University of Wisconsin-Madison AIDS Vaccine Research Laboratories | Gage Moreno, Katarina Braun, et al. AIDS Vaccine Research Laboratories |
| EPI_ISL_536658, EPI_ISL_536660, EPI_ISL_536661, EPI_ISL_536662, EPI_ISL_536663, EPI_ISL_536664, EPI_ISL_536665, EPI_ISL_536666, EPI_ISL_536667, EPI_ISL_536668, EPI_ISL_536669, EPI_ISL_536670, EPI_ISL_536671, EPI_ISL_536673, EPI_ISL_536674, EPI_ISL_536675, EPI_ISL_536676, EPI_ISL_536677, EPI_ISL_536678, EPI_ISL_536679, EPI_ISL_536680, EPI_ISL_536681, EPI_ISL_536682, EPI_ISL_536683, EPI_ISL_536685, EPI_ISL_536686, EPI_ISL_536687, EPI_ISL_536688, EPI_ISL_536689, EPI_ISL_536694, EPI_ISL_536697, EPI_ISL_536698, EPI_ISL_536699, EPI_ISL_536700, EPI_ISL_536701, EPI_ISL_536702, EPI_ISL_536703, EPI_ISL_536704, EPI_ISL_536705, EPI_ISL_536706, EPI_ISL_536707, EPI_ISL_536708, EPI_ISL_536709, EPI_ISL_536710, EPI_ISL_536711, EPI_ISL_536712, EPI_ISL_536713, EPI_ISL_536714, EPI_ISL_536715, EPI_ISL_536717, EPI_ISL_536718, EPI_ISL_536719, EPI_ISL_536720, EPI_ISL_536721, EPI_ISL_536722, EPI_ISL_536723, EPI_ISL_536724, EPI_ISL_536725, EPI_ISL_536727, EPI_ISL_536728, EPI_ISL_536729, EPI_ISL_536730, EPI_ISL_536731, EPI_ISL_536732, EPI_ISL_536733, EPI_ISL_536734, EPI_ISL_536735, EPI_ISL_536736, EPI_ISL_536737, EPI_ISL_536738, EPI_ISL_536739, EPI_ISL_536740, EPI_ISL_536741, EPI_ISL_536742, EPI_ISL_536743, EPI_ISL_536744, EPI_ISL_536745, EPI_ISL_536746, EPI_ISL_536747, EPI_ISL_536748, EPI_ISL_536749, EPI_ISL_536750, EPI_ISL_536752, EPI_ISL_536753, EPI_ISL_536754, EPI_ISL_536755, EPI_ISL_536756, EPI_ISL_536757, EPI_ISL_536758, EPI_ISL_536759, EPI_ISL_536760, EPI_ISL_536761, EPI_ISL_536762, EPI_ISL_536763, EPI_ISL_536764, EPI_ISL_536765, EPI_ISL_536766, EPI_ISL_536767, EPI_ISL_536768, EPI_ISL_536769, EPI_ISL_536772, EPI_ISL_536773, EPI_ISL_536774, EPI_ISL_536775, EPI_ISL_536776, EPI_ISL_536777, EPI_ISL_536778, EPI_ISL_536779, EPI_ISL_536780, EPI_ISL_536781, EPI_ISL_536782, EPI_ISL_536783, EPI_ISL_536784 | | | |
| see above | University of Wisconsin-Madison AIDS Vaccine Research Laboratories | University of Wisconsin-Madison AIDS Vaccine Research Laboratories | Gage Moreno, Katarina Braun, et al. AIDS Vaccine Research Laboratories |
| EPI_ISL_536785 | University of Wisconsin-Madison Campus AIDS Vaccine Research Laboratories | University of Wisconsin-Madison AIDS Vaccine Research Laboratories | Gage Moreno, Katarina Braun, et al. AIDS Vaccine Research Laboratories |

| | | | |
|---|---|---|---|
| EPI_ISL_536786 | University of Wisconsin-Madison AIDS Vaccine Research Laboratories | University of Wisconsin-Madison AIDS Vaccine Research Laboratories | Gage Moreno, Katarina Braun, et al. AIDS Vaccine Research Laboratories |
| EPI_ISL_536789 | Southern Community Labs Dunedin | Institute of Environmental Science and Research (ESR) | Xiaoyun Ren, Matt Storey, Nikki Freed, Muhammad Faisal, Jing Wang, Hermes Perez, Anja Werno, Antje van der Linden, Arlo Upton, Chris Mansell, David Hammer, Dragana Drinkovic, Gary McAuliffe, Hana Sofia Andersson, James Ussher, Jill Sherwood, Josh Freeman, Julia Howard, Juliet Elvy, Mary DeAlmeida, Matt Blakiston, Matthew Rogers, Max Bloomfield, Michael Addidle, Michelle Balm, Sally Roberts, Sarah Jefferies, Sharmini Muttaiyah, Susan Morpeth, Susan Taylor, Timothy Blackmore, Vani Sathyendran, Veronica Playle, Virginia Hope, Erasmus Smit, Lauren Jelly, Olin Silander, Joep de Ligt |
| EPI_ISL_536793 | Medtimes Molecular Laboratory | Medtimes Molecular Laboratory | Eric Chan, Winsome Wong, Jacqueline Tam, Isaac Chow |
| EPI_ISL_536819, EPI_ISL_536820, EPI_ISL_536821, EPI_ISL_536822, EPI_ISL_536825, EPI_ISL_536826, EPI_ISL_536827, EPI_ISL_536828, EPI_ISL_536829, EPI_ISL_536830, EPI_ISL_536831, EPI_ISL_536833, EPI_ISL_536835, EPI_ISL_536837, EPI_ISL_536838, EPI_ISL_536839, EPI_ISL_536840, EPI_ISL_536841, EPI_ISL_536842, EPI_ISL_536843, EPI_ISL_536845, EPI_ISL_536847, EPI_ISL_536848, EPI_ISL_536849, EPI_ISL_536851, EPI_ISL_536852, EPI_ISL_536853, EPI_ISL_536854, EPI_ISL_536855, EPI_ISL_536857, EPI_ISL_536860, EPI_ISL_536861, EPI_ISL_536862, EPI_ISL_536863, EPI_ISL_536864, EPI_ISL_536865, EPI_ISL_536866, EPI_ISL_536868, EPI_ISL_536869, EPI_ISL_536870, EPI_ISL_536871, EPI_ISL_536872, EPI_ISL_536873, EPI_ISL_536874, EPI_ISL_536875, EPI_ISL_536876, EPI_ISL_536877, EPI_ISL_536879, EPI_ISL_536880, EPI_ISL_536881, EPI_ISL_536882, EPI_ISL_536884, EPI_ISL_536885, EPI_ISL_536886, EPI_ISL_536887, EPI_ISL_536888, EPI_ISL_536890, EPI_ISL_536891, EPI_ISL_536892, EPI_ISL_536893, EPI_ISL_536894, EPI_ISL_536895, EPI_ISL_536896, EPI_ISL_536897, EPI_ISL_536898, EPI_ISL_536899, EPI_ISL_536904, EPI_ISL_536905, EPI_ISL_536906, EPI_ISL_536907, EPI_ISL_536908, EPI_ISL_536909, EPI_ISL_536910, EPI_ISL_536911, EPI_ISL_536912, EPI_ISL_536913, EPI_ISL_536914, EPI_ISL_536915, EPI_ISL_536917, EPI_ISL_536918, EPI_ISL_536919, EPI_ISL_536920, EPI_ISL_536921, EPI_ISL_536922, EPI_ISL_536923, EPI_ISL_536924, EPI_ISL_536925, EPI_ISL_536926, EPI_ISL_536927, EPI_ISL_536928, EPI_ISL_536929, EPI_ISL_536931, EPI_ISL_536932, EPI_ISL_536933, EPI_ISL_536934, EPI_ISL_536935, EPI_ISL_536937, EPI_ISL_536938, EPI_ISL_536939, EPI_ISL_536940, EPI_ISL_536943, EPI_ISL_536944, EPI_ISL_536945, EPI_ISL_536947, EPI_ISL_536948, EPI_ISL_536949, EPI_ISL_536950, EPI_ISL_536952, EPI_ISL_536953, EPI_ISL_536954, EPI_ISL_536958, EPI_ISL_536959, EPI_ISL_536960, EPI_ISL_536961, EPI_ISL_536962, EPI_ISL_536963, EPI_ISL_536964, EPI_ISL_536966, EPI_ISL_536967, EPI_ISL_536968, EPI_ISL_536969, EPI_ISL_536970, EPI_ISL_536971, EPI_ISL_536972, EPI_ISL_536973, EPI_ISL_536974, EPI_ISL_536975, EPI_ISL_536976, EPI_ISL_536977, EPI_ISL_536978, EPI_ISL_536979, EPI_ISL_536980, EPI_ISL_536981, EPI_ISL_536982, EPI_ISL_536983, EPI_ISL_536984, EPI_ISL_536985, EPI_ISL_536986, EPI_ISL_536987, EPI_ISL_536988, EPI_ISL_536989, EPI_ISL_536990, EPI_ISL_536991, EPI_ISL_536993, EPI_ISL_536994, EPI_ISL_536996, EPI_ISL_536997, EPI_ISL_536998, EPI_ISL_536999, EPI_ISL_537000, EPI_ISL_537001, EPI_ISL_537002, EPI_ISL_537003, EPI_ISL_537004, EPI_ISL_537005, EPI_ISL_537006, EPI_ISL_537007, EPI_ISL_537008, EPI_ISL_537009, EPI_ISL_537011, EPI_ISL_537012, EPI_ISL_537013, EPI_ISL_537014, EPI_ISL_537015, EPI_ISL_537016, EPI_ISL_537018, EPI_ISL_537019, EPI_ISL_537021, EPI_ISL_537022, EPI_ISL_537024, EPI_ISL_537025, EPI_ISL_537026, EPI_ISL_537027, EPI_ISL_537028, EPI_ISL_537029, EPI_ISL_537030, EPI_ISL_537031, EPI_ISL_537032, EPI_ISL_537033, EPI_ISL_537034, EPI_ISL_537036, EPI_ISL_537037, EPI_ISL_537038, EPI_ISL_537039, EPI_ISL_537042, EPI_ISL_537043, EPI_ISL_537044, EPI_ISL_537048, EPI_ISL_537049, EPI_ISL_537050, EPI_ISL_537051, EPI_ISL_537052, EPI_ISL_537053, EPI_ISL_537055, EPI_ISL_537056, EPI_ISL_537057, EPI_ISL_537058, EPI_ISL_537059, EPI_ISL_537060, EPI_ISL_537062, EPI_ISL_537064, EPI_ISL_537065, EPI_ISL_537066, EPI_ISL_537067, EPI_ISL_537068, EPI_ISL_537069, EPI_ISL_537070, EPI_ISL_537071, EPI_ISL_537072, EPI_ISL_537073, EPI_ISL_537075, EPI_ISL_537076, EPI_ISL_537078, EPI_ISL_537079, EPI_ISL_537080, EPI_ISL_537081, EPI_ISL_537082, EPI_ISL_537083, EPI_ISL_537084, EPI_ISL_537085, EPI_ISL_537086, EPI_ISL_537087, EPI_ISL_537088, EPI_ISL_537089, EPI_ISL_537090, EPI_ISL_537091, EPI_ISL_537092, EPI_ISL_537093, EPI_ISL_537094, EPI_ISL_537095, EPI_ISL_537096, EPI_ISL_537097, EPI_ISL_537098, EPI_ISL_537099, EPI_ISL_537100, EPI_ISL_537101, EPI_ISL_537102, EPI_ISL_537103, EPI_ISL_537104, EPI_ISL_537105, EPI_ISL_537106, EPI_ISL_537107, EPI_ISL_537108, EPI_ISL_537109, EPI_ISL_537110, EPI_ISL_537111, EPI_ISL_537112, EPI_ISL_537113, EPI_ISL_537115, EPI_ISL_537116, EPI_ISL_537117, EPI_ISL_537118, EPI_ISL_537119, EPI_ISL_537120, EPI_ISL_537121, EPI_ISL_537123, EPI_ISL_537124, EPI_ISL_537126, EPI_ISL_537127, EPI_ISL_537128, EPI_ISL_537129, EPI_ISL_537130, EPI_ISL_537131, EPI_ISL_537132, EPI_ISL_537133, EPI_ISL_537134, EPI_ISL_537136, EPI_ISL_537137, EPI_ISL_537138, EPI_ISL_537139, EPI_ISL_537140, EPI_ISL_537141, EPI_ISL_537142, EPI_ISL_537143, EPI_ISL_537144, EPI_ISL_537145, EPI_ISL_537146, EPI_ISL_537148, EPI_ISL_537149, EPI_ISL_537150, EPI_ISL_537151, EPI_ISL_537153, EPI_ISL_537154, EPI_ISL_537156, EPI_ISL_537157, EPI_ISL_537158, EPI_ISL_537159, EPI_ISL_537160, EPI_ISL_537161, EPI_ISL_537162, EPI_ISL_537163, EPI_ISL_537164, EPI_ISL_537165, EPI_ISL_537166, EPI_ISL_537167, EPI_ISL_537168, EPI_ISL_537169, EPI_ISL_537170, EPI_ISL_537171, EPI_ISL_537172, EPI_ISL_537173, EPI_ISL_537174, EPI_ISL_537175, EPI_ISL_537176, EPI_ISL_537177, EPI_ISL_537179, EPI_ISL_537180, EPI_ISL_537181, EPI_ISL_537182, EPI_ISL_537184, EPI_ISL_537185, EPI_ISL_537188, EPI_ISL_537189, EPI_ISL_537190, EPI_ISL_537191, EPI_ISL_537192, EPI_ISL_537193, EPI_ISL_537195, EPI_ISL_537196, EPI_ISL_537198, EPI_ISL_537199, EPI_ISL_537200, EPI_ISL_537201, EPI_ISL_537202, EPI_ISL_537204, EPI_ISL_537205 | | | |
| see above | Lighthouse Lab in Glasgow | Wellcome Sanger Institute for the COVID-19 Genomics UK Consortium | Harper VanSteenhouse, Yumi Kasai, David Gray, Carol Clugston, Anna Dominiczak and Alex Alderton, Roberto Amato, Sonia Goncalves, Ewan Harrison, David K. Jackson, Ian Johnston, Dominic Kwiatkowski, Cordelia Langford, John Sillitoe on behalf of the Wellcome Sanger Institute COVID-19 Surveillance Team |
| EPI_ISL_537206, EPI_ISL_537208, EPI_ISL_537209, EPI_ISL_537212, EPI_ISL_537214, EPI_ISL_537215, EPI_ISL_537216, EPI_ISL_537220, EPI_ISL_537221, EPI_ISL_537223, EPI_ISL_537224, EPI_ISL_537225, EPI_ISL_537226, EPI_ISL_537227, EPI_ISL_537229, EPI_ISL_537230, EPI_ISL_537231, EPI_ISL_537237, EPI_ISL_537238, EPI_ISL_537239, EPI_ISL_537241, EPI_ISL_537243, EPI_ISL_537244, EPI_ISL_537246, EPI_ISL_537248, EPI_ISL_537249, EPI_ISL_537251, EPI_ISL_537252, EPI_ISL_537253, EPI_ISL_537255, EPI_ISL_537256, EPI_ISL_537257, EPI_ISL_537258, EPI_ISL_537259, EPI_ISL_537261, EPI_ISL_537263, EPI_ISL_537264, EPI_ISL_537265, EPI_ISL_537266, EPI_ISL_537267, EPI_ISL_537269, EPI_ISL_537270, EPI_ISL_537272, EPI_ISL_537273, EPI_ISL_537274, EPI_ISL_537275, EPI_ISL_537277, EPI_ISL_537280, EPI_ISL_537282, EPI_ISL_537283, EPI_ISL_537284 | | | |
| see above | Virology Department, Sheffield Teaching Hospitals NHS Foundation Trust / Department of Infection, Immunity and Cardiovascular Disease, The Medical School, University of Sheffield | Wellcome Sanger Institute for the COVID-19 Genomics UK Consortium | Thushan de Silva, Matthew Parker,Adri Angyal, Rebecca Brown, Luke Green, Rachel Tucker, Paul Parsons, Danielle Groves, Alex Keeley, Dave Partridge, Matthew Wyles, Benjamin Lindsey, Mehmet Yavuz, Mohammad Raza, Cariad Evans and Alex Alderton, Roberto Amato, Sonia Goncalves, Ewan Harrison, David K. Jackson, Ian Johnston, Dominic Kwiatkowski, Cordelia Langford, John Sillitoe on behalf of the Wellcome Sanger Institute COVID-19 Surveillance Team |
| EPI_ISL_537286, EPI_ISL_537287 | Area of Virology, Serology and Virology Division (SAViD), New South Wales Health Pathology Randwick | Area of Virology, Serology and Virology Division (SAViD), New South Wales Health Pathology Randwick | Rawlinson, W., Deveson, I., Bull, R., Van Hal, S. |
| EPI_ISL_537288, EPI_ISL_537289, EPI_ISL_537290, EPI_ISL_537291, EPI_ISL_537293, EPI_ISL_537294, EPI_ISL_537295, EPI_ISL_537296, EPI_ISL_537297, EPI_ISL_537298, EPI_ISL_537299, EPI_ISL_537300, EPI_ISL_537301, EPI_ISL_537302, EPI_ISL_537303, EPI_ISL_537304, EPI_ISL_537305, EPI_ISL_537306, EPI_ISL_537307, EPI_ISL_537308, EPI_ISL_537309, EPI_ISL_537310, EPI_ISL_537311, EPI_ISL_537312, EPI_ISL_537313, EPI_ISL_537314, EPI_ISL_537315, EPI_ISL_537316, EPI_ISL_537317, EPI_ISL_537319, EPI_ISL_537320, EPI_ISL_537321, EPI_ISL_537322, EPI_ISL_537324, EPI_ISL_537326, EPI_ISL_537327, EPI_ISL_537330, EPI_ISL_537331, EPI_ISL_537332, EPI_ISL_537334, EPI_ISL_537335, EPI_ISL_537336, EPI_ISL_537337, EPI_ISL_537338, EPI_ISL_537339, EPI_ISL_537340, EPI_ISL_537341, EPI_ISL_537342, EPI_ISL_537343, EPI_ISL_537344, EPI_ISL_537346, EPI_ISL_537347, EPI_ISL_537348, EPI_ISL_537349, EPI_ISL_537350, EPI_ISL_537351, EPI_ISL_537352, EPI_ISL_537353, EPI_ISL_537354, EPI_ISL_537355, EPI_ISL_537357, EPI_ISL_537358, EPI_ISL_537359, EPI_ISL_537360, EPI_ISL_537361, EPI_ISL_537362, EPI_ISL_537364, EPI_ISL_537365, EPI_ISL_537366, EPI_ISL_537367, EPI_ISL_537368, EPI_ISL_537369, EPI_ISL_537370, EPI_ISL_537371, EPI_ISL_537372, EPI_ISL_537373, EPI_ISL_537374, EPI_ISL_537377 | | | |
| see above | Universidad de León | SeqCOVID-SPAIN consortium/IBV(CSIC) | Ana Carvajal, Vicente Martín, Héctor Argüello, Juan M. Fregeneda, Tania Fernández-Villa, Antonio J. Molina and SeqCOVID-SPAIN consortium |
| EPI_ISL_537380, EPI_ISL_537381 | Complejo Hospitalario Universitario de Vigo | SeqCOVID-SPAIN consortium/IBV(CSIC) | Benito Regueiro and SeqCOVID-SPAIN consortium |
| EPI_ISL_537382, EPI_ISL_537384, EPI_ISL_537385, EPI_ISL_537387, EPI_ISL_537390, EPI_ISL_537394, EPI_ISL_537395, EPI_ISL_537397, EPI_ISL_537398, EPI_ISL_537399, EPI_ISL_537403, EPI_ISL_537404, EPI_ISL_537406, EPI_ISL_537410, EPI_ISL_537411, EPI_ISL_537412, EPI_ISL_537414, EPI_ISL_537422, EPI_ISL_537424, EPI_ISL_537426, EPI_ISL_537427, EPI_ISL_537428, EPI_ISL_537429, EPI_ISL_537431, EPI_ISL_537432, EPI_ISL_537434, EPI_ISL_537436, EPI_ISL_537437, EPI_ISL_537438, EPI_ISL_537439, EPI_ISL_537440, EPI_ISL_537443, EPI_ISL_537445, EPI_ISL_537446, EPI_ISL_537447, EPI_ISL_537449, EPI_ISL_537451, EPI_ISL_537453, EPI_ISL_537460, EPI_ISL_537465, EPI_ISL_537466 | | | |
| see above | Centro de Investigación Biomédica de La Rioja - Hospital San Pedro Logroño | SeqCOVID-SPAIN consortium/IBV(CSIC) | María de Toro, José Manuel Azcona Gutiérrez, María Pilar Bea Escudero, Miriam Blasco Alberdi and SeqCOVID-SPAIN consortium |
| EPI_ISL_537467, EPI_ISL_537469, EPI_ISL_537470, EPI_ISL_537471, EPI_ISL_537475, EPI_ISL_537476, EPI_ISL_537477, EPI_ISL_537478, EPI_ISL_537479, EPI_ISL_537480, EPI_ISL_537481, EPI_ISL_537482, EPI_ISL_537483, EPI_ISL_537484, EPI_ISL_537485, EPI_ISL_537486, EPI_ISL_537487, EPI_ISL_537488, EPI_ISL_537489, EPI_ISL_537490, EPI_ISL_537491, EPI_ISL_537492, EPI_ISL_537493, EPI_ISL_537494, EPI_ISL_537495, EPI_ISL_537496, EPI_ISL_537497, EPI_ISL_537498, EPI_ISL_537499, EPI_ISL_537500, EPI_ISL_537501, EPI_ISL_537502, EPI_ISL_537503, EPI_ISL_537504, EPI_ISL_537505, EPI_ISL_537506, EPI_ISL_537507, EPI_ISL_537508, EPI_ISL_537509, EPI_ISL_537510, EPI_ISL_537512, EPI_ISL_537513, EPI_ISL_537514, EPI_ISL_537515, EPI_ISL_537517, EPI_ISL_537523, EPI_ISL_537525, EPI_ISL_537526, EPI_ISL_537527, EPI_ISL_537528, EPI_ISL_537529, EPI_ISL_537530, EPI_ISL_537531, EPI_ISL_537532, EPI_ISL_537533, EPI_ISL_537535, EPI_ISL_537537, EPI_ISL_537538, EPI_ISL_537541, EPI_ISL_537542, EPI_ISL_537545, EPI_ISL_537546, EPI_ISL_537547, EPI_ISL_537551, EPI_ISL_537554, EPI_ISL_537557, EPI_ISL_537558, EPI_ISL_537559, EPI_ISL_537560, EPI_ISL_537561, EPI_ISL_537562, EPI_ISL_537563, EPI_ISL_537564, EPI_ISL_537565, EPI_ISL_537566, EPI_ISL_537567, EPI_ISL_537568, EPI_ISL_537569, EPI_ISL_537570, EPI_ISL_537571, EPI_ISL_537573, EPI_ISL_537577, EPI_ISL_537578, EPI_ISL_537579, EPI_ISL_537582, EPI_ISL_537583, EPI_ISL_537584, EPI_ISL_537585, EPI_ISL_537586, EPI_ISL_537587, EPI_ISL_537590, EPI_ISL_537591, EPI_ISL_537592, EPI_ISL_537593, EPI_ISL_537594, EPI_ISL_537595, EPI_ISL_537596, EPI_ISL_537597, EPI_ISL_537598, EPI_ISL_537599, EPI_ISL_537602, EPI_ISL_537603, EPI_ISL_537604, EPI_ISL_537605, EPI_ISL_537606, EPI_ISL_537607 | | | |
| see above | UCLA Pathology Clinical Microbiology Lab | Kruglyak Lab | Guo et al. |
| EPI_ISL_537614, EPI_ISL_537615, EPI_ISL_537619, EPI_ISL_537621, EPI_ISL_537624, EPI_ISL_537626, EPI_ISL_537631, EPI_ISL_537638, EPI_ISL_537644, EPI_ISL_537645, EPI_ISL_537646, EPI_ISL_537651, EPI_ISL_537652, EPI_ISL_537663, EPI_ISL_537676 | | | |
| see above | Universidad de León | SeqCOVID-SPAIN consortium/IBV(CSIC) | Ana Carvajal, Vicente Martín, Héctor Argüello, Juan M. Fregeneda, Tania Fernández-Villa, Antonio J. Molina and SeqCOVID-SPAIN consortium |
| EPI_ISL_537693, EPI_ISL_537697, EPI_ISL_537700, EPI_ISL_537703, EPI_ISL_537706, EPI_ISL_537708, EPI_ISL_537709, EPI_ISL_537713, EPI_ISL_537715, EPI_ISL_537716, EPI_ISL_537718, EPI_ISL_537720, EPI_ISL_537721, EPI_ISL_537724 | | | |
| see above | Hospital Universitario de Gran Canaria Dr. Negrín | SeqCOVID-SPAIN consortium/IBV(CSIC) | M. Carmen Pérez González, Francisco J. Chamizo López, Ana Bordes Benítez and SeqCOVID-SPAIN consortium |
| EPI_ISL_537809, EPI_ISL_537810 | Hospital Universitario Marqués de Valdecilla (Santander), Servicio de Microbiología | SeqCOVID-SPAIN consortium/IBV(CSIC) | Mónica Gozalo Margüello, María Eliecer Cano García, Jose Manuel Méndez Legaza, Daniel Pablo Marcos, Jesús Rodríguez Lozano, María Siller Ruiz and SeqCOVID-SPAIN consortium |
| EPI_ISL_537811, EPI_ISL_537822, EPI_ISL_537828, EPI_ISL_537829, EPI_ISL_537831, EPI_ISL_537836, EPI_ISL_537837, EPI_ISL_537838, EPI_ISL_537839, EPI_ISL_537840, EPI_ISL_537841, EPI_ISL_537842, EPI_ISL_537844, EPI_ISL_537845, EPI_ISL_537847, EPI_ISL_537849, EPI_ISL_537850, EPI_ISL_537852, EPI_ISL_537853, EPI_ISL_537855, EPI_ISL_537856, EPI_ISL_537857, EPI_ISL_537858, EPI_ISL_537859, EPI_ISL_537863, EPI_ISL_537864, EPI_ISL_537865, EPI_ISL_537866, EPI_ISL_537870, EPI_ISL_537872, EPI_ISL_537873 | | | |
| see above | Centro de Investigación Biomédica de La Rioja - Hospital San Pedro Logroño | SeqCOVID-SPAIN consortium/IBV(CSIC) | María de Toro, José Manuel Azcona Gutiérrez, María Pilar Bea Escudero, Miriam Blasco Alberdi and SeqCOVID-SPAIN consortium |
| EPI_ISL_538012, EPI_ISL_538013, EPI_ISL_538014, EPI_ISL_538015, EPI_ISL_538016, EPI_ISL_538017, EPI_ISL_538018, EPI_ISL_538019, EPI_ISL_538020, EPI_ISL_538021, EPI_ISL_538022, EPI_ISL_538023, EPI_ISL_538024, EPI_ISL_538025, EPI_ISL_538026, EPI_ISL_538027, EPI_ISL_538028, EPI_ISL_538030, EPI_ISL_538033, EPI_ISL_538034, EPI_ISL_538035, EPI_ISL_538036, EPI_ISL_538037, EPI_ISL_538038, EPI_ISL_538039, EPI_ISL_538041, EPI_ISL_538042, EPI_ISL_538043, EPI_ISL_538044, EPI_ISL_538046, EPI_ISL_538047, EPI_ISL_538048, EPI_ISL_538049, EPI_ISL_538050, EPI_ISL_538051, EPI_ISL_538052, EPI_ISL_538054 | | | |
| see above | Hospital Universitari i Politècnic La Fe de València | SeqCOVID-SPAIN consortium/IBV(CSIC) | María Dolores Gómez Ruiz, Eva González Barbera, Ana Gil Brusola, Salvador Giner Almaraz, José Luis López Hontangas and SeqCOVID-SPAIN consortium |
| EPI_ISL_538119, EPI_ISL_538120, EPI_ISL_538121, EPI_ISL_538123, EPI_ISL_538124, EPI_ISL_538125, EPI_ISL_538127, EPI_ISL_538128, EPI_ISL_538129, EPI_ISL_538131, EPI_ISL_538134, EPI_ISL_538135, EPI_ISL_538137, EPI_ISL_538141, EPI_ISL_538147, EPI_ISL_538150, EPI_ISL_538151, EPI_ISL_538152, EPI_ISL_538153, EPI_ISL_538154, EPI_ISL_538156, EPI_ISL_538157, EPI_ISL_538160, EPI_ISL_538161, EPI_ISL_538162, EPI_ISL_538163, EPI_ISL_538165, EPI_ISL_538167 | | | |
| see above | Servicio de Microbiología y Parasitología clínica. UCEIMP. Hospital Universitario Virgen del Rocío/IBIS/CSIC/US | SeqCOVID-SPAIN consortium/IBV(CSIC) | Guillermo Martín Gutiérrez, Ángel Rodríguez Villodres, Lidia Gálvez Benitez, Verónica González Galán, Javier Aznar Martín and SeqCOVID-SPAIN consortium |
| EPI_ISL_538265, EPI_ISL_538266, EPI_ISL_538267, EPI_ISL_538268, EPI_ISL_538269, EPI_ISL_538270, EPI_ISL_538271, EPI_ISL_538272, EPI_ISL_538273, EPI_ISL_538274 | Maryland Public Health Laboratory | Maryland Public Health Laboratory | Maryland Department of Health Laboratories Administration |
| EPI_ISL_538280, EPI_ISL_538307 | Texas Department of State Health Services | Texas Department of State Health Services | Bonnie Oh, Rashmi Tuladhar, Jenny Zhang, Maliha Rahman, Anita Pokharel, Myong Koag, Chun Wang, Rachel Lee, Grace Kubin |
| EPI_ISL_538321, EPI_ISL_538322, EPI_ISL_538323, EPI_ISL_538324, EPI_ISL_538325, EPI_ISL_538326, EPI_ISL_538327, EPI_ISL_538328, EPI_ISL_538329, EPI_ISL_538330, EPI_ISL_538331, EPI_ISL_538332, EPI_ISL_538333, EPI_ISL_538335, EPI_ISL_538336, EPI_ISL_538338, EPI_ISL_538339, EPI_ISL_538340, EPI_ISL_538341, EPI_ISL_538342, EPI_ISL_538343, EPI_ISL_538345, EPI_ISL_538346, EPI_ISL_538347, EPI_ISL_538348, EPI_ISL_538349, EPI_ISL_538350, EPI_ISL_538351, EPI_ISL_538353, EPI_ISL_538355, EPI_ISL_538356, EPI_ISL_538358, EPI_ISL_538359, EPI_ISL_538360, EPI_ISL_538361, EPI_ISL_538362, EPI_ISL_538363, EPI_ISL_538364, EPI_ISL_538365, EPI_ISL_538367, EPI_ISL_538368, EPI_ISL_538369, EPI_ISL_538370, EPI_ISL_538371, EPI_ISL_538372, EPI_ISL_538373, EPI_ISL_538374, EPI_ISL_538375, EPI_ISL_538376, EPI_ISL_538377, EPI_ISL_538378, EPI_ISL_538379, EPI_ISL_538380, EPI_ISL_538381, EPI_ISL_538382, EPI_ISL_538383, EPI_ISL_538384, EPI_ISL_538385, EPI_ISL_538386 | | | |
| see above | Kingston Health Sciences Centre / Queen's University | Ontario Institute for Cancer Research | Prameet M. Sheth, Calvin Sjaarda, Robert Colautti, Katya Douchant, Ilinca Lungu, Bernard Lam, Paul Krzyzanowski, Michael Laszloffy, Lawrence E Heisler, Richard de Borja, Jared T. Simpson |
| EPI_ISL_538390, EPI_ISL_538391, EPI_ISL_538392, EPI_ISL_538393, EPI_ISL_538395, EPI_ISL_538396, EPI_ISL_538397, EPI_ISL_538399, EPI_ISL_538400, EPI_ISL_538401, EPI_ISL_538402, EPI_ISL_538404, EPI_ISL_538406, EPI_ISL_538407, EPI_ISL_538410, EPI_ISL_538411, EPI_ISL_538412, EPI_ISL_538413, EPI_ISL_538415, EPI_ISL_538417, EPI_ISL_538418, EPI_ISL_538420, EPI_ISL_538421, EPI_ISL_538422, EPI_ISL_538423, EPI_ISL_538425, EPI_ISL_538426, EPI_ISL_538427, EPI_ISL_538428, EPI_ISL_538430, EPI_ISL_538431, EPI_ISL_538432, EPI_ISL_538433, EPI_ISL_538434 | | | |
| see above | Microbiology Division, South Carolina Department of Health and Environmental Control | Microbiology Division, South Carolina Department of Health and Environmental Control | Flores,H. |
| EPI_ISL_538435, EPI_ISL_538436, EPI_ISL_538437, EPI_ISL_538438, EPI_ISL_538439, EPI_ISL_538440, EPI_ISL_538441, EPI_ISL_538442, EPI_ISL_538443, EPI_ISL_538444, EPI_ISL_538445, EPI_ISL_538446, EPI_ISL_538447, EPI_ISL_538448, EPI_ISL_538449, EPI_ISL_538450, EPI_ISL_538451, EPI_ISL_538452, EPI_ISL_538453, EPI_ISL_538454, EPI_ISL_538455, EPI_ISL_538456, EPI_ISL_538457, EPI_ISL_538458, EPI_ISL_538459, EPI_ISL_538460, EPI_ISL_538461, EPI_ISL_538462, EPI_ISL_538463, EPI_ISL_538464, EPI_ISL_538465, EPI_ISL_538466, EPI_ISL_538467, EPI_ISL_538468, EPI_ISL_538469, EPI_ISL_538470, EPI_ISL_538471, EPI_ISL_538472, EPI_ISL_538473, EPI_ISL_538474, EPI_ISL_538475, EPI_ISL_538476, EPI_ISL_538477, EPI_ISL_538478, EPI_ISL_538479, EPI_ISL_538480, EPI_ISL_538481, EPI_ISL_538482, EPI_ISL_538483, EPI_ISL_538484, EPI_ISL_538485, EPI_ISL_538486, EPI_ISL_538487, EPI_ISL_538488, EPI_ISL_538489, EPI_ISL_538490, EPI_ISL_538491, EPI_ISL_538492, EPI_ISL_538493, EPI_ISL_538494, EPI_ISL_538495, EPI_ISL_538496, EPI_ISL_538497 | | | |
| see above | Department of Laboratory Medicine, Tan Tock Seng Hospital | Department of Laboratory Medicine, Tan Tock Seng Hospital | Chen YYC, Zair X, Lim JX, Li C, Tang WY, Maurer-Stroh S, Barkham TMS, Nagarajan N, Sessions OM |
| EPI_ISL_538498 | RSUD Soediono Madiun East Java | National Institute of Health Research and Development | Pawestri, HA; Subangkit; Puspa, KD; Nugraha, AA; Ikawati, HD; Pangesti, KNA; Soekarso, T; Paisal; Setiawaty,V. |
| EPI_ISL_538499 | RS Lavallete Malang East Java | National Institute of Health Research and Development | Pawestri, HA; Subangkit; Puspa, KD; Nugraha, AA; Ikawati, HD; Pangesti, KNA; Soekarso, T; Paisal; Setiawaty,V. |
| EPI_ISL_538500, EPI_ISL_538501 | RS Prima Husada Cipta Medan North Sumatra | National Institute of Health Research and Development | Pawestri, HA; Subangkit; Puspa, KD; Nugraha, AA; Ikawati, HD; Pangesti, KNA; Soekarso, T; Paisal; Setiawaty,V. |
| EPI_ISL_538502 | RSUD Sultan Imanudin Pangkalan Bun Center Kalimantan | National Institute of Health Research and Development | Pawestri, HA; Subangkit; Puspa, KD; Nugraha, AA; Ikawati, HD; Pangesti, KNA; Soekarso, T; Paisal; Setiawaty,V. |

| | | | |
|---|---|---|---|
| EPI_ISL_538503 | RSUD Bali Mandara Denpasar Bali | National Institute of Health Research and Development | Pawestri, HA; Subangkit; Puspa, KD; Nugraha, AA; Ikawati, HD; Pangesti, KNA; Soekarso, T; Paisal; Setiawaty,V. |
| EPI_ISL_538504, EPI_ISL_538505 | National Institute of Health Research and Development | National Institute of Health Research and Development | Pawestri, HA; Subangkit; Puspa, KD; Nugraha, AA; Ikawati, HD; Pangesti, KNA; Soekarso, T; Susilarini, NK; Hariastuti, NI; Nikmah, UA; Mursinah; Febriyani, A; Herman, R; Susanti, N; Herna; Febriyanti, T; Nurhadi, M; Paisal; Ramadhany, R; Agustiningsih; Kurniawati, J; Kipuw, NL; Muna, F; Indalau, IL; Adam, K; Wibowo, HA; Rizki, A; Puspandary, N; Setiawaty,V. |
| EPI_ISL_538506, EPI_ISL_538507 | Balai Penelitian dan Pengembangan Biomedis Papua | National Institute of Health Research and Development | Pawestri, HA; Subangkit; Puspa, KD; Nugraha, AA; Ikawati, HD; Pangesti, KNA; Soekarso, T; Paisal; Oktavian, A; Hutapea, HML; Setiawaty,V. |
| EPI_ISL_538508, EPI_ISL_538509 | National Institute of Health Research and Development | National Institute of Health Research and Development | Pawestri, HA; Subangkit; Puspa, KD; Nugraha, AA; Ikawati, HD; Pangesti, KNA; Soekarso, T; Susilarini, NK; Hariastuti, NI; Nikmah, UA; Mursinah; Febriyani, A; Herman, R; Susanti, N; Herna; Febriyanti, T; Nurhadi, M; Paisal; Ramadhany, R; Agustiningsih; Kurniawati, J; Kipuw, NL; Muna, F; Indalau, IL; Adam, K; Wibowo, HA; Rizki, A; Puspandary, N; Setiawaty,V. |
| EPI_ISL_538510 | RSUD Ulin Banjarmasin South Kalimantan | National Institute of Health Research and Development | Pawestri, HA; Subangkit; Puspa, KD; Nugraha, AA; Ikawati, HD; Pangesti, KNA; Soekarso, T; Paisal; Pasaribu, M; Setiawaty,V. |
| EPI_ISL_538511 | RSUD Wahidin Sudirohusodo Mojokerto East Java | National Institute of Health Research and Development | Pawestri, HA; Subangkit; Puspa, KD; Nugraha, AA; Ikawati, HD; Pangesti, KNA; Soekarso, T; Paisal; Setiawaty,V. |
| EPI_ISL_538512 | Balai Penelitian dan Pengembangan Biomedis Papua | National Institute of Health Research and Development | Pawestri, HA; Subangkit; Puspa, KD; Nugraha, AA; Ikawati, HD; Pangesti, KNA; Soekarso, T; Paisal; Pasaribu, M; Setiawaty,V. |
| EPI_ISL_538513 | Provincial Health Laboratory Bekasi West Java | National Institute of Health Research and Development | Pawestri, HA; Subangkit; Puspa, KD; Nugraha, AA; Ikawati, HD; Pangesti, KNA; Soekarso, T; Paisal; Setiawaty,V. |
| EPI_ISL_538514, EPI_ISL_538515, EPI_ISL_538516, EPI_ISL_538517, EPI_ISL_538519, EPI_ISL_538520, EPI_ISL_538521 | Infectious Diseases, North Carolina State Laboratory of Public Health COVID-19 Response Team | Infectious Diseases, North Carolina State Laboratory of Public Health COVID-19 Response Team | Chase,K. |
| EPI_ISL_538522, EPI_ISL_538523 | Infectious Diseases, North Carolina State Laboratory of Public Health COVID-19 Response Team | North Carolina State Laboratory of Public Health | Chase,K. |
| EPI_ISL_538552 | Hospital Universitari Germans Trias i Pujol(HUGTiP)/Fundació Lluita contra la SIDA (FLSida) | IrsiCaixa AIDS Research Lab | Marc Noguera-Julian, Mariona Parera, Maria Pilar Armengol, Marta Massanella, Ester Ballana, Lidia Ruiz, Nuria Izquierdo, Jorge Carrillo, Roger Paredes, Julia Blanco, Joaquim Segalés, Bonaventura Clotet |
| EPI_ISL_538748, EPI_ISL_538749, EPI_ISL_538750, EPI_ISL_538751, EPI_ISL_538752, EPI_ISL_538753, EPI_ISL_538755, EPI_ISL_538756, EPI_ISL_538757, EPI_ISL_538758, EPI_ISL_538759, EPI_ISL_538760, EPI_ISL_538761, EPI_ISL_538762, EPI_ISL_538764, EPI_ISL_538765, EPI_ISL_538766, EPI_ISL_538767, EPI_ISL_538768, EPI_ISL_538769, EPI_ISL_538770, EPI_ISL_538772, EPI_ISL_538773, EPI_ISL_538774, EPI_ISL_538776, EPI_ISL_538777, EPI_ISL_538778, EPI_ISL_538779, EPI_ISL_538780, EPI_ISL_538781, EPI_ISL_538782, EPI_ISL_538783, EPI_ISL_538784, EPI_ISL_538786, EPI_ISL_538787, EPI_ISL_538788, EPI_ISL_538789, EPI_ISL_538790, EPI_ISL_538791, EPI_ISL_538792, EPI_ISL_538793, EPI_ISL_538794, EPI_ISL_538795, EPI_ISL_538796, EPI_ISL_538797, EPI_ISL_538798, EPI_ISL_538799, EPI_ISL_538800, EPI_ISL_538802, EPI_ISL_538803, EPI_ISL_538804, EPI_ISL_538805, EPI_ISL_538806, EPI_ISL_538807, EPI_ISL_538808, EPI_ISL_538809, EPI_ISL_538810, EPI_ISL_538812, EPI_ISL_538813, EPI_ISL_538814, EPI_ISL_538815, EPI_ISL_538816, EPI_ISL_538817, EPI_ISL_538819, EPI_ISL_538820, EPI_ISL_538821, EPI_ISL_538822, EPI_ISL_538823, EPI_ISL_538824, EPI_ISL_538825, EPI_ISL_538826, EPI_ISL_538827, EPI_ISL_538828, EPI_ISL_538829, EPI_ISL_538830, EPI_ISL_538831, EPI_ISL_538832, EPI_ISL_538833, EPI_ISL_538834, EPI_ISL_538835, EPI_ISL_538836, EPI_ISL_538837, EPI_ISL_538838, EPI_ISL_538839, EPI_ISL_538840, EPI_ISL_538842, EPI_ISL_538843, EPI_ISL_538844, EPI_ISL_538845, EPI_ISL_538846, EPI_ISL_538847, EPI_ISL_538848, EPI_ISL_538849, EPI_ISL_538852, EPI_ISL_538853, EPI_ISL_538854, EPI_ISL_538855, EPI_ISL_538856, EPI_ISL_538857, EPI_ISL_538858, EPI_ISL_538859, EPI_ISL_538860, EPI_ISL_538861, EPI_ISL_538862, EPI_ISL_538864, EPI_ISL_538865, EPI_ISL_538867, EPI_ISL_538868, EPI_ISL_538869, EPI_ISL_538870, EPI_ISL_538871, EPI_ISL_538872, EPI_ISL_538874, EPI_ISL_538877, EPI_ISL_538878, EPI_ISL_538879, EPI_ISL_538880, EPI_ISL_538881, EPI_ISL_538882, EPI_ISL_538883, EPI_ISL_538885, EPI_ISL_538886, EPI_ISL_538887, EPI_ISL_538888, EPI_ISL_538889, EPI_ISL_538890, EPI_ISL_538891, EPI_ISL_538892, EPI_ISL_538893, EPI_ISL_538894, EPI_ISL_538895, EPI_ISL_538896, EPI_ISL_538897, EPI_ISL_538900, EPI_ISL_538901, EPI_ISL_538902, EPI_ISL_538903, EPI_ISL_538904, EPI_ISL_538905, EPI_ISL_538906, EPI_ISL_538907, EPI_ISL_538908, EPI_ISL_538909, EPI_ISL_538910, EPI_ISL_538911, EPI_ISL_538912, EPI_ISL_538913, EPI_ISL_538914, EPI_ISL_538915, EPI_ISL_538916, EPI_ISL_538918, EPI_ISL_538919, EPI_ISL_538920, EPI_ISL_538921, EPI_ISL_538922, EPI_ISL_538923, EPI_ISL_538924, EPI_ISL_538925, EPI_ISL_538926, EPI_ISL_538928, EPI_ISL_538929, EPI_ISL_538930, EPI_ISL_538931, EPI_ISL_538932, EPI_ISL_538933, EPI_ISL_538934, EPI_ISL_538935, EPI_ISL_538936, EPI_ISL_538937, EPI_ISL_538938, EPI_ISL_538939, EPI_ISL_538940, EPI_ISL_538941, EPI_ISL_538942, EPI_ISL_538943, EPI_ISL_538944, EPI_ISL_538946, EPI_ISL_538947, EPI_ISL_538948, EPI_ISL_538949, EPI_ISL_538950, EPI_ISL_538951, EPI_ISL_538952, EPI_ISL_538953, EPI_ISL_538954, EPI_ISL_538955, EPI_ISL_538956, EPI_ISL_538957, EPI_ISL_538958, EPI_ISL_538959, EPI_ISL_538960, EPI_ISL_538961, EPI_ISL_538962, EPI_ISL_538963, EPI_ISL_538965, EPI_ISL_538966, EPI_ISL_538967, EPI_ISL_538968, EPI_ISL_538969, EPI_ISL_538970, EPI_ISL_538971, EPI_ISL_538972, EPI_ISL_538973, EPI_ISL_538974, EPI_ISL_538975, EPI_ISL_538976, EPI_ISL_538977, EPI_ISL_538978, EPI_ISL_538979, EPI_ISL_538980, EPI_ISL_538981, EPI_ISL_538982, EPI_ISL_538983, EPI_ISL_538984, EPI_ISL_538985, EPI_ISL_538986, EPI_ISL_538988, EPI_ISL_538989, EPI_ISL_538990, EPI_ISL_538991, EPI_ISL_538992, EPI_ISL_538993, EPI_ISL_538995, EPI_ISL_538996, EPI_ISL_538998, EPI_ISL_538999, EPI_ISL_539000, EPI_ISL_539001, EPI_ISL_539002, EPI_ISL_539003, EPI_ISL_539004, EPI_ISL_539007, EPI_ISL_539008, EPI_ISL_539010, EPI_ISL_539013, EPI_ISL_539014, EPI_ISL_539015, EPI_ISL_539016, EPI_ISL_539017, EPI_ISL_539018, EPI_ISL_539020, EPI_ISL_539021, EPI_ISL_539022, EPI_ISL_539023, EPI_ISL_539024, EPI_ISL_539026, EPI_ISL_539027, EPI_ISL_539028, EPI_ISL_539029, EPI_ISL_539031, EPI_ISL_539032, EPI_ISL_539033, EPI_ISL_539034, EPI_ISL_539036, EPI_ISL_539037, EPI_ISL_539038, EPI_ISL_539039, EPI_ISL_539040, EPI_ISL_539041, EPI_ISL_539043, EPI_ISL_539044, EPI_ISL_539045, EPI_ISL_539047, EPI_ISL_539051, EPI_ISL_539053, EPI_ISL_539054, EPI_ISL_539055, EPI_ISL_539058, EPI_ISL_539061, EPI_ISL_539063, EPI_ISL_539064, EPI_ISL_539065, EPI_ISL_539067, EPI_ISL_539068, EPI_ISL_539069, EPI_ISL_539071, EPI_ISL_539072, EPI_ISL_539073, EPI_ISL_539074, EPI_ISL_539075, EPI_ISL_539076, EPI_ISL_539077, EPI_ISL_539078, EPI_ISL_539079, EPI_ISL_539080, EPI_ISL_539081, EPI_ISL_539082, EPI_ISL_539083, EPI_ISL_539084, EPI_ISL_539085, EPI_ISL_539086, EPI_ISL_539087, EPI_ISL_539089, EPI_ISL_539090, EPI_ISL_539091, EPI_ISL_539092, EPI_ISL_539093, EPI_ISL_539096, EPI_ISL_539097, EPI_ISL_539099, EPI_ISL_539100, EPI_ISL_539102, EPI_ISL_539103, EPI_ISL_539104, EPI_ISL_539106, EPI_ISL_539107, EPI_ISL_539108, EPI_ISL_539109, EPI_ISL_539110, EPI_ISL_539111, EPI_ISL_539113, EPI_ISL_539114, EPI_ISL_539115, EPI_ISL_539118, EPI_ISL_539119, EPI_ISL_539121, EPI_ISL_539122, EPI_ISL_539123, EPI_ISL_539124, EPI_ISL_539126, EPI_ISL_539127, EPI_ISL_539128, EPI_ISL_539129, EPI_ISL_539130, EPI_ISL_539131, EPI_ISL_539132, EPI_ISL_539133, EPI_ISL_539134, EPI_ISL_539135, EPI_ISL_539137, EPI_ISL_539138, EPI_ISL_539142, EPI_ISL_539144, EPI_ISL_539145, EPI_ISL_539146, EPI_ISL_539147, EPI_ISL_539148, EPI_ISL_539153, EPI_ISL_539156, EPI_ISL_539158, EPI_ISL_539159, EPI_ISL_539160, EPI_ISL_539162, EPI_ISL_539163, EPI_ISL_539165, EPI_ISL_539166, EPI_ISL_539167, EPI_ISL_539169, EPI_ISL_539170, EPI_ISL_539171, EPI_ISL_539172, EPI_ISL_539173, EPI_ISL_539174, EPI_ISL_539175, EPI_ISL_539176, EPI_ISL_539178, EPI_ISL_539179, EPI_ISL_539180, EPI_ISL_539183, EPI_ISL_539184, EPI_ISL_539186, EPI_ISL_539187, EPI_ISL_539193, EPI_ISL_539194, EPI_ISL_539195, EPI_ISL_539196, EPI_ISL_539198, EPI_ISL_539199, EPI_ISL_539200, EPI_ISL_539202, EPI_ISL_539203, EPI_ISL_539205, EPI_ISL_539206, EPI_ISL_539207, EPI_ISL_539208, EPI_ISL_539209, EPI_ISL_539211, EPI_ISL_539212, EPI_ISL_539214, EPI_ISL_539215, EPI_ISL_539216, EPI_ISL_539217, EPI_ISL_539218, EPI_ISL_539219 | | | |
| see above | Leeds Teaching Hospitals NHS Trust and Public Health England, National Infection Service (Leeds laboratory) | Wellcome Sanger Institute for the COVID-19 Genomics UK Consortium | Louissa Macfarlane-Smith, Holli Carden, Katherine L. Harper, Antony Hale and Alex Alderton, Roberto Amato, Sonia Goncalves, Ewan Harrison, David K. Jackson, Ian Johnston, Dominic Kwiatkowski, Cordelia Langford, John Sillitoe on behalf of the Wellcome Sanger Institute COVID-19 Surveillance Team |
| EPI_ISL_539251, EPI_ISL_539252, EPI_ISL_539253, EPI_ISL_539254, EPI_ISL_539255, EPI_ISL_539256, EPI_ISL_539257, EPI_ISL_539258, EPI_ISL_539281, EPI_ISL_539282, EPI_ISL_539283 | | | |
| see above | Hospital Universitario de La Ribera (Alzira, València) | SeqCOVID-SPAIN consortium/IBV(CSIC) | Olalla Martínez Macias, Julia González and SeqCOVID-SPAIN consortium |
| EPI_ISL_539284, EPI_ISL_539285, EPI_ISL_539286, EPI_ISL_539287 | Servicio de Microbiología. Hospital General Universitario de Castellón | SeqCOVID-SPAIN consortium/IBV(CSIC) | Rosario Moreno, María Dolores Tirado and SeqCOVID-SPAIN consortium |
| EPI_ISL_539327 | Area of Virology, Serology and Virology Division (SAViD), New South Wales Health Pathology Randwick | Area of Virology, Serology and Virology Division (SAViD), New South Wales Health Pathology Randwick | Rawlinson, W., Bull, R., Deveson, I., Van Hal, S. |
| EPI_ISL_539333, EPI_ISL_539334, EPI_ISL_539335, EPI_ISL_539336, EPI_ISL_539337, EPI_ISL_539338, EPI_ISL_539339 | Institute of Disease Control and Prevention, People's Liberation Army | Institute of Disease Control and Prevention, People's Liberation Army | Qiu,S., Li,P. |
| EPI_ISL_539341, EPI_ISL_539342, EPI_ISL_539343, EPI_ISL_539344, EPI_ISL_539345, EPI_ISL_539346, EPI_ISL_539347, EPI_ISL_539348, EPI_ISL_539349, EPI_ISL_539350, EPI_ISL_539351, EPI_ISL_539353, EPI_ISL_539354, EPI_ISL_539356, EPI_ISL_539357, EPI_ISL_539358, EPI_ISL_539359, EPI_ISL_539360, EPI_ISL_539361, EPI_ISL_539362, EPI_ISL_539364, EPI_ISL_539365, EPI_ISL_539366, EPI_ISL_539367, EPI_ISL_539368, EPI_ISL_539369, EPI_ISL_539370, EPI_ISL_539372, EPI_ISL_539373, EPI_ISL_539375, EPI_ISL_539376, EPI_ISL_539377, EPI_ISL_539378, EPI_ISL_539379, EPI_ISL_539381, EPI_ISL_539382, EPI_ISL_539383, EPI_ISL_539384, EPI_ISL_539386, EPI_ISL_539387, EPI_ISL_539389, EPI_ISL_539392, EPI_ISL_539393, EPI_ISL_539397, EPI_ISL_539398, EPI_ISL_539399, EPI_ISL_539400, EPI_ISL_539401, EPI_ISL_539402, EPI_ISL_539403, EPI_ISL_539405, EPI_ISL_539406, EPI_ISL_539410, EPI_ISL_539411, EPI_ISL_539412, EPI_ISL_539415, EPI_ISL_539417, EPI_ISL_539418, EPI_ISL_539419, EPI_ISL_539421, EPI_ISL_539422, EPI_ISL_539423, EPI_ISL_539425, EPI_ISL_539426, EPI_ISL_539429, EPI_ISL_539430, EPI_ISL_539432, EPI_ISL_539433, EPI_ISL_539435, EPI_ISL_539436, EPI_ISL_539437, EPI_ISL_539438, EPI_ISL_539439, EPI_ISL_539440, EPI_ISL_539441, EPI_ISL_539442, EPI_ISL_539443, EPI_ISL_539444, EPI_ISL_539445, EPI_ISL_539446, EPI_ISL_539447, EPI_ISL_539448, EPI_ISL_539450, EPI_ISL_539452, EPI_ISL_539454, EPI_ISL_539455, EPI_ISL_539457, EPI_ISL_539458, EPI_ISL_539459, EPI_ISL_539460, EPI_ISL_539461, EPI_ISL_539462, EPI_ISL_539464, EPI_ISL_539465, EPI_ISL_539467, EPI_ISL_539468, EPI_ISL_539471, EPI_ISL_539472, EPI_ISL_539473, EPI_ISL_539474, EPI_ISL_539476, EPI_ISL_539477, EPI_ISL_539478, EPI_ISL_539480, EPI_ISL_539481, EPI_ISL_539482 | | | |
| see above | Viollier AG | Department of Biosystems Science and Engineering, ETH Zürich | Christian Beisel, Sarah Nadeau, Ivan Topolsky, Pedro Ferreira, Philipp Jablonski, Susana Posada-Céspedes, Tobias Schär, Ina Nissen, Natascha Santacroce, Elodie Burcklen, Christiane Beckmann, Maurice Redondo, Olivier Kobel, Christoph Noppen, Sophie Seidel, Noemie Santamaria de Souza, Niko Beerenwinkel, Tanja Stadler |
| EPI_ISL_539483, EPI_ISL_539486, EPI_ISL_539487 | Civil Hospital, Rupnagar | CSIR-Institute of Microbial Technology | Kanika Bansal, Sanjeet Kumar, Anu Singh, Debarghya Ghose, Amandeep Kaur, Rajesh Kumar Mishra, Poushali Chakraborty, Harsh Goar, Navin Baid, Ashwani Kumar, Dipak Dutta, Sanjeev Khosla, Prabhu B. Patil |
| EPI_ISL_539491 | IDSP unit, Dehradun | CSIR-Institute of Microbial Technology | Kanika Bansal, Sanjeet Kumar, Anu Singh, Debarghya Ghose, Amandeep Kaur, Rajesh Kumar Mishra, Poushali Chakraborty, Harsh Goar, Navin Baid, Ashwani Kumar, Dipak Dutta, Sanjeev Khosla, Prabhu B. Patil |
| EPI_ISL_539493, EPI_ISL_539494 | The National Institute of Public Health | State Veterinary Institute Prague | Nagy,A;Jirincova,H;Novakova,L;Trnka,D;Vecerova,J |
| EPI_ISL_539495 | Centers for Disease Control and Prevention, Dengue Branch | Centers for Disease Control and Prevention, Dengue Branch | Gilberto A. Santiago, Glenda Gonzalez, Betzabel Flores, Keyla Charriez, Jorge L. Munoz-Jordan, Gabriela Paz-Bailey, Janice Perez, Vanessa Rivera-Amill, Diego Sainz de la Peña, Jorge Bertran |
| EPI_ISL_539496 | Hospital Nostra Senyora de Meritxell | Instituto de Salud Carlos III | Iglesias-Caballero, M. Molinero Calamita, M. González-Esguevillas, M. Camarero, S. Pozo, F. Casas, I. Jiménez, P. Jiménez, M. Zaballos, A. Monzón, S. Varona, S. Juliá, M. Cuesta, I, F. Fernández |
| EPI_ISL_539497 | Hospital Virgen de las Nieves | Instituto de Salud Carlos III | Iglesias-Caballero, M. Molinero Calamita, M. González-Esguevillas, M. Camarero, S. Pozo, F. Casas, I. Jiménez, P. Jiménez, M. Zaballos, A. Monzón, S. Varona, S. Juliá, M. Cuesta, I, J. Lepe |
| EPI_ISL_539500, EPI_ISL_539501, EPI_ISL_539502 | Hospital Clínico Universitario Lozano Blesa | Instituto de Salud Carlos III | Iglesias-Caballero, M. Molinero Calamita, M. González-Esguevillas, M. Camarero, S. Pozo, F. Casas, I. Jiménez, P. Jiménez, M. Zaballos, A. Monzón, S. Varona, S. Juliá, M. Cuesta, I, R. Benito |
| EPI_ISL_539503 | Hospital Universitario Miguel Servet | Instituto de Salud Carlos III | Iglesias-Caballero, M. Molinero Calamita, M. González-Esguevillas, M. Camarero, S. Pozo, F. Casas, I. Jiménez, P. Jiménez, M. Zaballos, A. Monzón, S. Varona, S. Juliá, M. Cuesta, I, A. Rezusta |
| EPI_ISL_539504 | Hospital Universitario Miguel Servet | Instituto de Salud Carlos III | Iglesias-Caballero, M. Molinero Calamita, M. González-Esguevillas, M. Camarero, S. Pozo, F. Casas, I. Jiménez, P. Jiménez, M. Zaballos, A. Monzón, S. Varona, S. Juliá, M. Cuesta, I, R. Benito |
| EPI_ISL_539505, EPI_ISL_539506, EPI_ISL_539507, EPI_ISL_539508 | Hospital Universitario Miguel Servet | Instituto de Salud Carlos III | Iglesias-Caballero, M. Molinero Calamita, M. González-Esguevillas, M. Camarero, S. Pozo, F. Casas, I. Jiménez, P. Jiménez, M. Zaballos, A. Monzón, S. Varona, S. Juliá, M. Cuesta, I, A. Rezusta |
| EPI_ISL_539509, EPI_ISL_539510, EPI_ISL_539511, EPI_ISL_539512 | Hospital Clínico Universitario Lozano Blesa | Instituto de Salud Carlos III | Iglesias-Caballero, M. Molinero Calamita, M. González-Esguevillas, M. Camarero, S. Pozo, F. Casas, I. Jiménez, P. Jiménez, M. Zaballos, A. Monzón, S. Varona, S. Juliá, M. Cuesta, I, R. Benito |
| EPI_ISL_539513, EPI_ISL_539514, EPI_ISL_539515, EPI_ISL_539516, EPI_ISL_539517, EPI_ISL_539518, EPI_ISL_539519 | Hospital Universitario Miguel Servet | Instituto de Salud Carlos III | Iglesias-Caballero, M. Molinero Calamita, M. González-Esguevillas, M. Camarero, S. Pozo, F. Casas, I. Jiménez, P. Jiménez, M. Zaballos, A. Monzón, S. Varona, S. Juliá, M. Cuesta, I, A. Rezusta |
| EPI_ISL_539520, EPI_ISL_539521 | Hospital Universitario de Ceuta | Instituto de Salud Carlos III | Iglesias-Caballero, M. Molinero Calamita, M. González-Esguevillas, M. Camarero, S. Pozo, F. Casas, I. Jiménez, P. Jiménez, M. Zaballos, A. Monzón, S. Varona, S. Juliá, M. Cuesta, I, J. López |
| EPI_ISL_539522 | Hospital Universitario de Ceuta | Instituto de Salud Carlos III | Iglesias-Caballero, M. Molinero Calamita, M. González-Esguevillas, M. Camarero, S. Pozo, F. Casas, I. Jiménez, P. Jiménez, M. Zaballos, A. Monzón, S. Varona, S. Juliá, M. Cuesta, I, G. Sánchez |
| EPI_ISL_539523 | Hospital General de Segovia | Instituto de Salud Carlos III | Iglesias-Caballero, M. Molinero Calamita, M. González-Esguevillas, M. Camarero, S. Pozo, F. Casas, I. Jiménez, P. Jiménez, M. Zaballos, A. Monzón, S. Varona, S. Juliá, M. Cuesta, I, S. Hernando |
| EPI_ISL_539524 | Gerencia de Asistencia Sanitaria de Soria | Instituto de Salud Carlos III | Iglesias-Caballero, M. Molinero Calamita, M. González-Esguevillas, M. Camarero, S. Pozo, F. Casas, I. Jiménez, P. Jiménez, M. Zaballos, A. Monzón, S. Varona, S. Juliá, M. Cuesta, I, C. Aldea |
| EPI_ISL_539525 | Hospital Nuestra Señora de Sonsoles | Instituto de Salud Carlos III | Iglesias-Caballero, M. Molinero Calamita, M. González-Esguevillas, M. Camarero, S. Pozo, F. Casas, I. Jiménez, P. Jiménez, M. Zaballos, A. Monzón, S. Varona, S. Juliá, M. Cuesta, I, A. San Pedro |
| EPI_ISL_539526, EPI_ISL_539527, EPI_ISL_539528, EPI_ISL_539529, EPI_ISL_539530 | Consejeria de Sanidad y Asuntos Sociales | Instituto de Salud Carlos III | Iglesias-Caballero, M. Molinero Calamita, M. González-Esguevillas, M. Camarero, S. Pozo, F. Casas, I. Jiménez, P. Jiménez, M. Zaballos, A. Monzón, S. Varona, S. Juliá, M. Cuesta, I, G. Gutiérrez |
| EPI_ISL_539531 | C.H.U Nuestra Señora de Candelaria | Instituto de Salud Carlos III | Iglesias-Caballero, M. Molinero Calamita, M. González-Esguevillas, M. Camarero, S. Pozo, F. Casas, I. Jiménez, P. Jiménez, M. Zaballos, A. Monzón, S. Varona, S. Juliá, M. Cuesta, I, O. Díez |
| EPI_ISL_539532 | Hospital Universitario de Canarias | Instituto de Salud Carlos III | Iglesias-Caballero, M. Molinero Calamita, M. González-Esguevillas, M. Camarero, S. Pozo, F. Casas, I. Jiménez, P. Jiménez, M. Zaballos, A. Monzón, S. Varona, S. Juliá, M. Cuesta, I, B. Castro |
| EPI_ISL_539534, EPI_ISL_539535, EPI_ISL_539536, EPI_ISL_539537, EPI_ISL_539538, EPI_ISL_539539, EPI_ISL_539540, EPI_ISL_539541, EPI_ISL_539542, EPI_ISL_539543, EPI_ISL_539544, EPI_ISL_539545, EPI_ISL_539546, EPI_ISL_539547, EPI_ISL_539548, EPI_ISL_539549, EPI_ISL_539550, EPI_ISL_539551, EPI_ISL_539552, EPI_ISL_539553, EPI_ISL_539554, EPI_ISL_539555, EPI_ISL_539556 | | | |
| see above | Hospital Clínic | Instituto de Salud Carlos III | Iglesias-Caballero, M. Molinero Calamita, M. González-Esguevillas, M. Camarero, S. Pozo, F. Casas, I. Jiménez, P. Jiménez, M. Zaballos, A. Monzón, S. Varona, S. Juliá, M. Cuesta, I, M.A Marcos |
| EPI_ISL_539557 | Gerencia del área de salud de Badajoz, Llerena y Zafra | Instituto de Salud Carlos III | Iglesias-Caballero, M. Molinero Calamita, M. González-Esguevillas, M. Camarero, S. Pozo, F. Casas, I. Jiménez, P. Jiménez, M. Zaballos, A. Monzón, S. Varona, S. Juliá, M. Cuesta, I, C. Pazos |
| EPI_ISL_539558 | Hospital San Pedro de Alcántara | Instituto de Salud Carlos III | Iglesias-Caballero, M. Molinero Calamita, M. González-Esguevillas, M. Camarero, S. Pozo, F. Casas, I. Jiménez, P. Jiménez, M. Zaballos, A. Monzón, S. Varona, S. Juliá, M. Cuesta, I, E. Cerro |
| EPI_ISL_539559 | Hospital Campo Arañuelo | Instituto de Salud Carlos III | Iglesias-Caballero, M. Molinero Calamita, M. González-Esguevillas, M. Camarero, S. Pozo, F. Casas, I. Jiménez, P. Jiménez, M. Zaballos, A. Monzón, S. Varona, S. Juliá, M. Cuesta, I, J. López |
| EPI_ISL_539560 | Hospital Campo Arañuelo | Instituto de Salud Carlos III | Iglesias-Caballero, M. Molinero Calamita, M. González-Esguevillas, M. Camarero, S. Pozo, F. Casas, I. Jiménez, P. Jiménez, M. Zaballos, A. Monzón, S. Varona, S. Juliá, M. Cuesta, I, G. Rodríguez |

| | | | |
|---|---|---|---|
| EPI_ISL_539561 | Hospital San Pedro de Alcántara | Instituto de Salud Carlos III | Iglesias-Caballero, M. Molinero Calamita, M. González-Esguevillas, M. Camarero, S. Pozo, F. Casas, I. Jiménez, P. Jiménez, M. Zaballos, A. Monzón, S. Varona, S. Juliá, M. Cuesta, I, J. López |
| EPI_ISL_539562 | Hospital San Pedro de Alcántara | Instituto de Salud Carlos III | Iglesias-Caballero, M. Molinero Calamita, M. González-Esguevillas, M. Camarero, S. Pozo, F. Casas, I. Jiménez, P. Jiménez, M. Zaballos, A. Monzón, S. Varona, S. Juliá, M. Cuesta, I, M.A Cañizares |
| EPI_ISL_539567 | Hospital Comarcal de Melilla | Instituto de Salud Carlos III | Iglesias-Caballero, M. Molinero Calamita, M. González-Esguevillas, M. Camarero, S. Pozo, F. Casas, I. Jiménez, P. Jiménez, M. Zaballos, A. Monzón, S. Varona, S. Juliá, M. Cuesta, I, J. López |
| EPI_ISL_539568 | Hospital Comarcal de Melilla | Instituto de Salud Carlos III | Iglesias-Caballero, M. Molinero Calamita, M. González-Esguevillas, M. Camarero, S. Pozo, F. Casas, I. Jiménez, P. Jiménez, M. Zaballos, A. Monzón, S. Varona, S. Juliá, M. Cuesta, I, C. Ezpeleta |
| EPI_ISL_539569, EPI_ISL_539570, EPI_ISL_539571, EPI_ISL_539572 | Complejo Hospitalario de Navarra | Instituto de Salud Carlos III | Iglesias-Caballero, M. Molinero Calamita, M. González-Esguevillas, M. Camarero, S. Pozo, F. Casas, I. Jiménez, P. Jiménez, M. Zaballos, A. Monzón, S. Varona, S. Juliá, M. Cuesta, I, J. López |
| EPI_ISL_539573, EPI_ISL_539574, EPI_ISL_539575, EPI_ISL_539576 | Centre de Recherches Medicales de Lambarene (CERMEL) | Department of Emerging Infectious Diseases, Institute of Tropical Medicine, Nagasaki University | Haruka Abe, Yuri Ushijima, Rodrigue Bikangui, Akim A. Adegnika, Bertrand Lell, Jiro Yasuda |
| EPI_ISL_539577, EPI_ISL_539587, EPI_ISL_539591, EPI_ISL_539593, EPI_ISL_539596, EPI_ISL_539599, EPI_ISL_539600, EPI_ISL_539601, EPI_ISL_539602, EPI_ISL_539603, EPI_ISL_539604, EPI_ISL_539605, EPI_ISL_539606, EPI_ISL_539607, EPI_ISL_539609, EPI_ISL_539610, EPI_ISL_539611, EPI_ISL_539613, EPI_ISL_539614, EPI_ISL_539615 | | | |
| see above | ZOTZ KLIMAS MVZ Düsseldorf-Centrum GbR ÜBAG für Labormedizin, Genetik, Zytologie, Pathologie | Center of Medical Microbiology, Virology, and Hospital Hygiene, University of Duesseldorf | Maximilian Damagnez, Alexander Dilthey, Ashley-Jane Duplessis, Patrick Finzer, Katrin Hoffmann, Torsten Houwaart, Malte Kohns Vasconcelos, Marek Korencak, Nadine Lübke, Jessica Nicolai, Klaus Pfeffer, Daniel Strelow, Jörg Timm, Andreas Walker, Tobias Wienemann, Rainer Zotz |
| EPI_ISL_539616 | CSIR-Centre for Cellular and Molecular Biology | CSIR-Centre for Cellular and Molecular Biology | Lamuk Zaveri, Shagufta Khan, Namami Gaur, Sakshi Shambhavi, Nikhil Hajirnis, M Soujanya Reddy, Pratheusa Maccha, Tulasi Nagabandi, Purushotham Vodnala, Payel Mukherjee, Sofia Banu, Priya Singh, Onkar Kulkarni, Dhiviya Vedagiri, Divya Gupta, Vishal Sah, Santosh Kumar Kuncha, Krishnan Harinivas Harshan, Archana Bharadwaj Siva, Karthik Bharadwaj Tallapaka, Renu Sudhakar, Somesh Gorde, Gangumala Srinivas Reddy, Sujoy Deb, Swati Bayyana, Rakesh K Mishra, Divya Tej Sowpati |
| EPI_ISL_539617 | CSIR-Centre for Cellular and Molecular Biology | CSIR-Centre for Cellular and Molecular Biology | Lamuk Zaveri, Shagufta Khan,Nikhil Hajirnis, M Soujanya Reddy, Pratheusa Maccha, Namami Gaur, Sakshi Shambhavi, Tulasi Nagabandi, Purushotham Vodnala, Payel Mukherjee, Sofia Banu, Priya Singh, Onkar Kulkarni, Dhiviya Vedagiri, Divya Gupta, Vishal Sah, Santosh Kumar Kuncha, Krishnan Harinivas Harshan, Archana Bharadwaj Siva, Karthik Bharadwaj Tallapaka,Umesh Kumar, Unis Ahmad Bhat, Ajay Sarawagi, Priyanka Pant, Rajkanwar Nathawat, Rakesh K Mishra, Divya Tej Sowpati |
| EPI_ISL_539618 | CSIR-Centre for Cellular and Molecular Biology | CSIR-Centre for Cellular and Molecular Biology | Lamuk Zaveri, Shagufta Khan,Nikhil Hajirnis, M Soujanya Reddy, Pratheusa Maccha, Namami Gaur, Sakshi Shambhavi, Tulasi Nagabandi, Purushotham Vodnala, Payel Mukherjee, Sofia Banu, Priya Singh, Onkar Kulkarni, Dhiviya Vedagiri, Divya Gupta, Vishal Sah, Santosh Kumar Kuncha, Krishnan Harinivas Harshan, Archana Bharadwaj Siva, Karthik Bharadwaj Tallapaka,Zeba Rizvi, Zuberwasim Sayyad, Kakade Aishwarya Arun, Amrutha H C, Ananga Ghosh, Rakesh K Mishra, Divya Tej Sowpati |
| EPI_ISL_539619 | CSIR-Centre for Cellular and Molecular Biology | CSIR-Centre for Cellular and Molecular Biology | M Soujanya Reddy, Nikhil Hajirnis, Pratheusa Maccha, Namami Gaur, Sakshi Shambhavi, Lamuk Zaveri, Shagufta Khan, Tulasi Nagabandi, Purushotham Vodnala, Payel Mukherjee, Sofia Banu, Priya Singh, Onkar Kulkarni, Dhiviya Vedagiri, Divya Gupta, Vishal Sah, Santosh Kumar Kuncha, Krishnan Harinivas Harshan, Archana Bharadwaj Siva, Karthik Bharadwaj Tallapaka, Zeba Rizvi, Zuberwasim Sayyad, Kakade Aishwarya Arun, Amrutha H C, Ananga Ghosh, Rakesh K Mishra, Divya Tej Sowpati |
| EPI_ISL_539620 | CSIR-Centre for Cellular and Molecular Biology | CSIR-Centre for Cellular and Molecular Biology | M Soujanya Reddy, Nikhil Hajirnis, Pratheusa Maccha, Payel Mukherjee, Sofia Banu, Priya Singh, Onkar Kulkarni,Tulasi Nagabandi, Namami Gaur, Sakshi Shambhavi, Lamuk Zaveri, Shagufta Khan, Purushotham Vodnala, Dhiviya Vedagiri, Divya Gupta, Vishal Sah, Santosh Kumar Kuncha, Krishnan Harinivas Harshan, Archana Bharadwaj Siva, Karthik Bharadwaj Tallapaka,Kezia J Ann, Radhika Khandelwal, Roshan Maku Venkata, Shemin Mansuri, Sonu Uday, Rakesh K Mishra, Divya Tej Sowpati |
| EPI_ISL_539621 | CSIR-Centre for Cellular and Molecular Biology | CSIR-Centre for Cellular and Molecular Biology | M Soujanya Reddy, Nikhil Hajirnis, Pratheusa Maccha, Sakshi Shambhavi, Lamuk Zaveri, Shagufta Khan, Namami Gaur, Tulasi Nagabandi, Purushotham Vodnala, Payel Mukherjee, Sofia Banu, Priya Singh,Onkar Kulkarni, Dhiviya Vedagiri, Divya Gupta, Vishal Sah, Santosh Kumar Kuncha, Krishnan Harinivas Harshan, Archana Bharadwaj Siva, Karthik Bharadwaj Tallapaka, G. Aditya Kumar, Koushick Sivakumar, Pooja Ramesh Gupta, Rajan Kumar Jha, Shraddha Vijay Lahoti, Rakesh K Mishra, Divya Tej Sowpati |
| EPI_ISL_539622 | CSIR-Centre for Cellular and Molecular Biology | CSIR-Centre for Cellular and Molecular Biology | Namami Gaur, Sakshi Shambhavi, Lamuk Zaveri, Shagufta Khan, Nikhil Hajirnis, M Soujanya Reddy, Pratheusa Maccha, Tulasi Nagabandi, Purushotham Vodnala, Payel Mukherjee, Sofia Banu, Priya Singh, Onkar Kulkarni, Dhiviya Vedagiri, Divya Gupta, Vishal Sah, Santosh Kumar Kuncha, Krishnan Harinivas Harshan, Archana Bharadwaj Siva, Karthik Bharadwaj Tallapaka, Zeba Rizvi, Zuberwasim Sayyad, Kakade Aishwarya Arun, Amrutha H C, Ananga Ghosh, Rakesh K Mishra, Divya Tej Sowpati |
| EPI_ISL_539623 | CSIR-Centre for Cellular and Molecular Biology | CSIR-Centre for Cellular and Molecular Biology | Namami Gaur, Sakshi Shambhavi, Lamuk Zaveri, Shagufta Khan, Nikhil Hajirnis, M Soujanya Reddy, Pratheusa Maccha, Tulasi Nagabandi, Purushotham Vodnala, Payel Mukherjee, Sofia Banu, Priya Singh, Onkar Kulkarni, Dhiviya Vedagiri, Divya Gupta, Vishal Sah, Santosh Kumar Kuncha, Krishnan Harinivas Harshan, Archana Bharadwaj Siva, Karthik Bharadwaj Tallapaka,G. Aditya Kumar, Koushick Sivakumar, Rakesh K Mishra, Divya Tej Sowpati |
| EPI_ISL_539624 | CSIR-Centre for Cellular and Molecular Biology | CSIR-Centre for Cellular and Molecular Biology | Namami Gaur, Sakshi Shambhavi, Lamuk Zaveri, Shagufta Khan, Nikhil Hajirnis, M Soujanya Reddy, Pratheusa Maccha, Tulasi Nagabandi, Purushotham Vodnala, Payel Mukherjee, Sofia Banu, Priya Singh,Onkar Kulkarni, Dhiviya Vedagiri, Divya Gupta, Vishal Sah, Santosh Kumar Kuncha, Krishnan Harinivas Harshan, Archana Bharadwaj Siva, Karthik Bharadwaj Tallapaka, Zeba Rizvi, Zuberwasim Sayyad, Kakade Aishwarya Arun, Amrutha H C, Ananga Ghosh, Rakesh K Mishra, Divya Tej Sowpati |
| EPI_ISL_539625 | CSIR-Centre for Cellular and Molecular Biology | CSIR-Centre for Cellular and Molecular Biology | Nikhil Hajirnis, M Soujanya Reddy, Pratheusa Maccha, Lamuk Zaveri, Shagufta Khan, Namami Gaur, Sakshi Shambhavi, Tulasi Nagabandi, Purushotham Vodnala, Payel Mukherjee, Sofia Banu, Priya Singh, Onkar Kulkarni, Dhiviya Vedagiri, Divya Gupta, Vishal Sah, Santosh Kumar Kuncha, Krishnan Harinivas Harshan, Archana Bharadwaj Siva, Karthik Bharadwaj Tallapaka,Zeba Rizvi, Zuberwasim Sayyad, Kakade Aishwarya Arun, Amrutha H C, Ananga Ghosh, Rakesh K Mishra, Divya Tej Sowpati |
| EPI_ISL_539626 | CSIR-Centre for Cellular and Molecular Biology | CSIR-Centre for Cellular and Molecular Biology | Nikhil Hajirnis, M Soujanya Reddy, Pratheusa Maccha, Namami Gaur, Sakshi Shambhavi, Lamuk Zaveri, Shagufta Khan, Tulasi Nagabandi, Purushotham Vodnala, Payel Mukherjee, Sofia Banu, Priya Singh,Onkar Kulkarni, Dhiviya Vedagiri, Divya Gupta, Vishal Sah, Santosh Kumar Kuncha, Krishnan Harinivas Harshan, Archana Bharadwaj Siva, Karthik Bharadwaj Tallapaka,Kezia J Ann, Radhika Khandelwal, Roshan Maku Venkata, Shemin Mansuri, Sonu Uday, Rakesh K Mishra, Divya Tej Sowpati |
| EPI_ISL_539627 | CSIR-Centre for Cellular and Molecular Biology | CSIR-Centre for Cellular and Molecular Biology | Nikhil Hajirnis, M Soujanya Reddy, Pratheusa Maccha, Payel Mukherjee, Sofia Banu, Priya Singh,Onkar Kulkarni, Dhiviya Vedagiri, Divya Gupta, Vishal Sah, Santosh Kumar Kuncha, Krishnan Harinivas Harshan, Archana Bharadwaj Siva, Karthik Bharadwaj Tallapaka, Shagufta Khan, Lamuk Zaveri, Namami Gaur, Sakshi Shambhavi, Tulasi Nagabandi, Purushotham Vodnala,Deepak Kumar, Devi Prasad Vijayashankar, Disha Nanda, Divya Das, Jotin Gogoi, Manish Bhattacharjee, Rakesh K Mishra, Divya Tej Sowpati |
| EPI_ISL_539628 | CSIR-Centre for Cellular and Molecular Biology | CSIR-Centre for Cellular and Molecular Biology | Payel Mukherjee, Sofia Banu, Priya Singh, Onkar Kulkarni, Dhiviya Vedagiri, Divya Gupta, Vishal Sah, Santosh Kumar Kuncha, Krishnan Harinivas Harshan, Archana Bharadwaj Siva, Karthik Bharadwaj Tallapaka, Shagufta Khan, Lamuk Zaveri, Nikhil Hajirnis, M Soujanya Reddy, Pratheusa Maccha, Namami Gaur, Sakshi Shambhavi, Tulasi Nagabandi, Purushotham Vodnala, G. Aditya Kumar, Koushick Sivakumar, Pooja Ramesh Gupta, Rajan Kumar Jha, Shraddha Vijay Lahoti, Rakesh K Mishra, Divya Tej Sowpati |
| EPI_ISL_539629 | CSIR-Centre for Cellular and Molecular Biology | CSIR-Centre for Cellular and Molecular Biology | Payel Mukherjee, Sofia Banu, Priya Singh, Onkar Kulkarni, Dhiviya Vedagiri, Divya Gupta, Vishal Sah, Santosh Kumar Kuncha, Krishnan Harinivas Harshan, Archana Bharadwaj Siva, Karthik Bharadwaj Tallapaka, Shagufta Khan, Lamuk Zaveri, Nikhil Hajirnis, M Soujanya Reddy, Pratheusa Maccha, Namami Gaur, Sakshi Shambhavi, Tulasi Nagabandi, Purushotham Vodnala, Gokulan C G, Gunjan Purohit, Hanuman Tulashiram Kale, Pankaj Kumar, Prachand Issarapu, Rakesh K Mishra, Divya Tej Sowpati |
| EPI_ISL_539630 | CSIR-Centre for Cellular and Molecular Biology | CSIR-Centre for Cellular and Molecular Biology | Payel Mukherjee, Sofia Banu, Priya Singh, Onkar Kulkarni, Dhiviya Vedagiri, Divya Gupta, Vishal Sah, Santosh Kumar Kuncha, Krishnan Harinivas Harshan, Archana Bharadwaj Siva, Karthik Bharadwaj Tallapaka, Shagufta Khan, Lamuk Zaveri, Nikhil Hajirnis, M Soujanya Reddy, Pratheusa Maccha, Namami Gaur, Sakshi Shambhavi, Tulasi Nagabandi, Purushotham Vodnala, Rakesh K Mishra, Sonu Uday, Sudipta Mondal, Annapoorna P Karthyayani, Debabrata Jana, Debrya Saha, Divya Tej Sowpati |
| EPI_ISL_539631 | CSIR-Centre for Cellular and Molecular Biology | CSIR-Centre for Cellular and Molecular Biology | Pratheusa Maccha, Sakshi Shambhavi, Lamuk Zaveri, Shagufta Khan, Namami Gaur, Nikhil Hajirnis, M Soujanya Reddy, Tulasi Nagabandi, Purushotham Vodnala, Payel Mukherjee, Sofia Banu, Priya Singh,Onkar Kulkarni, Dhiviya Vedagiri, Divya Gupta, Vishal Sah, Santosh Kumar Kuncha, Krishnan Harinivas Harshan, Archana Bharadwaj Siva, Karthik Bharadwaj Tallapaka, G. Aditya Kumar, Koushick Sivakumar,Disha Nanda, Divya Das, Jotin Gogoi, Manish Bhattacharjee, Ravi Prasad Mukku, Rakesh K Mishra, Divya Tej Sowpati |
| EPI_ISL_539632 | CSIR-Centre for Cellular and Molecular Biology | CSIR-Centre for Cellular and Molecular Biology | Pratheusa Maccha, Shagufta Khan, Lamuk Zaveri, Namami Gaur, Sakshi Shambhavi, Tulasi Nagabandi, Nikhil Hajirnis, M Soujanya Reddy, Purushotham Vodnala, Payel Mukherjee, Sofia Banu, Priya Singh, Onkar Kulkarni, Dhiviya Vedagiri, Divya Gupta, Vishal Sah, Santosh Kumar Kuncha, Krishnan Harinivas Harshan, Archana Bharadwaj Siva, Karthik Bharadwaj Tallapaka, Disha Nanda, Divya Das, Jotin Gogoi, Manish Bhattacharjee, Ravi Prasad Mukku, Rakesh K Mishra, Divya Tej Sowpati |
| EPI_ISL_539633 | CSIR-Centre for Cellular and Molecular Biology | CSIR-Centre for Cellular and Molecular Biology | Pratheusa Maccha, Sofia Banu, Payel Mukherjee, Priya Singh, Onkar Kulkarni, Dhiviya Vedagiri, Divya Gupta, Vishal Sah, Santosh Kumar Kuncha, Krishnan Harinivas Harshan, Archana Bharadwaj Siva, Karthik Bharadwaj Tallapaka, Shagufta Khan, Lamuk Zaveri, Namami Gaur, Sakshi Shambhavi, Nikhil Hajirnis, M Soujanya Reddy, Tulasi Nagabandi, Purushotham Vodnala,Preethi Jampala, Sharada Ravi Iyer, Sulagana Mukherjee, Swetha Sundar, Peddapuvala Sai Uday Kiran, Rakesh K Mishra, Divya Tej Sowpati |
| EPI_ISL_539634 | CSIR-Centre for Cellular and Molecular Biology | CSIR-Centre for Cellular and Molecular Biology | Sakshi Shambhavi, Lamuk Zaveri, Shagufta Khan, Namami Gaur, Nikhil Hajirnis, M Soujanya Reddy, Pratheusa Maccha, Tulasi Nagabandi, Purushotham Vodnala, Payel Mukherjee, Sofia Banu, Priya Singh, Onkar Kulkarni, Dhiviya Vedagiri, Divya Gupta, Vishal Sah, Santosh Kumar Kuncha, Krishnan Harinivas Harshan, Archana Bharadwaj Siva, Karthik Bharadwaj Tallapaka, Deepak Kumar, Devi Prasad Vijayashankar, Disha Nanda, Divya Das, Jotin Gogoi, Manish Bhattacharjee, Rakesh K Mishra, Divya Tej Sowpati |
| EPI_ISL_539635 | CSIR-Centre for Cellular and Molecular Biology | CSIR-Centre for Cellular and Molecular Biology | Sakshi Shambhavi, Lamuk Zaveri, Shagufta Khan, Namami Gaur, Nikhil Hajirnis, M Soujanya Reddy, Pratheusa Maccha,Tulasi Nagabandi, Purushotham Vodnala, Payel Mukherjee, Sofia Banu, Priya Singh,Onkar Kulkarni, Dhiviya Vedagiri, Divya Gupta, Vishal Sah, Santosh Kumar Kuncha, Krishnan Harinivas Harshan, Archana Bharadwaj Siva, Karthik Bharadwaj Tallapaka, G. Aditya Kumar, Koushick Sivakumar, Pooja Ramesh Gupta, Rajan Kumar Jha, Shraddha Vijay Lahoti, Rakesh K Mishra, Divya Tej Sowpati |
| EPI_ISL_539636 | CSIR-Centre for Cellular and Molecular Biology | CSIR-Centre for Cellular and Molecular Biology | Sakshi Shambhavi, Lamuk Zaveri, Shagufta Khan, Nikhil Hajirnis, M Soujanya Reddy, Pratheusa Maccha, Namami Gaur, Tulasi Nagabandi, Purushotham Vodnala, Payel Mukherjee, Sofia Banu, Priya Singh, Onkar Kulkarni, Dhiviya Vedagiri, Divya Gupta, Vishal Sah, Santosh Kumar Kuncha, Krishnan Harinivas Harshan, Archana Bharadwaj Siva, Karthik Bharadwaj Tallapaka,G. Aditya Kumar, Koushick Sivakumar, Rakesh K Mishra, Divya Tej Sowpati |
| EPI_ISL_539637 | CSIR-Centre for Cellular and Molecular Biology | CSIR-Centre for Cellular and Molecular Biology | Shagufta Khan, Lamuk Zaveri, Namami Gaur, Sakshi Shambhavi, Nikhil Hajirnis, M Soujanya Reddy, Pratheusa Maccha, Tulasi Nagabandi, Purushotham Vodnala, Payel Mukherjee, Sofia Banu, Priya Singh, Onkar Kulkarni, Dhiviya Vedagiri, Divya Gupta, Vishal Sah, Santosh Kumar Kuncha, Krishnan Harinivas Harshan, Archana Bharadwaj Siva, Karthik Bharadwaj Tallapaka, Renu Sudhakar, Somesh Gorde, Gangumala Srinivas Reddy, Sujoy Deb, Swati Bayyana, Rakesh K Mishra, Divya Tej Sowpati |
| EPI_ISL_539638 | CSIR-Centre for Cellular and Molecular Biology | CSIR-Centre for Cellular and Molecular Biology | Shagufta Khan, Lamuk Zaveri, Namami Gaur, Sakshi Shambhavi, Nikhil Hajirnis, M Soujanya Reddy, Pratheusa Maccha, Tulasi Nagabandi, Purushotham Vodnala, Payel Mukherjee, Sofia Banu, Priya Singh, Onkar Kulkarni, Dhiviya Vedagiri, Divya Gupta, Vishal Sah, Santosh Kumar Kuncha, Krishnan Harinivas Harshan, Archana Bharadwaj Siva, Karthik Bharadwaj Tallapaka,Preethi Jampala, Sharada Ravi Iyer, Sulagana Mukherjee, Swetha Sundar, Peddapuvala Sai Uday Kiran Rakesh K Mishra, Divya Tej Sowpati |
| EPI_ISL_539639 | CSIR-Centre for Cellular and Molecular Biology | CSIR-Centre for Cellular and Molecular Biology | Shagufta Khan, Lamuk Zaveri, Namami Gaur, Sakshi Shambhavi, Nikhil Hajirnis, M Soujanya Reddy, Pratheusa Maccha,Tulasi Nagabandi, Purushotham Vodnala, Payel Mukherjee, Sofia Banu, Priya Singh, Onkar Kulkarni, Dhiviya Vedagiri, Divya Gupta, Vishal Sah, Santosh Kumar Kuncha, Krishnan Harinivas Harshan, Archana Bharadwaj Siva, Karthik Bharadwaj Tallapaka,Umesh Kumar, Unis Ahmad Bhat, Ajay Sarawagi, Priyanka Pant, Rajkanwar Nathawat, Rakesh K Mishra, Divya Tej Sowpati |
| EPI_ISL_539640 | CSIR-Centre for Cellular and Molecular Biology | CSIR-Centre for Cellular and Molecular Biology | Sofia Banu, Payel Mukherjee, Priya Singh,Onkar Kulkarni, Dhiviya Vedagiri, Divya Gupta, Vishal Sah, Santosh Kumar Kuncha, Krishnan Harinivas Harshan, Archana Bharadwaj Siva, Karthik Bharadwaj Tallapaka, Shagufta Khan, Lamuk Zaveri, Namami Gaur, Sakshi Shambhavi, Nikhil Hajirnis, M Soujanya Reddy, Pratheusa Maccha, Tulasi Nagabandi, Purushotham Vodnala, Disha Nanda, Divya Das, Jotin Gogoi, Manish Bhattacharjee, Ravi Prasad Mukku, Rakesh K Mishra, Divya Tej Sowpati |
| EPI_ISL_539641 | CSIR-Centre for Cellular and Molecular Biology | CSIR-Centre for Cellular and Molecular Biology | Sofia Banu, Payel Mukherjee, Priya Singh,Onkar Kulkarni, Dhiviya Vedagiri, Divya Gupta, Vishal Sah, Santosh Kumar Kuncha, Krishnan Harinivas Harshan, Archana Bharadwaj Siva, Karthik Bharadwaj Tallapaka, Shagufta Khan, Lamuk Zaveri, Namami Gaur, Sakshi Shambhavi, Nikhil Hajirnis, M Soujanya Reddy, Pratheusa Maccha,Tulasi Nagabandi, Purushotham Vodnala, Deepak Kumar, Devi Prasad Vijayashankar, Disha Nanda, Divya Das, Jotin Gogoi, Manish Bhattacharjee, Rakesh K Mishra, Divya Tej Sowpati |
| EPI_ISL_539642 | CSIR-Centre for Cellular and Molecular Biology | CSIR-Centre for Cellular and Molecular Biology | Sofia Banu, Payel Mukherjee, Priya Singh,Onkar Kulkarni, Dhiviya Vedagiri, Divya Gupta, Vishal Sah, Santosh Kumar Kuncha, Krishnan Harinivas Harshan, Archana Bharadwaj Siva, Karthik Bharadwaj Tallapaka, Shagufta Khan, Lamuk Zaveri, Namami Gaur, Sakshi Shambhavi, Tulasi Nagabandi, Nikhil Hajirnis, M Soujanya Reddy, Pratheusa Maccha, Purushotham Vodnala, Gokulan C G, Gunjan Purohit, Hanuman Tulashiram Kale, Pankaj Kumar, Prachand Issarapu, Rakesh K Mishra, Divya Tej Sowpati |
| EPI_ISL_539643 | CSIR-Centre for Cellular and Molecular Biology | CSIR-Centre for Cellular and Molecular Biology | Tulasi Nagabandi, Namami Gaur, Sakshi Shambhavi, Lamuk Zaveri, Shagufta Khan, Nikhil Hajirnis, M Soujanya Reddy, Pratheusa Maccha, Purushotham Vodnala, Payel Mukherjee, Sofia Banu, Priya Singh, Onkar Kulkarni, Dhiviya Vedagiri, Divya Gupta, Vishal Sah, Santosh Kumar Kuncha, Krishnan Harinivas Harshan, Archana Bharadwaj Siva, Karthik Bharadwaj Tallapaka,G. Aditya Kumar, Koushick Sivakumar, Pooja Ramesh Gupta, Rajan Kumar Jha, Shraddha Vijay Lahoti, Rakesh K Mishra, Divya Tej Sowpati |

| | | | |
|---|---|---|---|
| EPI_ISL_539644 | CSIR-Centre for Cellular and Molecular Biology | CSIR-Centre for Cellular and Molecular Biology | Tulasi Nagabandi, Namami Gaur, Sakshi Shambhavi, Lamuk Zaveri, Shagufta Khan, Nikhil Hajirnis, M Soujanya Reddy, Pratheusa Maccha, Purushotham Vodnala, Payel Mukherjee, Sofia Banu, Priya Singh,Onkar Kulkarni , Dhiviya Vedagiri, Divya Gupta, Vishal Sah, Santosh Kumar Kuncha, Krishnan Harinivas Harshan, Archana Bharadwaj Siva, Karthik Bharadwaj Tallapaka,G. Aditya Kumar, Koushick Sivakumar, Pooja Ramesh Gupta, Rajan Kumar Jha, Shraddha Vijay Lahoti, Rakesh K Mishra, Divya Tej Sowpati |
| EPI_ISL_539645 | CSIR-Centre for Cellular and Molecular Biology | CSIR-Centre for Cellular and Molecular Biology | Tulasi Nagabandi, Namami Gaur, Sakshi Shambhavi, Lamuk Zaveri, Shagufta Khan, Nikhil Hajirnis, M Soujanya Reddy, Pratheusa Maccha, Purushotham Vodnala, Payel Mukherjee, Sofia Banu, Priya Singh,Onkar Kulkarni, Dhiviya Vedagiri, Divya Gupta, Vishal Sah, Santosh Kumar Kuncha, Krishnan Harinivas Harshan, Archana Bharadwaj Siva, Karthik Bharadwaj Tallapaka,Kezia J Ann, Radhika Khandelwal, Roshan Maku Venkata, Shemin Mansuri, Sonu Uday, Rakesh K Mishra, Divya Tej Sowpati |
| EPI_ISL_539646 | CSIR-Centre for Cellular and Molecular Biology | CSIR-Centre for Cellular and Molecular Biology | Lamuk Zaveri, Shagufta Khan, Namami Gaur, Sakshi Shambhavi, Nikhil Hajirnis, M Soujanya Reddy, Pratheusa Maccha, Tulasi Nagabandi, Purushotham Vodnala, Payel Mukherjee, Sofia Banu, Priya Singh, Onkar Kulkarni, Dhiviya Vedagiri, Divya Gupta, Vishal Sah, Santosh Kumar Kuncha, Krishnan Harinivas Harshan, Archana Bharadwaj Siva, Karthik Bharadwaj Tallapaka, Renu Sudhakar, Somesh Gorde, Gangumala Srinivas Reddy, Sujoy Deb, Swati Bayyana, Rakesh K Mishra, Divya Tej Sowpati |
| EPI_ISL_539647 | CSIR-Centre for Cellular and Molecular Biology | CSIR-Centre for Cellular and Molecular Biology | Lamuk Zaveri, Shagufta Khan,Nikhil Hajirnis, M Soujanya Reddy, Pratheusa Maccha, Namami Gaur, Sakshi Shambhavi, Tulasi Nagabandi, Purushotham Vodnala, Payel Mukherjee, Sofia Banu, Priya Singh, Onkar Kulkarni, Dhiviya Vedagiri, Divya Gupta, Vishal Sah, Santosh Kumar Kuncha, Krishnan Harinivas Harshan, Archana Bharadwaj Siva, Karthik Bharadwaj Tallapaka,Umesh Kumar, Unis Ahmad Bhat, Ajay Sarawagi, Priyanka Pant, Rajkanwar Nathawat, Rakesh K Mishra, Divya Tej Sowpati |
| EPI_ISL_539648 | CSIR-Centre for Cellular and Molecular Biology | CSIR-Centre for Cellular and Molecular Biology | Lamuk Zaveri, Shagufta Khan,Nikhil Hajirnis, M Soujanya Reddy, Pratheusa Maccha, Namami Gaur, Sakshi Shambhavi, Tulasi Nagabandi, Purushotham Vodnala, Payel Mukherjee, Sofia Banu, Priya Singh, Onkar Kulkarni, Dhiviya Vedagiri, Divya Gupta, Vishal Sah, Santosh Kumar Kuncha, Krishnan Harinivas Harshan, Archana Bharadwaj Siva, Karthik Bharadwaj Tallapaka,Zeba Rizvi, Zuberwasim Sayyad, Kakade Aishwarya Arun, Amrutha H C, Ananga Ghosh, Rakesh K Mishra, Divya Tej Sowpati |
| EPI_ISL_539649 | CSIR-Centre for Cellular and Molecular Biology | CSIR-Centre for Cellular and Molecular Biology | M Soujanya Reddy, Nikhil Hajirnis, Pratheusa Maccha, Namami Gaur, Sakshi Shambhavi, Lamuk Zaveri, Shagufta Khan, Tulasi Nagabandi, Purushotham Vodnala, Payel Mukherjee, Sofia Banu, Priya Singh, Onkar Kulkarni, Dhiviya Vedagiri, Divya Gupta, Vishal Sah, Santosh Kumar Kuncha, Krishnan Harinivas Harshan, Archana Bharadwaj Siva, Karthik Bharadwaj Tallapaka, Zeba Rizvi, Zuberwasim Sayyad, Kakade Aishwarya Arun, Amrutha H C, Ananga Ghosh, Rakesh K Mishra, Divya Tej Sowpati |
| EPI_ISL_539650 | CSIR-Centre for Cellular and Molecular Biology | CSIR-Centre for Cellular and Molecular Biology | M Soujanya Reddy, Nikhil Hajirnis, Pratheusa Maccha, Payel Mukherjee, Sofia Banu, Priya Singh, Onkar Kulkarni,Tulasi Nagabandi, Namami Gaur, Sakshi Shambhavi, Lamuk Zaveri, Shagufta Khan, Purushotham Vodnala, Dhiviya Vedagiri, Divya Gupta, Vishal Sah, Santosh Kumar Kuncha, Krishnan Harinivas Harshan, Archana Bharadwaj Siva, Karthik Bharadwaj Tallapaka,Kezia J Ann, Radhika Khandelwal, Roshan Maku Venkata, Shemin Mansuri, Sonu Uday, Rakesh K Mishra, Divya Tej Sowpati |
| EPI_ISL_539651 | CSIR-Centre for Cellular and Molecular Biology | CSIR-Centre for Cellular and Molecular Biology | M Soujanya Reddy, Nikhil Hajirnis, Pratheusa Maccha, Sakshi Shambhavi, Lamuk Zaveri, Shagufta Khan, Namami Gaur, Tulasi Nagabandi, Purushotham Vodnala, Payel Mukherjee, Sofia Banu, Priya Singh,Onkar Kulkarni, Dhiviya Vedagiri, Divya Gupta, Vishal Sah, Santosh Kumar Kuncha, Krishnan Harinivas Harshan, Archana Bharadwaj Siva, Karthik Bharadwaj Tallapaka, G. Aditya Kumar, Koushick Sivakumar, Pooja Ramesh Gupta, Rajan Kumar Jha, Shraddha Vijay Lahoti, Rakesh K Mishra, Divya Tej Sowpati |
| EPI_ISL_539652 | CSIR-Centre for Cellular and Molecular Biology | CSIR-Centre for Cellular and Molecular Biology | Namami Gaur, Sakshi Shambhavi, Lamuk Zaveri, Shagufta Khan, Nikhil Hajirnis, M Soujanya Reddy, Pratheusa Maccha, Tulasi Nagabandi, Purushotham Vodnala, Payel Mukherjee, Sofia Banu, Priya Singh, Onkar Kulkarni, Dhiviya Vedagiri, Divya Gupta, Vishal Sah, Santosh Kumar Kuncha, Krishnan Harinivas Harshan, Archana Bharadwaj Siva, Karthik Bharadwaj Tallapaka, Zeba Rizvi, Zuberwasim Sayyad, Kakade Aishwarya Arun, Amrutha H C, Ananga Ghosh, Rakesh K Mishra, Divya Tej Sowpati |
| EPI_ISL_539653 | CSIR-Centre for Cellular and Molecular Biology | CSIR-Centre for Cellular and Molecular Biology | Namami Gaur, Sakshi Shambhavi, Lamuk Zaveri, Shagufta Khan, Nikhil Hajirnis, M Soujanya Reddy, Pratheusa Maccha, Tulasi Nagabandi, Purushotham Vodnala, Payel Mukherjee, Sofia Banu, Priya Singh, Onkar Kulkarni, Dhiviya Vedagiri, Divya Gupta, Vishal Sah, Santosh Kumar Kuncha, Krishnan Harinivas Harshan, Archana Bharadwaj Siva, Karthik Bharadwaj Tallapaka,G. Aditya Kumar, Koushick Sivakumar, Rakesh K Mishra, Divya Tej Sowpati |
| EPI_ISL_539654 | CSIR-Centre for Cellular and Molecular Biology | CSIR-Centre for Cellular and Molecular Biology | Namami Gaur, Sakshi Shambhavi, Lamuk Zaveri, Shagufta Khan, Nikhil Hajirnis, M Soujanya Reddy, Pratheusa Maccha, Tulasi Nagabandi, Purushotham Vodnala, Payel Mukherjee, Sofia Banu, Priya Singh,Onkar Kulkarni, Dhiviya Vedagiri, Divya Gupta, Vishal Sah, Santosh Kumar Kuncha, Krishnan Harinivas Harshan, Archana Bharadwaj Siva, Karthik Bharadwaj Tallapaka, Zeba Rizvi, Zuberwasim Sayyad, Kakade Aishwarya Arun, Amrutha H C, Ananga Ghosh, Rakesh K Mishra, Divya Tej Sowpati |
| EPI_ISL_539655 | CSIR-Centre for Cellular and Molecular Biology | CSIR-Centre for Cellular and Molecular Biology | Nikhil Hajirnis, M Soujanya Reddy, Pratheusa Maccha, Lamuk Zaveri, Shagufta Khan, Namami Gaur, Sakshi Shambhavi, Tulasi Nagabandi, Purushotham Vodnala, Payel Mukherjee, Sofia Banu, Priya Singh, Onkar Kulkarni, Dhiviya Vedagiri, Divya Gupta, Vishal Sah, Santosh Kumar Kuncha, Krishnan Harinivas Harshan, Archana Bharadwaj Siva, Karthik Bharadwaj Tallapaka,Zeba Rizvi, Zuberwasim Sayyad, Kakade Aishwarya Arun, Amrutha H C, Ananga Ghosh, Rakesh K Mishra, Divya Tej Sowpati |
| EPI_ISL_539656 | CSIR-Centre for Cellular and Molecular Biology | CSIR-Centre for Cellular and Molecular Biology | Nikhil Hajirnis, M Soujanya Reddy, Pratheusa Maccha, Namami Gaur, Sakshi Shambhavi, Lamuk Zaveri, Shagufta Khan, Tulasi Nagabandi, Purushotham Vodnala, Payel Mukherjee, Sofia Banu, Priya Singh,Onkar Kulkarni, Dhiviya Vedagiri, Divya Gupta, Vishal Sah, Santosh Kumar Kuncha, Krishnan Harinivas Harshan, Archana Bharadwaj Siva, Karthik Bharadwaj Tallapaka,Kezia J Ann, Radhika Khandelwal, Roshan Maku Venkata, Shemin Mansuri, Sonu Uday, Rakesh K Mishra, Divya Tej Sowpati |
| EPI_ISL_539657 | CSIR-Centre for Cellular and Molecular Biology | CSIR-Centre for Cellular and Molecular Biology | Nikhil Hajirnis, M Soujanya Reddy, Pratheusa Maccha, Payel Mukherjee, Sofia Banu, Priya Singh,Onkar Kulkarni, Dhiviya Vedagiri, Divya Gupta, Vishal Sah, Santosh Kumar Kuncha, Krishnan Harinivas Harshan, Archana Bharadwaj Siva, Karthik Bharadwaj Tallapaka, Shagufta Khan, Lamuk Zaveri, Namami Gaur, Sakshi Shambhavi, Tulasi Nagabandi, Purushotham Vodnala,Deepak Kumar, Devi Prasad Vijayashankar, Disha Nanda, Divya Das, Jotin Gogoi, Manish Bhattacharjee, Rakesh K Mishra, Divya Tej Sowpati |
| EPI_ISL_539658 | CSIR-Centre for Cellular and Molecular Biology | CSIR-Centre for Cellular and Molecular Biology | Payel Mukherjee, Sofia Banu, Priya Singh, Onkar Kulkarni, Dhiviya Vedagiri, Divya Gupta, Vishal Sah, Santosh Kumar Kuncha, Krishnan Harinivas Harshan, Archana Bharadwaj Siva, Karthik Bharadwaj Tallapaka, Shagufta Khan, Lamuk Zaveri, Nikhil Hajirnis, M Soujanya Reddy, Pratheusa Maccha, Namami Gaur, Sakshi Shambhavi, Tulasi Nagabandi, Purushotham Vodnala, G. Aditya Kumar, Koushick Sivakumar, Pooja Ramesh Gupta, Rajan Kumar Jha, Shraddha Vijay Lahoti, Rakesh K Mishra, Divya Tej Sowpati |
| EPI_ISL_539659 | CSIR-Centre for Cellular and Molecular Biology | CSIR-Centre for Cellular and Molecular Biology | Payel Mukherjee, Sofia Banu, Priya Singh, Onkar Kulkarni, Dhiviya Vedagiri, Divya Gupta, Vishal Sah, Santosh Kumar Kuncha, Krishnan Harinivas Harshan, Archana Bharadwaj Siva, Karthik Bharadwaj Tallapaka, Shagufta Khan, Lamuk Zaveri, Nikhil Hajirnis, M Soujanya Reddy, Pratheusa Maccha, Namami Gaur, Sakshi Shambhavi, Tulasi Nagabandi, Purushotham Vodnala, Gokulan C G, Gunjan Purohit, Hanuman Tulashiram Kale, Pankaj Kumar, Prachand Issarapu, Rakesh K Mishra, Divya Tej Sowpati |
| EPI_ISL_539660 | CSIR-Centre for Cellular and Molecular Biology | CSIR-Centre for Cellular and Molecular Biology | Payel Mukherjee, Sofia Banu, Priya Singh, Onkar Kulkarni, Dhiviya Vedagiri, Divya Gupta, Vishal Sah, Santosh Kumar Kuncha, Krishnan Harinivas Harshan, Archana Bharadwaj Siva, Karthik Bharadwaj Tallapaka, Shagufta Khan, Lamuk Zaveri, Nikhil Hajirnis, M Soujanya Reddy, Pratheusa Maccha, Namami Gaur, Sakshi Shambhavi, Tulasi Nagabandi, Purushotham Vodnala, Rakesh K Mishra, Sonu Uday, Sudipta Mondal, Annapoorna P Karthyayani, Debabrata Jana, Debrya Saha, Divya Tej Sowpati |
| EPI_ISL_539661 | CSIR-Centre for Cellular and Molecular Biology | CSIR-Centre for Cellular and Molecular Biology | Pratheusa Maccha, Sakshi Shambhavi, Lamuk Zaveri, Shagufta Khan, Namami Gaur, Nikhil Hajirnis, M Soujanya Reddy, Tulasi Nagabandi, Purushotham Vodnala, Payel Mukherjee, Sofia Banu, Priya Singh, Onkar Kulkarni, Dhiviya Vedagiri, Divya Gupta, Vishal Sah, Santosh Kumar Kuncha, Krishnan Harinivas Harshan, Archana Bharadwaj Siva, Karthik Bharadwaj Tallapaka, G. Aditya Kumar, Koushick Sivakumar,Disha Nanda, Divya Das, Jotin Gogoi, Manish Bhattacharjee, Ravi Prasad Mukku, Rakesh K Mishra, Divya Tej Sowpati |
| EPI_ISL_539662 | CSIR-Centre for Cellular and Molecular Biology | CSIR-Centre for Cellular and Molecular Biology | Pratheusa Maccha, Shagufta Khan, Lamuk Zaveri, Namami Gaur, Sakshi Shambhavi, Tulasi Nagabandi, Nikhil Hajirnis, M Soujanya Reddy, Purushotham Vodnala, Payel Mukherjee, Sofia Banu, Priya Singh, Onkar Kulkarni, Dhiviya Vedagiri, Divya Gupta, Vishal Sah, Santosh Kumar Kuncha, Krishnan Harinivas Harshan, Archana Bharadwaj Siva, Karthik Bharadwaj Tallapaka, Disha Nanda, Divya Das, Jotin Gogoi, Manish Bhattacharjee, Ravi Prasad Mukku, Rakesh K Mishra, Divya Tej Sowpati |
| EPI_ISL_539663 | CSIR-Centre for Cellular and Molecular Biology | CSIR-Centre for Cellular and Molecular Biology | Pratheusa Maccha, Sofia Banu, Payel Mukherjee, Priya Singh, Onkar Kulkarni, Dhiviya Vedagiri, Divya Gupta, Vishal Sah, Santosh Kumar Kuncha, Krishnan Harinivas Harshan, Archana Bharadwaj Siva, Karthik Bharadwaj Tallapaka, Shagufta Khan, Lamuk Zaveri, Namami Gaur, Sakshi Shambhavi, Nikhil Hajirnis, M Soujanya Reddy, Tulasi Nagabandi, Purushotham Vodnala,Preethi Jampala, Sharada Ravi Iyer, Sulagana Mukherjee, Swetha Sundar, Peddapuvala Sai Uday Kiran, Rakesh K Mishra, Divya Tej Sowpati |
| EPI_ISL_539664 | CSIR-Centre for Cellular and Molecular Biology | CSIR-Centre for Cellular and Molecular Biology | Sakshi Shambhavi, Lamuk Zaveri, Shagufta Khan, Namami Gaur, Nikhil Hajirnis, M Soujanya Reddy, Pratheusa Maccha, Tulasi Nagabandi, Purushotham Vodnala, Payel Mukherjee, Sofia Banu, Priya Singh, Onkar Kulkarni, Dhiviya Vedagiri, Divya Gupta, Vishal Sah, Santosh Kumar Kuncha, Krishnan Harinivas Harshan, Archana Bharadwaj Siva, Karthik Bharadwaj Tallapaka, Deepak Kumar, Devi Prasad Vijayashankar, Disha Nanda, Divya Das, Jotin Gogoi, Manish Bhattacharjee, Rakesh K Mishra, Divya Tej Sowpati |
| EPI_ISL_539665 | CSIR-Centre for Cellular and Molecular Biology | CSIR-Centre for Cellular and Molecular Biology | Sakshi Shambhavi, Lamuk Zaveri, Shagufta Khan, Namami Gaur, Nikhil Hajirnis, M Soujanya Reddy, Pratheusa Maccha,Tulasi Nagabandi, Purushotham Vodnala, Payel Mukherjee, Sofia Banu, Priya Singh,Onkar Kulkarni, Dhiviya Vedagiri, Divya Gupta, Vishal Sah, Santosh Kumar Kuncha, Krishnan Harinivas Harshan, Archana Bharadwaj Siva, Karthik Bharadwaj Tallapaka, G. Aditya Kumar, Koushick Sivakumar, Pooja Ramesh Gupta, Rajan Kumar Jha, Shraddha Vijay Lahoti, Rakesh K Mishra, Divya Tej Sowpati |
| EPI_ISL_539666 | CSIR-Centre for Cellular and Molecular Biology | CSIR-Centre for Cellular and Molecular Biology | Sakshi Shambhavi, Lamuk Zaveri, Shagufta Khan, Nikhil Hajirnis, M Soujanya Reddy, Pratheusa Maccha, Namami Gaur, Tulasi Nagabandi, Purushotham Vodnala, Payel Mukherjee, Sofia Banu, Priya Singh, Onkar Kulkarni, Dhiviya Vedagiri, Divya Gupta, Vishal Sah, Santosh Kumar Kuncha, Krishnan Harinivas Harshan, Archana Bharadwaj Siva, Karthik Bharadwaj Tallapaka,G. Aditya Kumar, Koushick Sivakumar, Rakesh K Mishra, Divya Tej Sowpati |
| EPI_ISL_539667 | CSIR-Centre for Cellular and Molecular Biology | CSIR-Centre for Cellular and Molecular Biology | Shagufta Khan, Lamuk Zaveri, Namami Gaur, Sakshi Shambhavi, Nikhil Hajirnis, M Soujanya Reddy, Pratheusa Maccha, Tulasi Nagabandi, Purushotham Vodnala, Payel Mukherjee, Sofia Banu, Priya Singh, Onkar Kulkarni, Dhiviya Vedagiri, Divya Gupta, Vishal Sah, Santosh Kumar Kuncha, Krishnan Harinivas Harshan, Archana Bharadwaj Siva, Karthik Bharadwaj Tallapaka, Renu Sudhakar, Somesh Gorde, Gangumala Srinivas Reddy, Sujoy Deb, Swati Bayyana, Rakesh K Mishra, Divya Tej Sowpati |
| EPI_ISL_539668 | CSIR-Centre for Cellular and Molecular Biology | CSIR-Centre for Cellular and Molecular Biology | Shagufta Khan, Lamuk Zaveri, Namami Gaur, Sakshi Shambhavi, Nikhil Hajirnis, M Soujanya Reddy, Pratheusa Maccha, Tulasi Nagabandi, Purushotham Vodnala, Payel Mukherjee, Sofia Banu, Priya Singh, Onkar Kulkarni, Dhiviya Vedagiri, Divya Gupta, Vishal Sah, Santosh Kumar Kuncha, Krishnan Harinivas Harshan, Archana Bharadwaj Siva, Karthik Bharadwaj Tallapaka,Preethi Jampala, Sharada Ravi Iyer, Sulagana Mukherjee, Swetha Sundar, Peddapuvala Sai Uday Kiran Rakesh K Mishra, Divya Tej Sowpati |
| EPI_ISL_539669 | CSIR-Centre for Cellular and Molecular Biology | CSIR-Centre for Cellular and Molecular Biology | Shagufta Khan, Lamuk Zaveri, Namami Gaur, Sakshi Shambhavi, Nikhil Hajirnis, M Soujanya Reddy, Pratheusa Maccha,Tulasi Nagabandi, Purushotham Vodnala, Payel Mukherjee, Sofia Banu, Priya Singh, Onkar Kulkarni, Dhiviya Vedagiri, Divya Gupta, Vishal Sah, Santosh Kumar Kuncha, Krishnan Harinivas Harshan, Archana Bharadwaj Siva, Karthik Bharadwaj Tallapaka,Umesh Kumar, Unis Ahmad Bhat, Ajay Sarawagi, Priyanka Pant, Rajkanwar Nathawat, Rakesh K Mishra, Divya Tej Sowpati |
| EPI_ISL_539670 | CSIR-Centre for Cellular and Molecular Biology | CSIR-Centre for Cellular and Molecular Biology | Sofia Banu, Payel Mukherjee, Priya Singh,Onkar Kulkarni, Dhiviya Vedagiri, Divya Gupta, Vishal Sah, Santosh Kumar Kuncha, Krishnan Harinivas Harshan, Archana Bharadwaj Siva, Karthik Bharadwaj Tallapaka, Shagufta Khan, Lamuk Zaveri, Namami Gaur, Sakshi Shambhavi, Nikhil Hajirnis, M Soujanya Reddy, Pratheusa Maccha, Tulasi Nagabandi, Purushotham Vodnala, Disha Nanda, Divya Das, Jotin Gogoi, Manish Bhattacharjee, Ravi Prasad Mukku, Rakesh K Mishra, Divya Tej Sowpati |
| EPI_ISL_539671 | CSIR-Centre for Cellular and Molecular Biology | CSIR-Centre for Cellular and Molecular Biology | Sofia Banu, Payel Mukherjee, Priya Singh,Onkar Kulkarni, Dhiviya Vedagiri, Divya Gupta, Vishal Sah, Santosh Kumar Kuncha, Krishnan Harinivas Harshan, Archana Bharadwaj Siva, Karthik Bharadwaj Tallapaka, Shagufta Khan, Lamuk Zaveri, Namami Gaur, Sakshi Shambhavi, Nikhil Hajirnis, M Soujanya Reddy, Pratheusa Maccha,Tulasi Nagabandi, Purushotham Vodnala, Deepak Kumar, Devi Prasad Vijayashankar, Disha Nanda, Divya Das, Jotin Gogoi, Manish Bhattacharjee, Rakesh K Mishra, Divya Tej Sowpati |
| EPI_ISL_539672 | CSIR-Centre for Cellular and Molecular Biology | CSIR-Centre for Cellular and Molecular Biology | Sofia Banu, Payel Mukherjee, Priya Singh,Onkar Kulkarni, Dhiviya Vedagiri, Divya Gupta, Vishal Sah, Santosh Kumar Kuncha, Krishnan Harinivas Harshan, Archana Bharadwaj Siva, Karthik Bharadwaj Tallapaka, Shagufta Khan, Lamuk Zaveri, Namami Gaur, Sakshi Shambhavi, Tulasi Nagabandi, Nikhil Hajirnis, M Soujanya Reddy, Pratheusa Maccha, Purushotham Vodnala, Gokulan C G, Gunjan Purohit, Hanuman Tulashiram Kale, Pankaj Kumar, Prachand Issarapu, Rakesh K Mishra, Divya Tej Sowpati |
| EPI_ISL_539673 | CSIR-Centre for Cellular and Molecular Biology | CSIR-Centre for Cellular and Molecular Biology | Tulasi Nagabandi, Namami Gaur, Sakshi Shambhavi, Lamuk Zaveri, Shagufta Khan, Nikhil Hajirnis, M Soujanya Reddy, Pratheusa Maccha, Purushotham Vodnala, Payel Mukherjee, Sofia Banu, Priya Singh, Onkar Kulkarni, Dhiviya Vedagiri, Divya Gupta, Vishal Sah, Santosh Kumar Kuncha, Krishnan Harinivas Harshan, Archana Bharadwaj Siva, Karthik Bharadwaj Tallapaka,G. Aditya Kumar, Koushick Sivakumar, Pooja Ramesh Gupta, Rajan Kumar Jha, Shraddha Vijay Lahoti, Rakesh K Mishra, Divya Tej Sowpati |
| EPI_ISL_539674 | CSIR-Centre for Cellular and Molecular Biology | CSIR-Centre for Cellular and Molecular Biology | Tulasi Nagabandi, Namami Gaur, Sakshi Shambhavi, Lamuk Zaveri, Shagufta Khan, Nikhil Hajirnis, M Soujanya Reddy, Pratheusa Maccha, Purushotham Vodnala, Payel Mukherjee, Sofia Banu, Priya Singh,Onkar Kulkarni , Dhiviya Vedagiri, Divya Gupta, Vishal Sah, Santosh Kumar Kuncha, Krishnan Harinivas Harshan, Archana Bharadwaj Siva, Karthik Bharadwaj Tallapaka,G. Aditya Kumar, Koushick Sivakumar, Pooja Ramesh Gupta, Rajan Kumar Jha, Shraddha Vijay Lahoti, Rakesh K Mishra, Divya Tej Sowpati |
| EPI_ISL_539675 | CSIR-Centre for Cellular and Molecular Biology | CSIR-Centre for Cellular and Molecular Biology | Tulasi Nagabandi, Namami Gaur, Sakshi Shambhavi, Lamuk Zaveri, Shagufta Khan, Nikhil Hajirnis, M Soujanya Reddy, Pratheusa Maccha, Purushotham Vodnala, Payel Mukherjee, Sofia Banu, Priya Singh,Onkar Kulkarni, Dhiviya Vedagiri, Divya Gupta, Vishal Sah, Santosh Kumar Kuncha, Krishnan Harinivas Harshan, Archana Bharadwaj Siva, Karthik Bharadwaj Tallapaka,Kezia J Ann, Radhika Khandelwal, Roshan Maku Venkata, Shemin Mansuri, Sonu Uday, Rakesh K Mishra, Divya Tej Sowpati |
| EPI_ISL_539676 | CSIR-Centre for Cellular and Molecular Biology | CSIR-Centre for Cellular and Molecular Biology | Lamuk Zaveri, Shagufta Khan, Namami Gaur, Sakshi Shambhavi, Nikhil Hajirnis, M Soujanya Reddy, Pratheusa Maccha, Tulasi Nagabandi, Purushotham Vodnala, Payel Mukherjee, Sofia Banu, Priya Singh, Onkar Kulkarni, Dhiviya Vedagiri, Divya Gupta, Vishal Sah, Santosh Kumar Kuncha, Krishnan Harinivas Harshan, Archana Bharadwaj Siva, Karthik Bharadwaj Tallapaka, Renu Sudhakar, Somesh Gorde, Gangumala Srinivas Reddy, Sujoy Deb, Swati Bayyana, |

| | | | |
|---|---|---|---|
| | | | Rakesh K Mishra, Divya Tej Sowpati |
| EPI_ISL_539677 | CSIR-Centre for Cellular and Molecular Biology | CSIR-Centre for Cellular and Molecular Biology | Lamuk Zaveri, Shagufta Khan,Nikhil Hajirnis, M Soujanya Reddy, Pratheusa Maccha, Namami Gaur, Sakshi Shambhavi, Tulasi Nagabandi, Purushotham Vodnala, Payel Mukherjee, Sofia Banu, Priya Singh, Onkar Kulkarni, Dhiviya Vedagiri, Divya Gupta, Vishal Sah, Santosh Kumar Kuncha, Krishnan Harinivas Harshan, Archana Bharadwaj Siva, Karthik Bharadwaj Tallapaka,Umesh Kumar, Unis Ahmad Bhat, Ajay Sarawagi, Priyanka Pant, Rajkanwar Nathawat, Rakesh K Mishra, Divya Tej Sowpati |
| EPI_ISL_539678 | CSIR-Centre for Cellular and Molecular Biology | CSIR-Centre for Cellular and Molecular Biology | Lamuk Zaveri, Shagufta Khan,Nikhil Hajirnis, M Soujanya Reddy, Pratheusa Maccha, Namami Gaur, Sakshi Shambhavi, Tulasi Nagabandi, Purushotham Vodnala, Payel Mukherjee, Sofia Banu, Priya Singh, Onkar Kulkarni, Dhiviya Vedagiri, Divya Gupta, Vishal Sah, Santosh Kumar Kuncha, Krishnan Harinivas Harshan, Archana Bharadwaj Siva, Karthik Bharadwaj Tallapaka,Zeba Rizvi, Zuberwasim Sayyad, Kakade Aishwarya Arun, Amrutha H C, Ananga Ghosh, Rakesh K Mishra, Divya Tej Sowpati |
| EPI_ISL_539679 | CSIR-Centre for Cellular and Molecular Biology | CSIR-Centre for Cellular and Molecular Biology | M Soujanya Reddy, Nikhil Hajirnis, Pratheusa Maccha, Namami Gaur, Sakshi Shambhavi, Lamuk Zaveri, Shagufta Khan, Tulasi Nagabandi, Purushotham Vodnala, Payel Mukherjee, Sofia Banu, Priya Singh, Onkar Kulkarni, Dhiviya Vedagiri, Divya Gupta, Vishal Sah, Santosh Kumar Kuncha, Krishnan Harinivas Harshan, Archana Bharadwaj Siva, Karthik Bharadwaj Tallapaka, Zeba Rizvi, Zuberwasim Sayyad, Kakade Aishwarya Arun, Amrutha H C, Ananga Ghosh, Rakesh K Mishra, Divya Tej Sowpati |
| EPI_ISL_539680 | CSIR-Centre for Cellular and Molecular Biology | CSIR-Centre for Cellular and Molecular Biology | M Soujanya Reddy, Nikhil Hajirnis, Pratheusa Maccha, Payel Mukherjee, Sofia Banu, Priya Singh, Onkar Kulkarni,Tulasi Nagabandi, Namami Gaur, Sakshi Shambhavi, Lamuk Zaveri, Shagufta Khan, Purushotham Vodnala, Dhiviya Vedagiri, Divya Gupta, Vishal Sah, Santosh Kumar Kuncha, Krishnan Harinivas Harshan, Archana Bharadwaj Siva, Karthik Bharadwaj Tallapaka,Kezia J Ann, Radhika Khandelwal, Roshan Maku Venkata, Shemin Mansuri, Sonu Uday, Rakesh K Mishra, Divya Tej Sowpati |
| EPI_ISL_539681 | CSIR-Centre for Cellular and Molecular Biology | CSIR-Centre for Cellular and Molecular Biology | M Soujanya Reddy, Nikhil Hajirnis, Pratheusa Maccha, Sakshi Shambhavi, Lamuk Zaveri, Shagufta Khan, Namami Gaur, Tulasi Nagabandi, Purushotham Vodnala, Payel Mukherjee, Sofia Banu, Priya Singh,Onkar Kulkarni, Dhiviya Vedagiri, Divya Gupta, Vishal Sah, Santosh Kumar Kuncha, Krishnan Harinivas Harshan, Archana Bharadwaj Siva, Karthik Bharadwaj Tallapaka, G. Aditya Kumar, Koushick Sivakumar, Pooja Ramesh Gupta, Rajan Kumar Jha, Shraddha Vijay Lahoti, Rakesh K Mishra, Divya Tej Sowpati |
| EPI_ISL_539682 | CSIR-Centre for Cellular and Molecular Biology | CSIR-Centre for Cellular and Molecular Biology | Namami Gaur, Sakshi Shambhavi, Lamuk Zaveri, Shagufta Khan, Nikhil Hajirnis, M Soujanya Reddy, Pratheusa Maccha, Tulasi Nagabandi, Purushotham Vodnala, Payel Mukherjee, Sofia Banu, Priya Singh, Onkar Kulkarni, Dhiviya Vedagiri, Divya Gupta, Vishal Sah, Santosh Kumar Kuncha, Krishnan Harinivas Harshan, Archana Bharadwaj Siva, Karthik Bharadwaj Tallapaka, Zeba Rizvi, Zuberwasim Sayyad, Kakade Aishwarya Arun, Amrutha H C, Ananga Ghosh, Rakesh K Mishra, Divya Tej Sowpati |
| EPI_ISL_539683 | CSIR-Centre for Cellular and Molecular Biology | CSIR-Centre for Cellular and Molecular Biology | Namami Gaur, Sakshi Shambhavi, Lamuk Zaveri, Shagufta Khan, Nikhil Hajirnis, M Soujanya Reddy, Pratheusa Maccha, Tulasi Nagabandi, Purushotham Vodnala, Payel Mukherjee, Sofia Banu, Priya Singh, Onkar Kulkarni, Dhiviya Vedagiri, Divya Gupta, Vishal Sah, Santosh Kumar Kuncha, Krishnan Harinivas Harshan, Archana Bharadwaj Siva, Karthik Bharadwaj Tallapaka,G. Aditya Kumar, Koushick Sivakumar, Rakesh K Mishra, Divya Tej Sowpati |
| EPI_ISL_539684 | CSIR-Centre for Cellular and Molecular Biology | CSIR-Centre for Cellular and Molecular Biology | Namami Gaur, Sakshi Shambhavi, Lamuk Zaveri, Shagufta Khan, Nikhil Hajirnis, M Soujanya Reddy, Pratheusa Maccha, Tulasi Nagabandi, Purushotham Vodnala, Payel Mukherjee, Sofia Banu, Priya Singh,Onkar Kulkarni, Dhiviya Vedagiri, Divya Gupta, Vishal Sah, Santosh Kumar Kuncha, Krishnan Harinivas Harshan, Archana Bharadwaj Siva, Karthik Bharadwaj Tallapaka, Zeba Rizvi, Zuberwasim Sayyad, Kakade Aishwarya Arun, Amrutha H C, Ananga Ghosh, Rakesh K Mishra, Divya Tej Sowpati |
| EPI_ISL_539685 | CSIR-Centre for Cellular and Molecular Biology | CSIR-Centre for Cellular and Molecular Biology | Nikhil Hajirnis, M Soujanya Reddy, Pratheusa Maccha, Lamuk Zaveri, Shagufta Khan, Namami Gaur, Sakshi Shambhavi, Tulasi Nagabandi, Purushotham Vodnala, Payel Mukherjee, Sofia Banu, Priya Singh, Onkar Kulkarni, Dhiviya Vedagiri, Divya Gupta, Vishal Sah, Santosh Kumar Kuncha, Krishnan Harinivas Harshan, Archana Bharadwaj Siva, Karthik Bharadwaj Tallapaka,Zeba Rizvi, Zuberwasim Sayyad, Kakade Aishwarya Arun, Amrutha H C, Ananga Ghosh, Rakesh K Mishra, Divya Tej Sowpati |
| EPI_ISL_539686 | CSIR-Centre for Cellular and Molecular Biology | CSIR-Centre for Cellular and Molecular Biology | Nikhil Hajirnis, M Soujanya Reddy, Pratheusa Maccha, Namami Gaur, Sakshi Shambhavi, Lamuk Zaveri, Shagufta Khan, Tulasi Nagabandi, Purushotham Vodnala, Payel Mukherjee, Sofia Banu, Priya Singh,Onkar Kulkarni, Dhiviya Vedagiri, Divya Gupta, Vishal Sah, Santosh Kumar Kuncha, Krishnan Harinivas Harshan, Archana Bharadwaj Siva, Karthik Bharadwaj Tallapaka,Kezia J Ann, Radhika Khandelwal, Roshan Maku Venkata, Shemin Mansuri, Sonu Uday, Rakesh K Mishra, Divya Tej Sowpati |
| EPI_ISL_539687 | CSIR-Centre for Cellular and Molecular Biology | CSIR-Centre for Cellular and Molecular Biology | Nikhil Hajirnis, M Soujanya Reddy, Pratheusa Maccha, Payel Mukherjee, Sofia Banu, Priya Singh,Onkar Kulkarni, Dhiviya Vedagiri, Divya Gupta, Vishal Sah, Santosh Kumar Kuncha, Krishnan Harinivas Harshan, Archana Bharadwaj Siva, Karthik Bharadwaj Tallapaka, Shagufta Khan, Lamuk Zaveri, Namami Gaur, Sakshi Shambhavi, Tulasi Nagabandi, Purushotham Vodnala,Deepak Kumar, Devi Prasad Vijayashankar, Disha Nanda, Divya Das, Jotin Gogoi, Manish Bhattacharjee, Rakesh K Mishra, Divya Tej Sowpati |
| EPI_ISL_539688 | CSIR-Centre for Cellular and Molecular Biology | CSIR-Centre for Cellular and Molecular Biology | Payel Mukherjee, Sofia Banu, Priya Singh, Onkar Kulkarni, Dhiviya Vedagiri, Divya Gupta, Vishal Sah, Santosh Kumar Kuncha, Krishnan Harinivas Harshan, Archana Bharadwaj Siva, Karthik Bharadwaj Tallapaka, Shagufta Khan, Lamuk Zaveri, Nikhil Hajirnis, M Soujanya Reddy, Pratheusa Maccha, Namami Gaur, Sakshi Shambhavi, Tulasi Nagabandi, Purushotham Vodnala, G. Aditya Kumar, Koushick Sivakumar, Pooja Ramesh Gupta, Rajan Kumar Jha, Shraddha Vijay Lahoti, Rakesh K Mishra, Divya Tej Sowpati |
| EPI_ISL_539689 | CSIR-Centre for Cellular and Molecular Biology | CSIR-Centre for Cellular and Molecular Biology | Payel Mukherjee, Sofia Banu, Priya Singh, Onkar Kulkarni, Dhiviya Vedagiri, Divya Gupta, Vishal Sah, Santosh Kumar Kuncha, Krishnan Harinivas Harshan, Archana Bharadwaj Siva, Karthik Bharadwaj Tallapaka, Shagufta Khan, Lamuk Zaveri, Nikhil Hajirnis, M Soujanya Reddy, Pratheusa Maccha, Namami Gaur, Sakshi Shambhavi, Tulasi Nagabandi, Purushotham Vodnala, Gokulan C G, Gunjan Purohit, Hanuman Tulashiram Kale, Pankaj Kumar, Prachand Issarapu, Rakesh K Mishra, Divya Tej Sowpati |
| EPI_ISL_539690 | CSIR-Centre for Cellular and Molecular Biology | CSIR-Centre for Cellular and Molecular Biology | Payel Mukherjee, Sofia Banu, Priya Singh, Onkar Kulkarni, Dhiviya Vedagiri, Divya Gupta, Vishal Sah, Santosh Kumar Kuncha, Krishnan Harinivas Harshan, Archana Bharadwaj Siva, Karthik Bharadwaj Tallapaka, Shagufta Khan, Lamuk Zaveri, Nikhil Hajirnis, M Soujanya Reddy, Pratheusa Maccha, Namami Gaur, Sakshi Shambhavi, Tulasi Nagabandi, Purushotham Vodnala, Rakesh K Mishra, Sonu Uday, Sudipta Mondal, Annapoorna P Karthyayani, Debabrata Jana, Debrya Saha, Divya Tej Sowpati |
| EPI_ISL_539691 | CSIR-Centre for Cellular and Molecular Biology | CSIR-Centre for Cellular and Molecular Biology | Pratheusa Maccha, Sakshi Shambhavi, Lamuk Zaveri, Shagufta Khan, Namami Gaur, Nikhil Hajirnis, M Soujanya Reddy, Tulasi Nagabandi, Purushotham Vodnala, Payel Mukherjee, Sofia Banu, Priya Singh,Onkar Kulkarni, Dhiviya Vedagiri, Divya Gupta, Vishal Sah, Santosh Kumar Kuncha, Krishnan Harinivas Harshan, Archana Bharadwaj Siva, Karthik Bharadwaj Tallapaka, G. Aditya Kumar, Koushick Sivakumar,Disha Nanda, Divya Das, Jotin Gogoi, Manish Bhattacharjee, Ravi Prasad Mukku, Rakesh K Mishra, Divya Tej Sowpati |
| EPI_ISL_539692 | CSIR-Centre for Cellular and Molecular Biology | CSIR-Centre for Cellular and Molecular Biology | Pratheusa Maccha, Shagufta Khan, Lamuk Zaveri, Namami Gaur, Sakshi Shambhavi, Tulasi Nagabandi, Nikhil Hajirnis, M Soujanya Reddy, Purushotham Vodnala, Payel Mukherjee, Sofia Banu, Priya Singh, Onkar Kulkarni, Dhiviya Vedagiri, Divya Gupta, Vishal Sah, Santosh Kumar Kuncha, Krishnan Harinivas Harshan, Archana Bharadwaj Siva, Karthik Bharadwaj Tallapaka, Disha Nanda, Divya Das, Jotin Gogoi, Manish Bhattacharjee, Ravi Prasad Mukku, Rakesh K Mishra, Divya Tej Sowpati |
| EPI_ISL_539693 | CSIR-Centre for Cellular and Molecular Biology | CSIR-Centre for Cellular and Molecular Biology | Pratheusa Maccha, Sofia Banu, Payel Mukherjee, Priya Singh, Onkar Kulkarni, Dhiviya Vedagiri, Divya Gupta, Vishal Sah, Santosh Kumar Kuncha, Krishnan Harinivas Harshan, Archana Bharadwaj Siva, Karthik Bharadwaj Tallapaka, Shagufta Khan, Lamuk Zaveri, Namami Gaur, Sakshi Shambhavi, Nikhil Hajirnis, M Soujanya Reddy, Tulasi Nagabandi, Purushotham Vodnala,Preethi Jampala, Sharada Ravi Iyer, Sulagana Mukherjee, Swetha Sundar, Peddapuvala Sai Uday Kiran, Rakesh K Mishra, Divya Tej Sowpati |
| EPI_ISL_539694 | CSIR-Centre for Cellular and Molecular Biology | CSIR-Centre for Cellular and Molecular Biology | Sakshi Shambhavi, Lamuk Zaveri, Shagufta Khan, Namami Gaur, Nikhil Hajirnis, M Soujanya Reddy, Pratheusa Maccha, Tulasi Nagabandi, Purushotham Vodnala, Payel Mukherjee, Sofia Banu, Priya Singh, Onkar Kulkarni, Dhiviya Vedagiri, Divya Gupta, Vishal Sah, Santosh Kumar Kuncha, Krishnan Harinivas Harshan, Archana Bharadwaj Siva, Karthik Bharadwaj Tallapaka, Deepak Kumar, Devi Prasad Vijayashankar, Disha Nanda, Divya Das, Jotin Gogoi, Manish Bhattacharjee, Rakesh K Mishra, Divya Tej Sowpati |
| EPI_ISL_539695 | CSIR-Centre for Cellular and Molecular Biology | CSIR-Centre for Cellular and Molecular Biology | Sakshi Shambhavi, Lamuk Zaveri, Shagufta Khan, Namami Gaur, Nikhil Hajirnis, M Soujanya Reddy, Pratheusa Maccha,Tulasi Nagabandi, Purushotham Vodnala, Payel Mukherjee, Sofia Banu, Priya Singh,Onkar Kulkarni, Dhiviya Vedagiri, Divya Gupta, Vishal Sah, Santosh Kumar Kuncha, Krishnan Harinivas Harshan, Archana Bharadwaj Siva, Karthik Bharadwaj Tallapaka, G. Aditya Kumar, Koushick Sivakumar, Pooja Ramesh Gupta, Rajan Kumar Jha, Shraddha Vijay Lahoti, Rakesh K Mishra, Divya Tej Sowpati |
| EPI_ISL_539696 | CSIR-Centre for Cellular and Molecular Biology | CSIR-Centre for Cellular and Molecular Biology | Sakshi Shambhavi, Lamuk Zaveri, Shagufta Khan, Nikhil Hajirnis, M Soujanya Reddy, Pratheusa Maccha, Namami Gaur, Tulasi Nagabandi, Purushotham Vodnala, Payel Mukherjee, Sofia Banu, Priya Singh, Onkar Kulkarni, Dhiviya Vedagiri, Divya Gupta, Vishal Sah, Santosh Kumar Kuncha, Krishnan Harinivas Harshan, Archana Bharadwaj Siva, Karthik Bharadwaj Tallapaka,G. Aditya Kumar, Koushick Sivakumar, Rakesh K Mishra, Divya Tej Sowpati |
| EPI_ISL_539697 | CSIR-Centre for Cellular and Molecular Biology | CSIR-Centre for Cellular and Molecular Biology | Shagufta Khan, Lamuk Zaveri, Namami Gaur, Sakshi Shambhavi, Nikhil Hajirnis, M Soujanya Reddy, Pratheusa Maccha, Tulasi Nagabandi, Purushotham Vodnala, Payel Mukherjee, Sofia Banu, Priya Singh, Onkar Kulkarni, Dhiviya Vedagiri, Divya Gupta, Vishal Sah, Santosh Kumar Kuncha, Krishnan Harinivas Harshan, Archana Bharadwaj Siva, Karthik Bharadwaj Tallapaka, Renu Sudhakar, Somesh Gorde, Gangumala Srinivas Reddy, Sujoy Deb, Swati Bayyana, Rakesh K Mishra, Divya Tej Sowpati |
| EPI_ISL_539698 | CSIR-Centre for Cellular and Molecular Biology | CSIR-Centre for Cellular and Molecular Biology | Shagufta Khan, Lamuk Zaveri, Namami Gaur, Sakshi Shambhavi, Nikhil Hajirnis, M Soujanya Reddy, Pratheusa Maccha, Tulasi Nagabandi, Purushotham Vodnala, Payel Mukherjee, Sofia Banu, Priya Singh, Onkar Kulkarni, Dhiviya Vedagiri, Divya Gupta, Vishal Sah, Santosh Kumar Kuncha, Krishnan Harinivas Harshan, Archana Bharadwaj Siva, Karthik Bharadwaj Tallapaka,Preethi Jampala, Sharada Ravi Iyer, Sulagana Mukherjee, Swetha Sundar, Peddapuvala Sai Uday Kiran Rakesh K Mishra, Divya Tej Sowpati |
| EPI_ISL_539699 | CSIR-Centre for Cellular and Molecular Biology | CSIR-Centre for Cellular and Molecular Biology | Shagufta Khan, Lamuk Zaveri, Namami Gaur, Sakshi Shambhavi, Nikhil Hajirnis, M Soujanya Reddy, Pratheusa Maccha,Tulasi Nagabandi, Purushotham Vodnala, Payel Mukherjee, Sofia Banu, Priya Singh, Onkar Kulkarni, Dhiviya Vedagiri, Divya Gupta, Vishal Sah, Santosh Kumar Kuncha, Krishnan Harinivas Harshan, Archana Bharadwaj Siva, Karthik Bharadwaj Tallapaka,Umesh Kumar, Unis Ahmad Bhat, Ajay Sarawagi, Priyanka Pant, Rajkanwar Nathawat, Rakesh K Mishra, Divya Tej Sowpati |
| EPI_ISL_539700 | CSIR-Centre for Cellular and Molecular Biology | CSIR-Centre for Cellular and Molecular Biology | Sofia Banu, Payel Mukherjee, Priya Singh,Onkar Kulkarni, Dhiviya Vedagiri, Divya Gupta, Vishal Sah, Santosh Kumar Kuncha, Krishnan Harinivas Harshan, Archana Bharadwaj Siva, Karthik Bharadwaj Tallapaka, Shagufta Khan, Lamuk Zaveri, Namami Gaur, Sakshi Shambhavi, Nikhil Hajirnis, M Soujanya Reddy, Pratheusa Maccha, Tulasi Nagabandi, Purushotham Vodnala, Disha Nanda, Divya Das, Jotin Gogoi, Manish Bhattacharjee, Ravi Prasad Mukku, Rakesh K Mishra, Divya Tej Sowpati |
| EPI_ISL_539701 | CSIR-Centre for Cellular and Molecular Biology | CSIR-Centre for Cellular and Molecular Biology | Sofia Banu, Payel Mukherjee, Priya Singh,Onkar Kulkarni, Dhiviya Vedagiri, Divya Gupta, Vishal Sah, Santosh Kumar Kuncha, Krishnan Harinivas Harshan, Archana Bharadwaj Siva, Karthik Bharadwaj Tallapaka, Shagufta Khan, Lamuk Zaveri, Namami Gaur, Sakshi Shambhavi, Nikhil Hajirnis, M Soujanya Reddy, Pratheusa Maccha,Tulasi Nagabandi, Purushotham Vodnala, Deepak Kumar, Devi Prasad Vijayashankar, Disha Nanda, Divya Das, Jotin Gogoi, Manish Bhattacharjee, Rakesh K Mishra, Divya Tej Sowpati |
| EPI_ISL_539702 | CSIR-Centre for Cellular and Molecular Biology | CSIR-Centre for Cellular and Molecular Biology | Sofia Banu, Payel Mukherjee, Priya Singh,Onkar Kulkarni, Dhiviya Vedagiri, Divya Gupta, Vishal Sah, Santosh Kumar Kuncha, Krishnan Harinivas Harshan, Archana Bharadwaj Siva, Karthik Bharadwaj Tallapaka, Shagufta Khan, Lamuk Zaveri, Namami Gaur, Sakshi Shambhavi, Tulasi Nagabandi, Nikhil Hajirnis, M Soujanya Reddy, Pratheusa Maccha, Purushotham Vodnala, Gokulan C G, Gunjan Purohit, Hanuman Tulashiram Kale, Pankaj Kumar, Prachand Issarapu, Rakesh K Mishra, Divya Tej Sowpati |
| EPI_ISL_539703 | CSIR-Centre for Cellular and Molecular Biology | CSIR-Centre for Cellular and Molecular Biology | Tulasi Nagabandi, Namami Gaur, Sakshi Shambhavi, Lamuk Zaveri, Shagufta Khan, Nikhil Hajirnis, M Soujanya Reddy, Pratheusa Maccha, Purushotham Vodnala, Payel Mukherjee, Sofia Banu, Priya Singh, Onkar Kulkarni, Dhiviya Vedagiri, Divya Gupta, Vishal Sah, Santosh Kumar Kuncha, Krishnan Harinivas Harshan, Archana Bharadwaj Siva, Karthik Bharadwaj Tallapaka,G. Aditya Kumar, Koushick Sivakumar, Pooja Ramesh Gupta, Rajan Kumar Jha, Shraddha Vijay Lahoti, Rakesh K Mishra, Divya Tej Sowpati |
| EPI_ISL_539704 | CSIR-Centre for Cellular and Molecular Biology | CSIR-Centre for Cellular and Molecular Biology | Tulasi Nagabandi, Namami Gaur, Sakshi Shambhavi, Lamuk Zaveri, Shagufta Khan, Nikhil Hajirnis, M Soujanya Reddy, Pratheusa Maccha, Purushotham Vodnala, Payel Mukherjee, Sofia Banu, Priya Singh,Onkar Kulkarni , Dhiviya Vedagiri, Divya Gupta, Vishal Sah, Santosh Kumar Kuncha, Krishnan Harinivas Harshan, Archana Bharadwaj Siva, Karthik Bharadwaj Tallapaka,G. Aditya Kumar, Koushick Sivakumar, Pooja Ramesh Gupta, Rajan Kumar Jha, Shraddha Vijay Lahoti, Rakesh K Mishra, Divya Tej Sowpati |
| EPI_ISL_539705 | CSIR-Centre for Cellular and Molecular Biology | CSIR-Centre for Cellular and Molecular Biology | Tulasi Nagabandi, Namami Gaur, Sakshi Shambhavi, Lamuk Zaveri, Shagufta Khan, Nikhil Hajirnis, M Soujanya Reddy, Pratheusa Maccha, Purushotham Vodnala, Payel Mukherjee, Sofia Banu, Priya Singh,Onkar Kulkarni, Dhiviya Vedagiri, Divya Gupta, Vishal Sah, Santosh Kumar Kuncha, Krishnan Harinivas Harshan, Archana Bharadwaj Siva, Karthik Bharadwaj Tallapaka,Kezia J Ann, Radhika Khandelwal, Roshan Maku Venkata, Shemin Mansuri, Sonu Uday, Rakesh K Mishra, Divya Tej Sowpati |
| EPI_ISL_539706 | CSIR-Centre for Cellular and Molecular Biology | CSIR-Centre for Cellular and Molecular Biology | Lamuk Zaveri, Shagufta Khan, Namami Gaur, Sakshi Shambhavi, Nikhil Hajirnis, M Soujanya Reddy, Pratheusa Maccha, Tulasi Nagabandi, Purushotham Vodnala, Payel Mukherjee, Sofia Banu, Priya Singh, Onkar Kulkarni, Dhiviya Vedagiri, Divya Gupta, Vishal Sah, Santosh Kumar Kuncha, Krishnan Harinivas Harshan, Archana Bharadwaj Siva, Karthik Bharadwaj Tallapaka, Renu Sudhakar, Somesh Gorde, Gangumala Srinivas Reddy, Sujoy Deb, Swati Bayyana, Rakesh K Mishra, Divya Tej Sowpati |
| EPI_ISL_539707 | CSIR-Centre for Cellular and Molecular Biology | CSIR-Centre for Cellular and Molecular Biology | Lamuk Zaveri, Shagufta Khan,Nikhil Hajirnis, M Soujanya Reddy, Pratheusa Maccha, Namami Gaur, Sakshi Shambhavi, Tulasi Nagabandi, Purushotham Vodnala, Payel Mukherjee, Sofia Banu, Priya Singh, Onkar Kulkarni, Dhiviya Vedagiri, Divya Gupta, Vishal Sah, Santosh Kumar Kuncha, Krishnan Harinivas Harshan, Archana Bharadwaj Siva, Karthik Bharadwaj Tallapaka,Umesh Kumar, Unis Ahmad Bhat, Ajay Sarawagi, Priyanka Pant, Rajkanwar Nathawat, Rakesh K Mishra, Divya Tej Sowpati |
| EPI_ISL_539708 | CSIR-Centre for Cellular and Molecular Biology | CSIR-Centre for Cellular and Molecular Biology | Lamuk Zaveri, Shagufta Khan,Nikhil Hajirnis, M Soujanya Reddy, Pratheusa Maccha, Namami Gaur, Sakshi Shambhavi, Tulasi Nagabandi, Purushotham Vodnala, Payel Mukherjee, Sofia Banu, Priya Singh, Onkar Kulkarni, Dhiviya Vedagiri, Divya Gupta, Vishal Sah, Santosh Kumar Kuncha, Krishnan Harinivas Harshan, Archana Bharadwaj Siva, Karthik Bharadwaj Tallapaka,Zeba Rizvi, Zuberwasim Sayyad, Kakade Aishwarya Arun, Amrutha H C, Ananga Ghosh, Rakesh K Mishra, Divya Tej Sowpati |

| | | | |
|---|---|---|---|
| EPI_ISL_539709 | CSIR-Centre for Cellular and Molecular Biology | CSIR-Centre for Cellular and Molecular Biology | M Soujanya Reddy, Nikhil Hajirnis, Pratheusa Maccha, Namami Gaur, Sakshi Shambhavi, Lamuk Zaveri, Shagufta Khan, Tulasi Nagabandi, Purushotham Vodnala, Payel Mukherjee, Sofia Banu, Priya Singh, Onkar Kulkarni, Dhiviya Vedagiri, Divya Gupta, Vishal Sah, Santosh Kumar Kuncha, Krishnan Harinivas Harshan, Archana Bharadwaj Siva, Karthik Bharadwaj Tallapaka, Zeba Rizvi, Zuberwasim Sayyad, Kakade Aishwarya Arun, Amrutha H C, Ananga Ghosh, Rakesh K Mishra, Divya Tej Sowpati |
| EPI_ISL_539710 | CSIR-Centre for Cellular and Molecular Biology | CSIR-Centre for Cellular and Molecular Biology | M Soujanya Reddy, Nikhil Hajirnis, Pratheusa Maccha, Payel Mukherjee, Sofia Banu, Priya Singh, Onkar Kulkarni,Tulasi Nagabandi, Namami Gaur, Sakshi Shambhavi, Lamuk Zaveri, Shagufta Khan, Purushotham Vodnala, Dhiviya Vedagiri, Divya Gupta, Vishal Sah, Santosh Kumar Kuncha, Krishnan Harinivas Harshan, Archana Bharadwaj Siva, Karthik Bharadwaj Tallapaka,Kezia J Ann, Radhika Khandelwal, Roshan Maku Venkata, Shemin Mansuri, Sonu Uday, Rakesh K Mishra, Divya Tej Sowpati |
| EPI_ISL_539711 | CSIR-Centre for Cellular and Molecular Biology | CSIR-Centre for Cellular and Molecular Biology | M Soujanya Reddy, Nikhil Hajirnis, Pratheusa Maccha, Sakshi Shambhavi, Lamuk Zaveri, Shagufta Khan, Namami Gaur, Tulasi Nagabandi, Purushotham Vodnala, Payel Mukherjee, Sofia Banu, Priya Singh,Onkar Kulkarni, Dhiviya Vedagiri, Divya Gupta, Vishal Sah, Santosh Kumar Kuncha, Krishnan Harinivas Harshan, Archana Bharadwaj Siva, Karthik Bharadwaj Tallapaka, G. Aditya Kumar, Koushick Sivakumar, Pooja Ramesh Gupta, Rajan Kumar Jha, Shraddha Vijay Lahoti, Rakesh K Mishra, Divya Tej Sowpati |
| EPI_ISL_539712 | CSIR-Centre for Cellular and Molecular Biology | CSIR-Centre for Cellular and Molecular Biology | Namami Gaur, Sakshi Shambhavi, Lamuk Zaveri, Shagufta Khan, Nikhil Hajirnis, M Soujanya Reddy, Pratheusa Maccha, Tulasi Nagabandi, Purushotham Vodnala, Payel Mukherjee, Sofia Banu, Priya Singh, Onkar Kulkarni, Dhiviya Vedagiri, Divya Gupta, Vishal Sah, Santosh Kumar Kuncha, Krishnan Harinivas Harshan, Archana Bharadwaj Siva, Karthik Bharadwaj Tallapaka, Zeba Rizvi, Zuberwasim Sayyad, Kakade Aishwarya Arun, Amrutha H C, Ananga Ghosh, Rakesh K Mishra, Divya Tej Sowpati |
| EPI_ISL_539713 | CSIR-Centre for Cellular and Molecular Biology | CSIR-Centre for Cellular and Molecular Biology | Namami Gaur, Sakshi Shambhavi, Lamuk Zaveri, Shagufta Khan, Nikhil Hajirnis, M Soujanya Reddy, Pratheusa Maccha, Tulasi Nagabandi, Purushotham Vodnala, Payel Mukherjee, Sofia Banu, Priya Singh, Onkar Kulkarni, Dhiviya Vedagiri, Divya Gupta, Vishal Sah, Santosh Kumar Kuncha, Krishnan Harinivas Harshan, Archana Bharadwaj Siva, Karthik Bharadwaj Tallapaka,G. Aditya Kumar, Koushick Sivakumar, Rakesh K Mishra, Divya Tej Sowpati |
| EPI_ISL_539714 | CSIR-Centre for Cellular and Molecular Biology | CSIR-Centre for Cellular and Molecular Biology | Namami Gaur, Sakshi Shambhavi, Lamuk Zaveri, Shagufta Khan, Nikhil Hajirnis, M Soujanya Reddy, Pratheusa Maccha, Tulasi Nagabandi, Purushotham Vodnala, Payel Mukherjee, Sofia Banu, Priya Singh,Onkar Kulkarni, Dhiviya Vedagiri, Divya Gupta, Vishal Sah, Santosh Kumar Kuncha, Krishnan Harinivas Harshan, Archana Bharadwaj Siva, Karthik Bharadwaj Tallapaka, Zeba Rizvi, Zuberwasim Sayyad, Kakade Aishwarya Arun, Amrutha H C, Ananga Ghosh, Rakesh K Mishra, Divya Tej Sowpati |
| EPI_ISL_539715 | CSIR-Centre for Cellular and Molecular Biology | CSIR-Centre for Cellular and Molecular Biology | Nikhil Hajirnis, M Soujanya Reddy, Pratheusa Maccha, Lamuk Zaveri, Shagufta Khan, Namami Gaur, Sakshi Shambhavi, Tulasi Nagabandi, Purushotham Vodnala, Payel Mukherjee, Sofia Banu, Priya Singh, Onkar Kulkarni, Dhiviya Vedagiri, Divya Gupta, Vishal Sah, Santosh Kumar Kuncha, Krishnan Harinivas Harshan, Archana Bharadwaj Siva, Karthik Bharadwaj Tallapaka,Zeba Rizvi, Zuberwasim Sayyad, Kakade Aishwarya Arun, Amrutha H C, Ananga Ghosh, Rakesh K Mishra, Divya Tej Sowpati |
| EPI_ISL_539716 | CSIR-Centre for Cellular and Molecular Biology | CSIR-Centre for Cellular and Molecular Biology | Nikhil Hajirnis, M Soujanya Reddy, Pratheusa Maccha, Namami Gaur, Sakshi Shambhavi, Lamuk Zaveri, Shagufta Khan, Tulasi Nagabandi, Purushotham Vodnala, Payel Mukherjee, Sofia Banu, Priya Singh,Onkar Kulkarni, Dhiviya Vedagiri, Divya Gupta, Vishal Sah, Santosh Kumar Kuncha, Krishnan Harinivas Harshan, Archana Bharadwaj Siva, Karthik Bharadwaj Tallapaka,Kezia J Ann, Radhika Khandelwal, Roshan Maku Venkata, Shemin Mansuri, Sonu Uday, Rakesh K Mishra, Divya Tej Sowpati |
| EPI_ISL_539717 | CSIR-Centre for Cellular and Molecular Biology | CSIR-Centre for Cellular and Molecular Biology | Nikhil Hajirnis, M Soujanya Reddy, Pratheusa Maccha, Payel Mukherjee, Sofia Banu, Priya Singh,Onkar Kulkarni, Dhiviya Vedagiri, Divya Gupta, Vishal Sah, Santosh Kumar Kuncha, Krishnan Harinivas Harshan, Archana Bharadwaj Siva, Karthik Bharadwaj Tallapaka, Shagufta Khan, Lamuk Zaveri, Namami Gaur, Sakshi Shambhavi, Tulasi Nagabandi, Purushotham Vodnala,Deepak Kumar, Devi Prasad Vijayashankar, Disha Nanda, Divya Das, Jotin Gogoi, Manish Bhattacharjee, Rakesh K Mishra, Divya Tej Sowpati |
| EPI_ISL_539718 | CSIR-Centre for Cellular and Molecular Biology | CSIR-Centre for Cellular and Molecular Biology | Payel Mukherjee, Sofia Banu, Priya Singh, Onkar Kulkarni, Dhiviya Vedagiri, Divya Gupta, Vishal Sah, Santosh Kumar Kuncha, Krishnan Harinivas Harshan, Archana Bharadwaj Siva, Karthik Bharadwaj Tallapaka, Shagufta Khan, Lamuk Zaveri, Nikhil Hajirnis, M Soujanya Reddy, Pratheusa Maccha, Namami Gaur, Sakshi Shambhavi, Tulasi Nagabandi, Purushotham Vodnala, G. Aditya Kumar, Koushick Sivakumar, Pooja Ramesh Gupta, Rajan Kumar Jha, Shraddha Vijay Lahoti, Rakesh K Mishra, Divya Tej Sowpati |
| EPI_ISL_539719 | CSIR-Centre for Cellular and Molecular Biology | CSIR-Centre for Cellular and Molecular Biology | Payel Mukherjee, Sofia Banu, Priya Singh, Onkar Kulkarni, Dhiviya Vedagiri, Divya Gupta, Vishal Sah, Santosh Kumar Kuncha, Krishnan Harinivas Harshan, Archana Bharadwaj Siva, Karthik Bharadwaj Tallapaka, Shagufta Khan, Lamuk Zaveri, Nikhil Hajirnis, M Soujanya Reddy, Pratheusa Maccha, Namami Gaur, Sakshi Shambhavi, Tulasi Nagabandi, Purushotham Vodnala, Gokulan C G, Gunjan Purohit, Hanuman Tulashiram Kale, Pankaj Kumar, Prachand Issarapu, Rakesh K Mishra, Divya Tej Sowpati |
| EPI_ISL_539720 | CSIR-Centre for Cellular and Molecular Biology | CSIR-Centre for Cellular and Molecular Biology | Payel Mukherjee, Sofia Banu, Priya Singh, Onkar Kulkarni, Dhiviya Vedagiri, Divya Gupta, Vishal Sah, Santosh Kumar Kuncha, Krishnan Harinivas Harshan, Archana Bharadwaj Siva, Karthik Bharadwaj Tallapaka, Shagufta Khan, Lamuk Zaveri, Nikhil Hajirnis, M Soujanya Reddy, Pratheusa Maccha, Namami Gaur, Sakshi Shambhavi, Tulasi Nagabandi, Purushotham Vodnala, Rakesh K Mishra, Sonu Uday, Sudipta Mondal, Annapoorna P Karthyayani, Debabrata Jana, Debrya Saha, Divya Tej Sowpati |
| EPI_ISL_539721 | CSIR-Centre for Cellular and Molecular Biology | CSIR-Centre for Cellular and Molecular Biology | Pratheusa Maccha, Sakshi Shambhavi, Lamuk Zaveri, Shagufta Khan, Namami Gaur, Nikhil Hajirnis, M Soujanya Reddy, Tulasi Nagabandi, Purushotham Vodnala, Payel Mukherjee, Sofia Banu, Priya Singh,Onkar Kulkarni, Dhiviya Vedagiri, Divya Gupta, Vishal Sah, Santosh Kumar Kuncha, Krishnan Harinivas Harshan, Archana Bharadwaj Siva, Karthik Bharadwaj Tallapaka, G. Aditya Kumar, Koushick Sivakumar,Disha Nanda, Divya Das, Jotin Gogoi, Manish Bhattacharjee, Ravi Prasad Mukku, Rakesh K Mishra, Divya Tej Sowpati |
| EPI_ISL_539722 | CSIR-Centre for Cellular and Molecular Biology | CSIR-Centre for Cellular and Molecular Biology | Pratheusa Maccha, Shagufta Khan, Lamuk Zaveri, Namami Gaur, Sakshi Shambhavi, Tulasi Nagabandi, Nikhil Hajirnis, M Soujanya Reddy, Purushotham Vodnala, Payel Mukherjee, Sofia Banu, Priya Singh, Onkar Kulkarni, Dhiviya Vedagiri, Divya Gupta, Vishal Sah, Santosh Kumar Kuncha, Krishnan Harinivas Harshan, Archana Bharadwaj Siva, Karthik Bharadwaj Tallapaka, Disha Nanda, Divya Das, Jotin Gogoi, Manish Bhattacharjee, Ravi Prasad Mukku, Rakesh K Mishra, Divya Tej Sowpati |
| EPI_ISL_539723 | CSIR-Centre for Cellular and Molecular Biology | CSIR-Centre for Cellular and Molecular Biology | Pratheusa Maccha, Sofia Banu, Payel Mukherjee, Priya Singh, Onkar Kulkarni, Dhiviya Vedagiri, Divya Gupta, Vishal Sah, Santosh Kumar Kuncha, Krishnan Harinivas Harshan, Archana Bharadwaj Siva, Karthik Bharadwaj Tallapaka, Shagufta Khan, Lamuk Zaveri, Namami Gaur, Sakshi Shambhavi, Nikhil Hajirnis, M Soujanya Reddy, Tulasi Nagabandi, Purushotham Vodnala,Preethi Jampala, Sharada Ravi Iyer, Sulagana Mukherjee, Swetha Sundar, Peddapuvala Sai Uday Kiran, Rakesh K Mishra, Divya Tej Sowpati |
| EPI_ISL_539724 | CSIR-Centre for Cellular and Molecular Biology | CSIR-Centre for Cellular and Molecular Biology | Sakshi Shambhavi, Lamuk Zaveri, Shagufta Khan, Namami Gaur, Nikhil Hajirnis, M Soujanya Reddy, Pratheusa Maccha, Tulasi Nagabandi, Purushotham Vodnala, Payel Mukherjee, Sofia Banu, Priya Singh, Onkar Kulkarni, Dhiviya Vedagiri, Divya Gupta, Vishal Sah, Santosh Kumar Kuncha, Krishnan Harinivas Harshan, Archana Bharadwaj Siva, Karthik Bharadwaj Tallapaka, Deepak Kumar, Devi Prasad Vijayashankar, Disha Nanda, Divya Das, Jotin Gogoi, Manish Bhattacharjee, Rakesh K Mishra, Divya Tej Sowpati |
| EPI_ISL_539725 | CSIR-Centre for Cellular and Molecular Biology | CSIR-Centre for Cellular and Molecular Biology | Sakshi Shambhavi, Lamuk Zaveri, Shagufta Khan, Namami Gaur, Nikhil Hajirnis, M Soujanya Reddy, Pratheusa Maccha,Tulasi Nagabandi, Purushotham Vodnala, Payel Mukherjee, Sofia Banu, Priya Singh,Onkar Kulkarni, Dhiviya Vedagiri, Divya Gupta, Vishal Sah, Santosh Kumar Kuncha, Krishnan Harinivas Harshan, Archana Bharadwaj Siva, Karthik Bharadwaj Tallapaka, G. Aditya Kumar, Koushick Sivakumar, Pooja Ramesh Gupta, Rajan Kumar Jha, Shraddha Vijay Lahoti, Rakesh K Mishra, Divya Tej Sowpati |
| EPI_ISL_539726 | CSIR-Centre for Cellular and Molecular Biology | CSIR-Centre for Cellular and Molecular Biology | Sakshi Shambhavi, Lamuk Zaveri, Shagufta Khan, Nikhil Hajirnis, M Soujanya Reddy, Pratheusa Maccha, Namami Gaur, Tulasi Nagabandi, Purushotham Vodnala, Payel Mukherjee, Sofia Banu, Priya Singh, Onkar Kulkarni, Dhiviya Vedagiri, Divya Gupta, Vishal Sah, Santosh Kumar Kuncha, Krishnan Harinivas Harshan, Archana Bharadwaj Siva, Karthik Bharadwaj Tallapaka,G. Aditya Kumar, Koushick Sivakumar, Rakesh K Mishra, Divya Tej Sowpati |
| EPI_ISL_539727 | CSIR-Centre for Cellular and Molecular Biology | CSIR-Centre for Cellular and Molecular Biology | Shagufta Khan, Lamuk Zaveri, Namami Gaur, Sakshi Shambhavi, Nikhil Hajirnis, M Soujanya Reddy, Pratheusa Maccha, Tulasi Nagabandi, Purushotham Vodnala, Payel Mukherjee, Sofia Banu, Priya Singh, Onkar Kulkarni, Dhiviya Vedagiri, Divya Gupta, Vishal Sah, Santosh Kumar Kuncha, Krishnan Harinivas Harshan, Archana Bharadwaj Siva, Karthik Bharadwaj Tallapaka, Renu Sudhakar, Somesh Gorde, Gangumala Srinivas Reddy, Sujoy Deb, Swati Bayyana, Rakesh K Mishra, Divya Tej Sowpati |
| EPI_ISL_539728 | CSIR-Centre for Cellular and Molecular Biology | CSIR-Centre for Cellular and Molecular Biology | Shagufta Khan, Lamuk Zaveri, Namami Gaur, Sakshi Shambhavi, Nikhil Hajirnis, M Soujanya Reddy, Pratheusa Maccha, Tulasi Nagabandi, Purushotham Vodnala, Payel Mukherjee, Sofia Banu, Priya Singh, Onkar Kulkarni, Dhiviya Vedagiri, Divya Gupta, Vishal Sah, Santosh Kumar Kuncha, Krishnan Harinivas Harshan, Archana Bharadwaj Siva, Karthik Bharadwaj Tallapaka,Preethi Jampala, Sharada Ravi Iyer, Sulagana Mukherjee, Swetha Sundar, Peddapuvala Sai Uday Kiran Rakesh K Mishra, Divya Tej Sowpati |
| EPI_ISL_539729 | CSIR-Centre for Cellular and Molecular Biology | CSIR-Centre for Cellular and Molecular Biology | Shagufta Khan, Lamuk Zaveri, Namami Gaur, Sakshi Shambhavi, Nikhil Hajirnis, M Soujanya Reddy, Pratheusa Maccha,Tulasi Nagabandi, Purushotham Vodnala, Payel Mukherjee, Sofia Banu, Priya Singh, Onkar Kulkarni, Dhiviya Vedagiri, Divya Gupta, Vishal Sah, Santosh Kumar Kuncha, Krishnan Harinivas Harshan, Archana Bharadwaj Siva, Karthik Bharadwaj Tallapaka,Umesh Kumar, Unis Ahmad Bhat, Ajay Sarawagi, Priyanka Pant, Rajkanwar Nathawat, Rakesh K Mishra, Divya Tej Sowpati |
| EPI_ISL_539730 | CSIR-Centre for Cellular and Molecular Biology | CSIR-Centre for Cellular and Molecular Biology | Sofia Banu, Payel Mukherjee, Priya Singh,Onkar Kulkarni, Dhiviya Vedagiri, Divya Gupta, Vishal Sah, Santosh Kumar Kuncha, Krishnan Harinivas Harshan, Archana Bharadwaj Siva, Karthik Bharadwaj Tallapaka, Shagufta Khan, Lamuk Zaveri, Namami Gaur, Sakshi Shambhavi, Nikhil Hajirnis, M Soujanya Reddy, Pratheusa Maccha, Tulasi Nagabandi, Purushotham Vodnala, Disha Nanda, Divya Das, Jotin Gogoi, Manish Bhattacharjee, Ravi Prasad Mukku, Rakesh K Mishra, Divya Tej Sowpati |
| EPI_ISL_539731 | CSIR-Centre for Cellular and Molecular Biology | CSIR-Centre for Cellular and Molecular Biology | Sofia Banu, Payel Mukherjee, Priya Singh,Onkar Kulkarni, Dhiviya Vedagiri, Divya Gupta, Vishal Sah, Santosh Kumar Kuncha, Krishnan Harinivas Harshan, Archana Bharadwaj Siva, Karthik Bharadwaj Tallapaka, Shagufta Khan, Lamuk Zaveri, Namami Gaur, Sakshi Shambhavi, Nikhil Hajirnis, M Soujanya Reddy, Pratheusa Maccha,Tulasi Nagabandi, Purushotham Vodnala, Deepak Kumar, Devi Prasad Vijayashankar, Disha Nanda, Divya Das, Jotin Gogoi, Manish Bhattacharjee, Rakesh K Mishra, Divya Tej Sowpati |
| EPI_ISL_539732 | CSIR-Centre for Cellular and Molecular Biology | CSIR-Centre for Cellular and Molecular Biology | Sofia Banu, Payel Mukherjee, Priya Singh,Onkar Kulkarni, Dhiviya Vedagiri, Divya Gupta, Vishal Sah, Santosh Kumar Kuncha, Krishnan Harinivas Harshan, Archana Bharadwaj Siva, Karthik Bharadwaj Tallapaka, Shagufta Khan, Lamuk Zaveri, Namami Gaur, Sakshi Shambhavi, Tulasi Nagabandi, Nikhil Hajirnis, M Soujanya Reddy, Pratheusa Maccha, Purushotham Vodnala, Gokulan C G, Gunjan Purohit, Hanuman Tulashiram Kale, Pankaj Kumar, Prachand Issarapu, Rakesh K Mishra, Divya Tej Sowpati |
| EPI_ISL_539733 | CSIR-Centre for Cellular and Molecular Biology | CSIR-Centre for Cellular and Molecular Biology | Tulasi Nagabandi, Namami Gaur, Sakshi Shambhavi, Lamuk Zaveri, Shagufta Khan, Nikhil Hajirnis, M Soujanya Reddy, Pratheusa Maccha, Purushotham Vodnala, Payel Mukherjee, Sofia Banu, Priya Singh, Onkar Kulkarni, Dhiviya Vedagiri, Divya Gupta, Vishal Sah, Santosh Kumar Kuncha, Krishnan Harinivas Harshan, Archana Bharadwaj Siva, Karthik Bharadwaj Tallapaka,G. Aditya Kumar, Koushick Sivakumar, Pooja Ramesh Gupta, Rajan Kumar Jha, Shraddha Vijay Lahoti, Rakesh K Mishra, Divya Tej Sowpati |
| EPI_ISL_539734 | CSIR-Centre for Cellular and Molecular Biology | CSIR-Centre for Cellular and Molecular Biology | Tulasi Nagabandi, Namami Gaur, Sakshi Shambhavi, Lamuk Zaveri, Shagufta Khan, Nikhil Hajirnis, M Soujanya Reddy, Pratheusa Maccha, Purushotham Vodnala, Payel Mukherjee, Sofia Banu, Priya Singh,Onkar Kulkarni , Dhiviya Vedagiri, Divya Gupta, Vishal Sah, Santosh Kumar Kuncha, Krishnan Harinivas Harshan, Archana Bharadwaj Siva, Karthik Bharadwaj Tallapaka,G. Aditya Kumar, Koushick Sivakumar, Pooja Ramesh Gupta, Rajan Kumar Jha, Shraddha Vijay Lahoti, Rakesh K Mishra, Divya Tej Sowpati |
| EPI_ISL_539735 | CSIR-Centre for Cellular and Molecular Biology | CSIR-Centre for Cellular and Molecular Biology | Tulasi Nagabandi, Namami Gaur, Sakshi Shambhavi, Lamuk Zaveri, Shagufta Khan, Nikhil Hajirnis, M Soujanya Reddy, Pratheusa Maccha, Purushotham Vodnala, Payel Mukherjee, Sofia Banu, Priya Singh,Onkar Kulkarni, Dhiviya Vedagiri, Divya Gupta, Vishal Sah, Santosh Kumar Kuncha, Krishnan Harinivas Harshan, Archana Bharadwaj Siva, Karthik Bharadwaj Tallapaka,Kezia J Ann, Radhika Khandelwal, Roshan Maku Venkata, Shemin Mansuri, Sonu Uday, Rakesh K Mishra, Divya Tej Sowpati |
| EPI_ISL_539736 | CSIR-Centre for Cellular and Molecular Biology | CSIR-Centre for Cellular and Molecular Biology | Lamuk Zaveri, Shagufta Khan, Namami Gaur, Sakshi Shambhavi, Nikhil Hajirnis, M Soujanya Reddy, Pratheusa Maccha, Tulasi Nagabandi, Purushotham Vodnala, Payel Mukherjee, Sofia Banu, Priya Singh, Onkar Kulkarni, Dhiviya Vedagiri, Divya Gupta, Vishal Sah, Santosh Kumar Kuncha, Krishnan Harinivas Harshan, Archana Bharadwaj Siva, Karthik Bharadwaj Tallapaka, Renu Sudhakar, Somesh Gorde, Gangumala Srinivas Reddy, Sujoy Deb, Swati Bayyana, Rakesh K Mishra, Divya Tej Sowpati |
| EPI_ISL_539737 | CSIR-Centre for Cellular and Molecular Biology | CSIR-Centre for Cellular and Molecular Biology | Lamuk Zaveri, Shagufta Khan,Nikhil Hajirnis, M Soujanya Reddy, Pratheusa Maccha, Namami Gaur, Sakshi Shambhavi, Tulasi Nagabandi, Purushotham Vodnala, Payel Mukherjee, Sofia Banu, Priya Singh, Onkar Kulkarni, Dhiviya Vedagiri, Divya Gupta, Vishal Sah, Santosh Kumar Kuncha, Krishnan Harinivas Harshan, Archana Bharadwaj Siva, Karthik Bharadwaj Tallapaka,Umesh Kumar, Unis Ahmad Bhat, Ajay Sarawagi, Priyanka Pant, Rajkanwar Nathawat, Rakesh K Mishra, Divya Tej Sowpati |
| EPI_ISL_539738 | CSIR-Centre for Cellular and Molecular Biology | CSIR-Centre for Cellular and Molecular Biology | Lamuk Zaveri, Shagufta Khan,Nikhil Hajirnis, M Soujanya Reddy, Pratheusa Maccha, Namami Gaur, Sakshi Shambhavi, Tulasi Nagabandi, Purushotham Vodnala, Payel Mukherjee, Sofia Banu, Priya Singh, Onkar Kulkarni, Dhiviya Vedagiri, Divya Gupta, Vishal Sah, Santosh Kumar Kuncha, Krishnan Harinivas Harshan, Archana Bharadwaj Siva, Karthik Bharadwaj Tallapaka,Zeba Rizvi, Zuberwasim Sayyad, Kakade Aishwarya Arun, Amrutha H C, Ananga Ghosh, Rakesh K Mishra, Divya Tej Sowpati |
| EPI_ISL_539739 | CSIR-Centre for Cellular and Molecular Biology | CSIR-Centre for Cellular and Molecular Biology | M Soujanya Reddy, Nikhil Hajirnis, Pratheusa Maccha, Namami Gaur, Sakshi Shambhavi, Lamuk Zaveri, Shagufta Khan, Tulasi Nagabandi, Purushotham Vodnala, Payel Mukherjee, Sofia Banu, Priya Singh, Onkar Kulkarni, Dhiviya Vedagiri, Divya Gupta, Vishal Sah, Santosh Kumar Kuncha, Krishnan Harinivas Harshan, Archana Bharadwaj Siva, Karthik Bharadwaj Tallapaka, Zeba Rizvi, Zuberwasim Sayyad, Kakade Aishwarya Arun, Amrutha H C, Ananga Ghosh, Rakesh K Mishra, Divya Tej Sowpati |
| EPI_ISL_539740 | CSIR-Centre for Cellular and Molecular Biology | CSIR-Centre for Cellular and Molecular Biology | M Soujanya Reddy, Nikhil Hajirnis, Pratheusa Maccha, Payel Mukherjee, Sofia Banu, Priya Singh, Onkar Kulkarni,Tulasi Nagabandi, Namami Gaur, Sakshi Shambhavi, Lamuk Zaveri, Shagufta Khan, Purushotham Vodnala, Dhiviya Vedagiri, Divya Gupta, Vishal Sah, Santosh Kumar Kuncha, Krishnan Harinivas Harshan, Archana Bharadwaj Siva, Karthik Bharadwaj Tallapaka,Kezia J Ann, Radhika Khandelwal, Roshan Maku Venkata, Shemin Mansuri, Sonu Uday, Rakesh K Mishra, Divya Tej Sowpati |
| EPI_ISL_539741 | CSIR-Centre for Cellular and Molecular Biology | CSIR-Centre for Cellular and Molecular | M Soujanya Reddy, Nikhil Hajirnis, Pratheusa Maccha, Sakshi Shambhavi, Lamuk Zaveri, Shagufta Khan, Namami Gaur, Tulasi Nagabandi, Purushotham Vodnala, Payel Mukherjee, Sofia Banu, Priya Singh,Onkar Kulkarni, Dhiviya |

| | | | |
|---|---|---|---|
| | | Biology | Vedagiri, Divya Gupta, Vishal Sah, Santosh Kumar Kuncha, Krishnan Harinivas Harshan, Archana Bharadwaj Siva, Karthik Bharadwaj Tallapaka, G. Aditya Kumar, Koushick Sivakumar, Pooja Ramesh Gupta, Rajan Kumar Jha, Shraddha Vijay Lahoti, Rakesh K Mishra, Divya Tej Sowpati |
| EPI_ISL_539742 | CSIR-Centre for Cellular and Molecular Biology | CSIR-Centre for Cellular and Molecular Biology | Namami Gaur, Sakshi Shambhavi, Lamuk Zaveri, Shagufta Khan, Nikhil Hajirnis, M Soujanya Reddy, Pratheusa Maccha, Tulasi Nagabandi, Purushotham Vodnala, Payel Mukherjee, Sofia Banu, Priya Singh, Onkar Kulkarni, Dhiviya Vedagiri, Divya Gupta, Vishal Sah, Santosh Kumar Kuncha, Krishnan Harinivas Harshan, Archana Bharadwaj Siva, Karthik Bharadwaj Tallapaka, Zeba Rizvi, Zuberwasim Sayyad, Kakade Aishwarya Arun, Amrutha H C, Ananga Ghosh, Rakesh K Mishra, Divya Tej Sowpati |
| EPI_ISL_539743 | CSIR-Centre for Cellular and Molecular Biology | CSIR-Centre for Cellular and Molecular Biology | Namami Gaur, Sakshi Shambhavi, Lamuk Zaveri, Shagufta Khan, Nikhil Hajirnis, M Soujanya Reddy, Pratheusa Maccha, Tulasi Nagabandi, Purushotham Vodnala, Payel Mukherjee, Sofia Banu, Priya Singh, Onkar Kulkarni, Dhiviya Vedagiri, Divya Gupta, Vishal Sah, Santosh Kumar Kuncha, Krishnan Harinivas Harshan, Archana Bharadwaj Siva, Karthik Bharadwaj Tallapaka,G. Aditya Kumar, Koushick Sivakumar, Rakesh K Mishra, Divya Tej Sowpati |
| EPI_ISL_539744 | CSIR-Centre for Cellular and Molecular Biology | CSIR-Centre for Cellular and Molecular Biology | Namami Gaur, Sakshi Shambhavi, Lamuk Zaveri, Shagufta Khan, Nikhil Hajirnis, M Soujanya Reddy, Pratheusa Maccha, Tulasi Nagabandi, Purushotham Vodnala, Payel Mukherjee, Sofia Banu, Priya Singh,Onkar Kulkarni, Dhiviya Vedagiri, Divya Gupta, Vishal Sah, Santosh Kumar Kuncha, Krishnan Harinivas Harshan, Archana Bharadwaj Siva, Karthik Bharadwaj Tallapaka, Zeba Rizvi, Zuberwasim Sayyad, Kakade Aishwarya Arun, Amrutha H C, Ananga Ghosh, Rakesh K Mishra, Divya Tej Sowpati |
| EPI_ISL_539745 | CSIR-Centre for Cellular and Molecular Biology | CSIR-Centre for Cellular and Molecular Biology | Nikhil Hajirnis, M Soujanya Reddy, Pratheusa Maccha, Lamuk Zaveri, Shagufta Khan, Namami Gaur, Sakshi Shambhavi, Tulasi Nagabandi, Purushotham Vodnala, Payel Mukherjee, Sofia Banu, Priya Singh, Onkar Kulkarni, Dhiviya Vedagiri, Divya Gupta, Vishal Sah, Santosh Kumar Kuncha, Krishnan Harinivas Harshan, Archana Bharadwaj Siva, Karthik Bharadwaj Tallapaka,Zeba Rizvi, Zuberwasim Sayyad, Kakade Aishwarya Arun, Amrutha H C, Ananga Ghosh, Rakesh K Mishra, Divya Tej Sowpati |
| EPI_ISL_539746 | CSIR-Centre for Cellular and Molecular Biology | CSIR-Centre for Cellular and Molecular Biology | Nikhil Hajirnis, M Soujanya Reddy, Pratheusa Maccha, Namami Gaur, Sakshi Shambhavi, Lamuk Zaveri, Shagufta Khan, Tulasi Nagabandi, Purushotham Vodnala, Payel Mukherjee, Sofia Banu, Priya Singh,Onkar Kulkarni, Dhiviya Vedagiri, Divya Gupta, Vishal Sah, Santosh Kumar Kuncha, Krishnan Harinivas Harshan, Archana Bharadwaj Siva, Karthik Bharadwaj Tallapaka,Kezia J Ann, Radhika Khandelwal, Roshan Maku Venkata, Shemin Mansuri, Sonu Uday, Rakesh K Mishra, Divya Tej Sowpati |
| EPI_ISL_539747 | CSIR-Centre for Cellular and Molecular Biology | CSIR-Centre for Cellular and Molecular Biology | Nikhil Hajirnis, M Soujanya Reddy, Pratheusa Maccha, Payel Mukherjee, Sofia Banu, Priya Singh,Onkar Kulkarni, Dhiviya Vedagiri, Divya Gupta, Vishal Sah, Santosh Kumar Kuncha, Krishnan Harinivas Harshan, Archana Bharadwaj Siva, Karthik Bharadwaj Tallapaka, Shagufta Khan, Lamuk Zaveri, Namami Gaur, Sakshi Shambhavi, Tulasi Nagabandi, Purushotham Vodnala,Deepak Kumar, Devi Prasad Vijayashankar, Disha Nanda, Divya Das, Jotin Gogoi, Manish Bhattacharjee, Rakesh K Mishra, Divya Tej Sowpati |
| EPI_ISL_539748 | CSIR-Centre for Cellular and Molecular Biology | CSIR-Centre for Cellular and Molecular Biology | Payel Mukherjee, Sofia Banu, Priya Singh, Onkar Kulkarni, Dhiviya Vedagiri, Divya Gupta, Vishal Sah, Santosh Kumar Kuncha, Krishnan Harinivas Harshan, Archana Bharadwaj Siva, Karthik Bharadwaj Tallapaka, Shagufta Khan, Lamuk Zaveri, Nikhil Hajirnis, M Soujanya Reddy, Pratheusa Maccha, Namami Gaur, Sakshi Shambhavi, Tulasi Nagabandi, Purushotham Vodnala, G. Aditya Kumar, Koushick Sivakumar, Pooja Ramesh Gupta, Rajan Kumar Jha, Shraddha Vijay Lahoti, Rakesh K Mishra, Divya Tej Sowpati |
| EPI_ISL_539749 | CSIR-Centre for Cellular and Molecular Biology | CSIR-Centre for Cellular and Molecular Biology | Payel Mukherjee, Sofia Banu, Priya Singh, Onkar Kulkarni, Dhiviya Vedagiri, Divya Gupta, Vishal Sah, Santosh Kumar Kuncha, Krishnan Harinivas Harshan, Archana Bharadwaj Siva, Karthik Bharadwaj Tallapaka, Shagufta Khan, Lamuk Zaveri, Nikhil Hajirnis, M Soujanya Reddy, Pratheusa Maccha, Namami Gaur, Sakshi Shambhavi, Tulasi Nagabandi, Purushotham Vodnala, Gokulan C G, Gunjan Purohit, Hanuman Tulashiram Kale, Pankaj Kumar, Prachand Issarapu, Rakesh K Mishra, Divya Tej Sowpati |
| EPI_ISL_539750 | CSIR-Centre for Cellular and Molecular Biology | CSIR-Centre for Cellular and Molecular Biology | Payel Mukherjee, Sofia Banu, Priya Singh, Onkar Kulkarni, Dhiviya Vedagiri, Divya Gupta, Vishal Sah, Santosh Kumar Kuncha, Krishnan Harinivas Harshan, Archana Bharadwaj Siva, Karthik Bharadwaj Tallapaka, Shagufta Khan, Lamuk Zaveri, Nikhil Hajirnis, M Soujanya Reddy, Pratheusa Maccha, Namami Gaur, Sakshi Shambhavi, Tulasi Nagabandi, Purushotham Vodnala, Rakesh K Mishra, Sonu Uday, Sudipta Mondal, Annapoorna P Karthyayani, Debabrata Jana, Debrya Saha, Divya Tej Sowpati |
| EPI_ISL_539751 | CSIR-Centre for Cellular and Molecular Biology | CSIR-Centre for Cellular and Molecular Biology | Pratheusa Maccha, Sakshi Shambhavi, Lamuk Zaveri, Shagufta Khan, Namami Gaur, Nikhil Hajirnis, M Soujanya Reddy, Tulasi Nagabandi, Purushotham Vodnala, Payel Mukherjee, Sofia Banu, Priya Singh,Onkar Kulkarni, Dhiviya Vedagiri, Divya Gupta, Vishal Sah, Santosh Kumar Kuncha, Krishnan Harinivas Harshan, Archana Bharadwaj Siva, Karthik Bharadwaj Tallapaka, G. Aditya Kumar, Koushick Sivakumar,Disha Nanda, Divya Das, Jotin Gogoi, Manish Bhattacharjee, Ravi Prasad Mukku, Rakesh K Mishra, Divya Tej Sowpati |
| EPI_ISL_539752 | CSIR-Centre for Cellular and Molecular Biology | CSIR-Centre for Cellular and Molecular Biology | Pratheusa Maccha, Shagufta Khan, Lamuk Zaveri, Namami Gaur, Sakshi Shambhavi, Tulasi Nagabandi, Nikhil Hajirnis, M Soujanya Reddy, Purushotham Vodnala, Payel Mukherjee, Sofia Banu, Priya Singh, Onkar Kulkarni, Dhiviya Vedagiri, Divya Gupta, Vishal Sah, Santosh Kumar Kuncha, Krishnan Harinivas Harshan, Archana Bharadwaj Siva, Karthik Bharadwaj Tallapaka, Disha Nanda, Divya Das, Jotin Gogoi, Manish Bhattacharjee, Ravi Prasad Mukku, Rakesh K Mishra, Divya Tej Sowpati |
| EPI_ISL_539753 | CSIR-Centre for Cellular and Molecular Biology | CSIR-Centre for Cellular and Molecular Biology | Pratheusa Maccha, Sofia Banu, Payel Mukherjee, Priya Singh, Onkar Kulkarni, Dhiviya Vedagiri, Divya Gupta, Vishal Sah, Santosh Kumar Kuncha, Krishnan Harinivas Harshan, Archana Bharadwaj Siva, Karthik Bharadwaj Tallapaka, Shagufta Khan, Lamuk Zaveri, Namami Gaur, Sakshi Shambhavi, Nikhil Hajirnis, M Soujanya Reddy, Tulasi Nagabandi, Purushotham Vodnala,Preethi Jampala, Sharada Ravi Iyer, Sulagana Mukherjee, Swetha Sundar, Peddapuvala Sai Uday Kiran, Rakesh K Mishra, Divya Tej Sowpati |
| EPI_ISL_539754 | CSIR-Centre for Cellular and Molecular Biology | CSIR-Centre for Cellular and Molecular Biology | Sakshi Shambhavi, Lamuk Zaveri, Shagufta Khan, Namami Gaur, Nikhil Hajirnis, M Soujanya Reddy, Pratheusa Maccha, Tulasi Nagabandi, Purushotham Vodnala, Payel Mukherjee, Sofia Banu, Priya Singh, Onkar Kulkarni, Dhiviya Vedagiri, Divya Gupta, Vishal Sah, Santosh Kumar Kuncha, Krishnan Harinivas Harshan, Archana Bharadwaj Siva, Karthik Bharadwaj Tallapaka, Deepak Kumar, Devi Prasad Vijayashankar, Disha Nanda, Divya Das, Jotin Gogoi, Manish Bhattacharjee, Rakesh K Mishra, Divya Tej Sowpati |
| EPI_ISL_539755 | CSIR-Centre for Cellular and Molecular Biology | CSIR-Centre for Cellular and Molecular Biology | Sakshi Shambhavi, Lamuk Zaveri, Shagufta Khan, Namami Gaur, Nikhil Hajirnis, M Soujanya Reddy, Pratheusa Maccha,Tulasi Nagabandi, Purushotham Vodnala, Payel Mukherjee, Sofia Banu, Priya Singh,Onkar Kulkarni, Dhiviya Vedagiri, Divya Gupta, Vishal Sah, Santosh Kumar Kuncha, Krishnan Harinivas Harshan, Archana Bharadwaj Siva, Karthik Bharadwaj Tallapaka, G. Aditya Kumar, Koushick Sivakumar, Pooja Ramesh Gupta, Rajan Kumar Jha, Shraddha Vijay Lahoti, Rakesh K Mishra, Divya Tej Sowpati |
| EPI_ISL_539756 | CSIR-Centre for Cellular and Molecular Biology | CSIR-Centre for Cellular and Molecular Biology | Sakshi Shambhavi, Lamuk Zaveri, Shagufta Khan, Nikhil Hajirnis, M Soujanya Reddy, Pratheusa Maccha, Namami Gaur, Tulasi Nagabandi, Purushotham Vodnala, Payel Mukherjee, Sofia Banu, Priya Singh, Onkar Kulkarni, Dhiviya Vedagiri, Divya Gupta, Vishal Sah, Santosh Kumar Kuncha, Krishnan Harinivas Harshan, Archana Bharadwaj Siva, Karthik Bharadwaj Tallapaka,G. Aditya Kumar, Koushick Sivakumar, Rakesh K Mishra, Divya Tej Sowpati |
| EPI_ISL_539757 | CSIR-Centre for Cellular and Molecular Biology | CSIR-Centre for Cellular and Molecular Biology | Shagufta Khan, Lamuk Zaveri, Namami Gaur, Sakshi Shambhavi, Nikhil Hajirnis, M Soujanya Reddy, Pratheusa Maccha, Tulasi Nagabandi, Purushotham Vodnala, Payel Mukherjee, Sofia Banu, Priya Singh, Onkar Kulkarni, Dhiviya Vedagiri, Divya Gupta, Vishal Sah, Santosh Kumar Kuncha, Krishnan Harinivas Harshan, Archana Bharadwaj Siva, Karthik Bharadwaj Tallapaka, Renu Sudhakar, Somesh Gorde, Gangumala Srinivas Reddy, Sujoy Deb, Swati Bayyana, Rakesh K Mishra, Divya Tej Sowpati |
| EPI_ISL_539758 | CSIR-Centre for Cellular and Molecular Biology | CSIR-Centre for Cellular and Molecular Biology | Shagufta Khan, Lamuk Zaveri, Namami Gaur, Sakshi Shambhavi, Nikhil Hajirnis, M Soujanya Reddy, Pratheusa Maccha, Tulasi Nagabandi, Purushotham Vodnala, Payel Mukherjee, Sofia Banu, Priya Singh, Onkar Kulkarni, Dhiviya Vedagiri, Divya Gupta, Vishal Sah, Santosh Kumar Kuncha, Krishnan Harinivas Harshan, Archana Bharadwaj Siva, Karthik Bharadwaj Tallapaka,Preethi Jampala, Sharada Ravi Iyer, Sulagana Mukherjee, Swetha Sundar, Peddapuvala Sai Uday Kiran Rakesh K Mishra, Divya Tej Sowpati |
| EPI_ISL_539759 | CSIR-Centre for Cellular and Molecular Biology | CSIR-Centre for Cellular and Molecular Biology | Shagufta Khan, Lamuk Zaveri, Namami Gaur, Sakshi Shambhavi, Nikhil Hajirnis, M Soujanya Reddy, Pratheusa Maccha,Tulasi Nagabandi, Purushotham Vodnala, Payel Mukherjee, Sofia Banu, Priya Singh, Onkar Kulkarni, Dhiviya Vedagiri, Divya Gupta, Vishal Sah, Santosh Kumar Kuncha, Krishnan Harinivas Harshan, Archana Bharadwaj Siva, Karthik Bharadwaj Tallapaka,Umesh Kumar, Unis Ahmad Bhat, Ajay Sarawagi, Priyanka Pant, Rajkanwar Nathawat, Rakesh K Mishra, Divya Tej Sowpati |
| EPI_ISL_539760 | CSIR-Centre for Cellular and Molecular Biology | CSIR-Centre for Cellular and Molecular Biology | Sofia Banu, Payel Mukherjee, Priya Singh,Onkar Kulkarni, Dhiviya Vedagiri, Divya Gupta, Vishal Sah, Santosh Kumar Kuncha, Krishnan Harinivas Harshan, Archana Bharadwaj Siva, Karthik Bharadwaj Tallapaka, Shagufta Khan, Lamuk Zaveri, Namami Gaur, Sakshi Shambhavi, Nikhil Hajirnis, M Soujanya Reddy, Pratheusa Maccha, Tulasi Nagabandi, Purushotham Vodnala, Disha Nanda, Divya Das, Jotin Gogoi, Manish Bhattacharjee, Ravi Prasad Mukku, Rakesh K Mishra, Divya Tej Sowpati |
| EPI_ISL_539761 | CSIR-Centre for Cellular and Molecular Biology | CSIR-Centre for Cellular and Molecular Biology | Sofia Banu, Payel Mukherjee, Priya Singh,Onkar Kulkarni, Dhiviya Vedagiri, Divya Gupta, Vishal Sah, Santosh Kumar Kuncha, Krishnan Harinivas Harshan, Archana Bharadwaj Siva, Karthik Bharadwaj Tallapaka, Shagufta Khan, Lamuk Zaveri, Namami Gaur, Sakshi Shambhavi, Nikhil Hajirnis, M Soujanya Reddy, Pratheusa Maccha,Tulasi Nagabandi, Purushotham Vodnala, Deepak Kumar, Devi Prasad Vijayashankar, Disha Nanda, Divya Das, Jotin Gogoi, Manish Bhattacharjee, Rakesh K Mishra, Divya Tej Sowpati |
| EPI_ISL_539762 | CSIR-Centre for Cellular and Molecular Biology | CSIR-Centre for Cellular and Molecular Biology | Sofia Banu, Payel Mukherjee, Priya Singh,Onkar Kulkarni, Dhiviya Vedagiri, Divya Gupta, Vishal Sah, Santosh Kumar Kuncha, Krishnan Harinivas Harshan, Archana Bharadwaj Siva, Karthik Bharadwaj Tallapaka, Shagufta Khan, Lamuk Zaveri, Namami Gaur, Sakshi Shambhavi, Tulasi Nagabandi, Nikhil Hajirnis, M Soujanya Reddy, Pratheusa Maccha, Purushotham Vodnala, Gokulan C G, Gunjan Purohit, Hanuman Tulashiram Kale, Pankaj Kumar, Prachand Issarapu, Rakesh K Mishra, Divya Tej Sowpati |
| EPI_ISL_539763 | CSIR-Centre for Cellular and Molecular Biology | CSIR-Centre for Cellular and Molecular Biology | Tulasi Nagabandi, Namami Gaur, Sakshi Shambhavi, Lamuk Zaveri, Shagufta Khan, Nikhil Hajirnis, M Soujanya Reddy, Pratheusa Maccha, Purushotham Vodnala, Payel Mukherjee, Sofia Banu, Priya Singh, Onkar Kulkarni, Dhiviya Vedagiri, Divya Gupta, Vishal Sah, Santosh Kumar Kuncha, Krishnan Harinivas Harshan, Archana Bharadwaj Siva, Karthik Bharadwaj Tallapaka,G. Aditya Kumar, Koushick Sivakumar, Pooja Ramesh Gupta, Rajan Kumar Jha, Shraddha Vijay Lahoti, Rakesh K Mishra, Divya Tej Sowpati |
| EPI_ISL_539764 | CSIR-Centre for Cellular and Molecular Biology | CSIR-Centre for Cellular and Molecular Biology | Tulasi Nagabandi, Namami Gaur, Sakshi Shambhavi, Lamuk Zaveri, Shagufta Khan, Nikhil Hajirnis, M Soujanya Reddy, Pratheusa Maccha, Purushotham Vodnala, Payel Mukherjee, Sofia Banu, Priya Singh,Onkar Kulkarni , Dhiviya Vedagiri, Divya Gupta, Vishal Sah, Santosh Kumar Kuncha, Krishnan Harinivas Harshan, Archana Bharadwaj Siva, Karthik Bharadwaj Tallapaka,G. Aditya Kumar, Koushick Sivakumar, Pooja Ramesh Gupta, Rajan Kumar Jha, Shraddha Vijay Lahoti, Rakesh K Mishra, Divya Tej Sowpati |
| EPI_ISL_539765 | CSIR-Centre for Cellular and Molecular Biology | CSIR-Centre for Cellular and Molecular Biology | Tulasi Nagabandi, Namami Gaur, Sakshi Shambhavi, Lamuk Zaveri, Shagufta Khan, Nikhil Hajirnis, M Soujanya Reddy, Pratheusa Maccha, Purushotham Vodnala, Payel Mukherjee, Sofia Banu, Priya Singh,Onkar Kulkarni, Dhiviya Vedagiri, Divya Gupta, Vishal Sah, Santosh Kumar Kuncha, Krishnan Harinivas Harshan, Archana Bharadwaj Siva, Karthik Bharadwaj Tallapaka,Kezia J Ann, Radhika Khandelwal, Roshan Maku Venkata, Shemin Mansuri, Sonu Uday, Rakesh K Mishra, Divya Tej Sowpati |
| EPI_ISL_539766 | CSIR-Centre for Cellular and Molecular Biology | CSIR-Centre for Cellular and Molecular Biology | Tulasi Nagabandi, Namami Gaur, Sakshi Shambhavi, Lamuk Zaveri, Shagufta Khan, Nikhil Hajirnis, M Soujanya Reddy, Pratheusa Maccha, Purushotham Vodnala, Payel Mukherjee, Sofia Banu, Priya Singh, Onkar Kulkarni, Dhiviya Vedagiri, Divya Gupta, Vishal Sah, Santosh Kumar Kuncha, Krishnan Harinivas Harshan, Archana Bharadwaj Siva, Karthik Bharadwaj Tallapaka,G. Aditya Kumar, Koushick Sivakumar, Pooja Ramesh Gupta, Rajan Kumar Jha, Shraddha Vijay Lahoti, Rakesh K Mishra, Divya Tej Sowpati |
| EPI_ISL_539767 | CSIR-Centre for Cellular and Molecular Biology | CSIR-Centre for Cellular and Molecular Biology | Tulasi Nagabandi, Namami Gaur, Sakshi Shambhavi, Lamuk Zaveri, Shagufta Khan, Nikhil Hajirnis, M Soujanya Reddy, Pratheusa Maccha, Purushotham Vodnala, Payel Mukherjee, Sofia Banu, Priya Singh,Onkar Kulkarni , Dhiviya Vedagiri, Divya Gupta, Vishal Sah, Santosh Kumar Kuncha, Krishnan Harinivas Harshan, Archana Bharadwaj Siva, Karthik Bharadwaj Tallapaka,G. Aditya Kumar, Koushick Sivakumar, Pooja Ramesh Gupta, Rajan Kumar Jha, Shraddha Vijay Lahoti, Rakesh K Mishra, Divya Tej Sowpati |
| EPI_ISL_539768 | CSIR-Centre for Cellular and Molecular Biology | CSIR-Centre for Cellular and Molecular Biology | Tulasi Nagabandi, Namami Gaur, Sakshi Shambhavi, Lamuk Zaveri, Shagufta Khan, Nikhil Hajirnis, M Soujanya Reddy, Pratheusa Maccha, Purushotham Vodnala, Payel Mukherjee, Sofia Banu, Priya Singh,Onkar Kulkarni, Dhiviya Vedagiri, Divya Gupta, Vishal Sah, Santosh Kumar Kuncha, Krishnan Harinivas Harshan, Archana Bharadwaj Siva, Karthik Bharadwaj Tallapaka,Kezia J Ann, Radhika Khandelwal, Roshan Maku Venkata, Shemin Mansuri, Sonu Uday, Rakesh K Mishra, Divya Tej Sowpati |
| EPI_ISL_539769 | CSIR-Centre for Cellular and Molecular Biology | CSIR-Centre for Cellular and Molecular Biology | Lamuk Zaveri, Shagufta Khan,Nikhil Hajirnis, M Soujanya Reddy, Pratheusa Maccha, Namami Gaur, Sakshi Shambhavi, Tulasi Nagabandi, Purushotham Vodnala, Payel Mukherjee, Sofia Banu, Priya Singh, Onkar Kulkarni, Dhiviya Vedagiri, Divya Gupta, Vishal Sah, Santosh Kumar Kuncha, Krishnan Harinivas Harshan, Archana Bharadwaj Siva, Karthik Bharadwaj Tallapaka,Umesh Kumar, Unis Ahmad Bhat, Ajay Sarawagi, Priyanka Pant, Rajkanwar Nathawat, Rakesh K Mishra, Divya Tej Sowpati |
| EPI_ISL_539770 | CSIR-Centre for Cellular and Molecular Biology | CSIR-Centre for Cellular and Molecular Biology | Lamuk Zaveri, Shagufta Khan,Nikhil Hajirnis, M Soujanya Reddy, Pratheusa Maccha, Namami Gaur, Sakshi Shambhavi, Tulasi Nagabandi, Purushotham Vodnala, Payel Mukherjee, Sofia Banu, Priya Singh, Onkar Kulkarni, Dhiviya Vedagiri, Divya Gupta, Vishal Sah, Santosh Kumar Kuncha, Krishnan Harinivas Harshan, Archana Bharadwaj Siva, Karthik Bharadwaj Tallapaka,Zeba Rizvi, Zuberwasim Sayyad, Kakade Aishwarya Arun, Amrutha H C, Ananga Ghosh, Rakesh K Mishra, Divya Tej Sowpati |
| EPI_ISL_539771 | CSIR-Centre for Cellular and Molecular Biology | CSIR-Centre for Cellular and Molecular Biology | Pratheusa Maccha, Sakshi Shambhavi, Lamuk Zaveri, Shagufta Khan, Namami Gaur, Nikhil Hajirnis, M Soujanya Reddy, Tulasi Nagabandi, Purushotham Vodnala, Payel Mukherjee, Sofia Banu, Priya Singh,Onkar Kulkarni, Dhiviya Vedagiri, Divya Gupta, Vishal Sah, Santosh Kumar Kuncha, Krishnan Harinivas Harshan, Archana Bharadwaj Siva, Karthik Bharadwaj Tallapaka, G. Aditya Kumar, Koushick Sivakumar,Disha Nanda, Divya Das, Jotin Gogoi, Manish Bhattacharjee, Ravi Prasad Mukku, Rakesh K Mishra, Divya Tej Sowpati |
| EPI_ISL_539772 | CSIR-Centre for Cellular and Molecular Biology | CSIR-Centre for Cellular and Molecular Biology | Pratheusa Maccha, Shagufta Khan, Lamuk Zaveri, Namami Gaur, Sakshi Shambhavi, Tulasi Nagabandi, Nikhil Hajirnis, M Soujanya Reddy, Purushotham Vodnala, Payel Mukherjee, Sofia Banu, Priya Singh, Onkar Kulkarni, Dhiviya Vedagiri, Divya Gupta, Vishal Sah, Santosh Kumar Kuncha, Krishnan Harinivas Harshan, Archana Bharadwaj Siva, Karthik Bharadwaj Tallapaka, Disha Nanda, Divya Das, Jotin Gogoi, Manish Bhattacharjee, Ravi Prasad Mukku, Rakesh K Mishra, Divya Tej Sowpati |
| EPI_ISL_539773 | CSIR-Centre for Cellular and Molecular Biology | CSIR-Centre for Cellular and Molecular Biology | Payel Mukherjee, Sofia Banu, Priya Singh, Onkar Kulkarni, Dhiviya Vedagiri, Divya Gupta, Vishal Sah, Santosh Kumar Kuncha, Krishnan Harinivas Harshan, Archana Bharadwaj Siva, Karthik Bharadwaj Tallapaka, Shagufta Khan, Lamuk Zaveri, Nikhil Hajirnis, M Soujanya Reddy, Pratheusa Maccha, Namami Gaur, Sakshi Shambhavi, Tulasi Nagabandi, Purushotham Vodnala, Gokulan C G, Gunjan Purohit, Hanuman Tulashiram Kale, Pankaj Kumar, Prachand Issarapu, Rakesh K Mishra, Divya Tej Sowpati |

| | | | |
|---|---|---|---|
| EPI_ISL_539774 | CSIR-Centre for Cellular and Molecular Biology | CSIR-Centre for Cellular and Molecular Biology | Payel Mukherjee, Sofia Banu, Priya Singh, Onkar Kulkarni, Dhiviya Vedagiri, Divya Gupta, Vishal Sah, Santosh Kumar Kuncha, Krishnan Harinivas Harshan, Archana Bharadwaj Siva, Karthik Bharadwaj Tallapaka, Shagufta Khan, Lamuk Zaveri, Nikhil Hajirnis, M Soujanya Reddy, Pratheusa Maccha, Namami Gaur, Sakshi Shambhavi, Tulasi Nagabandi, Purushotham Vodnala, Rakesh K Mishra, Sonu Uday, Sudipta Mondal, Annapoorna P Karthyayani, Debabrata Jana, Debrya Saha, Divya Tej Sowpati |
| EPI_ISL_539775 | CSIR-Centre for Cellular and Molecular Biology | CSIR-Centre for Cellular and Molecular Biology | Payel Mukherjee, Sofia Banu, Priya Singh, Onkar Kulkarni, Dhiviya Vedagiri, Divya Gupta, Vishal Sah, Santosh Kumar Kuncha, Krishnan Harinivas Harshan, Archana Bharadwaj Siva, Karthik Bharadwaj Tallapaka, Shagufta Khan, Lamuk Zaveri, Nikhil Hajirnis, M Soujanya Reddy, Pratheusa Maccha, Namami Gaur, Sakshi Shambhavi, Tulasi Nagabandi, Purushotham Vodnala, Gokulan C G, Gunjan Purohit, Hanuman Tulashiram Kale, Pankaj Kumar, Prachand Issarapu, Rakesh K Mishra, Divya Tej Sowpati |
| EPI_ISL_539776 | National Institute of Public Health (Czech Republic) | State Veterinary Institute Prague | Nagy, A; Jirincova, H; Novakova, L; Trnka, D; Vecerova, J. |
| EPI_ISL_539777 | The National Institute of Public Health | State Veterinary Institute Prague | Nagy,A;Jirincova,H;Novakova,L;Trnka,D;Vecerova,J |
| EPI_ISL_539778, EPI_ISL_539779, EPI_ISL_539780, EPI_ISL_539781, EPI_ISL_539782 | National Institute of Public Health (Czech Republic) | State Veterinary Institute Prague | Nagy, A; Jirincova, H; Novakova, L; Trnka, D; Vecerova, J. |
| EPI_ISL_539783, EPI_ISL_539784 | Universidad Regional Amazonica IKIAM | Institute of Microbiology, Universidad San Francisco de Quito | Fabian Aguilar, Katherine Apunte, Andrea Carrera, Nina Espinoza de los Monteros, Giovanna Moran, Marcelo Ortiz, Yeimy Rojas, Sonia Sislema, Carolina Proaño-Bolaños, Belén Prado-Vivar, Sully Márquez, Juan José Guadalupe, Monica Becerra-Wong, Bernardo Gutiérrez, Verónica Barragán, Patricio Rojas-Silva, Gabriel Trueba, Michelle Grunauer, Paúl Cárdenas |
| EPI_ISL_539785, EPI_ISL_539786, EPI_ISL_539787, EPI_ISL_539788 | Institute of Microbiology, Universidad San Francisco de Quito | Institute of Microbiology, Universidad San Francisco de Quito | Andrea Macias, Belén Prado-Vivar, Sully Márquez, Juan José Guadalupe, Monica Becerra-Wong, Bernardo Gutiérrez, Verónica Barragán, Patricio Rojas-Silva, Gabriel Trueba, Michelle Grunauer, Paúl Cárdenas |
| EPI_ISL_539789, EPI_ISL_539790, EPI_ISL_539791, EPI_ISL_539792, EPI_ISL_539793 | Institute of Microbiology, Universidad San Francisco de Quito | Institute of Microbiology, Universidad San Francisco de Quito | Belén Prado-Vivar, Sully Márquez, Juan José Guadalupe, Monica Becerra-Wong, Bernardo Gutiérrez, Ligia Briceño, Nabih Dahik, Verónica Barragán, Patricio Rojas-Silva, Gabriel Trueba, Michelle Grunauer, Paúl Cárdenas |
| EPI_ISL_539794, EPI_ISL_539795, EPI_ISL_539796, EPI_ISL_539797, EPI_ISL_539798, EPI_ISL_539799, EPI_ISL_539800, EPI_ISL_539801, EPI_ISL_539802 | Wyoming Public Health Laboratory | Wyoming Public Health Laboratory | Noah Hull, Rob Christensen, Jim Mildenberger, Joel Sevinsky, Cari Sloma, and Wanda Manley |
| EPI_ISL_539803 | Queen Mary Hospital | Hong Kong Department of Health | Alan K.L. Tsang, Peter C.W. Yip, Edman T.K. Lam, Rickjason C.W. Chan, Dominic N.C. Tsang |
| EPI_ISL_539804 | Yan Chai Hospital | Hong Kong Department of Health | Alan K.L. Tsang, Peter C.W. Yip, Edman T.K. Lam, Rickjason C.W. Chan, Dominic N.C. Tsang |
| EPI_ISL_539805 | Kwong Wah Hospital | Hong Kong Department of Health | Alan K.L. Tsang, Peter C.W. Yip, Edman T.K. Lam, Rickjason C.W. Chan, Dominic N.C. Tsang |
| EPI_ISL_539806, EPI_ISL_539807, EPI_ISL_539808 | Princess Margaret Hospital | Hong Kong Department of Health | Alan K.L. Tsang, Peter C.W. Yip, Edman T.K. Lam, Rickjason C.W. Chan, Dominic N.C. Tsang |
| EPI_ISL_539809, EPI_ISL_539810, EPI_ISL_539811, EPI_ISL_539812 | Asiaworld Expo Command Post | Hong Kong Department of Health | Alan K.L. Tsang, Peter C.W. Yip, Edman T.K. Lam, Rickjason C.W. Chan, Dominic N.C. Tsang |
| EPI_ISL_539813 | Prince of Wales Hospital | Hong Kong Department of Health | Alan K.L. Tsang, Peter C.W. Yip, Edman T.K. Lam, Rickjason C.W. Chan, Dominic N.C. Tsang |
| EPI_ISL_539814 | Tuen Mun Hospital | Hong Kong Department of Health | Alan K.L. Tsang, Peter C.W. Yip, Edman T.K. Lam, Rickjason C.W. Chan, Dominic N.C. Tsang |
| EPI_ISL_539815 | Queen Elizabeth Hospital | Hong Kong Department of Health | Alan K.L. Tsang, Peter C.W. Yip, Edman T.K. Lam, Rickjason C.W. Chan, Dominic N.C. Tsang |
| EPI_ISL_539816, EPI_ISL_539817 | Queen Mary Hospital | Hong Kong Department of Health | Alan K.L. Tsang, Peter C.W. Yip, Edman T.K. Lam, Rickjason C.W. Chan, Dominic N.C. Tsang |
| EPI_ISL_539818, EPI_ISL_539819 | Tuen Mun Hospital | Hong Kong Department of Health | Alan K.L. Tsang, Peter C.W. Yip, Edman T.K. Lam, Rickjason C.W. Chan, Dominic N.C. Tsang |
| EPI_ISL_539820 | Queen Mary Hospital | Hong Kong Department of Health | Alan K.L. Tsang, Peter C.W. Yip, Edman T.K. Lam, Rickjason C.W. Chan, Dominic N.C. Tsang |
| EPI_ISL_539821 | Asiaworld Expo Command Post | Hong Kong Department of Health | Alan K.L. Tsang, Peter C.W. Yip, Edman T.K. Lam, Rickjason C.W. Chan, Dominic N.C. Tsang |
| EPI_ISL_539822, EPI_ISL_539823 | Communicable Disease Branch | Hong Kong Department of Health | Alan K.L. Tsang, Peter C.W. Yip, Edman T.K. Lam, Rickjason C.W. Chan, Dominic N.C. Tsang |
| EPI_ISL_539824, EPI_ISL_539825, EPI_ISL_539826, EPI_ISL_539827, EPI_ISL_539828, EPI_ISL_539829, EPI_ISL_539830, EPI_ISL_539831, EPI_ISL_539832, EPI_ISL_539833, EPI_ISL_539834, EPI_ISL_539835, EPI_ISL_539836, EPI_ISL_539837, EPI_ISL_539838, EPI_ISL_539839, EPI_ISL_539840, EPI_ISL_539841 | | | |
| see above | Minnesota Department of Health, Public Health Laboratory | Minnesota Department of Health, Public Health Laboratory | Matt Plumb, Jacob Garfin, and Xiong Wang |
| EPI_ISL_539842, EPI_ISL_539843, EPI_ISL_539844, EPI_ISL_539845, EPI_ISL_539846, EPI_ISL_539847, EPI_ISL_539848, EPI_ISL_539849 | Mayo Clinic & Mayo Clinic Laboratories | Minnesota Department of Health, Public Health Laboratory | Matt Plumb, Jacob Garfin, and Xiong Wang |
| EPI_ISL_539850, EPI_ISL_539851 | Pok Oi Hospital | Hong Kong Department of Health | Alan K.L. Tsang, Peter C.W. Yip, Edman T.K. Lam, Rickjason C.W. Chan, Dominic N.C. Tsang |
| EPI_ISL_539879 | Microbiology Division, South Carolina Department of Health and Environmental Control | Microbiology Division, South Carolina Department of Health and Environmental Control | Flores,H. |
| EPI_ISL_539880 | Respiratory Virus Unit, Microbiology Services Colindale, Public Health England | Respiratory Virus Unit, Microbiology Services Colindale, Public Health England | PHE Covid Sequencing Team |
| EPI_ISL_539881 | Kungsbacka Narakut | The Public Health Agency of Sweden | Anna-Malin Linde, Maria Lind Karlberg, Oskar Karlsson Lindsjo, Olov Svartstrom, Mattias Haukland, Reza Advani, Sandra Broddesson, Anna Risberg, Theresa Enkirch, Mia Brytting, Karin Tegmark-Wisell |
| EPI_ISL_539882 | Narhalsan Olskroken VC | The Public Health Agency of Sweden | Anna-Malin Linde, Maria Lind Karlberg, Oskar Karlsson Lindsjo, Olov Svartstrom, Mattias Haukland, Reza Advani, Sandra Broddesson, Anna Risberg, Theresa Enkirch, Mia Brytting, Karin Tegmark-Wisell |
| EPI_ISL_539883, EPI_ISL_539884 | Omtanken Grimmered | The Public Health Agency of Sweden | Anna-Malin Linde, Maria Lind Karlberg, Oskar Karlsson Lindsjo, Olov Svartstrom, Mattias Haukland, Reza Advani, Sandra Broddesson, Anna Risberg, Theresa Enkirch, Mia Brytting, Karin Tegmark-Wisell |
| EPI_ISL_539885 | Narhalsan Fjallbacka VC | The Public Health Agency of Sweden | Anna-Malin Linde, Maria Lind Karlberg, Oskar Karlsson Lindsjo, Olov Svartstrom, Mattias Haukland, Reza Advani, Sandra Broddesson, Anna Risberg, Theresa Enkirch, Mia Brytting, Karin Tegmark-Wisell |
| EPI_ISL_539886 | Barnakuten | The Public Health Agency of Sweden | Anna-Malin Linde, Maria Lind Karlberg, Oskar Karlsson Lindsjo, Olov Svartstrom, Mattias Haukland, Reza Advani, Sandra Broddesson, Anna Risberg, Theresa Enkirch, Mia Brytting, Karin Tegmark-Wisell |
| EPI_ISL_539894, EPI_ISL_539896 | PHE South West Regional Laboratory, National Infection Service | Wellcome Sanger Institute for the COVID-19 Genomics UK Consortium | Stephanie Hutchings, Hannah Pymont, Dr Peter Muir, Barry Vipond, Rich Hopes; and Alex Alderton, Roberto Amato, Sonia Goncalves, Ewan Harrison, David K. Jackson, Ian Johnston, Dominic Kwiatkowski, Cordelia Langford, John Sillitoe on behalf of the Wellcome Sanger Institute COVID-19 Surveillance Team |
| EPI_ISL_539897, EPI_ISL_539898, EPI_ISL_539900, EPI_ISL_539901, EPI_ISL_539902, EPI_ISL_539903, EPI_ISL_539904, EPI_ISL_539905, EPI_ISL_539907, EPI_ISL_539908, EPI_ISL_539909, EPI_ISL_539910, EPI_ISL_539911, EPI_ISL_539912, EPI_ISL_539913, EPI_ISL_539914, EPI_ISL_539915, EPI_ISL_539916, EPI_ISL_539917, EPI_ISL_539918, EPI_ISL_539919, EPI_ISL_539921, EPI_ISL_539922, EPI_ISL_539923, EPI_ISL_539924, EPI_ISL_539925, EPI_ISL_539926, EPI_ISL_539927, EPI_ISL_539928, EPI_ISL_539929, EPI_ISL_539930, EPI_ISL_539931, EPI_ISL_539932, EPI_ISL_539933, EPI_ISL_539934, EPI_ISL_539935, EPI_ISL_539936, EPI_ISL_539937, EPI_ISL_539938, EPI_ISL_539939, EPI_ISL_539941, EPI_ISL_539942, EPI_ISL_539943, EPI_ISL_539944, EPI_ISL_539945, EPI_ISL_539946, EPI_ISL_539947, EPI_ISL_539948, EPI_ISL_539949, EPI_ISL_539950, EPI_ISL_539951, EPI_ISL_539952, EPI_ISL_539953, EPI_ISL_539954, EPI_ISL_539955, EPI_ISL_539956, EPI_ISL_539957, EPI_ISL_539958, EPI_ISL_539959, EPI_ISL_539960, EPI_ISL_539964, EPI_ISL_539965, EPI_ISL_539966, EPI_ISL_539967, EPI_ISL_539969, EPI_ISL_539970, EPI_ISL_539971, EPI_ISL_539972, EPI_ISL_539973, EPI_ISL_539974, EPI_ISL_539975, EPI_ISL_539976, EPI_ISL_539977, EPI_ISL_539978, EPI_ISL_539979, EPI_ISL_539980, EPI_ISL_539981, EPI_ISL_539983, EPI_ISL_539984, EPI_ISL_539985, EPI_ISL_539986, EPI_ISL_539987, EPI_ISL_539988, EPI_ISL_539990, EPI_ISL_539991, EPI_ISL_539992, EPI_ISL_539993, EPI_ISL_539994, EPI_ISL_539995, EPI_ISL_539996, EPI_ISL_539997, EPI_ISL_539998, EPI_ISL_539999, EPI_ISL_540000, EPI_ISL_540001, EPI_ISL_540002, EPI_ISL_540004, EPI_ISL_540005, EPI_ISL_540006, EPI_ISL_540007, EPI_ISL_540008, EPI_ISL_540009, EPI_ISL_540010, EPI_ISL_540011, EPI_ISL_540012, EPI_ISL_540013, EPI_ISL_540014, EPI_ISL_540016, EPI_ISL_540017, EPI_ISL_540018, EPI_ISL_540019, EPI_ISL_540022, EPI_ISL_540023, EPI_ISL_540025, EPI_ISL_540026, EPI_ISL_540027, EPI_ISL_540028, EPI_ISL_540029, EPI_ISL_540031, EPI_ISL_540032, EPI_ISL_540033, EPI_ISL_540034, EPI_ISL_540035, EPI_ISL_540036, EPI_ISL_540037, EPI_ISL_540038, EPI_ISL_540039, EPI_ISL_540040, EPI_ISL_540041, EPI_ISL_540042, EPI_ISL_540043, EPI_ISL_540044, EPI_ISL_540045, EPI_ISL_540046, EPI_ISL_540047, EPI_ISL_540048, EPI_ISL_540050, EPI_ISL_540051, EPI_ISL_540052, EPI_ISL_540054, EPI_ISL_540055, EPI_ISL_540056, EPI_ISL_540057, EPI_ISL_540058, EPI_ISL_540060, EPI_ISL_540061, EPI_ISL_540062, EPI_ISL_540063, EPI_ISL_540064, EPI_ISL_540065, EPI_ISL_540066, EPI_ISL_540067, EPI_ISL_540068, EPI_ISL_540069, EPI_ISL_540070, EPI_ISL_540071, EPI_ISL_540072, EPI_ISL_540074, EPI_ISL_540075, EPI_ISL_540076, EPI_ISL_540077, EPI_ISL_540078, EPI_ISL_540079, EPI_ISL_540080, EPI_ISL_540081, EPI_ISL_540082, EPI_ISL_540083, EPI_ISL_540084, EPI_ISL_540085, EPI_ISL_540086, EPI_ISL_540088, EPI_ISL_540089, EPI_ISL_540090, EPI_ISL_540091, EPI_ISL_540092, EPI_ISL_540093, EPI_ISL_540094, EPI_ISL_540095, EPI_ISL_540096, EPI_ISL_540097, EPI_ISL_540098, EPI_ISL_540101, EPI_ISL_540102, EPI_ISL_540103, EPI_ISL_540104, EPI_ISL_540105, EPI_ISL_540106, EPI_ISL_540107, EPI_ISL_540108, EPI_ISL_540109, EPI_ISL_540110, EPI_ISL_540112, EPI_ISL_540113, EPI_ISL_540115, EPI_ISL_540116, EPI_ISL_540117, EPI_ISL_540118, EPI_ISL_540119, EPI_ISL_540120, EPI_ISL_540121, EPI_ISL_540122, EPI_ISL_540123, EPI_ISL_540124, EPI_ISL_540125, EPI_ISL_540126, EPI_ISL_540127, EPI_ISL_540128, EPI_ISL_540129, EPI_ISL_540130, EPI_ISL_540131, EPI_ISL_540132, EPI_ISL_540134, EPI_ISL_540135, EPI_ISL_540136, EPI_ISL_540137, EPI_ISL_540138, EPI_ISL_540139, EPI_ISL_540140, EPI_ISL_540141, EPI_ISL_540142, EPI_ISL_540143, EPI_ISL_540144, EPI_ISL_540145, EPI_ISL_540146, EPI_ISL_540147, EPI_ISL_540148, EPI_ISL_540149, EPI_ISL_540150, EPI_ISL_540151, EPI_ISL_540152, EPI_ISL_540154, EPI_ISL_540155, EPI_ISL_540157, EPI_ISL_540159, EPI_ISL_540160, EPI_ISL_540162, EPI_ISL_540163, EPI_ISL_540164, EPI_ISL_540166, EPI_ISL_540167, EPI_ISL_540168, EPI_ISL_540169, EPI_ISL_540170, EPI_ISL_540171, EPI_ISL_540172, EPI_ISL_540173, EPI_ISL_540174, EPI_ISL_540175, EPI_ISL_540176, EPI_ISL_540177, EPI_ISL_540178, EPI_ISL_540179, EPI_ISL_540180, EPI_ISL_540181, EPI_ISL_540182, EPI_ISL_540183, EPI_ISL_540185, EPI_ISL_540186, EPI_ISL_540187, EPI_ISL_540188, EPI_ISL_540189, EPI_ISL_540191, EPI_ISL_540192, EPI_ISL_540193, EPI_ISL_540194, EPI_ISL_540195, EPI_ISL_540196, EPI_ISL_540197, EPI_ISL_540198, EPI_ISL_540199, EPI_ISL_540200, EPI_ISL_540201, EPI_ISL_540202, EPI_ISL_540203, EPI_ISL_540204, EPI_ISL_540205, EPI_ISL_540206, EPI_ISL_540207, EPI_ISL_540208, EPI_ISL_540209, EPI_ISL_540210, EPI_ISL_540211, EPI_ISL_540212, EPI_ISL_540213, EPI_ISL_540214, EPI_ISL_540215, EPI_ISL_540216, EPI_ISL_540217, EPI_ISL_540218, EPI_ISL_540219, EPI_ISL_540220, EPI_ISL_540221, EPI_ISL_540222, EPI_ISL_540223, EPI_ISL_540224, EPI_ISL_540225, EPI_ISL_540226, EPI_ISL_540227, EPI_ISL_540228, EPI_ISL_540230, EPI_ISL_540231, EPI_ISL_540233, EPI_ISL_540234, EPI_ISL_540235, EPI_ISL_540236, EPI_ISL_540239, EPI_ISL_540240, EPI_ISL_540242, EPI_ISL_540243, EPI_ISL_540244, EPI_ISL_540245, EPI_ISL_540246, EPI_ISL_540248, EPI_ISL_540249, EPI_ISL_540251, EPI_ISL_540252, EPI_ISL_540253, EPI_ISL_540254, EPI_ISL_540256, EPI_ISL_540257, EPI_ISL_540258, EPI_ISL_540259, EPI_ISL_540260, EPI_ISL_540261, EPI_ISL_540262, EPI_ISL_540263, EPI_ISL_540264, EPI_ISL_540265, EPI_ISL_540266, EPI_ISL_540267, EPI_ISL_540268, EPI_ISL_540269, EPI_ISL_540270, EPI_ISL_540271, EPI_ISL_540272, EPI_ISL_540274, EPI_ISL_540276, EPI_ISL_540277, EPI_ISL_540279, EPI_ISL_540280, EPI_ISL_540282, EPI_ISL_540283, EPI_ISL_540285, EPI_ISL_540286, EPI_ISL_540288, EPI_ISL_540289, EPI_ISL_540290, EPI_ISL_540291, EPI_ISL_540292, EPI_ISL_540293, EPI_ISL_540294, EPI_ISL_540296, EPI_ISL_540297, EPI_ISL_540298, EPI_ISL_540299, EPI_ISL_540300, EPI_ISL_540301, EPI_ISL_540302, EPI_ISL_540303, EPI_ISL_540304, EPI_ISL_540306, EPI_ISL_540307, EPI_ISL_540308, EPI_ISL_540310, EPI_ISL_540311, EPI_ISL_540312, EPI_ISL_540314, EPI_ISL_540315, EPI_ISL_540316, EPI_ISL_540317, EPI_ISL_540318, EPI_ISL_540319, EPI_ISL_540320, EPI_ISL_540322, EPI_ISL_540323, EPI_ISL_540324, EPI_ISL_540325, EPI_ISL_540326, EPI_ISL_540328, EPI_ISL_540330, EPI_ISL_540331, EPI_ISL_540335, EPI_ISL_540336, EPI_ISL_540337, EPI_ISL_540338, EPI_ISL_540339, EPI_ISL_540340, EPI_ISL_540341, EPI_ISL_540342, EPI_ISL_540343, EPI_ISL_540344, EPI_ISL_540345, EPI_ISL_540346, EPI_ISL_540347, EPI_ISL_540348, EPI_ISL_540349, EPI_ISL_540350, EPI_ISL_540352, EPI_ISL_540353, EPI_ISL_540354, EPI_ISL_540355, EPI_ISL_540356, EPI_ISL_540357, EPI_ISL_540358, EPI_ISL_540359, EPI_ISL_540360, EPI_ISL_540361, EPI_ISL_540362, EPI_ISL_540363, EPI_ISL_540364, EPI_ISL_540365, EPI_ISL_540366, EPI_ISL_540367, EPI_ISL_540368, EPI_ISL_540369, EPI_ISL_540370, EPI_ISL_540372, EPI_ISL_540373, EPI_ISL_540376, EPI_ISL_540377, EPI_ISL_540378, EPI_ISL_540379, EPI_ISL_540380, EPI_ISL_540381, EPI_ISL_540382, EPI_ISL_540383, EPI_ISL_540384, EPI_ISL_540385, EPI_ISL_540386, EPI_ISL_540387, EPI_ISL_540389, EPI_ISL_540390, EPI_ISL_540391, EPI_ISL_540392, EPI_ISL_540393, EPI_ISL_540394, EPI_ISL_540395, EPI_ISL_540396, EPI_ISL_540397, EPI_ISL_540398, EPI_ISL_540399, EPI_ISL_540401, EPI_ISL_540402, EPI_ISL_540403, EPI_ISL_540404, EPI_ISL_540405, EPI_ISL_540406, EPI_ISL_540407, EPI_ISL_540408, EPI_ISL_540409, EPI_ISL_540410, EPI_ISL_540411, EPI_ISL_540413, EPI_ISL_540414, EPI_ISL_540416, EPI_ISL_540419, EPI_ISL_540420 | | | |
| see above | Lighthouse Lab in Glasgow | Wellcome Sanger Institute for the COVID-19 Genomics UK Consortium | Harper VanSteenhouse, Yumi Kasai, David Gray, Carol Clugston, Anna Dominiczak and Alex Alderton, Roberto Amato, Sonia Goncalves, Ewan Harrison, David K. Jackson, Ian Johnston, Dominic Kwiatkowski, Cordelia Langford, John Sillitoe on behalf of the Wellcome Sanger Institute COVID-19 Surveillance Team |
| EPI_ISL_540421, EPI_ISL_540422 | Wyoming Public Health Laboratory | Wyoming Public Health Laboratory | Noah Hull, Rob Christensen, Jim Mildenberger, Joel Sevinsky, Cari Sloma, and Wanda Manley |
| EPI_ISL_540431 | IN State Department of Health Laboratory Services | Pathogen Discovery, Respiratory Viruses Branch, Division of Viral Diseases, Centers for Disease Control and Prevention | Yan Li, Jing Zhang, Anna Montmayeur, Krista Queen,Ying Tao, Anna Uehara, Clinton R. Paden, Rachel Marine, Haibin Wang, Suxiang Tong |
| EPI_ISL_540432, EPI_ISL_540433 | TN Division of Laboratory Services | Pathogen Discovery, Respiratory Viruses Branch, Division of Viral Diseases, Centers for Disease Control and Prevention | Yan Li, Jing Zhang, Anna Montmayeur, Krista Queen,Ying Tao, Anna Uehara, Clinton R. Paden, Rachel Marine, Haibin Wang, Suxiang Tong |
| EPI_ISL_540434 | TN Division of Laboratory Services | Pathogen Discovery, Respiratory Viruses Branch, Division of Viral Diseases, Centers for Disease Control and Prevention | Ying Tao, Yan Li, Jing Zhang, Krista Queen, Anna Uehara, Clinton R. Paden, Haibin Wang, Suxiang Tong |
| EPI_ISL_540435 | VA-Division of Consolidated Laboratory Services | Pathogen Discovery, Respiratory Viruses Branch, Division of Viral Diseases, Centers for Disease Control and Prevention | Krista Queen, Yan Li, Jing Zhang, Anna Montmayeur, Krista Queen,Ying Tao, Anna Uehara, Clinton R. Paden, Rachel Marine, Haibin Wang, Suxiang Tong |

| | | | |
|---|---|---|---|
| EPI_ISL_540436, EPI_ISL_540437, EPI_ISL_540438 | WI State Laboratory of Hygiene | Pathogen Discovery, Respiratory Viruses Branch, Division of Viral Diseases, Centers for Disease Control and Prevention | Yan Li, Jing Zhang, Anna Montmayeur, Krista Queen,Ying Tao, Anna Uehara, Clinton R. Paden, Rachel Marine, Haibin Wang, Suxiang Tong |
| EPI_ISL_540439 | WVDHHR - Office of Laboratory Services | Pathogen Discovery, Respiratory Viruses Branch, Division of Viral Diseases, Centers for Disease Control and Prevention | Ying Tao, Yan Li, Jing Zhang, Krista Queen, Anna Uehara, Clinton R. Paden, Haibin Wang, Suxiang Tong |
| EPI_ISL_540442, EPI_ISL_540443, EPI_ISL_540444, EPI_ISL_540445, EPI_ISL_540446, EPI_ISL_540447, EPI_ISL_540448, EPI_ISL_540449, EPI_ISL_540450, EPI_ISL_540452, EPI_ISL_540453, EPI_ISL_540454, EPI_ISL_540455, EPI_ISL_540457, EPI_ISL_540458, EPI_ISL_540459, EPI_ISL_540460, EPI_ISL_540461, EPI_ISL_540462, EPI_ISL_540463, EPI_ISL_540464, EPI_ISL_540465, EPI_ISL_540466, EPI_ISL_540467 | | | |
| see above | University of Liège COVID-19 testing center | GIGA Medical Genomics | Keith Durkin, Maria Artesi, Emmanuel André, Marc Van Ranst, Fabrice Bureau, Laurent Gillet, Wouter Coppieters, Vincent Bours |
| EPI_ISL_540469, EPI_ISL_540470, EPI_ISL_540471, EPI_ISL_540472, EPI_ISL_540473, EPI_ISL_540474, EPI_ISL_540475, EPI_ISL_540476, EPI_ISL_540477, EPI_ISL_540478, EPI_ISL_540479, EPI_ISL_540480, EPI_ISL_540481, EPI_ISL_540482, EPI_ISL_540484, EPI_ISL_540486, EPI_ISL_540487, EPI_ISL_540488, EPI_ISL_540489, EPI_ISL_540490, EPI_ISL_540491, EPI_ISL_540492, EPI_ISL_540493, EPI_ISL_540494, EPI_ISL_540495, EPI_ISL_540496, EPI_ISL_540497, EPI_ISL_540498, EPI_ISL_540499, EPI_ISL_540500, EPI_ISL_540501, EPI_ISL_540502, EPI_ISL_540503, EPI_ISL_540504, EPI_ISL_540505, EPI_ISL_540506, EPI_ISL_540507, EPI_ISL_540508, EPI_ISL_540510, EPI_ISL_540511, EPI_ISL_540512, EPI_ISL_540514, EPI_ISL_540515, EPI_ISL_540517, EPI_ISL_540523, EPI_ISL_540525, EPI_ISL_540527, EPI_ISL_540528, EPI_ISL_540531, EPI_ISL_540533, EPI_ISL_540534, EPI_ISL_540535, EPI_ISL_540537, EPI_ISL_540541, EPI_ISL_540542, EPI_ISL_540543, EPI_ISL_540544, EPI_ISL_540547, EPI_ISL_540548, EPI_ISL_540550, EPI_ISL_540551, EPI_ISL_540555, EPI_ISL_540558, EPI_ISL_540559, EPI_ISL_540560, EPI_ISL_540565, EPI_ISL_540566, EPI_ISL_540568, EPI_ISL_540569, EPI_ISL_540573, EPI_ISL_540574, EPI_ISL_540578 | | | |
| see above | Department of Clinical Microbiology | GIGA Medical Genomics | Keith Durkin, Maria Artesi, Sébastien Bontems, Raphaël Boreux, Bouchra Boujemla, Cécile Meex, Axelle Chaslain, Céline Fombellida-Lopez, Pierrette Melin, Marie-Pierre Hayette, Vincent Bours |
| EPI_ISL_540587, EPI_ISL_540588, EPI_ISL_540590, EPI_ISL_540591, EPI_ISL_540592, EPI_ISL_540593, EPI_ISL_540603, EPI_ISL_540604, EPI_ISL_540605, EPI_ISL_540610, EPI_ISL_540613, EPI_ISL_540614, EPI_ISL_540617, EPI_ISL_540618, EPI_ISL_540619, EPI_ISL_540620, EPI_ISL_540622, EPI_ISL_540623, EPI_ISL_540624, EPI_ISL_540625, EPI_ISL_540629 | | | |
| see above | Liverpool Clinical Laboratories | COVID-19 Genomics UK (COG-UK) Consortium | Sam Haldenby, Anita Lucaci, Steve Paterson, Julian Hiscox, Alistair Darby, M Almsaud, A Alrezaihi, Muhannad Alruwaili, Stuart D Armstrong, Jones Benjamin, Eleanor G Bentley, Anu Chawla, Jordan J Clark, Angela Cowell, Richard Eccles, Isabel García-Dorival, Matthew Gemmell, Alessandro Gerada, PKF Gilmore, Richard Gregory, Ximeng Han, Catherine Hartley, Margaret Hughes, Miren Iturriza-Gomara, James Johnson, L Luu, Jenifer Manson, Charlotte Nelson, Elaine O'Toole, Cassie Olateju, Rebekah Penrice-Randal , Lucille Rainbow, N.P Randle, Trevor Ian Robinson, Parul Sharma, Ghada T Shawli, James P Stewart, Neil Swainston, Ecaterina Vamos, Joanne Watts, Mark Whitehead |
| EPI_ISL_540640, EPI_ISL_540641, EPI_ISL_540642, EPI_ISL_540643, EPI_ISL_540645, EPI_ISL_540646, EPI_ISL_540647, EPI_ISL_540648, EPI_ISL_540649, EPI_ISL_540650, EPI_ISL_540651, EPI_ISL_540652, EPI_ISL_540656, EPI_ISL_540658, EPI_ISL_540659, EPI_ISL_540660, EPI_ISL_540661, EPI_ISL_540663, EPI_ISL_540664, EPI_ISL_540667, EPI_ISL_540668, EPI_ISL_540672, EPI_ISL_540673, EPI_ISL_540674, EPI_ISL_540676, EPI_ISL_540678, EPI_ISL_540679, EPI_ISL_540682, EPI_ISL_540683, EPI_ISL_540685, EPI_ISL_540686, EPI_ISL_540687, EPI_ISL_540688, EPI_ISL_540689 | | | |
| see above | Queens Medical Centre, Clinical Microbiology Department / DeepSeq Nottingham | COVID-19 Genomics UK (COG-UK) Consortium | Gemma Clark, Wendy Smith, Manjinder Khakh, Vicki M Fleming, Michelle M Lister, Hannah Howson-Wells, Jonathan Ball, Patrick McClure, Joseph Chappell, Theocharis Tsoleridis, Nadine Holmes, Matthew Carlisle, Christopher Moore, Fei Sang, Johnny Debebe, Victoria Wright, Matthew Loose |
| EPI_ISL_540691, EPI_ISL_540692, EPI_ISL_540693, EPI_ISL_540694, EPI_ISL_540695, EPI_ISL_540698 | Quadram Institute Bioscience | COVID-19 Genomics UK (COG-UK) Consortium | Dave J. Baker, Gemma L. Kay, Alp Aydin, Thanh Le-Viet, Steven Rudder, Ana P. Tedim, Anastasia Kolyva, Maria Diaz, Leonardo de Oliveira Martins, Nabil-Fareed Alikhan, Lizzie Meadows, Rachael Stanley, Ngozi Elumogo, Muhammed Yasir, Nicholas M. Thomson, Alexander J Trotter, Rachel Gilroy, Samuel Bloomfield, Claire Stuart, Andrew Bell, Reenesh Prakash, Samir Dervisevic, Alison E. Mather, John Wain, Mark Webber, Andrew J. Page, Justin O'Grady |
| EPI_ISL_540701, EPI_ISL_540702, EPI_ISL_540704, EPI_ISL_540705, EPI_ISL_540706, EPI_ISL_540708, EPI_ISL_540710, EPI_ISL_540711, EPI_ISL_540712, EPI_ISL_540713, EPI_ISL_540715, EPI_ISL_540717, EPI_ISL_540718 | | | |
| see above | Queens Medical Centre, Clinical Microbiology Department / DeepSeq Nottingham | COVID-19 Genomics UK (COG-UK) Consortium | Gemma Clark, Wendy Smith, Manjinder Khakh, Vicki M Fleming, Michelle M Lister, Hannah Howson-Wells, Jonathan Ball, Patrick McClure, Joseph Chappell, Theocharis Tsoleridis, Nadine Holmes, Matthew Carlisle, Christopher Moore, Fei Sang, Johnny Debebe, Victoria Wright, Matthew Loose |
| EPI_ISL_540719, EPI_ISL_540720, EPI_ISL_540721, EPI_ISL_540722, EPI_ISL_540728, EPI_ISL_540729, EPI_ISL_540732, EPI_ISL_540733, EPI_ISL_540734, EPI_ISL_540735, EPI_ISL_540736, EPI_ISL_540743, EPI_ISL_540744, EPI_ISL_540746, EPI_ISL_540751, EPI_ISL_540752, EPI_ISL_540753, EPI_ISL_540756, EPI_ISL_540758, EPI_ISL_540760, EPI_ISL_540767, EPI_ISL_540768, EPI_ISL_540770 | | | |
| see above | Virology Department, Sheffield Teaching Hospitals NHS Foundation Trust/Department of Infection, Immunity and Cardiovascular Disease, The Medical School, University of Sheffield | COVID-19 Genomics UK (COG-UK) Consortium | Thushan de Silva, Matthew Parker, Nikki Smith, Adri Angyal, Rebecca Brown, Luke Green, Rachel Tucker, Paul Parsons, Danielle Groves, Katie Johnson, Laura Carrilero, Alex Keeley, Dave Partridge, Matthew Wyles, Benjamin Lindsey, Mehmet Yavuz, Mohammad Raza, Cariad Evans |
| EPI_ISL_540775, EPI_ISL_540777, EPI_ISL_540778, EPI_ISL_540779, EPI_ISL_540780, EPI_ISL_540784, EPI_ISL_540785, EPI_ISL_540786, EPI_ISL_540787, EPI_ISL_540789, EPI_ISL_540791, EPI_ISL_540796, EPI_ISL_540810 | | | |
| see above | West of Scotland Specialist Virology Centre, NHSGGC / MRC-University of Glasgow Centre for Virus Research | COVID-19 Genomics UK (COG-UK) Consortium | Ana da Silva Filipe, Natasha Johnson, Kathy Smollett, Daniel Mair, Stephen Carmichael, Lily Tong, Jenna Nichols, Elihu Aranday-Cortes, Kyriaki Nomikou; Sarah McDonald, Marc Niebel, Patawee Asamaphan; Richard Orton, Joseph Hughes, Sreenu Vattipally, David L Robertson; Alasdair MacLean, Rory Gunson; Kathy Li, Igor Starinskij, Natasha Jesudason, Rajiv Shah, James Shepherd, Antonia Ho, Emma Thomson |
| EPI_ISL_540824, EPI_ISL_540825, EPI_ISL_540826, EPI_ISL_540827, EPI_ISL_540828, EPI_ISL_540829, EPI_ISL_540830, EPI_ISL_540831, EPI_ISL_540832, EPI_ISL_540833, EPI_ISL_540834, EPI_ISL_540835, EPI_ISL_540836, EPI_ISL_540839, EPI_ISL_540840, EPI_ISL_540841, EPI_ISL_540842, EPI_ISL_540843, EPI_ISL_540844, EPI_ISL_540846, EPI_ISL_540847, EPI_ISL_540849, EPI_ISL_540851, EPI_ISL_540852, EPI_ISL_540853, EPI_ISL_540855, EPI_ISL_540856, EPI_ISL_540857, EPI_ISL_540858, EPI_ISL_540859, EPI_ISL_540861, EPI_ISL_540863, EPI_ISL_540864, EPI_ISL_540866, EPI_ISL_540870 | | | |
| see above | Lighthouse Lab in Glasgow / MRC-University of Glasgow Centre for Virus Research | COVID-19 Genomics UK (COG-UK) Consortium | Ana da Silva Filipe, Natasha Johnson, Kathy Smollett, Daniel Mair, Stephen Carmichael, Lily Tong, Jenna Nichols, Elihu Aranday-Cortes, Kyriaki Nomikou; Sarah McDonald, Marc Niebel, Patawee Asamaphan; Harper VanSteenhouse, Yumi Kasai, David Gray, Carol Clugston, Anna Dominiczak; Alasdair MacLean, Rory Gunson; Richard Orton, Joseph Hughes, Sreenu Vattipally, David L Robertson; Sharif Shaaban, Matthew Holden; Kathy Li, Natasha Jesudason, Rajiv Shah, James Shepherd, Antonia Ho, Emma Thomson |
| EPI_ISL_540872, EPI_ISL_540880, EPI_ISL_540882, EPI_ISL_540883, EPI_ISL_540884, EPI_ISL_540885, EPI_ISL_540886, EPI_ISL_540889, EPI_ISL_540892, EPI_ISL_540893 | Virology Department, Royal Infirmary of Edinburgh, NHS Lothian / School of Biological Sciences, University of Edinburgh / Institute of Genetics and Molecular Medicine, University of Edinburgh | COVID-19 Genomics UK (COG-UK) Consortium | McHugh M, Dewar R, Rooke S, Gallagher M, Balcaza C, O'Toole Á, Scher E, Hill V, McCrone JT, Colquhoun R, Yu X, Jackson B, Rambaut A, Williams TC, Templeton K |
| EPI_ISL_540894, EPI_ISL_540895 | Queens Medical Centre, Clinical Microbiology Department / DeepSeq Nottingham | COVID-19 Genomics UK (COG-UK) Consortium | Gemma Clark, Wendy Smith, Manjinder Khakh, Vicki M Fleming, Michelle M Lister, Hannah Howson-Wells, Jonathan Ball, Patrick McClure, Joseph Chappell, Theocharis Tsoleridis, Nadine Holmes, Matthew Carlisle, Christopher Moore, Fei Sang, Johnny Debebe, Victoria Wright, Matthew Loose |
| EPI_ISL_540898, EPI_ISL_540899, EPI_ISL_540900, EPI_ISL_540901, EPI_ISL_540902, EPI_ISL_540903, EPI_ISL_540904, EPI_ISL_540905, EPI_ISL_540906, EPI_ISL_540907, EPI_ISL_540908, EPI_ISL_540909, EPI_ISL_540910, EPI_ISL_540911, EPI_ISL_540912, EPI_ISL_540913, EPI_ISL_540914, EPI_ISL_540915, EPI_ISL_540916, EPI_ISL_540917, EPI_ISL_540918, EPI_ISL_540919, EPI_ISL_540920 | | | |
| see above | Wales Specialist Virology Centre Sequencing lab: Pathogen Genomics Unit | COVID-19 Genomics UK (COG-UK) Consortium | Catherine Moore, Johnathan Evans, Laura Gifford, Malorie Perry, Simon Cottrell, Angela Marchbank, Alec Birchley, Alexander Adams, Amy Gaskin, Bree Gatica-Wilcox, Jason Coombes, Joel Southgate, Lauren Gilbert, Lee Graham, Nicole Pacchiarini, Sara Kumziene-Summerhayes, Sarah Taylor, Sophie Jones, Sara Rey, Matthew Bull, Joanne Watkins, Sally Corden, Tom Connor |
| EPI_ISL_540921 | Wyoming Public Health Laboratory | Wyoming Public Health Laboratory | Noah Hull, Rob Christensen, Jim Mildenberger, Joel Sevinsky, Cari Sloma, and Wanda Manley |
| EPI_ISL_540923, EPI_ISL_540924, EPI_ISL_540925, EPI_ISL_540926, EPI_ISL_540927, EPI_ISL_540928, EPI_ISL_540929, EPI_ISL_540930, EPI_ISL_540931, EPI_ISL_540932, EPI_ISL_540933, EPI_ISL_540934, EPI_ISL_540935, EPI_ISL_540936, EPI_ISL_540937, EPI_ISL_540938, EPI_ISL_540939, EPI_ISL_540940, EPI_ISL_540941, EPI_ISL_540942, EPI_ISL_540943, EPI_ISL_540944, EPI_ISL_540945, EPI_ISL_540946, EPI_ISL_540947, EPI_ISL_540948, EPI_ISL_540949, EPI_ISL_540950, EPI_ISL_540951, EPI_ISL_540952, EPI_ISL_540953, EPI_ISL_540954, EPI_ISL_540955, EPI_ISL_540956, EPI_ISL_540957, EPI_ISL_540958, EPI_ISL_540959, EPI_ISL_540960, EPI_ISL_540961, EPI_ISL_540962, EPI_ISL_540963, EPI_ISL_540964, EPI_ISL_540965, EPI_ISL_540966, EPI_ISL_540967, EPI_ISL_540968, EPI_ISL_540969, EPI_ISL_540970, EPI_ISL_540971, EPI_ISL_540972, EPI_ISL_540973, EPI_ISL_540974, EPI_ISL_540975, EPI_ISL_540976, EPI_ISL_540977, EPI_ISL_540978, EPI_ISL_540979, EPI_ISL_540980, EPI_ISL_540981, EPI_ISL_540982, EPI_ISL_540983, EPI_ISL_540984, EPI_ISL_540985, EPI_ISL_540986, EPI_ISL_540987, EPI_ISL_540988, EPI_ISL_540989, EPI_ISL_540990, EPI_ISL_540991, EPI_ISL_540992 | | | |
| see above | Laboratorio de Referencia Nacional de Virus Respiratorios, Instituto Nacional de Salud Peru | Laboratorio de Genómica Microbiana, Universidad Peruana Cayetano Heredia | Pablo Tsukayama, Alejandra Dávila-Barclay, Luis González, Pedro E. Romero, Brenda Ayzanoa, Janet Huancachoque, Pool Marcos, Maribel Huaringa, Camila Castillo-Vilcahuaman, Guillermo Salvatierra |
| EPI_ISL_540993, EPI_ISL_540994, EPI_ISL_540995, EPI_ISL_540996, EPI_ISL_540997, EPI_ISL_540998, EPI_ISL_540999, EPI_ISL_541000, EPI_ISL_541001, EPI_ISL_541002, EPI_ISL_541003, EPI_ISL_541004, EPI_ISL_541005, EPI_ISL_541006, EPI_ISL_541007 | | | |
| see above | Health and Environmental Research Institute of Gwangju Metropolitan city | Health and Environmental Research Institute of Gwangju Metropolitan city | Min Ji Kim, Ji-eun Lee |
| EPI_ISL_541082 | The National Institute of Public Health | State Veterinary Institute Prague | Nagy,A;Jirincova,H;Novakova,L;Trnka,D;Vecerova,J |
| EPI_ISL_541138 | The National Institute of Public Health | State Veterinary Institute Prague | Nagy,A; Jirincova,H; Novakova,L; Trnka,D; Vecerova,J |
| EPI_ISL_541144, EPI_ISL_541150, EPI_ISL_541154, EPI_ISL_541155, EPI_ISL_541158, EPI_ISL_541164, EPI_ISL_541168, EPI_ISL_541173, EPI_ISL_541174, EPI_ISL_541175, EPI_ISL_541176, EPI_ISL_541178, EPI_ISL_541179, EPI_ISL_541180, EPI_ISL_541181, EPI_ISL_541182, EPI_ISL_541185, EPI_ISL_541186, EPI_ISL_541187, EPI_ISL_541188, EPI_ISL_541189, EPI_ISL_541191, EPI_ISL_541192, EPI_ISL_541193, EPI_ISL_541195, EPI_ISL_541196, EPI_ISL_541198, EPI_ISL_541199, EPI_ISL_541200, EPI_ISL_541201, EPI_ISL_541204, EPI_ISL_541205, EPI_ISL_541209, EPI_ISL_541210, EPI_ISL_541213, EPI_ISL_541215, EPI_ISL_541217, EPI_ISL_541219, EPI_ISL_541220, EPI_ISL_541221, EPI_ISL_541222, EPI_ISL_541225, EPI_ISL_541226, EPI_ISL_541231, EPI_ISL_541232, EPI_ISL_541233, EPI_ISL_541234, EPI_ISL_541236, EPI_ISL_541237, EPI_ISL_541239, EPI_ISL_541240, EPI_ISL_541241, EPI_ISL_541244, EPI_ISL_541246, EPI_ISL_541250, EPI_ISL_541253, EPI_ISL_541255, EPI_ISL_541256, EPI_ISL_541257, EPI_ISL_541259, EPI_ISL_541261, EPI_ISL_541262, EPI_ISL_541263, EPI_ISL_541264, EPI_ISL_541265, EPI_ISL_541266, EPI_ISL_541267, EPI_ISL_541268, EPI_ISL_541269, EPI_ISL_541271, EPI_ISL_541272, EPI_ISL_541277, EPI_ISL_541278, EPI_ISL_541279, EPI_ISL_541281, EPI_ISL_541282, EPI_ISL_541285, EPI_ISL_541286, EPI_ISL_541287, EPI_ISL_541289, EPI_ISL_541291, EPI_ISL_541292, EPI_ISL_541294, EPI_ISL_541295, EPI_ISL_541297, EPI_ISL_541299, EPI_ISL_541301, EPI_ISL_541302, EPI_ISL_541303, EPI_ISL_541309, EPI_ISL_541315, EPI_ISL_541318, EPI_ISL_541319, EPI_ISL_541323, EPI_ISL_541331 | | | |
| see above | Florida Bureau of Public Health Laboratories, Florida Department of Health | Florida Bureau of Public Health Laboratories, Florida Department of Health | Schmedes,S., Blanton,J. |
| EPI_ISL_541332, EPI_ISL_541333, EPI_ISL_541334 | The National Institute of Public Health | State Veterinary Institute Prague | Nagy,A; Jirincova,H; Novakova,L; Trnka,D; Vecerova,J |
| EPI_ISL_541335 | The National Institute of Public Health | Sídlištní 136/24 165 03, Prague Czech Republic | Nagy,A; Jirincova,H; Novakova,L; Trnka,D; Vecerova,J |
| EPI_ISL_541336, EPI_ISL_541337 | The National Institute of Public Health | State Veterinary Institute Prague | Nagy,A; Jirincova,H; Novakova,L; Trnka,D; Vecerova,J |
| EPI_ISL_541340, EPI_ISL_541341, EPI_ISL_541342, EPI_ISL_541343, EPI_ISL_541344, EPI_ISL_541345, EPI_ISL_541346 | LACEN/PR | Laboratory of Respiratory Viruses and Measles, Oswaldo Cruz Institute, FIOCRUZ | Paola Resende, Luciana Appolinario, Fernando Motta, Anna Carolina Paixão, Ana Carolina Mendonça, Jonathan Lopes, Irina Riediger, Maria do Carmo Debur, Marilda Siqueira |
| EPI_ISL_541347, EPI_ISL_541348, EPI_ISL_541349, EPI_ISL_541350, EPI_ISL_541351, EPI_ISL_541352, EPI_ISL_541353, EPI_ISL_541354, EPI_ISL_541355, EPI_ISL_541356, EPI_ISL_541357, EPI_ISL_541358, EPI_ISL_541359, EPI_ISL_541360, EPI_ISL_541361, EPI_ISL_541362, EPI_ISL_541363, EPI_ISL_541364, EPI_ISL_541365, EPI_ISL_541366, EPI_ISL_541367, EPI_ISL_541368, EPI_ISL_541369 | | | |
| see above | Laboratory of Respiratory Viruses and Measles, Oswaldo Cruz Institute, FIOCRUZ | Laboratory of Respiratory Viruses and Measles, Oswaldo Cruz Institute, FIOCRUZ | Paola Resende, Luciana Appolinario, Fernando Motta, Anna Carolina Paixão, Ana Carolina Mendonça, Jonathan Lopes, Marilda Siqueira |
| EPI_ISL_541370 | LACEN/SC | Laboratory of Respiratory Viruses and Measles, Oswaldo Cruz Institute, FIOCRUZ | Paola Resende, Luciana Appolinario, Fernando Motta, Anna Carolina Paixão, Ana Carolina Mendonça, Jonathan Lopes, Sandra Bianchini, Marilda Siqueira |
| EPI_ISL_541372, EPI_ISL_541373, EPI_ISL_541374, EPI_ISL_541375, EPI_ISL_541376, EPI_ISL_541377, EPI_ISL_541378, EPI_ISL_541379, EPI_ISL_541380, EPI_ISL_541381, EPI_ISL_541382, EPI_ISL_541383, EPI_ISL_541384, EPI_ISL_541385, EPI_ISL_541386, EPI_ISL_541387, EPI_ISL_541388, EPI_ISL_541389, EPI_ISL_541390, EPI_ISL_541391, EPI_ISL_541393, EPI_ISL_541394, EPI_ISL_541395, EPI_ISL_541396 | | | |
| see above | LACEN/SE | Laboratory of Respiratory Viruses and Measles, Oswaldo Cruz Institute, FIOCRUZ | Paola Resende, Luciana Appolinario, Fernando Motta, Anna Carolina Paixão, Ana Carolina Mendonça, Jonathan Lopes, Clioma Santos, Marilda Siqueira |
| EPI_ISL_541401, EPI_ISL_541402, EPI_ISL_541403, EPI_ISL_541404, EPI_ISL_541405, EPI_ISL_541407, EPI_ISL_541408, EPI_ISL_541409, EPI_ISL_541410, EPI_ISL_541411, EPI_ISL_541412, EPI_ISL_541413, EPI_ISL_541414, EPI_ISL_541415, EPI_ISL_541416, EPI_ISL_541417, EPI_ISL_541418, EPI_ISL_541419, EPI_ISL_541420, EPI_ISL_541421, EPI_ISL_541422, EPI_ISL_541423, EPI_ISL_541424, EPI_ISL_541425, EPI_ISL_541426, EPI_ISL_541427, EPI_ISL_541428, EPI_ISL_541429, EPI_ISL_541430, EPI_ISL_541432, EPI_ISL_541433, EPI_ISL_541434, EPI_ISL_541435, EPI_ISL_541436, EPI_ISL_541437, EPI_ISL_541438, EPI_ISL_541439, EPI_ISL_541440, EPI_ISL_541441, EPI_ISL_541442, EPI_ISL_541443, EPI_ISL_541444, EPI_ISL_541445, EPI_ISL_541446, EPI_ISL_541447, EPI_ISL_541448, EPI_ISL_541449, EPI_ISL_541450, EPI_ISL_541451, EPI_ISL_541452, EPI_ISL_541453, EPI_ISL_541454, EPI_ISL_541455, EPI_ISL_541456, EPI_ISL_541457, EPI_ISL_541458, EPI_ISL_541459, EPI_ISL_541460, EPI_ISL_541461, EPI_ISL_541462, EPI_ISL_541463, EPI_ISL_541464, EPI_ISL_541465, EPI_ISL_541466, EPI_ISL_541467, EPI_ISL_541468, EPI_ISL_541469, EPI_ISL_541470, EPI_ISL_541471, EPI_ISL_541472, EPI_ISL_541473, EPI_ISL_541474, EPI_ISL_541475, EPI_ISL_541476, EPI_ISL_541477, EPI_ISL_541478, EPI_ISL_541479, EPI_ISL_541481, EPI_ISL_541482, EPI_ISL_541483, EPI_ISL_541484, EPI_ISL_541485, EPI_ISL_541486, EPI_ISL_541487, EPI_ISL_541488, EPI_ISL_541489, EPI_ISL_541490, EPI_ISL_541491, EPI_ISL_541492, EPI_ISL_541493, EPI_ISL_541494, EPI_ISL_541496, EPI_ISL_541498, EPI_ISL_541499, EPI_ISL_541500, EPI_ISL_541501, EPI_ISL_541503, EPI_ISL_541504, EPI_ISL_541505, EPI_ISL_541506, EPI_ISL_541507, EPI_ISL_541509, EPI_ISL_541510, EPI_ISL_541511, EPI_ISL_541512, EPI_ISL_541513, EPI_ISL_541514, EPI_ISL_541515, EPI_ISL_541516, EPI_ISL_541517, EPI_ISL_541518, EPI_ISL_541519, EPI_ISL_541520, EPI_ISL_541521, EPI_ISL_541522, EPI_ISL_541523, EPI_ISL_541524, EPI_ISL_541525, EPI_ISL_541526, EPI_ISL_541527, EPI_ISL_541528, EPI_ISL_541529, EPI_ISL_541530, EPI_ISL_541531, EPI_ISL_541532, EPI_ISL_541533, EPI_ISL_541534, EPI_ISL_541535, EPI_ISL_541536, EPI_ISL_541538, EPI_ISL_541539 | | | |
| see above | Viollier AG | Department of Biosystems Science and | Christian Beisel, Sarah Nadeau, Ivan Topolsky, Pedro Ferreira, Philipp Jablonski, Susana Posada-Céspedes, Tobias Schär, Ina Nissen, Natascha Santacroce, Elodie Burcklen, Christiane Beckmann, Maurice Redondo, Olivier Kobel, |

| | | | |
|---|---|---|---|
| | | Engineering, ETH Zürich | Christoph Noppen, Sophie Seidel, Noemie Santamaria de Souza, Niko Beerenwinkel, Tanja Stadler |
| EPI_ISL_541540, EPI_ISL_541541, EPI_ISL_541542, EPI_ISL_541543, EPI_ISL_541544, EPI_ISL_541545, EPI_ISL_541546, EPI_ISL_541547, EPI_ISL_541548, EPI_ISL_541549, EPI_ISL_541550 | | | |
| see above | University of Wisconsin-Madison AIDS Vaccine Research Laboratories | University of Wisconsin-Madison AIDS Vaccine Research Laboratories | Gage Moreno, Katarina Braun, et al. AIDS Vaccine Research Laboratories |
| EPI_ISL_541551, EPI_ISL_541552, EPI_ISL_541553, EPI_ISL_541554, EPI_ISL_541555, EPI_ISL_541556, EPI_ISL_541557, EPI_ISL_541558, EPI_ISL_541559, EPI_ISL_541560, EPI_ISL_541561, EPI_ISL_541562, EPI_ISL_541563, EPI_ISL_541564, EPI_ISL_541565 | | | |
| see above | University of Wisconsin-Madison Campus AIDS Vaccine Research Laboratories | University of Wisconsin-Madison AIDS Vaccine Research Laboratories | Gage Moreno, Katarina Braun, et al. AIDS Vaccine Research Laboratories |
| EPI_ISL_541566, EPI_ISL_541567, EPI_ISL_541569, EPI_ISL_541570, EPI_ISL_541571, EPI_ISL_541573, EPI_ISL_541574, EPI_ISL_541575, EPI_ISL_541576, EPI_ISL_541577, EPI_ISL_541578, EPI_ISL_541579, EPI_ISL_541580, EPI_ISL_541581, EPI_ISL_541582, EPI_ISL_541583, EPI_ISL_541584, EPI_ISL_541585, EPI_ISL_541586, EPI_ISL_541587, EPI_ISL_541588, EPI_ISL_541589, EPI_ISL_541590, EPI_ISL_541591, EPI_ISL_541592, EPI_ISL_541593, EPI_ISL_541594, EPI_ISL_541595, EPI_ISL_541596, EPI_ISL_541597, EPI_ISL_541598, EPI_ISL_541599, EPI_ISL_541600, EPI_ISL_541601, EPI_ISL_541602, EPI_ISL_541603, EPI_ISL_541604, EPI_ISL_541605, EPI_ISL_541606, EPI_ISL_541607, EPI_ISL_541608, EPI_ISL_541609, EPI_ISL_541610, EPI_ISL_541611, EPI_ISL_541612, EPI_ISL_541613, EPI_ISL_541614, EPI_ISL_541615, EPI_ISL_541616, EPI_ISL_541617, EPI_ISL_541618, EPI_ISL_541619, EPI_ISL_541620, EPI_ISL_541621, EPI_ISL_541623, EPI_ISL_541624, EPI_ISL_541625, EPI_ISL_541626, EPI_ISL_541627, EPI_ISL_541628, EPI_ISL_541629, EPI_ISL_541630, EPI_ISL_541631, EPI_ISL_541632, EPI_ISL_541633, EPI_ISL_541635, EPI_ISL_541636, EPI_ISL_541637, EPI_ISL_541638, EPI_ISL_541639, EPI_ISL_541640, EPI_ISL_541641, EPI_ISL_541642, EPI_ISL_541643, EPI_ISL_541644, EPI_ISL_541645, EPI_ISL_541646, EPI_ISL_541647, EPI_ISL_541648 | | | |
| see above | University of Wisconsin-Madison AIDS Vaccine Research Laboratories | University of Wisconsin-Madison AIDS Vaccine Research Laboratories | Gage Moreno, Katarina Braun, et al. AIDS Vaccine Research Laboratories |
| EPI_ISL_541649, EPI_ISL_541650, EPI_ISL_541651, EPI_ISL_541652, EPI_ISL_541653, EPI_ISL_541654, EPI_ISL_541655 | Laboratory Diagnostic, Veterinary Specialized Institute Kraljevo | Laboratory Diagnostic, Veterinary Specialized Institute Kraljevo | Vidanovic,D., Tesovic,B., Knezevic,A., Jovanovic,T., Jankovic,M., Sekler,M., Banovic Djeri,B., Volkening,J., Afonso,C., Petrovic,T. |
| EPI_ISL_541656 | Laboratory Diagnostic, Veterinary Specialized Institute Kraljevo | Laboratory Diagnostic, Veterinary Specialized Institute Kraljevo | Vidanovic,D., Tesovic,B., Knezevic,A., Jovanovic,T., Jankovic,M., Sekler,M., Banovic Djeri,B., Volkening,J., Afonso,C., Petrovic,T. |
| EPI_ISL_541657, EPI_ISL_541658, EPI_ISL_541659, EPI_ISL_541660, EPI_ISL_541661, EPI_ISL_541662 | Laboratory Diagnostic, Veterinary Specialized Institute Kraljevo | Laboratory Diagnostic, Veterinary Specialized Institute Kraljevo | Vidanovic,D., Tesovic,B., Knezevic,A., Jovanovic,T., Jankovic,M., Sekler,M., Banovic Djeri,B., Volkening,J., Afonso,C., Petrovic,T. |
| EPI_ISL_541663, EPI_ISL_541664, EPI_ISL_541665, EPI_ISL_541666, EPI_ISL_541667, EPI_ISL_541668, EPI_ISL_541669, EPI_ISL_541670, EPI_ISL_541672, EPI_ISL_541673, EPI_ISL_541674, EPI_ISL_541675, EPI_ISL_541677, EPI_ISL_541678, EPI_ISL_541679 | | | |
| see above | Microbiology Division, South Carolina Department of Health and Environmental Control | Microbiology Division, South Carolina Department of Health and Environmental Control | Flores,H. |
| EPI_ISL_541681, EPI_ISL_541682, EPI_ISL_541683, EPI_ISL_541684, EPI_ISL_541685, EPI_ISL_541687, EPI_ISL_541688, EPI_ISL_541689, EPI_ISL_541690, EPI_ISL_541691, EPI_ISL_541692, EPI_ISL_541693, EPI_ISL_541694, EPI_ISL_541695, EPI_ISL_541696, EPI_ISL_541697, EPI_ISL_541698, EPI_ISL_541699, EPI_ISL_541700, EPI_ISL_541701, EPI_ISL_541702, EPI_ISL_541704, EPI_ISL_541705, EPI_ISL_541706, EPI_ISL_541707, EPI_ISL_541708, EPI_ISL_541710, EPI_ISL_541711, EPI_ISL_541713, EPI_ISL_541714, EPI_ISL_541715, EPI_ISL_541716, EPI_ISL_541717, EPI_ISL_541718, EPI_ISL_541719, EPI_ISL_541720, EPI_ISL_541721, EPI_ISL_541723, EPI_ISL_541724, EPI_ISL_541725, EPI_ISL_541726, EPI_ISL_541727, EPI_ISL_541728, EPI_ISL_541729, EPI_ISL_541730, EPI_ISL_541731, EPI_ISL_541733, EPI_ISL_541734, EPI_ISL_541735, EPI_ISL_541736, EPI_ISL_541737, EPI_ISL_541738, EPI_ISL_541739, EPI_ISL_541740, EPI_ISL_541741, EPI_ISL_541742, EPI_ISL_541743, EPI_ISL_541744, EPI_ISL_541745, EPI_ISL_541746, EPI_ISL_541747, EPI_ISL_541748, EPI_ISL_541751 | | | |
| see above | National Institute of Virology, NIV Influenza | National Institute of Virology, NIV Influenza | Potdar V |
| EPI_ISL_541752, EPI_ISL_541754, EPI_ISL_541755 | Barts Health NHS Trust | Wellcome Sanger Institute for the COVID-19 Genomics UK Consortium | Teresa Cutino-Moguel, Mark Hopkins, Beatrix Kele, David Harrington and Alex Alderton, Roberto Amato, Sonia Goncalves, Ewan Harrison, David K. Jackson, Ian Johnston, Dominic Kwiatkowski, Cordelia Langford, John Sillitoe on behalf of the Wellcome Sanger Institute COVID-19 Surveillance Team |
| EPI_ISL_541756 | Microbiology Department, Barking Havering and Redbridge University Hospitals NHS trust | Wellcome Sanger Institute for the COVID-19 Genomics UK Consortium | Amy Ash, Fatima Ali, Cherian Koshy and Alex Alderton, Roberto Amato, Sonia Goncalves, Ewan Harrison, David K. Jackson, Ian Johnston, Dominic Kwiatkowski, Cordelia Langford, John Sillitoe on behalf of the Wellcome Sanger Institute COVID-19 Surveillance Team |
| EPI_ISL_541757, EPI_ISL_541759, EPI_ISL_541760, EPI_ISL_541761, EPI_ISL_541762, EPI_ISL_541763, EPI_ISL_541764, EPI_ISL_541765, EPI_ISL_541766 | Barts Health NHS Trust | Wellcome Sanger Institute for the COVID-19 Genomics UK Consortium | Teresa Cutino-Moguel, Mark Hopkins, Beatrix Kele, David Harrington and Alex Alderton, Roberto Amato, Sonia Goncalves, Ewan Harrison, David K. Jackson, Ian Johnston, Dominic Kwiatkowski, Cordelia Langford, John Sillitoe on behalf of the Wellcome Sanger Institute COVID-19 Surveillance Team |
| EPI_ISL_541767 | Microbiology Department, Barking Havering and Redbridge University Hospitals NHS trust | Wellcome Sanger Institute for the COVID-19 Genomics UK Consortium | Amy Ash, Fatima Ali, Cherian Koshy and Alex Alderton, Roberto Amato, Sonia Goncalves, Ewan Harrison, David K. Jackson, Ian Johnston, Dominic Kwiatkowski, Cordelia Langford, John Sillitoe on behalf of the Wellcome Sanger Institute COVID-19 Surveillance Team |
| EPI_ISL_541768, EPI_ISL_541769, EPI_ISL_541770, EPI_ISL_541771, EPI_ISL_541772 | Barts Health NHS Trust | Wellcome Sanger Institute for the COVID-19 Genomics UK Consortium | Teresa Cutino-Moguel, Mark Hopkins, Beatrix Kele, David Harrington and Alex Alderton, Roberto Amato, Sonia Goncalves, Ewan Harrison, David K. Jackson, Ian Johnston, Dominic Kwiatkowski, Cordelia Langford, John Sillitoe on behalf of the Wellcome Sanger Institute COVID-19 Surveillance Team |
| EPI_ISL_541773 | Microbiology Department, Barking Havering and Redbridge University Hospitals NHS trust | Wellcome Sanger Institute for the COVID-19 Genomics UK Consortium | Amy Ash, Fatima Ali, Cherian Koshy and Alex Alderton, Roberto Amato, Sonia Goncalves, Ewan Harrison, David K. Jackson, Ian Johnston, Dominic Kwiatkowski, Cordelia Langford, John Sillitoe on behalf of the Wellcome Sanger Institute COVID-19 Surveillance Team |
| EPI_ISL_541775, EPI_ISL_541776, EPI_ISL_541777, EPI_ISL_541778, EPI_ISL_541779, EPI_ISL_541780, EPI_ISL_541781, EPI_ISL_541782 | Barts Health NHS Trust | Wellcome Sanger Institute for the COVID-19 Genomics UK Consortium | Teresa Cutino-Moguel, Mark Hopkins, Beatrix Kele, David Harrington and Alex Alderton, Roberto Amato, Sonia Goncalves, Ewan Harrison, David K. Jackson, Ian Johnston, Dominic Kwiatkowski, Cordelia Langford, John Sillitoe on behalf of the Wellcome Sanger Institute COVID-19 Surveillance Team |
| EPI_ISL_541784, EPI_ISL_541785, EPI_ISL_541786, EPI_ISL_541787, EPI_ISL_541788, EPI_ISL_541789, EPI_ISL_541790, EPI_ISL_541791, EPI_ISL_541793, EPI_ISL_541795, EPI_ISL_541796, EPI_ISL_541797, EPI_ISL_541800, EPI_ISL_541801, EPI_ISL_541802, EPI_ISL_541803, EPI_ISL_541804, EPI_ISL_541805, EPI_ISL_541807, EPI_ISL_541808, EPI_ISL_541809, EPI_ISL_541810, EPI_ISL_541811, EPI_ISL_541812, EPI_ISL_541813, EPI_ISL_541814, EPI_ISL_541815, EPI_ISL_541816, EPI_ISL_541818, EPI_ISL_541819, EPI_ISL_541820, EPI_ISL_541821, EPI_ISL_541823, EPI_ISL_541824, EPI_ISL_541825, EPI_ISL_541826, EPI_ISL_541827, EPI_ISL_541828, EPI_ISL_541829, EPI_ISL_541830, EPI_ISL_541831, EPI_ISL_541832, EPI_ISL_541833, EPI_ISL_541834, EPI_ISL_541835, EPI_ISL_541836, EPI_ISL_541837, EPI_ISL_541838, EPI_ISL_541839, EPI_ISL_541840, EPI_ISL_541841, EPI_ISL_541842, EPI_ISL_541844, EPI_ISL_541846 | | | |
| see above | Lighthouse Lab in Glasgow | Wellcome Sanger Institute for the COVID-19 Genomics UK Consortium | Harper VanSteenhouse, Yumi Kasai, David Gray, Carol Clugston, Anna Dominiczak and Alex Alderton, Roberto Amato, Sonia Goncalves, Ewan Harrison, David K. Jackson, Ian Johnston, Dominic Kwiatkowski, Cordelia Langford, John Sillitoe on behalf of the Wellcome Sanger Institute COVID-19 Surveillance Team |
| EPI_ISL_541847, EPI_ISL_541848, EPI_ISL_541849, EPI_ISL_541850, EPI_ISL_541851, EPI_ISL_541852, EPI_ISL_541853, EPI_ISL_541854, EPI_ISL_541855, EPI_ISL_541856, EPI_ISL_541857, EPI_ISL_541858, EPI_ISL_541859, EPI_ISL_541860, EPI_ISL_541861, EPI_ISL_541862, EPI_ISL_541863, EPI_ISL_541864, EPI_ISL_541865, EPI_ISL_541866, EPI_ISL_541867, EPI_ISL_541868, EPI_ISL_541869, EPI_ISL_541870, EPI_ISL_541871, EPI_ISL_541872, EPI_ISL_541873, EPI_ISL_541874, EPI_ISL_541875, EPI_ISL_541876, EPI_ISL_541877 | | | |
| see above | Lithuanian University of Health Sciences Hospital, Department of Laboratory Medicine | Lithuanian University of Health Sciences, Laboratory of Molecular Cardiology | Lukas Zemaitis, Arnoldas Pautienius, Kamile Tamusauskaite, Dovydas Gecys, Laura Pareckaite, Vaiva Lesauskaite, Astra Vitkauskiene |
| EPI_ISL_541878, EPI_ISL_541884, EPI_ISL_541886, EPI_ISL_541887, EPI_ISL_541889, EPI_ISL_541890, EPI_ISL_541892, EPI_ISL_541894, EPI_ISL_541895, EPI_ISL_541896, EPI_ISL_541897, EPI_ISL_541898, EPI_ISL_541899, EPI_ISL_541900, EPI_ISL_541902, EPI_ISL_541903, EPI_ISL_541904, EPI_ISL_541905, EPI_ISL_541906, EPI_ISL_541907, EPI_ISL_541908, EPI_ISL_541911, EPI_ISL_541912, EPI_ISL_541913, EPI_ISL_541915, EPI_ISL_541916, EPI_ISL_541917, EPI_ISL_541918, EPI_ISL_541919, EPI_ISL_541920, EPI_ISL_541921, EPI_ISL_541922, EPI_ISL_541924, EPI_ISL_541925, EPI_ISL_541926, EPI_ISL_541927, EPI_ISL_541928, EPI_ISL_541929, EPI_ISL_541933, EPI_ISL_541934, EPI_ISL_541936, EPI_ISL_541938, EPI_ISL_541939, EPI_ISL_541941 | | | |
| see above | Hospital General Universitario Gregorio Marañón | SeqCOVID-SPAIN consortium/IBV(CSIC) | Laura Pérez-Lago, Marta Herranz, Jon Sicilia, Julia Suárez, Pilar Catalán, Patricia Muñoz, Darío García de Viedma and SeqCOVID-SPAIN consortium |
| EPI_ISL_541949, EPI_ISL_541950, EPI_ISL_541951, EPI_ISL_541953, EPI_ISL_541956, EPI_ISL_541958, EPI_ISL_541963, EPI_ISL_541964, EPI_ISL_541965 | Servicio de Microbiología, Hospital Universitario Son Espases | SeqCOVID-SPAIN consortium/IBV(CSIC) | Carla López-Causapé, Jordi Reina, Antonio Oliver and SeqCOVID-SPAIN consortium |
| EPI_ISL_541970 | Influenza Centre, University of Bergen | Norwegian Institute of Public Health, Department of Virology | Fan Zhou, Rebecca J Cox, Karl A Brokstad, Bjørn Blomberg, Kathrine Stene-Johansen, Kamilla Heddeland Instefjord, Hilde Elshaug, Rasmus Riis Kopperud, Hilde Synnøve Vollan, Karoline Bragstad, Olav Hungnes |
| EPI_ISL_542014, EPI_ISL_542016, EPI_ISL_542017, EPI_ISL_542018, EPI_ISL_542019, EPI_ISL_542020, EPI_ISL_542021, EPI_ISL_542022, EPI_ISL_542023, EPI_ISL_542024, EPI_ISL_542025, EPI_ISL_542026, EPI_ISL_542027, EPI_ISL_542029, EPI_ISL_542030, EPI_ISL_542031, EPI_ISL_542032, EPI_ISL_542033, EPI_ISL_542035, EPI_ISL_542038, EPI_ISL_542039, EPI_ISL_542040, EPI_ISL_542041, EPI_ISL_542042, EPI_ISL_542043, EPI_ISL_542044, EPI_ISL_542045, EPI_ISL_542046, EPI_ISL_542047, EPI_ISL_542048, EPI_ISL_542049, EPI_ISL_542050, EPI_ISL_542051, EPI_ISL_542052, EPI_ISL_542053, EPI_ISL_542054, EPI_ISL_542055, EPI_ISL_542056, EPI_ISL_542057, EPI_ISL_542058, EPI_ISL_542059, EPI_ISL_542060, EPI_ISL_542061, EPI_ISL_542062, EPI_ISL_542063, EPI_ISL_542065, EPI_ISL_542066, EPI_ISL_542067, EPI_ISL_542068, EPI_ISL_542069, EPI_ISL_542070, EPI_ISL_542071, EPI_ISL_542072, EPI_ISL_542073, EPI_ISL_542074, EPI_ISL_542077, EPI_ISL_542078, EPI_ISL_542079, EPI_ISL_542080, EPI_ISL_542082, EPI_ISL_542083, EPI_ISL_542084, EPI_ISL_542086, EPI_ISL_542087, EPI_ISL_542088, EPI_ISL_542089, EPI_ISL_542090, EPI_ISL_542091, EPI_ISL_542092, EPI_ISL_542093, EPI_ISL_542094, EPI_ISL_542095, EPI_ISL_542097 | | | |
| see above | New Mexico Department of Health Scientific Laboratory | New Mexico Department of Health Scientific Laboratory | Ellie Johnson, Anastacia Griego-Fisher, D'Eldra Malone |
| EPI_ISL_542098, EPI_ISL_542099, EPI_ISL_542100, EPI_ISL_542101, EPI_ISL_542102, EPI_ISL_542103, EPI_ISL_542104, EPI_ISL_542105, EPI_ISL_542106, EPI_ISL_542107, EPI_ISL_542108, EPI_ISL_542109, EPI_ISL_542110, EPI_ISL_542111, EPI_ISL_542112, EPI_ISL_542113, EPI_ISL_542114, EPI_ISL_542115, EPI_ISL_542116, EPI_ISL_542117, EPI_ISL_542118, EPI_ISL_542119, EPI_ISL_542120, EPI_ISL_542121, EPI_ISL_542122, EPI_ISL_542123, EPI_ISL_542124, EPI_ISL_542125, EPI_ISL_542126, EPI_ISL_542127, EPI_ISL_542128, EPI_ISL_542129, EPI_ISL_542130, EPI_ISL_542131, EPI_ISL_542132, EPI_ISL_542133, EPI_ISL_542134, EPI_ISL_542135, EPI_ISL_542136, EPI_ISL_542137, EPI_ISL_542138, EPI_ISL_542139, EPI_ISL_542140, EPI_ISL_542141, EPI_ISL_542142, EPI_ISL_542143, EPI_ISL_542144, EPI_ISL_542145, EPI_ISL_542146, EPI_ISL_542147, EPI_ISL_542148, EPI_ISL_542149, EPI_ISL_542150, EPI_ISL_542151, EPI_ISL_542152, EPI_ISL_542153, EPI_ISL_542154, EPI_ISL_542155, EPI_ISL_542156, EPI_ISL_542157, EPI_ISL_542158, EPI_ISL_542159, EPI_ISL_542160, EPI_ISL_542161, EPI_ISL_542162, EPI_ISL_542163, EPI_ISL_542164, EPI_ISL_542165, EPI_ISL_542166, EPI_ISL_542167, EPI_ISL_542168, EPI_ISL_542169, EPI_ISL_542170, EPI_ISL_542171, EPI_ISL_542172, EPI_ISL_542173, EPI_ISL_542174, EPI_ISL_542175, EPI_ISL_542176, EPI_ISL_542177, EPI_ISL_542178, EPI_ISL_542179, EPI_ISL_542180, EPI_ISL_542181, EPI_ISL_542182, EPI_ISL_542183, EPI_ISL_542184, EPI_ISL_542185, EPI_ISL_542186, EPI_ISL_542187, EPI_ISL_542188, EPI_ISL_542189, EPI_ISL_542190, EPI_ISL_542191, EPI_ISL_542192, EPI_ISL_542193, EPI_ISL_542194, EPI_ISL_542195, EPI_ISL_542196, EPI_ISL_542197, EPI_ISL_542198, EPI_ISL_542199, EPI_ISL_542200, EPI_ISL_542201, EPI_ISL_542202, EPI_ISL_542203, EPI_ISL_542204, EPI_ISL_542205, EPI_ISL_542206, EPI_ISL_542207, EPI_ISL_542208, EPI_ISL_542209, EPI_ISL_542210, EPI_ISL_542211, EPI_ISL_542212, EPI_ISL_542213, EPI_ISL_542214, EPI_ISL_542215, EPI_ISL_542216, EPI_ISL_542217, EPI_ISL_542218, EPI_ISL_542219, EPI_ISL_542220, EPI_ISL_542221, EPI_ISL_542222, EPI_ISL_542223, EPI_ISL_542224, EPI_ISL_542225, EPI_ISL_542226, EPI_ISL_542228, EPI_ISL_542229, EPI_ISL_542230, EPI_ISL_542231, EPI_ISL_542232, EPI_ISL_542233, EPI_ISL_542234, EPI_ISL_542235, EPI_ISL_542236, EPI_ISL_542237, EPI_ISL_542238, EPI_ISL_542239, EPI_ISL_542240, EPI_ISL_542241, EPI_ISL_542242, EPI_ISL_542243, EPI_ISL_542244, EPI_ISL_542245, EPI_ISL_542246, EPI_ISL_542247, EPI_ISL_542248, EPI_ISL_542249, EPI_ISL_542250, EPI_ISL_542251, EPI_ISL_542252, EPI_ISL_542253, EPI_ISL_542254, EPI_ISL_542255, EPI_ISL_542256, EPI_ISL_542257, EPI_ISL_542258, EPI_ISL_542259, EPI_ISL_542260, EPI_ISL_542261, EPI_ISL_542262, EPI_ISL_542263, EPI_ISL_542264, EPI_ISL_542265, EPI_ISL_542266, EPI_ISL_542267, EPI_ISL_542268, EPI_ISL_542269, EPI_ISL_542270, EPI_ISL_542271, EPI_ISL_542272, EPI_ISL_542273, EPI_ISL_542274, EPI_ISL_542275, EPI_ISL_542276, EPI_ISL_542277 | | | |
| see above | ASST GOM Niguarda | Dep. Of Oncology and Hemato-Oncology University of Milan | Claudia Alteri, Valeria Cento, Antonio Piralla, Valentino Costabile, Monica Tallarita, Luna Colagrossi, Silvia Renica, Federica Giardina, Federica Novazzi, Stefano Gaiarsa, Elisa Matarazzo, Maria Antonello, Chiara Vismara, Roberto Fumagalli, Oscar Massimiliano Epis, Massimo Puoti, Carlo Federico Perno, Fausto Baldanti |
| EPI_ISL_542278, EPI_ISL_542279, EPI_ISL_542280, EPI_ISL_542281, EPI_ISL_542282, EPI_ISL_542283, EPI_ISL_542284, EPI_ISL_542285, EPI_ISL_542286, EPI_ISL_542287, EPI_ISL_542288, EPI_ISL_542289, EPI_ISL_542290, EPI_ISL_542291, EPI_ISL_542292, EPI_ISL_542293, EPI_ISL_542294, EPI_ISL_542295, EPI_ISL_542296, EPI_ISL_542297, EPI_ISL_542298, EPI_ISL_542299, EPI_ISL_542300, EPI_ISL_542301, EPI_ISL_542302, EPI_ISL_542303, EPI_ISL_542304, EPI_ISL_542305, EPI_ISL_542306, EPI_ISL_542307, EPI_ISL_542308, EPI_ISL_542309, EPI_ISL_542310, EPI_ISL_542311, EPI_ISL_542312, EPI_ISL_542313, EPI_ISL_542314, EPI_ISL_542315, EPI_ISL_542316, EPI_ISL_542317, EPI_ISL_542318, EPI_ISL_542319, EPI_ISL_542320, EPI_ISL_542321, EPI_ISL_542322, EPI_ISL_542323, EPI_ISL_542324, EPI_ISL_542325, EPI_ISL_542326, EPI_ISL_542327, EPI_ISL_542328, EPI_ISL_542329, EPI_ISL_542330, EPI_ISL_542331, EPI_ISL_542332, EPI_ISL_542333, EPI_ISL_542334, EPI_ISL_542335, EPI_ISL_542336, EPI_ISL_542337, EPI_ISL_542338, EPI_ISL_542339, EPI_ISL_542340, EPI_ISL_542341, EPI_ISL_542342, EPI_ISL_542343, EPI_ISL_542344, EPI_ISL_542345, EPI_ISL_542346, EPI_ISL_542347, EPI_ISL_542348, EPI_ISL_542349, EPI_ISL_542350, EPI_ISL_542351, EPI_ISL_542352, EPI_ISL_542353, EPI_ISL_542354, EPI_ISL_542355, EPI_ISL_542356, EPI_ISL_542357, EPI_ISL_542358, EPI_ISL_542359, EPI_ISL_542360, EPI_ISL_542361, EPI_ISL_542362, EPI_ISL_542363, EPI_ISL_542364, EPI_ISL_542365, EPI_ISL_542366, EPI_ISL_542367, EPI_ISL_542368, EPI_ISL_542369, EPI_ISL_542370, EPI_ISL_542371, EPI_ISL_542372, EPI_ISL_542373, EPI_ISL_542374, EPI_ISL_542375, EPI_ISL_542376, EPI_ISL_542377, EPI_ISL_542378, EPI_ISL_542379, EPI_ISL_542380, EPI_ISL_542381, EPI_ISL_542382, EPI_ISL_542383, EPI_ISL_542384, EPI_ISL_542385, EPI_ISL_542386, EPI_ISL_542387, EPI_ISL_542388, EPI_ISL_542389, EPI_ISL_542390, EPI_ISL_542391, EPI_ISL_542392, EPI_ISL_542393, EPI_ISL_542394, EPI_ISL_542395, EPI_ISL_542396, EPI_ISL_542397, EPI_ISL_542398, EPI_ISL_542399 | | | |
| see above | San Matteo Hospital Pavia | Dep. Of Oncology and Hemato-Oncology University of Milan | Claudia Alteri, Valeria Cento, Antonio Piralla, Valentino Costabile, Monica Tallarita, Luna Colagrossi, Silvia Renica, Federica Giardina, Federica Novazzi, Stefano Gaiarsa, Elisa Matarazzo, Maria Antonello, Chiara Vismara, Roberto Fumagalli, Oscar Massimiliano Epis, Massimo Puoti, Carlo Federico Perno, Fausto Baldanti |
| EPI_ISL_542400, EPI_ISL_542401, EPI_ISL_542402, EPI_ISL_542403, EPI_ISL_542404, EPI_ISL_542405, EPI_ISL_542406, EPI_ISL_542407, EPI_ISL_542408, EPI_ISL_542409, EPI_ISL_542410, EPI_ISL_542411, EPI_ISL_542412, EPI_ISL_542413, EPI_ISL_542414, EPI_ISL_542415, EPI_ISL_542416, EPI_ISL_542417, EPI_ISL_542418, EPI_ISL_542419, EPI_ISL_542420, EPI_ISL_542421, EPI_ISL_542422, EPI_ISL_542423, EPI_ISL_542424, EPI_ISL_542425, EPI_ISL_542426, EPI_ISL_542427, EPI_ISL_542428, EPI_ISL_542429, EPI_ISL_542430, EPI_ISL_542431, EPI_ISL_542432, EPI_ISL_542433, EPI_ISL_542434, EPI_ISL_542435, EPI_ISL_542436, EPI_ISL_542437, EPI_ISL_542438, EPI_ISL_542439, EPI_ISL_542440, EPI_ISL_542441, EPI_ISL_542442, EPI_ISL_542443 | | | |
| see above | ASST GOM Niguarda | Dep. Of Oncology and Hemato-Oncology University of Milan | Claudia Alteri, Valeria Cento, Antonio Piralla, Valentino Costabile, Monica Tallarita, Luna Colagrossi, Silvia Renica, Federica Giardina, Federica Novazzi, Stefano Gaiarsa, Elisa Matarazzo, Maria Antonello, Chiara Vismara, Roberto Fumagalli, Oscar Massimiliano Epis, Massimo Puoti, Carlo Federico Perno, Fausto Baldanti |
| EPI_ISL_542464 | Texas Department of State Health Services | Texas Department of State Health Services | Rashmi Tuladhar, Bonnie Oh,Jenny Zhang, Maliha Rahman, Anita Pokharel, Myong Koag, Chun Wang, Rachel Lee, Grace Kubin |

| | | | |
|---|---|---|---|
| EPI_ISL_542487, EPI_ISL_542488, EPI_ISL_542491, EPI_ISL_542492, EPI_ISL_542493, EPI_ISL_542494, EPI_ISL_542495, EPI_ISL_542496, EPI_ISL_542497, EPI_ISL_542498, EPI_ISL_542500, EPI_ISL_542502, EPI_ISL_542503, EPI_ISL_542504, EPI_ISL_542505, EPI_ISL_542506, EPI_ISL_542507, EPI_ISL_542508, EPI_ISL_542509, EPI_ISL_542510, EPI_ISL_542511, EPI_ISL_542512, EPI_ISL_542513, EPI_ISL_542514, EPI_ISL_542515, EPI_ISL_542516, EPI_ISL_542517, EPI_ISL_542518, EPI_ISL_542519, EPI_ISL_542520, EPI_ISL_542521, EPI_ISL_542522, EPI_ISL_542523, EPI_ISL_542524, EPI_ISL_542526, EPI_ISL_542527, EPI_ISL_542528, EPI_ISL_542529, EPI_ISL_542530, EPI_ISL_542531, EPI_ISL_542532, EPI_ISL_542533, EPI_ISL_542534, EPI_ISL_542535, EPI_ISL_542536, EPI_ISL_542537, EPI_ISL_542538, EPI_ISL_542539, EPI_ISL_542540, EPI_ISL_542541, EPI_ISL_542542, EPI_ISL_542543, EPI_ISL_542544, EPI_ISL_542545, EPI_ISL_542546, EPI_ISL_542547, EPI_ISL_542548, EPI_ISL_542549, EPI_ISL_542550, EPI_ISL_542551, EPI_ISL_542552, EPI_ISL_542553, EPI_ISL_542556, EPI_ISL_542557, EPI_ISL_542558, EPI_ISL_542560, EPI_ISL_542561, EPI_ISL_542562, EPI_ISL_542565, EPI_ISL_542567, EPI_ISL_542568, EPI_ISL_542570, EPI_ISL_542571, EPI_ISL_542572, EPI_ISL_542573, EPI_ISL_542574, EPI_ISL_542575, EPI_ISL_542576, EPI_ISL_542577, EPI_ISL_542578, EPI_ISL_542579, EPI_ISL_542580, EPI_ISL_542581, EPI_ISL_542582, EPI_ISL_542583, EPI_ISL_542584, EPI_ISL_542585, EPI_ISL_542586, EPI_ISL_542587, EPI_ISL_542588, EPI_ISL_542589, EPI_ISL_542590, EPI_ISL_542591, EPI_ISL_542592, EPI_ISL_542593, EPI_ISL_542594, EPI_ISL_542595, EPI_ISL_542596, EPI_ISL_542597, EPI_ISL_542598, EPI_ISL_542599, EPI_ISL_542600, EPI_ISL_542601, EPI_ISL_542602, EPI_ISL_542603, EPI_ISL_542604, EPI_ISL_542605, EPI_ISL_542606, EPI_ISL_542607, EPI_ISL_542609, EPI_ISL_542610, EPI_ISL_542611, EPI_ISL_542612, EPI_ISL_542613, EPI_ISL_542614, EPI_ISL_542615, EPI_ISL_542616, EPI_ISL_542617, EPI_ISL_542618, EPI_ISL_542620, EPI_ISL_542621, EPI_ISL_542622, EPI_ISL_542623, EPI_ISL_542624, EPI_ISL_542625, EPI_ISL_542626, EPI_ISL_542628, EPI_ISL_542629, EPI_ISL_542630, EPI_ISL_542631, EPI_ISL_542634, EPI_ISL_542635, EPI_ISL_542637, EPI_ISL_542638, EPI_ISL_542639, EPI_ISL_542640, EPI_ISL_542641, EPI_ISL_542642, EPI_ISL_542643, EPI_ISL_542644, EPI_ISL_542645, EPI_ISL_542646, EPI_ISL_542647, EPI_ISL_542648, EPI_ISL_542649, EPI_ISL_542651, EPI_ISL_542652, EPI_ISL_542653, EPI_ISL_542654, EPI_ISL_542655, EPI_ISL_542657, EPI_ISL_542658, EPI_ISL_542659, EPI_ISL_542660, EPI_ISL_542663, EPI_ISL_542664, EPI_ISL_542665, EPI_ISL_542666, EPI_ISL_542667, EPI_ISL_542668, EPI_ISL_542669, EPI_ISL_542671, EPI_ISL_542673, EPI_ISL_542674, EPI_ISL_542675, EPI_ISL_542676, EPI_ISL_542677, EPI_ISL_542678, EPI_ISL_542679, EPI_ISL_542680, EPI_ISL_542681, EPI_ISL_542682, EPI_ISL_542683, EPI_ISL_542684, EPI_ISL_542685, EPI_ISL_542687, EPI_ISL_542688, EPI_ISL_542689, EPI_ISL_542690, EPI_ISL_542691, EPI_ISL_542692, EPI_ISL_542693, EPI_ISL_542694, EPI_ISL_542695, EPI_ISL_542696, EPI_ISL_542698, EPI_ISL_542699, EPI_ISL_542700, EPI_ISL_542701, EPI_ISL_542702, EPI_ISL_542703, EPI_ISL_542704, EPI_ISL_542706, EPI_ISL_542707, EPI_ISL_542708, EPI_ISL_542709, EPI_ISL_542711, EPI_ISL_542712, EPI_ISL_542713, EPI_ISL_542714, EPI_ISL_542715, EPI_ISL_542716, EPI_ISL_542717, EPI_ISL_542718, EPI_ISL_542719, EPI_ISL_542720, EPI_ISL_542721, EPI_ISL_542722, EPI_ISL_542723, EPI_ISL_542724, EPI_ISL_542725, EPI_ISL_542726, EPI_ISL_542727, EPI_ISL_542728, EPI_ISL_542729, EPI_ISL_542730, EPI_ISL_542731, EPI_ISL_542733, EPI_ISL_542734, EPI_ISL_542735, EPI_ISL_542736, EPI_ISL_542738, EPI_ISL_542739, EPI_ISL_542740, EPI_ISL_542742, EPI_ISL_542743, EPI_ISL_542744, EPI_ISL_542746, EPI_ISL_542748, EPI_ISL_542749, EPI_ISL_542751, EPI_ISL_542752, EPI_ISL_542753, EPI_ISL_542754, EPI_ISL_542755, EPI_ISL_542756, EPI_ISL_542757, EPI_ISL_542759, EPI_ISL_542760, EPI_ISL_542761, EPI_ISL_542763, EPI_ISL_542765, EPI_ISL_542766, EPI_ISL_542768, EPI_ISL_542769, EPI_ISL_542771, EPI_ISL_542772, EPI_ISL_542773, EPI_ISL_542775, EPI_ISL_542776, EPI_ISL_542777, EPI_ISL_542778, EPI_ISL_542779, EPI_ISL_542780, EPI_ISL_542782, EPI_ISL_542784, EPI_ISL_542785, EPI_ISL_542786, EPI_ISL_542787, EPI_ISL_542788, EPI_ISL_542789, EPI_ISL_542790, EPI_ISL_542791, EPI_ISL_542792, EPI_ISL_542794, EPI_ISL_542795, EPI_ISL_542796, EPI_ISL_542797, EPI_ISL_542798, EPI_ISL_542799, EPI_ISL_542800, EPI_ISL_542801, EPI_ISL_542802, EPI_ISL_542803, EPI_ISL_542804, EPI_ISL_542805, EPI_ISL_542806, EPI_ISL_542808, EPI_ISL_542809, EPI_ISL_542810, EPI_ISL_542811, EPI_ISL_542812, EPI_ISL_542813, EPI_ISL_542814, EPI_ISL_542815, EPI_ISL_542817, EPI_ISL_542818, EPI_ISL_542819, EPI_ISL_542820, EPI_ISL_542821, EPI_ISL_542822, EPI_ISL_542823, EPI_ISL_542824, EPI_ISL_542825, EPI_ISL_542826, EPI_ISL_542827, EPI_ISL_542828, EPI_ISL_542830, EPI_ISL_542831, EPI_ISL_542832, EPI_ISL_542833, EPI_ISL_542834, EPI_ISL_542835, EPI_ISL_542836, EPI_ISL_542838, EPI_ISL_542839, EPI_ISL_542840, EPI_ISL_542841, EPI_ISL_542842, EPI_ISL_542843, EPI_ISL_542844, EPI_ISL_542845, EPI_ISL_542846, EPI_ISL_542847, EPI_ISL_542848, EPI_ISL_542849, EPI_ISL_542850, EPI_ISL_542851, EPI_ISL_542852, EPI_ISL_542854, EPI_ISL_542855, EPI_ISL_542856, EPI_ISL_542857, EPI_ISL_542859, EPI_ISL_542860, EPI_ISL_542862, EPI_ISL_542863, EPI_ISL_542864, EPI_ISL_542865, EPI_ISL_542866, EPI_ISL_542868, EPI_ISL_542869, EPI_ISL_542871, EPI_ISL_542872, EPI_ISL_542873, EPI_ISL_542875, EPI_ISL_542876, EPI_ISL_542878, EPI_ISL_542880, EPI_ISL_542881, EPI_ISL_542882, EPI_ISL_542883, EPI_ISL_542885, EPI_ISL_542886, EPI_ISL_542887, EPI_ISL_542888, EPI_ISL_542889, EPI_ISL_542890, EPI_ISL_542891, EPI_ISL_542892, EPI_ISL_542893, EPI_ISL_542894, EPI_ISL_542895, EPI_ISL_542896, EPI_ISL_542897, EPI_ISL_542898, EPI_ISL_542899, EPI_ISL_542900, EPI_ISL_542901, EPI_ISL_542902, EPI_ISL_542904, EPI_ISL_542905, EPI_ISL_542906, EPI_ISL_542907, EPI_ISL_542908, EPI_ISL_542909, EPI_ISL_542910, EPI_ISL_542911, EPI_ISL_542913, EPI_ISL_542914, EPI_ISL_542915, EPI_ISL_542916, EPI_ISL_542917, EPI_ISL_542918, EPI_ISL_542920, EPI_ISL_542921, EPI_ISL_542922, EPI_ISL_542923, EPI_ISL_542924, EPI_ISL_542925, EPI_ISL_542926, EPI_ISL_542927, EPI_ISL_542929, EPI_ISL_542930, EPI_ISL_542931, EPI_ISL_542934 | | | |
| see above | Houston Methodist Hospital | Houston Methodist Hospital | S. Wesley Long, Randall J. Olsen, Paul A. Christensen, David W. Bernard, James J. Davis, Maulik Shukla, Marcus Nguyen, Matthew Ojeda Saavedra, Concepcion C. Cantu, Prasanti Yerramilli, Layne Pruitt, Sishir Subedi, Hung-Che Kuo, Heather Hendrickson, Ghazaleh Eskandari, Hoang A. T. Nguyen, J. Hunter Long, Muthiah Kumaraswami, Jule Goike, Daniel Boutz, Jimmy Gollihar, Jason S. McLellan, Chia-Wei Chou, Kamyab Javanmardi, Ilya J. Finkelstein, and James M. Musser |
| EPI_ISL_542951, EPI_ISL_542984, EPI_ISL_542988, EPI_ISL_542989, EPI_ISL_542997 | TriCore Reference Laboratories | Center for Global Health, University of New Mexico Health Sciences Center | Daryl Domman, Kurt Schwalm, Twila Kunde, Joseph Hicks, Michael Edwards, Darrell Dinwiddie |
| EPI_ISL_543007, EPI_ISL_543008, EPI_ISL_543009, EPI_ISL_543011, EPI_ISL_543012, EPI_ISL_543014, EPI_ISL_543015, EPI_ISL_543017, EPI_ISL_543018, EPI_ISL_543019, EPI_ISL_543020, EPI_ISL_543023, EPI_ISL_543024, EPI_ISL_543025, EPI_ISL_543027, EPI_ISL_543029, EPI_ISL_543030, EPI_ISL_543031, EPI_ISL_543033, EPI_ISL_543034, EPI_ISL_543035, EPI_ISL_543036, EPI_ISL_543037, EPI_ISL_543039, EPI_ISL_543040, EPI_ISL_543041, EPI_ISL_543042, EPI_ISL_543043, EPI_ISL_543044, EPI_ISL_543045, EPI_ISL_543047, EPI_ISL_543049, EPI_ISL_543050, EPI_ISL_543052, EPI_ISL_543053, EPI_ISL_543054, EPI_ISL_543055, EPI_ISL_543056, EPI_ISL_543057, EPI_ISL_543060, EPI_ISL_543061, EPI_ISL_543062, EPI_ISL_543063, EPI_ISL_543064, EPI_ISL_543065, EPI_ISL_543066, EPI_ISL_543067, EPI_ISL_543068, EPI_ISL_543070, EPI_ISL_543071, EPI_ISL_543072, EPI_ISL_543073, EPI_ISL_543074, EPI_ISL_543076, EPI_ISL_543077, EPI_ISL_543078, EPI_ISL_543079, EPI_ISL_543081, EPI_ISL_543082, EPI_ISL_543084, EPI_ISL_543085, EPI_ISL_543086, EPI_ISL_543087, EPI_ISL_543088, EPI_ISL_543089, EPI_ISL_543090, EPI_ISL_543091, EPI_ISL_543092, EPI_ISL_543093, EPI_ISL_543094, EPI_ISL_543095, EPI_ISL_543097, EPI_ISL_543099, EPI_ISL_543100, EPI_ISL_543101, EPI_ISL_543102, EPI_ISL_543104, EPI_ISL_543106, EPI_ISL_543107, EPI_ISL_543108, EPI_ISL_543109, EPI_ISL_543110, EPI_ISL_543111, EPI_ISL_543112, EPI_ISL_543114, EPI_ISL_543116, EPI_ISL_543117, EPI_ISL_543118, EPI_ISL_543119, EPI_ISL_543120, EPI_ISL_543122, EPI_ISL_543123, EPI_ISL_543124, EPI_ISL_543126, EPI_ISL_543130, EPI_ISL_543131, EPI_ISL_543132, EPI_ISL_543133, EPI_ISL_543135, EPI_ISL_543137, EPI_ISL_543138, EPI_ISL_543140, EPI_ISL_543141, EPI_ISL_543142, EPI_ISL_543143, EPI_ISL_543144, EPI_ISL_543145, EPI_ISL_543146, EPI_ISL_543147, EPI_ISL_543148, EPI_ISL_543149, EPI_ISL_543150, EPI_ISL_543154, EPI_ISL_543155, EPI_ISL_543156, EPI_ISL_543157, EPI_ISL_543158, EPI_ISL_543159, EPI_ISL_543160, EPI_ISL_543161, EPI_ISL_543164, EPI_ISL_543165, EPI_ISL_543166, EPI_ISL_543169, EPI_ISL_543170, EPI_ISL_543171, EPI_ISL_543173, EPI_ISL_543176, EPI_ISL_543177, EPI_ISL_543178, EPI_ISL_543179, EPI_ISL_543181, EPI_ISL_543182, EPI_ISL_543183, EPI_ISL_543184, EPI_ISL_543185, EPI_ISL_543186, EPI_ISL_543187, EPI_ISL_543188, EPI_ISL_543189, EPI_ISL_543190, EPI_ISL_543191, EPI_ISL_543193, EPI_ISL_543194, EPI_ISL_543196, EPI_ISL_543197, EPI_ISL_543198, EPI_ISL_543199, EPI_ISL_543200, EPI_ISL_543201, EPI_ISL_543202, EPI_ISL_543203, EPI_ISL_543204, EPI_ISL_543205, EPI_ISL_543207, EPI_ISL_543208, EPI_ISL_543209, EPI_ISL_543210, EPI_ISL_543211, EPI_ISL_543212, EPI_ISL_543213, EPI_ISL_543214, EPI_ISL_543215, EPI_ISL_543216, EPI_ISL_543217, EPI_ISL_543218, EPI_ISL_543219, EPI_ISL_543220, EPI_ISL_543221, EPI_ISL_543222, EPI_ISL_543223, EPI_ISL_543224, EPI_ISL_543225, EPI_ISL_543226, EPI_ISL_543227, EPI_ISL_543229, EPI_ISL_543231, EPI_ISL_543232, EPI_ISL_543233, EPI_ISL_543234, EPI_ISL_543235, EPI_ISL_543237, EPI_ISL_543239, EPI_ISL_543240, EPI_ISL_543241, EPI_ISL_543242, EPI_ISL_543243, EPI_ISL_543244, EPI_ISL_543245, EPI_ISL_543246, EPI_ISL_543247, EPI_ISL_543250, EPI_ISL_543251, EPI_ISL_543252, EPI_ISL_543253, EPI_ISL_543254, EPI_ISL_543255, EPI_ISL_543256, EPI_ISL_543259, EPI_ISL_543260, EPI_ISL_543261, EPI_ISL_543263, EPI_ISL_543264, EPI_ISL_543266, EPI_ISL_543267, EPI_ISL_543269, EPI_ISL_543270, EPI_ISL_543271, EPI_ISL_543272, EPI_ISL_543273, EPI_ISL_543275, EPI_ISL_543276, EPI_ISL_543277, EPI_ISL_543278, EPI_ISL_543280, EPI_ISL_543281, EPI_ISL_543282, EPI_ISL_543283, EPI_ISL_543285, EPI_ISL_543286, EPI_ISL_543288, EPI_ISL_543290, EPI_ISL_543291, EPI_ISL_543292, EPI_ISL_543293, EPI_ISL_543294, EPI_ISL_543298, EPI_ISL_543299, EPI_ISL_543300, EPI_ISL_543301, EPI_ISL_543303, EPI_ISL_543304, EPI_ISL_543305, EPI_ISL_543306, EPI_ISL_543308, EPI_ISL_543310, EPI_ISL_543311, EPI_ISL_543312, EPI_ISL_543313, EPI_ISL_543314, EPI_ISL_543316, EPI_ISL_543318, EPI_ISL_543319, EPI_ISL_543320, EPI_ISL_543321, EPI_ISL_543322, EPI_ISL_543323, EPI_ISL_543324, EPI_ISL_543325, EPI_ISL_543326, EPI_ISL_543327, EPI_ISL_543328, EPI_ISL_543329, EPI_ISL_543331, EPI_ISL_543332, EPI_ISL_543334, EPI_ISL_543335, EPI_ISL_543336, EPI_ISL_543338, EPI_ISL_543341, EPI_ISL_543342, EPI_ISL_543343, EPI_ISL_543345, EPI_ISL_543346, EPI_ISL_543348, EPI_ISL_543351, EPI_ISL_543353, EPI_ISL_543354, EPI_ISL_543356, EPI_ISL_543357, EPI_ISL_543358, EPI_ISL_543359, EPI_ISL_543360, EPI_ISL_543361, EPI_ISL_543363, EPI_ISL_543364, EPI_ISL_543367, EPI_ISL_543368, EPI_ISL_543369, EPI_ISL_543371, EPI_ISL_543372, EPI_ISL_543374, EPI_ISL_543375, EPI_ISL_543377, EPI_ISL_543378, EPI_ISL_543381, EPI_ISL_543382, EPI_ISL_543386, EPI_ISL_543387, EPI_ISL_543388, EPI_ISL_543389, EPI_ISL_543390, EPI_ISL_543391, EPI_ISL_543394, EPI_ISL_543395, EPI_ISL_543396, EPI_ISL_543397, EPI_ISL_543398, EPI_ISL_543399, EPI_ISL_543400, EPI_ISL_543401, EPI_ISL_543404, EPI_ISL_543405, EPI_ISL_543406, EPI_ISL_543407, EPI_ISL_543408, EPI_ISL_543409, EPI_ISL_543410, EPI_ISL_543411, EPI_ISL_543412, EPI_ISL_543413, EPI_ISL_543415, EPI_ISL_543416, EPI_ISL_543419, EPI_ISL_543420, EPI_ISL_543421, EPI_ISL_543422, EPI_ISL_543423, EPI_ISL_543425, EPI_ISL_543427, EPI_ISL_543429, EPI_ISL_543430, EPI_ISL_543431, EPI_ISL_543433, EPI_ISL_543434, EPI_ISL_543435, EPI_ISL_543436, EPI_ISL_543437, EPI_ISL_543438, EPI_ISL_543439, EPI_ISL_543440, EPI_ISL_543441, EPI_ISL_543442, EPI_ISL_543443, EPI_ISL_543444, EPI_ISL_543446, EPI_ISL_543447, EPI_ISL_543450, EPI_ISL_543453, EPI_ISL_543454, EPI_ISL_543455, EPI_ISL_543456, EPI_ISL_543458, EPI_ISL_543459, EPI_ISL_543460, EPI_ISL_543461, EPI_ISL_543464, EPI_ISL_543465, EPI_ISL_543466, EPI_ISL_543467, EPI_ISL_543469, EPI_ISL_543470, EPI_ISL_543471, EPI_ISL_543473, EPI_ISL_543474, EPI_ISL_543475, EPI_ISL_543476, EPI_ISL_543477, EPI_ISL_543478, EPI_ISL_543479, EPI_ISL_543481, EPI_ISL_543483, EPI_ISL_543485, EPI_ISL_543486, EPI_ISL_543487, EPI_ISL_543488, EPI_ISL_543489, EPI_ISL_543490, EPI_ISL_543491, EPI_ISL_543493, EPI_ISL_543494, EPI_ISL_543495, EPI_ISL_543497, EPI_ISL_543498, EPI_ISL_543500, EPI_ISL_543502, EPI_ISL_543503, EPI_ISL_543504, EPI_ISL_543506, EPI_ISL_543507, EPI_ISL_543509, EPI_ISL_543511, EPI_ISL_543512, EPI_ISL_543513, EPI_ISL_543514, EPI_ISL_543515, EPI_ISL_543518, EPI_ISL_543519, EPI_ISL_543520, EPI_ISL_543521, EPI_ISL_543523, EPI_ISL_543524, EPI_ISL_543525, EPI_ISL_543527, EPI_ISL_543528, EPI_ISL_543529, EPI_ISL_543530, EPI_ISL_543532, EPI_ISL_543533, EPI_ISL_543534, EPI_ISL_543535, EPI_ISL_543536, EPI_ISL_543537, EPI_ISL_543538, EPI_ISL_543540, EPI_ISL_543541, EPI_ISL_543543, EPI_ISL_543544, EPI_ISL_543545, EPI_ISL_543546, EPI_ISL_543547, EPI_ISL_543549, EPI_ISL_543550, EPI_ISL_543551, EPI_ISL_543552, EPI_ISL_543553, EPI_ISL_543554, EPI_ISL_543555, EPI_ISL_543556, EPI_ISL_543557, EPI_ISL_543558, EPI_ISL_543563, EPI_ISL_543565, EPI_ISL_543566, EPI_ISL_543567, EPI_ISL_543568, EPI_ISL_543569, EPI_ISL_543570, EPI_ISL_543571, EPI_ISL_543572, EPI_ISL_543574, EPI_ISL_543575, EPI_ISL_543576, EPI_ISL_543577, EPI_ISL_543579, EPI_ISL_543580, EPI_ISL_543581, EPI_ISL_543582, EPI_ISL_543583, EPI_ISL_543584, EPI_ISL_543585, EPI_ISL_543586, EPI_ISL_543587, EPI_ISL_543588, EPI_ISL_543589, EPI_ISL_543590, EPI_ISL_543591, EPI_ISL_543592, EPI_ISL_543593, EPI_ISL_543594, EPI_ISL_543596, EPI_ISL_543597, EPI_ISL_543598, EPI_ISL_543599, EPI_ISL_543600, EPI_ISL_543602, EPI_ISL_543603, EPI_ISL_543604, EPI_ISL_543605, EPI_ISL_543606, EPI_ISL_543607, EPI_ISL_543608, EPI_ISL_543609, EPI_ISL_543611, EPI_ISL_543612, EPI_ISL_543615, EPI_ISL_543619, EPI_ISL_543620, EPI_ISL_543621, EPI_ISL_543623, EPI_ISL_543625, EPI_ISL_543626, EPI_ISL_543627, EPI_ISL_543628, EPI_ISL_543629, EPI_ISL_543630, EPI_ISL_543631, EPI_ISL_543632, EPI_ISL_543633, EPI_ISL_543634, EPI_ISL_543635, EPI_ISL_543636, EPI_ISL_543637, EPI_ISL_543638, EPI_ISL_543639, EPI_ISL_543640, EPI_ISL_543641, EPI_ISL_543642, EPI_ISL_543643, EPI_ISL_543644, EPI_ISL_543645, EPI_ISL_543646, EPI_ISL_543650, EPI_ISL_543651, EPI_ISL_543652, EPI_ISL_543653, EPI_ISL_543654, EPI_ISL_543655, EPI_ISL_543656, EPI_ISL_543657, EPI_ISL_543658, EPI_ISL_543659, EPI_ISL_543662, EPI_ISL_543663, EPI_ISL_543665, EPI_ISL_543666, EPI_ISL_543667, EPI_ISL_543668, EPI_ISL_543670, EPI_ISL_543671, EPI_ISL_543672, EPI_ISL_543673, EPI_ISL_543674, EPI_ISL_543675, EPI_ISL_543676, EPI_ISL_543677, EPI_ISL_543678, EPI_ISL_543680, EPI_ISL_543682, EPI_ISL_543683, EPI_ISL_543684, EPI_ISL_543687, EPI_ISL_543688, EPI_ISL_543689, EPI_ISL_543691, EPI_ISL_543692, EPI_ISL_543693, EPI_ISL_543694, EPI_ISL_543695, EPI_ISL_543696, EPI_ISL_543697, EPI_ISL_543698, EPI_ISL_543699, EPI_ISL_543700, EPI_ISL_543701, EPI_ISL_543702, EPI_ISL_543703, EPI_ISL_543704, EPI_ISL_543705, EPI_ISL_543706, EPI_ISL_543707, EPI_ISL_543708, EPI_ISL_543709, EPI_ISL_543710, EPI_ISL_543711, EPI_ISL_543712, EPI_ISL_543715, EPI_ISL_543717, EPI_ISL_543719, EPI_ISL_543720, EPI_ISL_543721, EPI_ISL_543722, EPI_ISL_543723, EPI_ISL_543724, EPI_ISL_543725, EPI_ISL_543726, EPI_ISL_543727, EPI_ISL_543728, EPI_ISL_543729, EPI_ISL_543730, EPI_ISL_543731, EPI_ISL_543732, EPI_ISL_543733, EPI_ISL_543734, EPI_ISL_543735, EPI_ISL_543736, EPI_ISL_543737, EPI_ISL_543738, EPI_ISL_543739, EPI_ISL_543740, EPI_ISL_543742, EPI_ISL_543743, EPI_ISL_543744, EPI_ISL_543746, EPI_ISL_543748, EPI_ISL_543750, EPI_ISL_543751, EPI_ISL_543753, EPI_ISL_543756, EPI_ISL_543757, EPI_ISL_543758, EPI_ISL_543759, EPI_ISL_543760, EPI_ISL_543761, EPI_ISL_543762, EPI_ISL_543763, EPI_ISL_543764, EPI_ISL_543765, EPI_ISL_543767, EPI_ISL_543768, EPI_ISL_543769, EPI_ISL_543770, EPI_ISL_543772, EPI_ISL_543773, EPI_ISL_543774, EPI_ISL_543775, EPI_ISL_543776, EPI_ISL_543777, EPI_ISL_543781, EPI_ISL_543782, EPI_ISL_543783, EPI_ISL_543785, EPI_ISL_543786, EPI_ISL_543788 | | | |
| see above | Houston Methodist Hospital | Houston Methodist Hospital | S. Wesley Long, Randall J. Olsen, Paul A. Christensen, David W. Bernard, James J. Davis, Maulik Shukla, Marcus Nguyen, Matthew Ojeda Saavedra, Concepcion C. Cantu, Prasanti Yerramilli, Layne Pruitt, Sishir Subedi, Hung-Che Kuo, Heather Hendrickson, Ghazaleh Eskandari, Hoang A. T. Nguyen, J. Hunter Long, Muthiah Kumaraswami, Jule Goike, Daniel Boutz, Jimmy Gollihar, Jason S. McLellan, Chia-Wei Chou, Kamyab Javanmardi, Ilya J. Finkelstein, and James M. Musser |
| EPI_ISL_545955 | Laboratorio de Infecciones Respiratorias Agudas. Centro Nacional de Salud Publica,Instituto Nacional de Salud | Laboratorio de Infecciones Respiratorias Agudas. Centro Nacional de Salud Publica,Instituto Nacional de Salud | Juscamayta,E. |
| EPI_ISL_547878, EPI_ISL_547879, EPI_ISL_547880, EPI_ISL_547882, EPI_ISL_547883, EPI_ISL_547884, EPI_ISL_547885, EPI_ISL_547886, EPI_ISL_547887, EPI_ISL_547888, EPI_ISL_547891, EPI_ISL_547892, EPI_ISL_547893, EPI_ISL_547894, EPI_ISL_547895, EPI_ISL_547896, EPI_ISL_547897, EPI_ISL_547898, EPI_ISL_547899, EPI_ISL_547900, EPI_ISL_547901, EPI_ISL_547902, EPI_ISL_547903, EPI_ISL_547904, EPI_ISL_547905, EPI_ISL_547906, EPI_ISL_547907, EPI_ISL_547908, EPI_ISL_547910, EPI_ISL_547911, EPI_ISL_547912, EPI_ISL_547913, EPI_ISL_547914, EPI_ISL_547915, EPI_ISL_547916, EPI_ISL_547917, EPI_ISL_547918, EPI_ISL_547919, EPI_ISL_547920, EPI_ISL_547921, EPI_ISL_547922, EPI_ISL_547923, EPI_ISL_547924, EPI_ISL_547925, EPI_ISL_547926, EPI_ISL_547927, EPI_ISL_547928, EPI_ISL_547929, EPI_ISL_547930, EPI_ISL_547931, EPI_ISL_547932, EPI_ISL_547933, EPI_ISL_547934, EPI_ISL_547935, EPI_ISL_547936, EPI_ISL_547937, EPI_ISL_547938, EPI_ISL_547939, EPI_ISL_547940, EPI_ISL_547941, EPI_ISL_547942, EPI_ISL_547943, EPI_ISL_547944, EPI_ISL_547945, EPI_ISL_547946, EPI_ISL_547947, EPI_ISL_547948, EPI_ISL_547949, EPI_ISL_547950, EPI_ISL_547951, EPI_ISL_547952, EPI_ISL_547953, EPI_ISL_547954, EPI_ISL_547955, EPI_ISL_547956, EPI_ISL_547957, EPI_ISL_547958, EPI_ISL_547959, EPI_ISL_547960, EPI_ISL_547961, EPI_ISL_547962, EPI_ISL_547963, EPI_ISL_547964 | | | |
| see above | Laboratorio de Infecciones Respiratorias Agudas. Centro Nacional de Salud Publica, Instituto Nacional de Salud | Laboratorio de Infecciones Respiratorias Agudas. Centro Nacional de Salud Publica, Instituto Nacional de Salud | Juscamayta,E. |
| EPI_ISL_548942, EPI_ISL_548943, EPI_ISL_548944, EPI_ISL_548945, EPI_ISL_548946 | Institute of Microbiology, University of Veterinary and Animal sciences | Institute of Microbiology, University of Veterinary and Animal sciences | Yaqub,T., Nawaz,M., Ali,M.A., Altaf,I., Raza,S., Shabbir,M.A., Ashraf,M.A., Aziz,S.Z., Cheema,S.Q., Shah,M.B., Hassan,S., Rafique,S., Sardar,N., Mehmood,A., Aziz,M.W., Fazal,S., Khan,N., Khan,M.T., Attique,M.M., Asif,A., Anwar,M., Awan,N.A., Younis,M.U., Bhatti,M.A., Tahir,Z., Mukhtar,N., Sarwar,H., Rana,M.S., Shabbir,M.Z. |
