## Supplementary material for "Analysis of the Dynamics and Distribution of SARS-CoV-2 Mutations and its Possible Structural and Functional Implications": GISAID_acknowledgements_part_9

We gratefully acknowledge the following Authors from the Originating laboratories responsible for obtaining the specimens, as well as the Submitting laboratories where the genome data were generated and shared via GISAID, on which this research is based.

All Submitters of data may be contacted directly via www.gisaid.org

| Accession ID | Originating Laboratory | Submitting Laboratory | Authors |
|---|---|---|---|
| EPI_ISL_543789, EPI_ISL_543790, EPI_ISL_543791, EPI_ISL_543792, EPI_ISL_543794, EPI_ISL_543795, EPI_ISL_543796, EPI_ISL_543797, EPI_ISL_543798, EPI_ISL_543799, EPI_ISL_543801, EPI_ISL_543802, EPI_ISL_543803, EPI_ISL_543804, EPI_ISL_543805, EPI_ISL_543806, EPI_ISL_543807, EPI_ISL_543808, EPI_ISL_543809, EPI_ISL_543810, EPI_ISL_543811, EPI_ISL_543812, EPI_ISL_543813, EPI_ISL_543814, EPI_ISL_543815, EPI_ISL_543816, EPI_ISL_543817, EPI_ISL_543818, EPI_ISL_543819, EPI_ISL_543821, EPI_ISL_543822, EPI_ISL_543823, EPI_ISL_543825, EPI_ISL_543827, EPI_ISL_543828, EPI_ISL_543829, EPI_ISL_543830, EPI_ISL_543832, EPI_ISL_543834, EPI_ISL_543835, EPI_ISL_543836, EPI_ISL_543837, EPI_ISL_543838, EPI_ISL_543840, EPI_ISL_543841, EPI_ISL_543842, EPI_ISL_543843, EPI_ISL_543844, EPI_ISL_543845, EPI_ISL_543846, EPI_ISL_543847, EPI_ISL_543848, EPI_ISL_543849, EPI_ISL_543850, EPI_ISL_543851, EPI_ISL_543852, EPI_ISL_543853, EPI_ISL_543854, EPI_ISL_543855, EPI_ISL_543856, EPI_ISL_543857, EPI_ISL_543858, EPI_ISL_543859, EPI_ISL_543860, EPI_ISL_543861, EPI_ISL_543862, EPI_ISL_543864, EPI_ISL_543865, EPI_ISL_543866, EPI_ISL_543868, EPI_ISL_543869, EPI_ISL_543871, EPI_ISL_543872, EPI_ISL_543873, EPI_ISL_543876, EPI_ISL_543879, EPI_ISL_543880, EPI_ISL_543881, EPI_ISL_543882, EPI_ISL_543883, EPI_ISL_543886, EPI_ISL_543888, EPI_ISL_543892, EPI_ISL_543893, EPI_ISL_543894, EPI_ISL_543895, EPI_ISL_543897, EPI_ISL_543898, EPI_ISL_543899, EPI_ISL_543902, EPI_ISL_543903, EPI_ISL_543904, EPI_ISL_543906, EPI_ISL_543908, EPI_ISL_543909, EPI_ISL_543910, EPI_ISL_543914, EPI_ISL_543915, EPI_ISL_543916, EPI_ISL_543917, EPI_ISL_543919, EPI_ISL_543920, EPI_ISL_543921, EPI_ISL_543923, EPI_ISL_543925, EPI_ISL_543926, EPI_ISL_543927, EPI_ISL_543929, EPI_ISL_543930, EPI_ISL_543931, EPI_ISL_543932, EPI_ISL_543933, EPI_ISL_543935, EPI_ISL_543936, EPI_ISL_543939, EPI_ISL_543940, EPI_ISL_543943, EPI_ISL_543944, EPI_ISL_543946, EPI_ISL_543947, EPI_ISL_543948, EPI_ISL_543950, EPI_ISL_543952, EPI_ISL_543955, EPI_ISL_543956, EPI_ISL_543957, EPI_ISL_543959, EPI_ISL_543961, EPI_ISL_543962, EPI_ISL_543963, EPI_ISL_543966, EPI_ISL_543969, EPI_ISL_543970, EPI_ISL_543972, EPI_ISL_543973, EPI_ISL_543974, EPI_ISL_543977, EPI_ISL_543978, EPI_ISL_543979, EPI_ISL_543981, EPI_ISL_543982, EPI_ISL_543983, EPI_ISL_543984, EPI_ISL_543985, EPI_ISL_543987, EPI_ISL_543989, EPI_ISL_543990, EPI_ISL_543991, EPI_ISL_543992, EPI_ISL_543993, EPI_ISL_543997, EPI_ISL_543998, EPI_ISL_543999, EPI_ISL_544000, EPI_ISL_544001, EPI_ISL_544003, EPI_ISL_544004, EPI_ISL_544005, EPI_ISL_544006, EPI_ISL_544007, EPI_ISL_544008, EPI_ISL_544009, EPI_ISL_544010, EPI_ISL_544011, EPI_ISL_544012, EPI_ISL_544014, EPI_ISL_544016, EPI_ISL_544017, EPI_ISL_544018, EPI_ISL_544019, EPI_ISL_544020, EPI_ISL_544021, EPI_ISL_544022, EPI_ISL_544023, EPI_ISL_544024, EPI_ISL_544026, EPI_ISL_544027, EPI_ISL_544028, EPI_ISL_544030, EPI_ISL_544032, EPI_ISL_544033, EPI_ISL_544034, EPI_ISL_544035, EPI_ISL_544036, EPI_ISL_544037, EPI_ISL_544038, EPI_ISL_544039, EPI_ISL_544040, EPI_ISL_544041, EPI_ISL_544042, EPI_ISL_544044, EPI_ISL_544045, EPI_ISL_544046, EPI_ISL_544047, EPI_ISL_544048, EPI_ISL_544049, EPI_ISL_544050, EPI_ISL_544051, EPI_ISL_544053, EPI_ISL_544054, EPI_ISL_544055, EPI_ISL_544056, EPI_ISL_544057, EPI_ISL_544059, EPI_ISL_544060, EPI_ISL_544061, EPI_ISL_544062, EPI_ISL_544063, EPI_ISL_544064, EPI_ISL_544065, EPI_ISL_544066, EPI_ISL_544067, EPI_ISL_544068, EPI_ISL_544069, EPI_ISL_544070, EPI_ISL_544071, EPI_ISL_544074, EPI_ISL_544075, EPI_ISL_544077, EPI_ISL_544078, EPI_ISL_544079, EPI_ISL_544080, EPI_ISL_544081, EPI_ISL_544082, EPI_ISL_544083, EPI_ISL_544084, EPI_ISL_544085, EPI_ISL_544086, EPI_ISL_544087, EPI_ISL_544088, EPI_ISL_544089, EPI_ISL_544090, EPI_ISL_544091, EPI_ISL_544092, EPI_ISL_544093, EPI_ISL_544094, EPI_ISL_544095, EPI_ISL_544096, EPI_ISL_544097, EPI_ISL_544098, EPI_ISL_544100, EPI_ISL_544101, EPI_ISL_544102, EPI_ISL_544105, EPI_ISL_544106, EPI_ISL_544107, EPI_ISL_544109, EPI_ISL_544110, EPI_ISL_544112, EPI_ISL_544113, EPI_ISL_544114, EPI_ISL_544115, EPI_ISL_544117, EPI_ISL_544118, EPI_ISL_544120, EPI_ISL_544121, EPI_ISL_544122, EPI_ISL_544123, EPI_ISL_544124, EPI_ISL_544126, EPI_ISL_544127, EPI_ISL_544128, EPI_ISL_544129, EPI_ISL_544130, EPI_ISL_544131, EPI_ISL_544133, EPI_ISL_544134, EPI_ISL_544137, EPI_ISL_544138, EPI_ISL_544140, EPI_ISL_544142, EPI_ISL_544143, EPI_ISL_544144, EPI_ISL_544145, EPI_ISL_544146, EPI_ISL_544147, EPI_ISL_544148, EPI_ISL_544150, EPI_ISL_544151, EPI_ISL_544153, EPI_ISL_544154, EPI_ISL_544155, EPI_ISL_544156, EPI_ISL_544158, EPI_ISL_544159, EPI_ISL_544160, EPI_ISL_544161, EPI_ISL_544162, EPI_ISL_544163, EPI_ISL_544164, EPI_ISL_544165, EPI_ISL_544166, EPI_ISL_544167, EPI_ISL_544169, EPI_ISL_544170, EPI_ISL_544171, EPI_ISL_544172, EPI_ISL_544174, EPI_ISL_544175, EPI_ISL_544176, EPI_ISL_544179, EPI_ISL_544180, EPI_ISL_544182, EPI_ISL_544184, EPI_ISL_544185, EPI_ISL_544186, EPI_ISL_544188, EPI_ISL_544190, EPI_ISL_544191, EPI_ISL_544192, EPI_ISL_544194, EPI_ISL_544195, EPI_ISL_544196, EPI_ISL_544197, EPI_ISL_544199, EPI_ISL_544200, EPI_ISL_544203, EPI_ISL_544206, EPI_ISL_544207, EPI_ISL_544208, EPI_ISL_544209, EPI_ISL_544210, EPI_ISL_544211, EPI_ISL_544215, EPI_ISL_544216, EPI_ISL_544218, EPI_ISL_544219, EPI_ISL_544220, EPI_ISL_544221, EPI_ISL_544222, EPI_ISL_544223, EPI_ISL_544224, EPI_ISL_544225, EPI_ISL_544226, EPI_ISL_544227, EPI_ISL_544228, EPI_ISL_544229, EPI_ISL_544230, EPI_ISL_544231, EPI_ISL_544232, EPI_ISL_544233, EPI_ISL_544234, EPI_ISL_544235, EPI_ISL_544238, EPI_ISL_544240, EPI_ISL_544242, EPI_ISL_544243, EPI_ISL_544245, EPI_ISL_544246, EPI_ISL_544247, EPI_ISL_544249, EPI_ISL_544251, EPI_ISL_544254, EPI_ISL_544255, EPI_ISL_544257, EPI_ISL_544259, EPI_ISL_544261, EPI_ISL_544264, EPI_ISL_544265, EPI_ISL_544267, EPI_ISL_544268, EPI_ISL_544270, EPI_ISL_544271, EPI_ISL_544273, EPI_ISL_544275, EPI_ISL_544277, EPI_ISL_544278, EPI_ISL_544279, EPI_ISL_544281, EPI_ISL_544282, EPI_ISL_544283, EPI_ISL_544284, EPI_ISL_544285, EPI_ISL_544286, EPI_ISL_544288, EPI_ISL_544290, EPI_ISL_544294, EPI_ISL_544296, EPI_ISL_544297, EPI_ISL_544298, EPI_ISL_544299, EPI_ISL_544300, EPI_ISL_544302, EPI_ISL_544303, EPI_ISL_544304, EPI_ISL_544306, EPI_ISL_544307, EPI_ISL_544308, EPI_ISL_544309, EPI_ISL_544310, EPI_ISL_544311, EPI_ISL_544312, EPI_ISL_544314, EPI_ISL_544315, EPI_ISL_544316, EPI_ISL_544318, EPI_ISL_544319, EPI_ISL_544320, EPI_ISL_544321, EPI_ISL_544323, EPI_ISL_544324, EPI_ISL_544325, EPI_ISL_544326, EPI_ISL_544329, EPI_ISL_544330, EPI_ISL_544331, EPI_ISL_544332, EPI_ISL_544333, EPI_ISL_544334, EPI_ISL_544335, EPI_ISL_544337, EPI_ISL_544339, EPI_ISL_544342, EPI_ISL_544344, EPI_ISL_544345, EPI_ISL_544346, EPI_ISL_544348, EPI_ISL_544349, EPI_ISL_544351, EPI_ISL_544352, EPI_ISL_544353, EPI_ISL_544354, EPI_ISL_544355, EPI_ISL_544356, EPI_ISL_544357, EPI_ISL_544358, EPI_ISL_544359, EPI_ISL_544360, EPI_ISL_544362, EPI_ISL_544363, EPI_ISL_544364, EPI_ISL_544365, EPI_ISL_544366, EPI_ISL_544367, EPI_ISL_544368, EPI_ISL_544369, EPI_ISL_544370, EPI_ISL_544371, EPI_ISL_544372, EPI_ISL_544373, EPI_ISL_544374, EPI_ISL_544376, EPI_ISL_544377, EPI_ISL_544380, EPI_ISL_544381, EPI_ISL_544384, EPI_ISL_544386, EPI_ISL_544387, EPI_ISL_544389, EPI_ISL_544390, EPI_ISL_544391, EPI_ISL_544392, EPI_ISL_544393, EPI_ISL_544394, EPI_ISL_544396, EPI_ISL_544397, EPI_ISL_544398, EPI_ISL_544399, EPI_ISL_544400, EPI_ISL_544402, EPI_ISL_544403, EPI_ISL_544404, EPI_ISL_544406, EPI_ISL_544407, EPI_ISL_544408, EPI_ISL_544409, EPI_ISL_544410, EPI_ISL_544411, EPI_ISL_544412, EPI_ISL_544414, EPI_ISL_544415, EPI_ISL_544416, EPI_ISL_544421, EPI_ISL_544422, EPI_ISL_544423, EPI_ISL_544424, EPI_ISL_544425, EPI_ISL_544426, EPI_ISL_544428, EPI_ISL_544429, EPI_ISL_544430, EPI_ISL_544431, EPI_ISL_544432, EPI_ISL_544433, EPI_ISL_544434, EPI_ISL_544435, EPI_ISL_544436, EPI_ISL_544439, EPI_ISL_544440, EPI_ISL_544441, EPI_ISL_544443, EPI_ISL_544445, EPI_ISL_544446, EPI_ISL_544447, EPI_ISL_544452, EPI_ISL_544454, EPI_ISL_544456, EPI_ISL_544457, EPI_ISL_544458, EPI_ISL_544459, EPI_ISL_544460, EPI_ISL_544461, EPI_ISL_544462, EPI_ISL_544465, EPI_ISL_544466, EPI_ISL_544467, EPI_ISL_544468, EPI_ISL_544469, EPI_ISL_544470, EPI_ISL_544471, EPI_ISL_544473, EPI_ISL_544474, EPI_ISL_544475, EPI_ISL_544476, EPI_ISL_544478, EPI_ISL_544479, EPI_ISL_544480, EPI_ISL_544481, EPI_ISL_544482, EPI_ISL_544483, EPI_ISL_544484, EPI_ISL_544487, EPI_ISL_544489, EPI_ISL_544491, EPI_ISL_544492, EPI_ISL_544493, EPI_ISL_544495, EPI_ISL_544496, EPI_ISL_544497, EPI_ISL_544498, EPI_ISL_544501, EPI_ISL_544502, EPI_ISL_544504, EPI_ISL_544505, EPI_ISL_544506, EPI_ISL_544507, EPI_ISL_544508, EPI_ISL_544511, EPI_ISL_544512, EPI_ISL_544513, EPI_ISL_544514, EPI_ISL_544515, EPI_ISL_544517, EPI_ISL_544518, EPI_ISL_544519, EPI_ISL_544520, EPI_ISL_544521, EPI_ISL_544522, EPI_ISL_544523, EPI_ISL_544524, EPI_ISL_544527, EPI_ISL_544528, EPI_ISL_544529, EPI_ISL_544530, EPI_ISL_544532, EPI_ISL_544533, EPI_ISL_544534, EPI_ISL_544535, EPI_ISL_544536, EPI_ISL_544538, EPI_ISL_544541, EPI_ISL_544542, EPI_ISL_544543, EPI_ISL_544545, EPI_ISL_544546, EPI_ISL_544548, EPI_ISL_544549, EPI_ISL_544551, EPI_ISL_544552, EPI_ISL_544553, EPI_ISL_544556, EPI_ISL_544558, EPI_ISL_544560, EPI_ISL_544561, EPI_ISL_544562, EPI_ISL_544565, EPI_ISL_544566, EPI_ISL_544567, EPI_ISL_544568, EPI_ISL_544569, EPI_ISL_544570, EPI_ISL_544571, EPI_ISL_544572, EPI_ISL_544573, EPI_ISL_544574, EPI_ISL_544575, EPI_ISL_544576, EPI_ISL_544578, EPI_ISL_544579, EPI_ISL_544580, EPI_ISL_544581, EPI_ISL_544583, EPI_ISL_544584, EPI_ISL_544585, EPI_ISL_544586, EPI_ISL_544587, EPI_ISL_544588, EPI_ISL_544593, EPI_ISL_544594, EPI_ISL_544595, EPI_ISL_544596, EPI_ISL_544597, EPI_ISL_544598, EPI_ISL_544601, EPI_ISL_544602, EPI_ISL_544603, EPI_ISL_544604, EPI_ISL_544605, EPI_ISL_544607, EPI_ISL_544609, EPI_ISL_544611, EPI_ISL_544612, EPI_ISL_544613, EPI_ISL_544614, EPI_ISL_544615, EPI_ISL_544617, EPI_ISL_544619, EPI_ISL_544620, EPI_ISL_544622, EPI_ISL_544623, EPI_ISL_544624, EPI_ISL_544627, EPI_ISL_544629, EPI_ISL_544630, EPI_ISL_544633, EPI_ISL_544634, EPI_ISL_544635, EPI_ISL_544639, EPI_ISL_544640, EPI_ISL_544641, EPI_ISL_544642, EPI_ISL_544643, EPI_ISL_544645, EPI_ISL_544646, EPI_ISL_544647, EPI_ISL_544648, EPI_ISL_544649, EPI_ISL_544651, EPI_ISL_544652, EPI_ISL_544653, EPI_ISL_544656, EPI_ISL_544658, EPI_ISL_544659, EPI_ISL_544660, EPI_ISL_544661, EPI_ISL_544662, EPI_ISL_544664, EPI_ISL_544665, EPI_ISL_544666, EPI_ISL_544667, EPI_ISL_544668, EPI_ISL_544669, EPI_ISL_544670, EPI_ISL_544672, EPI_ISL_544673, EPI_ISL_544675, EPI_ISL_544676, EPI_ISL_544677, EPI_ISL_544679, EPI_ISL_544680, EPI_ISL_544681, EPI_ISL_544682, EPI_ISL_544683, EPI_ISL_544684, EPI_ISL_544687, EPI_ISL_544688, EPI_ISL_544690, EPI_ISL_544691, EPI_ISL_544692, EPI_ISL_544693, EPI_ISL_544694, EPI_ISL_544695, EPI_ISL_544696, EPI_ISL_544697, EPI_ISL_544698, EPI_ISL_544699, EPI_ISL_544700, EPI_ISL_544701, EPI_ISL_544702, EPI_ISL_544703, EPI_ISL_544705, EPI_ISL_544706, EPI_ISL_544707, EPI_ISL_544708, EPI_ISL_544709, EPI_ISL_544711, EPI_ISL_544712, EPI_ISL_544713, EPI_ISL_544714, EPI_ISL_544715, EPI_ISL_544716, EPI_ISL_544717, EPI_ISL_544718, EPI_ISL_544719, EPI_ISL_544720, EPI_ISL_544721, EPI_ISL_544722, EPI_ISL_544723, EPI_ISL_544724, EPI_ISL_544725, EPI_ISL_544726, EPI_ISL_544727, EPI_ISL_544728, EPI_ISL_544730, EPI_ISL_544733, EPI_ISL_544734, EPI_ISL_544737, EPI_ISL_544738, EPI_ISL_544739, EPI_ISL_544741, EPI_ISL_544742, EPI_ISL_544743, EPI_ISL_544744, EPI_ISL_544747, EPI_ISL_544748, EPI_ISL_544752, EPI_ISL_544756, EPI_ISL_544757, EPI_ISL_544759, EPI_ISL_544760, EPI_ISL_544762, EPI_ISL_544763, EPI_ISL_544764, EPI_ISL_544765, EPI_ISL_544766, EPI_ISL_544767, EPI_ISL_544768, EPI_ISL_544770, EPI_ISL_544772, EPI_ISL_544773, EPI_ISL_544775, EPI_ISL_544776, EPI_ISL_544777, EPI_ISL_544778, EPI_ISL_544779, EPI_ISL_544780, EPI_ISL_544781, EPI_ISL_544782, EPI_ISL_544783, EPI_ISL_544784, EPI_ISL_544788, EPI_ISL_544791, EPI_ISL_544792, EPI_ISL_544795, EPI_ISL_544796, EPI_ISL_544797, EPI_ISL_544798, EPI_ISL_544799, EPI_ISL_544800, EPI_ISL_544802, EPI_ISL_544803, EPI_ISL_544805, EPI_ISL_544806, EPI_ISL_544807, EPI_ISL_544812, EPI_ISL_544813, EPI_ISL_544815, EPI_ISL_544816, EPI_ISL_544819, EPI_ISL_544820, EPI_ISL_544821, EPI_ISL_544822, EPI_ISL_544823, EPI_ISL_544824, EPI_ISL_544825, EPI_ISL_544826, EPI_ISL_544828, EPI_ISL_544830, EPI_ISL_544831, EPI_ISL_544832, EPI_ISL_544833, EPI_ISL_544834, EPI_ISL_544835, EPI_ISL_544838, EPI_ISL_544839, EPI_ISL_544840, EPI_ISL_544841, EPI_ISL_544842, EPI_ISL_544843, EPI_ISL_544844, EPI_ISL_544845, EPI_ISL_544847, EPI_ISL_544848, EPI_ISL_544849, EPI_ISL_544850, EPI_ISL_544851, EPI_ISL_544852, EPI_ISL_544853, EPI_ISL_544854, EPI_ISL_544855, EPI_ISL_544857, EPI_ISL_544859, EPI_ISL_544860, EPI_ISL_544864, EPI_ISL_544866, EPI_ISL_544867, EPI_ISL_544868, EPI_ISL_544870, EPI_ISL_544871, EPI_ISL_544872, EPI_ISL_544873, EPI_ISL_544874, EPI_ISL_544875, EPI_ISL_544876, EPI_ISL_544878, EPI_ISL_544879, EPI_ISL_544882, EPI_ISL_544883, EPI_ISL_544884, EPI_ISL_544885, EPI_ISL_544886, EPI_ISL_544888, EPI_ISL_544889, EPI_ISL_544890, EPI_ISL_544891, EPI_ISL_544893, EPI_ISL_544895, EPI_ISL_544896, EPI_ISL_544899, EPI_ISL_544900, EPI_ISL_544902, EPI_ISL_544903, EPI_ISL_544904, EPI_ISL_544905, EPI_ISL_544906, EPI_ISL_544907, EPI_ISL_544910, EPI_ISL_544911, EPI_ISL_544913, EPI_ISL_544915, EPI_ISL_544916, EPI_ISL_544917, EPI_ISL_544921, EPI_ISL_544922, EPI_ISL_544923, EPI_ISL_544925, EPI_ISL_544928, EPI_ISL_544929, EPI_ISL_544930, EPI_ISL_544932, EPI_ISL_544933, EPI_ISL_544934, EPI_ISL_544935, EPI_ISL_544936, EPI_ISL_544937, EPI_ISL_544941, EPI_ISL_544944, EPI_ISL_544945, EPI_ISL_544946, EPI_ISL_544947, EPI_ISL_544948, EPI_ISL_544950, EPI_ISL_544953, EPI_ISL_544954 | | | |
| see above | Houston Methodist Hospital | Houston Methodist Hospital | S. Wesley Long, Randall J. Olsen, Paul A. Christensen, David W. Bernard, James J. Davis, Maulik Shukla, Marcus Nguyen, Matthew Ojeda Saavedra, Concepcion C. Cantu, Prasanti Yerramilli, Layne Pruitt, Sishir Subedi, Hung-Che Kuo, Heather Hendrickson, Ghazaleh Eskandari, Hoang A. T. Nguyen, J. Hunter Long, Muthiah Kumaraswami, Jule Goike, Daniel Boutz, Jimmy Gollihar, Jason S. McLellan, Chia-Wei Chou, Kamyab Javanmardi, Ilya J. Finkelstein, and James M. Musser |
| EPI_ISL_544955 | Pathology West - NSW Health Pathology | NSW Health Pathology - Institute of Clinical Pathology and Medical Research; Westmead Hospital; University of Sydney | CIDM-PH et al. |
| EPI_ISL_544956 | Sydney South West Pathology Service (SSWPS) - Concord Repatriation General Hospital - NSW Health Pathology | NSW Health Pathology - Institute of Clinical Pathology and Medical Research; Westmead Hospital; University of Sydney | CIDM-PH et al. |
| EPI_ISL_544957 | Laverty Pathology | NSW Health Pathology - Institute of Clinical Pathology and Medical Research; Westmead Hospital; University of Sydney | CIDM-PH et al. |
| EPI_ISL_544958, EPI_ISL_544959 | Sydney South West Pathology Service (SSWPS) - Liverpool Hospital - NSW Health Pathology | NSW Health Pathology - Institute of Clinical Pathology and Medical Research; Westmead Hospital; University of Sydney | CIDM-PH et al. |
| EPI_ISL_544960 | Histopath | NSW Health Pathology - Institute of Clinical Pathology and Medical Research; Westmead Hospital; University of Sydney | CIDM-PH et al. |
| EPI_ISL_544961 | St Vincent's Pathology (SydPath) | NSW Health Pathology - Institute of Clinical Pathology and Medical Research; Westmead Hospital; University of Sydney | CIDM-PH et al. |
| EPI_ISL_544962 | Laverty Pathology | NSW Health Pathology - Institute of Clinical Pathology and Medical Research; Westmead Hospital; University of Sydney | CIDM-PH et al. |
| EPI_ISL_544963 | The Children's Hospital at Westmead | NSW Health Pathology - Institute of Clinical Pathology and Medical Research; Westmead Hospital; University of Sydney | CIDM-PH et al. |
| EPI_ISL_544964, EPI_ISL_544965 | Sydney South West Pathology Service (SSWPS) - Liverpool Hospital - NSW Health Pathology | NSW Health Pathology - Institute of Clinical Pathology and Medical Research; Westmead Hospital; University of Sydney | CIDM-PH et al. |
| EPI_ISL_544966, EPI_ISL_544967 | Australian Clinical Labs | NSW Health Pathology - Institute of Clinical Pathology and Medical Research; Westmead Hospital; University of Sydney | CIDM-PH et al. |
| EPI_ISL_544968 | St Vincent's Pathology (SydPath) | NSW Health Pathology - Institute of Clinical Pathology and Medical Research; Westmead Hospital; University of Sydney | CIDM-PH et al. |
| EPI_ISL_544969 | Pathology North - Royal North Shore Hospital - NSW Health Pathology | NSW Health Pathology - Institute of Clinical Pathology and Medical Research; Westmead Hospital; University of Sydney | CIDM-PH et al. |
| EPI_ISL_544970 | South Eastern Area Laboratory Services (SEALS) | NSW Health Pathology - Institute of Clinical Pathology and Medical Research; Westmead Hospital; University of Sydney | CIDM-PH et al. |
| EPI_ISL_544971, EPI_ISL_544972, EPI_ISL_544973 | St Vincent's Pathology (SydPath) | NSW Health Pathology - Institute of Clinical Pathology and Medical Research; Westmead Hospital; University of Sydney | CIDM-PH et al. |
| EPI_ISL_544974 | Sydney South West Pathology Service (SSWPS) - Royal Prince Alfred Hospital - NSW Health Pathology | NSW Health Pathology - Institute of Clinical Pathology and Medical Research; Westmead Hospital; University of Sydney | CIDM-PH et al. |
| EPI_ISL_544975 | Histopath | NSW Health Pathology - Institute of Clinical Pathology and Medical Research; Westmead Hospital; University of Sydney | CIDM-PH et al. |
| EPI_ISL_544976 | Pathology North - Royal North Shore Hospital - NSW Health Pathology | NSW Health Pathology - Institute of Clinical Pathology and Medical Research; Westmead | CIDM-PH et al. |

| | | | |
|---|---|---|---|
| | | Hospital; University of Sydney | |
| EPI_ISL_544977, EPI_ISL_544978, EPI_ISL_544979, EPI_ISL_544980, EPI_ISL_544981 | St Vincent's Pathology (SydPath) | NSW Health Pathology - Institute of Clinical Pathology and Medical Research; Westmead Hospital; University of Sydney | CIDM-PH et al. |
| EPI_ISL_544982 | Pathology North - Royal North Shore Hospital - NSW Health Pathology | NSW Health Pathology - Institute of Clinical Pathology and Medical Research; Westmead Hospital; University of Sydney | CIDM-PH et al. |
| EPI_ISL_544983, EPI_ISL_544984 | South Eastern Area Laboratory Services (SEALS) | NSW Health Pathology - Institute of Clinical Pathology and Medical Research; Westmead Hospital; University of Sydney | CIDM-PH et al. |
| EPI_ISL_544985 | Sydney South West Pathology Service (SSWPS) - Liverpool Hospital - NSW Health Pathology | NSW Health Pathology - Institute of Clinical Pathology and Medical Research; Westmead Hospital; University of Sydney | CIDM-PH et al. |
| EPI_ISL_544986 | Pathology West - NSW Health Pathology | NSW Health Pathology - Institute of Clinical Pathology and Medical Research; Westmead Hospital; University of Sydney | CIDM-PH et al. |
| EPI_ISL_544987 | Histopath | NSW Health Pathology - Institute of Clinical Pathology and Medical Research; Westmead Hospital; University of Sydney | CIDM-PH et al. |
| EPI_ISL_544988 | Australian Clinical Labs | NSW Health Pathology - Institute of Clinical Pathology and Medical Research; Westmead Hospital; University of Sydney | CIDM-PH et al. |
| EPI_ISL_544989, EPI_ISL_544990 | Pathology West - NSW Health Pathology | NSW Health Pathology - Institute of Clinical Pathology and Medical Research; Westmead Hospital; University of Sydney | CIDM-PH et al. |
| EPI_ISL_544991 | Pathology North - Royal North Shore Hospital - NSW Health Pathology | NSW Health Pathology - Institute of Clinical Pathology and Medical Research; Westmead Hospital; University of Sydney | CIDM-PH et al. |
| EPI_ISL_544992, EPI_ISL_544993, EPI_ISL_544994, EPI_ISL_544995, EPI_ISL_544996, EPI_ISL_544997, EPI_ISL_544998, EPI_ISL_544999, EPI_ISL_545000, EPI_ISL_545001 | St Vincent's Pathology (SydPath) | NSW Health Pathology - Institute of Clinical Pathology and Medical Research; Westmead Hospital; University of Sydney | CIDM-PH et al. |
| EPI_ISL_545002 | The Children's Hospital at Westmead | NSW Health Pathology - Institute of Clinical Pathology and Medical Research; Westmead Hospital; University of Sydney | CIDM-PH et al. |
| EPI_ISL_545003 | Sydney South West Pathology Service (SSWPS) - Liverpool Hospital - NSW Health Pathology | NSW Health Pathology - Institute of Clinical Pathology and Medical Research; Westmead Hospital; University of Sydney | CIDM-PH et al. |
| EPI_ISL_545004, EPI_ISL_545005 | Sydney South West Pathology Service (SSWPS) - Royal Prince Alfred Hospital - NSW Health Pathology | NSW Health Pathology - Institute of Clinical Pathology and Medical Research; Westmead Hospital; University of Sydney | CIDM-PH et al. |
| EPI_ISL_545006 | Pathology West - NSW Health Pathology | NSW Health Pathology - Institute of Clinical Pathology and Medical Research; Westmead Hospital; University of Sydney | CIDM-PH et al. |
| EPI_ISL_545007 | Histopath | NSW Health Pathology - Institute of Clinical Pathology and Medical Research; Westmead Hospital; University of Sydney | CIDM-PH et al. |
| EPI_ISL_545008 | Pathology North - Royal North Shore Hospital - NSW Health Pathology | NSW Health Pathology - Institute of Clinical Pathology and Medical Research; Westmead Hospital; University of Sydney | CIDM-PH et al. |
| EPI_ISL_545009, EPI_ISL_545010, EPI_ISL_545011, EPI_ISL_545012, EPI_ISL_545013, EPI_ISL_545014, EPI_ISL_545015, EPI_ISL_545016 | South Eastern Area Laboratory Services (SEALS) | NSW Health Pathology - Institute of Clinical Pathology and Medical Research; Westmead Hospital; University of Sydney | CIDM-PH et al. |
| EPI_ISL_545017, EPI_ISL_545018, EPI_ISL_545019, EPI_ISL_545020, EPI_ISL_545021, EPI_ISL_545022, EPI_ISL_545023 | Sydney South West Pathology Service (SSWPS) - Royal Prince Alfred Hospital - NSW Health Pathology | NSW Health Pathology - Institute of Clinical Pathology and Medical Research; Westmead Hospital; University of Sydney | CIDM-PH et al. |
| EPI_ISL_545024 | Australian Clinical Labs | NSW Health Pathology - Institute of Clinical Pathology and Medical Research; Westmead Hospital; University of Sydney | CIDM-PH et al. |
| EPI_ISL_545025 | Sydney South West Pathology Service (SSWPS) - Liverpool Hospital - NSW Health Pathology | NSW Health Pathology - Institute of Clinical Pathology and Medical Research; Westmead Hospital; University of Sydney | CIDM-PH et al. |
| EPI_ISL_545026 | St Vincent's Pathology (SydPath) | NSW Health Pathology - Institute of Clinical Pathology and Medical Research; Westmead Hospital; University of Sydney | CIDM-PH et al. |
| EPI_ISL_545027 | Douglass Hanly Moir Pathology | NSW Health Pathology - Institute of Clinical Pathology and Medical Research; Westmead Hospital; University of Sydney | CIDM-PH et al. |
| EPI_ISL_545028 | Sydney South West Pathology Service (SSWPS) - Royal Prince Alfred Hospital - NSW Health Pathology | NSW Health Pathology - Institute of Clinical Pathology and Medical Research; Westmead Hospital; University of Sydney | CIDM-PH et al. |
| EPI_ISL_545029 | Pathology West - NSW Health Pathology | NSW Health Pathology - Institute of Clinical Pathology and Medical Research; Westmead Hospital; University of Sydney | CIDM-PH et al. |
| EPI_ISL_545030 | St Vincent's Pathology (SydPath) | NSW Health Pathology - Institute of Clinical Pathology and Medical Research; Westmead Hospital; University of Sydney | CIDM-PH et al. |
| EPI_ISL_545031 | South Eastern Area Laboratory Services (SEALS) | NSW Health Pathology - Institute of Clinical Pathology and Medical Research; Westmead Hospital; University of Sydney | CIDM-PH et al. |
| EPI_ISL_545032, EPI_ISL_545033 | Pathology West - NSW Health Pathology | NSW Health Pathology - Institute of Clinical Pathology and Medical Research; Westmead Hospital; University of Sydney | CIDM-PH et al. |
| EPI_ISL_545034, EPI_ISL_545035, EPI_ISL_545036 | Histopath | NSW Health Pathology - Institute of Clinical Pathology and Medical Research; Westmead Hospital; University of Sydney | CIDM-PH et al. |
| EPI_ISL_545037 | Sydney South West Pathology Service (SSWPS) - Concord Repatriation General Hospital - NSW Health Pathology | NSW Health Pathology - Institute of Clinical Pathology and Medical Research; Westmead Hospital; University of Sydney | CIDM-PH et al. |
| EPI_ISL_545038 | Pathology West - NSW Health Pathology | NSW Health Pathology - Institute of Clinical Pathology and Medical Research; Westmead Hospital; University of Sydney | CIDM-PH et al. |
| EPI_ISL_545039 | Sydney South West Pathology Service (SSWPS) - Royal Prince Alfred Hospital - NSW Health Pathology | NSW Health Pathology - Institute of Clinical Pathology and Medical Research; Westmead Hospital; University of Sydney | CIDM-PH et al. |
| EPI_ISL_545040 | Laverty Pathology | NSW Health Pathology - Institute of Clinical Pathology and Medical Research; Westmead | CIDM-PH et al. |

| | | | |
|---|---|---|---|
| | | Hospital; University of Sydney | |
| EPI_ISL_545041 | Sydney South West Pathology Service (SSWPS) - Royal Prince Alfred Hospital - NSW Health Pathology | NSW Health Pathology - Institute of Clinical Pathology and Medical Research; Westmead Hospital; University of Sydney | CIDM-PH et al. |
| EPI_ISL_545042 | Sydney South West Pathology Service (SSWPS) - Liverpool Hospital - NSW Health Pathology | NSW Health Pathology - Institute of Clinical Pathology and Medical Research; Westmead Hospital; University of Sydney | CIDM-PH et al. |
| EPI_ISL_545043 | Sydney South West Pathology Service (SSWPS) - Royal Prince Alfred Hospital - NSW Health Pathology | NSW Health Pathology - Institute of Clinical Pathology and Medical Research; Westmead Hospital; University of Sydney | CIDM-PH et al. |
| EPI_ISL_545044 | Sydney South West Pathology Service (SSWPS) - Concord Repatriation General Hospital - NSW Health Pathology | NSW Health Pathology - Institute of Clinical Pathology and Medical Research; Westmead Hospital; University of Sydney | CIDM-PH et al. |
| EPI_ISL_545045 | Sydney South West Pathology Service (SSWPS) - Royal Prince Alfred Hospital - NSW Health Pathology | NSW Health Pathology - Institute of Clinical Pathology and Medical Research; Westmead Hospital; University of Sydney | CIDM-PH et al. |
| EPI_ISL_545046, EPI_ISL_545047 | Pathology West - NSW Health Pathology | NSW Health Pathology - Institute of Clinical Pathology and Medical Research; Westmead Hospital; University of Sydney | CIDM-PH et al. |
| EPI_ISL_545048 | South Eastern Area Laboratory Services (SEALS) | NSW Health Pathology - Institute of Clinical Pathology and Medical Research; Westmead Hospital; University of Sydney | CIDM-PH et al. |
| EPI_ISL_545049 | Sydney South West Pathology Service (SSWPS) - Concord Repatriation General Hospital - NSW Health Pathology | NSW Health Pathology - Institute of Clinical Pathology and Medical Research; Westmead Hospital; University of Sydney | CIDM-PH et al. |
| EPI_ISL_545050 | Pathology West - NSW Health Pathology | NSW Health Pathology - Institute of Clinical Pathology and Medical Research; Westmead Hospital; University of Sydney | CIDM-PH et al. |
| EPI_ISL_545051, EPI_ISL_545052, EPI_ISL_545053, EPI_ISL_545055, EPI_ISL_545056, EPI_ISL_545057, EPI_ISL_545058, EPI_ISL_545059, EPI_ISL_545060, EPI_ISL_545061, EPI_ISL_545062, EPI_ISL_545063, EPI_ISL_545064, EPI_ISL_545065, EPI_ISL_545066, EPI_ISL_545068, EPI_ISL_545069, EPI_ISL_545070, EPI_ISL_545071, EPI_ISL_545072, EPI_ISL_545073, EPI_ISL_545075, EPI_ISL_545076, EPI_ISL_545078, EPI_ISL_545079, EPI_ISL_545080, EPI_ISL_545081, EPI_ISL_545082, EPI_ISL_545083, EPI_ISL_545084, EPI_ISL_545085, EPI_ISL_545087, EPI_ISL_545088, EPI_ISL_545090, EPI_ISL_545093, EPI_ISL_545094, EPI_ISL_545095, EPI_ISL_545096, EPI_ISL_545097, EPI_ISL_545098, EPI_ISL_545100, EPI_ISL_545101, EPI_ISL_545102, EPI_ISL_545103, EPI_ISL_545106, EPI_ISL_545108, EPI_ISL_545109, EPI_ISL_545110, EPI_ISL_545111, EPI_ISL_545112, EPI_ISL_545113, EPI_ISL_545114, EPI_ISL_545115, EPI_ISL_545116, EPI_ISL_545117, EPI_ISL_545118, EPI_ISL_545119, EPI_ISL_545120, EPI_ISL_545121, EPI_ISL_545122, EPI_ISL_545123, EPI_ISL_545125, EPI_ISL_545126, EPI_ISL_545127, EPI_ISL_545128, EPI_ISL_545129, EPI_ISL_545130, EPI_ISL_545131, EPI_ISL_545132, EPI_ISL_545133, EPI_ISL_545134, EPI_ISL_545135, EPI_ISL_545136, EPI_ISL_545137, EPI_ISL_545138, EPI_ISL_545139, EPI_ISL_545140, EPI_ISL_545142, EPI_ISL_545143, EPI_ISL_545144, EPI_ISL_545145, EPI_ISL_545146, EPI_ISL_545147, EPI_ISL_545149, EPI_ISL_545150, EPI_ISL_545152, EPI_ISL_545153, EPI_ISL_545155, EPI_ISL_545156, EPI_ISL_545157, EPI_ISL_545158, EPI_ISL_545159, EPI_ISL_545160, EPI_ISL_545161, EPI_ISL_545162, EPI_ISL_545163, EPI_ISL_545164, EPI_ISL_545165, EPI_ISL_545166, EPI_ISL_545168, EPI_ISL_545169, EPI_ISL_545170, EPI_ISL_545171, EPI_ISL_545172, EPI_ISL_545173, EPI_ISL_545175, EPI_ISL_545176, EPI_ISL_545177, EPI_ISL_545178, EPI_ISL_545179, EPI_ISL_545180, EPI_ISL_545181, EPI_ISL_545182, EPI_ISL_545183, EPI_ISL_545184, EPI_ISL_545185, EPI_ISL_545186, EPI_ISL_545187, EPI_ISL_545188, EPI_ISL_545189, EPI_ISL_545191, EPI_ISL_545193, EPI_ISL_545194, EPI_ISL_545195, EPI_ISL_545196, EPI_ISL_545198, EPI_ISL_545199, EPI_ISL_545200, EPI_ISL_545201, EPI_ISL_545204, EPI_ISL_545205, EPI_ISL_545206, EPI_ISL_545207, EPI_ISL_545208, EPI_ISL_545210, EPI_ISL_545214, EPI_ISL_545216, EPI_ISL_545217, EPI_ISL_545218, EPI_ISL_545219, EPI_ISL_545220, EPI_ISL_545221, EPI_ISL_545223, EPI_ISL_545224, EPI_ISL_545225, EPI_ISL_545226, EPI_ISL_545227, EPI_ISL_545228, EPI_ISL_545229, EPI_ISL_545230, EPI_ISL_545231, EPI_ISL_545233, EPI_ISL_545234, EPI_ISL_545235, EPI_ISL_545236, EPI_ISL_545237, EPI_ISL_545238, EPI_ISL_545239, EPI_ISL_545240, EPI_ISL_545241, EPI_ISL_545242, EPI_ISL_545243, EPI_ISL_545244, EPI_ISL_545246, EPI_ISL_545247, EPI_ISL_545249, EPI_ISL_545251, EPI_ISL_545255, EPI_ISL_545256, EPI_ISL_545258, EPI_ISL_545259, EPI_ISL_545260, EPI_ISL_545261, EPI_ISL_545262, EPI_ISL_545263, EPI_ISL_545264, EPI_ISL_545266, EPI_ISL_545267, EPI_ISL_545269, EPI_ISL_545270, EPI_ISL_545271, EPI_ISL_545272, EPI_ISL_545273, EPI_ISL_545274, EPI_ISL_545276, EPI_ISL_545277, EPI_ISL_545279, EPI_ISL_545281, EPI_ISL_545282, EPI_ISL_545283, EPI_ISL_545284, EPI_ISL_545285, EPI_ISL_545288, EPI_ISL_545289, EPI_ISL_545290, EPI_ISL_545291, EPI_ISL_545295, EPI_ISL_545296, EPI_ISL_545297, EPI_ISL_545298, EPI_ISL_545299, EPI_ISL_545300, EPI_ISL_545302, EPI_ISL_545303, EPI_ISL_545305, EPI_ISL_545306, EPI_ISL_545307, EPI_ISL_545308, EPI_ISL_545310, EPI_ISL_545311, EPI_ISL_545312, EPI_ISL_545313, EPI_ISL_545314, EPI_ISL_545315, EPI_ISL_545316, EPI_ISL_545317, EPI_ISL_545319, EPI_ISL_545320, EPI_ISL_545321, EPI_ISL_545322, EPI_ISL_545324, EPI_ISL_545325, EPI_ISL_545326, EPI_ISL_545327, EPI_ISL_545328, EPI_ISL_545329, EPI_ISL_545330, EPI_ISL_545331, EPI_ISL_545332, EPI_ISL_545333, EPI_ISL_545334, EPI_ISL_545335, EPI_ISL_545336, EPI_ISL_545337, EPI_ISL_545338, EPI_ISL_545339, EPI_ISL_545340, EPI_ISL_545342, EPI_ISL_545343, EPI_ISL_545344, EPI_ISL_545345, EPI_ISL_545346, EPI_ISL_545347, EPI_ISL_545348, EPI_ISL_545349, EPI_ISL_545351, EPI_ISL_545352, EPI_ISL_545353, EPI_ISL_545354, EPI_ISL_545355, EPI_ISL_545356, EPI_ISL_545357, EPI_ISL_545358, EPI_ISL_545359, EPI_ISL_545360, EPI_ISL_545361, EPI_ISL_545362, EPI_ISL_545363, EPI_ISL_545364, EPI_ISL_545366, EPI_ISL_545367, EPI_ISL_545368, EPI_ISL_545369, EPI_ISL_545371, EPI_ISL_545372, EPI_ISL_545373, EPI_ISL_545374, EPI_ISL_545375, EPI_ISL_545376, EPI_ISL_545377, EPI_ISL_545378, EPI_ISL_545381, EPI_ISL_545383, EPI_ISL_545384, EPI_ISL_545385, EPI_ISL_545387, EPI_ISL_545388, EPI_ISL_545389, EPI_ISL_545390, EPI_ISL_545391, EPI_ISL_545392, EPI_ISL_545393, EPI_ISL_545394, EPI_ISL_545395, EPI_ISL_545396, EPI_ISL_545397, EPI_ISL_545398, EPI_ISL_545399, EPI_ISL_545400, EPI_ISL_545401, EPI_ISL_545402, EPI_ISL_545405, EPI_ISL_545406, EPI_ISL_545407, EPI_ISL_545409, EPI_ISL_545411, EPI_ISL_545412, EPI_ISL_545413, EPI_ISL_545414, EPI_ISL_545416, EPI_ISL_545419, EPI_ISL_545420, EPI_ISL_545421, EPI_ISL_545422, EPI_ISL_545423, EPI_ISL_545424, EPI_ISL_545425, EPI_ISL_545426, EPI_ISL_545427, EPI_ISL_545429, EPI_ISL_545430, EPI_ISL_545431, EPI_ISL_545432, EPI_ISL_545433, EPI_ISL_545435, EPI_ISL_545436, EPI_ISL_545437, EPI_ISL_545438, EPI_ISL_545439, EPI_ISL_545440, EPI_ISL_545441, EPI_ISL_545442, EPI_ISL_545443, EPI_ISL_545444, EPI_ISL_545445, EPI_ISL_545446, EPI_ISL_545448, EPI_ISL_545449, EPI_ISL_545450, EPI_ISL_545451, EPI_ISL_545453, EPI_ISL_545454, EPI_ISL_545455, EPI_ISL_545456, EPI_ISL_545457, EPI_ISL_545458, EPI_ISL_545461, EPI_ISL_545462, EPI_ISL_545463, EPI_ISL_545464, EPI_ISL_545466, EPI_ISL_545467, EPI_ISL_545468, EPI_ISL_545469, EPI_ISL_545470, EPI_ISL_545471, EPI_ISL_545472, EPI_ISL_545475, EPI_ISL_545476, EPI_ISL_545477, EPI_ISL_545478, EPI_ISL_545479, EPI_ISL_545480, EPI_ISL_545482, EPI_ISL_545483, EPI_ISL_545484, EPI_ISL_545485, EPI_ISL_545487, EPI_ISL_545489, EPI_ISL_545490, EPI_ISL_545491, EPI_ISL_545492, EPI_ISL_545495, EPI_ISL_545496, EPI_ISL_545497, EPI_ISL_545498, EPI_ISL_545500, EPI_ISL_545501, EPI_ISL_545503, EPI_ISL_545504, EPI_ISL_545505, EPI_ISL_545506, EPI_ISL_545507, EPI_ISL_545509, EPI_ISL_545510, EPI_ISL_545511, EPI_ISL_545512, EPI_ISL_545513, EPI_ISL_545514, EPI_ISL_545515, EPI_ISL_545516, EPI_ISL_545518, EPI_ISL_545519, EPI_ISL_545520, EPI_ISL_545521, EPI_ISL_545522, EPI_ISL_545523, EPI_ISL_545524, EPI_ISL_545526, EPI_ISL_545527, EPI_ISL_545528, EPI_ISL_545529, EPI_ISL_545530, EPI_ISL_545531, EPI_ISL_545532, EPI_ISL_545533, EPI_ISL_545535, EPI_ISL_545537, EPI_ISL_545538, EPI_ISL_545539, EPI_ISL_545540, EPI_ISL_545541, EPI_ISL_545542, EPI_ISL_545543, EPI_ISL_545545, EPI_ISL_545546, EPI_ISL_545547, EPI_ISL_545548, EPI_ISL_545549, EPI_ISL_545550, EPI_ISL_545552, EPI_ISL_545553, EPI_ISL_545555, EPI_ISL_545556, EPI_ISL_545557, EPI_ISL_545558, EPI_ISL_545559, EPI_ISL_545560, EPI_ISL_545561, EPI_ISL_545562, EPI_ISL_545563, EPI_ISL_545564, EPI_ISL_545565, EPI_ISL_545566, EPI_ISL_545567, EPI_ISL_545568, EPI_ISL_545569, EPI_ISL_545570, EPI_ISL_545573, EPI_ISL_545574 | | | |
| see above | Houston Methodist Hospital | Houston Methodist Hospital | S. Wesley Long, Randall J. Olsen, Paul A. Christensen, David W. Bernard, James J. Davis, Maulik Shukla, Marcus Nguyen, Matthew Ojeda Saavedra, Concepcion C. Cantu, Prasanti Yerramilli, Layne Pruitt, Sishir Subedi, Hung-Che Kuo, Heather Hendrickson, Ghazaleh Eskandari, Hoang A. T. Nguyen, J. Hunter Long, Muthiah Kumaraswami, Jule Goike, Daniel Boutz, Jimmy Gollihar, Jason S. McLellan, Chia-Wei Chou, Kamyab Javanmardi, Ilya J. Finkelstein, and James M. Musser |
| EPI_ISL_545580 | The National Institute of Public Health | State Veterinary Institute Prague | Nagy,A;Jirincova,H;Novakova,L;Trnka,D;Vecerova,J |
| EPI_ISL_545582, EPI_ISL_545583, EPI_ISL_545584, EPI_ISL_545586, EPI_ISL_545587, EPI_ISL_545588, EPI_ISL_545589, EPI_ISL_545591, EPI_ISL_545593, EPI_ISL_545595, EPI_ISL_545596, EPI_ISL_545597, EPI_ISL_545599, EPI_ISL_545600, EPI_ISL_545601, EPI_ISL_545602, EPI_ISL_545606, EPI_ISL_545607, EPI_ISL_545608, EPI_ISL_545609, EPI_ISL_545610, EPI_ISL_545611, EPI_ISL_545612, EPI_ISL_545614, EPI_ISL_545615, EPI_ISL_545619, EPI_ISL_545621, EPI_ISL_545624, EPI_ISL_545625, EPI_ISL_545626, EPI_ISL_545627, EPI_ISL_545630, EPI_ISL_545633, EPI_ISL_545634, EPI_ISL_545636, EPI_ISL_545637, EPI_ISL_545638, EPI_ISL_545639, EPI_ISL_545640, EPI_ISL_545644, EPI_ISL_545645, EPI_ISL_545646, EPI_ISL_545647, EPI_ISL_545650, EPI_ISL_545651, EPI_ISL_545653, EPI_ISL_545656, EPI_ISL_545658, EPI_ISL_545659, EPI_ISL_545660, EPI_ISL_545661, EPI_ISL_545662, EPI_ISL_545663, EPI_ISL_545664, EPI_ISL_545666, EPI_ISL_545667, EPI_ISL_545668, EPI_ISL_545669, EPI_ISL_545670, EPI_ISL_545671, EPI_ISL_545672, EPI_ISL_545673, EPI_ISL_545674, EPI_ISL_545675, EPI_ISL_545676, EPI_ISL_545677, EPI_ISL_545678, EPI_ISL_545679, EPI_ISL_545680, EPI_ISL_545681, EPI_ISL_545682, EPI_ISL_545683, EPI_ISL_545684, EPI_ISL_545685, EPI_ISL_545686, EPI_ISL_545687, EPI_ISL_545688, EPI_ISL_545689, EPI_ISL_545691, EPI_ISL_545692, EPI_ISL_545693, EPI_ISL_545694, EPI_ISL_545695, EPI_ISL_545696, EPI_ISL_545697, EPI_ISL_545698, EPI_ISL_545700, EPI_ISL_545701, EPI_ISL_545702, EPI_ISL_545703, EPI_ISL_545704, EPI_ISL_545706, EPI_ISL_545707, EPI_ISL_545708, EPI_ISL_545709, EPI_ISL_545710, EPI_ISL_545712, EPI_ISL_545714, EPI_ISL_545715, EPI_ISL_545716, EPI_ISL_545717, EPI_ISL_545718, EPI_ISL_545719, EPI_ISL_545720, EPI_ISL_545721, EPI_ISL_545722, EPI_ISL_545723, EPI_ISL_545724, EPI_ISL_545725, EPI_ISL_545727, EPI_ISL_545728, EPI_ISL_545730, EPI_ISL_545731, EPI_ISL_545732, EPI_ISL_545733, EPI_ISL_545734, EPI_ISL_545735, EPI_ISL_545736, EPI_ISL_545737, EPI_ISL_545738, EPI_ISL_545739, EPI_ISL_545740, EPI_ISL_545744, EPI_ISL_545745, EPI_ISL_545746, EPI_ISL_545748, EPI_ISL_545749, EPI_ISL_545751, EPI_ISL_545752, EPI_ISL_545753, EPI_ISL_545754, EPI_ISL_545755, EPI_ISL_545756, EPI_ISL_545757, EPI_ISL_545758, EPI_ISL_545759, EPI_ISL_545760, EPI_ISL_545761, EPI_ISL_545762, EPI_ISL_545763, EPI_ISL_545764, EPI_ISL_545765, EPI_ISL_545766, EPI_ISL_545767, EPI_ISL_545769, EPI_ISL_545770, EPI_ISL_545771, EPI_ISL_545772, EPI_ISL_545776, EPI_ISL_545777, EPI_ISL_545778, EPI_ISL_545779, EPI_ISL_545780, EPI_ISL_545782, EPI_ISL_545783, EPI_ISL_545784, EPI_ISL_545785, EPI_ISL_545786, EPI_ISL_545787, EPI_ISL_545788, EPI_ISL_545789, EPI_ISL_545790, EPI_ISL_545791, EPI_ISL_545793, EPI_ISL_545794, EPI_ISL_545795, EPI_ISL_545796, EPI_ISL_545797, EPI_ISL_545798, EPI_ISL_545799, EPI_ISL_545800, EPI_ISL_545801, EPI_ISL_545802, EPI_ISL_545803, EPI_ISL_545804, EPI_ISL_545805, EPI_ISL_545807, EPI_ISL_545808, EPI_ISL_545809, EPI_ISL_545810, EPI_ISL_545811, EPI_ISL_545812, EPI_ISL_545814, EPI_ISL_545815, EPI_ISL_545816, EPI_ISL_545817, EPI_ISL_545818, EPI_ISL_545819, EPI_ISL_545820, EPI_ISL_545822, EPI_ISL_545823, EPI_ISL_545824, EPI_ISL_545825, EPI_ISL_545827, EPI_ISL_545828, EPI_ISL_545829, EPI_ISL_545830, EPI_ISL_545831, EPI_ISL_545832, EPI_ISL_545833, EPI_ISL_545834, EPI_ISL_545835, EPI_ISL_545836, EPI_ISL_545837, EPI_ISL_545838, EPI_ISL_545839, EPI_ISL_545840, EPI_ISL_545841, EPI_ISL_545842, EPI_ISL_545843, EPI_ISL_545844, EPI_ISL_545846, EPI_ISL_545847, EPI_ISL_545848, EPI_ISL_545849, EPI_ISL_545850, EPI_ISL_545852, EPI_ISL_545854, EPI_ISL_545855, EPI_ISL_545856, EPI_ISL_545857, EPI_ISL_545858, EPI_ISL_545859, EPI_ISL_545860, EPI_ISL_545861, EPI_ISL_545862, EPI_ISL_545863, EPI_ISL_545865, EPI_ISL_545867, EPI_ISL_545868, EPI_ISL_545869, EPI_ISL_545870, EPI_ISL_545871, EPI_ISL_545873, EPI_ISL_545874, EPI_ISL_545875, EPI_ISL_545876, EPI_ISL_545877, EPI_ISL_545878, EPI_ISL_545879, EPI_ISL_545880, EPI_ISL_545881, EPI_ISL_545882, EPI_ISL_545883, EPI_ISL_545884, EPI_ISL_545885, EPI_ISL_545886, EPI_ISL_545887, EPI_ISL_545888, EPI_ISL_545889, EPI_ISL_545890, EPI_ISL_545891, EPI_ISL_545892, EPI_ISL_545893, EPI_ISL_545894, EPI_ISL_545895, EPI_ISL_545897, EPI_ISL_545898, EPI_ISL_545899, EPI_ISL_545900, EPI_ISL_545901, EPI_ISL_545902, EPI_ISL_545903, EPI_ISL_545905, EPI_ISL_545906, EPI_ISL_545907, EPI_ISL_545908, EPI_ISL_545909, EPI_ISL_545911, EPI_ISL_545912, EPI_ISL_545913, EPI_ISL_545914, EPI_ISL_545915, EPI_ISL_545916, EPI_ISL_545917, EPI_ISL_545919, EPI_ISL_545920, EPI_ISL_545921, EPI_ISL_545922, EPI_ISL_545923, EPI_ISL_545924, EPI_ISL_545926, EPI_ISL_545927, EPI_ISL_545928, EPI_ISL_545929, EPI_ISL_545930, EPI_ISL_545931, EPI_ISL_545932, EPI_ISL_545933, EPI_ISL_545934, EPI_ISL_545935, EPI_ISL_545936, EPI_ISL_545937, EPI_ISL_545938, EPI_ISL_545939, EPI_ISL_545940, EPI_ISL_545941, EPI_ISL_545942, EPI_ISL_545943, EPI_ISL_545945, EPI_ISL_545946, EPI_ISL_545947, EPI_ISL_545948, EPI_ISL_545949, EPI_ISL_545951, EPI_ISL_545952, EPI_ISL_545953 | | | |
| see above | Houston Methodist Hospital | Houston Methodist Hospital | S. Wesley Long, Randall J. Olsen, Paul A. Christensen, David W. Bernard, James J. Davis, Maulik Shukla, Marcus Nguyen, Matthew Ojeda Saavedra, Concepcion C. Cantu, Prasanti Yerramilli, Layne Pruitt, Sishir Subedi, Hung-Che Kuo, Heather Hendrickson, Ghazaleh Eskandari, Hoang A. T. Nguyen, J. Hunter Long, Muthiah Kumaraswami, Jule Goike, Daniel Boutz, Jimmy Gollihar, Jason S. McLellan, Chia-Wei Chou, Kamyab Javanmardi, Ilya J. Finkelstein, and James M. Musser |
| EPI_ISL_545954, EPI_ISL_545956 | The National Institute of Public Health | State Veterinary Institute Prague | Nagy,A;Jirincova,H;Novakova,L;Trnka,D;Vecerova,J |
| EPI_ISL_545957 | The National Institute of Public Health | State Veterinary Institute Prague | Nagy,A; Jirincova,H; Novakova,L; Trnka,D; Vecerova,J |
| EPI_ISL_545958, EPI_ISL_545960, EPI_ISL_545961, EPI_ISL_545963, EPI_ISL_545964, EPI_ISL_545965, EPI_ISL_545967, EPI_ISL_545968, EPI_ISL_545970, EPI_ISL_545973, EPI_ISL_545974, EPI_ISL_545976, EPI_ISL_545977, EPI_ISL_545981, EPI_ISL_545982, EPI_ISL_545983, EPI_ISL_545985, EPI_ISL_545986, EPI_ISL_545987, EPI_ISL_545988, EPI_ISL_545989, EPI_ISL_545991, EPI_ISL_545992, EPI_ISL_545994, EPI_ISL_545995, EPI_ISL_545998, EPI_ISL_545999, EPI_ISL_546000, EPI_ISL_546001, EPI_ISL_546002, EPI_ISL_546004, EPI_ISL_546005, EPI_ISL_546006, EPI_ISL_546007, EPI_ISL_546008, EPI_ISL_546009, EPI_ISL_546010, EPI_ISL_546011, EPI_ISL_546013, EPI_ISL_546014, EPI_ISL_546017, EPI_ISL_546018, EPI_ISL_546019, EPI_ISL_546020, EPI_ISL_546021, EPI_ISL_546022, EPI_ISL_546024, EPI_ISL_546025, EPI_ISL_546026, EPI_ISL_546027, EPI_ISL_546028, EPI_ISL_546029, EPI_ISL_546031, EPI_ISL_546032, EPI_ISL_546033, EPI_ISL_546035, EPI_ISL_546036, EPI_ISL_546037, EPI_ISL_546038, EPI_ISL_546039, EPI_ISL_546040, EPI_ISL_546041, EPI_ISL_546042, EPI_ISL_546043, EPI_ISL_546044, EPI_ISL_546045, EPI_ISL_546046, EPI_ISL_546047, EPI_ISL_546048, EPI_ISL_546049, EPI_ISL_546051, EPI_ISL_546052, EPI_ISL_546053, EPI_ISL_546055, EPI_ISL_546056, EPI_ISL_546057, EPI_ISL_546058, EPI_ISL_546059, EPI_ISL_546060, EPI_ISL_546061, EPI_ISL_546062, EPI_ISL_546064, EPI_ISL_546065, EPI_ISL_546066, EPI_ISL_546067, EPI_ISL_546068, EPI_ISL_546069, EPI_ISL_546070, EPI_ISL_546071, EPI_ISL_546072, EPI_ISL_546073, EPI_ISL_546074, EPI_ISL_546075, EPI_ISL_546078, EPI_ISL_546080, EPI_ISL_546081, EPI_ISL_546082, EPI_ISL_546083, EPI_ISL_546084, EPI_ISL_546088, EPI_ISL_546090, EPI_ISL_546091, EPI_ISL_546093, EPI_ISL_546095, EPI_ISL_546096, EPI_ISL_546097, EPI_ISL_546098, EPI_ISL_546099, EPI_ISL_546100, EPI_ISL_546101, EPI_ISL_546102, EPI_ISL_546103, EPI_ISL_546105, EPI_ISL_546106, EPI_ISL_546108, EPI_ISL_546109, EPI_ISL_546110, EPI_ISL_546111, EPI_ISL_546113, EPI_ISL_546114, EPI_ISL_546116, EPI_ISL_546117, EPI_ISL_546119, EPI_ISL_546120, EPI_ISL_546121, EPI_ISL_546123, EPI_ISL_546127, EPI_ISL_546128, EPI_ISL_546129, EPI_ISL_546135, EPI_ISL_546136, EPI_ISL_546137, EPI_ISL_546138, EPI_ISL_546139, EPI_ISL_546140, EPI_ISL_546141, EPI_ISL_546144, EPI_ISL_546145, EPI_ISL_546146, EPI_ISL_546147, EPI_ISL_546148, EPI_ISL_546149, EPI_ISL_546150, EPI_ISL_546151, EPI_ISL_546154, EPI_ISL_546155, EPI_ISL_546156, EPI_ISL_546157, EPI_ISL_546158, EPI_ISL_546160, EPI_ISL_546162, EPI_ISL_546164, EPI_ISL_546165, EPI_ISL_546168, EPI_ISL_546170, EPI_ISL_546171, EPI_ISL_546172, EPI_ISL_546173, EPI_ISL_546177, EPI_ISL_546178, EPI_ISL_546179, EPI_ISL_546180, EPI_ISL_546181, EPI_ISL_546183, EPI_ISL_546184, EPI_ISL_546185, EPI_ISL_546188, EPI_ISL_546190, EPI_ISL_546191, EPI_ISL_546192, EPI_ISL_546193, EPI_ISL_546194, EPI_ISL_546195, EPI_ISL_546196, EPI_ISL_546197, EPI_ISL_546198, EPI_ISL_546199, EPI_ISL_546200, EPI_ISL_546201, EPI_ISL_546202, EPI_ISL_546206, EPI_ISL_546207, EPI_ISL_546208, EPI_ISL_546209, EPI_ISL_546210, EPI_ISL_546211, EPI_ISL_546212, EPI_ISL_546213, EPI_ISL_546214, EPI_ISL_546216, EPI_ISL_546217, EPI_ISL_546218, EPI_ISL_546219, EPI_ISL_546220, EPI_ISL_546221, EPI_ISL_546222, EPI_ISL_546224, EPI_ISL_546225, EPI_ISL_546226, EPI_ISL_546227, EPI_ISL_546228, EPI_ISL_546231, EPI_ISL_546232, EPI_ISL_546233, EPI_ISL_546234, EPI_ISL_546235, EPI_ISL_546236, EPI_ISL_546237, EPI_ISL_546239, EPI_ISL_546240, EPI_ISL_546241, EPI_ISL_546242, EPI_ISL_546243, EPI_ISL_546244, EPI_ISL_546245, EPI_ISL_546246, EPI_ISL_546247, EPI_ISL_546248, EPI_ISL_546250, EPI_ISL_546251, EPI_ISL_546252, EPI_ISL_546253, EPI_ISL_546254, EPI_ISL_546255, EPI_ISL_546256, EPI_ISL_546257, EPI_ISL_546258, EPI_ISL_546259, EPI_ISL_546260, EPI_ISL_546262, EPI_ISL_546263, EPI_ISL_546264, EPI_ISL_546265, EPI_ISL_546267, EPI_ISL_546269, EPI_ISL_546272, EPI_ISL_546277, EPI_ISL_546278, EPI_ISL_546279, EPI_ISL_546280, EPI_ISL_546282, EPI_ISL_546284, EPI_ISL_546287, EPI_ISL_546288, EPI_ISL_546290, EPI_ISL_546291, EPI_ISL_546292, EPI_ISL_546293, EPI_ISL_546294, EPI_ISL_546295, EPI_ISL_546296, EPI_ISL_546297, EPI_ISL_546299, EPI_ISL_546300, EPI_ISL_546301, EPI_ISL_546302, EPI_ISL_546303, EPI_ISL_546305, EPI_ISL_546306, EPI_ISL_546308, EPI_ISL_546309, EPI_ISL_546310, EPI_ISL_546311, EPI_ISL_546312, EPI_ISL_546313, EPI_ISL_546314, EPI_ISL_546315, EPI_ISL_546316, EPI_ISL_546317, EPI_ISL_546318, EPI_ISL_546319, EPI_ISL_546320, EPI_ISL_546322, EPI_ISL_546323, EPI_ISL_546324, EPI_ISL_546326, EPI_ISL_546327, EPI_ISL_546328, EPI_ISL_546329, EPI_ISL_546331, EPI_ISL_546332, EPI_ISL_546333, EPI_ISL_546334, EPI_ISL_546335, EPI_ISL_546336, EPI_ISL_546337, EPI_ISL_546338, EPI_ISL_546339, EPI_ISL_546340, EPI_ISL_546342, EPI_ISL_546343, EPI_ISL_546345, EPI_ISL_546348, EPI_ISL_546352, EPI_ISL_546353, EPI_ISL_546355, EPI_ISL_546356, EPI_ISL_546357, EPI_ISL_546360, EPI_ISL_546361, EPI_ISL_546362, EPI_ISL_546363, EPI_ISL_546365, EPI_ISL_546366, EPI_ISL_546368, EPI_ISL_546370, EPI_ISL_546372, EPI_ISL_546373, EPI_ISL_546376, EPI_ISL_546378, EPI_ISL_546379, EPI_ISL_546380, EPI_ISL_546384, EPI_ISL_546385, EPI_ISL_546386, EPI_ISL_546390, EPI_ISL_546391, EPI_ISL_546392, EPI_ISL_546396, EPI_ISL_546397, EPI_ISL_546399, EPI_ISL_546400, EPI_ISL_546403, EPI_ISL_546406, EPI_ISL_546408, EPI_ISL_546409, EPI_ISL_546410, EPI_ISL_546411, EPI_ISL_546413, EPI_ISL_546414, EPI_ISL_546419, EPI_ISL_546421, EPI_ISL_546423, EPI_ISL_546424, EPI_ISL_546425, EPI_ISL_546426, EPI_ISL_546427, EPI_ISL_546430, EPI_ISL_546432, EPI_ISL_546433, EPI_ISL_546434 | | | |
| see above | Houston Methodist Hospital | Houston Methodist Hospital | S. Wesley Long, Randall J. Olsen, Paul A. Christensen, David W. Bernard, James J. Davis, Maulik Shukla, Marcus Nguyen, Matthew Ojeda Saavedra, Concepcion C. Cantu, Prasanti Yerramilli, Layne Pruitt, Sishir Subedi, Hung-Che Kuo, Heather Hendrickson, Ghazaleh Eskandari, Hoang A. T. Nguyen, J. Hunter Long, Muthiah Kumaraswami, Jule Goike, Daniel Boutz, Jimmy Gollihar, Jason S. McLellan, Chia-Wei Chou, Kamyab Javanmardi, Ilya J. Finkelstein, and James M. Musser |
| EPI_ISL_546436 | The National Institute of Public Health | State Veterinary Institute Prague | Nagy,A; Jirincova,H; Novakova,L; Trnka,D; Vecerova,J |
| EPI_ISL_546437, EPI_ISL_546438, EPI_ISL_546439, EPI_ISL_546440, EPI_ISL_546443, EPI_ISL_546444, EPI_ISL_546446, EPI_ISL_546447, EPI_ISL_546448, EPI_ISL_546449, EPI_ISL_546450, EPI_ISL_546451, EPI_ISL_546454, EPI_ISL_546456, EPI_ISL_546457, EPI_ISL_546463, EPI_ISL_546464, EPI_ISL_546465, EPI_ISL_546466, EPI_ISL_546468, EPI_ISL_546469, EPI_ISL_546471, EPI_ISL_546473, EPI_ISL_546475, EPI_ISL_546476, EPI_ISL_546477, EPI_ISL_546478, EPI_ISL_546479, EPI_ISL_546480, EPI_ISL_546482, EPI_ISL_546483, EPI_ISL_546484, EPI_ISL_546486, EPI_ISL_546487, EPI_ISL_546488, EPI_ISL_546489, EPI_ISL_546490, EPI_ISL_546492, EPI_ISL_546494, EPI_ISL_546495, EPI_ISL_546498, EPI_ISL_546499, EPI_ISL_546500, EPI_ISL_546502, EPI_ISL_546503, EPI_ISL_546504, EPI_ISL_546505, EPI_ISL_546506, EPI_ISL_546507, EPI_ISL_546509, EPI_ISL_546510, EPI_ISL_546511, EPI_ISL_546512, EPI_ISL_546515, EPI_ISL_546517, EPI_ISL_546518, EPI_ISL_546519, EPI_ISL_546520, EPI_ISL_546521, EPI_ISL_546522, EPI_ISL_546523, EPI_ISL_546524, EPI_ISL_546525, EPI_ISL_546526, EPI_ISL_546527, EPI_ISL_546529, EPI_ISL_546531, EPI_ISL_546533, EPI_ISL_546536, EPI_ISL_546537, EPI_ISL_546538, EPI_ISL_546539, EPI_ISL_546541, EPI_ISL_546542, EPI_ISL_546545, EPI_ISL_546546, EPI_ISL_546547, EPI_ISL_546549, EPI_ISL_546550, EPI_ISL_546551, EPI_ISL_546552, EPI_ISL_546554, EPI_ISL_546555, EPI_ISL_546556, EPI_ISL_546557, EPI_ISL_546558, EPI_ISL_546563, EPI_ISL_546564, EPI_ISL_546565, EPI_ISL_546566, EPI_ISL_546567, EPI_ISL_546568, EPI_ISL_546569, EPI_ISL_546570, EPI_ISL_546571, EPI_ISL_546572, EPI_ISL_546574, EPI_ISL_546575, EPI_ISL_546579, EPI_ISL_546581, EPI_ISL_546582, EPI_ISL_546584, EPI_ISL_546585, EPI_ISL_546586, EPI_ISL_546587, EPI_ISL_546592, EPI_ISL_546593, EPI_ISL_546594, EPI_ISL_546598, EPI_ISL_546599, EPI_ISL_546600, EPI_ISL_546601, EPI_ISL_546603, EPI_ISL_546604, EPI_ISL_546605, EPI_ISL_546606, EPI_ISL_546607, EPI_ISL_546610, EPI_ISL_546611, EPI_ISL_546612, EPI_ISL_546613, EPI_ISL_546614, EPI_ISL_546615, EPI_ISL_546617, EPI_ISL_546618, EPI_ISL_546619, EPI_ISL_546621, EPI_ISL_546623, EPI_ISL_546624, EPI_ISL_546626, EPI_ISL_546627, EPI_ISL_546629, EPI_ISL_546631, EPI_ISL_546632, EPI_ISL_546636, EPI_ISL_546638, EPI_ISL_546639, EPI_ISL_546640, EPI_ISL_546641, EPI_ISL_546642, EPI_ISL_546645, EPI_ISL_546647, EPI_ISL_546650, EPI_ISL_546651, EPI_ISL_546652, EPI_ISL_546653, EPI_ISL_546656, EPI_ISL_546657, EPI_ISL_546658, EPI_ISL_546659, EPI_ISL_546660, EPI_ISL_546663, EPI_ISL_546664, EPI_ISL_546667, EPI_ISL_546668, EPI_ISL_546669, EPI_ISL_546670, EPI_ISL_546671, EPI_ISL_546672, EPI_ISL_546674, EPI_ISL_546675, EPI_ISL_546676, EPI_ISL_546677, EPI_ISL_546678, EPI_ISL_546679, EPI_ISL_546680, EPI_ISL_546681, EPI_ISL_546683, EPI_ISL_546684, EPI_ISL_546686, EPI_ISL_546687, EPI_ISL_546688, EPI_ISL_546689, EPI_ISL_546690, EPI_ISL_546691, EPI_ISL_546692, EPI_ISL_546693, EPI_ISL_546694, EPI_ISL_546697, EPI_ISL_546699, EPI_ISL_546700, EPI_ISL_546701, EPI_ISL_546702, EPI_ISL_546704, EPI_ISL_546706, EPI_ISL_546707, EPI_ISL_546709, EPI_ISL_546710, EPI_ISL_546711, EPI_ISL_546712, EPI_ISL_546713, EPI_ISL_546714, EPI_ISL_546715, EPI_ISL_546717, EPI_ISL_546718, EPI_ISL_546719, EPI_ISL_546721, EPI_ISL_546722, EPI_ISL_546723, EPI_ISL_546724, EPI_ISL_546725, EPI_ISL_546726, EPI_ISL_546728, EPI_ISL_546729, EPI_ISL_546730, EPI_ISL_546731, EPI_ISL_546733, EPI_ISL_546736, EPI_ISL_546738, EPI_ISL_546739, EPI_ISL_546740, EPI_ISL_546741, EPI_ISL_546742, EPI_ISL_546743, EPI_ISL_546744, EPI_ISL_546745, EPI_ISL_546746, EPI_ISL_546747, EPI_ISL_546749, EPI_ISL_546750, EPI_ISL_546752, EPI_ISL_546753, EPI_ISL_546754, EPI_ISL_546755, EPI_ISL_546756, EPI_ISL_546757, EPI_ISL_546758, EPI_ISL_546759, EPI_ISL_546760, EPI_ISL_546763, EPI_ISL_546764, EPI_ISL_546765, EPI_ISL_546766, EPI_ISL_546769, EPI_ISL_546770, EPI_ISL_546771, EPI_ISL_546774, EPI_ISL_546775, EPI_ISL_546776, EPI_ISL_546777, EPI_ISL_546778, EPI_ISL_546779, EPI_ISL_546780, EPI_ISL_546781, EPI_ISL_546782, EPI_ISL_546783, EPI_ISL_546785, EPI_ISL_546786, EPI_ISL_546787, EPI_ISL_546789, EPI_ISL_546790, EPI_ISL_546792, EPI_ISL_546793, EPI_ISL_546794, EPI_ISL_546795, EPI_ISL_546797, EPI_ISL_546800, EPI_ISL_546801, EPI_ISL_546802, EPI_ISL_546803, EPI_ISL_546804, EPI_ISL_546805, EPI_ISL_546807, EPI_ISL_546808, EPI_ISL_546810, EPI_ISL_546811, EPI_ISL_546812, EPI_ISL_546815, EPI_ISL_546816, EPI_ISL_546819, EPI_ISL_546820, EPI_ISL_546824, EPI_ISL_546825, EPI_ISL_546826, EPI_ISL_546827, EPI_ISL_546829, EPI_ISL_546831, EPI_ISL_546833, EPI_ISL_546834, EPI_ISL_546837, EPI_ISL_546838, EPI_ISL_546839, EPI_ISL_546840, EPI_ISL_546841, EPI_ISL_546842, EPI_ISL_546843, EPI_ISL_546844, EPI_ISL_546845, EPI_ISL_546847, EPI_ISL_546850, EPI_ISL_546852, EPI_ISL_546853, EPI_ISL_546854, EPI_ISL_546855, EPI_ISL_546856, EPI_ISL_546860, EPI_ISL_546861, EPI_ISL_546862, EPI_ISL_546863, EPI_ISL_546864, EPI_ISL_546865, EPI_ISL_546868, EPI_ISL_546869, EPI_ISL_546870, EPI_ISL_546871, EPI_ISL_546873, EPI_ISL_546879, EPI_ISL_546880, EPI_ISL_546881, EPI_ISL_546882, EPI_ISL_546884, EPI_ISL_546885, EPI_ISL_546886, EPI_ISL_546887, EPI_ISL_546888, EPI_ISL_546889, EPI_ISL_546890, EPI_ISL_546892, EPI_ISL_546893, EPI_ISL_546894, EPI_ISL_546895, EPI_ISL_546896, EPI_ISL_546897, EPI_ISL_546898, EPI_ISL_546899, EPI_ISL_546900, EPI_ISL_546902, EPI_ISL_546904, EPI_ISL_546905, EPI_ISL_546906, EPI_ISL_546907, EPI_ISL_546908, EPI_ISL_546909, EPI_ISL_546910, EPI_ISL_546911, EPI_ISL_546912, EPI_ISL_546913, EPI_ISL_546914, EPI_ISL_546916, EPI_ISL_546918, EPI_ISL_546920, EPI_ISL_546921, EPI_ISL_546922, EPI_ISL_546925, EPI_ISL_546927, EPI_ISL_546931, EPI_ISL_546932, EPI_ISL_546933 | | | |
| see above | Houston Methodist Hospital | Houston Methodist Hospital | S. Wesley Long, Randall J. Olsen, Paul A. Christensen, David W. Bernard, James J. Davis, Maulik Shukla, Marcus Nguyen, Matthew Ojeda Saavedra, Concepcion C. Cantu, Prasanti Yerramilli, Layne Pruitt, Sishir Subedi, Hung-Che Kuo, Heather |

| | | | |
|---|---|---|---|
| | | | Hendrickson, Ghazaleh Eskandari, Hoang A. T. Nguyen, J. Hunter Long, Muthiah Kumaraswami, Jule Goike, Daniel Boutz, Jimmy Gollihar, Jason S. McLellan, Chia-Wei Chou, Kamyab Javanmardi, Ilya J. Finkelstein, and James M. Musser |
| EPI_ISL_546935, EPI_ISL_546936 | The National Institute of Public Health | State Veterinary Institute Prague | Nagy,A; Jirincova,H; Novakova,L; Trnka,D; Vecerova,J |
| EPI_ISL_546937, EPI_ISL_546938, EPI_ISL_546940, EPI_ISL_546941, EPI_ISL_546945, EPI_ISL_546946, EPI_ISL_546948, EPI_ISL_546949, EPI_ISL_546950, EPI_ISL_546951, EPI_ISL_546952, EPI_ISL_546953, EPI_ISL_546954, EPI_ISL_546955, EPI_ISL_546956, EPI_ISL_546957, EPI_ISL_546958, EPI_ISL_546961, EPI_ISL_546962, EPI_ISL_546963, EPI_ISL_546964, EPI_ISL_546966, EPI_ISL_546967, EPI_ISL_546968, EPI_ISL_546973, EPI_ISL_546974, EPI_ISL_546975, EPI_ISL_546976, EPI_ISL_546977, EPI_ISL_546979, EPI_ISL_546980, EPI_ISL_546981, EPI_ISL_546983, EPI_ISL_546984, EPI_ISL_546985, EPI_ISL_546988, EPI_ISL_546989, EPI_ISL_546990, EPI_ISL_546991, EPI_ISL_546992, EPI_ISL_546995, EPI_ISL_546996, EPI_ISL_546997, EPI_ISL_546999, EPI_ISL_547000, EPI_ISL_547001, EPI_ISL_547002, EPI_ISL_547003, EPI_ISL_547004, EPI_ISL_547007, EPI_ISL_547008, EPI_ISL_547009, EPI_ISL_547010, EPI_ISL_547011, EPI_ISL_547012, EPI_ISL_547013, EPI_ISL_547014, EPI_ISL_547015, EPI_ISL_547016, EPI_ISL_547017, EPI_ISL_547020, EPI_ISL_547021, EPI_ISL_547022, EPI_ISL_547023, EPI_ISL_547025, EPI_ISL_547027, EPI_ISL_547028, EPI_ISL_547029, EPI_ISL_547031, EPI_ISL_547034, EPI_ISL_547035, EPI_ISL_547036, EPI_ISL_547037, EPI_ISL_547040, EPI_ISL_547041, EPI_ISL_547042, EPI_ISL_547044, EPI_ISL_547045, EPI_ISL_547046, EPI_ISL_547047, EPI_ISL_547048, EPI_ISL_547049, EPI_ISL_547050, EPI_ISL_547051, EPI_ISL_547054, EPI_ISL_547056, EPI_ISL_547057, EPI_ISL_547060, EPI_ISL_547062, EPI_ISL_547064, EPI_ISL_547066, EPI_ISL_547067, EPI_ISL_547068, EPI_ISL_547069, EPI_ISL_547071, EPI_ISL_547072, EPI_ISL_547073, EPI_ISL_547074, EPI_ISL_547076, EPI_ISL_547077, EPI_ISL_547079, EPI_ISL_547080, EPI_ISL_547081, EPI_ISL_547082, EPI_ISL_547083, EPI_ISL_547085, EPI_ISL_547086, EPI_ISL_547087, EPI_ISL_547088, EPI_ISL_547091, EPI_ISL_547092, EPI_ISL_547094, EPI_ISL_547095, EPI_ISL_547096, EPI_ISL_547097, EPI_ISL_547099, EPI_ISL_547100, EPI_ISL_547101, EPI_ISL_547102, EPI_ISL_547104, EPI_ISL_547105, EPI_ISL_547106, EPI_ISL_547107, EPI_ISL_547109, EPI_ISL_547110, EPI_ISL_547111, EPI_ISL_547112, EPI_ISL_547113, EPI_ISL_547118, EPI_ISL_547119, EPI_ISL_547121, EPI_ISL_547122, EPI_ISL_547123, EPI_ISL_547124, EPI_ISL_547125, EPI_ISL_547126, EPI_ISL_547127, EPI_ISL_547129, EPI_ISL_547130, EPI_ISL_547131, EPI_ISL_547132, EPI_ISL_547133, EPI_ISL_547135, EPI_ISL_547138, EPI_ISL_547139, EPI_ISL_547140, EPI_ISL_547143, EPI_ISL_547144, EPI_ISL_547146, EPI_ISL_547148, EPI_ISL_547149, EPI_ISL_547152, EPI_ISL_547155, EPI_ISL_547156, EPI_ISL_547158, EPI_ISL_547159, EPI_ISL_547160, EPI_ISL_547161, EPI_ISL_547162, EPI_ISL_547163, EPI_ISL_547165, EPI_ISL_547166, EPI_ISL_547168, EPI_ISL_547169, EPI_ISL_547170, EPI_ISL_547171, EPI_ISL_547172, EPI_ISL_547173, EPI_ISL_547174, EPI_ISL_547175, EPI_ISL_547176, EPI_ISL_547177, EPI_ISL_547178, EPI_ISL_547180, EPI_ISL_547181, EPI_ISL_547182, EPI_ISL_547183, EPI_ISL_547184, EPI_ISL_547185, EPI_ISL_547186, EPI_ISL_547187, EPI_ISL_547188, EPI_ISL_547190, EPI_ISL_547191, EPI_ISL_547194, EPI_ISL_547195, EPI_ISL_547197, EPI_ISL_547200, EPI_ISL_547201, EPI_ISL_547202, EPI_ISL_547203, EPI_ISL_547204, EPI_ISL_547205, EPI_ISL_547206, EPI_ISL_547207, EPI_ISL_547208, EPI_ISL_547209, EPI_ISL_547210, EPI_ISL_547211, EPI_ISL_547212, EPI_ISL_547213, EPI_ISL_547214, EPI_ISL_547215, EPI_ISL_547216, EPI_ISL_547217, EPI_ISL_547218, EPI_ISL_547219, EPI_ISL_547221, EPI_ISL_547222, EPI_ISL_547223, EPI_ISL_547224, EPI_ISL_547225, EPI_ISL_547226, EPI_ISL_547228, EPI_ISL_547230, EPI_ISL_547233, EPI_ISL_547234, EPI_ISL_547235, EPI_ISL_547236, EPI_ISL_547237, EPI_ISL_547238, EPI_ISL_547239, EPI_ISL_547240, EPI_ISL_547244, EPI_ISL_547245, EPI_ISL_547246, EPI_ISL_547248, EPI_ISL_547250, EPI_ISL_547251, EPI_ISL_547252, EPI_ISL_547253, EPI_ISL_547254, EPI_ISL_547255, EPI_ISL_547256, EPI_ISL_547259, EPI_ISL_547261, EPI_ISL_547262, EPI_ISL_547263, EPI_ISL_547264, EPI_ISL_547266, EPI_ISL_547268, EPI_ISL_547269, EPI_ISL_547270, EPI_ISL_547271, EPI_ISL_547272, EPI_ISL_547273, EPI_ISL_547274, EPI_ISL_547276, EPI_ISL_547277, EPI_ISL_547278, EPI_ISL_547279, EPI_ISL_547280, EPI_ISL_547283, EPI_ISL_547286, EPI_ISL_547287, EPI_ISL_547288, EPI_ISL_547289, EPI_ISL_547291, EPI_ISL_547293, EPI_ISL_547294, EPI_ISL_547295, EPI_ISL_547296, EPI_ISL_547297, EPI_ISL_547298, EPI_ISL_547299, EPI_ISL_547300, EPI_ISL_547301, EPI_ISL_547302, EPI_ISL_547303, EPI_ISL_547304, EPI_ISL_547306, EPI_ISL_547309, EPI_ISL_547310, EPI_ISL_547311, EPI_ISL_547312, EPI_ISL_547313, EPI_ISL_547314, EPI_ISL_547315, EPI_ISL_547316, EPI_ISL_547318, EPI_ISL_547320, EPI_ISL_547321, EPI_ISL_547322, EPI_ISL_547323, EPI_ISL_547324, EPI_ISL_547325, EPI_ISL_547327, EPI_ISL_547328, EPI_ISL_547331, EPI_ISL_547333, EPI_ISL_547334, EPI_ISL_547335, EPI_ISL_547336, EPI_ISL_547337, EPI_ISL_547339, EPI_ISL_547340, EPI_ISL_547341, EPI_ISL_547342, EPI_ISL_547343, EPI_ISL_547344, EPI_ISL_547345, EPI_ISL_547347, EPI_ISL_547348, EPI_ISL_547350, EPI_ISL_547352, EPI_ISL_547353, EPI_ISL_547354, EPI_ISL_547357, EPI_ISL_547359, EPI_ISL_547360, EPI_ISL_547362, EPI_ISL_547363, EPI_ISL_547365, EPI_ISL_547368, EPI_ISL_547369, EPI_ISL_547371, EPI_ISL_547372, EPI_ISL_547373, EPI_ISL_547374, EPI_ISL_547375, EPI_ISL_547376, EPI_ISL_547378, EPI_ISL_547379, EPI_ISL_547380, EPI_ISL_547381, EPI_ISL_547382, EPI_ISL_547383, EPI_ISL_547384, EPI_ISL_547387, EPI_ISL_547388, EPI_ISL_547390, EPI_ISL_547394, EPI_ISL_547395, EPI_ISL_547396, EPI_ISL_547397, EPI_ISL_547399, EPI_ISL_547402, EPI_ISL_547403, EPI_ISL_547404, EPI_ISL_547405, EPI_ISL_547408, EPI_ISL_547409, EPI_ISL_547410, EPI_ISL_547411, EPI_ISL_547412, EPI_ISL_547413, EPI_ISL_547414, EPI_ISL_547416, EPI_ISL_547418, EPI_ISL_547419, EPI_ISL_547420, EPI_ISL_547421, EPI_ISL_547423, EPI_ISL_547424, EPI_ISL_547426, EPI_ISL_547427, EPI_ISL_547431, EPI_ISL_547432 | | | |
| see above | Houston Methodist Hospital | Houston Methodist Hospital | S. Wesley Long, Randall J. Olsen, Paul A. Christensen, David W. Bernard, James J. Davis, Maulik Shukla, Marcus Nguyen, Matthew Ojeda Saavedra, Concepcion C. Cantu, Prasanti Yerramilli, Layne Pruitt, Sishir Subedi, Hung-Che Kuo, Heather Hendrickson, Ghazaleh Eskandari, Hoang A. T. Nguyen, J. Hunter Long, Muthiah Kumaraswami, Jule Goike, Daniel Boutz, Jimmy Gollihar, Jason S. McLellan, Chia-Wei Chou, Kamyab Javanmardi, Ilya J. Finkelstein, and James M. Musser |
| EPI_ISL_547433, EPI_ISL_547434, EPI_ISL_547435, EPI_ISL_547436, EPI_ISL_547437, EPI_ISL_547438, EPI_ISL_547439, EPI_ISL_547440, EPI_ISL_547441, EPI_ISL_547442, EPI_ISL_547443, EPI_ISL_547444 | | | |
| see above | Microbiology, Department of Pathology, St. Bernard's Hospital, Gibraltar Health Authority | Respiratory Virus Unit, Microbiology Services Colindale, Public Health England | PHE Covid Sequencing Team, Dr Nicholas Cortes (Gibraltar), Charlotte Gillborn-Jones (Gibraltar) |
| EPI_ISL_547445, EPI_ISL_547446, EPI_ISL_547447, EPI_ISL_547448, EPI_ISL_547449, EPI_ISL_547450, EPI_ISL_547451, EPI_ISL_547452, EPI_ISL_547453, EPI_ISL_547454, EPI_ISL_547455, EPI_ISL_547456, EPI_ISL_547457, EPI_ISL_547458, EPI_ISL_547459, EPI_ISL_547460, EPI_ISL_547461, EPI_ISL_547462, EPI_ISL_547463, EPI_ISL_547464, EPI_ISL_547465, EPI_ISL_547466, EPI_ISL_547467, EPI_ISL_547468, EPI_ISL_547469, EPI_ISL_547470, EPI_ISL_547471, EPI_ISL_547472, EPI_ISL_547473, EPI_ISL_547474, EPI_ISL_547475, EPI_ISL_547476, EPI_ISL_547477, EPI_ISL_547478, EPI_ISL_547479, EPI_ISL_547480, EPI_ISL_547481, EPI_ISL_547482, EPI_ISL_547483, EPI_ISL_547484, EPI_ISL_547485, EPI_ISL_547486, EPI_ISL_547487, EPI_ISL_547488, EPI_ISL_547489, EPI_ISL_547490, EPI_ISL_547491, EPI_ISL_547492, EPI_ISL_547493, EPI_ISL_547494, EPI_ISL_547495, EPI_ISL_547496, EPI_ISL_547497, EPI_ISL_547498, EPI_ISL_547499, EPI_ISL_547500, EPI_ISL_547501, EPI_ISL_547502, EPI_ISL_547503, EPI_ISL_547504, EPI_ISL_547505, EPI_ISL_547506, EPI_ISL_547507, EPI_ISL_547508, EPI_ISL_547509, EPI_ISL_547510, EPI_ISL_547511, EPI_ISL_547512, EPI_ISL_547513, EPI_ISL_547514, EPI_ISL_547515, EPI_ISL_547516, EPI_ISL_547517, EPI_ISL_547518, EPI_ISL_547519, EPI_ISL_547520, EPI_ISL_547521, EPI_ISL_547522, EPI_ISL_547523, EPI_ISL_547524, EPI_ISL_547525, EPI_ISL_547526, EPI_ISL_547527, EPI_ISL_547528, EPI_ISL_547529, EPI_ISL_547530, EPI_ISL_547531, EPI_ISL_547532, EPI_ISL_547533, EPI_ISL_547534, EPI_ISL_547535, EPI_ISL_547536, EPI_ISL_547537, EPI_ISL_547538, EPI_ISL_547539, EPI_ISL_547540, EPI_ISL_547541, EPI_ISL_547542, EPI_ISL_547543, EPI_ISL_547544, EPI_ISL_547545, EPI_ISL_547546, EPI_ISL_547547, EPI_ISL_547548, EPI_ISL_547549, EPI_ISL_547550, EPI_ISL_547551, EPI_ISL_547552, EPI_ISL_547553, EPI_ISL_547554, EPI_ISL_547555, EPI_ISL_547556, EPI_ISL_547557, EPI_ISL_547558, EPI_ISL_547559, EPI_ISL_547560, EPI_ISL_547561, EPI_ISL_547562, EPI_ISL_547563, EPI_ISL_547564, EPI_ISL_547565, EPI_ISL_547566, EPI_ISL_547567, EPI_ISL_547568 | | | |
| see above | Dutch COVID-19 response team | National Institute for Public Health and the Environment (RIVM) | Adam Meijer, Harry Vennema, Jeroen Cremer, Sharon van den Brink, Bas van der Veer, AnneMarie van den Brandt, Florian Zwagemaker, Dennis Schmitz, Chantal Reusken, on behalf of the national COVID-19 response team |
| EPI_ISL_547570 | PS Municipal Dr Augusto Gomes de Mattos | Instituto Adolfo Lutz, Interdiciplinary Procedures Center, Strategic Laboratory | Claudio Tavares Sacchi, Claudia Regina Gonçalves, Erica Valessa Ramos Gomes, Karoline Rodrigues Campos |
| EPI_ISL_547571 | Hospital Municipal Antônio Giglio | Instituto Adolfo Lutz, Interdiciplinary Procedures Center, Strategic Laboratory | Claudio Tavares Sacchi, Claudia Regina Gonçalves, Erica Valessa Ramos Gomes, Karoline Rodrigues Campos |
| EPI_ISL_547573 | Vigilância em Saúde de Cajamar | Instituto Adolfo Lutz, Interdiciplinary Procedures Center, Strategic Laboratory | Claudio Tavares Sacchi, Claudia Regina Gonçalves, Erica Valessa Ramos Gomes, Karoline Rodrigues Campos |
| EPI_ISL_547574 | Hospital Universitario da USP | Instituto Adolfo Lutz, Interdiciplinary Procedures Center, Strategic Laboratory | Claudio Tavares Sacchi, Claudia Regina Gonçalves, Erica Valessa Ramos Gomes, Karoline Rodrigues Campos |
| EPI_ISL_547575 | SVO Jundiaí | Instituto Adolfo Lutz, Interdiciplinary Procedures Center, Strategic Laboratory | Claudio Tavares Sacchi, Claudia Regina Gonçalves, Erica Valessa Ramos Gomes, Karoline Rodrigues Campos |
| EPI_ISL_547576 | Secretaria Municipal de Saúde | Instituto Adolfo Lutz, Interdiciplinary Procedures Center, Strategic Laboratory | Claudio Tavares Sacchi, Claudia Regina Gonçalves, Erica Valessa Ramos Gomes, Karoline Rodrigues Campos |
| EPI_ISL_547577 | Hospital e Maternidade Nossa Senhora das Graças | Instituto Adolfo Lutz, Interdiciplinary Procedures Center, Strategic Laboratory | Claudio Tavares Sacchi, Claudia Regina Gonçalves, Erica Valessa Ramos Gomes, Karoline Rodrigues Campos |
| EPI_ISL_547578 | Hospital Doutor Domingos Leonardo Cerávolo | Instituto Adolfo Lutz, Interdiciplinary Procedures Center, Strategic Laboratory | Claudio Tavares Sacchi, Claudia Regina Gonçalves, Erica Valessa Ramos Gomes, Karoline Rodrigues Campos |
| EPI_ISL_547579 | Santa Casa de Misericórdia de Araçatuba | Instituto Adolfo Lutz, Interdiciplinary Procedures Center, Strategic Laboratory | Claudio Tavares Sacchi, Claudia Regina Gonçalves, Erica Valessa Ramos Gomes, Karoline Rodrigues Campos |
| EPI_ISL_547580 | Santa Casa da Misericórdia de Presidente Prudente | Instituto Adolfo Lutz, Interdiciplinary Procedures Center, Strategic Laboratory | Claudio Tavares Sacchi, Claudia Regina Gonçalves, Erica Valessa Ramos Gomes, Karoline Rodrigues Campos |
| EPI_ISL_547584 | CMS, Roorkee | CSIR-Institute of Microbial Technology | Kanika Bansal, Sanjeet Kumar, Anu Singh, Debarghya Ghose, Rajesh Kumar Mishra, Dipak Dutta, Sanjeev Khosla, Prabhu B. Patil |
| EPI_ISL_547585, EPI_ISL_547586, EPI_ISL_547588, EPI_ISL_547589, EPI_ISL_547590, EPI_ISL_547591, EPI_ISL_547592, EPI_ISL_547593 | Civil Hospital, Panchkula | CSIR-Institute of Microbial Technology | Kanika Bansal, Sanjeet Kumar, Anu Singh, Debarghya Ghose, Rajesh Kumar Mishra, Dipak Dutta, Sanjeev Khosla, Prabhu B. Patil |
| EPI_ISL_547594 | Sydney South West Pathology Service (SSWPS) - Liverpool Hospital - NSW Health Pathology | NSW Health Pathology - Institute of Clinical Pathology and Medical Research; Westmead Hospital; University of Sydney | CIDM-PH et al. |
| EPI_ISL_547595, EPI_ISL_547596 | St Vincent's Pathology (SydPath) | NSW Health Pathology - Institute of Clinical Pathology and Medical Research; Westmead Hospital; University of Sydney | CIDM-PH et al. |
| EPI_ISL_547597 | Australian Clinical Labs | NSW Health Pathology - Institute of Clinical Pathology and Medical Research; Westmead Hospital; University of Sydney | CIDM-PH et al. |
| EPI_ISL_547598 | Sydney South West Pathology Service (SSWPS) - Concord Repatriation General Hospital - NSW Health Pathology | NSW Health Pathology - Institute of Clinical Pathology and Medical Research; Westmead Hospital; University of Sydney | CIDM-PH et al. |
| EPI_ISL_547599 | Sydney South West Pathology Service (SSWPS) - Royal Prince Alfred Hospital - NSW Health Pathology | NSW Health Pathology - Institute of Clinical Pathology and Medical Research; Westmead Hospital; University of Sydney | CIDM-PH et al. |
| EPI_ISL_547600 | Histopath | NSW Health Pathology - Institute of Clinical Pathology and Medical Research; Westmead Hospital; University of Sydney | CIDM-PH et al. |
| EPI_ISL_547601 | South Eastern Area Laboratory Services (SEALS) | NSW Health Pathology - Institute of Clinical Pathology and Medical Research; Westmead Hospital; University of Sydney | CIDM-PH et al. |
| EPI_ISL_547602, EPI_ISL_547603, EPI_ISL_547604, EPI_ISL_547605, EPI_ISL_547606, EPI_ISL_547607 | Gundersen Molecular Diagnostics Laboratory | Kabara Cancer Research Institute | Craig S. Richmond, Paraic A. Kenny |
| EPI_ISL_547608, EPI_ISL_547609, EPI_ISL_547610 | Gundersen Clinical Microbiology Laboratory | Kabara Cancer Research Institute | Craig S. Richmond, Paraic A. Kenny |
| EPI_ISL_547611, EPI_ISL_547612, EPI_ISL_547613, EPI_ISL_547614, EPI_ISL_547615, EPI_ISL_547616, EPI_ISL_547617, EPI_ISL_547618, EPI_ISL_547619, EPI_ISL_547620, EPI_ISL_547621, EPI_ISL_547622, EPI_ISL_547623, EPI_ISL_547624, EPI_ISL_547626, EPI_ISL_547627, EPI_ISL_547628, EPI_ISL_547629, EPI_ISL_547630, EPI_ISL_547631, EPI_ISL_547632, EPI_ISL_547633, EPI_ISL_547634, EPI_ISL_547635, EPI_ISL_547636, EPI_ISL_547637, EPI_ISL_547638, EPI_ISL_547639, EPI_ISL_547640, EPI_ISL_547641, EPI_ISL_547642, EPI_ISL_547643, EPI_ISL_547644, EPI_ISL_547645, EPI_ISL_547646, EPI_ISL_547647, EPI_ISL_547648, EPI_ISL_547649, EPI_ISL_547650, EPI_ISL_547651, EPI_ISL_547652, EPI_ISL_547653, EPI_ISL_547654, EPI_ISL_547655, EPI_ISL_547656, EPI_ISL_547657, EPI_ISL_547658, EPI_ISL_547659, EPI_ISL_547660, EPI_ISL_547661, EPI_ISL_547662, EPI_ISL_547664, EPI_ISL_547665 | | | |
| see above | Gundersen Molecular Diagnostics Laboratory | Kabara Cancer Research Institute | Craig S. Richmond, Paraic A. Kenny |
| EPI_ISL_547666, EPI_ISL_547667, EPI_ISL_547668, EPI_ISL_547669, EPI_ISL_547670, EPI_ISL_547671, EPI_ISL_547672, EPI_ISL_547673, EPI_ISL_547674, EPI_ISL_547675, EPI_ISL_547676 | | | |
| see above | Gundersen Clinical Microbiology Laboratory | Kabara Cancer Research Institute | Craig S. Richmond, Paraic A. Kenny |
| EPI_ISL_547678, EPI_ISL_547680, EPI_ISL_547681, EPI_ISL_547682 | Gundersen Molecular Diagnostics Laboratory | Kabara Cancer Research Institute | Craig S. Richmond, Paraic A. Kenny |
| EPI_ISL_547683 | Gundersen Clinical Microbiology Laboratory | Kabara Cancer Research Institute | Craig S. Richmond, Paraic A. Kenny |
| EPI_ISL_547685, EPI_ISL_547687, EPI_ISL_547688, EPI_ISL_547689, EPI_ISL_547690, EPI_ISL_547691, EPI_ISL_547692, EPI_ISL_547693, EPI_ISL_547694, EPI_ISL_547695, EPI_ISL_547696, EPI_ISL_547697, EPI_ISL_547698, EPI_ISL_547701, EPI_ISL_547702, EPI_ISL_547703, EPI_ISL_547704 | | | |
| see above | Gundersen Molecular Diagnostics Laboratory | Kabara Cancer Research Institute | Craig S. Richmond, Paraic A. Kenny |
| EPI_ISL_547705 | Gundersen Clinical Microbiology Laboratory | Kabara Cancer Research Institute | Craig S. Richmond, Paraic A. Kenny |
| EPI_ISL_547706, EPI_ISL_547708, EPI_ISL_547709, EPI_ISL_547710, EPI_ISL_547711, EPI_ISL_547712, EPI_ISL_547713, EPI_ISL_547714, EPI_ISL_547715, EPI_ISL_547717, EPI_ISL_547718, EPI_ISL_547719, EPI_ISL_547720, EPI_ISL_547721, EPI_ISL_547722, EPI_ISL_547723, EPI_ISL_547724, EPI_ISL_547725, EPI_ISL_547726, EPI_ISL_547727, EPI_ISL_547728, EPI_ISL_547729, EPI_ISL_547730, EPI_ISL_547731, EPI_ISL_547732, EPI_ISL_547733, EPI_ISL_547734, EPI_ISL_547735, EPI_ISL_547736, EPI_ISL_547740, EPI_ISL_547741, EPI_ISL_547742, EPI_ISL_547743, EPI_ISL_547744, EPI_ISL_547746, EPI_ISL_547747, EPI_ISL_547748, EPI_ISL_547749, EPI_ISL_547750, EPI_ISL_547751, EPI_ISL_547752, EPI_ISL_547753, EPI_ISL_547754, EPI_ISL_547755, EPI_ISL_547756, EPI_ISL_547757, EPI_ISL_547758, EPI_ISL_547759, EPI_ISL_547760, EPI_ISL_547761, EPI_ISL_547762, EPI_ISL_547763, EPI_ISL_547764, EPI_ISL_547765, EPI_ISL_547766, EPI_ISL_547767, EPI_ISL_547768, EPI_ISL_547769, EPI_ISL_547770, EPI_ISL_547771, EPI_ISL_547772, EPI_ISL_547773, EPI_ISL_547774, EPI_ISL_547775, EPI_ISL_547776, EPI_ISL_547777, EPI_ISL_547778, EPI_ISL_547779, EPI_ISL_547780, EPI_ISL_547781, EPI_ISL_547782, EPI_ISL_547783, EPI_ISL_547784, EPI_ISL_547785, EPI_ISL_547786, EPI_ISL_547787, EPI_ISL_547788, EPI_ISL_547789, EPI_ISL_547790, EPI_ISL_547791, EPI_ISL_547792, EPI_ISL_547793, EPI_ISL_547794, EPI_ISL_547795, EPI_ISL_547796, EPI_ISL_547797, EPI_ISL_547798, EPI_ISL_547802, EPI_ISL_547803, EPI_ISL_547804, EPI_ISL_547805, EPI_ISL_547806, EPI_ISL_547807, EPI_ISL_547808, EPI_ISL_547809, EPI_ISL_547810, | | | |

| | | | |
|---|---|---|---|
| EPI_ISL_547811, EPI_ISL_547812, EPI_ISL_547813, EPI_ISL_547814, EPI_ISL_547815, EPI_ISL_547816, EPI_ISL_547817, EPI_ISL_547818, EPI_ISL_547819, EPI_ISL_547820, EPI_ISL_547821, EPI_ISL_547822, EPI_ISL_547823, EPI_ISL_547824, EPI_ISL_547825, EPI_ISL_547826, EPI_ISL_547827, EPI_ISL_547828, EPI_ISL_547829, EPI_ISL_547831, EPI_ISL_547833, EPI_ISL_547834, EPI_ISL_547835, EPI_ISL_547836, EPI_ISL_547837, EPI_ISL_547838, EPI_ISL_547839, EPI_ISL_547840, EPI_ISL_547841, EPI_ISL_547842, EPI_ISL_547843, EPI_ISL_547844, EPI_ISL_547845, EPI_ISL_547846, EPI_ISL_547847, EPI_ISL_547848, EPI_ISL_547849, EPI_ISL_547850, EPI_ISL_547851, EPI_ISL_547853, EPI_ISL_547855, EPI_ISL_547856, EPI_ISL_547857, EPI_ISL_547858, EPI_ISL_547859, EPI_ISL_547860, EPI_ISL_547861, EPI_ISL_547862, EPI_ISL_547863, EPI_ISL_547864, EPI_ISL_547865, EPI_ISL_547866, EPI_ISL_547867, EPI_ISL_547868, EPI_ISL_547869 | | | |
| see above | Gundersen Molecular Diagnostics Laboratory | Kabara Cancer Research Institute | Craig S. Richmond, Paraic A. Kenny |
| EPI_ISL_547966, EPI_ISL_547967, EPI_ISL_547968 | The National Institute of Public Health | State Veterinary Institute Prague | Nagy,A;Jirincova,H;Novakova,L;Trnka,D;Vecerova,J |
| EPI_ISL_547969 | LabPLUS | Institute of Environmental Science and Research (ESR) | Xiaoyun Ren, Matt Storey, Nikki Freed, Muhammad Faisal, Jing Wang, Hermes Perez, Anja Werno, Antje van der Linden, Arlo Upton, Chris Mansell, David Hammer, Dragana Drinkovic, Gary McAuliffe, Hana Sofia Andersson, James Ussher, Jill Sherwood, Josh Freeman, Julia Howard, Juliet Elvy, Mary DeAlmeida, Matt Blakiston, Matthew Rogers, Max Bloomfield, Michael Addidle, Michelle Balm, Sally Roberts, Sarah Jefferies, Sharmini Muttaiyah, Susan Morpeth, Susan Taylor, Timothy Blackmore, Vani Sathyendran, Veronica Playle, Virginia Hope, Erasmus Smit, Lauren Jelly, Olin Silander, Joep de Ligt |
| EPI_ISL_547970, EPI_ISL_547971, EPI_ISL_547972, EPI_ISL_547973, EPI_ISL_547974, EPI_ISL_547975 | LabTests | Institute of Environmental Science and Research (ESR) | Xiaoyun Ren, Matt Storey, Nikki Freed, Muhammad Faisal, Jing Wang, Hermes Perez, Anja Werno, Antje van der Linden, Arlo Upton, Chris Mansell, David Hammer, Dragana Drinkovic, Gary McAuliffe, Hana Sofia Andersson, James Ussher, Jill Sherwood, Josh Freeman, Julia Howard, Juliet Elvy, Mary DeAlmeida, Matt Blakiston, Matthew Rogers, Max Bloomfield, Michael Addidle, Michelle Balm, Sally Roberts, Sarah Jefferies, Sharmini Muttaiyah, Susan Morpeth, Susan Taylor, Timothy Blackmore, Vani Sathyendran, Veronica Playle, Virginia Hope, Erasmus Smit, Lauren Jelly, Olin Silander, Joep de Ligt |
| EPI_ISL_547976 | LabPLUS | Institute of Environmental Science and Research (ESR) | Xiaoyun Ren, Matt Storey, Nikki Freed, Muhammad Faisal, Jing Wang, Hermes Perez, Anja Werno, Antje van der Linden, Arlo Upton, Chris Mansell, David Hammer, Dragana Drinkovic, Gary McAuliffe, Hana Sofia Andersson, James Ussher, Jill Sherwood, Josh Freeman, Julia Howard, Juliet Elvy, Mary DeAlmeida, Matt Blakiston, Matthew Rogers, Max Bloomfield, Michael Addidle, Michelle Balm, Sally Roberts, Sarah Jefferies, Sharmini Muttaiyah, Susan Morpeth, Susan Taylor, Timothy Blackmore, Vani Sathyendran, Veronica Playle, Virginia Hope, Erasmus Smit, Lauren Jelly, Olin Silander, Joep de Ligt |
| EPI_ISL_547977 | LabTests | Institute of Environmental Science and Research (ESR) | Xiaoyun Ren, Matt Storey, Nikki Freed, Muhammad Faisal, Jing Wang, Hermes Perez, Anja Werno, Antje van der Linden, Arlo Upton, Chris Mansell, David Hammer, Dragana Drinkovic, Gary McAuliffe, Hana Sofia Andersson, James Ussher, Jill Sherwood, Josh Freeman, Julia Howard, Juliet Elvy, Mary DeAlmeida, Matt Blakiston, Matthew Rogers, Max Bloomfield, Michael Addidle, Michelle Balm, Sally Roberts, Sarah Jefferies, Sharmini Muttaiyah, Susan Morpeth, Susan Taylor, Timothy Blackmore, Vani Sathyendran, Veronica Playle, Virginia Hope, Erasmus Smit, Lauren Jelly, Olin Silander, Joep de Ligt |
| EPI_ISL_547978, EPI_ISL_547979 | LabPLUS | Institute of Environmental Science and Research (ESR) | Xiaoyun Ren, Matt Storey, Nikki Freed, Muhammad Faisal, Jing Wang, Hermes Perez, Anja Werno, Antje van der Linden, Arlo Upton, Chris Mansell, David Hammer, Dragana Drinkovic, Gary McAuliffe, Hana Sofia Andersson, James Ussher, Jill Sherwood, Josh Freeman, Julia Howard, Juliet Elvy, Mary DeAlmeida, Matt Blakiston, Matthew Rogers, Max Bloomfield, Michael Addidle, Michelle Balm, Sally Roberts, Sarah Jefferies, Sharmini Muttaiyah, Susan Morpeth, Susan Taylor, Timothy Blackmore, Vani Sathyendran, Veronica Playle, Virginia Hope, Erasmus Smit, Lauren Jelly, Olin Silander, Joep de Ligt |
| EPI_ISL_547981, EPI_ISL_547982, EPI_ISL_547983, EPI_ISL_547984, EPI_ISL_547985 | Canterbury Health Laboratories | Institute of Environmental Science and Research (ESR) | Xiaoyun Ren, Matt Storey, Nikki Freed, Muhammad Faisal, Jing Wang, Hermes Perez, Anja Werno, Antje van der Linden, Arlo Upton, Chris Mansell, David Hammer, Dragana Drinkovic, Gary McAuliffe, Hana Sofia Andersson, James Ussher, Jill Sherwood, Josh Freeman, Julia Howard, Juliet Elvy, Mary DeAlmeida, Matt Blakiston, Matthew Rogers, Max Bloomfield, Michael Addidle, Michelle Balm, Sally Roberts, Sarah Jefferies, Sharmini Muttaiyah, Susan Morpeth, Susan Taylor, Timothy Blackmore, Vani Sathyendran, Veronica Playle, Virginia Hope, Erasmus Smit, Lauren Jelly, Olin Silander, Joep de Ligt |
| EPI_ISL_547993, EPI_ISL_547994, EPI_ISL_547995 | LabTests | Institute of Environmental Science and Research (ESR) | Xiaoyun Ren, Matt Storey, Nikki Freed, Muhammad Faisal, Jing Wang, Hermes Perez, Anja Werno, Antje van der Linden, Arlo Upton, Chris Mansell, David Hammer, Dragana Drinkovic, Gary McAuliffe, Hana Sofia Andersson, James Ussher, Jill Sherwood, Josh Freeman, Julia Howard, Juliet Elvy, Mary DeAlmeida, Matt Blakiston, Matthew Rogers, Max Bloomfield, Michael Addidle, Michelle Balm, Sally Roberts, Sarah Jefferies, Sharmini Muttaiyah, Susan Morpeth, Susan Taylor, Timothy Blackmore, Vani Sathyendran, Veronica Playle, Virginia Hope, Erasmus Smit, Lauren Jelly, Olin Silander, Joep de Ligt |
| EPI_ISL_547996, EPI_ISL_547997, EPI_ISL_547998, EPI_ISL_547999, EPI_ISL_548000 | LabPLUS | Institute of Environmental Science and Research (ESR) | Xiaoyun Ren, Matt Storey, Nikki Freed, Muhammad Faisal, Jing Wang, Hermes Perez, Anja Werno, Antje van der Linden, Arlo Upton, Chris Mansell, David Hammer, Dragana Drinkovic, Gary McAuliffe, Hana Sofia Andersson, James Ussher, Jill Sherwood, Josh Freeman, Julia Howard, Juliet Elvy, Mary DeAlmeida, Matt Blakiston, Matthew Rogers, Max Bloomfield, Michael Addidle, Michelle Balm, Sally Roberts, Sarah Jefferies, Sharmini Muttaiyah, Susan Morpeth, Susan Taylor, Timothy Blackmore, Vani Sathyendran, Veronica Playle, Virginia Hope, Erasmus Smit, Lauren Jelly, Olin Silander, Joep de Ligt |
| EPI_ISL_548002, EPI_ISL_548004, EPI_ISL_548005, EPI_ISL_548006, EPI_ISL_548008, EPI_ISL_548009, EPI_ISL_548010, EPI_ISL_548011 | Middlemore Hospital | Institute of Environmental Science and Research (ESR) | Xiaoyun Ren, Matt Storey, Nikki Freed, Muhammad Faisal, Jing Wang, Hermes Perez, Anja Werno, Antje van der Linden, Arlo Upton, Chris Mansell, David Hammer, Dragana Drinkovic, Gary McAuliffe, Hana Sofia Andersson, James Ussher, Jill Sherwood, Josh Freeman, Julia Howard, Juliet Elvy, Mary DeAlmeida, Matt Blakiston, Matthew Rogers, Max Bloomfield, Michael Addidle, Michelle Balm, Sally Roberts, Sarah Jefferies, Sharmini Muttaiyah, Susan Morpeth, Susan Taylor, Timothy Blackmore, Vani Sathyendran, Veronica Playle, Virginia Hope, Erasmus Smit, Lauren Jelly, Olin Silander, Joep de Ligt |
| EPI_ISL_548013, EPI_ISL_548014, EPI_ISL_548015, EPI_ISL_548016 | LabTests | Institute of Environmental Science and Research (ESR) | Xiaoyun Ren, Matt Storey, Nikki Freed, Muhammad Faisal, Jing Wang, Hermes Perez, Anja Werno, Antje van der Linden, Arlo Upton, Chris Mansell, David Hammer, Dragana Drinkovic, Gary McAuliffe, Hana Sofia Andersson, James Ussher, Jill Sherwood, Josh Freeman, Julia Howard, Juliet Elvy, Mary DeAlmeida, Matt Blakiston, Matthew Rogers, Max Bloomfield, Michael Addidle, Michelle Balm, Sally Roberts, Sarah Jefferies, Sharmini Muttaiyah, Susan Morpeth, Susan Taylor, Timothy Blackmore, Vani Sathyendran, Veronica Playle, Virginia Hope, Erasmus Smit, Lauren Jelly, Olin Silander, Joep de Ligt |
| EPI_ISL_548017, EPI_ISL_548021, EPI_ISL_548022, EPI_ISL_548023, EPI_ISL_548024, EPI_ISL_548025 | LabPLUS | Institute of Environmental Science and Research (ESR) | Xiaoyun Ren, Matt Storey, Nikki Freed, Muhammad Faisal, Jing Wang, Hermes Perez, Anja Werno, Antje van der Linden, Arlo Upton, Chris Mansell, David Hammer, Dragana Drinkovic, Gary McAuliffe, Hana Sofia Andersson, James Ussher, Jill Sherwood, Josh Freeman, Julia Howard, Juliet Elvy, Mary DeAlmeida, Matt Blakiston, Matthew Rogers, Max Bloomfield, Michael Addidle, Michelle Balm, Sally Roberts, Sarah Jefferies, Sharmini Muttaiyah, Susan Morpeth, Susan Taylor, Timothy Blackmore, Vani Sathyendran, Veronica Playle, Virginia Hope, Erasmus Smit, Lauren Jelly, Olin Silander, Joep de Ligt |
| EPI_ISL_548030 | North Shore Hospital | Institute of Environmental Science and Research (ESR) | Xiaoyun Ren, Matt Storey, Nikki Freed, Muhammad Faisal, Jing Wang, Hermes Perez, Anja Werno, Antje van der Linden, Arlo Upton, Chris Mansell, David Hammer, Dragana Drinkovic, Gary McAuliffe, Hana Sofia Andersson, James Ussher, Jill Sherwood, Josh Freeman, Julia Howard, Juliet Elvy, Mary DeAlmeida, Matt Blakiston, Matthew Rogers, Max Bloomfield, Michael Addidle, Michelle Balm, Sally Roberts, Sarah Jefferies, Sharmini Muttaiyah, Susan Morpeth, Susan Taylor, Timothy Blackmore, Vani Sathyendran, Veronica Playle, Virginia Hope, Erasmus Smit, Lauren Jelly, Olin Silander, Joep de Ligt |
| EPI_ISL_548040, EPI_ISL_548041 | LabPLUS | Institute of Environmental Science and Research (ESR) | Xiaoyun Ren, Matt Storey, Nikki Freed, Muhammad Faisal, Jing Wang, Hermes Perez, Anja Werno, Antje van der Linden, Arlo Upton, Chris Mansell, David Hammer, Dragana Drinkovic, Gary McAuliffe, Hana Sofia Andersson, James Ussher, Jill Sherwood, Josh Freeman, Julia Howard, Juliet Elvy, Mary DeAlmeida, Matt Blakiston, Matthew Rogers, Max Bloomfield, Michael Addidle, Michelle Balm, Sally Roberts, Sarah Jefferies, Sharmini Muttaiyah, Susan Morpeth, Susan Taylor, Timothy Blackmore, Vani Sathyendran, Veronica Playle, Virginia Hope, Erasmus Smit, Lauren Jelly, Olin Silander, Joep de Ligt |
| EPI_ISL_548045 | North Shore Hospital | Institute of Environmental Science and Research (ESR) | Xiaoyun Ren, Matt Storey, Nikki Freed, Muhammad Faisal, Jing Wang, Hermes Perez, Anja Werno, Antje van der Linden, Arlo Upton, Chris Mansell, David Hammer, Dragana Drinkovic, Gary McAuliffe, Hana Sofia Andersson, James Ussher, Jill Sherwood, Josh Freeman, Julia Howard, Juliet Elvy, Mary DeAlmeida, Matt Blakiston, Matthew Rogers, Max Bloomfield, Michael Addidle, Michelle Balm, Sally Roberts, Sarah Jefferies, Sharmini Muttaiyah, Susan Morpeth, Susan Taylor, Timothy Blackmore, Vani Sathyendran, Veronica Playle, Virginia Hope, Erasmus Smit, Lauren Jelly, Olin Silander, Joep de Ligt |
| EPI_ISL_548046 | LabTests | Institute of Environmental Science and Research (ESR) | Xiaoyun Ren, Matt Storey, Nikki Freed, Muhammad Faisal, Jing Wang, Hermes Perez, Anja Werno, Antje van der Linden, Arlo Upton, Chris Mansell, David Hammer, Dragana Drinkovic, Gary McAuliffe, Hana Sofia Andersson, James Ussher, Jill Sherwood, Josh Freeman, Julia Howard, Juliet Elvy, Mary DeAlmeida, Matt Blakiston, Matthew Rogers, Max Bloomfield, Michael Addidle, Michelle Balm, Sally Roberts, Sarah Jefferies, Sharmini Muttaiyah, Susan Morpeth, Susan Taylor, Timothy Blackmore, Vani Sathyendran, Veronica Playle, Virginia Hope, Erasmus Smit, Lauren Jelly, Olin Silander, Joep de Ligt |
| EPI_ISL_548047, EPI_ISL_548048 | North Shore Hospital | Institute of Environmental Science and Research (ESR) | Xiaoyun Ren, Matt Storey, Nikki Freed, Muhammad Faisal, Jing Wang, Hermes Perez, Anja Werno, Antje van der Linden, Arlo Upton, Chris Mansell, David Hammer, Dragana Drinkovic, Gary McAuliffe, Hana Sofia Andersson, James Ussher, Jill Sherwood, Josh Freeman, Julia Howard, Juliet Elvy, Mary DeAlmeida, Matt Blakiston, Matthew Rogers, Max Bloomfield, Michael Addidle, Michelle Balm, Sally Roberts, Sarah Jefferies, Sharmini Muttaiyah, Susan Morpeth, Susan Taylor, Timothy Blackmore, Vani Sathyendran, Veronica Playle, Virginia Hope, Erasmus Smit, Lauren Jelly, Olin Silander, Joep de Ligt |
| EPI_ISL_548049, EPI_ISL_548050 | LabPLUS | Institute of Environmental Science and Research (ESR) | Xiaoyun Ren, Matt Storey, Nikki Freed, Muhammad Faisal, Jing Wang, Hermes Perez, Anja Werno, Antje van der Linden, Arlo Upton, Chris Mansell, David Hammer, Dragana Drinkovic, Gary McAuliffe, Hana Sofia Andersson, James Ussher, Jill Sherwood, Josh Freeman, Julia Howard, Juliet Elvy, Mary DeAlmeida, Matt Blakiston, Matthew Rogers, Max Bloomfield, Michael Addidle, Michelle Balm, Sally Roberts, Sarah Jefferies, Sharmini Muttaiyah, Susan Morpeth, Susan Taylor, Timothy Blackmore, Vani Sathyendran, Veronica Playle, Virginia Hope, Erasmus Smit, Lauren Jelly, Olin Silander, Joep de Ligt |
| EPI_ISL_548057 | North Shore Hospital | Institute of Environmental Science and Research (ESR) | Xiaoyun Ren, Matt Storey, Nikki Freed, Muhammad Faisal, Jing Wang, Hermes Perez, Anja Werno, Antje van der Linden, Arlo Upton, Chris Mansell, David Hammer, Dragana Drinkovic, Gary McAuliffe, Hana Sofia Andersson, James Ussher, Jill Sherwood, Josh Freeman, Julia Howard, Juliet Elvy, Mary DeAlmeida, Matt Blakiston, Matthew Rogers, Max Bloomfield, Michael Addidle, Michelle Balm, Sally Roberts, Sarah Jefferies, Sharmini Muttaiyah, Susan Morpeth, Susan Taylor, Timothy Blackmore, Vani Sathyendran, Veronica Playle, Virginia Hope, Erasmus Smit, Lauren Jelly, Olin Silander, Joep de Ligt |
| EPI_ISL_548064 | Waikato Hospital | Institute of Environmental Science and Research (ESR) | Xiaoyun Ren, Matt Storey, Nikki Freed, Muhammad Faisal, Jing Wang, Hermes Perez, Anja Werno, Antje van der Linden, Arlo Upton, Chris Mansell, David Hammer, Dragana Drinkovic, Gary McAuliffe, Hana Sofia Andersson, James Ussher, Jill Sherwood, Josh Freeman, Julia Howard, Juliet Elvy, Mary DeAlmeida, Matt Blakiston, Matthew Rogers, Max Bloomfield, Michael Addidle, Michelle Balm, Sally Roberts, Sarah Jefferies, Sharmini Muttaiyah, Susan Morpeth, Susan Taylor, Timothy Blackmore, Vani Sathyendran, Veronica Playle, Virginia Hope, Erasmus Smit, Lauren Jelly, Olin Silander, Joep de Ligt |
| EPI_ISL_548067 | North Shore Hospital | Institute of Environmental Science and Research (ESR) | Xiaoyun Ren, Matt Storey, Nikki Freed, Muhammad Faisal, Jing Wang, Hermes Perez, Anja Werno, Antje van der Linden, Arlo Upton, Chris Mansell, David Hammer, Dragana Drinkovic, Gary McAuliffe, Hana Sofia Andersson, James Ussher, Jill Sherwood, Josh Freeman, Julia Howard, Juliet Elvy, Mary DeAlmeida, Matt Blakiston, Matthew Rogers, Max Bloomfield, Michael Addidle, Michelle Balm, Sally Roberts, Sarah Jefferies, Sharmini Muttaiyah, Susan Morpeth, Susan Taylor, Timothy Blackmore, Vani Sathyendran, Veronica Playle, Virginia Hope, Erasmus Smit, Lauren Jelly, Olin Silander, Joep de Ligt |
| EPI_ISL_548069 | LabPLUS | Institute of Environmental Science and Research (ESR) | Xiaoyun Ren, Matt Storey, Nikki Freed, Muhammad Faisal, Jing Wang, Hermes Perez, Anja Werno, Antje van der Linden, Arlo Upton, Chris Mansell, David Hammer, Dragana Drinkovic, Gary McAuliffe, Hana Sofia Andersson, James Ussher, Jill Sherwood, Josh Freeman, Julia Howard, Juliet Elvy, Mary DeAlmeida, Matt Blakiston, Matthew Rogers, Max Bloomfield, Michael Addidle, Michelle Balm, Sally Roberts, Sarah Jefferies, Sharmini Muttaiyah, Susan Morpeth, Susan Taylor, Timothy Blackmore, Vani Sathyendran, Veronica Playle, Virginia Hope, Erasmus Smit, Lauren Jelly, Olin Silander, Joep de Ligt |
| EPI_ISL_548075 | North Shore Hospital | Institute of Environmental Science and Research (ESR) | Xiaoyun Ren, Matt Storey, Nikki Freed, Muhammad Faisal, Jing Wang, Hermes Perez, Anja Werno, Antje van der Linden, Arlo Upton, Chris Mansell, David Hammer, Dragana Drinkovic, Gary McAuliffe, Hana Sofia Andersson, James Ussher, Jill Sherwood, Josh Freeman, Julia Howard, Juliet Elvy, Mary DeAlmeida, Matt Blakiston, Matthew Rogers, Max Bloomfield, Michael Addidle, Michelle Balm, Sally Roberts, Sarah Jefferies, Sharmini Muttaiyah, Susan Morpeth, Susan Taylor, Timothy Blackmore, Vani Sathyendran, Veronica Playle, Virginia Hope, Erasmus Smit, Lauren Jelly, Olin Silander, Joep de Ligt |
| EPI_ISL_548077 | LabPLUS | Institute of Environmental Science and Research (ESR) | Xiaoyun Ren, Matt Storey, Nikki Freed, Muhammad Faisal, Jing Wang, Hermes Perez, Anja Werno, Antje van der Linden, Arlo Upton, Chris Mansell, David Hammer, Dragana Drinkovic, Gary McAuliffe, Hana Sofia Andersson, James Ussher, Jill Sherwood, Josh Freeman, Julia Howard, Juliet Elvy, Mary DeAlmeida, Matt Blakiston, Matthew Rogers, Max Bloomfield, Michael Addidle, Michelle Balm, Sally Roberts, Sarah Jefferies, Sharmini Muttaiyah, Susan Morpeth, Susan Taylor, Timothy Blackmore, Vani Sathyendran, Veronica Playle, Virginia Hope, Erasmus Smit, Lauren Jelly, Olin Silander, Joep de Ligt |
| EPI_ISL_548081, EPI_ISL_548082 | Middlemore Hospital | Institute of Environmental Science and Research (ESR) | Xiaoyun Ren, Matt Storey, Nikki Freed, Muhammad Faisal, Jing Wang, Hermes Perez, Anja Werno, Antje van der Linden, Arlo Upton, Chris Mansell, David Hammer, Dragana Drinkovic, Gary McAuliffe, Hana Sofia Andersson, James Ussher, Jill Sherwood, Josh Freeman, Julia Howard, Juliet Elvy, Mary DeAlmeida, Matt Blakiston, Matthew Rogers, Max Bloomfield, Michael Addidle, Michelle Balm, Sally Roberts, Sarah Jefferies, Sharmini Muttaiyah, Susan Morpeth, Susan Taylor, Timothy Blackmore, Vani Sathyendran, Veronica Playle, Virginia Hope, Erasmus Smit, Lauren Jelly, Olin Silander, Joep de Ligt |
| EPI_ISL_548100 | North Shore Hospital | Institute of Environmental Science and Research (ESR) | Xiaoyun Ren, Matt Storey, Nikki Freed, Muhammad Faisal, Jing Wang, Hermes Perez, Anja Werno, Antje van der Linden, Arlo Upton, Chris Mansell, David Hammer, Dragana Drinkovic, Gary McAuliffe, Hana Sofia Andersson, James Ussher, Jill Sherwood, Josh Freeman, Julia Howard, Juliet Elvy, Mary DeAlmeida, Matt Blakiston, Matthew Rogers, Max Bloomfield, Michael Addidle, Michelle Balm, Sally Roberts, Sarah Jefferies, Sharmini Muttaiyah, Susan Morpeth, Susan Taylor, Timothy Blackmore, Vani Sathyendran, Veronica Playle, Virginia Hope, Erasmus Smit, Lauren Jelly, Olin Silander, Joep de Ligt |
| EPI_ISL_548101 | Waikato Hospital | Institute of Environmental Science and Research (ESR) | Xiaoyun Ren, Matt Storey, Nikki Freed, Muhammad Faisal, Jing Wang, Hermes Perez, Anja Werno, Antje van der Linden, Arlo Upton, Chris Mansell, David Hammer, Dragana Drinkovic, Gary McAuliffe, Hana Sofia Andersson, James Ussher, Jill Sherwood, Josh Freeman, Julia Howard, Juliet Elvy, Mary DeAlmeida, Matt Blakiston, Matthew Rogers, Max Bloomfield, Michael Addidle, Michelle Balm, Sally Roberts, Sarah Jefferies, Sharmini Muttaiyah, Susan Morpeth, Susan Taylor, Timothy Blackmore, Vani Sathyendran, Veronica Playle, Virginia Hope, Erasmus Smit, Lauren Jelly, Olin Silander, Joep de Ligt |
| EPI_ISL_548104 | LabTests | Institute of Environmental Science and Research (ESR) | Xiaoyun Ren, Matt Storey, Nikki Freed, Muhammad Faisal, Jing Wang, Hermes Perez, Anja Werno, Antje van der Linden, Arlo Upton, Chris Mansell, David Hammer, Dragana Drinkovic, Gary McAuliffe, Hana Sofia Andersson, James Ussher, Jill Sherwood, Josh Freeman, Julia Howard, Juliet Elvy, Mary DeAlmeida, Matt Blakiston, Matthew Rogers, Max Bloomfield, Michael Addidle, Michelle Balm, Sally Roberts, Sarah Jefferies, Sharmini Muttaiyah, Susan Morpeth, Susan Taylor, Timothy Blackmore, Vani Sathyendran, Veronica Playle, Virginia Hope, Erasmus Smit, Lauren Jelly, Olin Silander, Joep de Ligt |
| EPI_ISL_548106, EPI_ISL_548108, EPI_ISL_548110, EPI_ISL_548111, EPI_ISL_548120 | LabPLUS | Institute of Environmental Science and Research (ESR) | Xiaoyun Ren, Matt Storey, Nikki Freed, Muhammad Faisal, Jing Wang, Hermes Perez, Anja Werno, Antje van der Linden, Arlo Upton, Chris Mansell, David Hammer, Dragana Drinkovic, Gary McAuliffe, Hana Sofia Andersson, James Ussher, Jill Sherwood, Josh Freeman, Julia Howard, Juliet Elvy, Mary DeAlmeida, Matt Blakiston, Matthew Rogers, Max Bloomfield, Michael Addidle, Michelle Balm, Sally Roberts, Sarah Jefferies, Sharmini Muttaiyah, Susan Morpeth, Susan Taylor, Timothy Blackmore, Vani Sathyendran, Veronica Playle, Virginia Hope, Erasmus Smit, Lauren Jelly, Olin Silander, Joep de Ligt |
| EPI_ISL_548123, EPI_ISL_548124, EPI_ISL_548127 | Middlemore Hospital | Institute of Environmental Science and Research (ESR) | Xiaoyun Ren, Matt Storey, Nikki Freed, Muhammad Faisal, Jing Wang, Hermes Perez, Anja Werno, Antje van der Linden, Arlo Upton, Chris Mansell, David Hammer, Dragana Drinkovic, Gary McAuliffe, Hana Sofia Andersson, James Ussher, Jill Sherwood, Josh Freeman, Julia Howard, Juliet Elvy, Mary DeAlmeida, Matt Blakiston, Matthew Rogers, Max Bloomfield, Michael Addidle, Michelle Balm, Sally Roberts, Sarah Jefferies, Sharmini Muttaiyah, Susan Morpeth, Susan Taylor, Timothy Blackmore, Vani Sathyendran, Veronica Playle, Virginia Hope, Erasmus Smit, Lauren Jelly, Olin Silander, Joep de Ligt |
| EPI_ISL_548129 | Canterbury Health Laboratories | Institute of Environmental Science and Research (ESR) | Xiaoyun Ren, Matt Storey, Nikki Freed, Muhammad Faisal, Jing Wang, Hermes Perez, Anja Werno, Antje van der Linden, Arlo Upton, Chris Mansell, David Hammer, Dragana Drinkovic, Gary McAuliffe, Hana Sofia Andersson, James Ussher, Jill Sherwood, Josh Freeman, Julia Howard, Juliet Elvy, Mary DeAlmeida, Matt Blakiston, Matthew Rogers, Max Bloomfield, Michael Addidle, Michelle Balm, Sally Roberts, Sarah Jefferies, Sharmini Muttaiyah, Susan Morpeth, Susan Taylor, Timothy Blackmore, Vani Sathyendran, Veronica Playle, Virginia Hope, Erasmus Smit, Lauren Jelly, Olin Silander, Joep de Ligt |

| | | | |
|---|---|---|---|
| EPI_ISL_548132 | Middlemore Hospital | Institute of Environmental Science and Research (ESR) | Xiaoyun Ren, Matt Storey, Nikki Freed, Muhammad Faisal, Jing Wang, Hermes Perez, Anja Werno, Antje van der Linden, Arlo Upton, Chris Mansell, David Hammer, Dragana Drinkovic, Gary McAuliffe, Hana Sofia Andersson, James Ussher, Jill Sherwood, Josh Freeman, Julia Howard, Juliet Elvy, Mary DeAlmeida, Matt Blakiston, Matthew Rogers, Max Bloomfield, Michael Addidle, Michelle Balm, Sally Roberts, Sarah Jefferies, Sharmini Muttaiyah, Susan Morpeth, Susan Taylor, Timothy Blackmore, Vani Sathyendran, Veronica Playle, Virginia Hope, Erasmus Smit, Lauren Jelly, Olin Silander, Joep de Ligt |
| EPI_ISL_548139, EPI_ISL_548140 | Canterbury Health Laboratories | Institute of Environmental Science and Research (ESR) | Xiaoyun Ren, Matt Storey, Nikki Freed, Muhammad Faisal, Jing Wang, Hermes Perez, Anja Werno, Antje van der Linden, Arlo Upton, Chris Mansell, David Hammer, Dragana Drinkovic, Gary McAuliffe, Hana Sofia Andersson, James Ussher, Jill Sherwood, Josh Freeman, Julia Howard, Juliet Elvy, Mary DeAlmeida, Matt Blakiston, Matthew Rogers, Max Bloomfield, Michael Addidle, Michelle Balm, Sally Roberts, Sarah Jefferies, Sharmini Muttaiyah, Susan Morpeth, Susan Taylor, Timothy Blackmore, Vani Sathyendran, Veronica Playle, Virginia Hope, Erasmus Smit, Lauren Jelly, Olin Silander, Joep de Ligt |
| EPI_ISL_548141, EPI_ISL_548142 | LabPLUS | Institute of Environmental Science and Research (ESR) | Xiaoyun Ren, Matt Storey, Nikki Freed, Muhammad Faisal, Jing Wang, Hermes Perez, Anja Werno, Antje van der Linden, Arlo Upton, Chris Mansell, David Hammer, Dragana Drinkovic, Gary McAuliffe, Hana Sofia Andersson, James Ussher, Jill Sherwood, Josh Freeman, Julia Howard, Juliet Elvy, Mary DeAlmeida, Matt Blakiston, Matthew Rogers, Max Bloomfield, Michael Addidle, Michelle Balm, Sally Roberts, Sarah Jefferies, Sharmini Muttaiyah, Susan Morpeth, Susan Taylor, Timothy Blackmore, Vani Sathyendran, Veronica Playle, Virginia Hope, Erasmus Smit, Lauren Jelly, Olin Silander, Joep de Ligt |
| EPI_ISL_548143 | Middlemore Hospital | Institute of Environmental Science and Research (ESR) | Xiaoyun Ren, Matt Storey, Nikki Freed, Muhammad Faisal, Jing Wang, Hermes Perez, Anja Werno, Antje van der Linden, Arlo Upton, Chris Mansell, David Hammer, Dragana Drinkovic, Gary McAuliffe, Hana Sofia Andersson, James Ussher, Jill Sherwood, Josh Freeman, Julia Howard, Juliet Elvy, Mary DeAlmeida, Matt Blakiston, Matthew Rogers, Max Bloomfield, Michael Addidle, Michelle Balm, Sally Roberts, Sarah Jefferies, Sharmini Muttaiyah, Susan Morpeth, Susan Taylor, Timothy Blackmore, Vani Sathyendran, Veronica Playle, Virginia Hope, Erasmus Smit, Lauren Jelly, Olin Silander, Joep de Ligt |
| EPI_ISL_548147, EPI_ISL_548148, EPI_ISL_548149, EPI_ISL_548150, EPI_ISL_548151, EPI_ISL_548152, EPI_ISL_548153, EPI_ISL_548154, EPI_ISL_548155, EPI_ISL_548156, EPI_ISL_548157, EPI_ISL_548159, EPI_ISL_548160, EPI_ISL_548161, EPI_ISL_548162, EPI_ISL_548164, EPI_ISL_548165, EPI_ISL_548166, EPI_ISL_548167, EPI_ISL_548168, EPI_ISL_548170, EPI_ISL_548172, EPI_ISL_548173, EPI_ISL_548174, EPI_ISL_548175, EPI_ISL_548176, EPI_ISL_548177, EPI_ISL_548178, EPI_ISL_548179, EPI_ISL_548180, EPI_ISL_548181, EPI_ISL_548182, EPI_ISL_548183, EPI_ISL_548184, EPI_ISL_548185, EPI_ISL_548186, EPI_ISL_548187, EPI_ISL_548188, EPI_ISL_548189, EPI_ISL_548190, EPI_ISL_548191, EPI_ISL_548192, EPI_ISL_548193, EPI_ISL_548194, EPI_ISL_548195, EPI_ISL_548196, EPI_ISL_548197, EPI_ISL_548198, EPI_ISL_548199, EPI_ISL_548201, EPI_ISL_548202, EPI_ISL_548204, EPI_ISL_548205, EPI_ISL_548206, EPI_ISL_548207, EPI_ISL_548208, EPI_ISL_548209, EPI_ISL_548210, EPI_ISL_548211, EPI_ISL_548212, EPI_ISL_548213, EPI_ISL_548214, EPI_ISL_548215, EPI_ISL_548216, EPI_ISL_548217, EPI_ISL_548218, EPI_ISL_548220, EPI_ISL_548221, EPI_ISL_548222, EPI_ISL_548223, EPI_ISL_548224, EPI_ISL_548225, EPI_ISL_548226, EPI_ISL_548227, EPI_ISL_548228, EPI_ISL_548229, EPI_ISL_548230, EPI_ISL_548231, EPI_ISL_548232, EPI_ISL_548233, EPI_ISL_548234, EPI_ISL_548235, EPI_ISL_548237, EPI_ISL_548238, EPI_ISL_548240, EPI_ISL_548241, EPI_ISL_548242 | | | |
| see above | Laboratoire de Virologie, HUG | Swiss National Reference Centre for Influenza | LAUBSCHER F. |
| EPI_ISL_548243, EPI_ISL_548244 | Faith Laboratory, Immunology Institute, Icahn School of Medicine at Mount Sinai | van Bakel Laboratory, Genetics and Genomics Sciences, Icahn School of Medicine at Mount Sinai | Graham J. Britton, Alice Chen-Liaw, Francesca Cossarini, Alexandra Livanos, Matthew P. Spindler, Tamar Plitt, Joseph Eggers, Ilaria Mogno, Ana S. Gonzalez-Reiche, Sophia Sui, Michael Tankelevich, Lauren Tal Grinspan, Rebekah E. Dixon, Divya Jha, Gustavo Martinez-Delgado, Fatima Amanat, Daisy Hoagland, Benjamin R. tenOever, Marla C. Dubinsky, Miriam Merad, Harm Van Bakel, Florian Krammer, Gerold Bongers, Saurabh Mehandru and Jeremiah J. Faith |
| EPI_ISL_548245, EPI_ISL_548246 | Skovde/Unilabs | The Public Health Agency of Sweden | Anna-Malin Linde, Maria Lind Karlberg, Mattias Haukland, Reza Advani, Olov Svartstrom, Oskar Karlsson Lindsjo, Sandra Broddesson, Petra Edquist, Mia Brytting, Anna Risberg, Karin Tegmark-Wisell |
| EPI_ISL_548247 | Halmstad klinisk mikrobiologi | The Public Health Agency of Sweden | Anna-Malin Linde, Maria Lind Karlberg, Mattias Haukland, Reza Advani, Olov Svartstrom, Oskar Karlsson Lindsjo, Sandra Broddesson, Petra Edquist, Mia Brytting, Anna Risberg, Karin Tegmark-Wisell |
| EPI_ISL_548248 | Ostersund klinisk mikrobiologi | The Public Health Agency of Sweden | Anna-Malin Linde, Maria Lind Karlberg, Mattias Haukland, Reza Advani, Olov Svartstrom, Oskar Karlsson Lindsjo, Sandra Broddesson, Petra Edquist, Mia Brytting, Anna Risberg, Karin Tegmark-Wisell |
| EPI_ISL_548249 | Halmstad klinisk mikrobiologi | The Public Health Agency of Sweden | Anna-Malin Linde, Maria Lind Karlberg, Mattias Haukland, Reza Advani, Olov Svartstrom, Oskar Karlsson Lindsjo, Sandra Broddesson, Petra Edquist, Mia Brytting, Anna Risberg, Karin Tegmark-Wisell |
| EPI_ISL_548250, EPI_ISL_548251 | Klinisk mikrobiologi NAL Trollhattan | The Public Health Agency of Sweden | Anna-Malin Linde, Maria Lind Karlberg, Mattias Haukland, Reza Advani, Olov Svartstrom, Oskar Karlsson Lindsjo, Sandra Broddesson, Petra Edquist, Mia Brytting, Anna Risberg, Karin Tegmark-Wisell |
| EPI_ISL_548252 | Klinisk mikrobiologi centralsjukhuset Karlstad | The Public Health Agency of Sweden | Anna-Malin Linde, Maria Lind Karlberg, Mattias Haukland, Reza Advani, Olov Svartstrom, Oskar Karlsson Lindsjo, Sandra Broddesson, Petra Edquist, Mia Brytting, Anna Risberg, Karin Tegmark-Wisell |
| EPI_ISL_548253, EPI_ISL_548254 | Klinisk mikrobiologi NAL Trollhattan | The Public Health Agency of Sweden | Anna-Malin Linde, Maria Lind Karlberg, Mattias Haukland, Reza Advani, Olov Svartstrom, Oskar Karlsson Lindsjo, Sandra Broddesson, Petra Edquist, Mia Brytting, Anna Risberg, Karin Tegmark-Wisell |
| EPI_ISL_548255 | Lanssjukhuset Kalmar | The Public Health Agency of Sweden | Anna-Malin Linde, Maria Lind Karlberg, Mattias Haukland, Reza Advani, Olov Svartstrom, Oskar Karlsson Lindsjo, Sandra Broddesson, Petra Edquist, Mia Brytting, Anna Risberg, Karin Tegmark-Wisell |
| EPI_ISL_548256 | Centralsjukhuset | The Public Health Agency of Sweden | Anna-Malin Linde, Maria Lind Karlberg, Mattias Haukland, Reza Advani, Olov Svartstrom, Oskar Karlsson Lindsjo, Sandra Broddesson, Petra Edquist, Mia Brytting, Anna Risberg, Karin Tegmark-Wisell |
| EPI_ISL_548257, EPI_ISL_548258 | Karolinska universitetslaboratoriet | The Public Health Agency of Sweden | Anna-Malin Linde, Maria Lind Karlberg, Mattias Haukland, Reza Advani, Olov Svartstrom, Oskar Karlsson Lindsjo, Sandra Broddesson, Petra Edquist, Mia Brytting, Anna Risberg, Karin Tegmark-Wisell |
| EPI_ISL_548264 | County of Santa Clara Public Health Department | Chan-Zuckerberg Biohub | CZB Cliahub Consortium |
| EPI_ISL_548265 | County of San Luis Obispo Public Health Laboratory | Chan-Zuckerberg Biohub | CZB Cliahub Consortium |
| EPI_ISL_548266, EPI_ISL_548267 | County of Santa Clara Public Health Department | Chan-Zuckerberg Biohub | CZB Cliahub Consortium |
| EPI_ISL_548268, EPI_ISL_548269 | County of San Luis Obispo Public Health Laboratory | Chan-Zuckerberg Biohub | CZB Cliahub Consortium |
| EPI_ISL_548270, EPI_ISL_548271, EPI_ISL_548272, EPI_ISL_548273, EPI_ISL_548274, EPI_ISL_548275, EPI_ISL_548276 | Orange County Public Health Laboratory | Chan-Zuckerberg Biohub | CZB Cliahub Consortium |
| EPI_ISL_548279 | County of Santa Clara Public Health Department | Chan-Zuckerberg Biohub | CZB Cliahub Consortium |
| EPI_ISL_548280, EPI_ISL_548281 | County of San Luis Obispo Public Health Laboratory | Chan-Zuckerberg Biohub | CZB Cliahub Consortium |
| EPI_ISL_548282 | County of Santa Clara Public Health Department | Chan-Zuckerberg Biohub | CZB Cliahub Consortium |
| EPI_ISL_548283 | Orange County Public Health Laboratory | Chan-Zuckerberg Biohub | CZB Cliahub Consortium |
| EPI_ISL_548284 | County of San Luis Obispo Public Health Laboratory | Chan-Zuckerberg Biohub | CZB Cliahub Consortium |
| EPI_ISL_548285 | Orange County Public Health Laboratory | Chan-Zuckerberg Biohub | CZB Cliahub Consortium |
| EPI_ISL_548286, EPI_ISL_548288 | County of San Luis Obispo Public Health Laboratory | Chan-Zuckerberg Biohub | CZB Cliahub Consortium |
| EPI_ISL_548289 | Orange County Public Health Laboratory | Chan-Zuckerberg Biohub | CZB Cliahub Consortium |
| EPI_ISL_548292, EPI_ISL_548294, EPI_ISL_548295 | County of San Luis Obispo Public Health Laboratory | Chan-Zuckerberg Biohub | CZB Cliahub Consortium |
| EPI_ISL_548296, EPI_ISL_548298 | Orange County Public Health Laboratory | Chan-Zuckerberg Biohub | CZB Cliahub Consortium |
| EPI_ISL_548299 | County of Santa Clara Public Health Department | Chan-Zuckerberg Biohub | CZB Cliahub Consortium |
| EPI_ISL_548300, EPI_ISL_548301 | Orange County Public Health Laboratory | Chan-Zuckerberg Biohub | CZB Cliahub Consortium |
| EPI_ISL_548302, EPI_ISL_548303 | County of Santa Clara Public Health Department | Chan-Zuckerberg Biohub | CZB Cliahub Consortium |
| EPI_ISL_548304 | County of San Luis Obispo Public Health Laboratory | Chan-Zuckerberg Biohub | CZB Cliahub Consortium |
| EPI_ISL_548305, EPI_ISL_548306 | Orange County Public Health Laboratory | Chan-Zuckerberg Biohub | CZB Cliahub Consortium |
| EPI_ISL_548307 | County of Santa Clara Public Health Department | Chan-Zuckerberg Biohub | CZB Cliahub Consortium |
| EPI_ISL_548308, EPI_ISL_548309 | Orange County Public Health Laboratory | Chan-Zuckerberg Biohub | CZB Cliahub Consortium |
| EPI_ISL_548310 | County of Santa Clara Public Health Department | Chan-Zuckerberg Biohub | CZB Cliahub Consortium |
| EPI_ISL_548311 | County of San Luis Obispo Public Health Laboratory | Chan-Zuckerberg Biohub | CZB Cliahub Consortium |
| EPI_ISL_548312, EPI_ISL_548313 | Orange County Public Health Laboratory | Chan-Zuckerberg Biohub | CZB Cliahub Consortium |
| EPI_ISL_548314 | County of Santa Clara Public Health Department | Chan-Zuckerberg Biohub | CZB Cliahub Consortium |
| EPI_ISL_548315 | Orange County Public Health Laboratory | Chan-Zuckerberg Biohub | CZB Cliahub Consortium |
| EPI_ISL_548316 | County of San Luis Obispo Public Health Laboratory | Chan-Zuckerberg Biohub | CZB Cliahub Consortium |
| EPI_ISL_548317 | Orange County Public Health Laboratory | Chan-Zuckerberg Biohub | CZB Cliahub Consortium |
| EPI_ISL_548319 | County of Santa Clara Public Health Department | Chan-Zuckerberg Biohub | CZB Cliahub Consortium |
| EPI_ISL_548322 | County of San Luis Obispo Public Health Laboratory | Chan-Zuckerberg Biohub | CZB Cliahub Consortium |
| EPI_ISL_548323 | Orange County Public Health Laboratory | Chan-Zuckerberg Biohub | CZB Cliahub Consortium |
| EPI_ISL_548325 | County of San Luis Obispo Public Health Laboratory | Chan-Zuckerberg Biohub | CZB Cliahub Consortium |
| EPI_ISL_548326 | Orange County Public Health Laboratory | Chan-Zuckerberg Biohub | CZB Cliahub Consortium |

| | | | |
|---|---|---|---|
| EPI_ISL_548327 | County of Santa Clara Public Health Department | Chan-Zuckerberg Biohub | CZB Cliahub Consortium |
| EPI_ISL_548328 | Orange County Public Health Laboratory | Chan-Zuckerberg Biohub | CZB Cliahub Consortium |
| EPI_ISL_548329 | County of San Luis Obispo Public Health Laboratory | Chan-Zuckerberg Biohub | CZB Cliahub Consortium |
| EPI_ISL_548332, EPI_ISL_548334 | Orange County Public Health Laboratory | Chan-Zuckerberg Biohub | CZB Cliahub Consortium |
| EPI_ISL_548336 | County of San Luis Obispo Public Health Laboratory | Chan-Zuckerberg Biohub | CZB Cliahub Consortium |
| EPI_ISL_548337, EPI_ISL_548338, EPI_ISL_548339, EPI_ISL_548340, EPI_ISL_548341, EPI_ISL_548342, EPI_ISL_548343, EPI_ISL_548344, EPI_ISL_548345 | County of Santa Clara Public Health Department | Chan-Zuckerberg Biohub | CZB Cliahub Consortium |
| EPI_ISL_548346 | Ventura County Public Health Lab | Chan-Zuckerberg Biohub | CZB Cliahub Consortium |
| EPI_ISL_548347 | County of San Luis Obispo Public Health Laboratory | Chan-Zuckerberg Biohub | CZB Cliahub Consortium |
| EPI_ISL_548348, EPI_ISL_548349 | Ventura County Public Health Lab | Chan-Zuckerberg Biohub | CZB Cliahub Consortium |
| EPI_ISL_548350, EPI_ISL_548351 | County of San Luis Obispo Public Health Laboratory | Chan-Zuckerberg Biohub | CZB Cliahub Consortium |
| EPI_ISL_548352, EPI_ISL_548353, EPI_ISL_548354, EPI_ISL_548355, EPI_ISL_548356 | Ventura County Public Health Lab | Chan-Zuckerberg Biohub | CZB Cliahub Consortium |
| EPI_ISL_548357 | County of San Luis Obispo Public Health Laboratory | Chan-Zuckerberg Biohub | CZB Cliahub Consortium |
| EPI_ISL_548358 | Ventura County Public Health Lab | Chan-Zuckerberg Biohub | CZB Cliahub Consortium |
| EPI_ISL_548359, EPI_ISL_548360, EPI_ISL_548361 | County of San Luis Obispo Public Health Laboratory | Chan-Zuckerberg Biohub | CZB Cliahub Consortium |
| EPI_ISL_548362 | Ventura County Public Health Lab | Chan-Zuckerberg Biohub | CZB Cliahub Consortium |
| EPI_ISL_548363 | Tuolumne County Public Health | Chan-Zuckerberg Biohub | CZB Cliahub Consortium |
| EPI_ISL_548364 | Ventura County Public Health Lab | Chan-Zuckerberg Biohub | CZB Cliahub Consortium |
| EPI_ISL_548365, EPI_ISL_548366 | County of San Luis Obispo Public Health Laboratory | Chan-Zuckerberg Biohub | CZB Cliahub Consortium |
| EPI_ISL_548367, EPI_ISL_548368 | Ventura County Public Health Lab | Chan-Zuckerberg Biohub | CZB Cliahub Consortium |
| EPI_ISL_548369, EPI_ISL_548370 | County of San Luis Obispo Public Health Laboratory | Chan-Zuckerberg Biohub | CZB Cliahub Consortium |
| EPI_ISL_548373 | Orange County Public Health Laboratory | Chan-Zuckerberg Biohub | CZB Cliahub Consortium |
| EPI_ISL_548374 | University of California, Davis | Chan-Zuckerberg Biohub | CZB Cliahub Consortium |
| EPI_ISL_548375 | Orange County Public Health Laboratory | Chan-Zuckerberg Biohub | CZB Cliahub Consortium |
| EPI_ISL_548376 | Ventura County Public Health Lab | Chan-Zuckerberg Biohub | CZB Cliahub Consortium |
| EPI_ISL_548377, EPI_ISL_548378 | Orange County Public Health Laboratory | Chan-Zuckerberg Biohub | CZB Cliahub Consortium |
| EPI_ISL_548379 | County of Santa Clara Public Health Department | Chan-Zuckerberg Biohub | CZB Cliahub Consortium |
| EPI_ISL_548381 | Orange County Public Health Laboratory | Chan-Zuckerberg Biohub | CZB Cliahub Consortium |
| EPI_ISL_548382 | University of California, Davis | Chan-Zuckerberg Biohub | CZB Cliahub Consortium |
| EPI_ISL_548386, EPI_ISL_548387 | Orange County Public Health Laboratory | Chan-Zuckerberg Biohub | CZB Cliahub Consortium |
| EPI_ISL_548394, EPI_ISL_548395 | County of Santa Clara Public Health Department | Chan-Zuckerberg Biohub | CZB Cliahub Consortium |
| EPI_ISL_548398 | Ventura County Public Health Lab | Chan-Zuckerberg Biohub | CZB Cliahub Consortium |
| EPI_ISL_548401 | County of Santa Clara Public Health Department | Chan-Zuckerberg Biohub | CZB Cliahub Consortium |
| EPI_ISL_548402 | Ventura County Public Health Lab | Chan-Zuckerberg Biohub | CZB Cliahub Consortium |
| EPI_ISL_548404 | Orange County Public Health Laboratory | Chan-Zuckerberg Biohub | CZB Cliahub Consortium |
| EPI_ISL_548406 | Ventura County Public Health Lab | Chan-Zuckerberg Biohub | CZB Cliahub Consortium |
| EPI_ISL_548408 | Orange County Public Health Laboratory | Chan-Zuckerberg Biohub | CZB Cliahub Consortium |
| EPI_ISL_548409, EPI_ISL_548410 | Ventura County Public Health Lab | Chan-Zuckerberg Biohub | CZB Cliahub Consortium |
| EPI_ISL_548414 | University of California, Davis | Chan-Zuckerberg Biohub | CZB Cliahub Consortium |
| EPI_ISL_548417, EPI_ISL_548420, EPI_ISL_548423 | County of Santa Clara Public Health Department | Chan-Zuckerberg Biohub | CZB Cliahub Consortium |
| EPI_ISL_548425 | County of San Luis Obispo Public Health Laboratory | Chan-Zuckerberg Biohub | CZB Cliahub Consortium |
| EPI_ISL_548426 | County of Santa Clara Public Health Department | Chan-Zuckerberg Biohub | CZB Cliahub Consortium |
| EPI_ISL_548428 | County of San Luis Obispo Public Health Laboratory | Chan-Zuckerberg Biohub | CZB Cliahub Consortium |
| EPI_ISL_548429 | County of Santa Clara Public Health Department | Chan-Zuckerberg Biohub | CZB Cliahub Consortium |
| EPI_ISL_548430 | Ventura County Public Health Lab | Chan-Zuckerberg Biohub | CZB Cliahub Consortium |
| EPI_ISL_548431 | Orange County Public Health Laboratory | Chan-Zuckerberg Biohub | CZB Cliahub Consortium |
| EPI_ISL_548433, EPI_ISL_548434 | County of Santa Clara Public Health Department | Chan-Zuckerberg Biohub | CZB Cliahub Consortium |
| EPI_ISL_548439 | Orange County Public Health Laboratory | Chan-Zuckerberg Biohub | CZB Cliahub Consortium |
| EPI_ISL_548442, EPI_ISL_548443 | Ventura County Public Health Lab | Chan-Zuckerberg Biohub | CZB Cliahub Consortium |
| EPI_ISL_548444 | Orange County Public Health Laboratory | Chan-Zuckerberg Biohub | CZB Cliahub Consortium |
| EPI_ISL_548447 | University of California, Davis | Chan-Zuckerberg Biohub | CZB Cliahub Consortium |
| EPI_ISL_548450 | Ventura County Public Health Lab | Chan-Zuckerberg Biohub | CZB Cliahub Consortium |
| EPI_ISL_548452 | University of California, Davis | Chan-Zuckerberg Biohub | CZB Cliahub Consortium |
| EPI_ISL_548453 | County of San Luis Obispo Public Health Laboratory | Chan-Zuckerberg Biohub | CZB Cliahub Consortium |
| EPI_ISL_548455 | Ventura County Public Health Lab | Chan-Zuckerberg Biohub | CZB Cliahub Consortium |
| EPI_ISL_548457 | County of San Luis Obispo Public Health Laboratory | Chan-Zuckerberg Biohub | CZB Cliahub Consortium |
| EPI_ISL_548459 | Ventura County Public Health Lab | Chan-Zuckerberg Biohub | CZB Cliahub Consortium |
| EPI_ISL_548460 | Orange County Public Health Laboratory | Chan-Zuckerberg Biohub | CZB Cliahub Consortium |
| EPI_ISL_548461 | Ventura County Public Health Lab | Chan-Zuckerberg Biohub | CZB Cliahub Consortium |
| EPI_ISL_548463, EPI_ISL_548464 | Orange County Public Health Laboratory | Chan-Zuckerberg Biohub | CZB Cliahub Consortium |
| EPI_ISL_548470 | County of Santa Clara Public Health | Chan-Zuckerberg Biohub | CZB Cliahub Consortium |

| | | | |
|---|---|---|---|
| | Department | | |
| EPI_ISL_548471, EPI_ISL_548473 | Ventura County Public Health Lab | Chan-Zuckerberg Biohub | CZB Cliahub Consortium |
| EPI_ISL_548474 | University of California, Davis | Chan-Zuckerberg Biohub | CZB Cliahub Consortium |
| EPI_ISL_548475, EPI_ISL_548476 | Ventura County Public Health Lab | Chan-Zuckerberg Biohub | CZB Cliahub Consortium |
| EPI_ISL_548477 | University of California, Davis | Chan-Zuckerberg Biohub | CZB Cliahub Consortium |
| EPI_ISL_548478 | County of Santa Clara Public Health Department | Chan-Zuckerberg Biohub | CZB Cliahub Consortium |
| EPI_ISL_548479 | University of California, Davis | Chan-Zuckerberg Biohub | CZB Cliahub Consortium |
| EPI_ISL_548480, EPI_ISL_548481 | Ventura County Public Health Lab | Chan-Zuckerberg Biohub | CZB Cliahub Consortium |
| EPI_ISL_548482 | County of Santa Clara Public Health Department | Chan-Zuckerberg Biohub | CZB Cliahub Consortium |
| EPI_ISL_548483 | Ventura County Public Health Lab | Chan-Zuckerberg Biohub | CZB Cliahub Consortium |
| EPI_ISL_548484, EPI_ISL_548485 | Orange County Public Health Laboratory | Chan-Zuckerberg Biohub | CZB Cliahub Consortium |
| EPI_ISL_548486 | University of California, Davis | Chan-Zuckerberg Biohub | CZB Cliahub Consortium |
| EPI_ISL_548487 | Ventura County Public Health Lab | Chan-Zuckerberg Biohub | CZB Cliahub Consortium |
| EPI_ISL_548488 | Orange County Public Health Laboratory | Chan-Zuckerberg Biohub | CZB Cliahub Consortium |
| EPI_ISL_548489 | Ventura County Public Health Lab | Chan-Zuckerberg Biohub | CZB Cliahub Consortium |
| EPI_ISL_548490 | University of California, Davis | Chan-Zuckerberg Biohub | CZB Cliahub Consortium |
| EPI_ISL_548491, EPI_ISL_548492 | Contra Costa Public Health Lab | Chan-Zuckerberg Biohub | CZB Cliahub Consortium |
| EPI_ISL_548493 | County of San Luis Obispo Public Health Laboratory | Chan-Zuckerberg Biohub | CZB Cliahub Consortium |
| EPI_ISL_548494 | County of Santa Clara Public Health Department | Chan-Zuckerberg Biohub | CZB Cliahub Consortium |
| EPI_ISL_548495 | Contra Costa Public Health Lab | Chan-Zuckerberg Biohub | CZB Cliahub Consortium |
| EPI_ISL_548496, EPI_ISL_548497, EPI_ISL_548498 | County of Santa Clara Public Health Department | Chan-Zuckerberg Biohub | CZB Cliahub Consortium |
| EPI_ISL_548499 | Contra Costa Public Health Lab | Chan-Zuckerberg Biohub | CZB Cliahub Consortium |
| EPI_ISL_548500 | County of Santa Clara Public Health Department | Chan-Zuckerberg Biohub | CZB Cliahub Consortium |
| EPI_ISL_548501 | Contra Costa Public Health Lab | Chan-Zuckerberg Biohub | CZB Cliahub Consortium |
| EPI_ISL_548502, EPI_ISL_548504 | County of San Luis Obispo Public Health Laboratory | Chan-Zuckerberg Biohub | CZB Cliahub Consortium |
| EPI_ISL_548505 | UCSF Clinical Microbiology Laboratory | Chan-Zuckerberg Biohub | CZB Cliahub Consortium |
| EPI_ISL_548506, EPI_ISL_548507, EPI_ISL_548508 | Contra Costa Public Health Lab | Chan-Zuckerberg Biohub | CZB Cliahub Consortium |
| EPI_ISL_548509 | County of Santa Clara Public Health Department | Chan-Zuckerberg Biohub | CZB Cliahub Consortium |
| EPI_ISL_548510 | Contra Costa Public Health Lab | Chan-Zuckerberg Biohub | CZB Cliahub Consortium |
| EPI_ISL_548511, EPI_ISL_548512 | County of Santa Clara Public Health Department | Chan-Zuckerberg Biohub | CZB Cliahub Consortium |
| EPI_ISL_548513 | UCSF Clinical Microbiology Laboratory | Chan-Zuckerberg Biohub | CZB Cliahub Consortium |
| EPI_ISL_548514 | County of Santa Clara Public Health Department | Chan-Zuckerberg Biohub | CZB Cliahub Consortium |
| EPI_ISL_548515, EPI_ISL_548516 | Contra Costa Public Health Lab | Chan-Zuckerberg Biohub | CZB Cliahub Consortium |
| EPI_ISL_548517 | County of San Luis Obispo Public Health Laboratory | Chan-Zuckerberg Biohub | CZB Cliahub Consortium |
| EPI_ISL_548518 | County of Santa Clara Public Health Department | Chan-Zuckerberg Biohub | CZB Cliahub Consortium |
| EPI_ISL_548519 | County of San Luis Obispo Public Health Laboratory | Chan-Zuckerberg Biohub | CZB Cliahub Consortium |
| EPI_ISL_548520 | Contra Costa Public Health Lab | Chan-Zuckerberg Biohub | CZB Cliahub Consortium |
| EPI_ISL_548521 | County of San Luis Obispo Public Health Laboratory | Chan-Zuckerberg Biohub | CZB Cliahub Consortium |
| EPI_ISL_548522 | UCSF Clinical Microbiology Laboratory | Chan-Zuckerberg Biohub | CZB Cliahub Consortium |
| EPI_ISL_548524 | Contra Costa Public Health Lab | Chan-Zuckerberg Biohub | CZB Cliahub Consortium |
| EPI_ISL_548525 | County of Santa Clara Public Health Department | Chan-Zuckerberg Biohub | CZB Cliahub Consortium |
| EPI_ISL_548527, EPI_ISL_548528 | Contra Costa Public Health Lab | Chan-Zuckerberg Biohub | CZB Cliahub Consortium |
| EPI_ISL_548529 | County of Santa Clara Public Health Department | Chan-Zuckerberg Biohub | CZB Cliahub Consortium |
| EPI_ISL_548531 | Contra Costa Public Health Lab | Chan-Zuckerberg Biohub | CZB Cliahub Consortium |
| EPI_ISL_548532 | County of San Luis Obispo Public Health Laboratory | Chan-Zuckerberg Biohub | CZB Cliahub Consortium |
| EPI_ISL_548533 | County of Santa Clara Public Health Department | Chan-Zuckerberg Biohub | CZB Cliahub Consortium |
| EPI_ISL_548534, EPI_ISL_548535 | Contra Costa Public Health Lab | Chan-Zuckerberg Biohub | CZB Cliahub Consortium |
| EPI_ISL_548536 | County of Santa Clara Public Health Department | Chan-Zuckerberg Biohub | CZB Cliahub Consortium |
| EPI_ISL_548537, EPI_ISL_548538, EPI_ISL_548539, EPI_ISL_548540 | Contra Costa Public Health Lab | Chan-Zuckerberg Biohub | CZB Cliahub Consortium |
| EPI_ISL_548541 | County of Santa Clara Public Health Department | Chan-Zuckerberg Biohub | CZB Cliahub Consortium |
| EPI_ISL_548542 | County of San Luis Obispo Public Health Laboratory | Chan-Zuckerberg Biohub | CZB Cliahub Consortium |
| EPI_ISL_548543 | County of Santa Clara Public Health Department | Chan-Zuckerberg Biohub | CZB Cliahub Consortium |
| EPI_ISL_548544 | County of San Luis Obispo Public Health Laboratory | Chan-Zuckerberg Biohub | CZB Cliahub Consortium |
| EPI_ISL_548545 | Contra Costa Public Health Lab | Chan-Zuckerberg Biohub | CZB Cliahub Consortium |
| EPI_ISL_548546, EPI_ISL_548547 | County of Santa Clara Public Health Department | Chan-Zuckerberg Biohub | CZB Cliahub Consortium |
| EPI_ISL_548548, EPI_ISL_548549 | Contra Costa Public Health Lab | Chan-Zuckerberg Biohub | CZB Cliahub Consortium |
| EPI_ISL_548550, EPI_ISL_548551 | County of Santa Clara Public Health Department | Chan-Zuckerberg Biohub | CZB Cliahub Consortium |
| EPI_ISL_548552 | Contra Costa Public Health Lab | Chan-Zuckerberg Biohub | CZB Cliahub Consortium |
| EPI_ISL_548553 | County of Santa Clara Public Health | Chan-Zuckerberg Biohub | CZB Cliahub Consortium |

| | | | |
|---|---|---|---|
| | Department | | |
| EPI_ISL_548554 | Contra Costa Public Health Lab | Chan-Zuckerberg Biohub | CZB Cliahub Consortium |
| EPI_ISL_548555 | UCSF Clinical Microbiology Laboratory | Chan-Zuckerberg Biohub | CZB Cliahub Consortium |
| EPI_ISL_548556, EPI_ISL_548557, EPI_ISL_548558 | Contra Costa Public Health Lab | Chan-Zuckerberg Biohub | CZB Cliahub Consortium |
| EPI_ISL_548559 | County of Santa Clara Public Health Department | Chan-Zuckerberg Biohub | CZB Cliahub Consortium |
| EPI_ISL_548560 | County of San Luis Obispo Public Health Laboratory | Chan-Zuckerberg Biohub | CZB Cliahub Consortium |
| EPI_ISL_548561 | Contra Costa Public Health Lab | Chan-Zuckerberg Biohub | CZB Cliahub Consortium |
| EPI_ISL_548562, EPI_ISL_548563 | County of Santa Clara Public Health Department | Chan-Zuckerberg Biohub | CZB Cliahub Consortium |
| EPI_ISL_548564, EPI_ISL_548565, EPI_ISL_548566, EPI_ISL_548567 | Contra Costa Public Health Lab | Chan-Zuckerberg Biohub | CZB Cliahub Consortium |
| EPI_ISL_548568 | County of Santa Clara Public Health Department | Chan-Zuckerberg Biohub | CZB Cliahub Consortium |
| EPI_ISL_548569 | Contra Costa Public Health Lab | Chan-Zuckerberg Biohub | CZB Cliahub Consortium |
| EPI_ISL_548570 | County of Santa Clara Public Health Department | Chan-Zuckerberg Biohub | CZB Cliahub Consortium |
| EPI_ISL_548571 | UCSF Clinical Microbiology Laboratory | Chan-Zuckerberg Biohub | CZB Cliahub Consortium |
| EPI_ISL_548572, EPI_ISL_548573, EPI_ISL_548574, EPI_ISL_548575 | Contra Costa Public Health Lab | Chan-Zuckerberg Biohub | CZB Cliahub Consortium |
| EPI_ISL_548576 | County of Santa Clara Public Health Department | Chan-Zuckerberg Biohub | CZB Cliahub Consortium |
| EPI_ISL_548577 | Contra Costa Public Health Lab | Chan-Zuckerberg Biohub | CZB Cliahub Consortium |
| EPI_ISL_548578 | County of Santa Clara Public Health Department | Chan-Zuckerberg Biohub | CZB Cliahub Consortium |
| EPI_ISL_548579 | Contra Costa Public Health Lab | Chan-Zuckerberg Biohub | CZB Cliahub Consortium |
| EPI_ISL_548580, EPI_ISL_548581 | Orange County Public Health Laboratory | Chan-Zuckerberg Biohub | CZB Cliahub Consortium |
| EPI_ISL_548582, EPI_ISL_548583, EPI_ISL_548584 | County of Santa Clara Public Health Department | Chan-Zuckerberg Biohub | CZB Cliahub Consortium |
| EPI_ISL_548585, EPI_ISL_548586 | Orange County Public Health Laboratory | Chan-Zuckerberg Biohub | CZB Cliahub Consortium |
| EPI_ISL_548588 | County of Santa Clara Public Health Department | Chan-Zuckerberg Biohub | CZB Cliahub Consortium |
| EPI_ISL_548590, EPI_ISL_548591, EPI_ISL_548592 | Orange County Public Health Laboratory | Chan-Zuckerberg Biohub | CZB Cliahub Consortium |
| EPI_ISL_548593 | County of Santa Clara Public Health Department | Chan-Zuckerberg Biohub | CZB Cliahub Consortium |
| EPI_ISL_548594 | Orange County Public Health Laboratory | Chan-Zuckerberg Biohub | CZB Cliahub Consortium |
| EPI_ISL_548595, EPI_ISL_548596, EPI_ISL_548597, EPI_ISL_548598 | County of Santa Clara Public Health Department | Chan-Zuckerberg Biohub | CZB Cliahub Consortium |
| EPI_ISL_548599 | Tuolumne County Public Health | Chan-Zuckerberg Biohub | CZB Cliahub Consortium |
| EPI_ISL_548601 | Orange County Public Health Laboratory | Chan-Zuckerberg Biohub | CZB Cliahub Consortium |
| EPI_ISL_548602 | County of Santa Clara Public Health Department | Chan-Zuckerberg Biohub | CZB Cliahub Consortium |
| EPI_ISL_548603 | County of San Luis Obispo Public Health Laboratory | Chan-Zuckerberg Biohub | CZB Cliahub Consortium |
| EPI_ISL_548604, EPI_ISL_548605, EPI_ISL_548606, EPI_ISL_548607 | County of Santa Clara Public Health Department | Chan-Zuckerberg Biohub | CZB Cliahub Consortium |
| EPI_ISL_548608, EPI_ISL_548609 | Orange County Public Health Laboratory | Chan-Zuckerberg Biohub | CZB Cliahub Consortium |
| EPI_ISL_548610 | County of Santa Clara Public Health Department | Chan-Zuckerberg Biohub | CZB Cliahub Consortium |
| EPI_ISL_548611 | Orange County Public Health Laboratory | Chan-Zuckerberg Biohub | CZB Cliahub Consortium |
| EPI_ISL_548612 | County of Santa Clara Public Health Department | Chan-Zuckerberg Biohub | CZB Cliahub Consortium |
| EPI_ISL_548613, EPI_ISL_548614, EPI_ISL_548615 | Orange County Public Health Laboratory | Chan-Zuckerberg Biohub | CZB Cliahub Consortium |
| EPI_ISL_548616, EPI_ISL_548617, EPI_ISL_548618, EPI_ISL_548619 | County of Santa Clara Public Health Department | Chan-Zuckerberg Biohub | CZB Cliahub Consortium |
| EPI_ISL_548620 | Orange County Public Health Laboratory | Chan-Zuckerberg Biohub | CZB Cliahub Consortium |
| EPI_ISL_548621, EPI_ISL_548622 | County of Santa Clara Public Health Department | Chan-Zuckerberg Biohub | CZB Cliahub Consortium |
| EPI_ISL_548623 | Orange County Public Health Laboratory | Chan-Zuckerberg Biohub | CZB Cliahub Consortium |
| EPI_ISL_548624 | County of Santa Clara Public Health Department | Chan-Zuckerberg Biohub | CZB Cliahub Consortium |
| EPI_ISL_548625 | Orange County Public Health Laboratory | Chan-Zuckerberg Biohub | CZB Cliahub Consortium |
| EPI_ISL_548626 | Tuolumne County Public Health | Chan-Zuckerberg Biohub | CZB Cliahub Consortium |
| EPI_ISL_548627, EPI_ISL_548628 | Orange County Public Health Laboratory | Chan-Zuckerberg Biohub | CZB Cliahub Consortium |
| EPI_ISL_548629, EPI_ISL_548630 | County of Santa Clara Public Health Department | Chan-Zuckerberg Biohub | CZB Cliahub Consortium |
| EPI_ISL_548631 | County of San Luis Obispo Public Health Laboratory | Chan-Zuckerberg Biohub | CZB Cliahub Consortium |
| EPI_ISL_548632, EPI_ISL_548633 | County of Santa Clara Public Health Department | Chan-Zuckerberg Biohub | CZB Cliahub Consortium |
| EPI_ISL_548634, EPI_ISL_548635 | Orange County Public Health Laboratory | Chan-Zuckerberg Biohub | CZB Cliahub Consortium |
| EPI_ISL_548636 | County of Santa Clara Public Health Department | Chan-Zuckerberg Biohub | CZB Cliahub Consortium |
| EPI_ISL_548637, EPI_ISL_548638 | Orange County Public Health Laboratory | Chan-Zuckerberg Biohub | CZB Cliahub Consortium |
| EPI_ISL_548639 | County of Santa Clara Public Health Department | Chan-Zuckerberg Biohub | CZB Cliahub Consortium |
| EPI_ISL_548641 | Orange County Public Health Laboratory | Chan-Zuckerberg Biohub | CZB Cliahub Consortium |
| EPI_ISL_548642, EPI_ISL_548643 | County of Santa Clara Public Health Department | Chan-Zuckerberg Biohub | CZB Cliahub Consortium |
| EPI_ISL_548644 | County of San Luis Obispo Public Health Laboratory | Chan-Zuckerberg Biohub | CZB Cliahub Consortium |
| EPI_ISL_548645 | County of Santa Clara Public Health Department | Chan-Zuckerberg Biohub | CZB Cliahub Consortium |
| EPI_ISL_548646 | Orange County Public Health Laboratory | Chan-Zuckerberg Biohub | CZB Cliahub Consortium |
| EPI_ISL_548647 | County of Santa Clara Public Health Department | Chan-Zuckerberg Biohub | CZB Cliahub Consortium |

| | | | |
|---|---|---|---|
| EPI_ISL_548648 | Orange County Public Health Laboratory | Chan-Zuckerberg Biohub | CZB Cliahub Consortium |
| EPI_ISL_548649, EPI_ISL_548650 | County of Santa Clara Public Health Department | Chan-Zuckerberg Biohub | CZB Cliahub Consortium |
| EPI_ISL_548651 | Orange County Public Health Laboratory | Chan-Zuckerberg Biohub | CZB Cliahub Consortium |
| EPI_ISL_548652 | County of Santa Clara Public Health Department | Chan-Zuckerberg Biohub | CZB Cliahub Consortium |
| EPI_ISL_548653, EPI_ISL_548654 | Orange County Public Health Laboratory | Chan-Zuckerberg Biohub | CZB Cliahub Consortium |
| EPI_ISL_548655, EPI_ISL_548656, EPI_ISL_548657, EPI_ISL_548658, EPI_ISL_548659 | County of Santa Clara Public Health Department | Chan-Zuckerberg Biohub | CZB Cliahub Consortium |
| EPI_ISL_548660 | Orange County Public Health Laboratory | Chan-Zuckerberg Biohub | CZB Cliahub Consortium |
| EPI_ISL_548661 | County of Santa Clara Public Health Department | Chan-Zuckerberg Biohub | CZB Cliahub Consortium |
| EPI_ISL_548662 | Orange County Public Health Laboratory | Chan-Zuckerberg Biohub | CZB Cliahub Consortium |
| EPI_ISL_548663 | County of San Luis Obispo Public Health Laboratory | Chan-Zuckerberg Biohub | CZB Cliahub Consortium |
| EPI_ISL_548664 | County of Santa Clara Public Health Department | Chan-Zuckerberg Biohub | CZB Cliahub Consortium |
| EPI_ISL_548665, EPI_ISL_548666, EPI_ISL_548667, EPI_ISL_548668 | Orange County Public Health Laboratory | Chan-Zuckerberg Biohub | CZB Cliahub Consortium |
| EPI_ISL_548669, EPI_ISL_548670 | County of Santa Clara Public Health Department | Chan-Zuckerberg Biohub | CZB Cliahub Consortium |
| EPI_ISL_548671 | County of San Luis Obispo Public Health Laboratory | Chan-Zuckerberg Biohub | CZB Cliahub Consortium |
| EPI_ISL_548672, EPI_ISL_548673 | County of Santa Clara Public Health Department | Chan-Zuckerberg Biohub | CZB Cliahub Consortium |
| EPI_ISL_548674, EPI_ISL_548675 | Orange County Public Health Laboratory | Chan-Zuckerberg Biohub | CZB Cliahub Consortium |
| EPI_ISL_548676 | County of Santa Clara Public Health Department | Chan-Zuckerberg Biohub | CZB Cliahub Consortium |
| EPI_ISL_548677 | Orange County Public Health Laboratory | Chan-Zuckerberg Biohub | CZB Cliahub Consortium |
| EPI_ISL_548678, EPI_ISL_548679 | County of Santa Clara Public Health Department | Chan-Zuckerberg Biohub | CZB Cliahub Consortium |
| EPI_ISL_548687, EPI_ISL_548690, EPI_ISL_548691, EPI_ISL_548692, EPI_ISL_548693, EPI_ISL_548694, EPI_ISL_548695, EPI_ISL_548696, EPI_ISL_548697, EPI_ISL_548698, EPI_ISL_548699, EPI_ISL_548700, EPI_ISL_548701, EPI_ISL_548702, EPI_ISL_548703, EPI_ISL_548704, EPI_ISL_548705, EPI_ISL_548708, EPI_ISL_548709, EPI_ISL_548711, EPI_ISL_548712, EPI_ISL_548713, EPI_ISL_548714, EPI_ISL_548716, EPI_ISL_548717, EPI_ISL_548718, EPI_ISL_548720, EPI_ISL_548721, EPI_ISL_548722, EPI_ISL_548723, EPI_ISL_548724, EPI_ISL_548726, EPI_ISL_548727, EPI_ISL_548728, EPI_ISL_548729, EPI_ISL_548730, EPI_ISL_548731, EPI_ISL_548732, EPI_ISL_548733, EPI_ISL_548734, EPI_ISL_548735, EPI_ISL_548737, EPI_ISL_548738, EPI_ISL_548740, EPI_ISL_548741, EPI_ISL_548742, EPI_ISL_548743, EPI_ISL_548744, EPI_ISL_548745, EPI_ISL_548746, EPI_ISL_548747, EPI_ISL_548750, EPI_ISL_548751, EPI_ISL_548752, EPI_ISL_548753, EPI_ISL_548754, EPI_ISL_548755, EPI_ISL_548759, EPI_ISL_548760, EPI_ISL_548761, EPI_ISL_548762, EPI_ISL_548763, EPI_ISL_548764, EPI_ISL_548765, EPI_ISL_548766, EPI_ISL_548767, EPI_ISL_548768, EPI_ISL_548769, EPI_ISL_548770, EPI_ISL_548771, EPI_ISL_548772, EPI_ISL_548773, EPI_ISL_548774, EPI_ISL_548775, EPI_ISL_548776, EPI_ISL_548777, EPI_ISL_548778, EPI_ISL_548779, EPI_ISL_548780, EPI_ISL_548781, EPI_ISL_548782, EPI_ISL_548783, EPI_ISL_548784, EPI_ISL_548785, EPI_ISL_548786, EPI_ISL_548787, EPI_ISL_548788, EPI_ISL_548789, EPI_ISL_548790, EPI_ISL_548791, EPI_ISL_548792, EPI_ISL_548793, EPI_ISL_548794, EPI_ISL_548795, EPI_ISL_548796, EPI_ISL_548798, EPI_ISL_548799, EPI_ISL_548800, EPI_ISL_548801, EPI_ISL_548802, EPI_ISL_548803, EPI_ISL_548804, EPI_ISL_548805, EPI_ISL_548806, EPI_ISL_548807, EPI_ISL_548808, EPI_ISL_548809, EPI_ISL_548810, EPI_ISL_548811, EPI_ISL_548812, EPI_ISL_548813, EPI_ISL_548814, EPI_ISL_548815, EPI_ISL_548816, EPI_ISL_548817, EPI_ISL_548818, EPI_ISL_548819, EPI_ISL_548820, EPI_ISL_548821, EPI_ISL_548822, EPI_ISL_548823, EPI_ISL_548824, EPI_ISL_548825, EPI_ISL_548826, EPI_ISL_548827, EPI_ISL_548828, EPI_ISL_548829, EPI_ISL_548830, EPI_ISL_548831, EPI_ISL_548832, EPI_ISL_548833, EPI_ISL_548834, EPI_ISL_548835, EPI_ISL_548836, EPI_ISL_548837, EPI_ISL_548838, EPI_ISL_548839, EPI_ISL_548840, EPI_ISL_548841, EPI_ISL_548842, EPI_ISL_548843, EPI_ISL_548844, EPI_ISL_548845, EPI_ISL_548846, EPI_ISL_548847, EPI_ISL_548848, EPI_ISL_548849, EPI_ISL_548850, EPI_ISL_548851, EPI_ISL_548852, EPI_ISL_548853, EPI_ISL_548854, EPI_ISL_548855, EPI_ISL_548856, EPI_ISL_548857, EPI_ISL_548858, EPI_ISL_548859, EPI_ISL_548860, EPI_ISL_548861, EPI_ISL_548862, EPI_ISL_548864, EPI_ISL_548865, EPI_ISL_548866, EPI_ISL_548868, EPI_ISL_548869, EPI_ISL_548870, EPI_ISL_548871, EPI_ISL_548872, EPI_ISL_548873, EPI_ISL_548874, EPI_ISL_548875, EPI_ISL_548876, EPI_ISL_548877, EPI_ISL_548878, EPI_ISL_548879, EPI_ISL_548881, EPI_ISL_548884, EPI_ISL_548886, EPI_ISL_548887, EPI_ISL_548888, EPI_ISL_548889, EPI_ISL_548890, EPI_ISL_548891, EPI_ISL_548892, EPI_ISL_548893, EPI_ISL_548894, EPI_ISL_548895, EPI_ISL_548896, EPI_ISL_548898, EPI_ISL_548899, EPI_ISL_548900, EPI_ISL_548901, EPI_ISL_548902, EPI_ISL_548903, EPI_ISL_548904, EPI_ISL_548905, EPI_ISL_548906, EPI_ISL_548908, EPI_ISL_548909, EPI_ISL_548910, EPI_ISL_548911, EPI_ISL_548912, EPI_ISL_548913, EPI_ISL_548914, EPI_ISL_548915, EPI_ISL_548916, EPI_ISL_548917, EPI_ISL_548918, EPI_ISL_548919, EPI_ISL_548920, EPI_ISL_548921, EPI_ISL_548922, EPI_ISL_548923, EPI_ISL_548924, EPI_ISL_548925, EPI_ISL_548926, EPI_ISL_548927, EPI_ISL_548928, EPI_ISL_548929, EPI_ISL_548930, EPI_ISL_548931, EPI_ISL_548932, EPI_ISL_548933, EPI_ISL_548934, EPI_ISL_548935, EPI_ISL_548937, EPI_ISL_548938, EPI_ISL_548939, EPI_ISL_548940, EPI_ISL_548941 | | | |
| see above | Public Health Ontario Laboratory | Public Health Ontario Laboratory | Vanessa G Allen, Philip Banh, Richard de Borja, Yao Chen, Alireza Eshaghi, Nahuel Fittipaldi, Christine Frantz, Jonathan B Gubbay, Jennifer L Guthrie, Lawrence Heisler, Esha Joshi, Michael Laszloffy, Aimin Li, Michael CY Li, Dean Maxwell, Sandeep Nagra, Samir N Patel, Heather Rilkoff, Jared Simpson, Karthikeyan Sivaraman, Yogi Sundaravadanam, Sarah Teatero, Andre Villegas, Sandra Zittermann |
