## Supplementary material for "Analysis of the Dynamics and Distribution of SARS-CoV-2 Mutations and its Possible Structural and Functional Implications": GISAID_acknowledgements_part_11

We gratefully acknowledge the following Authors from the Originating laboratories responsible for obtaining the specimens, as well as the Submitting laboratories where the genome data were generated and shared via GISAID, on which this research is based.

All Submitters of data may be contacted directly via www.gisaid.org

| Accession ID | Originating Laboratory | Submitting Laboratory | Authors |
|---|---|---|---|
| EPI_ISL_560972 | Orebro klinisk mikrobiologi | The Public Health Agency of Sweden | Anna-Malin Linde, Maria Lind Karlberg, Mattias Haukland, Reza Advani, Olov Svartstrom, Oskar Karlsson Lindsjo, Sandra Broddesson, Petra Edquist, Mia Brytting, Anna Risberg, Karin Tegmark-Wisell |
| EPI_ISL_560973 | Klinisk mikrobiologi NAL Trollhattan | The Public Health Agency of Sweden | Anna-Malin Linde, Maria Lind Karlberg, Mattias Haukland, Reza Advani, Olov Svartstrom, Oskar Karlsson Lindsjo, Sandra Broddesson, Petra Edquist, Mia Brytting, Anna Risberg, Karin Tegmark-Wisell |
| EPI_ISL_560974 | Karolinska Universitetslaboratoriet | The Public Health Agency of Sweden | Anna-Malin Linde, Maria Lind Karlberg, Mattias Haukland, Reza Advani, Olov Svartstrom, Oskar Karlsson Lindsjo, Sandra Broddesson, Petra Edquist, Mia Brytting, Anna Risberg, Karin Tegmark-Wisell |
| EPI_ISL_560975 | Klinisk mikrobiologi centralsjukhuset Karlstad | The Public Health Agency of Sweden | Anna-Malin Linde, Maria Lind Karlberg, Mattias Haukland, Reza Advani, Olov Svartstrom, Oskar Karlsson Lindsjo, Sandra Broddesson, Petra Edquist, Mia Brytting, Anna Risberg, Karin Tegmark-Wisell |
| EPI_ISL_560976 | Klinsisk mikrobiologi Linkoping | The Public Health Agency of Sweden | Anna-Malin Linde, Maria Lind Karlberg, Mattias Haukland, Reza Advani, Olov Svartstrom, Oskar Karlsson Lindsjo, Sandra Broddesson, Petra Edquist, Mia Brytting, Anna Risberg, Karin Tegmark-Wisell |
| EPI_ISL_560977, EPI_ISL_560978, EPI_ISL_560979 | Universitetssjukhuset i Linkoping | The Public Health Agency of Sweden | Anna-Malin Linde, Maria Lind Karlberg, Mattias Haukland, Reza Advani, Olov Svartstrom, Oskar Karlsson Lindsjo, Sandra Broddesson, Petra Edquist, Mia Brytting, Anna Risberg, Karin Tegmark-Wisell |
| EPI_ISL_560980 | Skanes universitetssjukhus Lund | The Public Health Agency of Sweden | Anna-Malin Linde, Maria Lind Karlberg, Mattias Haukland, Reza Advani, Olov Svartstrom, Oskar Karlsson Lindsjo, Sandra Broddesson, Petra Edquist, Mia Brytting, Anna Risberg, Karin Tegmark-Wisell |
| EPI_ISL_560981 | Capio S:t Gorans sjukhus | The Public Health Agency of Sweden | Anna-Malin Linde, Maria Lind Karlberg, Mattias Haukland, Reza Advani, Olov Svartstrom, Oskar Karlsson Lindsjo, Sandra Broddesson, Petra Edquist, Mia Brytting, Anna Risberg, Karin Tegmark-Wisell |
| EPI_ISL_560982, EPI_ISL_560983, EPI_ISL_560984, EPI_ISL_560985 | Karolinska universitetslaboratoriet SOLNA | The Public Health Agency of Sweden | Anna-Malin Linde, Maria Lind Karlberg, Mattias Haukland, Reza Advani, Olov Svartstrom, Oskar Karlsson Lindsjo, Sandra Broddesson, Petra Edquist, Mia Brytting, Anna Risberg, Karin Tegmark-Wisell |
| EPI_ISL_560986 | Sundsvalls sjukhus | The Public Health Agency of Sweden | Anna-Malin Linde, Maria Lind Karlberg, Mattias Haukland, Reza Advani, Olov Svartstrom, Oskar Karlsson Lindsjo, Sandra Broddesson, Petra Edquist, Mia Brytting, Anna Risberg, Karin Tegmark-Wisell |
| EPI_ISL_560987 | Kliniskt mikrobiologiska laboratoriet | The Public Health Agency of Sweden | Anna-Malin Linde, Maria Lind Karlberg, Mattias Haukland, Reza Advani, Olov Svartstrom, Oskar Karlsson Lindsjo, Sandra Broddesson, Petra Edquist, Mia Brytting, Anna Risberg, Karin Tegmark-Wisell |
| EPI_ISL_560988 | Klinisk mikrobiologi, Laboratoriemedicin Gavleborg | The Public Health Agency of Sweden | Anna-Malin Linde, Maria Lind Karlberg, Mattias Haukland, Reza Advani, Olov Svartstrom, Oskar Karlsson Lindsjo, Sandra Broddesson, Petra Edquist, Mia Brytting, Anna Risberg, Karin Tegmark-Wisell |
| EPI_ISL_560989, EPI_ISL_560990 | Karolinska universitetslaboratoriet | The Public Health Agency of Sweden | Anna-Malin Linde, Maria Lind Karlberg, Mattias Haukland, Reza Advani, Olov Svartstrom, Oskar Karlsson Lindsjo, Sandra Broddesson, Petra Edquist, Mia Brytting, Anna Risberg, Karin Tegmark-Wisell |
| EPI_ISL_560991 | RSUD Dr. Soetomo | Institute of Tropical Disease, Universitas Airlangga | Rima R Prasetya, Krisnoadi Rahardjo, Aldise M Nastri, Jezzy R Dewantari, Joni Wahyuhadi, Gatot Soegiarto, Laksmi Wulandari, Retno A Setyoningrum, Resti Yudhawati, Yohko K Shimizu, Mitsuhiro Nishimura, Yasuko Mori, Soetjipto, Kazufumi Shimizu, Maria I Lusida |
| EPI_ISL_560992, EPI_ISL_560994, EPI_ISL_560995, EPI_ISL_560997, EPI_ISL_561000, EPI_ISL_561001, EPI_ISL_561003, EPI_ISL_561004, EPI_ISL_561006, EPI_ISL_561011, EPI_ISL_561012, EPI_ISL_561013, EPI_ISL_561021, EPI_ISL_561022, EPI_ISL_561034, EPI_ISL_561038, EPI_ISL_561040, EPI_ISL_561041, EPI_ISL_561042, EPI_ISL_561043, EPI_ISL_561044, EPI_ISL_561046, EPI_ISL_561047, EPI_ISL_561048, EPI_ISL_561049, EPI_ISL_561051, EPI_ISL_561052, EPI_ISL_561053, EPI_ISL_561054, EPI_ISL_561055, EPI_ISL_561056, EPI_ISL_561058, EPI_ISL_561059, EPI_ISL_561060, EPI_ISL_561061, EPI_ISL_561062, EPI_ISL_561063, EPI_ISL_561064, EPI_ISL_561065, EPI_ISL_561066, EPI_ISL_561067, EPI_ISL_561068, EPI_ISL_561069, EPI_ISL_561070, EPI_ISL_561071, EPI_ISL_561073, EPI_ISL_561074, EPI_ISL_561075, EPI_ISL_561076, EPI_ISL_561077, EPI_ISL_561079, EPI_ISL_561080, EPI_ISL_561081, EPI_ISL_561082, EPI_ISL_561083, EPI_ISL_561084, EPI_ISL_561085, EPI_ISL_561086, EPI_ISL_561087, EPI_ISL_561088, EPI_ISL_561089, EPI_ISL_561090, EPI_ISL_561091, EPI_ISL_561093, EPI_ISL_561094, EPI_ISL_561095, EPI_ISL_561096, EPI_ISL_561097, EPI_ISL_561098, EPI_ISL_561099, EPI_ISL_561100, EPI_ISL_561102, EPI_ISL_561105, EPI_ISL_561107, EPI_ISL_561108, EPI_ISL_561110, EPI_ISL_561111, EPI_ISL_561112, EPI_ISL_561113, EPI_ISL_561114, EPI_ISL_561115, EPI_ISL_561116, EPI_ISL_561117, EPI_ISL_561118, EPI_ISL_561119, EPI_ISL_561120, EPI_ISL_561121, EPI_ISL_561122, EPI_ISL_561123, EPI_ISL_561124, EPI_ISL_561125, EPI_ISL_561127, EPI_ISL_561129, EPI_ISL_561130, EPI_ISL_561131, EPI_ISL_561132, EPI_ISL_561133, EPI_ISL_561134, EPI_ISL_561135, EPI_ISL_561136, EPI_ISL_561137, EPI_ISL_561138, EPI_ISL_561139, EPI_ISL_561140, EPI_ISL_561141, EPI_ISL_561142, EPI_ISL_561143, EPI_ISL_561144, EPI_ISL_561145, EPI_ISL_561146, EPI_ISL_561148, EPI_ISL_561150, EPI_ISL_561151, EPI_ISL_561153, EPI_ISL_561154, EPI_ISL_561155, EPI_ISL_561156, EPI_ISL_561157, EPI_ISL_561158, EPI_ISL_561159, EPI_ISL_561160, EPI_ISL_561162, EPI_ISL_561163, EPI_ISL_561164, EPI_ISL_561165, EPI_ISL_561166, EPI_ISL_561167, EPI_ISL_561168, EPI_ISL_561169, EPI_ISL_561170, EPI_ISL_561171, EPI_ISL_561172, EPI_ISL_561173, EPI_ISL_561174, EPI_ISL_561176, EPI_ISL_561177, EPI_ISL_561178, EPI_ISL_561179, EPI_ISL_561180, EPI_ISL_561181, EPI_ISL_561182, EPI_ISL_561184, EPI_ISL_561185, EPI_ISL_561187, EPI_ISL_561188, EPI_ISL_561189, EPI_ISL_561190, EPI_ISL_561191, EPI_ISL_561193, EPI_ISL_561194, EPI_ISL_561195, EPI_ISL_561196, EPI_ISL_561198, EPI_ISL_561199, EPI_ISL_561200, EPI_ISL_561201, EPI_ISL_561202, EPI_ISL_561203, EPI_ISL_561204, EPI_ISL_561205, EPI_ISL_561206, EPI_ISL_561212, EPI_ISL_561213, EPI_ISL_561215, EPI_ISL_561218, EPI_ISL_561222, EPI_ISL_561223, EPI_ISL_561229, EPI_ISL_561230, EPI_ISL_561232, EPI_ISL_561236, EPI_ISL_561237, EPI_ISL_561238, EPI_ISL_561239, EPI_ISL_561240, EPI_ISL_561241, EPI_ISL_561242, EPI_ISL_561243, EPI_ISL_561244, EPI_ISL_561245, EPI_ISL_561247, EPI_ISL_561248, EPI_ISL_561249, EPI_ISL_561251, EPI_ISL_561257, EPI_ISL_561260, EPI_ISL_561263, EPI_ISL_561268, EPI_ISL_561270, EPI_ISL_561273, EPI_ISL_561277, EPI_ISL_561279, EPI_ISL_561282, EPI_ISL_561283, EPI_ISL_561284, EPI_ISL_561285, EPI_ISL_561287, EPI_ISL_561288, EPI_ISL_561289, EPI_ISL_561290, EPI_ISL_561291, EPI_ISL_561292, EPI_ISL_561293, EPI_ISL_561294, EPI_ISL_561295, EPI_ISL_561296, EPI_ISL_561297, EPI_ISL_561298, EPI_ISL_561299, EPI_ISL_561300, EPI_ISL_561303, EPI_ISL_561304, EPI_ISL_561305, EPI_ISL_561306, EPI_ISL_561307, EPI_ISL_561308, EPI_ISL_561309, EPI_ISL_561310, EPI_ISL_561311, EPI_ISL_561313, EPI_ISL_561314, EPI_ISL_561315, EPI_ISL_561324, EPI_ISL_561327, EPI_ISL_561330, EPI_ISL_561332, EPI_ISL_561333 | | | |
| see above | MRCG at LSHTM Genomics lab | MRCG at LSHTM Genomics lab | Abdul Karim sesay, Abdoulie Kanteh, Jarra Manneh, Mariama Kujabi, Bakary Sanyang |
| EPI_ISL_561335 | Delaware Public Health Lab | Delaware Public Health Lab | Gregory Hovan |
| EPI_ISL_561339, EPI_ISL_561340, EPI_ISL_561341, EPI_ISL_561343 | Civil Hospital, Panchkula | CSIR-Institute of Microbial Technology | Kanika Bansal, Sanjeet Kumar, Anu Singh, Debarghya Ghose, Rajesh Kumar Mishra, Dipak Dutta, Sanjeev Khosla, Prabhu B. Patil |
| EPI_ISL_561344 | Civil Hospital, Rupnagar | CSIR-Institute of Microbial Technology | Kanika Bansal, Sanjeet Kumar, Anu Singh, Debarghya Ghose, Amandeep Kaur, Rajesh Kumar Mishra, Poushali Chakraborty, Harsh Goar, Navin Baid, Ashwani Kumar, Dipak Dutta, Sanjeev Khosla, Prabhu B. Patil |
| EPI_ISL_561345, EPI_ISL_561346, EPI_ISL_561351, EPI_ISL_561352, EPI_ISL_561353, EPI_ISL_561356, EPI_ISL_561358 | Delaware Public Health Lab | Delaware Public Health Lab | Gregory Hovan |
| EPI_ISL_561370, EPI_ISL_561371, EPI_ISL_561372 | Hospital Universitario de Gran Canaria Dr. Negrín | SeqCOVID-SPAIN consortium/IBV(CSIC) | M. Carmen Pérez González, Francisco J. Chamizo López, Ana Bordes Benítez and SeqCOVID-SPAIN consortium |
| EPI_ISL_561373 | Centre for Clinical Infection and Diagnostics Research and Genomics Innovation Unit, Guy's and St. Thomas' NHS Trust | Centre for Clinical Infection and Diagnostics Research and Genomics Innovation Unit, Guy's and St. Thomas' NHS Trust | Chloe Fisher, Luke Snell, Rahul Batra, Jonathan Edgeworth, Ali Raza Awan |
| EPI_ISL_562168, EPI_ISL_562169, EPI_ISL_562170, EPI_ISL_562171, EPI_ISL_562172, EPI_ISL_562173, EPI_ISL_562174, EPI_ISL_562591, EPI_ISL_562592 | Microbiological Diagnostic Unit - Public Health Laboratory (MDU-PHL) | MDU-PHL | Seemann, T., Schultz M. B., Sait, M., Sherry, N. |
| EPI_ISL_562593 | Victorian Infectious Diseases Reference Laboratory (VIDRL) | VIDRL and MDU-PHL | Caly, L., Seemann, T., Sait, M., Schultz, M. B., Druce J., Sherry, N. |
| EPI_ISL_562594 | Microbiological Diagnostic Unit - Public Health Laboratory (MDU-PHL) | MDU-PHL | Seemann, T., Schultz M. B., Sait, M., Sherry, N. |
| EPI_ISL_562595 | Victorian Infectious Diseases Reference Laboratory (VIDRL) | VIDRL and MDU-PHL | Caly, L., Seemann, T., Sait, M., Schultz, M. B., Druce J., Sherry, N. |
| EPI_ISL_562596, EPI_ISL_562597, EPI_ISL_562598, EPI_ISL_562599, EPI_ISL_562600, EPI_ISL_562601, EPI_ISL_562602, EPI_ISL_562603, EPI_ISL_562604, EPI_ISL_562605, EPI_ISL_562606, EPI_ISL_562607, EPI_ISL_562608, EPI_ISL_562609, EPI_ISL_562610, EPI_ISL_562611, EPI_ISL_562612, EPI_ISL_562613, EPI_ISL_562614, EPI_ISL_562615, EPI_ISL_562616, EPI_ISL_562617, EPI_ISL_562618, EPI_ISL_562619, EPI_ISL_562620, EPI_ISL_562621, EPI_ISL_562622, EPI_ISL_562623, EPI_ISL_562624, EPI_ISL_562625, EPI_ISL_562626, EPI_ISL_562627 | | | |
| see above | Microbiological Diagnostic Unit - Public Health Laboratory (MDU-PHL) | MDU-PHL | Seemann, T., Schultz M. B., Sait, M., Sherry, N. |
| EPI_ISL_562628 | Victorian Infectious Diseases Reference Laboratory (VIDRL) | VIDRL and MDU-PHL | Caly, L., Seemann, T., Sait, M., Schultz, M. B., Druce J., Sherry, N. |
| EPI_ISL_562629, EPI_ISL_562630, EPI_ISL_562631, EPI_ISL_562632, EPI_ISL_562633, EPI_ISL_562634, EPI_ISL_562635, EPI_ISL_562636, EPI_ISL_562637, EPI_ISL_562638, EPI_ISL_562639 | | | |
| see above | Microbiological Diagnostic Unit - Public Health Laboratory (MDU-PHL) | MDU-PHL | Seemann, T., Schultz M. B., Sait, M., Sherry, N. |
| EPI_ISL_562640 | Victorian Infectious Diseases Reference Laboratory (VIDRL) | VIDRL and MDU-PHL | Caly, L., Seemann, T., Sait, M., Schultz, M. B., Druce J., Sherry, N. |
| EPI_ISL_562641, EPI_ISL_562642, EPI_ISL_562643, EPI_ISL_562644 | Microbiological Diagnostic Unit - Public Health Laboratory (MDU-PHL) | MDU-PHL | Seemann, T., Schultz M. B., Sait, M., Sherry, N. |
| EPI_ISL_562645, EPI_ISL_562646 | Victorian Infectious Diseases Reference Laboratory (VIDRL) | VIDRL and MDU-PHL | Caly, L., Seemann, T., Sait, M., Schultz, M. B., Druce J., Sherry, N. |
| EPI_ISL_562647, EPI_ISL_562648, EPI_ISL_562649, EPI_ISL_562650, EPI_ISL_562651, EPI_ISL_562652, EPI_ISL_562653, EPI_ISL_562654, EPI_ISL_562655, EPI_ISL_562656, EPI_ISL_562657 | | | |
| see above | Microbiological Diagnostic Unit - Public Health Laboratory (MDU-PHL) | MDU-PHL | Seemann, T., Schultz M. B., Sait, M., Sherry, N. |
| EPI_ISL_562658, EPI_ISL_562659 | Victorian Infectious Diseases Reference Laboratory (VIDRL) | VIDRL and MDU-PHL | Caly, L., Seemann, T., Sait, M., Schultz, M. B., Druce J., Sherry, N. |
| EPI_ISL_562660, EPI_ISL_562661, EPI_ISL_562662, EPI_ISL_562663, EPI_ISL_562664, EPI_ISL_562665, EPI_ISL_562666, EPI_ISL_562667, EPI_ISL_562668 | Microbiological Diagnostic Unit - Public Health Laboratory (MDU-PHL) | MDU-PHL | Seemann, T., Schultz M. B., Sait, M., Sherry, N. |
| EPI_ISL_562669 | Victorian Infectious Diseases Reference Laboratory (VIDRL) | VIDRL and MDU-PHL | Caly, L., Seemann, T., Sait, M., Schultz, M. B., Druce J., Sherry, N. |
| EPI_ISL_562670 | Microbiological Diagnostic Unit - Public Health Laboratory (MDU-PHL) | MDU-PHL | Seemann, T., Schultz M. B., Sait, M., Sherry, N. |
| EPI_ISL_562671 | Victorian Infectious Diseases Reference Laboratory (VIDRL) | VIDRL and MDU-PHL | Caly, L., Seemann, T., Sait, M., Schultz, M. B., Druce J., Sherry, N. |
| EPI_ISL_562672, EPI_ISL_562673, EPI_ISL_562674, EPI_ISL_562675, EPI_ISL_562676, EPI_ISL_562677, EPI_ISL_562678, EPI_ISL_562679, EPI_ISL_562680, EPI_ISL_562681, EPI_ISL_562682, EPI_ISL_562683, EPI_ISL_562684, EPI_ISL_562685, EPI_ISL_562686, EPI_ISL_562687, EPI_ISL_562688, EPI_ISL_562691, EPI_ISL_562692, EPI_ISL_562693, EPI_ISL_562694, EPI_ISL_562695 | | | |
| see above | Microbiological Diagnostic Unit - Public Health Laboratory (MDU-PHL) | MDU-PHL | Seemann, T., Schultz M. B., Sait, M., Sherry, N. |
| EPI_ISL_562696, EPI_ISL_562697 | Victorian Infectious Diseases Reference Laboratory (VIDRL) | VIDRL and MDU-PHL | Caly, L., Seemann, T., Sait, M., Schultz, M. B., Druce J., Sherry, N. |

| | | | |
|---|---|---|---|
| EPI_ISL_562698 | Microbiological Diagnostic Unit - Public Health Laboratory (MDU-PHL) | MDU-PHL | Seemann, T., Schultz M. B., Sait, M., Sherry, N. |
| EPI_ISL_562700 | Victorian Infectious Diseases Reference Laboratory (VIDRL) | VIDRL and MDU-PHL | Caly, L., Seemann, T., Sait, M., Schultz, M. B., Druce J., Sherry, N. |
| EPI_ISL_562701, EPI_ISL_562702, EPI_ISL_562703, EPI_ISL_562704 | Microbiological Diagnostic Unit - Public Health Laboratory (MDU-PHL) | MDU-PHL | Seemann, T., Schultz M. B., Sait, M., Sherry, N. |
| EPI_ISL_562705 | Victorian Infectious Diseases Reference Laboratory (VIDRL) | VIDRL and MDU-PHL | Caly, L., Seemann, T., Sait, M., Schultz, M. B., Druce J., Sherry, N. |
| EPI_ISL_562706, EPI_ISL_562707, EPI_ISL_562708, EPI_ISL_562728, EPI_ISL_562729, EPI_ISL_562730, EPI_ISL_562731, EPI_ISL_562732, EPI_ISL_562733, EPI_ISL_562734, EPI_ISL_562735, EPI_ISL_562736, EPI_ISL_562737, EPI_ISL_562738, EPI_ISL_562739, EPI_ISL_562740, EPI_ISL_562741, EPI_ISL_562742, EPI_ISL_562743, EPI_ISL_562744, EPI_ISL_562745, EPI_ISL_562746, EPI_ISL_562747, EPI_ISL_562748, EPI_ISL_562749, EPI_ISL_562750, EPI_ISL_562751, EPI_ISL_562752, EPI_ISL_562753, EPI_ISL_562761, EPI_ISL_562762, EPI_ISL_562763, EPI_ISL_562764, EPI_ISL_562765, EPI_ISL_562766, EPI_ISL_562767, EPI_ISL_562768, EPI_ISL_562769, EPI_ISL_562770, EPI_ISL_562771, EPI_ISL_562772, EPI_ISL_562773, EPI_ISL_562774, EPI_ISL_562775, EPI_ISL_562776, EPI_ISL_562777, EPI_ISL_562778, EPI_ISL_562779, EPI_ISL_562780, EPI_ISL_562781, EPI_ISL_562782, EPI_ISL_562783, EPI_ISL_562784, EPI_ISL_562785, EPI_ISL_562786, EPI_ISL_562787, EPI_ISL_562788, EPI_ISL_562789, EPI_ISL_562790 | | | |
| see above | Microbiological Diagnostic Unit - Public Health Laboratory (MDU-PHL) | MDU-PHL | Seemann, T., Schultz M. B., Sait, M., Sherry, N. |
| EPI_ISL_562791, EPI_ISL_562792 | Victorian Infectious Diseases Reference Laboratory (VIDRL) | VIDRL and MDU-PHL | Caly, L., Seemann, T., Sait, M., Schultz, M. B., Druce J., Sherry, N. |
| EPI_ISL_562793, EPI_ISL_562794, EPI_ISL_562795, EPI_ISL_562796, EPI_ISL_562797, EPI_ISL_562798, EPI_ISL_562799, EPI_ISL_562800, EPI_ISL_562801, EPI_ISL_562802 | Microbiological Diagnostic Unit - Public Health Laboratory (MDU-PHL) | MDU-PHL | Seemann, T., Schultz M. B., Sait, M., Sherry, N. |
| EPI_ISL_562803, EPI_ISL_562804, EPI_ISL_562805, EPI_ISL_562806, EPI_ISL_562807, EPI_ISL_562808, EPI_ISL_562809 | Victorian Infectious Diseases Reference Laboratory (VIDRL) | VIDRL and MDU-PHL | Caly, L., Seemann, T., Sait, M., Schultz, M. B., Druce J., Sherry, N. |
| EPI_ISL_562810, EPI_ISL_562811, EPI_ISL_562812, EPI_ISL_562813, EPI_ISL_562814, EPI_ISL_562815, EPI_ISL_562816, EPI_ISL_562817, EPI_ISL_562818, EPI_ISL_562819, EPI_ISL_562820, EPI_ISL_562821, EPI_ISL_562822, EPI_ISL_562823, EPI_ISL_562824, EPI_ISL_562825, EPI_ISL_562826, EPI_ISL_562827, EPI_ISL_562828, EPI_ISL_562829, EPI_ISL_562830, EPI_ISL_562831, EPI_ISL_562832, EPI_ISL_562833, EPI_ISL_562834, EPI_ISL_562835, EPI_ISL_562836, EPI_ISL_562837 | | | |
| see above | Microbiological Diagnostic Unit - Public Health Laboratory (MDU-PHL) | MDU-PHL | Seemann, T., Schultz M. B., Sait, M., Sherry, N. |
| EPI_ISL_562838, EPI_ISL_562839, EPI_ISL_562840, EPI_ISL_562841, EPI_ISL_562842, EPI_ISL_562843, EPI_ISL_562844, EPI_ISL_562845 | Victorian Infectious Diseases Reference Laboratory (VIDRL) | VIDRL and MDU-PHL | Caly, L., Seemann, T., Sait, M., Schultz, M. B., Druce J., Sherry, N. |
| EPI_ISL_562846, EPI_ISL_562847, EPI_ISL_562848, EPI_ISL_562849, EPI_ISL_562850, EPI_ISL_562851, EPI_ISL_562852, EPI_ISL_562853, EPI_ISL_562854, EPI_ISL_562855, EPI_ISL_562856, EPI_ISL_562857, EPI_ISL_562858, EPI_ISL_562859, EPI_ISL_562860, EPI_ISL_562861, EPI_ISL_562862, EPI_ISL_562863, EPI_ISL_562864, EPI_ISL_562865, EPI_ISL_562866, EPI_ISL_562867, EPI_ISL_562868, EPI_ISL_562869, EPI_ISL_562870, EPI_ISL_562871, EPI_ISL_562872, EPI_ISL_562873, EPI_ISL_562874, EPI_ISL_562875, EPI_ISL_562876, EPI_ISL_562877, EPI_ISL_562878, EPI_ISL_562879, EPI_ISL_562880, EPI_ISL_562881, EPI_ISL_562882, EPI_ISL_562883, EPI_ISL_562884, EPI_ISL_562885, EPI_ISL_562886, EPI_ISL_562887, EPI_ISL_562888, EPI_ISL_562889, EPI_ISL_562890, EPI_ISL_562891, EPI_ISL_562892, EPI_ISL_562893, EPI_ISL_562894, EPI_ISL_562895, EPI_ISL_562896, EPI_ISL_562897, EPI_ISL_562898, EPI_ISL_562899, EPI_ISL_562900, EPI_ISL_562901, EPI_ISL_562902, EPI_ISL_562903, EPI_ISL_562904, EPI_ISL_562905, EPI_ISL_562906, EPI_ISL_562907, EPI_ISL_562908, EPI_ISL_562909 | | | |
| see above | Microbiological Diagnostic Unit - Public Health Laboratory (MDU-PHL) | MDU-PHL | Seemann, T., Schultz M. B., Sait, M., Sherry, N. |
| EPI_ISL_562910, EPI_ISL_562911, EPI_ISL_562912, EPI_ISL_562913 | Victorian Infectious Diseases Reference Laboratory (VIDRL) | VIDRL and MDU-PHL | Caly, L., Seemann, T., Sait, M., Schultz, M. B., Druce J., Sherry, N. |
| EPI_ISL_562914, EPI_ISL_562915, EPI_ISL_562916, EPI_ISL_562917, EPI_ISL_562918, EPI_ISL_562919, EPI_ISL_562920, EPI_ISL_562921, EPI_ISL_562922, EPI_ISL_562923 | Microbiological Diagnostic Unit - Public Health Laboratory (MDU-PHL) | MDU-PHL | Seemann, T., Schultz M. B., Sait, M., Sherry, N. |
| EPI_ISL_562924 | Victorian Infectious Diseases Reference Laboratory (VIDRL) | VIDRL and MDU-PHL | Caly, L., Seemann, T., Sait, M., Schultz, M. B., Druce J., Sherry, N. |
| EPI_ISL_562925, EPI_ISL_562926, EPI_ISL_562927, EPI_ISL_562928 | Microbiological Diagnostic Unit - Public Health Laboratory (MDU-PHL) | MDU-PHL | Seemann, T., Schultz M. B., Sait, M., Sherry, N. |
| EPI_ISL_562929 | Victorian Infectious Diseases Reference Laboratory (VIDRL) | VIDRL and MDU-PHL | Caly, L., Seemann, T., Sait, M., Schultz, M. B., Druce J., Sherry, N. |
| EPI_ISL_562930, EPI_ISL_562931, EPI_ISL_562932, EPI_ISL_562933, EPI_ISL_562934, EPI_ISL_562935 | Microbiological Diagnostic Unit - Public Health Laboratory (MDU-PHL) | MDU-PHL | Seemann, T., Schultz M. B., Sait, M., Sherry, N. |
| EPI_ISL_562936 | Victorian Infectious Diseases Reference Laboratory (VIDRL) | VIDRL and MDU-PHL | Caly, L., Seemann, T., Sait, M., Schultz, M. B., Druce J., Sherry, N. |
| EPI_ISL_562937, EPI_ISL_562938, EPI_ISL_562939, EPI_ISL_562940, EPI_ISL_562941, EPI_ISL_562942, EPI_ISL_562943, EPI_ISL_562944, EPI_ISL_562945 | Microbiological Diagnostic Unit - Public Health Laboratory (MDU-PHL) | MDU-PHL | Seemann, T., Schultz M. B., Sait, M., Sherry, N. |
| EPI_ISL_562946 | Victorian Infectious Diseases Reference Laboratory (VIDRL) | VIDRL and MDU-PHL | Caly, L., Seemann, T., Sait, M., Schultz, M. B., Druce J., Sherry, N. |
| EPI_ISL_562947, EPI_ISL_562948, EPI_ISL_562949, EPI_ISL_562950 | Microbiological Diagnostic Unit - Public Health Laboratory (MDU-PHL) | MDU-PHL | Seemann, T., Schultz M. B., Sait, M., Sherry, N. |
| EPI_ISL_562951 | Victorian Infectious Diseases Reference Laboratory (VIDRL) | VIDRL and MDU-PHL | Caly, L., Seemann, T., Sait, M., Schultz, M. B., Druce J., Sherry, N. |
| EPI_ISL_562952, EPI_ISL_562953, EPI_ISL_562954, EPI_ISL_562955, EPI_ISL_562956, EPI_ISL_562957, EPI_ISL_562958, EPI_ISL_562959, EPI_ISL_562960 | Microbiological Diagnostic Unit - Public Health Laboratory (MDU-PHL) | MDU-PHL | Seemann, T., Schultz M. B., Sait, M., Sherry, N. |
| EPI_ISL_562961 | Victorian Infectious Diseases Reference Laboratory (VIDRL) | VIDRL and MDU-PHL | Caly, L., Seemann, T., Sait, M., Schultz, M. B., Druce J., Sherry, N. |
| EPI_ISL_562962, EPI_ISL_562963, EPI_ISL_562964, EPI_ISL_562965, EPI_ISL_562966, EPI_ISL_562967, EPI_ISL_562968, EPI_ISL_562969, EPI_ISL_562970, EPI_ISL_562971, EPI_ISL_562972, EPI_ISL_562973, EPI_ISL_562974, EPI_ISL_562975, EPI_ISL_562976, EPI_ISL_562977, EPI_ISL_562978, EPI_ISL_562979, EPI_ISL_562980, EPI_ISL_562981, EPI_ISL_562982, EPI_ISL_562983, EPI_ISL_562984, EPI_ISL_562985, EPI_ISL_562986, EPI_ISL_562987, EPI_ISL_562988, EPI_ISL_562989, EPI_ISL_562990, EPI_ISL_562991, EPI_ISL_562992, EPI_ISL_562993, EPI_ISL_562994, EPI_ISL_562995, EPI_ISL_562996, EPI_ISL_562997, EPI_ISL_562998, EPI_ISL_562999 | | | |
| see above | Microbiological Diagnostic Unit - Public Health Laboratory (MDU-PHL) | MDU-PHL | Seemann, T., Schultz M. B., Sait, M., Sherry, N. |
| EPI_ISL_563000 | Victorian Infectious Diseases Reference Laboratory (VIDRL) | VIDRL and MDU-PHL | Caly, L., Seemann, T., Sait, M., Schultz, M. B., Druce J., Sherry, N. |
| EPI_ISL_563001, EPI_ISL_563002, EPI_ISL_563003, EPI_ISL_563004, EPI_ISL_563005, EPI_ISL_563006, EPI_ISL_563007, EPI_ISL_563008, EPI_ISL_563009, EPI_ISL_563010, EPI_ISL_563011, EPI_ISL_563012, EPI_ISL_563013, EPI_ISL_563014, EPI_ISL_563015, EPI_ISL_563016, EPI_ISL_563017, EPI_ISL_563018, EPI_ISL_563019, EPI_ISL_563020, EPI_ISL_563021, EPI_ISL_563022, EPI_ISL_563023, EPI_ISL_563024, EPI_ISL_563025, EPI_ISL_563026, EPI_ISL_563027, EPI_ISL_563028, EPI_ISL_563029, EPI_ISL_563030, EPI_ISL_563031, EPI_ISL_563032, EPI_ISL_563033, EPI_ISL_563034, EPI_ISL_563035, EPI_ISL_563036, EPI_ISL_563037, EPI_ISL_563038, EPI_ISL_563039, EPI_ISL_563040, EPI_ISL_563041, EPI_ISL_563042, EPI_ISL_563043, EPI_ISL_563044, EPI_ISL_563045, EPI_ISL_563046, EPI_ISL_563047, EPI_ISL_563048, EPI_ISL_563049, EPI_ISL_563050, EPI_ISL_563051, EPI_ISL_563052, EPI_ISL_563053, EPI_ISL_563054, EPI_ISL_563055, EPI_ISL_563056, EPI_ISL_563057, EPI_ISL_563058, EPI_ISL_563059, EPI_ISL_563060, EPI_ISL_563061, EPI_ISL_563062, EPI_ISL_563063, EPI_ISL_563064, EPI_ISL_563065, EPI_ISL_563066, EPI_ISL_563067, EPI_ISL_563068, EPI_ISL_563069, EPI_ISL_563070, EPI_ISL_563071, EPI_ISL_563072, EPI_ISL_563073, EPI_ISL_563074, EPI_ISL_563075, EPI_ISL_563076, EPI_ISL_563077, EPI_ISL_563078, EPI_ISL_563079, EPI_ISL_563080, EPI_ISL_563081, EPI_ISL_563082, EPI_ISL_563083, EPI_ISL_563084, EPI_ISL_563085, EPI_ISL_563086, EPI_ISL_563087, EPI_ISL_563088, EPI_ISL_563089, EPI_ISL_563090, EPI_ISL_563091, EPI_ISL_563092, EPI_ISL_563093, EPI_ISL_563094, EPI_ISL_563095, EPI_ISL_563096, EPI_ISL_563097, EPI_ISL_563098, EPI_ISL_563099, EPI_ISL_563100, EPI_ISL_563101, EPI_ISL_563102, EPI_ISL_563103, EPI_ISL_563104, EPI_ISL_563105, EPI_ISL_563106, EPI_ISL_563107, EPI_ISL_563108, EPI_ISL_563109, EPI_ISL_563110, EPI_ISL_563111, EPI_ISL_563112, EPI_ISL_563113, EPI_ISL_563114, EPI_ISL_563115, EPI_ISL_563116, EPI_ISL_563117, EPI_ISL_563118, EPI_ISL_563119, EPI_ISL_563120, EPI_ISL_563121, EPI_ISL_563122, EPI_ISL_563123, EPI_ISL_563124, EPI_ISL_563125, EPI_ISL_563126, EPI_ISL_563127, EPI_ISL_563128, EPI_ISL_563129, EPI_ISL_563130 | | | |
| see above | Microbiological Diagnostic Unit - Public Health Laboratory (MDU-PHL) | MDU-PHL | Seemann, T., Schultz M. B., Sait, M., Sherry, N. |
| EPI_ISL_563131, EPI_ISL_563132, EPI_ISL_563133 | Victorian Infectious Diseases Reference Laboratory (VIDRL) | VIDRL and MDU-PHL | Caly, L., Seemann, T., Sait, M., Schultz, M. B., Druce J., Sherry, N. |
| EPI_ISL_563134, EPI_ISL_563135 | Microbiological Diagnostic Unit - Public Health Laboratory (MDU-PHL) | MDU-PHL | Seemann, T., Schultz M. B., Sait, M., Sherry, N. |
| EPI_ISL_563136, EPI_ISL_563137, EPI_ISL_563138, EPI_ISL_563139, EPI_ISL_563140, EPI_ISL_563141, EPI_ISL_563142, EPI_ISL_563143, EPI_ISL_563144 | Victorian Infectious Diseases Reference Laboratory (VIDRL) | VIDRL and MDU-PHL | Caly, L., Seemann, T., Sait, M., Schultz, M. B., Druce J., Sherry, N. |
| EPI_ISL_563145, EPI_ISL_563146, EPI_ISL_563147, EPI_ISL_563148, EPI_ISL_563149, EPI_ISL_563150, EPI_ISL_563151, EPI_ISL_563152, EPI_ISL_563153, EPI_ISL_563154, EPI_ISL_563155, EPI_ISL_563156, EPI_ISL_563157, EPI_ISL_563158, EPI_ISL_563159, EPI_ISL_563160, EPI_ISL_563161, EPI_ISL_563162, EPI_ISL_563163, EPI_ISL_563164, EPI_ISL_563165, EPI_ISL_563166, EPI_ISL_563167, EPI_ISL_563168, EPI_ISL_563169, EPI_ISL_563170, EPI_ISL_563171, EPI_ISL_563172, EPI_ISL_563173, EPI_ISL_563174, EPI_ISL_563175, EPI_ISL_563176, EPI_ISL_563177, EPI_ISL_563178, EPI_ISL_563179, EPI_ISL_563180, EPI_ISL_563181, EPI_ISL_563182, EPI_ISL_563183, EPI_ISL_563184, EPI_ISL_563185, EPI_ISL_563186, EPI_ISL_563187, EPI_ISL_563188, EPI_ISL_563189, EPI_ISL_563190, EPI_ISL_563191, EPI_ISL_563192, EPI_ISL_563193, EPI_ISL_563194, EPI_ISL_563195, EPI_ISL_563196, EPI_ISL_563197, EPI_ISL_563198, EPI_ISL_563199, EPI_ISL_563200, EPI_ISL_563201, EPI_ISL_563202, EPI_ISL_563203, EPI_ISL_563204, EPI_ISL_563205, EPI_ISL_563206, EPI_ISL_563207, EPI_ISL_563208, EPI_ISL_563209, EPI_ISL_563210, EPI_ISL_563211, EPI_ISL_563212, EPI_ISL_563213, EPI_ISL_563214, EPI_ISL_563215, EPI_ISL_563216, EPI_ISL_563217, EPI_ISL_563218, EPI_ISL_563219, EPI_ISL_563220, EPI_ISL_563221, EPI_ISL_563222, EPI_ISL_563223, EPI_ISL_563224, EPI_ISL_563225, EPI_ISL_563226, EPI_ISL_563227, EPI_ISL_563228, EPI_ISL_563229, EPI_ISL_563230, EPI_ISL_563231, EPI_ISL_563232, EPI_ISL_563233, EPI_ISL_563234, EPI_ISL_563235, EPI_ISL_563236, EPI_ISL_563237, EPI_ISL_563238, EPI_ISL_563239, EPI_ISL_563240, EPI_ISL_563241, EPI_ISL_563242, EPI_ISL_563243, EPI_ISL_563244, EPI_ISL_563245, EPI_ISL_563246, EPI_ISL_563247, EPI_ISL_563248, EPI_ISL_563249, EPI_ISL_563250, EPI_ISL_563251, EPI_ISL_563252, EPI_ISL_563253, EPI_ISL_563254, EPI_ISL_563255, EPI_ISL_563256, EPI_ISL_563257, EPI_ISL_563258, EPI_ISL_563259, EPI_ISL_563260, EPI_ISL_563261, EPI_ISL_563262, EPI_ISL_563263, EPI_ISL_563264, EPI_ISL_563265, EPI_ISL_563266, EPI_ISL_563267, EPI_ISL_563268, EPI_ISL_563269, EPI_ISL_563270, EPI_ISL_563271, EPI_ISL_563272, EPI_ISL_563273, EPI_ISL_563274, EPI_ISL_563275, EPI_ISL_563276, EPI_ISL_563277, EPI_ISL_563278, EPI_ISL_563279, EPI_ISL_563280, EPI_ISL_563281, EPI_ISL_563282, EPI_ISL_563283, EPI_ISL_563284, EPI_ISL_563285, EPI_ISL_563286, EPI_ISL_563287, EPI_ISL_563288, EPI_ISL_563289, EPI_ISL_563290, EPI_ISL_563291, EPI_ISL_563292, EPI_ISL_563293, EPI_ISL_563294, EPI_ISL_563295, EPI_ISL_563296, EPI_ISL_563297, EPI_ISL_563298, EPI_ISL_563299, EPI_ISL_563300, EPI_ISL_563301, EPI_ISL_563302, EPI_ISL_563303, EPI_ISL_563304, EPI_ISL_563305, EPI_ISL_563306, EPI_ISL_563307, EPI_ISL_563308, EPI_ISL_563309, EPI_ISL_563310, EPI_ISL_563311, EPI_ISL_563312, EPI_ISL_563313, EPI_ISL_563314, EPI_ISL_563315, EPI_ISL_563316, EPI_ISL_563317, EPI_ISL_563318, EPI_ISL_563319, EPI_ISL_563320, EPI_ISL_563321, EPI_ISL_563322, EPI_ISL_563323, EPI_ISL_563324, EPI_ISL_563325, EPI_ISL_563326, EPI_ISL_563327, EPI_ISL_563328, EPI_ISL_563329, EPI_ISL_563330, EPI_ISL_563331, EPI_ISL_563332, EPI_ISL_563333, EPI_ISL_563334, EPI_ISL_563335, EPI_ISL_563336, EPI_ISL_563337, EPI_ISL_563338, EPI_ISL_563339, EPI_ISL_563340, EPI_ISL_563341, EPI_ISL_563342, EPI_ISL_563343, EPI_ISL_563344, EPI_ISL_563345, EPI_ISL_563346, EPI_ISL_563347, EPI_ISL_563348, EPI_ISL_563349, EPI_ISL_563350, EPI_ISL_563351, EPI_ISL_563352, EPI_ISL_563353, EPI_ISL_563354, EPI_ISL_563355, EPI_ISL_563356, EPI_ISL_563357, EPI_ISL_563358, EPI_ISL_563359, EPI_ISL_563360, EPI_ISL_563361, EPI_ISL_563362, EPI_ISL_563363, EPI_ISL_563364, EPI_ISL_563365, EPI_ISL_563366, EPI_ISL_563367, EPI_ISL_563368, EPI_ISL_563369, EPI_ISL_563370, EPI_ISL_563371, EPI_ISL_563372, EPI_ISL_563373, EPI_ISL_563374, EPI_ISL_563375, EPI_ISL_563376, EPI_ISL_563377, EPI_ISL_563378, EPI_ISL_563379, EPI_ISL_563380, EPI_ISL_563381, EPI_ISL_563382, EPI_ISL_563383, EPI_ISL_563384, EPI_ISL_563385, EPI_ISL_563386, EPI_ISL_563387, EPI_ISL_563388, EPI_ISL_563389, EPI_ISL_563390, EPI_ISL_563391, EPI_ISL_563392, EPI_ISL_563393, EPI_ISL_563394, EPI_ISL_563395, EPI_ISL_563396, EPI_ISL_563397, EPI_ISL_563398, EPI_ISL_563399, EPI_ISL_563400, EPI_ISL_563401, EPI_ISL_563402, EPI_ISL_563403, EPI_ISL_563404, EPI_ISL_563405, EPI_ISL_563406 | | | |
| see above | Microbiological Diagnostic Unit - Public Health Laboratory (MDU-PHL) | MDU-PHL | Seemann, T., Schultz M. B., Sait, M., Sherry, N. |

| | | | |
|---|---|---|---|
| EPI_ISL_563407, EPI_ISL_563408, EPI_ISL_563409, EPI_ISL_563410, EPI_ISL_563411, EPI_ISL_563412, EPI_ISL_563413, EPI_ISL_563414, EPI_ISL_563415, EPI_ISL_563416, EPI_ISL_563417, EPI_ISL_563418, EPI_ISL_563419, EPI_ISL_563420, EPI_ISL_563421, EPI_ISL_563422, EPI_ISL_563423 | | | |
| see above | Victorian Infectious Diseases Reference Laboratory (VIDRL) | VIDRL and MDU-PHL | Caly, L., Seemann, T., Sait, M., Schultz, M. B., Druce J., Sherry, N. |
| EPI_ISL_563424, EPI_ISL_563425, EPI_ISL_563426, EPI_ISL_563427, EPI_ISL_563428, EPI_ISL_563429, EPI_ISL_563430, EPI_ISL_563431, EPI_ISL_563432, EPI_ISL_563433, EPI_ISL_563434, EPI_ISL_563435, EPI_ISL_563436, EPI_ISL_563437, EPI_ISL_563438, EPI_ISL_563439, EPI_ISL_563440, EPI_ISL_563441, EPI_ISL_563442, EPI_ISL_563443, EPI_ISL_563444, EPI_ISL_563445, EPI_ISL_563446, EPI_ISL_563447, EPI_ISL_563448, EPI_ISL_563449, EPI_ISL_563450, EPI_ISL_563451, EPI_ISL_563452, EPI_ISL_563453, EPI_ISL_563454, EPI_ISL_563455, EPI_ISL_563456, EPI_ISL_563457, EPI_ISL_563458, EPI_ISL_563459, EPI_ISL_563460, EPI_ISL_563461, EPI_ISL_563462, EPI_ISL_563463, EPI_ISL_563464, EPI_ISL_563465, EPI_ISL_563466, EPI_ISL_563467, EPI_ISL_563468, EPI_ISL_563469, EPI_ISL_563470, EPI_ISL_563471, EPI_ISL_563472, EPI_ISL_563473, EPI_ISL_563474, EPI_ISL_563475, EPI_ISL_563476, EPI_ISL_563477, EPI_ISL_563478, EPI_ISL_563479, EPI_ISL_563480, EPI_ISL_563481, EPI_ISL_563482, EPI_ISL_563483, EPI_ISL_563484, EPI_ISL_563485, EPI_ISL_563486, EPI_ISL_563487, EPI_ISL_563488, EPI_ISL_563489, EPI_ISL_563490, EPI_ISL_563491, EPI_ISL_563492, EPI_ISL_563493, EPI_ISL_563494, EPI_ISL_563495, EPI_ISL_563496 | | | |
| see above | Microbiological Diagnostic Unit - Public Health Laboratory (MDU-PHL) | MDU-PHL | Seemann, T., Schultz M. B., Sait, M., Sherry, N. |
| EPI_ISL_563497, EPI_ISL_563498, EPI_ISL_563499 | Victorian Infectious Diseases Reference Laboratory (VIDRL) | VIDRL and MDU-PHL | Caly, L., Seemann, T., Sait, M., Schultz, M. B., Druce J., Sherry, N. |
| EPI_ISL_563500, EPI_ISL_563501, EPI_ISL_563502, EPI_ISL_563503, EPI_ISL_563504, EPI_ISL_563505, EPI_ISL_563506, EPI_ISL_563507, EPI_ISL_563508, EPI_ISL_563509, EPI_ISL_563510, EPI_ISL_563511, EPI_ISL_563512, EPI_ISL_563513, EPI_ISL_563514, EPI_ISL_563515, EPI_ISL_563516, EPI_ISL_563517, EPI_ISL_563518, EPI_ISL_563519, EPI_ISL_563520, EPI_ISL_563521, EPI_ISL_563522, EPI_ISL_563523, EPI_ISL_563524, EPI_ISL_563525, EPI_ISL_563526, EPI_ISL_563527, EPI_ISL_563528, EPI_ISL_563529, EPI_ISL_563530, EPI_ISL_563531, EPI_ISL_563532, EPI_ISL_563533, EPI_ISL_563534, EPI_ISL_563535, EPI_ISL_563536, EPI_ISL_563537, EPI_ISL_563538, EPI_ISL_563539, EPI_ISL_563540, EPI_ISL_563541, EPI_ISL_563542, EPI_ISL_563543, EPI_ISL_563544, EPI_ISL_563545, EPI_ISL_563546, EPI_ISL_563547, EPI_ISL_563548, EPI_ISL_563549, EPI_ISL_563550 | | | |
| see above | Microbiological Diagnostic Unit - Public Health Laboratory (MDU-PHL) | MDU-PHL | Seemann, T., Schultz M. B., Sait, M., Sherry, N. |
| EPI_ISL_563551 | Victorian Infectious Diseases Reference Laboratory (VIDRL) | VIDRL and MDU-PHL | Caly, L., Seemann, T., Sait, M., Schultz, M. B., Druce J., Sherry, N. |
| EPI_ISL_563552, EPI_ISL_563553, EPI_ISL_563554, EPI_ISL_563555, EPI_ISL_563556, EPI_ISL_563557, EPI_ISL_563558, EPI_ISL_563559, EPI_ISL_563560, EPI_ISL_563561, EPI_ISL_563562, EPI_ISL_563563, EPI_ISL_563564, EPI_ISL_563565, EPI_ISL_563566, EPI_ISL_563567, EPI_ISL_563568, EPI_ISL_563569, EPI_ISL_563570, EPI_ISL_563571, EPI_ISL_563572, EPI_ISL_563573, EPI_ISL_563574, EPI_ISL_563575, EPI_ISL_563576, EPI_ISL_563577, EPI_ISL_563578, EPI_ISL_563579, EPI_ISL_563580, EPI_ISL_563581, EPI_ISL_563582, EPI_ISL_563583, EPI_ISL_563584, EPI_ISL_563585, EPI_ISL_563586, EPI_ISL_563587, EPI_ISL_563588, EPI_ISL_563589, EPI_ISL_563590, EPI_ISL_563591, EPI_ISL_563592, EPI_ISL_563593, EPI_ISL_563594, EPI_ISL_563595, EPI_ISL_563596, EPI_ISL_563597, EPI_ISL_563598, EPI_ISL_563599, EPI_ISL_563600, EPI_ISL_563601, EPI_ISL_563602, EPI_ISL_563603, EPI_ISL_563604, EPI_ISL_563605, EPI_ISL_563606, EPI_ISL_563607, EPI_ISL_563608 | | | |
| see above | Microbiological Diagnostic Unit - Public Health Laboratory (MDU-PHL) | MDU-PHL | Seemann, T., Schultz M. B., Sait, M., Sherry, N. |
| EPI_ISL_563609 | Victorian Infectious Diseases Reference Laboratory (VIDRL) | VIDRL and MDU-PHL | Caly, L., Seemann, T., Sait, M., Schultz, M. B., Druce J., Sherry, N. |
| EPI_ISL_563610, EPI_ISL_563611, EPI_ISL_563612, EPI_ISL_563613, EPI_ISL_563614, EPI_ISL_563615, EPI_ISL_563616, EPI_ISL_563617, EPI_ISL_563618, EPI_ISL_563619, EPI_ISL_563620, EPI_ISL_563621, EPI_ISL_563622, EPI_ISL_563623, EPI_ISL_563624, EPI_ISL_563625, EPI_ISL_563626, EPI_ISL_563627, EPI_ISL_563628, EPI_ISL_563629, EPI_ISL_563630, EPI_ISL_563631, EPI_ISL_563632, EPI_ISL_563633, EPI_ISL_563634, EPI_ISL_563635, EPI_ISL_563636, EPI_ISL_563637, EPI_ISL_563638, EPI_ISL_563639, EPI_ISL_563640, EPI_ISL_563641, EPI_ISL_563642, EPI_ISL_563643, EPI_ISL_563644, EPI_ISL_563645, EPI_ISL_563646, EPI_ISL_563647, EPI_ISL_563648, EPI_ISL_563649, EPI_ISL_563650, EPI_ISL_563651, EPI_ISL_563652, EPI_ISL_563653, EPI_ISL_563654, EPI_ISL_563655, EPI_ISL_563656, EPI_ISL_563657, EPI_ISL_563658, EPI_ISL_563659, EPI_ISL_563660, EPI_ISL_563661, EPI_ISL_563662, EPI_ISL_563663, EPI_ISL_563664, EPI_ISL_563665, EPI_ISL_563666, EPI_ISL_563667, EPI_ISL_563668, EPI_ISL_563669, EPI_ISL_563670, EPI_ISL_563671, EPI_ISL_563672, EPI_ISL_563673, EPI_ISL_563674, EPI_ISL_563675, EPI_ISL_563676, EPI_ISL_563677, EPI_ISL_563678, EPI_ISL_563679, EPI_ISL_563680, EPI_ISL_563681, EPI_ISL_563682, EPI_ISL_563683, EPI_ISL_563684, EPI_ISL_563685, EPI_ISL_563686, EPI_ISL_563687, EPI_ISL_563688, EPI_ISL_563689, EPI_ISL_563690, EPI_ISL_563691, EPI_ISL_563692, EPI_ISL_563693, EPI_ISL_563694, EPI_ISL_563695, EPI_ISL_563696, EPI_ISL_563697, EPI_ISL_563698, EPI_ISL_563699, EPI_ISL_563700, EPI_ISL_563701, EPI_ISL_563702, EPI_ISL_563703, EPI_ISL_563704, EPI_ISL_563705, EPI_ISL_563706, EPI_ISL_563707, EPI_ISL_563708, EPI_ISL_563709, EPI_ISL_563710, EPI_ISL_563711, EPI_ISL_563712, EPI_ISL_563713, EPI_ISL_563714, EPI_ISL_563715, EPI_ISL_563716, EPI_ISL_563717, EPI_ISL_563718, EPI_ISL_563719, EPI_ISL_563720 | | | |
| see above | Microbiological Diagnostic Unit - Public Health Laboratory (MDU-PHL) | MDU-PHL | Seemann, T., Schultz M. B., Sait, M., Sherry, N. |
| EPI_ISL_563721, EPI_ISL_563722, EPI_ISL_563723 | Victorian Infectious Diseases Reference Laboratory (VIDRL) | VIDRL and MDU-PHL | Caly, L., Seemann, T., Sait, M., Schultz, M. B., Druce J., Sherry, N. |
| EPI_ISL_563724, EPI_ISL_563725, EPI_ISL_563726, EPI_ISL_563727, EPI_ISL_563728, EPI_ISL_563729, EPI_ISL_563730, EPI_ISL_563731, EPI_ISL_563732, EPI_ISL_563733, EPI_ISL_563734, EPI_ISL_563735, EPI_ISL_563736, EPI_ISL_563737, EPI_ISL_563738, EPI_ISL_563739, EPI_ISL_563740, EPI_ISL_563741, EPI_ISL_563742, EPI_ISL_563743, EPI_ISL_563744, EPI_ISL_563745, EPI_ISL_563746, EPI_ISL_563747, EPI_ISL_563748, EPI_ISL_563749, EPI_ISL_563750, EPI_ISL_563751, EPI_ISL_563752, EPI_ISL_563753, EPI_ISL_563754, EPI_ISL_563755, EPI_ISL_563756, EPI_ISL_563757, EPI_ISL_563758, EPI_ISL_563759, EPI_ISL_563760, EPI_ISL_563761, EPI_ISL_563762, EPI_ISL_563763, EPI_ISL_563764, EPI_ISL_563765, EPI_ISL_563766, EPI_ISL_563767, EPI_ISL_563768, EPI_ISL_563769, EPI_ISL_563770, EPI_ISL_563771, EPI_ISL_563772, EPI_ISL_563773, EPI_ISL_563774, EPI_ISL_563775, EPI_ISL_563776, EPI_ISL_563777, EPI_ISL_563778, EPI_ISL_563779, EPI_ISL_563780, EPI_ISL_563781, EPI_ISL_563782, EPI_ISL_563783, EPI_ISL_563784, EPI_ISL_563785, EPI_ISL_563786, EPI_ISL_563787, EPI_ISL_563788, EPI_ISL_563789 | | | |
| see above | Microbiological Diagnostic Unit - Public Health Laboratory (MDU-PHL) | MDU-PHL | Seemann, T., Schultz M. B., Sait, M., Sherry, N. |
| EPI_ISL_563790 | Victorian Infectious Diseases Reference Laboratory (VIDRL) | VIDRL and MDU-PHL | Caly, L., Seemann, T., Sait, M., Schultz, M. B., Druce J., Sherry, N. |
| EPI_ISL_563791 | Microbiological Diagnostic Unit - Public Health Laboratory (MDU-PHL) | MDU-PHL | Seemann, T., Schultz M. B., Sait, M., Sherry, N. |
| EPI_ISL_563792 | Victorian Infectious Diseases Reference Laboratory (VIDRL) | VIDRL and MDU-PHL | Caly, L., Seemann, T., Sait, M., Schultz, M. B., Druce J., Sherry, N. |
| EPI_ISL_563793, EPI_ISL_563794, EPI_ISL_563795, EPI_ISL_563796, EPI_ISL_563797, EPI_ISL_563798, EPI_ISL_563799, EPI_ISL_563800, EPI_ISL_563801, EPI_ISL_563802, EPI_ISL_563803, EPI_ISL_563804, EPI_ISL_563805, EPI_ISL_563806, EPI_ISL_563807 | | | |
| see above | Microbiological Diagnostic Unit - Public Health Laboratory (MDU-PHL) | MDU-PHL | Seemann, T., Schultz M. B., Sait, M., Sherry, N. |
| EPI_ISL_563808 | Victorian Infectious Diseases Reference Laboratory (VIDRL) | VIDRL and MDU-PHL | Caly, L., Seemann, T., Sait, M., Schultz, M. B., Druce J., Sherry, N. |
| EPI_ISL_563809, EPI_ISL_563810 | Microbiological Diagnostic Unit - Public Health Laboratory (MDU-PHL) | MDU-PHL | Seemann, T., Schultz M. B., Sait, M., Sherry, N. |
| EPI_ISL_563811 | Victorian Infectious Diseases Reference Laboratory (VIDRL) | VIDRL and MDU-PHL | Caly, L., Seemann, T., Sait, M., Schultz, M. B., Druce J., Sherry, N. |
| EPI_ISL_563812, EPI_ISL_563813, EPI_ISL_563814, EPI_ISL_563815, EPI_ISL_563816 | Microbiological Diagnostic Unit - Public Health Laboratory (MDU-PHL) | MDU-PHL | Seemann, T., Schultz M. B., Sait, M., Sherry, N. |
| EPI_ISL_563817 | Victorian Infectious Diseases Reference Laboratory (VIDRL) | VIDRL and MDU-PHL | Caly, L., Seemann, T., Sait, M., Schultz, M. B., Druce J., Sherry, N. |
| EPI_ISL_563818, EPI_ISL_563819 | Microbiological Diagnostic Unit - Public Health Laboratory (MDU-PHL) | MDU-PHL | Seemann, T., Schultz M. B., Sait, M., Sherry, N. |
| EPI_ISL_563820 | Victorian Infectious Diseases Reference Laboratory (VIDRL) | VIDRL and MDU-PHL | Caly, L., Seemann, T., Sait, M., Schultz, M. B., Druce J., Sherry, N. |
| EPI_ISL_563821, EPI_ISL_563822, EPI_ISL_563823, EPI_ISL_563824, EPI_ISL_563825, EPI_ISL_563826, EPI_ISL_563827, EPI_ISL_563828, EPI_ISL_563829, EPI_ISL_563830, EPI_ISL_563831 | | | |
| see above | Microbiological Diagnostic Unit - Public Health Laboratory (MDU-PHL) | MDU-PHL | Seemann, T., Schultz M. B., Sait, M., Sherry, N. |
| EPI_ISL_563832 | Victorian Infectious Diseases Reference Laboratory (VIDRL) | VIDRL and MDU-PHL | Caly, L., Seemann, T., Sait, M., Schultz, M. B., Druce J., Sherry, N. |
| EPI_ISL_563833, EPI_ISL_563834, EPI_ISL_563835, EPI_ISL_563836, EPI_ISL_563837, EPI_ISL_563838, EPI_ISL_563839 | Microbiological Diagnostic Unit - Public Health Laboratory (MDU-PHL) | MDU-PHL | Seemann, T., Schultz M. B., Sait, M., Sherry, N. |
| EPI_ISL_563840 | Victorian Infectious Diseases Reference Laboratory (VIDRL) | VIDRL and MDU-PHL | Caly, L., Seemann, T., Sait, M., Schultz, M. B., Druce J., Sherry, N. |
| EPI_ISL_563841, EPI_ISL_563842, EPI_ISL_563843, EPI_ISL_563844, EPI_ISL_563845, EPI_ISL_563846, EPI_ISL_563847, EPI_ISL_563848, EPI_ISL_563849, EPI_ISL_563850, EPI_ISL_563851, EPI_ISL_563852, EPI_ISL_563853, EPI_ISL_563854, EPI_ISL_563855, EPI_ISL_563856, EPI_ISL_563857, EPI_ISL_563858, EPI_ISL_563859, EPI_ISL_563860, EPI_ISL_563861, EPI_ISL_563862, EPI_ISL_563863, EPI_ISL_563864, EPI_ISL_563865 | | | |
| see above | Microbiological Diagnostic Unit - Public Health Laboratory (MDU-PHL) | MDU-PHL | Seemann, T., Schultz M. B., Sait, M., Sherry, N. |
| EPI_ISL_563866 | Victorian Infectious Diseases Reference Laboratory (VIDRL) | VIDRL and MDU-PHL | Caly, L., Seemann, T., Sait, M., Schultz, M. B., Druce J., Sherry, N. |
| EPI_ISL_563867, EPI_ISL_563868, EPI_ISL_563869, EPI_ISL_563870, EPI_ISL_563871, EPI_ISL_563872, EPI_ISL_563873, EPI_ISL_563874, EPI_ISL_563875, EPI_ISL_563876, EPI_ISL_563877, EPI_ISL_563878, EPI_ISL_563879 | | | |
| see above | Microbiological Diagnostic Unit - Public Health Laboratory (MDU-PHL) | MDU-PHL | Seemann, T., Schultz M. B., Sait, M., Sherry, N. |
| EPI_ISL_563880 | Victorian Infectious Diseases Reference Laboratory (VIDRL) | VIDRL and MDU-PHL | Caly, L., Seemann, T., Sait, M., Schultz, M. B., Druce J., Sherry, N. |
| EPI_ISL_563881, EPI_ISL_563882, EPI_ISL_563883, EPI_ISL_563884, EPI_ISL_563885, EPI_ISL_563886, EPI_ISL_563887, EPI_ISL_563888, EPI_ISL_563889, EPI_ISL_563890, EPI_ISL_563891, EPI_ISL_563892, EPI_ISL_563893, EPI_ISL_563894, EPI_ISL_563895, EPI_ISL_563896, EPI_ISL_563897, EPI_ISL_563898, EPI_ISL_563899, EPI_ISL_563900, EPI_ISL_563901, EPI_ISL_563902, EPI_ISL_563903, EPI_ISL_563904, EPI_ISL_563905, EPI_ISL_563906, EPI_ISL_563907, EPI_ISL_563908, EPI_ISL_563909, EPI_ISL_563910, EPI_ISL_563911, EPI_ISL_563912, EPI_ISL_563913, EPI_ISL_563914, EPI_ISL_563915, EPI_ISL_563916, EPI_ISL_563917, EPI_ISL_563918, EPI_ISL_563919, EPI_ISL_563920, EPI_ISL_563921, EPI_ISL_563922, EPI_ISL_563923, EPI_ISL_563924 | | | |
| see above | Microbiological Diagnostic Unit - Public Health Laboratory (MDU-PHL) | MDU-PHL | Seemann, T., Schultz M. B., Sait, M., Sherry, N. |
| EPI_ISL_563925 | Victorian Infectious Diseases Reference Laboratory (VIDRL) | VIDRL and MDU-PHL | Caly, L., Seemann, T., Sait, M., Schultz, M. B., Druce J., Sherry, N. |
| EPI_ISL_563926, EPI_ISL_563927, EPI_ISL_563928, EPI_ISL_563929, EPI_ISL_563930, EPI_ISL_563931, EPI_ISL_563932, EPI_ISL_563933, EPI_ISL_563934, EPI_ISL_563935, EPI_ISL_563936, EPI_ISL_563937 | | | |
| see above | Microbiological Diagnostic Unit - Public Health Laboratory (MDU-PHL) | MDU-PHL | Seemann, T., Schultz M. B., Sait, M., Sherry, N. |

| | | | |
|---|---|---|---|
| EPI_ISL_563938 | Victorian Infectious Diseases Reference Laboratory (VIDRL) | VIDRL and MDU-PHL | Caly, L., Seemann, T., Sait, M., Schultz, M. B., Druce J., Sherry, N. |
| EPI_ISL_563939, EPI_ISL_563940, EPI_ISL_563941, EPI_ISL_563942, EPI_ISL_563943, EPI_ISL_563944, EPI_ISL_563945, EPI_ISL_563946, EPI_ISL_563947, EPI_ISL_563948, EPI_ISL_563949, EPI_ISL_563950, EPI_ISL_563951, EPI_ISL_563952, EPI_ISL_563953, EPI_ISL_563954, EPI_ISL_563955, EPI_ISL_563956, EPI_ISL_563957, EPI_ISL_563958, EPI_ISL_563959, EPI_ISL_563960, EPI_ISL_563961, EPI_ISL_563962, EPI_ISL_563963, EPI_ISL_563964, EPI_ISL_563965, EPI_ISL_563966, EPI_ISL_563967, EPI_ISL_563968, EPI_ISL_563969, EPI_ISL_563970, EPI_ISL_563971, EPI_ISL_563972 | | | |
| see above | Microbiological Diagnostic Unit - Public Health Laboratory (MDU-PHL) | MDU-PHL | Seemann, T., Schultz M. B., Sait, M., Sherry, N. |
| EPI_ISL_563973, EPI_ISL_563974, EPI_ISL_563975 | Victorian Infectious Diseases Reference Laboratory (VIDRL) | VIDRL and MDU-PHL | Caly, L., Seemann, T., Sait, M., Schultz, M. B., Druce J., Sherry, N. |
| EPI_ISL_563976 | Microbiological Diagnostic Unit - Public Health Laboratory (MDU-PHL) | MDU-PHL | Seemann, T., Schultz M. B., Sait, M., Sherry, N. |
| EPI_ISL_563977, EPI_ISL_563978 | Victorian Infectious Diseases Reference Laboratory (VIDRL) | VIDRL and MDU-PHL | Caly, L., Seemann, T., Sait, M., Schultz, M. B., Druce J., Sherry, N. |
| EPI_ISL_563979, EPI_ISL_563980, EPI_ISL_563981, EPI_ISL_563982, EPI_ISL_563983, EPI_ISL_563984, EPI_ISL_563985, EPI_ISL_563986 | Microbiological Diagnostic Unit - Public Health Laboratory (MDU-PHL) | MDU-PHL | Seemann, T., Schultz M. B., Sait, M., Sherry, N. |
| EPI_ISL_563987 | Victorian Infectious Diseases Reference Laboratory (VIDRL) | VIDRL and MDU-PHL | Caly, L., Seemann, T., Sait, M., Schultz, M. B., Druce J., Sherry, N. |
| EPI_ISL_563988, EPI_ISL_563989, EPI_ISL_563990, EPI_ISL_563991, EPI_ISL_563992, EPI_ISL_563993, EPI_ISL_563994, EPI_ISL_563995, EPI_ISL_563996 | Microbiological Diagnostic Unit - Public Health Laboratory (MDU-PHL) | MDU-PHL | Seemann, T., Schultz M. B., Sait, M., Sherry, N. |
| EPI_ISL_563997, EPI_ISL_563998, EPI_ISL_563999, EPI_ISL_564000, EPI_ISL_564001, EPI_ISL_564002, EPI_ISL_564003, EPI_ISL_564004, EPI_ISL_564005, EPI_ISL_564006, EPI_ISL_564007, EPI_ISL_564008, EPI_ISL_564009, EPI_ISL_564010, EPI_ISL_564011, EPI_ISL_564012, EPI_ISL_564013, EPI_ISL_564014, EPI_ISL_564015, EPI_ISL_564016, EPI_ISL_564017, EPI_ISL_564018, EPI_ISL_564019, EPI_ISL_564020, EPI_ISL_564021, EPI_ISL_564022, EPI_ISL_564023, EPI_ISL_564024, EPI_ISL_564025, EPI_ISL_564026, EPI_ISL_564027, EPI_ISL_564028, EPI_ISL_564029, EPI_ISL_564030, EPI_ISL_564031, EPI_ISL_564032, EPI_ISL_564033, EPI_ISL_564034, EPI_ISL_564035, EPI_ISL_564036, EPI_ISL_564037, EPI_ISL_564038, EPI_ISL_564039, EPI_ISL_564040, EPI_ISL_564041, EPI_ISL_564042, EPI_ISL_564043, EPI_ISL_564044, EPI_ISL_564045, EPI_ISL_564046, EPI_ISL_564047, EPI_ISL_564048, EPI_ISL_564049 | | | |
| see above | Victorian Infectious Diseases Reference Laboratory (VIDRL) | VIDRL and MDU-PHL | Caly, L., Seemann, T., Sait, M., Schultz, M. B., Druce J., Sherry, N. |
| EPI_ISL_564050, EPI_ISL_564051, EPI_ISL_564052, EPI_ISL_564053, EPI_ISL_564054 | Microbiological Diagnostic Unit - Public Health Laboratory (MDU-PHL) | MDU-PHL | Seemann, T., Schultz M. B., Sait, M., Sherry, N. |
| EPI_ISL_564055, EPI_ISL_564056, EPI_ISL_564057, EPI_ISL_564058, EPI_ISL_564059, EPI_ISL_564060, EPI_ISL_564061, EPI_ISL_564062, EPI_ISL_564063, EPI_ISL_564064, EPI_ISL_564065, EPI_ISL_564066, EPI_ISL_564067, EPI_ISL_564068, EPI_ISL_564069, EPI_ISL_564070, EPI_ISL_564071, EPI_ISL_564072, EPI_ISL_564073, EPI_ISL_564074, EPI_ISL_564075, EPI_ISL_564076, EPI_ISL_564077, EPI_ISL_564078, EPI_ISL_564079, EPI_ISL_564080, EPI_ISL_564081, EPI_ISL_564082, EPI_ISL_564083, EPI_ISL_564084, EPI_ISL_564085, EPI_ISL_564086, EPI_ISL_564087, EPI_ISL_564088, EPI_ISL_564089, EPI_ISL_564090, EPI_ISL_564091, EPI_ISL_564092, EPI_ISL_564093, EPI_ISL_564094, EPI_ISL_564095, EPI_ISL_564096, EPI_ISL_564097, EPI_ISL_564098, EPI_ISL_564099, EPI_ISL_564100, EPI_ISL_564101, EPI_ISL_564102, EPI_ISL_564103, EPI_ISL_564104, EPI_ISL_564105, EPI_ISL_564106, EPI_ISL_564107, EPI_ISL_564108, EPI_ISL_564109, EPI_ISL_564110 | | | |
| see above | Victorian Infectious Diseases Reference Laboratory (VIDRL) | VIDRL and MDU-PHL | Caly, L., Seemann, T., Sait, M., Schultz, M. B., Druce J., Sherry, N. |
| EPI_ISL_564111, EPI_ISL_564112, EPI_ISL_564113, EPI_ISL_564114, EPI_ISL_564115, EPI_ISL_564116, EPI_ISL_564117, EPI_ISL_564118, EPI_ISL_564119, EPI_ISL_564120, EPI_ISL_564121, EPI_ISL_564122, EPI_ISL_564123, EPI_ISL_564124, EPI_ISL_564125, EPI_ISL_564126, EPI_ISL_564127, EPI_ISL_564128, EPI_ISL_564129, EPI_ISL_564130, EPI_ISL_564131, EPI_ISL_564132, EPI_ISL_564133, EPI_ISL_564134, EPI_ISL_564135, EPI_ISL_564136, EPI_ISL_564137, EPI_ISL_564138, EPI_ISL_564139, EPI_ISL_564140, EPI_ISL_564141, EPI_ISL_564142, EPI_ISL_564143, EPI_ISL_564144, EPI_ISL_564145, EPI_ISL_564146, EPI_ISL_564147, EPI_ISL_564148, EPI_ISL_564149, EPI_ISL_564150, EPI_ISL_564151, EPI_ISL_564152, EPI_ISL_564153, EPI_ISL_564154, EPI_ISL_564155, EPI_ISL_564156, EPI_ISL_564157, EPI_ISL_564158, EPI_ISL_564159, EPI_ISL_564160, EPI_ISL_564161, EPI_ISL_564162, EPI_ISL_564163, EPI_ISL_564164, EPI_ISL_564165, EPI_ISL_564166, EPI_ISL_564167, EPI_ISL_564168, EPI_ISL_564169, EPI_ISL_564170, EPI_ISL_564171, EPI_ISL_564172, EPI_ISL_564173, EPI_ISL_564174, EPI_ISL_564175, EPI_ISL_564176, EPI_ISL_564177, EPI_ISL_564178, EPI_ISL_564179 | | | |
| see above | Microbiological Diagnostic Unit - Public Health Laboratory (MDU-PHL) | MDU-PHL | Seemann, T., Schultz M. B., Sait, M., Sherry, N. |
| EPI_ISL_564180, EPI_ISL_564181, EPI_ISL_564182, EPI_ISL_564183, EPI_ISL_564184, EPI_ISL_564185, EPI_ISL_564186, EPI_ISL_564187, EPI_ISL_564188, EPI_ISL_564189, EPI_ISL_564190, EPI_ISL_564191, EPI_ISL_564192, EPI_ISL_564193, EPI_ISL_564194, EPI_ISL_564195, EPI_ISL_564196, EPI_ISL_564197, EPI_ISL_564198, EPI_ISL_564199, EPI_ISL_564200, EPI_ISL_564201, EPI_ISL_564202, EPI_ISL_564203, EPI_ISL_564204, EPI_ISL_564205, EPI_ISL_564206, EPI_ISL_564207, EPI_ISL_564208, EPI_ISL_564209, EPI_ISL_564210, EPI_ISL_564211, EPI_ISL_564212, EPI_ISL_564213, EPI_ISL_564214, EPI_ISL_564215, EPI_ISL_564216, EPI_ISL_564217, EPI_ISL_564218, EPI_ISL_564219, EPI_ISL_564220, EPI_ISL_564221, EPI_ISL_564222, EPI_ISL_564223, EPI_ISL_564224, EPI_ISL_564225, EPI_ISL_564226, EPI_ISL_564227, EPI_ISL_564228, EPI_ISL_564229, EPI_ISL_564230, EPI_ISL_564231, EPI_ISL_564232, EPI_ISL_564233, EPI_ISL_564234, EPI_ISL_564235, EPI_ISL_564236, EPI_ISL_564237, EPI_ISL_564238, EPI_ISL_564239, EPI_ISL_564240, EPI_ISL_564241, EPI_ISL_564242, EPI_ISL_564243, EPI_ISL_564244, EPI_ISL_564245, EPI_ISL_564246, EPI_ISL_564247, EPI_ISL_564248, EPI_ISL_564249, EPI_ISL_564250, EPI_ISL_564251, EPI_ISL_564252, EPI_ISL_564253, EPI_ISL_564254 | | | |
| see above | Victorian Infectious Diseases Reference Laboratory (VIDRL) | VIDRL and MDU-PHL | Caly, L., Seemann, T., Sait, M., Schultz, M. B., Druce J., Sherry, N. |
| EPI_ISL_564255, EPI_ISL_564256, EPI_ISL_564257, EPI_ISL_564258, EPI_ISL_564259, EPI_ISL_564260, EPI_ISL_564261, EPI_ISL_564262, EPI_ISL_564263, EPI_ISL_564264, EPI_ISL_564265, EPI_ISL_564266, EPI_ISL_564267, EPI_ISL_564268, EPI_ISL_564269 | | | |
| see above | Microbiological Diagnostic Unit - Public Health Laboratory (MDU-PHL) | MDU-PHL | Seemann, T., Schultz M. B., Sait, M., Sherry, N. |
| EPI_ISL_564270 | Victorian Infectious Diseases Reference Laboratory (VIDRL) | VIDRL and MDU-PHL | Caly, L., Seemann, T., Sait, M., Schultz, M. B., Druce J., Sherry, N. |
| EPI_ISL_564271, EPI_ISL_564272, EPI_ISL_564273, EPI_ISL_564274, EPI_ISL_564275, EPI_ISL_564276, EPI_ISL_564277, EPI_ISL_564278, EPI_ISL_564279, EPI_ISL_564280, EPI_ISL_564281, EPI_ISL_564282, EPI_ISL_564283, EPI_ISL_564284, EPI_ISL_564285, EPI_ISL_564286, EPI_ISL_564287, EPI_ISL_564288, EPI_ISL_564289, EPI_ISL_564290, EPI_ISL_564291, EPI_ISL_564292, EPI_ISL_564293, EPI_ISL_564294, EPI_ISL_564295, EPI_ISL_564296, EPI_ISL_564297, EPI_ISL_564298, EPI_ISL_564299, EPI_ISL_564300, EPI_ISL_564301, EPI_ISL_564302, EPI_ISL_564303, EPI_ISL_564304, EPI_ISL_564305, EPI_ISL_564306, EPI_ISL_564307, EPI_ISL_564308, EPI_ISL_564309, EPI_ISL_564310, EPI_ISL_564311, EPI_ISL_564312, EPI_ISL_564313, EPI_ISL_564314, EPI_ISL_564315, EPI_ISL_564316, EPI_ISL_564317 | | | |
| see above | Microbiological Diagnostic Unit - Public Health Laboratory (MDU-PHL) | MDU-PHL | Seemann, T., Schultz M. B., Sait, M., Sherry, N. |
| EPI_ISL_564318, EPI_ISL_564319, EPI_ISL_564320, EPI_ISL_564321, EPI_ISL_564322, EPI_ISL_564323, EPI_ISL_564324, EPI_ISL_564325, EPI_ISL_564326, EPI_ISL_564327, EPI_ISL_564328, EPI_ISL_564329, EPI_ISL_564330, EPI_ISL_564331, EPI_ISL_564332, EPI_ISL_564333, EPI_ISL_564334, EPI_ISL_564335, EPI_ISL_564336, EPI_ISL_564337, EPI_ISL_564338, EPI_ISL_564339, EPI_ISL_564340, EPI_ISL_564341, EPI_ISL_564342, EPI_ISL_564343, EPI_ISL_564344, EPI_ISL_564345, EPI_ISL_564346, EPI_ISL_564347, EPI_ISL_564348, EPI_ISL_564349, EPI_ISL_564350 | | | |
| see above | Victorian Infectious Diseases Reference Laboratory (VIDRL) | VIDRL and MDU-PHL | Caly, L., Seemann, T., Sait, M., Schultz, M. B., Druce J., Sherry, N. |
| EPI_ISL_564351 | Microbiological Diagnostic Unit - Public Health Laboratory (MDU-PHL) | MDU-PHL | Seemann, T., Schultz M. B., Sait, M., Sherry, N. |
| EPI_ISL_564352, EPI_ISL_564353, EPI_ISL_564354, EPI_ISL_564355, EPI_ISL_564356, EPI_ISL_564357, EPI_ISL_564358, EPI_ISL_564359, EPI_ISL_564360, EPI_ISL_564361, EPI_ISL_564362, EPI_ISL_564363, EPI_ISL_564364, EPI_ISL_564365, EPI_ISL_564366, EPI_ISL_564367, EPI_ISL_564368, EPI_ISL_564369, EPI_ISL_564370, EPI_ISL_564371, EPI_ISL_564372, EPI_ISL_564373, EPI_ISL_564374, EPI_ISL_564375, EPI_ISL_564376, EPI_ISL_564377, EPI_ISL_564378, EPI_ISL_564379, EPI_ISL_564380, EPI_ISL_564381, EPI_ISL_564382, EPI_ISL_564383, EPI_ISL_564384, EPI_ISL_564385, EPI_ISL_564386, EPI_ISL_564387, EPI_ISL_564388, EPI_ISL_564389, EPI_ISL_564390, EPI_ISL_564391, EPI_ISL_564392, EPI_ISL_564393, EPI_ISL_564394, EPI_ISL_564395, EPI_ISL_564396, EPI_ISL_564397 | | | |
| see above | Victorian Infectious Diseases Reference Laboratory (VIDRL) | VIDRL and MDU-PHL | Caly, L., Seemann, T., Sait, M., Schultz, M. B., Druce J., Sherry, N. |
| EPI_ISL_564398, EPI_ISL_564399, EPI_ISL_564400, EPI_ISL_564401, EPI_ISL_564402, EPI_ISL_564403 | Microbiological Diagnostic Unit - Public Health Laboratory (MDU-PHL) | MDU-PHL | Seemann, T., Schultz M. B., Sait, M., Sherry, N. |
| EPI_ISL_564404 | Victorian Infectious Diseases Reference Laboratory (VIDRL) | VIDRL and MDU-PHL | Caly, L., Seemann, T., Sait, M., Schultz, M. B., Druce J., Sherry, N. |
| EPI_ISL_564405, EPI_ISL_564406, EPI_ISL_564407, EPI_ISL_564408, EPI_ISL_564409, EPI_ISL_564410, EPI_ISL_564411, EPI_ISL_564412, EPI_ISL_564413, EPI_ISL_564414, EPI_ISL_564415, EPI_ISL_564416, EPI_ISL_564417, EPI_ISL_564418, EPI_ISL_564419, EPI_ISL_564420, EPI_ISL_564421, EPI_ISL_564422, EPI_ISL_564423, EPI_ISL_564424, EPI_ISL_564425, EPI_ISL_564426, EPI_ISL_564427, EPI_ISL_564428, EPI_ISL_564429, EPI_ISL_564430, EPI_ISL_564431, EPI_ISL_564432, EPI_ISL_564433, EPI_ISL_564434, EPI_ISL_564435, EPI_ISL_564436, EPI_ISL_564437, EPI_ISL_564438, EPI_ISL_564439, EPI_ISL_564440, EPI_ISL_564441, EPI_ISL_564442, EPI_ISL_564443, EPI_ISL_564444, EPI_ISL_564445, EPI_ISL_564446, EPI_ISL_564447, EPI_ISL_564448, EPI_ISL_564449, EPI_ISL_564450, EPI_ISL_564451, EPI_ISL_564452, EPI_ISL_564453, EPI_ISL_564454, EPI_ISL_564455, EPI_ISL_564456, EPI_ISL_564457, EPI_ISL_564458, EPI_ISL_564459, EPI_ISL_564460, EPI_ISL_564461, EPI_ISL_564462, EPI_ISL_564463, EPI_ISL_564464, EPI_ISL_564465, EPI_ISL_564466, EPI_ISL_564467, EPI_ISL_564468, EPI_ISL_564469, EPI_ISL_564470, EPI_ISL_564471, EPI_ISL_564472, EPI_ISL_564473, EPI_ISL_564474, EPI_ISL_564475, EPI_ISL_564476, EPI_ISL_564477, EPI_ISL_564478, EPI_ISL_564479, EPI_ISL_564480, EPI_ISL_564481, EPI_ISL_564482, EPI_ISL_564483, EPI_ISL_564484, EPI_ISL_564485, EPI_ISL_564486, EPI_ISL_564487, EPI_ISL_564488, EPI_ISL_564489, EPI_ISL_564490, EPI_ISL_564491, EPI_ISL_564492, EPI_ISL_564493, EPI_ISL_564494, EPI_ISL_564495, EPI_ISL_564496, EPI_ISL_564497, EPI_ISL_564498, EPI_ISL_564499, EPI_ISL_564500, EPI_ISL_564501, EPI_ISL_564502, EPI_ISL_564503, EPI_ISL_564504, EPI_ISL_564505, EPI_ISL_564506, EPI_ISL_564507, EPI_ISL_564508, EPI_ISL_564509, EPI_ISL_564510, EPI_ISL_564511, EPI_ISL_564512, EPI_ISL_564513, EPI_ISL_564514, EPI_ISL_564515, EPI_ISL_564516, EPI_ISL_564517, EPI_ISL_564518, EPI_ISL_564519, EPI_ISL_564520, EPI_ISL_564521, EPI_ISL_564522, EPI_ISL_564523, EPI_ISL_564524, EPI_ISL_564525, EPI_ISL_564526, EPI_ISL_564527, EPI_ISL_564528, EPI_ISL_564529, EPI_ISL_564530, EPI_ISL_564531, EPI_ISL_564532, EPI_ISL_564533, EPI_ISL_564534, EPI_ISL_564535, EPI_ISL_564536, EPI_ISL_564537, EPI_ISL_564538, EPI_ISL_564539, EPI_ISL_564540, EPI_ISL_564541, EPI_ISL_564542, EPI_ISL_564543 | | | |
| see above | Microbiological Diagnostic Unit - Public Health Laboratory (MDU-PHL) | MDU-PHL | Seemann, T., Schultz M. B., Sait, M., Sherry, N. |
| EPI_ISL_564544 | Victorian Infectious Diseases Reference Laboratory (VIDRL) | VIDRL and MDU-PHL | Caly, L., Seemann, T., Sait, M., Schultz, M. B., Druce J., Sherry, N. |
| EPI_ISL_564545 | Microbiological Diagnostic Unit - Public Health Laboratory (MDU-PHL) | MDU-PHL | Seemann, T., Schultz M. B., Sait, M., Sherry, N. |
| EPI_ISL_564546, EPI_ISL_564547, EPI_ISL_564548, EPI_ISL_564549, EPI_ISL_564550, EPI_ISL_564551, EPI_ISL_564552, EPI_ISL_564553, EPI_ISL_564554, EPI_ISL_564555, EPI_ISL_564556, EPI_ISL_564557, EPI_ISL_564558, EPI_ISL_564559, EPI_ISL_564560, EPI_ISL_564561, EPI_ISL_564562, EPI_ISL_564563, EPI_ISL_564564 | | | |
| see above | Victorian Infectious Diseases Reference Laboratory (VIDRL) | VIDRL and MDU-PHL | Caly, L., Seemann, T., Sait, M., Schultz, M. B., Druce J., Sherry, N. |
| EPI_ISL_564565, EPI_ISL_564566, EPI_ISL_564567, EPI_ISL_564568, EPI_ISL_564569, EPI_ISL_564570, EPI_ISL_564571, EPI_ISL_564572, EPI_ISL_564573, EPI_ISL_564574, EPI_ISL_564575, EPI_ISL_564576, EPI_ISL_564577, EPI_ISL_564578, EPI_ISL_564579, EPI_ISL_564580, EPI_ISL_564581, EPI_ISL_564582, EPI_ISL_564583, EPI_ISL_564584, EPI_ISL_564585, EPI_ISL_564586, EPI_ISL_564587, EPI_ISL_564588, EPI_ISL_564589, EPI_ISL_564590, EPI_ISL_564591, EPI_ISL_564592, EPI_ISL_564593, EPI_ISL_564594, EPI_ISL_564595, EPI_ISL_564596, EPI_ISL_564597, EPI_ISL_564598, EPI_ISL_564599, EPI_ISL_564600, EPI_ISL_564601, EPI_ISL_564602, EPI_ISL_564603, EPI_ISL_564604, EPI_ISL_564605, EPI_ISL_564606, EPI_ISL_564607, EPI_ISL_564608, EPI_ISL_564609, EPI_ISL_564610, EPI_ISL_564611, EPI_ISL_564612, EPI_ISL_564613, EPI_ISL_564614, EPI_ISL_564615, EPI_ISL_564616, EPI_ISL_564617, EPI_ISL_564618, EPI_ISL_564619, EPI_ISL_564620, EPI_ISL_564621, EPI_ISL_564622, EPI_ISL_564623, EPI_ISL_564624, EPI_ISL_564625, EPI_ISL_564626, EPI_ISL_564627, EPI_ISL_564628, EPI_ISL_564629, EPI_ISL_564630, EPI_ISL_564631, EPI_ISL_564632, EPI_ISL_564633, EPI_ISL_564634, EPI_ISL_564635, EPI_ISL_564636, EPI_ISL_564637, EPI_ISL_564638, EPI_ISL_564639, EPI_ISL_564640, EPI_ISL_564641, EPI_ISL_564642, EPI_ISL_564643, EPI_ISL_564644, EPI_ISL_564645, EPI_ISL_564646, EPI_ISL_564647, EPI_ISL_564648, EPI_ISL_564649, EPI_ISL_564650, EPI_ISL_564651, EPI_ISL_564652, EPI_ISL_564653, EPI_ISL_564654, EPI_ISL_564655, EPI_ISL_564656, EPI_ISL_564657, EPI_ISL_564658, EPI_ISL_564659, EPI_ISL_564660, EPI_ISL_564661, EPI_ISL_564662, EPI_ISL_564663, EPI_ISL_564664, EPI_ISL_564665, EPI_ISL_564666, EPI_ISL_564667, EPI_ISL_564668, EPI_ISL_564669, EPI_ISL_564670, EPI_ISL_564671, EPI_ISL_564672, EPI_ISL_564673, EPI_ISL_564674, EPI_ISL_564675, EPI_ISL_564676, EPI_ISL_564677, EPI_ISL_564678, EPI_ISL_564679, EPI_ISL_564680, EPI_ISL_564681, EPI_ISL_564682, EPI_ISL_564683, EPI_ISL_564684, EPI_ISL_564685, EPI_ISL_564686, EPI_ISL_564687, EPI_ISL_564688, EPI_ISL_564689, EPI_ISL_564690, EPI_ISL_564691, EPI_ISL_564692, EPI_ISL_564693, EPI_ISL_564694, EPI_ISL_564695, EPI_ISL_564696, EPI_ISL_564697, EPI_ISL_564698, EPI_ISL_564699, EPI_ISL_564700, EPI_ISL_564701, EPI_ISL_564702, EPI_ISL_564703, EPI_ISL_564704, EPI_ISL_564705, EPI_ISL_564706, EPI_ISL_564707, EPI_ISL_564708, EPI_ISL_564709, EPI_ISL_564710, EPI_ISL_564711, EPI_ISL_564712, EPI_ISL_564713, EPI_ISL_564714, EPI_ISL_564715, EPI_ISL_564716, EPI_ISL_564717, EPI_ISL_564718, EPI_ISL_564719, EPI_ISL_564720, EPI_ISL_564721, EPI_ISL_564722, EPI_ISL_564723, EPI_ISL_564724, EPI_ISL_564725, EPI_ISL_564726, EPI_ISL_564727, EPI_ISL_564728, EPI_ISL_564729, EPI_ISL_564730, EPI_ISL_564731, EPI_ISL_564732, EPI_ISL_564733, EPI_ISL_564734, EPI_ISL_564735, EPI_ISL_564736, EPI_ISL_564737, EPI_ISL_564738, EPI_ISL_564739, EPI_ISL_564740, EPI_ISL_564741, EPI_ISL_564742, EPI_ISL_564743, EPI_ISL_564744, EPI_ISL_564745, EPI_ISL_564746, EPI_ISL_564747, EPI_ISL_564748, EPI_ISL_564749, EPI_ISL_564750, EPI_ISL_564751, EPI_ISL_564752, EPI_ISL_564753, EPI_ISL_564754, EPI_ISL_564755, EPI_ISL_564756, EPI_ISL_564757, EPI_ISL_564758, EPI_ISL_564759, EPI_ISL_564760, EPI_ISL_564761, EPI_ISL_564762, EPI_ISL_564763, EPI_ISL_564764, EPI_ISL_564765, EPI_ISL_564766, EPI_ISL_564767, EPI_ISL_564768, EPI_ISL_564769, EPI_ISL_564770, EPI_ISL_564771, EPI_ISL_564772, EPI_ISL_564773, EPI_ISL_564774, EPI_ISL_564775, EPI_ISL_564776, EPI_ISL_564777, EPI_ISL_564778, EPI_ISL_564779, EPI_ISL_564780, EPI_ISL_564781, EPI_ISL_564782, EPI_ISL_564783, EPI_ISL_564784, EPI_ISL_564785, EPI_ISL_564786, EPI_ISL_564787, EPI_ISL_564788, EPI_ISL_564789, EPI_ISL_564790, EPI_ISL_564791, EPI_ISL_564792, EPI_ISL_564793, EPI_ISL_564794, EPI_ISL_564795, EPI_ISL_564796, EPI_ISL_564797, EPI_ISL_564798, EPI_ISL_564799, EPI_ISL_564800, EPI_ISL_564801, EPI_ISL_564802, EPI_ISL_564803, EPI_ISL_564804, EPI_ISL_564805, EPI_ISL_564806, EPI_ISL_564807, EPI_ISL_564808, EPI_ISL_564809, EPI_ISL_564810, EPI_ISL_564811, EPI_ISL_564812, EPI_ISL_564813, EPI_ISL_564814, EPI_ISL_564815, EPI_ISL_564816, EPI_ISL_564817, EPI_ISL_564818, EPI_ISL_564819, EPI_ISL_564820, EPI_ISL_564821, EPI_ISL_564822, EPI_ISL_564823, EPI_ISL_564824, EPI_ISL_564825, EPI_ISL_564826, EPI_ISL_564827, EPI_ISL_564828, EPI_ISL_564829, EPI_ISL_564830, EPI_ISL_564831, EPI_ISL_564832, EPI_ISL_564833, EPI_ISL_564834, EPI_ISL_564835, EPI_ISL_564836, EPI_ISL_564837, EPI_ISL_564838, EPI_ISL_564839, EPI_ISL_564840, EPI_ISL_564841, EPI_ISL_564842, EPI_ISL_564843, EPI_ISL_564844, EPI_ISL_564845, EPI_ISL_564846, EPI_ISL_564847, EPI_ISL_564848, EPI_ISL_564849, EPI_ISL_564850, EPI_ISL_564851, EPI_ISL_564852, EPI_ISL_564853, EPI_ISL_564854, EPI_ISL_564855, EPI_ISL_564856, EPI_ISL_564857, EPI_ISL_564858, EPI_ISL_564859, EPI_ISL_564860, EPI_ISL_564861, EPI_ISL_564862, EPI_ISL_564863, EPI_ISL_564864, EPI_ISL_564865, EPI_ISL_564866, EPI_ISL_564867, EPI_ISL_564868, EPI_ISL_564869, EPI_ISL_564870, EPI_ISL_564871, EPI_ISL_564872, EPI_ISL_564873, EPI_ISL_564874, EPI_ISL_564875, EPI_ISL_564876, EPI_ISL_564877, EPI_ISL_564878, EPI_ISL_564879, EPI_ISL_564880, EPI_ISL_564881, EPI_ISL_564882, EPI_ISL_564883, EPI_ISL_564884, EPI_ISL_564885, EPI_ISL_564886, EPI_ISL_564887, EPI_ISL_564888, EPI_ISL_564889, EPI_ISL_564890, EPI_ISL_564891, EPI_ISL_564892, EPI_ISL_564893, EPI_ISL_564894, EPI_ISL_564895, EPI_ISL_564896, EPI_ISL_564897, EPI_ISL_564898, EPI_ISL_564899, EPI_ISL_564900, | | | |

| | | | |
|---|---|---|---|
| EPI_ISL_564901, EPI_ISL_564902, EPI_ISL_564903, EPI_ISL_564904, EPI_ISL_564905, EPI_ISL_564906, EPI_ISL_564907, EPI_ISL_564908, EPI_ISL_564909, EPI_ISL_564910, EPI_ISL_564911, EPI_ISL_564912, EPI_ISL_564913, EPI_ISL_564914, EPI_ISL_564915, EPI_ISL_564916, EPI_ISL_564917, EPI_ISL_564918, EPI_ISL_564919, EPI_ISL_564920, EPI_ISL_564921, EPI_ISL_564922, EPI_ISL_564923, EPI_ISL_564924, EPI_ISL_564925, EPI_ISL_564926, EPI_ISL_564927, EPI_ISL_564928, EPI_ISL_564929, EPI_ISL_564930, EPI_ISL_564931, EPI_ISL_564932, EPI_ISL_564933, EPI_ISL_564934, EPI_ISL_564935, EPI_ISL_564936, EPI_ISL_564937, EPI_ISL_564938, EPI_ISL_564939, EPI_ISL_564940, EPI_ISL_564941, EPI_ISL_564942, EPI_ISL_564943, EPI_ISL_564944, EPI_ISL_564945, EPI_ISL_564946, EPI_ISL_564947, EPI_ISL_564948, EPI_ISL_564949, EPI_ISL_564950, EPI_ISL_564951, EPI_ISL_564952, EPI_ISL_564953, EPI_ISL_564954, EPI_ISL_564955, EPI_ISL_564956, EPI_ISL_564957, EPI_ISL_564958, EPI_ISL_564959, EPI_ISL_564960, EPI_ISL_564961, EPI_ISL_564962, EPI_ISL_564963, EPI_ISL_564964, EPI_ISL_564965, EPI_ISL_564966, EPI_ISL_564967, EPI_ISL_564968, EPI_ISL_564969, EPI_ISL_564970, EPI_ISL_564971, EPI_ISL_564972, EPI_ISL_564973, EPI_ISL_564974, EPI_ISL_564975, EPI_ISL_564976, EPI_ISL_564977, EPI_ISL_564978, EPI_ISL_564979, EPI_ISL_564980, EPI_ISL_564981, EPI_ISL_564982, EPI_ISL_564983, EPI_ISL_564984, EPI_ISL_564985, EPI_ISL_564986, EPI_ISL_564987, EPI_ISL_564988, EPI_ISL_564989, EPI_ISL_564990, EPI_ISL_564991, EPI_ISL_564992, EPI_ISL_564993, EPI_ISL_564994, EPI_ISL_564995, EPI_ISL_564996, EPI_ISL_564997, EPI_ISL_564998, EPI_ISL_564999, EPI_ISL_565000, EPI_ISL_565001, EPI_ISL_565002, EPI_ISL_565003, EPI_ISL_565004, EPI_ISL_565005, EPI_ISL_565006, EPI_ISL_565007, EPI_ISL_565008, EPI_ISL_565009, EPI_ISL_565010, EPI_ISL_565011, EPI_ISL_565012, EPI_ISL_565013, EPI_ISL_565014, EPI_ISL_565015, EPI_ISL_565016, EPI_ISL_565017, EPI_ISL_565018, EPI_ISL_565019, EPI_ISL_565020, EPI_ISL_565021, EPI_ISL_565022, EPI_ISL_565023, EPI_ISL_565024, EPI_ISL_565025, EPI_ISL_565026, EPI_ISL_565027, EPI_ISL_565028, EPI_ISL_565029, EPI_ISL_565030, EPI_ISL_565031, EPI_ISL_565032, EPI_ISL_565033, EPI_ISL_565034, EPI_ISL_565035, EPI_ISL_565036, EPI_ISL_565037, EPI_ISL_565038, EPI_ISL_565039, EPI_ISL_565040, EPI_ISL_565041, EPI_ISL_565042, EPI_ISL_565043, EPI_ISL_565044, EPI_ISL_565045, EPI_ISL_565046, EPI_ISL_565047, EPI_ISL_565048, EPI_ISL_565049, EPI_ISL_565050, EPI_ISL_565051, EPI_ISL_565052, EPI_ISL_565053, EPI_ISL_565054, EPI_ISL_565055, EPI_ISL_565056, EPI_ISL_565057, EPI_ISL_565058, EPI_ISL_565059, EPI_ISL_565060, EPI_ISL_565061, EPI_ISL_565062, EPI_ISL_565063, EPI_ISL_565064, EPI_ISL_565065, EPI_ISL_565066, EPI_ISL_565067, EPI_ISL_565068, EPI_ISL_565069, EPI_ISL_565070, EPI_ISL_565071, EPI_ISL_565072, EPI_ISL_565073, EPI_ISL_565074, EPI_ISL_565075, EPI_ISL_565076, EPI_ISL_565077, EPI_ISL_565078, EPI_ISL_565079, EPI_ISL_565080, EPI_ISL_565081, EPI_ISL_565082, EPI_ISL_565083, EPI_ISL_565084, EPI_ISL_565085, EPI_ISL_565086, EPI_ISL_565087, EPI_ISL_565088, EPI_ISL_565089, EPI_ISL_565090, EPI_ISL_565091, EPI_ISL_565092, EPI_ISL_565093, EPI_ISL_565094, EPI_ISL_565095, EPI_ISL_565096, EPI_ISL_565097, EPI_ISL_565098, EPI_ISL_565099, EPI_ISL_565100, EPI_ISL_565101, EPI_ISL_565102, EPI_ISL_565103, EPI_ISL_565104, EPI_ISL_565105, EPI_ISL_565106, EPI_ISL_565107, EPI_ISL_565108, EPI_ISL_565109, EPI_ISL_565110, EPI_ISL_565111, EPI_ISL_565112, EPI_ISL_565113, EPI_ISL_565114, EPI_ISL_565115, EPI_ISL_565116, EPI_ISL_565117, EPI_ISL_565118, EPI_ISL_565119, EPI_ISL_565120, EPI_ISL_565121, EPI_ISL_565122, EPI_ISL_565123, EPI_ISL_565124, EPI_ISL_565125, EPI_ISL_565126, EPI_ISL_565127, EPI_ISL_565128, EPI_ISL_565129, EPI_ISL_565130, EPI_ISL_565131, EPI_ISL_565132, EPI_ISL_565133, EPI_ISL_565134, EPI_ISL_565135 | | | |
| see above | Microbiological Diagnostic Unit - Public Health Laboratory (MDU-PHL) | MDU-PHL | Seemann, T., Schultz M. B., Sait, M., Sherry, N. |
| EPI_ISL_565136, EPI_ISL_565137, EPI_ISL_565138, EPI_ISL_565139, EPI_ISL_565140, EPI_ISL_565141, EPI_ISL_565142, EPI_ISL_565143, EPI_ISL_565144, EPI_ISL_565145, EPI_ISL_565146, EPI_ISL_565147, EPI_ISL_565148, EPI_ISL_565149, EPI_ISL_565150, EPI_ISL_565151, EPI_ISL_565152, EPI_ISL_565153, EPI_ISL_565154, EPI_ISL_565155, EPI_ISL_565156, EPI_ISL_565157, EPI_ISL_565158, EPI_ISL_565159, EPI_ISL_565160, EPI_ISL_565161, EPI_ISL_565162, EPI_ISL_565163, EPI_ISL_565164, EPI_ISL_565165, EPI_ISL_565166, EPI_ISL_565167, EPI_ISL_565168, EPI_ISL_565169, EPI_ISL_565170, EPI_ISL_565171, EPI_ISL_565172, EPI_ISL_565173, EPI_ISL_565174, EPI_ISL_565175, EPI_ISL_565176, EPI_ISL_565177, EPI_ISL_565178, EPI_ISL_565179, EPI_ISL_565180, EPI_ISL_565181, EPI_ISL_565182, EPI_ISL_565183, EPI_ISL_565184, EPI_ISL_565185, EPI_ISL_565186, EPI_ISL_565187, EPI_ISL_565188, EPI_ISL_565189, EPI_ISL_565190, EPI_ISL_565191, EPI_ISL_565192, EPI_ISL_565193, EPI_ISL_565194, EPI_ISL_565195, EPI_ISL_565196, EPI_ISL_565197, EPI_ISL_565198, EPI_ISL_565199, EPI_ISL_565200, EPI_ISL_565201, EPI_ISL_565202, EPI_ISL_565203, EPI_ISL_565204, EPI_ISL_565205, EPI_ISL_565206, EPI_ISL_565207, EPI_ISL_565208, EPI_ISL_565209, EPI_ISL_565210, EPI_ISL_565211, EPI_ISL_565212, EPI_ISL_565213, EPI_ISL_565214, EPI_ISL_565215, EPI_ISL_565216, EPI_ISL_565217, EPI_ISL_565218, EPI_ISL_565219, EPI_ISL_565220, EPI_ISL_565221, EPI_ISL_565222 | | | |
| see above | Victorian Infectious Diseases Reference Laboratory (VIDRL) | VIDRL and MDU-PHL | Caly, L., Seemann, T., Sait, M., Schultz, M. B., Druce J., Sherry, N. |
| EPI_ISL_565223, EPI_ISL_565224, EPI_ISL_565225, EPI_ISL_565226, EPI_ISL_565227, EPI_ISL_565228, EPI_ISL_565229, EPI_ISL_565230, EPI_ISL_565231, EPI_ISL_565232, EPI_ISL_565233, EPI_ISL_565234, EPI_ISL_565235, EPI_ISL_565236, EPI_ISL_565237, EPI_ISL_565238, EPI_ISL_565239, EPI_ISL_565240, EPI_ISL_565241, EPI_ISL_565242, EPI_ISL_565243, EPI_ISL_565244, EPI_ISL_565245, EPI_ISL_565246, EPI_ISL_565247, EPI_ISL_565248, EPI_ISL_565249, EPI_ISL_565250, EPI_ISL_565251, EPI_ISL_565252, EPI_ISL_565253, EPI_ISL_565254, EPI_ISL_565255, EPI_ISL_565256, EPI_ISL_565257, EPI_ISL_565258, EPI_ISL_565259, EPI_ISL_565260, EPI_ISL_565261, EPI_ISL_565262, EPI_ISL_565263, EPI_ISL_565264, EPI_ISL_565265, EPI_ISL_565266, EPI_ISL_565267, EPI_ISL_565268, EPI_ISL_565269, EPI_ISL_565270, EPI_ISL_565271, EPI_ISL_565272, EPI_ISL_565273, EPI_ISL_565274, EPI_ISL_565275, EPI_ISL_565276, EPI_ISL_565277, EPI_ISL_565278, EPI_ISL_565279, EPI_ISL_565280, EPI_ISL_565281, EPI_ISL_565282, EPI_ISL_565283, EPI_ISL_565284, EPI_ISL_565285, EPI_ISL_565286, EPI_ISL_565287, EPI_ISL_565288, EPI_ISL_565289, EPI_ISL_565290, EPI_ISL_565291, EPI_ISL_565292, EPI_ISL_565293, EPI_ISL_565294, EPI_ISL_565295, EPI_ISL_565296, EPI_ISL_565297, EPI_ISL_565298, EPI_ISL_565299, EPI_ISL_565300, EPI_ISL_565301, EPI_ISL_565302, EPI_ISL_565303, EPI_ISL_565304, EPI_ISL_565305, EPI_ISL_565306, EPI_ISL_565307, EPI_ISL_565308, EPI_ISL_565309, EPI_ISL_565310, EPI_ISL_565311, EPI_ISL_565312, EPI_ISL_565313, EPI_ISL_565314, EPI_ISL_565315, EPI_ISL_565316, EPI_ISL_565317, EPI_ISL_565318, EPI_ISL_565319, EPI_ISL_565320, EPI_ISL_565321, EPI_ISL_565322, EPI_ISL_565323, EPI_ISL_565324, EPI_ISL_565325, EPI_ISL_565326, EPI_ISL_565327, EPI_ISL_565328, EPI_ISL_565329, EPI_ISL_565330, EPI_ISL_565331, EPI_ISL_565332, EPI_ISL_565333, EPI_ISL_565334, EPI_ISL_565335, EPI_ISL_565336, EPI_ISL_565337, EPI_ISL_565338, EPI_ISL_565339, EPI_ISL_565340, EPI_ISL_565341, EPI_ISL_565342, EPI_ISL_565343, EPI_ISL_565344, EPI_ISL_565345, EPI_ISL_565346, EPI_ISL_565347, EPI_ISL_565348, EPI_ISL_565349, EPI_ISL_565350, EPI_ISL_565351, EPI_ISL_565352, EPI_ISL_565353, EPI_ISL_565354, EPI_ISL_565355, EPI_ISL_565356, EPI_ISL_565357, EPI_ISL_565358, EPI_ISL_565359, EPI_ISL_565360, EPI_ISL_565361, EPI_ISL_565362, EPI_ISL_565363, EPI_ISL_565364, EPI_ISL_565365, EPI_ISL_565366, EPI_ISL_565367, EPI_ISL_565368, EPI_ISL_565369, EPI_ISL_565370, EPI_ISL_565371, EPI_ISL_565372, EPI_ISL_565373, EPI_ISL_565374, EPI_ISL_565375, EPI_ISL_565376, EPI_ISL_565377, EPI_ISL_565378, EPI_ISL_565379, EPI_ISL_565380, EPI_ISL_565381 | | | |
| see above | Microbiological Diagnostic Unit - Public Health Laboratory (MDU-PHL) | MDU-PHL | Seemann, T., Schultz M. B., Sait, M., Sherry, N. |
| EPI_ISL_565382 | Victorian Infectious Diseases Reference Laboratory (VIDRL) | VIDRL and MDU-PHL | Caly, L., Seemann, T., Sait, M., Schultz, M. B., Druce J., Sherry, N. |
| EPI_ISL_565383, EPI_ISL_565384 | Microbiological Diagnostic Unit - Public Health Laboratory (MDU-PHL) | MDU-PHL | Seemann, T., Schultz M. B., Sait, M., Sherry, N. |
| EPI_ISL_565385 | Victorian Infectious Diseases Reference Laboratory (VIDRL) | VIDRL and MDU-PHL | Caly, L., Seemann, T., Sait, M., Schultz, M. B., Druce J., Sherry, N. |
| EPI_ISL_565386, EPI_ISL_565387, EPI_ISL_565388, EPI_ISL_565389, EPI_ISL_565390, EPI_ISL_565391, EPI_ISL_565392, EPI_ISL_565393 | Microbiological Diagnostic Unit - Public Health Laboratory (MDU-PHL) | MDU-PHL | Seemann, T., Schultz M. B., Sait, M., Sherry, N. |
| EPI_ISL_565394, EPI_ISL_565395, EPI_ISL_565396 | Victorian Infectious Diseases Reference Laboratory (VIDRL) | VIDRL and MDU-PHL | Caly, L., Seemann, T., Sait, M., Schultz, M. B., Druce J., Sherry, N. |
| EPI_ISL_565397, EPI_ISL_565398 | Microbiological Diagnostic Unit - Public Health Laboratory (MDU-PHL) | MDU-PHL | Seemann, T., Schultz M. B., Sait, M., Sherry, N. |
| EPI_ISL_565399 | Victorian Infectious Diseases Reference Laboratory (VIDRL) | VIDRL and MDU-PHL | Caly, L., Seemann, T., Sait, M., Schultz, M. B., Druce J., Sherry, N. |
| EPI_ISL_565400 | Microbiological Diagnostic Unit - Public Health Laboratory (MDU-PHL) | MDU-PHL | Seemann, T., Schultz M. B., Sait, M., Sherry, N. |
| EPI_ISL_565401, EPI_ISL_565402, EPI_ISL_565403, EPI_ISL_565404 | Victorian Infectious Diseases Reference Laboratory (VIDRL) | VIDRL and MDU-PHL | Caly, L., Seemann, T., Sait, M., Schultz, M. B., Druce J., Sherry, N. |
| EPI_ISL_565405, EPI_ISL_565406, EPI_ISL_565407, EPI_ISL_565408 | Microbiological Diagnostic Unit - Public Health Laboratory (MDU-PHL) | MDU-PHL | Seemann, T., Schultz M. B., Sait, M., Sherry, N. |
| EPI_ISL_565409 | Victorian Infectious Diseases Reference Laboratory (VIDRL) | VIDRL and MDU-PHL | Caly, L., Seemann, T., Sait, M., Schultz, M. B., Druce J., Sherry, N. |
| EPI_ISL_565410, EPI_ISL_565411, EPI_ISL_565412, EPI_ISL_565413, EPI_ISL_565414 | Microbiological Diagnostic Unit - Public Health Laboratory (MDU-PHL) | MDU-PHL | Seemann, T., Schultz M. B., Sait, M., Sherry, N. |
| EPI_ISL_565415 | Victorian Infectious Diseases Reference Laboratory (VIDRL) | VIDRL and MDU-PHL | Caly, L., Seemann, T., Sait, M., Schultz, M. B., Druce J., Sherry, N. |
| EPI_ISL_565416, EPI_ISL_565417, EPI_ISL_565418, EPI_ISL_565419, EPI_ISL_565420, EPI_ISL_565421, EPI_ISL_565422, EPI_ISL_565423, EPI_ISL_565424, EPI_ISL_565425, EPI_ISL_565426, EPI_ISL_565427, EPI_ISL_565428, EPI_ISL_565429, EPI_ISL_565430, EPI_ISL_565431, EPI_ISL_565432, EPI_ISL_565433, EPI_ISL_565434, EPI_ISL_565435, EPI_ISL_565436, EPI_ISL_565437, EPI_ISL_565438, EPI_ISL_565439, EPI_ISL_565440, EPI_ISL_565441, EPI_ISL_565442, EPI_ISL_565443, EPI_ISL_565444, EPI_ISL_565445, EPI_ISL_565446, EPI_ISL_565447, EPI_ISL_565448, EPI_ISL_565449, EPI_ISL_565450, EPI_ISL_565451, EPI_ISL_565452, EPI_ISL_565453, EPI_ISL_565454, EPI_ISL_565455, EPI_ISL_565456, EPI_ISL_565457, EPI_ISL_565458 | | | |
| see above | Microbiological Diagnostic Unit - Public Health Laboratory (MDU-PHL) | MDU-PHL | Seemann, T., Schultz M. B., Sait, M., Sherry, N. |
| EPI_ISL_565459 | Victorian Infectious Diseases Reference Laboratory (VIDRL) | VIDRL and MDU-PHL | Caly, L., Seemann, T., Sait, M., Schultz, M. B., Druce J., Sherry, N. |
| EPI_ISL_565460 | Microbiological Diagnostic Unit - Public Health Laboratory (MDU-PHL) | MDU-PHL | Seemann, T., Schultz M. B., Sait, M., Sherry, N. |
| EPI_ISL_565461 | Victorian Infectious Diseases Reference Laboratory (VIDRL) | VIDRL and MDU-PHL | Caly, L., Seemann, T., Sait, M., Schultz, M. B., Druce J., Sherry, N. |
| EPI_ISL_565462 | Microbiological Diagnostic Unit - Public Health Laboratory (MDU-PHL) | MDU-PHL | Seemann, T., Schultz M. B., Sait, M., Sherry, N. |
| EPI_ISL_565463, EPI_ISL_565464, EPI_ISL_565465, EPI_ISL_565466, EPI_ISL_565467 | Victorian Infectious Diseases Reference Laboratory (VIDRL) | VIDRL and MDU-PHL | Caly, L., Seemann, T., Sait, M., Schultz, M. B., Druce J., Sherry, N. |
| EPI_ISL_565468 | Microbiological Diagnostic Unit - Public Health Laboratory (MDU-PHL) | MDU-PHL | Seemann, T., Schultz M. B., Sait, M., Sherry, N. |
| EPI_ISL_565469 | Victorian Infectious Diseases Reference Laboratory (VIDRL) | VIDRL and MDU-PHL | Caly, L., Seemann, T., Sait, M., Schultz, M. B., Druce J., Sherry, N. |
| EPI_ISL_565470, EPI_ISL_565471, EPI_ISL_565472, EPI_ISL_565473, EPI_ISL_565474, EPI_ISL_565475, EPI_ISL_565476, EPI_ISL_565477, EPI_ISL_565478, EPI_ISL_565479, EPI_ISL_565480, EPI_ISL_565481, EPI_ISL_565482, EPI_ISL_565483, EPI_ISL_565484, EPI_ISL_565485, EPI_ISL_565486, EPI_ISL_565487, EPI_ISL_565488, EPI_ISL_565489, EPI_ISL_565490, EPI_ISL_565491, EPI_ISL_565492, EPI_ISL_565493, EPI_ISL_565494, EPI_ISL_565495, EPI_ISL_565496, EPI_ISL_565497, EPI_ISL_565498, EPI_ISL_565499, EPI_ISL_565500, EPI_ISL_565501, EPI_ISL_565502, EPI_ISL_565503, EPI_ISL_565504, EPI_ISL_565505, EPI_ISL_565506, EPI_ISL_565507, EPI_ISL_565508 | | | |
| see above | Microbiological Diagnostic Unit - Public Health Laboratory (MDU-PHL) | MDU-PHL | Seemann, T., Schultz M. B., Sait, M., Sherry, N. |
| EPI_ISL_565509 | Victorian Infectious Diseases Reference Laboratory (VIDRL) | VIDRL and MDU-PHL | Caly, L., Seemann, T., Sait, M., Schultz, M. B., Druce J., Sherry, N. |
| EPI_ISL_565510, EPI_ISL_565511, EPI_ISL_565512, EPI_ISL_565513, EPI_ISL_565514, EPI_ISL_565515, EPI_ISL_565516, EPI_ISL_565517, EPI_ISL_565518, EPI_ISL_565519, EPI_ISL_565520, EPI_ISL_565521, EPI_ISL_565522, EPI_ISL_565523, EPI_ISL_565524, EPI_ISL_565525, EPI_ISL_565526, EPI_ISL_565527, EPI_ISL_565528, EPI_ISL_565529, EPI_ISL_565530, EPI_ISL_565531, EPI_ISL_565532, EPI_ISL_565533, EPI_ISL_565534, EPI_ISL_565535, EPI_ISL_565536, EPI_ISL_565537, EPI_ISL_565538, EPI_ISL_565539, EPI_ISL_565540, EPI_ISL_565541, EPI_ISL_565542, EPI_ISL_565543, EPI_ISL_565544, EPI_ISL_565545, EPI_ISL_565546, EPI_ISL_565547, EPI_ISL_565548, EPI_ISL_565549, EPI_ISL_565550, EPI_ISL_565551, EPI_ISL_565552, EPI_ISL_565553, EPI_ISL_565554, EPI_ISL_565555, EPI_ISL_565556, EPI_ISL_565557, EPI_ISL_565558, EPI_ISL_565559, EPI_ISL_565560, EPI_ISL_565561, EPI_ISL_565562, EPI_ISL_565563, EPI_ISL_565564, EPI_ISL_565565, EPI_ISL_565566, EPI_ISL_565567, EPI_ISL_565568, EPI_ISL_565569, EPI_ISL_565570, EPI_ISL_565571, EPI_ISL_565572, EPI_ISL_565573, EPI_ISL_565574, EPI_ISL_565575, EPI_ISL_565576, EPI_ISL_565577, EPI_ISL_565578, EPI_ISL_565579, EPI_ISL_565580, EPI_ISL_565581, EPI_ISL_565582, EPI_ISL_565583, EPI_ISL_565584, EPI_ISL_565585, EPI_ISL_565586, EPI_ISL_565587, EPI_ISL_565588, EPI_ISL_565589, EPI_ISL_565590, EPI_ISL_565591, EPI_ISL_565592, EPI_ISL_565593, EPI_ISL_565594, EPI_ISL_565595, EPI_ISL_565596, EPI_ISL_565597, EPI_ISL_565598, EPI_ISL_565599, EPI_ISL_565600, EPI_ISL_565601, EPI_ISL_565602, EPI_ISL_565603, EPI_ISL_565604, EPI_ISL_565605, EPI_ISL_565606, EPI_ISL_565607, EPI_ISL_565608, EPI_ISL_565609, EPI_ISL_565610, EPI_ISL_565611, EPI_ISL_565612, EPI_ISL_565613, EPI_ISL_565614, EPI_ISL_565615, EPI_ISL_565616, EPI_ISL_565617, EPI_ISL_565618, EPI_ISL_565619, EPI_ISL_565620, EPI_ISL_565621, EPI_ISL_565622, EPI_ISL_565623, EPI_ISL_565624, EPI_ISL_565625, EPI_ISL_565626, EPI_ISL_565627, EPI_ISL_565628, EPI_ISL_565629, EPI_ISL_565630, EPI_ISL_565631, EPI_ISL_565632, EPI_ISL_565633, EPI_ISL_565634, EPI_ISL_565635, EPI_ISL_565636, EPI_ISL_565637, EPI_ISL_565638, EPI_ISL_565639, EPI_ISL_565640, EPI_ISL_565641, EPI_ISL_565642, EPI_ISL_565643, EPI_ISL_565644, EPI_ISL_565645, EPI_ISL_565646, EPI_ISL_565647, EPI_ISL_565648, EPI_ISL_565649, EPI_ISL_565650, EPI_ISL_565651, EPI_ISL_565652, EPI_ISL_565653, EPI_ISL_565654, EPI_ISL_565655, EPI_ISL_565656, EPI_ISL_565657, EPI_ISL_565658, EPI_ISL_565659, EPI_ISL_565660, EPI_ISL_565661, EPI_ISL_565662, EPI_ISL_565663, EPI_ISL_565664, EPI_ISL_565665, EPI_ISL_565666, EPI_ISL_565667, EPI_ISL_565668, EPI_ISL_565669, EPI_ISL_565670, EPI_ISL_565671, EPI_ISL_565672, EPI_ISL_565673, EPI_ISL_565674, EPI_ISL_565675, EPI_ISL_565676, EPI_ISL_565677, EPI_ISL_565678, EPI_ISL_565679, EPI_ISL_565680, EPI_ISL_565681, EPI_ISL_565682, EPI_ISL_565683, EPI_ISL_565684, EPI_ISL_565685, EPI_ISL_565686, EPI_ISL_565687, EPI_ISL_565688, EPI_ISL_565689, EPI_ISL_565690, EPI_ISL_565691, EPI_ISL_565692, EPI_ISL_565693, EPI_ISL_565694, EPI_ISL_565695, EPI_ISL_565696, EPI_ISL_565697, EPI_ISL_565698, EPI_ISL_565699, EPI_ISL_565700, EPI_ISL_565701, EPI_ISL_565702, EPI_ISL_565703, EPI_ISL_565704, EPI_ISL_565705, EPI_ISL_565706, EPI_ISL_565707, EPI_ISL_565708, EPI_ISL_565709, EPI_ISL_565710, EPI_ISL_565711, EPI_ISL_565712, EPI_ISL_565713, EPI_ISL_565714, EPI_ISL_565715, EPI_ISL_565716, EPI_ISL_565717, EPI_ISL_565718, EPI_ISL_565719, EPI_ISL_565720, EPI_ISL_565721, EPI_ISL_565722, EPI_ISL_565723, EPI_ISL_565724, EPI_ISL_565725, EPI_ISL_565726, EPI_ISL_565727, EPI_ISL_565728, EPI_ISL_565729 | | | |
| see above | Microbiological Diagnostic Unit - Public Health Laboratory (MDU-PHL) | MDU-PHL | Seemann, T., Schultz M. B., Sait, M., Sherry, N. |
| EPI_ISL_565730, EPI_ISL_565731, EPI_ISL_565732, EPI_ISL_565733, EPI_ISL_565734, EPI_ISL_565735, EPI_ISL_565736, EPI_ISL_565737, EPI_ISL_565738, EPI_ISL_565739, EPI_ISL_565740, EPI_ISL_565741, EPI_ISL_565742 | | | |
| see above | Victorian Infectious Diseases Reference Laboratory (VIDRL) | VIDRL and MDU-PHL | Caly, L., Seemann, T., Sait, M., Schultz, M. B., Druce J., Sherry, N. |
| EPI_ISL_565743, EPI_ISL_565744, EPI_ISL_565745, EPI_ISL_565746, EPI_ISL_565747, EPI_ISL_565748, EPI_ISL_565749, EPI_ISL_565750, EPI_ISL_565751, EPI_ISL_565752, EPI_ISL_565753, EPI_ISL_565754, EPI_ISL_565755, EPI_ISL_565756, EPI_ISL_565757, EPI_ISL_565758, EPI_ISL_565759, EPI_ISL_565760, EPI_ISL_565761, EPI_ISL_565762, EPI_ISL_565763, EPI_ISL_565764, EPI_ISL_565765, EPI_ISL_565766, EPI_ISL_565767, EPI_ISL_565768, EPI_ISL_565769, EPI_ISL_565770, EPI_ISL_565771, EPI_ISL_565772, EPI_ISL_565773, EPI_ISL_565774, EPI_ISL_565775, EPI_ISL_565776, EPI_ISL_565777, EPI_ISL_565778, EPI_ISL_565779, EPI_ISL_565780, EPI_ISL_565781, EPI_ISL_565782, EPI_ISL_565783, EPI_ISL_565784, EPI_ISL_565785, EPI_ISL_565786, EPI_ISL_565787, EPI_ISL_565788, EPI_ISL_565789, EPI_ISL_565790, EPI_ISL_565791, EPI_ISL_565792, EPI_ISL_565793, EPI_ISL_565794, EPI_ISL_565795, EPI_ISL_565796, EPI_ISL_565797, EPI_ISL_565798, EPI_ISL_565799, EPI_ISL_565800, EPI_ISL_565801, EPI_ISL_565802, EPI_ISL_565803, EPI_ISL_565804, EPI_ISL_565805, EPI_ISL_565806, EPI_ISL_565807, EPI_ISL_565808, EPI_ISL_565809, EPI_ISL_565810, EPI_ISL_565811, EPI_ISL_565812, EPI_ISL_565813, EPI_ISL_565814, | | | |

| | | | |
|---|---|---|---|
| EPI_ISL_565815, EPI_ISL_565816, EPI_ISL_565817, EPI_ISL_565818, EPI_ISL_565819, EPI_ISL_565820, EPI_ISL_565821, EPI_ISL_565822, EPI_ISL_565823, EPI_ISL_565824, EPI_ISL_565825, EPI_ISL_565826, EPI_ISL_565827, EPI_ISL_565828, EPI_ISL_565829 | | | |
| see above | Microbiological Diagnostic Unit - Public Health Laboratory (MDU-PHL) | MDU-PHL | Seemann, T., Schultz M. B., Sait, M., Sherry, N. |
| EPI_ISL_565830, EPI_ISL_565832 | Delaware Public Health Lab | Delaware Public Health Lab | Gregory Hovan |
| EPI_ISL_565837, EPI_ISL_565838, EPI_ISL_565843, EPI_ISL_565846, EPI_ISL_565849, EPI_ISL_565853, EPI_ISL_565855, EPI_ISL_565873, EPI_ISL_565881, EPI_ISL_565884, EPI_ISL_565949, EPI_ISL_565951, EPI_ISL_565953, EPI_ISL_565954, EPI_ISL_565955, EPI_ISL_565956, EPI_ISL_565957, EPI_ISL_565958, EPI_ISL_565959, EPI_ISL_565960, EPI_ISL_565961, EPI_ISL_565962, EPI_ISL_565963, EPI_ISL_565964, EPI_ISL_565965, EPI_ISL_565966, EPI_ISL_565967, EPI_ISL_565968, EPI_ISL_565969, EPI_ISL_565970, EPI_ISL_565971, EPI_ISL_565972, EPI_ISL_565973, EPI_ISL_565974, EPI_ISL_565975, EPI_ISL_565976, EPI_ISL_565977, EPI_ISL_565978, EPI_ISL_565979, EPI_ISL_565980, EPI_ISL_565981, EPI_ISL_565982, EPI_ISL_565983, EPI_ISL_565984, EPI_ISL_565985, EPI_ISL_565986, EPI_ISL_565987, EPI_ISL_565988, EPI_ISL_565989, EPI_ISL_566019, EPI_ISL_566020, EPI_ISL_566021, EPI_ISL_566022, EPI_ISL_566023, EPI_ISL_566024, EPI_ISL_566025, EPI_ISL_566026, EPI_ISL_566027, EPI_ISL_566028, EPI_ISL_566029, EPI_ISL_566030, EPI_ISL_566031, EPI_ISL_566032, EPI_ISL_566033, EPI_ISL_566034, EPI_ISL_566035 | | | |
| see above | Michigan Department of Health and Human Services, Bureau of Laboratories | Michigan Department of Health and Human Services, Bureau of Laboratories | Blankenship HM, Riner D, Soehnlen MK |
| EPI_ISL_566047, EPI_ISL_566048, EPI_ISL_566049, EPI_ISL_566050, EPI_ISL_566051, EPI_ISL_566052, EPI_ISL_566053, EPI_ISL_566054, EPI_ISL_566055, EPI_ISL_566056, EPI_ISL_566057, EPI_ISL_566058, EPI_ISL_566059, EPI_ISL_566060, EPI_ISL_566061, EPI_ISL_566062, EPI_ISL_566063, EPI_ISL_566064, EPI_ISL_566065, EPI_ISL_566066, EPI_ISL_566067, EPI_ISL_566068, EPI_ISL_566069, EPI_ISL_566070, EPI_ISL_566071, EPI_ISL_566072, EPI_ISL_566073, EPI_ISL_566074 | | | |
| see above | Respiratory Virus Unit, Microbiology Services Colindale, Public Health England | Respiratory Virus Unit, Microbiology Services Colindale, Public Health England | PHE Covid Sequencing Team |
| EPI_ISL_566077, EPI_ISL_566078, EPI_ISL_566079, EPI_ISL_566080, EPI_ISL_566081, EPI_ISL_566082, EPI_ISL_566083 | Pathogenic Microorganisms Variability Laboratory | WHO National Influenza Centre Russian Federation | Andrey Komissarov, Artem Fadeev, Anna Ivanova, Kseniya Komissarova, Daria Danilenko, Dmitry Lioznov, Nadezhda Kuznetsova, Elena Shidlovskaya, Elizaveta Divisenko, Ekaterina Milashenko, Kirill Krasnoslobotsev, Evgeniya Mukasheva, Anna Ignatieva, Svetlana Trushakova, Alexey Shchetinin, Maria Nikiforova, Andrey Pochtovyy, Valeria Bacalin, Evgeny Usachev, Olga Burgasova, Ludmila Kolobukhina, Svetlana Smetanina, Elena Burtseva, Artem Tkachuk, Vladimir Gushchin, Alexander Gintsburg |
| EPI_ISL_566084 | Delaware Public Health Lab | Delaware Public Health Lab | Gregory Hovan |
| EPI_ISL_566085, EPI_ISL_566086, EPI_ISL_566087, EPI_ISL_566088, EPI_ISL_566089, EPI_ISL_566090, EPI_ISL_566091, EPI_ISL_566092, EPI_ISL_566093, EPI_ISL_566094, EPI_ISL_566095, EPI_ISL_566096, EPI_ISL_566097, EPI_ISL_566098, EPI_ISL_566099, EPI_ISL_566100, EPI_ISL_566101, EPI_ISL_566102, EPI_ISL_566103, EPI_ISL_566104, EPI_ISL_566105, EPI_ISL_566106, EPI_ISL_566107 | | | |
| see above | GA Department of Public Health Laboratory | Pathogen Discovery, Respiratory Viruses Branch, Division of Viral Diseases, Centers for Disease Control and Prevention | Jing Zhang, Brian Lynch, Yan Li, Anna Montmayeur, Krista Queen, Ying Tao, Anna Uehara, Clinton R. Paden, Rachel Marine, Haibin Wang, Suxiang Tong |
| EPI_ISL_566108 | Delaware Public Health Lab | Delaware Public Health Lab | Gregory Hovan |
| EPI_ISL_566111, EPI_ISL_566113, EPI_ISL_566114, EPI_ISL_566115, EPI_ISL_566116, EPI_ISL_566117, EPI_ISL_566118, EPI_ISL_566119, EPI_ISL_566120, EPI_ISL_566121, EPI_ISL_566122, EPI_ISL_566123, EPI_ISL_566124, EPI_ISL_566125, EPI_ISL_566126, EPI_ISL_566127, EPI_ISL_566128, EPI_ISL_566129, EPI_ISL_566130, EPI_ISL_566131, EPI_ISL_566132, EPI_ISL_566133, EPI_ISL_566134, EPI_ISL_566135, EPI_ISL_566136, EPI_ISL_566137, EPI_ISL_566138, EPI_ISL_566139, EPI_ISL_566140, EPI_ISL_566141, EPI_ISL_566142, EPI_ISL_566143, EPI_ISL_566144, EPI_ISL_566145, EPI_ISL_566146, EPI_ISL_566147, EPI_ISL_566148, EPI_ISL_566149, EPI_ISL_566151, EPI_ISL_566153, EPI_ISL_566154, EPI_ISL_566155, EPI_ISL_566156, EPI_ISL_566157, EPI_ISL_566159, EPI_ISL_566160, EPI_ISL_566161, EPI_ISL_566162, EPI_ISL_566163, EPI_ISL_566164, EPI_ISL_566166, EPI_ISL_566167, EPI_ISL_566168, EPI_ISL_566169, EPI_ISL_566170, EPI_ISL_566171, EPI_ISL_566174, EPI_ISL_566175, EPI_ISL_566176, EPI_ISL_566177, EPI_ISL_566179, EPI_ISL_566180, EPI_ISL_566181, EPI_ISL_566182, EPI_ISL_566183, EPI_ISL_566184, EPI_ISL_566185, EPI_ISL_566186, EPI_ISL_566187, EPI_ISL_566188, EPI_ISL_566189, EPI_ISL_566190, EPI_ISL_566191, EPI_ISL_566192, EPI_ISL_566194, EPI_ISL_566195, EPI_ISL_566196, EPI_ISL_566197, EPI_ISL_566198, EPI_ISL_566199, EPI_ISL_566200, EPI_ISL_566202, EPI_ISL_566203, EPI_ISL_566204, EPI_ISL_566205, EPI_ISL_566206, EPI_ISL_566207, EPI_ISL_566210, EPI_ISL_566211, EPI_ISL_566212, EPI_ISL_566213, EPI_ISL_566214, EPI_ISL_566215, EPI_ISL_566216, EPI_ISL_566217, EPI_ISL_566218, EPI_ISL_566219, EPI_ISL_566220, EPI_ISL_566221, EPI_ISL_566222, EPI_ISL_566223, EPI_ISL_566224, EPI_ISL_566225, EPI_ISL_566226, EPI_ISL_566227, EPI_ISL_566228, EPI_ISL_566229, EPI_ISL_566230, EPI_ISL_566231, EPI_ISL_566234, EPI_ISL_566235, EPI_ISL_566236, EPI_ISL_566237, EPI_ISL_566238, EPI_ISL_566239, EPI_ISL_566240, EPI_ISL_566241, EPI_ISL_566243, EPI_ISL_566244, EPI_ISL_566245, EPI_ISL_566246, EPI_ISL_566247, EPI_ISL_566248, EPI_ISL_566249, EPI_ISL_566250, EPI_ISL_566251, EPI_ISL_566252, EPI_ISL_566253, EPI_ISL_566255, EPI_ISL_566256, EPI_ISL_566257, EPI_ISL_566258, EPI_ISL_566259, EPI_ISL_566260, EPI_ISL_566261, EPI_ISL_566262, EPI_ISL_566263, EPI_ISL_566264, EPI_ISL_566265, EPI_ISL_566267, EPI_ISL_566268, EPI_ISL_566269, EPI_ISL_566270, EPI_ISL_566271, EPI_ISL_566272, EPI_ISL_566273, EPI_ISL_566275, EPI_ISL_566277, EPI_ISL_566278, EPI_ISL_566279, EPI_ISL_566280, EPI_ISL_566281, EPI_ISL_566282, EPI_ISL_566283, EPI_ISL_566284, EPI_ISL_566285, EPI_ISL_566286, EPI_ISL_566287, EPI_ISL_566288, EPI_ISL_566290, EPI_ISL_566291, EPI_ISL_566292, EPI_ISL_566294, EPI_ISL_566295, EPI_ISL_566296, EPI_ISL_566297, EPI_ISL_566298, EPI_ISL_566299, EPI_ISL_566300, EPI_ISL_566301, EPI_ISL_566302, EPI_ISL_566303, EPI_ISL_566304, EPI_ISL_566306, EPI_ISL_566308, EPI_ISL_566309, EPI_ISL_566310, EPI_ISL_566311, EPI_ISL_566312, EPI_ISL_566313, EPI_ISL_566314, EPI_ISL_566316, EPI_ISL_566317, EPI_ISL_566318, EPI_ISL_566319, EPI_ISL_566320, EPI_ISL_566321, EPI_ISL_566322, EPI_ISL_566323, EPI_ISL_566325, EPI_ISL_566326, EPI_ISL_566327, EPI_ISL_566328, EPI_ISL_566329, EPI_ISL_566330, EPI_ISL_566331, EPI_ISL_566333, EPI_ISL_566334, EPI_ISL_566335, EPI_ISL_566336, EPI_ISL_566337, EPI_ISL_566338, EPI_ISL_566339, EPI_ISL_566340, EPI_ISL_566341, EPI_ISL_566342, EPI_ISL_566343, EPI_ISL_566345, EPI_ISL_566346, EPI_ISL_566347, EPI_ISL_566348, EPI_ISL_566349, EPI_ISL_566350, EPI_ISL_566351, EPI_ISL_566352, EPI_ISL_566353, EPI_ISL_566354, EPI_ISL_566355, EPI_ISL_566356, EPI_ISL_566357, EPI_ISL_566358, EPI_ISL_566359, EPI_ISL_566360, EPI_ISL_566361, EPI_ISL_566362, EPI_ISL_566363, EPI_ISL_566364, EPI_ISL_566366, EPI_ISL_566367, EPI_ISL_566368, EPI_ISL_566369, EPI_ISL_566370, EPI_ISL_566372, EPI_ISL_566373, EPI_ISL_566374, EPI_ISL_566375, EPI_ISL_566376, EPI_ISL_566378, EPI_ISL_566380, EPI_ISL_566381, EPI_ISL_566382, EPI_ISL_566383, EPI_ISL_566384, EPI_ISL_566386, EPI_ISL_566387, EPI_ISL_566388, EPI_ISL_566389, EPI_ISL_566390, EPI_ISL_566391, EPI_ISL_566392, EPI_ISL_566393, EPI_ISL_566394, EPI_ISL_566395, EPI_ISL_566396, EPI_ISL_566397, EPI_ISL_566398, EPI_ISL_566399, EPI_ISL_566400, EPI_ISL_566402, EPI_ISL_566403, EPI_ISL_566404, EPI_ISL_566405, EPI_ISL_566407, EPI_ISL_566408, EPI_ISL_566409, EPI_ISL_566410, EPI_ISL_566411, EPI_ISL_566412, EPI_ISL_566413, EPI_ISL_566414, EPI_ISL_566415, EPI_ISL_566416, EPI_ISL_566417, EPI_ISL_566418, EPI_ISL_566419, EPI_ISL_566420 | | | |
| see above | Lighthouse Lab in Alderley Park | Wellcome Sanger Institute for the COVID-19 Genomics UK Consortium | Jacquelyn Wynn, Mairead Hyland, The Lighthouse Lab in Alderley Park and Alex Alderton, Roberto Amato, Sonia Goncalves, Ewan Harrison, David K. Jackson, Ian Johnston, Dominic Kwiatkowski, Cordelia Langford, John Sillitoe on behalf of the Wellcome Sanger Institute COVID-19 Surveillance Team |
| EPI_ISL_566421 | Lighthouse Lab in Milton Keynes | Wellcome Sanger Institute for the COVID-19 Genomics UK Consortium | The Lighthouse Lab in Milton Keynes and Alex Alderton, Roberto Amato, Sonia Goncalves, Ewan Harrison, David K. Jackson, Ian Johnston, Dominic Kwiatkowski, Cordelia Langford, John Sillitoe on behalf of the Wellcome Sanger Institute COVID-19 Surveillance Team |
| EPI_ISL_566423 | Lighthouse Lab in Cambridge | Wellcome Sanger Institute for the COVID-19 Genomics UK Consortium | Rob Howes, The Lighthouse Lab in Cambridge and Alex Alderton, Roberto Amato, Sonia Goncalves, Ewan Harrison, David K. Jackson, Ian Johnston, Dominic Kwiatkowski, Cordelia Langford, John Sillitoe on behalf of the Wellcome Sanger Institute COVID-19 Surveillance Team |
| EPI_ISL_566424, EPI_ISL_566425 | Lighthouse Lab in Milton Keynes | Wellcome Sanger Institute for the COVID-19 Genomics UK Consortium | The Lighthouse Lab in Milton Keynes and Alex Alderton, Roberto Amato, Sonia Goncalves, Ewan Harrison, David K. Jackson, Ian Johnston, Dominic Kwiatkowski, Cordelia Langford, John Sillitoe on behalf of the Wellcome Sanger Institute COVID-19 Surveillance Team |
| EPI_ISL_566429 | Lighthouse Lab in Cambridge | Wellcome Sanger Institute for the COVID-19 Genomics UK Consortium | Rob Howes, The Lighthouse Lab in Cambridge and Alex Alderton, Roberto Amato, Sonia Goncalves, Ewan Harrison, David K. Jackson, Ian Johnston, Dominic Kwiatkowski, Cordelia Langford, John Sillitoe on behalf of the Wellcome Sanger Institute COVID-19 Surveillance Team |
| EPI_ISL_566430, EPI_ISL_566431, EPI_ISL_566432 | Lighthouse Lab in Milton Keynes | Wellcome Sanger Institute for the COVID-19 Genomics UK Consortium | The Lighthouse Lab in Milton Keynes and Alex Alderton, Roberto Amato, Sonia Goncalves, Ewan Harrison, David K. Jackson, Ian Johnston, Dominic Kwiatkowski, Cordelia Langford, John Sillitoe on behalf of the Wellcome Sanger Institute COVID-19 Surveillance Team |
| EPI_ISL_566433, EPI_ISL_566434, EPI_ISL_566436, EPI_ISL_566437, EPI_ISL_566438, EPI_ISL_566439, EPI_ISL_566440, EPI_ISL_566441, EPI_ISL_566443 | Lighthouse Lab in Milton Keynes | Wellcome Sanger Institute for the COVID-19 Genomics UK Consortium | The Lighthouse Lab in Milton Keynes and Alex Alderton, Roberto Amato, Sonia Goncalves, Ewan Harrison, David K. Jackson, Ian Johnston, Dominic Kwiatkowski, Cordelia Langford, John Sillitoe on behalf of the Wellcome Sanger Institute COVID-19 Surveillance Team |
| EPI_ISL_566444, EPI_ISL_566445 | Lighthouse Lab in Cambridge | Wellcome Sanger Institute for the COVID-19 Genomics UK Consortium | Rob Howes, The Lighthouse Lab in Cambridge and Alex Alderton, Roberto Amato, Sonia Goncalves, Ewan Harrison, David K. Jackson, Ian Johnston, Dominic Kwiatkowski, Cordelia Langford, John Sillitoe on behalf of the Wellcome Sanger Institute COVID-19 Surveillance Team |
| EPI_ISL_566446, EPI_ISL_566447, EPI_ISL_566448, EPI_ISL_566449, EPI_ISL_566450, EPI_ISL_566451, EPI_ISL_566452 | Lighthouse Lab in Milton Keynes | Wellcome Sanger Institute for the COVID-19 Genomics UK Consortium | The Lighthouse Lab in Milton Keynes and Alex Alderton, Roberto Amato, Sonia Goncalves, Ewan Harrison, David K. Jackson, Ian Johnston, Dominic Kwiatkowski, Cordelia Langford, John Sillitoe on behalf of the Wellcome Sanger Institute COVID-19 Surveillance Team |
| EPI_ISL_566453 | Lighthouse Lab in Cambridge | Wellcome Sanger Institute for the COVID-19 Genomics UK Consortium | Rob Howes, The Lighthouse Lab in Cambridge and Alex Alderton, Roberto Amato, Sonia Goncalves, Ewan Harrison, David K. Jackson, Ian Johnston, Dominic Kwiatkowski, Cordelia Langford, John Sillitoe on behalf of the Wellcome Sanger Institute COVID-19 Surveillance Team |
| EPI_ISL_566454 | Lighthouse Lab in Milton Keynes | Wellcome Sanger Institute for the COVID-19 Genomics UK Consortium | The Lighthouse Lab in Milton Keynes and Alex Alderton, Roberto Amato, Sonia Goncalves, Ewan Harrison, David K. Jackson, Ian Johnston, Dominic Kwiatkowski, Cordelia Langford, John Sillitoe on behalf of the Wellcome Sanger Institute COVID-19 Surveillance Team |
| EPI_ISL_566457 | Lighthouse Lab in Cambridge | Wellcome Sanger Institute for the COVID-19 Genomics UK Consortium | Rob Howes, The Lighthouse Lab in Cambridge and Alex Alderton, Roberto Amato, Sonia Goncalves, Ewan Harrison, David K. Jackson, Ian Johnston, Dominic Kwiatkowski, Cordelia Langford, John Sillitoe on behalf of the Wellcome Sanger Institute COVID-19 Surveillance Team |
| EPI_ISL_566458, EPI_ISL_566459, EPI_ISL_566460, EPI_ISL_566461, EPI_ISL_566462, EPI_ISL_566463, EPI_ISL_566465, EPI_ISL_566466 | Lighthouse Lab in Milton Keynes | Wellcome Sanger Institute for the COVID-19 Genomics UK Consortium | The Lighthouse Lab in Milton Keynes and Alex Alderton, Roberto Amato, Sonia Goncalves, Ewan Harrison, David K. Jackson, Ian Johnston, Dominic Kwiatkowski, Cordelia Langford, John Sillitoe on behalf of the Wellcome Sanger Institute COVID-19 Surveillance Team |
| EPI_ISL_566467 | Lighthouse Lab in Cambridge | Wellcome Sanger Institute for the COVID-19 Genomics UK Consortium | Rob Howes, The Lighthouse Lab in Cambridge and Alex Alderton, Roberto Amato, Sonia Goncalves, Ewan Harrison, David K. Jackson, Ian Johnston, Dominic Kwiatkowski, Cordelia Langford, John Sillitoe on behalf of the Wellcome Sanger Institute COVID-19 Surveillance Team |
| EPI_ISL_566468, EPI_ISL_566469, EPI_ISL_566470, EPI_ISL_566471, EPI_ISL_566472, EPI_ISL_566475, EPI_ISL_566477, EPI_ISL_566478, EPI_ISL_566480, EPI_ISL_566481, EPI_ISL_566482, EPI_ISL_566483, EPI_ISL_566484, EPI_ISL_566485, EPI_ISL_566487, EPI_ISL_566488, EPI_ISL_566489, EPI_ISL_566490, EPI_ISL_566491 | | | |
| see above | Lighthouse Lab in Milton Keynes | Wellcome Sanger Institute for the COVID-19 Genomics UK Consortium | The Lighthouse Lab in Milton Keynes and Alex Alderton, Roberto Amato, Sonia Goncalves, Ewan Harrison, David K. Jackson, Ian Johnston, Dominic Kwiatkowski, Cordelia Langford, John Sillitoe on behalf of the Wellcome Sanger Institute COVID-19 Surveillance Team |
| EPI_ISL_566493, EPI_ISL_566495 | Lighthouse Lab in Cambridge | Wellcome Sanger Institute for the COVID-19 Genomics UK Consortium | Rob Howes, The Lighthouse Lab in Cambridge and Alex Alderton, Roberto Amato, Sonia Goncalves, Ewan Harrison, David K. Jackson, Ian Johnston, Dominic Kwiatkowski, Cordelia Langford, John Sillitoe on behalf of the Wellcome Sanger Institute COVID-19 Surveillance Team |
| EPI_ISL_566496, EPI_ISL_566497, EPI_ISL_566498, EPI_ISL_566499 | Lighthouse Lab in Milton Keynes | Wellcome Sanger Institute for the COVID-19 Genomics UK Consortium | The Lighthouse Lab in Milton Keynes and Alex Alderton, Roberto Amato, Sonia Goncalves, Ewan Harrison, David K. Jackson, Ian Johnston, Dominic Kwiatkowski, Cordelia Langford, John Sillitoe on behalf of the Wellcome Sanger Institute COVID-19 Surveillance Team |
| EPI_ISL_566500 | Lighthouse Lab in Cambridge | Wellcome Sanger Institute for the COVID-19 Genomics UK Consortium | Rob Howes, The Lighthouse Lab in Cambridge and Alex Alderton, Roberto Amato, Sonia Goncalves, Ewan Harrison, David K. Jackson, Ian Johnston, Dominic Kwiatkowski, Cordelia Langford, John Sillitoe on behalf of the Wellcome Sanger Institute COVID-19 Surveillance Team |
| EPI_ISL_566503, EPI_ISL_566506, EPI_ISL_566507 | Lighthouse Lab in Milton Keynes | Wellcome Sanger Institute for the COVID-19 Genomics UK Consortium | The Lighthouse Lab in Milton Keynes and Alex Alderton, Roberto Amato, Sonia Goncalves, Ewan Harrison, David K. Jackson, Ian Johnston, Dominic Kwiatkowski, Cordelia Langford, John Sillitoe on behalf of the Wellcome Sanger Institute COVID-19 Surveillance Team |
| EPI_ISL_566508 | Lighthouse Lab in Cambridge | Wellcome Sanger Institute for the COVID-19 Genomics UK Consortium | Rob Howes, The Lighthouse Lab in Cambridge and Alex Alderton, Roberto Amato, Sonia Goncalves, Ewan Harrison, David K. Jackson, Ian Johnston, Dominic Kwiatkowski, Cordelia Langford, John Sillitoe on behalf of the Wellcome Sanger Institute COVID-19 Surveillance Team |
| EPI_ISL_566509, EPI_ISL_566510, EPI_ISL_566511, EPI_ISL_566512, EPI_ISL_566514, EPI_ISL_566515, EPI_ISL_566516, EPI_ISL_566517 | Lighthouse Lab in Milton Keynes | Wellcome Sanger Institute for the COVID-19 Genomics UK Consortium | The Lighthouse Lab in Milton Keynes and Alex Alderton, Roberto Amato, Sonia Goncalves, Ewan Harrison, David K. Jackson, Ian Johnston, Dominic Kwiatkowski, Cordelia Langford, John Sillitoe on behalf of the Wellcome Sanger Institute COVID-19 Surveillance Team |
| EPI_ISL_566518, EPI_ISL_566519 | Lighthouse Lab in Cambridge | Wellcome Sanger Institute for the COVID-19 Genomics UK Consortium | Rob Howes, The Lighthouse Lab in Cambridge and Alex Alderton, Roberto Amato, Sonia Goncalves, Ewan Harrison, David K. Jackson, Ian Johnston, Dominic Kwiatkowski, Cordelia Langford, John Sillitoe on behalf of the Wellcome Sanger Institute COVID-19 Surveillance Team |
| EPI_ISL_566520, EPI_ISL_566521, EPI_ISL_566522, EPI_ISL_566523 | Lighthouse Lab in Milton Keynes | Wellcome Sanger Institute for the COVID-19 Genomics UK Consortium | The Lighthouse Lab in Milton Keynes and Alex Alderton, Roberto Amato, Sonia Goncalves, Ewan Harrison, David K. Jackson, Ian Johnston, Dominic Kwiatkowski, Cordelia Langford, John Sillitoe on behalf of the Wellcome Sanger Institute COVID-19 Surveillance Team |
| EPI_ISL_566524 | Lighthouse Lab in Cambridge | Wellcome Sanger Institute for the COVID-19 Genomics UK Consortium | Rob Howes, The Lighthouse Lab in Cambridge and Alex Alderton, Roberto Amato, Sonia Goncalves, Ewan Harrison, David K. Jackson, Ian Johnston, Dominic Kwiatkowski, Cordelia Langford, John Sillitoe on behalf of the Wellcome Sanger Institute COVID-19 Surveillance Team |
| EPI_ISL_566525, EPI_ISL_566526, EPI_ISL_566527 | Lighthouse Lab in Milton Keynes | Wellcome Sanger Institute for the COVID-19 Genomics UK Consortium | The Lighthouse Lab in Milton Keynes and Alex Alderton, Roberto Amato, Sonia Goncalves, Ewan Harrison, David K. Jackson, Ian Johnston, Dominic Kwiatkowski, Cordelia Langford, John Sillitoe on behalf of the Wellcome Sanger Institute COVID-19 Surveillance Team |
| EPI_ISL_566528 | Lighthouse Lab in Cambridge | Wellcome Sanger Institute for the COVID-19 Genomics UK Consortium | Rob Howes, The Lighthouse Lab in Cambridge and Alex Alderton, Roberto Amato, Sonia Goncalves, Ewan Harrison, David K. Jackson, Ian Johnston, Dominic Kwiatkowski, Cordelia Langford, John Sillitoe on behalf of the Wellcome Sanger Institute COVID-19 Surveillance Team |
| EPI_ISL_566530, EPI_ISL_566531, EPI_ISL_566532, EPI_ISL_566533 | Lighthouse Lab in Milton Keynes | Wellcome Sanger Institute for the COVID-19 Genomics UK Consortium | The Lighthouse Lab in Milton Keynes and Alex Alderton, Roberto Amato, Sonia Goncalves, Ewan Harrison, David K. Jackson, Ian Johnston, Dominic Kwiatkowski, Cordelia Langford, John Sillitoe on behalf of the Wellcome Sanger Institute COVID-19 Surveillance Team |
| EPI_ISL_566534 | Lighthouse Lab in Cambridge | Wellcome Sanger Institute for the COVID-19 Genomics UK Consortium | Rob Howes, The Lighthouse Lab in Cambridge and Alex Alderton, Roberto Amato, Sonia Goncalves, Ewan Harrison, David K. Jackson, Ian Johnston, Dominic Kwiatkowski, Cordelia Langford, John Sillitoe on behalf of the Wellcome Sanger Institute COVID-19 Surveillance Team |

| | | | |
|---|---|---|---|
| EPI_ISL_566535, EPI_ISL_566536, EPI_ISL_566539, EPI_ISL_566540, EPI_ISL_566541, EPI_ISL_566542 | Lighthouse Lab in Milton Keynes | Wellcome Sanger Institute for the COVID-19 Genomics UK Consortium | The Lighthouse Lab in Milton Keynes and Alex Alderton, Roberto Amato, Sonia Goncalves, Ewan Harrison, David K. Jackson, Ian Johnston, Dominic Kwiatkowski, Cordelia Langford, John Sillitoe on behalf of the Wellcome Sanger Institute COVID-19 Surveillance Team |
| EPI_ISL_566543, EPI_ISL_566544 | Lighthouse Lab in Cambridge | Wellcome Sanger Institute for the COVID-19 Genomics UK Consortium | Rob Howes, The Lighthouse Lab in Cambridge and Alex Alderton, Roberto Amato, Sonia Goncalves, Ewan Harrison, David K. Jackson, Ian Johnston, Dominic Kwiatkowski, Cordelia Langford, John Sillitoe on behalf of the Wellcome Sanger Institute COVID-19 Surveillance Team |
| EPI_ISL_566545, EPI_ISL_566546, EPI_ISL_566547, EPI_ISL_566548, EPI_ISL_566549, EPI_ISL_566550, EPI_ISL_566551, EPI_ISL_566552, EPI_ISL_566553, EPI_ISL_566555, EPI_ISL_566556, EPI_ISL_566557, EPI_ISL_566558, EPI_ISL_566559, EPI_ISL_566560, EPI_ISL_566561, EPI_ISL_566562, EPI_ISL_566564, EPI_ISL_566565, EPI_ISL_566566 | | | |
| see above | Lighthouse Lab in Milton Keynes | Wellcome Sanger Institute for the COVID-19 Genomics UK Consortium | The Lighthouse Lab in Milton Keynes and Alex Alderton, Roberto Amato, Sonia Goncalves, Ewan Harrison, David K. Jackson, Ian Johnston, Dominic Kwiatkowski, Cordelia Langford, John Sillitoe on behalf of the Wellcome Sanger Institute COVID-19 Surveillance Team |
| EPI_ISL_566567 | Lighthouse Lab in Cambridge | Wellcome Sanger Institute for the COVID-19 Genomics UK Consortium | Rob Howes, The Lighthouse Lab in Cambridge and Alex Alderton, Roberto Amato, Sonia Goncalves, Ewan Harrison, David K. Jackson, Ian Johnston, Dominic Kwiatkowski, Cordelia Langford, John Sillitoe on behalf of the Wellcome Sanger Institute COVID-19 Surveillance Team |
| EPI_ISL_566568, EPI_ISL_566569, EPI_ISL_566570, EPI_ISL_566571, EPI_ISL_566572, EPI_ISL_566573, EPI_ISL_566574 | Lighthouse Lab in Milton Keynes | Wellcome Sanger Institute for the COVID-19 Genomics UK Consortium | The Lighthouse Lab in Milton Keynes and Alex Alderton, Roberto Amato, Sonia Goncalves, Ewan Harrison, David K. Jackson, Ian Johnston, Dominic Kwiatkowski, Cordelia Langford, John Sillitoe on behalf of the Wellcome Sanger Institute COVID-19 Surveillance Team |
| EPI_ISL_566576 | Lighthouse Lab in Cambridge | Wellcome Sanger Institute for the COVID-19 Genomics UK Consortium | Rob Howes, The Lighthouse Lab in Cambridge and Alex Alderton, Roberto Amato, Sonia Goncalves, Ewan Harrison, David K. Jackson, Ian Johnston, Dominic Kwiatkowski, Cordelia Langford, John Sillitoe on behalf of the Wellcome Sanger Institute COVID-19 Surveillance Team |
| EPI_ISL_566577, EPI_ISL_566578, EPI_ISL_566579, EPI_ISL_566580, EPI_ISL_566581, EPI_ISL_566582, EPI_ISL_566583, EPI_ISL_566584, EPI_ISL_566585 | Lighthouse Lab in Milton Keynes | Wellcome Sanger Institute for the COVID-19 Genomics UK Consortium | The Lighthouse Lab in Milton Keynes and Alex Alderton, Roberto Amato, Sonia Goncalves, Ewan Harrison, David K. Jackson, Ian Johnston, Dominic Kwiatkowski, Cordelia Langford, John Sillitoe on behalf of the Wellcome Sanger Institute COVID-19 Surveillance Team |
| EPI_ISL_566586, EPI_ISL_566589 | Lighthouse Lab in Cambridge | Wellcome Sanger Institute for the COVID-19 Genomics UK Consortium | Rob Howes, The Lighthouse Lab in Cambridge and Alex Alderton, Roberto Amato, Sonia Goncalves, Ewan Harrison, David K. Jackson, Ian Johnston, Dominic Kwiatkowski, Cordelia Langford, John Sillitoe on behalf of the Wellcome Sanger Institute COVID-19 Surveillance Team |
| EPI_ISL_566590, EPI_ISL_566591, EPI_ISL_566592, EPI_ISL_566593, EPI_ISL_566594, EPI_ISL_566595, EPI_ISL_566596, EPI_ISL_566597, EPI_ISL_566598, EPI_ISL_566599, EPI_ISL_566600, EPI_ISL_566601, EPI_ISL_566602, EPI_ISL_566603, EPI_ISL_566604, EPI_ISL_566605, EPI_ISL_566606, EPI_ISL_566607, EPI_ISL_566608, EPI_ISL_566609 | | | |
| see above | Lighthouse Lab in Milton Keynes | Wellcome Sanger Institute for the COVID-19 Genomics UK Consortium | The Lighthouse Lab in Milton Keynes and Alex Alderton, Roberto Amato, Sonia Goncalves, Ewan Harrison, David K. Jackson, Ian Johnston, Dominic Kwiatkowski, Cordelia Langford, John Sillitoe on behalf of the Wellcome Sanger Institute COVID-19 Surveillance Team |
| EPI_ISL_566610, EPI_ISL_566611 | Lighthouse Lab in Cambridge | Wellcome Sanger Institute for the COVID-19 Genomics UK Consortium | Rob Howes, The Lighthouse Lab in Cambridge and Alex Alderton, Roberto Amato, Sonia Goncalves, Ewan Harrison, David K. Jackson, Ian Johnston, Dominic Kwiatkowski, Cordelia Langford, John Sillitoe on behalf of the Wellcome Sanger Institute COVID-19 Surveillance Team |
| EPI_ISL_566612, EPI_ISL_566613, EPI_ISL_566614, EPI_ISL_566615, EPI_ISL_566616, EPI_ISL_566617, EPI_ISL_566618, EPI_ISL_566620, EPI_ISL_566621, EPI_ISL_566622, EPI_ISL_566623, EPI_ISL_566624 | | | |
| see above | Lighthouse Lab in Milton Keynes | Wellcome Sanger Institute for the COVID-19 Genomics UK Consortium | The Lighthouse Lab in Milton Keynes and Alex Alderton, Roberto Amato, Sonia Goncalves, Ewan Harrison, David K. Jackson, Ian Johnston, Dominic Kwiatkowski, Cordelia Langford, John Sillitoe on behalf of the Wellcome Sanger Institute COVID-19 Surveillance Team |
| EPI_ISL_566625 | Lighthouse Lab in Cambridge | Wellcome Sanger Institute for the COVID-19 Genomics UK Consortium | Rob Howes, The Lighthouse Lab in Cambridge and Alex Alderton, Roberto Amato, Sonia Goncalves, Ewan Harrison, David K. Jackson, Ian Johnston, Dominic Kwiatkowski, Cordelia Langford, John Sillitoe on behalf of the Wellcome Sanger Institute COVID-19 Surveillance Team |
| EPI_ISL_566626, EPI_ISL_566627, EPI_ISL_566628, EPI_ISL_566629, EPI_ISL_566630, EPI_ISL_566632, EPI_ISL_566633, EPI_ISL_566634, EPI_ISL_566636 | Lighthouse Lab in Milton Keynes | Wellcome Sanger Institute for the COVID-19 Genomics UK Consortium | The Lighthouse Lab in Milton Keynes and Alex Alderton, Roberto Amato, Sonia Goncalves, Ewan Harrison, David K. Jackson, Ian Johnston, Dominic Kwiatkowski, Cordelia Langford, John Sillitoe on behalf of the Wellcome Sanger Institute COVID-19 Surveillance Team |
| EPI_ISL_566638 | Lighthouse Lab in Cambridge | Wellcome Sanger Institute for the COVID-19 Genomics UK Consortium | Rob Howes, The Lighthouse Lab in Cambridge and Alex Alderton, Roberto Amato, Sonia Goncalves, Ewan Harrison, David K. Jackson, Ian Johnston, Dominic Kwiatkowski, Cordelia Langford, John Sillitoe on behalf of the Wellcome Sanger Institute COVID-19 Surveillance Team |
| EPI_ISL_566639, EPI_ISL_566640, EPI_ISL_566641, EPI_ISL_566642, EPI_ISL_566643, EPI_ISL_566644, EPI_ISL_566645, EPI_ISL_566646, EPI_ISL_566649, EPI_ISL_566650, EPI_ISL_566651, EPI_ISL_566652, EPI_ISL_566653, EPI_ISL_566654, EPI_ISL_566655, EPI_ISL_566656, EPI_ISL_566657, EPI_ISL_566658 | | | |
| see above | Lighthouse Lab in Milton Keynes | Wellcome Sanger Institute for the COVID-19 Genomics UK Consortium | The Lighthouse Lab in Milton Keynes and Alex Alderton, Roberto Amato, Sonia Goncalves, Ewan Harrison, David K. Jackson, Ian Johnston, Dominic Kwiatkowski, Cordelia Langford, John Sillitoe on behalf of the Wellcome Sanger Institute COVID-19 Surveillance Team |
| EPI_ISL_566659 | Lighthouse Lab in Cambridge | Wellcome Sanger Institute for the COVID-19 Genomics UK Consortium | Rob Howes, The Lighthouse Lab in Cambridge and Alex Alderton, Roberto Amato, Sonia Goncalves, Ewan Harrison, David K. Jackson, Ian Johnston, Dominic Kwiatkowski, Cordelia Langford, John Sillitoe on behalf of the Wellcome Sanger Institute COVID-19 Surveillance Team |
| EPI_ISL_566660 | Lighthouse Lab in Milton Keynes | Wellcome Sanger Institute for the COVID-19 Genomics UK Consortium | The Lighthouse Lab in Milton Keynes and Alex Alderton, Roberto Amato, Sonia Goncalves, Ewan Harrison, David K. Jackson, Ian Johnston, Dominic Kwiatkowski, Cordelia Langford, John Sillitoe on behalf of the Wellcome Sanger Institute COVID-19 Surveillance Team |
| EPI_ISL_566661, EPI_ISL_566662 | Lighthouse Lab in Cambridge | Wellcome Sanger Institute for the COVID-19 Genomics UK Consortium | Rob Howes, The Lighthouse Lab in Cambridge and Alex Alderton, Roberto Amato, Sonia Goncalves, Ewan Harrison, David K. Jackson, Ian Johnston, Dominic Kwiatkowski, Cordelia Langford, John Sillitoe on behalf of the Wellcome Sanger Institute COVID-19 Surveillance Team |
| EPI_ISL_566663, EPI_ISL_566664, EPI_ISL_566665 | Lighthouse Lab in Milton Keynes | Wellcome Sanger Institute for the COVID-19 Genomics UK Consortium | The Lighthouse Lab in Milton Keynes and Alex Alderton, Roberto Amato, Sonia Goncalves, Ewan Harrison, David K. Jackson, Ian Johnston, Dominic Kwiatkowski, Cordelia Langford, John Sillitoe on behalf of the Wellcome Sanger Institute COVID-19 Surveillance Team |
| EPI_ISL_566666 | Lighthouse Lab in Cambridge | Wellcome Sanger Institute for the COVID-19 Genomics UK Consortium | Rob Howes, The Lighthouse Lab in Cambridge and Alex Alderton, Roberto Amato, Sonia Goncalves, Ewan Harrison, David K. Jackson, Ian Johnston, Dominic Kwiatkowski, Cordelia Langford, John Sillitoe on behalf of the Wellcome Sanger Institute COVID-19 Surveillance Team |
| EPI_ISL_566668, EPI_ISL_566669, EPI_ISL_566670, EPI_ISL_566671, EPI_ISL_566673 | Lighthouse Lab in Milton Keynes | Wellcome Sanger Institute for the COVID-19 Genomics UK Consortium | The Lighthouse Lab in Milton Keynes and Alex Alderton, Roberto Amato, Sonia Goncalves, Ewan Harrison, David K. Jackson, Ian Johnston, Dominic Kwiatkowski, Cordelia Langford, John Sillitoe on behalf of the Wellcome Sanger Institute COVID-19 Surveillance Team |
| EPI_ISL_566674 | Lighthouse Lab in Cambridge | Wellcome Sanger Institute for the COVID-19 Genomics UK Consortium | Rob Howes, The Lighthouse Lab in Cambridge and Alex Alderton, Roberto Amato, Sonia Goncalves, Ewan Harrison, David K. Jackson, Ian Johnston, Dominic Kwiatkowski, Cordelia Langford, John Sillitoe on behalf of the Wellcome Sanger Institute COVID-19 Surveillance Team |
| EPI_ISL_566675, EPI_ISL_566676, EPI_ISL_566677, EPI_ISL_566678, EPI_ISL_566679, EPI_ISL_566680, EPI_ISL_566681, EPI_ISL_566682, EPI_ISL_566683, EPI_ISL_566684, EPI_ISL_566685, EPI_ISL_566686, EPI_ISL_566687, EPI_ISL_566688, EPI_ISL_566689, EPI_ISL_566690, EPI_ISL_566691, EPI_ISL_566692, EPI_ISL_566693, EPI_ISL_566695, EPI_ISL_566696, EPI_ISL_566697, EPI_ISL_566698, EPI_ISL_566699, EPI_ISL_566700, EPI_ISL_566701, EPI_ISL_566702, EPI_ISL_566704, EPI_ISL_566706, EPI_ISL_566708, EPI_ISL_566709, EPI_ISL_566710, EPI_ISL_566711, EPI_ISL_566712, EPI_ISL_566713, EPI_ISL_566714, EPI_ISL_566715, EPI_ISL_566716, EPI_ISL_566717, EPI_ISL_566718, EPI_ISL_566719, EPI_ISL_566720, EPI_ISL_566721, EPI_ISL_566722, EPI_ISL_566723, EPI_ISL_566725, EPI_ISL_566726, EPI_ISL_566727, EPI_ISL_566728, EPI_ISL_566729, EPI_ISL_566730, EPI_ISL_566731, EPI_ISL_566732, EPI_ISL_566734, EPI_ISL_566735, EPI_ISL_566736, EPI_ISL_566737, EPI_ISL_566738, EPI_ISL_566739, EPI_ISL_566740, EPI_ISL_566741, EPI_ISL_566742, EPI_ISL_566743, EPI_ISL_566744, EPI_ISL_566745, EPI_ISL_566748, EPI_ISL_566749, EPI_ISL_566750, EPI_ISL_566751, EPI_ISL_566752, EPI_ISL_566753, EPI_ISL_566754, EPI_ISL_566756, EPI_ISL_566757, EPI_ISL_566758, EPI_ISL_566759, EPI_ISL_566760, EPI_ISL_566761, EPI_ISL_566762, EPI_ISL_566763, EPI_ISL_566764, EPI_ISL_566765, EPI_ISL_566766, EPI_ISL_566767, EPI_ISL_566768, EPI_ISL_566769, EPI_ISL_566770, EPI_ISL_566771, EPI_ISL_566772, EPI_ISL_566773, EPI_ISL_566774, EPI_ISL_566775, EPI_ISL_566776, EPI_ISL_566777, EPI_ISL_566778, EPI_ISL_566779, EPI_ISL_566780, EPI_ISL_566781, EPI_ISL_566782, EPI_ISL_566783, EPI_ISL_566784, EPI_ISL_566785, EPI_ISL_566787, EPI_ISL_566789, EPI_ISL_566790, EPI_ISL_566792, EPI_ISL_566793, EPI_ISL_566794, EPI_ISL_566796, EPI_ISL_566797, EPI_ISL_566798, EPI_ISL_566801, EPI_ISL_566802, EPI_ISL_566803, EPI_ISL_566804, EPI_ISL_566806, EPI_ISL_566807, EPI_ISL_566808, EPI_ISL_566809, EPI_ISL_566810, EPI_ISL_566812, EPI_ISL_566813, EPI_ISL_566814, EPI_ISL_566816, EPI_ISL_566817, EPI_ISL_566818, EPI_ISL_566819, EPI_ISL_566820, EPI_ISL_566821, EPI_ISL_566822, EPI_ISL_566824, EPI_ISL_566825, EPI_ISL_566826, EPI_ISL_566827, EPI_ISL_566828, EPI_ISL_566829, EPI_ISL_566830, EPI_ISL_566831, EPI_ISL_566832, EPI_ISL_566833, EPI_ISL_566834, EPI_ISL_566835, EPI_ISL_566836, EPI_ISL_566837, EPI_ISL_566839, EPI_ISL_566840, EPI_ISL_566842, EPI_ISL_566843, EPI_ISL_566844, EPI_ISL_566845, EPI_ISL_566846, EPI_ISL_566847, EPI_ISL_566848, EPI_ISL_566849, EPI_ISL_566850, EPI_ISL_566851, EPI_ISL_566852, EPI_ISL_566853, EPI_ISL_566854, EPI_ISL_566855, EPI_ISL_566856, EPI_ISL_566857, EPI_ISL_566858, EPI_ISL_566859, EPI_ISL_566860, EPI_ISL_566862, EPI_ISL_566864, EPI_ISL_566865, EPI_ISL_566867, EPI_ISL_566868, EPI_ISL_566869, EPI_ISL_566870, EPI_ISL_566871, EPI_ISL_566872, EPI_ISL_566874, EPI_ISL_566875, EPI_ISL_566877, EPI_ISL_566879, EPI_ISL_566880, EPI_ISL_566881, EPI_ISL_566882, EPI_ISL_566883, EPI_ISL_566884, EPI_ISL_566885, EPI_ISL_566886, EPI_ISL_566887, EPI_ISL_566888, EPI_ISL_566889, EPI_ISL_566890, EPI_ISL_566891, EPI_ISL_566892, EPI_ISL_566893, EPI_ISL_566894, EPI_ISL_566895, EPI_ISL_566896, EPI_ISL_566898, EPI_ISL_566899, EPI_ISL_566900, EPI_ISL_566901, EPI_ISL_566902, EPI_ISL_566903, EPI_ISL_566906, EPI_ISL_566907, EPI_ISL_566908, EPI_ISL_566909, EPI_ISL_566910, EPI_ISL_566911, EPI_ISL_566912, EPI_ISL_566913, EPI_ISL_566914, EPI_ISL_566915, EPI_ISL_566916, EPI_ISL_566917, EPI_ISL_566918, EPI_ISL_566919, EPI_ISL_566920, EPI_ISL_566921, EPI_ISL_566923, EPI_ISL_566924, EPI_ISL_566926, EPI_ISL_566928, EPI_ISL_566929, EPI_ISL_566930, EPI_ISL_566931, EPI_ISL_566933, EPI_ISL_566934, EPI_ISL_566935, EPI_ISL_566937, EPI_ISL_566938, EPI_ISL_566939, EPI_ISL_566940, EPI_ISL_566941, EPI_ISL_566942, EPI_ISL_566943, EPI_ISL_566944, EPI_ISL_566945, EPI_ISL_566946, EPI_ISL_566947, EPI_ISL_566948, EPI_ISL_566949, EPI_ISL_566950, EPI_ISL_566952, EPI_ISL_566953, EPI_ISL_566954, EPI_ISL_566955, EPI_ISL_566956, EPI_ISL_566957 | | | |
| see above | Lighthouse Lab in Milton Keynes | Wellcome Sanger Institute for the COVID-19 Genomics UK Consortium | The Lighthouse Lab in Milton Keynes and Alex Alderton, Roberto Amato, Sonia Goncalves, Ewan Harrison, David K. Jackson, Ian Johnston, Dominic Kwiatkowski, Cordelia Langford, John Sillitoe on behalf of the Wellcome Sanger Institute COVID-19 Surveillance Team |
| EPI_ISL_566958, EPI_ISL_566959, EPI_ISL_566960, EPI_ISL_566961, EPI_ISL_566962, EPI_ISL_566963, EPI_ISL_566965, EPI_ISL_566966, EPI_ISL_566967, EPI_ISL_566968, EPI_ISL_566969, EPI_ISL_566971, EPI_ISL_566972, EPI_ISL_566974, EPI_ISL_566975, EPI_ISL_566976, EPI_ISL_566977, EPI_ISL_566978, EPI_ISL_566979, EPI_ISL_566980, EPI_ISL_566981, EPI_ISL_566982, EPI_ISL_566983, EPI_ISL_566984, EPI_ISL_566985, EPI_ISL_566987, EPI_ISL_566988, EPI_ISL_566989, EPI_ISL_566991, EPI_ISL_566992, EPI_ISL_566993, EPI_ISL_566994, EPI_ISL_566995, EPI_ISL_566996, EPI_ISL_566997 | | | |
| see above | Lighthouse Lab in Milton Keynes | Wellcome Sanger Institute for the COVID-19 Genomics UK Consortium | The Lighthouse Lab in Milton Keynes and Alex Alderton, Roberto Amato, Sonia Goncalves, Ewan Harrison, David K. Jackson, Ian Johnston, Dominic Kwiatkowski, Cordelia Langford, John Sillitoe on behalf of the Wellcome Sanger Institute COVID-19 Surveillance Team |
| EPI_ISL_566998, EPI_ISL_566999, EPI_ISL_567001, EPI_ISL_567002, EPI_ISL_567003, EPI_ISL_567004, EPI_ISL_567005, EPI_ISL_567006, EPI_ISL_567007, EPI_ISL_567008, EPI_ISL_567009, EPI_ISL_567010, EPI_ISL_567011, EPI_ISL_567012, EPI_ISL_567013, EPI_ISL_567014, EPI_ISL_567015, EPI_ISL_567016, EPI_ISL_567017, EPI_ISL_567018, EPI_ISL_567019, EPI_ISL_567020, EPI_ISL_567021, EPI_ISL_567022, EPI_ISL_567023, EPI_ISL_567025, EPI_ISL_567026, EPI_ISL_567028, EPI_ISL_567029, EPI_ISL_567030, EPI_ISL_567031, EPI_ISL_567033, EPI_ISL_567035, EPI_ISL_567036, EPI_ISL_567038, EPI_ISL_567039, EPI_ISL_567040, EPI_ISL_567041, EPI_ISL_567042, EPI_ISL_567044, EPI_ISL_567045, EPI_ISL_567046, EPI_ISL_567047, EPI_ISL_567049, EPI_ISL_567050, EPI_ISL_567051, EPI_ISL_567052, EPI_ISL_567053, EPI_ISL_567054, EPI_ISL_567055, EPI_ISL_567056, EPI_ISL_567057, EPI_ISL_567058, EPI_ISL_567060, EPI_ISL_567061, EPI_ISL_567062, EPI_ISL_567063, EPI_ISL_567064, EPI_ISL_567065, EPI_ISL_567066, EPI_ISL_567067, EPI_ISL_567068, EPI_ISL_567069, EPI_ISL_567070, EPI_ISL_567072, EPI_ISL_567073, EPI_ISL_567074, EPI_ISL_567075, EPI_ISL_567076, EPI_ISL_567078, EPI_ISL_567079, EPI_ISL_567080, EPI_ISL_567081, EPI_ISL_567082, EPI_ISL_567083, EPI_ISL_567085 | | | |
| see above | Lighthouse Lab in Glasgow | Wellcome Sanger Institute for the COVID-19 Genomics UK Consortium | Harper VanSteenhouse, Yumi Kasai, David Gray, Carol Clugston, Anna Dominiczak and Alex Alderton, Roberto Amato, Sonia Goncalves, Ewan Harrison, David K. Jackson, Ian Johnston, Dominic Kwiatkowski, Cordelia Langford, John Sillitoe on behalf of the Wellcome Sanger Institute COVID-19 Surveillance Team |
| EPI_ISL_567086, EPI_ISL_567087, EPI_ISL_567088, EPI_ISL_567090, EPI_ISL_567091, EPI_ISL_567093, EPI_ISL_567094, EPI_ISL_567095, EPI_ISL_567096, EPI_ISL_567097, EPI_ISL_567098, EPI_ISL_567099, EPI_ISL_567100, EPI_ISL_567103, EPI_ISL_567104, EPI_ISL_567105, EPI_ISL_567106, EPI_ISL_567107, EPI_ISL_567108, EPI_ISL_567109, EPI_ISL_567111, EPI_ISL_567112, EPI_ISL_567115, EPI_ISL_567116, EPI_ISL_567117, EPI_ISL_567118, EPI_ISL_567119, EPI_ISL_567120, EPI_ISL_567121, EPI_ISL_567122, EPI_ISL_567123, EPI_ISL_567124, EPI_ISL_567125, EPI_ISL_567127, EPI_ISL_567129, EPI_ISL_567131, EPI_ISL_567132, EPI_ISL_567133, EPI_ISL_567135, EPI_ISL_567137, EPI_ISL_567140, EPI_ISL_567141, EPI_ISL_567142, EPI_ISL_567143, EPI_ISL_567144, EPI_ISL_567145, EPI_ISL_567146, EPI_ISL_567147, EPI_ISL_567148, EPI_ISL_567150, EPI_ISL_567151, EPI_ISL_567152, EPI_ISL_567153, EPI_ISL_567154, EPI_ISL_567155, EPI_ISL_567156, EPI_ISL_567157, EPI_ISL_567159, EPI_ISL_567160, EPI_ISL_567161, EPI_ISL_567162, EPI_ISL_567164, EPI_ISL_567165 | | | |
| see above | Lighthouse Lab in Glasgow | Wellcome Sanger Institute for the COVID-19 Genomics UK Consortium | Harper VanSteenhouse, Yumi Kasai, David Gray, Carol Clugston, Anna Dominiczak and Alex Alderton, Roberto Amato, Sonia Goncalves, Ewan Harrison, David K. Jackson, Ian Johnston, Dominic Kwiatkowski, Cordelia Langford, John Sillitoe on behalf of the Wellcome Sanger Institute COVID-19 Surveillance Team |
| EPI_ISL_567166 | Lighthouse Lab in Glasgow | Wellcome Sanger Institute for the COVID-19 Genomics UK Consortium | Harper VanSteenhouse, Yumi Kasai, David Gray, Carol Clugston, Anna Dominiczak and Alex Alderton, Roberto Amato, Sonia Goncalves, Ewan Harrison, David K. Jackson, Ian Johnston, Dominic Kwiatkowski, Cordelia Langford, John Sillitoe on behalf of the Wellcome Sanger Institute COVID-19 Surveillance Team |
| EPI_ISL_567167, EPI_ISL_567168, EPI_ISL_567169, EPI_ISL_567171, EPI_ISL_567172, EPI_ISL_567173, EPI_ISL_567174, EPI_ISL_567175, EPI_ISL_567176, EPI_ISL_567177, EPI_ISL_567178, EPI_ISL_567179, EPI_ISL_567180, EPI_ISL_567181, EPI_ISL_567182, EPI_ISL_567183, EPI_ISL_567185, EPI_ISL_567186, EPI_ISL_567187, EPI_ISL_567188, EPI_ISL_567189, EPI_ISL_567190, EPI_ISL_567191, EPI_ISL_567192, EPI_ISL_567193, EPI_ISL_567194, EPI_ISL_567195, EPI_ISL_567196, EPI_ISL_567197, EPI_ISL_567198, EPI_ISL_567199, EPI_ISL_567200, EPI_ISL_567201, EPI_ISL_567203, EPI_ISL_567204, EPI_ISL_567205, EPI_ISL_567206, EPI_ISL_567207, EPI_ISL_567208, EPI_ISL_567209, EPI_ISL_567210, EPI_ISL_567211, EPI_ISL_567212, EPI_ISL_567215, EPI_ISL_567216, EPI_ISL_567218, EPI_ISL_567219, EPI_ISL_567220, EPI_ISL_567222, EPI_ISL_567223, EPI_ISL_567225, EPI_ISL_567226, EPI_ISL_567228, EPI_ISL_567229, EPI_ISL_567230, EPI_ISL_567232, EPI_ISL_567233, EPI_ISL_567235, EPI_ISL_567236, EPI_ISL_567237, EPI_ISL_567238, EPI_ISL_567239, EPI_ISL_567240, EPI_ISL_567242, EPI_ISL_567243, EPI_ISL_567245, EPI_ISL_567246, EPI_ISL_567247, EPI_ISL_567248, EPI_ISL_567249, EPI_ISL_567250, EPI_ISL_567251, EPI_ISL_567252, EPI_ISL_567253, EPI_ISL_567254, EPI_ISL_567255, EPI_ISL_567256, EPI_ISL_567257, EPI_ISL_567258, EPI_ISL_567259, EPI_ISL_567261, EPI_ISL_567262, EPI_ISL_567264, EPI_ISL_567266, EPI_ISL_567267, EPI_ISL_567268, EPI_ISL_567269, EPI_ISL_567270, EPI_ISL_567271, EPI_ISL_567272, EPI_ISL_567273, EPI_ISL_567274, EPI_ISL_567275, EPI_ISL_567276, EPI_ISL_567277, EPI_ISL_567279, EPI_ISL_567281, EPI_ISL_567282, EPI_ISL_567283, EPI_ISL_567284, EPI_ISL_567285, EPI_ISL_567286, EPI_ISL_567287, EPI_ISL_567288, EPI_ISL_567290, EPI_ISL_567291, EPI_ISL_567292, EPI_ISL_567293, EPI_ISL_567294, EPI_ISL_567297, EPI_ISL_567299, EPI_ISL_567300, EPI_ISL_567301, EPI_ISL_567302, EPI_ISL_567303, EPI_ISL_567304, EPI_ISL_567306, EPI_ISL_567307, EPI_ISL_567308, EPI_ISL_567309, EPI_ISL_567310, EPI_ISL_567311, EPI_ISL_567312, EPI_ISL_567313, EPI_ISL_567314, EPI_ISL_567315, EPI_ISL_567316, EPI_ISL_567317, EPI_ISL_567320, EPI_ISL_567322, EPI_ISL_567323, EPI_ISL_567324, EPI_ISL_567325, EPI_ISL_567327, EPI_ISL_567329, EPI_ISL_567330, EPI_ISL_567331, EPI_ISL_567332, EPI_ISL_567333, EPI_ISL_567334, EPI_ISL_567335, EPI_ISL_567336, EPI_ISL_567337, EPI_ISL_567338, EPI_ISL_567339, EPI_ISL_567340, EPI_ISL_567341, EPI_ISL_567345, EPI_ISL_567346, EPI_ISL_567347, EPI_ISL_567348, EPI_ISL_567349, EPI_ISL_567350, EPI_ISL_567352, EPI_ISL_567353, EPI_ISL_567355, EPI_ISL_567356, EPI_ISL_567357, EPI_ISL_567358, EPI_ISL_567359, EPI_ISL_567360, EPI_ISL_567361, EPI_ISL_567362, EPI_ISL_567363, EPI_ISL_567364, EPI_ISL_567365, EPI_ISL_567366, EPI_ISL_567367, EPI_ISL_567368, EPI_ISL_567369, EPI_ISL_567370, EPI_ISL_567372, EPI_ISL_567373, EPI_ISL_567374, EPI_ISL_567376, EPI_ISL_567377, EPI_ISL_567378, EPI_ISL_567379, EPI_ISL_567380, EPI_ISL_567381, EPI_ISL_567382, EPI_ISL_567383, EPI_ISL_567384, EPI_ISL_567386, EPI_ISL_567388, EPI_ISL_567389, EPI_ISL_567391, EPI_ISL_567392, EPI_ISL_567393, EPI_ISL_567394, EPI_ISL_567396, EPI_ISL_567397, EPI_ISL_567398, EPI_ISL_567399, EPI_ISL_567400, EPI_ISL_567401, EPI_ISL_567402, EPI_ISL_567403, EPI_ISL_567404, EPI_ISL_567405, EPI_ISL_567406, EPI_ISL_567407, EPI_ISL_567408, EPI_ISL_567409, EPI_ISL_567410, EPI_ISL_567411, EPI_ISL_567412, EPI_ISL_567413, EPI_ISL_567414, EPI_ISL_567415, EPI_ISL_567416, EPI_ISL_567417, EPI_ISL_567418, EPI_ISL_567419, EPI_ISL_567420, EPI_ISL_567421, EPI_ISL_567422, EPI_ISL_567423, EPI_ISL_567424, EPI_ISL_567425, EPI_ISL_567426, EPI_ISL_567427, EPI_ISL_567428, EPI_ISL_567429, EPI_ISL_567430, EPI_ISL_567431, EPI_ISL_567433, EPI_ISL_567434, EPI_ISL_567435, EPI_ISL_567436, EPI_ISL_567437, EPI_ISL_567438, EPI_ISL_567439, EPI_ISL_567440, EPI_ISL_567441, EPI_ISL_567442, EPI_ISL_567443, EPI_ISL_567445, EPI_ISL_567446, EPI_ISL_567448, EPI_ISL_567449, EPI_ISL_567450, EPI_ISL_567451, EPI_ISL_567452, EPI_ISL_567453, EPI_ISL_567454, EPI_ISL_567455, EPI_ISL_567457, EPI_ISL_567458, EPI_ISL_567459, EPI_ISL_567460, EPI_ISL_567461, EPI_ISL_567462, EPI_ISL_567463, EPI_ISL_567465 | | | |
| see above | Lighthouse Lab in Glasgow | Wellcome Sanger Institute for the COVID-19 Genomics UK Consortium | Harper VanSteenhouse, Yumi Kasai, David Gray, Carol Clugston, Anna Dominiczak and Alex Alderton, Roberto Amato, Sonia Goncalves, Ewan Harrison, David K. Jackson, Ian Johnston, Dominic Kwiatkowski, Cordelia Langford, John Sillitoe on behalf of the Wellcome Sanger Institute COVID-19 Surveillance Team |
| EPI_ISL_567466 | Lighthouse Lab in Glasgow | Wellcome Sanger Institute for the COVID-19 | Harper VanSteenhouse, Yumi Kasai, David Gray, Carol Clugston, Anna Dominiczak and Alex Alderton, Roberto Amato, Sonia Goncalves, Ewan Harrison, David K. Jackson, Ian Johnston, Dominic Kwiatkowski, Cordelia |

| | | | |
|---|---|---|---|
| | | Genomics UK Consortium | Langford, John Sillitoe on behalf of the Wellcome Sanger Institute COVID-19 Surveillance Team |
| EPI_ISL_567467, EPI_ISL_567468, EPI_ISL_567470, EPI_ISL_567471, EPI_ISL_567472, EPI_ISL_567473, EPI_ISL_567474, EPI_ISL_567475, EPI_ISL_567477, EPI_ISL_567478, EPI_ISL_567479, EPI_ISL_567480, EPI_ISL_567481, EPI_ISL_567482, EPI_ISL_567483, EPI_ISL_567484, EPI_ISL_567485, EPI_ISL_567486, EPI_ISL_567488, EPI_ISL_567489, EPI_ISL_567490, EPI_ISL_567492, EPI_ISL_567493, EPI_ISL_567494, EPI_ISL_567495, EPI_ISL_567496, EPI_ISL_567497, EPI_ISL_567498, EPI_ISL_567499, EPI_ISL_567500, EPI_ISL_567501, EPI_ISL_567502, EPI_ISL_567503, EPI_ISL_567505, EPI_ISL_567506, EPI_ISL_567507, EPI_ISL_567508, EPI_ISL_567509, EPI_ISL_567510, EPI_ISL_567511, EPI_ISL_567512, EPI_ISL_567513, EPI_ISL_567514, EPI_ISL_567515, EPI_ISL_567517, EPI_ISL_567518, EPI_ISL_567520, EPI_ISL_567521, EPI_ISL_567522, EPI_ISL_567523, EPI_ISL_567524, EPI_ISL_567525, EPI_ISL_567526, EPI_ISL_567528, EPI_ISL_567529, EPI_ISL_567530, EPI_ISL_567531, EPI_ISL_567532 | | | |
| see above | Lighthouse Lab in Glasgow | Wellcome Sanger Institute for the COVID-19 Genomics UK Consortium | Harper VanSteenhouse, Yumi Kasai, David Gray, Carol Clugston, Anna Dominiczak and Alex Alderton, Roberto Amato, Sonia Goncalves, Ewan Harrison, David K. Jackson, Ian Johnston, Dominic Kwiatkowski, Cordelia Langford, John Sillitoe on behalf of the Wellcome Sanger Institute COVID-19 Surveillance Team |
| EPI_ISL_567533, EPI_ISL_567534, EPI_ISL_567535, EPI_ISL_567537, EPI_ISL_567538, EPI_ISL_567539, EPI_ISL_567540, EPI_ISL_567541, EPI_ISL_567542, EPI_ISL_567543, EPI_ISL_567544, EPI_ISL_567545, EPI_ISL_567546, EPI_ISL_567547, EPI_ISL_567548, EPI_ISL_567549, EPI_ISL_567550, EPI_ISL_567551, EPI_ISL_567552, EPI_ISL_567553, EPI_ISL_567554, EPI_ISL_567555, EPI_ISL_567556, EPI_ISL_567557, EPI_ISL_567558, EPI_ISL_567559, EPI_ISL_567560, EPI_ISL_567561, EPI_ISL_567562, EPI_ISL_567563, EPI_ISL_567564, EPI_ISL_567565, EPI_ISL_567566, EPI_ISL_567567, EPI_ISL_567568, EPI_ISL_567569, EPI_ISL_567570, EPI_ISL_567571, EPI_ISL_567572, EPI_ISL_567573, EPI_ISL_567574, EPI_ISL_567575, EPI_ISL_567576, EPI_ISL_567577, EPI_ISL_567578, EPI_ISL_567579, EPI_ISL_567580, EPI_ISL_567581, EPI_ISL_567582, EPI_ISL_567583, EPI_ISL_567584, EPI_ISL_567585, EPI_ISL_567586, EPI_ISL_567587, EPI_ISL_567589, EPI_ISL_567590, EPI_ISL_567591, EPI_ISL_567592, EPI_ISL_567595, EPI_ISL_567596, EPI_ISL_567597, EPI_ISL_567598, EPI_ISL_567599, EPI_ISL_567600, EPI_ISL_567601, EPI_ISL_567602, EPI_ISL_567604, EPI_ISL_567605, EPI_ISL_567606, EPI_ISL_567608, EPI_ISL_567609, EPI_ISL_567610, EPI_ISL_567611, EPI_ISL_567612, EPI_ISL_567613, EPI_ISL_567614, EPI_ISL_567615, EPI_ISL_567616, EPI_ISL_567617, EPI_ISL_567618, EPI_ISL_567619, EPI_ISL_567620, EPI_ISL_567621, EPI_ISL_567622, EPI_ISL_567624, EPI_ISL_567625, EPI_ISL_567626, EPI_ISL_567627, EPI_ISL_567628, EPI_ISL_567629, EPI_ISL_567630, EPI_ISL_567631, EPI_ISL_567632, EPI_ISL_567633, EPI_ISL_567634, EPI_ISL_567635, EPI_ISL_567636, EPI_ISL_567637, EPI_ISL_567638, EPI_ISL_567639, EPI_ISL_567640, EPI_ISL_567641, EPI_ISL_567642, EPI_ISL_567643, EPI_ISL_567644, EPI_ISL_567645, EPI_ISL_567646, EPI_ISL_567647, EPI_ISL_567648, EPI_ISL_567650, EPI_ISL_567651, EPI_ISL_567653, EPI_ISL_567654, EPI_ISL_567655, EPI_ISL_567656, EPI_ISL_567657, EPI_ISL_567658, EPI_ISL_567659, EPI_ISL_567660, EPI_ISL_567661, EPI_ISL_567662, EPI_ISL_567663, EPI_ISL_567664, EPI_ISL_567665, EPI_ISL_567666, EPI_ISL_567667, EPI_ISL_567669, EPI_ISL_567670, EPI_ISL_567671, EPI_ISL_567672, EPI_ISL_567673, EPI_ISL_567674, EPI_ISL_567675, EPI_ISL_567676, EPI_ISL_567677, EPI_ISL_567680, EPI_ISL_567681, EPI_ISL_567682, EPI_ISL_567683, EPI_ISL_567684, EPI_ISL_567685, EPI_ISL_567686, EPI_ISL_567687, EPI_ISL_567688, EPI_ISL_567689, EPI_ISL_567690, EPI_ISL_567691, EPI_ISL_567692, EPI_ISL_567693, EPI_ISL_567694, EPI_ISL_567695, EPI_ISL_567696, EPI_ISL_567697, EPI_ISL_567698, EPI_ISL_567700, EPI_ISL_567701, EPI_ISL_567702, EPI_ISL_567703, EPI_ISL_567704, EPI_ISL_567705, EPI_ISL_567706, EPI_ISL_567707, EPI_ISL_567708, EPI_ISL_567709, EPI_ISL_567710, EPI_ISL_567711, EPI_ISL_567712, EPI_ISL_567713, EPI_ISL_567714, EPI_ISL_567715, EPI_ISL_567716, EPI_ISL_567717, EPI_ISL_567718, EPI_ISL_567721, EPI_ISL_567722, EPI_ISL_567723, EPI_ISL_567724, EPI_ISL_567725, EPI_ISL_567726, EPI_ISL_567727, EPI_ISL_567728, EPI_ISL_567729, EPI_ISL_567730, EPI_ISL_567731, EPI_ISL_567732, EPI_ISL_567733, EPI_ISL_567734, EPI_ISL_567735, EPI_ISL_567736, EPI_ISL_567737, EPI_ISL_567739, EPI_ISL_567740, EPI_ISL_567741, EPI_ISL_567742, EPI_ISL_567743, EPI_ISL_567744, EPI_ISL_567745, EPI_ISL_567746, EPI_ISL_567747, EPI_ISL_567748, EPI_ISL_567749, EPI_ISL_567750, EPI_ISL_567751, EPI_ISL_567753, EPI_ISL_567754, EPI_ISL_567755, EPI_ISL_567756, EPI_ISL_567757, EPI_ISL_567758, EPI_ISL_567760, EPI_ISL_567761, EPI_ISL_567762, EPI_ISL_567763, EPI_ISL_567764, EPI_ISL_567766, EPI_ISL_567768, EPI_ISL_567769, EPI_ISL_567771, EPI_ISL_567772, EPI_ISL_567773, EPI_ISL_567774, EPI_ISL_567775, EPI_ISL_567776, EPI_ISL_567777, EPI_ISL_567778, EPI_ISL_567779, EPI_ISL_567780, EPI_ISL_567781, EPI_ISL_567782, EPI_ISL_567783, EPI_ISL_567784, EPI_ISL_567785, EPI_ISL_567786, EPI_ISL_567787, EPI_ISL_567788, EPI_ISL_567789, EPI_ISL_567790, EPI_ISL_567791, EPI_ISL_567792, EPI_ISL_567793, EPI_ISL_567794, EPI_ISL_567795, EPI_ISL_567796, EPI_ISL_567797, EPI_ISL_567798, EPI_ISL_567799, EPI_ISL_567800, EPI_ISL_567801, EPI_ISL_567802, EPI_ISL_567804, EPI_ISL_567805, EPI_ISL_567806, EPI_ISL_567807, EPI_ISL_567808, EPI_ISL_567809, EPI_ISL_567810, EPI_ISL_567813, EPI_ISL_567814, EPI_ISL_567815, EPI_ISL_567816, EPI_ISL_567817, EPI_ISL_567818, EPI_ISL_567819, EPI_ISL_567820, EPI_ISL_567821, EPI_ISL_567822, EPI_ISL_567823, EPI_ISL_567824, EPI_ISL_567825, EPI_ISL_567826, EPI_ISL_567827, EPI_ISL_567828, EPI_ISL_567829, EPI_ISL_567830, EPI_ISL_567832, EPI_ISL_567834, EPI_ISL_567835, EPI_ISL_567836, EPI_ISL_567837, EPI_ISL_567838, EPI_ISL_567839, EPI_ISL_567840, EPI_ISL_567841, EPI_ISL_567842, EPI_ISL_567843, EPI_ISL_567844, EPI_ISL_567846, EPI_ISL_567847, EPI_ISL_567848, EPI_ISL_567849, EPI_ISL_567850, EPI_ISL_567851, EPI_ISL_567852, EPI_ISL_567853, EPI_ISL_567854, EPI_ISL_567855, EPI_ISL_567856, EPI_ISL_567857, EPI_ISL_567858, EPI_ISL_567861, EPI_ISL_567863, EPI_ISL_567864, EPI_ISL_567865, EPI_ISL_567866, EPI_ISL_567867, EPI_ISL_567868, EPI_ISL_567869, EPI_ISL_567870, EPI_ISL_567871, EPI_ISL_567872, EPI_ISL_567873, EPI_ISL_567874, EPI_ISL_567875, EPI_ISL_567876, EPI_ISL_567877, EPI_ISL_567878, EPI_ISL_567879, EPI_ISL_567880, EPI_ISL_567881, EPI_ISL_567882, EPI_ISL_567883, EPI_ISL_567884, EPI_ISL_567885, EPI_ISL_567886, EPI_ISL_567888, EPI_ISL_567889, EPI_ISL_567890, EPI_ISL_567891, EPI_ISL_567892, EPI_ISL_567893, EPI_ISL_567894, EPI_ISL_567895, EPI_ISL_567896, EPI_ISL_567897, EPI_ISL_567898, EPI_ISL_567901, EPI_ISL_567902, EPI_ISL_567903, EPI_ISL_567904, EPI_ISL_567905, EPI_ISL_567906, EPI_ISL_567907, EPI_ISL_567908, EPI_ISL_567909, EPI_ISL_567910, EPI_ISL_567912, EPI_ISL_567913, EPI_ISL_567915, EPI_ISL_567916, EPI_ISL_567918, EPI_ISL_567919, EPI_ISL_567920, EPI_ISL_567921, EPI_ISL_567922, EPI_ISL_567923, EPI_ISL_567924, EPI_ISL_567925, EPI_ISL_567926, EPI_ISL_567927, EPI_ISL_567928, EPI_ISL_567929, EPI_ISL_567930, EPI_ISL_567931, EPI_ISL_567932, EPI_ISL_567933, EPI_ISL_567935, EPI_ISL_567936, EPI_ISL_567937, EPI_ISL_567938, EPI_ISL_567939, EPI_ISL_567940, EPI_ISL_567942, EPI_ISL_567943, EPI_ISL_567944, EPI_ISL_567945, EPI_ISL_567946, EPI_ISL_567947, EPI_ISL_567948, EPI_ISL_567949, EPI_ISL_567950, EPI_ISL_567951, EPI_ISL_567952, EPI_ISL_567953, EPI_ISL_567955, EPI_ISL_567957, EPI_ISL_567959, EPI_ISL_567960, EPI_ISL_567961, EPI_ISL_567962, EPI_ISL_567963, EPI_ISL_567964, EPI_ISL_567965, EPI_ISL_567966, EPI_ISL_567967, EPI_ISL_567968, EPI_ISL_567969, EPI_ISL_567971, EPI_ISL_567973, EPI_ISL_567974, EPI_ISL_567975, EPI_ISL_567976, EPI_ISL_567977, EPI_ISL_567978, EPI_ISL_567979, EPI_ISL_567980, EPI_ISL_567981, EPI_ISL_567982, EPI_ISL_567983, EPI_ISL_567985, EPI_ISL_567986, EPI_ISL_567987, EPI_ISL_567988, EPI_ISL_567989, EPI_ISL_567990, EPI_ISL_567992, EPI_ISL_567993, EPI_ISL_567994, EPI_ISL_567995, EPI_ISL_567996, EPI_ISL_567997, EPI_ISL_567998, EPI_ISL_567999, EPI_ISL_568000, EPI_ISL_568001, EPI_ISL_568002, EPI_ISL_568003, EPI_ISL_568004, EPI_ISL_568005, EPI_ISL_568006, EPI_ISL_568007, EPI_ISL_568008, EPI_ISL_568009, EPI_ISL_568010, EPI_ISL_568011, EPI_ISL_568013, EPI_ISL_568014, EPI_ISL_568016, EPI_ISL_568017, EPI_ISL_568018, EPI_ISL_568019, EPI_ISL_568020, EPI_ISL_568021, EPI_ISL_568022, EPI_ISL_568024, EPI_ISL_568025, EPI_ISL_568027, EPI_ISL_568030, EPI_ISL_568031, EPI_ISL_568033, EPI_ISL_568034, EPI_ISL_568035, EPI_ISL_568036, EPI_ISL_568038, EPI_ISL_568039, EPI_ISL_568041, EPI_ISL_568042, EPI_ISL_568043, EPI_ISL_568044, EPI_ISL_568045, EPI_ISL_568046, EPI_ISL_568047, EPI_ISL_568048, EPI_ISL_568049, EPI_ISL_568050, EPI_ISL_568051, EPI_ISL_568052, EPI_ISL_568053, EPI_ISL_568054, EPI_ISL_568055, EPI_ISL_568056, EPI_ISL_568057, EPI_ISL_568058, EPI_ISL_568059, EPI_ISL_568060, EPI_ISL_568061, EPI_ISL_568063, EPI_ISL_568064, EPI_ISL_568065, EPI_ISL_568066, EPI_ISL_568067, EPI_ISL_568069, EPI_ISL_568070, EPI_ISL_568071, EPI_ISL_568072, EPI_ISL_568073, EPI_ISL_568074, EPI_ISL_568075, EPI_ISL_568076, EPI_ISL_568079, EPI_ISL_568080, EPI_ISL_568081, EPI_ISL_568082, EPI_ISL_568083, EPI_ISL_568084, EPI_ISL_568085, EPI_ISL_568086, EPI_ISL_568087, EPI_ISL_568088, EPI_ISL_568089, EPI_ISL_568090, EPI_ISL_568091, EPI_ISL_568092, EPI_ISL_568093, EPI_ISL_568094, EPI_ISL_568095, EPI_ISL_568097, EPI_ISL_568098, EPI_ISL_568099, EPI_ISL_568100, EPI_ISL_568101, EPI_ISL_568102, EPI_ISL_568103, EPI_ISL_568105, EPI_ISL_568106, EPI_ISL_568107, EPI_ISL_568108, EPI_ISL_568109, EPI_ISL_568110, EPI_ISL_568111, EPI_ISL_568112, EPI_ISL_568113, EPI_ISL_568114, EPI_ISL_568115, EPI_ISL_568116, EPI_ISL_568118, EPI_ISL_568119, EPI_ISL_568120, EPI_ISL_568121, EPI_ISL_568123, EPI_ISL_568124, EPI_ISL_568125, EPI_ISL_568126, EPI_ISL_568127, EPI_ISL_568128, EPI_ISL_568129, EPI_ISL_568130, EPI_ISL_568131, EPI_ISL_568132, EPI_ISL_568133, EPI_ISL_568134, EPI_ISL_568135, EPI_ISL_568136, EPI_ISL_568137, EPI_ISL_568138, EPI_ISL_568139, EPI_ISL_568140, EPI_ISL_568141, EPI_ISL_568142, EPI_ISL_568143, EPI_ISL_568144, EPI_ISL_568146, EPI_ISL_568147, EPI_ISL_568149, EPI_ISL_568150, EPI_ISL_568151, EPI_ISL_568152, EPI_ISL_568153, EPI_ISL_568154, EPI_ISL_568155, EPI_ISL_568156, EPI_ISL_568157, EPI_ISL_568158, EPI_ISL_568159, EPI_ISL_568160, EPI_ISL_568161, EPI_ISL_568162, EPI_ISL_568163, EPI_ISL_568164, EPI_ISL_568165, EPI_ISL_568166, EPI_ISL_568167, EPI_ISL_568168, EPI_ISL_568169, EPI_ISL_568171, EPI_ISL_568172, EPI_ISL_568173, EPI_ISL_568174, EPI_ISL_568175, EPI_ISL_568176, EPI_ISL_568177, EPI_ISL_568178, EPI_ISL_568179, EPI_ISL_568180, EPI_ISL_568181, EPI_ISL_568183, EPI_ISL_568184, EPI_ISL_568185, EPI_ISL_568186, EPI_ISL_568188, EPI_ISL_568189, EPI_ISL_568190, EPI_ISL_568191, EPI_ISL_568193, EPI_ISL_568194, EPI_ISL_568195, EPI_ISL_568196, EPI_ISL_568197, EPI_ISL_568198, EPI_ISL_568199, EPI_ISL_568200, EPI_ISL_568201, EPI_ISL_568202, EPI_ISL_568203, EPI_ISL_568205, EPI_ISL_568206, EPI_ISL_568207, EPI_ISL_568208, EPI_ISL_568211, EPI_ISL_568212, EPI_ISL_568213, EPI_ISL_568214, EPI_ISL_568215, EPI_ISL_568216, EPI_ISL_568217, EPI_ISL_568218, EPI_ISL_568219, EPI_ISL_568220, EPI_ISL_568221, EPI_ISL_568222 | | | |
| see above | Lighthouse Lab in Alderley Park | Wellcome Sanger Institute for the COVID-19 Genomics UK Consortium | Jacquelyn Wynn, Mairead Hyland, The Lighthouse Lab in Alderley Park and Alex Alderton, Roberto Amato, Sonia Goncalves, Ewan Harrison, David K. Jackson, Ian Johnston, Dominic Kwiatkowski, Cordelia Langford, John Sillitoe on behalf of the Wellcome Sanger Institute COVID-19 Surveillance Team |
| EPI_ISL_568223, EPI_ISL_568224, EPI_ISL_568225, EPI_ISL_568226, EPI_ISL_568227, EPI_ISL_568228, EPI_ISL_568229, EPI_ISL_568230, EPI_ISL_568231, EPI_ISL_568232, EPI_ISL_568233, EPI_ISL_568234, EPI_ISL_568235, EPI_ISL_568236, EPI_ISL_568237, EPI_ISL_568238, EPI_ISL_568239, EPI_ISL_568240, EPI_ISL_568241, EPI_ISL_568242, EPI_ISL_568243, EPI_ISL_568244, EPI_ISL_568245, EPI_ISL_568246, EPI_ISL_568247, EPI_ISL_568248, EPI_ISL_568249, EPI_ISL_568250, EPI_ISL_568251, EPI_ISL_568252, EPI_ISL_568253, EPI_ISL_568254, EPI_ISL_568255, EPI_ISL_568256, EPI_ISL_568257, EPI_ISL_568258, EPI_ISL_568259, EPI_ISL_568260, EPI_ISL_568261, EPI_ISL_568262, EPI_ISL_568263, EPI_ISL_568264, EPI_ISL_568265, EPI_ISL_568266, EPI_ISL_568267, EPI_ISL_568268, EPI_ISL_568269, EPI_ISL_568270, EPI_ISL_568271, EPI_ISL_568272, EPI_ISL_568273, EPI_ISL_568274, EPI_ISL_568275, EPI_ISL_568276, EPI_ISL_568277, EPI_ISL_568278, EPI_ISL_568279, EPI_ISL_568281, EPI_ISL_568282, EPI_ISL_568283, EPI_ISL_568284, EPI_ISL_568285, EPI_ISL_568286, EPI_ISL_568287, EPI_ISL_568288 | | | |
| see above | Lighthouse Lab in Glasgow | Wellcome Sanger Institute for the COVID-19 Genomics UK Consortium | Harper VanSteenhouse, Yumi Kasai, David Gray, Carol Clugston, Anna Dominiczak and Alex Alderton, Roberto Amato, Sonia Goncalves, Ewan Harrison, David K. Jackson, Ian Johnston, Dominic Kwiatkowski, Cordelia Langford, John Sillitoe on behalf of the Wellcome Sanger Institute COVID-19 Surveillance Team |
| EPI_ISL_568289 | Lighthouse Lab in Milton Keynes | Wellcome Sanger Institute for the COVID-19 Genomics UK Consortium | The Lighthouse Lab in Milton Keynes and Alex Alderton, Roberto Amato, Sonia Goncalves, Ewan Harrison, David K. Jackson, Ian Johnston, Dominic Kwiatkowski, Cordelia Langford, John Sillitoe on behalf of the Wellcome Sanger Institute COVID-19 Surveillance Team |
| EPI_ISL_568291, EPI_ISL_568292, EPI_ISL_568293, EPI_ISL_568294, EPI_ISL_568295, EPI_ISL_568296, EPI_ISL_568297, EPI_ISL_568298, EPI_ISL_568299, EPI_ISL_568302, EPI_ISL_568304, EPI_ISL_568305, EPI_ISL_568306, EPI_ISL_568307, EPI_ISL_568308, EPI_ISL_568309, EPI_ISL_568310, EPI_ISL_568311, EPI_ISL_568312, EPI_ISL_568313, EPI_ISL_568314, EPI_ISL_568316, EPI_ISL_568318, EPI_ISL_568319, EPI_ISL_568320, EPI_ISL_568321, EPI_ISL_568322, EPI_ISL_568323, EPI_ISL_568324, EPI_ISL_568326, EPI_ISL_568327, EPI_ISL_568328, EPI_ISL_568329, EPI_ISL_568330, EPI_ISL_568331, EPI_ISL_568332, EPI_ISL_568333, EPI_ISL_568334, EPI_ISL_568335, EPI_ISL_568336, EPI_ISL_568337, EPI_ISL_568338, EPI_ISL_568339, EPI_ISL_568340, EPI_ISL_568341, EPI_ISL_568342, EPI_ISL_568343, EPI_ISL_568345, EPI_ISL_568346, EPI_ISL_568347, EPI_ISL_568348, EPI_ISL_568349, EPI_ISL_568350, EPI_ISL_568351, EPI_ISL_568352, EPI_ISL_568353, EPI_ISL_568354, EPI_ISL_568355, EPI_ISL_568357, EPI_ISL_568358, EPI_ISL_568359, EPI_ISL_568360, EPI_ISL_568361, EPI_ISL_568362, EPI_ISL_568363 | | | |
| see above | Lighthouse Lab in Glasgow | Wellcome Sanger Institute for the COVID-19 Genomics UK Consortium | Harper VanSteenhouse, Yumi Kasai, David Gray, Carol Clugston, Anna Dominiczak and Alex Alderton, Roberto Amato, Sonia Goncalves, Ewan Harrison, David K. Jackson, Ian Johnston, Dominic Kwiatkowski, Cordelia Langford, John Sillitoe on behalf of the Wellcome Sanger Institute COVID-19 Surveillance Team |
| EPI_ISL_568364, EPI_ISL_568365 | Lighthouse Lab in Milton Keynes | Wellcome Sanger Institute for the COVID-19 Genomics UK Consortium | The Lighthouse Lab in Milton Keynes and Alex Alderton, Roberto Amato, Sonia Goncalves, Ewan Harrison, David K. Jackson, Ian Johnston, Dominic Kwiatkowski, Cordelia Langford, John Sillitoe on behalf of the Wellcome Sanger Institute COVID-19 Surveillance Team |
| EPI_ISL_568366, EPI_ISL_568367, EPI_ISL_568368, EPI_ISL_568369, EPI_ISL_568370, EPI_ISL_568371, EPI_ISL_568372, EPI_ISL_568373, EPI_ISL_568374, EPI_ISL_568375, EPI_ISL_568376, EPI_ISL_568377, EPI_ISL_568378, EPI_ISL_568379, EPI_ISL_568380, EPI_ISL_568381, EPI_ISL_568382, EPI_ISL_568383, EPI_ISL_568384 | | | |
| see above | Lighthouse Lab in Glasgow | Wellcome Sanger Institute for the COVID-19 Genomics UK Consortium | Harper VanSteenhouse, Yumi Kasai, David Gray, Carol Clugston, Anna Dominiczak and Alex Alderton, Roberto Amato, Sonia Goncalves, Ewan Harrison, David K. Jackson, Ian Johnston, Dominic Kwiatkowski, Cordelia Langford, John Sillitoe on behalf of the Wellcome Sanger Institute COVID-19 Surveillance Team |
| EPI_ISL_568385 | Lighthouse Lab in Cambridge | Wellcome Sanger Institute for the COVID-19 Genomics UK Consortium | Rob Howes, The Lighthouse Lab in Cambridge and Alex Alderton, Roberto Amato, Sonia Goncalves, Ewan Harrison, David K. Jackson, Ian Johnston, Dominic Kwiatkowski, Cordelia Langford, John Sillitoe on behalf of the Wellcome Sanger Institute COVID-19 Surveillance Team |
| EPI_ISL_568386, EPI_ISL_568387 | Lighthouse Lab in Milton Keynes | Wellcome Sanger Institute for the COVID-19 Genomics UK Consortium | The Lighthouse Lab in Milton Keynes and Alex Alderton, Roberto Amato, Sonia Goncalves, Ewan Harrison, David K. Jackson, Ian Johnston, Dominic Kwiatkowski, Cordelia Langford, John Sillitoe on behalf of the Wellcome Sanger Institute COVID-19 Surveillance Team |
| EPI_ISL_568388, EPI_ISL_568389, EPI_ISL_568390, EPI_ISL_568391, EPI_ISL_568393, EPI_ISL_568394, EPI_ISL_568401, EPI_ISL_568402, EPI_ISL_568403, EPI_ISL_568404, EPI_ISL_568405, EPI_ISL_568406, EPI_ISL_568407, EPI_ISL_568408, EPI_ISL_568411, EPI_ISL_568412, EPI_ISL_568413, EPI_ISL_568414, EPI_ISL_568416, EPI_ISL_568417, EPI_ISL_568418, EPI_ISL_568420, EPI_ISL_568421, EPI_ISL_568423, EPI_ISL_568424, EPI_ISL_568425, EPI_ISL_568426, EPI_ISL_568427, EPI_ISL_568429, EPI_ISL_568430, EPI_ISL_568431, EPI_ISL_568432, EPI_ISL_568433, EPI_ISL_568434 | | | |
| see above | Lighthouse Lab in Glasgow | Wellcome Sanger Institute for the COVID-19 Genomics UK Consortium | Harper VanSteenhouse, Yumi Kasai, David Gray, Carol Clugston, Anna Dominiczak and Alex Alderton, Roberto Amato, Sonia Goncalves, Ewan Harrison, David K. Jackson, Ian Johnston, Dominic Kwiatkowski, Cordelia Langford, John Sillitoe on behalf of the Wellcome Sanger Institute COVID-19 Surveillance Team |
| EPI_ISL_568436 | Lighthouse Lab in Cambridge | Wellcome Sanger Institute for the COVID-19 Genomics UK Consortium | Rob Howes, The Lighthouse Lab in Cambridge and Alex Alderton, Roberto Amato, Sonia Goncalves, Ewan Harrison, David K. Jackson, Ian Johnston, Dominic Kwiatkowski, Cordelia Langford, John Sillitoe on behalf of the Wellcome Sanger Institute COVID-19 Surveillance Team |
| EPI_ISL_568437, EPI_ISL_568438, EPI_ISL_568439, EPI_ISL_568440, EPI_ISL_568441, EPI_ISL_568442, EPI_ISL_568443, EPI_ISL_568444, EPI_ISL_568445, EPI_ISL_568446, EPI_ISL_568447, EPI_ISL_568448, EPI_ISL_568449, EPI_ISL_568450, EPI_ISL_568451 | | | |
| see above | Lighthouse Lab in Glasgow | Wellcome Sanger Institute for the COVID-19 Genomics UK Consortium | Harper VanSteenhouse, Yumi Kasai, David Gray, Carol Clugston, Anna Dominiczak and Alex Alderton, Roberto Amato, Sonia Goncalves, Ewan Harrison, David K. Jackson, Ian Johnston, Dominic Kwiatkowski, Cordelia Langford, John Sillitoe on behalf of the Wellcome Sanger Institute COVID-19 Surveillance Team |
| EPI_ISL_568453, EPI_ISL_568454 | Lighthouse Lab in Cambridge | Wellcome Sanger Institute for the COVID-19 Genomics UK Consortium | Rob Howes, The Lighthouse Lab in Cambridge and Alex Alderton, Roberto Amato, Sonia Goncalves, Ewan Harrison, David K. Jackson, Ian Johnston, Dominic Kwiatkowski, Cordelia Langford, John Sillitoe on behalf of the Wellcome Sanger Institute COVID-19 Surveillance Team |
| EPI_ISL_568455, EPI_ISL_568456, EPI_ISL_568457 | Lighthouse Lab in Glasgow | Wellcome Sanger Institute for the COVID-19 Genomics UK Consortium | Harper VanSteenhouse, Yumi Kasai, David Gray, Carol Clugston, Anna Dominiczak and Alex Alderton, Roberto Amato, Sonia Goncalves, Ewan Harrison, David K. Jackson, Ian Johnston, Dominic Kwiatkowski, Cordelia Langford, John Sillitoe on behalf of the Wellcome Sanger Institute COVID-19 Surveillance Team |
| EPI_ISL_568458 | Lighthouse Lab in Cambridge | Wellcome Sanger Institute for the COVID-19 Genomics UK Consortium | Rob Howes, The Lighthouse Lab in Cambridge and Alex Alderton, Roberto Amato, Sonia Goncalves, Ewan Harrison, David K. Jackson, Ian Johnston, Dominic Kwiatkowski, Cordelia Langford, John Sillitoe on behalf of the Wellcome Sanger Institute COVID-19 Surveillance Team |
| EPI_ISL_568459 | Lighthouse Lab in Glasgow | Wellcome Sanger Institute for the COVID-19 Genomics UK Consortium | Harper VanSteenhouse, Yumi Kasai, David Gray, Carol Clugston, Anna Dominiczak and Alex Alderton, Roberto Amato, Sonia Goncalves, Ewan Harrison, David K. Jackson, Ian Johnston, Dominic Kwiatkowski, Cordelia Langford, John Sillitoe on behalf of the Wellcome Sanger Institute COVID-19 Surveillance Team |
| EPI_ISL_568461 | Lighthouse Lab in Cambridge | Wellcome Sanger Institute for the COVID-19 Genomics UK Consortium | Rob Howes, The Lighthouse Lab in Cambridge and Alex Alderton, Roberto Amato, Sonia Goncalves, Ewan Harrison, David K. Jackson, Ian Johnston, Dominic Kwiatkowski, Cordelia Langford, John Sillitoe on behalf of the Wellcome Sanger Institute COVID-19 Surveillance Team |
| EPI_ISL_568462, EPI_ISL_568463, EPI_ISL_568464 | Lighthouse Lab in Glasgow | Wellcome Sanger Institute for the COVID-19 Genomics UK Consortium | Harper VanSteenhouse, Yumi Kasai, David Gray, Carol Clugston, Anna Dominiczak and Alex Alderton, Roberto Amato, Sonia Goncalves, Ewan Harrison, David K. Jackson, Ian Johnston, Dominic Kwiatkowski, Cordelia Langford, John Sillitoe on behalf of the Wellcome Sanger Institute COVID-19 Surveillance Team |
| EPI_ISL_568467 | Lighthouse Lab in Alderley Park | Wellcome Sanger Institute for the COVID-19 Genomics UK Consortium | Jacquelyn Wynn, Mairead Hyland, The Lighthouse Lab in Alderley Park and Alex Alderton, Roberto Amato, Sonia Goncalves, Ewan Harrison, David K. Jackson, Ian Johnston, Dominic Kwiatkowski, Cordelia Langford, John Sillitoe on behalf of the Wellcome Sanger Institute COVID-19 Surveillance Team |
| EPI_ISL_568470, EPI_ISL_568471 | Lighthouse Lab in Milton Keynes | Wellcome Sanger Institute for the COVID-19 Genomics UK Consortium | The Lighthouse Lab in Milton Keynes and Alex Alderton, Roberto Amato, Sonia Goncalves, Ewan Harrison, David K. Jackson, Ian Johnston, Dominic Kwiatkowski, Cordelia Langford, John Sillitoe on behalf of the Wellcome Sanger Institute COVID-19 Surveillance Team |
| EPI_ISL_568473 | Lighthouse Lab in Cambridge | Wellcome Sanger Institute for the COVID-19 Genomics UK Consortium | Rob Howes, The Lighthouse Lab in Cambridge and Alex Alderton, Roberto Amato, Sonia Goncalves, Ewan Harrison, David K. Jackson, Ian Johnston, Dominic Kwiatkowski, Cordelia Langford, John Sillitoe on behalf of the Wellcome Sanger Institute COVID-19 Surveillance Team |
| EPI_ISL_568476 | Lighthouse Lab in Milton Keynes | Wellcome Sanger Institute for the COVID-19 Genomics UK Consortium | The Lighthouse Lab in Milton Keynes and Alex Alderton, Roberto Amato, Sonia Goncalves, Ewan Harrison, David K. Jackson, Ian Johnston, Dominic Kwiatkowski, Cordelia Langford, John Sillitoe on behalf of the Wellcome Sanger Institute COVID-19 Surveillance Team |
| EPI_ISL_568510, EPI_ISL_568511, EPI_ISL_568512, EPI_ISL_568513, EPI_ISL_568514, EPI_ISL_568515, EPI_ISL_568516, EPI_ISL_568517, EPI_ISL_568518, EPI_ISL_568519, EPI_ISL_568520, EPI_ISL_568521, EPI_ISL_568522, EPI_ISL_568523, EPI_ISL_568524, EPI_ISL_568525, EPI_ISL_568526, EPI_ISL_568527, EPI_ISL_568528, EPI_ISL_568529, EPI_ISL_568530, EPI_ISL_568531, EPI_ISL_568532, EPI_ISL_568533, EPI_ISL_568534, EPI_ISL_568535, EPI_ISL_568536, EPI_ISL_568537, EPI_ISL_568538, EPI_ISL_568539, EPI_ISL_568540, EPI_ISL_568541, EPI_ISL_568542, EPI_ISL_568543, EPI_ISL_568544, EPI_ISL_568545, EPI_ISL_568546, EPI_ISL_568547, EPI_ISL_568548, EPI_ISL_568549, EPI_ISL_568550, EPI_ISL_568551, EPI_ISL_568552, EPI_ISL_568553, EPI_ISL_568554, EPI_ISL_568555 | | | |
| see above | Laboratorio de Referencia Nacional de Virus Respiratorios, Instituto Nacional de Salud Peru | Laboratorio de Genómica Microbiana, Universidad Peruana Cayetano Heredia | Pablo Tsukayama, Alejandra Dávila-Barclay, Luis González, Pedro E. Romero, Brenda Ayzanoa, Janet Huancachoque, Pool Marcos, Maribel Huaringa, Camila Castillo-Vilcahuaman, Guillermo Salvatierra |
| EPI_ISL_568556, EPI_ISL_568557, EPI_ISL_568558, EPI_ISL_568559, EPI_ISL_568560, EPI_ISL_568561, EPI_ISL_568562, EPI_ISL_568563, EPI_ISL_568564, EPI_ISL_568565, EPI_ISL_568566, EPI_ISL_568567, EPI_ISL_568568, EPI_ISL_568569, EPI_ISL_568570, EPI_ISL_568571, EPI_ISL_568572, EPI_ISL_568573, EPI_ISL_568574, EPI_ISL_568575, EPI_ISL_568576 | | | |
| see above | Department of Infectious Diseases and Immunology, National Hospital Organization Nagoya Medical Center | Clinical Research Center, National Hospital Organization Nagoya Medical Center | Yoshihiro Nakata, Hirotaka Ode, Mai Kubota, Masakazu Matsuda, Kazuhiro Matsuoka, Nakasuji Miho, Mikiko Mori, Mayumi Imahashi, Yoshiyuki Yokomaku, Yasumasa Iwatani |
| EPI_ISL_568577 | The National Institute of Public Health | State Veterinary Institute Prague | Nagy, A; Jirincova, H; Novakova, L; Trnka,D; Vecerova, J |
| EPI_ISL_568579 | Virus Molecular Laboratory of the Microbiology and Virology Department | INMI Lazzaro Spallanzani IRCCS | Cesare E.M. Gruber, Martina Rueca, Barbara Bartolini, Francesco Messina, Silvia Meschi, Francesca Colavita, Concetta Castilletti, Elena Percivalle, Irene Cassaniti, Edoardo Vecchio Nepita, Fausto Baldanti, Maria R. Capobianchi, Antonino Di Caro |

| | | | |
|---|---|---|---|
| EPI_ISL_568580, EPI_ISL_568581, EPI_ISL_568582, EPI_ISL_568583, EPI_ISL_568584, EPI_ISL_568585, EPI_ISL_568586 | Ramathibodi Hospital | COVID-19 Network Investigations (CONI) Alliance | Elizabeth Batty, Wasun Chantratita, Thanat Chookajorn, Stefan Fernandez, Angkana Huang, Anthony R. Jones, Khajohn Joonsalak, Chonticha Klungtong, Theerarat Kochakarn, Namfon Kotanan, Krittikorn Kumpornsin, Wudtichai Manasatienkij, Bhakbhoom Panthan, Ekawat Pasomsub, Kingkan Rakmanee, Insee Sensorn, Janjira Thaipadungpanit, Arporn Wangwiwatsin,Treewat Watthanachockchai |
| EPI_ISL_568587, EPI_ISL_568590, EPI_ISL_568591, EPI_ISL_568595, EPI_ISL_568597, EPI_ISL_568598, EPI_ISL_568599, EPI_ISL_568600, EPI_ISL_568601, EPI_ISL_568602, EPI_ISL_568603, EPI_ISL_568604, EPI_ISL_568606, EPI_ISL_568607, EPI_ISL_568608, EPI_ISL_568610, EPI_ISL_568612, EPI_ISL_568616, EPI_ISL_568617, EPI_ISL_568618, EPI_ISL_568619, EPI_ISL_568620, EPI_ISL_568621, EPI_ISL_568622, EPI_ISL_568623, EPI_ISL_568624, EPI_ISL_568625, EPI_ISL_568626, EPI_ISL_568627, EPI_ISL_568628, EPI_ISL_568633, EPI_ISL_568637, EPI_ISL_568638, EPI_ISL_568639, EPI_ISL_568640, EPI_ISL_568644, EPI_ISL_568645, EPI_ISL_568648, EPI_ISL_568649, EPI_ISL_568651, EPI_ISL_568652, EPI_ISL_568654, EPI_ISL_568656, EPI_ISL_568657, EPI_ISL_568658, EPI_ISL_568659, EPI_ISL_568660, EPI_ISL_568661, EPI_ISL_568662, EPI_ISL_568663, EPI_ISL_568664, EPI_ISL_568665, EPI_ISL_568666, EPI_ISL_568667, EPI_ISL_568668, EPI_ISL_568669, EPI_ISL_568671, EPI_ISL_568675, EPI_ISL_568676, EPI_ISL_568677, EPI_ISL_568678, EPI_ISL_568679, EPI_ISL_568681 | | | |
| see above | Florida Bureau of Public Health Laboratories | Florida Bureau of Public Health Laboratories | Sarah Schmedes, Jason Blanton |
| EPI_ISL_568686 | RSUD Prof. DR. Margono Soekarjo | Eijkman Institute for Molecular Biology, Ministry of Research and Technology/National Agency for Research and Innovation | Frilasita A Yudhaputri, Edison Johar, Hidayat Trimarsanto, Iskandar A Adnan, Willy Agustine, David H Muljono, Safarina G Malik, Herawati Sudoyo, Khin Saw Myint, Amin Soebandrio |
| EPI_ISL_568687 | RSUP Fatmawati | Eijkman Institute for Molecular Biology, Ministry of Research and Technology/National Agency for Research and Innovation | Frilasita A Yudhaputri, Edison Johar, Hidayat Trimarsanto, Iskandar A Adnan, Willy Agustine, David H Muljono, Safarina G Malik, Herawati Sudoyo, Khin Saw Myint, Amin Soebandrio |
| EPI_ISL_568688 | RS Pelni | Eijkman Institute for Molecular Biology, Ministry of Research and Technology/National Agency for Research and Innovation | Frilasita A Yudhaputri, Edison Johar, Hidayat Trimarsanto, Iskandar A Adnan, Willy Agustine, David H Muljono, Safarina G Malik, Herawati Sudoyo, Khin Saw Myint, Amin Soebandrio |
| EPI_ISL_568689 | RS Pondok Indah Puri Indah | Eijkman Institute for Molecular Biology, Ministry of Research and Technology/National Agency for Research and Innovation | Frilasita A Yudhaputri, Edison Johar, Hidayat Trimarsanto, Iskandar A Adnan, Willy Agustine, David H Muljono, Safarina G Malik, Herawati Sudoyo, Khin Saw Myint, Amin Soebandrio |
| EPI_ISL_568690, EPI_ISL_568691 | RSUP Fatmawati | Eijkman Institute for Molecular Biology, Ministry of Research and Technology/National Agency for Research and Innovation | Frilasita A Yudhaputri, Edison Johar, Hidayat Trimarsanto, Iskandar A Adnan, Willy Agustine, David H Muljono, Safarina G Malik, Herawati Sudoyo, Khin Saw Myint, Amin Soebandrio |
| EPI_ISL_568692 | RS Kramat 128 | Eijkman Institute for Molecular Biology, Ministry of Research and Technology/National Agency for Research and Innovation | Frilasita A Yudhaputri, Edison Johar, Hidayat Trimarsanto, Iskandar A Adnan, Willy Agustine, David H Muljono, Safarina G Malik, Herawati Sudoyo, Khin Saw Myint, Amin Soebandrio |
| EPI_ISL_568693 | RS Freeport Tembagapura | Eijkman Institute for Molecular Biology, Ministry of Research and Technology/National Agency for Research and Innovation | Frilasita A Yudhaputri, Edison Johar, Hidayat Trimarsanto, Iskandar A Adnan, Willy Agustine, David H Muljono, Safarina G Malik, Herawati Sudoyo, Khin Saw Myint, Amin Soebandrio |
| EPI_ISL_568694 | RSUP Prof. Dr. R. Kandou Manado | Eijkman Institute for Molecular Biology, Ministry of Research and Technology/National Agency for Research and Innovation | Frilasita A Yudhaputri, Edison Johar, Hidayat Trimarsanto, Iskandar A Adnan, Willy Agustine, David H Muljono, Safarina G Malik, Herawati Sudoyo, Khin Saw Myint, Amin Soebandrio |
| EPI_ISL_568702, EPI_ISL_568725, EPI_ISL_568727, EPI_ISL_568735, EPI_ISL_568764, EPI_ISL_568847, EPI_ISL_568872 | KEMRI-Wellcome Trust Research Programme/KEMRI-CGMR-C Kilifi | KEMRI-Wellcome Trust Research Programme/KEMRI-CGMR-C Kilifi | Githinji et al 2020 |
| EPI_ISL_568873, EPI_ISL_568874, EPI_ISL_568875 | Malaysia Genome Institute | Malaysia Genome Institute | Mohd Noor Mat Isa, Irni Suhayu Sapian, Yusuf Muhammad Noor, Nurhezreen Md Iqbal, Mohd Faizal Abu Bakar, Enizza Kasim, Shamsidar Sopie, Siti Noraini Othman, Azrin Ahmad, Nor Azfa Johari, Shahrul Hisham Zainal Ariffin |
| EPI_ISL_568877, EPI_ISL_568878, EPI_ISL_568879, EPI_ISL_568880, EPI_ISL_568881, EPI_ISL_568882, EPI_ISL_568883, EPI_ISL_568884, EPI_ISL_568885, EPI_ISL_568886, EPI_ISL_568887, EPI_ISL_568888, EPI_ISL_568889, EPI_ISL_568890, EPI_ISL_568891, EPI_ISL_568892, EPI_ISL_568893, EPI_ISL_568894, EPI_ISL_568895, EPI_ISL_568896, EPI_ISL_568897, EPI_ISL_568898, EPI_ISL_568899, EPI_ISL_568901, EPI_ISL_568902, EPI_ISL_568903, EPI_ISL_568904, EPI_ISL_568905, EPI_ISL_568906, EPI_ISL_568907, EPI_ISL_568908, EPI_ISL_568909, EPI_ISL_568910, EPI_ISL_568911, EPI_ISL_568912, EPI_ISL_568914, EPI_ISL_568915, EPI_ISL_568916, EPI_ISL_568917, EPI_ISL_568918, EPI_ISL_568919, EPI_ISL_568920, EPI_ISL_568921, EPI_ISL_568922, EPI_ISL_568924, EPI_ISL_568925, EPI_ISL_568926, EPI_ISL_568927, EPI_ISL_568928, EPI_ISL_568929, EPI_ISL_568930, EPI_ISL_568931, EPI_ISL_568932, EPI_ISL_568933, EPI_ISL_568934, EPI_ISL_568935, EPI_ISL_568936, EPI_ISL_568939, EPI_ISL_568940, EPI_ISL_568941, EPI_ISL_568942, EPI_ISL_568944, EPI_ISL_568945, EPI_ISL_568947, EPI_ISL_568948, EPI_ISL_568949, EPI_ISL_568950, EPI_ISL_568951, EPI_ISL_568952, EPI_ISL_568953, EPI_ISL_568954, EPI_ISL_568955, EPI_ISL_568956, EPI_ISL_568958, EPI_ISL_568959, EPI_ISL_568960, EPI_ISL_568961, EPI_ISL_568962, EPI_ISL_568963, EPI_ISL_568964, EPI_ISL_568965, EPI_ISL_568966, EPI_ISL_568967, EPI_ISL_568968, EPI_ISL_568969, EPI_ISL_568970, EPI_ISL_568971, EPI_ISL_568972, EPI_ISL_568973, EPI_ISL_568974, EPI_ISL_568975, EPI_ISL_568976, EPI_ISL_568977, EPI_ISL_568978, EPI_ISL_568979, EPI_ISL_568980, EPI_ISL_568981, EPI_ISL_568982, EPI_ISL_568983, EPI_ISL_568984, EPI_ISL_568986, EPI_ISL_568992, EPI_ISL_568993, EPI_ISL_568994, EPI_ISL_568995, EPI_ISL_568996, EPI_ISL_568997, EPI_ISL_568998, EPI_ISL_569000, EPI_ISL_569002, EPI_ISL_569003, EPI_ISL_569004, EPI_ISL_569005, EPI_ISL_569006, EPI_ISL_569008, EPI_ISL_569009, EPI_ISL_569010, EPI_ISL_569011, EPI_ISL_569012, EPI_ISL_569013, EPI_ISL_569014, EPI_ISL_569015, EPI_ISL_569016, EPI_ISL_569017, EPI_ISL_569018, EPI_ISL_569019, EPI_ISL_569020, EPI_ISL_569021, EPI_ISL_569022, EPI_ISL_569023, EPI_ISL_569024, EPI_ISL_569025, EPI_ISL_569026, EPI_ISL_569027, EPI_ISL_569028, EPI_ISL_569029, EPI_ISL_569031, EPI_ISL_569032, EPI_ISL_569033, EPI_ISL_569034, EPI_ISL_569035, EPI_ISL_569036, EPI_ISL_569037, EPI_ISL_569038, EPI_ISL_569041, EPI_ISL_569043, EPI_ISL_569044, EPI_ISL_569045, EPI_ISL_569046, EPI_ISL_569047, EPI_ISL_569048, EPI_ISL_569049, EPI_ISL_569050, EPI_ISL_569051, EPI_ISL_569053, EPI_ISL_569054, EPI_ISL_569055, EPI_ISL_569056, EPI_ISL_569057, EPI_ISL_569058, EPI_ISL_569060, EPI_ISL_569061, EPI_ISL_569062, EPI_ISL_569063, EPI_ISL_569064, EPI_ISL_569067, EPI_ISL_569068, EPI_ISL_569070, EPI_ISL_569071, EPI_ISL_569072, EPI_ISL_569073, EPI_ISL_569074, EPI_ISL_569075, EPI_ISL_569076, EPI_ISL_569077, EPI_ISL_569078, EPI_ISL_569079, EPI_ISL_569080, EPI_ISL_569081, EPI_ISL_569082, EPI_ISL_569084, EPI_ISL_569085, EPI_ISL_569086, EPI_ISL_569087, EPI_ISL_569088, EPI_ISL_569089, EPI_ISL_569090, EPI_ISL_569091, EPI_ISL_569092, EPI_ISL_569093, EPI_ISL_569094, EPI_ISL_569096, EPI_ISL_569097, EPI_ISL_569098, EPI_ISL_569099, EPI_ISL_569100, EPI_ISL_569101, EPI_ISL_569102, EPI_ISL_569103, EPI_ISL_569104, EPI_ISL_569105, EPI_ISL_569106, EPI_ISL_569107, EPI_ISL_569108, EPI_ISL_569109, EPI_ISL_569110, EPI_ISL_569111, EPI_ISL_569113, EPI_ISL_569114, EPI_ISL_569115, EPI_ISL_569116, EPI_ISL_569117, EPI_ISL_569119, EPI_ISL_569120, EPI_ISL_569121, EPI_ISL_569123, EPI_ISL_569124, EPI_ISL_569125, EPI_ISL_569126, EPI_ISL_569127, EPI_ISL_569128, EPI_ISL_569129, EPI_ISL_569130, EPI_ISL_569132, EPI_ISL_569133, EPI_ISL_569134, EPI_ISL_569135, EPI_ISL_569136, EPI_ISL_569137, EPI_ISL_569138, EPI_ISL_569139, EPI_ISL_569140, EPI_ISL_569141, EPI_ISL_569142, EPI_ISL_569143, EPI_ISL_569144, EPI_ISL_569145, EPI_ISL_569146, EPI_ISL_569147, EPI_ISL_569149, EPI_ISL_569150, EPI_ISL_569151, EPI_ISL_569155, EPI_ISL_569156, EPI_ISL_569157, EPI_ISL_569158, EPI_ISL_569159, EPI_ISL_569160, EPI_ISL_569161, EPI_ISL_569162, EPI_ISL_569164, EPI_ISL_569165, EPI_ISL_569166, EPI_ISL_569168, EPI_ISL_569169, EPI_ISL_569170, EPI_ISL_569171, EPI_ISL_569172, EPI_ISL_569173, EPI_ISL_569174, EPI_ISL_569175, EPI_ISL_569176, EPI_ISL_569177, EPI_ISL_569178, EPI_ISL_569179, EPI_ISL_569180, EPI_ISL_569181, EPI_ISL_569182, EPI_ISL_569183, EPI_ISL_569184, EPI_ISL_569185, EPI_ISL_569186, EPI_ISL_569187, EPI_ISL_569188, EPI_ISL_569189, EPI_ISL_569190, EPI_ISL_569191, EPI_ISL_569192, EPI_ISL_569193, EPI_ISL_569195, EPI_ISL_569196, EPI_ISL_569197, EPI_ISL_569198, EPI_ISL_569199, EPI_ISL_569200, EPI_ISL_569201, EPI_ISL_569202, EPI_ISL_569203, EPI_ISL_569204, EPI_ISL_569205, EPI_ISL_569207, EPI_ISL_569208, EPI_ISL_569210, EPI_ISL_569211, EPI_ISL_569214, EPI_ISL_569216, EPI_ISL_569217, EPI_ISL_569219, EPI_ISL_569220, EPI_ISL_569221, EPI_ISL_569222, EPI_ISL_569223, EPI_ISL_569225, EPI_ISL_569226, EPI_ISL_569227, EPI_ISL_569228, EPI_ISL_569229, EPI_ISL_569230, EPI_ISL_569231, EPI_ISL_569232, EPI_ISL_569233, EPI_ISL_569234, EPI_ISL_569235, EPI_ISL_569236, EPI_ISL_569237, EPI_ISL_569238, EPI_ISL_569239, EPI_ISL_569240, EPI_ISL_569241, EPI_ISL_569243, EPI_ISL_569244, EPI_ISL_569246, EPI_ISL_569247, EPI_ISL_569248, EPI_ISL_569249, EPI_ISL_569250, EPI_ISL_569251, EPI_ISL_569252, EPI_ISL_569253, EPI_ISL_569254, EPI_ISL_569255, EPI_ISL_569256, EPI_ISL_569257, EPI_ISL_569258, EPI_ISL_569259, EPI_ISL_569261, EPI_ISL_569262, EPI_ISL_569263, EPI_ISL_569264, EPI_ISL_569265, EPI_ISL_569266, EPI_ISL_569267, EPI_ISL_569268, EPI_ISL_569269, EPI_ISL_569270, EPI_ISL_569271, EPI_ISL_569272, EPI_ISL_569273, EPI_ISL_569274, EPI_ISL_569275, EPI_ISL_569276, EPI_ISL_569277, EPI_ISL_569278, EPI_ISL_569279, EPI_ISL_569280, EPI_ISL_569281, EPI_ISL_569282, EPI_ISL_569284, EPI_ISL_569285, EPI_ISL_569288, EPI_ISL_569290, EPI_ISL_569291, EPI_ISL_569292, EPI_ISL_569293, EPI_ISL_569294, EPI_ISL_569295, EPI_ISL_569296, EPI_ISL_569297, EPI_ISL_569298, EPI_ISL_569299, EPI_ISL_569300, EPI_ISL_569301, EPI_ISL_569302, EPI_ISL_569303, EPI_ISL_569304, EPI_ISL_569305, EPI_ISL_569307, EPI_ISL_569308, EPI_ISL_569309, EPI_ISL_569310, EPI_ISL_569311, EPI_ISL_569312, EPI_ISL_569313, EPI_ISL_569314, EPI_ISL_569315, EPI_ISL_569316, EPI_ISL_569317, EPI_ISL_569318, EPI_ISL_569321, EPI_ISL_569323, EPI_ISL_569324, EPI_ISL_569325, EPI_ISL_569326, EPI_ISL_569327, EPI_ISL_569328, EPI_ISL_569329, EPI_ISL_569330, EPI_ISL_569331, EPI_ISL_569332, EPI_ISL_569333, EPI_ISL_569334, EPI_ISL_569335, EPI_ISL_569336, EPI_ISL_569337, EPI_ISL_569338, EPI_ISL_569339, EPI_ISL_569340, EPI_ISL_569341, EPI_ISL_569343, EPI_ISL_569344, EPI_ISL_569345, EPI_ISL_569346, EPI_ISL_569347, EPI_ISL_569348, EPI_ISL_569350, EPI_ISL_569351, EPI_ISL_569352, EPI_ISL_569353, EPI_ISL_569354, EPI_ISL_569355, EPI_ISL_569356, EPI_ISL_569357, EPI_ISL_569358, EPI_ISL_569359, EPI_ISL_569360, EPI_ISL_569361, EPI_ISL_569362, EPI_ISL_569363, EPI_ISL_569364, EPI_ISL_569365, EPI_ISL_569366, EPI_ISL_569367, EPI_ISL_569368, EPI_ISL_569369, EPI_ISL_569370, EPI_ISL_569371, EPI_ISL_569372, EPI_ISL_569373, EPI_ISL_569374, EPI_ISL_569375, EPI_ISL_569377, EPI_ISL_569378, EPI_ISL_569379, EPI_ISL_569380, EPI_ISL_569381, EPI_ISL_569382, EPI_ISL_569383, EPI_ISL_569384, EPI_ISL_569386, EPI_ISL_569387, EPI_ISL_569388, EPI_ISL_569389, EPI_ISL_569390, EPI_ISL_569391, EPI_ISL_569392, EPI_ISL_569393, EPI_ISL_569394, EPI_ISL_569395, EPI_ISL_569397, EPI_ISL_569398, EPI_ISL_569399, EPI_ISL_569400, EPI_ISL_569401, EPI_ISL_569402, EPI_ISL_569403, EPI_ISL_569404, EPI_ISL_569405, EPI_ISL_569406, EPI_ISL_569407, EPI_ISL_569408, EPI_ISL_569410, EPI_ISL_569411, EPI_ISL_569412, EPI_ISL_569413, EPI_ISL_569414, EPI_ISL_569415, EPI_ISL_569416, EPI_ISL_569417, EPI_ISL_569418, EPI_ISL_569419, EPI_ISL_569420, EPI_ISL_569421, EPI_ISL_569422, EPI_ISL_569423, EPI_ISL_569424, EPI_ISL_569425, EPI_ISL_569426, EPI_ISL_569427, EPI_ISL_569428, EPI_ISL_569429, EPI_ISL_569431, EPI_ISL_569432, EPI_ISL_569433, EPI_ISL_569434, EPI_ISL_569435, EPI_ISL_569436, EPI_ISL_569437, EPI_ISL_569438, EPI_ISL_569440, EPI_ISL_569441, EPI_ISL_569443, EPI_ISL_569444, EPI_ISL_569445, EPI_ISL_569446, EPI_ISL_569447, EPI_ISL_569448, EPI_ISL_569449, EPI_ISL_569450, EPI_ISL_569451, EPI_ISL_569456, EPI_ISL_569457, EPI_ISL_569458, EPI_ISL_569459, EPI_ISL_569460, EPI_ISL_569461, EPI_ISL_569462, EPI_ISL_569463, EPI_ISL_569464, EPI_ISL_569465, EPI_ISL_569466, EPI_ISL_569467, EPI_ISL_569468, EPI_ISL_569469, EPI_ISL_569470, EPI_ISL_569471, EPI_ISL_569472, EPI_ISL_569473, EPI_ISL_569474, EPI_ISL_569475, EPI_ISL_569476, EPI_ISL_569477, EPI_ISL_569478, EPI_ISL_569479, EPI_ISL_569481, EPI_ISL_569483, EPI_ISL_569484, EPI_ISL_569485, EPI_ISL_569489, EPI_ISL_569490, EPI_ISL_569491, EPI_ISL_569492, EPI_ISL_569493, EPI_ISL_569494, EPI_ISL_569495, EPI_ISL_569496, EPI_ISL_569497, EPI_ISL_569498, EPI_ISL_569499, EPI_ISL_569500, EPI_ISL_569501, EPI_ISL_569502, EPI_ISL_569503, EPI_ISL_569504, EPI_ISL_569505, EPI_ISL_569506, EPI_ISL_569507, EPI_ISL_569508, EPI_ISL_569509, EPI_ISL_569510, EPI_ISL_569511, EPI_ISL_569512, EPI_ISL_569513, EPI_ISL_569514, EPI_ISL_569515, EPI_ISL_569516, EPI_ISL_569517, EPI_ISL_569518, EPI_ISL_569519, EPI_ISL_569520, EPI_ISL_569521, EPI_ISL_569522, EPI_ISL_569523, EPI_ISL_569524, EPI_ISL_569525, EPI_ISL_569526, EPI_ISL_569527, EPI_ISL_569528, EPI_ISL_569529, EPI_ISL_569530, EPI_ISL_569531, EPI_ISL_569532, EPI_ISL_569533, EPI_ISL_569534, EPI_ISL_569535, EPI_ISL_569536, EPI_ISL_569537, EPI_ISL_569538, EPI_ISL_569540, EPI_ISL_569541, EPI_ISL_569542, EPI_ISL_569543, EPI_ISL_569544, EPI_ISL_569545, EPI_ISL_569546, EPI_ISL_569547, EPI_ISL_569548, EPI_ISL_569549, EPI_ISL_569550, EPI_ISL_569551, EPI_ISL_569552, EPI_ISL_569553, EPI_ISL_569554, EPI_ISL_569555, EPI_ISL_569556, EPI_ISL_569557, EPI_ISL_569558, EPI_ISL_569559, EPI_ISL_569560, EPI_ISL_569561, EPI_ISL_569562, EPI_ISL_569563, EPI_ISL_569564, EPI_ISL_569565, EPI_ISL_569566, EPI_ISL_569567, EPI_ISL_569568, EPI_ISL_569569, EPI_ISL_569570, EPI_ISL_569571, EPI_ISL_569572, EPI_ISL_569573, EPI_ISL_569574, EPI_ISL_569575, EPI_ISL_569576, EPI_ISL_569577, EPI_ISL_569578, EPI_ISL_569582, EPI_ISL_569583, EPI_ISL_569584, EPI_ISL_569585, EPI_ISL_569586, EPI_ISL_569587, EPI_ISL_569588, EPI_ISL_569589, EPI_ISL_569590, EPI_ISL_569592, EPI_ISL_569593, EPI_ISL_569594, EPI_ISL_569595, EPI_ISL_569597, EPI_ISL_569598, EPI_ISL_569599, EPI_ISL_569600, EPI_ISL_569601, EPI_ISL_569602, EPI_ISL_569603, EPI_ISL_569604, EPI_ISL_569605, EPI_ISL_569606, EPI_ISL_569607, EPI_ISL_569608 | | | |
| see above | MEPHI, Aix Marseille University | MEPHI, Aix Marseille University | Anthony LEVASSEUR |
| EPI_ISL_569609 | Bennet County Hospital | South Dakota Public Health Laboratory | Matt Plumb, Jacob Garfin, Xiong Wang, and Chris Carlson |
| EPI_ISL_569610 | Avera McKennan Hospital | South Dakota Public Health Laboratory | Matt Plumb, Jacob Garfin, Xiong Wang, and Chris Carlson |
| EPI_ISL_569611 | Sioux Falls Urgent Care | South Dakota Public Health Laboratory | Matt Plumb, Jacob Garfin, Xiong Wang, and Chris Carlson |
| EPI_ISL_569612, EPI_ISL_569613 | Cheyenne River Health Center | South Dakota Public Health Laboratory | Matt Plumb, Jacob Garfin, Xiong Wang, and Chris Carlson |
| EPI_ISL_569614 | IHS Fort Yates Hospital | South Dakota Public Health Laboratory | Matt Plumb, Jacob Garfin, Xiong Wang, and Chris Carlson |
| EPI_ISL_569616 | Aurora County Clinic | South Dakota Public Health Laboratory | Matt Plumb, Jacob Garfin, Xiong Wang, and Chris Carlson |
| EPI_ISL_569617 | SD Urban Indian Health Pierre | South Dakota Public Health Laboratory | Matt Plumb, Jacob Garfin, Xiong Wang, and Chris Carlson |
| EPI_ISL_569618, EPI_ISL_569619 | Bethel Lutheran Home | South Dakota Public Health Laboratory | Matt Plumb, Jacob Garfin, Xiong Wang, and Chris Carlson |
| EPI_ISL_569620 | McCrossan Boys Ranch | South Dakota Public Health Laboratory | Matt Plumb, Jacob Garfin, Xiong Wang, and Chris Carlson |
| EPI_ISL_569621 | Quick Care Watertown | South Dakota Public Health Laboratory | Matt Plumb, Jacob Garfin, Xiong Wang, and Chris Carlson |
| EPI_ISL_569622 | Woodrow Wilson Keeble Memorial Healthcare | South Dakota Public Health Laboratory | Matt Plumb, Jacob Garfin, Xiong Wang, and Chris Carlson |
| EPI_ISL_569623 | Quick Care Watertown | South Dakota Public Health Laboratory | Matt Plumb, Jacob Garfin, Xiong Wang, and Chris Carlson |
| EPI_ISL_569624 | Avera McKennan Hospital | South Dakota Public Health Laboratory | Matt Plumb, Jacob Garfin, Xiong Wang, and Chris Carlson |
| EPI_ISL_569632, EPI_ISL_569633, EPI_ISL_569634, EPI_ISL_569635, EPI_ISL_569636, EPI_ISL_569637, EPI_ISL_569638, EPI_ISL_569639, EPI_ISL_569640, EPI_ISL_569641, EPI_ISL_569642, EPI_ISL_569643, EPI_ISL_569644, EPI_ISL_569645, EPI_ISL_569646, EPI_ISL_569647, EPI_ISL_569648, EPI_ISL_569649, EPI_ISL_569650, EPI_ISL_569651, EPI_ISL_569652, EPI_ISL_569653, EPI_ISL_569654, EPI_ISL_569655, EPI_ISL_569656, EPI_ISL_569657, EPI_ISL_569658, EPI_ISL_569659, EPI_ISL_569660, EPI_ISL_569661, EPI_ISL_569662, EPI_ISL_569663 | | | |
| see above | New Mexico Department of Health Scientific Laboratory | New Mexico Department of Health Scientific Laboratory | Ellie Johnson, Anastacia Griego-Fisher, D'Eldra Malone |
| EPI_ISL_569720 | Texas Department of State Health Services (TXDSHS) | Texas Department of State Health Services (TXDSHS) | Rashmi Tuladhar, Bonnie Oh,Jenny Zhang, Maliha Rahman, Anita Pokharel, Myong Koag, Chun Wang, Rachel Lee, Grace Kubin |
| EPI_ISL_569735, EPI_ISL_569736, EPI_ISL_569737, EPI_ISL_569738, EPI_ISL_569739, EPI_ISL_569740, EPI_ISL_569741, EPI_ISL_569742, EPI_ISL_569743, EPI_ISL_569744, EPI_ISL_569745, EPI_ISL_569746, EPI_ISL_569747, EPI_ISL_569748, EPI_ISL_569749, EPI_ISL_569750, EPI_ISL_569751, EPI_ISL_569752, EPI_ISL_569753, EPI_ISL_569754, EPI_ISL_569755, EPI_ISL_569756, EPI_ISL_569757, EPI_ISL_569758, EPI_ISL_569759, EPI_ISL_569760, EPI_ISL_569761, EPI_ISL_569762, EPI_ISL_569763, EPI_ISL_569764, EPI_ISL_569765, EPI_ISL_569766, EPI_ISL_569767, EPI_ISL_569768, EPI_ISL_569769, EPI_ISL_569770, EPI_ISL_569771, EPI_ISL_569772, EPI_ISL_569773, EPI_ISL_569774, EPI_ISL_569775, EPI_ISL_569776, EPI_ISL_569777, EPI_ISL_569778, EPI_ISL_569779, EPI_ISL_569780, EPI_ISL_569781, EPI_ISL_569782, EPI_ISL_569783, EPI_ISL_569784, EPI_ISL_569785, EPI_ISL_569786, EPI_ISL_569787, EPI_ISL_569788, EPI_ISL_569789, EPI_ISL_569790, EPI_ISL_569791, EPI_ISL_569792, EPI_ISL_569793, EPI_ISL_569794, EPI_ISL_569795, EPI_ISL_569796, EPI_ISL_569797, EPI_ISL_569798, EPI_ISL_569799, EPI_ISL_569800, EPI_ISL_569801, EPI_ISL_569802, EPI_ISL_569803, EPI_ISL_569804, EPI_ISL_569805, EPI_ISL_569806, EPI_ISL_569807, EPI_ISL_569808, EPI_ISL_569809, EPI_ISL_569810, EPI_ISL_569811, EPI_ISL_569812, EPI_ISL_569813, EPI_ISL_569814, EPI_ISL_569815, EPI_ISL_569816, EPI_ISL_569817, EPI_ISL_569818, EPI_ISL_569819, EPI_ISL_569820, EPI_ISL_569821, EPI_ISL_569822, EPI_ISL_569823, EPI_ISL_569824, EPI_ISL_569825, EPI_ISL_569826, EPI_ISL_569827, EPI_ISL_569828, EPI_ISL_569829, EPI_ISL_569830, EPI_ISL_569831, EPI_ISL_569832, EPI_ISL_569833, EPI_ISL_569834, EPI_ISL_569835, EPI_ISL_569836, EPI_ISL_569837, EPI_ISL_569838, EPI_ISL_569839, EPI_ISL_569840, EPI_ISL_569841, EPI_ISL_569842, EPI_ISL_569843, EPI_ISL_569844, EPI_ISL_569845, EPI_ISL_569846, EPI_ISL_569847, EPI_ISL_569848, EPI_ISL_569849, EPI_ISL_569850, EPI_ISL_569851, EPI_ISL_569852, EPI_ISL_569853, EPI_ISL_569854, EPI_ISL_569855, EPI_ISL_569856, EPI_ISL_569857 | | | |
| see above | Omsk Research Institute of Natural Focal Infections | WHO National Influenza Centre Russian Federation | Artem Fadeev, Ekaterina Gradoboeva, Ekaterina Savkina, Daria Nashatyreva, Elena Poleshchuk, Aleksei Vasilenko, Valery Yakimenko, Andrey Komissarov |
| EPI_ISL_569858 | Area of Virology, Serology and Virology Division (SAViD), New South Wales Health Pathology Randwick | Area of Virology, Serology and Virology Division (SAViD), New South Wales Health Pathology Randwick | Rawlinson, W., Deveson, I., Bull, R., Van Hal, S. |
| EPI_ISL_569859, EPI_ISL_569860, EPI_ISL_569861, EPI_ISL_569862, EPI_ISL_569863 | CSIR-Indian Institute of Chemical Biology, MEDICA Supercpecialty Hospital Kolkata | CSIR-Indian Institute of Chemical Biology, MEDICA Supercpecialty Hospital Kolkata | Sujay Krishna Maity, Priyanka Mallick, Debaleena Bhowmik, Abhishake Lahiri, Dr. Aviral Roy, Dr. Soumen Saha, Dr. Arpita Ghosh Mitra, Dr. Rajesh Pandey, Dr. Sandip Paul, Dr. Partha Chakrabarti, Dr. Saikat Chakrabarti |

EPI_ISL_569865, EPI_ISL_569866, EPI_ISL_569867, EPI_ISL_569868, EPI_ISL_569869, EPI_ISL_569870, EPI_ISL_569871, EPI_ISL_569872, EPI_ISL_569873, EPI_ISL_569874, EPI_ISL_569875, EPI_ISL_569876, EPI_ISL_569877, EPI_ISL_569878, EPI_ISL_569879, EPI_ISL_569880, EPI_ISL_569881, EPI_ISL_569882, EPI_ISL_569883, EPI_ISL_569884, EPI_ISL_569885, EPI_ISL_569886

| | | | |
|---|---|---|---|
| see above | Amedeo di savoia | Crosetto lab, Karolinska Institutet, SciLifeLab | Michele Simonetti, Maria Grazia Milia, Luuk Harbers, Ning Zhang, Anna Sapino, Valeria Ghisetti, Nicola Crosetto |

EPI_ISL_569887, EPI_ISL_569888, EPI_ISL_569889, EPI_ISL_569890, EPI_ISL_569891, EPI_ISL_569892, EPI_ISL_569893, EPI_ISL_569894, EPI_ISL_569895, EPI_ISL_569896, EPI_ISL_569898, EPI_ISL_569899, EPI_ISL_569900, EPI_ISL_569901, EPI_ISL_569902, EPI_ISL_569904, EPI_ISL_569905, EPI_ISL_569906, EPI_ISL_569907, EPI_ISL_569909, EPI_ISL_569911, EPI_ISL_569913, EPI_ISL_569915, EPI_ISL_569932, EPI_ISL_569933, EPI_ISL_569934, EPI_ISL_569935, EPI_ISL_569939, EPI_ISL_569940, EPI_ISL_569941, EPI_ISL_569942, EPI_ISL_569943, EPI_ISL_569945, EPI_ISL_569947

| | | | |
|---|---|---|---|
| see above | Innovative Genomics Institute, UC Berkeley | Innovative Genomics Institute, UC Berkeley | Stacia Wyman, Haridha Shivram, Phil Frankino, Liana Lareau, Shana McDevitt, Justin Choi |

EPI_ISL_569949, EPI_ISL_569954, EPI_ISL_569955, EPI_ISL_569958, EPI_ISL_569961, EPI_ISL_569965, EPI_ISL_569966, EPI_ISL_569968, EPI_ISL_569969, EPI_ISL_569970, EPI_ISL_569971, EPI_ISL_569972, EPI_ISL_569973, EPI_ISL_569975, EPI_ISL_569976, EPI_ISL_569978, EPI_ISL_569980, EPI_ISL_569981, EPI_ISL_569982, EPI_ISL_569983, EPI_ISL_569986, EPI_ISL_569987, EPI_ISL_569991, EPI_ISL_569992, EPI_ISL_569993, EPI_ISL_569995, EPI_ISL_569996, EPI_ISL_569998, EPI_ISL_570000, EPI_ISL_570001, EPI_ISL_570002, EPI_ISL_570003, EPI_ISL_570004, EPI_ISL_570005, EPI_ISL_570006, EPI_ISL_570007, EPI_ISL_570008, EPI_ISL_570009, EPI_ISL_570011, EPI_ISL_570012, EPI_ISL_570014, EPI_ISL_570016, EPI_ISL_570017, EPI_ISL_570019, EPI_ISL_570020, EPI_ISL_570021, EPI_ISL_570023, EPI_ISL_570025

| | | | |
|---|---|---|---|
| see above | Unity Health Toronto | Ontario Institute for Cancer Research | Ramzi Fattouh, Larissa M. Matukas, Yan Chen,Mark Downing, Trina Otterman, Karel Boissinot, Wai Sum Siu, Zhi Cui, Le Luu, Samira Mubareka, TIBDN, Ilinca Lungu, Bernard Lam, Jeremy Johns, Paul Krzyzanowski, Richard de Borja, Felicia Vincelli, Philip Zuzarte, Jared T. Simpson |
