## Supplementary material for "Analysis of the Dynamics and Distribution of SARS-CoV-2 Mutations and its Possible Structural and Functional Implications": GISAID_acknowledgements_part_12

We gratefully acknowledge the following Authors from the Originating laboratories responsible for obtaining the specimens, as well as the Submitting laboratories where the genome data were generated and shared via GISAID, on which this research is based.

All Submitters of data may be contacted directly via www.gisaid.org

| Accession ID | Originating Laboratory | Submitting Laboratory | Authors |
|---|---|---|---|
| EPI_ISL_570028, EPI_ISL_570029, EPI_ISL_570030, EPI_ISL_570032, EPI_ISL_570033, EPI_ISL_570034, EPI_ISL_570037, EPI_ISL_570038, EPI_ISL_570040, EPI_ISL_570041, EPI_ISL_570042, EPI_ISL_570043, EPI_ISL_570044, EPI_ISL_570046, EPI_ISL_570047, EPI_ISL_570048, EPI_ISL_570049, EPI_ISL_570050, EPI_ISL_570051, EPI_ISL_570052, EPI_ISL_570053, EPI_ISL_570054, EPI_ISL_570056, EPI_ISL_570057, EPI_ISL_570059, EPI_ISL_570060, EPI_ISL_570061, EPI_ISL_570062, EPI_ISL_570063, EPI_ISL_570064, EPI_ISL_570065, EPI_ISL_570066, EPI_ISL_570067, EPI_ISL_570068, EPI_ISL_570069, EPI_ISL_570071, EPI_ISL_570072, EPI_ISL_570073, EPI_ISL_570074, EPI_ISL_570075, EPI_ISL_570076, EPI_ISL_570078, EPI_ISL_570079, EPI_ISL_570080, EPI_ISL_570081, EPI_ISL_570082, EPI_ISL_570083, EPI_ISL_570084, EPI_ISL_570086, EPI_ISL_570087, EPI_ISL_570088, EPI_ISL_570089, EPI_ISL_570090, EPI_ISL_570091, EPI_ISL_570092, EPI_ISL_570093, EPI_ISL_570094, EPI_ISL_570095, EPI_ISL_570096, EPI_ISL_570097, EPI_ISL_570098, EPI_ISL_570099, EPI_ISL_570100, EPI_ISL_570101, EPI_ISL_570102, EPI_ISL_570103, EPI_ISL_570104, EPI_ISL_570105, EPI_ISL_570106, EPI_ISL_570107, EPI_ISL_570108, EPI_ISL_570109, EPI_ISL_570110, EPI_ISL_570111, EPI_ISL_570112, EPI_ISL_570113, EPI_ISL_570114, EPI_ISL_570115, EPI_ISL_570116, EPI_ISL_570117, EPI_ISL_570118, EPI_ISL_570119, EPI_ISL_570120, EPI_ISL_570121, EPI_ISL_570122, EPI_ISL_570123, EPI_ISL_570124, EPI_ISL_570125, EPI_ISL_570126, EPI_ISL_570127, EPI_ISL_570128, EPI_ISL_570129, EPI_ISL_570130, EPI_ISL_570131, EPI_ISL_570132, EPI_ISL_570133, EPI_ISL_570135, EPI_ISL_570136, EPI_ISL_570137, EPI_ISL_570138, EPI_ISL_570139, EPI_ISL_570140, EPI_ISL_570141, EPI_ISL_570142, EPI_ISL_570143, EPI_ISL_570144, EPI_ISL_570145, EPI_ISL_570146, EPI_ISL_570147, EPI_ISL_570148, EPI_ISL_570150, EPI_ISL_570151, EPI_ISL_570152, EPI_ISL_570153, EPI_ISL_570154, EPI_ISL_570155, EPI_ISL_570156, EPI_ISL_570157, EPI_ISL_570158, EPI_ISL_570159, EPI_ISL_570161, EPI_ISL_570163, EPI_ISL_570164, EPI_ISL_570165, EPI_ISL_570166, EPI_ISL_570167, EPI_ISL_570168, EPI_ISL_570169, EPI_ISL_570170, EPI_ISL_570172, EPI_ISL_570173, EPI_ISL_570175, EPI_ISL_570176, EPI_ISL_570177, EPI_ISL_570178, EPI_ISL_570179, EPI_ISL_570180, EPI_ISL_570181, EPI_ISL_570182, EPI_ISL_570184, EPI_ISL_570185, EPI_ISL_570186, EPI_ISL_570187, EPI_ISL_570188, EPI_ISL_570190, EPI_ISL_570191, EPI_ISL_570192, EPI_ISL_570193, EPI_ISL_570194, EPI_ISL_570195, EPI_ISL_570196, EPI_ISL_570197, EPI_ISL_570198, EPI_ISL_570199, EPI_ISL_570200, EPI_ISL_570201, EPI_ISL_570202, EPI_ISL_570203, EPI_ISL_570205, EPI_ISL_570206, EPI_ISL_570207, EPI_ISL_570208, EPI_ISL_570211, EPI_ISL_570213, EPI_ISL_570214, EPI_ISL_570215, EPI_ISL_570217, EPI_ISL_570219, EPI_ISL_570221, EPI_ISL_570222, EPI_ISL_570223, EPI_ISL_570224, EPI_ISL_570225, EPI_ISL_570226, EPI_ISL_570227, EPI_ISL_570228, EPI_ISL_570229, EPI_ISL_570230, EPI_ISL_570231, EPI_ISL_570232, EPI_ISL_570233, EPI_ISL_570234, EPI_ISL_570235, EPI_ISL_570236, EPI_ISL_570237, EPI_ISL_570238, EPI_ISL_570239, EPI_ISL_570240, EPI_ISL_570241, EPI_ISL_570242, EPI_ISL_570243, EPI_ISL_570244, EPI_ISL_570245, EPI_ISL_570246, EPI_ISL_570247, EPI_ISL_570248, EPI_ISL_570249, EPI_ISL_570250, EPI_ISL_570252, EPI_ISL_570253, EPI_ISL_570254, EPI_ISL_570256, EPI_ISL_570257, EPI_ISL_570258, EPI_ISL_570259, EPI_ISL_570260, EPI_ISL_570261, EPI_ISL_570262, EPI_ISL_570264, EPI_ISL_570265, EPI_ISL_570266, EPI_ISL_570267, EPI_ISL_570268, EPI_ISL_570269, EPI_ISL_570270, EPI_ISL_570271, EPI_ISL_570272, EPI_ISL_570273, EPI_ISL_570274, EPI_ISL_570275, EPI_ISL_570276, EPI_ISL_570277, EPI_ISL_570278, EPI_ISL_570279, EPI_ISL_570280, EPI_ISL_570281, EPI_ISL_570282, EPI_ISL_570283, EPI_ISL_570284, EPI_ISL_570285, EPI_ISL_570286, EPI_ISL_570287, EPI_ISL_570288, EPI_ISL_570289, EPI_ISL_570290, EPI_ISL_570291, EPI_ISL_570292, EPI_ISL_570293, EPI_ISL_570294, EPI_ISL_570295, EPI_ISL_570297, EPI_ISL_570298, EPI_ISL_570299, EPI_ISL_570300, EPI_ISL_570301, EPI_ISL_570302, EPI_ISL_570303, EPI_ISL_570304, EPI_ISL_570306, EPI_ISL_570307, EPI_ISL_570308, EPI_ISL_570309, EPI_ISL_570310, EPI_ISL_570311, EPI_ISL_570312, EPI_ISL_570313, EPI_ISL_570314, EPI_ISL_570315, EPI_ISL_570316, EPI_ISL_570317, EPI_ISL_570318, EPI_ISL_570319, EPI_ISL_570320, EPI_ISL_570321, EPI_ISL_570322, EPI_ISL_570323, EPI_ISL_570324, EPI_ISL_570325, EPI_ISL_570326, EPI_ISL_570327, EPI_ISL_570328, EPI_ISL_570329, EPI_ISL_570330, EPI_ISL_570331, EPI_ISL_570332, EPI_ISL_570333, EPI_ISL_570334, EPI_ISL_570335, EPI_ISL_570336, EPI_ISL_570337, EPI_ISL_570338, EPI_ISL_570339, EPI_ISL_570340, EPI_ISL_570341, EPI_ISL_570342, EPI_ISL_570343, EPI_ISL_570344, EPI_ISL_570345, EPI_ISL_570346, EPI_ISL_570348, EPI_ISL_570349, EPI_ISL_570350, EPI_ISL_570351, EPI_ISL_570352, EPI_ISL_570353, EPI_ISL_570354, EPI_ISL_570355, EPI_ISL_570356, EPI_ISL_570358, EPI_ISL_570359, EPI_ISL_570360, EPI_ISL_570361, EPI_ISL_570362, EPI_ISL_570363, EPI_ISL_570364, EPI_ISL_570366, EPI_ISL_570367, EPI_ISL_570368, EPI_ISL_570369, EPI_ISL_570370, EPI_ISL_570371, EPI_ISL_570372, EPI_ISL_570373, EPI_ISL_570375, EPI_ISL_570376, EPI_ISL_570377, EPI_ISL_570378, EPI_ISL_570379, EPI_ISL_570380, EPI_ISL_570381, EPI_ISL_570382, EPI_ISL_570383, EPI_ISL_570384, EPI_ISL_570385, EPI_ISL_570386, EPI_ISL_570387, EPI_ISL_570388, EPI_ISL_570389, EPI_ISL_570390, EPI_ISL_570391, EPI_ISL_570392, EPI_ISL_570393, EPI_ISL_570394, EPI_ISL_570395, EPI_ISL_570396, EPI_ISL_570397, EPI_ISL_570398, EPI_ISL_570399, EPI_ISL_570400, EPI_ISL_570401, EPI_ISL_570402, EPI_ISL_570403, EPI_ISL_570404, EPI_ISL_570405, EPI_ISL_570406, EPI_ISL_570407, EPI_ISL_570408, EPI_ISL_570409, EPI_ISL_570410, EPI_ISL_570411, EPI_ISL_570412, EPI_ISL_570413, EPI_ISL_570414, EPI_ISL_570415, EPI_ISL_570416, EPI_ISL_570417, EPI_ISL_570418, EPI_ISL_570419, EPI_ISL_570420, EPI_ISL_570421, EPI_ISL_570422, EPI_ISL_570423, EPI_ISL_570424, EPI_ISL_570426, EPI_ISL_570427, EPI_ISL_570428, EPI_ISL_570429, EPI_ISL_570430, EPI_ISL_570431, EPI_ISL_570432, EPI_ISL_570433, EPI_ISL_570434, EPI_ISL_570435, EPI_ISL_570436, EPI_ISL_570437, EPI_ISL_570438, EPI_ISL_570439, EPI_ISL_570440, EPI_ISL_570441, EPI_ISL_570442, EPI_ISL_570444, EPI_ISL_570445, EPI_ISL_570446, EPI_ISL_570447, EPI_ISL_570449, EPI_ISL_570450, EPI_ISL_570451, EPI_ISL_570452, EPI_ISL_570453, EPI_ISL_570454, EPI_ISL_570455, EPI_ISL_570456, EPI_ISL_570457, EPI_ISL_570458, EPI_ISL_570459, EPI_ISL_570460, EPI_ISL_570461, EPI_ISL_570462, EPI_ISL_570463, EPI_ISL_570464, EPI_ISL_570465, EPI_ISL_570466, EPI_ISL_570467, EPI_ISL_570468, EPI_ISL_570469, EPI_ISL_570470, EPI_ISL_570471, EPI_ISL_570472, EPI_ISL_570473, EPI_ISL_570474, EPI_ISL_570475, EPI_ISL_570476, EPI_ISL_570478, EPI_ISL_570480, EPI_ISL_570481, EPI_ISL_570482, EPI_ISL_570483, EPI_ISL_570484, EPI_ISL_570485, EPI_ISL_570486, EPI_ISL_570487, EPI_ISL_570488, EPI_ISL_570489, EPI_ISL_570490, EPI_ISL_570491, EPI_ISL_570492, EPI_ISL_570493, EPI_ISL_570494, EPI_ISL_570496, EPI_ISL_570497, EPI_ISL_570498, EPI_ISL_570499, EPI_ISL_570500, EPI_ISL_570501, EPI_ISL_570502, EPI_ISL_570503, EPI_ISL_570504, EPI_ISL_570506, EPI_ISL_570507, EPI_ISL_570508, EPI_ISL_570509, EPI_ISL_570510, EPI_ISL_570511, EPI_ISL_570512, EPI_ISL_570513, EPI_ISL_570514, EPI_ISL_570515, EPI_ISL_570516, EPI_ISL_570517, EPI_ISL_570518, EPI_ISL_570519, EPI_ISL_570520, EPI_ISL_570521, EPI_ISL_570522, EPI_ISL_570523, EPI_ISL_570524, EPI_ISL_570525, EPI_ISL_570526, EPI_ISL_570527, EPI_ISL_570528, EPI_ISL_570529, EPI_ISL_570530, EPI_ISL_570531, EPI_ISL_570532, EPI_ISL_570533, EPI_ISL_570534, EPI_ISL_570535, EPI_ISL_570536, EPI_ISL_570537, EPI_ISL_570538, EPI_ISL_570539, EPI_ISL_570540, EPI_ISL_570541, EPI_ISL_570542, EPI_ISL_570543, EPI_ISL_570545, EPI_ISL_570546, EPI_ISL_570547, EPI_ISL_570558, EPI_ISL_570559, EPI_ISL_570560, EPI_ISL_570561, EPI_ISL_570562, EPI_ISL_570563, EPI_ISL_570564, EPI_ISL_570565, EPI_ISL_570566, EPI_ISL_570567, EPI_ISL_570568, EPI_ISL_570569, EPI_ISL_570570, EPI_ISL_570571, EPI_ISL_570572, EPI_ISL_570573, EPI_ISL_570574, EPI_ISL_570575, EPI_ISL_570576, EPI_ISL_570577, EPI_ISL_570578, EPI_ISL_570579, EPI_ISL_570580, EPI_ISL_570581, EPI_ISL_570582, EPI_ISL_570583, EPI_ISL_570584, EPI_ISL_570585, EPI_ISL_570586, EPI_ISL_570587, EPI_ISL_570588, EPI_ISL_570589, EPI_ISL_570590, EPI_ISL_570591, EPI_ISL_570592, EPI_ISL_570593, EPI_ISL_570594, EPI_ISL_570595, EPI_ISL_570596, EPI_ISL_570597, EPI_ISL_570598, EPI_ISL_570599, EPI_ISL_570600, EPI_ISL_570601, EPI_ISL_570602, EPI_ISL_570603, EPI_ISL_570604, EPI_ISL_570605, EPI_ISL_570606, EPI_ISL_570607, EPI_ISL_570608, EPI_ISL_570609, EPI_ISL_570610, EPI_ISL_570611, EPI_ISL_570612, EPI_ISL_570613, EPI_ISL_570614, EPI_ISL_570615, EPI_ISL_570616, EPI_ISL_570617, EPI_ISL_570618, EPI_ISL_570619, EPI_ISL_570620, EPI_ISL_570621, EPI_ISL_570622, EPI_ISL_570623, EPI_ISL_570624, EPI_ISL_570625, EPI_ISL_570626, EPI_ISL_570627, EPI_ISL_570628, EPI_ISL_570629, EPI_ISL_570630, EPI_ISL_570631, EPI_ISL_570632, EPI_ISL_570633, EPI_ISL_570634, EPI_ISL_570635, EPI_ISL_570636, EPI_ISL_570637, EPI_ISL_570638, EPI_ISL_570639, EPI_ISL_570640, EPI_ISL_570641, EPI_ISL_570642, EPI_ISL_570643, EPI_ISL_570644, EPI_ISL_570645, EPI_ISL_570646, EPI_ISL_570647, EPI_ISL_570648, EPI_ISL_570649, EPI_ISL_570650, EPI_ISL_570651, EPI_ISL_570652, EPI_ISL_570653, EPI_ISL_570654, EPI_ISL_570655, EPI_ISL_570656, EPI_ISL_570657, EPI_ISL_570658, EPI_ISL_570659, EPI_ISL_570660, EPI_ISL_570661, EPI_ISL_570662, EPI_ISL_570663, EPI_ISL_570664, EPI_ISL_570665, EPI_ISL_570666, EPI_ISL_570667, EPI_ISL_570668, EPI_ISL_570669, EPI_ISL_570670, EPI_ISL_570671, EPI_ISL_570672, EPI_ISL_570673, EPI_ISL_570674, EPI_ISL_570675, EPI_ISL_570676, EPI_ISL_570677, EPI_ISL_570678, EPI_ISL_570679, EPI_ISL_570680, EPI_ISL_570681, EPI_ISL_570682, EPI_ISL_570683, EPI_ISL_570684, EPI_ISL_570685, EPI_ISL_570686, EPI_ISL_570687, EPI_ISL_570688, EPI_ISL_570689, EPI_ISL_570690, EPI_ISL_570691, EPI_ISL_570692, EPI_ISL_570693, EPI_ISL_570694, EPI_ISL_570695, EPI_ISL_570696, EPI_ISL_570697, EPI_ISL_570698, EPI_ISL_570699, EPI_ISL_570700, EPI_ISL_570701, EPI_ISL_570702, EPI_ISL_570703, EPI_ISL_570704, EPI_ISL_570705, EPI_ISL_570706, EPI_ISL_570707, EPI_ISL_570708, EPI_ISL_570709, EPI_ISL_570710, EPI_ISL_570711, EPI_ISL_570712, EPI_ISL_570713, EPI_ISL_570714, EPI_ISL_570715, EPI_ISL_570716, EPI_ISL_570717, EPI_ISL_570718, EPI_ISL_570719, EPI_ISL_570720, EPI_ISL_570721, EPI_ISL_570722, EPI_ISL_570723, EPI_ISL_570724, EPI_ISL_570725, EPI_ISL_570726, EPI_ISL_570727, EPI_ISL_570728, EPI_ISL_570729, EPI_ISL_570730, EPI_ISL_570731, EPI_ISL_570732, EPI_ISL_570733, EPI_ISL_570734, EPI_ISL_570735, EPI_ISL_570736, EPI_ISL_570737, EPI_ISL_570738, EPI_ISL_570739, EPI_ISL_570740, EPI_ISL_570741, EPI_ISL_570742, EPI_ISL_570743, EPI_ISL_570744, EPI_ISL_570745, EPI_ISL_570746, EPI_ISL_570747, EPI_ISL_570748, EPI_ISL_570749, EPI_ISL_570750, EPI_ISL_570751, EPI_ISL_570752, EPI_ISL_570753, EPI_ISL_570754, EPI_ISL_570755, EPI_ISL_570756, EPI_ISL_570757, EPI_ISL_570758, EPI_ISL_570759, EPI_ISL_570760, EPI_ISL_570761, EPI_ISL_570762, EPI_ISL_570763, EPI_ISL_570764, EPI_ISL_570765, EPI_ISL_570766, EPI_ISL_570767, EPI_ISL_570768, EPI_ISL_570769, EPI_ISL_570770, EPI_ISL_570771, EPI_ISL_570772, EPI_ISL_570773, EPI_ISL_570774, EPI_ISL_570775, EPI_ISL_570776, EPI_ISL_570777, EPI_ISL_570778, EPI_ISL_570779, EPI_ISL_570780, EPI_ISL_570781, EPI_ISL_570782, EPI_ISL_570783, EPI_ISL_570784, EPI_ISL_570785, EPI_ISL_570786, EPI_ISL_570787, EPI_ISL_570788, EPI_ISL_570789, EPI_ISL_570790, EPI_ISL_570791, EPI_ISL_570792, EPI_ISL_570793, EPI_ISL_570794, EPI_ISL_570795, EPI_ISL_570796, EPI_ISL_570797, EPI_ISL_570818, EPI_ISL_570819, EPI_ISL_570820, EPI_ISL_570821, EPI_ISL_570822, EPI_ISL_570823, EPI_ISL_570824, EPI_ISL_570825, EPI_ISL_570826, EPI_ISL_570827, EPI_ISL_570828, EPI_ISL_570829, EPI_ISL_570830, EPI_ISL_570831, EPI_ISL_570832, EPI_ISL_570833, EPI_ISL_570834, EPI_ISL_570835, EPI_ISL_570836, EPI_ISL_570837, EPI_ISL_570838, EPI_ISL_570839, EPI_ISL_570840, EPI_ISL_570841, EPI_ISL_570842, EPI_ISL_570843, EPI_ISL_570844, EPI_ISL_570846, EPI_ISL_570847, EPI_ISL_570848, EPI_ISL_570849, EPI_ISL_570850, EPI_ISL_570851, EPI_ISL_570852, EPI_ISL_570853, EPI_ISL_570854, EPI_ISL_570855, EPI_ISL_570856, EPI_ISL_570857, EPI_ISL_570858, EPI_ISL_570859, EPI_ISL_570860, EPI_ISL_570861, EPI_ISL_570862, EPI_ISL_570863, EPI_ISL_570864, EPI_ISL_570865, EPI_ISL_570866, EPI_ISL_570867, EPI_ISL_570868, EPI_ISL_570869, EPI_ISL_570870, EPI_ISL_570871, EPI_ISL_570872, EPI_ISL_570873, EPI_ISL_570874, EPI_ISL_570875, EPI_ISL_570876, EPI_ISL_570877, EPI_ISL_570878, EPI_ISL_570879, EPI_ISL_570880, EPI_ISL_570881, EPI_ISL_570882, EPI_ISL_570883, EPI_ISL_570884, EPI_ISL_570885, EPI_ISL_570886, EPI_ISL_570887, EPI_ISL_570888, EPI_ISL_570889, EPI_ISL_570890, EPI_ISL_570891, EPI_ISL_570892, EPI_ISL_570893, EPI_ISL_570894, EPI_ISL_570895, EPI_ISL_570896, EPI_ISL_570897, EPI_ISL_570898, EPI_ISL_570899, EPI_ISL_570900, EPI_ISL_570901, EPI_ISL_570902, EPI_ISL_570903, EPI_ISL_570904, EPI_ISL_570905, EPI_ISL_570906, EPI_ISL_570907, EPI_ISL_570908, EPI_ISL_570909, EPI_ISL_570910, EPI_ISL_570911, EPI_ISL_570912, EPI_ISL_570913, EPI_ISL_570914, EPI_ISL_570915, EPI_ISL_570916, EPI_ISL_570917, EPI_ISL_570918, EPI_ISL_570919, EPI_ISL_570920, EPI_ISL_570921, EPI_ISL_570922, EPI_ISL_570923, EPI_ISL_570924, EPI_ISL_570925, EPI_ISL_570926, EPI_ISL_570927, EPI_ISL_570928, EPI_ISL_570929, EPI_ISL_570930, EPI_ISL_570931, EPI_ISL_570932, EPI_ISL_570933, EPI_ISL_570934, EPI_ISL_570935, EPI_ISL_570936, EPI_ISL_570937, EPI_ISL_570938, EPI_ISL_570939, EPI_ISL_570940, EPI_ISL_570941, EPI_ISL_570942, EPI_ISL_570943, EPI_ISL_570944, EPI_ISL_570945, EPI_ISL_570946, EPI_ISL_570947, EPI_ISL_570948, EPI_ISL_570949, EPI_ISL_570950, EPI_ISL_570951, EPI_ISL_570952, EPI_ISL_570953, EPI_ISL_570954, EPI_ISL_570955, EPI_ISL_570956, EPI_ISL_570957, EPI_ISL_570958, EPI_ISL_570959, EPI_ISL_570960, EPI_ISL_570961, EPI_ISL_570962, EPI_ISL_570963, EPI_ISL_570964, EPI_ISL_570965, EPI_ISL_570966, EPI_ISL_570967, EPI_ISL_570968, EPI_ISL_570969, EPI_ISL_570970, EPI_ISL_570971, EPI_ISL_570972, EPI_ISL_570973, EPI_ISL_570974, EPI_ISL_570975, EPI_ISL_570976, EPI_ISL_570977, EPI_ISL_570978, EPI_ISL_570979, EPI_ISL_570980, EPI_ISL_570981, EPI_ISL_570982, EPI_ISL_570983, EPI_ISL_570984, EPI_ISL_570985, EPI_ISL_570986, EPI_ISL_570987, EPI_ISL_570988, EPI_ISL_570989, EPI_ISL_570990, EPI_ISL_570991, EPI_ISL_570992, EPI_ISL_570993, EPI_ISL_570994, EPI_ISL_570995, EPI_ISL_570996, EPI_ISL_570997, EPI_ISL_570998, EPI_ISL_570999, EPI_ISL_571000, EPI_ISL_571001, EPI_ISL_571002, EPI_ISL_571003, EPI_ISL_571004, EPI_ISL_571005, EPI_ISL_571006, EPI_ISL_571007, EPI_ISL_571008, EPI_ISL_571009, EPI_ISL_571010, EPI_ISL_571011, EPI_ISL_571012, EPI_ISL_571013, EPI_ISL_571014, EPI_ISL_571015 | | | |
| see above | UW Virology Lab | UW Virology Lab | Pavitra Roychoudhury, Hong Xie, Lasata Shrestha, Amin Addetia, Victoria M Rachleff, Meei-Li Huang, Keith R Jerome, Alexander Greninger |
| EPI_ISL_571016, EPI_ISL_571017, EPI_ISL_571018, EPI_ISL_571019, EPI_ISL_571020, EPI_ISL_571021, EPI_ISL_571022, EPI_ISL_571023, EPI_ISL_571024, EPI_ISL_571025, EPI_ISL_571026, EPI_ISL_571027, EPI_ISL_571028, EPI_ISL_571029, EPI_ISL_571030, EPI_ISL_571031, EPI_ISL_571033, EPI_ISL_571034, EPI_ISL_571035, EPI_ISL_571036, EPI_ISL_571037, EPI_ISL_571038, EPI_ISL_571039, EPI_ISL_571040, EPI_ISL_571041, EPI_ISL_571042, EPI_ISL_571043, EPI_ISL_571044, EPI_ISL_571045, EPI_ISL_571046, EPI_ISL_571047, EPI_ISL_571048, EPI_ISL_571049, EPI_ISL_571050, EPI_ISL_571051, EPI_ISL_571052, EPI_ISL_571053, EPI_ISL_571054, EPI_ISL_571055, EPI_ISL_571056, EPI_ISL_571057, EPI_ISL_571058, EPI_ISL_571059, EPI_ISL_571060, EPI_ISL_571061, EPI_ISL_571062, EPI_ISL_571063, EPI_ISL_571064, EPI_ISL_571065, EPI_ISL_571066, EPI_ISL_571067, EPI_ISL_571068, EPI_ISL_571069, EPI_ISL_571070, EPI_ISL_571071, EPI_ISL_571072, EPI_ISL_571073, EPI_ISL_571074, EPI_ISL_571075, EPI_ISL_571076, EPI_ISL_571077, EPI_ISL_571078, EPI_ISL_571079, EPI_ISL_571081, EPI_ISL_571082, EPI_ISL_571083, EPI_ISL_571084, EPI_ISL_571085, EPI_ISL_571087, EPI_ISL_571088, EPI_ISL_571089, EPI_ISL_571090, EPI_ISL_571091, EPI_ISL_571092, EPI_ISL_571093, EPI_ISL_571094, EPI_ISL_571095, EPI_ISL_571096, EPI_ISL_571097, EPI_ISL_571098, EPI_ISL_571099, EPI_ISL_571100, EPI_ISL_571101, EPI_ISL_571102, EPI_ISL_571103, EPI_ISL_571104, EPI_ISL_571105, EPI_ISL_571106, EPI_ISL_571107, EPI_ISL_571108, EPI_ISL_571109, EPI_ISL_571110, EPI_ISL_571111, EPI_ISL_571112, EPI_ISL_571113, EPI_ISL_571114, EPI_ISL_571115, EPI_ISL_571116, EPI_ISL_571117, EPI_ISL_571118, EPI_ISL_571119, EPI_ISL_571120, EPI_ISL_571121, EPI_ISL_571122, EPI_ISL_571123, EPI_ISL_571124, EPI_ISL_571125, EPI_ISL_571126, EPI_ISL_571127, EPI_ISL_571128, EPI_ISL_571129, EPI_ISL_571130, EPI_ISL_571131, EPI_ISL_571132, EPI_ISL_571133, EPI_ISL_571134, EPI_ISL_571135, EPI_ISL_571136, EPI_ISL_571137, EPI_ISL_571138, EPI_ISL_571139, EPI_ISL_571140, EPI_ISL_571141, EPI_ISL_571142, EPI_ISL_571143, EPI_ISL_571144, EPI_ISL_571145, EPI_ISL_571146, EPI_ISL_571147, EPI_ISL_571148, EPI_ISL_571149, EPI_ISL_571150, EPI_ISL_571151, EPI_ISL_571152, EPI_ISL_571153, EPI_ISL_571154, EPI_ISL_571155, EPI_ISL_571156, EPI_ISL_571157, EPI_ISL_571158, EPI_ISL_571159, EPI_ISL_571160, EPI_ISL_571161, EPI_ISL_571162, EPI_ISL_571163, EPI_ISL_571164, EPI_ISL_571165, EPI_ISL_571166, EPI_ISL_571167, EPI_ISL_571168, EPI_ISL_571169, EPI_ISL_571170, EPI_ISL_571171, EPI_ISL_571172, EPI_ISL_571174, EPI_ISL_571175, EPI_ISL_571176, EPI_ISL_571177, EPI_ISL_571178, EPI_ISL_571179, EPI_ISL_571180, EPI_ISL_571181, EPI_ISL_571182, EPI_ISL_571183, EPI_ISL_571184, EPI_ISL_571185, EPI_ISL_571186, EPI_ISL_571187, EPI_ISL_571188, EPI_ISL_571189, EPI_ISL_571190, EPI_ISL_571191, EPI_ISL_571192, EPI_ISL_571193, EPI_ISL_571194, EPI_ISL_571195, EPI_ISL_571196, EPI_ISL_571197, EPI_ISL_571198, EPI_ISL_571199, EPI_ISL_571200, EPI_ISL_571201, EPI_ISL_571202, EPI_ISL_571203, EPI_ISL_571204, EPI_ISL_571205, EPI_ISL_571206, EPI_ISL_571207, EPI_ISL_571208, EPI_ISL_571209, EPI_ISL_571210, EPI_ISL_571211, EPI_ISL_571212, EPI_ISL_571213, EPI_ISL_571214, EPI_ISL_571215, EPI_ISL_571216, EPI_ISL_571217, EPI_ISL_571218, EPI_ISL_571219, EPI_ISL_571220, EPI_ISL_571221, EPI_ISL_571222, EPI_ISL_571223, EPI_ISL_571224, EPI_ISL_571225, EPI_ISL_571226, EPI_ISL_571227, EPI_ISL_571228, EPI_ISL_571229, EPI_ISL_571230, EPI_ISL_571231, EPI_ISL_571232, EPI_ISL_571233, EPI_ISL_571234, EPI_ISL_571235, EPI_ISL_571236, EPI_ISL_571237, EPI_ISL_571238, EPI_ISL_571239, EPI_ISL_571240, EPI_ISL_571241, EPI_ISL_571242, EPI_ISL_571243, EPI_ISL_571244, EPI_ISL_571245, EPI_ISL_571246, EPI_ISL_571247, EPI_ISL_571248, EPI_ISL_571249, EPI_ISL_571250, EPI_ISL_571251, EPI_ISL_571252, EPI_ISL_571254, EPI_ISL_571255, EPI_ISL_571256, EPI_ISL_571257, EPI_ISL_571258, EPI_ISL_571259, EPI_ISL_571260, EPI_ISL_571261, EPI_ISL_571262, EPI_ISL_571263, EPI_ISL_571264, EPI_ISL_571265, EPI_ISL_571266, EPI_ISL_571267, EPI_ISL_571268, EPI_ISL_571269, EPI_ISL_571270, EPI_ISL_571271, EPI_ISL_571272, EPI_ISL_571273, EPI_ISL_571274, EPI_ISL_571275, EPI_ISL_571276, EPI_ISL_571277, EPI_ISL_571278, EPI_ISL_571279, EPI_ISL_571280, EPI_ISL_571281, EPI_ISL_571283, EPI_ISL_571284, EPI_ISL_571285, EPI_ISL_571286, EPI_ISL_571287, EPI_ISL_571288, EPI_ISL_571289, EPI_ISL_571290, EPI_ISL_571291, EPI_ISL_571292, EPI_ISL_571293, EPI_ISL_571295, EPI_ISL_571296, EPI_ISL_571299, EPI_ISL_571300, EPI_ISL_571301, EPI_ISL_571302, EPI_ISL_571303, EPI_ISL_571304, EPI_ISL_571305, EPI_ISL_571306, EPI_ISL_571307, EPI_ISL_571308, EPI_ISL_571309, EPI_ISL_571310, EPI_ISL_571311, EPI_ISL_571312, EPI_ISL_571313, EPI_ISL_571314, EPI_ISL_571315, EPI_ISL_571316, EPI_ISL_571317, EPI_ISL_571318, EPI_ISL_571319, EPI_ISL_571320, EPI_ISL_571321, EPI_ISL_571322, EPI_ISL_571323, EPI_ISL_571324, EPI_ISL_571325, EPI_ISL_571326, EPI_ISL_571327, EPI_ISL_571328, EPI_ISL_571329, EPI_ISL_571330, EPI_ISL_571331, EPI_ISL_571332, EPI_ISL_571333, EPI_ISL_571334, EPI_ISL_571335, EPI_ISL_571336, EPI_ISL_571337, EPI_ISL_571338, EPI_ISL_571339, EPI_ISL_571340, EPI_ISL_571341, EPI_ISL_571342, EPI_ISL_571343, EPI_ISL_571344, EPI_ISL_571345, EPI_ISL_571346, EPI_ISL_571347, EPI_ISL_571348, EPI_ISL_571349, EPI_ISL_571350, EPI_ISL_571351, EPI_ISL_571352, EPI_ISL_571353, EPI_ISL_571354, EPI_ISL_571355, EPI_ISL_571356, EPI_ISL_571357, EPI_ISL_571358, EPI_ISL_571359, EPI_ISL_571360, EPI_ISL_571361, EPI_ISL_571362, EPI_ISL_571363, EPI_ISL_571364, EPI_ISL_571365, EPI_ISL_571366, EPI_ISL_571367, EPI_ISL_571368, EPI_ISL_571369, EPI_ISL_571370, EPI_ISL_571371, EPI_ISL_571372, EPI_ISL_571373, EPI_ISL_571374, EPI_ISL_571375, EPI_ISL_571376, EPI_ISL_571377, EPI_ISL_571378, EPI_ISL_571379, EPI_ISL_571380, EPI_ISL_571381, EPI_ISL_571382, EPI_ISL_571383, EPI_ISL_571384, EPI_ISL_571385, EPI_ISL_571386, EPI_ISL_571387, EPI_ISL_571389, EPI_ISL_571390, EPI_ISL_571391, EPI_ISL_571392, EPI_ISL_571393, EPI_ISL_571394, EPI_ISL_571395, EPI_ISL_571396, EPI_ISL_571398, EPI_ISL_571399, EPI_ISL_571401, EPI_ISL_571403, EPI_ISL_571404, EPI_ISL_571405, EPI_ISL_571406, EPI_ISL_571407, EPI_ISL_571408, EPI_ISL_571409, EPI_ISL_571410, EPI_ISL_571411, EPI_ISL_571412, EPI_ISL_571413, EPI_ISL_571414, EPI_ISL_571415, EPI_ISL_571416, EPI_ISL_571417, EPI_ISL_571418, EPI_ISL_571419, EPI_ISL_571420, EPI_ISL_571421, EPI_ISL_571422, EPI_ISL_571423, EPI_ISL_571424, EPI_ISL_571425, EPI_ISL_571426, EPI_ISL_571427, EPI_ISL_571428, EPI_ISL_571429, EPI_ISL_571430, EPI_ISL_571431, EPI_ISL_571432, EPI_ISL_571433, EPI_ISL_571434, EPI_ISL_571435, EPI_ISL_571436, EPI_ISL_571437, EPI_ISL_571438, EPI_ISL_571439, EPI_ISL_571440, EPI_ISL_571441, EPI_ISL_571442, EPI_ISL_571443, EPI_ISL_571444, EPI_ISL_571445, EPI_ISL_571446, EPI_ISL_571447, EPI_ISL_571448, EPI_ISL_571449, EPI_ISL_571450, EPI_ISL_571451, EPI_ISL_571452, EPI_ISL_571453, EPI_ISL_571454, EPI_ISL_571455, EPI_ISL_571456, EPI_ISL_571457, EPI_ISL_571458, EPI_ISL_571459, EPI_ISL_571460, EPI_ISL_571461, EPI_ISL_571462, EPI_ISL_571463, EPI_ISL_571464, EPI_ISL_571465, EPI_ISL_571466, EPI_ISL_571467, EPI_ISL_571468, EPI_ISL_571469, EPI_ISL_571470, EPI_ISL_571471, EPI_ISL_571472, EPI_ISL_571473, EPI_ISL_571474, EPI_ISL_571475, EPI_ISL_571476, EPI_ISL_571477, EPI_ISL_571478, EPI_ISL_571479, EPI_ISL_571480, EPI_ISL_571481, EPI_ISL_571482, EPI_ISL_571483, EPI_ISL_571484, EPI_ISL_571486, EPI_ISL_571487, EPI_ISL_571488, EPI_ISL_571489, EPI_ISL_571490, EPI_ISL_571491, EPI_ISL_571492, EPI_ISL_571493, EPI_ISL_571494, EPI_ISL_571495, EPI_ISL_571496, EPI_ISL_571497, EPI_ISL_571498, EPI_ISL_571499, EPI_ISL_571500, EPI_ISL_571501, EPI_ISL_571502, EPI_ISL_571503, EPI_ISL_571504, EPI_ISL_571505, EPI_ISL_571506, EPI_ISL_571507, EPI_ISL_571508, EPI_ISL_571509, EPI_ISL_571510, EPI_ISL_571511, EPI_ISL_571512, EPI_ISL_571514, EPI_ISL_571515, EPI_ISL_571516, EPI_ISL_571517, EPI_ISL_571518, EPI_ISL_571519, EPI_ISL_571520, EPI_ISL_571521, EPI_ISL_571522, EPI_ISL_571523, EPI_ISL_571524, EPI_ISL_571525, EPI_ISL_571526, EPI_ISL_571527, EPI_ISL_571528, EPI_ISL_571529, EPI_ISL_571530, EPI_ISL_571531, EPI_ISL_571532, EPI_ISL_571533, EPI_ISL_571534, EPI_ISL_571535, EPI_ISL_571536, EPI_ISL_571537, EPI_ISL_571538, EPI_ISL_571539, EPI_ISL_571540, EPI_ISL_571541, EPI_ISL_571543, EPI_ISL_571544, EPI_ISL_571545, EPI_ISL_571546, EPI_ISL_571547, EPI_ISL_571548, EPI_ISL_571549, EPI_ISL_571550, EPI_ISL_571551, EPI_ISL_571552, EPI_ISL_571553, EPI_ISL_571554, EPI_ISL_571555, EPI_ISL_571556, EPI_ISL_571557, EPI_ISL_571558, EPI_ISL_571559, EPI_ISL_571560, EPI_ISL_571561, EPI_ISL_571562, EPI_ISL_571563, EPI_ISL_571564, EPI_ISL_571565, EPI_ISL_571566, EPI_ISL_571567, EPI_ISL_571568, EPI_ISL_571569, EPI_ISL_571570, EPI_ISL_571571, EPI_ISL_571572, EPI_ISL_571573, EPI_ISL_571574, EPI_ISL_571575, EPI_ISL_571576, EPI_ISL_571577, EPI_ISL_571578, EPI_ISL_571579, EPI_ISL_571580, EPI_ISL_571581, EPI_ISL_571582, EPI_ISL_571583, EPI_ISL_571584, EPI_ISL_571585, EPI_ISL_571586, EPI_ISL_571587, EPI_ISL_571588, EPI_ISL_571589, EPI_ISL_571590, EPI_ISL_571591, EPI_ISL_571592, EPI_ISL_571593, EPI_ISL_571594, EPI_ISL_571595, EPI_ISL_571596, EPI_ISL_571597, EPI_ISL_571598, EPI_ISL_571599, EPI_ISL_571600, EPI_ISL_571601, EPI_ISL_571602, EPI_ISL_571603, EPI_ISL_571604, EPI_ISL_571605, EPI_ISL_571606, EPI_ISL_571607, EPI_ISL_571608, EPI_ISL_571609, EPI_ISL_571610, EPI_ISL_571611, EPI_ISL_571612, EPI_ISL_571613, EPI_ISL_571614, EPI_ISL_571615, EPI_ISL_571616, EPI_ISL_571617, EPI_ISL_571618, EPI_ISL_571619, EPI_ISL_571620, EPI_ISL_571621, EPI_ISL_571622, EPI_ISL_571623, EPI_ISL_571624, EPI_ISL_571625, EPI_ISL_571626, EPI_ISL_571627, EPI_ISL_571628, EPI_ISL_571629, EPI_ISL_571630, EPI_ISL_571631, EPI_ISL_571632, EPI_ISL_571633, EPI_ISL_571634, EPI_ISL_571635, EPI_ISL_571636, EPI_ISL_571637, EPI_ISL_571638, EPI_ISL_571639, EPI_ISL_571640, EPI_ISL_571641, EPI_ISL_571642, EPI_ISL_571643, EPI_ISL_571644, EPI_ISL_571645, EPI_ISL_571646, EPI_ISL_571647, EPI_ISL_571648, EPI_ISL_571649, EPI_ISL_571650, EPI_ISL_571651, EPI_ISL_571652, EPI_ISL_571653, EPI_ISL_571654, EPI_ISL_571655, EPI_ISL_571656, EPI_ISL_571657, EPI_ISL_571658, EPI_ISL_571659, EPI_ISL_571660, EPI_ISL_571661, EPI_ISL_571662, EPI_ISL_571663, EPI_ISL_571664, EPI_ISL_571665, EPI_ISL_571666, EPI_ISL_571667, EPI_ISL_571668, EPI_ISL_571669, EPI_ISL_571670, EPI_ISL_571671, EPI_ISL_571672, EPI_ISL_571673, EPI_ISL_571674, EPI_ISL_571675, EPI_ISL_571676, EPI_ISL_571677, EPI_ISL_571678, EPI_ISL_571679, EPI_ISL_571680, EPI_ISL_571681, EPI_ISL_571682, EPI_ISL_571683, EPI_ISL_571684, EPI_ISL_571685, EPI_ISL_571686, EPI_ISL_571687, EPI_ISL_571688, EPI_ISL_571689, EPI_ISL_571691, EPI_ISL_571692, EPI_ISL_571694, EPI_ISL_571695, EPI_ISL_571696, EPI_ISL_571697, EPI_ISL_571698, EPI_ISL_571699, EPI_ISL_571700, EPI_ISL_571701, EPI_ISL_571702, EPI_ISL_571703, EPI_ISL_571704, EPI_ISL_571705, EPI_ISL_571706, EPI_ISL_571707, EPI_ISL_571708, EPI_ISL_571709, EPI_ISL_571710, EPI_ISL_571711, EPI_ISL_571712, EPI_ISL_571713, EPI_ISL_571714, EPI_ISL_571715, EPI_ISL_571716, EPI_ISL_571717, EPI_ISL_571718, EPI_ISL_571719, EPI_ISL_571720, EPI_ISL_571721, EPI_ISL_571722, EPI_ISL_571723, EPI_ISL_571724, EPI_ISL_571725, EPI_ISL_571726, EPI_ISL_571727, EPI_ISL_571728, EPI_ISL_571729, EPI_ISL_571730, EPI_ISL_571731, EPI_ISL_571732, EPI_ISL_571733, EPI_ISL_571734, EPI_ISL_571735, EPI_ISL_571736, EPI_ISL_571737, EPI_ISL_571738, EPI_ISL_571739, EPI_ISL_571740, EPI_ISL_571741, EPI_ISL_571742, EPI_ISL_571743, EPI_ISL_571744, EPI_ISL_571745, EPI_ISL_571746, EPI_ISL_571748, EPI_ISL_571749, EPI_ISL_571750, EPI_ISL_571751, EPI_ISL_571752, EPI_ISL_571753, EPI_ISL_571754, EPI_ISL_571755, EPI_ISL_571756, EPI_ISL_571757, EPI_ISL_571758, EPI_ISL_571759, EPI_ISL_571760, EPI_ISL_571761, EPI_ISL_571762, EPI_ISL_571763, EPI_ISL_571764, EPI_ISL_571765, EPI_ISL_571766, EPI_ISL_571767, EPI_ISL_571768, EPI_ISL_571769, EPI_ISL_571770, EPI_ISL_571771, EPI_ISL_571772, EPI_ISL_571773, EPI_ISL_571774, EPI_ISL_571775, EPI_ISL_571776, EPI_ISL_571777, EPI_ISL_571778, EPI_ISL_571779, EPI_ISL_571780, EPI_ISL_571781, EPI_ISL_571782, EPI_ISL_571783, EPI_ISL_571784, EPI_ISL_571785, EPI_ISL_571786, EPI_ISL_571787, EPI_ISL_571788, EPI_ISL_571789, EPI_ISL_571790, EPI_ISL_571791, EPI_ISL_571792, EPI_ISL_571793, EPI_ISL_571794, EPI_ISL_571795, EPI_ISL_571796, EPI_ISL_571797, EPI_ISL_571798, EPI_ISL_571799, EPI_ISL_571800, EPI_ISL_571801, EPI_ISL_571802, EPI_ISL_571803, EPI_ISL_571804, EPI_ISL_571805, EPI_ISL_571806, EPI_ISL_571807, EPI_ISL_571808, EPI_ISL_571809, EPI_ISL_571810, EPI_ISL_571811, EPI_ISL_571812, EPI_ISL_571813, EPI_ISL_571814, EPI_ISL_571815, EPI_ISL_571816, EPI_ISL_571817, EPI_ISL_571818, EPI_ISL_571819, EPI_ISL_571820, EPI_ISL_571821, EPI_ISL_571822, EPI_ISL_571823, EPI_ISL_571824, EPI_ISL_571825, EPI_ISL_571826, EPI_ISL_571827, EPI_ISL_571828, EPI_ISL_571829, EPI_ISL_571830, EPI_ISL_571831, EPI_ISL_571832, EPI_ISL_571833, EPI_ISL_571834, EPI_ISL_571835, EPI_ISL_571836, EPI_ISL_571837, EPI_ISL_571838, EPI_ISL_571839, EPI_ISL_571840, EPI_ISL_571841, EPI_ISL_571842, EPI_ISL_571843, EPI_ISL_571844, EPI_ISL_571845, EPI_ISL_571846, EPI_ISL_571847, EPI_ISL_571848, EPI_ISL_571849, EPI_ISL_571850, EPI_ISL_571851, EPI_ISL_571852, EPI_ISL_571853, EPI_ISL_571854, EPI_ISL_571855, EPI_ISL_571856, EPI_ISL_571857, EPI_ISL_571858, EPI_ISL_571860, EPI_ISL_571862, EPI_ISL_571863, EPI_ISL_571864, EPI_ISL_571865, EPI_ISL_571866, EPI_ISL_571867, EPI_ISL_571868, EPI_ISL_571869, EPI_ISL_571870, EPI_ISL_571871, EPI_ISL_571872, EPI_ISL_571873, EPI_ISL_571874, EPI_ISL_571875, EPI_ISL_571876, EPI_ISL_571877, EPI_ISL_571878, EPI_ISL_571879, EPI_ISL_571880, EPI_ISL_571881, EPI_ISL_571882, EPI_ISL_571883, EPI_ISL_571884, EPI_ISL_571885, EPI_ISL_571886, EPI_ISL_571887, EPI_ISL_571888, EPI_ISL_571889, EPI_ISL_571890, EPI_ISL_571891, EPI_ISL_571892, EPI_ISL_571893, EPI_ISL_571894, EPI_ISL_571895, EPI_ISL_571897, EPI_ISL_571898, EPI_ISL_571899, EPI_ISL_571900, EPI_ISL_571901, EPI_ISL_571902, EPI_ISL_571903, EPI_ISL_571905, EPI_ISL_571906, EPI_ISL_571907, EPI_ISL_571908, EPI_ISL_571909, EPI_ISL_571910, EPI_ISL_571912, EPI_ISL_571913, EPI_ISL_571915, EPI_ISL_571916, EPI_ISL_571917, EPI_ISL_571918, EPI_ISL_571919, EPI_ISL_571920, EPI_ISL_571921, EPI_ISL_571922, EPI_ISL_571923, EPI_ISL_571924, EPI_ISL_571925, EPI_ISL_571926, EPI_ISL_571928, EPI_ISL_571929, EPI_ISL_571930, EPI_ISL_571931, EPI_ISL_571932, EPI_ISL_571933, EPI_ISL_571934, EPI_ISL_571936, EPI_ISL_571937, EPI_ISL_571938, EPI_ISL_571939, EPI_ISL_571940, EPI_ISL_571941, EPI_ISL_571942, EPI_ISL_571943, EPI_ISL_571944, EPI_ISL_571945, EPI_ISL_571946, EPI_ISL_571947, EPI_ISL_571949, EPI_ISL_571950, EPI_ISL_571951, EPI_ISL_571952, EPI_ISL_571953, EPI_ISL_571954, EPI_ISL_571955, EPI_ISL_571956, EPI_ISL_571957, EPI_ISL_571958, EPI_ISL_571959, EPI_ISL_571960, EPI_ISL_571961, EPI_ISL_571962, EPI_ISL_571963, EPI_ISL_571964, EPI_ISL_571965, EPI_ISL_571966, EPI_ISL_571967, EPI_ISL_571968, EPI_ISL_571969, EPI_ISL_571970, EPI_ISL_571971, EPI_ISL_571972, EPI_ISL_571973, EPI_ISL_571974, EPI_ISL_571975, EPI_ISL_571976, EPI_ISL_571977, EPI_ISL_571978, EPI_ISL_571979, EPI_ISL_571980, EPI_ISL_571981, EPI_ISL_571982, EPI_ISL_571983, EPI_ISL_571984, EPI_ISL_571985, EPI_ISL_571986, EPI_ISL_571987, EPI_ISL_571988, EPI_ISL_571989, EPI_ISL_571990, EPI_ISL_571991, EPI_ISL_571992, EPI_ISL_571993, EPI_ISL_571994, EPI_ISL_571995, EPI_ISL_571996, EPI_ISL_571997, EPI_ISL_571998, EPI_ISL_571999, EPI_ISL_572000, EPI_ISL_572001, EPI_ISL_572002, EPI_ISL_572003, EPI_ISL_572004, EPI_ISL_572005, EPI_ISL_572006, EPI_ISL_572007, EPI_ISL_572008, EPI_ISL_572009, EPI_ISL_572010, EPI_ISL_572011, EPI_ISL_572012, EPI_ISL_572013, EPI_ISL_572014, EPI_ISL_572015, EPI_ISL_572016, EPI_ISL_572017, EPI_ISL_572018, EPI_ISL_572019, EPI_ISL_572020, EPI_ISL_572021, EPI_ISL_572022, EPI_ISL_572023, EPI_ISL_572024, EPI_ISL_572025, EPI_ISL_572026, EPI_ISL_572027, EPI_ISL_572028, EPI_ISL_572029, EPI_ISL_572031, EPI_ISL_572032, EPI_ISL_572033, EPI_ISL_572034, EPI_ISL_572035, EPI_ISL_572036, EPI_ISL_572037, EPI_ISL_572038, EPI_ISL_572039, EPI_ISL_572040, EPI_ISL_572041, EPI_ISL_572042, EPI_ISL_572043, EPI_ISL_572044, EPI_ISL_572045, EPI_ISL_572046, EPI_ISL_572047, EPI_ISL_572048, EPI_ISL_572049, EPI_ISL_572050, EPI_ISL_572051, EPI_ISL_572052, EPI_ISL_572053, EPI_ISL_572054, EPI_ISL_572055, EPI_ISL_572056, EPI_ISL_572057, EPI_ISL_572058, EPI_ISL_572059, EPI_ISL_572060, EPI_ISL_572061, EPI_ISL_572062, EPI_ISL_572063, EPI_ISL_572064, EPI_ISL_572065, EPI_ISL_572066, EPI_ISL_572067, EPI_ISL_572068, EPI_ISL_572069, EPI_ISL_572070, EPI_ISL_572071, EPI_ISL_572072, EPI_ISL_572074, EPI_ISL_572075, EPI_ISL_572076, EPI_ISL_572077, EPI_ISL_572078, EPI_ISL_572079, EPI_ISL_572080, EPI_ISL_572081, EPI_ISL_572082, EPI_ISL_572083, EPI_ISL_572084, EPI_ISL_572085, EPI_ISL_572086, EPI_ISL_572087, EPI_ISL_572088, EPI_ISL_572089, EPI_ISL_572090, EPI_ISL_572091, EPI_ISL_572092, EPI_ISL_572093, EPI_ISL_572094, EPI_ISL_572095, EPI_ISL_572096, EPI_ISL_572097, EPI_ISL_572098, EPI_ISL_572099, EPI_ISL_572100, EPI_ISL_572101, EPI_ISL_572102, EPI_ISL_572103, EPI_ISL_572104, EPI_ISL_572105, EPI_ISL_572106, EPI_ISL_572107, EPI_ISL_572108, EPI_ISL_572109, EPI_ISL_572110, EPI_ISL_572111, EPI_ISL_572112, EPI_ISL_572113, EPI_ISL_572114, EPI_ISL_572115, EPI_ISL_572116, EPI_ISL_572117, EPI_ISL_572118, EPI_ISL_572119, EPI_ISL_572120, EPI_ISL_572121, EPI_ISL_572122, EPI_ISL_572123, EPI_ISL_572124, EPI_ISL_572125, EPI_ISL_572126, EPI_ISL_572127, EPI_ISL_572128, EPI_ISL_572129, EPI_ISL_572130, EPI_ISL_572131, EPI_ISL_572132, EPI_ISL_572133, EPI_ISL_572134, EPI_ISL_572135, EPI_ISL_572136, EPI_ISL_572137, EPI_ISL_572138, EPI_ISL_572139, EPI_ISL_572140, EPI_ISL_572141, EPI_ISL_572142, EPI_ISL_572143, EPI_ISL_572144, EPI_ISL_572145, EPI_ISL_572146, EPI_ISL_572147, EPI_ISL_572148, EPI_ISL_572149, EPI_ISL_572150, EPI_ISL_572151, EPI_ISL_572152, EPI_ISL_572153, EPI_ISL_572154, EPI_ISL_572155, EPI_ISL_572156, EPI_ISL_572157, EPI_ISL_572158, EPI_ISL_572159, EPI_ISL_572160, EPI_ISL_572161, EPI_ISL_572162, EPI_ISL_572163, EPI_ISL_572164, EPI_ISL_572165, EPI_ISL_572166, EPI_ISL_572167, EPI_ISL_572168, EPI_ISL_572169, EPI_ISL_572170, EPI_ISL_572171, EPI_ISL_572172, EPI_ISL_572173, EPI_ISL_572174, EPI_ISL_572175, EPI_ISL_572176, EPI_ISL_572177, EPI_ISL_572178, EPI_ISL_572179, EPI_ISL_572180, EPI_ISL_572181, EPI_ISL_572182, EPI_ISL_572183, EPI_ISL_572184, EPI_ISL_572185 | | | |
| see above | Quest Diagnostics | Quest Diagnostics | Rosenthal,S.H., Gerasimova,A., Kagan,R.M., Anderson, B., Grover, D., Livingston, K.E., Hua, M., Liu Y., Shalhout, D.F., Owen, R., Lacbawan, F. |
| EPI_ISL_572195, EPI_ISL_572196, EPI_ISL_572197, EPI_ISL_572198, EPI_ISL_572199, EPI_ISL_572200, EPI_ISL_572201, EPI_ISL_572202, EPI_ISL_572203, EPI_ISL_572204, EPI_ISL_572205, EPI_ISL_572206, EPI_ISL_572207, EPI_ISL_572208, EPI_ISL_572209, EPI_ISL_572210, EPI_ISL_572211, EPI_ISL_572212, EPI_ISL_572213, EPI_ISL_572214, EPI_ISL_572215, EPI_ISL_572216, EPI_ISL_572217, EPI_ISL_572218, EPI_ISL_572219, EPI_ISL_572220, EPI_ISL_572221, EPI_ISL_572222, EPI_ISL_572223, EPI_ISL_572224, EPI_ISL_572225, EPI_ISL_572226, EPI_ISL_572227, EPI_ISL_572228, EPI_ISL_572229, EPI_ISL_572230, EPI_ISL_572231, EPI_ISL_572232, EPI_ISL_572233, EPI_ISL_572234, EPI_ISL_572235, EPI_ISL_572236, EPI_ISL_572237, EPI_ISL_572238, EPI_ISL_572239, EPI_ISL_572240, EPI_ISL_572241, EPI_ISL_572242, EPI_ISL_572243, EPI_ISL_572244, EPI_ISL_572245, EPI_ISL_572248, EPI_ISL_572249, EPI_ISL_572250, EPI_ISL_572251, EPI_ISL_572252, EPI_ISL_572253, EPI_ISL_572254, EPI_ISL_572255, EPI_ISL_572256, EPI_ISL_572257, EPI_ISL_572258, EPI_ISL_572259, EPI_ISL_572260, EPI_ISL_572261, EPI_ISL_572262, EPI_ISL_572263, EPI_ISL_572264, EPI_ISL_572265, EPI_ISL_572266, EPI_ISL_572267, EPI_ISL_572268, EPI_ISL_572269, EPI_ISL_572270, EPI_ISL_572271, EPI_ISL_572272, EPI_ISL_572273, EPI_ISL_572274, EPI_ISL_572275, EPI_ISL_572276, EPI_ISL_572277, EPI_ISL_572278, EPI_ISL_572279, EPI_ISL_572280, EPI_ISL_572281, EPI_ISL_572282, EPI_ISL_572283, EPI_ISL_572284, EPI_ISL_572285, EPI_ISL_572287, EPI_ISL_572288, EPI_ISL_572289, EPI_ISL_572290, EPI_ISL_572291, EPI_ISL_572292, EPI_ISL_572293, EPI_ISL_572294, EPI_ISL_572295, EPI_ISL_572296, EPI_ISL_572297, EPI_ISL_572298, EPI_ISL_572299, EPI_ISL_572300, EPI_ISL_572301, EPI_ISL_572302, EPI_ISL_572303, EPI_ISL_572304, EPI_ISL_572305, EPI_ISL_572306, EPI_ISL_572307, EPI_ISL_572308, EPI_ISL_572309, EPI_ISL_572310, EPI_ISL_572311, EPI_ISL_572312, EPI_ISL_572313, EPI_ISL_572314, EPI_ISL_572315, EPI_ISL_572316, EPI_ISL_572317, EPI_ISL_572318 | | | |

| | | | |
|---|---|---|---|
| see above | Virginia DCLS | Virginia DCLS | Virginia DCLS |
| EPI_ISL_572320, EPI_ISL_572321, EPI_ISL_572322, EPI_ISL_572323, EPI_ISL_572324 | IZSM | IZSM | Maurizio Viscardi, Lorena Cardillo, Giovanna Fusco |
| EPI_ISL_572329 | Public Health Authority of the Slovak Republic, Bratislava | Faculty of Natural Sciences, Comenius University in Bratislava | Dominika Fričová, Viktória Hodorová, Kristína Boršová, Broňa Brejová, Viktória Čabanová, Sabina Fumačová Havlíková, Juraj Kopáček, Martina Ličková, Ľubomíra Lukáčiková, Martina Neboháčová, Monika Sláviková, Edita Staroňová, Elena Tichá, Tomáš Vinař, Boris Klempa, Jozef Nosek |
| EPI_ISL_572330, EPI_ISL_572331 | Institute for Virology, University Hospital Duesseldorf, Medical Faculty, Heinrich-Heine-University Duesseldorf | Institute for Virology, University Hospital Duesseldorf, Medical Faculty, Heinrich-Heine-University Duesseldorf | Maximilian Damagnez, Verena Keitel, Björn Jensen, Nadine Lübke, Lisa Müller, Philipp Ostermann, Tina Senff, Ortwin Adams, Philipp Albrecht, Gerald Antoch, Johannes Bode, Edwin Bölke, Saskia Elben, Torsten Feldt, Johannes C. Fischer, , Anselm Kunstein, Caroline Klindt, Alexander Killer, Tom Lüdde, Annemarie Mohring, Jennifer Neubert, Heiner Schaal, Ansgar Schulz, Jörg Timm, Andreas Walker |
| EPI_ISL_572334, EPI_ISL_572335, EPI_ISL_572351, EPI_ISL_572353, EPI_ISL_572366, EPI_ISL_572371, EPI_ISL_572386 | LACEN/PE | WallauLab, Aggeu Magalhaes Institute | Marcelo Henrique Santos Paiva, Duschinka Ribeiro Duarte Guedes, Cássia Docena, Matheus Filgueira Bezerra, Filipe Zimmer Dezordi, Laís Ceschini Machado, Larissa Krokovsky, Elisama Helvecio, Alexandre Freitas da Silva, Luydson Richardson Silva Vasconcelos, Antonio Mauro Rezende, Severino Jefferson Ribeiro da Silva, Kamila Gaudêncio da Silva Sales, Bruna Santos Lima Figueiredo de Sá, Derciliano Lopes da Cruz, Claudio Eduardo Cavalcanti, Armando de Menezes Neto, Caroline Targino Alves da Silva, Renata Pessôa Germano Mendes, Maria Almerice Lopes da Silva, Tiago Gräf, Paola Cristina Resende, Gonzalo Bello0, Michelle da Silva Barros, Wheverton Ricardo Correia do Nascimento,, Rodrigo Moraes Loyo Arcoverde, Luciane Caroline Albuquerque Bezerra, Sinval Pinto Brandão Filho, Constância Flávia Junqueira Ayres, Gabriel Luz Wallau |
| EPI_ISL_572397 | Institute for Virology, University Hospital Duesseldorf, Medical Faculty, Heinrich-Heine-University Duesseldorf | Institute for Virology, University Hospital Duesseldorf, Medical Faculty, Heinrich-Heine-University Duesseldorf | Maximilian Damagnez, Verena Keitel, Björn Jensen, Nadine Lübke, Lisa Müller, Philipp Ostermann, Tina Senff, Ortwin Adams, Philipp Albrecht, Gerald Antoch, Johannes Bode, Edwin Bölke, Saskia Elben, Torsten Feldt, Johannes C. Fischer, , Anselm Kunstein, Caroline Klindt, Alexander Killer, Tom Lüdde, Annemarie Mohring, Jennifer Neubert, Heiner Schaal, Ansgar Schulz, Jörg Timm, Andreas Walker |
| EPI_ISL_572398 | Laboratory of Biology and Identification of Arboviruses | Pathogenic Microorganisms Variability Laboratory | Alexey Shchetinin, Maria Nikiforova, Andrei Siniavin, Victor Larichev, Alina Kozlova, Muhammad Saifullin, Alexey Prilipov, Vladimir Gushchin, Alexander Gintsburg |
| EPI_ISL_572710 | Liverpool Clinical Laboratories | COVID-19 Genomics UK (COG-UK) Consortium | Sam Haldenby, Anita Lucaci, Steve Paterson, Julian Hiscox, Alistair Darby, M Almsaud, A Alrezaihi, Muhannad Alruwaili, Stuart D Armstrong, Jones Benjamin, Eleanor G Bentley, Anu Chawla, Jordan J Clark, Angela Cowell, Richard Eccles, Isabel García-Dorival, Matthew Gemmell, Alessandro Gerada, PKF Gilmore, Richard Gregory, Ximeng Han, Catherine Hartley, Margaret Hughes, Miren Iturriza-Gomara, James Johnson, L Luu, Jenifer Manson, Charlotte Nelson, Elaine O'Toole, Cassie Olateju, Rebekah Penrice-Randal , Lucille Rainbow, N.P Randle, Trevor Ian Robinson, Parul Sharma, Ghada T Shawli, James P Stewart, Neil Swainston, Ecaterina Vamos, Joanne Watts, Mark Whitehead |
| EPI_ISL_572711 | Quadram Institute Bioscience | COVID-19 Genomics UK (COG-UK) Consortium | Dave J. Baker, Gemma L. Kay, Alp Aydin, Thanh Le-Viet, Steven Rudder, Ana P. Tedim, Anastasia Kolyva, Maria Diaz, Leonardo de Oliveira Martins, Nabil-Fareed Alikhan, Lizzie Meadows, Rachael Stanley, Ngozi Elumogo, Muhammed Yasir, Nicholas M. Thomson, Alexander J Trotter, Rachel Gilroy, Samuel Bloomfield, Claire Stuart, Andrew Bell, Reenesh Prakash, Samir Dervisevic, Alison E. Mather, John Wain, Mark Webber, Andrew J. Page, Justin O'Grady |
| EPI_ISL_572712, EPI_ISL_572713 | Northumbria University / South Tees Hospitals NHS Foundation Trust / North Cumbria Integrated Care NHS Foundation Trust / North Tees and Hartlepool NHS Foundation Trust / Newcastle Hospitals NHS Foundation Trust | COVID-19 Genomics UK (COG-UK) Consortium | Darren L Smith,Andrew Nelson,Matthew Bashton,Greg R Young,Joshua Loh,John Allan,Mohammad A Tariq,Giles S Holt,Gary Black,Wen C Yew,Lynn Dover,Paul Baker,Steve Liggett,Sarah Essex,Jane Greenaway,Debra Padgett,Clive Graham,Garren Scott,Edward Barton,Emma Swindells,Brendan Payne,Jennifer Collins,Yusri Taha,Gary Eltringham |
| EPI_ISL_572714 | Oxford Viromics, NDM, University of Oxford; Oxford University Hospitals; Basingstoke and North Hampshire Hospital | COVID-19 Genomics UK (COG-UK) Consortium | Tanya Golubchik, David Bonsall, George Macintyre, Amy Trebes, Mariateresa de Cesare, Catrin Moore, Alex Mobbs, Anita Justice, Robert Shaw, Monique Andersson, Timothy Peto, Emma Wise, Nathan Moore, Jessica Lynch, Nick Cortes, Matilde Mori, Stephen Kidd, David Buck, John Todd, Christophe Fraser |
| EPI_ISL_572715, EPI_ISL_572716 | Centre for Enzyme Innovation, University of Portsmouth / Translational Research Laboratory, Portsmouth Hospitals NHS Trust | COVID-19 Genomics UK (COG-UK) Consortium | Angela Beckett,Yann Bourgeois,Garry Scarlett,Sharon Glaysher,Scott Elliott,Kelly Bicknell,Robert Impey,Allyson Lloyd,Sarah Wyllie,Ethan Butcher,Anoop Chauhan,Samuel Robson |
| EPI_ISL_572717, EPI_ISL_572718, EPI_ISL_572719, EPI_ISL_572720 | Quadram Institute Bioscience | COVID-19 Genomics UK (COG-UK) Consortium | Dave J. Baker, Gemma L. Kay, Alp Aydin, Thanh Le-Viet, Steven Rudder, Ana P. Tedim, Anastasia Kolyva, Maria Diaz, Leonardo de Oliveira Martins, Nabil-Fareed Alikhan, Lizzie Meadows, Rachael Stanley, Ngozi Elumogo, Muhammed Yasir, Nicholas M. Thomson, Alexander J Trotter, Rachel Gilroy, Samuel Bloomfield, Claire Stuart, Andrew Bell, Reenesh Prakash, Samir Dervisevic, Alison E. Mather, John Wain, Mark Webber, Andrew J. Page, Justin O'Grady |
| EPI_ISL_572721 | Centre for Enzyme Innovation, University of Portsmouth / Translational Research Laboratory, Portsmouth Hospitals NHS Trust | COVID-19 Genomics UK (COG-UK) Consortium | Angela Beckett,Yann Bourgeois,Garry Scarlett,Sharon Glaysher,Scott Elliott,Kelly Bicknell,Robert Impey,Allyson Lloyd,Sarah Wyllie,Ethan Butcher,Anoop Chauhan,Samuel Robson |
| EPI_ISL_572722 | Virology Department, Sheffield Teaching Hospitals NHS Foundation Trust/Department of Infection, Immunity and Cardiovascular Disease, The Medical School, University of Sheffield | COVID-19 Genomics UK (COG-UK) Consortium | Thushan de Silva, Matthew Parker, Nikki Smith, Adri Angyal, Rebecca Brown, Luke Green, Rachel Tucker, Paul Parsons, Danielle Groves, Katie Johnson, Laura Carrilero, Alex Keeley, Dave Partridge, Matthew Wyles, Benjamin Lindsey, Mehmet Yavuz, Mohammad Raza, Cariad Evans |
| EPI_ISL_572723, EPI_ISL_572724 | Oxford Viromics, NDM, University of Oxford; Oxford University Hospitals; Basingstoke and North Hampshire Hospital | COVID-19 Genomics UK (COG-UK) Consortium | Tanya Golubchik, David Bonsall, George Macintyre, Amy Trebes, Mariateresa de Cesare, Catrin Moore, Alex Mobbs, Anita Justice, Robert Shaw, Monique Andersson, Timothy Peto, Emma Wise, Nathan Moore, Jessica Lynch, Nick Cortes, Matilde Mori, Stephen Kidd, David Buck, John Todd, Christophe Fraser |
| EPI_ISL_572725 | University College London, Great Ormond Street Hospital for Children NHS Foundation Trust, Imperial College Healthcare NHS Trust | COVID-19 Genomics UK (COG-UK) Consortium | Sergi Castellano, Rachel Williams, Mark Kristiansen, Paola Resende Silva, Sunando Roy, Tony Brooks, Helena Tutill, Paola Niola, Patricia Dyal, Charlotte Williams, Leysa Forrest, Yasmin Panchbhaya, Jacqueline Findlay, Samuel Weeks, Julianne Brown, Kathryn Harris, Paul Randell, James Price, Alison Holmes, Judith Breuer |
| EPI_ISL_572726, EPI_ISL_572727, EPI_ISL_572728 | Oxford Viromics, NDM, University of Oxford; Oxford University Hospitals; Basingstoke and North Hampshire Hospital | COVID-19 Genomics UK (COG-UK) Consortium | Tanya Golubchik, David Bonsall, George Macintyre, Amy Trebes, Mariateresa de Cesare, Catrin Moore, Alex Mobbs, Anita Justice, Robert Shaw, Monique Andersson, Timothy Peto, Emma Wise, Nathan Moore, Jessica Lynch, Nick Cortes, Matilde Mori, Stephen Kidd, David Buck, John Todd, Christophe Fraser |
| EPI_ISL_572729 | Northumbria University / South Tees Hospitals NHS Foundation Trust / North Cumbria Integrated Care NHS Foundation Trust / North Tees and Hartlepool NHS Foundation Trust / Newcastle Hospitals NHS Foundation Trust | COVID-19 Genomics UK (COG-UK) Consortium | Darren L Smith,Andrew Nelson,Matthew Bashton,Greg R Young,Joshua Loh,John Allan,Mohammad A Tariq,Giles S Holt,Gary Black,Wen C Yew,Lynn Dover,Paul Baker,Steve Liggett,Sarah Essex,Jane Greenaway,Debra Padgett,Clive Graham,Garren Scott,Edward Barton,Emma Swindells,Brendan Payne,Jennifer Collins,Yusri Taha,Gary Eltringham |
| EPI_ISL_572846 | Oxford Viromics, NDM, University of Oxford; Oxford University Hospitals; Basingstoke and North Hampshire Hospital | COVID-19 Genomics UK (COG-UK) Consortium | Tanya Golubchik, David Bonsall, George Macintyre, Amy Trebes, Mariateresa de Cesare, Catrin Moore, Alex Mobbs, Anita Justice, Robert Shaw, Monique Andersson, Timothy Peto, Emma Wise, Nathan Moore, Jessica Lynch, Nick Cortes, Matilde Mori, Stephen Kidd, David Buck, John Todd, Christophe Fraser |
| EPI_ISL_572847 | Quadram Institute Bioscience | COVID-19 Genomics UK (COG-UK) Consortium | Dave J. Baker, Gemma L. Kay, Alp Aydin, Thanh Le-Viet, Steven Rudder, Ana P. Tedim, Anastasia Kolyva, Maria Diaz, Leonardo de Oliveira Martins, Nabil-Fareed Alikhan, Lizzie Meadows, Rachael Stanley, Ngozi Elumogo, Muhammed Yasir, Nicholas M. Thomson, Alexander J Trotter, Rachel Gilroy, Samuel Bloomfield, Claire Stuart, Andrew Bell, Reenesh Prakash, Samir Dervisevic, Alison E. Mather, John Wain, Mark Webber, Andrew J. Page, Justin O'Grady |
| EPI_ISL_572848 | Virology Department, Sheffield Teaching Hospitals NHS Foundation Trust/Department of Infection, Immunity and Cardiovascular Disease, The Medical School, University of Sheffield | COVID-19 Genomics UK (COG-UK) Consortium | Thushan de Silva, Matthew Parker, Nikki Smith, Adri Angyal, Rebecca Brown, Luke Green, Rachel Tucker, Paul Parsons, Danielle Groves, Katie Johnson, Laura Carrilero, Alex Keeley, Dave Partridge, Matthew Wyles, Benjamin Lindsey, Mehmet Yavuz, Mohammad Raza, Cariad Evans |
| EPI_ISL_572849, EPI_ISL_572850 | Oxford Viromics, NDM, University of Oxford; Oxford University Hospitals; Basingstoke and North Hampshire Hospital | COVID-19 Genomics UK (COG-UK) Consortium | Tanya Golubchik, David Bonsall, George Macintyre, Amy Trebes, Mariateresa de Cesare, Catrin Moore, Alex Mobbs, Anita Justice, Robert Shaw, Monique Andersson, Timothy Peto, Emma Wise, Nathan Moore, Jessica Lynch, Nick Cortes, Matilde Mori, Stephen Kidd, David Buck, John Todd, Christophe Fraser |
| EPI_ISL_572851, EPI_ISL_572852 | Department of Pathology, University of Cambridge | COVID-19 Genomics UK (COG-UK) Consortium | Aminu S. Jahun, Yasmin Chaudhry, Grant Hall, Iliana Georgana, Myra Hosmillo, Martin D. Curran, Malte Pinckert, Surendra Parmar, Ian Goodfellow |
| EPI_ISL_572853, EPI_ISL_572854, EPI_ISL_572855, EPI_ISL_572856 | Wales Specialist Virology Centre Sequencing lab: Pathogen Genomics Unit | COVID-19 Genomics UK (COG-UK) Consortium | Catherine Moore, Johnathan Evans, Laura Gifford, Malorie Perry, Simon Cottrell, Angela Marchbank, Alec Birchley, Alexander Adams, Amy Gaskin, Bree Gatica-Wilcox, Jason Coombes, Joel Southgate, Lauren Gilbert, Lee Graham, Nicole Pacchiarini, Sara Kumziene-Summerhayes, Sarah Taylor, Sophie Jones, Sara Rey, Matthew Bull, Joanne Watkins, Sally Corden, Tom Connor |
| EPI_ISL_572857, EPI_ISL_572858, EPI_ISL_572859, EPI_ISL_572860, EPI_ISL_572861 | Oxford Viromics, NDM, University of Oxford; Oxford University Hospitals; Basingstoke and North Hampshire Hospital | COVID-19 Genomics UK (COG-UK) Consortium | Tanya Golubchik, David Bonsall, George Macintyre, Amy Trebes, Mariateresa de Cesare, Catrin Moore, Alex Mobbs, Anita Justice, Robert Shaw, Monique Andersson, Timothy Peto, Emma Wise, Nathan Moore, Jessica Lynch, Nick Cortes, Matilde Mori, Stephen Kidd, David Buck, John Todd, Christophe Fraser |
| EPI_ISL_572862, EPI_ISL_572863, EPI_ISL_572864, EPI_ISL_572865, EPI_ISL_572866, EPI_ISL_572867, EPI_ISL_572868 | Department of Pathology, University of Cambridge | COVID-19 Genomics UK (COG-UK) Consortium | Aminu S. Jahun, Yasmin Chaudhry, Grant Hall, Iliana Georgana, Myra Hosmillo, Martin D. Curran, Malte Pinckert, Surendra Parmar, Ian Goodfellow |
| EPI_ISL_572869, EPI_ISL_572870 | Oxford Viromics, NDM, University of Oxford; Oxford University Hospitals; Basingstoke and North Hampshire Hospital | COVID-19 Genomics UK (COG-UK) Consortium | Tanya Golubchik, David Bonsall, George Macintyre, Amy Trebes, Mariateresa de Cesare, Catrin Moore, Alex Mobbs, Anita Justice, Robert Shaw, Monique Andersson, Timothy Peto, Emma Wise, Nathan Moore, Jessica Lynch, Nick Cortes, Matilde Mori, Stephen Kidd, David Buck, John Todd, Christophe Fraser |
| EPI_ISL_572871 | Virology Department, Royal Infirmary of Edinburgh, NHS Lothian / School of Biological Sciences, University of Edinburgh / Institute of Genetics and Molecular Medicine, University of Edinburgh | COVID-19 Genomics UK (COG-UK) Consortium | McHugh M, Dewar R, Rooke S, Gallagher M, Balcaza C, O'Toole Á, Scher E, Hill V, McCrone JT, Colquhoun R, Yu X, Jackson B, Rambaut A, Williams TC, Templeton K |
| EPI_ISL_572872 | Queens Medical Centre, Clinical Microbiology Department / DeepSeq Nottingham | COVID-19 Genomics UK (COG-UK) Consortium | Gemma Clark, Wendy Smith, Manjinder Khakh, Vicki M Fleming, Michelle M Lister, Hannah Howson-Wells, Jonathan Ball, Patrick McClure, Joseph Chappell, Theocharis Tsoleridis, Nadine Holmes, Matthew Carlisle, Christopher Moore, Fei Sang, Johnny Debebe, Victoria Wright, Matthew Loose |
| EPI_ISL_572873 | Oxford Viromics, NDM, University of Oxford; Oxford University Hospitals; Basingstoke and North Hampshire Hospital | COVID-19 Genomics UK (COG-UK) Consortium | Tanya Golubchik, David Bonsall, George Macintyre, Amy Trebes, Mariateresa de Cesare, Catrin Moore, Alex Mobbs, Anita Justice, Robert Shaw, Monique Andersson, Timothy Peto, Emma Wise, Nathan Moore, Jessica Lynch, Nick Cortes, Matilde Mori, Stephen Kidd, David Buck, John Todd, Christophe Fraser |
| EPI_ISL_572874 | Virology Department, Royal Infirmary of Edinburgh, NHS Lothian / School of Biological Sciences, University of Edinburgh / Institute of Genetics and Molecular Medicine, University of Edinburgh | COVID-19 Genomics UK (COG-UK) Consortium | McHugh M, Dewar R, Rooke S, Gallagher M, Balcaza C, O'Toole Á, Scher E, Hill V, McCrone JT, Colquhoun R, Yu X, Jackson B, Rambaut A, Williams TC, Templeton K |
| EPI_ISL_572875, EPI_ISL_572876, EPI_ISL_572877, EPI_ISL_572878, EPI_ISL_572879, EPI_ISL_572880, EPI_ISL_572881, EPI_ISL_572882, EPI_ISL_572883, EPI_ISL_572884, EPI_ISL_572885, EPI_ISL_572886, EPI_ISL_572887, EPI_ISL_572888, EPI_ISL_572889, EPI_ISL_572890 | | | |
| see above | Wales Specialist Virology Centre Sequencing lab: Pathogen Genomics Unit | COVID-19 Genomics UK (COG-UK) Consortium | Catherine Moore, Johnathan Evans, Laura Gifford, Malorie Perry, Simon Cottrell, Angela Marchbank, Alec Birchley, Alexander Adams, Amy Gaskin, Bree Gatica-Wilcox, Jason Coombes, Joel Southgate, Lauren Gilbert, Lee Graham, Nicole Pacchiarini, Sara Kumziene-Summerhayes, Sarah Taylor, Sophie Jones, Sara Rey, Matthew Bull, Joanne Watkins, Sally Corden, Tom Connor |
| EPI_ISL_572938 | Lincolnshire Hospitals and DeepSeq Nottingham | COVID-19 Genomics UK (COG-UK) Consortium | Nichola Duckworth, Tim Sloan, Sarah Walsh, Jonathan Ball, Patrick McClure, Joeseph Chappell, Nadine Holmes, Matthew Carlisle, Christopher Moore, Fei Sang, Johnny Debebe, Victoria Wright, Matthew Loose |
| EPI_ISL_572939, EPI_ISL_572940, EPI_ISL_572941 | Quadram Institute Bioscience | COVID-19 Genomics UK (COG-UK) Consortium | Dave J. Baker, Gemma L. Kay, Alp Aydin, Thanh Le-Viet, Steven Rudder, Ana P. Tedim, Anastasia Kolyva, Maria Diaz, Leonardo de Oliveira Martins, Nabil-Fareed Alikhan, Lizzie Meadows, Rachael Stanley, Ngozi Elumogo, Muhammed Yasir, Nicholas M. Thomson, Alexander J Trotter, Rachel Gilroy, Samuel Bloomfield, Claire Stuart, Andrew Bell, Reenesh Prakash, Samir Dervisevic, Alison E. Mather, John Wain, Mark Webber, Andrew J. Page, Justin O'Grady |
| EPI_ISL_572942 | Northumbria University / South Tees Hospitals NHS Foundation Trust / North Cumbria Integrated Care NHS Foundation Trust / | COVID-19 Genomics UK (COG-UK) Consortium | Darren L Smith,Andrew Nelson,Matthew Bashton,Greg R Young,Joshua Loh,John Allan,Mohammad A Tariq,Giles S Holt,Gary Black,Wen C Yew,Lynn Dover,Paul Baker,Steve Liggett,Sarah Essex,Jane Greenaway,Debra Padgett,Clive Graham,Garren Scott,Edward Barton,Emma Swindells,Brendan Payne,Jennifer Collins,Yusri Taha,Gary Eltringham |

| | | | |
|---|---|---|---|
| | North Tees and Hartlepool NHS Foundation Trust / Newcastle Hospitals NHS Foundation Trust | | |
| EPI_ISL_572943, EPI_ISL_572944 | Lincolnshire Hospitals and DeepSeq Nottingham | COVID-19 Genomics UK (COG-UK) Consortium | Nichola Duckworth, Tim Sloan, Sarah Walsh, Jonathan Ball, Patrick McClure, Joeseph Chappell, Nadine Holmes, Matthew Carlisle, Christopher Moore, Fei Sang, Johnny Debebe, Victoria Wright, Matthew Loose |
| EPI_ISL_572945, EPI_ISL_572946 | Virology Department, Sheffield Teaching Hospitals NHS Foundation Trust/Department of Infection, Immunity and Cardiovascular Disease, The Medical School, University of Sheffield | COVID-19 Genomics UK (COG-UK) Consortium | Thushan de Silva, Matthew Parker, Nikki Smith, Adri Angyal, Rebecca Brown, Luke Green, Rachel Tucker, Paul Parsons, Danielle Groves, Katie Johnson, Laura Carrilero, Alex Keeley, Dave Partridge, Matthew Wyles, Benjamin Lindsey, Mehmet Yavuz, Mohammad Raza, Cariad Evans |
| EPI_ISL_572947 | Oxford Viromics, NDM, University of Oxford; Oxford University Hospitals; Basingstoke and North Hampshire Hospital | COVID-19 Genomics UK (COG-UK) Consortium | Tanya Golubchik, David Bonsall, George Macintyre, Amy Trebes, Mariateresa de Cesare, Catrin Moore, Alex Mobbs, Anita Justice, Robert Shaw, Monique Andersson, Timothy Peto, Emma Wise, Nathan Moore, Jessica Lynch, Nick Cortes, Matilde Mori, Stephen Kidd, David Buck, John Todd, Christophe Fraser |
| EPI_ISL_572948, EPI_ISL_572949 | Virology Department, Sheffield Teaching Hospitals NHS Foundation Trust/Department of Infection, Immunity and Cardiovascular Disease, The Medical School, University of Sheffield | COVID-19 Genomics UK (COG-UK) Consortium | Thushan de Silva, Matthew Parker, Nikki Smith, Adri Angyal, Rebecca Brown, Luke Green, Rachel Tucker, Paul Parsons, Danielle Groves, Katie Johnson, Laura Carrilero, Alex Keeley, Dave Partridge, Matthew Wyles, Benjamin Lindsey, Mehmet Yavuz, Mohammad Raza, Cariad Evans |
| EPI_ISL_572950 | Northumbria University / South Tees Hospitals NHS Foundation Trust / North Cumbria Integrated Care NHS Foundation Trust / North Tees and Hartlepool NHS Foundation Trust / Newcastle Hospitals NHS Foundation Trust | COVID-19 Genomics UK (COG-UK) Consortium | Darren L Smith,Andrew Nelson,Matthew Bashton,Greg R Young,Joshua Loh,John Allan,Mohammad A Tariq,Giles S Holt,Gary Black,Wen C Yew,Lynn Dover,Paul Baker,Steve Liggett,Sarah Essex,Jane Greenaway,Debra Padgett,Clive Graham,Garren Scott,Edward Barton,Emma Swindells,Brendan Payne,Jennifer Collins,Yusri Taha,Gary Eltringham |
| EPI_ISL_572951 | Wales Specialist Virology Centre Sequencing lab: Pathogen Genomics Unit | COVID-19 Genomics UK (COG-UK) Consortium | Catherine Moore, Johnathan Evans, Laura Gifford, Malorie Perry, Simon Cottrell, Angela Marchbank, Alec Birchley, Alexander Adams, Amy Gaskin, Bree Gatica-Wilcox, Jason Coombes, Joel Southgate, Lauren Gilbert, Lee Graham, Nicole Pacchiarini, Sara Kumziene-Summerhayes, Sarah Taylor, Sophie Jones, Sara Rey, Matthew Bull, Joanne Watkins, Sally Corden, Tom Connor |
| EPI_ISL_572954 | Quadram Institute Bioscience | COVID-19 Genomics UK (COG-UK) Consortium | Dave J. Baker, Gemma L. Kay, Alp Aydin, Thanh Le-Viet, Steven Rudder, Ana P. Tedim, Anastasia Kolyva, Maria Diaz, Leonardo de Oliveira Martins, Nabil-Fareed Alikhan, Lizzie Meadows, Rachael Stanley, Ngozi Elumogo, Muhammed Yasir, Nicholas M. Thomson, Alexander J Trotter, Rachel Gilroy, Samuel Bloomfield, Claire Stuart, Andrew Bell, Reenesh Prakash, Samir Dervisevic, Alison E. Mather, John Wain, Mark Webber, Andrew J. Page, Justin O’Grady |
| EPI_ISL_572957 | University College London, Great Ormond Street Hospital for Children NHS Foundation Trust, Imperial College Healthcare NHS Trust | COVID-19 Genomics UK (COG-UK) Consortium | Sergi Castellano, Rachel Williams, Mark Kristiansen, Paola Resende Silva, Sunando Roy, Tony Brooks, Helena Tutill, Paola Niola, Patricia Dyal, Charlotte Williams, Leysa Forrest, Yasmin Panchbhaya, Jacqueline Findlay, Samuel Weeks, Julianne Brown, Kathryn Harris, Paul Randell, James Price, Alison Holmes, Judith Breuer |
| EPI_ISL_572958 | Oxford Viromics, NDM, University of Oxford; Oxford University Hospitals; Basingstoke and North Hampshire Hospital | COVID-19 Genomics UK (COG-UK) Consortium | Tanya Golubchik, David Bonsall, George Macintyre, Amy Trebes, Mariateresa de Cesare, Catrin Moore, Alex Mobbs, Anita Justice, Robert Shaw, Monique Andersson, Timothy Peto, Emma Wise, Nathan Moore, Jessica Lynch, Nick Cortes, Matilde Mori, Stephen Kidd, David Buck, John Todd, Christophe Fraser |
| EPI_ISL_572959 | Virology Department, Sheffield Teaching Hospitals NHS Foundation Trust/Department of Infection, Immunity and Cardiovascular Disease, The Medical School, University of Sheffield | COVID-19 Genomics UK (COG-UK) Consortium | Thushan de Silva, Matthew Parker, Nikki Smith, Adri Angyal, Rebecca Brown, Luke Green, Rachel Tucker, Paul Parsons, Danielle Groves, Katie Johnson, Laura Carrilero, Alex Keeley, Dave Partridge, Matthew Wyles, Benjamin Lindsey, Mehmet Yavuz, Mohammad Raza, Cariad Evans |
| EPI_ISL_572960 | Oxford Viromics, NDM, University of Oxford; Oxford University Hospitals; Basingstoke and North Hampshire Hospital | COVID-19 Genomics UK (COG-UK) Consortium | Tanya Golubchik, David Bonsall, George Macintyre, Amy Trebes, Mariateresa de Cesare, Catrin Moore, Alex Mobbs, Anita Justice, Robert Shaw, Monique Andersson, Timothy Peto, Emma Wise, Nathan Moore, Jessica Lynch, Nick Cortes, Matilde Mori, Stephen Kidd, David Buck, John Todd, Christophe Fraser |
| EPI_ISL_572961 | Virology Department, Sheffield Teaching Hospitals NHS Foundation Trust/Department of Infection, Immunity and Cardiovascular Disease, The Medical School, University of Sheffield | COVID-19 Genomics UK (COG-UK) Consortium | Thushan de Silva, Matthew Parker, Nikki Smith, Adri Angyal, Rebecca Brown, Luke Green, Rachel Tucker, Paul Parsons, Danielle Groves, Katie Johnson, Laura Carrilero, Alex Keeley, Dave Partridge, Matthew Wyles, Benjamin Lindsey, Mehmet Yavuz, Mohammad Raza, Cariad Evans |
| EPI_ISL_572962, EPI_ISL_572963, EPI_ISL_572964, EPI_ISL_572965, EPI_ISL_572966, EPI_ISL_572967, EPI_ISL_572968, EPI_ISL_572969, EPI_ISL_572970, EPI_ISL_572971, EPI_ISL_572972, EPI_ISL_572973, EPI_ISL_572974, EPI_ISL_572975, EPI_ISL_572976, EPI_ISL_572977, EPI_ISL_572978, EPI_ISL_572979, EPI_ISL_572980, EPI_ISL_572981, EPI_ISL_572982, EPI_ISL_572983, EPI_ISL_572984, EPI_ISL_572985, EPI_ISL_572986, EPI_ISL_572987, EPI_ISL_572988, EPI_ISL_572989, EPI_ISL_572990, EPI_ISL_572991, EPI_ISL_572992, EPI_ISL_572993, EPI_ISL_572994, EPI_ISL_572995, EPI_ISL_572996, EPI_ISL_572997, EPI_ISL_572998, EPI_ISL_572999, EPI_ISL_573000, EPI_ISL_573001, EPI_ISL_573002, EPI_ISL_573003, EPI_ISL_573004, EPI_ISL_573005, EPI_ISL_573006, EPI_ISL_573007, EPI_ISL_573008, EPI_ISL_573009, EPI_ISL_573010, EPI_ISL_573011, EPI_ISL_573012, EPI_ISL_573013, EPI_ISL_573014, EPI_ISL_573015, EPI_ISL_573016, EPI_ISL_573017, EPI_ISL_573018, EPI_ISL_573019, EPI_ISL_573020, EPI_ISL_573021, EPI_ISL_573022, EPI_ISL_573023, EPI_ISL_573024, EPI_ISL_573025, EPI_ISL_573026, EPI_ISL_573027, EPI_ISL_573028, EPI_ISL_573029, EPI_ISL_573030, EPI_ISL_573031, EPI_ISL_573032, EPI_ISL_573033, EPI_ISL_573034, EPI_ISL_573035, EPI_ISL_573036, EPI_ISL_573037, EPI_ISL_573038, EPI_ISL_573039, EPI_ISL_573040, EPI_ISL_573041, EPI_ISL_573042, EPI_ISL_573043, EPI_ISL_573044, EPI_ISL_573045, EPI_ISL_573046, EPI_ISL_573047, EPI_ISL_573048, EPI_ISL_573049, EPI_ISL_573050, EPI_ISL_573051, EPI_ISL_573052, EPI_ISL_573053, EPI_ISL_573054, EPI_ISL_573055, EPI_ISL_573056, EPI_ISL_573057, EPI_ISL_573058, EPI_ISL_573059, EPI_ISL_573060, EPI_ISL_573061, EPI_ISL_573062, EPI_ISL_573063, EPI_ISL_573064, EPI_ISL_573065, EPI_ISL_573066, EPI_ISL_573067, EPI_ISL_573068, EPI_ISL_573069, EPI_ISL_573070, EPI_ISL_573071, EPI_ISL_573072, EPI_ISL_573073, EPI_ISL_573074, EPI_ISL_573075, EPI_ISL_573076, EPI_ISL_573077, EPI_ISL_573078, EPI_ISL_573079, EPI_ISL_573080, EPI_ISL_573081, EPI_ISL_573082, EPI_ISL_573083, EPI_ISL_573084 | | | |
| see above | Oxford Viromics, NDM, University of Oxford; Oxford University Hospitals; Basingstoke and North Hampshire Hospital | COVID-19 Genomics UK (COG-UK) Consortium | Tanya Golubchik, David Bonsall, George Macintyre, Amy Trebes, Mariateresa de Cesare, Catrin Moore, Alex Mobbs, Anita Justice, Robert Shaw, Monique Andersson, Timothy Peto, Emma Wise, Nathan Moore, Jessica Lynch, Nick Cortes, Matilde Mori, Stephen Kidd, David Buck, John Todd, Christophe Fraser |
| EPI_ISL_573085, EPI_ISL_573086, EPI_ISL_573087, EPI_ISL_573088, EPI_ISL_573089, EPI_ISL_573090, EPI_ISL_573091, EPI_ISL_573092, EPI_ISL_573093, EPI_ISL_573094, EPI_ISL_573095, EPI_ISL_573096, EPI_ISL_573097, EPI_ISL_573098, EPI_ISL_573099, EPI_ISL_573100, EPI_ISL_573101, EPI_ISL_573102, EPI_ISL_573103, EPI_ISL_573104, EPI_ISL_573105, EPI_ISL_573106, EPI_ISL_573107, EPI_ISL_573108, EPI_ISL_573109, EPI_ISL_573110, EPI_ISL_573111, EPI_ISL_573112, EPI_ISL_573113, EPI_ISL_573114, EPI_ISL_573115, EPI_ISL_573116, EPI_ISL_573117, EPI_ISL_573118, EPI_ISL_573119, EPI_ISL_573120, EPI_ISL_573121, EPI_ISL_573122, EPI_ISL_573123, EPI_ISL_573124, EPI_ISL_573125, EPI_ISL_573126, EPI_ISL_573127, EPI_ISL_573128, EPI_ISL_573129, EPI_ISL_573130, EPI_ISL_573131 | | | |
| see above | Quadram Institute Bioscience | COVID-19 Genomics UK (COG-UK) Consortium | Dave J. Baker, Gemma L. Kay, Alp Aydin, Thanh Le-Viet, Steven Rudder, Ana P. Tedim, Anastasia Kolyva, Maria Diaz, Leonardo de Oliveira Martins, Nabil-Fareed Alikhan, Lizzie Meadows, Rachael Stanley, Ngozi Elumogo, Muhammed Yasir, Nicholas M. Thomson, Alexander J Trotter, Rachel Gilroy, Samuel Bloomfield, Claire Stuart, Andrew Bell, Reenesh Prakash, Samir Dervisevic, Alison E. Mather, John Wain, Mark Webber, Andrew J. Page, Justin O’Grady |
| EPI_ISL_573132, EPI_ISL_573133, EPI_ISL_573134, EPI_ISL_573135, EPI_ISL_573136, EPI_ISL_573137, EPI_ISL_573138, EPI_ISL_573139, EPI_ISL_573140, EPI_ISL_573141, EPI_ISL_573142, EPI_ISL_573143, EPI_ISL_573144, EPI_ISL_573145, EPI_ISL_573146, EPI_ISL_573147, EPI_ISL_573148, EPI_ISL_573149, EPI_ISL_573150, EPI_ISL_573151 | | | |
| see above | Department of Pathology, University of Cambridge | COVID-19 Genomics UK (COG-UK) Consortium | Aminu S. Jahun, Yasmin Chaudhry, Grant Hall, Iliana Georgana, Myra Hosmillo, Martin D. Curran, Malte Pinckert, Surendra Parmar, Ian Goodfellow |
| EPI_ISL_573152 | Institute for Virology, University Hospital Duesseldorf, Medical Faculty, Heinrich-Heine-University Duesseldorf | Institute for Virology, University Hospital Duesseldorf, Medical Faculty, Heinrich-Heine-University Duesseldorf | Maximilian Damagnez, Verena Keitel, Björn Jensen, Nadine Lübke, Lisa Müller, Philipp Ostermann, Tina Senff, Ortwin Adams, Philipp Albrecht, Gerald Antoch, Johannes Bode, Edwin Bölke, Saskia Elben, Torsten Feldt, Johannes C. Fischer, , Anselm Kunstein, Caroline Klindt, Alexander Killer, Tom Lüdde, Annemarie Mohring, Jennifer Neubert, Heiner Schaal, Ansgar Schulz, Jörg Timm, Andreas Walker |
| EPI_ISL_573153 | Department of Pathology, University of Cambridge | COVID-19 Genomics UK (COG-UK) Consortium | Aminu S. Jahun, Yasmin Chaudhry, Grant Hall, Iliana Georgana, Myra Hosmillo, Martin D. Curran, Malte Pinckert, Surendra Parmar, Ian Goodfellow |
| EPI_ISL_573154, EPI_ISL_573155, EPI_ISL_573156, EPI_ISL_573157, EPI_ISL_573158, EPI_ISL_573159, EPI_ISL_573160, EPI_ISL_573161, EPI_ISL_573162, EPI_ISL_573163, EPI_ISL_573164, EPI_ISL_573165, EPI_ISL_573166, EPI_ISL_573167, EPI_ISL_573168, EPI_ISL_573169, EPI_ISL_573170, EPI_ISL_573171, EPI_ISL_573172, EPI_ISL_573173, EPI_ISL_573174, EPI_ISL_573175, EPI_ISL_573176, EPI_ISL_573177, EPI_ISL_573178, EPI_ISL_573179, EPI_ISL_573180, EPI_ISL_573181, EPI_ISL_573182, EPI_ISL_573183, EPI_ISL_573184, EPI_ISL_573185, EPI_ISL_573186, EPI_ISL_573187, EPI_ISL_573188, EPI_ISL_573189, EPI_ISL_573190, EPI_ISL_573191, EPI_ISL_573192, EPI_ISL_573193, EPI_ISL_573194, EPI_ISL_573195, EPI_ISL_573196, EPI_ISL_573197, EPI_ISL_573198, EPI_ISL_573199, EPI_ISL_573200, EPI_ISL_573201, EPI_ISL_573202, EPI_ISL_573203, EPI_ISL_573204, EPI_ISL_573205, EPI_ISL_573206, EPI_ISL_573207, EPI_ISL_573208, EPI_ISL_573209, EPI_ISL_573210, EPI_ISL_573211, EPI_ISL_573212, EPI_ISL_573213, EPI_ISL_573214, EPI_ISL_573215, EPI_ISL_573216, EPI_ISL_573217, EPI_ISL_573218, EPI_ISL_573219, EPI_ISL_573220, EPI_ISL_573221, EPI_ISL_573222, EPI_ISL_573223, EPI_ISL_573224, EPI_ISL_573225, EPI_ISL_573226, EPI_ISL_573227, EPI_ISL_573228, EPI_ISL_573229, EPI_ISL_573230, EPI_ISL_573231, EPI_ISL_573232, EPI_ISL_573233, EPI_ISL_573234, EPI_ISL_573235 | | | |
| see above | Oxford Viromics, NDM, University of Oxford; Oxford University Hospitals; Basingstoke and North Hampshire Hospital | COVID-19 Genomics UK (COG-UK) Consortium | Tanya Golubchik, David Bonsall, George Macintyre, Amy Trebes, Mariateresa de Cesare, Catrin Moore, Alex Mobbs, Anita Justice, Robert Shaw, Monique Andersson, Timothy Peto, Emma Wise, Nathan Moore, Jessica Lynch, Nick Cortes, Matilde Mori, Stephen Kidd, David Buck, John Todd, Christophe Fraser |
| EPI_ISL_573236, EPI_ISL_573237, EPI_ISL_573238, EPI_ISL_573239, EPI_ISL_573240, EPI_ISL_573241, EPI_ISL_573242, EPI_ISL_573243, EPI_ISL_573244, EPI_ISL_573245, EPI_ISL_573246 | | | |
| see above | Liverpool Clinical Laboratories | COVID-19 Genomics UK (COG-UK) Consortium | Sam Haldenby, Anita Lucaci, Steve Paterson, Julian Hiscox, Alistair Darby, M Almsaud, A Alrezaihi, Muhannad Alruwaili, Stuart D Armstrong, Jones Benjamin, Eleanor G Bentley, Anu Chawla, Jordan J Clark, Angela Cowell, Richard Eccles, Isabel García-Dorival, Matthew Gemmell, Alessandro Gerada, PKF Gilmore, Richard Gregory, Ximeng Han, Catherine Hartley, Margaret Hughes, Miren Iturriza-Gomara, James Johnson, L Luu, Jenifer Manson, Charlotte Nelson, Elaine O’Toole, Cassie Olateju, Rebekah Penrice-Randal , Lucille Rainbow, N.P Randle, Trevor Ian Robinson, Parul Sharma, Ghada T Shawli, James P Stewart, Neil Swainston, Ecaterina Vamos, Joanne Watts, Mark Whitehead |
| EPI_ISL_573247, EPI_ISL_573248, EPI_ISL_573249, EPI_ISL_573250, EPI_ISL_573251, EPI_ISL_573252 | Virology Department, Royal Infirmary of Edinburgh, NHS Lothian / School of Biological Sciences, University of Edinburgh / Institute of Genetics and Molecular Medicine, University of Edinburgh | COVID-19 Genomics UK (COG-UK) Consortium | McHugh M, Dewar R, Rooke S, Gallagher M, Balcaza C, O’Toole Á, Scher E, Hill V, McCrone JT, Colquhoun R, Yu X, Jackson B, Rambaut A, Williams TC, Templeton K |
| EPI_ISL_573253, EPI_ISL_573254, EPI_ISL_573255, EPI_ISL_573256, EPI_ISL_573257, EPI_ISL_573258, EPI_ISL_573259, EPI_ISL_573260, EPI_ISL_573261, EPI_ISL_573262, EPI_ISL_573263, EPI_ISL_573264, EPI_ISL_573265 | | | |
| see above | Centre for Enzyme Innovation, University of Portsmouth / Translational Research Laboratory, Portsmouth Hospitals NHS Trust | COVID-19 Genomics UK (COG-UK) Consortium | Angela Beckett,Yann Bourgeois,Garry Scarlett,Sharon Glaysher,Scott Elliott,Kelly Bicknell,Robert Impey,Allyson Lloyd,Sarah Wyllie,Ethan Butcher,Anoop Chauhan,Samuel Robson |
| EPI_ISL_573266, EPI_ISL_573267, EPI_ISL_573268, EPI_ISL_573269, EPI_ISL_573270, EPI_ISL_573271, EPI_ISL_573272, EPI_ISL_573273, EPI_ISL_573274, EPI_ISL_573275, EPI_ISL_573276, EPI_ISL_573277, EPI_ISL_573278, EPI_ISL_573279, EPI_ISL_573280, EPI_ISL_573281, EPI_ISL_573282, EPI_ISL_573283, EPI_ISL_573284, EPI_ISL_573285, EPI_ISL_573286, EPI_ISL_573287, EPI_ISL_573288, EPI_ISL_573289, EPI_ISL_573290, EPI_ISL_573291, EPI_ISL_573292, EPI_ISL_573293, EPI_ISL_573294, EPI_ISL_573295, EPI_ISL_573296, EPI_ISL_573297, EPI_ISL_573298, EPI_ISL_573299, EPI_ISL_573300, EPI_ISL_573301, EPI_ISL_573302, EPI_ISL_573303, EPI_ISL_573304, EPI_ISL_573305, EPI_ISL_573306, EPI_ISL_573307, EPI_ISL_573308, EPI_ISL_573309, EPI_ISL_573310, EPI_ISL_573311, EPI_ISL_573312, EPI_ISL_573313, EPI_ISL_573314, EPI_ISL_573315, EPI_ISL_573316, EPI_ISL_573317, EPI_ISL_573318, EPI_ISL_573319, EPI_ISL_573320, EPI_ISL_573321, EPI_ISL_573322, EPI_ISL_573323, EPI_ISL_573324, EPI_ISL_573325, EPI_ISL_573326, EPI_ISL_573327, EPI_ISL_573328, EPI_ISL_573329, EPI_ISL_573330, EPI_ISL_573331, EPI_ISL_573332, EPI_ISL_573333, EPI_ISL_573334, EPI_ISL_573335, EPI_ISL_573336, EPI_ISL_573337, EPI_ISL_573338, EPI_ISL_573339, EPI_ISL_573340, EPI_ISL_573341, EPI_ISL_573342, EPI_ISL_573343, EPI_ISL_573344, EPI_ISL_573345, EPI_ISL_573346, EPI_ISL_573347, EPI_ISL_573348, EPI_ISL_573349, EPI_ISL_573350, EPI_ISL_573351, EPI_ISL_573352, EPI_ISL_573353, EPI_ISL_573354, EPI_ISL_573355, EPI_ISL_573356, EPI_ISL_573357, EPI_ISL_573358, EPI_ISL_573359, EPI_ISL_573360, EPI_ISL_573361, EPI_ISL_573362, EPI_ISL_573363, EPI_ISL_573364, EPI_ISL_573365, EPI_ISL_573366, EPI_ISL_573367, EPI_ISL_573368, EPI_ISL_573369, EPI_ISL_573370, EPI_ISL_573371, EPI_ISL_573372, EPI_ISL_573373, EPI_ISL_573374, EPI_ISL_573375, EPI_ISL_573376, EPI_ISL_573377, EPI_ISL_573378, EPI_ISL_573379, EPI_ISL_573380, EPI_ISL_573381, EPI_ISL_573382, EPI_ISL_573383, EPI_ISL_573384, EPI_ISL_573385, EPI_ISL_573386, EPI_ISL_573387, EPI_ISL_573388, EPI_ISL_573389, EPI_ISL_573390, EPI_ISL_573391, EPI_ISL_573392, EPI_ISL_573393, EPI_ISL_573394, EPI_ISL_573395, EPI_ISL_573396, EPI_ISL_573397, EPI_ISL_573398, EPI_ISL_573399, EPI_ISL_573400, EPI_ISL_573401, EPI_ISL_573402, EPI_ISL_573403, EPI_ISL_573404, EPI_ISL_573405, EPI_ISL_573406, EPI_ISL_573407, EPI_ISL_573408, EPI_ISL_573409, EPI_ISL_573410, EPI_ISL_573411, EPI_ISL_573412, EPI_ISL_573413 | | | |
| see above | Northumbria University / South Tees Hospitals NHS Foundation Trust / North Cumbria Integrated Care NHS Foundation Trust / North Tees and Hartlepool NHS Foundation Trust / Newcastle Hospitals NHS Foundation Trust | COVID-19 Genomics UK (COG-UK) Consortium | Darren L Smith,Andrew Nelson,Matthew Bashton,Greg R Young,Joshua Loh,John Allan,Mohammad A Tariq,Giles S Holt,Gary Black,Wen C Yew,Lynn Dover,Paul Baker,Steve Liggett,Sarah Essex,Jane Greenaway,Debra Padgett,Clive Graham,Garren Scott,Edward Barton,Emma Swindells,Brendan Payne,Jennifer Collins,Yusri Taha,Gary Eltringham |
| EPI_ISL_573414, EPI_ISL_573415, EPI_ISL_573416, EPI_ISL_573417 | Quadram Institute Bioscience | COVID-19 Genomics UK (COG-UK) Consortium | Dave J. Baker, Gemma L. Kay, Alp Aydin, Thanh Le-Viet, Steven Rudder, Ana P. Tedim, Anastasia Kolyva, Maria Diaz, Leonardo de Oliveira Martins, Nabil-Fareed Alikhan, Lizzie Meadows, Rachael Stanley, Ngozi Elumogo, Muhammed Yasir, Nicholas M. Thomson, Alexander J Trotter, Rachel Gilroy, Samuel Bloomfield, Claire Stuart, Andrew Bell, Reenesh Prakash, Samir Dervisevic, Alison E. Mather, John Wain, Mark Webber, Andrew J. Page, Justin O’Grady |
| EPI_ISL_573418, EPI_ISL_573419, EPI_ISL_573420, EPI_ISL_573421, EPI_ISL_573422, EPI_ISL_573423, EPI_ISL_573424, EPI_ISL_573425, EPI_ISL_573426, EPI_ISL_573427, EPI_ISL_573428, EPI_ISL_573429, EPI_ISL_573430, EPI_ISL_573431, EPI_ISL_573432, EPI_ISL_573433, EPI_ISL_573434, EPI_ISL_573435, EPI_ISL_573436 | | | |
| see above | Queens Medical Centre, Clinical Microbiology Department / DeepSeq Nottingham | COVID-19 Genomics UK (COG-UK) Consortium | Gemma Clark, Wendy Smith, Manjinder Khakh, Vicki M Fleming, Michelle M Lister, Hannah Howson-Wells, Jonathan Ball, Patrick McClure, Joseph Chappell, Theocharis Tsoleridis, Nadine Holmes, Matthew Carlisle, Christopher Moore, Fei Sang, Johnny Debebe, Victoria Wright, Matthew Loose |
| EPI_ISL_573437, EPI_ISL_573438, EPI_ISL_573439, EPI_ISL_573440, EPI_ISL_573441, EPI_ISL_573442, EPI_ISL_573443, EPI_ISL_573444, EPI_ISL_573445, EPI_ISL_573446, EPI_ISL_573447, EPI_ISL_573448, EPI_ISL_573449 | | | |
| see above | Lincolnshire Hospitals and DeepSeq Nottingham | COVID-19 Genomics UK (COG-UK) Consortium | Nichola Duckworth, Tim Sloan, Sarah Walsh, Jonathan Ball, Patrick McClure, Joeseph Chappell, Nadine Holmes, Matthew Carlisle, Christopher Moore, Fei Sang, Johnny Debebe, Victoria Wright, Matthew Loose |
| EPI_ISL_573450, EPI_ISL_573451, EPI_ISL_573452, EPI_ISL_573453, EPI_ISL_573454, EPI_ISL_573455, EPI_ISL_573456, EPI_ISL_573457, EPI_ISL_573458, EPI_ISL_573459, EPI_ISL_573460, EPI_ISL_573461, EPI_ISL_573462, EPI_ISL_573463, EPI_ISL_573464, EPI_ISL_573465, EPI_ISL_573466, EPI_ISL_573467, EPI_ISL_573468, EPI_ISL_573469, EPI_ISL_573470, EPI_ISL_573471, EPI_ISL_573472, EPI_ISL_573473, | | | |

| | | | |
|---|---|---|---|
| EPI_ISL_573474, EPI_ISL_573475, EPI_ISL_573476, EPI_ISL_573477 | | | |
| see above | Queens Medical Centre, Clinical Microbiology Department / DeepSeq Nottingham | COVID-19 Genomics UK (COG-UK) Consortium | Gemma Clark, Wendy Smith, Manjinder Khakh, Vicki M Fleming, Michelle M Lister, Hannah Howson-Wells, Jonathan Ball, Patrick McClure, Joseph Chappell, Theocharis Tsoleridis, Nadine Holmes, Matthew Carlisle, Christopher Moore, Fei Sang, Johnny Debebe, Victoria Wright, Matthew Loose |
| EPI_ISL_573478, EPI_ISL_573479, EPI_ISL_573480, EPI_ISL_573481, EPI_ISL_573482, EPI_ISL_573483, EPI_ISL_573484, EPI_ISL_573485, EPI_ISL_573486, EPI_ISL_573487, EPI_ISL_573488, EPI_ISL_573489, EPI_ISL_573490, EPI_ISL_573491, EPI_ISL_573492, EPI_ISL_573493, EPI_ISL_573494, EPI_ISL_573495, EPI_ISL_573496, EPI_ISL_573497, EPI_ISL_573498, EPI_ISL_573499, EPI_ISL_573500, EPI_ISL_573502, EPI_ISL_573503, EPI_ISL_573504, EPI_ISL_573505, EPI_ISL_573506, EPI_ISL_573507, EPI_ISL_573508, EPI_ISL_573509, EPI_ISL_573510, EPI_ISL_573511, EPI_ISL_573512, EPI_ISL_573513, EPI_ISL_573514, EPI_ISL_573515, EPI_ISL_573516, EPI_ISL_573517, EPI_ISL_573518, EPI_ISL_573519, EPI_ISL_573520, EPI_ISL_573521, EPI_ISL_573522, EPI_ISL_573523, EPI_ISL_573524, EPI_ISL_573525 | | | |
| see above | University College London, Great Ormond Street Hospital for Children NHS Foundation Trust, Imperial College Healthcare NHS Trust | COVID-19 Genomics UK (COG-UK) Consortium | Sergi Castellano, Rachel Williams, Mark Kristiansen, Paola Resende Silva, Sunando Roy, Tony Brooks, Helena Tutill, Paola Niola, Patricia Dyal, Charlotte Williams, Leysa Forrest, Yasmin Panchbhaya, Jacqueline Findlay, Samuel Weeks, Julianne Brown, Kathryn Harris, Paul Randell, James Price, Alison Holmes, Judith Breuer |
| EPI_ISL_573526, EPI_ISL_573527, EPI_ISL_573528, EPI_ISL_573529, EPI_ISL_573530, EPI_ISL_573531, EPI_ISL_573532, EPI_ISL_573533, EPI_ISL_573534, EPI_ISL_573535, EPI_ISL_573536, EPI_ISL_573537, EPI_ISL_573538, EPI_ISL_573539, EPI_ISL_573540, EPI_ISL_573541, EPI_ISL_573542, EPI_ISL_573543, EPI_ISL_573544, EPI_ISL_573545, EPI_ISL_573546, EPI_ISL_573547, EPI_ISL_573548, EPI_ISL_573549, EPI_ISL_573550, EPI_ISL_573551, EPI_ISL_573552, EPI_ISL_573553, EPI_ISL_573554, EPI_ISL_573555, EPI_ISL_573556, EPI_ISL_573557, EPI_ISL_573558, EPI_ISL_573559, EPI_ISL_573560, EPI_ISL_573561, EPI_ISL_573562, EPI_ISL_573563, EPI_ISL_573564, EPI_ISL_573565 | | | |
| see above | Virology Department, Sheffield Teaching Hospitals NHS Foundation Trust/Department of Infection, Immunity and Cardiovascular Disease, The Medical School, University of Sheffield | COVID-19 Genomics UK (COG-UK) Consortium | Thushan de Silva, Matthew Parker, Nikki Smith, Adri Angyal, Rebecca Brown, Luke Green, Rachel Tucker, Paul Parsons, Danielle Groves, Katie Johnson, Laura Carrilero, Alex Keeley, Dave Partridge, Matthew Wyles, Benjamin Lindsey, Mehmet Yavuz, Mohammad Raza, Cariad Evans |
| EPI_ISL_573566, EPI_ISL_573567, EPI_ISL_573568, EPI_ISL_573569, EPI_ISL_573570, EPI_ISL_573571, EPI_ISL_573572, EPI_ISL_573573, EPI_ISL_573574, EPI_ISL_573575, EPI_ISL_573576, EPI_ISL_573577, EPI_ISL_573578, EPI_ISL_573579, EPI_ISL_573580, EPI_ISL_573581, EPI_ISL_573582, EPI_ISL_573583, EPI_ISL_573584, EPI_ISL_573585, EPI_ISL_573586, EPI_ISL_573587, EPI_ISL_573588, EPI_ISL_573589, EPI_ISL_573590, EPI_ISL_573591, EPI_ISL_573592, EPI_ISL_573593, EPI_ISL_573594, EPI_ISL_573595, EPI_ISL_573596, EPI_ISL_573597, EPI_ISL_573598, EPI_ISL_573599, EPI_ISL_573600, EPI_ISL_573601, EPI_ISL_573602, EPI_ISL_573603, EPI_ISL_573604, EPI_ISL_573605, EPI_ISL_573606, EPI_ISL_573607, EPI_ISL_573608, EPI_ISL_573609, EPI_ISL_573610, EPI_ISL_573611, EPI_ISL_573612, EPI_ISL_573613, EPI_ISL_573614, EPI_ISL_573615, EPI_ISL_573616, EPI_ISL_573617, EPI_ISL_573618, EPI_ISL_573619, EPI_ISL_573620, EPI_ISL_573621, EPI_ISL_573622, EPI_ISL_573623, EPI_ISL_573624, EPI_ISL_573625, EPI_ISL_573626, EPI_ISL_573627, EPI_ISL_573628, EPI_ISL_573629, EPI_ISL_573630, EPI_ISL_573631, EPI_ISL_573632, EPI_ISL_573633, EPI_ISL_573634, EPI_ISL_573635, EPI_ISL_573636, EPI_ISL_573637, EPI_ISL_573638, EPI_ISL_573639, EPI_ISL_573640, EPI_ISL_573641, EPI_ISL_573642, EPI_ISL_573643, EPI_ISL_573644, EPI_ISL_573645, EPI_ISL_573646, EPI_ISL_573647, EPI_ISL_573648, EPI_ISL_573649, EPI_ISL_573650, EPI_ISL_573651 | | | |
| see above | Oxford Viromics, NDM, University of Oxford; Oxford University Hospitals; Basingstoke and North Hampshire Hospital | COVID-19 Genomics UK (COG-UK) Consortium | Tanya Golubchik, David Bonsall, George Macintyre, Amy Trebes, Mariateresa de Cesare, Catrin Moore, Alex Mobbs, Anita Justice, Robert Shaw, Monique Andersson, Timothy Peto, Emma Wise, Nathan Moore, Jessica Lynch, Nick Cortes, Matilde Mori, Stephen Kidd, David Buck, John Todd, Christophe Fraser |
| EPI_ISL_573652, EPI_ISL_573653, EPI_ISL_573654, EPI_ISL_573655, EPI_ISL_573656, EPI_ISL_573657, EPI_ISL_573658, EPI_ISL_573659, EPI_ISL_573660, EPI_ISL_573661, EPI_ISL_573662, EPI_ISL_573663, EPI_ISL_573664, EPI_ISL_573665, EPI_ISL_573666, EPI_ISL_573667, EPI_ISL_573668, EPI_ISL_573669, EPI_ISL_573670, EPI_ISL_573671, EPI_ISL_573672, EPI_ISL_573673, EPI_ISL_573674, EPI_ISL_573675, EPI_ISL_573676, EPI_ISL_573677, EPI_ISL_573678, EPI_ISL_573679, EPI_ISL_573680, EPI_ISL_573681, EPI_ISL_573682, EPI_ISL_573684, EPI_ISL_573685 | | | |
| see above | University College London, Great Ormond Street Hospital for Children NHS Foundation Trust, Imperial College Healthcare NHS Trust | COVID-19 Genomics UK (COG-UK) Consortium | Sergi Castellano, Rachel Williams, Mark Kristiansen, Paola Resende Silva, Sunando Roy, Tony Brooks, Helena Tutill, Paola Niola, Patricia Dyal, Charlotte Williams, Leysa Forrest, Yasmin Panchbhaya, Jacqueline Findlay, Samuel Weeks, Julianne Brown, Kathryn Harris, Paul Randell, James Price, Alison Holmes, Judith Breuer |
| EPI_ISL_573686, EPI_ISL_573687, EPI_ISL_573688, EPI_ISL_573689, EPI_ISL_573690, EPI_ISL_573691, EPI_ISL_573692, EPI_ISL_573693, EPI_ISL_573694, EPI_ISL_573695, EPI_ISL_573696, EPI_ISL_573697, EPI_ISL_573698, EPI_ISL_573699, EPI_ISL_573700, EPI_ISL_573701, EPI_ISL_573702, EPI_ISL_573703, EPI_ISL_573704, EPI_ISL_573705, EPI_ISL_573706, EPI_ISL_573707, EPI_ISL_573708, EPI_ISL_573709, EPI_ISL_573710, EPI_ISL_573711, EPI_ISL_573712, EPI_ISL_573713, EPI_ISL_573714, EPI_ISL_573715, EPI_ISL_573716, EPI_ISL_573717, EPI_ISL_573718, EPI_ISL_573719, EPI_ISL_573720, EPI_ISL_573721, EPI_ISL_573722, EPI_ISL_573723, EPI_ISL_573724, EPI_ISL_573725, EPI_ISL_573726, EPI_ISL_573727, EPI_ISL_573728, EPI_ISL_573729, EPI_ISL_573730, EPI_ISL_573731, EPI_ISL_573732, EPI_ISL_573733, EPI_ISL_573734, EPI_ISL_573735, EPI_ISL_573736, EPI_ISL_573737, EPI_ISL_573738, EPI_ISL_573739, EPI_ISL_573740, EPI_ISL_573741, EPI_ISL_573742, EPI_ISL_573743, EPI_ISL_573744, EPI_ISL_573745, EPI_ISL_573746, EPI_ISL_573747, EPI_ISL_573748, EPI_ISL_573749, EPI_ISL_573750, EPI_ISL_573751, EPI_ISL_573752, EPI_ISL_573753, EPI_ISL_573754, EPI_ISL_573755, EPI_ISL_573756, EPI_ISL_573757, EPI_ISL_573758 | | | |
| see above | Virology Department, Sheffield Teaching Hospitals NHS Foundation Trust/Department of Infection, Immunity and Cardiovascular Disease, The Medical School, University of Sheffield | COVID-19 Genomics UK (COG-UK) Consortium | Thushan de Silva, Matthew Parker, Nikki Smith, Adri Angyal, Rebecca Brown, Luke Green, Rachel Tucker, Paul Parsons, Danielle Groves, Katie Johnson, Laura Carrilero, Alex Keeley, Dave Partridge, Matthew Wyles, Benjamin Lindsey, Mehmet Yavuz, Mohammad Raza, Cariad Evans |
| EPI_ISL_573759 | Oxford Viromics, NDM, University of Oxford; Oxford University Hospitals; Basingstoke and North Hampshire Hospital | COVID-19 Genomics UK (COG-UK) Consortium | Tanya Golubchik, David Bonsall, George Macintyre, Amy Trebes, Mariateresa de Cesare, Catrin Moore, Alex Mobbs, Anita Justice, Robert Shaw, Monique Andersson, Timothy Peto, Emma Wise, Nathan Moore, Jessica Lynch, Nick Cortes, Matilde Mori, Stephen Kidd, David Buck, John Todd, Christophe Fraser |
| EPI_ISL_573760, EPI_ISL_573761 | Northumbria University / South Tees Hospitals NHS Foundation Trust / North Cumbria Integrated Care NHS Foundation Trust / North Tees and Hartlepool NHS Foundation Trust / Newcastle Hospitals NHS Foundation Trust | COVID-19 Genomics UK (COG-UK) Consortium | Darren L Smith,Andrew Nelson,Matthew Bashton,Greg R Young,Joshua Loh,John Allan,Mohammad A Tariq,Giles S Holt,Gary Black,Wen C Yew,Lynn Dover,Paul Baker,Steve Liggett,Sarah Essex,Jane Greenaway,Debra Padgett,Clive Graham,Garren Scott,Edward Barton,Emma Swindells,Brendan Payne,Jennifer Collins,Yusri Taha,Gary Eltringham |
| EPI_ISL_573762, EPI_ISL_573763 | Virology Department, Sheffield Teaching Hospitals NHS Foundation Trust/Department of Infection, Immunity and Cardiovascular Disease, The Medical School, University of Sheffield | COVID-19 Genomics UK (COG-UK) Consortium | Thushan de Silva, Matthew Parker, Nikki Smith, Adri Angyal, Rebecca Brown, Luke Green, Rachel Tucker, Paul Parsons, Danielle Groves, Katie Johnson, Laura Carrilero, Alex Keeley, Dave Partridge, Matthew Wyles, Benjamin Lindsey, Mehmet Yavuz, Mohammad Raza, Cariad Evans |
| EPI_ISL_573764 | University College London, Great Ormond Street Hospital for Children NHS Foundation Trust, Imperial College Healthcare NHS Trust | COVID-19 Genomics UK (COG-UK) Consortium | Sergi Castellano, Rachel Williams, Mark Kristiansen, Paola Resende Silva, Sunando Roy, Tony Brooks, Helena Tutill, Paola Niola, Patricia Dyal, Charlotte Williams, Leysa Forrest, Yasmin Panchbhaya, Jacqueline Findlay, Samuel Weeks, Julianne Brown, Kathryn Harris, Paul Randell, James Price, Alison Holmes, Judith Breuer |
| EPI_ISL_573765, EPI_ISL_573766, EPI_ISL_573767, EPI_ISL_573768 | West of Scotland Specialist Virology Centre, NHSGGC / MRC-University of Glasgow Centre for Virus Research | COVID-19 Genomics UK (COG-UK) Consortium | Ana da Silva Filipe, Natasha Johnson, Kathy Smollett, Daniel Mair, Stephen Carmichael, Lily Tong, Jenna Nichols, Elihu Aranday-Cortes, Kyriaki Nomikou; Sarah McDonald, Marc Niebel, Patawee Asamaphan; Richard Orton, Joseph Hughes, Sreenu Vattipally, David L Robertson; Alasdair MacLean, Rory Gunson; Kathy Li, Igor Starinskij, Natasha Jesudason, Rajiv Shah, James Shepherd, Antonia Ho, Emma Thomson |
| EPI_ISL_573769 | Lighthouse Lab in Glasgow / MRC-University of Glasgow Centre for Virus Research | COVID-19 Genomics UK (COG-UK) Consortium | Ana da Silva Filipe, Natasha Johnson, Kathy Smollett, Daniel Mair, Stephen Carmichael, Lily Tong, Jenna Nichols, Elihu Aranday-Cortes, Kyriaki Nomikou; Sarah McDonald, Marc Niebel, Patawee Asamaphan; Harper VanSteenhouse, Yumi Kasai, David Gray, Carol Clugston, Anna Dominiczak; Alasdair MacLean, Rory Gunson; Richard Orton, Joseph Hughes, Sreenu Vattipally, David L Robertson; Sharif Shaaban, Matthew Holden; Kathy Li, Natasha Jesudason, Rajiv Shah, James Shepherd, Antonia Ho, Emma Thomson |
| EPI_ISL_573770, EPI_ISL_573771, EPI_ISL_573772, EPI_ISL_573773, EPI_ISL_573774, EPI_ISL_573775, EPI_ISL_573776, EPI_ISL_573777, EPI_ISL_573778, EPI_ISL_573779, EPI_ISL_573780, EPI_ISL_573781, EPI_ISL_573782, EPI_ISL_573783, EPI_ISL_573784, EPI_ISL_573785, EPI_ISL_573786, EPI_ISL_573787, EPI_ISL_573788, EPI_ISL_573789, EPI_ISL_573790, EPI_ISL_573791, EPI_ISL_573792, EPI_ISL_573793 | | | |
| see above | West of Scotland Specialist Virology Centre, NHSGGC / MRC-University of Glasgow Centre for Virus Research | COVID-19 Genomics UK (COG-UK) Consortium | Ana da Silva Filipe, Natasha Johnson, Kathy Smollett, Daniel Mair, Stephen Carmichael, Lily Tong, Jenna Nichols, Elihu Aranday-Cortes, Kyriaki Nomikou; Sarah McDonald, Marc Niebel, Patawee Asamaphan; Richard Orton, Joseph Hughes, Sreenu Vattipally, David L Robertson; Alasdair MacLean, Rory Gunson; Kathy Li, Igor Starinskij, Natasha Jesudason, Rajiv Shah, James Shepherd, Antonia Ho, Emma Thomson |
| EPI_ISL_573794, EPI_ISL_573795, EPI_ISL_573796, EPI_ISL_573797, EPI_ISL_573798 | Virology Department, Royal Infirmary of Edinburgh, NHS Lothian / School of Biological Sciences, University of Edinburgh / Institute of Genetics and Molecular Medicine, University of Edinburgh | COVID-19 Genomics UK (COG-UK) Consortium | McHugh M, Dewar R, Rooke S, Gallagher M, Balcaza C, O'Toole Á, Scher E, Hill V, McCrone JT, Colquhoun R, Yu X, Jackson B, Rambaut A, Williams TC, Templeton K |
| EPI_ISL_573799 | Northumbria University / South Tees Hospitals NHS Foundation Trust / North Cumbria Integrated Care NHS Foundation Trust / North Tees and Hartlepool NHS Foundation Trust / Newcastle Hospitals NHS Foundation Trust | COVID-19 Genomics UK (COG-UK) Consortium | Darren L Smith,Andrew Nelson,Matthew Bashton,Greg R Young,Joshua Loh,John Allan,Mohammad A Tariq,Giles S Holt,Gary Black,Wen C Yew,Lynn Dover,Paul Baker,Steve Liggett,Sarah Essex,Jane Greenaway,Debra Padgett,Clive Graham,Garren Scott,Edward Barton,Emma Swindells,Brendan Payne,Jennifer Collins,Yusri Taha,Gary Eltringham |
| EPI_ISL_573800, EPI_ISL_573801, EPI_ISL_573802, EPI_ISL_573803, EPI_ISL_573804, EPI_ISL_573805 | University College London, Great Ormond Street Hospital for Children NHS Foundation Trust, Imperial College Healthcare NHS Trust | COVID-19 Genomics UK (COG-UK) Consortium | Sergi Castellano, Rachel Williams, Mark Kristiansen, Paola Resende Silva, Sunando Roy, Tony Brooks, Helena Tutill, Paola Niola, Patricia Dyal, Charlotte Williams, Leysa Forrest, Yasmin Panchbhaya, Jacqueline Findlay, Samuel Weeks, Julianne Brown, Kathryn Harris, Paul Randell, James Price, Alison Holmes, Judith Breuer |
| EPI_ISL_573806, EPI_ISL_573807, EPI_ISL_573808, EPI_ISL_573809, EPI_ISL_573810 | Virology Department, Sheffield Teaching Hospitals NHS Foundation Trust/Department of Infection, Immunity and Cardiovascular Disease, The Medical School, University of Sheffield | COVID-19 Genomics UK (COG-UK) Consortium | Thushan de Silva, Matthew Parker, Nikki Smith, Adri Angyal, Rebecca Brown, Luke Green, Rachel Tucker, Paul Parsons, Danielle Groves, Katie Johnson, Laura Carrilero, Alex Keeley, Dave Partridge, Matthew Wyles, Benjamin Lindsey, Mehmet Yavuz, Mohammad Raza, Cariad Evans |
| EPI_ISL_573811, EPI_ISL_573812, EPI_ISL_573813, EPI_ISL_573814, EPI_ISL_573815, EPI_ISL_573816, EPI_ISL_573817, EPI_ISL_573818, EPI_ISL_573819, EPI_ISL_573820, EPI_ISL_573821, EPI_ISL_573822, EPI_ISL_573823, EPI_ISL_573824, EPI_ISL_573825, EPI_ISL_573826, EPI_ISL_573827, EPI_ISL_573828, EPI_ISL_573829, EPI_ISL_573830, EPI_ISL_573831, EPI_ISL_573832, EPI_ISL_573833, EPI_ISL_573834, EPI_ISL_573835, EPI_ISL_573836, EPI_ISL_573837, EPI_ISL_573838, EPI_ISL_573839, EPI_ISL_573840, EPI_ISL_573841, EPI_ISL_573842, EPI_ISL_573843, EPI_ISL_573844, EPI_ISL_573845, EPI_ISL_573846, EPI_ISL_573847, EPI_ISL_573848, EPI_ISL_573849, EPI_ISL_573850, EPI_ISL_573851, EPI_ISL_573852, EPI_ISL_573853, EPI_ISL_573854, EPI_ISL_573855 | | | |
| see above | Oxford Viromics, NDM, University of Oxford; Oxford University Hospitals; Basingstoke and North Hampshire Hospital | COVID-19 Genomics UK (COG-UK) Consortium | Tanya Golubchik, David Bonsall, George Macintyre, Amy Trebes, Mariateresa de Cesare, Catrin Moore, Alex Mobbs, Anita Justice, Robert Shaw, Monique Andersson, Timothy Peto, Emma Wise, Nathan Moore, Jessica Lynch, Nick Cortes, Matilde Mori, Stephen Kidd, David Buck, John Todd, Christophe Fraser |
| EPI_ISL_573856, EPI_ISL_573857, EPI_ISL_573858, EPI_ISL_573859, EPI_ISL_573860, EPI_ISL_573861, EPI_ISL_573862, EPI_ISL_573863, EPI_ISL_573864, EPI_ISL_573865 | University College London, Great Ormond Street Hospital for Children NHS Foundation Trust, Imperial College Healthcare NHS Trust | COVID-19 Genomics UK (COG-UK) Consortium | Sergi Castellano, Rachel Williams, Mark Kristiansen, Paola Resende Silva, Sunando Roy, Tony Brooks, Helena Tutill, Paola Niola, Patricia Dyal, Charlotte Williams, Leysa Forrest, Yasmin Panchbhaya, Jacqueline Findlay, Samuel Weeks, Julianne Brown, Kathryn Harris, Paul Randell, James Price, Alison Holmes, Judith Breuer |
| EPI_ISL_573866, EPI_ISL_573867, EPI_ISL_573868, EPI_ISL_573869 | Oxford Viromics, NDM, University of Oxford; Oxford University Hospitals; Basingstoke and North Hampshire Hospital | COVID-19 Genomics UK (COG-UK) Consortium | Tanya Golubchik, David Bonsall, George Macintyre, Amy Trebes, Mariateresa de Cesare, Catrin Moore, Alex Mobbs, Anita Justice, Robert Shaw, Monique Andersson, Timothy Peto, Emma Wise, Nathan Moore, Jessica Lynch, Nick Cortes, Matilde Mori, Stephen Kidd, David Buck, John Todd, Christophe Fraser |
| EPI_ISL_573870 | Centre for Enzyme Innovation, University of Portsmouth / Translational Research Laboratory, Portsmouth Hospitals NHS Trust | COVID-19 Genomics UK (COG-UK) Consortium | Angela Beckett,Yann Bourgeois,Garry Scarlett,Sharon Glaysher,Scott Elliott,Kelly Bicknell,Robert Impey,Allyson Lloyd,Sarah Wyllie,Ethan Butcher,Anoop Chauhan,Samuel Robson |
| EPI_ISL_573871, EPI_ISL_573872 | Northumbria University / South Tees Hospitals NHS Foundation Trust / North Cumbria Integrated Care NHS Foundation Trust / North Tees and Hartlepool NHS Foundation Trust / Newcastle Hospitals NHS Foundation Trust | COVID-19 Genomics UK (COG-UK) Consortium | Darren L Smith,Andrew Nelson,Matthew Bashton,Greg R Young,Joshua Loh,John Allan,Mohammad A Tariq,Giles S Holt,Gary Black,Wen C Yew,Lynn Dover,Paul Baker,Steve Liggett,Sarah Essex,Jane Greenaway,Debra Padgett,Clive Graham,Garren Scott,Edward Barton,Emma Swindells,Brendan Payne,Jennifer Collins,Yusri Taha,Gary Eltringham |
| EPI_ISL_573873, EPI_ISL_573874, EPI_ISL_573875, EPI_ISL_573876, EPI_ISL_573877, EPI_ISL_573878, EPI_ISL_573879, EPI_ISL_573880, EPI_ISL_573881, EPI_ISL_573882, EPI_ISL_573883, EPI_ISL_573884, EPI_ISL_573885, EPI_ISL_573886, EPI_ISL_573887, EPI_ISL_573888, EPI_ISL_573889, EPI_ISL_573890, EPI_ISL_573891, EPI_ISL_573892, EPI_ISL_573893, EPI_ISL_573894, EPI_ISL_573895, EPI_ISL_573896, EPI_ISL_573897, EPI_ISL_573898, EPI_ISL_573899, EPI_ISL_573900, EPI_ISL_573901, EPI_ISL_573902, EPI_ISL_573903, EPI_ISL_573904, EPI_ISL_573905, EPI_ISL_573906, EPI_ISL_573907, EPI_ISL_573908, EPI_ISL_573909, EPI_ISL_573910, EPI_ISL_573911, EPI_ISL_573912, EPI_ISL_573913, EPI_ISL_573914, EPI_ISL_573915, EPI_ISL_573916, EPI_ISL_573917, EPI_ISL_573918, EPI_ISL_573919, EPI_ISL_573920, EPI_ISL_573921, EPI_ISL_573922, EPI_ISL_573923, EPI_ISL_573924, EPI_ISL_573925, EPI_ISL_573926, EPI_ISL_573927, EPI_ISL_573928, EPI_ISL_573929, EPI_ISL_573930, EPI_ISL_573931, EPI_ISL_573932, EPI_ISL_573933, EPI_ISL_573934, EPI_ISL_573935, EPI_ISL_573936, EPI_ISL_573937, EPI_ISL_573938, EPI_ISL_573939, EPI_ISL_573940, EPI_ISL_573941, EPI_ISL_573942, EPI_ISL_573943, EPI_ISL_573944, EPI_ISL_573945, EPI_ISL_573946, EPI_ISL_573947, EPI_ISL_573948, EPI_ISL_573949, EPI_ISL_573950, EPI_ISL_573951, EPI_ISL_573952, EPI_ISL_573953, EPI_ISL_573954, EPI_ISL_573955, EPI_ISL_573956, EPI_ISL_573957, EPI_ISL_573958, EPI_ISL_573959, EPI_ISL_573960, EPI_ISL_573961, EPI_ISL_573962, EPI_ISL_573963, EPI_ISL_573964, EPI_ISL_573965, EPI_ISL_573966, EPI_ISL_573967, EPI_ISL_573968, EPI_ISL_573969, EPI_ISL_573970, EPI_ISL_573971, EPI_ISL_573972, EPI_ISL_573973, EPI_ISL_573974, EPI_ISL_573975, EPI_ISL_573976, EPI_ISL_573977, EPI_ISL_573978, EPI_ISL_573979, EPI_ISL_573980, EPI_ISL_573981, EPI_ISL_573982, EPI_ISL_573983, EPI_ISL_573984, EPI_ISL_573985, EPI_ISL_573986, EPI_ISL_573987, EPI_ISL_573988, EPI_ISL_573989, EPI_ISL_573990, EPI_ISL_573991, EPI_ISL_573992, EPI_ISL_573993, EPI_ISL_573994, EPI_ISL_573995, EPI_ISL_573996, EPI_ISL_573997, EPI_ISL_573998, EPI_ISL_573999, EPI_ISL_574000, EPI_ISL_574001, EPI_ISL_574002, EPI_ISL_574003, EPI_ISL_574004, EPI_ISL_574005, EPI_ISL_574006, EPI_ISL_574007, EPI_ISL_574008, EPI_ISL_574009, EPI_ISL_574010, EPI_ISL_574011, EPI_ISL_574012, EPI_ISL_574013, EPI_ISL_574014, EPI_ISL_574015, EPI_ISL_574016, EPI_ISL_574017, EPI_ISL_574018, EPI_ISL_574019, EPI_ISL_574020, EPI_ISL_574021, EPI_ISL_574022, EPI_ISL_574023, EPI_ISL_574024, EPI_ISL_574025, EPI_ISL_574026, EPI_ISL_574027, EPI_ISL_574028, EPI_ISL_574029, EPI_ISL_574030, EPI_ISL_574031, EPI_ISL_574032, EPI_ISL_574033, EPI_ISL_574034, EPI_ISL_574035, EPI_ISL_574036, EPI_ISL_574037, EPI_ISL_574038, EPI_ISL_574039, EPI_ISL_574040, EPI_ISL_574041, EPI_ISL_574042, EPI_ISL_574043, EPI_ISL_574044, EPI_ISL_574045, EPI_ISL_574046, EPI_ISL_574047, EPI_ISL_574048, EPI_ISL_574049, EPI_ISL_574050, EPI_ISL_574051, EPI_ISL_574052, EPI_ISL_574053, EPI_ISL_574054, EPI_ISL_574055, EPI_ISL_574056, EPI_ISL_574057, EPI_ISL_574058, EPI_ISL_574059, EPI_ISL_574060, EPI_ISL_574061, EPI_ISL_574062, EPI_ISL_574063, EPI_ISL_574064, EPI_ISL_574065, EPI_ISL_574066, EPI_ISL_574067, EPI_ISL_574068, EPI_ISL_574069, EPI_ISL_574070, EPI_ISL_574071, EPI_ISL_574072, EPI_ISL_574073, EPI_ISL_574074, EPI_ISL_574075, EPI_ISL_574076, EPI_ISL_574077, EPI_ISL_574078, EPI_ISL_574079, EPI_ISL_574080, EPI_ISL_574081, EPI_ISL_574082, EPI_ISL_574083, EPI_ISL_574084, EPI_ISL_574085, EPI_ISL_574086, EPI_ISL_574087, EPI_ISL_574088, EPI_ISL_574089, EPI_ISL_574090, EPI_ISL_574091, EPI_ISL_574092, EPI_ISL_574093, EPI_ISL_574094, EPI_ISL_574095, EPI_ISL_574096, EPI_ISL_574097, EPI_ISL_574098, EPI_ISL_574099, EPI_ISL_574100, EPI_ISL_574101, EPI_ISL_574102, EPI_ISL_574103, EPI_ISL_574104, EPI_ISL_574105, EPI_ISL_574106, EPI_ISL_574107, EPI_ISL_574108, EPI_ISL_574109, EPI_ISL_574110, EPI_ISL_574111, EPI_ISL_574112, EPI_ISL_574113, EPI_ISL_574114, EPI_ISL_574115, EPI_ISL_574116, EPI_ISL_574117, EPI_ISL_574118, EPI_ISL_574119, EPI_ISL_574120, EPI_ISL_574121, EPI_ISL_574122, EPI_ISL_574123, EPI_ISL_574124, EPI_ISL_574125, EPI_ISL_574126, EPI_ISL_574127, EPI_ISL_574128, EPI_ISL_574129, EPI_ISL_574130, EPI_ISL_574131, EPI_ISL_574132, EPI_ISL_574133, EPI_ISL_574134, EPI_ISL_574135, EPI_ISL_574136, | | | |

| | | | |
|---|---|---|---|
| EPI_ISL_574137, EPI_ISL_574138, EPI_ISL_574139, EPI_ISL_574140, EPI_ISL_574141, EPI_ISL_574142, EPI_ISL_574143, EPI_ISL_574144, EPI_ISL_574145, EPI_ISL_574146, EPI_ISL_574147, EPI_ISL_574148, EPI_ISL_574149, EPI_ISL_574150, EPI_ISL_574151, EPI_ISL_574152, EPI_ISL_574153, EPI_ISL_574154, EPI_ISL_574155, EPI_ISL_574156, EPI_ISL_574157, EPI_ISL_574158, EPI_ISL_574159, EPI_ISL_574160, EPI_ISL_574161, EPI_ISL_574162, EPI_ISL_574163, EPI_ISL_574164, EPI_ISL_574165, EPI_ISL_574166, EPI_ISL_574167, EPI_ISL_574168, EPI_ISL_574169, EPI_ISL_574170, EPI_ISL_574171, EPI_ISL_574172, EPI_ISL_574173, EPI_ISL_574174, EPI_ISL_574175, EPI_ISL_574176, EPI_ISL_574177, EPI_ISL_574178, EPI_ISL_574179, EPI_ISL_574180, EPI_ISL_574181, EPI_ISL_574182, EPI_ISL_574183, EPI_ISL_574184, EPI_ISL_574185, EPI_ISL_574186, EPI_ISL_574187, EPI_ISL_574188, EPI_ISL_574189, EPI_ISL_574190, EPI_ISL_574191, EPI_ISL_574192, EPI_ISL_574193, EPI_ISL_574194, EPI_ISL_574195, EPI_ISL_574196, EPI_ISL_574197, EPI_ISL_574198, EPI_ISL_574199, EPI_ISL_574200, EPI_ISL_574201, EPI_ISL_574202, EPI_ISL_574203, EPI_ISL_574204, EPI_ISL_574205, EPI_ISL_574206, EPI_ISL_574207, EPI_ISL_574208, EPI_ISL_574209, EPI_ISL_574210, EPI_ISL_574211, EPI_ISL_574212, EPI_ISL_574213, EPI_ISL_574214, EPI_ISL_574215, EPI_ISL_574216, EPI_ISL_574217, EPI_ISL_574218, EPI_ISL_574219, EPI_ISL_574220, EPI_ISL_574221, EPI_ISL_574222, EPI_ISL_574223, EPI_ISL_574224, EPI_ISL_574225, EPI_ISL_574226, EPI_ISL_574227, EPI_ISL_574228, EPI_ISL_574229, EPI_ISL_574230, EPI_ISL_574231, EPI_ISL_574232, EPI_ISL_574233, EPI_ISL_574234, EPI_ISL_574235, EPI_ISL_574236, EPI_ISL_574237, EPI_ISL_574238, EPI_ISL_574239, EPI_ISL_574240, EPI_ISL_574241, EPI_ISL_574242, EPI_ISL_574243, EPI_ISL_574244, EPI_ISL_574245, EPI_ISL_574246, EPI_ISL_574247, EPI_ISL_574248, EPI_ISL_574249, EPI_ISL_574250, EPI_ISL_574251, EPI_ISL_574252, EPI_ISL_574253, EPI_ISL_574254, EPI_ISL_574255, EPI_ISL_574256 | | | |
| see above | Wales Specialist Virology Centre Sequencing lab: Pathogen Genomics Unit | COVID-19 Genomics UK (COG-UK) Consortium | Catherine Moore, Johnathan Evans, Laura Gifford, Malorie Perry, Simon Cottrell, Angela Marchbank, Alec Birchley, Alexander Adams, Amy Gaskin, Bree Gatica-Wilcox, Jason Coombes, Joel Southgate, Lauren Gilbert, Lee Graham, Nicole Pacchiarini, Sara Kumziene-Summerhayes, Sarah Taylor, Sophie Jones, Sara Rey, Matthew Bull, Joanne Watkins, Sally Corden, Tom Connor |
| EPI_ISL_574257 | Oxford Viromics, NDM, University of Oxford; Oxford University Hospitals; Basingstoke and North Hampshire Hospital | COVID-19 Genomics UK (COG-UK) Consortium | Tanya Golubchik, David Bonsall, George Macintyre, Amy Trebes, Mariateresa de Cesare, Catrin Moore, Alex Mobbs, Anita Justice, Robert Shaw, Monique Andersson, Timothy Peto, Emma Wise, Nathan Moore, Jessica Lynch, Nick Cortes, Matilde Mori, Stephen Kidd, David Buck, John Todd, Christophe Fraser |
| EPI_ISL_574258 | University College London, Great Ormond Street Hospital for Children NHS Foundation Trust, Imperial College Healthcare NHS Trust | COVID-19 Genomics UK (COG-UK) Consortium | Sergi Castellano, Rachel Williams, Mark Kristiansen, Paola Resende Silva, Sunando Roy, Tony Brooks, Helena Tutill, Paola Niola, Patricia Dyal, Charlotte Williams, Leysa Forrest, Yasmin Panchbhaya, Jacqueline Findlay, Samuel Weeks, Julianne Brown, Kathryn Harris, Paul Randell, James Price, Alison Holmes, Judith Breuer |
| EPI_ISL_574259 | Institute for Virology, University Hospital Duesseldorf, Medical Faculty, Heinrich-Heine-University Duesseldorf | Institute for Virology, University Hospital Duesseldorf, Medical Faculty, Heinrich-Heine-University Duesseldorf | Maximilian Damagnez, Verena Keitel, Björn Jensen, Nadine Lübke, Lisa Müller, Philipp Ostermann, Tina Senff, Ortwin Adams, Philipp Albrecht, Gerald Antoch, Johannes Bode, Edwin Bölke, Saskia Elben, Torsten Feldt, Johannes C. Fischer, , Anselm Kunstein, Caroline Klindt, Alexander Killer, Tom Lüdde, Annemarie Mohring, Jennifer Neubert, Heiner Schaal, Ansgar Schulz, Jörg Timm, Andreas Walker |
| EPI_ISL_574260, EPI_ISL_574261, EPI_ISL_574262, EPI_ISL_574263, EPI_ISL_574264, EPI_ISL_574265, EPI_ISL_574266, EPI_ISL_574267, EPI_ISL_574268, EPI_ISL_574269, EPI_ISL_574270, EPI_ISL_574271, EPI_ISL_574272, EPI_ISL_574273, EPI_ISL_574274, EPI_ISL_574275, EPI_ISL_574276, EPI_ISL_574277, EPI_ISL_574278, EPI_ISL_574279, EPI_ISL_574280, EPI_ISL_574281, EPI_ISL_574282, EPI_ISL_574283, EPI_ISL_574284, EPI_ISL_574285, EPI_ISL_574286, EPI_ISL_574287, EPI_ISL_574291 | | | |
| see above | New Mexico Department of Health Scientific Laboratory | New Mexico Department of Health Scientific Laboratory | Ellie Johnson, Anastacia Griego-Fisher, D'Eldra Malone |
| EPI_ISL_574295, EPI_ISL_574296, EPI_ISL_574297, EPI_ISL_574298, EPI_ISL_574299, EPI_ISL_574300 | LSUHS Emerging Viral Threat Laboratory | Microbial Genome Sequencing Center | Jeremy P. Kamil, Rona S. Scott, Maarten Van Diest, Malgorzata Bienkowska-Haba, Katarzyna Zwolinska, Andrew D. Yurochko, Christopher G. Kevil, Martin J. Sapp, Daniel J. Snyder, Vaughn S. Cooper, John A. Vanchiere |
| EPI_ISL_574301, EPI_ISL_574302, EPI_ISL_574303, EPI_ISL_574304, EPI_ISL_574305, EPI_ISL_574306, EPI_ISL_574307, EPI_ISL_574308, EPI_ISL_574309, EPI_ISL_574310, EPI_ISL_574311, EPI_ISL_574312, EPI_ISL_574313, EPI_ISL_574314, EPI_ISL_574315, EPI_ISL_574316, EPI_ISL_574317, EPI_ISL_574318, EPI_ISL_574319 | | | |
| see above | LSUHS Emerging Viral Threat Laboratory | Microbial Genome Sequencing Center | Rona S. Scott, Jeremy P. Kamil, Maarten Van Diest, Malgorzata Bienkowska-Haba, Katarzyna Zwolinska, Andrew D. Yurochko, Christopher G. Kevil, Martin J. Sapp, Daniel J. Snyder, Vaughn S. Cooper, John A. Vanchiere |
| EPI_ISL_574320, EPI_ISL_574321, EPI_ISL_574322, EPI_ISL_574323, EPI_ISL_574324, EPI_ISL_574325, EPI_ISL_574326, EPI_ISL_574327, EPI_ISL_574328, EPI_ISL_574329, EPI_ISL_574330, EPI_ISL_574331, EPI_ISL_574332, EPI_ISL_574333, EPI_ISL_574334, EPI_ISL_574335, EPI_ISL_574336, EPI_ISL_574337, EPI_ISL_574338, EPI_ISL_574339, EPI_ISL_574340, EPI_ISL_574341, EPI_ISL_574342, EPI_ISL_574343, EPI_ISL_574344, EPI_ISL_574345, EPI_ISL_574346, EPI_ISL_574347, EPI_ISL_574348, EPI_ISL_574349, EPI_ISL_574350, EPI_ISL_574351, EPI_ISL_574352, EPI_ISL_574353, EPI_ISL_574354, EPI_ISL_574355, EPI_ISL_574356, EPI_ISL_574357, EPI_ISL_574358, EPI_ISL_574359, EPI_ISL_574360, EPI_ISL_574361, EPI_ISL_574362, EPI_ISL_574363, EPI_ISL_574364, EPI_ISL_574365, EPI_ISL_574366, EPI_ISL_574367, EPI_ISL_574368, EPI_ISL_574369, EPI_ISL_574370, EPI_ISL_574371, EPI_ISL_574372, EPI_ISL_574373, EPI_ISL_574374, EPI_ISL_574375, EPI_ISL_574376, EPI_ISL_574377, EPI_ISL_574378, EPI_ISL_574379, EPI_ISL_574380, EPI_ISL_574381, EPI_ISL_574382, EPI_ISL_574383, EPI_ISL_574384, EPI_ISL_574385, EPI_ISL_574386, EPI_ISL_574387, EPI_ISL_574388, EPI_ISL_574389, EPI_ISL_574390, EPI_ISL_574391, EPI_ISL_574392, EPI_ISL_574393, EPI_ISL_574394, EPI_ISL_574395, EPI_ISL_574396, EPI_ISL_574397, EPI_ISL_574398, EPI_ISL_574399, EPI_ISL_574400, EPI_ISL_574401, EPI_ISL_574402, EPI_ISL_574403, EPI_ISL_574404, EPI_ISL_574405, EPI_ISL_574406, EPI_ISL_574407, EPI_ISL_574408, EPI_ISL_574409, EPI_ISL_574410, EPI_ISL_574411, EPI_ISL_574412, EPI_ISL_574413, EPI_ISL_574414, EPI_ISL_574415, EPI_ISL_574416, EPI_ISL_574417, EPI_ISL_574418, EPI_ISL_574419, EPI_ISL_574420, EPI_ISL_574421, EPI_ISL_574422, EPI_ISL_574423, EPI_ISL_574424, EPI_ISL_574425, EPI_ISL_574426, EPI_ISL_574427, EPI_ISL_574428, EPI_ISL_574429, EPI_ISL_574430 | | | |
| see above | LSUHS Emerging Viral Threat Laboratory | Microbial Genome Sequencing Center | Jeremy P. Kamil, Rona S. Scott, Maarten Van Diest, Malgorzata Bienkowska-Haba, Katarzyna Zwolinska, Andrew D. Yurochko, Christopher G. Kevil, Martin J. Sapp, Daniel J. Snyder, Vaughn S. Cooper, John A. Vanchiere |
| EPI_ISL_574543, EPI_ISL_574544, EPI_ISL_574545, EPI_ISL_574546, EPI_ISL_574547, EPI_ISL_574550, EPI_ISL_574551, EPI_ISL_574552, EPI_ISL_574553, EPI_ISL_574554, EPI_ISL_574555, EPI_ISL_574557, EPI_ISL_574558, EPI_ISL_574559, EPI_ISL_574560, EPI_ISL_574561, EPI_ISL_574562, EPI_ISL_574563, EPI_ISL_574564, EPI_ISL_574565, EPI_ISL_574566, EPI_ISL_574567, EPI_ISL_574568, EPI_ISL_574569, EPI_ISL_574570, EPI_ISL_574571, EPI_ISL_574573, EPI_ISL_574574, EPI_ISL_574575 | | | |
| see above | Microbiology Division, South Carolina Department of Health and Environmental Control | Microbiology Division, South Carolina Department of Health and Environmental Control | Flores,H. |
| EPI_ISL_574576 | Molecular Biology, New Mexico Department of Health Scientific Laboratory | Molecular Biology, New Mexico Department of Health Scientific Laboratory | Johnson,E.J., Griego-Fisher,A.M., Malone,D. |
